# Chemokine Receptor 1 and its associated immune pathway are downregulated in SF3B1^MT^ blood and non-blood cancers

**DOI:** 10.1101/2022.03.31.485699

**Authors:** Mahtab Dastpak, Hyunmin Kim, Christina L. Paraggio, Claudia E. Leonard, Shanye Yin, Binkai Chi, Kelsey W. Nassar, R. Grant Rowe, Zhonggang Hou, Jiuchun Zhang, Erin F. Cohn, Na Yang, William Marion, Yosra Zhang, George Q. Daley, Robin Reed

## Abstract

Mutation of the essential splicing factor SF3B1 is primarily associated with hematological cancers but also occurs in solid tumors. We edited the most common mutation, K700E, into human embryonic stem (ES) cells to determine the effects of this mutation alone in an undifferentiated/non-cancer background. Unexpectedly, >20% of the significantly upregulated genes in the SF3B1^K700E^ ES lines have immune functions. Thus, SF3B1 may have an additional role in proper expression of immune genes in appropriate cell types. In striking contrast, we found that published RNA-seq data from SF3B1 blood (MDS, CLL, AML) and non-blood (BRCA, UVM) cancers exhibited the opposite, downregulation of a multitude of immune pathways with 7 of the pathways shared among all 5 of the SF3B1 cancers. One of these pathways, “leukocyte migration”, is the 1^st^ reported pathway shared among all splicing factor cancers, including the 5 SF3B1 cancers and MDS associated with U2AF1, SRSF2 and ZRSR2. Importantly, we identified CCR1, which is in the leukocyte migration pathway as the only shared downregulated gene in the 5 SF3B1 cancers and in U2AF1^MT^ MDS. We conclude that downregulation of CCR1 and its associated immune pathway may play a key role in pathogenesis of these splicing factor cancers and are thus potential therapeutic targets.

## Introduction

SF3B1 is a component of the essential splicing factor U2 snRNP and is the most frequently mutated spliceosomal gene in cancer. For reasons that are not clear, SF3B1 mutations are primarily associated with hematological cancers. These mutations are most frequent in myelodysplastic syndromes (MDS) and are also found in acute myeloid leukemia (AML), chronic lymphocytic leukemia (CLL) and some solid tumors, such as breast cancer (BRCA) and uveal melanoma (UVM) (Harbour et al. 2013; Wan and Wu 2013; Anczuków and Krainer 2016; Wang et al. 2016; Jenkins and Kielkopf 2017; Pellagatti et al. 2018; Bamopoulos et al. 2020; Palomo and Solé 2020; Zhou et al. 2020; Liu et al. 2021; van der Werf et al. 2021; Niño et al. 2022). Three other splicing factors, U2AF1, SRSF2, and ZRSR2, are also frequently mutated in cancer. Thus, understanding the molecular mechanisms by which splicing factor mutations contribute to cancer is an area of intense investigation.

Previously, we cloned and characterized components of the U2 snRNP-associated complexes SF3a and SF3b (Bennett and Reed 1993; Champion-Arnaud and Reed 1994; Gozani et al. 1996, 1998; Wang et al. 1998; Das et al. 1999, 2000). Our studies, as well as recent cryo-EM and crystal work, revealed that the SF3B1 component of the SF3b complex binds to pre-mRNA on both sides of the branchpoint sequence (BPS) at the catalytic heart of the spliceosome (Gozani et al. 1996; 1998; Reed 2000; Will and Luhrmann 2011; Cretu et al. 2016; Finci et al. 2018). SF3B1 is recruited to the BPS region via direct interactions with U2AF1, which binds to the pyrimidine tract at the 3’ splice site (Gozani et al. 1998; Wang et al. 1998; Tholen et al. 2022). Additional SF3a/b components bind immediately upstream of the BPS, where they anchor U2 snRNP to the pre-mRNA during spliceosome assembly (Gozani et al. 1996, 1998; Wang et al. 1998; Will and Luhrmann 2011; Cretu et al. 2016; Finci et al. 2018). Consistent with these roles for the SF3a/b complexes, cancer-causing mutations in SF3B1 typically result in use of cryptic 3’ splice sites and are thought to use an incorrect BPS in these mis-splicing events (Darman et al. 2015; DeBoever et al. 2015; Alsafadi et al. 2016). In contrast, the most common effect of mutant U2AF1, which interacts directly with U2AF2 and binds to the AG at the 3’ splice site, is to alter 3’ splice site recognition. Mutant SRSF2, which binds to exonic splicing enhancers, alters exon inclusion (Kim et al. 2015; Ilagan et al. 2015; Zhang et al. 2015; Bonnal et al. 2020). ZRSR2 is mainly associated with the minor spliceosome, and its mutation primarily results in intron retention (Bonnal et al. 2020; Inoue et al. 2021). Notably, these 4 splicing factors have functions early in spliceosome assembly and specifically in the vicinity of the 3’ end of introns, suggesting a role for these parameters in contributing to splicing factor cancers. As the types of mis-splicing events differ among each splicing factor, the mis-splicing of specific genes is not shared among them. Instead, alterations in cellular pathways are common in different splicing factors cancers, and these altered pathways may contribute to pathogenesis (Pellagatti et al. 2018; Pollyea et al. 2019; Madan et al. 2020). For example, recent studies (Lee et al. 2018; Pollyea et al. 2019; Sallman and List 2019) revealed that mutant SF3B1 leads to mis-splicing of the NF-κB signaling inhibitor, MAP3K7, which was proposed to play a role in MDS pathogenesis. However, another study reported that MAP3K7 mis-splicing leads to downregulation of GATA1, a master regulator of erythroid differentiation, and that this in turn results in the severe anemia associated with MDS (Lieu et al. 2022). The discrepancy between these studies remains to be determined (Smith et al. 2019).

To date, studies of cancer-associated SF3B1 mutations have been carried out in blood cancer patient cells (MDS, AML, CLL), blood cells from mouse models, blood cancer cell lines (e.g. NALM and K562), as well as non-blood transformed cell lines (e.g. HeLa and HEK293 cells) (Darman et al. 2015; Alsafadi et al. 2016; Mupo et al. 2017; Hacken et al. 2018; Paolella et al. 2017; Saez et al. 2017; Lee et al. 2018; Yin et al. 2019; Alsafadi et al. 2021). In our study, we sought to determine the specific role of mutant SF3B1 itself and thus examined it in a non-blood/non-cancer cell type. Accordingly, we CRISPR-edited human ES cells to harbor a heterozygous K700E mutation and used RNA-seq to examine gene expression and splicing. Unexpectedly, we found that several master transcription regulators of hematopoiesis and a host of immune genes were upregulated in the SF3B1^K700E^ ES cells. These data suggest that wild type SF3B1 has a specific role in ensuring proper regulation of immune genes in the cell types that normally express them. In contrast, when mutant SF3B1 is present in blood/non-blood cancers, the opposite is observed: immune pathways constitute virtually all of the top downregulated pathways. Excitingly, we also identified a single gene, the chemokine receptor CCR1, that was downregulated in all SF3B1 blood/non-blood cancers as well as in U2AF1^MT^ MDS. Thus, loss of normal CCR1 levels and/or downregulation of specific immune pathways associated with CCR1 may be of central importance in splicing factor cancers.

## Results and Discussion

### Immune genes are upregulated in human SF3B1^K700E^ ES cells

We established 3 independent heterozygous SF3B1^K700E^ ES lines (Supplemental Fig. S1A). Sanger sequencing verified the presence of the mutation in these lines (Supplemental Fig. S1B). To determine how the K700E mutation affected gene expression and splicing in ES cells we carried out RNA-seq. These data confirmed that mutant and WT SF3B1 alleles were expressed at similar levels in the 3 SF3B1^K700E^ ES lines (Supplemental Fig. S1C). Consistent with previous studies, the predominant missplicing event was use of cryptic 3’ splice sites (Supplemental Fig. S2A,B; Supplemental Table S1). We also observed mis-splicing of many genes, including MAP3K7, DLST, DVL2, TMEM14C, DYNLL1, SNRPN, and BRD9, that were reported in studies of mutant SF3B1 cancers/cell lines (Supplemental Fig. S2C,D) (Darman et al. 2015; Obeng et al. 2016; Wang et al. 2016; Lee et al. 2018). Moreover, the mis-splicing events occurred at identical coordinates in SF3B1^K700E^ ES cells and SF3B1 cancers/transformed cell lines (Supplemental Fig. S2C,D; Supplemental Table S2). Representative mis-splicing events were validated by RT-PCR in SF3B1^K700E^ ES cells (Supplemental Fig. S2E). We conclude that the K700E mutation in ES cells shares the mis-splicing patterns observed in SF3B1 cancers/cell lines.

We next analyzed our RNA-seq data to determine how the K700E mutation affects gene expression in ES cells. The full lists of genes detected in 3 biological replicates of SF3B1^K700E^ and WT ES cell lines are shown in Supplemental Table S3. Significantly dysregulated genes in each line (fold change ≥1.5, p-value <0.05) are shown in Supplemental Table S4. Heatmap analysis for dysregulated genes revealed that ES cells with or without the SF3B1^K700E^ mutation clustered separately (Supplemental Fig. S3A). Thus, although the SF3B1 mutation does not have a major effect on splicing in ES cells, the mutation does have a major effect on gene expression in ES cells. As shown in Fig. 1A, we identified 1342 up- and 1177 down-regulated genes in SF3B1^K700E^ ES cells (line 3 data are shown, see Supplemental Table S4 for other lines). Surprisingly, several of the upregulated genes in the mutant ES lines are master regulators of hematopoiesis (Fig. 1B), with some playing key roles in hematopoietic stem cells (HSC) (de la Grange et al. 2006; Zhang et al. 2007; Parente et al. 2017; Vo et al. 2017). These transcriptional regulators are all associated with blood cancers and often with several different blood cancers (Fig. 1B). Whether these associations have any relationship to SF3B1 blood cancers remains to be determined. We also identified upregulated genes that play roles in the immune response (Fig. 1C). These include SERPINE genes, which are serine protease inhibitors, and GDF15, a well-known marker for ineffective erythropoiesis (Fig. 1C) (Gettins 2002; Ranjbaran et al. 2020). Upregulation of representative masters/immune genes in the 3 SF3B1^K700E^ ES cell lines was validated by qPCR (Fig. 1D; Supplemental Fig. S3B).

**Figure 1.**
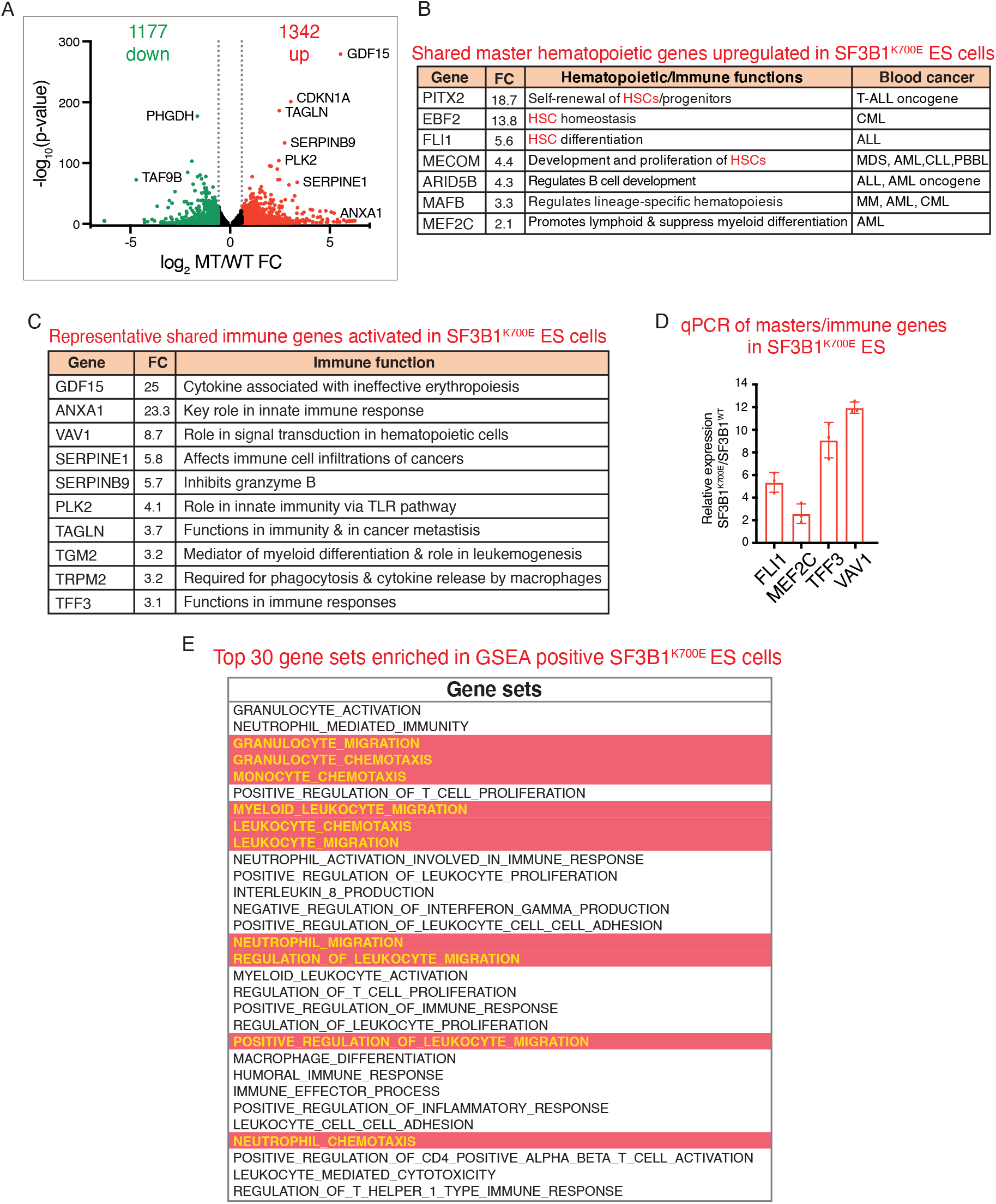
Upregulation of hematopoietic/immune genes in SF3B1^K700E^ ES cell lines. (A) Volcano plot of up- and downregulated genes in SF3B1^K700E^ versus WT ES cells (fold change ≥1.5, p-value <0.05, data for SF3B1^K700E^ ES line 3 are shown). Representative genes are labeled. (B, C) Representative shared master hematopoietic genes (B) and immune genes (C) upregulated in SF3B1^K700E^ ES lines are listed along with relevant functions. FC: Fold Change. (D) qPCR of representative hematopoietic/immune genes in SF3B1^K700E^ ES line 1 is shown. (E) Top 30 gene sets in GSEA positive in SF3B1^K700E^ ES lines with significant NES (NOM p-value < 0.05). Gene sets in red highlighting and yellow font are related to chemokine pathways described in Fig. 4. See also Fig. S1; Supplemental Tables S3-7.

To gain additional insight into the dysregulated genes in SF3B1^K700E^ ES lines, we carried out GSEA. In the GSEA negative data of each of the 3 mutant lines, the top processes were related to DNA replication and nucleosome/chromatin assembly (Supplemental Table S5). In marked contrast, immune gene sets were strongly enriched in the top 30 GSEA positive gene sets of the SF3B1^K700E^ ES lines (Fig. 1E). The data also revealed additional immune gene sets enriched in GSEA positive but not in GSEA negative of the SF3B1^K700E^ ES lines (Supplemental Table S5). To further investigate these findings, we identified the GSEA positive gene sets shared among the 3 SF3B1^K700E^ ES lines, revealing 63 common gene sets (Supplemental Table S6). A list of individual immune genes that were significantly upregulated in the 3 SF3B1^K700E^ ES lines is shown in Supplemental Table S7. In line 3, 296 of the upregulated genes, which is >20% of the total, play roles in the immune response, and similar data were obtained with the other 2 lines (Supplemental Table S7). Thus, both GSEA and direct examination of individual genes revealed that SF3B1 ^K700E^ mutation leads to upregulation of immune genes in ES cells. These data suggest that the essential splicing factor SF3B1 has a specific role in ensuring proper expression of immune genes only in the correct cell types (Matsunawa et al. 2013; Wang et al. 2014; De La Garza et al. 2016; Obeng et al. 2016; Pollyea et al. 2019). Specifically, WT SF3B1 may prevent inappropriate expression of master hematopoietic and immune genes in undifferentiated cell types, such as ES cells.

### Immune genes are robustly downregulated in SF3B1^MT^ MDS, AML and CLL

Considering our observation that immune genes are upregulated in SF3B1^K700E^ ES cells, we used GSEA to investigate gene expression in SF3B1^MT^ MDS using published RNA-seq data (Obeng et al. 2016; Zhang et al. 2019; Madan et al. 2020). MDS patients that were WT for SF3B1 were used as controls. In striking contrast to ES cells bearing SF3B1^K700E^ mutation, immune gene sets constituted almost all of the top 30 GSEA negative gene sets in SF3B1^MT^ MDS (Fig. 2A). This is readily apparent by looking at the large number of gene sets highlighted in green in Fig. 2A. Many additional immune gene sets can be seen in the full list shown in Supplemental Table S8. In sharp contrast, ~2 % of the gene sets enriched in GSEA positive for SF3B1^MT^ MDS were immune (Supplemental Table S8).

**Figure 2.**
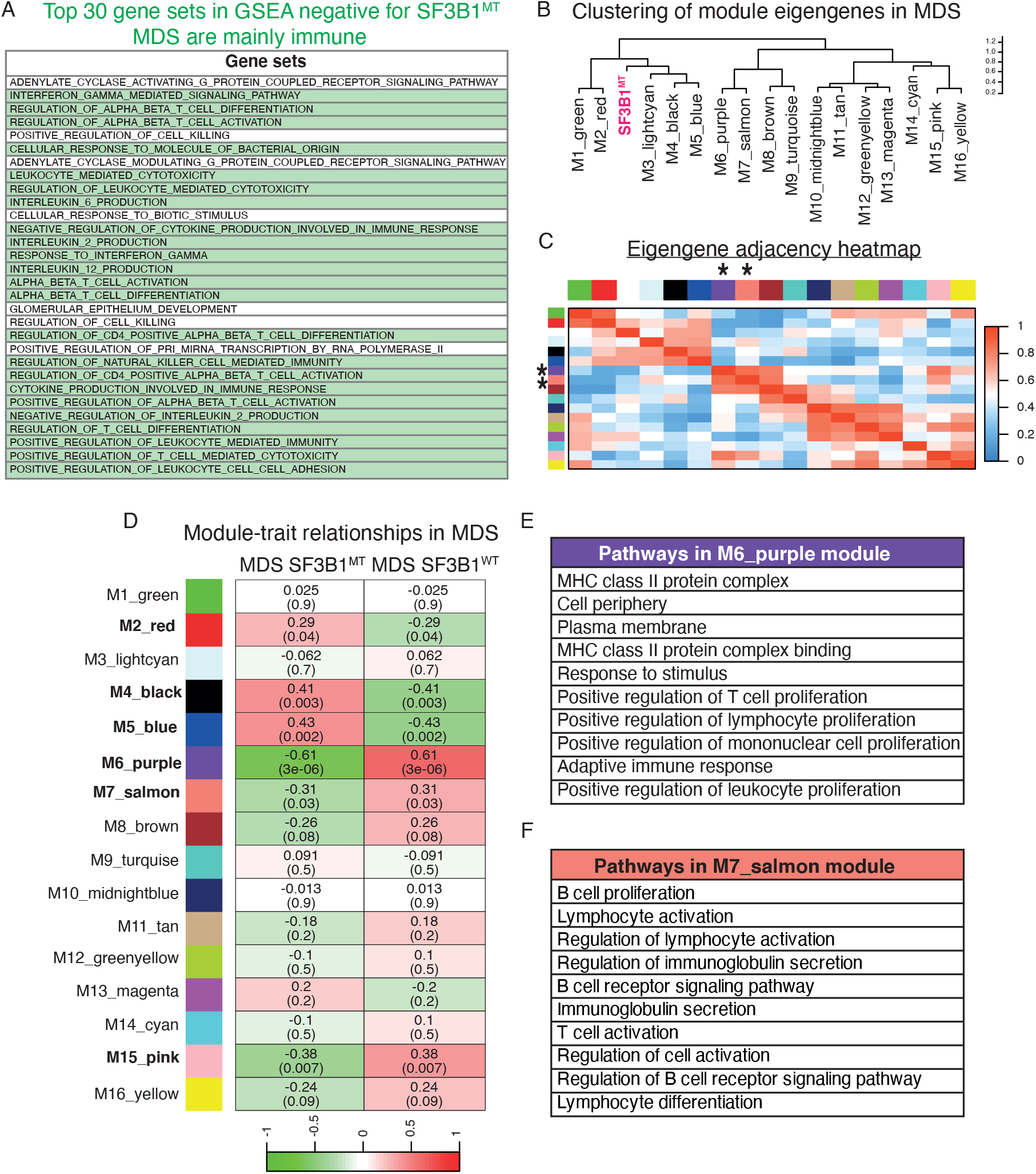
Hematopoietic/Immune gene sets are negatively enriched in SF3B1^MT^ MDS. (A) Top 30 significantly enriched gene sets in GSEA negative in SF3B1^MT^ MDS patients are listed. Hematopoietic/Immune pathways are highlighted in green. Gene sets with significant NES (NOM p-value < 0.05) are shown. (B) Gene co-expression network in SF3B1^MT^ patients using WGCNA shows clustering of 16 module eigengenes in MDS patients. SF3B1^MT^ module is labeled in pink. (C) Eigengene adjacency heatmap indicates the correlation between modules. Red and blue represent positive and negative correlations, respectively. Asterisks indicate modules negatively associated with immune pathways. (D) Heatmap of module-trait relationship displaying correlation coefficients between module eigengenes and SF3B1^MT^ phenotype. Colors indicate positive (red) or negative (green) correlations. Values without parentheses indicate correlation coefficient, corresponding p-values are in parentheses at the bottom of each row. Modules with significant p-values are in bold. (E, F) Enriched pathways in modules M6_purple and M7_salmon, respectively, are listed. See also Supplemental Fig. S4; Supplemental Tables S8-10.

To further examine these unexpected findings, we applied weighted gene correlation network analysis (WGCNA) (Langfelder and Horvath 2008) to identify gene networks correlated to SF3B1^MT^ MDS gene expression. Sixteen modules of densely interconnected genes were identified (Fig. 2B; Supplemental Fig. S4); the pathways within these modules are listed in Supplemental Table S9. High levels of co-expression in the SF3B1^MT^ MDS gene expression data are shown by the clustering of the 16 modules (Fig. 2B) and by the eigengene adjacency heat map (Fig. 2C). The M6_purple (r=0.61, p-value= 3e-06) and M7_salmon (r=0.31 and p-value= 0.03) modules, designated by the asterisks in Fig. 2D, were negatively associated with immune pathways (Fig. 2E,F). Among these are MHC class II pathways as well as pathways related to T and B cells (for list of gene interactions see Supplemental Table S10). Together, these results support the conclusion that immune genes are downregulated in SF3B1^MT^ MDS.

We also carried out GSEA of SF3B1^MT^ CLL and AML using published datasets with the corresponding WT splicing factor patient data as controls (Darman et al. 2015; Tyner et al. 2018). As observed with SF3B1^MT^ MDS, immune genes sets were strongly enriched only in the GSEA negative data for both AML and CLL bearing mutant SF3B1 (Fig. 3A; Supplemental Tables S11). We conclude that immune gene sets/genes are downregulated in 3 different SF3B1^MT^ blood cancers.

**Figure 3.**
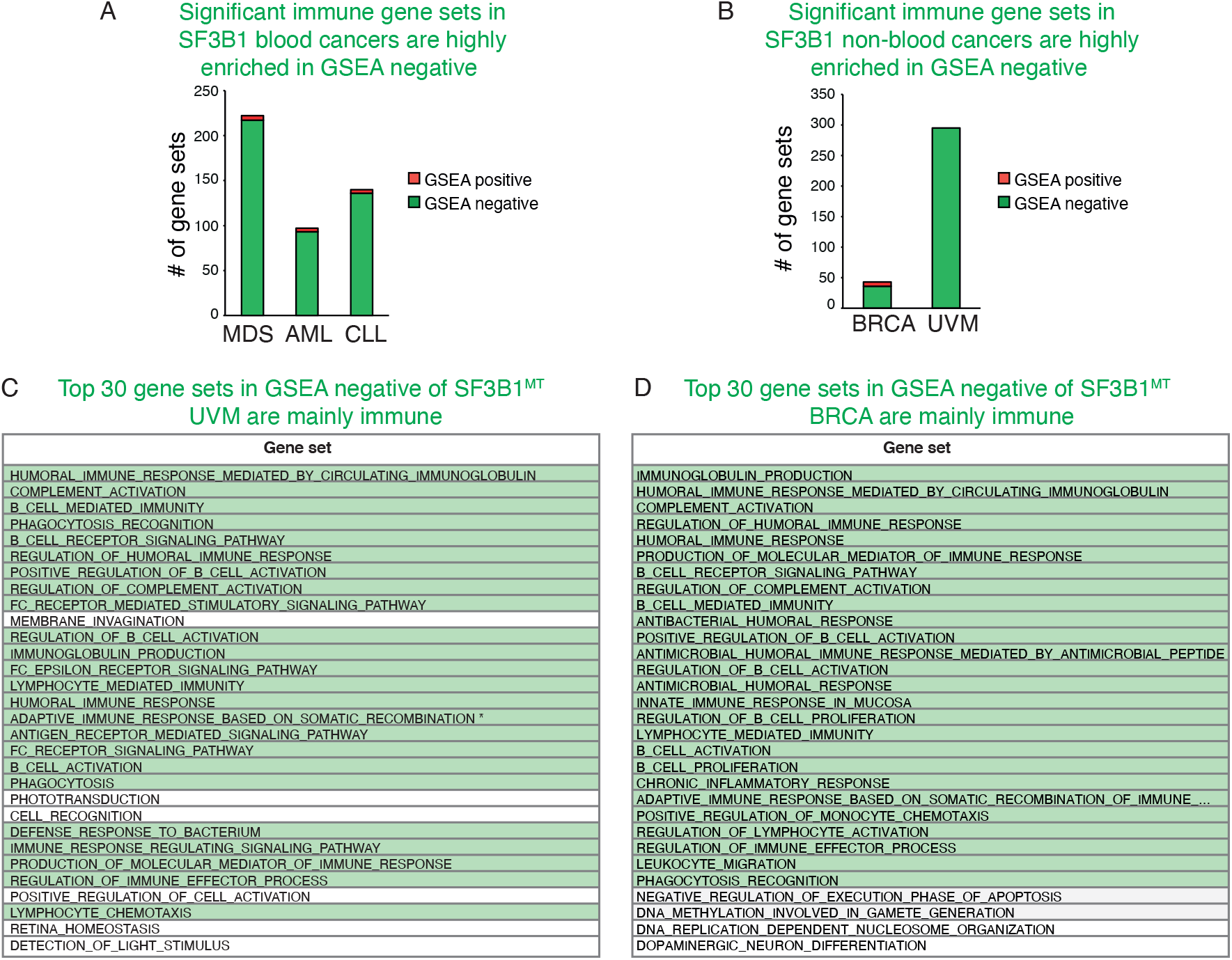
Hematopoietic/Immune gene sets are negatively enriched in SF3B1^MT^ blood and non-blood cancers. (A, B) Significant negative immune gene sets in blood and non-blood cancers, respectively. (C, D) Top 30 significantly enriched gene sets in GSEA negative of SF3B1^MT^ UVM patients and BRCA patients, respectively. Hematopoietic/immune pathways are highlighted in green. Gene sets with significant NES (NOM p-value < 0.05) are shown. See also Supplemental Tables S11,12.

### Immune gene sets are highly enriched in GSEA negative of SF3B1^MT^ UVM and BRCA

As SF3B1 mutation is also associated with non-blood cancers, we next asked whether the strong enrichment of immune gene sets in GSEA negative of MDS/CLL/AML was due to cell type. Accordingly, we carried out GSEA using 2 SF3B1 non-blood cancers, uveal melanoma (UVM) and breast cancer (BRCA). Strikingly, we obtained the same results with these 2 cell types as with the blood cancers. As shown in Fig. 3B, the significant immune gene sets in BRCA and UVM are extremely enriched in the GSEA negative data. Indeed, of the top 30 gene sets in the GSEA negative data for BRCA and UVM, most are immune, as indicated by the green highlighting (Fig. 3C,D). The high level of enrichment for these gene sets in BRCA and UVM is further evident in Supplemental Table S12.

We next sought to determine whether any of the GSEA negative gene sets overlapped in the blood cancers and non-blood cancers. This analysis revealed 30 overlapping gene sets in the blood cancers and 32 overlapping in the UVM and BRCA gene sets (Fig. 4A,B). In the blood cancers, 18 of the gene sets have functions in lymphocytes/leukocytes/T cells whereas in UVM and BRCA, 13 gene sets have roles in B cells and the humoral response (Supplemental Table S13). However, when the BRCA dataset is omitted from the analysis, there are important datasets shared between MDS, AML, CLL, and UVM (Supplemental Table S13). These are described in the section below.

**Figure 4.**
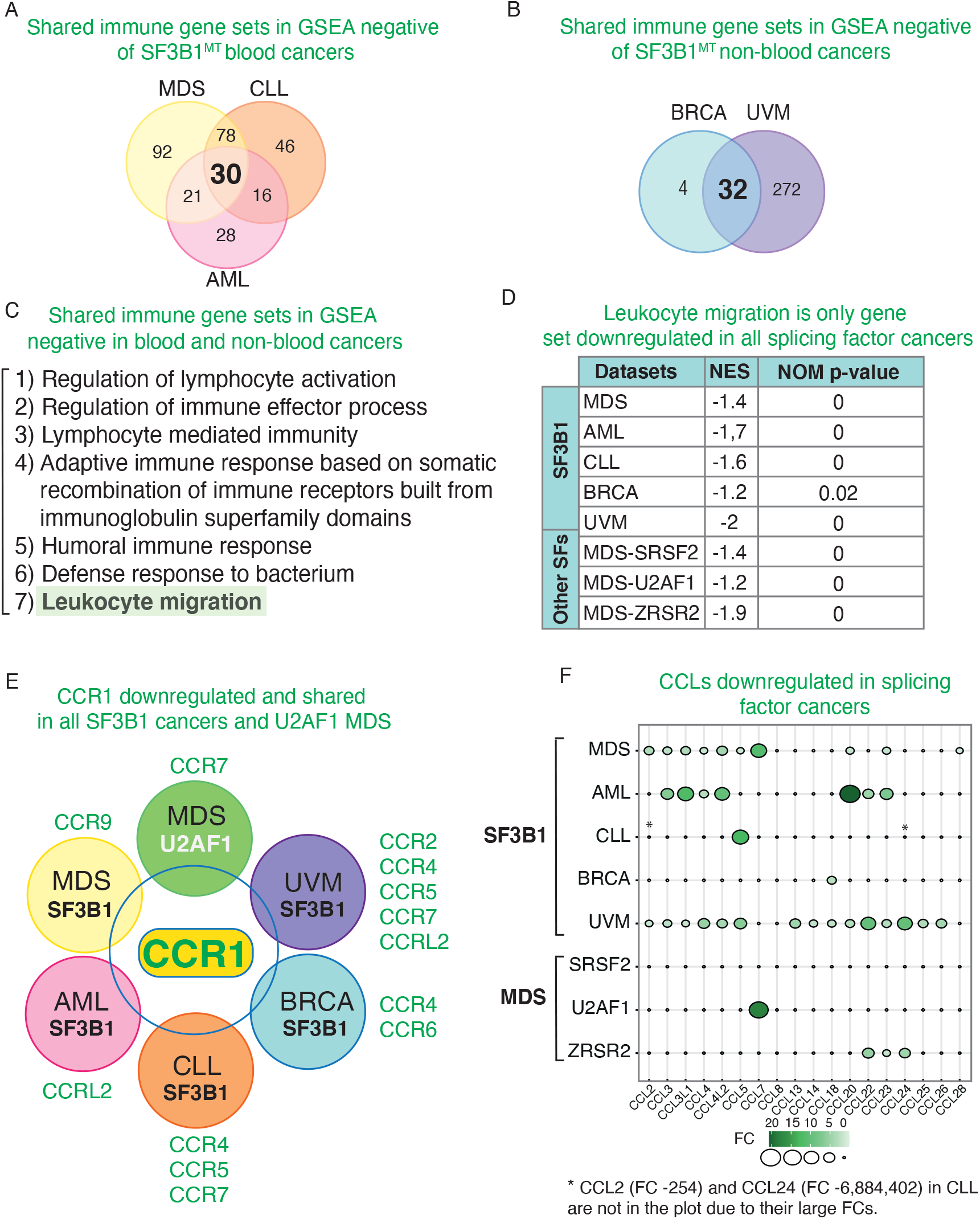
Downregulation of Chemokine Receptor 1 (CCR1) in 5 SF3B1 cancers and U2AF1 MDS. (A) Venn diagram depicting shared GSEA negative immune gene sets in blood cancers (MDS, CLL and AML). (B) Venn diagram showing shared GSEA negative immune gene sets in non-blood cancers (BRCA and UVM). (C) Shared immune GSEA negative gene sets in SF3B1^MT^ blood and non-blood cancers. Leukocyte migration gene set is highlighted in green. (D) Leukocyte migration is the only gene set shared among all splicing factors examined in this study. NES and NOM p-value are shown. (E) Venn diagram showing CCR1 as the only shared downregulated gene among the indicated SF3B1 splicing factor cancers and U2AF1 MDS. Additional CCRs present in the indicated cancers are shown. (F) Balloon plot showing downregulation of CCL genes in splicing factors (SF) cancers. See also Supplemental Tables S13-15.

Together, our data indicate that, although the mis-splicing patterns are similar between SF3B1^K700E^ ES cells and cancers with mutant SF3B1, there is a striking difference in expression of the immune gene sets in SF3B1^MT^ ES cells versus cancer cells. We conclude that upregulation of these genes in SF3B1^K700E^ ES cells versus their downregulation in the SF3B1^MT^ cancers indicate that the cancer cell background, not the cell type, explains the difference in expression of the immune genes.

### Downregulation of the LEUKOCYTE MIGRATION GSEA gene set shared among all 5 SF3B1 cancers and MDS associated with mutant U2AF1, SRSF2 and ZRSR2

To identify gene sets that have important roles in SF3B1 cancers, we compared the SF3B1^MT^ blood and non-blood cancers. This analysis revealed 7 gene sets in common. Among them were lymphocyte pathways and leukocyte migration (Fig. 4C). We then examined GSEA data for SRSF2^MT^ and ZRSR2^MT^ MDS. Remarkably, the leukocyte migration gene set was also downregulated in these splicing factor cancers (Fig. 4D). Moreover, this gene set was the only one in common among all the splicing factor cancers that we examined, SF3B1^MT^ MDS, AML, CLL, UVM and BRCA as well as U2AF1^MT^ MDS, SRSF2^MT^ MDS and ZRSR2^MT^ MDS (Fig. 4D). The significantly downregulated genes in each splicing factor cancer that overlap with the genes in the leukocyte migration pathway are listed in Supplemental Table S14.

### Downregulation of CCR1 shared among all SF3B1 cancers and U2AF1^MT^ MDS

At the level of individual genes, we identified one gene, CCR1, that is downregulated in all 5 SF3B1 cancers examined in this study. CCR1 is one of ~20 C-C motif chemokine receptors, which are 7 membrane proteins and couple to G proteins for signal transduction within cells (Allen et al. 2007). We observed downregulation of CCR1 by as much as 256-fold in CLL to ~2-3 fold in the other SF3B1 cancers (Supplemental Table S15). The potential importance of CCR1 in splicing factor cancers is further suggested by the observation that this gene is also downregulated in U2AF1^MT^ MDS (Fig. 4E). In general, CCRs may play an important role in splicing factor cancers. Specifically, additional CCRs are downregulated in SF3B1 blood and non-blood cancers and U2AF1 MDS, with downregulations ranging from 2-fold to as high as 50-fold (Fig. 4E; Supplemental Table S15). Moreover, in SRSF2 and ZRSR2 MDS, CCR2 is downregulated, and this CCR may substitute for downregulation of CCR1 as both receptors share a ligand (CCL7). Indeed, some CCRs have similar functions and/or have paralogs. Moreover, CCR1 knock out mice are not lethal indicating redundancy in the CCRs. Notably, the knockout mice show impaired hematopoiesis as well as impaired immune and inflammatory response (Gao et al. 1997), which supports our hypothesis that CCR1 downregulation plays a role in splicing factor cancers.

The CCRs have multiple ligands which are usually C-C motif chemokine ligands (CCLs). We found that many of the CCLs are likewise downregulated in the SF3B1 cancers, and many are shared and downregulated among the other splicing factor cancers (Fig. 4F; Supplemental Table S15). For example, the CCL ligands for CCR1 that are downregulated in SF3B1 MDS include CCL3, CCL4, CCL5, CCL7, CCL14 and CCL23 (Fig. 4F). Downregulations of the CCLs in the different splicing factor cancers vary from ~2-10 fold to 245-fold to as much as ~ 7 million-fold (Fig. 4F; Supplemental Table S15).

Regarding the mechanism for downregulation of these crucial immune factors, there are many possibilities. Although SF3B1 is best known for its essential role in splicing, we did not detect mis-splicing of the CCR/CCLs. However, our studies and those of others over the past 20 years have shown that the numerous steps in gene expression are extensively coupled to one another (Maniatis and Reed 2002; Reed and Hurt 2002; Keene 2007; Kornblihtt et al. 2013). For example, transcription promotes splicing, and reciprocally, splicing promotes transcription. Many other coupled steps in gene expression involve splicing, such as mRNA export, 3’ end formation, and mRNA degradation (Reed and Hurt 2002). Thus, any of these steps could be involved in downregulating the CCR/CCLs in splicing factor cancers.

The numerous CCRs and their CCL ligands have highly complex roles in both normal and cancer cells, as well as in cancer microenvironments (Korbecki et al. 2020a, 2020b; Morein et al. 2020). CCRs are key regulators of migration and chemotaxis of multiple blood cell types, including thymocytes, granulocytes, myeloid cells, and additional types of leukocytes. Notably, we observed shared downregulation of many of these chemokine-related pathways in SF3B1 MDS, AML, CLL and UVM (Supplemental Table S15). Moreover, as shown above in Fig. 1, we observed upregulation of immune pathways/genes in SF3B1^K700E^ ES lines, and among these are several chemokine-related pathways (Fig. 1E, highlighted in red with yellow font). Again, these data revealed the opposite results when mutant SF3B1 was expressed in ES cells versus cancer cells, indicating that the cancer background plays a key role in downregulating these crucial immune pathways. The pathways involved in chemokine function extend beyond cell migration and chemotaxis to include numerous pathways with key roles in normal and inflammatory conditions, and they play crucial roles in recruitment of effector immune cells to sites of inflammation (Morein et al. 2020). Which of these pathways may contribute to splicing factor cancers remains to be determined.

In contrast to the splicing factor cancers, CCRs and CCLs are upregulated in a multitude of non-splicing factor cancers, including breast, lung, pancreatic, prostate, and many others (Mollica Poeta et al. 2019; Korbecki et al. 2020a, 2020b; Morein et al. 2020). In this regard, inhibiting the chemokine system has been used as a therapy for these cancers. Thus, our observation that CCR/CCLs and their pathways are downregulated in splicing factor cancers may be a phenomenon relevant to these specific cancer types. There are some reports that cancers with downregulated CCR/CCLs have better prognoses (Korbecki et al. 2020a, 2020b). Thus, it is possible that CCR/CCLs are tumor suppressors, and their upregulation may be a therapy for splicing factor cancers. This therapeutic strategy has been used as part of a regimen for treatment of other cancers (Guiducci et al. 2004). Among the possible methods that could be used to upregulate CCR1 in SF3B1 and U2AF1 blood cancers include the exciting recent CRISPR activation technology, in which transcription activators are used to increase expression of gene(s) of interest.

## Materials and Methods

### Cell culture

H9 human ES cell lines obtained from the HMS Cell Biology Initiative for Genome Editing and Neurodegeneration were cultured using the mTeSR1^TM^1 kit (STEMCELL technology) and 1% penicillin-streptomycin (Sigma) on Matrigel-coated (CORNING) tissue culture plates and incubated at 37°C in humidified 5% (vol/vol) CO_2_.

### CRISPR editing SF3B1^K700E^ in ES cells

H9 ES cells (WA09) from WiCell Research Institute (Madison, WI) were cultured in E8 medium (Chen et al. 2011) on Matrigel-coated plates at 37 °C with 5% (vol/vol) CO2. CRISPR/CAS9 was used to generate ES cell lines containing an A>G point mutation resulting in K700E amino acid substitution in the SF3B1 gene. The CRISPR guide sequence was 5’-TGGATGAGCAGCAGAAAGTT-3’. The Ultramer sequence used to introduce the mutation was: 5’-TGTAACTTAGGTAATGTTGGGGCATAGTTAAAACCTGTGTTTGGTTTTGTAGGTCTTGTGGATGAGCAGCAGgAAGTTCGGACCATCAGTGCTTTGGCCATTGCTGCCTTGGCTGAAGCAGCAACTCCTTATGGTATCGAATCTTTTGAT-3 (point mutation lowercase; PAM region underscored) (Supplemental Fig. S1A). To create H9 cells harboring a heterozygous K700E mutation in SF3B1, 0.6 μg sgRNA was incubated with 3 μg SpCas9 protein for 10 minutes at room temperature and electroporated into 2×10^5^ H9 cells along with the Ultramer repair template. Mutants were identified by Illumina MiSeq. In addition, K700E mutation status was further confirmed by PCR amplification of genomic DNA followed by Sanger sequencing.

### RNA sequencing

Total RNA isolation from SF3B1^K700E^ and WT ES cells was performed using the RNeasy Kit (Qiagen) following manufacturer’s instructions. Bulk mRNA sequencing of triplicate samples was performed using NEBNext^®^ Ultra™ II RNA Library preparation (NEB #E7775) with PolyA selection on the Illumina HiSeq4000 using the standard protocol for Illumina. On average, ~31M paired-end reads were obtained per sample.

### Differential gene expression analysis

Raw FASTQ files for MDS and CLL patients, K562 and NALM-6 cell lines, were downloaded from NCBI-GEO datasets (accession numbers GSE85712, GSE128805, GSE128429, GSE72790, GSE95011) (Supplemental Table S16). Reads were aligned using STAR version 2.5.2b with the following parameters: --outSAMtype BAM – SortedByCoordinate --outSAMunmapped Within --outSAMattributes Standard --quantMode GeneCounts (Dobin et al. 2013). Ensembl database (human release 99) and Human assembly hg38 (GRCh38) were used as the gene annotation and reference genome, respectively. In addition, TCGA-BRCA, TCGA-UVM, and AML from the BEATAML1.0-COHORT, including aligned BAM files and STAR 2-Pass Genome counts, were downloaded from the GDC data portal (https://portal.gdc.cancer.gov/).

### Identification of differential splicing events

To estimate alternative splicing (AS) events, PSI-Sigma v1.9j was used to provide a comprehensive and accurate analysis of AS events (Lin and Krainer 2019). We used the default parameters of PSI-sigma, which required at least 10 supporting reads to detect splicing events with *Homo sapiens* GRCh38.99 as a reference genome. For GDC aligned BAM files, .SJ.out files were generated. To quantify changes in splicing patterns, percent-spliced-in (PSI) index and differential PSI (ΔPSI) values were calculated to find all isoforms in a specific gene region. Splicing events with ΔPSI ≥ 10% and p-value < 0.05 were included in analyses. Additionally, Integrative Genomic Viewer (IGV_2.8.9) was used for visualizing aberrant splicing events in SF3B1 mutant and wildtype samples.

### Gene set enrichment analysis (GSEA)

Gene ontology (GO) analysis was performed using GSEA (release v4.1.0) with gene set collection ‘c5.go.bp.v7.4.symbols.gmt’. The pre-ranked gene list was analyzed to identify biological processes positively or negatively associated with SF3B1 mutations. Gene sets with significant NES (NOM p-value < 0.05) were evaluated.

### Co-expression and weighted gene co-expression network analysis (WGCNA)

Weighted gene co-expression network analysis (WGCNA) was used to construct a transcriptional network from RNA expression profiles. WGCNA (R package, ‘WGCNA’ version 1.70-3) identifies the key modules and module-trait relationship using topological overlap measure (TOM) and detects similarly expressed gene sets across MDS samples (Langfelder and Horvath 2008).

### qPCR

Total RNA was isolated from cells using the RNeasy Kit (Qiagen) according to the manufacturer’s instructions. cDNA was synthesized using M-MLV reverse transcriptase (Invitrogen), and qPCR was performed with gene-specific primer sets (Supplemental Table S17) and SYBR Green PCR Master Mix (Thermo Fisher Scientific). All qPCR analyses were performed on an Applied Biosystems QuantStudio 7 Flex Cycler (Thermo Fisher Scientific), and relative expression values were calculated using the comparative CT method (Schmittgen and Livak 2008).

## Data availability

RNA sequencing data reported in this paper will be uploaded to Gene Expression Omnibus (GEO) database upon publication.

## Acknowledgements

We thank Dr. Thorsten M. Schlaeger for technical assistance with ES cell culture and characterization. We are grateful to Drs. John N. Hutchinson, Moein Farshchian, Kuan-Ting Lin, and Research Computing Consultant groups at HMS for assistance in RNA-seq analysis. We thank WiCell Research Institute (Madison, WI) for H9 ES cells (WA09). This work was supported by NIH grant NIGMS GM122524 to RR, NIDDK (1 K08 DK114527-01) and Boston Children’s Hospital Office of Faculty Development to RGR.

## Author contributions

RR conceived the project and interpretated the data in conjunction with MD. Computational analyses were performed on the O2 High Performance Compute Cluster, supported by the Research Computing Group, at HMS. HMK carried out RNA-seq, and HMK and MD performed bioinformatic analyses. KWN assisted with this work. ZH, JZ and EFC established the SF3B1^K700E^ lines. MD carried out all other experiments with assistance from RR, CLP, CEL, BC, and NY. SY performed experiments in the initial stages of the project. RGR carried out ES cell culture and quality control and WM and YZ taught ES cell culture to RR’s lab. The manuscript was written by RR and MD with assistance from CLP and input from GQD, and all authors contributed to the manuscript.

