## Supplemental_Figures for "Chemokine Receptor 1 and its associated immune pathway are downregulated in SF3B1^MT^ blood and non-blood cancers"

Figure S1

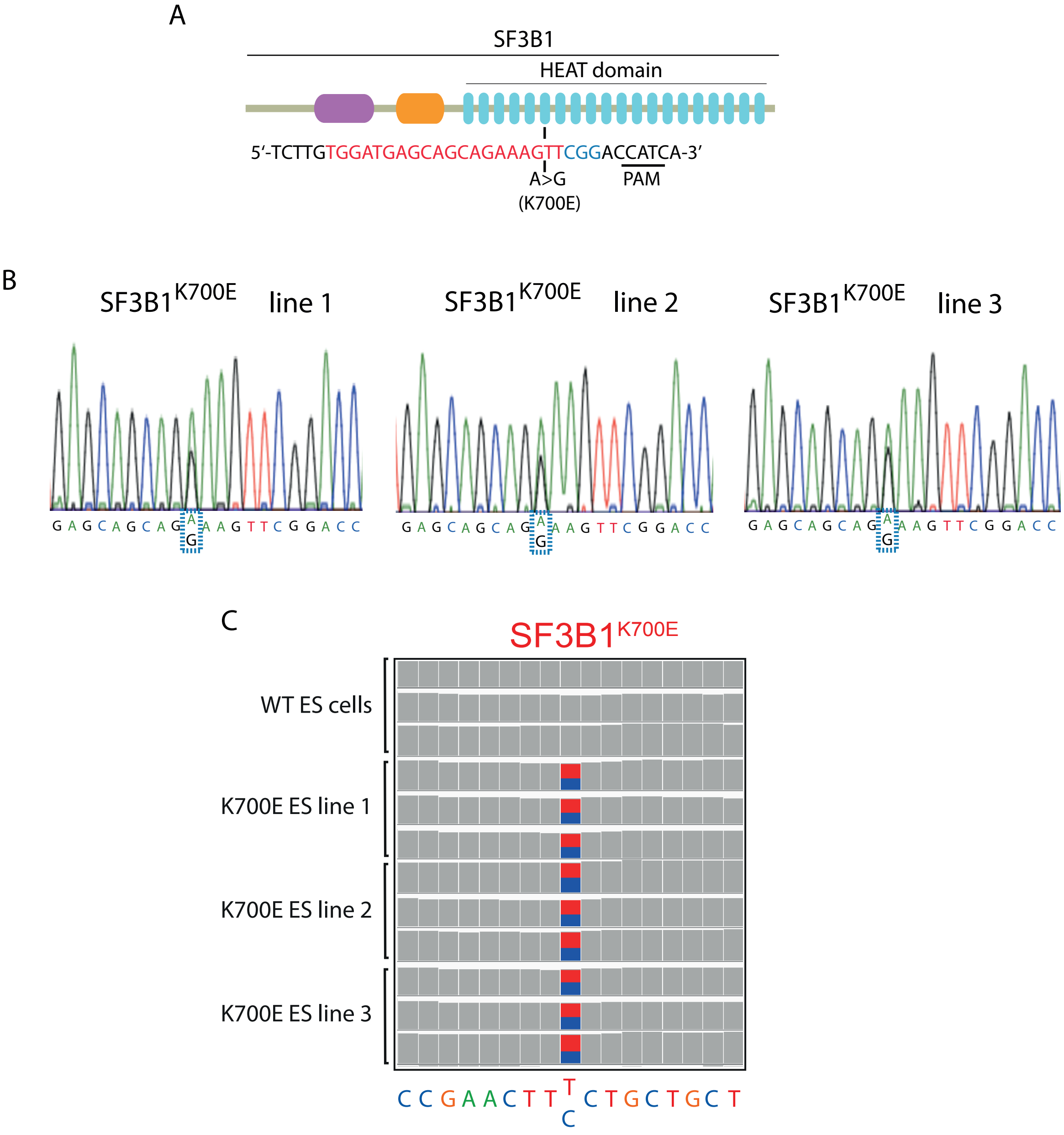


**Figure S1. CRISPR editing of K700E mutation into one allele of SF3B1 in human ES cells.**
(A) A schematic of SF3B1 gene showing domain structure and position of K700E mutation. Target sequence for sgRNA is labeled in red, followed by PAM sequence labeled in blue. (B) Sanger sequencing validation of K700E mutation in one allele of 3 independent ES cell lines. (C) IGV (integrated genome viewer) confirms that mutant and WT SF3B1 alleles were expressed at similar levels in SF3B1$^{K700E}$ ES cell lines.

Figure S2

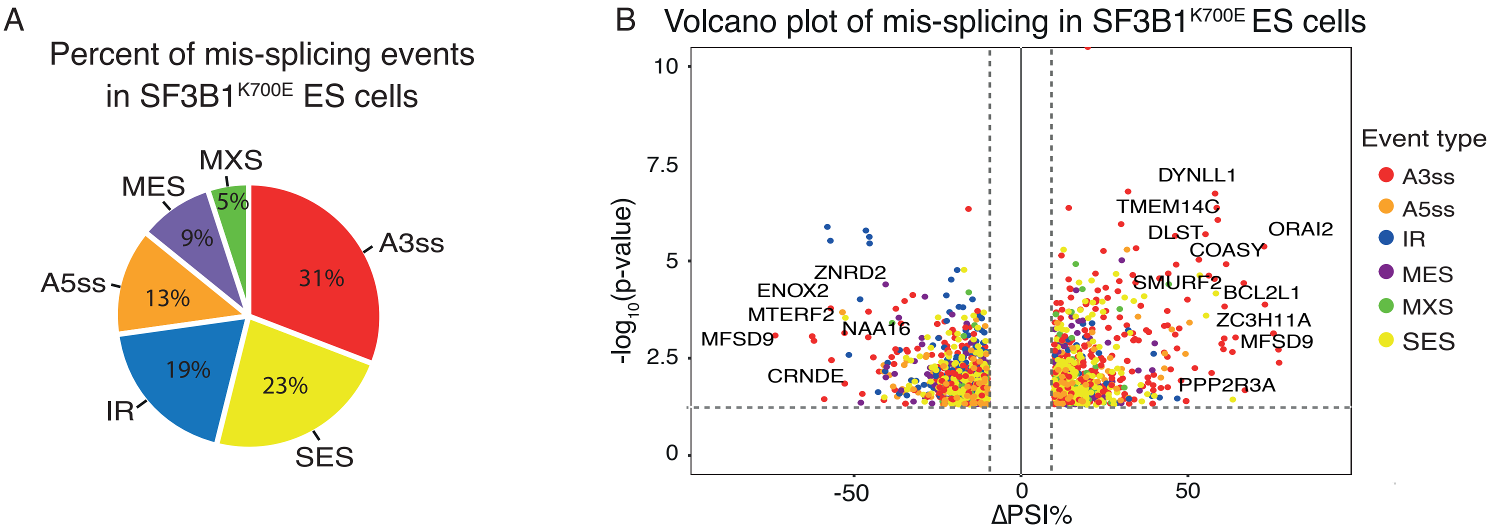


C

Shared mis-splicing events in SF3B1$^{K700E}$ ES cells and SF3B1$^{MT}$ cancers/cell lines

| Gene | Aberrant Junction | Event | Phenotype | Function |
|---|---|---|---|---|
| APPL2 | 12:105601825-105601935 | A3ss | 6 aa ins E 7-8 | Regulates TLR4 signaling, which functions in innate immunity |
| BRD9 | 5:869519-870587 | SES | NMD | BAF chromatin-remodeling complex factor; key regulator of AML |
| DLG1 | 3:196792335-196792578 | A3ss | NMD | Negative regulator of T cell activation |
| DLST | 14:75356053-75356581 | A3ss | NMD | Key role in MYC-mediated leukemogenesis |
| DVL2 | 17:7131031-7131295 | A3ss | 25 aa del E 10-11 | Mis-spliced DVL2 increases Notch signaling, which drives CLL |
| INPPL1 | 11:71939543-71939691 | A3ss | NMD | Suppresses a signaling pathway critical for myeloid cancers |
| MAP3K7 | 6:91269953-91271340 | A3ss | NMD | MAP3K7 mis-splicing causes anemia in MDS via P38 deactivation |
| PPOX | 1:161137025-161137129 | A3ss | NMD | Essential for heme biosynthesis |
| SMURF2 | 17:62574712-62576906 | A3ss | 6 aa ins E 8-9 | E3 ubiquitin ligase that regulates HSC self-renewal |
| TTI1 | 17:62574712-62576906 | A3ss | NMD | Role in mTOR pathway, which functions in hematopoiesis |

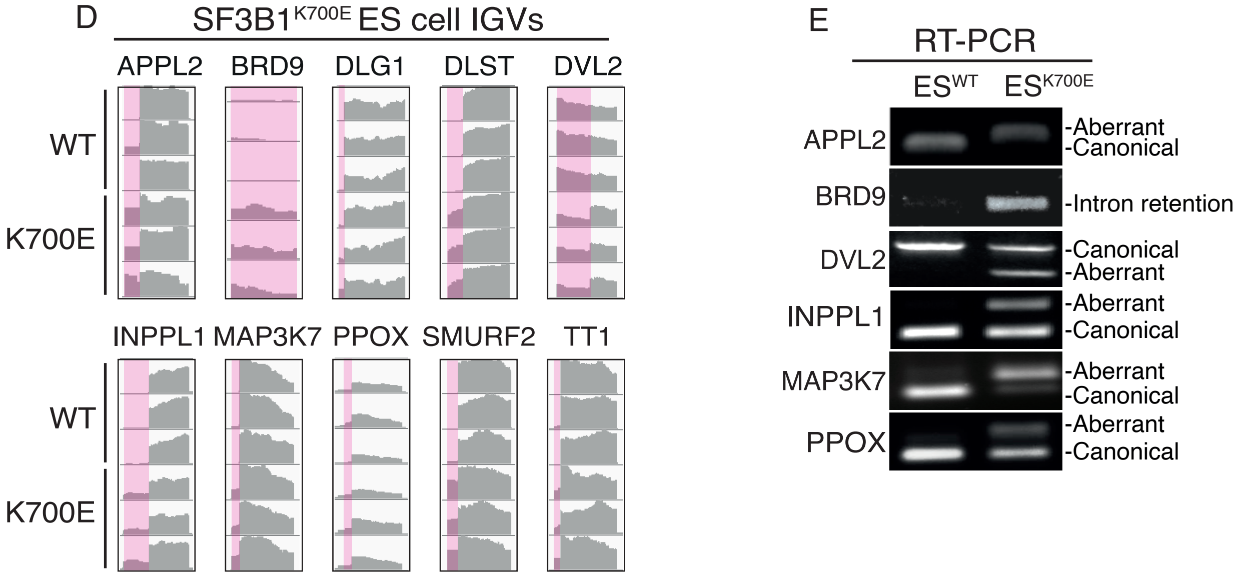


**Figure S2. Mis-splicing events in SF3B1$^{K700E}$ ES cells are shared with SF3B1$^{MT}$ cancers/cell lines.** (A) Distribution of mis-splicing events in SF3B1$^{K700E}$ ES line 2. (B) Volcano plot depicting mis-splicing events in SF3B1$^{K700E}$ ES line 2. Representative genes with top ΔPSI% are labeled. (C) Representative mis-spliced genes in SF3B1$^{K700E}$ ES cells and SF3B1$^{MT}$ blood cancers (MDS, AML and CLL), blood cell lines (K562 and NALM6) and non-blood cancers (BRCA and UVM) with immune functions. ΔPSI ≥ 10% and p-value < 0.05 were used as thresholds. SES, single-exon skipping; MES, multiple-exon skipping; MXS, mutually-exclusive splicing; A5ss, alternative 5′ splice site; A3ss, alternative 3′ splice site. (D) Integrated genome viewer (IGV) plots show mis-splicing in SF3B1$^{K700E}$ ES cell line 2 versus WT. Plots are from 3 biological replicates of ES lines. (E) Representative mis-spliced genes were validated by RT-PCR in SF3B1$^{K700E}$ ES line.

Figure S3

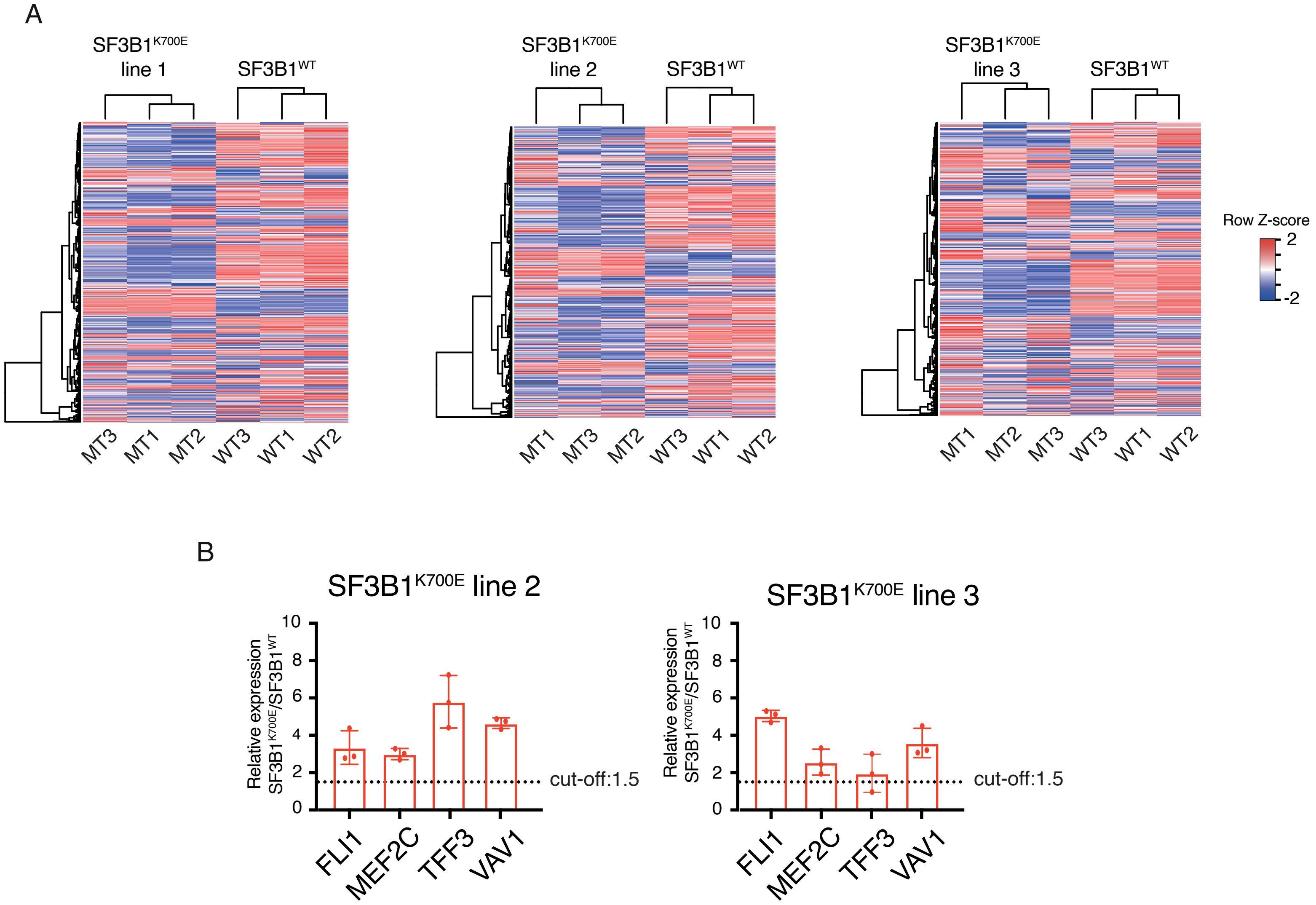


**Figure S3. Gene expression analysis in 3 independent SF3B1$^{K700E}$ ES lines.** (A) Heatmaps depicting differentially expressed genes in SF3B1$^{K700E}$ and WT ES cells in 3 biological replicates. Each row represents one gene and each column is one ES replicate. Z-scores in the matrix represent expression levels. (B) qPCR of upregulated immune genes in 2 independent ES cell lines (see Fig. 1 for SF3B1$^{K700E}$ line 1 data). Data in panel B are represented as mean of 3 biological replicates. See also Fig 1.

Figure S4

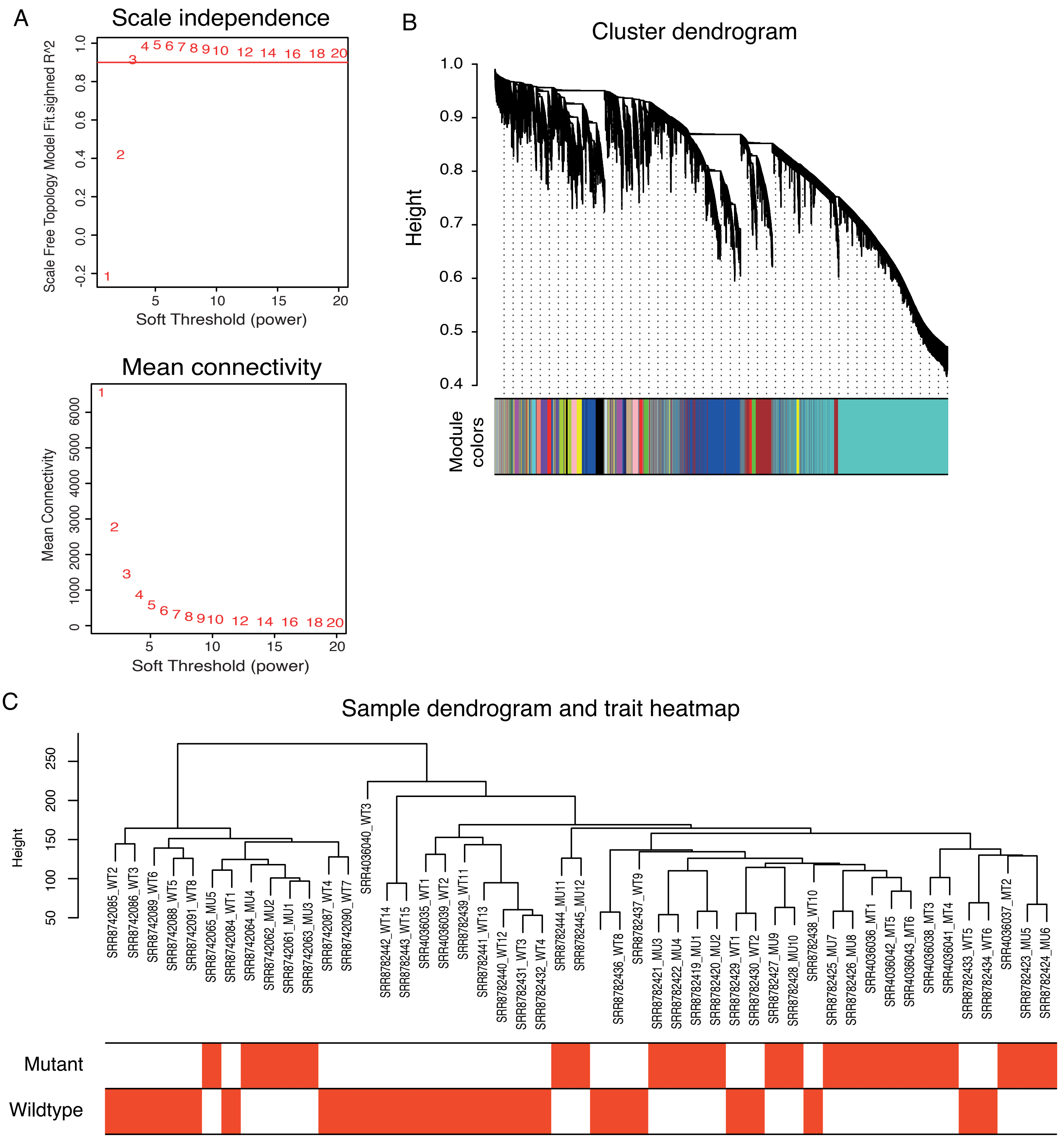


**Figure S4. Gene co-expression network analysis using WGCNA in SF3B1$^{MT}$ MDS patients.** (A) Determination of soft-thresholding power in WGCNA. Scale-free topology index and mean connectivity for each power are shown. In this study, the threshold was reached for a power of 3. (B) Dendrogram of differentially expressed genes clustered based on the difference metrics (1-TOM**).** (C) Sample dendrogram and trait heatmap based on expression data from 3 MDS datasets (see Table S16 for patient information).
