## Supplementary material for "Chemokine Receptor 1 and its associated immune pathway are downregulated in SF3B1^MT^ blood and non-blood cancers": Table_S1

**Table S1. Mis-splicing events in 3 biological replicates of SF3B1$^{K700E}$ (K700E ES line 1) compared to SF3B1$^{WT}$ ES cells.** The mis-splicing event code is as follows: SES, single-exon skipping; MES, multiple-exon skipping; MXS, mutually-exclusive splicing; A5ss, alternative 5′ splice site; A3ss, alternative 3′ splice site; TSS, transcription start site. Event Region: Genomic coordinates of splicing event; Target Exon: Genomic coordinates of the alternative exon. Events with p-values < 0.05 and ΔPSI (%)≥10% are listed.

| Gene Symbol | Event Region | Target Exon | Event | WT | K700E | Reference Transcript | ΔPSI (%) | p-value | FDR (BH) |
|---|---|---|---|---|---|---|---|---|---|
| GEN1 | 2:17753968-17759928 | 2:17759912-17759929 | A3ss | 2 | 3 | Ex.TSS.ENST00000317402 | 83.2 | 2.08E-03 | 7.44E-03 |
| APBB3 | 5:140561702-140561843 | 5:140561702-140561722 | A3ss | 2 | 3 | ENST00000354402 | 76.03 | 1.52E-02 | 1.03E-01 |
| HERC2P2 | 15:22561790-22562948 | 15:22561896-22562949 | A3ss | 3 | 3 | Ex.ENST00000613386 | 74.31 | 8.14E-05 | 1.36E-04 |
| VPS9D1-AS1 | 16:89712223-89712688 | 16:89712674-89712689 | A3ss | 3 | 3 | Ex.TSS.ENST00000562866 | 72.81 | 7.27E-05 | 1.42E-04 |
| PGBD1 | 6:28281919-28283775 | 6:28283756-28283775 | A3ss | 3 | 3 | Ex.TSS.ENST00000259883 | 71.67 | 5.65E-04 | 4.57E-03 |
| ORAI2 | 7:102433662-102436224 | 7:102436202-102436224 | TSSlA3ss | 3 | 3 | TSS.ENST00000356387 | 70.48 | 2.89E-05 | 2.67E-04 |
| DLST | 14:74889350-74889896 | 14:74889878-74889896 | A3ss | 3 | 3 | ENST00000555988 | 70.08 | 5.66E-05 | 9.35E-05 |
| GPR153 | 1:6255015-6260824 | 1:6255015-6255122 | A3ss | 2 | 3 | Ex.TSS.ENST00000377893 | 69.08 | 1.50E-02 | 4.33E-02 |
| NPR2 | 9:35801155-35801642 | 9:35801634-35801643 | A3ss | 3 | 3 | Ex.ENST00000342694 | 68.96 | 1.56E-03 | 4.71E-02 |
| CCDC88B | 11:64352387-64352743 | 11:64352727-64352744 | A3ss | 3 | 3 | Ex.ENST00000356786 | 66.86 | 1.65E-03 | 2.36E-03 |
| MFSD9 | 2:102732410-102736645 | 2:102732410-102732426 | A3ss | 3 | 3 | ENST00000462099 | 62.78 | 2.98E-03 | 1.04E-02 |
| MFSD9 | 2:102732393-102736645 | 2:102732393-102732426 | A3ss | 3 | 3 | ENST00000462099 | 62.78 | 2.98E-03 | 1.04E-02 |
| BCL2L1 | 20:31722331-31723863 | 20:31722331-31722348 | A3ss | 3 | 3 | ENST00000420488 | 62.33 | 9.21E-04 | 2.76E-03 |
| CCDC66 | 3:56615273-56615921 | 3:56615904-56615922 | A3ss | 3 | 3 | Ex.ENST00000326595 | 62.07 | 1.25E-02 | 7.12E-02 |
| ZNF91 | 19:23362726-23373741 | 19:23362726-23362739 | A3ss | 2 | 3 | Ex.TSS.ENST00000300619 | 61.28 | 6.06E-03 | 1.37E-02 |
| COASY | 17:42562220-42562611 | 17:42562356-42562611 | A3ss | 3 | 3 | ENST00000585811 | 61.07 | 2.17E-05 | 4.40E-05 |
| TMEM214 | 2:27037703-27038145 | 2:27037815-27037892 | SES | 3 | 3 | ENST00000425720 | 61.04 | 1.81E-04 | 6.77E-04 |
| AC007326.4 | 22:18939750-18946790 | 22:18945072-18946791 | A3ss | 3 | 3 | Ex.TSS.ENST00000638240 | 60.54 | 1.87E-03 | 6.16E-03 |
| SLC4A8 | 12:51440790-51450875 | 12:51450857-51450876 | A3ss | 3 | 3 | Ex.ENST00000319957 | 60.54 | 8.36E-03 | 1.28E-02 |
| ZC3H11A | 1:203801629-203802956 | 1:203802569-203802957 | A3ss | 3 | 3 | Ex.ENST00000367212 | 60.46 | 4.56E-03 | 1.18E-02 |
| ZC3H11A | 1:203801629-203802651 | 1:203802569-203802652 | A3ss | 3 | 3 | Ex.ENST00000332127 | 60.46 | 4.56E-03 | 1.18E-02 |
| TTC23 | 15:99228733-99249170 | 15:99245389-99245510 | MES | 3 | 2 | ENST00000394132 | 59.91 | 2.25E-02 | 3.89E-02 |
| GOLGA2P10 | 15:82475616-82476785 | 15:82475616-82475637 | A3ss | 3 | 3 | Ex.ENST00000614347 | 59.52 | 6.76E-05 | 1.16E-04 |
| HERC2P2 | 15:22561790-22562948 | 15:22561790-22562026 | A5ss | 3 | 3 | Ex.ENST00000613386 | 59.46 | 3.17E-04 | 5.29E-04 |
| DYNLL1 | 12:120496217-120496415 | 12:120496402-120496415 | TSSlA3ss | 3 | 3 | TSS.ENST00000392508 | 59.38 | 2.62E-07 | 3.89E-07 |
| TMEM14C | 6:10723242-10724569 | 6:10724556-10724570 | A3ss | 3 | 3 | Ex.TSS.ENST00000229563 | 58.49 | 8.80E-06 | 6.80E-05 |
| PPP2R3A | 3:136040963-136049258 | 3:136049244-136049259 | A3ss | 3 | 3 | Ex.ENST00000264977 | 57.26 | 2.21E-04 | 8.97E-04 |
| MFSD13A | 10:102466088-102468954 | 10:102468753-102468955 | A3ss | 3 | 3 | Ex.TSS.ENST00000238936 | 55.8 | 1.80E-03 | 2.37E-03 |
| TLCD5 | 11:120325369-120329976 | 11:120327452-120329977 | A3ss | 3 | 3 | Ex.ENST00000531346 | 53.34 | 1.16E-04 | 1.57E-04 |
| COASY | 17:42562220-42562611 | 17:42562356-42562487 | SES | 3 | 3 | ENST00000590958 | 53.33 | 5.64E-05 | 1.14E-04 |
| DZIP3 | 3:108684342-108687975 | 3:108686428-108687976 | A3ss | 3 | 2 | Ex.ENST00000495008 | 51.74 | 2.79E-03 | 1.11E-02 |
| DZIP3 | 3:108684342-108686444 | 3:108686428-108686445 | A3ss | 3 | 2 | Ex.ENST00000361582 | 51.74 | 2.79E-03 | 1.11E-02 |
| GCC2 | 2:108485909-108486510 | 2:108486499-108486511 | A3ss | 3 | 3 | Ex.ENST00000309863 | 51.46 | 4.71E-04 | 1.64E-03 |
| COASY | 17:42562220-42562611 | 17:42562220-42562487 | A5ss | 3 | 3 | ENST00000585909 | 51.31 | 7.78E-04 | 1.57E-03 |
| CCDC74A | 2:131530828-131531663 | 2:131531652-131531664 | A3ss | 3 | 3 | Ex.ENST00000295171 | 50.69 | 5.78E-03 | 2.04E-02 |
| BCL2L1 | 20:31722331-31723734 | 20:31722331-31722348 | A3ss | 3 | 3 | ENST00000456404 | 50.56 | 4.34E-03 | 1.30E-02 |
| ARMC9 | 2:231331898-231344974 | 2:231344949-231344975 | A3ss | 3 | 3 | Ex.TSS.ENST00000349938 | 50.33 | 2.24E-05 | 8.25E-05 |
| AC007326.4 | 22:18939750-18946790 | 22:18939750-18945166 | A5ss | 3 | 3 | Ex.TSS.ENST00000638240 | 49.27 | 2.48E-03 | 8.15E-03 |
| STAG3L5P | 7:100345969-100349798 | 7:100349717-100349798 | TSSlA3ss | 3 | 3 | TSS.ENST00000493499 | 49.24 | 2.47E-03 | 2.28E-02 |
| STAG3L5P | 7:100345969-100349730 | 7:100349717-100349730 | TSSlA3ss | 3 | 3 | TSS.ENST00000493499 | 49.24 | 2.47E-03 | 2.28E-02 |
| MAP3K7 | 6:90560215-90561621 | 6:90560215-90560234 | A3ss | 3 | 3 | Ex.ENST00000369325 | 48.8 | 1.28E-04 | 1.17E-03 |
| ZMYM5 | 13:19837822-19851354 | 13:19838700-19838985 | SES | 2 | 2 | ENST00000337963 | 48.64 | 2.41E-02 | 3.77E-02 |
| UXS1 | 2:106164785-106166055 | 2:106164785-106164799 | A3ss | 3 | 3 | ENST00000283148 | 48.1 | 1.62E-04 | 5.65E-04 |
| ENOSF1 | 18:683381-685920 | 18:683381-683395 | A3ss | 3 | 3 | ENST00000578647 | 47.59 | 1.06E-05 | 2.29E-05 |
| KIAA1217 | 10:24219910-24380868 | 10:24219910-24255596 | A5ss | 2 | 3 | Ex.ENST00000376452 | 47.42 | 2.16E-02 | 2.86E-02 |
| ZBED5 | 11:10855087-10856143 | 11:10855087-10855118 | TSSlA3ss | 3 | 3 | TSS.ENST00000432999 | 46.85 | 1.09E-03 | 1.46E-03 |
| ZBED5 | 11:10855082-10856143 | 11:10855082-10855118 | TSSlA3ss | 3 | 3 | TSS.ENST00000432999 | 46.85 | 1.09E-03 | 1.46E-03 |
| APPL2 | 12:105208030-105208157 | 12:105208030-105208047 | A3ss | 3 | 3 | ENST00000551662 | 46.36 | 6.23E-03 | 9.19E-03 |
| EMC3-AS1 | 3:9987265-10002716 | 3:9987815-9987905 | MES | 3 | 3 | ENST00000440568 | 45.99 | 1.12E-03 | 6.52E-03 |
| GLI1 | 12:57460202-57464672 | 12:57463665-57463791 | MES | 2 | 3 | ENST00000228682 | 45.94 | 4.43E-02 | 6.83E-02 |
| TMEM218 | 11:125102318-125102636 | 11:125102318-125102351 | A3ss | 3 | 3 | ENST00000279968 | 45.37 | 3.60E-03 | 4.87E-03 |
| GLI1 | 12:57460202-57464672 | 12:57463999-57464091 | MES | 2 | 3 | ENST00000228682 | 45.25 | 5.25E-04 | 8.12E-04 |
| MBD2 | 18:54203103-54204997 | 18:54203103-54203126 | TSSlA3ss | 3 | 3 | TSS.ENST00000398398 | 44.84 | 4.00E-04 | 8.61E-04 |
| ZNF337-AS1 | 20:25624136-25645530 | 20:25634049-25645531 | A3ss | 2 | 2 | Ex.TSS.ENST00000439498 | 44.25 | 7.32E-03 | 2.19E-02 |
| ZNF213-AS1 | 16:3132104-3134633 | 16:3132850-3132942 | SES | 2 | 3 | ENST00000572691 | 44.24 | 3.27E-03 | 6.15E-03 |
| CCDC74B | 2:130141297-130142132 | 2:130141297-130141308 | A3ss | 3 | 3 | Ex.ENST00000310463 | 43.21 | 1.28E-03 | 4.51E-03 |
| ZDHHC16 | 10:97454800-97455713 | 10:97455639-97455713 | A3ss | 3 | 3 | ENST00000487315 | 41.65 | 3.66E-04 | 4.91E-04 |
| ZDHHC16 | 10:97454800-97455659 | 10:97455639-97455659 | A3ss | 3 | 3 | ENST00000487315 | 41.65 | 3.66E-04 | 4.91E-04 |
| NDRG3 | 20:36653702-36656359 | 20:36653702-36653723 | A3ss | 3 | 3 | Ex.TSS.ENST00000349004 | 41.57 | 2.51E-03 | 7.55E-03 |
| LZTS1 | 8:20255316-20303739 | 8:20268195-20303740 | A5ss | 3 | 3 | Ex.TSS.ENST00000381569 | 41.41 | 3.99E-02 | 6.34E-01 |
| NMRK2 | 19:3936666-3937239 | 19:3937225-3937239 | A3ss | 3 | 3 | ENST00000593949 | 41.34 | 4.22E-06 | 9.72E-06 |
| AFG3L1P | 16:89972766-89978809 | 16:89977715-89978810 | A3ss | 3 | 2 | Ex.TSS.ENST00000429663 | 40.77 | 3.31E-02 | 6.46E-02 |
| SEPTIN6 | X:119625380-119629317 | X:119625380-119625396 | A3ss | 3 | 3 | ENST00000467310 | 40.63 | 7.20E-04 | 2.80E-02 |
| MTERF2 | 12:106978772-106986968 | 12:106985115-106985215 | SES | 3 | 3 | ENST00000392830 | 40.48 | 1.06E-04 | 1.56E-04 |
| ZNF213-AS1 | 16:3132104-3134633 | 16:3132850-3133367 | SES | 2 | 3 | ENST00000652946 | 40.29 | 1.87E-03 | 3.51E-03 |
| AC060780.1 | 17:43167251-43170125 | 17:43167251-43167262 | TSSlA3ss | 2 | 3 | TSS.ENST00000590740 | 39.77 | 9.42E-03 | 1.91E-02 |
| SPINT1-AS1 | 15:40838687-40844178 | 15:40838687-40838705 | A3ss | 3 | 3 | Ex.TSS.ENST00000564302 | 39.66 | 1.91E-03 | 3.22E-03 |
| SLC26A6 | 3:48633636-48635370 | 3:48634516-48635371 | A5ss | 3 | 3 | Ex.TSS.ENST00000307364 | 39.58 | 1.59E-03 | 7.10E-03 |
| ANKHD1 | 5:140436258-140438502 | 5:140438494-140438502 | A3ss | 3 | 3 | ENST00000394722 | 38.69 | 2.64E-05 | 1.80E-04 |
| THOC1 | 18:224180-224923 | 18:224180-224200 | A3ss | 3 | 3 | ENST00000579891 | 38.64 | 2.48E-05 | 5.31E-05 |
| RIMKLB | 12:8773467-8777589 | 12:8776673-8777590 | A3ss | 3 | 3 | Ex.TSS.ENST00000299673 | 38.54 | 3.35E-03 | 5.25E-03 |
| TRIM37 | 17:59001715-59012327 | 17:59001715-59001741 | A3ss | 3 | 3 | ENST00000585287 | 38.33 | 3.58E-03 | 7.39E-03 |
| DVL2 | 17:7227703-7227976 | 17:7227703-7227711 | A3ss | 3 | 3 | Ex.ENST00000575086 | 38.2 | 4.71E-04 | 9.84E-04 |
| SLC13A3 | 20:46596343-46610445 | 20:46599971-46600037 | SES | 3 | 3 | ENST00000279027 | 37.78 | 3.84E-02 | 1.23E-01 |
| SMURF2 | 17:64578577-64580788 | 17:64578577-64578594 | A3ss | 3 | 3 | Ex.ENST00000262435 | 36.4 | 2.36E-03 | 4.92E-03 |
| ZNF160 | 19:53086110-53086261 | 19:53086110-53086128 | A3ss | 3 | 3 | Ex.TSS.ENST00000596966 | 36.12 | 1.13E-03 | 2.64E-03 |
| ZNF160 | 19:53075184-53086261 | 19:53075184-53086128 | A3ss | 3 | 3 | Ex.ENST00000355147 | 36.12 | 1.13E-03 | 2.64E-03 |
| PIH1D1 | 19:49451579-49453375 | 19:49451579-49451591 | TSSlA3ss | 3 | 3 | TSS.ENST00000601825 | 35.85 | 4.13E-02 | 9.71E-02 |
| UBA1 | X:47199615-47200893 | X:47199615-47200544 | A5ss | 3 | 3 | Ex.ENST00000335972 | 35.84 | 6.29E-06 | 9.12E-04 |
| BRD9 | 5:869395-870472 | 5:869395-869404 | A3ss | 3 | 3 | ENST00000490814 | 35.22 | 1.47E-02 | 1.12E-01 |
| TRPM7 | 15:50583157-50586391 | 15:50583157-50583159 | A3ss | 3 | 3 | ENST00000646667 | 35.13 | 1.13E-02 | 1.91E-02 |
| MROH1 | 8:144258915-144259239 | 8:144259224-144259240 | A3ss | 3 | 3 | Ex.ENST00000326134 | 34.83 | 7.49E-03 | 1.14E-01 |
| PILRB | 7:100356884-100358263 | 7:100358227-100358264 | A3ss | 3 | 3 | Ex.ENST00000608825 | 34.77 | 1.41E-04 | 1.28E-03 |
| ZNF213-AS1 | 16:3132104-3134629 | 16:3132104-3132942 | A3ss | 3 | 3 | Ex.TSS.ENST00000571963 | 34.68 | 6.00E-04 | 1.13E-03 |
| PLA2G4B | 15:41842592-41843675 | 15:41842592-41843223 | A5ss | 3 | 3 | ENST00000483748 | 34.5 | 2.29E-02 | 3.85E-02 |
| SH3D19 | 4:151179398-151187422 | 4:151179398-151179416 | A3ss | 3 | 3 | Ex.ENST00000304527 | 34.33 | 2.69E-03 | 1.63E-02 |
| KANSL3 | 2:96619763-96631311 | 2:96619763-96619776 | A3ss | 3 | 3 | Ex.ENST00000354204 | 34.08 | 4.15E-04 | 1.65E-03 |
| KANSL3 | 2:96619545-96631311 | 2:96619545-96619776 | A3ss | 3 | 3 | Ex.ENST00000418735 | 34.08 | 4.15E-04 | 1.65E-03 |
| BRD9 | 5:865582-870472 | 5:869245-869404 | MXS | 3 | 3 | ENST00000519112 | 33.96 | 8.00E-03 | 6.11E-02 |
| DCAF16 | 4:17804757-17810446 | 4:17804757-17804771 | A3ss | 3 | 3 | Ex.ENST00000507768 | 33.96 | 2.00E-03 | 1.22E-02 |

| | | | | | | | | | |
|---|---|---|---|---|---|---|---|---|---|
| JMJD7-PLA2G4B | 15:41842592-41843675 | 15:41843126-41843223 | SES | 3 | 3 | ENST00000490848 | 33.81 | 2.02E-02 | 3.40E-02 |
| RACGAP1 | 12:50016716-50025372 | 12:50016716-50016719 | A3ss | 2 | 3 | Ex.TSS.ENST00000548824 | 33.8 | 7.24E-03 | 1.10E-02 |
| BCL2L1 | 20:31722331-31722617 | 20:31722331-31722348 | TSSIA3ss | 3 | 3 | TSS.ENST00000307677 | 33.62 | 2.73E-03 | 8.17E-03 |
| EMC3-AS1 | 3:9987265-9987905 | 3:9987815-9987905 | TSSIA3ss | 3 | 3 | TSS.ENST00000440568 | 33.56 | 1.67E-02 | 9.72E-02 |
| CAMTA2 | 17:4982161-4982756 | 17:4982161-4982175 | A3ss | 3 | 3 | ENST00000572543 | 33.54 | 1.46E-03 | 3.00E-03 |
| SARM1 | 17:28395905-28396820 | 17:28396027-28396156 | IR (overlapping region) | 3 | 3 | ENST00000579593 | 33.39 | 7.10E-03 | 1.41E-02 |
| ZNF771 | 16:30407665-30408044 | 16:30408032-30408045 | A3ss | 3 | 3 | Ex.TSS.ENST00000319296 | 33.39 | 2.21E-03 | 4.15E-03 |
| IGSF9B | 11:133912008-133919741 | 11:133918257-133919742 | A5ss | 2 | 3 | Ex.ENST00000533871 | 33.38 | 4.12E-02 | 5.59E-02 |
| NAA16 | 13:41373781-41374741 | 13:41373781-41373880 | A5ss | 3 | 3 | ENST00000477452 | 33.33 | 4.13E-03 | 6.51E-03 |
| PROSER3 | 19:35764937-35765066 | 19:35765034-35765066 | A3ss | 2 | 2 | ENST00000301165 | 33.08 | 2.52E-02 | 5.76E-02 |
| RECQL5 | 17:75628763-75628933 | 17:75628763-75628825 | TSSIA3ss | 3 | 3 | TSS.ENST00000582548 | 32.96 | 7.23E-03 | 1.52E-02 |
| PHKB | 16:47461427-47463898 | 16:47463882-47463898 | TSSIA3ss | 3 | 3 | TSS.ENST00000566044 | 32.91 | 7.88E-03 | 1.49E-02 |
| SLC7A3 | X:70929999-70930976 | X:70929999-70930022 | TSSIA3ss | 3 | 3 | TSS.ENST00000374299 | 32.84 | 1.34E-03 | 5.00E-01 |
| THTPA | 14:23556344-23556752 | 14:23556743-23556752 | TSSIA3ss | 3 | 3 | TSS.ENST00000404535 | 32.81 | 9.55E-03 | 1.56E-02 |
| TRIM16 | 17:15680954-15682860 | 17:15682854-15682860 | TSSIA5ss | 2 | 3 | TSS.ENST00000336708 | 32.75 | 1.52E-02 | 2.99E-02 |
| SLC26A6 | 3:48633636-48635370 | 3:48633636-48634590 | A3ss | 3 | 3 | Ex.TSS.ENST00000307364 | 32.67 | 1.07E-02 | 4.80E-02 |
| CCDC88A | 2:55296524-55301205 | 2:55297925-55301206 | A5ss | 3 | 3 | Ex.ENST00000444458 | 32.65 | 5.83E-03 | 2.21E-02 |
| CCDC88A | 2:55296524-55299838 | 2:55297925-55299839 | A5ss | 3 | 3 | Ex.ENST00000263630 | 32.65 | 5.83E-03 | 2.21E-02 |
| MSTO1 | 1:155611292-155611548 | 1:155611411-155611548 | TSSIA3ss | 3 | 3 | TSS.ENST00000490642 | 32.58 | 5.94E-03 | 1.50E-02 |
| MSTO1 | 1:155611292-155611426 | 1:155611411-155611426 | TSSIA3ss | 3 | 3 | TSS.ENST00000490642 | 32.58 | 5.94E-03 | 1.50E-02 |
| TMEM145 | 19:42323790-42324736 | 19:42324278-42324736 | TSSIA3ss | 2 | 3 | TSS.ENST00000673187 | 32.52 | 2.39E-02 | 5.53E-02 |
| ZBTB14 | 18:5293328-5295812 | 18:5293972-5294001 | SES | 3 | 2 | ENST00000357006 | 32.44 | 3.28E-02 | 6.03E-02 |
| RWDD4 | 4:183655962-183658928 | 4:183655962-183655974 | A3ss | 3 | 3 | Ex.TSS.ENST00000326397 | 32.41 | 1.20E-02 | 7.33E-02 |
| RWDD4 | 4:183651132-183658928 | 4:183651132-183655974 | A3ss | 3 | 3 | Ex.TSS.ENST00000510968 | 32.41 | 1.20E-02 | 7.33E-02 |
| COL23A1 | 5:178257568-178259720 | 5:178257568-178257582 | A3ss | 3 | 3 | Ex.ENST00000390654 | 32.33 | 8.27E-04 | 5.81E-03 |
| GLB1L3 | 11:134293210-134312348 | 11:134309626-134309763 | MES | 3 | 3 | ENST00000431683 | 32.17 | 2.77E-02 | 3.76E-02 |
| SMCO4 | 11:93479270-93499275 | 11:93479270-93479284 | TSSIA3ss | 3 | 3 | TSS.ENST00000529714 | 32.14 | 1.69E-05 | 2.48E-05 |
| CHTF18 | 16:793275-794053 | 16:794034-794054 | A3ss | 3 | 3 | Ex.ENST00000262315 | 32.06 | 5.60E-03 | 1.08E-02 |
| ZNF211 | 19:57633247-57634022 | 19:57633247-57633436 | TSSIA5ss | 3 | 3 | TSS.ENST00000240731 | 32.05 | 1.30E-02 | 3.11E-02 |
| ZNF814 | 19:57868580-57876915 | 19:57875156-57875226 | MXS | 2 | 3 | ENST00000597348 | 31.91 | 2.42E-02 | 5.81E-02 |
| TUG1 | 22:30969387-30972923 | 22:30971251-30972924 | A3ss | 3 | 3 | Ex.TSS.ENST00000643920 | 31.87 | 5.59E-03 | 1.90E-02 |
| TUG1 | 22:30969387-30972854 | 22:30971251-30972855 | A3ss | 3 | 3 | Ex.TSS.ENST00000569384 | 31.87 | 5.59E-03 | 1.90E-02 |
| TUG1 | 22:30969387-30972047 | 22:30971251-30972048 | A3ss | 3 | 3 | Ex.TSS.ENST00000602971 | 31.87 | 5.59E-03 | 1.90E-02 |
| ZNF219 | 14:21093291-21093585 | 14:21093291-21093304 | TSSIA3ss | 3 | 3 | TSS.ENST00000554478 | 31.55 | 6.01E-05 | 9.75E-05 |
| CAP1 | 1:40040784-40041428 | 1:40040784-40040801 | TSSIA5ss | 3 | 3 | TSS.ENST00000414281 | 31.44 | 7.14E-03 | 2.02E-02 |
| GANC | 15:42273445-42274167 | 15:42273445-42273611 | TSSIA5ss | 3 | 2 | TSS.ENST00000562859 | 31.43 | 1.45E-02 | 2.44E-02 |
| ACOXL | 2:110933643-110987107 | 2:110963624-110963713 | SES | 3 | 3 | ENST00000389811 | 31.33 | 3.24E-03 | 1.13E-02 |
| TCEA2 | 20:64070636-64071869 | 20:64071858-64071869 | A3ss | 3 | 3 | ENST00000465433 | 31.33 | 4.34E-04 | 1.40E-03 |
| DLG1 | 3:197065449-197065707 | 3:197065449-197065464 | A3ss | 3 | 3 | ENST00000661229 | 31.3 | 4.00E-03 | 1.68E-02 |
| B3GALT5 | 21:39613068-39659752 | 21:39646392-39659753 | A5ss | 3 | 3 | Ex.TSS.ENST00000615480 | 30.94 | 7.23E-03 | 2.36E-02 |
| CASTOR3 | 7:100211394-100211994 | 7:100211540-100211794 | IR (overlapping region) | 2 | 3 | ENST00000328453 | 30.86 | 3.88E-02 | 3.57E-01 |
| GLB1L3 | 11:134293210-134312348 | 11:134307124-134307208 | MES | 3 | 3 | ENST00000431683 | 30.67 | 4.09E-02 | 5.54E-02 |
| DPH5 | 1:100992741-100995109 | 1:100992741-100992754 | A3ss | 3 | 3 | Ex.ENST00000342173 | 30.61 | 4.03E-04 | 9.77E-04 |
| DPH5 | 1:100990632-100995109 | 1:100990632-100992754 | A3ss | 3 | 3 | Ex.ENST00000427040 | 30.61 | 4.03E-04 | 9.77E-04 |
| DPH5 | 1:100990629-100995109 | 1:100990629-100992754 | A3ss | 3 | 3 | Ex.TSS.ENST00000481871 | 30.61 | 4.03E-04 | 9.77E-04 |
| SERBP1 | 1:67424960-67425082 | 1:67424960-67424977 | A3ss | 3 | 3 | ENST00000361219 | 30.56 | 2.15E-06 | 6.20E-06 |
| THUMPD3-AS1 | 3:9390014-9391826 | 3:9390498-9391678 | IR | 3 | 3 | ENST00000518437 | 30.27 | 3.13E-03 | 1.82E-02 |
| ZFYVE27 | 10:97743165-97744728 | 10:97744712-97744729 | A3ss | 3 | 3 | Ex.ENST00000359980 | 30.24 | 1.62E-02 | 2.17E-02 |
| SEPTIN6 | X:119620052-119629317 | X:119625335-119625396 | SES | 3 | 3 | ENST00000467310 | 30.01 | 3.19E-04 | 1.24E-02 |
| AP5S1 | 20:3820703-3822101 | 20:3820703-3820758 | TSSIA5ss | 3 | 3 | TSS.ENST00000379573 | 29.94 | 6.29E-03 | 1.90E-02 |
| RABGAP1 | 9:122997362-122998596 | 9:122998576-122998597 | A3ss | 3 | 3 | Ex.ENST00000373647 | 29.87 | 5.60E-03 | 1.20E-01 |
| BAG6 | 6:31644195-31644524 | 6:31644307-31644414 | SES | 3 | 3 | ENST00000437771 | 29.66 | 7.75E-05 | 6.41E-04 |
| SNRPN | 15:24962210-24967931 | 15:24967029-24967152 | SES | 3 | 3 | ENST00000554227 | 29.5 | 1.61E-05 | 2.68E-05 |
| ECHDC2 | 1:52904834-52907867 | 1:52905034-52905090 | SES | 3 | 3 | ENST00000358358 | 29.46 | 1.58E-02 | 4.50E-02 |
| IMMP1L | 11:31433571-31509518 | 11:31460626-31460714 | MES | 3 | 3 | ENST00000278200 | 29.22 | 5.91E-03 | 8.06E-03 |
| CAMTA2 | 17:4981832-4982756 | 17:4982089-4982175 | SES | 3 | 3 | ENST00000572543 | 29.19 | 1.16E-02 | 2.38E-02 |
| MEGF6 | 1:3496784-3497232 | 1:3496988-3497119 | SES | 3 | 3 | ENST00000356575 | 29.1 | 1.99E-03 | 5.61E-03 |
| SNHG14 | 15:24982438-25019026 | 15:24998366-24998486 | MES | 3 | 3 | ENST00000551631 | 28.99 | 3.75E-02 | 6.26E-02 |
| BTBD10 | 11:13421839-13463091 | 11:13445024-13445181 | SES | 3 | 3 | ENST00000278174 | 28.96 | 1.46E-03 | 1.99E-03 |
| FDFT1 | 8:11803012-11803518 | 8:11803226-11803321 | IR | 3 | 3 | ENST00000528812 | 28.89 | 1.53E-02 | 2.19E-01 |
| ADGRG2 | X:19037491-19037597 | X:19037589-19037597 | A5ss | 3 | 2 | ENST00000356606 | 28.83 | 2.31E-02 | 1.00E+00 |
| NBEAL2 | 3:46991947-46993936 | 3:46992475-46992555 | SES | 3 | 3 | ENST00000450053 | 28.78 | 4.88E-03 | 2.12E-02 |
| TTC23 | 15:99241560-99249416 | 15:99245389-99245510 | MES | 2 | 3 | ENST00000490671 | 28.68 | 2.50E-02 | 4.31E-02 |
| IQSEC2 | X:53266328-53267064 | X:53266765-53266968 | IR | 2 | 2 | ENST00000639161 | 28.64 | 3.71E-02 | 1.00E+00 |
| TCF7L2 | 10:113146098-113150997 | 10:113150983-113150997 | A3ss | 3 | 3 | ENST00000543371 | 28.46 | 1.21E-02 | 1.60E-02 |
| TTC23 | 15:99241560-99249416 | 15:99245389-99245510 | MXS | 2 | 3 | ENST00000459771 | 28.2 | 2.61E-02 | 4.50E-02 |
| RMST | 12:97493286-97493894 | 12:97493891-97493894 | A3ss | 3 | 3 | ENST00000652810 | 28.05 | 1.17E-02 | 1.84E-02 |
| SUZ12P1 | 17:30709812-30759490 | 17:30734897-30734943 | MES | 3 | 3 | ENST00000579526 | 28.01 | 8.93E-03 | 1.78E-02 |
| ZNF213-AS1 | 16:3132104-3134629 | 16:3132850-3132945 | SES | 3 | 3 | ENST00000573447 | 27.96 | 1.12E-02 | 2.12E-02 |
| THUMPD3-AS1 | 3:9390014-9391826 | 3:9390498-9391704 | IR | 3 | 3 | ENST00000518437 | 27.95 | 4.90E-03 | 2.85E-02 |
| ZMYM1 | 1:35079443-35095818 | 1:35093914-35094083 | MES | 3 | 2 | ENST00000373330 | 27.91 | 1.09E-02 | 3.07E-02 |
| BTN2A2 | 6:26383245-26383791 | 6:26383245-26383299 | TSSIA5ss | 3 | 3 | TSS.ENST00000471116 | 27.8 | 2.45E-02 | 1.97E-01 |
| BTN2A2 | 6:26383182-26383791 | 6:26383182-26383299 | TSSIA5ss | 3 | 3 | TSS.ENST00000472507 | 27.8 | 2.45E-02 | 1.97E-01 |
| HINT2 | 9:35813146-35813265 | 9:35813146-35813156 | A3ss | 3 | 3 | Ex.TSS.ENST00000471774 | 27.75 | 1.14E-03 | 3.45E-02 |
| C1RL | 12:7097164-7101897 | 12:7099901-7100026 | MES | 3 | 3 | ENST00000266542 | 27.73 | 1.28E-02 | 1.99E-02 |
| CDK8 | 13:26396355-26397152 | 13:26397139-26397153 | A3ss | 3 | 3 | Ex.ENST00000381527 | 27.56 | 3.78E-04 | 5.94E-04 |
| STIM1 | 11:4082983-4083262 | 11:4083242-4083263 | A3ss | 3 | 3 | Ex.ENST00000300737 | 27.56 | 2.88E-03 | 3.94E-03 |
| EDA | X:70033523-70035357 | X:70033523-70035528 | TSSIA5ss | 3 | 3 | TSS.ENST00000374552 | 27.54 | 3.66E-02 | 1.00E+00 |
| THUMPD3-AS1 | 3:9390014-9391826 | 3:9390498-9391081 | IR | 3 | 3 | ENST00000518437 | 27.49 | 4.40E-03 | 2.56E-02 |
| NBPF9 | 1:149102848-149103300 | 1:149103271-149103301 | A5ss | 3 | 2 | Ex.TSS.ENST00000621645 | 27.44 | 4.50E-03 | 1.11E-02 |
| MIPOL1 | 14:37247241-37247828 | 14:37247799-37247828 | A3ss | 3 | 3 | ENST00000556615 | 27.32 | 2.77E-03 | 4.53E-03 |
| ACIN1 | 14:23081837-23090521 | 14:23089982-23090101 | SES | 3 | 3 | ENST00000262710 | 27.31 | 6.66E-04 | 1.09E-03 |
| ANO9 | 11:419582-420860 | 11:420616-420717 | IR | 3 | 3 | ENST00000532094 | 27.29 | 3.33E-03 | 4.56E-03 |
| THUMPD3-AS1 | 3:9390014-9391826 | 3:9390498-9391426 | IR | 3 | 3 | ENST00000518437 | 27.28 | 1.12E-03 | 6.49E-03 |
| TUT4 | 1:52436976-52438219 | 1:52436976-52436978 | A3ss | 3 | 3 | ENST00000257177 | 27.26 | 1.70E-02 | 4.26E-02 |
| TUT4 | 1:52436964-52438219 | 1:52436964-52436978 | A3ss | 3 | 3 | ENST00000257177 | 27.26 | 1.70E-02 | 4.26E-02 |
| UBA1 | X:47199615-47200893 | X:47200410-47200894 | A3ss | 3 | 3 | Ex.ENST00000335972 | 27.24 | 1.30E-05 | 1.88E-03 |
| RHBDD1 | 2:226836088-226864603 | 2:226839409-226839627 | MES | 3 | 3 | ENST00000392062 | 27.17 | 4.87E-02 | 1.79E-01 |
| MED6 | 14:70592989-70593295 | 14:70592989-70593005 | A3ss | 3 | 3 | ENST00000430055 | 27 | 2.78E-02 | 4.58E-02 |
| NFYA | 6:41073085-41079028 | 6:41079005-41079029 | A3ss | 3 | 3 | Ex.TSS.ENST00000341376 | 26.93 | 3.39E-04 | 2.88E-03 |
| ZNF532 | 18:58953800-58979054 | 18:58954234-58954334 | SES | 3 | 3 | ENST00000585662 | 26.92 | 1.56E-04 | 3.38E-04 |
| SKIV2L | 6:31969280-31969751 | 6:31969421-31969514 | IR | 3 | 3 | ENST00000465703 | 26.89 | 3.88E-03 | 3.22E-02 |
| TPRA1 | 3:127594934-127598195 | 3:127598007-127598196 | A5ss | 2 | 3 | Ex.TSS.ENST00000490643 | 26.85 | 4.47E-02 | 1.80E-01 |
| GLB1L3 | 11:134293210-134312348 | 11:134310571-134310651 | MES | 3 | 3 | ENST00000431683 | 26.78 | 2.32E-02 | 3.15E-02 |

| | | | | | | | | | |
|---|---|---|---|---|---|---|---|---|---|
| ZNF138 | 7:64815045-64831450 | 7:64830940-64831076 | MXS | 3 | 3 | ENST00000359735 | 26.76 | 2.26E-02 | 2.56E-01 |
| TOR1AIP2 | 1:179865855-179877238 | 1:179865855-179865869 | A3ss | 3 | 3 | Ex.TSS.ENST00000482587 | 26.74 | 1.64E-03 | 4.21E-03 |
| PHF20L1 | 8:132798861-132803818 | 8:132799083-132799172 | SES | 3 | 3 | ENST00000395376 | 26.73 | 2.38E-04 | 3.53E-03 |
| TTC14 | 3:180607648-180610974 | 3:180608811-180609629 | IR (overlapping region) | 3 | 3 | ENST00000412756 | 26.68 | 2.63E-02 | 1.09E-01 |
| SNHG14 | 15:25054206-25055230 | 15:25054867-25055073 | IR | 3 | 3 | ENST00000660717 | 26.56 | 1.07E-02 | 1.78E-02 |
| NEIL1 | 15:75352602-75354492 | 15:75352702-75353738 | IR | 3 | 2 | ENST00000561643 | 26.51 | 3.36E-03 | 5.72E-03 |
| CC2D2A | 4:15478807-15480703 | 4:15479223-15479328 | SES | 3 | 2 | ENST00000438599 | 26.49 | 3.25E-02 | 1.97E-01 |
| JAKMIP1 | 4:6085630-6112721 | 4:6105473-6105967 | SES | 3 | 3 | ENST00000282924 | 26.41 | 6.16E-03 | 3.91E-02 |
| MICAL1 | 6:109445863-109446135 | 6:109445863-109445875 | A3ss | 3 | 3 | Ex.ENST00000358577 | 26.29 | 4.76E-04 | 3.68E-03 |
| HEXIM2 | 17:45161940-45162749 | 17:45161940-45162031 | TSSlA5ss | 3 | 3 | TSS.ENST00000592695 | 26.06 | 2.09E-03 | 4.25E-03 |
| THUMPD3-AS1 | 3:9390014-9391826 | 3:9390498-9391106 | IR | 3 | 3 | ENST00000518437 | 26.01 | 2.11E-03 | 1.23E-02 |
| RSPO4 | 20:960467-967173 | 20:960600-967174 | A5ss | 3 | 3 | Ex.TSS.ENST00000400634 | 25.69 | 1.61E-02 | 5.21E-02 |
| INPPL1 | 11:72228499-72228726 | 11:72228647-72228727 | A3ss | 3 | 3 | Ex.ENST00000298229 | 25.63 | 3.61E-02 | 5.22E-02 |
| SNHG14 | 15:25054206-25055230 | 15:25054867-25055078 | IR | 3 | 3 | ENST00000660717 | 25.53 | 1.53E-02 | 2.55E-02 |
| CENPK | 5:65554899-65561496 | 5:65554899-65554946 | A3ss | 3 | 3 | ENST00000510354 | 25.39 | 5.86E-03 | 4.18E-02 |
| NEIL1 | 15:75352602-75354492 | 15:75354278-75354430 | IR | 3 | 3 | ENST00000561643 | 25.34 | 4.49E-02 | 7.54E-02 |
| STOX2 | 4:184011424-184017088 | 4:184011587-184011635 | SES | 3 | 3 | ENST00000506529 | 25.24 | 3.18E-02 | 1.79E-01 |
| THUMPD3-AS1 | 3:9390014-9391826 | 3:9390498-9390884 | IR | 3 | 3 | ENST00000518437 | 25.24 | 5.29E-03 | 3.08E-02 |
| STAG2 | X:124090765-124094017 | X:124090854-124090964 | SES | 3 | 3 | ENST00000218089 | 25.22 | 4.89E-05 | 1.91E-03 |
| PI4KAP2 | 22:21476161-21477789 | 22:21476644-21476764 | SES | 3 | 3 | ENST00000360806 | 25.18 | 1.21E-02 | 4.02E-02 |
| PRKCSH | 19:11447619-11447722 | 19:11447693-11447722 | A3ss | 3 | 3 | ENST00000585540 | 25.18 | 4.78E-04 | 1.04E-03 |
| FBXO21 | 12:117158064-117165484 | 12:117158064-117158084 | A3ss | 3 | 3 | ENST00000330622 | 25.17 | 3.75E-03 | 5.57E-03 |
| ZNF213-AS1 | 16:3132104-3134629 | 16:3132850-3133194 | SES | 3 | 3 | ENST00000571449 | 25.13 | 2.51E-02 | 4.72E-02 |
| KIAA0586 | 14:58458546-58460985 | 14:58459843-58459909 | SES | 3 | 3 | ENST00000651937 | 25.03 | 2.22E-03 | 3.65E-03 |
| DET1 | 15:88530623-88531473 | 15:88530977-88531296 | IR (overlapping region) | 3 | 3 | ENST00000557837 | 25.02 | 1.17E-02 | 2.02E-02 |
| UBXN2A | 2:23984832-23999671 | 2:23999655-23999672 | A3ss | 3 | 3 | Ex.TSS.ENST00000309033 | 24.9 | 5.42E-04 | 2.02E-03 |
| GOLGA2P7 | 15:84203021-84204425 | 15:84204177-84204331 | SES | 3 | 3 | ENST00000316967 | 24.83 | 9.58E-04 | 1.64E-03 |
| ATF7IP2 | 16:10386123-10428867 | 16:10419581-10419623 | MXS | 3 | 3 | ENST00000396559 | 24.75 | 2.23E-02 | 3.84E-02 |
| SNHG8 | 4:118278793-118279388 | 4:118278944-118279137 | SES | 3 | 3 | ENST00000602819 | 24.74 | 5.22E-04 | 3.12E-03 |
| TNS2 | 12:53060675-53060988 | 12:53060760-53060848 | IR | 3 | 2 | ENST00000551583 | 24.72 | 1.89E-02 | 2.89E-02 |
| TRIM37 | 17:58999460-59012327 | 17:59001598-59001741 | SES | 3 | 3 | ENST00000585287 | 24.66 | 5.58E-03 | 1.15E-02 |
| TTI1 | 20:38002777-38006196 | 20:38002777-38002793 | A3ss | 3 | 3 | Ex.ENST00000373447 | 24.65 | 4.70E-03 | 1.42E-02 |
| DNASE1 | 16:3656754-3657186 | 16:3656999-3657111 | SES | 3 | 3 | ENST00000246949 | 24.59 | 2.07E-02 | 3.92E-02 |
| RTN2 | 19:45489554-45494165 | 19:45493160-45493378 | SES | 3 | 3 | ENST00000245923 | 24.46 | 4.45E-03 | 1.03E-02 |
| CHD1L | 1:147242831-147265931 | 1:147255813-147255927 | MES | 3 | 3 | ENST00000369258 | 24.43 | 3.62E-03 | 8.88E-03 |
| PPOX | 1:161167235-161167370 | 1:161167339-161167370 | A3ss | 3 | 3 | ENST00000495483 | 24.36 | 1.04E-03 | 2.64E-03 |
| ECT2 | 3:172761684-172762415 | 3:172762398-172762416 | A3ss | 3 | 3 | Ex.ENST00000232458 | 24.3 | 2.21E-02 | 9.12E-02 |
| SNHG8 | 4:118278793-118279388 | 4:118278793-118279137 | A5ss | 3 | 3 | ENST00000652022 | 24.26 | 3.71E-04 | 2.22E-03 |
| GATD1 | 11:770399-771332 | 11:770679-770903 | MES | 3 | 3 | ENST00000354286 | 24.14 | 6.44E-03 | 9.41E-03 |
| VEZT | 12:95266633-95270050 | 12:95270039-95270050 | A3ss | 3 | 3 | ENST00000397792 | 24.1 | 6.58E-03 | 1.03E-02 |
| TM2D1 | 1:61683547-61694696 | 1:61686792-61686921 | SES | 3 | 3 | ENST00000488410 | 24.04 | 9.74E-03 | 2.80E-02 |
| SPART | 13:36335833-36346581 | 13:36335833-36336358 | A3ss | 3 | 3 | Ex.TSS.ENST00000451493 | 23.95 | 5.87E-04 | 9.25E-04 |
| SLTM | 15:58917000-58932355 | 15:58917000-58917020 | A3ss | 3 | 3 | Ex.ENST00000249736 | 23.94 | 6.89E-05 | 1.17E-04 |
| DPH7 | 9:137576403-137578509 | 9:137577470-137577603 | SES | 2 | 2 | ENST00000481839 | 23.87 | 2.90E-02 | 7.22E-01 |
| SLC25A29 | 14:100293378-100306198 | 14:100295626-100295697 | MES | 3 | 3 | ENST00000554224 | 23.85 | 1.36E-03 | 2.18E-03 |
| NOS1AP | 1:162287437-162300647 | 1:162300633-162300647 | A3ss | 3 | 3 | ENST00000361897 | 23.71 | 3.70E-03 | 9.41E-03 |
| THTPA | 14:23556344-23558694 | 14:23556743-23557304 | SES | 3 | 3 | ENST00000404535 | 23.58 | 5.68E-03 | 9.25E-03 |
| IQGAP3 | 1:156562666-156563133 | 1:156562666-156562674 | A3ss | 3 | 3 | Ex.ENST00000361170 | 23.57 | 8.81E-03 | 2.24E-02 |
| PPP1R16A | 8:144496726-144497453 | 8:144497294-144497409 | IR (overlapping region) | 3 | 3 | ENST00000435887 | 23.53 | 4.60E-02 | 6.96E-01 |
| ARAF | X:47561252-47562908 | X:47561252-47562762 | A5ss | 3 | 3 | Ex.TSS.ENST00000377039 | 23.46 | 6.25E-04 | 9.13E-02 |
| DNASE1 | 16:3656754-3657186 | 16:3656999-3657186 | A3ss | 3 | 3 | ENST00000570769 | 23.44 | 2.20E-02 | 4.16E-02 |
| ARL17A | 17:46554044-46557630 | 17:46555207-46555284 | IR (overlapping region) | 3 | 3 | ENST00000336125 | 23.42 | 4.44E-02 | 9.09E-02 |
| HELLS | 10:94545908-94546376 | 10:94546361-94546377 | A3ss | 3 | 3 | Ex.TSS.ENST00000419900 | 23.37 | 8.42E-05 | 1.13E-04 |
| GLB1L3 | 11:134293210-134312348 | 11:134311064-134311170 | MES | 3 | 3 | ENST00000431683 | 23.31 | 3.47E-02 | 4.71E-02 |
| SPIN2B | X:57119682-57120634 | X:57120102-57120404 | IR | 3 | 3 | ENST00000275988 | 23.28 | 9.53E-03 | 1.00E+00 |
| ZNF90 | 19:20105317-20126751 | 19:20125006-20126166 | MES | 3 | 3 | ENST00000469078 | 23.22 | 1.10E-02 | 2.49E-02 |
| GAS8 | 16:90027723-90031298 | 16:90031176-90031298 | A3ss | 3 | 2 | ENST00000566266 | 23.2 | 3.18E-02 | 6.21E-02 |
| GAS8 | 16:90027723-90031189 | 16:90031176-90031189 | TSSlA3ss | 3 | 2 | TSS.ENST00000563980 | 23.2 | 3.18E-02 | 6.21E-02 |
| TMEM214 | 2:27037703-27038145 | 2:27037790-27037892 | SES | 3 | 3 | ENST00000435172 | 23 | 1.24E-05 | 4.66E-05 |
| EDA | X:70030521-70035357 | X:70033398-70033528 | SES | 3 | 3 | ENST00000374552 | 22.93 | 3.02E-02 | 1.00E+00 |
| TP53TG1 | 7:87341685-87345304 | 7:87345043-87345233 | SES | 3 | 3 | ENST00000416560 | 22.83 | 4.17E-02 | 4.93E-01 |
| LRTOMT | 11:72080763-72088296 | 11:72080763-72080885 | TSSlA5ss | 2 | 3 | TSS.ENST00000289488 | 22.82 | 2.19E-02 | 3.19E-02 |
| GPR161 | 1:168114550-168137164 | 1:168136739-168137165 | A5ss | 3 | 3 | Ex.TSS.ENST00000367835 | 22.74 | 3.01E-03 | 7.69E-03 |
| UBR4 | 1:19153940-19154917 | 1:19153940-19153954 | A3ss | 3 | 3 | Ex.ENST00000375254 | 22.69 | 1.00E-03 | 2.57E-03 |
| AC009533.1 | 12:9298868-9299264 | 12:9299260-9299264 | A3ss | 3 | 3 | Ex.ENST00000539757 | 22.68 | 9.56E-03 | 1.50E-02 |
| ZNF567 | 19:36689498-36712385 | 19:36694802-36694876 | SES | 3 | 3 | ENST00000536254 | 22.65 | 1.29E-03 | 2.96E-03 |
| AC022140.1 | 5:25422856-25445819 | 5:25422856-25435273 | A3ss | 3 | 3 | Ex.TSS.ENST00000507887 | 22.59 | 5.18E-04 | 3.66E-03 |
| PPFIA1 | 11:70272437-70324401 | 11:70272437-70314314 | A5ss | 3 | 3 | Ex.ENST00000253925 | 22.57 | 2.18E-02 | 3.16E-02 |
| WDR53 | 3:196561492-196567190 | 3:196566890-196567190 | TSSlA5ss | 3 | 3 | TSS.ENST00000425888 | 22.5 | 3.49E-02 | 1.47E-01 |
| ZNF793 | 19:37533308-37536530 | 19:37533404-37533485 | IR | 3 | 2 | ENST00000587986 | 22.5 | 1.22E-03 | 2.81E-03 |
| EXO5 | 1:40508909-40514514 | 1:40509733-40509790 | SES | 3 | 3 | ENST00000443729 | 22.39 | 1.26E-02 | 3.56E-02 |
| ZFX | X:24149795-24172714 | X:24152720-24152830 | MXS | 3 | 3 | ENST00000419690 | 22.38 | 2.28E-02 | 1.00E+00 |
| LTBP4 | 19:40614447-40617099 | 19:40616889-40617099 | A3ss | 3 | 3 | ENST00000617753 | 22.37 | 4.29E-02 | 9.83E-02 |
| ZNF562 | 19:9656654-9660719 | 19:9659379-9659467 | MXS | 3 | 3 | ENST00000585688 | 22.37 | 4.09E-02 | 9.83E-02 |
| TRIM37 | 17:59070948-59075646 | 17:59070948-59070968 | A3ss | 3 | 3 | Ex.ENST00000262294 | 22.33 | 2.02E-03 | 4.17E-03 |
| BMS1P10 | 9:63380010-63385078 | 9:63380010-63380011 | A3ss | 3 | 3 | Ex.TSS.ENST00000647845 | 22.32 | 4.59E-02 | 9.38E-01 |
| SVIL | 10:29686652-29735754 | 10:29735751-29735754 | A5ss | 3 | 3 | Ex.TSS.ENST00000375400 | 22.27 | 1.36E-02 | 1.80E-02 |
| SLC25A29 | 14:100293378-100306198 | 14:100295626-100295993 | MES | 3 | 3 | ENST00000556868 | 22.22 | 8.47E-05 | 1.36E-04 |
| CAND2 | 3:12803632-12810058 | 3:12808210-12808333 | MES | 3 | 2 | ENST00000456430 | 22.17 | 4.39E-02 | 1.76E-01 |
| IMMT | 2:86171337-86173649 | 2:86171337-86171345 | A3ss | 3 | 3 | Ex.ENST00000410111 | 22.16 | 1.41E-02 | 5.55E-02 |
| IMMT | 2:86171313-86173649 | 2:86171313-86171345 | A3ss | 3 | 3 | ENST00000410111 | 22.16 | 1.41E-02 | 5.55E-02 |
| IMMT | 2:86171310-86173649 | 2:86171310-86171345 | A3ss | 3 | 3 | Ex.ENST00000410111 | 22.16 | 1.41E-02 | 5.55E-02 |
| ASB16-AS1 | 17:44176643-44181622 | 17:44176643-44177816 | TSSlA3ss | 2 | 3 | TSS.ENST00000585457 | 22.1 | 4.75E-02 | 9.66E-02 |
| FIP1L1 | 4:53399840-53425871 | 4:53414615-53414722 | SES | 3 | 3 | ENST00000337488 | 22.1 | 3.91E-03 | 2.48E-02 |
| FLYWCH1 | 16:2948688-2951208 | 16:2948714-2948843 | IR (overlapping region) | 3 | 3 | ENST00000253928 | 22.08 | 4.08E-02 | 7.65E-02 |
| R3HDM1 | 2:135605017-135621493 | 2:135616668-135616757 | SES | 3 | 3 | ENST00000456040 | 22.01 | 3.12E-02 | 1.10E-01 |
| TTC14 | 3:180609630-180611130 | 3:180610004-180610705 | IR (overlapping region) | 3 | 3 | ENST00000296015 | 22 | 3.04E-02 | 1.26E-01 |
| SKIV2L | 6:31969280-31969751 | 6:31969317-31969514 | IR | 3 | 3 | ENST00000465703 | 21.93 | 3.83E-03 | 3.18E-02 |
| FUT1 | 19:48752612-48753122 | 19:48752726-48752891 | IR | 2 | 3 | ENST00000601931 | 21.91 | 3.66E-02 | 8.60E-02 |
| SNHG14 | 15:24982438-25019026 | 15:25002725-25002888 | MES | 3 | 3 | ENST00000551631 | 21.9 | 1.84E-03 | 3.07E-03 |
| MTHFR | 1:11785723-11790898 | 1:11788048-11788148 | IR (overlapping region) | 3 | 3 | ENST00000376590 | 21.89 | 3.48E-02 | 8.45E-02 |
| DHX35 | 20:39006357-39010279 | 20:39010260-39010280 | A3ss | 3 | 3 | Ex.ENST00000252011 | 21.86 | 2.32E-02 | 7.02E-02 |
| PHKB | 16:47461427-47497398 | 16:47463882-47463993 | MXS | 3 | 3 | ENST00000566044 | 21.85 | 4.76E-03 | 9.03E-03 |
| TM2D3 | 15:101650162-101652270 | 15:101651696-101652270 | TSSlA5ss | 3 | 3 | TSS.ENST00000560013 | 21.85 | 2.48E-02 | 4.14E-02 |

| | | | | | | | | | |
|---|---|---|---|---|---|---|---|---|---|
| SPATA5L1 | 15:45415780-45417268 | 15:45415780-45415809 | A5ss | 3 | 3 | ENST00000531970 | 21.48 | 1.08E-03 | 1.81E-03 |
| SUZ12P1 | 17:30709812-30759490 | 17:30735033-30735097 | MES | 3 | 3 | ENST00000579526 | 21.48 | 1.09E-02 | 2.17E-02 |
| HTT | 4:3199940-3204006 | 4:3202977-3203066 | SES | 3 | 3 | ENST00000502820 | 21.43 | 3.82E-02 | 2.37E-01 |
| TM2D3 | 15:101650162-101652270 | 15:101651696-101651773 | SES | 3 | 3 | ENST00000333202 | 21.35 | 3.30E-02 | 5.50E-02 |
| FASTKD1 | 2:169537341-169540050 | 2:169538013-169538141 | SES | 3 | 3 | ENST00000453153 | 21.27 | 2.19E-03 | 7.79E-03 |
| DKK3 | 11:11967099-11968394 | 11:11967099-11967119 | A3ss | 3 | 3 | Ex.ENST00000326932 | 21.23 | 5.11E-04 | 6.90E-04 |
| CNOT2-DT | 12:70243066-70243360 | 12:70243121-70243189 | IR (overlapping region) | 3 | 3 | ENST00000549651 | 21.16 | 2.78E-02 | 4.33E-02 |
| DDX12P | 12:9437490-9438010 | 12:9437490-9437891 | A3ss | 3 | 3 | Ex.ENST00000432996 | 21.15 | 1.08E-02 | 1.70E-02 |
| SNED1 | 2:241091638-241095568 | 2:241094430-241094596 | IR (overlapping region) | 3 | 3 | ENST00000310397 | 21.15 | 2.05E-02 | 7.62E-02 |
| VEZT | 12:95256623-95257153 | 12:95257150-95257153 | A3ss | 3 | 3 | ENST00000356859 | 21.12 | 3.74E-02 | 5.85E-02 |
| PHF20L1 | 8:132798861-132803818 | 8:132799095-132799172 | SES | 3 | 3 | ENST00000315808 | 21.05 | 2.83E-03 | 4.21E-02 |
| SLC12A5 | 20:46021887-46023368 | 20:46022843-46023071 | SES | 3 | 3 | ENST00000626701 | 21.01 | 4.92E-02 | 1.58E-01 |
| ARID4A | 14:58365623-58366882 | 14:58366024-58366068 | SES | 2 | 2 | ENST00000395168 | 20.99 | 3.44E-02 | 5.65E-02 |
| FMNL1 | 17:45240476-45242146 | 17:45240605-45241110 | IR (overlapping region) | 3 | 2 | ENST00000587856 | 20.99 | 6.97E-03 | 1.42E-02 |
| ABHD18 | 4:127984424-128008919 | 4:127989721-127989821 | MXS | 3 | 3 | ENST00000398965 | 20.94 | 1.84E-03 | 1.10E-02 |
| LIPE | 19:42405562-42407173 | 19:42406161-42406388 | SES | 3 | 3 | ENST00000244289 | 20.91 | 3.52E-02 | 8.15E-02 |
| DTWD2 | 5:118944650-118988293 | 5:118952122-118988294 | A5ss | 3 | 3 | Ex.TSS.ENST00000506980 | 20.9 | 3.93E-02 | 2.59E-01 |
| DTWD2 | 5:118944650-118988129 | 5:118952122-118988130 | A5ss | 3 | 3 | Ex.TSS.ENST00000304058 | 20.9 | 3.93E-02 | 2.59E-01 |
| C11orf49 | 11:47161191-47161463 | 11:47161446-47161463 | TSSIA3ss | 3 | 3 | TSS.ENST00000378615 | 20.89 | 1.27E-02 | 1.74E-02 |
| SEPTIN8 | 5:132758825-132760801 | 5:132758825-132760013 | A3ss | 3 | 3 | Ex.TSS.ENST00000296873 | 20.83 | 1.28E-02 | 8.60E-02 |
| BIRC6 | 2:32471125-32473111 | 2:32471125-32471151 | A5ss | 3 | 3 | ENST00000648282 | 20.82 | 1.75E-03 | 6.61E-03 |
| PLEKHB1 | 11:73661466-73662819 | 11:73662423-73662510 | IR | 3 | 3 | ENST00000227214 | 20.8 | 1.11E-02 | 1.62E-02 |
| SLC12A5 | 20:46021887-46022929 | 20:46022843-46022929 | TSSIA3ss | 3 | 3 | TSS.ENST00000626701 | 20.78 | 2.99E-02 | 9.57E-02 |
| SLC12A5 | 20:46021887-46022889 | 20:46022843-46022889 | A3ss | 3 | 3 | Ex.TSS.ENST00000626701 | 20.78 | 2.99E-02 | 9.57E-02 |
| APOBEC3D | 22:39023015-39031693 | 22:39025070-39025349 | MES | 3 | 3 | ENST00000216099 | 20.75 | 3.00E-02 | 1.03E-01 |
| COQ8B | 19:40714636-40716861 | 19:40716587-40716861 | TSSIA5ss | 3 | 3 | TSS.ENST00000324464 | 20.73 | 2.66E-02 | 6.14E-02 |
| LINC02610 | 2:238225138-238227798 | 2:238225500-238227683 | IR | 2 | 2 | ENST00000659231 | 20.66 | 9.91E-03 | 3.67E-02 |
| PI4KAP2 | 22:21476161-21477789 | 22:21476161-21476764 | A3ss | 3 | 3 | ENST00000462560 | 20.62 | 8.46E-03 | 2.80E-02 |
| SNHG32 | 6:31835452-31837235 | 6:31836424-31836517 | SES | 3 | 3 | ENST00000375638 | 20.61 | 2.48E-03 | 2.06E-02 |
| MAPK12 | 22:50245737-50245909 | 22:50245737-50245812 | A3ss | 3 | 3 | Ex.ENST00000497036 | 20.57 | 6.86E-03 | 2.38E-02 |
| DNM1 | 9:128253101-128255246 | 9:128253138-128253784 | IR | 3 | 3 | ENST00000627543 | 20.55 | 4.99E-03 | 1.11E-01 |
| PRPF38A | 1:52414648-52414761 | 1:52414741-52414762 | A3ss | 3 | 3 | Ex.ENST00000257181 | 20.54 | 1.15E-03 | 3.29E-03 |
| SNHG11 | 20:38446654-38448730 | 20:38446968-38447465 | IR | 3 | 3 | ENST00000400436 | 20.54 | 3.20E-02 | 9.66E-02 |
| MACF1 | 1:39463687-39468614 | 1:39465095-39465112 | SES | 3 | 3 | ENST00000289893 | 20.51 | 1.51E-05 | 4.26E-05 |
| CLCC1 | 1:108950449-108963360 | 1:108962309-108962469 | SES | 3 | 3 | ENST00000369976 | 20.48 | 1.93E-02 | 4.68E-02 |
| AC105285.1 | 4:173164345-173166299 | 4:173165133-173165991 | IR | 2 | 2 | ENST00000510523 | 20.47 | 1.95E-02 | 1.18E-01 |
| RBM41 | X:107088840-107115351 | X:107113397-107113468 | SES | 3 | 3 | ENST00000495517 | 20.46 | 3.84E-02 | 7.91E-01 |
| GAS2L3 | 12:100573786-100591735 | 12:100574347-100591736 | A3ss | 3 | 3 | Ex.TSS.ENST00000547754 | 20.39 | 2.00E-02 | 2.94E-02 |
| PLEC | 8:143922396-143926783 | 8:143922504-143925884 | SES | 3 | 3 | ENST00000322810 | 20.35 | 4.18E-03 | 6.39E-02 |
| ZNF529 | 19:36548323-36554669 | 19:36551877-36551977 | SES | 3 | 3 | ENST00000334116 | 20.34 | 2.05E-03 | 4.68E-03 |
| FAM49B | 8:129871497-129903311 | 8:129879389-129879471 | MXS | 3 | 3 | ENST00000517654 | 20.29 | 2.31E-05 | 3.44E-04 |
| ORAI2 | 7:102433662-102438943 | 7:102436202-102436333 | SES | 3 | 3 | ENST00000356387 | 20.22 | 8.12E-03 | 7.38E-02 |
| ERCC3 | 2:127289502-127289688 | 2:127289502-127289519 | A3ss | 3 | 3 | ENST00000456257 | 20.2 | 4.54E-03 | 1.59E-02 |
| ZMYM4 | 1:35405469-35408007 | 1:35407987-35408008 | A3ss | 3 | 3 | Ex.ENST00000314607 | 20.18 | 4.41E-04 | 1.24E-03 |
| CTDSPL2 | 15:44502049-44514597 | 15:44502049-44503850 | A5ss | 3 | 3 | Ex.ENST00000560620 | 20.11 | 9.95E-04 | 1.68E-03 |
| CTDSPL2 | 15:44499814-44514597 | 15:44499814-44503850 | A5ss | 3 | 3 | Ex.ENST00000260327 | 20.11 | 9.95E-04 | 1.68E-03 |
| ZNF518A | 10:96133649-96155923 | 10:96155326-96155398 | MES | 2 | 3 | ENST00000488700 | 20.11 | 2.86E-02 | 3.77E-02 |
| GTF2H2 | 5:71037547-71041521 | 5:71037547-71038481 | A3ss | 3 | 3 | Ex.ENST00000274400 | 20.1 | 1.97E-02 | 1.42E-01 |
| DCAF10 | 9:37854983-37860047 | 9:37857241-37857351 | SES | 3 | 3 | ENST00000242323 | 20.09 | 2.85E-02 | 5.89E-01 |
| BCL9L | 11:118908656-118925237 | 11:118918826-118918879 | MES | 3 | 3 | ENST00000532899 | 20.08 | 1.12E-02 | 1.51E-02 |
| ESPL1 | 12:53272858-53274816 | 12:53274790-53274817 | A3ss | 3 | 3 | Ex.ENST00000257934 | 20.01 | 7.66E-03 | 1.17E-02 |
| HYI | 1:43452014-43452204 | 1:43452014-43452031 | A3ss | 3 | 3 | Ex.ENST00000372425 | 20.01 | 2.49E-02 | 7.05E-02 |
| DNAJC5 | 20:63931023-63931464 | 20:63931109-63931182 | SES | 3 | 3 | ENST00000470551 | 19.98 | 1.80E-02 | 5.78E-02 |
| AC119673.3 | 1:205891310-205894466 | 1:205894444-205894467 | A3ss | 3 | 3 | Ex.TSS.ENST00000652821 | 19.9 | 3.87E-05 | 1.00E-04 |
| SCAP | 3:47422537-47435007 | 3:47427447-47427667 | MES | 3 | 3 | ENST00000265565 | 19.84 | 4.26E-03 | 1.89E-02 |
| PRDM2 | 1:13731118-13742004 | 1:13732779-13732882 | SES | 3 | 3 | ENST00000235372 | 19.76 | 2.32E-02 | 5.70E-02 |
| ETV1 | 7:13989140-13989562 | 7:13989268-13989464 | SES | 3 | 3 | ENST00000430479 | 19.75 | 1.46E-02 | 1.43E-01 |
| NT5C3A | 7:33026916-33062567 | 7:33035934-33035988 | SES | 3 | 3 | ENST00000405342 | 19.71 | 1.77E-02 | 1.85E-01 |
| FOXN2 | 2:48314815-48346200 | 2:48328561-48328702 | SES | 3 | 3 | ENST00000340553 | 19.7 | 6.10E-03 | 2.32E-02 |
| PSME3IP1 | 16:57173870-57185544 | 16:57174169-57174286 | SES | 3 | 3 | ENST00000567044 | 19.66 | 4.63E-02 | 8.85E-02 |
| TMEM125 | 1:43271165-43272183 | 1:43271165-43271253 | A5ss | 3 | 3 | Ex.TSS.ENST00000456751 | 19.63 | 3.42E-02 | 9.70E-02 |
| TMEM125 | 1:43270794-43272183 | 1:43270794-43271253 | A5ss | 3 | 3 | Ex.ENST00000432792 | 19.63 | 3.42E-02 | 9.70E-02 |
| RFC5 | 12:118017690-118018053 | 12:118017816-118017994 | IR | 2 | 2 | ENST00000392542 | 19.6 | 2.10E-02 | 3.12E-02 |
| MIB2 | 1:1628273-1628722 | 1:1628400-1628488 | IR | 3 | 3 | ENST00000507229 | 19.59 | 3.55E-02 | 9.06E-02 |
| UQCC1 | 20:35346424-35347163 | 20:35346424-35346461 | A3ss | 3 | 3 | Ex.TSS.ENST00000491125 | 19.55 | 1.85E-04 | 5.56E-04 |
| UQCC1 | 20:35314766-35347163 | 20:35314766-35346461 | A3ss | 3 | 3 | Ex.ENST00000349714 | 19.55 | 1.85E-04 | 5.56E-04 |
| THTPA | 14:23557040-23558694 | 14:23557040-23557304 | A5ss | 3 | 3 | ENST00000288014 | 19.52 | 5.21E-03 | 8.49E-03 |
| THTPA | 14:23557035-23558694 | 14:23557035-23557304 | A5ss | 3 | 3 | ENST00000288014 | 19.52 | 5.21E-03 | 8.49E-03 |
| THTPA | 14:23556344-23558694 | 14:23556344-23557304 | A5ss | 3 | 3 | ENST00000288014 | 19.52 | 5.21E-03 | 8.49E-03 |
| DOCK6 | 19:11229391-11233202 | 19:11229391-11229852 | A3ss | 3 | 3 | Ex.ENST00000590680 | 19.5 | 2.72E-02 | 5.95E-02 |
| DOCK6 | 19:11229036-11233202 | 19:11229036-11229852 | A3ss | 3 | 3 | Ex.ENST00000294618 | 19.5 | 2.72E-02 | 5.95E-02 |
| TRPT1 | 11:64225581-64226049 | 11:64225786-64226049 | TSSIA5ss | 3 | 3 | TSS.ENST00000539595 | 19.47 | 3.65E-03 | 5.21E-03 |
| SOCS4 | 14:55027472-55042951 | 14:55031863-55031991 | SES | 3 | 3 | ENST00000555846 | 19.43 | 3.45E-03 | 5.66E-03 |
| KHNYN | 14:24429992-24430713 | 14:24429992-24430119 | TSSIA5ss | 3 | 3 | TSS.ENST00000553935 | 19.42 | 5.26E-03 | 8.60E-03 |
| DHX29 | 5:55270488-55270577 | 5:55270488-55270505 | A3ss | 3 | 3 | Ex.ENST00000251636 | 19.38 | 1.12E-03 | 7.99E-03 |
| STRN3 | 14:30913658-30935162 | 14:30918966-30919106 | MES | 3 | 3 | ENST00000357479 | 19.3 | 1.85E-02 | 3.02E-02 |
| INTS9 | 8:28768323-28769888 | 8:28768323-28768376 | TSSIA3ss | 3 | 3 | TSS.ENST00000517383 | 19.26 | 5.76E-03 | 7.34E-02 |
| LTBP4 | 19:40614447-40619346 | 19:40616889-40617020 | MES | 3 | 3 | ENST00000204005 | 19.24 | 4.93E-02 | 1.14E-01 |
| AKIP1 | 11:8911672-8914825 | 11:8912453-8912533 | SES | 3 | 3 | ENST00000309357 | 19.21 | 2.83E-02 | 4.16E-02 |
| OPA1 | 3:193618937-193631611 | 3:193626092-193626202 | MXS | 3 | 3 | ENST00000361150 | 19.13 | 2.52E-04 | 1.06E-03 |
| SOCS4 | 14:55027472-55031888 | 14:55031863-55031888 | TSSIA3ss | 3 | 3 | TSS.ENST00000555846 | 19.07 | 1.11E-02 | 1.83E-02 |
| ARHGEF4 | 2:130916965-130917498 | 2:130917062-130917396 | IR | 3 | 3 | ENST00000438985 | 19.05 | 4.40E-02 | 1.55E-01 |
| WDR24 | 16:689160-690398 | 16:690086-690369 | IR (overlapping region) | 3 | 3 | ENST00000293883 | 19.01 | 2.14E-03 | 4.14E-03 |
| PCBP1-AS1 | 2:70085625-70086123 | 2:70085816-70086123 | TSSIA5ss | 3 | 3 | TSS.ENST00000429599 | 18.91 | 3.73E-02 | 1.44E-01 |
| NCOA3 | 20:47649098-47650981 | 20:47649098-47649109 | A5ss | 2 | 3 | ENST00000371998 | 18.89 | 4.22E-02 | 1.35E-01 |
| RPAIN | 17:5422830-5432541 | 17:5425971-5426082 | MES | 3 | 3 | ENST00000381208 | 18.83 | 8.94E-05 | 1.84E-04 |
| UPRT | X:75293515-75296341 | X:75294563-75294620 | SES | 3 | 3 | ENST00000462237 | 18.81 | 1.63E-02 | 1.00E+00 |
| EXO5 | 1:40508909-40514514 | 1:40508909-40509790 | A5ss | 3 | 3 | ENST00000372703 | 18.78 | 1.88E-02 | 5.33E-02 |
| EXO5 | 1:40508890-40514514 | 1:40508890-40509790 | A5ss | 3 | 3 | ENST00000372703 | 18.78 | 1.88E-02 | 5.33E-02 |
| EXO5 | 1:40508878-40514514 | 1:40508878-40509790 | A5ss | 3 | 3 | ENST00000372703 | 18.78 | 1.88E-02 | 5.33E-02 |
| UQCC1 | 20:35344110-35347272 | 20:35346748-35347163 | IR | 3 | 3 | ENST00000397554 | 18.76 | 1.60E-03 | 4.81E-03 |
| HTT | 4:3140657-3142765 | 4:3142745-3142765 | A3ss | 3 | 3 | ENST00000649131 | 18.72 | 1.39E-02 | 8.62E-02 |
| AURKA | 20:56388203-56392167 | 20:56388687-56388784 | MES | 3 | 3 | ENST00000420474 | 18.7 | 8.18E-03 | 2.63E-02 |
| CHD3 | 17:7893936-7894126 | 17:7894115-7894126 | A3ss | 3 | 3 | ENST00000330494 | 18.7 | 2.03E-02 | 4.31E-02 |

| | | | | | | | | | |
|---|---|---|---|---|---|---|---|---|---|
| SLC29A4 | 7:5291822-5294864 | 7:5294860-5294864 | A3ss | 3 | 3 | ENST00000297195 | 18.67 | 2.12E-02 | 2.29E-01 |
| BRD9 | 5:865582-870472 | 5:869245-869404 | MES | 3 | 3 | ENST00000490814 | 18.66 | 3.42E-02 | 2.61E-01 |
| KIAA1217 | 10:24528120-24531832 | 10:24531830-24531832 | A3ss | 3 | 3 | ENST00000307544 | 18.65 | 2.31E-02 | 3.06E-02 |
| ADGRL2 | 1:81800513-81836884 | 1:81800513-81802229 | A5ss | 3 | 3 | Ex.TSS.ENST00000319517 | 18.6 | 9.28E-04 | 2.68E-03 |
| ATP2B4 | 1:203739546-203740319 | 1:203739645-203739764 | IR | 3 | 3 | ENST00000341360 | 18.6 | 2.94E-02 | 7.63E-02 |
| NFASC | 1:204944407-204952010 | 1:204950557-204950574 | SES | 3 | 3 | ENST00000339876 | 18.6 | 3.22E-02 | 8.36E-02 |
| MFSD14C | 9:96942753-96949554 | 9:96942753-96943154 | A3ss | 3 | 3 | Ex.TSS.ENST00000506067 | 18.56 | 2.94E-02 | 1.00E+00 |
| MFSD14C | 9:96918827-96949554 | 9:96918827-96943154 | A3ss | 3 | 3 | Ex.TSS.ENST00000637076 | 18.56 | 2.94E-02 | 1.00E+00 |
| WNK2 | 9:93298068-93308327 | 9:93299070-93299261 | MXS | 3 | 3 | ENST00000453718 | 18.55 | 8.48E-03 | 2.87E-01 |
| B4GALT7 | 5:177604542-177607301 | 5:177607283-177607301 | A3ss | 3 | 3 | ENST00000505433 | 18.54 | 2.01E-03 | 1.42E-02 |
| HDHD2 | 18:47137297-47150377 | 18:47137297-47137333 | A3ss | 3 | 3 | Ex.TSS.ENST00000586546 | 18.54 | 9.27E-05 | 1.99E-04 |
| HDHD2 | 18:47136450-47150377 | 18:47136450-47137333 | A3ss | 3 | 3 | Ex.TSS.ENST00000300605 | 18.54 | 9.27E-05 | 1.99E-04 |
| EHMT1 | 9:137728530-137743391 | 9:137743371-137743391 | A3ss | 3 | 3 | ENST00000371394 | 18.53 | 4.35E-04 | 1.17E-02 |
| TAF1C | 16:84184900-84186900 | 16:84184900-84184989 | TSSIA3ss | 3 | 3 | TSS.ENST00000564345 | 18.52 | 1.91E-03 | 3.71E-03 |
| C12orf73 | 12:103953535-103956630 | 12:103954875-103954968 | MES | 2 | 2 | ENST00000552460 | 18.51 | 4.65E-02 | 6.62E-02 |
| PHF20L1 | 8:132798861-132799094 | 8:132799083-132799094 | A3ss | 3 | 3 | ENST00000395376 | 18.48 | 7.60E-03 | 1.13E-01 |
| THUMPD3-AS1 | 3:9388800-9390497 | 3:9389797-9390289 | IR | 3 | 3 | ENST00000668063 | 18.47 | 1.65E-02 | 9.59E-02 |
| TBL1X | X:9640361-9654214 | X:9653545-9653689 | SES | 3 | 3 | ENST00000407597 | 18.45 | 9.79E-03 | 1.00E+00 |
| TMEM45A | 3:100492929-100519532 | 3:100519462-100519533 | A3ss | 3 | 3 | Ex.TSS.ENST00000449609 | 18.44 | 6.53E-03 | 2.59E-02 |
| ZNF783 | 7:149267223-149278398 | 7:149278136-149278399 | A3ss | 3 | 3 | Ex.ENST00000378052 | 18.44 | 2.24E-02 | 2.22E-01 |
| TRPT1 | 11:64225581-64226049 | 11:64225786-64225869 | SES | 3 | 3 | ENST00000317459 | 18.4 | 8.47E-03 | 1.21E-02 |
| FOSL1 | 11:65894117-65896808 | 11:65894117-65894121 | A3ss | 3 | 3 | ENST00000312562 | 18.38 | 3.15E-03 | 4.51E-03 |
| LRRCC1 | 8:85107400-85110114 | 8:85109576-85110115 | A3ss | 3 | 3 | Ex.TSS.ENST00000517875 | 18.38 | 5.78E-03 | 7.87E-02 |
| LRRCC1 | 8:85107400-85109594 | 8:85109576-85109595 | A3ss | 3 | 3 | Ex.TSS.ENST00000360375 | 18.38 | 5.78E-03 | 7.87E-02 |
| MRPL2 | 6:43054487-43058064 | 6:43056307-43056445 | MES | 3 | 3 | ENST00000230413 | 18.38 | 4.21E-02 | 3.61E-01 |
| PDLIM7 | 5:177491147-177491806 | 5:177491147-177491420 | TSSIA3ss | 3 | 3 | TSS.ENST00000504318 | 18.36 | 2.27E-02 | 1.60E-01 |
| TRAPPC2 | X:13719983-13734524 | X:13734044-13734185 | MES | 3 | 3 | ENST00000518847 | 18.34 | 3.71E-02 | 1.00E+00 |
| CXorf40A | X:149542067-149545367 | X:149542067-149542085 | A5ss | 3 | 3 | ENST00000393985 | 18.31 | 3.18E-04 | 1.33E-02 |
| PKIG | 20:44531979-44589796 | 20:44582585-44582731 | SES | 3 | 3 | ENST00000372887 | 18.31 | 6.72E-03 | 2.15E-02 |
| ACSF3 | 16:89093993-89102603 | 16:89100662-89101347 | SES | 3 | 3 | ENST00000537290 | 18.18 | 1.19E-02 | 2.31E-02 |
| NEK4 | 3:52746905-52751931 | 3:52749692-52749829 | SES | 3 | 3 | ENST00000233027 | 18.17 | 2.12E-03 | 1.20E-02 |
| SNRPN | 15:24962210-24967931 | 15:24967029-24967240 | SES | 3 | 3 | ENST00000584968 | 18.16 | 3.17E-05 | 5.29E-05 |
| LETMD1 | 12:51049186-51055834 | 12:51053812-51053860 | MXS | 3 | 3 | ENST00000549340 | 18.14 | 3.11E-02 | 4.75E-02 |
| PCNX3 | 11:65620967-65622244 | 11:65622224-65622245 | A3ss | 3 | 3 | Ex.ENST00000355703 | 18.11 | 6.27E-03 | 8.98E-03 |
| SCARF2 | 22:20430909-20431744 | 22:20431018-20431537 | SES | 3 | 3 | ENST00000622235 | 18.06 | 3.47E-02 | 1.15E-01 |
| TRIM16 | 17:15680954-15683035 | 17:15682854-15682964 | MXS | 3 | 3 | ENST00000580388 | 18.06 | 9.45E-03 | 1.86E-02 |
| CYB561D2 | 3:50350985-50351408 | 3:50351343-50351408 | TSSIA3ss | 3 | 3 | TSS.ENST00000232508 | 18.03 | 2.40E-02 | 1.09E-01 |
| SNHG14 | 15:25054206-25055230 | 15:25054867-25054969 | IR | 3 | 3 | ENST00000660717 | 17.96 | 1.83E-02 | 3.06E-02 |
| BCL2L12 | 19:49666800-49668850 | 19:49667019-49667161 | SES | 3 | 3 | ENST00000246784 | 17.94 | 1.51E-04 | 3.55E-04 |
| DERL3 | 22:23834933-23837063 | 22:23834933-23834979 | A3ss | 3 | 3 | Ex.TSS.ENST00000406855 | 17.91 | 1.80E-02 | 6.00E-02 |
| PUS7 | 7:105508521-105522051 | 7:105508521-105508544 | TSSIA3ss | 3 | 3 | TSS.ENST00000469408 | 17.81 | 1.07E-02 | 9.91E-02 |
| SHLD2 | 10:87170975-87180074 | 10:87175889-87176095 | SES | 3 | 3 | ENST00000298786 | 17.75 | 2.11E-02 | 2.82E-02 |
| RPSA | 3:39406765-39407620 | 3:39407603-39407620 | TSSIA3ss | 3 | 3 | TSS.ENST00000458478 | 17.74 | 4.15E-05 | 1.79E-04 |
| ZC3H13 | 13:46045517-46052403 | 13:46051932-46051989 | SES | 3 | 3 | ENST00000428921 | 17.73 | 4.26E-02 | 6.73E-02 |
| KANK1 | 9:504755-676889 | 9:539516-539633 | MES | 3 | 3 | ENST00000674102 | 17.72 | 1.70E-02 | 3.53E-01 |
| TNK1 | 17:7384266-7384754 | 17:7384377-7384483 | IR | 3 | 3 | ENST00000576136 | 17.69 | 3.89E-02 | 8.13E-02 |
| WASHC4 | 12:105120598-105121100 | 12:105121089-105121100 | A3ss | 3 | 3 | ENST00000547404 | 17.67 | 5.69E-03 | 8.40E-03 |
| AXIN1 | 16:289608-293487 | 16:291190-291297 | SES | 3 | 3 | ENST00000262320 | 17.66 | 2.48E-02 | 4.65E-02 |
| FMNL1 | 17:45240476-45242146 | 17:45240626-45241128 | IR (overlapping region) | 3 | 2 | ENST00000587856 | 17.65 | 6.70E-03 | 1.37E-02 |
| RNF2 | 1:185087641-185091578 | 1:185091565-185091579 | A3ss | 3 | 3 | Ex.ENST00000367509 | 17.63 | 3.34E-04 | 8.60E-04 |
| PTGR1 | 9:111563232-111570090 | 9:111563232-111564262 | TSSIA3ss | 3 | 3 | TSS.ENST00000374324 | 17.61 | 2.62E-02 | 3.74E-01 |
| RAB4A | 1:229271371-229295847 | 1:229286468-229295848 | A3ss | 3 | 3 | Ex.TSS.ENST00000473894 | 17.6 | 2.19E-04 | 6.02E-04 |
| RAB4A | 1:229271371-229286485 | 1:229286468-229286486 | A3ss | 3 | 3 | Ex.TSS.ENST00000366690 | 17.6 | 2.19E-04 | 6.02E-04 |
| ARHGAP12 | 10:31852598-31861394 | 10:31854066-31854206 | SES | 3 | 3 | ENST00000344936 | 17.59 | 2.99E-02 | 3.96E-02 |
| ZNF417 | 19:57910115-57916378 | 19:57912060-57912189 | SES | 3 | 3 | ENST00000312026 | 17.58 | 3.62E-02 | 8.66E-02 |
| ZNF839 | 14:102334647-102338815 | 14:102335689-102335838 | SES | 3 | 3 | ENST00000442396 | 17.54 | 4.74E-02 | 7.61E-02 |
| UBAP2L | 1:154268955-154270199 | 1:154269362-154269412 | SES | 3 | 3 | ENST00000428595 | 17.49 | 2.36E-02 | 5.92E-02 |
| ARRB2 | 17:4710745-4715972 | 17:4715013-4715043 | SES | 3 | 3 | ENST00000269260 | 17.48 | 9.83E-05 | 2.02E-04 |
| INSR | 19:7143091-7152725 | 19:7150497-7150532 | SES | 3 | 2 | ENST00000302850 | 17.48 | 1.15E-02 | 2.76E-02 |
| IKBKE | 1:206493120-206493919 | 1:206493266-206493378 | SES | 2 | 3 | ENST00000581977 | 17.42 | 7.48E-05 | 1.95E-04 |
| NINL | 20:25498347-25500839 | 20:25498347-25498361 | A3ss | 3 | 3 | Ex.ENST00000278886 | 17.38 | 2.63E-02 | 7.80E-02 |
| NSMCE2 | 8:125091959-125102050 | 8:125102014-125102050 | TSSIA3ss | 3 | 3 | TSS.ENST00000517532 | 17.37 | 5.11E-03 | 7.54E-02 |
| SLC25A36 | 3:140963228-140970926 | 3:140966934-140967036 | MXS | 3 | 3 | ENST00000512506 | 17.35 | 7.38E-03 | 2.99E-02 |
| MORC2 | 22:30926872-30928027 | 22:30928019-30928027 | TSSIA5ss | 3 | 3 | TSS.ENST00000215862 | 17.34 | 1.60E-02 | 5.42E-02 |
| EMSY | 11:76453389-76458182 | 11:76454749-76454790 | SES | 3 | 3 | ENST00000427574 | 17.33 | 5.84E-03 | 8.52E-03 |
| POLR3G | 5:90475021-90485534 | 5:90485525-90485534 | TSSIA3ss | 3 | 3 | TSS.ENST00000399107 | 17.32 | 1.59E-04 | 1.22E-03 |
| TNRC6A | 16:24794720-24797489 | 16:24795907-24795939 | SES | 3 | 3 | ENST00000315183 | 17.31 | 8.41E-03 | 1.57E-02 |
| BCOR | X:40071160-40072348 | X:40071637-40071690 | SES | 3 | 3 | ENST00000342274 | 17.29 | 7.76E-03 | 9.80E-01 |
| BRD9 | 5:865582-870472 | 5:869245-869394 | MXS | 3 | 3 | ENST00000489816 | 17.25 | 9.75E-03 | 7.45E-02 |
| ESPN | 1:6448803-6451602 | 1:6448893-6449091 | SES | 3 | 3 | ENST00000416731 | 17.25 | 1.14E-02 | 3.27E-02 |
| SNHG14 | 15:25065455-25070119 | 15:25067412-25067531 | SES | 3 | 3 | ENST00000653407 | 17.21 | 3.24E-02 | 5.40E-02 |
| ZMYM1 | 1:35079443-35093913 | 1:35093394-35093441 | SES | 3 | 3 | ENST00000373330 | 17.17 | 2.34E-02 | 6.58E-02 |
| PVT1 | 8:127989292-128096517 | 8:128010318-128010444 | MES | 3 | 3 | ENST00000522875 | 17.15 | 3.99E-02 | 5.90E-01 |
| ZNF814 | 19:57850562-57888766 | 19:57876916-57877042 | MXS | 3 | 3 | ENST00000596604 | 17.12 | 4.83E-02 | 1.16E-01 |
| BRPF3 | 6:36210529-36217916 | 6:36211258-36211560 | MES | 3 | 2 | ENST00000357641 | 17.1 | 3.23E-02 | 2.72E-01 |
| CD82 | 11:44618366-44619048 | 11:44618640-44618723 | SES | 3 | 3 | ENST00000227155 | 17.1 | 1.32E-02 | 1.81E-02 |
| BACE1-AS | 11:117288453-117291771 | 11:117291050-117291711 | IR (overlapping region) | 3 | 3 | ENST00000649580 | 17.06 | 3.51E-02 | 4.72E-02 |
| TMEM116 | 12:111938211-111991757 | 12:111943265-111943369 | MXS | 2 | 2 | ENST00000548283 | 17.04 | 3.78E-02 | 5.59E-02 |
| FKRP | 19:46747295-46748026 | 19:46747295-46747305 | A5ss | 3 | 3 | ENST00000600005 | 16.99 | 2.43E-02 | 5.64E-02 |
| ACD | 16:67658817-67658927 | 16:67658817-67658832 | TSSIA3ss | 3 | 3 | TSS.ENST00000602945 | 16.97 | 9.33E-03 | 1.80E-02 |
| TUBGCP4 | 15:43405202-43409771 | 15:43408089-43408896 | IR (overlapping region) | 3 | 3 | ENST00000564079 | 16.97 | 4.64E-03 | 7.82E-03 |
| FDPS | 1:155308966-155310063 | 1:155310043-155310063 | TSSIA3ss | 3 | 3 | TSS.ENST00000447866 | 16.96 | 5.65E-04 | 1.42E-03 |
| LTBP4 | 19:40614447-40619346 | 19:40617100-40617225 | MES | 3 | 3 | ENST00000204005 | 16.96 | 8.11E-03 | 1.87E-02 |
| FAM49B | 8:129903351-129970942 | 8:129904499-129904585 | MXS | 3 | 3 | ENST00000401979 | 16.93 | 2.45E-02 | 3.50E-01 |
| ARAF | X:47561252-47562908 | X:47562682-47562909 | A3ss | 3 | 3 | Ex.TSS.ENST00000377039 | 16.92 | 1.10E-03 | 1.61E-01 |
| DEPDC5 | 22:31754980-31758545 | 22:31758527-31758545 | A3ss | 3 | 3 | ENST00000456178 | 16.92 | 6.27E-03 | 2.13E-02 |
| ZFAND1 | 8:81715115-81718181 | 8:81717249-81717288 | SES | 3 | 3 | ENST00000220669 | 16.91 | 4.90E-02 | 6.63E-01 |
| LRRC14 | 8:144520911-144525172 | 8:144523410-144524440 | IR (overlapping region) | 3 | 3 | ENST00000292524 | 16.9 | 4.90E-02 | 7.58E-01 |
| DGCR2 | 22:19057163-19063201 | 22:19057163-19057186 | A3ss | 3 | 3 | Ex.ENST00000263196 | 16.89 | 2.25E-04 | 7.42E-04 |
| RAB4B | 19:40778392-40786664 | 19:40783778-40783840 | MES | 3 | 3 | ENST00000357052 | 16.86 | 2.79E-02 | 6.41E-02 |
| IMMP1L | 11:31433571-31509518 | 11:31456260-31456386 | MES | 3 | 3 | ENST00000278200 | 16.81 | 1.61E-02 | 2.19E-02 |
| PPFIBP1 | 12:27673828-27676427 | 12:27674192-27674221 | SES | 3 | 3 | ENST00000228425 | 16.79 | 1.64E-02 | 2.47E-02 |
| NUP210 | 3:13360492-13365945 | 3:13365669-13365946 | A5ss | 3 | 3 | Ex.ENST00000254508 | 16.77 | 6.89E-04 | 2.78E-03 |
| ESPN | 1:6448803-6451602 | 1:6448803-6449091 | A5ss | 3 | 3 | ENST00000636330 | 16.76 | 1.59E-02 | 4.59E-02 |

| | | | | | | | | | |
|---|---|---|---|---|---|---|---|---|---|
| PUF60 | 8:143824400-143828983 | 8:143827385-143827536 | MXS | 3 | 3 | ENST00000529999 | 16.76 | 3.28E-02 | 4.98E-01 |
| ZSWIM4 | 19:13799922-13804791 | 19:13804775-13804792 | A3ss | 3 | 3 | Ex.ENST00000254323 | 16.69 | 8.12E-03 | 1.79E-02 |
| TMCO4 | 1:19682242-19683444 | 1:19682715-19683287 | IR | 3 | 3 | ENST00000489814 | 16.67 | 3.38E-02 | 8.71E-02 |
| PPT2-EGFL8 | 6:32167006-32167146 | 6:32167103-32167146 | A3ss | 3 | 2 | ENST00000422437 | 16.6 | 2.57E-02 | 1.45E-01 |
| TTC14 | 3:180607648-180610974 | 3:180607766-180608700 | IR (overlapping region) | 3 | 3 | ENST00000412756 | 16.59 | 2.85E-02 | 1.18E-01 |
| PAM | 5:103019844-103028184 | 5:103025131-103025334 | SES | 3 | 3 | ENST00000348126 | 16.58 | 1.02E-02 | 6.70E-02 |
| BUB1B | 15:40200981-40202404 | 15:40202390-40202405 | A3ss | 3 | 3 | Ex.ENST00000287598 | 16.57 | 3.59E-04 | 6.03E-04 |
| CLK4 | 5:178618779-178623255 | 5:178619838-178619896 | MXS | 3 | 3 | ENST00000519583 | 16.57 | 1.90E-02 | 1.28E-01 |
| BBC3 | 19:47226755-47232514 | 19:47228158-47228446 | SES | 3 | 3 | ENST00000449228 | 16.54 | 3.75E-03 | 8.78E-03 |
| ANKRD54 | 22:37838594-37840186 | 22:37838594-37838598 | A3ss | 3 | 3 | ENST00000215941 | 16.52 | 3.51E-02 | 1.20E-01 |
| MICAL1 | 6:109445863-109446135 | 6:109445863-109445952 | A3ss | 3 | 3 | Ex.ENST00000358577 | 16.52 | 1.62E-02 | 1.24E-01 |
| ZNF211 | 19:57633247-57634628 | 19:57634023-57634061 | SES | 3 | 3 | ENST00000254182 | 16.49 | 3.89E-02 | 9.32E-02 |
| BABAM1 | 19:17267528-17268793 | 19:17268756-17268793 | TSSlA3ss | 3 | 3 | TSS.ENST00000359435 | 16.46 | 8.19E-03 | 1.82E-02 |
| LRRC14 | 8:144520911-144525172 | 8:144523810-144523923 | IR (overlapping region) | 3 | 3 | ENST00000292524 | 16.44 | 4.56E-02 | 7.07E-01 |
| WDR45 | X:49076751-49077836 | X:49077643-49077747 | SES | 3 | 3 | ENST00000322995 | 16.44 | 2.97E-02 | 1.00E+00 |
| PRKCQ-AS1 | 10:6580928-6583675 | 10:6582138-6582201 | SES | 3 | 3 | ENST00000455810 | 16.39 | 2.75E-02 | 3.62E-02 |
| SLC25A40 | 7:87856352-87858630 | 7:87856352-87856389 | A3ss | 2 | 3 | ENST00000429674 | 16.39 | 4.94E-02 | 5.84E-01 |
| EPOR | 19:11382106-11383096 | 19:11382847-11383096 | A5ss | 3 | 3 | ENST00000588681 | 16.36 | 1.81E-04 | 3.96E-04 |
| FOXRED2 | 22:36506424-36507008 | 22:36506912-36507008 | TSSlA5ss | 3 | 3 | TSS.ENST00000216187 | 16.36 | 3.24E-02 | 1.10E-01 |
| PLA2G2A | 1:19978485-19978733 | 1:19978485-19978524 | A3ss | 3 | 3 | ENST00000375111 | 16.36 | 3.49E-04 | 9.00E-04 |
| LRCH3 | 3:197854446-197865422 | 3:197858834-197858905 | SES | 3 | 3 | ENST00000334859 | 16.35 | 7.08E-03 | 2.98E-02 |
| UNC13A | 19:17648984-17649508 | 19:17649339-17649344 | SES | 3 | 3 | ENST00000519716 | 16.32 | 1.85E-02 | 4.10E-02 |
| TGFBRAP1 | 2:105308316-105329624 | 2:105308316-105308318 | A3ss | 3 | 3 | Ex.TSS.ENST00000393359 | 16.3 | 4.65E-03 | 1.62E-02 |
| FAM227B | 15:49326970-49328675 | 15:49328131-49328665 | IR (overlapping region) | 2 | 3 | ENST00000299338 | 16.29 | 4.77E-02 | 8.05E-02 |
| MIB2 | 1:1625429-1626649 | 1:1625546-1626649 | A3ss | 3 | 3 | ENST00000479659 | 16.16 | 5.00E-03 | 1.27E-02 |
| MIB2 | 1:1625429-1625557 | 1:1625546-1625557 | A3ss | 3 | 3 | ENST00000355826 | 16.16 | 5.00E-03 | 1.27E-02 |
| TNNT1 | 19:55145566-55146680 | 19:55146434-55146466 | SES | 3 | 3 | ENST00000291901 | 16.12 | 1.68E-03 | 4.00E-03 |
| PHF20 | 20:35801606-35842572 | 20:35823258-35842573 | A3ss | 3 | 3 | Ex.ENST00000339089 | 16.11 | 4.13E-03 | 1.24E-02 |
| FAM49B | 8:129903351-129939607 | 8:129904499-129904585 | SES | 3 | 3 | ENST00000519070 | 16.09 | 6.84E-03 | 9.80E-02 |
| VPS54 | 2:63972245-63983863 | 2:63981646-63981851 | SES | 3 | 3 | ENST00000409558 | 16.01 | 4.50E-02 | 1.70E-01 |
| ARL6IP4 | 12:122981267-122981594 | 12:122981267-122981299 | TSSlA5ss | 3 | 3 | TSS.ENST00000357866 | 15.98 | 2.95E-03 | 4.40E-03 |
| EI24 | 11:125572570-125575262 | 11:125575251-125575262 | A3ss | 3 | 3 | ENST00000618552 | 15.97 | 1.01E-06 | 1.36E-06 |
| HDAC7 | 12:47795768-47796206 | 12:47795906-47796016 | SES | 3 | 3 | ENST00000080059 | 15.91 | 2.70E-02 | 4.09E-02 |
| CENPK | 5:65554899-65561462 | 5:65554899-65554946 | A3ss | 3 | 3 | ENST00000396679 | 15.85 | 1.58E-02 | 1.13E-01 |
| PTPN21 | 14:88469499-88470050 | 14:88469734-88469921 | IR (overlapping region) | 2 | 3 | ENST00000554270 | 15.78 | 1.07E-02 | 1.77E-02 |
| TRIM16 | 17:15680954-15683035 | 17:15682854-15682951 | MXS | 3 | 3 | ENST00000460728 | 15.74 | 1.89E-02 | 3.71E-02 |
| ERGIC3 | 20:35556272-35556972 | 20:35556955-35556972 | A3ss | 3 | 3 | ENST00000451605 | 15.72 | 8.38E-05 | 2.52E-04 |
| ERGIC3 | 20:35556272-35556990 | 20:35556955-35556990 | A3ss | 3 | 3 | Ex.ENST00000451605 | 15.72 | 8.38E-05 | 2.52E-04 |
| MIS18BP1 | 14:45206171-45210379 | 14:45206738-45210380 | A5ss | 3 | 3 | Ex.ENST00000310806 | 15.72 | 3.04E-03 | 4.98E-03 |
| FAM149B1 | 10:73234108-73235318 | 10:73234941-73235192 | IR (overlapping region) | 3 | 3 | ENST00000468462 | 15.69 | 2.69E-02 | 3.59E-02 |
| HAGHL | 16:733469-735525 | 16:734348-734736 | IR (overlapping region) | 3 | 3 | ENST00000569604 | 15.69 | 1.08E-02 | 2.09E-02 |
| MIER2 | 19:334543-344773 | 19:336083-336173 | SES | 3 | 3 | ENST00000264819 | 15.68 | 5.03E-03 | 1.14E-02 |
| CLK2 | 1:155263401-155263949 | 1:155263573-155263950 | A5ss | 3 | 3 | Ex.TSS.ENST00000355560 | 15.67 | 1.60E-03 | 4.03E-03 |
| RPAIN | 17:5422830-5426235 | 17:5425971-5426235 | A3ss | 3 | 3 | ENST00000570883 | 15.66 | 4.15E-03 | 7.66E-03 |
| CEP95 | 17:64529428-64531889 | 17:64530926-64531018 | SES | 3 | 3 | ENST00000556440 | 15.63 | 1.54E-02 | 3.21E-02 |
| GCLC | 6:53516223-53520777 | 6:53516223-53516240 | A3ss | 3 | 3 | ENST00000643939 | 15.63 | 1.48E-03 | 1.29E-02 |
| MIB2 | 1:1627296-1627830 | 1:1627445-1627672 | IR | 3 | 3 | ENST00000505370 | 15.63 | 9.19E-03 | 2.34E-02 |
| OXNAD1 | 3:16269276-16270944 | 3:16270929-16270944 | A3ss | 3 | 3 | ENST00000435829 | 15.58 | 1.95E-03 | 8.04E-03 |
| PGPEP1 | 19:18356012-18363390 | 19:18357383-18357615 | SES | 3 | 3 | ENST00000252813 | 15.58 | 3.70E-02 | 8.24E-02 |
| LUC7L3 | 17:50750853-50752199 | 17:50751297-50751362 | SES | 3 | 3 | ENST00000503728 | 15.53 | 4.54E-02 | 9.36E-02 |
| SH3PXD2A | 10:103617315-103627088 | 10:103622470-103622553 | SES | 3 | 3 | ENST00000369774 | 15.53 | 3.62E-02 | 4.78E-02 |
| SKIV2L | 6:31968539-31968685 | 6:31968623-31968685 | A3ss | 3 | 3 | ENST00000465703 | 15.53 | 1.77E-02 | 1.47E-01 |
| ZFYVE19 | 15:40810758-40813337 | 15:40812699-40812902 | SES | 3 | 3 | ENST00000336455 | 15.51 | 8.74E-03 | 1.47E-02 |
| TAF1 | X:71458367-71459551 | X:71459150-71459251 | SES | 3 | 3 | ENST00000437147 | 15.49 | 4.34E-02 | 1.00E+00 |
| POLL | 10:101582892-101587245 | 10:101585316-101585478 | MES | 3 | 2 | ENST00000299206 | 15.43 | 1.37E-02 | 1.81E-02 |
| SH2D3A | 19:6754426-6754615 | 19:6754426-6754448 | A3ss | 3 | 3 | Ex.ENST00000245908 | 15.43 | 1.92E-03 | 4.61E-03 |
| STAU2 | 8:73688814-73738283 | 8:73709032-73709177 | SES | 3 | 3 | ENST00000521845 | 15.43 | 9.28E-07 | 1.25E-05 |
| DCAKD | 17:45035000-45051552 | 17:45051361-45051552 | TSSlA5ss | 3 | 3 | TSS.ENST00000593094 | 15.4 | 2.49E-02 | 5.07E-02 |
| SNHG11 | 20:38446654-38448730 | 20:38446949-38447465 | IR | 3 | 3 | ENST00000400436 | 15.4 | 3.63E-02 | 1.10E-01 |
| ICA1 | 7:8221399-8228600 | 7:8227638-8227851 | MXS | 3 | 3 | ENST00000455539 | 15.38 | 3.31E-03 | 3.91E-02 |
| CNTROB | 17:7932155-7933349 | 17:7932801-7932955 | IR | 3 | 3 | ENST00000380262 | 15.36 | 2.97E-02 | 6.31E-02 |
| AL117327.1 | X:103405024-103408157 | X:103405024-103405096 | A5ss | 3 | 3 | Ex.ENST00000628546 | 15.35 | 3.28E-03 | 1.21E-01 |
| SECISBP2 | 9:89332987-89334638 | 9:89334522-89334638 | A3ss | 3 | 3 | Ex.ENST00000339901 | 15.35 | 3.25E-02 | 6.65E-01 |
| SECISBP2 | 9:89332987-89334635 | 9:89334522-89334635 | A3ss | 3 | 3 | ENST00000339901 | 15.35 | 3.25E-02 | 6.65E-01 |
| SIDT2 | 11:117186637-117187377 | 11:117186990-117187001 | SES | 3 | 3 | ENST00000431081 | 15.31 | 2.46E-02 | 3.30E-02 |
| FNBP1 | 9:129923997-129924959 | 9:129923997-129924026 | A3ss | 3 | 3 | ENST00000420781 | 15.28 | 2.18E-02 | 5.44E-01 |
| FNBP1 | 9:129923940-129924959 | 9:129923940-129924026 | A3ss | 3 | 3 | ENST00000420781 | 15.28 | 2.18E-02 | 5.44E-01 |
| OGDHL | 10:49744123-49744649 | 10:49744123-49744140 | A3ss | 3 | 3 | Ex.ENST00000374103 | 15.24 | 7.14E-03 | 9.49E-03 |
| RAB3D | 19:11322068-11325585 | 19:11322983-11324116 | IR (overlapping region) | 3 | 3 | ENST00000222120 | 15.24 | 3.41E-02 | 7.46E-02 |
| RAD17 | 5:69371172-69372033 | 5:69371454-69371557 | SES | 3 | 3 | ENST00000354868 | 15.24 | 6.28E-03 | 4.52E-02 |
| UBAP2L | 1:154268955-154270880 | 1:154269362-154269412 | SES | 3 | 3 | ENST00000493867 | 15.17 | 3.43E-02 | 8.62E-02 |
| TRRAP | 7:98981981-98983263 | 7:98983240-98983264 | A3ss | 3 | 3 | Ex.ENST00000355540 | 15.16 | 6.92E-04 | 8.49E-03 |
| SNHG8 | 4:118278793-118279388 | 4:118278934-118279137 | SES | 3 | 3 | ENST00000602414 | 15.15 | 3.04E-03 | 1.81E-02 |
| FOXRED1 | 11:126273080-126273335 | 11:126273316-126273335 | A3ss | 3 | 3 | ENST00000524751 | 15.12 | 2.98E-04 | 4.03E-04 |
| FAM222B | 17:28766708-28842681 | 17:28790378-28790425 | MES | 3 | 3 | ENST00000583307 | 15.1 | 4.79E-05 | 9.53E-05 |
| MSTO1 | 1:155611216-155611603 | 1:155611356-155611548 | IR | 3 | 3 | ENST00000478756 | 15.1 | 2.25E-02 | 5.66E-02 |
| VARS2 | 6:30916960-30917104 | 6:30917095-30917105 | A3ss | 3 | 3 | Ex.ENST00000321897 | 15.09 | 2.01E-04 | 1.64E-03 |
| ECT2 | 3:172755375-172756982 | 3:172755483-172755575 | SES | 3 | 3 | ENST00000366090 | 15.05 | 9.52E-04 | 3.92E-03 |
| LRRC42 | 1:53946550-53951985 | 1:53947788-53947821 | MXS | 3 | 3 | ENST00000371370 | 15.01 | 1.05E-02 | 2.98E-02 |
| SMPD2 | 6:109442571-109442884 | 6:109442630-109442751 | IR | 3 | 3 | ENST00000458487 | 15 | 4.04E-02 | 3.08E-01 |
| SMPD2 | 6:109442571-109442884 | 6:109442626-109442751 | IR | 3 | 3 | ENST00000458487 | 15 | 4.04E-02 | 3.13E-01 |
| FAM131A | 3:184336124-184341723 | 3:184337311-184337371 | SES | 3 | 3 | ENST00000340957 | 14.96 | 1.13E-02 | 4.69E-02 |
| RNF20 | 9:101533915-101535397 | 9:101535376-101535398 | A3ss | 3 | 3 | Ex.TSS.ENST00000374819 | 14.96 | 2.11E-05 | 3.00E-04 |
| P2RX5 | 17:3689628-3690069 | 17:3689628-3689630 | A3ss | 3 | 3 | ENST00000225328 | 14.95 | 2.13E-02 | 4.27E-02 |
| PDLIM7 | 5:177491147-177491806 | 5:177491404-177491420 | SES | 3 | 3 | ENST00000493815 | 14.92 | 2.49E-02 | 1.75E-01 |
| B3GALNT2 | 1:235494829-235504140 | 1:235496186-235496308 | SES | 3 | 3 | ENST00000313984 | 14.9 | 6.70E-03 | 1.85E-02 |
| AURKA | 20:56388203-56392167 | 20:56390556-56390665 | MES | 3 | 3 | ENST00000420474 | 14.89 | 2.13E-02 | 6.84E-02 |
| INO80B | 2:74455175-74455405 | 2:74455391-74455405 | TSSlA3ss | 3 | 3 | TSS.ENST00000409493 | 14.89 | 7.55E-03 | 2.95E-02 |
| AC005670.3 | 17:47048163-47056753 | 17:47051401-47054517 | IR (overlapping region) | 3 | 3 | ENST00000658121 | 14.88 | 3.49E-02 | 7.15E-02 |
| THTPA | 14:23556344-23558694 | 14:23556743-23557034 | SES | 3 | 3 | ENST00000554970 | 14.88 | 2.00E-03 | 3.26E-03 |
| MADD | 11:47285066-47285450 | 11:47285066-47285194 | A5ss | 3 | 3 | ENST00000311027 | 14.87 | 5.10E-03 | 7.00E-03 |
| STRN3 | 14:30913658-30935162 | 14:30918966-30919106 | MXS | 3 | 3 | ENST00000554991 | 14.87 | 1.59E-02 | 2.59E-02 |
| BECN1 | 17:42810929-42814523 | 17:42811655-42811797 | MES | 3 | 3 | ENST00000361523 | 14.86 | 2.44E-02 | 4.93E-02 |
| ADGRL2 | 1:81936838-81942968 | 1:81942031-81942969 | A3ss | 3 | 3 | Ex.ENST00000319517 | 14.84 | 2.19E-03 | 6.30E-03 |

| | | | | | | | | | |
|---|---|---|---|---|---|---|---|---|---|
| THAP8 | 19:36039719-36054134 | 19:36039944-36040136 | SES | 3 | 3 | ENST00000292894 | 14.83 | 3.36E-03 | 7.68E-03 |
| CHTF18 | 16:794202-795684 | 16:795132-795356 | SES | 3 | 3 | ENST00000262315 | 14.78 | 5.79E-03 | 1.12E-02 |
| THUMPD3-AS1 | 3:9388800-9390497 | 3:9389797-9390279 | IR | 3 | 3 | ENST00000668063 | 14.71 | 1.21E-02 | 7.00E-02 |
| AC040160.1 | 16:67277510-67280806 | 16:67279277-67279875 | IR | 3 | 3 | ENST00000563969 | 14.7 | 1.49E-02 | 2.86E-02 |
| MBOAT2 | 2:8943265-9003539 | 2:8958497-8958642 | MXS | 3 | 3 | ENST00000305997 | 14.7 | 4.60E-02 | 1.82E-01 |
| TIA1 | 2:70216500-70216885 | 2:70216753-70216885 | A5ss | 3 | 3 | ENST00000454815 | 14.69 | 1.20E-02 | 4.66E-02 |
| UQCC1 | 20:35344110-35347272 | 20:35346424-35347163 | IR | 3 | 3 | ENST00000397554 | 14.69 | 1.70E-04 | 5.11E-04 |
| FBXW5 | 9:136942944-136943348 | 9:136942944-136942997 | A3ss | 3 | 3 | Ex.ENST00000325285 | 14.67 | 4.73E-04 | 1.25E-02 |
| HPS1 | 10:98429643-98430570 | 10:98429791-98429889 | SES | 3 | 3 | ENST00000325103 | 14.67 | 2.47E-03 | 3.31E-03 |
| ZNF124 | 1:247159814-247171847 | 1:247159814-247159859 | A3ss | 3 | 3 | Ex.TSS.ENST00000340684 | 14.67 | 3.54E-03 | 9.83E-03 |
| WDR90 | 16:662220-662736 | 16:662332-662672 | IR | 3 | 3 | ENST00000546923 | 14.65 | 5.24E-03 | 1.00E-02 |
| DAPK1 | 9:87642059-87645894 | 9:87643376-87643468 | SES | 3 | 3 | ENST00000358077 | 14.59 | 6.75E-03 | 2.17E-01 |
| TARBP1 | 1:234393838-234401180 | 1:234398382-234398553 | MXS | 3 | 3 | ENST00000040877 | 14.58 | 1.16E-03 | 3.18E-03 |
| GBAP1 | 1:155217240-155218049 | 1:155217847-155217963 | IR | 2 | 3 | ENST00000473223 | 14.57 | 4.48E-02 | 1.13E-01 |
| ZFAND6 | 15:80072648-80120327 | 15:80098416-80098578 | MES | 3 | 3 | ENST00000558272 | 14.57 | 4.52E-02 | 7.75E-02 |
| ZNF271P | 18:35290392-35305975 | 18:35291010-35291232 | SES | 3 | 3 | ENST00000639248 | 14.55 | 2.63E-02 | 5.64E-02 |
| INO80C | 18:35468743-35497718 | 18:35478282-35478349 | MES | 3 | 3 | ENST00000334598 | 14.5 | 3.61E-02 | 7.74E-02 |
| VPS52 | 6:33270284-33271690 | 6:33271586-33271690 | TSSlA5ss | 3 | 3 | TSS.ENST00000445902 | 14.5 | 3.41E-02 | 2.85E-01 |
| C1orf109 | 1:37689887-37692029 | 1:37690216-37690438 | SES | 3 | 3 | ENST00000488137 | 14.44 | 4.69E-02 | 1.32E-01 |
| SNHG11 | 20:38446654-38448730 | 20:38446941-38447465 | IR | 3 | 3 | ENST00000400436 | 14.43 | 3.82E-02 | 1.15E-01 |
| CNTROB | 17:7932155-7933349 | 17:7932801-7932927 | IR | 3 | 3 | ENST00000380262 | 14.41 | 3.84E-02 | 8.15E-02 |
| SLC37A4 | 11:119029565-119030848 | 11:119030548-119030735 | SES | 3 | 2 | ENST00000357590 | 14.39 | 3.03E-02 | 4.09E-02 |
| SRC | 20:37386175-37393894 | 20:37386460-37386477 | SES | 3 | 3 | ENST00000373558 | 14.36 | 2.36E-02 | 7.14E-02 |
| REST | 4:56929841-56935844 | 4:56931073-56931120 | IR | 3 | 3 | ENST00000309042 | 14.34 | 1.47E-02 | 9.36E-02 |
| TFR2 | 7:100627264-100627604 | 7:100627492-100627576 | IR | 3 | 3 | ENST00000461176 | 14.34 | 2.54E-02 | 1.43E-01 |
| STAU2 | 8:73688814-73746782 | 8:73709032-73709162 | MES | 3 | 3 | ENST00000519818 | 14.33 | 4.93E-04 | 6.62E-03 |
| THTPA | 14:23556344-23558694 | 14:23556743-23557039 | SES | 3 | 3 | ENST00000556015 | 14.31 | 6.02E-03 | 9.80E-03 |
| DCUN1D4 | 4:51886639-51891759 | 4:51886639-51887244 | A5ss | 3 | 3 | ENST00000477560 | 14.3 | 5.42E-03 | 3.42E-02 |
| NUB1 | 7:151368888-151375847 | 7:151374097-151374243 | SES | 3 | 3 | ENST00000470229 | 14.29 | 3.44E-02 | 3.44E-01 |
| TERF1 | 8:73027053-73032041 | 8:73030336-73030395 | SES | 3 | 3 | ENST00000276603 | 14.29 | 5.18E-04 | 6.95E-03 |
| PARD3 | 10:34696420-34814875 | 10:34696420-34696440 | A3ss | 3 | 3 | Ex.TSS.ENST00000340077 | 14.27 | 6.09E-03 | 8.08E-03 |
| ATG9A | 2:219228483-219229534 | 2:219228483-219228485 | TSSlA3ss | 3 | 3 | TSS.ENST00000361242 | 14.25 | 3.85E-03 | 1.41E-02 |
| MTERF2 | 12:106978772-106986968 | 12:106985115-106986968 | A5ss | 3 | 3 | ENST00000552029 | 14.23 | 7.05E-04 | 1.04E-03 |
| KLHL2 | 4:165220060-165238777 | 4:165228807-165228913 | SES | 3 | 3 | ENST00000226725 | 14.21 | 6.27E-03 | 3.81E-02 |
| PEX5 | 12:7201842-7202611 | 12:7202241-7202351 | SES | 3 | 3 | ENST00000266564 | 14.21 | 1.35E-03 | 2.10E-03 |
| TUBGCP4 | 15:43405202-43409771 | 15:43407571-43407942 | IR (overlapping region) | 3 | 3 | ENST00000564079 | 14.19 | 7.83E-03 | 1.32E-02 |
| ADAM23 | 2:206596163-206617578 | 2:206609910-206610000 | MXS | 3 | 3 | ENST00000264377 | 14.18 | 1.59E-03 | 5.79E-03 |
| NAV1 | 1:201786578-201789739 | 1:201788468-201788638 | SES | 3 | 3 | ENST00000367295 | 14.17 | 3.25E-03 | 8.39E-03 |
| FBXO7 | 22:32475399-32479132 | 22:32478981-32479132 | TSSlA3ss | 3 | 2 | TSS.ENST00000397426 | 14.16 | 1.51E-02 | 5.14E-02 |
| ZMYND8 | 20:47210898-47221313 | 20:47212642-47212725 | MES | 3 | 3 | ENST00000262975 | 14.15 | 4.65E-02 | 1.48E-01 |
| AGRN | 1:1050838-1051252 | 1:1051032-1051043 | SES | 3 | 3 | ENST00000620552 | 14.14 | 3.34E-03 | 8.09E-03 |
| FRMD4A | 10:13747820-13761646 | 10:13753056-13753100 | MXS | 3 | 3 | ENST00000650137 | 14.14 | 6.71E-03 | 8.89E-03 |
| HLA-F | 6:29723928-29724189 | 6:29724173-29724189 | A3ss | 3 | 3 | ENST00000259951 | 14.13 | 1.85E-03 | 1.50E-02 |
| PRDM15 | 21:41822038-41825959 | 21:41823318-41823449 | SES | 3 | 3 | ENST00000269844 | 14.11 | 2.79E-02 | 9.10E-02 |
| BCL2L12 | 19:49666800-49667021 | 19:49667019-49667021 | A3ss | 3 | 3 | ENST00000246784 | 14.09 | 2.52E-05 | 5.93E-05 |
| SUPT20H | 13:37047937-37051487 | 13:37048564-37048599 | SES | 3 | 3 | ENST00000350612 | 14.09 | 2.87E-02 | 4.52E-02 |
| NCAPG2 | 7:158646564-158652292 | 7:158650832-158650972 | SES | 3 | 3 | ENST00000356309 | 14.08 | 2.35E-03 | 2.41E-02 |
| VRK2 | 2:58046822-58048826 | 2:58046822-58046868 | TSSlA5ss | 3 | 3 | TSS.ENST00000340157 | 14.07 | 5.61E-03 | 2.15E-02 |
| FAM49B | 8:129903351-129970942 | 8:129904499-129904585 | MES | 3 | 3 | ENST00000519142 | 14.05 | 1.52E-02 | 2.18E-01 |
| RHBDD1 | 2:226836088-226864603 | 2:226839409-226839627 | MXS | 3 | 3 | ENST00000443477 | 14.03 | 3.78E-02 | 1.39E-01 |
| EIF2AK4 | 15:39943486-39949115 | 15:39945335-39949116 | A3ss | 3 | 3 | Ex.ENST00000263791 | 14.02 | 5.13E-03 | 8.60E-03 |
| LZTS2 | 10:100997176-101002496 | 10:100997176-100997234 | A5ss | 3 | 3 | Ex.TSS.ENST00000370223 | 14.02 | 5.17E-03 | 6.81E-03 |
| NET1 | 10:5455119-5456086 | 10:5456065-5456087 | A3ss | 3 | 3 | Ex.ENST00000355029 | 14.02 | 1.33E-02 | 1.77E-02 |
| FAM86C1 | 11:71796167-71799560 | 11:71796167-71796308 | TSSlA5ss | 3 | 3 | TSS.ENST00000528685 | 14.01 | 1.19E-02 | 1.72E-02 |
| TAZ | X:154413568-154419542 | X:154414101-154414190 | MXS | 3 | 3 | ENST00000426231 | 14 | 2.97E-02 | 1.00E+00 |
| TMEM63A | 1:225848555-225848896 | 1:225848555-225848614 | A3ss | 3 | 3 | Ex.ENST00000366835 | 14 | 6.79E-05 | 1.85E-04 |
| ZNF213-AS1 | 16:3132943-3134629 | 16:3132943-3132945 | A3ss | 3 | 3 | Ex.TSS.ENST00000573447 | 13.98 | 6.24E-03 | 1.17E-02 |
| HACE1 | 6:104811394-104843222 | 6:104833042-104833173 | SES | 2 | 3 | ENST00000262903 | 13.96 | 1.99E-02 | 1.53E-01 |
| MARK2 | 11:63903159-63904785 | 11:63903986-63904147 | SES | 3 | 3 | ENST00000402010 | 13.95 | 7.66E-03 | 1.09E-02 |
| SIGIRR | 11:408207-409867 | 11:408695-408893 | MXS | 3 | 3 | ENST00000332725 | 13.95 | 1.48E-02 | 2.02E-02 |
| AL357673.1 | 1:54184266-54195418 | 1:54187652-54187739 | MXS | 3 | 3 | ENST00000311841 | 13.93 | 4.17E-02 | 1.04E-01 |
| FAM221A | 7:23680284-23689268 | 7:23684499-23684672 | MXS | 3 | 3 | ENST00000344962 | 13.91 | 7.24E-04 | 7.50E-03 |
| FAM49B | 8:129903351-129970942 | 8:129904499-129904585 | MES | 3 | 3 | ENST00000518283 | 13.91 | 1.16E-02 | 1.66E-01 |
| IMMP1L | 11:31463306-31509518 | 11:31469740-31469797 | MES | 3 | 3 | ENST00000530023 | 13.91 | 1.58E-02 | 2.16E-02 |
| NPIPA1 | 16:14945982-14950049 | 16:14948930-14949037 | SES | 3 | 3 | ENST00000328085 | 13.87 | 2.30E-02 | 4.23E-02 |
| DOK4 | 16:57479442-57479874 | 16:57479686-57479860 | IR | 3 | 3 | ENST00000569548 | 13.85 | 2.39E-02 | 4.56E-02 |
| ATF7IP2 | 16:10386123-10428867 | 16:10419581-10419623 | MES | 3 | 2 | ENST00000561932 | 13.78 | 4.21E-02 | 7.26E-02 |
| CDC27 | 17:47151919-47154671 | 17:47151919-47151936 | A3ss | 3 | 3 | ENST00000525495 | 13.78 | 1.33E-02 | 2.73E-02 |
| OPLAH | 8:144052983-144053393 | 8:144053130-144053208 | IR | 3 | 3 | ENST00000527993 | 13.78 | 1.60E-02 | 2.44E-01 |
| TTC8 | 14:88841197-88852970 | 14:88841425-88841557 | MXS | 3 | 3 | ENST00000358622 | 13.75 | 5.01E-03 | 8.31E-03 |
| TTC8 | 14:88841197-88852970 | 14:88841425-88841514 | MXS | 3 | 3 | ENST00000556651 | 13.75 | 5.01E-03 | 8.31E-03 |
| TMEM191A | 22:20703716-20704040 | 22:20703884-20703967 | IR | 3 | 3 | ENST00000428752 | 13.74 | 4.53E-02 | 1.50E-01 |
| TXNRD2 | 22:19880622-19881347 | 22:19880946-19881035 | IR | 3 | 3 | ENST00000495655 | 13.7 | 4.50E-02 | 1.48E-01 |
| RCCD1 | 15:90955453-90956611 | 15:90955453-90955883 | A5ss | 3 | 3 | ENST00000554302 | 13.69 | 3.17E-02 | 5.46E-02 |
| AP1G2 | 14:23561636-23562287 | 14:23561962-23562066 | SES | 3 | 3 | ENST00000308724 | 13.66 | 8.06E-03 | 1.31E-02 |
| TMEM191A | 22:20703716-20704040 | 22:20703752-20703967 | IR | 3 | 2 | ENST00000428752 | 13.66 | 3.28E-03 | 1.09E-02 |
| AHSA2P | 2:61186617-61191203 | 2:61189070-61189697 | IR (overlapping region) | 3 | 3 | ENST00000394457 | 13.63 | 4.47E-02 | 1.69E-01 |
| TSC22D1 | 13:44539924-44573162 | 13:44539924-44539936 | TSSlA3ss | 3 | 3 | TSS.ENST00000486464 | 13.6 | 4.18E-02 | 6.60E-02 |
| IP6K2 | 3:48694241-48695089 | 3:48694702-48694824 | MXS | 3 | 3 | ENST00000412795 | 13.57 | 4.71E-02 | 2.08E-01 |
| STARD4 | 5:111501117-111506329 | 5:111501962-111502088 | MXS | 3 | 3 | ENST00000296632 | 13.55 | 4.09E-02 | 2.68E-01 |
| NEURL4 | 17:7318675-7319049 | 17:7318675-7318691 | A3ss | 3 | 3 | ENST00000573186 | 13.52 | 1.34E-04 | 2.80E-04 |
| STAU2 | 8:73709163-73738283 | 8:73709163-73709177 | A3ss | 3 | 3 | ENST00000521845 | 13.52 | 5.65E-03 | 7.59E-02 |
| RGL2 | 6:33293917-33294033 | 6:33293917-33293935 | A3ss | 3 | 3 | Ex.ENST00000437840 | 13.5 | 3.31E-04 | 2.76E-03 |
| CBARP | 19:1235001-1235342 | 19:1235146-1235336 | IR | 3 | 3 | ENST00000215376 | 13.49 | 4.27E-02 | 9.36E-02 |
| PWWP2A | 5:160078169-160093100 | 5:160089506-160089610 | MES | 3 | 3 | ENST00000521424 | 13.49 | 3.97E-02 | 2.76E-01 |
| BAHD1 | 15:40464548-40465343 | 15:40465335-40465343 | A3ss | 3 | 3 | ENST00000416165 | 13.48 | 9.09E-04 | 1.52E-03 |
| WDR90 | 16:662220-662736 | 16:662332-662678 | IR | 3 | 3 | ENST00000546923 | 13.47 | 5.02E-03 | 9.63E-03 |
| FAM49B | 8:129903351-129970942 | 8:129904499-129904585 | MES | 3 | 3 | ENST00000518879 | 13.44 | 1.06E-02 | 1.51E-01 |
| RNF125 | 18:32045733-32068297 | 18:32065902-32066009 | SES | 3 | 3 | ENST00000217740 | 13.38 | 1.31E-02 | 2.80E-02 |
| NAA40 | 11:63945936-63952237 | 11:63946951-63947003 | SES | 3 | 3 | ENST00000377793 | 13.37 | 2.61E-02 | 3.72E-02 |
| ZC3H4 | 19:47094609-47112147 | 19:47110895-47110956 | SES | 3 | 3 | ENST00000597069 | 13.35 | 1.11E-02 | 2.60E-02 |
| ADGRB2 | 1:31727606-31728181 | 1:31728025-31728081 | SES | 3 | 3 | ENST00000373658 | 13.33 | 4.98E-03 | 1.40E-02 |
| FIGNL1 | 7:50444129-50447297 | 7:50447017-50447131 | IR | 3 | 3 | ENST00000613602 | 13.32 | 6.82E-03 | 7.32E-02 |
| TSC1 | 9:132910690-132911001 | 9:132910690-132910692 | A3ss | 3 | 3 | ENST00000298552 | 13.31 | 2.86E-02 | 7.13E-01 |

| | | | | | | | | | |
|---|---|---|---|---|---|---|---|---|---|
| SHARPIN | 8:144098995-144099276 | 8:144099081-144099205 | SES | 3 | 3 | ENST00000398712 | 13.3 | 1.70E-02 | 2.60E-01 |
| B3GALNT1 | 3:161090063-161103426 | 3:161101139-161101233 | SES | 3 | 2 | ENST00000460353 | 13.29 | 2.42E-02 | 9.79E-02 |
| UBQLN1 | 9:83665146-83669184 | 9:83666350-83666184 | A5ss | 3 | 3 | ENST00000533705 | 13.28 | 1.00E-03 | 2.69E-02 |
| UHRF2 | 9:6486926-6493825 | 9:6492303-6492401 | SES | 3 | 3 | ENST00000468435 | 13.27 | 9.24E-03 | 1.92E-01 |
| WDR24 | 16:689160-690398 | 16:689643-689976 | IR (overlapping region) | 3 | 3 | ENST00000293883 | 13.25 | 4.60E-02 | 8.87E-02 |
| NXPE3 | 3:101779901-101782109 | 3:101779901-101779934 | A5ss | 3 | 3 | ENST00000474165 | 13.24 | 2.96E-02 | 1.18E-01 |
| RAD9A | 11:67396172-67396262 | 11:67396172-67396175 | A5ss | 3 | 3 | ENST00000307980 | 13.24 | 2.10E-02 | 3.05E-02 |
| FAM3A | X:154511872-154515759 | X:154512314-154512411 | MES | 3 | 3 | ENST00000322269 | 13.23 | 4.48E-03 | 2.61E-01 |
| NAA40 | 11:63945936-63952237 | 11:63945936-63947003 | A5ss | 3 | 3 | ENST00000338447 | 13.22 | 2.03E-02 | 2.90E-02 |
| TRPC4 | 13:37636641-37637625 | 13:37637229-37637483 | IR | 3 | 3 | ENST00000338947 | 13.22 | 4.97E-02 | 7.83E-02 |
| CRYBB2P1 | 22:25459516-25509182 | 22:25466059-25509183 | A3ss | 3 | 3 | Ex.ENST00000354451 | 13.21 | 2.89E-02 | 9.65E-02 |
| ASB16-AS1 | 17:44181721-44186564 | 17:44182930-44183093 | SES | 3 | 3 | ENST00000585457 | 13.17 | 4.59E-02 | 9.32E-02 |
| WDR27 | 6:169670694-169672254 | 6:169670694-169670957 | TSSIA3ss | 3 | 3 | TSS.ENST00000649806 | 13.17 | 1.96E-02 | 1.57E-01 |
| BLOC1S6 | 15:45587526-45605427 | 15:45603100-45603187 | MXS | 3 | 3 | ENST00000567461 | 13.16 | 8.99E-03 | 1.52E-02 |
| GSN | 9:121281563-121301962 | 9:121300056-121300126 | MXS | 3 | 3 | ENST00000373808 | 13.16 | 2.04E-03 | 4.26E-02 |
| APTX | 9:32987732-32988091 | 9:32988083-32988091 | A5ss | 3 | 3 | ENST00000464632 | 13.15 | 1.54E-02 | 4.32E-01 |
| ADGRB2 | 1:31727606-31728181 | 1:31727606-31728081 | TSSIA3ss | 3 | 3 | TSS.ENST00000465256 | 13.13 | 7.74E-03 | 2.17E-02 |
| PLXNB1 | 3:48416452-48417910 | 3:48416452-48416469 | A3ss | 3 | 3 | Ex.ENST00000296440 | 13.13 | 5.71E-05 | 2.54E-04 |
| CCT6P1 | 7:65751279-65755779 | 7:65754527-65755780 | A3ss | 3 | 3 | Ex.TSS.ENST00000443108 | 13.1 | 9.39E-03 | 1.06E-01 |
| LOXL3 | 2:74536472-74549368 | 2:74536709-74536928 | SES | 3 | 3 | ENST00000264094 | 13.1 | 1.53E-02 | 5.99E-02 |
| ZNF271P | 18:35290392-35305975 | 18:35290392-35291232 | A5ss | 3 | 3 | ENST00000639141 | 13.08 | 3.35E-02 | 7.18E-02 |
| MIB2 | 1:1626837-1627207 | 1:1627000-1627073 | IR | 3 | 3 | ENST00000506488 | 13.06 | 1.01E-02 | 2.58E-02 |
| AL109918.1 | 6:52666389-52669155 | 6:52666519-52666748 | IR | 3 | 3 | ENST00000602367 | 13.05 | 1.19E-02 | 1.03E-01 |
| FOXRED1 | 11:126274927-126275349 | 11:126275022-126275326 | IR | 3 | 3 | ENST00000524751 | 13.05 | 2.24E-03 | 3.03E-03 |
| GGT1 | 22:24608024-24610249 | 22:24609965-24610033 | SES | 3 | 3 | ENST00000248923 | 13.02 | 1.51E-02 | 5.02E-02 |
| NR1H3 | 11:47257979-47259259 | 11:47258130-47259179 | IR | 3 | 3 | ENST00000467728 | 13.02 | 1.72E-02 | 2.36E-02 |
| SLC25A39 | 17:44322529-44322807 | 17:44322529-44322552 | A3ss | 3 | 3 | ENST00000377095 | 13 | 1.83E-03 | 3.72E-03 |
| ZNF90 | 19:20105317-20126751 | 19:20121222-20121346 | MES | 3 | 3 | ENST00000469078 | 13 | 4.23E-02 | 9.56E-02 |
| ENTPD6 | 20:25195868-25207078 | 20:25207076-25207078 | TSSIA3ss | 3 | 3 | TSS.ENST00000418890 | 12.99 | 1.30E-02 | 3.88E-02 |
| LRRC14 | 8:144520911-144525172 | 8:144524106-144524260 | IR (overlapping region) | 3 | 3 | ENST00000292524 | 12.99 | 3.00E-02 | 4.64E-01 |
| DPP7 | 9:137114379-137114462 | 9:137114379-137114382 | A3ss | 3 | 3 | ENST00000371579 | 12.98 | 3.45E-02 | 9.24E-01 |
| ENTPD6 | 20:25195868-25209848 | 20:25207076-25207397 | MXS | 3 | 3 | ENST00000418890 | 12.98 | 7.95E-03 | 2.38E-02 |
| AP1S2 | X:15827373-15845378 | X:15831452-15831568 | MES | 3 | 3 | ENST00000671830 | 12.97 | 3.73E-03 | 2.23E-01 |
| SNHG8 | 4:118278793-118279388 | 4:118279107-118279137 | SES | 3 | 3 | ENST00000602483 | 12.97 | 8.50E-03 | 5.07E-02 |
| TOPBP1 | 3:133652615-133653344 | 3:133652615-133652629 | A3ss | 3 | 3 | ENST00000260810 | 12.96 | 4.71E-02 | 1.90E-01 |
| TOPBP1 | 3:133652548-133653344 | 3:133652548-133652629 | A3ss | 3 | 3 | ENST00000260810 | 12.96 | 4.71E-02 | 1.90E-01 |
| UBQLN1 | 9:83665146-83669184 | 9:83666350-83666433 | SES | 3 | 3 | ENST00000376395 | 12.96 | 3.41E-03 | 7.11E-02 |
| ARMC10 | 7:103075412-103083681 | 7:103075777-103075881 | MXS | 3 | 3 | ENST00000323716 | 12.95 | 5.50E-03 | 5.09E-02 |
| PIGX | 3:196722557-196727922 | 3:196726655-196726771 | SES | 3 | 3 | ENST00000415832 | 12.93 | 1.20E-02 | 5.05E-02 |
| TSTD1 | 1:161038076-161038879 | 1:161038551-161038673 | SES | 3 | 3 | ENST00000318289 | 12.91 | 9.72E-03 | 2.47E-02 |
| ATG9A | 2:219228054-219229534 | 2:219228434-219228485 | SES | 3 | 3 | ENST00000361242 | 12.9 | 2.60E-03 | 9.53E-03 |
| C1orf52 | 1:85258723-85259357 | 1:85258935-85259061 | SES | 3 | 3 | ENST00000294661 | 12.9 | 4.48E-02 | 1.29E-01 |
| SEMA6C | 1:151133319-151134400 | 1:151133319-151133517 | A3ss | 3 | 3 | Ex.TSS.ENST00000341697 | 12.88 | 6.21E-03 | 1.53E-02 |
| ARMC6 | 19:19033931-19042710 | 19:19033931-19034013 | TSSIA5ss | 3 | 3 | TSS.ENST00000269932 | 12.84 | 8.39E-03 | 1.87E-02 |
| RNF138 | 18:32092887-32123517 | 18:32113745-32113860 | MES | 3 | 3 | ENST00000261593 | 12.84 | 2.42E-02 | 5.19E-02 |
| TIA1 | 2:70224630-70227734 | 2:70225328-70225393 | SES | 3 | 3 | ENST00000474809 | 12.84 | 2.12E-04 | 8.20E-04 |
| TSPAN5 | 4:98482176-98507677 | 4:98486738-98486884 | SES | 3 | 3 | ENST00000305798 | 12.84 | 1.46E-02 | 9.51E-02 |
| LRR1 | 14:49599204-49614255 | 14:49607400-49608121 | MES | 3 | 3 | ENST00000298288 | 12.82 | 1.77E-02 | 2.90E-02 |
| TTK | 6:80040704-80042118 | 6:80042098-80042119 | A3ss | 3 | 3 | Ex.TSS.ENST00000230510 | 12.81 | 5.54E-03 | 4.98E-02 |
| PAM | 5:103019844-103025133 | 5:103025131-103025133 | A3ss | 3 | 3 | ENST00000304400 | 12.79 | 7.78E-03 | 5.09E-02 |
| UCKL1 | 20:63941209-63944396 | 20:63941383-63944397 | A5ss | 3 | 3 | Ex.ENST00000358711 | 12.78 | 1.19E-03 | 3.84E-03 |
| UCKL1 | 20:63941209-63943652 | 20:63941383-63943653 | A5ss | 3 | 3 | Ex.ENST00000354216 | 12.78 | 1.19E-03 | 3.84E-03 |
| SDR39U1 | 14:24441796-24442345 | 14:24442178-24442345 | TSSIA5ss | 3 | 3 | TSS.ENST00000554947 | 12.77 | 2.77E-04 | 4.52E-04 |
| SLC27A1 | 19:17486563-17487335 | 19:17486958-17487099 | IR | 3 | 2 | ENST00000600277 | 12.77 | 1.16E-03 | 2.57E-03 |
| NEK3 | 13:52153993-52156074 | 13:52153993-52154155 | A3ss | 3 | 3 | Ex.ENST00000551355 | 12.76 | 4.18E-04 | 6.62E-04 |
| C2CD5 | 12:22457162-22469708 | 12:22458484-22458585 | MES | 3 | 3 | ENST00000396028 | 12.75 | 3.17E-02 | 4.77E-02 |
| PNPLA8 | 7:108515575-108526028 | 7:108521476-108521521 | MXS | 3 | 3 | ENST00000257694 | 12.73 | 1.93E-02 | 1.80E-01 |
| PRPF40B | 12:49642676-49642929 | 12:49642927-49642929 | A3ss | 3 | 3 | ENST00000261897 | 12.73 | 4.80E-02 | 7.30E-02 |
| SCAP | 3:47422537-47435007 | 3:47428513-47428670 | MES | 3 | 3 | ENST00000265565 | 12.73 | 1.93E-02 | 8.53E-02 |
| ZNF394 | 7:99494632-99499637 | 7:99498716-99498842 | SES | 3 | 3 | ENST00000337673 | 12.73 | 9.02E-03 | 1.11E-01 |
| FKRP | 19:46746091-46748026 | 19:46747234-46747305 | MXS | 3 | 3 | ENST00000600005 | 12.71 | 2.83E-02 | 6.57E-02 |
| ZNF154 | 19:57697367-57701145 | 19:57697842-57700987 | IR (overlapping region) | 2 | 3 | ENST00000512439 | 12.69 | 7.67E-03 | 1.84E-02 |
| DOK4 | 16:57479442-57479874 | 16:57479689-57479844 | IR | 3 | 3 | ENST00000569548 | 12.67 | 4.33E-02 | 8.26E-02 |
| PPP1R16A | 8:144495757-144496460 | 8:144495757-144496183 | A5ss | 3 | 3 | Ex.TSS.ENST00000533829 | 12.65 | 8.44E-03 | 1.28E-01 |
| PPP1R16A | 8:144490213-144496460 | 8:144490213-144496183 | A5ss | 3 | 3 | Ex.TSS.ENST00000292539 | 12.65 | 8.44E-03 | 1.30E-01 |
| XRCC3 | 14:103711568-103715423 | 14:103712875-103712968 | SES | 3 | 3 | ENST00000554913 | 12.61 | 5.50E-03 | 8.85E-03 |
| CENPE | 4:103136141-103136359 | 4:103136172-103136279 | IR | 3 | 3 | ENST00000265148 | 12.6 | 4.99E-02 | 2.94E-01 |
| DENND10 | 10:119104198-119107967 | 10:119107948-119107967 | TSSIA3ss | 3 | 3 | TSS.ENST00000448258 | 12.6 | 2.29E-02 | 3.03E-02 |
| EXOC6B | 2:72179462-72184074 | 2:72182894-72182905 | SES | 3 | 3 | ENST00000634650 | 12.58 | 5.87E-04 | 2.28E-03 |
| MYH14 | 19:50292390-50293232 | 19:50292390-50292895 | A5ss | 3 | 3 | Ex.ENST00000262269 | 12.57 | 1.76E-02 | 4.10E-02 |
| ATP2C1 | 3:131001315-131016151 | 3:131001315-131001344 | TSSIA5ss | 3 | 3 | TSS.ENST00000359644 | 12.49 | 4.94E-02 | 1.98E-01 |
| MAP2K7 | 19:7911331-7911435 | 19:7911415-7911435 | A3ss | 3 | 3 | ENST00000397981 | 12.49 | 4.90E-03 | 1.18E-02 |
| DPP8 | 15:65452103-65463760 | 15:65456225-65456371 | MXS | 3 | 3 | ENST00000321147 | 12.47 | 2.11E-02 | 3.58E-02 |
| TOP2B | 3:25645456-25664228 | 3:25645456-25645470 | TSSIA3ss | 3 | 3 | TSS.ENST00000264331 | 12.47 | 1.49E-03 | 6.31E-03 |
| RGL3 | 19:11406865-11415936 | 19:11411253-11411444 | SES | 3 | 3 | ENST00000563726 | 12.45 | 2.35E-02 | 5.13E-02 |
| NSUN5P2 | 7:72950197-72954624 | 7:72953326-72953430 | MES | 3 | 3 | ENST00000388955 | 12.43 | 3.18E-02 | 3.63E-01 |
| CBSL | 21:6458056-6460186 | 21:6459456-6459638 | SES | 3 | 3 | ENST00000624921 | 12.4 | 4.78E-02 | 1.57E-01 |
| RBM25 | 14:73099751-73103191 | 14:73099751-73100362 | A5ss | 3 | 3 | Ex.ENST00000261973 | 12.39 | 5.65E-04 | 9.32E-04 |
| ARHGEF35-AS1 | 7:144207805-144239042 | 7:144239040-144239042 | A3ss | 3 | 3 | ENST00000493248 | 12.38 | 4.01E-02 | 3.94E-01 |
| LYRM1 | 16:20900536-20920121 | 16:20915556-20915714 | MES | 3 | 3 | ENST00000568820 | 12.36 | 1.87E-05 | 3.48E-05 |
| JPH4 | 14:23576457-23578179 | 14:23577869-23577994 | MES | 3 | 3 | ENST00000397118 | 12.35 | 2.17E-02 | 3.53E-02 |
| RPP38 | 10:15097410-15103304 | 10:15102218-15102336 | SES | 3 | 3 | ENST00000378201 | 12.34 | 3.74E-02 | 4.96E-02 |
| SAT2 | 17:7626794-7627142 | 17:7626943-7627044 | SES | 3 | 3 | ENST00000269298 | 12.27 | 4.05E-02 | 8.59E-02 |
| PFDN6 | 6:33296829-33298401 | 6:33297344-33297930 | IR (overlapping region) | 3 | 3 | ENST00000463584 | 12.25 | 2.92E-02 | 2.44E-01 |
| SUOX | 12:55997340-56002211 | 12:55997615-55997723 | MES | 3 | 3 | ENST00000394115 | 12.23 | 1.80E-03 | 2.78E-03 |
| CCDC107 | 9:35660557-35661187 | 9:35660648-35660765 | IR (overlapping region) | 3 | 3 | ENST00000378406 | 12.22 | 6.08E-03 | 1.83E-01 |
| HIP1 | 7:75563264-75568198 | 7:75563264-75563281 | A3ss | 3 | 3 | Ex.ENST00000336926 | 12.22 | 3.01E-03 | 3.48E-02 |
| SPG7 | 16:89554564-89556886 | 16:89554564-89556085 | TSSIA5ss | 3 | 3 | TSS.ENST00000644225 | 12.22 | 2.97E-02 | 5.80E-02 |
| ZNF227 | 19:44228573-44234701 | 19:44229733-44229816 | SES | 3 | 3 | ENST00000313040 | 12.2 | 3.33E-02 | 7.69E-02 |
| MIER1 | 1:66926243-66940027 | 1:66928890-66928963 | SES | 3 | 3 | ENST00000371012 | 12.19 | 3.41E-02 | 9.83E-02 |
| CHD1L | 1:147264429-147264584 | 1:147264533-147264533 | IR (overlapping region) | 3 | 3 | ENST00000361293 | 12.15 | 6.20E-03 | 1.52E-02 |
| CCDC107 | 9:35660557-35661187 | 9:35660648-35660773 | IR (overlapping region) | 3 | 3 | ENST00000378406 | 12.1 | 3.10E-03 | 9.34E-02 |
| CPNE1 | 20:35632924-35664759 | 20:35658930-35659014 | MXS | 3 | 3 | ENST00000434795 | 12.1 | 1.21E-02 | 3.61E-02 |
| HMGN1 | 21:39345830-39347292 | 21:39345959-39347246 | IR | 3 | 3 | ENST00000492280 | 12.1 | 1.59E-02 | 5.20E-02 |

| | | | | | | | | | |
|---|---|---|---|---|---|---|---|---|---|
| POLR1D | 13:27621105-27621336 | 13:27621105-27621140 | A5ss | 3 | 3 | Ex.ENST00000626064 | 12.09 | 4.56E-02 | 7.14E-02 |
| FDPS | 1:155308966-155309788 | 1:155308966-155309076 | TSSlA5ss | 3 | 3 | TSS.ENST00000356657 | 12.07 | 7.38E-03 | 1.86E-02 |
| ZFAND5 | 9:72363607-72364695 | 9:72363607-72363631 | A3ss | 3 | 3 | Ex.TSS.ENST00000376962 | 12.07 | 4.30E-02 | 1.00E+00 |
| ZFAND5 | 9:72360788-72364695 | 9:72360788-72363631 | A3ss | 3 | 3 | Ex.TSS.ENST00000376960 | 12.07 | 4.30E-02 | 1.00E+00 |
| MARK2 | 11:63905044-63908876 | 11:63906088-63906114 | MES | 3 | 3 | ENST00000402010 | 12.03 | 4.11E-02 | 5.86E-02 |
| RBM12 | 20:35658983-35664759 | 20:35658983-35659014 | TSSlA3ss | 3 | 3 | TSS.ENST00000374114 | 12.01 | 9.96E-03 | 2.97E-02 |
| RAB30-DT | 11:83072703-83073302 | 11:83072703-83072804 | TSSlA5ss | 3 | 3 | TSS.ENST00000501011 | 12 | 3.43E-02 | 5.03E-02 |
| IMMP2L | 7:110663722-110963499 | 7:110886593-110886695 | MXS | 3 | 3 | ENST00000331762 | 11.99 | 4.54E-02 | 4.25E-01 |
| XRCC3 | 14:103712932-103715423 | 14:103712932-103712941 | A3ss | 3 | 3 | Ex.TSS.ENST00000555055 | 11.97 | 3.37E-03 | 5.43E-03 |
| XRCC3 | 14:103711662-103715423 | 14:103711662-103712941 | A3ss | 3 | 3 | Ex.TSS.ENST00000553332 | 11.97 | 3.37E-03 | 5.43E-03 |
| XRCC3 | 14:103711568-103715423 | 14:103711568-103712941 | A3ss | 3 | 3 | Ex.TSS.ENST00000352127 | 11.97 | 3.37E-03 | 5.43E-03 |
| C16orf70 | 16:67147045-67148544 | 16:67147095-67147305 | IR | 3 | 3 | ENST00000219139 | 11.96 | 4.51E-03 | 8.67E-03 |
| ZNF202 | 11:123740128-123740536 | 11:123740233-123740392 | IR | 3 | 3 | ENST00000530944 | 11.96 | 4.27E-02 | 5.76E-02 |
| CARM1 | 19:10921128-10921614 | 19:10921375-10921443 | SES | 3 | 3 | ENST00000327064 | 11.93 | 1.23E-02 | 2.67E-02 |
| TSTD1 | 1:161038635-161038879 | 1:161038635-161038673 | A3ss | 3 | 3 | Ex.TSS.ENST00000318289 | 11.88 | 4.43E-02 | 1.13E-01 |
| TSTD1 | 1:161038076-161038879 | 1:161038076-161038673 | A3ss | 3 | 3 | ENST00000486084 | 11.88 | 4.43E-02 | 1.13E-01 |
| CARM1 | 19:10921128-10921614 | 19:10921375-10921614 | TSSlA3ss | 3 | 3 | TSS.ENST00000590699 | 11.85 | 1.63E-02 | 3.55E-02 |
| GALNS | 16:88837622-88837889 | 16:88837766-88837839 | IR | 3 | 3 | ENST00000566563 | 11.85 | 7.24E-03 | 1.41E-02 |
| AL354821.1 | 13:55554985-55583489 | 13:55554985-55581529 | A3ss | 3 | 3 | Ex.ENST00000617598 | 11.81 | 4.41E-02 | 6.99E-02 |
| KHDC4 | 1:155921525-155921860 | 1:155921525-155921527 | A3ss | 3 | 3 | Ex.TSS.ENST00000368319 | 11.8 | 4.21E-02 | 1.07E-01 |
| SKA3 | 13:21155812-21176374 | 13:21159902-21159987 | MES | 3 | 3 | ENST00000298260 | 11.8 | 2.39E-02 | 3.76E-02 |
| BLOC1S6 | 15:45587797-45592134 | 15:45587797-45587876 | TSSlA5ss | 3 | 3 | TSS.ENST00000565409 | 11.79 | 2.60E-02 | 4.38E-02 |
| BAIAP3 | 16:1339245-1340921 | 16:1339496-1339603 | SES | 2 | 3 | ENST00000324385 | 11.78 | 8.00E-03 | 1.38E-02 |
| WWP1 | 8:86342931-86374029 | 8:86368939-86369031 | SES | 3 | 3 | ENST00000517970 | 11.77 | 2.05E-03 | 2.81E-02 |
| PCYT1B | X:24619085-24646988 | X:24640342-24646989 | A5ss | 3 | 3 | Ex.TSS.ENST00000356768 | 11.76 | 6.40E-03 | 4.00E-01 |
| SMUG1 | 12:54187906-54188514 | 12:54188511-54188514 | A5ss | 3 | 3 | ENST00000513838 | 11.76 | 3.30E-02 | 5.08E-02 |
| AP1B1 | 22:29328896-29330377 | 22:29329712-29329720 | SES | 3 | 3 | ENST00000357586 | 11.75 | 1.20E-02 | 4.02E-02 |
| PLA2G2A | 1:19978122-19978733 | 1:19978380-19978524 | SES | 3 | 3 | ENST00000375111 | 11.75 | 1.08E-02 | 2.79E-02 |
| AKAP13 | 15:85578930-85582107 | 15:85579840-85580019 | IR (overlapping region) | 3 | 3 | ENST00000361243 | 11.72 | 1.22E-02 | 2.10E-02 |
| HS6ST2 | X:132957327-132958174 | X:132957446-132958175 | A5ss | 3 | 3 | Ex.TSS.ENST00000370833 | 11.72 | 3.86E-02 | 7.96E-01 |
| ZNF826P | 19:20409025-20424785 | 19:20409025-20409028 | A3ss | 3 | 3 | Ex.TSS.ENST00000615684 | 11.72 | 7.49E-04 | 1.69E-03 |
| CCDC107 | 9:35660557-35661187 | 9:35660648-35660745 | IR (overlapping region) | 3 | 3 | ENST00000378406 | 11.69 | 8.29E-03 | 2.50E-01 |
| PACRGL | 4:20713540-20727284 | 4:20724808-20724888 | MXS | 3 | 3 | ENST00000471979 | 11.66 | 2.76E-03 | 1.70E-02 |
| AMT | 3:49422258-49422360 | 3:49422258-49422271 | TSSlA3ss | 3 | 3 | TSS.ENST00000273588 | 11.65 | 3.54E-04 | 1.59E-03 |
| WDR45 | X:49075454-49075544 | X:49075454-49075465 | A3ss | 3 | 3 | ENST00000322995 | 11.65 | 3.61E-02 | 1.00E+00 |
| ALG13 | X:111681827-111682235 | X:111682132-111682235 | TSSlA3ss | 3 | 3 | TSS.ENST00000482742 | 11.61 | 3.31E-02 | 6.83E-01 |
| TNK2 | 3:195867017-195867360 | 3:195867169-195867264 | SES | 3 | 3 | ENST00000672548 | 11.61 | 1.68E-02 | 7.09E-02 |
| CREM | 10:35127194-35178888 | 10:35148368-35148491 | MXS | 3 | 3 | ENST00000374726 | 11.6 | 2.82E-02 | 3.74E-02 |
| PHF21B | 22:44885530-44885862 | 22:44885530-44885552 | A3ss | 3 | 3 | Ex.ENST00000313237 | 11.57 | 5.60E-03 | 1.93E-02 |
| TMEM222 | 1:27330805-27332069 | 1:27330985-27331137 | SES | 3 | 3 | ENST00000464720 | 11.55 | 2.24E-02 | 6.26E-02 |
| PSME3IP1 | 16:57173870-57185820 | 16:57174169-57174286 | MXS | 3 | 3 | ENST00000565760 | 11.54 | 4.36E-05 | 8.32E-05 |
| ATAD1 | 10:87756923-87767672 | 10:87756923-87759800 | A3ss | 3 | 3 | Ex.ENST00000308448 | 11.52 | 2.82E-02 | 3.77E-02 |
| CENPK | 5:65552550-65561496 | 5:65554797-65554946 | SES | 3 | 3 | ENST00000510354 | 11.51 | 9.41E-04 | 6.72E-03 |
| SCN5A | 3:38609965-38620842 | 3:38613975-38614066 | MXS | 2 | 2 | ENST00000413689 | 11.51 | 1.19E-03 | 5.10E-03 |
| SH3BP5-AS1 | 3:15258228-15260582 | 3:15259051-15259760 | IR (overlapping region) | 3 | 3 | ENST00000655760 | 11.5 | 5.80E-03 | 2.37E-02 |
| LARP7 | 4:112644769-112646350 | 4:112644769-112644871 | A5ss | 3 | 3 | ENST00000324052 | 11.47 | 4.56E-02 | 2.72E-01 |
| CD200 | 3:112333225-112344961 | 3:112340902-112340983 | MXS | 3 | 3 | ENST00000315711 | 11.44 | 8.96E-03 | 3.55E-02 |
| PKP4 | 2:158674001-158680428 | 2:158678581-158678654 | MXS | 3 | 3 | ENST00000389757 | 11.44 | 1.09E-02 | 3.86E-02 |
| WASF1 | 6:110127630-110179438 | 6:110160635-110160732 | MXS | 3 | 3 | ENST00000359451 | 11.44 | 3.26E-02 | 2.49E-01 |
| SLC26A6 | 3:48629642-48629978 | 3:48629712-48629871 | IR | 3 | 3 | ENST00000496469 | 11.43 | 4.02E-02 | 1.80E-01 |
| SSH3 | 11:67307437-67307845 | 11:67307549-67307737 | SES | 3 | 3 | ENST00000308127 | 11.43 | 4.45E-02 | 6.45E-02 |
| TCF12 | 15:57251424-57253261 | 15:57252421-57252492 | SES | 3 | 3 | ENST00000333725 | 11.43 | 6.24E-03 | 1.05E-02 |
| TMBIM4 | 12:66138166-66138723 | 12:66138166-66138166 | TSSlA3ss | 3 | 3 | TSS.ENST00000286424 | 11.43 | 3.71E-02 | 5.75E-02 |
| FAM222B | 17:28766708-28842681 | 17:28790378-28790425 | MXS | 3 | 3 | ENST00000581229 | 11.4 | 2.99E-03 | 5.52E-03 |
| NEK4 | 3:52746905-52751931 | 3:52749692-52749829 | MES | 3 | 3 | ENST00000487068 | 11.39 | 1.89E-02 | 1.07E-01 |
| PRRC2B | 9:131492261-131495739 | 9:131494405-131494486 | SES | 3 | 3 | ENST00000357304 | 11.39 | 9.92E-03 | 2.54E-01 |
| SETD1B | 12:121817370-121817704 | 12:121817519-121817596 | IR | 3 | 3 | ENST00000604567 | 11.36 | 1.55E-02 | 2.27E-02 |
| RAB17 | 2:237577383-237585997 | 2:237578004-237578155 | SES | 3 | 3 | ENST00000264601 | 11.35 | 4.61E-02 | 1.70E-01 |
| CHD1L | 1:147242831-147265931 | 1:147254870-147254976 | MES | 3 | 3 | ENST00000369258 | 11.33 | 1.49E-04 | 3.67E-04 |
| NR1H3 | 11:47258130-47259790 | 11:47258130-47259259 | TSSlA5ss | 3 | 3 | TSS.ENST00000467728 | 11.33 | 3.49E-02 | 4.79E-02 |
| GTF2H2 | 5:71037547-71041521 | 5:71038429-71041522 | A5ss | 3 | 3 | Ex.ENST00000274400 | 11.32 | 2.40E-02 | 1.73E-01 |
| NAP1L4 | 11:2972244-2976023 | 11:2972244-2972279 | A3ss | 3 | 3 | ENST00000448187 | 11.31 | 1.92E-02 | 2.61E-02 |
| SAFB2 | 19:5595498-5598792 | 19:5595498-5595510 | A3ss | 3 | 3 | Ex.ENST00000252542 | 11.31 | 4.69E-03 | 1.12E-02 |
| DALRD3 | 3:49015708-49015818 | 3:49015804-49015818 | TSSlA5ss | 3 | 3 | TSS.ENST00000313778 | 11.3 | 1.22E-02 | 5.39E-02 |
| CCNYL1 | 2:207737447-207742222 | 2:207740655-207740706 | SES | 3 | 3 | ENST00000295414 | 11.29 | 1.59E-02 | 5.79E-02 |
| C19orf12 | 19:29702978-29715124 | 19:29708254-29708423 | MXS | 3 | 3 | ENST00000323670 | 11.28 | 2.07E-02 | 4.71E-02 |
| CTSL | 9:87726309-87727593 | 9:87726309-87726398 | TSSlA5ss | 3 | 3 | TSS.ENST00000343150 | 11.28 | 1.98E-02 | 6.33E-01 |
| CTSL | 9:87726254-87727593 | 9:87726254-87726398 | TSSlA5ss | 3 | 3 | TSS.ENST00000343150 | 11.28 | 1.98E-02 | 6.33E-01 |
| RABGGTB | 1:75790058-75791284 | 1:75790446-75790517 | MES | 3 | 3 | ENST00000471759 | 11.24 | 1.63E-03 | 4.70E-03 |
| THOC3 | 5:175918173-175961050 | 5:175959920-175961051 | A5ss | 3 | 3 | Ex.TSS.ENST00000432305 | 11.23 | 3.14E-02 | 2.20E-01 |
| UQCC1 | 20:35344110-35347272 | 20:35346462-35347163 | IR | 3 | 3 | ENST00000397554 | 11.2 | 1.56E-02 | 4.64E-02 |
| AC092279.2 | 19:24046280-24049409 | 19:24049405-24049409 | A3ss | 3 | 3 | ENST00000594934 | 11.19 | 4.96E-02 | 1.13E-01 |
| FANCA | 16:89740867-89744958 | 16:89742800-89742938 | SES | 3 | 3 | ENST00000305699 | 11.19 | 6.02E-03 | 1.17E-02 |
| FASTKD1 | 2:169531491-169537226 | 2:169531491-169531508 | A3ss | 3 | 3 | Ex.ENST00000453153 | 11.17 | 3.97E-03 | 1.41E-02 |
| PPFIA1 | 11:70272437-70324401 | 11:70272437-70313132 | A5ss | 3 | 3 | Ex.ENST00000253925 | 11.17 | 4.45E-02 | 6.46E-02 |
| FAM120A | 9:93498887-93515669 | 9:93515667-93515669 | A3ss | 3 | 3 | Ex.ENST00000277165 | 11.16 | 1.15E-02 | 3.91E-01 |
| GIT2 | 12:109947505-109953091 | 12:109951167-109951316 | SES | 3 | 3 | ENST00000355312 | 11.14 | 2.95E-02 | 4.37E-02 |
| METTL26 | 16:634798-636093 | 16:634889-636093 | TSSlA5ss | 3 | 3 | TSS.ENST00000448973 | 11.14 | 8.59E-03 | 1.64E-02 |
| METTL26 | 16:634798-635611 | 16:634889-635611 | TSSlA5ss | 3 | 3 | TSS.ENST00000448973 | 11.14 | 8.59E-03 | 1.64E-02 |
| METTL26 | 16:634798-635280 | 16:634889-635280 | TSSlA5ss | 3 | 3 | TSS.ENST00000448973 | 11.14 | 8.59E-03 | 1.64E-02 |
| SETD1B | 12:121817370-121817704 | 12:121817516-121817593 | IR | 3 | 3 | ENST00000604567 | 11.14 | 1.40E-02 | 2.05E-02 |
| TNK1 | 17:7386656-7386989 | 17:7386975-7386989 | A3ss | 3 | 3 | ENST00000576812 | 11.14 | 1.50E-02 | 3.14E-02 |
| TUT4 | 1:52446668-52461133 | 1:52458336-52458449 | SES | 3 | 3 | ENST00000257177 | 11.14 | 2.19E-02 | 5.48E-02 |
| AMZ2 | 17:68248070-68249096 | 17:68248087-68249010 | IR | 3 | 3 | ENST00000578175 | 11.13 | 3.62E-02 | 7.55E-02 |
| RBM39 | 20:35739018-35740823 | 20:35740525-35740597 | MXS | 3 | 3 | ENST00000403542 | 11.13 | 3.23E-03 | 9.70E-03 |
| GPBP1 | 5:57214194-57230845 | 5:57221376-57221396 | SES | 3 | 3 | ENST00000264779 | 11.11 | 4.59E-02 | 2.58E-01 |
| RFX3 | 9:3293259-3330258 | 9:3301546-3301620 | SES | 3 | 3 | ENST00000302303 | 11.11 | 1.62E-02 | 4.58E-01 |
| LMCD1 | 3:8501981-8548567 | 3:8537185-8537440 | MXS | 3 | 3 | ENST00000454244 | 11.1 | 3.24E-02 | 1.88E-01 |
| EBLN3P | 9:37079957-37084292 | 9:37080031-37084198 | IR | 2 | 2 | ENST00000654889 | 11.09 | 2.18E-02 | 6.83E-01 |
| FCHSD1 | 5:141646087-141646722 | 5:141646192-141646602 | IR | 3 | 3 | ENST00000523856 | 11.06 | 1.40E-02 | 9.40E-02 |
| OS9 | 12:57719183-57720405 | 12:57719183-57720263 | A5ss | 3 | 3 | ENST00000549307 | 11.06 | 5.71E-03 | 8.84E-03 |
| ATG9A | 2:219228054-219229009 | 2:219228434-219228482 | SES | 3 | 3 | ENST00000443140 | 11.05 | 4.47E-02 | 1.64E-01 |
| SENP6 | 6:75621626-75623899 | 6:75622751-75622858 | SES | 3 | 3 | ENST00000493959 | 11.03 | 1.27E-03 | 1.13E-02 |
| ACAA2 | 18:49785352-49787290 | 18:49785352-49785372 | A3ss | 3 | 3 | Ex.ENST00000285093 | 11.02 | 3.51E-05 | 7.54E-05 |

| | | | | | | | | | |
|---|---|---|---|---|---|---|---|---|---|
| FAM234A | 16:234858-249548 | 16:248244-248397 | MXS | 3 | 3 | ENST00000420046 | 11.01 | 6.48E-04 | 1.21E-03 |
| MAPKAP1 | 9:125484584-125543058 | 9:125506310-125506417 | SES | 3 | 3 | ENST00000265960 | 11 | 2.58E-02 | 5.57E-01 |
| RPAIN | 17:5422830-5432541 | 17:5425971-5428211 | SES | 3 | 3 | ENST00000573126 | 11 | 2.18E-02 | 4.02E-02 |
| ZNF74 | 22:20394663-20400631 | 22:20395333-20395418 | MXS | 3 | 3 | ENST00000400451 | 10.99 | 4.41E-02 | 1.46E-01 |
| CLDN7 | 17:7262344-7263158 | 17:7262611-7263158 | A5ss | 3 | 3 | ENST00000573745 | 10.98 | 4.24E-02 | 7.79E-02 |
| PILRB | 7:100353124-100355142 | 7:100353210-100353270 | SES | 3 | 3 | ENST00000448382 | 10.98 | 2.73E-02 | 2.51E-01 |
| MAP3K12 | 12:53484257-53485214 | 12:53484366-53485055 | IR | 3 | 3 | ENST00000547020 | 10.97 | 7.07E-03 | 1.08E-02 |
| AEN | 15:88621664-88626145 | 15:88621664-88621797 | A5ss | 3 | 3 | ENST00000558327 | 10.95 | 2.00E-03 | 3.44E-03 |
| AEN | 15:88621383-88626145 | 15:88621383-88621797 | A5ss | 3 | 3 | ENST00000558327 | 10.95 | 2.00E-03 | 3.44E-03 |
| LARP4 | 12:50427910-50428934 | 12:50428917-50428934 | A3ss | 3 | 3 | ENST00000429001 | 10.95 | 4.23E-03 | 6.45E-03 |
| STAU2 | 8:73688814-73746782 | 8:73738284-73738349 | MES | 3 | 3 | ENST00000519818 | 10.95 | 7.39E-03 | 9.91E-02 |
| GEMIN2 | 14:39122544-39128279 | 14:39124992-39125036 | SES | 3 | 3 | ENST00000308317 | 10.94 | 1.31E-02 | 2.14E-02 |
| RBPJ | 4:26321049-26386352 | 4:26362446-26362628 | SES | 3 | 3 | ENST00000355476 | 10.92 | 3.10E-02 | 1.91E-01 |
| HMMR | 5:163464803-163469640 | 5:163467701-163467748 | SES | 3 | 3 | ENST00000393915 | 10.9 | 1.61E-02 | 1.12E-01 |
| STX16 | 20:58668128-58670511 | 20:58669291-58669453 | SES | 3 | 3 | ENST00000312283 | 10.9 | 2.45E-03 | 7.90E-03 |
| TRIB2 | 2:12717011-12717639 | 2:12717252-12717639 | A3ss | 3 | 3 | Ex.ENST00000483034 | 10.88 | 3.65E-02 | 1.28E-01 |
| CPNE1 | 20:35632924-35664759 | 20:35655202-35655344 | MES | 3 | 3 | ENST00000352393 | 10.87 | 9.86E-03 | 2.94E-02 |
| STX16 | 20:58652088-58659622 | 20:58652088-58652138 | A5ss | 3 | 3 | ENST00000359617 | 10.87 | 4.23E-02 | 1.35E-01 |
| MTF2 | 1:93079532-93114687 | 1:93110545-93110626 | MES | 3 | 3 | ENST00000370298 | 10.85 | 2.24E-02 | 6.66E-02 |
| RAB4B | 19:40778392-40786664 | 19:40780385-40780499 | MES | 3 | 3 | ENST00000357052 | 10.85 | 4.96E-02 | 1.14E-01 |
| RCAN3 | 1:24503118-24514313 | 1:24503118-24503150 | TSSlA5ss | 3 | 3 | TSS.ENST00000374395 | 10.85 | 1.81E-02 | 5.03E-02 |
| AC040160.1 | 16:67277510-67280806 | 16:67278883-67279875 | IR | 3 | 3 | ENST00000563969 | 10.81 | 4.39E-02 | 8.43E-02 |
| ARHGAP22 | 10:48459892-48479635 | 10:48479588-48479635 | A5ss | 3 | 3 | ENST00000374172 | 10.8 | 1.36E-02 | 1.81E-02 |
| CLN3 | 16:28484119-28487455 | 16:28486578-28486650 | MXS | 3 | 3 | ENST00000355477 | 10.8 | 1.09E-02 | 2.04E-02 |
| FRS2 | 12:69532057-69562179 | 12:69537916-69537983 | MES | 3 | 3 | ENST00000550937 | 10.8 | 9.40E-03 | 1.46E-02 |
| FMR1 | X:147937601-147940575 | X:147938099-147938161 | SES | 3 | 3 | ENST00000370470 | 10.77 | 1.75E-02 | 7.34E-01 |
| TOP3B | 22:21975808-21982729 | 22:21975981-21976187 | MXS | 3 | 3 | ENST00000449704 | 10.76 | 2.04E-03 | 6.76E-03 |
| C2CD5 | 12:22457162-22469708 | 12:22459492-22459542 | MES | 3 | 3 | ENST00000545552 | 10.75 | 3.50E-02 | 5.26E-02 |
| MIR302CHG | 4:112646969-112648653 | 4:112647415-112647467 | SES | 3 | 3 | ENST00000509938 | 10.75 | 1.11E-03 | 6.60E-03 |
| RNASEH1-AS1 | 2:3558999-3561316 | 2:3559448-3561317 | A3ss | 3 | 3 | Ex.ENST00000426725 | 10.72 | 3.65E-02 | 1.38E-01 |
| ANKRD11 | 16:89291778-89305205 | 16:89291778-89291799 | A3ss | 3 | 3 | ENST00000645278 | 10.71 | 1.42E-04 | 2.77E-04 |
| PELI1 | 2:64100500-64104700 | 2:64104554-64104597 | SES | 3 | 3 | ENST00000466177 | 10.71 | 1.12E-02 | 4.23E-02 |
| SUZ12P1 | 17:30709812-30759490 | 17:30743301-30743369 | MES | 3 | 3 | ENST00000579526 | 10.68 | 3.12E-03 | 6.21E-03 |
| ELMOD3 | 2:85354830-85356966 | 2:85355555-85355896 | MXS | 3 | 3 | ENST00000462891 | 10.67 | 4.27E-02 | 1.67E-01 |
| TM2D1 | 1:61683547-61694696 | 1:61686792-61686895 | SES | 3 | 3 | ENST00000468586 | 10.66 | 6.71E-03 | 1.93E-02 |
| ZNF74 | 22:20394663-20400631 | 22:20394663-20395418 | TSSlA5ss | 3 | 3 | TSS.ENST00000493734 | 10.66 | 1.70E-02 | 5.63E-02 |
| STAU1 | 20:49124588-49135832 | 20:49124588-49124605 | A3ss | 3 | 3 | ENST00000371802 | 10.65 | 2.71E-02 | 8.69E-02 |
| ZNF131 | 5:43174447-43175719 | 5:43174495-43175455 | IR | 3 | 3 | ENST00000306938 | 10.65 | 1.25E-02 | 8.87E-02 |
| AKT1S1 | 19:49873303-49877236 | 19:49876603-49877236 | TSSlA5ss | 3 | 3 | TSS.ENST00000391835 | 10.64 | 1.70E-02 | 3.90E-02 |
| AKT1S1 | 19:49873303-49877024 | 19:49876603-49877024 | TSSlA5ss | 3 | 3 | TSS.ENST00000391835 | 10.64 | 1.70E-02 | 3.90E-02 |
| DUS2 | 16:68023352-68038005 | 16:68025415-68025494 | SES | 3 | 3 | ENST00000432752 | 10.61 | 3.15E-02 | 6.08E-02 |
| SREBF1 | 17:17819538-17820864 | 17:17819726-17820089 | IR | 3 | 3 | ENST00000395751 | 10.61 | 4.87E-02 | 8.95E-02 |
| DLST | 14:74882017-74889094 | 14:74885586-74885634 | MXS | 3 | 3 | ENST00000555492 | 10.6 | 5.23E-04 | 8.65E-04 |
| ATM | 11:108223187-108227594 | 11:108224784-108224871 | MXS | 3 | 3 | ENST00000452508 | 10.59 | 1.86E-02 | 2.50E-02 |
| SIGIRR | 11:406539-407061 | 11:406843-406993 | SES | 3 | 3 | ENST00000332725 | 10.59 | 3.80E-02 | 5.20E-02 |
| PWWP3A | 19:1366382-1369264 | 19:1367160-1367220 | SES | 3 | 3 | ENST00000415183 | 10.58 | 3.25E-03 | 7.15E-03 |
| SLC25A36 | 3:140963228-140970926 | 3:140967000-140967036 | MXS | 3 | 3 | ENST00000502594 | 10.58 | 1.11E-02 | 4.50E-02 |
| GSN | 9:121268220-121301962 | 9:121281470-121281562 | MES | 3 | 3 | ENST00000373808 | 10.57 | 8.76E-03 | 1.82E-01 |
| PHF21A | 11:46084303-46120934 | 11:46092183-46092223 | SES | 3 | 3 | ENST00000323180 | 10.57 | 6.46E-03 | 8.86E-03 |
| MIB2 | 1:1625429-1626649 | 1:1625546-1625653 | SES | 3 | 3 | ENST00000355826 | 10.55 | 2.11E-02 | 5.37E-02 |
| TMEM179B | 11:62787528-62789022 | 11:62789010-62789023 | A3ss | 3 | 3 | Ex.TSS.ENST00000333449 | 10.55 | 1.87E-02 | 2.66E-02 |
| S100A16 | 1:153608178-153609177 | 1:153608899-153609177 | TSSlA5ss | 3 | 3 | TSS.ENST00000368705 | 10.54 | 3.49E-03 | 8.74E-03 |
| LMO3 | 12:16560539-16600654 | 12:16594137-16594334 | MES | 3 | 3 | ENST00000541764 | 10.53 | 1.44E-03 | 2.17E-03 |
| ATP9B | 18:79154556-79193182 | 18:79176813-79176907 | SES | 3 | 2 | ENST00000307671 | 10.5 | 1.96E-02 | 4.26E-02 |
| CPNE1 | 20:35658983-35664801 | 20:35658983-35659014 | A3ss | 3 | 3 | Ex.TSS.ENST00000414711 | 10.5 | 1.26E-02 | 3.76E-02 |
| CERS5 | 12:50144058-50167100 | 12:50144769-50144969 | MXS | 3 | 3 | ENST00000422340 | 10.49 | 1.31E-03 | 2.00E-03 |
| RPAIN | 17:5422830-5432541 | 17:5425971-5426082 | MXS | 3 | 3 | ENST00000327154 | 10.49 | 3.45E-03 | 7.12E-03 |
| SIPA1L2 | 1:232403572-232415493 | 1:232404125-232404178 | SES | 3 | 3 | ENST00000262861 | 10.49 | 1.74E-02 | 4.80E-02 |
| CHKA | 11:68074831-68097018 | 11:68081404-68081457 | SES | 3 | 3 | ENST00000265689 | 10.48 | 5.41E-03 | 7.85E-03 |
| OS9 | 12:57719183-57720405 | 12:57720099-57720263 | SES | 3 | 3 | ENST00000315970 | 10.46 | 2.27E-02 | 3.51E-02 |
| RBM5 | 3:50100052-50103082 | 3:50100532-50100605 | SES | 3 | 3 | ENST00000347869 | 10.45 | 4.69E-02 | 2.11E-01 |
| TRIM46 | 1:155174030-155175385 | 1:155174601-155174641 | SES | 3 | 3 | ENST00000392451 | 10.45 | 1.03E-02 | 2.58E-02 |
| SNHG14 | 15:25062089-25070119 | 15:25064715-25064855 | MES | 3 | 3 | ENST00000654223 | 10.43 | 6.24E-03 | 1.04E-02 |
| PICK1 | 22:38057437-38057752 | 22:38057437-38057489 | TSSlA5ss | 3 | 3 | TSS.ENST00000404072 | 10.42 | 1.89E-02 | 6.46E-02 |
| REC8 | 14:24172080-24172608 | 14:24172366-24172491 | IR | 3 | 3 | ENST00000611366 | 10.41 | 4.81E-02 | 7.81E-02 |
| MDM4 | 1:204542945-204546796 | 1:204544535-204544684 | SES | 2 | 3 | ENST00000367179 | 10.4 | 2.51E-02 | 6.51E-02 |
| NUB1 | 7:151374202-151375847 | 7:151374202-151374243 | A5ss | 3 | 3 | ENST00000470229 | 10.4 | 2.14E-02 | 2.13E-01 |
| CNNM3 | 2:96829135-96832551 | 2:96831969-96832068 | SES | 3 | 3 | ENST00000480035 | 10.39 | 2.20E-02 | 8.71E-02 |
| RPAIN | 17:5422830-5432541 | 17:5425971-5426299 | MXS | 3 | 3 | ENST00000571043 | 10.39 | 9.62E-04 | 1.99E-03 |
| APBB3 | 5:140562353-140562842 | 5:140562500-140562662 | IR (overlapping region) | 3 | 3 | ENST00000503979 | 10.38 | 4.66E-02 | 3.13E-01 |
| DCAF11 | 14:24115393-24115542 | 14:24115515-24115542 | TSSlA3ss | 3 | 3 | TSS.ENST00000559396 | 10.38 | 1.68E-02 | 2.73E-02 |
| L3MBTL1 | 20:43528657-43529368 | 20:43528746-43529263 | IR (overlapping region) | 3 | 3 | ENST00000648786 | 10.38 | 1.62E-02 | 5.17E-02 |
| UPF3A | 13:114282128-114286518 | 13:114286302-114286400 | MES | 3 | 3 | ENST00000375299 | 10.37 | 3.25E-02 | 5.11E-02 |
| VEGFA | 6:43778919-43784540 | 6:43780732-43780803 | MES | 3 | 3 | ENST00000425836 | 10.37 | 2.63E-02 | 2.28E-01 |
| DPH5 | 1:100995150-101021640 | 1:101001467-101001587 | MXS | 3 | 3 | ENST00000477293 | 10.36 | 9.73E-03 | 2.36E-02 |
| RPAIN | 17:5422830-5432541 | 17:5425971-5426082 | MES | 3 | 3 | ENST00000539417 | 10.33 | 1.77E-03 | 3.65E-03 |
| AC093752.1 | 4:119455134-119460182 | 4:119457971-119458077 | SES | 3 | 3 | ENST00000515251 | 10.32 | 3.61E-02 | 2.16E-01 |
| ACTR5 | 20:38756040-38765401 | 20:38761653-38765402 | A3ss | 3 | 3 | Ex.ENST00000243903 | 10.32 | 3.70E-02 | 1.12E-01 |
| ARFGAP2 | 11:47173883-47175014 | 11:47173883-47173915 | A3ss | 3 | 3 | ENST00000528444 | 10.3 | 2.50E-05 | 3.43E-05 |
| ARFGAP2 | 11:47173841-47175014 | 11:47173841-47173915 | A3ss | 3 | 3 | ENST00000528444 | 10.3 | 2.50E-05 | 3.43E-05 |
| CENPX | 17:82019382-82019857 | 17:82019641-82019694 | SES | 3 | 3 | ENST00000392359 | 10.3 | 4.32E-04 | 9.19E-04 |
| MALAT1 | 11:65497762-65506469 | 11:65502394-65504325 | IR | 3 | 3 | ENST00000534336 | 10.3 | 1.65E-02 | 2.36E-02 |
| THOC2 | X:123610958-123611439 | X:123610958-123610963 | A3ss | 3 | 3 | ENST00000245838 | 10.27 | 3.74E-03 | 1.46E-01 |
| ATXN3 | 14:92093319-92096092 | 14:92093746-92093850 | SES | 3 | 3 | ENST00000556374 | 10.26 | 4.47E-02 | 7.43E-02 |
| SGSM2 | 17:2371863-2374281 | 17:2372489-2372952 | IR | 3 | 3 | ENST00000573717 | 10.26 | 4.75E-02 | 9.44E-02 |
| NUMA1 | 11:72007189-72008845 | 11:72007436-72007925 | IR | 3 | 3 | ENST00000545721 | 10.25 | 1.00E-02 | 1.45E-02 |
| ST3GAL2 | 16:70399534-70438948 | 16:70400696-70401094 | SES | 3 | 3 | ENST00000567586 | 10.23 | 7.09E-03 | 1.37E-02 |
| ATRIP | 3:48460110-48460799 | 3:48460420-48460708 | IR | 3 | 3 | ENST00000320211 | 10.22 | 6.37E-03 | 2.83E-02 |
| TRPC4 | 13:37636641-37637625 | 13:37637232-37637483 | IR | 3 | 3 | ENST00000338947 | 10.21 | 4.78E-02 | 7.53E-02 |
| C8orf58 | 8:22603195-22604142 | 8:22603289-22603290 | IR | 3 | 3 | ENST00000289989 | 10.19 | 1.24E-02 | 1.56E-01 |
| AC058822.1 | 4:53377924-53379224 | 4:53379073-53379117 | SES | 3 | 3 | ENST00000507166 | 10.18 | 2.88E-02 | 1.83E-01 |
| BID | 22:17739489-17774380 | 22:17750105-17750174 | MXS | 3 | 3 | ENST00000614949 | 10.18 | 2.98E-02 | 9.80E-02 |
| TENT2 | 5:79649191-79668891 | 5:79656958-79657001 | SES | 3 | 3 | ENST00000296783 | 10.18 | 3.35E-03 | 2.46E-02 |
| NPEPL1 | 20:58712580-58713419 | 20:58713351-58713420 | A3ss | 3 | 3 | Ex.ENST00000356091 | 10.17 | 8.83E-03 | 2.84E-02 |

| | | | | | | | | | |
|---|---|---|---|---|---|---|---|---|---|
| NELFE | 6:31954878-31955058 | 6:31954878-31954892 | A3ss | 3 | 3 | ENST00000375425 | 10.16 | 2.23E-02 | 1.85E-01 |
| STK19 | 6:31972512-31978902 | 6:31972621-31972757 | SES | 3 | 3 | ENST00000375331 | 10.15 | 5.15E-03 | 3.66E-02 |
| AC084125.2 | 8:144498752-144499305 | 8:144498841-144498915 | IR (overlapping region) | 3 | 3 | ENST00000532766 | 10.14 | 7.15E-03 | 1.09E-01 |
| PLSCR1 | 3:146533551-146544466 | 3:146536540-146536565 | MXS | 3 | 3 | ENST00000342435 | 10.14 | 8.95E-04 | 3.65E-03 |
| PSMA3-AS1 | 14:58291708-58297954 | 14:58293766-58295811 | MXS | 3 | 3 | ENST00000557660 | 10.13 | 1.69E-02 | 2.77E-02 |
| SUOX | 12:56002272-56003617 | 12:56002272-56002720 | TSSlA5ss | 3 | 3 | TSS.ENST00000546712 | 10.13 | 2.56E-02 | 3.94E-02 |
| DOCK6 | 19:11250874-11252958 | 19:11252249-11252481 | IR | 3 | 3 | ENST00000585609 | 10.1 | 4.02E-02 | 8.78E-02 |
| PABPC4 | 1:39563923-39564470 | 1:39564423-39564470 | A5ss | 3 | 3 | ENST00000372858 | 10.1 | 4.91E-02 | 1.39E-01 |
| SREK1 | 5:66175042-66177516 | 5:66177514-66177516 | A3ss | 3 | 3 | ENST00000284041 | 10.1 | 7.97E-03 | 5.73E-02 |
| NPIPA9 | 16:18374832-18375069 | 16:18374885-18374952 | IR (overlapping region) | 3 | 3 | ENST00000427999 | 10.08 | 2.91E-02 | 5.36E-02 |
| INKA2 | 1:111722064-111727804 | 1:111723149-111727563 | IR | 3 | 3 | ENST00000357260 | 10.04 | 1.05E-03 | 2.56E-03 |
| AC002316.1 | 17:2985067-2991296 | 17:2985067-2985197 | A5ss | 3 | 3 | Ex.ENST00000254695 | 10.01 | 1.90E-02 | 3.50E-02 |
| BAZ2A | 12:56615608-56630124 | 12:56617395-56617532 | SES | 3 | 3 | ENST00000549884 | 10 | 3.81E-02 | 5.88E-02 |
| AMT | 3:49421573-49422360 | 3:49422104-49422257 | MXS | 3 | 3 | ENST00000427987 | -10 | 1.25E-02 | 5.55E-02 |
| BIRC2 | 11:102348598-102350749 | 11:102348765-102349877 | IR | 3 | 3 | ENST00000227758 | -10 | 4.66E-02 | 6.25E-02 |
| CIAPIN1 | 16:57440960-57447341 | 16:57440960-57440983 | TSSlA3ss | 3 | 3 | TSS.ENST00000394391 | -10 | 2.10E-03 | 4.02E-03 |
| CNST | 1:246591942-246631893 | 1:246621429-246621634 | SES | 3 | 3 | ENST00000366512 | -10.02 | 3.89E-02 | 1.07E-01 |
| ANKLE1 | 19:17283885-17284088 | 19:17283885-17283962 | A5ss | 3 | 3 | ENST00000394458 | -10.04 | 4.44E-02 | 9.83E-02 |
| DUSP23 | 1:159780994-159781098 | 1:159780994-159781000 | A5ss | 3 | 3 | Ex.TSS.ENST00000368108 | -10.06 | 3.79E-02 | 9.64E-02 |
| MRPS18C | 4:83456959-83459739 | 4:83458346-83458429 | SES | 3 | 3 | ENST00000295491 | -10.06 | 7.38E-04 | 4.79E-03 |
| CENPT | 16:67832109-67832454 | 16:67832228-67832354 | SES | 3 | 3 | ENST00000566067 | -10.07 | 9.06E-03 | 1.73E-02 |
| TPM1 | 15:63060940-63061712 | 15:63061198-63061273 | SES | 3 | 3 | ENST00000558264 | -10.07 | 5.65E-03 | 9.57E-03 |
| ALKBH4 | 7:102459711-102464713 | 7:102459711-102459801 | TSSlA3ss | 3 | 3 | TSS.ENST00000292566 | -10.12 | 2.77E-02 | 2.52E-01 |
| CDCA7 | 2:173358838-173359342 | 2:173359255-173359342 | A3ss | 3 | 3 | ENST00000306721 | -10.13 | 1.53E-02 | 5.48E-02 |
| DLG1 | 3:197075871-197081050 | 3:197076586-197076682 | SES | 3 | 3 | ENST00000469073 | -10.13 | 1.02E-02 | 4.31E-02 |
| ECPAS | 9:111451556-111484115 | 9:111451556-111473000 | A3ss | 3 | 3 | Ex.TSS.ENST00000338205 | -10.13 | 4.47E-02 | 6.38E-01 |
| AC093323.1 | 4:6673969-6676013 | 4:6674707-6674992 | IR | 3 | 3 | ENST00000635031 | -10.14 | 3.73E-03 | 2.37E-02 |
| NDRG2 | 14:21023322-21025028 | 14:21024030-21024139 | SES | 3 | 3 | ENST00000556008 | -10.16 | 1.15E-02 | 1.87E-02 |
| LYPLAL1 | 1:219173982-219179146 | 1:219173982-219174304 | TSSlA5ss | 3 | 3 | TSS.ENST00000483635 | -10.19 | 1.27E-03 | 3.45E-03 |
| AMDHD2 | 16:2529475-2531422 | 16:2529526-2530256 | IR | 3 | 3 | ENST00000302956 | -10.2 | 2.65E-02 | 4.95E-02 |
| ICMT | 1:6232120-6234885 | 1:6233474-6233643 | MXS | 3 | 3 | ENST00000343813 | -10.21 | 1.56E-02 | 4.47E-02 |
| MSI2 | 17:57652162-57674971 | 17:57674918-57674971 | A3ss | 3 | 3 | ENST00000416426 | -10.21 | 8.46E-04 | 1.75E-03 |
| SETD5 | 3:9433951-9434333 | 3:9434277-9434333 | A3ss | 3 | 3 | ENST00000407969 | -10.22 | 3.62E-02 | 2.10E-01 |
| HPAT5 | 6:167231958-167237240 | 6:167232479-167232560 | SES | 3 | 3 | ENST00000628362 | -10.23 | 1.03E-02 | 8.20E-02 |
| SUMF2 | 7:56073112-56074585 | 7:56074174-56074218 | SES | 3 | 3 | ENST00000275607 | -10.23 | 3.61E-02 | 4.03E-01 |
| TRIP12 | 2:229805803-229807707 | 2:229805803-229805883 | A3ss | 3 | 3 | ENST00000389045 | -10.23 | 2.83E-04 | 1.04E-03 |
| VPS9D1-AS1 | 16:89716471-89716904 | 16:89716625-89716729 | IR (overlapping region) | 3 | 3 | ENST00000562866 | -10.23 | 2.41E-03 | 4.71E-03 |
| PSMC3IP | 17:42577461-42577697 | 17:42577562-42577652 | IR | 3 | 3 | ENST00000587209 | -10.24 | 2.55E-02 | 5.18E-02 |
| PAXX | 9:136992640-136992974 | 9:136992705-136992924 | IR | 3 | 3 | ENST00000493968 | -10.26 | 4.76E-03 | 1.24E-01 |
| SLC39A11 | 17:73088776-73092610 | 17:73092495-73092610 | TSSlA5ss | 3 | 3 | TSS.ENST00000580557 | -10.26 | 3.92E-02 | 7.21E-02 |
| UPP1 | 7:48089419-48090187 | 7:48089509-48089636 | SES | 3 | 3 | ENST00000436673 | -10.27 | 4.12E-04 | 4.43E-03 |
| AC093323.1 | 4:6673969-6676013 | 4:6674638-6674992 | IR | 3 | 3 | ENST00000635031 | -10.28 | 6.22E-03 | 3.95E-02 |
| P2RX4 | 12:121228725-121229099 | 12:121228867-121228962 | IR | 3 | 3 | ENST00000541187 | -10.28 | 1.07E-02 | 1.59E-02 |
| ATG9A | 2:219228486-219229534 | 2:219229010-219229174 | SES | 3 | 3 | ENST00000436856 | -10.29 | 3.58E-02 | 1.31E-01 |
| SNTB2 | 16:69187734-69245601 | 16:69187734-69187746 | TSSlA5ss | 3 | 3 | TSS.ENST00000336278 | -10.29 | 3.89E-02 | 7.43E-02 |
| ABCD4 | 14:74292870-74295147 | 14:74292870-74293248 | A3ss | 3 | 3 | ENST00000496015 | -10.32 | 4.26E-02 | 7.04E-02 |
| AP1S2 | X:15831569-15845378 | X:15833444-15833524 | SES | 3 | 3 | ENST00000479184 | -10.33 | 2.67E-03 | 1.60E-01 |
| RNF44 | 5:176531882-176532516 | 5:176532194-176532365 | IR | 3 | 3 | ENST00000504434 | -10.34 | 3.08E-03 | 2.16E-02 |
| DROSHA | 5:31515220-31526078 | 5:31521123-31521215 | MES | 3 | 3 | ENST00000344624 | -10.35 | 4.95E-02 | 3.16E-01 |
| CYP2B6 | 19:41004447-41006936 | 19:41006905-41006936 | A3ss | 3 | 3 | Ex.ENST00000324071 | -10.36 | 4.11E-02 | 9.44E-02 |
| GGT1 | 22:24608024-24609964 | 22:24609962-24609964 | A3ss | 3 | 3 | ENST00000411974 | -10.36 | 2.69E-02 | 8.92E-02 |
| AHSA2P | 2:61185502-61186497 | 2:61185866-61186445 | IR | 3 | 3 | ENST00000471542 | -10.38 | 1.63E-02 | 6.17E-02 |
| PLPP5 | 8:38263407-38266140 | 8:38263407-38264604 | TSSlA3ss | 3 | 3 | TSS.ENST00000424479 | -10.38 | 2.93E-02 | 3.66E-01 |
| SECISBP2 | 9:89319798-89325547 | 9:89325427-89325547 | A3ss | 3 | 3 | ENST00000375807 | -10.39 | 4.36E-02 | 8.92E-01 |
| ZNF664 | 12:123974021-124011380 | 12:123988043-123988138 | MXS | 3 | 3 | ENST00000337815 | -10.42 | 4.19E-02 | 6.26E-02 |
| FOXH1 | 8:144475224-144475582 | 8:144475224-144475261 | TSSlA3ss | 3 | 3 | TSS.ENST00000377317 | -10.43 | 3.52E-02 | 5.41E-01 |
| FBXL6 | 8:144356808-144357973 | 8:144357122-144357456 | IR (overlapping region) | 3 | 3 | ENST00000524492 | -10.44 | 4.01E-02 | 6.09E-01 |
| BIRC2 | 11:102348598-102350749 | 11:102348765-102350000 | IR | 3 | 3 | ENST00000227758 | -10.45 | 4.94E-02 | 6.62E-02 |
| DDR1 | 6:30882984-30884537 | 6:30884058-30884145 | MES | 3 | 3 | ENST00000454612 | -10.47 | 4.35E-02 | 3.41E-01 |
| GNPDA1 | 5:142012042-142012994 | 5:142012288-142012339 | SES | 3 | 3 | ENST00000500692 | -10.47 | 4.23E-03 | 2.90E-02 |
| CTNND1 | 11:57762047-57794009 | 11:57789037-57789155 | MES | 3 | 3 | ENST00000534579 | -10.48 | 1.81E-02 | 2.50E-02 |
| SLC7A6 | 16:68297234-68301819 | 16:68297477-68298490 | IR (overlapping region) | 3 | 3 | ENST00000219343 | -10.48 | 4.46E-03 | 8.59E-03 |
| ASCC2 | 22:29825781-29834488 | 22:29832245-29832342 | MXS | 3 | 3 | ENST00000397771 | -10.49 | 1.59E-02 | 5.33E-02 |
| RUSC1 | 1:155324577-155324943 | 1:155324588-155324844 | IR | 3 | 3 | ENST00000292254 | -10.5 | 4.81E-02 | 1.21E-01 |
| CCDC82 | 11:96387614-96389843 | 11:96389365-96389523 | SES | 3 | 3 | ENST00000645366 | -10.53 | 9.10E-03 | 1.34E-02 |
| MAPK15 | 8:143721000-143721673 | 8:143721106-143721230 | IR | 3 | 3 | ENST00000461928 | -10.54 | 1.24E-02 | 1.88E-01 |
| PHF6 | X:134425201-134428789 | X:134425331-134425660 | IR | 3 | 3 | ENST00000332070 | -10.54 | 2.75E-03 | 1.11E-01 |
| ROBO3 | 11:124877519-124878410 | 11:124877659-124878297 | IR | 3 | 3 | ENST00000534598 | -10.59 | 2.54E-02 | 3.44E-02 |
| ZNF638 | 2:71331876-71355718 | 2:71348753-71350271 | SES | 3 | 3 | ENST00000264447 | -10.59 | 1.91E-02 | 7.43E-02 |
| BIRC2 | 11:102348598-102350749 | 11:102348720-102350345 | IR | 3 | 3 | ENST00000227758 | -10.6 | 1.32E-03 | 1.76E-03 |
| ZNF266 | 19:9418505-9419081 | 19:9418896-9419050 | IR | 3 | 3 | ENST00000591213 | -10.61 | 2.02E-02 | 4.86E-02 |
| POLM | 7:44074026-44074397 | 7:44074131-44074220 | SES | 3 | 3 | ENST00000430942 | -10.62 | 2.98E-02 | 3.14E-01 |
| RANGRF | 17:8289503-8290090 | 17:8289589-8289812 | IR (overlapping region) | 3 | 3 | ENST00000407006 | -10.62 | 2.87E-02 | 6.12E-02 |
| LPCAT4 | 15:34360210-34362195 | 15:34361169-34362196 | A5ss | 3 | 3 | Ex.ENST00000567507 | -10.63 | 8.43E-03 | 1.41E-02 |
| LPCAT4 | 15:34360210-34361399 | 15:34361169-34361400 | A5ss | 3 | 3 | Ex.ENST00000314891 | -10.63 | 8.43E-03 | 1.41E-02 |
| NEIL2 | 8:11769710-11770335 | 8:11769792-11770173 | IR | 3 | 3 | ENST00000284503 | -10.63 | 3.31E-02 | 4.75E-01 |
| ARHGEF19 | 1:16208226-16212501 | 1:16208643-16209083 | SES | 3 | 3 | ENST00000270747 | -10.66 | 2.09E-02 | 5.30E-02 |
| MAP3K10 | 19:40198556-40205120 | 19:40204485-40204633 | SES | 3 | 3 | ENST00000253055 | -10.66 | 4.64E-02 | 1.06E-01 |
| SCRIB | 8:143804826-143804954 | 8:143804934-143804954 | A5ss | 3 | 3 | ENST00000320476 | -10.67 | 3.46E-03 | 5.26E-02 |
| CLN3 | 16:28484119-28487455 | 16:28486347-28486490 | MXS | 3 | 3 | ENST00000567495 | -10.68 | 3.93E-02 | 7.31E-02 |
| TMEM25 | 11:118532150-118532461 | 11:118532200-118532334 | IR | 3 | 3 | ENST00000313236 | -10.69 | 1.03E-02 | 1.39E-02 |
| GSDMD | 8:143559777-143561423 | 8:143559970-143560602 | IR | 3 | 3 | ENST00000524846 | -10.7 | 1.27E-03 | 1.89E-02 |
| LUC7L3 | 17:50749345-50750746 | 17:50750501-50750746 | TSSlA3ss | 3 | 3 | TSS.ENST00000504065 | -10.7 | 2.87E-02 | 5.91E-02 |
| NDRG2 | 14:21023322-21025028 | 14:21024030-21024096 | SES | 3 | 3 | ENST00000397853 | -10.72 | 3.99E-02 | 6.48E-02 |
| NFRKB | 11:129892871-129894358 | 11:129893297-129893385 | MES | 3 | 3 | ENST00000526884 | -10.74 | 9.41E-03 | 1.28E-02 |
| NFRKB | 11:129892871-129894358 | 11:129893293-129893385 | MES | 3 | 3 | ENST00000532225 | -10.74 | 9.41E-03 | 1.28E-02 |
| NUTM2A-AS1 | 10:87238850-87246114 | 10:87238850-87238872 | TSSlA3ss | 3 | 3 | TSS.ENST00000433920 | -10.74 | 8.42E-06 | 1.13E-05 |
| PDE9A | 21:42653884-42698967 | 21:42686192-42686262 | MES | 3 | 3 | ENST00000335512 | -10.74 | 2.52E-02 | 8.23E-02 |
| TESMIN | 11:68736132-68738788 | 11:68737886-68738699 | IR | 3 | 3 | ENST00000443940 | -10.74 | 3.51E-02 | 5.09E-02 |
| ZNF43 | 19:21851028-21851904 | 19:21851028-21851162 | TSSlA3ss | 3 | 3 | TSS.ENST00000594312 | -10.74 | 8.42E-06 | 1.90E-05 |
| TESK1 | 9:35606972-35607409 | 9:35606984-35607326 | IR | 3 | 3 | ENST00000467424 | -10.8 | 1.39E-02 | 4.19E-01 |
| CYP2B6 | 19:41004447-41012297 | 19:41009219-41009395 | MES | 2 | 3 | ENST00000324071 | -10.82 | 3.42E-02 | 7.89E-02 |
| NEK3 | 13:52154156-52156074 | 13:52154156-52154173 | A3ss | 3 | 3 | Ex.TSS.ENST00000550841 | -10.83 | 1.71E-02 | 2.71E-02 |
| AACS | 12:125114558-125134002 | 12:125129335-125129460 | MES | 3 | 3 | ENST00000316519 | -10.84 | 3.00E-02 | 4.48E-02 |

| | | | | | | | | | |
|---|---|---|---|---|---|---|---|---|---|
| RETREG2 | 2:219179764-219180197 | 2:219180110-219180197 | A3ss | 3 | 3 | ENST00000430297 | -10.84 | 1.25E-02 | 4.57E-02 |
| RETREG2 | 2:219179764-219180166 | 2:219180110-219180166 | A3ss | 3 | 3 | Ex.ENST00000430297 | -10.84 | 1.25E-02 | 4.57E-02 |
| YWHAZ | 8:100948733-100952849 | 8:100948733-100948900 | TSSlA3ss | 3 | 3 | TSS.ENST00000457309 | -10.84 | 2.10E-02 | 2.61E-01 |
| PAN2 | 12:56323914-56324048 | 12:56324037-56324048 | A5ss | 3 | 3 | ENST00000257931 | -10.85 | 3.15E-02 | 4.86E-02 |
| ALAS1 | 3:52198256-52199220 | 3:52198672-52198848 | SES | 3 | 3 | ENST00000394965 | -10.86 | 1.42E-02 | 8.01E-02 |
| JPH4 | 14:23576457-23578179 | 14:23577075-23577624 | MXS | 3 | 3 | ENST00000356300 | -10.89 | 1.69E-02 | 2.75E-02 |
| USP21 | 1:161159679-161160619 | 1:161160366-161160506 | SES | 3 | 3 | ENST00000368002 | -10.9 | 4.17E-02 | 1.06E-01 |
| TTI2 | 8:33511967-33513109 | 8:33513005-33513081 | IR | 3 | 3 | ENST00000360742 | -10.91 | 1.01E-03 | 1.30E-02 |
| YDJC | 22:21629408-21629601 | 22:21629543-21629601 | A5ss | 3 | 3 | ENST00000415762 | -10.92 | 3.98E-02 | 1.32E-01 |
| MAPKAPK5-AS1 | 12:111839767-111840477 | 12:111840004-111840133 | IR | 3 | 3 | ENST00000443596 | -10.94 | 2.28E-02 | 3.37E-02 |
| TXNDC15 | 5:134899489-134901525 | 5:134900155-134901065 | IR | 3 | 3 | ENST00000358387 | -10.94 | 3.93E-02 | 2.65E-01 |
| METTL8 | 2:171360514-171433882 | 2:171414603-171414726 | MXS | 3 | 3 | ENST00000447486 | -10.96 | 3.05E-02 | 1.09E-01 |
| WDR37 | 10:1056565-1072115 | 10:1056565-1056968 | TSSlA5ss | 3 | 3 | TSS.ENST00000263150 | -10.96 | 1.83E-02 | 2.42E-02 |
| PILRB | 7:100353271-100355142 | 7:100353895-100354012 | MXS | 3 | 3 | ENST00000455145 | -10.97 | 3.91E-02 | 2.20E-01 |
| YDJC | 22:21629010-21629761 | 22:21629408-21629542 | IR | 3 | 3 | ENST00000464015 | -10.97 | 3.16E-02 | 1.05E-01 |
| GJC1 | 17:44798448-44805837 | 17:44804115-44804587 | IR | 3 | 3 | ENST00000590758 | -10.98 | 1.02E-02 | 2.07E-02 |
| FBXL6 | 8:144356808-144357973 | 8:144357122-144357438 | IR (overlapping region) | 3 | 3 | ENST00000524492 | -11.02 | 4.11E-02 | 6.23E-01 |
| LARP7 | 4:112650583-112653076 | 4:112650583-112652673 | A5ss | 3 | 3 | Ex.ENST00000324052 | -11.03 | 4.40E-02 | 2.62E-01 |
| AC113189.2 | 17:7442594-7444081 | 17:7442793-7443075 | IR (overlapping region) | 3 | 3 | ENST00000575310 | -11.05 | 2.82E-02 | 5.20E-02 |
| NEURL4 | 17:7318357-7319049 | 17:7318357-7318674 | A3ss | 3 | 3 | ENST00000572029 | -11.09 | 1.69E-02 | 3.54E-02 |
| DCXR | 17:82036859-82037547 | 17:82037089-82037449 | IR | 3 | 3 | ENST00000579334 | -11.11 | 6.19E-03 | 1.32E-02 |
| FKRP | 19:46748718-46755411 | 19:46748718-46748884 | TSSlA5ss | 3 | 3 | TSS.ENST00000598271 | -11.11 | 1.15E-02 | 2.67E-02 |
| FKRP | 19:46748666-46755411 | 19:46748666-46748884 | TSSlA5ss | 3 | 3 | TSS.ENST00000598271 | -11.11 | 1.15E-02 | 2.67E-02 |
| MRTFA | 22:40421101-40424205 | 22:40423536-40423685 | SES | 3 | 3 | ENST00000355630 | -11.11 | 6.77E-04 | 2.32E-03 |
| SGSH | 17:80209282-80214328 | 17:80211012-80212070 | IR | 3 | 3 | ENST00000575282 | -11.11 | 1.81E-02 | 3.33E-02 |
| MBD1 | 18:50268846-50271540 | 18:50269812-50271468 | IR | 3 | 3 | ENST00000591416 | -11.12 | 2.28E-03 | 4.89E-03 |
| PIDD1 | 11:799185-800244 | 11:799566-799814 | IR | 3 | 3 | ENST00000527357 | -11.12 | 3.89E-02 | 5.70E-02 |
| BEX3 | X:103377107-103377509 | X:103377463-103377509 | A3ss | 3 | 3 | ENST00000372645 | -11.14 | 5.88E-06 | 2.18E-04 |
| ATG9A | 2:219227814-219229534 | 2:219227922-219228053 | SES | 3 | 3 | ENST00000396761 | -11.16 | 4.91E-02 | 1.80E-01 |
| CCDC82 | 11:96359098-96364979 | 11:96359098-96359178 | A3ss | 3 | 3 | ENST00000278520 | -11.17 | 3.85E-02 | 5.66E-02 |
| AC008555.4 | 19:34860174-34860576 | 19:34860303-34860533 | IR (overlapping region) | 3 | 3 | ENST00000604749 | -11.2 | 1.55E-02 | 3.52E-02 |
| PARPBP | 12:102195951-102197520 | 12:102196702-102196972 | IR (overlapping region) | 3 | 3 | ENST00000327680 | -11.22 | 3.59E-02 | 5.11E-02 |
| ANKRD13D | 11:67299612-67299826 | 11:67299612-67299621 | A5ss | 3 | 3 | Ex.ENST00000308440 | -11.23 | 4.17E-02 | 6.04E-02 |
| ZNF197 | 3:44625144-44629073 | 3:44629065-44629073 | TSSlA3ss | 3 | 3 | TSS.ENST00000383745 | -11.24 | 3.35E-02 | 1.45E-01 |
| SEC31A | 4:82829059-82844385 | 4:82842140-82842481 | MXS | 3 | 3 | ENST00000348405 | -11.25 | 2.13E-02 | 1.37E-01 |
| SFI1 | 22:31614386-31615279 | 22:31614861-31615047 | IR | 3 | 3 | ENST00000357852 | -11.25 | 1.92E-02 | 6.55E-02 |
| CYBC1 | 17:82449293-82450479 | 17:82449431-82449480 | SES | 3 | 3 | ENST00000437807 | -11.28 | 3.88E-02 | 8.26E-02 |
| TCIRG1 | 11:68041347-68041752 | 11:68041347-68041388 | A5ss | 3 | 3 | ENST00000265686 | -11.28 | 1.45E-02 | 2.11E-02 |
| MFN1 | 3:179347811-179348847 | 3:179348845-179348847 | A3ss | 3 | 3 | Ex.TSS.ENST00000357390 | -11.29 | 3.91E-02 | 1.62E-01 |
| DERL3 | 22:23838711-23839128 | 22:23838774-23838894 | IR | 3 | 3 | ENST00000290730 | -11.3 | 2.30E-02 | 7.60E-02 |
| MFGE8 | 15:88899533-88901550 | 15:88899656-88899811 | SES | 3 | 3 | ENST00000542878 | -11.3 | 5.62E-06 | 9.65E-06 |
| ATG4B | 2:241651336-241653511 | 2:241653307-241653369 | MXS | 3 | 3 | ENST00000344376 | -11.32 | 1.46E-02 | 5.43E-02 |
| ECHDC2 | 1:52911791-52921552 | 1:52913895-52914095 | MES | 3 | 3 | ENST00000476477 | -11.32 | 4.22E-02 | 1.06E-01 |
| MIA2 | 14:39276956-39279336 | 14:39276956-39277065 | A5ss | 3 | 3 | ENST00000280083 | -11.32 | 3.93E-03 | 6.45E-03 |
| DELE1 | 5:141938521-141942047 | 5:141938905-141939505 | IR | 3 | 3 | ENST00000432126 | -11.34 | 3.06E-02 | 1.96E-01 |
| OSBPL5 | 11:3087116-3088343 | 11:3087784-3088177 | IR | 3 | 3 | ENST00000478260 | -11.36 | 4.00E-02 | 5.45E-02 |
| SNX22 | 15:64151744-64152742 | 15:64151851-64152242 | IR | 3 | 3 | ENST00000560945 | -11.37 | 1.27E-02 | 2.16E-02 |
| SLC17A9 | 20:62962624-62963369 | 20:62962755-62963272 | IR | 3 | 3 | ENST00000488738 | -11.38 | 3.14E-02 | 1.01E-01 |
| EPB41 | 1:29053313-29064981 | 1:29060422-29060484 | MES | 3 | 3 | ENST00000373797 | -11.42 | 4.92E-02 | 1.38E-01 |
| FAM193B | 5:177553168-177554541 | 5:177553351-177553496 | IR | 3 | 3 | ENST00000506955 | -11.42 | 4.11E-02 | 2.63E-01 |
| THOC5 | 22:29549159-29553670 | 22:29550728-29550843 | SES | 3 | 3 | ENST00000397873 | -11.43 | 3.72E-03 | 1.24E-02 |
| FCGRT | 19:49513195-49514486 | 19:49514134-49514210 | IR | 3 | 3 | ENST00000595881 | -11.45 | 6.70E-03 | 1.58E-02 |
| SYT7 | 11:61533125-61551383 | 11:61546031-61546255 | MXS | 3 | 3 | ENST00000540677 | -11.46 | 2.90E-02 | 4.00E-02 |
| UVSSA | 4:1383766-1388049 | 4:1383941-1385867 | IR | 3 | 3 | ENST00000503548 | -11.49 | 3.48E-02 | 2.10E-01 |
| AL669830.2 | 6:30913811-30915037 | 6:30914470-30914809 | IR (overlapping region) | 3 | 3 | ENST00000477288 | -11.5 | 2.18E-02 | 1.77E-01 |
| PHKG2 | 16:30751106-30751913 | 16:30751218-30751548 | IR (overlapping region) | 3 | 3 | ENST00000569762 | -11.5 | 2.25E-02 | 4.22E-02 |
| ZNF692 | 1:248857437-248858130 | 1:248857828-248858017 | SES | 3 | 3 | ENST00000533647 | -11.5 | 3.41E-03 | 9.46E-03 |
| DCTD | 4:182915576-182916672 | 4:182916286-182916539 | MES | 3 | 3 | ENST00000503988 | -11.52 | 2.10E-02 | 1.28E-01 |
| CIART | 1:150282554-150283633 | 1:150282824-150283461 | IR | 3 | 3 | ENST00000290363 | -11.53 | 5.41E-03 | 1.34E-02 |
| PKP3 | 11:394209-394524 | 11:394335-394508 | IR | 3 | 3 | ENST00000331563 | -11.53 | 4.52E-02 | 6.18E-02 |
| PLXNB2 | 22:50290598-50307552 | 22:50294719-50294778 | SES | 3 | 3 | ENST00000359337 | -11.53 | 2.01E-02 | 6.95E-02 |
| ZFYVE16 | 5:80445406-80449590 | 5:80448026-80448404 | SES | 3 | 3 | ENST00000338008 | -11.53 | 1.13E-04 | 8.34E-04 |
| NFRKB | 11:129892871-129895495 | 11:129894359-129894438 | MES | 3 | 3 | ENST00000526884 | -11.54 | 4.95E-02 | 6.70E-02 |
| PIGT | 20:45420694-45424215 | 20:45420694-45421583 | A5ss | 3 | 3 | ENST00000639250 | -11.54 | 3.49E-03 | 1.12E-02 |
| MPPE1 | 18:11885817-11886916 | 18:11886499-11886778 | SES | 3 | 3 | ENST00000592894 | -11.56 | 2.34E-02 | 4.98E-02 |
| CREBZF | 11:85661066-85665102 | 11:85662458-85663523 | IR | 3 | 3 | ENST00000527447 | -11.57 | 1.33E-02 | 1.95E-02 |
| HUWE1 | X:53542540-53543885 | X:53543841-53543885 | A5ss | 3 | 3 | ENST00000262854 | -11.57 | 4.86E-02 | 1.00E+00 |
| KCTD9 | 8:25440674-25458198 | 8:25446129-25446250 | MES | 3 | 3 | ENST00000221200 | -11.6 | 5.84E-03 | 7.39E-02 |
| TRIM26 | 6:30199154-30213304 | 6:30204656-30204765 | MES | 3 | 3 | ENST00000416596 | -11.6 | 3.08E-02 | 2.50E-01 |
| TRIM16 | 17:15683036-15684299 | 17:15683158-15684195 | IR | 3 | 3 | ENST00000416464 | -11.63 | 2.90E-02 | 5.70E-02 |
| ORC3 | 6:87594408-87601783 | 6:87594542-87601784 | A3ss | 3 | 3 | Ex.ENST00000257789 | -11.66 | 3.87E-03 | 3.52E-02 |
| TSPOAP1 | 17:58306342-58306867 | 17:58306414-58306799 | IR | 3 | 3 | ENST00000578511 | -11.66 | 2.38E-02 | 4.38E-02 |
| DERL3 | 22:23838711-23839128 | 22:23838777-23838894 | IR | 3 | 3 | ENST00000290730 | -11.67 | 2.19E-02 | 7.26E-02 |
| MYH14 | 19:50284764-50286694 | 19:50286536-50286536 | IR | 3 | 3 | ENST00000595016 | -11.67 | 5.57E-03 | 1.30E-02 |
| SLC37A4 | 11:119027628-119029034 | 11:119027873-119028193 | IR | 3 | 3 | ENST00000526275 | -11.67 | 2.41E-03 | 3.24E-03 |
| HSPA14 | 10:14840158-14843370 | 10:14842134-14842846 | SES | 3 | 3 | ENST00000640019 | -11.68 | 2.62E-02 | 3.47E-02 |
| SLC35C2 | 20:46355179-46355801 | 20:46355179-46355241 | A3ss | 3 | 3 | ENST00000243896 | -11.68 | 7.32E-03 | 2.34E-02 |
| POLQ | 3:121449427-121467518 | 3:121460050-121460234 | SES | 3 | 3 | ENST00000264233 | -11.69 | 1.67E-02 | 6.70E-02 |
| TARBP2 | 12:53503130-53503729 | 12:53503130-53503212 | A5ss | 3 | 3 | ENST00000549679 | -11.7 | 1.44E-02 | 2.21E-02 |
| KAT8 | 16:31130746-31131359 | 16:31130901-31131194 | IR | 3 | 3 | ENST00000448516 | -11.71 | 1.83E-02 | 3.45E-02 |
| AL117327.1 | X:103408300-103450496 | X:103449274-103450497 | A3ss | 3 | 3 | Ex.ENST00000629535 | -11.74 | 3.51E-02 | 7.24E-01 |
| NKAPD1 | 11:112074287-112074916 | 11:112074359-112074886 | IR | 3 | 3 | ENST00000532163 | -11.75 | 9.15E-03 | 1.23E-02 |
| ECHDC1 | 6:127314897-127327001 | 6:127316450-127316502 | MXS | 3 | 3 | ENST00000430841 | -11.77 | 2.82E-02 | 2.20E-01 |
| UPP1 | 7:48094828-48106872 | 7:48101824-48101982 | MES | 3 | 3 | ENST00000331803 | -11.78 | 2.03E-02 | 2.18E-01 |
| APOO | X:23833511-23840312 | X:23833511-23833612 | TSSlA3ss | 3 | 3 | TSS.ENST00000633432 | -11.79 | 3.86E-02 | 1.00E+00 |
| PIGO | 9:35091240-35092767 | 9:35091292-35092542 | IR | 3 | 3 | ENST00000378617 | -11.8 | 4.30E-03 | 1.28E-01 |
| RGL3 | 19:11406865-11416613 | 19:11415937-11416148 | SES | 3 | 3 | ENST00000380456 | -11.81 | 2.26E-02 | 4.93E-02 |
| TSPAN2 | 1:115050556-115057536 | 1:115053379-115053462 | SES | 3 | 3 | ENST00000369515 | -11.82 | 4.76E-02 | 1.16E-01 |
| STON1 | 2:48529880-48531917 | 2:48530217-48530712 | IR (overlapping region) | 3 | 3 | ENST00000484110 | -11.86 | 2.22E-02 | 8.46E-02 |
| CHRNE | 17:4899468-4900229 | 17:4899829-4899935 | IR (overlapping region) | 2 | 3 | ENST00000652550 | -11.87 | 1.31E-02 | 2.69E-02 |
| KRIT1 | 7:92245465-92246100 | 7:92245596-92245789 | IR (overlapping region) | 3 | 3 | ENST00000340022 | -11.87 | 4.76E-05 | 5.66E-04 |
| ATG16L1 | 2:233256196-233270001 | 2:233263130-233263235 | MES | 3 | 3 | ENST00000392017 | -11.9 | 1.98E-02 | 7.31E-02 |
| CHN1 | 2:174918566-174944887 | 2:174940816-174940894 | SES | 3 | 3 | ENST00000650770 | -11.92 | 3.56E-02 | 1.27E-01 |
| LDB2 | 4:16502874-16508534 | 4:16505840-16505982 | SES | 3 | 3 | ENST00000441778 | -11.93 | 3.01E-02 | 1.70E-01 |

| | | | | | | | | | |
|---|---|---|---|---|---|---|---|---|---|
| PATL1 | 11:59655741-59656498 | 11:59655956-59656045 | SES | 3 | 3 | ENST00000300146 | -11.94 | 4.60E-02 | 6.34E-02 |
| AL669830.2 | 6:30913811-30915037 | 6:30914470-30914804 | IR (overlapping region) | 3 | 3 | ENST00000477288 | -11.95 | 7.67E-03 | 6.27E-02 |
| PDCD4 | 10:110876071-110881274 | 10:110881233-110881274 | A3ss | 3 | 3 | ENST00000280154 | -11.97 | 4.95E-02 | 6.54E-02 |
| TP53I11 | 11:44938367-44949972 | 11:44939114-44939190 | MES | 3 | 3 | ENST00000533940 | -11.97 | 3.47E-02 | 4.76E-02 |
| GGT1 | 22:24608024-24610357 | 22:24609965-24610033 | MES | 3 | 3 | ENST00000400380 | -11.98 | 1.46E-02 | 4.86E-02 |
| MON2 | 12:62467319-62484169 | 12:62470701-62470778 | SES | 3 | 3 | ENST00000547095 | -11.99 | 5.84E-03 | 9.05E-03 |
| ZNF664 | 12:123974021-124011380 | 12:123988046-123988138 | MXS | 3 | 3 | ENST00000392404 | -11.99 | 1.28E-03 | 1.91E-03 |
| AP1S2 | X:15833008-15845378 | X:15833444-15833524 | SES | 3 | 3 | ENST00000673591 | -12 | 2.11E-02 | 1.00E+00 |
| APOO | X:23833511-23840312 | X:23833607-23833612 | SES | 3 | 3 | ENST00000634086 | -12 | 2.70E-02 | 1.00E+00 |
| DALRD3 | 3:49015525-49016340 | 3:49015873-49015972 | IR (overlapping region) | 3 | 3 | ENST00000467457 | -12.02 | 1.47E-02 | 6.50E-02 |
| LINC00869 | 1:149646599-149647610 | 1:149646946-149647380 | IR | 3 | 3 | ENST00000620190 | -12.02 | 2.50E-02 | 6.17E-02 |
| WDR90 | 16:662332-665678 | 16:662673-662844 | SES | 3 | 3 | ENST00000549024 | -12.02 | 4.56E-02 | 8.74E-02 |
| PHACTR2 | 6:143753913-143765260 | 6:143760401-143760640 | SES | 3 | 3 | ENST00000427704 | -12.04 | 1.15E-02 | 9.03E-02 |
| RALGDS | 9:133100383-133101519 | 9:133100496-133101519 | A5ss | 3 | 3 | ENST00000493438 | -12.04 | 6.50E-03 | 1.67E-01 |
| ZBTB8OS | 1:32634111-32650432 | 1:32634768-32634792 | SES | 3 | 3 | ENST00000473294 | -12.06 | 4.32E-04 | 1.22E-03 |
| MDC1 | 6:30711502-30711666 | 6:30711502-30711504 | A3ss | 3 | 3 | ENST00000376406 | -12.09 | 1.12E-02 | 9.13E-02 |
| ATRX | X:77688928-77696576 | X:77693824-77693937 | MES | 2 | 3 | ENST00000480283 | -12.1 | 3.03E-02 | 1.00E+00 |
| AP002856.2 | 11:131253592-131350308 | 11:131300358-131350309 | A3ss | 3 | 3 | Ex.ENST00000606885 | -12.11 | 3.54E-02 | 4.79E-02 |
| NDOR1 | 9:137215203-137215658 | 9:137215428-137215521 | SES | 3 | 3 | ENST00000458322 | -12.11 | 4.47E-02 | 1.00E+00 |
| APBB3 | 5:140561587-140562227 | 5:140561702-140561843 | IR | 3 | 3 | ENST00000510241 | -12.12 | 3.07E-02 | 2.06E-01 |
| MRRF | 9:122285288-122291748 | 9:122286039-122286166 | MES | 3 | 3 | ENST00000470366 | -12.14 | 4.59E-03 | 9.66E-02 |
| SNHG8 | 4:118278718-118279822 | 4:118278810-118278943 | IR | 3 | 3 | ENST00000667980 | -12.14 | 3.45E-03 | 2.06E-02 |
| CIART | 1:150282554-150283633 | 1:150282619-150283513 | IR | 3 | 3 | ENST00000290363 | -12.15 | 6.50E-03 | 1.60E-02 |
| SLC44A2 | 19:10642452-10643278 | 19:10642897-10643278 | TSSlA3ss | 3 | 3 | TSS.ENST00000586078 | -12.15 | 1.17E-02 | 2.55E-02 |
| MBD1 | 18:50268846-50271540 | 18:50269819-50271468 | IR | 3 | 3 | ENST00000591416 | -12.16 | 1.74E-03 | 3.75E-03 |
| KIF1B | 1:10278129-10282399 | 1:10282322-10282399 | A3ss | 3 | 3 | ENST00000620295 | -12.19 | 1.56E-02 | 3.77E-02 |
| SNHG11 | 20:38447466-38450447 | 20:38448094-38450335 | IR (overlapping region) | 3 | 3 | ENST00000659610 | -12.19 | 3.85E-04 | 1.17E-03 |
| ABCA2 | 9:137017562-137018075 | 9:137017693-137017786 | IR | 3 | 3 | ENST00000466707 | -12.2 | 1.83E-02 | 4.57E-01 |
| GDPD2 | X:70423383-70426051 | X:70425354-70425457 | MES | 3 | 3 | ENST00000374382 | -12.21 | 2.76E-02 | 1.00E+00 |
| ZNF131 | 5:43121598-43122177 | 5:43121829-43122043 | IR | 3 | 3 | ENST00000509634 | -12.24 | 1.71E-03 | 1.22E-02 |
| ACTR5 | 20:38765519-38766256 | 20:38766238-38766256 | A3ss | 3 | 3 | Ex.ENST00000243903 | -12.25 | 3.35E-02 | 1.01E-01 |
| JMJD7 | 15:41834894-41835223 | 15:41834894-41834969 | IR (overlapping region) | 3 | 3 | ENST00000408047 | -12.26 | 3.53E-02 | 5.94E-02 |
| GNB1L | 22:19806758-19854442 | 22:19821228-19821375 | MES | 3 | 3 | ENST00000329517 | -12.27 | 1.02E-02 | 3.36E-02 |
| STAT6 | 12:57108300-57111128 | 12:57109941-57110110 | SES | 3 | 3 | ENST00000553275 | -12.27 | 4.88E-02 | 7.54E-02 |
| ALG13 | X:111736880-111752789 | X:111744668-111744904 | SES | 3 | 3 | ENST00000394780 | -12.28 | 1.13E-02 | 4.32E-01 |
| RWDD4 | 4:183651132-183658928 | 4:183655881-183655961 | MXS | 3 | 3 | ENST00000514322 | -12.29 | 3.98E-02 | 2.43E-01 |
| ZNF326 | 1:90005245-90007611 | 1:90007345-90007611 | A3ss | 3 | 3 | ENST00000340281 | -12.3 | 2.59E-02 | 7.68E-02 |
| AFF1 | 4:86948572-87046165 | 4:86988438-87046166 | A3ss | 3 | 3 | Ex.ENST00000395146 | -12.31 | 4.26E-02 | 2.77E-01 |
| BBC3 | 19:47226755-47232514 | 19:47227432-47232515 | A5ss | 3 | 3 | Ex.TSS.ENST00000341983 | -12.31 | 4.12E-02 | 9.66E-02 |
| BBC3 | 19:47226755-47228157 | 19:47227432-47228158 | A5ss | 3 | 3 | Ex.ENST00000439096 | -12.31 | 4.12E-02 | 9.66E-02 |
| EHBP1 | 2:63045410-63046487 | 2:63046142-63046481 | IR (overlapping region) | 3 | 3 | ENST00000263991 | -12.33 | 4.69E-02 | 1.80E-01 |
| ZSCAN5A | 19:56313385-56314680 | 19:56314250-56314680 | TSSlA5ss | 3 | 3 | TSS.ENST00000592679 | -12.33 | 3.85E-03 | 9.16E-03 |
| ZSCAN5A | 19:56313385-56314619 | 19:56314250-56314619 | TSSlA5ss | 3 | 3 | TSS.ENST00000592679 | -12.33 | 3.85E-03 | 9.16E-03 |
| ZSCAN5A | 19:56313385-56314420 | 19:56314250-56314420 | TSSlA5ss | 3 | 3 | TSS.ENST00000592679 | -12.33 | 3.85E-03 | 9.16E-03 |
| MED6 | 14:70583589-70584943 | 14:70584131-70584816 | IR | 3 | 3 | ENST00000256379 | -12.35 | 2.12E-02 | 3.49E-02 |
| TMEM67 | 8:93754894-93755137 | 8:93754910-93755019 | IR | 3 | 3 | ENST00000453321 | -12.35 | 2.23E-02 | 3.09E-01 |
| SFXN2 | 10:102728530-102729722 | 10:102729319-102729394 | SES | 3 | 3 | ENST00000369893 | -12.36 | 3.61E-02 | 4.76E-02 |
| ZC3H13 | 13:46045514-46052403 | 13:46045514-46045516 | A3ss | 3 | 3 | Ex.TSS.ENST00000242848 | -12.38 | 3.07E-02 | 4.85E-02 |
| HTATSF1 | X:136497440-136497675 | X:136497624-136497675 | TSSlA3ss | 3 | 3 | TSS.ENST00000448450 | -12.41 | 6.70E-03 | 2.74E-01 |
| PVT1 | 8:127890999-127989161 | 8:127939508-127939676 | MXS | 3 | 3 | ENST00000517525 | -12.41 | 4.10E-02 | 6.05E-01 |
| DCXR | 17:82036859-82037309 | 17:82037014-82037162 | IR | 3 | 3 | ENST00000582613 | -12.42 | 3.08E-03 | 6.56E-03 |
| PEX26 | 22:18078008-18078606 | 22:18078048-18078295 | IR (overlapping region) | 3 | 3 | ENST00000399744 | -12.46 | 1.64E-02 | 5.39E-02 |
| STYXL1 | 7:76030525-76047811 | 7:76047662-76047811 | TSSlA5ss | 3 | 3 | TSS.ENST00000248600 | -12.46 | 4.47E-02 | 5.19E-01 |
| CEP83 | 12:94411848-94412516 | 12:94412318-94412516 | A5ss | 3 | 3 | Ex.ENST00000339839 | -12.48 | 3.74E-02 | 5.85E-02 |
| PEX26 | 22:18078008-18078606 | 22:18078128-18078295 | IR (overlapping region) | 3 | 3 | ENST00000399744 | -12.51 | 1.54E-02 | 5.08E-02 |
| SMPD5 | 8:144050795-144051522 | 8:144050859-144050941 | IR | 3 | 3 | ENST00000528912 | -12.56 | 6.22E-03 | 9.50E-02 |
| PXN | 12:120215274-120221622 | 12:120215560-120215661 | MXS | 3 | 3 | ENST00000228307 | -12.58 | 1.66E-02 | 2.46E-02 |
| MFSD11 | 17:76737387-76738527 | 17:76737999-76738281 | IR (overlapping region) | 3 | 2 | ENST00000590393 | -12.59 | 1.12E-02 | 2.36E-02 |
| CRELD1 | 3:9943516-9944364 | 3:9943994-9944095 | SES | 3 | 3 | ENST00000674057 | -12.6 | 2.02E-03 | 1.18E-02 |
| ZNF131 | 5:43121598-43122177 | 5:43121829-43122038 | IR | 3 | 3 | ENST00000509634 | -12.6 | 2.28E-03 | 1.62E-02 |
| SELENBP1 | 1:151368320-151369712 | 1:151369004-151369712 | A5ss | 3 | 3 | ENST00000470345 | -12.61 | 4.00E-02 | 9.87E-02 |
| SELENBP1 | 1:151368320-151369441 | 1:151369004-151369441 | A5ss | 3 | 3 | ENST00000463664 | -12.61 | 4.00E-02 | 9.89E-02 |
| LGMN | 14:92704362-92709671 | 14:92706483-92706653 | MES | 3 | 3 | ENST00000334869 | -12.65 | 6.89E-03 | 1.15E-02 |
| NRSN2 | 20:349364-353209 | 20:349638-349832 | SES | 3 | 3 | ENST00000382285 | -12.66 | 2.09E-02 | 6.28E-02 |
| SLC35C2 | 20:46355242-46356573 | 20:46355802-46355874 | SES | 3 | 3 | ENST00000243896 | -12.67 | 3.08E-02 | 9.81E-02 |
| WASHC5 | 8:125083259-125091614 | 8:125083713-125084022 | SES | 3 | 3 | ENST00000318410 | -12.76 | 3.73E-02 | 5.50E-01 |
| IFT20 | 17:28328326-28329810 | 17:28329069-28329172 | IR | 3 | 3 | ENST00000583796 | -12.77 | 4.74E-02 | 9.42E-02 |
| LDOC1 | X:141173235-141177129 | X:141176812-141176892 | IR | 3 | 3 | ENST00000370526 | -12.78 | 1.01E-02 | 4.20E-01 |
| PIGS | 17:28571049-28571536 | 17:28571189-28571462 | IR (overlapping region) | 3 | 3 | ENST00000395346 | -12.79 | 5.68E-03 | 1.13E-02 |
| SLC44A2 | 19:10642897-10644557 | 19:10643022-10643278 | IR | 3 | 3 | ENST00000586078 | -12.81 | 4.07E-03 | 8.84E-03 |
| CTNND1 | 11:57762047-57795576 | 11:57789037-57789155 | MES | 3 | 3 | ENST00000358694 | -12.82 | 8.92E-03 | 1.23E-02 |
| RNF220 | 1:44635589-44636029 | 1:44635589-44635666 | A5ss | 3 | 3 | ENST00000335497 | -12.84 | 5.64E-05 | 1.61E-04 |
| ZMYM1 | 1:35110448-35112086 | 1:35111776-35111912 | SES | 2 | 3 | ENST00000488455 | -12.84 | 4.61E-02 | 1.29E-01 |
| BRD3OS | 9:134026208-134031587 | 9:134027263-134028825 | IR | 3 | 3 | ENST00000603928 | -12.85 | 1.57E-03 | 4.08E-02 |
| MRPL35 | 2:86210480-86213790 | 2:86210614-86212425 | IR | 3 | 3 | ENST00000337109 | -12.87 | 1.72E-02 | 6.80E-02 |
| ATP11A | 13:112875942-112881875 | 13:112878217-112878303 | MXS | 3 | 3 | ENST00000375645 | -12.88 | 4.76E-03 | 7.48E-03 |
| LRFN4 | 11:66857064-66859093 | 11:66857297-66857734 | IR (overlapping region) | 3 | 3 | ENST00000309602 | -12.88 | 4.05E-02 | 5.86E-02 |
| SEC31A | 4:82829059-82842184 | 4:82842140-82842184 | A5ss | 3 | 3 | ENST00000348405 | -12.9 | 3.17E-02 | 2.04E-01 |
| MYO1B | 2:191400469-191408114 | 2:191402632-191402718 | MES | 3 | 3 | ENST00000304164 | -12.94 | 9.40E-03 | 3.38E-02 |
| SCAND1 | 20:35953629-35955102 | 20:35954340-35954447 | IR | 3 | 3 | ENST00000373991 | -12.94 | 2.06E-03 | 6.20E-03 |
| RFC4 | 3:186804725-186806424 | 3:186806290-186806424 | TSSlA5ss | 3 | 3 | TSS.ENST00000296273 | -12.96 | 2.42E-05 | 1.01E-04 |
| TMEM164 | X:110002773-110067346 | X:110003501-110067347 | A3ss | 3 | 3 | Ex.TSS.ENST00000372072 | -12.96 | 4.96E-02 | 1.00E+00 |
| TMEM164 | X:110002773-110003882 | X:110003501-110003883 | A3ss | 3 | 3 | Ex.TSS.ENST00000471255 | -12.96 | 4.96E-02 | 1.00E+00 |
| TSPOAP1 | 17:58306342-58306867 | 17:58306387-58306799 | IR | 3 | 3 | ENST00000578511 | -13.1 | 1.70E-02 | 3.14E-02 |
| DPH5 | 1:100995150-101021640 | 1:101013710-101013818 | MES | 3 | 3 | ENST00000490732 | -13.14 | 2.27E-03 | 5.51E-03 |
| SALL2 | 14:21522245-21525336 | 14:21522833-21524931 | SES | 3 | 3 | ENST00000450879 | -13.14 | 1.65E-03 | 2.68E-03 |
| BRD1 | 22:49787390-49787887 | 22:49787495-49787576 | IR | 3 | 3 | ENST00000404760 | -13.15 | 3.16E-03 | 1.09E-02 |
| DMAP1 | 1:44213471-44213662 | 1:44213528-44213662 | IR | 3 | 3 | ENST00000440641 | -13.15 | 4.65E-02 | 1.17E-01 |
| PLXNB2 | 22:50290598-50294725 | 22:50294719-50294725 | A5ss | 3 | 3 | ENST00000359337 | -13.27 | 4.60E-02 | 1.59E-01 |
| CPT1B | 22:50577035-50577355 | 22:50577324-50577355 | A5ss | 3 | 3 | ENST00000312108 | -13.28 | 3.36E-02 | 1.16E-01 |
| PEX19 | 1:160283110-160285054 | 1:160283530-160283639 | SES | 3 | 3 | ENST00000368072 | -13.28 | 3.27E-03 | 8.33E-03 |
| RWDD4 | 4:183651132-183658928 | 4:183655881-183655961 | MES | 3 | 3 | ENST00000510702 | -13.32 | 2.38E-02 | 1.45E-01 |
| ZNF263 | 16:3284206-3289392 | 16:3285059-3285239 | MES | 3 | 3 | ENST00000219069 | -13.32 | 6.69E-03 | 1.26E-02 |
| ARHGEF19 | 1:16208728-16212501 | 1:16208728-16209083 | A3ss | 3 | 3 | Ex.TSS.ENST00000270747 | -13.36 | 3.20E-02 | 8.12E-02 |

| | | | | | | | | | |
|---|---|---|---|---|---|---|---|---|---|
| PICK1 | 22:38057490-38057752 | 22:38057490-38057587 | TSSIA5ss | 3 | 3 | TSS.ENST00000356976 | -13.36 | 1.65E-02 | 5.62E-02 |
| PICK1 | 22:38057437-38057752 | 22:38057437-38057587 | TSSIA5ss | 3 | 3 | TSS.ENST00000356976 | -13.36 | 1.65E-02 | 5.62E-02 |
| OGFOD2 | 12:122975384-122976653 | 12:122975811-122975867 | SES | 3 | 3 | ENST00000228922 | -13.38 | 3.10E-02 | 4.61E-02 |
| ORAI2 | 7:102433662-102438943 | 7:102436230-102436333 | SES | 3 | 3 | ENST00000478730 | -13.38 | 1.50E-02 | 1.36E-01 |
| THEM4 | 1:151895195-151909359 | 1:151907321-151909360 | A5ss | 3 | 3 | Ex.TSS.ENST00000368814 | -13.38 | 1.62E-02 | 4.06E-02 |
| AC068587.4 | 8:12511258-12601157 | 8:12511258-12581044 | A3ss | 3 | 3 | Ex.ENST00000641839 | -13.4 | 2.26E-02 | 3.33E-01 |
| EME2 | 16:1776068-1780635 | 16:1777444-1777746 | IR (overlapping region) | 3 | 3 | ENST00000568449 | -13.4 | 2.62E-02 | 4.82E-02 |
| CHD7 | 8:60741259-60743097 | 8:60741886-60742819 | IR | 3 | 3 | ENST00000423902 | -13.47 | 3.84E-02 | 5.11E-01 |
| PRMT2 | 21:46635764-46636443 | 21:46636439-46636443 | A3ss | 3 | 3 | Ex.TSS.ENST00000355680 | -13.51 | 4.81E-03 | 1.58E-02 |
| WDR60 | 7:158902596-158905988 | 7:158902596-158905283 | A5ss | 3 | 3 | ENST00000467220 | -13.53 | 7.03E-04 | 7.21E-03 |
| CAMKV | 3:49860716-49861024 | 3:49860853-49860942 | IR | 3 | 3 | ENST00000478149 | -13.54 | 1.58E-04 | 7.13E-04 |
| IDH3B | 20:2658523-2659524 | 20:2658759-2658837 | SES | 3 | 3 | ENST00000474315 | -13.57 | 2.07E-03 | 6.20E-03 |
| PIGT | 20:45420694-45424215 | 20:45421383-45421583 | SES | 3 | 3 | ENST00000279035 | -13.58 | 2.67E-03 | 8.55E-03 |
| CAST | 5:96727531-96729611 | 5:96729153-96729209 | SES | 3 | 3 | ENST00000309190 | -13.6 | 3.89E-02 | 3.00E-01 |
| PPIP5K2 | 5:103173973-103177666 | 5:103176866-103176910 | SES | 3 | 3 | ENST00000613674 | -13.62 | 1.28E-02 | 8.38E-02 |
| WHRN | 9:114504184-114505473 | 9:114505052-114505194 | IR | 3 | 3 | ENST00000362057 | -13.63 | 2.99E-02 | 6.13E-01 |
| MPPE1 | 18:11886622-11886916 | 18:11886713-11886778 | SES | 3 | 3 | ENST00000317251 | -13.65 | 5.53E-03 | 1.18E-02 |
| USP33 | 1:77721902-77722023 | 1:77721902-77721925 | A3ss | 3 | 3 | ENST00000357428 | -13.66 | 9.88E-03 | 2.85E-02 |
| MINK1 | 17:4891125-4891515 | 17:4891456-4891515 | A3ss | 3 | 3 | ENST00000347992 | -13.67 | 5.17E-03 | 1.06E-02 |
| SNHG14 | 15:25053968-25056482 | 15:25053968-25055230 | TSSIA5ss | 3 | 3 | TSS.ENST00000657798 | -13.69 | 6.19E-03 | 1.03E-02 |
| ZSWIM8 | 10:73800760-73801084 | 10:73801017-73801084 | A3ss | 3 | 3 | ENST00000398706 | -13.69 | 4.99E-02 | 6.67E-02 |
| LINC01224 | 19:23405336-23415903 | 19:23415894-23415903 | TSSIA5ss | 3 | 3 | TSS.ENST00000593573 | -13.71 | 6.99E-03 | 1.58E-02 |
| SLC39A9 | 14:69458363-69462381 | 14:69458730-69461696 | IR | 3 | 3 | ENST00000031146 | -13.73 | 4.48E-02 | 7.40E-02 |
| ARHGEF26 | 3:154253184-154254784 | 3:154254720-154254784 | A3ss | 3 | 2 | ENST00000356448 | -13.75 | 6.75E-04 | 2.76E-03 |
| GSDMD | 8:143559777-143561423 | 8:143560772-143561001 | IR | 3 | 3 | ENST00000524846 | -13.75 | 9.03E-03 | 1.29E-01 |
| LRTOMT | 11:72095097-72096722 | 11:72095379-72096722 | TSSIA3ss | 2 | 2 | TSS.ENST00000289488 | -13.75 | 8.16E-03 | 1.19E-02 |
| ZNF513 | 2:27378616-27378883 | 2:27378616-27378777 | A3ss | 3 | 3 | Ex.TSS.ENST00000491924 | -13.75 | 3.05E-02 | 1.14E-01 |
| HLA-J | 6:30007318-30007593 | 6:30007397-30007523 | IR (overlapping region) | 3 | 2 | ENST00000494367 | -13.77 | 4.70E-02 | 3.83E-01 |
| TRIM16 | 17:15683036-15684299 | 17:15683372-15684195 | IR | 3 | 3 | ENST00000416464 | -13.78 | 4.04E-03 | 7.95E-03 |
| PRXL2A | 10:80408644-80420482 | 10:80420466-80420482 | TSSIA3ss | 3 | 3 | TSS.ENST00000606162 | -13.79 | 1.50E-02 | 2.00E-02 |
| FBXW2 | 9:120788187-120793148 | 9:120788187-120788278 | TSSIA3ss | 3 | 3 | TSS.ENST00000474117 | -13.82 | 2.11E-02 | 4.38E-01 |
| MMP17 | 12:131845450-131849938 | 12:131849802-131849938 | A3ss | 3 | 3 | ENST00000360564 | -13.85 | 5.67E-03 | 8.47E-03 |
| MPPE1 | 18:11886622-11886916 | 18:11886713-11886781 | SES | 3 | 3 | ENST00000587724 | -13.85 | 3.81E-02 | 8.12E-02 |
| RAB29 | 1:205774833-205775459 | 1:205775087-205775272 | IR | 3 | 3 | ENST00000414729 | -13.86 | 1.22E-02 | 3.15E-02 |
| PCNX2 | 1:233017155-233054267 | 1:233025146-233025399 | SES | 3 | 3 | ENST00000258229 | -13.89 | 1.26E-02 | 3.46E-02 |
| RGL3 | 19:11394521-11397243 | 19:11394606-11394678 | SES | 3 | 3 | ENST00000568628 | -13.9 | 3.95E-02 | 8.63E-02 |
| DHDDS | 1:26432493-26432890 | 1:26432493-26432518 | TSSIA5ss | 3 | 3 | TSS.ENST00000436153 | -13.92 | 3.51E-03 | 9.77E-03 |
| DHDDS | 1:26432416-26432890 | 1:26432416-26432518 | TSSIA5ss | 3 | 3 | TSS.ENST00000436153 | -13.92 | 3.51E-03 | 9.77E-03 |
| DHDDS | 1:26432377-26432890 | 1:26432377-26432518 | TSSIA5ss | 3 | 3 | TSS.ENST00000525165 | -13.92 | 3.51E-03 | 9.76E-03 |
| WIPI2 | 7:5190494-5214534 | 7:5193118-5193171 | MES | 3 | 3 | ENST00000288828 | -13.92 | 7.69E-04 | 8.30E-03 |
| KIF20B | 10:89725159-89726412 | 10:89726293-89726412 | A3ss | 3 | 3 | ENST00000371728 | -13.96 | 2.78E-02 | 3.72E-02 |
| MPI | 15:74897512-74902219 | 15:74901257-74901949 | IR (overlapping region) | 3 | 3 | ENST00000352410 | -13.98 | 3.57E-02 | 6.00E-02 |
| SGK3 | 8:66667702-66679720 | 8:66677642-66677954 | SES | 3 | 3 | ENST00000661636 | -13.99 | 9.64E-03 | 1.29E-01 |
| ZNF138 | 7:64815045-64815575 | 7:64815572-64815575 | A3ss | 3 | 3 | ENST00000440598 | -13.99 | 4.08E-02 | 4.62E-01 |
| EXOC7 | 17:76089321-76094413 | 17:76090329-76090397 | MES | 3 | 3 | ENST00000405575 | -14.01 | 2.74E-02 | 5.04E-02 |
| FBXL6 | 8:144356808-144357973 | 8:144357122-144357627 | IR (overlapping region) | 3 | 3 | ENST00000524492 | -14.04 | 2.08E-03 | 3.18E-02 |
| ALG3 | 3:184245813-184248781 | 3:184248745-184248781 | TSSIA5ss | 3 | 3 | TSS.ENST00000397676 | -14.05 | 1.63E-02 | 6.77E-02 |
| FUT11 | 10:73773042-73773193 | 10:73773190-73773193 | TSSIA3ss | 3 | 3 | TSS.ENST00000372841 | -14.06 | 3.65E-02 | 4.87E-02 |
| PRMT2 | 21:46635764-46636895 | 21:46636439-46636547 | SES | 3 | 3 | ENST00000355680 | -14.07 | 2.75E-03 | 9.03E-03 |
| DPH5 | 1:100995150-101021640 | 1:101013710-101013818 | MXS | 3 | 3 | ENST00000488789 | -14.1 | 8.91E-04 | 2.16E-03 |
| GALT | 9:34647541-34647849 | 9:34647568-34647656 | IR (overlapping region) | 3 | 3 | ENST00000554330 | -14.1 | 1.68E-02 | 4.92E-01 |
| SYT6 | 1:114103722-114137494 | 1:114103722-114137065 | A3ss | 3 | 3 | Ex.ENST00000369547 | -14.13 | 2.25E-02 | 5.49E-02 |
| POLL | 10:101587246-101588126 | 10:101587407-101587821 | IR (overlapping region) | 3 | 3 | ENST00000339310 | -14.16 | 1.92E-02 | 2.53E-02 |
| WDR19 | 4:39189782-39199477 | 4:39194544-39194659 | SES | 3 | 2 | ENST00000399820 | -14.16 | 1.29E-02 | 8.04E-02 |
| EME2 | 16:1776068-1780635 | 16:1777873-1777945 | IR (overlapping region) | 3 | 3 | ENST00000568449 | -14.18 | 4.60E-02 | 8.45E-02 |
| ZNF875 | 19:37318187-37323532 | 19:37322185-37322232 | SES | 3 | 3 | ENST00000544914 | -14.18 | 4.77E-02 | 1.09E-01 |
| SLC12A5-AS1 | 20:46013500-46014788 | 20:46014275-46014370 | IR (overlapping region) | 3 | 3 | ENST00000535913 | -14.22 | 3.98E-02 | 1.27E-01 |
| NCK1-DT | 3:136846587-136861746 | 3:136861615-136861746 | TSSIA5ss | 3 | 3 | TSS.ENST00000470236 | -14.23 | 3.90E-02 | 1.57E-01 |
| ZNF189 | 9:101399190-101399925 | 9:101399884-101399925 | TSSIA3ss | 3 | 3 | TSS.ENST00000339664 | -14.23 | 1.90E-02 | 2.68E-01 |
| NF2 | 22:29678324-29694751 | 22:29681439-29681601 | SES | 3 | 3 | ENST00000338641 | -14.27 | 4.80E-02 | 1.59E-01 |
| KRIT1 | 7:92245465-92246100 | 7:92245575-92245843 | IR (overlapping region) | 3 | 3 | ENST00000340022 | -14.28 | 1.14E-02 | 1.35E-01 |
| DDX11 | 12:31074328-31078389 | 12:31077945-31078288 | SES | 3 | 3 | ENST00000535317 | -14.33 | 1.93E-03 | 2.91E-03 |
| ENTR1 | 9:136407939-136410327 | 9:136410090-136410239 | MXS | 3 | 3 | ENST00000298537 | -14.34 | 4.49E-03 | 1.17E-01 |
| ARSL | X:2955538-2960377 | X:2958274-2958435 | MES | 3 | 3 | ENST00000545496 | -14.36 | 2.18E-03 | 1.36E-01 |
| ASB8 | 12:48149999-48151200 | 12:48149999-48150140 | TSSIA3ss | 2 | 3 | TSS.ENST00000539528 | -14.38 | 2.17E-02 | 3.28E-02 |
| ATP2A1-AS1 | 16:28879489-28879895 | 16:28879756-28879866 | IR (overlapping region) | 3 | 3 | ENST00000561547 | -14.39 | 3.15E-02 | 5.88E-02 |
| KHDC4 | 1:155921528-155921860 | 1:155921528-155921628 | A3ss | 3 | 3 | ENST00000368320 | -14.42 | 1.27E-02 | 3.23E-02 |
| KHDC4 | 1:155921525-155921860 | 1:155921525-155921628 | A3ss | 3 | 3 | Ex.ENST00000368320 | -14.42 | 1.27E-02 | 3.23E-02 |
| MTX3 | 5:79990237-79991157 | 5:79990594-79990663 | SES | 3 | 3 | ENST00000509852 | -14.43 | 3.85E-03 | 2.83E-02 |
| ZNF205 | 16:3112877-3113416 | 16:3112877-3112950 | A5ss | 3 | 3 | Ex.TSS.ENST00000414351 | -14.45 | 1.22E-04 | 2.29E-04 |
| GAS2L3 | 12:100573786-100594874 | 12:100591736-100591856 | SES | 2 | 3 | ENST00000547754 | -14.47 | 4.10E-02 | 6.04E-02 |
| ZNF84 | 12:133037546-133041284 | 12:133041100-133041284 | TSSIA3ss | 3 | 3 | TSS.ENST00000543758 | -14.5 | 3.59E-02 | 5.40E-02 |
| ZNF84 | 12:133037546-133041277 | 12:133041100-133041277 | TSSIA3ss | 3 | 3 | TSS.ENST00000543758 | -14.5 | 3.59E-02 | 5.40E-02 |
| DALRD3 | 3:49016061-49016157 | 3:49016061-49016086 | A3ss | 3 | 3 | ENST00000313778 | -14.55 | 1.34E-02 | 5.92E-02 |
| MRNIP | 5:179837266-179837885 | 5:179837853-179837853 | IR (overlapping region) | 3 | 3 | ENST00000376931 | -14.55 | 2.88E-02 | 2.03E-01 |
| METTL8 | 2:171360514-171433882 | 2:171392043-171392197 | MES | 3 | 3 | ENST00000442541 | -14.57 | 4.82E-02 | 1.72E-01 |
| PIAS1 | 15:68187542-68193846 | 15:68189271-68190492 | IR (overlapping region) | 3 | 3 | ENST00000249636 | -14.58 | 3.71E-02 | 6.28E-02 |
| POLL | 10:101587246-101588126 | 10:101587407-101587884 | IR (overlapping region) | 3 | 3 | ENST00000339310 | -14.59 | 4.14E-03 | 5.46E-03 |
| SORBS1 | 10:95422074-95432447 | 10:95422190-95422216 | SES | 3 | 3 | ENST00000277982 | -14.6 | 1.31E-03 | 1.75E-03 |
| TEAD2 | 19:49360082-49362372 | 19:49362333-49362372 | TSSIA5ss | 3 | 3 | TSS.ENST00000593945 | -14.6 | 2.65E-02 | 6.25E-02 |
| DSP | 6:7579773-7582641 | 6:7579773-7581569 | TSSIA5ss | 3 | 3 | TSS.ENST00000379802 | -14.62 | 3.95E-02 | 3.54E-01 |
| PRMT2 | 21:46635764-46636895 | 21:46635764-46636547 | TSSIA5ss | 3 | 3 | TSS.ENST00000397637 | -14.65 | 2.18E-03 | 7.14E-03 |
| ASB8 | 12:48149999-48151200 | 12:48150087-48150140 | SES | 2 | 3 | ENST00000535055 | -14.68 | 2.45E-02 | 3.70E-02 |
| GOLGA6L9 | 15:82430045-82432897 | 15:82431950-82432575 | IR (overlapping region) | 3 | 3 | ENST00000618706 | -14.68 | 1.56E-02 | 2.68E-02 |
| GOLGA6L9 | 15:82430045-82432897 | 15:82431950-82432571 | IR (overlapping region) | 3 | 3 | ENST00000618706 | -14.68 | 1.56E-02 | 2.68E-02 |
| ZC3H11A | 1:203801629-203816925 | 1:203802957-203803016 | SES | 3 | 3 | ENST00000367212 | -14.74 | 1.90E-02 | 4.93E-02 |
| FANCE | 6:35457601-35458296 | 6:35457916-35457984 | SES | 3 | 3 | ENST00000229769 | -14.76 | 1.97E-02 | 1.66E-01 |
| NR2C2AP | 19:19201426-19202575 | 19:19201564-19201930 | IR (overlapping region) | 3 | 3 | ENST00000537399 | -14.78 | 3.32E-02 | 7.39E-02 |
| AL137058.2 | 13:52627579-52637304 | 13:52632422-52632494 | SES | 3 | 3 | ENST00000398039 | -14.87 | 6.63E-03 | 1.05E-02 |
| DPP3 | 11:66480466-66482195 | 11:66482193-66482195 | TSSIA3ss | 3 | 3 | TSS.ENST00000526515 | -14.89 | 2.87E-02 | 4.12E-02 |
| GVQW3 | 11:76381305-76382904 | 11:76381375-76381788 | IR | 2 | 3 | ENST00000530460 | -14.89 | 1.36E-02 | 1.99E-02 |
| SEC16A | 9:136446950-136447226 | 9:136446950-136447015 | A3ss | 3 | 3 | ENST00000290037 | -14.89 | 3.62E-03 | 9.45E-02 |
| TPCN2 | 11:69054098-69057577 | 11:69055175-69055352 | MES | 3 | 3 | ENST00000294309 | -14.93 | 4.79E-02 | 6.96E-02 |
| ARSA | 22:50627556-50628152 | 22:50627793-50627990 | IR | 3 | 3 | ENST00000216124 | -14.98 | 4.97E-02 | 1.72E-01 |

| | | | | | | | | | |
|---|---|---|---|---|---|---|---|---|---|
| TMEM41B | 11:9280654-9283593 | 11:9281042-9283370 | IR | 3 | 3 | ENST00000524543 | -14.99 | 4.64E-02 | 6.82E-02 |
| GUSBP11 | 22:23683868-23686837 | 22:23686831-23686837 | A5ss | 3 | 3 | ENST00000421064 | -15 | 3.97E-02 | 1.32E-01 |
| SPIN3 | X:56990831-56994949 | X:56992584-56994722 | IR | 3 | 3 | ENST00000374919 | -15 | 1.93E-04 | 7.45E-02 |
| ATG9A | 2:219227814-219229534 | 2:219227814-219228053 | TSSIA3ss | 3 | 3 | TSS.ENST00000486766 | -15.02 | 4.59E-02 | 1.68E-01 |
| GIGYF1 | 7:100688719-100693936 | 7:100689302-100689555 | SES | 3 | 2 | ENST00000646601 | -15.09 | 5.44E-03 | 3.86E-02 |
| WDR90 | 16:655773-656537 | 16:655886-656301 | IR (overlapping region) | 3 | 3 | ENST00000552943 | -15.12 | 3.48E-02 | 6.68E-02 |
| ZFP62 | 5:180847615-180851493 | 5:180848083-180851380 | IR | 3 | 3 | ENST00000512132 | -15.12 | 2.98E-02 | 1.90E-01 |
| ZNF638 | 2:71331866-71348752 | 2:71331866-71331875 | TSSIA5ss | 3 | 3 | TSS.ENST00000264447 | -15.15 | 2.24E-02 | 8.67E-02 |
| ZFYVE16 | 5:80445406-80448282 | 5:80448026-80448282 | A3ss | 3 | 3 | ENST00000338008 | -15.2 | 1.80E-02 | 1.33E-01 |
| GDPD2 | X:70423383-70426051 | X:70425763-70425856 | MES | 3 | 3 | ENST00000374382 | -15.22 | 2.98E-02 | 1.00E+00 |
| APAF1 | 12:98727173-98727316 | 12:98727192-98727278 | IR (overlapping region) | 3 | 3 | ENST00000357310 | -15.23 | 2.56E-02 | 4.01E-02 |
| TPD52L2 | 20:63873817-63882718 | 20:63875816-63875875 | SES | 3 | 3 | ENST00000217121 | -15.24 | 3.31E-03 | 1.07E-02 |
| TRIM41 | 5:181232890-181233635 | 5:181233413-181233635 | A3ss | 3 | 3 | ENST00000508930 | -15.24 | 2.91E-03 | 2.06E-02 |
| PPP1R16A | 8:144500096-144500761 | 8:144500200-144500266 | IR (overlapping region) | 3 | 3 | ENST00000526183 | -15.26 | 1.78E-03 | 2.73E-02 |
| ATG16L2 | 11:72824843-72825348 | 11:72825302-72825348 | A3ss | 2 | 3 | ENST00000321297 | -15.28 | 3.40E-03 | 4.94E-03 |
| RSPO4 | 20:960467-967173 | 20:963935-967174 | A5ss | 3 | 3 | Ex.TSS.ENST00000400634 | -15.28 | 3.44E-02 | 1.11E-01 |
| LDOC1 | X:141173235-141177129 | X:141176409-141176892 | IR | 3 | 3 | ENST00000370526 | -15.31 | 2.87E-03 | 1.19E-01 |
| RREB1 | 6:7108061-7176654 | 6:7108061-7108064 | TSSIA5ss | 3 | 3 | TSS.ENST00000471433 | -15.32 | 2.30E-02 | 2.05E-01 |
| TTN-AS1 | 2:178523637-178535731 | 2:178531314-178531578 | MXS | 2 | 2 | ENST00000657210 | -15.33 | 2.67E-02 | 9.51E-02 |
| NSUN5P1 | 7:75412933-75414884 | 7:75414640-75414798 | SES | 3 | 3 | ENST00000464979 | -15.34 | 4.98E-02 | 5.75E-01 |
| STAT6 | 12:57108300-57110076 | 12:57109941-57110076 | A5ss | 3 | 3 | ENST00000553275 | -15.34 | 1.78E-02 | 2.75E-02 |
| STAT6 | 12:57108300-57110042 | 12:57109941-57110042 | A5ss | 3 | 3 | ENST00000553275 | -15.34 | 1.78E-02 | 2.75E-02 |
| LDOC1 | X:141173235-141177129 | X:141176427-141176892 | IR | 3 | 3 | ENST00000370526 | -15.38 | 3.22E-03 | 1.33E-01 |
| KANSL3 | 2:96619545-96631311 | 2:96619672-96619762 | SES | 3 | 3 | ENST00000354204 | -15.39 | 4.04E-02 | 1.60E-01 |
| TLCD5 | 11:120325369-120327465 | 11:120327122-120327465 | A3ss | 3 | 3 | Ex.TSS.ENST00000314475 | -15.45 | 6.35E-03 | 8.57E-03 |
| TLCD5 | 11:120325369-120327451 | 11:120327122-120327451 | A3ss | 3 | 3 | Ex.TSS.ENST00000314475 | -15.45 | 6.35E-03 | 8.57E-03 |
| TLCD5 | 11:120325369-120327440 | 11:120327122-120327440 | A3ss | 3 | 3 | Ex.TSS.ENST00000314475 | -15.45 | 6.35E-03 | 8.57E-03 |
| RIC3 | 11:8126805-8137377 | 11:8128219-8128354 | SES | 3 | 3 | ENST00000530060 | -15.48 | 2.15E-02 | 3.16E-02 |
| OSER1-DT | 20:44211078-44224820 | 20:44211078-44211356 | A5ss | 3 | 3 | Ex.ENST00000664132 | -15.49 | 3.75E-02 | 1.20E-01 |
| ANK3 | 10:60139088-60166590 | 10:60145968-60146021 | SES | 3 | 3 | ENST00000474360 | -15.51 | 2.87E-02 | 3.82E-02 |
| IL18BP | 11:71999063-72000012 | 11:71999371-71999616 | IR | 3 | 3 | ENST00000404792 | -15.52 | 2.81E-02 | 4.08E-02 |
| BCL9L | 11:118908656-118925237 | 11:118909914-118910015 | MES | 3 | 3 | ENST00000532899 | -15.55 | 7.53E-03 | 1.02E-02 |
| FAHD2B | 2:97091986-97094700 | 2:97091986-97091988 | A3ss | 3 | 3 | Ex.TSS.ENST00000414820 | -15.57 | 2.91E-02 | 1.15E-01 |
| CHKB | 22:50581749-50582820 | 22:50581863-50582248 | IR (overlapping region) | 3 | 3 | ENST00000479003 | -15.58 | 7.69E-03 | 2.67E-02 |
| HEPH | X:66208247-66258839 | X:66256105-66256330 | MES | 3 | 2 | ENST00000336279 | -15.6 | 5.67E-03 | 1.00E+00 |
| TACC1 | 8:38787744-38825307 | 8:38788704-38788819 | MES | 3 | 2 | ENST00000317827 | -15.67 | 6.28E-03 | 8.14E-02 |
| PQBP1 | X:48898015-48898491 | X:48898015-48898082 | TSSIA5ss | 3 | 3 | TSS.ENST00000376566 | -15.72 | 4.99E-03 | 8.52E-01 |
| PLD3 | 19:40348725-40365717 | 19:40365553-40365585 | SES | 3 | 3 | ENST00000392032 | -15.73 | 1.70E-02 | 3.91E-02 |
| STK39 | 2:168063571-168074981 | 2:168065319-168065381 | SES | 3 | 3 | ENST00000355999 | -15.74 | 3.23E-03 | 1.15E-02 |
| METTL8 | 2:171360514-171433882 | 2:171392043-171392197 | MES | 3 | 3 | ENST00000392604 | -15.75 | 8.67E-03 | 3.10E-02 |
| ADGRL2 | 1:81800513-81836884 | 1:81800513-81801068 | A5ss | 3 | 3 | Ex.TSS.ENST00000319517 | -15.76 | 8.18E-03 | 2.36E-02 |
| CDC27 | 17:47122537-47123885 | 17:47122537-47122600 | A3ss | 3 | 3 | ENST00000066544 | -15.76 | 4.78E-02 | 9.80E-02 |
| MICALL2 | 7:1436742-1437608 | 7:1436857-1437534 | IR | 3 | 3 | ENST00000496184 | -15.77 | 4.49E-02 | 2.52E-01 |
| STRADA | 17:63702835-63704908 | 17:63703752-63704340 | IR (overlapping region) | 3 | 3 | ENST00000245865 | -15.79 | 3.61E-02 | 7.48E-02 |
| ZBTB46 | 20:63743666-63747301 | 20:63745676-63745822 | IR (overlapping region) | 3 | 3 | ENST00000245663 | -15.79 | 9.95E-03 | 3.20E-02 |
| CHKB | 22:50581749-50582820 | 22:50582096-50582248 | IR (overlapping region) | 3 | 3 | ENST00000479003 | -15.83 | 5.00E-04 | 1.73E-03 |
| SNRPN | 15:24968083-24974386 | 15:24974311-24974386 | A3ss | 3 | 3 | ENST00000390687 | -15.83 | 1.39E-06 | 2.32E-06 |
| SUGP2 | 19:18990893-18993740 | 19:18991150-18993647 | IR | 3 | 3 | ENST00000452918 | -15.85 | 3.48E-03 | 7.77E-03 |
| ADSS1 | 14:104744719-104746385 | 14:104746135-104746235 | IR | 3 | 3 | ENST00000557271 | -15.86 | 2.79E-02 | 4.48E-02 |
| NSMCE2 | 8:125091959-125102316 | 8:125102051-125102146 | MXS | 3 | 3 | ENST00000287437 | -15.87 | 2.62E-02 | 3.87E-01 |
| SH3D21 | 1:36321056-36321778 | 1:36321210-36321723 | IR | 3 | 3 | ENST00000474766 | -15.9 | 1.23E-02 | 3.44E-02 |
| SLC25A29 | 14:100293378-100306198 | 14:100298842-100298885 | MXS | 3 | 3 | ENST00000359232 | -15.9 | 1.51E-04 | 2.42E-04 |
| ZMYM1 | 1:35079443-35095818 | 1:35093914-35094009 | SES | 3 | 3 | ENST00000466390 | -15.91 | 2.39E-02 | 6.72E-02 |
| PANK4 | 1:2510749-2511337 | 1:2510749-2510782 | A3ss | 3 | 3 | ENST00000378466 | -15.94 | 4.14E-02 | 1.14E-01 |
| ELMOD3 | 2:85354830-85356966 | 2:85355076-85355170 | MXS | 3 | 3 | ENST00000393852 | -15.96 | 4.04E-02 | 1.58E-01 |
| SLC12A5-AS1 | 20:46013500-46014788 | 20:46013797-46014123 | IR (overlapping region) | 2 | 3 | ENST00000535913 | -16.03 | 1.61E-02 | 5.15E-02 |
| STYXL1 | 7:76030528-76047811 | 7:76047662-76047811 | TSSIA5ss | 3 | 3 | TSS.ENST00000359697 | -16.07 | 4.84E-02 | 5.62E-01 |
| CCDC40 | 17:80036692-80038122 | 17:80036692-80036757 | TSSIA5ss | 3 | 3 | TSS.ENST00000574099 | -16.08 | 6.02E-03 | 1.11E-02 |
| FBXL18 | 7:5500488-5501919 | 7:5501001-5501182 | IR | 3 | 3 | ENST00000458142 | -16.08 | 3.09E-02 | 3.33E-01 |
| CHKB | 22:50581749-50582820 | 22:50581863-50582028 | IR (overlapping region) | 3 | 3 | ENST00000479003 | -16.09 | 1.64E-02 | 5.70E-02 |
| FNBP1 | 9:129923940-129924959 | 9:129923940-129923996 | A3ss | 3 | 3 | ENST00000449089 | -16.13 | 1.50E-02 | 3.75E-01 |
| ENTPD1-AS1 | 10:95897680-95907662 | 10:95904830-95904938 | SES | 3 | 3 | ENST00000661794 | -16.14 | 1.42E-02 | 1.91E-02 |
| GAK | 4:866502-866955 | 4:866502-866534 | A3ss | 3 | 3 | ENST00000314167 | -16.14 | 4.39E-02 | 2.85E-01 |
| CAMK2G | 10:73824085-73837467 | 10:73825321-73825347 | MXS | 3 | 3 | ENST00000474131 | -16.16 | 3.39E-02 | 4.53E-02 |
| CCDC88A | 2:55296524-55301205 | 2:55299839-55299919 | SES | 3 | 3 | ENST00000263630 | -16.16 | 4.35E-02 | 1.65E-01 |
| REC8 | 14:24179907-24180026 | 14:24180010-24180026 | A3ss | 3 | 3 | ENST00000558381 | -16.16 | 5.97E-05 | 9.74E-05 |
| AC005062.1 | 7:20045562-20125079 | 7:20122986-20125080 | A5ss | 3 | 3 | Ex.TSS.ENST00000415499 | -16.2 | 3.92E-02 | 4.03E-01 |
| SELENBP1 | 1:151368320-151369712 | 1:151369004-151369189 | MES | 3 | 3 | ENST00000368868 | -16.21 | 3.94E-02 | 9.73E-02 |
| CBWD1 | 9:175785-178815 | 9:177723-177820 | SES | 3 | 3 | ENST00000314367 | -16.22 | 4.12E-02 | 1.00E+00 |
| CDK10 | 16:89693274-89693495 | 16:89693327-89693397 | IR | 3 | 3 | ENST00000504473 | -16.22 | 2.15E-02 | 4.19E-02 |
| MICALL2 | 7:1439925-1440727 | 7:1440086-1440590 | IR | 3 | 2 | ENST00000487156 | -16.24 | 1.19E-02 | 1.17E-01 |
| SALL3 | 18:78992074-78995462 | 18:78994906-78995121 | IR | 3 | 3 | ENST00000537592 | -16.24 | 2.62E-02 | 5.66E-02 |
| TBRG1 | 11:124625904-124626903 | 11:124626473-124626609 | MXS | 3 | 3 | ENST00000441174 | -16.24 | 3.17E-02 | 4.28E-02 |
| BX890604.2 | X:3818501-3822373 | X:3818657-3818703 | MES | 3 | 3 | ENST00000669937 | -16.25 | 1.19E-02 | 1.00E+00 |
| MAPK15 | 8:143721000-143721673 | 8:143721412-143721548 | IR | 3 | 3 | ENST00000461928 | -16.29 | 2.13E-02 | 3.23E-01 |
| ADIPOR2 | 12:1691192-1754257 | 12:1691192-1691213 | A5ss | 3 | 3 | Ex.TSS.ENST00000357103 | -16.33 | 2.38E-02 | 3.57E-02 |
| MYL6 | 12:56159623-56160024 | 12:56159975-56160024 | A3ss | 3 | 3 | ENST00000548400 | -16.36 | 2.62E-02 | 4.04E-02 |
| IL4I1 | 19:49927816-49929416 | 19:49929326-49929416 | TSSIA5ss | 3 | 3 | TSS.ENST00000341114 | -16.41 | 4.28E-02 | 9.81E-02 |
| IL4I1 | 19:49927816-49929349 | 19:49929326-49929349 | TSSIA5ss | 3 | 3 | TSS.ENST00000341114 | -16.41 | 4.28E-02 | 9.81E-02 |
| UBR3 | 2:170007191-170008803 | 2:170007191-170007202 | A5ss | 2 | 3 | ENST00000392632 | -16.41 | 1.85E-02 | 6.59E-02 |
| BRD1 | 22:49787495-49794033 | 22:49787495-49787887 | A3ss | 3 | 3 | ENST00000404760 | -16.45 | 3.97E-02 | 1.37E-01 |
| HYI | 1:43451312-43451427 | 1:43451410-43451427 | TSSIA5ss | 3 | 2 | TSS.ENST00000372425 | -16.45 | 3.36E-02 | 9.54E-02 |
| ITGA7 | 12:55698918-55700261 | 12:55699870-55699989 | SES | 3 | 3 | ENST00000555728 | -16.45 | 3.02E-02 | 4.65E-02 |
| TRIM41 | 5:181232890-181233635 | 5:181233413-181233435 | SES | 3 | 3 | ENST00000315073 | -16.47 | 7.14E-03 | 5.04E-02 |
| HEPH | X:66208247-66258839 | X:66255035-66255141 | MES | 3 | 2 | ENST00000336279 | -16.5 | 3.39E-02 | 1.00E+00 |
| SPIN3 | X:56990831-56994949 | X:56992584-56993786 | IR | 3 | 3 | ENST00000374919 | -16.5 | 1.05E-03 | 3.97E-01 |
| ENTPD6 | 20:25224158-25225204 | 20:25225162-25225204 | A3ss | 3 | 3 | ENST00000376666 | -16.51 | 5.07E-03 | 1.52E-02 |
| ATXN2 | 12:111470743-111485264 | 12:111479070-111479102 | SES | 3 | 3 | ENST00000643669 | -16.55 | 3.82E-02 | 5.65E-02 |
| WDR90 | 16:655773-656537 | 16:655890-656301 | IR (overlapping region) | 3 | 3 | ENST00000552943 | -16.57 | 1.91E-02 | 3.66E-02 |
| SLC13A3 | 20:46613726-46684395 | 20:46613726-46670139 | A3ss | 3 | 3 | Ex.TSS.ENST00000372121 | -16.61 | 2.07E-02 | 6.64E-02 |
| CAMSAP3 | 19:7606572-7608125 | 19:7607821-7608126 | A3ss | 3 | 3 | Ex.ENST00000160298 | -16.63 | 3.48E-02 | 8.37E-02 |
| PCDHGC3 | 5:141475947-141478795 | 5:141476147-141478393 | IR (overlapping region) | 3 | 2 | ENST00000611950 | -16.68 | 7.94E-03 | 5.44E-02 |
| TMEM145 | 19:42323583-42325064 | 19:42323790-42324277 | IR | 3 | 3 | ENST00000673205 | -16.73 | 4.95E-02 | 1.14E-01 |
| VCAN | 5:83512397-83537006 | 5:83519349-83522309 | SES | 3 | 3 | ENST00000265077 | -16.73 | 4.06E-02 | 3.09E-01 |

| | | | | | | | | | |
|---|---|---|---|---|---|---|---|---|---|
| ZNF232 | 17:5109769-5111799 | 17:5109769-5109868 | A3ss | 3 | 3 | ENST00000250076 | -16.81 | 4.43E-02 | 9.13E-02 |
| HEXD | 17:82439631-82440532 | 17:82439714-82440472 | IR | 3 | 3 | ENST00000585077 | -16.82 | 1.61E-02 | 3.44E-02 |
| MTHFR | 1:11802881-11804532 | 1:11803130-11803798 | IR | 3 | 3 | ENST00000376592 | -16.83 | 3.43E-02 | 8.39E-02 |
| SPIN3 | X:56990831-56994949 | X:56992238-56994722 | IR | 3 | 3 | ENST00000374919 | -16.84 | 1.49E-03 | 5.49E-01 |
| IQCB1 | 3:121790216-121807343 | 3:121797118-121797227 | MES | 3 | 3 | ENST00000310864 | -16.85 | 1.89E-02 | 7.56E-02 |
| FNIP1 | 5:131671505-131672924 | 5:131672662-131672685 | IR (overlapping region) | 3 | 3 | ENST00000307954 | -16.87 | 3.11E-02 | 2.07E-01 |
| GUSBP11 | 22:23683868-23705428 | 22:23695173-23695517 | MXS | 3 | 3 | ENST00000422506 | -16.87 | 6.32E-04 | 2.10E-03 |
| STAG3L5P-PVRIG2P-PILRB | 7:100353484-100355142 | 7:100353484-100354012 | A5ss | 3 | 3 | ENST00000310771 | -16.87 | 1.84E-02 | 1.69E-01 |
| TULP1 | 6:35506174-35509208 | 6:35506274-35506279 | SES | 3 | 3 | ENST00000229771 | -16.93 | 1.74E-02 | 1.46E-01 |
| RELL2 | 5:141637749-141638794 | 5:141637749-141638409 | TSSIA5ss | 3 | 3 | TSS.ENST00000297164 | -16.94 | 3.17E-02 | 2.03E-01 |
| GGT1 | 22:24608024-24610357 | 22:24609962-24610033 | MXS | 3 | 3 | ENST00000411974 | -16.95 | 7.24E-03 | 2.41E-02 |
| DLG1 | 3:197142769-197194424 | 3:197149743-197149796 | SES | 3 | 2 | ENST00000357674 | -16.97 | 4.08E-02 | 1.73E-01 |
| SUGP2 | 19:18991150-18994365 | 19:18991150-18993740 | TSSIA3ss | 3 | 3 | TSS.ENST00000452918 | -16.98 | 3.27E-02 | 7.30E-02 |
| UPP1 | 7:48094828-48106872 | 7:48103297-48103411 | MES | 3 | 3 | ENST00000331803 | -16.99 | 8.20E-03 | 8.80E-02 |
| APBB3 | 5:140561587-140562227 | 5:140561723-140561843 | IR | 3 | 3 | ENST00000510241 | -17 | 1.75E-02 | 1.19E-01 |
| CAMSAP2 | 1:200844870-200847257 | 1:200847210-200847257 | A3ss | 3 | 3 | ENST00000236925 | -17.05 | 2.06E-02 | 5.16E-02 |
| TRIB2 | 2:12717011-12723259 | 2:12717640-12723260 | A3ss | 3 | 3 | Ex.TSS.ENST00000381465 | -17.05 | 4.80E-03 | 1.68E-02 |
| ZMYM5 | 13:19851355-19851796 | 13:19851449-19851688 | IR | 3 | 3 | ENST00000382909 | -17.12 | 9.04E-03 | 1.42E-02 |
| COQ4 | 9:128323148-128325778 | 9:128325143-128325239 | SES | 3 | 3 | ENST00000300452 | -17.13 | 4.91E-02 | 1.00E+00 |
| FAHD2B | 2:97091713-97094700 | 2:97091863-97091988 | SES | 3 | 3 | ENST00000414820 | -17.13 | 1.35E-02 | 5.35E-02 |
| CCDC88A | 2:55296521-55299838 | 2:55296521-55296523 | A3ss | 3 | 3 | ENST00000263630 | -17.23 | 3.38E-02 | 1.28E-01 |
| YIPF1 | 1:53852271-53860055 | 1:53855001-53860056 | A5ss | 3 | 3 | Ex.TSS.ENST00000072644 | -17.28 | 4.88E-02 | 1.40E-01 |
| PLA2G4B | 15:41842193-41843223 | 15:41842277-41842553 | IR (overlapping region) | 3 | 3 | ENST00000483748 | -17.3 | 9.04E-03 | 1.52E-02 |
| GOLGA6L9 | 15:82430045-82432897 | 15:82432632-82432816 | IR (overlapping region) | 3 | 3 | ENST00000618706 | -17.32 | 7.73E-03 | 1.33E-02 |
| SYTL1 | 1:27349398-27351336 | 1:27350486-27350793 | IR | 3 | 3 | ENST00000490170 | -17.34 | 4.56E-02 | 1.27E-01 |
| SKP2 | 5:36177293-36183839 | 5:36181818-36183839 | TSSIA3ss | 3 | 3 | TSS.ENST00000274255 | -17.38 | 1.70E-02 | 9.59E-02 |
| HDAC11 | 3:13483565-13501870 | 3:13496736-13496852 | MES | 3 | 3 | ENST00000295757 | -17.39 | 4.48E-02 | 1.81E-01 |
| NFRKB | 11:129892871-129894358 | 11:129893297-129893385 | MXS | 3 | 3 | ENST00000524746 | -17.41 | 2.05E-02 | 2.78E-02 |
| C1RL | 12:7097164-7101897 | 12:7099686-7099807 | MES | 3 | 3 | ENST00000544702 | -17.52 | 2.77E-02 | 4.31E-02 |
| HDAC10 | 22:50245813-50245909 | 22:50245813-50245827 | A3ss | 3 | 3 | Ex.ENST00000216271 | -17.58 | 2.36E-02 | 8.16E-02 |
| HDAC10 | 22:50245737-50245909 | 22:50245737-50245827 | A3ss | 3 | 3 | ENST00000216271 | -17.58 | 2.36E-02 | 8.16E-02 |
| SLC43A1 | 11:57494172-57500778 | 11:57496031-57496164 | SES | 3 | 3 | ENST00000533066 | -17.58 | 1.59E-02 | 2.19E-02 |
| THTPA | 14:23556294-23557039 | 14:23556344-23556752 | IR (overlapping region) | 3 | 3 | ENST00000554789 | -17.64 | 3.11E-04 | 5.06E-04 |
| ANAPC10 | 4:145097688-145098183 | 4:145097761-145097886 | IR (overlapping region) | 3 | 3 | ENST00000507134 | -17.65 | 2.69E-02 | 1.52E-01 |
| TANGO2 | 22:20036855-20043354 | 22:20036855-20037094 | A5ss | 3 | 3 | ENST00000399807 | -17.65 | 1.68E-03 | 5.55E-03 |
| TMEM170A | 16:75447689-75451737 | 16:75451669-75451737 | TSSIA5ss | 3 | 3 | TSS.ENST00000561878 | -17.65 | 1.07E-02 | 2.07E-02 |
| ANO8 | 19:17328162-17329541 | 19:17328265-17329052 | IR | 3 | 3 | ENST00000597643 | -17.7 | 7.71E-03 | 1.71E-02 |
| MAN2A2 | 15:90916257-90918193 | 15:90916586-90916660 | SES | 3 | 2 | ENST00000557990 | -17.82 | 3.24E-03 | 5.58E-03 |
| DYRK2 | 12:67649183-67657105 | 12:67649797-67649945 | SES | 3 | 3 | ENST00000344096 | -17.86 | 4.68E-02 | 7.22E-02 |
| ADAL | 15:43333435-43335695 | 15:43334945-43335170 | SES | 3 | 3 | ENST00000389651 | -17.88 | 1.30E-02 | 2.18E-02 |
| RNF44 | 5:176532133-176532365 | 5:176532133-176532193 | A3ss | 3 | 3 | ENST00000274811 | -17.88 | 2.60E-02 | 1.82E-01 |
| RNF44 | 5:176532130-176532365 | 5:176532130-176532193 | A3ss | 3 | 3 | ENST00000274811 | -17.88 | 2.60E-02 | 1.82E-01 |
| TAF1C | 16:84180329-84181042 | 16:84180329-84180344 | A3ss | 3 | 3 | Ex.ENST00000341690 | -17.98 | 2.69E-03 | 5.22E-03 |
| FGFR2 | 10:121551460-121593708 | 10:121565438-121565704 | MES | 3 | 2 | ENST00000346997 | -18 | 4.52E-02 | 5.97E-02 |
| AFG3L1P | 16:89972766-89980241 | 16:89977639-89977802 | MXS | 3 | 3 | ENST00000418696 | -18.01 | 3.38E-02 | 6.60E-02 |
| RFNG | 17:82049677-82050555 | 17:82049843-82049917 | IR | 3 | 3 | ENST00000580953 | -18.01 | 1.62E-02 | 3.46E-02 |
| EGFL8 | 6:32167006-32167146 | 6:32167087-32167146 | A3ss | 3 | 2 | ENST00000333845 | -18.02 | 1.57E-02 | 1.11E-01 |
| EGFL8 | 6:32167006-32167102 | 6:32167087-32167102 | A3ss | 3 | 2 | ENST00000333845 | -18.02 | 1.57E-02 | 1.11E-01 |
| LGMN | 14:92704362-92709671 | 14:92704640-92704707 | MES | 3 | 3 | ENST00000334869 | -18.03 | 2.44E-03 | 4.06E-03 |
| SMIM1 | 1:3772825-3773181 | 1:3772842-3773062 | IR | 2 | 3 | ENST00000452264 | -18.06 | 2.76E-02 | 7.79E-02 |
| SUGP2 | 19:18990893-18993740 | 19:18991150-18991339 | IR | 3 | 3 | ENST00000452918 | -18.1 | 1.71E-02 | 3.82E-02 |
| SUGP2 | 19:18990893-18993740 | 19:18991150-18993276 | IR | 3 | 3 | ENST00000452918 | -18.14 | 1.76E-03 | 3.92E-03 |
| GPRC5C | 17:74432142-74439744 | 17:74432142-74432523 | A5ss | 3 | 3 | ENST00000582473 | -18.16 | 2.48E-02 | 5.18E-02 |
| NEMP1 | 12:57064740-57070819 | 12:57069234-57069306 | SES | 3 | 3 | ENST00000554340 | -18.18 | 4.26E-02 | 6.59E-02 |
| ZNF778 | 16:89221153-89223156 | 16:89222092-89222183 | SES | 3 | 3 | ENST00000433976 | -18.18 | 3.38E-02 | 6.59E-02 |
| THTPA | 14:23556294-23557039 | 14:23556344-23556742 | IR (overlapping region) | 3 | 3 | ENST00000554789 | -18.19 | 2.59E-04 | 4.22E-04 |
| ALG13 | X:111681425-111682235 | X:111682132-111682235 | TSSIA3ss | 3 | 3 | TSS.ENST00000251943 | -18.21 | 2.83E-02 | 5.84E-01 |
| COASY | 17:42562148-42563322 | 17:42562220-42562932 | IR | 3 | 3 | ENST00000393818 | -18.21 | 2.51E-03 | 5.07E-03 |
| RECQL | 12:21501216-21501462 | 12:21501216-21501372 | TSSIA3ss | 3 | 3 | TSS.ENST00000314748 | -18.21 | 1.66E-03 | 2.49E-03 |
| ARRDC2 | 19:18008168-18008777 | 19:18008229-18008453 | IR | 2 | 3 | ENST00000593560 | -18.22 | 4.46E-02 | 9.91E-02 |
| CYB5RL | 1:54184266-54195418 | 1:54190738-54190896 | MXS | 3 | 3 | ENST00000493530 | -18.22 | 2.28E-02 | 5.71E-02 |
| AC008555.4 | 19:34860174-34860576 | 19:34860303-34860448 | IR (overlapping region) | 3 | 3 | ENST00000604749 | -18.25 | 1.87E-02 | 4.25E-02 |
| FAM126A | 7:22946164-22947032 | 7:22946952-22947032 | TSSIA5ss | 3 | 3 | TSS.ENST00000409923 | -18.25 | 6.45E-03 | 6.64E-02 |
| CCT6P1 | 7:65751279-65755779 | 7:65752164-65752202 | MES | 3 | 3 | ENST00000434500 | -18.34 | 4.14E-02 | 4.69E-01 |
| MAP3K12 | 12:53486947-53487428 | 12:53487156-53487254 | IR | 3 | 3 | ENST00000547035 | -18.41 | 4.87E-02 | 7.45E-02 |
| EP300 | 22:41158501-41160641 | 22:41160612-41160642 | A3ss | 3 | 3 | Ex.ENST00000263253 | -18.43 | 2.36E-03 | 8.11E-03 |
| RFNG | 17:82049677-82050555 | 17:82049843-82050401 | IR | 3 | 3 | ENST00000580953 | -18.44 | 1.38E-04 | 2.94E-04 |
| ASPHD1 | 16:29900486-29901920 | 16:29900921-29901743 | IR | 3 | 3 | ENST00000308748 | -18.49 | 7.76E-03 | 1.46E-02 |
| TMEM147-AS1 | 19:35541582-35545492 | 19:35541585-35541750 | IR (overlapping region) | 3 | 3 | ENST00000588286 | -18.63 | 1.60E-02 | 3.62E-02 |
| ENTR1 | 9:136410202-136410327 | 9:136410202-136410239 | A3ss | 3 | 3 | Ex.TSS.ENST00000298537 | -18.69 | 8.25E-03 | 2.15E-01 |
| MYO5C | 15:52239880-52244355 | 15:52242048-52242213 | SES | 3 | 2 | ENST00000261839 | -18.74 | 2.75E-02 | 4.65E-02 |
| PIF1 | 15:64821282-64822265 | 15:64821367-64821520 | SES | 3 | 3 | ENST00000268043 | -18.75 | 2.67E-02 | 4.54E-02 |
| KCTD17 | 22:37059439-37061538 | 22:37060823-37060922 | MES | 3 | 3 | ENST00000403888 | -18.76 | 4.66E-02 | 1.59E-01 |
| TTC31 | 2:74492446-74492675 | 2:74492646-74492675 | A3ss | 3 | 3 | ENST00000233623 | -18.77 | 8.28E-04 | 3.24E-03 |
| MAP4K1 | 19:38595485-38595729 | 19:38595556-38595639 | IR | 3 | 3 | ENST00000588938 | -18.81 | 8.87E-04 | 2.04E-03 |
| TMEM240 | 1:1540290-1540624 | 1:1540356-1540419 | IR | 3 | 3 | ENST00000378733 | -18.83 | 4.07E-02 | 1.02E-01 |
| CREBZF | 11:85663524-85665102 | 11:85663602-85663697 | IR | 3 | 3 | ENST00000490820 | -18.86 | 1.76E-02 | 2.58E-02 |
| LRSAM1 | 9:127451513-127452084 | 9:127451670-127451928 | IR | 3 | 3 | ENST00000323301 | -18.86 | 4.54E-02 | 9.93E-01 |
| ARHGAP5 | 14:32090502-32094386 | 14:32094090-32094090 | IR | 3 | 3 | ENST00000345122 | -18.88 | 1.08E-02 | 1.76E-02 |
| MICALL2 | 7:1437890-1439431 | 7:1438354-1438839 | IR | 2 | 3 | ENST00000496184 | -18.94 | 4.23E-02 | 2.37E-01 |
| AMMECR1L | 2:127866997-127869453 | 2:127866997-127867279 | A3ss | 3 | 3 | Ex.ENST00000272647 | -18.96 | 3.40E-02 | 1.20E-01 |
| CCDC78 | 16:724322-724806 | 16:724510-724680 | IR | 3 | 3 | ENST00000463539 | -19.01 | 3.31E-02 | 6.40E-02 |
| MIAT | 22:26663399-26665530 | 22:26665457-26665530 | TSSIA3ss | 3 | 3 | TSS.ENST00000418918 | -19.02 | 1.50E-02 | 4.99E-02 |
| ACSBG1 | 15:78167468-78171529 | 15:78169078-78171049 | IR (overlapping region) | 3 | 3 | ENST00000258873 | -19.08 | 3.57E-02 | 6.12E-02 |
| PPIP5K2 | 5:103184745-103190841 | 5:103187314-103187376 | MES | 3 | 3 | ENST00000358359 | -19.08 | 3.33E-02 | 2.18E-01 |
| MAP3K10 | 19:40213076-40213734 | 19:40213189-40213516 | IR | 3 | 3 | ENST00000594791 | -19.11 | 1.98E-02 | 4.57E-02 |
| HYAL3 | 3:50294854-50299212 | 3:50294854-50295619 | TSSIA3ss | 3 | 3 | TSS.ENST00000336307 | -19.12 | 2.18E-02 | 9.87E-02 |
| AMT | 3:49416794-49418372 | 3:49417368-49417817 | IR (overlapping region) | 3 | 3 | ENST00000636594 | -19.13 | 1.05E-03 | 4.72E-03 |
| ZNF142 | 2:218656464-218658979 | 2:218658701-218658876 | SES | 3 | 3 | ENST00000432460 | -19.15 | 8.79E-04 | 3.21E-03 |
| LRATD2 | 8:126552901-126557731 | 8:126553504-126556264 | IR | 3 | 3 | ENST00000652209 | -19.16 | 8.23E-04 | 1.22E-02 |
| EPB41L5 | 2:120174841-120179119 | 2:120175098-120175700 | IR (overlapping region) | 3 | 3 | ENST00000263713 | -19.18 | 1.35E-02 | 4.71E-02 |
| ZNF286A | 17:15699778-15700207 | 17:15700048-15700207 | TSSIA3ss | 3 | 3 | TSS.ENST00000395894 | -19.21 | 2.59E-02 | 5.10E-02 |
| ZNF286A | 17:15699778-15700134 | 17:15700048-15700134 | TSSIA3ss | 3 | 3 | TSS.ENST00000395894 | -19.21 | 2.59E-02 | 5.10E-02 |
| ZNF117 | 7:64974452-64979536 | 7:64977224-64978735 | IR | 3 | 3 | ENST00000620222 | -19.22 | 2.78E-02 | 3.15E-01 |

| | | | | | | | | | |
|---|---|---|---|---|---|---|---|---|---|
| CADM2 | 3:84959005-84959668 | 3:84959493-84959589 | IR | 3 | 3 | ENST00000485126 | -19.24 | 2.12E-02 | 1.23E-01 |
| DOCK6 | 19:11229036-11233202 | 19:11232185-11232289 | MXS | 3 | 3 | ENST00000587656 | -19.26 | 7.33E-04 | 1.60E-03 |
| PHKB | 16:47461427-47497398 | 16:47463899-47463993 | MXS | 3 | 3 | ENST00000566037 | -19.27 | 7.63E-03 | 1.45E-02 |
| MDC1 | 6:30707455-30711411 | 6:30707566-30708357 | SES | 3 | 3 | ENST00000376406 | -19.36 | 3.25E-02 | 2.55E-01 |
| RBM33 | 7:155707069-155711202 | 7:155711173-155711202 | A3ss | 3 | 3 | ENST00000440108 | -19.42 | 3.77E-03 | 3.85E-02 |
| NUDT12 | 5:103559469-103560096 | 5:103560043-103560096 | A5ss | 3 | 3 | ENST00000230792 | -19.43 | 3.50E-02 | 2.29E-01 |
| SH3KBP1 | X:19542194-19549973 | X:19545922-19546050 | SES | 3 | 3 | ENST00000379698 | -19.45 | 1.33E-02 | 8.12E-01 |
| GMPPB | 3:49719924-49722147 | 3:49720654-49720799 | IR (overlapping region) | 3 | 3 | ENST00000308375 | -19.46 | 4.30E-02 | 1.90E-01 |
| SEPTIN9 | 17:77450787-77482143 | 17:77451367-77451528 | SES | 3 | 3 | ENST00000586128 | -19.47 | 2.60E-03 | 5.52E-03 |
| GUSBP11 | 22:23683868-23690231 | 22:23686831-23686995 | SES | 3 | 3 | ENST00000421064 | -19.48 | 7.34E-03 | 2.44E-02 |
| ATG9B | 7:151012214-151013408 | 7:151012473-151013230 | IR (overlapping region) | 2 | 3 | ENST00000605952 | -19.5 | 3.01E-02 | 1.69E-01 |
| FANCI | 15:89244045-89247628 | 15:89244045-89244067 | TSSlA5ss | 3 | 3 | TSS.ENST00000563250 | -19.54 | 1.82E-02 | 3.13E-02 |
| FANCI | 15:89244034-89247628 | 15:89244034-89244067 | TSSlA5ss | 3 | 3 | TSS.ENST00000563250 | -19.54 | 1.82E-02 | 3.13E-02 |
| CEP83 | 12:94411848-94412516 | 12:94412318-94412429 | SES | 3 | 3 | ENST00000547232 | -19.56 | 7.76E-03 | 1.22E-02 |
| DLST | 14:74882017-74889094 | 14:74882591-74882624 | MXS | 3 | 3 | ENST00000554806 | -19.59 | 1.51E-08 | 2.49E-08 |
| CCDC88A | 2:55296524-55301205 | 2:55299839-55299919 | MES | 3 | 2 | ENST00000644825 | -19.67 | 4.90E-02 | 1.86E-01 |
| CRELD1 | 3:9943516-9943993 | 3:9943902-9943993 | A3ss | 3 | 3 | ENST00000326434 | -19.75 | 1.16E-05 | 6.77E-05 |
| TSC2 | 16:2076586-2079031 | 16:2077598-2077726 | SES | 3 | 3 | ENST00000219476 | -19.75 | 1.74E-02 | 3.24E-02 |
| AL645939.2 | 6:29722981-29723971 | 6:29723528-29723657 | IR (overlapping region) | 3 | 3 | ENST00000427340 | -19.79 | 2.07E-02 | 1.69E-01 |
| POLM | 7:44079742-44079860 | 7:44079742-44079756 | TSSlA3ss | 2 | 3 | TSS.ENST00000452049 | -19.85 | 5.21E-03 | 5.51E-02 |
| PRDM15 | 21:41798225-41801722 | 21:41798497-41798582 | IR | 3 | 3 | ENST00000441787 | -19.85 | 1.62E-02 | 5.31E-02 |
| FGFR2 | 10:121551460-121593708 | 10:121564502-121564579 | MES | 3 | 2 | ENST00000346997 | -19.87 | 4.98E-03 | 6.58E-03 |
| PLD3 | 19:40348769-40365717 | 19:40365553-40365717 | TSSlA3ss | 3 | 3 | TSS.ENST00000594085 | -19.91 | 1.15E-03 | 2.65E-03 |
| PLD3 | 19:40348769-40365661 | 19:40365553-40365661 | TSSlA3ss | 3 | 3 | TSS.ENST00000594085 | -19.91 | 1.15E-03 | 2.65E-03 |
| ZNF655 | 7:99560696-99563862 | 7:99561888-99561973 | MES | 3 | 2 | ENST00000423973 | -19.96 | 7.32E-03 | 9.02E-02 |
| GOLGB1 | 3:121730018-121730875 | 3:121730848-121730875 | A5ss | 3 | 3 | ENST00000482512 | -19.99 | 1.73E-02 | 6.92E-02 |
| CRELD1 | 3:9943516-9944364 | 3:9943902-9944095 | SES | 3 | 3 | ENST00000326434 | -20.05 | 1.03E-03 | 5.99E-03 |
| HDAC9 | 7:18495855-18496261 | 7:18495855-18496023 | TSSlA5ss | 3 | 3 | TSS.ENST00000413380 | -20.08 | 3.03E-02 | 3.12E-01 |
| MYO1B | 2:191400469-191408114 | 2:191400749-191400835 | MES | 3 | 3 | ENST00000304164 | -20.11 | 8.14E-03 | 2.93E-02 |
| GOLGA2P10 | 15:82477124-82477260 | 15:82477203-82477260 | A5ss | 3 | 3 | Ex.ENST00000550961 | -20.12 | 2.68E-02 | 4.59E-02 |
| ERGIC3 | 20:35556272-35556990 | 20:35556973-35556990 | A3ss | 3 | 3 | Ex.ENST00000348547 | -20.14 | 6.03E-06 | 1.81E-05 |
| AL645939.2 | 6:29722981-29723971 | 6:29723528-29723769 | IR (overlapping region) | 3 | 3 | ENST00000427340 | -20.3 | 2.06E-02 | 1.67E-01 |
| ZKSCAN5 | 7:99512592-99525812 | 7:99519827-99519909 | MXS | 3 | 3 | ENST00000454175 | -20.35 | 1.31E-03 | 1.62E-02 |
| TOR1AIP1 | 1:179884770-179889312 | 1:179889310-179889312 | A3ss | 3 | 3 | ENST00000271583 | -20.52 | 3.48E-02 | 8.93E-02 |
| STRADA | 17:63702835-63704908 | 17:63703752-63704004 | IR (overlapping region) | 3 | 3 | ENST00000245865 | -20.54 | 4.12E-02 | 8.54E-02 |
| ACAD10 | 12:111748476-111749172 | 12:111749011-111749172 | TSSlA3ss | 3 | 3 | TSS.ENST00000550198 | -20.58 | 3.44E-02 | 5.08E-02 |
| PTBP2 | 1:96804940-96806865 | 1:96806419-96806452 | SES | 3 | 3 | ENST00000370197 | -20.64 | 1.10E-02 | 3.28E-02 |
| DPP7 | 9:137114463-137114726 | 9:137114577-137114646 | IR | 3 | 3 | ENST00000478597 | -20.67 | 1.49E-02 | 4.00E-01 |
| LRRC14 | 8:144518042-144519700 | 8:144519615-144519701 | A3ss | 3 | 3 | Ex.TSS.ENST00000527730 | -20.67 | 3.68E-03 | 5.71E-02 |
| MPI | 15:74897512-74902219 | 15:74900138-74900495 | IR (overlapping region) | 3 | 3 | ENST00000352410 | -20.73 | 3.95E-02 | 6.73E-02 |
| SLC52A3 | 20:760080-761824 | 20:761239-761700 | IR | 3 | 3 | ENST00000381944 | -20.83 | 4.43E-02 | 1.41E-01 |
| CAPN7 | 3:15240854-15242177 | 3:15241453-15241588 | SES | 3 | 3 | ENST00000253693 | -20.89 | 2.91E-02 | 1.19E-01 |
| AC074117.1 | 2:27357808-27360256 | 2:27357903-27358884 | IR | 3 | 3 | ENST00000447070 | -20.9 | 2.54E-02 | 9.53E-02 |
| GPR108 | 19:6731055-6731198 | 19:6731055-6731111 | A3ss | 3 | 3 | ENST00000264080 | -20.99 | 5.23E-04 | 1.26E-03 |
| LMO3 | 12:16560539-16600713 | 12:16600655-16600713 | A5ss | 3 | 3 | ENST00000261169 | -21.02 | 2.02E-02 | 3.04E-02 |
| MPI | 15:74897512-74902219 | 15:74900600-74901949 | IR (overlapping region) | 3 | 3 | ENST00000352410 | -21.04 | 2.07E-02 | 3.53E-02 |
| MRPL4 | 19:10259172-10260025 | 19:10259276-10259823 | IR (overlapping region) | 3 | 3 | ENST00000393733 | -21.1 | 1.87E-02 | 4.02E-02 |
| CLCC1 | 1:108950439-108963360 | 1:108950439-108950448 | A3ss | 3 | 3 | Ex.TSS.ENST00000356970 | -21.16 | 1.85E-02 | 4.49E-02 |
| ZNF142 | 2:218658802-218658979 | 2:218658802-218658876 | TSSlA3ss | 3 | 3 | TSS.ENST00000432460 | -21.2 | 1.88E-02 | 6.87E-02 |
| GOLGA2P10 | 15:82477124-82477393 | 15:82477203-82477298 | SES | 3 | 3 | ENST00000550961 | -21.22 | 2.28E-02 | 3.91E-02 |
| ACAD11 | 3:132578882-132579491 | 3:132578882-132579056 | TSSlA3ss | 3 | 3 | TSS.ENST00000477604 | -21.25 | 2.65E-02 | 1.07E-01 |
| LETMD1 | 12:51048479-51056349 | 12:51049034-51049185 | MES | 3 | 3 | ENST00000262055 | -21.25 | 1.41E-04 | 2.15E-04 |
| C1RL | 12:7097164-7101897 | 12:7099686-7099760 | MXS | 3 | 3 | ENST00000543933 | -21.39 | 1.78E-02 | 2.77E-02 |
| ZFP62 | 5:180847615-180851493 | 5:180848083-180849153 | IR | 3 | 3 | ENST00000512132 | -21.4 | 6.70E-03 | 4.72E-02 |
| PPOX | 1:161167235-161170619 | 1:161167371-161167486 | MES | 3 | 3 | ENST00000352210 | -21.42 | 4.23E-02 | 1.07E-01 |
| CREBZF | 11:85663524-85665102 | 11:85663602-85663814 | IR | 3 | 3 | ENST00000490820 | -21.47 | 6.25E-03 | 9.18E-03 |
| MPI | 15:74897512-74902219 | 15:74900138-74901949 | IR (overlapping region) | 3 | 3 | ENST00000352410 | -21.47 | 5.02E-04 | 8.55E-04 |
| ZNF268 | 12:133192004-133202143 | 12:133193428-133193533 | SES | 3 | 3 | ENST00000534953 | -21.48 | 2.08E-02 | 3.13E-02 |
| HPN | 19:35041873-35049289 | 19:35041873-35042522 | TSSlA5ss | 3 | 3 | TSS.ENST00000262626 | -21.52 | 1.74E-02 | 3.95E-02 |
| SEPTIN4 | 17:58526314-58531960 | 17:58526932-58526978 | MXS | 3 | 3 | ENST00000578131 | -21.58 | 2.63E-02 | 4.84E-02 |
| LINC01145 | 1:145171186-145215638 | 1:145214677-145214731 | SES | 2 | 3 | ENST00000614510 | -21.62 | 3.84E-02 | 9.43E-02 |
| ATG16L2 | 11:72817856-72828358 | 11:72824060-72824122 | MES | 2 | 2 | ENST00000321297 | -21.63 | 3.06E-02 | 4.45E-02 |
| RIMKLB | 12:8777803-8780574 | 12:8780035-8780112 | SES | 2 | 3 | ENST00000542232 | -22 | 2.04E-02 | 3.20E-02 |
| ZMIZ1 | 10:79292357-79293380 | 10:79293363-79293380 | TSSlA3ss | 3 | 3 | TSS.ENST00000478357 | -22.03 | 2.50E-02 | 3.34E-02 |
| TRIM37 | 17:58999460-59012327 | 17:59001598-59001714 | SES | 3 | 3 | ENST00000262294 | -22.18 | 1.05E-02 | 2.16E-02 |
| DDHD2 | 8:38237628-38238106 | 8:38238089-38238106 | A3ss | 3 | 3 | Ex.ENST00000397166 | -22.19 | 1.16E-04 | 1.50E-03 |
| SEPTIN6 | X:119620052-119629317 | X:119625335-119625379 | SES | 3 | 3 | ENST00000394610 | -22.19 | 1.13E-02 | 4.40E-01 |
| AMT | 3:49417358-49417614 | 3:49417358-49417367 | TSSlA3ss | 3 | 3 | TSS.ENST00000395338 | -22.2 | 8.64E-03 | 3.83E-02 |
| AMT | 3:49417310-49417614 | 3:49417310-49417367 | TSSlA3ss | 3 | 3 | TSS.ENST00000395338 | -22.2 | 8.64E-03 | 3.83E-02 |
| LMO7 | 13:75834388-75835232 | 13:75835152-75835232 | A3ss | 2 | 3 | ENST00000485987 | -22.24 | 4.90E-02 | 7.77E-02 |
| RCC1 | 1:28508906-28529857 | 1:28516725-28516867 | SES | 3 | 3 | ENST00000649185 | -22.25 | 3.45E-04 | 9.66E-04 |
| TMEM147-AS1 | 19:35541582-35545492 | 19:35541585-35542910 | IR (overlapping region) | 3 | 3 | ENST00000588286 | -22.25 | 9.62E-03 | 2.17E-02 |
| EIF1AD | 11:66000482-66002039 | 11:66001910-66002039 | TSSlA5ss | 3 | 3 | TSS.ENST00000527249 | -22.4 | 4.11E-02 | 5.90E-02 |
| AL162258.2 | 1:153632034-153632197 | 1:153632064-153632154 | IR (overlapping region) | 2 | 3 | ENST00000607839 | -22.41 | 9.15E-03 | 2.29E-02 |
| HEXD | 17:82439631-82440532 | 17:82440335-82440472 | IR | 3 | 3 | ENST00000585077 | -22.43 | 1.65E-02 | 3.51E-02 |
| ANO8 | 19:17328162-17328983 | 19:17328265-17328539 | IR | 3 | 3 | ENST00000159087 | -22.51 | 8.03E-03 | 1.78E-02 |
| LRRC28 | 15:99287952-99352368 | 15:99334018-99334129 | SES | 3 | 3 | ENST00000558771 | -22.61 | 1.21E-02 | 2.09E-02 |
| ACOXL | 2:110996005-111092864 | 2:111031627-111031714 | MES | 3 | 2 | ENST00000389811 | -22.63 | 4.51E-02 | 1.57E-01 |
| CREBZF | 11:85660335-85661749 | 11:85660598-85661636 | IR | 3 | 3 | ENST00000525639 | -22.65 | 1.60E-02 | 2.36E-02 |
| FAM122B | X:134796108-134797155 | X:134796257-134797120 | IR | 3 | 3 | ENST00000343004 | -22.71 | 8.10E-04 | 3.29E-02 |
| UBE2D4 | 7:43938495-43948631 | 7:43942826-43942857 | MES | 3 | 2 | ENST00000222402 | -22.8 | 2.55E-02 | 2.68E-01 |
| AL139353.1 | 14:31453345-31457282 | 14:31457029-31457282 | TSSlA5ss | 3 | 3 | TSS.ENST00000547760 | -22.91 | 4.04E-02 | 6.61E-02 |
| BCAM | 19:44811347-44812225 | 19:44812163-44812225 | A3ss | 3 | 3 | ENST00000270233 | -22.95 | 1.10E-03 | 2.55E-03 |
| AP3B2 | 15:82685914-82688831 | 15:82688097-82688735 | IR (overlapping region) | 3 | 3 | ENST00000535513 | -23.01 | 3.73E-02 | 6.38E-02 |
| CNTROB | 17:7932801-7933092 | 17:7932928-7933092 | A3ss | 2 | 3 | ENST00000570782 | -23.01 | 1.15E-02 | 2.45E-02 |
| CNTROB | 17:7932801-7932955 | 17:7932928-7932955 | A3ss | 2 | 3 | ENST00000570782 | -23.01 | 1.15E-02 | 2.45E-02 |
| ZNF232 | 17:5106507-5108925 | 17:5106507-5106533 | A3ss | 3 | 3 | Ex.TSS.ENST00000250076 | -23.05 | 1.06E-02 | 2.18E-02 |
| MAST1 | 19:12865046-12865468 | 19:12865179-12865315 | IR (overlapping region) | 2 | 3 | ENST00000589040 | -23.08 | ####### | 0.00E+00 |
| SETD4 | 21:36058925-36060346 | 21:36058925-36058973 | TSSlA3ss | 3 | 3 | TSS.ENST00000399212 | -23.18 | 3.05E-02 | 9.90E-02 |
| TCIRG1 | 11:68042980-68043497 | 11:68043032-68043370 | IR | 3 | 3 | ENST00000532635 | -23.23 | 4.77E-02 | 6.91E-02 |
| ASPHD1 | 16:29900488-29901920 | 16:29900742-29901551 | IR | 3 | 3 | ENST00000308748 | -23.27 | 3.25E-03 | 6.11E-03 |
| NCOA2 | 8:70296801-70403699 | 8:70403334-70403700 | A5ss | 3 | 3 | Ex.TSS.ENST00000452400 | -23.41 | 3.56E-02 | 4.46E-01 |
| LINC02245 | 2:65048213-65055914 | 2:65048213-65055620 | A3ss | 3 | 3 | Ex.TSS.ENST00000653778 | -23.43 | 1.60E-02 | 6.08E-02 |
| TRIML2 | 4:188105588-188109242 | 4:188105588-188105590 | A3ss | 3 | 3 | Ex.ENST00000394461 | -23.46 | 2.83E-02 | 1.74E-01 |

| | | | | | | | | | |
|---|---|---|---|---|---|---|---|---|---|
| ADAM33 | 20:3673574-3673747 | 20:3673659-3673744 | IR | 2 | 3 | ENST00000466620 | -23.47 | 4.61E-02 | 1.39E-01 |
| CLEC11A | 19:50725022-50725708 | 19:50725082-50725315 | IR | 3 | 3 | ENST00000250340 | -23.49 | 6.04E-04 | 1.41E-03 |
| AHDC1 | 1:27553186-27558705 | 1:27558305-27558530 | SES | 3 | 3 | ENST00000642245 | -23.52 | 1.16E-02 | 3.26E-02 |
| MPI | 15:74897512-74902219 | 15:74900138-74901168 | IR (overlapping region) | 3 | 3 | ENST00000352410 | -23.53 | 3.17E-04 | 5.39E-04 |
| RIC3 | 11:8106056-8111137 | 11:8110408-8110658 | IR | 3 | 3 | ENST00000309737 | -23.62 | 3.25E-02 | 4.77E-02 |
| AGTPBP1 | 9:85655201-85657434 | 9:85655201-85655320 | A3ss | 3 | 3 | ENST00000337006 | -23.68 | 4.74E-03 | 1.52E-01 |
| MTERF2 | 12:106978772-106986968 | 12:106985115-106985225 | SES | 3 | 3 | ENST00000240050 | -23.69 | 7.73E-03 | 1.14E-02 |
| HPN | 19:35041873-35049289 | 19:35042453-35042522 | SES | 3 | 3 | ENST00000392226 | -23.72 | 1.58E-03 | 3.59E-03 |
| DCAF17 | 2:171458076-171468887 | 2:171458372-171458477 | SES | 3 | 3 | ENST00000375255 | -23.85 | 2.76E-02 | 9.86E-02 |
| PAQR5 | 15:69380011-69397467 | 15:69389654-69389780 | MES | 3 | 2 | ENST00000340965 | -23.89 | 2.53E-02 | 4.31E-02 |
| PARP11 | 12:3830019-3873211 | 12:3863836-3863957 | SES | 3 | 3 | ENST00000427057 | -23.93 | 3.78E-02 | 5.71E-02 |
| RFNG | 17:82049918-82050514 | 17:82050007-82050401 | IR | 3 | 3 | ENST00000580793 | -23.95 | 4.09E-03 | 8.69E-03 |
| BRSK2 | 11:1449837-1450586 | 11:1450221-1450587 | A3ss | 3 | 3 | Ex.ENST00000308219 | -23.97 | 1.16E-02 | 1.57E-02 |
| AC009005.1 | 19:567625-571439 | 19:567625-567648 | TSSIA3ss | 3 | 3 | TSS.ENST00000588908 | -24 | 2.39E-02 | 5.69E-02 |
| ARMCX3 | X:101623277-101623813 | X:101623786-101623814 | A3ss | 3 | 3 | Ex.TSS.ENST00000471229 | -24 | 1.43E-03 | 5.28E-02 |
| PCBP1-AS1 | 2:70085816-70086273 | 2:70085904-70086251 | IR | 3 | 3 | ENST00000429599 | -24.06 | 3.47E-02 | 1.34E-01 |
| EIF1AD | 11:66000477-66001909 | 11:66000477-66000481 | TSSIA3ss | 3 | 3 | TSS.ENST00000527249 | -24.09 | 3.96E-02 | 5.68E-02 |
| EIF1AD | 11:66000407-66001909 | 11:66000407-66000481 | TSSIA3ss | 3 | 3 | TSS.ENST00000527249 | -24.09 | 3.96E-02 | 5.68E-02 |
| SH2D3C | 9:127751301-127755084 | 9:127755079-127755084 | TSSIA5ss | 3 | 3 | TSS.ENST00000420366 | -24.16 | 3.26E-02 | 6.68E-01 |
| MPPE1 | 18:11885817-11886916 | 18:11886499-11886621 | MES | 3 | 2 | ENST00000317251 | -24.22 | 2.12E-04 | 4.53E-04 |
| DYSF | 2:71511922-71513715 | 2:71513240-71513332 | SES | 3 | 3 | ENST00000409582 | -24.34 | 2.67E-02 | 1.04E-01 |
| NETO2 | 16:47122785-47122888 | 16:47122868-47122888 | A5ss | 3 | 3 | ENST00000562435 | -24.42 | 4.47E-03 | 8.47E-03 |
| DPH5 | 1:100995150-101021640 | 1:101013710-101013818 | MES | 3 | 3 | ENST00000342173 | -24.44 | 6.06E-03 | 1.47E-02 |
| RIPOR2 | 6:24839600-24842861 | 6:24839600-24840775 | TSSIA3ss | 3 | 3 | TSS.ENST00000510784 | -24.51 | 3.87E-02 | 3.10E-01 |
| ASPHD1 | 16:29900488-29901920 | 16:29900567-29901551 | IR | 3 | 3 | ENST00000308748 | -24.61 | 7.52E-04 | 1.41E-03 |
| FIRRE | X:131689755-131692636 | X:131690016-131692437 | IR | 3 | 3 | ENST00000663405 | -24.91 | 1.39E-02 | 5.61E-01 |
| HDAC10 | 22:50250906-50251101 | 22:50250974-50251101 | TSSIA5ss | 3 | 3 | TSS.ENST00000216271 | -24.91 | 1.42E-03 | 4.90E-03 |
| ITGA7 | 12:55698918-55700261 | 12:55699870-55700261 | TSSIA5ss | 3 | 3 | TSS.ENST00000554543 | -24.91 | 2.46E-02 | 3.78E-02 |
| COASY | 17:42562148-42563322 | 17:42562237-42562611 | IR | 3 | 3 | ENST00000393818 | -24.94 | 1.43E-03 | 2.90E-03 |
| LETMD1 | 12:51049186-51056349 | 12:51052092-51052207 | MES | 3 | 3 | ENST00000262055 | -24.94 | 9.29E-03 | 1.41E-02 |
| SH2D3A | 19:6754933-6759593 | 19:6754933-6755315 | A3ss | 3 | 3 | ENST00000245908 | -24.98 | 2.61E-02 | 6.27E-02 |
| TOR1AIP2 | 1:179852812-179877238 | 1:179865436-179865854 | SES | 3 | 3 | ENST00000609928 | -24.99 | 8.25E-06 | 2.12E-05 |
| TMEM218 | 11:125101304-125102733 | 11:125102132-125102317 | SES | 3 | 3 | ENST00000526175 | -25.12 | 2.65E-03 | 3.58E-03 |
| FBF1 | 17:75933089-75937565 | 17:75935632-75935673 | SES | 2 | 3 | ENST00000592193 | -25.27 | 1.37E-02 | 2.53E-02 |
| PCBP1-AS1 | 2:70085816-70086273 | 2:70085904-70086054 | IR | 3 | 3 | ENST00000429599 | -25.32 | 3.31E-02 | 1.28E-01 |
| SLC12A8 | 3:125110336-125135668 | 3:125120599-125120686 | MES | 3 | 2 | ENST00000393469 | -25.38 | 4.35E-02 | 1.74E-01 |
| ZKSCAN5 | 7:99512592-99525812 | 7:99519827-99519909 | MES | 3 | 3 | ENST00000326775 | -25.39 | 1.45E-02 | 1.77E-01 |
| BBS1 | 11:66531951-66533598 | 11:66532206-66532295 | IR (overlapping region) | 3 | 3 | ENST00000318312 | -25.49 | 1.19E-02 | 1.71E-02 |
| CENPE | 4:103151378-103158299 | 4:103153047-103153250 | SES | 2 | 3 | ENST00000265148 | -25.52 | 4.63E-02 | 2.72E-01 |
| PLEKHA4 | 19:48854265-48858859 | 19:48857422-48857496 | SES | 2 | 3 | ENST00000263265 | -25.56 | 4.54E-02 | 1.07E-01 |
| GOLGA2P7 | 15:84203008-84204425 | 15:84203008-84203020 | TSSIA3ss | 3 | 3 | TSS.ENST00000557980 | -25.62 | 1.87E-04 | 3.21E-04 |
| DPPA4 | 3:109329082-109331733 | 3:109330524-109331734 | A5ss | 3 | 3 | Ex.ENST00000463966 | -25.83 | 6.91E-04 | 2.76E-03 |
| PRC1 | 15:90966654-90969446 | 15:90966654-90967202 | A3ss | 3 | 3 | Ex.TSS.ENST00000361188 | -25.84 | 4.18E-03 | 7.20E-03 |
| IMMT | 2:86171310-86173649 | 2:86171310-86171312 | A3ss | 3 | 3 | Ex.ENST00000254636 | -25.96 | 4.52E-03 | 1.78E-02 |
| OGFOD2 | 12:122975122-122976653 | 12:122975811-122975867 | SES | 3 | 3 | ENST00000454694 | -26.14 | 2.12E-02 | 3.16E-02 |
| DPH5 | 1:100990632-100995109 | 1:100992637-100992740 | SES | 3 | 3 | ENST00000464270 | -26.18 | 5.61E-03 | 1.36E-02 |
| INO80E | 16:30001531-30005367 | 16:30005221-30005367 | TSSIA3ss | 3 | 3 | TSS.ENST00000563197 | -26.26 | 3.63E-03 | 6.70E-03 |
| RPS6KL1 | 14:74908933-74911247 | 14:74909101-74909190 | SES | 2 | 2 | ENST00000553789 | -26.33 | 7.87E-03 | 1.30E-02 |
| TRIP12 | 2:229793131-229795178 | 2:229793131-229793145 | A3ss | 3 | 3 | Ex.ENST00000283943 | -26.41 | 7.56E-04 | 2.78E-03 |
| SPIB | 19:50422546-50423604 | 19:50422823-50423037 | SES | 3 | 3 | ENST00000270632 | -26.49 | 1.59E-02 | 3.71E-02 |
| NACC2 | 9:136012014-136013198 | 9:136012014-136012024 | A3ss | 3 | 3 | Ex.TSS.ENST00000371753 | -26.55 | 1.19E-02 | 3.09E-01 |
| ZNF142 | 2:218650527-218656149 | 2:218651701-218652300 | SES | 3 | 3 | ENST00000440934 | -26.57 | 3.81E-03 | 1.39E-02 |
| AFG3L1P | 16:89972766-89977714 | 16:89977639-89977714 | A3ss | 3 | 2 | Ex.TSS.ENST00000418696 | -26.59 | 1.18E-02 | 2.31E-02 |
| AFG3L1P | 16:89972766-89977677 | 16:89977639-89977677 | TSSIA3ss | 3 | 2 | TSS.ENST00000418696 | -26.59 | 1.18E-02 | 2.31E-02 |
| ZNF142 | 2:218651823-218656149 | 2:218651823-218652300 | TSSIA3ss | 3 | 3 | TSS.ENST00000440934 | -26.62 | 2.21E-03 | 8.08E-03 |
| SEPTIN4 | 17:58526314-58531960 | 17:58526682-58526978 | MXS | 3 | 3 | ENST00000317256 | -26.82 | 6.47E-04 | 1.34E-03 |
| DCAF8 | 1:160261326-160262275 | 1:160261326-160261358 | TSSIA3ss | 3 | 3 | TSS.ENST00000326837 | -26.86 | 3.23E-03 | 8.22E-03 |
| KRIT1 | 7:92245465-92246100 | 7:92245596-92245905 | IR (overlapping region) | 3 | 3 | ENST00000340022 | -26.86 | 7.48E-03 | 8.88E-02 |
| SENP1 | 12:48101517-48105456 | 12:48105402-48105456 | TSSIA5ss | 3 | 2 | TSS.ENST00000552189 | -26.88 | 2.98E-02 | 4.51E-02 |
| COG1 | 17:73201901-73203023 | 17:73203000-73203023 | A3ss | 3 | 3 | Ex.ENST00000299886 | -26.89 | 5.66E-03 | 1.18E-02 |
| GOLGA2P7 | 15:84204517-84204786 | 15:84204600-84204691 | SES | 3 | 3 | ENST00000400817 | -27.09 | 5.24E-04 | 8.99E-04 |
| C2orf27A | 2:131682600-131684708 | 2:131682600-131682988 | A5ss | 3 | 3 | Ex.TSS.ENST00000653458 | -27.22 | 1.12E-02 | 3.96E-02 |
| C2orf27A | 2:131682558-131684708 | 2:131682558-131682988 | A5ss | 3 | 3 | Ex.TSS.ENST00000657415 | -27.22 | 1.12E-02 | 3.96E-02 |
| FAM160A2 | 11:6218151-6218599 | 11:6218558-6218599 | A5ss | 3 | 3 | ENST00000265978 | -27.27 | 3.40E-02 | 4.69E-02 |
| SEPTIN6 | X:119616743-119629317 | X:119625335-119625379 | MES | 3 | 3 | ENST00000460411 | -27.3 | 2.90E-03 | 1.13E-01 |
| SPINT1-AS1 | 15:40836283-40844178 | 15:40838543-40838686 | SES | 3 | 3 | ENST00000564302 | -27.34 | 1.56E-02 | 2.62E-02 |
| RAD9A | 11:67393611-67393722 | 11:67393691-67393722 | A3ss | 3 | 3 | ENST00000307980 | -27.37 | 3.42E-03 | 4.96E-03 |
| MMP19 | 12:55835433-55837374 | 12:55835616-55836125 | IR (overlapping region) | 3 | 3 | ENST00000322569 | -27.38 | 1.93E-02 | 2.97E-02 |
| COASY | 17:42562148-42563322 | 17:42562214-42562611 | IR | 3 | 3 | ENST00000393818 | -27.41 | 1.14E-03 | 2.31E-03 |
| TSPOAP1 | 17:58306387-58306799 | 17:58306387-58306413 | A3ss | 3 | 3 | ENST00000268893 | -27.48 | 4.93E-02 | 9.05E-02 |
| ZNF655 | 7:99560696-99563862 | 7:99561888-99561973 | MES | 3 | 2 | ENST00000320583 | -27.65 | 4.35E-03 | 5.36E-02 |
| MFSD14C | 9:96918827-96949554 | 9:96942603-96942752 | MES | 3 | 3 | ENST00000506067 | -27.75 | 1.16E-03 | 3.99E-02 |
| KRIT1 | 7:92245465-92246100 | 7:92245596-92245843 | IR (overlapping region) | 3 | 3 | ENST00000340022 | -27.83 | 5.75E-03 | 6.83E-02 |
| MST1 | 3:49686168-49687067 | 3:49686193-49686312 | IR | 3 | 2 | ENST00000492370 | -27.88 | 6.23E-03 | 2.77E-02 |
| AC060780.1 | 17:43166980-43167849 | 17:43167251-43167733 | IR (overlapping region) | 2 | 3 | ENST00000635600 | -27.92 | 4.68E-03 | 9.50E-03 |
| VSIG10 | 12:118095679-118102988 | 12:118095679-118095814 | A3ss | 3 | 3 | ENST00000536905 | -27.97 | 2.11E-02 | 3.13E-02 |
| EXOC3L1 | 16:67184686-67185057 | 16:67184811-67184901 | IR | 2 | 3 | ENST00000545725 | -28.17 | 4.71E-02 | 9.00E-02 |
| MIOS | 7:7567073-7567606 | 7:7567073-7567484 | TSSIA5ss | 3 | 3 | TSS.ENST00000433056 | -28.2 | 6.83E-03 | 7.90E-02 |
| COASY | 17:42562148-42563322 | 17:42562220-42562611 | IR | 3 | 3 | ENST00000393818 | -28.25 | 1.86E-03 | 3.76E-03 |
| ANKHD1 | 5:140436258-140438502 | 5:140438461-140438502 | A3ss | 3 | 3 | ENST00000297183 | -28.36 | 8.06E-04 | 5.51E-03 |
| ANKHD1 | 5:140436258-140438493 | 5:140438461-140438493 | A3ss | 3 | 3 | ENST00000297183 | -28.36 | 8.06E-04 | 5.51E-03 |
| KRBOX4 | X:46463294-46472749 | X:46463294-46463308 | TSSIA5ss | 3 | 3 | TSS.ENST00000344302 | -28.69 | 3.80E-02 | 1.00E+00 |
| CCDC74B | 2:130141297-130143268 | 2:130142133-130142353 | SES | 3 | 3 | ENST00000423263 | -28.95 | 1.03E-02 | 3.62E-02 |
| MFSD9 | 2:102723879-102726844 | 2:102723879-102723909 | A3ss | 3 | 3 | ENST00000258436 | -29.08 | 3.44E-02 | 1.20E-01 |
| ZSCAN26 | 6:28272337-28272669 | 6:28272337-28272339 | A5ss | 3 | 3 | ENST00000421553 | -29.2 | 6.96E-03 | 5.62E-02 |
| ZSCAN26 | 6:28271935-28272669 | 6:28271935-28272339 | A5ss | 3 | 3 | ENST00000421553 | -29.2 | 6.96E-03 | 5.62E-02 |
| HEXIM2 | 17:45162032-45162749 | 17:45162488-45162635 | SES | 3 | 3 | ENST00000589230 | -29.32 | 4.37E-03 | 8.90E-03 |
| TAF1C | 16:84184990-84186900 | 16:84184990-84185060 | TSSIA3ss | 3 | 3 | TSS.ENST00000564774 | -29.41 | 4.86E-04 | 9.43E-04 |
| TAF1C | 16:84184900-84186900 | 16:84184900-84185060 | TSSIA3ss | 3 | 3 | TSS.ENST00000564774 | -29.41 | 4.86E-04 | 9.43E-04 |
| VEPH1 | 3:157459519-157460355 | 3:157459927-157460180 | IR | 2 | 3 | ENST00000494677 | -29.72 | 2.13E-02 | 8.73E-02 |
| TTLL4 | 2:218746232-218747029 | 2:218747003-218747029 | A3ss | 3 | 3 | ENST00000258398 | -29.78 | 6.33E-03 | 2.31E-02 |
| ORAI2 | 7:102433662-102438943 | 7:102436225-102436333 | SES | 3 | 3 | ENST00000495936 | -29.88 | 1.51E-03 | 1.38E-02 |
| WRAP73 | 1:3632318-3633397 | 1:3632318-3632338 | A3ss | 3 | 3 | Ex.ENST00000270708 | -29.97 | 2.35E-03 | 6.63E-03 |
| SEPTIN6 | X:119616743-119629317 | X:119625335-119625379 | MXS | 3 | 3 | ENST00000489216 | -30.13 | 2.57E-03 | 9.98E-02 |

| | | | | | | | | | |
|---|---|---|---|---|---|---|---|---|---|
| DOK1 | 2:74555669-74556307 | 2:74555894-74556078 | SES | 3 | 3 | ENST00000233668 | -30.24 | 6.04E-04 | 2.37E-03 |
| PPOX | 1:161167235-161170619 | 1:161168432-161168576 | MES | 3 | 3 | ENST00000352210 | -30.57 | 4.78E-02 | 1.21E-01 |
| CLIP1 | 12:122319349-122327946 | 12:122322114-122324381 | SES | 3 | 3 | ENST00000537123 | -30.62 | 9.70E-03 | 1.44E-02 |
| GK | X:30696684-30700413 | X:30697732-30697749 | SES | 3 | 2 | ENST00000378946 | -30.88 | 3.78E-02 | 1.00E+00 |
| HMGCR | 5:75338304-75342582 | 5:75340448-75342583 | A3ss | 2 | 2 | Ex.TSS.ENST00000507942 | -30.96 | 3.51E-02 | 2.55E-01 |
| CCDC88A | 2:55296524-55301205 | 2:55299839-55301205 | A5ss | 3 | 3 | ENST00000644456 | -31.35 | 1.79E-02 | 6.77E-02 |
| SMC5 | 9:70344270-70347065 | 9:70346605-70346649 | SES | 3 | 3 | ENST00000361138 | -31.37 | 4.02E-02 | 1.00E+00 |
| MXRA8 | 1:1355244-1355512 | 1:1355346-1355449 | IR | 3 | 3 | ENST00000460473 | -31.54 | 1.23E-02 | 3.02E-02 |
| RMST | 12:97493286-97493894 | 12:97493875-97493894 | A3ss | 3 | 3 | ENST00000538559 | -31.86 | 5.72E-03 | 8.97E-03 |
| RMST | 12:97493286-97493890 | 12:97493875-97493890 | A3ss | 3 | 3 | ENST00000538559 | -31.86 | 5.72E-03 | 8.97E-03 |
| CABYR | 18:24156043-24159471 | 18:24156043-24156999 | A5ss | 2 | 3 | ENST00000399481 | -31.91 | 1.97E-03 | 4.22E-03 |
| TTC23 | 15:99245511-99249416 | 15:99249171-99249416 | TSSlA5ss | 2 | 3 | TSS.ENST00000394132 | -31.91 | 6.03E-03 | 1.04E-02 |
| ALKBH3 | 11:43889829-43901515 | 11:43892041-43892129 | MXS | 3 | 3 | ENST00000302708 | -31.95 | 1.81E-02 | 2.48E-02 |
| ZNF681 | 19:23754898-23758746 | 19:23754898-23754918 | TSSlA3ss | 2 | 2 | TSS.ENST00000528059 | -32.05 | 4.29E-02 | 9.73E-02 |
| KCNJ11 | 11:17385859-17388659 | 11:17387847-17388017 | IR | 3 | 3 | ENST00000339994 | -32.08 | 1.09E-02 | 1.49E-02 |
| DLST | 14:74882017-74889094 | 14:74885586-74885634 | MES | 3 | 3 | ENST00000238671 | -32.11 | 8.19E-06 | 1.35E-05 |
| MRTFB | 16:14240237-14241425 | 16:14240485-14240614 | IR | 2 | 3 | ENST00000572567 | -32.44 | 3.12E-02 | 5.40E-02 |
| WDR55 | 5:140672378-140674124 | 5:140672907-140673270 | IR (overlapping region) | 3 | 3 | ENST00000504897 | -32.71 | 1.01E-02 | 6.89E-02 |
| NCOA6 | 20:34736759-34746806 | 20:34740363-34743341 | SES | 2 | 2 | ENST00000359003 | -32.78 | 3.73E-03 | 1.11E-02 |
| CELF5 | 19:3293487-3296757 | 19:3295044-3296757 | TSSlA3ss | 3 | 3 | TSS.ENST00000588350 | -33.27 | 3.20E-02 | 7.27E-02 |
| ZNF783 | 7:149262358-149266334 | 7:149262358-149264510 | A5ss | 2 | 3 | Ex.TSS.ENST00000434415 | -33.27 | 4.76E-03 | 4.71E-02 |
| RSKR | 17:28611393-28611656 | 17:28611482-28611566 | IR (overlapping region) | 3 | 3 | ENST00000494272 | -33.7 | 1.38E-02 | 2.75E-02 |
| HEXIM2 | 17:45161940-45162749 | 17:45162488-45162635 | SES | 3 | 3 | ENST00000591070 | -33.74 | 1.01E-02 | 2.06E-02 |
| DPH5 | 1:100990629-100995109 | 1:100992637-100992740 | SES | 3 | 3 | ENST00000342173 | -34.65 | 3.69E-03 | 8.95E-03 |
| ITGA7 | 12:55700394-55700898 | 12:55700566-55700899 | A5ss | 3 | 3 | Ex.ENST00000452168 | -34.97 | 8.28E-04 | 1.27E-03 |
| SEPTIN4 | 17:58526314-58531960 | 17:58526314-58526978 | TSSlA3ss | 3 | 3 | TSS.ENST00000584789 | -35.04 | 1.14E-02 | 2.35E-02 |
| NPIPA1 | 16:14950256-14951614 | 16:14950752-14950821 | SES | 3 | 3 | ENST00000472413 | -35.1 | 2.02E-02 | 3.73E-02 |
| ZNF613 | 19:51929825-51936027 | 19:51929825-51929896 | A5ss | 3 | 3 | ENST00000293471 | -35.11 | 4.22E-03 | 9.85E-03 |
| HSPBP1 | 19:55265379-55265885 | 19:55265379-55265389 | A3ss | 3 | 3 | Ex.ENST00000255631 | -35.2 | 7.08E-04 | 1.68E-03 |
| HSPBP1 | 19:55265368-55265885 | 19:55265368-55265389 | A3ss | 3 | 3 | Ex.ENST00000255631 | -35.2 | 7.08E-04 | 1.68E-03 |
| TLCD5 | 11:120325369-120329976 | 11:120327441-120329977 | A3ss | 3 | 3 | Ex.ENST00000531346 | -35.27 | 3.11E-03 | 4.21E-03 |
| AGO3 | 1:35973512-36004340 | 1:35982621-35982712 | MES | 3 | 3 | ENST00000634486 | -35.36 | 2.06E-02 | 5.75E-02 |
| CRELD1 | 3:9933853-9934419 | 3:9933853-9933920 | TSSlA5ss | 3 | 3 | TSS.ENST00000452070 | -35.38 | 1.72E-02 | 1.00E-01 |
| STAG3L5P-PVRIG2P-PILRB | 7:100345969-100349798 | 7:100349731-100349798 | A3ss | 3 | 3 | ENST00000444874 | -35.46 | 5.32E-03 | 4.91E-02 |
| GPR161 | 1:168114550-168137164 | 1:168136010-168137165 | A5ss | 3 | 3 | Ex.TSS.ENST00000367835 | -36.35 | 6.44E-03 | 1.64E-02 |
| AMT | 3:49416794-49418372 | 3:49417368-49417488 | IR (overlapping region) | 3 | 3 | ENST00000636594 | -37.24 | 3.54E-04 | 1.59E-03 |
| TPRA1 | 3:127594934-127598195 | 3:127598055-127598196 | A5ss | 2 | 3 | Ex.TSS.ENST00000490643 | -37.95 | 1.78E-02 | 7.15E-02 |
| ZDHHC16 | 10:97454800-97455713 | 10:97455660-97455713 | A3ss | 3 | 3 | ENST00000345745 | -38.59 | 1.57E-04 | 2.10E-04 |
| CCDC74B | 2:130141297-130143268 | 2:130142133-130142183 | SES | 3 | 3 | ENST00000409128 | -38.87 | 1.17E-03 | 4.13E-03 |
| ATG16L2 | 11:72817856-72828358 | 11:72822848-72822961 | MES | 2 | 2 | ENST00000321297 | -39.05 | 1.96E-02 | 2.86E-02 |
| RBM18 | 9:122261499-122264714 | 9:122261499-122261508 | A3ss | 3 | 3 | Ex.TSS.ENST00000417201 | -40.06 | 1.03E-07 | 2.18E-06 |
| UPP1 | 7:48101871-48106872 | 7:48103297-48103411 | SES | 3 | 3 | ENST00000444999 | -40.81 | 4.66E-03 | 5.01E-02 |
| ZBED5 | 11:10855082-10856143 | 11:10855082-10855086 | TSSlA3ss | 3 | 3 | TSS.ENST00000413761 | -42 | 1.40E-03 | 1.88E-03 |
| AC090114.3 | 7:128574792-128577963 | 7:128577953-128577964 | A3ss | 2 | 3 | Ex.ENST00000479267 | -42.38 | 5.72E-03 | 5.46E-02 |
| AC108010.1 | 7:128574792-128578889 | 7:128577953-128578890 | A3ss | 2 | 3 | Ex.TSS.ENST00000605862 | -42.38 | 5.72E-03 | 5.46E-02 |
| AP000459.3 | 21:23372427-23372947 | 21:23372427-23372448 | A3ss | 3 | 3 | ENST00000262354 | -42.75 | 1.58E-03 | 5.09E-03 |
| MGA | 15:41736699-41740052 | 15:41739906-41740053 | A3ss | 3 | 3 | Ex.ENST00000219905 | -43.47 | 4.80E-02 | 8.07E-02 |
| DVL2 | 17:7227712-7227976 | 17:7227712-7227783 | A3ss | 3 | 3 | ENST00000005340 | -44.23 | 1.22E-04 | 2.54E-04 |
| DVL2 | 17:7227703-7227976 | 17:7227703-7227783 | A3ss | 3 | 3 | Ex.ENST00000005340 | -44.23 | 1.22E-04 | 2.54E-04 |
| RHBDL1 | 16:677459-678267 | 16:677560-677638 | IR | 2 | 3 | ENST00000450775 | -44.43 | 5.67E-03 | 1.09E-02 |
| ZNF561 | 19:9618167-9619431 | 19:9618167-9618179 | A3ss | 3 | 3 | ENST00000302851 | -45.51 | 1.37E-03 | 3.29E-03 |
| COASY | 17:42562220-42562932 | 17:42562612-42562932 | TSSlA3ss | 3 | 3 | TSS.ENST00000421097 | -45.78 | 1.04E-03 | 2.10E-03 |
| AMT | 3:49416794-49418372 | 3:49417310-49417614 | IR (overlapping region) | 3 | 3 | ENST00000636594 | -46.1 | 1.17E-04 | 5.24E-04 |
| AMT | 3:49416794-49418372 | 3:49417358-49417614 | IR (overlapping region) | 3 | 3 | ENST00000636594 | -47.61 | 5.32E-05 | 2.39E-04 |
| AMT | 3:49416794-49418372 | 3:49417368-49417614 | IR (overlapping region) | 3 | 3 | ENST00000636594 | -47.73 | 4.65E-05 | 2.09E-04 |
| DLST | 14:74882017-74889094 | 14:74882591-74882624 | MES | 3 | 3 | ENST00000238671 | -48.35 | 4.92E-09 | 8.14E-09 |
| STAG3L5P-PVRIG2P-PILRB | 7:100356884-100358263 | 7:100358220-100358263 | A3ss | 3 | 3 | ENST00000310771 | -52.07 | 8.12E-04 | 7.33E-03 |
| STAG3L5P-PVRIG2P-PILRB | 7:100356884-100358226 | 7:100358220-100358226 | A3ss | 3 | 3 | Ex.ENST00000310771 | -52.07 | 8.12E-04 | 7.33E-03 |
| MGAT4B | 5:179798577-179799064 | 5:179798928-179799064 | A5ss | 2 | 2 | ENST00000520969 | -52.31 | 2.09E-02 | 1.41E-01 |
| MTERF2 | 12:106985216-106986968 | 12:106985216-106985225 | TSSlA3ss | 3 | 3 | TSS.ENST00000240050 | -52.64 | 4.42E-04 | 6.53E-04 |
| COASY | 17:42562356-42562894 | 17:42562488-42562611 | IR | 3 | 3 | ENST00000585811 | -53.37 | 4.34E-04 | 8.78E-04 |
| GLIDR | 9:39778945-39809730 | 9:39809486-39809615 | SES | 3 | 2 | ENST00000638724 | -53.59 | 1.84E-03 | 4.94E-02 |
| SLC36A4 | 11:93154217-93162705 | 11:93154217-93154277 | A3ss | 3 | 3 | Ex.ENST00000326402 | -54.81 | 4.87E-04 | 7.16E-04 |
| AC119673.3 | 1:205891310-205894466 | 1:205894168-205894466 | TSSlA3ss | 3 | 3 | TSS.ENST00000653632 | -57.84 | 3.56E-03 | 9.25E-03 |
| AC119673.3 | 1:205891310-205894443 | 1:205894168-205894443 | A3ss | 3 | 3 | Ex.TSS.ENST00000653632 | -57.84 | 3.56E-03 | 9.25E-03 |
| CRNDE | 16:54920339-54923584 | 16:54920339-54920410 | A3ss | 2 | 2 | ENST00000502066 | -58.68 | 1.71E-03 | 3.26E-03 |
| CRNDE | 16:54920328-54923584 | 16:54920328-54920410 | A3ss | 2 | 2 | ENST00000502066 | -58.68 | 1.71E-03 | 3.26E-03 |
| MFSD9 | 2:102732393-102736645 | 2:102732393-102732409 | TSSlA3ss | 3 | 3 | TSS.ENST00000258436 | -60.76 | 3.40E-03 | 1.18E-02 |
| COASY | 17:42562137-42562487 | 17:42562220-42562295 | IR | 3 | 3 | ENST00000585909 | -62.25 | 1.26E-06 | 2.54E-06 |
| COASY | 17:42562137-42562487 | 17:42562220-42562355 | IR | 3 | 3 | ENST00000585909 | -62.96 | 1.66E-06 | 3.36E-06 |
| ENOX2 | X:130637405-130656580 | X:130637405-130637410 | A3ss | 3 | 3 | Ex.ENST00000338144 | -63.87 | 3.82E-04 | 1.53E-02 |
| FAM122B | X:134796257-134797120 | X:134796257-134796355 | A3ss | 2 | 3 | Ex.TSS.ENST00000298090 | -64.33 | 5.51E-04 | 2.24E-02 |

**Table S1. Mis-splicing events in 3 biological replicates of SF3B1$^{K700E}$ (K700E ES line 2) compared to SF3B1$^{WT}$ ES cells.** The mis-splicing event code is as follows: SES, single-exon skipping; MES, multiple-exon skipping; MXS, mutually-exclusive splicing; A5ss, alternative 5′ splice site; A3ss, alternative 3′ splice site; TSS, transcription start site. Event Region: Genomic coordinates of splicing event; Target Exon: Genomic coordinates of the alternative exon. Events with p-values < 0.05 and ΔPSI (%)≥10% are listed.

| Gene Symbol | Event Region | Target Exon | Event | WT | K700E | Reference Transcript | ΔPSI (%) | p-value | FDR (BH) |
|---|---|---|---|---|---|---|---|---|---|
| GEN1 | 2:17753968-17759928 | 2:17759912-17759929 | A3ss | 2 | 3 | Ex.TSS.ENST00000317402 | 77.32 | 4.13E-03 | 1.77E-02 |
| ZNF91 | 19:23362726-23373741 | 19:23362726-23362739 | A3ss | 2 | 2 | Ex.TSS.ENST00000300619 | 77.14 | 1.90E-03 | 4.88E-03 |
| MFSD9 | 2:102732410-102736645 | 2:102732410-102732426 | A3ss | 3 | 3 | ENST00000462099 | 75.64 | 7.28E-04 | 3.03E-03 |
| MFSD9 | 2:102732393-102736645 | 2:102732393-102732426 | A3ss | 3 | 3 | ENST00000462099 | 75.64 | 7.28E-04 | 3.03E-03 |
| AL133352.1 | 10:100527075-100529365 | 10:100527075-100527094 | A3ss | 3 | 2 | Ex.ENST00000528174 | 73.08 | 1.32E-04 | 1.97E-04 |
| ORAI2 | 7:102433662-102436224 | 7:102436202-102436224 | TSSlA3ss | 3 | 3 | TSS.ENST00000356387 | 72.82 | 4.25E-06 | 4.94E-05 |
| APBB3 | 5:140561702-140561843 | 5:140561702-140561722 | A3ss | 2 | 3 | ENST00000354402 | 67.09 | 2.10E-02 | 1.71E-01 |
| VPS9D1-AS1 | 16:89712223-89712688 | 16:89712674-89712689 | A3ss | 3 | 3 | Ex.TSS.ENST00000562866 | 66.73 | 3.71E-05 | 8.35E-05 |
| PGBD1 | 6:28281919-28283775 | 6:28283756-28283775 | A3ss | 3 | 3 | Ex.TSS.ENST00000259883 | 64.27 | 9.25E-04 | 9.61E-03 |
| GUSBP11 | 22:23695518-23705428 | 22:23700726-23700845 | SES | 2 | 2 | ENST00000445682 | 63.45 | 3.65E-02 | 1.45E-01 |
| GPR153 | 1:6255015-6260824 | 1:6255015-6255122 | A3ss | 2 | 3 | Ex.TSS.ENST00000377893 | 63.34 | 2.19E-03 | 7.91E-03 |
| COASY | 17:42562220-42562611 | 17:42562356-42562611 | A3ss | 3 | 3 | ENST00000585811 | 61.45 | 1.21E-05 | 2.79E-05 |
| BCL2L1 | 20:31722331-31723734 | 20:31722331-31722348 | A3ss | 3 | 3 | ENST00000456404 | 60.99 | 1.48E-04 | 5.47E-04 |
| ZC3H11A | 1:203801629-203802956 | 1:203802569-203802957 | A3ss | 3 | 3 | Ex.ENST00000367212 | 60.87 | 9.83E-04 | 3.22E-03 |
| ZC3H11A | 1:203801629-203802651 | 1:203802569-203802652 | A3ss | 3 | 3 | Ex.ENST00000332127 | 60.87 | 9.83E-04 | 3.22E-03 |
| AC007326.4 | 22:18939750-18946790 | 22:18945072-18946791 | A3ss | 3 | 3 | Ex.TSS.ENST00000638240 | 60.65 | 1.86E-03 | 7.36E-03 |
| HERC2P2 | 15:22561790-22562948 | 15:22561896-22562949 | A3ss | 3 | 3 | Ex.ENST00000613386 | 60.1 | 1.35E-03 | 2.56E-03 |
| TLCD5 | 11:120325369-120329976 | 11:120327452-120329977 | A3ss | 3 | 3 | Ex.ENST00000531346 | 58.95 | 8.83E-07 | 1.37E-06 |
| TMEM14C | 6:10723242-10724569 | 6:10724556-10724570 | A3ss | 3 | 3 | Ex.TSS.ENST00000229563 | 58.63 | 4.29E-07 | 4.11E-06 |
| TMEM214 | 2:27037703-27038145 | 2:27037815-27037892 | SES | 3 | 3 | ENST00000425720 | 58.39 | 6.83E-05 | 3.17E-04 |
| DYNLL1 | 12:120496217-120496415 | 12:120496402-120496415 | TSSlA3ss | 3 | 3 | TSS.ENST00000392508 | 58.14 | 1.85E-07 | 3.12E-07 |
| GOLGA2P10 | 15:82475616-82476785 | 15:82475616-82475637 | A3ss | 3 | 3 | Ex.ENST00000614347 | 57.96 | 2.91E-05 | 5.71E-05 |
| PPP2R3A | 3:136040963-136049258 | 3:136049244-136049259 | A3ss | 3 | 3 | Ex.ENST00000264977 | 57.1 | 7.57E-03 | 4.18E-02 |
| SMURF2 | 17:64578577-64580788 | 17:64578577-64578594 | A3ss | 3 | 3 | Ex.ENST00000262435 | 56.27 | 2.38E-05 | 5.68E-05 |
| NAA16 | 13:41372831-41374741 | 13:41373768-41373880 | SES | 2 | 3 | ENST00000477452 | 55.45 | 2.54E-04 | 4.55E-04 |
| DLST | 14:74889350-74889896 | 14:74889878-74889896 | A3ss | 3 | 3 | ENST00000555988 | 55.27 | 2.05E-06 | 3.84E-06 |
| COASY | 17:42562220-42562611 | 17:42562356-42562487 | SES | 3 | 3 | ENST00000590958 | 53.56 | 2.34E-05 | 5.43E-05 |
| GCC2 | 2:108485909-108486510 | 2:108486499-108486511 | A3ss | 3 | 3 | Ex.ENST00000309863 | 53.29 | 9.28E-06 | 3.87E-05 |
| SLC4A8 | 12:51440790-51450875 | 12:51450857-51450876 | A3ss | 3 | 3 | Ex.ENST00000319957 | 52.09 | 5.68E-03 | 9.85E-03 |
| COASY | 17:42562220-42562611 | 17:42562220-42562487 | A5ss | 3 | 3 | ENST00000585909 | 50.48 | 3.84E-04 | 8.90E-04 |
| PILRB | 7:100356884-100358263 | 7:100358227-100358264 | A3ss | 3 | 3 | Ex.ENST00000608825 | 49.86 | 9.86E-05 | 1.14E-03 |
| MFAP3L | 4:170006737-170026233 | 4:170006737-170025842 | A3ss | 2 | 3 | Ex.TSS.ENST00000504999 | 49.47 | 4.00E-02 | 2.84E-01 |
| MFAP3L | 4:170006011-170026233 | 4:170006011-170025842 | A3ss | 2 | 3 | Ex.TSS.ENST00000361618 | 49.47 | 4.00E-02 | 2.84E-01 |
| MFAP3L | 4:170005892-170026233 | 4:170005892-170025842 | A3ss | 2 | 3 | Ex.ENST00000506764 | 49.47 | 4.00E-02 | 2.84E-01 |
| MFAP3L | 4:169992310-170026233 | 4:169992310-170025842 | A3ss | 2 | 3 | Ex.ENST00000512698 | 49.47 | 4.00E-02 | 2.84E-01 |
| AC007326.4 | 22:18939750-18946790 | 22:18939750-18945166 | A5ss | 3 | 3 | Ex.TSS.ENST00000638240 | 49.28 | 2.45E-03 | 9.70E-03 |
| APPL2 | 12:105208030-105208157 | 12:105208030-105208047 | A3ss | 3 | 3 | ENST00000551662 | 47.95 | 1.19E-02 | 1.98E-02 |
| ARMC9 | 2:231331898-231344974 | 2:231344949-231344975 | A3ss | 3 | 3 | Ex.TSS.ENST00000349938 | 47.4 | 5.42E-04 | 2.47E-03 |
| MRPL43 | 10:100977821-100979372 | 10:100979019-100979101 | IR (overlapping regio | 3 | 2 | ENST00000318325 | 46.81 | 3.43E-02 | 5.11E-02 |
| BCL2L1 | 20:31722331-31723863 | 20:31722331-31722348 | A3ss | 3 | 3 | ENST00000420488 | 46.74 | 1.77E-03 | 6.52E-03 |
| MIPOL1 | 14:37247241-37247828 | 14:37247799-37247828 | A3ss | 3 | 3 | ENST00000556615 | 46.48 | 1.24E-05 | 2.30E-05 |
| TMEM218 | 11:125102318-125102636 | 11:125102318-125102351 | A3ss | 3 | 3 | ENST00000279968 | 46.21 | 7.80E-04 | 1.21E-03 |
| NMRK2 | 19:3936666-3937239 | 19:3937225-3937239 | A3ss | 3 | 3 | ENST00000593949 | 46.2 | 2.26E-06 | 5.94E-06 |
| SLC26A6 | 3:48633636-48635370 | 3:48634516-48635371 | A5ss | 3 | 3 | Ex.TSS.ENST00000307364 | 45.18 | 5.76E-04 | 3.79E-03 |
| ENOSF1 | 18:683381-685920 | 18:683381-683395 | A3ss | 3 | 3 | ENST00000578647 | 44.84 | 6.11E-03 | 1.52E-02 |
| PKIB | 6:122482267-122585920 | 6:122581765-122581844 | MXS | 3 | 3 | ENST00000583007 | 44.26 | 3.96E-05 | 3.84E-04 |
| SH3D19 | 4:151179398-151187422 | 4:151179398-151179416 | A3ss | 3 | 3 | Ex.ENST00000304527 | 44.11 | 2.10E-05 | 1.47E-04 |
| IL17RC | 3:9918610-9920490 | 3:9920467-9920491 | A3ss | 3 | 3 | Ex.ENST00000295981 | 43.95 | 1.76E-02 | 1.21E-01 |
| SLC26A6 | 3:48633636-48635370 | 3:48633636-48634590 | A3ss | 3 | 3 | Ex.TSS.ENST00000307364 | 43.91 | 4.51E-04 | 2.97E-03 |
| ZNF771 | 16:30407665-30408044 | 16:30408032-30408045 | A3ss | 3 | 3 | Ex.TSS.ENST00000319296 | 43.86 | 2.92E-03 | 6.32E-03 |
| SEPTIN6 | X:119625380-119629317 | X:119625380-119625396 | A3ss | 3 | 3 | ENST00000467310 | 43.82 | 1.93E-04 | 8.77E-03 |
| HERC2P2 | 15:22561790-22562948 | 15:22561790-22562026 | A5ss | 3 | 3 | Ex.ENST00000613386 | 43.09 | 2.40E-03 | 4.54E-03 |
| AL009179.1 | 6:27694227-27702313 | 6:27694227-27694506 | A5ss | 2 | 3 | ENST00000625477 | 42.96 | 5.75E-03 | 5.96E-02 |
| MBD2 | 18:54203103-54204997 | 18:54203103-54203126 | TSSlA3ss | 3 | 3 | TSS.ENST00000398398 | 42.92 | 4.67E-04 | 1.16E-03 |
| UXS1 | 2:106164785-106166055 | 2:106164785-106164799 | A3ss | 3 | 3 | ENST00000283148 | 42.9 | 2.26E-03 | 9.43E-03 |
| MATN2 | 8:98032318-98033098 | 8:98033042-98033098 | A3ss | 3 | 3 | ENST00000254898 | 41.92 | 1.84E-02 | 4.70E-01 |
| NAA16 | 13:41373781-41374741 | 13:41373781-41373880 | A5ss | 3 | 3 | ENST00000477452 | 41.9 | 9.97E-03 | 1.79E-02 |
| ELF4 | X:130064928-130067525 | X:130067480-130067480 | IR | 2 | 2 | ENST00000308167 | 41.53 | 3.25E-02 | 1.00E+00 |
| ZDHHC16 | 10:97454800-97455713 | 10:97455639-97455713 | A3ss | 3 | 3 | ENST00000487315 | 41.5 | 2.80E-05 | 4.31E-05 |
| ZDHHC16 | 10:97454800-97455659 | 10:97455639-97455659 | A3ss | 3 | 3 | ENST00000487315 | 41.5 | 2.80E-05 | 4.31E-05 |
| SUZ12P1 | 17:30709812-30759490 | 17:30734897-30734943 | MES | 3 | 3 | ENST00000579526 | 40.06 | 1.31E-03 | 2.98E-03 |
| CCNT2-AS1 | 2:134869042-134918562 | 2:134877520-134877643 | SES | 2 | 3 | ENST00000428857 | 39.94 | 2.92E-02 | 1.23E-01 |
| NPR2 | 9:35801155-35801642 | 9:35801634-35801643 | A3ss | 3 | 3 | Ex.ENST00000342694 | 39.77 | 8.47E-03 | 3.10E-01 |
| FUT1 | 19:48752612-48753122 | 19:48752726-48752891 | IR | 2 | 3 | ENST00000601931 | 39.74 | 2.43E-03 | 6.53E-03 |
| PIH1D1 | 19:49451579-49453375 | 19:49451579-49451591 | TSSlA3ss | 3 | 3 | TSS.ENST00000601825 | 39.64 | 2.11E-02 | 5.68E-02 |
| ZSWIM8 | 10:73794331-73794555 | 10:73794541-73794555 | A3ss | 3 | 3 | ENST00000398706 | 39.59 | 1.56E-02 | 2.38E-02 |
| JPX | X:73998845-73999608 | X:73998954-73999608 | TSSlA3ss | 3 | 3 | TSS.ENST00000414209 | 39.43 | 4.66E-02 | 1.00E+00 |
| JPX | X:73998845-73999354 | X:73998954-73999354 | TSSlA3ss | 3 | 3 | TSS.ENST00000414209 | 39.43 | 4.66E-02 | 1.00E+00 |
| FGGY | 1:59296908-59346246 | 1:59321536-59321750 | MES | 3 | 3 | ENST00000413489 | 39.34 | 6.40E-03 | 2.31E-02 |
| CCDC189 | 16:30760284-30760668 | 16:30760454-30760605 | IR (overlapping regio | 3 | 2 | ENST00000433909 | 39.22 | 2.97E-02 | 6.45E-02 |
| ZBED5 | 11:10855087-10856143 | 11:10855087-10855118 | TSSlA3ss | 3 | 3 | TSS.ENST00000432999 | 38.27 | 6.72E-04 | 1.03E-03 |
| ZBED5 | 11:10855082-10856143 | 11:10855082-10855118 | TSSlA3ss | 3 | 3 | TSS.ENST00000432999 | 38.27 | 6.72E-04 | 1.03E-03 |
| CCDC74B | 2:130141297-130142132 | 2:130141297-130141308 | A3ss | 3 | 3 | Ex.ENST00000310463 | 38.14 | 1.34E-02 | 5.62E-02 |
| STAG3L5P | 7:100345969-100349798 | 7:100349717-100349798 | TSSlA3ss | 3 | 3 | TSS.ENST00000493499 | 37.99 | 9.89E-03 | 1.14E-01 |
| STAG3L5P | 7:100345969-100349730 | 7:100349717-100349730 | TSSlA3ss | 3 | 3 | TSS.ENST00000493499 | 37.99 | 9.89E-03 | 1.14E-01 |
| SLC13A3 | 20:46596343-46610445 | 20:46599971-46600037 | SES | 3 | 3 | ENST00000279027 | 37.86 | 2.65E-02 | 1.01E-01 |
| DVL2 | 17:7227703-7227976 | 17:7227703-7227711 | A3ss | 3 | 3 | Ex.ENST00000575086 | 37.54 | 5.99E-04 | 1.44E-03 |
| MTERF2 | 12:106978772-106986968 | 12:106985115-106985215 | SES | 3 | 3 | ENST00000392830 | 36.89 | 8.07E-05 | 1.35E-04 |
| ADGRV1 | 5:90558918-90614834 | 5:90592603-90614835 | A3ss | 2 | 3 | Ex.TSS.ENST00000405460 | 36.5 | 1.67E-02 | 1.58E-01 |
| ARHGAP27 | 17:45404097-45404444 | 17:45404269-45404334 | SES | 3 | 3 | ENST00000376922 | 36.19 | 4.07E-02 | 9.49E-02 |
| CCDC138 | 2:108856971-108876087 | 2:108873451-108873589 | SES | 2 | 3 | ENST00000295124 | 35.89 | 1.54E-02 | 6.42E-02 |
| KANSL3 | 2:96619763-96631311 | 2:96619763-96619776 | A3ss | 3 | 3 | Ex.ENST00000354204 | 35.87 | 1.42E-04 | 7.51E-04 |
| KANSL3 | 2:96619545-96631311 | 2:96619545-96619776 | A3ss | 3 | 3 | Ex.ENST00000418735 | 35.87 | 1.42E-04 | 7.51E-04 |
| FGGY | 1:59296908-59346246 | 1:59339958-59340069 | MES | 3 | 3 | ENST00000413489 | 35.52 | 2.17E-02 | 7.83E-02 |
| DCAF16 | 4:17804757-17810446 | 4:17804757-17804771 | A3ss | 3 | 3 | Ex.ENST00000507768 | 35.22 | 1.10E-03 | 7.86E-03 |
| SUZ12P1 | 17:30709812-30759490 | 17:30735033-30735097 | MES | 3 | 3 | ENST00000579526 | 35.17 | 6.53E-03 | 1.49E-02 |
| BIRC6 | 2:32471125-32473111 | 2:32471125-32471151 | A5ss | 3 | 3 | ENST00000648282 | 35.05 | 2.70E-04 | 1.26E-03 |
| ZNF219 | 14:21093291-21093585 | 14:21093291-21093304 | TSSlA3ss | 3 | 3 | TSS.ENST00000554478 | 34.47 | 4.70E-06 | 8.59E-06 |
| SPINT1-AS1 | 15:40838687-40844178 | 15:40838687-40838705 | A3ss | 3 | 3 | Ex.TSS.ENST00000564302 | 34.32 | 3.61E-05 | 6.87E-05 |
| THOC1 | 18:224180-224923 | 18:224180-224200 | A3ss | 3 | 3 | ENST00000579891 | 34.14 | 1.01E-03 | 2.47E-03 |
| ZNF26 | 12:132986874-133007041 | 12:132987665-133007042 | A3ss | 3 | 3 | Ex.TSS.ENST00000328654 | 33.88 | 3.63E-02 | 6.20E-02 |
| ZNF26 | 12:132986874-133006056 | 12:132987665-133006057 | A3ss | 3 | 3 | Ex.ENST00000545101 | 33.88 | 3.63E-02 | 6.20E-02 |
| SEPTIN6 | X:119620052-119629317 | X:119625335-119625396 | SES | 3 | 3 | ENST00000467310 | 33.78 | 1.07E-04 | 4.86E-03 |
| SLC7A3 | X:70929999-70930976 | X:70929999-70930022 | TSSlA3ss | 3 | 3 | TSS.ENST00000374299 | 33.69 | 1.10E-03 | 6.35E-01 |
| TOR1AIP2 | 1:179865855-179877238 | 1:179865855-179865869 | A3ss | 3 | 3 | Ex.TSS.ENST00000482587 | 33.5 | 7.19E-04 | 2.30E-03 |
| TRIM37 | 17:59001715-59012327 | 17:59001715-59001741 | A3ss | 3 | 3 | ENST00000585287 | 33.48 | 2.29E-05 | 5.43E-05 |

| | | | | | | | | | |
|---|---|---|---|---|---|---|---|---|---|
| TCF7L2 | 10:113146098-113150997 | 10:113150983-113150997 | A3ss | 3 | 3 | ENST00000543371 | 33.27 | 6.94E-03 | 1.04E-02 |
| AC007405.4 | 2:170771428-170777376 | 2:170777147-170777177 | SES | 3 | 3 | ENST00000653024 | 32.66 | 7.06E-04 | 3.01E-03 |
| PLXND1 | 3:129566018-129566526 | 3:129566018-129566029 | A3ss | 3 | 3 | Ex.ENST00000324093 | 32.41 | 4.67E-02 | 2.56E-01 |
| DNMBP | 10:99955214-99957205 | 10:99955833-99956481 | IR (overlapping regio | 3 | 2 | ENST00000324109 | 32.33 | 3.49E-02 | 5.37E-02 |
| CRAMP1 | 16:1612423-1614638 | 16:1612423-1612657 | A5ss | 3 | 3 | Ex.TSS.ENST00000397412 | 32.23 | 3.28E-02 | 6.93E-02 |
| ZNF706 | 8:101202333-101205287 | 8:101204780-101205210 | IR | 2 | 3 | ENST00000520347 | 32.16 | 6.03E-04 | 9.91E-03 |
| NDRG3 | 20:36653702-36656359 | 20:36653702-36653723 | A3ss | 3 | 3 | Ex.TSS.ENST00000349004 | 32.07 | 1.64E-07 | 6.11E-07 |
| CASD1 | 7:94528251-94533204 | 7:94533189-94533205 | A3ss | 3 | 2 | Ex.ENST00000297273 | 31.85 | 3.74E-03 | 5.52E-02 |
| BCL2L1 | 20:31722331-31722617 | 20:31722331-31722348 | TSSIA3ss | 3 | 3 | TSS.ENST00000307677 | 31.8 | 3.67E-03 | 1.35E-02 |
| UBA1 | X:47199615-47200893 | X:47199615-47200544 | A5ss | 3 | 3 | Ex.ENST00000335972 | 31.75 | 5.11E-06 | 5.85E-04 |
| CHTF18 | 16:793275-794053 | 16:794034-794054 | A3ss | 3 | 3 | Ex.ENST00000262315 | 31.7 | 4.51E-04 | 1.00E-03 |
| BAG6 | 6:31644195-31644524 | 6:31644307-31644414 | SES | 3 | 3 | ENST00000437771 | 31.06 | 8.67E-05 | 9.11E-04 |
| RWDD4 | 4:183655962-183658928 | 4:183655962-183655974 | A3ss | 3 | 3 | Ex.TSS.ENST00000326397 | 31.05 | 2.64E-02 | 1.88E-01 |
| RWDD4 | 4:183651132-183658928 | 4:183651132-183655974 | A3ss | 3 | 3 | Ex.TSS.ENST00000510968 | 31.05 | 2.64E-02 | 8.23E-02 |
| ZNF300 | 5:150898555-150903140 | 5:150898555-150902566 | A3ss | 3 | 3 | Ex.ENST00000274599 | 31.03 | 3.51E-02 | 2.91E-01 |
| SMCO4 | 11:93479270-93499275 | 11:93479270-93479284 | TSSIA3ss | 3 | 3 | TSS.ENST00000529714 | 31.01 | 7.81E-04 | 1.30E-03 |
| RTF2 | 20:56468767-56472325 | 20:56471614-56471669 | MXS | 2 | 3 | ENST00000487211 | 30.95 | 1.80E-02 | 6.87E-02 |
| KLHDC10 | 7:130070810-130116444 | 7:130096921-130116445 | A3ss | 3 | 2 | Ex.TSS.ENST00000463413 | 30.9 | 3.42E-02 | 4.14E-01 |
| CAMTA2 | 17:4982161-4982756 | 17:4982161-4982175 | A3ss | 3 | 3 | ENST00000572543 | 30.45 | 1.04E-02 | 2.45E-02 |
| PKIB | 6:122482267-122585920 | 6:122581765-122581844 | MES | 3 | 3 | ENST00000618977 | 30.19 | 9.59E-06 | 9.32E-05 |
| BAZ2A | 12:56630866-56636181 | 12:56634945-56636182 | A5ss | 3 | 3 | Ex.TSS.ENST00000549506 | 30.17 | 5.97E-03 | 1.05E-02 |
| PHF20L1 | 8:132798861-132803818 | 8:132799083-132799172 | SES | 3 | 3 | ENST00000395376 | 30.04 | 7.62E-03 | 1.31E-01 |
| FIRRE | X:131794467-131825221 | X:131804972-131805037 | MES | 3 | 3 | ENST00000427391 | 30.02 | 4.53E-02 | 1.00E+00 |
| SERBP1 | 1:67424960-67425082 | 1:67424960-67424977 | A3ss | 3 | 3 | ENST00000361219 | 30 | 1.13E-06 | 4.08E-06 |
| ZSCAN30 | 18:35264456-35290130 | 18:35290084-35290130 | TSSIA5ss | 3 | 2 | TSS.ENST00000333206 | 29.85 | 3.84E-02 | 8.04E-02 |
| MAP3K7 | 6:90560215-90561621 | 6:90560215-90560234 | A3ss | 3 | 3 | Ex.ENST00000369325 | 29.76 | 1.16E-02 | 1.33E-01 |
| DERL3 | 22:23834933-23837063 | 22:23834933-23834979 | A3ss | 3 | 3 | Ex.TSS.ENST00000406855 | 29.69 | 1.79E-02 | 7.14E-02 |
| MED6 | 14:70592989-70593295 | 14:70592989-70593009 | A3ss | 3 | 3 | ENST00000430055 | 29.6 | 2.81E-04 | 5.26E-04 |
| BRD9 | 5:865582-870472 | 5:869245-869404 | MXS | 3 | 3 | ENST00000519112 | 29.58 | 1.72E-04 | 1.61E-03 |
| ANKHD1 | 5:140436258-140438502 | 5:140438494-140438502 | A3ss | 3 | 3 | ENST00000394722 | 29.45 | 3.30E-05 | 2.70E-04 |
| SKIV2L | 6:31969280-31969751 | 6:31969421-31969514 | IR | 3 | 3 | ENST00000465703 | 29.04 | 4.46E-02 | 1.39E-01 |
| TTI1 | 20:38002777-38006196 | 20:38002777-38002793 | A3ss | 3 | 3 | Ex.ENST00000373447 | 29.02 | 5.99E-05 | 2.23E-04 |
| STOX2 | 4:184011424-184017088 | 4:184011587-184011635 | SES | 3 | 3 | ENST00000506529 | 28.9 | 5.82E-03 | 4.15E-02 |
| GRIP1 | 12:66445509-66462923 | 12:66455409-66455561 | SES | 2 | 3 | ENST00000543172 | 28.8 | 2.40E-02 | 4.23E-02 |
| ZNF160 | 19:53086110-53086261 | 19:53086110-53086128 | A3ss | 3 | 3 | Ex.TSS.ENST00000596966 | 28.78 | 9.61E-04 | 2.63E-03 |
| ZNF160 | 19:53086089-53086261 | 19:53086089-53086128 | A3ss | 3 | 3 | Ex.TSS.ENST00000594760 | 28.78 | 9.61E-04 | 2.63E-03 |
| ZNF160 | 19:53075184-53086261 | 19:53075184-53086128 | A3ss | 3 | 3 | Ex.ENST00000355147 | 28.78 | 9.61E-04 | 2.63E-03 |
| JAKMIP1 | 4:6085630-6112721 | 4:6105473-6105967 | SES | 3 | 3 | ENST00000282924 | 28.67 | 1.61E-03 | 1.21E-02 |
| BTN2A2 | 6:26383245-26383791 | 6:26383245-26383299 | TSSIA5ss | 3 | 3 | TSS.ENST00000471116 | 28.63 | 1.92E-02 | 1.97E-01 |
| BTN2A2 | 6:26383182-26383791 | 6:26383182-26383299 | TSSIA5ss | 3 | 3 | TSS.ENST00000472507 | 28.63 | 1.92E-02 | 1.97E-01 |
| ZC3H13 | 13:46045517-46052403 | 13:46051932-46051989 | SES | 3 | 3 | ENST00000428921 | 28.5 | 1.62E-03 | 2.91E-03 |
| SNRPN | 15:24962210-24967931 | 15:24967029-24967152 | SES | 3 | 3 | ENST00000554227 | 28.48 | 2.15E-05 | 4.08E-05 |
| GRIP1 | 12:66445509-66462923 | 12:66455409-66455564 | SES | 2 | 3 | ENST00000398016 | 27.85 | 4.86E-02 | 8.57E-02 |
| ECHDC2 | 1:52904834-52907867 | 1:52905034-52905090 | SES | 3 | 3 | ENST00000358358 | 27.67 | 3.88E-02 | 1.39E-01 |
| WASHC4 | 12:105120598-105121100 | 12:105121089-105121100 | A3ss | 3 | 3 | ENST00000547404 | 27.54 | 7.76E-03 | 1.30E-02 |
| DPH7 | 9:137576403-137578509 | 9:137577470-137577603 | SES | 2 | 2 | ENST00000481839 | 27.31 | 7.11E-03 | 2.41E-01 |
| TMEM39B | 1:32075823-32091674 | 1:32077164-32077318 | MES | 3 | 3 | ENST00000336294 | 27.31 | 1.27E-02 | 4.37E-02 |
| SETD4 | 21:36043957-36045581 | 21:36043957-36043969 | A3ss | 3 | 3 | Ex.ENST00000332131 | 27.27 | 2.72E-03 | 1.06E-02 |
| NBPF15 | 1:144450803-144456536 | 1:144450803-144450870 | TSSIA3ss | 2 | 3 | TSS.ENST00000578953 | 27.25 | 1.03E-02 | 2.90E-02 |
| ESCO1 | 18:21584441-21600622 | 18:21600415-21600623 | A5ss | 3 | 2 | Ex.TSS.ENST00000269214 | 27.24 | 1.15E-02 | 2.82E-02 |
| CAMTA2 | 17:4981832-4982756 | 17:4982089-4982175 | SES | 3 | 3 | ENST00000572543 | 27.23 | 1.52E-02 | 3.57E-02 |
| EMC3-AS1 | 3:9987265-9987905 | 3:9987815-9987905 | TSSIA3ss | 3 | 3 | TSS.ENST00000440568 | 27.14 | 1.55E-02 | 1.04E-01 |
| ANKRD12 | 18:9195699-9208656 | 18:9204476-9204544 | MXS | 3 | 2 | ENST00000262126 | 27.1 | 4.39E-02 | 1.10E-01 |
| PHF20L1 | 8:132798861-132799094 | 8:132799083-132799094 | A3ss | 3 | 3 | ENST00000395376 | 26.96 | 1.65E-02 | 2.83E-01 |
| MICAL1 | 6:109445863-109446135 | 6:109445863-109445875 | A3ss | 3 | 3 | Ex.ENST00000358577 | 26.93 | 8.02E-03 | 7.71E-02 |
| CCDC74A | 2:131530828-131531663 | 2:131531638-131531664 | A3ss | 3 | 3 | Ex.ENST00000295171 | 26.76 | 4.59E-02 | 1.93E-01 |
| SOS1 | 2:38987592-38995122 | 2:38989270-38989314 | SES | 3 | 3 | ENST00000402219 | 26.72 | 1.77E-02 | 8.33E-02 |
| CAP1 | 1:40040784-40041428 | 1:40040784-40040801 | TSSIA5ss | 3 | 3 | TSS.ENST00000414281 | 26.71 | 3.11E-03 | 1.09E-02 |
| MAPK12 | 22:50245737-50245909 | 22:50245737-50245812 | A3ss | 3 | 3 | Ex.ENST00000497036 | 26.62 | 1.75E-02 | 7.19E-02 |
| CCDC189 | 16:30761198-30762221 | 16:30761230-30761544 | IR | 3 | 2 | ENST00000543128 | 26.29 | 1.58E-02 | 3.43E-02 |
| DNASE1 | 16:3656754-3657186 | 16:3656999-3657111 | SES | 3 | 2 | ENST00000246949 | 26.28 | 3.25E-02 | 7.08E-02 |
| ZNF91 | 19:23362726-23395324 | 19:23374638-23374764 | SES | 3 | 3 | ENST00000397082 | 26.23 | 2.70E-02 | 6.93E-02 |
| SGSH | 17:80214766-80215032 | 17:80214766-80214783 | A3ss | 3 | 3 | ENST00000570427 | 26.19 | 5.23E-04 | 1.27E-03 |
| SGSH | 17:80214680-80215032 | 17:80214680-80214783 | A3ss | 3 | 3 | ENST00000570427 | 26.19 | 5.23E-04 | 1.27E-03 |
| ZNF100 | 19:21744113-21744940 | 19:21744113-21744115 | A3ss | 3 | 3 | ENST00000358296 | 26.18 | 2.47E-02 | 6.34E-02 |
| GPR161 | 1:168114550-168137164 | 1:168136739-168137165 | A5ss | 3 | 3 | Ex.TSS.ENST00000367835 | 26.11 | 6.78E-03 | 2.16E-02 |
| ARHGAP12 | 10:31831801-31843460 | 10:31839637-31839711 | MES | 3 | 3 | ENST00000344936 | 26.06 | 1.26E-03 | 1.92E-03 |
| NFYA | 6:41073085-41079028 | 6:41079005-41079029 | A3ss | 3 | 3 | Ex.TSS.ENST00000341376 | 25.96 | 8.90E-05 | 9.64E-04 |
| CAV1 | 7:116526293-116526689 | 7:116526389-116526524 | IR | 3 | 3 | ENST00000393467 | 25.61 | 1.40E-02 | 1.66E-01 |
| UBA1 | X:47199615-47200893 | X:47200410-47200894 | A3ss | 3 | 3 | Ex.ENST00000335972 | 25.45 | 1.93E-04 | 2.21E-02 |
| PRKCSH | 19:11447619-11447722 | 19:11447693-11447722 | A3ss | 3 | 3 | ENST00000585540 | 25.38 | 4.87E-05 | 1.23E-04 |
| DDX12P | 12:9421492-9421817 | 12:9421492-9421546 | A3ss | 3 | 3 | Ex.ENST00000432996 | 25.37 | 1.82E-02 | 3.25E-02 |
| DICER1 | 14:95133504-95157229 | 14:95141683-95141748 | MES | 3 | 3 | ENST00000531162 | 25.1 | 3.10E-02 | 5.68E-02 |
| ZMYM1 | 1:35094010-35095818 | 1:35094010-35094083 | A5ss | 3 | 3 | ENST00000359858 | 25.06 | 1.15E-02 | 3.97E-02 |
| AFG3L1P | 16:89972766-89978809 | 16:89977715-89978810 | A3ss | 3 | 3 | Ex.TSS.ENST00000429663 | 24.92 | 1.61E-02 | 3.63E-02 |
| PRSS16 | 6:27252041-27254985 | 6:27254693-27254872 | MXS | 3 | 3 | ENST00000471463 | 24.75 | 4.38E-02 | 4.51E-01 |
| ZSWIM4 | 19:13799922-13804791 | 19:13804775-13804792 | A3ss | 3 | 3 | Ex.ENST00000254323 | 24.73 | 2.79E-04 | 7.08E-04 |
| HTT | 4:3140657-3142765 | 4:3142745-3142765 | A3ss | 3 | 3 | ENST00000649131 | 24.71 | 5.53E-06 | 4.07E-05 |
| DAB1 | 1:57026044-57062889 | 1:57062884-57062889 | A5ss | 3 | 3 | ENST00000371236 | 24.65 | 3.78E-02 | 1.36E-01 |
| IQGAP3 | 1:156562666-156563133 | 1:156562666-156562674 | A3ss | 3 | 3 | Ex.ENST00000361170 | 24.62 | 3.58E-03 | 1.13E-02 |
| PI4KAP2 | 22:21476161-21477789 | 22:21476644-21476764 | SES | 3 | 3 | ENST00000360806 | 24.6 | 2.04E-04 | 8.09E-04 |
| CU634019.6 | 21:6995314-6997591 | 21:6995314-6995324 | TSSIA3ss | 3 | 3 | TSS.ENST00000624728 | 24.6 | 2.28E-02 | 8.98E-02 |
| KIF13A | 6:17764947-17771907 | 6:17771114-17771218 | SES | 3 | 3 | ENST00000259711 | 24.56 | 4.29E-02 | 4.40E-01 |
| DYNC2LI1 | 2:43774147-43776781 | 2:43774403-43776782 | A3ss | 3 | 3 | Ex.TSS.ENST00000398823 | 24.43 | 2.72E-02 | 1.28E-01 |
| ORAI2 | 7:102433662-102438943 | 7:102436202-102436333 | SES | 3 | 3 | ENST00000356387 | 24.35 | 1.11E-04 | 1.28E-03 |
| ADGRG2 | X:19037491-19037597 | X:19037589-19037597 | A5ss | 3 | 3 | ENST00000356606 | 24.18 | 9.21E-03 | 7.49E-01 |
| HINT2 | 9:35813146-35813265 | 9:35813146-35813156 | A3ss | 3 | 3 | Ex.TSS.ENST00000471774 | 24.14 | 3.12E-02 | 1.00E+00 |
| ARAF | X:47561252-47562908 | X:47561252-47562762 | A5ss | 3 | 3 | Ex.TSS.ENST00000377039 | 24.09 | 5.24E-03 | 6.06E-01 |
| NR2C2 | 3:14947907-15003875 | 3:14959157-14959196 | MES | 3 | 3 | ENST00000435454 | 24.07 | 2.13E-02 | 1.19E-01 |
| PHKB | 16:47461427-47463898 | 16:47463882-47463898 | TSSIA3ss | 3 | 3 | TSS.ENST00000566044 | 24.01 | 4.79E-03 | 1.04E-02 |
| DPH5 | 1:100992741-100995109 | 1:100992741-100992754 | A3ss | 3 | 3 | Ex.ENST00000342173 | 23.98 | 3.11E-03 | 8.62E-03 |
| DPH5 | 1:100990632-100995109 | 1:100990632-100992754 | A3ss | 3 | 3 | Ex.ENST00000427040 | 23.98 | 3.11E-03 | 8.62E-03 |
| ADARB1 | 21:45183511-45184922 | 21:45184474-45184593 | SES | 3 | 2 | ENST00000360697 | 23.93 | 3.77E-02 | 1.48E-01 |
| PLEKHA8P1 | 12:45176434-45215939 | 12:45190794-45190961 | SES | 3 | 3 | ENST00000545609 | 23.55 | 9.06E-03 | 1.56E-02 |
| ZNF778 | 16:89222184-89224718 | 16:89223157-89223283 | SES | 3 | 2 | ENST00000433976 | 23.47 | 1.85E-02 | 4.13E-02 |
| TMEM214 | 2:27037703-27038145 | 2:27037790-27037892 | SES | 3 | 3 | ENST00000435172 | 23.46 | 1.62E-03 | 7.50E-03 |
| RHBDD1 | 2:226836088-226864603 | 2:226838073-226838154 | MES | 3 | 3 | ENST00000392062 | 23.4 | 3.56E-02 | 1.61E-01 |
| TTC14 | 3:180607648-180610974 | 3:180607766-180608700 | IR (overlapping regio | 3 | 3 | ENST00000412756 | 23.34 | 2.33E-03 | 1.34E-02 |
| CU638689.4 | 21:6668244-6670521 | 21:6668244-6668254 | TSSIA3ss | 3 | 3 | TSS.ENST00000623324 | 23.28 | 4.28E-02 | 1.69E-01 |
| TRPC1 | 3:142736534-142748257 | 3:142743485-142743586 | SES | 2 | 2 | ENST00000476941 | 23.27 | 4.17E-02 | 2.31E-01 |
| PRDM2 | 1:13731118-13742004 | 1:13732779-13732882 | SES | 3 | 2 | ENST00000235372 | 23.21 | 2.52E-02 | 7.04E-02 |

| | | | | | | | | | |
|---|---|---|---|---|---|---|---|---|---|
| LRRC37B | 17:32008133-32024710 | 17:32017972-32018054 | MES | 3 | 3 | ENST00000543378 | 23.2 | 1.53E-02 | 3.49E-02 |
| SLC16A5 | 17:75088045-75089150 | 17:75089114-75089150 | TSSIA3ss | 3 | 3 | TSS.ENST00000578376 | 23.09 | 2.88E-02 | 6.76E-02 |
| GTF3C2 | 2:27343579-27356738 | 2:27350314-27350632 | MXS | 3 | 3 | ENST00000423998 | 23.04 | 1.93E-02 | 8.97E-02 |
| XRCC3 | 14:103712932-103715423 | 14:103715337-103715424 | A5ss | 3 | 3 | Ex.TSS.ENST00000555055 | 23.04 | 3.02E-02 | 5.49E-02 |
| DHX29 | 5:55270488-55270577 | 5:55270488-55270505 | A3ss | 3 | 3 | Ex.ENST00000251636 | 22.99 | 5.70E-05 | 5.16E-04 |
| ZNF250 | 8:144890404-144901398 | 8:144901337-144901398 | TSSIA5ss | 3 | 3 | TSS.ENST00000533622 | 22.96 | 1.26E-02 | 2.28E-01 |
| MAP11 | 7:100155165-100155261 | 7:100155165-100155181 | A3ss | 3 | 3 | Ex.TSS.ENST00000316937 | 22.81 | 4.03E-02 | 4.61E-01 |
| EHMT1 | 9:137728530-137743391 | 9:137743371-137743391 | A3ss | 3 | 3 | ENST00000371394 | 22.56 | 7.97E-03 | 2.70E-01 |
| INPPL1 | 11:72228499-72228726 | 11:72228647-72228727 | A3ss | 3 | 3 | Ex.ENST00000298229 | 22.54 | 9.83E-05 | 1.61E-04 |
| OCEL1 | 19:17228310-17229008 | 19:17228803-17228915 | SES | 3 | 3 | ENST00000595573 | 22.53 | 4.97E-02 | 1.26E-01 |
| ACIN1 | 14:23081837-23090521 | 14:23089982-23090101 | SES | 3 | 3 | ENST00000262710 | 22.46 | 3.52E-03 | 6.43E-03 |
| PACRGL | 4:20713540-20727284 | 4:20724808-20724888 | MXS | 3 | 3 | ENST00000471979 | 22.42 | 5.13E-03 | 3.74E-02 |
| HELLS | 10:94545908-94546376 | 10:94546361-94546377 | A3ss | 3 | 3 | Ex.TSS.ENST00000419900 | 22.4 | 1.13E-04 | 1.73E-04 |
| TRIM37 | 17:59070948-59075646 | 17:59070948-59070968 | A3ss | 3 | 3 | Ex.ENST00000262294 | 22.33 | 1.71E-04 | 4.04E-04 |
| SLTM | 15:58917000-58932355 | 15:58917000-58917020 | A3ss | 3 | 3 | Ex.ENST00000249736 | 22.32 | 2.11E-05 | 4.06E-05 |
| DCLK1 | 13:35822876-35827634 | 13:35822876-35822893 | A3ss | 3 | 3 | Ex.ENST00000255448 | 22.32 | 1.67E-03 | 2.99E-03 |
| CCDC189 | 16:30761198-30762221 | 16:30761336-30761544 | IR | 3 | 2 | ENST00000543128 | 22.29 | 2.82E-02 | 6.11E-02 |
| EMSY | 11:76453389-76458182 | 11:76454749-76454790 | SES | 3 | 3 | ENST00000427574 | 22.27 | 1.28E-02 | 2.10E-02 |
| VEZT | 12:95266633-95270050 | 12:95270039-95270050 | A3ss | 3 | 3 | ENST00000397792 | 22.21 | 6.06E-03 | 1.08E-02 |
| LUC7L3 | 17:50750853-50752199 | 17:50751297-50751362 | SES | 3 | 3 | ENST00000503728 | 22.12 | 1.74E-02 | 4.11E-02 |
| PDE3B | 11:14771988-14789105 | 11:14786437-14786532 | SES | 3 | 3 | ENST00000455098 | 21.99 | 1.63E-02 | 2.55E-02 |
| ZFYVE27 | 10:97750471-97751390 | 10:97751376-97751390 | A3ss | 3 | 3 | ENST00000370610 | 21.95 | 1.43E-02 | 2.19E-02 |
| ATP2C1 | 3:131001315-131016151 | 3:131001315-131001344 | TSSIA5ss | 3 | 3 | TSS.ENST00000359644 | 21.88 | 1.86E-03 | 1.02E-02 |
| TRIM37 | 17:58999460-59012327 | 17:59001598-59001741 | SES | 3 | 3 | ENST00000585287 | 21.87 | 4.01E-03 | 9.49E-03 |
| CDPF1 | 22:46245239-46250255 | 22:46248172-46248284 | MES | 3 | 3 | ENST00000314567 | 21.7 | 3.33E-02 | 1.38E-01 |
| THAP8 | 19:36039719-36054134 | 19:36039944-36040136 | SES | 3 | 3 | ENST00000292894 | 21.55 | 1.17E-03 | 3.06E-03 |
| SNHG8 | 4:118278793-118279388 | 4:118278944-118279137 | SES | 3 | 3 | ENST00000602819 | 21.52 | 1.17E-04 | 8.10E-04 |
| FBXO21 | 12:117158064-117165484 | 12:117158064-117158084 | A3ss | 3 | 3 | ENST00000330622 | 21.51 | 8.52E-03 | 1.44E-02 |
| ZNF76 | 6:35293045-35294455 | 6:35293751-35293915 | SES | 3 | 3 | ENST00000373953 | 21.43 | 9.05E-03 | 9.77E-02 |
| BTBD10 | 11:13421839-13463091 | 11:13445024-13445181 | SES | 3 | 3 | ENST00000278174 | 21.38 | 2.38E-04 | 3.73E-04 |
| ARHGAP12 | 10:31831801-31843460 | 10:31839637-31839711 | MXS | 3 | 3 | ENST00000311380 | 21.38 | 2.18E-02 | 3.31E-02 |
| CC2D1A | 19:13920923-13923332 | 19:13923318-13923333 | A3ss | 3 | 3 | Ex.ENST00000318003 | 21.35 | 5.65E-04 | 1.43E-03 |
| SNHG8 | 4:118278793-118279388 | 4:118278793-118279137 | A5ss | 3 | 3 | ENST00000652022 | 21.33 | 6.84E-04 | 4.75E-03 |
| IMMT | 2:86171313-86173649 | 2:86171313-86171345 | A3ss | 3 | 3 | ENST00000410111 | 21.3 | 2.93E-03 | 1.51E-02 |
| IMMT | 2:86171310-86173649 | 2:86171310-86171345 | A3ss | 3 | 3 | Ex.ENST00000410111 | 21.3 | 2.93E-03 | 1.51E-02 |
| WDR90 | 16:662220-662736 | 16:662332-662672 | IR | 3 | 3 | ENST00000546923 | 21.29 | 3.99E-04 | 8.78E-04 |
| ESPL1 | 12:53272858-53274816 | 12:53274790-53274817 | A3ss | 3 | 3 | Ex.ENST00000257934 | 21.2 | 2.06E-03 | 3.57E-03 |
| ZNF532 | 18:58953800-58979054 | 18:58954234-58954334 | SES | 3 | 3 | ENST00000585662 | 21.15 | 3.18E-05 | 7.89E-05 |
| DKK3 | 11:11967099-11968394 | 11:11967099-11967119 | A3ss | 3 | 3 | Ex.ENST00000326932 | 21.09 | 6.62E-03 | 1.03E-02 |
| TTC32 | 2:19898036-19901705 | 2:19901476-19901706 | A5ss | 3 | 3 | Ex.ENST00000333610 | 21.06 | 2.88E-02 | 1.24E-01 |
| CD27-AS1 | 12:6450341-6451517 | 12:6450709-6450892 | IR (overlapping regio | 2 | 3 | ENST00000535639 | 21.01 | 2.64E-02 | 4.66E-02 |
| STAP2 | 19:4324198-4325215 | 19:4324455-4324664 | SES | 3 | 3 | ENST00000599736 | 20.96 | 2.42E-02 | 6.42E-02 |
| PCNX3 | 11:65620967-65622244 | 11:65622224-65622245 | A3ss | 3 | 3 | Ex.ENST00000355703 | 20.9 | 2.65E-03 | 4.27E-03 |
| TOR1AIP1 | 1:179884770-179889312 | 1:179889310-179889312 | A3ss | 3 | 3 | ENST00000271583 | 20.9 | 1.48E-02 | 4.75E-02 |
| ZNF875 | 19:37334783-37335199 | 19:37335169-37335199 | TSSIA3ss | 3 | 3 | TSS.ENST00000324411 | 20.83 | 1.47E-02 | 3.09E-02 |
| PREX2 | 8:68083389-68087723 | 8:68084746-68087724 | A3ss | 3 | 3 | Ex.ENST00000288368 | 20.77 | 3.53E-02 | 8.47E-01 |
| KIAA1217 | 10:24528120-24531832 | 10:24531830-24531832 | A3ss | 3 | 3 | ENST00000307544 | 20.76 | 1.13E-02 | 1.70E-02 |
| DNASE1 | 16:3656754-3657186 | 16:3656999-3657186 | A3ss | 3 | 2 | ENST00000570769 | 20.66 | 4.15E-02 | 9.04E-02 |
| FAM222B | 17:28766708-28842681 | 17:28790378-28790425 | MES | 3 | 3 | ENST00000583307 | 20.58 | 9.70E-04 | 2.21E-03 |
| ECT2 | 3:172761684-172762415 | 3:172762398-172762416 | A3ss | 3 | 3 | Ex.ENST00000232458 | 20.48 | 2.96E-02 | 1.70E-01 |
| C2orf74 | 2:61145197-61162841 | 2:61157862-61158012 | MES | 3 | 3 | ENST00000398622 | 20.44 | 1.94E-02 | 9.35E-02 |
| ENTPD1-AS1 | 10:95876661-95907777 | 10:95904830-95904935 | SES | 2 | 3 | ENST00000661375 | 20.41 | 1.39E-02 | 2.14E-02 |
| PRPF38A | 1:52414648-52414761 | 1:52414741-52414762 | A3ss | 3 | 3 | Ex.ENST00000257181 | 20.33 | 1.37E-03 | 4.90E-03 |
| SEPTIN8 | 5:132758825-132760801 | 5:132759937-132760802 | A5ss | 3 | 3 | Ex.TSS.ENST00000296873 | 20.32 | 1.41E-03 | 1.12E-02 |
| ZNF562 | 19:9656654-9660719 | 19:9659379-9659467 | MXS | 3 | 3 | ENST00000585688 | 20.31 | 3.68E-02 | 1.02E-01 |
| KLHL2 | 4:165317970-165322031 | 4:165321312-165321375 | SES | 3 | 3 | ENST00000509028 | 20.3 | 8.70E-03 | 6.15E-02 |
| SKIV2L | 6:31968539-31968685 | 6:31968623-31968685 | A3ss | 3 | 3 | ENST00000465703 | 20.25 | 1.98E-02 | 1.32E-01 |
| ARRB2 | 17:4710745-4715972 | 17:4715013-4715043 | SES | 3 | 3 | ENST00000269260 | 20.12 | 2.95E-04 | 6.89E-04 |
| LETMD1 | 12:51049186-51055834 | 12:51053812-51053860 | MXS | 3 | 3 | ENST00000549340 | 20.12 | 1.54E-02 | 2.66E-02 |
| GATD1 | 11:770399-771332 | 11:770679-770903 | MES | 3 | 3 | ENST00000354286 | 20.03 | 9.54E-03 | 1.57E-02 |
| LTBP4 | 19:40614447-40617099 | 19:40616889-40617099 | A3ss | 3 | 3 | ENST00000617753 | 20.02 | 2.25E-02 | 5.97E-02 |
| ZNF382 | 19:36610743-36626132 | 19:36626130-36626132 | TSSIA3ss | 2 | 3 | TSS.ENST00000292928 | 20 | 0.00E+00 | 0.00E+00 |
| ZMYM1 | 1:35079443-35093913 | 1:35093320-35093441 | SES | 3 | 3 | ENST00000488455 | 19.85 | 4.49E-02 | 1.55E-01 |
| ANO9 | 11:419582-420860 | 11:420616-420717 | IR | 3 | 3 | ENST00000532094 | 19.83 | 1.49E-02 | 2.35E-02 |
| AFG3L1P | 16:89984600-89990865 | 16:89988852-89988951 | SES | 2 | 2 | ENST00000388970 | 19.81 | 1.45E-02 | 3.28E-02 |
| ZCCHC8 | 12:122482696-122489385 | 12:122483460-122483563 | MES | 3 | 3 | ENST00000633063 | 19.79 | 1.71E-02 | 2.90E-02 |
| FRMD4A | 10:13675045-13693897 | 10:13693482-13693517 | SES | 3 | 3 | ENST00000632570 | 19.79 | 4.96E-02 | 7.49E-02 |
| FOSL1 | 11:65894117-65896808 | 11:65894117-65894121 | A3ss | 3 | 3 | ENST00000312562 | 19.77 | 1.07E-04 | 1.73E-04 |
| THTPA | 14:23556344-23556752 | 14:23556743-23556752 | TSSIA3ss | 3 | 3 | TSS.ENST00000404535 | 19.77 | 8.23E-04 | 1.51E-03 |
| ZNF79 | 9:127424804-127428831 | 9:127424804-127424910 | A5ss | 2 | 3 | Ex.TSS.ENST00000342483 | 19.71 | 4.64E-02 | 1.00E+00 |
| SH2D3A | 19:6754426-6754615 | 19:6754426-6754448 | A3ss | 3 | 3 | Ex.ENST00000245908 | 19.7 | 9.78E-03 | 2.71E-02 |
| SEPTIN8 | 5:132758825-132760801 | 5:132758825-132760013 | A3ss | 3 | 3 | Ex.TSS.ENST00000296873 | 19.67 | 3.22E-02 | 2.55E-01 |
| FIRRE | X:131794467-131825221 | X:131804972-131805037 | MXS | 3 | 3 | ENST00000658376 | 19.66 | 7.49E-03 | 3.93E-01 |
| R3HCC1L | 10:98134707-98163292 | 10:98156093-98156147 | MES | 3 | 3 | ENST00000298999 | 19.65 | 1.46E-02 | 2.25E-02 |
| ELAC1 | 18:50974562-50984486 | 18:50984096-50984486 | A3ss | 2 | 3 | ENST00000269466 | 19.61 | 2.92E-02 | 7.23E-02 |
| TRA2A | 7:23522433-23531202 | 7:23531181-23531202 | A5ss | 3 | 3 | Ex.ENST00000392502 | 19.58 | 2.60E-02 | 3.39E-01 |
| ZFP1 | 16:75148644-75166769 | 16:75152909-75152966 | SES | 2 | 3 | ENST00000464850 | 19.58 | 3.41E-02 | 7.14E-02 |
| MORC2 | 22:30926872-30928027 | 22:30928019-30928027 | TSSIA5ss | 3 | 3 | TSS.ENST00000215862 | 19.54 | 9.71E-03 | 3.95E-02 |
| PI4KAP2 | 22:21476742-21477789 | 22:21476742-21476764 | A3ss | 3 | 3 | ENST00000360806 | 19.49 | 1.29E-04 | 5.14E-04 |
| PI4KAP2 | 22:21476161-21477789 | 22:21476161-21476764 | A3ss | 3 | 3 | ENST00000462560 | 19.49 | 1.29E-04 | 5.14E-04 |
| CHD6 | 20:41533571-41618339 | 20:41551305-41551360 | MXS | 2 | 3 | ENST00000373233 | 19.47 | 1.92E-02 | 7.15E-02 |
| PHF20L1 | 8:132798861-132803818 | 8:132799095-132799172 | SES | 3 | 3 | ENST00000315808 | 19.46 | 2.77E-02 | 4.76E-01 |
| CAMTA2 | 17:4969428-4969701 | 17:4969521-4969629 | IR | 3 | 3 | ENST00000576872 | 19.46 | 4.91E-02 | 1.15E-01 |
| HELQ | 4:83453946-83455507 | 4:83455397-83455507 | TSSIA5ss | 3 | 3 | TSS.ENST00000295488 | 19.44 | 1.52E-02 | 1.16E-01 |
| PRH1 | 12:11047189-11171421 | 12:11121010-11121180 | SES | 3 | 3 | ENST00000546265 | 19.42 | 3.13E-02 | 5.25E-02 |
| VSIG10 | 12:118095815-118103592 | 12:118102989-118103170 | SES | 3 | 3 | ENST00000536905 | 19.38 | 1.88E-02 | 3.17E-02 |
| WDR90 | 16:662220-662736 | 16:662332-662678 | IR | 3 | 3 | ENST00000546923 | 19.27 | 9.03E-04 | 1.98E-03 |
| STAG2 | X:124090765-124094017 | X:124090854-124090964 | SES | 3 | 3 | ENST00000218089 | 19.26 | 1.09E-04 | 5.75E-03 |
| TMEM161B-AS1 | 5:88270586-88410072 | 5:88270586-88282106 | A5ss | 3 | 3 | Ex.ENST00000501869 | 19.25 | 1.52E-02 | 1.43E-01 |
| AKIP1 | 11:8911672-8914825 | 11:8912453-8912533 | SES | 3 | 3 | ENST00000309357 | 19.23 | 2.12E-02 | 3.52E-02 |
| LINC01002 | 19:201814-203946 | 19:203871-203946 | A5ss | 2 | 2 | ENST00000633344 | 19.23 | 3.77E-02 | 9.68E-02 |
| LINC01002 | 19:201814-203874 | 19:203871-203874 | A5ss | 2 | 2 | ENST00000633344 | 19.23 | 3.77E-02 | 9.68E-02 |
| FOXP1 | 3:70966057-70976940 | 3:70970736-70970805 | SES | 3 | 2 | ENST00000650402 | 19.2 | 1.49E-02 | 1.02E-01 |
| PPOX | 1:161167235-161167370 | 1:161167339-161167370 | A3ss | 3 | 3 | ENST00000495483 | 19.13 | 1.60E-03 | 5.06E-03 |
| PPOX | 1:161167235-161167354 | 1:161167339-161167354 | A3ss | 3 | 3 | ENST00000495483 | 19.13 | 1.60E-03 | 5.06E-03 |
| NEPRO | 3:113008431-113019541 | 3:113017448-113017602 | MES | 3 | 3 | ENST00000314400 | 19.13 | 3.02E-02 | 1.59E-01 |
| HDHD2 | 18:47137297-47150377 | 18:47137297-47137333 | A3ss | 3 | 3 | Ex.TSS.ENST00000586546 | 19.12 | 6.00E-04 | 1.48E-03 |
| HDHD2 | 18:47136450-47150377 | 18:47136450-47137333 | A3ss | 3 | 3 | Ex.TSS.ENST00000300605 | 19.12 | 6.00E-04 | 1.48E-03 |
| CENPK | 5:65554899-65561496 | 5:65554899-65554946 | A3ss | 3 | 3 | ENST00000510354 | 19.1 | 1.96E-02 | 1.31E-01 |
| DDIAS | 11:82914852-82931731 | 11:82928777-82928938 | SES | 3 | 3 | ENST00000524921 | 19.1 | 2.05E-02 | 3.39E-02 |
| ZNF232 | 17:5109869-5111799 | 17:5111076-5111219 | SES | 3 | 3 | ENST00000570486 | 19.1 | 4.52E-02 | 1.07E-01 |

| | | | | | | | | | |
|---|---|---|---|---|---|---|---|---|---|
| FIP1L1 | 4:53399840-53425871 | 4:53414615-53414722 | SES | 3 | 3 | ENST00000337488 | 19.09 | 2.15E-02 | 1.60E-01 |
| MFSD14C | 9:96942753-96949554 | 9:96942753-96943154 | A3ss | 3 | 3 | Ex.TSS.ENST00000506067 | 19.02 | 3.77E-02 | 1.00E+00 |
| MFSD14C | 9:96918827-96949554 | 9:96918827-96943154 | A3ss | 3 | 3 | Ex.TSS.ENST00000637076 | 19.02 | 3.77E-02 | 1.00E+00 |
| FDPS | 1:155308966-155310063 | 1:155310043-155310063 | TSSIA3ss | 3 | 3 | TSS.ENST00000447866 | 19.01 | 1.48E-03 | 4.64E-03 |
| CUEDC1 | 17:57885229-57885811 | 17:57885442-57885645 | IR | 3 | 3 | ENST00000407144 | 19.01 | 3.83E-02 | 9.04E-02 |
| RNF2 | 1:185087641-185091578 | 1:185091565-185091579 | A3ss | 3 | 3 | Ex.ENST00000367509 | 18.84 | 7.85E-03 | 2.52E-02 |
| INO80B | 2:74455175-74455405 | 2:74455391-74455405 | TSSIA3ss | 3 | 3 | TSS.ENST00000409493 | 18.84 | 1.29E-02 | 6.61E-02 |
| THUMPD3-AS1 | 3:9390280-9391904 | 3:9390498-9391678 | IR | 3 | 3 | ENST00000665171 | 18.84 | 2.60E-02 | 1.77E-01 |
| NOS1AP | 1:162287437-162300647 | 1:162300633-162300647 | A3ss | 3 | 3 | ENST00000361897 | 18.77 | 1.27E-02 | 4.03E-02 |
| PNCK | X:153671499-153672200 | X:153672019-153672125 | IR | 3 | 3 | ENST00000488168 | 18.72 | 1.11E-02 | 7.71E-01 |
| RPSA | 3:39406765-39407620 | 3:39407603-39407620 | TSSIA3ss | 3 | 3 | TSS.ENST00000458478 | 18.69 | 4.90E-03 | 3.12E-02 |
| DENND10 | 10:119104198-119107967 | 10:119107948-119107967 | TSSIA3ss | 3 | 3 | TSS.ENST00000448258 | 18.66 | 4.05E-03 | 6.06E-03 |
| PLEKHH1 | 14:67575473-67575822 | 14:67575610-67575823 | A3ss | 3 | 3 | Ex.ENST00000329153 | 18.62 | 1.94E-02 | 3.63E-02 |
| ERMP1 | 9:5801329-5810010 | 9:5805027-5805217 | MES | 3 | 3 | ENST00000214893 | 18.59 | 3.18E-02 | 1.00E+00 |
| TUG1 | 22:30970182-30973587 | 22:30971770-30972923 | IR | 3 | 3 | ENST00000644027 | 18.56 | 4.14E-04 | 1.68E-03 |
| TRRAP | 7:98981961-98983263 | 7:98983240-98983264 | A3ss | 3 | 3 | Ex.ENST00000355540 | 18.5 | 2.51E-02 | 3.86E-01 |
| RPAIN | 17:5422830-5432541 | 17:5425971-5426082 | MES | 3 | 3 | ENST00000381208 | 18.49 | 3.38E-04 | 7.99E-04 |
| ZNF250 | 8:144890060-144890307 | 8:144890060-144890074 | A3ss | 3 | 3 | ENST00000292579 | 18.48 | 2.14E-02 | 3.88E-01 |
| GGT1 | 22:24603336-24607953 | 22:24603336-24603527 | TSSIA5ss | 3 | 3 | TSS.ENST00000400382 | 18.46 | 4.07E-02 | 1.63E-01 |
| GGT1 | 22:24603322-24607953 | 22:24603322-24603527 | TSSIA5ss | 3 | 3 | TSS.ENST00000400382 | 18.46 | 4.07E-02 | 1.63E-01 |
| NR2C2 | 3:14947907-15003875 | 3:14959536-14959641 | MES | 3 | 3 | ENST00000435454 | 18.45 | 1.71E-02 | 9.49E-02 |
| DDR1 | 6:30883166-30888687 | 6:30884538-30884710 | MES | 3 | 3 | ENST00000504651 | 18.44 | 3.41E-03 | 3.56E-02 |
| FAM49B | 8:129871497-129903311 | 8:129879389-129879471 | MXS | 3 | 3 | ENST00000517654 | 18.4 | 5.29E-05 | 9.06E-04 |
| PKIA | 8:78516469-78598357 | 8:78572811-78572939 | SES | 3 | 3 | ENST00000396418 | 18.4 | 4.27E-02 | 1.00E+00 |
| SZT2 | 1:43450337-43452650 | 1:43451545-43451647 | IR (overlapping regio | 3 | 3 | ENST00000460536 | 18.38 | 2.83E-03 | 9.93E-03 |
| TEX30 | 13:102769542-102773681 | 13:102770012-102770086 | SES | 3 | 3 | ENST00000376029 | 18.31 | 3.89E-03 | 6.94E-03 |
| ZNF124 | 1:247159814-247171847 | 1:247159814-247159859 | A3ss | 3 | 3 | Ex.TSS.ENST00000340684 | 18.31 | 2.65E-02 | 8.87E-02 |
| PIGH | 14:67593743-67593952 | 14:67593814-67593909 | IR | 3 | 3 | ENST00000216452 | 18.31 | 2.70E-02 | 5.03E-02 |
| RSRP1 | 1:25244225-25245149 | 1:25244225-25244498 | A3ss | 3 | 3 | ENST00000491378 | 18.31 | 3.76E-02 | 1.26E-01 |
| ZNF875 | 19:37334783-37347189 | 19:37335169-37335257 | MXS | 3 | 3 | ENST00000392153 | 18.21 | 1.29E-02 | 2.70E-02 |
| RAPGEF2 | 4:159241369-159304341 | 4:159243774-159243791 | SES | 3 | 3 | ENST00000504604 | 18.2 | 4.74E-02 | 3.35E-01 |
| TNRC6A | 16:24794720-24797489 | 16:24795907-24795939 | SES | 3 | 3 | ENST00000315183 | 18.13 | 3.13E-03 | 6.69E-03 |
| PPFIBP1 | 12:27673828-27676427 | 12:27674192-27674221 | SES | 3 | 3 | ENST00000228425 | 18.1 | 1.07E-02 | 1.84E-02 |
| PLEC | 8:143922396-143926783 | 8:143922504-143925884 | SES | 3 | 3 | ENST00000322810 | 18.07 | 1.85E-02 | 3.28E-01 |
| TMCO6 | 5:140639609-140639738 | 5:140639609-140639612 | TSSIA5ss | 3 | 3 | TSS.ENST00000252100 | 18.04 | 3.88E-02 | 3.16E-01 |
| ZNF76 | 6:35293045-35294455 | 6:35293751-35294455 | TSSIA3ss | 3 | 3 | TSS.ENST00000229405 | 17.99 | 1.09E-02 | 1.18E-01 |
| DTD2 | 14:31453345-31457282 | 14:31454839-31457283 | A5ss | 3 | 3 | Ex.TSS.ENST00000310850 | 17.98 | 3.12E-03 | 5.75E-03 |
| AL139353.1 | 14:31453345-31457028 | 14:31454839-31457029 | A5ss | 3 | 3 | Ex.TSS.ENST00000547760 | 17.98 | 3.12E-03 | 5.75E-03 |
| ZNF730 | 19:23075388-23134079 | 19:23095012-23134080 | A3ss | 3 | 3 | Ex.TSS.ENST00000593635 | 17.97 | 2.61E-02 | 6.70E-02 |
| AC074135.1 | 19:23075388-23098339 | 19:23095012-23098340 | A3ss | 3 | 3 | Ex.TSS.ENST00000600223 | 17.97 | 2.61E-02 | 6.70E-02 |
| GLT1D1 | 12:128853650-128888638 | 12:128875914-128876062 | SES | 3 | 3 | ENST00000281703 | 17.94 | 2.57E-02 | 4.38E-02 |
| TESMIN | 11:68738789-68747207 | 11:68742318-68742394 | MES | 3 | 3 | ENST00000255087 | 17.85 | 3.53E-02 | 5.77E-02 |
| NEK3 | 13:52153993-52156074 | 13:52153993-52154155 | A3ss | 3 | 3 | Ex.ENST00000551355 | 17.8 | 1.77E-04 | 3.19E-04 |
| MIER2 | 19:334543-344773 | 19:336083-336173 | SES | 3 | 3 | ENST00000264819 | 17.79 | 2.95E-03 | 7.61E-03 |
| MACF1 | 1:39463687-39468614 | 1:39465095-39465112 | SES | 3 | 3 | ENST00000289893 | 17.79 | 1.83E-02 | 6.39E-02 |
| BCL2L12 | 19:49666800-49668850 | 19:49667019-49667161 | SES | 3 | 3 | ENST00000246784 | 17.77 | 1.82E-04 | 4.92E-04 |
| VRK2 | 2:58146475-58154758 | 2:58154754-58154758 | A3ss | 3 | 3 | ENST00000417641 | 17.76 | 2.24E-02 | 1.07E-01 |
| TAF4 | 20:62064451-62065810 | 20:62065459-62065747 | IR | 3 | 3 | ENST00000252996 | 17.65 | 2.83E-02 | 1.09E-01 |
| FAM131A | 3:184336124-184341723 | 3:184337311-184337371 | SES | 3 | 3 | ENST00000340957 | 17.64 | 2.55E-03 | 1.53E-02 |
| ZNF202 | 11:123730986-123740416 | 11:123740117-123740232 | SES | 3 | 3 | ENST00000529250 | 17.56 | 2.91E-02 | 4.51E-02 |
| TMEM125 | 1:43271165-43272183 | 1:43271165-43271253 | A5ss | 3 | 3 | Ex.TSS.ENST00000456751 | 17.5 | 2.87E-03 | 1.01E-02 |
| TMEM125 | 1:43270794-43272183 | 1:43270794-43271253 | A5ss | 3 | 3 | Ex.ENST00000432792 | 17.5 | 2.87E-03 | 1.01E-02 |
| DDR1 | 6:30881126-30888687 | 6:30885625-30885680 | MES | 3 | 3 | ENST00000324771 | 17.49 | 4.57E-02 | 4.59E-01 |
| TNK2 | 3:195867259-195867360 | 3:195867259-195867264 | A3ss | 3 | 3 | ENST00000672548 | 17.45 | 4.85E-03 | 2.93E-02 |
| MIER1 | 1:66926243-66940027 | 1:66928887-66928963 | SES | 3 | 3 | ENST00000357692 | 17.43 | 2.76E-02 | 9.99E-02 |
| DGCR2 | 22:19057163-19063201 | 22:19057163-19057186 | A3ss | 3 | 3 | Ex.ENST00000263196 | 17.42 | 2.02E-03 | 7.97E-03 |
| CYB561 | 17:63434595-63435085 | 17:63434595-63434620 | A3ss | 3 | 3 | ENST00000585153 | 17.42 | 1.20E-02 | 2.85E-02 |
| LYSMD3 | 5:90519426-90525034 | 5:90519426-90519484 | TSSIA3ss | 3 | 3 | TSS.ENST00000315948 | 17.4 | 3.68E-02 | 3.48E-01 |
| SNRPN | 15:24962210-24967931 | 15:24967029-24967240 | SES | 3 | 3 | ENST00000584968 | 17.39 | 3.58E-05 | 6.78E-05 |
| ARAF | X:47561252-47562908 | X:47562682-47562909 | A3ss | 3 | 3 | Ex.TSS.ENST00000377039 | 17.39 | 3.27E-02 | 1.00E+00 |
| PXMP2 | 12:132690377-132704618 | 12:132701250-132701369 | MES | 3 | 3 | ENST00000317479 | 17.38 | 4.57E-02 | 7.80E-02 |
| FIGNL1 | 7:50444129-50447297 | 7:50447017-50447131 | IR | 3 | 3 | ENST00000613602 | 17.3 | 7.83E-03 | 1.07E-01 |
| SIDT2 | 11:117186637-117187377 | 11:117186990-117187001 | SES | 3 | 3 | ENST00000431081 | 17.29 | 2.08E-02 | 3.21E-02 |
| ZMYND8 | 20:47210898-47221313 | 20:47212642-47212725 | MES | 3 | 3 | ENST00000262975 | 17.25 | 3.49E-04 | 1.32E-03 |
| UCKL1 | 20:63941209-63944396 | 20:63941383-63944397 | A5ss | 3 | 3 | Ex.ENST00000358711 | 17.19 | 4.02E-03 | 1.55E-02 |
| UCKL1 | 20:63941209-63943652 | 20:63941383-63943653 | A5ss | 3 | 3 | Ex.ENST00000354216 | 17.19 | 4.02E-03 | 1.55E-02 |
| CD82 | 11:44618366-44619048 | 11:44618640-44618723 | SES | 3 | 2 | ENST00000227155 | 17.1 | 4.61E-02 | 7.30E-02 |
| FASTKD1 | 2:169537341-169540050 | 2:169538013-169538141 | SES | 3 | 3 | ENST00000453153 | 17.09 | 1.30E-02 | 5.52E-02 |
| FTSJ1 | X:48481716-48482408 | X:48482403-48482408 | A3ss | 3 | 3 | ENST00000348411 | 17.07 | 2.88E-03 | 3.37E-01 |
| AC119673.3 | 1:205891310-205894466 | 1:205894444-205894467 | A3ss | 3 | 3 | Ex.TSS.ENST00000652821 | 16.98 | 5.92E-03 | 1.94E-02 |
| USP28 | 11:113841769-113875444 | 11:113854258-113854335 | MES | 3 | 3 | ENST00000003302 | 16.88 | 7.07E-04 | 1.09E-03 |
| TRPT1 | 11:64225581-64226049 | 11:64225786-64225869 | SES | 3 | 3 | ENST00000317459 | 16.79 | 1.66E-03 | 2.67E-03 |
| TCEA2 | 20:64070636-64071869 | 20:64071858-64071869 | A3ss | 3 | 3 | ENST00000465433 | 16.71 | 3.51E-02 | 1.35E-01 |
| GTF3C2 | 2:27343579-27356738 | 2:27343579-27350389 | A3ss | 3 | 3 | Ex.TSS.ENST00000264720 | 16.69 | 1.90E-02 | 8.83E-02 |
| TTC14 | 3:180607648-180610974 | 3:180608811-180609629 | IR (overlapping regio | 3 | 3 | ENST00000412756 | 16.62 | 1.45E-03 | 8.34E-03 |
| AC009533.1 | 12:9294519-9297780 | 12:9294817-9294974 | SES | 3 | 3 | ENST00000539757 | 16.61 | 1.99E-02 | 3.54E-02 |
| ALMS1 | 2:73520017-73557219 | 2:73550267-73550437 | MXS | 3 | 3 | ENST00000651057 | 16.6 | 8.04E-03 | 4.09E-02 |
| KLHL24 | 3:183635794-183650295 | 3:183643480-183643542 | MXS | 3 | 3 | ENST00000242810 | 16.6 | 3.53E-02 | 2.10E-01 |
| LYRM1 | 16:20900536-20920121 | 16:20915556-20915714 | MES | 3 | 3 | ENST00000568820 | 16.53 | 5.66E-05 | 1.20E-04 |
| CBSL | 21:6458056-6460186 | 21:6459456-6459638 | SES | 3 | 3 | ENST00000624921 | 16.51 | 2.58E-02 | 1.02E-01 |
| OPLAH | 8:144052983-144053393 | 8:144053130-144053208 | IR | 3 | 3 | ENST00000527993 | 16.49 | 4.64E-02 | 8.21E-01 |
| MIER1 | 1:66926243-66928889 | 1:66928887-66928889 | A3ss | 3 | 3 | ENST00000357692 | 16.46 | 4.60E-02 | 1.66E-01 |
| BRD9 | 5:865582-870472 | 5:869245-869394 | MXS | 3 | 3 | ENST00000489816 | 16.4 | 1.20E-05 | 1.13E-04 |
| AP1B1 | 22:29328896-29330377 | 22:29329712-29329720 | SES | 3 | 3 | ENST00000357586 | 16.4 | 2.68E-04 | 1.08E-03 |
| CLASP1 | 2:121425307-121430072 | 2:121427404-121427430 | SES | 3 | 3 | ENST00000263710 | 16.38 | 4.41E-02 | 1.85E-01 |
| ZW10 | 11:113748421-113758553 | 11:113757662-113757853 | SES | 3 | 3 | ENST00000200135 | 16.37 | 3.48E-02 | 5.36E-02 |
| DICER1 | 14:95133504-95157229 | 14:95140659-95140759 | MES | 3 | 3 | ENST00000531162 | 16.36 | 3.45E-02 | 6.52E-02 |
| ZNF213-AS1 | 16:3132104-3134633 | 16:3132850-3132942 | SES | 2 | 3 | ENST00000572691 | 16.33 | 7.12E-03 | 1.55E-02 |
| SPAG9 | 17:51021366-51041500 | 17:51031681-51031722 | SES | 3 | 3 | ENST00000262013 | 16.32 | 2.15E-02 | 5.08E-02 |
| C12orf73 | 12:103951631-103965515 | 12:103953414-103953534 | MXS | 3 | 3 | ENST00000543740 | 16.29 | 1.71E-02 | 2.86E-02 |
| CPT1B | 22:50576626-50577034 | 22:50576638-50576856 | IR (overlapping regio | 3 | 3 | ENST00000460853 | 16.29 | 3.02E-02 | 1.22E-01 |
| MRPL2 | 6:43054487-43058064 | 6:43056307-43056445 | MES | 3 | 3 | ENST00000230413 | 16.24 | 4.84E-02 | 1.50E-01 |
| CEP57L1 | 6:109095576-109145218 | 6:109140421-109140675 | MXS | 3 | 3 | ENST00000518329 | 16.18 | 4.54E-02 | 4.36E-01 |
| SLC25A14 | X:130349346-130358639 | X:130350646-130350731 | MXS | 3 | 3 | ENST00000218197 | 16.15 | 1.51E-02 | 8.14E-01 |
| RTN2 | 19:45489554-45494165 | 19:45493160-45493378 | SES | 3 | 3 | ENST00000245923 | 16.13 | 3.48E-02 | 9.30E-02 |
| ZNF154 | 19:57697367-57701145 | 19:57697842-57700987 | IR (overlapping regio | 2 | 2 | ENST00000512439 | 16.11 | 2.85E-02 | 7.81E-02 |
| UBR4 | 1:19153940-19154917 | 1:19153940-19153954 | A3ss | 3 | 3 | Ex.ENST00000375254 | 16.1 | 1.57E-02 | 5.04E-02 |
| CCDC77 | 12:405565-411746 | 12:409368-409421 | SES | 3 | 3 | ENST00000239830 | 16.04 | 8.10E-03 | 1.39E-02 |
| TRDMT1 | 10:17160375-17174550 | 10:17168841-17168917 | MXS | 3 | 3 | ENST00000313936 | 15.97 | 1.37E-02 | 2.06E-02 |
| TNNT1 | 19:55145566-55146680 | 19:55146434-55146466 | SES | 3 | 3 | ENST00000291901 | 15.94 | 4.20E-03 | 1.15E-02 |
| ZFAND1 | 8:81715115-81718181 | 8:81717249-81717288 | SES | 3 | 3 | ENST00000220669 | 15.93 | 9.32E-03 | 2.29E-01 |

| | | | | | | | | | |
|---|---|---|---|---|---|---|---|---|---|
| VARS2 | 6:30916960-30917104 | 6:30917095-30917105 | A3ss | 3 | 3 | Ex.ENST00000321897 | 15.87 | 5.37E-03 | 5.44E-02 |
| LTBP3 | 11:65539882-65540244 | 11:65540013-65540153 | SES | 3 | 3 | ENST00000301873 | 15.86 | 2.78E-02 | 4.47E-02 |
| THUMPD3-AS1 | 3:9390280-9391904 | 3:9390498-9391081 | IR | 3 | 3 | ENST00000665171 | 15.86 | 3.21E-02 | 2.19E-01 |
| THUMPD3-AS1 | 3:9390280-9391904 | 3:9390498-9391426 | IR | 3 | 3 | ENST00000665171 | 15.86 | 4.04E-02 | 2.75E-01 |
| ANKRD13A | 12:110025824-110027707 | 12:110027705-110027707 | A3ss | 3 | 3 | ENST00000261739 | 15.85 | 4.78E-02 | 8.01E-02 |
| CHKA | 11:68074831-68097018 | 11:68081404-68081457 | SES | 3 | 3 | ENST00000265689 | 15.82 | 2.08E-03 | 3.40E-03 |
| BRICD5 | 16:2209979-2210281 | 16:2210052-2210125 | IR | 3 | 3 | ENST00000566018 | 15.76 | 4.19E-02 | 8.92E-02 |
| LTBP4 | 19:40619494-40623603 | 19:40622401-40622667 | MES | 3 | 3 | ENST00000204005 | 15.72 | 4.64E-02 | 1.23E-01 |
| MSTO1 | 1:155611216-155611603 | 1:155611356-155611548 | IR | 3 | 3 | ENST00000478756 | 15.7 | 2.96E-02 | 9.30E-02 |
| XRCC3 | 14:103711568-103715423 | 14:103712875-103712968 | SES | 3 | 3 | ENST00000554913 | 15.69 | 7.42E-03 | 1.34E-02 |
| ZNF468 | 19:52849214-52854257 | 19:52853872-52854126 | SES | 3 | 3 | ENST00000601980 | 15.54 | 3.69E-02 | 1.01E-01 |
| TRPT1 | 11:64225581-64226049 | 11:64225786-64226049 | TSSIA5ss | 3 | 3 | TSS.ENST00000539595 | 15.51 | 1.73E-03 | 2.77E-03 |
| NBPF15 | 1:144456806-144461380 | 1:144459366-144459483 | MXS | 3 | 3 | ENST00000579734 | 15.51 | 3.13E-02 | 8.75E-02 |
| RAB11FIP5 | 2:73076183-73088049 | 2:73079651-73081663 | SES | 3 | 3 | ENST00000486777 | 15.47 | 3.20E-02 | 1.62E-01 |
| PPOX | 1:161167235-161168992 | 1:161167339-161167486 | MES | 3 | 3 | ENST00000495483 | 15.37 | 2.62E-04 | 8.29E-04 |
| FAM222B | 17:28766708-28842681 | 17:28790378-28790425 | MXS | 3 | 3 | ENST00000581229 | 15.37 | 3.32E-03 | 7.56E-03 |
| KHNYN | 14:24429992-24430713 | 14:24429992-24430119 | TSSIA5ss | 3 | 3 | TSS.ENST00000553935 | 15.36 | 1.38E-02 | 2.53E-02 |
| WNK2 | 9:93298068-93308327 | 9:93299070-93299261 | MXS | 3 | 3 | ENST00000453718 | 15.36 | 1.88E-02 | 7.63E-01 |
| NAP1L4 | 11:2972244-2976023 | 11:2972244-2972279 | A3ss | 3 | 3 | ENST00000448187 | 15.31 | 5.89E-03 | 9.23E-03 |
| RABGAP1L | 1:174957550-174969276 | 1:174957891-174958173 | SES | 3 | 3 | ENST00000469553 | 15.31 | 6.01E-03 | 1.92E-02 |
| DEPDC5 | 22:31754980-31758545 | 22:31758527-31758545 | A3ss | 3 | 3 | ENST00000456178 | 15.28 | 1.08E-03 | 4.41E-03 |
| PIP5KL1 | 9:127925261-127927152 | 9:127925958-127925979 | SES | 3 | 3 | ENST00000495448 | 15.28 | 1.93E-02 | 5.29E-01 |
| ABCA11P | 4:426974-472574 | 4:437446-437511 | SES | 3 | 3 | ENST00000514396 | 15.26 | 1.30E-02 | 9.66E-02 |
| AC010326.1 | 19:57942648-57947441 | 19:57943934-57947442 | A5ss | 3 | 3 | Ex.ENST00000598928 | 15.24 | 5.98E-03 | 1.65E-02 |
| ATG13 | 11:46617891-46644278 | 11:46630045-46630100 | MXS | 3 | 3 | ENST00000524625 | 15.2 | 9.47E-04 | 1.50E-03 |
| ZNF227 | 19:44213245-44228445 | 19:44217791-44217852 | SES | 3 | 3 | ENST00000313040 | 15.2 | 9.23E-03 | 2.45E-02 |
| PSPH | 7:56034107-56051137 | 7:56034107-56034154 | TSSIA3ss | 3 | 3 | TSS.ENST00000395471 | 15.18 | 4.26E-02 | 5.95E-01 |
| PSPH | 7:56034104-56051137 | 7:56034104-56034154 | TSSIA3ss | 3 | 3 | TSS.ENST00000395471 | 15.18 | 4.26E-02 | 5.95E-01 |
| BRD9 | 5:865582-870472 | 5:869245-869404 | MES | 3 | 3 | ENST00000490814 | 15.17 | 4.63E-04 | 4.35E-03 |
| SEMA6C | 1:151133319-151134400 | 1:151133319-151133517 | A3ss | 3 | 3 | Ex.TSS.ENST00000341697 | 15.17 | 4.82E-03 | 1.37E-02 |
| SLC25A39 | 17:44322529-44322807 | 17:44322529-44322552 | A3ss | 3 | 3 | ENST00000377095 | 15.15 | 2.74E-04 | 6.38E-04 |
| GSN | 9:121281563-121301962 | 9:121300056-121300126 | MXS | 3 | 3 | ENST00000373808 | 15.13 | 2.04E-04 | 5.37E-03 |
| CNDP2 | 18:74497596-74499936 | 18:74499882-74499936 | A3ss | 3 | 3 | ENST00000584613 | 15.11 | 4.48E-02 | 1.12E-01 |
| MED24 | 17:40053636-40054360 | 17:40053636-40053657 | TSSIA3ss | 3 | 3 | TSS.ENST00000394127 | 15.07 | 2.53E-02 | 5.85E-02 |
| ZNF213-AS1 | 16:3132104-3134633 | 16:3132850-3133367 | SES | 2 | 3 | ENST00000652946 | 15.05 | 3.64E-03 | 7.91E-03 |
| LOXL2 | 8:23328566-23333400 | 8:23328566-23328583 | A3ss | 3 | 3 | Ex.ENST00000389131 | 14.97 | 8.71E-05 | 1.91E-03 |
| ABTB1 | 3:127676536-127677286 | 3:127677084-127677167 | IR | 3 | 3 | ENST00000464431 | 14.96 | 4.51E-02 | 2.45E-01 |
| SCAP | 3:47422537-47435007 | 3:47427447-47427667 | MES | 3 | 3 | ENST00000265565 | 14.87 | 2.90E-02 | 1.86E-01 |
| TEX30 | 13:102770087-102773681 | 13:102770087-102770100 | A3ss | 3 | 3 | Ex.TSS.ENST00000376029 | 14.84 | 5.28E-05 | 9.42E-05 |
| TEX30 | 13:102769542-102773681 | 13:102769542-102770100 | A3ss | 3 | 3 | Ex.TSS.ENST00000376021 | 14.84 | 5.28E-05 | 9.42E-05 |
| EML3 | 11:62608806-62608961 | 11:62608806-62608826 | A3ss | 3 | 3 | Ex.ENST00000278845 | 14.79 | 1.02E-03 | 1.64E-03 |
| NCOA2 | 8:70296801-70403699 | 8:70403198-70403700 | A5ss | 3 | 2 | Ex.TSS.ENST00000452400 | 14.79 | 1.00E-02 | 2.43E-01 |
| MICAL1 | 6:109445863-109446135 | 6:109445863-109445962 | A3ss | 3 | 3 | Ex.ENST00000358577 | 14.76 | 7.34E-03 | 7.05E-02 |
| PPFIBP2 | 11:7577162-7593131 | 11:7577162-7577638 | TSSIA5ss | 2 | 3 | TSS.ENST00000532926 | 14.76 | 3.06E-02 | 5.02E-02 |
| RPGR | X:38276773-38290958 | X:38287094-38287245 | MXS | 3 | 3 | ENST00000645124 | 14.74 | 4.81E-02 | 1.00E+00 |
| ST6GALNAC4 | 9:127908582-127912267 | 9:127909951-127910058 | SES | 3 | 3 | ENST00000361444 | 14.7 | 4.45E-02 | 1.00E+00 |
| MAP3K10 | 19:40205297-40209102 | 19:40205488-40209103 | A3ss | 3 | 3 | Ex.ENST00000593502 | 14.63 | 4.50E-02 | 1.19E-01 |
| MAP3K10 | 19:40205297-40205910 | 19:40205488-40205911 | A3ss | 3 | 3 | Ex.ENST00000253055 | 14.63 | 4.50E-02 | 1.19E-01 |
| PCYT1B | X:24619085-24646988 | X:24619085-24640461 | A3ss | 3 | 3 | Ex.TSS.ENST00000356768 | 14.6 | 5.07E-05 | 4.24E-03 |
| RAB4A | 1:229271371-229295847 | 1:229286468-229295848 | A3ss | 3 | 3 | Ex.TSS.ENST00000473894 | 14.59 | 2.07E-04 | 7.06E-04 |
| RAB4A | 1:229271371-229286485 | 1:229286468-229286486 | A3ss | 3 | 3 | Ex.TSS.ENST00000366690 | 14.59 | 2.07E-04 | 7.06E-04 |
| IFT122 | 3:129451999-129463559 | 3:129461228-129461304 | MES | 3 | 3 | ENST00000347300 | 14.59 | 2.50E-02 | 1.37E-01 |
| GBGT1 | 9:133162533-133163753 | 9:133162533-133162545 | A3ss | 3 | 3 | Ex.TSS.ENST00000372036 | 14.56 | 1.26E-03 | 3.92E-02 |
| AC105339.2 | 15:82752884-82757208 | 15:82754785-82756729 | IR (overlapping regio | 3 | 2 | ENST00000607520 | 14.55 | 3.73E-02 | 7.27E-02 |
| TFR2 | 7:100627264-100627604 | 7:100627492-100627576 | IR | 3 | 3 | ENST00000461176 | 14.53 | 1.19E-02 | 1.37E-01 |
| HLA-DPB2 | 6:33117221-33127817 | 6:33117221-33117587 | A5ss | 3 | 3 | Ex.ENST00000470997 | 14.52 | 2.87E-02 | 3.07E-01 |
| TBC1D1 | 4:38049860-38054198 | 4:38049860-38049898 | A5ss | 3 | 3 | Ex.ENST00000261439 | 14.49 | 3.98E-02 | 2.94E-01 |
| CHTF18 | 16:794202-795684 | 16:795132-795356 | SES | 3 | 3 | ENST00000262315 | 14.48 | 8.65E-03 | 1.92E-02 |
| FOXRED1 | 11:126274927-126275349 | 11:126275022-126275326 | IR | 3 | 3 | ENST00000524751 | 14.47 | 8.91E-03 | 1.38E-02 |
| DIS3 | 13:72778381-72781288 | 13:72780846-72781003 | SES | 3 | 3 | ENST00000545453 | 14.47 | 2.04E-02 | 3.67E-02 |
| KMT2D | 12:49022876-49024577 | 12:49024053-49024100 | SES | 3 | 3 | ENST00000526209 | 14.47 | 3.87E-02 | 6.69E-02 |
| NINL | 20:25498347-25500839 | 20:25498347-25498361 | A3ss | 3 | 3 | Ex.ENST00000278886 | 14.45 | 1.37E-02 | 5.05E-02 |
| PTGR1 | 9:111563232-111570090 | 9:111564157-111564262 | SES | 3 | 3 | ENST00000466771 | 14.44 | 3.02E-02 | 7.83E-01 |
| POLR3G | 5:90475021-90485534 | 5:90485525-90485534 | TSSIA3ss | 3 | 3 | TSS.ENST00000399107 | 14.37 | 9.44E-03 | 8.97E-02 |
| AXIN1 | 16:289608-293487 | 16:291190-291297 | SES | 3 | 3 | ENST00000262320 | 14.36 | 3.45E-03 | 7.45E-03 |
| ERGIC3 | 20:35556272-35556972 | 20:35556955-35556972 | A3ss | 3 | 3 | ENST00000451605 | 14.34 | 2.00E-04 | 7.40E-04 |
| ZNF692 | 1:248857437-248857827 | 1:248857437-248857497 | A3ss | 3 | 3 | ENST00000306601 | 14.34 | 3.19E-02 | 1.07E-01 |
| EI24 | 11:125572570-125575262 | 11:125575251-125575262 | A3ss | 3 | 3 | ENST00000618552 | 14.3 | 4.32E-07 | 6.71E-07 |
| IRF3 | 19:49664674-49665671 | 19:49665635-49665671 | A5ss | 3 | 3 | Ex.TSS.ENST00000593818 | 14.3 | 2.18E-02 | 5.89E-02 |
| CRYBG3 | 3:97943226-97944984 | 3:97943571-97944833 | IR (overlapping regio | 3 | 3 | ENST00000389622 | 14.3 | 3.69E-02 | 1.15E-01 |
| STIM1 | 11:4082983-4083262 | 11:4083242-4083263 | A3ss | 3 | 3 | Ex.ENST00000300737 | 14.29 | 6.64E-03 | 1.05E-02 |
| BCL2L12 | 19:49666800-49667021 | 19:49667019-49667021 | A3ss | 3 | 3 | ENST00000246784 | 14.27 | 2.97E-05 | 8.02E-05 |
| SLC25A29 | 14:100293378-100298841 | 14:100295626-100295697 | SES | 3 | 3 | ENST00000554224 | 14.26 | 5.15E-03 | 9.29E-03 |
| PKIB | 6:122482267-122585920 | 6:122585493-122585553 | MES | 3 | 3 | ENST00000618977 | 14.26 | 6.10E-03 | 5.92E-02 |
| CHRNA5 | 15:78593092-78595269 | 15:78593192-78593913 | IR (overlapping regio | 3 | 3 | ENST00000299565 | 14.23 | 3.81E-02 | 7.47E-02 |
| STAU2 | 8:73688814-73738283 | 8:73709032-73709177 | SES | 3 | 3 | ENST00000521845 | 14.22 | 2.01E-04 | 4.88E-03 |
| AC007731.5 | 22:20429341-20429535 | 22:20429341-20429355 | TSSIA3ss | 3 | 3 | TSS.ENST00000429594 | 14.15 | 4.24E-02 | 1.59E-01 |
| EIF2AK4 | 15:39943486-39949115 | 15:39945335-39949116 | A3ss | 3 | 3 | Ex.ENST00000263791 | 14.14 | 8.83E-03 | 1.68E-02 |
| USHBP1 | 19:17264348-17264670 | 19:17264561-17264670 | TSSIA5ss | 2 | 3 | TSS.ENST00000594190 | 14.14 | 3.49E-02 | 8.88E-02 |
| HLA-F | 6:29723928-29724189 | 6:29724173-29724189 | A3ss | 3 | 3 | ENST00000259951 | 14.13 | 1.85E-03 | 1.87E-02 |
| FOXRED1 | 11:126273080-126273335 | 11:126273316-126273335 | A3ss | 3 | 3 | ENST00000524751 | 14.13 | 4.72E-02 | 7.02E-02 |
| SLC39A13 | 11:47414862-47415159 | 11:47414910-47415038 | IR | 3 | 3 | ENST00000524886 | 14.12 | 1.96E-03 | 3.12E-03 |
| CCDC136 | 7:128801474-128804649 | 7:128801474-128801509 | A5ss | 3 | 3 | ENST00000297788 | 14.12 | 2.77E-02 | 3.33E-01 |
| PHKB | 16:47461427-47497398 | 16:47463882-47463993 | MXS | 3 | 3 | ENST00000566044 | 14.12 | 3.76E-02 | 8.18E-02 |
| PRDM15 | 21:41822038-41825959 | 21:41823318-41823449 | SES | 3 | 3 | ENST00000269844 | 14.11 | 2.79E-02 | 1.05E-01 |
| ZNF202 | 11:123740193-123740416 | 11:123740193-123740232 | A3ss | 3 | 3 | ENST00000525391 | 14.1 | 2.01E-02 | 2.99E-02 |
| ZNF202 | 11:123740190-123740416 | 11:123740190-123740232 | A3ss | 3 | 3 | ENST00000525391 | 14.1 | 2.01E-02 | 2.99E-02 |
| KCTD2 | 17:75047225-75047589 | 17:75047300-75047418 | IR | 3 | 3 | ENST00000322444 | 14.08 | 4.38E-02 | 1.03E-01 |
| TMEM94 | 17:75492560-75493102 | 17:75492790-75492928 | IR | 3 | 3 | ENST00000578624 | 14.07 | 1.05E-03 | 2.53E-03 |
| SPECC1L | 22:24321288-24321688 | 22:24321324-24321404 | IR (overlapping regio | 3 | 3 | ENST00000440893 | 14.07 | 4.50E-02 | 1.81E-01 |
| ACD | 16:67658817-67658927 | 16:67658817-67658832 | TSSIA3ss | 3 | 3 | TSS.ENST00000602945 | 14.06 | 1.82E-02 | 4.02E-02 |
| ZNF143 | 11:9474025-9474552 | 11:9474550-9474552 | A3ss | 3 | 3 | ENST00000396597 | 14.06 | 2.46E-02 | 4.11E-02 |
| WDR45 | X:49076751-49077836 | X:49077643-49077747 | SES | 3 | 3 | ENST00000322995 | 14.05 | 1.52E-02 | 1.00E+00 |
| BAIAP3 | 16:1339245-1340921 | 16:1339496-1339603 | SES | 2 | 3 | ENST00000324385 | 14.02 | 8.41E-03 | 1.66E-02 |
| TUG1 | 22:30970182-30973587 | 22:30972173-30972923 | IR | 3 | 3 | ENST00000644027 | 14.01 | 5.81E-03 | 2.36E-02 |
| CCDC24 | 1:43992356-43992639 | 1:43992388-43992522 | IR (overlapping regio | 3 | 3 | ENST00000479055 | 13.94 | 1.53E-02 | 4.76E-02 |
| ZMYM1 | 1:35079443-35093913 | 1:35093394-35093441 | SES | 3 | 3 | ENST00000373330 | 13.93 | 1.19E-02 | 4.12E-02 |
| BUB1B | 15:40200981-40202404 | 15:40202390-40202405 | A3ss | 3 | 3 | Ex.ENST00000287598 | 13.9 | 2.64E-03 | 5.01E-03 |
| STAU2 | 8:73709163-73738283 | 8:73709163-73709177 | A3ss | 3 | 3 | ENST00000521845 | 13.88 | 3.48E-03 | 8.47E-02 |
| ACOXL | 2:110933643-110987107 | 2:110963624-110963713 | SES | 3 | 3 | ENST00000389811 | 13.85 | 5.10E-03 | 2.13E-02 |
| SLC7A6 | 16:68264577-68266592 | 16:68264577-68264741 | TSSIA5ss | 3 | 3 | TSS.ENST00000568088 | 13.83 | 1.52E-02 | 3.36E-02 |

| | | | | | | | | | |
|---|---|---|---|---|---|---|---|---|---|
| UQCC1 | 20:35344110-35347272 | 20:35346748-35347163 | IR | 3 | 3 | ENST00000397554 | 13.83 | 2.11E-02 | 7.73E-02 |
| INPPL1 | 11:72228727-72228847 | 11:72228787-72228788 | IR | 3 | 3 | ENST00000298229 | 13.82 | 5.66E-03 | 9.27E-03 |
| BTBD6 | 14:105248626-105249247 | 14:105248896-105249017 | IR (overlapping regio | 3 | 3 | ENST00000463376 | 13.82 | 4.98E-02 | 9.07E-02 |
| CREM | 10:35127194-35178888 | 10:35148368-35148491 | MXS | 3 | 3 | ENST00000374726 | 13.79 | 1.30E-02 | 1.97E-02 |
| DPP7 | 9:137114379-137114462 | 9:137114379-137114382 | A3ss | 3 | 3 | ENST00000371579 | 13.72 | 2.98E-02 | 9.79E-01 |
| SNHG8 | 4:118278793-118279388 | 4:118278934-118279137 | SES | 3 | 3 | ENST00000602414 | 13.7 | 8.37E-03 | 5.82E-02 |
| CNOT9 | 2:218594108-218597080 | 2:218596299-218596334 | IR | 3 | 3 | ENST00000273064 | 13.69 | 4.12E-02 | 1.85E-01 |
| LZTS2 | 10:100997176-101002496 | 10:100997176-100997234 | A5ss | 3 | 3 | Ex.TSS.ENST00000370223 | 13.68 | 5.75E-03 | 8.56E-03 |
| TUBG2 | 17:42660708-42663376 | 17:42662973-42663052 | SES | 3 | 3 | ENST00000251412 | 13.68 | 1.43E-02 | 3.31E-02 |
| PARD3 | 10:34696420-34814875 | 10:34696420-34696440 | A3ss | 3 | 3 | Ex.TSS.ENST00000340077 | 13.66 | 2.73E-03 | 4.14E-03 |
| TM2D3 | 15:101650162-101652270 | 15:101651696-101651773 | SES | 3 | 3 | ENST00000333202 | 13.65 | 2.22E-02 | 4.20E-02 |
| FGGY | 1:59296908-59346246 | 1:59321536-59321750 | MES | 3 | 3 | ENST00000462744 | 13.63 | 1.16E-02 | 4.17E-02 |
| MTA1 | 14:105445512-105450057 | 14:105449359-105449409 | SES | 3 | 3 | ENST00000331320 | 13.62 | 5.96E-04 | 1.08E-03 |
| UQCC1 | 20:35346424-35347163 | 20:35346424-35346461 | A3ss | 3 | 3 | Ex.TSS.ENST00000491125 | 13.61 | 7.70E-03 | 2.83E-02 |
| UQCC1 | 20:35314766-35347163 | 20:35314766-35346461 | A3ss | 3 | 3 | Ex.ENST00000349714 | 13.61 | 7.70E-03 | 2.83E-02 |
| ZNF131 | 5:43174447-43175719 | 5:43174495-43175455 | IR | 3 | 3 | ENST00000306938 | 13.61 | 3.01E-02 | 2.68E-01 |
| SNHG14 | 15:25100891-25103858 | 15:25101823-25103858 | A3ss | 3 | 3 | ENST00000657285 | 13.6 | 1.57E-02 | 2.97E-02 |
| GPBP1 | 5:57236033-57246299 | 5:57237113-57237172 | SES | 3 | 3 | ENST00000424459 | 13.59 | 1.95E-02 | 1.77E-01 |
| MIB2 | 1:1625429-1626649 | 1:1625546-1626649 | A3ss | 3 | 3 | ENST00000479659 | 13.55 | 8.26E-03 | 2.62E-02 |
| MIB2 | 1:1625429-1625557 | 1:1625546-1625557 | A3ss | 3 | 3 | ENST00000355826 | 13.55 | 8.26E-03 | 2.62E-02 |
| XRCC3 | 14:103711568-103715423 | 14:103712875-103712931 | SES | 3 | 3 | ENST00000555055 | 13.52 | 5.32E-03 | 9.63E-03 |
| C12orf73 | 12:103951631-103956630 | 12:103953414-103953534 | MES | 3 | 3 | ENST00000549960 | 13.5 | 4.72E-02 | 7.88E-02 |
| C20orf96 | 20:284082-290258 | 20:289559-289676 | SES | 3 | 3 | ENST00000360321 | 13.43 | 3.70E-02 | 1.36E-01 |
| ARMC6 | 19:19033931-19042710 | 19:19033931-19034013 | TSSIA5ss | 3 | 3 | TSS.ENST00000269932 | 13.41 | 6.34E-03 | 1.62E-02 |
| NSRP1 | 17:30116864-30172541 | 17:30117302-30117332 | MES | 3 | 2 | ENST00000475652 | 13.4 | 7.29E-03 | 1.66E-02 |
| MARK2 | 11:63905044-63908876 | 11:63906088-63906114 | MES | 3 | 3 | ENST00000402010 | 13.39 | 5.57E-03 | 8.92E-03 |
| PARD3 | 10:34360260-34374873 | 10:34372498-34372536 | SES | 3 | 3 | ENST00000340077 | 13.39 | 3.10E-02 | 4.70E-02 |
| ATP11A | 13:112875942-112881875 | 13:112881816-112881875 | TSSIA3ss | 3 | 3 | TSS.ENST00000375630 | 13.37 | 3.38E-02 | 6.04E-02 |
| CSTF1 | 20:56392539-56395520 | 20:56392539-56392713 | A5ss | 3 | 3 | Ex.TSS.ENST00000613138 | 13.36 | 1.05E-02 | 4.01E-02 |
| SPART | 13:36335833-36346581 | 13:36335833-36336358 | A3ss | 3 | 3 | Ex.TSS.ENST00000451493 | 13.35 | 4.25E-03 | 7.61E-03 |
| UBE3B | 12:109507736-109510343 | 12:109509596-109509714 | SES | 3 | 3 | ENST00000342494 | 13.35 | 3.72E-02 | 6.23E-02 |
| ZNF419 | 19:57487984-57491467 | 19:57490131-57490185 | SES | 3 | 3 | ENST00000519310 | 13.34 | 3.17E-02 | 8.70E-02 |
| CMTM7 | 3:32442014-32454240 | 3:32452392-32452473 | SES | 3 | 3 | ENST00000349718 | 13.32 | 2.36E-02 | 1.48E-01 |
| THTPA | 14:23556344-23558694 | 14:23556743-23557304 | SES | 3 | 3 | ENST00000404535 | 13.3 | 4.06E-03 | 7.43E-03 |
| VPS33A | 12:122232969-122235785 | 12:122232969-122232995 | A3ss | 3 | 3 | Ex.ENST00000267199 | 13.26 | 3.58E-03 | 6.06E-03 |
| TNK2 | 3:195867017-195867360 | 3:195867169-195867264 | SES | 3 | 3 | ENST00000672548 | 13.26 | 5.53E-03 | 3.35E-02 |
| WWP1 | 8:86342931-86374029 | 8:86368939-86369031 | SES | 3 | 3 | ENST00000517970 | 13.23 | 3.48E-02 | 8.58E-01 |
| ATF1 | 12:50780239-50795908 | 12:50795898-50795909 | A3ss | 3 | 3 | Ex.ENST00000262053 | 13.16 | 1.40E-02 | 2.42E-02 |
| RABGGTB | 1:75790058-75791284 | 1:75790446-75790517 | MES | 3 | 3 | ENST00000471759 | 13.15 | 2.83E-02 | 1.02E-01 |
| RIC3 | 11:8111138-8137377 | 11:8126659-8126804 | SES | 3 | 3 | ENST00000343202 | 13.13 | 2.64E-02 | 4.36E-02 |
| LHFPL4 | 3:9552845-9553684 | 3:9552845-9552847 | A3ss | 3 | 3 | Ex.TSS.ENST00000287585 | 13.1 | 8.61E-04 | 5.88E-03 |
| LHFPL4 | 3:9552763-9553684 | 3:9552763-9552847 | TSSIA3ss | 3 | 3 | TSS.ENST00000287585 | 13.1 | 8.61E-04 | 5.88E-03 |
| SZT2 | 1:43450337-43452650 | 1:43451718-43451934 | IR (overlapping regio | 3 | 3 | ENST00000460536 | 13.06 | 1.50E-03 | 5.25E-03 |
| ECT2 | 3:172755375-172756982 | 3:172755483-172755575 | SES | 3 | 3 | ENST00000366090 | 13.06 | 2.40E-02 | 1.38E-01 |
| ARL6IP4 | 12:122981267-122981594 | 12:122981267-122981299 | TSSIA5ss | 3 | 3 | TSS.ENST00000357866 | 13.05 | 1.39E-02 | 2.35E-02 |
| THOC5 | 22:29549159-29553447 | 22:29550728-29550843 | SES | 3 | 3 | ENST00000397872 | 13.03 | 2.69E-02 | 1.08E-01 |
| USP9X | X:41151058-41152947 | X:41152933-41152948 | A3ss | 3 | 3 | Ex.ENST00000324545 | 13.03 | 4.44E-02 | 1.00E+00 |
| ZNF721 | 4:435880-472574 | 4:437446-437511 | SES | 3 | 3 | ENST00000515578 | 13.02 | 1.31E-02 | 9.75E-02 |
| CARD8 | 19:48231811-48234402 | 19:48232453-48234402 | A5ss | 2 | 3 | ENST00000522051 | 13.02 | 4.34E-02 | 1.16E-01 |
| NNT | 5:43648183-43649146 | 5:43648183-43648440 | A5ss | 3 | 2 | ENST00000657973 | 13 | 2.26E-02 | 2.04E-01 |
| GPBP1L1 | 1:45661226-45686992 | 1:45685576-45685620 | SES | 3 | 3 | ENST00000488278 | 13 | 3.72E-02 | 1.32E-01 |
| AGRN | 1:1050838-1051252 | 1:1051032-1051043 | SES | 3 | 3 | ENST00000620552 | 12.98 | 5.80E-03 | 1.62E-02 |
| HMMR | 5:163464803-163469640 | 5:163467701-163467748 | SES | 3 | 3 | ENST00000393915 | 12.98 | 7.41E-03 | 6.22E-02 |
| BABAM1 | 19:17267528-17268793 | 19:17268756-17268793 | TSSIA3ss | 3 | 3 | TSS.ENST00000359435 | 12.98 | 7.95E-03 | 2.02E-02 |
| AC084125.2 | 8:144495618-144498161 | 8:144497294-144497409 | IR (overlapping regio | 2 | 3 | ENST00000528207 | 12.98 | 4.67E-02 | 7.66E-01 |
| MAPKAP1 | 9:125484584-125543058 | 9:125506310-125506417 | SES | 3 | 3 | ENST00000265960 | 12.96 | 2.41E-02 | 6.55E-01 |
| RAD17 | 5:69371172-69373829 | 5:69371454-69371557 | MES | 3 | 3 | ENST00000354868 | 12.96 | 2.44E-02 | 7.60E-02 |
| FIP1L1 | 4:53428184-53442652 | 4:53440582-53440608 | SES | 3 | 3 | ENST00000358575 | 12.94 | 4.09E-03 | 3.05E-02 |
| RPAIN | 17:5422830-5426235 | 17:5425971-5426235 | A3ss | 3 | 3 | ENST00000570883 | 12.93 | 2.70E-02 | 6.38E-02 |
| TMEM125 | 1:43270794-43272183 | 1:43270901-43272184 | A3ss | 3 | 3 | Ex.ENST00000432792 | 12.92 | 1.17E-03 | 4.09E-03 |
| JOSD1 | 22:38700619-38700903 | 22:38700800-38700903 | A5ss | 3 | 3 | ENST00000493939 | 12.91 | 4.11E-02 | 1.69E-01 |
| JOSD1 | 22:38700619-38700861 | 22:38700800-38700861 | A5ss | 3 | 3 | Ex.ENST00000493939 | 12.91 | 4.11E-02 | 1.69E-01 |
| RBPJ | 4:26321049-26386352 | 4:26362446-26362628 | SES | 3 | 3 | ENST00000355476 | 12.9 | 1.51E-02 | 1.10E-01 |
| SLC38A9 | 5:55626660-55627890 | 5:55626660-55626673 | A3ss | 3 | 3 | Ex.TSS.ENST00000318672 | 12.89 | 2.65E-02 | 2.39E-01 |
| LTBP4 | 19:40614447-40619346 | 19:40616889-40617020 | MES | 3 | 3 | ENST00000204005 | 12.87 | 1.49E-02 | 3.95E-02 |
| GLB1L3 | 11:134292178-134293144 | 11:134292178-134292213 | A5ss | 3 | 3 | Ex.ENST00000389887 | 12.87 | 2.28E-02 | 3.56E-02 |
| WNK2 | 9:93298068-93308327 | 9:93299070-93299261 | MES | 3 | 3 | ENST00000297954 | 12.86 | 1.56E-03 | 6.34E-02 |
| UBQLN1 | 9:83665146-83669184 | 9:83666350-83669184 | A5ss | 3 | 3 | ENST00000533705 | 12.83 | 1.06E-03 | 4.19E-02 |
| CEP95 | 17:64529428-64531889 | 17:64530926-64531018 | SES | 3 | 3 | ENST00000556440 | 12.83 | 1.01E-02 | 2.42E-02 |
| PTGR2 | 14:73852117-73858815 | 14:73852117-73852120 | TSSIA5ss | 3 | 3 | TSS.ENST00000267568 | 12.82 | 2.59E-02 | 4.75E-02 |
| PTGR2 | 14:73852005-73858815 | 14:73852005-73852120 | TSSIA5ss | 3 | 3 | TSS.ENST00000267568 | 12.82 | 2.59E-02 | 4.75E-02 |
| HNRNPD | 4:82373722-82374118 | 4:82373722-82373737 | A3ss | 3 | 3 | Ex.TSS.ENST00000507010 | 12.82 | 3.80E-02 | 2.89E-01 |
| HNRNPD | 4:82373664-82374118 | 4:82373664-82373737 | A3ss | 3 | 3 | Ex.TSS.ENST00000507010 | 12.82 | 3.80E-02 | 2.89E-01 |
| ARHGEF9 | X:63724712-63754920 | X:63754315-63754676 | MES | 3 | 2 | ENST00000635967 | 12.81 | 9.30E-03 | 1.00E+00 |
| ARHGEF9 | X:63724712-63754920 | X:63754304-63754676 | MES | 3 | 2 | ENST00000637417 | 12.81 | 9.30E-03 | 1.00E+00 |
| ALG13 | X:111685104-111687886 | X:111686137-111686174 | SES | 3 | 3 | ENST00000473389 | 12.8 | 7.11E-03 | 3.20E-01 |
| BIN1 | 2:127050913-127059010 | 2:127051154-127051243 | MES | 3 | 3 | ENST00000259238 | 12.8 | 1.61E-02 | 6.77E-02 |
| PWWP3A | 19:1366382-1369264 | 19:1367160-1367220 | SES | 3 | 3 | ENST00000415183 | 12.77 | 3.99E-03 | 1.01E-02 |
| TBL1XR1 | 3:177065023-177196339 | 3:177098466-177098541 | SES | 3 | 3 | ENST00000430069 | 12.76 | 5.12E-06 | 2.94E-05 |
| PRDM2 | 1:13731118-13742004 | 1:13732779-13732855 | SES | 3 | 2 | ENST00000484063 | 12.75 | 4.29E-02 | 1.20E-01 |
| GOLGA2P7 | 15:84203021-84204425 | 15:84204177-84204331 | SES | 3 | 3 | ENST00000316967 | 12.73 | 2.03E-02 | 3.99E-02 |
| AGER | 6:32181240-32181523 | 6:32181351-32181523 | A5ss | 2 | 2 | ENST00000469940 | 12.7 | 1.53E-02 | 1.61E-01 |
| SSH3 | 11:67307437-67307845 | 11:67307549-67307737 | SES | 3 | 3 | ENST00000308127 | 12.7 | 3.08E-02 | 5.04E-02 |
| MAP4K1 | 19:38587818-38593281 | 19:38589178-38589275 | MXS | 3 | 3 | ENST00000591517 | 12.67 | 6.71E-03 | 1.76E-02 |
| RBMXL1 | 1:88984067-88992584 | 1:88988252-88988351 | SES | 3 | 3 | ENST00000399794 | 12.67 | 4.40E-02 | 1.61E-01 |
| MTO1 | 6:73480127-73480674 | 6:73480127-73480367 | A5ss | 3 | 3 | ENST00000462039 | 12.65 | 4.79E-02 | 5.39E-01 |
| AC022140.1 | 5:25422856-25445819 | 5:25422856-25435273 | A3ss | 3 | 3 | Ex.TSS.ENST00000507887 | 12.61 | 2.49E-03 | 2.19E-02 |
| PDLIM7 | 5:177491147-177491806 | 5:177491147-177491420 | TSSIA3ss | 3 | 3 | TSS.ENST00000504318 | 12.61 | 1.16E-02 | 9.84E-02 |
| MARK2 | 11:63905044-63908876 | 11:63908260-63908304 | MES | 3 | 3 | ENST00000402010 | 12.6 | 1.89E-02 | 3.03E-02 |
| SHARPIN | 8:144098995-144099276 | 8:144099081-144099205 | SES | 3 | 3 | ENST00000398712 | 12.59 | 2.41E-02 | 4.28E-01 |
| RAB17 | 2:237577383-237585997 | 2:237578004-237578155 | SES | 3 | 3 | ENST00000264601 | 12.59 | 3.78E-02 | 1.73E-01 |
| LRCH3 | 3:197854446-197865422 | 3:197858834-197858905 | SES | 3 | 3 | ENST00000334859 | 12.56 | 1.74E-02 | 1.06E-01 |
| ATRIP | 3:48460110-48460799 | 3:48460420-48460708 | IR | 3 | 3 | ENST00000320211 | 12.55 | 1.71E-02 | 1.12E-01 |
| LRRC8B | 1:89582625-89584789 | 1:89583329-89583329 | IR (overlapping regio | 3 | 3 | ENST00000330947 | 12.55 | 4.31E-02 | 1.58E-01 |
| ANKS6 | 9:98784877-98796132 | 9:98790104-98790606 | SES | 3 | 3 | ENST00000353234 | 12.52 | 1.25E-02 | 5.22E-01 |
| SLC5A2 | 16:31489123-31490769 | 16:31489339-31490103 | IR | 3 | 3 | ENST00000568188 | 12.51 | 2.72E-02 | 5.91E-02 |
| ACP6 | 1:147669830-147670524 | 1:147669950-147670392 | IR | 3 | 3 | ENST00000583509 | 12.5 | 5.23E-03 | 1.48E-02 |
| THTPA | 14:23556344-23558694 | 14:23556743-23557034 | SES | 3 | 3 | ENST00000554970 | 12.48 | 5.78E-04 | 1.06E-03 |
| KLF7 | 2:207165467-207165948 | 2:207165707-207165912 | IR | 3 | 3 | ENST00000309446 | 12.47 | 1.45E-02 | 6.50E-02 |
| SH3BP1 | 22:37639847-37641373 | 22:37641126-37641168 | SES | 3 | 3 | ENST00000417536 | 12.47 | 3.38E-02 | 1.38E-01 |
| PBRM1 | 3:52550818-52561766 | 3:52558249-52558413 | MES | 3 | 3 | ENST00000296302 | 12.46 | 4.50E-03 | 3.02E-02 |

| | | | | | | | | | |
|---|---|---|---|---|---|---|---|---|---|
| SZT2 | 1:43450337-43452650 | 1:43451848-43451934 | IR (overlapping regio | 3 | 3 | ENST00000460536 | 12.46 | 1.92E-02 | 6.73E-02 |
| GLB1L3 | 11:134293210-134312348 | 11:134311064-134311170 | MES | 3 | 3 | ENST00000431683 | 12.46 | 3.91E-02 | 6.11E-02 |
| SLC39A13 | 11:47413597-47416494 | 11:47414910-47415287 | IR | 3 | 3 | ENST00000524928 | 12.45 | 8.37E-03 | 1.33E-02 |
| SYVN1 | 11:65127281-65129915 | 11:65128489-65128562 | IR | 3 | 3 | ENST00000449943 | 12.44 | 4.20E-02 | 6.76E-02 |
| FCHSD1 | 5:141639302-141641563 | 5:141640003-141640176 | IR (overlapping regio | 3 | 3 | ENST00000435817 | 12.43 | 9.93E-03 | 8.16E-02 |
| BSCL2 | 11:62705618-62706258 | 11:62706203-62706258 | TSSIA5ss | 3 | 3 | TSS.ENST00000278893 | 12.43 | 4.64E-02 | 7.43E-02 |
| CAPN15 | 16:533999-546816 | 16:536029-536142 | SES | 3 | 3 | ENST00000219611 | 12.41 | 5.45E-03 | 1.19E-02 |
| UBQLN1 | 9:83665146-83669184 | 9:83666350-83666433 | SES | 3 | 3 | ENST00000376395 | 12.39 | 3.77E-03 | 1.48E-01 |
| SCAP | 3:47422537-47435007 | 3:47428513-47428670 | MES | 3 | 3 | ENST00000265565 | 12.34 | 1.18E-02 | 7.57E-02 |
| LETMD1 | 12:51048479-51049033 | 12:51049013-51049033 | TSSIA3ss | 3 | 3 | TSS.ENST00000548401 | 12.33 | 1.10E-03 | 1.91E-03 |
| MSANTD4 | 11:106010456-106011067 | 11:106010555-106010556 | IR | 3 | 3 | ENST00000301919 | 12.32 | 4.64E-02 | 7.12E-02 |
| SDR39U1 | 14:24441796-24442345 | 14:24442178-24442345 | TSSIA5ss | 3 | 3 | TSS.ENST00000554947 | 12.31 | 9.43E-03 | 1.73E-02 |
| ACAA2 | 18:49785352-49787290 | 18:49785352-49785372 | A3ss | 3 | 3 | Ex.ENST00000285093 | 12.3 | 1.79E-04 | 4.43E-04 |
| SNHG32 | 6:31835452-31837235 | 6:31836424-31836517 | SES | 3 | 3 | ENST00000375638 | 12.3 | 1.12E-03 | 1.18E-02 |
| TFDP1 | 13:113623287-113633119 | 13:113631623-113631744 | SES | 3 | 3 | ENST00000375370 | 12.3 | 5.82E-03 | 1.04E-02 |
| MADD | 11:47285066-47285450 | 11:47285066-47285194 | A5ss | 3 | 3 | ENST00000311027 | 12.3 | 1.07E-02 | 1.70E-02 |
| LRRC42 | 1:53946550-53951985 | 1:53947788-53947821 | MXS | 3 | 3 | ENST00000371370 | 12.29 | 4.56E-03 | 1.64E-02 |
| SCRIB | 8:143791441-143791875 | 8:143791666-143791740 | SES | 3 | 3 | ENST00000356994 | 12.27 | 3.26E-02 | 5.76E-01 |
| BAHD1 | 15:40464548-40465343 | 15:40465335-40465343 | A3ss | 3 | 3 | ENST00000416165 | 12.26 | 1.89E-02 | 3.60E-02 |
| OXNAD1 | 3:16269276-16270944 | 3:16270929-16270944 | A3ss | 3 | 3 | ENST00000435829 | 12.24 | 3.73E-03 | 2.12E-02 |
| CREB1 | 2:207596914-207605988 | 2:207597484-207599079 | IR (overlapping regio | 3 | 3 | ENST00000353267 | 12.23 | 3.96E-03 | 1.77E-02 |
| MYH14 | 19:50292390-50293232 | 19:50292390-50292895 | A5ss | 3 | 3 | Ex.ENST00000262269 | 12.21 | 2.24E-03 | 6.07E-03 |
| MTR | 1:236795738-236806143 | 1:236803428-236803642 | SES | 3 | 3 | ENST00000463959 | 12.21 | 4.32E-03 | 1.44E-02 |
| REC8 | 14:24172366-24172491 | 14:24172467-24172492 | A3ss | 3 | 3 | Ex.TSS.ENST00000557806 | 12.21 | 2.34E-02 | 4.28E-02 |
| DBP | 19:48633656-48634507 | 19:48634349-48634507 | TSSIA5ss | 2 | 3 | TSS.ENST00000593500 | 12.17 | 1.28E-02 | 3.45E-02 |
| AC091230.1 | 15:74840234-74840641 | 15:74840317-74840497 | IR | 3 | 3 | ENST00000561725 | 12.16 | 2.71E-02 | 5.27E-02 |
| ZMYM4 | 1:35325406-35370028 | 1:35361619-35361789 | MES | 3 | 3 | ENST00000314607 | 12.16 | 3.30E-02 | 1.14E-01 |
| INSIG2 | 2:118088542-118103196 | 2:118096419-118096426 | SES | 2 | 2 | ENST00000485520 | 12.14 | 3.69E-02 | 1.55E-01 |
| URI1 | 19:29942665-29985222 | 19:29971193-29971227 | SES | 3 | 3 | ENST00000392271 | 12.14 | 4.65E-02 | 1.19E-01 |
| PROSER1 | 13:39031632-39034130 | 13:39031632-39031645 | A3ss | 3 | 3 | Ex.ENST00000352251 | 12.13 | 7.24E-06 | 1.30E-05 |
| HDAC7 | 12:47795768-47796206 | 12:47795906-47796016 | SES | 3 | 3 | ENST00000080059 | 12.13 | 3.97E-02 | 6.84E-02 |
| SLC39A13 | 11:47414862-47415159 | 11:47414910-47415059 | IR | 3 | 3 | ENST00000524886 | 12.07 | 3.12E-02 | 4.96E-02 |
| SLC39A13 | 11:47414862-47415159 | 11:47414906-47415059 | IR | 3 | 3 | ENST00000524886 | 12.07 | 3.12E-02 | 4.96E-02 |
| EIF1AD | 11:66000482-66002039 | 11:66000482-66000505 | TSSIA3ss | 3 | 3 | TSS.ENST00000530462 | 12.07 | 4.28E-02 | 6.88E-02 |
| EIF1AD | 11:66000407-66002039 | 11:66000407-66000505 | TSSIA3ss | 3 | 3 | TSS.ENST00000530462 | 12.07 | 4.28E-02 | 6.91E-02 |
| GPBP1 | 5:57176401-57230845 | 5:57221376-57221396 | MES | 3 | 3 | ENST00000511209 | 12.01 | 2.38E-02 | 2.15E-01 |
| BCOR | X:40071160-40072348 | X:40071637-40071690 | SES | 3 | 3 | ENST00000342274 | 11.98 | 7.32E-03 | 7.06E-01 |
| ZNF786 | 7:149074539-149090622 | 7:149080591-149080717 | SES | 3 | 3 | ENST00000491431 | 11.98 | 4.69E-02 | 5.89E-01 |
| ABCA7 | 19:1057430-1058145 | 19:1057915-1058059 | SES | 2 | 3 | ENST00000263094 | 11.97 | 4.30E-04 | 1.08E-03 |
| SZT2 | 1:43450337-43452650 | 1:43451718-43451827 | IR (overlapping regio | 3 | 3 | ENST00000460536 | 11.96 | 4.59E-02 | 1.61E-01 |
| TRIB2 | 2:12717011-12717639 | 2:12717252-12717639 | A3ss | 3 | 3 | Ex.ENST00000483034 | 11.94 | 1.07E-03 | 4.50E-03 |
| TMUB2 | 17:44188935-44189588 | 17:44189092-44189206 | IR | 3 | 3 | ENST00000319511 | 11.93 | 4.22E-02 | 9.82E-02 |
| CRACD | 4:56179431-56272320 | 4:56196344-56196427 | SES | 3 | 2 | ENST00000646253 | 11.92 | 1.58E-02 | 1.18E-01 |
| GLT1D1 | 12:128899288-128945325 | 12:128926359-128926454 | MES | 3 | 3 | ENST00000442111 | 11.92 | 4.19E-02 | 7.14E-02 |
| MAP2K7 | 19:7911331-7911435 | 19:7911415-7911435 | A3ss | 3 | 3 | ENST00000397981 | 11.9 | 2.95E-04 | 8.20E-04 |
| PHF21A | 11:45946004-45948885 | 11:45946004-45946098 | A3ss | 3 | 3 | ENST00000534766 | 11.9 | 4.30E-02 | 6.79E-02 |
| ZNF740 | 12:53181993-53184890 | 12:53181993-53182190 | A5ss | 3 | 3 | ENST00000552593 | 11.88 | 3.62E-02 | 6.28E-02 |
| KANSL3 | 2:96595632-96601642 | 2:96598081-96598160 | SES | 3 | 3 | ENST00000670907 | 11.87 | 3.17E-02 | 1.67E-01 |
| AC091230.1 | 15:74838746-74839009 | 15:74838746-74838763 | A3ss | 3 | 3 | Ex.ENST00000440863 | 11.85 | 1.20E-04 | 2.33E-04 |
| SCAMP4 | 19:1918991-1923069 | 19:1918991-1921009 | A5ss | 3 | 3 | Ex.ENST00000316097 | 11.81 | 5.38E-03 | 1.37E-02 |
| SCAMP4 | 19:1918284-1923069 | 19:1918284-1921009 | A5ss | 3 | 3 | Ex.ENST00000409472 | 11.81 | 5.38E-03 | 1.37E-02 |
| PPP6R3 | 11:68505250-68519500 | 11:68505250-68505271 | A5ss | 3 | 3 | ENST00000529907 | 11.81 | 2.02E-02 | 3.30E-02 |
| PIGQ | 16:579181-580857 | 16:580183-580263 | SES | 3 | 3 | ENST00000026218 | 11.8 | 1.43E-02 | 3.01E-02 |
| ABHD18 | 4:127984424-128008919 | 4:127989721-127989821 | MXS | 3 | 3 | ENST00000398965 | 11.8 | 2.64E-02 | 8.22E-02 |
| TM2D3 | 15:101650162-101652270 | 15:101651696-101652270 | TSSIA5ss | 3 | 3 | TSS.ENST00000560013 | 11.8 | 2.95E-02 | 5.57E-02 |
| WDR34 | 9:128635768-128636280 | 9:128635768-128635784 | A3ss | 3 | 3 | Ex.ENST00000372715 | 11.78 | 6.61E-03 | 1.89E-01 |
| OPA1 | 3:193618937-193631611 | 3:193626092-193626202 | MXS | 3 | 3 | ENST00000361150 | 11.77 | 2.65E-02 | 1.60E-01 |
| NSMCE2 | 8:125091959-125102050 | 8:125102014-125102050 | TSSIA3ss | 3 | 3 | TSS.ENST00000517532 | 11.75 | 3.91E-02 | 6.56E-01 |
| RAD51D | 17:35106482-35119110 | 17:35107366-35107447 | MXS | 3 | 3 | ENST00000588594 | 11.74 | 2.31E-02 | 5.31E-02 |
| ANKRD17 | 4:73135263-73141740 | 4:73139531-73140283 | SES | 3 | 3 | ENST00000558247 | 11.74 | 3.03E-02 | 2.28E-01 |
| RPARP-AS1 | 10:102450331-102455336 | 10:102452090-102455337 | A3ss | 3 | 3 | Ex.ENST00000473970 | 11.73 | 1.18E-02 | 1.76E-02 |
| TBC1D5 | 3:17508606-17623848 | 3:17622648-17622765 | MES | 3 | 3 | ENST00000481396 | 11.73 | 1.81E-02 | 1.04E-01 |
| CREM | 10:35127194-35206894 | 10:35148368-35148491 | MES | 3 | 3 | ENST00000495960 | 11.7 | 1.34E-03 | 2.03E-03 |
| PCNX2 | 1:233259345-233261284 | 1:233259345-233259774 | A3ss | 3 | 3 | Ex.ENST00000258229 | 11.69 | 8.51E-04 | 2.90E-03 |
| FAM234A | 16:234858-249548 | 16:248244-248397 | MXS | 3 | 3 | ENST00000420046 | 11.66 | 7.63E-04 | 1.63E-03 |
| SUPT20H | 13:37047937-37051487 | 13:37048564-37048599 | SES | 3 | 3 | ENST00000350612 | 11.65 | 1.83E-02 | 3.28E-02 |
| WDR45 | X:49075454-49075544 | X:49075454-49075465 | A3ss | 3 | 3 | ENST00000322995 | 11.65 | 3.61E-02 | 1.00E+00 |
| MFF | 2:227340381-227355676 | 2:227352514-227352573 | MXS | 3 | 3 | ENST00000349901 | 11.61 | 1.82E-02 | 8.22E-02 |
| BRD9 | 5:886705-887360 | 5:886705-886707 | A3ss | 3 | 3 | Ex.ENST00000466684 | 11.61 | 3.84E-02 | 3.64E-01 |
| PLXNB1 | 3:48416452-48417910 | 3:48416452-48416469 | A3ss | 3 | 3 | Ex.ENST00000296440 | 11.6 | 3.14E-03 | 2.07E-02 |
| TBC1D5 | 3:17508606-17623848 | 3:17622648-17622765 | MES | 3 | 3 | ENST00000497531 | 11.59 | 1.83E-02 | 1.05E-01 |
| TBC1D5 | 3:17508606-17623848 | 3:17622648-17622765 | MES | 3 | 3 | ENST00000444471 | 11.59 | 1.83E-02 | 1.05E-01 |
| STAG3L5P-PVRIG2P-PILRB | 7:100358367-100358932 | 7:100358690-100358932 | A3ss | 3 | 3 | ENST00000310771 | 11.58 | 1.30E-02 | 1.48E-01 |
| STAG3L5P-PVRIG2P-PILRB | 7:100358367-100358898 | 7:100358690-100358898 | A3ss | 3 | 3 | ENST00000310771 | 11.58 | 1.30E-02 | 1.48E-01 |
| AC005261.1 | 19:57351326-57351697 | 19:57351418-57351491 | IR (overlapping regio | 3 | 3 | ENST00000443917 | 11.58 | 4.18E-02 | 8.75E-02 |
| FAM222B | 17:28766708-28842681 | 17:28834293-28834326 | MES | 3 | 3 | ENST00000583307 | 11.56 | 3.78E-02 | 7.92E-02 |
| POLG | 15:89327064-89327166 | 15:89327064-89327079 | A3ss | 3 | 3 | ENST00000636774 | 11.55 | 3.32E-03 | 6.53E-03 |
| TRIM39 | 6:30326996-30327651 | 6:30327647-30327651 | TSSIA3ss | 3 | 3 | TSS.ENST00000428404 | 11.53 | 2.07E-02 | 2.08E-01 |
| MORC4 | X:106941464-106942221 | X:106941633-106941975 | IR | 3 | 2 | ENST00000478924 | 11.49 | 3.88E-02 | 1.00E+00 |
| CARMIL2 | 16:67656646-67656833 | 16:67656801-67656833 | A3ss | 3 | 3 | ENST00000334583 | 11.48 | 1.11E-03 | 2.46E-03 |
| PICALM | 11:85983974-85996825 | 11:85990250-85990399 | SES | 3 | 3 | ENST00000356360 | 11.38 | 2.32E-02 | 3.84E-02 |
| RSPRY1 | 16:57186786-57204503 | 16:57186786-57187050 | TSSIA5ss | 3 | 3 | TSS.ENST00000537866 | 11.38 | 4.73E-02 | 1.04E-01 |
| RSPRY1 | 16:57186452-57204503 | 16:57186452-57187050 | TSSIA5ss | 3 | 3 | TSS.ENST00000537866 | 11.38 | 4.73E-02 | 1.04E-01 |
| RSPRY1 | 16:57186443-57204503 | 16:57186443-57187050 | TSSIA5ss | 3 | 3 | TSS.ENST00000537866 | 11.38 | 4.73E-02 | 1.04E-01 |
| RSPRY1 | 16:57186430-57204503 | 16:57186430-57187050 | TSSIA5ss | 3 | 3 | TSS.ENST00000537866 | 11.38 | 4.73E-02 | 1.04E-01 |
| FBXW5 | 9:136942944-136943348 | 9:136942944-136942997 | A3ss | 3 | 3 | Ex.ENST00000325285 | 11.37 | 3.92E-04 | 1.27E-02 |
| CLYBL | 13:99606758-99772823 | 13:99657893-99772824 | A3ss | 3 | 3 | Ex.TSS.ENST00000339105 | 11.37 | 4.46E-02 | 8.05E-02 |
| CHD8 | 14:21430801-21431858 | 14:21431725-21431810 | IR | 3 | 3 | ENST00000557364 | 11.36 | 1.04E-02 | 1.90E-02 |
| ZFYVE19 | 15:40810758-40813337 | 15:40812699-40812902 | SES | 3 | 3 | ENST00000336455 | 11.34 | 3.12E-02 | 5.71E-02 |
| CIZ1 | 9:128180800-128185546 | 9:128180800-128180814 | A3ss | 3 | 3 | ENST00000277465 | 11.33 | 1.46E-02 | 4.02E-01 |
| LYRM1 | 16:20900536-20920121 | 16:20915556-20915714 | MES | 3 | 3 | ENST00000569023 | 11.33 | 2.89E-02 | 6.15E-02 |
| HAUS4 | 14:22950411-22952327 | 14:22951555-22951689 | SES | 3 | 3 | ENST00000206474 | 11.32 | 2.98E-02 | 5.43E-02 |
| AC138969.1 | 16:16318080-16318922 | 16:16318551-16318734 | SES | 3 | 3 | ENST00000536260 | 11.3 | 4.84E-03 | 1.02E-02 |
| STAU1 | 20:49124588-49135832 | 20:49124588-49124605 | A3ss | 3 | 3 | ENST00000371802 | 11.27 | 3.28E-02 | 1.25E-01 |
| NSUN5P1 | 7:75415235-75415980 | 7:75415592-75415770 | SES | 3 | 3 | ENST00000428392 | 11.25 | 1.80E-02 | 2.58E-01 |
| SETD5 | 3:9434486-9434823 | 3:9434590-9434630 | SES | 3 | 3 | ENST00000490791 | 11.25 | 3.21E-02 | 2.20E-01 |
| METTL5 | 2:169812507-169815476 | 2:169812507-169812524 | A3ss | 3 | 3 | ENST00000442181 | 11.19 | 5.16E-03 | 2.20E-02 |
| FAM131A | 3:184336124-184342065 | 3:184337311-184337371 | MXS | 3 | 3 | ENST00000418281 | 11.18 | 1.05E-03 | 6.33E-03 |
| BACE1-AS | 11:117288453-117291771 | 11:117291050-117291711 | IR (overlapping regio | 3 | 3 | ENST00000649580 | 11.18 | 7.56E-03 | 1.17E-02 |
| GUCD1 | 22:24549002-24555614 | 22:24554949-24555136 | SES | 3 | 3 | ENST00000407973 | 11.17 | 4.38E-02 | 1.75E-01 |
| NSRP1 | 17:30117333-30172541 | 17:30118080-30118173 | MES | 3 | 2 | ENST00000467446 | 11.16 | 2.17E-03 | 4.94E-03 |

| | | | | | | | | | |
|---|---|---|---|---|---|---|---|---|---|
| GRAMD1B | 11:123560528-123577366 | 11:123560528-123560652 | TSSIA5ss | 3 | 3 | TSS.ENST00000534764 | 11.14 | 6.29E-03 | 9.75E-03 |
| ANKRD13A | 12:110025824-110028511 | 12:110027705-110027766 | SES | 3 | 3 | ENST00000261739 | 11.11 | 1.86E-02 | 3.11E-02 |
| STRN3 | 14:30913658-30935162 | 14:30918966-30919106 | MES | 3 | 3 | ENST00000357479 | 11.11 | 2.47E-02 | 4.56E-02 |
| RAD51D | 17:35106482-35119110 | 17:35107366-35107447 | MES | 3 | 3 | ENST00000460118 | 11.1 | 3.21E-02 | 7.38E-02 |
| THTPA | 14:23556344-23558694 | 14:23556743-23557039 | SES | 3 | 3 | ENST00000556015 | 11.09 | 3.49E-03 | 6.38E-03 |
| AP1G2 | 14:23561636-23562287 | 14:23561962-23562066 | SES | 3 | 3 | ENST00000308724 | 11.09 | 2.63E-02 | 4.80E-02 |
| RETREG2 | 2:219179764-219180197 | 2:219180167-219180198 | A3ss | 3 | 3 | Ex.ENST00000273048 | 11.09 | 2.71E-02 | 1.22E-01 |
| ZNF217 | 20:53583169-53597630 | 20:53594107-53597631 | A5ss | 3 | 3 | Ex.TSS.ENST00000540425 | 11.09 | 4.13E-02 | 1.57E-01 |
| AC008105.1 | 17:45238028-45239241 | 17:45238639-45238954 | IR (overlapping regio | 3 | 3 | ENST00000587534 | 11.08 | 2.46E-02 | 5.73E-02 |
| ANKFY1 | 17:4182350-4183397 | 17:4182350-4182352 | A3ss | 3 | 3 | ENST00000572412 | 11.07 | 1.19E-03 | 2.75E-03 |
| SLC25A23 | 19:6444302-6454322 | 19:6444302-6452479 | A3ss | 3 | 3 | Ex.ENST00000598908 | 11.07 | 1.10E-02 | 3.05E-02 |
| XRCC3 | 14:103712932-103715423 | 14:103712932-103712941 | A3ss | 3 | 3 | Ex.TSS.ENST00000555055 | 11.05 | 9.25E-04 | 1.67E-03 |
| XRCC3 | 14:103711662-103715423 | 14:103711662-103712941 | A3ss | 3 | 3 | Ex.TSS.ENST00000553332 | 11.05 | 9.25E-04 | 1.67E-03 |
| XRCC3 | 14:103711568-103715423 | 14:103711568-103712941 | A3ss | 3 | 3 | Ex.TSS.ENST00000352127 | 11.05 | 9.25E-04 | 1.67E-03 |
| DGKA | 12:55940082-55940722 | 12:55940434-55940623 | IR | 3 | 3 | ENST00000552652 | 11.05 | 9.02E-03 | 1.57E-02 |
| CPZ | 4:8592922-8601122 | 4:8599453-8599485 | MXS | 3 | 3 | ENST00000360986 | 11.04 | 1.75E-02 | 1.33E-01 |
| HDHD2 | 18:47136450-47150377 | 18:47137235-47137338 | MXS | 3 | 3 | ENST00000590815 | 11.01 | 1.68E-02 | 4.14E-02 |
| RRN3 | 16:15086259-15091314 | 16:15086365-15086454 | SES | 3 | 3 | ENST00000198767 | 11.01 | 2.83E-02 | 5.97E-02 |
| RPP14 | 3:58306418-58310308 | 3:58306418-58306651 | TSSIA5ss | 3 | 3 | TSS.ENST00000445193 | 11 | 2.73E-02 | 1.84E-01 |
| ADGRL2 | 1:81800513-81836884 | 1:81800513-81802229 | A5ss | 3 | 3 | Ex.TSS.ENST00000319517 | 10.99 | 1.12E-02 | 4.07E-02 |
| RBPJ | 4:26320816-26386352 | 4:26362446-26362628 | SES | 3 | 3 | ENST00000514807 | 10.98 | 1.63E-03 | 1.19E-02 |
| PLCG2 | 16:81956695-81956879 | 16:81956848-81956848 | IR | 3 | 3 | ENST00000564138 | 10.98 | 3.76E-02 | 8.35E-02 |
| PRICKLE3 | X:49179807-49183733 | X:49179807-49179836 | TSSIA3ss | 3 | 3 | TSS.ENST00000376317 | 10.93 | 4.69E-02 | 1.00E+00 |
| MRPL55 | 1:228109022-228109260 | 1:228109149-228109260 | TSSIA5ss | 3 | 3 | TSS.ENST00000336520 | 10.89 | 3.90E-04 | 1.32E-03 |
| AP2A1 | 19:49802142-49802948 | 19:49802534-49802599 | SES | 3 | 3 | ENST00000359032 | 10.89 | 5.51E-03 | 1.49E-02 |
| MIR302CHG | 4:112646969-112648653 | 4:112647415-112647467 | SES | 3 | 3 | ENST00000509938 | 10.88 | 5.41E-04 | 3.75E-03 |
| SYVN1 | 11:65133253-65133469 | 11:65133253-65133269 | A3ss | 3 | 3 | Ex.ENST00000294256 | 10.88 | 6.35E-03 | 1.02E-02 |
| VPS51 | 11:65109924-65110481 | 11:65110463-65110482 | A3ss | 3 | 3 | Ex.ENST00000279281 | 10.86 | 1.01E-02 | 1.63E-02 |
| PXN | 12:120215274-120221622 | 12:120215274-120215417 | A3ss | 3 | 3 | ENST00000267257 | 10.8 | 2.18E-02 | 3.69E-02 |
| ARSJ | 4:113978437-113979647 | 4:113979093-113979455 | IR | 3 | 3 | ENST00000315366 | 10.8 | 4.83E-02 | 3.36E-01 |
| LTBP4 | 19:40614447-40619346 | 19:40616889-40617020 | MXS | 3 | 3 | ENST00000593463 | 10.79 | 3.47E-02 | 9.21E-02 |
| SYNGAP1 | 6:33447933-33451759 | 6:33447933-33447933 | TSSIA5ss | 3 | 3 | TSS.ENST00000428982 | 10.79 | 4.50E-02 | 4.83E-01 |
| DPH5 | 1:101001545-101021640 | 1:101001545-101001587 | A3ss | 3 | 3 | ENST00000477293 | 10.76 | 7.43E-03 | 2.06E-02 |
| XYLT2 | 17:50356774-50358206 | 17:50357118-50357252 | SES | 3 | 3 | ENST00000511654 | 10.75 | 1.03E-03 | 2.43E-03 |
| ABHD12 | 20:25295031-25302218 | 20:25300699-25300884 | SES | 3 | 3 | ENST00000465694 | 10.74 | 6.01E-03 | 2.22E-02 |
| RAB40C | 16:590066-590433 | 16:590120-590260 | IR | 3 | 3 | ENST00000248139 | 10.73 | 1.16E-02 | 2.54E-02 |
| MIS18BP1 | 14:45206171-45210379 | 14:45206171-45206872 | A3ss | 3 | 3 | Ex.ENST00000310806 | 10.72 | 3.22E-02 | 5.96E-02 |
| LTBP4 | 19:40614447-40619346 | 19:40616889-40617042 | MXS | 3 | 3 | ENST00000594448 | 10.71 | 1.19E-02 | 3.16E-02 |
| CENPK | 5:65554899-65561462 | 5:65554899-65554946 | A3ss | 3 | 3 | ENST00000396679 | 10.71 | 4.80E-02 | 1.49E-01 |
| PIGQ | 16:582883-584109 | 16:583667-583753 | IR | 3 | 3 | ENST00000026218 | 10.7 | 4.15E-03 | 9.12E-03 |
| PXN | 12:120215274-120215559 | 12:120215274-120215417 | A3ss | 3 | 3 | ENST00000637624 | 10.7 | 1.72E-02 | 2.91E-02 |
| LAMTOR4 | 7:100149034-100149498 | 7:100149034-100149053 | A5ss | 3 | 3 | Ex.TSS.ENST00000468582 | 10.7 | 3.82E-02 | 4.37E-01 |
| SAFB2 | 19:5595498-5598792 | 19:5595498-5595510 | A3ss | 3 | 3 | Ex.ENST00000252542 | 10.68 | 1.20E-04 | 3.29E-04 |
| B3GALNT2 | 1:235494829-235504140 | 1:235496186-235496308 | SES | 3 | 3 | ENST00000313984 | 10.67 | 6.40E-03 | 2.13E-02 |
| HDHD2 | 18:47136450-47150377 | 18:47137235-47137296 | MXS | 3 | 3 | ENST00000586546 | 10.67 | 9.73E-03 | 2.41E-02 |
| PILRB | 7:100355143-100358015 | 7:100355241-100356393 | IR (overlapping regio | 3 | 3 | ENST00000493091 | 10.65 | 9.81E-03 | 1.13E-01 |
| NELFE | 6:31954878-31955058 | 6:31954878-31954892 | A3ss | 3 | 3 | ENST00000375425 | 10.65 | 1.29E-02 | 1.36E-01 |
| SMURF1 | 7:99033113-99035514 | 7:99033113-99033121 | A3ss | 3 | 3 | Ex.ENST00000361125 | 10.63 | 3.63E-02 | 5.57E-01 |
| SDR39U1 | 14:24440927-24442345 | 14:24442178-24442260 | MXS | 3 | 3 | ENST00000538105 | 10.62 | 7.14E-03 | 1.31E-02 |
| LARP4 | 12:50427910-50428934 | 12:50428917-50428934 | A3ss | 3 | 3 | ENST00000429001 | 10.61 | 8.95E-03 | 1.54E-02 |
| AC009533.1 | 12:9298868-9299264 | 12:9299260-9299264 | A3ss | 3 | 3 | Ex.ENST00000539757 | 10.61 | 4.05E-02 | 7.22E-02 |
| AL355987.1 | 9:136740049-136742786 | 9:136741881-136742020 | MES | 3 | 3 | ENST00000435202 | 10.59 | 8.40E-03 | 2.69E-01 |
| SORBS2 | 4:185678823-185734098 | 4:185690562-185690611 | SES | 3 | 3 | ENST00000430503 | 10.58 | 8.04E-03 | 5.75E-02 |
| TRIM32 | 9:116687382-116697661 | 9:116697373-116697433 | SES | 3 | 3 | ENST00000411410 | 10.58 | 2.28E-02 | 5.95E-01 |
| ZSCAN25 | 7:99617017-99618524 | 7:99618135-99618209 | SES | 3 | 3 | ENST00000431485 | 10.57 | 4.59E-02 | 7.11E-01 |
| AL138720.1 | 6:14790099-14997415 | 6:14790099-14876270 | A3ss | 3 | 3 | Ex.ENST00000629853 | 10.56 | 2.09E-04 | 2.08E-03 |
| HMGN1 | 21:39345830-39347292 | 21:39345959-39347113 | IR | 3 | 3 | ENST00000492280 | 10.56 | 1.14E-02 | 4.43E-02 |
| IDI1 | 10:1043394-1048863 | 10:1043394-1044171 | TSSIA3ss | 3 | 3 | TSS.ENST00000482091 | 10.55 | 1.54E-02 | 2.29E-02 |
| OGFOD3 | 17:82392535-82398195 | 17:82394331-82394543 | SES | 3 | 3 | ENST00000329197 | 10.54 | 6.12E-03 | 1.49E-02 |
| PHTF1 | 1:113713439-113726417 | 1:113724759-113724893 | SES | 3 | 3 | ENST00000357783 | 10.53 | 3.70E-02 | 1.04E-01 |
| PLA2G12A | 4:109717714-109729601 | 4:109718683-109718759 | SES | 3 | 3 | ENST00000243501 | 10.49 | 7.87E-03 | 5.45E-02 |
| ZFAS1 | 20:49279209-49289044 | 20:49280485-49280570 | MXS | 3 | 3 | ENST00000326677 | 10.49 | 2.69E-02 | 1.02E-01 |
| TMEM145 | 19:42314347-42314612 | 19:42314421-42314528 | SES | 3 | 2 | ENST00000598766 | 10.47 | 2.21E-02 | 5.87E-02 |
| DDR1 | 6:30894672-30896620 | 6:30895404-30895514 | SES | 3 | 3 | ENST00000324771 | 10.46 | 3.04E-02 | 3.18E-01 |
| ZSCAN21 | 7:100063788-100065040 | 7:100063980-100064082 | IR | 3 | 3 | ENST00000292450 | 10.45 | 2.67E-02 | 3.08E-01 |
| PVT1 | 8:128010445-128096517 | 8:128070160-128070272 | MXS | 3 | 3 | ENST00000523190 | 10.44 | 2.35E-04 | 4.00E-03 |
| CKAP2 | 13:52455627-52456522 | 13:52456503-52456523 | A3ss | 3 | 3 | Ex.TSS.ENST00000258607 | 10.42 | 2.44E-02 | 4.38E-02 |
| SNHG8 | 4:118278793-118279388 | 4:118279107-118279137 | SES | 3 | 3 | ENST00000602483 | 10.42 | 2.64E-02 | 1.84E-01 |
| RPAIN | 17:5422830-5432541 | 17:5425971-5428211 | SES | 3 | 3 | ENST00000573126 | 10.42 | 3.68E-02 | 8.68E-02 |
| SLC25A26 | 3:66221128-66243202 | 3:66236544-66236700 | SES | 3 | 3 | ENST00000354883 | 10.41 | 1.38E-02 | 9.34E-02 |
| U2AF1L4 | 19:35742804-35743808 | 19:35743155-35743345 | SES | 3 | 3 | ENST00000587987 | 10.41 | 1.52E-02 | 3.98E-02 |
| CREM | 10:35127194-35206894 | 10:35148368-35148491 | MXS | 3 | 3 | ENST00000490460 | 10.4 | 9.96E-03 | 1.51E-02 |
| TIA1 | 2:70224630-70227734 | 2:70225328-70225393 | SES | 3 | 3 | ENST00000474809 | 10.38 | 7.45E-03 | 3.76E-02 |
| RCAN3 | 1:24503118-24514313 | 1:24503118-24503150 | TSSIA5ss | 3 | 3 | TSS.ENST00000374395 | 10.37 | 2.74E-02 | 9.17E-02 |
| FAM222B | 17:28766708-28812208 | 17:28766708-28790425 | A3ss | 3 | 3 | Ex.TSS.ENST00000452648 | 10.36 | 5.43E-03 | 1.24E-02 |
| KCTD10 | 12:109458079-109460635 | 12:109458079-109458081 | A3ss | 3 | 3 | ENST00000537165 | 10.34 | 3.11E-03 | 5.21E-03 |
| TAF1C | 16:84184900-84186900 | 16:84184900-84184989 | TSSIA3ss | 3 | 3 | TSS.ENST00000564345 | 10.34 | 7.32E-03 | 1.63E-02 |
| DDR1 | 6:30881126-30888687 | 6:30884538-30884710 | MES | 3 | 3 | ENST00000324771 | 10.33 | 1.40E-02 | 1.46E-01 |
| ARID4A | 14:58365623-58366882 | 14:58366024-58366068 | SES | 2 | 3 | ENST00000395168 | 10.33 | 3.33E-02 | 6.19E-02 |
| MIB2 | 1:1628273-1628722 | 1:1628400-1628488 | IR | 3 | 3 | ENST00000507229 | 10.3 | 2.89E-02 | 9.17E-02 |
| PHF12 | 17:28911238-28912481 | 17:28911238-28911384 | A3ss | 3 | 3 | ENST00000579563 | 10.3 | 3.81E-02 | 7.98E-02 |
| THOC2 | X:123619437-123620906 | X:123619437-123619517 | A3ss | 3 | 3 | ENST00000441692 | 10.25 | 3.72E-02 | 1.00E+00 |
| ANKRD13D | 11:67301282-67301823 | 11:67301652-67301731 | IR | 3 | 3 | ENST00000507915 | 10.24 | 1.80E-02 | 2.95E-02 |
| FMNL3 | 12:49636499-49645903 | 12:49642979-49643222 | IR (overlapping regio | 3 | 3 | ENST00000335154 | 10.22 | 2.61E-02 | 4.50E-02 |
| FCHSD1 | 5:141639302-141641563 | 5:141640446-141640670 | IR (overlapping regio | 3 | 3 | ENST00000435817 | 10.2 | 2.63E-03 | 2.16E-02 |
| TMEM108 | 3:133046021-133229269 | 3:133229266-133229269 | A3ss | 3 | 3 | ENST00000321871 | 10.2 | 3.76E-02 | 2.06E-01 |
| ANO8 | 19:17325387-17327234 | 19:17325387-17325414 | A3ss | 3 | 3 | Ex.ENST00000159087 | 10.18 | 3.82E-04 | 9.72E-04 |
| RHNO1 | 12:2877283-2885282 | 12:2885263-2885282 | TSSIA3ss | 3 | 3 | TSS.ENST00000538636 | 10.16 | 2.03E-03 | 3.49E-03 |
| IMMP2L | 7:110663722-110963499 | 7:110886593-110886695 | MXS | 3 | 3 | ENST00000331762 | 10.16 | 2.14E-02 | 2.53E-01 |
| CHD1L | 1:147242831-147265931 | 1:147254870-147254976 | MES | 3 | 3 | ENST00000369258 | 10.15 | 5.71E-04 | 1.62E-03 |
| ECHDC2 | 1:52897437-52899224 | 1:52897485-52899173 | IR | 3 | 3 | ENST00000479183 | 10.13 | 2.05E-02 | 7.33E-02 |
| C11orf95 | 11:63763870-63765806 | 11:63763870-63764598 | A3ss | 3 | 3 | Ex.ENST00000338498 | 10.12 | 7.05E-03 | 1.13E-02 |
| RTF1 | 15:41470393-41471171 | 15:41471152-41471172 | A3ss | 3 | 3 | Ex.ENST00000389629 | 10.11 | 2.76E-03 | 5.26E-03 |
| DBF4B | 17:44708840-44722879 | 17:44709304-44709366 | SES | 3 | 3 | ENST00000315005 | 10.09 | 1.14E-02 | 2.66E-02 |
| ACOT9 | X:23733218-23743124 | X:23735919-23736016 | MES | 3 | 3 | ENST00000379303 | 10.08 | 4.04E-03 | 3.36E-01 |
| CBLL1 | 7:107749048-107753410 | 7:107752545-107752617 | SES | 3 | 3 | ENST00000432748 | 10.07 | 1.67E-02 | 1.97E-01 |
| TP73-AS1 | 1:3740388-3742803 | 1:3740388-3740402 | A3ss | 3 | 3 | Ex.ENST00000418088 | 10.07 | 1.69E-02 | 5.86E-02 |
| LYRM1 | 16:20900536-20920121 | 16:20915556-20915714 | MXS | 3 | 3 | ENST00000439021 | 10.07 | 4.78E-02 | 1.02E-01 |
| GMPPA | 2:219505556-219505714 | 2:219505695-219505714 | A3ss | 3 | 3 | ENST00000443704 | 10.05 | 8.59E-03 | 3.88E-02 |
| CARMIL2 | 16:67656646-67657238 | 16:67656801-67656881 | SES | 3 | 3 | ENST00000334583 | 10.05 | 2.04E-02 | 4.49E-02 |
| CCDC106 | 19:55648278-55649001 | 19:55648946-55649001 | TSSIA3ss | 3 | 3 | TSS.ENST00000588740 | 10.05 | 3.12E-02 | 8.57E-02 |

| | | | | | | | | | |
|---|---|---|---|---|---|---|---|---|---|
| CLCN6 | 1:11840143-11843143 | 1:11840372-11841057 | IR | 3 | 3 | ENST00000312413 | 10.04 | 1.52E-02 | 4.27E-02 |
| DICER1 | 14:95133504-95157229 | 14:95141683-95141748 | MES | 3 | 3 | ENST00000529720 | 10.03 | 4.37E-02 | 8.00E-02 |
| RAB34 | 17:28717292-28718121 | 17:28717292-28717425 | TSSIA3ss | 3 | 3 | TSS.ENST00000301043 | 10.02 | 4.84E-02 | 1.01E-01 |
| PIGQ | 16:579181-580857 | 16:580183-580857 | A3ss | 3 | 3 | ENST00000480424 | 10.01 | 2.14E-02 | 4.49E-02 |
| ZNF738 | 19:21375965-21378628 | 19:21377389-21378628 | TSSIA3ss | 3 | 3 | TSS.ENST00000311015 | 10 | 1.41E-03 | 3.62E-03 |
| MAGI1 | 3:65383624-65391141 | 3:65387143-65387226 | SES | 3 | 3 | ENST00000330909 | 10 | 2.17E-02 | 1.46E-01 |
| FHL2 | 2:105386541-105398969 | 2:105398842-105398969 | TSSIA5ss | 3 | 3 | TSS.ENST00000393353 | -10 | 2.44E-02 | 1.02E-01 |
| PPIL3 | 2:200882383-200885697 | 2:200882383-200882435 | A3ss | 3 | 3 | ENST00000392283 | -10.02 | 1.07E-02 | 4.64E-02 |
| CDKN2AIP | 4:183446088-183448198 | 4:183446348-183447789 | IR | 3 | 3 | ENST00000302350 | -10.02 | 2.93E-02 | 9.13E-02 |
| PLXNB2 | 22:50290598-50307552 | 22:50294719-50294778 | SES | 3 | 3 | ENST00000359337 | -10.04 | 3.47E-02 | 1.41E-01 |
| ALG13 | X:111736880-111752789 | X:111744668-111744904 | SES | 3 | 3 | ENST00000394780 | -10.05 | 1.64E-02 | 7.37E-01 |
| NCOR2 | 12:124402559-124419956 | 12:124402559-124402561 | A3ss | 3 | 3 | Ex.ENST00000356219 | -10.07 | 4.13E-02 | 7.03E-02 |
| UVSSA | 4:1354848-1355116 | 4:1355007-1355116 | A3ss | 3 | 3 | ENST00000505716 | -10.09 | 2.30E-02 | 1.54E-01 |
| DCAF8 | 1:160261359-160262448 | 1:160262117-160262448 | TSSIA5ss | 3 | 3 | TSS.ENST00000368073 | -10.09 | 4.77E-02 | 1.48E-01 |
| DCAF8 | 1:160261359-160262275 | 1:160262117-160262275 | TSSIA5ss | 3 | 3 | TSS.ENST00000368073 | -10.09 | 4.77E-02 | 1.48E-01 |
| CEP131 | 17:81220074-81222768 | 17:81222646-81222769 | A5ss | 3 | 3 | Ex.TSS.ENST00000269392 | -10.1 | 1.48E-02 | 3.11E-02 |
| DENND4B | 1:153930858-153931022 | 1:153930947-153931022 | A5ss | 3 | 3 | Ex.ENST00000361217 | -10.11 | 8.78E-03 | 2.74E-02 |
| ARHGAP23 | 17:38497827-38500596 | 17:38498414-38498510 | MES | 3 | 3 | ENST00000620417 | -10.11 | 4.09E-02 | 9.41E-02 |
| PIGT | 20:45420694-45424215 | 20:45421383-45421583 | SES | 3 | 3 | ENST00000279035 | -10.12 | 6.72E-03 | 2.53E-02 |
| NKD2 | 5:1036385-1037804 | 5:1037520-1037698 | SES | 3 | 3 | ENST00000274150 | -10.12 | 3.67E-02 | 2.80E-01 |
| NEK3 | 13:52153993-52156074 | 13:52154080-52154173 | SES | 3 | 3 | ENST00000552973 | -10.14 | 1.49E-02 | 2.67E-02 |
| UBAP2L | 1:154220976-154225083 | 1:154220976-154221040 | TSSIA5ss | 3 | 3 | TSS.ENST00000412596 | -10.14 | 1.81E-02 | 5.64E-02 |
| RHNO1 | 12:2877283-2887910 | 12:2885283-2885534 | MXS | 3 | 3 | ENST00000489288 | -10.15 | 5.19E-03 | 8.90E-03 |
| BRD1 | 22:49787495-49794033 | 22:49787577-49794034 | A5ss | 3 | 3 | Ex.ENST00000216267 | -10.15 | 4.36E-02 | 1.77E-01 |
| MYO9B | 19:17211651-17213284 | 19:17211775-17211894 | IR | 3 | 3 | ENST00000597073 | -10.2 | 4.23E-02 | 1.06E-01 |
| TP53I11 | 11:44938367-44949972 | 11:44939114-44939190 | MES | 3 | 3 | ENST00000533940 | -10.24 | 5.23E-03 | 8.27E-03 |
| PDPR | 16:70156475-70162537 | 16:70160312-70160734 | IR (overlapping regio | 3 | 3 | ENST00000288050 | -10.24 | 3.14E-02 | 6.94E-02 |
| IFI27L1 | 14:94100772-94102476 | 14:94101814-94102059 | SES | 3 | 3 | ENST00000554562 | -10.25 | 2.31E-02 | 4.36E-02 |
| TMPO | 12:98537573-98544950 | 12:98544230-98544349 | MES | 3 | 3 | ENST00000556029 | -10.29 | 4.04E-03 | 7.21E-03 |
| SNRPB | 20:2461794-2462635 | 20:2461794-2461939 | A3ss | 3 | 3 | Ex.TSS.ENST00000438552 | -10.3 | 6.43E-03 | 2.36E-02 |
| ZNF559 | 19:9341102-9342084 | 19:9341185-9341616 | IR (overlapping regio | 3 | 3 | ENST00000592504 | -10.31 | 2.19E-03 | 6.07E-03 |
| AC084018.2 | 12:121800797-121803403 | 12:121801809-121802878 | IR (overlapping regio | 3 | 3 | ENST00000613093 | -10.31 | 2.06E-02 | 3.44E-02 |
| APTX | 9:32989896-33001566 | 9:32997243-32997348 | SES | 3 | 3 | ENST00000379819 | -10.32 | 4.08E-02 | 1.00E+00 |
| SATB1 | 3:18423627-18425339 | 3:18423727-18424689 | IR (overlapping regio | 3 | 3 | ENST00000338745 | -10.33 | 2.88E-02 | 1.72E-01 |
| RLIM | X:74596001-74614421 | X:74612582-74612643 | SES | 3 | 3 | ENST00000349225 | -10.34 | 2.12E-02 | 1.00E+00 |
| LINC00665 | 19:36324924-36327170 | 19:36326754-36326996 | IR | 3 | 3 | ENST00000666458 | -10.37 | 2.31E-02 | 6.06E-02 |
| PARPBP | 12:102195951-102197520 | 12:102196702-102196972 | IR (overlapping regio | 3 | 3 | ENST00000327680 | -10.37 | 4.35E-02 | 7.27E-02 |
| ST7L | 1:112610455-112611003 | 1:112610691-112610840 | IR | 3 | 3 | ENST00000470683 | -10.38 | 2.87E-02 | 8.04E-02 |
| MAPKAPK5-AS1 | 12:111840478-111841683 | 12:111841327-111841683 | TSSIA5ss | 3 | 3 | TSS.ENST00000456429 | -10.4 | 2.50E-03 | 4.19E-03 |
| AL669830.2 | 6:30913811-30915037 | 6:30914345-30914809 | IR (overlapping regio | 3 | 3 | ENST00000477288 | -10.4 | 4.49E-02 | 4.51E-01 |
| EIF3C | 16:28711519-28711656 | 16:28711519-28711567 | TSSIA5ss | 3 | 3 | TSS.ENST00000395587 | -10.41 | 5.76E-03 | 1.24E-02 |
| GPBP1 | 5:57176401-57230845 | 5:57214074-57214193 | MXS | 3 | 3 | ENST00000424459 | -10.41 | 1.04E-02 | 9.45E-02 |
| NDUFAF5 | 20:13788653-13793179 | 20:13793160-13793179 | A3ss | 3 | 3 | ENST00000463598 | -10.41 | 3.64E-02 | 1.34E-01 |
| SPP1 | 4:87980127-87981474 | 4:87980393-87980434 | SES | 3 | 3 | ENST00000395080 | -10.43 | 4.97E-02 | 3.80E-01 |
| NFKBIB | 19:38907399-38908893 | 19:38907660-38908730 | IR | 3 | 3 | ENST00000509705 | -10.44 | 1.54E-02 | 3.23E-02 |
| MYO9B | 19:17211651-17213284 | 19:17211787-17211894 | IR | 3 | 3 | ENST00000597073 | -10.44 | 2.30E-02 | 5.79E-02 |
| ARHGAP21 | 10:24600789-24601977 | 10:24600789-24600930 | A3ss | 3 | 2 | ENST00000376410 | -10.48 | 3.45E-02 | 5.23E-02 |
| CHMP2A | 19:58554237-58554987 | 19:58554865-58554987 | TSSIA5ss | 3 | 3 | TSS.ENST00000597209 | -10.51 | 4.81E-02 | 1.33E-01 |
| AKR1A1 | 1:45568178-45568926 | 1:45568485-45568684 | SES | 3 | 3 | ENST00000351829 | -10.52 | 5.31E-03 | 1.88E-02 |
| CERS4 | 19:8209495-8241340 | 19:8210706-8210862 | MES | 3 | 3 | ENST00000559490 | -10.53 | 4.40E-03 | 1.22E-02 |
| SEPTIN9 | 17:77450787-77482143 | 17:77451367-77451528 | SES | 3 | 3 | ENST00000586128 | -10.54 | 2.36E-03 | 5.71E-03 |
| PGGHG | 11:290658-291113 | 11:290782-291001 | IR | 3 | 3 | ENST00000409655 | -10.58 | 1.11E-02 | 1.75E-02 |
| RMST | 12:97530860-97560536 | 12:97551832-97552082 | MXS | 3 | 3 | ENST00000665975 | -10.63 | 1.47E-02 | 2.63E-02 |
| CIAO2A | 15:64075538-64088686 | 15:64081102-64081151 | SES | 3 | 3 | ENST00000300030 | -10.64 | 1.14E-03 | 2.19E-03 |
| COPS2 | 15:49137438-49139527 | 15:49137438-49137458 | A3ss | 3 | 3 | ENST00000299259 | -10.65 | 2.16E-02 | 4.13E-02 |
| TMEM63A | 1:225859350-225860947 | 1:225860860-225860947 | A5ss | 3 | 3 | Ex.ENST00000366835 | -10.66 | 5.13E-03 | 1.73E-02 |
| ATP2A1-AS1 | 16:28879489-28879895 | 16:28879756-28879866 | IR (overlapping regio | 3 | 3 | ENST00000561547 | -10.72 | 9.39E-03 | 2.02E-02 |
| AC026412.1 | 5:1632644-1632906 | 5:1632752-1632844 | IR | 3 | 3 | ENST00000507290 | -10.72 | 4.64E-02 | 1.44E-01 |
| RFC4 | 3:186804725-186806424 | 3:186806290-186806424 | TSSIA5ss | 3 | 3 | TSS.ENST00000296273 | -10.73 | 4.70E-04 | 2.83E-03 |
| SPIN3 | X:56990942-56995555 | X:56993889-56994722 | IR | 3 | 3 | ENST00000638289 | -10.74 | 4.68E-03 | 1.00E+00 |
| PIBF1 | 13:72782350-72792446 | 13:72783423-72783721 | SES | 3 | 3 | ENST00000617689 | -10.74 | 1.24E-02 | 2.23E-02 |
| PDZD4 | X:153806841-153830238 | X:153808342-153808595 | MES | 3 | 3 | ENST00000393758 | -10.74 | 1.66E-02 | 1.00E+00 |
| FAM133B | 7:92577677-92578149 | 7:92577677-92577717 | A3ss | 3 | 3 | ENST00000427372 | -10.75 | 1.61E-03 | 2.37E-02 |
| COP1 | 1:176135087-176149005 | 1:176136488-176136547 | SES | 3 | 3 | ENST00000367667 | -10.75 | 1.89E-02 | 6.04E-02 |
| METTL26 | 16:634798-636093 | 16:635612-635774 | MES | 3 | 3 | ENST00000301686 | -10.76 | 2.22E-02 | 4.66E-02 |
| ZNF211 | 19:57633437-57634628 | 19:57633713-57633905 | MES | 3 | 2 | ENST00000535785 | -10.76 | 3.03E-02 | 8.32E-02 |
| R3HDM2 | 12:57284057-57295398 | 12:57292572-57292667 | MXS | 3 | 3 | ENST00000547019 | -10.76 | 4.23E-02 | 7.44E-02 |
| GTPBP8 | 3:112993125-112996891 | 3:112995137-112995265 | SES | 3 | 3 | ENST00000619116 | -10.77 | 3.71E-02 | 1.98E-01 |
| SHPRH | 6:145955355-145964156 | 6:145963731-145964156 | TSSIA5ss | 3 | 3 | TSS.ENST00000275233 | -10.79 | 1.06E-02 | 1.06E-01 |
| SLC26A6 | 3:48633477-48633846 | 3:48633636-48633723 | IR (overlapping regio | 3 | 3 | ENST00000358747 | -10.83 | 3.36E-02 | 2.15E-01 |
| DOK1 | 2:74555669-74556307 | 2:74555894-74556078 | SES | 3 | 3 | ENST00000233668 | -10.86 | 2.29E-02 | 1.17E-01 |
| TRPT1 | 11:64223968-64224284 | 11:64224100-64224162 | SES | 3 | 3 | ENST00000537907 | -10.88 | 2.56E-02 | 4.09E-02 |
| ALAS1 | 3:52198256-52199220 | 3:52198672-52198848 | SES | 3 | 3 | ENST00000394965 | -10.89 | 1.42E-02 | 9.48E-02 |
| GALNT16 | 14:69352031-69354466 | 14:69352171-69353331 | IR | 3 | 2 | ENST00000448469 | -10.91 | 1.47E-02 | 2.75E-02 |
| TPM1 | 15:63060940-63061712 | 15:63061198-63061273 | SES | 3 | 3 | ENST00000558264 | -10.92 | 5.38E-03 | 1.04E-02 |
| RAP1GDS1 | 4:98352602-98391951 | 4:98379017-98379163 | MES | 3 | 3 | ENST00000503745 | -10.95 | 2.53E-02 | 1.93E-01 |
| KRTCAP3 | 2:27443949-27444291 | 2:27444062-27444188 | IR | 3 | 3 | ENST00000494572 | -10.98 | 6.28E-03 | 2.93E-02 |
| KRTCAP3 | 2:27443949-27444291 | 2:27444062-27444185 | IR | 3 | 3 | ENST00000494572 | -10.98 | 6.28E-03 | 2.93E-02 |
| PHACTR2 | 6:143753913-143765260 | 6:143760401-143760640 | SES | 3 | 3 | ENST00000427704 | -10.99 | 4.39E-03 | 4.36E-02 |
| LRFN4 | 11:66857735-66858876 | 11:66858270-66858335 | IR (overlapping regio | 3 | 3 | ENST00000393952 | -11 | 6.92E-04 | 1.13E-03 |
| SLC5A6 | 2:27211534-27211766 | 2:27211711-27211766 | TSSIA5ss | 3 | 3 | TSS.ENST00000401463 | -11 | 1.28E-02 | 5.74E-02 |
| FBXL6 | 8:144356808-144357973 | 8:144357122-144357456 | IR (overlapping regio | 3 | 3 | ENST00000524492 | -11.01 | 6.89E-03 | 1.23E-01 |
| SELENBP1 | 1:151369507-151369712 | 1:151369507-151369554 | A3ss | 3 | 3 | ENST00000368868 | -11.01 | 1.25E-02 | 3.54E-02 |
| LGALS8 | 1:236523599-236524061 | 1:236523625-236523969 | IR (overlapping regio | 3 | 3 | ENST00000366584 | -11.01 | 3.85E-02 | 1.28E-01 |
| PROSER1 | 13:39012234-39017499 | 13:39012691-39014476 | SES | 3 | 3 | ENST00000352251 | -11.02 | 4.01E-02 | 7.19E-02 |
| CENPO | 2:24814494-24815597 | 2:24815497-24815597 | A3ss | 3 | 3 | ENST00000260662 | -11.05 | 8.71E-03 | 4.03E-02 |
| C2CD3 | 11:74129843-74132843 | 11:74129843-74129972 | TSSIA3ss | 3 | 2 | TSS.ENST00000436679 | -11.05 | 8.91E-03 | 1.46E-02 |
| PLEKHH3 | 17:42670706-42671002 | 17:42670994-42671002 | A5ss | 3 | 3 | ENST00000591022 | -11.08 | 4.62E-02 | 1.07E-01 |
| VAMP1 | 12:6462237-6464941 | 12:6462843-6464889 | IR (overlapping regio | 3 | 3 | ENST00000361716 | -11.1 | 1.06E-02 | 1.88E-02 |
| WIPF1 | 2:174559572-174562602 | 2:174562126-174562550 | IR (overlapping regio | 3 | 3 | ENST00000392547 | -11.1 | 1.14E-02 | 4.85E-02 |
| PRRC2C | 1:171587222-171588378 | 1:171587648-171587751 | SES | 3 | 3 | ENST00000338920 | -11.18 | 2.11E-04 | 6.73E-04 |
| CDCA7 | 2:173358838-173359342 | 2:173359255-173359342 | A3ss | 3 | 3 | ENST00000306721 | -11.19 | 1.89E-02 | 8.08E-02 |
| CHRNE | 17:4899468-4900229 | 17:4899829-4899935 | IR (overlapping regio | 2 | 2 | ENST00000652550 | -11.19 | 3.85E-02 | 9.04E-02 |
| PI4KB | 1:151316510-151327270 | 1:151326172-151326373 | MXS | 3 | 3 | ENST00000368875 | -11.2 | 1.58E-02 | 4.48E-02 |
| CYLD | 16:50742842-50749575 | 16:50744730-50744866 | SES | 3 | 3 | ENST00000311559 | -11.21 | 1.96E-02 | 4.12E-02 |
| TWNK | 10:100989454-100989705 | 10:100989644-100989705 | TSSIA3ss | 3 | 3 | TSS.ENST00000311916 | -11.22 | 1.81E-02 | 2.70E-02 |
| PDE9A | 21:42699012-42732069 | 21:42731770-42731949 | SES | 3 | 3 | ENST00000291539 | -11.24 | 5.10E-03 | 1.99E-02 |
| CDC27 | 17:47158304-47181561 | 17:47169917-47170042 | MES | 3 | 3 | ENST00000066544 | -11.26 | 1.12E-02 | 2.61E-02 |
| MBD1 | 18:50268846-50271540 | 18:50269812-50271468 | IR | 3 | 3 | ENST00000591416 | -11.27 | 1.26E-03 | 3.10E-03 |
| SLC15A3 | 11:60939639-60942034 | 11:60941122-60941290 | SES | 3 | 3 | ENST00000227880 | -11.3 | 5.15E-03 | 8.22E-03 |
| PRSS16 | 6:27252041-27254692 | 6:27252808-27252949 | SES | 3 | 3 | ENST00000230582 | -11.32 | 1.07E-02 | 1.11E-01 |

| | | | | | | | | | |
|---|---|---|---|---|---|---|---|---|---|
| LGMN | 14:92704362-92709671 | 14:92706483-92706653 | MES | 3 | 3 | ENST00000334869 | -11.32 | 1.13E-02 | 2.13E-02 |
| INPP5F | 10:119781772-119791868 | 10:119791517-119791645 | SES | 2 | 2 | ENST00000369081 | -11.32 | 3.03E-02 | 4.54E-02 |
| SLC30A5 | 5:69094339-69103061 | 5:69100807-69100929 | SES | 3 | 3 | ENST00000380860 | -11.34 | 3.78E-02 | 1.18E-01 |
| ECHDC1 | 6:127314897-127327001 | 6:127316450-127316502 | MES | 3 | 3 | ENST00000525745 | -11.35 | 3.24E-03 | 3.17E-02 |
| ZNF644 | 1:90982371-91021989 | 1:91021689-91021990 | A5ss | 3 | 3 | Ex.TSS.ENST00000337393 | -11.36 | 2.15E-02 | 7.87E-02 |
| CABIN1 | 22:24098106-24113565 | 22:24098106-24098192 | A5ss | 3 | 3 | ENST00000263119 | -11.37 | 2.61E-02 | 1.05E-01 |
| LPCAT4 | 15:34360210-34362195 | 15:34361169-34362196 | A5ss | 3 | 3 | Ex.ENST00000567507 | -11.41 | 4.39E-03 | 8.34E-03 |
| LPCAT4 | 15:34360210-34361399 | 15:34361169-34361400 | A5ss | 3 | 3 | Ex.ENST00000314891 | -11.41 | 4.39E-03 | 8.34E-03 |
| FCGRT | 19:49513195-49514486 | 19:49514134-49514210 | IR | 3 | 3 | ENST00000595881 | -11.41 | 8.68E-03 | 2.34E-02 |
| RIC8B | 12:106860413-106885903 | 12:106870823-106870942 | MES | 3 | 3 | ENST00000355478 | -11.45 | 3.26E-02 | 5.45E-02 |
| CTC1 | 17:8230268-8230468 | 17:8230349-8230446 | IR | 3 | 3 | ENST00000584439 | -11.48 | 7.51E-03 | 1.83E-02 |
| SFI1 | 22:31614386-31615279 | 22:31614861-31615047 | IR | 3 | 3 | ENST00000357852 | -11.48 | 1.39E-02 | 5.67E-02 |
| DLG1 | 3:197075871-197081050 | 3:197076586-197076682 | SES | 3 | 3 | ENST00000469073 | -11.52 | 1.51E-02 | 9.18E-02 |
| SLC2A6 | 9:133475109-133475399 | 9:133475109-133475113 | A3ss | 3 | 3 | Ex.ENST00000371897 | -11.54 | 2.06E-02 | 6.01E-01 |
| SEMA4G | 10:100983305-100985615 | 10:100983809-100984489 | IR (overlapping regio | 3 | 3 | ENST00000210633 | -11.59 | 8.68E-03 | 1.29E-02 |
| EPN2 | 17:19312145-19328710 | 17:19313105-19313279 | SES | 3 | 3 | ENST00000314728 | -11.61 | 8.65E-03 | 1.82E-02 |
| C11orf95 | 11:63764599-63765806 | 11:63764868-63765254 | SES | 3 | 3 | ENST00000433688 | -11.62 | 1.74E-02 | 2.78E-02 |
| PLXNB2 | 22:50290598-50294725 | 22:50294719-50294725 | A5ss | 3 | 3 | ENST00000359337 | -11.63 | 4.12E-02 | 1.67E-01 |
| CDH11 | 16:64998857-65053803 | 16:65004642-65005041 | SES | 3 | 3 | ENST00000268603 | -11.63 | 4.27E-02 | 8.94E-02 |
| JAKMIP2 | 5:147661446-147782455 | 5:147671678-147671954 | SES | 3 | 3 | ENST00000265272 | -11.65 | 4.31E-02 | 3.55E-01 |
| LRRC20 | 10:70301509-70376451 | 10:70340553-70340702 | MXS | 3 | 3 | ENST00000357631 | -11.66 | 4.06E-02 | 6.18E-02 |
| TTC3 | 21:37156907-37159698 | 21:37157161-37157214 | MXS | 3 | 3 | ENST00000411496 | -11.69 | 3.35E-02 | 1.30E-01 |
| STIL | 1:47280241-47281209 | 1:47280364-47280945 | IR | 3 | 3 | ENST00000337817 | -11.71 | 1.42E-02 | 5.06E-02 |
| AC138028.2 | 16:88736800-88738405 | 16:88737647-88737727 | IR (overlapping regio | 3 | 3 | ENST00000567968 | -11.72 | 1.94E-03 | 4.32E-03 |
| ZNF875 | 19:37334783-37347231 | 19:37347190-37347231 | TSSIA3ss | 3 | 3 | TSS.ENST00000591417 | -11.73 | 2.87E-02 | 7.52E-02 |
| MBD1 | 18:50268846-50271540 | 18:50269819-50271468 | IR | 3 | 3 | ENST00000591416 | -11.76 | 2.99E-04 | 7.39E-04 |
| PKD1P1 | 16:16327736-16328516 | 16:16327907-16328004 | IR (overlapping regio | 3 | 3 | ENST00000458669 | -11.76 | 6.07E-04 | 1.28E-03 |
| FBXO27 | 19:39024021-39025554 | 19:39024243-39024399 | IR | 3 | 3 | ENST00000292853 | -11.78 | 3.00E-03 | 7.88E-03 |
| WDR90 | 16:655773-656537 | 16:655890-656301 | IR (overlapping regio | 3 | 3 | ENST00000552943 | -11.81 | 3.73E-02 | 7.82E-02 |
| CDK5RAP3 | 17:47974400-47974583 | 17:47974449-47974580 | IR (overlapping regio | 3 | 3 | ENST00000581662 | -11.82 | 2.13E-02 | 4.99E-02 |
| RIOX2 | 3:97950013-97950785 | 3:97950013-97950015 | A3ss | 3 | 3 | ENST00000333396 | -11.83 | 2.46E-02 | 1.68E-01 |
| ZBTB44 | 11:130234274-130237009 | 11:130236793-130237009 | A5ss | 3 | 3 | ENST00000357899 | -11.83 | 4.96E-02 | 7.71E-02 |
| TMEM132A | 11:60927049-60927418 | 11:60927071-60927203 | IR | 3 | 3 | ENST00000534983 | -11.87 | 3.11E-02 | 4.97E-02 |
| LRATD2 | 8:126552901-126557731 | 8:126553504-126556264 | IR | 3 | 3 | ENST00000652209 | -11.88 | 4.57E-03 | 7.57E-02 |
| AC068587.4 | 8:12511258-12601157 | 8:12511258-12581044 | A3ss | 3 | 3 | Ex.ENST00000641839 | -11.9 | 4.28E-02 | 7.09E-01 |
| ERBIN | 5:66054952-66076315 | 5:66075024-66075230 | MES | 3 | 2 | ENST00000506030 | -11.91 | 4.36E-03 | 3.84E-02 |
| GTPBP3 | 19:17339600-17341106 | 19:17341044-17341106 | A3ss | 3 | 3 | ENST00000324894 | -11.96 | 1.65E-02 | 4.15E-02 |
| MPI | 15:74897512-74902219 | 15:74901257-74901949 | IR (overlapping regio | 3 | 3 | ENST00000352410 | -11.96 | 2.75E-02 | 5.36E-02 |
| PHF6 | X:134425201-134428789 | X:134425331-134425660 | IR | 3 | 3 | ENST00000332070 | -11.97 | 2.29E-03 | 1.24E-01 |
| SORBS2 | 4:185678823-185775226 | 4:185684754-185684846 | MXS | 3 | 3 | ENST00000319471 | -11.98 | 1.22E-02 | 8.70E-02 |
| STYXL1 | 7:76030525-76047811 | 7:76047662-76047811 | TSSIA5ss | 3 | 3 | TSS.ENST00000248600 | -11.98 | 4.66E-02 | 6.78E-01 |
| ZER1 | 9:128742746-128750621 | 9:128750616-128750621 | A5ss | 3 | 3 | Ex.ENST00000291900 | -11.99 | 4.06E-02 | 1.00E+00 |
| FBXL6 | 8:144356808-144357973 | 8:144357122-144357438 | IR (overlapping regio | 3 | 3 | ENST00000524492 | -12.01 | 3.88E-03 | 6.94E-02 |
| SPIN3 | X:56990831-56994949 | X:56992238-56994722 | IR | 3 | 3 | ENST00000374919 | -12.02 | 5.31E-03 | 1.00E+00 |
| RASSF7 | 11:563189-564012 | 11:563318-563557 | IR | 3 | 3 | ENST00000397582 | -12.04 | 7.26E-03 | 1.16E-02 |
| DNAJC4 | 11:64233894-64234281 | 11:64233981-64234072 | IR (overlapping regio | 3 | 3 | ENST00000321460 | -12.05 | 1.86E-02 | 2.98E-02 |
| SELENBP1 | 1:151368320-151369712 | 1:151369004-151369554 | SES | 3 | 3 | ENST00000463664 | -12.05 | 2.86E-02 | 8.11E-02 |
| FAM200B | 4:15681902-15686235 | 4:15681902-15681968 | TSSIA5ss | 3 | 3 | TSS.ENST00000503600 | -12.05 | 4.37E-02 | 3.07E-01 |
| FAM200B | 4:15681785-15686235 | 4:15681785-15681968 | A5ss | 3 | 3 | ENST00000503617 | -12.05 | 4.37E-02 | 3.07E-01 |
| PLSCR3 | 17:7392953-7393466 | 17:7393144-7393466 | A5ss | 3 | 3 | ENST00000571541 | -12.08 | 1.67E-02 | 3.93E-02 |
| ZNF616 | 19:52117025-52123922 | 19:52122360-52122457 | SES | 3 | 3 | ENST00000330123 | -12.18 | 3.05E-02 | 8.33E-02 |
| FGGY | 1:59667404-59757930 | 1:59674039-59674133 | SES | 2 | 3 | ENST00000303721 | -12.21 | 2.55E-03 | 9.18E-03 |
| FBXL6 | 8:144356808-144357973 | 8:144357122-144357627 | IR (overlapping regio | 3 | 3 | ENST00000524492 | -12.21 | 2.62E-03 | 4.70E-02 |
| RANGRF | 17:8289503-8290090 | 17:8289589-8289812 | IR (overlapping regio | 3 | 3 | ENST00000407006 | -12.21 | 1.42E-02 | 3.47E-02 |
| RNF167 | 17:4940643-4940872 | 17:4940643-4940726 | A5ss | 3 | 3 | ENST00000571816 | -12.21 | 3.13E-02 | 7.34E-02 |
| HP1BP3 | 1:20779912-20787194 | 1:20780345-20780540 | SES | 3 | 3 | ENST00000438032 | -12.23 | 1.77E-02 | 5.75E-02 |
| SLC37A3 | 7:140348768-140355667 | 7:140351273-140351451 | MES | 3 | 3 | ENST00000326232 | -12.25 | 1.06E-02 | 1.30E-01 |
| CNOT2-DT | 12:70242894-70243311 | 12:70243066-70243204 | SES | 3 | 2 | ENST00000550216 | -12.25 | 1.49E-02 | 2.64E-02 |
| SECISBP2 | 9:89319798-89325547 | 9:89325427-89325547 | A3ss | 3 | 3 | ENST00000375807 | -12.3 | 1.94E-03 | 6.58E-02 |
| CREBZF | 11:85663524-85665102 | 11:85663602-85663814 | IR | 3 | 3 | ENST00000490820 | -12.3 | 4.61E-02 | 7.62E-02 |
| LGMN | 14:92704362-92709671 | 14:92704640-92704707 | MES | 3 | 3 | ENST00000334869 | -12.31 | 1.25E-02 | 2.37E-02 |
| PCNX2 | 1:233139856-233161270 | 1:233160283-233160433 | SES | 3 | 3 | ENST00000258229 | -12.31 | 2.76E-02 | 9.39E-02 |
| ZSCAN5A | 19:56313385-56314680 | 19:56314250-56314680 | TSSIA5ss | 3 | 3 | TSS.ENST00000592679 | -12.33 | 3.85E-03 | 1.06E-02 |
| ZSCAN5A | 19:56313385-56314619 | 19:56314250-56314619 | TSSIA5ss | 3 | 3 | TSS.ENST00000592679 | -12.33 | 3.85E-03 | 1.06E-02 |
| ZSCAN5A | 19:56313385-56314420 | 19:56314250-56314420 | TSSIA5ss | 3 | 3 | TSS.ENST00000592679 | -12.33 | 3.85E-03 | 1.06E-02 |
| CHTF8 | 16:69120650-69121435 | 16:69121053-69121170 | SES | 3 | 3 | ENST00000398235 | -12.36 | 2.08E-03 | 4.61E-03 |
| STK33 | 11:8475067-8594082 | 11:8480534-8480614 | SES | 3 | 3 | ENST00000454443 | -12.37 | 3.82E-02 | 6.33E-02 |
| CAMKV | 3:49860716-49861024 | 3:49860853-49860942 | IR | 3 | 3 | ENST00000478149 | -12.41 | 2.49E-04 | 1.62E-03 |
| ZGPAT | 20:63708170-63708552 | 20:63708170-63708188 | TSSIA5ss | 3 | 3 | TSS.ENST00000328969 | -12.45 | 1.39E-02 | 5.37E-02 |
| ZGPAT | 20:63708119-63708552 | 20:63708119-63708188 | TSSIA5ss | 3 | 3 | TSS.ENST00000328969 | -12.45 | 1.39E-02 | 5.37E-02 |
| CCDC77 | 12:389487-411746 | 12:405511-405564 | SES | 3 | 3 | ENST00000422000 | -12.45 | 3.45E-02 | 5.94E-02 |
| STAT6 | 12:57108300-57110076 | 12:57109941-57110076 | A5ss | 3 | 3 | ENST00000553275 | -12.47 | 2.61E-02 | 4.58E-02 |
| STAT6 | 12:57108300-57110042 | 12:57109941-57110042 | A5ss | 3 | 3 | ENST00000553275 | -12.47 | 2.61E-02 | 4.58E-02 |
| WDR60 | 7:158902596-158905988 | 7:158902596-158905283 | A5ss | 3 | 3 | ENST00000467220 | -12.48 | 4.51E-03 | 5.84E-02 |
| GABPB1-AS1 | 15:50356100-50372202 | 15:50356258-50370329 | IR | 2 | 2 | ENST00000499624 | -12.52 | 3.28E-02 | 6.29E-02 |
| DEDD2 | 19:42215253-42217631 | 19:42216680-42217045 | SES | 3 | 3 | ENST00000336034 | -12.53 | 1.19E-02 | 3.17E-02 |
| ARID5A | 2:96551099-96552634 | 2:96551327-96552210 | IR | 3 | 3 | ENST00000357485 | -12.57 | 5.42E-03 | 2.85E-02 |
| PPIL3 | 2:200881489-200885697 | 2:200882342-200882435 | MXS | 3 | 3 | ENST00000392283 | -12.6 | 1.66E-02 | 7.17E-02 |
| FOXH1 | 8:144475224-144475582 | 8:144475224-144475261 | TSSIA3ss | 3 | 3 | TSS.ENST00000377317 | -12.64 | 6.11E-03 | 1.09E-01 |
| LSM14B | 20:62126440-62130218 | 20:62129785-62129952 | MXS | 3 | 3 | ENST00000279068 | -12.64 | 1.34E-02 | 5.14E-02 |
| ERBIN | 5:66054952-66076315 | 5:66075024-66075230 | MES | 3 | 2 | ENST00000284037 | -12.7 | 3.50E-02 | 1.09E-01 |
| CEP170 | 1:243172847-243186258 | 1:243185779-243186072 | SES | 3 | 3 | ENST00000366542 | -12.71 | 2.35E-02 | 7.84E-02 |
| GSDMD | 8:143559777-143561423 | 8:143560772-143561001 | IR | 3 | 3 | ENST00000524846 | -12.73 | 9.22E-03 | 1.61E-01 |
| TRIT1 | 1:39850262-39883317 | 1:39852731-39852876 | MXS | 3 | 3 | ENST00000441669 | -12.74 | 6.81E-03 | 2.38E-02 |
| SLC37A4 | 11:119027628-119029034 | 11:119027873-119028193 | IR | 3 | 3 | ENST00000526275 | -12.81 | 4.92E-03 | 7.62E-03 |
| BRD8 | 5:138168079-138169221 | 5:138168470-138168688 | SES | 3 | 3 | ENST00000230901 | -12.81 | 1.95E-02 | 1.58E-01 |
| HUWE1 | X:53542540-53543885 | X:53543841-53543885 | A5ss | 3 | 3 | ENST00000262854 | -12.81 | 3.04E-02 | 1.00E+00 |
| TAF2 | 8:119744394-119746704 | 8:119744881-119745036 | SES | 3 | 3 | ENST00000529653 | -12.86 | 1.55E-03 | 2.57E-02 |
| EMC9 | 14:24141107-24141585 | 14:24141317-24141437 | IR | 3 | 3 | ENST00000419198 | -12.87 | 1.06E-02 | 1.95E-02 |
| HTATSF1 | X:136497440-136497675 | X:136497624-136497675 | TSSIA3ss | 3 | 3 | TSS.ENST00000448450 | -12.87 | 4.22E-02 | 1.00E+00 |
| TRDMT1 | 10:17160375-17174550 | 10:17161483-17161548 | MXS | 3 | 3 | ENST00000424636 | -12.89 | 3.89E-02 | 5.87E-02 |
| TRMU | 22:46331733-46337778 | 22:46333082-46337779 | A3ss | 3 | 3 | Ex.TSS.ENST00000486620 | -12.91 | 4.78E-02 | 1.98E-01 |
| SLC44A2 | 19:10642452-10643278 | 19:10642897-10643021 | SES | 3 | 3 | ENST00000586549 | -12.92 | 5.19E-04 | 1.30E-03 |
| SLC9A6 | X:135994890-135998107 | X:135994890-135994985 | A5ss | 3 | 3 | ENST00000370695 | -13 | 1.91E-02 | 9.93E-01 |
| PTP4A3 | 8:141392085-141421387 | 8:141392085-141418018 | A5ss | 3 | 3 | Ex.TSS.ENST00000520105 | -13 | 3.55E-02 | 6.10E-01 |
| TNFRSF25 | 1:6462863-6464719 | 1:6463148-6464374 | IR | 3 | 3 | ENST00000513135 | -13.01 | 4.39E-02 | 1.59E-01 |
| TMEM127 | 2:96265490-96265868 | 2:96265490-96265512 | TSSIA3ss | 3 | 3 | TSS.ENST00000258439 | -13.02 | 1.93E-04 | 1.02E-03 |
| PTAR1 | 9:69741859-69759852 | 9:69750781-69750950 | SES | 3 | 3 | ENST00000340434 | -13.05 | 2.70E-02 | 1.00E+00 |
| TTLL5 | 14:75752956-75764614 | 14:75757831-75757869 | SES | 3 | 3 | ENST00000554510 | -13.06 | 1.17E-02 | 2.19E-02 |
| ARVCF | 22:19982092-20016588 | 22:19990585-19990812 | MES | 3 | 3 | ENST00000263207 | -13.06 | 4.70E-02 | 1.76E-01 |
| RAB29 | 1:205774833-205775459 | 1:205775087-205775272 | IR | 3 | 3 | ENST00000414729 | -13.09 | 9.03E-03 | 2.93E-02 |

| | | | | | | | | | |
|---|---|---|---|---|---|---|---|---|---|
| HCFC1R1 | 16:3023847-3024246 | 16:3024087-3024192 | IR (overlapping regio | 3 | 3 | ENST00000248089 | -13.1 | 5.50E-04 | 1.19E-03 |
| GLI4 | 8:143274704-143275552 | 8:143274803-143274977 | IR (overlapping regio | 3 | 3 | ENST00000344692 | -13.12 | 1.92E-02 | 3.27E-01 |
| GZF1 | 20:23368930-23370090 | 20:23369584-23369741 | SES | 3 | 3 | ENST00000338121 | -13.16 | 3.42E-02 | 1.25E-01 |
| RECQL | 12:21501170-21501635 | 12:21501216-21501462 | IR | 3 | 3 | ENST00000444129 | -13.17 | 3.77E-02 | 6.46E-02 |
| CHFR | 12:132861635-132870723 | 12:132869619-132869834 | SES | 3 | 3 | ENST00000432561 | -13.2 | 8.21E-03 | 1.40E-02 |
| PLAT | 8:42182891-42188933 | 8:42185081-42185158 | MES | 3 | 3 | ENST00000519510 | -13.21 | 3.92E-02 | 9.08E-01 |
| AL138720.1 | 6:14997476-15021820 | 6:15008925-15021821 | A5ss | 3 | 3 | Ex.TSS.ENST00000629853 | -13.22 | 1.94E-02 | 1.93E-01 |
| ZNF562 | 19:9660822-9675014 | 19:9660822-9660874 | TSSIA3ss | 2 | 3 | TSS.ENST00000293648 | -13.23 | 1.48E-02 | 4.11E-02 |
| HEPH | X:66208247-66258839 | X:66255035-66255141 | MES | 3 | 3 | ENST00000336279 | -13.25 | 3.02E-02 | 1.00E+00 |
| RESF1 | 12:31970357-31987238 | 12:31980878-31985957 | SES | 3 | 3 | ENST00000312561 | -13.25 | 3.14E-02 | 5.36E-02 |
| NKAPD1 | 11:112074287-112074916 | 11:112074359-112074886 | IR | 3 | 3 | ENST00000532163 | -13.26 | 3.42E-02 | 5.26E-02 |
| EPN2 | 17:19312145-19313125 | 17:19313105-19313125 | A3ss | 3 | 3 | ENST00000314728 | -13.31 | 2.14E-02 | 4.50E-02 |
| EPN2 | 17:19312145-19313122 | 17:19313105-19313122 | A3ss | 3 | 3 | ENST00000314728 | -13.31 | 2.14E-02 | 4.50E-02 |
| RNF220 | 1:44635589-44636029 | 1:44635589-44635666 | A5ss | 3 | 3 | ENST00000335497 | -13.36 | 3.21E-04 | 1.13E-03 |
| PLPPR2 | 19:11357740-11359606 | 19:11359532-11359606 | A3ss | 3 | 3 | ENST00000251473 | -13.37 | 4.00E-03 | 1.00E-02 |
| ZNF589 | 3:48260940-48286245 | 3:48267915-48270969 | SES | 3 | 3 | ENST00000448461 | -13.37 | 1.71E-02 | 1.10E-01 |
| ZNF142 | 2:218650527-218656149 | 2:218651701-218651822 | SES | 3 | 3 | ENST00000433921 | -13.4 | 1.46E-02 | 6.55E-02 |
| ZNF473 | 19:50044670-50048776 | 19:50044788-50047958 | IR (overlapping regio | 3 | 3 | ENST00000270617 | -13.4 | 2.02E-02 | 5.48E-02 |
| SLC37A4 | 11:119027628-119029034 | 11:119027771-119028193 | IR | 3 | 3 | ENST00000526275 | -13.41 | 3.12E-02 | 4.65E-02 |
| RECQL | 12:21501170-21501635 | 12:21501216-21501480 | IR | 3 | 3 | ENST00000444129 | -13.41 | 4.45E-02 | 7.61E-02 |
| RAB29 | 1:205774833-205775459 | 1:205775092-205775373 | IR | 3 | 3 | ENST00000414729 | -13.42 | 4.44E-02 | 1.44E-01 |
| FNBP4 | 11:47731840-47732670 | 11:47732220-47732536 | IR | 3 | 3 | ENST00000534003 | -13.44 | 1.16E-03 | 1.86E-03 |
| ZNF131 | 5:43121598-43122177 | 5:43121829-43122043 | IR | 3 | 3 | ENST00000509634 | -13.45 | 2.36E-03 | 2.10E-02 |
| CCDC78 | 16:724106-725880 | 16:725581-725793 | IR | 3 | 3 | ENST00000481804 | -13.46 | 2.96E-02 | 6.56E-02 |
| RASSF7 | 11:563189-564012 | 11:563318-563395 | IR | 3 | 3 | ENST00000397582 | -13.47 | 1.44E-03 | 2.29E-03 |
| ZFP62 | 5:180847615-180851493 | 5:180848083-180851380 | IR | 3 | 3 | ENST00000512132 | -13.47 | 1.17E-02 | 1.03E-01 |
| ZNF266 | 19:9418505-9419081 | 19:9418632-9419050 | IR | 3 | 3 | ENST00000591213 | -13.48 | 4.79E-02 | 1.33E-01 |
| GLI4 | 8:143274704-143275552 | 8:143274803-143275194 | IR (overlapping regio | 3 | 3 | ENST00000344692 | -13.52 | 2.47E-02 | 4.21E-01 |
| DPH5 | 1:101001588-101021640 | 1:101013710-101013818 | SES | 3 | 3 | ENST00000342173 | -13.53 | 2.28E-03 | 6.33E-03 |
| APH1A | 1:150267992-150268697 | 1:150267992-150268127 | TSSIA3ss | 3 | 3 | TSS.ENST00000360244 | -13.54 | 4.22E-03 | 1.19E-02 |
| BIRC2 | 11:102348598-102350749 | 11:102348720-102350345 | IR | 3 | 3 | ENST00000227758 | -13.56 | 1.63E-03 | 2.50E-03 |
| NOL3 | 16:67174162-67174944 | 16:67174341-67174620 | IR | 3 | 3 | ENST00000568503 | -13.56 | 4.29E-02 | 9.44E-02 |
| ASPHD1 | 16:29900488-29901920 | 16:29900921-29901743 | IR | 3 | 3 | ENST00000308748 | -13.61 | 2.45E-02 | 5.14E-02 |
| PPIP5K2 | 5:103173973-103177666 | 5:103176866-103176910 | SES | 3 | 2 | ENST00000613674 | -13.62 | 4.51E-02 | 3.44E-01 |
| SH3D21 | 1:36321056-36321778 | 1:36321210-36321723 | IR | 3 | 3 | ENST00000474766 | -13.63 | 2.60E-02 | 9.03E-02 |
| ZNF692 | 1:248857975-248858385 | 1:248858035-248858130 | IR | 3 | 3 | ENST00000533976 | -13.67 | 3.37E-02 | 1.13E-01 |
| DERL3 | 22:23834503-23837217 | 22:23834933-23837063 | IR (overlapping regio | 3 | 3 | ENST00000404056 | -13.71 | 1.14E-03 | 4.56E-03 |
| SUGP2 | 19:18990893-18993740 | 19:18991150-18993647 | IR | 3 | 3 | ENST00000452918 | -13.71 | 1.93E-02 | 4.94E-02 |
| ATP6V0D1 | 16:67443179-67480956 | 16:67453544-67453715 | MES | 3 | 3 | ENST00000290949 | -13.75 | 7.96E-03 | 1.75E-02 |
| VPS54 | 2:63972245-63983863 | 2:63981646-63981887 | SES | 3 | 3 | ENST00000272322 | -13.75 | 3.78E-02 | 1.84E-01 |
| AL669830.2 | 6:30913811-30915037 | 6:30914470-30914809 | IR (overlapping regio | 3 | 3 | ENST00000477288 | -13.79 | 8.11E-03 | 8.47E-02 |
| AL669830.2 | 6:30913811-30915037 | 6:30914470-30914804 | IR (overlapping regio | 3 | 3 | ENST00000477288 | -13.79 | 8.11E-03 | 8.47E-02 |
| SEMA4C | 2:96867924-96869875 | 2:96869187-96869876 | A5ss | 3 | 3 | Ex.TSS.ENST00000305476 | -13.8 | 1.98E-03 | 1.05E-02 |
| C19orf54 | 19:40749481-40750477 | 19:40749608-40750284 | IR | 3 | 3 | ENST00000378313 | -13.84 | 4.18E-02 | 1.11E-01 |
| ZNF131 | 5:43121598-43122177 | 5:43121829-43122038 | IR | 3 | 3 | ENST00000509634 | -13.88 | 4.19E-03 | 3.74E-02 |
| IFT81 | 12:110124444-110127359 | 12:110124444-110124861 | TSSIA5ss | 3 | 3 | TSS.ENST00000242591 | -13.9 | 4.46E-02 | 7.48E-02 |
| FAM131A | 3:184336124-184342065 | 3:184341724-184341817 | MXS | 3 | 3 | ENST00000433578 | -13.91 | 4.47E-04 | 2.69E-03 |
| DELE1 | 5:141938521-141942047 | 5:141938905-141939505 | IR | 3 | 3 | ENST00000432126 | -13.94 | 3.29E-02 | 2.70E-01 |
| LRFN4 | 11:66857064-66859093 | 11:66857297-66857734 | IR (overlapping regio | 3 | 3 | ENST00000309602 | -14 | 1.17E-02 | 1.91E-02 |
| ELP5 | 17:7252286-7252596 | 17:7252338-7252425 | IR (overlapping regio | 3 | 3 | ENST00000396628 | -14.03 | 8.52E-03 | 2.00E-02 |
| ARHGAP23 | 17:38497827-38500596 | 17:38498414-38498510 | MXS | 3 | 3 | ENST00000614693 | -14.04 | 1.79E-02 | 4.12E-02 |
| TJAP1 | 6:43503707-43505623 | 6:43504761-43505623 | A3ss | 3 | 3 | Ex.TSS.ENST00000259751 | -14.05 | 1.30E-03 | 1.42E-02 |
| CELSR2 | 1:109273166-109275607 | 1:109273671-109274160 | IR | 3 | 3 | ENST00000498157 | -14.05 | 1.56E-02 | 4.37E-02 |
| THOC5 | 22:29549159-29553670 | 22:29553448-29553670 | TSSIA5ss | 3 | 3 | TSS.ENST00000428374 | -14.05 | 3.04E-02 | 1.22E-01 |
| ANKRD13D | 11:67299612-67299826 | 11:67299612-67299621 | A5ss | 3 | 3 | Ex.ENST00000308440 | -14.05 | 4.14E-02 | 6.77E-02 |
| ATP6V0D1 | 16:67443179-67480956 | 16:67444528-67444706 | MES | 3 | 3 | ENST00000290949 | -14.08 | 1.16E-03 | 2.55E-03 |
| DDX11 | 12:31073970-31078389 | 12:31073970-31074091 | TSSIA5ss | 3 | 3 | TSS.ENST00000350437 | -14.09 | 5.65E-03 | 9.71E-03 |
| GPSM2 | 1:108877229-108896863 | 1:108885275-108885578 | SES | 3 | 3 | ENST00000264126 | -14.09 | 3.92E-02 | 1.08E-01 |
| AC069281.2 | 7:100573297-100575782 | 7:100573380-100573596 | IR (overlapping regio | 3 | 3 | ENST00000485071 | -14.12 | 2.91E-02 | 3.33E-01 |
| CHTF8 | 16:69120650-69121435 | 16:69120650-69121170 | TSSIA3ss | 3 | 3 | TSS.ENST00000522091 | -14.13 | 7.52E-04 | 1.66E-03 |
| ZSCAN2 | 15:84601136-84603819 | 15:84601136-84601188 | TSSIA5ss | 3 | 3 | TSS.ENST00000448803 | -14.13 | 1.84E-02 | 3.61E-02 |
| MPDZ | 9:13140150-13150510 | 9:13147548-13147658 | MES | 3 | 3 | ENST00000319217 | -14.14 | 4.55E-02 | 1.00E+00 |
| AC069281.2 | 7:100573297-100575782 | 7:100573401-100573596 | IR (overlapping regio | 3 | 3 | ENST00000485071 | -14.16 | 4.03E-02 | 4.60E-01 |
| ECHDC2 | 1:52896598-52899173 | 1:52897437-52899173 | TSSIA5ss | 3 | 3 | TSS.ENST00000479183 | -14.17 | 4.55E-02 | 1.63E-01 |
| MFGE8 | 15:88899533-88901550 | 15:88899656-88899811 | SES | 3 | 3 | ENST00000542878 | -14.19 | 2.53E-04 | 4.98E-04 |
| MAPKAPK5-AS1 | 12:111840478-111841683 | 12:111841327-111841470 | SES | 3 | 3 | ENST00000590479 | -14.19 | 1.04E-03 | 1.75E-03 |
| TRIM26 | 6:30199154-30213304 | 6:30201035-30201138 | MES | 3 | 3 | ENST00000416596 | -14.21 | 1.15E-02 | 1.16E-01 |
| SERPINI1 | 3:167735813-167789110 | 3:167735813-167735823 | TSSIA5ss | 3 | 3 | TSS.ENST00000446050 | -14.27 | 3.30E-03 | 1.88E-02 |
| HDAC10 | 22:50250906-50251101 | 22:50250974-50251101 | TSSIA5ss | 3 | 3 | TSS.ENST00000216271 | -14.33 | 4.76E-02 | 1.93E-01 |
| BRD1 | 22:49787495-49794033 | 22:49787495-49787887 | A3ss | 3 | 3 | ENST00000404760 | -14.35 | 2.90E-03 | 1.20E-02 |
| AC132008.2 | 3:197624198-197627779 | 3:197627572-197627779 | TSSIA5ss | 3 | 3 | TSS.ENST00000437008 | -14.42 | 3.26E-02 | 1.98E-01 |
| ELP5 | 17:7252286-7252596 | 17:7252346-7252425 | IR (overlapping regio | 3 | 3 | ENST00000396628 | -14.43 | 1.03E-02 | 2.42E-02 |
| AMBRA1 | 11:46548501-46593931 | 11:46593828-46593931 | A5ss | 3 | 3 | Ex.TSS.ENST00000531542 | -14.45 | 2.89E-02 | 4.57E-02 |
| PRMT2 | 21:46635764-46636443 | 21:46636439-46636443 | A3ss | 3 | 3 | Ex.TSS.ENST00000355680 | -14.47 | 1.40E-02 | 5.25E-02 |
| PIGO | 9:35095567-35096154 | 9:35095879-35095976 | SES | 2 | 2 | ENST00000298004 | -14.47 | 3.20E-02 | 1.00E+00 |
| DAG1 | 3:49470275-49510418 | 3:49470275-49470433 | TSSIA5ss | 3 | 3 | TSS.ENST00000308775 | -14.48 | 4.42E-02 | 2.63E-01 |
| MDC1 | 6:30713803-30714183 | 6:30713933-30714030 | IR | 3 | 3 | ENST00000376406 | -14.51 | 2.22E-02 | 2.31E-01 |
| RALGDS | 9:133100383-133101519 | 9:133100496-133101519 | A5ss | 3 | 3 | ENST00000493438 | -14.52 | 1.13E-03 | 3.52E-02 |
| COASY | 17:42562148-42563322 | 17:42562220-42562932 | IR | 3 | 3 | ENST00000393818 | -14.53 | 9.69E-05 | 2.25E-04 |
| ANO8 | 19:17328162-17329541 | 19:17328265-17329052 | IR | 3 | 3 | ENST00000597643 | -14.55 | 1.88E-02 | 4.75E-02 |
| TRIM16 | 17:15683036-15684299 | 17:15683158-15684195 | IR | 3 | 3 | ENST00000416464 | -14.55 | 2.06E-02 | 4.66E-02 |
| ZNF692 | 1:248857975-248858385 | 1:248858018-248858130 | IR | 3 | 3 | ENST00000533976 | -14.57 | 2.20E-02 | 7.37E-02 |
| TOP3B | 22:21970407-21971876 | 22:21970980-21971061 | SES | 3 | 3 | ENST00000444502 | -14.59 | 4.50E-03 | 1.79E-02 |
| SALL2 | 14:21522245-21525336 | 14:21522833-21524931 | SES | 3 | 3 | ENST00000450879 | -14.65 | 9.37E-03 | 1.71E-02 |
| PRMT2 | 21:46635764-46636895 | 21:46636439-46636547 | SES | 3 | 3 | ENST00000355680 | -14.67 | 5.39E-03 | 2.10E-02 |
| LDOC1 | X:141173235-141177129 | X:141176812-141176892 | IR | 3 | 3 | ENST00000370526 | -14.7 | 2.68E-03 | 1.48E-01 |
| TNFRSF25 | 1:6462863-6464719 | 1:6463128-6464374 | IR | 3 | 3 | ENST00000513135 | -14.7 | 1.81E-02 | 6.54E-02 |
| STK39 | 2:168063571-168074981 | 2:168065319-168065381 | SES | 3 | 3 | ENST00000355999 | -14.76 | 1.06E-02 | 4.50E-02 |
| OSBPL7 | 17:47819721-47820096 | 17:47819783-47819970 | IR | 3 | 3 | ENST00000585051 | -14.79 | 1.58E-02 | 3.69E-02 |
| ABHD16A | 6:31701029-31703149 | 6:31701274-31701340 | MES | 3 | 3 | ENST00000395952 | -14.8 | 3.49E-02 | 3.51E-01 |
| FGGY | 1:59626050-59660218 | 1:59638228-59638375 | MES | 3 | 3 | ENST00000371218 | -14.81 | 9.36E-03 | 3.38E-02 |
| U2AF1L4 | 19:35742620-35743808 | 19:35742620-35742803 | TSSIA3ss | 3 | 3 | TSS.ENST00000292879 | -14.82 | 3.10E-03 | 8.11E-03 |
| TANGO2 | 22:20052585-20063337 | 22:20061530-20061683 | MXS | 3 | 3 | ENST00000398042 | -14.82 | 3.38E-02 | 1.27E-01 |
| PPP1R37 | 19:45145094-45148077 | 19:45145261-45145352 | IR (overlapping regio | 3 | 3 | ENST00000422370 | -14.86 | 1.94E-02 | 5.18E-02 |
| CHST4 | 16:71526313-71536659 | 16:71526313-71526495 | A5ss | 3 | 3 | ENST00000539698 | -14.86 | 4.50E-02 | 9.97E-02 |
| STAG3L5P-PVRIG2P-PILRB | 7:100336221-100338178 | 7:100336221-100336337 | TSSIA5ss | 3 | 3 | TSS.ENST00000310771 | -14.88 | 1.08E-02 | 1.24E-01 |
| CHML | 1:241635798-241639881 | 1:241635798-241636073 | A3ss | 3 | 3 | ENST00000366553 | -14.9 | 1.05E-02 | 3.50E-02 |
| CHML | 1:241635795-241639881 | 1:241635795-241636073 | A3ss | 3 | 3 | Ex.ENST00000366553 | -14.9 | 1.05E-02 | 3.50E-02 |
| WIPI2 | 7:5190494-5214534 | 7:5193118-5193171 | MES | 3 | 3 | ENST00000288828 | -14.93 | 1.27E-03 | 1.76E-02 |
| PRMT2 | 21:46635764-46636895 | 21:46635764-46636547 | TSSIA5ss | 3 | 3 | TSS.ENST00000397637 | -14.93 | 1.98E-03 | 7.81E-03 |
| CYP2B6 | 19:41004447-41012297 | 19:41009219-41009395 | MES | 2 | 3 | ENST00000324071 | -14.95 | 2.25E-02 | 5.97E-02 |

| | | | | | | | | | |
|---|---|---|---|---|---|---|---|---|---|
| LINC00869 | 1:149646599-149647610 | 1:149646946-149647380 | IR | 3 | 3 | ENST00000620190 | -14.96 | 5.50E-03 | 1.55E-02 |
| GUSBP11 | 22:23683868-23705428 | 22:23695173-23695517 | MXS | 3 | 3 | ENST00000422506 | -15.04 | 2.47E-02 | 9.84E-02 |
| TTC31 | 2:74492446-74492675 | 2:74492646-74492675 | A3ss | 3 | 3 | ENST00000233623 | -15.05 | 5.37E-03 | 2.71E-02 |
| POM121 | 7:72948328-72951440 | 7:72948598-72948667 | IR (overlapping regio | 3 | 3 | ENST00000395270 | -15.06 | 4.09E-02 | 5.80E-01 |
| FAHD2B | 2:97091986-97094700 | 2:97091986-97091988 | A3ss | 3 | 3 | Ex.TSS.ENST00000414820 | -15.09 | 2.24E-02 | 1.18E-01 |
| HYAL3 | 3:50294854-50299212 | 3:50294854-50295619 | TSSIA3ss | 3 | 3 | TSS.ENST00000336307 | -15.1 | 6.81E-03 | 4.46E-02 |
| ENTPD6 | 20:25224158-25225204 | 20:25225162-25225204 | A3ss | 3 | 3 | ENST00000376666 | -15.12 | 7.48E-03 | 2.76E-02 |
| FBXL19 | 16:30928500-30930072 | 16:30928500-30928628 | A5ss | 3 | 3 | ENST00000338343 | -15.13 | 6.49E-03 | 1.41E-02 |
| CAMK1 | 3:9757347-9757846 | 3:9757622-9757728 | IR (overlapping regio | 3 | 3 | ENST00000496534 | -15.17 | 1.13E-02 | 7.56E-02 |
| SUMF1 | 3:4362255-4410864 | 3:4376330-4376389 | SES | 3 | 3 | ENST00000272902 | -15.22 | 8.46E-03 | 5.40E-02 |
| SORBS2 | 4:185684843-185690561 | 4:185684843-185684846 | A3ss | 3 | 3 | ENST00000355634 | -15.22 | 2.16E-02 | 1.54E-01 |
| KRIT1 | 7:92245465-92246100 | 7:92245596-92245905 | IR (overlapping regio | 3 | 3 | ENST00000340022 | -15.25 | 1.01E-02 | 1.48E-01 |
| RFNG | 17:82049677-82050555 | 17:82049843-82050401 | IR | 3 | 3 | ENST00000580953 | -15.25 | 1.32E-02 | 3.22E-02 |
| AFG3L1P | 16:89972766-89980241 | 16:89977639-89977802 | MXS | 3 | 3 | ENST00000418696 | -15.25 | 3.13E-02 | 7.07E-02 |
| TSC2 | 16:2076586-2079031 | 16:2077598-2077726 | SES | 3 | 3 | ENST00000219476 | -15.26 | 1.78E-02 | 3.79E-02 |
| RAD52 | 12:929764-930144 | 12:929887-930050 | IR | 3 | 3 | ENST00000481052 | -15.26 | 4.31E-02 | 7.67E-02 |
| RAD52 | 12:929764-930144 | 12:929887-929972 | IR | 3 | 3 | ENST00000481052 | -15.26 | 4.31E-02 | 7.67E-02 |
| AC006504.5 | 19:27805605-27806785 | 19:27806186-27806322 | IR (overlapping regio | 3 | 3 | ENST00000585917 | -15.27 | 3.43E-02 | 8.83E-02 |
| BIRC2 | 11:102348598-102350749 | 11:102348765-102350000 | IR | 3 | 3 | ENST00000227758 | -15.28 | 7.64E-03 | 1.17E-02 |
| GUSBP11 | 22:23683868-23690231 | 22:23686831-23686995 | SES | 3 | 3 | ENST00000421064 | -15.32 | 4.94E-02 | 1.97E-01 |
| C15orf41 | 15:36808324-36810231 | 15:36808490-36808903 | IR (overlapping regio | 3 | 3 | ENST00000338183 | -15.36 | 1.81E-02 | 3.44E-02 |
| SELENBP1 | 1:151368320-151369712 | 1:151369004-151369712 | A5ss | 3 | 3 | ENST00000470345 | -15.36 | 3.49E-02 | 9.89E-02 |
| GUSBP2 | 6:26925276-26956388 | 6:26951086-26951159 | SES | 3 | 3 | ENST00000479900 | -15.39 | 1.90E-02 | 1.96E-01 |
| ALKBH6 | 19:36010646-36010893 | 19:36010646-36010683 | A3ss | 3 | 3 | ENST00000252984 | -15.41 | 4.50E-02 | 1.18E-01 |
| ZBTB44 | 11:130238557-130239811 | 11:130238557-130238607 | A3ss | 3 | 3 | ENST00000357899 | -15.43 | 1.20E-02 | 1.86E-02 |
| ZBTB44 | 11:130238554-130239811 | 11:130238554-130238607 | A3ss | 3 | 3 | Ex.ENST00000357899 | -15.43 | 1.20E-02 | 1.86E-02 |
| CLINT1 | 5:157789514-157791702 | 5:157789514-157789567 | A3ss | 3 | 3 | ENST00000523094 | -15.43 | 3.80E-02 | 3.18E-01 |
| NFKB1 | 4:102529915-102537857 | 4:102533845-102533885 | SES | 3 | 2 | ENST00000226574 | -15.45 | 9.27E-03 | 6.21E-02 |
| RBM33 | 7:155707069-155711202 | 7:155711173-155711202 | A3ss | 3 | 3 | ENST00000440108 | -15.48 | 5.06E-04 | 6.51E-03 |
| GIT2 | 12:109947505-109951166 | 12:109947505-109948846 | TSSIA3ss | 3 | 3 | TSS.ENST00000547815 | -15.48 | 1.78E-02 | 2.98E-02 |
| STAU2 | 8:73688814-73747386 | 8:73738284-73738349 | SES | 3 | 3 | ENST00000522509 | -15.5 | 1.22E-02 | 2.97E-01 |
| LYRM7 | 5:131199531-131205428 | 5:131202814-131203590 | IR | 3 | 3 | ENST00000379380 | -15.5 | 4.82E-02 | 3.78E-01 |
| COX11 | 17:54961327-54963305 | 17:54961327-54962915 | TSSIA3ss | 3 | 3 | TSS.ENST00000299335 | -15.52 | 1.57E-02 | 3.70E-02 |
| COX11 | 17:54961313-54963305 | 17:54961313-54962915 | TSSIA3ss | 3 | 3 | TSS.ENST00000299335 | -15.52 | 1.57E-02 | 3.70E-02 |
| SCAND1 | 20:35953629-35955102 | 20:35954340-35954447 | IR | 3 | 3 | ENST00000373991 | -15.61 | 7.23E-03 | 2.68E-02 |
| CHD2 | 15:92900825-92924320 | 15:92901167-92901299 | SES | 3 | 3 | ENST00000394196 | -15.62 | 2.95E-02 | 5.82E-02 |
| ZNF84 | 12:133037546-133041284 | 12:133041100-133041284 | TSSIA3ss | 3 | 3 | TSS.ENST00000543758 | -15.63 | 2.46E-02 | 4.21E-02 |
| ZNF84 | 12:133037546-133041277 | 12:133041100-133041277 | TSSIA3ss | 3 | 3 | TSS.ENST00000543758 | -15.63 | 2.46E-02 | 4.21E-02 |
| DLST | 14:74882017-74889094 | 14:74882591-74882624 | MXS | 3 | 3 | ENST00000554806 | -15.65 | 6.40E-05 | 1.20E-04 |
| GOLGB1 | 3:121730018-121730875 | 3:121730848-121730875 | A5ss | 3 | 3 | ENST00000482512 | -15.66 | 4.31E-02 | 2.31E-01 |
| C3orf18 | 3:50561748-50567462 | 3:50565466-50565861 | MES | 3 | 3 | ENST00000430746 | -15.7 | 2.63E-02 | 1.73E-01 |
| TMEM164 | X:110002773-110067346 | X:110003501-110067347 | A3ss | 3 | 3 | Ex.TSS.ENST00000372072 | -15.7 | 3.26E-02 | 1.00E+00 |
| TMEM164 | X:110002773-110003882 | X:110003501-110003883 | A3ss | 3 | 3 | Ex.TSS.ENST00000471255 | -15.7 | 3.26E-02 | 1.00E+00 |
| SNRPN | 15:24968083-24974386 | 15:24974311-24974386 | A3ss | 3 | 3 | ENST00000390687 | -15.73 | 4.60E-07 | 8.72E-07 |
| PIGG | 4:530840-533981 | 4:532821-533100 | IR | 3 | 3 | ENST00000513239 | -15.79 | 8.14E-04 | 6.07E-03 |
| ARMCX3 | X:101623277-101623813 | X:101623786-101623814 | A3ss | 3 | 3 | Ex.TSS.ENST00000471229 | -15.8 | 1.45E-02 | 6.16E-01 |
| CD4 | 12:6818543-6818846 | 12:6818543-6818650 | A5ss | 3 | 3 | Ex.ENST00000011653 | -15.83 | 2.90E-02 | 5.08E-02 |
| VRK3 | 19:50020648-50025345 | 19:50025267-50025345 | TSSIA5ss | 3 | 3 | TSS.ENST00000316763 | -16.01 | 1.70E-02 | 4.60E-02 |
| CREBZF | 11:85663524-85665102 | 11:85663602-85663697 | IR | 3 | 3 | ENST00000490820 | -16.01 | 2.10E-02 | 3.48E-02 |
| C1RL | 12:7097164-7101897 | 12:7099686-7099760 | SES | 3 | 3 | ENST00000543933 | -16.04 | 1.30E-02 | 2.31E-02 |
| FANCI | 15:89244045-89247628 | 15:89244045-89244067 | TSSIA5ss | 3 | 3 | TSS.ENST00000563250 | -16.08 | 2.93E-02 | 5.70E-02 |
| FANCI | 15:89244034-89247628 | 15:89244034-89244067 | TSSIA5ss | 3 | 3 | TSS.ENST00000563250 | -16.08 | 2.93E-02 | 5.70E-02 |
| TLCD5 | 11:120325369-120327451 | 11:120327122-120327451 | A3ss | 3 | 3 | Ex.TSS.ENST00000314475 | -16.09 | 6.90E-03 | 1.07E-02 |
| TLCD5 | 11:120325369-120327440 | 11:120327122-120327440 | A3ss | 3 | 3 | Ex.TSS.ENST00000314475 | -16.09 | 6.90E-03 | 1.07E-02 |
| ADCY3 | 2:24828024-24830708 | 2:24828024-24828161 | A3ss | 3 | 3 | ENST00000260600 | -16.1 | 1.50E-03 | 6.96E-03 |
| AP002856.2 | 11:131253592-131350308 | 11:131300358-131350309 | A3ss | 3 | 3 | Ex.ENST00000606885 | -16.17 | 5.39E-03 | 8.38E-03 |
| SUGP2 | 19:18990893-18993740 | 19:18991150-18991339 | IR | 3 | 3 | ENST00000452918 | -16.17 | 2.33E-02 | 5.95E-02 |
| ECHDC2 | 1:52896598-52899173 | 1:52897437-52897484 | SES | 3 | 3 | ENST00000358358 | -16.21 | 3.35E-02 | 1.20E-01 |
| ZC3H11A | 1:203795776-203796450 | 1:203796293-203796450 | TSSIA3ss | 3 | 3 | TSS.ENST00000367212 | -16.23 | 4.66E-02 | 1.53E-01 |
| MTHFR | 1:11785723-11790898 | 1:11788379-11788651 | IR (overlapping regio | 3 | 3 | ENST00000376590 | -16.24 | 2.54E-02 | 7.17E-02 |
| DDB2 | 11:47217050-47237836 | 11:47232814-47232959 | MES | 3 | 3 | ENST00000256996 | -16.27 | 4.06E-04 | 6.44E-04 |
| MAPK15 | 8:143721000-143721673 | 8:143721412-143721548 | IR | 3 | 3 | ENST00000461928 | -16.29 | 2.13E-02 | 3.77E-01 |
| FMNL1 | 17:45240476-45242146 | 17:45241891-45242079 | IR | 3 | 3 | ENST00000587856 | -16.34 | 4.02E-02 | 9.37E-02 |
| MAPKBP1 | 15:41814740-41815623 | 15:41815259-41815623 | A3ss | 2 | 3 | ENST00000505061 | -16.36 | 4.40E-02 | 8.05E-02 |
| SPEG | 2:219444985-219447973 | 2:219444985-219445161 | A5ss | 3 | 3 | ENST00000312358 | -16.38 | 4.60E-02 | 2.08E-01 |
| UPP1 | 7:48094828-48106872 | 7:48103297-48103411 | MES | 3 | 3 | ENST00000331803 | -16.42 | 2.37E-03 | 3.24E-02 |
| MPI | 15:74897512-74902219 | 15:74900600-74901949 | IR (overlapping regio | 3 | 3 | ENST00000352410 | -16.48 | 1.99E-02 | 3.88E-02 |
| ZNF266 | 19:9418762-9419299 | 19:9418896-9419050 | IR | 3 | 3 | ENST00000588221 | -16.49 | 2.16E-03 | 5.99E-03 |
| SLC12A4 | 16:67950494-67951139 | 16:67950654-67950711 | MES | 3 | 2 | ENST00000316341 | -16.5 | 1.24E-02 | 2.74E-02 |
| ZBTB46 | 20:63743666-63747301 | 20:63745312-63746871 | IR (overlapping regio | 3 | 3 | ENST00000245663 | -16.53 | 4.07E-03 | 1.57E-02 |
| RB1CC1 | 8:52684014-52686852 | 8:52685399-52685520 | MES | 3 | 2 | ENST00000518211 | -16.54 | 4.71E-02 | 1.00E+00 |
| MPI | 15:74897512-74902219 | 15:74900201-74900483 | IR (overlapping regio | 3 | 3 | ENST00000352410 | -16.55 | 6.07E-04 | 1.18E-03 |
| SUGP2 | 19:18990893-18993740 | 19:18991150-18993276 | IR | 3 | 3 | ENST00000452918 | -16.56 | 1.27E-02 | 3.24E-02 |
| RNF19A | 8:100303366-100309866 | 8:100303366-100303401 | TSSIA3ss | 2 | 2 | TSS.ENST00000523481 | -16.59 | 9.07E-03 | 1.41E-01 |
| SLC44A2 | 19:10642452-10643278 | 19:10642897-10643278 | TSSIA3ss | 3 | 3 | TSS.ENST00000586078 | -16.6 | 5.11E-04 | 1.28E-03 |
| CAMSAP2 | 1:200844870-200847257 | 1:200847210-200847257 | A3ss | 3 | 3 | ENST00000236925 | -16.71 | 9.36E-04 | 3.04E-03 |
| DPP7 | 9:137114463-137114726 | 9:137114577-137114646 | IR | 3 | 3 | ENST00000478597 | -16.71 | 1.67E-02 | 5.48E-01 |
| LDOC1 | X:141173235-141177129 | X:141176409-141176892 | IR | 3 | 3 | ENST00000370526 | -16.73 | 1.85E-03 | 1.02E-01 |
| RFNG | 17:82049677-82050555 | 17:82049843-82049917 | IR | 3 | 3 | ENST00000580953 | -16.75 | 2.51E-02 | 6.11E-02 |
| FBXO36 | 2:229922610-229976240 | 2:229932771-229932886 | SES | 2 | 3 | ENST00000373652 | -16.76 | 8.13E-03 | 3.68E-02 |
| CPT1B | 22:50577035-50577355 | 22:50577324-50577355 | A5ss | 3 | 3 | ENST00000312108 | -16.76 | 2.05E-02 | 8.42E-02 |
| GALNT13 | 2:153872177-153900935 | 2:153872177-153872303 | A5ss | 3 | 3 | Ex.TSS.ENST00000434213 | -16.78 | 1.45E-02 | 6.16E-02 |
| HPN | 19:35041873-35049289 | 19:35041873-35042522 | TSSIA5ss | 3 | 3 | TSS.ENST00000262626 | -16.83 | 4.67E-02 | 1.21E-01 |
| CHD1 | 5:98868636-98869753 | 5:98869014-98869277 | SES | 3 | 3 | ENST00000511067 | -16.86 | 2.88E-02 | 2.75E-01 |
| SALL3 | 18:78992074-78995462 | 18:78994906-78995121 | IR | 3 | 3 | ENST00000537592 | -16.9 | 2.10E-02 | 5.25E-02 |
| ZSCAN30 | 18:35264456-35290130 | 18:35267476-35267679 | SES | 2 | 2 | ENST00000592211 | -16.95 | 2.87E-04 | 7.08E-04 |
| FAHD2B | 2:97091713-97094700 | 2:97091863-97091988 | SES | 3 | 3 | ENST00000414820 | -16.96 | 1.17E-02 | 6.17E-02 |
| CYB5RL | 1:54184266-54195418 | 1:54190738-54190896 | MXS | 3 | 3 | ENST00000493530 | -16.99 | 1.58E-02 | 5.69E-02 |
| FOXP1 | 3:70970806-70972554 | 3:70972011-70972180 | SES | 3 | 2 | ENST00000327590 | -17 | 4.05E-02 | 2.76E-01 |
| TPD52L2 | 20:63873817-63882718 | 20:63875816-63875875 | SES | 3 | 3 | ENST00000217121 | -17.06 | 1.69E-05 | 6.51E-05 |
| SRPK2 | 7:105125875-105126992 | 7:105126248-105126340 | SES | 3 | 3 | ENST00000465072 | -17.08 | 2.35E-02 | 2.75E-01 |
| SYT7 | 11:61533125-61551383 | 11:61546031-61546255 | MXS | 3 | 3 | ENST00000540677 | -17.13 | 8.45E-04 | 1.35E-03 |
| KCTD17 | 22:37060823-37061408 | 22:37060923-37061103 | IR | 3 | 3 | ENST00000462640 | -17.13 | 2.28E-02 | 9.31E-02 |
| AMT | 3:49416794-49418372 | 3:49417368-49417817 | IR (overlapping regio | 3 | 3 | ENST00000636594 | -17.17 | 2.85E-03 | 1.84E-02 |
| GOLGA2P10 | 15:82477124-82477260 | 15:82477203-82477260 | A5ss | 3 | 3 | Ex.ENST00000550961 | -17.17 | 3.03E-03 | 5.94E-03 |
| CLEC11A | 19:50725022-50725708 | 19:50725082-50725315 | IR | 3 | 3 | ENST00000250340 | -17.22 | 1.43E-02 | 3.89E-02 |
| NDE1 | 16:15696861-15724190 | 16:15699729-15699854 | SES | 3 | 3 | ENST00000572967 | -17.26 | 2.00E-02 | 4.23E-02 |
| FAM122C | X:134807169-134807708 | X:134807342-134807539 | IR | 3 | 3 | ENST00000646004 | -17.27 | 1.32E-02 | 6.88E-01 |
| SLC12A5-AS1 | 20:46013500-46014788 | 20:46013797-46014123 | IR (overlapping regio | 2 | 3 | ENST00000535913 | -17.3 | 2.73E-02 | 1.03E-01 |
| GPSM2 | 1:108877229-108885506 | 1:108885275-108885506 | TSSIA3ss | 3 | 3 | TSS.ENST00000264126 | -17.32 | 4.86E-03 | 1.36E-02 |
| MTFR1L | 1:25826827-25829686 | 1:25829509-25829686 | A3ss | 3 | 3 | ENST00000374300 | -17.36 | 8.82E-03 | 3.01E-02 |

| | | | | | | | | | |
|---|---|---|---|---|---|---|---|---|---|
| MTFR1L | 1:25826827-25829617 | 1:25829509-25829617 | A3ss | 3 | 3 | ENST00000374300 | -17.36 | 8.82E-03 | 3.01E-02 |
| MTFR1L | 1:25826827-25829544 | 1:25829509-25829544 | A3ss | 3 | 3 | ENST00000374300 | -17.36 | 8.82E-03 | 3.01E-02 |
| ZNF263 | 16:3284206-3289392 | 16:3285059-3285239 | MES | 3 | 3 | ENST00000219069 | -17.36 | 2.44E-02 | 5.29E-02 |
| KREMEN2 | 16:2967320-2968380 | 16:2967446-2967809 | IR | 3 | 3 | ENST00000572045 | -17.36 | 3.38E-02 | 7.29E-02 |
| SPEG | 2:219451280-219461881 | 2:219451625-219451807 | SES | 3 | 3 | ENST00000312358 | -17.37 | 1.10E-02 | 4.98E-02 |
| SLC44A2 | 19:10642897-10644557 | 19:10643022-10643278 | IR | 3 | 3 | ENST00000586078 | -17.38 | 1.55E-04 | 3.90E-04 |
| PDCD4 | 10:110876071-110881274 | 10:110881233-110881274 | A3ss | 3 | 3 | ENST00000280154 | -17.48 | 4.02E-03 | 6.00E-03 |
| ZNF266 | 19:9418505-9419081 | 19:9418632-9418765 | IR | 3 | 3 | ENST00000591213 | -17.53 | 9.94E-03 | 2.75E-02 |
| ZNF266 | 19:9418505-9419081 | 19:9418632-9418761 | IR | 3 | 3 | ENST00000591213 | -17.53 | 9.94E-03 | 2.75E-02 |
| ARSL | X:2955538-2960377 | X:2958274-2958435 | MES | 3 | 3 | ENST00000545496 | -17.54 | 1.11E-02 | 9.21E-01 |
| NEK3 | 13:52154156-52156074 | 13:52154156-52154173 | A3ss | 3 | 3 | Ex.TSS.ENST00000550841 | -17.6 | 1.66E-03 | 2.99E-03 |
| TMEM218 | 11:125101304-125102733 | 11:125102132-125102317 | SES | 3 | 3 | ENST00000526175 | -17.62 | 4.68E-03 | 7.27E-03 |
| ASPHD1 | 16:29900488-29901920 | 16:29900567-29901551 | IR | 3 | 3 | ENST00000308748 | -17.64 | 5.46E-03 | 1.18E-02 |
| ARMCX5 | X:102599520-102604159 | X:102599573-102599705 | IR (overlapping regio | 3 | 3 | ENST00000604957 | -17.64 | 6.61E-03 | 1.70E-01 |
| RNF44 | 5:176532133-176532365 | 5:176532133-176532193 | A3ss | 3 | 3 | ENST00000274811 | -17.65 | 1.51E-03 | 1.27E-02 |
| RNF44 | 5:176532130-176532365 | 5:176532130-176532193 | A3ss | 3 | 3 | ENST00000274811 | -17.65 | 1.51E-03 | 1.27E-02 |
| GOLGA2P10 | 15:82477124-82477393 | 15:82477203-82477298 | SES | 3 | 3 | ENST00000550961 | -17.69 | 2.90E-03 | 5.69E-03 |
| CCPG1 | 15:55355250-55360944 | 15:55356410-55359538 | IR (overlapping regio | 3 | 2 | ENST00000310958 | -17.69 | 4.03E-02 | 7.75E-02 |
| MFSD8 | 4:127917925-127921771 | 4:127920837-127921117 | IR | 2 | 3 | ENST00000641482 | -17.73 | 3.84E-02 | 2.68E-01 |
| KIF20B | 10:89725159-89726412 | 10:89726293-89726412 | A3ss | 3 | 3 | ENST00000371728 | -17.74 | 8.86E-03 | 1.36E-02 |
| SLC16A5 | 17:75088045-75093588 | 17:75089151-75089304 | SES | 3 | 3 | ENST00000329783 | -17.77 | 3.26E-03 | 7.86E-03 |
| AMMECR1L | 2:127866997-127869453 | 2:127866997-127867279 | A3ss | 3 | 3 | Ex.ENST00000272647 | -17.77 | 1.15E-02 | 4.84E-02 |
| CDK10 | 16:89693274-89693495 | 16:89693327-89693397 | IR | 3 | 3 | ENST00000504473 | -17.79 | 1.65E-02 | 3.72E-02 |
| MPI | 15:74897512-74902219 | 15:74900138-74900508 | IR (overlapping regio | 3 | 3 | ENST00000352410 | -17.82 | 8.71E-03 | 1.70E-02 |
| ZNF232 | 17:5109769-5111799 | 17:5109769-5109868 | A3ss | 3 | 3 | ENST00000250076 | -17.88 | 2.30E-02 | 5.43E-02 |
| C9orf24 | 9:34381369-34381531 | 9:34381445-34381525 | IR | 3 | 3 | ENST00000379124 | -17.89 | 4.94E-02 | 1.00E+00 |
| MPI | 15:74897512-74902219 | 15:74900138-74900495 | IR (overlapping regio | 3 | 3 | ENST00000352410 | -17.94 | 4.44E-03 | 8.65E-03 |
| PHKB | 16:47461427-47497398 | 16:47463899-47463993 | MXS | 3 | 3 | ENST00000566037 | -18.07 | 2.53E-02 | 5.51E-02 |
| RB1CC1 | 8:52624788-52628040 | 8:52628032-52628040 | A5ss | 3 | 3 | ENST00000025008 | -18.11 | 3.07E-02 | 6.70E-01 |
| ANO8 | 19:17328162-17328983 | 19:17328265-17328539 | IR | 3 | 3 | ENST00000159087 | -18.23 | 2.36E-02 | 5.94E-02 |
| BAG4 | 8:38176588-38177139 | 8:38176590-38177004 | IR (overlapping regio | 3 | 3 | ENST00000432471 | -18.25 | 3.34E-03 | 7.55E-02 |
| AL645939.2 | 6:29722981-29723971 | 6:29723528-29723657 | IR (overlapping regio | 3 | 3 | ENST00000427340 | -18.26 | 3.55E-02 | 3.57E-01 |
| TMEM52 | 1:1918253-1918934 | 1:1918432-1918892 | IR | 3 | 3 | ENST00000602604 | -18.29 | 2.56E-02 | 8.25E-02 |
| FRAS1 | 4:78464584-78469977 | 4:78466208-78466435 | SES | 3 | 3 | ENST00000512123 | -18.29 | 3.74E-02 | 1.16E-01 |
| LNP1 | 3:100429886-100455776 | 3:100451736-100451949 | SES | 3 | 3 | ENST00000466996 | -18.33 | 2.39E-02 | 1.26E-01 |
| LNP1 | 3:100429886-100455776 | 3:100451680-100451949 | SES | 3 | 3 | ENST00000489752 | -18.33 | 2.39E-02 | 1.26E-01 |
| ARMCX5 | X:102599520-102604159 | X:102599573-102599711 | IR (overlapping regio | 3 | 3 | ENST00000604957 | -18.36 | 6.56E-03 | 1.69E-01 |
| TTC13 | 1:230933862-230954332 | 1:230940440-230940556 | MES | 3 | 3 | ENST00000366661 | -18.39 | 1.55E-02 | 5.26E-02 |
| PLD3 | 19:40348769-40365717 | 19:40365553-40365717 | TSSlA3ss | 3 | 3 | TSS.ENST00000594085 | -18.45 | 9.51E-03 | 2.52E-02 |
| PLD3 | 19:40348769-40365661 | 19:40365553-40365661 | TSSlA3ss | 3 | 3 | TSS.ENST00000594085 | -18.45 | 9.51E-03 | 2.52E-02 |
| PLD3 | 19:40348725-40365717 | 19:40365553-40365585 | SES | 3 | 3 | ENST00000392032 | -18.49 | 6.73E-03 | 1.79E-02 |
| INO80E | 16:30004658-30005367 | 16:30005221-30005367 | A3ss | 3 | 3 | Ex.TSS.ENST00000540562 | -18.54 | 4.15E-02 | 8.69E-02 |
| DNASE1 | 16:3657720-3658207 | 16:3657817-3657905 | IR (overlapping regio | 3 | 3 | ENST00000576792 | -18.56 | 8.12E-03 | 1.77E-02 |
| MPI | 15:74897512-74902219 | 15:74900138-74901949 | IR (overlapping regio | 3 | 3 | ENST00000352410 | -18.59 | 1.51E-04 | 2.94E-04 |
| ODF2L | 1:86384802-86386914 | 1:86385458-86385590 | SES | 3 | 3 | ENST00000294678 | -18.73 | 1.05E-03 | 3.84E-03 |
| ANK2 | 4:113369806-113373089 | 4:113372533-113372625 | SES | 2 | 3 | ENST00000510275 | -18.75 | 4.86E-02 | 3.34E-01 |
| SYNE4 | 19:36005438-36008553 | 19:36007125-36007268 | MES | 3 | 3 | ENST00000324444 | -18.78 | 9.46E-03 | 2.47E-02 |
| MTHFR | 1:11802881-11804532 | 1:11803130-11803798 | IR | 3 | 3 | ENST00000376592 | -18.78 | 1.62E-02 | 4.56E-02 |
| CCDC78 | 16:724542-725145 | 16:724807-724910 | IR | 3 | 3 | ENST00000538176 | -18.79 | 1.09E-02 | 2.42E-02 |
| AMMECR1L | 2:127866997-127869453 | 2:127867145-127869454 | A5ss | 3 | 3 | Ex.ENST00000272647 | -18.8 | 2.16E-02 | 9.10E-02 |
| ALMS1 | 2:73520017-73557219 | 2:73534824-73534949 | MXS | 3 | 3 | ENST00000476650 | -18.82 | 1.54E-02 | 7.83E-02 |
| DAB1 | 1:57026044-57062883 | 1:57033523-57033573 | MES | 3 | 3 | ENST00000371231 | -18.87 | 1.24E-02 | 4.48E-02 |
| ZNF513 | 2:27378616-27378883 | 2:27378616-27378777 | A3ss | 3 | 3 | Ex.TSS.ENST00000491924 | -18.88 | 6.55E-03 | 3.06E-02 |
| USP30 | 12:109052762-109067523 | 12:109057926-109058108 | MES | 3 | 3 | ENST00000257548 | -18.95 | 9.58E-03 | 1.60E-02 |
| TRIB2 | 2:12717011-12723259 | 2:12717640-12723260 | A3ss | 3 | 3 | Ex.TSS.ENST00000381465 | -18.97 | 2.31E-03 | 9.69E-03 |
| ASPHD1 | 16:29900488-29901920 | 16:29900742-29901551 | IR | 3 | 3 | ENST00000308748 | -19 | 4.72E-03 | 1.02E-02 |
| AGAP1 | 2:235908907-235968461 | 2:235930765-235930923 | SES | 3 | 3 | ENST00000304032 | -19.01 | 7.43E-04 | 3.39E-03 |
| SFI1 | 22:31508377-31546860 | 22:31528690-31528863 | MXS | 3 | 3 | ENST00000540643 | -19.05 | 1.04E-02 | 4.25E-02 |
| SPIB | 19:50422546-50423604 | 19:50422823-50423037 | SES | 3 | 3 | ENST00000270632 | -19.1 | 2.10E-02 | 5.70E-02 |
| GALT | 9:34647541-34647849 | 9:34647568-34647656 | IR (overlapping regio | 3 | 3 | ENST00000554330 | -19.13 | 1.71E-05 | 6.25E-04 |
| THAP9 | 4:82907936-82916943 | 4:82910697-82910811 | SES | 3 | 3 | ENST00000506208 | -19.18 | 4.21E-02 | 1.31E-01 |
| RSPRY1 | 16:57186443-57204503 | 16:57186443-57186451 | TSSlA5ss | 3 | 3 | TSS.ENST00000394420 | -19.2 | 3.30E-02 | 7.23E-02 |
| RSPRY1 | 16:57186430-57204503 | 16:57186430-57186451 | TSSlA5ss | 3 | 3 | TSS.ENST00000394420 | -19.2 | 3.30E-02 | 7.23E-02 |
| LRSAM1 | 9:127451513-127452084 | 9:127451670-127451928 | IR | 3 | 3 | ENST00000323301 | -19.2 | 4.42E-02 | 1.00E+00 |
| ZNF461 | 19:36656544-36658335 | 19:36658299-36658335 | A5ss | 2 | 3 | ENST00000360357 | -19.22 | 4.17E-02 | 1.09E-01 |
| CLUHP3 | 16:31707907-31711986 | 16:31708208-31708460 | IR | 3 | 3 | ENST00000562354 | -19.26 | 2.89E-03 | 6.28E-03 |
| AL645939.2 | 6:29722981-29723971 | 6:29723528-29723769 | IR (overlapping regio | 3 | 3 | ENST00000427340 | -19.35 | 3.03E-02 | 3.05E-01 |
| ZFP62 | 5:180847615-180851493 | 5:180848083-180849153 | IR | 3 | 3 | ENST00000512132 | -19.37 | 3.30E-03 | 2.90E-02 |
| AC084125.2 | 8:144497977-144500280 | 8:144500200-144500266 | IR (overlapping regio | 3 | 3 | ENST00000569326 | -19.5 | 2.12E-02 | 3.77E-01 |
| FLVCR2 | 14:75635014-75640954 | 14:75639352-75639462 | SES | 3 | 2 | ENST00000238667 | -19.53 | 1.37E-02 | 2.56E-02 |
| SLC13A3 | 20:46613726-46684395 | 20:46613726-46670139 | A3ss | 3 | 3 | Ex.TSS.ENST00000372121 | -19.55 | 5.93E-03 | 2.25E-02 |
| TRIM37 | 17:58999460-59012327 | 17:59001598-59001714 | SES | 3 | 3 | ENST00000262294 | -19.58 | 4.18E-03 | 9.89E-03 |
| ARHGAP26 | 5:143147382-143213996 | 5:143207198-143207308 | MXS | 3 | 3 | ENST00000645722 | -19.58 | 3.62E-02 | 2.99E-01 |
| NFKB1 | 4:102529915-102533847 | 4:102533845-102533847 | A3ss | 3 | 2 | ENST00000226574 | -19.62 | 2.50E-02 | 7.80E-02 |
| MXRA8 | 1:1354193-1354509 | 1:1354233-1354353 | IR | 3 | 3 | ENST00000476718 | -19.65 | 2.23E-02 | 6.24E-02 |
| MAPKBP1 | 15:41814740-41815623 | 15:41815259-41815405 | SES | 2 | 2 | ENST00000456763 | -19.68 | 1.62E-02 | 3.09E-02 |
| AFG3L1P | 16:89972766-89977714 | 16:89977639-89977714 | A3ss | 3 | 3 | Ex.TSS.ENST00000418696 | -19.7 | 1.85E-02 | 4.17E-02 |
| AFG3L1P | 16:89972766-89977677 | 16:89977639-89977677 | TSSlA3ss | 3 | 3 | TSS.ENST00000418696 | -19.7 | 1.85E-02 | 4.17E-02 |
| BRD3OS | 9:134026208-134031587 | 9:134027263-134028825 | IR | 3 | 3 | ENST00000603928 | -19.74 | 8.66E-03 | 2.72E-01 |
| CLK4 | 5:178616949-178618555 | 5:178617344-178618555 | A5ss | 3 | 3 | ENST00000520878 | -19.74 | 1.56E-02 | 1.36E-01 |
| PRXL2A | 10:80408644-80420482 | 10:80420466-80420482 | TSSlA3ss | 3 | 3 | TSS.ENST00000606162 | -19.77 | 8.03E-03 | 1.23E-02 |
| ZNF326 | 1:90005245-90007611 | 1:90007345-90007611 | A3ss | 3 | 3 | ENST00000340281 | -19.8 | 7.25E-04 | 2.65E-03 |
| ZNF326 | 1:90005245-90007608 | 1:90007345-90007608 | A3ss | 3 | 3 | Ex.ENST00000340281 | -19.8 | 7.25E-04 | 2.65E-03 |
| SYT6 | 1:114103722-114137494 | 1:114103722-114137065 | A3ss | 3 | 3 | Ex.ENST00000369547 | -19.85 | 9.53E-04 | 2.68E-03 |
| ZNF142 | 2:218656464-218658979 | 2:218658701-218658876 | SES | 3 | 3 | ENST00000432460 | -19.85 | 2.51E-02 | 1.13E-01 |
| NSUN5P1 | 7:75412933-75414884 | 7:75414640-75414798 | SES | 3 | 3 | ENST00000464979 | -19.87 | 4.43E-03 | 6.35E-02 |
| DOCK7 | 1:62477826-62488933 | 1:62477826-62477834 | A3ss | 3 | 3 | ENST00000454575 | -19.88 | 3.97E-02 | 1.43E-01 |
| AL138720.1 | 6:14997476-15021820 | 6:14997476-15009084 | A3ss | 3 | 3 | Ex.TSS.ENST00000629853 | -20.02 | 4.42E-02 | 4.40E-01 |
| DNAJC1 | 10:21806100-21882281 | 10:21807191-21882282 | A5ss | 3 | 3 | Ex.ENST00000376980 | -20.02 | 4.83E-02 | 7.28E-02 |
| ZSCAN30 | 18:35264456-35290083 | 18:35267476-35267679 | SES | 2 | 2 | ENST00000420878 | -20.03 | 1.60E-03 | 3.94E-03 |
| CCDC74A | 2:131529692-131531663 | 2:131530777-131530827 | SES | 3 | 3 | ENST00000409856 | -20.06 | 1.11E-02 | 4.67E-02 |
| APBB3 | 5:140561587-140562227 | 5:140561702-140561843 | IR | 3 | 3 | ENST00000510241 | -20.08 | 8.40E-03 | 6.86E-02 |
| NFKBIZ | 3:101855903-101857291 | 3:101857073-101857183 | SES | 2 | 3 | ENST00000326151 | -20.12 | 1.02E-02 | 5.40E-02 |
| TAZ | X:154419624-154420031 | X:154419624-154419746 | A5ss | 3 | 3 | ENST00000426231 | -20.13 | 4.77E-03 | 3.54E-01 |
| STEAP2 | 7:90232337-90237631 | 7:90232485-90236890 | IR | 3 | 3 | ENST00000287908 | -20.15 | 4.51E-02 | 6.54E-01 |
| PRRT2 | 16:29812990-29815880 | 16:29813934-29814627 | IR (overlapping regio | 3 | 3 | ENST00000300797 | -20.26 | 2.78E-02 | 5.83E-02 |
| PANK4 | 1:2510749-2511337 | 1:2510749-2510782 | A3ss | 3 | 3 | ENST00000378466 | -20.28 | 4.67E-02 | 1.56E-01 |
| AC009005.1 | 19:567625-571439 | 19:567625-567648 | TSSlA3ss | 3 | 3 | TSS.ENST00000588908 | -20.41 | 3.63E-02 | 7.61E-02 |
| ZC3H11A | 1:203801629-203816925 | 1:203802957-203803016 | SES | 3 | 3 | ENST00000367212 | -20.42 | 4.56E-02 | 1.50E-01 |
| KCTD21-AS1 | 11:78140073-78141594 | 11:78141506-78141594 | TSSlA3ss | 3 | 2 | TSS.ENST00000667996 | -20.45 | 7.49E-03 | 1.24E-02 |
| LRRC14 | 8:144518042-144519700 | 8:144519615-144519701 | A3ss | 3 | 3 | Ex.TSS.ENST00000527730 | -20.51 | 5.87E-04 | 1.06E-02 |

| | | | | | | | | | |
|---|---|---|---|---|---|---|---|---|---|
| RFNG | 17:82049918-82050514 | 17:82050007-82050401 | IR | 3 | 3 | ENST00000580793 | -20.57 | 2.58E-02 | 6.30E-02 |
| ACSBG1 | 15:78167468-78171529 | 15:78169078-78171049 | IR (overlapping regio | 3 | 3 | ENST00000258873 | -20.58 | 2.09E-02 | 4.08E-02 |
| SYT7 | 11:61533125-61551383 | 11:61546031-61546255 | MES | 3 | 3 | ENST00000535826 | -20.6 | 2.38E-04 | 3.80E-04 |
| B3GNT8 | 19:41426440-41427283 | 19:41426440-41426810 | A3ss | 2 | 3 | Ex.ENST00000601379 | -20.63 | 2.75E-02 | 5.77E-02 |
| C3orf18 | 3:50561748-50567462 | 3:50565466-50565861 | MXS | 3 | 3 | ENST00000426034 | -20.63 | 3.21E-02 | 2.11E-01 |
| FAM122B | X:134796108-134797155 | X:134796257-134797120 | IR | 3 | 3 | ENST00000343004 | -20.64 | 6.38E-03 | 3.36E-01 |
| ZNF189 | 9:101399190-101399925 | 9:101399884-101399925 | TSSIA3ss | 3 | 3 | TSS.ENST00000339664 | -20.65 | 3.51E-02 | 8.94E-01 |
| ZC3H13 | 13:46045517-46052697 | 13:46052404-46052698 | A5ss | 3 | 3 | Ex.TSS.ENST00000282007 | -20.76 | 3.47E-02 | 6.21E-02 |
| ELF2 | 4:139072004-139073453 | 4:139072004-139072039 | A3ss | 3 | 3 | ENST00000358635 | -20.8 | 3.72E-02 | 1.16E-01 |
| ACAD11 | 3:132578882-132579491 | 3:132578882-132579056 | TSSIA3ss | 3 | 3 | TSS.ENST00000477604 | -20.81 | 3.48E-02 | 1.91E-01 |
| BANP | 16:87984260-88006089 | 16:88004295-88004411 | SES | 3 | 3 | ENST00000286122 | -20.82 | 4.72E-02 | 1.05E-01 |
| GALNT10 | 5:154329739-154380447 | 5:154376277-154376462 | MES | 3 | 3 | ENST00000520647 | -20.88 | 4.35E-03 | 3.64E-02 |
| MPI | 15:74897512-74902219 | 15:74900138-74901168 | IR (overlapping regio | 3 | 3 | ENST00000352410 | -20.9 | 3.01E-05 | 5.86E-05 |
| MIAT | 22:26663399-26665530 | 22:26665457-26665530 | TSSIA3ss | 3 | 3 | TSS.ENST00000418918 | -20.9 | 1.42E-02 | 5.71E-02 |
| ZNF382 | 19:36626130-36634114 | 19:36627489-36628373 | IR | 2 | 3 | ENST00000292928 | -21.14 | 3.36E-02 | 8.81E-02 |
| TNS2 | 12:53059047-53060258 | 12:53059972-53060166 | IR | 2 | 3 | ENST00000314250 | -21.18 | 2.41E-02 | 4.15E-02 |
| VCAN | 5:83512397-83537006 | 5:83519349-83522309 | SES | 3 | 3 | ENST00000265077 | -21.22 | 2.01E-02 | 1.89E-01 |
| LETMD1 | 12:51048479-51056349 | 12:51049034-51049185 | MES | 3 | 3 | ENST00000262055 | -21.26 | 2.22E-02 | 3.84E-02 |
| SPG11 | 15:44570659-44583813 | 15:44572683-44572820 | SES | 3 | 3 | ENST00000535302 | -21.26 | 3.89E-02 | 7.43E-02 |
| ANKHD1 | 5:140436258-140438502 | 5:140438461-140438502 | A3ss | 3 | 3 | ENST00000297183 | -21.27 | 6.86E-03 | 5.60E-02 |
| ANKHD1 | 5:140436258-140438493 | 5:140438461-140438493 | A3ss | 3 | 3 | ENST00000297183 | -21.27 | 6.86E-03 | 5.60E-02 |
| PPP4R4 | 14:94208567-94230586 | 14:94227275-94227378 | SES | 3 | 2 | ENST00000555690 | -21.33 | 2.40E-02 | 4.52E-02 |
| TRIM16L | 17:18734877-18736118 | 17:18735202-18736016 | IR | 3 | 3 | ENST00000395671 | -21.34 | 4.15E-02 | 9.41E-02 |
| THTPA | 14:23556294-23557039 | 14:23556344-23556742 | IR (overlapping regio | 3 | 3 | ENST00000554789 | -21.4 | 2.06E-04 | 3.77E-04 |
| ZNF620 | 3:40506126-40506303 | 3:40506126-40506138 | TSSIA5ss | 3 | 3 | TSS.ENST00000314529 | -21.49 | 2.28E-03 | 1.45E-02 |
| SARDH | 9:133685287-133702915 | 9:133696223-133696361 | MES | 3 | 3 | ENST00000371868 | -21.56 | 9.35E-03 | 2.93E-01 |
| TAZ | X:154419624-154420031 | X:154419705-154419746 | SES | 3 | 3 | ENST00000439735 | -21.58 | 2.87E-03 | 2.13E-01 |
| RCC1 | 1:28508906-28529857 | 1:28516725-28516867 | SES | 3 | 3 | ENST00000649185 | -21.59 | 7.03E-04 | 2.41E-03 |
| KANSL3 | 2:96619545-96631311 | 2:96619672-96619762 | SES | 3 | 3 | ENST00000354204 | -21.7 | 2.09E-02 | 1.10E-01 |
| DPPA4 | 3:109329082-109331733 | 3:109330524-109331734 | A5ss | 3 | 3 | Ex.ENST00000463966 | -21.78 | 4.52E-03 | 2.40E-02 |
| INPP5B | 1:37860697-37862430 | 1:37861092-37861494 | IR (overlapping regio | 3 | 3 | ENST00000373024 | -21.82 | 6.43E-03 | 2.24E-02 |
| ZNF142 | 2:218651823-218656149 | 2:218651823-218652300 | TSSIA3ss | 3 | 3 | TSS.ENST00000440934 | -21.83 | 7.48E-03 | 3.37E-02 |
| ZC3H13 | 13:46045514-46052403 | 13:46045514-46045516 | A3ss | 3 | 3 | Ex.TSS.ENST00000242848 | -21.92 | 7.34E-03 | 1.32E-02 |
| POLM | 7:44079742-44079860 | 7:44079742-44079756 | TSSIA3ss | 2 | 3 | TSS.ENST00000452049 | -21.93 | 1.94E-03 | 2.59E-02 |
| TRIP12 | 2:229793131-229795178 | 2:229793131-229793145 | A3ss | 3 | 3 | Ex.ENST00000283943 | -21.96 | 2.11E-03 | 9.55E-03 |
| MICALL2 | 7:1437890-1439431 | 7:1438354-1438839 | IR | 2 | 3 | ENST00000496184 | -21.97 | 1.15E-02 | 1.43E-01 |
| OBSCN | 1:228359597-228361250 | 1:228360313-228360889 | IR (overlapping regio | 3 | 3 | ENST00000284548 | -21.99 | 4.79E-03 | 1.63E-02 |
| COASY | 17:42562148-42563322 | 17:42562237-42562611 | IR | 3 | 3 | ENST00000393818 | -22.04 | 3.38E-03 | 7.81E-03 |
| PCNX2 | 1:233261322-233262957 | 1:233262045-233262165 | SES | 3 | 3 | ENST00000258229 | -22.05 | 2.38E-03 | 8.13E-03 |
| MRPL4 | 19:10259172-10260025 | 19:10259242-10259823 | IR (overlapping regio | 3 | 3 | ENST00000393733 | -22.08 | 8.28E-03 | 2.07E-02 |
| DOP1A | 6:83162920-83167861 | 6:83164660-83164719 | SES | 2 | 3 | ENST00000237163 | -22.13 | 2.04E-02 | 2.32E-01 |
| IMMT | 2:86171310-86173649 | 2:86171310-86171312 | A3ss | 3 | 3 | Ex.ENST00000254636 | -22.15 | 1.54E-03 | 7.91E-03 |
| HEXIM2 | 17:45161940-45162749 | 17:45162488-45162635 | SES | 3 | 3 | ENST00000591070 | -22.17 | 3.64E-02 | 8.48E-02 |
| THTPA | 14:23556294-23557039 | 14:23556344-23556752 | IR (overlapping regio | 3 | 3 | ENST00000554789 | -22.26 | 3.47E-04 | 6.36E-04 |
| MYO1B | 2:191400469-191408114 | 2:191400749-191400835 | MES | 3 | 3 | ENST00000304164 | -22.29 | 3.42E-02 | 1.48E-01 |
| TRIM16 | 17:15683036-15684299 | 17:15683372-15684195 | IR | 3 | 3 | ENST00000416464 | -22.36 | 7.67E-05 | 1.74E-04 |
| C3orf18 | 3:50565701-50567462 | 3:50565701-50565861 | TSSIA3ss | 3 | 2 | TSS.ENST00000426034 | -22.38 | 1.59E-02 | 1.04E-01 |
| SH2D3A | 19:6754933-6759593 | 19:6754933-6755315 | A3ss | 3 | 3 | ENST00000245908 | -22.38 | 1.82E-02 | 5.03E-02 |
| HAGHL | 16:727141-727411 | 16:727141-727175 | TSSIA5ss | 3 | 3 | TSS.ENST00000564537 | -22.39 | 1.44E-02 | 3.20E-02 |
| MTERF2 | 12:106978772-106986968 | 12:106985115-106985225 | SES | 3 | 3 | ENST00000240050 | -22.58 | 1.03E-02 | 1.72E-02 |
| HPN | 19:35041873-35049289 | 19:35042453-35042522 | SES | 3 | 3 | ENST00000392226 | -22.67 | 5.50E-03 | 1.42E-02 |
| PRSS16 | 6:27251265-27252807 | 6:27251750-27252040 | SES | 3 | 3 | ENST00000230582 | -22.69 | 4.30E-03 | 4.43E-02 |
| ZNF10 | 12:133130755-133144433 | 12:133130755-133131014 | TSSIA5ss | 2 | 2 | TSS.ENST00000426665 | -22.73 | 3.77E-02 | 6.46E-02 |
| APBB3 | 5:140563752-140564469 | 5:140563916-140564196 | IR (overlapping regio | 3 | 3 | ENST00000506958 | -22.8 | 3.62E-02 | 2.95E-01 |
| SNHG14 | 15:25051677-25053265 | 15:25051677-25052140 | TSSIA5ss | 3 | 3 | TSS.ENST00000657798 | -22.83 | 4.14E-02 | 7.84E-02 |
| TTLL4 | 2:218746232-218747029 | 2:218747003-218747029 | A3ss | 3 | 3 | ENST00000258398 | -22.87 | 3.61E-04 | 1.63E-03 |
| ZSCAN9 | 6:28225331-28227011 | 6:28225331-28225366 | TSSIA5ss | 3 | 3 | TSS.ENST00000252207 | -22.87 | 4.81E-02 | 4.99E-01 |
| BCAM | 19:44811347-44812225 | 19:44812163-44812225 | A3ss | 3 | 3 | ENST00000270233 | -22.9 | 1.26E-03 | 3.34E-03 |
| COASY | 17:42562148-42563322 | 17:42562214-42562611 | IR | 3 | 3 | ENST00000393818 | -22.99 | 1.67E-03 | 3.88E-03 |
| RIPOR2 | 6:24839600-24842861 | 6:24839600-24840775 | TSSIA3ss | 3 | 3 | TSS.ENST00000510784 | -23.03 | 4.74E-02 | 4.87E-01 |
| ZNF268 | 12:133192004-133202143 | 12:133193428-133193533 | SES | 3 | 3 | ENST00000534953 | -23.04 | 1.05E-02 | 1.80E-02 |
| SCUBE1 | 22:43238955-43258218 | 22:43255488-43255577 | SES | 2 | 3 | ENST00000290460 | -23.06 | 1.88E-02 | 7.78E-02 |
| FST | 5:53485228-53485950 | 5:53485687-53485950 | TSSIA3ss | 3 | 2 | TSS.ENST00000396947 | -23.09 | 2.08E-02 | 1.88E-01 |
| AC068279.2 | 2:87323697-87348658 | 2:87323697-87345262 | A3ss | 3 | 3 | Ex.ENST00000664547 | -23.12 | 5.72E-03 | 2.95E-02 |
| ZNF696 | 8:143291611-143292971 | 8:143291611-143291767 | TSSIA5ss | 3 | 3 | TSS.ENST00000330143 | -23.18 | 5.02E-03 | 8.72E-02 |
| CD47 | 3:108047293-108057476 | 3:108051939-108051970 | MES | 3 | 3 | ENST00000361309 | -23.18 | 1.12E-02 | 5.88E-02 |
| NEDD1 | 12:96907625-96909751 | 12:96907625-96907856 | A5ss | 3 | 3 | ENST00000266742 | -23.2 | 1.88E-02 | 3.35E-02 |
| VSIG10 | 12:118095679-118102988 | 12:118095679-118095814 | A3ss | 3 | 3 | ENST00000536905 | -23.28 | 3.75E-02 | 6.33E-02 |
| CNTROB | 17:7948620-7949084 | 17:7949082-7949084 | A3ss | 3 | 3 | ENST00000565740 | -23.29 | 2.61E-02 | 5.47E-02 |
| C3orf18 | 3:50561748-50565525 | 3:50565466-50565525 | A5ss | 2 | 3 | ENST00000357203 | -23.3 | 4.14E-02 | 2.72E-01 |
| BBS9 | 7:33351324-33352858 | 7:33351324-33352049 | A5ss | 3 | 3 | Ex.ENST00000242067 | -23.33 | 2.58E-02 | 3.41E-01 |
| NCOA2 | 8:70296801-70403699 | 8:70403334-70403700 | A5ss | 3 | 2 | Ex.TSS.ENST00000452400 | -23.34 | 4.82E-02 | 1.00E+00 |
| RPS6KL1 | 14:74922443-74923179 | 14:74922443-74922485 | TSSIA3ss | 3 | 3 | TSS.ENST00000553894 | -23.43 | 3.34E-02 | 6.26E-02 |
| ZNF562 | 19:9656654-9660719 | 19:9658009-9658135 | MXS | 3 | 3 | ENST00000588653 | -23.48 | 4.05E-02 | 1.13E-01 |
| MAP6D1 | 3:183818108-183825146 | 3:183818108-183818111 | TSSIA3ss | 2 | 2 | TSS.ENST00000318631 | -23.68 | 1.21E-02 | 7.23E-02 |
| C11orf74 | 11:36610240-36648015 | 11:36636051-36636117 | MES | 3 | 3 | ENST00000334307 | -23.68 | 2.12E-02 | 3.35E-02 |
| WASHC2C | 10:45785632-45787034 | 10:45786612-45786674 | SES | 3 | 3 | ENST00000336378 | -23.7 | 1.88E-02 | 2.85E-02 |
| AC060780.1 | 17:43166980-43167849 | 17:43167251-43167733 | IR (overlapping regio | 2 | 3 | ENST00000635600 | -23.73 | 1.69E-02 | 3.93E-02 |
| VPS54 | 2:63981852-63983863 | 2:63981852-63981887 | A3ss | 3 | 3 | ENST00000272322 | -23.76 | 9.24E-03 | 4.49E-02 |
| APBB3 | 5:140561587-140562227 | 5:140561702-140562099 | IR | 3 | 3 | ENST00000510241 | -23.81 | 3.14E-02 | 2.56E-01 |
| ZNF142 | 2:218650527-218656149 | 2:218651701-218652300 | SES | 3 | 3 | ENST00000440934 | -23.82 | 3.99E-03 | 1.80E-02 |
| TAZ | X:154419624-154420031 | X:154419705-154420031 | A3ss | 3 | 3 | ENST00000470127 | -23.83 | 2.66E-03 | 1.98E-01 |
| APOL2 | 22:36231435-36233152 | 22:36231435-36231466 | A3ss | 3 | 3 | Ex.ENST00000249066 | -23.99 | 8.22E-03 | 3.36E-02 |
| SLC8A2 | 19:47457595-47471788 | 19:47465729-47466147 | SES | 2 | 3 | ENST00000594353 | -24.08 | 1.21E-02 | 3.23E-02 |
| CMTM7 | 3:32442014-32452391 | 3:32449454-32449552 | SES | 3 | 3 | ENST00000334983 | -24.14 | 1.32E-02 | 8.27E-02 |
| STAG3L5P-PVRIG2P-PILRB | 7:100345969-100349798 | 7:100349731-100349798 | A3ss | 3 | 3 | ENST00000444874 | -24.25 | 3.17E-02 | 3.63E-01 |
| EFHC1 | 6:52438304-52438454 | 6:52438376-52438454 | IR | 3 | 3 | ENST00000636311 | -24.34 | 2.96E-02 | 3.24E-01 |
| INO80E | 16:30001531-30005367 | 16:30005221-30005367 | TSSIA3ss | 3 | 3 | TSS.ENST00000563197 | -24.36 | 2.49E-02 | 5.22E-02 |
| YIPF2 | 19:10923175-10923271 | 19:10923175-10923191 | TSSIA3ss | 3 | 3 | TSS.ENST00000592646 | -24.37 | 1.76E-02 | 3.69E-02 |
| GPR108 | 19:6731055-6731198 | 19:6731055-6731111 | A3ss | 3 | 3 | ENST00000264080 | -24.58 | 9.44E-04 | 2.62E-03 |
| HDAC10 | 22:50245813-50245909 | 22:50245813-50245827 | A3ss | 3 | 3 | Ex.ENST00000216271 | -24.61 | 2.80E-02 | 1.13E-01 |
| HDAC10 | 22:50245737-50245909 | 22:50245737-50245827 | A3ss | 3 | 3 | ENST00000216271 | -24.61 | 2.80E-02 | 1.13E-01 |
| WASHC2A | 10:50126180-50127582 | 10:50127160-50127222 | SES | 3 | 3 | ENST00000282633 | -24.71 | 1.81E-02 | 2.75E-02 |
| COASY | 17:42562148-42563322 | 17:42562220-42562611 | IR | 3 | 3 | ENST00000393818 | -24.72 | 9.58E-04 | 2.22E-03 |
| R3HDM2 | 12:57292650-57295398 | 12:57292650-57292667 | A3ss | 3 | 3 | ENST00000402412 | -24.8 | 1.33E-02 | 2.33E-02 |
| ORAI2 | 7:102433662-102438943 | 7:102436225-102436333 | SES | 3 | 3 | ENST00000495936 | -24.83 | 2.82E-04 | 3.28E-03 |
| IQCJ-SCHIP1 | 3:159866282-159892096 | 3:159887708-159887905 | MES | 3 | 3 | ENST00000412423 | -24.92 | 2.08E-02 | 1.18E-01 |
| APBB3 | 5:140561587-140562227 | 5:140561723-140561843 | IR | 3 | 3 | ENST00000510241 | -24.96 | 6.23E-03 | 5.09E-02 |
| BRSK2 | 11:1449837-1450586 | 11:1450221-1450587 | A3ss | 3 | 3 | Ex.ENST00000308219 | -25 | 5.28E-03 | 8.26E-03 |
| FAM111A | 11:59144901-59145826 | 11:59144901-59145419 | A5ss | 3 | 3 | Ex.TSS.ENST00000529358 | -25.04 | 2.09E-02 | 3.34E-02 |
| FAM111A | 11:59144897-59145826 | 11:59144897-59145419 | A5ss | 3 | 3 | Ex.TSS.ENST00000361723 | -25.04 | 2.09E-02 | 3.34E-02 |

| | | | | | | | | | |
|---|---|---|---|---|---|---|---|---|---|
| FAM111A | 11:59143305-59145826 | 11:59143305-59145419 | A5ss | 3 | 3 | Ex.ENST00000420244 | -25.04 | 2.09E-02 | 3.34E-02 |
| TAF1C | 16:84184990-84186900 | 16:84184990-84185060 | TSSIA3ss | 3 | 3 | TSS.ENST00000564774 | -25.07 | 4.16E-04 | 9.25E-04 |
| TAF1C | 16:84184900-84186900 | 16:84184900-84185060 | TSSIA3ss | 3 | 3 | TSS.ENST00000564774 | -25.07 | 4.16E-04 | 9.25E-04 |
| PPP4R4 | 14:94208567-94227285 | 14:94227275-94227285 | A3ss | 3 | 2 | ENST00000555690 | -25.29 | 1.11E-02 | 2.09E-02 |
| AHDC1 | 1:27553186-27558705 | 1:27558305-27558530 | SES | 3 | 3 | ENST00000642245 | -25.45 | 6.43E-03 | 2.20E-02 |
| PRRT2 | 16:29812990-29815880 | 16:29813934-29814343 | IR (overlapping regio | 3 | 3 | ENST00000300797 | -25.45 | 7.21E-03 | 1.52E-02 |
| PRRT2 | 16:29812990-29815880 | 16:29813934-29814332 | IR (overlapping regio | 3 | 3 | ENST00000300797 | -25.58 | 9.13E-03 | 1.97E-02 |
| DPH5 | 1:100990632-100995109 | 1:100992637-100992740 | SES | 3 | 3 | ENST00000464270 | -25.83 | 5.11E-03 | 1.42E-02 |
| SLC29A3 | 10:71323055-71356080 | 10:71351562-71351788 | MES | 3 | 2 | ENST00000373189 | -25.85 | 1.56E-02 | 2.38E-02 |
| ANO7 | 2:241223186-241223955 | 2:241223782-241223904 | IR | 3 | 3 | ENST00000459928 | -26.16 | 1.84E-02 | 8.43E-02 |
| FAM160A2 | 11:6218151-6218599 | 11:6218558-6218599 | A5ss | 3 | 3 | ENST00000265978 | -26.3 | 7.42E-03 | 1.19E-02 |
| AATK | 17:81120201-81122823 | 17:81122427-81122534 | IR | 2 | 2 | ENST00000326724 | -26.4 | 1.47E-03 | 3.59E-03 |
| MKS1 | 17:58206381-58207893 | 17:58207085-58207218 | MES | 3 | 2 | ENST00000393119 | -26.65 | 4.59E-02 | 1.08E-01 |
| SGSH | 17:80214680-80215032 | 17:80214680-80214765 | A3ss | 3 | 3 | ENST00000326317 | -26.7 | 2.59E-03 | 5.44E-03 |
| CLK4 | 5:178616949-178618555 | 5:178617344-178617434 | SES | 3 | 3 | ENST00000316308 | -26.73 | 5.45E-03 | 4.74E-02 |
| SPINT1-AS1 | 15:40836283-40844178 | 15:40838543-40838686 | SES | 3 | 3 | ENST00000564302 | -26.85 | 2.54E-02 | 4.83E-02 |
| EML2 | 19:45640930-45642352 | 19:45641767-45641878 | IR (overlapping regio | 3 | 3 | ENST00000585386 | -26.87 | 1.55E-02 | 4.13E-02 |
| CTIF | 18:48758406-48761389 | 18:48758406-48758411 | A5ss | 3 | 3 | ENST00000382998 | -26.95 | 3.12E-02 | 7.72E-02 |
| GOLGA2P7 | 15:84204517-84204786 | 15:84204600-84204691 | SES | 3 | 3 | ENST00000400817 | -26.99 | 1.70E-03 | 3.35E-03 |
| EML2 | 19:45640930-45642352 | 19:45641749-45641917 | IR (overlapping regio | 3 | 3 | ENST00000585386 | -27.05 | 1.57E-02 | 4.20E-02 |
| EML2 | 19:45640930-45642352 | 19:45641746-45641917 | IR (overlapping regio | 3 | 3 | ENST00000585386 | -27.05 | 1.57E-02 | 4.20E-02 |
| FAM234A | 16:249655-254328 | 16:253833-253929 | SES | 3 | 3 | ENST00000417499 | -27.06 | 1.26E-02 | 2.70E-02 |
| MFSD14C | 9:96918827-96949554 | 9:96942603-96942752 | MES | 3 | 3 | ENST00000506067 | -27.17 | 9.38E-03 | 3.89E-01 |
| LETMD1 | 12:51049186-51056349 | 12:51052092-51052207 | MES | 3 | 3 | ENST00000262055 | -27.21 | 5.57E-03 | 9.64E-03 |
| CABYR | 18:24156043-24159471 | 18:24156043-24156999 | A5ss | 2 | 3 | ENST00000399481 | -27.55 | 1.46E-02 | 3.60E-02 |
| AL139353.1 | 14:31453345-31457282 | 14:31457029-31457282 | TSSIA5ss | 3 | 3 | TSS.ENST00000547760 | -27.69 | 2.78E-02 | 5.12E-02 |
| ZNF606 | 19:58002780-58003349 | 19:58003047-58003210 | IR (overlapping regio | 3 | 3 | ENST00000341164 | -27.78 | 2.71E-02 | 7.47E-02 |
| PLD3 | 19:40348725-40365717 | 19:40365553-40365717 | TSSIA3ss | 3 | 3 | TSS.ENST00000598962 | -27.96 | 3.10E-02 | 8.21E-02 |
| SEPTIN6 | X:119620052-119629317 | X:119625335-119625379 | SES | 3 | 3 | ENST00000394610 | -28.02 | 9.11E-04 | 4.13E-02 |
| COL13A1 | 10:69930088-69935349 | 10:69930400-69930552 | MES | 2 | 3 | ENST00000354547 | -28.05 | 8.08E-03 | 1.23E-02 |
| NEPRO | 3:113011407-113013952 | 3:113012734-113012811 | MES | 3 | 3 | ENST00000314400 | -28.18 | 3.08E-02 | 1.62E-01 |
| AC007405.4 | 2:170771188-170777376 | 2:170771188-170771427 | TSSIA5ss | 3 | 3 | TSS.ENST00000664870 | -28.28 | 2.61E-03 | 1.11E-02 |
| HDAC9 | 7:18495855-18496261 | 7:18495855-18496023 | TSSIA5ss | 3 | 3 | TSS.ENST00000413380 | -28.47 | 2.95E-03 | 3.83E-02 |
| ADGRB2 | 1:31756816-31763883 | 1:31757201-31757281 | MES | 3 | 3 | ENST00000373655 | -28.63 | 3.35E-02 | 1.15E-01 |
| DLST | 14:74882017-74889094 | 14:74885586-74885634 | MES | 3 | 3 | ENST00000238671 | -28.78 | 8.29E-05 | 1.55E-04 |
| GABBR1 | 6:29629108-29631395 | 6:29630458-29630643 | SES | 3 | 3 | ENST00000376977 | -28.86 | 4.15E-02 | 4.32E-01 |
| TCIRG1 | 11:68042946-68043497 | 11:68043032-68043370 | IR | 3 | 3 | ENST00000533947 | -29.05 | 1.53E-02 | 2.51E-02 |
| RPS6KL1 | 14:74921562-74923179 | 14:74922341-74922485 | SES | 3 | 2 | ENST00000555647 | -29.23 | 2.73E-02 | 5.13E-02 |
| COL13A1 | 10:69930088-69935349 | 10:69930400-69930552 | MES | 2 | 3 | ENST00000479733 | -29.63 | 1.39E-03 | 2.12E-03 |
| EP400P1 | 12:132090944-132108571 | 12:132104375-132105304 | MES | 2 | 2 | ENST00000332441 | -29.7 | 9.49E-04 | 1.62E-03 |
| TOR1AIP2 | 1:179852812-179877238 | 1:179865436-179865854 | SES | 3 | 3 | ENST00000609928 | -30.28 | 2.83E-03 | 9.06E-03 |
| SLC1A6 | 19:14972918-14979512 | 19:14979309-14979512 | A5ss | 3 | 3 | ENST00000594383 | -30.78 | 2.42E-02 | 6.10E-02 |
| SYTL1 | 1:27349398-27351336 | 1:27350486-27350793 | IR | 3 | 3 | ENST00000490170 | -30.81 | 1.79E-02 | 6.11E-02 |
| WRAP73 | 1:3632318-3633397 | 1:3632318-3632338 | A3ss | 3 | 3 | Ex.ENST00000270708 | -30.99 | 1.79E-03 | 6.20E-03 |
| GRK6 | 5:177403204-177403526 | 5:177403398-177403480 | IR (overlapping regio | 2 | 3 | ENST00000502598 | -31.41 | 1.20E-02 | 1.01E-01 |
| FGFR1OP | 6:167010894-167022408 | 6:167013508-167013567 | SES | 3 | 3 | ENST00000366847 | -31.83 | 1.26E-02 | 1.28E-01 |
| AGO3 | 1:35973512-36004340 | 1:35982621-35982712 | MES | 3 | 3 | ENST00000634486 | -31.88 | 3.76E-03 | 1.30E-02 |
| ZNF160 | 19:53086089-53086261 | 19:53086089-53086109 | TSSIA3ss | 3 | 3 | TSS.ENST00000596966 | -31.99 | 2.10E-02 | 5.77E-02 |
| ZNF23 | 16:71456829-71462209 | 16:71461659-71462209 | TSSIA5ss | 3 | 3 | TSS.ENST00000393539 | -32.18 | 1.99E-02 | 4.41E-02 |
| TSGA10IP | 11:65945823-65953566 | 11:65946880-65947016 | MES | 2 | 3 | ENST00000608857 | -32.19 | 4.11E-02 | 6.60E-02 |
| HSPBP1 | 19:55265379-55265885 | 19:55265379-55265389 | A3ss | 3 | 3 | Ex.ENST00000255631 | -32.42 | 7.52E-05 | 2.07E-04 |
| HSPBP1 | 19:55265368-55265885 | 19:55265368-55265389 | A3ss | 3 | 3 | Ex.ENST00000255631 | -32.42 | 7.52E-05 | 2.07E-04 |
| INTS6L | X:135581528-135582509 | X:135581565-135582240 | IR | 3 | 3 | ENST00000370752 | -32.85 | 1.17E-02 | 6.41E-01 |
| GPR161 | 1:168114550-168137164 | 1:168136010-168137165 | A5ss | 3 | 3 | Ex.TSS.ENST00000367835 | -32.86 | 1.16E-02 | 3.70E-02 |
| ABCG1 | 21:42288038-42288210 | 21:42288038-42288073 | A5ss | 3 | 3 | ENST00000361802 | -32.93 | 1.93E-02 | 7.55E-02 |
| CLIP1 | 12:122319349-122327946 | 12:122322114-122324381 | SES | 3 | 3 | ENST00000537123 | -32.98 | 3.62E-03 | 6.12E-03 |
| RGS19 | 20:64076647-64079293 | 20:64076857-64076954 | SES | 3 | 2 | ENST00000395042 | -33.52 | 1.60E-02 | 6.14E-02 |
| RBM18 | 9:122261499-122264714 | 9:122261499-122261508 | A3ss | 3 | 3 | Ex.TSS.ENST00000417201 | -33.97 | 2.15E-03 | 5.66E-02 |
| NEDD1 | 12:96907625-96909751 | 12:96907702-96907856 | SES | 3 | 3 | ENST00000557092 | -34.07 | 2.02E-02 | 3.59E-02 |
| FAM111A | 11:59143083-59143547 | 11:59143305-59143496 | IR | 2 | 2 | ENST00000531676 | -34.19 | 2.86E-02 | 4.57E-02 |
| NETO2 | 16:47122785-47122888 | 16:47122868-47122888 | A5ss | 3 | 3 | ENST00000562435 | -34.21 | 3.66E-03 | 7.98E-03 |
| TLCD5 | 11:120325369-120329976 | 11:120327441-120329977 | A3ss | 3 | 3 | Ex.ENST00000531346 | -34.31 | 1.92E-03 | 2.98E-03 |
| DPH5 | 1:100990629-100995109 | 1:100992637-100992740 | SES | 3 | 3 | ENST00000342173 | -34.4 | 3.07E-03 | 8.52E-03 |
| LINC01503 | 9:129338918-129341831 | 9:129341736-129341831 | A3ss | 2 | 3 | ENST00000654735 | -34.6 | 4.66E-02 | 1.00E+00 |
| LINC01503 | 9:129338918-129341751 | 9:129341736-129341751 | A3ss | 2 | 3 | ENST00000654735 | -34.6 | 4.66E-02 | 1.00E+00 |
| RAD9A | 11:67393611-67393722 | 11:67393691-67393722 | A3ss | 3 | 3 | ENST00000307980 | -34.87 | 1.07E-04 | 1.76E-04 |
| C2orf27A | 2:131682600-131684708 | 2:131682600-131682988 | A5ss | 3 | 3 | Ex.TSS.ENST00000653458 | -34.98 | 4.72E-03 | 1.99E-02 |
| C2orf27A | 2:131682558-131684708 | 2:131682558-131682988 | A5ss | 3 | 3 | Ex.TSS.ENST00000657415 | -34.98 | 4.72E-03 | 1.99E-02 |
| GABBR1 | 6:29629108-29630521 | 6:29630458-29630521 | A5ss | 3 | 3 | ENST00000376977 | -35.22 | 2.34E-02 | 2.44E-01 |
| AC243919.2 | 15:82418793-82419979 | 15:82418793-82419465 | A5ss | 3 | 2 | Ex.TSS.ENST00000613086 | -35.57 | 1.96E-02 | 3.85E-02 |
| SARDH | 9:133685287-133702915 | 9:133694258-133694371 | MES | 3 | 3 | ENST00000371868 | -35.9 | 2.11E-02 | 6.59E-01 |
| AP000459.3 | 21:23372427-23372947 | 21:23372427-23372448 | A3ss | 3 | 3 | ENST00000262354 | -35.93 | 4.04E-04 | 1.56E-03 |
| AMT | 3:49416794-49418372 | 3:49417368-49417488 | IR (overlapping regio | 3 | 3 | ENST00000636594 | -36.34 | 1.95E-03 | 1.26E-02 |
| INTS6L | X:135581528-135582509 | X:135581632-135582240 | IR | 3 | 2 | ENST00000370752 | -36.43 | 2.58E-02 | 1.00E+00 |
| SEPTIN6 | X:119616743-119629317 | X:119625335-119625379 | MES | 3 | 3 | ENST00000460411 | -36.6 | 2.87E-04 | 1.30E-02 |
| ZSCAN26 | 6:28272337-28272669 | 6:28272337-28272339 | A5ss | 3 | 3 | ENST00000421553 | -37.06 | 3.06E-02 | 3.18E-01 |
| ZSCAN26 | 6:28271935-28272669 | 6:28271935-28272339 | A5ss | 3 | 3 | ENST00000421553 | -37.06 | 3.06E-02 | 3.18E-01 |
| ZNF561 | 19:9618167-9619431 | 19:9618167-9618179 | A3ss | 3 | 3 | ENST00000302851 | -37.11 | 2.21E-03 | 6.13E-03 |
| ZDHHC16 | 10:97454800-97455713 | 10:97455660-97455713 | A3ss | 3 | 3 | ENST00000345745 | -37.49 | 1.69E-04 | 2.59E-04 |
| ZBED5 | 11:10855082-10856143 | 11:10855082-10855086 | TSSIA3ss | 3 | 3 | TSS.ENST00000413761 | -37.58 | 1.85E-03 | 2.85E-03 |
| STAG3L5P-PVRIG2P-PILRB | 7:100351411-100352261 | 7:100352163-100352261 | A3ss | 2 | 2 | ENST00000310771 | -37.78 | 2.07E-02 | 2.36E-01 |
| RTF2 | 20:56468767-56472325 | 20:56470600-56470941 | MES | 2 | 3 | ENST00000484084 | -37.85 | 7.99E-03 | 3.05E-02 |
| CNTROB | 17:7932801-7933092 | 17:7932928-7933092 | A3ss | 2 | 3 | ENST00000570782 | -38.05 | 4.45E-03 | 1.08E-02 |
| CNTROB | 17:7932801-7932955 | 17:7932928-7932955 | A3ss | 2 | 3 | ENST00000570782 | -38.05 | 4.45E-03 | 1.08E-02 |
| ABTB1 | 3:127674419-127674544 | 3:127674419-127674453 | A5ss | 3 | 2 | ENST00000232744 | -38.3 | 1.68E-02 | 9.12E-02 |
| SEPTIN6 | X:119616743-119629317 | X:119625335-119625379 | MXS | 3 | 3 | ENST00000489216 | -38.59 | 3.91E-04 | 1.77E-02 |
| SLC36A4 | 11:93154217-93162705 | 11:93154217-93154277 | A3ss | 3 | 3 | Ex.ENST00000326402 | -38.74 | 2.95E-03 | 4.92E-03 |
| ZNF783 | 7:149262358-149266334 | 7:149262358-149264510 | A5ss | 2 | 3 | Ex.TSS.ENST00000434415 | -39.42 | 8.61E-03 | 1.09E-01 |
| MXRA8 | 1:1355244-1355512 | 1:1355346-1355449 | IR | 3 | 3 | ENST00000460473 | -39.94 | 4.13E-03 | 1.17E-02 |
| ST3GAL6 | 3:98732533-98770878 | 3:98768430-98768529 | MES | 2 | 2 | ENST00000489112 | -40.25 | 5.28E-03 | 3.60E-02 |
| SARDH | 9:133685287-133702915 | 9:133690380-133690527 | MES | 3 | 3 | ENST00000371868 | -40.5 | 9.80E-03 | 3.07E-01 |
| DLST | 14:74882017-74889094 | 14:74882591-74882624 | MES | 3 | 3 | ENST00000238671 | -40.55 | 4.00E-05 | 7.50E-05 |
| AL162258.2 | 1:153632034-153632197 | 1:153632064-153632154 | IR (overlapping regio | 2 | 2 | ENST00000607839 | -40.68 | 6.83E-03 | 2.13E-02 |
| NPIPA1 | 16:14950256-14951614 | 16:14950752-14950821 | SES | 3 | 3 | ENST00000472413 | -40.94 | 3.20E-02 | 6.74E-02 |
| PXK | 3:58410160-58412903 | 3:58412901-58412903 | A3ss | 3 | 2 | ENST00000356151 | -41.07 | 3.76E-02 | 2.53E-01 |
| BUD31 | 7:99409005-99410036 | 7:99409005-99409245 | A5ss | 2 | 3 | Ex.TSS.ENST00000496696 | -41.09 | 4.76E-03 | 7.33E-02 |
| GRID2 | 4:92304745-92590130 | 4:92304745-92442310 | A5ss | 3 | 3 | Ex.TSS.ENST00000282020 | -41.29 | 6.92E-03 | 5.30E-02 |
| TSPOAP1 | 17:58306387-58306799 | 17:58306387-58306413 | A3ss | 3 | 3 | ENST00000268893 | -42.52 | 6.97E-03 | 1.64E-02 |
| VEPH1 | 3:157459519-157460355 | 3:157459927-157460180 | IR | 2 | 3 | ENST00000494677 | -42.81 | 2.35E-02 | 1.32E-01 |
| AC119673.3 | 1:205891310-205894466 | 1:205894168-205894466 | TSSIA3ss | 3 | 3 | TSS.ENST00000653632 | -44.23 | 3.00E-03 | 9.84E-03 |
| AC119673.3 | 1:205891310-205894443 | 1:205894168-205894443 | A3ss | 3 | 3 | Ex.TSS.ENST00000653632 | -44.23 | 3.00E-03 | 9.84E-03 |

| | | | | | | | | | |
|---|---|---|---|---|---|---|---|---|---|
| AMT | 3:49416794-49418372 | 3:49417310-49417614 | IR (overlapping regio | 3 | 3 | ENST00000636594 | -45.35 | 3.54E-06 | 2.29E-05 |
| AMT | 3:49416794-49418372 | 3:49417358-49417614 | IR (overlapping regio | 3 | 3 | ENST00000636594 | -45.43 | 2.40E-06 | 1.55E-05 |
| DVL2 | 17:7227712-7227976 | 17:7227712-7227783 | A3ss | 3 | 3 | ENST00000005340 | -45.68 | 2.01E-04 | 4.83E-04 |
| DVL2 | 17:7227703-7227976 | 17:7227703-7227783 | A3ss | 3 | 3 | Ex.ENST00000005340 | -45.68 | 2.01E-04 | 4.83E-04 |
| COASY | 17:42562220-42562932 | 17:42562612-42562932 | TSSIA3ss | 3 | 3 | TSS.ENST00000421097 | -45.77 | 9.16E-04 | 2.12E-03 |
| AMT | 3:49416794-49418372 | 3:49417368-49417614 | IR (overlapping regio | 3 | 3 | ENST00000636594 | -46.46 | 1.65E-06 | 1.07E-05 |
| ARHGEF35-AS1 | 7:144207805-144215858 | 7:144214014-144215858 | TSSIA3ss | 3 | 3 | TSS.ENST00000650250 | -47.53 | 2.61E-02 | 3.25E-01 |
| ZBTB49 | 4:4313115-4320639 | 4:4315809-4315970 | MES | 2 | 2 | ENST00000337872 | -47.97 | 4.39E-02 | 3.26E-01 |
| COASY | 17:42562356-42562894 | 17:42562488-42562611 | IR | 3 | 3 | ENST00000585811 | -48.2 | 9.67E-05 | 2.24E-04 |
| RHBDL1 | 16:677459-678267 | 16:677560-677638 | IR | 2 | 3 | ENST00000450775 | -51.56 | 2.60E-03 | 5.73E-03 |
| NAA16 | 13:41372831-41374741 | 13:41373637-41373780 | SES | 2 | 3 | ENST00000379406 | -52.6 | 2.89E-04 | 5.18E-04 |
| CRNDE | 16:54920339-54923584 | 16:54920339-54920410 | A3ss | 2 | 2 | ENST00000502066 | -52.8 | 1.42E-02 | 3.11E-02 |
| CRNDE | 16:54920328-54923584 | 16:54920328-54920410 | A3ss | 2 | 2 | ENST00000502066 | -52.8 | 1.42E-02 | 3.11E-02 |
| MTERF2 | 12:106985216-106986968 | 12:106985216-106985225 | TSSIA3ss | 3 | 3 | TSS.ENST00000240050 | -52.86 | 7.12E-04 | 1.19E-03 |
| ZNRD2 | 11:65570742-65570885 | 11:65570742-65570755 | A5ss | 3 | 3 | Ex.ENST00000309328 | -53.48 | 2.08E-04 | 3.35E-04 |
| ZNRD2 | 11:65570651-65570885 | 11:65570651-65570755 | A5ss | 3 | 3 | Ex.ENST00000309328 | -53.48 | 2.08E-04 | 3.35E-04 |
| AC108010.1 | 7:128574792-128578889 | 7:128577953-128578890 | A3ss | 2 | 3 | Ex.TSS.ENST00000605862 | -56.77 | 3.54E-03 | 4.26E-02 |
| AC090114.3 | 7:128574792-128577963 | 7:128577953-128577964 | A3ss | 2 | 3 | Ex.ENST00000479267 | -56.77 | 3.54E-03 | 4.26E-02 |
| ENOX2 | X:130637405-130656580 | X:130637405-130637410 | A3ss | 3 | 3 | Ex.ENST00000338144 | -57.01 | 1.65E-04 | 8.71E-03 |
| COASY | 17:42562137-42562487 | 17:42562220-42562295 | IR | 3 | 3 | ENST00000585909 | -57.09 | 3.02E-06 | 7.00E-06 |
| COASY | 17:42562137-42562487 | 17:42562220-42562355 | IR | 3 | 3 | ENST00000585909 | -58 | 1.33E-06 | 3.08E-06 |
| NAA16 | 13:41372831-41373767 | 13:41373637-41373767 | A3ss | 2 | 3 | ENST00000379406 | -58.95 | 3.56E-02 | 6.38E-02 |
| FAM122B | X:134796257-134797120 | X:134796257-134796355 | A3ss | 2 | 3 | Ex.TSS.ENST00000298090 | -62.01 | 1.13E-03 | 5.95E-02 |
| STAG3L5P-PVRIG2P-PILRB | 7:100356884-100358263 | 7:100358220-100358263 | A3ss | 3 | 3 | ENST00000310771 | -62.55 | 8.64E-04 | 9.99E-03 |
| STAG3L5P-PVRIG2P-PILRB | 7:100356884-100358226 | 7:100358220-100358226 | A3ss | 3 | 3 | Ex.ENST00000310771 | -62.55 | 8.64E-04 | 9.99E-03 |
| MFSD9 | 2:102732393-102736645 | 2:102732393-102732409 | TSSIA3ss | 3 | 3 | TSS.ENST00000258436 | -73.62 | 8.23E-04 | 3.43E-03 |

**Table S1. Mis-splicing events in 3 biological replicates of SF3B1$^{K700E}$ (K700E ES line 3) compared to SF3B1$^{WT}$ ES cells.** The mis-splicing event code is as follows: SES, single-exon skipping; MES, multiple-exon skipping; MXS, mutually-exclusive splicing; A5ss, alternative 5′ splice site; A3ss, alternative 3′ splice site; TSS, transcription start site. Event Region: Genomic coordinates of splicing event; Target Exon: Genomic coordinates of the alternative exon. Events with p-values < 0.05 and ΔPSI (%)≥10% are listed.

| Gene Symbol | Event Region | Target Exon | Event | WT | K700E | Reference Transcript | ΔPSI (%) | p-value | FDR (BH) |
|---|---|---|---|---|---|---|---|---|---|
| GPR153 | 1:6255015-6260824 | 1:6255015-6255122 | A3ss | 2 | 2 | Ex.TSS.ENST00000377893 | 80.62 | 6.01E-05 | 1.13E-04 |
| GEN1 | 2:17753968-17759928 | 2:17759912-17759929 | A3ss | 2 | 3 | Ex.TSS.ENST00000317402 | 79.53 | 4.50E-04 | 9.53E-04 |
| APBB3 | 5:140561702-140561843 | 5:140561702-140561722 | A3ss | 2 | 3 | ENST00000354402 | 78.55 | 1.36E-02 | 3.96E-02 |
| BCL2L1 | 20:31722331-31723863 | 20:31722331-31722348 | A3ss | 3 | 3 | ENST00000420488 | 76.69 | 1.12E-05 | 2.13E-05 |
| MFSD9 | 2:102732410-102736645 | 2:102732410-102732426 | A3ss | 3 | 3 | ENST00000462099 | 75.83 | 5.81E-05 | 1.21E-04 |
| MFSD9 | 2:102732393-102736645 | 2:102732393-102732426 | A3ss | 3 | 3 | ENST00000462099 | 75.83 | 5.81E-05 | 1.21E-04 |
| ZNF91 | 19:23362726-23373741 | 19:23362726-23362739 | A3ss | 2 | 3 | Ex.TSS.ENST00000300619 | 74.48 | 5.88E-04 | 3.24E-03 |
| ORAI2 | 7:102433662-102436224 | 7:102436202-102436224 | TSSIA3ss | 3 | 3 | TSS.ENST00000356387 | 73.46 | 1.13E-05 | 1.32E-04 |
| DLST | 14:74889350-74889896 | 14:74889878-74889896 | A3ss | 3 | 3 | ENST00000555988 | 70.97 | 4.77E-07 | 1.52E-06 |
| CCDC74A | 2:131530828-131531663 | 2:131531652-131531664 | A3ss | 3 | 2 | Ex.ENST00000295171 | 70.4 | 1.45E-07 | 3.05E-07 |
| HERC2P2 | 15:22561790-22562948 | 15:22561896-22562949 | A3ss | 3 | 2 | Ex.ENST00000613386 | 68.18 | 1.35E-03 | 4.39E-03 |
| PGBD1 | 6:28281919-28283775 | 6:28283756-28283775 | A3ss | 3 | 3 | Ex.TSS.ENST00000259883 | 67.59 | 7.24E-04 | 6.52E-03 |
| BCL2L1 | 20:31722331-31723734 | 20:31722331-31722348 | A3ss | 3 | 3 | ENST00000456404 | 65.87 | 8.22E-04 | 1.56E-03 |
| ZC3H11A | 1:203801629-203802956 | 1:203802569-203802957 | A3ss | 3 | 2 | Ex.ENST00000367212 | 65.54 | 4.73E-03 | 8.42E-03 |
| ZC3H11A | 1:203801629-203802651 | 1:203802569-203802652 | A3ss | 3 | 2 | Ex.ENST00000332127 | 65.54 | 4.73E-03 | 8.42E-03 |
| APPL2 | 12:105208030-105208157 | 12:105208030-105208047 | A3ss | 3 | 3 | ENST00000551662 | 65.05 | 2.44E-03 | 3.78E-03 |
| KIAA1217 | 10:24219910-24380868 | 10:24219910-24255596 | A5ss | 2 | 3 | Ex.ENST00000376452 | 64.82 | 1.02E-02 | 1.44E-02 |
| MAP3K7 | 6:90560215-90561621 | 6:90560215-90560234 | A3ss | 3 | 3 | Ex.ENST00000369325 | 64.48 | 3.61E-05 | 4.11E-04 |
| TMEM214 | 2:27037703-27038145 | 2:27037815-27037892 | SES | 3 | 3 | ENST00000425720 | 63.8 | 1.42E-04 | 3.11E-04 |
| GOLGA2P10 | 15:82475616-82476785 | 15:82475616-82475637 | A3ss | 3 | 3 | Ex.ENST00000614347 | 63.76 | 2.76E-05 | 9.42E-05 |
| COASY | 17:42562220-42562611 | 17:42562356-42562611 | A3ss | 3 | 3 | ENST00000585811 | 61.86 | 1.46E-05 | 6.19E-05 |
| VPS9D1-AS1 | 16:89712223-89712688 | 16:89712674-89712689 | A3ss | 3 | 3 | Ex.TSS.ENST00000562866 | 61.86 | 1.11E-03 | 4.37E-03 |
| TMEM14C | 6:10723242-10724569 | 6:10724556-10724570 | A3ss | 3 | 3 | Ex.TSS.ENST00000229563 | 60.46 | 1.04E-06 | 3.18E-06 |
| DYNLL1 | 12:120496217-120496415 | 12:120496402-120496415 | TSSIA3ss | 3 | 3 | TSS.ENST00000392508 | 60.15 | 4.45E-07 | 6.95E-07 |
| AC007326.4 | 22:18939750-18946790 | 22:18945072-18946791 | A3ss | 3 | 3 | Ex.TSS.ENST00000638240 | 59.79 | 1.97E-03 | 3.84E-03 |
| GCC2 | 2:108485909-108486510 | 2:108486499-108486511 | A3ss | 3 | 3 | Ex.ENST00000309863 | 58.5 | 2.63E-04 | 5.47E-04 |
| TMCC2 | 1:205271256-205271812 | 1:205271796-205271813 | A3ss | 3 | 2 | Ex.TSS.ENST00000329800 | 58.15 | 2.24E-03 | 3.98E-03 |
| ARMC9 | 2:231331898-231344974 | 2:231344949-231344975 | A3ss | 3 | 3 | Ex.TSS.ENST00000349938 | 56.33 | 5.01E-03 | 1.09E-02 |
| GAS8 | 16:90027723-90031298 | 16:90031176-90031298 | A3ss | 3 | 3 | ENST00000566266 | 54.53 | 3.94E-03 | 1.56E-02 |
| GAS8 | 16:90027723-90031189 | 16:90031176-90031189 | TSSIA3ss | 3 | 3 | TSS.ENST00000563980 | 54.53 | 3.94E-03 | 1.56E-02 |
| TMEM218 | 11:125102318-125102636 | 11:125102318-125102351 | A3ss | 3 | 3 | ENST00000279968 | 54.18 | 1.61E-04 | 2.32E-04 |
| TLCD5 | 11:120325369-120329976 | 11:120327452-120329977 | A3ss | 3 | 3 | Ex.ENST00000531346 | 53.88 | 5.55E-05 | 8.00E-05 |
| COASY | 17:42562220-42562611 | 17:42562356-42562487 | SES | 3 | 3 | ENST00000590958 | 53.46 | 3.50E-05 | 1.48E-04 |
| COASY | 17:42562220-42562611 | 17:42562220-42562487 | A5ss | 3 | 3 | ENST00000585909 | 52.11 | 3.73E-04 | 1.58E-03 |
| ENOSF1 | 18:683381-685920 | 18:683381-683395 | A3ss | 3 | 3 | ENST00000578647 | 50.95 | 2.36E-03 | 1.20E-02 |
| UXS1 | 2:106164785-106166055 | 2:106164785-106164799 | A3ss | 3 | 3 | ENST00000283148 | 50.73 | 6.14E-04 | 1.28E-03 |
| AC007326.4 | 22:18939750-18946790 | 22:18939750-18945166 | A5ss | 3 | 3 | Ex.TSS.ENST00000638240 | 49 | 2.48E-03 | 4.84E-03 |
| NMRK2 | 19:3936666-3937239 | 19:3937225-3937239 | A3ss | 3 | 3 | ENST00000593949 | 47.94 | 4.61E-05 | 3.03E-04 |
| SMURF2 | 17:64578577-64580788 | 17:64578577-64578594 | A3ss | 3 | 3 | Ex.ENST00000262435 | 47.13 | 1.13E-02 | 5.14E-02 |
| HERC2P2 | 15:22561790-22562948 | 15:22561790-22562026 | A5ss | 3 | 2 | Ex.ENST00000613386 | 46.18 | 1.08E-02 | 3.52E-02 |
| TRIM37 | 17:59001715-59012327 | 17:59001715-59001741 | A3ss | 3 | 3 | ENST00000585287 | 45.61 | 3.12E-05 | 1.39E-04 |
| THOC1 | 18:224180-224923 | 18:224180-224200 | A3ss | 3 | 3 | ENST00000579891 | 45.59 | 4.04E-05 | 2.00E-04 |
| MFAP3L | 4:170006737-170026233 | 4:170006737-170025842 | A3ss | 2 | 3 | Ex.TSS.ENST00000504999 | 45.44 | 1.46E-02 | 3.93E-02 |
| MFAP3L | 4:170006011-170026233 | 4:170006011-170025842 | A3ss | 2 | 3 | Ex.TSS.ENST00000361618 | 45.44 | 1.46E-02 | 3.93E-02 |
| MFAP3L | 4:170005892-170026233 | 4:170005892-170025842 | A3ss | 2 | 3 | Ex.ENST00000506764 | 45.44 | 1.46E-02 | 3.93E-02 |
| MFAP3L | 4:169992310-170026233 | 4:169992310-170025842 | A3ss | 2 | 3 | Ex.ENST00000512698 | 45.44 | 1.46E-02 | 3.93E-02 |
| SH3D19 | 4:151179398-151187422 | 4:151179398-151179416 | A3ss | 3 | 3 | Ex.ENST00000304527 | 44.41 | 6.08E-04 | 1.62E-03 |
| TAC1 | 7:97733820-97734825 | 7:97734248-97734292 | SES | 2 | 3 | ENST00000319273 | 43.89 | 9.01E-03 | 1.42E-01 |
| SLC13A3 | 20:46596343-46610445 | 20:46599971-46600037 | SES | 3 | 3 | ENST00000279027 | 43.83 | 1.94E-02 | 3.70E-02 |
| PILRB | 7:100356884-100358263 | 7:100358227-100358264 | A3ss | 3 | 3 | Ex.ENST00000608825 | 43.65 | 1.36E-02 | 1.57E-01 |
| ZNF213-AS1 | 16:3132104-3134633 | 16:3132850-3132942 | SES | 2 | 3 | ENST00000572691 | 42.66 | 1.58E-03 | 5.80E-03 |
| MBD2 | 18:54203103-54204997 | 18:54203103-54203126 | TSSIA3ss | 3 | 3 | TSS.ENST00000398398 | 42.64 | 6.94E-04 | 3.50E-03 |
| PIH1D1 | 19:49451579-49453375 | 19:49451579-49451591 | TSSIA3ss | 3 | 3 | TSS.ENST00000601825 | 42.64 | 1.02E-02 | 7.06E-02 |
| ZBED5 | 11:10855087-10856143 | 11:10855087-10855118 | TSSIA3ss | 3 | 3 | TSS.ENST00000432999 | 42.62 | 2.18E-03 | 3.12E-03 |
| ZBED5 | 11:10855082-10856143 | 11:10855082-10855118 | TSSIA3ss | 3 | 3 | TSS.ENST00000432999 | 42.62 | 2.18E-03 | 3.12E-03 |
| DDX12P | 12:9437490-9438010 | 12:9437490-9437891 | A3ss | 3 | 2 | Ex.ENST00000432996 | 41.89 | 4.40E-04 | 6.99E-04 |
| MTERF2 | 12:106978772-106986968 | 12:106985115-106985215 | SES | 3 | 3 | ENST00000392830 | 41.59 | 1.14E-04 | 1.76E-04 |
| CHTF18 | 16:793275-794053 | 16:794034-794054 | A3ss | 3 | 3 | Ex.ENST00000262315 | 41.58 | 5.89E-04 | 2.27E-03 |
| STAG3L5P | 7:100345969-100349798 | 7:100349717-100349798 | TSSIA3ss | 3 | 3 | TSS.ENST00000493499 | 41.47 | 4.50E-03 | 5.20E-02 |
| STAG3L5P | 7:100345969-100349730 | 7:100349717-100349730 | TSSIA3ss | 3 | 3 | TSS.ENST00000493499 | 41.47 | 4.50E-03 | 5.20E-02 |
| SPINT1-AS1 | 15:40838687-40844178 | 15:40838687-40838705 | A3ss | 3 | 3 | Ex.TSS.ENST00000564302 | 40.79 | 6.26E-05 | 2.05E-04 |
| DCAF16 | 4:17804757-17810446 | 4:17804757-17804771 | A3ss | 3 | 3 | Ex.ENST00000507768 | 40.54 | 7.07E-04 | 1.90E-03 |
| BRD9 | 5:865582-870472 | 5:869245-869404 | MXS | 3 | 3 | ENST00000519112 | 39.9 | 1.37E-06 | 4.16E-06 |
| ZNF213-AS1 | 16:3132104-3134633 | 16:3132850-3133367 | SES | 2 | 3 | ENST00000652946 | 39.72 | 1.74E-03 | 6.36E-03 |
| ACIN1 | 14:23081837-23090521 | 14:23089982-23090101 | SES | 3 | 3 | ENST00000262710 | 39.47 | 2.23E-04 | 3.78E-04 |
| BCL2L1 | 20:31722331-31722617 | 20:31722331-31722348 | TSSIA3ss | 3 | 3 | TSS.ENST00000307677 | 39 | 1.91E-03 | 3.62E-03 |
| KANSL3 | 2:96619763-96631311 | 2:96619763-96619776 | A3ss | 3 | 3 | Ex.ENST00000354204 | 38.96 | 6.68E-04 | 1.52E-03 |
| KANSL3 | 2:96619545-96631311 | 2:96619545-96619776 | A3ss | 3 | 3 | Ex.ENST00000418735 | 38.96 | 6.68E-04 | 1.52E-03 |
| SNHG14 | 15:24982438-25019026 | 15:24998366-24998486 | MES | 3 | 3 | ENST00000551631 | 38.66 | 6.99E-03 | 2.27E-02 |
| NDRG3 | 20:36653702-36656359 | 20:36653702-36653723 | A3ss | 3 | 3 | Ex.TSS.ENST00000349004 | 38.55 | 1.63E-05 | 3.11E-05 |
| MFSD13A | 10:102466088-102468954 | 10:102468753-102468955 | A3ss | 3 | 3 | Ex.TSS.ENST00000238936 | 38.45 | 2.81E-03 | 3.93E-03 |
| ZNF771 | 16:30407665-30408044 | 16:30408032-30408045 | A3ss | 3 | 3 | Ex.TSS.ENST00000319296 | 38.43 | 2.93E-03 | 1.07E-02 |
| RGS3 | 9:113565361-113583449 | 9:113565361-113565730 | A5ss | 3 | 3 | Ex.TSS.ENST00000488620 | 38.41 | 5.91E-03 | 1.52E-01 |
| RGS3 | 9:113536919-113583449 | 9:113536919-113565730 | A5ss | 3 | 3 | Ex.ENST00000343817 | 38.41 | 5.91E-03 | 1.52E-01 |
| SEPTIN6 | X:119625380-119629317 | X:119625380-119625396 | A3ss | 3 | 3 | ENST00000467310 | 38.38 | 2.13E-03 | 1.05E-01 |
| SLC16A5 | 17:75088045-75089150 | 17:75089114-75089150 | TSSIA3ss | 3 | 3 | TSS.ENST00000578376 | 38.37 | 7.13E-03 | 3.28E-02 |
| TTI1 | 20:38002777-38006196 | 20:38002777-38002793 | A3ss | 3 | 3 | Ex.ENST00000373447 | 38.21 | 7.22E-05 | 1.38E-04 |
| RACGAP1 | 12:50016716-50025372 | 12:50016716-50016719 | A3ss | 2 | 3 | Ex.TSS.ENST00000548824 | 36.87 | 2.13E-02 | 3.50E-02 |
| LRP4 | 11:46889534-46889943 | 11:46889534-46889591 | A3ss | 3 | 3 | Ex.ENST00000378623 | 36.67 | 1.79E-02 | 2.63E-02 |
| SLC4A8 | 12:51440790-51450875 | 12:51450857-51450876 | A3ss | 3 | 3 | Ex.ENST00000319957 | 36.53 | 8.06E-04 | 1.33E-03 |
| HINT2 | 9:35813146-35813265 | 9:35813146-35813156 | A3ss | 3 | 3 | Ex.TSS.ENST00000471774 | 35.81 | 2.13E-03 | 6.44E-02 |
| ANKHD1 | 5:140436258-140438502 | 5:140438494-140438502 | A3ss | 3 | 3 | ENST00000394722 | 35.7 | 8.69E-06 | 2.53E-05 |
| CAMTA2 | 17:4982161-4982756 | 17:4982161-4982175 | A3ss | 3 | 3 | ENST00000572543 | 35.64 | 1.06E-02 | 4.63E-02 |
| GOLGA2P7 | 15:84203021-84204425 | 15:84204177-84204331 | SES | 3 | 3 | ENST00000316967 | 35.59 | 4.78E-02 | 1.65E-01 |
| STX1A | 7:73702983-73703754 | 7:73702983-73703642 | A3ss | 3 | 3 | Ex.ENST00000222812 | 35.47 | 4.07E-02 | 5.68E-01 |
| SLC7A3 | X:70929999-70930976 | X:70929999-70930022 | TSSIA3ss | 3 | 3 | TSS.ENST00000374299 | 34.89 | 1.19E-03 | 5.45E-01 |
| TMEM214 | 2:27037703-27038145 | 2:27037790-27037892 | SES | 3 | 3 | ENST00000435172 | 34.16 | 2.40E-03 | 5.25E-03 |
| DVL2 | 17:7227703-7227976 | 17:7227703-7227711 | A3ss | 3 | 3 | Ex.ENST00000575086 | 34.14 | 2.33E-05 | 1.06E-04 |
| ZBTB14 | 18:5293328-5295812 | 18:5293972-5294001 | SES | 3 | 2 | ENST00000357006 | 34.04 | 5.33E-03 | 2.69E-02 |
| RIMKLB | 12:8773467-8777589 | 12:8776673-8777590 | A3ss | 3 | 3 | Ex.TSS.ENST00000299673 | 34 | 3.72E-02 | 6.28E-02 |

| | | | | | | | | | |
|---|---|---|---|---|---|---|---|---|---|
| DLG1 | 3:197065449-197065707 | 3:197065449-197065464 | A3ss | 3 | 3 | ENST00000661229 | 33.94 | 4.12E-03 | 9.75E-03 |
| WASHC4 | 12:105120598-105121100 | 12:105121089-105121100 | A3ss | 3 | 3 | ENST00000547404 | 33.38 | 4.04E-03 | 6.27E-03 |
| CAMTA2 | 17:4981832-4982756 | 17:4982089-4982175 | SES | 3 | 3 | ENST00000572543 | 33.34 | 1.59E-02 | 5.60E-02 |
| ZNF532 | 18:58953800-58979054 | 18:58954234-58954334 | SES | 3 | 3 | ENST00000585662 | 33.3 | 1.81E-04 | 9.16E-04 |
| SLC26A6 | 3:48633636-48635370 | 3:48633636-48634590 | A3ss | 3 | 3 | Ex.TSS.ENST00000307364 | 33.15 | 5.71E-05 | 1.38E-04 |
| UBA1 | X:47199615-47200893 | X:47199615-47200544 | A5ss | 3 | 3 | Ex.ENST00000335972 | 32.84 | 4.41E-06 | 4.66E-04 |
| PCNX3 | 11:65635804-65636173 | 11:65635804-65636031 | A5ss | 3 | 3 | Ex.ENST00000355703 | 32.77 | 2.16E-02 | 3.28E-02 |
| ATF7IP2 | 16:10386123-10428867 | 16:10419581-10419623 | MXS | 3 | 3 | ENST00000396559 | 32.72 | 6.77E-03 | 2.37E-02 |
| EMC3-AS1 | 3:9987265-9987905 | 3:9987815-9987905 | TSSlA3ss | 3 | 3 | TSS.ENST00000440568 | 32.58 | 2.02E-02 | 5.26E-02 |
| PHKB | 16:47461427-47463898 | 16:47463882-47463898 | TSSlA3ss | 3 | 3 | TSS.ENST00000566044 | 32.43 | 5.18E-03 | 1.91E-02 |
| ZDHHC16 | 10:97454800-97455713 | 10:97455639-97455713 | A3ss | 3 | 3 | ENST00000487315 | 32.34 | 8.93E-04 | 1.28E-03 |
| ZDHHC16 | 10:97454800-97455659 | 10:97455639-97455659 | A3ss | 3 | 3 | ENST00000487315 | 32.34 | 8.93E-04 | 1.28E-03 |
| PHF20L1 | 8:132798861-132803818 | 8:132799083-132799172 | SES | 3 | 3 | ENST00000395376 | 32.31 | 4.88E-03 | 9.60E-02 |
| HERC1 | 15:63775650-63833826 | 15:63833822-63833826 | A5ss | 2 | 2 | ENST00000558532 | 32.06 | 6.50E-03 | 2.17E-02 |
| AC026412.1 | 5:1602736-1632844 | 5:1626786-1626926 | MES | 3 | 2 | ENST00000507841 | 32.02 | 2.79E-02 | 8.23E-02 |
| TMEM241 | 18:23370262-23373711 | 18:23371422-23371470 | SES | 2 | 2 | ENST00000383233 | 31.82 | 1.93E-02 | 9.57E-02 |
| SERBP1 | 1:67424960-67425082 | 1:67424960-67424977 | A3ss | 3 | 3 | ENST00000361219 | 31.59 | 4.40E-07 | 8.28E-07 |
| SLC12A5 | 20:46021887-46023368 | 20:46022843-46023071 | SES | 3 | 2 | ENST00000626701 | 31.56 | 1.06E-02 | 2.03E-02 |
| GRIP1 | 12:66445509-66462923 | 12:66455409-66455561 | SES | 2 | 2 | ENST00000543172 | 31.54 | 3.37E-02 | 5.64E-02 |
| ZNF160 | 19:53086110-53086261 | 19:53086110-53086128 | A3ss | 3 | 3 | Ex.TSS.ENST00000596966 | 31.46 | 2.79E-03 | 1.96E-02 |
| ZNF160 | 19:53086089-53086261 | 19:53086089-53086128 | A3ss | 3 | 3 | Ex.TSS.ENST00000594760 | 31.46 | 2.79E-03 | 1.96E-02 |
| ZNF160 | 19:53075184-53086261 | 19:53075184-53086128 | A3ss | 3 | 3 | Ex.ENST00000355147 | 31.46 | 2.79E-03 | 1.96E-02 |
| ANO9 | 11:419582-420860 | 11:420616-420717 | IR | 3 | 3 | ENST00000532094 | 31.36 | 3.93E-03 | 5.78E-03 |
| RNF2 | 1:185087641-185091578 | 1:185091565-185091579 | A3ss | 3 | 3 | Ex.ENST00000367509 | 31.28 | 8.80E-04 | 1.55E-03 |
| R3HDM1 | 2:135605017-135621493 | 2:135616668-135616757 | SES | 3 | 3 | ENST00000456040 | 31.18 | 7.30E-03 | 1.53E-02 |
| SLC26A6 | 3:48633636-48635370 | 3:48634516-48635371 | A5ss | 3 | 3 | Ex.TSS.ENST00000307364 | 30.96 | 6.43E-04 | 1.55E-03 |
| FGGY | 1:59296908-59346246 | 1:59321536-59321750 | MES | 3 | 3 | ENST00000413489 | 30.9 | 1.00E-02 | 1.88E-02 |
| ECHDC2 | 1:52904834-52907867 | 1:52905034-52905090 | SES | 3 | 3 | ENST00000358358 | 30.9 | 2.07E-02 | 3.87E-02 |
| INPPL1 | 11:72228499-72228726 | 11:72228647-72228727 | A3ss | 3 | 3 | Ex.ENST00000298229 | 30.79 | 4.57E-05 | 7.00E-05 |
| DPH5 | 1:100992741-100995109 | 1:100992741-100992754 | A3ss | 3 | 3 | Ex.ENST00000342173 | 30.76 | 1.48E-07 | 1.21E-06 |
| DPH5 | 1:100990632-100995109 | 1:100990632-100992754 | A3ss | 3 | 3 | Ex.ENST00000427040 | 30.76 | 1.48E-07 | 1.21E-06 |
| ITPA | 20:3209536-3213257 | 20:3209536-3209617 | TSSlA5ss | 3 | 3 | TSS.ENST00000609835 | 30.47 | 4.02E-02 | 7.63E-02 |
| STAP2 | 19:4324665-4325215 | 19:4324665-4324667 | TSSlA3ss | 3 | 3 | TSS.ENST00000597593 | 30.24 | 2.07E-02 | 1.37E-01 |
| STAP2 | 19:4324530-4325215 | 19:4324530-4324667 | A3ss | 3 | 3 | ENST00000600324 | 30.24 | 2.07E-02 | 1.37E-01 |
| GIT2 | 12:109939248-109947255 | 12:109945260-109945349 | SES | 3 | 3 | ENST00000355312 | 30.23 | 1.25E-02 | 1.95E-02 |
| TCEA2 | 20:64070636-64071869 | 20:64071858-64071869 | A3ss | 3 | 3 | ENST00000465433 | 30.03 | 1.17E-02 | 2.25E-02 |
| SNRPN | 15:24962210-24967931 | 15:24967029-24967152 | SES | 3 | 3 | ENST00000554227 | 30.02 | 1.26E-05 | 4.09E-05 |
| AL049834.1 | 14:88391456-88396155 | 14:88393393-88393488 | MXS | 3 | 3 | ENST00000045347 | 30.02 | 2.36E-02 | 7.50E-02 |
| MEGF6 | 1:3496784-3497232 | 1:3496988-3497119 | SES | 3 | 3 | ENST00000356575 | 29.95 | 3.87E-03 | 7.16E-03 |
| SUZ12P1 | 17:30709812-30759490 | 17:30734897-30734943 | MES | 3 | 3 | ENST00000579526 | 29.91 | 3.26E-02 | 1.31E-01 |
| ARL17A | 17:46554044-46557630 | 17:46555207-46555284 | IR (overlapping | 3 | 3 | ENST00000336125 | 29.64 | 1.80E-02 | 7.80E-02 |
| RWDD4 | 4:183655962-183658928 | 4:183655962-183655974 | A3ss | 3 | 3 | Ex.TSS.ENST00000326397 | 29.57 | 1.83E-03 | 4.92E-03 |
| RWDD4 | 4:183651132-183658928 | 4:183651132-183655974 | A3ss | 3 | 3 | Ex.TSS.ENST00000510968 | 29.57 | 1.83E-03 | 4.92E-03 |
| RECQL5 | 17:75628763-75628933 | 17:75628763-75628825 | TSSlA3ss | 3 | 3 | TSS.ENST00000582548 | 29.57 | 2.21E-02 | 1.02E-01 |
| SEPTIN6 | X:119620052-119629317 | X:119625335-119625396 | SES | 3 | 3 | ENST00000467310 | 29.51 | 8.35E-04 | 4.12E-02 |
| SETD4 | 21:36043957-36045581 | 21:36043957-36043969 | A3ss | 3 | 3 | Ex.ENST00000332131 | 29.38 | 9.73E-04 | 1.88E-03 |
| AC022140.1 | 5:25422856-25445819 | 5:25422856-25435273 | A3ss | 3 | 3 | Ex.TSS.ENST00000507887 | 29 | 6.01E-03 | 1.55E-02 |
| DENND2A | 7:140602543-140640503 | 7:140605705-140605806 | SES | 2 | 2 | ENST00000496613 | 28.66 | 4.84E-02 | 5.89E-01 |
| NSMCE2 | 8:125091959-125102050 | 8:125102014-125102050 | TSSlA3ss | 3 | 3 | TSS.ENST00000517532 | 28.56 | 2.31E-04 | 4.41E-03 |
| TRIM37 | 17:58999460-59012327 | 17:59001598-59001741 | SES | 3 | 3 | ENST00000585287 | 28.52 | 1.50E-03 | 6.71E-03 |
| BRPF1 | 3:9743254-9743580 | 3:9743578-9743580 | A3ss | 3 | 3 | ENST00000383829 | 28.52 | 4.13E-02 | 1.07E-01 |
| ZNF219 | 14:21093291-21093585 | 14:21093291-21093304 | TSSlA3ss | 3 | 3 | TSS.ENST00000554478 | 28.37 | 8.15E-05 | 1.38E-04 |
| GBAP1 | 1:155217240-155218049 | 1:155217847-155217963 | IR | 2 | 3 | ENST00000473223 | 28.3 | 2.42E-02 | 4.19E-02 |
| E2F7 | 12:77064636-77065344 | 12:77064636-77064653 | TSSlA3ss | 3 | 3 | TSS.ENST00000547316 | 28.25 | 1.94E-02 | 3.27E-02 |
| CDK8 | 13:26396355-26397152 | 13:26397139-26397153 | A3ss | 3 | 3 | Ex.ENST00000381527 | 28.18 | 4.80E-02 | 7.75E-02 |
| DDX12P | 12:9421492-9421817 | 12:9421492-9421546 | A3ss | 3 | 3 | Ex.ENST00000432996 | 28.01 | 2.30E-04 | 3.65E-04 |
| TTC32 | 2:19898036-19901705 | 2:19901476-19901706 | A5ss | 3 | 3 | Ex.ENST00000333610 | 27.89 | 5.34E-03 | 1.13E-02 |
| MFSD14C | 9:96942753-96949554 | 9:96942753-96943154 | A3ss | 3 | 3 | Ex.TSS.ENST00000506067 | 27.88 | 3.26E-02 | 8.89E-01 |
| MFSD14C | 9:96918827-96949554 | 9:96918827-96943154 | A3ss | 3 | 3 | Ex.TSS.ENST00000637076 | 27.88 | 3.26E-02 | 1.00E+00 |
| TRAPPC9 | 8:140405699-140435111 | 8:140426615-140426641 | SES | 3 | 3 | ENST00000389328 | 27.83 | 4.86E-02 | 9.62E-01 |
| EHMT1 | 9:137728530-137743391 | 9:137743371-137743391 | A3ss | 3 | 3 | ENST00000371394 | 27.72 | 1.08E-03 | 2.96E-02 |
| FBXO21 | 12:117158064-117165484 | 12:117158064-117158084 | A3ss | 3 | 3 | ENST00000330622 | 27.62 | 7.89E-04 | 1.23E-03 |
| NOS1AP | 1:162287437-162300647 | 1:162300633-162300647 | A3ss | 3 | 3 | ENST00000361897 | 27.42 | 9.27E-03 | 1.59E-02 |
| PHF20L1 | 8:132798861-132803818 | 8:132799095-132799172 | SES | 3 | 3 | ENST00000315808 | 27.37 | 2.46E-03 | 4.84E-02 |
| ZNF138 | 7:64815045-64831450 | 7:64830940-64831076 | MXS | 3 | 3 | ENST00000359735 | 27.19 | 1.84E-03 | 2.66E-02 |
| MICAL1 | 6:109445863-109446135 | 6:109445863-109445875 | A3ss | 3 | 3 | Ex.ENST00000358577 | 27.09 | 1.43E-04 | 4.39E-04 |
| ARAF | X:47561252-47562908 | X:47561252-47562762 | A5ss | 3 | 3 | Ex.TSS.ENST00000377039 | 27.08 | 5.76E-04 | 6.14E-02 |
| TMEM125 | 1:43271165-43272183 | 1:43271165-43271253 | A5ss | 3 | 3 | Ex.TSS.ENST00000456751 | 27.01 | 4.49E-04 | 8.35E-04 |
| TMEM125 | 1:43270794-43272183 | 1:43270794-43271253 | A5ss | 3 | 3 | Ex.ENST00000432792 | 27.01 | 4.49E-04 | 8.35E-04 |
| ZNF213-AS1 | 16:3132104-3134629 | 16:3132104-3132942 | A3ss | 3 | 3 | Ex.TSS.ENST00000571963 | 26.98 | 2.91E-02 | 1.07E-01 |
| TUBGCP4 | 15:43405202-43409771 | 15:43407571-43407942 | IR (overlapping | 3 | 3 | ENST00000564079 | 26.77 | 7.80E-04 | 2.58E-03 |
| THTPA | 14:23556344-23556752 | 14:23556743-23556752 | TSSlA3ss | 3 | 3 | TSS.ENST00000404535 | 26.73 | 9.36E-03 | 1.51E-02 |
| HLTF | 3:149041669-149042165 | 3:149041669-149041680 | A3ss | 3 | 3 | ENST00000467858 | 26.72 | 3.50E-03 | 8.09E-03 |
| MIER1 | 1:66926243-66928889 | 1:66928887-66928889 | A3ss | 3 | 3 | ENST00000357692 | 26.59 | 1.70E-02 | 3.19E-02 |
| ZNF410 | 14:73892209-73893796 | 14:73893776-73893796 | A3ss | 3 | 3 | ENST00000557495 | 26.5 | 1.70E-04 | 5.41E-04 |
| CNDP2 | 18:74497596-74499936 | 18:74499882-74499936 | A3ss | 3 | 3 | ENST00000584613 | 26.47 | 3.83E-02 | 1.95E-01 |
| ORAI2 | 7:102433662-102438943 | 7:102436202-102436333 | SES | 3 | 3 | ENST00000356387 | 26.46 | 1.56E-04 | 1.82E-03 |
| OCEL1 | 19:17228310-17229008 | 19:17228803-17228915 | SES | 3 | 3 | ENST00000595573 | 26.36 | 4.46E-02 | 2.39E-01 |
| MCEE | 2:71124544-71130179 | 2:71129717-71130179 | A5ss | 3 | 3 | ENST00000494660 | 26.34 | 1.64E-02 | 3.68E-02 |
| HELLS | 10:94545908-94546376 | 10:94546361-94546377 | A3ss | 3 | 3 | Ex.TSS.ENST00000419900 | 26.3 | 4.54E-04 | 6.49E-04 |
| SNHG8 | 4:118278793-118279388 | 4:118278944-118279137 | SES | 3 | 3 | ENST00000602819 | 26 | 5.53E-04 | 1.45E-03 |
| NINL | 20:25498347-25500839 | 20:25498347-25498361 | A3ss | 3 | 3 | Ex.ENST00000278886 | 25.87 | 5.06E-03 | 9.58E-03 |
| SLC35C1 | 11:45805149-45806336 | 11:45805317-45805809 | IR | 3 | 3 | ENST00000314134 | 25.84 | 3.58E-02 | 5.27E-02 |
| BAG6 | 6:31644195-31644524 | 6:31644307-31644414 | SES | 3 | 3 | ENST00000437771 | 25.81 | 1.33E-04 | 1.22E-03 |
| ADPRM | 17:10705528-10706437 | 17:10705528-10705615 | A5ss | 3 | 3 | ENST00000468843 | 25.56 | 3.52E-02 | 1.37E-01 |
| SLC25A36 | 3:140963228-140966999 | 3:140966934-140966999 | TSSlA3ss | 3 | 3 | TSS.ENST00000393015 | 25.55 | 2.13E-02 | 4.89E-02 |
| EME2 | 16:1776068-1780635 | 16:1777406-1778454 | IR (overlapping | 3 | 3 | ENST00000568449 | 25.53 | 3.51E-02 | 1.23E-01 |
| PDLIM7 | 5:177491147-177491806 | 5:177491147-177491420 | TSSlA3ss | 3 | 3 | TSS.ENST00000504318 | 25.44 | 4.02E-03 | 1.19E-02 |
| OCIAD1 | 4:48831053-48831226 | 4:48831053-48831141 | TSSlA5ss | 2 | 2 | TSS.ENST00000514981 | 25.44 | 4.87E-02 | 1.33E-01 |
| NCOA1 | 2:24762777-24768223 | 2:24768079-24768135 | SES | 3 | 3 | ENST00000407230 | 25.41 | 5.47E-03 | 1.20E-02 |
| ARHGAP5 | 14:32094387-32117142 | 14:32117140-32117142 | A3ss | 3 | 3 | ENST00000345122 | 25.39 | 4.10E-02 | 7.01E-02 |
| SMCO4 | 11:93479270-93499275 | 11:93479270-93479284 | TSSlA3ss | 3 | 3 | TSS.ENST00000529714 | 25.38 | 9.04E-05 | 1.40E-04 |
| TUT4 | 1:52436976-52438219 | 1:52436976-52436978 | A3ss | 3 | 3 | ENST00000257177 | 25.32 | 3.62E-02 | 6.77E-02 |
| TUT4 | 1:52436964-52438219 | 1:52436964-52436978 | A3ss | 3 | 3 | ENST00000257177 | 25.32 | 3.62E-02 | 6.77E-02 |

| | | | | | | | | | |
|---|---|---|---|---|---|---|---|---|---|
| ZNF213-AS1 | 16:3132104-3134629 | 16:3132850-3132945 | SES | 3 | 3 | ENST00000573447 | 25.31 | 1.36E-02 | 4.99E-02 |
| VPS54 | 2:63972245-63983863 | 2:63981646-63981851 | SES | 3 | 3 | ENST00000409558 | 25.23 | 5.30E-03 | 1.18E-02 |
| BRD9 | 5:865582-870472 | 5:869245-869404 | MES | 3 | 3 | ENST00000490814 | 25.21 | 7.73E-04 | 2.35E-03 |
| SLC25A36 | 3:140963228-140970926 | 3:140966934-140967036 | MXS | 3 | 3 | ENST00000512506 | 25.18 | 2.22E-02 | 5.11E-02 |
| SNHG8 | 4:118278793-118279388 | 4:118278793-118279137 | A5ss | 3 | 3 | ENST00000652022 | 25.16 | 6.53E-04 | 1.72E-03 |
| NFYA | 6:41073085-41079028 | 6:41079005-41079029 | A3ss | 3 | 3 | Ex.TSS.ENST00000341376 | 25.02 | 1.86E-03 | 1.76E-02 |
| MED6 | 14:70592989-70593295 | 14:70592989-70593009 | A3ss | 3 | 3 | ENST00000430055 | 24.93 | 2.96E-02 | 9.30E-02 |
| PRKCSH | 19:11447619-11447722 | 19:11447693-11447722 | A3ss | 3 | 3 | ENST00000585540 | 24.89 | 2.90E-04 | 1.50E-03 |
| ENTPD1-AS1 | 10:95876661-95907777 | 10:95904830-95904938 | SES | 2 | 3 | ENST00000669099 | 24.74 | 1.83E-02 | 2.60E-02 |
| MTHFR | 1:11785723-11790898 | 1:11788048-11788242 | IR (overlapping | 3 | 3 | ENST00000376590 | 24.67 | 3.56E-03 | 2.90E-02 |
| MIER1 | 1:66926243-66940027 | 1:66928887-66928963 | SES | 3 | 3 | ENST00000357692 | 24.66 | 6.34E-03 | 1.19E-02 |
| ZBED3 | 5:77077896-77087110 | 5:77078594-77078728 | SES | 3 | 3 | ENST00000255198 | 24.61 | 2.40E-02 | 7.27E-02 |
| ZNF493 | 19:21397268-21422912 | 19:21405129-21405255 | MES | 3 | 3 | ENST00000392288 | 24.57 | 4.65E-02 | 2.49E-01 |
| PNCK | X:153671499-153672200 | X:153672019-153672125 | IR | 3 | 2 | ENST00000488168 | 24.55 | 1.71E-02 | 1.00E+00 |
| C1orf52 | 1:85258723-85259357 | 1:85258935-85259061 | SES | 3 | 3 | ENST00000294661 | 24.4 | 6.23E-03 | 1.18E-02 |
| HTT | 4:3140657-3142765 | 4:3142745-3142765 | A3ss | 3 | 3 | ENST00000649131 | 24.22 | 5.17E-03 | 1.40E-02 |
| MTHFR | 1:11785723-11790898 | 1:11788048-11788148 | IR (overlapping | 3 | 3 | ENST00000376590 | 24.21 | 5.66E-03 | 4.61E-02 |
| DDX11 | 12:31074328-31078389 | 12:31078178-31078390 | A3ss | 3 | 3 | Ex.TSS.ENST00000542244 | 24.02 | 8.38E-03 | 1.34E-02 |
| TRIM46 | 1:155174030-155175385 | 1:155174601-155174641 | SES | 3 | 3 | ENST00000392451 | 23.97 | 4.37E-03 | 7.56E-03 |
| ZSCAN26 | 6:28267454-28272669 | 6:28271854-28271934 | SES | 2 | 3 | ENST00000614088 | 23.92 | 5.74E-03 | 5.17E-02 |
| AC119673.3 | 1:205891310-205894466 | 1:205894444-205894467 | A3ss | 3 | 3 | Ex.TSS.ENST00000652821 | 23.86 | 7.43E-04 | 1.32E-03 |
| FASTKD1 | 2:169531491-169537226 | 2:169531491-169531508 | A3ss | 3 | 3 | Ex.ENST00000453153 | 23.74 | 4.36E-03 | 9.17E-03 |
| UBE3A | 15:25405503-25438488 | 15:25411908-25411971 | MES | 3 | 3 | ENST00000428984 | 23.72 | 1.17E-02 | 3.82E-02 |
| ARFIP1 | 4:152863715-152872451 | 4:152870753-152870848 | SES | 3 | 3 | ENST00000353617 | 23.69 | 2.18E-03 | 5.83E-03 |
| NOVA1 | 14:26480144-26595409 | 14:26549680-26549781 | MXS | 3 | 2 | ENST00000483536 | 23.67 | 1.15E-02 | 1.96E-02 |
| FOXRED1 | 11:126274927-126275349 | 11:126275022-126275326 | IR | 3 | 3 | ENST00000524751 | 23.47 | 3.43E-02 | 4.96E-02 |
| RBM41 | X:107088840-107115351 | X:107113397-107113468 | SES | 3 | 3 | ENST00000495517 | 23.4 | 4.19E-02 | 1.00E+00 |
| COL23A1 | 5:178257568-178259720 | 5:178257568-178257582 | A3ss | 3 | 3 | Ex.ENST00000390654 | 23.39 | 4.06E-02 | 1.21E-01 |
| AC005062.1 | 7:19919261-19960921 | 7:19919866-19960922 | A5ss | 2 | 3 | Ex.TSS.ENST00000435968 | 23.28 | 4.97E-02 | 6.36E-01 |
| TRA2A | 7:23522433-23531202 | 7:23531181-23531202 | A5ss | 3 | 3 | Ex.ENST00000392502 | 23.25 | 8.39E-03 | 1.08E-01 |
| TUBGCP4 | 15:43405202-43409771 | 15:43409097-43409646 | IR (overlapping | 3 | 3 | ENST00000564079 | 23.19 | 8.63E-03 | 2.86E-02 |
| TTC14 | 3:180607648-180610974 | 3:180608811-180609629 | IR (overlapping | 3 | 3 | ENST00000412756 | 23.1 | 2.39E-02 | 5.59E-02 |
| UBA1 | X:47199615-47200893 | X:47200410-47200894 | A3ss | 3 | 3 | Ex.ENST00000335972 | 22.98 | 2.61E-04 | 2.74E-02 |
| DYNC2LI1 | 2:43774147-43776781 | 2:43774403-43776782 | A3ss | 3 | 3 | Ex.TSS.ENST00000398823 | 22.91 | 4.31E-02 | 9.48E-02 |
| IMMT | 2:86171313-86173649 | 2:86171313-86171345 | A3ss | 3 | 3 | ENST00000410111 | 22.9 | 1.11E-03 | 2.52E-03 |
| IMMT | 2:86171310-86173649 | 2:86171310-86171345 | A3ss | 3 | 3 | Ex.ENST00000410111 | 22.9 | 1.11E-03 | 2.49E-03 |
| P2RX5 | 17:3690681-3691643 | 17:3690956-3691027 | SES | 3 | 3 | ENST00000225328 | 22.9 | 3.64E-02 | 1.48E-01 |
| MTHFR | 1:11785723-11790898 | 1:11788048-11788163 | IR (overlapping | 3 | 3 | ENST00000376590 | 22.87 | 7.61E-03 | 6.20E-02 |
| HDAC7 | 12:47795768-47796206 | 12:47795906-47796016 | SES | 3 | 3 | ENST00000080059 | 22.84 | 8.80E-04 | 1.44E-03 |
| EIF2AK4 | 15:39943486-39949115 | 15:39945335-39949116 | A3ss | 3 | 3 | Ex.ENST00000263791 | 22.82 | 2.14E-03 | 7.00E-03 |
| UQCC1 | 20:35346424-35347163 | 20:35346424-35346461 | A3ss | 3 | 3 | Ex.TSS.ENST00000491125 | 22.81 | 5.96E-04 | 1.13E-03 |
| UQCC1 | 20:35314766-35347163 | 20:35314766-35346461 | A3ss | 3 | 3 | Ex.ENST00000349714 | 22.81 | 5.96E-04 | 1.13E-03 |
| SLTM | 15:58917000-58932355 | 15:58917000-58917020 | A3ss | 3 | 3 | Ex.ENST00000249736 | 22.77 | 4.02E-03 | 1.34E-02 |
| TRIM37 | 17:59070948-59075646 | 17:59070948-59070968 | A3ss | 3 | 3 | Ex.ENST00000262294 | 22.76 | 1.34E-04 | 6.00E-04 |
| CYB561D2 | 3:50350985-50351408 | 3:50351343-50351408 | TSSIA3ss | 3 | 3 | TSS.ENST00000232508 | 22.73 | 1.86E-02 | 4.54E-02 |
| RTF2 | 20:56468767-56472325 | 20:56471614-56471669 | MXS | 2 | 3 | ENST00000487211 | 22.65 | 3.17E-02 | 4.41E-02 |
| ADARB1 | 21:45222018-45226560 | 21:45222108-45225546 | IR | 3 | 3 | ENST00000348831 | 22.6 | 7.60E-03 | 1.48E-02 |
| PACRGL | 4:20700707-20704465 | 4:20702141-20702221 | SES | 3 | 3 | ENST00000504630 | 22.58 | 1.75E-02 | 4.74E-02 |
| TCF7L2 | 10:113158718-113165554 | 10:113159920-113159992 | SES | 3 | 3 | ENST00000355995 | 22.51 | 2.74E-02 | 3.84E-02 |
| PHF20L1 | 8:132798861-132799094 | 8:132799083-132799094 | A3ss | 3 | 3 | ENST00000395376 | 22.51 | 3.52E-02 | 6.93E-01 |
| SGSH | 17:80214766-80215032 | 17:80214766-80214783 | A3ss | 3 | 3 | ENST00000570427 | 22.49 | 5.75E-05 | 2.78E-04 |
| SGSH | 17:80214680-80215032 | 17:80214680-80214783 | A3ss | 3 | 3 | ENST00000570427 | 22.49 | 5.75E-05 | 2.78E-04 |
| TCF12 | 15:57251424-57253261 | 15:57252421-57252492 | SES | 3 | 3 | ENST00000333725 | 22.42 | 1.31E-02 | 4.36E-02 |
| DTWD2 | 5:118944650-118988293 | 5:118952122-118988294 | A5ss | 3 | 3 | Ex.TSS.ENST00000506980 | 22.17 | 2.86E-02 | 8.18E-02 |
| TCF7 | 5:134146224-134148210 | 5:134146245-134146369 | IR | 3 | 3 | ENST00000342854 | 22.13 | 3.01E-02 | 8.67E-02 |
| VEZT | 12:95266633-95270050 | 12:95270039-95270050 | A3ss | 3 | 3 | ENST00000397792 | 22.11 | 1.40E-02 | 2.23E-02 |
| TMEM161B-AS1 | 5:88282603-88287627 | 5:88285852-88287438 | IR | 2 | 3 | ENST00000669353 | 22.06 | 3.02E-02 | 9.17E-02 |
| PCNX3 | 11:65620967-65622244 | 11:65622224-65622245 | A3ss | 3 | 3 | Ex.ENST00000355703 | 22.01 | 3.86E-03 | 5.86E-03 |
| DLG2 | 11:83472778-83532707 | 11:83484129-83484228 | MXS | 3 | 3 | ENST00000280241 | 21.94 | 2.32E-02 | 3.58E-02 |
| ARHGEF9 | X:63724712-63754920 | X:63754315-63754676 | MES | 3 | 2 | ENST00000635967 | 21.89 | 3.72E-02 | 1.00E+00 |
| TRRAP | 7:98981961-98983263 | 7:98983240-98983264 | A3ss | 3 | 3 | Ex.ENST00000355540 | 21.86 | 6.81E-04 | 1.08E-02 |
| ATXN2 | 12:111457360-111470107 | 12:111464662-111464715 | MXS | 3 | 3 | ENST00000389153 | 21.83 | 5.63E-04 | 8.76E-04 |
| VEZT | 12:95256623-95257153 | 12:95257150-95257153 | A3ss | 3 | 3 | ENST00000356859 | 21.83 | 3.17E-02 | 4.42E-02 |
| TOR1AIP2 | 1:179865855-179877238 | 1:179865855-179865869 | A3ss | 3 | 3 | Ex.TSS.ENST00000482587 | 21.82 | 3.25E-03 | 5.71E-03 |
| ERCC3 | 2:127289502-127289688 | 2:127289502-127289519 | A3ss | 3 | 3 | ENST00000456257 | 21.8 | 1.01E-02 | 2.11E-02 |
| ZNF79 | 9:127424804-127428831 | 9:127424804-127424910 | A5ss | 2 | 3 | Ex.TSS.ENST00000342483 | 21.77 | 5.16E-03 | 1.81E-01 |
| ADCY5 | 3:123303220-123314234 | 3:123304067-123304183 | SES | 3 | 3 | ENST00000309879 | 21.77 | 3.02E-02 | 6.91E-02 |
| KLHL7 | 7:23106795-23123776 | 7:23117834-23117964 | SES | 3 | 2 | ENST00000410047 | 21.71 | 2.13E-02 | 2.74E-01 |
| EME1 | 17:50376194-50378599 | 17:50378595-50378599 | A3ss | 2 | 3 | ENST00000338165 | 21.7 | 1.36E-02 | 5.95E-02 |
| CNOT2-DT | 12:70243066-70243360 | 12:70243121-70243189 | IR (overlapping | 3 | 3 | ENST00000549651 | 21.67 | 4.64E-02 | 7.80E-02 |
| CYB561 | 17:63434595-63435085 | 17:63434595-63434620 | A3ss | 3 | 3 | ENST00000585153 | 21.62 | 6.91E-03 | 3.11E-02 |
| STIM1 | 11:4082983-4083262 | 11:4083242-4083263 | A3ss | 3 | 3 | Ex.ENST00000300737 | 21.62 | 4.84E-02 | 7.10E-02 |
| DCAF10 | 9:37854983-37860047 | 9:37857241-37857351 | SES | 3 | 3 | ENST00000242323 | 21.59 | 1.58E-02 | 4.33E-01 |
| FIP1L1 | 4:53428184-53442652 | 4:53440582-53440608 | SES | 3 | 3 | ENST00000358575 | 21.51 | 1.81E-02 | 5.08E-02 |
| CAND2 | 3:12803632-12810058 | 3:12808210-12808333 | MES | 3 | 2 | ENST00000456430 | 21.45 | 4.13E-02 | 9.47E-02 |
| SLC25A29 | 14:100293378-100298841 | 14:100295626-100295697 | SES | 3 | 3 | ENST00000554224 | 21.39 | 7.26E-04 | 1.23E-03 |
| PSPH | 7:56034104-56051137 | 7:56034104-56034154 | TSSIA3ss | 3 | 3 | TSS.ENST00000395471 | 21.36 | 2.79E-02 | 3.97E-01 |
| TNK1 | 17:7384266-7384754 | 17:7384377-7384483 | IR | 3 | 3 | ENST00000576136 | 21.35 | 1.09E-02 | 4.99E-02 |
| PRKD3 | 2:37293272-37324680 | 2:37293272-37317179 | A3ss | 3 | 3 | Ex.TSS.ENST00000443187 | 21.31 | 4.58E-02 | 1.01E-01 |
| PPOX | 1:161167235-161168992 | 1:161167339-161167486 | MES | 3 | 3 | ENST00000495483 | 21.27 | 1.78E-03 | 3.10E-03 |
| SLC2A11 | 22:23862203-23868480 | 22:23867735-23867899 | SES | 3 | 3 | ENST00000418102 | 21.18 | 1.01E-02 | 1.98E-02 |
| PDLIM7 | 5:177491147-177491806 | 5:177491404-177491420 | SES | 3 | 3 | ENST00000493815 | 21.1 | 1.09E-02 | 3.22E-02 |
| STAG2 | X:124090765-124094017 | X:124090854-124090964 | SES | 3 | 3 | ENST00000218089 | 21.02 | 5.12E-03 | 2.82E-01 |
| TRAF2 | 9:136890543-136898712 | 9:136890543-136898599 | A5ss | 3 | 3 | Ex.ENST00000414589 | 21 | 2.46E-02 | 1.00E+00 |
| TRAF2 | 9:136886542-136898712 | 9:136886542-136898599 | A5ss | 3 | 3 | Ex.TSS.ENST00000247668 | 21 | 2.46E-02 | 1.00E+00 |
| PACRGL | 4:20700618-20704465 | 4:20702141-20702221 | MXS | 3 | 3 | ENST00000295290 | 20.91 | 1.52E-02 | 4.11E-02 |
| THTPA | 14:23556344-23558694 | 14:23556743-23557304 | SES | 3 | 3 | ENST00000404535 | 20.89 | 2.73E-03 | 4.44E-03 |
| APTX | 9:32987732-32988091 | 9:32988083-32988091 | A5ss | 3 | 3 | ENST00000464632 | 20.86 | 1.88E-03 | 5.35E-02 |
| ZNF793 | 19:37536897-37543335 | 19:37537514-37537514 | IR | 3 | 2 | ENST00000445217 | 20.8 | 1.75E-02 | 1.13E-01 |
| INTS9 | 8:28768323-28769888 | 8:28768323-28768376 | TSSIA3ss | 3 | 3 | TSS.ENST00000517383 | 20.79 | 7.95E-03 | 1.71E-01 |
| SPART | 13:36335833-36346581 | 13:36335833-36336358 | A3ss | 3 | 3 | Ex.TSS.ENST00000451493 | 20.77 | 4.27E-04 | 6.90E-04 |
| TUBGCP4 | 15:43405202-43409771 | 15:43408089-43408896 | IR (overlapping | 3 | 3 | ENST00000564079 | 20.75 | 3.07E-02 | 1.02E-01 |
| ZMAT5 | 22:29748572-29766948 | 22:29766872-29766948 | TSSIA5ss | 3 | 3 | TSS.ENST00000344318 | 20.64 | 2.33E-02 | 4.73E-02 |
| PI4KAP2 | 22:21476161-21477789 | 22:21476644-21476764 | SES | 3 | 2 | ENST00000360806 | 20.5 | 8.08E-03 | 1.58E-02 |

| | | | | | | | | | |
|---|---|---|---|---|---|---|---|---|---|
| DCAF11 | 14:24115393-24115542 | 14:24115515-24115542 | TSSlA3ss | 3 | 3 | TSS.ENST00000559396 | 20.5 | 1.41E-02 | 2.28E-02 |
| SKIV2L | 6:31968539-31968685 | 6:31968623-31968685 | A3ss | 3 | 3 | ENST00000465703 | 20.45 | 2.32E-02 | 2.13E-01 |
| DGCR2 | 22:19057163-19063201 | 22:19057163-19057186 | A3ss | 3 | 3 | Ex.ENST00000263196 | 20.43 | 2.06E-03 | 4.02E-03 |
| GTF3C2 | 2:27343579-27356738 | 2:27350314-27350632 | MXS | 3 | 3 | ENST00000423998 | 20.38 | 2.04E-02 | 4.47E-02 |
| TMEM143 | 19:48345355-48363897 | 19:48360072-48360183 | MXS | 3 | 3 | ENST00000598258 | 20.38 | 2.38E-02 | 1.63E-01 |
| ARAF | X:47561252-47562908 | X:47562682-47562909 | A3ss | 3 | 3 | Ex.TSS.ENST00000377039 | 20.34 | 4.86E-03 | 5.18E-01 |
| PTOV1-AS2 | 19:49858843-49859287 | 19:49859098-49859205 | IR (overlapping | 3 | 3 | ENST00000601893 | 20.33 | 5.52E-03 | 3.83E-02 |
| LINC01224 | 19:23403748-23415893 | 19:23405258-23405335 | MES | 3 | 3 | ENST00000593573 | 20.32 | 3.46E-02 | 1.91E-01 |
| PHKB | 16:47461427-47497398 | 16:47463882-47463993 | MXS | 3 | 3 | ENST00000566044 | 20.28 | 1.68E-02 | 6.20E-02 |
| ZNF250 | 8:144890404-144901398 | 8:144901337-144901398 | TSSlA5ss | 3 | 3 | TSS.ENST00000533622 | 20.28 | 2.55E-02 | 5.22E-01 |
| AKT1S1 | 19:49873303-49877236 | 19:49876603-49877236 | TSSlA5ss | 3 | 3 | TSS.ENST00000391835 | 20.24 | 2.27E-03 | 1.57E-02 |
| AKT1S1 | 19:49873303-49877024 | 19:49876603-49877024 | TSSlA5ss | 3 | 3 | TSS.ENST00000391835 | 20.24 | 2.27E-03 | 1.57E-02 |
| AL162411.1 | 9:6669729-6670424 | 9:6669999-6670425 | A3ss | 3 | 3 | Ex.ENST00000413145 | 20.23 | 7.89E-03 | 2.42E-01 |
| PPOX | 1:161167235-161167370 | 1:161167339-161167370 | A3ss | 3 | 3 | ENST00000495483 | 20.2 | 8.44E-04 | 1.47E-03 |
| AC058822.1 | 4:53377924-53379224 | 4:53379073-53379117 | SES | 3 | 3 | ENST00000507166 | 20.19 | 1.38E-03 | 3.88E-03 |
| SREBF1 | 17:17817458-17817695 | 17:17817548-17817695 | A5ss | 3 | 3 | ENST00000395756 | 20.17 | 9.01E-03 | 3.18E-02 |
| SRGAP2B | 1:144955602-144995007 | 1:144960338-144995008 | A5ss | 3 | 3 | Ex.ENST00000612199 | 20.06 | 4.12E-02 | 3.35E-01 |
| POLR3G | 5:90475021-90485534 | 5:90485525-90485534 | TSSlA3ss | 3 | 3 | TSS.ENST00000399107 | 20.02 | 1.04E-03 | 3.17E-03 |
| SREBF1 | 17:17817458-17817695 | 17:17817548-17817591 | SES | 3 | 3 | ENST00000395751 | 20.02 | 4.06E-03 | 1.61E-02 |
| STOX2 | 4:184011424-184017088 | 4:184011587-184011635 | SES | 3 | 3 | ENST00000506529 | 20.02 | 1.73E-02 | 4.67E-02 |
| MLLT10 | 10:21681377-21713771 | 10:21682225-21682257 | MXS | 3 | 3 | ENST00000307729 | 20.02 | 4.97E-02 | 6.99E-02 |
| RFFL | 17:35017607-35026373 | 17:35021371-35021781 | SES | 3 | 3 | ENST00000315249 | 19.96 | 4.93E-02 | 2.00E-01 |
| PI4KAP2 | 22:21476161-21477789 | 22:21476161-21476764 | A3ss | 3 | 3 | ENST00000462560 | 19.9 | 5.65E-04 | 1.11E-03 |
| FBXL3 | 13:77018723-77021656 | 13:77021513-77021656 | A5ss | 2 | 3 | ENST00000355619 | 19.86 | 2.45E-02 | 4.01E-02 |
| IQGAP3 | 1:156562666-156563133 | 1:156562666-156562674 | A3ss | 3 | 3 | Ex.ENST00000361170 | 19.85 | 1.94E-03 | 3.37E-03 |
| LUC7L3 | 17:50750853-50752199 | 17:50751297-50751362 | SES | 3 | 3 | ENST00000503728 | 19.85 | 9.49E-03 | 4.21E-02 |
| ZNF213-AS1 | 16:3132104-3134629 | 16:3132850-3133194 | SES | 3 | 3 | ENST00000571449 | 19.78 | 3.84E-02 | 1.41E-01 |
| ENTPD1-AS1 | 10:95876661-95907662 | 10:95904830-95904935 | MXS | 3 | 3 | ENST00000449419 | 19.75 | 4.48E-02 | 6.38E-02 |
| TRPT1 | 11:64225581-64226049 | 11:64225786-64226049 | TSSlA5ss | 3 | 3 | TSS.ENST00000539595 | 19.72 | 8.21E-03 | 1.23E-02 |
| TRAPPC2 | X:13719983-13734524 | X:13734044-13734185 | MES | 3 | 3 | ENST00000518847 | 19.71 | 2.07E-03 | 1.24E-01 |
| TCEAL1 | X:103629026-103629884 | X:103629494-103629561 | SES | 3 | 3 | ENST00000372624 | 19.7 | 1.04E-02 | 2.63E-01 |
| ADGRL2 | 1:81800513-81836884 | 1:81800513-81802229 | A5ss | 3 | 3 | Ex.TSS.ENST00000319517 | 19.66 | 2.12E-04 | 4.00E-04 |
| EZH2 | 7:148818118-148819595 | 7:148818978-148819059 | SES | 3 | 3 | ENST00000483012 | 19.66 | 2.91E-02 | 3.58E-01 |
| FAM49B | 8:129871497-129903311 | 8:129879389-129879471 | MXS | 3 | 3 | ENST00000517654 | 19.6 | 1.00E-03 | 1.96E-02 |
| ECT2 | 3:172761684-172762415 | 3:172762398-172762416 | A3ss | 3 | 3 | Ex.ENST00000232458 | 19.53 | 1.18E-06 | 2.75E-06 |
| MCEE | 2:71124544-71130179 | 2:71129717-71129817 | SES | 3 | 3 | ENST00000486135 | 19.51 | 1.10E-02 | 2.46E-02 |
| MAP3K7 | 6:90536402-90547257 | 6:90544552-90544632 | SES | 3 | 3 | ENST00000369325 | 19.49 | 1.25E-02 | 1.42E-01 |
| AC007161.3 | 7:7640822-7673308 | 7:7641504-7641705 | SES | 3 | 2 | ENST00000469183 | 19.43 | 1.29E-02 | 1.94E-01 |
| KDM4C | 9:7103871-7128065 | 9:7105402-7105500 | SES | 3 | 3 | ENST00000420847 | 19.31 | 4.07E-03 | 1.25E-01 |
| RTN2 | 19:45495140-45496791 | 19:45495140-45495214 | TSSlA3ss | 3 | 3 | TSS.ENST00000590526 | 19.3 | 3.27E-03 | 2.19E-02 |
| ARHGEF9 | X:63724712-63754920 | X:63754304-63754676 | MES | 3 | 2 | ENST00000637417 | 19.26 | 2.11E-02 | 1.00E+00 |
| DZIP3 | 3:108647943-108648962 | 3:108648113-108648917 | IR | 3 | 2 | ENST00000495008 | 19.25 | 3.04E-02 | 6.92E-02 |
| NEURL4 | 17:7318675-7319049 | 17:7318675-7318691 | A3ss | 3 | 3 | ENST00000573186 | 19.23 | 3.81E-02 | 1.74E-01 |
| AC138969.1 | 16:16318080-16318922 | 16:16318551-16318734 | SES | 3 | 3 | ENST00000536260 | 19.21 | 3.31E-02 | 1.16E-01 |
| ORMDL1 | 2:189783124-189784268 | 2:189783879-189784269 | A5ss | 3 | 3 | Ex.TSS.ENST00000392349 | 19.17 | 3.93E-02 | 8.31E-02 |
| NBEAL2 | 3:46991947-46993936 | 3:46992475-46992555 | SES | 3 | 3 | ENST00000450053 | 19.16 | 1.13E-03 | 2.71E-03 |
| MYO6 | 6:75892691-75907604 | 6:75894814-75894840 | MES | 3 | 3 | ENST00000430435 | 19.15 | 2.91E-02 | 2.97E-01 |
| CC2D2A | 4:15478807-15480703 | 4:15479223-15479328 | SES | 3 | 3 | ENST00000438599 | 19.15 | 3.23E-02 | 8.62E-02 |
| AP1S2 | X:15827373-15845378 | X:15831452-15831568 | MES | 3 | 3 | ENST00000671830 | 19.14 | 4.57E-03 | 3.31E-01 |
| PDXDC2P | 16:70036579-70042350 | 16:70038905-70038985 | SES | 3 | 3 | ENST00000527016 | 19.1 | 1.22E-02 | 4.69E-02 |
| FDPS | 1:155308966-155309788 | 1:155308966-155309076 | TSSlA5ss | 3 | 3 | TSS.ENST00000356657 | 19.06 | 2.83E-03 | 4.89E-03 |
| LIG3 | 17:35004273-35009743 | 17:35007455-35007871 | IR (overlapping | 3 | 3 | ENST00000378526 | 19.04 | 3.53E-02 | 1.43E-01 |
| HDDC3 | 15:90931406-90931703 | 15:90931406-90931556 | TSSlA3ss | 3 | 3 | TSS.ENST00000330334 | 19.01 | 3.01E-03 | 1.05E-02 |
| TMEM161B-AS1 | 5:88270586-88436272 | 5:88270586-88282106 | A5ss | 3 | 3 | Ex.TSS.ENST00000504922 | 19.01 | 1.43E-02 | 4.37E-02 |
| EME2 | 16:1776068-1780635 | 16:1776340-1776860 | IR | 3 | 3 | ENST00000568449 | 18.99 | 4.56E-02 | 1.60E-01 |
| SLC25A29 | 14:100293378-100306198 | 14:100295626-100295993 | MES | 3 | 3 | ENST00000556868 | 18.85 | 2.05E-02 | 3.46E-02 |
| GABARAPL1 | 12:10213429-10214187 | 12:10213715-10214073 | IR | 3 | 2 | ENST00000546017 | 18.85 | 4.79E-02 | 7.16E-02 |
| MORC2 | 22:30926872-30928027 | 22:30928019-30928027 | TSSlA5ss | 3 | 3 | TSS.ENST00000215862 | 18.82 | 6.14E-03 | 1.25E-02 |
| FDPS | 1:155308966-155310063 | 1:155310043-155310063 | TSSlA3ss | 3 | 3 | TSS.ENST00000447866 | 18.81 | 3.71E-04 | 6.42E-04 |
| JAKMIP1 | 4:6085630-6112721 | 4:6105473-6105967 | SES | 3 | 3 | ENST00000282924 | 18.81 | 3.35E-02 | 9.43E-02 |
| STAU2 | 8:73709163-73738283 | 8:73709163-73709177 | A3ss | 3 | 3 | ENST00000521845 | 18.8 | 1.87E-03 | 4.38E-02 |
| ATF2 | 2:175118370-175130137 | 2:175121444-175121540 | SES | 3 | 3 | ENST00000264110 | 18.78 | 8.86E-03 | 1.87E-02 |
| ICA1 | 7:8221399-8228600 | 7:8227638-8227851 | MXS | 3 | 3 | ENST00000455539 | 18.69 | 2.52E-02 | 3.83E-01 |
| AP5S1 | 20:3820703-3822101 | 20:3820703-3820758 | TSSlA5ss | 3 | 3 | TSS.ENST00000379573 | 18.64 | 2.33E-02 | 4.45E-02 |
| CNTROB | 17:7932155-7933349 | 17:7932801-7933092 | IR | 3 | 3 | ENST00000380262 | 18.61 | 1.16E-02 | 5.61E-02 |
| ZNF468 | 19:52849214-52854257 | 19:52853872-52854126 | SES | 3 | 3 | ENST00000601980 | 18.61 | 1.57E-02 | 1.11E-01 |
| UBR4 | 1:19153940-19154917 | 1:19153940-19153954 | A3ss | 3 | 3 | Ex.ENST00000375254 | 18.6 | 9.08E-03 | 1.61E-02 |
| TRAPPC2 | X:13719983-13734524 | X:13734050-13734185 | MXS | 3 | 3 | ENST00000458511 | 18.6 | 1.79E-02 | 9.72E-01 |
| DHX34 | 19:47376561-47377099 | 19:47376915-47376989 | SES | 3 | 3 | ENST00000460681 | 18.57 | 8.60E-03 | 5.78E-02 |
| SNRPN | 15:24962210-24967931 | 15:24967029-24967240 | SES | 3 | 3 | ENST00000584968 | 18.55 | 1.83E-05 | 5.94E-05 |
| PACRGL | 4:20700788-20704465 | 4:20702141-20702221 | MXS | 3 | 3 | ENST00000506745 | 18.52 | 3.49E-03 | 9.45E-03 |
| METTL5 | 2:169812507-169815476 | 2:169812507-169812524 | A3ss | 3 | 3 | ENST00000442181 | 18.5 | 1.48E-03 | 3.13E-03 |
| AC022384.1 | 3:10278463-10278884 | 3:10278664-10278760 | SES | 3 | 3 | ENST00000437082 | 18.49 | 7.66E-03 | 1.75E-02 |
| SNTG2 | 2:1083656-1209102 | 2:1165548-1209103 | A3ss | 3 | 3 | Ex.ENST00000407292 | 18.48 | 4.77E-03 | 9.91E-03 |
| POMT1 | 9:131503074-131506113 | 9:131504189-131504340 | SES | 3 | 3 | ENST00000372228 | 18.43 | 1.70E-02 | 6.73E-01 |
| TRPT1 | 11:64225581-64226049 | 11:64225786-64225869 | SES | 3 | 3 | ENST00000317459 | 18.42 | 1.73E-02 | 2.59E-02 |
| VARS2 | 6:30916960-30917104 | 6:30917095-30917105 | A3ss | 3 | 3 | Ex.ENST00000321897 | 18.39 | 1.90E-02 | 1.71E-01 |
| FBXO27 | 19:39031318-39031863 | 19:39031318-39031320 | A3ss | 3 | 3 | ENST00000292853 | 18.31 | 3.13E-02 | 2.04E-01 |
| ZNF85 | 19:20923404-20948743 | 19:20934024-20934150 | MXS | 3 | 3 | ENST00000345030 | 18.29 | 6.71E-03 | 3.69E-02 |
| DERL3 | 22:23834933-23837063 | 22:23834933-23834979 | A3ss | 3 | 3 | Ex.TSS.ENST00000406855 | 18.28 | 1.12E-02 | 2.17E-02 |
| HDAC10 | 22:50250161-50250770 | 22:50250427-50250523 | SES | 3 | 3 | ENST00000216271 | 18.22 | 3.35E-02 | 6.93E-02 |
| MIS18BP1 | 14:45206171-45210379 | 14:45206738-45210380 | A5ss | 3 | 3 | Ex.ENST00000310806 | 18.18 | 3.63E-02 | 1.14E-01 |
| PATJ | 1:62160908-62163920 | 1:62162865-62162867 | IR | 3 | 3 | ENST00000371158 | 18.18 | 3.66E-02 | 6.88E-02 |
| AC091230.1 | 15:74842421-74843003 | 15:74842535-74842674 | SES | 3 | 3 | ENST00000568679 | 18.16 | 4.62E-02 | 1.56E-01 |
| B4GALT7 | 5:177604542-177607301 | 5:177607283-177607301 | A3ss | 3 | 3 | ENST00000505433 | 18.1 | 1.30E-02 | 3.86E-02 |
| NT5C3A | 7:33026916-33062567 | 7:33035934-33035988 | SES | 3 | 3 | ENST00000405342 | 18.06 | 1.81E-02 | 2.43E-01 |
| MACF1 | 1:39463687-39468614 | 1:39465095-39465112 | SES | 3 | 3 | ENST00000289893 | 18.04 | 1.98E-03 | 3.67E-03 |
| DKK3 | 11:11967099-11968394 | 11:11967099-11967119 | A3ss | 3 | 3 | Ex.ENST00000326932 | 18.03 | 2.00E-03 | 2.88E-03 |
| ZMYM4 | 1:35405469-35408007 | 1:35407987-35408008 | A5ss | 3 | 3 | Ex.ENST00000314607 | 17.95 | 2.63E-02 | 4.86E-02 |
| BACE1 | 11:117293189-117295130 | 11:117293871-117293933 | SES | 3 | 3 | ENST00000510630 | 17.94 | 2.06E-02 | 2.96E-02 |
| MIER1 | 1:66926243-66940027 | 1:66928890-66928963 | SES | 3 | 3 | ENST00000371012 | 17.93 | 1.45E-03 | 2.73E-03 |
| CEP104 | 1:3839777-3844906 | 1:3839777-3842067 | A3ss | 3 | 3 | Ex.ENST00000378230 | 17.92 | 3.89E-02 | 7.21E-02 |
| EFEMP2 | 11:65870536-65872019 | 11:65871410-65871969 | IR | 3 | 2 | ENST00000531005 | 17.89 | 1.03E-02 | 1.56E-02 |
| RCCD1 | 15:90955453-90956611 | 15:90955453-90955883 | A5ss | 3 | 3 | ENST00000554302 | 17.89 | 1.20E-02 | 4.20E-02 |

| | | | | | | | | | |
|---|---|---|---|---|---|---|---|---|---|
| ANAPC4 | 4:25405620-25406828 | 4:25405620-25405622 | A5ss | 3 | 3 | ENST00000510092 | 17.89 | 4.26E-02 | 1.15E-01 |
| SNX13 | 7:17821649-17826021 | 7:17821649-17821681 | A3ss | 3 | 3 | ENST00000611725 | 17.87 | 2.20E-02 | 2.82E-01 |
| RBM15 | 1:110342182-110346307 | 1:110342182-110342268 | A5ss | 3 | 3 | ENST00000487146 | 17.87 | 3.64E-02 | 3.03E-01 |
| CHD1L | 1:147242831-147265931 | 1:147256531-147256562 | MES | 3 | 3 | ENST00000369258 | 17.87 | 4.37E-02 | 3.75E-01 |
| TAF1C | 16:84184900-84186900 | 16:84184900-84184989 | TSSlA3ss | 3 | 3 | TSS.ENST00000564345 | 17.85 | 7.31E-04 | 2.83E-03 |
| XRCC3 | 14:103712932-103715423 | 14:103715337-103715424 | A5ss | 3 | 3 | Ex.TSS.ENST00000555055 | 17.83 | 4.91E-02 | 7.96E-02 |
| DENND10 | 10:119104198-119107967 | 10:119107948-119107967 | TSSlA3ss | 3 | 3 | TSS.ENST00000448258 | 17.81 | 8.68E-03 | 1.22E-02 |
| B3GALT5-AS1 | 21:39597990-39609582 | 21:39601419-39601543 | MES | 3 | 3 | ENST00000670883 | 17.81 | 4.65E-02 | 9.03E-02 |
| PRPF38A | 1:52414648-52414761 | 1:52414741-52414762 | A3ss | 3 | 3 | Ex.ENST00000257181 | 17.77 | 1.64E-03 | 3.07E-03 |
| CARMIL2 | 16:67656646-67657238 | 16:67656801-67656881 | SES | 3 | 3 | ENST00000334583 | 17.76 | 1.34E-03 | 5.08E-03 |
| BRD9 | 5:865582-870472 | 5:869245-869394 | MXS | 3 | 3 | ENST00000489816 | 17.72 | 4.18E-03 | 1.27E-02 |
| RWDD3 | 1:95234316-95246541 | 1:95244211-95244698 | MXS | 3 | 3 | ENST00000370202 | 17.71 | 6.11E-03 | 1.15E-02 |
| TNRC6A | 16:24794720-24797489 | 16:24795907-24795939 | SES | 3 | 3 | ENST00000315183 | 17.7 | 1.27E-02 | 4.57E-02 |
| JPH4 | 14:23576457-23578179 | 14:23577869-23577994 | MES | 3 | 3 | ENST00000397118 | 17.68 | 2.77E-07 | 4.69E-07 |
| AGRN | 1:1050838-1051252 | 1:1051032-1051043 | SES | 3 | 3 | ENST00000620552 | 17.63 | 1.13E-03 | 9.24E-03 |
| CHD3 | 17:7907489-7907893 | 17:7907601-7907702 | SES | 3 | 3 | ENST00000330494 | 17.62 | 2.60E-02 | 1.25E-01 |
| ZW10 | 11:113748421-113758553 | 11:113757662-113757853 | SES | 3 | 3 | ENST00000200135 | 17.61 | 2.55E-02 | 3.65E-02 |
| SEPTIN8 | 5:132758825-132760801 | 5:132759937-132760802 | A5ss | 3 | 3 | Ex.TSS.ENST00000296873 | 17.59 | 4.25E-04 | 1.22E-03 |
| TBC1D16 | 17:79950576-79951449 | 17:79950576-79950578 | A3ss | 3 | 3 | ENST00000310924 | 17.55 | 5.11E-03 | 2.47E-02 |
| SCMH1 | 1:41143113-41160874 | 1:41151614-41151684 | MES | 3 | 3 | ENST00000326197 | 17.51 | 1.79E-03 | 3.32E-03 |
| STAU2 | 8:73688814-73738283 | 8:73709032-73709177 | SES | 3 | 3 | ENST00000521845 | 17.5 | 2.88E-04 | 6.75E-03 |
| ZSCAN30 | 18:35264456-35290130 | 18:35267476-35267679 | SES | 2 | 2 | ENST00000592211 | 17.49 | 4.90E-02 | 2.45E-01 |
| SH2D3A | 19:6754426-6754615 | 19:6754426-6754448 | A3ss | 3 | 3 | Ex.ENST00000245908 | 17.48 | 5.02E-03 | 3.74E-02 |
| POLL | 10:101582892-101587245 | 10:101583508-101583681 | MES | 3 | 3 | ENST00000299206 | 17.48 | 4.12E-02 | 5.74E-02 |
| ARHGAP12 | 10:31831801-31843460 | 10:31839637-31839711 | MES | 3 | 3 | ENST00000344936 | 17.47 | 2.35E-02 | 3.32E-02 |
| CCDC150 | 2:196658861-196669832 | 2:196666719-196666848 | MES | 2 | 3 | ENST00000389175 | 17.47 | 2.66E-02 | 5.63E-02 |
| THTPA | 14:23557040-23558694 | 14:23557040-23557304 | A5ss | 3 | 3 | ENST00000288014 | 17.44 | 1.99E-03 | 3.23E-03 |
| THTPA | 14:23557035-23558694 | 14:23557035-23557304 | A5ss | 3 | 3 | ENST00000288014 | 17.44 | 1.99E-03 | 3.23E-03 |
| THTPA | 14:23556344-23558694 | 14:23556344-23557304 | A5ss | 3 | 3 | ENST00000288014 | 17.44 | 1.99E-03 | 3.23E-03 |
| FIP1L1 | 4:53399840-53425871 | 4:53414615-53414722 | SES | 3 | 3 | ENST00000337488 | 17.39 | 1.57E-02 | 4.40E-02 |
| MYH14 | 19:50292390-50293232 | 19:50292390-50292895 | A5ss | 3 | 3 | Ex.ENST00000262269 | 17.39 | 2.03E-02 | 1.41E-01 |
| RPSA | 3:39406765-39407620 | 3:39407603-39407620 | TSSlA3ss | 3 | 3 | TSS.ENST00000458478 | 17.38 | 3.93E-04 | 9.39E-04 |
| CCDC78 | 16:725294-725445 | 16:725413-725445 | A5ss | 3 | 2 | ENST00000293889 | 17.36 | 2.63E-02 | 1.01E-01 |
| MSL3 | X:11765467-11767977 | X:11767170-11767818 | IR | 3 | 3 | ENST00000337339 | 17.33 | 4.53E-02 | 1.00E+00 |
| PHF21B | 22:44885530-44885862 | 22:44885530-44885552 | A3ss | 3 | 3 | Ex.ENST00000313237 | 17.28 | 1.68E-03 | 3.46E-03 |
| WASF1 | 6:110127630-110179438 | 6:110160635-110160732 | MXS | 3 | 3 | ENST00000359451 | 17.28 | 3.98E-02 | 1.22E-01 |
| HDAC2 | 6:113960019-113970479 | 6:113968634-113968696 | MXS | 3 | 3 | ENST00000518690 | 17.22 | 4.25E-02 | 1.30E-01 |
| SFI1 | 22:31602786-31604868 | 22:31604309-31604404 | MXS | 3 | 3 | ENST00000382162 | 17.17 | 8.27E-03 | 1.69E-02 |
| OPA1 | 3:193618937-193631611 | 3:193626092-193626202 | MXS | 3 | 3 | ENST00000361150 | 17.16 | 2.97E-04 | 7.01E-04 |
| PLXND1 | 3:129566018-129566526 | 3:129566018-129566029 | A3ss | 3 | 3 | Ex.ENST00000324093 | 17.14 | 2.00E-03 | 4.59E-03 |
| ANK2 | 4:113373174-113381456 | 4:113373285-113373449 | SES | 3 | 3 | ENST00000264366 | 17.14 | 2.74E-02 | 7.15E-02 |
| RPP14 | 3:58306418-58310308 | 3:58306418-58306651 | TSSlA5ss | 3 | 3 | TSS.ENST00000445193 | 17.13 | 1.27E-03 | 3.28E-03 |
| ECT2 | 3:172755375-172756982 | 3:172755483-172755575 | SES | 3 | 3 | ENST00000366090 | 17.13 | 6.15E-03 | 1.43E-02 |
| ANKRD11 | 16:89275192-89288527 | 16:89286039-89286186 | MES | 3 | 2 | ENST00000301030 | 17.12 | 2.46E-02 | 9.47E-02 |
| CD82 | 11:44618366-44619048 | 11:44618640-44618723 | SES | 3 | 3 | ENST00000227155 | 17.1 | 1.32E-02 | 1.94E-02 |
| THUMPD3-AS1 | 3:9390014-9391826 | 3:9390498-9391106 | IR | 3 | 3 | ENST00000518437 | 17.09 | 4.20E-02 | 1.08E-01 |
| FAM3A | X:154507862-154508288 | X:154507862-154507882 | A3ss | 3 | 3 | ENST00000369641 | 17.06 | 4.10E-02 | 1.00E+00 |
| TPM1 | 15:63061274-63062214 | 15:63061274-63061788 | A5ss | 3 | 3 | ENST00000561242 | 17.05 | 9.16E-05 | 3.06E-04 |
| PWWP2A | 5:160094066-160118804 | 5:160108545-160108614 | SES | 3 | 3 | ENST00000520662 | 17.04 | 5.32E-03 | 1.58E-02 |
| STAU2 | 8:73688814-73746782 | 8:73738284-73738349 | MES | 3 | 3 | ENST00000519818 | 17.01 | 4.46E-03 | 1.04E-01 |
| DNAJC5 | 20:63931023-63931464 | 20:63931109-63931182 | SES | 3 | 3 | ENST00000470551 | 17 | 3.84E-02 | 5.35E-02 |
| ARHGAP12 | 10:31831801-31843460 | 10:31839637-31839711 | MXS | 3 | 3 | ENST00000311380 | 16.93 | 1.66E-02 | 2.35E-02 |
| ELMOD3 | 2:85354830-85356966 | 2:85355555-85355598 | MXS | 3 | 3 | ENST00000409331 | 16.9 | 2.54E-02 | 5.75E-02 |
| EMC3-AS1 | 3:9987906-9988711 | 3:9987906-9988007 | A5ss | 3 | 3 | ENST00000366264 | 16.9 | 2.86E-02 | 7.42E-02 |
| ATF7IP2 | 16:10386123-10428867 | 16:10419581-10419623 | MES | 3 | 3 | ENST00000561932 | 16.88 | 1.77E-02 | 6.21E-02 |
| TRPM4 | 19:49210843-49211163 | 19:49211015-49211087 | SES | 3 | 3 | ENST00000252826 | 16.83 | 1.52E-04 | 1.05E-03 |
| BPTF | 17:67875021-67891843 | 17:67875556-67875744 | MES | 3 | 3 | ENST00000321892 | 16.82 | 9.25E-04 | 4.23E-03 |
| PIGG | 4:521660-522842 | 4:521942-522026 | IR | 3 | 3 | ENST00000503111 | 16.8 | 4.17E-02 | 1.14E-01 |
| SLC12A9 | 7:100855838-100858834 | 7:100855838-100857176 | A5ss | 3 | 3 | Ex.TSS.ENST00000475687 | 16.79 | 3.95E-02 | 4.56E-01 |
| ZNF875 | 19:37334783-37347189 | 19:37335169-37335257 | MES | 3 | 3 | ENST00000324411 | 16.67 | 4.86E-02 | 3.14E-01 |
| TTC3 | 21:37198026-37200231 | 21:37200214-37200232 | A3ss | 3 | 3 | Ex.ENST00000354749 | 16.62 | 2.16E-02 | 4.20E-02 |
| MKI67 | 10:128113603-128116490 | 10:128114928-128116007 | SES | 3 | 3 | ENST00000368654 | 16.61 | 3.80E-02 | 5.35E-02 |
| BCL2L12 | 19:49666800-49668850 | 19:49667019-49667161 | SES | 3 | 3 | ENST00000246784 | 16.6 | 3.21E-04 | 2.22E-03 |
| WASF1 | 6:110178743-110179840 | 6:110179439-110179840 | TSSlA5ss | 3 | 3 | TSS.ENST00000392588 | 16.57 | 1.20E-02 | 3.68E-02 |
| PSMA3-AS1 | 14:58267394-58291634 | 14:58273979-58274135 | SES | 3 | 3 | ENST00000555275 | 16.56 | 1.58E-02 | 4.98E-02 |
| BUB1B | 15:40200981-40202404 | 15:40202390-40202405 | A3ss | 3 | 3 | Ex.ENST00000287598 | 16.56 | 2.18E-02 | 7.15E-02 |
| PATJ | 1:62160908-62163920 | 1:62162880-62162884 | IR | 3 | 3 | ENST00000371158 | 16.56 | 3.89E-02 | 7.31E-02 |
| LTBP4 | 19:40614447-40617099 | 19:40616889-40617099 | A3ss | 3 | 3 | ENST00000617753 | 16.5 | 4.01E-02 | 2.58E-01 |
| KIAA0895L | 16:67180667-67183261 | 16:67181606-67181697 | SES | 3 | 3 | ENST00000290881 | 16.49 | 4.98E-03 | 1.89E-02 |
| METTL26 | 16:634798-636093 | 16:634889-636093 | TSSlA5ss | 3 | 3 | TSS.ENST00000448973 | 16.47 | 5.03E-03 | 1.90E-02 |
| METTL26 | 16:634798-635611 | 16:634889-635611 | TSSlA5ss | 3 | 3 | TSS.ENST00000448973 | 16.47 | 5.03E-03 | 1.90E-02 |
| METTL26 | 16:634798-635280 | 16:634889-635280 | TSSlA5ss | 3 | 3 | TSS.ENST00000448973 | 16.47 | 5.03E-03 | 1.90E-02 |
| SLC44A3 | 1:94820479-94824492 | 1:94820949-94821056 | SES | 2 | 2 | ENST00000271227 | 16.42 | 4.15E-02 | 7.84E-02 |
| PWWP2A | 5:160078169-160093100 | 5:160089506-160089610 | MES | 3 | 3 | ENST00000521424 | 16.42 | 4.26E-02 | 1.26E-01 |
| ZNF875 | 19:37334783-37347189 | 19:37335169-37335257 | MES | 3 | 3 | ENST00000587437 | 16.39 | 3.74E-02 | 2.42E-01 |
| PCM1 | 8:17972688-17985446 | 8:17980591-17980755 | MXS | 3 | 3 | ENST00000325083 | 16.38 | 3.16E-03 | 6.55E-02 |
| BIRC6 | 2:32471125-32473111 | 2:32471125-32471151 | A5ss | 3 | 3 | ENST00000648282 | 16.37 | 9.48E-03 | 2.08E-02 |
| ARID1B | 6:157174118-157180968 | 6:157174847-157175005 | SES | 3 | 3 | ENST00000319584 | 16.36 | 3.28E-02 | 2.87E-01 |
| ITGA7 | 12:55699990-55700898 | 12:55700262-55700393 | SES | 3 | 3 | ENST00000555728 | 16.34 | 1.11E-02 | 1.84E-02 |
| KCNMB2-AS1 | 3:178859624-178860293 | 3:178860209-178860294 | A5ss | 3 | 3 | Ex.TSS.ENST00000437488 | 16.31 | 5.54E-03 | 1.29E-02 |
| RCCD1 | 15:90954949-90956611 | 15:90955800-90955883 | SES | 3 | 3 | ENST00000555737 | 16.28 | 1.69E-02 | 5.89E-02 |
| SLC25A39 | 17:44322529-44322807 | 17:44322529-44322552 | A3ss | 3 | 3 | ENST00000377095 | 16.24 | 6.10E-05 | 2.60E-04 |
| ETV1 | 7:13989140-13989562 | 7:13989268-13989464 | SES | 3 | 3 | ENST00000430479 | 16.19 | 1.46E-03 | 1.78E-02 |
| CCDC78 | 16:724106-725880 | 16:724206-724321 | IR | 3 | 2 | ENST00000481804 | 16.18 | 1.55E-02 | 5.97E-02 |
| DEF8 | 16:89963444-89964220 | 16:89964170-89964220 | A3ss | 3 | 3 | ENST00000268676 | 16.14 | 1.26E-02 | 4.99E-02 |
| MBP | 18:77016832-77017261 | 18:77016914-77016979 | IR | 3 | 3 | ENST00000354542 | 16.13 | 3.78E-02 | 1.92E-01 |
| PACRGL | 4:20700579-20704465 | 4:20702141-20702221 | MES | 3 | 3 | ENST00000510051 | 16.13 | 4.03E-02 | 1.09E-01 |
| SAT2 | 17:7626794-7627142 | 17:7626943-7627044 | SES | 3 | 3 | ENST00000269298 | 16.12 | 1.56E-02 | 7.36E-02 |
| PLEKHA8P1 | 12:45176434-45215939 | 12:45190794-45190961 | SES | 3 | 3 | ENST00000545609 | 16.08 | 3.31E-02 | 5.32E-02 |
| UBE3A | 15:25405503-25438488 | 15:25409088-25409207 | MES | 3 | 3 | ENST00000428984 | 16.07 | 6.36E-03 | 2.07E-02 |
| B3GALNT2 | 1:235494829-235504140 | 1:235496186-235496308 | SES | 3 | 3 | ENST00000313984 | 16.07 | 1.08E-02 | 1.94E-02 |
| ZNF232 | 17:5109769-5111799 | 17:5111076-5111219 | SES | 3 | 3 | ENST00000571076 | 16.06 | 4.09E-02 | 1.82E-01 |
| AMIGO1 | 1:109508978-109509419 | 1:109508978-109508999 | A3ss | 3 | 2 | Ex.ENST00000369862 | 16.04 | 6.05E-03 | 5.00E-02 |
| ARID4A | 14:58365623-58366882 | 14:58366024-58366230 | SES | 2 | 2 | ENST00000355431 | 16.04 | 1.75E-02 | 5.50E-02 |

| | | | | | | | | | |
|---|---|---|---|---|---|---|---|---|---|
| UPRT | X:75293515-75296341 | X:75294563-75294620 | SES | 3 | 3 | ENST00000462237 | 15.99 | 6.69E-04 | 1.00E+00 |
| ADAM23 | 2:206596163-206617578 | 2:206609910-206610000 | MXS | 3 | 3 | ENST00000264377 | 15.99 | 2.71E-02 | 5.81E-02 |
| NSMCE2 | 8:125091959-125102316 | 8:125102014-125102146 | MXS | 3 | 3 | ENST00000517532 | 15.95 | 2.74E-03 | 5.25E-02 |
| FOXRED1 | 11:126273080-126273335 | 11:126273316-126273335 | A3ss | 3 | 3 | ENST00000524751 | 15.95 | 1.74E-02 | 2.52E-02 |
| ERGIC3 | 20:35556272-35556972 | 20:35556955-35556972 | A3ss | 3 | 3 | ENST00000451605 | 15.92 | 6.96E-05 | 1.32E-04 |
| RIMKLB | 12:8777803-8780577 | 12:8780575-8780577 | A3ss | 3 | 3 | Ex.ENST00000544257 | 15.92 | 1.95E-02 | 3.29E-02 |
| SENP6 | 6:75621626-75623899 | 6:75622751-75622858 | SES | 3 | 3 | ENST00000493959 | 15.91 | 5.50E-03 | 5.61E-02 |
| ZNF124 | 1:247159814-247171847 | 1:247159814-247159859 | A3ss | 3 | 3 | Ex.TSS.ENST00000340684 | 15.9 | 1.50E-02 | 2.70E-02 |
| KHNYN | 14:24429992-24430713 | 14:24429992-24430119 | TSSIA5ss | 3 | 3 | TSS.ENST00000553935 | 15.88 | 1.08E-02 | 1.84E-02 |
| NEIL2 | 8:11769792-11771445 | 8:11770174-11770335 | SES | 3 | 3 | ENST00000455213 | 15.86 | 2.66E-02 | 5.01E-01 |
| IMMP1L | 11:31433571-31509518 | 11:31456260-31456386 | MES | 3 | 3 | ENST00000278200 | 15.85 | 2.51E-02 | 3.64E-02 |
| EI24 | 11:125572570-125575262 | 11:125575251-125575262 | A3ss | 3 | 3 | ENST00000618552 | 15.79 | 4.89E-05 | 7.06E-05 |
| RAB4A | 1:229271371-229295847 | 1:229286468-229295848 | A3ss | 3 | 3 | Ex.TSS.ENST00000473894 | 15.74 | 1.41E-03 | 2.54E-03 |
| RAB4A | 1:229271371-229286485 | 1:229286468-229286486 | A3ss | 3 | 3 | Ex.TSS.ENST00000366690 | 15.74 | 1.41E-03 | 2.54E-03 |
| MTMR2 | 11:95888262-95923691 | 11:95914242-95914312 | SES | 3 | 3 | ENST00000469574 | 15.74 | 6.58E-03 | 1.02E-02 |
| RIC3 | 11:8111138-8137377 | 11:8126659-8126804 | SES | 3 | 3 | ENST00000343202 | 15.74 | 9.77E-03 | 1.50E-02 |
| TNNT1 | 19:55145566-55146680 | 19:55146434-55146466 | SES | 3 | 3 | ENST00000291901 | 15.7 | 5.44E-03 | 3.89E-02 |
| ADGRL2 | 1:81936838-81942968 | 1:81942031-81942969 | A3ss | 3 | 3 | Ex.ENST00000319517 | 15.69 | 2.69E-03 | 5.06E-03 |
| TUBE1 | 6:112081208-112086555 | 6:112084189-112084246 | SES | 3 | 3 | ENST00000368657 | 15.68 | 3.21E-03 | 9.86E-03 |
| PRSS16 | 6:27252041-27254985 | 6:27254693-27254872 | MXS | 3 | 3 | ENST00000471463 | 15.68 | 8.67E-03 | 7.78E-02 |
| ARL6IP4 | 12:122981267-122981594 | 12:122981267-122981299 | TSSIA5ss | 3 | 3 | TSS.ENST00000357866 | 15.65 | 3.45E-03 | 5.49E-03 |
| RUBCN | 3:197705176-197736654 | 3:197717977-197718130 | SES | 3 | 3 | ENST00000296343 | 15.65 | 2.02E-02 | 4.78E-02 |
| PTPRD | 9:8492980-8499646 | 9:8497242-8497268 | SES | 3 | 3 | ENST00000356435 | 15.64 | 1.39E-02 | 4.51E-01 |
| SNHG14 | 15:24982438-25019026 | 15:25002725-25002888 | MES | 3 | 3 | ENST00000551631 | 15.62 | 3.58E-03 | 1.16E-02 |
| MED24 | 17:40053636-40054360 | 17:40053636-40053657 | TSSIA3ss | 3 | 3 | TSS.ENST00000394127 | 15.62 | 1.50E-02 | 6.28E-02 |
| CC2D1A | 19:13920923-13923332 | 19:13923318-13923333 | A3ss | 3 | 3 | Ex.ENST00000318003 | 15.61 | 3.44E-02 | 1.84E-01 |
| VPS52 | 6:33270284-33271690 | 6:33271586-33271690 | TSSIA5ss | 3 | 3 | TSS.ENST00000445902 | 15.58 | 2.77E-02 | 2.57E-01 |
| SEMA6C | 1:151133319-151134400 | 1:151133319-151133517 | A3ss | 3 | 3 | Ex.TSS.ENST00000341697 | 15.56 | 1.44E-03 | 2.46E-03 |
| BABAM1 | 19:17267528-17268793 | 19:17268756-17268793 | TSSIA3ss | 3 | 3 | TSS.ENST00000359435 | 15.56 | 2.32E-03 | 1.26E-02 |
| SLC39A13 | 11:47414862-47415159 | 11:47414910-47415038 | IR | 3 | 3 | ENST00000524886 | 15.54 | 3.68E-04 | 5.44E-04 |
| TSC22D1 | 13:44539924-44573162 | 13:44539924-44539936 | TSSIA3ss | 3 | 3 | TSS.ENST00000486464 | 15.54 | 6.58E-03 | 1.07E-02 |
| ADCYAP1R1 | 7:31078099-31084140 | 7:31081713-31081754 | MES | 3 | 3 | ENST00000304166 | 15.54 | 3.34E-02 | 4.48E-01 |
| ZFYVE19 | 15:40810758-40813337 | 15:40812699-40812902 | SES | 3 | 3 | ENST00000336455 | 15.51 | 8.06E-03 | 2.65E-02 |
| OSGEP | 14:20448964-20449398 | 14:20449014-20449170 | IR | 3 | 3 | ENST00000554249 | 15.5 | 4.28E-02 | 5.95E-02 |
| IMMP1L | 11:31463306-31509518 | 11:31473724-31473786 | MES | 3 | 3 | ENST00000530023 | 15.48 | 8.16E-03 | 1.19E-02 |
| ANKRD17 | 4:73135263-73141740 | 4:73139531-73140283 | SES | 3 | 3 | ENST00000558247 | 15.46 | 3.75E-02 | 1.05E-01 |
| ANO8 | 19:17325387-17327234 | 19:17325387-17325414 | A3ss | 3 | 3 | Ex.ENST00000159087 | 15.45 | 1.14E-04 | 6.18E-04 |
| PXN | 12:120215274-120215559 | 12:120215274-120215417 | A3ss | 3 | 3 | ENST00000637624 | 15.36 | 4.50E-02 | 7.03E-02 |
| PPFIBP1 | 12:27673828-27676427 | 12:27674192-27674221 | SES | 3 | 3 | ENST00000228425 | 15.3 | 2.29E-02 | 3.67E-02 |
| NET1 | 10:5455119-5456086 | 10:5456065-5456087 | A3ss | 3 | 3 | Ex.ENST00000355029 | 15.28 | 1.61E-05 | 2.28E-05 |
| IMMP1L | 11:31463306-31509518 | 11:31469740-31469797 | MES | 3 | 3 | ENST00000530023 | 15.28 | 1.48E-02 | 2.15E-02 |
| PACRGL | 4:20700579-20704465 | 4:20702141-20702221 | MXS | 3 | 3 | ENST00000511089 | 15.28 | 2.60E-02 | 7.02E-02 |
| ARMCX5-GPRASP2 | X:102599848-102605481 | X:102601464-102601509 | MXS | 3 | 3 | ENST00000475738 | 15.23 | 2.38E-02 | 6.00E-01 |
| SHARPIN | 8:144098995-144099276 | 8:144099081-144099205 | SES | 3 | 3 | ENST00000398712 | 15.16 | 1.20E-02 | 2.43E-01 |
| PCNX3 | 11:65635804-65636173 | 11:65635951-65636174 | A3ss | 3 | 3 | Ex.ENST00000355703 | 15.16 | 3.09E-02 | 4.68E-02 |
| CEP95 | 17:64529428-64531889 | 17:64530926-64531018 | SES | 3 | 3 | ENST00000556440 | 15.15 | 5.31E-04 | 2.42E-03 |
| TP53BP1 | 15:43403061-43407570 | 15:43404553-43406600 | IR (overlapping | 3 | 3 | ENST00000382044 | 15.1 | 1.41E-02 | 4.65E-02 |
| HDHD2 | 18:47137297-47150377 | 18:47137297-47137333 | A3ss | 3 | 3 | Ex.TSS.ENST00000586546 | 15.09 | 4.99E-03 | 2.50E-02 |
| HDHD2 | 18:47136450-47150377 | 18:47136450-47137333 | A3ss | 3 | 3 | Ex.TSS.ENST00000300605 | 15.09 | 4.99E-03 | 2.50E-02 |
| ZNF76 | 6:35293045-35294455 | 6:35293751-35294455 | TSSIA3ss | 3 | 3 | TSS.ENST00000229405 | 15.09 | 2.91E-02 | 2.73E-01 |
| GAS8 | 16:90043196-90044960 | 16:90043990-90044113 | IR (overlapping | 3 | 3 | ENST00000268699 | 15.08 | 5.77E-03 | 2.29E-02 |
| SNHG8 | 4:118278793-118279388 | 4:118278934-118279137 | SES | 3 | 3 | ENST00000602414 | 15.07 | 5.83E-03 | 1.53E-02 |
| BSCL2 | 11:62705618-62706258 | 11:62706203-62706258 | TSSIA5ss | 3 | 3 | TSS.ENST00000278893 | 15.05 | 2.10E-02 | 3.11E-02 |
| SLC25A36 | 3:140963228-140970926 | 3:140967000-140967036 | MXS | 3 | 3 | ENST00000502594 | 15.04 | 3.73E-02 | 8.60E-02 |
| ANKRD13A | 12:110025824-110027707 | 12:110027705-110027707 | A3ss | 3 | 3 | ENST00000261739 | 15.01 | 1.85E-02 | 2.87E-02 |
| BTBD10 | 11:13421839-13463091 | 11:13445024-13445181 | SES | 3 | 3 | ENST00000278174 | 14.99 | 1.66E-02 | 2.40E-02 |
| LIPT1 | 2:99155081-99155544 | 2:99155369-99155481 | IR (overlapping | 3 | 3 | ENST00000436234 | 14.99 | 4.52E-02 | 1.03E-01 |
| NSD3 | 8:38279682-38281499 | 8:38281467-38281499 | A5ss | 3 | 3 | ENST00000317025 | 14.97 | 1.56E-02 | 3.31E-01 |
| MYO6 | 6:75892691-75907604 | 6:75895231-75895260 | MES | 3 | 3 | ENST00000430435 | 14.96 | 8.74E-03 | 8.93E-02 |
| RBPJ | 4:26321049-26386352 | 4:26362446-26362628 | SES | 3 | 3 | ENST00000355476 | 14.96 | 1.64E-02 | 4.44E-02 |
| CNTROB | 17:7932155-7933349 | 17:7932801-7932955 | IR | 3 | 3 | ENST00000380262 | 14.96 | 2.94E-02 | 1.42E-01 |
| GPBP1 | 5:57176401-57230845 | 5:57221376-57221396 | MES | 3 | 3 | ENST00000511209 | 14.91 | 2.67E-03 | 8.05E-03 |
| ZNF821 | 16:71860673-71864137 | 16:71861776-71861942 | SES | 3 | 3 | ENST00000313565 | 14.9 | 3.21E-02 | 1.23E-01 |
| SLC39A9 | 14:69455867-69461696 | 14:69458363-69458729 | SES | 2 | 3 | ENST00000555840 | 14.89 | 8.26E-03 | 2.62E-02 |
| AC005562.1 | 17:30631896-30633097 | 17:30631896-30631927 | A5ss | 3 | 3 | ENST00000398849 | 14.89 | 2.61E-02 | 1.05E-01 |
| ACOX1 | 17:75957567-75978533 | 17:75960215-75960375 | MXS | 3 | 3 | ENST00000293217 | 14.89 | 3.63E-02 | 1.72E-01 |
| MLH3 | 14:75038413-75041614 | 14:75039911-75040015 | SES | 3 | 3 | ENST00000355774 | 14.83 | 4.45E-02 | 1.42E-01 |
| YPEL3 | 16:30095336-30096096 | 16:30095336-30095434 | TSSIA3ss | 3 | 3 | TSS.ENST00000398838 | 14.81 | 4.84E-03 | 1.77E-02 |
| AL096870.10 | 14:24271311-24298613 | 14:24292198-24298614 | A3ss | 3 | 3 | Ex.ENST00000669726 | 14.81 | 4.78E-02 | 8.13E-02 |
| RPAIN | 17:5422830-5432541 | 17:5425971-5426082 | MES | 3 | 3 | ENST00000381208 | 14.8 | 1.31E-02 | 5.82E-02 |
| QTRT2 | 3:114056865-114070625 | 3:114066228-114066283 | MES | 3 | 3 | ENST00000466050 | 14.78 | 1.24E-02 | 2.82E-02 |
| MSH5 | 6:31743904-31744299 | 6:31744026-31744138 | IR (overlapping | 2 | 3 | ENST00000650702 | 14.77 | 3.22E-02 | 1.00E-01 |
| SIDT2 | 11:117186637-117187377 | 11:117186990-117187001 | SES | 3 | 3 | ENST00000431081 | 14.74 | 1.59E-02 | 2.28E-02 |
| ABCA7 | 19:1057430-1058145 | 19:1057915-1058059 | SES | 2 | 3 | ENST00000263094 | 14.73 | 3.66E-06 | 1.88E-05 |
| SNHG32 | 6:31835452-31837235 | 6:31836424-31836517 | SES | 3 | 3 | ENST00000375638 | 14.71 | 3.56E-04 | 3.26E-03 |
| USP47 | 11:11842225-11884466 | 11:11880177-11880380 | MES | 3 | 3 | ENST00000525257 | 14.69 | 4.94E-02 | 7.12E-02 |
| ETV1 | 7:13989140-13989562 | 7:13989268-13989562 | TSSIA5ss | 3 | 3 | TSS.ENST00000433547 | 14.62 | 1.84E-02 | 2.23E-01 |
| CLK4 | 5:178618779-178623255 | 5:178619838-178619896 | MXS | 3 | 3 | ENST00000519583 | 14.6 | 2.36E-02 | 7.02E-02 |
| FAM49B | 8:129903351-129970942 | 8:129904499-129904585 | MXS | 3 | 3 | ENST00000401979 | 14.59 | 5.79E-03 | 1.13E-01 |
| SUPT20H | 13:37009810-37012191 | 13:37010552-37010655 | SES | 3 | 3 | ENST00000350612 | 14.58 | 5.43E-03 | 8.77E-03 |
| TMEM63A | 1:225848555-225848896 | 1:225848555-225848614 | A3ss | 3 | 3 | Ex.ENST00000366835 | 14.55 | 7.69E-04 | 1.38E-03 |
| RABGAP1L | 1:174811961-174957456 | 1:174811961-174874506 | A5ss | 3 | 3 | Ex.ENST00000251507 | 14.53 | 3.36E-03 | 5.89E-03 |
| MAP2K7 | 19:7911331-7911435 | 19:7911415-7911435 | A3ss | 3 | 3 | ENST00000397981 | 14.51 | 3.56E-02 | 2.66E-01 |
| UBQLN1 | 9:83665146-83669184 | 9:83666350-83669184 | A5ss | 3 | 3 | ENST00000533705 | 14.5 | 7.01E-04 | 2.25E-02 |
| ADGRB2 | 1:31727606-31728181 | 1:31728025-31728081 | SES | 3 | 3 | ENST00000373658 | 14.5 | 2.71E-03 | 5.02E-03 |
| LRRC20 | 10:70301509-70376451 | 10:70323863-70324030 | MXS | 3 | 3 | ENST00000358141 | 14.49 | 2.59E-02 | 3.69E-02 |
| LDLRAD3 | 11:35944145-36081652 | 11:36036103-36036249 | MXS | 3 | 3 | ENST00000315571 | 14.48 | 2.15E-02 | 3.16E-02 |
| DRAM2 | 1:111137587-111140037 | 1:111139588-111139711 | SES | 3 | 3 | ENST00000463682 | 14.46 | 4.63E-03 | 3.86E-02 |
| ZNF138 | 7:64794572-64831450 | 7:64830940-64831076 | MES | 3 | 3 | ENST00000494380 | 14.46 | 4.91E-02 | 7.11E-01 |
| SPPL2A | 15:50725324-50726320 | 15:50726073-50726320 | TSSIA5ss | 3 | 3 | TSS.ENST00000559293 | 14.44 | 6.07E-04 | 2.02E-03 |
| SLC39A13 | 11:47414862-47415159 | 11:47414906-47415059 | IR | 3 | 3 | ENST00000524886 | 14.44 | 1.27E-03 | 1.87E-03 |
| GRK6 | 5:177441054-177441736 | 5:177441735-177441736 | TSSIA3ss | 3 | 3 | TSS.ENST00000528793 | 14.44 | 4.62E-03 | 1.37E-02 |
| YY1AP1 | 1:155668718-155670319 | 1:155668718-155668777 | A3ss | 3 | 3 | ENST00000347088 | 14.44 | 1.01E-02 | 1.75E-02 |
| ADGRB2 | 1:31727606-31728181 | 1:31727606-31728081 | TSSIA3ss | 3 | 3 | TSS.ENST00000465256 | 14.41 | 3.65E-03 | 6.75E-03 |

| | | | | | | | | | |
|---|---|---|---|---|---|---|---|---|---|
| ARID3A | 19:926060-929261 | 19:926060-926197 | A5ss | 3 | 3 | Ex.TSS.ENST00000263620 | 14.41 | 6.71E-03 | 5.13E-02 |
| SUPT20H | 13:37047937-37051487 | 13:37048564-37048599 | SES | 3 | 3 | ENST00000350612 | 14.38 | 1.86E-02 | 3.01E-02 |
| ZNF207 | 17:32365488-32367771 | 17:32366665-32366757 | SES | 3 | 3 | ENST00000321233 | 14.37 | 4.78E-04 | 1.94E-03 |
| FOSL1 | 11:65894117-65896808 | 11:65894117-65894121 | A3ss | 3 | 3 | ENST00000312562 | 14.36 | 1.54E-04 | 2.33E-04 |
| TSC1 | 9:132910690-132911001 | 9:132910690-132910692 | A3ss | 3 | 3 | ENST00000298552 | 14.36 | 2.02E-02 | 7.98E-01 |
| SMARCA1 | X:129492094-129496749 | X:129493040-129493075 | SES | 3 | 3 | ENST00000371121 | 14.35 | 1.16E-02 | 6.50E-01 |
| DNASE1L1 | X:154405656-154409111 | X:154408558-154409111 | TSSIA5ss | 3 | 3 | TSS.ENST00000369808 | 14.34 | 2.60E-03 | 1.82E-01 |
| PEX26 | 22:18078607-18079873 | 22:18078607-18079256 | A5ss | 3 | 3 | Ex.ENST00000329627 | 14.34 | 1.93E-02 | 3.76E-02 |
| TMEM125 | 1:43270794-43272183 | 1:43270901-43272184 | A3ss | 3 | 3 | Ex.ENST00000432792 | 14.29 | 1.00E-03 | 1.86E-03 |
| CNTROB | 17:7932155-7933349 | 17:7932801-7932927 | IR | 3 | 3 | ENST00000380262 | 14.29 | 3.74E-02 | 1.80E-01 |
| ATG9A | 2:219228483-219229534 | 2:219228483-219228485 | TSSIA3ss | 3 | 3 | TSS.ENST00000361242 | 14.28 | 4.35E-03 | 9.40E-03 |
| PEX2 | 8:76986297-76999989 | 8:76988307-76988338 | SES | 3 | 3 | ENST00000357039 | 14.26 | 1.08E-02 | 2.52E-01 |
| ACD | 16:67658817-67658927 | 16:67658817-67658832 | TSSIA3ss | 3 | 3 | TSS.ENST00000602945 | 14.26 | 1.76E-02 | 6.70E-02 |
| TBC1D1 | 4:38049860-38054198 | 4:38049860-38049898 | A5ss | 3 | 3 | Ex.ENST00000261439 | 14.23 | 4.61E-03 | 1.25E-02 |
| CARMIL2 | 16:67656646-67657238 | 16:67656834-67656881 | SES | 3 | 3 | ENST00000602368 | 14.22 | 4.02E-02 | 1.53E-01 |
| CHERP | 19:16530881-16532597 | 19:16530881-16530913 | A3ss | 3 | 3 | ENST00000198939 | 14.21 | 1.98E-03 | 1.07E-02 |
| NPRL3 | 16:112776-138149 | 16:119126-119255 | MES | 3 | 3 | ENST00000399953 | 14.17 | 1.40E-02 | 4.91E-02 |
| DCTD | 4:182915576-182917310 | 4:182917288-182917310 | TSSIA5ss | 3 | 3 | TSS.ENST00000357067 | 14.17 | 2.14E-02 | 5.76E-02 |
| PLEKHB1 | 11:73661466-73662819 | 11:73662423-73662510 | IR | 3 | 3 | ENST00000227214 | 14.16 | 1.72E-02 | 2.64E-02 |
| KLC1 | 14:103679546-103700654 | 14:103687081-103687211 | SES | 3 | 3 | ENST00000452929 | 14.14 | 3.57E-02 | 4.98E-02 |
| LTBP4 | 19:40619494-40623603 | 19:40622401-40622667 | MES | 3 | 3 | ENST00000204005 | 14.14 | 3.84E-02 | 2.47E-01 |
| HLA-F | 6:29723928-29724189 | 6:29724173-29724189 | A3ss | 3 | 3 | ENST00000259951 | 14.13 | 1.85E-03 | 1.67E-02 |
| ZNF551 | 19:57682245-57686480 | 19:57685262-57685385 | SES | 3 | 3 | ENST00000282296 | 14.11 | 3.92E-02 | 2.87E-01 |
| ANKFY1 | 17:4181373-4183397 | 17:4182181-4182352 | SES | 3 | 3 | ENST00000572412 | 14.08 | 1.86E-02 | 7.84E-02 |
| KIAA0586 | 14:58458546-58460985 | 14:58459843-58459909 | SES | 3 | 2 | ENST00000651937 | 14.08 | 4.78E-02 | 1.50E-01 |
| TAZ | X:154413568-154420031 | X:154419543-154419623 | SES | 3 | 3 | ENST00000369776 | 14.06 | 7.65E-04 | 5.58E-02 |
| REPIN1 | 7:150368942-150371230 | 7:150371228-150371230 | A3ss | 3 | 3 | Ex.ENST00000444957 | 14.02 | 2.25E-04 | 2.80E-03 |
| AL049834.1 | 14:88391456-88396155 | 14:88393393-88393488 | MES | 3 | 3 | ENST00000555715 | 14.02 | 3.38E-02 | 1.07E-01 |
| NAXD | 13:110635589-110638377 | 13:110637129-110637249 | SES | 3 | 3 | ENST00000424185 | 13.99 | 1.03E-03 | 1.67E-03 |
| FCHO1 | 19:17754362-17754591 | 19:17754362-17754466 | A5ss | 3 | 3 | ENST00000593385 | 13.95 | 9.33E-03 | 5.08E-02 |
| PCM1 | 8:17972688-17985446 | 8:17980591-17980755 | MES | 3 | 3 | ENST00000522275 | 13.92 | 2.85E-02 | 5.84E-01 |
| ASIC1 | 12:50081545-50083611 | 12:50081779-50082715 | IR | 3 | 3 | ENST00000228468 | 13.87 | 3.37E-02 | 5.53E-02 |
| TUT4 | 1:52446668-52461133 | 1:52458336-52458449 | SES | 3 | 3 | ENST00000257177 | 13.86 | 7.01E-03 | 1.31E-02 |
| HOOK2 | 19:12765725-12765926 | 19:12765829-12765834 | SES | 3 | 3 | ENST00000397668 | 13.85 | 4.58E-02 | 2.45E-01 |
| MSL3 | X:11763780-11768999 | X:11763939-11765192 | IR | 3 | 3 | ENST00000473380 | 13.83 | 1.31E-02 | 6.39E-01 |
| LIN9 | 1:226287798-226297718 | 1:226295842-226295946 | SES | 3 | 3 | ENST00000328205 | 13.78 | 6.28E-03 | 1.08E-02 |
| NEK3 | 13:52153993-52156074 | 13:52153993-52154155 | A3ss | 3 | 3 | Ex.ENST00000551355 | 13.78 | 1.04E-02 | 1.69E-02 |
| USP35 | 11:78200809-78207529 | 11:78205842-78206035 | SES | 2 | 3 | ENST00000526425 | 13.77 | 3.69E-03 | 5.68E-03 |
| CENPX | 17:82019382-82019857 | 17:82019641-82019694 | SES | 3 | 3 | ENST00000392359 | 13.76 | 9.78E-04 | 4.78E-03 |
| TRIM65 | 17:75892501-75896523 | 17:75892755-75892850 | SES | 3 | 3 | ENST00000269383 | 13.74 | 2.58E-03 | 1.22E-02 |
| ZNF875 | 19:37324266-37347776 | 19:37347190-37347316 | MES | 3 | 3 | ENST00000652615 | 13.74 | 1.08E-02 | 6.95E-02 |
| BAIAP3 | 16:1339245-1340921 | 16:1339496-1339603 | SES | 2 | 3 | ENST00000324385 | 13.73 | 3.99E-02 | 1.40E-01 |
| AC005670.3 | 17:47056754-47099834 | 17:47072472-47072559 | SES | 3 | 3 | ENST00000573341 | 13.7 | 1.13E-02 | 4.88E-02 |
| FAM49B | 8:129903351-129939607 | 8:129904499-129904585 | SES | 3 | 3 | ENST00000519070 | 13.68 | 1.83E-03 | 3.57E-02 |
| UBQLN1 | 9:83665146-83669184 | 9:83666350-83666433 | SES | 3 | 3 | ENST00000376395 | 13.64 | 2.59E-03 | 8.32E-02 |
| ACAA2 | 18:49785352-49787290 | 18:49785352-49785372 | A3ss | 3 | 3 | Ex.ENST00000285093 | 13.59 | 5.41E-06 | 2.72E-05 |
| STAU1 | 20:49124588-49135832 | 20:49124588-49124605 | A3ss | 3 | 3 | ENST00000371802 | 13.58 | 2.51E-02 | 4.30E-02 |
| ZNF706 | 8:101202333-101205287 | 8:101202448-101204649 | IR | 2 | 3 | ENST00000520347 | 13.56 | 2.51E-02 | 4.70E-01 |
| HSCB | 22:28744705-28745863 | 22:28744705-28744709 | A5ss | 3 | 3 | ENST00000420442 | 13.56 | 4.31E-02 | 8.47E-02 |
| LRIF1 | 1:110950124-110963620 | 1:110951288-110952815 | SES | 3 | 3 | ENST00000369763 | 13.54 | 3.82E-03 | 3.18E-02 |
| STAU2 | 8:73688814-73746782 | 8:73739756-73739868 | MES | 3 | 3 | ENST00000519818 | 13.51 | 2.81E-02 | 6.56E-01 |
| ATAD1 | 10:87756923-87767672 | 10:87756923-87759800 | A3ss | 3 | 3 | Ex.ENST00000308448 | 13.47 | 2.75E-02 | 3.93E-02 |
| ENDOV | 17:80415315-80415821 | 17:80415519-80415649 | IR | 3 | 3 | ENST00000522751 | 13.44 | 8.67E-03 | 4.18E-02 |
| SAFB2 | 19:5595498-5598792 | 19:5595498-5595510 | A3ss | 3 | 3 | Ex.ENST00000252542 | 13.41 | 2.54E-05 | 1.82E-04 |
| NEK8 | 17:28734772-28735309 | 17:28735005-28735239 | IR (overlapping | 3 | 3 | ENST00000584342 | 13.39 | 2.50E-02 | 1.00E-01 |
| ANO9 | 11:428827-429569 | 11:429474-429569 | A5ss | 3 | 2 | ENST00000534161 | 13.38 | 1.83E-02 | 2.69E-02 |
| SPTAN1 | 9:128609285-128611713 | 9:128609651-128609665 | SES | 3 | 3 | ENST00000372739 | 13.36 | 3.47E-05 | 1.24E-03 |
| SLC39A13 | 11:47414862-47415159 | 11:47414910-47415059 | IR | 3 | 3 | ENST00000524886 | 13.35 | 2.08E-03 | 3.06E-03 |
| NEK8 | 17:28734772-28735309 | 17:28734955-28735239 | IR (overlapping | 3 | 3 | ENST00000584342 | 13.35 | 2.28E-02 | 9.17E-02 |
| LOXL2 | 8:23328566-23333400 | 8:23328566-23328583 | A3ss | 3 | 3 | Ex.ENST00000389131 | 13.31 | 3.85E-04 | 8.25E-03 |
| CHFR | 12:132869676-132870723 | 12:132869676-132869798 | A3ss | 3 | 3 | ENST00000450056 | 13.31 | 2.35E-02 | 3.76E-02 |
| STAU2 | 8:73688814-73746782 | 8:73739756-73739868 | MES | 3 | 3 | ENST00000355780 | 13.29 | 5.92E-03 | 1.39E-01 |
| HAUS2 | 15:42566607-42569994 | 15:42569466-42569600 | IR (overlapping | 3 | 3 | ENST00000260372 | 13.29 | 8.71E-03 | 2.88E-02 |
| PPFIBP2 | 11:7653024-7653756 | 11:7653201-7653501 | IR | 3 | 3 | ENST00000299492 | 13.25 | 1.66E-02 | 2.55E-02 |
| DDR1 | 6:30881126-30888687 | 6:30885625-30885680 | MES | 3 | 3 | ENST00000324771 | 13.25 | 4.45E-02 | 1.38E-01 |
| DLG5 | 10:77854359-77856729 | 10:77854359-77854370 | A3ss | 3 | 3 | Ex.ENST00000372391 | 13.24 | 2.67E-02 | 3.80E-02 |
| RNF125 | 18:32045733-32068297 | 18:32065902-32066009 | SES | 3 | 2 | ENST00000217740 | 13.24 | 3.28E-02 | 1.64E-01 |
| ZSCAN21 | 7:100063788-100065040 | 7:100063980-100064082 | IR | 3 | 3 | ENST00000292450 | 13.24 | 3.51E-02 | 4.04E-01 |
| WDR90 | 16:662220-662736 | 16:662332-662672 | IR | 3 | 3 | ENST00000546923 | 13.23 | 4.31E-02 | 1.63E-01 |
| TOP2B | 3:25645456-25664228 | 3:25645456-25645470 | TSSIA3ss | 3 | 3 | TSS.ENST00000264331 | 13.22 | 8.91E-03 | 2.11E-02 |
| DOC2A | 16:30009961-30010340 | 16:30010027-30010272 | IR | 2 | 3 | ENST00000565273 | 13.22 | 1.81E-02 | 6.58E-02 |
| GBAP1 | 1:155217240-155218049 | 1:155217594-155217963 | IR | 2 | 3 | ENST00000473223 | 13.22 | 2.92E-02 | 5.05E-02 |
| KLRG2 | 7:139479773-139482885 | 7:139480146-139480247 | SES | 3 | 3 | ENST00000340940 | 13.21 | 4.15E-02 | 5.03E-01 |
| VPS33A | 12:122232969-122235785 | 12:122232969-122232995 | A3ss | 3 | 3 | Ex.ENST00000267199 | 13.16 | 1.46E-03 | 2.33E-03 |
| ANKRD17 | 4:73135263-73139530 | 4:73135263-73135265 | A3ss | 3 | 3 | ENST00000358602 | 13.16 | 1.05E-02 | 2.94E-02 |
| ZCCHC9 | 5:81301671-81304740 | 5:81301671-81301698 | A5ss | 3 | 3 | Ex.TSS.ENST00000380199 | 13.16 | 3.61E-02 | 1.10E-01 |
| BCL2L12 | 19:49666800-49667021 | 19:49667019-49667021 | A3ss | 3 | 3 | ENST00000246784 | 13.12 | 3.52E-05 | 2.44E-04 |
| RNF175 | 4:153723459-153751437 | 4:153745669-153745727 | MES | 3 | 3 | ENST00000506505 | 13.09 | 7.16E-05 | 1.91E-04 |
| LINC01503 | 9:129338918-129341831 | 9:129341752-129341831 | A3ss | 2 | 3 | ENST00000436710 | 13.08 | 1.91E-02 | 7.12E-01 |
| AGO1 | 1:35893279-35893673 | 1:35893592-35893673 | A3ss | 3 | 3 | ENST00000635259 | 13.06 | 1.87E-02 | 3.46E-02 |
| TNK2 | 3:195867259-195867360 | 3:195867259-195867264 | A3ss | 3 | 3 | ENST00000672548 | 13.03 | 4.39E-02 | 1.04E-01 |
| DAPK1 | 9:87642059-87645894 | 9:87643376-87643468 | SES | 3 | 3 | ENST00000358077 | 13.01 | 1.86E-02 | 6.06E-01 |
| RNASEH1-AS1 | 2:3558999-3559447 | 2:3559404-3559447 | A3ss | 3 | 3 | Ex.TSS.ENST00000438436 | 13.01 | 1.96E-02 | 4.30E-02 |
| PRDM15 | 21:41836468-41836894 | 21:41836650-41836875 | IR | 2 | 3 | ENST00000489661 | 12.99 | 1.27E-02 | 2.46E-02 |
| PLXNB2 | 22:50294779-50307552 | 22:50300277-50307553 | A5ss | 3 | 3 | Ex.TSS.ENST00000359337 | 12.99 | 1.28E-02 | 2.66E-02 |
| INO80B | 2:74455175-74455405 | 2:74455391-74455405 | TSSIA3ss | 3 | 3 | TSS.ENST00000409493 | 12.98 | 4.92E-03 | 1.11E-02 |
| PUS7 | 7:105508521-105522051 | 7:105508521-105508544 | TSSIA3ss | 3 | 3 | TSS.ENST00000469408 | 12.98 | 1.73E-02 | 2.04E-01 |
| ZNF875 | 19:37334783-37335199 | 19:37335169-37335199 | TSSIA3ss | 3 | 3 | TSS.ENST00000324411 | 12.98 | 4.13E-02 | 2.67E-01 |
| C18orf25 | 18:46216596-46253696 | 18:46240005-46240187 | SES | 3 | 3 | ENST00000615052 | 12.96 | 3.99E-02 | 2.00E-01 |
| ANK3 | 10:60139088-60166823 | 10:60166591-60166653 | MXS | 3 | 3 | ENST00000280772 | 12.94 | 1.96E-02 | 2.79E-02 |
| NSRP1 | 17:30117333-30172541 | 17:30118080-30118173 | MES | 3 | 2 | ENST00000467446 | 12.92 | 5.43E-04 | 2.19E-03 |
| FTSJ1 | X:48481716-48482408 | X:48482403-48482408 | A3ss | 3 | 3 | ENST00000348411 | 12.91 | 1.04E-02 | 1.00E+00 |
| CHD2 | 15:92997404-92998498 | 15:92998149-92998261 | SES | 3 | 3 | ENST00000625662 | 12.89 | 4.16E-03 | 1.45E-02 |
| MIB2 | 1:1623680-1623773 | 1:1623680-1623699 | A5ss | 3 | 3 | ENST00000355826 | 12.89 | 1.54E-02 | 2.67E-02 |

| | | | | | | | | | |
|---|---|---|---|---|---|---|---|---|---|
| MIB2 | 1:1623483-1623773 | 1:1623483-1623699 | A5ss | 3 | 3 | ENST00000355826 | 12.89 | 1.54E-02 | 2.63E-02 |
| TERF1 | 8:73027053-73032041 | 8:73030336-73030395 | SES | 3 | 3 | ENST00000276603 | 12.85 | 6.27E-04 | 1.44E-02 |
| OS9 | 12:57719183-57720405 | 12:57719183-57720263 | A5ss | 3 | 3 | ENST00000549307 | 12.85 | 5.25E-03 | 8.76E-03 |
| ARMC10 | 7:103075412-103083681 | 7:103075777-103075881 | MXS | 3 | 3 | ENST00000323716 | 12.84 | 2.08E-03 | 2.44E-02 |
| PSRC1 | 1:109282790-109283073 | 1:109283063-109283074 | A5ss | 3 | 3 | Ex.TSS.ENST00000369903 | 12.82 | 3.70E-02 | 3.05E-01 |
| ATXN2 | 12:111457360-111470107 | 12:111464662-111464715 | MES | 3 | 3 | ENST00000673273 | 12.76 | 2.32E-03 | 3.62E-03 |
| TMEM251 | 14:93185033-93186259 | 14:93185033-93185485 | A5ss | 3 | 3 | ENST00000415050 | 12.76 | 3.83E-02 | 1.24E-01 |
| SPPL2A | 15:50725324-50726320 | 15:50726073-50726126 | SES | 3 | 3 | ENST00000558934 | 12.75 | 2.87E-03 | 9.55E-03 |
| PCYT1B | X:24619085-24646988 | X:24619085-24640461 | A3ss | 3 | 3 | Ex.TSS.ENST00000356768 | 12.74 | 2.88E-02 | 1.00E+00 |
| MYH10 | 17:8569813-8577235 | 17:8576643-8576672 | SES | 3 | 3 | ENST00000360416 | 12.71 | 3.51E-03 | 1.72E-02 |
| TAF1C | 16:84183779-84186900 | 16:84184851-84184989 | MXS | 3 | 3 | ENST00000564345 | 12.68 | 7.00E-04 | 2.71E-03 |
| NAP1L4 | 11:2972244-2976023 | 11:2972244-2972279 | A3ss | 3 | 3 | ENST00000448187 | 12.68 | 1.77E-02 | 2.57E-02 |
| SLC43A1 | 11:57514125-57515443 | 11:57514582-57515444 | A5ss | 3 | 3 | Ex.TSS.ENST00000278426 | 12.65 | 4.69E-02 | 6.92E-02 |
| SLC43A1 | 11:57514125-57515099 | 11:57514582-57515100 | A5ss | 3 | 3 | Ex.ENST00000533263 | 12.65 | 4.69E-02 | 6.92E-02 |
| SLC43A1 | 11:57514125-57514780 | 11:57514582-57514781 | A5ss | 3 | 3 | Ex.TSS.ENST00000528450 | 12.65 | 4.69E-02 | 6.92E-02 |
| PCYT1B | X:24619085-24646988 | X:24640342-24646989 | A5ss | 3 | 3 | Ex.TSS.ENST00000356768 | 12.59 | 5.59E-03 | 4.71E-01 |
| POLR1D | 13:27621105-27621336 | 13:27621105-27621140 | A5ss | 3 | 3 | Ex.ENST00000626064 | 12.57 | 8.38E-03 | 1.35E-02 |
| PELI1 | 2:64100500-64104700 | 2:64104554-64104597 | SES | 3 | 3 | ENST00000466177 | 12.55 | 3.38E-04 | 7.53E-04 |
| CACFD1 | 9:133463556-133467920 | 9:133465322-133465447 | MXS | 3 | 3 | ENST00000316948 | 12.55 | 3.78E-02 | 1.00E+00 |
| SEMA4D | 9:89405700-89455887 | 9:89418056-89418118 | MES | 3 | 3 | ENST00000356444 | 12.53 | 2.37E-02 | 6.46E-01 |
| SSBP3 | 1:54251861-54281437 | 1:54257127-54257186 | MXS | 3 | 3 | ENST00000326956 | 12.52 | 2.90E-03 | 5.43E-03 |
| OARD1 | 6:41067438-41071131 | 6:41068841-41068953 | MES | 3 | 3 | ENST00000424266 | 12.52 | 2.06E-02 | 1.95E-01 |
| UPF3A | 13:114282128-114286518 | 13:114286302-114286400 | MES | 3 | 3 | ENST00000375299 | 12.5 | 8.13E-03 | 1.31E-02 |
| MAGI2-AS3 | 7:79454086-79455374 | 7:79454245-79454865 | IR | 3 | 2 | ENST00000653779 | 12.48 | 3.11E-02 | 4.69E-01 |
| AC139769.1 | 19:23763135-23831596 | 19:23807826-23831597 | A3ss | 3 | 3 | Ex.ENST00000475499 | 12.44 | 4.58E-02 | 2.54E-01 |
| TNFSF11 | 13:42574152-42574522 | 13:42574253-42574468 | IR | 3 | 3 | ENST00000398795 | 12.36 | 3.38E-03 | 5.49E-03 |
| PCNX3 | 11:65619068-65620338 | 11:65619754-65619932 | MES | 3 | 2 | ENST00000355703 | 12.36 | 3.29E-02 | 4.99E-02 |
| SREBF1 | 17:17819538-17820864 | 17:17819726-17820089 | IR | 3 | 3 | ENST00000395751 | 12.34 | 1.16E-02 | 4.11E-02 |
| MAN2C1 | 15:75362454-75364487 | 15:75362642-75362748 | MXS | 3 | 3 | ENST00000421803 | 12.34 | 4.15E-02 | 1.40E-01 |
| ACTR5 | 20:38756040-38765401 | 20:38761653-38765402 | A3ss | 3 | 3 | Ex.ENST00000243903 | 12.3 | 4.08E-03 | 7.79E-03 |
| MAGI1 | 3:65383624-65391141 | 3:65387143-65387226 | SES | 3 | 3 | ENST00000330909 | 12.3 | 5.87E-03 | 1.52E-02 |
| UHRF2 | 9:6413199-6413643 | 9:6413356-6413482 | IR | 3 | 3 | ENST00000276893 | 12.26 | 1.08E-02 | 2.98E-01 |
| C18orf32 | 18:49483772-49487206 | 18:49487043-49487206 | TSSIA5ss | 3 | 3 | TSS.ENST00000318240 | 12.24 | 2.76E-02 | 1.39E-01 |
| DNAJC21 | 5:34945804-34950169 | 5:34949543-34949677 | SES | 3 | 3 | ENST00000382021 | 12.24 | 4.22E-02 | 1.26E-01 |
| TSPAN5 | 4:98482176-98507677 | 4:98486738-98486884 | SES | 3 | 3 | ENST00000305798 | 12.23 | 4.05E-02 | 1.15E-01 |
| FKRP | 19:46747295-46748026 | 19:46747295-46747305 | A5ss | 3 | 3 | ENST00000600005 | 12.2 | 2.91E-02 | 1.93E-01 |
| ZNF875 | 19:37334783-37347189 | 19:37335169-37335257 | MXS | 3 | 3 | ENST00000392153 | 12.2 | 4.52E-02 | 2.92E-01 |
| STON1 | 2:48530217-48580586 | 2:48530713-48530829 | SES | 3 | 3 | ENST00000406226 | 12.16 | 1.69E-02 | 3.73E-02 |
| RERE | 1:8352404-8355120 | 1:8353179-8354942 | IR | 3 | 3 | ENST00000400908 | 12.15 | 1.43E-02 | 2.70E-02 |
| TARBP1 | 1:234393838-234401180 | 1:234398382-234398553 | MXS | 3 | 3 | ENST00000040877 | 12.09 | 4.90E-02 | 6.82E-02 |
| OS9 | 12:57719183-57720405 | 12:57720099-57720263 | SES | 3 | 3 | ENST00000315970 | 12.08 | 1.66E-02 | 2.77E-02 |
| SCAMP4 | 19:1918991-1923069 | 19:1918991-1921009 | A5ss | 3 | 3 | Ex.ENST00000316097 | 12.08 | 2.38E-02 | 1.28E-01 |
| TPM1 | 15:63061274-63062214 | 15:63061713-63061788 | SES | 3 | 3 | ENST00000558264 | 12.07 | 2.62E-04 | 8.75E-04 |
| ERI3 | 1:44339317-44352849 | 1:44339317-44339322 | A3ss | 3 | 3 | ENST00000372257 | 12.07 | 2.47E-02 | 4.24E-02 |
| DPM3 | 1:155140246-155140490 | 1:155140246-155140271 | A3ss | 3 | 3 | Ex.ENST00000341298 | 12.01 | 1.63E-04 | 2.81E-04 |
| CD200 | 3:112333225-112344961 | 3:112340902-112340983 | MXS | 3 | 3 | ENST00000315711 | 12.01 | 3.57E-03 | 8.15E-03 |
| DPP7 | 9:137114379-137114462 | 9:137114379-137114382 | A3ss | 3 | 3 | ENST00000371579 | 12.01 | 4.03E-02 | 1.00E+00 |
| STX16 | 20:58668128-58670511 | 20:58669291-58669453 | SES | 3 | 3 | ENST00000312283 | 11.97 | 2.31E-02 | 4.43E-02 |
| PICALM | 11:85976683-85981128 | 11:85978070-85978093 | SES | 3 | 3 | ENST00000526961 | 11.97 | 4.76E-02 | 7.35E-02 |
| DTX2 | 7:76461837-76480420 | 7:76463654-76480421 | A3ss | 3 | 3 | Ex.TSS.ENST00000413936 | 11.95 | 2.19E-02 | 3.28E-01 |
| DTX2 | 7:76461837-76477113 | 7:76463654-76477114 | A3ss | 3 | 3 | Ex.TSS.ENST00000438930 | 11.95 | 2.19E-02 | 3.28E-01 |
| DTX2 | 7:76461837-76476942 | 7:76463654-76476943 | A3ss | 3 | 3 | Ex.TSS.ENST00000451769 | 11.95 | 2.19E-02 | 3.28E-01 |
| R3HDM1 | 2:135605017-135621493 | 2:135616668-135616757 | MES | 3 | 3 | ENST00000264160 | 11.93 | 3.64E-02 | 7.62E-02 |
| KANK1 | 9:504755-676889 | 9:539516-539633 | MES | 3 | 3 | ENST00000674102 | 11.91 | 9.18E-03 | 2.52E-01 |
| CCDC91 | 12:28190642-28305648 | 12:28257202-28257245 | MES | 3 | 3 | ENST00000536442 | 11.9 | 4.16E-02 | 6.66E-02 |
| RNASEH1-AS1 | 2:3558999-3561316 | 2:3559404-3559729 | SES | 3 | 3 | ENST00000438436 | 11.89 | 2.75E-02 | 6.05E-02 |
| CARM1 | 19:10921128-10921614 | 19:10921375-10921443 | SES | 3 | 3 | ENST00000327064 | 11.88 | 9.07E-03 | 4.68E-02 |
| ZNF615 | 19:52002282-52008140 | 19:52007293-52007330 | MXS | 3 | 3 | ENST00000593650 | 11.88 | 4.58E-02 | 3.21E-01 |
| PKIG | 20:44531979-44589796 | 20:44582585-44582731 | SES | 3 | 3 | ENST00000372887 | 11.87 | 4.99E-04 | 9.54E-04 |
| ACSL4 | X:109683667-109733138 | X:109696144-109696196 | MES | 3 | 3 | ENST00000340800 | 11.86 | 4.38E-02 | 1.00E+00 |
| MAZ | 16:29808742-29810076 | 16:29809540-29809764 | SES | 3 | 3 | ENST00000219782 | 11.85 | 4.85E-03 | 1.77E-02 |
| APTX | 9:32989896-33025022 | 9:33025000-33025022 | TSSIA5ss | 3 | 3 | TSS.ENST00000460940 | 11.85 | 1.24E-02 | 3.53E-01 |
| TRIM26 | 6:30201124-30213304 | 6:30201124-30201138 | A3ss | 3 | 3 | Ex.TSS.ENST00000453195 | 11.84 | 8.66E-04 | 7.82E-03 |
| MBD1 | 18:50274351-50274976 | 18:50274351-50274353 | A3ss | 3 | 3 | ENST00000269468 | 11.84 | 3.44E-02 | 1.72E-01 |
| FAM131A | 3:184336124-184341723 | 3:184337311-184337371 | SES | 3 | 3 | ENST00000340957 | 11.84 | 3.78E-02 | 8.91E-02 |
| SIGIRR | 11:406539-407061 | 11:406843-406993 | SES | 3 | 3 | ENST00000332725 | 11.83 | 1.85E-02 | 2.72E-02 |
| REC8 | 14:24172366-24172491 | 14:24172467-24172492 | A3ss | 3 | 3 | Ex.TSS.ENST00000557806 | 11.81 | 1.15E-02 | 1.95E-02 |
| AP1G2 | 14:23561636-23562287 | 14:23561962-23562066 | SES | 3 | 3 | ENST00000308724 | 11.78 | 1.91E-02 | 3.08E-02 |
| NSRP1 | 17:30116864-30172541 | 17:30117302-30117332 | MES | 3 | 3 | ENST00000475652 | 11.78 | 2.73E-02 | 1.10E-01 |
| RBPJ | 4:26320816-26386352 | 4:26362446-26362628 | SES | 3 | 3 | ENST00000514807 | 11.77 | 1.61E-03 | 4.36E-03 |
| GBGT1 | 9:133162533-133163753 | 9:133162533-133162545 | A3ss | 3 | 3 | Ex.TSS.ENST00000372036 | 11.76 | 7.00E-03 | 2.84E-01 |
| PRPF40A | 2:152663968-152664137 | 2:152663968-152663979 | A3ss | 3 | 3 | Ex.ENST00000410080 | 11.76 | 1.87E-02 | 3.93E-02 |
| ZNF473 | 19:50026123-50039160 | 19:50031014-50031091 | SES | 3 | 3 | ENST00000391821 | 11.73 | 1.87E-03 | 1.30E-02 |
| LAMP2 | X:120431463-120441729 | X:120439076-120439293 | SES | 3 | 3 | ENST00000486593 | 11.73 | 3.45E-02 | 1.00E+00 |
| ZNF644 | 1:90918155-90937484 | 1:90918155-90935530 | A3ss | 3 | 3 | Ex.ENST00000337393 | 11.72 | 4.31E-03 | 8.13E-03 |
| LTBP4 | 19:40614447-40619346 | 19:40616889-40617020 | MES | 3 | 3 | ENST00000204005 | 11.72 | 1.18E-02 | 7.76E-02 |
| NUP210 | 3:13360492-13365945 | 3:13365669-13365946 | A5ss | 3 | 3 | Ex.ENST00000254508 | 11.72 | 3.51E-02 | 8.05E-02 |
| TBC1D5 | 3:17508606-17623848 | 3:17622648-17622765 | MES | 3 | 3 | ENST00000481396 | 11.71 | 4.27E-03 | 9.96E-03 |
| SCOC | 4:140379193-140380961 | 4:140379569-140379652 | SES | 3 | 3 | ENST00000338517 | 11.7 | 1.31E-03 | 3.50E-03 |
| RPS24 | 10:78037305-78040614 | 10:78040204-78040225 | SES | 3 | 3 | ENST00000372360 | 11.69 | 6.16E-04 | 8.77E-04 |
| SNHG14 | 15:25064856-25067411 | 15:25067409-25067411 | A3ss | 3 | 3 | ENST00000654223 | 11.68 | 2.23E-03 | 7.25E-03 |
| TUG1 | 22:30971780-30972854 | 22:30971780-30972172 | TSSIA5ss | 3 | 3 | TSS.ENST00000563812 | 11.68 | 1.60E-02 | 3.26E-02 |
| MTERF2 | 12:106978772-106986968 | 12:106985115-106986968 | A5ss | 3 | 3 | ENST00000552029 | 11.67 | 1.72E-02 | 2.67E-02 |
| QTRT2 | 3:114056865-114070625 | 3:114066228-114066283 | MES | 3 | 3 | ENST00000281273 | 11.66 | 4.79E-02 | 1.09E-01 |
| PANK1 | 10:89612049-89644599 | 10:89612049-89626632 | A3ss | 3 | 3 | Ex.TSS.ENST00000307534 | 11.65 | 4.57E-04 | 6.54E-04 |
| CLCC1 | 1:108950449-108963360 | 1:108962309-108962469 | SES | 3 | 3 | ENST00000369976 | 11.65 | 1.48E-02 | 1.21E-01 |
| ZMIZ2 | 7:44756300-44756424 | 7:44756410-44756425 | A3ss | 3 | 3 | Ex.TSS.ENST00000265346 | 11.62 | 7.28E-05 | 1.00E-03 |
| ZNF143 | 11:9474025-9474552 | 11:9474550-9474552 | A3ss | 3 | 3 | ENST00000396597 | 11.59 | 4.39E-03 | 6.78E-03 |
| TIA1 | 2:70224630-70227734 | 2:70225328-70225393 | SES | 3 | 3 | ENST00000474809 | 11.57 | 2.95E-02 | 6.58E-02 |
| NADK2 | 5:36207266-36211843 | 5:36208625-36208690 | SES | 3 | 3 | ENST00000506945 | 11.53 | 4.96E-02 | 1.48E-01 |
| PPP2R5C | 14:101762971-101856685 | 14:101786018-101786183 | MXS | 3 | 3 | ENST00000328724 | 11.5 | 5.03E-03 | 8.15E-03 |
| KMT5A | 12:123407493-123409353 | 12:123408807-123408870 | IR | 3 | 3 | ENST00000402868 | 11.48 | 4.46E-02 | 7.10E-02 |
| SVBP | 1:42816581-42817189 | 1:42816581-42816605 | A3ss | 3 | 3 | Ex.TSS.ENST00000372521 | 11.47 | 5.74E-03 | 1.07E-02 |
| SNHG8 | 4:118278793-118279388 | 4:118279107-118279137 | SES | 3 | 3 | ENST00000602483 | 11.47 | 2.26E-02 | 5.94E-02 |

| | | | | | | | | | |
|---|---|---|---|---|---|---|---|---|---|
| SH3BP1 | 22:37639847-37641373 | 22:37641126-37641168 | SES | 3 | 3 | ENST00000417536 | 11.46 | 3.65E-02 | 7.47E-02 |
| FAM49B | 8:129903351-129970942 | 8:129904499-129904585 | MES | 3 | 3 | ENST00000519142 | 11.45 | 4.24E-03 | 8.29E-02 |
| MLLT10 | 10:21534495-21534644 | 10:21534495-21534520 | TSSIA5ss | 3 | 3 | TSS.ENST00000307729 | 11.45 | 4.36E-02 | 6.14E-02 |
| POLL | 10:101582892-101587245 | 10:101583508-101583681 | MXS | 3 | 3 | ENST00000339310 | 11.44 | 4.24E-02 | 5.92E-02 |
| LYSMD4 | 15:99732008-99733344 | 15:99733285-99733344 | TSSIA5ss | 3 | 3 | TSS.ENST00000493256 | 11.42 | 7.18E-03 | 2.51E-02 |
| RFX3 | 9:3293259-3330258 | 9:3301546-3301620 | SES | 3 | 3 | ENST00000302303 | 11.41 | 4.25E-02 | 1.00E+00 |
| FAM49B | 8:129903351-129970942 | 8:129904499-129904585 | MES | 3 | 3 | ENST00000518283 | 11.39 | 4.13E-03 | 8.06E-02 |
| RBM39 | 20:35739018-35740823 | 20:35740525-35740597 | MXS | 3 | 3 | ENST00000403542 | 11.39 | 3.11E-02 | 5.91E-02 |
| CARMIL2 | 16:67656646-67656833 | 16:67656801-67656833 | A3ss | 3 | 3 | ENST00000334583 | 11.36 | 1.26E-02 | 4.81E-02 |
| CARM1 | 19:10921128-10921614 | 19:10921375-10921614 | TSSIA3ss | 3 | 3 | TSS.ENST00000590699 | 11.35 | 1.25E-02 | 6.44E-02 |
| CARD8 | 19:48231811-48234402 | 19:48232453-48232789 | SES | 2 | 3 | ENST00000519646 | 11.35 | 1.58E-02 | 1.06E-01 |
| CARD8 | 19:48231811-48234402 | 19:48232453-48232493 | MES | 2 | 3 | ENST00000522773 | 11.35 | 1.58E-02 | 1.06E-01 |
| CBSL | 21:6458056-6460186 | 21:6459456-6459638 | SES | 3 | 3 | ENST00000624921 | 11.35 | 2.51E-02 | 4.89E-02 |
| CDK11B | 1:1655485-1657374 | 1:1657256-1657302 | SES | 3 | 3 | ENST00000340677 | 11.34 | 5.10E-03 | 8.90E-03 |
| SLC2A11 | 22:23877172-23877720 | 22:23877385-23877720 | A3ss | 3 | 2 | ENST00000405340 | 11.33 | 2.51E-02 | 4.93E-02 |
| ATG9A | 2:219228054-219229534 | 2:219228434-219228485 | SES | 3 | 3 | ENST00000361242 | 11.32 | 7.80E-04 | 1.69E-03 |
| MBD1 | 18:50273864-50274976 | 18:50274186-50274353 | SES | 3 | 3 | ENST00000269468 | 11.31 | 1.91E-03 | 9.61E-03 |
| COASY | 17:42562220-42562611 | 17:42562296-42562487 | SES | 3 | 3 | ENST00000587858 | 11.31 | 2.45E-02 | 1.03E-01 |
| AP1B1 | 22:29328896-29330377 | 22:29329712-29329720 | SES | 3 | 3 | ENST00000357586 | 11.26 | 2.58E-04 | 5.07E-04 |
| VEZT | 12:95252072-95266356 | 12:95256550-95256622 | MES | 3 | 3 | ENST00000356859 | 11.26 | 2.04E-02 | 3.25E-02 |
| CEP70 | 3:138572933-138591853 | 3:138572933-138572984 | A3ss | 3 | 3 | ENST00000464035 | 11.25 | 3.49E-02 | 8.04E-02 |
| LRRC20 | 10:70301509-70340552 | 10:70323863-70324030 | SES | 3 | 3 | ENST00000355790 | 11.22 | 1.60E-02 | 2.29E-02 |
| AMT | 3:49422258-49422360 | 3:49422258-49422271 | TSSIA3ss | 3 | 3 | TSS.ENST00000273588 | 11.21 | 1.74E-02 | 4.22E-02 |
| BPTF | 17:67875021-67891843 | 17:67875556-67875744 | MXS | 3 | 3 | ENST00000544778 | 11.19 | 2.28E-02 | 1.04E-01 |
| FAM49B | 8:129903351-129970942 | 8:129904499-129904585 | MES | 3 | 3 | ENST00000518879 | 11.17 | 5.71E-03 | 1.12E-01 |
| RAB30-DT | 11:83072703-83073302 | 11:83072703-83072804 | TSSIA5ss | 3 | 3 | TSS.ENST00000501011 | 11.17 | 4.96E-02 | 7.64E-02 |
| YY1AP1 | 1:155661424-155670319 | 1:155668627-155668777 | SES | 3 | 3 | ENST00000347088 | 11.15 | 2.50E-03 | 4.33E-03 |
| AC093752.1 | 4:119455134-119460182 | 4:119457971-119458077 | SES | 3 | 3 | ENST00000515251 | 11.14 | 7.72E-03 | 2.03E-02 |
| IQSEC2 | X:53235185-53236321 | X:53235783-53235832 | SES | 3 | 3 | ENST00000642864 | 11.13 | 3.44E-02 | 1.00E+00 |
| ATP11A | 13:112875942-112881875 | 13:112881816-112881875 | TSSIA3ss | 3 | 3 | TSS.ENST00000375630 | 11.12 | 1.46E-02 | 2.36E-02 |
| TNK2 | 3:195867017-195867360 | 3:195867169-195867264 | SES | 3 | 3 | ENST00000672548 | 11.1 | 2.92E-02 | 6.90E-02 |
| PROSER3 | 19:35758199-35759991 | 19:35758227-35759373 | IR | 3 | 3 | ENST00000600988 | 11.08 | 2.96E-03 | 1.69E-02 |
| MAPKAP1 | 9:125484584-125543058 | 9:125506310-125506417 | SES | 3 | 3 | ENST00000265960 | 11.08 | 1.59E-02 | 5.48E-01 |
| OTX2 | 14:56804340-56805359 | 14:56804340-56804363 | TSSIA3ss | 3 | 3 | TSS.ENST00000554845 | 11.07 | 4.72E-02 | 1.48E-01 |
| NELFE | 6:31954878-31955058 | 6:31954878-31954892 | A3ss | 3 | 3 | ENST00000375425 | 11.03 | 2.00E-02 | 5.15E-02 |
| PLA2G12A | 4:109717714-109729601 | 4:109718683-109718759 | SES | 3 | 3 | ENST00000243501 | 11.02 | 1.55E-02 | 4.05E-02 |
| ACSL4 | X:109683667-109733138 | X:109696144-109696196 | MXS | 3 | 3 | ENST00000469796 | 11.01 | 4.27E-02 | 1.00E+00 |
| ATF1 | 12:50780239-50795908 | 12:50795898-50795909 | A3ss | 3 | 3 | Ex.ENST00000262053 | 10.99 | 4.91E-02 | 8.05E-02 |
| JPX | X:73944663-74004269 | X:73994840-73994961 | MXS | 3 | 3 | ENST00000660836 | 10.96 | 5.69E-03 | 1.00E+00 |
| ATXN2L | 16:28831073-28832204 | 16:28832187-28832204 | A3ss | 3 | 3 | ENST00000564304 | 10.96 | 2.00E-02 | 7.29E-02 |
| PLXNB1 | 3:48416452-48417910 | 3:48416452-48416469 | A3ss | 3 | 3 | Ex.ENST00000296440 | 10.95 | 4.00E-03 | 9.48E-03 |
| CEP89 | 19:32918340-32926189 | 19:32923439-32923542 | SES | 2 | 3 | ENST00000305768 | 10.95 | 2.20E-02 | 1.23E-01 |
| MIAT | 22:26666938-26668157 | 22:26667185-26667310 | SES | 3 | 3 | ENST00000613780 | 10.95 | 3.65E-02 | 7.18E-02 |
| NCAPG2 | 7:158646564-158652292 | 7:158650832-158650972 | SES | 3 | 3 | ENST00000356309 | 10.93 | 3.86E-03 | 4.93E-02 |
| NR1H3 | 11:47261450-47261918 | 11:47261547-47261726 | SES | 3 | 3 | ENST00000395397 | 10.93 | 1.75E-02 | 2.59E-02 |
| RPL13P5 | 12:6883704-6884616 | 12:6883920-6884370 | IR (overlapping | 3 | 3 | ENST00000412023 | 10.93 | 2.97E-02 | 4.98E-02 |
| PLSCR1 | 3:146533551-146544466 | 3:146536540-146536565 | MXS | 3 | 3 | ENST00000342435 | 10.92 | 2.36E-03 | 5.45E-03 |
| PABPC4 | 1:39563923-39564470 | 1:39564423-39564470 | A5ss | 3 | 3 | ENST00000372858 | 10.92 | 1.43E-02 | 2.65E-02 |
| EPS8L2 | 11:720972-721352 | 11:721188-721284 | IR (overlapping | 3 | 3 | ENST00000531471 | 10.92 | 3.70E-02 | 5.66E-02 |
| LPCAT3 | 12:6977367-6978507 | 12:6977526-6977597 | IR (overlapping | 3 | 3 | ENST00000535479 | 10.91 | 2.08E-02 | 3.48E-02 |
| MOCS1 | 6:39907118-39909834 | 6:39909055-39909102 | SES | 3 | 3 | ENST00000340692 | 10.91 | 2.70E-02 | 2.55E-01 |
| SNHG14 | 15:25062089-25070119 | 15:25064715-25064855 | MES | 3 | 3 | ENST00000654223 | 10.9 | 4.70E-03 | 1.53E-02 |
| PHRF1 | 11:605301-605607 | 11:605605-605607 | A3ss | 3 | 3 | ENST00000264555 | 10.9 | 4.40E-02 | 6.50E-02 |
| MARK2 | 11:63905044-63908876 | 11:63908260-63908304 | MES | 3 | 3 | ENST00000402010 | 10.89 | 2.44E-02 | 3.66E-02 |
| ZNF350 | 19:51969132-51974345 | 19:51969132-51969145 | A3ss | 3 | 2 | ENST00000601430 | 10.88 | 4.80E-04 | 3.37E-03 |
| PRRC2C | 1:171512201-171513000 | 1:171512995-171513000 | A3ss | 3 | 3 | ENST00000367742 | 10.88 | 2.41E-02 | 4.23E-02 |
| FAM193B | 5:177525206-177532442 | 5:177531282-177531521 | MXS | 3 | 3 | ENST00000524677 | 10.88 | 4.63E-02 | 1.30E-01 |
| PBRM1 | 3:52550818-52561766 | 3:52558249-52558413 | MES | 3 | 3 | ENST00000296302 | 10.87 | 8.63E-03 | 2.23E-02 |
| FKRP | 19:46746091-46748026 | 19:46747234-46747305 | MXS | 3 | 3 | ENST00000600005 | 10.86 | 5.79E-03 | 3.88E-02 |
| FOXD3-AS1 | 1:63321464-63322315 | 1:63321464-63321944 | A3ss | 3 | 3 | Ex.ENST00000449386 | 10.82 | 2.59E-02 | 4.86E-02 |
| FASTK | 7:151078562-151079049 | 7:151078702-151078922 | IR | 3 | 3 | ENST00000478883 | 10.79 | 4.49E-03 | 5.66E-02 |
| TUSC3 | 8:15748466-15764202 | 8:15757791-15757855 | SES | 3 | 3 | ENST00000503731 | 10.77 | 4.73E-02 | 9.66E-01 |
| PPP1R26 | 9:135482816-135483949 | 9:135483352-135483553 | SES | 3 | 3 | ENST00000652373 | 10.76 | 9.21E-03 | 3.82E-01 |
| ZBTB17 | 1:15946295-15946934 | 1:15946295-15946316 | TSSIA3ss | 3 | 3 | TSS.ENST00000490899 | 10.76 | 2.59E-02 | 4.50E-02 |
| THTPA | 14:23556344-23558694 | 14:23556743-23557034 | SES | 3 | 3 | ENST00000554970 | 10.75 | 4.35E-02 | 6.05E-02 |
| METTL26 | 16:634798-635280 | 16:634889-635063 | SES | 3 | 3 | ENST00000456420 | 10.72 | 3.48E-03 | 1.32E-02 |
| SMUG1 | 12:54187906-54188514 | 12:54188511-54188514 | A5ss | 3 | 3 | ENST00000513838 | 10.71 | 2.48E-04 | 4.10E-04 |
| RPARP-AS1 | 10:102450331-102455336 | 10:102451399-102452187 | SES | 3 | 3 | ENST00000494270 | 10.71 | 3.82E-02 | 5.32E-02 |
| MTERF4 | 2:241094433-241102252 | 2:241099396-241099894 | MXS | 3 | 3 | ENST00000401626 | 10.69 | 4.50E-02 | 9.78E-02 |
| DALRD3 | 3:49015708-49015818 | 3:49015804-49015818 | TSSIA5ss | 3 | 3 | TSS.ENST00000313778 | 10.67 | 6.53E-03 | 1.57E-02 |
| MAGI2-AS3 | 7:79454086-79455374 | 7:79454245-79454885 | IR | 3 | 2 | ENST00000653779 | 10.65 | 9.89E-03 | 1.49E-01 |
| BNIP1 | 5:173146959-173154321 | 5:173151566-173151694 | SES | 3 | 3 | ENST00000231668 | 10.64 | 2.06E-02 | 6.00E-02 |
| GMPPA | 2:219505556-219505714 | 2:219505695-219505714 | A3ss | 3 | 3 | ENST00000443704 | 10.63 | 6.32E-04 | 1.37E-03 |
| SYVN1 | 11:65133253-65133469 | 11:65133253-65133269 | A3ss | 3 | 3 | Ex.ENST00000294256 | 10.63 | 2.70E-02 | 4.09E-02 |
| GALNT7 | 4:173295844-173298114 | 4:173297795-173298114 | TSSIA3ss | 3 | 3 | TSS.ENST00000512285 | 10.61 | 9.21E-03 | 2.47E-02 |
| DDB2 | 11:47234673-47235269 | 11:47234757-47235269 | A3ss | 3 | 3 | ENST00000612309 | 10.6 | 2.64E-02 | 3.87E-02 |
| PEX2 | 8:76984196-76999989 | 8:76986187-76986296 | MES | 3 | 3 | ENST00000519956 | 10.6 | 3.02E-02 | 7.05E-01 |
| RNF20 | 9:101533915-101535397 | 9:101535376-101535398 | A3ss | 3 | 3 | Ex.TSS.ENST00000374819 | 10.58 | 1.93E-02 | 4.78E-01 |
| WIZ | 19:15427534-15429585 | 19:15428110-15428508 | SES | 3 | 3 | ENST00000545156 | 10.56 | 4.71E-02 | 2.52E-01 |
| GNPTG | 16:1352307-1361871 | 16:1361686-1361872 | A3ss | 3 | 3 | Ex.ENST00000527168 | 10.52 | 7.89E-05 | 2.77E-04 |
| GNPTG | 16:1352307-1361742 | 16:1361686-1361743 | A3ss | 3 | 3 | Ex.ENST00000204679 | 10.52 | 7.89E-05 | 2.77E-04 |
| GCC2 | 2:108449281-108449632 | 2:108449281-108449289 | TSSIA5ss | 3 | 3 | TSS.ENST00000409821 | 10.52 | 2.94E-02 | 6.12E-02 |
| MVK | 12:109579947-109586021 | 12:109581395-109581550 | SES | 3 | 3 | ENST00000228510 | 10.47 | 2.54E-02 | 3.94E-02 |
| CCNL1 | 3:157153157-157158865 | 3:157156936-157157083 | SES | 3 | 3 | ENST00000477127 | 10.47 | 4.64E-02 | 1.07E-01 |
| STAU2 | 8:73688814-73746782 | 8:73709032-73709162 | MES | 3 | 3 | ENST00000519818 | 10.46 | 1.45E-02 | 3.39E-01 |
| TRA2A | 7:23521841-23531788 | 7:23531181-23531270 | MES | 3 | 3 | ENST00000392502 | 10.45 | 1.44E-04 | 1.86E-03 |
| BIN1 | 2:127050913-127059010 | 2:127051154-127051243 | SES | 3 | 3 | ENST00000352848 | 10.43 | 4.55E-02 | 9.47E-02 |
| SLC29A2 | 11:66368672-66369368 | 11:66369060-66369199 | SES | 3 | 3 | ENST00000311161 | 10.4 | 2.96E-02 | 4.50E-02 |
| SPIB | 19:50423038-50428037 | 19:50423640-50423755 | SES | 3 | 3 | ENST00000597855 | 10.39 | 2.51E-02 | 1.76E-01 |
| PACRGL | 4:20713540-20727284 | 4:20724808-20724888 | MXS | 3 | 3 | ENST00000471979 | 10.37 | 4.39E-03 | 1.19E-02 |
| ANKZF1 | 2:219229901-219232489 | 2:219230228-219230405 | MES | 3 | 3 | ENST00000323348 | 10.37 | 4.79E-02 | 1.03E-01 |
| UPP1 | 7:48094828-48106872 | 7:48101824-48101870 | MXS | 3 | 3 | ENST00000482015 | 10.36 | 4.52E-04 | 6.31E-03 |
| PEAK1 | 15:77252527-77286422 | 15:77283883-77284030 | MES | 2 | 3 | ENST00000567337 | 10.36 | 3.01E-02 | 1.02E-01 |
| AKAP8L | 19:15399411-15400294 | 19:15399411-15399967 | A3ss | 3 | 3 | Ex.ENST00000397410 | 10.34 | 3.40E-03 | 1.84E-02 |

| | | | | | | | | | |
|---|---|---|---|---|---|---|---|---|---|
| EGFL8 | 6:32167006-32167146 | 6:32167087-32167146 | A3ss | 3 | 3 | ENST00000333845 | 10.34 | 4.93E-02 | 1.27E-01 |
| EGFL8 | 6:32167006-32167102 | 6:32167087-32167102 | A3ss | 3 | 3 | ENST00000333845 | 10.34 | 4.93E-02 | 1.27E-01 |
| APTX | 9:32987732-32989758 | 9:32988083-32988124 | MXS | 3 | 3 | ENST00000464632 | 10.33 | 1.14E-03 | 3.25E-02 |
| AURKA | 20:56388203-56392167 | 20:56388687-56388784 | MES | 3 | 3 | ENST00000420474 | 10.33 | 3.11E-03 | 5.94E-03 |
| FASTK | 7:151078562-151079049 | 7:151078700-151078922 | IR | 3 | 3 | ENST00000478883 | 10.3 | 5.14E-03 | 6.48E-02 |
| CHD1L | 1:147242831-147265931 | 1:147255813-147255927 | MES | 3 | 3 | ENST00000369258 | 10.3 | 4.80E-02 | 4.12E-01 |
| AL357673.1 | 1:54184266-54195418 | 1:54187652-54187739 | MXS | 3 | 3 | ENST00000311841 | 10.29 | 2.47E-02 | 4.63E-02 |
| CPNE1 | 20:35632924-35664759 | 20:35658930-35659014 | MXS | 3 | 3 | ENST00000434795 | 10.26 | 2.83E-03 | 5.38E-03 |
| MAP3K6 | 1:27355470-27356025 | 1:27355669-27355745 | SES | 3 | 3 | ENST00000357582 | 10.26 | 1.11E-02 | 2.05E-02 |
| TMEM94 | 17:75492560-75493102 | 17:75492790-75492928 | IR | 3 | 3 | ENST00000578624 | 10.2 | 9.67E-03 | 4.46E-02 |
| APTX | 9:32987732-32989758 | 9:32988083-32988129 | MXS | 3 | 3 | ENST00000478279 | 10.17 | 1.80E-02 | 4.92E-01 |
| VPS51 | 11:65109924-65110481 | 11:65110463-65110482 | A3ss | 3 | 3 | Ex.ENST00000279281 | 10.15 | 3.13E-03 | 4.74E-03 |
| PAM | 5:103019844-103025133 | 5:103025131-103025133 | A3ss | 3 | 3 | ENST00000304400 | 10.14 | 2.18E-02 | 6.21E-02 |
| FCHSD1 | 5:141646087-141646722 | 5:141646192-141646602 | IR | 3 | 3 | ENST00000523856 | 10.09 | 2.80E-02 | 8.18E-02 |
| SIKE1 | 1:114779285-114780448 | 1:114780110-114780448 | A5ss | 3 | 3 | ENST00000471969 | 10.08 | 2.78E-03 | 2.34E-02 |
| ARFGAP2 | 11:47173883-47175014 | 11:47173883-47173915 | A3ss | 3 | 3 | ENST00000528444 | 10.08 | 1.38E-02 | 2.03E-02 |
| ARFGAP2 | 11:47173841-47175014 | 11:47173841-47173915 | A3ss | 3 | 3 | ENST00000528444 | 10.08 | 1.38E-02 | 2.03E-02 |
| HDHD2 | 18:47136450-47150377 | 18:47137235-47137296 | MXS | 3 | 3 | ENST00000586546 | 10.06 | 1.46E-02 | 7.30E-02 |
| MSTO1 | 1:155611828-155611982 | 1:155611921-155611982 | A3ss | 3 | 2 | ENST00000483832 | 10.05 | 4.32E-02 | 6.02E-02 |
| RPL17 | 18:49491585-49492257 | 18:49491736-49491833 | SES | 3 | 3 | ENST00000618613 | 10.02 | 5.18E-04 | 2.61E-03 |
| SIGIRR | 11:406539-407061 | 11:406539-406993 | A3ss | 3 | 3 | ENST00000534145 | 10.02 | 8.83E-03 | 1.30E-02 |
| TSSC2 | 11:3384285-3401758 | 11:3397117-3397406 | SES | 3 | 3 | ENST00000529482 | 10.02 | 3.14E-02 | 4.61E-02 |
| MOCS1 | 6:39925846-39934294 | 6:39927329-39927455 | SES | 3 | 3 | ENST00000340692 | 10.02 | 3.67E-02 | 3.47E-01 |
| CYB5D1 | 17:7858285-7858479 | 17:7858295-7858402 | IR (overlapping | 3 | 3 | ENST00000574196 | 10.01 | 2.35E-02 | 1.12E-01 |
| SNHG14 | 15:25062089-25070119 | 15:25067409-25067531 | MES | 3 | 3 | ENST00000654223 | 10.01 | 2.40E-02 | 7.81E-02 |
| MED15 | 22:20553175-20564449 | 22:20554936-20555148 | SES | 3 | 3 | ENST00000263205 | 10 | 1.31E-03 | 2.56E-03 |
| JPX | X:73944663-73998767 | X:73994840-73994961 | MXS | 3 | 3 | ENST00000602546 | 10 | 2.08E-03 | 1.00E+00 |
| ANKRD13A | 12:110025824-110028511 | 12:110027705-110027766 | SES | 3 | 3 | ENST00000261739 | 10 | 3.36E-03 | 5.23E-03 |
| DMAP1 | 1:44213471-44213858 | 1:44213528-44213748 | IR | 3 | 3 | ENST00000372289 | -10 | 4.45E-03 | 8.29E-03 |
| ALKBH6 | 19:36013090-36014174 | 19:36013090-36013422 | TSSIA3ss | 3 | 3 | TSS.ENST00000471323 | -10 | 1.57E-02 | 8.93E-02 |
| MAP7 | 6:136377869-136388392 | 6:136383671-136383781 | SES | 3 | 3 | ENST00000354570 | -10.01 | 1.38E-04 | 4.30E-04 |
| MAPK7 | 17:19378529-19382466 | 17:19379948-19380607 | IR | 3 | 3 | ENST00000570306 | -10.01 | 5.15E-03 | 1.82E-02 |
| RNF123 | 3:49689607-49691129 | 3:49689607-49689822 | TSSIA5ss | 3 | 3 | TSS.ENST00000454491 | -10.01 | 4.05E-02 | 9.82E-02 |
| DMAP1 | 1:44213471-44213858 | 1:44213663-44213748 | IR | 3 | 3 | ENST00000372289 | -10.02 | 2.28E-03 | 4.25E-03 |
| FAM177A1 | 14:35046403-35046628 | 14:35046407-35046504 | IR | 3 | 3 | ENST00000280987 | -10.03 | 3.53E-02 | 6.04E-02 |
| GOSR2 | 17:46938599-46941600 | 17:46938705-46940554 | IR (overlapping | 3 | 3 | ENST00000638216 | -10.05 | 8.52E-04 | 3.69E-03 |
| AC138028.2 | 16:88736800-88738405 | 16:88737647-88737727 | IR (overlapping | 3 | 3 | ENST00000567968 | -10.05 | 3.05E-03 | 1.19E-02 |
| TMEM129 | 4:1717429-1718151 | 4:1717516-1717675 | SES | 3 | 3 | ENST00000382936 | -10.06 | 3.04E-03 | 8.15E-03 |
| SEPTIN4 | 17:58520743-58522101 | 17:58521854-58521966 | IR (overlapping | 3 | 3 | ENST00000582270 | -10.08 | 5.61E-03 | 2.50E-02 |
| PIEZO1 | 16:88717212-88719573 | 16:88717537-88717615 | SES | 3 | 3 | ENST00000419505 | -10.08 | 3.15E-02 | 1.21E-01 |
| MAPKAPK5-AS1 | 12:111839767-111840477 | 12:111840004-111840133 | IR | 3 | 3 | ENST00000443596 | -10.08 | 3.86E-02 | 6.00E-02 |
| AL162231.1 | 9:34665613-34666045 | 9:34665681-34665977 | IR (overlapping | 3 | 3 | ENST00000421828 | -10.1 | 4.06E-02 | 1.00E+00 |
| GLE1 | 9:128504905-128508875 | 9:128507216-128508876 | A3ss | 3 | 3 | Ex.TSS.ENST00000372770 | -10.11 | 5.20E-03 | 1.85E-01 |
| ARVCF | 22:19982092-19987072 | 22:19982092-19983819 | A3ss | 3 | 3 | Ex.TSS.ENST00000401994 | -10.11 | 7.74E-03 | 1.51E-02 |
| DALRD3 | 3:49015491-49017513 | 3:49015708-49015972 | IR (overlapping | 3 | 3 | ENST00000498794 | -10.11 | 1.62E-02 | 3.91E-02 |
| PAQR4 | 16:2971515-2973482 | 16:2972755-2973137 | IR (overlapping | 3 | 3 | ENST00000293978 | -10.11 | 2.28E-02 | 8.31E-02 |
| RABGAP1L | 1:174811961-174957456 | 1:174877392-174877605 | SES | 3 | 3 | ENST00000469553 | -10.11 | 4.20E-02 | 5.85E-02 |
| CXXC1 | 18:50285248-50285748 | 18:50285325-50285351 | SES | 3 | 3 | ENST00000285106 | -10.12 | 9.06E-03 | 4.56E-02 |
| MTERF4 | 2:241094433-241102252 | 2:241097243-241097427 | MXS | 3 | 3 | ENST00000455202 | -10.12 | 1.92E-02 | 4.18E-02 |
| CNTN1 | 12:40692593-40908356 | 12:40895950-40896109 | SES | 3 | 3 | ENST00000551424 | -10.13 | 6.70E-03 | 1.08E-02 |
| TP53I11 | 11:44938367-44949972 | 11:44939114-44939190 | MES | 3 | 3 | ENST00000533940 | -10.13 | 1.03E-02 | 1.51E-02 |
| CENPT | 16:67832109-67832454 | 16:67832228-67832354 | SES | 3 | 3 | ENST00000566067 | -10.13 | 2.65E-02 | 1.01E-01 |
| SPIB | 19:50423038-50428037 | 19:50423605-50423751 | SES | 3 | 3 | ENST00000270632 | -10.17 | 3.89E-02 | 2.72E-01 |
| MBD1 | 18:50268846-50271540 | 18:50269812-50271468 | IR | 3 | 3 | ENST00000591416 | -10.18 | 1.60E-02 | 8.02E-02 |
| UPP1 | 7:48094828-48106872 | 7:48101824-48101982 | MES | 3 | 3 | ENST00000331803 | -10.19 | 1.52E-02 | 2.12E-01 |
| MYO5C | 15:52239880-52244355 | 15:52242048-52242213 | SES | 3 | 3 | ENST00000261839 | -10.24 | 2.73E-02 | 9.06E-02 |
| RAD17 | 5:69372034-69372217 | 5:69372055-69372156 | IR | 3 | 3 | ENST00000282891 | -10.26 | 1.75E-02 | 5.27E-02 |
| TTLL5 | 14:75902225-75954423 | 14:75906555-75906677 | MXS | 3 | 3 | ENST00000554972 | -10.28 | 3.04E-02 | 9.68E-02 |
| SPIN3 | X:56990831-56994949 | X:56993889-56994722 | IR | 3 | 3 | ENST00000374919 | -10.29 | 6.69E-04 | 1.57E-01 |
| CENPV | 17:16348686-16349930 | 17:16349140-16349333 | SES | 3 | 3 | ENST00000482983 | -10.29 | 1.94E-03 | 7.67E-03 |
| TSEN2 | 3:12503785-12516610 | 3:12505154-12505231 | SES | 3 | 3 | ENST00000284995 | -10.3 | 8.46E-03 | 1.89E-02 |
| CCND1 | 11:69651118-69654474 | 11:69653320-69654157 | IR (overlapping | 3 | 3 | ENST00000227507 | -10.3 | 3.60E-02 | 5.48E-02 |
| NDRG2 | 14:21018240-21018762 | 14:21018457-21018504 | SES | 3 | 3 | ENST00000298684 | -10.31 | 4.61E-02 | 7.48E-02 |
| PCBP2 | 12:53467294-53468779 | 12:53467805-53467843 | SES | 3 | 3 | ENST00000439930 | -10.33 | 9.23E-05 | 1.52E-04 |
| AL669830.2 | 6:30913811-30915037 | 6:30914470-30914809 | IR (overlapping | 3 | 3 | ENST00000477288 | -10.35 | 2.51E-02 | 7.80E-02 |
| CCND1 | 11:69651118-69654474 | 11:69653317-69654157 | IR (overlapping | 3 | 3 | ENST00000227507 | -10.35 | 3.57E-02 | 5.43E-02 |
| CTC1 | 17:8229891-8230468 | 17:8230349-8230446 | IR | 3 | 3 | ENST00000578240 | -10.36 | 1.90E-02 | 9.18E-02 |
| DERL3 | 22:23838711-23839128 | 22:23838774-23838894 | IR | 3 | 3 | ENST00000290730 | -10.37 | 1.83E-02 | 3.55E-02 |
| TMEM131L | 4:153621850-153622900 | 4:153622898-153622900 | A3ss | 3 | 3 | Ex.ENST00000240487 | -10.38 | 3.26E-02 | 8.69E-02 |
| DDX11 | 12:31073970-31078389 | 12:31073970-31074091 | TSSIA5ss | 3 | 3 | TSS.ENST00000350437 | -10.4 | 8.67E-03 | 1.39E-02 |
| DERL3 | 22:23838711-23839128 | 22:23838777-23838894 | IR | 3 | 3 | ENST00000290730 | -10.41 | 1.60E-02 | 3.11E-02 |
| TULP3 | 12:2939311-2941138 | 12:2939425-2940194 | IR | 3 | 3 | ENST00000448120 | -10.43 | 2.43E-02 | 3.90E-02 |
| PAQR4 | 16:2971515-2973482 | 16:2972104-2973417 | IR (overlapping | 3 | 3 | ENST00000293978 | -10.44 | 7.00E-04 | 2.55E-03 |
| GIPC1 | 19:14480779-14496036 | 19:14482689-14483006 | MES | 3 | 3 | ENST00000393033 | -10.45 | 1.59E-02 | 8.59E-02 |
| CEP250 | 20:35474053-35476448 | 20:35475502-35475646 | SES | 3 | 3 | ENST00000397527 | -10.45 | 3.83E-02 | 7.27E-02 |
| UNC5D | 8:35722357-35726151 | 8:35722357-35722395 | A5ss | 3 | 3 | ENST00000404895 | -10.46 | 4.81E-02 | 8.86E-01 |
| STAU2 | 8:73688814-73746782 | 8:73738284-73738349 | MXS | 3 | 3 | ENST00000521447 | -10.47 | 2.15E-02 | 5.03E-01 |
| EBP | X:48521908-48523982 | X:48523699-48523982 | A3ss | 3 | 3 | Ex.TSS.ENST00000495186 | -10.48 | 4.41E-04 | 4.81E-02 |
| EBP | X:48521908-48523922 | X:48523699-48523922 | A3ss | 3 | 3 | Ex.TSS.ENST00000495186 | -10.48 | 4.41E-04 | 4.81E-02 |
| EBP | X:48521908-48523753 | X:48523699-48523753 | TSSIA3ss | 3 | 3 | TSS.ENST00000495186 | -10.48 | 4.41E-04 | 4.81E-02 |
| MINK1 | 17:4891125-4891515 | 17:4891456-4891515 | A3ss | 3 | 3 | ENST00000347992 | -10.49 | 3.27E-02 | 1.43E-01 |
| MORF4L2 | X:103678621-103684680 | X:103678621-103678651 | A3ss | 3 | 3 | ENST00000434230 | -10.5 | 1.07E-02 | 2.70E-01 |
| MORF4L2 | X:103678601-103684680 | X:103678601-103678651 | A3ss | 3 | 3 | ENST00000434230 | -10.5 | 1.07E-02 | 2.70E-01 |
| LINC00339 | 1:22030280-22031223 | 1:22030692-22030693 | IR (overlapping | 3 | 3 | ENST00000404210 | -10.5 | 2.17E-02 | 3.88E-02 |
| SIRT6 | 19:4175112-4175679 | 19:4175112-4175151 | A3ss | 3 | 3 | ENST00000337491 | -10.51 | 2.86E-02 | 1.89E-01 |
| ULK4 | 3:41935941-41938097 | 3:41937229-41937340 | SES | 3 | 2 | ENST00000453224 | -10.52 | 4.28E-02 | 1.02E-01 |
| CDK5RAP1 | 20:33396937-33401427 | 20:33396937-33397084 | TSSIA3ss | 3 | 3 | TSS.ENST00000339269 | -10.53 | 1.27E-02 | 2.40E-02 |
| CDK5RAP1 | 20:33396853-33401427 | 20:33396853-33397084 | A3ss | 3 | 3 | Ex.TSS.ENST00000339269 | -10.53 | 1.27E-02 | 2.40E-02 |
| SLC43A3 | 11:57426317-57426587 | 11:57426317-57426352 | A3ss | 3 | 3 | ENST00000352187 | -10.54 | 5.18E-03 | 7.64E-03 |
| AC093323.1 | 4:6673969-6676013 | 4:6674638-6674992 | IR | 3 | 3 | ENST00000635031 | -10.54 | 8.19E-03 | 2.30E-02 |
| RABGAP1L | 1:174874507-174957456 | 1:174877392-174877605 | SES | 3 | 3 | ENST00000465412 | -10.55 | 1.44E-02 | 2.52E-02 |
| U2SURP | 3:143021383-143021472 | 3:143021383-143021385 | A5ss | 3 | 3 | ENST00000461591 | -10.55 | 3.64E-02 | 8.40E-02 |
| AL669830.2 | 6:30913811-30915037 | 6:30914470-30914804 | IR (overlapping | 3 | 3 | ENST00000477288 | -10.56 | 2.56E-02 | 7.97E-02 |
| ZNF589 | 3:48260940-48267917 | 3:48267915-48267917 | TSSIA3ss | 3 | 3 | TSS.ENST00000354698 | -10.57 | 4.03E-02 | 9.54E-02 |

| | | | | | | | | | |
|---|---|---|---|---|---|---|---|---|---|
| ZNF544 | 19:58246795-58261555 | 19:58260851-58261555 | A3ss | 3 | 3 | Ex.TSS.ENST00000269829 | -10.58 | 1.45E-02 | 1.07E-01 |
| TMEM127 | 2:96265490-96265868 | 2:96265490-96265512 | TSSlA3ss | 3 | 3 | TSS.ENST00000258439 | -10.6 | 2.88E-02 | 6.53E-02 |
| PPOX | 1:161167235-161170619 | 1:161170409-161170519 | MES | 3 | 3 | ENST00000539753 | -10.6 | 4.72E-02 | 8.21E-02 |
| CCDC78 | 16:724911-725145 | 16:724990-725077 | IR | 3 | 3 | ENST00000463539 | -10.62 | 2.60E-02 | 1.00E-01 |
| MAP3K9 | 14:70738399-70740041 | 14:70739898-70740041 | A5ss | 3 | 3 | ENST00000553414 | -10.63 | 3.42E-02 | 1.08E-01 |
| ZNF692 | 1:248857975-248858385 | 1:248858018-248858130 | IR | 3 | 3 | ENST00000533976 | -10.65 | 2.04E-02 | 3.68E-02 |
| CSNK1D | 17:82244832-82248874 | 17:82245976-82246039 | SES | 3 | 3 | ENST00000269361 | -10.66 | 9.43E-03 | 4.62E-02 |
| MBD1 | 18:50268846-50271540 | 18:50269819-50271468 | IR | 3 | 3 | ENST00000591416 | -10.67 | 1.05E-02 | 5.26E-02 |
| RMND5B | 5:178131025-178131376 | 5:178131055-178131236 | IR | 3 | 3 | ENST00000507457 | -10.68 | 7.49E-03 | 2.23E-02 |
| ANGEL1 | 14:76803913-76806415 | 14:76804375-76804415 | SES | 3 | 3 | ENST00000555079 | -10.69 | 1.07E-02 | 3.41E-02 |
| MRPL20-AS1 | 1:1399530-1400303 | 1:1399690-1400157 | IR | 3 | 3 | ENST00000418833 | -10.7 | 9.95E-03 | 8.09E-02 |
| MRPL20-AS1 | 1:1399530-1400303 | 1:1399686-1400157 | IR | 3 | 3 | ENST00000418833 | -10.7 | 9.95E-03 | 8.09E-02 |
| DROSHA | 5:31466282-31467962 | 5:31467939-31467962 | A5ss | 3 | 3 | Ex.ENST00000344624 | -10.7 | 1.84E-02 | 5.50E-02 |
| GMPR2 | 14:24232901-24233340 | 14:24232978-24233218 | IR | 3 | 3 | ENST00000420554 | -10.72 | 3.18E-03 | 5.40E-03 |
| KBTBD2 | 7:32875158-32879700 | 7:32879435-32879700 | A5ss | 3 | 3 | ENST00000304056 | -10.73 | 9.62E-03 | 1.30E-01 |
| ZNF43 | 19:21851028-21851904 | 19:21851028-21851162 | TSSlA3ss | 3 | 3 | TSS.ENST00000594312 | -10.74 | 8.42E-06 | 4.63E-05 |
| CHST8 | 19:33689392-33771412 | 19:33752518-33771413 | A3ss | 3 | 3 | Ex.ENST00000262622 | -10.75 | 4.13E-03 | 2.36E-02 |
| OGT | X:71538073-71547906 | X:71544567-71544635 | SES | 3 | 3 | ENST00000373701 | -10.77 | 1.76E-02 | 1.00E+00 |
| ACTN1 | 14:68877241-68879961 | 14:68878989-68879069 | MXS | 3 | 3 | ENST00000193403 | -10.78 | 9.10E-04 | 2.89E-03 |
| ACBD4 | 17:45136500-45136776 | 17:45136621-45136691 | IR | 3 | 3 | ENST00000589752 | -10.78 | 4.66E-02 | 1.99E-01 |
| OGT | X:71538073-71547906 | X:71544567-71547906 | A3ss | 3 | 3 | Ex.ENST00000488174 | -10.81 | 1.04E-02 | 1.00E+00 |
| AC064802.1 | 8:114284515-114295465 | 8:114284515-114284533 | TSSlA5ss | 3 | 3 | TSS.ENST00000507929 | -10.81 | 1.62E-02 | 3.05E-01 |
| CIART | 1:150282554-150283633 | 1:150282824-150283461 | IR | 3 | 3 | ENST00000290363 | -10.82 | 5.41E-03 | 4.66E-02 |
| PUM2 | 2:20263461-20282946 | 2:20278583-20278819 | SES | 3 | 3 | ENST00000338086 | -10.82 | 4.04E-02 | 8.58E-02 |
| SIRT7 | 17:81914087-81914504 | 17:81914168-81914293 | IR | 3 | 3 | ENST00000572976 | -10.83 | 9.42E-03 | 4.60E-02 |
| NFRKB | 11:129892871-129895495 | 11:129894359-129894438 | MES | 3 | 3 | ENST00000526884 | -10.86 | 4.25E-02 | 6.15E-02 |
| DOC2A | 16:30007031-30009105 | 16:30007300-30008995 | IR | 3 | 3 | ENST00000564233 | -10.87 | 1.07E-03 | 3.90E-03 |
| HUWE1 | X:53542540-53543885 | X:53543841-53543885 | A5ss | 3 | 3 | ENST00000262854 | -10.87 | 4.92E-02 | 1.00E+00 |
| TMEM131L | 4:153589008-153593798 | 4:153591053-153591194 | MES | 3 | 2 | ENST00000409663 | -10.89 | 7.48E-05 | 2.00E-04 |
| TTLL5 | 14:75752956-75764614 | 14:75757831-75757869 | SES | 3 | 3 | ENST00000554510 | -10.89 | 1.27E-02 | 4.06E-02 |
| NEO1 | 15:73178428-73236346 | 15:73236287-73236346 | A3ss | 3 | 3 | ENST00000560328 | -10.91 | 3.46E-02 | 1.16E-01 |
| ECI2 | 6:4133702-4135510 | 6:4133702-4133711 | TSSlA3ss | 3 | 3 | TSS.ENST00000380118 | -10.93 | 2.65E-02 | 2.61E-01 |
| HPAT5 | 6:167231958-167237240 | 6:167232479-167232560 | SES | 3 | 3 | ENST00000628362 | -10.94 | 1.50E-02 | 1.33E-01 |
| MRTFA | 22:40421101-40424205 | 22:40423536-40423685 | SES | 3 | 3 | ENST00000355630 | -10.96 | 3.89E-02 | 7.99E-02 |
| CAST | 5:96727531-96729611 | 5:96729153-96729209 | SES | 3 | 3 | ENST00000309190 | -10.97 | 1.22E-02 | 3.73E-02 |
| CAPN1 | 11:65182462-65182968 | 11:65182563-65182700 | IR | 3 | 3 | ENST00000527323 | -10.98 | 1.97E-03 | 2.99E-03 |
| IDH3B | 20:2658523-2659524 | 20:2658523-2658837 | TSSlA3ss | 3 | 3 | TSS.ENST00000488299 | -10.98 | 7.69E-03 | 1.46E-02 |
| UNKL | 16:1414010-1414614 | 16:1414010-1414055 | TSSlA3ss | 3 | 2 | TSS.ENST00000301712 | -10.99 | 2.08E-02 | 7.33E-02 |
| AP3M2 | 8:42153106-42157940 | 8:42154616-42154960 | SES | 3 | 3 | ENST00000396926 | -11.01 | 1.05E-02 | 2.32E-01 |
| SLC2A8 | 9:127400007-127403659 | 9:127402557-127402753 | SES | 3 | 3 | ENST00000373360 | -11.01 | 3.35E-02 | 1.00E+00 |
| PPP4R3A | 14:91509506-91510480 | 14:91509705-91510074 | IR | 3 | 3 | ENST00000554390 | -11.02 | 2.22E-03 | 7.12E-03 |
| RECQL5 | 17:75666428-75667088 | 17:75666572-75667043 | IR | 3 | 3 | ENST00000578201 | -11.02 | 1.91E-02 | 8.80E-02 |
| SLC35B1 | 17:49705897-49706964 | 17:49706204-49706334 | SES | 3 | 3 | ENST00000240333 | -11.03 | 6.98E-03 | 3.05E-02 |
| SETD6 | 16:58516929-58518400 | 16:58518051-58518231 | SES | 3 | 3 | ENST00000219315 | -11.03 | 3.19E-02 | 1.20E-01 |
| CDC25A | 3:48187778-48188417 | 3:48188216-48188369 | IR | 3 | 3 | ENST00000302506 | -11.06 | 4.65E-02 | 1.10E-01 |
| COBL | 7:51017569-51026545 | 7:51025109-51025231 | SES | 3 | 3 | ENST00000395542 | -11.09 | 3.92E-02 | 5.47E-01 |
| ZNF197 | 3:44632600-44646490 | 3:44641900-44646490 | TSSlA3ss | 3 | 3 | TSS.ENST00000344387 | -11.1 | 9.67E-03 | 2.32E-02 |
| NDRG4 | 16:58500270-58503797 | 16:58500988-58501077 | MES | 3 | 3 | ENST00000568464 | -11.1 | 2.39E-02 | 8.99E-02 |
| TAZ | X:154413568-154420031 | X:154419543-154419746 | SES | 3 | 3 | ENST00000483674 | -11.11 | 3.63E-02 | 1.00E+00 |
| SLC12A9 | 7:100854379-100858834 | 7:100854379-100855837 | A5ss | 3 | 3 | Ex.ENST00000416675 | -11.12 | 8.45E-03 | 9.77E-02 |
| FAM204A | 10:118336424-118342269 | 10:118341727-118341927 | SES | 3 | 3 | ENST00000369183 | -11.12 | 1.85E-02 | 2.59E-02 |
| PIGT | 20:45420694-45424215 | 20:45421383-45421583 | SES | 3 | 3 | ENST00000279035 | -11.13 | 3.97E-03 | 7.60E-03 |
| LRRC20 | 10:70324031-70376451 | 10:70340553-70340702 | SES | 3 | 3 | ENST00000355790 | -11.13 | 4.51E-02 | 6.43E-02 |
| CHID1 | 11:867860-869956 | 11:868672-869857 | IR | 3 | 3 | ENST00000449825 | -11.13 | 4.88E-02 | 7.52E-02 |
| ENTPD1-AS1 | 10:95897680-95907662 | 10:95904830-95904938 | SES | 3 | 3 | ENST00000661794 | -11.18 | 3.36E-02 | 4.77E-02 |
| NCAPG | 4:17840210-17842309 | 4:17840607-17840693 | SES | 3 | 3 | ENST00000251496 | -11.19 | 1.65E-02 | 4.43E-02 |
| CNST | 1:246591942-246631893 | 1:246621429-246621634 | SES | 3 | 3 | ENST00000366512 | -11.19 | 2.08E-02 | 3.76E-02 |
| ZNF638 | 2:71331866-71355718 | 2:71348427-71350271 | SES | 3 | 3 | ENST00000409544 | -11.19 | 2.62E-02 | 5.87E-02 |
| SNHG14 | 15:25041974-25042515 | 15:25042201-25042475 | IR | 3 | 3 | ENST00000656959 | -11.26 | 6.76E-04 | 2.20E-03 |
| RNF220 | 1:44635589-44636029 | 1:44635589-44635666 | A5ss | 3 | 3 | ENST00000335497 | -11.28 | 3.41E-03 | 6.35E-03 |
| SYT6 | 1:114103722-114137494 | 1:114103722-114137065 | A3ss | 3 | 3 | Ex.ENST00000369547 | -11.29 | 3.68E-03 | 3.09E-02 |
| MED6 | 14:70583589-70584943 | 14:70584131-70584816 | IR | 3 | 3 | ENST00000256379 | -11.29 | 2.78E-02 | 8.75E-02 |
| ZNF513 | 2:27378616-27378883 | 2:27378616-27378777 | A3ss | 3 | 3 | Ex.TSS.ENST00000491924 | -11.29 | 4.24E-02 | 9.29E-02 |
| DSN1 | 20:36771194-36773661 | 20:36771194-36771473 | TSSlA3ss | 3 | 3 | TSS.ENST00000480153 | -11.33 | 2.53E-02 | 4.81E-02 |
| MIER1 | 1:66984572-66988030 | 1:66984593-66986402 | IR | 3 | 3 | ENST00000355356 | -11.35 | 4.26E-02 | 7.97E-02 |
| JMJD7 | 15:41834894-41835223 | 15:41834894-41834969 | IR (overlapping | 3 | 3 | ENST00000408047 | -11.35 | 4.39E-02 | 1.45E-01 |
| ZBTB44 | 11:130238557-130239811 | 11:130238557-130238607 | A3ss | 3 | 3 | ENST00000357899 | -11.38 | 2.35E-02 | 3.41E-02 |
| ZBTB44 | 11:130238554-130239811 | 11:130238554-130238607 | A3ss | 3 | 3 | Ex.ENST00000357899 | -11.38 | 2.35E-02 | 3.41E-02 |
| PDPN | 1:13614406-13615904 | 1:13614406-13614411 | TSSlA5ss | 3 | 3 | TSS.ENST00000294489 | -11.41 | 1.19E-02 | 1.01E-01 |
| EHMT2 | 6:31888748-31889227 | 6:31888969-31889070 | SES | 3 | 3 | ENST00000375537 | -11.42 | 9.16E-03 | 2.36E-02 |
| MLXIPL | 7:73593202-73594403 | 7:73593984-73594273 | IR | 2 | 2 | ENST00000434326 | -11.44 | 8.15E-04 | 1.20E-02 |
| BAG4 | 8:38176588-38177139 | 8:38176590-38177004 | IR (overlapping | 3 | 3 | ENST00000432471 | -11.44 | 6.86E-03 | 1.50E-01 |
| AAR2 | 20:36236783-36239820 | 20:36236783-36237037 | A5ss | 3 | 3 | Ex.TSS.ENST00000373932 | -11.45 | 2.95E-02 | 5.63E-02 |
| AAR2 | 20:36236504-36239820 | 20:36236504-36237037 | A5ss | 3 | 3 | Ex.TSS.ENST00000320849 | -11.45 | 2.95E-02 | 5.63E-02 |
| GLE1 | 9:128504905-128508875 | 9:128504905-128507340 | A5ss | 3 | 3 | Ex.TSS.ENST00000372770 | -11.48 | 2.54E-03 | 9.05E-02 |
| PLXNB2 | 22:50290598-50307552 | 22:50294719-50294778 | SES | 3 | 3 | ENST00000359337 | -11.48 | 7.07E-03 | 1.46E-02 |
| PIGQ | 16:575839-577217 | 16:575971-576133 | IR (overlapping | 3 | 3 | ENST00000470411 | -11.49 | 2.16E-02 | 8.14E-02 |
| CHN1 | 2:174824519-174915057 | 2:174877840-174878128 | MXS | 3 | 3 | ENST00000409156 | -11.49 | 2.82E-02 | 5.93E-02 |
| AJUBA | 14:22971177-22973568 | 14:22972181-22972924 | IR (overlapping | 3 | 3 | ENST00000262713 | -11.5 | 1.27E-02 | 2.16E-02 |
| ZNF692 | 1:248857975-248858385 | 1:248858035-248858130 | IR | 3 | 3 | ENST00000533976 | -11.51 | 3.05E-03 | 5.50E-03 |
| KLC1 | 14:103677524-103679407 | 14:103679309-103679407 | A3ss | 3 | 3 | Ex.ENST00000380038 | -11.55 | 5.88E-04 | 9.52E-04 |
| KLC1 | 14:103677524-103679383 | 14:103679309-103679383 | A3ss | 3 | 3 | ENST00000380038 | -11.55 | 5.88E-04 | 9.52E-04 |
| TRIM16 | 17:15683036-15684299 | 17:15683158-15684195 | IR | 3 | 3 | ENST00000416464 | -11.56 | 3.57E-02 | 1.40E-01 |
| AMT | 3:49421573-49422360 | 3:49422104-49422257 | MXS | 3 | 3 | ENST00000427987 | -11.57 | 2.16E-02 | 5.22E-02 |
| CAPN1 | 11:65182462-65182968 | 11:65182580-65182700 | IR | 3 | 3 | ENST00000527323 | -11.58 | 1.40E-03 | 2.12E-03 |
| GSDMD | 8:143559777-143561423 | 8:143560772-143561001 | IR | 3 | 3 | ENST00000524846 | -11.58 | 1.93E-02 | 3.84E-01 |
| TELO2 | 16:1493606-1494280 | 16:1494246-1494280 | A3ss | 3 | 3 | Ex.TSS.ENST00000262319 | -11.58 | 4.64E-02 | 1.63E-01 |
| ZFYVE21 | 14:103729183-103732619 | 14:103729793-103729846 | SES | 3 | 3 | ENST00000216602 | -11.61 | 2.87E-03 | 4.64E-03 |
| ALPK3 | 15:84827606-84856391 | 15:84838980-84839097 | MES | 3 | 3 | ENST00000258888 | -11.62 | 2.86E-02 | 9.88E-02 |
| ICMT | 1:6232120-6234885 | 1:6233474-6233643 | MXS | 3 | 3 | ENST00000343813 | -11.63 | 5.31E-03 | 9.98E-03 |
| DUSP23 | 1:159780962-159781367 | 1:159780994-159781098 | IR | 3 | 3 | ENST00000368107 | -11.63 | 8.70E-03 | 1.51E-02 |
| LARGE2 | 11:45922778-45923167 | 11:45922965-45923057 | IR | 3 | 3 | ENST00000325468 | -11.65 | 3.73E-02 | 5.49E-02 |
| RAB6B | 3:133824235-133828852 | 3:133827346-133827427 | IR | 3 | 3 | ENST00000285208 | -11.67 | 2.44E-02 | 5.60E-02 |
| ALG3 | 3:184245813-184248781 | 3:184248745-184248781 | TSSlA5ss | 3 | 3 | TSS.ENST00000397676 | -11.67 | 3.92E-02 | 9.23E-02 |

| | | | | | | | | | |
|---|---|---|---|---|---|---|---|---|---|
| CEP131 | 17:81220074-81222768 | 17:81222646-81222769 | A5ss | 3 | 3 | Ex.TSS.ENST00000269392 | -11.68 | 9.12E-03 | 4.43E-02 |
| APIP | 11:34890553-34916227 | 11:34895010-34895110 | SES | 3 | 3 | ENST00000395787 | -11.68 | 4.44E-02 | 6.52E-02 |
| CREBZF | 11:85661066-85665102 | 11:85662458-85663523 | IR | 3 | 3 | ENST00000527447 | -11.69 | 1.94E-02 | 3.00E-02 |
| PARD3 | 10:34336244-34337364 | 10:34337275-34337364 | A5ss | 3 | 3 | ENST00000340077 | -11.7 | 1.61E-02 | 2.27E-02 |
| PARD3 | 10:34336244-34337277 | 10:34337275-34337277 | A5ss | 3 | 3 | ENST00000340077 | -11.7 | 1.61E-02 | 2.27E-02 |
| STYXL1 | 7:76030525-76047811 | 7:76047662-76047811 | TSSlA5ss | 3 | 3 | TSS.ENST00000248600 | -11.7 | 4.69E-02 | 7.01E-01 |
| SNHG14 | 15:25041974-25042515 | 15:25042126-25042475 | IR | 3 | 3 | ENST00000656959 | -11.72 | 6.53E-04 | 2.12E-03 |
| SFI1 | 22:31614371-31616877 | 22:31614861-31615047 | IR | 3 | 3 | ENST00000464333 | -11.74 | 1.13E-02 | 2.31E-02 |
| CDCA7 | 2:173358838-173363225 | 2:173359255-173359491 | SES | 3 | 3 | ENST00000306721 | -11.74 | 4.44E-02 | 9.36E-02 |
| ZFP62 | 5:180847615-180851493 | 5:180848083-180851380 | IR | 3 | 3 | ENST00000512132 | -11.76 | 3.70E-02 | 1.04E-01 |
| GOLGA2P10 | 15:82477124-82477260 | 15:82477203-82477260 | A5ss | 3 | 3 | Ex.ENST00000550961 | -11.8 | 1.69E-02 | 5.78E-02 |
| METTL26 | 16:634798-635611 | 16:635281-635340 | SES | 3 | 3 | ENST00000397666 | -11.82 | 2.02E-02 | 7.63E-02 |
| MSI2 | 17:57652162-57674971 | 17:57674918-57674971 | A3ss | 3 | 3 | ENST00000416426 | -11.82 | 2.77E-02 | 1.23E-01 |
| AC026412.1 | 5:1632644-1632906 | 5:1632752-1632844 | IR | 3 | 3 | ENST00000507290 | -11.83 | 2.03E-02 | 5.98E-02 |
| STK33 | 11:8475067-8594082 | 11:8480534-8480614 | SES | 3 | 3 | ENST00000454443 | -11.83 | 4.34E-02 | 6.70E-02 |
| ABHD14B | 3:51970185-51973964 | 3:51971460-51971892 | SES | 3 | 3 | ENST00000395008 | -11.84 | 3.47E-03 | 8.94E-03 |
| UBXN11 | 1:26294332-26306591 | 1:26296919-26296995 | MES | 3 | 2 | ENST00000357089 | -11.84 | 9.21E-03 | 1.67E-02 |
| TMEM182 | 2:102737014-102762586 | 2:102758389-102758473 | MXS | 3 | 3 | ENST00000409173 | -11.84 | 1.62E-02 | 3.36E-02 |
| NCOR1 | 17:16086443-16091862 | 17:16087172-16087345 | SES | 3 | 3 | ENST00000436068 | -11.84 | 4.43E-02 | 1.76E-01 |
| GPR161 | 1:168104895-168137389 | 1:168136010-168137390 | A5ss | 3 | 3 | Ex.TSS.ENST00000271357 | -11.86 | 3.57E-02 | 6.25E-02 |
| DUSP23 | 1:159780962-159781367 | 1:159781001-159781098 | IR | 3 | 3 | ENST00000368107 | -11.87 | 3.24E-03 | 5.63E-03 |
| TRIM16 | 17:15683036-15684299 | 17:15683372-15684195 | IR | 3 | 3 | ENST00000416464 | -11.88 | 1.17E-03 | 4.62E-03 |
| CHID1 | 11:867860-869956 | 11:868672-869213 | IR | 3 | 3 | ENST00000449825 | -11.88 | 4.37E-02 | 6.74E-02 |
| NFKBIB | 19:38907399-38908893 | 19:38907660-38908730 | IR | 3 | 3 | ENST00000509705 | -11.88 | 4.40E-02 | 2.83E-01 |
| THUMPD3 | 3:9374647-9383198 | 3:9380503-9380618 | MES | 3 | 3 | ENST00000345094 | -11.89 | 3.24E-02 | 8.34E-02 |
| EXOC7 | 17:76089321-76094413 | 17:76090329-76090397 | MES | 3 | 3 | ENST00000405575 | -11.91 | 4.59E-02 | 2.17E-01 |
| UFD1 | 22:19468004-19479082 | 22:19475470-19475602 | MES | 3 | 3 | ENST00000263202 | -11.92 | 1.95E-02 | 3.80E-02 |
| ZBTB44 | 11:130226677-130231715 | 11:130228357-130229828 | IR | 3 | 3 | ENST00000357899 | -11.92 | 4.56E-02 | 6.59E-02 |
| RECQL5 | 17:75666428-75667088 | 17:75666621-75666900 | IR | 3 | 3 | ENST00000578201 | -11.93 | 3.13E-02 | 1.45E-01 |
| CAPN1 | 11:65182462-65182968 | 11:65182542-65182700 | IR | 3 | 3 | ENST00000527323 | -11.94 | 7.78E-04 | 1.18E-03 |
| ATG4B | 2:241651336-241653511 | 2:241653307-241653369 | MXS | 3 | 3 | ENST00000344376 | -11.96 | 1.35E-02 | 2.94E-02 |
| SLC26A6 | 3:48632338-48632973 | 3:48632338-48632396 | A3ss | 3 | 3 | ENST00000307364 | -11.97 | 1.97E-03 | 4.74E-03 |
| NAGK | 2:71068713-71070740 | 2:71070502-71070740 | A3ss | 3 | 3 | ENST00000465105 | -11.97 | 7.98E-03 | 1.79E-02 |
| PLEKHA4 | 19:48838130-48845369 | 19:48841149-48841310 | SES | 3 | 3 | ENST00000597129 | -11.98 | 1.56E-02 | 1.07E-01 |
| ZSCAN2 | 15:84601136-84603819 | 15:84601136-84601188 | TSSlA5ss | 3 | 3 | TSS.ENST00000448803 | -11.99 | 1.35E-02 | 4.68E-02 |
| B3GAT1 | 11:134387548-134387940 | 11:134387695-134387826 | IR | 3 | 3 | ENST00000312527 | -12 | 2.32E-02 | 3.36E-02 |
| SEMA4C | 2:96867924-96869875 | 2:96867924-96867926 | A3ss | 3 | 3 | Ex.TSS.ENST00000305476 | -12 | 3.15E-02 | 7.03E-02 |
| DSN1 | 20:36771194-36773661 | 20:36771425-36771473 | SES | 3 | 3 | ENST00000373750 | -12.01 | 4.35E-03 | 8.28E-03 |
| PDCD4 | 10:110876071-110881274 | 10:110881233-110881274 | A3ss | 3 | 3 | ENST00000280154 | -12.02 | 1.80E-02 | 2.53E-02 |
| ZNF286A | 17:15700135-15700677 | 17:15700169-15700272 | IR (overlapping | 3 | 3 | ENST00000579516 | -12.05 | 2.87E-02 | 1.13E-01 |
| FOXH1 | 8:144475224-144475582 | 8:144475224-144475261 | TSSlA3ss | 3 | 3 | TSS.ENST00000377317 | -12.06 | 5.92E-03 | 1.21E-01 |
| MRPS12 | 19:38930999-38931275 | 19:38930999-38931171 | TSSlA5ss | 3 | 3 | TSS.ENST00000402029 | -12.06 | 8.17E-03 | 5.34E-02 |
| SAC3D1 | 11:65040970-65041928 | 11:65041107-65041285 | IR | 3 | 3 | ENST00000398846 | -12.06 | 4.28E-02 | 6.43E-02 |
| WIPI2 | 7:5190494-5214534 | 7:5193118-5193171 | MES | 3 | 3 | ENST00000288828 | -12.07 | 2.56E-02 | 3.58E-01 |
| CNKSR1 | 1:26183415-26183728 | 1:26183415-26183435 | A5ss | 3 | 3 | ENST00000374253 | -12.07 | 4.97E-02 | 8.98E-02 |
| XIAP | X:123860294-123888618 | X:123885631-123886539 | SES | 3 | 3 | ENST00000371199 | -12.12 | 2.33E-02 | 1.00E+00 |
| FAM126A | 7:22946164-22947032 | 7:22946952-22947032 | TSSlA5ss | 3 | 3 | TSS.ENST00000409923 | -12.12 | 4.60E-02 | 5.93E-01 |
| CNOT2 | 12:70242994-70243480 | 12:70243238-70243332 | IR (overlapping | 3 | 3 | ENST00000229195 | -12.12 | 4.83E-02 | 8.13E-02 |
| FBXL6 | 8:144356808-144357973 | 8:144357122-144357627 | IR (overlapping | 3 | 3 | ENST00000524492 | -12.14 | 2.92E-03 | 5.95E-02 |
| B3GAT1 | 11:134387548-134387940 | 11:134387695-134387843 | IR | 3 | 3 | ENST00000312527 | -12.15 | 1.37E-02 | 1.98E-02 |
| TLCD5 | 11:120325369-120329976 | 11:120327122-120327640 | SES | 3 | 3 | ENST00000314475 | -12.15 | 4.77E-02 | 6.88E-02 |
| AC093323.1 | 4:6673969-6676013 | 4:6674707-6674992 | IR | 3 | 3 | ENST00000635031 | -12.16 | 1.27E-03 | 3.56E-03 |
| PLAUR | 19:43667692-43668043 | 19:43667777-43668043 | TSSlA5ss | 3 | 3 | TSS.ENST00000593939 | -12.16 | 3.46E-03 | 2.28E-02 |
| PLAUR | 19:43667692-43667778 | 19:43667777-43667778 | TSSlA5ss | 3 | 3 | TSS.ENST00000593939 | -12.16 | 3.46E-03 | 2.28E-02 |
| RBPJ | 4:26320816-26386352 | 4:26320816-26321048 | TSSlA5ss | 3 | 3 | TSS.ENST00000348160 | -12.16 | 4.14E-02 | 1.12E-01 |
| ZDHHC15 | X:75478986-75505820 | X:75478986-75502268 | A3ss | 3 | 3 | Ex.ENST00000373367 | -12.17 | 2.05E-02 | 1.00E+00 |
| GNB1L | 22:19812448-19854442 | 22:19821228-19821375 | MXS | 3 | 3 | ENST00000453108 | -12.18 | 9.39E-03 | 1.84E-02 |
| NBR2 | 17:43138657-43140283 | 17:43138923-43139815 | IR (overlapping | 3 | 3 | ENST00000657841 | -12.18 | 3.13E-02 | 1.34E-01 |
| CDR2 | 16:22349850-22374230 | 16:22364902-22365014 | SES | 3 | 3 | ENST00000268383 | -12.18 | 4.84E-02 | 1.74E-01 |
| MRPL20-AS1 | 1:1399530-1400303 | 1:1399690-1399980 | IR | 3 | 3 | ENST00000418833 | -12.2 | 2.05E-03 | 1.74E-02 |
| ABCA2 | 9:137017562-137018075 | 9:137017693-137017786 | IR | 3 | 3 | ENST00000466707 | -12.2 | 1.83E-02 | 7.55E-01 |
| SPIN3 | X:56990831-56994949 | X:56992238-56994722 | IR | 3 | 3 | ENST00000374919 | -12.25 | 7.17E-03 | 1.00E+00 |
| TMEM132A | 11:60930660-60931688 | 11:60931686-60931688 | A3ss | 3 | 3 | ENST00000005286 | -12.26 | 3.70E-02 | 5.47E-02 |
| CAP2 | 6:17426691-17507168 | 6:17462996-17463073 | SES | 3 | 3 | ENST00000229922 | -12.28 | 4.26E-03 | 3.79E-02 |
| SYVN1 | 11:65130173-65130250 | 11:65130173-65130175 | A3ss | 2 | 2 | ENST00000377190 | -12.29 | 1.70E-03 | 2.57E-03 |
| PSMC3IP | 17:42577461-42577697 | 17:42577562-42577652 | IR | 3 | 3 | ENST00000587209 | -12.31 | 1.93E-02 | 8.17E-02 |
| ECHDC2 | 1:52911791-52921552 | 1:52913895-52914095 | MES | 3 | 3 | ENST00000476477 | -12.32 | 2.72E-02 | 5.08E-02 |
| RAP1GDS1 | 4:98352602-98391951 | 4:98379017-98379163 | MXS | 3 | 3 | ENST00000339360 | -12.34 | 3.32E-03 | 9.44E-03 |
| PPM1N | 19:45500562-45500680 | 19:45500650-45500680 | A3ss | 3 | 3 | ENST00000396735 | -12.36 | 1.75E-02 | 1.17E-01 |
| FLII | 17:18251814-18251998 | 17:18251814-18251816 | A3ss | 3 | 3 | ENST00000327031 | -12.4 | 2.12E-02 | 7.46E-02 |
| BRD1 | 22:49787390-49787887 | 22:49787495-49787576 | IR | 3 | 3 | ENST00000404760 | -12.41 | 4.00E-03 | 8.29E-03 |
| ZNF275 | X:153334286-153345519 | X:153336634-153336710 | SES | 3 | 3 | ENST00000370251 | -12.43 | 2.78E-03 | 1.84E-01 |
| DPP3 | 11:66480466-66482195 | 11:66482193-66482195 | TSSlA3ss | 3 | 3 | TSS.ENST00000526515 | -12.47 | 2.82E-02 | 4.28E-02 |
| CUTA | 6:33417481-33417921 | 6:33417696-33417870 | IR | 3 | 3 | ENST00000440279 | -12.53 | 2.23E-03 | 2.07E-02 |
| CUTA | 6:33417481-33417921 | 6:33417696-33417868 | IR | 3 | 3 | ENST00000440279 | -12.53 | 2.23E-03 | 2.07E-02 |
| IL18BP | 11:71999063-72000012 | 11:71999371-71999926 | IR | 3 | 3 | ENST00000404792 | -12.53 | 6.59E-03 | 1.01E-02 |
| PRSS16 | 6:27252041-27254985 | 6:27252808-27252949 | MXS | 3 | 3 | ENST00000481125 | -12.54 | 2.54E-03 | 2.28E-02 |
| ZNF780A | 19:40084799-40090905 | 19:40090166-40090235 | MES | 2 | 3 | ENST00000414720 | -12.54 | 1.07E-02 | 6.98E-02 |
| ALKBH4 | 7:102459711-102464713 | 7:102459711-102459801 | TSSlA3ss | 3 | 3 | TSS.ENST00000292566 | -12.54 | 1.08E-02 | 1.26E-01 |
| EMC9 | 14:24141107-24141585 | 14:24141317-24141437 | IR | 3 | 3 | ENST00000419198 | -12.54 | 1.45E-02 | 2.34E-02 |
| MAP3K10 | 19:40198556-40205120 | 19:40204485-40204633 | SES | 3 | 3 | ENST00000253055 | -12.54 | 3.86E-02 | 2.52E-01 |
| ANKRD11 | 16:89286039-89288670 | 16:89287114-89288527 | IR | 3 | 3 | ENST00000645212 | -12.55 | 7.20E-03 | 2.78E-02 |
| PTP4A3 | 8:141392085-141421387 | 8:141392085-141418018 | A5ss | 3 | 3 | Ex.TSS.ENST00000520105 | -12.55 | 2.06E-02 | 4.01E-01 |
| MAPK7 | 17:19378529-19382466 | 17:19379948-19380483 | IR | 3 | 3 | ENST00000570306 | -12.56 | 1.14E-03 | 4.51E-03 |
| TRIM26 | 6:30199154-30213304 | 6:30201035-30201138 | MES | 3 | 3 | ENST00000416596 | -12.56 | 1.79E-02 | 1.60E-01 |
| PHKA2 | X:18897334-18901484 | X:18900670-18900699 | MES | 3 | 3 | ENST00000379942 | -12.58 | 2.29E-02 | 1.00E+00 |
| TSPAN9 | 12:3083720-3278420 | 12:3201177-3201256 | MES | 3 | 3 | ENST00000444315 | -12.59 | 4.33E-03 | 6.94E-03 |
| PREPL | 2:44359497-44360674 | 2:44359751-44360618 | IR | 3 | 3 | ENST00000260648 | -12.6 | 4.59E-02 | 1.01E-01 |
| ADAL | 15:43333435-43335695 | 15:43334945-43335170 | SES | 3 | 3 | ENST00000389651 | -12.62 | 2.41E-02 | 7.96E-02 |
| MPPE1 | 18:11886622-11886916 | 18:11886622-11886778 | A3ss | 3 | 3 | ENST00000592894 | -12.62 | 2.98E-02 | 1.47E-01 |
| ZNF577 | 19:51873530-51877281 | 19:51873530-51873706 | TSSlA3ss | 3 | 3 | TSS.ENST00000301399 | -12.63 | 3.87E-02 | 2.71E-01 |
| PKP3 | 11:394209-394524 | 11:394335-394508 | IR | 3 | 3 | ENST00000331563 | -12.64 | 1.46E-02 | 2.14E-02 |
| ZNF142 | 2:218658980-218659539 | 2:218659131-218659368 | IR (overlapping | 3 | 3 | ENST00000432460 | -12.64 | 4.05E-02 | 8.69E-02 |
| CLCC1 | 1:108950439-108963360 | 1:108950439-108950448 | A3ss | 3 | 3 | Ex.TSS.ENST00000356970 | -12.69 | 2.34E-02 | 1.92E-01 |

| | | | | | | | | | |
|---|---|---|---|---|---|---|---|---|---|
| PACRGL | 4:20700438-20700706 | 4:20700579-20700670 | IR | 3 | 3 | ENST00000508753 | -12.71 | 2.39E-02 | 6.45E-02 |
| PHKA2 | X:18897334-18901484 | X:18899173-18899226 | MES | 3 | 3 | ENST00000379942 | -12.72 | 1.01E-02 | 8.08E-01 |
| WDR60 | 7:158902596-158905988 | 7:158902596-158905283 | A5ss | 3 | 3 | ENST00000467220 | -12.72 | 2.01E-02 | 2.57E-01 |
| REPIN1 | 7:150368942-150371227 | 7:150369671-150369868 | MXS | 3 | 3 | ENST00000489432 | -12.73 | 5.44E-04 | 6.76E-03 |
| REPIN1 | 7:150368942-150371227 | 7:150369671-150369868 | MES | 3 | 3 | ENST00000397281 | -12.74 | 1.61E-04 | 2.00E-03 |
| LNP1 | 3:100429886-100455776 | 3:100451736-100451949 | SES | 3 | 3 | ENST00000466996 | -12.74 | 2.17E-03 | 4.94E-03 |
| LNP1 | 3:100429886-100455776 | 3:100451680-100451949 | SES | 3 | 3 | ENST00000489752 | -12.74 | 2.17E-03 | 4.94E-03 |
| PARP2 | 14:20354248-20354834 | 14:20354809-20354834 | A3ss | 3 | 3 | ENST00000250416 | -12.74 | 8.72E-03 | 1.41E-02 |
| MAPK7 | 17:19378529-19382466 | 17:19379948-19380394 | IR | 3 | 3 | ENST00000570306 | -12.75 | 1.24E-02 | 4.37E-02 |
| MCM3AP | 21:46285395-46286150 | 21:46285699-46286150 | A5ss | 3 | 3 | ENST00000426537 | -12.75 | 4.23E-02 | 8.22E-02 |
| MCM3AP | 21:46285395-46285802 | 21:46285699-46285802 | A5ss | 3 | 3 | ENST00000426537 | -12.75 | 4.23E-02 | 8.22E-02 |
| TTF1 | 9:132400259-132406789 | 9:132401455-132402828 | SES | 3 | 3 | ENST00000334270 | -12.77 | 8.43E-04 | 3.34E-02 |
| PAXX | 9:136992442-136992974 | 9:136992701-136992924 | IR | 3 | 3 | ENST00000481187 | -12.84 | 4.50E-03 | 1.93E-01 |
| UBAP2L | 1:154220976-154225083 | 1:154220976-154221040 | TSSIA5ss | 3 | 3 | TSS.ENST00000412596 | -12.84 | 2.77E-02 | 4.78E-02 |
| TMEM63A | 1:225859350-225860947 | 1:225860860-225860947 | A5ss | 3 | 3 | Ex.ENST00000366835 | -12.84 | 3.22E-02 | 4.49E-02 |
| TAF1C | 16:84180170-84181186 | 16:84180345-84180867 | IR | 3 | 3 | ENST00000564774 | -12.84 | 4.46E-02 | 1.72E-01 |
| RSRP1 | 1:25242249-25242705 | 1:25242427-25242538 | IR | 3 | 3 | ENST00000498238 | -12.87 | 5.60E-03 | 1.01E-02 |
| PGAP2 | 11:3808316-3824269 | 11:3823883-3824135 | MES | 3 | 3 | ENST00000396991 | -12.9 | 3.78E-02 | 5.55E-02 |
| GGT1 | 22:24608024-24610357 | 22:24609962-24610033 | SES | 3 | 3 | ENST00000411974 | -12.94 | 2.34E-02 | 4.54E-02 |
| PPP1R35 | 7:100435631-100436136 | 7:100435768-100435847 | IR (overlapping | 3 | 3 | ENST00000470295 | -12.95 | 9.56E-04 | 1.10E-02 |
| UBTF | 17:44212007-44212818 | 17:44212344-44212454 | SES | 3 | 3 | ENST00000302904 | -12.95 | 3.55E-03 | 1.51E-02 |
| SLC17A9 | 20:62962624-62963369 | 20:62962735-62963272 | IR | 3 | 3 | ENST00000488738 | -12.95 | 4.84E-02 | 6.73E-02 |
| PAXX | 9:136992442-136992974 | 9:136992705-136992924 | IR | 3 | 3 | ENST00000481187 | -12.96 | 3.84E-03 | 1.64E-01 |
| GALT | 9:34647541-34647849 | 9:34647568-34647656 | IR (overlapping | 3 | 3 | ENST00000554330 | -12.97 | 2.56E-03 | 7.04E-02 |
| CNOT2 | 12:70242994-70243480 | 12:70243205-70243311 | IR (overlapping | 3 | 3 | ENST00000229195 | -12.97 | 2.97E-02 | 5.00E-02 |
| PIGO | 9:35091240-35092767 | 9:35091292-35092542 | IR | 3 | 3 | ENST00000378617 | -12.98 | 2.43E-03 | 7.28E-02 |
| PPOX | 1:161167235-161168992 | 1:161168432-161168576 | MXS | 3 | 3 | ENST00000652103 | -13 | 3.48E-02 | 6.06E-02 |
| ATP6V1H | 8:53801897-53814661 | 8:53811164-53811217 | MES | 3 | 3 | ENST00000522849 | -13 | 4.89E-02 | 8.99E-01 |
| CPSF7 | 11:61416520-61419948 | 11:61419922-61419948 | A5ss | 3 | 3 | ENST00000340437 | -13.02 | 4.39E-03 | 6.50E-03 |
| CDC42BPA | 1:226989866-226994399 | 1:226992611-226993069 | IR (overlapping | 3 | 3 | ENST00000366764 | -13.02 | 2.15E-02 | 3.68E-02 |
| TMEM164 | X:110003157-110003882 | X:110003501-110003882 | A3ss | 3 | 3 | Ex.TSS.ENST00000372073 | -13.02 | 3.46E-02 | 1.00E+00 |
| CHID1 | 11:867860-869956 | 11:869346-869857 | IR | 3 | 3 | ENST00000449825 | -13.03 | 3.11E-04 | 4.81E-04 |
| EP400 | 12:131979794-131982092 | 12:131981489-131981596 | SES | 3 | 3 | ENST00000333577 | -13.03 | 2.24E-02 | 3.57E-02 |
| MPPE1 | 18:11886622-11886916 | 18:11886713-11886778 | SES | 3 | 3 | ENST00000317251 | -13.06 | 1.59E-02 | 7.86E-02 |
| DNAJC10 | 2:182732543-182740298 | 2:182736249-182736386 | MXS | 3 | 3 | ENST00000264065 | -13.07 | 2.35E-02 | 4.98E-02 |
| SNHG17 | 20:38421099-38426418 | 20:38422092-38422241 | MES | 3 | 3 | ENST00000654092 | -13.1 | 9.98E-03 | 1.90E-02 |
| ATG13 | 11:46617891-46618324 | 11:46618260-46618324 | TSSIA3ss | 3 | 3 | TSS.ENST00000395549 | -13.1 | 4.29E-02 | 6.32E-02 |
| SFI1 | 22:31602786-31604868 | 22:31603744-31603819 | MXS | 3 | 3 | ENST00000417682 | -13.13 | 4.99E-03 | 1.02E-02 |
| FBXO7 | 22:32475399-32479132 | 22:32478981-32479132 | TSSIA3ss | 3 | 2 | TSS.ENST00000397426 | -13.13 | 2.24E-02 | 4.58E-02 |
| SLTM | 15:58898853-58901259 | 15:58899469-58899883 | SES | 3 | 3 | ENST00000249736 | -13.17 | 3.32E-02 | 4.62E-02 |
| BAZ2A | 12:56617533-56636181 | 12:56630776-56630865 | SES | 3 | 3 | ENST00000549506 | -13.2 | 1.91E-02 | 3.18E-02 |
| PPM1N | 19:45500562-45502016 | 19:45500650-45500710 | SES | 3 | 3 | ENST00000396735 | -13.22 | 2.04E-02 | 1.36E-01 |
| CLEC16A | 16:10982992-11020192 | 16:11003074-11003305 | SES | 3 | 3 | ENST00000409790 | -13.22 | 2.25E-02 | 7.88E-02 |
| AC022140.1 | 5:25413845-25445819 | 5:25422730-25422855 | SES | 3 | 3 | ENST00000507887 | -13.22 | 3.76E-02 | 9.69E-02 |
| ACAP1 | 17:7345331-7350228 | 17:7348222-7348305 | IR | 3 | 3 | ENST00000570439 | -13.23 | 3.27E-02 | 1.50E-01 |
| AP1S2 | X:15830059-15833007 | X:15831569-15832917 | IR | 3 | 3 | ENST00000380291 | -13.24 | 8.18E-04 | 5.95E-02 |
| ANK2 | 4:113373450-113381456 | 4:113374764-113374946 | MXS | 3 | 3 | ENST00000505342 | -13.24 | 3.11E-02 | 8.13E-02 |
| RBM12B | 8:93729356-93736438 | 8:93733785-93733895 | IR (overlapping | 3 | 3 | ENST00000399300 | -13.24 | 4.35E-02 | 1.00E+00 |
| TSEN2 | 3:12503446-12503784 | 3:12503470-12503646 | IR | 3 | 3 | ENST00000490981 | -13.25 | 2.66E-02 | 5.95E-02 |
| STAG3L5P-PVRIG2P-PILRB | 7:100353484-100355142 | 7:100353484-100354012 | A5ss | 3 | 3 | ENST00000310771 | -13.26 | 2.00E-02 | 2.31E-01 |
| CSNK1D | 17:82244832-82248874 | 17:82245976-82248874 | TSSIA5ss | 3 | 3 | TSS.ENST00000581241 | -13.27 | 2.50E-02 | 1.22E-01 |
| LPCAT4 | 15:34360210-34362195 | 15:34361169-34362196 | A5ss | 3 | 3 | Ex.ENST00000567507 | -13.29 | 1.07E-03 | 3.47E-03 |
| LPCAT4 | 15:34360210-34361399 | 15:34361169-34361400 | A5ss | 3 | 3 | Ex.ENST00000314891 | -13.29 | 1.07E-03 | 3.47E-03 |
| PLD3 | 19:40348769-40366418 | 19:40365662-40365930 | MXS | 3 | 3 | ENST00000600948 | -13.33 | 2.34E-03 | 1.53E-02 |
| SSC4D | 7:76398798-76409413 | 7:76401008-76401043 | MES | 3 | 3 | ENST00000275560 | -13.36 | 4.19E-02 | 6.28E-01 |
| MFSD13A | 10:102470076-102471272 | 10:102470665-102471117 | SES | 3 | 3 | ENST00000238936 | -13.4 | 1.10E-02 | 1.53E-02 |
| ARHGEF19 | 1:16208728-16212501 | 1:16208728-16209083 | A3ss | 3 | 3 | Ex.TSS.ENST00000270747 | -13.4 | 1.30E-02 | 2.23E-02 |
| UCKL1 | 20:63946641-63956259 | 20:63946641-63946643 | A3ss | 3 | 3 | Ex.TSS.ENST00000354216 | -13.4 | 3.51E-02 | 4.89E-02 |
| FCGRT | 19:49513154-49513473 | 19:49513244-49513386 | IR | 3 | 3 | ENST00000426395 | -13.41 | 3.28E-02 | 2.27E-01 |
| NEK3 | 13:52154156-52156074 | 13:52154156-52154173 | A3ss | 3 | 3 | Ex.TSS.ENST00000550841 | -13.46 | 9.72E-03 | 1.59E-02 |
| IDH3B | 20:2658523-2659524 | 20:2658638-2658837 | SES | 3 | 3 | ENST00000466494 | -13.46 | 1.40E-02 | 2.66E-02 |
| USP21 | 1:161159679-161160619 | 1:161160366-161160506 | SES | 3 | 3 | ENST00000368002 | -13.47 | 3.09E-02 | 5.38E-02 |
| NSMCE2 | 8:125091959-125102316 | 8:125102051-125102146 | MXS | 3 | 3 | ENST00000287437 | -13.47 | 3.99E-02 | 7.62E-01 |
| NBPF3 | 1:21445220-21470631 | 1:21468688-21468897 | MXS | 3 | 3 | ENST00000318249 | -13.49 | 3.42E-02 | 6.11E-02 |
| NUBP2 | 16:1786656-1787676 | 16:1786757-1786955 | SES | 3 | 3 | ENST00000262302 | -13.55 | 2.81E-02 | 9.92E-02 |
| DOCK6 | 19:11202114-11202393 | 19:11202114-11202125 | A3ss | 3 | 3 | Ex.ENST00000294618 | -13.56 | 8.58E-04 | 4.44E-03 |
| LGMN | 14:92704362-92709671 | 14:92704640-92704707 | MES | 3 | 3 | ENST00000334869 | -13.57 | 1.74E-02 | 5.62E-02 |
| AC069281.2 | 7:100573297-100575782 | 7:100573380-100575596 | IR (overlapping | 3 | 3 | ENST00000485071 | -13.59 | 2.27E-02 | 2.62E-01 |
| SLC37A4 | 11:119027628-119030863 | 11:119029565-119030848 | IR | 3 | 3 | ENST00000532085 | -13.6 | 3.20E-03 | 4.61E-03 |
| DBF4 | 7:87878134-87884978 | 7:87878134-87878225 | A5ss | 3 | 3 | ENST00000265728 | -13.61 | 2.19E-03 | 3.35E-02 |
| UPF3A | 13:114282128-114286518 | 13:114282837-114282943 | MXS | 3 | 3 | ENST00000351487 | -13.61 | 3.99E-02 | 6.43E-02 |
| KCND1 | X:48969151-48971844 | X:48970394-48971740 | IR | 3 | 3 | ENST00000218176 | -13.62 | 2.92E-02 | 1.00E+00 |
| DNASE1L1 | X:154411891-154412112 | X:154411953-154412085 | IR (overlapping | 3 | 3 | ENST00000309585 | -13.64 | 8.48E-03 | 5.92E-01 |
| AC132008.2 | 3:197624198-197627779 | 3:197627572-197627779 | TSSIA5ss | 3 | 3 | TSS.ENST00000437008 | -13.64 | 4.12E-02 | 9.77E-02 |
| PACRGL | 4:20700748-20704465 | 4:20700748-20700787 | TSSIA5ss | 3 | 3 | TSS.ENST00000360916 | -13.65 | 1.09E-02 | 2.96E-02 |
| PACRGL | 4:20700707-20704465 | 4:20700707-20700787 | TSSIA5ss | 3 | 3 | TSS.ENST00000360916 | -13.65 | 1.09E-02 | 2.96E-02 |
| PACRGL | 4:20700618-20704465 | 4:20700618-20700787 | TSSIA5ss | 3 | 3 | TSS.ENST00000360916 | -13.65 | 1.09E-02 | 2.96E-02 |
| PACRGL | 4:20700612-20704465 | 4:20700612-20700787 | TSSIA5ss | 3 | 3 | TSS.ENST00000360916 | -13.65 | 1.09E-02 | 2.96E-02 |
| PACRGL | 4:20700579-20704465 | 4:20700579-20700787 | TSSIA5ss | 3 | 3 | TSS.ENST00000360916 | -13.65 | 1.09E-02 | 2.96E-02 |
| DTNB | 2:25451636-25482795 | 2:25455405-25455494 | SES | 3 | 3 | ENST00000288642 | -13.66 | 5.14E-03 | 1.12E-02 |
| SSNA1 | 9:137188699-137189151 | 9:137188779-137189065 | IR | 3 | 3 | ENST00000463511 | -13.69 | 2.62E-02 | 1.00E+00 |
| ZKSCAN5 | 7:99512592-99525812 | 7:99519827-99519909 | MXS | 3 | 3 | ENST00000454175 | -13.7 | 9.76E-03 | 1.55E-01 |
| C8orf82 | 8:144525733-144527787 | 8:144526640-144526959 | IR (overlapping | 3 | 3 | ENST00000524821 | -13.73 | 1.00E-02 | 2.06E-01 |
| CCPG1 | 15:55355250-55360944 | 15:55356410-55359538 | IR (overlapping | 3 | 3 | ENST00000310958 | -13.74 | 3.97E-02 | 1.32E-01 |
| HTATSF1 | X:136497440-136497675 | X:136497624-136497675 | TSSIA3ss | 3 | 3 | TSS.ENST00000448450 | -13.75 | 4.29E-03 | 2.56E-01 |
| CEP95 | 17:64531019-64531889 | 17:64531019-64531022 | A5ss | 3 | 3 | ENST00000553412 | -13.75 | 2.36E-02 | 1.08E-01 |
| ZNF708 | 19:21309342-21329209 | 19:21327960-21328141 | MXS | 3 | 3 | ENST00000598046 | -13.76 | 2.21E-02 | 1.21E-01 |
| STAT6 | 12:57108300-57111128 | 12:57109941-57110110 | SES | 3 | 3 | ENST00000553275 | -13.77 | 3.14E-02 | 5.23E-02 |
| ZNF561-AS1 | 19:9621646-9632227 | 19:9626784-9626860 | MXS | 3 | 3 | ENST00000587536 | -13.77 | 3.59E-02 | 2.73E-01 |
| EME2 | 16:1776068-1780635 | 16:1778049-1778133 | IR (overlapping | 3 | 3 | ENST00000568449 | -13.78 | 4.99E-03 | 1.76E-02 |
| ST3GAL3 | 1:43899728-43930131 | 1:43920404-43920550 | MXS | 3 | 3 | ENST00000372368 | -13.78 | 1.73E-02 | 3.22E-02 |
| CPS1 | 2:210477767-210556718 | 2:210519551-210519784 | SES | 3 | 3 | ENST00000673630 | -13.78 | 3.86E-02 | 8.29E-02 |
| TMEM64 | 8:90625863-90645110 | 8:90631552-90631707 | SES | 3 | 3 | ENST00000458549 | -13.81 | 7.53E-03 | 1.81E-01 |
| GGT1 | 22:24608024-24609964 | 22:24609962-24609964 | A3ss | 3 | 3 | ENST00000411974 | -13.81 | 2.02E-02 | 3.93E-02 |

| | | | | | | | | | |
|---|---|---|---|---|---|---|---|---|---|
| ST6GALNAC6 | 9:127896333-127899502 | 9:127897956-127898010 | SES | 3 | 3 | ENST00000373142 | -13.81 | 3.77E-02 | 1.00E+00 |
| CDCA7 | 2:173358838-173359342 | 2:173359255-173359342 | A3ss | 3 | 3 | ENST00000306721 | -13.86 | 1.20E-02 | 2.53E-02 |
| PHF6 | X:134425201-134428789 | X:134425331-134425660 | IR | 3 | 3 | ENST00000332070 | -13.88 | 1.51E-03 | 8.91E-02 |
| GLIDR | 9:39807361-39809615 | 9:39808762-39809485 | IR | 3 | 3 | ENST00000625350 | -13.88 | 1.41E-02 | 4.31E-01 |
| STON1 | 2:48529880-48531917 | 2:48530217-48530712 | IR (overlapping | 3 | 3 | ENST00000484110 | -13.89 | 1.11E-02 | 2.45E-02 |
| LGMN | 14:92704362-92709671 | 14:92706483-92706653 | MES | 3 | 3 | ENST00000334869 | -13.89 | 1.95E-02 | 6.28E-02 |
| POLR3GL | 1:145975437-145977482 | 1:145977084-145977152 | SES | 2 | 3 | ENST00000369314 | -13.91 | 1.50E-02 | 1.28E-01 |
| WDR27 | 6:169667136-169668776 | 6:169667188-169667981 | IR (overlapping | 3 | 2 | ENST00000647790 | -13.91 | 2.12E-02 | 1.88E-01 |
| ENTPD6 | 20:25224158-25225204 | 20:25225162-25225204 | A3ss | 3 | 3 | ENST00000376666 | -13.95 | 1.28E-02 | 2.42E-02 |
| SLC17A9 | 20:62962624-62963369 | 20:62962755-62963272 | IR | 3 | 3 | ENST00000488738 | -13.97 | 4.47E-02 | 6.22E-02 |
| KANSL3 | 2:96601777-96602752 | 2:96602116-96602338 | SES | 3 | 3 | ENST00000354204 | -13.98 | 2.93E-02 | 6.66E-02 |
| CCDC150 | 2:196729193-196729817 | 2:196729388-196729792 | IR | 3 | 2 | ENST00000487663 | -13.99 | 2.07E-02 | 4.39E-02 |
| JPH4 | 14:23576457-23578179 | 14:23577075-23577624 | MXS | 3 | 3 | ENST00000356300 | -14.01 | 3.74E-07 | 6.33E-07 |
| C18orf54 | 18:54378174-54382032 | 18:54380929-54381675 | IR | 3 | 3 | ENST00000300091 | -14.02 | 3.70E-02 | 1.85E-01 |
| TMEM106C | 12:47963634-47964208 | 12:47963634-47963704 | TSSIA5ss | 3 | 3 | TSS.ENST00000429772 | -14.02 | 3.74E-02 | 6.10E-02 |
| C5orf22 | 5:31534326-31535743 | 5:31534326-31534417 | A5ss | 3 | 3 | ENST00000325366 | -14.03 | 1.22E-02 | 3.64E-02 |
| NUDT1 | 7:2242976-2244562 | 7:2242976-2243048 | TSSIA5ss | 3 | 3 | TSS.ENST00000343985 | -14.08 | 1.74E-02 | 2.24E-01 |
| SHPRH | 6:145954690-145955256 | 6:145954854-145955186 | IR | 3 | 3 | ENST00000433355 | -14.11 | 1.75E-02 | 5.47E-02 |
| RSRP1 | 1:25242249-25242705 | 1:25242414-25242538 | IR | 3 | 3 | ENST00000498238 | -14.12 | 1.74E-03 | 3.15E-03 |
| RCC1 | 1:28508906-28529857 | 1:28516725-28516867 | SES | 3 | 3 | ENST00000649185 | -14.12 | 1.59E-02 | 2.92E-02 |
| BRD1 | 22:49787495-49794033 | 22:49787495-49787887 | A3ss | 3 | 3 | ENST00000404760 | -14.16 | 7.77E-03 | 1.61E-02 |
| DCAF8 | 1:160261359-160262448 | 1:160262117-160262448 | TSSIA5ss | 3 | 3 | TSS.ENST00000368073 | -14.17 | 2.63E-02 | 4.50E-02 |
| DCAF8 | 1:160261359-160262275 | 1:160262117-160262275 | TSSIA5ss | 3 | 3 | TSS.ENST00000368073 | -14.17 | 2.63E-02 | 4.50E-02 |
| TLCD5 | 11:120325369-120327451 | 11:120327122-120327451 | A3ss | 3 | 3 | Ex.TSS.ENST00000314475 | -14.19 | 1.45E-02 | 2.09E-02 |
| TLCD5 | 11:120325369-120327440 | 11:120327122-120327440 | A3ss | 3 | 3 | Ex.TSS.ENST00000314475 | -14.19 | 1.45E-02 | 2.09E-02 |
| PPOX | 1:161169721-161170408 | 1:161169721-161170024 | TSSIA5ss | 3 | 3 | TSS.ENST00000541818 | -14.21 | 3.52E-02 | 6.12E-02 |
| FBXW2 | 9:120788187-120793148 | 9:120788187-120788278 | TSSIA3ss | 3 | 3 | TSS.ENST00000474117 | -14.24 | 2.96E-02 | 7.73E-01 |
| CENPV | 17:16348686-16349930 | 17:16348686-16349333 | A3ss | 3 | 3 | ENST00000476243 | -14.25 | 3.35E-03 | 1.32E-02 |
| BICD1 | 12:32106835-32108796 | 12:32107545-32107661 | IR | 3 | 3 | ENST00000551848 | -14.25 | 1.33E-02 | 2.13E-02 |
| CCDC82 | 11:96383962-96389887 | 11:96389524-96389843 | IR | 3 | 3 | ENST00000645439 | -14.3 | 3.97E-02 | 6.15E-02 |
| LIAS | 4:39465343-39470018 | 4:39467518-39467646 | MES | 3 | 3 | ENST00000638422 | -14.31 | 3.85E-02 | 1.05E-01 |
| MPHOSPH9 | 12:123202320-123202690 | 12:123202624-123202690 | A5ss | 3 | 3 | ENST00000302373 | -14.33 | 4.86E-02 | 7.74E-02 |
| THRA | 17:40089334-40093136 | 17:40093020-40093136 | TSSIA3ss | 3 | 3 | TSS.ENST00000264637 | -14.37 | 1.96E-03 | 8.22E-03 |
| DDX11 | 12:31074328-31078389 | 12:31077945-31078288 | SES | 3 | 3 | ENST00000535317 | -14.38 | 4.25E-02 | 6.82E-02 |
| AC009022.1 | 16:70036579-70042350 | 16:70038905-70039100 | SES | 3 | 3 | ENST00000530079 | -14.43 | 1.70E-02 | 6.52E-02 |
| ZNF263 | 16:3284206-3289392 | 16:3285059-3285239 | MES | 3 | 3 | ENST00000219069 | -14.44 | 8.88E-03 | 3.25E-02 |
| ST3GAL3 | 1:43899264-43930131 | 1:43920782-43920928 | MES | 3 | 3 | ENST00000262915 | -14.44 | 1.58E-02 | 2.94E-02 |
| SNHG14 | 15:25041689-25042910 | 15:25042516-25042602 | IR | 3 | 3 | ENST00000640631 | -14.46 | 1.31E-02 | 4.27E-02 |
| SLC2A8 | 9:127397287-127399906 | 9:127397905-127398111 | MES | 3 | 3 | ENST00000373360 | -14.51 | 5.79E-03 | 2.03E-01 |
| TMEM170A | 16:75447689-75451737 | 16:75451669-75451737 | TSSIA5ss | 3 | 3 | TSS.ENST00000561878 | -14.52 | 1.21E-02 | 4.67E-02 |
| CEP83 | 12:94411848-94412516 | 12:94412318-94412516 | A5ss | 3 | 3 | Ex.ENST00000339839 | -14.53 | 7.46E-03 | 1.19E-02 |
| TMPO | 12:98537573-98544950 | 12:98544230-98544349 | MES | 3 | 3 | ENST00000556029 | -14.55 | 2.96E-03 | 4.76E-03 |
| SCLY | 2:238064470-238068068 | 2:238068065-238068068 | A3ss | 3 | 3 | ENST00000254663 | -14.55 | 6.39E-03 | 1.39E-02 |
| SSH3 | 11:67303692-67304117 | 11:67304049-67304118 | A3ss | 3 | 3 | Ex.TSS.ENST00000308127 | -14.6 | 4.16E-03 | 6.34E-03 |
| SNHG14 | 15:25072864-25075734 | 15:25073892-25074051 | SES | 3 | 3 | ENST00000549804 | -14.62 | 1.02E-04 | 3.32E-04 |
| PIGS | 17:28571049-28571536 | 17:28571189-28571462 | IR (overlapping | 3 | 3 | ENST00000395346 | -14.65 | 7.78E-05 | 3.11E-04 |
| TPD52L1 | 6:125248384-125262833 | 6:125257098-125257158 | MXS | 3 | 3 | ENST00000527711 | -14.65 | 2.93E-03 | 9.01E-03 |
| KLC1 | 14:103701201-103714249 | 14:103712942-103714104 | IR (overlapping | 3 | 3 | ENST00000348520 | -14.66 | 7.04E-04 | 1.14E-03 |
| CYFIP2 | 5:157266196-157285338 | 5:157266196-157266584 | TSSIA5ss | 3 | 3 | TSS.ENST00000611075 | -14.66 | 2.12E-02 | 6.29E-02 |
| CABYR | 18:24156043-24159471 | 18:24156043-24156999 | A5ss | 2 | 3 | ENST00000399481 | -14.66 | 3.44E-02 | 1.71E-01 |
| AFF1 | 4:86948572-87046165 | 4:86988438-87046166 | A3ss | 3 | 3 | Ex.ENST00000395146 | -14.67 | 1.82E-02 | 5.16E-02 |
| SALL3 | 18:78992074-78995462 | 18:78994906-78995121 | IR | 3 | 3 | ENST00000537592 | -14.67 | 3.47E-02 | 1.73E-01 |
| TPD52L2 | 20:63873817-63882718 | 20:63875816-63875875 | SES | 3 | 3 | ENST00000217121 | -14.68 | 5.97E-05 | 1.15E-04 |
| RAB29 | 1:205774833-205775459 | 1:205775087-205775373 | IR | 3 | 3 | ENST00000414729 | -14.68 | 2.70E-02 | 4.81E-02 |
| PACRGL | 4:20700707-20704465 | 4:20700707-20700747 | TSSIA5ss | 3 | 3 | TSS.ENST00000513459 | -14.68 | 3.49E-02 | 9.43E-02 |
| PACRGL | 4:20700618-20704465 | 4:20700618-20700747 | TSSIA5ss | 3 | 3 | TSS.ENST00000513459 | -14.68 | 3.49E-02 | 9.43E-02 |
| PACRGL | 4:20700612-20704465 | 4:20700612-20700747 | TSSIA5ss | 3 | 3 | TSS.ENST00000513459 | -14.68 | 3.49E-02 | 9.43E-02 |
| PACRGL | 4:20700579-20704465 | 4:20700579-20700747 | TSSIA5ss | 3 | 3 | TSS.ENST00000513459 | -14.68 | 3.49E-02 | 9.43E-02 |
| PEX19 | 1:160283110-160285054 | 1:160283530-160283639 | SES | 3 | 3 | ENST00000368072 | -14.72 | 2.09E-04 | 3.62E-04 |
| METTL23 | 17:76727179-76729689 | 17:76727179-76727213 | A5ss | 3 | 3 | ENST00000586738 | -14.72 | 1.38E-02 | 6.53E-02 |
| METTL23 | 17:76726880-76729689 | 17:76726880-76727213 | TSSIA5ss | 3 | 3 | TSS.ENST00000592849 | -14.72 | 1.38E-02 | 6.53E-02 |
| AMZ2 | 17:68248043-68248705 | 17:68248111-68248639 | IR | 3 | 3 | ENST00000359904 | -14.72 | 1.48E-02 | 6.71E-02 |
| AP3M2 | 8:42153106-42154615 | 8:42154478-42154536 | SES | 3 | 3 | ENST00000518421 | -14.75 | 2.15E-02 | 4.77E-01 |
| PAQR4 | 16:2971515-2973482 | 16:2973065-2973137 | IR (overlapping | 3 | 3 | ENST00000293978 | -14.75 | 2.55E-02 | 9.29E-02 |
| PLXNB2 | 22:50290598-50294725 | 22:50294719-50294725 | A5ss | 3 | 3 | ENST00000359337 | -14.77 | 1.31E-03 | 2.71E-03 |
| PLD3 | 19:40348769-40366418 | 19:40365553-40365930 | SES | 3 | 3 | ENST00000594085 | -14.8 | 4.33E-04 | 2.84E-03 |
| POC1A | 3:52075986-52122378 | 3:52096569-52096712 | SES | 3 | 3 | ENST00000474012 | -14.8 | 1.37E-02 | 3.52E-02 |
| TAF1C | 16:84180329-84181042 | 16:84180329-84180344 | A3ss | 3 | 3 | Ex.ENST00000341690 | -14.83 | 2.20E-02 | 8.53E-02 |
| TRIB2 | 2:12717011-12723259 | 2:12717640-12723260 | A3ss | 3 | 3 | Ex.TSS.ENST00000381465 | -14.84 | 4.92E-03 | 1.03E-02 |
| PPOX | 1:161167235-161168992 | 1:161168445-161168576 | MES | 3 | 3 | ENST00000651150 | -14.86 | 3.22E-02 | 5.60E-02 |
| LMNA | 1:156139080-156140089 | 1:156139103-156139769 | IR | 3 | 3 | ENST00000347559 | -14.86 | 4.45E-02 | 6.19E-02 |
| CDK5RAP1 | 20:33392243-33401427 | 20:33396761-33397084 | MES | 3 | 3 | ENST00000339269 | -14.87 | 1.47E-03 | 2.79E-03 |
| GPS1 | 17:82050691-82051143 | 17:82050765-82051011 | IR (overlapping | 3 | 3 | ENST00000583983 | -14.87 | 1.47E-02 | 7.10E-02 |
| SIRT2 | 19:38893868-38899505 | 19:38898379-38898425 | MXS | 3 | 3 | ENST00000249396 | -14.87 | 4.11E-02 | 2.64E-01 |
| DNASE1L1 | X:154411891-154412112 | X:154412007-154412085 | IR (overlapping | 3 | 3 | ENST00000309585 | -14.88 | 7.58E-03 | 5.29E-01 |
| LINC02245 | 2:65048213-65055914 | 2:65051279-65055915 | A5ss | 3 | 3 | Ex.TSS.ENST00000653778 | -14.95 | 4.32E-02 | 9.61E-02 |
| RAD52 | 12:916533-916638 | 12:916533-916535 | A3ss | 3 | 3 | ENST00000461568 | -14.96 | 4.16E-02 | 6.61E-02 |
| RAD52 | 12:916484-916638 | 12:916484-916535 | A3ss | 3 | 3 | ENST00000461568 | -14.96 | 4.16E-02 | 6.61E-02 |
| WNK2 | 9:93317520-93319208 | 9:93317632-93318027 | IR | 3 | 3 | ENST00000411624 | -14.96 | 4.74E-02 | 1.00E+00 |
| GUSBP11 | 22:23683868-23705428 | 22:23695173-23695517 | MXS | 3 | 3 | ENST00000422506 | -14.99 | 1.10E-03 | 2.15E-03 |
| SETD5 | 3:9433951-9434333 | 3:9434277-9434333 | A3ss | 3 | 3 | ENST00000407969 | -15 | 7.41E-03 | 1.92E-02 |
| LINC00869 | 1:149646599-149647610 | 1:149646655-149647380 | IR | 3 | 3 | ENST00000620190 | -15.07 | 2.83E-02 | 2.44E-01 |
| RGL3 | 19:11406865-11416613 | 19:11415937-11416148 | MXS | 3 | 3 | ENST00000380456 | -15.12 | 2.03E-02 | 1.05E-01 |
| RBSN | 3:15096711-15099034 | 3:15098119-15098274 | SES | 3 | 3 | ENST00000253699 | -15.12 | 4.40E-02 | 1.02E-01 |
| DALRD3 | 3:49015525-49016340 | 3:49015873-49015972 | IR (overlapping | 3 | 3 | ENST00000467457 | -15.13 | 2.59E-03 | 6.24E-03 |
| ADSS1 | 14:104744719-104746385 | 14:104746135-104746235 | IR | 3 | 3 | ENST00000557271 | -15.15 | 3.00E-02 | 4.76E-02 |
| UPP1 | 7:48094828-48106872 | 7:48103297-48103411 | MES | 3 | 3 | ENST00000331803 | -15.16 | 3.92E-03 | 5.47E-02 |
| AMZ2 | 17:68248043-68248705 | 17:68248163-68248639 | IR | 3 | 3 | ENST00000359904 | -15.16 | 1.11E-02 | 5.04E-02 |
| ADGRL2 | 1:81800513-81836884 | 1:81800513-81801068 | A5ss | 3 | 3 | Ex.TSS.ENST00000319517 | -15.16 | 1.39E-02 | 2.60E-02 |
| AL049834.1 | 14:88385838-88396155 | 14:88391381-88391455 | SES | 3 | 3 | ENST00000356583 | -15.17 | 3.70E-02 | 1.18E-01 |
| FBRSL1 | 12:132570541-132571306 | 12:132571068-132571136 | SES | 3 | 3 | ENST00000650108 | -15.18 | 1.19E-02 | 1.90E-02 |
| PDE2A | 11:72608752-72674136 | 11:72642254-72642326 | MXS | 3 | 2 | ENST00000334456 | -15.19 | 2.17E-02 | 3.33E-02 |
| SNRPN | 15:24968083-24974386 | 15:24974311-24974386 | A3ss | 3 | 3 | ENST00000390687 | -15.2 | 3.45E-09 | 1.12E-08 |
| ADIPOR2 | 12:1691192-1754257 | 12:1691192-1691213 | A5ss | 3 | 3 | Ex.TSS.ENST00000357103 | -15.22 | 2.25E-02 | 3.60E-02 |

| | | | | | | | | | |
|---|---|---|---|---|---|---|---|---|---|
| UPP1 | 7:48089419-48090187 | 7:48089509-48089636 | SES | 3 | 3 | ENST00000436673 | -15.27 | 1.29E-05 | 1.80E-04 |
| TBC1D22A | 22:46793842-46878652 | 22:46797444-46797620 | SES | 3 | 3 | ENST00000337137 | -15.28 | 6.59E-03 | 1.37E-02 |
| POLL | 10:101582892-101587245 | 10:101585862-101586156 | MES | 3 | 3 | ENST00000299206 | -15.33 | 7.61E-03 | 1.06E-02 |
| MFAP3L | 4:170005892-170026233 | 4:170005892-170006010 | TSSlA3ss | 2 | 3 | TSS.ENST00000361618 | -15.33 | 3.34E-02 | 8.96E-02 |
| ZBTB8OS | 1:32634111-32650432 | 1:32634768-32634792 | SES | 3 | 3 | ENST00000473294 | -15.34 | 2.20E-02 | 4.07E-02 |
| TCAIM | 3:44361519-44367531 | 3:44367456-44367531 | A3ss | 3 | 2 | ENST00000342649 | -15.35 | 5.08E-03 | 1.22E-02 |
| KLC1 | 14:103701201-103714249 | 14:103712932-103714104 | IR (overlapping | 3 | 3 | ENST00000348520 | -15.36 | 8.43E-04 | 1.37E-03 |
| DYRK2 | 12:67649183-67657105 | 12:67649797-67649945 | SES | 3 | 3 | ENST00000344096 | -15.36 | 2.55E-03 | 4.27E-03 |
| SYNRG | 17:37577614-37584647 | 17:37579132-37579434 | SES | 3 | 3 | ENST00000614000 | -15.39 | 1.88E-02 | 7.64E-02 |
| FAHD2B | 2:97091713-97094700 | 2:97091863-97091988 | SES | 3 | 3 | ENST00000414820 | -15.4 | 2.33E-02 | 5.20E-02 |
| CDCA8 | 1:37692516-37692784 | 1:37692579-37692675 | IR | 3 | 3 | ENST00000373055 | -15.41 | 4.30E-03 | 7.97E-03 |
| KRIT1 | 7:92245465-92246100 | 7:92245596-92245789 | IR (overlapping | 3 | 3 | ENST00000340022 | -15.42 | 1.39E-03 | 2.14E-02 |
| CHID1 | 11:884170-899339 | 11:893427-893519 | SES | 3 | 3 | ENST00000323578 | -15.42 | 1.56E-03 | 2.41E-03 |
| MBD4 | 3:129434137-129436460 | 3:129436443-129436460 | A5ss | 3 | 3 | ENST00000249910 | -15.43 | 2.24E-02 | 5.00E-02 |
| DTX3 | 12:57604859-57606189 | 12:57605344-57605731 | MXS | 3 | 3 | ENST00000549583 | -15.43 | 4.25E-02 | 7.09E-02 |
| LETMD1 | 12:51048479-51056349 | 12:51049034-51049185 | MES | 3 | 3 | ENST00000262055 | -15.46 | 4.14E-02 | 6.81E-02 |
| WDR19 | 4:39189782-39199477 | 4:39194544-39194659 | SES | 3 | 3 | ENST00000399820 | -15.47 | 3.60E-02 | 9.79E-02 |
| AMZ2 | 17:68248043-68248705 | 17:68248158-68248639 | IR | 3 | 3 | ENST00000359904 | -15.55 | 1.52E-02 | 6.88E-02 |
| FANCI | 15:89244045-89247628 | 15:89244045-89244067 | TSSlA5ss | 3 | 3 | TSS.ENST00000563250 | -15.55 | 2.50E-02 | 8.68E-02 |
| FANCI | 15:89244034-89247628 | 15:89244034-89244067 | TSSlA5ss | 3 | 3 | TSS.ENST00000563250 | -15.55 | 2.50E-02 | 8.68E-02 |
| GOLGA2P10 | 15:82477124-82477393 | 15:82477203-82477298 | SES | 3 | 3 | ENST00000550961 | -15.57 | 3.30E-03 | 1.13E-02 |
| SPIN3 | X:56990942-56995555 | X:56992584-56994722 | IR | 3 | 3 | ENST00000638289 | -15.57 | 6.40E-03 | 1.00E+00 |
| THTPA | 14:23556294-23557039 | 14:23556344-23556742 | IR (overlapping | 3 | 3 | ENST00000554789 | -15.58 | 1.68E-03 | 2.73E-03 |
| CLUHP3 | 16:31707907-31711986 | 16:31708208-31708460 | IR | 3 | 2 | ENST00000562354 | -15.62 | 6.76E-03 | 2.47E-02 |
| RNF111 | 15:59067084-59075956 | 15:59075954-59075956 | A3ss | 3 | 3 | Ex.ENST00000348370 | -15.62 | 2.33E-02 | 7.63E-02 |
| PWWP3A | 19:1358465-1360135 | 19:1358588-1358693 | SES | 3 | 3 | ENST00000588810 | -15.66 | 2.31E-02 | 1.24E-01 |
| MED12 | X:71137726-71137943 | X:71137781-71137855 | IR | 3 | 3 | ENST00000333646 | -15.66 | 4.87E-02 | 1.00E+00 |
| CCNE1 | 19:29821818-29822239 | 19:29821996-29822130 | SES | 3 | 3 | ENST00000262643 | -15.68 | 6.78E-03 | 3.77E-02 |
| NUP214 | 9:131140711-131144567 | 9:131144280-131144567 | A3ss | 3 | 3 | ENST00000359428 | -15.7 | 8.59E-04 | 3.31E-02 |
| KLC1 | 14:103679546-103687080 | 14:103684986-103685036 | SES | 3 | 3 | ENST00000554228 | -15.7 | 4.98E-02 | 8.06E-02 |
| POLL | 10:101582892-101587245 | 10:101585316-101585478 | MES | 3 | 3 | ENST00000299206 | -15.78 | 1.01E-02 | 1.41E-02 |
| AP1S2 | X:15831569-15845378 | X:15833444-15833524 | SES | 3 | 3 | ENST00000479184 | -15.79 | 1.60E-03 | 1.16E-01 |
| TCF3 | 19:1610501-1611703 | 19:1610501-1611369 | TSSlA3ss | 3 | 3 | TSS.ENST00000588136 | -15.79 | 2.14E-02 | 1.16E-01 |
| TCF3 | 19:1609600-1611703 | 19:1609600-1611369 | TSSlA3ss | 3 | 3 | TSS.ENST00000588136 | -15.79 | 2.14E-02 | 1.16E-01 |
| PLPPR2 | 19:11357740-11359606 | 19:11359532-11359606 | A3ss | 3 | 3 | ENST00000251473 | -15.83 | 8.78E-03 | 4.54E-02 |
| TNNT2 | 1:201367804-201368161 | 1:201367804-201367806 | A3ss | 3 | 3 | ENST00000236918 | -15.83 | 1.50E-02 | 2.66E-02 |
| PQBP1 | X:48898015-48898491 | X:48898015-48898082 | TSSlA5ss | 3 | 3 | TSS.ENST00000376566 | -15.85 | 3.11E-03 | 3.52E-01 |
| KLHL3 | 5:137735633-137736089 | 5:137735743-137735817 | IR | 3 | 3 | ENST00000309755 | -15.85 | 7.38E-03 | 2.13E-02 |
| FAM200B | 4:15681902-15686235 | 4:15681902-15681968 | TSSlA5ss | 3 | 3 | TSS.ENST00000503600 | -15.88 | 1.30E-02 | 3.49E-02 |
| FAM200B | 4:15681785-15686235 | 4:15681785-15681968 | A5ss | 3 | 3 | ENST00000503617 | -15.88 | 1.30E-02 | 3.49E-02 |
| ANKS3 | 16:4724832-4726978 | 16:4726659-4726780 | SES | 3 | 3 | ENST00000304283 | -15.94 | 6.67E-03 | 2.46E-02 |
| TMEM147-AS1 | 19:35541582-35545492 | 19:35542075-35545318 | IR (overlapping | 3 | 3 | ENST00000588286 | -15.95 | 2.22E-02 | 1.29E-01 |
| OTUD7B | 1:149967388-149971062 | 1:149967388-149967521 | A3ss | 3 | 3 | ENST00000417191 | -15.95 | 3.09E-02 | 2.67E-01 |
| CYP2B6 | 19:41004447-41012297 | 19:41009219-41009395 | MES | 2 | 3 | ENST00000324071 | -15.97 | 4.19E-02 | 2.77E-01 |
| SPIB | 19:50422546-50423604 | 19:50422823-50423037 | SES | 3 | 3 | ENST00000270632 | -15.98 | 2.63E-02 | 1.84E-01 |
| AP1S2 | X:15833008-15845378 | X:15833444-15833524 | SES | 3 | 3 | ENST00000673591 | -15.99 | 2.03E-02 | 1.00E+00 |
| ABHD14B | 3:51970185-51973964 | 3:51971460-51971698 | MES | 3 | 3 | ENST00000525795 | -16 | 4.52E-03 | 1.17E-02 |
| HSD11B1L | 19:5681272-5686415 | 19:5684819-5684905 | MXS | 3 | 3 | ENST00000582346 | -16.02 | 2.90E-02 | 2.12E-01 |
| KLHL4 | X:87666475-87670050 | X:87666881-87669328 | IR | 3 | 3 | ENST00000373119 | -16.06 | 8.56E-04 | 1.00E+00 |
| SLC9A7 | X:46605318-46607203 | X:46606371-46606907 | IR | 3 | 3 | ENST00000328306 | -16.06 | 6.27E-03 | 6.53E-01 |
| AMH | 19:2249744-2250760 | 19:2249745-2250336 | IR | 3 | 3 | ENST00000589313 | -16.06 | 1.99E-02 | 1.10E-01 |
| AGO3 | 1:35973512-36004340 | 1:35984470-35984548 | MES | 3 | 3 | ENST00000634486 | -16.07 | 1.60E-02 | 2.94E-02 |
| METTL23 | 17:76726880-76732977 | 17:76729690-76729794 | MES | 3 | 3 | ENST00000586738 | -16.14 | 1.62E-02 | 7.69E-02 |
| MAGI2 | 7:78255943-78256581 | 7:78256159-78256247 | IR | 3 | 3 | ENST00000354212 | -16.2 | 4.30E-02 | 5.99E-01 |
| MGAT1 | 5:180809077-180809633 | 5:180809364-180809633 | A5ss | 3 | 3 | ENST00000506708 | -16.22 | 4.15E-02 | 1.16E-01 |
| RAB29 | 1:205774833-205775459 | 1:205775087-205775272 | IR | 3 | 3 | ENST00000414729 | -16.23 | 5.13E-03 | 9.14E-03 |
| ACIN1 | 14:23069618-23071426 | 14:23071125-23071186 | SES | 3 | 3 | ENST00000357481 | -16.25 | 1.53E-03 | 2.49E-03 |
| MYO1B | 2:191400469-191408114 | 2:191400749-191400835 | MES | 3 | 3 | ENST00000304164 | -16.25 | 2.95E-02 | 6.25E-02 |
| AHDC1 | 1:27553186-27558705 | 1:27558305-27558530 | SES | 3 | 3 | ENST00000642245 | -16.26 | 4.72E-02 | 8.68E-02 |
| HPN | 19:35041873-35049289 | 19:35041873-35042522 | TSSlA5ss | 3 | 3 | TSS.ENST00000262626 | -16.3 | 2.61E-02 | 1.50E-01 |
| CYP27C1 | 2:127203572-127219988 | 2:127205900-127206090 | SES | 2 | 3 | ENST00000664447 | -16.3 | 4.76E-02 | 9.89E-02 |
| STAT6 | 12:57108300-57110076 | 12:57109941-57110076 | A5ss | 3 | 3 | ENST00000553275 | -16.31 | 1.01E-02 | 1.68E-02 |
| STAT6 | 12:57108300-57110042 | 12:57109941-57110042 | A5ss | 3 | 3 | ENST00000553275 | -16.31 | 1.01E-02 | 1.68E-02 |
| ATP11A | 13:112875942-112881875 | 13:112878217-112878303 | MXS | 3 | 3 | ENST00000375645 | -16.32 | 1.26E-02 | 2.03E-02 |
| RASSF8 | 12:26064498-26065387 | 12:26064713-26064714 | IR | 3 | 3 | ENST00000282884 | -16.32 | 2.14E-02 | 3.42E-02 |
| SLC25A30 | 13:45405978-45411361 | 13:45408927-45409074 | SES | 3 | 3 | ENST00000519676 | -16.33 | 3.54E-02 | 5.75E-02 |
| SSNA1 | 9:137188699-137189151 | 9:137188779-137189000 | IR | 3 | 3 | ENST00000463511 | -16.36 | 2.73E-02 | 1.00E+00 |
| APBB3 | 5:140561587-140562227 | 5:140561702-140561843 | IR | 3 | 3 | ENST00000510241 | -16.38 | 1.29E-02 | 3.75E-02 |
| RAD54L | 1:46247763-46248408 | 1:46247802-46248398 | IR | 3 | 3 | ENST00000371975 | -16.41 | 2.52E-03 | 4.70E-03 |
| CUTC | 10:99747391-99754528 | 10:99750369-99750396 | SES | 3 | 3 | ENST00000370472 | -16.41 | 1.60E-02 | 2.30E-02 |
| MYCL | 1:39900939-39901917 | 1:39901444-39901807 | IR | 3 | 3 | ENST00000372816 | -16.46 | 1.74E-03 | 3.22E-03 |
| HYAL3 | 3:50294854-50299212 | 3:50294854-50295619 | TSSlA3ss | 3 | 3 | TSS.ENST00000336307 | -16.49 | 6.42E-04 | 1.57E-03 |
| AP3B2 | 15:82685914-82688831 | 15:82687020-82688735 | IR (overlapping | 3 | 3 | ENST00000535513 | -16.53 | 2.30E-02 | 7.84E-02 |
| SUPT20H | 13:37022081-37024034 | 13:37022081-37022317 | A3ss | 3 | 3 | ENST00000475892 | -16.56 | 2.72E-02 | 4.32E-02 |
| RNF19A | 8:100303366-100309866 | 8:100303366-100303401 | TSSlA3ss | 2 | 3 | TSS.ENST00000523481 | -16.59 | 7.91E-04 | 1.46E-02 |
| SLC35C2 | 20:46355242-46356573 | 20:46355802-46355874 | SES | 3 | 3 | ENST00000243896 | -16.59 | 2.94E-03 | 5.61E-03 |
| SLC9A3 | 5:470456-473382 | 5:471824-472678 | IR (overlapping | 3 | 3 | ENST00000264938 | -16.65 | 3.61E-02 | 1.08E-01 |
| STYXL1 | 7:76030528-76047811 | 7:76047662-76047811 | TSSlA5ss | 3 | 3 | TSS.ENST00000359697 | -16.67 | 4.07E-02 | 6.07E-01 |
| ARTN | 1:43936069-43936345 | 1:43936232-43936301 | IR | 3 | 3 | ENST00000472435 | -16.67 | 4.66E-02 | 6.49E-02 |
| SYT7 | 11:61533125-61551383 | 11:61547177-61547308 | MES | 3 | 3 | ENST00000535826 | -16.7 | 4.10E-02 | 6.06E-02 |
| CAMK2G | 10:73824085-73837467 | 10:73825321-73825347 | MXS | 3 | 3 | ENST00000474131 | -16.78 | 2.91E-02 | 4.14E-02 |
| WDR90 | 16:655773-656537 | 16:655886-656301 | IR (overlapping | 3 | 3 | ENST00000552943 | -16.8 | 3.97E-02 | 1.50E-01 |
| CDK10 | 16:89693274-89693495 | 16:89693327-89693397 | IR | 3 | 3 | ENST00000504473 | -16.81 | 2.36E-02 | 9.31E-02 |
| SMPD5 | 8:144050795-144051522 | 8:144050859-144050941 | IR | 3 | 3 | ENST00000528912 | -16.85 | 9.78E-03 | 1.98E-01 |
| TSC22D1 | 13:44574412-44576565 | 13:44576247-44576434 | IR (overlapping | 3 | 3 | ENST00000501704 | -16.88 | 7.27E-03 | 1.18E-02 |
| SLC25A29 | 14:100293378-100306198 | 14:100298842-100298885 | MXS | 3 | 3 | ENST00000359232 | -16.89 | 2.28E-03 | 3.86E-03 |
| LDOC1 | X:141173235-141177129 | X:141176409-141176892 | IR | 3 | 3 | ENST00000370526 | -16.9 | 4.22E-03 | 2.55E-01 |
| MAP3K10 | 19:40213076-40213734 | 19:40213189-40213516 | IR | 3 | 3 | ENST00000594791 | -16.91 | 3.28E-02 | 2.15E-01 |
| MIA2 | 14:39276956-39279336 | 14:39276956-39277065 | A5ss | 3 | 3 | ENST00000280083 | -16.94 | 1.08E-03 | 3.39E-03 |
| SLTM | 15:58899710-58901259 | 15:58899710-58899883 | A3ss | 3 | 3 | ENST00000249736 | -16.99 | 1.87E-03 | 6.25E-03 |
| SUGP2 | 19:18991150-18994365 | 19:18993648-18993740 | MXS | 3 | 3 | ENST00000594773 | -17.03 | 1.39E-02 | 7.61E-02 |
| OXR1 | 8:106658086-106679209 | 8:106658086-106658185 | TSSlA5ss | 3 | 3 | TSS.ENST00000312046 | -17.06 | 9.17E-03 | 1.72E-01 |
| AC073111.4 | 7:150408489-150410159 | 7:150408489-150408784 | A5ss | 3 | 3 | Ex.TSS.ENST00000498682 | -17.06 | 4.51E-02 | 5.60E-01 |
| AC073111.4 | 7:150405318-150410159 | 7:150405318-150408784 | A5ss | 3 | 3 | Ex.ENST00000641234 | -17.06 | 4.51E-02 | 5.60E-01 |

| | | | | | | | | | |
|---|---|---|---|---|---|---|---|---|---|
| AC073111.3 | 7:150388502-150410159 | 7:150388502-150408784 | A5ss | 3 | 3 | Ex.TSS.ENST00000483664 | -17.06 | 4.51E-02 | 5.60E-01 |
| NUBP2 | 16:1786656-1787676 | 16:1786656-1786955 | A5ss | 3 | 3 | ENST00000565603 | -17.1 | 3.52E-02 | 1.24E-01 |
| SLC13A3 | 20:46613726-46684395 | 20:46613726-46670139 | A3ss | 3 | 3 | Ex.TSS.ENST00000372121 | -17.11 | 2.09E-03 | 4.01E-03 |
| RNF44 | 5:176532133-176532365 | 5:176532133-176532193 | A3ss | 3 | 3 | ENST00000274811 | -17.12 | 1.14E-03 | 3.37E-03 |
| RNF44 | 5:176532130-176532365 | 5:176532130-176532193 | A3ss | 3 | 3 | ENST00000274811 | -17.12 | 1.14E-03 | 3.37E-03 |
| KANSL3 | 2:96619545-96631311 | 2:96619672-96619762 | SES | 3 | 3 | ENST00000354204 | -17.14 | 9.21E-03 | 2.09E-02 |
| CEP170 | 1:243140108-243142366 | 1:243142316-243142366 | A5ss | 3 | 3 | ENST00000366542 | -17.18 | 1.41E-02 | 2.54E-02 |
| TPM1 | 15:63060940-63061712 | 15:63061198-63061712 | A3ss | 3 | 3 | ENST00000560975 | -17.21 | 1.56E-04 | 5.22E-04 |
| CYB5RL | 1:54187740-54195418 | 1:54190738-54190896 | SES | 3 | 3 | ENST00000528287 | -17.22 | 2.61E-02 | 4.89E-02 |
| USP37 | 2:218560884-218562717 | 2:218562673-218562717 | A5ss | 3 | 2 | ENST00000258399 | -17.22 | 4.22E-02 | 9.12E-02 |
| AP5S1 | 20:3820759-3822101 | 20:3820759-3820854 | TSSIA5ss | 3 | 3 | TSS.ENST00000246041 | -17.24 | 2.71E-02 | 5.16E-02 |
| AP5S1 | 20:3820703-3822101 | 20:3820703-3820854 | TSSIA5ss | 3 | 3 | TSS.ENST00000246041 | -17.24 | 2.71E-02 | 5.16E-02 |
| ABCB8 | 7:151028611-151034272 | 7:151033605-151033917 | MXS | 3 | 3 | ENST00000358849 | -17.26 | 1.98E-02 | 2.46E-01 |
| ZSCAN18 | 19:58090387-58098173 | 19:58098115-58098173 | TSSIA5ss | 3 | 3 | TSS.ENST00000600845 | -17.27 | 1.13E-02 | 8.31E-02 |
| FIRRE | X:131689755-131692636 | X:131689841-131689945 | IR | 3 | 3 | ENST00000663405 | -17.29 | 4.53E-02 | 1.00E+00 |
| GOLGA6L9 | 15:82430045-82432897 | 15:82432632-82432816 | IR (overlapping | 3 | 3 | ENST00000618706 | -17.3 | 2.18E-02 | 7.43E-02 |
| UBR3 | 2:170007191-170008803 | 2:170007191-170007202 | A5ss | 2 | 2 | ENST00000392632 | -17.31 | 8.17E-03 | 1.72E-02 |
| PPIP5K2 | 5:103184745-103190841 | 5:103187314-103187376 | MES | 3 | 3 | ENST00000358359 | -17.32 | 9.46E-04 | 2.70E-03 |
| CYLD | 16:50742842-50749575 | 16:50744730-50744866 | SES | 3 | 3 | ENST00000311559 | -17.33 | 1.60E-02 | 5.90E-02 |
| SNHG14 | 15:25041689-25042910 | 15:25042516-25042797 | IR | 3 | 3 | ENST00000640631 | -17.37 | 4.59E-03 | 1.49E-02 |
| SNHG1 | 11:62855424-62855885 | 11:62855746-62855864 | IR | 3 | 3 | ENST00000535689 | -17.37 | 2.17E-02 | 3.24E-02 |
| FKRP | 19:46746087-46748026 | 19:46746087-46746090 | TSSIA5ss | 3 | 3 | TSS.ENST00000318584 | -17.4 | 9.44E-03 | 6.32E-02 |
| BIRC2 | 11:102348598-102350749 | 11:102348720-102350345 | IR | 3 | 3 | ENST00000227758 | -17.42 | 4.71E-03 | 6.75E-03 |
| RGL3 | 19:11394521-11397243 | 19:11394606-11394678 | SES | 3 | 3 | ENST00000568628 | -17.44 | 4.15E-02 | 2.13E-01 |
| IFT122 | 3:129488398-129495445 | 3:129492141-129492194 | SES | 3 | 3 | ENST00000296266 | -17.45 | 6.61E-04 | 1.52E-03 |
| TCF25 | 16:89893859-89895082 | 16:89895038-89895082 | A3ss | 3 | 3 | ENST00000263346 | -17.5 | 4.03E-03 | 1.59E-02 |
| THOC5 | 22:29549159-29553670 | 22:29553448-29553670 | TSSIA5ss | 3 | 3 | TSS.ENST00000428374 | -17.51 | 3.89E-03 | 7.65E-03 |
| SPIN3 | X:56990831-56994949 | X:56992584-56993786 | IR | 3 | 3 | ENST00000374919 | -17.54 | 8.80E-04 | 2.04E-01 |
| ZNF84 | 12:133037546-133041284 | 12:133041100-133041284 | TSSIA3ss | 3 | 3 | TSS.ENST00000543758 | -17.54 | 1.57E-02 | 2.50E-02 |
| ZNF84 | 12:133037546-133041277 | 12:133041100-133041277 | TSSIA3ss | 3 | 3 | TSS.ENST00000543758 | -17.54 | 1.57E-02 | 2.50E-02 |
| HEXD | 17:82439631-82440532 | 17:82440335-82440472 | IR | 3 | 3 | ENST00000585077 | -17.6 | 3.34E-02 | 1.64E-01 |
| EXO5 | 1:40508747-40509790 | 1:40509626-40509732 | IR | 2 | 3 | ENST00000372703 | -17.61 | 1.91E-02 | 3.54E-02 |
| NME6 | 3:48298524-48301352 | 3:48301296-48301352 | A5ss | 3 | 3 | ENST00000302378 | -17.61 | 4.53E-02 | 1.07E-01 |
| IDH3B | 20:2658523-2659524 | 20:2658759-2658837 | SES | 3 | 3 | ENST00000474315 | -17.69 | 3.98E-03 | 7.54E-03 |
| AMZ2 | 17:68248043-68248705 | 17:68248087-68248643 | IR | 3 | 3 | ENST00000359904 | -17.69 | 1.32E-02 | 5.99E-02 |
| LRRC20 | 10:70301509-70376451 | 10:70340553-70340702 | MXS | 3 | 3 | ENST00000357631 | -17.7 | 1.29E-02 | 1.84E-02 |
| SPRY4 | 5:142315156-142323889 | 5:142319733-142319794 | SES | 2 | 3 | ENST00000511815 | -17.7 | 1.59E-02 | 4.63E-02 |
| TRIML2 | 4:188105588-188109242 | 4:188105588-188105590 | A3ss | 3 | 3 | Ex.ENST00000394461 | -17.72 | 3.00E-02 | 8.12E-02 |
| RABGGTA | 14:24271113-24271599 | 14:24271249-24271486 | IR (overlapping | 3 | 3 | ENST00000399409 | -17.74 | 3.50E-02 | 5.95E-02 |
| TMEM143 | 19:48345355-48363897 | 19:48363291-48363531 | MXS | 3 | 3 | ENST00000435956 | -17.76 | 1.29E-02 | 8.86E-02 |
| ZNF749 | 19:57435594-57443290 | 19:57441885-57442011 | SES | 3 | 3 | ENST00000334181 | -17.76 | 4.26E-02 | 3.12E-01 |
| SLC9A7 | X:46605318-46607203 | X:46606371-46606923 | IR | 3 | 3 | ENST00000328306 | -17.77 | 9.45E-03 | 9.82E-01 |
| ANKRD13D | 11:67299612-67299826 | 11:67299612-67299621 | A5ss | 3 | 3 | Ex.ENST00000308440 | -17.85 | 3.23E-02 | 4.93E-02 |
| AMZ2 | 17:68248043-68248705 | 17:68248087-68248639 | IR | 3 | 3 | ENST00000359904 | -17.86 | 1.26E-02 | 5.69E-02 |
| RBM33 | 7:155707069-155711202 | 7:155711173-155711202 | A3ss | 3 | 3 | ENST00000440108 | -17.88 | 2.19E-03 | 2.80E-02 |
| BIRC2 | 11:102348598-102350749 | 11:102348765-102350000 | IR | 3 | 3 | ENST00000227758 | -17.88 | 6.99E-03 | 1.00E-02 |
| THEM4 | 1:151895195-151909359 | 1:151907321-151909360 | A5ss | 3 | 3 | Ex.TSS.ENST00000368814 | -17.93 | 1.13E-02 | 1.94E-02 |
| C16orf70 | 16:67109990-67110144 | 16:67110119-67110144 | TSSIA3ss | 2 | 2 | TSS.ENST00000569600 | -17.97 | 4.38E-02 | 1.66E-01 |
| PANK2 | 20:3908136-3910576 | 20:3908136-3908278 | A5ss | 3 | 3 | ENST00000316562 | -18.03 | 1.96E-02 | 3.74E-02 |
| ZC3H13 | 13:46045514-46052403 | 13:46045514-46045516 | A3ss | 3 | 3 | Ex.TSS.ENST00000242848 | -18.04 | 2.03E-02 | 3.30E-02 |
| TMEM231 | 16:75545955-75555803 | 16:75545955-75546077 | A3ss | 2 | 3 | ENST00000562410 | -18.09 | 1.48E-02 | 5.71E-02 |
| TYK2 | 19:10353099-10354041 | 19:10353528-10354041 | TSSIA5ss | 3 | 2 | TSS.ENST00000529739 | -18.12 | 4.84E-02 | 2.42E-01 |
| FAAP24 | 19:32976431-32978229 | 19:32977317-32977496 | IR | 3 | 3 | ENST00000588258 | -18.13 | 4.10E-03 | 2.28E-02 |
| CCDC82 | 11:96387614-96389843 | 11:96389365-96389523 | SES | 3 | 3 | ENST00000645366 | -18.14 | 1.21E-02 | 1.88E-02 |
| AL645939.2 | 6:29722981-29723971 | 6:29723528-29723657 | IR (overlapping | 3 | 3 | ENST00000427340 | -18.17 | 3.75E-02 | 1.17E-01 |
| EME2 | 16:1776068-1780635 | 16:1777444-1777746 | IR (overlapping | 3 | 3 | ENST00000568449 | -18.18 | 4.62E-02 | 1.62E-01 |
| PRKAR1A | 17:68512971-68515393 | 17:68514987-68515119 | SES | 3 | 3 | ENST00000589480 | -18.19 | 3.27E-02 | 1.48E-01 |
| AMZ2 | 17:68248043-68248705 | 17:68248059-68248643 | IR | 3 | 3 | ENST00000359904 | -18.2 | 1.19E-02 | 5.40E-02 |
| ZNF43 | 19:21851163-21851904 | 19:21851163-21851360 | TSSIA3ss | 3 | 3 | TSS.ENST00000599414 | -18.2 | 2.02E-02 | 1.11E-01 |
| ACAD10 | 12:111748476-111749172 | 12:111749011-111749172 | TSSIA3ss | 3 | 3 | TSS.ENST00000550198 | -18.2 | 2.41E-02 | 3.74E-02 |
| WDR90 | 16:655773-656537 | 16:655890-656301 | IR (overlapping | 3 | 3 | ENST00000552943 | -18.25 | 2.47E-02 | 9.35E-02 |
| AMT | 3:49416794-49418372 | 3:49417368-49417817 | IR (overlapping | 3 | 3 | ENST00000636594 | -18.27 | 1.67E-03 | 4.06E-03 |
| MINDY1 | 1:151001783-151006311 | 1:151002165-151002323 | SES | 2 | 3 | ENST00000312210 | -18.32 | 1.49E-02 | 1.29E-01 |
| DPH5 | 1:100995150-101021640 | 1:101013710-101013818 | MES | 3 | 3 | ENST00000342173 | -18.34 | 3.60E-02 | 2.92E-01 |
| APBB3 | 5:140561587-140562227 | 5:140561702-140562099 | IR | 3 | 3 | ENST00000510241 | -18.35 | 3.51E-02 | 1.02E-01 |
| TANGO2 | 22:20052585-20063337 | 22:20061530-20061683 | MXS | 3 | 3 | ENST00000398042 | -18.37 | 8.23E-03 | 1.60E-02 |
| TRPC4 | 13:37636641-37637625 | 13:37637232-37637483 | IR | 3 | 2 | ENST00000338947 | -18.37 | 2.28E-02 | 3.62E-02 |
| TAZ | X:154419624-154420031 | X:154419705-154420031 | A3ss | 3 | 3 | ENST00000470127 | -18.41 | 1.44E-02 | 1.00E+00 |
| AC004967.2 | 7:97966978-97969769 | 7:97969005-97969769 | A5ss | 3 | 3 | Ex.TSS.ENST00000650204 | -18.42 | 3.33E-02 | 5.12E-01 |
| NPAS1 | 19:47035359-47035963 | 19:47035860-47035963 | TSSIA3ss | 3 | 3 | TSS.ENST00000594670 | -18.42 | 3.37E-02 | 2.26E-01 |
| NDRG2 | 14:21023322-21025676 | 14:21024030-21024096 | SES | 3 | 3 | ENST00000298687 | -18.42 | 4.12E-02 | 6.69E-02 |
| ACAD11 | 3:132578882-132579491 | 3:132578882-132579056 | TSSIA3ss | 3 | 3 | TSS.ENST00000477604 | -18.43 | 3.79E-02 | 8.69E-02 |
| PRELID3A | 18:12427324-12431149 | 18:12429350-12429436 | SES | 3 | 2 | ENST00000440960 | -18.48 | 2.14E-02 | 1.06E-01 |
| KLC4 | 6:43060160-43061310 | 6:43060160-43060605 | TSSIA5ss | 3 | 3 | TSS.ENST00000481499 | -18.49 | 2.43E-03 | 2.41E-02 |
| P2RX5 | 17:3689628-3690069 | 17:3689628-3689630 | A3ss | 3 | 2 | ENST00000225328 | -18.55 | 1.70E-03 | 6.91E-03 |
| BAD | 11:64284182-64284639 | 11:64284377-64284630 | IR (overlapping | 3 | 3 | ENST00000394532 | -18.61 | 9.48E-03 | 1.42E-02 |
| RFNG | 17:82049677-82050555 | 17:82049843-82049917 | IR | 3 | 3 | ENST00000580953 | -18.63 | 1.46E-02 | 7.03E-02 |
| SLC26A11 | 17:80220497-80221547 | 17:80220916-80221115 | SES | 3 | 3 | ENST00000361193 | -18.63 | 3.68E-02 | 1.77E-01 |
| RAB29 | 1:205775087-205775373 | 1:205775273-205775373 | TSSIA5ss | 3 | 3 | TSS.ENST00000367139 | -18.63 | 4.26E-02 | 7.58E-02 |
| TUG1 | 22:30969387-30972854 | 22:30969387-30971779 | TSSIA5ss | 3 | 3 | TSS.ENST00000644773 | -18.66 | 1.10E-02 | 2.23E-02 |
| TAZ | X:154419624-154420031 | X:154419705-154419746 | SES | 3 | 3 | ENST00000439735 | -18.66 | 1.14E-02 | 8.26E-01 |
| BBS1 | 11:66531951-66533598 | 11:66532206-66532295 | IR (overlapping | 3 | 3 | ENST00000318312 | -18.66 | 1.86E-02 | 2.83E-02 |
| KLC1 | 14:103679519-103684985 | 14:103679519-103679545 | A5ss | 3 | 3 | ENST00000389744 | -18.68 | 3.71E-03 | 6.01E-03 |
| RGL3 | 19:11416115-11416613 | 19:11416115-11416148 | A3ss | 3 | 3 | ENST00000380456 | -18.68 | 1.81E-02 | 9.34E-02 |
| TRAPPC10 | 21:44086961-44087821 | 21:44087699-44087821 | A3ss | 3 | 3 | Ex.ENST00000291574 | -18.7 | 1.97E-02 | 3.82E-02 |
| ZNF266 | 19:9418505-9419081 | 19:9418632-9418765 | IR | 3 | 3 | ENST00000591213 | -18.71 | 2.36E-03 | 1.80E-02 |
| ZNF266 | 19:9418505-9419081 | 19:9418632-9418761 | IR | 3 | 3 | ENST00000591213 | -18.71 | 2.36E-03 | 1.80E-02 |
| ATXN2 | 12:111470743-111485264 | 12:111479070-111479102 | SES | 3 | 3 | ENST00000643669 | -18.72 | 3.47E-02 | 5.39E-02 |
| KRIT1 | 7:92245465-92246100 | 7:92245677-92245789 | IR (overlapping | 3 | 3 | ENST00000340022 | -18.74 | 5.68E-03 | 8.74E-02 |
| ZNF232 | 17:5106507-5108925 | 17:5106507-5106533 | A3ss | 3 | 3 | Ex.TSS.ENST00000250076 | -18.74 | 6.44E-03 | 2.86E-02 |
| ZNF232 | 17:5106498-5108925 | 17:5106498-5106533 | A3ss | 3 | 3 | Ex.TSS.ENST00000250076 | -18.74 | 6.44E-03 | 2.86E-02 |
| AMMECR1L | 2:127866997-127869453 | 2:127866997-127867279 | A3ss | 3 | 3 | Ex.ENST00000272647 | -18.77 | 4.37E-03 | 9.15E-03 |
| TRPT1 | 11:64224971-64226049 | 11:64224971-64225580 | A3ss | 3 | 3 | ENST00000542040 | -18.81 | 2.62E-02 | 3.93E-02 |
| AGTPBP1 | 9:85655201-85657434 | 9:85655201-85655320 | A3ss | 3 | 3 | ENST00000337006 | -18.82 | 2.39E-03 | 7.79E-02 |

| | | | | | | | | | |
|---|---|---|---|---|---|---|---|---|---|
| ANO7 | 2:241223186-241223955 | 2:241223782-241223904 | IR | 3 | 3 | ENST00000459928 | -18.85 | 4.13E-02 | 8.98E-02 |
| SNTG2 | 2:1083656-1165547 | 2:1098196-1098252 | MES | 3 | 3 | ENST00000308624 | -18.89 | 1.66E-02 | 3.45E-02 |
| MXD3 | 5:177311761-177312309 | 5:177311858-177312120 | IR | 3 | 3 | ENST00000439742 | -18.91 | 4.07E-02 | 1.21E-01 |
| ZNF267 | 16:31885257-31914475 | 16:31890207-31890250 | SES | 3 | 3 | ENST00000394846 | -18.95 | 2.88E-02 | 1.05E-01 |
| GNAS-AS1 | 20:58842546-58850529 | 20:58848879-58848917 | SES | 3 | 3 | ENST00000424094 | -18.95 | 4.50E-02 | 6.27E-02 |
| BIRC2 | 11:102348598-102350749 | 11:102348765-102349877 | IR | 3 | 3 | ENST00000227758 | -18.98 | 8.79E-03 | 1.26E-02 |
| WDR37 | 10:1056880-1072115 | 10:1056880-1056968 | A5ss | 3 | 3 | Ex.TSS.ENST00000263150 | -18.99 | 4.56E-02 | 6.37E-02 |
| WDR37 | 10:1056565-1072115 | 10:1056565-1056968 | TSSIA5ss | 3 | 3 | TSS.ENST00000263150 | -18.99 | 4.56E-02 | 6.37E-02 |
| MAPK8IP3 | 16:1743332-1745103 | 16:1744190-1744977 | IR | 3 | 3 | ENST00000561765 | -19.02 | 4.42E-02 | 1.55E-01 |
| ARMCX3 | X:101623277-101623813 | X:101623786-101623814 | A3ss | 3 | 3 | Ex.TSS.ENST00000471229 | -19.05 | 8.89E-03 | 3.94E-01 |
| TMEM41B | 11:9280654-9283593 | 11:9281042-9283370 | IR | 3 | 3 | ENST00000524543 | -19.16 | 3.68E-02 | 5.68E-02 |
| THTPA | 14:23556271-23557304 | 14:23556344-23556752 | IR (overlapping | 3 | 3 | ENST00000288014 | -19.19 | 1.80E-03 | 2.92E-03 |
| ABCD4 | 14:74292870-74295147 | 14:74292870-74293248 | A3ss | 3 | 3 | ENST00000496015 | -19.19 | 1.78E-02 | 5.65E-02 |
| SEMA6C | 1:151144435-151146432 | 1:151145024-151146432 | TSSIA5ss | 3 | 3 | TSS.ENST00000341697 | -19.19 | 1.78E-02 | 3.05E-02 |
| RIC3 | 11:8126805-8137377 | 11:8128219-8128354 | SES | 3 | 3 | ENST00000530060 | -19.2 | 6.76E-03 | 1.04E-02 |
| SUGP2 | 19:18990893-18993740 | 19:18991150-18993647 | IR | 3 | 3 | ENST00000452918 | -19.23 | 2.99E-03 | 1.64E-02 |
| AL645939.2 | 6:29722981-29723971 | 6:29723528-29723769 | IR (overlapping | 3 | 3 | ENST00000427340 | -19.26 | 3.20E-02 | 9.97E-02 |
| MTHFD2L | 4:74158282-74174505 | 4:74160055-74160088 | SES | 3 | 3 | ENST00000359107 | -19.28 | 1.62E-02 | 4.56E-02 |
| SLC5A6 | 2:27211534-27211766 | 2:27211711-27211766 | TSSIA5ss | 3 | 3 | TSS.ENST00000401463 | -19.37 | 7.11E-04 | 1.56E-03 |
| MMP19 | 12:55835433-55837374 | 12:55835616-55836125 | IR (overlapping | 3 | 3 | ENST00000322569 | -19.42 | 1.11E-02 | 1.84E-02 |
| FKBP7 | 2:178465932-178478278 | 2:178469652-178469785 | MXS | 3 | 2 | ENST00000419184 | -19.56 | 3.37E-02 | 7.13E-02 |
| FAM122C | X:134807169-134807708 | X:134807342-134807539 | IR | 3 | 3 | ENST00000646004 | -19.57 | 7.15E-03 | 3.91E-01 |
| FIRRE | X:131768697-131783886 | X:131777623-131777744 | SES | 3 | 3 | ENST00000647966 | -19.58 | 1.68E-02 | 9.80E-01 |
| LINC00869 | 1:149646599-149647610 | 1:149646946-149647380 | IR | 3 | 3 | ENST00000620190 | -19.58 | 3.77E-02 | 3.25E-01 |
| DOCK6 | 19:11229036-11233202 | 19:11232185-11232289 | MXS | 3 | 3 | ENST00000587656 | -19.6 | 3.39E-02 | 1.74E-01 |
| BRD8 | 5:138168079-138169221 | 5:138168470-138168688 | SES | 3 | 3 | ENST00000230901 | -19.65 | 8.25E-04 | 2.39E-03 |
| TOR1AIP2 | 1:179852812-179877238 | 1:179865436-179865854 | SES | 3 | 3 | ENST00000609928 | -19.66 | 1.87E-04 | 3.29E-04 |
| ZNF639 | 3:179323292-179327560 | 3:179325042-179325322 | MXS | 3 | 3 | ENST00000491818 | -19.68 | 3.34E-02 | 7.79E-02 |
| LDOC1 | X:141173235-141177129 | X:141176812-141176892 | IR | 3 | 3 | ENST00000370526 | -19.69 | 1.22E-03 | 7.42E-02 |
| FIRRE | X:131689755-131692636 | X:131690016-131692437 | IR | 3 | 3 | ENST00000663405 | -19.73 | 4.75E-02 | 1.00E+00 |
| AL137058.2 | 13:52627579-52637304 | 13:52632422-52632494 | SES | 3 | 3 | ENST00000398039 | -19.8 | 3.93E-02 | 6.40E-02 |
| PIEZO2 | 18:10731522-10736603 | 18:10731522-10735330 | A3ss | 3 | 3 | Ex.ENST00000302079 | -19.81 | 1.13E-02 | 5.59E-02 |
| CLUHP3 | 16:31700739-31703155 | 16:31702535-31702681 | SES | 2 | 2 | ENST00000532304 | -19.82 | 1.11E-02 | 4.05E-02 |
| FAM122B | X:134796108-134797155 | X:134796257-134797120 | IR | 3 | 3 | ENST00000343004 | -19.85 | 3.07E-03 | 1.81E-01 |
| CREBZF | 11:85661066-85665102 | 11:85661750-85662388 | IR | 3 | 3 | ENST00000527447 | -19.85 | 4.09E-02 | 6.31E-02 |
| CCNE1 | 19:29821818-29822239 | 19:29821818-29822130 | TSSIA5ss | 3 | 3 | TSS.ENST00000574121 | -19.88 | 1.48E-02 | 8.21E-02 |
| TAZ | X:154419624-154420031 | X:154419624-154419746 | A5ss | 3 | 3 | ENST00000426231 | -19.9 | 2.00E-02 | 1.00E+00 |
| ZNF43 | 19:21819222-21851904 | 19:21850938-21851027 | MES | 3 | 3 | ENST00000594012 | -19.97 | 2.55E-02 | 1.40E-01 |
| ACOX1 | 17:75960376-75978533 | 17:75973625-75973785 | SES | 3 | 3 | ENST00000573078 | -20 | 1.47E-02 | 6.96E-02 |
| TRMU | 22:46337945-46346421 | 22:46343262-46343368 | SES | 3 | 3 | ENST00000381019 | -20.07 | 2.05E-02 | 4.24E-02 |
| SUGP2 | 19:18990893-18993740 | 19:18991150-18991339 | IR | 3 | 3 | ENST00000452918 | -20.07 | 3.83E-02 | 2.10E-01 |
| BUB1B | 15:40200981-40202449 | 15:40202405-40202449 | A3ss | 3 | 3 | Ex.ENST00000287598 | -20.09 | 9.26E-03 | 3.03E-02 |
| LETMD1 | 12:51049186-51056349 | 12:51052092-51052207 | MES | 3 | 3 | ENST00000262055 | -20.09 | 3.02E-02 | 4.95E-02 |
| BCAM | 19:44811347-44812225 | 19:44812163-44812225 | A3ss | 3 | 3 | ENST00000270233 | -20.12 | 2.64E-03 | 1.76E-02 |
| MTFR2 | 6:136249154-136250263 | 6:136249976-136250263 | TSSIA5ss | 3 | 3 | TSS.ENST00000420702 | -20.12 | 2.20E-02 | 6.87E-02 |
| CREBZF | 11:85660335-85661749 | 11:85660598-85661636 | IR | 3 | 3 | ENST00000525639 | -20.17 | 3.60E-02 | 5.56E-02 |
| BORA | 13:72729094-72737961 | 13:72731281-72731387 | MES | 3 | 3 | ENST00000377815 | -20.18 | 1.47E-02 | 2.40E-02 |
| SLC17A9 | 20:62966703-62968597 | 20:62966733-62967336 | IR | 3 | 3 | ENST00000488738 | -20.19 | 3.81E-02 | 5.31E-02 |
| ATP2A1-AS1 | 16:28879489-28879895 | 16:28879756-28879866 | IR (overlapping | 3 | 3 | ENST00000561547 | -20.22 | 1.14E-04 | 4.15E-04 |
| CCDC82 | 11:96359098-96364979 | 11:96359098-96359178 | A3ss | 3 | 3 | ENST00000278520 | -20.32 | 2.17E-02 | 3.37E-02 |
| TMEM147-AS1 | 19:35541582-35545492 | 19:35541585-35542910 | IR (overlapping | 3 | 3 | ENST00000588286 | -20.41 | 3.11E-02 | 1.81E-01 |
| SLC9A1 | 1:27098815-27100644 | 1:27099005-27099211 | IR | 3 | 3 | ENST00000374089 | -20.42 | 1.61E-02 | 2.97E-02 |
| SPINT1-AS1 | 15:40836283-40844178 | 15:40838543-40838686 | SES | 3 | 3 | ENST00000564302 | -20.43 | 1.09E-02 | 3.57E-02 |
| APBB3 | 5:140561587-140562227 | 5:140561723-140561843 | IR | 3 | 3 | ENST00000510241 | -20.43 | 1.27E-02 | 3.70E-02 |
| TAF1C | 16:84183779-84186900 | 16:84184851-84185060 | MXS | 3 | 3 | ENST00000564774 | -20.46 | 9.62E-04 | 3.72E-03 |
| EME2 | 16:1778455-1781708 | 16:1778613-1781192 | IR (overlapping | 3 | 3 | ENST00000561903 | -20.48 | 5.59E-03 | 1.97E-02 |
| AC091230.1 | 15:74842418-74843003 | 15:74842418-74842420 | TSSIA3ss | 3 | 3 | TSS.ENST00000440863 | -20.48 | 4.54E-02 | 6.31E-02 |
| ORAI2 | 7:102433662-102438943 | 7:102436225-102436333 | SES | 3 | 3 | ENST00000495936 | -20.56 | 1.68E-02 | 1.96E-02 |
| PCSK4 | 19:1486853-1487313 | 19:1487066-1487140 | IR | 3 | 3 | ENST00000586616 | -20.56 | 2.91E-02 | 1.57E-01 |
| SYT7 | 11:61533125-61551383 | 11:61546031-61546255 | MXS | 3 | 3 | ENST00000540677 | -20.59 | 1.04E-02 | 1.54E-02 |
| TPM1 | 15:63060940-63061712 | 15:63061198-63061273 | SES | 3 | 3 | ENST00000558264 | -20.61 | 2.75E-04 | 9.22E-04 |
| ASPHD1 | 16:29900488-29901920 | 16:29900742-29901551 | IR | 3 | 3 | ENST00000308748 | -20.62 | 2.16E-02 | 7.88E-02 |
| TRNT1 | 3:3127020-3128018 | 3:3127290-3127945 | IR | 3 | 3 | ENST00000402675 | -20.68 | 2.88E-02 | 6.85E-02 |
| BAZ2B | 2:159429261-159432756 | 2:159430863-159431156 | SES | 2 | 2 | ENST00000392782 | -20.74 | 1.88E-02 | 3.95E-02 |
| COASY | 17:42562148-42563322 | 17:42562220-42562932 | IR | 3 | 3 | ENST00000393818 | -20.78 | 8.97E-06 | 3.79E-05 |
| SEPTIN9 | 17:77450787-77482143 | 17:77451367-77451528 | SES | 3 | 3 | ENST00000586128 | -20.81 | 2.72E-04 | 1.30E-03 |
| MTHFR | 1:11802881-11804532 | 1:11803130-11803798 | IR | 3 | 3 | ENST00000376592 | -20.81 | 1.23E-02 | 1.04E-01 |
| EPM2A | 6:145686297-145735244 | 6:145735198-145735244 | TSSIA5ss | 3 | 3 | TSS.ENST00000367519 | -20.87 | 2.75E-02 | 8.56E-02 |
| KCTD17 | 22:37059439-37061538 | 22:37060823-37060922 | MES | 3 | 3 | ENST00000403888 | -20.9 | 6.16E-03 | 1.26E-02 |
| PHKB | 16:47461427-47497398 | 16:47463899-47463993 | MXS | 3 | 3 | ENST00000566037 | -21.08 | 5.35E-04 | 1.97E-03 |
| PACRGL | 4:20704815-20713431 | 4:20707803-20707870 | MES | 3 | 3 | ENST00000295290 | -21.1 | 7.27E-03 | 1.97E-02 |
| FAM122B | X:134796108-134797155 | X:134796356-134797120 | IR | 3 | 3 | ENST00000343004 | -21.26 | 1.21E-02 | 6.60E-01 |
| ZFP62 | 5:180847615-180851493 | 5:180848083-180849153 | IR | 3 | 3 | ENST00000512132 | -21.28 | 8.61E-04 | 2.57E-03 |
| TANGO2 | 22:20036855-20043354 | 22:20036855-20037094 | A5ss | 3 | 3 | ENST00000399807 | -21.3 | 2.79E-03 | 5.45E-03 |
| PCNX1 | 14:71011550-71012999 | 14:71012985-71012999 | A3ss | 3 | 3 | ENST00000304743 | -21.3 | 2.40E-02 | 7.56E-02 |
| CYB5RL | 1:54184266-54195418 | 1:54190738-54190896 | MXS | 3 | 3 | ENST00000493530 | -21.33 | 3.22E-02 | 6.03E-02 |
| TSPAN9 | 12:3077454-3201176 | 12:3083653-3083719 | SES | 3 | 3 | ENST00000011898 | -21.34 | 3.33E-02 | 5.34E-02 |
| SH2D3C | 9:127751301-127755084 | 9:127755079-127755084 | TSSIA5ss | 3 | 3 | TSS.ENST00000420366 | -21.35 | 1.66E-03 | 5.88E-02 |
| STK39 | 2:168063571-168074981 | 2:168065319-168065381 | SES | 3 | 3 | ENST00000355999 | -21.36 | 2.52E-03 | 5.30E-03 |
| LRATD2 | 8:126552901-126557731 | 8:126553504-126556264 | IR | 3 | 3 | ENST00000652209 | -21.38 | 6.12E-03 | 1.17E-01 |
| ELN | 7:74056714-74059885 | 7:74057640-74057696 | MXS | 3 | 2 | ENST00000252034 | -21.4 | 6.59E-03 | 9.78E-02 |
| ZNF652 | 17:49294565-49298924 | 17:49298480-49298631 | IR (overlapping | 3 | 3 | ENST00000362063 | -21.45 | 1.30E-02 | 5.66E-02 |
| AC016876.2 | 17:7582161-7583917 | 17:7582161-7582163 | A3ss | 3 | 3 | ENST00000417897 | -21.47 | 2.45E-02 | 1.16E-01 |
| ZBTB38 | 3:141404032-141438267 | 3:141438243-141438267 | A3ss | 3 | 3 | ENST00000504673 | -21.48 | 2.81E-02 | 6.47E-02 |
| SLC35D2 | 9:96321342-96324090 | 9:96321998-96322080 | SES | 3 | 3 | ENST00000253270 | -21.51 | 7.04E-05 | 2.36E-03 |
| ZNF268 | 12:133192004-133202143 | 12:133193428-133193533 | SES | 3 | 3 | ENST00000534953 | -21.51 | 1.60E-02 | 2.56E-02 |
| GLYCTK | 3:52291112-52292259 | 3:52291747-52291922 | SES | 2 | 2 | ENST00000436784 | -21.69 | 4.34E-02 | 1.12E-01 |
| PWWP3A | 19:1358465-1360135 | 19:1358588-1358700 | SES | 3 | 3 | ENST00000587460 | -21.7 | 3.42E-02 | 1.83E-01 |
| EME2 | 16:1778455-1781708 | 16:1778613-1781357 | IR (overlapping | 3 | 3 | ENST00000561903 | -21.71 | 1.02E-02 | 3.60E-02 |
| ZNF100 | 19:21726193-21727989 | 19:21726605-21726607 | IR | 3 | 3 | ENST00000305570 | -21.73 | 1.41E-02 | 7.73E-02 |
| GPR108 | 19:6731055-6731198 | 19:6731055-6731111 | A3ss | 3 | 3 | ENST00000264080 | -21.74 | 2.86E-02 | 2.03E-01 |
| SLC16A5 | 17:75088045-75093588 | 17:75089151-75089304 | SES | 3 | 3 | ENST00000329783 | -21.8 | 9.88E-03 | 4.55E-02 |
| RFNG | 17:82049677-82050555 | 17:82049843-82050401 | IR | 3 | 3 | ENST00000580953 | -21.92 | 1.48E-04 | 7.25E-04 |
| MPI | 15:74897512-74902219 | 15:74901257-74901949 | IR (overlapping | 3 | 3 | ENST00000352410 | -21.98 | 2.59E-02 | 8.77E-02 |

| | | | | | | | | | |
|---|---|---|---|---|---|---|---|---|---|
| EP300 | 22:41158501-41160641 | 22:41160612-41160642 | A3ss | 3 | 3 | Ex.ENST00000263253 | -22.02 | 1.92E-03 | 3.95E-03 |
| FRG1HP | 9:41011731-41032833 | 9:41013781-41032834 | A3ss | 2 | 3 | Ex.TSS.ENST00000611606 | -22.08 | 8.53E-03 | 2.34E-01 |
| IQCB1 | 3:121790216-121807343 | 3:121797118-121797227 | MES | 3 | 3 | ENST00000310864 | -22.08 | 3.74E-02 | 8.54E-02 |
| CREBZF | 11:85663524-85665102 | 11:85663602-85663814 | IR | 3 | 3 | ENST00000490820 | -22.11 | 3.80E-03 | 5.86E-03 |
| LMO7 | 13:75834388-75835232 | 13:75835152-75835232 | A3ss | 2 | 3 | ENST00000485987 | -22.21 | 2.29E-02 | 3.74E-02 |
| FAM193B | 5:177536487-177537872 | 5:177536487-177536745 | A3ss | 3 | 3 | ENST00000506955 | -22.21 | 2.82E-02 | 8.36E-02 |
| DPH5 | 1:100990632-100995109 | 1:100992637-100992740 | SES | 3 | 3 | ENST00000464270 | -22.26 | 2.42E-03 | 1.97E-02 |
| SLC27A1 | 19:17470708-17487459 | 19:17486563-17486957 | MES | 3 | 3 | ENST00000252595 | -22.26 | 2.66E-02 | 1.44E-01 |
| HDAC11 | 3:13483565-13501870 | 3:13498513-13498555 | MES | 3 | 3 | ENST00000295757 | -22.26 | 4.82E-02 | 1.11E-01 |
| SGSH | 17:80214680-80215032 | 17:80214680-80214765 | A3ss | 3 | 3 | ENST00000326317 | -22.29 | 8.03E-04 | 3.88E-03 |
| MRPS31P4 | 13:52627579-52637304 | 13:52632422-52632494 | MES | 3 | 2 | ENST00000509685 | -22.3 | 3.63E-02 | 5.93E-02 |
| MORN1 | 1:2388338-2391457 | 1:2389925-2389996 | SES | 3 | 3 | ENST00000378529 | -22.33 | 3.85E-02 | 6.94E-02 |
| TMEM147-AS1 | 19:35541582-35545492 | 19:35541585-35541750 | IR (overlapping | 3 | 3 | ENST00000588286 | -22.41 | 1.54E-03 | 8.93E-03 |
| PLD3 | 19:40348725-40365717 | 19:40365553-40365585 | SES | 3 | 3 | ENST00000392032 | -22.41 | 2.44E-03 | 1.60E-02 |
| EMID1 | 22:29243490-29258816 | 22:29254203-29254287 | MXS | 2 | 3 | ENST00000334018 | -22.42 | 3.09E-02 | 6.08E-02 |
| LRSAM1 | 9:127451513-127452084 | 9:127451670-127451928 | IR | 3 | 3 | ENST00000323301 | -22.43 | 2.81E-02 | 9.84E-01 |
| ENTPD1-AS1 | 10:95904939-95907777 | 10:95907663-95907777 | TSSIA5ss | 3 | 3 | TSS.ENST00000657446 | -22.43 | 3.62E-02 | 5.15E-02 |
| SGK3 | 8:66667702-66679720 | 8:66677642-66677954 | SES | 3 | 3 | ENST00000661636 | -22.45 | 6.65E-03 | 1.52E-01 |
| ENTPD4 | 8:23439916-23441592 | 8:23441569-23441592 | A5ss | 3 | 3 | ENST00000358689 | -22.57 | 2.39E-02 | 5.10E-01 |
| MAN2A2 | 15:90916257-90918193 | 15:90916586-90916660 | SES | 3 | 3 | ENST00000557990 | -22.58 | 1.78E-04 | 6.20E-04 |
| MEF2A | 15:99566105-99632977 | 15:99598430-99598511 | MXS | 3 | 3 | ENST00000354410 | -22.6 | 3.37E-02 | 1.18E-01 |
| LGALS8 | 1:236542788-236543559 | 1:236542915-236543040 | SES | 2 | 3 | ENST00000352231 | -22.67 | 5.92E-03 | 1.06E-02 |
| PLPP5 | 8:38263407-38266140 | 8:38263407-38264604 | TSSIA3ss | 3 | 3 | TSS.ENST00000424479 | -22.67 | 1.83E-02 | 3.87E-01 |
| KIF20B | 10:89725159-89726412 | 10:89726293-89726412 | A3ss | 3 | 3 | ENST00000371728 | -22.69 | 6.77E-03 | 9.68E-03 |
| C11orf74 | 11:36610240-36648015 | 11:36636051-36636117 | MES | 3 | 3 | ENST00000334307 | -22.84 | 3.44E-02 | 5.05E-02 |
| FBN3 | 19:8124009-8126296 | 19:8125892-8126017 | SES | 3 | 3 | ENST00000270509 | -22.87 | 2.69E-02 | 2.04E-01 |
| EML4 | 2:42256631-42263177 | 2:42261121-42261294 | SES | 3 | 3 | ENST00000318522 | -22.92 | 4.37E-03 | 9.62E-03 |
| DLST | 14:74882017-74889094 | 14:74882591-74882624 | MXS | 3 | 3 | ENST00000554806 | -22.93 | 3.29E-07 | 1.05E-06 |
| RFNG | 17:82049918-82050514 | 17:82050007-82050401 | IR | 3 | 3 | ENST00000580793 | -22.97 | 1.46E-02 | 7.02E-02 |
| MFSD8 | 4:127917925-127921771 | 4:127920837-127921117 | IR | 2 | 3 | ENST00000641482 | -22.98 | 7.64E-03 | 2.03E-02 |
| GNB1L | 22:19812448-19854442 | 22:19821228-19821375 | MES | 3 | 3 | ENST00000329517 | -23.02 | 1.37E-02 | 2.68E-02 |
| TMEM218 | 11:125101304-125102733 | 11:125102132-125102317 | SES | 3 | 3 | ENST00000526175 | -23.03 | 6.82E-04 | 9.86E-04 |
| AC074117.1 | 2:27357808-27360256 | 2:27357903-27358884 | IR | 3 | 3 | ENST00000447070 | -23.03 | 1.89E-02 | 4.14E-02 |
| SEPTIN6 | X:119620052-119629317 | X:119625335-119625379 | SES | 3 | 3 | ENST00000394610 | -23.07 | 2.06E-02 | 1.00E+00 |
| CEP83 | 12:94411848-94412516 | 12:94412318-94412429 | SES | 3 | 3 | ENST00000547232 | -23.08 | 4.08E-03 | 6.49E-03 |
| DTX3 | 12:57604859-57606189 | 12:57605571-57605731 | MXS | 3 | 3 | ENST00000337737 | -23.19 | 1.04E-02 | 1.73E-02 |
| CREM | 10:35207052-35211655 | 10:35211254-35211655 | TSSIA3ss | 3 | 3 | TSS.ENST00000344351 | -23.23 | 2.39E-02 | 3.39E-02 |
| AC068279.2 | 2:87323697-87348658 | 2:87323697-87345262 | A3ss | 3 | 3 | Ex.ENST00000664547 | -23.28 | 4.69E-03 | 1.06E-02 |
| ZDHHC19 | 3:196198870-196207397 | 3:196198870-196198874 | A3ss | 3 | 3 | ENST00000296326 | -23.35 | 2.60E-02 | 6.16E-02 |
| MRE11 | 11:94492907-94493790 | 11:94493684-94493790 | TSSIA5ss | 3 | 3 | TSS.ENST00000323977 | -23.37 | 2.18E-03 | 3.37E-03 |
| PPOX | 1:161167235-161170619 | 1:161170409-161170519 | MES | 3 | 3 | ENST00000352210 | -23.4 | 1.87E-02 | 3.26E-02 |
| UMAD1 | 7:7640822-7673321 | 7:7673309-7673321 | TSSIA3ss | 3 | 2 | TSS.ENST00000463725 | -23.44 | 4.79E-02 | 7.17E-01 |
| GLI2 | 2:120972064-120974974 | 2:120974924-120974974 | A3ss | 3 | 3 | ENST00000361492 | -23.47 | 2.35E-02 | 4.88E-02 |
| DNASE1 | 16:3657720-3658207 | 16:3657817-3657905 | IR (overlapping | 3 | 3 | ENST00000576792 | -23.49 | 4.21E-02 | 1.54E-01 |
| MERTK | 2:111898797-111944959 | 2:111929120-111929304 | SES | 3 | 3 | ENST00000439966 | -23.5 | 2.50E-02 | 5.21E-02 |
| LINC00665 | 19:36325888-36330252 | 19:36326997-36327225 | MES | 3 | 3 | ENST00000658301 | -23.51 | 3.62E-02 | 2.33E-01 |
| ASPHD1 | 16:29900488-29901920 | 16:29900567-29901551 | IR | 3 | 3 | ENST00000308748 | -23.52 | 7.52E-03 | 2.74E-02 |
| LINC00667 | 18:5240648-5243619 | 18:5241890-5243593 | IR | 2 | 3 | ENST00000657129 | -23.52 | 2.80E-02 | 1.41E-01 |
| AMT | 3:49417358-49417614 | 3:49417358-49417367 | TSSIA3ss | 3 | 3 | TSS.ENST00000395338 | -23.53 | 8.49E-03 | 2.05E-02 |
| AMT | 3:49417310-49417614 | 3:49417310-49417367 | TSSIA3ss | 3 | 3 | TSS.ENST00000395338 | -23.53 | 8.49E-03 | 2.05E-02 |
| ZNF142 | 2:218651823-218656149 | 2:218651823-218652300 | TSSIA3ss | 3 | 3 | TSS.ENST00000440934 | -23.63 | 2.27E-02 | 4.91E-02 |
| CSPP1 | 8:67172556-67175295 | 8:67172769-67172973 | SES | 3 | 2 | ENST00000521324 | -23.73 | 2.03E-02 | 4.65E-01 |
| OBSCN | 1:228359597-228361250 | 1:228360313-228360889 | IR (overlapping | 3 | 3 | ENST00000284548 | -23.74 | 6.54E-03 | 1.17E-02 |
| IMMT | 2:86171310-86173649 | 2:86171310-86171312 | A3ss | 3 | 3 | Ex.ENST00000254636 | -23.8 | 9.03E-04 | 2.04E-03 |
| ZNF100 | 19:21726193-21727989 | 19:21726635-21726636 | IR | 3 | 3 | ENST00000305570 | -23.85 | 7.23E-03 | 3.98E-02 |
| CREBZF | 11:85663524-85665102 | 11:85663602-85663697 | IR | 3 | 3 | ENST00000490820 | -23.86 | 2.18E-03 | 3.36E-03 |
| LIN54 | 4:82924603-82928303 | 4:82924851-82928055 | IR | 3 | 3 | ENST00000340417 | -23.86 | 4.68E-02 | 1.33E-01 |
| SUGP2 | 19:18990893-18993740 | 19:18991150-18993276 | IR | 3 | 3 | ENST00000452918 | -23.89 | 5.38E-04 | 2.95E-03 |
| ZSCAN26 | 6:28272337-28272669 | 6:28272337-28272339 | A5ss | 3 | 3 | ENST00000421553 | -23.92 | 1.72E-02 | 1.55E-01 |
| ZSCAN26 | 6:28271935-28272669 | 6:28271935-28272339 | A5ss | 3 | 3 | ENST00000421553 | -23.92 | 1.72E-02 | 1.55E-01 |
| HPN | 19:35041873-35049289 | 19:35042453-35042522 | SES | 3 | 3 | ENST00000392226 | -24.01 | 3.20E-03 | 1.84E-02 |
| SH2D3A | 19:6754933-6759593 | 19:6754933-6755315 | A3ss | 3 | 3 | ENST00000245908 | -24.02 | 3.72E-02 | 2.63E-01 |
| TCEANC | X:13655374-13662500 | X:13659652-13662501 | A3ss | 2 | 2 | Ex.ENST00000380600 | -24.02 | 4.53E-02 | 1.00E+00 |
| LRRC14 | 8:144518042-144519700 | 8:144519615-144519701 | A3ss | 3 | 3 | Ex.TSS.ENST00000527730 | -24.07 | 2.87E-02 | 5.90E-01 |
| RIMS2 | 8:103766538-103885297 | 8:103809280-103809321 | SES | 3 | 3 | ENST00000668113 | -24.09 | 1.40E-02 | 2.63E-01 |
| AP3B2 | 15:82685914-82688831 | 15:82688097-82688735 | IR (overlapping | 3 | 3 | ENST00000535513 | -24.12 | 6.28E-03 | 2.14E-02 |
| STPG1 | 1:24384004-24401318 | 1:24391561-24391679 | MXS | 3 | 3 | ENST00000337248 | -24.16 | 1.29E-02 | 2.33E-02 |
| ZNF142 | 2:218650527-218656149 | 2:218651701-218652300 | SES | 3 | 3 | ENST00000440934 | -24.19 | 3.05E-04 | 6.58E-04 |
| SUGP2 | 19:18991150-18994365 | 19:18991150-18993740 | TSSIA3ss | 3 | 3 | TSS.ENST00000452918 | -24.2 | 9.13E-03 | 5.00E-02 |
| PVT1 | 8:127890999-127989161 | 8:127939508-127939676 | MXS | 3 | 3 | ENST00000517525 | -24.24 | 7.91E-03 | 1.52E-01 |
| C2orf27A | 2:131682600-131684708 | 2:131682600-131682988 | A5ss | 3 | 3 | Ex.TSS.ENST00000653458 | -24.29 | 2.73E-02 | 5.71E-02 |
| C2orf27A | 2:131682558-131684708 | 2:131682558-131682988 | A5ss | 3 | 3 | Ex.TSS.ENST00000657415 | -24.29 | 2.73E-02 | 5.71E-02 |
| STIL | 1:47299777-47300152 | 1:47299969-47300109 | IR | 3 | 3 | ENST00000433827 | -24.29 | 2.75E-02 | 5.13E-02 |
| ADGRB1 | 8:142543709-142544219 | 8:142544217-142544220 | A3ss | 3 | 3 | Ex.TSS.ENST00000323289 | -24.43 | 4.99E-02 | 9.74E-01 |
| NT5M | 17:17344909-17346804 | 17:17344909-17344926 | TSSIA5ss | 3 | 3 | TSS.ENST00000470418 | -24.5 | 7.92E-03 | 2.80E-02 |
| RSPRY1 | 16:57186430-57204503 | 16:57186430-57186451 | TSSIA5ss | 3 | 3 | TSS.ENST00000394420 | -24.52 | 1.87E-02 | 6.97E-02 |
| AL162258.2 | 1:153632034-153632240 | 1:153632064-153632154 | IR (overlapping | 3 | 3 | ENST00000469931 | -24.52 | 3.75E-02 | 6.45E-02 |
| KNTC1 | 12:122594386-122602568 | 12:122601536-122601625 | MES | 3 | 3 | ENST00000333479 | -24.81 | 4.48E-02 | 7.13E-02 |
| TMEM52 | 1:1918253-1918934 | 1:1918432-1918892 | IR | 3 | 3 | ENST00000602604 | -25.25 | 6.94E-03 | 1.23E-02 |
| ZNF681 | 19:23754898-23758746 | 19:23754898-23754918 | TSSIA3ss | 2 | 3 | TSS.ENST00000528059 | -25.25 | 4.77E-02 | 2.64E-01 |
| CLIP1 | 12:122319349-122327946 | 12:122322114-122324381 | SES | 3 | 3 | ENST00000537123 | -25.36 | 6.37E-03 | 1.01E-02 |
| MPHOSPH9 | 12:123203085-123210055 | 12:123203250-123203375 | SES | 3 | 3 | ENST00000539336 | -25.36 | 1.76E-02 | 2.80E-02 |
| PLD3 | 19:40348769-40365717 | 19:40365553-40365717 | TSSIA3ss | 3 | 3 | TSS.ENST00000594085 | -25.42 | 3.46E-03 | 2.27E-02 |
| PLD3 | 19:40348769-40365661 | 19:40365553-40365661 | TSSIA3ss | 3 | 3 | TSS.ENST00000594085 | -25.42 | 3.46E-03 | 2.27E-02 |
| ATG10 | 5:81972307-81987558 | 5:81976193-81976327 | MXS | 3 | 3 | ENST00000458350 | -25.42 | 1.73E-02 | 5.25E-02 |
| DPPA4 | 3:109329082-109331733 | 3:109330524-109331734 | A5ss | 3 | 2 | Ex.ENST00000463966 | -25.48 | 2.41E-03 | 5.50E-03 |
| NANOGP1 | 12:7898352-7898534 | 12:7898490-7898534 | TSSIA3ss | 3 | 3 | TSS.ENST00000525030 | -25.51 | 4.48E-03 | 7.55E-03 |
| ZNF160 | 19:53086089-53086261 | 19:53086089-53086109 | TSSIA3ss | 3 | 3 | TSS.ENST00000596966 | -25.62 | 2.44E-02 | 1.71E-01 |
| TRIM37 | 17:58999460-59012327 | 17:59001598-59001714 | SES | 3 | 3 | ENST00000262294 | -25.65 | 7.49E-03 | 3.34E-02 |
| MADD | 11:47269751-47275898 | 11:47273827-47273976 | MES | 3 | 3 | ENST00000311027 | -25.72 | 1.54E-02 | 2.27E-02 |
| SLC52A3 | 20:760080-761824 | 20:761239-761700 | IR | 3 | 3 | ENST00000381944 | -25.88 | 2.74E-02 | 5.27E-02 |
| STEAP2 | 7:90232337-90237631 | 7:90232485-90236890 | IR | 3 | 2 | ENST00000287908 | -26.04 | 3.62E-02 | 5.56E-01 |
| MPI | 15:74897512-74902219 | 15:74900600-74901949 | IR (overlapping | 3 | 3 | ENST00000352410 | -26.33 | 2.16E-02 | 7.31E-02 |
| CMTM7 | 3:32442014-32452391 | 3:32449454-32449552 | SES | 3 | 3 | ENST00000334983 | -26.46 | 9.38E-03 | 2.23E-02 |

| | | | | | | | | | |
|---|---|---|---|---|---|---|---|---|---|
| LINC01315 | 22:42364726-42368714 | 22:42364726-42365144 | A3ss | 3 | 3 | ENST00000424852 | -26.46 | 1.69E-02 | 3.48E-02 |
| CPT1B | 22:50577035-50577355 | 22:50577324-50577355 | A5ss | 3 | 3 | ENST00000312108 | -26.61 | 7.33E-03 | 1.52E-02 |
| SKP2 | 5:36177293-36183839 | 5:36181818-36183839 | TSSlA3ss | 3 | 3 | TSS.ENST00000274255 | -26.65 | 2.48E-02 | 7.43E-02 |
| SCAND1 | 20:35953629-35955102 | 20:35954340-35954447 | IR | 3 | 3 | ENST00000373991 | -26.73 | 9.41E-04 | 1.79E-03 |
| IKBKG | X:154556377-154560406 | X:154558535-154558650 | SES | 3 | 3 | ENST00000615874 | -26.75 | 3.30E-02 | 1.00E+00 |
| NEMP1 | 12:57064740-57070819 | 12:57069234-57069306 | SES | 3 | 3 | ENST00000554340 | -26.86 | 1.60E-02 | 2.66E-02 |
| OGG1 | 3:9757348-9762324 | 3:9759576-9759671 | IR (overlapping | 3 | 3 | ENST00000602976 | -26.91 | 4.43E-02 | 1.15E-01 |
| COASY | 17:42562148-42563322 | 17:42562237-42562611 | IR | 3 | 3 | ENST00000393818 | -26.99 | 9.17E-04 | 3.87E-03 |
| AC060780.1 | 17:43166980-43167849 | 17:43167251-43167733 | IR (overlapping | 2 | 3 | ENST00000635600 | -27.11 | 3.57E-03 | 1.53E-02 |
| PRKCG | 19:53892644-53893361 | 19:53892988-53893075 | SES | 3 | 3 | ENST00000263431 | -27.13 | 3.92E-02 | 2.77E-01 |
| WRAP73 | 1:3632318-3633397 | 1:3632318-3632338 | A3ss | 3 | 3 | Ex.ENST00000270708 | -27.14 | 4.98E-02 | 9.14E-02 |
| MDC1 | 6:30707455-30708135 | 6:30707566-30708135 | A5ss | 3 | 3 | Ex.ENST00000376406 | -27.18 | 1.40E-02 | 1.28E-01 |
| ANKHD1 | 5:140436258-140438502 | 5:140438461-140438502 | A3ss | 3 | 3 | ENST00000297183 | -27.21 | 1.66E-03 | 4.83E-03 |
| ANKHD1 | 5:140436258-140438493 | 5:140438461-140438493 | A3ss | 3 | 3 | ENST00000297183 | -27.21 | 1.66E-03 | 4.83E-03 |
| CYP2R1 | 11:14880360-14885775 | 11:14880360-14880768 | A3ss | 3 | 3 | ENST00000334636 | -27.34 | 3.01E-02 | 4.35E-02 |
| WDR27 | 6:169634526-169658275 | 6:169649198-169649275 | MES | 3 | 3 | ENST00000423258 | -27.36 | 2.23E-02 | 1.99E-01 |
| AC016027.1 | 22:18077186-18078884 | 22:18078128-18078295 | IR (overlapping | 3 | 3 | ENST00000607927 | -27.62 | 1.07E-02 | 2.08E-02 |
| AC009022.1 | 16:70038986-70042350 | 16:70038986-70039100 | A3ss | 3 | 3 | ENST00000530079 | -27.66 | 3.32E-03 | 1.27E-02 |
| UBXN11 | 1:26294332-26306591 | 1:26297962-26298062 | MES | 3 | 2 | ENST00000357089 | -27.71 | 1.21E-04 | 2.20E-04 |
| STAG3L5P-PVRIG2P-PILRB | 7:100345969-100349798 | 7:100349731-100349798 | A3ss | 3 | 3 | ENST00000444874 | -27.86 | 1.17E-02 | 1.35E-01 |
| AC016027.1 | 22:18077186-18078884 | 22:18078048-18078295 | IR (overlapping | 3 | 3 | ENST00000607927 | -27.98 | 1.09E-02 | 2.12E-02 |
| TRIP12 | 2:229793131-229795178 | 2:229793131-229793145 | A3ss | 3 | 3 | Ex.ENST00000283943 | -28.19 | 1.60E-02 | 3.47E-02 |
| SYT7 | 11:61533125-61551383 | 11:61546031-61546255 | MES | 3 | 3 | ENST00000535826 | -28.25 | 3.50E-05 | 5.17E-05 |
| TTLL4 | 2:218746232-218747029 | 2:218747003-218747029 | A3ss | 3 | 3 | ENST00000258398 | -28.28 | 2.17E-03 | 4.69E-03 |
| SEC22C | 3:42548658-42555929 | 3:42548658-42553448 | TSSlA3ss | 3 | 3 | TSS.ENST00000451653 | -28.39 | 2.85E-02 | 6.83E-02 |
| COASY | 17:42562148-42563322 | 17:42562214-42562611 | IR | 3 | 3 | ENST00000393818 | -28.51 | 6.70E-04 | 2.83E-03 |
| ITGA7 | 12:55698918-55700261 | 12:55699870-55699989 | SES | 3 | 3 | ENST00000555728 | -28.52 | 6.54E-03 | 1.08E-02 |
| BORA | 13:72729094-72737961 | 13:72734960-72735005 | MES | 3 | 3 | ENST00000377815 | -28.54 | 4.86E-02 | 7.93E-02 |
| SLC29A4 | 7:5300663-5303215 | 7:5302797-5303215 | TSSlA3ss | 3 | 2 | TSS.ENST00000297195 | -28.59 | 7.85E-03 | 1.10E-01 |
| KRIT1 | 7:92245465-92246100 | 7:92245575-92245843 | IR (overlapping | 3 | 2 | ENST00000340022 | -29.01 | 8.89E-03 | 1.37E-01 |
| INO80E | 16:30001531-30005367 | 16:30005221-30005367 | TSSlA3ss | 3 | 3 | TSS.ENST00000563197 | -29.24 | 1.22E-03 | 4.45E-03 |
| MFSD14C | 9:96918827-96949554 | 9:96942603-96942752 | MES | 3 | 3 | ENST00000506067 | -29.26 | 1.48E-02 | 4.96E-01 |
| CAMK2G | 10:73824085-73825320 | 10:73825279-73825320 | A5ss | 3 | 3 | ENST00000305762 | -29.28 | 3.19E-02 | 4.54E-02 |
| ATG9B | 7:151012214-151013408 | 7:151012473-151013230 | IR (overlapping | 2 | 3 | ENST00000605952 | -29.44 | 7.88E-04 | 9.79E-03 |
| SLC1A6 | 19:14972918-14979512 | 19:14979309-14979512 | A5ss | 3 | 2 | ENST00000594383 | -29.47 | 1.29E-02 | 7.00E-02 |
| ECHDC2 | 1:52905091-52907867 | 1:52906519-52906611 | SES | 3 | 3 | ENST00000371522 | -29.5 | 1.72E-02 | 3.21E-02 |
| COASY | 17:42562148-42563322 | 17:42562220-42562611 | IR | 3 | 3 | ENST00000393818 | -29.53 | 4.86E-04 | 2.05E-03 |
| SEPTIN6 | X:119616743-119629317 | X:119625335-119625379 | MES | 3 | 3 | ENST00000460411 | -29.58 | 1.36E-02 | 6.71E-01 |
| PLEKHA2 | 8:38969421-38973912 | 8:38971851-38972231 | IR (overlapping | 3 | 3 | ENST00000617275 | -29.64 | 4.29E-03 | 9.44E-02 |
| CAMK2G | 10:73824085-73837467 | 10:73825279-73825347 | MXS | 3 | 3 | ENST00000322680 | -29.65 | 1.50E-02 | 2.13E-02 |
| MPI | 15:74897512-74902219 | 15:74900138-74901949 | IR (overlapping | 3 | 3 | ENST00000352410 | -29.83 | 1.27E-02 | 4.30E-02 |
| MRTFB | 16:14240237-14241425 | 16:14240485-14240614 | IR | 2 | 3 | ENST00000572567 | -29.85 | 4.10E-02 | 1.44E-01 |
| TAF1C | 16:84184990-84186900 | 16:84184990-84185060 | TSSlA3ss | 3 | 3 | TSS.ENST00000564774 | -30.01 | 3.85E-04 | 1.49E-03 |
| TAF1C | 16:84184900-84186900 | 16:84184900-84185060 | TSSlA3ss | 3 | 3 | TSS.ENST00000564774 | -30.01 | 3.85E-04 | 1.49E-03 |
| AC108010.1 | 7:128574792-128578889 | 7:128577953-128578890 | A3ss | 2 | 3 | Ex.TSS.ENST00000605862 | -30.01 | 2.65E-02 | 3.18E-01 |
| AC090114.3 | 7:128574792-128577963 | 7:128577953-128577964 | A3ss | 2 | 3 | Ex.ENST00000479267 | -30.01 | 2.65E-02 | 3.18E-01 |
| AP3M2 | 8:42153106-42157940 | 8:42154616-42154960 | MES | 3 | 3 | ENST00000518421 | -30.02 | 5.20E-03 | 1.15E-01 |
| MPI | 15:74897512-74902219 | 15:74900201-74900483 | IR (overlapping | 3 | 3 | ENST00000352410 | -30.02 | 4.06E-02 | 1.37E-01 |
| TCIRG1 | 11:68042946-68043497 | 11:68043032-68043370 | IR | 3 | 3 | ENST00000533947 | -30.06 | 1.51E-02 | 2.30E-02 |
| GALNT13 | 2:153872177-153900935 | 2:153872177-153872303 | A5ss | 3 | 3 | Ex.TSS.ENST00000434213 | -30.21 | 3.25E-02 | 6.82E-02 |
| TMEM260 | 14:56618764-56625381 | 14:56621531-56621702 | SES | 3 | 3 | ENST00000261556 | -30.32 | 5.17E-03 | 1.63E-02 |
| EIF4G2 | 11:10807320-10808848 | 11:10808705-10808848 | A5ss | 3 | 3 | Ex.TSS.ENST00000530211 | -30.51 | 9.19E-03 | 1.31E-02 |
| DYSF | 2:71511922-71513715 | 2:71513240-71513332 | SES | 3 | 3 | ENST00000409582 | -30.55 | 4.75E-03 | 1.06E-02 |
| IFT81 | 12:110124444-110127359 | 12:110124444-110124861 | TSSlA5ss | 3 | 2 | TSS.ENST00000242591 | -30.64 | 2.42E-02 | 3.76E-02 |
| CLEC16A | 16:11003258-11020192 | 16:11003258-11003305 | A5ss | 3 | 3 | ENST00000409790 | -30.65 | 1.03E-02 | 3.60E-02 |
| TLCD5 | 11:120325369-120329976 | 11:120327441-120329977 | A3ss | 3 | 3 | Ex.ENST00000531346 | -30.67 | 6.27E-03 | 9.04E-03 |
| HSPBP1 | 19:55265379-55265885 | 19:55265379-55265389 | A3ss | 3 | 3 | Ex.ENST00000255631 | -30.75 | 8.76E-06 | 6.27E-05 |
| HSPBP1 | 19:55265368-55265885 | 19:55265368-55265389 | A3ss | 3 | 3 | Ex.ENST00000255631 | -30.75 | 8.76E-06 | 6.27E-05 |
| RAD9A | 11:67393611-67393722 | 11:67393691-67393722 | A3ss | 3 | 3 | ENST00000307980 | -30.78 | 5.89E-03 | 8.97E-03 |
| VPS54 | 2:63981852-63983863 | 2:63981852-63981887 | A3ss | 3 | 3 | ENST00000272322 | -30.88 | 3.10E-02 | 6.88E-02 |
| ITGA7 | 12:55698918-55700261 | 12:55699870-55700261 | TSSlA5ss | 3 | 3 | TSS.ENST00000554543 | -30.94 | 2.31E-02 | 3.82E-02 |
| ACOT2 | 14:73569397-73573387 | 14:73569397-73569883 | TSSlA5ss | 3 | 3 | TSS.ENST00000238651 | -31.03 | 3.99E-02 | 1.26E-01 |
| INVS | 9:100292326-100293043 | 9:100292437-100292946 | IR | 3 | 3 | ENST00000262457 | -31.15 | 3.44E-02 | 8.51E-01 |
| TBCEL | 11:121047668-121053629 | 11:121053551-121053629 | A3ss | 3 | 3 | ENST00000422003 | -31.18 | 2.25E-02 | 3.24E-02 |
| DCAF17 | 2:171458076-171468887 | 2:171458372-171458477 | SES | 3 | 3 | ENST00000375255 | -31.25 | 4.96E-03 | 1.05E-02 |
| CAPN7 | 3:15240854-15242177 | 3:15241453-15241588 | SES | 3 | 3 | ENST00000253693 | -31.54 | 3.26E-03 | 7.54E-03 |
| ZNF142 | 2:218658802-218658979 | 2:218658802-218658876 | TSSlA3ss | 3 | 3 | TSS.ENST00000432460 | -31.55 | 4.26E-02 | 9.15E-02 |
| CRELD1 | 3:9933853-9934419 | 3:9933853-9933920 | TSSlA5ss | 3 | 3 | TSS.ENST00000452070 | -31.64 | 1.21E-02 | 3.15E-02 |
| SEPTIN6 | X:119616743-119629317 | X:119625335-119625379 | MXS | 3 | 3 | ENST00000489216 | -31.73 | 6.60E-03 | 3.26E-01 |
| GOLGA2P7 | 15:84203008-84204425 | 15:84203008-84203020 | TSSlA3ss | 3 | 3 | TSS.ENST00000557980 | -31.75 | 4.60E-02 | 1.59E-01 |
| TPCN2 | 11:69054098-69057577 | 11:69055175-69055352 | MES | 3 | 3 | ENST00000294309 | -32.08 | 2.43E-02 | 3.70E-02 |
| FZD3 | 8:28494344-28502669 | 8:28499933-28499978 | SES | 3 | 3 | ENST00000240093 | -32.66 | 2.86E-03 | 6.17E-02 |
| MDC1 | 6:30707455-30711411 | 6:30707566-30708357 | SES | 3 | 3 | ENST00000376406 | -32.86 | 5.14E-03 | 4.68E-02 |
| AP000459.3 | 21:23372427-23372947 | 21:23372427-23372448 | A3ss | 3 | 3 | ENST00000262354 | -33.06 | 5.29E-06 | 1.02E-05 |
| ZDHHC16 | 10:97454800-97455713 | 10:97455660-97455713 | A3ss | 3 | 3 | ENST00000345745 | -33.35 | 1.87E-04 | 2.67E-04 |
| MTERF2 | 12:106978772-106986968 | 12:106985115-106985225 | SES | 3 | 3 | ENST00000240050 | -33.38 | 8.63E-03 | 1.34E-02 |
| KRIT1 | 7:92245465-92246100 | 7:92245596-92245905 | IR (overlapping | 3 | 3 | ENST00000340022 | -33.41 | 1.04E-02 | 1.59E-01 |
| ZNF445 | 3:44450963-44455120 | 3:44451314-44451482 | SES | 3 | 3 | ENST00000396077 | -34.02 | 4.48E-03 | 1.08E-02 |
| MPI | 15:74897512-74902219 | 15:74900138-74901168 | IR (overlapping | 3 | 3 | ENST00000352410 | -34.38 | 1.53E-02 | 5.16E-02 |
| MISP3 | 19:14072876-14075062 | 19:14073878-14074389 | IR | 2 | 2 | ENST00000591982 | -34.39 | 4.84E-02 | 2.59E-01 |
| INSIG2 | 2:118108281-118110997 | 2:118108305-118108305 | IR | 3 | 2 | ENST00000245787 | -34.52 | 4.17E-02 | 8.69E-02 |
| ACOXL | 2:110996005-111092864 | 2:111031627-111031714 | MES | 3 | 2 | ENST00000389811 | -34.86 | 3.12E-02 | 6.49E-02 |
| CAPN7 | 3:15242254-15246731 | 3:15245526-15245671 | SES | 3 | 3 | ENST00000253693 | -35.13 | 2.68E-03 | 6.21E-03 |
| PPOX | 1:161167235-161170619 | 1:161167371-161167486 | MES | 3 | 3 | ENST00000352210 | -35.16 | 6.17E-03 | 1.07E-02 |
| KRIT1 | 7:92245465-92246100 | 7:92245596-92245843 | IR (overlapping | 3 | 3 | ENST00000340022 | -35.21 | 1.03E-02 | 1.58E-01 |
| DPH5 | 1:100990629-100995109 | 1:100992637-100992740 | SES | 3 | 3 | ENST00000342173 | -35.23 | 5.10E-05 | 4.16E-04 |
| TANGO2 | 22:20061682-20063337 | 22:20061682-20061683 | A5ss | 3 | 3 | ENST00000327374 | -35.36 | 1.56E-02 | 3.03E-02 |
| PPOX | 1:161167235-161170619 | 1:161168993-161169183 | MES | 3 | 3 | ENST00000352210 | -36.01 | 1.65E-02 | 2.87E-02 |
| ELN | 7:74046768-74048141 | 7:74047675-74047716 | SES | 3 | 3 | ENST00000252034 | -36.16 | 2.72E-02 | 4.04E-01 |
| PLD3 | 19:40348725-40365717 | 19:40365553-40365717 | TSSlA3ss | 3 | 3 | TSS.ENST00000598962 | -37.27 | 1.15E-02 | 7.54E-02 |
| ZBED5 | 11:10855082-10856143 | 11:10855082-10855086 | TSSlA3ss | 3 | 3 | TSS.ENST00000413761 | -37.58 | 8.02E-03 | 1.15E-02 |
| MXRA8 | 1:1355244-1355512 | 1:1355346-1355449 | IR | 3 | 3 | ENST00000460473 | -37.79 | 9.12E-03 | 7.78E-02 |
| DLST | 14:74882017-74889094 | 14:74885586-74885634 | MES | 3 | 3 | ENST00000238671 | -37.93 | 2.62E-05 | 8.35E-05 |
| ZNF606 | 19:58002780-58003349 | 19:58003047-58003210 | IR (overlapping | 3 | 3 | ENST00000341164 | -38.66 | 1.05E-02 | 7.71E-02 |

| | | | | | | | | | |
|---|---|---|---|---|---|---|---|---|---|
| RBM18 | 9:122261499-122264714 | 9:122261499-122261508 | A3ss | 3 | 3 | Ex.TSS.ENST00000417201 | -39.41 | 1.53E-03 | 5.19E-02 |
| FGFR1OP | 6:167010894-167022408 | 6:167013508-167013567 | SES | 3 | 3 | ENST00000366847 | -39.77 | 9.58E-03 | 8.49E-02 |
| INO80E | 16:30004658-30005367 | 16:30005221-30005367 | A3ss | 3 | 3 | Ex.TSS.ENST00000540562 | -39.79 | 9.99E-03 | 3.64E-02 |
| GPR161 | 1:168114550-168137164 | 1:168136010-168137165 | A5ss | 3 | 2 | Ex.TSS.ENST00000367835 | -39.87 | 3.20E-02 | 5.59E-02 |
| ZNF510 | 9:96776246-96778033 | 9:96777582-96778033 | TSSIA5ss | 3 | 3 | TSS.ENST00000375231 | -39.99 | 2.47E-02 | 8.25E-01 |
| GABBR1 | 6:29629108-29630521 | 6:29630458-29630521 | A5ss | 3 | 2 | ENST00000376977 | -40.48 | 4.93E-02 | 1.53E-01 |
| COL27A1 | 9:114267558-114270727 | 9:114269241-114269294 | SES | 3 | 2 | ENST00000356083 | -40.97 | 2.10E-02 | 5.42E-01 |
| AMT | 3:49416794-49418372 | 3:49417368-49417488 | IR (overlapping | 3 | 3 | ENST00000636594 | -41.41 | 2.66E-03 | 6.46E-03 |
| UPP1 | 7:48101871-48106872 | 7:48103297-48103411 | SES | 3 | 3 | ENST00000444999 | -41.69 | 3.04E-03 | 4.23E-02 |
| DVL2 | 17:7227712-7227976 | 17:7227712-7227783 | A3ss | 3 | 3 | ENST00000005340 | -41.7 | 3.25E-06 | 1.49E-05 |
| DVL2 | 17:7227703-7227976 | 17:7227703-7227783 | A3ss | 3 | 3 | Ex.ENST00000005340 | -41.7 | 3.25E-06 | 1.49E-05 |
| ZNF561 | 19:9618167-9619431 | 19:9618167-9618179 | A3ss | 3 | 3 | ENST00000302851 | -42.42 | 5.40E-05 | 4.13E-04 |
| ZNF445 | 3:44450963-44455120 | 3:44451314-44451431 | MES | 3 | 3 | ENST00000617032 | -42.52 | 3.22E-03 | 7.74E-03 |
| AGO3 | 1:35973512-36004340 | 1:35982621-35982712 | MES | 3 | 3 | ENST00000634486 | -42.86 | 7.03E-03 | 1.30E-02 |
| TSPOAP1 | 17:58319295-58320530 | 17:58320109-58320129 | SES | 3 | 2 | ENST00000268893 | -43.18 | 1.93E-02 | 8.58E-02 |
| PPOX | 1:161167235-161170619 | 1:161167995-161168127 | MES | 3 | 3 | ENST00000352210 | -44.34 | 2.63E-02 | 4.57E-02 |
| C18orf54 | 18:54361949-54362770 | 18:54361949-54362431 | A5ss | 2 | 3 | ENST00000382911 | -44.86 | 2.12E-02 | 1.06E-01 |
| C18orf54 | 18:54361892-54362770 | 18:54361892-54362431 | A5ss | 2 | 3 | ENST00000382911 | -44.86 | 2.12E-02 | 1.06E-01 |
| TSC22D3 | X:107775100-107775787 | X:107775526-107775779 | IR (overlapping | 2 | 3 | ENST00000372383 | -44.9 | 3.35E-02 | 8.59E-01 |
| PPOX | 1:161167235-161170619 | 1:161168432-161168576 | MES | 3 | 3 | ENST00000352210 | -45.02 | 1.46E-02 | 2.54E-02 |
| ITGA7 | 12:55700394-55700898 | 12:55700566-55700899 | A5ss | 3 | 3 | Ex.ENST00000452168 | -45.48 | 1.54E-03 | 2.55E-03 |
| COASY | 17:42562220-42562932 | 17:42562612-42562932 | TSSIA3ss | 3 | 3 | TSS.ENST00000421097 | -45.7 | 9.48E-04 | 4.01E-03 |
| ATP9B | 18:79374102-79377246 | 18:79375394-79375426 | SES | 3 | 2 | ENST00000426216 | -46.01 | 9.40E-03 | 4.79E-02 |
| NPIPA1 | 16:14950256-14951614 | 16:14950752-14950821 | SES | 3 | 3 | ENST00000472413 | -46.53 | 7.31E-03 | 2.58E-02 |
| AMT | 3:49416794-49418372 | 3:49417310-49417614 | IR (overlapping | 3 | 3 | ENST00000636594 | -47.6 | 5.42E-05 | 1.31E-04 |
| AMT | 3:49416794-49418372 | 3:49417368-49417614 | IR (overlapping | 3 | 3 | ENST00000636594 | -48.92 | 1.69E-05 | 4.09E-05 |
| AMT | 3:49416794-49418372 | 3:49417358-49417614 | IR (overlapping | 3 | 3 | ENST00000636594 | -49 | 1.55E-05 | 3.75E-05 |
| CRNDE | 16:54920339-54923584 | 16:54920339-54920410 | A3ss | 2 | 3 | ENST00000502066 | -50.35 | 1.91E-02 | 7.10E-02 |
| CRNDE | 16:54920328-54923584 | 16:54920328-54920410 | A3ss | 2 | 3 | ENST00000502066 | -50.35 | 1.91E-02 | 7.10E-02 |
| AC119673.3 | 1:205891310-205894466 | 1:205894168-205894466 | TSSIA3ss | 3 | 3 | TSS.ENST00000653632 | -51.48 | 5.60E-03 | 9.97E-03 |
| AC119673.3 | 1:205891310-205894443 | 1:205894168-205894443 | A3ss | 3 | 3 | Ex.TSS.ENST00000653632 | -51.48 | 5.60E-03 | 9.97E-03 |
| ENOX2 | X:130637405-130656580 | X:130637405-130637410 | A3ss | 3 | 3 | Ex.ENST00000338144 | -51.63 | 2.78E-03 | 1.63E-01 |
| SLC36A4 | 11:93154217-93162705 | 11:93154217-93154277 | A3ss | 3 | 3 | Ex.ENST00000326402 | -52.09 | 1.20E-03 | 1.85E-03 |
| COASY | 17:42562356-42562894 | 17:42562488-42562611 | IR | 3 | 3 | ENST00000585811 | -53.19 | 4.30E-05 | 1.82E-04 |
| DLST | 14:74882017-74889094 | 14:74882591-74882624 | MES | 3 | 3 | ENST00000238671 | -53.72 | 1.83E-06 | 5.83E-06 |
| GLIDR | 9:39778945-39809730 | 9:39809486-39809615 | SES | 3 | 3 | ENST00000638724 | -56.78 | 3.69E-04 | 1.13E-02 |
| MTERF2 | 12:106985216-106986968 | 12:106985216-106985225 | TSSIA3ss | 3 | 3 | TSS.ENST00000240050 | -58.8 | 1.85E-04 | 2.87E-04 |
| MGA | 15:41736699-41740052 | 15:41739906-41740053 | A3ss | 3 | 3 | Ex.ENST00000219905 | -59.13 | 9.70E-03 | 3.18E-02 |
| STAG3L5P-PVRIG2P-PILRB | 7:100356884-100358263 | 7:100358220-100358263 | A3ss | 3 | 3 | ENST00000310771 | -60.15 | 1.23E-02 | 1.42E-01 |
| STAG3L5P-PVRIG2P-PILRB | 7:100356884-100358226 | 7:100358220-100358226 | A3ss | 3 | 3 | Ex.ENST00000310771 | -60.15 | 1.23E-02 | 1.42E-01 |
| FAM122B | X:134796257-134797120 | X:134796257-134796355 | A3ss | 2 | 3 | Ex.TSS.ENST00000298090 | -62.01 | 1.77E-04 | 1.05E-02 |
| COASY | 17:42562137-42562487 | 17:42562220-42562355 | IR | 3 | 3 | ENST00000585909 | -64.85 | 6.92E-06 | 2.93E-05 |
| COASY | 17:42562137-42562487 | 17:42562220-42562295 | IR | 3 | 3 | ENST00000585909 | -65.07 | 1.08E-05 | 4.58E-05 |
| MFSD9 | 2:102732393-102736645 | 2:102732393-102732409 | TSSIA3ss | 3 | 3 | TSS.ENST00000258436 | -73.81 | 6.91E-05 | 1.43E-04 |
