## Supplementary material for "Chemokine Receptor 1 and its associated immune pathway are downregulated in SF3B1^MT^ blood and non-blood cancers": Table_S2

**Table S2. ΔPSI% and p-values for shared mis-splicing events in SF3B1$^{K700E}$ ES cells and SF3B1$^{MT}$ cancers/cell lines.** Mis-splicing event codes are as follows: SES, single-exon skipping; MES, multiple-exon skipping; MXS, mutually-exclusive splicing; A5ss, alternative 5′ splice site; A3ss, alternative 3′ splice site; TSS, transcription start site. Event region: Genomic coordinates of the splicing event; Target exon: Genomic coordinates of the alternative exon. Events with p-values < 0.05 and ΔPSI (%)≥10% are listed. Signatures in Figure S3C are shown in blue. * and ** indicate mis-splicing events with ΔPSI% < 10% and p-value > 0.05, repectively. NA is not available. See also Table S16 for list of patients.

**ΔPSI%**

| Gene Symbol | Event Region | Target Exon | Event Type | ES line 1 | ES line 2 | ES line 3 | MDS-O | MDS-Z | MDS-M | CLL | AML | K562 | NALM6 | BRCA | UVM |
|---|---|---|---|---|---|---|---|---|---|---|---|---|---|---|---|
| APPL2 | 12:105208030-105208157 | 12:105208030-105208047 | A3ss | 46.36 | 47.95 | 65.05 | 34.64 | 35.15 | 19.88 | 30.74 | 21.82 | 41.98 | 32.29 | 16.26 | 40.01 |
| BRD9 | 5:865582-870472 | 5:869245-869404 | MXS | 33.96 | 29.58 | 39.9 | 46.1 | 43.83 | 32.87 | 37.81 | 26.64 | 55.11 | 45.09 | 11.76 | 36.75 |
| DLG1 | 3:197065449-197065707 | 3:197065449-197065464 | A3ss | 31.3 | 27.75 | 33.94 | 37.34 | 33.96 | 34.47 | 45.39 | 29.68 | 29.13 | 29.44 | 10.47 | 26.04 |
| DLST | 14:74889350-74889896 | 14:74889878-74889896 | A3ss | 70.08 | 55.27 | 70.97 | 42.9 | 42.18 | 25.99 | 52.8 | 16.5 | 51.41 | 55.02 | 11.54 | 48.69 |
| DVL2 | 17:7227712-7227976 | 17:7227712-7227783 | A3ss | -44.23 | -45.68 | -41.7 | -31.01 | -30.57 | -26.12 | -25.68 | -21.3 | -37.04 | -33.51 | -18.1 | -30.91 |
| INPPL1 | 11:72228499-72228726 | 11:72228647-72228727 | A3ss | 25.63 | 22.54 | 30.79 | 20.68 | 21.63 | 18.88 | 34.96 | 19.22 | 22.2 | 20.13 | 15.33 | 9.35 * |
| MAP3K7 | 6:90560215-90561621 | 6:90560215-90560234 | A3ss | 48.8 | 29.76 | 64.48 | 43.51 | 51.51 | 40.59 | 60.08 | 46.84 | 64.59 | 55.79 | 16.15 | 66.7 |
| PPOX | 1:161167235-161167370 | 1:161167339-161167370 | A3ss | 24.36 | 19.13 | 20.2 | 24.62 | 21.52 | 21.94 | 29.43 | 27.59 | 36.17 | 31.15 | 16.39 | 35.16 |
| SMURF2 | 17:64578577-64580788 | 17:64578577-64578594 | A3ss | 36.4 | 56.27 | 47.13 | 30.66 | 22.28 | 18.48 | 24.38 | 16.16 | 35.59 | NA | 15.81 | 56.55 |
| TTI1 | 20:38002777-38006196 | 20:38002777-38002793 | A3ss | 24.65 | 29.02 | 38.21 | 39.2 | 22.44 | 32.52 | 44.01 | 50.57 | 33.11 | 41.88 | 21.45 | 39.13 |
| ANKHD1 | 5:140436258-140438493 | 5:140438461-140438493 | A3ss | -28.36 | -21.27 | -27.21 | -37.8 | -28.63 | -26.61 | -30.98 | -28.74 | -31.57 | -28.84 | -13.65 | -48.94 |
| ANKHD1 | 5:140436258-140438502 | 5:140438461-140438502 | A3ss | -28.36 | -21.27 | -27.21 | -37.8 | -28.63 | -26.61 | -30.98 | -28.74 | -31.57 | -28.84 | -13.65 | -48.94 |
| ANKHD1 | 5:140436258-140438502 | 5:140438494-140438502 | A3ss | 38.69 | 29.45 | 35.7 | 39.46 | 37.53 | 27.87 | 33.81 | 34.46 | 27.02 | 31.16 | 21.75 | 61 |
| BAG6 | 6:31644195-31644524 | 6:31644307-31644414 | SES | 29.66 | 31.06 | 25.81 | 23.07 | 26.46 | 36.2 | 17.81 | 28.35 | 22.36 | 19.61 | 17.32 | 13.23 |
| BCL2L1 | 20:31722331-31722617 | 20:31722331-31722348 | TSSIA3ss | 33.62 | 60.99 | 76.69 | 14.88 | 19.38 | 16.26 | 29.37 | 14.9 | 16.96 | 14.72 | 11.44 | 27.63 |
| CHTF18 | 16:793275-794053 | 16:794034-794054 | A3ss | 32.06 | 31.7 | 41.58 | 46.86 | 45.86 | 41.39 | 27.52 | 45.48 | 49.03 | 48.44 | 25.3 | 45.44 |
| COASY | 17:42562137-42562487 | 17:42562220-42562295 | IR | -62.25 | -57.09 | -65.07 | -27.25 | -38.5 | -17.04 | -25.82 | -11.23 | -32.36 | -36.57 | -25.25 | -14.23 |
| COASY | 17:42562137-42562487 | 17:42562220-42562355 | IR | -62.96 | -58 | -64.85 | -26.13 | -38.31 | -16.97 | -24.37 | -10.51 | -30.33 | -35.34 | -26.04 | -13.92 |
| COASY | 17:42562220-42562611 | 17:42562356-42562611 | A3ss | 61.07 | 61.45 | 61.86 | 68.45 | 58.65 | 58.34 | 62.8 | 43.02 | 59.95 | 68.79 | 36.85 | 55.34 |
| COASY | 17:42562220-42562611 | 17:42562356-42562487 | SES | 53.33 | 53.56 | 53.46 | 59.88 | 43.6 | 53.54 | 59.32 | 40.23 | 46.22 | 61.73 | 30.06 | 45.01 |
| DPH5 | 1:100990629-100995109 | 1:100992637-100992740 | SES | -34.65 | -34.4 | -35.23 | -32.21 | -19.6 | -20.55 | -24.23 | -34.93 | -34.24 | -34.24 | -18.17 | -55.72 |
| DPH5 | 1:100990632-100995109 | 1:100990632-100992754 | A3ss | 30.61 | 23.98 | 30.76 | 30.4 | 24.45 | 22.96 | 24.5 | 36.55 | 33.33 | 28.5 | 19.84 | 68.2 |
| DPH5 | 1:100992741-100995109 | 1:100992741-100992754 | A3ss | 30.61 | 23.98 | 30.76 | 30.4 | 24.45 | 22.96 | 24.5 | 36.55 | 33.33 | 28.5 | 19.84 | 68.2 |
| DYNLL1 | 12:120496217-120496415 | 12:120496402-120496415 | TSSIA3ss | 59.38 | 58.14 | 60.15 | 46.7 | 48.84 | 38.86 | 51.54 | 35.28 | 46.6 | 43.93 | 26.03 | 56 |
| ENOX2 | X:130637405-130656580 | X:130637405-130637410 | A3ss | -63.87 | -57.01 | -51.63 | -47.04 | -54 | -43.99 | -62.97 | -55.17 | -56.42 | -32.51 | -31.76 | -65.31 |
| ERGIC3 | 20:35556272-35556972 | 20:35556955-35556972 | A3ss | 15.72 | 14.34 | 15.92 | 23.86 | 26.93 | 24.43 | 22.22 | 19.93 | 20.45 | 21.49 | 16.15 | 32.53 |
| FDPS | 1:155308966-155310063 | 1:155310043-155310063 | TSSIA3ss | 16.96 | 19.01 | 18.81 | 25.34 | 24.94 | 17.86 | 26.1 | 21.08 | 11.18 | 17.13 | 17 | 18.69 |
| GCC2 | 2:108485909-108486510 | 2:108486499-108486511 | A3ss | 51.46 | 53.29 | 58.5 | 55.95 | 55.02 | 41.23 | 67.47 | 41.96 | 64.74 | 64.82 | 38.62 | 82.29 |
| GPR108 | 19:6731055-6731198 | 19:6731055-6731111 | A3ss | -20.99 | -24.58 | -21.74 | -17.16 | -15.65 | -16.39 | -14.31 | -23.03 | -22.77 | -14.13 | -18.39 | -22.94 |
| HINT2 | 9:35813146-35813265 | 9:35813146-35813156 | A3ss | 27.75 | 24.14 | 35.81 | 33.05 | 34.27 | 33.38 | 21 | 26.83 | 31.99 | 19.6 | 24.44 | 53.41 |
| HSPBP1 | 19:55265368-55265885 | 19:55265368-55265389 | A3ss | -35.2 | -32.42 | -30.75 | -30.1 | -27.48 | -26.52 | -20 | -18.91 | -26.89 | -25.08 | -16.67 | -35.79 |
| HSPBP1 | 19:55265379-55265885 | 19:55265379-55265389 | A3ss | -35.2 | -32.42 | -30.75 | -30.1 | -27.48 | -26.52 | -20 | -18.91 | -26.89 | -25.08 | -16.67 | -35.79 |
| KANSL3 | 2:96619763-96631311 | 2:96619763-96619776 | A3ss | 34.08 | 35.87 | 38.96 | 36.08 | 39.32 | 25.2 | 47.6 | 24.3 | 29.7 | 32.64 | 25.98 | 50.66 |
| KANSL3 | 2:96619545-96631311 | 2:96619545-96619776 | A3ss | 34.08 | 35.87 | 38.96 | 36.08 | 39.32 | 25.2 | 47.6 | 24.3 | 29.7 | 32.64 | 25.98 | 50.66 |
| LRSAM1 | 9:127451513-127452084 | 9:127451670-127451928 | IR | -18.86 | -19.2 | -22.43 | -23 | -13.36 | -10.31 | -13.79 | -15.7 | -18.78 | -11.74 | -10.65 | -16.16 |
| MED6 | 14:70592989-70593295 | 14:70592989-70593009 | A3ss | 27 | 29.6 | 24.93 | 19.04 | 19.72 | 12.09 | 12.19 | 10.48 | 25.65 | 22.09 | 11.67 | 38.86 |
| MTERF2 | 12:106985216-106986968 | 12:106985216-106985225 | TSSIA3ss | -52.64 | -52.86 | -58.8 | -69.25 | -59.2 | -42.89 | -45.19 | -62.76 | -68.39 | -59.96 | -43.91 | -71.39 |
| NDRG3 | 20:36653702-36656359 | 20:36653702-36653723 | A3ss | 41.57 | 32.07 | 38.55 | 19.03 | 15.79 | 12.77 | 28.77 | 22.11 | 29.03 | 27.84 | 15.02 | 15 |
| NFYA | 6:41073085-41079028 | 6:41079005-41079029 | A3ss | 26.93 | 25.96 | 25.02 | 23.32 | 28.14 | 15.83 | 23.7 | 23.34 | 21.91 | 19.03 | 19.53 | 53.66 |
| ORAI2 | 7:102433662-102436224 | 7:102436202-102436224 | TSSIA3ss | 70.48 | 72.82 | 73.46 | 60.51 | 59.62 | 25.14 | 59.62 | 37.19 | 70.92 | 60.61 | 36.79 | 73.71 |
| PRPF38A | 1:52414648-52414761 | 1:52414741-52414762 | A3ss | 20.54 | 20.33 | 17.77 | 20.24 | 18.46 | 12.53 | 19.9 | 10.11 | 28.24 | 17.29 | 15.97 | 23.02 |
| RBM18 | 9:122261499-122264714 | 9:122261499-122261508 | A3ss | -40.06 | -33.97 | -39.41 | -23.8 | -24.98 | -20.48 | -33.66 | -23.36 | -29.3 | -31.61 | -18.18 | -30.96 |
| RNF2 | 1:185087641-185091578 | 1:185091565-185091579 | A3ss | 17.63 | 18.84 | 31.28 | 14.59 | 22 | 12.86 | 26.53 | 15.23 | 20.64 | 17.75 | 12.37 | 21.25 |
| RWDD4 | 4:183651132-183658928 | 4:183651132-183655974 | A3ss | 32.41 | 31.05 | 29.57 | 26.4 | 33.41 | 24.99 | 25.37 | 25.67 | 35.53 | 22.4 | 30.63 | 68.37 |
| RWDD4 | 4:183655962-183658928 | 4:183655962-183655974 | A3ss | 32.41 | 31.05 | 29.57 | 26.4 | 33.41 | 24.99 | 25.37 | 25.67 | 35.53 | 22.4 | 30.63 | 68.37 |
| SEPTIN6 | X:119625380-119629317 | X:119625380-119625396 | A3ss | 40.63 | 43.82 | 38.38 | 36.64 | 42.03 | 32.16 | 39.19 | 34.24 | 56.39 | 52.4 | 13.85 | 21.74 |
| SERBP1 | 1:67424960-67425082 | 1:67424960-67424977 | A3ss | 30.56 | 30 | 31.59 | 26.12 | 24.65 | 21.27 | 29.64 | 21.77 | 28.73 | 25.85 | 18.82 | 30.08 |
| SNRPN | 15:24962210-24967931 | 15:24967029-24967152 | SES | 29.5 | 28.48 | 30.02 | 15.4 | 15.88 | 19.23 | 12.4 | 22.37 | NA | 28.2 | 12.2 | 55.21 |
| THOC1 | 18:224180-224923 | 18:224180-224200 | A3ss | 38.64 | 34.14 | 45.59 | 23.59 | 26.38 | 15.49 | 17.85 | 10.38 | 32.35 | 24.04 | 13.74 | 33.63 |
| TMEM14C | 6:10723242-10724569 | 6:10724556-10724570 | A3ss | 58.49 | 58.63 | 60.46 | 59.01 | 62.4 | 44.35 | 53.64 | 44.49 | 59.9 | 53.21 | 33.37 | 70.16 |
| TMEM214 | 2:27037703-27038145 | 2:27037815-27037892 | SES | 61.04 | 58.39 | 63.8 | 28.47 | 16.71 | 26.63 | 20.76 | 28.33 | 44.87 | 36.52 | 12 | 32.17 |
| UXS1 | 2:106164785-106166055 | 2:106164785-106164799 | A3ss | 48.1 | 42.9 | 50.73 | 36.91 | 36.75 | 28.87 | 49.51 | 33.52 | 43.88 | 43.65 | 19.51 | 56.5 |
| VEZT | 12:95266633-95270050 | 12:95270039-95270050 | A3ss | 24.1 | 22.21 | 22.11 | 16.21 | 22.83 | 21.26 | 21.87 | 26.16 | 20.63 | 21.5 | 20.99 | 26.25 |
| WRAP73 | 1:3632318-3633397 | 1:3632318-3632338 | A3ss | -29.97 | -30.99 | -27.14 | -25.82 | -27.52 | -13.72 | -11.26 | -16.66 | -22.51 | -22.09 | -23.83 | -40.23 |
| ZBED5 | 11:10855087-10856143 | 11:10855087-10855118 | TSSIA3ss | 46.85 | 38.27 | 42.62 | 12.55 | 22.38 | 31.21 | 42.37 | 42.55 | 37.78 | 36.75 | 17.52 | 51.89 |
| ZBED5 | 11:10855082-10856143 | 11:10855082-10855118 | TSSIA3ss | 46.85 | 38.27 | 42.62 | 12.55 | 22.38 | 31.21 | 42.37 | 42.55 | 37.78 | 36.75 | 17.52 | 51.89 |
| ZDHHC16 | 10:97454800-97455659 | 10:97455639-97455659 | A3ss | 41.65 | 41.5 | 32.34 | 40.27 | 42.73 | 30.55 | 37.37 | 24.2 | 53.24 | 51.02 | 29.18 | 52.47 |
| ZDHHC16 | 10:97454800-97455713 | 10:97455639-97455713 | A3ss | 41.65 | 41.5 | 32.34 | 40.27 | 42.73 | 30.55 | 37.37 | 24.2 | 53.24 | 51.02 | 29.18 | 52.47 |
| ZDHHC16 | 10:97454800-97455713 | 10:97455660-97455713 | A3ss | -38.59 | -37.49 | -33.35 | -33.61 | -37.11 | -31.08 | -28.79 | -19.12 | -45.34 | -38.8 | -22.41 | -42.18 |

**p-values**

| Gene Symbol | ES line 1 | ES line 2 | ES line 3 | MDS-O | MDS-Z | MDS-M | CLL | AML | K562 | NALM6 | BRCA | UVM |
|---|---|---|---|---|---|---|---|---|---|---|---|---|
| APPL2 | 6.23E-03 | 1.19E-02 | 2.44E-03 | 4.56E-05 | 4.41E-09 | 5.40E-04 | 2.25E-06 | 5.87E-05 | 2.01E-03 | 2.73E-02 | 8.24E-06 | 3.46E-35 |
| BRD9 | 8.00E-03 | 1.72E-04 | 1.37E-06 | 2.00E-05 | 5.68E-09 | 8.12E-07 | 5.50E-08 | 2.25E-07 | 3.35E-06 | 1.94E-05 | 2.39E-06 | 1.30E-32 |
| DLG1 | 4.00E-03 | 0.0576465 ** | 4.12E-03 | 7.55E-05 | 1.10E-06 | 1.38E-05 | 1.57E-07 | 3.05E-04 | 3.61E-05 | 3.18E-04 | 6.02E-06 | 1.17E-24 |
| DLST | 5.66E-05 | 2.05E-06 | 4.77E-07 | 3.87E-04 | 4.25E-12 | 8.01E-04 | 9.90E-10 | 2.73E-02 | 1.26E-06 | 1.47E-04 | 8.05E-06 | 3.24E-43 |
| DVL2 | 1.22E-04 | 2.01E-04 | 3.25E-06 | 9.64E-04 | 3.02E-08 | 3.76E-06 | 3.30E-07 | 8.29E-06 | 1.18E-03 | 2.04E-05 | 8.11E-07 | 1.23E-35 |
| INPPL1 | 3.61E-02 | 9.83E-05 | 4.57E-05 | 1.88E-04 | 9.98E-09 | 1.45E-03 | 2.46E-05 | 1.20E-03 | 3.05E-03 | 3.98E-03 | 1.43E-06 | 2.18E-20 |
| MAP3K7 | 1.28E-04 | 1.16E-02 | 3.61E-05 | 8.21E-05 | 6.74E-09 | 1.25E-05 | 9.94E-08 | 2.90E-06 | 4.12E-06 | 2.57E-04 | 3.50E-03 | 3.94E-16 |
| PPOX | 1.04E-03 | 1.60E-03 | 8.44E-04 | 5.83E-03 | 4.22E-08 | 6.10E-08 | 1.21E-06 | 9.35E-04 | 4.22E-04 | 1.20E-02 | 1.39E-05 | 2.11E-24 |
| SMURF2 | 2.36E-03 | 2.38E-05 | 1.13E-02 | 5.18E-05 | 7.25E-05 | 6.19E-04 | 7.25E-04 | 7.30E-04 | 2.77E-04 | NA | 9.18E-05 | 2.95E-19 |
| TTI1 | 4.70E-03 | 5.99E-05 | 7.22E-05 | 1.82E-03 | 5.21E-04 | 4.12E-05 | 9.58E-07 | 1.03E-05 | 1.39E-03 | 7.43E-05 | 1.96E-05 | 1.47E-26 |
| ANKHD1 | 8.06E-04 | 6.86E-03 | 1.66E-03 | 1.02E-06 | 2.23E-04 | 1.66E-06 | 2.18E-06 | 2.89E-05 | 1.08E-04 | 1.50E-04 | 2.09E-03 | 1.89E-22 |
| ANKHD1 | 8.06E-04 | 6.86E-03 | 1.66E-03 | 1.02E-06 | 2.23E-04 | 1.66E-06 | 2.18E-06 | 2.89E-05 | 1.08E-04 | 1.50E-04 | 2.09E-03 | 1.89E-22 |
| ANKHD1 | 2.64E-05 | 3.30E-05 | 8.69E-06 | 1.35E-07 | 4.50E-11 | 1.46E-06 | 1.76E-06 | 2.09E-07 | 5.18E-04 | 4.30E-04 | 9.99E-06 | 2.33E-38 |
| BAG6 | 7.75E-05 | 8.67E-05 | 1.33E-04 | 2.10E-02 | 1.01E-05 | 2.10E-05 | 5.33E-04 | 1.32E-06 | 1.45E-04 | 5.20E-04 | 4.88E-05 | 2.57E-07 |
| BCL2L1 | 2.73E-03 | 1.48E-04 | 1.12E-05 | 1.03E-05 | 3.37E-06 | 4.35E-08 | 9.61E-04 | 2.51E-02 | 3.12E-03 | 2.79E-02 | 8.90E-04 | 1.55E-23 |
| CHTF18 | 5.60E-03 | 4.51E-04 | 5.89E-04 | 4.62E-05 | 1.76E-10 | 1.18E-06 | 3.59E-02 | 1.59E-04 | 5.38E-05 | 5.66E-05 | 3.07E-05 | 1.63E-22 |
| COASY | 1.26E-06 | 3.02E-06 | 1.08E-05 | 1.23E-02 | 4.60E-05 | 2.10E-03 | 3.25E-03 | 2.27E-02 | 4.49E-04 | 2.22E-04 | 1.28E-05 | 1.38E-05 |
| COASY | 1.66E-06 | 1.33E-06 | 6.92E-06 | 1.14E-02 | 1.73E-05 | 1.47E-03 | 4.00E-03 | 1.74E-02 | 4.59E-04 | 2.29E-04 | 4.68E-06 | 1.57E-05 |
| COASY | 2.17E-05 | 1.21E-05 | 1.46E-05 | 1.98E-06 | 4.02E-09 | 2.10E-08 | 1.10E-05 | 1.02E-03 | 3.90E-05 | 8.66E-05 | 7.47E-06 | 2.47E-32 |
| COASY | 5.64E-05 | 2.34E-05 | 3.50E-05 | 5.19E-06 | 3.20E-04 | 4.08E-06 | 2.78E-05 | 2.33E-03 | 5.80E-05 | 3.67E-05 | 1.34E-04 | 1.25E-23 |
| DPH5 | 3.69E-03 | 3.07E-03 | 5.10E-05 | 2.39E-03 | 2.11E-04 | 6.37E-04 | 3.42E-04 | 7.39E-06 | 4.66E-05 | 2.13E-03 | 6.23E-06 | 1.04E-47 |
| DPH5 | 4.03E-04 | 3.11E-03 | 1.48E-07 | 7.71E-05 | 2.14E-07 | 2.36E-05 | 3.89E-04 | 7.27E-06 | 1.90E-04 | 3.09E-05 | 4.46E-06 | 1.24E-52 |
| DPH5 | 4.03E-04 | 3.11E-03 | 1.48E-07 | 7.71E-05 | 2.14E-07 | 2.36E-05 | 3.89E-04 | 7.27E-06 | 1.90E-04 | 3.09E-05 | 4.46E-06 | 1.24E-52 |
| DYNLL1 | 2.62E-07 | 1.85E-07 | 4.45E-07 | 3.35E-08 | 7.85E-11 | 3.37E-07 | 1.72E-10 | 1.02E-04 | 2.66E-06 | 4.47E-06 | 1.45E-08 | 7.67E-77 |
| ENOX2 | 3.82E-04 | 1.65E-04 | 2.78E-03 | 3.18E-03 | 7.18E-07 | 2.69E-06 | 2.05E-09 | 7.99E-08 | 1.72E-04 | 2.11E-02 | 1.99E-07 | 8.77E-26 |
| ERGIC3 | 8.38E-05 | 2.00E-04 | 6.96E-05 | 2.72E-06 | 8.10E-10 | 2.94E-06 | 1.91E-05 | 3.10E-05 | 9.40E-07 | 1.79E-05 | 2.07E-07 | 2.85E-54 |
| FDPS | 5.65E-04 | 1.48E-03 | 3.71E-04 | 9.23E-03 | 1.99E-07 | 4.74E-05 | 1.83E-04 | 1.07E-06 | 5.59E-03 | 2.49E-04 | 1.69E-04 | 1.47E-14 |
| GCC2 | 4.71E-04 | 9.28E-06 | 2.63E-04 | 1.78E-04 | 1.10E-07 | 2.39E-04 | 8.54E-10 | 1.26E-04 | 3.27E-04 | 5.37E-05 | 1.32E-06 | 1.52E-30 |
| GPR108 | 5.23E-04 | 9.44E-04 | 2.86E-02 | 4.39E-04 | 9.80E-06 | 1.08E-04 | 2.41E-03 | 2.01E-06 | 2.77E-04 | 4.93E-05 | 2.70E-06 | 1.36E-24 |
| HINT2 | 1.14E-03 | 3.12E-02 | 2.13E-03 | 3.31E-05 | 1.07E-08 | 3.32E-06 | 4.47E-03 | 2.50E-04 | 7.70E-06 | 1.38E-05 | 2.00E-07 | 4.25E-54 |
| HSPBP1 | 7.08E-04 | 7.52E-05 | 8.76E-06 | 2.62E-05 | 3.39E-07 | 1.47E-06 | 8.57E-04 | 2.52E-02 | 1.24E-04 | 5.27E-04 | 9.13E-08 | 1.01E-48 |
| HSPBP1 | 7.08E-04 | 7.52E-05 | 8.76E-06 | 2.62E-05 | 3.39E-07 | 1.47E-06 | 8.57E-04 | 2.52E-02 | 1.24E-04 | 5.27E-04 | 9.13E-08 | 1.01E-48 |
| KANSL3 | 4.15E-04 | 1.42E-04 | 6.68E-04 | 3.30E-05 | 2.53E-07 | 6.01E-04 | 9.06E-08 | 3.11E-03 | 3.31E-03 | 8.67E-04 | 8.15E-06 | 1.00E-41 |
| KANSL3 | 4.15E-04 | 1.42E-04 | 6.68E-04 | 3.30E-05 | 2.53E-07 | 6.01E-04 | 9.06E-08 | 3.11E-03 | 3.31E-03 | 8.67E-04 | 8.15E-06 | 1.00E-41 |
| LRSAM1 | 4.54E-02 | 4.42E-02 | 2.81E-02 | 8.18E-06 | 1.08E-03 | 2.08E-02 | 3.12E-02 | 5.21E-03 | 6.95E-03 | 4.43E-03 | 4.18E-05 | 4.21E-19 |
| MED6 | 2.78E-02 | 2.81E-04 | 2.96E-02 | 2.54E-06 | 1.73E-09 | 9.19E-03 | 4.62E-02 | 4.53E-02 | 7.95E-05 | 1.38E-04 | 1.23E-05 | 1.36E-34 |
| MTERF2 | 4.42E-04 | 7.12E-04 | 1.85E-04 | 5.77E-07 | 3.18E-07 | 4.18E-04 | 4.30E-05 | 1.30E-02 | 8.81E-07 | 3.59E-04 | 9.38E-07 | 1.83E-35 |
| NDRG3 | 2.51E-03 | 1.64E-07 | 1.63E-05 | 4.10E-02 | 1.31E-04 | 4.91E-06 | 3.50E-06 | 5.60E-05 | 1.09E-04 | 1.86E-04 | 1.88E-05 | 2.56E-23 |
| NFYA | 3.39E-04 | 8.90E-05 | 1.86E-03 | 7.26E-04 | 1.20E-09 | 1.21E-04 | 7.78E-04 | 2.38E-04 | 7.31E-04 | 2.78E-06 | 5.94E-05 | 1.82E-54 |
| ORAI2 | 2.89E-05 | 4.25E-06 | 1.13E-05 | 1.43E-08 | 2.98E-14 | 1.93E-03 | 6.23E-11 | 4.27E-03 | 6.68E-05 | 2.82E-04 | 8.76E-06 | 1.79E-30 |
| PRPF38A | 1.15E-03 | 1.37E-03 | 1.64E-03 | 3.11E-04 | 2.78E-09 | 5.70E-04 | 1.60E-07 | 8.12E-03 | 4.31E-06 | 3.31E-04 | 7.55E-07 | 2.98E-43 |
| RBM18 | 1.03E-07 | 2.15E-03 | 1.53E-03 | 1.14E-03 | 3.24E-08 | 6.60E-06 | 1.16E-10 | 7.50E-07 | 3.19E-05 | 1.41E-04 | 3.08E-07 | 4.34E-27 |
| RNF2 | 3.34E-04 | 7.85E-03 | 8.80E-04 | 1.02E-03 | 2.02E-08 | 2.50E-04 | 2.52E-06 | 9.62E-04 | 3.79E-03 | 5.41E-03 | 7.72E-05 | 6.61E-18 |
| RWDD4 | 1.20E-02 | 2.64E-02 | 1.83E-03 | 1.29E-05 | 5.28E-07 | 5.42E-05 | 1.43E-04 | 2.16E-03 | 2.91E-04 | 1.74E-04 | 1.08E-07 | 2.01E-17 |
| RWDD4 | 1.20E-02 | 2.64E-02 | 1.83E-03 | 1.29E-05 | 5.28E-07 | 5.42E-05 | 1.43E-04 | 2.16E-03 | 2.91E-04 | 1.74E-04 | 1.08E-07 | 2.01E-17 |
| SEPTIN6 | 7.20E-04 | 1.93E-04 | 2.13E-03 | 2.15E-06 | 7.84E-08 | 2.10E-07 | 3.24E-07 | 6.69E-08 | 2.40E-05 | 7.71E-06 | 1.64E-06 | 2.12E-19 |
| SERBP1 | 2.15E-06 | 1.13E-06 | 4.40E-07 | 3.32E-07 | 2.26E-06 | 3.26E-06 | 5.02E-09 | 3.35E-06 | 9.07E-06 | 9.90E-06 | 1.10E-07 | 3.55E-45 |
| SNRPN | 1.61E-05 | 2.15E-05 | 1.26E-05 | 1.83E-04 | 2.46E-08 | 9.73E-08 | 8.50E-04 | 1.05E-03 | NA | 1.98E-04 | 2.77E-07 | 5.32E-42 |
| THOC1 | 2.48E-05 | 1.01E-03 | 4.04E-05 | 7.75E-04 | 6.32E-08 | 4.22E-04 | 7.47E-04 | 2.65E-02 | 8.92E-06 | 7.65E-04 | 7.62E-06 | 2.27E-20 |
| TMEM14C | 8.80E-06 | 4.29E-07 | 1.04E-06 | 5.41E-07 | 8.62E-12 | 3.41E-06 | 6.35E-08 | 4.71E-06 | 3.59E-07 | 2.95E-05 | 2.69E-07 | 4.51E-78 |
| TMEM214 | 1.81E-04 | 6.83E-05 | 1.42E-04 | 4.13E-03 | 1.64E-03 | 3.78E-09 | 5.21E-03 | 4.52E-04 | 7.68E-06 | 4.68E-06 | 2.74E-07 | 5.65E-36 |
| UXS1 | 1.62E-04 | 2.26E-03 | 6.14E-04 | 7.54E-04 | 6.77E-07 | 9.19E-06 | 7.98E-09 | 2.32E-06 | 7.85E-05 | 2.57E-04 | 5.09E-06 | 1.50E-39 |
| VEZT | 6.58E-03 | 6.06E-03 | 1.40E-02 | 1.12E-03 | 1.59E-08 | 5.44E-04 | 1.22E-07 | 2.98E-08 | 2.80E-03 | 1.06E-02 | 1.06E-05 | 1.44E-06 |
| WRAP73 | 2.35E-03 | 1.79E-03 | 4.98E-02 | 1.07E-02 | 6.45E-07 | 4.13E-02 | 5.57E-03 | 3.43E-02 | 5.96E-04 | 1.82E-03 | 5.96E-04 | 1.07E-24 |
| ZBED5 | 1.09E-03 | 6.72E-04 | 2.18E-03 | 3.98E-02 | 2.81E-02 | 8.42E-04 | 2.27E-08 | 1.51E-07 | 5.13E-04 | 3.64E-04 | 2.18E-05 | 1.42E-25 |
| ZBED5 | 1.09E-03 | 6.72E-04 | 2.18E-03 | 3.98E-02 | 2.81E-02 | 8.42E-04 | 2.27E-08 | 1.51E-07 | 5.13E-04 | 3.64E-04 | 2.18E-05 | 1.42E-25 |
| ZDHHC16 | 3.66E-04 | 2.80E-05 | 8.93E-04 | 1.17E-02 | 2.24E-09 | 2.97E-05 | 2.20E-06 | 1.99E-03 | 1.61E-04 | 2.23E-04 | 4.92E-07 | 2.63E-55 |
| ZDHHC16 | 3.66E-04 | 2.80E-05 | 8.93E-04 | 1.17E-02 | 2.24E-09 | 2.97E-05 | 2.20E-06 | 1.99E-03 | 1.61E-04 | 2.23E-04 | 4.92E-07 | 2.63E-55 |
| ZDHHC16 | 1.57E-04 | 1.69E-04 | 1.87E-04 | 1.87E-02 | 7.56E-10 | 3.97E-04 | 7.91E-05 | 1.73E-02 | 6.91E-04 | 5.65E-04 | 1.25E-05 | 5.37E-38 |
