## Supplementary material for "Chemokine Receptor 1 and its associated immune pathway are downregulated in SF3B1^MT^ blood and non-blood cancers": Table_S3

| **Table S3. Total gene expression using RNA-seq data in 3 biological replicates of $SF3B1^{K700E}$ versus $SF3B1^{WT}$ ES cells (K700E ES line 1).** Average baseMean, log2fold change, fold change (FC) and p-values are shown. | | | | | | | |
|---|---|---|---|---|---|---|---|
| **Gene Symbol** | **log2 FC** | **FC** | **p-value** | **padj** | **WT baseMean** | **MT baseMean** | **Ensembl ID** |
| HORMAD1 | 7.709351052 | 209.2887643 | 2.44E-10 | 3.22E-09 | 0.323808629 | 75.24437789 | ENSG00000143452 |
| NA | 7.532265103 | 185.1133462 | 2.70E-09 | 3.10E-08 | 0 | 34.16857361 | ENSG00000233309 |
| KHDC3L | 7.238165959 | 150.97501 | 1.96E-08 | 1.99E-07 | 0 | 27.85557538 | ENSG00000203908 |
| LOC101928254 | 7.034596426 | 131.106593 | 3.68E-11 | 5.28E-10 | 0.653199063 | 87.60284232 | ENSG00000219445 |
| NA | 6.920385247 | 121.1277195 | 2.92E-08 | 2.89E-07 | 0.323808629 | 43.51614836 | ENSG00000265206 |
| NA | 6.682479089 | 102.7132926 | 5.62E-07 | 4.59E-06 | 0 | 18.93928567 | ENSG00000272068 |
| NA | 6.672410795 | 101.9989715 | 1.27E-07 | 1.15E-06 | 0.326599532 | 36.64441137 | ENSG00000272405 |
| PRAM1 | 6.507107875 | 90.95669163 | 1.44E-06 | 1.10E-05 | 0 | 16.79604861 | ENSG00000133246 |
| MAGEA4 | 6.464545728 | 88.31249863 | 1.23E-67 | 3.48E-65 | 5.655543462 | 501.7202351 | ENSG00000147381 |
| NA | 6.220438534 | 74.56561169 | 1.14E-05 | 7.39E-05 | 0 | 13.75691047 | ENSG00000257275 |
| MEG3 | 6.200776765 | 73.5562877 | 1.64E-06 | 1.24E-05 | 0.326599532 | 26.41062234 | ENSG00000214548 |
| LINC01366 | 6.179765936 | 72.49280641 | 1.02E-05 | 6.69E-05 | 0 | 13.36300922 | ENSG00000235172 |
| CRYGD | 6.002812812 | 64.12490206 | 2.24E-05 | 0.000138212 | 0 | 11.83819395 | ENSG00000118231 |
| NPAP1 | 5.970279957 | 62.69506392 | 5.17E-06 | 3.58E-05 | 0.353832839 | 22.51295481 | ENSG00000185823 |
| NA | 5.965071008 | 62.46910729 | 2.87E-05 | 0.000173707 | 0 | 11.53731883 | ENSG00000233064 |
| TDRD1 | 5.888482557 | 59.23929461 | 1.33E-10 | 1.81E-09 | 0.979798595 | 59.25309434 | ENSG00000095627 |
| MIR205HG | 5.8340645 | 57.0464221 | 7.52E-85 | 3.92E-82 | 9.606877233 | 552.5740447 | ENSG00000230937 |
| CD48 | 5.71803415 | 52.63805037 | 0.000207086 | 0.001041153 | 0 | 9.745504575 | ENSG00000117091 |
| MIR7-3HG | 5.716492653 | 52.58183749 | 0.000153041 | 0.000789811 | 0 | 9.736902281 | ENSG00000176840 |
| DMRTB1 | 5.68093887 | 51.30184727 | 2.00E-05 | 0.000124772 | 0.353832839 | 18.45368483 | ENSG00000143006 |
| IL6 | 5.639623035 | 49.85350505 | 0.000138494 | 0.000722859 | 0 | 9.202643856 | ENSG00000136244 |
| CYTOR | 5.567756627 | 47.43094235 | 8.13E-07 | 6.50E-06 | 0.677641468 | 31.78737952 | ENSG00000222041 |
| LOC105369438 | 5.490179679 | 44.94783367 | 0.000317857 | 0.001532943 | 0 | 8.312921933 | ENSG00000255929 |
| MOV10L1 | 5.457889884 | 43.95300455 | 0.00042085 | 0.001969584 | 0 | 8.094059608 | ENSG00000073146 |
| MROH3P | 5.415333516 | 42.67542205 | 0.000434134 | 0.002025365 | 0 | 7.875457747 | ENSG00000233217 |
| NA | 5.413691767 | 42.62688617 | 0.000390072 | 0.001845688 | 0 | 7.864704878 | ENSG00000275160 |
| LCN1 | 5.350583359 | 40.80243534 | 0.000521267 | 0.002378936 | 0 | 7.526185506 | ENSG00000160349 |
| CPA4 | 5.332015031 | 40.28064923 | 0.001374929 | 0.005669893 | 0 | 7.465145408 | ENSG00000128510 |
| MAS1 | 5.291909685 | 39.17631199 | 3.63E-06 | 2.58E-05 | 0.677641468 | 26.2108426 | ENSG00000130368 |
| TH | 5.282015601 | 38.90855781 | 0.000744063 | 0.00325629 | 0 | 7.172612117 | ENSG00000180176 |
| LGALS12 | 5.25966229 | 38.31034971 | 1.88E-08 | 1.93E-07 | 1.007031902 | 38.43003436 | ENSG00000133317 |
| KRT80 | 5.250022216 | 38.05521369 | 0.001484866 | 0.006071332 | 0 | 7.044352774 | ENSG00000167767 |
| DPEP3 | 5.23372785 | 37.62782122 | 7.70E-108 | 7.82E-105 | 16.04062268 | 603.5524554 | ENSG00000141096 |
| NA | 5.229639078 | 37.52133041 | 5.72E-06 | 3.94E-05 | 0.647617257 | 25.00357401 | ENSG00000286717 |
| NA | 5.224830737 | 37.39648419 | 0.000860242 | 0.003716669 | 0 | 6.903462563 | ENSG00000261435 |
| C1GALT1C1L | 5.212732984 | 37.08420623 | 0.000978606 | 0.004182162 | 0 | 6.834092744 | ENSG00000223658 |
| LOC101927314 | 5.161489326 | 35.79011628 | 5.52E-19 | 1.69E-17 | 2.716147677 | 95.92760162 | ENSG00000271860 |
| TCL1A | 5.159910435 | 35.75096886 | 0.001211165 | 0.005070439 | 0 | 6.604738015 | ENSG00000100721 |
| NA | 5.151088328 | 35.53301808 | 0.001207601 | 0.005059912 | 0 | 6.556340895 | ENSG00000256540 |
| LOC100506178 | 5.137405929 | 35.19761889 | 0.001486326 | 0.006074728 | 0 | 6.482669929 | ENSG00000232759 |
| NA | 5.135461305 | 35.15020761 | 3.06E-12 | 4.94E-11 | 1.714697581 | 58.94871157 | ENSG00000267279 |
| NA | 5.092767788 | 34.12525185 | 0.001600056 | 0.006484596 | 0 | 6.310314614 | ENSG00000232949 |
| FOXC2 | 5.092767788 | 34.12525185 | 0.001600056 | 0.006484596 | 0 | 6.310314614 | ENSG00000176692 |
| LOC105373085 | 5.08941074 | 34.04593713 | 7.09E-10 | 8.76E-09 | 1.360864742 | 45.60641459 | ENSG00000286381 |
| NA | 5.083894275 | 33.91600352 | 0.001507795 | 0.006155952 | 0 | 6.264068068 | ENSG00000229967 |
| C11orf16 | 5.03551052 | 32.79742217 | 0.000353655 | 0.001687869 | 0.323808629 | 11.78334511 | ENSG00000176029 |
| LINC01487 | 4.866190736 | 29.16549664 | 0.000671357 | 0.002971811 | 0.326599532 | 10.47928227 | ENSG00000241336 |
| LINC00707 | 4.671646886 | 25.48624431 | 8.71E-15 | 1.85E-13 | 2.686123467 | 68.23362869 | ENSG00000238266 |
| NA | 4.670770157 | 25.47076096 | 0.001423894 | 0.005851798 | 0.326599532 | 9.147795015 | ENSG00000285634 |
| OR2W3 | 4.668332813 | 25.42776602 | 0.001473478 | 0.006032443 | 0.323808629 | 9.12897289 | ENSG00000238243 |
| NA | 4.666159652 | 25.38949249 | 0.001803454 | 0.007213441 | 0.323808629 | 9.113918874 | ENSG00000271662 |
| NA | 4.666035831 | 25.3873135 | 0.001712396 | 0.00689066 | 0.326599532 | 9.112301338 | ENSG00000287264 |
| NA | 4.665285941 | 25.37412101 | 0.001563884 | 0.006347334 | 0.353832839 | 9.105849617 | ENSG00000233854 |
| PLA2G2C | 4.603287774 | 24.30679503 | 0.002792873 | 0.010549814 | 0.353832839 | 8.773509391 | ENSG00000187980 |
| NECTIN4 | 4.571522879 | 23.77746289 | 6.40E-11 | 8.96E-10 | 2.011272902 | 47.72087016 | ENSG00000143217 |
| TBX3 | 4.518753338 | 22.92346678 | 0.000244422 | 0.001210562 | 0.65040816 | 15.28575431 | ENSG00000135111 |
| NA | 4.516762327 | 22.89185275 | 4.64E-13 | 8.18E-12 | 2.691705273 | 61.2551502 | ENSG00000287839 |
| PRAME | 4.502780945 | 22.67107577 | 0.000310944 | 0.001504493 | 0.707665678 | 15.35942528 | ENSG00000185686 |
| NA | 4.469003652 | 22.14645151 | 1.38E-09 | 1.65E-08 | 2.038506209 | 44.48573251 | ENSG00000226539 |
| LOC283299 | 4.402350888 | 21.14655707 | 4.43E-07 | 3.68E-06 | 1.330840531 | 28.24947664 | ENSG00000254951 |
| TCL1B | 4.402024718 | 21.14177672 | 1.21E-09 | 1.45E-08 | 2.032924404 | 42.46241295 | ENSG00000213231 |
| NA | 4.395381785 | 21.04465253 | 0.003664986 | 0.013370128 | 0.323808629 | 7.568130904 | ENSG00000248356 |
| PRSS21 | 4.384836363 | 20.89138686 | 2.46E-08 | 2.46E-07 | 1.739139985 | 35.16961075 | ENSG00000007038 |
| C5orf63 | 4.37132788 | 20.6966861 | 0.005321128 | 0.018502141 | 0.323808629 | 7.418638398 | ENSG00000164241 |
| ICAM2 | 4.329727043 | 20.10840914 | 0.004505441 | 0.016029963 | 0.353832839 | 7.229611531 | ENSG00000108622 |
| NA | 4.267302209 | 19.25688183 | 0.005460269 | 0.018934066 | 0.353832839 | 6.928736409 | ENSG00000228714 |
| LINC01139 | 4.25268631 | 19.06277586 | 1.13E-93 | 8.04E-91 | 29.38762507 | 556.1092518 | ENSG00000215808 |
| CALML5 | 4.229934169 | 18.76450293 | 0.009435311 | 0.029956648 | 0.353832839 | 6.777353794 | ENSG00000178372 |
| EBF2 | 4.22617039 | 18.7156129 | 0.000929662 | 0.003990678 | 0.647617257 | 12.4227396 | ENSG00000221818 |
| PF4 | 4.221940043 | 18.66081438 | 3.19E-06 | 2.29E-05 | 4.000894302 | 74.7345877 | ENSG00000163737 |
| LOC101927793 | 4.194623845 | 18.31081176 | 0.006787189 | 0.022814368 | 0.353832839 | 6.585915889 | ENSG00000259881 |
| NA | 4.190127007 | 18.25382635 | 0.006857214 | 0.023009658 | 0.353832839 | 6.560642042 | ENSG00000259162 |
| SERPINB5 | 4.18711038 | 18.21569809 | 0.00778498 | 0.025545168 | 0.323808629 | 6.543970491 | ENSG00000206075 |

| | | | | | | | |
|---|---|---|---|---|---|---|---|
| NA | 4.186744378 | 18.21107748 | 0.007422261 | 0.02453734 | 0.323808629 | 6.541819917 | ENSG00000248243 |
| DPPA5 | 4.17031804 | 18.00490447 | 3.54E-34 | 2.77E-32 | 10.27056419 | 186.0106071 | ENSG00000203909 |
| NA | 4.140828673 | 17.64061158 | 0.009490247 | 0.03010629 | 0.323808629 | 6.363012881 | ENSG00000233936 |
| PCDHGB5 | 4.138211679 | 17.60864118 | 8.90E-22 | 3.43E-20 | 6.074330806 | 106.0931174 | ENSG00000276547 |
| CH25H | 4.131105927 | 17.52212601 | 0.008698943 | 0.028019923 | 0.323808629 | 6.312465188 | ENSG00000138135 |
| NA | 4.131095743 | 17.52200232 | 0.008644816 | 0.0278735 | 0.353832839 | 6.312465188 | ENSG00000287862 |
| NA | 4.121214263 | 17.40239859 | 0.008839102 | 0.0283719 | 0.353832839 | 6.259766921 | ENSG00000270127 |
| NA | 4.116872622 | 17.35010661 | 0.008709276 | 0.02804384 | 0.353832839 | 6.236643648 | ENSG00000242607 |
| F7 | 4.086801334 | 16.99220691 | 8.92E-05 | 0.00048721 | 1.007031902 | 17.04637604 | ENSG00000057593 |
| GDF15 | 4.063999448 | 16.72575521 | 4.51E-133 | 7.91E-130 | 43.62260166 | 731.6650691 | ENSG00000130513 |
| SDR42E1 | 4.051002407 | 16.57575188 | 0.010908114 | 0.033940069 | 0.326599532 | 5.969644667 | ENSG00000184860 |
| CHI3L1 | 4.035098312 | 16.3940263 | 0.011556399 | 0.035624292 | 0.326599532 | 5.88952198 | ENSG00000133048 |
| AGBL4 | 4.016627476 | 16.18547136 | 0.000175529 | 0.000895091 | 1.004240999 | 16.26366818 | ENSG00000186094 |
| KCNIP1-AS1 | 3.966226076 | 15.62978552 | 0.014093276 | 0.042047856 | 0.353832839 | 5.626824147 | ENSG00000253591 |
| ANXA1 | 3.95626295 | 15.52221943 | 4.33E-09 | 4.87E-08 | 2.628865949 | 41.38285855 | ENSG00000135046 |
| NPIPA5 | 3.952946233 | 15.48657526 | 0.000214549 | 0.001073633 | 0.974216789 | 15.43068521 | ENSG00000183793 |
| IZUMO2 | 3.92758043 | 15.21666641 | 3.67E-07 | 3.11E-06 | 1.986830497 | 30.42820738 | ENSG00000161652 |
| LINC02236 | 3.909363529 | 15.02573365 | 0.003144911 | 0.011667014 | 0.65040816 | 10.01035298 | ENSG00000251365 |
| TRIM58 | 3.881638408 | 14.7397322 | 9.51E-37 | 8.74E-35 | 12.41309754 | 182.3874626 | ENSG00000162722 |
| NA | 3.830828548 | 14.22965272 | 9.70E-06 | 6.40E-05 | 1.65464916 | 23.66752514 | ENSG00000272141 |
| NA | 3.816610741 | 14.09010777 | 0.000460604 | 0.002130819 | 1.03426521 | 14.25944333 | ENSG00000257869 |
| AADACL3 | 3.808707343 | 14.01313013 | 9.48E-06 | 6.28E-05 | 1.709115775 | 23.60082682 | ENSG00000188984 |
| SLC8A3 | 3.761170692 | 13.55892309 | 6.49E-11 | 9.07E-10 | 3.774855292 | 50.16713292 | ENSG00000100678 |
| CYP2B6 | 3.761143052 | 13.55866332 | 6.81E-150 | 1.88E-146 | 61.10925564 | 828.9639954 | ENSG00000197408 |
| NA | 3.73740257 | 13.33737246 | 0.004967386 | 0.017438734 | 0.65040816 | 8.857672762 | ENSG00000286186 |
| LINC01414 | 3.711398515 | 13.09912477 | 0.001125431 | 0.004739289 | 0.979798595 | 13.03524271 | ENSG00000253554 |
| NA | 3.704644516 | 13.03794434 | 2.08E-29 | 1.29E-27 | 11.51499887 | 148.6251607 | ENSG00000280623 |
| BCO1 | 3.681035694 | 12.82632261 | 0.006130797 | 0.020903005 | 0.680432371 | 8.628318033 | ENSG00000135697 |
| SIX3 | 3.670681853 | 12.734601 | 0.000854839 | 0.00369581 | 1.001450097 | 12.75318972 | ENSG00000138083 |
| NA | 3.668357854 | 12.71410369 | 0.005938629 | 0.020362975 | 0.680432371 | 8.533674367 | ENSG00000230650 |
| TMEM105 | 3.646491493 | 12.52285403 | 0.007342714 | 0.024325655 | 0.707665678 | 8.489578395 | ENSG00000185332 |
| MYBPHL | 3.646094208 | 12.519406 | 0.00880127 | 0.028278697 | 0.647617257 | 8.315072506 | ENSG00000221986 |
| COL8A1 | 3.645819131 | 12.51701916 | 1.55E-08 | 1.61E-07 | 3.015513901 | 37.72744919 | ENSG00000144810 |
| ADTRP | 3.644992323 | 12.50984772 | 0.008252182 | 0.026810415 | 0.647617257 | 8.310771359 | ENSG00000111863 |
| CER1 | 3.641423747 | 12.47894225 | 3.34E-17 | 8.95E-16 | 6.887938652 | 87.08284441 | ENSG00000147869 |
| NA | 3.630735789 | 12.38683575 | 0.007962909 | 0.026009562 | 0.647617257 | 8.21397712 | ENSG00000185168 |
| NA | 3.613470698 | 12.23948286 | 0.007788102 | 0.025551066 | 0.680432371 | 8.220428841 | ENSG00000270110 |
| KRT81 | 3.611101273 | 12.21939772 | 0.000124557 | 0.000657948 | 2.008481999 | 24.58061869 | ENSG00000205426 |
| SCG2 | 3.608959547 | 12.2012711 | 1.35E-10 | 1.83E-09 | 4.079803322 | 49.13652079 | ENSG00000171951 |
| GPRC5D-AS1 | 3.572624753 | 11.89781506 | 7.73E-05 | 0.000429005 | 1.632997658 | 19.75774767 | ENSG00000247498 |
| NA | 3.560747367 | 11.80026511 | 0.001805395 | 0.007218211 | 1.007031902 | 11.85109739 | ENSG00000254186 |
| SPDYE2 | 3.560266724 | 11.79633443 | 0.009234859 | 0.029417052 | 0.65040816 | 7.818458333 | ENSG00000205238 |
| VAV1 | 3.549641049 | 11.70977174 | 1.06E-57 | 2.27E-55 | 28.08122695 | 326.0017978 | ENSG00000141968 |
| PRPH2 | 3.539320095 | 11.62629968 | 7.42E-06 | 5.01E-05 | 2.035715307 | 23.42606047 | ENSG00000112619 |
| LINC-ROR | 3.536734161 | 11.60547897 | 4.59E-33 | 3.49E-31 | 14.73143365 | 170.8065688 | ENSG00000258609 |
| MPO | 3.524388994 | 11.50659437 | 0.012088703 | 0.036840942 | 0.707665678 | 7.809856036 | ENSG00000005381 |
| NA | 3.52068142 | 11.4770616 | 0.012017681 | 0.036670845 | 0.647617257 | 7.616528024 | ENSG00000284070 |
| POT1-AS1 | 3.511627644 | 11.40526169 | 0.000550386 | 0.002497061 | 1.295234515 | 15.1029187 | ENSG00000224897 |
| NA | 3.511021701 | 11.4004724 | 0.012489557 | 0.037936711 | 0.647617257 | 7.553609926 | ENSG00000277349 |
| CELP | 3.49696534 | 11.28993551 | 0.011505549 | 0.035523934 | 0.677641468 | 7.578883773 | ENSG00000170827 |
| FOXL1 | 3.496275548 | 11.28453877 | 0.002047551 | 0.008056152 | 0.974216789 | 11.24289543 | ENSG00000176678 |
| PITX2 | 3.476115717 | 11.1279483 | 6.00E-10 | 7.50E-09 | 4.324702931 | 48.28444311 | ENSG00000164093 |
| SP5 | 3.469673202 | 11.078366 | 0.002568122 | 0.009814664 | 1.007031902 | 11.11222505 | ENSG00000204335 |
| LCP2 | 3.459119213 | 10.99761829 | 0.015345636 | 0.045106302 | 0.65040816 | 7.307583645 | ENSG00000043462 |
| NA | 3.449714766 | 10.92616164 | 0.01668698 | 0.048305602 | 0.647617257 | 7.236063252 | ENSG00000205930 |
| LEFTY2 | 3.44596579 | 10.89780586 | 2.79E-63 | 6.90E-61 | 135.7222926 | 1476.540506 | ENSG00000143768 |
| NA | 3.431173427 | 10.78663844 | 0.000769575 | 0.003355 | 1.385307146 | 14.60871557 | ENSG00000256417 |
| NA | 3.404008565 | 10.58543436 | 0.019400336 | 0.054795298 | 0.677641468 | 7.082269599 | ENSG00000240219 |
| NA | 3.403888943 | 10.5845567 | 0.002953034 | 0.011046534 | 0.974216789 | 10.53413111 | ENSG00000237346 |
| NA | 3.398586628 | 10.54572684 | 0.003030706 | 0.011299837 | 0.974216789 | 10.48573399 | ENSG00000173727 |
| PKP1 | 3.376722505 | 10.38711072 | 7.80E-21 | 2.80E-19 | 10.04240999 | 104.2090706 | ENSG00000081277 |
| NA | 3.371934702 | 10.35269665 | 0.02491658 | 0.067433467 | 0.65040816 | 6.84914676 | ENSG00000286654 |
| KIF25 | 3.368397745 | 10.32734674 | 3.44E-05 | 0.000205379 | 2.038506209 | 20.83460635 | ENSG00000125337 |
| LOC157273 | 3.366569077 | 10.31426475 | 5.51E-10 | 6.91E-09 | 5.249359189 | 54.72328964 | ENSG00000248538 |
| TEX11 | 3.351548526 | 10.20743536 | 0.018086922 | 0.051622254 | 0.707665678 | 6.951859682 | ENSG00000120498 |
| CTCFL | 3.351414341 | 10.20648601 | 1.52E-16 | 3.84E-15 | 9.621031907 | 96.51052974 | ENSG00000124092 |
| SIGLEC17P | 3.349327983 | 10.19173654 | 0.001335707 | 0.005525867 | 1.355282936 | 13.72921348 | ENSG00000171101 |
| NA | 3.340253362 | 10.12783121 | 0.018337669 | 0.052168114 | 0.707665678 | 6.884640437 | ENSG00000263786 |
| LINC00304 | 3.334471294 | 10.08732188 | 8.93E-06 | 5.93E-05 | 2.362314838 | 23.6677856 | ENSG00000180422 |
| TRDN | 3.334204556 | 10.08545702 | 4.82E-07 | 3.98E-06 | 2.985489691 | 30.28757763 | ENSG00000186439 |
| LINC02078 | 3.327166128 | 10.03637327 | 2.12E-06 | 1.58E-05 | 2.68891437 | 26.86314054 | ENSG00000267719 |
| NA | 3.309560413 | 9.914640166 | 0.02084767 | 0.058145812 | 0.680432371 | 6.684860702 | ENSG00000224892 |
| HCG20 | 3.304152622 | 9.877545741 | 0.022852991 | 0.062778697 | 0.653199063 | 6.554190321 | ENSG00000228022 |
| PTPRB | 3.303855818 | 9.875513855 | 6.16E-15 | 1.32E-13 | 7.800191998 | 76.26261962 | ENSG00000127329 |
| NA | 3.289252118 | 9.776053079 | 0.005060984 | 0.01772217 | 1.001450097 | 9.785038936 | ENSG00000287150 |
| RGS4 | 3.280316748 | 9.715691949 | 6.32E-05 | 0.000356498 | 1.978457788 | 19.41492715 | ENSG00000117152 |
| NA | 3.274917643 | 9.679400143 | 0.023164432 | 0.063421099 | 0.677641468 | 6.480519355 | ENSG00000269514 |

| | | | | | | | |
|---|---|---|---|---|---|---|---|
| NA | 3.267806493 | 9.63180706 | 0.028219116 | 0.074745979 | 0.677641468 | 6.447176252 | ENSG00000274751 |
| TMEM82 | 3.26650214 | 9.623102793 | 4.47E-07 | 3.72E-06 | 3.336531627 | 32.13477376 | ENSG00000162460 |
| LINC00989 | 3.257482906 | 9.563130124 | 0.000160531 | 0.000825817 | 1.972875983 | 19.07640777 | ENSG00000250334 |
| PLAAT4 | 3.249189433 | 9.508313245 | 9.57E-05 | 0.000519717 | 1.984039594 | 18.99628509 | ENSG00000133321 |
| NA | 3.241602016 | 9.458438422 | 0.015021119 | 0.044328042 | 0.979798595 | 9.376616707 | ENSG00000270948 |
| ZNF385D-AS1 | 3.240076515 | 9.44844239 | 0.001589596 | 0.006446266 | 1.298025418 | 12.52813614 | ENSG00000225542 |
| FGF17 | 3.237723815 | 9.433046734 | 0.000123428 | 0.000652701 | 2.062948614 | 19.09738047 | ENSG00000158815 |
| GIP | 3.233672871 | 9.406596828 | 0.023072434 | 0.063205102 | 0.65040816 | 6.238794221 | ENSG00000159224 |
| MCCC2 | 3.230398919 | 9.385274353 | 1.73E-06 | 1.30E-05 | 3.015513901 | 28.25324475 | ENSG00000131844 |
| KRT17 | 3.228281938 | 9.371512697 | 7.49E-10 | 9.23E-09 | 5.32894393 | 50.0872707 | ENSG00000128422 |
| NA | 3.225912643 | 9.356134773 | 0.00025157 | 0.001242774 | 2.092972825 | 19.05973622 | ENSG00000248587 |
| CBY2 | 3.213209356 | 9.274113323 | 0.008165738 | 0.026573087 | 0.971425886 | 9.20479443 | ENSG00000174015 |
| LOC101926892 | 3.202367296 | 9.204678268 | 0.028473967 | 0.075327933 | 0.680432371 | 6.175876124 | ENSG00000253967 |
| NA | 3.19794358 | 9.176497318 | 0.011620848 | 0.035748329 | 0.979798595 | 9.1117683 | ENSG00000242880 |
| NA | 3.196007913 | 9.164193449 | 0.02843781 | 0.075242599 | 0.680432371 | 6.141999982 | ENSG00000233754 |
| MIR4435-2HG | 3.175204461 | 9.032995315 | 8.34E-14 | 1.59E-12 | 8.51064858 | 75.97464794 | ENSG00000172965 |
| NA | 3.16632821 | 8.977589997 | 0.032810321 | 0.084482634 | 0.680432371 | 6.064288333 | ENSG00000241547 |
| LINC00917 | 3.164987664 | 8.969251933 | 0.041210524 | 0.102105346 | 0.647617257 | 5.929849843 | ENSG00000168367 |
| ST6GALNAC6 | 3.154415713 | 8.903766217 | 0.029953547 | 0.078446086 | 0.65040816 | 5.900274849 | ENSG00000160408 |
| EVPL | 3.15145577 | 8.885517298 | 6.64E-22 | 2.60E-20 | 15.56178692 | 138.9842591 | ENSG00000167880 |
| NA | 3.139025195 | 8.809286626 | 0.043249943 | 0.106244066 | 0.647617257 | 5.803480609 | ENSG00000248187 |
| LOC107984525 | 3.105834364 | 8.608932549 | 7.82E-13 | 1.35E-11 | 8.154024838 | 69.40299257 | ENSG00000286069 |
| NA | 3.100297713 | 8.575957244 | 0.001091665 | 0.004611189 | 1.709115775 | 14.4398679 | ENSG00000256654 |
| PIWIL2 | 3.099197064 | 8.56941704 | 1.17E-13 | 2.20E-12 | 8.643148334 | 74.34873729 | ENSG00000197181 |
| LAPTM5 | 3.099163248 | 8.569216181 | 7.97E-05 | 0.00044003 | 2.30505732 | 19.93440413 | ENSG00000162511 |
| NA | 3.098320659 | 8.56421289 | 0.037078445 | 0.093586325 | 0.65040816 | 5.696193966 | ENSG00000215481 |
| NA | 3.094421341 | 8.541096781 | 0.001338638 | 0.005536807 | 1.681882467 | 14.27423688 | ENSG00000236212 |
| MYCT1 | 3.088252613 | 8.504654411 | 0.010238993 | 0.032080626 | 1.03426521 | 8.63261918 | ENSG00000120279 |
| CCDC60 | 3.086961143 | 8.497044632 | 0.013666952 | 0.040952029 | 0.977007692 | 8.42235915 | ENSG00000183273 |
| ZNF648 | 3.074770211 | 8.425546177 | 0.000825052 | 0.003575842 | 1.681882467 | 14.1034991 | ENSG00000179930 |
| GLIS2-AS1 | 3.074038071 | 8.42127146 | 0.036383505 | 0.092085305 | 0.677641468 | 5.670920119 | ENSG00000262686 |
| OR52N2 | 3.068403113 | 8.388443349 | 0.036683434 | 0.092747069 | 0.677641468 | 5.643495699 | ENSG00000180988 |
| S100A6 | 3.062310864 | 8.353095107 | 3.22E-46 | 4.78E-44 | 35.15872271 | 295.2482229 | ENSG00000197956 |
| NA | 3.061991406 | 8.351245672 | 2.26E-05 | 0.000139166 | 3.10279563 | 25.43081836 | ENSG00000282024 |
| NA | 3.054339966 | 8.307071466 | 0.035600177 | 0.090411139 | 0.677641468 | 5.57412588 | ENSG00000264548 |
| NA | 3.053788544 | 8.30389697 | 0.016020468 | 0.046746253 | 1.007031902 | 8.3656202 | ENSG00000222220 |
| MYL7 | 3.043729029 | 8.246197582 | 4.61E-05 | 0.000266859 | 2.634447755 | 21.92625858 | ENSG00000106631 |
| LINC02709 | 3.043494972 | 8.244859861 | 0.01183416 | 0.03623695 | 0.974216789 | 8.186552699 | ENSG00000245522 |
| NA | 3.040082505 | 8.225380993 | 0.012321877 | 0.037480501 | 0.974216789 | 8.161278853 | ENSG00000254683 |
| NA | 3.037425533 | 8.210246475 | 0.000354467 | 0.001690487 | 1.981248691 | 16.39622867 | ENSG00000259345 |
| CECR3 | 3.035189724 | 8.197532541 | 0.042468255 | 0.104670386 | 0.680432371 | 5.496153767 | ENSG00000241832 |
| NA | 3.034505235 | 8.193644135 | 0.001790206 | 0.007167883 | 1.687464273 | 13.72948605 | ENSG00000248213 |
| LEFTY1 | 3.030830244 | 8.172798945 | 1.55E-154 | 4.99E-151 | 136.1337463 | 1110.347549 | ENSG00000243709 |
| FAM90A1 | 3.017583944 | 8.098102728 | 4.59E-05 | 0.000266008 | 2.691705273 | 21.70550615 | ENSG00000171847 |
| NA | 3.011047551 | 8.061495784 | 0.003580225 | 0.013108013 | 1.358073839 | 10.82855451 | ENSG00000228157 |
| ODF3L2 | 3.004576951 | 8.025420309 | 0.004395247 | 0.015689991 | 1.360864742 | 10.78230797 | ENSG00000181781 |
| GZMM | 3.003289259 | 8.018260332 | 0.000567633 | 0.002565652 | 2.014063805 | 16.10610642 | ENSG00000197540 |
| LOC284865 | 2.999522616 | 7.997353256 | 0.004441533 | 0.015837635 | 1.358073839 | 10.72530855 | ENSG00000249923 |
| LINC01162 | 2.992000885 | 7.955766227 | 0.001645976 | 0.00665253 | 1.651858257 | 13.26862602 | ENSG00000232790 |
| GSTT2 | 2.991995276 | 7.955735299 | 0.014075713 | 0.042007205 | 0.977007692 | 7.917403145 | ENSG00000099984 |
| ALOX15B | 2.989806011 | 7.943671763 | 0.000209124 | 0.001049755 | 2.307848223 | 18.52520522 | ENSG00000179593 |
| LOC100505774 | 2.986598422 | 7.926029966 | 0.003238679 | 0.011978021 | 1.687464273 | 13.32724297 | ENSG00000226833 |
| SLC16A3 | 2.983863439 | 7.911018463 | 1.27E-100 | 1.07E-97 | 117.5315276 | 926.4134256 | ENSG00000141526 |
| SMC1B | 2.981177075 | 7.896301491 | 0.005228462 | 0.018232514 | 1.333631434 | 10.55725439 | ENSG00000077935 |
| LINC00607 | 2.981124178 | 7.896011975 | 6.99E-05 | 0.000390814 | 2.631656852 | 20.99674183 | ENSG00000235770 |
| C1orf87 | 2.965078506 | 7.808678994 | 0.017754129 | 0.050898337 | 1.03426521 | 7.92600544 | ENSG00000162598 |
| NA | 2.964828201 | 7.80732432 | 0.013938553 | 0.04166878 | 1.004240999 | 7.818458333 | ENSG00000282206 |
| NA | 2.962354201 | 7.793947434 | 4.17E-05 | 0.000243785 | 3.075562322 | 23.63014135 | ENSG00000236975 |
| LINC02188 | 2.959423983 | 7.778133425 | 8.46E-60 | 1.97E-57 | 46.7714912 | 363.8773945 | ENSG00000261175 |
| SLC25A3P1 | 2.95350784 | 7.746302494 | 0.008493721 | 0.027441363 | 1.306398126 | 10.28380368 | ENSG00000236253 |
| DUSP27 | 2.940705907 | 7.677868797 | 2.13E-07 | 1.86E-06 | 4.667372158 | 35.91870293 | ENSG00000198842 |
| SYT4 | 2.937606972 | 7.661394296 | 7.35E-34 | 5.72E-32 | 27.97699964 | 214.8768637 | ENSG00000132872 |
| PRDM1 | 2.927750325 | 7.609229233 | 6.36E-05 | 0.000358675 | 3.100004727 | 23.18218476 | ENSG00000057657 |
| PLA2G4C | 2.925155979 | 7.595558128 | 1.00E-24 | 4.87E-23 | 19.42113304 | 147.3044263 | ENSG00000105499 |
| NA | 2.907338324 | 7.502327925 | 0.003018001 | 0.011256818 | 1.649067354 | 12.47543787 | ENSG00000280164 |
| KRT42P | 2.905355061 | 7.492021612 | 0.028916073 | 0.076215324 | 1.061498518 | 7.721924557 | ENSG00000214514 |
| EPHA3 | 2.89781722 | 7.452979131 | 9.86E-05 | 0.0005338 | 3.012722998 | 22.38228443 | ENSG00000044524 |
| CDKN1A | 2.881340427 | 7.36834405 | 7.52E-180 | 3.62E-176 | 238.6018974 | 1761.026169 | ENSG00000124762 |
| GALR1 | 2.865794114 | 7.289369911 | 0.000264851 | 0.001300062 | 2.716147677 | 19.65691274 | ENSG00000166573 |
| H1-4 | 2.861344684 | 7.266923324 | 0.022140856 | 0.061119554 | 1.001450097 | 7.296830776 | ENSG00000168298 |
| SCARA5 | 2.856648154 | 7.243305143 | 0.000257035 | 0.001266529 | 2.694496175 | 19.43374927 | ENSG00000168079 |
| LINC01679 | 2.854301301 | 7.231531932 | 0.021611497 | 0.059952169 | 1.001450097 | 7.252734804 | ENSG00000237989 |
| NA | 2.853991525 | 7.229979342 | 0.01718365 | 0.049550043 | 1.295234515 | 9.535244545 | ENSG00000286458 |
| NA | 2.853592955 | 7.227982207 | 6.83E-13 | 1.19E-11 | 9.658552945 | 69.97542828 | ENSG00000286699 |
| SRRM5 | 2.85337481 | 7.226889371 | 0.000667618 | 0.002958654 | 2.367896644 | 16.96410277 | ENSG00000226763 |
| NA | 2.839600819 | 7.158219678 | 0.022887066 | 0.062831134 | 1.007031902 | 7.179063838 | ENSG00000272320 |
| SPAAR | 2.832474973 | 7.122950544 | 0.023279403 | 0.063706281 | 1.03426521 | 7.221009236 | ENSG00000235387 |

| | | | | | | | |
|---|---|---|---|---|---|---|---|
| LOC100420587 | 2.831757454 | 7.11940885 | 0.033331615 | 0.08552809 | 0.977007692 | 7.099201615 | ENSG00000283403 |
| LOC101927447 | 2.816690217 | 7.045441998 | 0.015469035 | 0.045406748 | 1.295234515 | 9.31826022 | ENSG00000203527 |
| NTSR2 | 2.812820666 | 7.026570253 | 0.033149712 | 0.085129271 | 0.979798595 | 7.000256802 | ENSG00000169006 |
| MIR2052HG | 2.802570548 | 6.976824521 | 0.00170442 | 0.00686286 | 2.065739517 | 14.14974565 | ENSG00000254349 |
| NA | 2.80149512 | 6.971625725 | 0.013508221 | 0.040539374 | 1.303607223 | 9.238137533 | ENSG00000261118 |
| NA | 2.800050673 | 6.964649129 | 0.000409369 | 0.001921914 | 2.944301869 | 20.76953768 | ENSG00000189229 |
| ITPRID1 | 2.79875885 | 6.958415608 | 4.89E-07 | 4.03E-06 | 5.336640917 | 37.19534134 | ENSG00000180347 |
| MOGAT1 | 2.798452594 | 6.956938629 | 0.026098631 | 0.070130756 | 0.974216789 | 6.90776371 | ENSG00000124003 |
| LINC02231 | 2.795505846 | 6.942743374 | 0.00991822 | 0.031227744 | 1.328049628 | 9.276314822 | ENSG00000248995 |
| KRT16P3 | 2.793779655 | 6.93444132 | 0.003828008 | 0.013906988 | 1.65464916 | 11.56259267 | ENSG00000214822 |
| CYTH4 | 2.792264881 | 6.927164244 | 0.030097367 | 0.07874434 | 0.974216789 | 6.865818312 | ENSG00000100055 |
| NA | 2.789315949 | 6.91301928 | 0.026648407 | 0.071336459 | 1.007031902 | 6.954010256 | ENSG00000279267 |
| PAX8-AS1 | 2.784392059 | 6.889465498 | 0.004310812 | 0.015428549 | 1.681882467 | 11.55022227 | ENSG00000189223 |
| USH2A | 2.78148893 | 6.875615792 | 0.008106494 | 0.026411479 | 1.621834046 | 11.3184565 | ENSG00000042781 |
| LNCTAM34A | 2.765166598 | 6.798264989 | 2.39E-05 | 0.000146826 | 3.926891366 | 27.06909942 | ENSG00000234546 |
| MIR210HG | 2.747674218 | 6.716335124 | 1.62E-22 | 6.60E-21 | 22.99160082 | 153.3719497 | ENSG00000247095 |
| HCLS1 | 2.739709869 | 6.679359978 | 1.57E-16 | 3.95E-15 | 15.76079278 | 105.0520129 | ENSG00000180353 |
| C14orf39 | 2.734259354 | 6.654172903 | 8.38E-09 | 9.05E-08 | 7.386310738 | 49.02761661 | ENSG00000179008 |
| GRAMD1C | 2.7293319 | 6.631484671 | 7.66E-08 | 7.16E-07 | 6.768717691 | 44.59541808 | ENSG00000178075 |
| KIF25-AS1 | 2.725582125 | 6.614270854 | 4.23E-08 | 4.12E-07 | 7.277377508 | 48.45867643 | ENSG00000229921 |
| ANGPT4 | 2.724800429 | 6.61068801 | 6.49E-07 | 5.26E-06 | 5.698846465 | 37.56370828 | ENSG00000101280 |
| CD163 | 2.721424315 | 6.59523614 | 0.045183168 | 0.110026318 | 0.979798595 | 6.546121064 | ENSG00000177575 |
| MEIOB | 2.717584023 | 6.577703705 | 2.59E-34 | 2.04E-32 | 37.02932331 | 242.5791785 | ENSG00000162039 |
| LINC02473 | 2.705581378 | 6.523206857 | 0.001622703 | 0.006566722 | 2.422363259 | 15.48553405 | ENSG00000261121 |
| NA | 2.68545194 | 6.432822763 | 0.003038577 | 0.011322619 | 2.005691096 | 12.92365492 | ENSG00000260492 |
| NA | 2.683180144 | 6.422701036 | 1.70E-08 | 1.75E-07 | 7.266213896 | 47.0287774 | ENSG00000277883 |
| NA | 2.6809834 | 6.412928847 | 1.57E-09 | 1.88E-08 | 9.068193206 | 57.95063061 | ENSG00000254275 |
| SP140L | 2.680839568 | 6.412289532 | 1.09E-10 | 1.50E-09 | 9.628528734 | 61.97413434 | ENSG00000185404 |
| NA | 2.668539825 | 6.357853725 | 0.004736426 | 0.016749843 | 2.038506209 | 12.8419147 | ENSG00000276975 |
| NA | 2.662631932 | 6.331871299 | 0.003352053 | 0.012349973 | 2.038506209 | 12.7677107 | ENSG00000287527 |
| PLBD1-AS1 | 2.656445789 | 6.304778948 | 0.047855098 | 0.115367744 | 0.979798595 | 6.268369215 | ENSG00000256751 |
| FAM215A | 2.653274373 | 6.290934637 | 0.045498115 | 0.110667525 | 1.001450097 | 6.318916909 | ENSG00000267496 |
| NR1I2 | 2.651859895 | 6.284769767 | 3.40E-20 | 1.18E-18 | 20.05758668 | 126.1017135 | ENSG00000144852 |
| NA | 2.646332043 | 6.260735024 | 0.009586747 | 0.03036249 | 1.942851772 | 12.35121921 | ENSG00000267069 |
| LINC01239 | 2.646148327 | 6.259937819 | 0.04544575 | 0.110568956 | 0.977007692 | 6.207068654 | ENSG00000234840 |
| DNAJC22 | 2.645157162 | 6.255638576 | 0.042828902 | 0.10541112 | 1.007031902 | 6.293643062 | ENSG00000178401 |
| NA | 2.638858552 | 6.228386839 | 0.023302119 | 0.063752827 | 1.415331357 | 8.531523793 | ENSG00000157306 |
| NA | 2.629775949 | 6.1892987 | 3.07E-05 | 0.000184954 | 4.702978174 | 29.04911651 | ENSG00000227712 |
| GDA | 2.628490033 | 6.183784457 | 0.018278809 | 0.052054399 | 1.328049628 | 8.255922518 | ENSG00000119125 |
| NA | 2.6190028 | 6.143253004 | 0.002460498 | 0.009466587 | 2.356733032 | 14.40437422 | ENSG00000258636 |
| NA | 2.615668855 | 6.12907286 | 0.018132238 | 0.051720981 | 1.328049628 | 8.163429427 | ENSG00000280202 |
| MUC4 | 2.613954767 | 6.12179514 | 7.13E-16 | 1.69E-14 | 18.63388262 | 114.3595018 | ENSG00000145113 |
| IL1A | 2.612756185 | 6.116711302 | 0.00039852 | 0.001879211 | 3.300925611 | 20.33207349 | ENSG00000115008 |
| NA | 2.610698547 | 6.107993584 | 0.000305981 | 0.001482341 | 3.681991758 | 22.49654372 | ENSG00000235888 |
| PTPN7 | 2.60171894 | 6.070094347 | 0.021472672 | 0.059597672 | 1.385307146 | 8.234949819 | ENSG00000143851 |
| ASB2 | 2.599200165 | 6.059505926 | 0.000547532 | 0.002486418 | 2.985489691 | 18.2033574 | ENSG00000100628 |
| CTSLP2 | 2.599137942 | 6.059244586 | 0.001512624 | 0.006173054 | 2.748962791 | 16.38170769 | ENSG00000280913 |
| S100A2 | 2.580854862 | 5.982941109 | 0.003821561 | 0.013888796 | 2.800638503 | 16.35805138 | ENSG00000196754 |
| PTPRH | 2.580458169 | 5.981296225 | 0.002510316 | 0.009625658 | 2.392339049 | 14.11425197 | ENSG00000080031 |
| ATP1A4 | 2.571304311 | 5.943465218 | 9.75E-05 | 0.000529027 | 3.95970648 | 23.73553788 | ENSG00000132681 |
| CLPSL1 | 2.567191934 | 5.926547597 | 0.051895639 | 0.122896977 | 1.007031902 | 5.963192946 | ENSG00000204140 |
| NA | 2.56649945 | 5.923703578 | 0.002157102 | 0.00843215 | 2.65051745 | 15.78210802 | ENSG00000267055 |
| NA | 2.562039501 | 5.905419311 | 0.059870206 | 0.137684004 | 0.974216789 | 5.836823713 | ENSG00000286859 |
| NA | 2.559691945 | 5.895817816 | 0.01016711 | 0.031907227 | 1.68467337 | 9.89043547 | ENSG00000233058 |
| DARS-AS1 | 2.555059131 | 5.876915403 | 0.052635328 | 0.124160933 | 1.031474307 | 5.971795241 | ENSG00000231890 |
| NA | 2.550709007 | 5.859221568 | 0.046891593 | 0.113513078 | 1.004240999 | 5.868549281 | ENSG00000249655 |
| TTC22 | 2.550623473 | 5.8588742 | 1.59E-11 | 2.38E-10 | 13.0662966 | 76.50298768 | ENSG00000006555 |
| TNFRSF10C | 2.548111886 | 5.84868336 | 4.04E-17 | 1.07E-15 | 18.69393104 | 109.4237044 | ENSG00000173535 |
| DLGAP2 | 2.544437737 | 5.833807318 | 1.13E-11 | 1.72E-10 | 12.70130015 | 74.05943907 | ENSG00000198010 |
| GJC2 | 2.542920056 | 5.827673519 | 9.69E-25 | 4.71E-23 | 28.46141721 | 165.7094414 | ENSG00000198835 |
| GPAT2 | 2.542774811 | 5.827086839 | 4.71E-05 | 0.000272008 | 4.340772627 | 25.31305142 | ENSG00000186281 |
| NA | 2.541285296 | 5.821073749 | 0.053235123 | 0.125361177 | 1.031474307 | 5.900274849 | ENSG00000233929 |
| CCDC80 | 2.539990112 | 5.81585021 | 2.80E-37 | 2.67E-35 | 52.2524712 | 304.714116 | ENSG00000091986 |
| NA | 2.538720296 | 5.810733526 | 6.40E-57 | 1.33E-54 | 112.0886563 | 649.5021625 | ENSG00000253507 |
| NA | 2.536260858 | 5.800836102 | 0.005926627 | 0.020339934 | 2.296684611 | 13.45765288 | ENSG00000233750 |
| RRAD | 2.536116967 | 5.800257572 | 0.052758439 | 0.124390498 | 1.031474307 | 5.872850428 | ENSG00000166592 |
| ARSL | 2.534581524 | 5.794087708 | 1.02E-65 | 2.68E-63 | 89.72549983 | 520.6957959 | ENSG00000157399 |
| STMN2 | 2.534260157 | 5.792797195 | 5.52E-24 | 2.52E-22 | 30.05410293 | 173.0651617 | ENSG00000104435 |
| IQSEC3 | 2.531613022 | 5.782178001 | 5.03E-27 | 2.74E-25 | 31.79025185 | 183.6135534 | ENSG00000120645 |
| NEAT1 | 2.530895436 | 5.779302701 | 3.43E-24 | 1.59E-22 | 29.59691867 | 170.4901066 | ENSG00000245532 |
| NA | 2.52973136 | 5.774641402 | 0.061411735 | 0.140536268 | 1.061498518 | 5.940069673 | ENSG00000267114 |
| CMKLR1 | 2.520336874 | 5.737160482 | 1.36E-15 | 3.13E-14 | 21.47359376 | 122.9101387 | ENSG00000174600 |
| NA | 2.509273084 | 5.693331421 | 0.034447112 | 0.087922631 | 1.360864742 | 7.696650711 | ENSG00000233695 |
| OTOF | 2.509136082 | 5.692790791 | 7.95E-05 | 0.000438923 | 4.319121125 | 24.68817791 | ENSG00000115155 |
| HAO2 | 2.50721825 | 5.685228167 | 0.005195098 | 0.018129277 | 2.367896644 | 13.34848824 | ENSG00000116882 |
| UNC93A | 2.507014364 | 5.684424769 | 0.000122011 | 0.000646272 | 4.422472549 | 24.86483437 | ENSG00000112494 |
| NA | 2.504239622 | 5.673502396 | 0.075718853 | 0.166006637 | 0.979798595 | 5.641345125 | ENSG00000250920 |

| | | | | | | | |
|---|---|---|---|---|---|---|---|
| NA | 2.503241247 | 5.669577573 | 0.025801462 | 0.069477387 | 1.330840531 | 7.570281478 | ENSG00000243155 |
| GUCY1B2 | 2.502240651 | 5.665646741 | 0.002555879 | 0.009773133 | 2.746171888 | 15.29409614 | ENSG00000123201 |
| MACC1-AS1 | 2.497318168 | 5.646348466 | 0.060174023 | 0.138241881 | 0.974216789 | 5.6015503 | ENSG00000228598 |
| NODAL | 2.496970809 | 5.644989154 | 1.78E-36 | 1.60E-34 | 46.58731573 | 262.5781183 | ENSG00000156574 |
| RGN | 2.495114013 | 5.637728539 | 2.65E-05 | 0.000161902 | 5.456737759 | 30.38196083 | ENSG00000130988 |
| NA | 2.491544422 | 5.623796621 | 0.034990882 | 0.08916897 | 1.360864742 | 7.581034346 | ENSG00000229282 |
| NLRP1 | 2.490439673 | 5.619491828 | 1.79E-07 | 1.58E-06 | 8.082812805 | 45.19045614 | ENSG00000091592 |
| NA | 2.487904826 | 5.609626924 | 0.066256765 | 0.14937002 | 1.004240999 | 5.639194551 | ENSG00000283405 |
| NA | 2.487552275 | 5.608256272 | 0.062147708 | 0.141880644 | 0.974216789 | 5.553153181 | ENSG00000256258 |
| NA | 2.484739451 | 5.597332499 | 0.055288666 | 0.129469975 | 1.004240999 | 5.624673573 | ENSG00000247903 |
| GSTT2B | 2.483989281 | 5.594422764 | 8.47E-17 | 2.19E-15 | 21.21473964 | 118.158243 | ENSG00000133433 |
| CHL1 | 2.46748711 | 5.530795922 | 0.037268216 | 0.093966952 | 1.388098049 | 7.50736338 | ENSG00000134121 |
| LINC01451 | 2.465040259 | 5.521423488 | 0.06849887 | 0.153313806 | 1.03426521 | 5.616071279 | ENSG00000279141 |
| KLHDC8A | 2.464306491 | 5.518615956 | 1.88E-38 | 1.94E-36 | 51.90422017 | 287.2848457 | ENSG00000162873 |
| PTGDR2 | 2.454461828 | 5.481086215 | 0.017257564 | 0.049711164 | 1.65464916 | 9.103699043 | ENSG00000183134 |
| FOXA2 | 2.45427706 | 5.480384292 | 1.60E-05 | 0.00010095 | 5.677194963 | 31.10954727 | ENSG00000125798 |
| SOX15 | 2.453598122 | 5.477805808 | 3.46E-24 | 1.60E-22 | 31.39802209 | 171.9525489 | ENSG00000129194 |
| HLA-DQA1 | 2.450904366 | 5.467587357 | 2.74E-07 | 2.36E-06 | 8.346573013 | 45.663414 | ENSG00000196735 |
| NA | 2.449442705 | 5.462050697 | 0.032055335 | 0.082927766 | 1.328049628 | 7.29898135 | ENSG00000213514 |
| CHADL | 2.448644217 | 5.459028455 | 0.003022792 | 0.011272509 | 2.626075046 | 14.49901789 | ENSG00000100399 |
| NA | 2.442396202 | 5.435437646 | 0.106878251 | 0.217763872 | 1.007031902 | 5.441304926 | ENSG00000254739 |
| NA | 2.441360411 | 5.431536645 | 0.032565076 | 0.083974452 | 1.330840531 | 7.254885378 | ENSG00000229953 |
| SIGLEC15 | 2.440188021 | 5.427124562 | 0.000332175 | 0.001595885 | 3.986939787 | 21.72056016 | ENSG00000197046 |
| C6orf223 | 2.439542251 | 5.424695849 | 2.32E-17 | 6.30E-16 | 22.00881116 | 119.6314139 | ENSG00000181577 |
| NA | 2.43069781 | 5.391541483 | 0.031266012 | 0.081242353 | 1.632997658 | 8.922741433 | ENSG00000285712 |
| ADGRD1 | 2.428978891 | 5.385121484 | 7.34E-05 | 0.000408884 | 5.276592496 | 28.63424255 | ENSG00000111452 |
| NA | 2.427312104 | 5.378903491 | 0.036072487 | 0.091454112 | 1.360864742 | 7.248433657 | ENSG00000125514 |
| NA | 2.420579682 | 5.353860995 | 0.03761807 | 0.094712893 | 1.360864742 | 7.206488258 | ENSG00000226942 |
| MMP9 | 2.418413645 | 5.345828833 | 9.70E-44 | 1.29E-41 | 67.13335019 | 359.0082833 | ENSG00000100985 |
| NA | 2.416637883 | 5.339252889 | 0.000449361 | 0.002086313 | 4.098663921 | 21.60226019 | ENSG00000260725 |
| HHLA1 | 2.415433285 | 5.334796669 | 2.25E-57 | 4.77E-55 | 88.49330479 | 471.337735 | ENSG00000132297 |
| NA | 2.408354548 | 5.308685034 | 0.025861361 | 0.069619246 | 1.709115775 | 8.981891421 | ENSG00000256424 |
| FAT4 | 2.40733059 | 5.304918511 | 2.77E-33 | 2.12E-31 | 48.81278831 | 258.9880381 | ENSG00000196159 |
| NA | 2.407160392 | 5.304292713 | 2.65E-09 | 3.05E-08 | 16.07642886 | 84.44781525 | ENSG00000287492 |
| TDRD5 | 2.406803219 | 5.302979675 | 7.74E-08 | 7.22E-07 | 10.47515185 | 55.30431554 | ENSG00000162782 |
| LOC101928517 | 2.4013525 | 5.282982021 | 1.39E-06 | 1.07E-05 | 7.759004176 | 40.78406449 | ENSG00000269072 |
| NA | 2.395730066 | 5.262433384 | 0.077743188 | 0.169424293 | 0.974216789 | 5.20388094 | ENSG00000225387 |
| VSIR | 2.388513001 | 5.236173862 | 0.000890573 | 0.003835688 | 3.663131159 | 19.18101081 | ENSG00000107738 |
| NA | 2.388296914 | 5.235389645 | 0.036773477 | 0.09290167 | 1.619043143 | 8.63261918 | ENSG00000224786 |
| FMO1 | 2.3857394 | 5.226116915 | 0.107056955 | 0.218081914 | 1.03426521 | 5.33455132 | ENSG00000010932 |
| HR | 2.381113158 | 5.209385343 | 0.025846977 | 0.069590235 | 1.956806287 | 10.34672178 | ENSG00000168453 |
| PADI1 | 2.378028378 | 5.198258499 | 8.54E-07 | 6.80E-06 | 8.02834619 | 41.71089763 | ENSG00000142623 |
| VSIG1 | 2.377742716 | 5.197229314 | 7.62E-19 | 2.32E-17 | 25.6363363 | 133.583803 | ENSG00000101842 |
| NA | 2.377472333 | 5.196255367 | 0.073533262 | 0.162303221 | 1.355282936 | 7.013160244 | ENSG00000253328 |
| LOC400499 | 2.369032655 | 5.16594633 | 9.51E-13 | 1.61E-11 | 16.41889793 | 84.6742259 | ENSG00000188897 |
| TRPM2 | 2.367066925 | 5.158912312 | 0.000139707 | 0.000728011 | 4.936714172 | 25.71018774 | ENSG00000142185 |
| NA | 2.364356472 | 5.149229143 | 0.001779859 | 0.00713386 | 3.276483206 | 17.06089701 | ENSG00000276627 |
| PCDHGB3 | 2.364034967 | 5.148081762 | 0.000225316 | 0.001124486 | 4.340772627 | 22.36587334 | ENSG00000262209 |
| IFIT3 | 2.35951169 | 5.131966277 | 0.001014098 | 0.004315677 | 3.630316045 | 18.77768323 | ENSG00000119917 |
| NA | 2.358555735 | 5.128566873 | 0.005068489 | 0.017738793 | 2.713356774 | 13.8004734 | ENSG00000286496 |
| SLC52A1 | 2.352794301 | 5.108126668 | 0.000364238 | 0.001733229 | 4.400821048 | 22.31155754 | ENSG00000132517 |
| LPAR5 | 2.35086389 | 5.101296265 | 0.004024913 | 0.014537466 | 2.979907885 | 15.31963046 | ENSG00000184574 |
| PCDHGA11 | 2.348611914 | 5.093339606 | 1.34E-24 | 6.39E-23 | 36.6424752 | 187.0054408 | ENSG00000253873 |
| SEMA3C | 2.346941466 | 5.087445613 | 0.000159104 | 0.000818692 | 5.018414094 | 25.57170857 | ENSG00000075223 |
| NA | 2.344367123 | 5.078375677 | 1.65E-07 | 1.47E-06 | 10.15692503 | 51.29049861 | ENSG00000285689 |
| MFAP5 | 2.336611288 | 5.051147908 | 0.107874101 | 0.219237177 | 0.979798595 | 5.029375051 | ENSG00000197614 |
| SNORD99 | 2.333070656 | 5.038766689 | 0.051778279 | 0.12272083 | 1.385307146 | 6.829791597 | ENSG00000221539 |
| PAPPA2 | 2.324513596 | 5.008968697 | 1.38E-08 | 1.44E-07 | 12.7904969 | 64.69787638 | ENSG00000116183 |
| LINC01508 | 2.32440117 | 5.008578374 | 0.067156956 | 0.150958383 | 1.385307146 | 6.796448493 | ENSG00000231107 |
| BIRC7 | 2.322897938 | 5.003362349 | 0.035690935 | 0.090593926 | 1.95959719 | 9.924311612 | ENSG00000101197 |
| LOC101929759 | 2.318662353 | 4.9886946 | 0.06865558 | 0.153611087 | 1.385307146 | 6.764722925 | ENSG00000246528 |
| DENND2D | 2.318582795 | 4.988419505 | 7.16E-08 | 6.72E-07 | 10.00680398 | 49.98832588 | ENSG00000162777 |
| ARID5B | 2.31634748 | 4.980696422 | 6.68E-06 | 4.55E-05 | 7.465219758 | 36.8880145 | ENSG00000150347 |
| NA | 2.310250661 | 4.959692447 | 0.05416062 | 0.127183509 | 1.358073839 | 6.703682827 | ENSG00000265055 |
| NA | 2.30758714 | 4.950544251 | 0.096826375 | 0.202063781 | 1.007031902 | 4.985279079 | ENSG00000244124 |
| MLKL | 2.307481132 | 4.950180503 | 0.005073655 | 0.017753653 | 3.591919126 | 17.95922123 | ENSG00000168404 |
| RNASEL | 2.30635339 | 4.946312501 | 4.95E-09 | 5.49E-08 | 12.0809162 | 59.66015949 | ENSG00000135828 |
| BTG2 | 2.304645055 | 4.940458901 | 1.16E-34 | 9.22E-33 | 63.3019131 | 313.2811087 | ENSG00000159388 |
| KCNE3 | 2.304344979 | 4.93943141 | 2.12E-15 | 4.79E-14 | 26.75038642 | 131.1765537 | ENSG00000175538 |
| PCDH11Y | 2.303711888 | 4.937264339 | 0.000253414 | 0.001249964 | 5.012832289 | 24.77852042 | ENSG00000099715 |
| LOC343052 | 2.302547889 | 4.933282446 | 0.002938539 | 0.011005117 | 3.426604259 | 16.65945954 | ENSG00000231827 |
| NA | 2.302449482 | 4.932945956 | 0.060745819 | 0.139389509 | 1.333631434 | 6.59236761 | ENSG00000270084 |
| SIM2 | 2.302202646 | 4.932102034 | 0.057998824 | 0.134364756 | 1.303607223 | 6.510094349 | ENSG00000159263 |
| FLJ22447 | 2.300625643 | 4.926713724 | 0.020483785 | 0.057376607 | 2.011272902 | 9.902805874 | ENSG00000232774 |
| TBX1 | 2.300080091 | 4.92485105 | 0.026495584 | 0.071009491 | 1.956806287 | 9.743093538 | ENSG00000184058 |
| NA | 2.296585151 | 4.912934997 | 0.02719907 | 0.072643855 | 1.681882467 | 8.237100393 | ENSG00000273243 |
| ROPN1B | 2.292461922 | 4.898913854 | 0.058530215 | 0.135254717 | 1.325258725 | 6.512244923 | ENSG00000114547 |

| | | | | | | | |
|---|---|---|---|---|---|---|---|
| GLIS1 | 2.28495169 | 4.873477861 | 0.054577001 | 0.128021028 | 1.330840531 | 6.484820503 | ENSG00000174332 |
| NA | 2.284326576 | 4.871366657 | 0.093431501 | 0.196231583 | 1.03426521 | 4.955704085 | ENSG00000267733 |
| IL32 | 2.283034679 | 4.867006426 | 3.14E-05 | 0.000188883 | 7.939149439 | 38.032365 | ENSG00000008517 |
| NA | 2.278757463 | 4.852598385 | 2.05E-39 | 2.30E-37 | 95.63491608 | 462.7991838 | ENSG00000251095 |
| GJB5 | 2.270608747 | 4.825266909 | 3.07E-06 | 2.21E-05 | 8.972538768 | 43.43896975 | ENSG00000189280 |
| FLI1 | 2.265810307 | 4.80924461 | 8.53E-06 | 5.69E-05 | 7.389101641 | 35.4263899 | ENSG00000151702 |
| AGMO | 2.265165099 | 4.807094279 | 0.027870804 | 0.074057257 | 2.014063805 | 9.690655735 | ENSG00000187546 |
| NA | 2.26321722 | 4.800608279 | 0.006740522 | 0.022685173 | 2.656099256 | 12.79728569 | ENSG00000225603 |
| NA | 2.262555655 | 4.798407409 | 0.000136501 | 0.000713613 | 5.375037836 | 25.74191331 | ENSG00000249626 |
| LINC00504 | 2.261946235 | 4.796380904 | 0.010798224 | 0.033641559 | 3.157262245 | 14.8084953 | ENSG00000248360 |
| NA | 2.26148063 | 4.794833204 | 0.021713534 | 0.060148733 | 2.340663336 | 11.23214256 | ENSG00000227242 |
| TFF3 | 2.2584256 | 4.78469048 | 2.81E-08 | 2.79E-07 | 11.71871066 | 56.05906594 | ENSG00000160180 |
| NA | 2.258264751 | 4.784157054 | 0.000659318 | 0.002925903 | 4.760235692 | 22.5425298 | ENSG00000270977 |
| NLRC4 | 2.258061736 | 4.783483879 | 0.074095731 | 0.163264505 | 1.415331357 | 6.579464168 | ENSG00000091106 |
| GATA4 | 2.257106249 | 4.780316859 | 0.047962587 | 0.115569107 | 1.649067354 | 7.949128713 | ENSG00000136574 |
| KCNMB1 | 2.252521526 | 4.765149659 | 1.35E-07 | 1.22E-06 | 10.60832733 | 50.749528 | ENSG00000145936 |
| CFHR3 | 2.248227006 | 4.750986168 | 0.010891234 | 0.033893014 | 2.721729483 | 12.81018913 | ENSG00000116785 |
| TMC8 | 2.246112035 | 4.744026389 | 5.03E-16 | 1.21E-14 | 25.71245442 | 121.9994441 | ENSG00000167895 |
| TRIM4 | 2.243651556 | 4.735942469 | 1.21E-23 | 5.38E-22 | 38.01661873 | 180.2587465 | ENSG00000146833 |
| PRSS30P | 2.242944256 | 4.733621183 | 0.000397382 | 0.001875679 | 5.07288071 | 23.82777051 | ENSG00000172460 |
| NA | 2.241393473 | 4.728535649 | 0.023659608 | 0.064602619 | 2.032924404 | 9.543313802 | ENSG00000225511 |
| RAB17 | 2.240501204 | 4.725612076 | 6.39E-190 | 4.43E-186 | 412.6543296 | 1948.649129 | ENSG00000124839 |
| NA | 2.240243682 | 4.724768626 | 0.045929985 | 0.111549207 | 1.624624949 | 7.772211787 | ENSG00000286813 |
| LINC02348 | 2.240099737 | 4.724297235 | 0.003087043 | 0.011478827 | 3.363764935 | 15.85147784 | ENSG00000286733 |
| NA | 2.238441983 | 4.718871819 | 1.93E-08 | 1.97E-07 | 11.99642538 | 56.67292612 | ENSG00000246263 |
| SERPINB9 | 2.236615779 | 4.712902314 | 3.70E-104 | 3.40E-101 | 231.1099568 | 1088.956536 | ENSG00000170542 |
| ANGPTL6 | 2.2358425 | 4.710376895 | 0.027011276 | 0.072221001 | 2.06853042 | 9.598162643 | ENSG00000130812 |
| CD93 | 2.231201355 | 4.695247978 | 0.026266276 | 0.070473258 | 2.367896644 | 11.03210236 | ENSG00000125810 |
| LINC00458 | 2.224658585 | 4.674002755 | 2.87E-84 | 1.42E-81 | 209.4393091 | 977.3898425 | ENSG00000234787 |
| MAGEB17 | 2.223474101 | 4.670166873 | 5.57E-05 | 0.000317795 | 6.327603124 | 29.62505986 | ENSG00000182798 |
| LGR5 | 2.222795024 | 4.667969142 | 0.017758697 | 0.050903866 | 2.277824012 | 10.77800682 | ENSG00000139292 |
| NA | 2.221950175 | 4.665236358 | 0.041679114 | 0.103067785 | 1.709115775 | 7.873307173 | ENSG00000245146 |
| IP6K3 | 2.220739567 | 4.66132326 | 0.004124783 | 0.014845198 | 3.284855915 | 15.46671192 | ENSG00000161896 |
| NA | 2.218303044 | 4.65345754 | 0.050041578 | 0.119400382 | 1.709115775 | 7.856102584 | ENSG00000288065 |
| LOC100132215 | 2.212333156 | 4.63424127 | 0.077387407 | 0.168853295 | 1.360864742 | 6.24954709 | ENSG00000231609 |
| LINC01686 | 2.210988875 | 4.629925166 | 0.03145968 | 0.081614939 | 1.951224481 | 9.160698457 | ENSG00000261504 |
| KCNJ9 | 2.210973816 | 4.629876839 | 0.021067879 | 0.058646531 | 2.299475514 | 10.72100741 | ENSG00000162728 |
| NA | 2.204642461 | 4.609602862 | 0.051783088 | 0.12272083 | 1.651858257 | 7.685897842 | ENSG00000279443 |
| NA | 2.20236364 | 4.602327471 | 0.015203814 | 0.044781591 | 2.389548146 | 10.87480106 | ENSG00000273812 |
| TDRD10 | 2.196822963 | 4.584686109 | 0.006718535 | 0.022619067 | 3.037165403 | 13.86392454 | ENSG00000163239 |
| NA | 2.191796243 | 4.568739682 | 0.031371738 | 0.081441555 | 2.310639126 | 10.64167822 | ENSG00000267662 |
| ACTA1 | 2.191793667 | 4.568731524 | 3.97E-100 | 3.19E-97 | 201.4160322 | 920.4551158 | ENSG00000143632 |
| NA | 2.188124749 | 4.557127544 | 0.017459555 | 0.050203051 | 2.392339049 | 10.7505824 | ENSG00000244349 |
| TLE6 | 2.181237782 | 4.53542511 | 3.90E-10 | 5.00E-09 | 17.40986013 | 78.82262075 | ENSG00000104953 |
| HAND2-AS1 | 2.180198901 | 4.532160336 | 0.03699097 | 0.093426658 | 1.954015384 | 9.002864121 | ENSG00000237125 |
| GPER1 | 2.17516706 | 4.516380572 | 0.053559483 | 0.126017318 | 1.660230966 | 7.517583211 | ENSG00000164850 |
| NA | 2.175051285 | 4.516018152 | 0.044501504 | 0.108695685 | 1.68467337 | 7.578883773 | ENSG00000266217 |
| NA | 2.174028735 | 4.512818433 | 0.079608435 | 0.172805933 | 1.300816321 | 5.971795241 | ENSG00000234311 |
| NA | 2.172443702 | 4.507863097 | 3.43E-05 | 0.000204679 | 8.54067279 | 38.08721384 | ENSG00000258417 |
| PPFIA4 | 2.171477652 | 4.504845577 | 5.23E-58 | 1.15E-55 | 140.8864852 | 632.7567348 | ENSG00000143847 |
| TGFB2 | 2.165140453 | 4.485100936 | 0.00035125 | 0.00167722 | 5.88090675 | 26.70960735 | ENSG00000092969 |
| EBF1 | 2.164451756 | 4.482960402 | 0.055110872 | 0.129100648 | 1.651858257 | 7.425090119 | ENSG00000164330 |
| INMT | 2.164448939 | 4.482951649 | 0.008332232 | 0.027029615 | 3.045538112 | 13.58617269 | ENSG00000241644 |
| NA | 2.163710286 | 4.480656985 | 3.26E-58 | 7.31E-56 | 180.3751292 | 806.1544545 | ENSG00000255666 |
| TNRC6C-AS1 | 2.160246571 | 4.46991244 | 1.33E-06 | 1.02E-05 | 9.930685861 | 44.61613031 | ENSG00000204282 |
| PRF1 | 2.157451172 | 4.461259822 | 0.080840046 | 0.174830195 | 1.303607223 | 5.893823127 | ENSG00000180644 |
| PTGS2 | 2.156071346 | 4.456995012 | 0.029592167 | 0.077636679 | 1.981248691 | 8.897467587 | ENSG00000073756 |
| PLD6 | 2.1557813 | 4.456099047 | 0.000225641 | 0.001125636 | 5.970979382 | 26.74563406 | ENSG00000179598 |
| NA | 2.152791841 | 4.446874966 | 0.022273806 | 0.061427303 | 2.332290628 | 10.35721419 | ENSG00000227355 |
| NRK | 2.151601574 | 4.443207674 | 1.38E-49 | 2.25E-47 | 102.6935512 | 456.2834802 | ENSG00000123572 |
| NA | 2.150122809 | 4.438655711 | 0.025944631 | 0.06980794 | 2.272242207 | 10.25637926 | ENSG00000283696 |
| TSPYL5 | 2.146913713 | 4.428793446 | 9.85E-32 | 6.86E-30 | 59.16572814 | 261.9820263 | ENSG00000180543 |
| NA | 2.141853154 | 4.413285727 | 0.029566991 | 0.077602288 | 2.038506209 | 8.927042581 | ENSG00000225087 |
| NA | 2.140025305 | 4.407697775 | 0.009498809 | 0.030128497 | 3.072771419 | 13.36946094 | ENSG00000237227 |
| NA | 2.139617555 | 4.406452198 | 0.059266822 | 0.136564294 | 1.741930888 | 7.505212806 | ENSG00000266573 |
| HAPLN1 | 2.139376617 | 4.405716359 | 1.05E-14 | 2.21E-13 | 27.30426426 | 120.5294777 | ENSG00000145681 |
| CREB5 | 2.135680276 | 4.394442886 | 1.68E-06 | 1.27E-05 | 10.92376325 | 48.312128 | ENSG00000146592 |
| LOC105274304 | 2.134805977 | 4.391780581 | 0.086277718 | 0.184419219 | 1.388098049 | 5.99276794 | ENSG00000261335 |
| LOC349160 | 2.132393677 | 4.384443312 | 8.33E-05 | 0.000458493 | 7.287865398 | 32.07992492 | ENSG00000234352 |
| CFAP300 | 2.13037924 | 4.378325582 | 0.000199986 | 0.001007819 | 6.754763177 | 29.45862323 | ENSG00000137691 |
| BCAS1 | 2.129166669 | 4.374647185 | 0.00249067 | 0.009569312 | 3.98973069 | 17.53223734 | ENSG00000064787 |
| SLC7A14 | 2.122891904 | 4.355661694 | 3.90E-05 | 0.000229618 | 8.055579497 | 35.05587239 | ENSG00000013293 |
| CD8A | 2.122691179 | 4.355055726 | 0.005954687 | 0.020407154 | 3.336531627 | 14.57053828 | ENSG00000153563 |
| NA | 2.121933409 | 4.352768851 | 0.048523954 | 0.116572303 | 1.65464916 | 7.248433657 | ENSG00000230863 |
| LINC01460 | 2.121889247 | 4.35263561 | 0.002815434 | 0.010620361 | 4.392448339 | 19.00220377 | ENSG00000205334 |
| CFAP100 | 2.121746555 | 4.352205127 | 0.052505204 | 0.123929756 | 1.68467337 | 7.319954049 | ENSG00000163885 |
| NA | 2.118817344 | 4.343377487 | 0.036739074 | 0.092860277 | 2.038506209 | 8.752276228 | ENSG00000255710 |

| | | | | | | | |
|---|---|---|---|---|---|---|---|
| SKAP1 | 2.118330732 | 4.341912739 | 0.092878878 | 0.195292334 | 1.355282936 | 5.824453309 | ENSG00000141293 |
| NA | 2.117613254 | 4.339753965 | 0.085125218 | 0.182416068 | 1.358073839 | 5.824453309 | ENSG00000287117 |
| LAIR1 | 2.112900472 | 4.325600632 | 0.087871888 | 0.187101169 | 1.385307146 | 5.870699855 | ENSG00000167613 |
| NA | 2.106134709 | 4.305362484 | 0.057049406 | 0.132659125 | 1.687464273 | 7.227460957 | ENSG00000232293 |
| LCN15 | 2.102856117 | 4.295589457 | 0.026491348 | 0.071008002 | 2.974326079 | 12.84406527 | ENSG00000177984 |
| PRSS42P | 2.101359102 | 4.291134443 | 0.059536828 | 0.137022781 | 1.687464273 | 7.197885963 | ENSG00000178055 |
| CSMD1 | 2.099966564 | 4.286994495 | 0.000453713 | 0.002103564 | 5.666031352 | 24.33245394 | ENSG00000183117 |
| LAMA2 | 2.099604821 | 4.285919703 | 9.78E-26 | 5.03E-24 | 71.49887752 | 305.3650506 | ENSG00000196569 |
| NA | 2.098399149 | 4.282339421 | 0.007832759 | 0.025671381 | 3.943636784 | 17.01249989 | ENSG00000251396 |
| RASSF9 | 2.09074801 | 4.259688724 | 0.000335568 | 0.001609908 | 7.155365644 | 30.19723511 | ENSG00000198774 |
| LOC105373958 | 2.085598864 | 4.2445125 | 0.010984274 | 0.034111017 | 3.009932095 | 12.75964144 | ENSG00000234949 |
| ARHGDIB | 2.085162509 | 4.243228906 | 0.03710294 | 0.0936359 | 2.011272902 | 8.514852242 | ENSG00000111348 |
| LOC101928978 | 2.083727553 | 4.239010538 | 0.094124255 | 0.19734279 | 1.300816321 | 5.6015503 | ENSG00000250546 |
| NA | 2.083650303 | 4.238783565 | 0.115233687 | 0.231150366 | 1.325258725 | 5.654248567 | ENSG00000168852 |
| ERVH48-1 | 2.082459598 | 4.235286596 | 1.25E-27 | 7.06E-26 | 534.3344308 | 2262.065906 | ENSG00000233056 |
| DPPA2 | 2.080326485 | 4.229029093 | 1.03E-39 | 1.19E-37 | 87.8121967 | 370.8104321 | ENSG00000163530 |
| COL4A4 | 2.080273953 | 4.228875105 | 7.54E-07 | 6.05E-06 | 11.66982585 | 49.36372494 | ENSG00000081052 |
| FZD10-AS1 | 2.078534815 | 4.223780359 | 0.094111625 | 0.19734279 | 1.300816321 | 5.576276454 | ENSG00000250208 |
| CR1L | 2.077466878 | 4.220654916 | 3.98E-07 | 3.33E-06 | 12.2930007 | 52.09417916 | ENSG00000197721 |
| NA | 2.075339222 | 4.214434969 | 0.10424373 | 0.213864276 | 1.358073839 | 5.700495113 | ENSG00000279035 |
| NA | 2.072797318 | 4.207016037 | 8.15E-15 | 1.73E-13 | 36.13381538 | 152.1079116 | ENSG00000223745 |
| CFP | 2.072115624 | 4.205028632 | 0.008424534 | 0.027254349 | 3.698737175 | 15.5231783 | ENSG00000126759 |
| RASAL3 | 2.071026427 | 4.201855145 | 1.30E-05 | 8.37E-05 | 9.122659821 | 38.10603596 | ENSG00000105122 |
| NA | 2.0709846 | 4.201733324 | 0.005305483 | 0.018464503 | 3.978567079 | 16.79389803 | ENSG00000257951 |
| NA | 2.067692428 | 4.192156069 | 0.073031668 | 0.161454722 | 1.627415852 | 6.922284688 | ENSG00000276968 |
| FAM238C | 2.067542737 | 4.191721123 | 0.100003254 | 0.207169541 | 1.333631434 | 5.6015503 | ENSG00000283709 |
| PCDHGA7 | 2.064316057 | 4.182356545 | 0.009313145 | 0.029627291 | 3.421022453 | 14.13522467 | ENSG00000253537 |
| ZAP70 | 2.062387482 | 4.176769364 | 0.099998684 | 0.207169541 | 1.333631434 | 5.576276454 | ENSG00000115085 |
| MS4A15 | 2.062359793 | 4.176689201 | 0.002114098 | 0.008287558 | 5.135720033 | 21.23442628 | ENSG00000166961 |
| NA | 2.0619722 | 4.175567248 | 0.10272861 | 0.21145301 | 1.328049628 | 5.559604902 | ENSG00000259726 |
| NA | 2.061917316 | 4.175408402 | 0.070579062 | 0.157131067 | 1.68467337 | 7.03198237 | ENSG00000235072 |
| NA | 2.060692098 | 4.171863914 | 0.027026321 | 0.072241202 | 2.365105741 | 9.795791805 | ENSG00000253252 |
| NA | 2.060501756 | 4.171313535 | 0.027166037 | 0.072574438 | 2.392339049 | 9.860860476 | ENSG00000198857 |
| PADI3 | 2.060368897 | 4.170929413 | 3.45E-13 | 6.20E-12 | 29.6127882 | 123.9167915 | ENSG00000142619 |
| GOLGA6C | 2.057742284 | 4.163342606 | 0.04809003 | 0.115774962 | 1.984039594 | 8.331744058 | ENSG00000167195 |
| ADGRG3 | 2.052631405 | 4.148619677 | 0.020610798 | 0.057657034 | 3.130028937 | 12.72791587 | ENSG00000182885 |
| NA | 2.052438677 | 4.148065506 | 0.059123748 | 0.13629974 | 1.65464916 | 6.903462563 | ENSG00000268401 |
| ABCC3 | 2.051235106 | 4.144606416 | 0.006584358 | 0.022225522 | 3.660340256 | 15.23305605 | ENSG00000108846 |
| LCP1 | 2.049931734 | 4.140863753 | 1.15E-14 | 2.42E-13 | 30.27926606 | 125.6435492 | ENSG00000136167 |
| NA | 2.046761531 | 4.131774538 | 0.00847988 | 0.027401235 | 4.242327286 | 17.71534552 | ENSG00000180066 |
| COL6A3 | 2.043003258 | 4.12102512 | 2.46E-42 | 3.08E-40 | 106.4860278 | 437.8849167 | ENSG00000163359 |
| NA | 2.042984005 | 4.120970124 | 4.66E-13 | 8.21E-12 | 29.55553068 | 122.1795961 | ENSG00000262152 |
| TMEM232 | 2.042609546 | 4.119900645 | 0.000113999 | 0.000608845 | 7.326262317 | 30.24993338 | ENSG00000186952 |
| EBI3 | 2.039200334 | 4.110176461 | 0.000166757 | 0.000855334 | 7.620046736 | 31.52253111 | ENSG00000105246 |
| PCDH17 | 2.039034674 | 4.109704529 | 2.27E-08 | 2.29E-07 | 17.97318673 | 73.36652227 | ENSG00000118946 |
| CHRNE | 2.035681111 | 4.100162565 | 0.009846157 | 0.031026169 | 3.336531627 | 13.66388434 | ENSG00000108556 |
| SCARF1 | 2.034385777 | 4.096482856 | 1.44E-36 | 1.30E-34 | 88.02842354 | 360.2712063 | ENSG00000074660 |
| NA | 2.033866025 | 4.095007303 | 0.080160842 | 0.173692328 | 1.687464273 | 6.871736995 | ENSG00000284128 |
| SERPINA11 | 2.033485311 | 4.093926812 | 0.042677909 | 0.105106541 | 2.005691096 | 8.209675972 | ENSG00000186910 |
| SGCA | 2.032400201 | 4.090848759 | 0.073715717 | 0.162586414 | 1.956806287 | 8.073086909 | ENSG00000108823 |
| TNN | 2.032396307 | 4.090837718 | 4.94E-10 | 6.26E-09 | 20.45539825 | 83.56375155 | ENSG00000120332 |
| LINC00622 | 2.031400107 | 4.088013915 | 0.020754891 | 0.057919738 | 2.628865949 | 10.84952721 | ENSG00000260941 |
| MYOM3 | 2.031051696 | 4.087026777 | 0.030637525 | 0.079933526 | 2.329499725 | 9.54761495 | ENSG00000142661 |
| NA | 2.029771104 | 4.083400584 | 0.019157275 | 0.054211969 | 2.92544127 | 12.10949408 | ENSG00000272195 |
| CALHM3 | 2.028840851 | 4.080768448 | 0.05214937 | 0.123285967 | 2.005691096 | 8.176332869 | ENSG00000183128 |
| LINC00858 | 2.028465891 | 4.079707984 | 0.00020427 | 0.001028067 | 7.356286528 | 30.03348209 | ENSG00000229404 |
| AKAP14 | 2.028080006 | 4.078616909 | 0.152726128 | 0.286710756 | 1.385307146 | 5.561222438 | ENSG00000186471 |
| HSD52 | 2.025316585 | 4.070811969 | 0.040680169 | 0.101011861 | 2.335081531 | 9.512121272 | ENSG00000224609 |
| CFAP44 | 2.024696489 | 4.069062639 | 0.004950136 | 0.017394003 | 4.92555056 | 20.23769029 | ENSG00000206530 |
| COL13A1 | 2.024048425 | 4.067235211 | 1.50E-47 | 2.31E-45 | 120.8365586 | 492.2968631 | ENSG00000197467 |
| NA | 2.023750159 | 4.066394429 | 0.04359926 | 0.106979562 | 2.038506209 | 8.216127694 | ENSG00000286570 |
| DNAH2 | 2.023637823 | 4.06607781 | 0.000314963 | 0.001520608 | 7.726189063 | 31.29533907 | ENSG00000183914 |
| C20orf144 | 2.022991442 | 4.064256462 | 0.032274567 | 0.083370203 | 2.332290628 | 9.486314388 | ENSG00000149609 |
| NA | 2.022895462 | 4.063986085 | 0.072831006 | 0.161066484 | 1.709115775 | 6.855065443 | ENSG00000253966 |
| LINC02714 | 2.021110873 | 4.058962112 | 0.020189793 | 0.056660067 | 2.661681062 | 10.82855451 | ENSG00000251226 |
| NHLRC1 | 2.017830941 | 4.049744641 | 4.57E-06 | 3.20E-05 | 10.66000304 | 43.21527324 | ENSG00000187566 |
| PTP4A1 | 2.017416648 | 4.048581858 | 0.050294467 | 0.119814577 | 2.038506209 | 8.174182295 | ENSG00000112245 |
| HTR2C | 2.015909087 | 4.044353455 | 1.32E-14 | 2.72E-13 | 33.80643084 | 136.6178281 | ENSG00000147246 |
| PKN2-AS1 | 2.014342363 | 4.039963793 | 0.117105841 | 0.234175265 | 1.300816321 | 5.351222872 | ENSG00000237505 |
| APOBEC3H | 2.014267107 | 4.039753058 | 0.025481333 | 0.068711267 | 2.683332564 | 10.81403353 | ENSG00000100298 |
| NA | 2.004641748 | 4.012890384 | 1.10E-05 | 7.17E-05 | 10.67395756 | 42.92326088 | ENSG00000225205 |
| LOC100130691 | 2.000277348 | 4.000769046 | 0.000268769 | 0.001317283 | 7.66402546 | 30.74816509 | ENSG00000213963 |
| TAFA2 | 1.999934391 | 3.999818097 | 0.00031426 | 0.001517875 | 6.645829947 | 26.6843335 | ENSG00000198673 |
| NA | 1.998035022 | 3.994555633 | 0.000237674 | 0.001181389 | 7.473592467 | 29.614307 | ENSG00000203362 |
| GABRP | 1.997592233 | 3.993329819 | 4.64E-16 | 1.12E-14 | 49.55027804 | 198.4642119 | ENSG00000094755 |
| PDE4B | 1.997299131 | 3.992518607 | 9.42E-59 | 2.14E-56 | 153.9363998 | 614.357844 | ENSG00000184588 |
| PDCD1 | 1.994301426 | 3.984231362 | 7.15E-10 | 8.84E-09 | 21.67383892 | 86.38968558 | ENSG00000188389 |

| | | | | | | | |
|---|---|---|---|---|---|---|---|
| OTOGL | 1.994269749 | 3.98414388 | 0.002898307 | 0.01087967 | 4.751862984 | 18.7922042 | ENSG00000165899 |
| NA | 1.99276313 | 3.979985378 | 1.20E-09 | 1.45E-08 | 20.50428306 | 81.34174886 | ENSG00000256969 |
| HLA-DQB1-AS1 | 1.989720008 | 3.971599116 | 0.108482419 | 0.220187521 | 1.328049628 | 5.304976326 | ENSG00000223534 |
| TOGARAM2 | 1.987961872 | 3.966762088 | 6.53E-05 | 0.000367714 | 8.387760835 | 33.2110994 | ENSG00000189350 |
| CXCL1 | 1.987379005 | 3.965159788 | 0.008071274 | 0.026320394 | 3.968079188 | 15.83480629 | ENSG00000163739 |
| NA | 1.987182734 | 3.964620385 | 5.08E-06 | 3.52E-05 | 11.57975322 | 46.10760248 | ENSG00000251271 |
| TXNIP | 1.983673506 | 3.954988517 | 3.38E-27 | 1.85E-25 | 113.2875606 | 447.4201002 | ENSG00000265972 |
| MYRFL | 1.983323004 | 3.954027773 | 0.055997287 | 0.130685175 | 2.008481999 | 7.949128713 | ENSG00000166268 |
| SLC25A45 | 1.982812779 | 3.952629633 | 0.00543539 | 0.018856732 | 4.291887817 | 17.07164988 | ENSG00000162241 |
| LPO | 1.98179893 | 3.949852913 | 0.002322665 | 0.008984667 | 5.345013626 | 21.14140015 | ENSG00000167419 |
| NA | 1.980531064 | 3.946383237 | 0.028374508 | 0.075126631 | 2.653308353 | 10.51315841 | ENSG00000253649 |
| NA | 1.978441601 | 3.940671805 | 0.01917766 | 0.054261696 | 3.018304804 | 11.88659107 | ENSG00000258623 |
| NA | 1.977363823 | 3.937728992 | 1.36E-06 | 1.05E-05 | 13.27000839 | 52.51282122 | ENSG00000257474 |
| NA | 1.977320256 | 3.93761008 | 0.132753693 | 0.256865746 | 1.303607223 | 5.185058814 | ENSG00000250762 |
| SPTY2D1OS | 1.977231367 | 3.937367479 | 0.000777804 | 0.003386281 | 6.812020695 | 26.56441599 | ENSG00000247595 |
| NA | 1.975358445 | 3.932259262 | 0.117317427 | 0.234538813 | 1.355282936 | 5.300675179 | ENSG00000229334 |
| NA | 1.974211721 | 3.929134953 | 0.125425732 | 0.246218357 | 1.355282936 | 5.296374031 | ENSG00000279133 |
| CEACAM1 | 1.973881455 | 3.928235587 | 2.79E-06 | 2.02E-05 | 12.48921566 | 48.77487814 | ENSG00000079385 |
| SLC39A4 | 1.97311471 | 3.926148415 | 5.93E-12 | 9.31E-11 | 26.55417146 | 103.8977151 | ENSG00000147804 |
| LINC01356 | 1.970754642 | 3.91973098 | 1.50E-64 | 3.80E-62 | 165.6753225 | 648.8974987 | ENSG00000215866 |
| NA | 1.969829257 | 3.917217561 | 0.108737008 | 0.22050306 | 1.358073839 | 5.279702479 | ENSG00000241057 |
| LOC101928335 | 1.969053688 | 3.915112294 | 0.000147522 | 0.000763782 | 8.85610871 | 34.39980634 | ENSG00000236064 |
| FALEC | 1.968174978 | 3.912728422 | 0.020683853 | 0.057777616 | 3.306507417 | 13.05002416 | ENSG00000228126 |
| NA | 1.967558803 | 3.911057652 | 0.117036045 | 0.234059957 | 1.360864742 | 5.277551906 | ENSG00000262434 |
| NA | 1.967200557 | 3.910086592 | 0.000195305 | 0.000987066 | 7.655652752 | 30.03079848 | ENSG00000175772 |
| KCNJ2 | 1.966055314 | 3.906983908 | 3.53E-05 | 0.000210064 | 9.721392269 | 37.8742702 | ENSG00000123700 |
| PANCR | 1.964720116 | 3.903369713 | 0.096497266 | 0.20144235 | 1.739139985 | 6.680559554 | ENSG00000250103 |
| WNT3 | 1.964395036 | 3.902490273 | 3.43E-35 | 2.84E-33 | 88.05988721 | 343.2635406 | ENSG00000108379 |
| HSPA2 | 1.961860178 | 3.895639502 | 1.56E-140 | 3.34E-137 | 659.0975074 | 2566.760206 | ENSG00000126803 |
| LINC01152 | 1.961392567 | 3.894377038 | 0.067542272 | 0.151664005 | 2.06853042 | 7.946978139 | ENSG00000256124 |
| ATP8B3 | 1.960292232 | 3.891407952 | 6.77E-20 | 2.26E-18 | 54.53483788 | 211.5739617 | ENSG00000130270 |
| GDF3 | 1.959273262 | 3.888660435 | 3.46E-39 | 3.77E-37 | 115.430933 | 449.4332489 | ENSG00000184344 |
| C4orf51 | 1.957662574 | 3.884321389 | 0.01564866 | 0.045857253 | 3.948542868 | 15.44735676 | ENSG00000237136 |
| NA | 1.957436615 | 3.883713062 | 0.04570798 | 0.111037985 | 2.386757243 | 9.170918288 | ENSG00000273002 |
| ADGRE5 | 1.954700748 | 3.876355128 | 1.42E-14 | 2.91E-13 | 33.91324889 | 131.1663217 | ENSG00000123146 |
| PLA2G2A | 1.95215738 | 3.869527413 | 1.38E-13 | 2.56E-12 | 34.14419398 | 132.0933969 | ENSG00000188257 |
| NA | 1.950455302 | 3.864964872 | 0.000426554 | 0.001994341 | 7.06250211 | 27.19144008 | ENSG00000277270 |
| DCC | 1.950151667 | 3.864151523 | 0.000146928 | 0.000760909 | 8.752757286 | 33.66765829 | ENSG00000187323 |
| BGN | 1.949702689 | 3.862949157 | 2.01E-27 | 1.11E-25 | 76.65517302 | 295.9645171 | ENSG00000182492 |
| UNC5C | 1.947176283 | 3.85619039 | 7.16E-07 | 5.77E-06 | 14.47883706 | 55.65064371 | ENSG00000182168 |
| SNTG1 | 1.944799367 | 3.849842344 | 0.015568384 | 0.045649748 | 3.36934674 | 12.9467782 | ENSG00000147481 |
| NA | 1.94456899 | 3.84922763 | 0.154762193 | 0.289687314 | 1.415331357 | 5.321647878 | ENSG00000241439 |
| NA | 1.941668669 | 3.841497112 | 0.14619851 | 0.276738684 | 1.415331357 | 5.302825752 | ENSG00000270019 |
| NA | 1.938812618 | 3.833899762 | 0.195303813 | 0.345619467 | 1.415331357 | 5.31734673 | ENSG00000265010 |
| NA | 1.937417768 | 3.830194799 | 0.181071108 | 0.326300116 | 1.355282936 | 5.143113416 | ENSG00000265916 |
| NA | 1.936363477 | 3.827396796 | 0.101965655 | 0.210174105 | 1.651858257 | 6.388286727 | ENSG00000264176 |
| COL5A2 | 1.935541441 | 3.825216599 | 9.58E-141 | 2.31E-137 | 469.9839871 | 1798.740606 | ENSG00000204262 |
| BCL2L14 | 1.934870779 | 3.823438794 | 0.000599362 | 0.00268953 | 6.91517196 | 26.65475851 | ENSG00000121380 |
| NA | 1.933197257 | 3.819006187 | 0.079265144 | 0.172215777 | 2.06853042 | 7.763609492 | ENSG00000267159 |
| SATB2-AS1 | 1.931334255 | 3.814077754 | 0.023768026 | 0.064834417 | 3.007141193 | 11.46364786 | ENSG00000225953 |
| NA | 1.928567755 | 3.806770919 | 0.097396031 | 0.2029233 | 1.627415852 | 6.291492488 | ENSG00000286451 |
| CFD | 1.927637999 | 3.804318405 | 4.87E-11 | 6.91E-10 | 26.12633568 | 99.21492873 | ENSG00000197766 |
| NA | 1.927411995 | 3.803722491 | 0.000522762 | 0.002385194 | 7.495243969 | 28.23442262 | ENSG00000252051 |
| GDNF | 1.927345444 | 3.80354703 | 0.025009277 | 0.067617934 | 3.744831081 | 14.13145656 | ENSG00000168621 |
| CAV1 | 1.923699906 | 3.793948001 | 1.55E-30 | 1.01E-28 | 79.12766039 | 300.4091044 | ENSG00000105974 |
| NR2F1-AS1 | 1.92286334 | 3.791748668 | 9.87E-05 | 0.000534266 | 9.443677547 | 35.60304636 | ENSG00000237187 |
| SERPINE1 | 1.922231384 | 3.790088101 | 4.62E-18 | 1.32E-16 | 44.62492764 | 168.7778941 | ENSG00000106366 |
| ABHD1 | 1.920668354 | 3.785984104 | 0.00463161 | 0.016439445 | 4.664581255 | 17.67501766 | ENSG00000143994 |
| NA | 1.920586225 | 3.785768584 | 0.062909225 | 0.143229301 | 2.011272902 | 7.616528024 | ENSG00000237713 |
| FRMPD4 | 1.920132619 | 3.784578466 | 1.69E-08 | 1.75E-07 | 21.15748212 | 79.82849207 | ENSG00000169933 |
| NA | 1.912826651 | 3.765461386 | 0.002373776 | 0.009165836 | 5.668822255 | 21.37558907 | ENSG00000280322 |
| NA | 1.912638487 | 3.764970308 | 0.007600227 | 0.025019792 | 5.012832289 | 18.90620303 | ENSG00000260412 |
| RTL4 | 1.912075275 | 3.763500792 | 0.021725833 | 0.060174163 | 3.300925611 | 12.50501286 | ENSG00000187823 |
| LINC01270 | 1.910316063 | 3.758914403 | 0.022390371 | 0.061704664 | 3.772064389 | 13.99218388 | ENSG00000203999 |
| PLK2 | 1.909319633 | 3.756319121 | 9.22E-66 | 2.47E-63 | 206.0195259 | 773.3135758 | ENSG00000145632 |
| LINC01140 | 1.909169213 | 3.755927495 | 0.000453839 | 0.002103564 | 7.685676962 | 28.86091367 | ENSG00000267272 |
| NA | 1.908467756 | 3.754101758 | 7.34E-45 | 1.01E-42 | 132.9672355 | 499.4979294 | ENSG00000243004 |
| ENPP6 | 1.907986517 | 3.752849714 | 0.029892246 | 0.078323977 | 3.069980516 | 11.38352517 | ENSG00000164303 |
| BANK1 | 1.907544906 | 3.751701136 | 8.37E-05 | 0.000460295 | 9.326371608 | 35.02441939 | ENSG00000153064 |
| MOGAT3 | 1.905130687 | 3.745428255 | 0.021293897 | 0.059181756 | 3.344904336 | 12.53458786 | ENSG00000106384 |
| NA | 1.904601464 | 3.744054573 | 0.077393646 | 0.168853295 | 1.984039594 | 7.446062818 | ENSG00000260907 |
| HJV | 1.901702878 | 3.736539771 | 0.055778005 | 0.130362672 | 2.773405195 | 10.20206346 | ENSG00000168509 |
| ANKRD33 | 1.898447396 | 3.728117668 | 0.001456653 | 0.005972431 | 6.78757829 | 25.07483393 | ENSG00000167612 |
| NA | 1.897870362 | 3.726626834 | 0.144953202 | 0.275144369 | 1.300816321 | 4.913758686 | ENSG00000228577 |
| NA | 1.896965855 | 3.724291135 | 0.051730105 | 0.122658033 | 2.389548146 | 8.786152369 | ENSG00000228528 |
| LOC107984827 | 1.896818119 | 3.723909775 | 0.130261256 | 0.253247231 | 1.330840531 | 4.980977931 | ENSG00000261170 |
| ITGA11 | 1.894814694 | 3.718742089 | 0.028171449 | 0.07465047 | 3.018304804 | 11.21547101 | ENSG00000137809 |

| | | | | | | | |
|---|---|---|---|---|---|---|---|
| IER3 | 1.894700614 | 3.718448045 | 1.47E-76 | 6.05E-74 | 249.1957066 | 927.3018331 | ENSG00000137331 |
| GP1BA | 1.893598073 | 3.715607404 | 0.00323551 | 0.011970889 | 5.660449546 | 21.0827832 | ENSG00000185245 |
| RSPH6A | 1.892280585 | 3.712215813 | 0.089834733 | 0.190460691 | 1.681882467 | 6.215670948 | ENSG00000104941 |
| NA | 1.892237699 | 3.712105465 | 0.075417196 | 0.165512963 | 2.041297112 | 7.498761085 | ENSG00000267547 |
| CD79B | 1.892022899 | 3.711552816 | 0.00010425 | 0.000561756 | 10.48910637 | 38.69434973 | ENSG00000007312 |
| NA | 1.891068939 | 3.709099421 | 0.004940148 | 0.017365232 | 5.075671612 | 18.7056298 | ENSG00000284630 |
| NA | 1.890417967 | 3.70742618 | 0.14814114 | 0.279736759 | 1.325258725 | 4.936881959 | ENSG00000232140 |
| NA | 1.888213142 | 3.701764566 | 0.018449684 | 0.05246357 | 3.690364466 | 13.59451452 | ENSG00000230982 |
| TUSC8 | 1.887956329 | 3.701105676 | 0.021625843 | 0.059974718 | 3.804879502 | 13.87629494 | ENSG00000237361 |
| NA | 1.886501576 | 3.697375518 | 0.130838918 | 0.254001567 | 1.330840531 | 4.934731385 | ENSG00000251520 |
| LOC100507642 | 1.884745865 | 3.692878675 | 0.001438605 | 0.005905965 | 6.890729555 | 25.68706447 | ENSG00000240859 |
| NA | 1.881374316 | 3.684258567 | 0.133730374 | 0.258496258 | 1.330840531 | 4.911608113 | ENSG00000284740 |
| EID3 | 1.880905498 | 3.683061526 | 0.000355359 | 0.001694323 | 7.998321979 | 29.48416965 | ENSG00000255150 |
| NA | 1.879906991 | 3.680513317 | 0.134062285 | 0.258982129 | 1.330840531 | 4.905156391 | ENSG00000285367 |
| KCNAB2 | 1.879482874 | 3.679431495 | 1.20E-55 | 2.42E-53 | 189.9218459 | 697.4349834 | ENSG00000069424 |
| RBPJL | 1.879328965 | 3.679038987 | 0.005933124 | 0.020350659 | 4.637347948 | 17.15984183 | ENSG00000124232 |
| EZHIP | 1.878906963 | 3.677962991 | 0.141955276 | 0.270786041 | 1.355282936 | 4.955704085 | ENSG00000187690 |
| DPPA3 | 1.875426948 | 3.669101839 | 1.91E-12 | 3.15E-11 | 35.27814383 | 129.6826584 | ENSG00000187569 |
| MIR3681HG | 1.871876258 | 3.660082729 | 0.159193851 | 0.295881257 | 1.325258725 | 4.856759272 | ENSG00000224184 |
| UPK1A | 1.871347664 | 3.658741943 | 0.002813415 | 0.010615999 | 7.612549908 | 27.4390839 | ENSG00000105668 |
| TMCC1-AS1 | 1.866459284 | 3.646365764 | 9.06E-05 | 0.000494324 | 10.92934505 | 40.09008162 | ENSG00000271270 |
| LOC101929457 | 1.864907762 | 3.642446449 | 0.061906733 | 0.141464438 | 2.302266417 | 8.430428407 | ENSG00000259362 |
| NA | 1.862467744 | 3.636291216 | 0.001291217 | 0.005370678 | 6.651411752 | 24.23189159 | ENSG00000277287 |
| TLX2 | 1.853491446 | 3.61373683 | 0.104952484 | 0.214861336 | 1.956806287 | 7.134967866 | ENSG00000115297 |
| MYOM1 | 1.853163964 | 3.612916629 | 0.000371968 | 0.001766527 | 8.662684654 | 31.38379148 | ENSG00000101605 |
| NA | 1.852770335 | 3.611931006 | 0.074443158 | 0.163946146 | 2.367896644 | 8.497647652 | ENSG00000259347 |
| NA | 1.851738798 | 3.609349373 | 0.075390963 | 0.165494614 | 2.011272902 | 7.254885378 | ENSG00000261757 |
| NA | 1.850989794 | 3.607475993 | 0.012419911 | 0.037742991 | 4.256281801 | 15.5317806 | ENSG00000237118 |
| NA | 1.85040119 | 3.606004481 | 0.078090098 | 0.170084237 | 1.978457788 | 7.172612117 | ENSG00000258768 |
| NA | 1.849305248 | 3.603266224 | 0.089922854 | 0.190584682 | 2.011272902 | 7.239831362 | ENSG00000287464 |
| NA | 1.848156127 | 3.60039733 | 5.94E-06 | 4.07E-05 | 13.95323166 | 50.42417253 | ENSG00000287535 |
| LINC01630 | 1.847465768 | 3.598674879 | 0.090141437 | 0.190859232 | 1.975666886 | 7.145187696 | ENSG00000227115 |
| NA | 1.84632932 | 3.595841225 | 0.061609582 | 0.140918953 | 2.419572356 | 8.561098787 | ENSG00000256139 |
| OR2H2 | 1.845568617 | 3.593945713 | 0.017244474 | 0.049680878 | 4.223466687 | 15.36561654 | ENSG00000204657 |
| LOC105372273 | 1.844422917 | 3.591092754 | 0.010928992 | 0.033980369 | 5.032368609 | 18.03666031 | ENSG00000267082 |
| NA | 1.84420835 | 3.590558704 | 0.00762524 | 0.025073458 | 5.002344398 | 17.95922123 | ENSG00000279481 |
| SGCD | 1.842380079 | 3.586011411 | 0.115862779 | 0.232146746 | 1.627415852 | 5.9169464 | ENSG00000170624 |
| ANKDD1B | 1.840644156 | 3.581699137 | 0.112792002 | 0.227271734 | 1.651858257 | 5.963192946 | ENSG00000189045 |
| NA | 1.839531703 | 3.578938375 | 0.007425358 | 0.024540446 | 5.10290492 | 18.11678299 | ENSG00000242861 |
| RIN1 | 1.839088189 | 3.577838307 | 3.43E-13 | 6.16E-12 | 39.22756258 | 140.1912246 | ENSG00000174791 |
| ZFP57 | 1.838835678 | 3.577212143 | 0.104270772 | 0.213897007 | 9.498819883 | 33.92631544 | ENSG00000204644 |
| NA | 1.838482308 | 3.576336056 | 0.032129414 | 0.083061984 | 3.015513901 | 10.7505824 | ENSG00000259723 |
| C4orf47 | 1.836683834 | 3.571880549 | 2.59E-05 | 0.000158031 | 13.45226884 | 47.90021023 | ENSG00000205129 |
| ADGRL4 | 1.836680343 | 3.571871905 | 0.000126511 | 0.000667012 | 9.963500975 | 35.67456676 | ENSG00000162618 |
| NTN1 | 1.836427659 | 3.571246357 | 1.51E-27 | 8.46E-26 | 88.68796815 | 316.5759784 | ENSG00000065320 |
| ADAMTS4 | 1.8348418 | 3.567322878 | 7.32E-56 | 1.50E-53 | 187.9045909 | 670.4607333 | ENSG00000158859 |
| NA | 1.830625083 | 3.55691151 | 0.014484766 | 0.043001787 | 4.40361195 | 15.59523173 | ENSG00000261068 |
| NA | 1.829256981 | 3.553540105 | 0.01584818 | 0.046350402 | 4.547275316 | 16.35643385 | ENSG00000267317 |
| ESPNL | 1.82831362 | 3.551217245 | 4.72E-10 | 5.99E-09 | 25.64191811 | 91.20744353 | ENSG00000144488 |
| NA | 1.827201807 | 3.548481554 | 0.01856846 | 0.052754663 | 3.968079188 | 14.16264909 | ENSG00000287830 |
| LINC00540 | 1.826300995 | 3.546266592 | 4.82E-10 | 6.11E-09 | 30.54043552 | 108.1072711 | ENSG00000276476 |
| RORC | 1.824104365 | 3.540871196 | 0.000493214 | 0.002262684 | 8.618705929 | 30.66589183 | ENSG00000143365 |
| NA | 1.82231939 | 3.536492962 | 0.090975345 | 0.192266147 | 2.06853042 | 7.200036537 | ENSG00000234353 |
| FOXJ1 | 1.821028446 | 3.533329874 | 5.92E-36 | 5.14E-34 | 121.7486118 | 430.9959994 | ENSG00000129654 |
| DNM3 | 1.818659892 | 3.527533765 | 6.09E-15 | 1.31E-13 | 43.50529572 | 153.9193841 | ENSG00000197959 |
| DDX60L | 1.818170914 | 3.526338367 | 0.000353797 | 0.001688128 | 8.662684654 | 30.61157603 | ENSG00000181381 |
| PAX7 | 1.817454644 | 3.524588044 | 0.066522219 | 0.149845882 | 2.776196098 | 9.646559763 | ENSG00000009709 |
| NA | 1.816652702 | 3.522629397 | 0.023941567 | 0.06519997 | 3.780437098 | 13.14010621 | ENSG00000254013 |
| BHLHE41 | 1.814373594 | 3.51706889 | 3.39E-06 | 2.42E-05 | 16.97173663 | 59.22458542 | ENSG00000123095 |
| LOC105369306 | 1.81339779 | 3.514690834 | 1.12E-11 | 1.71E-10 | 31.1642861 | 109.3855271 | ENSG00000233393 |
| LOC101926913 | 1.809020209 | 3.504042345 | 0.019837608 | 0.055826011 | 3.660340256 | 12.86880608 | ENSG00000234350 |
| OBSCN-AS1 | 1.808673892 | 3.503201305 | 0.035770732 | 0.090760645 | 2.979907885 | 10.48358342 | ENSG00000162913 |
| NA | 1.807706517 | 3.500853078 | 0.010933177 | 0.033980369 | 5.241662201 | 18.54402735 | ENSG00000261326 |
| PTAFR | 1.807181296 | 3.499578805 | 2.30E-12 | 3.78E-11 | 43.43910192 | 152.0934212 | ENSG00000169403 |
| TRIM10 | 1.806778809 | 3.498602619 | 0.103385072 | 0.212518042 | 2.005691096 | 7.046503348 | ENSG00000204613 |
| PTPRVP | 1.806094525 | 3.496943593 | 0.009657035 | 0.03053497 | 4.982808078 | 17.49244252 | ENSG00000243323 |
| SNCG | 1.804395521 | 3.492827807 | 1.08E-28 | 6.43E-27 | 93.31012228 | 325.1860194 | ENSG00000173267 |
| PMFBP1 | 1.803962786 | 3.491780295 | 0.000421634 | 0.001972772 | 9.027005384 | 31.50936721 | ENSG00000118557 |
| RBM43 | 1.803580929 | 3.4908562 | 9.37E-07 | 7.40E-06 | 18.23204085 | 63.32444372 | ENSG00000184898 |
| PTPRR | 1.802710408 | 3.488750457 | 0.007045749 | 0.02350736 | 5.366665128 | 18.68088899 | ENSG00000153233 |
| OR52A1 | 1.796280061 | 3.473235082 | 0.000341288 | 0.001634349 | 9.010935688 | 31.35421648 | ENSG00000182070 |
| NA | 1.794197139 | 3.468224142 | 0.158058963 | 0.294287758 | 1.330840531 | 4.636006837 | ENSG00000249669 |
| TUBB8B | 1.793959423 | 3.467652722 | 0.020015872 | 0.05624562 | 3.720388677 | 12.82471011 | ENSG00000173213 |
| NA | 1.793185886 | 3.465793951 | 0.071487523 | 0.158657333 | 3.130028937 | 10.64948701 | ENSG00000228863 |
| CYSRT1 | 1.792466593 | 3.46406642 | 8.35E-05 | 0.000459339 | 10.71726056 | 37.07542383 | ENSG00000197191 |
| GPA33 | 1.79236502 | 3.463822542 | 0.012973015 | 0.03920765 | 4.395239242 | 15.14271353 | ENSG00000143167 |
| PENK | 1.790122568 | 3.458442733 | 0.085195714 | 0.182510886 | 2.008481999 | 6.954010256 | ENSG00000181195 |

| | | | | | | | |
|---|---|---|---|---|---|---|---|
| COLEC11 | 1.788567837 | 3.454717726 | 0.00257979 | 0.009850895 | 6.906799251 | 24.06330439 | ENSG00000118004 |
| NA | 1.78574586 | 3.447966745 | 0.09777417 | 0.203578308 | 1.981248691 | 6.865818312 | ENSG00000223476 |
| NOS3 | 1.784608786 | 3.445250267 | 3.93E-16 | 9.54E-15 | 53.40057573 | 184.0526351 | ENSG00000164867 |
| LAMP3 | 1.783811377 | 3.443346529 | 0.019612721 | 0.055289932 | 4.049779111 | 13.90156879 | ENSG00000078081 |
| S100B | 1.783733718 | 3.443161182 | 0.136704312 | 0.262770316 | 1.741930888 | 5.872850428 | ENSG00000160307 |
| C2orf91 | 1.782736765 | 3.440782659 | 0.040776737 | 0.101212562 | 3.276483206 | 11.35018207 | ENSG00000205086 |
| PDE6G | 1.782447352 | 3.440092485 | 7.69E-18 | 2.18E-16 | 53.83678421 | 185.1267917 | ENSG00000185527 |
| NA | 1.781787061 | 3.43851839 | 0.015834844 | 0.046320885 | 4.337981724 | 14.96820764 | ENSG00000283312 |
| NA | 1.7811838 | 3.437080877 | 0.00069173 | 0.003050791 | 8.003903785 | 27.52565831 | ENSG00000287238 |
| NA | 1.779957412 | 3.434160369 | 9.88E-06 | 6.51E-05 | 15.65676563 | 53.7641979 | ENSG00000277011 |
| NA | 1.77921623 | 3.432396528 | 0.052592661 | 0.124075458 | 2.661681062 | 9.145644441 | ENSG00000276412 |
| LOC101927531 | 1.779014495 | 3.431916602 | 0.053476771 | 0.125868764 | 2.656099256 | 9.131123463 | ENSG00000255595 |
| DNAH3 | 1.776752295 | 3.426539445 | 5.99E-05 | 0.000339757 | 12.41588844 | 42.48579669 | ENSG00000158486 |
| TNFSF9 | 1.775849313 | 3.424395446 | 9.93E-14 | 1.88E-12 | 40.68878769 | 139.4477907 | ENSG00000125657 |
| OAZ3 | 1.774955585 | 3.422274743 | 0.035903267 | 0.091060994 | 3.578640333 | 12.3807942 | ENSG00000143450 |
| ZNF843 | 1.771360038 | 3.41375622 | 0.015419118 | 0.045294687 | 4.307957513 | 14.76601687 | ENSG00000176723 |
| C12orf56 | 1.769727398 | 3.409895195 | 0.002195118 | 0.00855648 | 7.492453066 | 25.32972297 | ENSG00000185306 |
| TBCE | 1.769170805 | 3.408579908 | 0.008414477 | 0.027238549 | 5.426713549 | 18.35904117 | ENSG00000284770 |
| AMDHD1 | 1.767862446 | 3.405490118 | 0.004879745 | 0.017181085 | 5.698846465 | 19.37674986 | ENSG00000139344 |
| TEX37 | 1.766860704 | 3.403126323 | 0.072980886 | 0.161360948 | 2.307848223 | 7.917403145 | ENSG00000172073 |
| NA | 1.766503959 | 3.402284912 | 0.126779256 | 0.248243872 | 1.981248691 | 6.749668909 | ENSG00000229259 |
| DMTN | 1.76569032 | 3.400366661 | 7.47E-08 | 6.99E-07 | 23.78846325 | 80.71284677 | ENSG00000158856 |
| LIPA | 1.765349768 | 3.399564089 | 8.05E-90 | 4.85E-87 | 454.174761 | 1545.823216 | ENSG00000107798 |
| ATP1A1-AS1 | 1.764514962 | 3.397597524 | 0.014727306 | 0.043607745 | 4.58009043 | 15.71273821 | ENSG00000203865 |
| NA | 1.764115634 | 3.396657222 | 0.226641704 | 0.385868664 | 1.385307146 | 4.644609132 | ENSG00000286962 |
| KCNK7 | 1.7624809 | 3.392810611 | 0.013566054 | 0.040673332 | 4.343563529 | 14.72622204 | ENSG00000173338 |
| PRKAR2A-AS1 | 1.761039591 | 3.389422753 | 0.03233446 | 0.08346903 | 3.747621984 | 12.61632808 | ENSG00000224424 |
| HAPLN2 | 1.760704729 | 3.388636129 | 0.003434907 | 0.012633516 | 6.637457238 | 22.5425298 | ENSG00000132702 |
| SATB2 | 1.760066163 | 3.387136583 | 3.38E-15 | 7.41E-14 | 48.33048592 | 163.9195173 | ENSG00000119042 |
| SLC22A2 | 1.758155395 | 3.382653481 | 0.0395012 | 0.098706949 | 3.012722998 | 10.1827083 | ENSG00000112499 |
| ADCY8 | 1.757625116 | 3.381410376 | 2.14E-11 | 3.15E-10 | 34.56365705 | 116.6544004 | ENSG00000155897 |
| ZIC1 | 1.753852056 | 3.372578575 | 0.028488145 | 0.075344777 | 5.130138228 | 17.10741613 | ENSG00000152977 |
| GREM2 | 1.753837422 | 3.372544365 | 0.027357502 | 0.072944515 | 4.077012419 | 13.6090355 | ENSG00000180875 |
| MYO1F | 1.753772857 | 3.372393439 | 7.81E-22 | 3.04E-20 | 88.58481689 | 298.1220452 | ENSG00000142347 |
| NA | 1.753319095 | 3.371332907 | 0.148245744 | 0.279866931 | 1.68467337 | 5.67737184 | ENSG00000262147 |
| PLAT | 1.752983149 | 3.37054795 | 1.96E-34 | 1.56E-32 | 130.5815543 | 439.1449078 | ENSG00000104368 |
| RAC2 | 1.752949536 | 3.370469421 | 9.29E-11 | 1.29E-09 | 44.23508847 | 149.7993466 | ENSG00000128340 |
| OR2A1 | 1.751603881 | 3.367327125 | 0.154731403 | 0.289657788 | 1.621834046 | 5.527879334 | ENSG00000221970 |
| NA | 1.750834312 | 3.365531387 | 0.168499007 | 0.308918706 | 1.630206755 | 5.540249739 | ENSG00000229325 |
| NA | 1.749915144 | 3.363387827 | 0.043301192 | 0.106342877 | 3.015513901 | 10.11333848 | ENSG00000267505 |
| NA | 1.748074353 | 3.359099088 | 0.078505761 | 0.17085454 | 2.362314838 | 7.900731594 | ENSG00000253854 |
| SLCO6A1 | 1.746449782 | 3.355318646 | 0.072721695 | 0.160885601 | 2.362314838 | 7.887828151 | ENSG00000205359 |
| MANEA-DT | 1.746071637 | 3.354439297 | 0.161592325 | 0.299019429 | 1.630206755 | 5.519277039 | ENSG00000261366 |
| LINC01814 | 1.745687337 | 3.353545873 | 0.146943203 | 0.277800993 | 1.679091564 | 5.626824147 | ENSG00000236008 |
| TNS4 | 1.744898987 | 3.351713853 | 0.000124854 | 0.000659156 | 12.57928829 | 41.86307375 | ENSG00000131746 |
| GJA5 | 1.744398849 | 3.35055212 | 1.88E-09 | 2.22E-08 | 32.86570488 | 110.08621 | ENSG00000265107 |
| ACOXL-AS1 | 1.743979549 | 3.349578468 | 0.015532723 | 0.045563174 | 5.108486726 | 16.93936197 | ENSG00000204581 |
| PARP15 | 1.743053203 | 3.347428414 | 0.086616928 | 0.184924588 | 2.335081531 | 7.807705464 | ENSG00000173200 |
| SYTL2 | 1.740359107 | 3.341183242 | 4.35E-15 | 9.44E-14 | 49.95040495 | 166.6809036 | ENSG00000137501 |
| NA | 1.739766979 | 3.339812195 | 0.004984095 | 0.01749421 | 5.98983998 | 20.05217107 | ENSG00000232057 |
| POLN | 1.739612552 | 3.339454718 | 0.030961849 | 0.080587021 | 3.747621984 | 12.37649306 | ENSG00000130997 |
| CD19 | 1.737484842 | 3.334533266 | 0.004624987 | 0.016421982 | 6.022655094 | 20.08174606 | ENSG00000177455 |
| CDKN2A | 1.736183185 | 3.331526075 | 0.079846017 | 0.173146303 | 2.365105741 | 7.82275948 | ENSG00000147889 |
| NA | 1.735038323 | 3.328883365 | 0.003901563 | 0.014131628 | 6.648620849 | 22.24191514 | ENSG00000265975 |
| MT1G | 1.733137548 | 3.324500393 | 5.24E-90 | 3.26E-87 | 455.2158842 | 1514.065127 | ENSG00000125144 |
| EMP1 | 1.732541347 | 3.32312681 | 0.008417479 | 0.027240663 | 5.312198513 | 17.70459266 | ENSG00000134531 |
| DBH | 1.730774643 | 3.319059847 | 0.000319817 | 0.001540088 | 10.10245842 | 33.37538546 | ENSG00000123454 |
| CD38 | 1.724774297 | 3.305284135 | 0.151716552 | 0.285120702 | 1.95959719 | 6.535368196 | ENSG00000004468 |
| APOBEC3D | 1.724074214 | 3.303680602 | 0.007988316 | 0.026079302 | 5.619937445 | 18.65131399 | ENSG00000243811 |
| XKR9 | 1.723948486 | 3.303392704 | 0.145969078 | 0.276419896 | 1.986830497 | 6.594518184 | ENSG00000221947 |
| KCNG2 | 1.720496236 | 3.295497409 | 0.110283579 | 0.223029723 | 2.283405818 | 7.620829171 | ENSG00000178342 |
| NXPH1 | 1.719710901 | 3.293703983 | 0.069370921 | 0.154924108 | 2.71893858 | 8.887247756 | ENSG00000122584 |
| CLC | 1.718199099 | 3.290254314 | 0.000933408 | 0.004003196 | 8.689917962 | 28.5965983 | ENSG00000105205 |
| NDUFV2-AS1 | 1.716964835 | 3.287440619 | 0.014046508 | 0.041946004 | 4.667372158 | 15.34034269 | ENSG00000266053 |
| NA | 1.716351522 | 3.286043372 | 0.139766864 | 0.267564015 | 1.711906678 | 5.548852033 | ENSG00000260278 |
| RGMB-AS1 | 1.715591059 | 3.284311712 | 0.026137338 | 0.07020543 | 4.041406403 | 13.24765332 | ENSG00000246763 |
| PPP1R3B | 1.715559222 | 3.284239235 | 8.28E-77 | 3.47E-74 | 405.2136403 | 1330.689519 | ENSG00000173281 |
| S100A4 | 1.714713898 | 3.282315451 | 6.26E-26 | 3.23E-24 | 94.42867816 | 309.8233349 | ENSG00000196154 |
| DLK1 | 1.712532728 | 3.277356762 | 1.44E-10 | 1.96E-09 | 34.27669374 | 112.5427047 | ENSG00000185559 |
| LOC101927483 | 1.709412662 | 3.270276595 | 0.011641721 | 0.035782032 | 5.767267595 | 18.73735536 | ENSG00000286595 |
| FMO4 | 1.708324694 | 3.267811339 | 4.31E-05 | 0.00025074 | 14.25259789 | 46.79135341 | ENSG00000076258 |
| NA | 1.707575767 | 3.266115403 | 0.109520865 | 0.221905998 | 2.011272902 | 6.556340895 | ENSG00000259868 |
| NA | 1.707284312 | 3.265455645 | 0.000532164 | 0.002424079 | 9.982361574 | 32.61607346 | ENSG00000227012 |
| NA | 1.707112431 | 3.265066625 | 0.021404869 | 0.059447355 | 4.628975239 | 15.20993277 | ENSG00000253553 |
| FMN1 | 1.70645732 | 3.263584334 | 4.97E-11 | 7.04E-10 | 37.05655662 | 120.4665596 | ENSG00000248905 |
| GTSF1 | 1.706431792 | 3.263526585 | 0.013917917 | 0.041619988 | 5.345013626 | 17.45694884 | ENSG00000170627 |
| PI15 | 1.706046121 | 3.262654274 | 0.078863667 | 0.171478697 | 2.740590082 | 8.832398915 | ENSG00000137558 |

| | | | | | | | |
|---|---|---|---|---|---|---|---|
| NA | 1.705696543 | 3.261863799 | 0.06970217 | 0.155519844 | 2.743380985 | 8.828097768 | ENSG00000235351 |
| NLRP10 | 1.705579059 | 3.261598185 | 0.122525601 | 0.242250832 | 1.984039594 | 6.486971076 | ENSG00000182261 |
| BTBD19 | 1.704857033 | 3.259966261 | 0.009973603 | 0.031356059 | 5.369456031 | 17.48599079 | ENSG00000222009 |
| DEPP1 | 1.703176032 | 3.256170021 | 0.053766282 | 0.126422789 | 3.611455446 | 11.85539854 | ENSG00000165507 |
| MARCHF11 | 1.701767111 | 3.252991632 | 0.007484365 | 0.024718521 | 6.019864191 | 19.53888534 | ENSG00000183654 |
| LOXHD1 | 1.701021728 | 3.251311374 | 0.000132527 | 0.000694155 | 12.14096462 | 39.26705802 | ENSG00000167210 |
| CLEC6A | 1.700577551 | 3.250310515 | 0.116974482 | 0.233985352 | 2.065739517 | 6.632162435 | ENSG00000205846 |
| NA | 1.696392255 | 3.240894943 | 0.116706301 | 0.233594236 | 2.065739517 | 6.606888588 | ENSG00000278000 |
| LYST | 1.69564422 | 3.23921498 | 0.039260648 | 0.098284086 | 3.366555838 | 10.87265049 | ENSG00000143669 |
| NA | 1.695470355 | 3.238824634 | 2.56E-19 | 8.08E-18 | 79.17096339 | 256.3132872 | ENSG00000234261 |
| PKDREJ | 1.694034732 | 3.235603288 | 0.073806508 | 0.162737218 | 2.977116982 | 9.713779008 | ENSG00000130943 |
| PRTN3 | 1.692419081 | 3.231981817 | 0.033016123 | 0.084899232 | 3.693155369 | 11.93498819 | ENSG00000196415 |
| NKAIN2 | 1.691632712 | 3.23022064 | 0.034631205 | 0.088345752 | 3.687573563 | 11.91401549 | ENSG00000188580 |
| TEX19 | 1.690021377 | 3.226614847 | 0.018255596 | 0.052042071 | 5.227707687 | 17.02917145 | ENSG00000182459 |
| MECOM | 1.688559867 | 3.223347806 | 0.003755088 | 0.013678137 | 7.375147127 | 23.76242927 | ENSG00000085276 |
| NA | 1.685493547 | 3.216504144 | 0.028796667 | 0.075983681 | 4.016963998 | 12.94032648 | ENSG00000259349 |
| LOC574538 | 1.685369717 | 3.216228074 | 0.165662357 | 0.305087593 | 8.959460135 | 28.71840593 | ENSG00000249695 |
| NA | 1.685221864 | 3.215898479 | 0.132073895 | 0.255806978 | 2.062948614 | 6.541819917 | ENSG00000229835 |
| RASGRP4 | 1.684464624 | 3.214210967 | 1.72E-13 | 3.17E-12 | 52.57907074 | 169.2019155 | ENSG00000171777 |
| LOC105373159 | 1.684376247 | 3.214014075 | 3.61E-17 | 9.66E-16 | 63.06905299 | 202.3522353 | ENSG00000177788 |
| AHSG | 1.683650539 | 3.212397761 | 0.193962154 | 0.343788016 | 1.769164196 | 5.544550886 | ENSG00000145192 |
| HLA-DQB1 | 1.681758915 | 3.208188508 | 1.28E-50 | 2.20E-48 | 223.2996772 | 715.4947243 | ENSG00000179344 |
| IL6R | 1.680494407 | 3.205377795 | 1.37E-26 | 7.36E-25 | 104.4805 | 335.55478 | ENSG00000160712 |
| LINC02572 | 1.67913582 | 3.202360708 | 0.0913222 | 0.192914648 | 2.365105741 | 7.526185506 | ENSG00000229536 |
| GRIN2D | 1.676831546 | 3.197249978 | 1.54E-111 | 1.65E-108 | 568.8535598 | 1816.977319 | ENSG00000105464 |
| NA | 1.676470759 | 3.196450514 | 0.048218434 | 0.115968406 | 3.712015968 | 11.81076953 | ENSG00000249863 |
| LOC107986412 | 1.674873409 | 3.192913368 | 0.184949444 | 0.331246841 | 1.649067354 | 5.3071269 | ENSG00000250155 |
| CHRNA9 | 1.673972707 | 3.190920592 | 6.93E-07 | 5.60E-06 | 21.32000608 | 68.08602629 | ENSG00000174343 |
| LOC105378909 | 1.673499964 | 3.189875163 | 0.108809785 | 0.220782676 | 2.386757243 | 7.540706484 | ENSG00000231246 |
| ITGAX | 1.673449507 | 3.189763601 | 0.165996072 | 0.305643812 | 1.624624949 | 5.254428633 | ENSG00000140678 |
| NA | 1.672555828 | 3.18778831 | 1.37E-07 | 1.23E-06 | 27.05724947 | 86.16597064 | ENSG00000287064 |
| LGALS1 | 1.670876218 | 3.184079192 | 1.01E-33 | 7.78E-32 | 149.5161546 | 475.6467751 | ENSG00000100097 |
| LINC00334 | 1.670747154 | 3.183794355 | 0.157411499 | 0.293335389 | 1.651858257 | 5.300675179 | ENSG00000182586 |
| NA | 1.670295548 | 3.18279789 | 4.55E-06 | 3.19E-05 | 18.41410113 | 58.58706892 | ENSG00000242759 |
| SLC44A5 | 1.667062624 | 3.175673569 | 6.44E-10 | 8.00E-09 | 33.04708945 | 105.0775472 | ENSG00000137968 |
| NA | 1.663489887 | 3.167818958 | 0.006749686 | 0.022704131 | 6.349254625 | 20.13821244 | ENSG00000225465 |
| RGS5 | 1.663210844 | 3.167206304 | 0.105961013 | 0.216374924 | 2.392339049 | 7.482089533 | ENSG00000232995 |
| LUCAT1 | 1.663075364 | 3.166908892 | 0.157048163 | 0.292801402 | 1.681882467 | 5.325949025 | ENSG00000248323 |
| MYO15B | 1.663037665 | 3.16682614 | 2.70E-27 | 1.48E-25 | 126.2146996 | 399.3605418 | ENSG00000266714 |
| CABYR | 1.663007505 | 3.166759936 | 4.35E-19 | 1.35E-17 | 82.27164384 | 260.685457 | ENSG00000154040 |
| LOC285095 | 1.662991365 | 3.166724509 | 0.019498006 | 0.055030875 | 4.642929753 | 14.75364646 | ENSG00000216921 |
| TBC1D10C | 1.662270122 | 3.165141773 | 0.001025018 | 0.004356381 | 9.010935688 | 28.53099659 | ENSG00000175463 |
| NA | 1.662174886 | 3.164932841 | 0.164052024 | 0.302738833 | 1.651858257 | 5.25872978 | ENSG00000196796 |
| LINC01063 | 1.662095505 | 3.164758702 | 0.178812768 | 0.32310576 | 1.649067354 | 5.250127485 | ENSG00000232065 |
| ADAMTS14 | 1.660485987 | 3.161229961 | 2.17E-15 | 4.89E-14 | 61.15901633 | 192.8032938 | ENSG00000138316 |
| ETV3L | 1.660320372 | 3.160867088 | 0.003540561 | 0.012987443 | 8.124000627 | 25.50799697 | ENSG00000253831 |
| NA | 1.658731502 | 3.157387875 | 0.143879241 | 0.273390153 | 1.948433578 | 6.234493074 | ENSG00000283031 |
| FXYD2 | 1.657378862 | 3.154428962 | 0.158696953 | 0.295133802 | 1.681882467 | 5.298524605 | ENSG00000137731 |
| ARSD | 1.655565741 | 3.150467094 | 8.92E-23 | 3.71E-21 | 88.42276848 | 278.2249943 | ENSG00000006756 |
| NA | 1.654179016 | 3.147440304 | 0.18196944 | 0.327460118 | 1.687464273 | 5.296374031 | ENSG00000231464 |
| IL15 | 1.652540948 | 3.14386866 | 0.001200165 | 0.005030941 | 10.95187244 | 34.06747822 | ENSG00000164136 |
| ATXN1 | 1.652281226 | 3.143302734 | 3.41E-07 | 2.89E-06 | 23.50584244 | 73.75989048 | ENSG00000124788 |
| NA | 1.651928666 | 3.142534679 | 0.124810853 | 0.245424828 | 1.978457788 | 6.266218642 | ENSG00000227117 |
| MX2 | 1.649852702 | 3.138015987 | 1.29E-13 | 2.41E-12 | 57.47423367 | 179.8333923 | ENSG00000183486 |
| NA | 1.644053826 | 3.125428125 | 0.150408455 | 0.282996346 | 2.014063805 | 6.293643062 | ENSG00000287783 |
| NA | 1.643747918 | 3.124765484 | 0.006159568 | 0.020989971 | 7.310192621 | 22.88104917 | ENSG00000260193 |
| MLNR | 1.643276025 | 3.123743567 | 0.103317617 | 0.212416087 | 2.631656852 | 8.296250381 | ENSG00000102539 |
| CAPN11 | 1.641225814 | 3.119307574 | 6.84E-08 | 6.44E-07 | 28.89204389 | 89.80847656 | ENSG00000137225 |
| PCDH10 | 1.64092468 | 3.118656547 | 3.50E-14 | 6.97E-13 | 72.88004232 | 226.5948676 | ENSG00000138650 |
| CD7 | 1.64069396 | 3.118157843 | 0.00028714 | 0.001398086 | 13.83034392 | 43.40939475 | ENSG00000173762 |
| RAD51AP2 | 1.638948322 | 3.114387202 | 0.18224077 | 0.327734379 | 1.65464916 | 5.15548382 | ENSG00000214842 |
| ZNF37A | 1.636097879 | 3.108239946 | 2.27E-16 | 5.64E-15 | 61.97107257 | 192.9401433 | ENSG00000075407 |
| EYS | 1.633830091 | 3.103357906 | 0.182055941 | 0.327554669 | 1.714697581 | 5.25872978 | ENSG00000188107 |
| CATSPER2 | 1.633686399 | 3.10304883 | 0.003342452 | 0.012319306 | 8.542587812 | 26.74133292 | ENSG00000166762 |
| PSTPIP1 | 1.631881244 | 3.099168605 | 0.00508916 | 0.017801447 | 7.296238107 | 22.70492575 | ENSG00000140368 |
| NA | 1.629970666 | 3.095067055 | 0.112829046 | 0.227272124 | 2.335081531 | 7.236063252 | ENSG00000229646 |
| CEMIP | 1.626906865 | 3.088501144 | 2.21E-08 | 2.23E-07 | 30.41052651 | 94.01586986 | ENSG00000103888 |
| PIWIL3 | 1.622666676 | 3.079437134 | 0.109685359 | 0.222076015 | 2.329499725 | 7.176913264 | ENSG00000184571 |
| NA | 1.622634028 | 3.079367447 | 0.107257454 | 0.218328958 | 2.362314838 | 7.244132509 | ENSG00000284747 |
| NA | 1.620064469 | 3.07388772 | 0.051754829 | 0.122699078 | 3.690364466 | 11.36711409 | ENSG00000279989 |
| PLCB2 | 1.619234559 | 3.072119975 | 4.53E-55 | 8.83E-53 | 257.6741406 | 790.5855932 | ENSG00000137841 |
| LINC00954 | 1.619075674 | 3.07178166 | 0.016227574 | 0.047237985 | 5.317780318 | 16.36665368 | ENSG00000228784 |
| NA | 1.618247787 | 3.070019432 | 0.015704456 | 0.046006783 | 5.726079773 | 17.54514078 | ENSG00000273759 |
| NA | 1.617216957 | 3.067826634 | 0.070776109 | 0.157460721 | 2.985489691 | 9.191890987 | ENSG00000280077 |
| GRAP | 1.61529759 | 3.063747899 | 0.086528571 | 0.184811199 | 2.661681062 | 8.161278853 | ENSG00000154016 |
| CPA6 | 1.613941639 | 3.060869716 | 0.20696269 | 0.361379187 | 1.942851772 | 6.04116506 | ENSG00000165078 |
| DNAH1 | 1.612370664 | 3.057538497 | 5.28E-05 | 0.000302481 | 16.84952461 | 51.35153871 | ENSG00000114841 |

| | | | | | | | |
|---|---|---|---|---|---|---|---|
| LOC101929563 | 1.609191522 | 3.050808282 | 0.123420683 | 0.243471562 | 2.395129951 | 7.229611531 | ENSG00000283982 |
| NA | 1.608289927 | 3.048902312 | 0.075476537 | 0.16560713 | 3.742040179 | 11.36281294 | ENSG00000246203 |
| ARHGEF33 | 1.607604994 | 3.047455161 | 0.055302143 | 0.129471165 | 3.33932253 | 10.18055772 | ENSG00000214694 |
| NA | 1.605440424 | 3.042886294 | 0.010757526 | 0.033525596 | 6.613014833 | 20.27103339 | ENSG00000204054 |
| EMP3 | 1.605230575 | 3.042443718 | 1.66E-10 | 2.24E-09 | 46.28603449 | 140.1737596 | ENSG00000142227 |
| STPG3 | 1.603971772 | 3.039790236 | 0.216007369 | 0.373119462 | 1.769164196 | 5.252278059 | ENSG00000197768 |
| POLR2J3 | 1.60269126 | 3.037093366 | 0.014067845 | 0.041996719 | 5.731661579 | 17.31122444 | ENSG00000285437 |
| LTB | 1.601967064 | 3.035569205 | 0.136366361 | 0.262303666 | 2.367896644 | 7.130666718 | ENSG00000227507 |
| NA | 1.599874237 | 3.031168888 | 0.232081615 | 0.392692433 | 1.619043143 | 4.983128505 | ENSG00000231907 |
| SNORA65 | 1.599214953 | 3.029784018 | 0.128265897 | 0.250341706 | 2.419572356 | 7.218858663 | ENSG00000201302 |
| CDA | 1.597998772 | 3.027231009 | 4.84E-20 | 1.65E-18 | 91.11870412 | 275.7155288 | ENSG00000158825 |
| CCDC9B | 1.597233138 | 3.025624893 | 6.55E-05 | 0.000368589 | 15.30014189 | 46.42702715 | ENSG00000188549 |
| NA | 1.59559114 | 3.022183247 | 0.058928388 | 0.135963107 | 3.995312496 | 12.08045212 | ENSG00000244055 |
| PTK6 | 1.595532992 | 3.022061441 | 0.017466381 | 0.050215196 | 6.017073288 | 18.2119597 | ENSG00000101213 |
| NA | 1.594544177 | 3.019990846 | 0.033963373 | 0.086860324 | 4.398030145 | 13.22453005 | ENSG00000270441 |
| SLFN13 | 1.593572932 | 3.017958425 | 1.26E-38 | 1.33E-36 | 189.1059227 | 569.9458251 | ENSG00000154760 |
| ZYG11A | 1.592586047 | 3.015894678 | 2.30E-08 | 2.32E-07 | 29.69169722 | 89.61325844 | ENSG00000203995 |
| NA | 1.590505155 | 3.011547794 | 1.05E-11 | 1.61E-10 | 53.81389341 | 161.573829 | ENSG00000235152 |
| SLC12A5-AS1 | 1.589561484 | 3.009578577 | 0.047018923 | 0.113721421 | 4.022545804 | 12.07184982 | ENSG00000204044 |
| NA | 1.589475227 | 3.009398642 | 6.49E-10 | 8.06E-09 | 37.25448644 | 112.3827198 | ENSG00000250519 |
| ANKRD31 | 1.589442421 | 3.00933021 | 0.141335032 | 0.269843043 | 2.422363259 | 7.170461543 | ENSG00000145700 |
| FAM163B | 1.587501936 | 3.005285258 | 8.30E-12 | 1.28E-10 | 51.80365965 | 156.2271556 | ENSG00000196990 |
| LOC101927021 | 1.587380712 | 3.005032745 | 0.000397979 | 0.001878037 | 13.62105033 | 41.12016072 | ENSG00000229043 |
| DEGS2 | 1.586761865 | 3.003744007 | 0.000508051 | 0.002326327 | 13.01115427 | 39.11944351 | ENSG00000168350 |
| NA | 1.576522549 | 2.982500851 | 0.014098378 | 0.042055079 | 6.953568879 | 20.77357836 | ENSG00000261172 |
| RGS7 | 1.576260602 | 2.981959372 | 9.73E-08 | 8.99E-07 | 28.86481058 | 85.95327536 | ENSG00000182901 |
| CTHRC1 | 1.573791806 | 2.976860892 | 1.34E-05 | 8.62E-05 | 20.1309139 | 59.83198177 | ENSG00000164932 |
| ABI3 | 1.573787932 | 2.976852897 | 1.16E-08 | 1.23E-07 | 37.49380424 | 111.8686406 | ENSG00000108798 |
| GABRG2 | 1.573347021 | 2.975943262 | 0.007166857 | 0.023828996 | 9.96003435 | 29.67749767 | ENSG00000113327 |
| NA | 1.573256303 | 2.975756137 | 0.126741446 | 0.248220228 | 2.30505732 | 6.912064858 | ENSG00000251414 |
| DUSP10 | 1.571941122 | 2.973044633 | 0.000249482 | 0.001233198 | 15.27378447 | 45.13586776 | ENSG00000143507 |
| MSH5 | 1.571939236 | 2.973040747 | 0.008352266 | 0.027079505 | 7.347913819 | 21.80849164 | ENSG00000204410 |
| BVES-AS1 | 1.571848209 | 2.972853168 | 0.103267674 | 0.212336037 | 2.686123467 | 7.993224685 | ENSG00000203808 |
| SPARCL1 | 1.571690121 | 2.972527425 | 0.002898839 | 0.01087967 | 10.14085533 | 30.0248798 | ENSG00000152583 |
| NA | 1.570830033 | 2.970755829 | 0.20862927 | 0.363532404 | 1.65464916 | 4.941183107 | ENSG00000223599 |
| NA | 1.569771304 | 2.968576524 | 0.119662156 | 0.237979654 | 2.335081531 | 6.951859682 | ENSG00000255621 |
| HPSE2 | 1.566778032 | 2.962423773 | 0.002240672 | 0.008712922 | 10.92463913 | 32.04389821 | ENSG00000172987 |
| LINC01618 | 1.566218717 | 2.9612755 | 0.041689465 | 0.103072807 | 4.349145335 | 12.89192936 | ENSG00000250302 |
| NA | 1.565901919 | 2.960625312 | 0.077373249 | 0.168853295 | 3.012722998 | 8.927042581 | ENSG00000250131 |
| NA | 1.565673717 | 2.960157045 | 0.028948676 | 0.076269995 | 4.637347948 | 13.78595243 | ENSG00000272720 |
| PLAC8L1 | 1.565212 | 2.959209834 | 0.038323763 | 0.096250971 | 4.628975239 | 13.75637743 | ENSG00000173261 |
| GGT1 | 1.564716322 | 2.95819329 | 1.08E-05 | 7.05E-05 | 21.1568064 | 62.46161318 | ENSG00000100031 |
| LINC01780 | 1.564241512 | 2.957219869 | 0.001240107 | 0.005181484 | 12.04810109 | 35.56003489 | ENSG00000232650 |
| NA | 1.563606642 | 2.955918806 | 0.179783659 | 0.324485231 | 1.986830497 | 5.893823127 | ENSG00000288568 |
| NA | 1.562318132 | 2.953279973 | 0.154582087 | 0.289490636 | 2.395129951 | 7.029831796 | ENSG00000233332 |
| TMEM249 | 1.560859254 | 2.950295077 | 0.087436133 | 0.186358463 | 3.045538112 | 8.952316427 | ENSG00000261587 |
| SNORD15B | 1.560679091 | 2.949926668 | 0.040240899 | 0.100101506 | 4.316330222 | 12.76179201 | ENSG00000207445 |
| KCNH3 | 1.559977606 | 2.948492668 | 0.009283762 | 0.02955331 | 9.680880168 | 28.61514786 | ENSG00000135519 |
| TAC4 | 1.559645729 | 2.947814474 | 0.139628425 | 0.267378594 | 2.356733032 | 6.930886983 | ENSG00000176358 |
| SPAG17 | 1.55900393 | 2.946503399 | 0.055735269 | 0.130310152 | 3.665922062 | 10.80973239 | ENSG00000155761 |
| CAPN9 | 1.55750817 | 2.943450101 | 0.048288998 | 0.116094599 | 3.984148884 | 11.74570086 | ENSG00000135773 |
| NA | 1.557316301 | 2.943058666 | 0.13758534 | 0.264226859 | 2.389548146 | 6.98358525 | ENSG00000287037 |
| OR10A3 | 1.556868371 | 2.942145043 | 0.267889941 | 0.434173333 | 1.632997658 | 4.831485425 | ENSG00000170683 |
| C3P1 | 1.556312125 | 2.941010887 | 0.158343735 | 0.294710049 | 2.392339049 | 6.987886397 | ENSG00000167798 |
| DCST2 | 1.556186555 | 2.940754918 | 0.049717379 | 0.118759034 | 4.041406403 | 11.84679624 | ENSG00000163354 |
| CDH5 | 1.555318636 | 2.938986304 | 0.204805388 | 0.358358343 | 9.451850096 | 27.84535555 | ENSG00000179776 |
| NA | 1.553946633 | 2.936192657 | 0.075560531 | 0.165753718 | 3.312089222 | 9.740942964 | ENSG00000272654 |
| PWWP3B | 1.553579325 | 2.935445201 | 3.39E-05 | 0.0002024 | 17.60519921 | 51.8118657 | ENSG00000157502 |
| TGM2 | 1.552682931 | 2.93362188 | 4.34E-12 | 6.89E-11 | 51.22956977 | 150.0784556 | ENSG00000198959 |
| VAC14-AS1 | 1.551877031 | 2.931983595 | 0.155202596 | 0.290201927 | 2.008481999 | 5.875001002 | ENSG00000214353 |
| NMI | 1.550728928 | 2.929651237 | 3.72E-08 | 3.64E-07 | 37.09878358 | 109.1779386 | ENSG00000123609 |
| NA | 1.549726612 | 2.927616561 | 0.190638574 | 0.339043857 | 1.68467337 | 4.907306965 | ENSG00000242088 |
| DRD3 | 1.54871934 | 2.925573248 | 0.088199723 | 0.187695627 | 3.012722998 | 8.788302943 | ENSG00000151577 |
| MMRN2 | 1.546230191 | 2.920529973 | 1.97E-09 | 2.32E-08 | 39.01527793 | 113.9954666 | ENSG00000173269 |
| NA | 1.545592332 | 2.919239004 | 0.085962454 | 0.183887968 | 3.069980516 | 8.876494887 | ENSG00000287392 |
| LOC101927825 | 1.545049151 | 2.918140105 | 0.027636945 | 0.073602233 | 5.282174302 | 15.51941019 | ENSG00000282556 |
| ALOX5 | 1.544253057 | 2.916530289 | 1.28E-07 | 1.16E-06 | 31.95086079 | 92.83094315 | ENSG00000012779 |
| LIPH | 1.543895466 | 2.915807478 | 0.220817626 | 0.379251472 | 1.711906678 | 4.941183107 | ENSG00000163898 |
| DPEP1 | 1.543741933 | 2.915497191 | 1.61E-13 | 2.98E-12 | 62.57646286 | 181.8937925 | ENSG00000015413 |
| NA | 1.542153972 | 2.912289896 | 0.122814413 | 0.242647772 | 2.748962791 | 7.915252572 | ENSG00000231407 |
| NA | 1.542029878 | 2.912039405 | 3.13E-05 | 0.000188022 | 20.83367349 | 60.4711158 | ENSG00000260528 |
| GADL1 | 1.540799009 | 2.90955599 | 0.190414801 | 0.338739573 | 2.272242207 | 6.712285122 | ENSG00000144644 |
| NA | 1.539338821 | 2.906612644 | 0.039272485 | 0.098288208 | 4.61011464 | 13.50013132 | ENSG00000286214 |
| NA | 1.538652693 | 2.905230625 | 0.054475817 | 0.127845862 | 4.488102776 | 12.87310723 | ENSG00000270069 |
| AIPL1 | 1.537485954 | 2.902882052 | 0.000611987 | 0.002740921 | 13.60218973 | 39.58083658 | ENSG00000129221 |
| WDR66 | 1.536808166 | 2.901518578 | 0.000506249 | 0.002318624 | 15.65888082 | 45.45315397 | ENSG00000158023 |
| KCTD19 | 1.536158279 | 2.900211832 | 0.021877479 | 0.060489958 | 6.412093949 | 18.55209661 | ENSG00000168676 |

| | | | | | | | |
|---|---|---|---|---|---|---|---|
| TSPAN32 | 1.53578564 | 2.899462824 | 0.033137898 | 0.085110265 | 5.43787716 | 15.62454626 | ENSG00000064201 |
| ARRB1 | 1.535483099 | 2.898854853 | 3.57E-58 | 7.92E-56 | 472.4525181 | 1367.80266 | ENSG00000137486 |
| ENPP2 | 1.535254414 | 2.898395386 | 1.00E-06 | 7.88E-06 | 34.59260522 | 100.7846335 | ENSG00000136960 |
| C5AR2 | 1.534736748 | 2.897355573 | 0.025397697 | 0.068514469 | 5.284965205 | 15.38658924 | ENSG00000134830 |
| CRYZL2P | 1.534299951 | 2.896478489 | 8.77E-05 | 0.00048057 | 17.14330902 | 49.50461515 | ENSG00000242193 |
| LINC02525 | 1.530885159 | 2.889630767 | 0.005926638 | 0.020339934 | 9.152684031 | 26.28934775 | ENSG00000230269 |
| SMIM10 | 1.530880372 | 2.88962118 | 2.89E-07 | 2.48E-06 | 32.82750812 | 94.38585434 | ENSG00000184785 |
| LINC00698 | 1.530737971 | 2.889335973 | 2.32E-15 | 5.21E-14 | 78.60831252 | 227.4082655 | ENSG00000244342 |
| FGF8 | 1.530198507 | 2.888255773 | 1.21E-05 | 7.84E-05 | 23.84013896 | 68.65820154 | ENSG00000107831 |
| NA | 1.527189615 | 2.882238288 | 0.08054389 | 0.174346071 | 3.279274109 | 9.541163229 | ENSG00000272129 |
| NA | 1.524451356 | 2.876772942 | 0.107224688 | 0.218285296 | 2.952674577 | 8.573469192 | ENSG00000228626 |
| NA | 1.524215317 | 2.876302311 | 0.123510947 | 0.243574902 | 2.656099256 | 7.694500137 | ENSG00000197358 |
| NA | 1.524133261 | 2.876138722 | 0.043817985 | 0.10735807 | 4.37358774 | 12.55771113 | ENSG00000286592 |
| PPARGC1A | 1.522216466 | 2.87231996 | 1.31E-05 | 8.39E-05 | 22.8693888 | 65.54914844 | ENSG00000109819 |
| SLC22A20P | 1.5221025 | 2.87209307 | 3.02E-20 | 1.06E-18 | 115.0839581 | 329.6195629 | ENSG00000197847 |
| NA | 1.521437942 | 2.870770384 | 0.025776707 | 0.069420417 | 5.345013626 | 15.31936999 | ENSG00000223784 |
| RHBDL2 | 1.520444489 | 2.868794224 | 0.064888109 | 0.146739719 | 4.743490275 | 13.55067901 | ENSG00000158315 |
| ITGB2 | 1.518632603 | 2.865193556 | 0.004700931 | 0.01664568 | 9.019308396 | 25.81962496 | ENSG00000160255 |
| LDHD | 1.516836206 | 2.861628131 | 3.17E-22 | 1.26E-20 | 106.4390581 | 304.4487282 | ENSG00000166816 |
| MOG | 1.516361503 | 2.860686699 | 0.017145793 | 0.049448271 | 6.313648609 | 18.12323472 | ENSG00000204655 |
| MT1H | 1.510583472 | 2.849252487 | 8.01E-11 | 1.11E-09 | 48.93567606 | 139.3369964 | ENSG00000205358 |
| C1orf74 | 1.510514247 | 2.849115774 | 2.85E-22 | 1.13E-20 | 112.7827309 | 321.0243091 | ENSG00000162757 |
| NA | 1.508275978 | 2.844698941 | 0.030553468 | 0.079768162 | 5.647170753 | 16.1120251 | ENSG00000273314 |
| NA | 1.505894846 | 2.84000771 | 0.016788284 | 0.048576989 | 7.808564707 | 22.04617609 | ENSG00000253706 |
| NA | 1.505474742 | 2.839180837 | 0.055444008 | 0.129739247 | 4.044197305 | 11.41740131 | ENSG00000270112 |
| CNIH3-AS2 | 1.505256246 | 2.838750875 | 0.001657321 | 0.006691371 | 11.25315368 | 32.06056976 | ENSG00000233384 |
| CERS3 | 1.504925209 | 2.838099578 | 0.033949489 | 0.08683633 | 5.489552873 | 15.39089038 | ENSG00000154227 |
| CFAP45 | 1.504607675 | 2.837474988 | 0.111500644 | 0.225042783 | 2.958256383 | 8.434729554 | ENSG00000213085 |
| LCN2 | 1.503974271 | 2.836229469 | 0.007321356 | 0.024267401 | 8.292106398 | 23.58604538 | ENSG00000148346 |
| PCOLCE-AS1 | 1.503549571 | 2.835394684 | 0.000495406 | 0.00227112 | 13.2972417 | 37.80946199 | ENSG00000224729 |
| CCDC169 | 1.502183996 | 2.832712127 | 8.57E-14 | 1.63E-12 | 64.17086344 | 181.8182315 | ENSG00000242715 |
| LINC01772 | 1.501707421 | 2.831776534 | 0.037450396 | 0.094364628 | 5.581540526 | 15.9633261 | ENSG00000226029 |
| NA | 1.500413192 | 2.829237311 | 0.004488324 | 0.015977903 | 10.18136744 | 28.6261733 | ENSG00000234232 |
| NA | 1.500024132 | 2.828474436 | 0.023399559 | 0.064001262 | 5.67440406 | 16.049107 | ENSG00000285847 |
| PPFIBP2 | 1.499522052 | 2.827490256 | 1.08E-30 | 7.06E-29 | 170.240019 | 481.3782023 | ENSG00000166387 |
| AMY2B | 1.497577999 | 2.823682736 | 0.05824007 | 0.134737379 | 4.814702308 | 13.43022846 | ENSG00000240038 |
| SULT1C2 | 1.496579809 | 2.821729727 | 0.000585814 | 0.002636094 | 14.0677467 | 39.63138427 | ENSG00000198203 |
| NA | 1.496053218 | 2.820699968 | 0.001410832 | 0.005806776 | 11.60419562 | 32.85805905 | ENSG00000234537 |
| BNIP3 | 1.49535659 | 2.819338278 | 5.16E-61 | 1.22E-58 | 815.7135219 | 2298.686115 | ENSG00000176171 |
| NA | 1.495067645 | 2.818773673 | 0.206229875 | 0.360393198 | 1.956806287 | 5.57412588 | ENSG00000223026 |
| SCAT2 | 1.493189335 | 2.815106173 | 0.13635769 | 0.262303666 | 3.252040801 | 9.261793844 | ENSG00000257596 |
| TCAM1P | 1.492283584 | 2.813339351 | 0.006216811 | 0.021158875 | 8.417785045 | 23.58174423 | ENSG00000240280 |
| LOC728755 | 1.491952101 | 2.812693015 | 0.076018884 | 0.166456333 | 4.329609015 | 12.20413774 | ENSG00000258932 |
| FOXS1 | 1.491845124 | 2.812484458 | 0.177948486 | 0.321875691 | 2.335081531 | 6.573545485 | ENSG00000179772 |
| NA | 1.491248804 | 2.811322193 | 5.05E-05 | 0.000289839 | 19.40717852 | 54.53237267 | ENSG00000244041 |
| MUC19 | 1.490571172 | 2.810002029 | 0.004297773 | 0.015387592 | 9.873428343 | 27.94214979 | ENSG00000205592 |
| LHX5 | 1.489211252 | 2.807354499 | 0.222155766 | 0.38057072 | 1.956806287 | 5.542400312 | ENSG00000089116 |
| TBX10 | 1.489157607 | 2.807250113 | 0.002091315 | 0.008208251 | 11.48218376 | 32.05196746 | ENSG00000167800 |
| LINC00886 | 1.488098513 | 2.805190045 | 0.080713626 | 0.174615516 | 4.161503245 | 11.49967458 | ENSG00000240875 |
| NA | 1.486581803 | 2.802242489 | 0.052001638 | 0.123036053 | 4.730211482 | 13.20732546 | ENSG00000231680 |
| NA | 1.485567123 | 2.800272302 | 0.126091288 | 0.247222993 | 2.713356774 | 7.542857057 | ENSG00000273576 |
| NA | 1.484735462 | 2.798658511 | 0.050148372 | 0.119566462 | 4.316330222 | 12.1073435 | ENSG00000255130 |
| NINJ2 | 1.484087066 | 2.797400983 | 0.236052787 | 0.397481288 | 1.627415852 | 4.612883564 | ENSG00000171840 |
| C22orf15 | 1.482872688 | 2.79504728 | 5.62E-07 | 4.59E-06 | 31.27301917 | 87.55741981 | ENSG00000169314 |
| RERG | 1.481261343 | 2.791927238 | 0.126013406 | 0.247160465 | 2.71893858 | 7.524034932 | ENSG00000134533 |
| C1orf220 | 1.480800045 | 2.791034669 | 0.00794718 | 0.02596698 | 9.054238691 | 25.21787472 | ENSG00000213057 |
| NA | 1.478847005 | 2.787258879 | 0.175818212 | 0.31917817 | 2.008481999 | 5.597249153 | ENSG00000259659 |
| NA | 1.478477389 | 2.786544878 | 0.155145962 | 0.290152278 | 2.359523935 | 6.541819917 | ENSG00000234807 |
| ANXA3 | 1.477332684 | 2.784334773 | 5.94E-37 | 5.56E-35 | 198.1940157 | 551.9373827 | ENSG00000138772 |
| TMPRSS9 | 1.477241629 | 2.784159047 | 0.000598393 | 0.002686433 | 13.62105033 | 38.00924173 | ENSG00000178297 |
| NA | 1.476495271 | 2.782719075 | 0.009617545 | 0.030430056 | 9.054238691 | 25.12591466 | ENSG00000287808 |
| LMX1B | 1.476104898 | 2.781966213 | 0.09195927 | 0.193857095 | 3.336531627 | 9.31826022 | ENSG00000136944 |
| NA | 1.475214351 | 2.780249489 | 0.013402681 | 0.040291589 | 8.129582433 | 22.5110647 | ENSG00000280099 |
| NEK7 | 1.474628581 | 2.779120869 | 1.31E-23 | 5.75E-22 | 129.3734014 | 359.1287154 | ENSG00000151414 |
| RASGRF2-AS1 | 1.473951218 | 2.777816344 | 0.059386889 | 0.136742954 | 4.016963998 | 11.14395061 | ENSG00000251450 |
| RNASE1 | 1.473860277 | 2.777641247 | 0.033788806 | 0.086494142 | 6.134379227 | 16.93908939 | ENSG00000129538 |
| IFI35 | 1.473537687 | 2.777020229 | 3.61E-05 | 0.000214152 | 23.089846 | 64.01574298 | ENSG00000068079 |
| ADGRG1 | 1.473316717 | 2.776594921 | 1.12E-06 | 8.77E-06 | 32.04496363 | 89.02226106 | ENSG00000205336 |
| CALHM5 | 1.47323103 | 2.776430014 | 0.184298698 | 0.330424947 | 2.035715307 | 5.620372426 | ENSG00000178033 |
| FMO5 | 1.472834298 | 2.775666618 | 0.002213522 | 0.008616032 | 11.97967996 | 33.35252265 | ENSG00000131781 |
| LOC105370802 | 1.472405992 | 2.774842704 | 0.254190973 | 0.418967708 | 2.313430029 | 6.421902405 | ENSG00000259200 |
| PYY | 1.471964249 | 2.773993196 | 0.000456782 | 0.00211466 | 16.53899477 | 45.61313888 | ENSG00000131096 |
| H2AC14 | 1.471781579 | 2.773641983 | 0.066371261 | 0.149540762 | 4.253490898 | 11.93283761 | ENSG00000276368 |
| ALG1L | 1.46975744 | 2.769753219 | 1.38E-06 | 1.06E-05 | 27.84661507 | 77.03053392 | ENSG00000189366 |
| LRFN5 | 1.469674451 | 2.769593897 | 0.007376845 | 0.024417969 | 8.632660444 | 23.98371474 | ENSG00000165379 |
| NA | 1.469218352 | 2.768718446 | 0.142374173 | 0.271424071 | 2.933813978 | 8.207525399 | ENSG00000224963 |
| XCL1 | 1.467300001 | 2.765039329 | 0.153513104 | 0.28790796 | 2.770614292 | 7.545007631 | ENSG00000143184 |

| | | | | | | | |
|---|---|---|---|---|---|---|---|
| NA | 1.466378 | 2.763272805 | 0.254265925 | 0.419055443 | 1.657440063 | 4.596212013 | ENSG00000266456 |
| SLC44A4 | 1.465579165 | 2.761743177 | 5.89E-07 | 4.80E-06 | 33.27591936 | 92.18587832 | ENSG00000204385 |
| CR2 | 1.465021699 | 2.760676229 | 5.47E-26 | 2.83E-24 | 135.4580199 | 373.841501 | ENSG00000117322 |
| EXOC8 | 1.464473183 | 2.759626812 | 1.99E-19 | 6.37E-18 | 104.2336854 | 287.6873614 | ENSG00000116903 |
| SGCG | 1.464164901 | 2.759037186 | 0.154426984 | 0.289368717 | 2.966629092 | 8.234949819 | ENSG00000102683 |
| NLRP14 | 1.464158832 | 2.759025579 | 0.163189196 | 0.301482741 | 2.389548146 | 6.51654607 | ENSG00000158077 |
| GABRE | 1.463385038 | 2.757546163 | 1.04E-13 | 1.97E-12 | 74.36359465 | 204.4298706 | ENSG00000102287 |
| GAL3ST2 | 1.462899615 | 2.75661849 | 0.05818787 | 0.134705652 | 4.634557045 | 12.83492994 | ENSG00000154252 |
| APOBEC3G | 1.461673011 | 2.754275761 | 5.95E-05 | 0.00033806 | 20.48474673 | 56.33843532 | ENSG00000239713 |
| AIF1 | 1.46093036 | 2.752858317 | 2.51E-07 | 2.17E-06 | 35.82799147 | 98.96217815 | ENSG00000204472 |
| SLC4A1 | 1.459866018 | 2.750828158 | 0.179594162 | 0.324305062 | 2.389548146 | 6.497723945 | ENSG00000004939 |
| LOC105370854 | 1.459635237 | 2.750388156 | 0.207865892 | 0.36256247 | 2.06853042 | 5.620372426 | ENSG00000259370 |
| SYCP2 | 1.458748325 | 2.748697846 | 0.106211202 | 0.216718026 | 3.303716514 | 9.10800019 | ENSG00000196074 |
| MYOZ1 | 1.45869673 | 2.748599546 | 0.000600816 | 0.002695428 | 16.73154294 | 45.95890348 | ENSG00000177791 |
| B3GNT3 | 1.456435023 | 2.744293953 | 0.000127465 | 0.000671287 | 17.92621694 | 49.37420524 | ENSG00000179913 |
| DBH-AS1 | 1.456208737 | 2.743863546 | 0.006677715 | 0.022497345 | 8.986493283 | 24.72797273 | ENSG00000225756 |
| MYH2 | 1.455923239 | 2.74332061 | 2.28E-08 | 2.29E-07 | 40.34141253 | 110.3881818 | ENSG00000125414 |
| PEX5L | 1.455922847 | 2.743319864 | 2.47E-07 | 2.14E-06 | 35.22088631 | 96.85900848 | ENSG00000114757 |
| NR4A2 | 1.455425518 | 2.742374345 | 0.035426862 | 0.090053972 | 6.335300111 | 17.38489541 | ENSG00000153234 |
| NA | 1.45423212 | 2.74010679 | 0.237427146 | 0.399146586 | 1.65464916 | 4.541363172 | ENSG00000248780 |
| NA | 1.45420306 | 2.740051596 | 0.133860879 | 0.258696676 | 3.018304804 | 8.270976534 | ENSG00000234273 |
| NA | 1.454097295 | 2.739850728 | 2.19E-05 | 0.000135612 | 22.39545912 | 61.33689042 | ENSG00000268970 |
| NAALADL1 | 1.453913834 | 2.739502336 | 2.43E-07 | 2.11E-06 | 34.47916622 | 94.42511612 | ENSG00000168060 |
| ISLR | 1.453001774 | 2.737770992 | 0.007970619 | 0.02603034 | 11.49546255 | 31.36658689 | ENSG00000129009 |
| NA | 1.452338253 | 2.736512133 | 0.050872122 | 0.121007905 | 5.326153027 | 14.63775752 | ENSG00000232093 |
| OCA2 | 1.452307761 | 2.736454295 | 0.00435864 | 0.015570837 | 9.718601366 | 26.54559387 | ENSG00000104044 |
| NKPD1 | 1.452060534 | 2.735985404 | 0.002208127 | 0.008600237 | 11.8443893 | 32.23318554 | ENSG00000179846 |
| LINC01844 | 1.450394704 | 2.732828081 | 5.26E-10 | 6.61E-09 | 56.96681315 | 155.7100897 | ENSG00000236714 |
| MRGPRG-AS1 | 1.44957284 | 2.731271707 | 0.113092741 | 0.227684416 | 4.028127609 | 11.01569127 | ENSG00000236301 |
| SLC14A2 | 1.448847344 | 2.729898562 | 0.252256403 | 0.416740453 | 1.687464273 | 4.58330857 | ENSG00000132874 |
| PCED1B | 1.445802507 | 2.724143132 | 1.19E-14 | 2.48E-13 | 78.18047675 | 213.3859552 | ENSG00000179715 |
| NA | 1.44559437 | 2.723750149 | 0.258413426 | 0.42358906 | 1.657440063 | 4.511788178 | ENSG00000271945 |
| COLCA1 | 1.445251793 | 2.723103453 | 0.049978455 | 0.119279273 | 5.056811014 | 13.69560991 | ENSG00000196167 |
| HPGD | 1.444188143 | 2.721096542 | 0.000111282 | 0.000595323 | 18.99608816 | 51.7156045 | ENSG00000164120 |
| FAM157A | 1.443986127 | 2.720715541 | 8.94E-13 | 1.53E-11 | 78.51740091 | 212.889874 | ENSG00000236438 |
| NA | 1.443123946 | 2.719090079 | 0.212748048 | 0.368744168 | 2.065739517 | 5.532180482 | ENSG00000229931 |
| AZU1 | 1.442199627 | 2.717348544 | 0.013321377 | 0.040072148 | 8.423366851 | 22.76113166 | ENSG00000172232 |
| CFAP70 | 1.440970669 | 2.715034761 | 0.114665169 | 0.230249379 | 3.360974032 | 9.076274623 | ENSG00000156042 |
| CCDC162P | 1.440573772 | 2.714287936 | 5.80E-05 | 0.000329852 | 20.1337048 | 54.55281233 | ENSG00000203799 |
| NA | 1.440296688 | 2.713766679 | 0.241307374 | 0.403664634 | 1.711906678 | 4.591910865 | ENSG00000272473 |
| NRROS | 1.439551962 | 2.712366182 | 1.01E-14 | 2.14E-13 | 81.17541519 | 219.7648461 | ENSG00000174004 |
| NA | 1.439515483 | 2.7122976 | 0.250748785 | 0.414889339 | 1.709115775 | 4.585459144 | ENSG00000286257 |
| MYBPC2 | 1.438400488 | 2.710202195 | 2.77E-17 | 7.48E-16 | 102.1332157 | 276.5001573 | ENSG00000086967 |
| ABHD12B | 1.437289874 | 2.708116633 | 3.33E-10 | 4.32E-09 | 52.35390761 | 141.6017685 | ENSG00000131969 |
| WDR31 | 1.436912955 | 2.707409202 | 0.020647445 | 0.057709341 | 7.068083915 | 19.08447703 | ENSG00000148225 |
| HCN1 | 1.436584501 | 2.706792884 | 1.63E-07 | 1.45E-06 | 42.4004942 | 115.2922415 | ENSG00000164588 |
| NA | 1.435975382 | 2.705650294 | 0.035410664 | 0.090036527 | 6.033818705 | 16.27201001 | ENSG00000253348 |
| MUC5B | 1.435423716 | 2.704615888 | 0.097006264 | 0.202351626 | 3.611455446 | 9.844188924 | ENSG00000117983 |
| OXCT2 | 1.435348639 | 2.704475146 | 0.00052753 | 0.002404104 | 17.76301725 | 47.9101696 | ENSG00000198754 |
| HGC6.3 | 1.434979429 | 2.703783113 | 0.14861319 | 0.28031294 | 2.68891437 | 7.259186525 | ENSG00000235994 |
| TRADD | 1.434420841 | 2.702736455 | 0.121040246 | 0.240083025 | 4.071430613 | 10.96567661 | ENSG00000102871 |
| SMIM38 | 1.434283214 | 2.702478638 | 0.138108023 | 0.26496687 | 2.658890159 | 7.197885963 | ENSG00000284713 |
| PHLDA3 | 1.434118063 | 2.702169293 | 1.17E-74 | 4.51E-72 | 914.1972132 | 2472.52064 | ENSG00000174307 |
| IL18 | 1.434059725 | 2.702060028 | 0.099766475 | 0.206878972 | 3.608664543 | 9.825366799 | ENSG00000150782 |
| TAGLN | 1.4335246 | 2.701057963 | 1.48E-70 | 4.77E-68 | 541.4533288 | 1461.936618 | ENSG00000149591 |
| TMEM213 | 1.432702475 | 2.699519194 | 0.175066042 | 0.318093531 | 2.302266417 | 6.268369215 | ENSG00000214128 |
| PRCD | 1.432490876 | 2.699123287 | 0.022677882 | 0.062367057 | 6.684226866 | 18.03504277 | ENSG00000214140 |
| TRG-AS1 | 1.432393511 | 2.698941134 | 0.009344767 | 0.029713187 | 9.147102226 | 24.51341155 | ENSG00000281103 |
| FOXB1 | 1.432101357 | 2.698394639 | 7.76E-10 | 9.57E-09 | 68.32734846 | 184.4116062 | ENSG00000171956 |
| SH2D2A | 1.431753899 | 2.697744836 | 8.03E-18 | 2.27E-16 | 95.38614953 | 257.1543089 | ENSG00000027869 |
| PYROXD2 | 1.43163507 | 2.697522643 | 5.21E-07 | 4.28E-06 | 32.55796595 | 87.64184365 | ENSG00000119943 |
| CAV2 | 1.431210702 | 2.696729285 | 1.95E-05 | 0.000121809 | 26.57370778 | 71.52362729 | ENSG00000105971 |
| FAM124B | 1.430280491 | 2.694991068 | 0.00031254 | 0.001511457 | 16.64214603 | 44.91700544 | ENSG00000124019 |
| TMEM204 | 1.429602543 | 2.693724941 | 0.073822684 | 0.162737218 | 3.98973069 | 10.75273297 | ENSG00000131634 |
| C5orf47 | 1.428423579 | 2.691524539 | 0.082470791 | 0.177785896 | 4.343563529 | 11.65750891 | ENSG00000185056 |
| RALYL | 1.427365196 | 2.689550721 | 0.141140292 | 0.269551269 | 3.478279971 | 9.217164834 | ENSG00000184672 |
| KLHL30 | 1.427223828 | 2.689287187 | 0.284866204 | 0.453748984 | 1.681882467 | 4.488664905 | ENSG00000168427 |
| FAM107A | 1.426176389 | 2.687335395 | 0.035171169 | 0.089502693 | 6.801532805 | 18.17378241 | ENSG00000168309 |
| F10 | 1.42607742 | 2.687151049 | 5.29E-08 | 5.07E-07 | 37.42067719 | 100.5891549 | ENSG00000126218 |
| MISP | 1.424706429 | 2.684598667 | 0.017356981 | 0.049967695 | 7.595604331 | 20.54018295 | ENSG00000099812 |
| RASSF10-DT | 1.423836643 | 2.682980637 | 0.088515446 | 0.188263668 | 4.419681646 | 11.76452298 | ENSG00000254670 |
| RNF157-AS1 | 1.423235905 | 2.681863678 | 0.027209204 | 0.072659572 | 6.585781526 | 17.79116706 | ENSG00000267128 |
| KCNJ6 | 1.420909314 | 2.677542201 | 0.000105079 | 0.000565749 | 19.95423526 | 53.55177519 | ENSG00000157542 |
| SLC9A7P1 | 1.416853752 | 2.670025928 | 0.003612296 | 0.013210389 | 11.13672363 | 29.61645757 | ENSG00000227825 |
| MASP1 | 1.415547943 | 2.667610333 | 2.79E-07 | 2.40E-06 | 42.94707538 | 114.8321871 | ENSG00000127241 |
| TNFAIP6 | 1.415382939 | 2.667305252 | 0.043960883 | 0.107659944 | 5.701637368 | 15.20616466 | ENSG00000123610 |
| LOC102723566 | 1.414942769 | 2.666491574 | 0.04897091 | 0.117392478 | 5.655543462 | 15.11098796 | ENSG00000225032 |

| | | | | | | | |
|---|---|---|---|---|---|---|---|
| FGF4 | 1.414521394 | 2.665712872 | 2.30E-05 | 0.000141926 | 25.54626367 | 68.31295787 | ENSG00000075388 |
| LGR6 | 1.413540472 | 2.663901009 | 6.74E-05 | 0.000378002 | 24.42905924 | 65.04714862 | ENSG00000133067 |
| NA | 1.413229041 | 2.663326021 | 0.10501229 | 0.214960959 | 3.309298319 | 8.851221041 | ENSG00000274373 |
| NA | 1.412623847 | 2.66220902 | 0.159754964 | 0.296762585 | 2.928232173 | 7.883527004 | ENSG00000277369 |
| SBK3 | 1.412503078 | 2.661986175 | 0.003196494 | 0.011840163 | 11.37045962 | 30.27252362 | ENSG00000231274 |
| PCDH11X | 1.41187231 | 2.660822567 | 9.28E-18 | 2.61E-16 | 112.8869582 | 299.8635296 | ENSG00000102290 |
| NA | 1.411681468 | 2.660470613 | 0.06823817 | 0.152854452 | 4.71904787 | 12.51576573 | ENSG00000285907 |
| WEE2-AS1 | 1.410681628 | 2.65862745 | 0.222604218 | 0.381026377 | 1.975666886 | 5.298524605 | ENSG00000228775 |
| CAVIN3 | 1.410319681 | 2.657960531 | 1.60E-21 | 6.01E-20 | 171.4677593 | 456.5908312 | ENSG00000170955 |
| NA | 1.409782307 | 2.656970679 | 0.219722498 | 0.377715976 | 2.005691096 | 5.351222872 | ENSG00000260455 |
| NA | 1.408504767 | 2.654618911 | 0.002294966 | 0.008888211 | 12.32581581 | 32.7462108 | ENSG00000241634 |
| PARVG | 1.408447356 | 2.654513274 | 0.064277761 | 0.145657498 | 4.343563529 | 11.54377055 | ENSG00000138964 |
| NA | 1.408373109 | 2.654376665 | 0.118503169 | 0.236180365 | 3.698737175 | 9.778587215 | ENSG00000231720 |
| WDR72 | 1.407879135 | 2.653467971 | 4.10E-06 | 2.90E-05 | 30.37982658 | 80.69937976 | ENSG00000166415 |
| GBGT1 | 1.406776838 | 2.651441354 | 8.60E-17 | 2.22E-15 | 110.1589712 | 291.6382665 | ENSG00000148288 |
| MAFB | 1.40540266 | 2.648917038 | 6.49E-07 | 5.26E-06 | 32.38061159 | 85.85217997 | ENSG00000204103 |
| NA | 1.40351468 | 2.645452804 | 0.211104848 | 0.366752848 | 1.981248691 | 5.277551906 | ENSG00000275880 |
| CD79A | 1.403471978 | 2.645374503 | 0.066118617 | 0.149110854 | 4.670163061 | 12.33024651 | ENSG00000105369 |
| WFDC1 | 1.401372417 | 2.641527477 | 0.006236135 | 0.021214178 | 11.46332316 | 30.16067536 | ENSG00000103175 |
| PRDM16 | 1.400927715 | 2.640713367 | 0.000570289 | 0.002575244 | 15.70844135 | 41.51165093 | ENSG00000142611 |
| FGR | 1.400593805 | 2.64010225 | 0.056039129 | 0.130719569 | 5.810570598 | 15.22713736 | ENSG00000000938 |
| ALDH1A1 | 1.400536442 | 2.639997278 | 0.000154561 | 0.000796805 | 20.81971898 | 54.94429043 | ENSG00000165092 |
| GJB7 | 1.399997021 | 2.639010371 | 0.004757403 | 0.01680554 | 11.89048321 | 31.15202571 | ENSG00000164411 |
| TPRG1 | 1.399729413 | 2.638520903 | 0.128577023 | 0.250796698 | 2.985489691 | 7.913101998 | ENSG00000188001 |
| FAM86EP | 1.399479516 | 2.63806391 | 1.04E-07 | 9.54E-07 | 45.25109356 | 118.8804438 | ENSG00000251669 |
| NDRG1 | 1.399244203 | 2.63763366 | 1.31E-08 | 1.37E-07 | 46.35637064 | 122.0438006 | ENSG00000104419 |
| NA | 1.399153854 | 2.637468483 | 0.21463093 | 0.371173937 | 2.005691096 | 5.300675179 | ENSG00000278467 |
| ARIH2OS | 1.398229128 | 2.635778483 | 0.086038189 | 0.18402957 | 3.962497383 | 10.50025497 | ENSG00000221883 |
| GNGT1 | 1.397808431 | 2.63500989 | 0.000395143 | 0.001866479 | 20.09809878 | 52.91721488 | ENSG00000127928 |
| NA | 1.397779671 | 2.634957462 | 0.069646173 | 0.15544884 | 4.37358774 | 11.48677113 | ENSG00000224272 |
| NA | 1.396232005 | 2.632132299 | 0.016393918 | 0.047614434 | 7.734561772 | 20.27937522 | ENSG00000282221 |
| SCTR | 1.395910946 | 2.631546606 | 0.009290537 | 0.029565118 | 9.699740767 | 25.48755731 | ENSG00000080293 |
| LINC01138 | 1.395883387 | 2.631496337 | 3.22E-07 | 2.75E-06 | 36.40036649 | 95.89587606 | ENSG00000274020 |
| NA | 1.395601174 | 2.630981628 | 0.130380794 | 0.253432654 | 3.015513901 | 7.938375845 | ENSG00000266904 |
| SLFNL1 | 1.395417437 | 2.630646575 | 0.118530088 | 0.236180365 | 3.614246349 | 9.591710922 | ENSG00000171790 |
| LINC02617 | 1.395293332 | 2.630420288 | 0.000129553 | 0.000680611 | 20.52872546 | 53.82254228 | ENSG00000256288 |
| FRK | 1.39471913 | 2.629373572 | 0.135903135 | 0.26155786 | 2.985489691 | 7.875457747 | ENSG00000111816 |
| NA | 1.394556748 | 2.629077641 | 0.125738708 | 0.246732319 | 3.360974032 | 8.798522774 | ENSG00000224728 |
| NA | 1.393511847 | 2.627174169 | 0.023263228 | 0.063673515 | 8.814720729 | 23.3733622 | ENSG00000198580 |
| INPP5J | 1.393449283 | 2.62706024 | 5.73E-20 | 1.92E-18 | 129.7992852 | 341.490288 | ENSG00000185133 |
| FAM20A | 1.392095491 | 2.624596224 | 0.012000168 | 0.036623198 | 8.684336156 | 22.84151481 | ENSG00000108950 |
| GPR141 | 1.391898035 | 2.624237029 | 0.311624531 | 0.482622738 | 1.95959719 | 5.170004798 | ENSG00000187037 |
| SORBS2 | 1.391878268 | 2.624201075 | 4.00E-47 | 6.07E-45 | 403.797196 | 1061.096175 | ENSG00000154556 |
| TFAP2C | 1.390171078 | 2.621097605 | 2.75E-15 | 6.10E-14 | 102.6008879 | 268.7206392 | ENSG00000087510 |
| SH3TC2 | 1.3896554 | 2.620160886 | 0.119474636 | 0.237655723 | 3.399370951 | 8.830248342 | ENSG00000169247 |
| NPFFR2 | 1.389329453 | 2.619568982 | 3.56E-14 | 7.06E-13 | 90.08252389 | 235.4915417 | ENSG00000056291 |
| SLA2 | 1.389289168 | 2.619495834 | 0.197753071 | 0.348898343 | 2.362314838 | 6.137698835 | ENSG00000101082 |
| NA | 1.388604924 | 2.61825375 | 0.210717818 | 0.366278377 | 2.008481999 | 5.254428633 | ENSG00000270557 |
| CA4 | 1.388400946 | 2.61788359 | 2.12E-09 | 2.47E-08 | 50.91325796 | 133.5012693 | ENSG00000167434 |
| PTPN14 | 1.388183461 | 2.617488975 | 7.22E-87 | 3.98E-84 | 1146.724876 | 3000.574179 | ENSG00000152104 |
| CTD-2297D10.2 | 1.387280285 | 2.615850852 | 2.68E-23 | 1.15E-21 | 138.5196277 | 362.1267322 | ENSG00000250579 |
| NTSR1 | 1.387250692 | 2.615797196 | 0.000381291 | 0.001807245 | 19.07778809 | 49.85927304 | ENSG00000101188 |
| QPCT | 1.385753182 | 2.613083421 | 5.06E-12 | 7.97E-11 | 70.38136079 | 183.7138247 | ENSG00000115828 |
| MARCHF10 | 1.384901508 | 2.611541279 | 0.099523348 | 0.206508002 | 3.660340256 | 9.596012069 | ENSG00000173838 |
| NA | 1.384226214 | 2.61031916 | 0.000749849 | 0.003280124 | 17.91226242 | 46.8819685 | ENSG00000276855 |
| POLH | 1.383964319 | 2.609845348 | 3.62E-19 | 1.13E-17 | 134.0761794 | 350.1129236 | ENSG00000170734 |
| TENM1 | 1.383692864 | 2.60935433 | 9.19E-06 | 6.10E-05 | 38.99816909 | 101.2223702 | ENSG00000009694 |
| NA | 1.383556773 | 2.609108198 | 0.157036575 | 0.292801402 | 2.979907885 | 7.784582191 | ENSG00000227953 |
| NA | 1.383161772 | 2.608393938 | 0.295978772 | 0.466214297 | 2.062948614 | 5.342620577 | ENSG00000267301 |
| MGARP | 1.38306259 | 2.608214623 | 0.07814952 | 0.170161338 | 4.988389884 | 13.09412013 | ENSG00000137463 |
| NPIPB6 | 1.382713907 | 2.607584324 | 0.001161783 | 0.004881716 | 16.11115899 | 41.99536166 | ENSG00000198156 |
| CCDC144A | 1.382413986 | 2.607042291 | 0.135549946 | 0.261019565 | 3.453837566 | 8.885097182 | ENSG00000170160 |
| ZMAT3 | 1.381376969 | 2.605169007 | 6.68E-23 | 2.79E-21 | 201.7079016 | 525.6525355 | ENSG00000172667 |
| NA | 1.380243039 | 2.603122201 | 0.141806781 | 0.270583047 | 4.593369223 | 12.06135742 | ENSG00000234694 |
| SMIM10L2A | 1.380057691 | 2.602787791 | 1.03E-25 | 5.31E-24 | 170.4625913 | 443.4684932 | ENSG00000178947 |
| NA | 1.379719518 | 2.602177758 | 0.097366636 | 0.202905884 | 4.48252097 | 11.52494842 | ENSG00000271815 |
| FGF16 | 1.376470963 | 2.596324957 | 2.48E-06 | 1.81E-05 | 30.96528023 | 80.55580594 | ENSG00000196468 |
| PPP1R3C | 1.376435275 | 2.596260733 | 0.000134638 | 0.000704259 | 22.26419866 | 57.98208361 | ENSG00000119938 |
| TOR4A | 1.376426019 | 2.596244075 | 4.94E-28 | 2.84E-26 | 222.4400635 | 576.6895569 | ENSG00000198113 |
| FZD4-DT | 1.375421699 | 2.594437351 | 0.240638138 | 0.402824568 | 2.005691096 | 5.187209388 | ENSG00000246523 |
| TRPM6 | 1.374627554 | 2.593009612 | 0.002123206 | 0.00831988 | 15.19208454 | 39.18478475 | ENSG00000119121 |
| NA | 1.374015676 | 2.591910093 | 0.017094927 | 0.049323696 | 8.450600159 | 21.8297248 | ENSG00000239407 |
| NA | 1.373630139 | 2.59121754 | 0.000361785 | 0.001722839 | 18.10078039 | 46.79969524 | ENSG00000258053 |
| NA | 1.372715092 | 2.58957455 | 0.226468506 | 0.385664191 | 2.038506209 | 5.233455934 | ENSG00000224520 |
| NA | 1.372376182 | 2.588966293 | 0.039487016 | 0.098697076 | 6.360418237 | 16.46559849 | ENSG00000269842 |
| NPNT | 1.371013217 | 2.58652156 | 0.032529224 | 0.083904433 | 6.975220381 | 18.09634333 | ENSG00000168743 |
| SPOCD1 | 1.369414154 | 2.583656283 | 0.000633768 | 0.002828264 | 17.13214541 | 44.10929631 | ENSG00000134668 |

| | | | | | | | |
|---|---|---|---|---|---|---|---|
| PROX1 | 1.36841155 | 2.58186139 | 0.004083543 | 0.014721664 | 11.78434088 | 30.35694745 | ENSG00000117707 |
| PRELP | 1.366735477 | 2.578863614 | 0.148996555 | 0.280932124 | 3.072771419 | 7.839431032 | ENSG00000188783 |
| NA | 1.366608205 | 2.578636121 | 0.18019927 | 0.325063037 | 3.048329015 | 7.803937354 | ENSG00000272328 |
| FRMPD2 | 1.365926359 | 2.577417695 | 0.132436577 | 0.256380733 | 3.728761385 | 9.564286502 | ENSG00000170324 |
| NYX | 1.365779639 | 2.577155589 | 0.079686529 | 0.172917078 | 4.316330222 | 11.15470348 | ENSG00000188937 |
| CD27-AS1 | 1.36523452 | 2.576182002 | 4.64E-05 | 0.000268272 | 24.44166231 | 62.86977495 | ENSG00000215039 |
| PCDHGA5 | 1.364206117 | 2.574346264 | 0.032856945 | 0.084563706 | 7.260632091 | 18.80725822 | ENSG00000253485 |
| NABP1 | 1.362734085 | 2.571720909 | 7.38E-06 | 4.98E-05 | 31.43920992 | 80.92846192 | ENSG00000173559 |
| NA | 1.362732483 | 2.571718053 | 0.017625578 | 0.050612537 | 9.857558807 | 25.23964092 | ENSG00000235493 |
| C5orf58 | 1.361521431 | 2.569560163 | 0.075971128 | 0.166389536 | 4.781887194 | 12.17671332 | ENSG00000234511 |
| LOC100507336 | 1.360992462 | 2.568618197 | 0.134796004 | 0.259982974 | 3.36934674 | 8.649290732 | ENSG00000228793 |
| PYGM | 1.36005674 | 2.566952749 | 1.73E-27 | 9.68E-26 | 209.7902389 | 537.79358 | ENSG00000068976 |
| GAPLINC | 1.359936381 | 2.566738607 | 0.264327232 | 0.430125402 | 2.041297112 | 5.187209388 | ENSG00000266835 |
| INPP5D | 1.359273671 | 2.565559831 | 2.97E-35 | 2.48E-33 | 256.1992364 | 657.1493621 | ENSG00000168918 |
| NA | 1.359107823 | 2.565264919 | 0.23607918 | 0.397481288 | 2.296684611 | 5.942220247 | ENSG00000260578 |
| LINC01376 | 1.35887763 | 2.564855645 | 0.025009081 | 0.067617934 | 7.468010661 | 19.01295664 | ENSG00000236204 |
| NA | 1.358141799 | 2.563547801 | 0.06526706 | 0.147431716 | 5.758894886 | 14.74315406 | ENSG00000267364 |
| IRF2BP2 | 1.356690585 | 2.560970412 | 1.54E-51 | 2.69E-49 | 462.4178945 | 1185.747265 | ENSG00000168264 |
| NA | 1.356618963 | 2.560843276 | 0.009340793 | 0.029705447 | 9.743043771 | 24.86698494 | ENSG00000286909 |
| MVP | 1.356127474 | 2.559971012 | 7.79E-05 | 0.00043133 | 28.63318977 | 72.97263313 | ENSG00000013364 |
| NA | 1.355681184 | 2.559179222 | 0.289759404 | 0.459192366 | 1.679091564 | 4.293186317 | ENSG00000235776 |
| NA | 1.355660674 | 2.559142839 | 0.307915849 | 0.479032632 | 1.687464273 | 4.309857869 | ENSG00000286373 |
| PCDHGB8P | 1.353874741 | 2.555976799 | 0.203683552 | 0.356783747 | 2.332290628 | 5.988466793 | ENSG00000248449 |
| PKP3 | 1.352041533 | 2.552731026 | 2.03E-39 | 2.29E-37 | 353.2691465 | 902.7075175 | ENSG00000184363 |
| KLK8 | 1.351886769 | 2.552457199 | 0.027289214 | 0.072812756 | 7.389101641 | 18.84920362 | ENSG00000129455 |
| ACTG2 | 1.351847801 | 2.552388256 | 3.88E-06 | 2.75E-05 | 32.05122116 | 81.76281407 | ENSG00000163017 |
| NA | 1.349013646 | 2.547379045 | 0.326041876 | 0.497852606 | 1.739139985 | 4.385679409 | ENSG00000259635 |
| ERVMER34-1 | 1.348886792 | 2.547155068 | 1.55E-31 | 1.07E-29 | 219.7716366 | 559.5619921 | ENSG00000226887 |
| SERHL | 1.347541093 | 2.544780273 | 0.084703362 | 0.181698271 | 4.428054355 | 11.17352561 | ENSG00000172250 |
| KIAA1257 | 1.34661322 | 2.543144116 | 1.53E-06 | 1.16E-05 | 35.26207414 | 89.92031271 | ENSG00000114656 |
| SERPINB1 | 1.345842043 | 2.54178507 | 1.85E-24 | 8.72E-23 | 194.0243399 | 493.2182558 | ENSG00000021355 |
| DCST1 | 1.345058086 | 2.540404245 | 0.108658452 | 0.220442921 | 3.965288286 | 10.10688676 | ENSG00000163357 |
| UBE2D3-AS1 | 1.344914235 | 2.540150955 | 0.031842338 | 0.082474563 | 7.008035494 | 17.83741361 | ENSG00000246560 |
| NA | 1.34428887 | 2.539050114 | 0.216229137 | 0.373335387 | 2.310639126 | 5.895973701 | ENSG00000204529 |
| LINC00471 | 1.343953003 | 2.538459078 | 0.001983173 | 0.007834813 | 14.81592448 | 37.49836704 | ENSG00000181798 |
| POU6F2 | 1.343880749 | 2.538331949 | 0.004425473 | 0.015786204 | 15.66322332 | 39.50904361 | ENSG00000106536 |
| LINC02615 | 1.343183158 | 2.537104877 | 0.010511016 | 0.032836925 | 10.47794276 | 26.41277291 | ENSG00000251432 |
| ARHGAP24 | 1.342655869 | 2.536177764 | 0.082127046 | 0.177236027 | 4.988389884 | 12.64134146 | ENSG00000138639 |
| TRAF1 | 1.342416317 | 2.53575668 | 1.45E-05 | 9.28E-05 | 29.95912421 | 75.69663563 | ENSG00000056558 |
| NA | 1.341698166 | 2.534494733 | 0.124590463 | 0.245202471 | 3.603082738 | 9.219315408 | ENSG00000237438 |
| NA | 1.341479214 | 2.534110113 | 0.198619678 | 0.350037407 | 2.332290628 | 5.921247548 | ENSG00000245904 |
| CD200R1 | 1.340824807 | 2.5329609 | 0.292664496 | 0.462315001 | 1.709115775 | 4.291035743 | ENSG00000163606 |
| MIR34AHG | 1.338620697 | 2.529094066 | 5.12E-12 | 8.06E-11 | 79.07195447 | 199.7046058 | ENSG00000228526 |
| MAMLD1 | 1.338609386 | 2.529074237 | 0.016832451 | 0.048690182 | 9.321465523 | 23.6048675 | ENSG00000013619 |
| HES4 | 1.338543185 | 2.528958188 | 0.000491309 | 0.002254481 | 20.06528367 | 50.68015818 | ENSG00000188290 |
| MAP1LC3C | 1.338170883 | 2.528305649 | 0.015422773 | 0.045298524 | 11.55618669 | 29.07223978 | ENSG00000197769 |
| NA | 1.338165319 | 2.528295899 | 0.223371646 | 0.381907508 | 2.623284143 | 6.710134549 | ENSG00000227107 |
| SOX3 | 1.33772018 | 2.527515921 | 8.02E-52 | 1.42E-49 | 385.999961 | 975.9422606 | ENSG00000134595 |
| CLIC5 | 1.336768127 | 2.525848531 | 0.206075619 | 0.360156258 | 2.335081531 | 5.900274849 | ENSG00000112782 |
| HACD4 | 1.335638538 | 2.523871638 | 4.80E-05 | 0.000277319 | 25.47871842 | 64.26874191 | ENSG00000188921 |
| NA | 1.335181558 | 2.523072318 | 0.000431864 | 0.002016235 | 19.3254786 | 48.79101665 | ENSG00000287315 |
| TACR1 | 1.33484203 | 2.5224786 | 0.063359995 | 0.144068559 | 5.323362124 | 13.4517342 | ENSG00000115353 |
| ATP10B | 1.334605971 | 2.522065897 | 0.070548591 | 0.157081355 | 5.818943307 | 14.56031845 | ENSG00000118322 |
| LMCD1 | 1.334593869 | 2.522044741 | 1.23E-23 | 5.45E-22 | 172.0584314 | 434.2943461 | ENSG00000071282 |
| NR5A1 | 1.334013618 | 2.52103058 | 0.06377905 | 0.144816578 | 5.287756108 | 13.37591266 | ENSG00000136931 |
| FZD10 | 1.332710898 | 2.518755175 | 0.021830625 | 0.060386377 | 10.91626642 | 27.24172731 | ENSG00000111432 |
| LOC107986163 | 1.331588159 | 2.516795783 | 0.052280988 | 0.123445844 | 6.324812221 | 15.99451863 | ENSG00000272970 |
| ZNF124 | 1.329126517 | 2.512505087 | 3.77E-19 | 1.17E-17 | 161.4944462 | 406.6476553 | ENSG00000196418 |
| APCDD1L-DT | 1.328025566 | 2.510588472 | 0.160842984 | 0.298090307 | 3.042747209 | 7.61437745 | ENSG00000231290 |
| S100A14 | 1.327519885 | 2.509708636 | 0.000257995 | 0.001270285 | 24.01382654 | 60.28477255 | ENSG00000189334 |
| MEIS3 | 1.327428827 | 2.509550237 | 3.04E-72 | 1.03E-69 | 640.1458977 | 1605.749073 | ENSG00000105419 |
| LOC101929551 | 1.327178774 | 2.509115312 | 0.061691516 | 0.14105476 | 5.399480241 | 13.49583017 | ENSG00000226506 |
| LOC101928222 | 1.326616974 | 2.508138426 | 0.096169846 | 0.20095456 | 4.68902366 | 11.7290293 | ENSG00000227210 |
| TAS2R5 | 1.324873084 | 2.50510849 | 0.125261107 | 0.246045414 | 3.638688754 | 9.145644441 | ENSG00000127366 |
| ARHGAP9 | 1.324586215 | 2.504610417 | 4.90E-06 | 3.41E-05 | 32.30660866 | 81.01398236 | ENSG00000123329 |
| LIF | 1.324229717 | 2.50399159 | 1.66E-13 | 3.06E-12 | 85.05812767 | 213.173805 | ENSG00000128342 |
| PLIN4 | 1.323638207 | 2.502965156 | 0.000118114 | 0.000628043 | 27.57918808 | 69.07443256 | ENSG00000167676 |
| NA | 1.323461006 | 2.502657744 | 1.71E-06 | 1.29E-05 | 38.27076693 | 95.60549334 | ENSG00000266869 |
| RTL9 | 1.322715315 | 2.50136452 | 0.089047937 | 0.189187637 | 4.340772627 | 10.87049991 | ENSG00000243978 |
| NA | 1.321988381 | 2.500104469 | 0.014834181 | 0.043882399 | 9.070984109 | 22.61189962 | ENSG00000228192 |
| LOC100129434 | 1.321959335 | 2.500054136 | 0.171274166 | 0.313019775 | 3.045538112 | 7.578883773 | ENSG00000233251 |
| NA | 1.321633304 | 2.499489219 | 0.020849055 | 0.058145812 | 8.382179029 | 20.88919472 | ENSG00000286207 |
| FAM183A | 1.321366822 | 2.499027578 | 0.044882139 | 0.109472922 | 6.300369816 | 15.81168302 | ENSG00000186973 |
| DANT2 | 1.32076466 | 2.497984733 | 0.060860699 | 0.139553517 | 6.368115224 | 15.92084766 | ENSG00000235244 |
| COLQ | 1.320657826 | 2.497799761 | 0.047987675 | 0.115577979 | 6.585781526 | 16.58981715 | ENSG00000206561 |
| WNT9B | 1.320620694 | 2.497735473 | 0.0280544 | 0.074432325 | 7.590022525 | 19.06995605 | ENSG00000158955 |
| NA | 1.320338294 | 2.497246603 | 2.02E-07 | 1.78E-06 | 42.79850593 | 106.8914002 | ENSG00000259820 |

| | | | | | | | |
|---|---|---|---|---|---|---|---|
| PPT2-EGFL8 | 1.319935439 | 2.496549374 | 0.096355787 | 0.201255898 | 4.307957513 | 10.78230797 | ENSG00000258388 |
| BBC3 | 1.319123465 | 2.495144668 | 1.09E-37 | 1.06E-35 | 282.3703145 | 704.3539904 | ENSG00000105327 |
| GRHL3 | 1.318474823 | 2.494023093 | 0.011394161 | 0.035230769 | 10.97332378 | 27.38046695 | ENSG00000158055 |
| PTH1R | 1.31846018 | 2.493997779 | 0.014110722 | 0.042079136 | 10.34177622 | 25.77391145 | ENSG00000160801 |
| PDGFB | 1.318443446 | 2.49396885 | 1.69E-18 | 5.03E-17 | 127.8840671 | 318.838842 | ENSG00000100311 |
| VIM | 1.318303998 | 2.4937278 | 1.47E-23 | 6.44E-22 | 212.9997276 | 531.8287147 | ENSG00000026025 |
| OVGP1 | 1.318215708 | 2.493575193 | 0.01741026 | 0.050091174 | 9.005353882 | 22.51966699 | ENSG00000085465 |
| SPATA12 | 1.318140549 | 2.493445291 | 0.124670078 | 0.245295666 | 3.72317958 | 9.242438681 | ENSG00000186451 |
| S100A11 | 1.31730013 | 2.491993197 | 3.02E-82 | 1.36E-79 | 1194.75104 | 2976.996805 | ENSG00000163191 |
| MEF2C-AS1 | 1.317186535 | 2.491796989 | 0.030382937 | 0.07936591 | 9.977655649 | 24.61073883 | ENSG00000248309 |
| KIAA1107 | 1.317027864 | 2.491522952 | 7.11E-06 | 4.82E-05 | 37.63276168 | 94.04302171 | ENSG00000189195 |
| CEBPZOS | 1.315692924 | 2.489218586 | 0.007814609 | 0.025629312 | 11.35159902 | 28.27690106 | ENSG00000218739 |
| NA | 1.315506538 | 2.488897018 | 1.75E-05 | 0.000109813 | 30.13002089 | 74.85074921 | ENSG00000237886 |
| RAB37 | 1.315101252 | 2.488197928 | 2.07E-09 | 2.42E-08 | 69.63653749 | 173.4784366 | ENSG00000172794 |
| NA | 1.314124174 | 2.486513345 | 0.076756738 | 0.167748298 | 5.380619642 | 13.31031095 | ENSG00000285130 |
| PNLIPRP2 | 1.313727347 | 2.485829499 | 0.073790471 | 0.162721991 | 6.820393404 | 16.78556831 | ENSG00000266200 |
| NA | 1.313709223 | 2.485798272 | 0.20403699 | 0.357240658 | 3.00435029 | 7.450363966 | ENSG00000249894 |
| NA | 1.313430569 | 2.48531819 | 0.036744525 | 0.092860277 | 7.051338498 | 17.51341521 | ENSG00000287750 |
| SAMSN1 | 1.312595852 | 2.483880647 | 0.039438307 | 0.098588101 | 6.790369193 | 16.74765149 | ENSG00000155307 |
| STAT6 | 1.311854995 | 2.482605444 | 1.70E-88 | 9.92E-86 | 1011.431859 | 2509.530845 | ENSG00000166888 |
| KCNN4 | 1.311240761 | 2.481548689 | 0.013108458 | 0.039561212 | 10.0912948 | 24.97399901 | ENSG00000104783 |
| PADI4 | 1.310631946 | 2.4805017 | 0.197554072 | 0.348611015 | 2.634447755 | 6.581614742 | ENSG00000159339 |
| SOX14 | 1.309972804 | 2.479368661 | 0.279536268 | 0.447438943 | 2.356733032 | 5.807781757 | ENSG00000168875 |
| NA | 1.308339187 | 2.476562769 | 0.275138857 | 0.441940357 | 2.299475514 | 5.767714358 | ENSG00000249199 |
| LOC101927708 | 1.307825009 | 2.475680276 | 0.251423481 | 0.415649188 | 2.008481999 | 4.983128505 | ENSG00000255367 |
| WASIR2 | 1.307502206 | 2.475126406 | 0.035671618 | 0.090556808 | 7.026896093 | 17.37467558 | ENSG00000231439 |
| C11orf86 | 1.307220857 | 2.474643762 | 0.008271737 | 0.026859127 | 11.62094104 | 28.82111885 | ENSG00000173237 |
| NA | 1.307178991 | 2.474571951 | 0.040991638 | 0.101641347 | 6.67864506 | 16.56884445 | ENSG00000259972 |
| NA | 1.304607746 | 2.47016557 | 0.211301016 | 0.366994498 | 3.007141193 | 7.397665699 | ENSG00000271959 |
| TTC39C | 1.304119939 | 2.469330493 | 4.39E-43 | 5.69E-41 | 339.3529972 | 838.017383 | ENSG00000168234 |
| LINC00173 | 1.302714025 | 2.466925291 | 0.002635398 | 0.010035423 | 16.57739169 | 40.68727025 | ENSG00000196668 |
| ADCY10P1 | 1.30237696 | 2.466348996 | 1.66E-09 | 1.97E-08 | 66.11180019 | 162.9270703 | ENSG00000161912 |
| NA | 1.302289013 | 2.466198651 | 0.210345304 | 0.365927609 | 2.919859464 | 7.301131924 | ENSG00000287595 |
| NA | 1.301876584 | 2.46549373 | 0.008676614 | 0.027961734 | 12.54069121 | 31.0444786 | ENSG00000287625 |
| TRPC6 | 1.301705452 | 2.465201292 | 0.11904809 | 0.236978299 | 5.144092742 | 12.58513555 | ENSG00000137672 |
| NA | 1.300751687 | 2.463572086 | 0.000207289 | 0.001041901 | 23.02633095 | 56.74497955 | ENSG00000224063 |
| NA | 1.300714194 | 2.463508064 | 0.262133105 | 0.427637187 | 2.038506209 | 5.004101204 | ENSG00000236349 |
| OVCH2 | 1.29987349 | 2.462072918 | 0.114524211 | 0.230086082 | 4.277933303 | 10.62877478 | ENSG00000183378 |
| KLK7 | 1.298335458 | 2.459449543 | 0.00249843 | 0.009591487 | 15.17813003 | 37.19964249 | ENSG00000169035 |
| FOSL1 | 1.298004316 | 2.45888509 | 3.21E-20 | 1.12E-18 | 162.026346 | 397.8451039 | ENSG00000175592 |
| CSRP1 | 1.297785502 | 2.458512179 | 0.033060097 | 0.084967003 | 7.389101641 | 18.14635799 | ENSG00000159176 |
| CRYBB1 | 1.297557507 | 2.458123681 | 6.38E-06 | 4.36E-05 | 32.46510242 | 79.73089222 | ENSG00000100122 |
| ARMCX3 | 1.296523586 | 2.456362675 | 4.71E-45 | 6.54E-43 | 380.1253118 | 934.1334785 | ENSG00000102401 |
| NA | 1.29628953 | 2.455964198 | 0.087024604 | 0.185645431 | 4.939505075 | 12.22726101 | ENSG00000226421 |
| KLHDC7A | 1.295650809 | 2.454877116 | 4.47E-53 | 8.13E-51 | 536.6664986 | 1317.669773 | ENSG00000179023 |
| GPR179 | 1.294871914 | 2.453552112 | 0.160364863 | 0.29766175 | 3.360974032 | 8.230648672 | ENSG00000277399 |
| C16orf89 | 1.293815293 | 2.451755805 | 5.21E-07 | 4.29E-06 | 45.59423835 | 111.6373958 | ENSG00000153446 |
| KANK3 | 1.293011336 | 2.450389918 | 2.31E-07 | 2.01E-06 | 47.04083321 | 115.468898 | ENSG00000186994 |
| DEUP1 | 1.292609404 | 2.449707339 | 0.192084034 | 0.341237048 | 2.686123467 | 6.56494319 | ENSG00000165325 |
| NA | 1.291046637 | 2.447055185 | 0.28157068 | 0.449985916 | 1.978457788 | 4.852458124 | ENSG00000213433 |
| NA | 1.290742377 | 2.446539164 | 2.18E-22 | 8.73E-21 | 219.3057162 | 535.9339829 | ENSG00000227671 |
| NA | 1.290570559 | 2.44624781 | 0.002187774 | 0.008534319 | 18.05747739 | 44.2025829 | ENSG00000282718 |
| CPZ | 1.289978195 | 2.445243597 | 0.01272423 | 0.038570571 | 10.96495107 | 26.8626075 | ENSG00000109625 |
| TAPBPL | 1.288520601 | 2.442774349 | 0.026313782 | 0.070581087 | 10.3033793 | 25.27890271 | ENSG00000139192 |
| SLC30A1 | 1.287751938 | 2.441473194 | 3.94E-32 | 2.84E-30 | 508.027578 | 1242.075147 | ENSG00000170385 |
| NFASC | 1.287345687 | 2.440785792 | 1.48E-39 | 1.67E-37 | 329.2546811 | 803.8162782 | ENSG00000163531 |
| NA | 1.286838131 | 2.439927248 | 0.185692405 | 0.332114982 | 2.982698788 | 7.328556344 | ENSG00000286602 |
| TLR3 | 1.285160653 | 2.437091898 | 0.006289119 | 0.021371041 | 12.71246377 | 30.99849252 | ENSG00000164342 |
| FAXDC2 | 1.285130895 | 2.437041629 | 4.01E-06 | 2.84E-05 | 38.6781905 | 94.4700056 | ENSG00000170271 |
| GCC2-AS1 | 1.284925505 | 2.436694703 | 0.00985975 | 0.031058857 | 11.77038637 | 28.61326986 | ENSG00000214184 |
| AKR1D1 | 1.284818671 | 2.436514267 | 0.260662633 | 0.426030394 | 2.038506209 | 4.930430238 | ENSG00000122787 |
| NA | 1.284584417 | 2.436118678 | 0.000648822 | 0.002884708 | 18.96885486 | 46.29688981 | ENSG00000282012 |
| ICAM3 | 1.28434309 | 2.43571121 | 6.42E-10 | 7.98E-09 | 65.85207019 | 160.3122446 | ENSG00000076662 |
| NA | 1.28334437 | 2.434025648 | 0.101277869 | 0.209091543 | 6.202800357 | 14.90933023 | ENSG00000224846 |
| NA | 1.282716981 | 2.432967386 | 0.19474426 | 0.344851433 | 3.012722998 | 7.353830191 | ENSG00000288271 |
| TUBA3E | 1.282692976 | 2.432926903 | 0.215804368 | 0.372835579 | 2.661681062 | 6.463847803 | ENSG00000152086 |
| INPP4B | 1.282632729 | 2.432825306 | 0.071947323 | 0.159459537 | 5.734452481 | 13.92684264 | ENSG00000109452 |
| NA | 1.27961927 | 2.427748997 | 0.049001333 | 0.117441261 | 6.966847672 | 16.93291024 | ENSG00000263503 |
| NA | 1.278830475 | 2.426421986 | 0.084702442 | 0.181698271 | 5.459528662 | 13.13203695 | ENSG00000228242 |
| ART3 | 1.27817709 | 2.425323326 | 0.180266463 | 0.325153841 | 3.012722998 | 7.32640577 | ENSG00000156219 |
| ATP2A3 | 1.277635513 | 2.424413049 | 9.87E-83 | 4.65E-80 | 1039.182006 | 2519.846883 | ENSG00000074370 |
| NA | 1.277439847 | 2.42408426 | 0.01179858 | 0.036150954 | 10.95936927 | 26.6509904 | ENSG00000257681 |
| UHMK1 | 1.27552723 | 2.420872721 | 8.89E-32 | 6.26E-30 | 422.6941247 | 1023.021264 | ENSG00000152332 |
| TMPRSS4 | 1.274542759 | 2.41922132 | 0.2994019 | 0.469954021 | 2.065739517 | 4.941183107 | ENSG00000137648 |
| C4orf50 | 1.273718569 | 2.417839649 | 0.019237432 | 0.054414853 | 11.0089298 | 26.64615621 | ENSG00000181215 |
| NA | 1.273256091 | 2.417064698 | 0.002987049 | 0.011158631 | 14.67447614 | 35.51243127 | ENSG00000154608 |
| LOC105375431 | 1.272435927 | 2.415691002 | 0.063893114 | 0.14497319 | 5.95981577 | 14.45707249 | ENSG00000274993 |

| | | | | | | | |
|---|---|---|---|---|---|---|---|
| OXCT1-AS1 | 1.272266719 | 2.415407692 | 0.211737844 | 0.367551955 | 3.0343745 | 7.324255197 | ENSG00000248668 |
| S100Z | 1.270940985 | 2.413189123 | 0.270065858 | 0.436554421 | 2.623284143 | 6.404958279 | ENSG00000171643 |
| NA | 1.270708212 | 2.412799796 | 2.59E-51 | 4.50E-49 | 956.2019684 | 2305.354632 | ENSG00000254339 |
| KLHL31 | 1.270564149 | 2.412558873 | 0.075519875 | 0.165683373 | 6.13717013 | 14.66330395 | ENSG00000124743 |
| CD300A | 1.270350053 | 2.412200876 | 0.011568443 | 0.035638269 | 11.62863802 | 28.08277954 | ENSG00000167851 |
| NMNAT2 | 1.270140106 | 2.411849869 | 4.31E-42 | 5.37E-40 | 370.2857746 | 892.4884672 | ENSG00000157064 |
| MIR3936HG | 1.269769615 | 2.411230574 | 0.171594705 | 0.313498765 | 3.777646195 | 9.05126124 | ENSG00000233006 |
| MRGPRF-AS1 | 1.269695951 | 2.411107459 | 0.234295856 | 0.395260284 | 2.332290628 | 5.647796846 | ENSG00000256508 |
| NA | 1.269646843 | 2.411025388 | 0.018306622 | 0.052087469 | 9.449259352 | 22.68987173 | ENSG00000286138 |
| CBX7 | 1.269570994 | 2.410898634 | 1.23E-11 | 1.87E-10 | 87.27630358 | 210.0833136 | ENSG00000100307 |
| MYMX | 1.268654218 | 2.409367089 | 0.000320685 | 0.001543881 | 28.58833516 | 68.94455568 | ENSG00000262179 |
| CRABP2 | 1.268542459 | 2.409180454 | 6.50E-102 | 5.70E-99 | 1498.208663 | 3607.715721 | ENSG00000143320 |
| COLCA2 | 1.268017774 | 2.408304433 | 0.203552575 | 0.356608822 | 3.366555838 | 8.054264783 | ENSG00000214290 |
| RASEF | 1.267849219 | 2.408023079 | 1.70E-08 | 1.75E-07 | 86.69220137 | 208.6821413 | ENSG00000165105 |
| NA | 1.267746537 | 2.407851697 | 0.183822677 | 0.329796678 | 2.985489691 | 7.208638832 | ENSG00000245025 |
| NA | 1.265044119 | 2.403345594 | 0.055905591 | 0.130551884 | 7.20145955 | 17.14532085 | ENSG00000246334 |
| RYR2 | 1.263580113 | 2.400907985 | 2.78E-12 | 4.51E-11 | 109.384436 | 262.0841756 | ENSG00000198626 |
| ZP3 | 1.262420909 | 2.398979632 | 0.080296266 | 0.173907631 | 6.828766112 | 16.29539375 | ENSG00000188372 |
| ITPR2 | 1.261686582 | 2.39775887 | 8.89E-54 | 1.67E-51 | 850.575506 | 2038.371152 | ENSG00000123104 |
| LURAP1L | 1.261648664 | 2.397695852 | 0.146117538 | 0.27664672 | 4.104245726 | 9.719970265 | ENSG00000153714 |
| LRRC7 | 1.260417155 | 2.395650011 | 0.022393625 | 0.061704818 | 9.032587189 | 21.61301305 | ENSG00000033122 |
| RASIP1 | 1.25802544 | 2.391681767 | 1.32E-20 | 4.69E-19 | 172.3618647 | 411.6318804 | ENSG00000105538 |
| NA | 1.257083301 | 2.390120412 | 1.24E-05 | 7.98E-05 | 36.71647813 | 87.83543213 | ENSG00000188013 |
| MXRA8 | 1.257036922 | 2.390043575 | 1.17E-21 | 4.45E-20 | 170.478661 | 407.3666515 | ENSG00000162576 |
| ROCK1P1 | 1.256199626 | 2.38865687 | 0.13482748 | 0.260017687 | 3.995312496 | 9.570738223 | ENSG00000263006 |
| STARD13 | 1.255639555 | 2.387729746 | 2.32E-08 | 2.33E-07 | 57.48105477 | 137.3419253 | ENSG00000133121 |
| IFIT1 | 1.254078082 | 2.38514683 | 0.000139834 | 0.000728279 | 28.96325592 | 68.84372076 | ENSG00000185745 |
| ATOH1 | 1.253453892 | 2.384115106 | 0.061169905 | 0.140096005 | 7.198668648 | 16.9705545 | ENSG00000172238 |
| NA | 1.253429924 | 2.3840755 | 0.259491277 | 0.424728171 | 2.329499725 | 5.557454328 | ENSG00000271216 |
| PARD6G-AS1 | 1.253142109 | 2.383599929 | 0.005508284 | 0.019073831 | 15.3085146 | 36.57745258 | ENSG00000267270 |
| PADI2 | 1.252335467 | 2.382267578 | 1.44E-15 | 3.28E-14 | 146.2643139 | 347.5930262 | ENSG00000117115 |
| LOC102724057 | 1.25230708 | 2.382220705 | 0.127691571 | 0.249548992 | 4.776305388 | 11.31871697 | ENSG00000230303 |
| CYP26B1 | 1.252151683 | 2.381964123 | 0.059148337 | 0.136340133 | 8.260167165 | 19.49504983 | ENSG00000003137 |
| DOC2A | 1.252053743 | 2.381802424 | 1.40E-17 | 3.84E-16 | 130.6649691 | 311.1857663 | ENSG00000149927 |
| SNORD14E | 1.251062348 | 2.380166253 | 0.235514114 | 0.396795001 | 3.072771419 | 7.263487673 | ENSG00000200879 |
| NA | 1.250877224 | 2.379860853 | 0.239219678 | 0.401250918 | 2.362314838 | 5.6015503 | ENSG00000255237 |
| SOHLH2 | 1.249912831 | 2.378270528 | 0.097067709 | 0.202393807 | 4.697396369 | 11.14610119 | ENSG00000120669 |
| FLNB-AS1 | 1.249712402 | 2.377940146 | 0.130388217 | 0.253432654 | 4.016963998 | 9.551916097 | ENSG00000244161 |
| SULF1 | 1.24916211 | 2.377033292 | 4.70E-19 | 1.45E-17 | 303.1521859 | 719.3065988 | ENSG00000137573 |
| PPP5D1 | 1.248480501 | 2.375910517 | 0.011273229 | 0.0349016 | 11.39769293 | 27.07985229 | ENSG00000230510 |
| NA | 1.247837892 | 2.374852467 | 0.035424112 | 0.090053972 | 9.23996576 | 21.76035499 | ENSG00000227799 |
| LPL | 1.247309763 | 2.373983261 | 1.52E-14 | 3.12E-13 | 113.898696 | 270.2701953 | ENSG00000175445 |
| KIFAP3 | 1.246887709 | 2.373288865 | 2.40E-08 | 2.41E-07 | 57.2492338 | 136.1521338 | ENSG00000075945 |
| OLFML3 | 1.246816651 | 2.373171974 | 5.87E-76 | 2.36E-73 | 811.3709095 | 1924.605689 | ENSG00000116774 |
| LRRC4C | 1.246178459 | 2.372122409 | 0.028409615 | 0.075188623 | 9.315883717 | 22.17899705 | ENSG00000148948 |
| NA | 1.24525083 | 2.370597663 | 0.144274141 | 0.2739902 | 4.071430613 | 9.610533047 | ENSG00000277715 |
| NA | 1.244370023 | 2.369150787 | 0.070172634 | 0.156424808 | 6.109936822 | 14.3559771 | ENSG00000262703 |
| SRMS | 1.244202893 | 2.368876347 | 0.042276304 | 0.104250565 | 8.384969932 | 19.86959592 | ENSG00000125508 |
| ELK4 | 1.243729157 | 2.368098609 | 1.22E-28 | 7.21E-27 | 318.6530116 | 754.1438446 | ENSG00000158711 |
| NEB | 1.242703119 | 2.366415027 | 0.000224425 | 0.001120441 | 27.87943018 | 65.79436279 | ENSG00000183091 |
| LOC101929494 | 1.24261748 | 2.366274561 | 2.41E-10 | 3.19E-09 | 77.45885655 | 183.7235115 | ENSG00000276170 |
| MSS51 | 1.241709337 | 2.364785515 | 0.020266989 | 0.056851896 | 10.34177622 | 24.52604242 | ENSG00000166343 |
| MIR302C | 1.241571084 | 2.364558909 | 0.244722105 | 0.407856081 | 2.362314838 | 5.553153181 | ENSG00000199102 |
| EFCAB13 | 1.241566896 | 2.364552044 | 0.041242682 | 0.10215878 | 8.003903785 | 18.91024372 | ENSG00000178852 |
| GNMT | 1.238726547 | 2.359901342 | 0.062284568 | 0.142125809 | 6.403721241 | 15.03946757 | ENSG00000124713 |
| CALCR | 1.238333997 | 2.359259311 | 0.000168596 | 0.000863845 | 26.64336821 | 63.02167849 | ENSG00000004948 |
| LEPR | 1.237568478 | 2.358007779 | 4.04E-05 | 0.000236955 | 32.20046633 | 76.13033171 | ENSG00000116678 |
| LINC00908 | 1.237106753 | 2.357253235 | 0.207711533 | 0.362412219 | 3.347695239 | 7.887828151 | ENSG00000266256 |
| NA | 1.235690857 | 2.354940904 | 0.011944786 | 0.036488814 | 13.21083585 | 30.96676695 | ENSG00000285184 |
| PAX5 | 1.235645503 | 2.354866873 | 0.008855918 | 0.028411694 | 12.40193393 | 29.14511724 | ENSG00000196092 |
| PCDHGC5 | 1.234521928 | 2.353033611 | 0.160765755 | 0.297982515 | 3.660340256 | 8.657893027 | ENSG00000240764 |
| TLR5 | 1.234075895 | 2.352306244 | 2.32E-07 | 2.02E-06 | 65.1969561 | 153.2184044 | ENSG00000187554 |
| PPP1R15B | 1.233691423 | 2.351679447 | 3.52E-68 | 1.03E-65 | 1020.432819 | 2398.773568 | ENSG00000158615 |
| CGB2 | 1.232893681 | 2.35037944 | 0.223738863 | 0.382230464 | 2.656099256 | 6.268369215 | ENSG00000104818 |
| NA | 1.232052669 | 2.349009697 | 0.26514199 | 0.431160257 | 2.359523935 | 5.502605488 | ENSG00000249311 |
| METTL27 | 1.231792524 | 2.348586164 | 2.19E-09 | 2.55E-08 | 77.7669958 | 182.3016938 | ENSG00000165171 |
| NA | 1.23138016 | 2.347914967 | 0.00616683 | 0.021011006 | 13.76000777 | 32.26706168 | ENSG00000279393 |
| LOXL4 | 1.231233242 | 2.347675877 | 0.156018952 | 0.291397471 | 4.555648025 | 10.7925278 | ENSG00000138131 |
| ITGAM | 1.230669284 | 2.346758335 | 0.066537853 | 0.149863599 | 6.052679304 | 14.17932064 | ENSG00000169896 |
| LINC02175 | 1.230505838 | 2.346492482 | 0.015414086 | 0.045286802 | 13.19409043 | 30.88743777 | ENSG00000262155 |
| NME9 | 1.230470704 | 2.346435338 | 0.26606359 | 0.432258103 | 2.419572356 | 5.6015503 | ENSG00000181322 |
| NA | 1.227277577 | 2.3412477 | 4.17E-06 | 2.94E-05 | 42.50962759 | 99.54159862 | ENSG00000253320 |
| TTN | 1.225919081 | 2.33904413 | 1.48E-10 | 2.00E-09 | 90.67076844 | 211.6562471 | ENSG00000155657 |
| RNF212 | 1.225379896 | 2.338170112 | 4.32E-08 | 4.19E-07 | 62.90131063 | 147.3686893 | ENSG00000178222 |
| LOC101929705 | 1.225061654 | 2.337654396 | 0.01547993 | 0.045424906 | 10.95936927 | 25.71448889 | ENSG00000237686 |
| NA | 1.224729242 | 2.337115837 | 0.230253544 | 0.390533268 | 3.097213824 | 7.137118439 | ENSG00000235092 |
| PPARG | 1.224315669 | 2.33644596 | 0.176602623 | 0.320161752 | 3.630316045 | 8.522921499 | ENSG00000132170 |

| | | | | | | | |
|---|---|---|---|---|---|---|---|
| CRLF1 | 1.223937997 | 2.335834399 | 4.96E-15 | 1.07E-13 | 131.3579165 | 306.835841 | ENSG00000006016 |
| VLDLR-AS1 | 1.223729083 | 2.335496176 | 1.73E-05 | 0.000109118 | 38.2239973 | 89.10265632 | ENSG00000236404 |
| GABRB2 | 1.223483154 | 2.335098089 | 0.027621123 | 0.073586394 | 8.953678169 | 20.96716684 | ENSG00000145864 |
| EWSAT1 | 1.22329496 | 2.334793505 | 0.127631729 | 0.249491771 | 4.631766142 | 10.86619876 | ENSG00000212766 |
| RHBDL1 | 1.222602154 | 2.333672568 | 1.47E-07 | 1.32E-06 | 63.65662182 | 148.2495606 | ENSG00000103269 |
| FLVCR1-DT | 1.221698322 | 2.332211007 | 1.31E-22 | 5.37E-21 | 186.2820811 | 434.2561504 | ENSG00000198468 |
| NA | 1.221652839 | 2.332137482 | 0.013269491 | 0.039959686 | 13.47738696 | 31.28190259 | ENSG00000181097 |
| NA | 1.221310554 | 2.331584238 | 0.357038541 | 0.530388919 | 1.95959719 | 4.610732991 | ENSG00000203392 |
| MYCBPAP | 1.220297843 | 2.329948139 | 0.008832226 | 0.028357663 | 13.11239051 | 30.44756254 | ENSG00000136449 |
| LINC01291 | 1.2197563 | 2.329073713 | 0.152617733 | 0.286563041 | 4.101454824 | 9.44436899 | ENSG00000204792 |
| NA | 1.219041835 | 2.327920573 | 0.052018146 | 0.123036053 | 7.590022525 | 17.77019436 | ENSG00000272316 |
| NEU4 | 1.218368994 | 2.326835135 | 0.065111438 | 0.147131868 | 6.406512144 | 14.8714134 | ENSG00000204099 |
| LRRN4 | 1.218366592 | 2.32683126 | 0.017773758 | 0.050939468 | 11.62584712 | 27.07744125 | ENSG00000125872 |
| ADAMTSL2 | 1.218007371 | 2.326251968 | 3.52E-05 | 0.000209611 | 38.1618337 | 88.81629007 | ENSG00000197859 |
| NA | 1.216639571 | 2.324047525 | 0.121359168 | 0.24048174 | 5.045647402 | 11.65912645 | ENSG00000276603 |
| PDZD2 | 1.216297204 | 2.323496068 | 4.28E-29 | 2.60E-27 | 278.3173811 | 645.8774432 | ENSG00000133401 |
| ACAP1 | 1.215664668 | 2.322477577 | 4.90E-49 | 7.68E-47 | 557.5110092 | 1293.649977 | ENSG00000072818 |
| ACER2 | 1.215348848 | 2.321969219 | 4.44E-11 | 6.33E-10 | 83.49076024 | 193.973439 | ENSG00000177076 |
| HMCN1 | 1.215303325 | 2.321895953 | 3.05E-15 | 6.72E-14 | 243.4954704 | 564.4251542 | ENSG00000143341 |
| AATBC | 1.214953102 | 2.321332367 | 0.069304705 | 0.154806963 | 6.733111675 | 15.62642426 | ENSG00000215458 |
| OXCT2P1 | 1.214106297 | 2.319970237 | 0.015871498 | 0.046411561 | 12.18705853 | 28.09138184 | ENSG00000237624 |
| NA | 1.213833719 | 2.31953195 | 0.32734489 | 0.499415336 | 1.984039594 | 4.629555116 | ENSG00000260459 |
| HEPHL1 | 1.212839779 | 2.317934468 | 0.256378243 | 0.421525713 | 2.95546548 | 6.895393306 | ENSG00000181333 |
| YPEL4 | 1.211968895 | 2.316535666 | 0.002284452 | 0.00885471 | 20.64583124 | 47.88032203 | ENSG00000166793 |
| TIGD6 | 1.2117913 | 2.316250519 | 6.64E-05 | 0.000372975 | 30.49992342 | 70.53685066 | ENSG00000164296 |
| TNNC1 | 1.21124532 | 2.315374113 | 0.000133534 | 0.000699053 | 29.02242846 | 67.2264124 | ENSG00000114854 |
| CPEB4 | 1.21052822 | 2.314223529 | 1.08E-17 | 3.00E-16 | 152.6060348 | 353.1542487 | ENSG00000113742 |
| EDA2R | 1.210025184 | 2.313416751 | 1.32E-14 | 2.73E-13 | 121.7962573 | 281.9941299 | ENSG00000131080 |
| NA | 1.209906308 | 2.313226137 | 0.172372928 | 0.314592825 | 3.725970483 | 8.579920913 | ENSG00000226823 |
| ADARB2 | 1.209896576 | 2.313210532 | 1.73E-06 | 1.30E-05 | 63.5425071 | 146.3913206 | ENSG00000185736 |
| LINC01275 | 1.209224883 | 2.312133794 | 0.200451878 | 0.352338695 | 3.396580048 | 7.776512934 | ENSG00000237595 |
| ASCL5 | 1.208887868 | 2.31159374 | 0.203071471 | 0.355905492 | 4.191527455 | 9.549765524 | ENSG00000232237 |
| MKLN1-AS | 1.2086622 | 2.311232187 | 0.000486787 | 0.00223799 | 24.19935345 | 56.11365432 | ENSG00000236753 |
| TRIM22 | 1.208649247 | 2.311211436 | 3.09E-32 | 2.24E-30 | 457.3160514 | 1055.805974 | ENSG00000132274 |
| PLD1 | 1.207094471 | 2.308722011 | 0.000824297 | 0.003574175 | 22.74650105 | 52.58004047 | ENSG00000075651 |
| ELAVL2 | 1.206581875 | 2.307901857 | 9.19E-25 | 4.49E-23 | 274.7126218 | 634.6095159 | ENSG00000107105 |
| SPATA32 | 1.204955634 | 2.305301799 | 0.21170899 | 0.367551955 | 3.33932253 | 7.742897257 | ENSG00000184361 |
| MIR1281 | 1.20440381 | 2.304420201 | 0.314420429 | 0.485549511 | 1.984039594 | 4.591910865 | ENSG00000284015 |
| LOC100507053 | 1.203964103 | 2.303717962 | 0.000183739 | 0.000933009 | 29.56689445 | 68.05697222 | ENSG00000246090 |
| SUCLG2 | 1.203418048 | 2.30284618 | 4.66E-50 | 7.75E-48 | 675.7860838 | 1555.482946 | ENSG00000172340 |
| NA | 1.202511777 | 2.301400033 | 0.308132149 | 0.479175919 | 2.446805664 | 5.525728761 | ENSG00000251161 |
| SPATA31E1 | 1.202330504 | 2.301110883 | 0.315297192 | 0.486545151 | 1.984039594 | 4.58330857 | ENSG00000177992 |
| NA | 1.201634708 | 2.30000135 | 0.072524318 | 0.160491096 | 6.724738966 | 15.42476653 | ENSG00000255145 |
| HTR1D | 1.199860164 | 2.297174041 | 0.000131658 | 0.00068998 | 29.09017387 | 66.74862035 | ENSG00000179546 |
| SPSB2 | 1.199518456 | 2.29663001 | 2.63E-12 | 4.29E-11 | 114.998079 | 264.638016 | ENSG00000111671 |
| CACNA2D4 | 1.197489647 | 2.293402614 | 0.141093235 | 0.26948808 | 5.396689338 | 12.30120455 | ENSG00000151062 |
| SDCBP2-AS1 | 1.197405194 | 2.293268366 | 0.020689758 | 0.057785741 | 10.95099656 | 25.15710719 | ENSG00000234684 |
| NEFH | 1.197184873 | 2.292918177 | 3.80E-13 | 6.78E-12 | 119.6164031 | 274.1114148 | ENSG00000100285 |
| NA | 1.19674447 | 2.292218339 | 0.027678042 | 0.073663641 | 10.99497528 | 25.30686016 | ENSG00000251615 |
| C1QTNF12 | 1.195954051 | 2.29096283 | 0.001275542 | 0.005312281 | 23.95935992 | 54.92709795 | ENSG00000184163 |
| CST1 | 1.194113951 | 2.28804266 | 0.000120267 | 0.00063773 | 30.81815024 | 70.49301515 | ENSG00000170373 |
| RRNAD1 | 1.193850373 | 2.287624676 | 1.40E-24 | 6.63E-23 | 245.4255942 | 561.1593817 | ENSG00000143303 |
| MSH2-OT1 | 1.193761921 | 2.287484425 | 0.072724195 | 0.160885601 | 6.324812221 | 14.50762018 | ENSG00000235760 |
| YOD1 | 1.193123493 | 2.286472382 | 3.57E-17 | 9.55E-16 | 154.1517508 | 352.780484 | ENSG00000180667 |
| INTS6-AS1 | 1.19292075 | 2.286151085 | 0.109680946 | 0.222076015 | 5.559889024 | 12.82686068 | ENSG00000236778 |
| NA | 1.192529437 | 2.285531079 | 0.223042342 | 0.381547381 | 3.774855292 | 8.548728383 | ENSG00000286216 |
| C10orf55 | 1.192520718 | 2.285517267 | 0.333212871 | 0.505846687 | 2.280614915 | 5.263030928 | ENSG00000222047 |
| SPDYC | 1.191552324 | 2.283983651 | 0.223024831 | 0.381547381 | 3.007141193 | 6.859366591 | ENSG00000204710 |
| CDIPTOSP | 1.191347058 | 2.28365871 | 0.310771295 | 0.481688017 | 2.011272902 | 4.58330857 | ENSG00000214725 |
| NA | 1.189518106 | 2.280765476 | 0.19911807 | 0.350591233 | 3.309298319 | 7.589103603 | ENSG00000226070 |
| NA | 1.188995633 | 2.279939644 | 0.059874092 | 0.137684004 | 7.086944514 | 16.08728429 | ENSG00000225518 |
| PHYHD1 | 1.188990119 | 2.27993093 | 2.25E-11 | 3.31E-10 | 95.27530128 | 216.9120207 | ENSG00000175287 |
| LINC00346 | 1.188976558 | 2.2799095 | 0.285391261 | 0.454435231 | 2.332290628 | 5.346921724 | ENSG00000255874 |
| NA | 1.188313809 | 2.278862389 | 3.24E-17 | 8.69E-16 | 189.1922164 | 431.2333807 | ENSG00000258649 |
| LACTB2 | 1.187558063 | 2.277668935 | 2.19E-06 | 1.62E-05 | 47.27803583 | 107.7932321 | ENSG00000147592 |
| C2orf80 | 1.187512898 | 2.277597631 | 0.320048537 | 0.491508852 | 2.065739517 | 4.661280684 | ENSG00000188674 |
| METTL11B | 1.187177236 | 2.277067781 | 0.098282674 | 0.204351267 | 5.375037836 | 12.23156216 | ENSG00000203740 |
| TUBA3D | 1.186797788 | 2.27646896 | 0.149506311 | 0.281659566 | 5.462319565 | 12.38750639 | ENSG00000075886 |
| GRIK2 | 1.18668559 | 2.276291926 | 0.000252898 | 0.001248056 | 29.33997956 | 66.80561976 | ENSG00000164418 |
| ALOX12P2 | 1.186349241 | 2.275761294 | 4.48E-07 | 3.73E-06 | 56.88857985 | 129.4922744 | ENSG00000262943 |
| NA | 1.185934518 | 2.27510719 | 0.022767995 | 0.062584409 | 16.75925181 | 38.35394025 | ENSG00000253878 |
| PAH | 1.185849037 | 2.274972391 | 0.000612272 | 0.002741087 | 24.79617087 | 56.30483176 | ENSG00000171759 |
| CDK10 | 1.18511802 | 2.273819949 | 1.11E-25 | 5.68E-24 | 265.9576741 | 604.2297788 | ENSG00000185324 |
| PLAAT5 | 1.18486438 | 2.273420225 | 3.89E-09 | 4.39E-08 | 73.06748425 | 165.8229193 | ENSG00000168004 |
| POLD4 | 1.183163033 | 2.270740797 | 0.00026368 | 0.001295301 | 28.4069506 | 64.44352037 | ENSG00000175482 |
| NA | 1.181528619 | 2.268169756 | 0.001031507 | 0.004380102 | 23.19810351 | 52.46631421 | ENSG00000228956 |
| LOC105369519 | 1.18146476 | 2.268069359 | 0.328842264 | 0.501083221 | 2.005691096 | 4.532760877 | ENSG00000255422 |

| | | | | | | | |
|---|---|---|---|---|---|---|---|
| ANXA2 | 1.181122453 | 2.26753128 | 3.81E-72 | 1.27E-69 | 859.2970382 | 1947.686833 | ENSG00000182718 |
| FAM95C | 1.181090282 | 2.267480716 | 0.104512776 | 0.214188458 | 5.317780318 | 12.0653981 | ENSG00000283486 |
| LMO7 | 1.181076883 | 2.267459658 | 9.25E-25 | 4.51E-23 | 236.0315652 | 534.9399368 | ENSG00000136153 |
| LOC105370941 | 1.180950216 | 2.267260587 | 0.130038168 | 0.252905162 | 4.642929753 | 10.58037766 | ENSG00000261136 |
| VCX3A | 1.180388827 | 2.266378509 | 0.308733468 | 0.479762956 | 2.299475514 | 5.245826338 | ENSG00000169059 |
| GPR20 | 1.179484589 | 2.264958456 | 0.002896514 | 0.010875175 | 19.95164451 | 44.9697037 | ENSG00000204882 |
| NA | 1.179250234 | 2.264590562 | 0.091925371 | 0.193849184 | 6.324812221 | 14.31564924 | ENSG00000259209 |
| NA | 1.177950694 | 2.262551599 | 0.09164619 | 0.193386117 | 7.236189686 | 16.4779689 | ENSG00000231940 |
| ACHE | 1.177076661 | 2.261181284 | 4.85E-08 | 4.67E-07 | 81.31955411 | 184.1558505 | ENSG00000087085 |
| NA | 1.175726651 | 2.259066361 | 0.000195727 | 0.000988682 | 31.03390152 | 69.80926422 | ENSG00000250903 |
| DHRS12 | 1.175656337 | 2.258956261 | 0.000156662 | 0.000806773 | 29.04128906 | 65.6209414 | ENSG00000102796 |
| NA | 1.175280641 | 2.258368078 | 0.09614278 | 0.200941536 | 6.877650922 | 15.35109556 | ENSG00000256967 |
| FAM178B | 1.175189367 | 2.258225204 | 0.199590835 | 0.351049125 | 3.36934674 | 7.591254177 | ENSG00000168754 |
| FZD5 | 1.173878101 | 2.256173634 | 2.20E-112 | 2.50E-109 | 1651.262559 | 3725.686592 | ENSG00000163251 |
| NA | 1.172534996 | 2.254074183 | 0.020671834 | 0.057760768 | 12.49758837 | 27.99888874 | ENSG00000266389 |
| MAP4K3-DT | 1.17221978 | 2.253581742 | 2.62E-07 | 2.26E-06 | 62.97128337 | 141.762559 | ENSG00000231312 |
| NA | 1.172114842 | 2.253417828 | 0.178620684 | 0.322849397 | 3.633106948 | 8.237100393 | ENSG00000234975 |
| PCDHA13 | 1.171997656 | 2.253234797 | 0.18207529 | 0.327558932 | 4.86637802 | 10.81403353 | ENSG00000239389 |
| NA | 1.169671995 | 2.249605451 | 0.206763486 | 0.36117962 | 3.981357982 | 9.009315842 | ENSG00000267745 |
| CLTRN | 1.169407158 | 2.249192526 | 0.047102691 | 0.113909741 | 8.564239314 | 19.38105101 | ENSG00000147003 |
| EIF2AK3-DT | 1.168913814 | 2.248423524 | 0.016148436 | 0.047057324 | 12.38586423 | 27.80125958 | ENSG00000234028 |
| KRBA2 | 1.16869716 | 2.248085896 | 0.02187455 | 0.060489958 | 10.95936927 | 24.7301233 | ENSG00000184619 |
| SFR1 | 1.16855174 | 2.247859306 | 3.20E-06 | 2.29E-05 | 47.77361702 | 107.5095495 | ENSG00000156384 |
| EXOC3L2 | 1.167690103 | 2.246517193 | 8.51E-07 | 6.78E-06 | 51.52682081 | 115.7291848 | ENSG00000283632 |
| AFDN-DT | 1.167623622 | 2.246413674 | 0.001486247 | 0.006074728 | 27.94294523 | 62.59040556 | ENSG00000198221 |
| RBMS2 | 1.167072843 | 2.245556221 | 7.23E-17 | 1.88E-15 | 165.4810226 | 372.046458 | ENSG00000076067 |
| OPCML | 1.166631887 | 2.244869978 | 0.000139201 | 0.000725962 | 37.93280363 | 85.34398889 | ENSG00000183715 |
| MIXL1 | 1.166329381 | 2.24439932 | 0.010322957 | 0.032286017 | 22.02535642 | 49.68368265 | ENSG00000185155 |
| TNNI1 | 1.166048314 | 2.243962107 | 0.000450352 | 0.002089405 | 28.13740842 | 63.38306067 | ENSG00000159173 |
| NA | 1.163912665 | 2.240642785 | 0.261505162 | 0.426881063 | 2.631656852 | 5.946521394 | ENSG00000233337 |
| CD22 | 1.163768496 | 2.240418888 | 0.199623847 | 0.351075178 | 4.079803322 | 9.051000776 | ENSG00000012124 |
| P4HA2 | 1.163436587 | 2.239903512 | 1.64E-14 | 3.35E-13 | 146.1570956 | 327.5050647 | ENSG00000072682 |
| NA | 1.162961705 | 2.239166341 | 0.062175212 | 0.141926638 | 8.913366228 | 19.78947323 | ENSG00000286811 |
| ARHGAP31-AS1 | 1.162254043 | 2.238068267 | 0.040503845 | 0.100694695 | 8.953678169 | 20.13014318 | ENSG00000241155 |
| LOC100506071 | 1.159599986 | 2.233954785 | 0.319008953 | 0.490475947 | 2.302266417 | 5.153333246 | ENSG00000258525 |
| ELAVL4 | 1.157380927 | 2.230521303 | 0.066214474 | 0.149309576 | 7.728979966 | 17.23781394 | ENSG00000162374 |
| ACP5 | 1.156766885 | 2.229572147 | 1.65E-09 | 1.96E-08 | 96.45478146 | 214.4977866 | ENSG00000102575 |
| CA9 | 1.154831706 | 2.226583484 | 0.077369478 | 0.168853295 | 7.410753143 | 16.4569962 | ENSG00000107159 |
| NA | 1.154241823 | 2.225673276 | 4.91E-05 | 0.000282653 | 42.70305165 | 94.80585347 | ENSG00000237978 |
| DACT1 | 1.15409348 | 2.225444435 | 2.90E-35 | 2.43E-33 | 401.3378241 | 893.4943754 | ENSG00000165617 |
| SNORA71B | 1.153155834 | 2.223998529 | 0.270746355 | 0.437418346 | 2.634447755 | 5.891672554 | ENSG00000235408 |
| SNORA33 | 1.153124514 | 2.223950248 | 0.038314373 | 0.096250971 | 11.59390789 | 25.59510442 | ENSG00000200534 |
| NA | 1.152554992 | 2.223072489 | 0.262749784 | 0.428353425 | 3.344904336 | 7.40411742 | ENSG00000285668 |
| SPATA17 | 1.152052364 | 2.222298116 | 0.16747947 | 0.307523935 | 4.675744867 | 10.36151533 | ENSG00000162814 |
| GBP1P1 | 1.152025775 | 2.222257159 | 0.200211888 | 0.351948927 | 3.725970483 | 8.24140154 | ENSG00000225492 |
| HPX | 1.151667582 | 2.221705485 | 0.298920534 | 0.469353412 | 3.021095707 | 6.733257821 | ENSG00000110169 |
| HPD | 1.151135288 | 2.22088592 | 0.358249329 | 0.531369199 | 2.092972825 | 4.564486445 | ENSG00000158104 |
| SYT5 | 1.15088033 | 2.220493472 | 0.000152323 | 0.000786507 | 36.62784496 | 81.52752855 | ENSG00000129990 |
| NA | 1.150722403 | 2.220250415 | 0.000351584 | 0.001678397 | 31.47673095 | 70.07573017 | ENSG00000268903 |
| LINC01771 | 1.150369512 | 2.219707397 | 0.306975179 | 0.477916083 | 2.604423544 | 5.820152161 | ENSG00000225675 |
| TENT5C | 1.149776072 | 2.218794527 | 0.00106019 | 0.004492013 | 23.9197237 | 52.89732668 | ENSG00000183508 |
| FKBP6 | 1.149092479 | 2.217743444 | 0.286694655 | 0.455908547 | 2.637238657 | 5.872850428 | ENSG00000077800 |
| NA | 1.148941198 | 2.217510902 | 0.298243339 | 0.468671574 | 2.365105741 | 5.208182087 | ENSG00000224334 |
| NA | 1.148537602 | 2.216890638 | 0.002061012 | 0.008104154 | 23.66645138 | 52.23480891 | ENSG00000213062 |
| NA | 1.148435565 | 2.216733851 | 0.054962975 | 0.128785467 | 8.006694688 | 17.7718119 | ENSG00000279602 |
| IL2RB | 1.148248938 | 2.216447112 | 7.93E-05 | 0.000438099 | 38.64346037 | 85.4937724 | ENSG00000100385 |
| CD81-AS1 | 1.148221864 | 2.216405518 | 0.126708562 | 0.24818102 | 5.075671612 | 11.19664888 | ENSG00000238184 |
| NA | 1.147905439 | 2.215919449 | 0.289892153 | 0.459365002 | 2.637238657 | 5.866398707 | ENSG00000280399 |
| PCSK1 | 1.147354314 | 2.215073106 | 0.000686414 | 0.003031507 | 26.88826782 | 59.70882919 | ENSG00000175426 |
| POMK | 1.145991117 | 2.212981081 | 3.70E-05 | 0.000218756 | 38.32435767 | 84.97106034 | ENSG00000185900 |
| ASH1L | 1.145649388 | 2.212456957 | 1.74E-30 | 1.13E-28 | 448.4632972 | 991.6971499 | ENSG00000116539 |
| TRPV2 | 1.145319347 | 2.211950877 | 0.000493494 | 0.00226343 | 28.48932624 | 63.00877505 | ENSG00000187688 |
| NA | 1.145289158 | 2.211904593 | 0.260397011 | 0.425741057 | 2.68891437 | 5.940069673 | ENSG00000272945 |
| LINC01484 | 1.144194649 | 2.210227155 | 3.14E-09 | 3.59E-08 | 83.50336331 | 184.7081376 | ENSG00000253686 |
| ZFHX3 | 1.142878111 | 2.208211121 | 1.06E-05 | 6.92E-05 | 45.81334411 | 101.1339057 | ENSG00000140836 |
| RAD51-AS1 | 1.142670899 | 2.207893983 | 3.65E-05 | 0.000216242 | 43.98077701 | 96.95365215 | ENSG00000245849 |
| NA | 1.141820479 | 2.206592887 | 2.45E-05 | 0.000150058 | 346.2426734 | 763.6411908 | ENSG00000274090 |
| DDR2 | 1.141600011 | 2.206255709 | 0.016555315 | 0.048003694 | 15.09922101 | 33.20115215 | ENSG00000162733 |
| LINC00958 | 1.14134695 | 2.205868746 | 0.271624354 | 0.438522505 | 2.691705273 | 5.923398121 | ENSG00000251381 |
| SPART-AS1 | 1.140464859 | 2.204520448 | 0.237470429 | 0.399184563 | 2.982698788 | 6.611189736 | ENSG00000120664 |
| NA | 1.140151761 | 2.204042069 | 0.016245243 | 0.047268022 | 13.43061733 | 29.53901849 | ENSG00000224616 |
| VPREB3 | 1.139069865 | 2.202389848 | 0.243511323 | 0.406491998 | 2.988280594 | 6.611189736 | ENSG00000128218 |
| NA | 1.13823857 | 2.201121176 | 0.008196637 | 0.026660145 | 16.51646739 | 36.5516457 | ENSG00000249602 |
| SEMA3G | 1.138178293 | 2.201029213 | 2.49E-09 | 2.88E-08 | 87.78408751 | 193.2719505 | ENSG00000010319 |
| RASAL2 | 1.137661246 | 2.200240527 | 1.02E-43 | 1.35E-41 | 603.0068902 | 1325.921812 | ENSG00000075391 |
| BMPER | 1.13730099 | 2.199691173 | 0.25834455 | 0.423534012 | 2.949883674 | 6.533217622 | ENSG00000164619 |
| MYOF | 1.136896328 | 2.199074267 | 0.000126398 | 0.000666761 | 34.09251827 | 74.89107708 | ENSG00000138119 |

| | | | | | | | |
|---|---|---|---|---|---|---|---|
| NA | 1.136670205 | 2.19872962 | 0.023813113 | 0.064920685 | 11.67261675 | 25.71072078 | ENSG00000262468 |
| LINC01748 | 1.136614039 | 2.198644021 | 0.16617775 | 0.305861557 | 4.343563529 | 9.559985354 | ENSG00000226476 |
| NA | 1.136597311 | 2.198618528 | 0.281784433 | 0.450277013 | 2.746171888 | 5.990617367 | ENSG00000203709 |
| LINC00548 | 1.136539589 | 2.198530564 | 0.161475601 | 0.2988321 | 4.337981724 | 9.549765524 | ENSG00000215483 |
| DISC1 | 1.136465902 | 2.198418275 | 1.72E-05 | 0.000108268 | 41.63365598 | 91.67233213 | ENSG00000162946 |
| HES2 | 1.135719647 | 2.197281405 | 0.01862096 | 0.052872672 | 13.90155595 | 30.69546682 | ENSG00000069812 |
| NA | 1.135517168 | 2.196973043 | 0.031361671 | 0.08142638 | 13.97854995 | 30.50026081 | ENSG00000285804 |
| FAM50B | 1.134404294 | 2.195278984 | 3.80E-08 | 3.71E-07 | 69.54590128 | 152.6190894 | ENSG00000145945 |
| NA | 1.134201143 | 2.194969881 | 0.132822848 | 0.256948011 | 5.426713549 | 11.84464567 | ENSG00000250993 |
| PLXDC2 | 1.133307336 | 2.193610431 | 7.70E-28 | 4.38E-26 | 394.0820981 | 865.0165185 | ENSG00000120594 |
| NCMAP | 1.133148908 | 2.193369554 | 0.105158192 | 0.215168294 | 6.850417614 | 14.91550937 | ENSG00000184454 |
| MMP23B | 1.131799335 | 2.191318721 | 0.000665125 | 0.002948284 | 28.26886904 | 61.78080633 | ENSG00000189409 |
| PLA2G4A | 1.131784102 | 2.191295584 | 0.000477347 | 0.002198782 | 28.95958914 | 63.5172387 | ENSG00000116711 |
| ZBTB41 | 1.131570415 | 2.19097104 | 2.75E-07 | 2.37E-06 | 65.25401346 | 143.0015411 | ENSG00000177888 |
| GAS5 | 1.130806065 | 2.189810556 | 8.56E-100 | 6.61E-97 | 3880.315331 | 8494.623767 | ENSG00000234741 |
| NA | 1.130506156 | 2.189355382 | 0.225552278 | 0.384613189 | 3.627525142 | 8.014197384 | ENSG00000224358 |
| PPP1R17 | 1.130148256 | 2.18881232 | 0.002045076 | 0.008048055 | 21.99764755 | 48.17904658 | ENSG00000106341 |
| NA | 1.130057079 | 2.188673994 | 0.293966505 | 0.463764135 | 2.710565871 | 5.895973701 | ENSG00000231443 |
| ELAVL3 | 1.129863692 | 2.188380632 | 0.004069403 | 0.014678918 | 20.80644018 | 45.49804344 | ENSG00000196361 |
| PCDHGA4 | 1.129827214 | 2.1883253 | 0.048725082 | 0.116968088 | 9.01372659 | 19.74322669 | ENSG00000262576 |
| NA | 1.129480639 | 2.187799667 | 0.206251553 | 0.360398434 | 4.277933303 | 9.383601466 | ENSG00000228748 |
| NA | 1.128357999 | 2.186097883 | 0.237002336 | 0.398536617 | 3.012722998 | 6.585915889 | ENSG00000236372 |
| NA | 1.128217266 | 2.185884642 | 0.292224005 | 0.461985166 | 2.691705273 | 5.854028303 | ENSG00000225742 |
| SNORA79B | 1.12790772 | 2.185415686 | 0.265198195 | 0.431215305 | 2.958256383 | 6.491272224 | ENSG00000222489 |
| NA | 1.126341916 | 2.183045071 | 9.27E-05 | 0.000504803 | 40.68579662 | 89.20724724 | ENSG00000260912 |
| NA | 1.126323515 | 2.183017227 | 0.160402374 | 0.297702731 | 4.37358774 | 9.518039956 | ENSG00000261474 |
| CEACAM19 | 1.126313337 | 2.183001826 | 0.016472287 | 0.047806061 | 14.88067883 | 32.64590891 | ENSG00000186567 |
| NA | 1.126257934 | 2.182917994 | 0.114974344 | 0.230726107 | 6.828766112 | 14.79344129 | ENSG00000242242 |
| PLOD2 | 1.126200459 | 2.182831033 | 4.16E-35 | 3.41E-33 | 422.3638825 | 921.697817 | ENSG00000152952 |
| LINC00472 | 1.124845252 | 2.180781535 | 0.125274976 | 0.246047604 | 7.282959314 | 15.96977782 | ENSG00000233237 |
| CCL25 | 1.124518304 | 2.180287376 | 0.088933133 | 0.189006177 | 7.592813428 | 16.69091254 | ENSG00000131142 |
| VASH2 | 1.124014656 | 2.179526366 | 1.01E-82 | 4.65E-80 | 1463.45764 | 3190.432561 | ENSG00000143494 |
| NA | 1.123708299 | 2.179063591 | 4.88E-05 | 0.000281327 | 37.36621057 | 81.49607556 | ENSG00000267278 |
| NDUFA4L2 | 1.122773261 | 2.177651757 | 0.177584884 | 0.321360098 | 5.208847088 | 11.43837401 | ENSG00000185633 |
| ACOT11 | 1.122182473 | 2.176760185 | 8.64E-10 | 1.06E-08 | 98.63627531 | 214.3671041 | ENSG00000162390 |
| NA | 1.121514921 | 2.175753204 | 0.048208599 | 0.115968406 | 9.854767904 | 21.28471351 | ENSG00000226121 |
| NA | 1.121360672 | 2.175520591 | 0.088075858 | 0.187514776 | 6.673063254 | 14.55386673 | ENSG00000223374 |
| RAG1 | 1.120450584 | 2.174148651 | 0.012731458 | 0.03858642 | 15.48865986 | 33.59639836 | ENSG00000166349 |
| SLC5A4-AS1 | 1.120441276 | 2.174134625 | 0.11101396 | 0.224295171 | 8.84562082 | 19.14577759 | ENSG00000242082 |
| NA | 1.120225674 | 2.173809737 | 0.373614435 | 0.547087239 | 2.422363259 | 5.193128071 | ENSG00000255176 |
| NA | 1.119586758 | 2.172847251 | 0.303417034 | 0.474136735 | 2.71893858 | 5.856178876 | ENSG00000223653 |
| GOLGA6B | 1.118260234 | 2.170850287 | 0.25701284 | 0.422172504 | 3.037165403 | 6.56494319 | ENSG00000215186 |
| NA | 1.117709922 | 2.170022381 | 0.142057057 | 0.270926612 | 5.048438305 | 10.9506226 | ENSG00000229970 |
| NA | 1.11726422 | 2.169352084 | 0.183792011 | 0.329796678 | 4.398030145 | 9.484163814 | ENSG00000265458 |
| C17orf50 | 1.117028772 | 2.168998074 | 0.22268139 | 0.381065085 | 3.611455446 | 7.894279872 | ENSG00000270806 |
| LAMC3 | 1.116825784 | 2.168692917 | 8.53E-67 | 2.35E-64 | 1040.687386 | 2257.145066 | ENSG00000050555 |
| NA | 1.116043347 | 2.167517057 | 0.056876568 | 0.132321016 | 9.073775011 | 19.59803533 | ENSG00000262903 |
| UCMA | 1.115743107 | 2.167066021 | 0.006761633 | 0.022740352 | 17.28697239 | 37.58225783 | ENSG00000165623 |
| HNMT | 1.115559543 | 2.166790308 | 0.00387088 | 0.014041585 | 21.14631851 | 45.66745468 | ENSG00000150540 |
| TIRAP | 1.114989045 | 2.165933644 | 0.007259009 | 0.024089719 | 17.54323577 | 37.87453066 | ENSG00000150455 |
| CORO2A | 1.113694516 | 2.163991026 | 8.07E-73 | 2.83E-70 | 1397.217769 | 3022.351036 | ENSG00000106789 |
| NPC1L1 | 1.113629403 | 2.16389336 | 0.354054903 | 0.527135897 | 2.272242207 | 4.985279079 | ENSG00000015520 |
| MTMR11 | 1.113332417 | 2.163447958 | 1.53E-05 | 9.68E-05 | 46.07307411 | 99.38969508 | ENSG00000014914 |
| CNNM3-DT | 1.112699993 | 2.162499791 | 0.367315994 | 0.541114382 | 1.951224481 | 4.26791247 | ENSG00000273265 |
| KC6 | 1.111440623 | 2.160612905 | 1.69E-06 | 1.28E-05 | 55.40136074 | 119.8419465 | ENSG00000267313 |
| MAML2 | 1.111239208 | 2.160311284 | 9.24E-10 | 1.13E-08 | 98.81554469 | 213.414985 | ENSG00000184384 |
| NA | 1.111114073 | 2.160123913 | 0.411795295 | 0.583920707 | 1.95959719 | 4.26791247 | ENSG00000232043 |
| RNF152 | 1.111023836 | 2.159988806 | 0.004492186 | 0.015988701 | 23.24611244 | 50.34404985 | ENSG00000176641 |
| ELOVL2 | 1.11024105 | 2.158817144 | 7.21E-05 | 0.000402468 | 43.27175989 | 93.19741998 | ENSG00000197977 |
| NA | 1.110066394 | 2.15855581 | 0.140719055 | 0.269017681 | 5.0239959 | 10.83285566 | ENSG00000273449 |
| ANO9 | 1.109320607 | 2.157440253 | 3.23E-20 | 1.13E-18 | 278.3511473 | 600.176155 | ENSG00000185101 |
| KLHL32 | 1.107226688 | 2.154311229 | 0.159414313 | 0.296239429 | 4.721838773 | 10.11548905 | ENSG00000186231 |
| NA | 1.106610121 | 2.153390734 | 0.199309815 | 0.35068884 | 4.016963998 | 8.678865726 | ENSG00000231066 |
| NA | 1.106470496 | 2.153182337 | 0.14403714 | 0.273593981 | 5.0239959 | 10.79682895 | ENSG00000287252 |
| NA | 1.106309722 | 2.1529424 | 0.226651048 | 0.385868664 | 3.663131159 | 7.902882167 | ENSG00000286366 |
| HSH2D | 1.105470649 | 2.15169061 | 0.004946064 | 0.017382859 | 24.8429405 | 53.41329602 | ENSG00000196684 |
| NA | 1.105285336 | 2.151414245 | 0.257145716 | 0.422286371 | 3.330949821 | 7.160241712 | ENSG00000206147 |
| RBM15 | 1.104682574 | 2.150515566 | 2.13E-28 | 1.24E-26 | 380.3111638 | 816.8375572 | ENSG00000162775 |
| SERTM2 | 1.10417198 | 2.149754597 | 0.004671754 | 0.016560606 | 20.61022522 | 44.4228023 | ENSG00000260802 |
| H2BC18 | 1.103246985 | 2.148376707 | 0.243854916 | 0.40676223 | 3.390998242 | 7.246283083 | ENSG00000203814 |
| RGS2 | 1.103169577 | 2.14826144 | 1.59E-15 | 3.62E-14 | 190.8803564 | 410.2984788 | ENSG00000116741 |
| FHIT | 1.1007358 | 2.144640451 | 0.007617164 | 0.025062914 | 18.27725887 | 39.24985343 | ENSG00000189283 |
| HHIP-AS1 | 1.099168243 | 2.142311462 | 0.372179277 | 0.54561377 | 2.27503311 | 4.923978517 | ENSG00000248890 |
| NA | 1.097765119 | 2.140228925 | 0.004136485 | 0.014876386 | 25.84072381 | 55.46242245 | ENSG00000260400 |
| TEX41 | 1.096741174 | 2.138710446 | 0.154461091 | 0.289376411 | 5.372246934 | 11.45987975 | ENSG00000226674 |
| NA | 1.095732541 | 2.137215731 | 0.203193579 | 0.356087149 | 3.981357982 | 8.533674367 | ENSG00000274922 |
| CBLC | 1.094541 | 2.135451304 | 9.80E-18 | 2.74E-16 | 202.5007339 | 432.3565044 | ENSG00000142273 |

| | | | | | | | |
|---|---|---|---|---|---|---|---|
| NOTUM | 1.09446348 | 2.135336564 | 0.07092128 | 0.157656421 | 7.710119367 | 16.44892694 | ENSG00000185269 |
| MIR7111 | 1.094238967 | 2.135004288 | 0.380767762 | 0.554160007 | 2.283405818 | 4.91590926 | ENSG00000276712 |
| JSRP1 | 1.09339231 | 2.13375171 | 0.129995561 | 0.252847788 | 5.756103983 | 12.23586331 | ENSG00000167476 |
| CNTNAP5 | 1.092434948 | 2.132336238 | 0.22530292 | 0.384357856 | 3.951333771 | 8.455702254 | ENSG00000155052 |
| UCN | 1.092215191 | 2.132011458 | 0.367116929 | 0.540947951 | 1.984039594 | 4.242638624 | ENSG00000163794 |
| TAGLN2 | 1.091969678 | 2.131648669 | 6.97E-93 | 4.80E-90 | 1656.738334 | 3531.330022 | ENSG00000158710 |
| OR2A7 | 1.091803097 | 2.131402552 | 0.11759882 | 0.234917557 | 6.733111675 | 14.26940269 | ENSG00000243896 |
| GPR37L1 | 1.091153709 | 2.130443378 | 0.135016362 | 0.260225862 | 5.679985866 | 12.0863708 | ENSG00000170075 |
| SYTL1 | 1.09108878 | 2.130347499 | 2.52E-11 | 3.69E-10 | 118.5833772 | 252.604616 | ENSG00000142765 |
| NA | 1.090612592 | 2.129644455 | 0.026037893 | 0.069996794 | 12.0753344 | 25.69566677 | ENSG00000251611 |
| ZMYND12 | 1.090370264 | 2.129286771 | 0.120060958 | 0.238600591 | 6.082703515 | 12.91935378 | ENSG00000066185 |
| ADAMTSL4 | 1.089994914 | 2.12873286 | 2.98E-10 | 3.89E-09 | 137.1453209 | 291.2284993 | ENSG00000143382 |
| CACNA1S | 1.089784221 | 2.128422001 | 0.357136405 | 0.530452601 | 2.005691096 | 4.26791247 | ENSG00000081248 |
| MIR4479 | 1.089518815 | 2.128030481 | 0.012886639 | 0.038995474 | 16.0762287 | 34.18901327 | ENSG00000266507 |
| NA | 1.089228099 | 2.127601707 | 0.020632425 | 0.057691964 | 13.40338403 | 28.45517505 | ENSG00000284620 |
| NA | 1.089067084 | 2.127364265 | 0.007125642 | 0.023724672 | 18.63388262 | 39.70290467 | ENSG00000267325 |
| LGALS8 | 1.08903559 | 2.127317824 | 4.90E-13 | 8.59E-12 | 151.9685421 | 322.7400292 | ENSG00000116977 |
| VAMP4 | 1.08827901 | 2.126202506 | 6.69E-10 | 8.29E-09 | 134.9989206 | 287.0533099 | ENSG00000117533 |
| CNTD2 | 1.087633987 | 2.125252103 | 0.00143646 | 0.005899669 | 24.75777395 | 52.60720443 | ENSG00000105219 |
| NA | 1.087621882 | 2.12523427 | 0.035235333 | 0.089627831 | 13.85267114 | 29.58096389 | ENSG00000236809 |
| NA | 1.087493274 | 2.125044825 | 0.334212055 | 0.50690186 | 2.596050835 | 5.578427028 | ENSG00000283897 |
| CHRNA10 | 1.087482705 | 2.125029259 | 0.128734853 | 0.251028408 | 5.731661579 | 12.1283162 | ENSG00000129749 |
| PRODH | 1.087337817 | 2.124815855 | 0.00053769 | 0.002445786 | 29.41956431 | 62.53743472 | ENSG00000100033 |
| GAL3ST1 | 1.087190014 | 2.12459818 | 2.30E-19 | 7.31E-18 | 220.2141906 | 467.4852784 | ENSG00000128242 |
| NA | 1.08555273 | 2.122188387 | 0.161393926 | 0.298738267 | 5.663240449 | 11.99979639 | ENSG00000250999 |
| LINC02582 | 1.08530164 | 2.121819068 | 0.007140208 | 0.0237591 | 19.82105978 | 41.95047219 | ENSG00000261780 |
| NA | 1.085215294 | 2.12169208 | 0.401947182 | 0.574301425 | 1.986830497 | 4.219515351 | ENSG00000273998 |
| APOBEC2 | 1.085011415 | 2.121392268 | 0.374900591 | 0.548268312 | 2.005691096 | 4.249090345 | ENSG00000124701 |
| LOC101929536 | 1.084879469 | 2.121198259 | 0.282227935 | 0.450625649 | 3.273692303 | 7.027681222 | ENSG00000261798 |
| PTPN6 | 1.083969857 | 2.119861275 | 3.95E-28 | 2.29E-26 | 345.2056315 | 731.4139545 | ENSG00000111679 |
| GCSAM | 1.083494154 | 2.119162404 | 0.013361208 | 0.04017317 | 16.18707695 | 34.45869586 | ENSG00000174500 |
| INHBC | 1.083401747 | 2.119026672 | 0.118024566 | 0.235548511 | 6.433745451 | 13.59047384 | ENSG00000175189 |
| NA | 1.082638915 | 2.117906522 | 0.23603421 | 0.397481288 | 3.714806871 | 7.818458333 | ENSG00000280401 |
| PHF11 | 1.082476312 | 2.117667832 | 7.73E-07 | 6.19E-06 | 77.34071163 | 163.4812291 | ENSG00000136147 |
| CAPN6 | 1.082332033 | 2.117456061 | 1.68E-13 | 3.10E-12 | 173.6889504 | 367.470431 | ENSG00000077274 |
| NA | 1.08226171 | 2.117352851 | 0.385161896 | 0.558658507 | 1.981248691 | 4.198542652 | ENSG00000228217 |
| TRIML2 | 1.081828807 | 2.116717601 | 3.75E-32 | 2.71E-30 | 520.3906493 | 1101.243498 | ENSG00000179046 |
| TP73 | 1.081719734 | 2.116557575 | 3.62E-09 | 4.11E-08 | 93.52758843 | 197.8818715 | ENSG00000078900 |
| NA | 1.081337209 | 2.115996453 | 0.086488501 | 0.184767405 | 10.13160675 | 21.64661662 | ENSG00000273654 |
| VWA5B1 | 1.081272146 | 2.115901027 | 0.002055119 | 0.008082632 | 32.56095701 | 68.55038186 | ENSG00000158816 |
| NA | 1.081129851 | 2.115692343 | 0.105433798 | 0.215595018 | 6.730320772 | 14.22341661 | ENSG00000272669 |
| MYO3B | 1.080980585 | 2.115473458 | 0.063396916 | 0.144135522 | 9.089844707 | 19.21084626 | ENSG00000071909 |
| NA | 1.080637292 | 2.114970136 | 0.112713792 | 0.227206036 | 6.319230415 | 13.36731037 | ENSG00000281852 |
| ESRRG | 1.080515321 | 2.114791336 | 0.000400796 | 0.001887635 | 31.88802147 | 67.30600205 | ENSG00000196482 |
| NA | 1.080497404 | 2.114765071 | 0.340429664 | 0.512984584 | 2.307848223 | 4.903005818 | ENSG00000260808 |
| STAC3 | 1.080429148 | 2.114665021 | 0.001331677 | 0.005510376 | 28.48865052 | 60.19792556 | ENSG00000185482 |
| NA | 1.079900868 | 2.113890824 | 0.408415032 | 0.580707327 | 2.002900193 | 4.225967072 | ENSG00000253939 |
| EMILIN1 | 1.079804547 | 2.113749696 | 1.92E-13 | 3.53E-12 | 150.565977 | 318.6912396 | ENSG00000138080 |
| NA | 1.079767273 | 2.113695084 | 0.001900207 | 0.00755034 | 29.469325 | 62.05183388 | ENSG00000260776 |
| NA | 1.079016607 | 2.112595569 | 0.340643502 | 0.513186553 | 2.65051745 | 5.626824147 | ENSG00000231822 |
| WNT8B | 1.078927968 | 2.112465775 | 0.063714876 | 0.144711586 | 9.509307773 | 19.93870528 | ENSG00000075290 |
| MIR17HG | 1.077239152 | 2.109994373 | 2.91E-10 | 3.81E-09 | 114.5333979 | 241.9048539 | ENSG00000215417 |
| ILDR1 | 1.076861914 | 2.109442721 | 0.000488597 | 0.002245242 | 31.64571261 | 66.904292 | ENSG00000145103 |
| MYL3 | 1.076755399 | 2.109286985 | 0.412922191 | 0.585186558 | 1.978457788 | 4.173268805 | ENSG00000160808 |
| CBR3 | 1.076702409 | 2.109209513 | 0.325069053 | 0.496744859 | 2.335081531 | 4.930430238 | ENSG00000159231 |
| SNORD116-5 | 1.076127664 | 2.108369408 | 0.285214248 | 0.454190859 | 2.952674577 | 6.285040767 | ENSG00000207191 |
| A2ML1 | 1.076111295 | 2.108345486 | 9.92E-13 | 1.68E-11 | 159.8306975 | 336.8429527 | ENSG00000166535 |
| OTOA | 1.074309004 | 2.105713274 | 0.347555802 | 0.52027691 | 2.389548146 | 5.008402352 | ENSG00000155719 |
| NA | 1.074212911 | 2.105573023 | 0.331514735 | 0.504198262 | 2.359523935 | 4.957854658 | ENSG00000225970 |
| LRRC37A6P | 1.074191803 | 2.105542219 | 0.003689817 | 0.013450539 | 23.10523997 | 48.61059208 | ENSG00000230445 |
| BOLA1 | 1.074123504 | 2.105442541 | 3.54E-11 | 5.09E-10 | 126.7773505 | 266.7148152 | ENSG00000178096 |
| TP53INP1 | 1.073504452 | 2.104539302 | 3.66E-10 | 4.71E-09 | 110.8748094 | 233.2122486 | ENSG00000164938 |
| LINC01237 | 1.07300349 | 2.103808647 | 0.14909099 | 0.281062669 | 5.919303669 | 12.54695826 | ENSG00000233806 |
| SPAG4 | 1.072281484 | 2.102756047 | 0.000147607 | 0.000764015 | 40.60305756 | 85.29720931 | ENSG00000061656 |
| TBILA | 1.071539867 | 2.101675404 | 0.12832711 | 0.250433996 | 7.015732482 | 14.73697491 | ENSG00000261488 |
| NA | 1.071263555 | 2.101272919 | 0.275240413 | 0.442061033 | 3.597500932 | 7.637500723 | ENSG00000275223 |
| ADA2 | 1.071020842 | 2.100919439 | 7.96E-43 | 1.02E-40 | 948.7988753 | 1992.396946 | ENSG00000093072 |
| APOBEC3B | 1.071009334 | 2.100902681 | 3.17E-08 | 3.12E-07 | 86.69112533 | 182.3998209 | ENSG00000179750 |
| LINC01694 | 1.070939909 | 2.100801584 | 0.002268172 | 0.008803883 | 30.13771787 | 63.1580192 | ENSG00000233922 |
| LINC01836 | 1.070535004 | 2.100212058 | 0.016015006 | 0.046739105 | 15.95613186 | 33.6093018 | ENSG00000267530 |
| NA | 1.070467692 | 2.100114071 | 0.310213941 | 0.481279598 | 2.628865949 | 5.553153181 | ENSG00000248588 |
| NA | 1.070247932 | 2.099794192 | 0.367568167 | 0.541279199 | 2.038506209 | 4.242638624 | ENSG00000240970 |
| NA | 1.069957146 | 2.099371006 | 0.187433704 | 0.334546964 | 5.551516316 | 11.73117988 | ENSG00000278997 |
| NA | 1.068869338 | 2.097788654 | 0.37384709 | 0.547183931 | 2.038506209 | 4.238337476 | ENSG00000213684 |
| LSMEM2 | 1.06842475 | 2.097142289 | 0.156367381 | 0.29192714 | 5.0239959 | 10.53198054 | ENSG00000179564 |
| S100A16 | 1.067917938 | 2.096405702 | 6.67E-20 | 2.23E-18 | 251.5970189 | 527.1118101 | ENSG00000188643 |
| ZMAT1 | 1.067428422 | 2.095694499 | 0.00798203 | 0.026063194 | 18.67507044 | 39.20224981 | ENSG00000166432 |

| | | | | | | | |
|---|---|---|---|---|---|---|---|
| NA | 1.066982064 | 2.095046207 | 0.031656992 | 0.082060606 | 12.3363037 | 25.84489881 | ENSG00000287858 |
| KBTBD12 | 1.066704671 | 2.094643424 | 0.249285046 | 0.413353775 | 4.095873018 | 8.506249947 | ENSG00000187715 |
| NA | 1.066314761 | 2.094077391 | 0.349391305 | 0.522003381 | 2.392339049 | 4.976676784 | ENSG00000273355 |
| MIR6835 | 1.066288138 | 2.094038748 | 0.402814759 | 0.575138739 | 2.032924404 | 4.223816498 | ENSG00000276404 |
| VIL1 | 1.066286983 | 2.094037071 | 9.30E-13 | 1.58E-11 | 144.5862615 | 302.428668 | ENSG00000127831 |
| MC1R | 1.065576732 | 2.093006413 | 0.039874491 | 0.099377141 | 14.17109813 | 29.48605976 | ENSG00000258839 |
| TLE2 | 1.065517226 | 2.092920086 | 3.47E-26 | 1.82E-24 | 379.7976362 | 795.2678587 | ENSG00000065717 |
| NUTM2F | 1.065400377 | 2.092750579 | 0.306523165 | 0.477520663 | 2.963838189 | 6.238794221 | ENSG00000130950 |
| NA | 1.064873942 | 2.091987081 | 0.004078703 | 0.014709714 | 21.38842721 | 44.77207454 | ENSG00000232325 |
| LINC00939 | 1.064651829 | 2.09166503 | 0.340521135 | 0.51304227 | 2.993862399 | 6.28073962 | ENSG00000249267 |
| TNFRSF14 | 1.063943845 | 2.090638823 | 0.255330624 | 0.420307454 | 3.630316045 | 7.650404165 | ENSG00000157873 |
| C1orf167 | 1.06354132 | 2.090055597 | 0.02688731 | 0.071919452 | 12.73132436 | 26.57301829 | ENSG00000215910 |
| MINAR1 | 1.063127353 | 2.089455963 | 0.020974314 | 0.058419793 | 16.30525877 | 33.87415021 | ENSG00000169330 |
| VPS13B-DT | 1.062834119 | 2.089031316 | 0.039701373 | 0.099078809 | 11.68589554 | 24.45882318 | ENSG00000253948 |
| RALY-AS1 | 1.062402763 | 2.088406803 | 0.029806608 | 0.078124903 | 13.52627177 | 28.07901143 | ENSG00000285230 |
| HCP5 | 1.062161995 | 2.088058303 | 8.29E-05 | 0.000456567 | 43.51925024 | 91.12381319 | ENSG00000206337 |
| WDR88 | 1.062126997 | 2.088007649 | 0.075855093 | 0.166210882 | 9.40807153 | 19.57437902 | ENSG00000166359 |
| HLA-DMB | 1.060178472 | 2.085189459 | 0.09373414 | 0.196674495 | 7.965506866 | 16.6255834 | ENSG00000242574 |
| IQCH | 1.060075347 | 2.085040414 | 0.096413926 | 0.201343946 | 7.437986451 | 15.43283578 | ENSG00000103599 |
| HEATR4 | 1.059798136 | 2.084639816 | 0.070664091 | 0.157265919 | 8.711569464 | 18.16733069 | ENSG00000187105 |
| CPB2-AS1 | 1.059604313 | 2.084359767 | 0.240588152 | 0.402775843 | 4.223466687 | 8.903919308 | ENSG00000235903 |
| CEP350 | 1.059273243 | 2.083881503 | 4.79E-32 | 3.42E-30 | 557.9010996 | 1162.154198 | ENSG00000135837 |
| NA | 1.057896162 | 2.081893346 | 0.155175839 | 0.290180021 | 5.707219174 | 11.87207009 | ENSG00000287608 |
| IFNL4 | 1.057630512 | 2.081510033 | 0.293167459 | 0.462882097 | 2.979907885 | 6.22212267 | ENSG00000272395 |
| DUOX1 | 1.056761416 | 2.080256485 | 0.005974237 | 0.020459615 | 21.22993345 | 44.28352962 | ENSG00000137857 |
| BEST1 | 1.056611426 | 2.080040222 | 0.256733987 | 0.42189755 | 3.600291835 | 7.551459352 | ENSG00000167995 |
| BAIAP3 | 1.056600242 | 2.080024097 | 9.66E-18 | 2.70E-16 | 222.0262575 | 461.7726855 | ENSG00000007516 |
| ZFP92 | 1.056527571 | 2.079919326 | 0.013280286 | 0.039973473 | 18.74349157 | 39.02290973 | ENSG00000189420 |
| GCNT4 | 1.056480306 | 2.079851184 | 0.000388218 | 0.001837818 | 34.98224423 | 72.83119777 | ENSG00000176928 |
| KCNS1 | 1.055975333 | 2.079123322 | 8.58E-06 | 5.73E-05 | 60.01079966 | 124.9049373 | ENSG00000124134 |
| NA | 1.055816863 | 2.078894956 | 0.126371788 | 0.247722607 | 6.297578913 | 13.13795564 | ENSG00000262619 |
| TRPV4 | 1.055438386 | 2.07834965 | 0.000120126 | 0.000637159 | 43.49588387 | 90.11794187 | ENSG00000111199 |
| ITGA10 | 1.055357236 | 2.078232749 | 0.259947124 | 0.425271432 | 3.33932253 | 6.958311403 | ENSG00000143127 |
| SNHG8 | 1.055158809 | 2.07794693 | 1.04E-34 | 8.39E-33 | 568.3507714 | 1182.055865 | ENSG00000269893 |
| NA | 1.055135112 | 2.077912799 | 0.071535491 | 0.158710886 | 8.675963447 | 18.03074162 | ENSG00000269399 |
| NUDT18 | 1.054693557 | 2.077276926 | 0.000555377 | 0.002517927 | 32.71490811 | 67.99002555 | ENSG00000275074 |
| FGF18 | 1.054629662 | 2.077184927 | 0.033448347 | 0.085781981 | 17.0572666 | 35.67214361 | ENSG00000156427 |
| NA | 1.0545191 | 2.077025748 | 0.31555162 | 0.486789789 | 2.661681062 | 5.523578187 | ENSG00000205583 |
| ATF6 | 1.054445155 | 2.076919292 | 1.75E-36 | 1.58E-34 | 480.1202245 | 997.4119482 | ENSG00000118217 |
| RGS16 | 1.054407639 | 2.076865285 | 2.21E-23 | 9.63E-22 | 327.7983252 | 681.6646701 | ENSG00000143333 |
| CARD14 | 1.05437736 | 2.076821695 | 0.226318595 | 0.385443502 | 4.691814563 | 9.702765675 | ENSG00000141527 |
| HBQ1 | 1.054275458 | 2.076675008 | 0.000742173 | 0.003249492 | 30.6791928 | 63.78718175 | ENSG00000086506 |
| LINC02482 | 1.053998782 | 2.076276788 | 0.373757219 | 0.547176938 | 2.359523935 | 4.869662714 | ENSG00000251580 |
| LINC01503 | 1.053743206 | 2.075909005 | 0.000108795 | 0.000583801 | 42.03358273 | 87.41489996 | ENSG00000233901 |
| NA | 1.052890164 | 2.074681917 | 0.022565766 | 0.062134768 | 16.0573681 | 33.24686566 | ENSG00000273145 |
| NA | 1.052853058 | 2.074628557 | 0.287564215 | 0.45693058 | 3.273692303 | 6.83194217 | ENSG00000215452 |
| HYPK | 1.051918299 | 2.073284787 | 0.15345613 | 0.28785708 | 5.375037836 | 11.10200521 | ENSG00000242028 |
| NA | 1.051331333 | 2.072441435 | 0.286968314 | 0.4562309 | 3.009932095 | 6.234493074 | ENSG00000204778 |
| LOC101929054 | 1.050070135 | 2.070630506 | 0.030056894 | 0.078663347 | 12.64962444 | 26.22374604 | ENSG00000243224 |
| RD3 | 1.049211602 | 2.069398662 | 0.245924118 | 0.409293604 | 3.687573563 | 7.643952444 | ENSG00000198570 |
| COA6-AS1 | 1.048865756 | 2.068902641 | 0.145671475 | 0.276046013 | 5.728870676 | 11.80646838 | ENSG00000231663 |
| STK32A | 1.047699736 | 2.067231181 | 0.054850562 | 0.128553298 | 9.955128266 | 20.62434632 | ENSG00000169302 |
| NA | 1.04724361 | 2.066577703 | 0.099220225 | 0.205989815 | 8.670381641 | 17.8957701 | ENSG00000227598 |
| PYCR2 | 1.046840474 | 2.066000315 | 7.18E-31 | 4.75E-29 | 444.3984506 | 918.61571 | ENSG00000143811 |
| RNF213 | 1.04554523 | 2.064146303 | 1.41E-31 | 9.73E-30 | 1053.382352 | 2173.365728 | ENSG00000173821 |
| LAMC2 | 1.045273251 | 2.063757203 | 5.41E-22 | 2.12E-20 | 277.1621431 | 571.5892129 | ENSG00000058085 |
| VSTM5 | 1.045223268 | 2.063685704 | 0.104900906 | 0.214778536 | 7.800191998 | 15.97758661 | ENSG00000214376 |
| NEURL1 | 1.04504612 | 2.063432321 | 0.011513817 | 0.035538088 | 19.3254786 | 39.88064563 | ENSG00000107954 |
| IRAK1 | 1.044249064 | 2.062292636 | 1.00E-54 | 1.93E-52 | 1083.033414 | 2234.123158 | ENSG00000184216 |
| SH2D3A | 1.043830244 | 2.061694032 | 6.10E-38 | 6.09E-36 | 539.7738369 | 1112.581916 | ENSG00000125731 |
| CTSK | 1.041765732 | 2.058745835 | 1.01E-09 | 1.22E-08 | 113.4545904 | 233.8718102 | ENSG00000143387 |
| CCDC172 | 1.041754209 | 2.058729393 | 0.002247158 | 0.008734622 | 25.61747571 | 52.82795686 | ENSG00000182645 |
| LINC01873 | 1.041414975 | 2.058245361 | 0.366766971 | 0.540553262 | 2.392339049 | 4.865361567 | ENSG00000232164 |
| LENG9 | 1.040784422 | 2.057345969 | 1.67E-07 | 1.49E-06 | 74.85251799 | 153.9952057 | ENSG00000275183 |
| ELN | 1.040552091 | 2.057014681 | 1.34E-14 | 2.77E-13 | 168.3144762 | 346.2752544 | ENSG00000049540 |
| OPN3 | 1.0401977 | 2.056509448 | 0.011266967 | 0.034893411 | 17.70364455 | 36.40215319 | ENSG00000054277 |
| NA | 1.040115182 | 2.056391825 | 0.248247927 | 0.411917319 | 3.693155369 | 7.589103603 | ENSG00000215003 |
| NA | 1.040000528 | 2.056228407 | 0.252409796 | 0.416958159 | 3.638688754 | 7.500911659 | ENSG00000260495 |
| CEBPB-AS1 | 1.039763691 | 2.055890877 | 0.109623711 | 0.222044416 | 8.218779183 | 16.97862375 | ENSG00000277449 |
| FOXP3 | 1.039354586 | 2.055307971 | 0.205732997 | 0.35965521 | 4.343563529 | 8.950165854 | ENSG00000049768 |
| GIPC3 | 1.039227282 | 2.055126617 | 1.91E-06 | 1.43E-05 | 73.66111046 | 151.4776951 | ENSG00000179855 |
| CARF | 1.039024344 | 2.054837551 | 0.006508458 | 0.022003971 | 25.99575095 | 53.53159599 | ENSG00000138380 |
| TMSB15B-AS1 | 1.038666186 | 2.054327489 | 0.267953503 | 0.434194962 | 3.665922062 | 7.5347878 | ENSG00000231728 |
| NA | 1.038039109 | 2.053434755 | 0.268742547 | 0.434962033 | 3.393789145 | 6.933037557 | ENSG00000283491 |
| MESP2 | 1.037750689 | 2.05302428 | 0.050275297 | 0.119795054 | 11.46611406 | 23.49973143 | ENSG00000188095 |
| TDRD6 | 1.037634679 | 2.052859197 | 0.224952604 | 0.383862068 | 4.316330222 | 8.88939833 | ENSG00000180113 |
| CNGA4 | 1.037588019 | 2.052792805 | 0.320353967 | 0.491681787 | 3.075562322 | 6.266218642 | ENSG00000132259 |

| | | | | | | | |
|---|---|---|---|---|---|---|---|
| AKT3 | 1.037559229 | 2.052751841 | 9.40E-19 | 2.84E-17 | 284.4122873 | 583.9000311 | ENSG00000117020 |
| NA | 1.037006519 | 2.051965563 | 0.436418162 | 0.607295878 | 2.095763727 | 4.225967072 | ENSG00000235660 |
| OBSCN | 1.035520124 | 2.04985253 | 1.55E-68 | 4.60E-66 | 1252.834707 | 2567.052249 | ENSG00000154358 |
| NA | 1.035502821 | 2.049827945 | 0.045988855 | 0.111621917 | 12.0536829 | 24.70753307 | ENSG00000231711 |
| C22orf42 | 1.034754089 | 2.048764399 | 0.265550069 | 0.431605559 | 3.633106948 | 7.454665113 | ENSG00000205856 |
| LTBR | 1.03405134 | 2.047766672 | 0.000208116 | 0.001045239 | 51.02740958 | 104.1144269 | ENSG00000111321 |
| CSTF3-DT | 1.033870159 | 2.047509519 | 0.288581908 | 0.457852915 | 3.641479657 | 7.46273437 | ENSG00000247151 |
| NA | 1.033685308 | 2.04724719 | 0.024226235 | 0.065825543 | 14.98470597 | 30.77989066 | ENSG00000282572 |
| MIR100HG | 1.033548333 | 2.047052826 | 0.171045731 | 0.312673477 | 6.131588324 | 12.45876632 | ENSG00000255248 |
| DUSP19 | 1.033471738 | 2.046944148 | 0.060869522 | 0.139557161 | 10.63556064 | 21.86145037 | ENSG00000162999 |
| PYROXD1 | 1.03305802 | 2.046357235 | 1.16E-05 | 7.49E-05 | 58.03389347 | 118.6911443 | ENSG00000121350 |
| LY75 | 1.032987993 | 2.046257909 | 0.002494064 | 0.009577505 | 29.26262215 | 59.7249677 | ENSG00000054219 |
| ADGRA2 | 1.032048672 | 2.04492605 | 3.86E-07 | 3.25E-06 | 100.699187 | 206.4064106 | ENSG00000020181 |
| SPIB | 1.031036237 | 2.043491492 | 1.80E-08 | 1.85E-07 | 98.24085433 | 200.9414493 | ENSG00000269404 |
| TUBB6 | 1.030399242 | 2.042589426 | 1.43E-84 | 7.28E-82 | 3084.244985 | 6300.897145 | ENSG00000176014 |
| MCHR1 | 1.03002673 | 2.042062087 | 0.000199468 | 0.001005472 | 47.50706591 | 96.82729502 | ENSG00000128285 |
| NLRP6 | 1.029797938 | 2.041738268 | 0.155951329 | 0.291319559 | 6.387651545 | 12.97609273 | ENSG00000174885 |
| MEF2D | 1.029611582 | 2.041474549 | 9.05E-30 | 5.71E-28 | 492.5829706 | 1005.306022 | ENSG00000116604 |
| NA | 1.029374912 | 2.041139679 | 0.220835197 | 0.379251472 | 4.310748416 | 8.81357679 | ENSG00000217624 |
| NA | 1.02872576 | 2.040221458 | 0.225926027 | 0.384982474 | 4.019754901 | 8.186552699 | ENSG00000255471 |
| NA | 1.028483775 | 2.039879279 | 0.00038111 | 0.001807245 | 36.4303907 | 74.34175253 | ENSG00000281490 |
| EIF1B-AS1 | 1.027960031 | 2.039138872 | 0.098152582 | 0.204168801 | 8.042300704 | 16.36880425 | ENSG00000280739 |
| GRIP2 | 1.027039858 | 2.037838693 | 2.46E-15 | 5.50E-14 | 200.7152998 | 409.1097961 | ENSG00000144596 |
| NAXE | 1.025477607 | 2.035633173 | 1.95E-74 | 7.24E-72 | 1427.384374 | 2905.21086 | ENSG00000163382 |
| NA | 1.025011425 | 2.0349755 | 0.198987894 | 0.350403968 | 5.078462515 | 10.31337868 | ENSG00000286757 |
| SEZ6L | 1.024659082 | 2.034478568 | 0.127899815 | 0.249829419 | 6.648620849 | 13.58832326 | ENSG00000100095 |
| TRPV1 | 1.023939987 | 2.033464757 | 0.000199088 | 0.001004118 | 48.9747487 | 99.36254323 | ENSG00000196689 |
| SLC17A9 | 1.023542121 | 2.032904046 | 2.29E-39 | 2.55E-37 | 749.4157448 | 1523.805144 | ENSG00000101194 |
| USP46-AS1 | 1.023063819 | 2.032230188 | 0.079941336 | 0.173275091 | 9.36755943 | 19.02317647 | ENSG00000248866 |
| NA | 1.022654273 | 2.031653363 | 0.057395463 | 0.133238974 | 10.60274552 | 21.59795904 | ENSG00000262583 |
| SIT1 | 1.022540472 | 2.031493111 | 6.72E-06 | 4.57E-05 | 87.74895706 | 178.3271508 | ENSG00000137078 |
| ISLR2 | 1.022513917 | 2.031455718 | 0.001903305 | 0.007559534 | 29.30504927 | 59.5668729 | ENSG00000167178 |
| NA | 1.022113681 | 2.030892225 | 0.269334561 | 0.435664348 | 3.693155369 | 7.464884944 | ENSG00000279368 |
| ADCY7 | 1.021801777 | 2.030453203 | 5.61E-08 | 5.34E-07 | 92.20021452 | 187.2525516 | ENSG00000121281 |
| RENBP | 1.021426057 | 2.029924482 | 7.06E-17 | 1.84E-15 | 233.8223255 | 474.8135743 | ENSG00000102032 |
| KLK5 | 1.021023648 | 2.029358357 | 0.166172676 | 0.305861557 | 6.893520458 | 14.06532181 | ENSG00000167754 |
| SSPN | 1.020972507 | 2.029286422 | 0.056416228 | 0.131456149 | 11.85188613 | 24.15391948 | ENSG00000123096 |
| RIMBP3 | 1.020852498 | 2.029117625 | 0.057734807 | 0.133881737 | 10.31175201 | 20.94619414 | ENSG00000275793 |
| ZCCHC12 | 1.019708455 | 2.027509192 | 0.000741656 | 0.003248702 | 36.34889094 | 73.47595949 | ENSG00000174460 |
| APOBEC3C | 1.019453233 | 2.027150545 | 1.62E-25 | 8.12E-24 | 375.7254547 | 761.8458994 | ENSG00000244509 |
| LOC112268389 | 1.019400261 | 2.027076114 | 0.012996345 | 0.039253562 | 17.63243252 | 35.81545697 | ENSG00000255052 |
| NA | 1.019262922 | 2.026883153 | 0.001830955 | 0.007305264 | 29.9451697 | 60.56493543 | ENSG00000123870 |
| SPN | 1.01885032 | 2.02630356 | 0.000661548 | 0.002935124 | 35.96503389 | 72.64674462 | ENSG00000197471 |
| CDK18 | 1.018203122 | 2.025394757 | 3.99E-70 | 1.26E-67 | 1265.421731 | 2563.304371 | ENSG00000117266 |
| ZNF687-AS1 | 1.018193744 | 2.025381591 | 0.391073656 | 0.564219438 | 2.598841738 | 5.332400746 | ENSG00000232671 |
| MYO7A | 1.018061122 | 2.025195414 | 4.12E-09 | 4.64E-08 | 107.4885171 | 217.6928689 | ENSG00000137474 |
| MSX2 | 1.01767764 | 2.024657169 | 0.166120025 | 0.305813667 | 5.97935209 | 12.13046678 | ENSG00000120149 |
| NA | 1.017619674 | 2.024575822 | 0.127819697 | 0.249698206 | 7.056920304 | 14.29923815 | ENSG00000270728 |
| MRVI1 | 1.017365779 | 2.024219555 | 2.22E-06 | 1.64E-05 | 72.43294563 | 146.8500059 | ENSG00000072952 |
| ACRBP | 1.017162883 | 2.023934895 | 0.059353747 | 0.136715599 | 11.95802846 | 24.24372896 | ENSG00000111644 |
| VMO1 | 1.016877527 | 2.023534613 | 0.313620337 | 0.484779641 | 3.57584943 | 7.328556344 | ENSG00000182853 |
| ZNF57 | 1.01686615 | 2.023518656 | 0.215673043 | 0.372642067 | 4.370796837 | 8.804974495 | ENSG00000171970 |
| NA | 1.016155528 | 2.022522186 | 0.269714189 | 0.436059036 | 3.750412887 | 7.515432637 | ENSG00000175730 |
| NA | 1.015992772 | 2.02229403 | 0.032997495 | 0.084862642 | 13.59660792 | 27.58265772 | ENSG00000179066 |
| CHRM3 | 1.015161661 | 2.021129357 | 1.57E-12 | 2.62E-11 | 155.4752946 | 313.992042 | ENSG00000133019 |
| RASGRF2 | 1.015107059 | 2.021052865 | 5.26E-19 | 1.62E-17 | 361.0345331 | 729.5804426 | ENSG00000113319 |
| NA | 1.014788141 | 2.020606146 | 0.184651347 | 0.330866537 | 5.048438305 | 10.16603675 | ENSG00000237017 |
| NA | 1.014445813 | 2.020126746 | 0.171840694 | 0.313823289 | 5.638798044 | 11.40288034 | ENSG00000214578 |
| FER1L5 | 1.014282329 | 2.019897842 | 0.107669123 | 0.218958997 | 8.64873014 | 17.52631866 | ENSG00000249715 |
| NA | 1.014086754 | 2.019624038 | 0.000237887 | 0.001182143 | 40.66713618 | 82.2570051 | ENSG00000224592 |
| NA | 1.012790288 | 2.017809935 | 0.34603417 | 0.518513684 | 3.124447132 | 6.213520375 | ENSG00000260329 |
| RCSD1 | 1.012407044 | 2.017273987 | 0.175596052 | 0.318936393 | 7.119759628 | 14.25864983 | ENSG00000198771 |
| PIFO | 1.012333212 | 2.017170752 | 3.76E-05 | 0.000221931 | 52.37065302 | 105.456667 | ENSG00000173947 |
| SBSN | 1.012249747 | 2.017054055 | 0.400069464 | 0.572585667 | 3.184495553 | 6.335588461 | ENSG00000189001 |
| BATF2 | 1.011940866 | 2.016622249 | 0.003264249 | 0.01205872 | 30.65965648 | 61.92923276 | ENSG00000168062 |
| NA | 1.01184395 | 2.016486783 | 0.200961383 | 0.353137734 | 5.042856499 | 10.1321606 | ENSG00000245768 |
| SBF2-AS1 | 1.011613992 | 2.016165392 | 0.056580432 | 0.131791007 | 11.34601722 | 22.85630837 | ENSG00000246273 |
| CDH11 | 1.011347514 | 2.015793024 | 2.85E-15 | 6.31E-14 | 197.9265887 | 399.3268956 | ENSG00000140937 |
| IRF6 | 1.011034877 | 2.015356241 | 5.18E-24 | 2.38E-22 | 407.1729493 | 820.8540399 | ENSG00000117595 |
| RNVU1-15 | 1.01087634 | 2.015134787 | 0.476468466 | 0.644795032 | 2.122997035 | 4.189940357 | ENSG00000207205 |
| DPY19L2P4 | 1.010718394 | 2.014914182 | 0.349869688 | 0.522556222 | 3.363764935 | 6.730846784 | ENSG00000235436 |
| STAB1 | 1.010533149 | 2.01465548 | 0.015318156 | 0.045052988 | 21.22869415 | 42.63287816 | ENSG00000010327 |
| MAG | 1.009880581 | 2.013744406 | 0.190103597 | 0.338248339 | 5.366665128 | 10.77585625 | ENSG00000105695 |
| SH3TC1 | 1.00868975 | 2.012082905 | 0.257293279 | 0.422420822 | 4.555648025 | 9.269863101 | ENSG00000125089 |
| DNAJA4 | 1.007664672 | 2.010653767 | 5.53E-08 | 5.28E-07 | 127.1015595 | 254.8900699 | ENSG00000140403 |
| NA | 1.006893605 | 2.009579434 | 0.008406508 | 0.027223426 | 23.19040652 | 46.46441094 | ENSG00000263826 |
| NPL | 1.006888046 | 2.009571691 | 2.12E-05 | 0.000131678 | 60.40996267 | 121.5409951 | ENSG00000135838 |

| | | | | | | | |
|---|---|---|---|---|---|---|---|
| BSND | 1.006684134 | 2.009287676 | 0.173921979 | 0.316700858 | 6.2647638 | 12.70075192 | ENSG00000162399 |
| GADD45G | 1.005671817 | 2.007878284 | 6.88E-13 | 1.19E-11 | 162.9331808 | 327.2198193 | ENSG00000130222 |
| AMIGO2 | 1.005492859 | 2.007629233 | 2.71E-10 | 3.56E-09 | 132.9250085 | 266.4376085 | ENSG00000139211 |
| TRHDE | 1.005201814 | 2.007224262 | 0.002739003 | 0.010374652 | 31.45998554 | 63.44032055 | ENSG00000072657 |
| ZDHHC21 | 1.005175526 | 2.007187687 | 2.86E-06 | 2.07E-05 | 103.1396471 | 207.5085126 | ENSG00000175893 |
| SPINK5 | 1.005149387 | 2.007151322 | 9.05E-06 | 6.01E-05 | 63.83656692 | 128.140087 | ENSG00000133710 |
| CLDN18 | 1.003238431 | 2.004494461 | 0.398525291 | 0.570915742 | 2.27503311 | 4.615034138 | ENSG00000066405 |
| SARDH | 1.00286203 | 2.003971555 | 1.35E-21 | 5.10E-20 | 311.4941794 | 623.8121724 | ENSG00000123453 |
| SULT1C4 | 1.002569714 | 2.003565554 | 0.014719521 | 0.043591385 | 21.29767886 | 42.74634395 | ENSG00000198075 |
| RRM2B | 1.002540463 | 2.003524933 | 4.94E-46 | 7.23E-44 | 1218.219113 | 2441.336512 | ENSG00000048392 |
| MIRLET7BHG | 1.002240423 | 2.003108299 | 0.310284854 | 0.481279598 | 3.8020886 | 7.519733784 | ENSG00000197182 |
| LAMC1 | 1.001004419 | 2.001392905 | 5.69E-116 | 7.31E-113 | 4192.538256 | 8389.687545 | ENSG00000135862 |
| NA | 1.000293194 | 2.000406495 | 0.005355745 | 0.01860574 | 27.19688263 | 54.34711391 | ENSG00000259692 |
| PLD5 | 0.999949149 | 1.999929507 | 2.19E-05 | 0.000135499 | 55.10267024 | 110.1951263 | ENSG00000180287 |
| STEAP2 | 0.998330856 | 1.997687413 | 1.22E-12 | 2.06E-11 | 198.417427 | 397.1121503 | ENSG00000157214 |
| NA | 0.998298835 | 1.997643074 | 0.010594435 | 0.033065672 | 22.05769597 | 44.07836424 | ENSG00000285517 |
| RAB20 | 0.997981609 | 1.997203873 | 1.39E-22 | 5.67E-21 | 326.8603149 | 652.8567273 | ENSG00000139832 |
| TIGD4 | 0.997261614 | 1.996207391 | 0.018462409 | 0.052492016 | 25.27624594 | 50.45054299 | ENSG00000169989 |
| KCNQ5 | 0.995312158 | 1.993511818 | 0.005503512 | 0.019060727 | 26.78790745 | 53.41007937 | ENSG00000185760 |
| MARCHF1 | 0.994585701 | 1.992508254 | 0.003448522 | 0.012676342 | 29.23471312 | 58.22783732 | ENSG00000145416 |
| RNF180 | 0.994466144 | 1.992343141 | 0.007116441 | 0.023706315 | 24.23515962 | 48.21318319 | ENSG00000164197 |
| SH3BP5-AS1 | 0.994286075 | 1.992094483 | 0.08126524 | 0.175621423 | 9.666925654 | 19.24446194 | ENSG00000224660 |
| RNASE10 | 0.994101766 | 1.991840003 | 0.404666038 | 0.576913252 | 2.302266417 | 4.619335286 | ENSG00000182545 |
| NA | 0.993537819 | 1.991061548 | 0.180995933 | 0.326195118 | 5.786128194 | 11.41740131 | ENSG00000277968 |
| PDK1 | 0.993149532 | 1.990525746 | 2.67E-14 | 5.40E-13 | 218.5127718 | 434.6513903 | ENSG00000152256 |
| TMEM63A | 0.992824765 | 1.990077706 | 7.56E-47 | 1.14E-44 | 1673.066554 | 3327.767314 | ENSG00000196187 |
| CSPG5 | 0.992438545 | 1.98954502 | 9.12E-09 | 9.79E-08 | 110.8409182 | 220.816964 | ENSG00000114646 |
| ABO | 0.991793575 | 1.988655774 | 0.023037729 | 0.06312796 | 15.71123225 | 31.25527167 | ENSG00000175164 |
| REXO5 | 0.991647934 | 1.988455028 | 1.57E-06 | 1.20E-05 | 83.71313246 | 166.9089012 | ENSG00000005189 |
| ASIC4 | 0.991532965 | 1.988296574 | 0.160670335 | 0.297917416 | 6.433745451 | 12.69834088 | ENSG00000072182 |
| SESN1 | 0.991371197 | 1.98807364 | 1.37E-21 | 5.19E-20 | 322.8172689 | 642.2252383 | ENSG00000080546 |
| SLC5A3 | 0.991215985 | 1.987859765 | 6.12E-22 | 2.40E-20 | 533.7151371 | 1060.564224 | ENSG00000198743 |
| NA | 0.990266724 | 1.986552229 | 0.34884131 | 0.521545563 | 3.018304804 | 6.011590066 | ENSG00000269694 |
| FGF11 | 0.990160704 | 1.986406247 | 0.019328101 | 0.054635819 | 17.71413244 | 35.20080328 | ENSG00000161958 |
| USH1C | 0.99012207 | 1.986353053 | 0.003611537 | 0.013210119 | 32.6010688 | 64.98636898 | ENSG00000006611 |
| NAV3 | 0.989693505 | 1.985763078 | 0.002320888 | 0.008979595 | 31.64782779 | 62.97973309 | ENSG00000067798 |
| NA | 0.989269951 | 1.985180173 | 0.368622027 | 0.542409011 | 2.691705273 | 5.353373446 | ENSG00000230552 |
| ASB16 | 0.988512484 | 1.984138156 | 0.036539512 | 0.092443766 | 13.43899004 | 26.60636139 | ENSG00000161664 |
| ADAM28 | 0.988284088 | 1.983824068 | 0.085219362 | 0.182541273 | 9.571271216 | 19.0952299 | ENSG00000042980 |
| RORA | 0.988162377 | 1.983656712 | 0.000288546 | 0.001403871 | 47.5825083 | 94.39848521 | ENSG00000069667 |
| OR2A42 | 0.988051692 | 1.983504529 | 0.317398325 | 0.488583712 | 4.164294147 | 8.148908448 | ENSG00000212807 |
| NA | 0.987913882 | 1.98331507 | 8.26E-11 | 1.15E-09 | 215.8435059 | 427.9637912 | ENSG00000244879 |
| LNCPRESS1 | 0.987899173 | 1.983294848 | 9.01E-16 | 2.11E-14 | 251.3264376 | 497.8217741 | ENSG00000232301 |
| SLC4A9 | 0.987108492 | 1.982208185 | 0.415508593 | 0.587747169 | 2.008481999 | 3.992311195 | ENSG00000113073 |
| NFAM1 | 0.986837944 | 1.981836497 | 0.010307098 | 0.032252106 | 24.23862625 | 47.90129473 | ENSG00000235568 |
| NA | 0.985997334 | 1.980682084 | 0.406443528 | 0.578761733 | 2.337872433 | 4.642458558 | ENSG00000269102 |
| NA | 0.985864382 | 1.980499561 | 0.219554019 | 0.377622733 | 4.931132366 | 9.823216225 | ENSG00000277406 |
| NEK5 | 0.985561773 | 1.98008419 | 0.135629808 | 0.261147282 | 6.999662786 | 13.90371936 | ENSG00000197168 |
| NA | 0.985105492 | 1.979458049 | 0.436525968 | 0.6073972 | 2.395129951 | 4.730650503 | ENSG00000233231 |
| CCDC146 | 0.984869467 | 1.979134236 | 0.313764945 | 0.48496431 | 3.284855915 | 6.539669343 | ENSG00000135205 |
| LOC107985206 | 0.983755038 | 1.977606017 | 0.17286111 | 0.315215397 | 6.659784461 | 13.20355735 | ENSG00000286391 |
| NA | 0.983720304 | 1.977558406 | 0.275311032 | 0.442106479 | 4.26465451 | 8.478825526 | ENSG00000230847 |
| PLB1 | 0.98369038 | 1.977517388 | 0.278934162 | 0.446709166 | 3.638688754 | 7.227460957 | ENSG00000163803 |
| CRYAB | 0.983399717 | 1.977119013 | 0.026486009 | 0.071003557 | 16.59537641 | 32.91720904 | ENSG00000109846 |
| ZNF91 | 0.982758822 | 1.976240903 | 1.95E-18 | 5.74E-17 | 283.0559284 | 559.8168506 | ENSG00000167232 |
| ZBTB18 | 0.982741905 | 1.976217731 | 9.85E-16 | 2.30E-14 | 271.0105429 | 535.4327708 | ENSG00000179456 |
| LHFPL1 | 0.982617532 | 1.976047371 | 0.000296017 | 0.001436956 | 46.79680949 | 92.29531554 | ENSG00000182508 |
| APOLD1 | 0.982136303 | 1.975388345 | 0.244968041 | 0.408089679 | 4.335190821 | 8.582071487 | ENSG00000178878 |
| LOC105370295 | 0.981943187 | 1.975123942 | 0.057473215 | 0.133387367 | 11.94474967 | 23.66563503 | ENSG00000287376 |
| NA | 0.980623761 | 1.973318405 | 2.94E-08 | 2.91E-07 | 103.5448803 | 204.50568 | ENSG00000177822 |
| LPAR6 | 0.980539547 | 1.973203221 | 0.001320888 | 0.005472778 | 34.91469898 | 68.7071317 | ENSG00000139679 |
| SYCP1 | 0.980015433 | 1.972486509 | 0.367505731 | 0.541239183 | 2.716147677 | 5.346921724 | ENSG00000198765 |
| NA | 0.979917049 | 1.972352 | 0.282061659 | 0.45039744 | 3.668712964 | 7.248433657 | ENSG00000278112 |
| NA | 0.979732275 | 1.972099406 | 0.263020169 | 0.428597094 | 3.95970648 | 7.839431032 | ENSG00000178660 |
| ZBBX | 0.979719658 | 1.97208216 | 0.182773973 | 0.328448319 | 6.05826111 | 11.94359048 | ENSG00000169064 |
| C3 | 0.979249344 | 1.971439373 | 9.14E-17 | 2.35E-15 | 270.6338193 | 533.0351898 | ENSG00000125730 |
| TMEM220-AS1 | 0.978867994 | 1.970918328 | 0.090770357 | 0.191917027 | 10.13248263 | 19.947035 | ENSG00000263400 |
| SLC6A16 | 0.977819747 | 1.969486799 | 0.164606819 | 0.303578407 | 7.470801564 | 14.68023596 | ENSG00000063127 |
| NA | 0.977462608 | 1.968999313 | 0.458262963 | 0.628178129 | 2.014063805 | 3.964886775 | ENSG00000249889 |
| LOC91370 | 0.977103632 | 1.968509442 | 0.39397758 | 0.566676241 | 2.329499725 | 4.591910865 | ENSG00000268812 |
| PPL | 0.97689428 | 1.968223809 | 1.36E-44 | 1.84E-42 | 1015.712095 | 1998.206231 | ENSG00000118898 |
| SVOPL | 0.975932215 | 1.966911731 | 0.001894369 | 0.007531796 | 33.14553479 | 65.18642129 | ENSG00000157703 |
| RCN3 | 0.975828815 | 1.966770764 | 1.91E-17 | 5.20E-16 | 292.8537511 | 575.8509036 | ENSG00000142552 |
| NA | 0.975630481 | 1.966500402 | 0.031905311 | 0.082615407 | 18.39803144 | 36.17978322 | ENSG00000261799 |
| PURPL | 0.975093099 | 1.965768046 | 1.14E-07 | 1.04E-06 | 139.34336 | 273.7448891 | ENSG00000250337 |
| NA | 0.974716419 | 1.965254862 | 0.351786123 | 0.524484589 | 3.360974032 | 6.598286293 | ENSG00000269984 |
| NA | 0.974646061 | 1.965159022 | 0.342509525 | 0.515313621 | 3.039956306 | 5.961042373 | ENSG00000228285 |

| | | | | | | | |
|---|---|---|---|---|---|---|---|
| COQ8A | 0.974500253 | 1.96496042 | 1.14E-46 | 1.71E-44 | 972.1954496 | 1911.34784 | ENSG00000163050 |
| IPP | 0.974139788 | 1.964469526 | 1.39E-06 | 1.07E-05 | 84.64377085 | 166.2192316 | ENSG00000197429 |
| IER5 | 0.974023624 | 1.964311356 | 1.81E-30 | 1.16E-28 | 523.884011 | 1029.279971 | ENSG00000162783 |
| TRAF3IP3 | 0.974009691 | 1.964292386 | 0.461981194 | 0.631715787 | 2.014063805 | 3.95251637 | ENSG00000009790 |
| ADAMTS16 | 0.973902331 | 1.964146215 | 5.78E-13 | 1.01E-11 | 215.7143237 | 424.3880205 | ENSG00000145536 |
| ATP1B1 | 0.973328632 | 1.963365313 | 6.33E-38 | 6.30E-36 | 801.3925043 | 1574.690328 | ENSG00000143153 |
| DLEC1 | 0.973306595 | 1.963335322 | 0.15863872 | 0.295053947 | 7.933367474 | 15.60142299 | ENSG00000008226 |
| TSKS | 0.973095203 | 1.963047664 | 0.387123219 | 0.560491896 | 2.710565871 | 5.304976326 | ENSG00000126467 |
| NA | 0.972945841 | 1.962844441 | 8.60E-06 | 5.74E-05 | 68.37846059 | 134.0806897 | ENSG00000250320 |
| LDLRAD2 | 0.972387096 | 1.962084393 | 0.163853172 | 0.30247766 | 6.947987073 | 13.65151394 | ENSG00000187942 |
| NA | 0.972363769 | 1.962052669 | 0.142687676 | 0.271870922 | 9.0032387 | 17.66479783 | ENSG00000284968 |
| C1orf94 | 0.972237678 | 1.961881193 | 2.04E-07 | 1.79E-06 | 110.2331374 | 216.6243279 | ENSG00000142698 |
| SULT2B1 | 0.971674517 | 1.961115515 | 0.134305266 | 0.259347637 | 7.628419444 | 14.97224832 | ENSG00000088002 |
| SCN11A | 0.971657133 | 1.961091885 | 0.183228203 | 0.329111291 | 5.965397576 | 11.70967414 | ENSG00000168356 |
| ZMAT4 | 0.970451117 | 1.959453201 | 1.68E-13 | 3.10E-12 | 198.4059 | 389.1616645 | ENSG00000165061 |
| NA | 0.97044274 | 1.959441824 | 0.066366251 | 0.149540762 | 11.43608985 | 22.35727104 | ENSG00000188459 |
| MSRB3 | 0.969990319 | 1.95882745 | 0.023716324 | 0.064729996 | 19.48030558 | 38.29774644 | ENSG00000174099 |
| HPCAL4 | 0.969836589 | 1.958618734 | 0.000519003 | 0.002369725 | 38.99083552 | 76.43497494 | ENSG00000116983 |
| HLA-DPA1 | 0.969708719 | 1.958445145 | 0.001582989 | 0.006420824 | 33.27611952 | 65.01327248 | ENSG00000231389 |
| ELFN1 | 0.969566803 | 1.958252503 | 6.52E-08 | 6.15E-07 | 112.4415011 | 220.2751451 | ENSG00000225968 |
| EBF4 | 0.968590801 | 1.956928168 | 0.001815957 | 0.007255929 | 37.90876154 | 73.83921967 | ENSG00000088881 |
| NA | 0.967883787 | 1.955969381 | 0.24844786 | 0.412178161 | 4.716256968 | 9.196192135 | ENSG00000254428 |
| TOMM20 | 0.967109871 | 1.954920407 | 9.97E-84 | 4.81E-81 | 4486.619348 | 8770.783287 | ENSG00000173726 |
| BAHCC1 | 0.966758853 | 1.95444482 | 0.004123703 | 0.014845198 | 29.3706795 | 57.52284111 | ENSG00000266074 |
| ACTN3 | 0.965297064 | 1.952465511 | 2.13E-20 | 7.47E-19 | 359.1393268 | 700.8475342 | ENSG00000248746 |
| COX20 | 0.964820792 | 1.951821056 | 0.071229995 | 0.158233284 | 12.49479746 | 24.29427665 | ENSG00000203667 |
| KDELR3 | 0.964627263 | 1.951559248 | 5.63E-09 | 6.20E-08 | 122.2396872 | 238.445723 | ENSG00000100196 |
| NA | 0.962743979 | 1.949013359 | 4.70E-10 | 5.97E-09 | 157.2912285 | 306.0415805 | ENSG00000235688 |
| NA | 0.962655403 | 1.9488937 | 0.385806204 | 0.559340709 | 2.710565871 | 5.254428633 | ENSG00000280138 |
| HLA-DRA | 0.961885693 | 1.9478542 | 0.000524332 | 0.002391228 | 44.27456143 | 86.14660337 | ENSG00000204287 |
| SCX | 0.961873184 | 1.94783731 | 0.000858855 | 0.003711507 | 39.90567961 | 77.89283143 | ENSG00000260428 |
| CALB1 | 0.961414732 | 1.947218436 | 3.90E-22 | 1.54E-20 | 356.4032425 | 694.367833 | ENSG00000104327 |
| CCDC191 | 0.961351196 | 1.947132682 | 0.104226328 | 0.213851319 | 9.645274152 | 18.84490247 | ENSG00000163617 |
| MDM2 | 0.960301614 | 1.945716629 | 5.93E-19 | 1.81E-17 | 404.1945689 | 786.4490957 | ENSG00000135679 |
| NA | 0.960220936 | 1.945607825 | 0.065675595 | 0.148250397 | 12.40472483 | 24.0442218 | ENSG00000280435 |
| PCDHGA6 | 0.959744212 | 1.944965025 | 0.045612369 | 0.110847594 | 17.65475974 | 34.34791368 | ENSG00000253731 |
| PHYHIPL | 0.959601098 | 1.944772095 | 0.223340693 | 0.381888434 | 5.396689338 | 10.44163802 | ENSG00000165443 |
| NA | 0.959197408 | 1.944227992 | 0.39149548 | 0.564598636 | 2.362314838 | 4.564486445 | ENSG00000231198 |
| LACC1 | 0.958598138 | 1.943420562 | 0.000258283 | 0.001271055 | 45.53139902 | 88.45008581 | ENSG00000179630 |
| FAM71E1 | 0.958454922 | 1.943227648 | 1.15E-07 | 1.05E-06 | 98.65984183 | 191.7253385 | ENSG00000142530 |
| ZNF808 | 0.958334714 | 1.943065742 | 0.000150996 | 0.000780299 | 57.9195786 | 112.3098424 | ENSG00000198482 |
| IAH1 | 0.958152784 | 1.942820728 | 2.88E-13 | 5.22E-12 | 190.6340173 | 370.4068319 | ENSG00000134330 |
| C1RL-AS1 | 0.958069267 | 1.942708262 | 0.007360892 | 0.024376984 | 25.42425181 | 49.30027381 | ENSG00000205885 |
| STMN4 | 0.958013824 | 1.942633605 | 0.417277604 | 0.589333206 | 2.335081531 | 4.520390473 | ENSG00000015592 |
| PLIN5 | 0.957856932 | 1.942422358 | 0.050709923 | 0.120666723 | 13.94764986 | 27.12637141 | ENSG00000214456 |
| ARMC12 | 0.957579086 | 1.942048306 | 0.121581437 | 0.240778925 | 10.17857653 | 19.65718582 | ENSG00000157343 |
| SLC9A9 | 0.957556974 | 1.94201854 | 0.321794175 | 0.493264318 | 3.36934674 | 6.505793202 | ENSG00000181804 |
| LINC01600 | 0.957489596 | 1.941927845 | 0.155703974 | 0.290970198 | 7.331844123 | 14.294937 | ENSG00000164385 |
| DMBX1 | 0.956325685 | 1.940361804 | 0.107381647 | 0.2185587 | 8.959259975 | 17.4440454 | ENSG00000197587 |
| DARS2 | 0.956113081 | 1.940075881 | 7.80E-34 | 6.05E-32 | 764.6372531 | 1484.624042 | ENSG00000117593 |
| PLEKHA2 | 0.955948395 | 1.93985443 | 1.30E-07 | 1.17E-06 | 148.0537782 | 286.8441344 | ENSG00000169499 |
| FAAH2 | 0.954938296 | 1.938496723 | 1.13E-05 | 7.31E-05 | 71.19564436 | 138.0262214 | ENSG00000165591 |
| MUC12 | 0.954537127 | 1.937957762 | 0.032249113 | 0.083326767 | 16.16351043 | 31.24021765 | ENSG00000205277 |
| PSD4 | 0.954261065 | 1.937586965 | 0.002188548 | 0.008534319 | 40.36266372 | 78.46500668 | ENSG00000125637 |
| PLAAT3 | 0.953741829 | 1.93688974 | 4.31E-40 | 5.07E-38 | 1089.4194 | 2109.787281 | ENSG00000176485 |
| NA | 0.952475484 | 1.935190355 | 0.41410308 | 0.58635221 | 2.770614292 | 5.3071269 | ENSG00000269907 |
| PLS3-AS1 | 0.951821249 | 1.934312982 | 0.153486719 | 0.287886466 | 7.846285905 | 15.04806986 | ENSG00000271826 |
| NA | 0.951243371 | 1.933538339 | 0.398850673 | 0.571339447 | 2.691705273 | 5.172155372 | ENSG00000254207 |
| IL23A | 0.950405189 | 1.932415311 | 0.022903493 | 0.062867284 | 20.28382585 | 39.01726362 | ENSG00000110944 |
| ZBTB7B | 0.950393157 | 1.932399195 | 5.96E-41 | 7.33E-39 | 750.4902467 | 1450.431564 | ENSG00000160685 |
| S100A10 | 0.949367739 | 1.9310262 | 1.98E-35 | 1.68E-33 | 621.1083094 | 1199.124832 | ENSG00000197747 |
| NA | 0.94902118 | 1.930562393 | 0.004917184 | 0.017293963 | 28.51568367 | 55.23845337 | ENSG00000273329 |
| PKD1L2 | 0.948745479 | 1.930193494 | 0.002937642 | 0.011005117 | 31.32190398 | 60.56278486 | ENSG00000166473 |
| DNAJC6 | 0.948627016 | 1.930035007 | 1.84E-07 | 1.63E-06 | 95.61218854 | 184.4873972 | ENSG00000116675 |
| SLFN5 | 0.948486676 | 1.929847271 | 4.10E-07 | 3.43E-06 | 93.53057949 | 180.272183 | ENSG00000166750 |
| MIR4258 | 0.948355764 | 1.929672161 | 0.256471863 | 0.421579483 | 4.939505075 | 9.60246379 | ENSG00000264349 |
| DGKH | 0.947671429 | 1.928757049 | 7.66E-07 | 6.14E-06 | 87.82894212 | 169.2121232 | ENSG00000102780 |
| ARMCX6 | 0.947627772 | 1.928698684 | 3.18E-17 | 8.56E-16 | 279.4059127 | 538.3442374 | ENSG00000198960 |
| NA | 0.947131631 | 1.928035521 | 0.253563008 | 0.418254295 | 5.141301839 | 9.812463357 | ENSG00000287516 |
| LOC100507551 | 0.946663231 | 1.927409647 | 0.218538256 | 0.376244676 | 5.818943307 | 11.09985464 | ENSG00000269439 |
| NA | 0.946212462 | 1.926807524 | 0.243142733 | 0.40599554 | 6.109936822 | 11.76693402 | ENSG00000229666 |
| ALS2CL | 0.946042739 | 1.926580862 | 1.60E-10 | 2.16E-09 | 165.0451264 | 317.7388905 | ENSG00000178038 |
| NA | 0.945852331 | 1.926326606 | 0.036316474 | 0.091988066 | 15.59671721 | 30.10044087 | ENSG00000267424 |
| LINC01876 | 0.945692252 | 1.926112876 | 0.010931655 | 0.033980369 | 25.26451875 | 48.49444268 | ENSG00000226383 |
| TWIST1 | 0.945541404 | 1.925911493 | 0.353323804 | 0.526248417 | 3.069980516 | 5.847576582 | ENSG00000122691 |
| WNK4 | 0.945107093 | 1.9253318 | 0.051938434 | 0.122953041 | 13.67830785 | 26.37969027 | ENSG00000126562 |
| NA | 0.944700804 | 1.924789668 | 0.150356173 | 0.282950048 | 10.28539458 | 19.63567958 | ENSG00000225913 |

| | | | | | | | |
|---|---|---|---|---|---|---|---|
| DIAPH2 | 0.943940045 | 1.92377496 | 5.07E-44 | 6.84E-42 | 965.0770399 | 1855.629947 | ENSG00000147202 |
| MT1X | 0.943899907 | 1.923721439 | 0.000207675 | 0.001043566 | 58.4142839 | 112.162773 | ENSG00000187193 |
| NA | 0.943879975 | 1.923694861 | 0.435375177 | 0.606135187 | 2.694496175 | 5.149032099 | ENSG00000230010 |
| SYNPR-AS1 | 0.94384243 | 1.923644799 | 0.122465118 | 0.242220162 | 9.024214481 | 17.34295001 | ENSG00000241359 |
| TMIGD2 | 0.94281413 | 1.922274184 | 0.080236444 | 0.173817097 | 13.49816258 | 26.02747395 | ENSG00000167664 |
| COL25A1 | 0.942425558 | 1.921756513 | 2.91E-08 | 2.88E-07 | 117.6231149 | 226.1842948 | ENSG00000188517 |
| IGSF21 | 0.94152898 | 1.920562589 | 1.09E-17 | 3.03E-16 | 407.7119087 | 783.6586801 | ENSG00000117154 |
| TCIRG1 | 0.940614972 | 1.919346216 | 2.19E-17 | 5.96E-16 | 287.6543158 | 552.2008862 | ENSG00000110719 |
| DICER1-AS1 | 0.940585428 | 1.919306913 | 0.002806476 | 0.010596931 | 31.91226372 | 61.37023352 | ENSG00000235706 |
| BFSP1 | 0.940002465 | 1.918531517 | 0.000238708 | 0.001185315 | 61.83319117 | 118.4626136 | ENSG00000125864 |
| TRANK1 | 0.939344378 | 1.917656576 | 2.72E-14 | 5.47E-13 | 259.5908351 | 497.0339532 | ENSG00000168016 |
| ITLN2 | 0.938757639 | 1.91687683 | 1.16E-06 | 9.06E-06 | 91.23042825 | 174.6109497 | ENSG00000158764 |
| TENM2 | 0.938150027 | 1.91606968 | 0.001379524 | 0.005687627 | 42.62760925 | 81.67325295 | ENSG00000145934 |
| BHLHE23 | 0.937997699 | 1.91586738 | 0.149834112 | 0.282160275 | 7.655652752 | 14.66115337 | ENSG00000125533 |
| MDM4 | 0.937692371 | 1.915461955 | 3.54E-20 | 1.22E-18 | 371.9105997 | 711.9546403 | ENSG00000198625 |
| ADCY10 | 0.937354837 | 1.915013865 | 0.162039421 | 0.299559412 | 7.002453689 | 13.40710519 | ENSG00000143199 |
| AKAP7 | 0.937273311 | 1.914905651 | 1.41E-14 | 2.91E-13 | 261.3051323 | 500.2524072 | ENSG00000118507 |
| CATSPER4 | 0.937257145 | 1.914884194 | 0.164869872 | 0.303889231 | 7.756213273 | 14.81011284 | ENSG00000188782 |
| KLHL12 | 0.937082983 | 1.914653044 | 6.43E-41 | 7.80E-39 | 933.5606728 | 1786.241512 | ENSG00000117153 |
| SCYL3 | 0.936386765 | 1.91372929 | 5.89E-08 | 5.59E-07 | 109.309357 | 209.0462618 | ENSG00000000457 |
| DLG2 | 0.936182017 | 1.913457712 | 4.54E-08 | 4.39E-07 | 119.7714302 | 228.9292821 | ENSG00000150672 |
| ALPG | 0.936162628 | 1.913431995 | 0.024989897 | 0.067603437 | 21.25784248 | 40.77384466 | ENSG00000163286 |
| SELENBP1 | 0.935767315 | 1.912907768 | 4.38E-20 | 1.49E-18 | 406.1244541 | 776.0661783 | ENSG00000143416 |
| MKRN2OS | 0.935261703 | 1.912237481 | 0.24995161 | 0.414103101 | 5.054020111 | 9.669683036 | ENSG00000225526 |
| NA | 0.935156502 | 1.912098046 | 0.129171315 | 0.25157434 | 9.465329048 | 18.0119195 | ENSG00000214992 |
| SORCS2 | 0.934990621 | 1.911878206 | 8.24E-05 | 0.00045414 | 65.09687114 | 124.5174877 | ENSG00000184985 |
| CCL26 | 0.934792214 | 1.911615293 | 0.000104476 | 0.000562818 | 56.95564954 | 108.7907616 | ENSG00000006606 |
| B3GALT4 | 0.934349121 | 1.911028271 | 0.008904117 | 0.028547336 | 24.41721991 | 46.64831263 | ENSG00000235863 |
| NA | 0.934043565 | 1.910623567 | 0.3279406 | 0.500102933 | 3.638688754 | 7.006708523 | ENSG00000254595 |
| NA | 0.934043366 | 1.910623303 | 0.175844984 | 0.31917817 | 6.354836431 | 12.13261735 | ENSG00000249406 |
| FSTL5 | 0.934031313 | 1.910607341 | 0.296658118 | 0.466954055 | 4.332399918 | 8.308620785 | ENSG00000168843 |
| EPHA1-AS1 | 0.934013008 | 1.910583099 | 0.00015052 | 0.000778677 | 58.55180188 | 111.6634631 | ENSG00000229153 |
| KCNA7 | 0.933894679 | 1.9104264 | 0.118255293 | 0.235813816 | 9.710228657 | 18.5816716 | ENSG00000104848 |
| IFITM1 | 0.933862816 | 1.910384208 | 2.87E-08 | 2.85E-07 | 832.045351 | 1589.161133 | ENSG00000185885 |
| INSL3 | 0.933669946 | 1.910128831 | 0.336679074 | 0.50947557 | 3.668712964 | 7.048653921 | ENSG00000248099 |
| ADRB1 | 0.933301251 | 1.909640741 | 0.276625042 | 0.443810187 | 4.289096914 | 8.24140154 | ENSG00000043591 |
| ZNF365 | 0.933244019 | 1.909564987 | 0.204654845 | 0.358159904 | 5.647170753 | 10.80113009 | ENSG00000138311 |
| CD46 | 0.932629649 | 1.908751974 | 5.90E-38 | 5.93E-36 | 1558.452396 | 2973.30696 | ENSG00000117335 |
| KIAA1211L | 0.932240338 | 1.908236967 | 0.146793505 | 0.277599598 | 7.410753143 | 14.08467697 | ENSG00000196872 |
| NA | 0.931716337 | 1.907544003 | 0.058980995 | 0.13605194 | 15.63164751 | 29.86086631 | ENSG00000283341 |
| PCDHB17P | 0.931618021 | 1.907414013 | 0.243140962 | 0.40599554 | 5.326153027 | 10.17248847 | ENSG00000255622 |
| TUBB2A | 0.93145384 | 1.907196958 | 1.22E-92 | 8.09E-90 | 2824.085074 | 5385.091255 | ENSG00000137267 |
| NA | 0.930523226 | 1.905967113 | 0.0100963 | 0.031700475 | 29.53475506 | 56.22658707 | ENSG00000267432 |
| ADAMTS5 | 0.929906253 | 1.905152195 | 0.002094938 | 0.008219127 | 34.72618101 | 66.12589741 | ENSG00000154736 |
| C3orf35 | 0.929040151 | 1.904008806 | 0.241023382 | 0.403358803 | 5.655543462 | 10.7715551 | ENSG00000198590 |
| NMUR1 | 0.928923312 | 1.903854613 | 0.07079803 | 0.157473583 | 13.04117848 | 24.78041053 | ENSG00000171596 |
| NA | 0.928920286 | 1.90385062 | 0.123437657 | 0.243480146 | 9.290765591 | 17.73631822 | ENSG00000285852 |
| NA | 0.928068563 | 1.902726977 | 0.078886112 | 0.171508169 | 11.69705915 | 22.23090181 | ENSG00000213513 |
| NA | 0.927968981 | 1.902595645 | 0.000265924 | 0.001304997 | 58.3143991 | 110.7068066 | ENSG00000274652 |
| CGN | 0.927812567 | 1.90238938 | 1.44E-68 | 4.33E-66 | 1907.074123 | 3628.497243 | ENSG00000143375 |
| RIMBP2 | 0.926908254 | 1.901197295 | 0.013886461 | 0.041545242 | 24.49333802 | 46.45472415 | ENSG00000060709 |
| LOC400622 | 0.926773436 | 1.901019639 | 0.313584407 | 0.484762945 | 3.633106948 | 6.95616083 | ENSG00000267665 |
| P4HA1 | 0.926742757 | 1.900979214 | 1.03E-23 | 4.61E-22 | 529.6917907 | 1007.2023 | ENSG00000122884 |
| BRCA2 | 0.926699504 | 1.900922222 | 2.65E-10 | 3.48E-09 | 219.9736846 | 418.6092438 | ENSG00000139618 |
| NA | 0.926424001 | 1.900559249 | 0.01652951 | 0.047948804 | 26.34679289 | 50.15555602 | ENSG00000224411 |
| LOC729867 | 0.926341765 | 1.900450916 | 0.238531424 | 0.400479522 | 5.410643853 | 10.25422869 | ENSG00000227741 |
| RNF227 | 0.926055212 | 1.900073481 | 0.031001059 | 0.080674683 | 18.79516728 | 35.61164866 | ENSG00000179859 |
| OSCAR | 0.925721145 | 1.899633555 | 0.000436796 | 0.002037288 | 51.48264192 | 97.96518169 | ENSG00000170909 |
| NAALADL2 | 0.925298558 | 1.899077205 | 0.31969216 | 0.491252306 | 4.259072704 | 8.127402711 | ENSG00000177694 |
| CTNNA3 | 0.924917087 | 1.898575126 | 0.230355299 | 0.390662336 | 6.093867126 | 11.52279785 | ENSG00000183230 |
| NA | 0.924754091 | 1.898360637 | 0.153787998 | 0.288339417 | 7.598395234 | 14.50546961 | ENSG00000253540 |
| NA | 0.924132152 | 1.897542438 | 0.048215137 | 0.115968406 | 14.77820328 | 28.01151961 | ENSG00000189212 |
| QRICH2 | 0.923491207 | 1.896699606 | 0.001479837 | 0.006054622 | 38.74661163 | 73.48617932 | ENSG00000129646 |
| TRIM6 | 0.923426551 | 1.896614605 | 1.22E-13 | 2.29E-12 | 223.0530259 | 423.1258846 | ENSG00000121236 |
| LINC-PINT | 0.92324478 | 1.896375659 | 0.289492511 | 0.4588825 | 4.291887817 | 8.167730521 | ENSG00000231721 |
| LOC105376199 | 0.922881954 | 1.895898795 | 0.357571946 | 0.530813421 | 3.273692303 | 6.261917494 | ENSG00000228317 |
| CACNG6 | 0.922437563 | 1.895314895 | 0.018903465 | 0.053580184 | 21.1030155 | 39.92527464 | ENSG00000130433 |
| NA | 0.922380964 | 1.89524054 | 0.018619086 | 0.052872672 | 22.76603737 | 43.14831446 | ENSG00000265749 |
| NA | 0.921723873 | 1.894377529 | 0.269549748 | 0.435935678 | 4.313539319 | 8.190853847 | ENSG00000242611 |
| NA | 0.920816954 | 1.893187044 | 0.040645021 | 0.100963572 | 19.73445377 | 37.32951937 | ENSG00000233290 |
| NA | 0.92072058 | 1.89306058 | 0.279582612 | 0.447475995 | 4.398030145 | 8.310771359 | ENSG00000271727 |
| NA | 0.920659939 | 1.892981011 | 0.015575156 | 0.045657223 | 29.82718804 | 56.32124284 | ENSG00000236333 |
| NA | 0.920573294 | 1.892867326 | 0.048218474 | 0.115968406 | 14.42157954 | 27.25865933 | ENSG00000279249 |
| NA | 0.920379266 | 1.892612772 | 0.370266024 | 0.544051977 | 3.57584943 | 6.815270618 | ENSG00000286970 |
| APOBEC3F | 0.920171658 | 1.892340439 | 1.72E-05 | 0.000108306 | 78.21887367 | 148.3646013 | ENSG00000128394 |
| NA | 0.920112679 | 1.892263079 | 0.006402796 | 0.021694419 | 28.68187442 | 54.25005921 | ENSG00000260025 |
| LMO3 | 0.919706569 | 1.891730493 | 3.21E-07 | 2.74E-06 | 140.910127 | 266.9842252 | ENSG00000048540 |

| | | | | | | | |
|---|---|---|---|---|---|---|---|
| TPO | 0.919138908 | 1.890986295 | 0.043379368 | 0.10650775 | 18.78467939 | 35.4807057 | ENSG00000115705 |
| CASC9 | 0.918750878 | 1.890477761 | 0.004385676 | 0.01566162 | 35.7557403 | 67.38880835 | ENSG00000249395 |
| LOC103611081 | 0.918707322 | 1.890420687 | 0.068270178 | 0.152908396 | 13.85566221 | 26.12882981 | ENSG00000255455 |
| LINC02751 | 0.918517369 | 1.8901718 | 0.084109191 | 0.180564257 | 11.67261675 | 22.10694362 | ENSG00000254946 |
| DYNC1I1 | 0.918290592 | 1.889874708 | 0.027904504 | 0.074126383 | 18.05189559 | 34.09652018 | ENSG00000158560 |
| OXSM | 0.918187197 | 1.889739269 | 0.043610807 | 0.106994286 | 19.56567228 | 36.80117963 | ENSG00000151093 |
| ZBTB37 | 0.917950119 | 1.889428754 | 2.23E-07 | 1.95E-06 | 165.7443441 | 312.6118667 | ENSG00000185278 |
| EPCAM-DT | 0.917899207 | 1.889362079 | 0.029572009 | 0.077604901 | 19.67863572 | 37.23513617 | ENSG00000234690 |
| HMGB3 | 0.916453269 | 1.887469418 | 6.43E-124 | 9.53E-121 | 6576.241435 | 12413.56698 | ENSG00000029993 |
| NA | 0.91624196 | 1.887192983 | 0.008422863 | 0.027253514 | 26.89875571 | 50.88449953 | ENSG00000205918 |
| RAP2C-AS1 | 0.916195946 | 1.887132793 | 0.147974468 | 0.279531478 | 8.697614949 | 16.40268039 | ENSG00000232160 |
| LINC00857 | 0.915634947 | 1.886399116 | 0.320069268 | 0.491508852 | 3.660340256 | 6.924435262 | ENSG00000237523 |
| XPR1 | 0.915457562 | 1.886167189 | 3.25E-55 | 6.46E-53 | 1443.101576 | 2722.263768 | ENSG00000143324 |
| SAMD4A | 0.91508264 | 1.885677084 | 5.10E-06 | 3.54E-05 | 87.04707335 | 164.3836003 | ENSG00000020577 |
| NA | 0.914756733 | 1.885251154 | 0.281896044 | 0.450277013 | 4.395239242 | 8.260223666 | ENSG00000213862 |
| LINC02076 | 0.914007235 | 1.884271997 | 0.102859964 | 0.211621887 | 10.55386071 | 19.97419895 | ENSG00000220161 |
| CKM | 0.913947648 | 1.884194174 | 0.337413181 | 0.510173133 | 3.630316045 | 6.865818312 | ENSG00000104879 |
| NA | 0.91379072 | 1.883989233 | 0.064644065 | 0.146291764 | 12.37749153 | 23.30614296 | ENSG00000287562 |
| NPIPA1 | 0.913377871 | 1.883450178 | 1.85E-07 | 1.63E-06 | 115.1184881 | 216.6020223 | ENSG00000183426 |
| LOC100507071 | 0.913016236 | 1.88297812 | 0.221683741 | 0.38009964 | 5.913721863 | 11.2240733 | ENSG00000254064 |
| MEF2C | 0.912948612 | 1.88288986 | 0.05142521 | 0.12203761 | 14.64973373 | 27.63750656 | ENSG00000081189 |
| PLBD1 | 0.912701765 | 1.882567723 | 7.46E-11 | 1.04E-09 | 179.888084 | 338.5874664 | ENSG00000121316 |
| PURA | 0.912579574 | 1.882408283 | 4.95E-06 | 3.44E-05 | 92.43172319 | 174.2842677 | ENSG00000185129 |
| NA | 0.911829014 | 1.881429218 | 0.278659814 | 0.446406451 | 4.343563529 | 8.159128279 | ENSG00000279800 |
| ASPM | 0.911447007 | 1.880931105 | 3.00E-29 | 1.86E-27 | 1074.123801 | 2019.579232 | ENSG00000066279 |
| GCNA | 0.910949749 | 1.880282911 | 0.12891875 | 0.25123463 | 9.862264732 | 18.63625998 | ENSG00000147174 |
| GOLGA6D | 0.910750264 | 1.880022938 | 0.223897776 | 0.382400354 | 6.169985243 | 11.48892171 | ENSG00000140478 |
| NA | 0.910215815 | 1.879326609 | 0.310399417 | 0.481304972 | 4.313539319 | 8.102661903 | ENSG00000281501 |
| SNORA66 | 0.909290152 | 1.878121181 | 0.336727672 | 0.50947557 | 3.665922062 | 6.891092158 | ENSG00000207523 |
| ENG | 0.909023951 | 1.877774668 | 1.20E-05 | 7.75E-05 | 82.96024876 | 155.9424311 | ENSG00000106991 |
| UEVLD | 0.908103499 | 1.876577013 | 3.03E-05 | 0.000182416 | 80.12006205 | 150.3126445 | ENSG00000151116 |
| NA | 0.908085827 | 1.876554027 | 0.221201025 | 0.379575607 | 6.022655094 | 11.33968966 | ENSG00000285103 |
| LRRK1 | 0.907646999 | 1.875983318 | 1.98E-05 | 0.000123711 | 107.5403049 | 201.7321838 | ENSG00000154237 |
| CYTL1 | 0.907646483 | 1.875982648 | 1.39E-07 | 1.25E-06 | 115.7668691 | 217.0773912 | ENSG00000170891 |
| NA | 0.907628441 | 1.875959187 | 0.306968409 | 0.477916083 | 4.044197305 | 7.563829757 | ENSG00000280407 |
| PIPOX | 0.907606154 | 1.875930206 | 2.17E-27 | 1.19E-25 | 826.5124824 | 1549.959938 | ENSG00000179761 |
| NA | 0.907505628 | 1.875799498 | 0.196621512 | 0.347314861 | 8.618705929 | 16.15801118 | ENSG00000250731 |
| NA | 0.907048265 | 1.875204927 | 0.008252566 | 0.026810415 | 44.06016159 | 82.62967319 | ENSG00000261786 |
| MAOB | 0.906484657 | 1.874472495 | 0.000568506 | 0.002568997 | 47.26986329 | 88.54688005 | ENSG00000069535 |
| NA | 0.906209927 | 1.874115577 | 0.136547163 | 0.262625257 | 8.736011868 | 16.30803673 | ENSG00000253170 |
| LOC105378663 | 0.905869109 | 1.873672894 | 0.422025967 | 0.593804067 | 2.637238657 | 4.980977931 | ENSG00000228436 |
| NA | 0.905493285 | 1.873184863 | 0.289115064 | 0.458547949 | 4.787469 | 8.89961816 | ENSG00000273001 |
| SWT1 | 0.905298158 | 1.872931529 | 3.66E-07 | 3.10E-06 | 107.3699718 | 201.1484622 | ENSG00000116668 |
| SNORA73B | 0.905197338 | 1.872800647 | 0.022932492 | 0.062911078 | 21.07299129 | 39.46656518 | ENSG00000200087 |
| RNF32 | 0.905032527 | 1.872586713 | 0.05936645 | 0.136728533 | 15.31967821 | 28.77083162 | ENSG00000105982 |
| KITLG | 0.904909667 | 1.872427251 | 5.83E-06 | 4.00E-05 | 87.94046609 | 164.7575974 | ENSG00000049130 |
| TMEM38A | 0.904826661 | 1.872319524 | 4.54E-11 | 6.46E-10 | 189.1979104 | 353.9649019 | ENSG00000072954 |
| NOSTRIN | 0.904596223 | 1.872020486 | 0.236038816 | 0.397481288 | 5.619937445 | 10.58467881 | ENSG00000163072 |
| DPH6 | 0.904459354 | 1.871842895 | 1.96E-08 | 1.99E-07 | 129.4081315 | 242.3280455 | ENSG00000134146 |
| LRRC8C-DT | 0.904089091 | 1.871362555 | 0.441481231 | 0.612573863 | 3.538328392 | 6.510094349 | ENSG00000231999 |
| RHBG | 0.904053375 | 1.871316227 | 0.019760501 | 0.055649608 | 20.37860441 | 38.15658366 | ENSG00000132677 |
| NA | 0.904022624 | 1.871276341 | 0.321012598 | 0.49234012 | 3.665922062 | 6.857216017 | ENSG00000287917 |
| GPSM3 | 0.903866407 | 1.871073727 | 3.92E-05 | 0.000230399 | 84.43303779 | 157.6584359 | ENSG00000213654 |
| NA | 0.903473068 | 1.870563664 | 0.02618475 | 0.070303411 | 23.75373311 | 44.17221441 | ENSG00000259048 |
| ADGRE2 | 0.903451644 | 1.870535886 | 0.009084164 | 0.029008873 | 28.3475779 | 53.12909245 | ENSG00000127507 |
| INKA2 | 0.90290072 | 1.869821718 | 1.58E-12 | 2.63E-11 | 308.1555695 | 575.2725676 | ENSG00000197852 |
| NA | 0.902315604 | 1.869063526 | 0.028747707 | 0.075885607 | 20.31664096 | 37.79279044 | ENSG00000263301 |
| MME | 0.901864335 | 1.868478981 | 2.69E-11 | 3.92E-10 | 238.6110707 | 445.7152003 | ENSG00000196549 |
| MIR573 | 0.901818547 | 1.868419681 | 0.303671865 | 0.474419659 | 4.346354432 | 8.08975846 | ENSG00000207697 |
| H2BC14 | 0.900477208 | 1.866683334 | 0.431524445 | 0.602593258 | 2.604423544 | 4.905156391 | ENSG00000273703 |
| PTAR1 | 0.90014645 | 1.86625542 | 3.92E-26 | 2.05E-24 | 709.5969963 | 1324.161432 | ENSG00000188647 |
| NA | 0.899214146 | 1.86504979 | 0.373974945 | 0.547214396 | 3.012722998 | 5.626824147 | ENSG00000279863 |
| CAPG | 0.899067142 | 1.86485976 | 4.30E-31 | 2.89E-29 | 693.1475106 | 1291.77724 | ENSG00000042493 |
| MMP23A | 0.898890546 | 1.864631502 | 0.402624942 | 0.574952828 | 2.656099256 | 4.980977931 | ENSG00000215914 |
| NA | 0.897801616 | 1.863224631 | 0.336764629 | 0.509491527 | 3.663131159 | 6.813120045 | ENSG00000230289 |
| NA | 0.897782042 | 1.863199351 | 0.018349069 | 0.052192846 | 22.01718387 | 41.05268101 | ENSG00000264015 |
| NDUFS2 | 0.897524562 | 1.862866853 | 4.46E-74 | 1.62E-71 | 2181.738259 | 4063.36283 | ENSG00000158864 |
| NA | 0.897065713 | 1.862274462 | 0.046964771 | 0.113618936 | 18.76147629 | 35.11529495 | ENSG00000239415 |
| PTGER3 | 0.896619687 | 1.861698807 | 0.129522305 | 0.252105214 | 10.29568231 | 19.25924338 | ENSG00000050628 |
| UBE2D4 | 0.8965725 | 1.861637916 | 0.010261859 | 0.032136776 | 27.383849 | 50.91381406 | ENSG00000078967 |
| CSF2RA | 0.896173842 | 1.861123563 | 0.014862076 | 0.043944166 | 24.57571367 | 45.59457722 | ENSG00000198223 |
| FTCDNL1 | 0.895791736 | 1.860630699 | 0.083410727 | 0.179464235 | 11.69705915 | 21.79584866 | ENSG00000226124 |
| LINC00551 | 0.895766019 | 1.860597533 | 0.107092679 | 0.218108623 | 11.95611344 | 22.09457321 | ENSG00000272274 |
| GNAS-AS1 | 0.895717865 | 1.860535432 | 0.123696198 | 0.243865442 | 10.32012472 | 19.18316138 | ENSG00000235590 |
| PDCD4 | 0.895383403 | 1.860104151 | 4.94E-14 | 9.61E-13 | 255.8938128 | 475.8185974 | ENSG00000150593 |
| NA | 0.894403953 | 1.858841749 | 0.35691442 | 0.530336733 | 3.698737175 | 6.848613722 | ENSG00000224614 |
| FCSK | 0.893866398 | 1.858149266 | 6.54E-08 | 6.17E-07 | 151.1645462 | 280.9048766 | ENSG00000157353 |

| | | | | | | | |
|---|---|---|---|---|---|---|---|
| NA | 0.892805854 | 1.856783817 | 0.24837238 | 0.412088379 | 5.314989415 | 9.907107022 | ENSG00000271474 |
| NA | 0.892576272 | 1.856488363 | 0.431125243 | 0.602337652 | 2.634447755 | 4.91590926 | ENSG00000273568 |
| NA | 0.892012781 | 1.855763393 | 2.73E-69 | 8.51E-67 | 2370.909925 | 4399.569243 | ENSG00000249790 |
| NA | 0.891646677 | 1.855292527 | 0.403838733 | 0.576115175 | 2.661681062 | 4.955704085 | ENSG00000230131 |
| NA | 0.891590843 | 1.855220727 | 0.156321357 | 0.291869464 | 9.334744316 | 17.40612857 | ENSG00000250138 |
| NLGN4Y | 0.890810745 | 1.854217838 | 0.375135892 | 0.548487684 | 3.039956306 | 5.622523 | ENSG00000165246 |
| C1QTNF6 | 0.890324848 | 1.853593446 | 0.000118565 | 0.000630265 | 61.96078484 | 114.6875593 | ENSG00000133466 |
| CDC42BPA | 0.890220101 | 1.85345887 | 4.30E-33 | 3.28E-31 | 824.489921 | 1528.198201 | ENSG00000143776 |
| NA | 0.890137154 | 1.853352309 | 0.00051177 | 0.002340579 | 52.87911268 | 98.17812533 | ENSG00000260077 |
| NA | 0.889686717 | 1.852773748 | 0.386014443 | 0.559523944 | 3.015513901 | 5.580577601 | ENSG00000287836 |
| ALDH1A3 | 0.889381318 | 1.852381582 | 0.002510325 | 0.009625658 | 39.26663522 | 72.63356861 | ENSG00000184254 |
| JAK1 | 0.889374553 | 1.852372896 | 1.15E-28 | 6.84E-27 | 664.2869545 | 1230.074339 | ENSG00000162434 |
| ABCC2 | 0.889109837 | 1.852033041 | 0.139072475 | 0.266499168 | 8.687127059 | 16.08943487 | ENSG00000023839 |
| NA | 0.888762573 | 1.851587301 | 0.102764423 | 0.211487786 | 10.69281815 | 19.83356921 | ENSG00000223551 |
| MGST2 | 0.888256953 | 1.850938491 | 6.57E-05 | 0.000369085 | 69.97027043 | 129.4344815 | ENSG00000085871 |
| SNHG31 | 0.888256689 | 1.850938152 | 0.426702739 | 0.598221114 | 2.949883674 | 5.473030494 | ENSG00000229267 |
| FAHD2CP | 0.887989957 | 1.850595973 | 0.002194676 | 0.00855648 | 43.16928434 | 79.574948 | ENSG00000231584 |
| MFSD4A | 0.887862543 | 1.850432542 | 0.222266198 | 0.380692287 | 6.079912612 | 11.21547101 | ENSG00000174514 |
| HLA-G | 0.887758123 | 1.850298616 | 0.000719658 | 0.003160236 | 48.35859511 | 89.30163045 | ENSG00000204632 |
| CHRFAM7A | 0.887467777 | 1.849926276 | 0.071815562 | 0.159204091 | 14.62250042 | 27.15944194 | ENSG00000166664 |
| SIM1 | 0.88726323 | 1.84966401 | 0.268372362 | 0.43470554 | 4.966738382 | 9.230068276 | ENSG00000112246 |
| NA | 0.886609107 | 1.848825555 | 0.193346747 | 0.343069546 | 7.410753143 | 13.61763779 | ENSG00000269378 |
| TSEN15 | 0.886385286 | 1.84853875 | 4.58E-26 | 2.37E-24 | 666.1274698 | 1231.605848 | ENSG00000198860 |
| CHODL | 0.886247187 | 1.848361811 | 0.023562364 | 0.064382658 | 23.71880282 | 43.60864145 | ENSG00000154645 |
| DYDC2 | 0.886056927 | 1.848118068 | 4.67E-05 | 0.000270069 | 67.88499459 | 125.415812 | ENSG00000133665 |
| RASAL2-AS1 | 0.885936751 | 1.847964127 | 0.021811375 | 0.060341781 | 24.9428253 | 46.0425217 | ENSG00000224687 |
| NA | 0.885900164 | 1.847917262 | 0.054573718 | 0.128021028 | 15.29176918 | 28.34196973 | ENSG00000279756 |
| NA | 0.884990562 | 1.84675254 | 0.266262334 | 0.432475357 | 4.958365673 | 9.198342708 | ENSG00000215861 |
| MGMT | 0.884969914 | 1.84672611 | 1.78E-15 | 4.04E-14 | 274.5315254 | 507.1166506 | ENSG00000170430 |
| FBXO4 | 0.884810251 | 1.846521744 | 0.07252923 | 0.160491096 | 13.81447438 | 25.47491433 | ENSG00000151876 |
| NA | 0.884190211 | 1.845728318 | 0.037392249 | 0.094255048 | 18.27167707 | 33.79402752 | ENSG00000283662 |
| RIPPLY2 | 0.884036249 | 1.845531355 | 0.438242124 | 0.609218766 | 2.691705273 | 4.964306379 | ENSG00000203877 |
| NA | 0.883584842 | 1.844953995 | 0.006420636 | 0.021745171 | 31.09586497 | 57.43249859 | ENSG00000280142 |
| ZDHHC1 | 0.883584183 | 1.844953152 | 0.000485434 | 0.002232834 | 53.60304822 | 98.7874118 | ENSG00000159714 |
| KRCC1 | 0.88326298 | 1.844542435 | 6.42E-09 | 7.02E-08 | 138.0596525 | 254.6661008 | ENSG00000172086 |
| CYP2W1 | 0.882953653 | 1.844146991 | 0.369190008 | 0.542962907 | 3.426604259 | 6.264068068 | ENSG00000073067 |
| NA | 0.882464239 | 1.843521496 | 0.121218029 | 0.240355329 | 10.25170359 | 18.95218911 | ENSG00000273295 |
| CD74 | 0.881993438 | 1.84291999 | 1.29E-29 | 8.01E-28 | 613.1882207 | 1129.909236 | ENSG00000019582 |
| UCP3 | 0.880495943 | 1.84100806 | 0.189998836 | 0.338093124 | 7.331844123 | 13.52917328 | ENSG00000175564 |
| GMPR | 0.880492751 | 1.841003987 | 0.049296771 | 0.11794418 | 16.53811889 | 30.57769989 | ENSG00000137198 |
| NA | 0.880375865 | 1.840854836 | 0.006548824 | 0.022117187 | 35.57280413 | 65.3969539 | ENSG00000249898 |
| NA | 0.880116825 | 1.840524335 | 0.188169524 | 0.335597049 | 7.650070946 | 14.09973099 | ENSG00000287406 |
| RNF112 | 0.879930545 | 1.840286703 | 0.001429892 | 0.005875198 | 45.59268674 | 84.12543439 | ENSG00000128482 |
| SSUH2 | 0.879771419 | 1.840083735 | 0.224364996 | 0.382927111 | 6.05826111 | 11.09770407 | ENSG00000125046 |
| DNER | 0.879598887 | 1.839863692 | 0.000178023 | 0.000906611 | 60.63062003 | 111.4913804 | ENSG00000187957 |
| SCAPER | 0.878986807 | 1.839083274 | 1.05E-05 | 6.85E-05 | 107.6555837 | 198.0213805 | ENSG00000140386 |
| TSHZ1 | 0.878704449 | 1.838723372 | 0.224281239 | 0.382851905 | 6.654202655 | 12.28856158 | ENSG00000179981 |
| NA | 0.878703228 | 1.838721816 | 0.392948479 | 0.565783632 | 3.578640333 | 6.661737429 | ENSG00000268047 |
| PAQR7 | 0.87852852 | 1.838499164 | 3.84E-07 | 3.23E-06 | 127.8414398 | 235.1358056 | ENSG00000182749 |
| NA | 0.87852203 | 1.838490893 | 0.307513349 | 0.478560827 | 4.286306011 | 7.938375845 | ENSG00000183022 |
| C16orf74 | 0.878483695 | 1.838442042 | 0.020718869 | 0.057850296 | 26.79839534 | 49.21717651 | ENSG00000154102 |
| NA | 0.878481437 | 1.838439164 | 0.096941713 | 0.202238844 | 12.40751574 | 22.82000908 | ENSG00000270012 |
| GDPGP1 | 0.878259082 | 1.838155837 | 0.316308787 | 0.487378373 | 5.361759043 | 9.846339498 | ENSG00000183208 |
| NA | 0.878195093 | 1.838074309 | 0.386197024 | 0.559606508 | 3.57584943 | 6.657436282 | ENSG00000286918 |
| SDCCAG8 | 0.877943652 | 1.837753988 | 1.14E-05 | 7.41E-05 | 93.75402775 | 171.9323881 | ENSG00000054282 |
| NA | 0.877904868 | 1.837704584 | 0.414472784 | 0.586746494 | 3.048329015 | 5.571975306 | ENSG00000243289 |
| LAMA3 | 0.877893409 | 1.837689988 | 0.011921645 | 0.036446982 | 28.90532268 | 53.04843672 | ENSG00000053747 |
| PGLYRP1 | 0.877739347 | 1.837493756 | 0.251727588 | 0.416047886 | 6.001003592 | 10.99660868 | ENSG00000008438 |
| SP100 | 0.877636729 | 1.83736306 | 0.41621427 | 0.588305208 | 3.0343745 | 5.548852033 | ENSG00000067066 |
| MYH7 | 0.876502679 | 1.835919344 | 0.39266515 | 0.565589811 | 3.072771419 | 5.597249153 | ENSG00000092054 |
| GNGT2 | 0.876424178 | 1.835819449 | 0.271160527 | 0.437993614 | 4.988389884 | 9.168767714 | ENSG00000167083 |
| ADAMTS10 | 0.876278499 | 1.835634082 | 2.60E-10 | 3.42E-09 | 179.9242536 | 330.321615 | ENSG00000142303 |
| TEN1-CDK3 | 0.875802593 | 1.835028657 | 0.154692048 | 0.289612219 | 8.384969932 | 15.33819212 | ENSG00000261408 |
| BTN3A2 | 0.87565733 | 1.8348439 | 0.000153407 | 0.000791278 | 74.78776364 | 137.2671577 | ENSG00000186470 |
| LINC02204 | 0.875582672 | 1.834748951 | 0.454152546 | 0.624130737 | 2.740590082 | 5.00195063 | ENSG00000280639 |
| TM4SF1 | 0.87554332 | 1.834698906 | 0.182157998 | 0.327646614 | 7.628419444 | 14.00885543 | ENSG00000169908 |
| NA | 0.875521705 | 1.834671418 | 0.164576044 | 0.303555397 | 8.790478483 | 16.04695643 | ENSG00000262333 |
| GORAB | 0.875430109 | 1.834554938 | 9.91E-09 | 1.06E-07 | 207.6936745 | 381.4185436 | ENSG00000120370 |
| SPART | 0.87535391 | 1.834458045 | 2.24E-40 | 2.67E-38 | 931.3623597 | 1708.764602 | ENSG00000133104 |
| FAM167B | 0.875291346 | 1.834378494 | 0.034036714 | 0.087024818 | 21.69337524 | 39.73975541 | ENSG00000183615 |
| ATRNL1 | 0.875029152 | 1.834045145 | 5.24E-09 | 5.80E-08 | 156.7981259 | 287.7766378 | ENSG00000107518 |
| DOCK2 | 0.874710838 | 1.833640529 | 2.59E-07 | 2.24E-06 | 119.5987818 | 219.3125883 | ENSG00000134516 |
| SPI1 | 0.874309545 | 1.833130563 | 0.288773808 | 0.458119699 | 4.996762593 | 9.162849031 | ENSG00000066336 |
| NA | 0.873988437 | 1.832722601 | 3.64E-05 | 0.000215348 | 80.03692267 | 146.8976095 | ENSG00000282508 |
| PRKN | 0.873584563 | 1.832209612 | 0.227518817 | 0.386835108 | 7.149783838 | 12.98039387 | ENSG00000185345 |
| JMJD4 | 0.873501206 | 1.832103753 | 0.005054709 | 0.017706625 | 34.62562049 | 63.46884158 | ENSG00000081692 |
| NA | 0.873157482 | 1.831667304 | 0.239444395 | 0.401523107 | 5.758894886 | 10.48358342 | ENSG00000231381 |

| | | | | | | | |
|---|---|---|---|---|---|---|---|
| ARRDC2 | 0.872890869 | 1.831328839 | 7.75E-06 | 5.21E-05 | 96.80793857 | 176.8966945 | ENSG00000105643 |
| SPATA18 | 0.872715551 | 1.831106307 | 0.025448575 | 0.068632526 | 20.84974319 | 38.07646097 | ENSG00000163071 |
| MRPL24 | 0.872455842 | 1.830776708 | 1.12E-21 | 4.28E-20 | 436.7229862 | 799.4529344 | ENSG00000143314 |
| SPATS2L | 0.871624941 | 1.829722599 | 1.84E-26 | 9.82E-25 | 943.9901055 | 1726.814748 | ENSG00000196141 |
| TMEM154 | 0.871068854 | 1.829017468 | 0.089994909 | 0.190674551 | 15.56968407 | 28.39036685 | ENSG00000170006 |
| BIK | 0.870821839 | 1.828704336 | 0.075745735 | 0.166046701 | 13.26721749 | 24.34750796 | ENSG00000100290 |
| RO60 | 0.870072983 | 1.82775536 | 5.27E-11 | 7.45E-10 | 319.3112432 | 584.4547229 | ENSG00000116747 |
| MEIOC | 0.870008051 | 1.827673099 | 7.60E-06 | 5.12E-05 | 104.3833309 | 190.5971265 | ENSG00000180336 |
| NA | 0.869900557 | 1.827536926 | 0.095267738 | 0.199436781 | 11.72150156 | 21.42345315 | ENSG00000258815 |
| PHF24 | 0.869748244 | 1.827343994 | 0.143620012 | 0.273205172 | 9.380838223 | 17.1410197 | ENSG00000122733 |
| DCAF6 | 0.869734297 | 1.827326328 | 1.40E-14 | 2.88E-13 | 285.5928067 | 521.8003879 | ENSG00000143164 |
| MATN4 | 0.869082058 | 1.826500386 | 0.045105643 | 0.109879943 | 20.76054644 | 37.66642121 | ENSG00000124159 |
| LINC01091 | 0.868733283 | 1.826058878 | 0.459235319 | 0.629108688 | 2.30505732 | 4.221665925 | ENSG00000249464 |
| HES7 | 0.868475867 | 1.825733088 | 0.235801776 | 0.397216745 | 6.330394027 | 11.59646881 | ENSG00000179111 |
| XG | 0.868438483 | 1.82568578 | 0.07610836 | 0.166557728 | 14.14107392 | 25.78600928 | ENSG00000124343 |
| PRKG2 | 0.868321877 | 1.825538225 | 0.031360119 | 0.08142638 | 30.12878159 | 54.80313976 | ENSG00000138669 |
| SDE2 | 0.867913227 | 1.825021205 | 1.01E-17 | 2.82E-16 | 404.5167748 | 737.3304041 | ENSG00000143751 |
| C10orf105 | 0.867572791 | 1.824590601 | 0.184382755 | 0.330530317 | 8.121209724 | 14.73912548 | ENSG00000214688 |
| AP3B2 | 0.865543069 | 1.822025397 | 1.75E-12 | 2.90E-11 | 264.0785375 | 480.607647 | ENSG00000103723 |
| PRELID2 | 0.865427066 | 1.8218789 | 0.000942328 | 0.004038756 | 50.28450131 | 91.76750884 | ENSG00000186314 |
| COL6A2 | 0.865323205 | 1.821747746 | 9.28E-68 | 2.67E-65 | 3325.788053 | 6059.551733 | ENSG00000142173 |
| LYSMD1 | 0.864979383 | 1.82131364 | 8.41E-07 | 6.71E-06 | 106.1089919 | 193.1442242 | ENSG00000163155 |
| GDPD5 | 0.86482037 | 1.821112906 | 1.22E-06 | 9.52E-06 | 115.048152 | 209.1581101 | ENSG00000158555 |
| RPH3AL | 0.86453701 | 1.820755256 | 0.016721075 | 0.048397038 | 27.86894229 | 50.58336394 | ENSG00000181031 |
| GNG12 | 0.864493199 | 1.820699965 | 5.60E-20 | 1.89E-18 | 524.9044849 | 956.461386 | ENSG00000172380 |
| NA | 0.864426437 | 1.820615713 | 0.452195533 | 0.622381851 | 2.365105741 | 4.293186317 | ENSG00000259374 |
| KTI12 | 0.864122611 | 1.820232338 | 0.028900308 | 0.076184181 | 21.66267531 | 39.49237206 | ENSG00000198841 |
| GRIN2A | 0.863664108 | 1.819653943 | 0.000212936 | 0.001066991 | 75.87948653 | 137.6078519 | ENSG00000183454 |
| ZADH2 | 0.863577774 | 1.819545054 | 3.87E-05 | 0.00022753 | 80.58226454 | 146.4434737 | ENSG00000180011 |
| ACVRL1 | 0.863436712 | 1.819367154 | 0.153906489 | 0.288533535 | 9.076565914 | 16.49517348 | ENSG00000139567 |
| KLHL11 | 0.863430665 | 1.819359528 | 0.000188231 | 0.000953679 | 68.71363299 | 124.7239797 | ENSG00000178502 |
| PGPEP1 | 0.863408639 | 1.819331751 | 1.41E-28 | 8.24E-27 | 615.593275 | 1119.619241 | ENSG00000130517 |
| NA | 0.862439581 | 1.818110116 | 0.2279172 | 0.387370537 | 6.768717691 | 12.27995928 | ENSG00000243926 |
| PSMB9 | 0.862343695 | 1.817989283 | 0.002062504 | 0.00810837 | 42.08804935 | 76.53230221 | ENSG00000240065 |
| SENP8 | 0.861364862 | 1.81675624 | 0.00110652 | 0.004665766 | 46.52168551 | 84.49165076 | ENSG00000166192 |
| LINC02453 | 0.861190767 | 1.81653702 | 0.3664199 | 0.540190067 | 3.603082738 | 6.606888588 | ENSG00000245017 |
| NA | 0.860910756 | 1.816184484 | 0.34273271 | 0.515435532 | 4.022545804 | 7.322104623 | ENSG00000251661 |
| MGLL | 0.86084037 | 1.816095878 | 0.094690552 | 0.198357633 | 14.6177945 | 26.42808739 | ENSG00000074416 |
| CACNB4 | 0.860458029 | 1.815614643 | 0.009371095 | 0.029782174 | 32.9186199 | 59.90671881 | ENSG00000182389 |
| NA | 0.860217175 | 1.815311556 | 0.472994602 | 0.641759661 | 2.661681062 | 4.814813873 | ENSG00000228373 |
| C1orf116 | 0.860040666 | 1.815089473 | 9.01E-06 | 5.99E-05 | 104.7203303 | 189.7093068 | ENSG00000182795 |
| LINC00605 | 0.859675881 | 1.814630586 | 0.009983187 | 0.031375962 | 31.9941638 | 57.92240058 | ENSG00000251533 |
| NA | 0.859311806 | 1.814172707 | 0.363513414 | 0.536988841 | 3.630316045 | 6.636463582 | ENSG00000233916 |
| NA | 0.859289533 | 1.8141447 | 0.123151446 | 0.243114464 | 12.56321859 | 22.67320018 | ENSG00000262580 |
| NA | 0.859104452 | 1.813911982 | 0.073508678 | 0.162286096 | 13.76838048 | 24.91323149 | ENSG00000272145 |
| NA | 0.858392164 | 1.813016637 | 0.308668503 | 0.479700644 | 4.37358774 | 7.917403145 | ENSG00000272084 |
| SLC66A1L | 0.858041724 | 1.812576297 | 0.130134998 | 0.253042459 | 10.32291562 | 18.74380708 | ENSG00000174899 |
| DES | 0.857927673 | 1.812433011 | 0.001235757 | 0.005167141 | 47.33740853 | 85.82072698 | ENSG00000175084 |
| CYS1 | 0.857797747 | 1.812269795 | 0.249724607 | 0.413833639 | 6.670272351 | 12.08045212 | ENSG00000205795 |
| GRM1 | 0.857785995 | 1.812255032 | 0.102984839 | 0.211822213 | 13.42782643 | 24.29427665 | ENSG00000152822 |
| ZNF814 | 0.857682317 | 1.812124801 | 1.75E-14 | 3.58E-13 | 273.0596123 | 494.9574144 | ENSG00000204514 |
| EFNA1 | 0.857569772 | 1.811983442 | 0.00042901 | 0.002004852 | 56.67302873 | 102.4732018 | ENSG00000169242 |
| ACAD10 | 0.857134338 | 1.811436632 | 1.34E-11 | 2.03E-10 | 303.6894688 | 550.0452787 | ENSG00000111271 |
| LINC00910 | 0.857004244 | 1.811273294 | 0.007276186 | 0.024138419 | 37.00871095 | 67.28717992 | ENSG00000188825 |
| NA | 0.856944459 | 1.811198236 | 0.357631664 | 0.530817142 | 4.046988208 | 7.330706918 | ENSG00000242553 |
| NA | 0.856738487 | 1.810939673 | 0.031070945 | 0.080823841 | 24.6441348 | 44.42630995 | ENSG00000240086 |
| EDIL3 | 0.856478868 | 1.810613816 | 1.76E-18 | 5.21E-17 | 772.6093411 | 1398.034497 | ENSG00000164176 |
| TMCO4 | 0.856403116 | 1.810518748 | 7.18E-06 | 4.86E-05 | 109.1888967 | 197.4220655 | ENSG00000162542 |
| TMEM161B-AS1 | 0.856180549 | 1.810239458 | 8.63E-06 | 5.75E-05 | 108.9398548 | 196.7525751 | ENSG00000247828 |
| PKLR | 0.856013187 | 1.81002947 | 0.000114913 | 0.000613175 | 66.97341697 | 121.4275293 | ENSG00000143627 |
| LINC01238 | 0.855794 | 1.809754495 | 0.110310015 | 0.223059802 | 11.78992269 | 21.24706926 | ENSG00000237940 |
| NA | 0.855272833 | 1.809100848 | 0.306165838 | 0.477156663 | 5.054020111 | 9.090795601 | ENSG00000259088 |
| NA | 0.85514277 | 1.80893776 | 0.419633407 | 0.591664088 | 3.097213824 | 5.527879334 | ENSG00000287445 |
| CALML6 | 0.85501617 | 1.808779028 | 0.291932158 | 0.46168743 | 5.257731897 | 9.585259201 | ENSG00000169885 |
| TNRC18 | 0.854586746 | 1.808240717 | 8.43E-09 | 9.08E-08 | 158.8505866 | 287.5029146 | ENSG00000182095 |
| NA | 0.854114966 | 1.807649496 | 2.17E-16 | 5.41E-15 | 332.6332765 | 600.9913698 | ENSG00000229191 |
| COX6B2 | 0.85400429 | 1.807510828 | 0.026246016 | 0.070438494 | 28.12779641 | 50.78717225 | ENSG00000160471 |
| NA | 0.853946476 | 1.807438397 | 0.282343369 | 0.450660759 | 5.905349155 | 10.71508872 | ENSG00000279406 |
| NA | 0.853686041 | 1.807112147 | 0.117336275 | 0.234538813 | 11.02220859 | 19.88385644 | ENSG00000231154 |
| ARNT | 0.853618986 | 1.807028156 | 2.63E-32 | 1.92E-30 | 807.2431725 | 1458.817297 | ENSG00000143437 |
| MSC-AS1 | 0.853412391 | 1.806769407 | 0.435426956 | 0.606135187 | 2.716147677 | 4.858909846 | ENSG00000235531 |
| LOC100506606 | 0.853374919 | 1.80672248 | 0.443198862 | 0.614029679 | 3.130028937 | 5.578427028 | ENSG00000257176 |
| CD70 | 0.853351335 | 1.806692945 | 0.020323788 | 0.056986364 | 26.30069898 | 47.67488408 | ENSG00000125726 |
| KLF5 | 0.852879849 | 1.806102597 | 0.026373923 | 0.070732567 | 23.01026126 | 41.66329401 | ENSG00000102554 |
| NA | 0.852676527 | 1.805848078 | 0.454685064 | 0.624382799 | 2.362314838 | 4.24048805 | ENSG00000249916 |
| IL13RA1 | 0.852522656 | 1.805655484 | 1.56E-24 | 7.39E-23 | 555.9691233 | 1003.547224 | ENSG00000131724 |
| NA | 0.852173673 | 1.805218755 | 0.337402552 | 0.510173133 | 4.376378643 | 7.877608321 | ENSG00000265574 |

| | | | | | | | |
|---|---|---|---|---|---|---|---|
| TRIM56 | 0.85185592 | 1.8048212 | 1.14E-30 | 7.44E-29 | 745.759784 | 1345.789821 | ENSG00000169871 |
| CCDC121 | 0.851801478 | 1.804753095 | 0.043158322 | 0.106073027 | 18.45461323 | 33.2826198 | ENSG00000176714 |
| GVQW2 | 0.851395211 | 1.804244943 | 0.221912754 | 0.380289503 | 6.727529869 | 12.09067195 | ENSG00000279968 |
| NA | 0.850798683 | 1.803499075 | 0.317628592 | 0.488860154 | 4.398030145 | 7.896430446 | ENSG00000224543 |
| MELTF | 0.850653284 | 1.803317322 | 7.75E-05 | 0.000429631 | 86.5457471 | 155.7692518 | ENSG00000163975 |
| HS3ST6 | 0.850643467 | 1.803305052 | 0.482276264 | 0.650237029 | 2.367896644 | 4.244789198 | ENSG00000162040 |
| TCTEX1D2 | 0.850490645 | 1.803114041 | 0.476240516 | 0.644667447 | 2.332290628 | 4.189940357 | ENSG00000213123 |
| NA | 0.850447569 | 1.803060203 | 0.099398613 | 0.206279864 | 15.08997242 | 27.40385068 | ENSG00000267284 |
| PCAT6 | 0.850383372 | 1.802979973 | 0.000636185 | 0.002836294 | 65.94318196 | 119.1348119 | ENSG00000228288 |
| RASSF10 | 0.850328113 | 1.802910916 | 0.000199096 | 0.001004118 | 63.54489769 | 114.6165599 | ENSG00000189431 |
| MPPE1 | 0.850266472 | 1.802833886 | 2.19E-05 | 0.000135755 | 102.0581736 | 183.6000863 | ENSG00000154889 |
| METAP1D | 0.850034514 | 1.802544047 | 0.000136728 | 0.000714609 | 67.04203826 | 120.6496678 | ENSG00000172878 |
| GJA4 | 0.849795109 | 1.802244953 | 0.00034724 | 0.001660126 | 69.10307185 | 124.0991183 | ENSG00000187513 |
| NA | 0.849792371 | 1.802241532 | 0.346309391 | 0.518845485 | 3.95970648 | 7.153789991 | ENSG00000279148 |
| CEL | 0.849652078 | 1.802066285 | 0.002636929 | 0.010039271 | 39.79136473 | 71.7255576 | ENSG00000170835 |
| NA | 0.849297222 | 1.801623089 | 3.30E-06 | 2.36E-05 | 101.6948921 | 183.1102149 | ENSG00000232098 |
| CYP2A6 | 0.848604969 | 1.800758818 | 0.424637499 | 0.596335198 | 3.282065012 | 5.958891799 | ENSG00000255974 |
| NA | 0.848171806 | 1.800218229 | 0.210484964 | 0.365939568 | 7.356286528 | 13.27507774 | ENSG00000261235 |
| SLC2A1-AS1 | 0.848083882 | 1.800108519 | 0.208023691 | 0.362785266 | 7.383519835 | 13.31487256 | ENSG00000227533 |
| NA | 0.847112918 | 1.798897416 | 0.029894741 | 0.078323977 | 21.30114548 | 38.36496569 | ENSG00000228613 |
| DAGLA | 0.846400332 | 1.798009111 | 3.30E-11 | 4.78E-10 | 260.3080759 | 468.1375948 | ENSG00000134780 |
| NLRP7 | 0.846346123 | 1.797941553 | 0.024782322 | 0.06712885 | 27.77627892 | 49.66029892 | ENSG00000167634 |
| FCHO2 | 0.845374299 | 1.796730836 | 1.25E-06 | 9.71E-06 | 129.5551493 | 233.0928826 | ENSG00000157107 |
| NMRK1 | 0.845247343 | 1.796572733 | 0.000838486 | 0.003629988 | 53.2322698 | 95.87490336 | ENSG00000106733 |
| NUAK2 | 0.844675474 | 1.795860731 | 1.21E-09 | 1.46E-08 | 222.1308482 | 399.1932996 | ENSG00000163545 |
| FGF13-AS1 | 0.84449671 | 1.795638221 | 0.005152077 | 0.018000995 | 40.482285 | 72.78547215 | ENSG00000226031 |
| NA | 0.844392413 | 1.795508413 | 0.323931466 | 0.495477475 | 5.271686412 | 9.507820125 | ENSG00000286540 |
| SPOCK3 | 0.84370271 | 1.794650247 | 0.32369169 | 0.495307083 | 4.400821048 | 7.848033327 | ENSG00000196104 |
| NA | 0.841929669 | 1.792446016 | 0.489763288 | 0.656203889 | 2.422363259 | 4.293186317 | ENSG00000226191 |
| HLA-F | 0.841362449 | 1.791741424 | 0.00234728 | 0.00907079 | 43.85509836 | 78.53652707 | ENSG00000204642 |
| SEMG1 | 0.841055383 | 1.791360106 | 0.019792565 | 0.055715508 | 24.36275329 | 43.66133972 | ENSG00000124233 |
| IL17C | 0.840785209 | 1.79102467 | 0.030430147 | 0.079478469 | 20.70799484 | 37.09209538 | ENSG00000124391 |
| NA | 0.840235096 | 1.790341866 | 0.016331745 | 0.047476747 | 27.41945502 | 49.01660327 | ENSG00000280058 |
| MYOCD | 0.840030375 | 1.79008783 | 0.065119346 | 0.147132503 | 16.85913661 | 30.32952303 | ENSG00000141052 |
| NA | 0.839650559 | 1.78961662 | 0.245138872 | 0.408339002 | 7.582525697 | 13.42215921 | ENSG00000267466 |
| EXD3 | 0.839546949 | 1.789488099 | 1.48E-06 | 1.13E-05 | 126.4023417 | 226.113871 | ENSG00000187609 |
| NA | 0.839057053 | 1.788880545 | 0.047229501 | 0.114159165 | 20.08761089 | 35.84423846 | ENSG00000261490 |
| ARHGAP25 | 0.838690627 | 1.788426251 | 0.056482349 | 0.131578431 | 16.63377333 | 29.77214133 | ENSG00000163219 |
| NEBL | 0.838635956 | 1.788358479 | 6.04E-05 | 0.000341997 | 78.89651513 | 141.3302079 | ENSG00000078114 |
| VENTX | 0.838412 | 1.788080886 | 1.73E-09 | 2.05E-08 | 194.1843852 | 346.6882503 | ENSG00000151650 |
| CASP8 | 0.838369546 | 1.788028269 | 0.001302541 | 0.005411869 | 49.63775993 | 88.61760694 | ENSG00000064012 |
| NA | 0.837957173 | 1.78751726 | 0.257882095 | 0.422919621 | 6.270345606 | 11.29344312 | ENSG00000259033 |
| ITPRIPL2 | 0.83753482 | 1.786994036 | 3.26E-06 | 2.34E-05 | 107.4042264 | 191.860601 | ENSG00000205730 |
| NA | 0.83748851 | 1.786936676 | 0.378389974 | 0.551850761 | 3.72317958 | 6.638614156 | ENSG00000235297 |
| MEIS1 | 0.837442623 | 1.786879841 | 0.500747387 | 0.666686425 | 2.386757243 | 4.223816498 | ENSG00000143995 |
| TRIM72 | 0.836999486 | 1.786331068 | 0.007617374 | 0.025062914 | 37.29375924 | 66.43967597 | ENSG00000177238 |
| USP53 | 0.836961793 | 1.786284398 | 1.02E-13 | 1.93E-12 | 330.7624005 | 590.8427555 | ENSG00000145390 |
| KIRREL1 | 0.836829347 | 1.786120416 | 4.25E-46 | 6.26E-44 | 1562.48134 | 2790.156667 | ENSG00000183853 |
| GUCY2C | 0.836758826 | 1.78603311 | 0.006949947 | 0.023260137 | 34.8705201 | 62.44279105 | ENSG00000070019 |
| SUCLG2-AS1 | 0.836025285 | 1.785125229 | 0.032395139 | 0.083614481 | 24.19867772 | 43.31206748 | ENSG00000241316 |
| HIVEP2 | 0.835768234 | 1.784807194 | 2.56E-08 | 2.56E-07 | 194.8302876 | 347.0638688 | ENSG00000010818 |
| EGFLAM | 0.835711206 | 1.784736644 | 0.393379121 | 0.566303538 | 3.698737175 | 6.59236761 | ENSG00000164318 |
| NA | 0.83531424 | 1.784245632 | 0.465092159 | 0.634608349 | 3.039956306 | 5.445866537 | ENSG00000214185 |
| EFR3A | 0.834998395 | 1.783855055 | 3.16E-26 | 1.66E-24 | 689.0264825 | 1229.690918 | ENSG00000132294 |
| NA | 0.834962979 | 1.783811264 | 0.028966884 | 0.076307547 | 23.59033327 | 41.91766212 | ENSG00000247735 |
| PDP1 | 0.834692445 | 1.783476795 | 3.64E-13 | 6.52E-12 | 282.7099927 | 504.3407128 | ENSG00000164951 |
| WNT9A | 0.833961025 | 1.782572834 | 0.000258172 | 0.001270834 | 62.79325328 | 111.9457887 | ENSG00000143816 |
| NA | 0.833694161 | 1.782243132 | 0.279316558 | 0.447167916 | 6.575293635 | 11.77312528 | ENSG00000253210 |
| NA | 0.832545576 | 1.780824784 | 0.224118657 | 0.382642093 | 8.513439483 | 15.05667216 | ENSG00000233013 |
| TMEM119 | 0.832519562 | 1.780792674 | 0.444650763 | 0.615289998 | 3.252040801 | 5.826603882 | ENSG00000183160 |
| DAP3 | 0.831991716 | 1.780141245 | 1.98E-38 | 2.04E-36 | 1487.769993 | 2647.273565 | ENSG00000132676 |
| APOL6 | 0.831765373 | 1.779861982 | 0.369028694 | 0.542833495 | 3.663131159 | 6.514395497 | ENSG00000221963 |
| PLK3 | 0.831564961 | 1.779614749 | 5.52E-07 | 4.51E-06 | 121.8509241 | 216.804201 | ENSG00000173846 |
| MST1L | 0.831476995 | 1.779506244 | 0.001254108 | 0.005234313 | 50.30981959 | 89.42584911 | ENSG00000186715 |
| KBTBD3 | 0.831449615 | 1.779472272 | 0.033119773 | 0.0850977 | 25.93311179 | 46.0406437 | ENSG00000182359 |
| NA | 0.831448972 | 1.779471679 | 0.411417058 | 0.58361986 | 3.284855915 | 5.875001002 | ENSG00000288573 |
| JPX | 0.831355915 | 1.779356903 | 0.2742796 | 0.441074338 | 6.70866927 | 11.9500422 | ENSG00000225470 |
| C17orf113 | 0.831271882 | 1.779253264 | 0.002168784 | 0.008474381 | 47.45559035 | 84.25799488 | ENSG00000267221 |
| MPL | 0.831209724 | 1.779176607 | 0.395258929 | 0.567965735 | 3.333740724 | 5.946521394 | ENSG00000117400 |
| NA | 0.830832527 | 1.778711496 | 0.306477605 | 0.477495365 | 4.996762593 | 8.910371029 | ENSG00000275318 |
| CPNE7 | 0.83012157 | 1.777835167 | 1.27E-13 | 2.38E-12 | 366.7215275 | 652.2550197 | ENSG00000178773 |
| NA | 0.830079525 | 1.777783356 | 0.27409123 | 0.440918439 | 5.696055562 | 10.09236578 | ENSG00000267506 |
| ABCG2 | 0.829453884 | 1.777012567 | 0.006077752 | 0.02077723 | 40.43810611 | 72.11085655 | ENSG00000118777 |
| LOC100130449 | 0.829398034 | 1.776943776 | 0.45792628 | 0.627850452 | 3.007141193 | 5.370044997 | ENSG00000218416 |
| NA | 0.829078514 | 1.776550272 | 0.21795865 | 0.375514899 | 7.375147127 | 13.06858582 | ENSG00000205740 |
| NA | 0.828715689 | 1.776103542 | 0.016858094 | 0.048757046 | 28.23605392 | 50.07920144 | ENSG00000264112 |
| ADORA2A | 0.828312235 | 1.775606919 | 0.313229933 | 0.484370232 | 4.955574771 | 8.828097768 | ENSG00000128271 |

| | | | | | | | |
|---|---|---|---|---|---|---|---|
| MYH11 | 0.828182445 | 1.775447186 | 0.037934922 | 0.095411028 | 20.78132206 | 36.87779467 | ENSG00000133392 |
| TGM1 | 0.827981288 | 1.775199649 | 0.067549486 | 0.151664005 | 19.88090804 | 35.37100802 | ENSG00000092295 |
| PLIN1 | 0.827933655 | 1.775141039 | 0.492683439 | 0.659017905 | 2.392339049 | 4.192090931 | ENSG00000166819 |
| TRIM50 | 0.827489471 | 1.774594584 | 0.460005699 | 0.629716856 | 2.933813978 | 5.252278059 | ENSG00000146755 |
| NA | 0.827034286 | 1.77403477 | 0.38912985 | 0.562490145 | 3.725970483 | 6.579464168 | ENSG00000260645 |
| SOSTDC1 | 0.82681702 | 1.773767626 | 0.393270105 | 0.566204462 | 3.366555838 | 5.969644667 | ENSG00000171243 |
| SMYD3 | 0.826397852 | 1.773252341 | 9.12E-13 | 1.56E-11 | 311.950651 | 553.6415075 | ENSG00000185420 |
| NA | 0.825675242 | 1.772364385 | 0.501719254 | 0.667627423 | 3.2709014 | 5.814233478 | ENSG00000272750 |
| MGAT3 | 0.825636033 | 1.772316218 | 6.30E-20 | 2.11E-18 | 446.495854 | 791.4470667 | ENSG00000128268 |
| CYP1B1-AS1 | 0.825536063 | 1.772193411 | 0.388716383 | 0.562039255 | 3.336531627 | 5.919096974 | ENSG00000232973 |
| CPO | 0.825286162 | 1.771886461 | 0.081621339 | 0.176302458 | 17.48876915 | 30.84497144 | ENSG00000144410 |
| CEP170 | 0.825084285 | 1.771638537 | 1.86E-31 | 1.27E-29 | 1011.13228 | 1791.613616 | ENSG00000143702 |
| PDGFRB | 0.824794625 | 1.771282869 | 4.62E-13 | 8.15E-12 | 300.5109313 | 532.1387557 | ENSG00000113721 |
| SUGP1 | 0.824728076 | 1.771201165 | 1.79E-30 | 1.15E-28 | 885.337807 | 1567.982708 | ENSG00000105705 |
| TATDN3 | 0.82465695 | 1.771113846 | 5.30E-11 | 7.48E-10 | 217.1375522 | 384.8418778 | ENSG00000203705 |
| SCNN1B | 0.824610223 | 1.771056482 | 0.330078512 | 0.502489312 | 4.939505075 | 8.773248927 | ENSG00000168447 |
| MPI | 0.824594963 | 1.771037749 | 5.49E-21 | 2.00E-19 | 545.0428204 | 965.5731849 | ENSG00000178802 |
| FZD9 | 0.824369621 | 1.770761143 | 1.07E-06 | 8.40E-06 | 131.5370737 | 233.3149615 | ENSG00000188763 |
| ARL11 | 0.824185558 | 1.770535239 | 0.10382838 | 0.21321623 | 13.69226236 | 24.23189159 | ENSG00000152213 |
| KIAA1614 | 0.823832122 | 1.770101541 | 0.002619996 | 0.009988604 | 46.16717695 | 81.657678 | ENSG00000135835 |
| WDR26 | 0.823731238 | 1.769977767 | 5.97E-46 | 8.67E-44 | 1561.224727 | 2762.559924 | ENSG00000162923 |
| EOMES | 0.823318254 | 1.769471167 | 0.337258084 | 0.510118057 | 4.618487349 | 8.21397712 | ENSG00000163508 |
| NA | 0.823293548 | 1.769440866 | 0.233314664 | 0.394121995 | 7.827425306 | 13.75260932 | ENSG00000280195 |
| GRIN3A | 0.822562385 | 1.768544333 | 0.393397631 | 0.566303538 | 3.663131159 | 6.459546656 | ENSG00000198785 |
| NXNL2 | 0.822521286 | 1.768493952 | 0.060440447 | 0.138787843 | 21.26573963 | 37.49836704 | ENSG00000130045 |
| LMOD1 | 0.822173849 | 1.768068106 | 0.093485208 | 0.196301637 | 14.42157954 | 25.52331145 | ENSG00000163431 |
| LCTL | 0.822066113 | 1.767936077 | 1.27E-06 | 9.85E-06 | 125.9195638 | 222.5880539 | ENSG00000188501 |
| NA | 0.821642511 | 1.767417055 | 0.395647035 | 0.568227141 | 3.36934674 | 5.944370821 | ENSG00000287790 |
| TRPS1 | 0.82164124 | 1.767415498 | 6.54E-05 | 0.000368168 | 78.16739811 | 138.2639058 | ENSG00000104447 |
| P3H2 | 0.821446529 | 1.767176978 | 2.21E-15 | 4.97E-14 | 415.6259159 | 734.8644647 | ENSG00000090530 |
| NA | 0.820682512 | 1.76624137 | 0.092838599 | 0.195262692 | 13.75442596 | 24.23996085 | ENSG00000260267 |
| NA | 0.820657991 | 1.766211351 | 0.289667005 | 0.459083651 | 5.927000657 | 10.52982996 | ENSG00000235001 |
| NA | 0.820453718 | 1.765961288 | 0.431434771 | 0.602534436 | 2.95546548 | 5.256579206 | ENSG00000268201 |
| MYH7B | 0.820290394 | 1.765761379 | 0.10338924 | 0.212518042 | 13.34891741 | 23.56238907 | ENSG00000078814 |
| ZNF654 | 0.820214122 | 1.76566803 | 2.79E-05 | 0.000169307 | 93.55482173 | 165.0958659 | ENSG00000175105 |
| RPL23P8 | 0.819394613 | 1.764665345 | 0.400708571 | 0.573192128 | 3.363764935 | 5.923398121 | ENSG00000219451 |
| ALG10 | 0.818969872 | 1.764145889 | 0.032626403 | 0.084110108 | 24.24054127 | 42.91895973 | ENSG00000139133 |
| PPIAP46 | 0.818475998 | 1.763542077 | 0.308356077 | 0.479446852 | 5.0239959 | 8.846919893 | ENSG00000260266 |
| UAP1L1 | 0.81822981 | 1.763241163 | 3.21E-10 | 4.17E-09 | 205.0152849 | 361.1364479 | ENSG00000197355 |
| THPO | 0.818229069 | 1.763240258 | 0.320918822 | 0.492320611 | 5.679985866 | 10.06305125 | ENSG00000090534 |
| NA | 0.817972992 | 1.762927312 | 0.480009976 | 0.648175374 | 2.631656852 | 4.688705104 | ENSG00000287804 |
| HLA-DRB1 | 0.817151596 | 1.761923878 | 3.08E-11 | 4.47E-10 | 281.2283923 | 494.8382904 | ENSG00000196126 |
| TDRKH | 0.817138508 | 1.761907895 | 6.48E-21 | 2.34E-19 | 499.6741696 | 880.2340423 | ENSG00000182134 |
| LMNA | 0.817105557 | 1.761867654 | 8.99E-33 | 6.67E-31 | 1249.406828 | 2202.659964 | ENSG00000160789 |
| NA | 0.817009539 | 1.761750396 | 0.282549907 | 0.450766644 | 6.74706619 | 11.85701607 | ENSG00000261570 |
| F8A1 | 0.816972655 | 1.761705357 | 0.000108532 | 0.000582552 | 80.79817908 | 142.5232281 | ENSG00000277203 |
| EFCAB10 | 0.816916841 | 1.761637202 | 0.146465489 | 0.277087943 | 10.44791855 | 18.3273156 | ENSG00000185055 |
| ALDH1A2 | 0.816661315 | 1.761325214 | 0.47281339 | 0.641649181 | 2.922650367 | 5.18290824 | ENSG00000128918 |
| LOC107985075 | 0.816437753 | 1.761052297 | 0.213371859 | 0.369659321 | 7.957809879 | 14.03412928 | ENSG00000214999 |
| ZNF79 | 0.816434635 | 1.761048492 | 7.08E-06 | 4.80E-05 | 107.3789081 | 189.3796381 | ENSG00000196152 |
| ORAI3 | 0.816100583 | 1.760640772 | 0.011934177 | 0.036473734 | 36.63756911 | 64.64113743 | ENSG00000175938 |
| ADORA2A-AS1 | 0.816007487 | 1.760527164 | 0.350838768 | 0.523517225 | 4.656208547 | 8.211826546 | ENSG00000178803 |
| ENO3 | 0.815877842 | 1.760368964 | 2.63E-12 | 4.29E-11 | 272.3378288 | 478.8736378 | ENSG00000108515 |
| WDR49 | 0.815804705 | 1.760279725 | 0.259023927 | 0.424164675 | 6.406512144 | 11.26386812 | ENSG00000174776 |
| FAM166B | 0.815565739 | 1.759988179 | 0.219897832 | 0.37791083 | 7.715701173 | 13.59262441 | ENSG00000215187 |
| KLF11 | 0.81521141 | 1.759555976 | 4.26E-10 | 5.43E-09 | 215.3279128 | 378.4119061 | ENSG00000172059 |
| EPS8L2 | 0.814490671 | 1.758677159 | 2.53E-23 | 1.09E-21 | 591.5311272 | 1040.3844 | ENSG00000177106 |
| TUSC1 | 0.81436617 | 1.758525396 | 0.000537549 | 0.002445721 | 64.52940221 | 113.5795081 | ENSG00000198680 |
| NOS2 | 0.81419674 | 1.758318887 | 0.000270769 | 0.001326746 | 67.92618242 | 119.3214156 | ENSG00000007171 |
| TMEM59L | 0.814086152 | 1.758184111 | 6.47E-16 | 1.54E-14 | 375.9090665 | 660.9346547 | ENSG00000105696 |
| PPIC | 0.813847033 | 1.757892725 | 0.00018359 | 0.0009325 | 72.13146423 | 126.8591475 | ENSG00000168938 |
| OGFOD2 | 0.813621366 | 1.757617776 | 0.401520756 | 0.573951189 | 4.270236315 | 7.547158205 | ENSG00000111325 |
| ZC3HAV1L | 0.8128569 | 1.756686683 | 0.000252215 | 0.001245324 | 70.23961244 | 123.3051245 | ENSG00000146858 |
| NA | 0.812398404 | 1.756128488 | 0.369305245 | 0.543049583 | 4.343563529 | 7.652554739 | ENSG00000230224 |
| FTX | 0.81238821 | 1.756116079 | 6.01E-08 | 5.70E-07 | 154.2570172 | 271.1825195 | ENSG00000230590 |
| GGACT | 0.811415098 | 1.754931961 | 0.183692964 | 0.329731177 | 10.49102139 | 18.5693012 | ENSG00000134864 |
| SOX6 | 0.810843085 | 1.754236287 | 0.315116912 | 0.486430371 | 5.105695823 | 8.901768734 | ENSG00000110693 |
| SLC35D3 | 0.810754584 | 1.754128679 | 0.134442309 | 0.259560306 | 13.47950214 | 23.56023849 | ENSG00000182747 |
| TMC1 | 0.810673356 | 1.754029919 | 0.451175401 | 0.621421153 | 3.567476721 | 6.337739034 | ENSG00000165091 |
| FNDC1 | 0.810131941 | 1.753371789 | 0.118125717 | 0.235652866 | 16.61299771 | 28.95152876 | ENSG00000164694 |
| NA | 0.810009264 | 1.7532227 | 0.100529582 | 0.207924966 | 14.17667993 | 24.80594484 | ENSG00000268575 |
| ARMCX2 | 0.809620641 | 1.752750493 | 4.41E-52 | 7.88E-50 | 2813.533928 | 4930.499149 | ENSG00000184867 |
| GEMIN8 | 0.809367117 | 1.752442511 | 1.31E-08 | 1.37E-07 | 171.918635 | 301.0076196 | ENSG00000046647 |
| EFCAB12 | 0.809255674 | 1.752307145 | 0.471094519 | 0.639991856 | 3.402161854 | 5.933617952 | ENSG00000172771 |
| BLZF1 | 0.809247263 | 1.75229693 | 1.41E-08 | 1.48E-07 | 184.2482808 | 323.0424914 | ENSG00000117475 |
| GPRIN3 | 0.80916055 | 1.752191612 | 0.001550592 | 0.006300014 | 50.17365305 | 87.83355413 | ENSG00000185477 |
| CAHM | 0.808789322 | 1.751740803 | 0.217644118 | 0.375241101 | 8.278827604 | 14.58344172 | ENSG00000270419 |

| | | | | | | | |
|---|---|---|---|---|---|---|---|
| CENPF | 0.80858814 | 1.751496542 | 6.39E-41 | 7.80E-39 | 2956.018067 | 5176.503737 | ENSG00000117724 |
| CREBRF | 0.808049397 | 1.750842606 | 0.013757377 | 0.041184599 | 32.9201715 | 57.54785449 | ENSG00000164463 |
| NA | 0.807387475 | 1.750039487 | 0.022925815 | 0.062910655 | 33.2000014 | 58.09823301 | ENSG00000234614 |
| TMEM256 | 0.807302103 | 1.749935931 | 0.041239544 | 0.10215878 | 23.99564166 | 42.12818262 | ENSG00000205544 |
| NA | 0.807145379 | 1.749745841 | 0.01416747 | 0.042222256 | 32.79381713 | 57.31285366 | ENSG00000257261 |
| CTSF | 0.807061324 | 1.7496439 | 0.478919091 | 0.647138101 | 2.947092772 | 5.174305945 | ENSG00000174080 |
| PSMB10 | 0.806542019 | 1.749014221 | 0.275926214 | 0.442873166 | 7.056920304 | 12.36815123 | ENSG00000205220 |
| NEDD9 | 0.806114927 | 1.748496523 | 6.30E-11 | 8.84E-10 | 229.7863377 | 401.4698722 | ENSG00000111859 |
| TNFRSF1B | 0.805010656 | 1.747158697 | 1.37E-05 | 8.75E-05 | 97.36440716 | 170.3099546 | ENSG00000028137 |
| ELFN2 | 0.803908485 | 1.745824435 | 0.000139771 | 0.000728144 | 79.26749371 | 138.194003 | ENSG00000166897 |
| PLEKHA8P1 | 0.803579213 | 1.745426024 | 2.52E-06 | 1.84E-05 | 112.9892336 | 197.2440825 | ENSG00000134297 |
| NA | 0.80295142 | 1.744666663 | 0.257057227 | 0.422176994 | 7.340216832 | 12.81610782 | ENSG00000233483 |
| GSTM3 | 0.802158857 | 1.74370847 | 0.003925902 | 0.014214449 | 46.08403756 | 80.35361517 | ENSG00000134202 |
| IL1RAP | 0.801925447 | 1.743426383 | 1.77E-05 | 0.000111121 | 114.6035708 | 199.7908771 | ENSG00000196083 |
| NA | 0.801801189 | 1.74327623 | 0.431543715 | 0.602593258 | 3.363764935 | 5.828754456 | ENSG00000286974 |
| NA | 0.801799992 | 1.743274783 | 0.219665032 | 0.377710335 | 8.142861226 | 14.11210139 | ENSG00000241741 |
| HHLA3 | 0.801300153 | 1.74267091 | 0.000117693 | 0.000626147 | 77.59589897 | 135.2532766 | ENSG00000197568 |
| SMIM10L2B | 0.801138106 | 1.74247518 | 9.57E-07 | 7.55E-06 | 128.7221173 | 224.1260573 | ENSG00000196972 |
| NA | 0.801025465 | 1.742339138 | 0.179668998 | 0.324379471 | 9.933476764 | 17.35370288 | ENSG00000259845 |
| SNORD101 | 0.800466552 | 1.741664271 | 0.481233735 | 0.649239806 | 2.664471965 | 4.663431258 | ENSG00000206754 |
| RDM1 | 0.800383143 | 1.741563581 | 0.001209314 | 0.00506599 | 73.5428534 | 128.0056364 | ENSG00000278023 |
| RFPL2 | 0.800214944 | 1.741360549 | 0.46395124 | 0.633634221 | 3.358183129 | 5.818001588 | ENSG00000128253 |
| TMEM184A | 0.800152173 | 1.741284785 | 0.04194317 | 0.103574718 | 21.06182768 | 36.62720678 | ENSG00000164855 |
| NA | 0.799569741 | 1.740581951 | 0.311983908 | 0.48298544 | 5.293337914 | 9.269863101 | ENSG00000213598 |
| NA | 0.798569967 | 1.739376162 | 0.336617859 | 0.509469254 | 4.751862984 | 8.216127694 | ENSG00000262136 |
| NA | 0.797881849 | 1.738546734 | 0.353999421 | 0.527132599 | 4.664581255 | 8.087607887 | ENSG00000285706 |
| NA | 0.797332823 | 1.737885246 | 0.000756743 | 0.003305779 | 62.25245408 | 108.0435595 | ENSG00000230707 |
| MBTPS2 | 0.796984904 | 1.73746619 | 2.60E-06 | 1.90E-05 | 156.1086451 | 271.4067491 | ENSG00000012174 |
| LRRC75A | 0.796525124 | 1.736912556 | 4.59E-14 | 8.95E-13 | 334.0048292 | 579.6329853 | ENSG00000181350 |
| RBMS3 | 0.79616755 | 1.736482113 | 0.015079147 | 0.044462907 | 31.52003396 | 54.899934 | ENSG00000144642 |
| ZNF552 | 0.796164681 | 1.73647866 | 8.20E-06 | 5.48E-05 | 115.5275513 | 200.4053008 | ENSG00000178935 |
| NA | 0.795449743 | 1.735618349 | 0.284985347 | 0.453901282 | 6.259181994 | 10.9506226 | ENSG00000228981 |
| NA | 0.795071574 | 1.735163456 | 0.363800403 | 0.537371636 | 4.400821048 | 7.61437745 | ENSG00000260633 |
| NA | 0.795003638 | 1.73508175 | 0.466534781 | 0.635722431 | 2.991071497 | 5.18290824 | ENSG00000257790 |
| LOC100507250 | 0.794711273 | 1.734730168 | 0.418938626 | 0.590975423 | 3.600291835 | 6.312465188 | ENSG00000247363 |
| PRRC2C | 0.79458857 | 1.734582633 | 3.76E-37 | 3.57E-35 | 4575.798097 | 7935.076245 | ENSG00000117523 |
| LOC285097 | 0.794536695 | 1.734520263 | 0.387606411 | 0.56093888 | 4.362424128 | 7.5557605 | ENSG00000280119 |
| NA | 0.79439562 | 1.734350661 | 0.063539898 | 0.144359422 | 17.98068355 | 31.24021765 | ENSG00000275765 |
| NA | 0.79435495 | 1.734301769 | 0.450766472 | 0.62116837 | 3.41823155 | 5.872850428 | ENSG00000267413 |
| NA | 0.794336241 | 1.734279279 | 0.137975777 | 0.264792577 | 12.19264034 | 21.04728953 | ENSG00000258072 |
| LINC00106 | 0.794325292 | 1.734266117 | 0.041962089 | 0.103608173 | 25.26172784 | 43.69845093 | ENSG00000236871 |
| ECHDC3 | 0.793879042 | 1.733729762 | 8.21E-16 | 1.93E-14 | 395.5770142 | 685.8363519 | ENSG00000134463 |
| SLC39A8 | 0.793594233 | 1.733387533 | 7.10E-20 | 2.37E-18 | 535.3890855 | 928.3673695 | ENSG00000138821 |
| FGD2 | 0.79330449 | 1.733039444 | 0.448403823 | 0.618752371 | 3.8020886 | 6.505793202 | ENSG00000146192 |
| NA | 0.792991615 | 1.732663642 | 0.217932966 | 0.375504184 | 8.69829067 | 15.05237101 | ENSG00000248740 |
| XK | 0.792841518 | 1.732483387 | 0.026148748 | 0.070226299 | 28.94005282 | 50.31473532 | ENSG00000047597 |
| GLRA4 | 0.792191852 | 1.7317034 | 0.18786906 | 0.335137973 | 10.53500012 | 18.32301445 | ENSG00000188828 |
| P2RY1 | 0.792119979 | 1.731617131 | 7.35E-18 | 2.09E-16 | 703.741757 | 1217.298323 | ENSG00000169860 |
| COL28A1 | 0.792091645 | 1.731583123 | 0.234483462 | 0.395473025 | 7.389101641 | 12.75534029 | ENSG00000215018 |
| COL11A1 | 0.791940881 | 1.73140218 | 5.02E-12 | 7.91E-11 | 319.0357174 | 551.7021214 | ENSG00000060718 |
| YY1AP1 | 0.791886384 | 1.731336779 | 1.13E-19 | 3.73E-18 | 549.6276409 | 951.8044491 | ENSG00000163374 |
| NA | 0.791882823 | 1.731332505 | 0.465701409 | 0.635040226 | 3.747621984 | 6.421902405 | ENSG00000281741 |
| PARVB | 0.791811809 | 1.731247286 | 3.11E-10 | 4.06E-09 | 247.3440916 | 428.1066205 | ENSG00000188677 |
| ARID4B | 0.791505679 | 1.730879966 | 5.92E-19 | 1.81E-17 | 534.309439 | 924.2140316 | ENSG00000054267 |
| SUSD2 | 0.79099981 | 1.730273154 | 5.56E-10 | 6.97E-09 | 217.3211641 | 376.115148 | ENSG00000099994 |
| CRYGB | 0.7904924 | 1.729664706 | 0.331823724 | 0.5045091 | 6.142751936 | 10.56532364 | ENSG00000182187 |
| SLC16A8 | 0.790346446 | 1.729489729 | 0.099934193 | 0.207093191 | 16.89474263 | 29.23358176 | ENSG00000100156 |
| NID1 | 0.790257737 | 1.729383389 | 1.17E-22 | 4.80E-21 | 1308.242462 | 2262.379648 | ENSG00000116962 |
| PPP1R12B | 0.790058723 | 1.729144843 | 1.18E-11 | 1.79E-10 | 261.66068 | 452.4240141 | ENSG00000077157 |
| TDRG1 | 0.789391208 | 1.728344977 | 0.10118762 | 0.208930903 | 14.12221332 | 24.36633008 | ENSG00000204091 |
| GRID2IP | 0.789115291 | 1.728014461 | 0.019326844 | 0.054635819 | 31.1265649 | 53.7241305 | ENSG00000215045 |
| CIART | 0.788826722 | 1.727668857 | 2.39E-05 | 0.000146747 | 122.77518 | 212.218233 | ENSG00000159208 |
| WIPF1 | 0.788784795 | 1.727618649 | 1.07E-06 | 8.41E-06 | 155.0552311 | 268.3329292 | ENSG00000115935 |
| EPRS1 | 0.788036917 | 1.726723302 | 2.29E-78 | 9.84E-76 | 4536.270925 | 7833.361612 | ENSG00000136628 |
| ABRAXAS1 | 0.787985423 | 1.726661671 | 1.42E-09 | 1.69E-08 | 299.4324872 | 517.3702731 | ENSG00000163322 |
| NA | 0.787561206 | 1.72615403 | 0.472885811 | 0.641657184 | 3.048329015 | 5.23130536 | ENSG00000270460 |
| NA | 0.787523794 | 1.726109268 | 0.081835735 | 0.176666647 | 17.55439938 | 30.1821811 | ENSG00000263412 |
| P2RX4 | 0.787326758 | 1.725873541 | 1.72E-05 | 0.0001084 | 103.9247071 | 179.0953135 | ENSG00000135124 |
| LRRC15 | 0.786832431 | 1.725282286 | 0.228490625 | 0.388180888 | 7.985043186 | 13.80262398 | ENSG00000172061 |
| NA | 0.786777204 | 1.725216242 | 0.466519251 | 0.635722431 | 2.686123467 | 4.636006837 | ENSG00000276538 |
| LINC01405 | 0.786717728 | 1.725145121 | 6.97E-07 | 5.62E-06 | 169.3248995 | 291.6269927 | ENSG00000185847 |
| NA | 0.786647169 | 1.72506075 | 0.47340186 | 0.641990664 | 3.048329015 | 5.227004212 | ENSG00000206762 |
| BDKRB2 | 0.786582461 | 1.724983379 | 0.122903924 | 0.242725185 | 14.07891031 | 24.24641257 | ENSG00000168398 |
| NA | 0.785762383 | 1.724003118 | 0.427766774 | 0.599149617 | 3.657549353 | 6.342040182 | ENSG00000234685 |
| PAG1 | 0.785702368 | 1.723931402 | 0.260035087 | 0.425329563 | 6.703087465 | 11.5604421 | ENSG00000076641 |
| IRF1-AS1 | 0.785700815 | 1.723929546 | 0.54483548 | 0.703179317 | 2.307848223 | 4.013283894 | ENSG00000197536 |
| CRYGS | 0.78507551 | 1.723182508 | 0.370663379 | 0.54446564 | 4.730211482 | 8.08975846 | ENSG00000213139 |

| | | | | | | | |
|---|---|---|---|---|---|---|---|
| NA | 0.784592054 | 1.722605156 | 0.393962059 | 0.566676241 | 4.319121125 | 7.425090119 | ENSG00000263766 |
| IFI27L2 | 0.784280027 | 1.72223263 | 2.66E-06 | 1.94E-05 | 129.4966525 | 223.2769115 | ENSG00000119632 |
| CLDN9 | 0.784279359 | 1.722231832 | 0.219619396 | 0.377701504 | 8.909699444 | 15.40756194 | ENSG00000213937 |
| TBXA2R | 0.783935345 | 1.72182121 | 0.108490093 | 0.220187521 | 14.28473728 | 24.65860291 | ENSG00000006638 |
| ETV3 | 0.783912602 | 1.721794068 | 3.12E-20 | 1.09E-18 | 728.8940534 | 1254.996519 | ENSG00000117036 |
| DCAF8 | 0.783888045 | 1.721764761 | 1.56E-11 | 2.34E-10 | 303.4764332 | 521.9670913 | ENSG00000132716 |
| LOC101928063 | 0.783827118 | 1.721692049 | 0.529370883 | 0.690519555 | 2.277824012 | 3.967037349 | ENSG00000266921 |
| ANXA9 | 0.783705612 | 1.721547052 | 0.000933283 | 0.004003196 | 66.14673049 | 113.8031925 | ENSG00000143412 |
| CLDN4 | 0.783702859 | 1.721543767 | 3.72E-12 | 5.96E-11 | 306.9492946 | 528.3094225 | ENSG00000189143 |
| NA | 0.783660419 | 1.721493125 | 0.203440049 | 0.356462052 | 9.266323187 | 15.98188776 | ENSG00000225173 |
| FOXL2NB | 0.783477248 | 1.72127457 | 0.018938286 | 0.053655231 | 31.26763752 | 53.7555835 | ENSG00000206262 |
| SDS | 0.783360789 | 1.721135628 | 0.566161823 | 0.72000935 | 2.286196721 | 3.969187922 | ENSG00000135094 |
| TPCN2 | 0.783354563 | 1.721128201 | 0.271559947 | 0.438491917 | 7.532765007 | 13.03094157 | ENSG00000287725 |
| JUNB | 0.783268851 | 1.72102595 | 4.35E-05 | 0.000253094 | 95.22641647 | 163.6167642 | ENSG00000171223 |
| WWC3 | 0.783186911 | 1.720928205 | 2.19E-33 | 1.68E-31 | 1220.013369 | 2099.038414 | ENSG00000047644 |
| NA | 0.783168207 | 1.720905893 | 0.040297145 | 0.100215576 | 22.30817739 | 38.45530821 | ENSG00000258592 |
| RIMKLA | 0.782013394 | 1.719528937 | 4.86E-09 | 5.41E-08 | 247.977591 | 425.7270377 | ENSG00000177181 |
| HFE | 0.781863001 | 1.719349694 | 0.066993162 | 0.150677986 | 17.75532026 | 30.48090565 | ENSG00000010704 |
| IVNS1ABP | 0.781678799 | 1.719130184 | 3.80E-67 | 1.06E-64 | 5769.322553 | 9917.285426 | ENSG00000116679 |
| CREB3L4 | 0.78146192 | 1.718871767 | 1.28E-10 | 1.74E-09 | 297.7799021 | 511.0116466 | ENSG00000143578 |
| SPRY3 | 0.781206025 | 1.718566914 | 0.000619426 | 0.002771186 | 64.78778077 | 111.3854508 | ENSG00000168939 |
| TOB1 | 0.781088128 | 1.718426478 | 7.56E-14 | 1.44E-12 | 347.0041073 | 596.6978925 | ENSG00000141232 |
| LINC02018 | 0.78102115 | 1.718346701 | 0.135210291 | 0.260527315 | 13.04396938 | 22.43147505 | ENSG00000272690 |
| EML2 | 0.780905359 | 1.718208791 | 3.70E-14 | 7.33E-13 | 375.3267161 | 644.2905338 | ENSG00000125746 |
| ANKRD29 | 0.780314814 | 1.717505613 | 0.00154514 | 0.006284483 | 59.84962714 | 102.7684187 | ENSG00000154065 |
| NA | 0.780137361 | 1.717294372 | 0.379550555 | 0.552847309 | 4.044197305 | 6.930886983 | ENSG00000223486 |
| NA | 0.780126323 | 1.717281233 | 0.514446286 | 0.678391094 | 2.302266417 | 3.992311195 | ENSG00000286705 |
| RNVU1-1 | 0.779700883 | 1.716774894 | 0.472544321 | 0.641419399 | 2.68891437 | 4.608582417 | ENSG00000207340 |
| MDN1 | 0.779341068 | 1.716346775 | 4.92E-09 | 5.46E-08 | 5207.823077 | 8938.105752 | ENSG00000112159 |
| CGAS | 0.778918313 | 1.715843905 | 0.003858952 | 0.014003582 | 51.6143027 | 88.24789504 | ENSG00000164430 |
| BDNF | 0.778853548 | 1.71576688 | 0.000471133 | 0.002173274 | 67.48506785 | 115.8749092 | ENSG00000176697 |
| LINC01108 | 0.778779436 | 1.715678742 | 4.24E-12 | 6.73E-11 | 302.6019011 | 518.7131194 | ENSG00000226673 |
| HECW1 | 0.778762548 | 1.715658659 | 1.46E-05 | 9.28E-05 | 124.6238168 | 214.2248568 | ENSG00000002746 |
| ONECUT2 | 0.778489327 | 1.715333774 | 0.030763107 | 0.080174425 | 28.43119284 | 48.96416547 | ENSG00000119547 |
| COL23A1 | 0.778350894 | 1.715169188 | 9.15E-13 | 1.56E-11 | 333.488948 | 571.6824995 | ENSG00000050767 |
| NOTCH4 | 0.778175896 | 1.714961152 | 0.009450052 | 0.029988643 | 36.76747812 | 63.04830941 | ENSG00000204301 |
| GAL | 0.77812174 | 1.714896777 | 7.21E-20 | 2.39E-18 | 829.6928001 | 1423.669543 | ENSG00000069482 |
| MTR | 0.777943625 | 1.714685069 | 3.87E-19 | 1.20E-17 | 683.8645541 | 1172.464869 | ENSG00000116984 |
| ITGA9-AS1 | 0.777682122 | 1.714374293 | 0.049918732 | 0.119166224 | 21.32279699 | 36.65059051 | ENSG00000235257 |
| CDC42SE1 | 0.777640092 | 1.714324349 | 9.68E-39 | 1.03E-36 | 1434.582116 | 2458.872585 | ENSG00000197622 |
| NA | 0.777472545 | 1.714125268 | 0.494510625 | 0.660880488 | 3.290437721 | 5.689742245 | ENSG00000279711 |
| KCNT2 | 0.777254823 | 1.713866602 | 1.68E-06 | 1.27E-05 | 147.5901362 | 252.6532736 | ENSG00000162687 |
| NA | 0.777046719 | 1.713619401 | 0.158446656 | 0.294810421 | 13.11709644 | 22.60759847 | ENSG00000236200 |
| NA | 0.776746798 | 1.713263195 | 0.201266308 | 0.353416015 | 9.361977624 | 16.01738144 | ENSG00000176933 |
| NA | 0.776665397 | 1.71316653 | 0.390703289 | 0.563982127 | 4.022545804 | 6.878188716 | ENSG00000279529 |
| NA | 0.776027277 | 1.712408945 | 0.483696628 | 0.651423601 | 2.716147677 | 4.63170569 | ENSG00000243696 |
| NA | 0.775797401 | 1.712136115 | 0.544724233 | 0.703082777 | 2.269451304 | 3.922941376 | ENSG00000260781 |
| PHETA2 | 0.775792427 | 1.712130212 | 0.000101655 | 0.000549308 | 93.97484838 | 161.0965579 | ENSG00000177096 |
| STC1 | 0.775742594 | 1.712071074 | 1.48E-10 | 2.00E-09 | 330.386953 | 564.8010757 | ENSG00000159167 |
| CA7 | 0.775416866 | 1.71168457 | 0.027842223 | 0.073991504 | 28.51867473 | 48.67970144 | ENSG00000168748 |
| ACTN2 | 0.775156119 | 1.711375235 | 0.00385425 | 0.013989152 | 48.10300745 | 82.30299118 | ENSG00000077522 |
| DCP1B | 0.774916348 | 1.711090834 | 4.84E-13 | 8.50E-12 | 323.512932 | 553.1755834 | ENSG00000151065 |
| TMEM88B | 0.774902976 | 1.711074974 | 0.085425291 | 0.182880845 | 17.81728371 | 30.655672 | ENSG00000205116 |
| FES | 0.774868887 | 1.711034545 | 2.35E-05 | 0.000144436 | 106.0187191 | 181.4345621 | ENSG00000182511 |
| TM7SF3 | 0.774616895 | 1.710735709 | 1.61E-19 | 5.19E-18 | 1058.603866 | 1812.207757 | ENSG00000064115 |
| RAB4A | 0.774445751 | 1.710532779 | 1.28E-19 | 4.19E-18 | 654.3049322 | 1119.244132 | ENSG00000168118 |
| RP1L1 | 0.774283729 | 1.710340689 | 0.233916246 | 0.39477762 | 7.652861849 | 13.10246196 | ENSG00000183638 |
| POU2F2 | 0.77378584 | 1.709750534 | 4.40E-05 | 0.00025547 | 100.0207066 | 171.004731 | ENSG00000028277 |
| NRSN2-AS1 | 0.773649223 | 1.709588636 | 0.001578776 | 0.006406432 | 79.48072949 | 135.9695587 | ENSG00000225377 |
| KIAA1217 | 0.77360292 | 1.709533768 | 4.44E-14 | 8.72E-13 | 368.4099556 | 630.2821809 | ENSG00000120549 |
| PCDHA8 | 0.773504474 | 1.709417117 | 0.14245248 | 0.271519691 | 12.93224525 | 22.22309302 | ENSG00000204962 |
| AFF3 | 0.773453105 | 1.709356253 | 0.077250973 | 0.168675553 | 23.2317945 | 39.46064649 | ENSG00000144218 |
| NA | 0.773214195 | 1.709073206 | 0.41077357 | 0.583050028 | 4.025336707 | 6.861517164 | ENSG00000277687 |
| FLRT3 | 0.773118297 | 1.708959606 | 0.315238711 | 0.486505618 | 6.615805736 | 11.39668908 | ENSG00000125848 |
| LOC651337 | 0.772592518 | 1.708336903 | 0.27971978 | 0.447621259 | 6.727529869 | 11.45342803 | ENSG00000255585 |
| NTPCR | 0.772537048 | 1.70827122 | 3.77E-29 | 2.30E-27 | 1017.599366 | 1737.286724 | ENSG00000135778 |
| ADSS2 | 0.772519223 | 1.708250114 | 1.54E-40 | 1.85E-38 | 1526.261645 | 2607.887631 | ENSG00000035687 |
| OSGEPL1 | 0.772187915 | 1.707857867 | 0.000230146 | 0.001146914 | 82.82918846 | 141.4724368 | ENSG00000128694 |
| CLMN | 0.772084487 | 1.707735434 | 2.69E-05 | 0.00016372 | 114.8898584 | 195.9870597 | ENSG00000165959 |
| CA3 | 0.771995112 | 1.707629643 | 0.23077444 | 0.391098105 | 8.997656894 | 15.36346596 | ENSG00000164879 |
| ZNF280D | 0.771911714 | 1.707530933 | 0.006135378 | 0.020911233 | 42.13760988 | 71.93555716 | ENSG00000137871 |
| BATF3 | 0.771850928 | 1.70745899 | 0.00337357 | 0.012424502 | 48.64101576 | 83.17415144 | ENSG00000123685 |
| NA | 0.771602761 | 1.707165304 | 0.539524931 | 0.698803239 | 2.299475514 | 3.95251637 | ENSG00000263606 |
| DYRK3 | 0.771463432 | 1.707000442 | 3.82E-09 | 4.31E-08 | 213.6608239 | 364.9550467 | ENSG00000143479 |
| MAPK13 | 0.771306011 | 1.706814191 | 2.49E-11 | 3.65E-10 | 306.553435 | 522.4704419 | ENSG00000156711 |
| NA | 0.771147562 | 1.706626745 | 0.509881459 | 0.674379942 | 2.748962791 | 4.661280684 | ENSG00000260884 |
| LIN28B-AS1 | 0.771011239 | 1.706465489 | 0.199176932 | 0.350616517 | 9.098217416 | 15.4898352 | ENSG00000203809 |

| | | | | | | | |
|---|---|---|---|---|---|---|---|
| TIMP4 | 0.770655362 | 1.7060446 | 0.000690642 | 0.003047385 | 73.64994685 | 125.7704698 | ENSG00000157150 |
| NA | 0.770328607 | 1.705658242 | 0.016145177 | 0.047054934 | 40.00257335 | 68.51191357 | ENSG00000259865 |
| SLC25A5-AS1 | 0.770062449 | 1.705343599 | 0.084828406 | 0.181865391 | 17.54044487 | 29.78236116 | ENSG00000224281 |
| NA | 0.769646605 | 1.704852121 | 0.005546094 | 0.019187536 | 47.11782721 | 80.12049233 | ENSG00000239213 |
| KRTCAP2 | 0.769443271 | 1.704611855 | 0.002830351 | 0.010670375 | 52.62671624 | 89.59013516 | ENSG00000163463 |
| ZSCAN31 | 0.769102207 | 1.704208919 | 0.000768575 | 0.003351397 | 72.44881517 | 123.674818 | ENSG00000235109 |
| CDC42EP5 | 0.768987357 | 1.704073257 | 9.50E-05 | 0.000516876 | 117.6236273 | 200.0097699 | ENSG00000167617 |
| NA | 0.767948468 | 1.70284659 | 0.309520404 | 0.480618909 | 6.687017769 | 11.35233264 | ENSG00000256742 |
| SEC14L5 | 0.767798588 | 1.702669692 | 1.43E-10 | 1.95E-09 | 243.5703875 | 414.7441564 | ENSG00000103184 |
| CNIH3 | 0.767246296 | 1.702018002 | 8.84E-05 | 0.000483783 | 100.9426841 | 171.5567575 | ENSG00000143786 |
| NA | 0.766645125 | 1.701308918 | 0.345100896 | 0.517566141 | 4.988389884 | 8.489578395 | ENSG00000273381 |
| RBM46 | 0.766473534 | 1.70110658 | 0.347819514 | 0.520501609 | 5.807779696 | 9.791490657 | ENSG00000151962 |
| NIT1 | 0.766243898 | 1.700835835 | 8.67E-17 | 2.23E-15 | 501.3119101 | 852.0168426 | ENSG00000158793 |
| ABL2 | 0.766011463 | 1.700561832 | 2.75E-18 | 7.98E-17 | 543.0093324 | 923.2890517 | ENSG00000143322 |
| NTRK1 | 0.765994713 | 1.700542088 | 0.419703178 | 0.591719214 | 3.633106948 | 6.194698249 | ENSG00000198400 |
| NA | 0.765902743 | 1.700433684 | 0.178167094 | 0.322150287 | 11.06060551 | 18.81102633 | ENSG00000261087 |
| LOC286059 | 0.765492106 | 1.699949756 | 0.152210094 | 0.285853284 | 12.0753344 | 20.54663467 | ENSG00000246130 |
| NA | 0.765399315 | 1.699840422 | 0.011182966 | 0.034661086 | 39.30714732 | 66.69241444 | ENSG00000286388 |
| DDX60 | 0.765182716 | 1.699585236 | 0.005896524 | 0.020254602 | 49.18615747 | 83.70490222 | ENSG00000137628 |
| PLEKHN1 | 0.764783121 | 1.699114553 | 0.024379706 | 0.066203393 | 34.52505997 | 58.70054682 | ENSG00000187583 |
| IQCN | 0.764471018 | 1.698747018 | 0.425005684 | 0.596709217 | 3.665922062 | 6.2323425 | ENSG00000130518 |
| DENND3 | 0.764091113 | 1.698299745 | 0.000231796 | 0.001153655 | 118.8635946 | 202.0172235 | ENSG00000105339 |
| NA | 0.763854981 | 1.6980218 | 0.368349637 | 0.542264412 | 4.664581255 | 7.946978139 | ENSG00000272572 |
| REL | 0.76384723 | 1.698012677 | 0.007057165 | 0.023541375 | 43.81044391 | 74.42025768 | ENSG00000162924 |
| SDR42E2 | 0.763793161 | 1.697949041 | 0.432452554 | 0.603512723 | 3.668712964 | 6.2323425 | ENSG00000183921 |
| NA | 0.763302963 | 1.69737221 | 0.522886544 | 0.685570353 | 2.773405195 | 4.665581831 | ENSG00000260136 |
| CDH8 | 0.763155725 | 1.69719899 | 0.000895173 | 0.003852058 | 67.7879007 | 115.1285006 | ENSG00000150394 |
| TRIM5 | 0.763023588 | 1.69704355 | 2.38E-08 | 2.39E-07 | 252.8512657 | 429.001134 | ENSG00000132256 |
| LGALS3BP | 0.762851438 | 1.696841061 | 5.03E-36 | 4.39E-34 | 1259.122274 | 2136.221324 | ENSG00000108679 |
| OGFRL1 | 0.762716502 | 1.696682363 | 2.06E-06 | 1.54E-05 | 140.0374346 | 237.5301878 | ENSG00000119900 |
| CALHM6 | 0.762392201 | 1.696301012 | 0.303496903 | 0.474223129 | 6.379278836 | 10.79037722 | ENSG00000188820 |
| OMA1 | 0.762289404 | 1.696180148 | 1.78E-05 | 0.000111682 | 113.9908838 | 193.1270196 | ENSG00000162600 |
| NA | 0.761893267 | 1.695714473 | 0.572683423 | 0.725196003 | 2.310639126 | 3.931543671 | ENSG00000229512 |
| THORLNC | 0.76172647 | 1.695518435 | 0.350365038 | 0.52297214 | 4.996762593 | 8.462153975 | ENSG00000226856 |
| IGFBP7 | 0.761588651 | 1.695356471 | 0.298676674 | 0.469123247 | 6.357627334 | 10.7505824 | ENSG00000163453 |
| OLFM3 | 0.761158566 | 1.694851139 | 0.371006158 | 0.544720283 | 4.607323737 | 7.845882753 | ENSG00000118733 |
| SLFN12 | 0.761119572 | 1.694805332 | 0.000339101 | 0.001625645 | 99.52456181 | 168.6348228 | ENSG00000172123 |
| BRINP3 | 0.761010055 | 1.694676681 | 0.557791603 | 0.713430492 | 2.326708822 | 3.95251637 | ENSG00000162670 |
| MFSD6L | 0.760916089 | 1.694566307 | 0.029868849 | 0.078277402 | 29.8830061 | 50.54759769 | ENSG00000185156 |
| HPN | 0.760395824 | 1.693955322 | 4.13E-08 | 4.03E-07 | 224.4964664 | 379.9791814 | ENSG00000105707 |
| FN1 | 0.76036573 | 1.693919987 | 1.97E-26 | 1.05E-24 | 2693.31661 | 4560.816455 | ENSG00000115414 |
| NA | 0.760201003 | 1.693726586 | 0.541325597 | 0.700278174 | 2.340663336 | 3.971338496 | ENSG00000223935 |
| NA | 0.760173623 | 1.693694443 | 0.227000127 | 0.386228563 | 8.684336156 | 14.68804475 | ENSG00000133519 |
| ZNF292 | 0.760033955 | 1.693530483 | 1.20E-11 | 1.82E-10 | 372.1999295 | 630.5096518 | ENSG00000188994 |
| LMCD1-AS1 | 0.759933687 | 1.693412786 | 0.226318933 | 0.385443502 | 9.590131815 | 16.26179018 | ENSG00000227110 |
| TMEM196 | 0.759863223 | 1.693330079 | 0.43665286 | 0.60747357 | 3.671503867 | 6.213520375 | ENSG00000173452 |
| TMEM185A | 0.759634057 | 1.693061122 | 5.93E-13 | 1.03E-11 | 340.223387 | 575.8390968 | ENSG00000269556 |
| CASTOR1 | 0.759049122 | 1.692374815 | 0.546383492 | 0.704470255 | 2.340663336 | 3.967037349 | ENSG00000239282 |
| NCF1C | 0.758949486 | 1.69225794 | 0.540132501 | 0.699308461 | 2.392339049 | 4.040708315 | ENSG00000165178 |
| NOD2 | 0.758763427 | 1.69203971 | 0.160769375 | 0.297982515 | 11.68589554 | 19.80614479 | ENSG00000167207 |
| OSBPL5 | 0.758620143 | 1.69187167 | 5.98E-12 | 9.37E-11 | 317.5348192 | 537.0162457 | ENSG00000021762 |
| NA | 0.758476723 | 1.691703488 | 0.294391955 | 0.464283454 | 6.403721241 | 10.78230797 | ENSG00000176700 |
| ARHGAP4 | 0.758430361 | 1.691649124 | 8.51E-22 | 3.28E-20 | 730.8979657 | 1235.752875 | ENSG00000089820 |
| TTC32 | 0.758395551 | 1.691608308 | 0.001159632 | 0.004873739 | 65.77615223 | 111.1270541 | ENSG00000183891 |
| ZFHX4 | 0.758252873 | 1.691441022 | 0.020008052 | 0.056240031 | 32.71211721 | 55.41752087 | ENSG00000091656 |
| NA | 0.757777824 | 1.690884158 | 6.42E-07 | 5.21E-06 | 150.6773377 | 254.5039774 | ENSG00000225889 |
| RIN2 | 0.757567476 | 1.690637641 | 0.096912411 | 0.202221452 | 20.35906809 | 34.50682041 | ENSG00000132669 |
| ACTL10 | 0.757325455 | 1.69035405 | 0.135717658 | 0.261264278 | 13.39289614 | 22.60974905 | ENSG00000182584 |
| CALHM1 | 0.757202722 | 1.690210254 | 0.260512314 | 0.425893479 | 7.889388749 | 13.39903593 | ENSG00000185933 |
| ZNRD1ASP | 0.757181151 | 1.690184983 | 0.003887253 | 0.014093028 | 49.5099661 | 83.73206618 | ENSG00000204623 |
| RALGPS2 | 0.757135372 | 1.690131351 | 2.36E-15 | 5.28E-14 | 588.9175672 | 995.5071401 | ENSG00000116191 |
| BROX | 0.756843742 | 1.689789739 | 4.24E-10 | 5.42E-09 | 259.0667444 | 437.7117616 | ENSG00000162819 |
| SGIP1 | 0.756820456 | 1.689762465 | 0.189424 | 0.337474915 | 10.72005146 | 18.11033127 | ENSG00000118473 |
| ANKRD65 | 0.756661223 | 1.689575972 | 0.137445029 | 0.264036253 | 13.45505974 | 22.68342001 | ENSG00000235098 |
| ELF3 | 0.756407453 | 1.689278802 | 0.00214107 | 0.008380749 | 58.93111626 | 99.6373389 | ENSG00000163435 |
| NA | 0.755874465 | 1.688654832 | 0.514285954 | 0.678226028 | 2.335081531 | 3.948215223 | ENSG00000280122 |
| NA | 0.755628417 | 1.68836686 | 0.062261577 | 0.142090155 | 21.19520332 | 35.65951274 | ENSG00000246792 |
| NA | 0.755050753 | 1.687690963 | 0.304374761 | 0.475094548 | 6.025445997 | 10.15528388 | ENSG00000226200 |
| NA | 0.755038636 | 1.687676788 | 0.351193791 | 0.523781345 | 5.290547011 | 8.998562973 | ENSG00000251292 |
| RAB4B | 0.754709802 | 1.687292159 | 0.347549524 | 0.52027691 | 5.938164268 | 10.02891464 | ENSG00000167578 |
| VEPH1 | 0.754620928 | 1.687188221 | 0.005703205 | 0.019660565 | 58.4321806 | 98.63709526 | ENSG00000197415 |
| PLEKHH1 | 0.754615508 | 1.687181882 | 6.40E-18 | 1.82E-16 | 681.4840303 | 1149.686836 | ENSG00000054690 |
| TP53I11 | 0.754417196 | 1.68694998 | 1.74E-27 | 9.72E-26 | 1184.476572 | 1998.834018 | ENSG00000175274 |
| NA | 0.754246875 | 1.686750834 | 0.505294061 | 0.670320137 | 2.713356774 | 4.534911451 | ENSG00000259182 |
| AMT | 0.75401159 | 1.686475768 | 0.338981233 | 0.51152927 | 5.587122332 | 9.467492263 | ENSG00000145020 |
| NA | 0.754009353 | 1.686473154 | 0.029218899 | 0.076855982 | 34.14630916 | 57.57689645 | ENSG00000261167 |
| NA | 0.753880087 | 1.686322052 | 0.236033823 | 0.397481288 | 7.976670478 | 13.48722788 | ENSG00000272989 |

| | | | | | | | |
|---|---|---|---|---|---|---|---|
| NA | 0.753639631 | 1.686041014 | 0.463463254 | 0.6331919 | 3.282065012 | 5.567674159 | ENSG00000286415 |
| C2orf72 | 0.753596798 | 1.685990957 | 0.003187335 | 0.01181304 | 54.9853643 | 92.75484903 | ENSG00000204128 |
| CLDN19 | 0.753490996 | 1.685867317 | 5.20E-06 | 3.60E-05 | 153.972206 | 259.3937767 | ENSG00000164007 |
| FBXO48 | 0.753454455 | 1.685824617 | 0.126851351 | 0.248334627 | 18.4050527 | 31.04957325 | ENSG00000204923 |
| IL12A | 0.753360188 | 1.685714468 | 0.371738561 | 0.545131782 | 6.011491482 | 10.12355831 | ENSG00000168811 |
| EMC1-AS1 | 0.751975968 | 1.684097853 | 0.427688813 | 0.599124602 | 4.515336084 | 7.496610512 | ENSG00000230424 |
| ZC3H6 | 0.751768469 | 1.683855652 | 0.000418147 | 0.001958833 | 81.27422395 | 137.140528 | ENSG00000188177 |
| MYO15A | 0.751527505 | 1.683574431 | 0.124344175 | 0.24481618 | 14.0705376 | 23.67423732 | ENSG00000091536 |
| FAM162A | 0.75125509 | 1.683256563 | 2.63E-21 | 9.78E-20 | 760.0103916 | 1278.239446 | ENSG00000114023 |
| NA | 0.751048742 | 1.683015825 | 0.598554809 | 0.74523821 | 2.283405818 | 3.853571558 | ENSG00000232901 |
| NA | 0.751039009 | 1.683004471 | 0.446027841 | 0.616426443 | 3.309298319 | 5.595098579 | ENSG00000282390 |
| STEAP1B | 0.750492557 | 1.682367117 | 0.001181648 | 0.004958706 | 76.61733968 | 129.1128941 | ENSG00000105889 |
| INE1 | 0.749914891 | 1.681693619 | 0.063224826 | 0.143845987 | 20.14486841 | 33.81069907 | ENSG00000224975 |
| NPAS1 | 0.749781159 | 1.68153774 | 2.82E-05 | 0.000171266 | 121.4934245 | 204.2177326 | ENSG00000130751 |
| NA | 0.749762164 | 1.681515601 | 0.416761801 | 0.588807116 | 4.2751424 | 7.231762105 | ENSG00000264281 |
| MIR1244-4 | 0.749551155 | 1.68126968 | 0.461191825 | 0.630982375 | 3.284855915 | 5.551002607 | ENSG00000283475 |
| HMGCL | 0.749132007 | 1.68078129 | 9.82E-06 | 6.48E-05 | 215.9937519 | 362.313578 | ENSG00000117305 |
| COL6A4P1 | 0.748815601 | 1.680412708 | 0.451574848 | 0.621749368 | 3.306507417 | 5.578427028 | ENSG00000230524 |
| TNXB | 0.748547697 | 1.68010069 | 2.98E-07 | 2.55E-06 | 162.3255121 | 272.5640214 | ENSG00000168477 |
| NA | 0.748541952 | 1.680093999 | 0.245840472 | 0.409196866 | 7.998321979 | 13.44743305 | ENSG00000287200 |
| MYO1G | 0.748536251 | 1.68008736 | 0.491084168 | 0.65747135 | 3.2709014 | 5.521427613 | ENSG00000136286 |
| ACOT4 | 0.748343006 | 1.679862332 | 0.062380977 | 0.142278485 | 21.71993283 | 36.49894743 | ENSG00000177465 |
| NA | 0.74831928 | 1.679834705 | 0.310333156 | 0.481279598 | 6.055470207 | 10.1321606 | ENSG00000271714 |
| NA | 0.748129007 | 1.679613172 | 0.521720554 | 0.684660626 | 2.335081531 | 3.918640229 | ENSG00000252202 |
| LOC101929140 | 0.748027129 | 1.679494568 | 0.322565549 | 0.494054149 | 5.698846465 | 9.572888796 | ENSG00000278390 |
| EMBP1 | 0.747476882 | 1.678854127 | 0.016425972 | 0.047685994 | 40.67060281 | 68.44173814 | ENSG00000231752 |
| KLF7 | 0.747400759 | 1.678765545 | 1.23E-10 | 1.69E-09 | 275.4845663 | 462.7220783 | ENSG00000118263 |
| RGS5 | 0.747273792 | 1.678617809 | 1.96E-08 | 1.99E-07 | 221.911467 | 372.4685893 | ENSG00000143248 |
| VWDE | 0.747133451 | 1.678454526 | 1.55E-07 | 1.39E-06 | 209.1909429 | 350.72974 | ENSG00000146530 |
| NPTXR | 0.747113327 | 1.678431114 | 4.16E-15 | 9.05E-14 | 448.4546874 | 752.7653784 | ENSG00000221890 |
| NA | 0.747096757 | 1.678411836 | 0.233987688 | 0.394843989 | 8.44222745 | 14.10780025 | ENSG00000228420 |
| NA | 0.74685713 | 1.67813308 | 0.024703183 | 0.066949861 | 32.96327434 | 55.35434231 | ENSG00000268858 |
| LOC150051 | 0.746729499 | 1.677984628 | 0.01628649 | 0.047367028 | 39.81214036 | 67.0623868 | ENSG00000237594 |
| NA | 0.746621067 | 1.677858516 | 0.429712266 | 0.601216952 | 4.681326673 | 7.845882753 | ENSG00000274356 |
| NA | 0.746338895 | 1.677530381 | 0.315636898 | 0.48684348 | 6.316439512 | 10.65404862 | ENSG00000260908 |
| IQCH-AS1 | 0.745904915 | 1.677025836 | 0.259396049 | 0.424608331 | 8.270454896 | 13.96286935 | ENSG00000259673 |
| EDEM3 | 0.745178089 | 1.676181167 | 4.49E-10 | 5.72E-09 | 301.495636 | 505.1688737 | ENSG00000116406 |
| LOC257396 | 0.745170979 | 1.676172907 | 0.058172945 | 0.134687264 | 27.01203144 | 45.07699035 | ENSG00000247796 |
| RINL | 0.745114619 | 1.676107427 | 1.19E-07 | 1.08E-06 | 191.7171674 | 321.006844 | ENSG00000187994 |
| SLC52A3 | 0.744991259 | 1.675964115 | 6.04E-08 | 5.72E-07 | 187.2213676 | 313.8745356 | ENSG00000101276 |
| NA | 0.744803647 | 1.675746182 | 0.00093296 | 0.004003054 | 77.80431668 | 130.5702113 | ENSG00000267640 |
| NA | 0.744666424 | 1.6755868 | 0.439533173 | 0.610661466 | 4.626184336 | 7.751239088 | ENSG00000287460 |
| NA | 0.744632871 | 1.675547831 | 0.212674272 | 0.36871568 | 9.454841158 | 15.7842586 | ENSG00000276900 |
| RYR1 | 0.744595071 | 1.67550393 | 1.22E-25 | 6.16E-24 | 1055.920383 | 1768.812606 | ENSG00000196218 |
| SMIM14 | 0.744334646 | 1.675201508 | 4.54E-07 | 3.77E-06 | 204.5015189 | 342.3969236 | ENSG00000163683 |
| PROK2 | 0.74422023 | 1.675068657 | 0.060369629 | 0.138641728 | 25.6672364 | 42.79501364 | ENSG00000163421 |
| SUMF2 | 0.743970904 | 1.674779198 | 2.79E-22 | 1.11E-20 | 1112.575901 | 1862.350786 | ENSG00000129103 |
| NA | 0.743904892 | 1.674702569 | 0.422306304 | 0.593870163 | 4.289096914 | 7.212939979 | ENSG00000234134 |
| GADD45B | 0.743771606 | 1.674547855 | 4.55E-05 | 0.000263924 | 111.4335933 | 186.6591432 | ENSG00000099860 |
| EIF4EBP3 | 0.743569996 | 1.674313861 | 0.566287547 | 0.720074687 | 2.326708822 | 3.891215809 | ENSG00000243056 |
| NA | 0.743119063 | 1.673790614 | 0.327056438 | 0.499070221 | 6.591363331 | 11.06167735 | ENSG00000236409 |
| JUN | 0.743114934 | 1.673785824 | 9.26E-10 | 1.13E-08 | 367.6645815 | 615.1174712 | ENSG00000177606 |
| NFE2 | 0.742911541 | 1.673549868 | 0.061527613 | 0.140764845 | 23.11919449 | 38.73063691 | ENSG00000123405 |
| HSPG2 | 0.742276376 | 1.672813229 | 3.18E-39 | 3.49E-37 | 2831.48077 | 4735.312908 | ENSG00000142798 |
| NA | 0.741695887 | 1.672140284 | 0.433548556 | 0.604567384 | 3.962497383 | 6.684860702 | ENSG00000255062 |
| ATP6V0A4 | 0.741580442 | 1.672006484 | 0.156162917 | 0.291630089 | 14.60383998 | 24.28243929 | ENSG00000105929 |
| C1orf61 | 0.741546778 | 1.671967469 | 0.001098095 | 0.004634292 | 67.67761602 | 113.2390982 | ENSG00000125462 |
| CCDC113 | 0.741416339 | 1.671816308 | 1.88E-06 | 1.41E-05 | 148.2989532 | 247.8244718 | ENSG00000103021 |
| RGPD1 | 0.741240287 | 1.671612309 | 0.355928047 | 0.529105953 | 5.375037836 | 9.002864121 | ENSG00000187627 |
| CCDC114 | 0.741209019 | 1.67157608 | 0.040014467 | 0.099628173 | 31.21576165 | 52.21519328 | ENSG00000105479 |
| NA | 0.740707746 | 1.670995382 | 0.014894634 | 0.044011354 | 38.58253607 | 64.54461576 | ENSG00000248092 |
| PLA2G6 | 0.740347307 | 1.670577957 | 8.42E-06 | 5.63E-05 | 128.080282 | 213.6760774 | ENSG00000184381 |
| CHGB | 0.740311867 | 1.670536919 | 0.000118733 | 0.000630812 | 96.77982939 | 161.4307882 | ENSG00000089199 |
| NA | 0.739776685 | 1.669917333 | 0.040908901 | 0.101488377 | 26.89684068 | 44.86671821 | ENSG00000233396 |
| DPP6 | 0.739768814 | 1.669908221 | 1.31E-06 | 1.01E-05 | 148.0287353 | 247.2899408 | ENSG00000130226 |
| NA | 0.739654584 | 1.669776006 | 0.435325224 | 0.606135187 | 4.261863607 | 7.145187696 | ENSG00000265460 |
| TTLL10-AS1 | 0.738987587 | 1.669004202 | 0.448590335 | 0.618920522 | 3.33932253 | 5.571975306 | ENSG00000205231 |
| ZNF678 | 0.738762377 | 1.668743685 | 7.67E-08 | 7.16E-07 | 190.0830937 | 317.4019703 | ENSG00000181450 |
| NA | 0.738349265 | 1.668265912 | 0.40371214 | 0.576115175 | 4.286306011 | 7.174762691 | ENSG00000246982 |
| TTC17 | 0.738292025 | 1.668199724 | 1.04E-24 | 5.03E-23 | 985.8975468 | 1644.134442 | ENSG00000052841 |
| NA | 0.738228908 | 1.668126743 | 0.065008366 | 0.146957838 | 25.50296067 | 42.75090556 | ENSG00000255495 |
| CST3 | 0.737597624 | 1.667396976 | 1.88E-12 | 3.11E-11 | 358.6778753 | 598.6279133 | ENSG00000101439 |
| NA | 0.737566897 | 1.667361464 | 0.505969549 | 0.670789539 | 2.947092772 | 4.957854658 | ENSG00000235689 |
| NA | 0.737460954 | 1.667239027 | 0.3774018 | 0.550884256 | 6.234739589 | 10.43895441 | ENSG00000280332 |
| IGIP | 0.737326252 | 1.667083367 | 0.348666044 | 0.521366113 | 6.631875432 | 11.06005982 | ENSG00000182700 |
| TMEM176B | 0.737301965 | 1.667055302 | 0.109854359 | 0.222336876 | 15.36577212 | 25.6219958 | ENSG00000106565 |
| DECR2 | 0.736995562 | 1.666701287 | 0.000240509 | 0.001193634 | 99.81834623 | 166.1668243 | ENSG00000242612 |

| | | | | | | | |
|---|---|---|---|---|---|---|---|
| NA | 0.73669955 | 1.666359349 | 0.540188412 | 0.699333914 | 2.332290628 | 3.872393683 | ENSG00000270571 |
| RPL39L | 0.736585672 | 1.666227822 | 6.27E-16 | 1.50E-14 | 511.0911235 | 851.2981069 | ENSG00000163923 |
| NA | 0.736193448 | 1.665774887 | 0.543943748 | 0.702451389 | 2.419572356 | 3.994461769 | ENSG00000248180 |
| NA | 0.736057746 | 1.665618209 | 0.485490422 | 0.653129864 | 2.982698788 | 5.006251778 | ENSG00000278876 |
| PFKFB2 | 0.736032117 | 1.66558862 | 3.34E-14 | 6.65E-13 | 478.4594669 | 796.1586409 | ENSG00000123836 |
| TSACC | 0.735024797 | 1.664426077 | 0.065360288 | 0.147625023 | 22.23205927 | 37.0985471 | ENSG00000163467 |
| TSTD1 | 0.734864731 | 1.66424142 | 7.24E-06 | 4.90E-05 | 145.0848338 | 241.0498201 | ENSG00000215845 |
| TMC4 | 0.73432563 | 1.663619649 | 0.002266422 | 0.008800635 | 68.29975173 | 113.4851128 | ENSG00000167608 |
| NA | 0.734250547 | 1.663533071 | 0.397628864 | 0.570131354 | 4.721838773 | 7.809856038 | ENSG00000217950 |
| LHX4 | 0.733886031 | 1.66311281 | 0.013706842 | 0.041052422 | 41.47392292 | 68.9528854 | ENSG00000121454 |
| ASAH2B | 0.733780964 | 1.662991695 | 0.004879504 | 0.017181085 | 63.0261134 | 104.9560243 | ENSG00000204147 |
| GOLGA6L7 | 0.733773275 | 1.662982832 | 0.539328123 | 0.698681435 | 2.389548146 | 3.941763502 | ENSG00000261649 |
| HYLS1 | 0.733761977 | 1.662969809 | 5.12E-07 | 4.22E-06 | 170.7814939 | 283.9733535 | ENSG00000198331 |
| NA | 0.733072576 | 1.662175339 | 0.421845332 | 0.593784627 | 4.307957513 | 7.168310969 | ENSG00000223751 |
| NUFIP2 | 0.733026483 | 1.662122234 | 6.71E-33 | 5.02E-31 | 2150.516601 | 3573.191666 | ENSG00000108256 |
| NA | 0.732920759 | 1.662000434 | 0.086855187 | 0.185366016 | 24.05656596 | 39.83681012 | ENSG00000285796 |
| LINC02293 | 0.732735241 | 1.661786729 | 0.55134457 | 0.708216014 | 2.395129951 | 3.948215223 | ENSG00000257185 |
| GUSBP11 | 0.732688271 | 1.661732628 | 0.046082736 | 0.111807579 | 25.02990687 | 41.58076029 | ENSG00000228315 |
| CSRNP3 | 0.732612065 | 1.661644853 | 0.000213785 | 0.001070365 | 99.13568654 | 164.9732406 | ENSG00000178662 |
| ENTPD1-AS1 | 0.732353027 | 1.66134653 | 0.254144292 | 0.41892656 | 8.725523978 | 14.48234634 | ENSG00000226688 |
| RGS6 | 0.73223499 | 1.661210608 | 0.064247241 | 0.145622551 | 23.79472078 | 39.5149623 | ENSG00000182732 |
| NA | 0.731405997 | 1.660256328 | 0.284185357 | 0.452851455 | 8.44222745 | 13.92066349 | ENSG00000286403 |
| LOC100505797 | 0.731138291 | 1.659948279 | 0.267935169 | 0.434194962 | 7.761795079 | 12.84568281 | ENSG00000263958 |
| THSD4 | 0.730994813 | 1.659783204 | 0.000142911 | 0.000742899 | 96.99538051 | 160.8376402 | ENSG00000187720 |
| ARHGEF11 | 0.730842103 | 1.659607524 | 1.10E-25 | 5.65E-24 | 993.5222994 | 1647.97843 | ENSG00000132694 |
| PHLDA2 | 0.730809531 | 1.659570055 | 0.000127594 | 0.0006715 | 99.51272248 | 165.0350863 | ENSG00000181649 |
| CNTNAP2 | 0.730765822 | 1.659519776 | 1.28E-36 | 1.17E-34 | 1507.404715 | 2501.958024 | ENSG00000174469 |
| NIT2 | 0.730625173 | 1.659357997 | 7.29E-09 | 7.91E-08 | 298.6477764 | 494.8996152 | ENSG00000114021 |
| PTPDC1 | 0.730386652 | 1.659083678 | 6.41E-06 | 4.38E-05 | 134.3289761 | 222.6649536 | ENSG00000158079 |
| RGL3 | 0.730359067 | 1.659051955 | 3.69E-18 | 1.06E-16 | 613.8580403 | 1018.758811 | ENSG00000205517 |
| NA | 0.730162734 | 1.658826194 | 0.477773046 | 0.646081043 | 4.40361195 | 7.288761519 | ENSG00000272800 |
| PCNX2 | 0.73013639 | 1.658795904 | 3.15E-31 | 2.13E-29 | 1365.144572 | 2264.251973 | ENSG00000135749 |
| NA | 0.729944022 | 1.658574736 | 0.303947605 | 0.47465824 | 7.119759628 | 11.72042701 | ENSG00000242798 |
| SCG5 | 0.729909739 | 1.658535324 | 5.77E-10 | 7.22E-09 | 344.0884271 | 570.5351439 | ENSG00000166922 |
| NA | 0.729755971 | 1.65835856 | 0.092790804 | 0.195183448 | 18.13080461 | 30.06037348 | ENSG00000224699 |
| C22orf23 | 0.729626816 | 1.658210105 | 0.244895195 | 0.408074044 | 9.133823432 | 15.06904256 | ENSG00000128346 |
| LINC01184 | 0.729357772 | 1.657900898 | 0.002689797 | 0.010212337 | 59.16293724 | 98.15364499 | ENSG00000245937 |
| C12orf66 | 0.72830975 | 1.656696982 | 0.000200155 | 0.001008409 | 96.20657848 | 159.6032077 | ENSG00000174206 |
| ARL8A | 0.727267079 | 1.655500078 | 6.06E-21 | 2.21E-19 | 775.2086726 | 1283.356165 | ENSG00000143862 |
| NA | 0.726915545 | 1.65509674 | 0.123231672 | 0.243185399 | 16.17399832 | 26.71820964 | ENSG00000279347 |
| NA | 0.726587182 | 1.654720077 | 0.308381804 | 0.47944821 | 6.705878368 | 11.05145752 | ENSG00000227214 |
| NA | 0.726355628 | 1.654454514 | 0.371655493 | 0.5450514 | 5.611564737 | 9.36665734 | ENSG00000249675 |
| SLC8A2 | 0.725611978 | 1.65360193 | 4.04E-06 | 2.86E-05 | 208.6990796 | 345.0422094 | ENSG00000118160 |
| MCMDC2 | 0.725574698 | 1.653559201 | 0.198605063 | 0.350037407 | 11.48497466 | 18.874217 | ENSG00000178460 |
| NA | 0.725463292 | 1.653431517 | 0.3706487 | 0.54446564 | 5.326153027 | 8.802823921 | ENSG00000230415 |
| NA | 0.725200276 | 1.653130109 | 5.22E-10 | 6.59E-09 | 297.2038603 | 490.6803787 | ENSG00000281406 |
| KLK6 | 0.725156297 | 1.653079716 | 0.194324451 | 0.34432929 | 11.13114182 | 18.31656273 | ENSG00000167755 |
| NA | 0.725132134 | 1.65305203 | 0.134141864 | 0.25908397 | 17.34231489 | 28.48690062 | ENSG00000268030 |
| NA | 0.724046247 | 1.651808279 | 0.202532501 | 0.355122204 | 13.7879168 | 22.7511723 | ENSG00000232022 |
| ITGB1 | 0.723943677 | 1.651690846 | 4.44E-55 | 8.73E-53 | 5979.88533 | 9877.58802 | ENSG00000150093 |
| CILP2 | 0.723536142 | 1.651224339 | 1.57E-12 | 2.62E-11 | 362.0922768 | 598.1861359 | ENSG00000160161 |
| MCEE | 0.723439297 | 1.6511135 | 0.000144941 | 0.000752034 | 97.27541057 | 160.4340401 | ENSG00000124370 |
| RAB26 | 0.722820621 | 1.650405599 | 0.001346399 | 0.005564136 | 74.92373002 | 123.6605759 | ENSG00000167964 |
| TRMT2B | 0.722161296 | 1.649651521 | 3.63E-15 | 7.94E-14 | 460.9970183 | 760.381646 | ENSG00000188917 |
| LOH12CR2 | 0.721768069 | 1.649201947 | 0.471922089 | 0.640800241 | 3.807670405 | 6.192547676 | ENSG00000205791 |
| THEM4 | 0.72172622 | 1.649154108 | 3.44E-07 | 2.92E-06 | 188.4709085 | 310.3608526 | ENSG00000159445 |
| RCOR3 | 0.721429012 | 1.648814403 | 7.00E-09 | 7.62E-08 | 248.2231295 | 409.556644 | ENSG00000117625 |
| MDGA2 | 0.720195979 | 1.647405806 | 0.020520876 | 0.057472157 | 35.61957376 | 58.59352064 | ENSG00000139915 |
| GPATCH2 | 0.720071297 | 1.647263439 | 8.39E-09 | 9.05E-08 | 278.9691918 | 459.6447445 | ENSG00000092978 |
| NA | 0.719977255 | 1.647156066 | 0.073262184 | 0.161797458 | 20.69683123 | 34.08630035 | ENSG00000241860 |
| SNAPC5 | 0.719919505 | 1.647090133 | 2.89E-08 | 2.87E-07 | 205.3729478 | 338.3979065 | ENSG00000174446 |
| LOC101059948 | 0.719279086 | 1.646359144 | 0.501212646 | 0.667096257 | 3.007141193 | 4.960005232 | ENSG00000268655 |
| INHA | 0.719245155 | 1.646320423 | 0.422616789 | 0.594133544 | 3.986939787 | 6.583765315 | ENSG00000123999 |
| NA | 0.719141612 | 1.64620227 | 0.347994109 | 0.520682104 | 6.047097499 | 9.964106436 | ENSG00000261251 |
| NA | 0.719121497 | 1.646179318 | 0.281838358 | 0.450277013 | 8.061161303 | 13.283147 | ENSG00000235499 |
| KCNA6 | 0.718620476 | 1.64560773 | 0.008086438 | 0.026363948 | 51.18259998 | 84.48922761 | ENSG00000130035 |
| TGFB1I1 | 0.718594208 | 1.645577768 | 0.000100687 | 0.000544686 | 101.959328 | 167.7429808 | ENSG00000140682 |
| IBA57 | 0.718452423 | 1.645416053 | 4.87E-09 | 5.42E-08 | 245.24167 | 403.7921768 | ENSG00000181873 |
| APLNR | 0.718054757 | 1.64496257 | 0.320298644 | 0.491635992 | 7.479174273 | 12.25683601 | ENSG00000134817 |
| CRISPLD2 | 0.717975001 | 1.644871634 | 6.71E-05 | 0.000376613 | 106.7969211 | 175.7760003 | ENSG00000103196 |
| NA | 0.716898157 | 1.643644342 | 0.415730373 | 0.587836592 | 4.398030145 | 7.183364985 | ENSG00000261441 |
| EXPH5 | 0.716865498 | 1.643607135 | 0.016810766 | 0.048634745 | 38.2051367 | 62.72699462 | ENSG00000110723 |
| NA | 0.71656227 | 1.643261714 | 0.339291298 | 0.511748899 | 6.022655094 | 9.907107022 | ENSG00000164616 |
| ZNF160 | 0.716134132 | 1.642774128 | 5.19E-12 | 8.17E-11 | 401.4702728 | 659.1345044 | ENSG00000170949 |
| ZBTB44-DT | 0.716095391 | 1.642730015 | 0.478937881 | 0.647138101 | 3.426604259 | 5.57412588 | ENSG00000175773 |
| NA | 0.71602038 | 1.642644606 | 0.322874861 | 0.494449388 | 7.549510424 | 12.47543787 | ENSG00000264456 |
| SIX4 | 0.715909075 | 1.642517879 | 3.02E-14 | 6.06E-13 | 485.7882831 | 798.3117642 | ENSG00000100625 |

| | | | | | | | |
|---|---|---|---|---|---|---|---|
| NA | 0.715813844 | 1.642409462 | 0.116281714 | 0.232889384 | 18.60174322 | 30.63846741 | ENSG00000233871 |
| DAW1 | 0.715737404 | 1.642322442 | 0.29071379 | 0.46047785 | 8.879675234 | 14.62512666 | ENSG00000123977 |
| ZFP36 | 0.715724223 | 1.642307438 | 0.000440731 | 0.002052667 | 96.11295121 | 158.0464064 | ENSG00000128016 |
| NA | 0.715628036 | 1.642197946 | 0.049597959 | 0.118488449 | 25.64470901 | 42.21826467 | ENSG00000280120 |
| C15orf40 | 0.715397479 | 1.641935527 | 4.46E-06 | 3.13E-05 | 163.6541253 | 268.6023819 | ENSG00000169609 |
| DEPDC4 | 0.715228645 | 1.641743388 | 0.048316409 | 0.116146023 | 28.16272671 | 46.25090373 | ENSG00000166153 |
| FAM241A | 0.714454919 | 1.640863147 | 0.031452297 | 0.08160676 | 35.04420768 | 57.68901728 | ENSG00000174749 |
| ADAMTS20 | 0.713727196 | 1.640035673 | 1.53E-05 | 9.71E-05 | 141.751093 | 232.1343238 | ENSG00000173157 |
| NA | 0.713589722 | 1.639879402 | 0.436671815 | 0.60747357 | 4.922759657 | 8.094059608 | ENSG00000271978 |
| AR | 0.713494681 | 1.639771375 | 2.35E-09 | 2.73E-08 | 266.8210426 | 437.0917528 | ENSG00000169083 |
| TFPI | 0.713013795 | 1.63922489 | 0.000754013 | 0.003294602 | 78.01727706 | 128.0892668 | ENSG00000003436 |
| HLA-V | 0.712857123 | 1.639046885 | 0.334204606 | 0.50690186 | 6.052679304 | 9.913558743 | ENSG00000181126 |
| ZNF490 | 0.712838511 | 1.63902574 | 0.055538745 | 0.129945171 | 24.64046801 | 40.4600661 | ENSG00000188033 |
| NA | 0.71280042 | 1.638982465 | 0.341103278 | 0.51373985 | 6.049888401 | 9.909257596 | ENSG00000267141 |
| TRPV3 | 0.712645303 | 1.638806254 | 0.149584863 | 0.281773419 | 14.01048918 | 22.97192473 | ENSG00000167723 |
| SP8 | 0.71240578 | 1.638534194 | 9.60E-05 | 0.000521279 | 113.0257155 | 184.880505 | ENSG00000164651 |
| ZNF8 | 0.712310939 | 1.638426482 | 1.10E-05 | 7.14E-05 | 171.9998224 | 282.2473893 | ENSG00000278129 |
| CLEC11A | 0.711892153 | 1.637950948 | 5.72E-06 | 3.93E-05 | 149.4348181 | 244.9092919 | ENSG00000105472 |
| TMC7 | 0.711731951 | 1.637769074 | 0.002199578 | 0.008570402 | 69.62548602 | 113.9683026 | ENSG00000170537 |
| PDZD3 | 0.711542083 | 1.637553548 | 0.566679057 | 0.7202656 | 2.392339049 | 3.868092536 | ENSG00000172367 |
| UROC1 | 0.711498867 | 1.637504495 | 0.439294 | 0.610417097 | 3.992521593 | 6.541819917 | ENSG00000159650 |
| SPRY4-AS1 | 0.71145946 | 1.637459767 | 0.13200943 | 0.255759156 | 15.53196287 | 25.35069567 | ENSG00000231185 |
| TMEM179B | 0.71040185 | 1.636259819 | 0.017459377 | 0.050203051 | 45.08729339 | 73.99301332 | ENSG00000185475 |
| ASPDH | 0.710272099 | 1.636112667 | 0.486379948 | 0.653850563 | 3.396580048 | 5.502605488 | ENSG00000204653 |
| TFR2 | 0.709969973 | 1.635770071 | 4.09E-05 | 0.000239686 | 122.5369383 | 200.1934232 | ENSG00000106327 |
| ZNF669 | 0.709856208 | 1.635641086 | 2.60E-08 | 2.59E-07 | 225.10585 | 368.5760469 | ENSG00000188295 |
| STX6 | 0.70950516 | 1.635243138 | 4.04E-23 | 1.70E-21 | 977.0283368 | 1598.778094 | ENSG00000135823 |
| NDUFAF1 | 0.709499518 | 1.635236743 | 1.62E-10 | 2.19E-09 | 339.6071454 | 555.4311591 | ENSG00000137806 |
| PNPLA7 | 0.708683259 | 1.634311808 | 0.095833243 | 0.200403153 | 19.0945335 | 31.17084783 | ENSG00000130653 |
| GCOM1 | 0.708167375 | 1.63372751 | 0.565812096 | 0.719812768 | 2.419572356 | 3.893366382 | ENSG00000137878 |
| NA | 0.707860919 | 1.633380511 | 0.435822781 | 0.606598649 | 4.041406403 | 6.585915889 | ENSG00000255200 |
| LOC107985177 | 0.707491892 | 1.632962762 | 0.535553287 | 0.695480206 | 3.097213824 | 5.033676198 | ENSG00000287281 |
| LINC00649 | 0.707058359 | 1.632472127 | 0.000267844 | 0.001313416 | 111.5985448 | 181.9472964 | ENSG00000237945 |
| TMEM37 | 0.706882469 | 1.632273111 | 4.83E-08 | 4.65E-07 | 234.0031465 | 381.9501121 | ENSG00000171227 |
| NA | 0.706792693 | 1.632171542 | 0.421115204 | 0.593059818 | 4.628975239 | 7.595555324 | ENSG00000228952 |
| NA | 0.706514022 | 1.631856302 | 0.382709326 | 0.556045981 | 5.67440406 | 9.286534653 | ENSG00000278200 |
| NA | 0.705904942 | 1.631167507 | 0.487874972 | 0.654792398 | 3.665922062 | 6.009439492 | ENSG00000260711 |
| DDHD2 | 0.705840569 | 1.631094725 | 2.73E-09 | 3.13E-08 | 287.6793587 | 468.846589 | ENSG00000085788 |
| NA | 0.705727262 | 1.630966627 | 0.112016337 | 0.22605996 | 16.71268235 | 27.25435818 | ENSG00000228648 |
| NA | 0.705607974 | 1.630831777 | 0.496063258 | 0.662298865 | 3.015513901 | 4.907306965 | ENSG00000244425 |
| SFT2D2 | 0.704714369 | 1.629821954 | 8.15E-07 | 6.51E-06 | 187.001386 | 304.9958901 | ENSG00000213064 |
| ALDH3B1 | 0.704613174 | 1.629707638 | 0.086260103 | 0.184419219 | 24.19588682 | 39.57411228 | ENSG00000006534 |
| CCDC122 | 0.70460255 | 1.629695636 | 0.011719911 | 0.035961319 | 43.7371167 | 71.31766841 | ENSG00000151773 |
| IKZF3 | 0.704243057 | 1.629289596 | 0.229041205 | 0.388858804 | 10.78847259 | 17.56181233 | ENSG00000161405 |
| IRAK4 | 0.704164573 | 1.629200964 | 0.028816597 | 0.076015449 | 35.57539487 | 58.15416635 | ENSG00000198001 |
| SEC14L6 | 0.704061242 | 1.629084278 | 0.035448628 | 0.09008556 | 30.95132572 | 50.47392673 | ENSG00000214491 |
| PBXIP1 | 0.703887147 | 1.628887704 | 1.28E-18 | 3.83E-17 | 667.8073493 | 1087.524293 | ENSG00000163346 |
| AKR7A3 | 0.703734912 | 1.62871583 | 0.355715949 | 0.52883141 | 5.995421786 | 9.749545259 | ENSG00000162482 |
| NANOS1 | 0.703726602 | 1.628706448 | 0.006870504 | 0.023050242 | 75.50427759 | 123.0206484 | ENSG00000188613 |
| ASB9 | 0.703459453 | 1.628404883 | 0.444119838 | 0.614996468 | 4.022545804 | 6.533217622 | ENSG00000102048 |
| NA | 0.703161701 | 1.628068839 | 0.389099896 | 0.562490145 | 5.078462515 | 8.232799245 | ENSG00000255992 |
| SLC5A12 | 0.702949487 | 1.627829374 | 0.008918345 | 0.028583452 | 49.61331752 | 80.62759891 | ENSG00000148942 |
| RPL32P3 | 0.702620719 | 1.62745846 | 0.000190959 | 0.000966872 | 103.9812889 | 169.0671923 | ENSG00000251474 |
| NA | 0.702263646 | 1.627055706 | 0.360304491 | 0.533392214 | 6.950777976 | 11.30178495 | ENSG00000275494 |
| ATP7A | 0.701997414 | 1.62675548 | 0.009014584 | 0.028829631 | 58.08365416 | 94.24522459 | ENSG00000165240 |
| POLR3GL | 0.701683354 | 1.62640139 | 0.21191446 | 0.367761952 | 14.69582764 | 23.88745354 | ENSG00000121851 |
| DDTL | 0.70159559 | 1.626302454 | 0.055571966 | 0.129991367 | 30.57305048 | 49.78130093 | ENSG00000099974 |
| PEAK1 | 0.701564616 | 1.626267539 | 7.57E-15 | 1.61E-13 | 877.3205108 | 1425.927475 | ENSG00000173517 |
| CXCL5 | 0.701093902 | 1.625737016 | 0.004054164 | 0.014632158 | 73.19843241 | 119.1458252 | ENSG00000163735 |
| NA | 0.700837504 | 1.625448114 | 0.28931271 | 0.458748272 | 7.685676962 | 12.49856114 | ENSG00000253930 |
| INSM1 | 0.700529286 | 1.625100889 | 0.090410955 | 0.191324893 | 20.01639886 | 32.56363565 | ENSG00000173404 |
| DRAM1 | 0.700517993 | 1.625088168 | 3.07E-05 | 0.00018506 | 129.0429349 | 209.9937646 | ENSG00000136048 |
| FKBP2 | 0.700235754 | 1.624770279 | 0.42258802 | 0.594133544 | 4.658999449 | 7.591254177 | ENSG00000173486 |
| ZNF687 | 0.700139232 | 1.624661578 | 1.68E-32 | 1.24E-30 | 1843.470891 | 2995.318437 | ENSG00000143373 |
| NA | 0.699835936 | 1.624320064 | 0.005552168 | 0.019205105 | 55.29609429 | 89.74931446 | ENSG00000286279 |
| LOC100631378 | 0.699817454 | 1.624299256 | 0.235805617 | 0.397216745 | 10.36063682 | 16.85519859 | ENSG00000225868 |
| PKNOX2-AS1 | 0.699737182 | 1.624208881 | 0.598229352 | 0.745025334 | 2.601632641 | 4.257159602 | ENSG00000254880 |
| ARHGAP31 | 0.699179756 | 1.623581444 | 1.45E-06 | 1.11E-05 | 195.6435832 | 317.243082 | ENSG00000031081 |
| TNMD | 0.699170226 | 1.623570718 | 0.007517447 | 0.024815034 | 51.08849714 | 82.93001527 | ENSG00000000005 |
| CUL9 | 0.698971927 | 1.623347573 | 1.92E-05 | 0.000119675 | 186.4291231 | 302.7072012 | ENSG00000112659 |
| ACBD3 | 0.698674052 | 1.623012433 | 5.83E-20 | 1.95E-18 | 748.1075197 | 1214.703622 | ENSG00000182827 |
| PAQR6 | 0.698655694 | 1.622991781 | 0.000432896 | 0.002020078 | 90.02295103 | 146.107638 | ENSG00000160781 |
| RPSAP52 | 0.698124523 | 1.622394339 | 0.458259858 | 0.628178129 | 4.077012419 | 6.577313594 | ENSG00000241749 |
| CERS2 | 0.698006123 | 1.622261196 | 1.39E-38 | 1.45E-36 | 2206.766077 | 3580.016393 | ENSG00000143418 |
| NA | 0.697449587 | 1.621635511 | 0.117830295 | 0.235282498 | 19.82385068 | 32.06003672 | ENSG00000260000 |
| ADGRD2 | 0.696865215 | 1.620978792 | 0.074839424 | 0.164621133 | 25.24199136 | 40.98922987 | ENSG00000180264 |
| RICTOR | 0.696734252 | 1.620831651 | 1.26E-06 | 9.77E-06 | 190.9091413 | 309.2500872 | ENSG00000164327 |

| | | | | | | | |
|---|---|---|---|---|---|---|---|
| NUF2 | 0.696692997 | 1.620785303 | 1.43E-12 | 2.41E-11 | 421.1395477 | 682.8592652 | ENSG00000143228 |
| TRIM32 | 0.696398474 | 1.620454456 | 1.72E-11 | 2.56E-10 | 437.647279 | 709.313953 | ENSG00000119401 |
| MADCAM1 | 0.696050144 | 1.620063255 | 0.08405132 | 0.180460102 | 24.27547156 | 39.48833138 | ENSG00000099866 |
| DET1 | 0.695804085 | 1.619786969 | 0.323910177 | 0.495477475 | 6.70866927 | 10.85597893 | ENSG00000140543 |
| NA | 0.695386388 | 1.619318067 | 0.305974918 | 0.477011056 | 8.673848266 | 14.03412928 | ENSG00000278192 |
| MDFI | 0.695156306 | 1.619059838 | 1.61E-25 | 8.11E-24 | 2129.106865 | 3447.820176 | ENSG00000112559 |
| HMBOX1 | 0.695020962 | 1.618907956 | 6.01E-05 | 0.000340584 | 125.3648101 | 202.88192 | ENSG00000147421 |
| ZNF34 | 0.695020368 | 1.618907289 | 0.000640497 | 0.002853538 | 88.4163108 | 143.1217617 | ENSG00000196378 |
| NA | 0.694911747 | 1.618785405 | 0.276431659 | 0.443545848 | 8.384969932 | 13.58832326 | ENSG00000230438 |
| NA | 0.694826718 | 1.618690001 | 0.010453433 | 0.032672907 | 52.90778545 | 85.67930373 | ENSG00000225077 |
| NA | 0.694455922 | 1.618274025 | 0.448443147 | 0.618762373 | 4.310748416 | 7.029831796 | ENSG00000266680 |
| SSTR1 | 0.694314768 | 1.6181157 | 0.175819397 | 0.31917817 | 13.95044076 | 22.61028208 | ENSG00000139874 |
| PDGFRA | 0.6942019 | 1.617989113 | 0.212517778 | 0.368510602 | 14.20303736 | 22.99478754 | ENSG00000134853 |
| CUTALP | 0.694052314 | 1.61782136 | 0.011894152 | 0.036380226 | 45.16640257 | 73.08877042 | ENSG00000226752 |
| KLF2 | 0.693717948 | 1.61744645 | 0.559465306 | 0.714744716 | 2.623284143 | 4.272213618 | ENSG00000127528 |
| NKX2-6 | 0.693452062 | 1.617148384 | 0.556173076 | 0.71233135 | 2.691705273 | 4.366857283 | ENSG00000180053 |
| NA | 0.693336817 | 1.617019209 | 0.302490289 | 0.473186844 | 7.296238107 | 11.83819395 | ENSG00000267249 |
| STRADA | 0.692908452 | 1.616539155 | 0.053588142 | 0.126069372 | 28.25000844 | 45.53327666 | ENSG00000266173 |
| SLCO1A2 | 0.69267436 | 1.616276876 | 0.018018813 | 0.051496432 | 44.07854663 | 71.11788867 | ENSG00000084453 |
| CDKL3 | 0.692632541 | 1.616230027 | 0.399370972 | 0.571872433 | 6.017073288 | 9.686094123 | ENSG00000006837 |
| C1orf43 | 0.692372422 | 1.615938645 | 2.52E-37 | 2.42E-35 | 2154.112301 | 3480.599473 | ENSG00000143612 |
| HELZ | 0.692080625 | 1.615611841 | 2.20E-12 | 3.62E-11 | 420.7383817 | 679.7935082 | ENSG00000198265 |
| INO80D | 0.691964685 | 1.61548201 | 5.24E-07 | 4.30E-06 | 229.1860536 | 370.1960205 | ENSG00000114933 |
| PABPC5 | 0.691818603 | 1.61531844 | 0.011520468 | 0.035552928 | 45.55584143 | 73.50500144 | ENSG00000174740 |
| LOXL2 | 0.691445218 | 1.614900433 | 1.16E-23 | 5.15E-22 | 952.3070305 | 1537.221845 | ENSG00000134013 |
| GLRX2 | 0.691321324 | 1.614761756 | 6.83E-05 | 0.000382525 | 176.8027606 | 286.0302582 | ENSG00000023572 |
| DBIL5P | 0.690888508 | 1.614277392 | 0.340140993 | 0.51274562 | 6.934032559 | 11.26816927 | ENSG00000231784 |
| NA | 0.690365969 | 1.613692813 | 0.032799953 | 0.084467214 | 33.64390687 | 54.14277256 | ENSG00000279059 |
| MORN1 | 0.689993721 | 1.613276497 | 0.001236292 | 0.005167373 | 95.59141292 | 153.8715322 | ENSG00000116151 |
| TARBP1 | 0.689912339 | 1.613185495 | 2.11E-13 | 3.86E-12 | 542.1372278 | 874.5340375 | ENSG00000059588 |
| LOC100130357 | 0.689770287 | 1.613026664 | 0.37115213 | 0.544810207 | 5.728870676 | 9.217164834 | ENSG00000215022 |
| NA | 0.689724002 | 1.612974915 | 0.192549577 | 0.341906654 | 13.52818679 | 21.89023186 | ENSG00000203644 |
| NA | 0.689717813 | 1.612967995 | 0.042672373 | 0.105106326 | 30.89494408 | 49.68127162 | ENSG00000259802 |
| NA | 0.688821856 | 1.611966605 | 0.542900938 | 0.701668363 | 3.064398711 | 4.903005818 | ENSG00000224376 |
| AKR7L | 0.688759376 | 1.611896796 | 0.442566934 | 0.613506664 | 4.615696446 | 7.450363966 | ENSG00000211454 |
| AGPAT3 | 0.688705919 | 1.61183707 | 7.99E-06 | 5.37E-05 | 160.154306 | 258.0902774 | ENSG00000160216 |
| GAB3 | 0.688699352 | 1.611829733 | 0.358130241 | 0.531369199 | 6.376487933 | 10.2902554 | ENSG00000160219 |
| NA | 0.688527532 | 1.611637781 | 0.053665787 | 0.12623664 | 30.61443846 | 49.31961686 | ENSG00000185495 |
| VSTM2A | 0.688234777 | 1.611310777 | 0.253322047 | 0.417928299 | 9.356395818 | 15.06474142 | ENSG00000170419 |
| RAB40A | 0.687904792 | 1.610942268 | 0.054256403 | 0.127392924 | 27.02443435 | 43.51829893 | ENSG00000172476 |
| PAPPA | 0.687660592 | 1.610669612 | 0.443189661 | 0.614029679 | 4.92555056 | 8.009896237 | ENSG00000182752 |
| HTR6 | 0.687316713 | 1.61028574 | 0.012042891 | 0.03673615 | 49.31183612 | 79.23911224 | ENSG00000158748 |
| LOC101928673 | 0.686648908 | 1.609540532 | 0.009955231 | 0.031308505 | 49.5476873 | 79.65560372 | ENSG00000203739 |
| SUCO | 0.686449569 | 1.609318156 | 1.19E-20 | 4.26E-19 | 868.0248775 | 1396.799048 | ENSG00000094975 |
| CEP85L | 0.68632548 | 1.609179741 | 0.001447708 | 0.005938282 | 84.68371937 | 136.0327494 | ENSG00000111860 |
| HLTF | 0.685897396 | 1.608702328 | 2.70E-19 | 8.48E-18 | 890.5555904 | 1433.276992 | ENSG00000071794 |
| STX17 | 0.685654881 | 1.60843193 | 3.20E-09 | 3.65E-08 | 297.5618355 | 478.1809815 | ENSG00000136874 |
| KAT2B | 0.685140498 | 1.607858557 | 0.177458742 | 0.321211875 | 13.07678449 | 20.99029011 | ENSG00000114166 |
| SH3RF2 | 0.685070536 | 1.607780586 | 0.193525767 | 0.343260903 | 14.639446 | 23.41745817 | ENSG00000156463 |
| NA | 0.684949831 | 1.607646076 | 0.467871297 | 0.63706857 | 3.663131159 | 5.891672554 | ENSG00000235957 |
| FBXO24 | 0.684586552 | 1.607241312 | 0.225406108 | 0.384465892 | 11.24757188 | 18.17378241 | ENSG00000106336 |
| TOR1AIP2 | 0.68450202 | 1.607147141 | 3.77E-13 | 6.74E-12 | 526.4592621 | 845.9719216 | ENSG00000169905 |
| ACTRT3 | 0.684371833 | 1.60700212 | 0.006334651 | 0.021499253 | 55.39577893 | 89.09378145 | ENSG00000184378 |
| PTP4A3 | 0.684326054 | 1.606951129 | 1.77E-16 | 4.46E-15 | 849.8022343 | 1364.291976 | ENSG00000184489 |
| OR1F1 | 0.684099991 | 1.606699347 | 0.544062943 | 0.702558286 | 2.628865949 | 4.244789198 | ENSG00000262668 |
| NA | 0.684096739 | 1.606695726 | 0.238014262 | 0.399944836 | 10.43184885 | 16.70570609 | ENSG00000269947 |
| CCDC27 | 0.68407377 | 1.606670146 | 0.316358714 | 0.487378373 | 8.505066774 | 13.62004883 | ENSG00000162592 |
| GLP1R | 0.683796465 | 1.606361352 | 0.001515992 | 0.006182877 | 86.89244653 | 139.3477614 | ENSG00000112164 |
| PHYH | 0.683725728 | 1.606282593 | 1.03E-07 | 9.50E-07 | 267.8268352 | 430.0521979 | ENSG00000107537 |
| LOC100134368 | 0.6835923 | 1.606134042 | 0.099402707 | 0.206279864 | 25.08907941 | 40.3753818 | ENSG00000236829 |
| PPIL6 | 0.683427594 | 1.605950688 | 0.325049419 | 0.496744859 | 7.389101641 | 11.85271493 | ENSG00000185250 |
| NA | 0.683236396 | 1.605737868 | 0.119248789 | 0.237353321 | 18.6010675 | 29.95282637 | ENSG00000278974 |
| ZBTB5 | 0.683136073 | 1.605626211 | 1.60E-10 | 2.16E-09 | 339.9429935 | 545.4915451 | ENSG00000168795 |
| PKIA-AS1 | 0.682974826 | 1.605446764 | 0.555375259 | 0.711498481 | 2.653308353 | 4.272213618 | ENSG00000254266 |
| NA | 0.682454428 | 1.604867763 | 0.35510887 | 0.528213834 | 6.289206204 | 10.1111879 | ENSG00000259032 |
| MYEF2 | 0.682291284 | 1.604686291 | 1.93E-19 | 6.19E-18 | 1002.397781 | 1607.743865 | ENSG00000104177 |
| DHTKD1 | 0.681907701 | 1.604259694 | 7.79E-29 | 4.65E-27 | 1503.336261 | 2411.199307 | ENSG00000181192 |
| RPS27L | 0.681853397 | 1.60419931 | 9.43E-18 | 2.64E-16 | 690.8720402 | 1108.64979 | ENSG00000185088 |
| ZNF670 | 0.681793925 | 1.604133182 | 3.15E-06 | 2.26E-05 | 216.5467922 | 347.6895784 | ENSG00000277462 |
| NA | 0.681552442 | 1.603864699 | 0.536871119 | 0.696488282 | 3.069980516 | 4.875581397 | ENSG00000268564 |
| ZNF793-AS1 | 0.681372949 | 1.603665166 | 0.227537088 | 0.386835108 | 10.72005146 | 17.21254009 | ENSG00000266916 |
| NA | 0.681330555 | 1.603618043 | 0.621415983 | 0.761915061 | 2.476829874 | 3.899818104 | ENSG00000260273 |
| STUM | 0.681121589 | 1.603385785 | 0.027199552 | 0.072643855 | 41.88501328 | 67.03014031 | ENSG00000203685 |
| ZNF81 | 0.680271315 | 1.602441084 | 0.000274269 | 0.001341886 | 114.7381857 | 183.9657576 | ENSG00000197779 |
| GABRG1 | 0.680195973 | 1.602357402 | 0.448622399 | 0.618920522 | 4.289096914 | 6.909914284 | ENSG00000163285 |
| NA | 0.679983145 | 1.602121038 | 0.397613184 | 0.570131354 | 5.810570598 | 9.248890402 | ENSG00000255320 |
| SYNPO2L | 0.679770676 | 1.601885106 | 0.549066755 | 0.706513259 | 3.127238035 | 4.951402937 | ENSG00000166317 |

| | | | | | | | |
|---|---|---|---|---|---|---|---|
| APOA1 | 0.679769399 | 1.601883689 | 0.532310425 | 0.693135363 | 3.069980516 | 4.865361567 | ENSG00000118137 |
| FCGBP | 0.679567911 | 1.601659984 | 0.203727549 | 0.356817101 | 12.16540703 | 19.41277657 | ENSG00000275395 |
| TRAF3IP2 | 0.679221376 | 1.601275312 | 6.67E-14 | 1.29E-12 | 470.0752238 | 752.7766401 | ENSG00000056972 |
| NA | 0.679143348 | 1.601188709 | 0.619868827 | 0.76111575 | 2.332290628 | 3.771558761 | ENSG00000281195 |
| SMIM29 | 0.678948982 | 1.600973005 | 0.001984621 | 0.007838928 | 82.45669519 | 132.0361249 | ENSG00000186577 |
| TIE1 | 0.678855231 | 1.600868972 | 0.002651033 | 0.010086999 | 83.09150921 | 133.1944634 | ENSG00000066056 |
| NUDT7 | 0.678657266 | 1.600649318 | 0.211168353 | 0.3668051 | 11.74594396 | 18.79005363 | ENSG00000140876 |
| SDHC | 0.67862727 | 1.600616038 | 1.05E-37 | 1.03E-35 | 1925.120147 | 3080.787087 | ENSG00000143252 |
| NA | 0.67861471 | 1.600602103 | 0.401470874 | 0.573942095 | 5.284965205 | 8.4809761 | ENSG00000260015 |
| CYP11A1 | 0.678532952 | 1.600511399 | 0.041663714 | 0.103042911 | 34.52872675 | 55.2153301 | ENSG00000140459 |
| ADAD2 | 0.678432182 | 1.60039961 | 0.472272047 | 0.641196734 | 4.637347948 | 7.410569141 | ENSG00000140955 |
| SRGAP2 | 0.678344595 | 1.600302452 | 2.08E-19 | 6.63E-18 | 887.3757865 | 1419.43301 | ENSG00000266028 |
| NES | 0.677866096 | 1.599771767 | 2.75E-52 | 4.96E-50 | 6401.071849 | 10240.50488 | ENSG00000132688 |
| ZNF547 | 0.6774699 | 1.599332494 | 0.174939402 | 0.317910521 | 13.38173252 | 21.41108274 | ENSG00000152433 |
| EPSTI1 | 0.677358964 | 1.599209518 | 0.174024689 | 0.316857978 | 15.52271428 | 24.91323149 | ENSG00000133106 |
| BCYRN1 | 0.677244583 | 1.599082734 | 0.551229757 | 0.708115685 | 2.71893858 | 4.339432863 | ENSG00000236824 |
| SIPA1L2 | 0.676956927 | 1.598763927 | 1.63E-11 | 2.43E-10 | 413.9202156 | 661.4167537 | ENSG00000116991 |
| TMEM139 | 0.676943598 | 1.598749157 | 0.539134305 | 0.698545187 | 2.686123467 | 4.293186317 | ENSG00000178826 |
| LINC00847 | 0.676785566 | 1.59857404 | 0.385881282 | 0.559407515 | 5.905349155 | 9.497067257 | ENSG00000245060 |
| MYH3 | 0.676484543 | 1.598240527 | 0.06450294 | 0.146064822 | 28.57294119 | 45.68842738 | ENSG00000109063 |
| ALPK3 | 0.676316791 | 1.5980547 | 4.81E-32 | 3.42E-30 | 1746.329656 | 2789.750843 | ENSG00000136383 |
| COLEC12 | 0.676268251 | 1.598000933 | 0.016232766 | 0.04724597 | 50.55519476 | 80.95509284 | ENSG00000158270 |
| ATG16L2 | 0.676191353 | 1.597915759 | 4.59E-08 | 4.43E-07 | 243.8708298 | 389.3074194 | ENSG00000168010 |
| NA | 0.67604633 | 1.597755142 | 0.042392039 | 0.1045226 | 31.75464584 | 50.78958329 | ENSG00000286358 |
| GAP43 | 0.67590347 | 1.597596935 | 3.32E-11 | 4.80E-10 | 403.1391646 | 643.4279574 | ENSG00000172020 |
| NA | 0.675873059 | 1.597563259 | 0.039859784 | 0.099377141 | 35.37188325 | 56.4147899 | ENSG00000237863 |
| GOLGA6A | 0.675598443 | 1.597259193 | 0.370373934 | 0.544164779 | 5.962606673 | 9.568587649 | ENSG00000159289 |
| ITPKB | 0.675566165 | 1.597223457 | 8.62E-09 | 9.28E-08 | 297.4544169 | 475.374421 | ENSG00000143772 |
| TLR4 | 0.675552524 | 1.597208355 | 0.553072393 | 0.709584965 | 2.691705273 | 4.295336891 | ENSG00000136869 |
| ZCCHC4 | 0.675440131 | 1.59708393 | 0.000164978 | 0.000846883 | 131.2583199 | 209.6791804 | ENSG00000168228 |
| ZNF486 | 0.675385681 | 1.597023654 | 1.82E-10 | 2.44E-09 | 366.0226348 | 584.8994812 | ENSG00000256229 |
| PTPN3 | 0.675128108 | 1.596738553 | 1.67E-05 | 0.000105233 | 155.653488 | 248.9236482 | ENSG00000070159 |
| NA | 0.675127735 | 1.596738141 | 0.059666624 | 0.137272372 | 26.71458024 | 42.65976954 | ENSG00000223611 |
| NA | 0.674846691 | 1.596427118 | 0.449338598 | 0.619599063 | 4.316330222 | 6.912064858 | ENSG00000275665 |
| IGFBP2 | 0.674466591 | 1.596006571 | 1.81E-08 | 1.86E-07 | 5889.046162 | 9398.797282 | ENSG00000115457 |
| IKBIP | 0.674308761 | 1.595831978 | 1.95E-07 | 1.72E-06 | 230.7146238 | 368.4762902 | ENSG00000166130 |
| SOCS1 | 0.674274371 | 1.595793938 | 3.91E-07 | 3.29E-06 | 395.766922 | 632.1345448 | ENSG00000185338 |
| CPM | 0.673647892 | 1.595101127 | 0.166943858 | 0.306883541 | 15.45863565 | 24.6016035 | ENSG00000135678 |
| LINC02731 | 0.67349539 | 1.594932524 | 0.293314491 | 0.46300056 | 8.033927996 | 12.80373741 | ENSG00000204241 |
| IGSF9B | 0.67266794 | 1.594018022 | 2.80E-21 | 1.04E-19 | 1505.846819 | 2399.633442 | ENSG00000080854 |
| AK5 | 0.672591217 | 1.593933253 | 0.002772087 | 0.010487612 | 79.39999346 | 126.5569153 | ENSG00000154027 |
| VSNL1 | 0.672194017 | 1.593494475 | 1.12E-22 | 4.64E-21 | 1314.357458 | 2094.902982 | ENSG00000163032 |
| STARD8 | 0.672055216 | 1.593341173 | 4.59E-05 | 0.000265657 | 137.0430086 | 218.0706317 | ENSG00000130052 |
| PGF | 0.6718585 | 1.59312393 | 0.000295482 | 0.00143472 | 103.5935649 | 165.2526221 | ENSG00000119630 |
| NUP133 | 0.671766805 | 1.593022678 | 6.49E-30 | 4.13E-28 | 1856.210813 | 2958.070433 | ENSG00000069248 |
| NA | 0.671692402 | 1.592940524 | 0.014675845 | 0.043482065 | 47.58298386 | 75.93647065 | ENSG00000273599 |
| RAMP2-AS1 | 0.671588778 | 1.592826113 | 0.032045356 | 0.082922464 | 39.10168377 | 62.48526949 | ENSG00000197291 |
| NA | 0.671268849 | 1.59247293 | 0.15375524 | 0.28830602 | 21.1435276 | 33.74346772 | ENSG00000205959 |
| NA | 0.671192365 | 1.592388509 | 0.403013325 | 0.575331266 | 6.550175509 | 10.49380325 | ENSG00000227848 |
| NA | 0.671059336 | 1.592241683 | 0.507074526 | 0.67183892 | 4.13706084 | 6.510094349 | ENSG00000261542 |
| MCAM | 0.670835437 | 1.591994594 | 1.39E-14 | 2.87E-13 | 529.4616093 | 842.6531599 | ENSG00000076706 |
| ZNF19 | 0.669622365 | 1.590656548 | 0.076072007 | 0.166519539 | 24.27826247 | 38.66530777 | ENSG00000157429 |
| NA | 0.669537565 | 1.590563053 | 0.273513563 | 0.440282889 | 10.36621862 | 16.50539332 | ENSG00000220323 |
| NA | 0.669413647 | 1.590426441 | 0.552363456 | 0.708958311 | 2.691705273 | 4.270063044 | ENSG00000273369 |
| VWF | 0.669117051 | 1.590099507 | 0.08333912 | 0.179346704 | 24.52124705 | 38.86293693 | ENSG00000110799 |
| KDM3A | 0.669079553 | 1.590058179 | 4.52E-14 | 8.85E-13 | 595.6100035 | 946.3034854 | ENSG00000115548 |
| NA | 0.668887038 | 1.589846013 | 0.119590531 | 0.237861731 | 18.38128602 | 29.22093878 | ENSG00000254297 |
| PSMD4 | 0.668691977 | 1.589631071 | 2.66E-49 | 4.27E-47 | 4373.307834 | 6951.244747 | ENSG00000159352 |
| SLC16A5 | 0.668308348 | 1.589208426 | 9.25E-08 | 8.57E-07 | 278.9842976 | 443.9099617 | ENSG00000170190 |
| PRR13 | 0.667987152 | 1.58885465 | 6.31E-11 | 8.84E-10 | 442.540402 | 703.0935026 | ENSG00000205352 |
| NA | 0.667956824 | 1.58882125 | 0.332084924 | 0.504747107 | 7.661234558 | 12.19983659 | ENSG00000257027 |
| DDX28 | 0.667655662 | 1.588489619 | 1.11E-08 | 1.18E-07 | 345.2446672 | 548.3683421 | ENSG00000182810 |
| RUNX1T1 | 0.667623809 | 1.588454547 | 1.08E-05 | 7.02E-05 | 188.4506085 | 299.1741388 | ENSG00000079102 |
| FBLN2 | 0.666930294 | 1.587691148 | 5.66E-27 | 3.08E-25 | 1178.977857 | 1871.312583 | ENSG00000163520 |
| MACROD1 | 0.666843758 | 1.587595918 | 1.35E-09 | 1.61E-08 | 372.8960681 | 591.6671603 | ENSG00000133315 |
| KLHL28 | 0.666813169 | 1.587562257 | 0.001155432 | 0.004858205 | 110.6073824 | 175.4092567 | ENSG00000179454 |
| ZNF619 | 0.666588908 | 1.587315496 | 0.000723749 | 0.003175311 | 94.03856358 | 149.2244878 | ENSG00000177873 |
| NFIA | 0.666390392 | 1.587097095 | 0.182559643 | 0.32815487 | 14.17109813 | 22.43954431 | ENSG00000162599 |
| KCTD13 | 0.666328262 | 1.587028748 | 4.80E-06 | 3.34E-05 | 172.7395395 | 274.3991076 | ENSG00000174943 |
| GRIN2C | 0.666263159 | 1.586957134 | 0.519154032 | 0.682639249 | 3.342113433 | 5.321647878 | ENSG00000161509 |
| C8orf31 | 0.666207024 | 1.586895387 | 0.336712744 | 0.50947557 | 8.387760835 | 13.24094113 | ENSG00000177335 |
| CD1D | 0.666172224 | 1.586857109 | 0.038760805 | 0.097209398 | 43.46151716 | 69.04133781 | ENSG00000158473 |
| LINC00205 | 0.665765144 | 1.586409414 | 5.54E-08 | 5.29E-07 | 259.1405472 | 411.1132274 | ENSG00000223768 |
| SH3PXD2A | 0.665634811 | 1.586266105 | 2.00E-15 | 4.52E-14 | 573.2393502 | 909.0591897 | ENSG00000107957 |
| NA | 0.665348553 | 1.585951391 | 0.293507802 | 0.463192001 | 9.22022928 | 14.67352378 | ENSG00000231073 |
| SLC19A2 | 0.665023012 | 1.585593565 | 3.30E-14 | 6.58E-13 | 625.1813796 | 991.0663941 | ENSG00000117479 |
| NA | 0.664861913 | 1.585416519 | 0.150675592 | 0.283385282 | 18.17757423 | 28.80470776 | ENSG00000284606 |

| | | | | | | | |
|---|---|---|---|---|---|---|---|
| KDM5B | 0.664850228 | 1.585403677 | 8.66E-44 | 1.16E-41 | 5058.809699 | 8018.454128 | ENSG00000117139 |
| SCART1 | 0.664804588 | 1.585353524 | 0.392775746 | 0.565659377 | 5.976561187 | 9.488464962 | ENSG00000214279 |
| NA | 0.664798404 | 1.585346728 | 0.507445038 | 0.672145167 | 3.312089222 | 5.275401332 | ENSG00000259594 |
| NCF1B | 0.664684617 | 1.585221695 | 0.401052986 | 0.573557206 | 5.372246934 | 8.485277248 | ENSG00000182487 |
| PIPSL | 0.664675129 | 1.58521127 | 0.214329505 | 0.370852137 | 11.99084357 | 19.02586008 | ENSG00000180764 |
| NA | 0.664294325 | 1.584792903 | 0.236864871 | 0.398390103 | 11.47939285 | 18.14635799 | ENSG00000230793 |
| C18orf25 | 0.663896476 | 1.584355929 | 1.10E-10 | 1.52E-09 | 379.2589891 | 600.6213853 | ENSG00000152242 |
| NA | 0.663877527 | 1.584335119 | 0.448104033 | 0.618471411 | 4.346354432 | 6.88033929 | ENSG00000236519 |
| CTBS | 0.663858726 | 1.584314472 | 0.346665723 | 0.519309753 | 7.470801564 | 11.750002 | ENSG00000117151 |
| NSG2 | 0.663615556 | 1.584047454 | 0.413485055 | 0.58564906 | 5.410643853 | 8.531523793 | ENSG00000170091 |
| ASH1L-AS1 | 0.662824754 | 1.583179409 | 0.009216419 | 0.029372863 | 52.67193427 | 83.48793001 | ENSG00000235919 |
| CSGALNACT1 | 0.662603009 | 1.58293609 | 0.421378684 | 0.593344248 | 5.846176615 | 9.15639731 | ENSG00000147408 |
| DCX | 0.662503865 | 1.582827312 | 0.005043396 | 0.017670205 | 103.4985862 | 164.2523542 | ENSG00000077279 |
| LINC00909 | 0.662488919 | 1.582810914 | 0.001661109 | 0.006703351 | 81.6784932 | 129.2519064 | ENSG00000264247 |
| UPK1B | 0.66239319 | 1.582705892 | 0.546069068 | 0.704338851 | 3.2709014 | 5.201730366 | ENSG00000114638 |
| C2CD4D | 0.66178943 | 1.582043677 | 0.057664681 | 0.133736452 | 32.26262993 | 51.1915538 | ENSG00000225556 |
| GLUD2 | 0.661449132 | 1.581670554 | 0.067448269 | 0.151489644 | 27.00557376 | 42.77323533 | ENSG00000182890 |
| CKMT2 | 0.661095053 | 1.581282413 | 0.06464866 | 0.146291764 | 27.52559734 | 43.49113498 | ENSG00000131730 |
| ARL4D | 0.660903887 | 1.581072898 | 4.15E-06 | 2.93E-05 | 230.8915395 | 364.5337211 | ENSG00000175906 |
| ARL4A | 0.660800233 | 1.580959306 | 1.57E-08 | 1.63E-07 | 326.3155607 | 515.9337471 | ENSG00000122644 |
| NA | 0.660685535 | 1.58083362 | 0.142008653 | 0.270861076 | 18.56075556 | 29.24406206 | ENSG00000205041 |
| NA | 0.660594837 | 1.580734242 | 0.319244922 | 0.490669175 | 8.121209724 | 12.7677107 | ENSG00000279583 |
| NLRC3 | 0.660579244 | 1.580717156 | 0.538555941 | 0.698157561 | 3.965288286 | 6.301712319 | ENSG00000167984 |
| SPTSSB | 0.660575773 | 1.580713354 | 5.23E-05 | 0.000299974 | 157.1005953 | 248.0118933 | ENSG00000196542 |
| POGK | 0.660458508 | 1.580584875 | 7.93E-30 | 5.03E-28 | 1360.126133 | 2149.393731 | ENSG00000143157 |
| NA | 0.659796013 | 1.579859227 | 0.35247757 | 0.525312484 | 8.26487309 | 13.09224213 | ENSG00000228589 |
| SMIM24 | 0.659706614 | 1.579761331 | 0.005185557 | 0.018099254 | 73.57382873 | 116.4094402 | ENSG00000095932 |
| NA | 0.659641189 | 1.579689693 | 0.312116186 | 0.483126914 | 8.319339705 | 13.17183178 | ENSG00000254910 |
| ZNF425 | 0.659601935 | 1.579646711 | 4.80E-06 | 3.34E-05 | 186.5058928 | 294.8376622 | ENSG00000204947 |
| C17orf97 | 0.659210045 | 1.579217679 | 0.381477508 | 0.554983606 | 6.025445997 | 9.503518978 | ENSG00000187624 |
| NA | 0.658748037 | 1.578712031 | 4.49E-05 | 0.000260553 | 150.8220402 | 238.3992159 | ENSG00000197462 |
| SERPINB6 | 0.658475457 | 1.57841378 | 2.02E-11 | 2.99E-10 | 449.9555104 | 709.5301864 | ENSG00000124570 |
| PRNP | 0.658444491 | 1.578379902 | 7.61E-12 | 1.18E-10 | 472.8897047 | 746.3641761 | ENSG00000171867 |
| ARHGEF37 | 0.65785714 | 1.577737442 | 0.351043026 | 0.523659992 | 7.666140642 | 12.09067195 | ENSG00000183111 |
| MIAT | 0.657842939 | 1.577721912 | 4.07E-13 | 7.22E-12 | 1070.831109 | 1688.028839 | ENSG00000225783 |
| NA | 0.657206376 | 1.577025924 | 0.235288724 | 0.396484519 | 11.01662678 | 17.37037443 | ENSG00000275088 |
| SPRTN | 0.656926707 | 1.576720244 | 4.73E-07 | 3.91E-06 | 209.0877547 | 329.7426971 | ENSG00000010072 |
| MIR31HG | 0.65688412 | 1.576673702 | 0.588950106 | 0.738474911 | 2.710565871 | 4.249090345 | ENSG00000171889 |
| HDAC4-AS1 | 0.656845212 | 1.576631181 | 0.479420661 | 0.647563644 | 4.392448339 | 6.899161415 | ENSG00000222020 |
| THBS3 | 0.656713878 | 1.57648766 | 0.007124776 | 0.023724672 | 60.92631947 | 96.16691572 | ENSG00000169231 |
| NA | 0.656620625 | 1.576385763 | 0.535349207 | 0.695403425 | 3.742040179 | 5.851877729 | ENSG00000227973 |
| CD37 | 0.656435199 | 1.576183167 | 0.154447697 | 0.289376411 | 18.94461261 | 29.69416922 | ENSG00000104894 |
| LPXN | 0.656296643 | 1.576031798 | 0.19372692 | 0.343459793 | 13.2972417 | 20.99244068 | ENSG00000110031 |
| LINC02367 | 0.656187537 | 1.575912612 | 0.361293955 | 0.534290242 | 6.621387542 | 10.45830957 | ENSG00000260423 |
| NA | 0.656046331 | 1.575758376 | 0.584718297 | 0.734696846 | 2.740590082 | 4.286734596 | ENSG00000204053 |
| METTL23 | 0.655825548 | 1.575517248 | 6.88E-05 | 0.000385228 | 152.8256162 | 241.2632846 | ENSG00000181038 |
| IARS2 | 0.65578194 | 1.575469625 | 4.90E-40 | 5.73E-38 | 2589.382161 | 4080.326842 | ENSG00000067704 |
| C2orf92 | 0.655408289 | 1.57506164 | 0.177328451 | 0.32103567 | 15.49077505 | 24.34105624 | ENSG00000228486 |
| LOC112267895 | 0.655167272 | 1.574798531 | 0.292029188 | 0.461727323 | 9.669716557 | 15.28575431 | ENSG00000261241 |
| HLA-DRB5 | 0.655022149 | 1.574640128 | 0.015995769 | 0.04669709 | 52.77249479 | 83.01579618 | ENSG00000198502 |
| ERN2 | 0.65498849 | 1.574603391 | 0.119047403 | 0.236978299 | 19.79382647 | 31.09287572 | ENSG00000134398 |
| NEDD4 | 0.654700868 | 1.574289502 | 2.59E-06 | 1.89E-05 | 193.4121285 | 304.451424 | ENSG00000069869 |
| TNFRSF10B | 0.654268211 | 1.573817451 | 4.46E-22 | 1.76E-20 | 1923.223399 | 3025.83357 | ENSG00000120889 |
| PGM1 | 0.653954043 | 1.573474766 | 9.07E-15 | 1.92E-13 | 1344.153479 | 2114.115615 | ENSG00000079739 |
| RAB24 | 0.653747128 | 1.573249111 | 0.006397373 | 0.021681609 | 63.91460006 | 100.7655388 | ENSG00000169228 |
| RASGRP2 | 0.653365472 | 1.572832973 | 1.20E-14 | 2.50E-13 | 566.292002 | 890.9230821 | ENSG00000068831 |
| KIZ-AS1 | 0.653194427 | 1.57264651 | 0.569572007 | 0.722631607 | 3.617037252 | 5.746741659 | ENSG00000232712 |
| SLC50A1 | 0.653136629 | 1.572583507 | 6.35E-16 | 1.52E-14 | 721.8099608 | 1134.405468 | ENSG00000169241 |
| CAPS | 0.652707625 | 1.572115949 | 0.310846361 | 0.481765659 | 8.306060912 | 13.06858582 | ENSG00000105519 |
| TNFSF4 | 0.652605229 | 1.572004371 | 0.341940689 | 0.514819155 | 7.269004799 | 11.47009958 | ENSG00000117586 |
| TSHZ3 | 0.652481017 | 1.571869031 | 5.05E-06 | 3.50E-05 | 202.7869846 | 319.1948385 | ENSG00000121297 |
| NTF4 | 0.652374288 | 1.571752751 | 0.151998583 | 0.285511653 | 17.52070839 | 27.66869909 | ENSG00000225950 |
| NA | 0.651967818 | 1.571309982 | 0.518087336 | 0.681701302 | 3.309298319 | 5.206031513 | ENSG00000266473 |
| NA | 0.651846388 | 1.571177732 | 0.197121662 | 0.348070844 | 14.24422518 | 22.49439315 | ENSG00000287670 |
| LPP | 0.650989435 | 1.570244739 | 6.03E-08 | 5.71E-07 | 407.06135 | 639.0470453 | ENSG00000145012 |
| NLGN1 | 0.650781089 | 1.570017989 | 0.000994625 | 0.004243103 | 116.7820736 | 183.5653862 | ENSG00000169760 |
| TSPAN10 | 0.650778936 | 1.570015645 | 0.175838113 | 0.31917817 | 18.79361568 | 29.64818314 | ENSG00000182612 |
| HASPIN | 0.650594839 | 1.569815314 | 3.47E-09 | 3.94E-08 | 316.1154576 | 495.9971498 | ENSG00000177602 |
| NA | 0.650585292 | 1.569804926 | 0.011479128 | 0.035461704 | 55.16012792 | 86.39478023 | ENSG00000279407 |
| DST | 0.650545014 | 1.5697611 | 3.07E-13 | 5.55E-12 | 1327.706084 | 2083.383018 | ENSG00000151914 |
| BHLHE40-AS1 | 0.650505129 | 1.569717702 | 0.446702675 | 0.617067046 | 6.611099811 | 10.28165311 | ENSG00000235831 |
| C5orf24 | 0.650483697 | 1.569694384 | 0.000143729 | 0.000746147 | 172.2149732 | 270.8176418 | ENSG00000181904 |
| FGF19 | 0.650103951 | 1.569281264 | 5.11E-10 | 6.46E-09 | 357.4923747 | 560.6186837 | ENSG00000162344 |
| TTPAL | 0.650088705 | 1.569264681 | 8.64E-09 | 9.29E-08 | 304.7456368 | 478.5310777 | ENSG00000124120 |
| NA | 0.649771877 | 1.568920094 | 0.123237803 | 0.243185399 | 21.46733623 | 33.68297277 | ENSG00000228106 |
| DPYD | 0.649753771 | 1.568900404 | 0.292967034 | 0.462717133 | 9.383629126 | 14.6799755 | ENSG00000188641 |
| MRTFB | 0.649561061 | 1.56869085 | 1.51E-07 | 1.36E-06 | 290.5069182 | 455.5072298 | ENSG00000186260 |

| | | | | | | | |
|---|---|---|---|---|---|---|---|
| GADD45A | 0.649261597 | 1.568365266 | 5.91E-10 | 7.39E-09 | 408.3888872 | 640.6703029 | ENSG00000116717 |
| SYNPO | 0.648680585 | 1.567733771 | 0.001423691 | 0.005851798 | 94.06523331 | 147.2304827 | ENSG00000171992 |
| RNF2 | 0.648419297 | 1.567449863 | 1.22E-11 | 1.85E-10 | 640.8377833 | 1005.276659 | ENSG00000121481 |
| SLFNL1-AS1 | 0.648086666 | 1.56708851 | 0.238129321 | 0.4000133 | 11.71871066 | 18.33591789 | ENSG00000281207 |
| NA | 0.647977282 | 1.566969699 | 0.573582245 | 0.72591498 | 3.012722998 | 4.755924349 | ENSG00000221184 |
| POGZ | 0.647933519 | 1.566922167 | 2.05E-28 | 1.20E-26 | 1341.570559 | 2102.2176 | ENSG00000143442 |
| LAMA5 | 0.647835548 | 1.566815763 | 2.03E-32 | 1.48E-30 | 3617.258085 | 5667.219763 | ENSG00000130702 |
| PLEKHA4 | 0.647689599 | 1.566657266 | 3.77E-08 | 3.69E-07 | 358.9640365 | 561.5810105 | ENSG00000105559 |
| KIZ | 0.646970061 | 1.565876097 | 9.38E-13 | 1.59E-11 | 470.5503048 | 736.8781222 | ENSG00000088970 |
| LOC100129129 | 0.646596135 | 1.565470297 | 0.384725074 | 0.558192796 | 6.319230415 | 9.932380869 | ENSG00000255020 |
| JAG2 | 0.64642809 | 1.565287962 | 1.12E-11 | 1.71E-10 | 478.7068082 | 749.3245607 | ENSG00000184916 |
| NA | 0.645993617 | 1.56481664 | 0.488750389 | 0.655460281 | 3.986939787 | 6.257616347 | ENSG00000274220 |
| NA | 0.645938211 | 1.564756545 | 0.222581225 | 0.381026377 | 12.8241879 | 19.98065068 | ENSG00000280173 |
| MFNG | 0.645670288 | 1.564465981 | 0.004325696 | 0.015467471 | 70.56274535 | 110.3698927 | ENSG00000100060 |
| NPTN-IT1 | 0.645657265 | 1.564451859 | 0.20064541 | 0.352646738 | 14.10056181 | 22.02950454 | ENSG00000281183 |
| TINCR | 0.645374095 | 1.564144821 | 0.186919419 | 0.333907075 | 17.85568063 | 27.95909392 | ENSG00000223573 |
| MAP3K15 | 0.644979645 | 1.563717224 | 0.023756146 | 0.064811174 | 48.91314867 | 76.51752077 | ENSG00000180815 |
| GAL3ST4 | 0.644877414 | 1.563606421 | 0.00089263 | 0.003841971 | 95.14663157 | 148.7281462 | ENSG00000197093 |
| NA | 0.644847694 | 1.56357421 | 0.473739295 | 0.642182893 | 4.40361195 | 6.834092744 | ENSG00000232389 |
| SNHG5 | 0.644833003 | 1.563558289 | 1.62E-38 | 1.68E-36 | 3786.568189 | 5918.649787 | ENSG00000203875 |
| LOC100268168 | 0.644644028 | 1.563353496 | 0.558080043 | 0.713636623 | 2.985489691 | 4.709677803 | ENSG00000204758 |
| NA | 0.644562862 | 1.563265544 | 0.28187013 | 0.450277013 | 10.70956357 | 16.72882936 | ENSG00000214772 |
| CD2BP2-DT | 0.644464262 | 1.563158707 | 0.122112669 | 0.241657313 | 20.45472252 | 31.96188541 | ENSG00000260219 |
| AK7 | 0.644431116 | 1.563122794 | 0.01059795 | 0.033065672 | 67.71708898 | 105.6292828 | ENSG00000140057 |
| RIPOR2 | 0.644000595 | 1.562656405 | 1.25E-14 | 2.61E-13 | 623.1256397 | 973.5369198 | ENSG00000111913 |
| PLXNB3 | 0.643862552 | 1.56250689 | 0.019738584 | 0.055602347 | 48.14563473 | 75.21695347 | ENSG00000198753 |
| REPS2 | 0.643672422 | 1.562300984 | 0.001946416 | 0.0077038 | 104.1369918 | 162.409514 | ENSG00000169891 |
| IGSF10 | 0.642198193 | 1.56070535 | 0.094101603 | 0.19734279 | 23.90556903 | 37.38651878 | ENSG00000152580 |
| SYT14 | 0.641519317 | 1.559971117 | 0.000100721 | 0.000544716 | 207.4739811 | 324.0093614 | ENSG00000143469 |
| EIF4E1B | 0.641206848 | 1.559633283 | 0.04440716 | 0.10849272 | 38.77663584 | 60.38130632 | ENSG00000175766 |
| VWA5A | 0.640967078 | 1.5593741 | 6.31E-14 | 1.22E-12 | 540.7852113 | 843.087389 | ENSG00000110002 |
| PDGFC | 0.640166547 | 1.558509066 | 0.107908732 | 0.21924634 | 26.76924701 | 41.4882672 | ENSG00000145431 |
| THNSL1 | 0.639778702 | 1.558090142 | 4.95E-05 | 0.000284984 | 151.679827 | 236.6754144 | ENSG00000185875 |
| H19 | 0.639497526 | 1.557786506 | 0.03408256 | 0.087118944 | 42.85018164 | 66.61956751 | ENSG00000130600 |
| SLC45A3 | 0.639235141 | 1.557503214 | 2.51E-09 | 2.90E-08 | 471.3060546 | 734.2458435 | ENSG00000158715 |
| NA | 0.638933989 | 1.557178131 | 0.311562328 | 0.482599863 | 9.345907928 | 14.5979627 | ENSG00000284655 |
| DTNA | 0.638653577 | 1.556875497 | 2.26E-05 | 0.000139352 | 181.243455 | 282.1153861 | ENSG00000134769 |
| PCSK2 | 0.638611436 | 1.556830021 | 0.261505482 | 0.426881063 | 10.75565748 | 16.69280265 | ENSG00000125851 |
| LINC02328 | 0.638602791 | 1.556820693 | 0.431326606 | 0.602514233 | 5.976561187 | 9.36665734 | ENSG00000258733 |
| PDLIM3 | 0.637947217 | 1.55611342 | 0.005327473 | 0.018520866 | 72.09518249 | 112.3205952 | ENSG00000154553 |
| NA | 0.63791844 | 1.55608238 | 0.351263531 | 0.523826901 | 7.350704722 | 11.41740131 | ENSG00000279145 |
| FZD6 | 0.637908107 | 1.556071236 | 0.000153331 | 0.000791094 | 148.2193685 | 230.6509025 | ENSG00000164930 |
| TPCN1 | 0.637674756 | 1.555819567 | 4.27E-10 | 5.45E-09 | 524.343935 | 815.2138034 | ENSG00000186815 |
| GPR78 | 0.637594986 | 1.555733544 | 0.248570128 | 0.412345545 | 12.09207982 | 18.79865593 | ENSG00000155269 |
| NA | 0.637296382 | 1.555411577 | 0.600294669 | 0.746440922 | 2.71893858 | 4.185639209 | ENSG00000243810 |
| RAB42 | 0.637082218 | 1.555180698 | 8.06E-06 | 5.40E-05 | 177.4894874 | 276.0599384 | ENSG00000188060 |
| EFCAB2 | 0.636837889 | 1.554917341 | 0.000300561 | 0.00145828 | 117.8358751 | 183.4331167 | ENSG00000203666 |
| DSEL | 0.636708173 | 1.554777541 | 5.03E-05 | 0.000288981 | 182.9741342 | 284.5791018 | ENSG00000171451 |
| SNAPIN | 0.636692242 | 1.554760373 | 4.21E-09 | 4.73E-08 | 354.237543 | 550.2227713 | ENSG00000143553 |
| NA | 0.63668963 | 1.554757558 | 0.428106028 | 0.599404368 | 6.261972897 | 9.758147554 | ENSG00000272054 |
| NA | 0.636508161 | 1.554562005 | 0.099034151 | 0.205647773 | 24.96985845 | 38.90997698 | ENSG00000224152 |
| MYL9 | 0.636436 | 1.554484251 | 2.58E-27 | 1.42E-25 | 1444.761458 | 2246.499038 | ENSG00000101335 |
| PTCD2 | 0.636386046 | 1.554430427 | 2.18E-05 | 0.000134988 | 157.9240153 | 245.390319 | ENSG00000049883 |
| RBM34 | 0.63614307 | 1.554168655 | 0.493992777 | 0.660402956 | 3.984148884 | 6.194698249 | ENSG00000188739 |
| NA | 0.63609105 | 1.554112616 | 0.29643996 | 0.466750186 | 9.304720106 | 14.50762018 | ENSG00000265194 |
| PLEKHF2 | 0.635523854 | 1.553501736 | 0.002074879 | 0.008151888 | 112.5935372 | 175.0569856 | ENSG00000175895 |
| SNORD104 | 0.635451392 | 1.553423711 | 0.044722142 | 0.109165488 | 41.02635067 | 63.90898938 | ENSG00000199753 |
| AHCTF1 | 0.635204429 | 1.553157816 | 2.96E-24 | 1.38E-22 | 1539.404885 | 2390.919628 | ENSG00000153207 |
| ZNF18 | 0.634825386 | 1.552749804 | 1.56E-05 | 9.88E-05 | 166.7415269 | 258.8450278 | ENSG00000154957 |
| NA | 0.634621885 | 1.552530795 | 0.117851085 | 0.235299657 | 25.01124643 | 38.7688142 | ENSG00000272077 |
| FGF2 | 0.634599876 | 1.552507111 | 6.50E-16 | 1.55E-14 | 920.9376954 | 1430.360231 | ENSG00000138685 |
| NA | 0.634477494 | 1.552375419 | 0.016344392 | 0.047499196 | 54.54935598 | 84.60700666 | ENSG00000187534 |
| CKS1B | 0.634337104 | 1.552224363 | 7.46E-25 | 3.66E-23 | 3390.992942 | 5262.788057 | ENSG00000173207 |
| NA | 0.633964995 | 1.551824055 | 0.009538374 | 0.030234108 | 68.29500891 | 106.213568 | ENSG00000179277 |
| NA | 0.633909406 | 1.551764263 | 0.547654733 | 0.705307928 | 3.366555838 | 5.195278645 | ENSG00000231482 |
| SBK2 | 0.633802052 | 1.551648797 | 0.001126538 | 0.004742918 | 93.71802142 | 145.541654 | ENSG00000187550 |
| SERPINA10 | 0.633708735 | 1.551548435 | 0.461760076 | 0.63158066 | 4.700187272 | 7.301131924 | ENSG00000140093 |
| SLC5A5 | 0.633614677 | 1.551447285 | 0.053354073 | 0.12561062 | 39.36372912 | 60.99379732 | ENSG00000105641 |
| NA | 0.633415623 | 1.55123324 | 0.410186954 | 0.582345933 | 6.58019972 | 10.28810483 | ENSG00000227627 |
| NA | 0.633381818 | 1.551196892 | 0.192962916 | 0.342514502 | 14.40762503 | 22.32392794 | ENSG00000286989 |
| NSUN7 | 0.63326354 | 1.551069724 | 2.98E-05 | 0.000180008 | 165.8775948 | 257.2494856 | ENSG00000179299 |
| PLEKHG4 | 0.633164895 | 1.550963672 | 5.32E-10 | 6.69E-09 | 407.7610064 | 632.1242824 | ENSG00000196155 |
| CATSPER3 | 0.632986057 | 1.550771425 | 0.548752975 | 0.706250826 | 3.64427056 | 5.694043392 | ENSG00000152705 |
| USP51 | 0.632878503 | 1.550655818 | 0.001402124 | 0.005773397 | 91.55403672 | 142.0126139 | ENSG00000247746 |
| STPG1 | 0.63271901 | 1.550484399 | 3.51E-05 | 0.000209076 | 158.2506148 | 245.3599384 | ENSG00000001460 |
| BHLHB9 | 0.632604029 | 1.550360832 | 0.599310321 | 0.745793794 | 2.776196098 | 4.244789198 | ENSG00000198908 |
| FAM78B | 0.632319864 | 1.550055491 | 0.007239409 | 0.024032943 | 73.53898645 | 114.1056973 | ENSG00000188859 |

| | | | | | | | |
|---|---|---|---|---|---|---|---|
| LINC02139 | 0.632226546 | 1.549955232 | 0.447367868 | 0.617720531 | 5.257731897 | 8.195154994 | ENSG00000278214 |
| APELA | 0.632106803 | 1.549826591 | 8.78E-06 | 5.85E-05 | 230.8755082 | 358.283362 | ENSG00000248329 |
| CENPL | 0.632024339 | 1.549738007 | 1.43E-05 | 9.14E-05 | 193.9316765 | 300.3666076 | ENSG00000120334 |
| VPS72 | 0.631505475 | 1.549180745 | 1.55E-20 | 5.49E-19 | 1755.252395 | 2718.333302 | ENSG00000163159 |
| NA | 0.631465813 | 1.549138156 | 0.194874693 | 0.345019062 | 15.09643011 | 23.36691048 | ENSG00000282386 |
| RPS27 | 0.631253825 | 1.548910544 | 3.45E-34 | 2.71E-32 | 10243.62012 | 15865.26323 | ENSG00000177954 |
| ZNF888 | 0.631184192 | 1.548835786 | 0.000406327 | 0.001909953 | 119.8864208 | 185.4397768 | ENSG00000213793 |
| DPF3 | 0.631183699 | 1.548835257 | 0.5793772 | 0.730794139 | 2.952674577 | 4.60428127 | ENSG00000205683 |
| EFHC2 | 0.63113359 | 1.548781463 | 0.0326593 | 0.084174469 | 40.07398554 | 62.08706709 | ENSG00000183690 |
| IDI2-AS1 | 0.631021296 | 1.548660916 | 0.245251477 | 0.408420774 | 12.0753344 | 18.65776571 | ENSG00000232656 |
| LOC101927752 | 0.630873201 | 1.548501951 | 0.002937991 | 0.011005117 | 94.50220553 | 146.0154175 | ENSG00000253982 |
| NCSTN | 0.630534282 | 1.548138219 | 8.64E-24 | 3.90E-22 | 1482.951462 | 2295.838296 | ENSG00000162736 |
| CPQ | 0.630230199 | 1.547811946 | 0.176116085 | 0.319519929 | 16.66937934 | 25.77983014 | ENSG00000104324 |
| HSD3B7 | 0.63020454 | 1.547784417 | 0.228622995 | 0.38827064 | 13.63432912 | 21.18172802 | ENSG00000099377 |
| SVEP1 | 0.630192922 | 1.547771953 | 0.117392461 | 0.234555102 | 22.46108934 | 34.69396928 | ENSG00000165124 |
| LINC00491 | 0.630100066 | 1.547672337 | 0.026533978 | 0.071092635 | 51.86602341 | 80.3665065 | ENSG00000250682 |
| NA | 0.62956974 | 1.547103527 | 0.502449999 | 0.668228848 | 3.992521593 | 6.167273829 | ENSG00000228252 |
| NXPH4 | 0.629351727 | 1.546869754 | 0.013445525 | 0.040395207 | 65.07578322 | 100.7892072 | ENSG00000182379 |
| TTC9B | 0.629139992 | 1.546642746 | 0.592459342 | 0.741062844 | 3.021095707 | 4.686554531 | ENSG00000174521 |
| ANKRA2 | 0.629137569 | 1.546640148 | 0.002262054 | 0.008788424 | 84.06225938 | 130.1211886 | ENSG00000164331 |
| POU2F1 | 0.62907032 | 1.546568056 | 2.85E-14 | 5.74E-13 | 1028.461343 | 1591.150758 | ENSG00000143190 |
| GIPR | 0.628847877 | 1.546329616 | 0.282511623 | 0.450766644 | 10.44512764 | 16.10610642 | ENSG00000010310 |
| ZNF548 | 0.628767924 | 1.546243922 | 0.002170471 | 0.008479252 | 88.82181935 | 137.4147843 | ENSG00000188785 |
| PYY2 | 0.628596845 | 1.546060574 | 0.388476298 | 0.561818447 | 7.09252632 | 10.95492375 | ENSG00000237575 |
| SMTNL1 | 0.628556171 | 1.546016987 | 0.345098975 | 0.517566141 | 9.735346783 | 15.00827504 | ENSG00000214872 |
| STK17A | 0.628530648 | 1.545989637 | 1.02E-07 | 9.36E-07 | 282.8052468 | 437.6499159 | ENSG00000164543 |
| NA | 0.628100298 | 1.545528543 | 0.310174725 | 0.481279598 | 11.21543249 | 17.40425057 | ENSG00000272153 |
| NA | 0.627917279 | 1.54533249 | 0.576175442 | 0.728231701 | 3.363764935 | 5.167854224 | ENSG00000227638 |
| KCTD18 | 0.627736832 | 1.545139219 | 0.004171414 | 0.014990823 | 81.42609676 | 125.5459493 | ENSG00000155729 |
| CALB2 | 0.627726803 | 1.545128477 | 0.045948139 | 0.111569713 | 35.12120167 | 54.27129237 | ENSG00000172137 |
| ANP32E | 0.62770118 | 1.545101035 | 7.56E-38 | 7.48E-36 | 4036.870044 | 6238.77006 | ENSG00000143401 |
| NEK2 | 0.627272656 | 1.544642161 | 2.61E-12 | 4.27E-11 | 589.9687028 | 910.5315793 | ENSG00000117650 |
| C11orf54 | 0.627238597 | 1.544605696 | 0.024447735 | 0.066341433 | 55.43138495 | 85.61771217 | ENSG00000182919 |
| NA | 0.626937492 | 1.544283355 | 0.415988577 | 0.588072375 | 5.998212689 | 9.301588669 | ENSG00000259553 |
| LIN37 | 0.626914855 | 1.544259125 | 0.530316149 | 0.691238435 | 3.627525142 | 5.645646273 | ENSG00000267796 |
| FAM160B2 | 0.626815258 | 1.544152519 | 3.44E-09 | 3.91E-08 | 347.0100014 | 535.5992622 | ENSG00000158863 |
| FOLR1 | 0.626672569 | 1.543999803 | 0.301682835 | 0.472498464 | 10.21888847 | 15.88535398 | ENSG00000110195 |
| NA | 0.625879861 | 1.543151665 | 0.201235393 | 0.353393898 | 14.05658309 | 21.6802323 | ENSG00000230699 |
| KCNJ3 | 0.625803415 | 1.543069898 | 0.463670647 | 0.633340682 | 5.135720033 | 7.848033327 | ENSG00000162989 |
| LINC00672 | 0.624870957 | 1.542072887 | 0.193227789 | 0.342921553 | 17.61847801 | 27.20219295 | ENSG00000263874 |
| OAS3 | 0.624569346 | 1.541750534 | 0.164962346 | 0.304022294 | 22.56376504 | 34.76790071 | ENSG00000111331 |
| FAM20B | 0.624386609 | 1.541555262 | 2.85E-15 | 6.31E-14 | 945.9205302 | 1458.46986 | ENSG00000116199 |
| DUSP5 | 0.624152489 | 1.54130512 | 1.49E-17 | 4.08E-16 | 913.1787891 | 1406.906268 | ENSG00000138166 |
| GAN | 0.624146441 | 1.541298659 | 0.000489644 | 0.002248446 | 123.9897906 | 191.3023832 | ENSG00000261609 |
| RILP | 0.623510881 | 1.54061981 | 0.00014606 | 0.00075682 | 155.0434287 | 238.5761145 | ENSG00000167705 |
| DDX25 | 0.623333699 | 1.540430612 | 4.70E-06 | 3.28E-05 | 204.0316194 | 314.4114897 | ENSG00000109832 |
| FCRL5 | 0.623276562 | 1.540369606 | 0.111112018 | 0.224422777 | 22.77931616 | 35.02441939 | ENSG00000143297 |
| IPMK | 0.623268836 | 1.540361357 | 1.62E-05 | 0.0001025 | 187.5455396 | 289.2662199 | ENSG00000151151 |
| SIL1 | 0.623054832 | 1.540132883 | 0.008339205 | 0.027046334 | 73.85298291 | 113.997605 | ENSG00000120725 |
| BCAN | 0.621842853 | 1.538839592 | 0.001388657 | 0.005721611 | 109.4974746 | 168.7991273 | ENSG00000132692 |
| TNFRSF10D | 0.621777396 | 1.538769774 | 4.24E-08 | 4.13E-07 | 337.3979058 | 519.3533011 | ENSG00000173530 |
| NA | 0.621767641 | 1.538759369 | 0.500352313 | 0.666503001 | 4.044197305 | 6.188246528 | ENSG00000231414 |
| SLC22A3 | 0.621762588 | 1.53875398 | 0.070625252 | 0.157215756 | 40.23842453 | 62.16718978 | ENSG00000146477 |
| NA | 0.621284565 | 1.538244213 | 0.517733994 | 0.681561791 | 4.014173095 | 6.148451703 | ENSG00000262904 |
| NA | 0.621227707 | 1.53818359 | 0.505621678 | 0.670442219 | 4.256281801 | 6.604738015 | ENSG00000287310 |
| DCAF12L1 | 0.621033295 | 1.537976324 | 0.000161254 | 0.000828651 | 136.8487087 | 210.4643235 | ENSG00000198889 |
| ZBTB4 | 0.62030184 | 1.537196759 | 2.82E-10 | 3.70E-09 | 473.6887575 | 728.5872696 | ENSG00000174282 |
| TBX19 | 0.620055602 | 1.536934414 | 0.233703537 | 0.394571585 | 13.09073901 | 20.10701991 | ENSG00000143178 |
| GATAD2B | 0.619668779 | 1.536522379 | 5.35E-10 | 6.72E-09 | 437.1537761 | 672.4017796 | ENSG00000143614 |
| NA | 0.619526502 | 1.536370856 | 0.564886535 | 0.719098994 | 3.015513901 | 4.640307985 | ENSG00000227518 |
| MORN3 | 0.619358711 | 1.536192181 | 0.288901263 | 0.458284209 | 10.99497528 | 16.96436324 | ENSG00000139714 |
| NA | 0.618926458 | 1.535731983 | 0.196011882 | 0.346491819 | 16.97173663 | 25.91856978 | ENSG00000273702 |
| GUCA1B | 0.618411493 | 1.535183906 | 0.176358132 | 0.319868817 | 18.11473491 | 27.83701372 | ENSG00000112599 |
| SYDE2 | 0.618345045 | 1.5351132 | 0.00220966 | 0.008602736 | 115.5160243 | 177.0870358 | ENSG00000097096 |
| AIM2 | 0.618335627 | 1.535103179 | 0.177513534 | 0.321269687 | 18.11473491 | 27.83486315 | ENSG00000163568 |
| GALNT9 | 0.618323898 | 1.535090699 | 0.166387833 | 0.306102202 | 19.86157188 | 30.47901554 | ENSG00000182870 |
| ZBTB34 | 0.618282265 | 1.5350464 | 8.36E-08 | 7.78E-07 | 370.5466687 | 568.2806221 | ENSG00000177125 |
| MACF1 | 0.618277346 | 1.535041166 | 5.84E-24 | 2.67E-22 | 4749.831584 | 7289.978958 | ENSG00000127603 |
| MUC3A | 0.618044916 | 1.534793879 | 3.93E-07 | 3.30E-06 | 271.5823175 | 416.7101977 | ENSG00000169894 |
| INAVA | 0.618030302 | 1.534778331 | 1.12E-15 | 2.62E-14 | 889.773909 | 1366.058504 | ENSG00000163362 |
| NA | 0.617886044 | 1.534624873 | 0.453837056 | 0.623840007 | 6.502166581 | 9.922161038 | ENSG00000266903 |
| LBX2 | 0.617877675 | 1.534615971 | 0.585448936 | 0.735192012 | 3.069980516 | 4.705376656 | ENSG00000179528 |
| SPEGNB | 0.617844781 | 1.534580982 | 0.601197358 | 0.747033702 | 2.95546548 | 4.549965467 | ENSG00000286095 |
| NA | 0.617736271 | 1.534465565 | 0.195589403 | 0.346030371 | 14.97912417 | 23.03269225 | ENSG00000258744 |
| RALGAPA1P1 | 0.617680179 | 1.534405906 | 0.45212755 | 0.622332683 | 5.756103983 | 8.767330244 | ENSG00000229419 |
| PRKAA1 | 0.617630318 | 1.534352876 | 1.87E-07 | 1.65E-06 | 361.5592626 | 555.0162667 | ENSG00000132356 |
| NA | 0.617359542 | 1.534064925 | 0.45146653 | 0.621688974 | 6.571827011 | 10.1197902 | ENSG00000273783 |

| | | | | | | | |
|---|---|---|---|---|---|---|---|
| TADA2A | 0.617182595 | 1.533876782 | 1.51E-06 | 1.15E-05 | 291.947768 | 447.3693531 | ENSG00000276234 |
| PIP5K1A | 0.616823728 | 1.533495283 | 1.09E-27 | 6.17E-26 | 1932.650335 | 2964.397777 | ENSG00000143398 |
| VCX3B | 0.61668676 | 1.5333497 | 0.475474982 | 0.643857075 | 4.928341463 | 7.620829171 | ENSG00000205642 |
| NA | 0.616627213 | 1.533286413 | 0.077926877 | 0.169767068 | 28.72585314 | 44.07137948 | ENSG00000267519 |
| ZFPM2 | 0.616504035 | 1.533155506 | 0.607632821 | 0.751721564 | 3.309298319 | 5.128319864 | ENSG00000169946 |
| LOC105375624 | 0.61632701 | 1.532967392 | 0.45556522 | 0.625368864 | 5.314989415 | 8.148908448 | ENSG00000254231 |
| NCALD | 0.615980337 | 1.532599072 | 0.068202519 | 0.15282783 | 30.67149582 | 47.08362624 | ENSG00000104490 |
| F8 | 0.615638884 | 1.532236383 | 0.085884452 | 0.183741485 | 28.3692294 | 43.5395321 | ENSG00000185010 |
| SERPINF2 | 0.615407709 | 1.53199088 | 0.028796677 | 0.075983681 | 47.2201026 | 72.42814276 | ENSG00000167711 |
| SNORD14A | 0.615159243 | 1.531727058 | 0.268493087 | 0.434752592 | 12.53510941 | 19.25683235 | ENSG00000272034 |
| ATP2B3 | 0.615055314 | 1.531616719 | 0.22523141 | 0.384269844 | 14.93861207 | 22.91492532 | ENSG00000067842 |
| GALNT2 | 0.614882129 | 1.531432871 | 1.04E-31 | 7.22E-30 | 3307.197896 | 5063.496649 | ENSG00000143641 |
| MARVELD2 | 0.614707241 | 1.531247237 | 4.61E-08 | 4.45E-07 | 305.2641457 | 467.2349205 | ENSG00000152939 |
| UFSP1 | 0.614540775 | 1.531070564 | 0.048117315 | 0.115797296 | 41.48911674 | 63.75786722 | ENSG00000176125 |
| CD177 | 0.614092331 | 1.530594723 | 0.060246064 | 0.138374428 | 35.57906166 | 54.32534771 | ENSG00000204936 |
| TTC39A | 0.614037456 | 1.530536506 | 0.001271741 | 0.00529874 | 111.2979392 | 169.9625725 | ENSG00000085831 |
| TMEM54 | 0.613902359 | 1.530393189 | 0.035024902 | 0.089232091 | 51.72726613 | 79.00760693 | ENSG00000121900 |
| CHROMR | 0.613576229 | 1.530047274 | 0.000950545 | 0.004070357 | 105.3384869 | 161.2105567 | ENSG00000223960 |
| IDH1-AS1 | 0.613189651 | 1.529637345 | 0.287312899 | 0.456665827 | 10.71446966 | 16.36073499 | ENSG00000231908 |
| CAPS2 | 0.613072864 | 1.529513525 | 0.154572811 | 0.289490636 | 20.69404033 | 31.5999823 | ENSG00000180881 |
| LOC105369779 | 0.612694582 | 1.529112531 | 0.007905172 | 0.025851615 | 74.74936672 | 114.0441363 | ENSG00000258056 |
| NA | 0.612639087 | 1.529053713 | 0.51314688 | 0.677418494 | 4.104245726 | 6.215670948 | ENSG00000258704 |
| EPPK1 | 0.612600755 | 1.529013087 | 4.87E-07 | 4.02E-06 | 283.2357102 | 432.7149362 | ENSG00000261150 |
| NA | 0.612198035 | 1.528586332 | 0.485966427 | 0.653385722 | 4.985598981 | 7.662774569 | ENSG00000108958 |
| SSC5D | 0.612145169 | 1.528530319 | 0.100231977 | 0.207487399 | 27.32467646 | 41.60173299 | ENSG00000179954 |
| ICAM1 | 0.612041609 | 1.528420601 | 0.009039519 | 0.028895005 | 66.65818121 | 101.7905048 | ENSG00000090339 |
| TTLL6 | 0.612037266 | 1.528416001 | 0.005933487 | 0.020350659 | 80.13277727 | 122.3962895 | ENSG00000170703 |
| PCDHA7 | 0.61194764 | 1.528321052 | 0.139259462 | 0.266804477 | 20.94731264 | 32.03314534 | ENSG00000204963 |
| NA | 0.61185059 | 1.528218245 | 0.564012162 | 0.718460025 | 3.039956306 | 4.638157411 | ENSG00000230333 |
| CXCR2 | 0.611504855 | 1.527852059 | 0.265013481 | 0.430987613 | 12.97901488 | 19.90939075 | ENSG00000180871 |
| PIGM | 0.611306546 | 1.527642059 | 2.73E-06 | 1.98E-05 | 238.4862311 | 364.5539003 | ENSG00000143315 |
| ASCL2 | 0.611290963 | 1.527625559 | 0.254717859 | 0.41951355 | 12.58678512 | 19.25898292 | ENSG00000183734 |
| C8orf34-AS1 | 0.611133878 | 1.527459235 | 0.249414397 | 0.413448377 | 12.67406685 | 19.37244871 | ENSG00000248801 |
| NA | 0.610978221 | 1.527294441 | 0.50257924 | 0.668279176 | 4.310748416 | 6.615490883 | ENSG00000232909 |
| NA | 0.610603363 | 1.526897654 | 0.565414519 | 0.719486246 | 3.725970483 | 5.689742245 | ENSG00000214252 |
| RGL1 | 0.610396458 | 1.526678688 | 0.004195665 | 0.01505834 | 91.91480281 | 140.2455525 | ENSG00000143344 |
| SSBP3-AS1 | 0.610321709 | 1.52659959 | 0.397649701 | 0.570131354 | 7.247353297 | 11.10415579 | ENSG00000198711 |
| NAT9 | 0.610263386 | 1.526537876 | 1.13E-15 | 2.62E-14 | 756.3987871 | 1154.578313 | ENSG00000109065 |
| SPATC1 | 0.609175806 | 1.525387525 | 0.554468749 | 0.710667506 | 3.624734239 | 5.54670146 | ENSG00000186583 |
| TSTD3 | 0.609019686 | 1.525222465 | 0.296758734 | 0.467034501 | 12.37470062 | 18.81560005 | ENSG00000279170 |
| CRIM1 | 0.609005425 | 1.525207388 | 8.72E-09 | 9.38E-08 | 371.9937022 | 567.6103077 | ENSG00000150938 |
| TRMT1L | 0.608938272 | 1.525136397 | 3.72E-13 | 6.66E-12 | 611.5760739 | 933.356683 | ENSG00000121486 |
| LOC112268411 | 0.608541516 | 1.524717026 | 0.345671256 | 0.518130863 | 9.119868918 | 13.87414437 | ENSG00000236106 |
| RELL1 | 0.608444809 | 1.524614824 | 0.005628575 | 0.019434551 | 75.06181158 | 114.5961202 | ENSG00000181826 |
| COQ5 | 0.608321783 | 1.524484818 | 6.43E-10 | 8.00E-09 | 513.8095865 | 783.001548 | ENSG00000110871 |
| PHF7 | 0.608211837 | 1.524368643 | 0.001289461 | 0.005365999 | 117.6958785 | 179.3889007 | ENSG00000010318 |
| TRIM68 | 0.608188995 | 1.524344509 | 0.000249503 | 0.001233198 | 168.7721729 | 256.7354064 | ENSG00000167333 |
| BCL11B | 0.608050441 | 1.52419812 | 0.010195301 | 0.031974892 | 74.74637566 | 114.0900981 | ENSG00000127152 |
| ANKRD50 | 0.607641857 | 1.523766515 | 4.51E-11 | 6.42E-10 | 639.2549083 | 974.0498293 | ENSG00000151458 |
| NA | 0.607441727 | 1.523555154 | 0.600204916 | 0.746377427 | 2.95546548 | 4.50748703 | ENSG00000229337 |
| TMEM9B-AS1 | 0.607436018 | 1.523549124 | 0.076008621 | 0.166452754 | 32.88456548 | 49.95687289 | ENSG00000254860 |
| SMIM1 | 0.606989112 | 1.523077245 | 0.218445353 | 0.376172171 | 15.84719863 | 24.21683758 | ENSG00000235169 |
| C1orf127 | 0.606967807 | 1.523054754 | 0.558241641 | 0.71379595 | 3.600291835 | 5.502605488 | ENSG00000175262 |
| COL16A1 | 0.606796777 | 1.522874207 | 6.60E-07 | 5.34E-06 | 305.4149425 | 464.5875941 | ENSG00000084636 |
| ZNF467 | 0.606537791 | 1.522600852 | 0.392268184 | 0.565211211 | 7.389101641 | 11.25364829 | ENSG00000181444 |
| ANK1 | 0.606480022 | 1.522539885 | 6.03E-07 | 4.90E-06 | 280.5465957 | 427.0034156 | ENSG00000029534 |
| NA | 0.606156959 | 1.52219898 | 0.454210327 | 0.624130737 | 5.372246934 | 8.144607301 | ENSG00000262921 |
| NELL1 | 0.606115661 | 1.522155408 | 0.408027355 | 0.580390481 | 6.670272351 | 10.1531333 | ENSG00000165973 |
| ZBED8 | 0.605960489 | 1.521991698 | 0.002171667 | 0.008480491 | 104.9427025 | 159.7532332 | ENSG00000221886 |
| SYT1 | 0.605921333 | 1.52195039 | 1.06E-12 | 1.80E-11 | 611.6028317 | 931.3193755 | ENSG00000067715 |
| ANKRD19P | 0.605873295 | 1.521899714 | 0.530640712 | 0.691614754 | 3.660340256 | 5.578427028 | ENSG00000187984 |
| RNPEP | 0.605825971 | 1.521849793 | 1.85E-19 | 5.95E-18 | 1085.414226 | 1652.26514 | ENSG00000176393 |
| NA | 0.605682725 | 1.521698694 | 0.170452821 | 0.31182602 | 19.06450929 | 29.03836364 | ENSG00000223813 |
| ZSWIM6 | 0.605509295 | 1.521515779 | 1.95E-06 | 1.46E-05 | 280.9347711 | 427.101579 | ENSG00000130449 |
| NA | 0.605453482 | 1.521456918 | 0.630992732 | 0.76847862 | 2.386757243 | 3.603244129 | ENSG00000273374 |
| FGD6 | 0.605403259 | 1.521403953 | 3.81E-12 | 6.09E-11 | 561.9891507 | 855.3197142 | ENSG00000180263 |
| PIGC | 0.605253955 | 1.521246512 | 4.44E-10 | 5.65E-09 | 473.4940431 | 719.9801968 | ENSG00000135845 |
| LRRC27 | 0.605195319 | 1.521184685 | 0.037599246 | 0.094677856 | 48.0110198 | 72.82340109 | ENSG00000148814 |
| JAZF1 | 0.604434496 | 1.520382681 | 0.0010109 | 0.004304918 | 131.8017098 | 200.5211592 | ENSG00000153814 |
| FLNC | 0.603288011 | 1.519174939 | 3.19E-16 | 7.83E-15 | 763.354183 | 1159.950515 | ENSG00000128591 |
| SFI1 | 0.603219574 | 1.519102875 | 2.05E-07 | 1.80E-06 | 299.2707511 | 454.3190743 | ENSG00000198089 |
| PARS2 | 0.603078008 | 1.518953819 | 1.60E-06 | 1.21E-05 | 281.0698616 | 427.2972633 | ENSG00000162396 |
| PRKG1 | 0.603031043 | 1.518904372 | 0.00011647 | 0.000620325 | 155.5445547 | 236.5600707 | ENSG00000185532 |
| PTCHD4 | 0.60296508 | 1.518834926 | 0.185843772 | 0.33235489 | 19.62504498 | 29.72615525 | ENSG00000244694 |
| GALNT12 | 0.602955958 | 1.518825323 | 2.36E-06 | 1.73E-05 | 332.9766966 | 505.9117867 | ENSG00000119514 |
| NA | 0.602935886 | 1.518804192 | 0.241519772 | 0.40384484 | 17.00993339 | 25.82445915 | ENSG00000279716 |
| SETDB1 | 0.602902899 | 1.518769466 | 2.48E-13 | 4.51E-12 | 648.1626799 | 984.0726501 | ENSG00000143379 |

| | | | | | | | |
|---|---|---|---|---|---|---|---|
| NA | 0.602844606 | 1.5187081 | 0.45315139 | 0.623386113 | 5.405062047 | 8.161278853 | ENSG00000278918 |
| NA | 0.602840266 | 1.518703531 | 0.479917295 | 0.648098323 | 5.423922646 | 8.193004421 | ENSG00000219470 |
| DOK7 | 0.602539081 | 1.518386511 | 0.350238388 | 0.52286401 | 9.729089256 | 14.74504417 | ENSG00000175920 |
| NA | 0.60252939 | 1.518376312 | 0.504935077 | 0.670016685 | 4.340772627 | 6.602587441 | ENSG00000213070 |
| H6PD | 0.602361768 | 1.518199907 | 1.49E-07 | 1.33E-06 | 293.8292071 | 446.1852078 | ENSG00000049239 |
| CD163L1 | 0.602193724 | 1.518023078 | 0.626795269 | 0.765264913 | 2.653308353 | 4.068132735 | ENSG00000177675 |
| SCN8A | 0.601928517 | 1.51774405 | 1.49E-10 | 2.02E-09 | 507.8960279 | 770.8874923 | ENSG00000196876 |
| CLHC1 | 0.601763674 | 1.517570641 | 0.056659581 | 0.131896656 | 40.55752723 | 61.62701267 | ENSG00000162994 |
| PARP1 | 0.601318042 | 1.517101953 | 6.38E-62 | 1.54E-59 | 18831.54173 | 28568.52438 | ENSG00000143799 |
| C6orf163 | 0.601280644 | 1.517062627 | 0.281042619 | 0.449400772 | 12.18984943 | 18.41335697 | ENSG00000203872 |
| NA | 0.601197438 | 1.516975135 | 0.26778799 | 0.434109085 | 11.66424404 | 17.70459266 | ENSG00000233577 |
| IMPDH2 | 0.600875982 | 1.516637166 | 2.44E-39 | 2.70E-37 | 4184.24869 | 6345.868291 | ENSG00000178035 |
| FHDC1 | 0.600496037 | 1.5162378 | 1.56E-17 | 4.27E-16 | 883.5607825 | 1339.320878 | ENSG00000137460 |
| NA | 0.600319275 | 1.516052038 | 0.409524369 | 0.581790604 | 8.758339091 | 13.27722831 | ENSG00000227001 |
| CPT1A | 0.600315158 | 1.516047712 | 0.000647397 | 0.002879781 | 138.3558644 | 209.5057527 | ENSG00000110090 |
| GAS2L1 | 0.600040053 | 1.515758647 | 2.25E-06 | 1.66E-05 | 294.939078 | 447.2658404 | ENSG00000185340 |
| THBS2 | 0.599889312 | 1.51560028 | 2.59E-10 | 3.41E-09 | 553.85818 | 840.145324 | ENSG00000186340 |
| LOC101927556 | 0.599635378 | 1.515333538 | 0.427871324 | 0.599249725 | 6.085494418 | 9.175219436 | ENSG00000260927 |
| GABRD | 0.599628396 | 1.515326204 | 0.065543974 | 0.147970602 | 37.72071913 | 57.19319661 | ENSG00000187730 |
| NA | 0.599533222 | 1.515226242 | 0.072372294 | 0.160254123 | 32.46231151 | 49.12415038 | ENSG00000265519 |
| NA | 0.599476776 | 1.51516696 | 0.328119396 | 0.500336041 | 10.18694924 | 15.31936999 | ENSG00000215086 |
| DGAT2 | 0.599387257 | 1.515072947 | 0.005901908 | 0.020269246 | 89.2449492 | 135.16211 | ENSG00000062282 |
| NSMCE1 | 0.598863714 | 1.514523238 | 9.58E-06 | 6.34E-05 | 203.8281078 | 308.4727771 | ENSG00000169189 |
| NA | 0.598707707 | 1.514359473 | 0.593250101 | 0.741562369 | 2.988280594 | 4.513938752 | ENSG00000269887 |
| PRR5-ARHGAP8 | 0.598360977 | 1.513995563 | 0.5080862 | 0.672809644 | 4.370796837 | 6.615490883 | ENSG00000248405 |
| KCNIP1 | 0.598164191 | 1.513789065 | 0.132373581 | 0.256310222 | 22.26978047 | 33.76230195 | ENSG00000182132 |
| KDM6B | 0.598038496 | 1.513657182 | 2.77E-05 | 0.000168496 | 188.0014846 | 284.5017054 | ENSG00000132510 |
| NA | 0.597884172 | 1.513495275 | 0.622801105 | 0.762596559 | 2.389548146 | 3.577970282 | ENSG00000282021 |
| ATP6V1G2 | 0.597563477 | 1.51315888 | 0.215048885 | 0.371796742 | 16.94450332 | 25.48540674 | ENSG00000213760 |
| PCLO | 0.597361778 | 1.512947345 | 0.003582493 | 0.013113826 | 93.72727001 | 141.6759846 | ENSG00000186472 |
| FAM86B3P | 0.597332113 | 1.512916236 | 0.007858716 | 0.025738963 | 76.51342467 | 115.7122528 | ENSG00000173295 |
| UPK1A-AS1 | 0.597091721 | 1.512664164 | 0.432526133 | 0.603534295 | 6.267554703 | 9.526642251 | ENSG00000226510 |
| CORO7 | 0.596769756 | 1.512326621 | 0.022795202 | 0.06265027 | 52.30223189 | 78.94845694 | ENSG00000262246 |
| SRP14-AS1 | 0.596550792 | 1.512097107 | 0.062666692 | 0.142795082 | 37.39152886 | 56.34731019 | ENSG00000248508 |
| FAM27C | 0.596117954 | 1.511643514 | 0.539692806 | 0.698943569 | 3.660340256 | 5.525728761 | ENSG00000231527 |
| NBEAL2 | 0.595845353 | 1.511357913 | 1.21E-23 | 5.36E-22 | 1929.003987 | 2914.630635 | ENSG00000160796 |
| CYBA | 0.595434972 | 1.510928061 | 1.79E-27 | 9.93E-26 | 3063.985223 | 4628.599753 | ENSG00000051523 |
| CEP112 | 0.595421791 | 1.510914257 | 0.035499839 | 0.090190424 | 45.87319237 | 69.4978967 | ENSG00000154240 |
| IKBKE | 0.59532288 | 1.510810672 | 3.67E-06 | 2.61E-05 | 230.0874188 | 347.6793465 | ENSG00000263528 |
| ASPG | 0.595226309 | 1.510709545 | 0.071356446 | 0.158459446 | 35.41307107 | 53.37296815 | ENSG00000166183 |
| NA | 0.595211764 | 1.510694314 | 0.282525926 | 0.450766644 | 12.93591203 | 19.43589985 | ENSG00000280152 |
| NA | 0.594993097 | 1.510465358 | 0.519517625 | 0.682977682 | 4.749072081 | 7.109694019 | ENSG00000267605 |
| MYSM1 | 0.594675385 | 1.510132758 | 3.21E-14 | 6.42E-13 | 1042.464545 | 1573.757557 | ENSG00000162601 |
| ADAM19 | 0.594622903 | 1.510077825 | 3.29E-15 | 7.22E-14 | 826.5124327 | 1248.108922 | ENSG00000135074 |
| ATP6AP1L | 0.594194881 | 1.509629878 | 0.278536708 | 0.446283358 | 12.90309692 | 19.37083117 | ENSG00000205464 |
| VIM-AS1 | 0.594045039 | 1.509473092 | 0.450669435 | 0.621079016 | 6.330394027 | 9.579340518 | ENSG00000229124 |
| ACVR1C | 0.593962865 | 1.509387116 | 0.230666841 | 0.39098445 | 15.80688669 | 23.7497984 | ENSG00000123612 |
| ZNF132 | 0.593916262 | 1.50933836 | 0.128933605 | 0.2512382 | 25.82138765 | 38.87584038 | ENSG00000131849 |
| ARHGAP40 | 0.593788696 | 1.509204907 | 0.414056415 | 0.586329171 | 6.763135886 | 10.15528388 | ENSG00000124143 |
| SPTBN5 | 0.593631525 | 1.509040499 | 0.009536259 | 0.030232371 | 73.87618602 | 111.4083136 | ENSG00000137877 |
| CSRNP1 | 0.593599027 | 1.509006507 | 0.000242644 | 0.001202685 | 142.1979527 | 214.6244163 | ENSG00000144655 |
| PWRN1 | 0.593376904 | 1.508774193 | 0.42477345 | 0.596469983 | 7.819052597 | 11.70805661 | ENSG00000259905 |
| MRPS9-AS1 | 0.593361002 | 1.508757562 | 0.596757347 | 0.74400866 | 3.037165403 | 4.55588415 | ENSG00000231851 |
| NA | 0.593354856 | 1.508751135 | 0.634986461 | 0.771283454 | 2.419572356 | 3.605394703 | ENSG00000232952 |
| GNPAT | 0.593312502 | 1.508706842 | 7.06E-21 | 2.55E-19 | 1098.210705 | 1656.948193 | ENSG00000116906 |
| PYCARD | 0.593251345 | 1.508642888 | 0.002375983 | 0.00917252 | 107.4831354 | 162.1984178 | ENSG00000103490 |
| ADAMTS6 | 0.593040615 | 1.508422542 | 0.246518869 | 0.410000475 | 14.12221332 | 21.22179541 | ENSG00000049192 |
| NA | 0.592917927 | 1.508294269 | 0.011533797 | 0.035582677 | 72.46835149 | 109.5430888 | ENSG00000246695 |
| SMAD3 | 0.592542366 | 1.507901682 | 2.52E-09 | 2.91E-08 | 456.8337121 | 688.8125414 | ENSG00000166949 |
| FLNB | 0.59253191 | 1.507890754 | 3.86E-36 | 3.40E-34 | 4207.811939 | 6343.296404 | ENSG00000136068 |
| NA | 0.592502287 | 1.507859793 | 0.486800522 | 0.654233584 | 5.056811014 | 7.618678597 | ENSG00000213236 |
| XIAP | 0.592487472 | 1.507844309 | 1.15E-11 | 1.75E-10 | 738.4677152 | 1113.935381 | ENSG00000101966 |
| ENAH | 0.591993019 | 1.507327615 | 3.85E-50 | 6.52E-48 | 6554.125418 | 9879.978272 | ENSG00000154380 |
| NA | 0.591966375 | 1.507299778 | 0.526445781 | 0.688566382 | 4.392448339 | 6.604738015 | ENSG00000273828 |
| NA | 0.59190524 | 1.507235907 | 0.347170096 | 0.519933105 | 8.717151269 | 13.11698294 | ENSG00000287104 |
| TBX22 | 0.591790635 | 1.507116179 | 0.333007992 | 0.505711815 | 9.370350333 | 14.11210139 | ENSG00000122145 |
| FRMD6 | 0.591477382 | 1.506788975 | 3.01E-06 | 2.17E-05 | 261.9292342 | 394.8995434 | ENSG00000139926 |
| LINC00599 | 0.591193232 | 1.506492229 | 1.89E-08 | 1.93E-07 | 559.149025 | 841.4268451 | ENSG00000253230 |
| XPC | 0.5910042 | 1.506294852 | 7.52E-13 | 1.30E-11 | 695.9329553 | 1047.72188 | ENSG00000154767 |
| RPL3L | 0.590993719 | 1.506283908 | 0.241740172 | 0.404108289 | 13.92041655 | 21.03868723 | ENSG00000140986 |
| NA | 0.590916945 | 1.506203753 | 0.605862408 | 0.750413593 | 3.064398711 | 4.58330857 | ENSG00000268333 |
| COCH | 0.590838828 | 1.506122199 | 4.99E-05 | 0.00028681 | 232.3077099 | 349.8787653 | ENSG00000100473 |
| MAGI3 | 0.590737618 | 1.506016544 | 0.000163696 | 0.000840608 | 213.0134948 | 321.0199716 | ENSG00000081026 |
| ADCK5 | 0.590550378 | 1.505821098 | 0.000137076 | 0.000716235 | 153.228534 | 230.9249047 | ENSG00000173137 |
| CCDC141 | 0.590544366 | 1.505814822 | 0.002516729 | 0.009642546 | 105.0462541 | 158.0598308 | ENSG00000163492 |
| DCAF4L1 | 0.590113624 | 1.505365302 | 0.629721007 | 0.767545836 | 2.988280594 | 4.482213184 | ENSG00000182308 |
| PRDM5 | 0.59000671 | 1.505253748 | 2.32E-07 | 2.01E-06 | 313.248313 | 471.7292194 | ENSG00000138738 |

| | | | | | | | |
|---|---|---|---|---|---|---|---|
| NA | 0.589898115 | 1.505140449 | 0.289431511 | 0.458867272 | 11.39769293 | 17.15769125 | ENSG00000246877 |
| EHHADH | 0.589662706 | 1.50489487 | 0.085094219 | 0.182374467 | 32.96818043 | 49.59039606 | ENSG00000113790 |
| ARPC5 | 0.589650284 | 1.504881913 | 3.76E-23 | 1.59E-21 | 2358.653722 | 3550.472394 | ENSG00000162704 |
| QSOX1 | 0.589380018 | 1.504600023 | 6.21E-35 | 5.05E-33 | 3331.081328 | 5012.421645 | ENSG00000116260 |
| NUCKS1 | 0.589283207 | 1.504499062 | 4.43E-39 | 4.80E-37 | 12165.91395 | 18304.8125 | ENSG00000069275 |
| NDUFA13 | 0.589157026 | 1.504367481 | 0.074517056 | 0.164053485 | 33.28728313 | 49.98187416 | ENSG00000186010 |
| LOC102724301 | 0.589098985 | 1.50430696 | 0.239906894 | 0.40194927 | 13.73277446 | 20.62004517 | ENSG00000254854 |
| LRATD1 | 0.588976571 | 1.504179324 | 0.000925524 | 0.003974686 | 116.0703777 | 174.4641408 | ENSG00000162981 |
| PUDP | 0.588785432 | 1.503980052 | 3.06E-13 | 5.53E-12 | 812.0742951 | 1221.81872 | ENSG00000130021 |
| SLC12A4 | 0.588750131 | 1.503943252 | 3.85E-20 | 1.32E-18 | 1201.360713 | 1806.561984 | ENSG00000124067 |
| CTC1 | 0.58871333 | 1.503904889 | 1.97E-10 | 2.63E-09 | 503.4705643 | 757.0147476 | ENSG00000178971 |
| HIPK3 | 0.588328621 | 1.503503912 | 2.91E-05 | 0.00017587 | 207.5954662 | 311.893434 | ENSG00000110422 |
| FNIP1 | 0.588295126 | 1.503469005 | 0.000645954 | 0.002876451 | 166.4532122 | 250.492541 | ENSG00000217128 |
| ABCB10 | 0.588120878 | 1.503287427 | 4.76E-06 | 3.32E-05 | 271.3208357 | 408.1937463 | ENSG00000135776 |
| C3orf62 | 0.587699314 | 1.502848222 | 0.129412392 | 0.251916696 | 24.89306461 | 37.52364088 | ENSG00000188315 |
| VANGL2 | 0.587631837 | 1.502777933 | 2.15E-24 | 1.01E-22 | 2048.115685 | 3077.406783 | ENSG00000162738 |
| NA | 0.587573037 | 1.502716686 | 0.243039292 | 0.405893081 | 14.0649558 | 21.15860474 | ENSG00000271936 |
| KIRREL3 | 0.58729071 | 1.502422642 | 0.245494199 | 0.408789694 | 14.45160375 | 21.66571132 | ENSG00000149571 |
| POU3F2 | 0.587283749 | 1.502415393 | 0.608060304 | 0.751928931 | 3.075562322 | 4.58330857 | ENSG00000184486 |
| TSPEAR | 0.587145766 | 1.502271705 | 0.132649475 | 0.256689841 | 29.40716139 | 44.0332022 | ENSG00000175894 |
| RGS9 | 0.586941933 | 1.502059469 | 5.89E-05 | 0.000334571 | 171.376084 | 257.5629916 | ENSG00000108370 |
| LOC100507291 | 0.586326065 | 1.501418396 | 0.108580025 | 0.220323684 | 28.16763279 | 42.31290834 | ENSG00000248932 |
| MRPS14 | 0.586167392 | 1.501253273 | 5.23E-10 | 6.59E-09 | 475.11022 | 712.6766112 | ENSG00000120333 |
| ZNF860 | 0.586072849 | 1.501154896 | 0.013195434 | 0.03978014 | 69.83390373 | 104.9810255 | ENSG00000197385 |
| WDR78 | 0.585888001 | 1.500962571 | 0.520716027 | 0.683672188 | 5.114068532 | 7.650404165 | ENSG00000152763 |
| CX3CL1 | 0.585582105 | 1.500644354 | 0.032272447 | 0.083370203 | 48.45963119 | 72.77284128 | ENSG00000006210 |
| NA | 0.585404877 | 1.500460018 | 0.415935668 | 0.588047687 | 7.465219758 | 11.13373078 | ENSG00000189423 |
| HIF3A | 0.585202022 | 1.500249056 | 3.22E-06 | 2.31E-05 | 270.6454585 | 405.9845924 | ENSG00000124440 |
| INA | 0.585082241 | 1.500124501 | 5.84E-08 | 5.55E-07 | 409.0446401 | 614.2833611 | ENSG00000148798 |
| RTBDN | 0.584996207 | 1.500035046 | 0.358176723 | 0.531369199 | 8.766036079 | 13.09385966 | ENSG00000132026 |
| TENT5A | 0.584416084 | 1.499431987 | 0.013902778 | 0.041587609 | 66.45937551 | 99.49079046 | ENSG00000112773 |
| ZNF391 | 0.584312929 | 1.499324779 | 0.000751582 | 0.003286212 | 134.0058432 | 200.783324 | ENSG00000124613 |
| AHNAK2 | 0.583367863 | 1.498342938 | 0.018100603 | 0.051646019 | 60.89928632 | 91.29212782 | ENSG00000185567 |
| KATNAL2 | 0.583113383 | 1.498078666 | 0.000635528 | 0.002834021 | 135.8811497 | 203.2909058 | ENSG00000167216 |
| MTMR10 | 0.583010898 | 1.49797225 | 2.10E-05 | 0.000130319 | 205.5382627 | 308.1702724 | ENSG00000166912 |
| GALNT3 | 0.582736108 | 1.497686959 | 1.25E-05 | 8.04E-05 | 364.6186431 | 546.3572892 | ENSG00000115339 |
| HIVEP1 | 0.582543319 | 1.497486835 | 2.10E-07 | 1.84E-06 | 332.9242331 | 499.0365605 | ENSG00000095951 |
| NIPSNAP2 | 0.581704472 | 1.496616382 | 4.34E-09 | 4.87E-08 | 577.6663286 | 864.8342159 | ENSG00000146729 |
| NA | 0.581506215 | 1.496410729 | 0.014100474 | 0.042055079 | 69.70842524 | 104.2526456 | ENSG00000182912 |
| NA | 0.581184449 | 1.49607702 | 0.024985788 | 0.0676018 | 59.3415309 | 88.91014023 | ENSG00000244627 |
| CALD1 | 0.580962748 | 1.495847134 | 1.98E-37 | 1.92E-35 | 3782.872046 | 5658.836133 | ENSG00000122786 |
| FDXR | 0.580855927 | 1.495736381 | 9.35E-15 | 1.98E-13 | 854.039982 | 1277.101505 | ENSG00000161513 |
| LCAT | 0.580823232 | 1.495702484 | 0.001481492 | 0.006060107 | 109.87595 | 164.3233779 | ENSG00000213398 |
| NA | 0.580770214 | 1.495647519 | 0.557918566 | 0.713477431 | 4.022545804 | 6.043315634 | ENSG00000232906 |
| LOC107985911 | 0.580746558 | 1.495622995 | 0.606647294 | 0.751144606 | 3.336531627 | 5.012703499 | ENSG00000283196 |
| JAKMIP3 | 0.580574112 | 1.495444233 | 0.025680883 | 0.069200995 | 56.76848301 | 84.98181321 | ENSG00000188385 |
| ACSL4 | 0.580502023 | 1.495369511 | 1.30E-19 | 4.25E-18 | 1809.84821 | 2706.991414 | ENSG00000068366 |
| LTC4S | 0.580306637 | 1.495167005 | 0.280275934 | 0.448374094 | 12.04810109 | 17.98019393 | ENSG00000213316 |
| NA | 0.58022926 | 1.495086815 | 0.632534534 | 0.769419491 | 3.10279563 | 4.596212013 | ENSG00000186543 |
| NA | 0.580109065 | 1.494962261 | 0.281134189 | 0.449457198 | 14.72027004 | 21.99616144 | ENSG00000267939 |
| IPCEF1 | 0.579912791 | 1.49475889 | 0.454641874 | 0.624367936 | 7.546919681 | 11.20095003 | ENSG00000074706 |
| TMPRSS11E | 0.579618246 | 1.494453746 | 0.309973198 | 0.481146843 | 14.30514948 | 21.23846697 | ENSG00000087128 |
| MYO6 | 0.57951266 | 1.494344377 | 9.85E-13 | 1.67E-11 | 813.6695091 | 1216.007097 | ENSG00000196586 |
| NA | 0.579317471 | 1.494142213 | 0.443993071 | 0.614865064 | 6.34367282 | 9.465341689 | ENSG00000270083 |
| ADAM12 | 0.579306362 | 1.494130708 | 0.471249098 | 0.640156774 | 7.195677585 | 10.81403353 | ENSG00000148848 |
| ZNF839 | 0.579286642 | 1.494110285 | 0.001190314 | 0.004992187 | 121.8996087 | 182.1382013 | ENSG00000022976 |
| SIK1 | 0.579228376 | 1.494049944 | 0.180465147 | 0.325420935 | 22.03692035 | 32.74836137 | ENSG00000142178 |
| HEYL | 0.579100391 | 1.493917408 | 0.000574734 | 0.002589862 | 137.2533414 | 205.1571966 | ENSG00000163909 |
| BEST4 | 0.578850084 | 1.493658237 | 0.071068278 | 0.157928597 | 34.50639953 | 51.46689461 | ENSG00000142959 |
| NA | 0.578595958 | 1.493395158 | 0.399269132 | 0.571769047 | 8.439436547 | 12.58728612 | ENSG00000279531 |
| ZNF587B | 0.578592856 | 1.493391946 | 1.43E-06 | 1.10E-05 | 324.0686879 | 484.5351316 | ENSG00000269343 |
| NAT1 | 0.578569672 | 1.493367948 | 0.029373532 | 0.077168025 | 57.02407067 | 85.15766406 | ENSG00000171428 |
| TULP1 | 0.578565938 | 1.493364082 | 0.00466913 | 0.016554343 | 95.0042075 | 141.9268635 | ENSG00000112041 |
| NA | 0.57846429 | 1.493258869 | 0.006852889 | 0.022999144 | 83.29127881 | 124.285722 | ENSG00000250986 |
| NA | 0.577887686 | 1.492662175 | 0.580342207 | 0.731378652 | 3.687573563 | 5.47087992 | ENSG00000270171 |
| PTGR2 | 0.57770981 | 1.49247815 | 0.334498455 | 0.507176742 | 9.694158961 | 14.47374404 | ENSG00000140043 |
| RAB3GAP2 | 0.577682697 | 1.492450101 | 1.26E-19 | 4.16E-18 | 1640.704294 | 2448.326635 | ENSG00000118873 |
| FAM162B | 0.577442961 | 1.492202118 | 0.100035385 | 0.207191605 | 28.79638945 | 42.90579582 | ENSG00000183807 |
| STON1 | 0.577377224 | 1.492134126 | 0.000145613 | 0.000754705 | 165.5189439 | 247.0261769 | ENSG00000243244 |
| AGMAT | 0.577144202 | 1.491893139 | 9.75E-05 | 0.000529027 | 213.6869811 | 319.3317064 | ENSG00000116771 |
| KIFC3 | 0.576972729 | 1.491715829 | 0.000118884 | 0.000631091 | 177.7791295 | 265.3639728 | ENSG00000140859 |
| HECA | 0.576788402 | 1.491525251 | 0.016633013 | 0.048178296 | 72.29801839 | 108.1131898 | ENSG00000112406 |
| GCNT2 | 0.57635073 | 1.491072834 | 2.12E-16 | 5.29E-15 | 997.0614299 | 1487.345318 | ENSG00000111846 |
| BCL2L13 | 0.576045427 | 1.490757326 | 4.09E-12 | 6.50E-11 | 785.7285158 | 1171.017517 | ENSG00000099968 |
| ST8SIA3 | 0.575685456 | 1.49038541 | 0.001404567 | 0.005782224 | 136.1553609 | 203.4159179 | ENSG00000177511 |
| LRRC9 | 0.575639568 | 1.490338005 | 0.23871523 | 0.400648643 | 16.78321866 | 24.98045073 | ENSG00000131951 |
| NA | 0.575590495 | 1.490287313 | 0.091479782 | 0.193120647 | 32.17969071 | 47.88945736 | ENSG00000254469 |

| | | | | | | | |
|---|---|---|---|---|---|---|---|
| ZBTB3 | 0.575337583 | 1.49002608 | 1.85E-09 | 2.18E-08 | 476.9128382 | 710.576695 | ENSG00000185670 |
| LHX1 | 0.574992001 | 1.489669203 | 0.431901086 | 0.60292379 | 7.026896093 | 10.44593917 | ENSG00000273706 |
| TRIM61 | 0.574569888 | 1.489233411 | 0.000366727 | 0.001743783 | 171.2293174 | 255.3557704 | ENSG00000183439 |
| SPDYE1 | 0.574556875 | 1.489219977 | 0.614094272 | 0.75677096 | 3.26041351 | 4.884183692 | ENSG00000136206 |
| USHBP1 | 0.574502657 | 1.489164012 | 0.001457004 | 0.005972603 | 116.416038 | 173.1426008 | ENSG00000130307 |
| NA | 0.574380634 | 1.489038065 | 0.624923308 | 0.763774412 | 3.744831081 | 5.540782776 | ENSG00000267892 |
| FAM160A1 | 0.574203546 | 1.488855299 | 5.18E-11 | 7.33E-10 | 631.2220436 | 939.9533275 | ENSG00000164142 |
| BRWD3 | 0.574168216 | 1.488818839 | 4.94E-12 | 7.81E-11 | 856.7654294 | 1276.189132 | ENSG00000165288 |
| NEK3 | 0.574162686 | 1.488813133 | 6.59E-08 | 6.22E-07 | 543.0145749 | 808.0441475 | ENSG00000136098 |
| COLGALT2 | 0.573935911 | 1.488579126 | 0.001986725 | 0.007845633 | 109.5870717 | 163.2330706 | ENSG00000198756 |
| CCNG2 | 0.573847891 | 1.488488309 | 9.85E-05 | 0.000533553 | 183.4325578 | 273.2114547 | ENSG00000138764 |
| LINC02026 | 0.573697942 | 1.48833361 | 0.623907261 | 0.763014769 | 2.653308353 | 3.967037349 | ENSG00000214146 |
| PARD6B | 0.573104611 | 1.487721634 | 0.003829286 | 0.01390901 | 126.4234808 | 188.5958457 | ENSG00000124171 |
| TRPM4 | 0.573089147 | 1.487705688 | 2.33E-05 | 0.000143724 | 254.5322354 | 379.1857086 | ENSG00000130529 |
| SLC46A1 | 0.572429337 | 1.487025448 | 0.060108932 | 0.138146165 | 41.78098613 | 62.14568404 | ENSG00000076351 |
| SOX10 | 0.572313368 | 1.486905921 | 0.000988394 | 0.00422119 | 125.4496643 | 186.6739247 | ENSG00000100146 |
| TMEM45B | 0.572083287 | 1.486668808 | 0.596652 | 0.74400866 | 3.728761385 | 5.500454914 | ENSG00000151715 |
| CNST | 0.571919898 | 1.486500448 | 3.14E-08 | 3.10E-07 | 429.9692621 | 639.4297644 | ENSG00000162852 |
| SMG7-AS1 | 0.571672655 | 1.48624572 | 0.274754792 | 0.441507263 | 14.25538879 | 21.21104255 | ENSG00000232860 |
| NDUFA6-DT | 0.571536648 | 1.486105613 | 0.164212206 | 0.302995473 | 21.08906099 | 31.36873746 | ENSG00000237037 |
| KDR | 0.571515503 | 1.486083833 | 3.32E-20 | 1.16E-18 | 3330.349489 | 4947.92359 | ENSG00000128052 |
| NT5DC1 | 0.571188207 | 1.485746731 | 4.03E-12 | 6.42E-11 | 673.142714 | 1000.743056 | ENSG00000178425 |
| LINC02035 | 0.570945721 | 1.48549703 | 0.210458872 | 0.365939568 | 17.44479043 | 25.85135053 | ENSG00000273033 |
| CENPE | 0.570828861 | 1.485376709 | 2.19E-12 | 3.61E-11 | 778.2237351 | 1155.966745 | ENSG00000138778 |
| NA | 0.570588471 | 1.485129227 | 0.181964688 | 0.327460118 | 20.33462568 | 30.23057822 | ENSG00000253981 |
| LINC01123 | 0.570484994 | 1.48502271 | 0.444498523 | 0.615255867 | 7.625628541 | 11.30985421 | ENSG00000204588 |
| NA | 0.57037102 | 1.484905396 | 0.054586391 | 0.128027485 | 43.05389343 | 63.93049511 | ENSG00000261655 |
| ELF1 | 0.570269596 | 1.484801009 | 0.00014541 | 0.000754261 | 198.8270027 | 295.2162305 | ENSG00000120690 |
| SNX27 | 0.570264811 | 1.484796084 | 9.82E-19 | 2.95E-17 | 1094.509013 | 1624.938141 | ENSG00000143376 |
| ZBTB20 | 0.568867409 | 1.4833586 | 0.435390839 | 0.606135187 | 7.680095156 | 11.36308551 | ENSG00000181722 |
| SCAMP1-AS1 | 0.568329856 | 1.482805998 | 0.026894217 | 0.071927955 | 60.98656805 | 90.36073308 | ENSG00000245556 |
| CDC73 | 0.568199657 | 1.482672185 | 2.94E-19 | 9.21E-18 | 1160.766956 | 1720.82413 | ENSG00000134371 |
| SPATA6L | 0.568133425 | 1.482604119 | 0.352974703 | 0.525931232 | 9.691368058 | 14.33715498 | ENSG00000106686 |
| ZMYM1 | 0.568011484 | 1.482478811 | 3.15E-05 | 0.000189235 | 220.2257176 | 326.8110939 | ENSG00000197056 |
| HEIH | 0.56782481 | 1.482287001 | 0.434460721 | 0.605532965 | 6.648620849 | 9.883983749 | ENSG00000278970 |
| NA | 0.567662294 | 1.482120034 | 0.302547932 | 0.473215828 | 15.06361499 | 22.40162748 | ENSG00000254365 |
| FILIP1 | 0.567635626 | 1.482092638 | 0.422265099 | 0.593855509 | 7.350704722 | 10.91674646 | ENSG00000118407 |
| CFLAR | 0.567579228 | 1.482034701 | 0.009924647 | 0.031235521 | 97.05862015 | 143.4027302 | ENSG00000003402 |
| RGS10 | 0.567052093 | 1.481493291 | 0.003311433 | 0.012218984 | 99.45478924 | 147.2724281 | ENSG00000148908 |
| HELB | 0.566680599 | 1.481111856 | 0.001281476 | 0.005334692 | 162.1127519 | 240.2546992 | ENSG00000127311 |
| MIR9-3HG | 0.566636321 | 1.481066399 | 0.367190043 | 0.541011466 | 9.79192858 | 14.45062077 | ENSG00000255571 |
| TPSG1 | 0.566566906 | 1.48099514 | 0.067692229 | 0.151913772 | 40.46833048 | 59.9669533 | ENSG00000116176 |
| AKAP3 | 0.566010067 | 1.480423629 | 0.339170022 | 0.511645988 | 9.955128266 | 14.77676974 | ENSG00000111254 |
| SPTLC3 | 0.565956323 | 1.48036848 | 0.51411825 | 0.678170569 | 4.694605466 | 6.958311403 | ENSG00000172296 |
| FUT1 | 0.565837599 | 1.480246661 | 7.29E-06 | 4.93E-05 | 250.2399842 | 370.3377468 | ENSG00000174951 |
| NA | 0.565687929 | 1.480093103 | 0.480063074 | 0.648175374 | 5.317780318 | 7.889978725 | ENSG00000232842 |
| RBMX2 | 0.565276705 | 1.479671279 | 6.10E-09 | 6.68E-08 | 478.0027341 | 706.991982 | ENSG00000134597 |
| INKA2-AS1 | 0.564952462 | 1.479338763 | 0.593195407 | 0.741562369 | 3.948542868 | 5.856178876 | ENSG00000227811 |
| PSEN2 | 0.564614601 | 1.478992361 | 5.34E-08 | 5.11E-07 | 432.2418535 | 638.7129308 | ENSG00000143801 |
| STRC | 0.564480541 | 1.478854934 | 0.621751229 | 0.762165785 | 2.683332564 | 3.971338496 | ENSG00000242866 |
| CRTC1 | 0.564423116 | 1.478796071 | 1.31E-07 | 1.19E-06 | 368.6294617 | 544.9976451 | ENSG00000105662 |
| TARS2 | 0.564326102 | 1.478696634 | 2.86E-17 | 7.71E-16 | 1222.747701 | 1807.382955 | ENSG00000143374 |
| NA | 0.564291701 | 1.478661374 | 0.442363833 | 0.613376392 | 7.285074495 | 10.79467837 | ENSG00000261812 |
| DENND4B | 0.564291579 | 1.478661249 | 1.70E-18 | 5.04E-17 | 1763.305323 | 2607.501424 | ENSG00000198837 |
| COL1A2 | 0.564144205 | 1.478510209 | 4.08E-24 | 1.88E-22 | 2350.516551 | 3475.303216 | ENSG00000164692 |
| FBXO32 | 0.563859772 | 1.478218744 | 0.440269183 | 0.611204314 | 8.082812805 | 11.94789163 | ENSG00000156804 |
| CDX4 | 0.563758769 | 1.478115257 | 0.352161027 | 0.524921833 | 11.13114182 | 16.39676171 | ENSG00000131264 |
| NA | 0.563506725 | 1.477857048 | 0.621333545 | 0.761915061 | 3.725970483 | 5.460660089 | ENSG00000231249 |
| HS1BP3 | 0.563393013 | 1.477740569 | 1.81E-08 | 1.86E-07 | 390.2627005 | 576.6446674 | ENSG00000118960 |
| NRAV | 0.563354919 | 1.47770155 | 0.001316832 | 0.005459491 | 124.4967356 | 183.7466348 | ENSG00000248008 |
| ST20 | 0.563326183 | 1.477672117 | 0.360202763 | 0.533288445 | 9.370350333 | 13.85747282 | ENSG00000180953 |
| TCEAL7 | 0.563228747 | 1.477572322 | 0.009683063 | 0.030607236 | 84.42311348 | 124.9138 | ENSG00000182916 |
| KLKB1 | 0.563156311 | 1.477498137 | 0.00098983 | 0.004225449 | 149.2314697 | 220.9102505 | ENSG00000164344 |
| NA | 0.563130085 | 1.477471278 | 0.617051407 | 0.759048006 | 3.388207339 | 5.00195063 | ENSG00000285257 |
| NA | 0.56305906 | 1.477398543 | 0.631385503 | 0.768762048 | 2.664471965 | 3.941763502 | ENSG00000240898 |
| PTER | 0.562425239 | 1.476749619 | 0.001496758 | 0.006113479 | 119.3936305 | 176.2352428 | ENSG00000165983 |
| TBX6 | 0.561970367 | 1.476284082 | 0.100106162 | 0.20731594 | 39.58331044 | 58.4405326 | ENSG00000149922 |
| RPS6KC1 | 0.561854203 | 1.476165218 | 2.34E-08 | 2.35E-07 | 475.2866105 | 701.0702135 | ENSG00000136643 |
| ZNF575 | 0.56159479 | 1.475899811 | 0.546628852 | 0.704645318 | 4.044197305 | 5.96534352 | ENSG00000176472 |
| FXYD1 | 0.561258309 | 1.475555626 | 0.64267086 | 0.776758449 | 2.710565871 | 3.994461769 | ENSG00000266964 |
| NA | 0.561118632 | 1.475412774 | 0.266546473 | 0.432732536 | 16.47404026 | 24.24641257 | ENSG00000268649 |
| PFDN2 | 0.561107144 | 1.475401026 | 3.95E-13 | 7.02E-12 | 1248.060544 | 1841.47166 | ENSG00000143256 |
| ARHGAP30 | 0.560947748 | 1.475238026 | 0.224310459 | 0.382867904 | 16.96037286 | 25.10681997 | ENSG00000186517 |
| NOCT | 0.560876924 | 1.475165606 | 1.94E-08 | 1.97E-07 | 403.5562754 | 595.5914772 | ENSG00000151014 |
| NA | 0.560141841 | 1.47441417 | 0.314885356 | 0.486150732 | 12.72853346 | 18.76316225 | ENSG00000249684 |
| ARHGAP5 | 0.560024319 | 1.474294068 | 5.19E-06 | 3.59E-05 | 408.2439845 | 602.0721902 | ENSG00000100852 |
| RGCC | 0.56001896 | 1.474288592 | 0.625003518 | 0.763774412 | 2.716147677 | 3.996612343 | ENSG00000102760 |

| | | | | | | | |
|---|---|---|---|---|---|---|---|
| GALNT6 | 0.559771301 | 1.474035531 | 7.71E-05 | 0.000427981 | 217.2002283 | 319.667839 | ENSG00000139629 |
| MEX3A | 0.559425228 | 1.473681983 | 3.10E-24 | 1.45E-22 | 2105.75895 | 3103.467817 | ENSG00000254726 |
| CHTF8 | 0.559189232 | 1.473440938 | 0.357508694 | 0.530760367 | 10.53520027 | 15.43283578 | ENSG00000168802 |
| LACTB | 0.558885178 | 1.473130436 | 1.76E-06 | 1.32E-05 | 293.3425622 | 432.159172 | ENSG00000103642 |
| NA | 0.558736434 | 1.472978562 | 0.005377614 | 0.018671627 | 92.60995343 | 136.3054066 | ENSG00000279088 |
| SUGCT | 0.558510625 | 1.472748032 | 0.049271187 | 0.117921892 | 53.24295785 | 78.42332174 | ENSG00000175600 |
| PPP1R3D | 0.558274182 | 1.472506683 | 0.018302312 | 0.052082889 | 78.89237279 | 116.2268956 | ENSG00000132825 |
| NA | 0.558094501 | 1.472323301 | 0.62417337 | 0.763058842 | 2.686123467 | 3.950365797 | ENSG00000231767 |
| NA | 0.557940636 | 1.472166284 | 0.52492056 | 0.687247537 | 4.700187272 | 6.912064858 | ENSG00000287935 |
| GALNT10 | 0.557798076 | 1.47202082 | 2.46E-10 | 3.24E-09 | 623.2344977 | 917.3691433 | ENSG00000164574 |
| LINC00639 | 0.557730731 | 1.471952107 | 0.548480833 | 0.705994779 | 4.046988208 | 5.946521394 | ENSG00000259070 |
| C1RL | 0.557655609 | 1.471875464 | 0.000447213 | 0.002079339 | 190.0865972 | 279.4569732 | ENSG00000139178 |
| PTH2 | 0.5576201 | 1.471839236 | 0.568159493 | 0.721408859 | 3.965288286 | 5.84112486 | ENSG00000142538 |
| AZIN2 | 0.557613122 | 1.471832118 | 0.195321312 | 0.345619467 | 18.75118856 | 27.57405543 | ENSG00000142920 |
| TBC1D2 | 0.557431564 | 1.471646904 | 0.000230203 | 0.001146914 | 163.405671 | 240.512587 | ENSG00000095383 |
| NA | 0.557062751 | 1.471270739 | 0.445858974 | 0.616326889 | 7.786237484 | 11.37975706 | ENSG00000272894 |
| NA | 0.55699317 | 1.471199781 | 0.60450632 | 0.749600272 | 3.317671028 | 4.882033118 | ENSG00000274528 |
| NA | 0.556916358 | 1.471121454 | 0.222434525 | 0.380946771 | 21.6097603 | 31.64622884 | ENSG00000266651 |
| SIRT4 | 0.556461813 | 1.470658025 | 0.143793386 | 0.273351275 | 27.20158855 | 40.11886311 | ENSG00000089163 |
| NA | 0.556445686 | 1.470641586 | 0.620331463 | 0.76139303 | 2.686123467 | 3.943914076 | ENSG00000248268 |
| TMEM168 | 0.556395559 | 1.470590488 | 4.08E-05 | 0.000239233 | 229.2390808 | 336.7642113 | ENSG00000146802 |
| LRRC75B | 0.556349995 | 1.470544044 | 0.000172401 | 0.000881006 | 206.8933455 | 304.456828 | ENSG00000178026 |
| NA | 0.556261575 | 1.470453921 | 0.601436973 | 0.74723518 | 3.344904336 | 4.913758686 | ENSG00000256824 |
| NA | 0.556088366 | 1.470277389 | 0.076077805 | 0.166519539 | 44.53533057 | 65.56447503 | ENSG00000269981 |
| UBR4 | 0.556012888 | 1.47020047 | 2.43E-05 | 0.000148858 | 4441.684948 | 6529.722924 | ENSG00000127481 |
| HS3ST4 | 0.555979206 | 1.470166146 | 0.00732548 | 0.0242769 | 90.52602904 | 133.0293959 | ENSG00000182601 |
| NA | 0.555708377 | 1.469890186 | 0.464488386 | 0.634087662 | 6.055470207 | 8.882946609 | ENSG00000272462 |
| PATL2 | 0.555560905 | 1.469739942 | 0.647443807 | 0.78051734 | 2.710565871 | 3.97348907 | ENSG00000229474 |
| MTX1 | 0.55535585 | 1.469531058 | 1.49E-08 | 1.55E-07 | 609.9487077 | 895.9782519 | ENSG00000173171 |
| KMT2C | 0.55532445 | 1.469499074 | 2.58E-06 | 1.89E-05 | 491.2805885 | 721.3516599 | ENSG00000055609 |
| NA | 0.555282161 | 1.469456 | 0.659446214 | 0.789485573 | 2.694496175 | 3.95251637 | ENSG00000233613 |
| DCXR | 0.554862909 | 1.469029033 | 6.70E-15 | 1.43E-13 | 963.1977174 | 1415.311695 | ENSG00000169738 |
| PRMT8 | 0.554490904 | 1.468650287 | 0.241343598 | 0.403690223 | 18.21038935 | 26.68648408 | ENSG00000111218 |
| MICOS13 | 0.553922266 | 1.468071532 | 7.71E-06 | 5.19E-05 | 294.5916276 | 432.3686143 | ENSG00000174917 |
| CACNA1C | 0.553824383 | 1.467971931 | 0.022261077 | 0.061400978 | 63.40614041 | 92.95165416 | ENSG00000151067 |
| CCSAP | 0.553667157 | 1.467811959 | 3.00E-06 | 2.16E-05 | 416.781442 | 611.9147979 | ENSG00000154429 |
| B3GNT4 | 0.55363821 | 1.467782509 | 0.174181757 | 0.316985609 | 22.78980405 | 33.38613833 | ENSG00000176383 |
| EFHB | 0.553527525 | 1.467669904 | 0.494767347 | 0.661163542 | 5.377828739 | 7.873307173 | ENSG00000163576 |
| NA | 0.55350245 | 1.467644395 | 0.575012627 | 0.727190809 | 4.340772627 | 6.413560574 | ENSG00000271387 |
| RAMP2 | 0.55332287 | 1.46746172 | 0.011142997 | 0.034553859 | 79.27479038 | 116.4258513 | ENSG00000131477 |
| STX16 | 0.552663842 | 1.466791531 | 9.82E-10 | 1.19E-08 | 967.8957994 | 1419.317345 | ENSG00000124222 |
| ATP2C2 | 0.552598684 | 1.466725287 | 0.017642006 | 0.050651174 | 67.90923684 | 99.70939233 | ENSG00000064270 |
| NOXA1 | 0.5525795 | 1.466705783 | 0.131439062 | 0.254858875 | 29.6653398 | 43.39434074 | ENSG00000188747 |
| NA | 0.551896694 | 1.466011777 | 0.297471562 | 0.467725479 | 15.84807451 | 23.22413016 | ENSG00000188383 |
| NA | 0.551743872 | 1.465856495 | 0.642943549 | 0.776937109 | 2.922650367 | 4.341583437 | ENSG00000281100 |
| RC3H1 | 0.551736384 | 1.465848886 | 0.000114421 | 0.00061093 | 212.3110475 | 311.0177 | ENSG00000135870 |
| KLK14 | 0.551669318 | 1.465780746 | 0.371804839 | 0.54518753 | 9.571271216 | 14.0825264 | ENSG00000129437 |
| GLMP | 0.551594794 | 1.465705031 | 1.16E-07 | 1.06E-06 | 367.6028055 | 538.7558947 | ENSG00000198715 |
| COP1 | 0.551534121 | 1.465643391 | 6.85E-13 | 1.19E-11 | 1046.870887 | 1534.62598 | ENSG00000143207 |
| LHX6 | 0.551383124 | 1.465490001 | 0.069557254 | 0.155322257 | 44.03070096 | 64.7376712 | ENSG00000106852 |
| NA | 0.551159537 | 1.465262899 | 0.527802247 | 0.689499096 | 4.730211482 | 6.905613136 | ENSG00000287262 |
| CCDC110 | 0.551139067 | 1.465242109 | 0.315619923 | 0.48684348 | 13.36498711 | 19.54318649 | ENSG00000168491 |
| NA | 0.551025955 | 1.465127233 | 0.595927253 | 0.743598489 | 3.336531627 | 4.877731971 | ENSG00000251485 |
| GALE | 0.550960622 | 1.465060886 | 1.04E-07 | 9.53E-07 | 521.9851507 | 764.0615357 | ENSG00000117308 |
| NBEAL1 | 0.550951112 | 1.465051229 | 0.039877583 | 0.099377141 | 53.78877528 | 78.578212 | ENSG00000144426 |
| ZKSCAN7-AS1 | 0.550942542 | 1.465042525 | 0.591440713 | 0.740202594 | 3.342113433 | 4.884183692 | ENSG00000236869 |
| NA | 0.550908924 | 1.465008387 | 0.150140421 | 0.282571597 | 24.72774974 | 36.26744213 | ENSG00000249456 |
| NA | 0.550827935 | 1.464926148 | 0.312257769 | 0.483254282 | 12.61401843 | 18.57145177 | ENSG00000237753 |
| LINC00926 | 0.550799084 | 1.464896852 | 0.289435331 | 0.458867272 | 14.39925232 | 21.09945475 | ENSG00000247982 |
| NA | 0.550676509 | 1.464772397 | 0.174839774 | 0.317802136 | 27.38173382 | 40.039013 | ENSG00000203635 |
| NA | 0.550646079 | 1.464741501 | 0.618229038 | 0.759924581 | 3.695946272 | 5.448017111 | ENSG00000248161 |
| GPSM1 | 0.550269637 | 1.464359356 | 5.35E-11 | 7.55E-10 | 599.2176432 | 877.6445088 | ENSG00000160360 |
| HMMR | 0.550140844 | 1.464228635 | 2.75E-12 | 4.47E-11 | 848.2007017 | 1242.387084 | ENSG00000072571 |
| NA | 0.550110105 | 1.464197437 | 0.642482791 | 0.776672235 | 2.664471965 | 3.893366382 | ENSG00000253541 |
| HSPB6 | 0.550094737 | 1.464181841 | 0.413209314 | 0.585516445 | 8.237639782 | 12.11164465 | ENSG00000004776 |
| FZD7 | 0.550087491 | 1.464174487 | 9.01E-22 | 3.46E-20 | 6674.939155 | 9774.374886 | ENSG00000155760 |
| PCYOX1L | 0.549742838 | 1.463824745 | 4.05E-09 | 4.56E-08 | 482.7929059 | 706.1676135 | ENSG00000145882 |
| NIPSNAP1 | 0.549322641 | 1.463398456 | 7.35E-27 | 3.96E-25 | 3709.957609 | 5428.76402 | ENSG00000184117 |
| TCAF1 | 0.549295203 | 1.463370623 | 1.45E-06 | 1.11E-05 | 365.7131058 | 534.7186514 | ENSG00000198420 |
| NA | 0.549224452 | 1.463298861 | 0.184302132 | 0.330424947 | 21.91315673 | 32.12348786 | ENSG00000267023 |
| SLC37A2 | 0.549159696 | 1.463233181 | 0.051153386 | 0.121542059 | 48.42681608 | 70.8538643 | ENSG00000134955 |
| NA | 0.549089846 | 1.463162338 | 0.568956791 | 0.722088517 | 4.049779111 | 5.902425422 | ENSG00000287202 |
| NA | 0.548994543 | 1.463065687 | 0.631462201 | 0.768793772 | 2.691705273 | 3.922941376 | ENSG00000238260 |
| CCDC87 | 0.548819118 | 1.462887796 | 0.505556602 | 0.670442219 | 5.375037836 | 7.835129884 | ENSG00000182791 |
| IL18BP | 0.548125871 | 1.462185015 | 0.002150091 | 0.008411561 | 116.8427977 | 170.8377734 | ENSG00000137496 |
| LOC100506207 | 0.547843044 | 1.461898394 | 0.520009314 | 0.683268285 | 5.551516316 | 8.167730574 | ENSG00000285219 |
| ITGAV | 0.546843615 | 1.460886014 | 1.54E-11 | 2.31E-10 | 759.2542925 | 1108.885845 | ENSG00000138448 |

| | | | | | | | |
|---|---|---|---|---|---|---|---|
| CHML | 0.546727493 | 1.460768432 | 4.92E-11 | 6.97E-10 | 1095.342471 | 1600.848429 | ENSG00000203668 |
| FAM49A | 0.546667386 | 1.460707573 | 0.000785029 | 0.003414656 | 150.6070016 | 219.8336523 | ENSG00000197872 |
| NA | 0.546617066 | 1.460656626 | 0.396575713 | 0.569264237 | 10.00401308 | 14.57242839 | ENSG00000223749 |
| TNNT2 | 0.546482618 | 1.46052051 | 0.455212526 | 0.62492918 | 6.790369193 | 9.86301105 | ENSG00000118194 |
| NDUFB1 | 0.546476696 | 1.460514514 | 3.17E-08 | 3.12E-07 | 447.450259 | 653.2138204 | ENSG00000183648 |
| ACADVL | 0.546444752 | 1.460482177 | 2.31E-23 | 1.00E-21 | 2596.103256 | 3790.260153 | ENSG00000072778 |
| CTAGE7P | 0.546368702 | 1.460405191 | 0.459823311 | 0.629627933 | 8.022764384 | 11.68278276 | ENSG00000233122 |
| FAM72A | 0.546361782 | 1.460398186 | 0.00653947 | 0.022090699 | 102.4724183 | 149.5871965 | ENSG00000196550 |
| NA | 0.546267621 | 1.460302873 | 0.595494317 | 0.743298665 | 3.393789145 | 4.928279664 | ENSG00000258626 |
| SLC16A7 | 0.546010694 | 1.460042833 | 0.075595459 | 0.165811483 | 39.54558924 | 57.87400346 | ENSG00000118596 |
| RPRD2 | 0.545902257 | 1.459933097 | 4.99E-12 | 7.88E-11 | 802.5207701 | 1171.521989 | ENSG00000163125 |
| B4GALT6 | 0.545853869 | 1.459884131 | 4.43E-06 | 3.11E-05 | 413.1691348 | 603.4429266 | ENSG00000118276 |
| EGLN1 | 0.545817462 | 1.459847291 | 0.007826051 | 0.025658116 | 91.27767344 | 133.1917919 | ENSG00000135766 |
| GFY | 0.545622395 | 1.459649918 | 0.160731614 | 0.297969783 | 27.97353302 | 40.82144828 | ENSG00000261949 |
| RNPC3 | 0.545569711 | 1.459596616 | 0.001987959 | 0.007848627 | 134.9605237 | 197.0015333 | ENSG00000185946 |
| BASP1-AS1 | 0.545302775 | 1.459326577 | 0.504930835 | 0.670016685 | 5.633216238 | 8.258073092 | ENSG00000215196 |
| OCEL1 | 0.545004894 | 1.459025294 | 2.30E-05 | 0.000141926 | 231.3376354 | 337.3280326 | ENSG00000099330 |
| EME1 | 0.54483028 | 1.458848714 | 0.000718391 | 0.003156111 | 157.3776343 | 229.292827 | ENSG00000154920 |
| NA | 0.544797388 | 1.458815454 | 0.102865504 | 0.211621887 | 32.13850289 | 46.80991507 | ENSG00000272977 |
| TMEM51-AS1 | 0.544507171 | 1.458522024 | 0.417044307 | 0.589133108 | 8.393342641 | 12.24823371 | ENSG00000175147 |
| NA | 0.54443465 | 1.458448709 | 0.439667961 | 0.610760757 | 7.668931545 | 11.2031006 | ENSG00000187952 |
| NA | 0.544215598 | 1.458227281 | 0.382810144 | 0.556125499 | 9.606877233 | 14.02767756 | ENSG00000231064 |
| NA | 0.544195324 | 1.45820679 | 0.552490272 | 0.709073903 | 4.697396369 | 6.821189302 | ENSG00000259562 |
| PSMB4 | 0.544170659 | 1.45818186 | 4.80E-33 | 3.62E-31 | 5757.458208 | 8394.728166 | ENSG00000159377 |
| MIA3 | 0.544096092 | 1.458106494 | 1.58E-14 | 3.23E-13 | 1011.143695 | 1475.002751 | ENSG00000154305 |
| MSTO1 | 0.544083738 | 1.458094008 | 1.36E-07 | 1.23E-06 | 404.8885923 | 590.2184455 | ENSG00000125459 |
| ANKRD37 | 0.5440588 | 1.458068804 | 0.295492144 | 0.465561823 | 14.0893982 | 20.57190851 | ENSG00000186352 |
| WSCD2 | 0.543902866 | 1.457911217 | 0.047140452 | 0.113986771 | 50.61265244 | 73.64563119 | ENSG00000075035 |
| SLC32A1 | 0.543781693 | 1.457788772 | 0.462872202 | 0.632628496 | 6.78757829 | 9.835586629 | ENSG00000101438 |
| CLDN11 | 0.543707406 | 1.457713709 | 0.007927897 | 0.025912751 | 86.77282525 | 126.5959166 | ENSG00000013297 |
| C8G | 0.543242775 | 1.457244316 | 0.486871438 | 0.654258193 | 6.387651545 | 9.322561368 | ENSG00000176919 |
| CD86 | 0.543191219 | 1.457192242 | 0.638394433 | 0.773863705 | 2.746171888 | 3.971338496 | ENSG00000114013 |
| ZNF236 | 0.542662569 | 1.456658377 | 3.61E-05 | 0.000214 | 230.8598388 | 336.3173392 | ENSG00000130856 |
| NA | 0.542619143 | 1.456614531 | 0.207635911 | 0.362390696 | 20.63187672 | 30.05581186 | ENSG00000259605 |
| PLCG2 | 0.542553009 | 1.456547761 | 5.17E-06 | 3.58E-05 | 280.7887924 | 409.0834441 | ENSG00000197943 |
| CCDC50 | 0.542509391 | 1.456503725 | 6.61E-07 | 5.35E-06 | 407.2454006 | 593.5750998 | ENSG00000152492 |
| CXCL14 | 0.542386223 | 1.456379383 | 0.476715454 | 0.644993534 | 8.045091607 | 11.76478345 | ENSG00000145824 |
| NA | 0.542329387 | 1.45632201 | 0.586300312 | 0.735915815 | 4.667372158 | 6.775475794 | ENSG00000277475 |
| SYN1 | 0.542084449 | 1.456074779 | 0.006949742 | 0.023260137 | 89.70875441 | 130.7288575 | ENSG00000008056 |
| NA | 0.54172043 | 1.45570743 | 0.624797646 | 0.763716347 | 4.409193756 | 6.430232126 | ENSG00000240291 |
| CRYZL1 | 0.541537249 | 1.455522608 | 0.000114968 | 0.000613175 | 222.186503 | 323.304644 | ENSG00000205758 |
| LINC00634 | 0.541289886 | 1.455273068 | 0.473803266 | 0.642224481 | 6.382069739 | 9.299438095 | ENSG00000205704 |
| CDH2 | 0.541286223 | 1.455269373 | 7.24E-09 | 7.86E-08 | 580.3502247 | 844.9218811 | ENSG00000170558 |
| MRC1 | 0.541255014 | 1.455237892 | 0.636185609 | 0.772253749 | 2.713356774 | 3.920790803 | ENSG00000260314 |
| CEP83-DT | 0.541249622 | 1.455232453 | 0.319860847 | 0.491386438 | 13.5891111 | 19.65450171 | ENSG00000278916 |
| NA | 0.541241193 | 1.455223951 | 0.05639172 | 0.131414916 | 47.27524493 | 68.85877478 | ENSG00000272734 |
| NA | 0.541147714 | 1.455129664 | 0.415671875 | 0.58779696 | 8.396133544 | 12.21274004 | ENSG00000224738 |
| LIFR | 0.540836283 | 1.454815581 | 7.01E-05 | 0.000391387 | 207.2915574 | 301.6531932 | ENSG00000113594 |
| AKAP1 | 0.540717317 | 1.454695621 | 2.12E-26 | 1.12E-24 | 2329.810484 | 3388.418848 | ENSG00000121057 |
| LIN9 | 0.540700356 | 1.454678519 | 1.11E-05 | 7.23E-05 | 299.559117 | 436.0506177 | ENSG00000183814 |
| NA | 0.540475023 | 1.454451332 | 1.61E-05 | 0.000101538 | 268.8436793 | 390.4767833 | ENSG00000282048 |
| ANGEL2 | 0.540129241 | 1.454102775 | 8.53E-10 | 1.04E-08 | 625.8534776 | 910.8703591 | ENSG00000174606 |
| REV3L | 0.540020529 | 1.453993207 | 0.007310497 | 0.024239739 | 93.99382112 | 136.6232137 | ENSG00000009413 |
| NA | 0.540001604 | 1.453974134 | 0.566136878 | 0.72000935 | 4.046988208 | 5.845426008 | ENSG00000225579 |
| NA | 0.539953383 | 1.453925536 | 0.477104968 | 0.645339499 | 6.37369703 | 9.276314822 | ENSG00000261098 |
| UBE2W | 0.539527486 | 1.453496388 | 0.001808764 | 0.007230184 | 153.7822117 | 223.9103995 | ENSG00000104343 |
| NA | 0.539411349 | 1.453379386 | 0.542993413 | 0.701682367 | 4.958365673 | 7.236063252 | ENSG00000285872 |
| LYRM1 | 0.539268852 | 1.45323584 | 3.53E-05 | 0.000209792 | 236.7268279 | 343.6738409 | ENSG00000102897 |
| PRKAB2 | 0.539072581 | 1.453038149 | 9.34E-09 | 1.00E-07 | 729.0624572 | 1058.757398 | ENSG00000131791 |
| FADD | 0.538674843 | 1.452637614 | 2.34E-08 | 2.35E-07 | 419.8776039 | 609.9692326 | ENSG00000168040 |
| TNS1 | 0.538587041 | 1.452549211 | 0.000344218 | 0.00164731 | 166.701215 | 242.104882 | ENSG00000079308 |
| KCNK2 | 0.538373097 | 1.452333821 | 0.159866412 | 0.296907957 | 26.58977747 | 38.55802113 | ENSG00000082482 |
| NA | 0.538297745 | 1.452257967 | 0.373841061 | 0.547183931 | 10.42347614 | 15.10453624 | ENSG00000164845 |
| TEX22 | 0.53828272 | 1.452242843 | 0.329429001 | 0.501738107 | 16.07535282 | 23.41934828 | ENSG00000226174 |
| GOLPH3L | 0.537952955 | 1.451910934 | 1.51E-11 | 2.26E-10 | 820.2099094 | 1190.35423 | ENSG00000143457 |
| PLPPR4 | 0.537687452 | 1.451643759 | 0.082914895 | 0.178656455 | 42.29542792 | 61.38233135 | ENSG00000117600 |
| KCNH8 | 0.537669449 | 1.451625644 | 0.093421543 | 0.196231583 | 41.08468423 | 59.40554303 | ENSG00000183960 |
| SLC26A1 | 0.537574607 | 1.451530218 | 0.332019707 | 0.50469379 | 15.60875671 | 22.53230997 | ENSG00000145217 |
| CCDC183 | 0.536689492 | 1.450639956 | 0.375954464 | 0.54935143 | 9.773067981 | 14.13737524 | ENSG00000213213 |
| NR5A2 | 0.536327512 | 1.450276028 | 0.000372205 | 0.001767218 | 167.1594384 | 242.6345788 | ENSG00000116833 |
| NA | 0.53610224 | 1.450049591 | 0.452423043 | 0.622561732 | 7.927785668 | 11.55614095 | ENSG00000213700 |
| BTN3A1 | 0.535893948 | 1.449840252 | 0.017644284 | 0.050651174 | 77.42009621 | 112.1893918 | ENSG00000026950 |
| NA | 0.535563893 | 1.4495086 | 0.03508695 | 0.089354765 | 89.6462785 | 130.2295413 | ENSG00000248131 |
| LYL1 | 0.535497433 | 1.449441828 | 0.332165441 | 0.504789943 | 13.28396291 | 19.26758521 | ENSG00000104903 |
| CCDC33 | 0.535329526 | 1.449273146 | 0.268599472 | 0.434766942 | 17.82027477 | 25.82633715 | ENSG00000140481 |
| UNC93B1 | 0.535313775 | 1.449257323 | 6.54E-06 | 4.46E-05 | 437.885772 | 634.8513205 | ENSG00000110057 |
| TPBGL | 0.535138599 | 1.44908136 | 0.232601236 | 0.393337074 | 19.52360858 | 28.43661339 | ENSG00000261594 |

| | | | | | | | |
|---|---|---|---|---|---|---|---|
| MKNK1 | 0.534992454 | 1.448934576 | 0.000588894 | 0.002648717 | 204.161165 | 295.8093542 | ENSG00000079277 |
| USP21 | 0.534966647 | 1.448908657 | 1.18E-08 | 1.24E-07 | 542.3794117 | 785.6758874 | ENSG00000143258 |
| ILDR2 | 0.534608827 | 1.448549341 | 0.16103445 | 0.298341927 | 25.12488559 | 36.32633165 | ENSG00000143195 |
| NA | 0.534422123 | 1.448361891 | 0.497738167 | 0.664121384 | 6.621387542 | 9.65086091 | ENSG00000239911 |
| TJP3 | 0.534400316 | 1.448339999 | 1.63E-06 | 1.24E-05 | 354.7015483 | 513.6804972 | ENSG00000105289 |
| NA | 0.534350267 | 1.448289755 | 0.489538363 | 0.656003026 | 6.382069739 | 9.238137533 | ENSG00000273343 |
| DESI2 | 0.534076345 | 1.448014797 | 9.17E-21 | 3.29E-19 | 2768.366201 | 4009.329282 | ENSG00000121644 |
| LTBP4 | 0.533635651 | 1.447572546 | 3.30E-21 | 1.22E-19 | 1799.342258 | 2605.068706 | ENSG00000090006 |
| BAMBI | 0.533593911 | 1.447530665 | 3.64E-10 | 4.69E-09 | 620.9378883 | 899.1867586 | ENSG00000095739 |
| TK2 | 0.533459742 | 1.447396053 | 0.076648575 | 0.167549876 | 40.28694592 | 58.24639898 | ENSG00000166548 |
| MST1R | 0.533246156 | 1.447181786 | 0.058440017 | 0.135094831 | 56.56497138 | 81.7434468 | ENSG00000164078 |
| MROH5 | 0.533171107 | 1.447106505 | 0.588199762 | 0.737773837 | 3.660340256 | 5.325949025 | ENSG00000226807 |
| AMZ1 | 0.532759655 | 1.446693855 | 0.073079181 | 0.161530855 | 41.6664711 | 60.29929352 | ENSG00000174945 |
| TMEM141 | 0.532603301 | 1.446537076 | 0.032950205 | 0.08475232 | 61.14610095 | 88.60738711 | ENSG00000244187 |
| UNC13D | 0.532335458 | 1.446268544 | 0.000734152 | 0.003217294 | 161.2233893 | 233.2926623 | ENSG00000092929 |
| LOC105372233 | 0.532310753 | 1.446243778 | 0.499089011 | 0.665399753 | 5.677194963 | 8.21397712 | ENSG00000267751 |
| NA | 0.532047118 | 1.445979519 | 0.562933757 | 0.717512725 | 4.942295977 | 7.16454286 | ENSG00000268496 |
| CACNA1B | 0.53181017 | 1.445742051 | 3.02E-07 | 2.58E-06 | 380.6909537 | 550.0133048 | ENSG00000148408 |
| SLC16A12 | 0.531540648 | 1.445471985 | 0.020902448 | 0.058260511 | 76.15178271 | 110.2247013 | ENSG00000152779 |
| TNNC2 | 0.53129906 | 1.445229952 | 0.040514346 | 0.100694695 | 54.32466841 | 78.38273342 | ENSG00000101470 |
| CHTOP | 0.531056359 | 1.444986845 | 2.64E-19 | 8.30E-18 | 1762.150823 | 2545.921948 | ENSG00000160679 |
| TRIM52 | 0.531040435 | 1.444970896 | 0.016571578 | 0.048021978 | 89.51237619 | 129.0679862 | ENSG00000183718 |
| RNF170 | 0.531001794 | 1.444932194 | 0.000535832 | 0.002439061 | 205.2941268 | 296.7714327 | ENSG00000120925 |
| LYPD5 | 0.530947265 | 1.444877582 | 0.477951414 | 0.646167266 | 6.37369703 | 9.196192135 | ENSG00000159871 |
| ZNF701 | 0.530871121 | 1.444801325 | 0.02194823 | 0.060650809 | 75.89024982 | 109.6132643 | ENSG00000167562 |
| TP53BP2 | 0.530843504 | 1.444773668 | 2.54E-14 | 5.16E-13 | 1188.278747 | 1715.968214 | ENSG00000143514 |
| PPP1R35 | 0.530745325 | 1.44467535 | 0.000257886 | 0.001270076 | 200.0296491 | 289.4657271 | ENSG00000160813 |
| PRKCQ-AS1 | 0.530644965 | 1.444574856 | 2.36E-05 | 0.000145109 | 261.1689658 | 376.9403527 | ENSG00000237943 |
| CHCHD10 | 0.530599357 | 1.44452919 | 0.030858678 | 0.080347517 | 557.003353 | 804.7671129 | ENSG00000250479 |
| CA5B | 0.53044679 | 1.444376437 | 0.008762972 | 0.02818851 | 98.69911463 | 142.3893105 | ENSG00000169239 |
| NA | 0.530094771 | 1.444024051 | 0.323511446 | 0.495149104 | 13.60709581 | 19.69214596 | ENSG00000271643 |
| NAV1 | 0.52855788 | 1.442486563 | 1.37E-16 | 3.48E-15 | 1252.454768 | 1806.051982 | ENSG00000134369 |
| CGREF1 | 0.528453312 | 1.442382014 | 0.039605687 | 0.098879359 | 59.3186401 | 85.39294958 | ENSG00000138028 |
| NA | 0.528399981 | 1.442328695 | 0.660879252 | 0.790735077 | 2.922650367 | 4.249090345 | ENSG00000259250 |
| NA | 0.528363673 | 1.442292397 | 0.47068214 | 0.639611804 | 7.048547595 | 10.13861232 | ENSG00000280434 |
| HAS2 | 0.528271767 | 1.44220052 | 0.006908097 | 0.023148185 | 95.39173134 | 137.4516351 | ENSG00000170961 |
| PEG10 | 0.528014595 | 1.441943459 | 1.68E-22 | 6.81E-21 | 2768.31379 | 3993.034156 | ENSG00000242265 |
| HAX1 | 0.528013779 | 1.441942643 | 7.58E-17 | 1.97E-15 | 1292.478178 | 1862.792921 | ENSG00000143575 |
| NAIP | 0.52794075 | 1.441869655 | 0.547832923 | 0.705490314 | 4.961156576 | 7.162392286 | ENSG00000249437 |
| KLK10 | 0.527580457 | 1.441509612 | 0.036180757 | 0.091680414 | 61.52804298 | 88.67622389 | ENSG00000129451 |
| KCNF1 | 0.527511215 | 1.441440429 | 0.058214072 | 0.134727659 | 47.2649572 | 68.08198561 | ENSG00000162975 |
| MCTP2 | 0.52725273 | 1.441182191 | 0.066777102 | 0.150262111 | 50.26440141 | 72.31146032 | ENSG00000140563 |
| DBT | 0.527219546 | 1.441149043 | 3.24E-09 | 3.69E-08 | 587.9135775 | 847.7548183 | ENSG00000137992 |
| C21orf91 | 0.527053012 | 1.440982697 | 1.01E-05 | 6.65E-05 | 338.9068644 | 488.5061791 | ENSG00000154642 |
| METTL18 | 0.52686117 | 1.440791095 | 0.018605967 | 0.052845658 | 77.61667459 | 112.0568313 | ENSG00000171806 |
| KDM4B | 0.526414502 | 1.440345086 | 3.72E-16 | 9.05E-15 | 1104.460163 | 1590.894089 | ENSG00000127663 |
| VMAC | 0.525966711 | 1.439898093 | 0.29178997 | 0.461500395 | 14.99519386 | 21.58988978 | ENSG00000187650 |
| LYPD1 | 0.525707688 | 1.439639596 | 0.218457147 | 0.376172171 | 20.12321691 | 28.94533751 | ENSG00000150551 |
| GMFG | 0.525596087 | 1.439528236 | 0.534243958 | 0.694668387 | 4.988389884 | 7.181214412 | ENSG00000130755 |
| ENDOG | 0.525197881 | 1.439130958 | 0.002493596 | 0.009577505 | 178.3097162 | 256.7009909 | ENSG00000167136 |
| NA | 0.525145004 | 1.439078213 | 0.586421608 | 0.735974187 | 4.585672235 | 6.663888003 | ENSG00000268938 |
| NA | 0.525013088 | 1.438946634 | 0.397561103 | 0.570131354 | 11.783465 | 17.05659587 | ENSG00000283877 |
| NA | 0.524615871 | 1.438550504 | 0.277116112 | 0.444339367 | 22.98003689 | 33.21808416 | ENSG00000259877 |
| COL6A1 | 0.524585282 | 1.438520003 | 1.14E-38 | 1.21E-36 | 6768.100873 | 9734.990083 | ENSG00000142156 |
| BLVRB | 0.52419205 | 1.438127962 | 3.05E-06 | 2.20E-05 | 387.5556013 | 557.7612604 | ENSG00000090013 |
| NA | 0.523339206 | 1.437278069 | 0.52350897 | 0.686069036 | 6.744275287 | 9.70947786 | ENSG00000258433 |
| PUS7L | 0.523241552 | 1.437180785 | 0.01207163 | 0.036801783 | 106.9494327 | 153.8777113 | ENSG00000129317 |
| SEMA3E | 0.523109005 | 1.437048751 | 0.000242233 | 0.001200954 | 226.9636843 | 325.6917689 | ENSG00000170381 |
| RHBDF1 | 0.523064972 | 1.437004891 | 3.39E-09 | 3.86E-08 | 576.5829784 | 828.3810978 | ENSG00000007384 |
| DECR1 | 0.522915015 | 1.436855533 | 6.89E-06 | 4.68E-05 | 407.6202092 | 586.2021628 | ENSG00000104325 |
| CAP2 | 0.522777185 | 1.436718267 | 3.61E-06 | 2.57E-05 | 309.565082 | 444.4942406 | ENSG00000112186 |
| PLXNA2 | 0.52264532 | 1.436586955 | 0.046563269 | 0.112803217 | 65.22167391 | 93.88304891 | ENSG00000076356 |
| PLSCR1 | 0.522617144 | 1.436558899 | 1.19E-09 | 1.44E-08 | 575.1424537 | 826.0046953 | ENSG00000188313 |
| USP32 | 0.5225837 | 1.436525597 | 6.13E-21 | 2.23E-19 | 1661.966741 | 2387.491496 | ENSG00000170832 |
| MIR5587 | 0.522535749 | 1.436477852 | 0.520161018 | 0.683357817 | 5.728870676 | 8.203224251 | ENSG00000266124 |
| LINC01970 | 0.52240761 | 1.436350271 | 0.464864275 | 0.634475556 | 7.315098706 | 10.53198054 | ENSG00000265692 |
| LAD1 | 0.521765173 | 1.435710801 | 4.30E-09 | 4.83E-08 | 507.3953405 | 728.42431 | ENSG00000159166 |
| NA | 0.521538087 | 1.435484833 | 0.509971209 | 0.674379942 | 5.946536977 | 8.58422206 | ENSG00000276997 |
| TMEM42 | 0.521403399 | 1.435350824 | 0.002694554 | 0.01022637 | 139.9218804 | 200.4448352 | ENSG00000169964 |
| NA | 0.521117336 | 1.435066246 | 0.320759513 | 0.492186746 | 14.81871538 | 21.27637168 | ENSG00000177855 |
| CASP10 | 0.520887186 | 1.434837331 | 0.077482172 | 0.168969995 | 45.4308385 | 65.25525807 | ENSG00000003400 |
| MYEOV | 0.520715802 | 1.434666891 | 0.337426727 | 0.510173133 | 12.85142121 | 18.3482883 | ENSG00000172927 |
| SEC14L4 | 0.520690863 | 1.43464209 | 0.110824548 | 0.223959392 | 34.47079351 | 49.49197218 | ENSG00000133488 |
| C15orf56 | 0.520561542 | 1.434513497 | 0.602455741 | 0.748190409 | 3.638688754 | 5.233455934 | ENSG00000176753 |
| ZDHHC24 | 0.520492078 | 1.434444428 | 0.001506595 | 0.006152355 | 167.0630202 | 239.6247673 | ENSG00000174165 |
| KLHL20 | 0.520488377 | 1.434440749 | 6.00E-05 | 0.000340065 | 254.7583113 | 365.4540904 | ENSG00000076321 |
| ZNF469 | 0.520381141 | 1.43433413 | 0.049019223 | 0.117469545 | 56.29878369 | 80.85453049 | ENSG00000225614 |

| | | | | | | | |
|---|---|---|---|---|---|---|---|
| NA | 0.52027514 | 1.434228747 | 0.130748513 | 0.253877157 | 38.34341842 | 54.80449683 | ENSG00000272455 |
| NA | 0.520259051 | 1.434212753 | 0.412677551 | 0.585020771 | 11.626723 | 16.60487117 | ENSG00000274525 |
| MB | 0.519731851 | 1.433688749 | 0.426850898 | 0.59834186 | 9.952337363 | 14.35838814 | ENSG00000198125 |
| KLC3 | 0.519706561 | 1.433663616 | 0.026604007 | 0.07126047 | 86.54523464 | 124.1464309 | ENSG00000104892 |
| TRAPPC6A | 0.519447469 | 1.433406169 | 2.22E-09 | 2.58E-08 | 687.7366297 | 986.1010032 | ENSG00000007255 |
| SIX2 | 0.51941463 | 1.433373543 | 0.402016201 | 0.574301425 | 13.51702318 | 19.38079054 | ENSG00000170577 |
| TMEM267 | 0.519141574 | 1.433102277 | 0.021031559 | 0.058570782 | 76.50995805 | 109.6428393 | ENSG00000151881 |
| LINC00339 | 0.519137468 | 1.433098197 | 0.241095007 | 0.403358803 | 20.63486779 | 29.41641737 | ENSG00000218510 |
| ZNF197 | 0.519119184 | 1.433080035 | 7.36E-06 | 4.97E-05 | 321.2502649 | 460.7880168 | ENSG00000186448 |
| CCNA1 | 0.518795318 | 1.432758364 | 0.143877557 | 0.273390153 | 30.60953237 | 43.78555838 | ENSG00000133101 |
| NA | 0.518691418 | 1.432655183 | 0.244231223 | 0.40730705 | 21.40449691 | 30.58980983 | ENSG00000271533 |
| RBP5 | 0.518500751 | 1.432465855 | 0.217080052 | 0.374502887 | 21.50852405 | 30.70380865 | ENSG00000139194 |
| PTGES | 0.518130231 | 1.43209801 | 0.025112265 | 0.067829838 | 76.24185534 | 109.4169801 | ENSG00000148344 |
| ZNF285 | 0.518113543 | 1.432081444 | 0.121369044 | 0.24048174 | 39.10343554 | 55.95608044 | ENSG00000267508 |
| ITSN2 | 0.518013307 | 1.431981949 | 0.000216254 | 0.001081325 | 235.607196 | 337.510032 | ENSG00000198399 |
| FAM151A | 0.517719534 | 1.431690387 | 0.457433688 | 0.627327439 | 7.386310738 | 10.57607651 | ENSG00000162391 |
| NA | 0.517696549 | 1.431667578 | 5.16E-06 | 3.58E-05 | 305.276824 | 437.3654519 | ENSG00000234072 |
| ANKRD49 | 0.517535905 | 1.431508171 | 0.032736651 | 0.084349236 | 72.57102719 | 103.982672 | ENSG00000168876 |
| AZIN1 | 0.517202271 | 1.431177162 | 4.62E-31 | 3.09E-29 | 4797.486624 | 6866.755015 | ENSG00000155096 |
| LINC00894 | 0.517166303 | 1.431141482 | 0.039810496 | 0.099286915 | 63.86400039 | 91.30879938 | ENSG00000235703 |
| APOOL | 0.516991182 | 1.430967774 | 0.000871402 | 0.003758994 | 155.1862161 | 222.0002852 | ENSG00000155008 |
| RNF128 | 0.516984261 | 1.430960909 | 0.145064022 | 0.275292987 | 33.2132802 | 47.52377404 | ENSG00000133135 |
| NLRC5 | 0.516954602 | 1.430931492 | 0.17905138 | 0.323459841 | 29.97364231 | 42.9079464 | ENSG00000140853 |
| NA | 0.516745069 | 1.430723682 | 0.644360415 | 0.777772291 | 2.949883674 | 4.242638624 | ENSG00000214192 |
| RAB13 | 0.516694269 | 1.430673305 | 5.07E-05 | 0.000291117 | 263.6956075 | 377.5202941 | ENSG00000143545 |
| CBX5 | 0.516475283 | 1.43045616 | 1.46E-16 | 3.68E-15 | 3340.987783 | 4779.564118 | ENSG00000094916 |
| ABCA11P | 0.516430986 | 1.430412239 | 0.573144936 | 0.725447081 | 4.313539319 | 6.165123255 | ENSG00000251595 |
| ZNF658 | 0.51557524 | 1.429564031 | 0.138679251 | 0.265798453 | 31.31353127 | 44.7486908 | ENSG00000274349 |
| NA | 0.515405483 | 1.429395828 | 0.513714361 | 0.677859608 | 5.940955171 | 8.527222646 | ENSG00000232024 |
| SUSD6 | 0.515124946 | 1.429117905 | 0.005814581 | 0.019998055 | 106.9262665 | 152.6196224 | ENSG00000100647 |
| AMZ2P1 | 0.515095213 | 1.429088452 | 0.009721097 | 0.030692255 | 124.3440238 | 177.6159086 | ENSG00000214174 |
| KCTD12 | 0.514850695 | 1.42884626 | 7.58E-05 | 0.000421301 | 265.6550414 | 379.3916796 | ENSG00000178695 |
| ALPK1 | 0.514497257 | 1.428496258 | 0.601074949 | 0.747006465 | 4.283515108 | 6.116726136 | ENSG00000073331 |
| STK17B | 0.51444124 | 1.428440794 | 0.001420213 | 0.005840403 | 155.366837 | 222.0398075 | ENSG00000081320 |
| NA | 0.514393481 | 1.428393506 | 0.28951811 | 0.458885372 | 18.90669125 | 27.10942728 | ENSG00000255306 |
| ABHD14B | 0.514389967 | 1.428390028 | 0.028484615 | 0.075344777 | 75.3354961 | 107.5931798 | ENSG00000114779 |
| RABGAP1L | 0.51427539 | 1.428276591 | 3.24E-15 | 7.11E-14 | 1242.139513 | 1774.285775 | ENSG00000152061 |
| F11R | 0.513968172 | 1.427972477 | 2.25E-18 | 6.57E-17 | 1689.620683 | 2413.204113 | ENSG00000158769 |
| NA | 0.513661121 | 1.427668591 | 0.269303443 | 0.435650543 | 18.57471008 | 26.47810205 | ENSG00000263571 |
| MYT1L | 0.513625871 | 1.427633709 | 0.587357299 | 0.736908796 | 4.362424128 | 6.211369801 | ENSG00000186487 |
| NA | 0.513600504 | 1.427608607 | 0.56082265 | 0.715862486 | 4.99397169 | 7.107543445 | ENSG00000241061 |
| RNF43 | 0.513477026 | 1.427486425 | 0.427020806 | 0.598536537 | 9.337535219 | 13.2936394 | ENSG00000108375 |
| NA | 0.513430644 | 1.427440533 | 0.518164713 | 0.681756614 | 5.756103983 | 8.163429427 | ENSG00000263004 |
| BCKDHB | 0.513328252 | 1.427339227 | 1.01E-06 | 7.95E-06 | 437.6572913 | 624.5109589 | ENSG00000083123 |
| NAGLU | 0.513162337 | 1.427175087 | 2.93E-05 | 0.000176716 | 301.7482198 | 430.866674 | ENSG00000108784 |
| TSSK4 | 0.513157835 | 1.427170634 | 0.223493083 | 0.382047414 | 25.04964335 | 35.55464924 | ENSG00000139908 |
| RIMS1 | 0.51305441 | 1.427068326 | 0.085632807 | 0.18328442 | 45.12712977 | 64.61586358 | ENSG00000079841 |
| CD55 | 0.512756667 | 1.426773838 | 1.49E-05 | 9.48E-05 | 315.8591573 | 451.1223808 | ENSG00000196352 |
| NA | 0.512735525 | 1.426752929 | 0.300876575 | 0.471503656 | 16.48173725 | 23.40267673 | ENSG00000283638 |
| NA | 0.512593449 | 1.42661243 | 0.679293316 | 0.803872602 | 2.746171888 | 3.865941962 | ENSG00000270000 |
| BAX | 0.512406394 | 1.426427472 | 5.17E-24 | 2.38E-22 | 2886.934258 | 4117.471647 | ENSG00000087088 |
| NA | 0.512264634 | 1.426287317 | 0.306844567 | 0.477828426 | 14.78590027 | 21.05750936 | ENSG00000232648 |
| CMYA5 | 0.51202735 | 1.426052752 | 0.239978929 | 0.402035042 | 19.14620922 | 27.25005703 | ENSG00000164309 |
| MIR589 | 0.512000445 | 1.426026157 | 0.505486878 | 0.670442219 | 5.965397576 | 8.531523793 | ENSG00000207973 |
| SLA | 0.511969181 | 1.425995255 | 0.261507219 | 0.426881063 | 19.1294638 | 27.23285244 | ENSG00000155926 |
| OSER1-DT | 0.511867919 | 1.425895168 | 0.030577575 | 0.079818309 | 66.22264845 | 94.32697692 | ENSG00000223891 |
| HBP1 | 0.511820937 | 1.425848734 | 3.65E-06 | 2.59E-05 | 377.460201 | 537.9277033 | ENSG00000105856 |
| GEM | 0.511727271 | 1.425756165 | 0.014590164 | 0.043254779 | 85.48317254 | 121.9410876 | ENSG00000164949 |
| RPL12 | 0.511720444 | 1.425749418 | 8.09E-37 | 7.51E-35 | 14295.77571 | 20380.28418 | ENSG00000197958 |
| TMEM8B | 0.511690017 | 1.425719349 | 0.000131372 | 0.000688665 | 233.6620289 | 332.9240946 | ENSG00000137103 |
| NALCN | 0.511661887 | 1.42569155 | 0.378345033 | 0.551850761 | 12.79137279 | 18.13372712 | ENSG00000102452 |
| CNTNAP3 | 0.511422798 | 1.425455299 | 0.03011611 | 0.078769221 | 67.96737024 | 96.92056951 | ENSG00000106714 |
| NA | 0.51132628 | 1.425359937 | 0.337700219 | 0.510343519 | 13.26442659 | 18.92476469 | ENSG00000265750 |
| AKAP9 | 0.511319794 | 1.425353529 | 1.63E-06 | 1.23E-05 | 511.0046055 | 727.9916432 | ENSG00000127914 |
| PCDHB3 | 0.511268568 | 1.42530292 | 0.267608154 | 0.433939628 | 17.91851995 | 25.59887253 | ENSG00000113205 |
| GRB7 | 0.511266877 | 1.425301249 | 3.19E-08 | 3.14E-07 | 557.7678745 | 795.3858925 | ENSG00000141738 |
| ZNF582 | 0.511202112 | 1.425237266 | 0.300235187 | 0.470881057 | 17.39099954 | 24.82906812 | ENSG00000018869 |
| IGFBPL1 | 0.511141303 | 1.425177195 | 5.70E-13 | 9.97E-12 | 1343.093296 | 1914.52533 | ENSG00000137142 |
| USF3 | 0.510981195 | 1.425019039 | 0.000685403 | 0.003028887 | 221.65081 | 315.6442805 | ENSG00000176542 |
| CNKSR1 | 0.510939638 | 1.424977993 | 4.85E-05 | 0.00027988 | 268.5955885 | 382.8540398 | ENSG00000142675 |
| SIRT5 | 0.510585325 | 1.424628074 | 8.09E-06 | 5.42E-05 | 328.9996939 | 468.5422063 | ENSG00000124523 |
| SLC27A3 | 0.510190564 | 1.424238309 | 4.61E-07 | 3.82E-06 | 633.0850494 | 901.756265 | ENSG00000143554 |
| MOB1B | 0.510147317 | 1.424195616 | 0.00011562 | 0.000616142 | 303.1723753 | 432.4863507 | ENSG00000173542 |
| LFNG | 0.509996506 | 1.424046747 | 2.31E-07 | 2.01E-06 | 540.3252361 | 768.9180281 | ENSG00000106003 |
| KCNJ14 | 0.509890101 | 1.423941721 | 0.463012115 | 0.632709974 | 7.353495625 | 10.45400842 | ENSG00000182324 |
| FIGNL1 | 0.509880185 | 1.423931934 | 0.001466216 | 0.006007809 | 171.5877682 | 244.7127472 | ENSG00000132436 |
| PGBD4 | 0.509420558 | 1.423478357 | 0.233310462 | 0.394121995 | 20.83646439 | 29.61860814 | ENSG00000182405 |

| | | | | | | | |
|---|---|---|---|---|---|---|---|
| NA | 0.50921092 | 1.423271526 | 0.57647645 | 0.728564413 | 4.672953964 | 6.684860702 | ENSG00000261821 |
| NHSL2 | 0.509208101 | 1.423268746 | 0.000210743 | 0.001056784 | 214.2385053 | 304.6404387 | ENSG00000204131 |
| RPRML | 0.508926436 | 1.4229909 | 0.236350805 | 0.397787766 | 31.58710364 | 45.00196231 | ENSG00000179673 |
| PPFIA1 | 0.508481878 | 1.422552481 | 1.37E-17 | 3.77E-16 | 1353.063119 | 1925.000123 | ENSG00000131626 |
| FOXN2 | 0.508164739 | 1.422239804 | 0.002228405 | 0.008668716 | 236.3162643 | 336.3380025 | ENSG00000170802 |
| NBPF26 | 0.508105197 | 1.422181108 | 0.02924339 | 0.076899432 | 70.72749664 | 100.4756891 | ENSG00000273136 |
| L1CAM | 0.508030241 | 1.42210722 | 1.38E-10 | 1.87E-09 | 1132.023291 | 1608.840994 | ENSG00000198910 |
| NA | 0.507833505 | 1.421913304 | 0.118923813 | 0.236804219 | 48.56019172 | 69.18302153 | ENSG00000232593 |
| ODR4 | 0.507778176 | 1.421858774 | 1.10E-05 | 7.14E-05 | 292.2240064 | 415.5284818 | ENSG00000157181 |
| ATM | 0.507712329 | 1.421793879 | 4.33E-08 | 4.21E-07 | 646.4254678 | 919.155814 | ENSG00000149311 |
| MOSPD1 | 0.507635235 | 1.421717904 | 9.05E-07 | 7.17E-06 | 441.7971062 | 628.6070554 | ENSG00000101928 |
| NA | 0.507503378 | 1.42158797 | 0.649542997 | 0.781978702 | 3.012722998 | 4.284584022 | ENSG00000223668 |
| KLHL6 | 0.507487273 | 1.421572101 | 0.705587005 | 0.822866524 | 2.830662713 | 3.969187922 | ENSG00000172578 |
| CCDC125 | 0.507393937 | 1.421480134 | 0.03327033 | 0.08541628 | 70.66099053 | 100.6587852 | ENSG00000183323 |
| LGI4 | 0.507204134 | 1.421293135 | 0.039218119 | 0.098203107 | 64.3642875 | 91.29322443 | ENSG00000153902 |
| ZNF468 | 0.507151454 | 1.421241238 | 0.000243431 | 0.001206274 | 206.6935006 | 293.7091897 | ENSG00000204604 |
| ZBTB6 | 0.507042559 | 1.421133966 | 0.000320832 | 0.001544206 | 191.2593074 | 271.685325 | ENSG00000186130 |
| ANKZF1 | 0.506771851 | 1.420867328 | 2.45E-12 | 4.01E-11 | 896.5565428 | 1273.386443 | ENSG00000163516 |
| SULT1A1 | 0.506120104 | 1.420225587 | 0.024786626 | 0.06712885 | 78.27932241 | 110.9646582 | ENSG00000196502 |
| NA | 0.505999519 | 1.420106885 | 0.606892469 | 0.751272767 | 3.695946272 | 5.23130536 | ENSG00000240036 |
| OXER1 | 0.505772396 | 1.419883336 | 0.374009736 | 0.547214396 | 12.55676091 | 17.9508794 | ENSG00000162881 |
| PBK | 0.505663676 | 1.419776339 | 0.002921438 | 0.010955353 | 161.7967524 | 229.4718824 | ENSG00000168078 |
| CPNE4 | 0.505468402 | 1.41958418 | 0.14034621 | 0.268486587 | 34.38929375 | 48.69878403 | ENSG00000196353 |
| NA | 0.505439408 | 1.419555651 | 0.549647963 | 0.706883924 | 5.263313703 | 7.498761085 | ENSG00000202058 |
| NA | 0.504467729 | 1.418599879 | 0.003688605 | 0.013448662 | 131.8559763 | 187.1958248 | ENSG00000247134 |
| PLEKHG5 | 0.504274151 | 1.418409547 | 1.68E-10 | 2.26E-09 | 808.6857003 | 1146.947065 | ENSG00000171680 |
| TNFRSF13C | 0.504244535 | 1.418380429 | 0.210798982 | 0.366369311 | 23.24957906 | 33.00298995 | ENSG00000159958 |
| NPBWR1 | 0.503936377 | 1.418077497 | 0.527900986 | 0.689522636 | 6.871193235 | 9.793641231 | ENSG00000183729 |
| AGRN | 0.50382732 | 1.417970305 | 1.16E-32 | 8.59E-31 | 8425.336985 | 11947.77791 | ENSG00000188157 |
| POLR2A | 0.503765265 | 1.417909314 | 6.60E-27 | 3.57E-25 | 5874.282307 | 8328.521443 | ENSG00000181222 |
| GBAP1 | 0.503637495 | 1.417783745 | 0.079644679 | 0.172865155 | 49.99206833 | 70.94312232 | ENSG00000160766 |
| NA | 0.503575175 | 1.417722502 | 0.378408681 | 0.551850761 | 12.20850987 | 17.37467558 | ENSG00000238058 |
| HNF4G | 0.503091807 | 1.417247581 | 0.187415664 | 0.334545718 | 27.99653596 | 39.70505524 | ENSG00000164749 |
| GLI1 | 0.503046249 | 1.417202827 | 1.85E-05 | 0.000116149 | 288.8380264 | 409.210631 | ENSG00000111087 |
| EPHA2 | 0.502945312 | 1.417103677 | 3.58E-16 | 8.75E-15 | 1239.744495 | 1757.025928 | ENSG00000142627 |
| ZNFX1 | 0.502936666 | 1.417095185 | 7.20E-05 | 0.00040178 | 352.6880353 | 500.0080716 | ENSG00000124201 |
| G2E3-AS1 | 0.50278755 | 1.416948723 | 0.576543603 | 0.728601549 | 5.100114017 | 7.172612117 | ENSG00000257636 |
| NA | 0.502751591 | 1.416913406 | 0.654820411 | 0.785826654 | 3.018304804 | 4.272213618 | ENSG00000243746 |
| MUCL3 | 0.502613363 | 1.416777654 | 0.620855662 | 0.761842541 | 4.624069155 | 6.575163021 | ENSG00000168631 |
| ETS1 | 0.502536989 | 1.416702654 | 1.24E-17 | 3.42E-16 | 1556.552149 | 2204.741591 | ENSG00000134954 |
| TPD52 | 0.502425606 | 1.416593283 | 0.00147157 | 0.006027188 | 248.3856165 | 352.4809354 | ENSG00000076554 |
| NA | 0.5024059 | 1.416573933 | 0.672978606 | 0.799651441 | 2.993862399 | 4.24048805 | ENSG00000254477 |
| FOXO3 | 0.502213562 | 1.41638509 | 8.27E-06 | 5.53E-05 | 367.968878 | 520.9985548 | ENSG00000118689 |
| NA | 0.502128602 | 1.416301682 | 0.395350413 | 0.568012573 | 10.67395756 | 15.13626181 | ENSG00000214770 |
| SERINC3 | 0.501685474 | 1.415866728 | 3.22E-10 | 4.19E-09 | 782.7115625 | 1107.753592 | ENSG00000132824 |
| ALDOC | 0.501361151 | 1.415548471 | 1.26E-10 | 1.73E-09 | 875.39557 | 1238.714283 | ENSG00000109107 |
| OTUD1 | 0.501282109 | 1.415470919 | 0.007694525 | 0.025282759 | 116.8820705 | 165.3741571 | ENSG00000165312 |
| MTERF1 | 0.501141621 | 1.415333088 | 0.000573441 | 0.002586595 | 187.8501243 | 266.0275566 | ENSG00000127989 |
| PPM1N | 0.501063556 | 1.415256506 | 0.000792694 | 0.003447218 | 181.0273403 | 255.8505187 | ENSG00000213889 |
| PTPRG | 0.500991028 | 1.415185359 | 1.26E-13 | 2.36E-12 | 1780.788498 | 2519.642667 | ENSG00000144724 |
| RTL10 | 0.500707747 | 1.414907507 | 1.24E-08 | 1.30E-07 | 537.4820583 | 760.2700582 | ENSG00000215012 |
| DDIT4 | 0.500700443 | 1.414900344 | 0.057378052 | 0.133238974 | 725.4575461 | 1026.188165 | ENSG00000168209 |
| SMPD3 | 0.500650893 | 1.414851749 | 0.169344696 | 0.310210666 | 36.36631208 | 51.37466198 | ENSG00000103056 |
| KIAA2026 | 0.500338757 | 1.414545671 | 0.000568928 | 0.002570303 | 201.1047387 | 284.646612 | ENSG00000183354 |
| RUNX1 | 0.499867089 | 1.414083281 | 0.482085957 | 0.650071312 | 7.410753143 | 10.426584 | ENSG00000159216 |
| SH2D4A | 0.499706694 | 1.413926076 | 0.001631404 | 0.006599162 | 166.9949994 | 236.08094 | ENSG00000104611 |
| APLN | 0.499506326 | 1.413729717 | 0.01083892 | 0.033746548 | 116.5256469 | 164.3828005 | ENSG00000171388 |
| RIBC2 | 0.499494503 | 1.413718132 | 0.222840816 | 0.38127026 | 21.42124233 | 30.28757763 | ENSG00000128408 |
| LOC653653 | 0.499016814 | 1.413250115 | 0.370730444 | 0.544522689 | 11.37045962 | 16.05340815 | ENSG00000280852 |
| BRWD1 | 0.49897239 | 1.413206597 | 2.03E-07 | 1.79E-06 | 645.4220146 | 912.106869 | ENSG00000185658 |
| MUC16 | 0.498772588 | 1.413010894 | 0.670990885 | 0.798249756 | 3.048329015 | 4.295336891 | ENSG00000181143 |
| GPR156 | 0.498747156 | 1.412985985 | 0.073656767 | 0.162495856 | 51.85553552 | 73.13234547 | ENSG00000175697 |
| JUND | 0.498707626 | 1.412947269 | 3.47E-11 | 5.01E-10 | 1062.728561 | 1502.191428 | ENSG00000130522 |
| IFI27 | 0.498542568 | 1.412785625 | 0.521007181 | 0.683957097 | 6.019864191 | 8.487427821 | ENSG00000165949 |
| NBPF14 | 0.498473605 | 1.412718093 | 0.26626457 | 0.432475357 | 18.28563158 | 25.85565168 | ENSG00000270629 |
| NEDD4L | 0.498285828 | 1.41253423 | 7.28E-16 | 1.72E-14 | 1330.651427 | 1879.33428 | ENSG00000049759 |
| CLIC6 | 0.49796818 | 1.412223256 | 0.036999059 | 0.093434854 | 69.47488941 | 98.00630306 | ENSG00000159212 |
| NA | 0.497821774 | 1.41207995 | 0.665516647 | 0.794116512 | 3.048329015 | 4.291035743 | ENSG00000232936 |
| ERFE | 0.497480206 | 1.41174567 | 0.161291537 | 0.298653172 | 33.36041019 | 47.06722726 | ENSG00000178752 |
| ZNF600 | 0.497431169 | 1.411697686 | 0.018663825 | 0.052968651 | 86.92852811 | 122.6501245 | ENSG00000189190 |
| MIR635 | 0.497416166 | 1.411683005 | 0.556620597 | 0.712762558 | 5.320571221 | 7.513282063 | ENSG00000207561 |
| FTCD | 0.497194566 | 1.411466185 | 0.129809916 | 0.252614052 | 43.33180831 | 60.97177066 | ENSG00000160282 |
| M1AP | 0.497021665 | 1.411297037 | 0.025439322 | 0.068617166 | 91.25574654 | 128.5200188 | ENSG00000159374 |
| TULP4 | 0.496978777 | 1.411255083 | 0.00063376 | 0.002828264 | 240.4486561 | 339.2055851 | ENSG00000130338 |
| TMEM74B | 0.496753214 | 1.411034453 | 0.158517767 | 0.294864946 | 32.70182948 | 46.01913796 | ENSG00000125895 |
| SLC6A15 | 0.496590876 | 1.410875686 | 2.26E-07 | 1.97E-06 | 472.1370964 | 666.4942596 | ENSG00000072041 |
| ABHD3 | 0.496551092 | 1.41083678 | 0.008223739 | 0.026734774 | 118.2085685 | 166.8102474 | ENSG00000158201 |

| | | | | | | | |
|---|---|---|---|---|---|---|---|
| NA | 0.496479673 | 1.41076694 | 0.519314148 | 0.682803246 | 6.291997107 | 8.92059086 | ENSG00000280383 |
| SEC1P | 0.496354995 | 1.410645026 | 0.383536849 | 0.556806306 | 12.82909398 | 18.02859105 | ENSG00000232871 |
| BCL6 | 0.496080788 | 1.410376937 | 0.074951178 | 0.164810576 | 52.30569852 | 73.62035735 | ENSG00000113916 |
| PDLIM5 | 0.496047274 | 1.410344173 | 3.72E-09 | 4.20E-08 | 823.228765 | 1161.01602 | ENSG00000163110 |
| DYSF | 0.495674137 | 1.409979451 | 3.37E-10 | 4.36E-09 | 866.7192303 | 1221.680586 | ENSG00000135636 |
| LOC283731 | 0.495474667 | 1.409784517 | 0.520345872 | 0.68346886 | 6.346463723 | 8.979740848 | ENSG00000248540 |
| LOC112267871 | 0.495384392 | 1.409696305 | 0.25021342 | 0.414430782 | 19.78266286 | 27.828684 | ENSG00000228549 |
| CCDC14 | 0.495265682 | 1.409580315 | 1.52E-08 | 1.58E-07 | 738.707006 | 1041.253403 | ENSG00000175455 |
| LENG8-AS1 | 0.495049944 | 1.409369545 | 0.223523943 | 0.382066311 | 26.31206275 | 37.03588947 | ENSG00000226696 |
| ZNF83 | 0.494796845 | 1.409122314 | 0.000474653 | 0.002187942 | 213.3485792 | 300.2929487 | ENSG00000167766 |
| TACR2 | 0.494792615 | 1.409118182 | 0.662387536 | 0.791698422 | 2.985489691 | 4.198542652 | ENSG00000075073 |
| VCPIP1 | 0.49476007 | 1.409086395 | 1.35E-05 | 8.65E-05 | 321.8198859 | 453.2599412 | ENSG00000175073 |
| JPH2 | 0.49455208 | 1.408883265 | 0.116827163 | 0.233763386 | 38.51499082 | 54.23473262 | ENSG00000149596 |
| CCND2-AS1 | 0.494512622 | 1.408844732 | 0.607029103 | 0.751272767 | 3.984148884 | 5.643495699 | ENSG00000255920 |
| INTU | 0.494494207 | 1.408826749 | 5.70E-05 | 0.000324656 | 266.3640585 | 375.0124703 | ENSG00000164066 |
| NA | 0.49444876 | 1.40878237 | 0.514146944 | 0.678170569 | 6.909590154 | 9.761915663 | ENSG00000280279 |
| NEK11 | 0.494445305 | 1.408778996 | 0.379013886 | 0.552436638 | 12.45081874 | 17.53223734 | ENSG00000114670 |
| CENPBD1 | 0.494328499 | 1.408664941 | 0.00226798 | 0.008803883 | 153.1534919 | 215.4571388 | ENSG00000177946 |
| RELA-DT | 0.494064658 | 1.408407347 | 0.331935859 | 0.504639818 | 15.75242007 | 22.10721619 | ENSG00000285533 |
| NRP2 | 0.493942174 | 1.408287779 | 2.98E-09 | 3.41E-08 | 696.4346322 | 980.9735563 | ENSG00000118257 |
| NA | 0.493737607 | 1.408088105 | 7.99E-07 | 6.39E-06 | 547.5790599 | 770.2831069 | ENSG00000230623 |
| FKBP9 | 0.493621924 | 1.407975202 | 3.83E-05 | 0.000225482 | 274.5639771 | 386.3631854 | ENSG00000122642 |
| NA | 0.493603617 | 1.407957335 | 0.651399772 | 0.783266107 | 3.045538112 | 4.26791247 | ENSG00000287265 |
| GFPT1 | 0.493417035 | 1.407775257 | 1.35E-15 | 3.10E-14 | 1902.494782 | 2678.369541 | ENSG00000198380 |
| NA | 0.492888412 | 1.407259524 | 0.674350876 | 0.800199707 | 3.018304804 | 4.234036329 | ENSG00000285407 |
| NA | 0.492884602 | 1.407255807 | 0.459100007 | 0.628967988 | 8.923653959 | 12.65020422 | ENSG00000230207 |
| NCAN | 0.492713828 | 1.407089238 | 0.000354203 | 0.001689646 | 218.8346285 | 308.3162877 | ENSG00000130287 |
| EVA1C | 0.492677109 | 1.407053426 | 0.156445405 | 0.29203511 | 33.50215853 | 47.03335112 | ENSG00000166979 |
| GSTO2 | 0.492663275 | 1.407039934 | 0.004816516 | 0.01698947 | 142.0181197 | 199.6854926 | ENSG00000065621 |
| FAM214A | 0.492468601 | 1.406850084 | 0.036087002 | 0.091466872 | 70.45448784 | 99.25577752 | ENSG00000047346 |
| CADM3 | 0.492363055 | 1.406747164 | 0.061849589 | 0.141350601 | 54.88412806 | 77.39432714 | ENSG00000162706 |
| RHBDD1 | 0.492284842 | 1.406670902 | 1.05E-05 | 6.87E-05 | 349.2590774 | 490.8967874 | ENSG00000144468 |
| PLPP2 | 0.49218735 | 1.406575848 | 2.44E-15 | 5.47E-14 | 1615.311893 | 2271.964478 | ENSG00000141934 |
| GSG1L | 0.492179952 | 1.406568635 | 0.398933681 | 0.571415922 | 10.75007567 | 15.08571411 | ENSG00000169181 |
| MAPT | 0.492139714 | 1.406529405 | 0.109842971 | 0.222336876 | 40.96929331 | 57.77101797 | ENSG00000186868 |
| IER5L | 0.491964525 | 1.406358618 | 0.276800702 | 0.443981238 | 17.94017145 | 25.28347643 | ENSG00000188483 |
| HLA-E | 0.491797226 | 1.406195542 | 3.46E-12 | 5.56E-11 | 1004.525647 | 1411.874355 | ENSG00000204592 |
| NAPEPLD | 0.491494215 | 1.405900228 | 0.005866476 | 0.020158565 | 135.7445076 | 190.5307129 | ENSG00000161048 |
| NA | 0.491411671 | 1.405819792 | 0.267682698 | 0.434011341 | 19.95722632 | 27.94860151 | ENSG00000234338 |
| LOC100506302 | 0.491216693 | 1.40562981 | 0.269202085 | 0.435576741 | 20.18884714 | 28.33766858 | ENSG00000216895 |
| NA | 0.491209065 | 1.405622378 | 0.584765978 | 0.73470878 | 4.71904787 | 6.630011861 | ENSG00000237493 |
| HM13-AS1 | 0.490886618 | 1.405308252 | 0.578096953 | 0.729895171 | 4.700187272 | 6.604738015 | ENSG00000230613 |
| NA | 0.490790701 | 1.405214824 | 0.627386156 | 0.765619074 | 3.924100463 | 5.544550886 | ENSG00000231991 |
| CD109 | 0.490636253 | 1.405064396 | 0.007887614 | 0.02580732 | 119.9543664 | 168.2734711 | ENSG00000156535 |
| F11 | 0.490596644 | 1.405025821 | 0.613745044 | 0.756749457 | 5.894185543 | 8.34841561 | ENSG00000088926 |
| SERTAD1 | 0.4905112 | 1.40494261 | 0.001236462 | 0.005167373 | 187.2568615 | 263.1150908 | ENSG00000197019 |
| CTNNA2 | 0.490271336 | 1.404709043 | 0.000692994 | 0.003055669 | 202.2963464 | 284.1454241 | ENSG00000066032 |
| MGC32805 | 0.490131973 | 1.404573355 | 0.288150745 | 0.457432194 | 17.12656361 | 24.02781071 | ENSG00000250328 |
| NA | 0.489883594 | 1.40433156 | 0.53481874 | 0.695114833 | 6.088285321 | 8.504099373 | ENSG00000280310 |
| ADPRM | 0.489704666 | 1.404157401 | 0.018276979 | 0.052054399 | 85.38492736 | 119.7825361 | ENSG00000170222 |
| MSX1 | 0.489437828 | 1.403897715 | 0.625863866 | 0.764553217 | 4.253490898 | 6.043315634 | ENSG00000163132 |
| ZNF674 | 0.48930845 | 1.403771822 | 0.129826983 | 0.25262178 | 37.35592284 | 52.34665717 | ENSG00000251192 |
| NA | 0.489147309 | 1.403615037 | 0.000695807 | 0.003066669 | 197.1440811 | 276.4915244 | ENSG00000244198 |
| TAF10 | 0.489114688 | 1.403583301 | 0.011880095 | 0.036348756 | 99.66408283 | 139.8261049 | ENSG00000166337 |
| C19orf71 | 0.488436558 | 1.402923709 | 0.164852175 | 0.303885647 | 29.66725482 | 41.63991027 | ENSG00000183397 |
| NA | 0.488414166 | 1.402901935 | 0.239057493 | 0.401048622 | 21.68221163 | 30.39701485 | ENSG00000278600 |
| ALKAL2 | 0.488383174 | 1.402871799 | 0.185617978 | 0.332012648 | 35.87707644 | 50.20451671 | ENSG00000189292 |
| ARHGEF39 | 0.488001289 | 1.402500504 | 0.000487345 | 0.00224002 | 215.7441847 | 302.3966698 | ENSG00000137135 |
| NA | 0.487854016 | 1.402357341 | 0.519151658 | 0.682639249 | 6.322021318 | 8.885097182 | ENSG00000279880 |
| NA | 0.487679938 | 1.40218814 | 0.331584119 | 0.504264032 | 16.04341359 | 22.55139256 | ENSG00000235726 |
| TMEM147-AS1 | 0.487230022 | 1.401750925 | 0.000141316 | 0.000735597 | 273.7133238 | 383.2299672 | ENSG00000236144 |
| RGS14 | 0.487212866 | 1.401734256 | 2.14E-06 | 1.59E-05 | 384.0671216 | 538.2194673 | ENSG00000169220 |
| SKIL | 0.486994284 | 1.401521896 | 2.76E-09 | 3.16E-08 | 1608.807803 | 2255.999988 | ENSG00000136603 |
| NA | 0.486189759 | 1.400740549 | 0.654493117 | 0.785579064 | 3.284855915 | 4.640307985 | ENSG00000268093 |
| TENT2 | 0.486101334 | 1.400654699 | 4.12E-07 | 3.45E-06 | 457.8165898 | 641.1290245 | ENSG00000164329 |
| INTS12 | 0.485928994 | 1.400487391 | 0.00881323 | 0.028309709 | 106.835318 | 149.6412518 | ENSG00000138785 |
| TPM3P9 | 0.485837928 | 1.400398992 | 0.000558129 | 0.002528866 | 205.5684871 | 287.8005851 | ENSG00000241015 |
| NA | 0.485815028 | 1.400376763 | 0.608766476 | 0.752512757 | 4.455287663 | 6.167273829 | ENSG00000206549 |
| NA | 0.485771995 | 1.400334993 | 0.441074654 | 0.612097774 | 10.05847969 | 14.05080083 | ENSG00000244171 |
| ZFAS1 | 0.485746943 | 1.400310677 | 0.004893281 | 0.017222455 | 131.811434 | 184.7073441 | ENSG00000177410 |
| PHF19 | 0.485739099 | 1.400303064 | 6.53E-06 | 4.45E-05 | 346.629699 | 485.3304703 | ENSG00000119403 |
| PLCD4 | 0.485680366 | 1.400246057 | 0.08394554 | 0.180273118 | 53.18079425 | 74.52028699 | ENSG00000115556 |
| INTS7 | 0.485105714 | 1.399688424 | 3.18E-08 | 3.13E-07 | 562.3929075 | 787.6787188 | ENSG00000143493 |
| FAM181B | 0.485070351 | 1.399654116 | 0.170507094 | 0.311879408 | 30.87819867 | 43.20263026 | ENSG00000182103 |
| HELZ2 | 0.485012462 | 1.399597956 | 6.46E-09 | 7.05E-08 | 678.0629581 | 949.3999139 | ENSG00000130589 |
| PI4K2B | 0.484727134 | 1.399321178 | 2.10E-05 | 0.000130487 | 422.1191973 | 591.270315 | ENSG00000038210 |
| ZNF169 | 0.484711892 | 1.399306395 | 0.044317116 | 0.108313883 | 66.24342407 | 92.77635477 | ENSG00000175787 |

| | | | | | | | |
|---|---|---|---|---|---|---|---|
| TESMIN | 0.484644361 | 1.399240896 | 2.46E-08 | 2.46E-07 | 549.7135712 | 769.3188657 | ENSG00000132749 |
| C7orf25 | 0.484471716 | 1.399073461 | 0.559787351 | 0.715014089 | 5.581540526 | 7.848033327 | ENSG00000136197 |
| HIBADH | 0.484076949 | 1.398690683 | 4.29E-08 | 4.17E-07 | 538.8257774 | 754.0688655 | ENSG00000106049 |
| NA | 0.484044113 | 1.398658849 | 0.690946346 | 0.812548522 | 3.075562322 | 4.274364192 | ENSG00000277782 |
| NA | 0.483737124 | 1.398361262 | 0.154493721 | 0.289381335 | 37.08271388 | 51.94012505 | ENSG00000275764 |
| SMUG1 | 0.483656384 | 1.398283004 | 9.87E-06 | 6.51E-05 | 349.3883859 | 488.7293489 | ENSG00000123415 |
| MMP19 | 0.483569968 | 1.398199252 | 0.315229772 | 0.486505618 | 17.87753229 | 24.90677977 | ENSG00000123342 |
| NA | 0.483413246 | 1.398047372 | 0.657116088 | 0.787600276 | 3.276483206 | 4.617184712 | ENSG00000279434 |
| PLAC8 | 0.483088714 | 1.397732919 | 0.514199326 | 0.678170569 | 6.978011284 | 9.743093538 | ENSG00000145287 |
| MINDY1 | 0.482946179 | 1.397594832 | 0.001360612 | 0.005615656 | 172.1425588 | 240.4109586 | ENSG00000143409 |
| SNORD117 | 0.482749402 | 1.397404219 | 0.665819474 | 0.794428716 | 3.100004727 | 4.293186317 | ENSG00000201785 |
| ZNF483 | 0.482556679 | 1.397217559 | 0.000403114 | 0.001897163 | 361.0067873 | 504.883543 | ENSG00000173258 |
| NA | 0.482533791 | 1.397195393 | 0.660597072 | 0.790446478 | 3.039956306 | 4.215214203 | ENSG00000239528 |
| NA | 0.482472457 | 1.397135995 | 0.5199538 | 0.683268285 | 6.376487933 | 8.906069881 | ENSG00000280422 |
| CHMP3 | 0.482463589 | 1.397127407 | 0.028842202 | 0.076047524 | 95.45417034 | 133.7055985 | ENSG00000115561 |
| NUDT19 | 0.48220533 | 1.396877327 | 0.099880796 | 0.207027015 | 45.60731698 | 63.82267543 | ENSG00000213965 |
| NFATC1 | 0.482134411 | 1.396808662 | 0.004795011 | 0.016922896 | 158.1920811 | 221.2229751 | ENSG00000131196 |
| JOSD2 | 0.482084767 | 1.396760598 | 0.134987582 | 0.260196387 | 46.83046358 | 65.64621525 | ENSG00000161677 |
| IGFBP3 | 0.48206401 | 1.396740502 | 0.064331384 | 0.145744771 | 55.7314269 | 77.89928315 | ENSG00000146674 |
| NA | 0.481727763 | 1.396415003 | 0.032060301 | 0.082927766 | 80.71005837 | 112.52361 | ENSG00000237094 |
| NID2 | 0.481487943 | 1.396182896 | 0.00041037 | 0.001926144 | 216.7310045 | 302.5270492 | ENSG00000087303 |
| LPAR1 | 0.481483824 | 1.39617891 | 0.049457107 | 0.118225162 | 69.09382326 | 96.39599788 | ENSG00000198121 |
| FBXW12 | 0.481366558 | 1.396065429 | 0.541400131 | 0.700278174 | 6.784787387 | 9.393821296 | ENSG00000164049 |
| PITPNM3 | 0.481333527 | 1.396033466 | 0.018533306 | 0.052682057 | 106.2901763 | 148.6434377 | ENSG00000091622 |
| SIDT2 | 0.48132436 | 1.396024596 | 1.58E-05 | 0.000100402 | 405.5685364 | 566.0045883 | ENSG00000149577 |
| EXD1 | 0.481030853 | 1.395740613 | 0.662861213 | 0.791976692 | 3.069980516 | 4.246939771 | ENSG00000178997 |
| UBAP2L | 0.480870361 | 1.395585353 | 1.19E-23 | 5.28E-22 | 5603.621294 | 7820.47816 | ENSG00000143569 |
| ZDHHC11B | 0.480586747 | 1.395311026 | 0.513401308 | 0.677642351 | 7.002453689 | 9.749545259 | ENSG00000206077 |
| NA | 0.48057898 | 1.395303515 | 0.121100663 | 0.240171983 | 43.63308955 | 61.00885134 | ENSG00000272335 |
| AQP11 | 0.480246784 | 1.394982269 | 0.131870719 | 0.255593091 | 37.66278589 | 52.60343632 | ENSG00000178301 |
| NA | 0.480141929 | 1.394880884 | 0.607875767 | 0.751797117 | 4.749072081 | 6.600436867 | ENSG00000286788 |
| NFAT5 | 0.479998074 | 1.394741804 | 0.000274617 | 0.001342874 | 229.7591044 | 320.2187384 | ENSG00000102908 |
| GAU1 | 0.479985025 | 1.394729189 | 0.711950794 | 0.826368397 | 3.00435029 | 4.171118231 | ENSG00000255474 |
| PMEPA1 | 0.479754639 | 1.394506481 | 9.20E-07 | 7.28E-06 | 524.8001696 | 732.4537019 | ENSG00000124225 |
| NA | 0.479321552 | 1.394087923 | 0.562565032 | 0.717227624 | 5.435086257 | 7.524034932 | ENSG00000240087 |
| NA | 0.479284834 | 1.394052442 | 0.620609485 | 0.761637354 | 4.074221516 | 5.673070693 | ENSG00000236861 |
| CDHR2 | 0.478967412 | 1.393745756 | 0.436897652 | 0.607641853 | 10.27335509 | 14.32640211 | ENSG00000074276 |
| NA | 0.478859046 | 1.393641071 | 0.487857524 | 0.654792398 | 8.243221588 | 11.55829153 | ENSG00000205267 |
| FGF1 | 0.478695985 | 1.393483563 | 0.520022029 | 0.683268285 | 6.379278836 | 8.876494887 | ENSG00000113578 |
| LRRIQ1 | 0.47863218 | 1.393421936 | 0.610418376 | 0.754119814 | 4.901108155 | 6.871736995 | ENSG00000133640 |
| GBX2 | 0.478368067 | 1.393166867 | 0.106835293 | 0.217722337 | 46.8135549 | 65.06328713 | ENSG00000168505 |
| KNL1 | 0.478294309 | 1.393095642 | 1.71E-05 | 0.00010809 | 369.5961817 | 514.8807626 | ENSG00000137812 |
| LOC101927468 | 0.477733298 | 1.392554024 | 0.063823457 | 0.144866255 | 61.03824376 | 84.95762386 | ENSG00000227733 |
| DUSP2 | 0.477616934 | 1.392441709 | 0.003419416 | 0.012581338 | 142.9775061 | 199.0254164 | ENSG00000158050 |
| BEX4 | 0.477065296 | 1.391909388 | 5.06E-10 | 6.40E-09 | 993.1307213 | 1382.347603 | ENSG00000102409 |
| DISP1 | 0.476954857 | 1.39180284 | 0.026199899 | 0.070334295 | 89.43518203 | 124.5578156 | ENSG00000154309 |
| NA | 0.476941755 | 1.391790201 | 0.466002339 | 0.635311034 | 8.774408787 | 12.17671332 | ENSG00000279619 |
| PIK3CB | 0.476893652 | 1.391743796 | 2.30E-06 | 1.70E-05 | 732.1152409 | 1018.237649 | ENSG00000051382 |
| NA | 0.476486373 | 1.391350956 | 0.565238997 | 0.719310343 | 5.728870676 | 7.972251986 | ENSG00000271013 |
| SMG1P1 | 0.4764012 | 1.391268817 | 0.262478872 | 0.428056464 | 22.95387962 | 31.82261273 | ENSG00000237296 |
| ERO1B | 0.476399774 | 1.391267441 | 0.05190592 | 0.122906235 | 65.82292186 | 91.46878429 | ENSG00000086619 |
| GAREM1 | 0.476160353 | 1.391036575 | 0.033724134 | 0.086374438 | 82.888361 | 115.3936216 | ENSG00000141441 |
| RAB25 | 0.476013477 | 1.390894965 | 2.88E-06 | 2.08E-05 | 441.8606581 | 613.9432669 | ENSG00000132698 |
| NA | 0.475582509 | 1.390479533 | 0.505601829 | 0.670442219 | 7.962715963 | 11.07458079 | ENSG00000219102 |
| NDFIP2 | 0.475512109 | 1.390411682 | 3.73E-05 | 0.000220104 | 413.8435609 | 575.6387961 | ENSG00000102471 |
| GJD2 | 0.475429392 | 1.390331965 | 0.59930429 | 0.745793794 | 4.901108155 | 6.855065443 | ENSG00000159248 |
| AEN | 0.475033904 | 1.389950884 | 2.22E-23 | 9.65E-22 | 3086.273675 | 4288.841964 | ENSG00000181026 |
| ERMP1 | 0.47498468 | 1.389903461 | 9.61E-07 | 7.57E-06 | 852.7587531 | 1185.802004 | ENSG00000099219 |
| FH | 0.474940066 | 1.389860479 | 1.35E-13 | 2.52E-12 | 1505.288224 | 2092.406439 | ENSG00000091483 |
| GABBR2 | 0.474879417 | 1.389802053 | 0.14622999 | 0.276739389 | 45.02812085 | 62.67508986 | ENSG00000136928 |
| NA | 0.474763659 | 1.389690543 | 0.669458411 | 0.797003432 | 3.605873641 | 5.075621597 | ENSG00000279400 |
| PRPF40B | 0.474748384 | 1.389675829 | 0.000839129 | 0.003631957 | 192.8942201 | 268.1401648 | ENSG00000110844 |
| BCHE | 0.474443276 | 1.389381965 | 0.089462437 | 0.189817408 | 51.39296961 | 71.28594284 | ENSG00000114200 |
| MFSD2B | 0.474162121 | 1.389111227 | 0.487348263 | 0.654508508 | 9.966291878 | 13.89081592 | ENSG00000205639 |
| NA | 0.473784515 | 1.388747693 | 0.039193676 | 0.098167386 | 87.98895058 | 122.5022374 | ENSG00000176654 |
| ZNF397 | 0.473627106 | 1.388596179 | 2.25E-07 | 1.96E-06 | 731.4931052 | 1014.89391 | ENSG00000186812 |
| NA | 0.473597001 | 1.388567203 | 0.051224657 | 0.121696412 | 62.97339855 | 87.37322713 | ENSG00000287356 |
| NOL3 | 0.473191052 | 1.388176539 | 0.075102666 | 0.165049616 | 57.27945817 | 79.55879737 | ENSG00000140939 |
| TMC6 | 0.472819583 | 1.387819154 | 8.29E-12 | 1.28E-10 | 1419.436766 | 1970.710136 | ENSG00000141524 |
| VPS13D | 0.472600539 | 1.387608457 | 2.07E-06 | 1.55E-05 | 759.4356019 | 1053.28736 | ENSG00000048707 |
| AKR1E2 | 0.47254737 | 1.387557319 | 0.131316882 | 0.254707725 | 46.39668258 | 64.51019447 | ENSG00000165568 |
| CSDC2 | 0.472520197 | 1.387531185 | 0.168511303 | 0.308918706 | 33.54267063 | 46.42676669 | ENSG00000172346 |
| NA | 0.472266307 | 1.387287024 | 0.597836793 | 0.74472876 | 4.700187272 | 6.486971076 | ENSG00000279253 |
| NA | 0.472036606 | 1.387066163 | 0.587533193 | 0.737081539 | 6.091076224 | 8.382031287 | ENSG00000224126 |
| NA | 0.472003292 | 1.387034134 | 0.298727018 | 0.469164122 | 17.78813538 | 24.63547964 | ENSG00000234287 |
| ADAMTS1 | 0.471929702 | 1.386963385 | 5.14E-07 | 4.23E-06 | 501.7012368 | 695.3135125 | ENSG00000154734 |
| PRUNE1 | 0.471762985 | 1.386803118 | 1.76E-07 | 1.56E-06 | 696.2875518 | 965.2989222 | ENSG00000143363 |

| | | | | | | | |
|---|---|---|---|---|---|---|---|
| NA | 0.471701171 | 1.3867437 | 0.277787395 | 0.445267689 | 20.96146732 | 28.97276193 | ENSG00000272419 |
| SPIRE2 | 0.47158163 | 1.3866288 | 0.000237313 | 0.001179896 | 232.9662906 | 323.0892894 | ENSG00000204991 |
| NA | 0.47135711 | 1.386413022 | 0.53742516 | 0.69697269 | 6.594154234 | 9.181671157 | ENSG00000273486 |
| GRM4 | 0.471218245 | 1.38627958 | 1.63E-09 | 1.94E-08 | 770.5032911 | 1067.860129 | ENSG00000124493 |
| IDH3G | 0.471213788 | 1.386275298 | 2.14E-07 | 1.87E-06 | 529.5385664 | 734.1742989 | ENSG00000067829 |
| NDUFV2 | 0.471045166 | 1.386113279 | 0.557806541 | 0.713430492 | 6.488212066 | 8.952316427 | ENSG00000178127 |
| LOC283387 | 0.470188528 | 1.385290483 | 0.65804503 | 0.788517714 | 3.279274109 | 4.564486445 | ENSG00000245651 |
| CLYBL | 0.470183525 | 1.385285679 | 0.010061011 | 0.031605107 | 115.1827669 | 159.5174268 | ENSG00000125246 |
| GVQW3 | 0.47011486 | 1.385219748 | 0.007222962 | 0.023990731 | 133.9052827 | 185.4392316 | ENSG00000179240 |
| PGK1 | 0.47003336 | 1.385141497 | 3.66E-16 | 8.94E-15 | 9832.778147 | 13618.50378 | ENSG00000102144 |
| AURKC | 0.469945529 | 1.385057173 | 0.22543451 | 0.384480341 | 26.18339304 | 36.33708452 | ENSG00000105146 |
| IRS4 | 0.469618686 | 1.384743423 | 0.001305405 | 0.005421434 | 218.6296022 | 302.5326953 | ENSG00000133124 |
| MASP2 | 0.469520031 | 1.384648734 | 0.391087163 | 0.564219438 | 14.0488861 | 19.47139352 | ENSG00000009724 |
| BTBD11 | 0.469425063 | 1.38455759 | 0.00130747 | 0.005428841 | 202.4872549 | 280.2316423 | ENSG00000151136 |
| PXN | 0.469421569 | 1.384554236 | 6.01E-10 | 7.50E-09 | 1024.833716 | 1418.301401 | ENSG00000089159 |
| NA | 0.469414818 | 1.384547757 | 0.617719953 | 0.759443988 | 4.019754901 | 5.551002607 | ENSG00000221930 |
| ERCC1 | 0.469167966 | 1.384310876 | 6.03E-08 | 5.72E-07 | 564.3349203 | 780.9411841 | ENSG00000012061 |
| RFX5 | 0.469137063 | 1.384281223 | 2.19E-11 | 3.22E-10 | 1070.053521 | 1480.461278 | ENSG00000143390 |
| AKAP13 | 0.469098067 | 1.384243807 | 3.56E-06 | 2.54E-05 | 729.3361319 | 1009.746726 | ENSG00000170776 |
| PAX3 | 0.469067596 | 1.384214571 | 0.544462175 | 0.702938966 | 6.687017769 | 9.276314822 | ENSG00000135903 |
| NA | 0.469032169 | 1.38418058 | 0.670387593 | 0.797758553 | 3.399370951 | 4.70752723 | ENSG00000238018 |
| KCTD3 | 0.469020235 | 1.38416913 | 2.41E-18 | 7.01E-17 | 1893.959854 | 2621.853996 | ENSG00000136636 |
| FEM1C | 0.468945584 | 1.384097509 | 0.000258365 | 0.001271137 | 343.3454435 | 475.0498591 | ENSG00000145780 |
| CA10 | 0.468936904 | 1.384089182 | 0.483296151 | 0.651157009 | 8.063952206 | 11.12727906 | ENSG00000154975 |
| ALDH9A1 | 0.468834261 | 1.383990712 | 1.15E-18 | 3.46E-17 | 1890.12659 | 2615.806179 | ENSG00000143149 |
| EEF1A2 | 0.468642726 | 1.383806983 | 1.65E-12 | 2.75E-11 | 1291.617227 | 1787.796944 | ENSG00000101210 |
| CD72 | 0.468550729 | 1.383718744 | 0.117935198 | 0.235406782 | 44.69825486 | 61.72810806 | ENSG00000137101 |
| NA | 0.468491617 | 1.38366205 | 0.473725993 | 0.642182893 | 10.31733381 | 14.24466188 | ENSG00000225507 |
| SOD3 | 0.468270917 | 1.383450397 | 0.02442511 | 0.066289363 | 85.59422095 | 118.4599116 | ENSG00000109610 |
| PAQR9-AS1 | 0.467931601 | 1.383125053 | 0.61088282 | 0.754500319 | 5.012832289 | 6.945407961 | ENSG00000241570 |
| TSNAX | 0.467823262 | 1.383021191 | 0.003003031 | 0.011209657 | 222.2204084 | 307.8903699 | ENSG00000116918 |
| LHPP | 0.467713381 | 1.382915859 | 5.31E-08 | 5.08E-07 | 834.8868154 | 1154.655177 | ENSG00000107902 |
| ELOA-AS1 | 0.46763018 | 1.382836108 | 0.453342246 | 0.623559734 | 9.952337363 | 13.79885587 | ENSG00000236810 |
| NMRAL1 | 0.467606991 | 1.382813881 | 1.19E-08 | 1.26E-07 | 868.7819546 | 1201.183966 | ENSG00000153406 |
| RHOU | 0.467340761 | 1.382558725 | 0.004708327 | 0.016665751 | 199.2742626 | 275.7547721 | ENSG00000116574 |
| OTUD7B | 0.467315069 | 1.382534104 | 7.70E-08 | 7.19E-07 | 607.3874623 | 839.8291039 | ENSG00000264522 |
| LINC01637 | 0.46721554 | 1.382438729 | 0.630609083 | 0.768239964 | 4.104245726 | 5.645646273 | ENSG00000237476 |
| PABPC1L2B | 0.466967243 | 1.382200823 | 0.500560524 | 0.666686425 | 8.154024838 | 11.22192273 | ENSG00000184388 |
| BPHL | 0.466795713 | 1.382036495 | 4.13E-05 | 0.000241719 | 366.7786345 | 506.3430902 | ENSG00000137274 |
| CHRNB2 | 0.466748525 | 1.381991292 | 0.006081759 | 0.020787247 | 133.9611888 | 184.9073964 | ENSG00000160716 |
| NT5C2 | 0.466415158 | 1.381671988 | 6.93E-08 | 6.51E-07 | 647.9913957 | 895.1699729 | ENSG00000076685 |
| ITGB5 | 0.466302978 | 1.381564558 | 5.39E-31 | 3.57E-29 | 7161.269414 | 9893.088235 | ENSG00000082781 |
| HPS6 | 0.466199783 | 1.381465739 | 3.38E-07 | 2.87E-06 | 532.8098681 | 735.9131239 | ENSG00000166189 |
| VIPAS39 | 0.466074005 | 1.381345305 | 6.87E-06 | 4.66E-05 | 405.0528312 | 559.5772882 | ENSG00000151445 |
| NA | 0.465891705 | 1.381170767 | 0.705425214 | 0.822754402 | 2.653308353 | 3.695737221 | ENSG00000213772 |
| PYGO2 | 0.465854975 | 1.381135605 | 3.48E-11 | 5.02E-10 | 1047.584998 | 1446.630963 | ENSG00000163348 |
| DOK6 | 0.465544314 | 1.380838231 | 0.339123623 | 0.511622821 | 15.89050163 | 22.02090225 | ENSG00000206052 |
| PCP4L1 | 0.465469599 | 1.380766721 | 0.317779781 | 0.489053829 | 24.050784 | 33.14146912 | ENSG00000248485 |
| CARD11 | 0.465305175 | 1.380609364 | 2.77E-12 | 4.50E-11 | 1216.610182 | 1679.668234 | ENSG00000198286 |
| PRR14L | 0.464965884 | 1.380284713 | 8.42E-08 | 7.83E-07 | 913.4814346 | 1260.817064 | ENSG00000183530 |
| LOC400684 | 0.46491579 | 1.380236787 | 0.298575013 | 0.46906992 | 19.89418684 | 27.51275486 | ENSG00000267213 |
| ZNF430 | 0.464719864 | 1.380049356 | 0.006387191 | 0.021650904 | 139.8032983 | 193.2200457 | ENSG00000118620 |
| DUSP23 | 0.464519228 | 1.379857445 | 1.48E-07 | 1.33E-06 | 604.4847869 | 834.60777 | ENSG00000158716 |
| TUBB2B | 0.464343988 | 1.379689848 | 3.96E-23 | 1.67E-21 | 10675.81519 | 14729.34654 | ENSG00000137285 |
| OBSL1 | 0.46432147 | 1.379668313 | 9.02E-22 | 3.46E-20 | 3333.209632 | 4598.426598 | ENSG00000124006 |
| KCNA3 | 0.464315389 | 1.379662498 | 0.419559915 | 0.591664088 | 12.69013654 | 17.54729136 | ENSG00000177272 |
| CPT1C | 0.464297712 | 1.379645593 | 7.13E-14 | 1.37E-12 | 1265.722774 | 1745.694059 | ENSG00000169169 |
| GBE1 | 0.46397249 | 1.37933462 | 0.00265425 | 0.010097248 | 193.1136012 | 266.8320612 | ENSG00000114480 |
| NA | 0.463910918 | 1.379275753 | 0.477854145 | 0.646081043 | 8.357736624 | 11.5394694 | ENSG00000231881 |
| SPP1 | 0.463881871 | 1.379247984 | 0.000271513 | 0.001330056 | 291.0590684 | 401.4704174 | ENSG00000118785 |
| NA | 0.463745555 | 1.379117668 | 0.627307507 | 0.765592317 | 4.261863607 | 5.919096974 | ENSG00000261348 |
| CYB5RL | 0.463705448 | 1.379079329 | 0.000743658 | 0.003255252 | 213.4534453 | 294.6760597 | ENSG00000215883 |
| PKDCC | 0.463656594 | 1.379032631 | 6.42E-06 | 4.38E-05 | 926.1058174 | 1277.227791 | ENSG00000162878 |
| NA | 0.463614299 | 1.378992202 | 0.649649921 | 0.781978702 | 4.052570014 | 5.565523585 | ENSG00000279212 |
| CCDC102B | 0.463550221 | 1.378930955 | 0.449601106 | 0.619828123 | 9.748625576 | 13.39258421 | ENSG00000150636 |
| SEC22A | 0.463547137 | 1.378928008 | 0.091944231 | 0.193851956 | 57.01529764 | 78.82234817 | ENSG00000121542 |
| TBC1D8-AS1 | 0.463289023 | 1.378681324 | 0.655094766 | 0.786056673 | 3.363764935 | 4.638157411 | ENSG00000272902 |
| POLI | 0.463060926 | 1.378463365 | 5.33E-05 | 0.000304957 | 323.9388158 | 446.4151382 | ENSG00000101751 |
| GPRC5A | 0.463016009 | 1.378420448 | 0.397943042 | 0.570346957 | 13.16406622 | 18.09150915 | ENSG00000013588 |
| LLPH-DT | 0.462893553 | 1.378303454 | 0.376211418 | 0.549602004 | 14.92744845 | 20.60284058 | ENSG00000239335 |
| CDH26 | 0.462834494 | 1.378247032 | 0.21523966 | 0.372093224 | 25.75643315 | 35.49360915 | ENSG00000124215 |
| PGGHG | 0.462601121 | 1.378024102 | 1.17E-08 | 1.24E-07 | 1490.764005 | 2053.345292 | ENSG00000142102 |
| NA | 0.462527206 | 1.377953501 | 0.671098877 | 0.798289611 | 3.284855915 | 4.537062025 | ENSG00000228487 |
| RFX3-AS1 | 0.462507137 | 1.377934334 | 0.600023263 | 0.746343979 | 6.423257561 | 8.811426216 | ENSG00000232104 |
| ACSS3 | 0.461994143 | 1.377444454 | 3.74E-07 | 3.16E-06 | 560.968001 | 772.5176978 | ENSG00000111058 |
| PHOSPHO2 | 0.461989921 | 1.377440423 | 0.269212663 | 0.435576741 | 23.88882361 | 33.03014179 | ENSG00000144362 |
| NA | 0.461958792 | 1.377410702 | 0.139277246 | 0.266812051 | 56.71669515 | 78.29103383 | ENSG00000229152 |

| | | | | | | | |
|---|---|---|---|---|---|---|---|
| COPA | 0.461833921 | 1.377291487 | 2.41E-25 | 1.20E-23 | 4408.348208 | 6070.687996 | ENSG00000122218 |
| MIR4326 | 0.461564594 | 1.377034394 | 0.674203771 | 0.800127035 | 3.303716514 | 4.55588415 | ENSG00000266104 |
| DEGS1 | 0.461400569 | 1.376877844 | 6.37E-09 | 6.96E-08 | 773.8806982 | 1065.682966 | ENSG00000143753 |
| CNTN2 | 0.461336784 | 1.376816969 | 0.012375636 | 0.037626223 | 132.6465812 | 182.9131188 | ENSG00000184144 |
| F12 | 0.461304633 | 1.376786286 | 0.119299543 | 0.237429845 | 48.29296488 | 66.32782772 | ENSG00000131187 |
| NA | 0.461036636 | 1.376530557 | 0.073090118 | 0.161530855 | 60.39688403 | 83.0020687 | ENSG00000260853 |
| NA | 0.460931304 | 1.376430059 | 0.007421256 | 0.02453734 | 140.398601 | 192.9826217 | ENSG00000204789 |
| NA | 0.460436286 | 1.375957859 | 0.624158178 | 0.763058842 | 4.277933303 | 5.921247548 | ENSG00000249119 |
| FCGRT | 0.460247834 | 1.375778137 | 1.60E-05 | 0.000101045 | 442.3297201 | 609.1717007 | ENSG00000104870 |
| C1orf131 | 0.459971016 | 1.375514184 | 0.000454996 | 0.002108416 | 222.5651417 | 306.1663385 | ENSG00000143633 |
| LOC148709 | 0.459827545 | 1.375377401 | 0.103811518 | 0.213204301 | 51.8724811 | 71.12354689 | ENSG00000234996 |
| LINC01670 | 0.45977553 | 1.375327813 | 0.339216329 | 0.511675831 | 18.53910406 | 25.36548922 | ENSG00000279094 |
| TMEM68 | 0.459647869 | 1.375206119 | 0.000887363 | 0.003822715 | 282.5196718 | 388.3219509 | ENSG00000167904 |
| LNPEP | 0.459572592 | 1.375134365 | 0.112294726 | 0.226564806 | 58.95942561 | 81.01288575 | ENSG00000113441 |
| NA | 0.459517041 | 1.375081417 | 0.462022604 | 0.631715787 | 9.364768527 | 12.90053165 | ENSG00000254198 |
| XRN1 | 0.459238999 | 1.374816431 | 0.002788193 | 0.010540277 | 180.3779713 | 248.1530318 | ENSG00000114127 |
| PKM | 0.459114574 | 1.374697865 | 1.13E-24 | 5.43E-23 | 20452.90681 | 28114.5728 | ENSG00000067225 |
| NA | 0.459043028 | 1.374629693 | 0.170619222 | 0.312012078 | 32.49512663 | 44.61397974 | ENSG00000255026 |
| FGD3 | 0.459032824 | 1.37461997 | 0.076382461 | 0.167043878 | 60.10413876 | 82.81035822 | ENSG00000127084 |
| C16orf58 | 0.459029092 | 1.374616414 | 2.00E-05 | 0.000124807 | 372.2534322 | 511.7321994 | ENSG00000140688 |
| NA | 0.458876749 | 1.374471268 | 0.169681964 | 0.310769463 | 33.61579769 | 46.25709499 | ENSG00000229939 |
| CLPSL2 | 0.458795935 | 1.374394278 | 0.34954178 | 0.522187755 | 15.85298059 | 21.73293057 | ENSG00000196748 |
| NA | 0.45850795 | 1.374119954 | 0.358554971 | 0.531627402 | 17.76281709 | 24.49458943 | ENSG00000229358 |
| MAP4K1 | 0.458384937 | 1.374002793 | 0.016127714 | 0.047018246 | 104.4983215 | 143.4481711 | ENSG00000104814 |
| NA | 0.457946954 | 1.373585728 | 0.348864626 | 0.521545563 | 15.77407157 | 21.6173142 | ENSG00000268218 |
| FOXD3-AS1 | 0.457838793 | 1.373482752 | 2.15E-11 | 3.16E-10 | 1579.58818 | 2168.73589 | ENSG00000230798 |
| VLDLR | 0.457287329 | 1.372957844 | 4.57E-05 | 0.000264669 | 490.3276229 | 672.8099046 | ENSG00000147852 |
| SPDYA | 0.457160747 | 1.372837386 | 0.632388791 | 0.769387751 | 4.071430613 | 5.548852033 | ENSG00000163806 |
| RNF207 | 0.457035071 | 1.372717801 | 0.068905018 | 0.154090265 | 62.29854798 | 85.47710084 | ENSG00000158286 |
| SEMA3A | 0.457000341 | 1.372684756 | 1.36E-09 | 1.63E-08 | 905.859311 | 1243.96029 | ENSG00000075213 |
| EBF3 | 0.456994494 | 1.372679192 | 0.634362202 | 0.771010663 | 4.29467872 | 5.921247548 | ENSG00000108001 |
| LINC00467 | 0.456915821 | 1.37260434 | 0.106736244 | 0.217543463 | 47.93893189 | 65.84222688 | ENSG00000153363 |
| ATRX | 0.456723756 | 1.372421618 | 3.94E-07 | 3.30E-06 | 1250.551639 | 1715.842479 | ENSG00000085224 |
| FOXH1 | 0.45665945 | 1.372360446 | 4.55E-08 | 4.40E-07 | 1154.777681 | 1585.2563 | ENSG00000160973 |
| TRPT1 | 0.456366982 | 1.372082265 | 0.00051375 | 0.002348517 | 223.0795834 | 306.0913105 | ENSG00000149743 |
| CFAP97 | 0.456200622 | 1.371924056 | 4.68E-07 | 3.87E-06 | 608.8749568 | 835.8942825 | ENSG00000164323 |
| AKAP11 | 0.455204495 | 1.37097712 | 1.49E-05 | 9.49E-05 | 487.7346895 | 668.9920871 | ENSG00000023516 |
| RPS6KA5 | 0.455168327 | 1.370942751 | 0.172469192 | 0.314619686 | 34.32154834 | 46.92983259 | ENSG00000100784 |
| NA | 0.454953854 | 1.37073896 | 0.435367662 | 0.606135187 | 13.62105033 | 18.69809358 | ENSG00000251141 |
| NA | 0.454753534 | 1.370548645 | 0.724864708 | 0.834977314 | 2.604423544 | 3.594641834 | ENSG00000237301 |
| GGCX | 0.454578996 | 1.370382845 | 4.87E-08 | 4.68E-07 | 604.765805 | 828.9715195 | ENSG00000115486 |
| DHRS4L1 | 0.454461314 | 1.370271065 | 0.717424286 | 0.830622489 | 2.607214447 | 3.598942982 | ENSG00000225766 |
| ZNF221 | 0.454460741 | 1.370270522 | 0.303730761 | 0.474434827 | 21.16864573 | 28.9749125 | ENSG00000159905 |
| ANP32B | 0.454424131 | 1.37023575 | 1.50E-19 | 4.86E-18 | 2607.24749 | 3572.653223 | ENSG00000136938 |
| APP | 0.454299731 | 1.370117603 | 3.37E-29 | 2.07E-27 | 20588.67017 | 28209.37412 | ENSG00000142192 |
| LINC01547 | 0.454213367 | 1.370035586 | 0.331640791 | 0.504310461 | 18.90968232 | 25.77552899 | ENSG00000183250 |
| LOC102546294 | 0.454065907 | 1.369895559 | 0.642793735 | 0.776853398 | 4.817493211 | 6.535368196 | ENSG00000247199 |
| NA | 0.4537943 | 1.369637683 | 0.600111439 | 0.746371046 | 5.056811014 | 6.90776371 | ENSG00000279300 |
| LDHA | 0.453676284 | 1.369525648 | 1.94E-18 | 5.73E-17 | 9560.921539 | 13094.15829 | ENSG00000134333 |
| SCN1B | 0.453548052 | 1.369403925 | 0.124930106 | 0.24549523 | 46.71790047 | 63.78853883 | ENSG00000105711 |
| RNF113A | 0.453522562 | 1.369379729 | 0.001088801 | 0.00460175 | 223.1823713 | 305.3328041 | ENSG00000125352 |
| CXXC4 | 0.453450761 | 1.369311579 | 0.20825468 | 0.363053507 | 29.71334872 | 40.6915714 | ENSG00000168772 |
| HLA-DOA | 0.45316842 | 1.369043626 | 6.28E-06 | 4.29E-05 | 460.9115763 | 631.1256626 | ENSG00000204252 |
| PDE4A | 0.453117772 | 1.368995564 | 0.006103757 | 0.020846797 | 163.6786798 | 224.1690445 | ENSG00000065989 |
| NA | 0.452718603 | 1.368616839 | 0.029307676 | 0.077047476 | 93.92627587 | 128.3796616 | ENSG00000203761 |
| GRIA4 | 0.452138909 | 1.368067021 | 0.001190403 | 0.004992187 | 201.4902353 | 275.6163235 | ENSG00000152578 |
| LINC00667 | 0.452109755 | 1.368039376 | 0.017897392 | 0.051240516 | 130.8301959 | 179.0681495 | ENSG00000263753 |
| TMEM64 | 0.451994394 | 1.367929989 | 1.53E-06 | 1.16E-05 | 515.0959231 | 704.7621642 | ENSG00000180694 |
| ZNF805 | 0.451656135 | 1.367609296 | 0.020877677 | 0.058209462 | 96.94302907 | 132.5190543 | ENSG00000204524 |
| LCA10 | 0.451633537 | 1.367587875 | 0.70726122 | 0.823594399 | 3.130028937 | 4.217364777 | ENSG00000273769 |
| ARID3A | 0.451458091 | 1.367421573 | 4.01E-15 | 8.73E-14 | 2149.374094 | 2938.954716 | ENSG00000116017 |
| NA | 0.450978207 | 1.366966803 | 0.702298983 | 0.820646295 | 2.634447755 | 3.624216828 | ENSG00000241170 |
| PTGES3L | 0.450760332 | 1.366760379 | 0.485145536 | 0.652828023 | 8.681545253 | 11.88874164 | ENSG00000267060 |
| PROSER3 | 0.450745614 | 1.366746437 | 0.008001986 | 0.026119511 | 145.0627819 | 198.1037085 | ENSG00000167595 |
| SDAD1P1 | 0.450708436 | 1.366711216 | 0.447127828 | 0.617477482 | 10.30896111 | 14.11210139 | ENSG00000228451 |
| NA | 0.450485585 | 1.366500118 | 0.09556443 | 0.199945829 | 56.59065309 | 77.2004782 | ENSG00000238278 |
| NA | 0.450359123 | 1.366380341 | 0.161056079 | 0.298341927 | 37.35592284 | 50.92214378 | ENSG00000273230 |
| C1orf53 | 0.450150708 | 1.366182965 | 0.589435769 | 0.738795752 | 5.756103983 | 7.809856038 | ENSG00000203724 |
| AOC1 | 0.450064843 | 1.366101655 | 0.649276775 | 0.781798781 | 4.079803322 | 5.525728761 | ENSG00000002726 |
| NA | 0.450046189 | 1.366083993 | 0.512143872 | 0.676434317 | 7.710119367 | 10.52552882 | ENSG00000270055 |
| SLC25A40 | 0.450029952 | 1.366068618 | 0.007626927 | 0.025073458 | 142.6500307 | 195.0301306 | ENSG00000075303 |
| XRCC3 | 0.449802964 | 1.365853703 | 1.13E-05 | 7.32E-05 | 443.992979 | 606.4038627 | ENSG00000126215 |
| UTRN | 0.449791405 | 1.365842759 | 9.50E-08 | 8.79E-07 | 906.1785755 | 1236.989614 | ENSG00000152818 |
| NA | 0.449650497 | 1.365709364 | 0.70599228 | 0.823224556 | 2.656099256 | 3.645189528 | ENSG00000257497 |
| NA | 0.449640653 | 1.365700045 | 0.664474854 | 0.793258787 | 4.095873018 | 5.548852033 | ENSG00000230537 |
| TXNRD2 | 0.449635134 | 1.365694821 | 0.010813782 | 0.033684592 | 131.8633099 | 180.1237629 | ENSG00000184470 |
| CAPN13 | 0.44960433 | 1.365665661 | 0.12181462 | 0.241191197 | 47.64044154 | 65.07645104 | ENSG00000162949 |

| | | | | | | | |
|---|---|---|---|---|---|---|---|
| C16orf96 | 0.44957237 | 1.365635408 | 0.217241474 | 0.374680848 | 29.90398188 | 40.77707973 | ENSG00000205832 |
| ARFGEF3 | 0.449527886 | 1.365593301 | 1.20E-06 | 9.36E-06 | 866.6990679 | 1183.236951 | ENSG00000112379 |
| ACCS | 0.449503 | 1.365569745 | 0.033039969 | 0.084949226 | 96.40031484 | 131.4521428 | ENSG00000110455 |
| BMP1 | 0.449481285 | 1.365549191 | 0.001412967 | 0.005814321 | 224.9618232 | 306.8183881 | ENSG00000168487 |
| DUS2 | 0.449322448 | 1.365398856 | 0.000204144 | 0.0010277 | 289.2239986 | 395.0904361 | ENSG00000167264 |
| CREG1 | 0.449143729 | 1.365229723 | 4.49E-06 | 3.15E-05 | 518.3016457 | 708.2997824 | ENSG00000143162 |
| LBR | 0.449084669 | 1.365173835 | 5.19E-15 | 1.12E-13 | 1940.40114 | 2648.927105 | ENSG00000143815 |
| MTHFR | 0.449043361 | 1.365134747 | 0.00045654 | 0.002114043 | 263.2708749 | 359.1782276 | ENSG00000177000 |
| NA | 0.44898365 | 1.365078248 | 0.653180735 | 0.784504704 | 4.098663921 | 5.544550886 | ENSG00000287576 |
| NA | 0.44853381 | 1.364652675 | 0.256885344 | 0.422038424 | 27.45438531 | 37.44405123 | ENSG00000274828 |
| PJA2 | 0.448443324 | 1.364567087 | 1.05E-09 | 1.27E-08 | 1289.981625 | 1760.144019 | ENSG00000198961 |
| IGF1R | 0.44817332 | 1.364311729 | 4.21E-13 | 7.45E-12 | 2026.674543 | 2764.725623 | ENSG00000140443 |
| NA | 0.447816207 | 1.36397406 | 0.64403017 | 0.77751962 | 4.640138851 | 6.379684433 | ENSG00000227620 |
| NA | 0.447813857 | 1.363971838 | 0.508883931 | 0.673520474 | 8.439436547 | 11.46149729 | ENSG00000286966 |
| NDUFAF3 | 0.447756899 | 1.363917989 | 0.006338379 | 0.021504337 | 141.1971783 | 192.7430472 | ENSG00000178057 |
| LINC01137 | 0.447381696 | 1.363563319 | 0.527648078 | 0.689424922 | 9.528168372 | 12.9591486 | ENSG00000233621 |
| CCNG1 | 0.447187619 | 1.3633799 | 4.58E-13 | 8.10E-12 | 3734.675534 | 5091.934005 | ENSG00000113328 |
| RAB30 | 0.447130612 | 1.363326028 | 0.044973045 | 0.109666921 | 84.95198534 | 115.9343318 | ENSG00000137502 |
| PPP1R21 | 0.446599943 | 1.362824646 | 6.63E-05 | 0.000372553 | 363.1481935 | 495.0627804 | ENSG00000162869 |
| HSD11B1L | 0.446567602 | 1.362794095 | 0.094734868 | 0.198407376 | 79.4189662 | 108.1263537 | ENSG00000167733 |
| AFAP1L1 | 0.446388885 | 1.362625287 | 0.00789162 | 0.025811671 | 162.2588427 | 220.8728668 | ENSG00000157510 |
| NA | 0.446336728 | 1.362576025 | 0.217429652 | 0.374971877 | 29.12298898 | 39.62708313 | ENSG00000260086 |
| KIF12 | 0.446317294 | 1.362557671 | 0.173852767 | 0.316664501 | 36.04653365 | 49.0897412 | ENSG00000136883 |
| SNHG12 | 0.445873561 | 1.362138651 | 0.002592786 | 0.009892687 | 323.2530259 | 439.9975491 | ENSG00000197989 |
| RSKR | 0.445663722 | 1.361940543 | 0.047012521 | 0.113720195 | 84.07553817 | 114.439122 | ENSG00000167524 |
| PCSK9 | 0.445599507 | 1.361879924 | 2.06E-13 | 3.76E-12 | 2280.69588 | 3106.076882 | ENSG00000169174 |
| GYG1 | 0.445529359 | 1.361813707 | 2.58E-07 | 2.23E-06 | 570.1742758 | 776.1951764 | ENSG00000163754 |
| ZC3H10 | 0.445440969 | 1.361730275 | 0.019542683 | 0.055140836 | 103.0152447 | 140.2388282 | ENSG00000135482 |
| LOC105371814 | 0.445298661 | 1.36159596 | 0.702889263 | 0.821286298 | 2.631656852 | 3.601093555 | ENSG00000248278 |
| ONECUT1 | 0.445210029 | 1.361512313 | 0.273353098 | 0.440191783 | 32.34049981 | 43.85922934 | ENSG00000169856 |
| BCDIN3D-AS1 | 0.445194845 | 1.361497983 | 0.629922028 | 0.767672076 | 4.310748416 | 5.875001002 | ENSG00000258057 |
| UNC5D | 0.445046906 | 1.361358378 | 8.48E-10 | 1.04E-08 | 1309.373372 | 1782.762352 | ENSG00000156687 |
| SHISA4 | 0.444983746 | 1.36129878 | 0.029463463 | 0.077362142 | 108.3919222 | 147.33992 | ENSG00000198892 |
| RRAS | 0.444845903 | 1.36116872 | 0.000323934 | 0.001558355 | 277.3593972 | 377.7127981 | ENSG00000126458 |
| ZNF506 | 0.444550188 | 1.360889744 | 0.608199382 | 0.752015913 | 5.32894393 | 7.273707503 | ENSG00000081665 |
| YBX2 | 0.444425122 | 1.360771775 | 1.39E-06 | 1.07E-05 | 660.0070324 | 898.1303817 | ENSG00000006047 |
| PDLIM1 | 0.444395194 | 1.360743546 | 7.10E-21 | 2.56E-19 | 2999.20141 | 4081.373077 | ENSG00000107438 |
| ZNF37BP | 0.444316197 | 1.360669039 | 0.008608423 | 0.027774728 | 140.8291908 | 191.890709 | ENSG00000234420 |
| TMOD1 | 0.444303221 | 1.360656801 | 0.397752858 | 0.570147896 | 15.34412062 | 20.95882501 | ENSG00000136842 |
| B4GALT3 | 0.444270912 | 1.360626329 | 5.99E-08 | 5.69E-07 | 700.6939419 | 953.377922 | ENSG00000158850 |
| PFKP | 0.4442631 | 1.360618962 | 7.74E-21 | 2.79E-19 | 3722.672378 | 5064.585648 | ENSG00000067057 |
| NA | 0.444228928 | 1.360586734 | 0.58960503 | 0.738863885 | 8.246888372 | 11.13964947 | ENSG00000226912 |
| LYRM9 | 0.443753225 | 1.360138179 | 0.491030524 | 0.657452395 | 8.950887267 | 12.23371274 | ENSG00000232859 |
| SYT2 | 0.443417994 | 1.359822168 | 3.12E-07 | 2.67E-06 | 843.8860991 | 1147.576513 | ENSG00000143858 |
| PIGX | 0.443230969 | 1.359645898 | 0.001356515 | 0.005601143 | 215.1374429 | 292.8390834 | ENSG00000163964 |
| RASSF5 | 0.443211457 | 1.359627509 | 0.161982152 | 0.299539656 | 44.85220595 | 60.98897525 | ENSG00000266094 |
| DDAH1 | 0.442909051 | 1.359342545 | 5.02E-08 | 4.82E-07 | 785.6218483 | 1068.166831 | ENSG00000153904 |
| SSBP4 | 0.442901437 | 1.359335371 | 2.35E-07 | 2.04E-06 | 806.0844552 | 1095.794209 | ENSG00000130511 |
| STARD4-AS1 | 0.44283809 | 1.359275685 | 0.089355643 | 0.189653395 | 70.01771578 | 95.16290397 | ENSG00000246859 |
| NA | 0.442732219 | 1.35917594 | 0.689464893 | 0.811401055 | 3.690364466 | 5.035826772 | ENSG00000263884 |
| PIDD1 | 0.442494778 | 1.358952262 | 1.11E-05 | 7.20E-05 | 562.8331703 | 764.8837658 | ENSG00000177595 |
| NEMP2 | 0.44217443 | 1.358650543 | 0.061255074 | 0.140241114 | 76.58396099 | 104.1114707 | ENSG00000189362 |
| ATP2B4 | 0.442154305 | 1.358631591 | 2.95E-06 | 2.13E-05 | 526.5372441 | 715.0714901 | ENSG00000058668 |
| ARF1 | 0.441841218 | 1.358336779 | 1.03E-29 | 6.47E-28 | 6589.011244 | 8950.701173 | ENSG00000143761 |
| NA | 0.441827575 | 1.358323933 | 0.634508667 | 0.771091515 | 4.368005934 | 5.925548695 | ENSG00000261220 |
| RYK | 0.441741475 | 1.358242871 | 5.71E-10 | 7.15E-09 | 924.8325964 | 1255.868999 | ENSG00000163785 |
| NA | 0.441741326 | 1.358242731 | 0.395406201 | 0.568050419 | 15.22613896 | 20.76523653 | ENSG00000231770 |
| NA | 0.441638102 | 1.358145552 | 0.601860642 | 0.747617106 | 5.786128194 | 7.786732765 | ENSG00000227288 |
| LIAS | 0.441592797 | 1.358102904 | 0.003055244 | 0.011371546 | 218.0745234 | 296.3678752 | ENSG00000121897 |
| HARS2 | 0.441367499 | 1.357890832 | 1.73E-05 | 0.000109118 | 431.7763846 | 585.7919294 | ENSG00000112855 |
| THEM5 | 0.440629352 | 1.357196253 | 0.101893024 | 0.210069288 | 50.74515219 | 68.86684403 | ENSG00000196407 |
| JHY | 0.44051874 | 1.3570922 | 0.365460137 | 0.539162703 | 16.56901898 | 22.41857161 | ENSG00000109944 |
| IL20RB | 0.440195397 | 1.356788077 | 0.234320531 | 0.395267345 | 27.3677793 | 37.16576635 | ENSG00000174564 |
| NA | 0.440190656 | 1.356783618 | 0.757636613 | 0.856114219 | 2.612796253 | 3.55484701 | ENSG00000272622 |
| AMACR | 0.440110796 | 1.356708516 | 0.302280921 | 0.473007588 | 20.81971898 | 28.17957378 | ENSG00000242110 |
| MALL | 0.440055931 | 1.356656922 | 0.385621342 | 0.559240814 | 14.82987899 | 20.06884262 | ENSG00000144063 |
| DNAH10 | 0.439779876 | 1.356397355 | 0.555202654 | 0.711324593 | 6.964056769 | 9.450820711 | ENSG00000197653 |
| KIAA0825 | 0.439748871 | 1.356368204 | 0.221076638 | 0.379429664 | 32.12367249 | 43.70758627 | ENSG00000185261 |
| LINC00562 | 0.439738603 | 1.356358551 | 0.73713573 | 0.843705033 | 2.607214447 | 3.548395288 | ENSG00000260388 |
| NA | 0.439711198 | 1.356332786 | 0.220440451 | 0.37870841 | 31.14195887 | 42.27311351 | ENSG00000280832 |
| NA | 0.43962318 | 1.35625004 | 0.674098013 | 0.800127035 | 4.13706084 | 5.551002607 | ENSG00000270804 |
| MRI1 | 0.439576353 | 1.35620602 | 0.004714777 | 0.016682459 | 197.7056559 | 268.0667358 | ENSG00000037757 |
| TTC3 | 0.439529104 | 1.356161604 | 5.54E-07 | 4.53E-06 | 766.6131939 | 1039.60489 | ENSG00000182670 |
| TAFA3 | 0.439528233 | 1.356160785 | 0.59480594 | 0.742733338 | 7.306725997 | 9.945284311 | ENSG00000184599 |
| ALG13 | 0.439485115 | 1.356120254 | 4.49E-10 | 5.72E-09 | 1105.988131 | 1499.448639 | ENSG00000101901 |
| PLCB1 | 0.439185599 | 1.355838741 | 0.019047435 | 0.05392487 | 138.7809973 | 187.9707117 | ENSG00000182621 |
| LRCH4 | 0.439076082 | 1.355735821 | 0.000691064 | 0.00304855 | 271.6144569 | 367.9998673 | ENSG00000077454 |

| | | | | | | | |
|---|---|---|---|---|---|---|---|
| SHTN1 | 0.439059905 | 1.355720619 | 0.000467448 | 0.002158678 | 323.4478142 | 437.9478469 | ENSG00000187164 |
| ALX1 | 0.439044643 | 1.355706277 | 0.292288059 | 0.461985166 | 25.13863994 | 34.20191671 | ENSG00000180318 |
| CCS | 0.438773121 | 1.355451151 | 1.52E-07 | 1.36E-06 | 660.4769077 | 895.5945641 | ENSG00000173992 |
| PRDX4 | 0.438591158 | 1.355280203 | 4.43E-17 | 1.17E-15 | 2218.668624 | 3006.892443 | ENSG00000123131 |
| INKA1 | 0.438439083 | 1.355137349 | 0.086619594 | 0.184924588 | 59.47904888 | 80.53457278 | ENSG00000185614 |
| SHISA8 | 0.438388298 | 1.355089648 | 0.40974355 | 0.581930563 | 13.28607809 | 18.03504277 | ENSG00000234965 |
| WIPI1 | 0.438308191 | 1.355014407 | 0.0003398 | 0.001628187 | 348.7768871 | 471.9688303 | ENSG00000070540 |
| CPNE8 | 0.438202584 | 1.354915222 | 0.01615674 | 0.047060195 | 114.2413652 | 154.6367507 | ENSG00000139117 |
| OGDHL | 0.437747077 | 1.354487498 | 4.70E-06 | 3.28E-05 | 624.4521235 | 845.1924367 | ENSG00000197444 |
| PLCB4 | 0.43772944 | 1.354470939 | 0.00222091 | 0.008643043 | 230.7727941 | 312.2400226 | ENSG00000101333 |
| GOLGA8A | 0.437682557 | 1.354426924 | 0.00023213 | 0.001155021 | 328.0294193 | 443.9059574 | ENSG00000175265 |
| TECPR2 | 0.437631116 | 1.354378631 | 1.70E-08 | 1.76E-07 | 741.9260443 | 1004.460081 | ENSG00000196663 |
| STOM | 0.437367339 | 1.354131024 | 3.15E-07 | 2.69E-06 | 649.0693274 | 879.0009668 | ENSG00000148175 |
| PPP1R15A | 0.437100062 | 1.353880178 | 3.63E-05 | 0.000214922 | 372.7762467 | 504.5728001 | ENSG00000087074 |
| PRKAG1 | 0.43695453 | 1.353743612 | 1.06E-05 | 6.92E-05 | 422.6004094 | 571.8941166 | ENSG00000181929 |
| EPHX1 | 0.436837542 | 1.353633841 | 1.74E-12 | 2.88E-11 | 1382.306354 | 1870.726959 | ENSG00000143819 |
| SOX4 | 0.436664512 | 1.353471503 | 3.75E-22 | 1.49E-20 | 5562.248753 | 7527.731489 | ENSG00000124766 |
| ADAM8 | 0.436660374 | 1.353467621 | 0.03940369 | 0.098570579 | 89.85329366 | 121.5571458 | ENSG00000151651 |
| RPRD1A | 0.436251994 | 1.353084553 | 7.08E-17 | 1.85E-15 | 2435.523206 | 3296.148939 | ENSG00000141425 |
| DMD | 0.43589552 | 1.352750261 | 0.00907303 | 0.028982915 | 137.7742408 | 186.3784168 | ENSG00000198947 |
| NMB | 0.435457451 | 1.352339566 | 0.004661737 | 0.01653117 | 179.9855413 | 243.5778046 | ENSG00000197696 |
| SENP7 | 0.43544253 | 1.35232558 | 0.044189514 | 0.108056777 | 86.89224637 | 117.5981471 | ENSG00000138468 |
| TMEM241 | 0.435192939 | 1.352091643 | 0.028014985 | 0.074337974 | 112.9373577 | 152.99684 | ENSG00000134490 |
| NT5E | 0.435147004 | 1.352048593 | 0.484223996 | 0.652042797 | 12.61401843 | 17.08993897 | ENSG00000135318 |
| NSL1 | 0.435108553 | 1.352012559 | 6.30E-09 | 6.89E-08 | 951.8266062 | 1287.427763 | ENSG00000117697 |
| NA | 0.435058297 | 1.351965463 | 0.505226652 | 0.670313776 | 8.948096364 | 12.12616563 | ENSG00000259994 |
| NA | 0.434793902 | 1.351717718 | 0.202464942 | 0.355036015 | 40.03857968 | 54.05860919 | ENSG00000260265 |
| ATAD5 | 0.434521383 | 1.351462408 | 0.000317743 | 0.001532779 | 334.1754505 | 451.1782957 | ENSG00000176208 |
| NA | 0.43415705 | 1.351121158 | 0.580980408 | 0.731628608 | 6.303160719 | 8.537975515 | ENSG00000235008 |
| NA | 0.434124201 | 1.351090394 | 0.117941348 | 0.235406782 | 52.37045286 | 70.75168441 | ENSG00000271369 |
| C2orf50 | 0.434043292 | 1.351014625 | 0.489158719 | 0.655667096 | 10.64672425 | 14.38932021 | ENSG00000150873 |
| NA | 0.433835554 | 1.350820102 | 0.541838493 | 0.700670758 | 7.761795079 | 10.426584 | ENSG00000260123 |
| TBCE | 0.433735369 | 1.350726301 | 0.155254459 | 0.290270766 | 41.7447044 | 56.36398174 | ENSG00000285053 |
| ARHGEF2 | 0.433629686 | 1.350627358 | 9.30E-13 | 1.58E-11 | 1479.938514 | 1999.108257 | ENSG00000116584 |
| SYTL5 | 0.433538547 | 1.350542038 | 0.529273178 | 0.690485485 | 8.782781496 | 11.81937182 | ENSG00000147041 |
| NA | 0.433536435 | 1.350540061 | 0.63374331 | 0.770452621 | 5.878115847 | 8.020649106 | ENSG00000261549 |
| GEN1 | 0.433257975 | 1.350279413 | 0.001339441 | 0.005538942 | 222.5838021 | 300.644099 | ENSG00000178295 |
| NA | 0.433196992 | 1.350222338 | 0.751008604 | 0.852669471 | 2.680541661 | 3.617765107 | ENSG00000231536 |
| MATN3 | 0.433159484 | 1.350187234 | 0.018267755 | 0.052049988 | 130.6555203 | 176.5880227 | ENSG00000132031 |
| NA | 0.433121014 | 1.350151232 | 0.303251692 | 0.473955148 | 24.76268004 | 33.39367455 | ENSG00000259363 |
| EPB41L4B | 0.433119099 | 1.35014944 | 2.74E-08 | 2.72E-07 | 733.5094105 | 990.160019 | ENSG00000095203 |
| NA | 0.433102693 | 1.350134086 | 0.338092365 | 0.510699225 | 22.88266759 | 30.82104255 | ENSG00000275557 |
| MALAT1 | 0.433009684 | 1.350047048 | 9.95E-06 | 6.56E-05 | 1073.891832 | 1450.921283 | ENSG00000251562 |
| TMPRSS13 | 0.43298586 | 1.350024754 | 0.637734007 | 0.773520244 | 5.982142993 | 8.055882319 | ENSG00000137747 |
| LRRC38 | 0.432888411 | 1.349933567 | 0.585993861 | 0.735676661 | 6.942405268 | 9.364246302 | ENSG00000162494 |
| ZNF627 | 0.432745886 | 1.349800213 | 0.000401825 | 0.001892019 | 273.8205053 | 369.6970016 | ENSG00000198551 |
| RCAN3 | 0.432677561 | 1.349736289 | 0.000518966 | 0.002369725 | 256.7784325 | 346.5451974 | ENSG00000117602 |
| TRMT5 | 0.432547164 | 1.349614299 | 6.79E-06 | 4.62E-05 | 542.5582424 | 731.657836 | ENSG00000126814 |
| TNFAIP8L1 | 0.432438627 | 1.349512769 | 0.002979155 | 0.011133452 | 215.8298027 | 291.5554418 | ENSG00000185361 |
| NA | 0.432413254 | 1.349489035 | 0.718170693 | 0.831037781 | 2.922650367 | 3.994461769 | ENSG00000230773 |
| NA | 0.432376526 | 1.34945468 | 0.629922025 | 0.767672076 | 4.612905543 | 6.264068066 | ENSG00000243759 |
| NA | 0.432283929 | 1.34936807 | 0.421045393 | 0.593046306 | 16.33787373 | 22.07790166 | ENSG00000256673 |
| NA | 0.432138601 | 1.34923215 | 0.42290921 | 0.594457999 | 14.75376088 | 19.83545932 | ENSG00000260805 |
| CXorf38 | 0.431979077 | 1.349082969 | 0.002067028 | 0.008124497 | 196.2530789 | 264.9477539 | ENSG00000185753 |
| GSG1 | 0.431914686 | 1.349022757 | 0.603506491 | 0.748841816 | 5.671613158 | 7.694500137 | ENSG00000111305 |
| ARL17A | 0.431486383 | 1.348622323 | 0.40153674 | 0.573951189 | 14.88713651 | 19.97634953 | ENSG00000228696 |
| SLC6A1 | 0.431293859 | 1.348442365 | 0.427190885 | 0.598608428 | 13.91291972 | 18.64701285 | ENSG00000157103 |
| NA | 0.430902691 | 1.348076802 | 0.006640185 | 0.022390459 | 152.4587047 | 205.6766615 | ENSG00000263072 |
| DCLK1 | 0.430709574 | 1.347896362 | 3.92E-07 | 3.30E-06 | 725.9022423 | 978.6966011 | ENSG00000133083 |
| SLC2A14 | 0.430313125 | 1.347526014 | 0.023035685 | 0.06312796 | 108.9047243 | 146.5424307 | ENSG00000173262 |
| NA | 0.430130973 | 1.347355889 | 0.692986684 | 0.814253022 | 4.191527455 | 5.578427028 | ENSG00000237976 |
| MLF1 | 0.42986671 | 1.347109112 | 8.52E-06 | 5.69E-05 | 490.5304588 | 660.3334311 | ENSG00000178053 |
| NA | 0.429359138 | 1.346635253 | 0.445323057 | 0.615843321 | 14.23951926 | 19.10598277 | ENSG00000285979 |
| SLC45A1 | 0.429114092 | 1.346406542 | 0.072889693 | 0.161177794 | 75.09809332 | 101.2643035 | ENSG00000162426 |
| CYP2F1 | 0.428851022 | 1.346161052 | 0.66101513 | 0.790750519 | 4.406402853 | 5.902425422 | ENSG00000197446 |
| KMT2A | 0.42874826 | 1.34606517 | 1.25E-06 | 9.71E-06 | 758.2781607 | 1020.456701 | ENSG00000118058 |
| MED13 | 0.428737325 | 1.346054967 | 3.29E-05 | 0.000197159 | 527.5872667 | 710.2620314 | ENSG00000108510 |
| STXBP3 | 0.428636891 | 1.345961265 | 1.10E-06 | 8.62E-06 | 636.9043079 | 857.8259267 | ENSG00000116266 |
| CFAP58 | 0.428553131 | 1.345883122 | 0.459703757 | 0.629539253 | 12.51086716 | 16.78314516 | ENSG00000120051 |
| PPFIBP1 | 0.428519886 | 1.345852109 | 3.60E-06 | 2.57E-05 | 548.4096885 | 738.1627872 | ENSG00000110841 |
| NA | 0.428410212 | 1.345749801 | 0.554657945 | 0.71086277 | 7.830216209 | 10.46046015 | ENSG00000198416 |
| C6orf62 | 0.428292557 | 1.345640056 | 7.03E-09 | 7.65E-08 | 971.0890453 | 1307.273454 | ENSG00000112308 |
| SH3YL1 | 0.428284976 | 1.345632985 | 5.70E-05 | 0.000324663 | 446.2779123 | 600.5299278 | ENSG00000035115 |
| AFF1 | 0.42827643 | 1.345625015 | 2.70E-09 | 3.10E-08 | 943.6849459 | 1269.371529 | ENSG00000172493 |
| CHP2 | 0.428186327 | 1.345540976 | 0.180434537 | 0.325396154 | 38.25681242 | 51.46715507 | ENSG00000166869 |
| LOC152048 | 0.428041389 | 1.345405806 | 0.676545274 | 0.801771402 | 4.37358774 | 5.860480024 | ENSG00000270194 |
| TBC1D8 | 0.427965672 | 1.345335196 | 8.21E-07 | 6.55E-06 | 708.3554888 | 952.7616692 | ENSG00000204634 |

| | | | | | | | |
|---|---|---|---|---|---|---|---|
| FGF12-AS2 | 0.427911489 | 1.345284671 | 0.469867218 | 0.638774384 | 11.85746794 | 16.00501103 | ENSG00000230126 |
| ZNF273 | 0.427867685 | 1.345243825 | 0.001353685 | 0.005591849 | 243.2846635 | 327.7110357 | ENSG00000198039 |
| PSD | 0.427865056 | 1.345241374 | 0.016951849 | 0.048976805 | 145.1555333 | 195.2382706 | ENSG00000059915 |
| NA | 0.427380104 | 1.344789256 | 0.00220965 | 0.008602736 | 238.737225 | 320.9713446 | ENSG00000214796 |
| SNAP47 | 0.42727816 | 1.344694234 | 2.18E-09 | 2.54E-08 | 1002.721767 | 1348.706667 | ENSG00000143740 |
| ZNF398 | 0.427037298 | 1.344469752 | 5.25E-12 | 8.25E-11 | 1544.548401 | 2076.578653 | ENSG00000197024 |
| ALG6 | 0.427036627 | 1.344469127 | 0.000145499 | 0.000754318 | 403.966815 | 542.9299265 | ENSG00000088035 |
| PSD2 | 0.426855504 | 1.344300346 | 0.602735518 | 0.748318373 | 5.372246934 | 7.202187111 | ENSG00000146005 |
| GABARAPL1 | 0.42654891 | 1.344014693 | 1.52E-08 | 1.58E-07 | 1207.296212 | 1622.513596 | ENSG00000139112 |
| CDHR4 | 0.426430898 | 1.343904758 | 0.525842637 | 0.688057405 | 8.798851192 | 11.75592069 | ENSG00000187492 |
| NA | 0.426370855 | 1.343848827 | 0.009191019 | 0.029301594 | 151.0440491 | 203.1992062 | ENSG00000272168 |
| NA | 0.42635946 | 1.343838213 | 0.688469349 | 0.810722879 | 4.316330222 | 5.78250791 | ENSG00000205871 |
| SMG1P5 | 0.426338927 | 1.343819087 | 0.266402359 | 0.432626307 | 27.66993643 | 37.25180772 | ENSG00000183604 |
| TGDS | 0.426264571 | 1.343749828 | 0.002844659 | 0.010715938 | 231.1510325 | 310.9327126 | ENSG00000088451 |
| SNAI2 | 0.426202366 | 1.343691892 | 0.509840932 | 0.674379942 | 8.98370238 | 12.0653981 | ENSG00000019549 |
| CCDC65 | 0.426132162 | 1.343626506 | 0.278405117 | 0.446146628 | 25.71871195 | 34.59313435 | ENSG00000139537 |
| NA | 0.426012722 | 1.343515273 | 0.703966822 | 0.82190951 | 3.647061463 | 4.907306965 | ENSG00000273062 |
| NA | 0.425934272 | 1.343442218 | 0.724045887 | 0.834405189 | 3.478279971 | 4.629555116 | ENSG00000271830 |
| NA | 0.42581939 | 1.343335244 | 0.487378296 | 0.654508508 | 10.09966751 | 13.55874827 | ENSG00000213080 |
| MMAA | 0.425768908 | 1.343288239 | 0.04408002 | 0.107843711 | 93.21725875 | 125.1786484 | ENSG00000151611 |
| SRD5A3 | 0.425591217 | 1.343122803 | 9.85E-05 | 0.0005338 | 361.7827861 | 486.2207437 | ENSG00000128039 |
| MOSPD2 | 0.425276799 | 1.342830116 | 0.006996111 | 0.023370039 | 176.6400734 | 237.3037772 | ENSG00000130150 |
| SNHG29 | 0.425258486 | 1.342813072 | 1.29E-15 | 2.97E-14 | 3438.731001 | 4616.201754 | ENSG00000175061 |
| SPIN3 | 0.425126908 | 1.342690609 | 0.000472849 | 0.002180667 | 270.7563067 | 363.2877542 | ENSG00000204271 |
| NA | 0.424947724 | 1.342523855 | 0.553770891 | 0.710086055 | 8.317224524 | 11.18804658 | ENSG00000273004 |
| NA | 0.424772245 | 1.342360571 | 0.68885181 | 0.81092575 | 3.611455446 | 4.858909846 | ENSG00000259045 |
| OPRD1 | 0.42474838 | 1.342338366 | 0.105206066 | 0.215220596 | 67.15779259 | 90.10424492 | ENSG00000116329 |
| CATSPER2P1 | 0.424728071 | 1.342319469 | 0.410523538 | 0.582738011 | 13.72998356 | 18.42626041 | ENSG00000205771 |
| GPC4 | 0.424592485 | 1.342193323 | 8.18E-24 | 3.71E-22 | 16400.46541 | 22014.69023 | ENSG00000076716 |
| NA | 0.424590192 | 1.342191189 | 0.574299258 | 0.726479147 | 6.382069739 | 8.554647066 | ENSG00000231409 |
| OXLD1 | 0.424525589 | 1.342131088 | 5.99E-06 | 4.11E-05 | 487.1846417 | 653.8623502 | ENSG00000204237 |
| CCHCR1 | 0.424460894 | 1.342070904 | 7.69E-07 | 6.17E-06 | 869.3701097 | 1165.768287 | ENSG00000204536 |
| NA | 0.424433447 | 1.342045372 | 0.710547831 | 0.825733627 | 2.982698788 | 4.042858888 | ENSG00000286024 |
| SLC9A8 | 0.424142941 | 1.34177516 | 0.003845746 | 0.013963539 | 206.3202436 | 276.68647 | ENSG00000197818 |
| B3GALNT2 | 0.424121945 | 1.341755633 | 5.84E-06 | 4.01E-05 | 546.6108124 | 733.0277857 | ENSG00000162885 |
| CCDC8 | 0.423959313 | 1.341604388 | 1.05E-06 | 8.29E-06 | 715.6351206 | 960.1343826 | ENSG00000169515 |
| OTOG | 0.423912337 | 1.341560704 | 0.172195914 | 0.314358984 | 43.74078348 | 58.47119209 | ENSG00000188162 |
| LOC112268124 | 0.423789608 | 1.341446584 | 0.534883607 | 0.695125281 | 8.085603708 | 10.80543124 | ENSG00000258738 |
| ECH1 | 0.423720759 | 1.341382568 | 1.10E-05 | 7.14E-05 | 602.8291738 | 809.0304337 | ENSG00000104823 |
| PDE12 | 0.423715756 | 1.341377917 | 0.000127066 | 0.00066937 | 381.75597 | 512.4538855 | ENSG00000174840 |
| CIAO2B | 0.423642135 | 1.341309468 | 8.41E-09 | 9.06E-08 | 1127.698903 | 1512.494938 | ENSG00000166595 |
| SGO1 | 0.423399853 | 1.341084231 | 0.020062486 | 0.05634378 | 133.5511376 | 179.3891733 | ENSG00000129810 |
| NA | 0.423370879 | 1.341057298 | 0.689877701 | 0.811588653 | 3.899658059 | 5.281853053 | ENSG00000285589 |
| FAM83B | 0.423352996 | 1.341040675 | 0.151742306 | 0.285141324 | 44.30091885 | 59.49184486 | ENSG00000168143 |
| CCDC192 | 0.423332535 | 1.341021656 | 0.730930662 | 0.839058816 | 2.694496175 | 3.601093555 | ENSG00000230561 |
| SEMA6A | 0.423258581 | 1.340952915 | 3.21E-19 | 1.00E-17 | 4855.259558 | 6510.657101 | ENSG00000092421 |
| NA | 0.423234793 | 1.340930805 | 0.713265824 | 0.827546221 | 2.716147677 | 3.624216828 | ENSG00000237276 |
| MYO5A | 0.423226603 | 1.340923193 | 1.19E-06 | 9.26E-06 | 720.2043966 | 965.3099477 | ENSG00000197535 |
| NA | 0.423033695 | 1.340743905 | 0.451086595 | 0.621387568 | 12.79695459 | 17.12649872 | ENSG00000228274 |
| NA | 0.422991461 | 1.340704657 | 0.491820852 | 0.658000975 | 10.90211175 | 14.63372895 | ENSG00000230979 |
| ELMO3 | 0.422948233 | 1.340664485 | 0.002199472 | 0.008570402 | 259.7489285 | 347.8213333 | ENSG00000102890 |
| TMEM185B | 0.422754657 | 1.340484611 | 1.28E-05 | 8.25E-05 | 430.1406713 | 576.6548752 | ENSG00000226479 |
| ABCC10 | 0.422748484 | 1.340478875 | 6.94E-05 | 0.000388276 | 466.2098075 | 624.8863838 | ENSG00000124574 |
| NHS | 0.42247602 | 1.340225739 | 0.000766073 | 0.003342754 | 322.3488458 | 431.9932258 | ENSG00000188158 |
| RLF | 0.422451827 | 1.340203265 | 1.22E-05 | 7.85E-05 | 512.6536728 | 686.8467543 | ENSG00000117000 |
| NSUN5P2 | 0.422415618 | 1.340169629 | 0.22013547 | 0.37828553 | 36.7278419 | 49.03972655 | ENSG00000106133 |
| FAM114A1 | 0.422196946 | 1.339966512 | 0.005843773 | 0.020087709 | 182.9689527 | 244.8660078 | ENSG00000197712 |
| FAM83E | 0.422016328 | 1.339798766 | 0.069097713 | 0.154439112 | 78.23282818 | 105.010328 | ENSG00000105523 |
| TCAIM | 0.421890647 | 1.339682054 | 0.009477316 | 0.030070213 | 193.3831434 | 259.4997062 | ENSG00000179152 |
| NA | 0.421667636 | 1.339474982 | 0.698462957 | 0.817989588 | 3.451046663 | 4.564486445 | ENSG00000246250 |
| COL5A3 | 0.421617583 | 1.339428512 | 0.092905383 | 0.195317976 | 69.77038869 | 93.52141837 | ENSG00000080573 |
| WDR5B | 0.421521151 | 1.339338985 | 0.140977546 | 0.269347117 | 53.6472271 | 71.65350417 | ENSG00000196981 |
| NA | 0.421303459 | 1.339136904 | 0.100201913 | 0.207448393 | 74.91986308 | 100.4786637 | ENSG00000261963 |
| ADAR | 0.421266387 | 1.339102493 | 9.33E-24 | 4.19E-22 | 7947.214295 | 10641.95301 | ENSG00000160710 |
| UNC13A | 0.421183584 | 1.339025638 | 5.47E-14 | 1.06E-12 | 1842.417764 | 2466.269809 | ENSG00000130477 |
| SMG1 | 0.421022103 | 1.33887577 | 0.004956582 | 0.01740714 | 1777.516117 | 2379.57371 | ENSG00000157106 |
| LOC100130950 | 0.420887803 | 1.33875114 | 0.612966835 | 0.756106094 | 5.701637368 | 7.650404165 | ENSG00000261879 |
| CDK11A | 0.420838425 | 1.33870532 | 0.503841718 | 0.669027801 | 10.40740645 | 13.95426706 | ENSG00000008128 |
| ELF2 | 0.420824168 | 1.338692091 | 1.99E-05 | 0.000124194 | 623.8981335 | 834.8577701 | ENSG00000109381 |
| CHD1L | 0.420801636 | 1.338671183 | 0.640449296 | 0.775088292 | 4.631766142 | 6.215670948 | ENSG00000131778 |
| ATP12A | 0.420734648 | 1.338609026 | 0.207580759 | 0.362327219 | 41.06514791 | 54.95666084 | ENSG00000075673 |
| FAT3 | 0.42055244 | 1.338439975 | 0.020959647 | 0.058387372 | 906.0464761 | 1212.627736 | ENSG00000165323 |
| NA | 0.420537232 | 1.338425866 | 0.639041412 | 0.774453315 | 4.697396369 | 6.293643062 | ENSG00000233885 |
| NA | 0.420446669 | 1.338341851 | 0.474157976 | 0.642434437 | 11.8227378 | 15.76758705 | ENSG00000281344 |
| PIEZO2 | 0.420388073 | 1.338287494 | 3.17E-05 | 0.000190398 | 503.8103915 | 673.9034532 | ENSG00000154864 |
| SCCPDH | 0.420273469 | 1.338181188 | 3.33E-06 | 2.38E-05 | 576.5659207 | 771.7433928 | ENSG00000143653 |
| EIF3J-DT | 0.420096295 | 1.33801686 | 0.011554282 | 0.035624292 | 180.4995076 | 241.6155131 | ENSG00000179523 |

| | | | | | | | |
|---|---|---|---|---|---|---|---|
| SBNO2 | 0.419951396 | 1.337882481 | 7.16E-10 | 8.85E-09 | 1401.880852 | 1875.901623 | ENSG00000064932 |
| ZNF311 | 0.41994823 | 1.337879545 | 0.094709774 | 0.198376357 | 71.36395029 | 95.55815019 | ENSG00000197935 |
| ADARB1 | 0.419891391 | 1.337826837 | 0.000118833 | 0.000630996 | 339.9165992 | 455.0068538 | ENSG00000197381 |
| DDX50 | 0.41975528 | 1.337700626 | 4.58E-09 | 5.12E-08 | 909.5539058 | 1217.039902 | ENSG00000107625 |
| KDF1 | 0.41955624 | 1.337516083 | 8.77E-05 | 0.00048057 | 352.4598059 | 471.3318347 | ENSG00000175707 |
| KIAA0040 | 0.419469807 | 1.337435954 | 6.34E-05 | 0.000357659 | 392.7957488 | 525.0707466 | ENSG00000235750 |
| UROS | 0.419321494 | 1.337298469 | 1.90E-07 | 1.68E-06 | 708.8266787 | 947.8372003 | ENSG00000188690 |
| MFAP4 | 0.419289674 | 1.337268974 | 0.056949916 | 0.132443741 | 78.74168816 | 105.3340659 | ENSG00000166482 |
| SYCE1L | 0.419213578 | 1.337198441 | 0.07992605 | 0.173261426 | 68.61498749 | 91.60485242 | ENSG00000205078 |
| BTK | 0.419144753 | 1.33713465 | 0.613774284 | 0.756749457 | 5.605982931 | 7.521884358 | ENSG00000010671 |
| RPP25L | 0.418950172 | 1.336954318 | 7.71E-05 | 0.00042797 | 398.0322806 | 532.1761395 | ENSG00000164967 |
| MYO3A | 0.418855924 | 1.336866981 | 0.037064437 | 0.093563211 | 94.21850869 | 126.0425635 | ENSG00000095777 |
| DSG2-AS1 | 0.418844354 | 1.33685626 | 0.692886979 | 0.814185461 | 3.896867156 | 5.256579206 | ENSG00000264859 |
| RAPGEF6 | 0.418808168 | 1.336822729 | 0.12716121 | 0.248764517 | 53.19387288 | 71.22302474 | ENSG00000158987 |
| TDRD7 | 0.418691539 | 1.336714664 | 0.097679467 | 0.203403994 | 58.47413216 | 78.1684327 | ENSG00000196116 |
| TGFBR2 | 0.418532922 | 1.336567706 | 0.002781794 | 0.010520208 | 228.5507513 | 305.7178183 | ENSG00000163513 |
| CCDC71L | 0.418230127 | 1.336287215 | 3.02E-05 | 0.00018202 | 428.5979095 | 573.1892862 | ENSG00000253276 |
| DUS4L | 0.418226239 | 1.336283614 | 0.22384262 | 0.382340003 | 35.82607645 | 47.80771714 | ENSG00000105865 |
| SFMBT2 | 0.418187777 | 1.336247989 | 0.021647417 | 0.060008667 | 144.6138951 | 192.826417 | ENSG00000198879 |
| NA | 0.418134166 | 1.336198334 | 0.087729094 | 0.186858999 | 68.50605426 | 91.62472851 | ENSG00000228327 |
| SKOR2 | 0.418115997 | 1.336181507 | 0.628034938 | 0.766071576 | 5.64437985 | 7.559528609 | ENSG00000215474 |
| NA | 0.417931052 | 1.336010228 | 0.246985765 | 0.410493875 | 29.45796122 | 39.30255169 | ENSG00000274925 |
| SV2B | 0.41784539 | 1.335930902 | 0.429402485 | 0.600827108 | 16.15234682 | 21.59230082 | ENSG00000185518 |
| PGM2 | 0.417595502 | 1.335699527 | 4.91E-07 | 4.05E-06 | 710.7965636 | 949.9424473 | ENSG00000169299 |
| PLEKHM3 | 0.417313473 | 1.335438439 | 0.019206074 | 0.05433412 | 130.5309177 | 174.2528147 | ENSG00000178385 |
| CD58 | 0.417224441 | 1.335356029 | 0.123850224 | 0.244094265 | 55.8300724 | 74.33987453 | ENSG00000116815 |
| NA | 0.417141181 | 1.335278966 | 0.675350679 | 0.800828482 | 3.72317958 | 4.957854658 | ENSG00000181359 |
| CPEB1 | 0.416921386 | 1.335075552 | 0.332023684 | 0.50469379 | 20.65073732 | 27.6095491 | ENSG00000214575 |
| PPDPF | 0.416874274 | 1.335031955 | 8.95E-07 | 7.10E-06 | 832.8283229 | 1112.444242 | ENSG00000125534 |
| NA | 0.416700059 | 1.334870751 | 0.725152905 | 0.835021231 | 2.947092772 | 3.967037349 | ENSG00000233538 |
| NA | 0.416625147 | 1.334801439 | 0.579739656 | 0.731024514 | 7.416334949 | 9.878065066 | ENSG00000272721 |
| ZNF281 | 0.416496171 | 1.334682114 | 2.02E-12 | 3.33E-11 | 1732.252923 | 2312.382029 | ENSG00000162702 |
| UGP2 | 0.416419248 | 1.334610952 | 6.38E-18 | 1.81E-16 | 8025.957763 | 10712.26684 | ENSG00000169764 |
| ZNF75A | 0.416330921 | 1.334529245 | 0.005907411 | 0.020281166 | 186.5156169 | 248.8435497 | ENSG00000162086 |
| HFM1 | 0.416015201 | 1.334237228 | 0.402280616 | 0.574546188 | 16.0978802 | 21.49282297 | ENSG00000162669 |
| SYCP2L | 0.415842978 | 1.334077962 | 0.300842386 | 0.47148838 | 25.46264873 | 33.86313688 | ENSG00000153157 |
| BCL3 | 0.415766996 | 1.334007702 | 0.131983308 | 0.255759156 | 50.90297023 | 67.80450633 | ENSG00000069399 |
| RDH16 | 0.415711901 | 1.333956759 | 0.609506048 | 0.753313613 | 5.728870676 | 7.646103018 | ENSG00000139547 |
| FASTKD3 | 0.415509222 | 1.333769369 | 0.012577355 | 0.038179347 | 136.7260211 | 182.3683679 | ENSG00000124279 |
| FAS | 0.415492292 | 1.333753718 | 0.330564406 | 0.502990847 | 23.59224829 | 31.52065311 | ENSG00000026103 |
| ZNF691 | 0.415298771 | 1.333574821 | 0.000109296 | 0.000586164 | 372.5552259 | 496.4666427 | ENSG00000164011 |
| ARL6IP4 | 0.415244406 | 1.33352457 | 0.514208746 | 0.678170569 | 9.392001834 | 12.51953384 | ENSG00000182196 |
| CCDC90B | 0.415116191 | 1.333406061 | 7.96E-06 | 5.35E-05 | 498.4192471 | 664.6967992 | ENSG00000137500 |
| NA | 0.415053822 | 1.333348419 | 0.673992492 | 0.800111161 | 4.395239242 | 5.81370044 | ENSG00000228878 |
| TIPRL | 0.414857868 | 1.33316733 | 4.27E-05 | 0.000248862 | 544.2468339 | 725.6144656 | ENSG00000143155 |
| MRPL38 | 0.414439621 | 1.332780891 | 0.287582863 | 0.45693058 | 31.35838587 | 41.68132264 | ENSG00000204316 |
| ROBO2 | 0.414416745 | 1.332759758 | 0.049363846 | 0.118031477 | 93.83082159 | 124.7952396 | ENSG00000185008 |
| RPUSD3 | 0.414412049 | 1.33275542 | 7.06E-11 | 9.84E-10 | 1311.241658 | 1747.130689 | ENSG00000156990 |
| POT1 | 0.414339511 | 1.332688411 | 0.001669896 | 0.006735094 | 309.9954332 | 413.0206033 | ENSG00000128513 |
| NA | 0.414337877 | 1.332686901 | 2.42E-05 | 0.000148711 | 550.8690462 | 733.4106552 | ENSG00000254277 |
| MED12L | 0.414175945 | 1.332537326 | 0.050183995 | 0.119621828 | 165.0787421 | 219.6964997 | ENSG00000144893 |
| NA | 0.41313756 | 1.331578573 | 0.637944932 | 0.773642659 | 7.544128778 | 10.01465413 | ENSG00000108785 |
| AKR1C1 | 0.413095977 | 1.331540193 | 0.397047283 | 0.569559721 | 16.1167408 | 21.45517872 | ENSG00000187134 |
| NIFK-AS1 | 0.413045306 | 1.331493427 | 0.080502627 | 0.174295868 | 71.19150201 | 94.86311335 | ENSG00000236859 |
| ATF7 | 0.412965531 | 1.331419803 | 0.045507424 | 0.110676213 | 105.3110534 | 140.4670864 | ENSG00000170653 |
| MTRES1 | 0.412685546 | 1.331161438 | 0.006936867 | 0.023224419 | 181.4864396 | 241.5644323 | ENSG00000130349 |
| POLM | 0.412361395 | 1.33086238 | 0.001438513 | 0.005905965 | 239.7328931 | 318.9122646 | ENSG00000122678 |
| PLGRKT | 0.412131744 | 1.330650548 | 0.008304799 | 0.026948339 | 181.3162186 | 241.0519522 | ENSG00000107020 |
| STING1 | 0.412121425 | 1.330641031 | 0.619424536 | 0.760753231 | 6.025445997 | 8.06044393 | ENSG00000184584 |
| RLN1 | 0.412097637 | 1.33061909 | 0.604869896 | 0.749761946 | 5.668822255 | 7.547158205 | ENSG00000107018 |
| CEP44 | 0.41203548 | 1.330561763 | 0.008037907 | 0.026225711 | 195.7090132 | 260.582211 | ENSG00000164118 |
| NT5DC3 | 0.411996105 | 1.330525449 | 0.002509244 | 0.009625339 | 321.2618529 | 427.6672357 | ENSG00000111696 |
| RILPL1 | 0.411767126 | 1.330314289 | 0.006479352 | 0.021927997 | 173.7958565 | 231.1053292 | ENSG00000188026 |
| DDX59 | 0.411717863 | 1.330268865 | 0.000286807 | 0.001396818 | 334.2223834 | 444.4826821 | ENSG00000118197 |
| KIF27 | 0.411677629 | 1.330231767 | 0.067866306 | 0.152198198 | 78.94906673 | 104.9433813 | ENSG00000165115 |
| TMEM106B | 0.411575251 | 1.330137373 | 0.000932545 | 0.004002164 | 364.7008938 | 485.738329 | ENSG00000106460 |
| CCDC7 | 0.411565705 | 1.330128572 | 0.575822653 | 0.727976591 | 7.326262317 | 9.719970265 | ENSG00000216937 |
| LGALS9 | 0.411550004 | 1.330114096 | 0.599026891 | 0.745585377 | 6.469351467 | 8.554647066 | ENSG00000168961 |
| CLN5 | 0.411535353 | 1.330100589 | 0.43071892 | 0.602014105 | 14.36990383 | 19.13932587 | ENSG00000102805 |
| FOXO1 | 0.411383131 | 1.329960254 | 1.50E-06 | 1.15E-05 | 661.5088576 | 880.148013 | ENSG00000150907 |
| CRYBG3 | 0.410924228 | 1.329537278 | 0.021694675 | 0.060113758 | 153.8320092 | 204.4139805 | ENSG00000080200 |
| FRAT1 | 0.41070288 | 1.329333306 | 0.053049752 | 0.124973473 | 92.38438998 | 122.7961399 | ENSG00000165879 |
| KANSL1-AS1 | 0.410681905 | 1.32931398 | 0.517699867 | 0.681561791 | 10.51633968 | 13.95426706 | ENSG00000214401 |
| NA | 0.410622984 | 1.32925969 | 0.723795743 | 0.83439367 | 3.456628469 | 4.534911451 | ENSG00000255741 |
| PGBD5 | 0.410520949 | 1.329165681 | 1.14E-06 | 8.93E-06 | 830.866196 | 1104.270903 | ENSG00000177614 |
| ZKSCAN1 | 0.410453004 | 1.329103084 | 3.23E-07 | 2.75E-06 | 779.8462449 | 1036.374605 | ENSG00000106261 |
| PCDH12 | 0.410320421 | 1.328980947 | 0.415594772 | 0.587774102 | 18.13080461 | 23.98722238 | ENSG00000113555 |

| | | | | | | | |
|---|---|---|---|---|---|---|---|
| SNORD19 | 0.410314173 | 1.328975191 | 0.687228792 | 0.809904736 | 3.665922062 | 4.856759272 | ENSG00000212493 |
| CD47 | 0.410275571 | 1.328939632 | 0.000992227 | 0.004233809 | 289.4056955 | 384.4678526 | ENSG00000196776 |
| PEAR1 | 0.410176576 | 1.328848446 | 0.35868257 | 0.531725617 | 21.15469122 | 28.006958 | ENSG00000187800 |
| EVC2 | 0.410137595 | 1.328812542 | 0.005109034 | 0.017858013 | 199.6915225 | 265.5508491 | ENSG00000173040 |
| NA | 0.409996468 | 1.328682561 | 0.536391032 | 0.696038255 | 9.89037392 | 13.03094157 | ENSG00000243829 |
| PBX1 | 0.409869445 | 1.328565582 | 3.38E-18 | 9.74E-17 | 3404.321535 | 4521.990545 | ENSG00000185630 |
| SERINC5 | 0.409864049 | 1.328560613 | 3.61E-10 | 4.66E-09 | 1763.964665 | 2342.643018 | ENSG00000164300 |
| CAMKK1 | 0.409401683 | 1.328134893 | 0.004111806 | 0.014812321 | 190.8563896 | 253.6543108 | ENSG00000004660 |
| FBXO28 | 0.409381251 | 1.328116084 | 1.13E-06 | 8.85E-06 | 732.7090077 | 973.9447358 | ENSG00000143756 |
| SORCS3 | 0.409368698 | 1.328104528 | 0.210381793 | 0.365939568 | 40.86123596 | 54.13444284 | ENSG00000156395 |
| ZNF786 | 0.409198657 | 1.327948002 | 0.00012981 | 0.000681108 | 396.378107 | 526.2928393 | ENSG00000197362 |
| LYRM7 | 0.409162167 | 1.327914415 | 0.011112217 | 0.034463952 | 175.0402912 | 232.4120636 | ENSG00000186687 |
| DIO3 | 0.40906139 | 1.327821659 | 0.627670765 | 0.765821057 | 6.939614365 | 9.256959659 | ENSG00000197406 |
| KCNE5 | 0.408933403 | 1.327703868 | 0.192038682 | 0.3411879 | 52.69493721 | 69.92730374 | ENSG00000176076 |
| COL1A1 | 0.40890757 | 1.327680094 | 1.53E-08 | 1.59E-07 | 1738.711602 | 2309.689856 | ENSG00000108821 |
| DNASE1L1 | 0.408867488 | 1.327643208 | 0.211412532 | 0.367122076 | 35.47224361 | 47.11535181 | ENSG00000013563 |
| IFT80 | 0.408799129 | 1.327580302 | 0.072140044 | 0.159813228 | 91.05478875 | 120.9481319 | ENSG00000068885 |
| ZNF28 | 0.408700415 | 1.327489468 | 0.002632014 | 0.010026496 | 283.6448486 | 376.9351849 | ENSG00000198538 |
| SLC48A1 | 0.408671512 | 1.327462874 | 0.001253434 | 0.005232633 | 253.7296279 | 336.7550517 | ENSG00000211584 |
| MYLIP | 0.408649134 | 1.327442283 | 2.82E-06 | 2.05E-05 | 572.1413698 | 759.5738821 | ENSG00000007944 |
| NA | 0.408641701 | 1.327435444 | 0.721047316 | 0.832444884 | 3.015513901 | 4.021886189 | ENSG00000264589 |
| NA | 0.408411737 | 1.327223868 | 0.533532948 | 0.694024552 | 10.55944252 | 14.05133387 | ENSG00000219201 |
| UCK2 | 0.408184012 | 1.327014387 | 3.60E-11 | 5.18E-10 | 1359.536537 | 1803.944735 | ENSG00000143179 |
| NA | 0.408024473 | 1.326867649 | 0.590992318 | 0.739910841 | 6.697505659 | 8.885097182 | ENSG00000230295 |
| ERCC6 | 0.407835539 | 1.326693894 | 0.006236359 | 0.021214178 | 200.5160186 | 265.8442185 | ENSG00000225830 |
| NA | 0.407773733 | 1.32663706 | 0.37834286 | 0.551850761 | 18.05468649 | 23.93746819 | ENSG00000259953 |
| NHLH2 | 0.40759241 | 1.326470333 | 0.046286451 | 0.112231263 | 96.57157493 | 128.3143204 | ENSG00000177551 |
| NA | 0.407379412 | 1.326274509 | 0.530795599 | 0.691676432 | 9.685786253 | 12.8628874 | ENSG00000286504 |
| ETNK2 | 0.407267139 | 1.3261713 | 6.85E-05 | 0.000383248 | 435.2633127 | 576.835033 | ENSG00000143845 |
| ZNF223 | 0.407117868 | 1.326034093 | 0.541629943 | 0.700448027 | 9.315883717 | 12.33239708 | ENSG00000178386 |
| TAPT1-AS1 | 0.407102129 | 1.326019626 | 0.379065932 | 0.552436638 | 18.74839765 | 24.80191627 | ENSG00000263327 |
| ZNF525 | 0.407068916 | 1.3259891 | 6.34E-06 | 4.33E-05 | 587.089706 | 778.6806768 | ENSG00000203326 |
| GPNMB | 0.406921026 | 1.325853181 | 0.504428407 | 0.66957465 | 10.72563327 | 14.23847063 | ENSG00000136235 |
| ARHGEF1 | 0.406901065 | 1.325834836 | 1.14E-09 | 1.37E-08 | 1356.422164 | 1797.584697 | ENSG00000076928 |
| NA | 0.406749646 | 1.325695689 | 0.655434499 | 0.786295949 | 4.781887194 | 6.312465188 | ENSG00000261613 |
| LETMD1 | 0.406679557 | 1.325631286 | 0.000117744 | 0.000626246 | 371.2258249 | 491.7954026 | ENSG00000050426 |
| H3-3A | 0.406679146 | 1.325630909 | 6.43E-25 | 3.16E-23 | 8340.358888 | 11056.22003 | ENSG00000163041 |
| PCDHAC1 | 0.406514744 | 1.325479855 | 0.098556306 | 0.204809834 | 66.19453926 | 87.94943095 | ENSG00000248383 |
| TRPA1 | 0.406424939 | 1.325397349 | 0.471417993 | 0.640296029 | 12.3090704 | 16.30803673 | ENSG00000104321 |
| CAMSAP2 | 0.406156808 | 1.325151042 | 1.67E-06 | 1.26E-05 | 685.4251516 | 908.3754082 | ENSG00000118200 |
| CFAP99 | 0.406112252 | 1.325110116 | 0.705368313 | 0.822737779 | 3.908030767 | 5.208182087 | ENSG00000206113 |
| FAM110C | 0.40610302 | 1.325101636 | 0.011471143 | 0.035451749 | 162.772772 | 215.5310823 | ENSG00000184731 |
| ZNF718 | 0.406063644 | 1.325065471 | 0.00539188 | 0.018714423 | 192.585442 | 255.0546348 | ENSG00000250312 |
| FRS2 | 0.406061578 | 1.325063573 | 0.000336186 | 0.001612473 | 336.979831 | 446.1481271 | ENSG00000166225 |
| MTFR2 | 0.4060117 | 1.325017763 | 0.019488327 | 0.055017989 | 126.4470842 | 167.4829666 | ENSG00000146410 |
| PLEKHA5 | 0.405858233 | 1.324876821 | 9.40E-10 | 1.14E-08 | 1256.613505 | 1664.629729 | ENSG00000052126 |
| MPZL3 | 0.405684005 | 1.324716831 | 0.122454979 | 0.242220162 | 54.11728984 | 71.74437972 | ENSG00000160588 |
| ZNF337-AS1 | 0.405604059 | 1.324643425 | 0.310519763 | 0.481452882 | 24.327823 | 32.2858838 | ENSG00000213742 |
| COQ9 | 0.405520551 | 1.324566753 | 6.90E-07 | 5.57E-06 | 653.179267 | 864.9372746 | ENSG00000088682 |
| NINJ2-AS1 | 0.40548205 | 1.324531404 | 0.220866224 | 0.379270992 | 36.20136063 | 48.0911151 | ENSG00000177406 |
| MXD1 | 0.405410796 | 1.324465988 | 0.285448275 | 0.454462067 | 28.4286021 | 37.59651835 | ENSG00000059728 |
| BMPR1B | 0.405369522 | 1.324428097 | 0.495356058 | 0.66162942 | 12.46198235 | 16.48011947 | ENSG00000138696 |
| SAMD11 | 0.405362206 | 1.324421381 | 0.681445066 | 0.80560396 | 4.488102776 | 5.875001002 | ENSG00000187634 |
| UAP1 | 0.405341252 | 1.324402145 | 2.43E-08 | 2.44E-07 | 885.7136547 | 1173.498219 | ENSG00000117143 |
| TFCP2L1 | 0.405316706 | 1.324379612 | 0.036467206 | 0.092272938 | 125.2686801 | 165.6070011 | ENSG00000115112 |
| ZNF681 | 0.405264133 | 1.324331351 | 0.035481723 | 0.090157788 | 122.2439928 | 162.1688549 | ENSG00000196172 |
| NA | 0.405177569 | 1.324251891 | 0.745520078 | 0.849098159 | 2.721729483 | 3.573669135 | ENSG00000176857 |
| HPAT5 | 0.405168098 | 1.324243198 | 1.30E-08 | 1.36E-07 | 974.8993434 | 1290.785084 | ENSG00000280707 |
| ZNF217 | 0.404958289 | 1.324050629 | 1.81E-09 | 2.14E-08 | 1259.095729 | 1666.705753 | ENSG00000171940 |
| NA | 0.404858505 | 1.323959054 | 0.675561036 | 0.800938489 | 3.962497383 | 5.275401332 | ENSG00000279794 |
| NA | 0.404788198 | 1.323894535 | 0.446074741 | 0.616426443 | 13.59102612 | 18.03719334 | ENSG00000276805 |
| MTFMT | 0.404667769 | 1.323784028 | 0.00189641 | 0.007538357 | 349.5687697 | 463.4587759 | ENSG00000103707 |
| PFKL | 0.404403566 | 1.323541623 | 2.39E-19 | 7.58E-18 | 4071.521369 | 5389.136028 | ENSG00000141959 |
| RCCD1 | 0.404277804 | 1.323426253 | 1.35E-05 | 8.65E-05 | 496.8416799 | 657.3346393 | ENSG00000166965 |
| SEC14L1 | 0.404010559 | 1.323181124 | 1.73E-10 | 2.32E-09 | 1935.321726 | 2560.930368 | ENSG00000129657 |
| NA | 0.403932135 | 1.323109199 | 0.305720456 | 0.476693608 | 24.40673202 | 32.32997978 | ENSG00000278730 |
| IQSEC2 | 0.40392113 | 1.323099106 | 8.34E-10 | 1.02E-08 | 1523.127237 | 2015.794347 | ENSG00000124313 |
| LINC02610 | 0.403744647 | 1.322937263 | 0.277433635 | 0.444811524 | 36.73207227 | 48.54094969 | ENSG00000186235 |
| VAMP2 | 0.403575881 | 1.322782515 | 0.01233324 | 0.037509149 | 160.442033 | 211.9235371 | ENSG00000220205 |
| PIGV | 0.403038401 | 1.322289801 | 0.009441567 | 0.029971575 | 168.655706 | 223.0852132 | ENSG00000060642 |
| CLK2 | 0.403028591 | 1.322280809 | 7.36E-10 | 9.08E-09 | 1261.02917 | 1667.399561 | ENSG00000176444 |
| MMACHC | 0.402914256 | 1.322176022 | 0.027378351 | 0.072990017 | 133.603489 | 176.6366924 | ENSG00000132763 |
| CYTH2 | 0.402610547 | 1.321897713 | 2.07E-08 | 2.09E-07 | 1534.067303 | 2026.710306 | ENSG00000105443 |
| CLU | 0.402434932 | 1.321736812 | 1.25E-16 | 3.19E-15 | 3916.716757 | 5178.410776 | ENSG00000120885 |
| MAVS | 0.402347193 | 1.321656431 | 4.99E-09 | 5.53E-08 | 1088.659282 | 1438.954406 | ENSG00000088888 |
| TLCD4 | 0.402144468 | 1.321470728 | 0.255309779 | 0.420307454 | 31.29678585 | 41.42292595 | ENSG00000152078 |
| AGBL2 | 0.401564282 | 1.3209394 | 0.367512937 | 0.541239183 | 20.99129137 | 27.81604102 | ENSG00000165923 |

| | | | | | | | |
|---|---|---|---|---|---|---|---|
| NA | 0.401530832 | 1.320908773 | 0.566139454 | 0.72000935 | 10.13997945 | 13.47647501 | ENSG00000277476 |
| HOMEZ | 0.40150968 | 1.320889406 | 0.013043363 | 0.039377075 | 185.0338916 | 244.451679 | ENSG00000215271 |
| ART5 | 0.401413543 | 1.320801389 | 0.02377507 | 0.064844462 | 122.4788049 | 161.8238838 | ENSG00000167311 |
| COX15 | 0.401302719 | 1.320699933 | 0.001725859 | 0.006941942 | 296.903418 | 392.332527 | ENSG00000014919 |
| IRAK3 | 0.401139627 | 1.32055064 | 0.168275557 | 0.308580934 | 48.94963057 | 64.52579363 | ENSG00000090376 |
| MYO1D | 0.401133609 | 1.320545132 | 1.40E-05 | 8.93E-05 | 644.0469824 | 850.6638497 | ENSG00000176658 |
| NA | 0.401093897 | 1.320508783 | 0.618217171 | 0.759924581 | 5.698846465 | 7.500911659 | ENSG00000275198 |
| ANKRD33B | 0.40104021 | 1.320459644 | 0.000151944 | 0.000784876 | 388.0275038 | 512.1473948 | ENSG00000164236 |
| GPR37 | 0.400648993 | 1.320101622 | 0.074943837 | 0.164810576 | 117.6776184 | 154.9755306 | ENSG00000170775 |
| SLC23A1 | 0.400631031 | 1.320085186 | 0.618641694 | 0.760227954 | 5.701637368 | 7.500911659 | ENSG00000170482 |
| CNTNAP1 | 0.400251104 | 1.319737594 | 0.00154205 | 0.006277209 | 356.8801122 | 470.4300334 | ENSG00000108797 |
| GRAMD1A | 0.400069394 | 1.319571381 | 4.58E-17 | 1.21E-15 | 3757.357083 | 4958.781562 | ENSG00000089351 |
| RBM4B | 0.399985802 | 1.319494925 | 5.99E-05 | 0.000339723 | 427.7073445 | 564.3595893 | ENSG00000173914 |
| NA | 0.399954488 | 1.319466285 | 0.131814552 | 0.2555099 | 53.89683263 | 71.12595793 | ENSG00000233230 |
| NA | 0.399898891 | 1.319415438 | 0.721997368 | 0.833266601 | 2.958256383 | 3.920790803 | ENSG00000253361 |
| GTF2A1 | 0.399843328 | 1.319364625 | 2.65E-05 | 0.000161894 | 686.858594 | 906.9619203 | ENSG00000165417 |
| NUDT6 | 0.399803362 | 1.319328075 | 0.173875268 | 0.316675586 | 46.4511492 | 61.23579503 | ENSG00000170917 |
| PSPC1 | 0.399664112 | 1.319200739 | 5.94E-16 | 1.42E-14 | 2575.99875 | 3398.26778 | ENSG00000121390 |
| FUT2 | 0.399540828 | 1.319088012 | 0.029764916 | 0.078027022 | 113.3018786 | 149.4565261 | ENSG00000176920 |
| NA | 0.399470916 | 1.319024092 | 0.549940083 | 0.707071057 | 8.966956963 | 11.86346779 | ENSG00000258526 |
| NA | 0.398935224 | 1.318534412 | 0.261952839 | 0.427415395 | 32.02889393 | 42.21208552 | ENSG00000279672 |
| CCDC200 | 0.398724373 | 1.318341721 | 0.69996095 | 0.818956009 | 4.817493211 | 6.318916909 | ENSG00000236383 |
| NA | 0.398617981 | 1.318244503 | 0.409624859 | 0.581847666 | 15.68958075 | 20.69156556 | ENSG00000274565 |
| FEZ2 | 0.398614089 | 1.318240947 | 0.000139641 | 0.000727862 | 453.8692862 | 598.5781349 | ENSG00000171055 |
| LOC389831 | 0.398557664 | 1.31818939 | 0.027725948 | 0.073733288 | 120.0028509 | 158.2531588 | ENSG00000276256 |
| GSDMB | 0.398435535 | 1.318077806 | 0.03038134 | 0.07936591 | 128.9383442 | 169.9270788 | ENSG00000073605 |
| BSPRY | 0.398432046 | 1.318074618 | 0.009218572 | 0.029374873 | 168.1937036 | 221.8440684 | ENSG00000119411 |
| NA | 0.39789078 | 1.3175802 | 0.623040203 | 0.762604966 | 5.938164268 | 7.8501839 | ENSG00000178429 |
| C18orf54 | 0.397848936 | 1.317541985 | 0.001789169 | 0.007165218 | 380.7780112 | 501.8686432 | ENSG00000166845 |
| NA | 0.397685433 | 1.317392674 | 0.4705111 | 0.63942442 | 14.70487607 | 19.37083117 | ENSG00000254978 |
| NA | 0.397580894 | 1.317297218 | 0.408218256 | 0.580576406 | 16.50617966 | 21.68453345 | ENSG00000259366 |
| PHPT1 | 0.397528829 | 1.31724968 | 5.51E-10 | 6.91E-09 | 1346.522043 | 1774.440096 | ENSG00000054148 |
| LOC101927741 | 0.397328592 | 1.317066866 | 0.525182007 | 0.687472762 | 9.751416479 | 12.81825839 | ENSG00000287299 |
| NA | 0.397130929 | 1.316886428 | 0.543440503 | 0.70198947 | 9.068193206 | 11.96026203 | ENSG00000269427 |
| TMEM169 | 0.397090494 | 1.31684952 | 0.168279064 | 0.308580934 | 51.0265337 | 67.1320292 | ENSG00000163449 |
| ALDH1L2 | 0.396950399 | 1.316721651 | 0.03685847 | 0.093104196 | 146.945422 | 193.2805222 | ENSG00000136010 |
| PCDHB8 | 0.396862889 | 1.316641785 | 0.537307907 | 0.69691433 | 9.710228657 | 12.76394259 | ENSG00000120322 |
| MRS2 | 0.39645667 | 1.316271111 | 6.84E-09 | 7.45E-08 | 1389.719786 | 1829.126225 | ENSG00000124532 |
| NA | 0.396455328 | 1.316269886 | 0.182885844 | 0.328618742 | 44.9562331 | 59.16973658 | ENSG00000185834 |
| NA | 0.396390032 | 1.316210313 | 0.7178851 | 0.830806975 | 3.009932095 | 3.969187922 | ENSG00000157021 |
| LOC101926933 | 0.396383834 | 1.316204659 | 0.48028507 | 0.648392687 | 16.85789731 | 22.08786103 | ENSG00000257285 |
| ADIPOR1 | 0.396286327 | 1.316115705 | 9.15E-11 | 1.27E-09 | 1492.857354 | 1963.910707 | ENSG00000159346 |
| TRIM62 | 0.396146804 | 1.315988429 | 0.001652774 | 0.006674412 | 321.9569667 | 424.1543583 | ENSG00000116525 |
| ZNF516 | 0.396041212 | 1.315892114 | 4.09E-05 | 0.000239538 | 542.663472 | 714.4791725 | ENSG00000101493 |
| PIBF1 | 0.395864124 | 1.315730601 | 0.016984585 | 0.049049347 | 153.8426604 | 202.5770163 | ENSG00000083535 |
| XIRP2 | 0.395835035 | 1.315704072 | 0.714837274 | 0.828621912 | 3.698737175 | 4.833635999 | ENSG00000163092 |
| NIPAL1 | 0.395791422 | 1.315664299 | 0.05733868 | 0.133187295 | 108.367555 | 142.8773224 | ENSG00000163293 |
| LINC01285 | 0.395206742 | 1.315131208 | 0.672859055 | 0.799564127 | 4.969529285 | 6.548271638 | ENSG00000203650 |
| MTFP1 | 0.394910134 | 1.314860855 | 0.345106696 | 0.517566141 | 23.82877519 | 31.39590142 | ENSG00000242114 |
| JMJD1C | 0.394873524 | 1.314827489 | 2.22E-10 | 2.96E-09 | 2632.28558 | 3461.214836 | ENSG00000171988 |
| RLIM | 0.394513115 | 1.314499064 | 1.33E-06 | 1.03E-05 | 773.4949631 | 1016.814202 | ENSG00000131263 |
| NADK | 0.394485648 | 1.314474038 | 1.60E-11 | 2.39E-10 | 1841.373248 | 2419.317665 | ENSG00000008130 |
| PTPRN2 | 0.394351909 | 1.314352191 | 0.006922536 | 0.023184485 | 209.4580576 | 275.1202424 | ENSG00000155093 |
| NRXN1 | 0.394206674 | 1.314219883 | 0.006842886 | 0.02296957 | 211.011946 | 277.4979167 | ENSG00000179915 |
| NA | 0.393964077 | 1.313998908 | 0.683857846 | 0.807164813 | 3.986939787 | 5.247976912 | ENSG00000272505 |
| LOC339803 | 0.393797638 | 1.313847325 | 0.154958749 | 0.289942699 | 61.02532839 | 80.32242264 | ENSG00000212978 |
| CSPG4 | 0.393641217 | 1.313704882 | 0.015230266 | 0.044830976 | 161.7573674 | 212.3841245 | ENSG00000173546 |
| PMVK | 0.393640247 | 1.313703999 | 6.49E-05 | 0.000365904 | 547.8455372 | 720.0078633 | ENSG00000163344 |
| BMP8B | 0.393610243 | 1.313676678 | 0.256678258 | 0.421841904 | 33.92931858 | 44.55347268 | ENSG00000116985 |
| STARD4 | 0.3935827 | 1.313651598 | 9.97E-06 | 6.57E-05 | 628.0322927 | 825.6624078 | ENSG00000164211 |
| ACSBG1 | 0.39352368 | 1.313597859 | 0.066927121 | 0.150547001 | 95.80214597 | 125.8530036 | ENSG00000103740 |
| FSTL4 | 0.393370606 | 1.313458489 | 0.049352476 | 0.118018919 | 97.86297941 | 128.4108541 | ENSG00000053108 |
| CLN3 | 0.393045279 | 1.313162339 | 0.709663546 | 0.825105495 | 3.970870091 | 5.222703065 | ENSG00000188603 |
| DGAT1 | 0.392968588 | 1.313092535 | 5.42E-05 | 0.000309905 | 514.4305724 | 675.8684773 | ENSG00000185000 |
| MTHFD2 | 0.392900763 | 1.313030804 | 2.34E-13 | 4.27E-12 | 2474.020687 | 3248.253588 | ENSG00000065911 |
| RHOXF1-AS1 | 0.392757606 | 1.31290052 | 0.164736704 | 0.303783539 | 47.00311202 | 61.77597214 | ENSG00000258545 |
| TSLP | 0.392739764 | 1.312884284 | 0.634061455 | 0.770742252 | 5.761685789 | 7.517583211 | ENSG00000145777 |
| OVOL1 | 0.392627725 | 1.312782329 | 0.146209148 | 0.276738684 | 54.74903756 | 71.80944839 | ENSG00000172818 |
| KLHL24 | 0.392437427 | 1.312609179 | 0.001178284 | 0.004946742 | 335.478294 | 439.9138761 | ENSG00000114796 |
| NA | 0.392160262 | 1.31235703 | 0.580986919 | 0.731628608 | 7.8218435 | 10.21228329 | ENSG00000226478 |
| PRR3 | 0.392111558 | 1.312312727 | 0.010122467 | 0.031777464 | 169.2998566 | 222.1936133 | ENSG00000204576 |
| SHROOM3 | 0.391974888 | 1.312188414 | 3.91E-10 | 5.01E-09 | 1308.494885 | 1717.288293 | ENSG00000138771 |
| NA | 0.391945 | 1.31216123 | 0.663282729 | 0.792185928 | 5.981467272 | 7.854485048 | ENSG00000274248 |
| ZSWIM9 | 0.39192318 | 1.312141385 | 0.000332231 | 0.001595885 | 386.879236 | 507.4659107 | ENSG00000185453 |
| SNORD83A | 0.391920424 | 1.312138877 | 0.531416702 | 0.692345485 | 9.966291878 | 13.09170909 | ENSG00000209482 |
| RND3 | 0.391818886 | 1.312046532 | 2.71E-06 | 1.97E-05 | 681.4185874 | 894.210336 | ENSG00000115963 |
| ACBD6 | 0.391793568 | 1.312023507 | 2.02E-07 | 1.77E-06 | 1139.344086 | 1495.161342 | ENSG00000230124 |

| | | | | | | | |
|---|---|---|---|---|---|---|---|
| NA | 0.39153902 | 1.311792035 | 0.718668294 | 0.831214704 | 3.755994693 | 4.884183692 | ENSG00000233799 |
| WDR92 | 0.391438272 | 1.311700432 | 0.128089627 | 0.250090753 | 60.92285285 | 79.97046678 | ENSG00000243667 |
| LAMB3 | 0.391329365 | 1.311601417 | 0.317933275 | 0.489211998 | 24.03894466 | 31.54539392 | ENSG00000196878 |
| NA | 0.391242786 | 1.311522707 | 0.17074952 | 0.312191166 | 60.78042878 | 79.95646673 | ENSG00000268205 |
| SSH1 | 0.391226215 | 1.311507643 | 1.41E-06 | 1.08E-05 | 686.4786649 | 900.228693 | ENSG00000084112 |
| PHOSPHO1 | 0.390939182 | 1.311246736 | 0.481875195 | 0.649877964 | 12.91059375 | 16.94743122 | ENSG00000173868 |
| ENTPD8 | 0.390850089 | 1.311165763 | 0.558949364 | 0.714356707 | 9.672507459 | 12.64994376 | ENSG00000188833 |
| RBMS1 | 0.390770783 | 1.311093689 | 8.57E-08 | 7.96E-07 | 1137.642166 | 1492.308801 | ENSG00000153250 |
| PLEKHH3 | 0.390693687 | 1.311023628 | 2.79E-05 | 0.000169307 | 559.1189398 | 732.7126864 | ENSG00000068137 |
| IMMP2L | 0.390457068 | 1.310808622 | 0.048277568 | 0.116094599 | 130.4970635 | 171.0891427 | ENSG00000184903 |
| LDB3 | 0.390409756 | 1.310765636 | 0.524507259 | 0.686869156 | 11.23987489 | 14.78698957 | ENSG00000122367 |
| KIF17 | 0.390341457 | 1.310703584 | 0.001039308 | 0.004408375 | 297.708453 | 390.5638665 | ENSG00000117245 |
| XKR5 | 0.390125394 | 1.310507303 | 0.058940914 | 0.135975744 | 98.72479634 | 129.2919738 | ENSG00000275591 |
| TMBIM4 | 0.390006213 | 1.310399047 | 0.223296699 | 0.381847055 | 49.79940801 | 65.3759812 | ENSG00000155957 |
| NA | 0.389896456 | 1.310299359 | 0.633085975 | 0.769822396 | 6.245903201 | 8.234949819 | ENSG00000213225 |
| PLXND1 | 0.389831545 | 1.310240406 | 1.49E-10 | 2.02E-09 | 1478.210412 | 1936.307021 | ENSG00000004399 |
| PPM1D | 0.389790475 | 1.310203107 | 0.000106172 | 0.000571313 | 415.292332 | 543.9995399 | ENSG00000170836 |
| RGMB | 0.38970626 | 1.310126628 | 0.018080676 | 0.051612064 | 141.3801145 | 185.2128331 | ENSG00000174136 |
| IQGAP3 | 0.389639152 | 1.310065688 | 1.71E-08 | 1.76E-07 | 1076.312178 | 1409.859941 | ENSG00000183856 |
| OSGIN1 | 0.389617659 | 1.310046172 | 0.406319417 | 0.57872917 | 18.07075618 | 23.66831864 | ENSG00000140961 |
| NA | 0.389527341 | 1.30996416 | 0.624220185 | 0.763058842 | 5.968188479 | 7.82275948 | ENSG00000250461 |
| SNED1 | 0.389496457 | 1.309936118 | 0.318659845 | 0.490017303 | 37.19858036 | 48.77057699 | ENSG00000162804 |
| ITPR3 | 0.389424143 | 1.30987046 | 1.16E-10 | 1.59E-09 | 2206.709656 | 2889.695625 | ENSG00000096433 |
| CDON | 0.38871689 | 1.309228479 | 0.000374813 | 0.001778726 | 353.7235767 | 462.9121713 | ENSG00000064309 |
| PIK3IP1 | 0.388456809 | 1.30899248 | 0.26845752 | 0.434752592 | 33.40717981 | 43.75598338 | ENSG00000100100 |
| CADM2 | 0.388349009 | 1.308894674 | 0.047540195 | 0.11475199 | 107.5946594 | 140.9365671 | ENSG00000175161 |
| NA | 0.388297712 | 1.308848135 | 0.348645059 | 0.521366113 | 25.30503085 | 33.08284006 | ENSG00000280088 |
| AVPR1B | 0.388138824 | 1.308703996 | 0.489816753 | 0.656229945 | 14.5002884 | 19.05920318 | ENSG00000198049 |
| SSC4D | 0.38809629 | 1.308665413 | 0.001560973 | 0.006336854 | 288.5868692 | 377.4431397 | ENSG00000146700 |
| SMAD1 | 0.388094436 | 1.308663732 | 0.003006958 | 0.011222138 | 247.317534 | 323.3950292 | ENSG00000170365 |
| JADE1 | 0.388010176 | 1.308587302 | 1.19E-14 | 2.49E-13 | 2822.914591 | 3693.386956 | ENSG00000077684 |
| PEX19 | 0.388001061 | 1.308579034 | 9.59E-06 | 6.34E-05 | 647.3971932 | 847.1299753 | ENSG00000162735 |
| CRABP1 | 0.387920216 | 1.308505706 | 3.82E-06 | 2.71E-05 | 798.0089007 | 1044.38747 | ENSG00000166426 |
| ESD | 0.387911601 | 1.308497893 | 1.25E-07 | 1.14E-06 | 814.5539287 | 1065.808809 | ENSG00000139684 |
| PEA15 | 0.387886617 | 1.308475233 | 1.57E-12 | 2.62E-11 | 1760.30728 | 2303.119539 | ENSG00000162734 |
| LENG1 | 0.387455976 | 1.308084715 | 0.024077055 | 0.065493442 | 147.2586306 | 192.4943373 | ENSG00000105617 |
| KLHL35 | 0.387285489 | 1.307930144 | 0.421913016 | 0.593797713 | 16.10346201 | 21.03868723 | ENSG00000149243 |
| SPATA25 | 0.387218495 | 1.307869409 | 0.520065658 | 0.683279072 | 11.91751636 | 15.58851955 | ENSG00000149634 |
| DENND4A | 0.387157519 | 1.307814133 | 0.05272466 | 0.124341249 | 119.0385584 | 155.5875007 | ENSG00000174485 |
| ELK3 | 0.387095334 | 1.307757763 | 0.004587824 | 0.016299027 | 278.720174 | 364.7515052 | ENSG00000111145 |
| TOR3A | 0.387047661 | 1.307714549 | 5.06E-07 | 4.17E-06 | 772.3254825 | 1009.795469 | ENSG00000186283 |
| ZNF518A | 0.386904749 | 1.307585015 | 0.00013434 | 0.000703081 | 418.9968142 | 548.2402884 | ENSG00000177853 |
| SH2D3C | 0.386859199 | 1.307543731 | 0.080768599 | 0.174714851 | 91.63206986 | 119.8279891 | ENSG00000095370 |
| NA | 0.386804832 | 1.307494459 | 0.697151719 | 0.817153648 | 3.95970648 | 5.178607093 | ENSG00000249465 |
| UBE2Q1 | 0.386786873 | 1.307478183 | 1.17E-09 | 1.41E-08 | 1361.982567 | 1780.801533 | ENSG00000160714 |
| LOC100288637 | 0.386678884 | 1.307380318 | 0.020202532 | 0.056687569 | 175.8462021 | 229.9020708 | ENSG00000187951 |
| NA | 0.386540214 | 1.307254661 | 0.651152751 | 0.783195701 | 5.021204997 | 6.556340895 | ENSG00000267096 |
| MRPS27 | 0.386496911 | 1.307215424 | 5.99E-10 | 7.48E-09 | 1803.500717 | 2356.950604 | ENSG00000113048 |
| NA | 0.386412391 | 1.307138843 | 0.295120163 | 0.465089703 | 34.4030481 | 45.07321013 | ENSG00000229852 |
| ZNF215 | 0.386321488 | 1.307056484 | 0.029182857 | 0.076782119 | 142.5257035 | 186.5843756 | ENSG00000149054 |
| ZNF77 | 0.386274152 | 1.3070136 | 0.108039687 | 0.219481213 | 73.90341932 | 96.69551593 | ENSG00000175691 |
| PIM1 | 0.386238452 | 1.306981257 | 1.11E-09 | 1.34E-08 | 1233.146898 | 1611.49908 | ENSG00000137193 |
| CETN3 | 0.386137821 | 1.306890096 | 0.001258372 | 0.00524984 | 349.0299352 | 456.1003657 | ENSG00000153140 |
| BNIP3L | 0.385759031 | 1.306547007 | 6.03E-05 | 0.000341379 | 598.033494 | 781.5183871 | ENSG00000104765 |
| NA | 0.385194595 | 1.306035937 | 0.625458758 | 0.764233905 | 6.05826111 | 7.896430446 | ENSG00000287721 |
| PRR35 | 0.385159935 | 1.306004561 | 0.350327369 | 0.522956378 | 25.24968835 | 33.10596333 | ENSG00000161992 |
| TMEM130 | 0.385128559 | 1.305976158 | 0.115398813 | 0.231337265 | 65.95609734 | 86.09767321 | ENSG00000166448 |
| LINC02361 | 0.385032775 | 1.305889454 | 0.272534945 | 0.439294086 | 36.12197605 | 47.1145583 | ENSG00000256576 |
| NA | 0.384614442 | 1.305510844 | 0.630393072 | 0.76802531 | 5.992630883 | 7.814157185 | ENSG00000230896 |
| FGF23 | 0.384424806 | 1.305339252 | 0.460125489 | 0.629836146 | 14.68813065 | 19.23181896 | ENSG00000118972 |
| CTF1 | 0.384363042 | 1.30528337 | 0.495069405 | 0.661429731 | 14.09777091 | 18.37194461 | ENSG00000150281 |
| LRFN2 | 0.384154464 | 1.305094671 | 0.191102145 | 0.339805648 | 45.32613563 | 59.24796916 | ENSG00000156564 |
| CLDN23 | 0.384025355 | 1.304977882 | 0.026163303 | 0.070255608 | 135.7830678 | 177.2744572 | ENSG00000253958 |
| FRYL | 0.38394982 | 1.30490956 | 0.000560169 | 0.002536185 | 539.6207986 | 703.5599719 | ENSG00000075539 |
| ESRG | 0.383790766 | 1.304765704 | 4.07E-13 | 7.22E-12 | 8124.781747 | 10600.45131 | ENSG00000265992 |
| NA | 0.383674975 | 1.304660988 | 0.178007913 | 0.321952996 | 48.07509842 | 62.74958486 | ENSG00000227081 |
| NA | 0.383547568 | 1.304545775 | 0.340758144 | 0.513319175 | 27.52492162 | 35.87837506 | ENSG00000279821 |
| ZNF587 | 0.383295685 | 1.304318032 | 0.002295824 | 0.008889752 | 306.8513249 | 400.4244481 | ENSG00000198466 |
| LOC442497 | 0.383209908 | 1.304240485 | 0.452378744 | 0.622561732 | 15.55552939 | 20.35089562 | ENSG00000249574 |
| NFATC2 | 0.382824514 | 1.303892123 | 0.273542104 | 0.440292093 | 30.96528023 | 40.44286151 | ENSG00000101096 |
| FBLN7 | 0.382764614 | 1.303837988 | 0.695212693 | 0.815874684 | 4.014173095 | 5.222703065 | ENSG00000144152 |
| NA | 0.382750867 | 1.303825564 | 0.141209326 | 0.269656414 | 56.60161654 | 73.89621908 | ENSG00000280385 |
| NA | 0.382615966 | 1.303703653 | 1.25E-06 | 9.71E-06 | 882.5440776 | 1151.005946 | ENSG00000235655 |
| FBXO22 | 0.382546299 | 1.3036407 | 1.71E-08 | 1.76E-07 | 1541.506094 | 2009.910101 | ENSG00000167196 |
| CES2 | 0.382452545 | 1.303555985 | 0.000723076 | 0.003173079 | 380.9791564 | 496.2883623 | ENSG00000172831 |
| IDS | 0.381994489 | 1.303142171 | 0.020824162 | 0.058093842 | 153.1325161 | 199.5101811 | ENSG00000010404 |
| NA | 0.381957625 | 1.303108874 | 0.217600957 | 0.375200219 | 40.04117043 | 52.20093277 | ENSG00000236017 |

| | | | | | | | |
|---|---|---|---|---|---|---|---|
| LINC02614 | 0.38187334 | 1.303032745 | 0.593257584 | 0.741562369 | 7.320680512 | 9.566437075 | ENSG00000241288 |
| SOX13 | 0.38178667 | 1.302954468 | 1.89E-12 | 3.12E-11 | 2548.709756 | 3321.318296 | ENSG00000143842 |
| SQSTM1 | 0.381720711 | 1.302894899 | 3.64E-08 | 3.56E-07 | 1443.128736 | 1880.394068 | ENSG00000161011 |
| TMEM198B | 0.381718872 | 1.302893239 | 0.026287855 | 0.070521349 | 146.3423471 | 190.4938805 | ENSG00000182796 |
| NOP2 | 0.381713846 | 1.302888699 | 0.592511177 | 0.741062844 | 8.142861226 | 10.57392594 | ENSG00000111641 |
| PELI1 | 0.380935237 | 1.302185733 | 8.19E-09 | 8.84E-08 | 1222.153838 | 1591.87866 | ENSG00000197329 |
| ERV3-1 | 0.38086498 | 1.30212232 | 0.186842233 | 0.333800104 | 46.65697616 | 60.83355195 | ENSG00000213462 |
| HDHD5 | 0.38082329 | 1.302084693 | 4.70E-13 | 8.27E-12 | 2125.307614 | 2767.348597 | ENSG00000069998 |
| PILRB | 0.380813573 | 1.302075923 | 0.241567014 | 0.403888826 | 36.90364466 | 48.00077258 | ENSG00000121716 |
| NA | 0.380607196 | 1.301889674 | 0.284860407 | 0.453748984 | 31.29399495 | 40.82170874 | ENSG00000254602 |
| NA | 0.380600944 | 1.301884033 | 0.143250458 | 0.272699892 | 59.32980371 | 77.15610966 | ENSG00000275131 |
| RUSC1-AS1 | 0.380531577 | 1.301821437 | 0.153636636 | 0.28811163 | 61.32903712 | 79.8263415 | ENSG00000225855 |
| NA | 0.380488713 | 1.301782759 | 0.408959994 | 0.581202832 | 18.65274322 | 24.31148124 | ENSG00000217165 |
| AMPD3 | 0.380401805 | 1.301704343 | 0.05376835 | 0.126422789 | 115.3962909 | 150.197016 | ENSG00000133805 |
| FAM86C2P | 0.380389087 | 1.301692868 | 0.126661646 | 0.248114319 | 71.07957772 | 92.61825997 | ENSG00000160172 |
| TMEM238 | 0.380382289 | 1.301686734 | 0.161113754 | 0.298391459 | 53.36353025 | 69.46591067 | ENSG00000233493 |
| KIF26B | 0.380319246 | 1.301629854 | 0.388347689 | 0.561716673 | 24.64134389 | 31.92129708 | ENSG00000162849 |
| COL24A1 | 0.38031217 | 1.30162347 | 0.391627927 | 0.564598636 | 22.16172312 | 28.88267987 | ENSG00000171502 |
| TAB1 | 0.380265513 | 1.301581376 | 1.31E-06 | 1.01E-05 | 965.4171937 | 1255.855091 | ENSG00000100324 |
| SLC44A3-AS1 | 0.380245094 | 1.301562954 | 0.324352986 | 0.495925612 | 29.39599778 | 38.14152964 | ENSG00000224081 |
| NA | 0.38024436 | 1.301562292 | 0.666570799 | 0.795139722 | 5.551516316 | 7.296830776 | ENSG00000237877 |
| C5orf34 | 0.380234473 | 1.301553372 | 0.070507033 | 0.157006946 | 89.9140178 | 117.0612051 | ENSG00000172244 |
| KANSL1L | 0.380214069 | 1.301534965 | 0.013844438 | 0.041428161 | 167.4406198 | 218.0730427 | ENSG00000144445 |
| LRP6 | 0.38008619 | 1.301419603 | 1.87E-06 | 1.41E-05 | 1173.005539 | 1526.423871 | ENSG00000070018 |
| OPTC | 0.379939741 | 1.301287502 | 0.462951359 | 0.632671767 | 19.06016679 | 24.90247862 | ENSG00000188770 |
| GTF2IP12 | 0.37992453 | 1.301273782 | 0.053473869 | 0.125868764 | 101.2940894 | 131.9151534 | ENSG00000283050 |
| SELENOH | 0.379921685 | 1.301271216 | 0.026070654 | 0.070065337 | 180.9170556 | 235.4721744 | ENSG00000211450 |
| EGFL7 | 0.379714731 | 1.301084562 | 1.07E-06 | 8.40E-06 | 941.0357061 | 1225.224001 | ENSG00000172889 |
| NA | 0.379682292 | 1.301055308 | 0.691695898 | 0.81298395 | 4.280724206 | 5.603700874 | ENSG00000230593 |
| GPR39 | 0.379541278 | 1.300928144 | 0.622916544 | 0.762604966 | 6.354836431 | 8.287648086 | ENSG00000183840 |
| SURF1 | 0.379442436 | 1.300839018 | 0.000456007 | 0.002112204 | 456.6680977 | 593.6808421 | ENSG00000148290 |
| SMG7 | 0.379423476 | 1.300821922 | 8.93E-12 | 1.38E-10 | 1750.205497 | 2276.963509 | ENSG00000116698 |
| ZDHHC19 | 0.379055371 | 1.300490059 | 0.131047355 | 0.254278267 | 61.52804298 | 79.94519294 | ENSG00000163958 |
| NA | 0.37876978 | 1.300232644 | 0.497312547 | 0.663645293 | 13.28049628 | 17.26712847 | ENSG00000237350 |
| DAB1 | 0.378488245 | 1.299978934 | 0.00014146 | 0.000735951 | 444.7074658 | 578.2377438 | ENSG00000173406 |
| RAB3IP | 0.378282099 | 1.299793194 | 0.001693233 | 0.006823513 | 411.2048317 | 534.6301379 | ENSG00000127328 |
| LIPT1 | 0.378139844 | 1.299665036 | 0.215605739 | 0.372592521 | 45.39543264 | 58.79734106 | ENSG00000144182 |
| SKA3 | 0.377890076 | 1.29944005 | 1.06E-07 | 9.73E-07 | 971.0681576 | 1262.277459 | ENSG00000165480 |
| IL34 | 0.377757836 | 1.299320946 | 0.062088543 | 0.141812702 | 101.7249163 | 132.2713983 | ENSG00000157368 |
| ISG15 | 0.377688359 | 1.299258375 | 0.000814817 | 0.003534658 | 332.6407733 | 431.9948859 | ENSG00000187608 |
| LINC01023 | 0.377527976 | 1.299113946 | 0.57950766 | 0.730863859 | 9.375256417 | 12.17886389 | ENSG00000272523 |
| LINC01852 | 0.377384268 | 1.298984547 | 0.734357537 | 0.841740419 | 4.000894302 | 5.185058814 | ENSG00000236914 |
| NA | 0.37729751 | 1.298906434 | 0.491566204 | 0.657856494 | 12.37749153 | 16.06201045 | ENSG00000213881 |
| PTPRS | 0.377167182 | 1.2987891 | 2.87E-15 | 6.35E-14 | 3848.260963 | 4998.763382 | ENSG00000105426 |
| AP1G2 | 0.376965295 | 1.298607363 | 0.003890098 | 0.014097729 | 293.8048016 | 381.0641278 | ENSG00000213983 |
| HENMT1 | 0.376909823 | 1.298557432 | 0.03391908 | 0.086770054 | 145.9423834 | 189.7007045 | ENSG00000162639 |
| TDGF1 | 0.376851158 | 1.29850463 | 1.79E-18 | 5.28E-17 | 5715.509838 | 7422.127164 | ENSG00000241186 |
| MED7 | 0.376674265 | 1.298345426 | 0.044021502 | 0.107731619 | 117.6342785 | 152.7341543 | ENSG00000155868 |
| TMCO6 | 0.376667466 | 1.298339308 | 0.007165279 | 0.023827856 | 315.8844245 | 409.7080997 | ENSG00000113119 |
| ARMC4 | 0.376554675 | 1.298237806 | 0.033130571 | 0.085107861 | 137.2151446 | 178.2617853 | ENSG00000169126 |
| ROR2 | 0.376525538 | 1.298211587 | 0.058610004 | 0.135385952 | 129.9652006 | 168.8356749 | ENSG00000169071 |
| NA | 0.376484588 | 1.298174739 | 0.639575349 | 0.774741394 | 6.055470207 | 7.827060627 | ENSG00000278356 |
| AHSA2P | 0.37643912 | 1.298133826 | 0.00034134 | 0.001634349 | 428.9819156 | 556.4048266 | ENSG00000173209 |
| PSD3 | 0.376348449 | 1.298052243 | 0.002993062 | 0.011176763 | 360.9119208 | 469.010281 | ENSG00000156011 |
| ZNF765 | 0.376340007 | 1.298044648 | 0.004015881 | 0.014510274 | 247.5090061 | 321.3321874 | ENSG00000196417 |
| IDUA | 0.376289923 | 1.297999586 | 0.157500557 | 0.293417985 | 59.78419707 | 77.5814639 | ENSG00000127415 |
| FHL3 | 0.376255359 | 1.297968489 | 0.001291997 | 0.005372691 | 395.6427325 | 513.2613768 | ENSG00000183386 |
| DPP10 | 0.376243618 | 1.297957926 | 0.012908632 | 0.039043655 | 198.3421848 | 257.8579481 | ENSG00000175497 |
| PLAAT1 | 0.376156454 | 1.297879508 | 0.079752061 | 0.173000901 | 92.22801141 | 119.9489606 | ENSG00000127252 |
| PANK3 | 0.375950887 | 1.29769459 | 1.48E-08 | 1.54E-07 | 1541.724587 | 2001.593392 | ENSG00000120137 |
| MMP24OS | 0.375800857 | 1.297559645 | 0.009858079 | 0.031058664 | 242.0728438 | 314.4978037 | ENSG00000126005 |
| FAM86B1 | 0.375754914 | 1.297518325 | 0.26620531 | 0.432451928 | 41.12778707 | 53.40362765 | ENSG00000186523 |
| FLJ20021 | 0.375697176 | 1.297466398 | 0.232977715 | 0.393794093 | 43.27993244 | 56.12844787 | ENSG00000254531 |
| HLCS | 0.375635319 | 1.297410769 | 0.000722305 | 0.00317114 | 332.4180888 | 431.2433706 | ENSG00000159267 |
| CR1 | 0.375540181 | 1.297325214 | 0.566736382 | 0.7202656 | 9.403165446 | 12.18961676 | ENSG00000203710 |
| AHRR | 0.37544442 | 1.297239106 | 0.547104927 | 0.704929277 | 13.09756012 | 17.08617086 | ENSG00000286169 |
| SLC39A5 | 0.375102932 | 1.296932083 | 0.732445214 | 0.84020366 | 3.039956306 | 3.920790803 | ENSG00000139540 |
| BBS4 | 0.37486342 | 1.296716789 | 4.55E-05 | 0.000263924 | 638.2518555 | 827.2434048 | ENSG00000140463 |
| GHR | 0.374733044 | 1.29659961 | 0.693638589 | 0.81467166 | 4.277933303 | 5.57412588 | ENSG00000112964 |
| NA | 0.37467746 | 1.296549655 | 0.363474886 | 0.536988841 | 23.23562455 | 30.18863282 | ENSG00000245205 |
| ZNF284 | 0.374625297 | 1.296502778 | 0.478453721 | 0.646665083 | 15.26541176 | 19.69429653 | ENSG00000186026 |
| UBALD2 | 0.374224006 | 1.296142201 | 0.000447917 | 0.002082111 | 371.4897483 | 481.7796519 | ENSG00000185262 |
| FAM66C | 0.374211354 | 1.296130834 | 0.305690005 | 0.476684656 | 30.04553006 | 39.03581318 | ENSG00000226711 |
| C11orf71 | 0.374155334 | 1.296080506 | 0.430658978 | 0.60197394 | 17.01204857 | 22.08812149 | ENSG00000180425 |
| TVP23B | 0.374149699 | 1.296075444 | 0.00816186 | 0.02656943 | 243.2749394 | 315.5316509 | ENSG00000171928 |
| NA | 0.373990772 | 1.295932676 | 0.639435797 | 0.774736588 | 6.34367282 | 8.228498098 | ENSG00000246100 |
| HCN2 | 0.373981901 | 1.295924707 | 0.078938858 | 0.171603506 | 101.1216412 | 131.1821814 | ENSG00000099822 |

| | | | | | | | |
|---|---|---|---|---|---|---|---|
| CASK | 0.373875159 | 1.295828829 | 2.17E-06 | 1.61E-05 | 1073.507822 | 1391.403686 | ENSG00000147044 |
| VPS45 | 0.373817265 | 1.295776829 | 0.00011716 | 0.000623657 | 480.3587786 | 622.3572842 | ENSG00000136631 |
| DRAM2 | 0.373759231 | 1.295724706 | 0.001651558 | 0.006670898 | 282.9424894 | 366.5193964 | ENSG00000156171 |
| SLC35G2 | 0.373753738 | 1.295719773 | 0.134793144 | 0.259982974 | 62.88890772 | 81.46273245 | ENSG00000168917 |
| CXCR4 | 0.373535655 | 1.295523921 | 0.362236759 | 0.535430949 | 22.80934038 | 29.49223891 | ENSG00000121966 |
| OGT | 0.37348503 | 1.295478462 | 3.07E-09 | 3.51E-08 | 2092.444523 | 2709.561471 | ENSG00000147162 |
| NA | 0.373484323 | 1.295477827 | 0.430787292 | 0.602050472 | 17.03370007 | 22.09887436 | ENSG00000272221 |
| LOC155060 | 0.373466044 | 1.295461413 | 0.354905228 | 0.528150684 | 23.1275672 | 29.94690769 | ENSG00000244560 |
| TNFRSF10A | 0.373258133 | 1.295274734 | 0.019269389 | 0.054489267 | 202.0359406 | 261.8636836 | ENSG00000104689 |
| MAN2B2 | 0.373224174 | 1.295244245 | 0.006659714 | 0.022448462 | 207.9105018 | 269.263803 | ENSG00000013288 |
| NA | 0.373224128 | 1.295244204 | 0.643100857 | 0.776981188 | 6.433745451 | 8.329593484 | ENSG00000226329 |
| EPHA8 | 0.373206741 | 1.295228594 | 0.002847928 | 0.010721972 | 269.8808987 | 349.7996902 | ENSG00000070886 |
| PPP1R1C | 0.373086129 | 1.295120315 | 0.257692849 | 0.422821821 | 35.61678286 | 46.06538451 | ENSG00000150722 |
| ZDBF2 | 0.372989652 | 1.29503371 | 0.001811086 | 0.007237968 | 423.0176821 | 547.6094657 | ENSG00000204186 |
| TAF5L | 0.372745316 | 1.294814401 | 3.99E-06 | 2.82E-05 | 701.3967526 | 908.6908227 | ENSG00000135801 |
| BCL2L11 | 0.372605812 | 1.294689202 | 0.010185327 | 0.031948805 | 243.7239383 | 315.9879372 | ENSG00000153094 |
| NA | 0.37224452 | 1.294365015 | 0.689683193 | 0.811485516 | 4.307957513 | 5.597249153 | ENSG00000261625 |
| GTPBP10 | 0.372093301 | 1.294229351 | 0.008913148 | 0.028571541 | 209.6562628 | 271.5597492 | ENSG00000105793 |
| DNAH14 | 0.372032086 | 1.294174437 | 0.003548826 | 0.013015285 | 270.844152 | 350.2460719 | ENSG00000185842 |
| DPY19L2P2 | 0.37199629 | 1.294142327 | 0.408168707 | 0.580548738 | 19.27380289 | 25.00572458 | ENSG00000170629 |
| GATA2 | 0.371888795 | 1.294045904 | 0.595049512 | 0.742935741 | 8.722733075 | 11.31226524 | ENSG00000179348 |
| ZNF784 | 0.371841567 | 1.294003543 | 0.005313873 | 0.018483473 | 288.5119152 | 372.9673966 | ENSG00000179922 |
| THRB | 0.371777695 | 1.293946255 | 0.152633774 | 0.286565269 | 57.13838554 | 73.95536907 | ENSG00000151090 |
| ARHGEF3 | 0.371516144 | 1.293711692 | 0.055806425 | 0.130413294 | 106.4458792 | 137.7202516 | ENSG00000163947 |
| NA | 0.371464866 | 1.29366571 | 0.490874827 | 0.65741185 | 14.20949504 | 18.2767679 | ENSG00000272631 |
| CBFA2T2 | 0.371335939 | 1.293550106 | 2.46E-05 | 0.000150497 | 538.0703029 | 696.0424255 | ENSG00000078699 |
| B9D1 | 0.371324907 | 1.293540215 | 0.001553497 | 0.006309161 | 317.3338983 | 410.1831592 | ENSG00000108641 |
| SNAPC3 | 0.370902534 | 1.293161565 | 9.20E-06 | 6.10E-05 | 801.9669562 | 1036.846509 | ENSG00000164975 |
| SLC16A10 | 0.370887878 | 1.293148429 | 0.009920699 | 0.03123045 | 217.5145882 | 281.4593079 | ENSG00000112394 |
| NA | 0.370882835 | 1.293143908 | 0.001022841 | 0.004350006 | 358.9437876 | 464.0156303 | ENSG00000260834 |
| ANKEF1 | 0.370794347 | 1.293064595 | 0.015210756 | 0.044791742 | 188.1925933 | 243.6095181 | ENSG00000132623 |
| CCDC186 | 0.370746315 | 1.293021546 | 0.009182053 | 0.029282688 | 222.303772 | 287.674458 | ENSG00000165813 |
| SPDYE5 | 0.370379009 | 1.292692388 | 0.696570289 | 0.816821923 | 5.168535147 | 6.638614156 | ENSG00000170092 |
| LINC02210 | 0.370317738 | 1.292637488 | 0.000782452 | 0.003404981 | 389.7386467 | 503.8548515 | ENSG00000204650 |
| AEBP2 | 0.369763566 | 1.292141052 | 2.20E-05 | 0.000136159 | 824.7395776 | 1066.097265 | ENSG00000139154 |
| ZNF641 | 0.36950798 | 1.291912159 | 0.032859341 | 0.084563706 | 135.340714 | 174.9002479 | ENSG00000167528 |
| CFAP157 | 0.369438999 | 1.291850389 | 0.286056164 | 0.455118305 | 36.82174458 | 47.69721385 | ENSG00000160401 |
| AMZ2 | 0.36894531 | 1.291408394 | 1.04E-05 | 6.80E-05 | 821.981115 | 1061.699488 | ENSG00000196704 |
| PATJ | 0.368803399 | 1.291281371 | 6.64E-06 | 4.53E-05 | 1071.516698 | 1383.341364 | ENSG00000132849 |
| TOR1AIP1 | 0.368770864 | 1.291252251 | 3.47E-05 | 0.000206865 | 530.2709868 | 684.6771894 | ENSG00000143337 |
| PLSCR4 | 0.36868878 | 1.291178786 | 0.411779786 | 0.583920707 | 19.29266349 | 24.94710763 | ENSG00000114698 |
| GAA | 0.36866738 | 1.291159633 | 9.13E-07 | 7.23E-06 | 1060.065402 | 1368.46568 | ENSG00000171298 |
| NA | 0.368355501 | 1.290880543 | 0.56665719 | 0.7202656 | 9.865055635 | 12.80158684 | ENSG00000279453 |
| RDH12 | 0.368137055 | 1.290685099 | 0.566594693 | 0.7202656 | 12.09120393 | 15.68531378 | ENSG00000139988 |
| CRACR2A | 0.368027173 | 1.290586798 | 0.122455045 | 0.242220162 | 66.98757164 | 86.38832851 | ENSG00000130038 |
| NR2F2 | 0.368007734 | 1.290569409 | 0.700325826 | 0.819184181 | 4.664581255 | 6.066438906 | ENSG00000185551 |
| MTBP | 0.367990565 | 1.29055405 | 0.010517258 | 0.032851104 | 202.7929667 | 261.7349155 | ENSG00000172167 |
| NA | 0.367989627 | 1.290553212 | 0.710738122 | 0.825855265 | 4.041406403 | 5.180757667 | ENSG00000287077 |
| NA | 0.367766108 | 1.290353279 | 0.644357333 | 0.777772291 | 6.29478801 | 8.121484028 | ENSG00000272702 |
| FERMT2 | 0.367765131 | 1.290352406 | 9.47E-13 | 1.61E-11 | 2540.859527 | 3279.068935 | ENSG00000073712 |
| NA | 0.36771894 | 1.290311093 | 0.696184286 | 0.816684112 | 4.58009043 | 5.967494094 | ENSG00000242338 |
| RNF213-AS1 | 0.367661561 | 1.290259776 | 0.668376957 | 0.796192093 | 6.10714592 | 7.830828737 | ENSG00000263069 |
| TSSK6 | 0.36754245 | 1.290153255 | 0.222531372 | 0.381026377 | 42.02241912 | 54.2815122 | ENSG00000178093 |
| MARCHF3 | 0.367424673 | 1.290047935 | 0.025765845 | 0.069400854 | 164.8460084 | 212.7347417 | ENSG00000173926 |
| TMEM182 | 0.367170891 | 1.289821024 | 0.22667571 | 0.385868664 | 60.38552026 | 77.85814336 | ENSG00000170417 |
| COPS6 | 0.367125098 | 1.289780084 | 3.37E-11 | 4.87E-10 | 2236.648171 | 2885.040371 | ENSG00000168090 |
| C1orf35 | 0.366823052 | 1.289510081 | 4.52E-06 | 3.16E-05 | 877.3863057 | 1131.380324 | ENSG00000143793 |
| E2F7 | 0.366687756 | 1.289389157 | 0.000182268 | 0.000926026 | 474.3082145 | 611.9228672 | ENSG00000165891 |
| THBS1 | 0.3666448 | 1.289350766 | 0.000573887 | 0.002587856 | 431.9617866 | 556.9662305 | ENSG00000137801 |
| ZBTB25 | 0.36660968 | 1.289319379 | 0.074036022 | 0.163170215 | 121.4030764 | 156.2989607 | ENSG00000089775 |
| GPC6 | 0.366546977 | 1.289263343 | 0.0026744 | 0.010161877 | 357.7536563 | 461.0396587 | ENSG00000183098 |
| RNF169 | 0.36654409 | 1.289260763 | 0.011868274 | 0.036318348 | 203.9623593 | 262.7270777 | ENSG00000166439 |
| MYH14 | 0.366259543 | 1.289006504 | 2.60E-08 | 2.60E-07 | 1477.112245 | 1904.288749 | ENSG00000105357 |
| JMJD1C-AS1 | 0.36601128 | 1.288784707 | 0.755633513 | 0.855091825 | 3.924100463 | 5.12616929 | ENSG00000272767 |
| SERTAD4 | 0.365937644 | 1.288718929 | 0.154488605 | 0.289381335 | 57.65003642 | 74.21376576 | ENSG00000082497 |
| UQCC1 | 0.365848564 | 1.288639358 | 3.99E-07 | 3.34E-06 | 919.4306375 | 1184.619059 | ENSG00000101019 |
| NUDT12 | 0.365743509 | 1.288545525 | 0.012047542 | 0.036744526 | 210.1403169 | 270.796657 | ENSG00000112874 |
| RAB19 | 0.365717647 | 1.288522426 | 0.323013324 | 0.494582906 | 30.06785728 | 38.74543046 | ENSG00000146955 |
| BTBD1 | 0.365565912 | 1.288386914 | 0.000115006 | 0.000613206 | 727.7719812 | 938.0494651 | ENSG00000064726 |
| APH1B | 0.365524879 | 1.288350269 | 0.001342215 | 0.005549224 | 346.5315659 | 446.3777118 | ENSG00000138613 |
| SLC13A3 | 0.365359952 | 1.288202996 | 0.007344759 | 0.024328254 | 235.3485052 | 303.5100762 | ENSG00000158296 |
| BDP1 | 0.36529931 | 1.288148849 | 0.001101862 | 0.004647714 | 525.0460317 | 675.7398965 | ENSG00000145734 |
| KTN1-AS1 | 0.365279738 | 1.288131374 | 0.149007179 | 0.280932124 | 66.6674298 | 85.65429035 | ENSG00000186615 |
| NR1H3 | 0.365233734 | 1.288090298 | 0.002273117 | 0.008817756 | 308.8363284 | 397.591863 | ENSG00000025434 |
| DGKB | 0.364885718 | 1.287779615 | 0.272802282 | 0.439614804 | 42.72816978 | 54.93864432 | ENSG00000136267 |
| ZNF524 | 0.364763463 | 1.287670492 | 0.242926613 | 0.405751522 | 45.6457139 | 58.84761618 | ENSG00000171443 |
| CBX4 | 0.364670216 | 1.287587268 | 0.072137182 | 0.159813228 | 108.3508096 | 139.6921994 | ENSG00000141582 |

| | | | | | | | |
|---|---|---|---|---|---|---|---|
| JAK2 | 0.36458689 | 1.287512903 | 0.255392823 | 0.420373972 | 42.07265537 | 54.18256739 | ENSG00000096968 |
| IFT74 | 0.364564761 | 1.287493154 | 0.01489796 | 0.044011723 | 187.3903493 | 241.3194905 | ENSG00000096872 |
| SHARPIN | 0.364494069 | 1.287430068 | 6.50E-06 | 4.43E-05 | 724.5262348 | 933.1832921 | ENSG00000179526 |
| UGDH-AS1 | 0.364143017 | 1.287116835 | 0.620303018 | 0.76139303 | 7.859364539 | 10.1827083 | ENSG00000249348 |
| USO1 | 0.364140696 | 1.287114764 | 1.24E-12 | 2.09E-11 | 3454.276122 | 4447.484015 | ENSG00000138768 |
| NA | 0.36412391 | 1.287099789 | 0.069228993 | 0.154678771 | 111.4016541 | 143.3865796 | ENSG00000251363 |
| CEP19 | 0.363953611 | 1.286947866 | 0.371562732 | 0.545038336 | 28.96748629 | 37.22061519 | ENSG00000174007 |
| HHIPL1 | 0.363772437 | 1.28678626 | 0.050635372 | 0.12053393 | 122.9763011 | 158.4158457 | ENSG00000182218 |
| CMBL | 0.363760192 | 1.286775339 | 0.000236579 | 0.001176855 | 529.3937149 | 681.7015024 | ENSG00000164237 |
| NA | 0.363722552 | 1.286741768 | 0.128864075 | 0.25117883 | 84.09682625 | 107.9975734 | ENSG00000273151 |
| INTS5 | 0.363519117 | 1.286560336 | 9.94E-05 | 0.000538043 | 628.3509199 | 808.4972598 | ENSG00000185085 |
| NCOA3 | 0.363319932 | 1.28638272 | 0.000155834 | 0.000802938 | 517.1750663 | 665.2452876 | ENSG00000124151 |
| CRTAC1 | 0.363263143 | 1.286332085 | 0.156681262 | 0.292343382 | 60.48608078 | 77.79550415 | ENSG00000095713 |
| FAR2 | 0.363252553 | 1.286322643 | 0.000386032 | 0.001827917 | 449.583744 | 578.1839916 | ENSG00000064763 |
| SZT2 | 0.363204004 | 1.286279357 | 0.004475486 | 0.015941026 | 268.5656523 | 345.3166956 | ENSG00000198198 |
| PRPF3 | 0.362936418 | 1.286040804 | 4.54E-08 | 4.39E-07 | 1236.260018 | 1589.210106 | ENSG00000117360 |
| ERC1 | 0.362839406 | 1.285954329 | 1.42E-06 | 1.09E-05 | 1087.4481 | 1398.089656 | ENSG00000082805 |
| SGSM1 | 0.36262291 | 1.285761369 | 0.051435152 | 0.122046189 | 121.0128738 | 155.3533238 | ENSG00000167037 |
| SLC6A6 | 0.362342954 | 1.28551189 | 1.41E-12 | 2.38E-11 | 3417.87913 | 4392.303265 | ENSG00000131389 |
| PRRG3 | 0.362255867 | 1.285434293 | 0.049308994 | 0.11794418 | 125.0696742 | 160.594025 | ENSG00000130032 |
| CA14 | 0.362147795 | 1.285338005 | 1.75E-06 | 1.32E-05 | 908.3666023 | 1166.859357 | ENSG00000118298 |
| LOC440173 | 0.362144057 | 1.285334675 | 0.266548183 | 0.432732536 | 38.77942675 | 49.80925838 | ENSG00000269994 |
| SGO2 | 0.362112024 | 1.285306136 | 0.000102236 | 0.00055183 | 517.7339255 | 665.7776801 | ENSG00000163535 |
| ILF2 | 0.361808511 | 1.285035763 | 3.51E-16 | 8.62E-15 | 9185.152932 | 11802.17049 | ENSG00000143621 |
| VPS13C | 0.361757033 | 1.284989912 | 0.059380426 | 0.136742954 | 218.5543599 | 280.8491853 | ENSG00000129003 |
| ITGA5 | 0.361708539 | 1.284946719 | 1.17E-05 | 7.57E-05 | 827.1415769 | 1063.107366 | ENSG00000161638 |
| RAD51B | 0.361685689 | 1.284926368 | 0.203739587 | 0.356817101 | 51.90509605 | 66.76366226 | ENSG00000182185 |
| NSUN3 | 0.361640193 | 1.284885848 | 0.079702885 | 0.172922287 | 98.93843244 | 126.9336425 | ENSG00000178694 |
| CNN1 | 0.361510961 | 1.284770757 | 0.001444566 | 0.005929176 | 337.2338302 | 433.410264 | ENSG00000130176 |
| PNRC1 | 0.361097346 | 1.284402471 | 0.015565272 | 0.045647563 | 191.5663706 | 245.907918 | ENSG00000146278 |
| DENND5B | 0.36102709 | 1.284339925 | 0.011383275 | 0.035202751 | 233.9539863 | 300.4953878 | ENSG00000170456 |
| EIF5A2 | 0.360891458 | 1.284219186 | 0.10228852 | 0.210727022 | 84.23056531 | 108.2062159 | ENSG00000163577 |
| TMEM192 | 0.360636166 | 1.283991958 | 0.011470269 | 0.035451749 | 251.9014913 | 323.3823436 | ENSG00000170088 |
| NA | 0.360469135 | 1.283843309 | 0.291392518 | 0.461107051 | 32.69883841 | 41.98918252 | ENSG00000287763 |
| SF3B4 | 0.360375941 | 1.283760379 | 3.03E-08 | 3.00E-07 | 2151.482223 | 2762.975628 | ENSG00000143368 |
| CMTR2 | 0.360360295 | 1.283746457 | 0.033743217 | 0.086400371 | 185.3877614 | 237.6657108 | ENSG00000180917 |
| GAMT | 0.360177179 | 1.283583526 | 0.000295107 | 0.001433622 | 434.2070836 | 557.3582601 | ENSG00000130005 |
| CCDC159 | 0.360135117 | 1.283546103 | 0.301017208 | 0.471685724 | 32.63687497 | 41.82704703 | ENSG00000183401 |
| ZFAND6 | 0.360060304 | 1.283479546 | 1.83E-08 | 1.87E-07 | 2226.976962 | 2858.167707 | ENSG00000086666 |
| DEDD | 0.360013472 | 1.283437882 | 0.001316254 | 0.005458267 | 355.1477564 | 455.7395776 | ENSG00000158796 |
| FAM13C | 0.359988724 | 1.283415867 | 0.579753717 | 0.731024514 | 8.711569464 | 11.17137503 | ENSG00000148541 |
| NA | 0.359659557 | 1.283123074 | 0.582915492 | 0.733248614 | 9.470910854 | 12.08422023 | ENSG00000225218 |
| NA | 0.359658272 | 1.283121932 | 0.638454748 | 0.77388819 | 6.681435963 | 8.588523208 | ENSG00000213280 |
| NA | 0.359517516 | 1.28299675 | 0.218862936 | 0.376635584 | 49.73812029 | 63.76218048 | ENSG00000265415 |
| RBM33 | 0.359512439 | 1.282992235 | 1.61E-08 | 1.66E-07 | 1387.67601 | 1779.45959 | ENSG00000184863 |
| EBPL | 0.359511209 | 1.282991141 | 7.98E-09 | 8.63E-08 | 1535.486017 | 1969.927706 | ENSG00000123179 |
| NUMBL | 0.35928783 | 1.282792505 | 0.000237269 | 0.001179896 | 513.9164798 | 658.9057316 | ENSG00000105245 |
| PVT1 | 0.359212953 | 1.282725929 | 0.0166612 | 0.048252698 | 195.6080524 | 251.2112647 | ENSG00000249859 |
| RAD23A | 0.35890563 | 1.282452712 | 4.40E-08 | 4.26E-07 | 1296.567832 | 1662.871021 | ENSG00000179262 |
| SPON2 | 0.358814192 | 1.282371433 | 0.288089643 | 0.457410478 | 35.53141615 | 45.66609761 | ENSG00000159674 |
| ZNF462 | 0.358686358 | 1.282257809 | 1.98E-08 | 2.01E-07 | 1303.462754 | 1670.980129 | ENSG00000148143 |
| MAMSTR | 0.358685171 | 1.282256755 | 0.330621943 | 0.503038718 | 31.84384259 | 41.00024321 | ENSG00000176909 |
| EPM2A | 0.358353094 | 1.281961641 | 0.106366525 | 0.216904506 | 80.13748319 | 102.7495965 | ENSG00000112425 |
| NA | 0.358347503 | 1.281956673 | 0.000833786 | 0.003610454 | 412.3743875 | 528.8640595 | ENSG00000224032 |
| ZNF70 | 0.358010103 | 1.2816569 | 0.221032465 | 0.379404031 | 54.15991712 | 69.33524029 | ENSG00000187792 |
| FBXL4 | 0.358008671 | 1.281655627 | 0.015966616 | 0.046647279 | 223.79004 | 287.0436415 | ENSG00000112234 |
| XDH | 0.357831226 | 1.281497999 | 0.23987257 | 0.40194927 | 43.38607476 | 55.54336918 | ENSG00000158125 |
| IGSF1 | 0.357714037 | 1.281393909 | 6.52E-07 | 5.29E-06 | 1083.305057 | 1387.761303 | ENSG00000147255 |
| GATM | 0.357572824 | 1.281268489 | 0.005427083 | 0.018833219 | 334.1101326 | 427.5153443 | ENSG00000171766 |
| TMEM209 | 0.357340747 | 1.281062397 | 0.003568417 | 0.013074706 | 280.6560925 | 359.6629738 | ENSG00000146842 |
| GATA3 | 0.357141514 | 1.280885497 | 0.742358574 | 0.84758755 | 3.344904336 | 4.28888517 | ENSG00000107485 |
| NUP50-DT | 0.35708504 | 1.280835358 | 0.247996876 | 0.41164238 | 54.49421364 | 69.74419555 | ENSG00000226328 |
| HPN-AS1 | 0.356957453 | 1.28072209 | 0.368567047 | 0.542405084 | 25.1793522 | 32.20145997 | ENSG00000227392 |
| TEX15 | 0.356585743 | 1.280392156 | 0.004510568 | 0.016045247 | 284.8458682 | 365.1688023 | ENSG00000133863 |
| NA | 0.356558248 | 1.280367754 | 0.037637931 | 0.094750531 | 158.7992232 | 203.345191 | ENSG00000214425 |
| NA | 0.356532255 | 1.280344686 | 0.476815768 | 0.645084016 | 15.5883445 | 20.00162337 | ENSG00000258011 |
| SMOC2 | 0.356425078 | 1.280249573 | 0.105128971 | 0.215131323 | 101.0562111 | 129.3670018 | ENSG00000112562 |
| DYRK1A | 0.355926219 | 1.279806961 | 3.20E-05 | 0.000191987 | 777.5382093 | 994.8941529 | ENSG00000157540 |
| HES3 | 0.355613055 | 1.279529185 | 0.003549738 | 0.013016155 | 310.775113 | 397.9236396 | ENSG00000173673 |
| SSPO | 0.355601835 | 1.279519234 | 0.528936047 | 0.690185693 | 13.99374377 | 17.98475554 | ENSG00000197558 |
| TMEM9 | 0.355523536 | 1.279449793 | 8.75E-11 | 1.21E-09 | 2361.331384 | 3021.138844 | ENSG00000116857 |
| BAIAP2L1 | 0.355443866 | 1.27937914 | 3.30E-05 | 0.00019768 | 745.5636811 | 953.7990056 | ENSG00000006453 |
| HYAL1 | 0.355374609 | 1.279317725 | 0.124757562 | 0.245385568 | 74.45817305 | 95.30246764 | ENSG00000114378 |
| NA | 0.355180907 | 1.27914597 | 0.631396293 | 0.768762048 | 7.386310738 | 9.398122444 | ENSG00000250896 |
| SPR | 0.355103473 | 1.279077316 | 0.001051305 | 0.004456137 | 400.9535668 | 513.1271683 | ENSG00000116096 |
| NA | 0.355028956 | 1.279011252 | 0.018665745 | 0.052968651 | 194.3463087 | 248.8163736 | ENSG00000189343 |
| TIGAR | 0.355012853 | 1.278996976 | 0.094559403 | 0.198125931 | 99.70718568 | 127.7800498 | ENSG00000078237 |

| | | | | | | | |
|---|---|---|---|---|---|---|---|
| PDE1C | 0.35492471 | 1.278918837 | 0.707918283 | 0.823963032 | 5.894185543 | 7.584802456 | ENSG00000154678 |
| TAOK1 | 0.35482297 | 1.27882865 | 0.000158513 | 0.000815955 | 1277.693655 | 1633.679802 | ENSG00000160551 |
| PGAP1 | 0.354767697 | 1.278779655 | 0.00078458 | 0.003413473 | 426.2438041 | 545.373754 | ENSG00000197121 |
| NA | 0.354763796 | 1.278776198 | 0.5087692 | 0.673436712 | 13.10959961 | 16.72022707 | ENSG00000267750 |
| C4A | 0.354687954 | 1.278708974 | 0.646309957 | 0.77973517 | 6.697505659 | 8.565399935 | ENSG00000244731 |
| SLC25A36 | 0.354489948 | 1.278533487 | 0.000188254 | 0.000953679 | 815.5794194 | 1042.133166 | ENSG00000114120 |
| NA | 0.35444613 | 1.278494655 | 0.685706408 | 0.808505734 | 5.707219174 | 7.307583645 | ENSG00000255458 |
| ASPH | 0.354350787 | 1.278410167 | 1.10E-08 | 1.16E-07 | 1471.614217 | 1881.064521 | ENSG00000198363 |
| MTMR1 | 0.354289702 | 1.278356039 | 0.000251787 | 0.001243529 | 549.567229 | 702.5288514 | ENSG00000063601 |
| NA | 0.354235082 | 1.278307642 | 0.663083201 | 0.792102411 | 5.698846465 | 7.29898135 | ENSG00000217241 |
| NA | 0.353918817 | 1.278027444 | 0.263032554 | 0.428597094 | 51.54444211 | 65.74892817 | ENSG00000259768 |
| LOC100506691 | 0.353836566 | 1.277954584 | 0.138359789 | 0.265317962 | 76.87447894 | 98.18483752 | ENSG00000256546 |
| HCG23 | 0.353825528 | 1.277944806 | 0.30413345 | 0.474871581 | 33.35482838 | 42.63717931 | ENSG00000225914 |
| ACAT1 | 0.353765729 | 1.277891837 | 0.000707355 | 0.003111166 | 457.7488444 | 585.0836434 | ENSG00000075239 |
| ACSF2 | 0.35330096 | 1.277480226 | 0.006404523 | 0.021694419 | 256.3977667 | 327.2883651 | ENSG00000167107 |
| TST | 0.353059317 | 1.277266273 | 0.000264064 | 0.001296857 | 442.8571652 | 565.7233473 | ENSG00000128311 |
| DOCK3 | 0.352881204 | 1.277108594 | 0.008071715 | 0.026320394 | 263.1127815 | 335.7580553 | ENSG00000088538 |
| EPB41L2 | 0.352827648 | 1.277061185 | 1.24E-13 | 2.32E-12 | 3068.215034 | 3917.373838 | ENSG00000079819 |
| TADA1 | 0.352572118 | 1.276835013 | 0.000126515 | 0.000667012 | 533.1359041 | 680.7977804 | ENSG00000152382 |
| NA | 0.352443738 | 1.276721397 | 0.631889797 | 0.768926138 | 7.802982901 | 9.936682016 | ENSG00000229119 |
| SRGAP3 | 0.352147609 | 1.276459362 | 0.043121248 | 0.105995412 | 134.3690879 | 171.7317844 | ENSG00000196220 |
| FBF1 | 0.351976156 | 1.276307675 | 0.204140014 | 0.357388602 | 51.96446875 | 66.21408935 | ENSG00000188878 |
| RPAP2 | 0.351852566 | 1.276198343 | 0.005342047 | 0.01856484 | 279.8625845 | 357.0900935 | ENSG00000122484 |
| LINC00240 | 0.351807762 | 1.27615871 | 0.573852193 | 0.726147603 | 10.33898532 | 13.22883119 | ENSG00000224843 |
| SHANK1 | 0.351807476 | 1.276158457 | 0.119930746 | 0.238390934 | 74.86300588 | 95.51945197 | ENSG00000161681 |
| SEPTIN10 | 0.351770883 | 1.276126089 | 2.06E-08 | 2.09E-07 | 1472.496898 | 1879.25761 | ENSG00000186522 |
| PDK3 | 0.35162219 | 1.275994571 | 8.19E-07 | 6.54E-06 | 1105.667952 | 1411.184588 | ENSG00000067992 |
| KLHL36 | 0.351579925 | 1.275957189 | 2.11E-06 | 1.57E-05 | 928.6272251 | 1184.30313 | ENSG00000135686 |
| CMC1 | 0.351512482 | 1.275897542 | 0.00426864 | 0.015288958 | 302.5998739 | 385.8361906 | ENSG00000187118 |
| MCF2L2 | 0.35139853 | 1.275796769 | 0.12781189 | 0.249698206 | 71.60047719 | 91.31284006 | ENSG00000053524 |
| NA | 0.351121055 | 1.275551417 | 0.655700198 | 0.786538522 | 6.44211816 | 8.163429427 | ENSG00000214719 |
| NA | 0.351004926 | 1.275448747 | 0.684785613 | 0.807913843 | 5.896976446 | 7.572432051 | ENSG00000268584 |
| ACKR3 | 0.350921339 | 1.275374852 | 0.42396192 | 0.595650938 | 21.94876274 | 28.08142247 | ENSG00000144476 |
| NA | 0.350744873 | 1.275218861 | 0.763554645 | 0.85991903 | 3.072771419 | 3.872393683 | ENSG00000258539 |
| KLHDC9 | 0.350726252 | 1.275202402 | 0.107828231 | 0.219167044 | 97.8830793 | 124.8431037 | ENSG00000162755 |
| NANP | 0.35069499 | 1.27517477 | 0.023334296 | 0.063831806 | 179.7464237 | 229.3393219 | ENSG00000170191 |
| NOP53-AS1 | 0.350691729 | 1.275171888 | 0.746340772 | 0.849747228 | 3.624734239 | 4.65913011 | ENSG00000269656 |
| GIT2 | 0.350669288 | 1.275152052 | 0.039997017 | 0.099618566 | 139.0996997 | 177.1373351 | ENSG00000139436 |
| MPHOSPH9 | 0.350653451 | 1.275138054 | 0.000448636 | 0.002083947 | 459.6910205 | 586.1016368 | ENSG00000051825 |
| KCNH5 | 0.3506504 | 1.275135358 | 0.29482603 | 0.464765403 | 33.96963052 | 43.37713615 | ENSG00000140015 |
| THOC2 | 0.35050566 | 1.275007435 | 1.60E-06 | 1.22E-05 | 1187.817896 | 1514.649921 | ENSG00000125676 |
| UCHL3 | 0.350370608 | 1.274888086 | 0.688305387 | 0.810627756 | 5.345013626 | 6.783545051 | ENSG00000118939 |
| POLA2 | 0.350338294 | 1.274859531 | 0.616757344 | 0.758837926 | 7.737352675 | 9.835586629 | ENSG00000285816 |
| NHLRC4 | 0.350259002 | 1.274789466 | 0.628161157 | 0.766129882 | 7.8218435 | 9.936682016 | ENSG00000257108 |
| SLC9A5 | 0.35016135 | 1.274703181 | 0.0169605 | 0.04899446 | 261.6882257 | 333.200762 | ENSG00000135740 |
| ACTG1 | 0.349660995 | 1.274261166 | 5.21E-25 | 2.59E-23 | 55733.89505 | 71019.87947 | ENSG00000184009 |
| TNKS | 0.349499178 | 1.274118249 | 1.32E-05 | 8.46E-05 | 856.0651342 | 1090.404984 | ENSG00000173273 |
| MPZL1 | 0.349481348 | 1.274102502 | 1.43E-08 | 1.49E-07 | 1869.111234 | 2382.465065 | ENSG00000197965 |
| PROCR | 0.349280637 | 1.273925259 | 0.071507609 | 0.158667275 | 133.9190371 | 170.6893349 | ENSG00000101000 |
| RUSC1 | 0.349230524 | 1.273881009 | 3.61E-07 | 3.06E-06 | 1324.471267 | 1687.670854 | ENSG00000160753 |
| RGP1 | 0.349055994 | 1.273726911 | 0.000131153 | 0.000687707 | 540.178982 | 687.7370214 | ENSG00000107185 |
| SHC2 | 0.348874344 | 1.273566546 | 0.0582417 | 0.134737379 | 113.0409093 | 143.9598393 | ENSG00000129946 |
| FGGY | 0.348635573 | 1.273355783 | 0.059086954 | 0.136247482 | 151.0280163 | 192.0315992 | ENSG00000172456 |
| NA | 0.348484585 | 1.273222525 | 0.66178804 | 0.791282576 | 6.763135886 | 8.592824355 | ENSG00000279278 |
| DENND6A | 0.348425657 | 1.273170519 | 0.011768281 | 0.036075306 | 231.0659781 | 294.312527 | ENSG00000174839 |
| NA | 0.348422311 | 1.273167567 | 0.715834342 | 0.829478634 | 4.337981724 | 5.502605488 | ENSG00000285399 |
| SV2A | 0.34836113 | 1.273113577 | 2.65E-09 | 3.05E-08 | 2647.329922 | 3369.351704 | ENSG00000159164 |
| MRPL55 | 0.348310855 | 1.273069212 | 0.000591811 | 0.002660597 | 429.796926 | 547.2785373 | ENSG00000162910 |
| LYNX1 | 0.348288879 | 1.273049819 | 0.105757351 | 0.216027642 | 90.56095933 | 115.2949373 | ENSG00000180155 |
| CD99L2 | 0.348095003 | 1.272878753 | 0.000634889 | 0.002831827 | 507.6222923 | 645.6523712 | ENSG00000102181 |
| NPTX2 | 0.347976969 | 1.272774617 | 4.11E-05 | 0.00024074 | 778.681209 | 991.8913446 | ENSG00000106236 |
| RBFOX3 | 0.34778843 | 1.272608295 | 0.001075407 | 0.004549491 | 461.1767391 | 587.2387605 | ENSG00000167281 |
| NA | 0.347562799 | 1.27240928 | 2.25E-16 | 5.60E-15 | 648937.3474 | 825712.7293 | N_noFeature |
| SMYD2 | 0.347554574 | 1.272402026 | 0.043522313 | 0.106831523 | 141.9076349 | 180.6091033 | ENSG00000143499 |
| STXBP5 | 0.34752495 | 1.272375899 | 0.051407689 | 0.122026054 | 149.6252511 | 190.6535744 | ENSG00000164506 |
| FHL1 | 0.347412399 | 1.272276639 | 1.22E-06 | 9.47E-06 | 1223.974953 | 1557.088763 | ENSG00000022267 |
| NA | 0.347306752 | 1.272183475 | 0.577973666 | 0.729835046 | 10.08080691 | 12.79728569 | ENSG00000271646 |
| NA | 0.347292891 | 1.272171252 | 0.641532883 | 0.775864262 | 6.705878368 | 8.512701668 | ENSG00000215156 |
| NA | 0.347012237 | 1.271923795 | 0.752608809 | 0.853582018 | 3.344904336 | 4.246939771 | ENSG00000213399 |
| POPDC3 | 0.346688494 | 1.271638406 | 0.193369999 | 0.343079248 | 53.42569386 | 67.94754712 | ENSG00000132429 |
| ITGA3 | 0.346634752 | 1.271591036 | 5.14E-06 | 3.56E-05 | 821.903282 | 1045.191726 | ENSG00000005884 |
| EFCC1 | 0.346615469 | 1.27157404 | 0.547360302 | 0.705176488 | 13.5714898 | 17.27358019 | ENSG00000114654 |
| MBD5 | 0.346423939 | 1.27140524 | 0.031164523 | 0.081045406 | 160.9656865 | 204.9278297 | ENSG00000204406 |
| NA | 0.346201029 | 1.271208811 | 0.352081761 | 0.524844236 | 31.37078879 | 39.95134198 | ENSG00000235174 |
| NA | 0.346191375 | 1.271200304 | 0.669040519 | 0.796744284 | 5.962606673 | 7.622979745 | ENSG00000235280 |
| PHYKPL | 0.346099784 | 1.271119603 | 0.005983717 | 0.020484806 | 255.0255382 | 323.9513264 | ENSG00000175309 |
| TCERG1L | 0.345892511 | 1.270936994 | 0.00963371 | 0.030476205 | 228.3505941 | 290.1191216 | ENSG00000176769 |

| | | | | | | | |
|---|---|---|---|---|---|---|---|
| PTCH2 | 0.345841679 | 1.270892214 | 0.658115752 | 0.788553477 | 7.315098706 | 9.349985788 | ENSG00000117425 |
| NDUFB5 | 0.345835991 | 1.270887204 | 0.000774886 | 0.003375193 | 737.6284768 | 938.0813843 | ENSG00000136521 |
| ABHD15 | 0.345833391 | 1.270884914 | 0.401273396 | 0.573764151 | 24.69772553 | 31.47628457 | ENSG00000168792 |
| DOCK1 | 0.345797582 | 1.270853369 | 2.21E-06 | 1.63E-05 | 1400.411526 | 1778.806965 | ENSG00000150760 |
| VRK3 | 0.345779442 | 1.27083739 | 0.000302401 | 0.001466469 | 530.1612898 | 673.5362255 | ENSG00000105053 |
| LINC00466 | 0.345735163 | 1.270798386 | 0.757577055 | 0.856114219 | 3.423813356 | 4.335131715 | ENSG00000224209 |
| RBAK | 0.345733266 | 1.270796716 | 0.009809308 | 0.03094038 | 261.1097564 | 332.1292463 | ENSG00000146587 |
| THUMPD2 | 0.345439752 | 1.2705382 | 0.00294098 | 0.011012119 | 328.7650452 | 418.0860293 | ENSG00000138050 |
| NA | 0.345434823 | 1.270533859 | 0.67725927 | 0.802386422 | 6.721948064 | 8.521303963 | ENSG00000254419 |
| ZNF644 | 0.345399791 | 1.270503008 | 7.77E-06 | 5.22E-05 | 956.2703811 | 1215.777003 | ENSG00000122482 |
| LIN7C | 0.345257604 | 1.270377798 | 0.002722598 | 0.010321435 | 306.75731 | 389.6604598 | ENSG00000148943 |
| PABPC4L | 0.345162529 | 1.270294081 | 0.188660026 | 0.336289317 | 57.75318769 | 73.4520306 | ENSG00000254535 |
| LINC01607 | 0.345104394 | 1.270242895 | 0.776579974 | 0.868967765 | 3.124447132 | 3.918640229 | ENSG00000272138 |
| NFKBIZ | 0.344926605 | 1.270086368 | 0.055583804 | 0.130003292 | 125.4546825 | 159.2813599 | ENSG00000144802 |
| SYT6 | 0.344899281 | 1.270062313 | 4.07E-05 | 0.000238894 | 711.3576117 | 902.8822838 | ENSG00000134207 |
| LINC02323 | 0.344797937 | 1.269973098 | 0.616513036 | 0.758685441 | 8.033927996 | 10.203681 | ENSG00000259230 |
| SLC35B2 | 0.344675893 | 1.26986567 | 0.000161129 | 0.000828638 | 670.3020886 | 850.925705 | ENSG00000157593 |
| BOK | 0.344550103 | 1.269754955 | 9.49E-07 | 7.49E-06 | 1372.29586 | 1743.646472 | ENSG00000176720 |
| NA | 0.344362627 | 1.269589962 | 0.15834688 | 0.294710049 | 73.51330475 | 93.24260045 | ENSG00000204055 |
| UVSSA | 0.344312501 | 1.269545851 | 0.008464432 | 0.027365067 | 339.6169817 | 430.8682915 | ENSG00000163945 |
| TLCD5 | 0.344195477 | 1.269442876 | 0.006813205 | 0.022886735 | 251.9928785 | 320.0834943 | ENSG00000181264 |
| PTPN4 | 0.344132797 | 1.269387725 | 0.000443186 | 0.002063106 | 486.4229097 | 617.4830172 | ENSG00000088179 |
| TICAM1 | 0.343860962 | 1.269148568 | 0.333713276 | 0.50622496 | 35.80095832 | 45.4528814 | ENSG00000127666 |
| PCIF1 | 0.343065746 | 1.268449203 | 7.44E-05 | 0.000414365 | 639.4386323 | 810.4632584 | ENSG00000100982 |
| RAB11B-AS1 | 0.343056354 | 1.268440946 | 0.594238063 | 0.742276808 | 10.29847322 | 13.07073639 | ENSG00000269386 |
| NA | 0.343051427 | 1.268436613 | 0.726973787 | 0.83605886 | 4.686232757 | 5.950822542 | ENSG00000259687 |
| ERMARD | 0.342894598 | 1.268298735 | 0.04875868 | 0.117005062 | 130.3767665 | 165.2614969 | ENSG00000130023 |
| EDARADD | 0.342687661 | 1.268116826 | 0.093429152 | 0.196231583 | 91.86735746 | 116.4823177 | ENSG00000186197 |
| HAGHL | 0.342660941 | 1.268093339 | 0.000213338 | 0.001068682 | 573.8503607 | 728.0164209 | ENSG00000103253 |
| NCOA4 | 0.342616627 | 1.268054389 | 3.32E-09 | 3.78E-08 | 1715.477714 | 2175.19855 | ENSG00000266412 |
| DTL | 0.342590587 | 1.268031502 | 1.10E-07 | 1.01E-06 | 1254.753355 | 1590.955674 | ENSG00000143476 |
| ZFPM1 | 0.342574643 | 1.268017489 | 0.057408545 | 0.133253308 | 136.5570394 | 173.0243129 | ENSG00000179588 |
| SUGP2 | 0.342399415 | 1.267863485 | 1.66E-07 | 1.48E-06 | 3435.902399 | 4355.007913 | ENSG00000064607 |
| ETFRF1 | 0.342028492 | 1.267537555 | 0.256356375 | 0.421525713 | 49.5504782 | 62.89772029 | ENSG00000205707 |
| C21orf58 | 0.341985968 | 1.267500194 | 0.010262232 | 0.032136776 | 329.2644422 | 417.0320397 | ENSG00000160298 |
| NA | 0.341726134 | 1.267271934 | 0.716601889 | 0.829919376 | 4.658999449 | 5.912645253 | ENSG00000204092 |
| FKBP9P1 | 0.341624915 | 1.267183026 | 0.707340833 | 0.823594399 | 4.727420579 | 5.99276794 | ENSG00000176826 |
| PITRM1 | 0.341378318 | 1.266966447 | 5.31E-05 | 0.000304121 | 693.0034469 | 877.5009286 | ENSG00000107959 |
| ZNF526 | 0.341263658 | 1.266865757 | 0.000599068 | 0.002688839 | 477.2962949 | 604.7666903 | ENSG00000167625 |
| CLCN1 | 0.341245532 | 1.26684984 | 0.520355479 | 0.68346886 | 15.51501729 | 19.74054308 | ENSG00000188037 |
| IL27RA | 0.34094303 | 1.266584237 | 0.000519281 | 0.002370433 | 459.3412931 | 582.1789864 | ENSG00000104998 |
| ZFYVE16 | 0.340738331 | 1.266404538 | 0.000886312 | 0.003819893 | 399.4267483 | 505.7547643 | ENSG00000039319 |
| TUBGCP4 | 0.340718543 | 1.266387169 | 0.004627031 | 0.016426216 | 426.5802399 | 540.2612937 | ENSG00000137822 |
| ITM2B | 0.340164436 | 1.265900871 | 2.04E-08 | 2.07E-07 | 1798.668609 | 2278.110761 | ENSG00000136156 |
| NA | 0.340163359 | 1.265899926 | 0.679928738 | 0.804446813 | 5.957024867 | 7.570281478 | ENSG00000236989 |
| APRT | 0.340028859 | 1.265781914 | 4.54E-09 | 5.07E-08 | 2050.096408 | 2594.469398 | ENSG00000198931 |
| CHD5 | 0.340019103 | 1.265773355 | 0.136038947 | 0.261778286 | 73.8971618 | 93.48350155 | ENSG00000116254 |
| MAP10 | 0.339688359 | 1.265483203 | 0.337860364 | 0.510468693 | 35.03970191 | 44.19156957 | ENSG00000212916 |
| BBS12 | 0.339503297 | 1.265320884 | 0.208031238 | 0.362785266 | 54.18224434 | 68.49659909 | ENSG00000181004 |
| LNX1 | 0.339327786 | 1.265166961 | 0.11739306 | 0.234555102 | 83.4139664 | 105.6604753 | ENSG00000072201 |
| SH3PXD2B | 0.33927415 | 1.265119926 | 6.27E-11 | 8.81E-10 | 2551.538629 | 3228.959129 | ENSG00000174705 |
| KDM7A-DT | 0.339080832 | 1.264950414 | 0.198243307 | 0.349620469 | 57.72615454 | 72.91615464 | ENSG00000260231 |
| DHX58 | 0.339029274 | 1.264905209 | 0.170128041 | 0.311409055 | 65.75729163 | 83.0563845 | ENSG00000108771 |
| LINC02593 | 0.338358529 | 1.264317259 | 0.592722734 | 0.741158069 | 9.364768527 | 11.8424951 | ENSG00000223764 |
| ALG10B | 0.33796849 | 1.263975491 | 0.123597327 | 0.243720336 | 90.36426881 | 114.2360951 | ENSG00000175548 |
| PLCD3 | 0.337823045 | 1.26384807 | 0.000162795 | 0.000836346 | 681.7441862 | 861.8659084 | ENSG00000161714 |
| NA | 0.337808139 | 1.263835012 | 0.056237533 | 0.131103111 | 153.7447275 | 194.2517118 | ENSG00000241743 |
| MFSD14C | 0.337653156 | 1.263699251 | 0.000346611 | 0.001657939 | 534.5746387 | 674.9542872 | ENSG00000196312 |
| SLC25A11 | 0.337645398 | 1.263692456 | 3.19E-06 | 2.29E-05 | 1436.440639 | 1814.118333 | ENSG00000108528 |
| TNFSF15 | 0.337537462 | 1.263597915 | 0.765277643 | 0.861065676 | 3.929682269 | 5.012703499 | ENSG00000181634 |
| ZNF106 | 0.337295543 | 1.263386046 | 2.07E-07 | 1.81E-06 | 1616.31975 | 2041.384259 | ENSG00000103994 |
| AQP4 | 0.337197085 | 1.263299828 | 0.421366993 | 0.593344248 | 25.62393339 | 32.31760937 | ENSG00000171885 |
| PCCA-DT | 0.337027706 | 1.26315152 | 0.024537137 | 0.066555949 | 209.8470081 | 265.0932178 | ENSG00000274605 |
| NOD1 | 0.336759567 | 1.262916772 | 0.122640025 | 0.242433689 | 86.96413412 | 109.8380453 | ENSG00000106100 |
| DLL3 | 0.336754741 | 1.262912548 | 0.010287822 | 0.03220746 | 304.3964716 | 384.106743 | ENSG00000090932 |
| DNAH7 | 0.336449086 | 1.262645011 | 0.753836313 | 0.85412054 | 3.399370951 | 4.265761897 | ENSG00000118997 |
| GPR143 | 0.336429047 | 1.262627472 | 0.001244234 | 0.005196474 | 407.8382389 | 515.4029416 | ENSG00000101850 |
| ZNF141 | 0.336400409 | 1.262602409 | 0.007395999 | 0.024468566 | 303.6723359 | 383.0914696 | ENSG00000131127 |
| SRR | 0.336298239 | 1.262512997 | 0.32922687 | 0.501507307 | 34.22222712 | 43.26582093 | ENSG00000167720 |
| SMTN | 0.336193689 | 1.262421507 | 0.005475482 | 0.01897483 | 274.0919625 | 346.0628316 | ENSG00000183963 |
| MARCHF7 | 0.336129075 | 1.262364968 | 4.16E-07 | 3.47E-06 | 1526.324172 | 1927.456284 | ENSG00000136536 |
| UBE2S | 0.336054305 | 1.262299546 | 1.96E-14 | 3.99E-13 | 7123.350712 | 8993.19648 | ENSG00000108106 |
| SUSD1 | 0.335886285 | 1.262152544 | 0.093711756 | 0.196674495 | 109.9793015 | 138.985301 | ENSG00000106868 |
| ANO7 | 0.335876403 | 1.262143899 | 0.172442316 | 0.314600408 | 66.28548777 | 83.5575724 | ENSG00000146205 |
| NA | 0.335691704 | 1.261982324 | 0.603101994 | 0.748628825 | 9.906243457 | 12.57868383 | ENSG00000232630 |
| PLAU | 0.335571995 | 1.261877614 | 0.024087309 | 0.065510823 | 209.1067274 | 263.6910342 | ENSG00000122861 |
| NA | 0.335431643 | 1.26175486 | 0.653576168 | 0.78472259 | 7.952228073 | 10.01869481 | ENSG00000286379 |

| | | | | | | | |
|---|---|---|---|---|---|---|---|
| PFKFB3 | 0.335421185 | 1.261745713 | 3.55E-05 | 0.000210496 | 1084.420322 | 1367.703819 | ENSG00000170525 |
| KIF21B | 0.335382419 | 1.26171181 | 1.73E-08 | 1.78E-07 | 2002.922238 | 2527.476345 | ENSG00000116852 |
| AIG1 | 0.335102213 | 1.261466779 | 0.000852632 | 0.003687092 | 448.7204507 | 566.4944661 | ENSG00000146416 |
| SMG1P6 | 0.335047397 | 1.26141885 | 0.757255887 | 0.855942967 | 3.725970483 | 4.709677803 | ENSG00000254634 |
| NA | 0.334952931 | 1.261336257 | 0.024195009 | 0.06576665 | 195.4773924 | 246.4811656 | ENSG00000213553 |
| C2orf16 | 0.334761661 | 1.261169041 | 0.541129643 | 0.700176533 | 13.34333561 | 16.87402072 | ENSG00000221843 |
| ZNF623 | 0.33470244 | 1.261117273 | 7.92E-05 | 0.00043778 | 654.0016479 | 824.5304277 | ENSG00000183309 |
| USP45 | 0.334596269 | 1.261024468 | 0.001908343 | 0.007573305 | 406.1703479 | 511.9171976 | ENSG00000123552 |
| YBEY | 0.334113905 | 1.260602916 | 0.002877571 | 0.010823011 | 334.7126582 | 421.868111 | ENSG00000182362 |
| NA | 0.334009002 | 1.260511257 | 0.694927414 | 0.815746479 | 5.668822255 | 7.122064424 | ENSG00000263745 |
| WHRN | 0.333893267 | 1.260410141 | 0.012536743 | 0.038068047 | 277.8915356 | 350.4321305 | ENSG00000095397 |
| NA | 0.333806517 | 1.260334354 | 0.604614602 | 0.749638171 | 9.337535219 | 11.75807126 | ENSG00000235636 |
| DTNB | 0.333720116 | 1.260258877 | 8.14E-05 | 0.000449015 | 655.3148671 | 825.8336849 | ENSG00000138101 |
| RNF215 | 0.333707412 | 1.260247779 | 0.013181792 | 0.039751438 | 244.9574465 | 308.3560947 | ENSG00000099999 |
| NA | 0.333606535 | 1.260159662 | 0.338358942 | 0.510981865 | 37.26218343 | 47.03791273 | ENSG00000228502 |
| MBOAT1 | 0.333565215 | 1.260123571 | 0.008332648 | 0.027029615 | 260.9394602 | 328.6193193 | ENSG00000172197 |
| FRAS1 | 0.333447247 | 1.260020536 | 5.68E-06 | 3.91E-05 | 2812.595018 | 3543.14305 | ENSG00000138759 |
| MCL1 | 0.3334454 | 1.260018923 | 1.71E-18 | 5.06E-17 | 7061.198376 | 8897.6933 | ENSG00000143384 |
| RGS3 | 0.33335998 | 1.259944321 | 0.002415847 | 0.009314702 | 432.5783533 | 544.6752337 | ENSG00000138835 |
| EID2B | 0.333022447 | 1.259649578 | 0.582582561 | 0.73296812 | 11.0522328 | 13.95856821 | ENSG00000176401 |
| NA | 0.333014914 | 1.259643001 | 0.500617859 | 0.666686425 | 16.40773432 | 20.6388673 | ENSG00000213189 |
| FBXL3 | 0.332910254 | 1.259551625 | 0.103622196 | 0.212928823 | 99.22344383 | 125.1294578 | ENSG00000005812 |
| RFX7 | 0.332864562 | 1.259511733 | 3.20E-05 | 0.000191987 | 704.8977601 | 887.8126656 | ENSG00000181827 |
| ALG1L9P | 0.332848657 | 1.259497848 | 0.35762665 | 0.530817142 | 30.06718156 | 37.9170091 | ENSG00000248671 |
| SP2-AS1 | 0.33284657 | 1.259496026 | 0.75298333 | 0.853786015 | 3.393789145 | 4.242638624 | ENSG00000234494 |
| ERMAP | 0.332573574 | 1.259257718 | 0.305422917 | 0.476345177 | 37.42625899 | 47.1992426 | ENSG00000164010 |
| IDO1 | 0.332560893 | 1.25924665 | 0.002444 | 0.009408743 | 404.1322803 | 509.3330072 | ENSG00000131203 |
| RNF125 | 0.332395157 | 1.259101996 | 0.009951093 | 0.031300591 | 262.6314302 | 330.9211721 | ENSG00000101695 |
| GXYLT2 | 0.332365875 | 1.259076442 | 0.209984388 | 0.365464529 | 56.58084092 | 71.16602533 | ENSG00000172986 |
| AIDA | 0.332285427 | 1.259006234 | 7.31E-06 | 4.94E-05 | 1794.654952 | 2260.498389 | ENSG00000186063 |
| TDGF1P3 | 0.332200912 | 1.258932483 | 0.000278857 | 0.001362916 | 528.754219 | 665.7935766 | ENSG00000225366 |
| ABCC4 | 0.33220044 | 1.25893207 | 0.003862992 | 0.014015608 | 367.0751729 | 462.4270977 | ENSG00000125257 |
| AMER2 | 0.332172246 | 1.258907467 | 0.604329757 | 0.749477682 | 9.688577155 | 12.21704118 | ENSG00000165566 |
| TCP11L2 | 0.332090661 | 1.258836278 | 0.483623625 | 0.651370756 | 16.68544904 | 21.01341338 | ENSG00000166046 |
| NA | 0.331813882 | 1.258594796 | 0.775342273 | 0.868388748 | 3.573058527 | 4.513938752 | ENSG00000223393 |
| LOC107984660 | 0.331772289 | 1.25855851 | 0.721833424 | 0.833127234 | 5.054020111 | 6.381835006 | ENSG00000258457 |
| NA | 0.331760541 | 1.258548261 | 0.729186986 | 0.837555591 | 4.455287663 | 5.551002607 | ENSG00000237821 |
| ITGB4 | 0.331602403 | 1.258410316 | 0.075043276 | 0.164975481 | 120.8662194 | 152.1127763 | ENSG00000132470 |
| NPIPB2 | 0.331576284 | 1.258387534 | 0.677654834 | 0.802741136 | 6.055470207 | 7.622979745 | ENSG00000234719 |
| NA | 0.331485572 | 1.258308413 | 0.493375527 | 0.659806398 | 19.27121214 | 24.11574219 | ENSG00000230805 |
| GLUL | 0.331448354 | 1.258275952 | 1.40E-13 | 2.61E-12 | 5784.324385 | 7278.019342 | ENSG00000135821 |
| DNAH17 | 0.331334201 | 1.258176395 | 0.24980831 | 0.413901238 | 52.7564251 | 66.29771968 | ENSG00000187775 |
| PRKCI | 0.331327691 | 1.258170718 | 2.07E-09 | 2.42E-08 | 2519.289052 | 3169.643619 | ENSG00000163558 |
| LINC00937 | 0.331300598 | 1.258147091 | 0.364416686 | 0.537974382 | 28.46699902 | 35.7837314 | ENSG00000226091 |
| NA | 0.33129954 | 1.258146168 | 0.748484185 | 0.851005388 | 3.630316045 | 4.58330857 | ENSG00000254896 |
| IDNK | 0.331061796 | 1.257938853 | 0.689978563 | 0.811657812 | 5.75331308 | 7.197885963 | ENSG00000148057 |
| NA | 0.33097106 | 1.257859739 | 0.545787509 | 0.704108719 | 13.99374377 | 17.57607285 | ENSG00000269054 |
| MAP3K20 | 0.3308424 | 1.257747568 | 0.010763168 | 0.03353776 | 283.5513463 | 356.5442524 | ENSG00000091436 |
| NA | 0.330792626 | 1.257704176 | 0.540863974 | 0.700109467 | 15.8424927 | 19.82255587 | ENSG00000267102 |
| MMP11 | 0.330738272 | 1.257656792 | 0.027635797 | 0.073602233 | 199.1446659 | 250.4796375 | ENSG00000099953 |
| TRIM16 | 0.330578897 | 1.257517866 | 0.091511779 | 0.193167057 | 119.5394091 | 150.4502875 | ENSG00000221926 |
| SCML2 | 0.330530534 | 1.257475711 | 0.000276641 | 0.001352429 | 642.0607283 | 807.5907626 | ENSG00000102098 |
| DHRS7B | 0.33053036 | 1.25747556 | 0.01169786 | 0.035922224 | 244.334635 | 307.160645 | ENSG00000109016 |
| RETSAT | 0.330250911 | 1.257232011 | 0.004735879 | 0.016749843 | 325.2091324 | 408.7172335 | ENSG00000042445 |
| RXYLT1 | 0.330118785 | 1.257116876 | 0.090993515 | 0.192283481 | 115.0813305 | 144.6099988 | ENSG00000118600 |
| CDKL5 | 0.329820623 | 1.256857094 | 0.007201754 | 0.023928529 | 293.0924812 | 368.3832519 | ENSG00000008086 |
| UTP25 | 0.329640674 | 1.256700334 | 0.000982618 | 0.004198378 | 515.1245958 | 647.2253841 | ENSG00000117597 |
| TBC1D8B | 0.329530755 | 1.25660459 | 0.019508868 | 0.055053477 | 213.825463 | 268.8042816 | ENSG00000133138 |
| NA | 0.329518176 | 1.256593633 | 0.711095331 | 0.825872368 | 4.985598981 | 6.291492488 | ENSG00000240950 |
| FGF13 | 0.329453533 | 1.256537331 | 9.62E-06 | 6.36E-05 | 1207.511688 | 1516.531854 | ENSG00000129682 |
| MFAP3L | 0.329388147 | 1.256480383 | 0.126577464 | 0.248024973 | 89.89535736 | 112.7532373 | ENSG00000198948 |
| RAB33B | 0.329262196 | 1.256370693 | 0.015080649 | 0.044462907 | 230.1909704 | 289.1258627 | ENSG00000172007 |
| HMBS | 0.329097187 | 1.256227004 | 1.27E-05 | 8.19E-05 | 1070.094346 | 1343.646621 | ENSG00000256269 |
| CCT3 | 0.329081947 | 1.256213734 | 2.17E-19 | 6.91E-18 | 13264.04657 | 16662.54151 | ENSG00000163468 |
| TMEM177 | 0.328955287 | 1.25610345 | 4.09E-05 | 0.000239684 | 964.6445877 | 1211.839887 | ENSG00000144120 |
| ZNF41 | 0.328897606 | 1.256053231 | 0.113999588 | 0.229203656 | 100.0327461 | 125.4733444 | ENSG00000147124 |
| LMLN | 0.328880059 | 1.256037954 | 0.193649486 | 0.343354066 | 64.3466662 | 80.92121669 | ENSG00000185621 |
| TMEM178A | 0.328877595 | 1.256035777 | 0.00360025 | 0.013173829 | 368.304489 | 462.8718497 | ENSG00000152154 |
| NPHS1 | 0.328856185 | 1.256017169 | 0.386019603 | 0.559523944 | 27.58006396 | 34.5974355 | ENSG00000161270 |
| ROBO3 | 0.328828564 | 1.255993122 | 0.039951804 | 0.099523535 | 169.2771659 | 212.3789993 | ENSG00000154134 |
| LPCAT4 | 0.328818946 | 1.255984749 | 4.31E-05 | 0.000250838 | 870.0632588 | 1092.384305 | ENSG00000176454 |
| SLC9B2 | 0.328808625 | 1.255975764 | 0.19746642 | 0.34855199 | 59.16851905 | 74.34228556 | ENSG00000164038 |
| TRAF5 | 0.328694963 | 1.255876816 | 0.002847619 | 0.010721972 | 339.0647946 | 425.7818865 | ENSG00000082512 |
| VSIG10L | 0.328558658 | 1.255758167 | 0.004742882 | 0.016763394 | 350.050358 | 439.544964 | ENSG00000186806 |
| DIS3L2 | 0.328455195 | 1.255668114 | 0.004415117 | 0.015755092 | 327.8667832 | 411.6791809 | ENSG00000144535 |
| DSTYK | 0.328381023 | 1.255603559 | 0.000482743 | 0.002220984 | 487.0963208 | 611.649447 | ENSG00000133059 |
| DPP8 | 0.32829971 | 1.255532793 | 0.001817827 | 0.007261899 | 427.449805 | 537.26627 | ENSG00000074603 |

| | | | | | | | |
|---|---|---|---|---|---|---|---|
| LINC00877 | 0.328285788 | 1.255520677 | 0.469705567 | 0.638689651 | 20.06037758 | 25.14205318 | ENSG00000241163 |
| ADRB2 | 0.328283368 | 1.25551857 | 0.466065856 | 0.635352726 | 19.07220628 | 23.91811303 | ENSG00000169252 |
| MRM2 | 0.328190442 | 1.255437703 | 0.621793947 | 0.762169718 | 10.95446318 | 13.77519956 | ENSG00000122687 |
| LINC00869 | 0.32801663 | 1.255286461 | 0.238151619 | 0.400015923 | 58.38772631 | 73.07749663 | ENSG00000277147 |
| FIGN | 0.328010645 | 1.255281253 | 0.01163409 | 0.035777656 | 257.4309558 | 323.0841763 | ENSG00000182263 |
| FAM172A | 0.327941335 | 1.255220949 | 0.080413265 | 0.174141479 | 114.6871857 | 143.9646735 | ENSG00000113391 |
| NA | 0.327941258 | 1.255220882 | 0.533305277 | 0.693924025 | 14.1110497 | 17.65619554 | ENSG00000285128 |
| NA | 0.327932863 | 1.255213578 | 0.519861804 | 0.683212997 | 17.92968356 | 22.51295481 | ENSG00000234405 |
| ZNF710-AS1 | 0.327745004 | 1.255050142 | 0.662777937 | 0.791973727 | 7.038059705 | 8.807125069 | ENSG00000259291 |
| LMNTD2 | 0.327650781 | 1.254968177 | 0.176255514 | 0.319712752 | 84.03502607 | 105.3572013 | ENSG00000185522 |
| CACNA1D | 0.327351135 | 1.254707549 | 0.420513444 | 0.592558426 | 25.94060861 | 32.62870432 | ENSG00000157388 |
| ENTPD5 | 0.327230075 | 1.254602267 | 0.058268458 | 0.134766618 | 134.4188486 | 168.6630712 | ENSG00000187097 |
| NA | 0.327198364 | 1.254574691 | 0.621428208 | 0.761915061 | 10.17279457 | 12.84353224 | ENSG00000257557 |
| NA | 0.327186705 | 1.254564552 | 0.353001793 | 0.525931232 | 31.07142256 | 38.94305962 | ENSG00000279198 |
| SEMA3F | 0.327075903 | 1.254468203 | 2.41E-06 | 1.76E-05 | 1322.398668 | 1658.372997 | ENSG00000001617 |
| FDXACB1 | 0.32692297 | 1.25433523 | 0.486564163 | 0.654052629 | 18.87878222 | 23.78985369 | ENSG00000255561 |
| ATF7IP | 0.326855317 | 1.254276411 | 4.32E-07 | 3.60E-06 | 1940.632974 | 2434.030262 | ENSG00000171681 |
| ALS2 | 0.326768523 | 1.254200955 | 4.92E-06 | 3.42E-05 | 1194.17297 | 1497.045685 | ENSG00000003393 |
| MPDZ | 0.326718505 | 1.254157472 | 1.87E-08 | 1.91E-07 | 1713.273593 | 2148.499423 | ENSG00000107186 |
| CAPN7 | 0.326652634 | 1.254100211 | 0.013558308 | 0.040658063 | 306.0370044 | 384.4401556 | ENSG00000131375 |
| TUNAR | 0.326589755 | 1.254045553 | 1.09E-05 | 7.12E-05 | 1069.792152 | 1341.284999 | ENSG00000250366 |
| NA | 0.326556696 | 1.254016817 | 0.759655935 | 0.857270428 | 3.638688754 | 4.570938166 | ENSG00000288546 |
| UBR3 | 0.326546853 | 1.254008261 | 0.002671909 | 0.010156412 | 430.0122386 | 538.5854232 | ENSG00000144357 |
| RBPJ | 0.326412385 | 1.253891386 | 6.13E-08 | 5.80E-07 | 2380.838419 | 2985.045664 | ENSG00000168214 |
| NGFR | 0.326257995 | 1.253757208 | 0.011381151 | 0.035201826 | 256.5240841 | 321.9132133 | ENSG00000064300 |
| NA | 0.326140352 | 1.253654976 | 0.687604106 | 0.810099598 | 6.088285321 | 7.620829171 | ENSG00000226564 |
| H2AC20 | 0.326131783 | 1.253647529 | 0.389093082 | 0.562490145 | 28.95891342 | 36.31181068 | ENSG00000184260 |
| OSBPL10 | 0.32611081 | 1.253629305 | 3.38E-07 | 2.87E-06 | 1444.957495 | 1811.057615 | ENSG00000144645 |
| LOC101930085 | 0.326103847 | 1.253623255 | 0.350971065 | 0.523640184 | 40.69572093 | 51.08615726 | ENSG00000271122 |
| TBC1D5 | 0.326083394 | 1.253605483 | 0.000327623 | 0.001575319 | 626.4159894 | 785.2760248 | ENSG00000131374 |
| SP4 | 0.325989835 | 1.253524189 | 0.003295666 | 0.012165458 | 363.9134801 | 456.6416273 | ENSG00000105866 |
| NHLRC2 | 0.325974684 | 1.253511024 | 0.011642933 | 0.035782032 | 335.3465963 | 420.7696855 | ENSG00000196865 |
| ALDH5A1 | 0.325967083 | 1.25350442 | 0.09883404 | 0.205276428 | 104.7716937 | 131.412348 | ENSG00000112294 |
| RELL2 | 0.325930628 | 1.253472746 | 0.011941244 | 0.03648764 | 351.3577953 | 439.978739 | ENSG00000164620 |
| SPARC | 0.325788444 | 1.253349216 | 1.58E-10 | 2.13E-09 | 4101.187905 | 5139.975865 | ENSG00000113140 |
| VCL | 0.325774105 | 1.25333676 | 7.28E-14 | 1.40E-12 | 10709.24593 | 13422.30654 | ENSG00000035403 |
| MGST3 | 0.325727403 | 1.253296188 | 2.22E-06 | 1.64E-05 | 1205.571604 | 1511.015855 | ENSG00000143198 |
| RBFOX1 | 0.325709469 | 1.253280609 | 0.338764956 | 0.511352842 | 35.39265887 | 44.38247444 | ENSG00000078328 |
| NA | 0.325410794 | 1.253021174 | 0.389387325 | 0.562629833 | 28.28473857 | 35.50221144 | ENSG00000272556 |
| NA | 0.325375793 | 1.252990775 | 0.602892332 | 0.748416729 | 10.00680398 | 12.57438268 | ENSG00000213326 |
| FOXO4 | 0.325271954 | 1.252900593 | 0.000286658 | 0.001396661 | 625.9332994 | 784.2242042 | ENSG00000184481 |
| RTN4IP1 | 0.325235171 | 1.25286865 | 0.007929739 | 0.025914381 | 303.4151966 | 380.4276652 | ENSG00000130347 |
| LCK | 0.324960909 | 1.252630497 | 6.20E-08 | 5.86E-07 | 1891.65098 | 2368.714183 | ENSG00000182866 |
| HAAO | 0.324895699 | 1.25257388 | 0.373091784 | 0.546534559 | 30.11116029 | 37.6750235 | ENSG00000162882 |
| KSR1 | 0.324746841 | 1.252444645 | 0.000106598 | 0.00057329 | 885.7601347 | 1108.312125 | ENSG00000141068 |
| STX12 | 0.324727811 | 1.252428125 | 0.051761644 | 0.122700158 | 176.5928282 | 221.2463283 | ENSG00000117758 |
| SCARA3 | 0.324579283 | 1.252299191 | 0.006558162 | 0.02214097 | 396.7279849 | 497.1737957 | ENSG00000168077 |
| NA | 0.323834726 | 1.251653062 | 0.001797217 | 0.007192774 | 690.1181813 | 863.7596115 | ENSG00000197582 |
| ASH2L | 0.323791568 | 1.25161562 | 2.12E-06 | 1.58E-05 | 1367.846665 | 1711.982226 | ENSG00000129691 |
| CENPJ | 0.323719681 | 1.251553255 | 0.000898012 | 0.003860469 | 605.9812163 | 758.3894521 | ENSG00000151849 |
| NAA80 | 0.32364462 | 1.25148814 | 0.204247809 | 0.357512429 | 79.19205131 | 98.85302562 | ENSG00000243477 |
| FIRRE | 0.323560522 | 1.251415191 | 0.001090988 | 0.004609341 | 451.4667347 | 565.1928448 | ENSG00000213468 |
| NBR1 | 0.323511399 | 1.251372581 | 4.13E-05 | 0.000241495 | 941.2346225 | 1177.264702 | ENSG00000188554 |
| SPRED1 | 0.323496112 | 1.251359322 | 9.81E-05 | 0.000531879 | 661.0191705 | 827.4730442 | ENSG00000166068 |
| LUZP1 | 0.323480185 | 1.251345507 | 8.45E-09 | 9.10E-08 | 1751.430032 | 2191.468682 | ENSG00000169641 |
| TRIM11 | 0.323119998 | 1.251033132 | 0.001515793 | 0.006182877 | 541.2287164 | 677.048873 | ENSG00000154370 |
| NA | 0.323002138 | 1.250930934 | 0.31721401 | 0.48837793 | 43.96826195 | 54.95693341 | ENSG00000269001 |
| ADCY9 | 0.322996958 | 1.250926443 | 0.00296536 | 0.011088343 | 440.2633417 | 550.4859838 | ENSG00000162104 |
| COG2 | 0.322829751 | 1.250781469 | 5.82E-05 | 0.000330878 | 703.9816767 | 880.4986181 | ENSG00000135775 |
| MAGEA8 | 0.322823189 | 1.25077578 | 0.787936405 | 0.876337918 | 2.658890159 | 3.352916701 | ENSG00000156009 |
| SEC24B-AS1 | 0.322598266 | 1.250580793 | 0.505302755 | 0.670320137 | 18.39244963 | 22.99934915 | ENSG00000247950 |
| OPRK1 | 0.322575325 | 1.250560908 | 0.635679033 | 0.771784519 | 9.038168995 | 11.34614139 | ENSG00000082556 |
| HECTD1 | 0.322476927 | 1.250475617 | 9.05E-08 | 8.40E-07 | 1921.060969 | 2401.679661 | ENSG00000092148 |
| RGS17 | 0.322232675 | 1.250263926 | 0.12701077 | 0.248545827 | 93.81300013 | 117.3604627 | ENSG00000091844 |
| FAM160B1 | 0.32197538 | 1.25004097 | 0.041566205 | 0.102814933 | 211.9057774 | 265.3959709 | ENSG00000151553 |
| TMEM186 | 0.321909876 | 1.249984215 | 0.021765436 | 0.060247816 | 233.3038905 | 291.2110343 | ENSG00000184857 |
| CBWD6 | 0.321536648 | 1.249660884 | 0.391962793 | 0.56495474 | 34.04151828 | 42.51457818 | ENSG00000215126 |
| SESN2 | 0.321525616 | 1.249651327 | 1.28E-05 | 8.23E-05 | 978.1550637 | 1222.157227 | ENSG00000130766 |
| SDHAF4 | 0.321432687 | 1.249570835 | 0.038155429 | 0.095865641 | 177.9704384 | 222.1809824 | ENSG00000154079 |
| MLH3 | 0.321350361 | 1.249499532 | 0.011292292 | 0.034955008 | 257.4122954 | 321.5660916 | ENSG00000119684 |
| ANXA7 | 0.321103876 | 1.249286073 | 4.16E-06 | 2.94E-05 | 1002.95554 | 1253.118608 | ENSG00000138279 |
| ZNF75D | 0.321061592 | 1.249249458 | 0.056012134 | 0.130704014 | 143.1304181 | 178.8059847 | ENSG00000186376 |
| OXTR | 0.321034548 | 1.24922604 | 0.217129238 | 0.374554245 | 59.5579579 | 74.29792913 | ENSG00000180914 |
| TTC30A | 0.320969972 | 1.249170125 | 0.349080293 | 0.521661052 | 35.1286985 | 44.00200967 | ENSG00000197557 |
| RASL12 | 0.32048387 | 1.248749301 | 0.006931919 | 0.023211882 | 299.4930112 | 374.2383647 | ENSG00000103710 |
| NHLH1 | 0.320479336 | 1.248745376 | 0.732919416 | 0.840491526 | 4.939505075 | 6.181794807 | ENSG00000171786 |
| ZNF225 | 0.320414508 | 1.248689264 | 0.119343229 | 0.237443299 | 101.8406706 | 127.1178167 | ENSG00000256294 |

| | | | | | | | |
|---|---|---|---|---|---|---|---|
| RUFY3 | 0.320390083 | 1.248668124 | 0.000697165 | 0.003071251 | 525.9168985 | 657.1827236 | ENSG00000018189 |
| PDS5B | 0.320152383 | 1.248462409 | 0.000115404 | 0.000615161 | 842.8856002 | 1053.075713 | ENSG00000083642 |
| ZNF555 | 0.319911597 | 1.248254058 | 0.10376013 | 0.213121452 | 104.710406 | 130.7694459 | ENSG00000186300 |
| NA | 0.319215291 | 1.247651743 | 0.256769396 | 0.421919796 | 58.08680849 | 72.57521212 | ENSG00000223459 |
| GH2 | 0.319105035 | 1.247556396 | 0.678380738 | 0.803241041 | 7.557207412 | 9.461040541 | ENSG00000136487 |
| NA | 0.319041738 | 1.247501663 | 0.369250104 | 0.543009892 | 36.86659919 | 45.94384946 | ENSG00000251687 |
| PRRG1 | 0.318932876 | 1.247407532 | 0.143786991 | 0.273351275 | 97.03713191 | 121.2099936 | ENSG00000130962 |
| SYNGR1 | 0.318871381 | 1.247354363 | 0.000831303 | 0.003601318 | 608.9117652 | 760.0234988 | ENSG00000100321 |
| U2AF1 | 0.318845696 | 1.247332156 | 0.423608272 | 0.595310496 | 27.9150362 | 34.72837846 | ENSG00000160201 |
| RPP38 | 0.318801611 | 1.247294041 | 0.090267208 | 0.19106262 | 120.0264174 | 149.9673887 | ENSG00000152464 |
| IQGAP2 | 0.318785584 | 1.247280185 | 1.96E-08 | 2.00E-07 | 2080.012229 | 2594.992418 | ENSG00000145703 |
| PRORSD1P | 0.318766478 | 1.247263667 | 0.633105195 | 0.769822396 | 8.962050878 | 11.21977215 | ENSG00000162997 |
| LOC107987373 | 0.318682233 | 1.247190836 | 0.05288199 | 0.124651327 | 156.1962391 | 194.5679441 | ENSG00000276345 |
| NCR3LG1 | 0.318522568 | 1.247052816 | 0.316364263 | 0.487378373 | 42.37991874 | 52.81316331 | ENSG00000188211 |
| NA | 0.318366545 | 1.246917958 | 0.720818257 | 0.832304124 | 5.042856499 | 6.289341915 | ENSG00000227775 |
| AFF4 | 0.318355836 | 1.246908702 | 0.000118717 | 0.000630812 | 1008.756438 | 1258.458484 | ENSG00000072364 |
| NIPAL3 | 0.318120505 | 1.246705325 | 0.033771054 | 0.086460173 | 178.2181289 | 222.0645604 | ENSG00000001461 |
| NA | 0.317875902 | 1.246493968 | 0.594810395 | 0.742733338 | 11.27480519 | 14.07822525 | ENSG00000280734 |
| BBIP1 | 0.31781509 | 1.246441428 | 0.083604986 | 0.179801982 | 130.279197 | 162.288221 | ENSG00000214413 |
| RBM12B | 0.317562339 | 1.246223078 | 0.000406962 | 0.001912036 | 680.4744829 | 848.3068023 | ENSG00000183808 |
| MCM9 | 0.317517692 | 1.246184512 | 0.053811543 | 0.126456143 | 151.102746 | 188.512779 | ENSG00000111877 |
| PCDHA10 | 0.317438406 | 1.246116028 | 0.023281804 | 0.063706281 | 222.8009928 | 277.8512296 | ENSG00000250120 |
| ZNF816 | 0.317408727 | 1.246090393 | 0.122092557 | 0.241642301 | 117.7147391 | 146.6596525 | ENSG00000180257 |
| SNHG32 | 0.317369183 | 1.246056238 | 0.00025755 | 0.001268743 | 614.7740853 | 765.7788307 | ENSG00000204387 |
| NA | 0.316992486 | 1.245730927 | 0.526891912 | 0.688817103 | 15.85298059 | 19.67762498 | ENSG00000234678 |
| NENF | 0.316831483 | 1.245591913 | 0.007621578 | 0.02506869 | 309.7345392 | 385.6245856 | ENSG00000117691 |
| LIMD2 | 0.316592592 | 1.245385677 | 0.000230937 | 0.001150273 | 1488.35311 | 1853.723771 | ENSG00000136490 |
| TCF7L2 | 0.316505916 | 1.245310858 | 0.002896331 | 0.010875175 | 441.0950223 | 549.4179757 | ENSG00000148737 |
| APOC1 | 0.316432576 | 1.245247553 | 0.145012585 | 0.275230006 | 100.9990657 | 125.625018 | ENSG00000130208 |
| SIRT1 | 0.316356131 | 1.245181572 | 1.02E-05 | 6.73E-05 | 1413.452208 | 1760.681846 | ENSG00000096717 |
| TMEM263 | 0.31620523 | 1.245051337 | 0.019969835 | 0.056173533 | 235.8119469 | 293.8398601 | ENSG00000151135 |
| TUBB4A | 0.316173079 | 1.245023591 | 0.000175177 | 0.000893532 | 682.4258947 | 849.5861912 | ENSG00000104833 |
| ETV7 | 0.316135047 | 1.244990771 | 0.560861934 | 0.715865261 | 14.27704029 | 17.84171476 | ENSG00000010030 |
| ZBTB42 | 0.316125825 | 1.244982812 | 0.141561196 | 0.27019462 | 90.06836921 | 111.9638173 | ENSG00000179627 |
| NA | 0.315785951 | 1.24468955 | 0.649709949 | 0.781978702 | 9.54491379 | 11.80646838 | ENSG00000228010 |
| PRR15 | 0.315734773 | 1.244645397 | 0.046482916 | 0.112636846 | 163.5975435 | 203.5813006 | ENSG00000176532 |
| MRPS31 | 0.315679306 | 1.244597546 | 0.001747271 | 0.007017824 | 483.0091328 | 600.8451124 | ENSG00000102738 |
| LINC02289 | 0.315486968 | 1.244431629 | 0.532535524 | 0.693288082 | 15.79293217 | 19.58083074 | ENSG00000258819 |
| SOX2 | 0.315476549 | 1.244422642 | 1.58E-09 | 1.88E-08 | 5605.491271 | 6975.832907 | ENSG00000181449 |
| CCDC151 | 0.315432276 | 1.244384453 | 0.251392528 | 0.415633636 | 55.14039144 | 68.60738127 | ENSG00000198003 |
| RAD50 | 0.315400136 | 1.244356732 | 0.141882144 | 0.270700073 | 115.8872783 | 143.8326766 | ENSG00000113522 |
| ZNF850 | 0.31539759 | 1.244354535 | 0.126820653 | 0.248299728 | 107.5665502 | 134.1003238 | ENSG00000267041 |
| NA | 0.315335522 | 1.244301002 | 0.778977208 | 0.870236815 | 3.67429477 | 4.564486445 | ENSG00000286407 |
| ORMDL2 | 0.315183384 | 1.244169792 | 0.038526314 | 0.096671583 | 186.7874745 | 232.3002274 | ENSG00000123353 |
| STAU2 | 0.315029664 | 1.244037233 | 0.000180523 | 0.000918133 | 742.6198307 | 924.2868359 | ENSG00000040341 |
| SLC7A3 | 0.314959092 | 1.24397638 | 5.66E-09 | 6.22E-08 | 2889.353966 | 3593.591893 | ENSG00000165349 |
| ABCA7 | 0.314706733 | 1.2437588 | 0.000850858 | 0.003681072 | 526.1629621 | 654.7359094 | ENSG00000064687 |
| NA | 0.31455246 | 1.243625807 | 0.075853176 | 0.166210882 | 149.1234493 | 185.2722435 | ENSG00000267809 |
| NA | 0.314500832 | 1.243581303 | 0.183418458 | 0.329361023 | 71.46730171 | 88.78645461 | ENSG00000227838 |
| COG4 | 0.314485745 | 1.243568299 | 8.35E-06 | 5.59E-05 | 1145.379593 | 1424.017749 | ENSG00000103051 |
| NUTM1 | 0.314472152 | 1.243556582 | 0.479295522 | 0.647439949 | 25.42983362 | 31.54029927 | ENSG00000184507 |
| HVCN1 | 0.314266501 | 1.24337933 | 0.125055529 | 0.245691639 | 94.72525348 | 117.7737375 | ENSG00000122986 |
| CFAP74 | 0.314142851 | 1.243272767 | 0.67161756 | 0.798777703 | 7.413544046 | 9.189740414 | ENSG00000142609 |
| CLCN3 | 0.314103116 | 1.243238525 | 0.000700524 | 0.003084639 | 738.4266125 | 917.9956281 | ENSG00000109572 |
| LINC00571 | 0.314019427 | 1.243166409 | 0.792247686 | 0.879072498 | 2.661681062 | 3.327642854 | ENSG00000223685 |
| ASCC3 | 0.31395331 | 1.243109437 | 1.09E-07 | 1.00E-06 | 1945.148636 | 2418.145756 | ENSG00000112249 |
| FAN1 | 0.313831895 | 1.243004823 | 0.001490621 | 0.006090993 | 685.4153252 | 851.388722 | ENSG00000198690 |
| PLAA | 0.31382112 | 1.24299554 | 1.08E-06 | 8.43E-06 | 1376.609749 | 1711.767386 | ENSG00000137055 |
| ZFC3H1 | 0.313652994 | 1.242850695 | 2.93E-05 | 0.000176839 | 859.5162561 | 1067.863303 | ENSG00000133858 |
| ENSA | 0.313621571 | 1.242823625 | 4.28E-14 | 8.41E-13 | 5222.365695 | 6490.058429 | ENSG00000143420 |
| HDAC2-AS2 | 0.313542311 | 1.242755348 | 3.40E-05 | 0.000202975 | 885.7407248 | 1100.326164 | ENSG00000228624 |
| SLCO1C1 | 0.313490032 | 1.242710314 | 0.770359178 | 0.864741191 | 3.638688754 | 4.513938752 | ENSG00000139155 |
| ICE2 | 0.313262919 | 1.242514699 | 2.30E-05 | 0.000141926 | 892.3843274 | 1108.971081 | ENSG00000128915 |
| SCUBE1 | 0.312803975 | 1.242119498 | 0.024965746 | 0.06755705 | 233.1934056 | 289.4600994 | ENSG00000159307 |
| TSPAN11 | 0.312784083 | 1.242102372 | 0.050297862 | 0.119814577 | 169.9729923 | 211.0555692 | ENSG00000110900 |
| 1-Mar | 0.312778443 | 1.242097516 | 0.00960403 | 0.030396591 | 333.7940593 | 414.9350492 | ENSG00000186205 |
| TIGD2 | 0.31260899 | 1.241951634 | 0.090120929 | 0.190857706 | 122.198975 | 151.8449838 | ENSG00000180346 |
| NA | 0.312550413 | 1.241901208 | 0.594021468 | 0.742132513 | 11.38162323 | 14.12877295 | ENSG00000236364 |
| NIPA1 | 0.312233756 | 1.241628653 | 0.002658071 | 0.010109789 | 448.0385788 | 556.6430135 | ENSG00000170113 |
| KLHL8 | 0.312232666 | 1.241627715 | 0.027020445 | 0.072235504 | 219.9478396 | 272.8925815 | ENSG00000145332 |
| NA | 0.312227595 | 1.241623351 | 0.728529881 | 0.837199615 | 4.985598981 | 6.186095954 | ENSG00000226790 |
| TRIM71 | 0.312156224 | 1.241561928 | 1.29E-08 | 1.35E-07 | 4342.203234 | 5392.077257 | ENSG00000206557 |
| PLEKHG4B | 0.312127337 | 1.241537069 | 8.05E-07 | 6.44E-06 | 1295.689595 | 1608.071408 | ENSG00000153404 |
| KIAA0319L | 0.312070162 | 1.241487867 | 2.43E-06 | 1.78E-05 | 1556.379762 | 1931.718352 | ENSG00000142687 |
| TMEM125 | 0.312042613 | 1.24146416 | 0.006744163 | 0.022691254 | 367.3350151 | 455.6030006 | ENSG00000179178 |
| NA | 0.311981021 | 1.24141116 | 0.650280927 | 0.782519454 | 8.662684654 | 10.73606142 | ENSG00000241081 |
| ZNF185 | 0.311949716 | 1.241384223 | 0.011863679 | 0.036310049 | 296.8730304 | 368.2700771 | ENSG00000147394 |

| | | | | | | | |
|---|---|---|---|---|---|---|---|
| PPP1R2 | 0.311895626 | 1.241337682 | 0.003252838 | 0.012021167 | 432.7552704 | 537.684912 | ENSG00000184203 |
| NA | 0.311704716 | 1.241173428 | 0.720597384 | 0.832149671 | 5.018414094 | 6.219972096 | ENSG00000205578 |
| VPS13B | 0.31137583 | 1.240890514 | 0.006036162 | 0.02064969 | 339.2848884 | 420.7272376 | ENSG00000132549 |
| CARNMT1 | 0.311321493 | 1.240843779 | 0.001117435 | 0.00470973 | 549.3943563 | 681.8864823 | ENSG00000156017 |
| LYSMD3 | 0.311251677 | 1.240783733 | 0.058218328 | 0.134727659 | 195.9022002 | 243.4256285 | ENSG00000176018 |
| NA | 0.311220829 | 1.240757202 | 0.722508873 | 0.833657436 | 4.996762593 | 6.192547676 | ENSG00000217716 |
| NA | 0.311105006 | 1.240657595 | 0.269390255 | 0.435717901 | 51.01537009 | 63.25910248 | ENSG00000246451 |
| ZFAND1 | 0.311060877 | 1.240619647 | 0.008269173 | 0.026858393 | 392.6417366 | 487.103202 | ENSG00000104231 |
| CYP1B1 | 0.311012721 | 1.240578236 | 0.446150405 | 0.616436575 | 26.47717746 | 32.98389525 | ENSG00000138061 |
| DOC2B | 0.310997713 | 1.240565332 | 0.065530279 | 0.147957 | 143.3416267 | 177.9622489 | ENSG00000272636 |
| KLHL42 | 0.31073455 | 1.24033906 | 0.006638383 | 0.022388295 | 357.3721529 | 443.6558783 | ENSG00000087448 |
| LOC105371730 | 0.310653025 | 1.240268972 | 0.770983085 | 0.86506338 | 4.615696446 | 5.790837631 | ENSG00000214708 |
| PDE6B | 0.310576169 | 1.240202901 | 0.06756053 | 0.151671149 | 149.2517187 | 184.8850666 | ENSG00000133256 |
| NA | 0.310158278 | 1.239843716 | 0.581180119 | 0.731773991 | 14.44044014 | 17.81402987 | ENSG00000237977 |
| UBQLN4 | 0.310103511 | 1.23979665 | 1.88E-07 | 1.66E-06 | 1812.247321 | 2247.409218 | ENSG00000160803 |
| P2RX3 | 0.309926026 | 1.239644135 | 0.695862174 | 0.816487878 | 6.379278836 | 7.900731594 | ENSG00000109991 |
| NA | 0.30977308 | 1.239512723 | 0.68011076 | 0.804514145 | 7.867737247 | 9.795791805 | ENSG00000235734 |
| PPCS | 0.309742533 | 1.239486479 | 0.004840038 | 0.017063078 | 357.547517 | 443.0030289 | ENSG00000127125 |
| SYT15 | 0.309665915 | 1.239420654 | 0.762721573 | 0.859242109 | 3.693155369 | 4.558034724 | ENSG00000204176 |
| GNA13 | 0.309598274 | 1.239362545 | 2.39E-05 | 0.000146826 | 1336.99778 | 1658.126868 | ENSG00000120063 |
| CBWD1 | 0.309553024 | 1.239323673 | 0.014093693 | 0.042047856 | 265.7406835 | 329.2200034 | ENSG00000172785 |
| ARHGAP27 | 0.309294368 | 1.239101499 | 0.014116159 | 0.042088843 | 316.8058896 | 392.5874586 | ENSG00000159314 |
| NA | 0.308942553 | 1.238799369 | 0.666582906 | 0.795139722 | 8.031137093 | 9.980777988 | ENSG00000284707 |
| NA | 0.308883921 | 1.238749024 | 0.767990486 | 0.863060962 | 3.744831081 | 4.615034138 | ENSG00000231424 |
| SERPINI1 | 0.308870664 | 1.238737641 | 0.189412488 | 0.337474915 | 84.49133445 | 104.8178056 | ENSG00000163536 |
| HTATIP2 | 0.308734406 | 1.238620652 | 3.31E-05 | 0.000197864 | 855.1644831 | 1059.472936 | ENSG00000109854 |
| SNX9 | 0.308676725 | 1.238571131 | 8.40E-05 | 0.000461798 | 818.3066569 | 1013.39194 | ENSG00000130340 |
| AKAP10 | 0.308574569 | 1.238483432 | 0.002930356 | 0.010982999 | 399.2954879 | 494.5858003 | ENSG00000108599 |
| MOCS3 | 0.308509123 | 1.238427251 | 0.013976676 | 0.04176333 | 321.4751526 | 398.1516678 | ENSG00000124217 |
| FAM126A | 0.308358871 | 1.23829828 | 4.00E-07 | 3.35E-06 | 1927.238638 | 2386.145693 | ENSG00000122591 |
| NIPAL2 | 0.308281779 | 1.238232112 | 0.335146317 | 0.507799842 | 40.33216394 | 50.00768105 | ENSG00000104361 |
| CBL | 0.308257352 | 1.238211147 | 1.13E-05 | 7.31E-05 | 1762.894869 | 2182.751499 | ENSG00000110395 |
| SULT4A1 | 0.307892263 | 1.237897844 | 0.039064986 | 0.097870474 | 216.1340238 | 267.0883195 | ENSG00000130540 |
| NA | 0.307879781 | 1.237887134 | 0.393798809 | 0.566676241 | 29.73220932 | 36.82079525 | ENSG00000224934 |
| KCNMB4 | 0.307833026 | 1.237847017 | 0.006214527 | 0.021154835 | 381.8086337 | 473.0184887 | ENSG00000135643 |
| IRF9 | 0.307798586 | 1.237817468 | 0.670388466 | 0.797758553 | 7.658443655 | 9.478245131 | ENSG00000213928 |
| SOWAHC | 0.307575106 | 1.237625739 | 0.001508822 | 0.006158841 | 472.8309197 | 585.3079156 | ENSG00000198142 |
| NA | 0.307511073 | 1.237570809 | 0.03210691 | 0.083014932 | 196.8078326 | 243.726231 | ENSG00000196810 |
| NA | 0.307463278 | 1.237529811 | 0.725249118 | 0.835021231 | 5.081253418 | 6.266218642 | ENSG00000285859 |
| AARD | 0.307441594 | 1.23751121 | 0.596414779 | 0.743899395 | 15.10604211 | 18.8008065 | ENSG00000205002 |
| USP34 | 0.307440823 | 1.237510549 | 0.000251414 | 0.001242321 | 1352.736793 | 1673.823951 | ENSG00000115464 |
| C1orf56 | 0.307405586 | 1.237480323 | 0.220691698 | 0.379072527 | 67.71880384 | 83.8861324 | ENSG00000143443 |
| SERF1A | 0.307398949 | 1.23747463 | 0.693513122 | 0.814573894 | 6.324812221 | 7.818458333 | ENSG00000172058 |
| UGCG | 0.30739069 | 1.237467547 | 0.001841075 | 0.007341086 | 527.5183203 | 653.2070777 | ENSG00000148154 |
| RHOH | 0.30737037 | 1.237450117 | 0.023076255 | 0.063206593 | 250.4470504 | 310.0266343 | ENSG00000168421 |
| ACVR1 | 0.307218674 | 1.237320009 | 0.003095036 | 0.01149968 | 399.8364872 | 494.5099909 | ENSG00000115170 |
| ZCCHC14 | 0.306935574 | 1.237077233 | 3.70E-05 | 0.000218538 | 855.7598611 | 1058.469772 | ENSG00000140948 |
| ZCCHC7 | 0.306891262 | 1.237039237 | 0.002945163 | 0.011023503 | 422.3169127 | 522.4972906 | ENSG00000147905 |
| CDKL1 | 0.306804777 | 1.236965083 | 0.488902492 | 0.655460281 | 22.51834686 | 27.7657659 | ENSG00000100490 |
| RIOX2 | 0.306790553 | 1.236952888 | 0.015995247 | 0.04669709 | 283.2629435 | 350.3250859 | ENSG00000170854 |
| UNC80 | 0.306785872 | 1.236948874 | 0.638335767 | 0.773863705 | 9.008144785 | 11.14180004 | ENSG00000144406 |
| NA | 0.306763957 | 1.236930085 | 0.083172358 | 0.179031238 | 131.7830494 | 162.9835488 | ENSG00000261409 |
| ACBD5 | 0.306711531 | 1.236885137 | 0.004013251 | 0.014503489 | 366.3678195 | 453.0125884 | ENSG00000107897 |
| SERPINA5 | 0.306175168 | 1.236425375 | 0.31155908 | 0.482599863 | 46.35916154 | 57.25343109 | ENSG00000188488 |
| SALL4 | 0.306164707 | 1.23641641 | 1.02E-09 | 1.23E-08 | 3196.128598 | 3951.303788 | ENSG00000101115 |
| NA | 0.306040961 | 1.236310362 | 0.765253946 | 0.861065676 | 3.693155369 | 4.541363172 | ENSG00000285813 |
| PRAG1 | 0.305979858 | 1.236258001 | 2.95E-05 | 0.00017809 | 1114.713181 | 1378.330958 | ENSG00000275342 |
| FUOM | 0.305855128 | 1.236151123 | 0.018022671 | 0.051499829 | 270.572895 | 334.3520914 | ENSG00000148803 |
| C11orf65 | 0.305826606 | 1.236126685 | 0.775687823 | 0.868533369 | 4.283515108 | 5.33455132 | ENSG00000166323 |
| ACAP2 | 0.305744568 | 1.236056396 | 0.004956384 | 0.01740714 | 586.9556178 | 725.8112949 | ENSG00000114331 |
| ALMS1 | 0.305490622 | 1.235838842 | 4.33E-05 | 0.000251704 | 1043.08498 | 1289.265006 | ENSG00000116127 |
| RRP15 | 0.305157376 | 1.23555341 | 0.000553324 | 0.002509209 | 913.4923385 | 1129.469894 | ENSG00000067533 |
| MYRF | 0.305134945 | 1.2355342 | 0.023618665 | 0.064516589 | 268.1157136 | 330.8819529 | ENSG00000124920 |
| NA | 0.305113736 | 1.235516036 | 0.7568754 | 0.855780331 | 3.984148884 | 4.936881959 | ENSG00000269978 |
| ZNF264 | 0.305040226 | 1.235453084 | 0.087864564 | 0.187101169 | 194.4325272 | 240.4133575 | ENSG00000083844 |
| SH3RF1 | 0.30498234 | 1.235403514 | 0.020670647 | 0.057760768 | 261.1581656 | 322.7668901 | ENSG00000154447 |
| ABI1 | 0.304954591 | 1.235379753 | 0.000362328 | 0.001724789 | 752.2519652 | 929.8117953 | ENSG00000136754 |
| NA | 0.304843299 | 1.235284457 | 0.734855021 | 0.842023481 | 5.029577706 | 6.192547676 | ENSG00000214263 |
| ZSCAN20 | 0.304799419 | 1.235246886 | 0.247145518 | 0.410688619 | 64.1534423 | 79.28455317 | ENSG00000121903 |
| ALDOA | 0.304782368 | 1.235232287 | 0.308984643 | 0.48007593 | 48.92431229 | 60.54074608 | ENSG00000149925 |
| ITM2C | 0.304739105 | 1.235195246 | 4.91E-16 | 1.19E-14 | 13272.53762 | 16393.95941 | ENSG00000135916 |
| OSBPL6 | 0.304642331 | 1.235112394 | 0.045999623 | 0.111634007 | 191.8171643 | 236.7364545 | ENSG00000079156 |
| RFX3 | 0.30443479 | 1.234934727 | 0.028158029 | 0.074635414 | 238.5001856 | 294.7295636 | ENSG00000080298 |
| KLF4 | 0.304426338 | 1.234927493 | 0.263824184 | 0.42959672 | 55.43485158 | 68.41002468 | ENSG00000136826 |
| HSPA4L | 0.304387514 | 1.23489426 | 0.06455397 | 0.146128917 | 155.1403224 | 191.5212576 | ENSG00000164070 |
| NA | 0.3042939 | 1.234814133 | 0.7383291 | 0.844486343 | 5.559889024 | 6.914215431 | ENSG00000248923 |
| ORAI1 | 0.304206251 | 1.234739116 | 0.002722813 | 0.010321435 | 546.9725808 | 675.2715671 | ENSG00000276045 |

| | | | | | | | |
|---|---|---|---|---|---|---|---|
| IL4I1 | 0.304110119 | 1.234656843 | 0.676675028 | 0.801875912 | 7.718492076 | 9.513738808 | ENSG00000104951 |
| MBL1P | 0.304031144 | 1.234589258 | 0.800952667 | 0.885251398 | 2.68891437 | 3.32549228 | ENSG00000242600 |
| NA | 0.304031144 | 1.234589258 | 0.800952667 | 0.885251398 | 2.68891437 | 3.32549228 | ENSG00000197813 |
| NA | 0.303693969 | 1.234300754 | 0.344971522 | 0.517524352 | 49.34800571 | 60.90642941 | ENSG00000249896 |
| RALGDS | 0.303513443 | 1.234146314 | 0.33474498 | 0.50740055 | 48.13370738 | 59.38698137 | ENSG00000160271 |
| NA | 0.303288219 | 1.233953662 | 0.671815838 | 0.798838096 | 7.770167788 | 9.564286502 | ENSG00000228305 |
| PPA2 | 0.303183626 | 1.233864206 | 0.001324649 | 0.005483648 | 538.2582573 | 664.1154702 | ENSG00000138777 |
| MGA | 0.303182405 | 1.233863161 | 0.000794653 | 0.003454179 | 1001.743407 | 1235.495713 | ENSG00000174197 |
| MFAP3 | 0.303006383 | 1.233712628 | 0.03155231 | 0.081833237 | 242.4839711 | 299.3663702 | ENSG00000037749 |
| PLLP | 0.302526999 | 1.233302754 | 0.206916738 | 0.361331654 | 88.98290385 | 109.7689665 | ENSG00000102934 |
| SLC7A2 | 0.302200496 | 1.233023671 | 0.001354742 | 0.005595018 | 523.4460876 | 645.070025 | ENSG00000003989 |
| ATP2B2 | 0.302044675 | 1.232890503 | 0.421488997 | 0.59341295 | 30.01271495 | 36.99583418 | ENSG00000157087 |
| MBNL1 | 0.301946315 | 1.23280645 | 0.109772867 | 0.222229864 | 133.705489 | 164.9630329 | ENSG00000152601 |
| ZNF702P | 0.30190789 | 1.232773615 | 0.167192258 | 0.307259774 | 86.71923451 | 106.8784968 | ENSG00000242779 |
| EHD2 | 0.301893779 | 1.232761558 | 0.00517578 | 0.018074941 | 385.3998771 | 475.0765142 | ENSG00000024422 |
| PDLIM2 | 0.301888946 | 1.232757428 | 0.143524037 | 0.273049527 | 171.9171714 | 211.7847974 | ENSG00000120913 |
| SOS1 | 0.301813226 | 1.232692728 | 0.001889015 | 0.007512057 | 478.8757899 | 590.5290011 | ENSG00000115904 |
| CDC37L1 | 0.301711608 | 1.232605906 | 0.004521912 | 0.016082633 | 414.857438 | 511.5625766 | ENSG00000106993 |
| NA | 0.301689903 | 1.232587361 | 0.681957993 | 0.805942337 | 8.689917962 | 10.66884218 | ENSG00000280211 |
| HACE1 | 0.301667333 | 1.232568078 | 0.01395841 | 0.041721676 | 369.3911426 | 455.3724945 | ENSG00000085382 |
| LOC105374727 | 0.301637097 | 1.232542246 | 0.774537617 | 0.867840226 | 3.744831081 | 4.58330857 | ENSG00000286193 |
| SGSM3 | 0.301608544 | 1.232517853 | 0.70526902 | 0.822671706 | 6.901217446 | 8.554647066 | ENSG00000100359 |
| NA | 0.3007698 | 1.231801508 | 0.811412794 | 0.891703555 | 2.680541661 | 3.304519581 | ENSG00000266538 |
| ZNF415 | 0.300663667 | 1.231710893 | 0.071009771 | 0.157816762 | 167.4401442 | 206.3367377 | ENSG00000170954 |
| FSCN1 | 0.30055899 | 1.231621527 | 7.92E-12 | 1.23E-10 | 13202.39521 | 16261.39272 | ENSG00000075618 |
| NA | 0.300488451 | 1.23156131 | 0.589951312 | 0.739181092 | 13.66714423 | 16.88208998 | ENSG00000234129 |
| CCND1 | 0.300450923 | 1.231529275 | 2.82E-05 | 0.000170856 | 2041.795645 | 2515.4329 | ENSG00000110092 |
| CYP2R1 | 0.300416559 | 1.231499942 | 0.069809296 | 0.155686836 | 186.4419263 | 229.4883119 | ENSG00000186104 |
| ZNF714 | 0.29999137 | 1.231137049 | 0.003532758 | 0.012963702 | 461.8985836 | 568.6363703 | ENSG00000160352 |
| DDB2 | 0.299783884 | 1.230960002 | 0.002438704 | 0.0093921 | 604.5765971 | 743.7210593 | ENSG00000134574 |
| PRB3 | 0.29971271 | 1.230899274 | 0.487242362 | 0.654462637 | 22.26698956 | 27.40789137 | ENSG00000197870 |
| KCTD20 | 0.299659726 | 1.23085407 | 9.51E-05 | 0.000516932 | 909.6291097 | 1119.008077 | ENSG00000112078 |
| ANKRD18A | 0.299437269 | 1.230664293 | 0.25023533 | 0.414430782 | 59.13291303 | 72.72686731 | ENSG00000180071 |
| DTWD2 | 0.29935161 | 1.230591225 | 0.09223986 | 0.194292233 | 145.4519085 | 179.2816262 | ENSG00000169570 |
| TJP1 | 0.299309051 | 1.230554924 | 1.38E-08 | 1.45E-07 | 2629.068269 | 3234.64434 | ENSG00000104067 |
| PABPC1L2A | 0.299277704 | 1.230528186 | 0.749456272 | 0.851859589 | 5.059601916 | 6.196848823 | ENSG00000186288 |
| NA | 0.299192117 | 1.230455188 | 0.355604595 | 0.528706609 | 45.57490218 | 55.94076596 | ENSG00000279369 |
| RIT1 | 0.299169493 | 1.230435892 | 0.005296285 | 0.018445682 | 454.2336439 | 558.247982 | ENSG00000143622 |
| BICDL2 | 0.299102179 | 1.230378484 | 0.017995978 | 0.051454042 | 291.0087569 | 357.8935078 | ENSG00000162069 |
| REEP3 | 0.299099023 | 1.230375792 | 0.02176938 | 0.060247816 | 370.3479524 | 456.3445203 | ENSG00000165476 |
| MPG | 0.299089926 | 1.230368034 | 0.010643031 | 0.033200955 | 383.5712522 | 471.8263409 | ENSG00000103152 |
| TDRP | 0.299065471 | 1.230347178 | 2.59E-05 | 0.000158223 | 1124.385562 | 1383.463196 | ENSG00000180190 |
| MAGI2 | 0.298885737 | 1.230193908 | 0.002408472 | 0.009289858 | 467.1959517 | 574.8525927 | ENSG00000187391 |
| SMAD5 | 0.298821788 | 1.23013938 | 0.000108521 | 0.000582552 | 951.1606167 | 1170.175665 | ENSG00000113658 |
| NA | 0.298771196 | 1.230096242 | 0.080581259 | 0.174387824 | 195.6052984 | 240.543749 | ENSG00000248605 |
| GSTA4 | 0.29870957 | 1.230043699 | 0.041081974 | 0.101826077 | 240.5652494 | 296.2600129 | ENSG00000170899 |
| TTC41P | 0.298611304 | 1.22995992 | 0.798820503 | 0.883907026 | 2.686123467 | 3.304519581 | ENSG00000214198 |
| CEP68 | 0.298552062 | 1.229909415 | 0.000729549 | 0.003199304 | 585.6215108 | 720.7115087 | ENSG00000011523 |
| HSD17B11 | 0.298520027 | 1.229882104 | 0.096184183 | 0.20096275 | 131.3207589 | 161.6735673 | ENSG00000198189 |
| COL4A5 | 0.298408759 | 1.229787254 | 7.80E-05 | 0.000432092 | 901.5674601 | 1108.098085 | ENSG00000188153 |
| CACNA1F | 0.298339589 | 1.229728293 | 0.561851969 | 0.716559985 | 17.98155943 | 22.01928471 | ENSG00000102001 |
| WDR17 | 0.298328244 | 1.229718623 | 0.317274584 | 0.488432207 | 46.93277586 | 57.72530446 | ENSG00000150627 |
| PKHD1L1 | 0.298199874 | 1.229609208 | 0.769342126 | 0.864048241 | 4.289096914 | 5.309277473 | ENSG00000205038 |
| BPNT1 | 0.298069951 | 1.22949848 | 0.017861971 | 0.051169482 | 306.1274021 | 376.3294366 | ENSG00000162813 |
| FBXO41 | 0.298053918 | 1.229484816 | 0.007819283 | 0.025640284 | 511.9474707 | 628.68567 | ENSG00000163013 |
| RFTN2 | 0.297680445 | 1.229166579 | 0.549479603 | 0.706799555 | 22.79941606 | 28.15025925 | ENSG00000162944 |
| CGGBP1 | 0.29763918 | 1.229131421 | 8.84E-05 | 0.000483696 | 1139.706643 | 1400.941687 | ENSG00000163320 |
| EPHA6 | 0.297293617 | 1.228837047 | 0.461149917 | 0.630969788 | 23.81079047 | 29.22093878 | ENSG00000080224 |
| KIAA0355 | 0.297158441 | 1.228721915 | 0.000452158 | 0.002097276 | 651.2881662 | 800.2902185 | ENSG00000166398 |
| NA | 0.297105726 | 1.228677019 | 0.689646738 | 0.811485516 | 7.704537561 | 9.434149159 | ENSG00000287825 |
| ZNF766 | 0.296804131 | 1.228420192 | 0.002542815 | 0.009728966 | 517.1810115 | 634.9736244 | ENSG00000196214 |
| TCHP | 0.296790582 | 1.228408655 | 0.015747915 | 0.046120089 | 278.7057439 | 342.2423427 | ENSG00000139437 |
| ARCN1 | 0.29675658 | 1.228379704 | 4.35E-09 | 4.88E-08 | 2933.471451 | 3604.057586 | ENSG00000095139 |
| RBBP6 | 0.296735675 | 1.228361904 | 5.34E-06 | 3.69E-05 | 1382.89281 | 1698.34533 | ENSG00000122257 |
| BTNL9 | 0.296733496 | 1.228360049 | 0.460657849 | 0.630430658 | 25.98546322 | 31.80217307 | ENSG00000165810 |
| SNRNP48 | 0.296710659 | 1.228340605 | 0.017025915 | 0.049153984 | 516.6893115 | 635.4126271 | ENSG00000168566 |
| HSD17B8 | 0.296446739 | 1.228115919 | 0.225846329 | 0.384910463 | 75.42633247 | 92.51849112 | ENSG00000204228 |
| NA | 0.296370206 | 1.22805077 | 0.782081562 | 0.872240007 | 3.935264075 | 4.837937146 | ENSG00000273148 |
| TMEM200B | 0.296197083 | 1.227903413 | 0.050128709 | 0.119541141 | 200.3107552 | 246.0009504 | ENSG00000253304 |
| AFF2 | 0.295905182 | 1.227654996 | 0.027663427 | 0.07363808 | 247.4652275 | 304.0072234 | ENSG00000155966 |
| ZDHHC11 | 0.295709649 | 1.227488619 | 0.122680922 | 0.242433689 | 115.0640215 | 141.1914683 | ENSG00000188818 |
| MMP16 | 0.295571996 | 1.227371506 | 0.004961032 | 0.017419598 | 443.3453376 | 544.1396366 | ENSG00000156103 |
| FIBIN | 0.29549183 | 1.227303306 | 0.740201305 | 0.84572929 | 5.32894393 | 6.539669343 | ENSG00000176971 |
| ALG9 | 0.295286218 | 1.227128405 | 0.411810752 | 0.583920707 | 33.81480355 | 41.50466617 | ENSG00000086848 |
| COQ6 | 0.295234544 | 1.227084452 | 0.659399491 | 0.789485573 | 12.94707564 | 15.82673703 | ENSG00000119723 |
| PICALM | 0.295209242 | 1.227062932 | 0.000111472 | 0.000596174 | 1168.764464 | 1434.947574 | ENSG00000073921 |
| NA | 0.295197284 | 1.227052762 | 0.637491138 | 0.773400512 | 11.2454567 | 13.86177397 | ENSG00000237015 |

| | | | | | | | |
|---|---|---|---|---|---|---|---|
| EXOC1 | 0.29507793 | 1.226951252 | 0.019995673 | 0.056221619 | 255.727622 | 313.5812088 | ENSG00000090989 |
| KHDC4 | 0.295073713 | 1.226947666 | 3.13E-06 | 2.25E-05 | 1273.098396 | 1561.775572 | ENSG00000132680 |
| ATP5PD | 0.295070905 | 1.226945277 | 4.06E-06 | 2.87E-05 | 1395.621556 | 1711.945745 | ENSG00000167863 |
| PLAG1 | 0.294995466 | 1.226881122 | 0.057323991 | 0.133169209 | 198.2409854 | 243.0884357 | ENSG00000181690 |
| GPX1 | 0.29484011 | 1.226749012 | 2.70E-06 | 1.97E-05 | 4282.410203 | 5253.080211 | ENSG00000233276 |
| NA | 0.294750519 | 1.226672834 | 0.760542028 | 0.857655235 | 4.046988208 | 4.957854658 | ENSG00000286001 |
| DYRK2 | 0.294680215 | 1.226613058 | 0.001997391 | 0.007879689 | 748.899689 | 918.5785683 | ENSG00000127334 |
| ZNF608 | 0.294472081 | 1.226436111 | 0.000103329 | 0.000557262 | 901.2566549 | 1104.754813 | ENSG00000168916 |
| SLC25A44 | 0.294469625 | 1.226434023 | 0.000315003 | 0.001520608 | 760.0793394 | 931.7741295 | ENSG00000160785 |
| COQ10B | 0.294244405 | 1.226242579 | 0.030800394 | 0.080228249 | 271.5374998 | 332.882367 | ENSG00000115520 |
| GLIPR1 | 0.294163065 | 1.226173444 | 0.771826554 | 0.865657328 | 4.286306011 | 5.286154201 | ENSG00000139278 |
| IFI16 | 0.294111756 | 1.226129836 | 0.651260018 | 0.783257918 | 10.12410992 | 12.33454766 | ENSG00000163565 |
| PDHB | 0.293931225 | 1.225976415 | 2.82E-06 | 2.04E-05 | 2227.649837 | 2730.533812 | ENSG00000168291 |
| HERC4 | 0.293909813 | 1.225958219 | 0.000809027 | 0.003511914 | 655.5675759 | 803.721362 | ENSG00000148634 |
| BIRC2 | 0.293815266 | 1.225877879 | 0.000824605 | 0.003574709 | 591.4564486 | 724.9920642 | ENSG00000110330 |
| THAP8 | 0.293409904 | 1.225533485 | 0.103210633 | 0.212241373 | 131.5216798 | 161.2960771 | ENSG00000161277 |
| EIF2D | 0.293350371 | 1.225482915 | 0.000303913 | 0.001472693 | 822.5512484 | 1007.47529 | ENSG00000143486 |
| BICRAL | 0.293301664 | 1.225441542 | 0.029683459 | 0.077855012 | 265.7546011 | 325.8461262 | ENSG00000112624 |
| NIBAN1 | 0.293218467 | 1.225370875 | 0.004346204 | 0.015535041 | 571.0290715 | 699.1770861 | ENSG00000135842 |
| ZNF333 | 0.293147102 | 1.225310263 | 0.129257594 | 0.251691557 | 126.4191752 | 155.0354741 | ENSG00000160961 |
| TRAPPC2B | 0.293106517 | 1.225275793 | 0.444614453 | 0.615289998 | 27.1256706 | 33.23045457 | ENSG00000256060 |
| FUT9 | 0.292977236 | 1.225166 | 0.273776597 | 0.440559254 | 76.05421325 | 93.11836336 | ENSG00000172461 |
| LINC00659 | 0.292949595 | 1.225142527 | 0.722858353 | 0.833877469 | 5.671613158 | 6.977133529 | ENSG00000228705 |
| CCPG1 | 0.292931848 | 1.225127456 | 0.46405308 | 0.633683583 | 24.48005923 | 29.98455194 | ENSG00000260916 |
| CTSB | 0.292432772 | 1.224703717 | 4.40E-05 | 0.00025547 | 1082.703058 | 1325.719094 | ENSG00000164733 |
| EMILIN2 | 0.292385019 | 1.224663181 | 0.039492541 | 0.098698096 | 209.9129265 | 256.8327337 | ENSG00000132205 |
| NA | 0.292267594 | 1.224563506 | 0.640640903 | 0.775170981 | 10.99497528 | 13.48130919 | ENSG00000254610 |
| PTPRE | 0.292046638 | 1.224375973 | 0.49653694 | 0.662747811 | 26.22881123 | 32.1697344 | ENSG00000132334 |
| VPS35L | 0.291997518 | 1.224334286 | 0.000175077 | 0.000893257 | 989.5839194 | 1211.973131 | ENSG00000103544 |
| CNTRL | 0.29193676 | 1.224282725 | 0.021923117 | 0.060598775 | 254.7429173 | 311.7178741 | ENSG00000119397 |
| ELAC1 | 0.291803378 | 1.224169542 | 0.475384458 | 0.643857075 | 23.39411831 | 28.66596812 | ENSG00000141642 |
| GCNT1 | 0.291780684 | 1.224150285 | 0.051417873 | 0.122035211 | 189.9261515 | 232.4879093 | ENSG00000187210 |
| NA | 0.291667028 | 1.22405385 | 0.722903352 | 0.833877469 | 7.144202032 | 8.752536692 | ENSG00000224418 |
| C6orf120 | 0.291590171 | 1.223988643 | 0.001607583 | 0.006508267 | 522.2768952 | 639.2907032 | ENSG00000185127 |
| RGPD5 | 0.29146831 | 1.22388526 | 0.646154551 | 0.779596437 | 9.68299535 | 11.86776894 | ENSG00000015568 |
| URB2 | 0.291356301 | 1.223790243 | 5.64E-06 | 3.88E-05 | 1189.313139 | 1455.518471 | ENSG00000135763 |
| RMND5B | 0.291310412 | 1.223751316 | 2.54E-06 | 1.86E-05 | 1646.817158 | 2014.717458 | ENSG00000145916 |
| ZNF529-AS1 | 0.291265303 | 1.223713054 | 0.403806833 | 0.576115175 | 34.92797778 | 42.66460373 | ENSG00000233527 |
| IFITM10 | 0.291252679 | 1.223702346 | 0.81168893 | 0.891854596 | 3.478279971 | 4.194241504 | ENSG00000244242 |
| TGFBR3L | 0.291160871 | 1.223624477 | 0.433456608 | 0.604482861 | 33.18026492 | 40.67489985 | ENSG00000260001 |
| TMTC3 | 0.291026544 | 1.223510552 | 0.010315406 | 0.032272867 | 483.482401 | 592.3008845 | ENSG00000139324 |
| WDR44 | 0.290838066 | 1.223350719 | 0.000161166 | 0.000828638 | 850.659872 | 1040.369085 | ENSG00000131725 |
| DRP2 | 0.290818689 | 1.223334289 | 0.493547167 | 0.659990185 | 23.56309996 | 28.70335191 | ENSG00000102385 |
| CLCN5 | 0.290804323 | 1.223322107 | 0.012716666 | 0.038559761 | 339.8653238 | 415.7077914 | ENSG00000171365 |
| GFPT2 | 0.29078742 | 1.223307774 | 1.48E-08 | 1.54E-07 | 2856.641976 | 3494.2307 | ENSG00000131459 |
| NA | 0.290745742 | 1.223272435 | 0.804065731 | 0.887065768 | 2.68891437 | 3.281396308 | ENSG00000224415 |
| CYC1 | 0.290660603 | 1.223200247 | 1.01E-10 | 1.39E-09 | 4800.706089 | 5872.588291 | ENSG00000179091 |
| MIA2 | 0.290635139 | 1.223178657 | 0.018034839 | 0.051519335 | 275.5930239 | 337.2175293 | ENSG00000150527 |
| BAZ2B | 0.290597012 | 1.223146332 | 0.565097755 | 0.719287468 | 17.5723841 | 21.49954726 | ENSG00000226266 |
| SOCS4 | 0.290501207 | 1.223065109 | 0.008602759 | 0.027761094 | 421.3062141 | 515.5193331 | ENSG00000180008 |
| CEP295 | 0.29036301 | 1.222947956 | 0.052559532 | 0.124027636 | 182.88251 | 223.6694984 | ENSG00000166004 |
| HERC5 | 0.290339424 | 1.222927963 | 0.010068661 | 0.031623989 | 347.8833454 | 425.8321616 | ENSG00000138646 |
| NFE2L2 | 0.290236507 | 1.222840727 | 0.006975733 | 0.023314079 | 472.8469525 | 578.897287 | ENSG00000116044 |
| HPS4 | 0.28999009 | 1.222631879 | 0.002082724 | 0.008177861 | 770.4495612 | 941.6827809 | ENSG00000100099 |
| TPM1 | 0.289978479 | 1.22262204 | 6.15E-12 | 9.62E-11 | 4164.54868 | 5091.486947 | ENSG00000140416 |
| DNHD1 | 0.289853068 | 1.222515763 | 0.145106713 | 0.275327393 | 103.9596374 | 127.0145587 | ENSG00000179532 |
| DMXL2 | 0.289674559 | 1.222364507 | 0.107894588 | 0.21924634 | 181.7327786 | 222.2390237 | ENSG00000104093 |
| BBS2 | 0.289657929 | 1.222350417 | 0.015375242 | 0.045186441 | 348.4987721 | 425.7283947 | ENSG00000125124 |
| NA | 0.289568768 | 1.222274876 | 0.064012525 | 0.145160428 | 174.7403245 | 213.7013513 | ENSG00000254635 |
| TRIOBP | 0.289455755 | 1.222179134 | 2.56E-05 | 0.000156652 | 1362.469901 | 1665.788135 | ENSG00000100106 |
| SATL1 | 0.28926679 | 1.222019063 | 0.629230455 | 0.76709332 | 10.72284237 | 13.07073639 | ENSG00000184788 |
| NETO1 | 0.289141476 | 1.221912921 | 0.032600743 | 0.084055191 | 270.9157644 | 330.7340658 | ENSG00000166342 |
| PAM | 0.289093999 | 1.221872711 | 0.000149578 | 0.000774009 | 999.9161347 | 1222.305065 | ENSG00000145730 |
| PIGK | 0.289079033 | 1.221860035 | 0.025013418 | 0.067619653 | 303.2362907 | 370.9617905 | ENSG00000142892 |
| CARD8 | 0.288927281 | 1.221731519 | 0.04262785 | 0.105010069 | 214.816262 | 262.2664903 | ENSG00000105483 |
| SERPINF1 | 0.288863496 | 1.221677504 | 0.005005154 | 0.017557731 | 412.6192872 | 503.8309653 | ENSG00000132386 |
| BBS10 | 0.288813965 | 1.221635562 | 0.031909564 | 0.082615407 | 229.3158746 | 280.0463714 | ENSG00000179941 |
| MAP4K2 | 0.288762452 | 1.221591943 | 0.002622317 | 0.009995478 | 545.5143467 | 666.5034254 | ENSG00000168067 |
| TMEM245 | 0.288634529 | 1.22148363 | 4.37E-06 | 3.07E-05 | 1446.206962 | 1766.94669 | ENSG00000106771 |
| BCS1L | 0.28834312 | 1.221236928 | 0.001765279 | 0.00707836 | 541.1598197 | 660.6421456 | ENSG00000074582 |
| GOLGA1 | 0.288150233 | 1.221073661 | 0.131321499 | 0.254707725 | 114.6168496 | 139.921542 | ENSG00000136935 |
| GPBP1L1 | 0.28808372 | 1.221017366 | 1.06E-06 | 8.33E-06 | 1824.312978 | 2226.760379 | ENSG00000159592 |
| ERBIN | 0.28801783 | 1.220961602 | 0.004237907 | 0.01519581 | 463.9324719 | 566.7049866 | ENSG00000112851 |
| LOC100133091 | 0.287958327 | 1.220911245 | 0.315016399 | 0.486314129 | 50.18952259 | 61.37668524 | ENSG00000205485 |
| LYRM2 | 0.287952947 | 1.220906692 | 0.029677846 | 0.077850875 | 255.8104732 | 312.3575291 | ENSG00000083099 |
| LOC100129917 | 0.287920944 | 1.220879609 | 0.227142192 | 0.386402119 | 73.2064417 | 89.19461638 | ENSG00000249592 |
| JMJD7-PLA2G4B | 0.287759251 | 1.220742785 | 0.251019508 | 0.415194833 | 70.23000043 | 85.53867398 | ENSG00000168970 |

| | | | | | | | |
|---|---|---|---|---|---|---|---|
| MYCBP2 | 0.287748273 | 1.220733496 | 0.004650555 | 0.016494549 | 704.6214465 | 859.5836102 | ENSG00000005810 |
| SYNPR | 0.287693843 | 1.220687441 | 0.639278715 | 0.774692235 | 12.55205498 | 15.30457644 | ENSG00000163630 |
| NA | 0.287666228 | 1.220664075 | 0.617456831 | 0.759372316 | 12.73690617 | 15.56995788 | ENSG00000253816 |
| ZNF780A | 0.287483661 | 1.220509615 | 0.189864432 | 0.337972785 | 101.7564921 | 124.0894194 | ENSG00000197782 |
| PCDHGB2 | 0.287140574 | 1.220219401 | 0.496156843 | 0.66237797 | 28.26954476 | 34.41244932 | ENSG00000253910 |
| AGL | 0.287060425 | 1.220151614 | 0.003118608 | 0.011580571 | 535.875455 | 653.5235278 | ENSG00000162688 |
| ADAP1 | 0.28703845 | 1.220133028 | 0.343907049 | 0.51665064 | 43.60932287 | 53.31166759 | ENSG00000105963 |
| ZBTB10 | 0.287006188 | 1.220105744 | 0.051731157 | 0.122658033 | 255.4155037 | 312.1001985 | ENSG00000205189 |
| PRKCQ | 0.286974176 | 1.220078671 | 0.001988297 | 0.007848627 | 512.1080428 | 624.9212896 | ENSG00000065675 |
| DLEU1 | 0.286829823 | 1.219956599 | 0.307809819 | 0.4789063 | 61.252919 | 74.68081705 | ENSG00000176124 |
| THTPA | 0.286807835 | 1.219938005 | 0.080869945 | 0.17487525 | 164.8485991 | 201.4391175 | ENSG00000259431 |
| MIR5581 | 0.28679257 | 1.219925098 | 0.821668785 | 0.898470272 | 2.740590082 | 3.329793428 | ENSG00000263675 |
| CITED2 | 0.286706241 | 1.219852101 | 0.000760551 | 0.003320159 | 862.3669946 | 1051.571999 | ENSG00000164442 |
| FGFR1OP2 | 0.286690806 | 1.21983905 | 0.004141408 | 0.014888538 | 444.3691902 | 542.3948745 | ENSG00000111790 |
| DIPK2A | 0.286613295 | 1.219773514 | 0.015077755 | 0.044462907 | 419.2693106 | 512.0892441 | ENSG00000181744 |
| CLDN1 | 0.286579939 | 1.219745313 | 0.253207648 | 0.417775294 | 66.51296624 | 81.12583062 | ENSG00000163347 |
| NA | 0.285852157 | 1.219130155 | 0.725562861 | 0.835289284 | 5.671613158 | 6.928736409 | ENSG00000255050 |
| BLOC1S3 | 0.285758164 | 1.21905073 | 0.124204903 | 0.244643331 | 122.6685621 | 149.75121 | ENSG00000189114 |
| RAD51C | 0.28564201 | 1.218952586 | 0.056285041 | 0.131198009 | 191.6644156 | 233.8175065 | ENSG00000108384 |
| ZFP2 | 0.285582504 | 1.21890231 | 0.649669304 | 0.781978702 | 11.3048294 | 13.80315702 | ENSG00000198939 |
| ADA | 0.28558095 | 1.218900996 | 0.020905047 | 0.058260511 | 328.8705487 | 400.9113818 | ENSG00000196839 |
| RMI2 | 0.285569298 | 1.218891152 | 1.68E-06 | 1.27E-05 | 1506.499419 | 1836.606231 | ENSG00000175643 |
| LITAF | 0.285545967 | 1.218871441 | 0.000166644 | 0.000854982 | 1993.144204 | 2430.685499 | ENSG00000189067 |
| NA | 0.2855427 | 1.21886868 | 0.706804849 | 0.823594399 | 8.045091607 | 9.778587215 | ENSG00000285722 |
| CRYGN | 0.285368041 | 1.218721127 | 0.606224037 | 0.75080146 | 14.73980636 | 18.02670094 | ENSG00000127377 |
| ZNF551 | 0.285327233 | 1.218686655 | 0.002034581 | 0.008013293 | 542.9369563 | 661.8582464 | ENSG00000204519 |
| COX10-AS1 | 0.285249816 | 1.218621261 | 0.201160233 | 0.353294068 | 82.82380682 | 100.8136754 | ENSG00000236088 |
| ANKAR | 0.285052298 | 1.218454432 | 0.647349702 | 0.780452665 | 12.99999066 | 15.87675169 | ENSG00000151687 |
| NA | 0.285004508 | 1.218414071 | 0.75642819 | 0.855549731 | 5.35896814 | 6.510094349 | ENSG00000237938 |
| YTHDC1 | 0.284982592 | 1.218395562 | 3.95E-07 | 3.31E-06 | 2046.39871 | 2493.946523 | ENSG00000083896 |
| RMDN1 | 0.284760463 | 1.218207982 | 0.012788987 | 0.038739036 | 409.5116223 | 498.5550067 | ENSG00000176623 |
| MAPK12 | 0.284700103 | 1.218157016 | 0.004936607 | 0.017355948 | 574.5802272 | 699.4297819 | ENSG00000188130 |
| NA | 0.284598566 | 1.218071285 | 0.669865057 | 0.797264277 | 9.971197962 | 12.17079464 | ENSG00000285761 |
| LOC729870 | 0.284569662 | 1.218046881 | 0.576611316 | 0.728639388 | 17.71480816 | 21.5431102 | ENSG00000287642 |
| TRIP11 | 0.284545079 | 1.218026127 | 0.007877134 | 0.025787258 | 422.25958 | 513.8883583 | ENSG00000100815 |
| SMG1P3 | 0.284424291 | 1.217924152 | 0.348667473 | 0.521366113 | 61.55846751 | 74.8050236 | ENSG00000180747 |
| STK3 | 0.284324222 | 1.217839677 | 0.040660674 | 0.100989451 | 216.1258144 | 263.314307 | ENSG00000104375 |
| HK1 | 0.284101254 | 1.217651476 | 2.79E-12 | 4.53E-11 | 9549.961843 | 11627.63019 | ENSG00000156515 |
| CLVS2 | 0.284088815 | 1.217640977 | 0.583491314 | 0.733585176 | 15.88858661 | 19.23585965 | ENSG00000146352 |
| ADAT2 | 0.28377916 | 1.217379654 | 0.045604837 | 0.110847594 | 227.9202798 | 277.1169431 | ENSG00000189007 |
| ZC3H12A | 0.283735737 | 1.217343013 | 0.045653332 | 0.110919197 | 275.3602391 | 335.2679233 | ENSG00000163874 |
| NA | 0.283683922 | 1.217299293 | 0.660003132 | 0.789931779 | 9.669716557 | 11.76237241 | ENSG00000283959 |
| SYNM | 0.283608703 | 1.217235827 | 0.229269791 | 0.389197778 | 87.97647243 | 106.8569911 | ENSG00000182253 |
| BTN2A2 | 0.283295571 | 1.216971659 | 0.053966684 | 0.126789832 | 213.858315 | 259.9441857 | ENSG00000124508 |
| EPHA7 | 0.283277196 | 1.216956159 | 0.074074555 | 0.163236492 | 163.6891677 | 199.1208536 | ENSG00000135333 |
| WIF1 | 0.283175113 | 1.216870052 | 0.266914035 | 0.43316553 | 60.97241338 | 74.29388845 | ENSG00000156076 |
| EEF2K | 0.282960322 | 1.216688896 | 0.003046061 | 0.011348314 | 611.8275561 | 744.5757963 | ENSG00000103319 |
| RNF141 | 0.282939507 | 1.216671342 | 0.009651785 | 0.030523374 | 455.4755773 | 554.3707663 | ENSG00000110315 |
| ALG11 | 0.282780666 | 1.216537394 | 0.098710852 | 0.205070404 | 184.7163533 | 224.4107939 | ENSG00000253710 |
| PACSIN3 | 0.282746662 | 1.216508721 | 4.85E-05 | 0.000279717 | 1165.99853 | 1417.84323 | ENSG00000165912 |
| GJD3 | 0.28259651 | 1.216382115 | 0.440600165 | 0.611527288 | 30.55439004 | 37.10499882 | ENSG00000183153 |
| ITGA2B | 0.282547503 | 1.216340797 | 0.045583706 | 0.110836793 | 223.9616124 | 272.205626 | ENSG00000005961 |
| RPL3 | 0.282513627 | 1.216312236 | 8.68E-16 | 2.04E-14 | 55430.70596 | 67419.88091 | ENSG00000100316 |
| SUCLA2 | 0.282500638 | 1.216301286 | 4.69E-05 | 0.000270677 | 1114.001834 | 1355.106984 | ENSG00000136143 |
| ORC5 | 0.282329428 | 1.216156951 | 0.007307407 | 0.024233657 | 377.3947341 | 458.9838506 | ENSG00000164815 |
| GXYLT1 | 0.282194352 | 1.21604309 | 0.026001051 | 0.06991724 | 360.4286545 | 438.8539373 | ENSG00000151233 |
| GZF1 | 0.282130406 | 1.215989192 | 0.005105285 | 0.017851378 | 427.7085838 | 520.066385 | ENSG00000125812 |
| ITM2A | 0.282078547 | 1.215945483 | 0.004012728 | 0.014503489 | 530.1941929 | 645.0065007 | ENSG00000078596 |
| DNM1 | 0.281962806 | 1.215847937 | 0.000445785 | 0.002073702 | 793.4541044 | 964.1428196 | ENSG00000106976 |
| LINC01006 | 0.28195395 | 1.215840473 | 0.138135008 | 0.264992287 | 112.7904279 | 136.9781442 | ENSG00000182648 |
| NA | 0.281896283 | 1.215791875 | 0.516845145 | 0.68071685 | 21.8651478 | 26.49504617 | ENSG00000260563 |
| GPCPD1 | 0.281732169 | 1.21565358 | 0.061698205 | 0.14105476 | 190.5439447 | 231.7648724 | ENSG00000125772 |
| CSNK1G1 | 0.281694006 | 1.215621424 | 0.068065379 | 0.15255597 | 218.4226623 | 265.8468837 | ENSG00000169118 |
| NA | 0.281520633 | 1.215475348 | 0.574446137 | 0.726582986 | 16.89195173 | 20.55282593 | ENSG00000234176 |
| COL27A1 | 0.281492424 | 1.215451582 | 0.001476391 | 0.006041804 | 792.1118121 | 962.2168884 | ENSG00000196739 |
| NA | 0.281397061 | 1.215371242 | 0.074509565 | 0.164053485 | 181.2971578 | 220.1348306 | ENSG00000249673 |
| MAML3 | 0.28132941 | 1.215314253 | 0.070234413 | 0.156526347 | 180.2804018 | 219.1533969 | ENSG00000196782 |
| HSCB | 0.281238728 | 1.215237865 | 0.31101506 | 0.481988393 | 57.78620296 | 70.04615517 | ENSG00000100209 |
| SREBF2-AS1 | 0.281166999 | 1.215177446 | 0.328937413 | 0.501187239 | 54.20581087 | 65.95300905 | ENSG00000184068 |
| B3GNT9 | 0.28115064 | 1.215163667 | 0.089944318 | 0.190588295 | 150.7105163 | 183.2817462 | ENSG00000237172 |
| NA | 0.281044135 | 1.215073963 | 0.316177369 | 0.487356879 | 53.67426025 | 65.17996957 | ENSG00000239665 |
| CPSF2 | 0.281040405 | 1.215070822 | 1.10E-05 | 7.14E-05 | 1907.523648 | 2319.097295 | ENSG00000165934 |
| LRRC57 | 0.281001824 | 1.215038328 | 0.244365734 | 0.407473366 | 81.65771758 | 99.12564017 | ENSG00000180979 |
| UFC1 | 0.280901789 | 1.214954081 | 0.004204671 | 0.015085049 | 517.7669776 | 628.7176198 | ENSG00000143222 |
| ECI1 | 0.280839989 | 1.214902038 | 2.68E-06 | 1.95E-05 | 1765.274092 | 2143.735981 | ENSG00000167969 |
| PSTPIP2 | 0.280795572 | 1.214864635 | 0.296500253 | 0.466807017 | 61.33290406 | 74.3525054 | ENSG00000152229 |
| CMTM1 | 0.280700839 | 1.214784864 | 0.656155694 | 0.786791366 | 9.990734282 | 12.15143947 | ENSG00000089505 |

| | | | | | | | |
|---|---|---|---|---|---|---|---|
| CEP63 | 0.280352264 | 1.214491391 | 0.017576935 | 0.050495406 | 334.5445524 | 406.5298946 | ENSG00000182923 |
| CIITA | 0.280344742 | 1.214485059 | 0.643076127 | 0.776981188 | 11.11228122 | 13.45980346 | ENSG00000179583 |
| CNNM1 | 0.280214378 | 1.214375321 | 0.331052145 | 0.503574116 | 61.35802219 | 74.36809245 | ENSG00000119946 |
| STOX2 | 0.280133932 | 1.214307609 | 0.000458915 | 0.002123515 | 826.4919084 | 1004.182559 | ENSG00000173320 |
| SHB | 0.279989614 | 1.214186144 | 0.221934187 | 0.38029245 | 84.37634385 | 102.5299286 | ENSG00000107338 |
| BORCS6 | 0.279954486 | 1.214156579 | 0.367442762 | 0.541239183 | 42.37067015 | 51.43731962 | ENSG00000196544 |
| NLRX1 | 0.279891636 | 1.214103687 | 0.00295088 | 0.011040617 | 565.7002028 | 687.2333982 | ENSG00000160703 |
| SEC24A | 0.279849218 | 1.21406799 | 0.00613367 | 0.020909107 | 427.3838113 | 519.1745004 | ENSG00000113615 |
| PHIP | 0.279807071 | 1.214032523 | 0.001415523 | 0.005822359 | 1075.15022 | 1305.484008 | ENSG00000146247 |
| TUBGCP5 | 0.279738623 | 1.213974926 | 0.009997956 | 0.031412146 | 392.3898399 | 476.3773483 | ENSG00000275835 |
| NA | 0.279688354 | 1.213932626 | 0.711030741 | 0.825872368 | 8.673848266 | 10.54273341 | ENSG00000263535 |
| ZNF146 | 0.279639203 | 1.21389127 | 9.79E-06 | 6.46E-05 | 2308.145785 | 2802.932585 | ENSG00000167635 |
| FSD1L | 0.27947744 | 1.213755169 | 0.386208222 | 0.559606508 | 41.73354079 | 50.70354192 | ENSG00000106701 |
| SLC25A34 | 0.279398674 | 1.213688904 | 0.720183889 | 0.832069677 | 6.784787387 | 8.186552699 | ENSG00000162461 |
| NA | 0.279214045 | 1.213533592 | 0.462484067 | 0.632212312 | 29.13606762 | 35.46403415 | ENSG00000247627 |
| NCEH1 | 0.279076675 | 1.213418048 | 0.26665494 | 0.432781462 | 74.89390597 | 90.77937514 | ENSG00000144959 |
| SLC5A11 | 0.278801681 | 1.213186779 | 0.819598833 | 0.897020372 | 2.716147677 | 3.274944587 | ENSG00000158865 |
| RFX1 | 0.278605995 | 1.213022235 | 0.011530239 | 0.035577391 | 528.3527918 | 641.2145876 | ENSG00000132005 |
| NA | 0.278553288 | 1.212977919 | 0.779775564 | 0.870674909 | 4.289096914 | 5.212483234 | ENSG00000226986 |
| LRIG2 | 0.278537666 | 1.212964784 | 0.00923045 | 0.029407865 | 420.8701177 | 510.7007695 | ENSG00000198799 |
| STRCP1 | 0.278223135 | 1.212700367 | 0.66989045 | 0.797264277 | 11.74661969 | 14.2615939 | ENSG00000166763 |
| S100PBP | 0.277996804 | 1.212510133 | 0.000899853 | 0.003866585 | 772.6564614 | 936.5168231 | ENSG00000116497 |
| SEMA6A-AS1 | 0.277840246 | 1.212378561 | 0.579771446 | 0.731024514 | 15.72518676 | 19.06135376 | ENSG00000248445 |
| RIPOR1 | 0.277822618 | 1.212363748 | 0.075281275 | 0.165310311 | 197.1233807 | 239.0829669 | ENSG00000039523 |
| RPL13AP5 | 0.277819131 | 1.212360817 | 0.047990235 | 0.115577979 | 208.9657428 | 253.2058333 | ENSG00000236552 |
| ZBTB32 | 0.277793176 | 1.212339006 | 0.623874345 | 0.763014769 | 15.92698353 | 19.29311953 | ENSG00000011590 |
| LNPK | 0.277731436 | 1.212287126 | 0.104379955 | 0.214026449 | 140.9829786 | 170.8587461 | ENSG00000144320 |
| SKIDA1 | 0.277730706 | 1.212286512 | 0.297842017 | 0.468117187 | 59.40225504 | 72.09203443 | ENSG00000180592 |
| MED13L | 0.277635634 | 1.212206626 | 0.001962326 | 0.007757218 | 696.4282483 | 843.9345836 | ENSG00000123066 |
| NA | 0.277414725 | 1.212021025 | 0.590994772 | 0.739910841 | 16.78321866 | 20.29604678 | ENSG00000274840 |
| KBTBD7 | 0.277342171 | 1.211960073 | 0.009682706 | 0.030607236 | 397.3277934 | 481.5220913 | ENSG00000120696 |
| SH3GLB2 | 0.277029692 | 1.211697598 | 0.000129704 | 0.000680849 | 1375.610236 | 1665.71828 | ENSG00000148341 |
| NA | 0.276808591 | 1.211511913 | 0.36477024 | 0.538433139 | 43.33998086 | 52.55073805 | ENSG00000271576 |
| EIF4ENIF1 | 0.276805952 | 1.211509696 | 0.020565063 | 0.057545784 | 316.1627028 | 382.7755589 | ENSG00000184708 |
| NA | 0.276709821 | 1.211428973 | 0.261644228 | 0.427020203 | 82.35381932 | 99.92904815 | ENSG00000224546 |
| E2F6 | 0.276526514 | 1.21127506 | 0.013328528 | 0.040087408 | 453.7077262 | 549.6072573 | ENSG00000169016 |
| GTF2IRD2B | 0.27649218 | 1.211246234 | 0.245805603 | 0.409196866 | 70.45727875 | 85.35394826 | ENSG00000174428 |
| CDKAL1 | 0.276478128 | 1.211234436 | 0.011577186 | 0.035653816 | 389.3684688 | 471.613603 | ENSG00000145996 |
| NA | 0.276443774 | 1.211205594 | 0.77092253 | 0.86504575 | 4.316330222 | 5.233455934 | ENSG00000287005 |
| PPP3CA | 0.276400389 | 1.211169171 | 0.000102199 | 0.000551785 | 1244.192412 | 1507.763427 | ENSG00000138814 |
| PDZK1 | 0.276338603 | 1.211117302 | 0.735660258 | 0.842640381 | 7.933567634 | 9.530943398 | ENSG00000174827 |
| NA | 0.276283987 | 1.211071453 | 0.712038157 | 0.826420076 | 7.065293013 | 8.535824941 | ENSG00000282978 |
| MRPS33 | 0.276242495 | 1.211036624 | 0.003105371 | 0.011535857 | 549.8323535 | 665.3636302 | ENSG00000090263 |
| CHGA | 0.275574758 | 1.210476237 | 8.11E-05 | 0.000447642 | 1776.625156 | 2150.67641 | ENSG00000100604 |
| NA | 0.275497155 | 1.210411127 | 0.349016885 | 0.521661052 | 51.74122064 | 62.56916029 | ENSG00000259943 |
| TAB2 | 0.275244528 | 1.210199193 | 8.68E-05 | 0.000475929 | 1245.071762 | 1507.467398 | ENSG00000055208 |
| PNMA5 | 0.275166451 | 1.2101337 | 0.723299745 | 0.834108096 | 7.282283592 | 8.828097768 | ENSG00000198883 |
| TNRC6B | 0.275154955 | 1.210124057 | 0.00210167 | 0.008242185 | 672.7528365 | 814.1487211 | ENSG00000100354 |
| C17orf67 | 0.275032337 | 1.210021211 | 0.296885037 | 0.467034501 | 63.05912867 | 76.39544058 | ENSG00000214226 |
| KAZN | 0.275004125 | 1.209997549 | 0.010216769 | 0.032031803 | 403.9710326 | 489.1001268 | ENSG00000189337 |
| KMT2E | 0.274839671 | 1.209859628 | 0.012816258 | 0.038800758 | 500.4457507 | 605.5872545 | ENSG00000005483 |
| AGO4 | 0.274811286 | 1.209835824 | 0.039266635 | 0.098286317 | 285.8542884 | 345.8001217 | ENSG00000134698 |
| EIF4B | 0.274628963 | 1.209682939 | 5.43E-09 | 6.00E-08 | 12680.29712 | 15338.66359 | ENSG00000063046 |
| RPS6KA6 | 0.274585113 | 1.209646172 | 0.041465508 | 0.102618487 | 255.687073 | 309.6190056 | ENSG00000072133 |
| FGD5 | 0.274534543 | 1.209603771 | 0.00061804 | 0.002765625 | 758.0171034 | 916.4533783 | ENSG00000154783 |
| MIR570 | 0.274496908 | 1.209572218 | 0.748151822 | 0.850784433 | 6.21587899 | 7.572432051 | ENSG00000207650 |
| ZNF410 | 0.274461166 | 1.209542251 | 0.825886717 | 0.901037344 | 2.773405195 | 3.327642854 | ENSG00000119725 |
| IGDCC4 | 0.27437814 | 1.209472645 | 0.002259135 | 0.008779404 | 552.7891335 | 668.5347958 | ENSG00000103742 |
| NUDT13 | 0.274149419 | 1.209280914 | 0.442495337 | 0.613451495 | 30.396572 | 36.75572658 | ENSG00000166321 |
| BHLHA15 | 0.274073273 | 1.209217089 | 0.40690221 | 0.579174392 | 36.2099335 | 43.76028453 | ENSG00000180535 |
| GOLGA4 | 0.273963128 | 1.209124773 | 0.000238478 | 0.001184773 | 919.2447103 | 1111.716608 | ENSG00000144674 |
| BTG1 | 0.273898244 | 1.209070395 | 0.098079961 | 0.204039742 | 234.1950942 | 283.4027895 | ENSG00000133639 |
| CLEC2D | 0.273786058 | 1.20897638 | 0.541557293 | 0.700401028 | 20.47146794 | 24.74894543 | ENSG00000069493 |
| SLC34A2 | 0.273549486 | 1.208778148 | 0.358621646 | 0.531676158 | 47.33270261 | 57.12491129 | ENSG00000157765 |
| APLF | 0.273545857 | 1.208775109 | 0.274736275 | 0.441507263 | 65.11784693 | 78.69651198 | ENSG00000169621 |
| SAMD10 | 0.273146069 | 1.208440188 | 0.200206591 | 0.351948927 | 102.4154362 | 123.8816009 | ENSG00000130590 |
| RUNX2 | 0.27313284 | 1.208429108 | 0.613865337 | 0.756749457 | 15.46055068 | 18.7922042 | ENSG00000124813 |
| FNTA | 0.272734753 | 1.208095708 | 0.006715187 | 0.022612858 | 462.0998679 | 558.7612556 | ENSG00000168522 |
| NAPRT | 0.272731449 | 1.208092941 | 0.008108569 | 0.02641378 | 505.3961582 | 610.8376909 | ENSG00000147813 |
| PCDHAC2 | 0.272528741 | 1.207923208 | 0.140594666 | 0.268935222 | 130.9268143 | 158.2018055 | ENSG00000243232 |
| CAMK2D | 0.272460579 | 1.20786614 | 0.070846486 | 0.157544619 | 184.2156659 | 222.3993117 | ENSG00000145349 |
| DARS1 | 0.272289767 | 1.20772314 | 7.33E-05 | 0.000408573 | 1099.978993 | 1328.973048 | ENSG00000115866 |
| ZNF320 | 0.272262198 | 1.207700061 | 0.000870857 | 0.003757481 | 732.2004728 | 884.5877235 | ENSG00000182986 |
| MUTYH | 0.272225374 | 1.207669236 | 0.548028553 | 0.705648033 | 19.78545377 | 23.82131879 | ENSG00000132781 |
| RALGAPA1 | 0.272193551 | 1.207642597 | 0.030117764 | 0.078769221 | 327.8262342 | 396.2724614 | ENSG00000174373 |
| GON4L | 0.272116901 | 1.207578437 | 0.00031843 | 0.001534953 | 1017.646499 | 1228.163722 | ENSG00000116580 |
| COA6 | 0.272092223 | 1.207557781 | 0.010238468 | 0.032080626 | 375.4098186 | 453.4903804 | ENSG00000168275 |

| | | | | | | | |
|---|---|---|---|---|---|---|---|
| LOC100505549 | 0.272008048 | 1.207487327 | 0.61116552 | 0.754801156 | 14.63924584 | 17.69007168 | ENSG00000267040 |
| DYRK4 | 0.271983575 | 1.207466845 | 0.451517166 | 0.621714322 | 32.62080528 | 39.26920859 | ENSG00000010219 |
| GALK2 | 0.271879035 | 1.207379353 | 0.166098199 | 0.305802667 | 118.5450924 | 142.9892012 | ENSG00000156958 |
| SLC41A1 | 0.27155283 | 1.207106385 | 0.000252972 | 0.001248101 | 949.6024851 | 1146.31248 | ENSG00000133065 |
| MRNIP | 0.27127212 | 1.206871537 | 0.038390631 | 0.096402985 | 305.7950447 | 368.7938674 | ENSG00000161010 |
| SBNO1 | 0.271267315 | 1.206867518 | 4.93E-05 | 0.000283739 | 1162.181248 | 1402.593405 | ENSG00000139697 |
| FNDC3B | 0.271177173 | 1.206792113 | 0.00176375 | 0.007073699 | 707.6300655 | 853.8215725 | ENSG00000075420 |
| TEX30 | 0.271079776 | 1.206710645 | 0.210324954 | 0.365925206 | 84.85932197 | 102.3868878 | ENSG00000151287 |
| SLC25A14 | 0.270934655 | 1.206589268 | 0.079593146 | 0.172792187 | 197.7937255 | 238.3279681 | ENSG00000102078 |
| SYDE1 | 0.270931253 | 1.206586422 | 0.016389841 | 0.047609762 | 444.9007408 | 536.8855811 | ENSG00000105137 |
| G2E3 | 0.270720589 | 1.206410248 | 0.007078781 | 0.023601238 | 940.5721153 | 1135.639736 | ENSG00000092140 |
| CACNG8 | 0.270541391 | 1.206260409 | 0.048537817 | 0.116591089 | 240.0191807 | 289.2727021 | ENSG00000142408 |
| TNS2 | 0.270330233 | 1.206083869 | 0.156760141 | 0.29246227 | 125.7677279 | 151.401601 | ENSG00000111077 |
| GTF2H2C | 0.270308853 | 1.206065995 | 0.049348216 | 0.118018919 | 259.2273164 | 313.0337743 | ENSG00000183474 |
| BACH1 | 0.270121101 | 1.205909048 | 0.002116842 | 0.008296627 | 613.9225561 | 740.035554 | ENSG00000156273 |
| NA | 0.269978472 | 1.205789834 | 0.580829122 | 0.731628608 | 17.40918441 | 20.96286569 | ENSG00000172974 |
| NA | 0.269967228 | 1.205780437 | 0.748312847 | 0.850910883 | 5.957024867 | 7.197885963 | ENSG00000258655 |
| SLC17A7 | 0.269881182 | 1.205708523 | 0.320961134 | 0.492323626 | 62.94548952 | 75.95932135 | ENSG00000104888 |
| DCUN1D1 | 0.269657535 | 1.205521629 | 0.009055271 | 0.028930976 | 478.4121479 | 576.9898806 | ENSG00000043093 |
| SYNE4 | 0.269627258 | 1.205496329 | 0.046903622 | 0.113513708 | 271.820396 | 327.3932528 | ENSG00000181392 |
| AK4 | 0.269627075 | 1.205496176 | 6.58E-07 | 5.33E-06 | 3420.433742 | 4122.391628 | ENSG00000162433 |
| ADCK1 | 0.269581909 | 1.205458437 | 0.0350622 | 0.089303525 | 250.2755902 | 301.6650427 | ENSG00000063761 |
| LMBRD2 | 0.269561666 | 1.205441523 | 0.223674218 | 0.382177042 | 84.31015005 | 101.5055077 | ENSG00000164187 |
| MIB1 | 0.269420235 | 1.205323356 | 0.000166301 | 0.000853449 | 1310.404208 | 1578.885914 | ENSG00000101752 |
| ZBTB7A | 0.26941813 | 1.205321598 | 0.008792833 | 0.028256295 | 465.4571751 | 561.5739952 | ENSG00000178951 |
| RHD | 0.269411864 | 1.205316362 | 0.783286874 | 0.87312996 | 4.321912028 | 5.20388094 | ENSG00000187010 |
| BLOC1S4 | 0.269376608 | 1.205286908 | 0.004952383 | 0.017398731 | 464.7418493 | 560.1556731 | ENSG00000186222 |
| JRKL | 0.269169118 | 1.205113575 | 0.124653017 | 0.245295666 | 134.4872697 | 162.0578002 | ENSG00000183340 |
| C2CD4C | 0.26912851 | 1.205079655 | 0.020903841 | 0.058260511 | 382.4419452 | 460.972773 | ENSG00000183186 |
| TIAM1 | 0.269057006 | 1.205019929 | 0.005029938 | 0.017632661 | 610.699225 | 735.4213197 | ENSG00000156299 |
| DOCK6 | 0.268596863 | 1.204635653 | 0.000752574 | 0.003289806 | 1025.705132 | 1235.234736 | ENSG00000130158 |
| PDXK | 0.268531075 | 1.204580721 | 1.08E-06 | 8.47E-06 | 2128.138954 | 2564.077926 | ENSG00000160209 |
| GOLGA6L9 | 0.268503096 | 1.204557361 | 0.14325551 | 0.272699892 | 119.3489761 | 143.7006491 | ENSG00000197978 |
| LINC01234 | 0.268137897 | 1.204252482 | 0.439752013 | 0.610833531 | 37.44021351 | 44.99848519 | ENSG00000249550 |
| PABPC3 | 0.268122364 | 1.204239516 | 0.659517531 | 0.789485573 | 10.36900953 | 12.48404016 | ENSG00000151846 |
| BARD1 | 0.267867483 | 1.204026782 | 0.015985445 | 0.046681081 | 345.0750468 | 415.3332637 | ENSG00000138376 |
| CAND2 | 0.267837116 | 1.204001439 | 0.039413622 | 0.098577489 | 297.7186286 | 358.7480086 | ENSG00000144712 |
| NLRP12 | 0.267765335 | 1.203941535 | 0.075651677 | 0.165897067 | 187.0892313 | 225.3077794 | ENSG00000142405 |
| AP1M2 | 0.267701837 | 1.203888547 | 4.70E-07 | 3.89E-06 | 2462.354939 | 2963.59386 | ENSG00000129354 |
| THOC7 | 0.267697819 | 1.203885194 | 0.004180978 | 0.015014007 | 622.2681283 | 749.5336752 | ENSG00000163634 |
| ZNF724 | 0.267614459 | 1.203815634 | 0.25073773 | 0.414889339 | 79.48149323 | 95.75175077 | ENSG00000196081 |
| TMEM101 | 0.267457177 | 1.203684403 | 0.009303442 | 0.029601302 | 421.5150321 | 507.4215359 | ENSG00000091947 |
| SLC35E2A | 0.267253677 | 1.203514628 | 0.073817556 | 0.162737218 | 206.120562 | 247.8685983 | ENSG00000215790 |
| C12orf76 | 0.267211151 | 1.203479153 | 0.124759818 | 0.245385568 | 163.1193082 | 196.1777041 | ENSG00000174456 |
| NA | 0.266964179 | 1.203273149 | 0.468428289 | 0.637287157 | 30.97086204 | 37.25799898 | ENSG00000257270 |
| STAM2 | 0.266911463 | 1.203229183 | 0.106380642 | 0.216910364 | 157.0063803 | 188.796189 | ENSG00000115145 |
| KLF6 | 0.266885619 | 1.203207629 | 0.008189967 | 0.026642943 | 447.0925081 | 537.589487 | ENSG00000067082 |
| PARP4 | 0.266827447 | 1.203159114 | 0.00010945 | 0.000586826 | 1147.454032 | 1380.963981 | ENSG00000102699 |
| CLEC2L | 0.266621985 | 1.202987778 | 0.596981585 | 0.744192035 | 16.15792862 | 19.39610502 | ENSG00000236279 |
| POLR3D | 0.266491394 | 1.20287889 | 3.63E-05 | 0.000214922 | 1393.324658 | 1676.020263 | ENSG00000168495 |
| ENPP1 | 0.266190009 | 1.20262763 | 0.002743145 | 0.010384225 | 666.8544958 | 802.1973339 | ENSG00000197594 |
| PPP3CC | 0.266171838 | 1.202612482 | 0.368644681 | 0.542409011 | 46.69066716 | 56.01040836 | ENSG00000120910 |
| SACS | 0.266162308 | 1.202604538 | 0.003567675 | 0.013074473 | 1121.563009 | 1348.507589 | ENSG00000151835 |
| DENND1B | 0.26615647 | 1.202599672 | 0.404715517 | 0.576913252 | 54.04157204 | 64.84790192 | ENSG00000213047 |
| MYO18B | 0.266140592 | 1.202586436 | 0.565991955 | 0.719983567 | 19.77496587 | 23.80275713 | ENSG00000133454 |
| ZBTB43 | 0.266029895 | 1.202494166 | 0.043767746 | 0.107274683 | 270.4254768 | 325.1241553 | ENSG00000169155 |
| RBM26 | 0.266015851 | 1.20248246 | 0.000105008 | 0.000565525 | 1242.81709 | 1494.767719 | ENSG00000139746 |
| GHRLOS | 0.2659854 | 1.20245708 | 0.639640341 | 0.774741394 | 13.37335982 | 16.07276331 | ENSG00000240288 |
| DEPDC1 | 0.265951579 | 1.202428891 | 0.01978012 | 0.055696731 | 561.0295769 | 674.6589666 | ENSG00000024526 |
| NA | 0.265879385 | 1.202368722 | 0.732664727 | 0.84029936 | 8.04788251 | 9.713779008 | ENSG00000229976 |
| TANC2 | 0.265854919 | 1.202348332 | 0.05497307 | 0.128793478 | 233.9828465 | 281.4276129 | ENSG00000170921 |
| HOMER1 | 0.26585316 | 1.202346866 | 0.000565534 | 0.002556763 | 813.3551124 | 977.7727852 | ENSG00000152413 |
| CBR4 | 0.265780778 | 1.202286544 | 0.019892684 | 0.055972835 | 376.9962468 | 453.5686009 | ENSG00000145439 |
| APAF1 | 0.265730734 | 1.20224484 | 0.014346552 | 0.042670264 | 383.5339313 | 461.1112096 | ENSG00000120868 |
| MAP4K3 | 0.265705925 | 1.202224166 | 0.000111018 | 0.000594079 | 1161.213452 | 1396.412937 | ENSG00000011566 |
| DSG2 | 0.265596607 | 1.202133073 | 1.79E-09 | 2.12E-08 | 4111.373781 | 4943.156428 | ENSG00000046604 |
| UST | 0.265416781 | 1.201983241 | 0.00177184 | 0.007103193 | 779.3192369 | 935.9965769 | ENSG00000111962 |
| NA | 0.265260588 | 1.201853116 | 0.826518598 | 0.90121921 | 3.83211281 | 4.547814893 | ENSG00000286799 |
| LRRC58 | 0.265206655 | 1.201808187 | 0.021153987 | 0.058843773 | 459.3608053 | 552.2906893 | ENSG00000163428 |
| S100A13 | 0.265020618 | 1.201653223 | 0.44941173 | 0.619627613 | 34.2175212 | 41.07150313 | ENSG00000189171 |
| PCDHA9 | 0.264868176 | 1.201526257 | 0.505644934 | 0.670442219 | 28.11296602 | 33.90964389 | ENSG00000204961 |
| ABHD18 | 0.264797354 | 1.201467276 | 0.055775391 | 0.130362672 | 233.3424506 | 280.1499021 | ENSG00000164074 |
| CDT1 | 0.264663768 | 1.201356031 | 2.51E-05 | 0.000153683 | 2341.207141 | 2813.419428 | ENSG00000167513 |
| KIAA0556 | 0.264615794 | 1.201316083 | 0.024165229 | 0.06569496 | 320.1935092 | 384.6598541 | ENSG00000047578 |
| NA | 0.264544834 | 1.201256997 | 0.463160415 | 0.632822974 | 31.55863104 | 37.88528353 | ENSG00000259612 |
| PBLD | 0.264539534 | 1.201252583 | 0.434356878 | 0.605519477 | 36.10311545 | 43.36906689 | ENSG00000108187 |
| MRPL23 | 0.26442366 | 1.201156106 | 0.056037364 | 0.130719569 | 220.6668322 | 265.1878615 | ENSG00000214026 |

| | | | | | | | |
|---|---|---|---|---|---|---|---|
| RGPD3 | 0.264224621 | 1.200990402 | 0.752694998 | 0.853629583 | 6.036609608 | 7.248433657 | ENSG00000153165 |
| POM121L9P | 0.264205414 | 1.200974413 | 0.740093057 | 0.84572929 | 6.322021318 | 7.625130318 | ENSG00000128262 |
| C2orf69 | 0.264195114 | 1.200965838 | 0.032011091 | 0.082844909 | 449.404601 | 540.2620993 | ENSG00000178074 |
| USP9X | 0.26415886 | 1.200935659 | 2.11E-05 | 0.000130812 | 4902.097023 | 5887.280138 | ENSG00000124486 |
| MOB3A | 0.264133674 | 1.200914694 | 0.164790712 | 0.303801379 | 134.2847972 | 161.4025461 | ENSG00000172081 |
| TATDN1 | 0.263675747 | 1.200533571 | 0.016254524 | 0.047287895 | 358.1842461 | 429.9634971 | ENSG00000147687 |
| ZNF837 | 0.263654923 | 1.200516242 | 0.260285025 | 0.425662591 | 98.43898431 | 118.4462025 | ENSG00000152475 |
| NA | 0.263623912 | 1.200490438 | 0.287995127 | 0.457335693 | 69.76847367 | 83.67454584 | ENSG00000234160 |
| LOC101928674 | 0.263621964 | 1.200488817 | 0.406271375 | 0.578703471 | 48.91143381 | 58.49216479 | ENSG00000266970 |
| CACYBP | 0.263570896 | 1.200446323 | 2.38E-09 | 2.75E-08 | 4589.870971 | 5509.516274 | ENSG00000116161 |
| TRIM21 | 0.263546862 | 1.200426325 | 0.038892387 | 0.097463372 | 256.5950208 | 307.8575904 | ENSG00000132109 |
| NSD1 | 0.263503966 | 1.200390632 | 4.35E-08 | 4.22E-07 | 3847.166196 | 4618.463478 | ENSG00000165671 |
| HACD1 | 0.263503885 | 1.200390565 | 0.074540303 | 0.164056702 | 192.2193695 | 230.9625489 | ENSG00000165996 |
| LINC00678 | 0.263424168 | 1.200324239 | 1.73E-06 | 1.30E-05 | 3395.392484 | 4074.441669 | ENSG00000254934 |
| GOLT1A | 0.263356884 | 1.20026826 | 0.267895142 | 0.434173333 | 69.49634075 | 83.42071076 | ENSG00000174567 |
| TFB2M | 0.263203955 | 1.200141035 | 0.008098043 | 0.026390304 | 528.8997974 | 634.6687391 | ENSG00000162851 |
| RAB11FIP1 | 0.263120082 | 1.200071266 | 0.002489437 | 0.009566479 | 611.4556506 | 733.5074494 | ENSG00000156675 |
| ZSWIM4 | 0.263054707 | 1.200016886 | 0.012789846 | 0.038739036 | 421.9436557 | 506.1731459 | ENSG00000132003 |
| ISYNA1 | 0.263047929 | 1.200011248 | 1.03E-08 | 1.10E-07 | 3991.227294 | 4789.887864 | ENSG00000105655 |
| MINK1 | 0.263013865 | 1.199982915 | 2.26E-05 | 0.000139583 | 2513.499007 | 3015.647082 | ENSG00000141503 |
| MXI1 | 0.263001589 | 1.199972704 | 0.153093142 | 0.287315868 | 159.1541321 | 190.6909582 | ENSG00000119950 |
| MAP3K21 | 0.262692454 | 1.199715607 | 0.001755446 | 0.007047721 | 802.7058726 | 963.4320799 | ENSG00000143674 |
| SMIM19 | 0.262620585 | 1.199655843 | 0.056659975 | 0.131896656 | 220.0121937 | 264.1467569 | ENSG00000176209 |
| SPRY4 | 0.262560565 | 1.199605935 | 6.84E-09 | 7.45E-08 | 4133.591084 | 4959.167098 | ENSG00000187678 |
| ZNF426-DT | 0.262551701 | 1.199598565 | 0.568280809 | 0.721515406 | 18.66390683 | 22.44384545 | ENSG00000278611 |
| RBM48 | 0.26248987 | 1.199547153 | 0.186958789 | 0.333946481 | 106.0733859 | 127.0898593 | ENSG00000127993 |
| NA | 0.262467508 | 1.19952856 | 0.708710261 | 0.824392766 | 8.725523978 | 10.43088515 | ENSG00000279307 |
| SCN1A | 0.262267201 | 1.199362027 | 0.486920579 | 0.654258193 | 30.21451171 | 36.20829214 | ENSG00000144285 |
| ZNF280C | 0.262216585 | 1.199319949 | 0.097460554 | 0.20301388 | 226.2337666 | 271.5218203 | ENSG00000056277 |
| NA | 0.262211315 | 1.199315568 | 0.511500241 | 0.6757534 | 24.33831089 | 29.2064178 | ENSG00000269973 |
| DCUN1D3 | 0.262106934 | 1.199228799 | 0.111356982 | 0.224823395 | 180.8370705 | 216.9246821 | ENSG00000188215 |
| ASB12 | 0.262092298 | 1.199216633 | 0.686822641 | 0.809524993 | 9.035378092 | 10.82640394 | ENSG00000198881 |
| NA | 0.262026906 | 1.199162278 | 0.324055738 | 0.495543064 | 65.47100404 | 78.64541914 | ENSG00000268362 |
| USP33 | 0.261894849 | 1.199052518 | 9.07E-05 | 0.000494853 | 1595.462309 | 1912.739743 | ENSG00000077254 |
| NA | 0.26186203 | 1.199025242 | 0.846746522 | 0.912319442 | 2.949883674 | 3.538175458 | ENSG00000250068 |
| ZC3HAV1 | 0.261704843 | 1.19889461 | 0.000527197 | 0.002403154 | 989.4581414 | 1185.765324 | ENSG00000105939 |
| CLPX | 0.261670875 | 1.198866383 | 6.28E-05 | 0.000354506 | 1169.869479 | 1402.912048 | ENSG00000166855 |
| MED17 | 0.26161923 | 1.198823467 | 0.477207242 | 0.645432581 | 30.856347 | 37.06682154 | ENSG00000042429 |
| LRRC2 | 0.261613199 | 1.198818456 | 0.187984482 | 0.335312866 | 105.6790409 | 126.6160958 | ENSG00000163827 |
| NA | 0.26160638 | 1.198812789 | 0.385526813 | 0.559145759 | 44.37703697 | 53.23852966 | ENSG00000244479 |
| TNK1 | 0.261538379 | 1.198756284 | 0.079154939 | 0.172015087 | 200.1253035 | 240.2052602 | ENSG00000174292 |
| LOC105376805 | 0.261512902 | 1.198735116 | 0.744591972 | 0.848722265 | 6.052679304 | 7.246283083 | ENSG00000238142 |
| LHX1-DT | 0.261444511 | 1.198678291 | 0.583930574 | 0.733964094 | 17.75252936 | 21.2277141 | ENSG00000277268 |
| NA | 0.261074448 | 1.19837086 | 0.389144859 | 0.562490145 | 44.59402755 | 53.50767922 | ENSG00000213025 |
| ZNF154 | 0.261051402 | 1.198351717 | 0.045584942 | 0.110836793 | 248.9700679 | 298.3594265 | ENSG00000179909 |
| SORBS1 | 0.260916505 | 1.198239672 | 0.003518514 | 0.012918851 | 693.8375038 | 830.9931973 | ENSG00000095637 |
| NA | 0.260867359 | 1.198198855 | 0.716468965 | 0.829914904 | 9.321465523 | 11.18642905 | ENSG00000228906 |
| NA | 0.260840192 | 1.198176291 | 0.485089837 | 0.652828023 | 34.57482066 | 41.48018583 | ENSG00000236255 |
| THAP2 | 0.260834643 | 1.198171683 | 0.316225884 | 0.487356879 | 61.8812001 | 74.0809448 | ENSG00000173451 |
| LINC00294 | 0.26057537 | 1.197956374 | 0.210933814 | 0.366521726 | 98.95988378 | 118.5236353 | ENSG00000280798 |
| GRIK4 | 0.260562767 | 1.197945909 | 0.191834444 | 0.340895698 | 113.7333811 | 136.1279445 | ENSG00000149403 |
| NA | 0.260442975 | 1.197846443 | 0.585467371 | 0.735192012 | 18.26677098 | 21.89453301 | ENSG00000272293 |
| SOCS2-AS1 | 0.260358262 | 1.197776109 | 0.522937434 | 0.685570353 | 25.25056423 | 30.15313914 | ENSG00000246985 |
| FAH | 0.260331877 | 1.197754204 | 0.021661233 | 0.060029716 | 335.7401023 | 402.2036378 | ENSG00000103876 |
| LMF1 | 0.260265791 | 1.197699339 | 0.177334306 | 0.32103567 | 115.977314 | 139.0697369 | ENSG00000103227 |
| MED23 | 0.260185726 | 1.197632872 | 0.045009671 | 0.109714628 | 333.8181383 | 399.9737958 | ENSG00000112282 |
| COA5 | 0.260157705 | 1.197609612 | 0.01159934 | 0.035704388 | 490.4096862 | 586.6746787 | ENSG00000183513 |
| CYREN | 0.260111544 | 1.197571293 | 0.083628463 | 0.179832424 | 182.5979742 | 218.5790832 | ENSG00000122783 |
| INPP5B | 0.260053991 | 1.197523519 | 0.011685986 | 0.035891474 | 482.0694091 | 577.7618419 | ENSG00000204084 |
| CUX1 | 0.260031759 | 1.197505066 | 0.00154948 | 0.006296824 | 1119.288188 | 1340.577277 | ENSG00000257923 |
| PTPRN | 0.259996292 | 1.197475627 | 0.173569441 | 0.316267884 | 117.7732359 | 141.0099776 | ENSG00000054356 |
| C6orf141 | 0.259888416 | 1.197386091 | 0.246159429 | 0.409579226 | 87.5175364 | 104.6987121 | ENSG00000197261 |
| PGA5 | 0.259774355 | 1.197291427 | 0.841694344 | 0.909490034 | 3.347695239 | 4.024036763 | ENSG00000256713 |
| LINC01535 | 0.259639305 | 1.197179355 | 0.71840539 | 0.831109947 | 8.505066774 | 10.10473618 | ENSG00000226686 |
| LCLAT1 | 0.259615076 | 1.19715925 | 0.090473649 | 0.191432663 | 182.7006499 | 218.7218514 | ENSG00000172954 |
| CSTB | 0.259605654 | 1.197151431 | 0.00012278 | 0.000650164 | 1452.535944 | 1739.208191 | ENSG00000160213 |
| C6orf99 | 0.259525449 | 1.197084878 | 0.564145632 | 0.718582594 | 21.54816027 | 25.82822726 | ENSG00000203711 |
| SSR2 | 0.259518294 | 1.197078941 | 0.000466995 | 0.002157282 | 1224.705771 | 1465.700095 | ENSG00000163479 |
| CPLX1 | 0.259404913 | 1.196984867 | 0.536022913 | 0.695818206 | 26.72419225 | 32.08369303 | ENSG00000168993 |
| LCOR | 0.259389829 | 1.196972352 | 0.027035626 | 0.072256062 | 316.8215717 | 379.0069442 | ENSG00000196233 |
| SMARCD3 | 0.259331635 | 1.196924071 | 0.140845625 | 0.269221702 | 133.9674463 | 160.2189459 | ENSG00000082014 |
| PLPP3 | 0.259318798 | 1.196913421 | 0.017573979 | 0.050494434 | 426.2572831 | 509.8261748 | ENSG00000162407 |
| SATB1 | 0.25931837 | 1.196913066 | 0.004440301 | 0.01583617 | 612.0430192 | 732.2128066 | ENSG00000182568 |
| FAM185A | 0.259303569 | 1.196900786 | 0.184006481 | 0.329986895 | 105.6057137 | 126.3808224 | ENSG00000222011 |
| TGFA | 0.259296275 | 1.196894735 | 0.16163875 | 0.299076647 | 130.3525242 | 155.8450854 | ENSG00000163235 |
| DUSP12 | 0.259097518 | 1.196729852 | 0.011774933 | 0.036084301 | 420.815539 | 503.664256 | ENSG00000081721 |
| BRMS1L | 0.259079384 | 1.196714811 | 0.156470628 | 0.29203511 | 150.0946381 | 180.0275017 | ENSG00000100916 |

| | | | | | | | |
|---|---|---|---|---|---|---|---|
| JAK3 | 0.258973728 | 1.196627172 | 0.253616724 | 0.418307132 | 77.39968401 | 92.69864312 | ENSG00000105639 |
| PRICKLE2 | 0.25896529 | 1.196620173 | 0.35460221 | 0.52778576 | 58.77840447 | 70.37148011 | ENSG00000163637 |
| OIP5-AS1 | 0.258950294 | 1.196607735 | 0.000350574 | 0.001674403 | 1528.772605 | 1830.097396 | ENSG00000247556 |
| NA | 0.258857046 | 1.196530395 | 0.825012911 | 0.900593514 | 2.988280594 | 3.571518561 | ENSG00000214243 |
| PDZD4 | 0.258743213 | 1.196435989 | 3.95E-08 | 3.86E-07 | 7137.177244 | 8538.087995 | ENSG00000067840 |
| FAM217B | 0.258641634 | 1.196351752 | 0.018041622 | 0.051531083 | 382.1411785 | 457.5897057 | ENSG00000196227 |
| ASB16-AS1 | 0.258582912 | 1.196303058 | 0.281972697 | 0.450329937 | 78.73957297 | 94.24012994 | ENSG00000267080 |
| NKAPP1 | 0.258521985 | 1.196252537 | 0.595635108 | 0.743377944 | 19.072882 | 22.86033694 | ENSG00000233382 |
| TSGA10 | 0.258456485 | 1.196198228 | 0.413397241 | 0.585608724 | 39.82417985 | 47.61170552 | ENSG00000135951 |
| FLOT1 | 0.258419414 | 1.196167491 | 7.20E-08 | 6.75E-07 | 3055.764578 | 3655.319897 | ENSG00000137312 |
| MDGA1 | 0.258350088 | 1.196110013 | 0.509596181 | 0.674253767 | 31.00455303 | 36.95550632 | ENSG00000112139 |
| NA | 0.258219704 | 1.196001918 | 0.818141572 | 0.89613724 | 3.015513901 | 3.598942982 | ENSG00000257605 |
| NA | 0.258176723 | 1.195966288 | 0.627938903 | 0.766051311 | 14.75866696 | 17.58682572 | ENSG00000277142 |
| NA | 0.258162575 | 1.195954559 | 2.25E-10 | 2.99E-09 | 11118.573 | 13298.2384 | ENSG00000224078 |
| SRPK2 | 0.258119856 | 1.195919147 | 9.58E-05 | 0.000519981 | 1169.864334 | 1398.748206 | ENSG00000135250 |
| SHC1 | 0.258049458 | 1.195860793 | 2.69E-09 | 3.09E-08 | 6349.081507 | 7591.51119 | ENSG00000160691 |
| XPO1 | 0.258004958 | 1.195823906 | 3.13E-08 | 3.08E-07 | 6550.80926 | 7834.138769 | ENSG00000082898 |
| NA | 0.257951003 | 1.195779185 | 0.718588559 | 0.831214704 | 8.502275871 | 10.08376348 | ENSG00000286447 |
| SEMA4C | 0.257880242 | 1.195720536 | 1.62E-06 | 1.23E-05 | 2431.058955 | 2906.703181 | ENSG00000168758 |
| GTF2H2 | 0.257720439 | 1.195588097 | 0.040738835 | 0.101144513 | 292.675809 | 349.730072 | ENSG00000145736 |
| LRRC56 | 0.257272174 | 1.195216669 | 0.383939074 | 0.557261957 | 53.69695089 | 64.32279725 | ENSG00000161328 |
| NA | 0.257239822 | 1.195189867 | 0.376758998 | 0.550235282 | 57.51474577 | 68.50063978 | ENSG00000281091 |
| KMT2D | 0.257152782 | 1.195117762 | 0.00012677 | 0.000668176 | 2014.205641 | 2407.346249 | ENSG00000167548 |
| BTD | 0.257070013 | 1.195049198 | 0.043588414 | 0.106979562 | 283.0723104 | 338.2379216 | ENSG00000169814 |
| XPNPEP2 | 0.257015363 | 1.19500393 | 0.748054312 | 0.850782425 | 6.384860642 | 7.64180187 | ENSG00000122121 |
| LRG1 | 0.256967597 | 1.194964366 | 0.570145164 | 0.723168543 | 20.03593518 | 23.93746819 | ENSG00000171236 |
| C16orf95 | 0.256957776 | 1.194956231 | 0.606936595 | 0.751272767 | 19.07587306 | 22.67104961 | ENSG00000260456 |
| SH3D21 | 0.256950699 | 1.19495037 | 0.056788214 | 0.132147338 | 285.0893652 | 340.3090989 | ENSG00000214193 |
| CLCC1 | 0.256930996 | 1.19493405 | 0.041698625 | 0.103076393 | 282.9877074 | 338.3806898 | ENSG00000121940 |
| MCOLN3 | 0.256831501 | 1.194851644 | 0.503121524 | 0.668630645 | 28.04195415 | 33.52487797 | ENSG00000055732 |
| PYGL | 0.256575771 | 1.194639865 | 0.000143104 | 0.000743502 | 1274.71421 | 1522.344216 | ENSG00000100504 |
| UBA3 | 0.256395372 | 1.194490494 | 0.000308891 | 0.00149531 | 1120.889548 | 1339.333739 | ENSG00000144744 |
| ACACB | 0.256372253 | 1.194471352 | 0.032541669 | 0.083925312 | 413.282923 | 493.4487135 | ENSG00000076555 |
| SLC35D2 | 0.256259863 | 1.194378303 | 0.300710825 | 0.471320485 | 72.87214518 | 86.8610259 | ENSG00000130958 |
| NA | 0.256097565 | 1.194243948 | 0.703327283 | 0.821599044 | 9.029796286 | 10.75703412 | ENSG00000261559 |
| TCTE3 | 0.256032685 | 1.194190242 | 0.599000322 | 0.745585377 | 18.3400982 | 21.88378014 | ENSG00000184786 |
| NA | 0.256004245 | 1.194166701 | 0.781238432 | 0.871562348 | 4.661790352 | 5.569824733 | ENSG00000179978 |
| FAM169A | 0.255877516 | 1.194061808 | 0.007608936 | 0.025044188 | 586.3840803 | 700.1583797 | ENSG00000198780 |
| LINC01574 | 0.255416861 | 1.193680603 | 0.725621436 | 0.835289284 | 7.718492076 | 9.202643856 | ENSG00000248859 |
| SENP2 | 0.255349036 | 1.193624486 | 0.000801194 | 0.003479476 | 842.6181732 | 1006.250538 | ENSG00000163904 |
| NEURL1B | 0.255282424 | 1.193569375 | 0.030673871 | 0.079985083 | 390.8664512 | 466.4998347 | ENSG00000214357 |
| OGA | 0.255275114 | 1.193563327 | 7.80E-07 | 6.25E-06 | 2790.533807 | 3330.678641 | ENSG00000198408 |
| CLCN4 | 0.255268068 | 1.193557498 | 0.06352707 | 0.144359422 | 281.4001648 | 335.4813695 | ENSG00000073464 |
| ADAMTS13 | 0.255053881 | 1.193380312 | 0.148441013 | 0.280056291 | 145.4241116 | 173.5004874 | ENSG00000160323 |
| NPM2 | 0.255048514 | 1.193375873 | 0.502586402 | 0.668279176 | 27.77886966 | 33.11860631 | ENSG00000158806 |
| TMEM108 | 0.254856388 | 1.193216959 | 0.060804812 | 0.139508282 | 229.2813078 | 273.620713 | ENSG00000144868 |
| NA | 0.254792625 | 1.193164224 | 0.316719082 | 0.487849554 | 72.35635195 | 86.23560093 | ENSG00000216866 |
| NA | 0.254754937 | 1.193133055 | 0.704993346 | 0.822564008 | 9.152684031 | 10.88125278 | ENSG00000271869 |
| FAM216A | 0.254750052 | 1.193129014 | 0.100137649 | 0.207358887 | 173.5076539 | 206.8403671 | ENSG00000204856 |
| GCC2 | 0.254659453 | 1.19305409 | 0.001211157 | 0.005070439 | 730.8781171 | 872.0985887 | ENSG00000135968 |
| CAPRIN2 | 0.254565219 | 1.192976165 | 0.012466475 | 0.037872563 | 794.6133456 | 947.3652716 | ENSG00000110888 |
| EGF | 0.254390659 | 1.192831829 | 0.285587149 | 0.454522043 | 78.861785 | 94.12368954 | ENSG00000138798 |
| LOC100507283 | 0.254201165 | 1.192675164 | 0.557360138 | 0.713357435 | 21.83791449 | 26.02181574 | ENSG00000245498 |
| SGK3 | 0.254173927 | 1.192652646 | 0.097994897 | 0.203928756 | 179.2710545 | 213.782831 | ENSG00000104205 |
| NA | 0.254054353 | 1.192553801 | 0.316953334 | 0.488132425 | 89.47908551 | 106.4523369 | ENSG00000234648 |
| IQCG | 0.253909356 | 1.19243395 | 0.35427713 | 0.527383295 | 71.34664129 | 84.88260793 | ENSG00000114473 |
| LOC283856 | 0.253751968 | 1.192303871 | 0.789027976 | 0.876995775 | 4.901108155 | 5.875001002 | ENSG00000246379 |
| C12orf4 | 0.253727133 | 1.192283346 | 0.142095594 | 0.270973319 | 135.9911221 | 162.110771 | ENSG00000047621 |
| LY6E | 0.253704273 | 1.192264454 | 3.21E-05 | 0.000192071 | 1654.489733 | 1972.3866 | ENSG00000160932 |
| LATS1 | 0.253535534 | 1.192125014 | 0.028384053 | 0.07514159 | 336.1768744 | 400.9242973 | ENSG00000131023 |
| FKTN | 0.253527296 | 1.192118207 | 0.075093579 | 0.165048449 | 224.9532503 | 268.2135447 | ENSG00000106692 |
| GPAA1 | 0.253441217 | 1.19204708 | 0.000350528 | 0.001674403 | 1202.984389 | 1434.167666 | ENSG00000197858 |
| APOE | 0.253381641 | 1.191997856 | 2.00E-06 | 1.49E-05 | 16927.87041 | 20179.18967 | ENSG00000130203 |
| CTNND1 | 0.253160039 | 1.191814776 | 0.205260614 | 0.359037764 | 108.6347066 | 129.311862 | ENSG00000198561 |
| SERPINB8 | 0.253118053 | 1.191780091 | 0.720283037 | 0.832084584 | 8.2948973 | 9.932380869 | ENSG00000166401 |
| KRTCAP3 | 0.253101256 | 1.191766216 | 0.008251683 | 0.026810415 | 473.9217667 | 564.8774487 | ENSG00000157992 |
| SLC25A15 | 0.253096713 | 1.191762463 | 0.005249496 | 0.018292634 | 666.1409488 | 793.732539 | ENSG00000102743 |
| ABHD11 | 0.253039724 | 1.191715387 | 0.040836844 | 0.101348714 | 288.9535805 | 344.3992888 | ENSG00000106077 |
| NOTCH1 | 0.252934396 | 1.191628386 | 2.52E-07 | 2.18E-06 | 3030.370728 | 3611.623082 | ENSG00000148400 |
| GBA | 0.252840175 | 1.191550565 | 0.106464123 | 0.21705764 | 189.6955698 | 226.2082542 | ENSG00000177628 |
| COL5A1 | 0.252794789 | 1.19151308 | 0.003685859 | 0.013441193 | 721.9061661 | 859.872709 | ENSG00000130635 |
| IRF5 | 0.252764871 | 1.191488371 | 0.699345632 | 0.818434639 | 10.02075849 | 11.98768645 | ENSG00000128604 |
| CEPT1 | 0.252588299 | 1.191342553 | 0.081703053 | 0.1764197 | 266.9090512 | 318.3752799 | ENSG00000134255 |
| PGBD2 | 0.252540298 | 1.191302916 | 0.174064475 | 0.316900508 | 136.4495457 | 162.7684425 | ENSG00000185220 |
| COX18 | 0.252437752 | 1.191218242 | 0.003923132 | 0.014207083 | 604.2710244 | 719.6690592 | ENSG00000163626 |
| NA | 0.25239527 | 1.191183166 | 0.701688493 | 0.820229901 | 9.454841158 | 11.24719657 | ENSG00000259424 |
| NA | 0.252360212 | 1.19115422 | 0.784860386 | 0.874043418 | 4.700187272 | 5.592948006 | ENSG00000256995 |

| | | | | | | | |
|---|---|---|---|---|---|---|---|
| NA | 0.252276053 | 1.191084736 | 0.000510262 | 0.002334788 | 1209.217863 | 1439.583115 | ENSG00000196205 |
| USP24 | 0.25225289 | 1.191065613 | 0.000667919 | 0.002959306 | 1571.421677 | 1871.484314 | ENSG00000162402 |
| IGF2BP3 | 0.252186049 | 1.191010431 | 0.000106245 | 0.000571548 | 3329.713924 | 3964.701988 | ENSG00000136231 |
| CPVL | 0.251753311 | 1.19065324 | 0.006957751 | 0.023278178 | 557.3518397 | 663.3249841 | ENSG00000106066 |
| CAPN12 | 0.251728143 | 1.190632469 | 0.318537034 | 0.489867501 | 65.96638507 | 78.54056775 | ENSG00000182472 |
| NA | 0.251599794 | 1.190526549 | 0.777016885 | 0.869299514 | 4.988389884 | 5.96534352 | ENSG00000250412 |
| CUEDC2 | 0.251589525 | 1.190518076 | 0.002890606 | 0.010857219 | 924.4690522 | 1100.337426 | ENSG00000107874 |
| MTMR3 | 0.251307945 | 1.190285737 | 0.334360283 | 0.507046947 | 68.79425688 | 82.0596364 | ENSG00000100330 |
| TMEM151A | 0.251248198 | 1.190236445 | 0.483195736 | 0.65106719 | 31.96952123 | 38.09581613 | ENSG00000179292 |
| ZNF783 | 0.251191595 | 1.190189747 | 0.064812784 | 0.146594375 | 250.7716469 | 298.4269489 | ENSG00000204946 |
| NA | 0.251141145 | 1.190148128 | 0.61919522 | 0.760607426 | 16.57930671 | 19.81259651 | ENSG00000278784 |
| KIF13B | 0.251133045 | 1.190141446 | 0.054339369 | 0.127541156 | 283.5289069 | 337.3809913 | ENSG00000197892 |
| NA | 0.250916605 | 1.189962908 | 0.091086945 | 0.192438752 | 226.0539848 | 269.3780623 | ENSG00000196922 |
| NA | 0.250881156 | 1.18993367 | 0.432479389 | 0.603512723 | 44.46623372 | 52.96213489 | ENSG00000286431 |
| DMTF1 | 0.250879988 | 1.189932706 | 0.000895735 | 0.003853618 | 1023.163911 | 1217.587361 | ENSG00000135164 |
| ST3GAL3 | 0.250806966 | 1.18987248 | 0.56481662 | 0.719057442 | 21.87005388 | 25.97126804 | ENSG00000126091 |
| ZNF24 | 0.250719885 | 1.189800661 | 0.000249456 | 0.001233198 | 1464.405238 | 1743.152541 | ENSG00000172466 |
| UBAP1L | 0.250657678 | 1.189749359 | 0.649023843 | 0.781685316 | 14.37201901 | 17.10552602 | ENSG00000246922 |
| IPO9 | 0.250608079 | 1.189708457 | 2.58E-08 | 2.57E-07 | 3719.565514 | 4425.199713 | ENSG00000198700 |
| UCKL1 | 0.250510635 | 1.189628104 | 0.004240584 | 0.015202583 | 647.9977285 | 770.8364663 | ENSG00000198276 |
| SLC19A3 | 0.250301141 | 1.18945537 | 0.19567465 | 0.346149444 | 129.0944473 | 153.3727432 | ENSG00000135917 |
| TMEM128 | 0.250298718 | 1.189453373 | 0.058633506 | 0.135396094 | 232.169228 | 276.1866207 | ENSG00000132406 |
| ZCRB1 | 0.250166443 | 1.189344321 | 0.032166148 | 0.083134669 | 436.143641 | 519.0019088 | ENSG00000139168 |
| ZNF101 | 0.250157895 | 1.189337274 | 0.005902888 | 0.020269246 | 633.5452858 | 753.5786726 | ENSG00000181896 |
| MAP3K9 | 0.250057139 | 1.189254215 | 0.006625927 | 0.022350193 | 825.9535352 | 981.5206755 | ENSG00000006432 |
| SEC31B | 0.2498875 | 1.189114385 | 0.353126277 | 0.526035445 | 55.9896053 | 66.62843026 | ENSG00000075826 |
| LRCH3 | 0.249746805 | 1.188998426 | 0.001092795 | 0.004614953 | 865.6624746 | 1028.795231 | ENSG00000186001 |
| ACBD4 | 0.248993173 | 1.188377481 | 0.093117501 | 0.195721258 | 198.282849 | 235.4643898 | ENSG00000181513 |
| ETFB | 0.248987122 | 1.188372497 | 0.330210891 | 0.502574863 | 70.14328229 | 83.46721777 | ENSG00000105379 |
| HCFC1R1 | 0.248936794 | 1.188331042 | 0.020635247 | 0.057691964 | 391.9256102 | 466.0709422 | ENSG00000103145 |
| EIF5AL1 | 0.24892163 | 1.188318552 | 0.473285232 | 0.64197338 | 37.85832513 | 44.93797813 | ENSG00000253626 |
| CRIP3 | 0.248889622 | 1.188292187 | 0.176805221 | 0.320438699 | 173.4661027 | 205.6522238 | ENSG00000146215 |
| ZNF562 | 0.248719533 | 1.1881521 | 0.052195282 | 0.123350208 | 311.7783518 | 370.32322 | ENSG00000171466 |
| CHKA | 0.248637512 | 1.188084552 | 0.007192864 | 0.023907226 | 678.1103524 | 805.7371574 | ENSG00000110721 |
| SOX5 | 0.248484359 | 1.187958435 | 0.605299246 | 0.7500773 | 17.15937872 | 20.32562177 | ENSG00000134532 |
| FAM86JP | 0.248263202 | 1.187776342 | 0.560909646 | 0.715878794 | 27.18591918 | 32.12563843 | ENSG00000171084 |
| LOC105374042 | 0.248224623 | 1.18774458 | 0.754356473 | 0.854310609 | 6.733111675 | 8.016347958 | ENSG00000239482 |
| SNX19 | 0.248153928 | 1.187686379 | 0.002512433 | 0.009627999 | 696.1227878 | 826.4362834 | ENSG00000120451 |
| TMPRSS2 | 0.247996407 | 1.187556708 | 0.059778205 | 0.137496287 | 236.5728399 | 281.0221347 | ENSG00000184012 |
| MRPS35 | 0.247861751 | 1.187445872 | 0.003012953 | 0.011240161 | 835.0290776 | 992.06101 | ENSG00000061794 |
| CREB3L1 | 0.247691686 | 1.187305904 | 0.058520053 | 0.135247436 | 250.2342022 | 297.1470754 | ENSG00000157613 |
| MECR | 0.247665828 | 1.187284623 | 0.008772765 | 0.028201201 | 573.7490492 | 681.1449632 | ENSG00000116353 |
| TLE5 | 0.247649621 | 1.187271285 | 1.74E-06 | 1.31E-05 | 3212.901467 | 3814.340553 | ENSG00000104964 |
| UBN2 | 0.247585614 | 1.187218612 | 0.017990379 | 0.051445656 | 439.1625704 | 521.6855408 | ENSG00000157741 |
| TRIP10 | 0.247460002 | 1.187115248 | 1.07E-05 | 6.99E-05 | 2064.031594 | 2450.709211 | ENSG00000125733 |
| MARS2 | 0.247350677 | 1.187025294 | 0.001546752 | 0.006289712 | 789.2424771 | 936.507421 | ENSG00000247626 |
| ZNF565 | 0.247304007 | 1.186986895 | 0.510910734 | 0.67539087 | 31.16129503 | 37.07112268 | ENSG00000196357 |
| REST | 0.247272522 | 1.186960991 | 0.007009464 | 0.023406537 | 629.1653284 | 746.8833316 | ENSG00000084093 |
| ZDHHC17 | 0.247182535 | 1.186886957 | 0.022230854 | 0.061335155 | 413.5582613 | 491.0234292 | ENSG00000186908 |
| SRP9 | 0.247174046 | 1.186879974 | 8.33E-07 | 6.64E-06 | 5117.920757 | 6075.604328 | ENSG00000143742 |
| HROB | 0.24704091 | 1.18677045 | 0.029451695 | 0.077341771 | 341.6061786 | 405.3487297 | ENSG00000125319 |
| SIKE1 | 0.246988125 | 1.18672703 | 0.001885181 | 0.007499904 | 749.3328936 | 889.643469 | ENSG00000052723 |
| UXT | 0.246789011 | 1.186563254 | 0.210808115 | 0.366369311 | 117.1517759 | 139.2740783 | ENSG00000126756 |
| IQCA1 | 0.246590636 | 1.18640011 | 0.165610974 | 0.305051208 | 140.8658728 | 166.8979 | ENSG00000132321 |
| TOX3 | 0.24643308 | 1.18627055 | 0.073969995 | 0.163043325 | 282.2724569 | 334.9164395 | ENSG00000103460 |
| SBF2 | 0.246389021 | 1.186234323 | 0.004175233 | 0.01499617 | 767.7768926 | 910.7587471 | ENSG00000133812 |
| B4GALT5 | 0.246365859 | 1.186215279 | 1.36E-05 | 8.71E-05 | 2459.215135 | 2916.475788 | ENSG00000158470 |
| MTMR6 | 0.246322202 | 1.186179383 | 0.075145822 | 0.165121253 | 265.7663283 | 315.7631441 | ENSG00000139505 |
| GUSBP2 | 0.24626372 | 1.186131301 | 0.479039537 | 0.647184796 | 35.66845857 | 42.15508611 | ENSG00000241549 |
| ORC3 | 0.246017125 | 1.185928577 | 0.002463908 | 0.009477818 | 873.5471205 | 1036.263811 | ENSG00000135336 |
| PRDM2 | 0.24590345 | 1.185835137 | 0.009038468 | 0.028895005 | 537.6213281 | 637.3531646 | ENSG00000116731 |
| EEF1AKNMT | 0.245860211 | 1.185799596 | 5.93E-06 | 4.07E-05 | 2621.529414 | 3109.607382 | ENSG00000010165 |
| CDC42SE2 | 0.245733004 | 1.185695046 | 0.006537658 | 0.022090699 | 586.4024284 | 695.4137718 | ENSG00000158985 |
| CHST4 | 0.245703142 | 1.185670503 | 0.108935585 | 0.220836105 | 243.6090599 | 289.2204943 | ENSG00000140835 |
| LANCL1 | 0.2454477 | 1.185460588 | 0.0005741 | 0.002588215 | 947.8885881 | 1123.483566 | ENSG00000115365 |
| RPAP1 | 0.245381857 | 1.185406487 | 7.59E-05 | 0.000421413 | 1808.265623 | 2143.36364 | ENSG00000103932 |
| NA | 0.24533128 | 1.18536493 | 0.640678354 | 0.775170981 | 17.44267525 | 20.64155091 | ENSG00000272078 |
| CYB5R1 | 0.245249321 | 1.185297592 | 0.073720562 | 0.162586414 | 242.5706523 | 287.8062128 | ENSG00000159348 |
| ZBTB8B | 0.24518246 | 1.185242661 | 0.265451189 | 0.431481201 | 94.71129897 | 112.282951 | ENSG00000273274 |
| HEATR3 | 0.245004485 | 1.185096455 | 0.010943303 | 0.03400568 | 472.7985049 | 560.1414126 | ENSG00000155393 |
| CHEK2 | 0.244888185 | 1.185000925 | 0.000311782 | 0.001508171 | 1388.135421 | 1644.697209 | ENSG00000183765 |
| DDX6 | 0.244671432 | 1.184822901 | 7.46E-05 | 0.000415159 | 2040.335631 | 2418.179335 | ENSG00000110367 |
| LOC106660606 | 0.244612393 | 1.184774417 | 0.748856082 | 0.851227545 | 6.643039044 | 7.889978725 | ENSG00000260778 |
| MOB4 | 0.244547797 | 1.18472137 | 0.144670592 | 0.274689015 | 344.4349647 | 408.4052961 | ENSG00000115540 |
| NEIL1 | 0.244525047 | 1.184702688 | 0.272372459 | 0.439134203 | 105.8221407 | 125.1552768 | ENSG00000140398 |
| ATG10 | 0.244499824 | 1.184681975 | 0.134765962 | 0.259977018 | 173.6085778 | 205.7777568 | ENSG00000152348 |
| TMTC4 | 0.244386658 | 1.184589052 | 0.001897842 | 0.007542495 | 832.240101 | 985.3425336 | ENSG00000125247 |

| | | | | | | | |
|---|---|---|---|---|---|---|---|
| NA | 0.244328046 | 1.184540927 | 0.828901496 | 0.902644017 | 3.039956306 | 3.575819709 | ENSG00000261557 |
| PTCH1 | 0.244291851 | 1.184511209 | 0.034341708 | 0.087688404 | 354.5341183 | 419.6210949 | ENSG00000185920 |
| SCHIP1 | 0.24426695 | 1.184490764 | 0.714635448 | 0.828587115 | 9.389210931 | 11.08533366 | ENSG00000151967 |
| LOC105371083 | 0.244214023 | 1.184447311 | 0.753343052 | 0.853912734 | 6.996871883 | 8.329593484 | ENSG00000263080 |
| FAM193B | 0.244197013 | 1.184433346 | 0.006097685 | 0.020830607 | 956.5927232 | 1132.99762 | ENSG00000146067 |
| ZFP91 | 0.244137728 | 1.184384675 | 0.020304545 | 0.056940687 | 456.3125004 | 540.9824469 | ENSG00000186660 |
| NA | 0.244021409 | 1.184289186 | 0.653719359 | 0.784845666 | 13.0579239 | 15.46241078 | ENSG00000225850 |
| PIK3R5 | 0.243841859 | 1.184141805 | 0.07876255 | 0.171316761 | 217.1476767 | 257.1169252 | ENSG00000141506 |
| ACTR1B | 0.243839694 | 1.184140028 | 0.001067037 | 0.004517055 | 951.8885313 | 1126.492087 | ENSG00000115073 |
| CHSY1 | 0.24382486 | 1.184127852 | 0.000346796 | 0.001658415 | 1291.349685 | 1529.210313 | ENSG00000131873 |
| TRMO | 0.243677281 | 1.18400673 | 0.387519796 | 0.560897688 | 61.63119425 | 73.07910205 | ENSG00000136932 |
| LINC01842 | 0.243624211 | 1.183963176 | 0.817825899 | 0.895882273 | 4.713466065 | 5.565523585 | ENSG00000267147 |
| PLEC | 0.243595767 | 1.183939834 | 7.99E-06 | 5.37E-05 | 4833.807885 | 5722.965961 | ENSG00000178209 |
| CFAP77 | 0.243555465 | 1.18390676 | 0.683302574 | 0.806854989 | 11.63142893 | 13.79670529 | ENSG00000188523 |
| UBE2E2 | 0.243513504 | 1.183872327 | 0.012396836 | 0.03768474 | 600.5187846 | 710.8100294 | ENSG00000182247 |
| GPSM2 | 0.24343774 | 1.183810157 | 0.121281414 | 0.24040688 | 227.8761009 | 269.7757132 | ENSG00000121957 |
| SMAD6 | 0.243430785 | 1.18380445 | 0.087458869 | 0.186386328 | 200.3386642 | 237.0876169 | ENSG00000137834 |
| GTF2H5 | 0.243429017 | 1.183802999 | 0.022811783 | 0.062686909 | 416.0951396 | 492.9171387 | ENSG00000272047 |
| NA | 0.243349469 | 1.183737728 | 0.006715518 | 0.022612858 | 1325.149318 | 1567.785538 | ENSG00000236673 |
| MAST4 | 0.243270209 | 1.183672697 | 0.212737042 | 0.368744168 | 114.9900186 | 136.1505163 | ENSG00000069020 |
| MORC2 | 0.243247677 | 1.18365421 | 4.66E-05 | 0.000269279 | 2283.374828 | 2702.213124 | ENSG00000133422 |
| ZNF581 | 0.243208412 | 1.183621996 | 0.002864414 | 0.010777727 | 773.4217622 | 915.9277221 | ENSG00000171425 |
| NA | 0.243166331 | 1.183587472 | 0.75187086 | 0.85314633 | 6.651411752 | 7.887828151 | ENSG00000244363 |
| NA | 0.243137351 | 1.183563697 | 0.503690492 | 0.66897596 | 36.59658145 | 43.22440857 | ENSG00000254536 |
| LOC441204 | 0.24311511 | 1.183545451 | 0.828122612 | 0.902152321 | 3.284855915 | 3.89766753 | ENSG00000214870 |
| TMEM220 | 0.24308174 | 1.183518076 | 0.431590152 | 0.602593258 | 41.14106586 | 48.70738632 | ENSG00000187824 |
| SNAI3-AS1 | 0.243001415 | 1.183452183 | 0.343223113 | 0.516025024 | 66.48660881 | 78.59137591 | ENSG00000260630 |
| SGCB | 0.242957863 | 1.183416457 | 0.032287797 | 0.08339321 | 660.3124829 | 782.2406 | ENSG00000163069 |
| CLIP2 | 0.242885826 | 1.183357368 | 5.98E-05 | 0.000339606 | 1931.473095 | 2285.700216 | ENSG00000106665 |
| EGFR | 0.24288403 | 1.183355894 | 0.128062643 | 0.250071511 | 160.873899 | 190.2500169 | ENSG00000146648 |
| RBM26-AS1 | 0.242588327 | 1.183113373 | 0.822856796 | 0.899004953 | 4.490893679 | 5.271100185 | ENSG00000227354 |
| COMMD6 | 0.242572018 | 1.183099997 | 0.01909895 | 0.054057702 | 441.5185156 | 522.701923 | ENSG00000188243 |
| NA | 0.242502862 | 1.183043287 | 0.73854887 | 0.844637554 | 8.09676732 | 9.559985354 | ENSG00000226964 |
| UHRF1BP1L | 0.242366905 | 1.182931805 | 0.024910829 | 0.067427361 | 455.6180624 | 538.795138 | ENSG00000111647 |
| SLC22A17 | 0.24236265 | 1.182928315 | 0.000405276 | 0.001906041 | 1159.999805 | 1372.234368 | ENSG00000092096 |
| TMEM181 | 0.242331012 | 1.182902374 | 0.011601011 | 0.035704388 | 567.2896973 | 671.3182209 | ENSG00000146433 |
| PARP10 | 0.242250358 | 1.182836246 | 0.306088029 | 0.477112489 | 86.29909573 | 102.2403273 | ENSG00000178685 |
| MAP3K2 | 0.242195757 | 1.18279148 | 0.126592 | 0.248028262 | 211.1786123 | 249.8431992 | ENSG00000169967 |
| INSC | 0.24204489 | 1.182667799 | 0.631004165 | 0.76847862 | 16.16071953 | 19.07210663 | ENSG00000188487 |
| RHOBTB1 | 0.241935715 | 1.182578305 | 0.05482577 | 0.128510805 | 285.9690547 | 338.3115925 | ENSG00000072422 |
| SLC36A4 | 0.241901973 | 1.182550646 | 0.033422494 | 0.085738474 | 353.765128 | 418.5511905 | ENSG00000180773 |
| SHC3 | 0.241865105 | 1.182520427 | 0.065021522 | 0.146957838 | 252.8843562 | 298.9872747 | ENSG00000148082 |
| HELLS | 0.2418524 | 1.182510013 | 1.28E-05 | 8.24E-05 | 2035.683076 | 2407.923562 | ENSG00000119969 |
| CRTC2 | 0.24164899 | 1.182343299 | 0.009802708 | 0.030924617 | 542.6731961 | 641.6568738 | ENSG00000160741 |
| NA | 0.241335171 | 1.182086141 | 0.831594721 | 0.904389657 | 3.069980516 | 3.598942982 | ENSG00000286608 |
| ZSCAN26 | 0.241312175 | 1.182067299 | 0.236220655 | 0.397603408 | 110.0297379 | 129.9821764 | ENSG00000197062 |
| DZIP1L | 0.241308544 | 1.182064323 | 0.169884456 | 0.311022211 | 155.1088587 | 183.094882 | ENSG00000158163 |
| KCNK6 | 0.24117656 | 1.181956188 | 0.023912964 | 0.065137672 | 437.8062994 | 517.854057 | ENSG00000099337 |
| SNHG16 | 0.241022838 | 1.181830256 | 0.002416192 | 0.009314702 | 918.1569793 | 1084.887895 | ENSG00000163597 |
| BCAR1 | 0.240902949 | 1.181732049 | 0.000176154 | 0.000897565 | 2872.56302 | 3395.136635 | ENSG00000050820 |
| PTPN21 | 0.240727279 | 1.181588164 | 0.071803973 | 0.159196696 | 235.8381411 | 278.468337 | ENSG00000070778 |
| ARFGEF1 | 0.240713419 | 1.181576812 | 5.32E-05 | 0.000304516 | 2542.544312 | 3004.684269 | ENSG00000066777 |
| MATK | 0.240689679 | 1.181557369 | 0.006424455 | 0.021754284 | 677.0448506 | 800.2017845 | ENSG00000007264 |
| LOC100506411 | 0.240540809 | 1.181435452 | 0.227082088 | 0.38633394 | 132.0553825 | 155.8617754 | ENSG00000259153 |
| SPC24 | 0.240470897 | 1.181378201 | 0.01059692 | 0.033065672 | 619.9631207 | 731.9802416 | ENSG00000161888 |
| ZNF611 | 0.240374463 | 1.181299237 | 0.248847866 | 0.412735291 | 99.02974437 | 116.9498899 | ENSG00000213020 |
| LPIN2 | 0.240370186 | 1.181295735 | 0.033694869 | 0.086310943 | 368.9046978 | 435.5965188 | ENSG00000101577 |
| MAPK11 | 0.240345327 | 1.18127538 | 0.427768787 | 0.599149617 | 51.11764547 | 60.41733304 | ENSG00000185386 |
| NA | 0.240088319 | 1.181064961 | 0.772991491 | 0.866695002 | 6.357627334 | 7.488008217 | ENSG00000154898 |
| SLC26A2 | 0.240075793 | 1.181054707 | 0.09593441 | 0.200571226 | 243.5816392 | 287.8997599 | ENSG00000155850 |
| KCNA1 | 0.24003566 | 1.181021853 | 0.800174329 | 0.884896069 | 5.363874225 | 6.358711733 | ENSG00000111262 |
| OLFML2A | 0.23993522 | 1.180939633 | 0.080477326 | 0.174260647 | 249.4586279 | 294.4813867 | ENSG00000185585 |
| PCNT | 0.239732974 | 1.180774093 | 0.001603823 | 0.0064985 | 1125.496494 | 1327.923971 | ENSG00000160299 |
| DUSP14 | 0.239589421 | 1.180656609 | 0.022915362 | 0.062890915 | 437.169207 | 515.8178398 | ENSG00000276023 |
| TMEM183A | 0.239556224 | 1.180629442 | 3.97E-06 | 2.81E-05 | 2426.476738 | 2864.184822 | ENSG00000163444 |
| NA | 0.239416589 | 1.180515177 | 0.682728225 | 0.806522364 | 11.75431667 | 13.87844552 | ENSG00000213885 |
| URI1 | 0.239293 | 1.180414052 | 0.000690341 | 0.003046754 | 1373.013785 | 1619.841583 | ENSG00000105176 |
| INTS2 | 0.239244555 | 1.180374415 | 0.028729661 | 0.075866446 | 421.1262689 | 497.4342819 | ENSG00000108506 |
| SLC13A5 | 0.239203765 | 1.180341042 | 0.546961344 | 0.70479907 | 28.88088028 | 34.11560277 | ENSG00000141485 |
| POGLUT3 | 0.239107144 | 1.180261994 | 0.062763112 | 0.142997897 | 327.7661858 | 387.4494278 | ENSG00000178202 |
| FBXL18 | 0.23903704 | 1.180204644 | 0.021748138 | 0.06021865 | 442.1156453 | 521.5070674 | ENSG00000155034 |
| NA | 0.23890266 | 1.180094718 | 0.504918093 | 0.670016685 | 36.61524189 | 43.22926118 | ENSG00000214820 |
| ETFBKMT | 0.238714129 | 1.179940514 | 0.679147643 | 0.803872602 | 15.75541113 | 18.49347966 | ENSG00000139160 |
| NA | 0.23871288 | 1.179939492 | 0.526527816 | 0.688570564 | 29.48798544 | 34.74289943 | ENSG00000269825 |
| PPP1R9A | 0.23869944 | 1.179928501 | 0.072080698 | 0.159736789 | 293.581917 | 346.235429 | ENSG00000158528 |
| NA | 0.23852066 | 1.179782292 | 0.832843014 | 0.905198861 | 3.663131159 | 4.345884584 | ENSG00000250049 |
| SYT7 | 0.238479932 | 1.179748986 | 0.014533948 | 0.043101368 | 552.2487346 | 651.3523701 | ENSG00000011347 |

| | | | | | | | |
|---|---|---|---|---|---|---|---|
| HEATR5A | 0.238452212 | 1.179726319 | 0.102410068 | 0.210887342 | 211.0721946 | 248.8857434 | ENSG00000129493 |
| ZNF710 | 0.238404839 | 1.179687582 | 0.022674525 | 0.062367057 | 643.6527525 | 759.5997437 | ENSG00000140548 |
| NA | 0.238397961 | 1.179681958 | 0.838044129 | 0.907681168 | 3.388207339 | 3.994461769 | ENSG00000235215 |
| RGPD2 | 0.238377268 | 1.179665037 | 0.54289425 | 0.701668363 | 33.88621574 | 39.85239717 | ENSG00000185304 |
| PDE3A | 0.23830383 | 1.17960499 | 0.10587638 | 0.216247882 | 197.670162 | 232.9060184 | ENSG00000172572 |
| NA | 0.238289486 | 1.179593262 | 0.733898367 | 0.841314084 | 8.341666928 | 9.842038351 | ENSG00000275029 |
| NQO2 | 0.238142717 | 1.179473265 | 0.00041154 | 0.001930228 | 1215.985729 | 1433.875841 | ENSG00000124588 |
| ACP4 | 0.238021072 | 1.179373818 | 0.793222568 | 0.879731894 | 5.081253418 | 5.988466793 | ENSG00000142513 |
| TYW5 | 0.237998725 | 1.17935555 | 0.11630079 | 0.23290341 | 200.4161338 | 236.0946611 | ENSG00000162971 |
| NA | 0.237931171 | 1.179300328 | 0.831503061 | 0.904389657 | 3.393789145 | 3.998762916 | ENSG00000259376 |
| BBX | 0.237904186 | 1.17927827 | 0.005352114 | 0.018596476 | 790.6239542 | 932.2139195 | ENSG00000114439 |
| C6orf52 | 0.23789202 | 1.179268325 | 0.611328833 | 0.754893995 | 18.66111592 | 22.0375738 | ENSG00000137434 |
| SPOCK1 | 0.237873641 | 1.179253302 | 0.039601875 | 0.098879359 | 376.2925602 | 443.329723 | ENSG00000152377 |
| RHOB | 0.237850408 | 1.179234312 | 0.068485572 | 0.153302654 | 365.8717259 | 431.2705588 | ENSG00000143878 |
| NA | 0.237729842 | 1.179135767 | 0.744251563 | 0.848593706 | 8.102349125 | 9.522341103 | ENSG00000230170 |
| NFU1 | 0.23761079 | 1.179038468 | 0.043168914 | 0.106085543 | 416.8571087 | 491.5963737 | ENSG00000169599 |
| NARF | 0.237602928 | 1.179032044 | 0.00068759 | 0.003036003 | 1212.19568 | 1428.555338 | ENSG00000141562 |
| GPR161 | 0.237590164 | 1.179021612 | 0.018008759 | 0.051475324 | 591.6047298 | 697.8380021 | ENSG00000143147 |
| TCP11L1 | 0.237570368 | 1.179005434 | 0.138756843 | 0.26592075 | 183.1643671 | 216.0446285 | ENSG00000176148 |
| ROBO1 | 0.237547426 | 1.178986685 | 1.53E-05 | 9.70E-05 | 2546.567196 | 3001.852308 | ENSG00000169855 |
| PROCA1 | 0.237519883 | 1.178964177 | 0.083049933 | 0.178833574 | 239.6541842 | 282.2724269 | ENSG00000167525 |
| KCNN3 | 0.237481911 | 1.178933148 | 0.327892659 | 0.500098897 | 74.97752091 | 88.47508708 | ENSG00000143603 |
| ZNF519 | 0.237420196 | 1.178882716 | 0.112624963 | 0.227098166 | 187.6888396 | 221.4273044 | ENSG00000175322 |
| USP20 | 0.237416611 | 1.178879787 | 0.027093472 | 0.072390606 | 448.5629818 | 528.4696679 | ENSG00000136878 |
| STMN1 | 0.237211244 | 1.178711986 | 4.45E-07 | 3.70E-06 | 5973.878512 | 7040.332325 | ENSG00000117632 |
| IL15RA | 0.237154164 | 1.178665351 | 0.597992348 | 0.744849761 | 21.87264462 | 25.81155571 | ENSG00000134470 |
| FAM135A | 0.237112841 | 1.178631591 | 0.029945331 | 0.078435217 | 534.3986741 | 630.4934949 | ENSG00000082269 |
| MYO9A | 0.236919313 | 1.178473496 | 0.049863427 | 0.119063667 | 317.6780822 | 374.7048525 | ENSG00000066933 |
| TRIM46 | 0.236859685 | 1.178424789 | 0.026836615 | 0.071793808 | 480.7336625 | 566.7676869 | ENSG00000163462 |
| NA | 0.236549823 | 1.178171715 | 0.837699826 | 0.907563032 | 3.344904336 | 3.939612928 | ENSG00000238230 |
| TUBA4A | 0.236485548 | 1.178119226 | 0.001694974 | 0.006827675 | 981.6214916 | 1156.148358 | ENSG00000127824 |
| DNAJB14 | 0.236482952 | 1.178117107 | 0.043229585 | 0.106207581 | 369.8588658 | 435.8882218 | ENSG00000164031 |
| ANTKMT | 0.236308829 | 1.177974924 | 0.041916298 | 0.103521611 | 387.1822832 | 456.5674354 | ENSG00000103254 |
| MAP3K1 | 0.23622502 | 1.177906495 | 0.051291373 | 0.121824905 | 304.9988339 | 359.2594042 | ENSG00000095015 |
| NA | 0.235968307 | 1.177696918 | 0.84008755 | 0.908905319 | 3.069980516 | 3.58227143 | ENSG00000259169 |
| TSG101 | 0.235836023 | 1.177588937 | 0.005811513 | 0.019991065 | 972.4240538 | 1145.164288 | ENSG00000074319 |
| HOMER3 | 0.235784476 | 1.177546863 | 0.029234697 | 0.076887055 | 382.0558487 | 449.8309609 | ENSG00000051128 |
| NRBP2 | 0.235676708 | 1.177458904 | 0.096600929 | 0.201636931 | 206.8436217 | 243.4543979 | ENSG00000185189 |
| SLC38A6 | 0.235674904 | 1.177457432 | 0.41910785 | 0.591095886 | 50.00255622 | 58.9788196 | ENSG00000139974 |
| MUC1 | 0.235441009 | 1.177266554 | 0.582498382 | 0.732910011 | 22.53633158 | 26.60636139 | ENSG00000185499 |
| PAAF1 | 0.23536205 | 1.177202123 | 0.032831735 | 0.08452309 | 388.6555847 | 457.7657986 | ENSG00000175575 |
| GLTP | 0.235123034 | 1.177007109 | 0.016286636 | 0.047367028 | 505.6708307 | 594.8558457 | ENSG00000139433 |
| GALNTL6 | 0.23470896 | 1.176669339 | 0.714021877 | 0.828174365 | 10.86092393 | 12.85213453 | ENSG00000174473 |
| USP8 | 0.234690375 | 1.176654182 | 0.002090767 | 0.008207772 | 847.0611828 | 996.6424164 | ENSG00000138592 |
| FAM76A | 0.234687941 | 1.176652197 | 0.118361932 | 0.236002071 | 195.0462022 | 229.1957602 | ENSG00000009780 |
| DCDC2C | 0.234658379 | 1.176628086 | 0.821741828 | 0.898499212 | 3.95970648 | 4.709677803 | ENSG00000214866 |
| NA | 0.234402591 | 1.17641949 | 0.118537053 | 0.236180365 | 219.4440859 | 257.7751539 | ENSG00000231698 |
| PSAP | 0.234401847 | 1.176418883 | 8.75E-09 | 9.40E-08 | 13218.41539 | 15549.77579 | ENSG00000197746 |
| MAFA-AS1 | 0.23437587 | 1.176397701 | 0.710935944 | 0.825872368 | 10.25728539 | 12.08852138 | ENSG00000254338 |
| PAN2 | 0.234359691 | 1.176384509 | 0.031036368 | 0.080755674 | 511.9964677 | 601.9746388 | ENSG00000135473 |
| PELI2 | 0.234354181 | 1.176380016 | 0.075802839 | 0.166134124 | 256.5369995 | 301.7895218 | ENSG00000139946 |
| ANAPC16 | 0.23420322 | 1.176256928 | 0.011024043 | 0.034229005 | 591.3477155 | 695.5603018 | ENSG00000166295 |
| SLC35B3 | 0.23411497 | 1.176184978 | 0.130579627 | 0.253676897 | 183.8581662 | 216.4490101 | ENSG00000124786 |
| ZNF417 | 0.234111747 | 1.17618235 | 0.220479532 | 0.378741821 | 143.3463326 | 169.0166325 | ENSG00000173480 |
| FOXG1 | 0.234078423 | 1.176155182 | 0.792404592 | 0.879178523 | 5.369456031 | 6.333437887 | ENSG00000176165 |
| ZBED4 | 0.233972908 | 1.176069165 | 9.54E-05 | 0.000518272 | 1541.877723 | 1813.708795 | ENSG00000100426 |
| SLX4IP | 0.233958538 | 1.176057451 | 0.192735174 | 0.342204716 | 140.890991 | 165.7828398 | ENSG00000149346 |
| PIP4P2 | 0.233879074 | 1.175992675 | 0.058804773 | 0.135742836 | 375.6386116 | 442.4394438 | ENSG00000155099 |
| BCDIN3D | 0.23383537 | 1.175957051 | 0.613405707 | 0.7565507 | 24.08483841 | 28.47479068 | ENSG00000186666 |
| NA | 0.233404506 | 1.175605901 | 0.825110969 | 0.900593514 | 3.693155369 | 4.362556136 | ENSG00000274080 |
| TNFAIP1 | 0.233367086 | 1.175575409 | 0.031406593 | 0.081521068 | 433.2097007 | 509.2181906 | ENSG00000109079 |
| RAB12 | 0.233284086 | 1.175507779 | 0.04894101 | 0.117340421 | 329.341087 | 386.9663048 | ENSG00000206418 |
| CXADR | 0.233166266 | 1.175411783 | 0.003645161 | 0.013310392 | 1778.864923 | 2091.726339 | ENSG00000154639 |
| NA | 0.233155687 | 1.175403164 | 0.732494176 | 0.84020366 | 8.624287735 | 10.15528388 | ENSG00000267053 |
| OLMALINC | 0.23314429 | 1.175393879 | 0.007800997 | 0.02558902 | 676.3792345 | 794.6055772 | ENSG00000235823 |
| PLCD1 | 0.232953591 | 1.175238522 | 0.033052552 | 0.084967003 | 387.163972 | 454.8936485 | ENSG00000187091 |
| MEF2A | 0.232730434 | 1.17505675 | 0.074948365 | 0.164810576 | 303.4958205 | 356.7631205 | ENSG00000068305 |
| MCRIP2 | 0.232646908 | 1.174988721 | 0.004881492 | 0.0171841 | 829.4928673 | 975.0708583 | ENSG00000172366 |
| INAFM2 | 0.232582062 | 1.174935909 | 0.00215343 | 0.008421209 | 830.6833478 | 976.221097 | ENSG00000259330 |
| MTSS2 | 0.232420549 | 1.174804379 | 0.070228453 | 0.156526347 | 353.0295774 | 414.7180891 | ENSG00000132613 |
| NA | 0.232257887 | 1.17467193 | 0.840979896 | 0.90907448 | 3.388207339 | 3.971338496 | ENSG00000236534 |
| FANCF | 0.232232643 | 1.174651375 | 0.05040529 | 0.120045489 | 335.8259957 | 394.8586825 | ENSG00000183161 |
| C19orf84 | 0.232175578 | 1.174604914 | 0.344967008 | 0.517524352 | 72.38705188 | 85.03451148 | ENSG00000262874 |
| SDK1 | 0.232089581 | 1.174534899 | 0.048108057 | 0.115789461 | 417.8846649 | 490.3273079 | ENSG00000146555 |
| NA | 0.231724663 | 1.174237848 | 0.444312489 | 0.615130776 | 47.52505063 | 55.93107917 | ENSG00000178458 |
| DCPS | 0.231577029 | 1.174117691 | 0.017286686 | 0.049787618 | 490.3089255 | 575.3223339 | ENSG00000110063 |
| ARL14EP | 0.231529651 | 1.174079134 | 0.034078002 | 0.087118836 | 467.0109131 | 548.2034619 | ENSG00000152219 |

| | | | | | | | |
|---|---|---|---|---|---|---|---|
| EP400 | 0.23149675 | 1.17405236 | 3.12E-05 | 0.000187539 | 2395.215684 | 2811.504164 | ENSG00000183495 |
| TRIQK | 0.231468928 | 1.174029718 | 0.148431016 | 0.280056291 | 190.978126 | 224.2179868 | ENSG00000205133 |
| FAM72B | 0.231353236 | 1.173935575 | 0.055314635 | 0.129482212 | 312.3053597 | 366.8622048 | ENSG00000188610 |
| FUT11 | 0.231345171 | 1.173929012 | 0.179153126 | 0.323569231 | 147.4846696 | 173.0573529 | ENSG00000196968 |
| ROMO1 | 0.231308342 | 1.173899045 | 0.006786887 | 0.022814368 | 1012.68747 | 1188.096497 | ENSG00000125995 |
| GNPTG | 0.231037033 | 1.173678306 | 0.028524575 | 0.075420447 | 401.4658181 | 470.979891 | ENSG00000090581 |
| REXO1 | 0.230745066 | 1.173440805 | 0.001763058 | 0.007072398 | 1016.799401 | 1193.56201 | ENSG00000079313 |
| EMID1 | 0.230646323 | 1.173360493 | 0.206773507 | 0.36117962 | 128.1431213 | 150.2088534 | ENSG00000186998 |
| NA | 0.230403432 | 1.173162964 | 0.644817172 | 0.778274921 | 16.85443069 | 19.72172095 | ENSG00000233830 |
| RPS6KB2 | 0.230316151 | 1.173091991 | 0.01156161 | 0.035628596 | 569.7582794 | 667.9873735 | ENSG00000175634 |
| VAMP1 | 0.230300943 | 1.173079625 | 0.282262536 | 0.450635717 | 134.5719607 | 157.4917388 | ENSG00000139190 |
| NA | 0.230207741 | 1.173003844 | 0.401810515 | 0.574214879 | 55.01538851 | 64.53386289 | ENSG00000214176 |
| TBL1XR1 | 0.230159126 | 1.172964317 | 0.0172239 | 0.049636428 | 675.4437042 | 792.8196448 | ENSG00000177565 |
| NA | 0.230096419 | 1.172913336 | 0.765156598 | 0.861065676 | 7.334635026 | 8.636920327 | ENSG00000280145 |
| CNIH4 | 0.230050639 | 1.172876117 | 0.002724632 | 0.010326304 | 1520.664448 | 1783.873638 | ENSG00000143771 |
| COG7 | 0.229879471 | 1.172736969 | 0.165656505 | 0.305087593 | 163.4564709 | 191.6775049 | ENSG00000168434 |
| FBXL22 | 0.229858544 | 1.172719958 | 0.764991409 | 0.860961539 | 6.727529869 | 7.866855452 | ENSG00000197361 |
| FBXL2 | 0.229631039 | 1.172535042 | 0.36443156 | 0.537974382 | 69.98230992 | 81.92708802 | ENSG00000153558 |
| CEP164 | 0.229562275 | 1.172479156 | 0.086456985 | 0.184739617 | 347.0950189 | 407.2935504 | ENSG00000110274 |
| POLR2J3 | 0.229534724 | 1.172456765 | 0.772637594 | 0.866415848 | 7.0136173 | 8.230648672 | ENSG00000168255 |
| LIMS2 | 0.229500569 | 1.172429008 | 0.415571362 | 0.587774102 | 49.86658984 | 58.42520601 | ENSG00000072163 |
| MPV17 | 0.229441948 | 1.17238137 | 0.005980244 | 0.02047655 | 786.0947516 | 921.3509863 | ENSG00000115204 |
| RNF13 | 0.229225182 | 1.172205232 | 0.063113444 | 0.143643399 | 330.4524342 | 387.180042 | ENSG00000082996 |
| NA | 0.229192366 | 1.172178569 | 0.222484678 | 0.380998839 | 115.3918603 | 135.3734667 | ENSG00000250462 |
| EDNRB | 0.229103186 | 1.172106113 | 0.097892922 | 0.203760505 | 254.7656449 | 298.3594265 | ENSG00000136160 |
| NMD3 | 0.229098014 | 1.172101911 | 0.001658423 | 0.006694419 | 936.7136308 | 1098.078003 | ENSG00000169251 |
| GDF9 | 0.229091492 | 1.172096612 | 0.833408131 | 0.905506971 | 3.396580048 | 3.967037349 | ENSG00000164404 |
| APH1A | 0.229071057 | 1.172080011 | 8.14E-05 | 0.000449015 | 1891.80727 | 2216.691985 | ENSG00000117362 |
| TOMM7 | 0.228989502 | 1.172013756 | 0.000406968 | 0.001912036 | 1767.095296 | 2070.218174 | ENSG00000196683 |
| AP4E1 | 0.2288928 | 1.171935199 | 0.112790865 | 0.227271734 | 222.2333237 | 260.7621147 | ENSG00000081014 |
| PIR | 0.228875019 | 1.171920756 | 0.004495043 | 0.01599592 | 854.7649936 | 1001.185118 | ENSG00000087842 |
| LOC102606465 | 0.228828756 | 1.171883176 | 0.597141542 | 0.744295244 | 21.05345497 | 24.69032848 | ENSG00000233184 |
| PCOLCE | 0.228799379 | 1.171859314 | 0.036679201 | 0.092747069 | 457.6655049 | 535.9813382 | ENSG00000106333 |
| MCPH1 | 0.22875528 | 1.171823494 | 0.083883241 | 0.180179445 | 305.5929726 | 357.91176 | ENSG00000147316 |
| SH3BP2 | 0.228693693 | 1.171773472 | 0.010184931 | 0.031948805 | 842.9353466 | 986.8418572 | ENSG00000087266 |
| CDK1 | 0.228628239 | 1.17172031 | 0.002633113 | 0.010028702 | 1201.552586 | 1407.677877 | ENSG00000170312 |
| SLC25A24 | 0.22860374 | 1.171700412 | 0.012828669 | 0.038826906 | 537.461446 | 629.8450198 | ENSG00000085491 |
| NA | 0.228459857 | 1.171583562 | 0.808787366 | 0.890136757 | 4.988389884 | 5.83090503 | ENSG00000232732 |
| PMF1 | 0.228457425 | 1.171581587 | 0.063244951 | 0.143874805 | 365.4636507 | 428.1068868 | ENSG00000160783 |
| FGF14-AS2 | 0.228286142 | 1.171442501 | 0.499671818 | 0.666010436 | 38.31127903 | 44.85219723 | ENSG00000272143 |
| NA | 0.228203225 | 1.171375175 | 0.502695931 | 0.668378745 | 35.37447399 | 41.48852766 | ENSG00000246067 |
| SRXN1 | 0.228067376 | 1.17126488 | 0.367908238 | 0.541738632 | 76.91690606 | 89.99131217 | ENSG00000271303 |
| RAP1GAP2 | 0.227728688 | 1.170989945 | 0.000560324 | 0.002536185 | 1593.535314 | 1865.671796 | ENSG00000132359 |
| LOC648987 | 0.22759497 | 1.170881415 | 0.650106475 | 0.782358329 | 16.04062268 | 18.80887576 | ENSG00000177738 |
| PTTG1 | 0.227351756 | 1.170684042 | 0.000370525 | 0.001760975 | 1514.432673 | 1772.311683 | ENSG00000164611 |
| TTC21B | 0.227170132 | 1.170536671 | 0.044289795 | 0.108260826 | 359.9109831 | 421.3451142 | ENSG00000123607 |
| SSX2IP | 0.227018647 | 1.170413769 | 0.04541239 | 0.110514749 | 490.2969982 | 573.9579634 | ENSG00000117155 |
| HRH2 | 0.226937225 | 1.170347716 | 0.061238803 | 0.140220504 | 314.504587 | 367.8748125 | ENSG00000113749 |
| ZSCAN12 | 0.226733565 | 1.170182514 | 0.129068291 | 0.251475245 | 214.3110688 | 250.5033244 | ENSG00000158691 |
| ABCB6 | 0.226609492 | 1.170081881 | 0.732200602 | 0.840066719 | 10.21418255 | 11.90541319 | ENSG00000115657 |
| VWCE | 0.226500168 | 1.169993218 | 0.09557309 | 0.199945829 | 288.0165214 | 336.8120938 | ENSG00000167992 |
| BIRC6 | 0.226478675 | 1.169975789 | 0.005584462 | 0.019292594 | 1390.859055 | 1626.894452 | ENSG00000115760 |
| KIF2C | 0.226163374 | 1.169720118 | 0.002419498 | 0.009325581 | 1478.534694 | 1728.480944 | ENSG00000142945 |
| ADAMTS12 | 0.226104747 | 1.169672586 | 0.08220242 | 0.177339172 | 419.8083807 | 490.6719759 | ENSG00000151388 |
| NA | 0.226060051 | 1.169636348 | 0.779412947 | 0.870480335 | 6.439327257 | 7.473487239 | ENSG00000236439 |
| LOC101928059 | 0.22605887 | 1.169635391 | 0.81101231 | 0.891568183 | 5.864161333 | 6.90776371 | ENSG00000284738 |
| ACTA2 | 0.225922914 | 1.169525172 | 0.770034207 | 0.864544733 | 7.114177822 | 8.312921933 | ENSG00000107796 |
| NA | 0.225892487 | 1.169500507 | 0.840782632 | 0.909051234 | 3.393789145 | 3.95251637 | ENSG00000280193 |
| CHD6 | 0.225773473 | 1.169404034 | 0.001905632 | 0.007564104 | 1298.847494 | 1518.092181 | ENSG00000124177 |
| NA | 0.225721358 | 1.169361792 | 0.794092519 | 0.880342439 | 8.295773181 | 9.653011484 | ENSG00000230084 |
| SLC7A7 | 0.225714108 | 1.169355915 | 0.291626036 | 0.461392427 | 101.381171 | 118.8065002 | ENSG00000155465 |
| NCOA5 | 0.225504508 | 1.16918604 | 0.005385724 | 0.018696418 | 944.1296535 | 1103.786659 | ENSG00000124160 |
| GLB1L | 0.225448533 | 1.169140677 | 0.448157997 | 0.61850164 | 49.05489702 | 57.34136257 | ENSG00000163521 |
| NA | 0.225416931 | 1.169115068 | 0.57103333 | 0.723801675 | 27.63086379 | 32.40365074 | ENSG00000180581 |
| CSNK1G3 | 0.225405937 | 1.169106159 | 0.008869035 | 0.028445151 | 602.0584303 | 703.9245954 | ENSG00000151292 |
| RNF139 | 0.225297823 | 1.16901855 | 0.027693012 | 0.07368633 | 446.347648 | 522.0211466 | ENSG00000170881 |
| FSTL3 | 0.225244475 | 1.168975323 | 0.118532216 | 0.236180365 | 217.0086071 | 253.8312277 | ENSG00000070404 |
| CNFN | 0.224854379 | 1.168659282 | 0.29455456 | 0.464501922 | 92.86390147 | 108.4761895 | ENSG00000105427 |
| ITPRID2 | 0.224714415 | 1.168545909 | 0.168075451 | 0.308295477 | 168.1409888 | 196.1989615 | ENSG00000138434 |
| SNTB2 | 0.224629119 | 1.168476824 | 0.098261187 | 0.204328614 | 283.035465 | 330.4608877 | ENSG00000168807 |
| URB1-AS1 | 0.224582548 | 1.168439105 | 0.797867531 | 0.883157134 | 5.693264659 | 6.678408981 | ENSG00000256073 |
| COIL | 0.224453037 | 1.168334219 | 0.010522048 | 0.032860746 | 594.8152194 | 694.9222644 | ENSG00000121058 |
| CHIC1 | 0.224380772 | 1.168275698 | 0.233531171 | 0.394315082 | 121.3838155 | 141.9179702 | ENSG00000204116 |
| LINC00624 | 0.224380691 | 1.168275633 | 0.760274243 | 0.857637004 | 7.32905322 | 8.586372634 | ENSG00000278811 |
| NA | 0.224342244 | 1.168244499 | 0.60748635 | 0.751604616 | 21.17634272 | 24.71560233 | ENSG00000224261 |
| ZXDB | 0.224304618 | 1.168214031 | 0.131935817 | 0.255693575 | 198.3523093 | 231.8684094 | ENSG00000198455 |
| SAMD8 | 0.224188223 | 1.168119785 | 0.044060438 | 0.107809476 | 431.9295719 | 504.6673827 | ENSG00000156671 |

| | | | | | | | |
|---|---|---|---|---|---|---|---|
| STOML1 | 0.223978665 | 1.167950122 | 0.487895181 | 0.654792398 | 50.9331946 | 59.40983207 | ENSG00000067221 |
| FZD3 | 0.223956869 | 1.167932478 | 0.026739186 | 0.071553008 | 600.2374414 | 701.7848234 | ENSG00000104290 |
| PPIE | 0.223891335 | 1.167879426 | 0.013267342 | 0.039959453 | 688.2505448 | 803.6797076 | ENSG00000084072 |
| NA | 0.223834585 | 1.167833486 | 0.755201605 | 0.854964446 | 8.398924446 | 9.768367384 | ENSG00000220157 |
| FRS3 | 0.223816954 | 1.167819215 | 0.391539563 | 0.564598636 | 74.56479094 | 87.11403694 | ENSG00000137218 |
| ZNF761 | 0.22377797 | 1.167787659 | 0.021771164 | 0.060247816 | 491.7276866 | 574.6533218 | ENSG00000160336 |
| ETS2 | 0.223764675 | 1.167776897 | 0.004834295 | 0.017045949 | 779.4204874 | 910.1611044 | ENSG00000157557 |
| CARNS1 | 0.223712558 | 1.167734713 | 0.578294689 | 0.730049267 | 25.9309966 | 30.19938569 | ENSG00000172508 |
| ZBTB39 | 0.223466088 | 1.167535233 | 0.001543074 | 0.006279544 | 1052.084552 | 1228.466718 | ENSG00000166860 |
| C1orf210 | 0.223460432 | 1.167530657 | 0.199285897 | 0.35068884 | 136.3318763 | 159.0858813 | ENSG00000253313 |
| PLPP1 | 0.223326698 | 1.167422435 | 0.023751619 | 0.064807988 | 473.2048524 | 552.4815942 | ENSG00000067113 |
| ARL5B | 0.223276838 | 1.167382089 | 0.005544523 | 0.01918554 | 1008.768152 | 1178.326451 | ENSG00000165997 |
| TLE4 | 0.223144484 | 1.167274997 | 0.019652219 | 0.055393186 | 542.4389846 | 633.3600294 | ENSG00000106829 |
| ZBTB47 | 0.223135077 | 1.167267386 | 0.27194279 | 0.438889681 | 101.0584384 | 117.9143673 | ENSG00000114853 |
| C1orf21 | 0.223083283 | 1.167225481 | 0.005307851 | 0.018469294 | 740.7995544 | 864.6310444 | ENSG00000116667 |
| NA | 0.222997016 | 1.167155688 | 0.285561734 | 0.454519093 | 103.5273711 | 120.8593948 | ENSG00000267002 |
| PLCH1 | 0.222951271 | 1.16711868 | 0.013279123 | 0.039973473 | 583.9946954 | 681.6684382 | ENSG00000114805 |
| CLOCK | 0.222709888 | 1.166923421 | 0.104082459 | 0.213624293 | 301.6454688 | 351.7208541 | ENSG00000134852 |
| HERC1 | 0.222642425 | 1.166868855 | 0.017349865 | 0.049954664 | 740.9117157 | 864.280354 | ENSG00000103657 |
| NFS1 | 0.222585867 | 1.166823112 | 0.103741904 | 0.213106705 | 242.7385948 | 283.2255879 | ENSG00000244005 |
| ABCA2 | 0.222568685 | 1.166809215 | 0.018664119 | 0.052968651 | 771.4034922 | 900.1359884 | ENSG00000107331 |
| NA | 0.222517023 | 1.166767433 | 0.826697624 | 0.901230101 | 3.984148884 | 4.684403957 | ENSG00000258611 |
| NA | 0.222395247 | 1.166668952 | 0.827287883 | 0.901548444 | 3.95970648 | 4.656979537 | ENSG00000267778 |
| NA | 0.222346988 | 1.166629927 | 0.711093676 | 0.825872368 | 14.74538817 | 17.12570522 | ENSG00000178464 |
| VSTM2L | 0.222339512 | 1.166623881 | 0.40231305 | 0.574549974 | 62.9043017 | 73.40201595 | ENSG00000132821 |
| PIEZO1 | 0.222301166 | 1.166592873 | 0.000560218 | 0.002536185 | 1690.039121 | 1971.701783 | ENSG00000103335 |
| REX1BD | 0.221802574 | 1.166189772 | 0.003933934 | 0.014232844 | 856.2719634 | 998.5253909 | ENSG00000006015 |
| DDHD1 | 0.221790397 | 1.166179928 | 0.041859244 | 0.103393941 | 412.9921807 | 481.6252946 | ENSG00000100523 |
| PHC3 | 0.221781612 | 1.166172827 | 0.10815227 | 0.219663659 | 240.9869398 | 281.2170498 | ENSG00000173889 |
| DNAJC27 | 0.221765159 | 1.166159527 | 0.44611302 | 0.616429072 | 50.53565844 | 58.99738126 | ENSG00000115137 |
| TERF2 | 0.221670523 | 1.166083034 | 0.007439881 | 0.02458423 | 679.9644588 | 793.0189405 | ENSG00000132604 |
| ADAM22 | 0.221641426 | 1.166059516 | 0.041150356 | 0.101982466 | 430.4795985 | 501.6011538 | ENSG00000008277 |
| C8orf34 | 0.221604674 | 1.166029812 | 0.67269197 | 0.799543055 | 15.40907512 | 17.90867354 | ENSG00000165084 |
| ANKRD13C | 0.221603658 | 1.166028991 | 0.034415571 | 0.087863123 | 393.9448924 | 459.1666983 | ENSG00000118454 |
| PMAIP1 | 0.221601072 | 1.166026901 | 0.000409028 | 0.001920777 | 2409.096152 | 2809.039509 | ENSG00000141682 |
| SULT1A2 | 0.221475504 | 1.165925417 | 0.798762651 | 0.883907026 | 6.474933273 | 7.492309364 | ENSG00000197165 |
| MICB | 0.221448473 | 1.165903572 | 0.056177102 | 0.130993892 | 389.9847104 | 454.7624146 | ENSG00000204516 |
| LOC100287896 | 0.221446038 | 1.165901604 | 0.573004063 | 0.725403106 | 27.20091283 | 31.76399578 | ENSG00000254837 |
| FGFR4 | 0.221419941 | 1.165880514 | 0.000128918 | 0.000677646 | 2601.521687 | 3033.725852 | ENSG00000160867 |
| CHMP1B2P | 0.221395318 | 1.165860616 | 0.43518421 | 0.606135187 | 53.71756325 | 62.45758461 | ENSG00000278530 |
| MRPL35 | 0.22133186 | 1.165809336 | 0.008163859 | 0.026571452 | 726.3223569 | 846.6557031 | ENSG00000132313 |
| ZNF426 | 0.22104733 | 1.165579436 | 0.063822931 | 0.144866255 | 329.6666105 | 384.6471927 | ENSG00000130818 |
| SNORD20 | 0.220931185 | 1.165485604 | 0.841787687 | 0.90951788 | 3.33932253 | 3.872393683 | ENSG00000207280 |
| XPNPEP1 | 0.220918517 | 1.16547537 | 0.003240504 | 0.011982472 | 946.6446659 | 1102.811646 | ENSG00000108039 |
| TRIM69 | 0.22083803 | 1.165410351 | 0.233808669 | 0.394679999 | 135.462926 | 158.0127786 | ENSG00000185880 |
| GDPD1 | 0.220710739 | 1.16530753 | 0.340904796 | 0.513499994 | 78.45580088 | 91.33758087 | ENSG00000153982 |
| TRIM54 | 0.220186097 | 1.164883838 | 0.287984323 | 0.457335693 | 102.131464 | 119.1119491 | ENSG00000138100 |
| ZNF404 | 0.220003435 | 1.16473636 | 0.622814203 | 0.762596559 | 22.58609227 | 26.22321301 | ENSG00000176222 |
| PALM | 0.219900189 | 1.164653009 | 0.032322872 | 0.083450284 | 468.2789015 | 545.5267904 | ENSG00000099864 |
| ENTPD4 | 0.2198778 | 1.164634935 | 0.00379723 | 0.013810776 | 818.5209318 | 953.4992271 | ENSG00000197217 |
| NA | 0.219701896 | 1.164492942 | 0.488988363 | 0.655529848 | 42.96102989 | 50.01764041 | ENSG00000225031 |
| IL11 | 0.21962017 | 1.164426978 | 0.601097991 | 0.747006465 | 28.83199547 | 33.6617275 | ENSG00000095752 |
| C2orf74 | 0.219532286 | 1.164356047 | 0.839041874 | 0.908201842 | 3.366555838 | 3.895516956 | ENSG00000237651 |
| SEMA3D | 0.219491646 | 1.164323248 | 0.301377885 | 0.472097506 | 92.80684411 | 107.9314387 | ENSG00000153993 |
| EAF2 | 0.219389727 | 1.164240998 | 0.371360939 | 0.54495085 | 82.96706986 | 96.8538833 | ENSG00000145088 |
| CCM2 | 0.219195444 | 1.164084224 | 0.01150856 | 0.035527545 | 634.5415175 | 738.8250751 | ENSG00000136280 |
| ATP13A3 | 0.219171636 | 1.164065014 | 0.000898128 | 0.003860469 | 1310.893118 | 1525.858989 | ENSG00000133657 |
| ATF3 | 0.219137764 | 1.164037684 | 0.032834795 | 0.08452309 | 437.0058071 | 508.8092169 | ENSG00000162772 |
| RNASEH1-AS1 | 0.218889663 | 1.163837521 | 0.060835052 | 0.139527878 | 367.7624264 | 428.2579847 | ENSG00000234171 |
| COX10 | 0.218872437 | 1.163823625 | 0.034208458 | 0.087406007 | 551.0016964 | 641.2637661 | ENSG00000006695 |
| IL17RE | 0.218770462 | 1.163741364 | 0.388092132 | 0.561526829 | 69.46699226 | 80.75909332 | ENSG00000163701 |
| GRIK3 | 0.218761439 | 1.163734086 | 0.240453331 | 0.402654967 | 123.0511799 | 143.1661181 | ENSG00000163873 |
| NA | 0.218428666 | 1.163465689 | 0.752181148 | 0.853347831 | 9.800301289 | 11.3184565 | ENSG00000279861 |
| ATE1 | 0.2182254 | 1.163301777 | 0.041512614 | 0.102708714 | 418.2128303 | 486.5979855 | ENSG00000107669 |
| WDR53 | 0.218215319 | 1.163293648 | 0.117888495 | 0.235349991 | 212.4410317 | 247.1165315 | ENSG00000185798 |
| NA | 0.218164936 | 1.163253022 | 0.117708915 | 0.235088797 | 245.4606877 | 285.8070826 | ENSG00000231205 |
| SLC17A5 | 0.218069314 | 1.163175925 | 0.082294227 | 0.17751738 | 266.5180096 | 310.0416763 | ENSG00000119899 |
| PPP3CB-AS1 | 0.218065382 | 1.163172754 | 0.807678147 | 0.889275996 | 5.30940761 | 6.169424403 | ENSG00000221817 |
| NA | 0.217979526 | 1.163103535 | 0.844401861 | 0.910884992 | 3.372137643 | 3.89766753 | ENSG00000287778 |
| PRRG4 | 0.217904458 | 1.163043017 | 0.512530612 | 0.676808081 | 44.01271624 | 51.3286759 | ENSG00000135378 |
| COX19 | 0.217843102 | 1.162993555 | 0.087724343 | 0.186858999 | 291.0226745 | 338.8539202 | ENSG00000240230 |
| OSGIN2 | 0.217804813 | 1.16296269 | 0.124170664 | 0.244606897 | 244.7852736 | 284.6237734 | ENSG00000164823 |
| SLC2A10 | 0.217719725 | 1.162894102 | 0.146026365 | 0.276501237 | 197.8899435 | 230.1448804 | ENSG00000197496 |
| ELL2 | 0.217461751 | 1.162686179 | 0.113780162 | 0.228925022 | 261.8892346 | 304.3766564 | ENSG00000118985 |
| SHANK2 | 0.217370158 | 1.162612366 | 0.009567289 | 0.030307458 | 927.6400177 | 1078.154104 | ENSG00000162105 |
| ZNF629 | 0.217294021 | 1.162551011 | 0.006295002 | 0.021387263 | 877.5832205 | 1020.254863 | ENSG00000102870 |
| C19orf81 | 0.217176493 | 1.162456309 | 0.321948321 | 0.493428784 | 93.73409111 | 108.8967217 | ENSG00000235034 |

| | | | | | | | |
|---|---|---|---|---|---|---|---|
| ZNF652 | 0.217044848 | 1.16235024 | 0.167536259 | 0.307569659 | 223.6018855 | 260.2692381 | ENSG00000198740 |
| GGH | 0.217042959 | 1.162348718 | 0.017739115 | 0.050870412 | 622.7619194 | 723.9361355 | ENSG00000137563 |
| C15orf61 | 0.216997197 | 1.16231185 | 0.616076125 | 0.758292893 | 22.97832203 | 26.61711426 | ENSG00000189227 |
| C2CD4B | 0.216952249 | 1.162275637 | 0.835335428 | 0.906737673 | 4.863587117 | 5.599399727 | ENSG00000205502 |
| FAM122C | 0.216843863 | 1.162188322 | 0.437192095 | 0.607978137 | 53.13190944 | 61.7533819 | ENSG00000156500 |
| ZFAND4 | 0.216753774 | 1.162115751 | 0.136656328 | 0.262756625 | 214.2275049 | 249.1161642 | ENSG00000172671 |
| RHBDF2 | 0.216592197 | 1.161985606 | 0.050912849 | 0.12108985 | 509.5542942 | 592.6205939 | ENSG00000129667 |
| LOC100506688 | 0.216584068 | 1.161979058 | 0.836348924 | 0.907067358 | 3.665922062 | 4.261460749 | ENSG00000215246 |
| RAPGEF5 | 0.216573008 | 1.16197015 | 0.122672349 | 0.242433689 | 241.6298142 | 281.0933947 | ENSG00000136237 |
| SDHAP2 | 0.216568788 | 1.161966751 | 0.31271755 | 0.483810591 | 117.3834337 | 136.0365296 | ENSG00000242086 |
| GOLGA3 | 0.216491942 | 1.16190486 | 0.005617341 | 0.019399235 | 868.3873343 | 1009.034515 | ENSG00000090615 |
| NA | 0.216410187 | 1.161839019 | 0.008353643 | 0.027079505 | 817.8402241 | 949.8140249 | ENSG00000281566 |
| AIMP1 | 0.216349496 | 1.161790144 | 0.013179867 | 0.039751438 | 667.5115249 | 775.7649575 | ENSG00000164022 |
| IDH2 | 0.216313962 | 1.161761529 | 0.000653006 | 0.002901898 | 1548.723596 | 1799.164288 | ENSG00000182054 |
| AMMECR1 | 0.216282205 | 1.161735957 | 0.003065034 | 0.01140358 | 1075.577042 | 1249.903916 | ENSG00000101935 |
| CDK19 | 0.216187547 | 1.161659736 | 0.167707635 | 0.307814424 | 176.1322895 | 204.4250364 | ENSG00000155111 |
| GPATCH4 | 0.216093594 | 1.161584087 | 0.001000155 | 0.004262921 | 1843.674867 | 2141.850123 | ENSG00000160818 |
| ZNF675 | 0.216059 | 1.161556234 | 0.149993089 | 0.282349403 | 205.4285657 | 238.770527 | ENSG00000197372 |
| PLPP7 | 0.216039686 | 1.161540683 | 0.381749983 | 0.555170669 | 76.45673074 | 89.04511176 | ENSG00000160539 |
| ME2 | 0.215912333 | 1.161438153 | 0.000273368 | 0.001338121 | 1688.815261 | 1961.825456 | ENSG00000082212 |
| HPSE | 0.215909421 | 1.16143581 | 0.293709374 | 0.46339638 | 98.17542426 | 114.0925213 | ENSG00000173083 |
| ZNF653 | 0.215883103 | 1.161414622 | 0.338790144 | 0.511352842 | 87.17466702 | 101.3465889 | ENSG00000161914 |
| ZSCAN23 | 0.215831757 | 1.161373288 | 0.51466472 | 0.678586366 | 38.53385142 | 44.73227972 | ENSG00000187987 |
| USP27X-AS1 | 0.215808915 | 1.1613549 | 0.689263275 | 0.811311113 | 14.0838164 | 16.30158501 | ENSG00000234390 |
| FRMD8 | 0.215705268 | 1.161271468 | 0.021470567 | 0.059597672 | 569.9715663 | 662.2099418 | ENSG00000126391 |
| UCP2 | 0.215431871 | 1.161051423 | 0.009605392 | 0.030396591 | 747.0708639 | 867.2612272 | ENSG00000175567 |
| FAM120B | 0.21536946 | 1.161001197 | 0.021799797 | 0.060318399 | 513.2109293 | 595.7151444 | ENSG00000112584 |
| TBC1D12 | 0.215342983 | 1.16097989 | 0.434947515 | 0.605992526 | 60.60617763 | 70.21689295 | ENSG00000108239 |
| SCRT2 | 0.215333916 | 1.160972594 | 0.709621563 | 0.825105495 | 13.16685713 | 15.28790489 | ENSG00000215397 |
| PHF1 | 0.215281459 | 1.160930382 | 0.005577546 | 0.019272155 | 801.4497107 | 930.3267228 | ENSG00000112511 |
| TEX264 | 0.215274732 | 1.160924968 | 0.138420272 | 0.26540756 | 215.4036306 | 250.150805 | ENSG00000164081 |
| GAS2 | 0.215251543 | 1.160906308 | 0.773880655 | 0.867313818 | 7.056920304 | 8.161278853 | ENSG00000148935 |
| PAK1 | 0.215242681 | 1.160899177 | 4.70E-06 | 3.28E-05 | 4289.620204 | 4979.339143 | ENSG00000149269 |
| B4GALT4 | 0.215190678 | 1.160857333 | 0.032456805 | 0.083751234 | 497.7475877 | 577.826347 | ENSG00000121578 |
| BZW2 | 0.215118986 | 1.160799647 | 0.002699966 | 0.010242879 | 1328.498342 | 1542.13235 | ENSG00000136261 |
| SAR1B | 0.214933276 | 1.160650234 | 0.106126572 | 0.216620957 | 299.2337936 | 346.9689583 | ENSG00000152700 |
| RHBDD2 | 0.21473181 | 1.160488165 | 0.013708968 | 0.041052422 | 659.6446394 | 765.5473191 | ENSG00000005486 |
| PAPSS2 | 0.214714243 | 1.160474035 | 0.029764956 | 0.078027022 | 512.4874693 | 594.6786441 | ENSG00000198682 |
| IGLON5 | 0.21470849 | 1.160469407 | 0.000674541 | 0.002984536 | 1528.138091 | 1774.115576 | ENSG00000142549 |
| NA | 0.214699754 | 1.16046238 | 0.815600635 | 0.894616739 | 5.054020111 | 5.824453309 | ENSG00000261033 |
| NOVA2 | 0.214677539 | 1.160444511 | 0.028590246 | 0.075552662 | 582.4953119 | 676.6777429 | ENSG00000104967 |
| MED18 | 0.214671769 | 1.16043987 | 0.064116164 | 0.145342528 | 325.2040261 | 377.4111415 | ENSG00000130772 |
| WWC2 | 0.214668583 | 1.160437307 | 6.55E-05 | 0.000368589 | 2503.437759 | 2905.013146 | ENSG00000151718 |
| CPAMD8 | 0.214511925 | 1.160311305 | 0.114660761 | 0.230249379 | 230.9563691 | 267.9613577 | ENSG00000160111 |
| NSUN5P1 | 0.214457091 | 1.160267205 | 0.031707435 | 0.082180321 | 617.8959658 | 716.2968116 | ENSG00000223705 |
| GNE | 0.214337385 | 1.160170938 | 0.058896895 | 0.135922958 | 388.9482051 | 451.5517573 | ENSG00000159921 |
| TYSND1 | 0.214289397 | 1.160132347 | 0.01568812 | 0.045965904 | 775.0641703 | 898.5250949 | ENSG00000156521 |
| TTC21A | 0.214287015 | 1.160130432 | 0.550362454 | 0.707422466 | 32.91112307 | 38.12915924 | ENSG00000168026 |
| NFKBIA | 0.214207356 | 1.160066377 | 0.095617862 | 0.200017802 | 292.869145 | 339.376105 | ENSG00000100906 |
| NA | 0.21403948 | 1.159931396 | 0.113895157 | 0.229060829 | 255.5258253 | 296.2336787 | ENSG00000233836 |
| CCND2 | 0.213992361 | 1.159893513 | 0.003284908 | 0.012128071 | 1357.078056 | 1574.740839 | ENSG00000118971 |
| MANEA | 0.213671883 | 1.159635884 | 0.332832286 | 0.50556438 | 93.85283651 | 108.906378 | ENSG00000172469 |
| SMC5 | 0.213652913 | 1.159620636 | 0.019099958 | 0.054057702 | 626.5559107 | 726.5811056 | ENSG00000198887 |
| TTLL2 | 0.213648663 | 1.159617221 | 0.838985775 | 0.908201842 | 4.817493211 | 5.532180482 | ENSG00000120440 |
| SERPINE2 | 0.213592534 | 1.159572105 | 0.001101961 | 0.004647714 | 1352.741962 | 1568.166573 | ENSG00000135919 |
| NA | 0.213575318 | 1.159558269 | 0.779581387 | 0.870508482 | 7.698955755 | 8.943714132 | ENSG00000250220 |
| RBL2 | 0.213524558 | 1.159517471 | 0.097265481 | 0.202738886 | 338.1121413 | 392.2999958 | ENSG00000103479 |
| NA | 0.213449873 | 1.159457447 | 0.566543681 | 0.720257951 | 29.35393408 | 34.01262938 | ENSG00000287126 |
| CEP250 | 0.213429712 | 1.159441245 | 0.005180068 | 0.018086642 | 821.1482831 | 951.8934956 | ENSG00000126001 |
| TMEFF1 | 0.213377352 | 1.159399166 | 0.158413403 | 0.29477698 | 254.2744815 | 294.5870864 | ENSG00000241697 |
| ZNF853 | 0.213323451 | 1.159355849 | 0.099963296 | 0.207131251 | 254.490484 | 295.2227007 | ENSG00000236609 |
| SLC22A7 | 0.213320579 | 1.159353542 | 0.801035537 | 0.885292269 | 5.64437985 | 6.554190321 | ENSG00000137204 |
| KDSR | 0.213081013 | 1.159161042 | 0.011743459 | 0.036010661 | 700.2593731 | 811.6414424 | ENSG00000119537 |
| IMMP1L | 0.21304497 | 1.159132082 | 0.360206751 | 0.533288445 | 84.77136452 | 98.2165752 | ENSG00000148950 |
| MAP11 | 0.212924985 | 1.159035685 | 0.116055843 | 0.232485283 | 260.8346324 | 302.639982 | ENSG00000146826 |
| TMEM92 | 0.212923438 | 1.159034442 | 0.107460382 | 0.218672809 | 273.9984232 | 317.3600554 | ENSG00000167105 |
| LRCH1 | 0.212862262 | 1.158985295 | 0.09052767 | 0.191487859 | 270.2921892 | 313.2348806 | ENSG00000136141 |
| FA2H | 0.212784215 | 1.158922598 | 0.635961615 | 0.772030427 | 26.08929021 | 30.23326183 | ENSG00000103089 |
| ZKSCAN8 | 0.212755338 | 1.158899402 | 0.033462141 | 0.085794547 | 476.5901056 | 552.3111048 | ENSG00000198315 |
| ATP11A | 0.212736062 | 1.158883917 | 0.000795275 | 0.003456104 | 1561.970589 | 1809.881285 | ENSG00000068650 |
| SYK | 0.21270826 | 1.158861585 | 0.258967523 | 0.424158812 | 116.7480192 | 135.2245072 | ENSG00000165025 |
| PCDHGA8 | 0.212652771 | 1.158817013 | 0.673805241 | 0.800111161 | 16.3316162 | 18.89734027 | ENSG00000253767 |
| RCOR1 | 0.212578036 | 1.158756986 | 0.007140908 | 0.0237591 | 796.9860504 | 923.8916018 | ENSG00000089902 |
| ZNF776 | 0.212455276 | 1.15865839 | 0.112540781 | 0.226975871 | 272.3383555 | 315.8658691 | ENSG00000152443 |
| NPIPB13 | 0.212452243 | 1.158655954 | 0.835886087 | 0.906924912 | 3.690364466 | 4.270063044 | ENSG00000198064 |
| PRDX6 | 0.212377765 | 1.158596141 | 3.47E-06 | 2.48E-05 | 4813.759681 | 5577.604767 | ENSG00000117592 |
| FAM3C | 0.212369745 | 1.158589701 | 0.015899535 | 0.046472413 | 1157.158593 | 1341.434988 | ENSG00000196937 |

| | | | | | | | |
|---|---|---|---|---|---|---|---|
| EEF1A1 | 0.21229962 | 1.158533387 | 1.48E-06 | 1.13E-05 | 206688.9791 | 239454.7595 | ENSG00000156508 |
| OCIAD2 | 0.212186493 | 1.158442545 | 0.004730012 | 0.016733294 | 1631.593925 | 1890.370369 | ENSG00000145247 |
| MAPK6-DT | 0.212175182 | 1.158433462 | 0.850064784 | 0.914641071 | 3.956915577 | 4.602130696 | ENSG00000259438 |
| PIGP | 0.212130659 | 1.158397713 | 0.324822024 | 0.496485355 | 88.7177922 | 102.8566106 | ENSG00000185808 |
| ACOT9 | 0.212093116 | 1.158367569 | 0.014658207 | 0.043443152 | 822.9971811 | 953.0759687 | ENSG00000123130 |
| MSRB2 | 0.212001802 | 1.158294253 | 0.152827667 | 0.286873457 | 233.3326385 | 270.2314786 | ENSG00000148450 |
| RPARP-AS1 | 0.21190102 | 1.158213341 | 0.231891937 | 0.392474763 | 155.7414454 | 180.3308121 | ENSG00000269609 |
| GIHCG | 0.211802331 | 1.158134115 | 0.583980081 | 0.733964094 | 29.38395829 | 34.12205449 | ENSG00000257698 |
| GAS6 | 0.21178329 | 1.15811883 | 0.020159164 | 0.056582342 | 590.4706167 | 683.6543555 | ENSG00000183087 |
| TMEM91 | 0.211713973 | 1.158063187 | 0.709533334 | 0.825101425 | 12.69571835 | 14.6713732 | ENSG00000142046 |
| WEE1 | 0.21163991 | 1.158003738 | 0.000655226 | 0.002909084 | 1424.439347 | 1649.951965 | ENSG00000166483 |
| LEPROT | 0.211625254 | 1.157991974 | 0.110606359 | 0.223557696 | 281.6699455 | 326.4607372 | ENSG00000213625 |
| CCDC167 | 0.211483045 | 1.157877834 | 0.185096735 | 0.331387575 | 180.9085949 | 209.4742876 | ENSG00000198937 |
| UBE4A | 0.211406345 | 1.157816278 | 0.02647846 | 0.070993185 | 607.1805082 | 702.7896897 | ENSG00000110344 |
| NOTCH3 | 0.211335581 | 1.157759489 | 9.84E-05 | 0.000533167 | 3635.773852 | 4209.888438 | ENSG00000074181 |
| NA | 0.211302953 | 1.157733305 | 0.426488937 | 0.598095238 | 59.41553384 | 68.7490771 | ENSG00000260469 |
| NA | 0.211297586 | 1.157728998 | 0.406406143 | 0.578761733 | 68.08100939 | 78.73819691 | ENSG00000229335 |
| BAZ2B | 0.211254943 | 1.157694779 | 0.047530847 | 0.114743784 | 406.9759833 | 470.9000166 | ENSG00000123636 |
| LINC01176 | 0.21119413 | 1.15764598 | 0.865245851 | 0.924580829 | 3.896867156 | 4.526842194 | ENSG00000281404 |
| DHFR2 | 0.211193436 | 1.157645424 | 0.436048191 | 0.606868599 | 57.5046213 | 66.69403197 | ENSG00000178700 |
| SNRNP35 | 0.211119064 | 1.157585748 | 0.132493583 | 0.256465353 | 262.9068296 | 304.378286 | ENSG00000184209 |
| HLA-DMA | 0.211046109 | 1.157527211 | 0.509131653 | 0.673777782 | 40.83611783 | 47.22263845 | ENSG00000204257 |
| FREM2 | 0.210966802 | 1.157463582 | 0.052228431 | 0.123382152 | 922.1193631 | 1066.693617 | ENSG00000150893 |
| NRG1 | 0.210908025 | 1.157416427 | 0.508235513 | 0.67296117 | 47.59135657 | 55.14112609 | ENSG00000157168 |
| ZNF574 | 0.210851858 | 1.157371367 | 0.00334401 | 0.012322692 | 990.1831913 | 1146.442575 | ENSG00000105732 |
| SNX30 | 0.210833383 | 1.157356546 | 0.068875382 | 0.154049274 | 405.5871216 | 469.4813793 | ENSG00000148158 |
| ABCB4 | 0.210739012 | 1.157280842 | 0.677787369 | 0.802799542 | 19.14274259 | 22.16179246 | ENSG00000005471 |
| NEK1 | 0.210460814 | 1.157057703 | 0.140707189 | 0.269017681 | 280.3797292 | 324.7598048 | ENSG00000137601 |
| NA | 0.210389631 | 1.157000615 | 0.68073925 | 0.805060132 | 18.26888617 | 21.11424831 | ENSG00000188599 |
| CCDC88C | 0.210342995 | 1.156963214 | 0.001820227 | 0.007269981 | 1387.104762 | 1604.63623 | ENSG00000015133 |
| ADAM10 | 0.210337922 | 1.156959147 | 0.000244204 | 0.001209794 | 2282.906408 | 2641.817659 | ENSG00000137845 |
| KLHL29 | 0.210321237 | 1.156945766 | 0.101440219 | 0.209337105 | 265.1882082 | 307.0837453 | ENSG00000119771 |
| WDR13 | 0.210241714 | 1.156881996 | 0.03077076 | 0.080183539 | 471.4477277 | 545.2926015 | ENSG00000101940 |
| SIAE | 0.210237241 | 1.156878409 | 0.338986584 | 0.51152927 | 97.98395213 | 113.1506341 | ENSG00000110013 |
| NA | 0.210154311 | 1.15681191 | 0.778342309 | 0.869988497 | 7.383519835 | 8.525072072 | ENSG00000260196 |
| NA | 0.210141706 | 1.156801803 | 0.6062534 | 0.75080146 | 27.36239766 | 31.55614679 | ENSG00000237506 |
| ZNF845 | 0.210084462 | 1.156755903 | 0.060928469 | 0.13964252 | 401.4048938 | 464.7464276 | ENSG00000213799 |
| TBC1D23 | 0.210040034 | 1.156720282 | 0.052570968 | 0.124039451 | 429.4763966 | 496.9366138 | ENSG00000036054 |
| KDM2A | 0.20990456 | 1.156611667 | 0.000682993 | 0.003020151 | 1653.454528 | 1912.342456 | ENSG00000173120 |
| ACVR2B | 0.209888024 | 1.15659841 | 0.002958796 | 0.011065944 | 1882.289016 | 2177.750608 | ENSG00000114739 |
| ANXA6 | 0.209728197 | 1.156470285 | 4.03E-05 | 0.000236582 | 2733.113776 | 3160.715875 | ENSG00000197043 |
| SCAMP3 | 0.209640851 | 1.156400271 | 0.007882407 | 0.025797461 | 975.7957783 | 1128.3861 | ENSG00000116521 |
| PPP2R2A | 0.209565745 | 1.15634007 | 0.000865536 | 0.003735359 | 1649.793525 | 1907.931108 | ENSG00000221914 |
| ACYP1 | 0.209555469 | 1.156331833 | 0.222676403 | 0.381065085 | 168.0242464 | 194.3845449 | ENSG00000119640 |
| OPHN1 | 0.209548866 | 1.156326542 | 0.292481418 | 0.462252911 | 109.7813716 | 126.7532179 | ENSG00000079482 |
| INF2 | 0.209500794 | 1.156288012 | 0.000463571 | 0.002143002 | 1830.44821 | 2116.440544 | ENSG00000203485 |
| CCDC62 | 0.209354696 | 1.156170924 | 0.859623671 | 0.92112427 | 3.366555838 | 3.863791388 | ENSG00000130783 |
| NDUFA10 | 0.209320451 | 1.15614348 | 0.001118543 | 0.004713371 | 1578.973639 | 1824.855436 | ENSG00000130414 |
| DAZAP2 | 0.209121762 | 1.155984266 | 0.099727273 | 0.206819912 | 341.4532156 | 394.3569552 | ENSG00000183283 |
| TAF1 | 0.20903174 | 1.155912137 | 0.006790754 | 0.022818399 | 876.7910257 | 1013.167165 | ENSG00000147133 |
| DCAF16 | 0.208980961 | 1.155871453 | 0.012026259 | 0.036691217 | 850.155955 | 981.9307274 | ENSG00000163257 |
| FXR1 | 0.208858151 | 1.155773063 | 1.20E-05 | 7.75E-05 | 5047.703565 | 5833.871025 | ENSG00000114416 |
| NA | 0.208663045 | 1.15561677 | 0.650492031 | 0.782675842 | 19.25705747 | 22.2900518 | ENSG00000198134 |
| GLS2 | 0.208574098 | 1.155545525 | 0.777473135 | 0.869419662 | 7.685676962 | 8.887247756 | ENSG00000135423 |
| NA | 0.208556856 | 1.155531714 | 0.806271036 | 0.888381001 | 5.638798044 | 6.51654607 | ENSG00000273387 |
| EPB41L4A-DT | 0.208464041 | 1.155457376 | 0.691655489 | 0.81298395 | 15.69727773 | 18.17163183 | ENSG00000278921 |
| INTS13 | 0.208287286 | 1.155315822 | 0.004418701 | 0.015764964 | 1123.51123 | 1298.407161 | ENSG00000064102 |
| NBPF3 | 0.208273902 | 1.155305104 | 0.026242676 | 0.070438494 | 538.9792898 | 622.6398644 | ENSG00000142794 |
| LRP12 | 0.208196864 | 1.155243413 | 0.164057424 | 0.302738833 | 204.8309093 | 236.4243056 | ENSG00000147650 |
| NA | 0.208180033 | 1.155229936 | 0.600018106 | 0.746343979 | 32.47714191 | 37.38006706 | ENSG00000244625 |
| WDR18 | 0.208128843 | 1.155188947 | 0.000475227 | 0.002189541 | 1785.366274 | 2062.776884 | ENSG00000065268 |
| PLEKHH2 | 0.207985242 | 1.155073968 | 0.31610171 | 0.487287685 | 100.6747815 | 116.211278 | ENSG00000152527 |
| NA | 0.207889857 | 1.154997603 | 0.6915843 | 0.81298395 | 17.79304146 | 20.57836024 | ENSG00000279159 |
| WDR54 | 0.207752129 | 1.154887345 | 0.015030088 | 0.044340942 | 741.1786303 | 855.833012 | ENSG00000005448 |
| LRRTM1 | 0.207708242 | 1.154852214 | 0.605527683 | 0.750277421 | 27.33987027 | 31.622833 | ENSG00000162951 |
| NA | 0.207579215 | 1.154748935 | 0.704758419 | 0.822374465 | 13.37056891 | 15.4392875 | ENSG00000237742 |
| NA | 0.207578491 | 1.154748355 | 0.774880237 | 0.86807286 | 8.28440941 | 9.619135342 | ENSG00000279500 |
| KIF5B | 0.207555075 | 1.154729613 | 0.000448102 | 0.002082469 | 1983.359281 | 2290.974447 | ENSG00000170759 |
| EMB | 0.207317786 | 1.154539703 | 0.022305845 | 0.061494773 | 691.9698957 | 799.2496775 | ENSG00000170571 |
| ITPR1 | 0.207294565 | 1.15452112 | 0.236086289 | 0.397481288 | 177.9507388 | 205.2477875 | ENSG00000150995 |
| C1GALT1 | 0.207047545 | 1.154323458 | 0.011706297 | 0.035925819 | 1025.419835 | 1184.319207 | ENSG00000106392 |
| N4BP2L2 | 0.20697004 | 1.154261447 | 0.009158666 | 0.029217766 | 746.9239582 | 861.9623995 | ENSG00000244754 |
| MMGT1 | 0.206921334 | 1.15422248 | 0.036079489 | 0.091459846 | 657.139212 | 759.1600437 | ENSG00000169446 |
| CCNK | 0.20682214 | 1.154143122 | 0.047459523 | 0.11461463 | 537.4029108 | 619.5746526 | ENSG00000090061 |
| MRPL10 | 0.206796889 | 1.154122922 | 0.000733544 | 0.003215358 | 1650.794849 | 1904.393315 | ENSG00000159111 |
| NA | 0.206748519 | 1.154084228 | 0.498704428 | 0.66504266 | 45.07632994 | 51.96109775 | ENSG00000226824 |
| MB21D2 | 0.206701079 | 1.154046278 | 0.070713974 | 0.157346302 | 402.6291164 | 464.859924 | ENSG00000180611 |

| | | | | | | | |
|---|---|---|---|---|---|---|---|
| ZSCAN10 | 0.206666114 | 1.15401831 | 0.000141402 | 0.000735846 | 5183.24319 | 5982.058541 | ENSG00000130182 |
| C10orf25 | 0.20659849 | 1.153964218 | 0.59758404 | 0.744461977 | 26.43686552 | 30.47068582 | ENSG00000165511 |
| TMEM251 | 0.206590356 | 1.153957712 | 0.353117946 | 0.526035445 | 113.8588596 | 131.3101865 | ENSG00000153485 |
| FNDC10 | 0.206569586 | 1.153941099 | 0.086510871 | 0.184794731 | 545.7496215 | 629.8587473 | ENSG00000228594 |
| NA | 0.206507693 | 1.153891595 | 0.462211737 | 0.631884815 | 55.70698449 | 64.29643891 | ENSG00000228782 |
| TMEM235 | 0.20646819 | 1.15386 | 0.850803647 | 0.915101063 | 3.638688754 | 4.189940357 | ENSG00000204278 |
| TRIM13 | 0.206415269 | 1.153817675 | 0.142204824 | 0.271128015 | 247.4902576 | 285.6764122 | ENSG00000204977 |
| MRPS18B | 0.206249787 | 1.153685336 | 0.008039959 | 0.026225711 | 964.5875303 | 1113.581802 | ENSG00000204568 |
| LOC102724330 | 0.206216086 | 1.153658386 | 0.485311063 | 0.652959668 | 50.76209777 | 58.4510129 | ENSG00000228801 |
| NKIRAS1 | 0.206053878 | 1.153528683 | 0.387244692 | 0.560605674 | 81.15201192 | 93.67951317 | ENSG00000197885 |
| METTL4 | 0.206023896 | 1.153504711 | 0.199223916 | 0.350627549 | 164.7321691 | 190.1518656 | ENSG00000101574 |
| NA | 0.205948277 | 1.153444252 | 0.69623542 | 0.816684112 | 17.20459674 | 19.85643202 | ENSG00000286536 |
| NA | 0.205908343 | 1.153412325 | 0.679312068 | 0.803872602 | 16.72384596 | 19.24446194 | ENSG00000230091 |
| FAM86C1 | 0.205898409 | 1.153404382 | 0.306935643 | 0.477916083 | 107.6463351 | 124.016845 | ENSG00000158483 |
| RIPK1 | 0.205810204 | 1.153333866 | 0.048058659 | 0.115713878 | 450.2005733 | 518.8378953 | ENSG00000137275 |
| IRX4 | 0.205752928 | 1.153288079 | 0.495283638 | 0.661578498 | 46.96192419 | 54.23149755 | ENSG00000113430 |
| NA | 0.205690775 | 1.153238395 | 0.759322503 | 0.857014416 | 10.26219148 | 11.88444049 | ENSG00000285077 |
| CLCNKB | 0.205599343 | 1.15316531 | 0.550409237 | 0.707422466 | 37.76960394 | 43.50377796 | ENSG00000184908 |
| SPEN | 0.20559256 | 1.153159888 | 0.001272348 | 0.005300122 | 1731.495607 | 1995.812315 | ENSG00000065526 |
| TSPAN31 | 0.205367031 | 1.152979635 | 0.125631994 | 0.246573079 | 247.648476 | 285.3723205 | ENSG00000135452 |
| CPSF6 | 0.205333611 | 1.152952927 | 3.50E-05 | 0.000208168 | 3130.384124 | 3609.03187 | ENSG00000111605 |
| FAM114A2 | 0.205261182 | 1.152895045 | 0.21044011 | 0.365939568 | 181.2594366 | 209.0626487 | ENSG00000055147 |
| LMBRD1 | 0.205249967 | 1.152886083 | 0.109966018 | 0.222504139 | 283.8404759 | 327.3808582 | ENSG00000168216 |
| GLMN | 0.2049002 | 1.152606611 | 0.058423088 | 0.135071882 | 390.5072878 | 450.4356552 | ENSG00000174842 |
| SAYSD1 | 0.204882554 | 1.152592513 | 0.298775663 | 0.46917245 | 144.8212368 | 166.7454634 | ENSG00000112167 |
| NA | 0.204866942 | 1.152580041 | 0.172224204 | 0.314380878 | 189.9675395 | 218.8331788 | ENSG00000274265 |
| FRA10AC1 | 0.204760684 | 1.152495154 | 0.079536168 | 0.172687925 | 349.1951251 | 402.7301117 | ENSG00000148690 |
| CACNA2D1 | 0.20470484 | 1.152450544 | 0.077441544 | 0.168900489 | 348.9602747 | 401.8775009 | ENSG00000153956 |
| NA | 0.204432871 | 1.15223331 | 0.562388343 | 0.71705443 | 37.48929848 | 43.07411045 | ENSG00000257135 |
| SDHB | 0.20435758 | 1.152173179 | 0.005247811 | 0.018290067 | 925.4356713 | 1065.940849 | ENSG00000117118 |
| ACVR2A | 0.204207615 | 1.15205342 | 0.203908665 | 0.357048385 | 169.4829929 | 195.1247743 | ENSG00000121989 |
| SEPHS2 | 0.204201257 | 1.152048343 | 0.000822914 | 0.003568981 | 1742.66386 | 2007.489779 | ENSG00000179918 |
| CAMLG | 0.204195995 | 1.152044141 | 0.016127468 | 0.047018246 | 991.8509319 | 1142.670458 | ENSG00000164615 |
| CACNA1H | 0.203957251 | 1.151853511 | 0.001065466 | 0.00451207 | 1941.47862 | 2235.547126 | ENSG00000196557 |
| CACNA1A | 0.203925792 | 1.151828394 | 0.129216676 | 0.251637281 | 296.0398493 | 340.8664927 | ENSG00000141837 |
| POMZP3 | 0.203854545 | 1.151771513 | 0.71019667 | 0.825424989 | 13.88760144 | 16.03458603 | ENSG00000146707 |
| PLIN3 | 0.203767056 | 1.151701668 | 0.000551841 | 0.002503071 | 1709.245904 | 1968.236715 | ENSG00000105355 |
| SURF6 | 0.203756902 | 1.151693563 | 0.00062228 | 0.002783307 | 3130.79891 | 3605.842684 | ENSG00000148296 |
| SEMA3F-AS1 | 0.203368014 | 1.151383158 | 0.805039743 | 0.887405972 | 6.291997107 | 7.282309798 | ENSG00000235016 |
| VAMP7 | 0.203352625 | 1.151370876 | 0.091042441 | 0.192365797 | 364.035492 | 419.4205096 | ENSG00000124333 |
| TXNRD3 | 0.203349353 | 1.151368265 | 0.258460068 | 0.423615393 | 152.2047567 | 175.2772172 | ENSG00000197763 |
| PILRA | 0.203331451 | 1.151353978 | 0.7925548 | 0.879294608 | 7.146992935 | 8.167730574 | ENSG00000085514 |
| U2AF1L5 | 0.203317771 | 1.151343061 | 0.603454139 | 0.748838988 | 27.90108168 | 32.01217264 | ENSG00000275895 |
| ZSCAN2 | 0.203310735 | 1.151337446 | 0.096335801 | 0.201235944 | 437.4947162 | 503.035354 | ENSG00000176371 |
| LINC00032 | 0.203183078 | 1.151235574 | 0.801335832 | 0.885522687 | 7.198668648 | 8.230648672 | ENSG00000283945 |
| FAM227B | 0.203085057 | 1.151157358 | 0.516120945 | 0.680227448 | 48.80875811 | 56.03729974 | ENSG00000166262 |
| ZNF546 | 0.203039902 | 1.151121329 | 0.71510491 | 0.828882343 | 16.90100016 | 19.53699523 | ENSG00000187187 |
| H1-0 | 0.202872555 | 1.150987811 | 0.001482143 | 0.006061483 | 3406.525966 | 3919.845804 | ENSG00000189060 |
| RSAD1 | 0.20280781 | 1.150936158 | 0.00788951 | 0.025809145 | 899.8751409 | 1035.38735 | ENSG00000136444 |
| YES1 | 0.202734586 | 1.150877744 | 0.001243119 | 0.005192944 | 1743.27208 | 2007.07611 | ENSG00000176105 |
| NUDT9 | 0.20268887 | 1.150841275 | 0.007169585 | 0.023833958 | 894.5023303 | 1029.708573 | ENSG00000170502 |
| EEF2 | 0.202660866 | 1.150818937 | 1.11E-08 | 1.17E-07 | 83645.68943 | 96260.39332 | ENSG00000167658 |
| CHST9 | 0.202596164 | 1.150767326 | 0.121788583 | 0.241164396 | 296.4692367 | 341.4464283 | ENSG00000154080 |
| GLUD1 | 0.202586004 | 1.150759222 | 0.000449653 | 0.002087163 | 2158.044812 | 2483.737655 | ENSG00000148672 |
| EPPIN | 0.202162236 | 1.150421255 | 0.856583853 | 0.919296751 | 3.426604259 | 3.89766753 | ENSG00000101448 |
| MAP1B | 0.202016364 | 1.150304941 | 0.004792437 | 0.016920004 | 1383.192626 | 1590.815238 | ENSG00000131711 |
| ATRIP | 0.201974195 | 1.150271319 | 0.071260533 | 0.158282895 | 349.2892649 | 401.6624067 | ENSG00000164053 |
| CCDC77 | 0.201947899 | 1.150250353 | 0.0374466 | 0.094364628 | 528.6881501 | 607.8372694 | ENSG00000120647 |
| NA | 0.201685889 | 1.150041473 | 0.090299984 | 0.191111024 | 320.7352964 | 368.7172097 | ENSG00000249152 |
| TMEM191C | 0.201546294 | 1.149930201 | 0.749866015 | 0.852024107 | 11.44446256 | 13.12988638 | ENSG00000206140 |
| MELTF-AS1 | 0.201497711 | 1.149891478 | 0.706102927 | 0.823231454 | 15.46421746 | 17.75514035 | ENSG00000228109 |
| CSMD2 | 0.201474266 | 1.149872791 | 0.062843593 | 0.143096761 | 474.1929357 | 545.6077008 | ENSG00000121904 |
| MNT | 0.201402277 | 1.149815415 | 0.184843079 | 0.33112866 | 265.5348197 | 305.3083848 | ENSG00000070444 |
| HSPA1A | 0.201347803 | 1.149772 | 0.020454308 | 0.057302362 | 809.3204149 | 930.6157853 | ENSG00000204389 |
| PROKR2 | 0.20126054 | 1.149702457 | 0.590543972 | 0.739752192 | 31.57170967 | 36.32471412 | ENSG00000101292 |
| USE1 | 0.201243739 | 1.149689068 | 0.202423682 | 0.354995932 | 172.6672883 | 198.3231038 | ENSG00000053501 |
| DEPDC5 | 0.201161618 | 1.149623627 | 0.235108942 | 0.396216178 | 194.2120572 | 223.4226786 | ENSG00000100150 |
| C8orf82 | 0.201042519 | 1.149528726 | 0.04837242 | 0.116266178 | 626.6496643 | 720.516339 | ENSG00000213563 |
| ZBTB38 | 0.201020307 | 1.149511028 | 0.185191306 | 0.331433852 | 186.2325206 | 214.1396089 | ENSG00000177311 |
| IMPACT | 0.201009242 | 1.149502212 | 0.011919685 | 0.036446767 | 807.4461717 | 928.6666208 | ENSG00000154059 |
| LRP8 | 0.20092996 | 1.149439044 | 0.010597682 | 0.033065672 | 1315.991103 | 1512.262337 | ENSG00000157193 |
| IFNGR2 | 0.200905368 | 1.149419451 | 0.089515204 | 0.189895943 | 318.0851424 | 365.6764724 | ENSG00000159128 |
| NA | 0.200647694 | 1.149214176 | 0.334891545 | 0.507544432 | 135.5343382 | 155.8617327 | ENSG00000255135 |
| NFIL3 | 0.200509123 | 1.149103799 | 0.214463653 | 0.370951163 | 226.5743207 | 260.5141983 | ENSG00000165030 |
| ZNF195 | 0.200506183 | 1.149101457 | 0.020533909 | 0.057491966 | 899.9607702 | 1034.384944 | ENSG00000005801 |
| NUMB | 0.200478278 | 1.149079231 | 0.028364179 | 0.075109592 | 681.3339717 | 783.0101987 | ENSG00000133961 |
| SPOPL | 0.200368206 | 1.148991564 | 0.272170118 | 0.439080963 | 131.203453 | 150.9918217 | ENSG00000144228 |

| | | | | | | | |
|---|---|---|---|---|---|---|---|
| SLC39A9 | 0.200329848 | 1.148961016 | 0.02934405 | 0.077111576 | 540.4963329 | 621.0257727 | ENSG00000029364 |
| SIK1B | 0.200320176 | 1.148953313 | 0.384593817 | 0.558044327 | 79.61765976 | 91.36957901 | ENSG00000275993 |
| TMEM79 | 0.200314301 | 1.148948634 | 0.586133445 | 0.735756135 | 32.4727994 | 37.22895702 | ENSG00000163472 |
| RASSF8 | 0.200204261 | 1.148861003 | 0.08100535 | 0.175109168 | 389.3030018 | 447.1749648 | ENSG00000123094 |
| REEP5 | 0.19983534 | 1.148567257 | 0.004263067 | 0.015271835 | 1131.464684 | 1299.662354 | ENSG00000129625 |
| EIF3D | 0.199798131 | 1.148537635 | 9.19E-06 | 6.10E-05 | 4196.199549 | 4818.961091 | ENSG00000100353 |
| CKAP2L | 0.199744537 | 1.148494969 | 0.140271316 | 0.268396545 | 238.258189 | 273.6742232 | ENSG00000169607 |
| ZBED1 | 0.19969879 | 1.148458552 | 0.002026377 | 0.00798424 | 1506.088804 | 1730.060905 | ENSG00000214717 |
| PTBP3 | 0.199578964 | 1.148363168 | 0.006363321 | 0.02158137 | 1553.816415 | 1784.937036 | ENSG00000119314 |
| TET2 | 0.199495652 | 1.148296855 | 0.559611345 | 0.714836608 | 34.86023237 | 39.95484963 | ENSG00000168769 |
| COBL | 0.19939427 | 1.148216164 | 0.016559778 | 0.04800942 | 913.4561915 | 1049.040645 | ENSG00000106078 |
| NA | 0.19936549 | 1.148193259 | 0.839530313 | 0.908525708 | 3.98973069 | 4.585459144 | ENSG00000258427 |
| RASL11A | 0.199310425 | 1.148149435 | 0.540968934 | 0.700109467 | 44.59402755 | 51.15768977 | ENSG00000122035 |
| WDR37 | 0.199270659 | 1.148117788 | 0.081684981 | 0.176400422 | 355.8411552 | 408.7696776 | ENSG00000047056 |
| BDH1 | 0.199170261 | 1.148037893 | 0.041351813 | 0.102376517 | 501.1490738 | 575.2712106 | ENSG00000161267 |
| SLC25A6 | 0.199049468 | 1.147941775 | 3.73E-06 | 2.65E-05 | 16203.14634 | 18599.89366 | ENSG00000169100 |
| INTS8 | 0.199040239 | 1.147934431 | 0.002864171 | 0.010777727 | 1336.045088 | 1534.354759 | ENSG00000164941 |
| INTS4P2 | 0.198918783 | 1.147837794 | 0.841586344 | 0.909490034 | 3.98973069 | 4.58330857 | ENSG00000273024 |
| FAR1 | 0.198846344 | 1.147780162 | 0.01213821 | 0.036968456 | 953.9772127 | 1095.07892 | ENSG00000197601 |
| PATL1 | 0.19872838 | 1.147686316 | 0.010329789 | 0.032302148 | 895.8846833 | 1028.071097 | ENSG00000166889 |
| MIB2 | 0.198676953 | 1.147645405 | 0.005304802 | 0.018464503 | 1302.019491 | 1494.556478 | ENSG00000197530 |
| C18orf21 | 0.19862393 | 1.147603227 | 0.336430178 | 0.509225152 | 107.3997959 | 123.3881913 | ENSG00000141428 |
| CIRBP | 0.198599316 | 1.147583648 | 0.00022947 | 0.001143851 | 2775.373308 | 3183.852928 | ENSG00000099622 |
| TRIAP1 | 0.19857145 | 1.147561482 | 0.004590589 | 0.016305846 | 1115.449193 | 1280.383689 | ENSG00000170855 |
| LOC100128770 | 0.198297487 | 1.147343585 | 0.538723091 | 0.698233482 | 38.07945806 | 43.70543569 | ENSG00000205890 |
| GNG4 | 0.198168801 | 1.147241248 | 0.003951911 | 0.014295203 | 1321.333891 | 1515.827173 | ENSG00000168243 |
| CDNF | 0.198162921 | 1.147236572 | 0.803065978 | 0.886394088 | 6.319230415 | 7.278008651 | ENSG00000185267 |
| NME3 | 0.198077035 | 1.147168277 | 0.007877472 | 0.025787258 | 1254.105565 | 1439.038383 | ENSG00000103024 |
| NA | 0.198054455 | 1.147150323 | 0.851285335 | 0.915341698 | 3.693155369 | 4.21306363 | ENSG00000177173 |
| NAPSA | 0.198021723 | 1.147124297 | 0.805428543 | 0.887728253 | 5.995421786 | 6.88033929 | ENSG00000131400 |
| ATG9B | 0.197985783 | 1.14709572 | 0.603709369 | 0.749045371 | 36.72955677 | 42.20510077 | ENSG00000181652 |
| IFFO2 | 0.197955076 | 1.147071305 | 0.091814866 | 0.193658492 | 342.291554 | 392.3113243 | ENSG00000169991 |
| NA | 0.197837416 | 1.146977759 | 0.717226678 | 0.830493388 | 13.67830785 | 15.66864223 | ENSG00000260066 |
| NA | 0.197837024 | 1.146977447 | 0.843120029 | 0.910214813 | 4.577299527 | 5.296374031 | ENSG00000253520 |
| ATP11B | 0.197827416 | 1.146969809 | 0.068955142 | 0.154174043 | 400.0503987 | 458.7324507 | ENSG00000058063 |
| RPP14 | 0.197653396 | 1.146831468 | 0.179650619 | 0.324376646 | 206.4977612 | 237.0082877 | ENSG00000163684 |
| NUDT1 | 0.197547792 | 1.146747523 | 0.010299891 | 0.032237919 | 904.3851833 | 1036.923979 | ENSG00000106268 |
| CORO1B | 0.197031924 | 1.146337551 | 0.000176014 | 0.000897092 | 2432.877632 | 2788.567274 | ENSG00000172725 |
| SPIN2A | 0.197017229 | 1.146325875 | 0.557844878 | 0.713430492 | 49.92018058 | 57.2294959 | ENSG00000147059 |
| ADCY5 | 0.196930337 | 1.146256836 | 0.035346551 | 0.089897207 | 555.1546026 | 636.4583785 | ENSG00000173175 |
| NDUFV3 | 0.19692239 | 1.146250521 | 0.014408754 | 0.042809068 | 725.7042003 | 831.6173016 | ENSG00000160194 |
| VASN | 0.196692202 | 1.146067647 | 0.130776056 | 0.253905082 | 341.51886 | 391.8010132 | ENSG00000168140 |
| CLN8 | 0.196675324 | 1.146054239 | 0.011891576 | 0.036378116 | 793.9390728 | 910.033366 | ENSG00000182372 |
| SUPT20H | 0.196653132 | 1.146036611 | 0.005767487 | 0.019854308 | 1125.247328 | 1289.201628 | ENSG00000102710 |
| PIAS3 | 0.196637058 | 1.146023842 | 0.016313321 | 0.047430336 | 791.0052957 | 905.9456297 | ENSG00000131788 |
| FAM66B | 0.196600489 | 1.145994793 | 0.734115033 | 0.841512455 | 13.17714486 | 15.19326122 | ENSG00000215374 |
| NA | 0.196343435 | 1.145790622 | 0.606885291 | 0.751272767 | 29.36230679 | 33.7265357 | ENSG00000256943 |
| RIC1 | 0.196342574 | 1.145789938 | 0.034595813 | 0.088267139 | 534.5523867 | 612.269771 | ENSG00000107036 |
| CD4 | 0.196227856 | 1.145698833 | 0.426827136 | 0.59834186 | 81.43917539 | 93.32701217 | ENSG00000010610 |
| DFFA | 0.196222047 | 1.145694219 | 0.002139742 | 0.008378243 | 1568.821758 | 1797.337598 | ENSG00000160049 |
| TMEM80 | 0.196070964 | 1.145574245 | 0.19580975 | 0.346229687 | 227.8482671 | 261.1872206 | ENSG00000177042 |
| NA | 0.196063631 | 1.145568423 | 0.76313393 | 0.85955606 | 10.38228832 | 11.90971434 | ENSG00000242477 |
| FAHD2B | 0.195990581 | 1.145510419 | 0.045643738 | 0.110909855 | 511.2734592 | 585.4625638 | ENSG00000144199 |
| ZNF267 | 0.195881375 | 1.145423712 | 0.132779966 | 0.256890816 | 244.8432948 | 280.5029303 | ENSG00000185947 |
| NT5C | 0.195829231 | 1.145382313 | 0.176492833 | 0.319992786 | 221.2579798 | 253.3725303 | ENSG00000125458 |
| CHMP4C | 0.195813397 | 1.145369742 | 0.656445318 | 0.7869973 | 22.16730492 | 25.37381895 | ENSG00000164695 |
| MATN2 | 0.195784197 | 1.145346561 | 0.088373259 | 0.188002706 | 326.7302057 | 374.3733484 | ENSG00000132561 |
| TAF1A-AS1 | 0.195768754 | 1.1453343 | 0.751471473 | 0.852957914 | 10.63556064 | 12.22726101 | ENSG00000225265 |
| B3GNT8 | 0.195578262 | 1.145183082 | 0.727992991 | 0.836782029 | 14.01539527 | 16.04318832 | ENSG00000177191 |
| HERC6 | 0.195567238 | 1.145174331 | 0.319224361 | 0.490669175 | 106.5087185 | 122.0798273 | ENSG00000138642 |
| TTC9 | 0.195346423 | 1.144999067 | 0.007765005 | 0.025492632 | 1026.188449 | 1175.054561 | ENSG00000133985 |
| PFKM | 0.195309823 | 1.14497002 | 0.000385292 | 0.001824863 | 2757.772927 | 3156.55823 | ENSG00000152556 |
| NA | 0.19528132 | 1.144947399 | 0.473395958 | 0.641990664 | 64.36727856 | 73.50016726 | ENSG00000137970 |
| PCDHB9 | 0.19509489 | 1.144799455 | 0.609531501 | 0.753313613 | 29.12019808 | 33.38156461 | ENSG00000177839 |
| SEC23A | 0.194968702 | 1.144699327 | 0.006539836 | 0.022090699 | 1123.840844 | 1286.281032 | ENSG00000100934 |
| NA | 0.194950475 | 1.144684865 | 0.64370968 | 0.777327289 | 22.41990152 | 25.66824235 | ENSG00000231160 |
| DYNC1H1 | 0.19489221 | 1.144638636 | 0.079528034 | 0.172687925 | 6808.568134 | 7792.895564 | ENSG00000197102 |
| SLC25A4 | 0.19449704 | 1.144325151 | 0.001745453 | 0.007011982 | 1469.706835 | 1681.406768 | ENSG00000151729 |
| MLXIPL | 0.194249485 | 1.14412881 | 0.391548505 | 0.564598636 | 112.7313306 | 129.1529615 | ENSG00000009950 |
| SELENOO | 0.194237392 | 1.14411922 | 0.057267323 | 0.133085643 | 508.0285518 | 580.8418836 | ENSG00000073169 |
| IFT27 | 0.194120755 | 1.144026726 | 0.088154901 | 0.187641642 | 371.7353605 | 424.7988901 | ENSG00000100360 |
| ZNF460 | 0.194090274 | 1.144002555 | 0.600628127 | 0.746663049 | 34.61812366 | 39.46630471 | ENSG00000197714 |
| EML6 | 0.194047729 | 1.143968818 | 0.627530696 | 0.765698587 | 31.50907051 | 35.97759245 | ENSG00000214595 |
| TMOD3 | 0.193911726 | 1.143860982 | 0.015960942 | 0.046637767 | 732.5247939 | 837.7617005 | ENSG00000138594 |
| LARP4 | 0.19390776 | 1.143857838 | 0.006188445 | 0.021073485 | 1139.658446 | 1303.785312 | ENSG00000161813 |
| FBXO46 | 0.193809195 | 1.143779691 | 0.042891269 | 0.105524228 | 536.9574921 | 614.1570345 | ENSG00000177051 |
| ZBED9 | 0.193674467 | 1.143672883 | 0.419614859 | 0.591664088 | 80.05319253 | 91.56236188 | ENSG00000232040 |

| | | | | | | | |
|---|---|---|---|---|---|---|---|
| PON1 | 0.193663519 | 1.143664204 | 0.781897407 | 0.87209434 | 8.615915026 | 9.892586044 | ENSG00000005421 |
| RPL22 | 0.193590339 | 1.143606194 | 0.011347381 | 0.035103004 | 1367.476238 | 1564.566739 | ENSG00000116251 |
| MRPL19 | 0.193567483 | 1.143588077 | 0.020851834 | 0.058145812 | 750.7280129 | 858.8702721 | ENSG00000115364 |
| RYR3 | 0.193512081 | 1.143544162 | 0.55901347 | 0.714356707 | 46.19875276 | 52.74971217 | ENSG00000198838 |
| RPLP0 | 0.193502327 | 1.14353643 | 8.75E-08 | 8.12E-07 | 42348.7618 | 48426.30596 | ENSG00000089157 |
| DRD4 | 0.193483579 | 1.14352157 | 0.485523942 | 0.653129864 | 54.61757695 | 62.58153069 | ENSG00000069696 |
| NGLY1 | 0.193360639 | 1.143424128 | 0.061220377 | 0.140194953 | 420.2195974 | 480.5855777 | ENSG00000151092 |
| TMEM143 | 0.193181902 | 1.143282477 | 0.154646578 | 0.289555192 | 254.3794725 | 290.5544352 | ENSG00000161558 |
| KLF1 | 0.193141807 | 1.143250703 | 0.76500657 | 0.860961539 | 12.57562151 | 14.40275669 | ENSG00000105610 |
| BTBD9 | 0.193128668 | 1.143240292 | 0.266362277 | 0.432597633 | 142.4276585 | 162.7673701 | ENSG00000183826 |
| ZSCAN22 | 0.19309185 | 1.143211116 | 0.491761834 | 0.657967647 | 59.58778195 | 68.2782577 | ENSG00000182318 |
| LIMD1 | 0.193057657 | 1.143184021 | 0.121448632 | 0.240590014 | 348.2557506 | 398.3196914 | ENSG00000144791 |
| IRF2BP1 | 0.192956829 | 1.143104129 | 0.029324665 | 0.077081633 | 913.5770294 | 1044.863451 | ENSG00000170604 |
| PEX16 | 0.192858792 | 1.143026453 | 0.07516604 | 0.165121253 | 466.4147955 | 532.9763614 | ENSG00000121680 |
| PURB | 0.192689822 | 1.142892588 | 0.016681455 | 0.048304106 | 816.6425079 | 933.2448226 | ENSG00000146676 |
| LOC105375421 | 0.192660338 | 1.142869232 | 0.849476075 | 0.914224265 | 4.014173095 | 4.581157997 | ENSG00000284523 |
| NA | 0.192654117 | 1.142864303 | 0.442174118 | 0.61325164 | 71.34229878 | 81.36299414 | ENSG00000223804 |
| TRIM14 | 0.192590103 | 1.142813594 | 0.035235989 | 0.089627831 | 529.6902391 | 605.3582271 | ENSG00000106785 |
| SDHA | 0.192517095 | 1.142755763 | 0.594576109 | 0.742536915 | 31.45248871 | 35.9837716 | ENSG00000073578 |
| IL11RA | 0.192464502 | 1.142714105 | 0.588869953 | 0.738474911 | 35.33974386 | 40.25572475 | ENSG00000137070 |
| SLC2A3 | 0.192403947 | 1.142666142 | 0.000151963 | 0.000784876 | 13003.05527 | 14857.5334 | ENSG00000059804 |
| LOC100130548 | 0.192359819 | 1.142631192 | 0.846819342 | 0.912319442 | 4.365215031 | 5.00195063 | ENSG00000235138 |
| NA | 0.192170028 | 1.142480885 | 0.636507143 | 0.77256924 | 25.69638473 | 29.42958128 | ENSG00000196741 |
| MGME1 | 0.192104302 | 1.142428838 | 0.014299502 | 0.042550006 | 1168.530376 | 1334.695315 | ENSG00000125871 |
| BRSK1 | 0.192097098 | 1.142423133 | 0.04249514 | 0.104723269 | 608.7385901 | 695.3140698 | ENSG00000160469 |
| ZNF92 | 0.192059696 | 1.142393516 | 0.095924273 | 0.200571226 | 363.14086 | 415.3321671 | ENSG00000146757 |
| DAB2 | 0.191990718 | 1.142338897 | 0.364266109 | 0.537935965 | 115.2338421 | 131.9137964 | ENSG00000153071 |
| FBXO16 | 0.191961198 | 1.142315523 | 0.46211862 | 0.631802289 | 71.61308026 | 81.7700593 | ENSG00000214050 |
| NA | 0.19193789 | 1.142297068 | 0.591015818 | 0.739910841 | 39.04406284 | 44.46018609 | ENSG00000248927 |
| SEC31A | 0.191866154 | 1.142240271 | 0.001049878 | 0.004452229 | 1927.768387 | 2202.510447 | ENSG00000138674 |
| FANCL | 0.191758439 | 1.142154991 | 0.03345602 | 0.085790255 | 561.977787 | 642.1341023 | ENSG00000115392 |
| NTS | 0.191755073 | 1.142152327 | 0.212032091 | 0.367899899 | 186.5501838 | 212.8344921 | ENSG00000133636 |
| SPTBN4 | 0.191564558 | 1.14200151 | 0.389220114 | 0.562514625 | 99.18679867 | 113.2571152 | ENSG00000160460 |
| TPM3 | 0.191417118 | 1.141884806 | 1.20E-05 | 7.78E-05 | 13982.84494 | 15966.77223 | ENSG00000143549 |
| UBA6 | 0.191398102 | 1.141869755 | 0.028980837 | 0.076323459 | 736.4319119 | 841.4461028 | ENSG00000033178 |
| GSDMD | 0.191145955 | 1.141670202 | 0.01960567 | 0.055278132 | 711.1758027 | 811.8254052 | ENSG00000104518 |
| ZNF296 | 0.190973426 | 1.14153368 | 0.302224082 | 0.472989952 | 127.265398 | 145.3926945 | ENSG00000170684 |
| NA | 0.190857412 | 1.141441888 | 0.287734387 | 0.457109781 | 149.6746483 | 170.7049403 | ENSG00000280511 |
| POC1B | 0.190819773 | 1.141412109 | 0.170612586 | 0.312012078 | 220.3508327 | 251.5594708 | ENSG00000139323 |
| PIK3CA | 0.190792963 | 1.141390898 | 0.216297011 | 0.373375092 | 179.2768365 | 204.6882552 | ENSG00000121879 |
| GCN1 | 0.19078192 | 1.141382161 | 9.15E-06 | 6.07E-05 | 7323.406259 | 8357.569893 | ENSG00000089154 |
| STK10 | 0.190778724 | 1.141379633 | 0.030963195 | 0.080587021 | 690.5330378 | 788.0694582 | ENSG00000072786 |
| ZFP1 | 0.190769588 | 1.141372405 | 0.336161449 | 0.508938198 | 114.01022 | 130.2580381 | ENSG00000184517 |
| MAFA | 0.190761109 | 1.141365697 | 0.389186837 | 0.562508674 | 87.06381876 | 99.53299633 | ENSG00000182759 |
| POLL | 0.190718016 | 1.141331605 | 0.063512401 | 0.144347046 | 430.2101684 | 490.7709696 | ENSG00000166169 |
| NKD1 | 0.190705085 | 1.141321375 | 0.181256299 | 0.326450869 | 221.5835913 | 253.093161 | ENSG00000140807 |
| ASPHD1 | 0.190507127 | 1.14116478 | 0.09136144 | 0.192955281 | 450.4409302 | 513.8120463 | ENSG00000174939 |
| PRMT2 | 0.190489448 | 1.141150797 | 0.003064562 | 0.01140358 | 1328.334894 | 1515.965064 | ENSG00000160310 |
| PRPSAP2 | 0.190484353 | 1.141146767 | 0.013285516 | 0.039976743 | 867.1022995 | 989.2076942 | ENSG00000141127 |
| PRIMPOL | 0.190475314 | 1.141139617 | 0.482721852 | 0.650701367 | 72.75483924 | 83.11013674 | ENSG00000164306 |
| NA | 0.190147143 | 1.14088007 | 0.551990187 | 0.708573509 | 42.05244333 | 48.0158266 | ENSG00000230724 |
| SNX14 | 0.190057618 | 1.140809276 | 0.126762277 | 0.248235822 | 305.802993 | 348.9492543 | ENSG00000135317 |
| BMPR1A | 0.190015299 | 1.140775813 | 0.004393684 | 0.015687315 | 1649.458026 | 1881.853602 | ENSG00000107779 |
| RBPMS-AS1 | 0.189978427 | 1.140746658 | 0.847012464 | 0.912404925 | 4.658999449 | 5.346921724 | ENSG00000254109 |
| MSANTD4 | 0.18996492 | 1.140735978 | 0.052271416 | 0.12343835 | 483.134387 | 551.0667676 | ENSG00000170903 |
| SACM1L | 0.189921741 | 1.140701837 | 0.006487026 | 0.021943051 | 1419.82166 | 1619.949032 | ENSG00000211456 |
| NA | 0.189867248 | 1.140658751 | 0.362133609 | 0.535401553 | 96.21949386 | 109.6127312 | ENSG00000188185 |
| LIFR-AS1 | 0.189746002 | 1.140562893 | 0.862292923 | 0.922713518 | 4.028127609 | 4.585459144 | ENSG00000244968 |
| MFSD3 | 0.189744421 | 1.140561643 | 0.012423675 | 0.037748482 | 1275.423255 | 1455.000927 | ENSG00000167700 |
| RGS20 | 0.189735197 | 1.140554351 | 0.858117624 | 0.920622544 | 3.95970648 | 4.509637604 | ENSG00000147509 |
| ERF | 0.18956878 | 1.140422793 | 0.012098156 | 0.036858107 | 1431.686564 | 1632.698885 | ENSG00000105722 |
| NUSAP1 | 0.189561388 | 1.140416951 | 0.843702958 | 0.910436526 | 4.337981724 | 4.960005232 | ENSG00000137804 |
| EFNA4 | 0.189536643 | 1.14039739 | 0.155128949 | 0.290152278 | 234.9698665 | 267.925864 | ENSG00000243364 |
| NA | 0.189472039 | 1.140346325 | 0.845618818 | 0.91184084 | 5.497925581 | 6.217821522 | ENSG00000260288 |
| SNCA | 0.189413066 | 1.140299711 | 0.194603256 | 0.344728321 | 202.3932401 | 230.7759452 | ENSG00000145335 |
| DDIT3 | 0.189298218 | 1.14020894 | 0.637140768 | 0.773121308 | 26.25392935 | 30.03294906 | ENSG00000175197 |
| NA | 0.189227237 | 1.140152843 | 0.741269696 | 0.84669511 | 13.69995935 | 15.56539627 | ENSG00000239377 |
| INTS4 | 0.189131211 | 1.140076957 | 0.040152267 | 0.099906796 | 570.6680172 | 650.4021469 | ENSG00000149262 |
| NA | 0.189087506 | 1.140042419 | 0.651395764 | 0.783266107 | 24.97476453 | 28.47668079 | ENSG00000228506 |
| TEAD3 | 0.188881254 | 1.139879447 | 0.017122197 | 0.049387606 | 775.051042 | 883.7318899 | ENSG00000007866 |
| TUFT1 | 0.188856747 | 1.139860084 | 0.130717954 | 0.253868927 | 342.8313141 | 390.3345239 | ENSG00000143367 |
| NA | 0.188835698 | 1.139843454 | 0.469069633 | 0.637959835 | 63.86994562 | 72.90755235 | ENSG00000256008 |
| TAPBP | 0.188629636 | 1.13968066 | 0.040036528 | 0.099657376 | 740.2671151 | 844.0746987 | ENSG00000231925 |
| NA | 0.18850591 | 1.139582925 | 0.687587327 | 0.810099598 | 21.66893284 | 24.74034313 | ENSG00000243517 |
| FOXK1 | 0.188468961 | 1.139553739 | 0.100393556 | 0.207732713 | 389.6193505 | 443.7669025 | ENSG00000164916 |
| SRPX | 0.188430059 | 1.139523012 | 0.287597901 | 0.45693058 | 138.7547151 | 158.1391599 | ENSG00000101955 |
| TBCC | 0.188272957 | 1.13939893 | 0.073233991 | 0.161772234 | 457.0944572 | 520.8550299 | ENSG00000124659 |

| | | | | | | | |
|---|---|---|---|---|---|---|---|
| PNMA2 | 0.188156933 | 1.139307302 | 0.308518577 | 0.479622199 | 119.5338273 | 136.1959694 | ENSG00000240694 |
| PCDHGA9 | 0.187800899 | 1.139026173 | 0.790196084 | 0.87763484 | 8.882466137 | 10.1617356 | ENSG00000261934 |
| AFAP1L2 | 0.187658216 | 1.138913529 | 0.342011757 | 0.514835246 | 110.5966944 | 126.0804803 | ENSG00000169129 |
| FOXA3 | 0.187640228 | 1.138899329 | 0.058774793 | 0.135689868 | 520.7709787 | 593.4611252 | ENSG00000170608 |
| SRRM3 | 0.18759944 | 1.13886713 | 0.076666971 | 0.167571101 | 458.841094 | 522.8431042 | ENSG00000177679 |
| DNMBP | 0.187480254 | 1.138773048 | 0.045136548 | 0.109940549 | 676.6331825 | 770.1887179 | ENSG00000107554 |
| ALMS1-IT1 | 0.187361562 | 1.138679364 | 0.677990639 | 0.802991001 | 21.70107223 | 24.75162904 | ENSG00000230002 |
| RAP1GAP | 0.187073939 | 1.138452374 | 0.10211582 | 0.210416181 | 422.5699608 | 481.1494232 | ENSG00000076864 |
| FOXJ2 | 0.187016328 | 1.138406914 | 0.099388988 | 0.206279864 | 343.1879009 | 390.4786613 | ENSG00000065970 |
| NA | 0.186982701 | 1.138380379 | 0.802894681 | 0.886331404 | 7.851867711 | 8.882946609 | ENSG00000260448 |
| CACNA1I | 0.186964089 | 1.138365693 | 0.44835988 | 0.618735992 | 88.78581301 | 101.2449605 | ENSG00000100346 |
| GPR63 | 0.186884711 | 1.138303061 | 0.167394474 | 0.30749961 | 290.6996538 | 331.2096768 | ENSG00000112218 |
| HIC1 | 0.186791872 | 1.138229812 | 0.81415043 | 0.893693863 | 6.961265867 | 7.967950839 | ENSG00000177374 |
| PTPRM | 0.186778437 | 1.138219213 | 0.2664602 | 0.432683814 | 166.7293242 | 189.6998684 | ENSG00000173482 |
| SNORD6 | 0.186762714 | 1.138206808 | 0.849568478 | 0.914260042 | 4.044197305 | 4.585459144 | ENSG00000202314 |
| CKMT1A | 0.186739534 | 1.138188521 | 0.294797746 | 0.464765403 | 146.1373591 | 166.377066 | ENSG00000223572 |
| LSM11 | 0.186506909 | 1.13800501 | 0.273106208 | 0.439957566 | 144.7166461 | 164.744431 | ENSG00000155858 |
| PPARGC1B | 0.186500152 | 1.13799968 | 0.129126088 | 0.251537046 | 311.2295433 | 354.1496339 | ENSG00000155846 |
| NA | 0.186361654 | 1.137890438 | 0.846309148 | 0.912075406 | 4.346354432 | 4.953553511 | ENSG00000279573 |
| SH3GLB1 | 0.186353851 | 1.137884284 | 0.026915916 | 0.071976009 | 828.7014986 | 943.5004021 | ENSG00000097033 |
| ELL | 0.186286325 | 1.137831026 | 0.062307453 | 0.142161216 | 494.726343 | 563.066564 | ENSG00000105656 |
| SLC22A23 | 0.186134746 | 1.137711484 | 0.012366517 | 0.037604425 | 1194.171806 | 1358.263666 | ENSG00000137266 |
| RHOQ | 0.186125365 | 1.137704086 | 0.072387121 | 0.160268559 | 576.6495214 | 655.7837077 | ENSG00000119729 |
| SNX2 | 0.186109351 | 1.137691458 | 0.005200457 | 0.018141414 | 1437.428387 | 1635.77474 | ENSG00000205302 |
| TNRC6A | 0.185880248 | 1.137510804 | 0.000340857 | 0.001632846 | 2355.320666 | 2679.07976 | ENSG00000090905 |
| METTL7B | 0.185843539 | 1.137481861 | 0.63775022 | 0.773520244 | 27.33987027 | 31.08910761 | ENSG00000170439 |
| NA | 0.185695698 | 1.137365302 | 0.878215249 | 0.933167192 | 3.807670405 | 4.299638038 | ENSG00000272482 |
| ZNF597 | 0.185654283 | 1.137332653 | 0.465704944 | 0.635040226 | 72.98470829 | 83.22254856 | ENSG00000167981 |
| NA | 0.185494547 | 1.137206734 | 0.747545772 | 0.850422569 | 12.45360964 | 14.13092352 | ENSG00000261420 |
| NECAP1 | 0.185432237 | 1.137157619 | 0.174948841 | 0.317910521 | 223.5337766 | 254.211432 | ENSG00000089818 |
| LYPLAL1 | 0.185426118 | 1.137152796 | 0.437555381 | 0.608395621 | 79.45146902 | 90.27604878 | ENSG00000143353 |
| NA | 0.185414976 | 1.137144013 | 0.865709559 | 0.924922635 | 4.245118189 | 4.835786573 | ENSG00000278383 |
| STX8 | 0.185383979 | 1.137119582 | 0.147278896 | 0.278353795 | 348.0361309 | 395.2023086 | ENSG00000170310 |
| LRRC71 | 0.185299266 | 1.137052814 | 0.814529546 | 0.893906558 | 6.354836431 | 7.22315981 | ENSG00000160838 |
| MTPAP | 0.18513654 | 1.136924569 | 0.037235231 | 0.093896057 | 591.3666513 | 672.7029273 | ENSG00000107951 |
| MPC2 | 0.185065009 | 1.1368682 | 0.309833973 | 0.480969435 | 123.3798946 | 140.2447469 | ENSG00000143158 |
| CEP120 | 0.185018729 | 1.136831731 | 0.141272468 | 0.269750291 | 351.93947 | 400.5368357 | ENSG00000168944 |
| HSPB1 | 0.184923796 | 1.136756928 | 0.000439744 | 0.00204906 | 2714.355481 | 3085.518821 | ENSG00000106211 |
| NA | 0.184908557 | 1.13674492 | 0.698586472 | 0.818040309 | 17.63522342 | 20.09626704 | ENSG00000285976 |
| LOC101927322 | 0.184766079 | 1.136632663 | 0.862205702 | 0.922712204 | 4.019754901 | 4.553733576 | ENSG00000267226 |
| NA | 0.184741358 | 1.136613187 | 0.74677985 | 0.849969325 | 13.03627239 | 14.83161858 | ENSG00000228204 |
| TNFRSF11B | 0.184724936 | 1.136600248 | 0.758292197 | 0.856402715 | 12.68734564 | 14.41459405 | ENSG00000164761 |
| NA | 0.184698834 | 1.136579684 | 0.861418313 | 0.922330059 | 3.750412887 | 4.219515351 | ENSG00000233589 |
| LOC374443 | 0.184690756 | 1.136573321 | 0.787753146 | 0.876235135 | 11.93426178 | 13.62811809 | ENSG00000256594 |
| XYLT1 | 0.184671837 | 1.136558416 | 0.016151189 | 0.047058236 | 1001.767711 | 1138.188821 | ENSG00000103489 |
| EXOC4 | 0.18462133 | 1.136518628 | 0.005933983 | 0.020350659 | 1295.860016 | 1472.709984 | ENSG00000131558 |
| COX7A2L | 0.184588359 | 1.136492654 | 0.007782436 | 0.025541165 | 1544.346906 | 1755.445355 | ENSG00000115944 |
| PREPL | 0.184493087 | 1.136417605 | 0.034831328 | 0.088821038 | 624.9812466 | 709.8351022 | ENSG00000138078 |
| GLI3 | 0.184452281 | 1.136385463 | 0.058907243 | 0.135930576 | 462.1150986 | 525.1726661 | ENSG00000106571 |
| NA | 0.184439807 | 1.136375637 | 0.77745642 | 0.869419662 | 11.37045962 | 12.95699803 | ENSG00000227347 |
| WWTR1 | 0.184322521 | 1.136283258 | 0.185604082 | 0.332012648 | 267.5693837 | 304.3954664 | ENSG00000018408 |
| EMC4 | 0.184295777 | 1.136262194 | 0.015108939 | 0.044539509 | 1023.069997 | 1162.252944 | ENSG00000128463 |
| KIAA1522 | 0.184234636 | 1.13621404 | 0.0017828 | 0.007144161 | 1687.440316 | 1917.234158 | ENSG00000162522 |
| TTC31 | 0.184144156 | 1.136142784 | 0.066732995 | 0.150226399 | 503.0682342 | 571.5784601 | ENSG00000115282 |
| SMURF1 | 0.184078124 | 1.136090784 | 0.025576949 | 0.068959457 | 701.6569951 | 796.9332795 | ENSG00000198742 |
| MSN | 0.183945451 | 1.135986312 | 4.56E-06 | 3.19E-05 | 8010.755087 | 9099.153203 | ENSG00000147065 |
| EVI5 | 0.183780205 | 1.135856203 | 0.408686635 | 0.580985547 | 101.993019 | 115.872486 | ENSG00000067208 |
| PMS2 | 0.183747784 | 1.135830678 | 0.128711143 | 0.251019351 | 288.0745426 | 327.2415855 | ENSG00000122512 |
| ZNF646 | 0.183722352 | 1.135810656 | 0.028076176 | 0.074474127 | 933.7612444 | 1060.27858 | ENSG00000167395 |
| TBC1D9B | 0.183715253 | 1.135805067 | 0.002879134 | 0.010826777 | 1689.004054 | 1917.998353 | ENSG00000197226 |
| ZFYVE19 | 0.183516027 | 1.135648231 | 0.158936865 | 0.295503619 | 305.8860813 | 346.9361851 | ENSG00000166140 |
| ATF6B | 0.1835155 | 1.135647816 | 0.046752769 | 0.113233857 | 605.7808462 | 687.8346581 | ENSG00000213676 |
| ATL3 | 0.183446182 | 1.135593252 | 0.14538006 | 0.275618051 | 311.5184216 | 353.8285612 | ENSG00000184743 |
| LMBR1L | 0.183436663 | 1.13558576 | 0.051978073 | 0.123001596 | 592.7460501 | 672.4502131 | ENSG00000139636 |
| LPIN1 | 0.183397754 | 1.135555134 | 0.112796569 | 0.227271734 | 395.500933 | 449.0135408 | ENSG00000134324 |
| CHST15 | 0.183352617 | 1.135519607 | 0.108252903 | 0.219821763 | 357.7729555 | 406.4815096 | ENSG00000182022 |
| NAGA | 0.183333537 | 1.13550459 | 0.042405325 | 0.104541997 | 555.2925209 | 630.5505369 | ENSG00000198951 |
| ZNF347 | 0.183256245 | 1.135443757 | 0.263777085 | 0.429556284 | 198.2806969 | 225.536298 | ENSG00000197937 |
| SPG11 | 0.183210354 | 1.13540764 | 0.067271294 | 0.15114495 | 739.6264297 | 839.4889364 | ENSG00000104133 |
| NA | 0.183098151 | 1.135319338 | 0.001001914 | 0.004269478 | 8890.833417 | 10093.31417 | ENSG00000240342 |
| PRSS46P | 0.183053301 | 1.135284045 | 0.837628626 | 0.907563032 | 6.039400511 | 6.83194217 | ENSG00000261603 |
| PELI3 | 0.182973338 | 1.135221122 | 0.342641113 | 0.515408609 | 141.800929 | 160.8494776 | ENSG00000174516 |
| TINAGL1 | 0.182929171 | 1.135186369 | 0.231085142 | 0.391383974 | 193.1633619 | 219.5572576 | ENSG00000142910 |
| SFN | 0.182851629 | 1.135125357 | 0.293600705 | 0.463300711 | 144.457792 | 164.1026134 | ENSG00000175793 |
| TMPPE | 0.182834246 | 1.13511168 | 0.675075306 | 0.800668782 | 22.3409925 | 25.41576434 | ENSG00000188167 |
| MRPS31P5 | 0.182820142 | 1.135100583 | 0.807667002 | 0.889275996 | 7.764585982 | 8.752276228 | ENSG00000243406 |
| JAG1 | 0.182817668 | 1.135098636 | 0.092382809 | 0.1944949 | 643.5513388 | 730.432484 | ENSG00000101384 |

| | | | | | | | |
|---|---|---|---|---|---|---|---|
| FTL | 0.182580162 | 1.134911784 | 2.19E-06 | 1.62E-05 | 12722.81811 | 14438.58284 | ENSG00000087086 |
| HP1BP3 | 0.182544234 | 1.134883521 | 0.002271792 | 0.008814389 | 1500.437416 | 1702.803287 | ENSG00000127483 |
| NA | 0.182539194 | 1.134879556 | 0.78464724 | 0.873939371 | 10.49468817 | 11.88013935 | ENSG00000224790 |
| JARID2 | 0.182423723 | 1.134788726 | 1.04E-06 | 8.20E-06 | 8322.059291 | 9443.707642 | ENSG00000008083 |
| RPL28 | 0.182415961 | 1.13478262 | 6.56E-05 | 0.000368589 | 13730.02451 | 15580.26914 | ENSG00000108107 |
| MYLK-AS1 | 0.182315527 | 1.134703625 | 0.81411064 | 0.893693863 | 6.917962863 | 7.889978725 | ENSG00000239523 |
| YLPM1 | 0.182205825 | 1.134617346 | 0.000318797 | 0.001535656 | 2506.42987 | 2843.328146 | ENSG00000119596 |
| NA | 0.182079294 | 1.134517839 | 0.847769908 | 0.912986619 | 4.670163061 | 5.321647878 | ENSG00000279019 |
| ASB13 | 0.181916666 | 1.134389957 | 0.037295048 | 0.094022317 | 643.9622673 | 730.7522544 | ENSG00000196372 |
| MED1 | 0.181804812 | 1.13430201 | 0.004745377 | 0.016766129 | 1428.408064 | 1620.467226 | ENSG00000125686 |
| MRC2 | 0.18169754 | 1.134217672 | 0.032401857 | 0.083620632 | 736.8737391 | 835.3283959 | ENSG00000011028 |
| ST6GALNAC5 | 0.18156048 | 1.134109923 | 0.360021447 | 0.533095927 | 139.4800532 | 158.4787454 | ENSG00000117069 |
| TRAFD1 | 0.181451001 | 1.134023865 | 0.055927108 | 0.130568781 | 749.9695715 | 850.0919954 | ENSG00000135148 |
| ZNF2 | 0.181353749 | 1.133947423 | 0.503527908 | 0.668887233 | 58.48462005 | 66.30711548 | ENSG00000275111 |
| TWF1 | 0.181308381 | 1.133911765 | 0.120178578 | 0.238711382 | 396.4617972 | 450.1307394 | ENSG00000151239 |
| RBM18 | 0.18130255 | 1.133907181 | 0.274334841 | 0.441089633 | 191.7717092 | 217.7482387 | ENSG00000119446 |
| SLC30A5 | 0.181279119 | 1.133888766 | 0.052688501 | 0.124271166 | 548.1032642 | 621.9265079 | ENSG00000145740 |
| CIT | 0.181277818 | 1.133887743 | 0.0154765 | 0.04542175 | 1148.442765 | 1301.628105 | ENSG00000122966 |
| MYRIP | 0.181264439 | 1.133877228 | 0.600178393 | 0.746377427 | 37.4884226 | 42.53204324 | ENSG00000170011 |
| LOC100996842 | 0.181241345 | 1.133859077 | 0.309477695 | 0.480609742 | 131.8596431 | 149.4810065 | ENSG00000233223 |
| WBP2 | 0.181151791 | 1.133788696 | 0.007641899 | 0.025118396 | 1580.955014 | 1792.014412 | ENSG00000132471 |
| CERS6 | 0.181142124 | 1.133781099 | 0.064773227 | 0.146539267 | 474.6200205 | 537.947622 | ENSG00000172292 |
| PRICKLE3 | 0.181009822 | 1.133677131 | 0.247406867 | 0.410981302 | 195.3846041 | 221.6733428 | ENSG00000012211 |
| LOC171391 | 0.180998035 | 1.133667869 | 0.509439792 | 0.67409308 | 59.47904888 | 67.47378364 | ENSG00000255284 |
| FKRP | 0.180951969 | 1.133631671 | 0.051644632 | 0.122513107 | 653.6484539 | 740.6795648 | ENSG00000181027 |
| EIF4A2 | 0.180939551 | 1.133621913 | 0.000249193 | 0.001232609 | 6953.284257 | 7881.753219 | ENSG00000156976 |
| VCAN | 0.180891787 | 1.133584382 | 4.25E-05 | 0.000248012 | 5269.915825 | 5972.86723 | ENSG00000038427 |
| SECISBP2 | 0.180864245 | 1.133562742 | 0.008720478 | 0.028064881 | 1501.463257 | 1702.2859 | ENSG00000187742 |
| WDR47 | 0.18062084 | 1.133371508 | 0.290442962 | 0.46012443 | 149.9033561 | 169.8412979 | ENSG00000085433 |
| NA | 0.180586287 | 1.133344364 | 0.788396078 | 0.876551964 | 8.989284186 | 10.21013272 | ENSG00000231551 |
| TRIM38 | 0.180480023 | 1.133260888 | 0.55331703 | 0.709662844 | 46.96539082 | 53.31166759 | ENSG00000112343 |
| SDCBP | 0.180470343 | 1.133253285 | 0.00165101 | 0.00667008 | 1843.16543 | 2088.79272 | ENSG00000137575 |
| AMOT | 0.18023253 | 1.133066495 | 0.148253599 | 0.279866931 | 358.4951762 | 406.0622977 | ENSG00000126016 |
| CTSV | 0.180203367 | 1.133043591 | 0.003431352 | 0.012622849 | 1563.245024 | 1770.573972 | ENSG00000136943 |
| DCAF17 | 0.180150169 | 1.133001812 | 0.077147828 | 0.168469407 | 416.6201815 | 471.9540246 | ENSG00000115827 |
| YIPF2 | 0.180062592 | 1.132933037 | 0.030791526 | 0.080215981 | 870.233819 | 986.3500104 | ENSG00000130733 |
| ADAT1 | 0.18002093 | 1.132900321 | 0.160530434 | 0.29785444 | 299.3260936 | 338.8945207 | ENSG00000065457 |
| SERINC1 | 0.180016231 | 1.132896631 | 0.042927508 | 0.105581979 | 797.079804 | 903.634308 | ENSG00000111897 |
| CEBPA | 0.180007819 | 1.132890025 | 0.653191362 | 0.784504704 | 26.89963159 | 30.36098813 | ENSG00000245848 |
| SLC25A5 | 0.17990662 | 1.13281056 | 0.000123111 | 0.000651381 | 6539.913241 | 7408.449424 | ENSG00000005022 |
| RASL11B | 0.17984529 | 1.132762405 | 0.659548468 | 0.789485573 | 33.90870622 | 38.49699314 | ENSG00000128045 |
| DALRD3 | 0.179804734 | 1.132730565 | 0.104803249 | 0.214624147 | 414.9208778 | 469.9180558 | ENSG00000178149 |
| MAP2K1 | 0.179796513 | 1.132724107 | 0.002904527 | 0.010898899 | 1527.961425 | 1731.031289 | ENSG00000169032 |
| MORC3 | 0.179744653 | 1.13268339 | 0.169250831 | 0.310127069 | 310.970401 | 352.4166787 | ENSG00000159256 |
| MOB1A | 0.179729712 | 1.13267166 | 0.009713877 | 0.030679502 | 1571.888387 | 1781.196155 | ENSG00000114978 |
| CDHR3 | 0.179699412 | 1.132647871 | 0.361298854 | 0.534290242 | 133.8604281 | 151.7441305 | ENSG00000128536 |
| MRPL17 | 0.179625449 | 1.132589805 | 0.013439556 | 0.040389854 | 1137.171316 | 1288.681545 | ENSG00000158042 |
| NA | 0.179613548 | 1.132580462 | 0.762202347 | 0.858807632 | 12.40193393 | 13.99648503 | ENSG00000237758 |
| NDUFB3 | 0.179557815 | 1.132536711 | 0.097383382 | 0.202918862 | 416.2783512 | 471.5112053 | ENSG00000119013 |
| ABTB2 | 0.179538733 | 1.132521731 | 0.046661333 | 0.113026592 | 658.2395958 | 745.4537662 | ENSG00000166016 |
| MARK1 | 0.179383085 | 1.132399553 | 0.046496031 | 0.112654474 | 558.9668157 | 633.0422406 | ENSG00000116141 |
| THAP5 | 0.179358179 | 1.132380004 | 0.333344471 | 0.505943996 | 142.9068577 | 161.991114 | ENSG00000177683 |
| ITGB3BP | 0.179198295 | 1.132254517 | 0.08231689 | 0.177535026 | 440.1019449 | 498.3829176 | ENSG00000142856 |
| C6orf89 | 0.179131692 | 1.132202247 | 0.119332422 | 0.237443299 | 392.2055892 | 443.8959738 | ENSG00000198663 |
| KANSL1 | 0.179113185 | 1.132187723 | 0.019740836 | 0.055602347 | 1044.950875 | 1182.814899 | ENSG00000120071 |
| ARHGEF12 | 0.179046321 | 1.132135251 | 0.01222879 | 0.037220826 | 1063.406891 | 1204.12969 | ENSG00000196914 |
| NA | 0.17897002 | 1.132075377 | 0.620379069 | 0.761403017 | 35.80462511 | 40.40736783 | ENSG00000214199 |
| ZNF232 | 0.178957222 | 1.132065334 | 0.088800699 | 0.188766311 | 457.292299 | 517.8664395 | ENSG00000167840 |
| KLRG2 | 0.178724743 | 1.131882925 | 0.110060048 | 0.222671042 | 480.9159229 | 544.3647329 | ENSG00000188883 |
| PRSS22 | 0.178718194 | 1.131877787 | 0.718817623 | 0.831337575 | 18.86012179 | 21.27879483 | ENSG00000005001 |
| VRK1 | 0.178513579 | 1.131717266 | 0.009700087 | 0.030650571 | 1094.655943 | 1238.983638 | ENSG00000100749 |
| ANKRD63 | 0.178411384 | 1.131637103 | 0.786699637 | 0.875421825 | 9.337535219 | 10.60350093 | ENSG00000230778 |
| NA | 0.178178931 | 1.131454783 | 0.445892029 | 0.616326889 | 80.66117356 | 91.15285515 | ENSG00000230551 |
| RSU1 | 0.178149344 | 1.13143158 | 0.006297932 | 0.021393451 | 1301.38231 | 1472.711935 | ENSG00000148484 |
| FKBP10 | 0.178139328 | 1.131423725 | 0.000364633 | 0.001734255 | 4620.797502 | 5227.396255 | ENSG00000141756 |
| ERLEC1 | 0.177828911 | 1.131180308 | 0.106132676 | 0.216620957 | 490.2833319 | 554.3932954 | ENSG00000068912 |
| THAP9 | 0.177821883 | 1.131174798 | 0.262097723 | 0.427615627 | 171.8632925 | 194.4872578 | ENSG00000168152 |
| ARFGAP3 | 0.177629583 | 1.131024032 | 0.128055407 | 0.250071511 | 344.2702872 | 389.2711017 | ENSG00000242247 |
| NA | 0.177611549 | 1.131009893 | 0.621074764 | 0.761915061 | 32.28583304 | 36.43172819 | ENSG00000136149 |
| HPS3 | 0.177584252 | 1.130988494 | 0.017701655 | 0.050785638 | 1080.593191 | 1222.616687 | ENSG00000163755 |
| RALB | 0.177575746 | 1.130981826 | 0.042068735 | 0.103824141 | 742.1111837 | 838.9294468 | ENSG00000144118 |
| DMAC2L | 0.177545026 | 1.130957743 | 0.372372141 | 0.545833134 | 106.5408579 | 120.5106556 | ENSG00000125375 |
| NA | 0.177497808 | 1.130920729 | 0.423974031 | 0.595650938 | 91.11870412 | 103.0378287 | ENSG00000267886 |
| FAM222B | 0.177393304 | 1.130838811 | 0.005036335 | 0.01765188 | 1908.944422 | 2159.150515 | ENSG00000173065 |
| NA | 0.177372887 | 1.130822808 | 0.444987976 | 0.615568217 | 85.64992687 | 96.95365215 | ENSG00000272163 |
| NA | 0.177363869 | 1.13081574 | 0.780560043 | 0.871147457 | 11.91684064 | 13.52917328 | ENSG00000279227 |
| S1PR2 | 0.177344102 | 1.130800246 | 0.205917249 | 0.359912083 | 256.5075389 | 290.1430568 | ENSG00000267534 |

| | | | | | | | |
|---|---|---|---|---|---|---|---|
| SLC22A15 | 0.177180564 | 1.13067207 | 0.318138001 | 0.489370877 | 130.0885639 | 147.0180721 | ENSG00000163393 |
| LINC02649 | 0.17716149 | 1.130657121 | 0.871951532 | 0.928865039 | 4.125897228 | 4.638157411 | ENSG00000215244 |
| RAI1 | 0.177153297 | 1.130650701 | 0.350572105 | 0.523200254 | 121.619979 | 137.314513 | ENSG00000108557 |
| TMEM60 | 0.177053127 | 1.130572199 | 0.22213806 | 0.38057072 | 230.4206762 | 260.5427072 | ENSG00000135211 |
| MKLN1 | 0.177042832 | 1.130564132 | 0.060834963 | 0.139527878 | 623.527118 | 705.2641093 | ENSG00000128585 |
| FABP3 | 0.176875912 | 1.130433333 | 0.213864435 | 0.370310868 | 281.3859732 | 317.8857236 | ENSG00000121769 |
| NA | 0.176803266 | 1.130376412 | 0.548142339 | 0.705700338 | 49.62639616 | 56.15613275 | ENSG00000212802 |
| APC | 0.17675757 | 1.130340609 | 0.074196868 | 0.163468682 | 495.573003 | 559.8416341 | ENSG00000134982 |
| ZNF260 | 0.176704678 | 1.130299169 | 0.178495325 | 0.322683283 | 387.0517865 | 437.8056118 | ENSG00000254004 |
| LPGAT1 | 0.176230087 | 1.129927406 | 0.185147697 | 0.331433852 | 267.1192207 | 301.9626828 | ENSG00000123684 |
| PPP1R16B | 0.176229861 | 1.129927229 | 0.053787287 | 0.126422789 | 669.695283 | 756.261057 | ENSG00000101445 |
| C9orf72 | 0.176211205 | 1.129912618 | 0.244237935 | 0.40730705 | 256.1574239 | 289.6805729 | ENSG00000147894 |
| TPGS1 | 0.176175435 | 1.129884603 | 0.522557937 | 0.685293349 | 54.13335953 | 61.20164632 | ENSG00000141933 |
| ZSCAN30 | 0.17614844 | 1.129863461 | 0.230436685 | 0.39073166 | 217.4605219 | 245.5720701 | ENSG00000186814 |
| ETV4 | 0.17607647 | 1.129807098 | 0.000193059 | 0.000976739 | 3571.014491 | 4034.600432 | ENSG00000175832 |
| DNASE1 | 0.176054997 | 1.129790282 | 0.161435836 | 0.298787173 | 269.552184 | 304.6479749 | ENSG00000213918 |
| FUZ | 0.175988594 | 1.129738283 | 0.132432774 | 0.256380733 | 337.5454361 | 381.2227861 | ENSG00000010361 |
| ATXN3 | 0.175762173 | 1.129560992 | 0.140172014 | 0.268259754 | 306.797622 | 346.6008703 | ENSG00000066427 |
| RAPGEF3 | 0.175722123 | 1.129529635 | 0.177061848 | 0.320692903 | 264.530979 | 298.8415503 | ENSG00000079337 |
| MANSC1 | 0.175611083 | 1.129442702 | 0.473840136 | 0.64222933 | 91.41770692 | 103.0674037 | ENSG00000111261 |
| SLC37A1 | 0.17557451 | 1.129414071 | 0.051866567 | 0.122873594 | 767.3145652 | 866.2013369 | ENSG00000160190 |
| IKZF1 | 0.175482312 | 1.129341896 | 0.592099685 | 0.740787017 | 44.7645608 | 50.52258431 | ENSG00000185811 |
| TMEM63C | 0.17533404 | 1.129225834 | 0.405216071 | 0.5774561 | 116.6650799 | 131.6094441 | ENSG00000165548 |
| FAM120AOS | 0.175308606 | 1.129205927 | 0.030895023 | 0.080431294 | 775.1042693 | 875.4493364 | ENSG00000188938 |
| JCAD | 0.175265423 | 1.129172127 | 0.226688589 | 0.385868664 | 222.8327688 | 251.5995261 | ENSG00000165757 |
| HOOK1 | 0.175210978 | 1.129129515 | 0.047931152 | 0.115507788 | 699.8649401 | 790.3685542 | ENSG00000134709 |
| JAGN1 | 0.175139403 | 1.129073498 | 0.051322654 | 0.121884196 | 589.1988366 | 664.9197264 | ENSG00000171135 |
| MIEF1 | 0.17506588 | 1.12901596 | 0.0075123 | 0.02480229 | 1228.511465 | 1386.919693 | ENSG00000100335 |
| NA | 0.175035818 | 1.128992434 | 0.866509753 | 0.925470028 | 4.041406403 | 4.532760877 | ENSG00000273091 |
| POTEI | 0.175006289 | 1.128969326 | 0.892034325 | 0.940288616 | 2.952674577 | 3.3738894 | ENSG00000196834 |
| NA | 0.174823389 | 1.128826208 | 0.359046196 | 0.532109371 | 142.4536525 | 160.4907912 | ENSG00000258634 |
| PIWIL4 | 0.17463912 | 1.128682038 | 0.830187011 | 0.903443473 | 7.043641511 | 7.963649691 | ENSG00000134627 |
| GGNBP2 | 0.174523862 | 1.12859187 | 0.03214645 | 0.083094891 | 785.8159849 | 886.3483879 | ENSG00000278311 |
| FAM91A1 | 0.174404108 | 1.128498193 | 0.07542504 | 0.165512963 | 489.3067997 | 551.8900822 | ENSG00000176853 |
| NA | 0.174318994 | 1.128431618 | 0.480303328 | 0.648392687 | 74.02822209 | 83.53229855 | ENSG00000204253 |
| ERICH2 | 0.174306459 | 1.128421813 | 0.506753635 | 0.671592013 | 60.10992072 | 67.78568421 | ENSG00000239467 |
| F2RL1 | 0.174239194 | 1.128369202 | 0.127647217 | 0.249491771 | 442.2639024 | 499.3562272 | ENSG00000164251 |
| FKBP11 | 0.174228852 | 1.128361113 | 0.09248452 | 0.194649969 | 455.0080542 | 513.5974183 | ENSG00000134285 |
| ARHGAP12 | 0.174203798 | 1.128341518 | 0.027719081 | 0.073725189 | 921.9480548 | 1040.528246 | ENSG00000165322 |
| GPD2 | 0.1741598 | 1.128307107 | 0.03492262 | 0.089030298 | 801.4946022 | 903.8437561 | ENSG00000115159 |
| B3GNT2 | 0.174155237 | 1.128303539 | 0.063717807 | 0.144711586 | 721.9874273 | 814.4963758 | ENSG00000170340 |
| NGF | 0.173901785 | 1.128105337 | 0.746985041 | 0.850102556 | 13.93927715 | 15.76113532 | ENSG00000134259 |
| NA | 0.173819905 | 1.128041312 | 0.478407341 | 0.646647703 | 77.16574786 | 87.22184452 | ENSG00000229321 |
| ZNF620 | 0.173779059 | 1.128009376 | 0.198819428 | 0.350299167 | 245.5415487 | 276.8633869 | ENSG00000177842 |
| CARD10 | 0.173727799 | 1.127969297 | 0.022165149 | 0.061162622 | 1154.926487 | 1303.00484 | ENSG00000100065 |
| HSPB7 | 0.173599249 | 1.127868795 | 0.859107154 | 0.920821105 | 4.716256968 | 5.325949025 | ENSG00000173641 |
| RNF187 | 0.173580504 | 1.127854141 | 0.004185377 | 0.015027007 | 2426.717858 | 2736.558118 | ENSG00000168159 |
| NA | 0.173534261 | 1.12781799 | 0.775037518 | 0.868148227 | 20.64064975 | 23.12277431 | ENSG00000230615 |
| TTC23 | 0.173482296 | 1.127777367 | 0.176624913 | 0.320172073 | 291.4334015 | 328.5171515 | ENSG00000103852 |
| RBBP5 | 0.173431857 | 1.127737939 | 0.04438164 | 0.108444105 | 678.3455406 | 764.9622162 | ENSG00000117222 |
| TTC8 | 0.17339929 | 1.127712482 | 0.391174645 | 0.56427397 | 152.8875044 | 172.6050831 | ENSG00000165533 |
| GUK1 | 0.173236015 | 1.127584862 | 0.004314216 | 0.015437868 | 1503.613063 | 1695.013913 | ENSG00000143774 |
| ARSG | 0.173110795 | 1.127486997 | 0.507147617 | 0.671889614 | 64.21907253 | 72.44024059 | ENSG00000141337 |
| PDE8A | 0.173090852 | 1.127471411 | 0.369999996 | 0.543818897 | 121.7891239 | 137.451102 | ENSG00000073417 |
| SPIN4 | 0.17306982 | 1.127454975 | 0.227326486 | 0.386637444 | 254.3681598 | 286.9799542 | ENSG00000186767 |
| C1orf226 | 0.173001581 | 1.127401648 | 0.115027667 | 0.230809104 | 466.8261982 | 526.0970576 | ENSG00000239887 |
| AHR | 0.172996374 | 1.127397579 | 0.579270588 | 0.730755226 | 44.14820706 | 49.82887401 | ENSG00000106546 |
| CYP2D6 | 0.172957182 | 1.127366952 | 0.691494804 | 0.812945716 | 23.38363042 | 26.40228051 | ENSG00000100197 |
| NEMP1 | 0.172936299 | 1.127350634 | 0.010834848 | 0.033739314 | 1076.100094 | 1213.231796 | ENSG00000166881 |
| SAXO2 | 0.172869791 | 1.127298664 | 0.696459178 | 0.816741277 | 22.128908 | 24.90301166 | ENSG00000188659 |
| RHOD | 0.172763976 | 1.127215985 | 0.211873003 | 0.367723085 | 251.7959878 | 284.1981224 | ENSG00000173156 |
| TESC | 0.172687835 | 1.127156496 | 0.265827357 | 0.431947064 | 184.0855567 | 207.657763 | ENSG00000088992 |
| BCL11A | 0.172687008 | 1.12715585 | 0.124675963 | 0.245295666 | 374.2895109 | 422.2520408 | ENSG00000119866 |
| ZCCHC10 | 0.172542923 | 1.127043284 | 0.183714319 | 0.329738829 | 303.5569081 | 342.5886404 | ENSG00000155329 |
| VPS4B | 0.172526404 | 1.127030379 | 0.06163741 | 0.14096589 | 635.4285278 | 716.5594668 | ENSG00000119541 |
| NA | 0.172495023 | 1.127005865 | 0.817362506 | 0.895589009 | 7.320680512 | 8.285497512 | ENSG00000270964 |
| DUSP13 | 0.1724481 | 1.12696921 | 0.703981708 | 0.82190951 | 27.08814956 | 30.39324674 | ENSG00000079393 |
| CDH20 | 0.172432928 | 1.126957358 | 0.82269338 | 0.898928235 | 6.942405268 | 7.839431032 | ENSG00000101542 |
| CHST13 | 0.172120524 | 1.126713351 | 0.488524059 | 0.655408416 | 79.38842953 | 89.69687666 | ENSG00000180767 |
| SNAI3 | 0.172074462 | 1.126677378 | 0.338524531 | 0.511151904 | 138.1451825 | 155.8991286 | ENSG00000185669 |
| FHL2 | 0.171993719 | 1.126614323 | 0.057521074 | 0.133456692 | 665.2298836 | 749.3008497 | ENSG00000115641 |
| NRBF2 | 0.171926684 | 1.126561977 | 0.090186472 | 0.190933629 | 527.2651218 | 593.7859539 | ENSG00000148572 |
| PRXL2C | 0.171794947 | 1.126459111 | 0.438461113 | 0.609435358 | 93.78086074 | 105.7809259 | ENSG00000158122 |
| HEXIM1 | 0.171556782 | 1.126273167 | 0.010471749 | 0.032724853 | 1298.42916 | 1462.557736 | ENSG00000186834 |
| GLG1 | 0.171410418 | 1.126158911 | 0.000896841 | 0.003857515 | 5174.17122 | 5826.033811 | ENSG00000090863 |
| ATXN1L | 0.171291478 | 1.12606607 | 0.022543109 | 0.062081243 | 922.6404514 | 1038.598547 | ENSG00000224470 |
| TNFRSF21 | 0.171274118 | 1.126052521 | 0.006473429 | 0.021912428 | 1431.27765 | 1611.702882 | ENSG00000146072 |

| | | | | | | | |
|---|---|---|---|---|---|---|---|
| NA | 0.171190874 | 1.125987549 | 0.825557546 | 0.900739697 | 6.738693481 | 7.572432051 | ENSG00000251468 |
| RPS2 | 0.171168471 | 1.125970064 | 5.24E-06 | 3.62E-05 | 57927.27799 | 65224.70851 | ENSG00000140988 |
| KLK1 | 0.171164812 | 1.125967208 | 0.666749917 | 0.795243764 | 25.6146848 | 28.88026884 | ENSG00000167748 |
| R3HCC1L | 0.171146173 | 1.125952661 | 0.143804559 | 0.273351275 | 401.186901 | 451.2713218 | ENSG00000166024 |
| SYCE2 | 0.171118106 | 1.125930756 | 0.738521646 | 0.844637554 | 18.32961031 | 20.64155091 | ENSG00000161860 |
| ARFIP1 | 0.170914967 | 1.125772231 | 0.057203623 | 0.132969645 | 718.6096852 | 809.5165371 | ENSG00000164144 |
| FAM102B | 0.170853779 | 1.125724485 | 0.138282996 | 0.265197065 | 392.3716309 | 442.0197173 | ENSG00000162636 |
| ATP5IF1 | 0.170730613 | 1.125628384 | 0.01488188 | 0.043984426 | 2140.63911 | 2409.572439 | ENSG00000130770 |
| NA | 0.170687928 | 1.12559508 | 0.871493775 | 0.928633805 | 4.681326673 | 5.273250758 | ENSG00000275263 |
| RC3H2 | 0.170428498 | 1.12539269 | 0.095033157 | 0.198988888 | 544.9332867 | 613.8800214 | ENSG00000056586 |
| TEPP | 0.170385293 | 1.125358989 | 0.778720116 | 0.870151168 | 12.33418852 | 13.92899321 | ENSG00000159648 |
| NA | 0.170267855 | 1.125267386 | 0.790714924 | 0.878010798 | 11.82465282 | 13.33988595 | ENSG00000226332 |
| VPS37A | 0.16999946 | 1.125058064 | 0.123293003 | 0.243244562 | 433.7150812 | 487.9530985 | ENSG00000155975 |
| ZNF671 | 0.169982359 | 1.125044728 | 0.288125616 | 0.457429945 | 166.1043976 | 186.8694032 | ENSG00000083814 |
| DSCAM | 0.169823928 | 1.124921186 | 0.845429101 | 0.911738193 | 6.425372742 | 7.19573539 | ENSG00000171587 |
| CARS2 | 0.169751702 | 1.124864871 | 0.009756259 | 0.030793192 | 1292.529268 | 1454.474411 | ENSG00000134905 |
| NA | 0.169549836 | 1.124707487 | 0.802032362 | 0.885936266 | 9.269114089 | 10.4351863 | ENSG00000213104 |
| DTX3L | 0.169456706 | 1.124634887 | 0.337638665 | 0.510343519 | 161.9839189 | 181.9833294 | ENSG00000163840 |
| CHST3 | 0.169296814 | 1.124510252 | 0.121491991 | 0.240651196 | 371.0830005 | 417.4617072 | ENSG00000122863 |
| EIF2AK2 | 0.169189455 | 1.124426574 | 0.027268898 | 0.072778682 | 926.3223693 | 1042.05039 | ENSG00000055332 |
| OSTF1 | 0.169128996 | 1.124379454 | 0.167411876 | 0.30749961 | 285.3088344 | 320.8092028 | ENSG00000134996 |
| NA | 0.169128287 | 1.124378901 | 0.783977155 | 0.873614559 | 17.25012707 | 19.33910561 | ENSG00000271043 |
| A1BG-AS1 | 0.169105238 | 1.124360938 | 0.858910748 | 0.920821105 | 6.253600188 | 7.103502762 | ENSG00000268895 |
| USP28 | 0.16884056 | 1.12415468 | 0.002874963 | 0.010815311 | 2158.027779 | 2426.585605 | ENSG00000048028 |
| DISP2 | 0.168809447 | 1.124130438 | 0.579370646 | 0.730794139 | 48.56856442 | 54.57540256 | ENSG00000140323 |
| ACTL8 | 0.168770722 | 1.124100264 | 0.634702723 | 0.771175881 | 40.60776348 | 45.60615412 | ENSG00000117148 |
| SNX18 | 0.168754709 | 1.124087787 | 0.084286471 | 0.180904574 | 491.4695962 | 552.5579672 | ENSG00000178996 |
| SMC4 | 0.168714858 | 1.124056737 | 0.012618079 | 0.038296942 | 1527.820701 | 1717.563585 | ENSG00000113810 |
| SLK | 0.168639184 | 1.123997779 | 0.022608832 | 0.06223692 | 907.1426309 | 1019.456246 | ENSG00000065613 |
| SNORD100 | 0.168591324 | 1.123960491 | 0.757022816 | 0.855820859 | 15.1124998 | 16.91355508 | ENSG00000221500 |
| PI4KAP2 | 0.168572349 | 1.123945708 | 0.159117555 | 0.29577344 | 363.0084724 | 408.0340219 | ENSG00000183506 |
| THAP6 | 0.168570407 | 1.123944196 | 0.534496074 | 0.694855702 | 64.20435428 | 72.11221363 | ENSG00000174796 |
| MPND | 0.168547206 | 1.123926121 | 0.21616403 | 0.373289795 | 297.6609324 | 334.4978158 | ENSG00000008382 |
| UBE2Q2 | 0.168422638 | 1.123829081 | 0.120677311 | 0.239603335 | 426.4162524 | 479.3201947 | ENSG00000140367 |
| GTF2B | 0.16840421 | 1.123814726 | 0.250560224 | 0.414755216 | 232.249213 | 260.9207609 | ENSG00000137947 |
| CDC42BPG | 0.168304156 | 1.12373679 | 0.068543464 | 0.153378026 | 590.5370859 | 663.3355007 | ENSG00000171219 |
| TMEM131 | 0.168201622 | 1.123656927 | 0.06719836 | 0.151033854 | 618.0340105 | 694.5732526 | ENSG00000075568 |
| SLF2 | 0.168189499 | 1.123647485 | 0.087547347 | 0.186554277 | 507.015789 | 570.1929054 | ENSG00000119906 |
| CPT2 | 0.168068678 | 1.123553387 | 0.201658478 | 0.353975769 | 268.4271704 | 301.5039854 | ENSG00000157184 |
| PHACTR4 | 0.16806733 | 1.123552337 | 0.017925723 | 0.051306398 | 981.1990375 | 1102.39192 | ENSG00000204138 |
| RNF123 | 0.167999736 | 1.123499698 | 0.044700509 | 0.10912649 | 711.3677873 | 798.8538983 | ENSG00000164068 |
| G6PC3 | 0.167976835 | 1.123481864 | 0.012911118 | 0.039045051 | 2138.696684 | 2403.256503 | ENSG00000141349 |
| CLDND1 | 0.167960282 | 1.123468973 | 0.257032609 | 0.422172504 | 399.3462139 | 449.1119399 | ENSG00000080822 |
| AMOTL2 | 0.16795187 | 1.123462423 | 0.022156174 | 0.061146604 | 1107.930358 | 1244.856507 | ENSG00000114019 |
| KLHL26 | 0.167857275 | 1.123388761 | 0.14153842 | 0.270177881 | 381.5855858 | 428.4674818 | ENSG00000167487 |
| KIAA0586 | 0.167654482 | 1.123230863 | 0.100251905 | 0.207506384 | 438.0043783 | 492.0048386 | ENSG00000100578 |
| NIN | 0.167426401 | 1.123053301 | 0.143434556 | 0.272973975 | 437.5340402 | 491.0726198 | ENSG00000100503 |
| GPS2 | 0.167378795 | 1.123016243 | 0.705267846 | 0.822671706 | 25.03269777 | 28.19435522 | ENSG00000132522 |
| GOLGA5 | 0.167315185 | 1.122966729 | 0.104503861 | 0.214188458 | 562.944744 | 632.092551 | ENSG00000066455 |
| TNNT1 | 0.167211336 | 1.122885898 | 0.008620531 | 0.027804491 | 1586.243772 | 1781.953093 | ENSG00000105048 |
| PRRX2 | 0.167175821 | 1.122858256 | 0.412839976 | 0.585167134 | 121.1304552 | 136.1037367 | ENSG00000167157 |
| SNCAIP | 0.167086691 | 1.122788888 | 0.500051574 | 0.666286429 | 88.17507797 | 98.73955982 | ENSG00000064692 |
| SDSL | 0.167082595 | 1.1227857 | 0.77499909 | 0.868148227 | 11.75431667 | 13.19280448 | ENSG00000139410 |
| ZNF394 | 0.167055335 | 1.122764485 | 0.092486735 | 0.194649969 | 493.325991 | 553.6006772 | ENSG00000160908 |
| AARS1 | 0.16680933 | 1.12257305 | 8.82E-05 | 0.000482808 | 11194.38324 | 12565.09231 | ENSG00000090861 |
| PLCH2 | 0.166790527 | 1.12255842 | 0.809312088 | 0.890310719 | 9.32916251 | 10.47121301 | ENSG00000149527 |
| BACH2 | 0.166761004 | 1.122535448 | 0.552753057 | 0.709222429 | 55.88278725 | 62.85981558 | ENSG00000112182 |
| ASAH1 | 0.166665437 | 1.122461092 | 0.029588782 | 0.077636679 | 971.2079525 | 1089.83874 | ENSG00000104763 |
| PRIM2 | 0.16665546 | 1.122453329 | 0.049035849 | 0.11749479 | 649.3380178 | 728.7251852 | ENSG00000146143 |
| AHI1 | 0.166557105 | 1.122376809 | 0.341117295 | 0.51373985 | 155.453443 | 174.6017838 | ENSG00000135541 |
| GPRIN1 | 0.166548653 | 1.122370234 | 0.128214589 | 0.250292213 | 521.1754354 | 585.1649237 | ENSG00000169258 |
| ZNF337 | 0.166510512 | 1.122340561 | 0.707041272 | 0.823594399 | 20.99898836 | 23.56292211 | ENSG00000130684 |
| OSBPL8 | 0.166475743 | 1.122313514 | 0.055405134 | 0.129664012 | 1026.072283 | 1151.982503 | ENSG00000091039 |
| BTNL2 | 0.166450137 | 1.122293594 | 0.86258903 | 0.922764098 | 5.040065596 | 5.670920119 | ENSG00000204290 |
| LOC102724788 | 0.166436798 | 1.122283217 | 0.250072939 | 0.414268533 | 206.6960914 | 231.9641312 | ENSG00000277196 |
| ABRAXAS2 | 0.166431344 | 1.122278975 | 0.134207499 | 0.25918479 | 359.6381745 | 403.4189878 | ENSG00000165660 |
| EXO1 | 0.166328045 | 1.122198621 | 0.031802478 | 0.082393447 | 825.036302 | 925.6116657 | ENSG00000174371 |
| YTHDF3 | 0.166203574 | 1.122101805 | 0.014699089 | 0.04353756 | 1873.504808 | 2102.584827 | ENSG00000185728 |
| TERT | 0.166156015 | 1.122064816 | 0.110434677 | 0.223241679 | 530.0647595 | 594.5009458 | ENSG00000164362 |
| PLEKHG2 | 0.166124551 | 1.122040344 | 0.023621412 | 0.064516589 | 1194.141508 | 1339.554068 | ENSG00000090924 |
| NA | 0.166110081 | 1.122029091 | 0.430132449 | 0.60158668 | 98.40513006 | 110.3629079 | ENSG00000197258 |
| TMEM44 | 0.166007412 | 1.121949244 | 0.288290794 | 0.457579208 | 208.6651373 | 233.9054564 | ENSG00000145014 |
| MIR600HG | 0.165831813 | 1.121812694 | 0.187309435 | 0.334387036 | 378.7138233 | 424.6926695 | ENSG00000236901 |
| SGF29 | 0.165812279 | 1.121797504 | 0.326592797 | 0.498481023 | 163.4067102 | 183.5186371 | ENSG00000176476 |
| NA | 0.16577574 | 1.121769093 | 0.428810448 | 0.600166354 | 95.56888554 | 107.271865 | ENSG00000229638 |
| C15orf41 | 0.165769998 | 1.121764629 | 0.264319321 | 0.430125402 | 209.8110898 | 235.528665 | ENSG00000186073 |
| TBCD | 0.16575749 | 1.121754904 | 0.000490815 | 0.002252749 | 4200.526507 | 4711.856863 | ENSG00000141556 |

| | | | | | | | |
|---|---|---|---|---|---|---|---|
| FAM72C | 0.165682662 | 1.121696723 | 0.541999717 | 0.700832263 | 61.67046705 | 69.17013019 | ENSG00000263513 |
| KDM1B | 0.165652782 | 1.121673492 | 0.258714166 | 0.423923807 | 206.8230093 | 232.2198322 | ENSG00000165097 |
| GRM5 | 0.165638248 | 1.121662191 | 0.898611124 | 0.94422938 | 2.919859464 | 3.308820728 | ENSG00000168959 |
| EGLN2 | 0.165530554 | 1.121578465 | 0.485572988 | 0.653129864 | 92.47621438 | 103.4710038 | ENSG00000269858 |
| SLC15A4 | 0.165492815 | 1.121549127 | 0.083049194 | 0.178833574 | 704.404155 | 789.7582074 | ENSG00000139370 |
| ARHGEF15 | 0.165389023 | 1.121468441 | 0.652164281 | 0.783856769 | 34.39399967 | 38.63867685 | ENSG00000198844 |
| NA | 0.165360751 | 1.121446465 | 0.574456674 | 0.726582986 | 60.76208064 | 67.9523813 | ENSG00000170919 |
| SPG21 | 0.165305541 | 1.121403549 | 0.01103976 | 0.03427229 | 1260.729916 | 1414.161656 | ENSG00000090487 |
| GDAP1L1 | 0.165255027 | 1.121364285 | 0.407601399 | 0.579908483 | 144.2949557 | 161.7340865 | ENSG00000124194 |
| NA | 0.165227157 | 1.121342623 | 0.627149178 | 0.765475153 | 42.47537302 | 47.61924174 | ENSG00000260261 |
| ANLN | 0.165226324 | 1.121341976 | 0.010726809 | 0.033440672 | 1466.907773 | 1644.919076 | ENSG00000011426 |
| NA | 0.165152991 | 1.121284979 | 0.324610673 | 0.496240945 | 160.5822298 | 180.1350488 | ENSG00000174028 |
| NA | 0.165152823 | 1.121284848 | 0.558679693 | 0.714203884 | 53.36373041 | 59.74378983 | ENSG00000261512 |
| ARHGEF5 | 0.16492407 | 1.121107072 | 0.102371127 | 0.210829659 | 563.4013647 | 631.2450592 | ENSG00000050327 |
| CAMTA2 | 0.164855256 | 1.121053598 | 0.370553891 | 0.544387717 | 126.7427836 | 142.1618459 | ENSG00000108509 |
| ZNF385A | 0.164767877 | 1.120985702 | 0.187604208 | 0.33478934 | 327.440262 | 367.235709 | ENSG00000161642 |
| ARHGAP19 | 0.164764369 | 1.120982976 | 0.05950251 | 0.136960138 | 683.7716168 | 766.8995249 | ENSG00000213390 |
| SLC25A43 | 0.164696529 | 1.120930266 | 0.33308916 | 0.505795261 | 156.2218328 | 175.0656185 | ENSG00000077713 |
| NA | 0.164691546 | 1.120926393 | 0.745091824 | 0.848999786 | 16.63935513 | 18.65938325 | ENSG00000237356 |
| NA | 0.164594097 | 1.120850682 | 0.76475648 | 0.860830742 | 13.98325588 | 15.66219051 | ENSG00000235454 |
| ATG4D | 0.164562054 | 1.120825787 | 0.06454622 | 0.146128521 | 605.5883491 | 679.0344993 | ENSG00000130734 |
| C3orf33 | 0.164554907 | 1.120820235 | 0.822348979 | 0.898735651 | 8.556742486 | 9.520190529 | ENSG00000174928 |
| HMCN2 | 0.164496297 | 1.120774702 | 0.122798703 | 0.242641584 | 403.3545156 | 451.946743 | ENSG00000148357 |
| RBIS | 0.164335408 | 1.12064972 | 0.531515057 | 0.692377304 | 61.89862124 | 69.35862402 | ENSG00000176731 |
| MRPL32 | 0.164311814 | 1.120631393 | 0.006106674 | 0.020849112 | 1622.22754 | 1817.549482 | ENSG00000106591 |
| SNRPE | 0.164304603 | 1.120625792 | 0.008942785 | 0.028637993 | 1960.788906 | 2196.689216 | ENSG00000182004 |
| CXCL3 | 0.164295997 | 1.120619107 | 0.590165943 | 0.739374672 | 43.74548941 | 49.07978184 | ENSG00000163734 |
| IRX1 | 0.164268405 | 1.120597675 | 0.787463294 | 0.876013748 | 11.93079515 | 13.38021381 | ENSG00000170549 |
| USP38 | 0.164249236 | 1.120582787 | 0.218538261 | 0.376244676 | 269.7162595 | 302.3797136 | ENSG00000170185 |
| IL6ST | 0.164247777 | 1.120581653 | 0.437976059 | 0.608936659 | 103.2006595 | 115.8431836 | ENSG00000134352 |
| ARL17A | 0.164234416 | 1.120571275 | 0.561175567 | 0.716063622 | 52.7403554 | 59.12536803 | ENSG00000185829 |
| BSN-DT | 0.1641566 | 1.120510835 | 0.82797675 | 0.902095306 | 7.389101641 | 8.28979866 | ENSG00000226913 |
| NA | 0.164123708 | 1.120485289 | 0.858333 | 0.920712962 | 4.933923269 | 5.54670146 | ENSG00000241255 |
| SUSD3 | 0.163983807 | 1.120376639 | 0.446863152 | 0.617244527 | 100.3166062 | 112.3897046 | ENSG00000157303 |
| FADS1 | 0.163970527 | 1.120366326 | 0.002524657 | 0.009667161 | 3359.561839 | 3763.292276 | ENSG00000149485 |
| HEXD | 0.163857869 | 1.120278842 | 0.151679854 | 0.285079508 | 381.6006916 | 427.2591408 | ENSG00000169660 |
| NVL | 0.163855838 | 1.120277264 | 0.023889904 | 0.06509324 | 942.5784679 | 1056.298275 | ENSG00000143748 |
| CTSH | 0.163677963 | 1.12013915 | 0.163821886 | 0.30247766 | 360.9163002 | 403.9655982 | ENSG00000103811 |
| ZNF444 | 0.16359233 | 1.120072665 | 0.038479539 | 0.096566774 | 903.9561466 | 1012.905322 | ENSG00000167685 |
| SLC30A6 | 0.163553259 | 1.120042331 | 0.083753007 | 0.180019974 | 581.6888375 | 651.7325496 | ENSG00000152683 |
| GEMIN4 | 0.163529245 | 1.120023688 | 0.010912281 | 0.033947557 | 1656.563581 | 1855.170226 | ENSG00000179409 |
| PODXL2 | 0.163447655 | 1.119960348 | 0.00973652 | 0.030735921 | 1443.919653 | 1617.472232 | ENSG00000114631 |
| DCP1A | 0.163288352 | 1.119836689 | 0.122850151 | 0.242681852 | 425.2032814 | 476.0847966 | ENSG00000272886 |
| OSBPL9 | 0.163134233 | 1.119717066 | 0.020755749 | 0.057919738 | 1378.555891 | 1543.554743 | ENSG00000117859 |
| LOC103344931 | 0.163087245 | 1.119680598 | 0.609691845 | 0.753415245 | 40.91291167 | 45.80268668 | ENSG00000260917 |
| NUAK1 | 0.16301246 | 1.119622558 | 0.073563863 | 0.162335441 | 547.9634677 | 613.4119588 | ENSG00000074590 |
| CHRNA4 | 0.162943903 | 1.119569355 | 0.511365242 | 0.675720602 | 85.65582097 | 95.82972289 | ENSG00000101204 |
| GOLGB1 | 0.162927141 | 1.119556347 | 0.100940442 | 0.208528814 | 1022.270917 | 1144.008348 | ENSG00000173230 |
| NA | 0.162894896 | 1.119531325 | 0.623780589 | 0.763014769 | 39.54279834 | 44.35505002 | ENSG00000227097 |
| CISD3 | 0.162850704 | 1.119497032 | 0.24581772 | 0.409196866 | 271.6825657 | 304.2115889 | ENSG00000277972 |
| LOC105372482 | 0.162621852 | 1.119319463 | 0.642277331 | 0.776527012 | 34.60887507 | 38.80834856 | ENSG00000268516 |
| TMEM132D | 0.162592102 | 1.119296382 | 0.11532237 | 0.231256117 | 464.7219496 | 519.9141904 | ENSG00000151952 |
| MYO1C | 0.162445103 | 1.11918234 | 0.00324401 | 0.011993139 | 2074.452816 | 2322.017122 | ENSG00000197879 |
| MICU3 | 0.162407891 | 1.119153473 | 0.525114744 | 0.687431361 | 63.52536137 | 71.14693063 | ENSG00000155970 |
| CCDC18 | 0.162394209 | 1.11914286 | 0.276785805 | 0.443981238 | 196.0764003 | 219.538708 | ENSG00000122483 |
| CRYBA4 | 0.162308059 | 1.119076032 | 0.700219194 | 0.819158791 | 25.48218505 | 28.45517505 | ENSG00000196431 |
| NA | 0.162278472 | 1.119053082 | 0.774855631 | 0.86807286 | 12.94340886 | 14.54741501 | ENSG00000260233 |
| HOOK3 | 0.162166978 | 1.118966603 | 0.175521333 | 0.3188307 | 335.3653817 | 375.0213573 | ENSG00000168172 |
| KREMEN2 | 0.162136665 | 1.118943092 | 0.279627839 | 0.447511253 | 204.9278031 | 229.419996 | ENSG00000131650 |
| NA | 0.161979641 | 1.118821312 | 0.31406057 | 0.485226851 | 190.5118053 | 212.9124764 | ENSG00000286619 |
| GSK3B | 0.161967335 | 1.118811769 | 0.043331021 | 0.106402588 | 781.8511481 | 874.800001 | ENSG00000082701 |
| NA | 0.161959223 | 1.118805478 | 0.843116567 | 0.910214813 | 6.439327257 | 7.149488844 | ENSG00000261349 |
| ISCA2 | 0.161755711 | 1.118647666 | 0.213452274 | 0.36976543 | 300.7498857 | 336.2893391 | ENSG00000165898 |
| JAKMIP2-AS1 | 0.161754866 | 1.118647011 | 0.218087254 | 0.375669366 | 259.0236415 | 289.4843008 | ENSG00000280780 |
| LOC101930370 | 0.161732348 | 1.118629551 | 0.604059816 | 0.749287417 | 47.12253314 | 52.77390152 | ENSG00000245213 |
| NARS2 | 0.161706914 | 1.11860983 | 0.024383364 | 0.066204008 | 1058.758418 | 1184.957306 | ENSG00000137513 |
| NUP58 | 0.16155808 | 1.118494436 | 0.030729457 | 0.080107647 | 1091.939158 | 1221.763604 | ENSG00000139496 |
| FER1L4 | 0.161426849 | 1.1183927 | 0.594977398 | 0.742893785 | 46.62503693 | 52.02454888 | ENSG00000088340 |
| SMURF2 | 0.161383756 | 1.118359294 | 0.075169501 | 0.165121253 | 590.0927929 | 659.8728621 | ENSG00000108854 |
| CPNE2 | 0.161214488 | 1.118228087 | 0.107712659 | 0.21900136 | 703.2284665 | 785.7194445 | ENSG00000140848 |
| FAM227A | 0.161097788 | 1.118137638 | 0.623066274 | 0.762604966 | 38.53106051 | 43.05528833 | ENSG00000184949 |
| NA | 0.161093272 | 1.118134137 | 0.896503767 | 0.942785335 | 3.015513901 | 3.40131382 | ENSG00000260274 |
| SLC4A11 | 0.161019974 | 1.11807733 | 0.091918824 | 0.193849184 | 591.8813174 | 661.6334838 | ENSG00000088836 |
| TENT4B | 0.160930102 | 1.118007683 | 0.184195805 | 0.330295713 | 338.6266342 | 378.7160248 | ENSG00000121274 |
| OLFML2B | 0.160830568 | 1.117930552 | 0.340377884 | 0.512946626 | 186.1595568 | 208.2291448 | ENSG00000162745 |
| NA | 0.160702455 | 1.117831283 | 0.599509036 | 0.74594484 | 44.89897558 | 50.27844814 | ENSG00000232573 |
| KCNH6 | 0.160697876 | 1.117827735 | 0.593873279 | 0.741995429 | 54.67994071 | 61.04623513 | ENSG00000173826 |

| | | | | | | | |
|---|---|---|---|---|---|---|---|
| ISG20L2 | 0.160650545 | 1.117791063 | 0.009600346 | 0.030395539 | 1441.712677 | 1611.786779 | ENSG00000143319 |
| ZNF395 | 0.160588701 | 1.117743148 | 0.169709738 | 0.310790826 | 358.9161909 | 400.807627 | ENSG00000186918 |
| SCNM1 | 0.160436841 | 1.117625498 | 0.070888869 | 0.157602532 | 666.8224813 | 744.6059896 | ENSG00000163156 |
| HOOK2 | 0.160397913 | 1.117595342 | 0.00803944 | 0.026225711 | 1671.515261 | 1868.047095 | ENSG00000095066 |
| RAD54L2 | 0.160255609 | 1.117485111 | 0.052012837 | 0.123036053 | 814.5325143 | 909.9142114 | ENSG00000164080 |
| FARP1 | 0.160252077 | 1.117482375 | 0.008930825 | 0.028604516 | 2077.009642 | 2321.287075 | ENSG00000152767 |
| PLXNB2 | 0.160220369 | 1.117457815 | 0.003670585 | 0.013388023 | 3801.718079 | 4249.362207 | ENSG00000196576 |
| NA | 0.160191149 | 1.117435182 | 0.497230658 | 0.663581918 | 84.82862203 | 94.76982675 | ENSG00000267106 |
| ASB1 | 0.16002004 | 1.117302658 | 0.036335364 | 0.091991812 | 837.2740723 | 935.5668667 | ENSG00000065802 |
| INHBA | 0.159860535 | 1.117179136 | 0.803410536 | 0.886596456 | 10.89932084 | 12.24823371 | ENSG00000122641 |
| EIF3F | 0.159793635 | 1.117127331 | 0.000371245 | 0.001763961 | 4859.062928 | 5428.584378 | ENSG00000175390 |
| UBE2H | 0.159788846 | 1.117123623 | 0.006966987 | 0.023292921 | 2300.920821 | 2571.341297 | ENSG00000186591 |
| NA | 0.159775277 | 1.117113116 | 0.651510443 | 0.783266107 | 41.97209485 | 46.91746218 | ENSG00000205534 |
| SRGAP1 | 0.159665641 | 1.117028226 | 0.34403108 | 0.516676027 | 154.2233631 | 172.0958502 | ENSG00000196935 |
| RMDN2 | 0.15945082 | 1.11686191 | 0.667304913 | 0.795634837 | 32.45740543 | 36.28034557 | ENSG00000115841 |
| C12orf60 | 0.159360121 | 1.116791697 | 0.69760609 | 0.817418609 | 25.40260031 | 28.41994184 | ENSG00000182993 |
| NA | 0.159209397 | 1.116675028 | 0.779036372 | 0.870252513 | 13.54792327 | 15.04591929 | ENSG00000261884 |
| SAV1 | 0.159143839 | 1.116624286 | 0.027577161 | 0.073479421 | 1528.06076 | 1707.425809 | ENSG00000151748 |
| PKN3 | 0.159130449 | 1.116613922 | 0.032660095 | 0.084174469 | 977.2122226 | 1090.594084 | ENSG00000160447 |
| STK38L | 0.159080259 | 1.116575076 | 0.059563523 | 0.137067867 | 1084.264984 | 1211.581635 | ENSG00000211455 |
| PPP1R3G | 0.159036804 | 1.116541445 | 0.842297773 | 0.909743682 | 7.638907334 | 8.546044772 | ENSG00000219607 |
| TRIM2 | 0.158941564 | 1.116467739 | 0.061401328 | 0.140536268 | 764.3496637 | 853.5798047 | ENSG00000109654 |
| SNX6 | 0.158798362 | 1.116356924 | 0.026662013 | 0.071356396 | 991.9607651 | 1107.159663 | ENSG00000129515 |
| ALOX12 | 0.15867536 | 1.116261749 | 0.59237764 | 0.741062844 | 61.63889123 | 68.72407582 | ENSG00000108839 |
| SYN2 | 0.158564849 | 1.116176246 | 0.344014068 | 0.516676027 | 170.9541423 | 190.7065637 | ENSG00000157152 |
| ECSIT | 0.158449979 | 1.116087378 | 0.03510363 | 0.089385445 | 1121.504314 | 1251.832368 | ENSG00000130159 |
| SNX21 | 0.158420001 | 1.116064187 | 0.261456114 | 0.426881063 | 226.0307064 | 252.3814405 | ENSG00000124104 |
| RBM12 | 0.158291907 | 1.115965098 | 0.00232323 | 0.008985052 | 2472.158483 | 2758.062421 | ENSG00000244462 |
| WBP1 | 0.158151131 | 1.115856209 | 0.550005732 | 0.707108336 | 82.46650735 | 91.85626435 | ENSG00000239779 |
| ARHGEF6 | 0.15800935 | 1.115746553 | 0.155817092 | 0.29109699 | 403.3699095 | 450.0401059 | ENSG00000129675 |
| CTSZ | 0.157948552 | 1.115699535 | 0.102367206 | 0.210829659 | 736.7526925 | 822.3963807 | ENSG00000101160 |
| NA | 0.157928048 | 1.115683678 | 0.896482736 | 0.942785335 | 3.8020886 | 4.187789783 | ENSG00000252311 |
| RAB22A | 0.157850511 | 1.115623718 | 0.254331723 | 0.419128076 | 227.3935983 | 253.705131 | ENSG00000124209 |
| NA | 0.157798542 | 1.115583531 | 0.8739691 | 0.929987198 | 4.433636161 | 4.909457539 | ENSG00000212195 |
| BUB1B | 0.157739658 | 1.115538 | 0.002915153 | 0.010934514 | 3309.640384 | 3690.796665 | ENSG00000156970 |
| ADAM17 | 0.157698795 | 1.115506403 | 0.114869505 | 0.23056369 | 541.1962264 | 603.471224 | ENSG00000151694 |
| ZFAT | 0.157667962 | 1.115482563 | 0.32222184 | 0.493684499 | 196.1276004 | 219.0345334 | ENSG00000066827 |
| SNHG1 | 0.157645431 | 1.115465143 | 0.049522248 | 0.118351547 | 1675.323791 | 1868.389407 | ENSG00000255717 |
| NA | 0.157615657 | 1.115442122 | 0.591281576 | 0.74009946 | 47.39745696 | 52.89732668 | ENSG00000245248 |
| CADM4 | 0.157599567 | 1.115429682 | 0.115172077 | 0.231050805 | 590.0549596 | 657.9342632 | ENSG00000105767 |
| KCTD14 | 0.157587585 | 1.115420418 | 0.555187227 | 0.711324593 | 58.52369269 | 65.22327204 | ENSG00000151364 |
| NA | 0.157562838 | 1.115401285 | 0.794710655 | 0.880471663 | 13.78425001 | 15.45810963 | ENSG00000226450 |
| GPR139 | 0.157453205 | 1.115316527 | 0.823211349 | 0.899239532 | 9.204359744 | 10.21228329 | ENSG00000180269 |
| NA | 0.15742059 | 1.115291313 | 0.866737682 | 0.925559735 | 4.721838773 | 5.247976912 | ENSG00000270060 |
| FBLIM1 | 0.157416942 | 1.115288493 | 0.005253554 | 0.018303471 | 2335.205511 | 2604.606623 | ENSG00000162458 |
| NA | 0.157398237 | 1.115274033 | 0.711448343 | 0.826133147 | 22.32703799 | 24.90893034 | ENSG00000185641 |
| RWDD2B | 0.157356227 | 1.115241558 | 0.48711852 | 0.654341855 | 84.09998057 | 93.79406346 | ENSG00000156253 |
| SSTR5 | 0.157334732 | 1.115224942 | 0.723486748 | 0.834137153 | 25.61556068 | 28.5861059 | ENSG00000162009 |
| HAGH | 0.157257019 | 1.11516487 | 0.119363476 | 0.237459092 | 454.6383518 | 506.9929529 | ENSG00000063854 |
| EBLN3P | 0.157202548 | 1.115122766 | 0.061736707 | 0.14112606 | 839.3216781 | 936.5616821 | ENSG00000281649 |
| GALNT13 | 0.157067117 | 1.11501809 | 0.26101556 | 0.426318784 | 217.6291032 | 242.6047434 | ENSG00000144278 |
| UTP23 | 0.156969389 | 1.114942561 | 0.409400249 | 0.581657107 | 172.4229154 | 192.6112923 | ENSG00000147679 |
| EXOC5 | 0.156876968 | 1.114871139 | 0.011539195 | 0.035593637 | 1389.402561 | 1549.342619 | ENSG00000070367 |
| MRPL44 | 0.156743355 | 1.114767892 | 0.027733231 | 0.073742489 | 1018.240688 | 1134.628886 | ENSG00000135900 |
| NA | 0.156569745 | 1.114633752 | 0.536935025 | 0.696524348 | 71.14628398 | 79.09338784 | ENSG00000242071 |
| SRSF6 | 0.156496567 | 1.114577216 | 0.025321129 | 0.068339533 | 1439.674871 | 1604.66526 | ENSG00000124193 |
| LINC02701 | 0.156175377 | 1.114329103 | 0.807682584 | 0.889275996 | 10.97332378 | 12.28426023 | ENSG00000250508 |
| GPATCH2L | 0.156041232 | 1.114225495 | 0.026610072 | 0.071266819 | 1264.551791 | 1408.698573 | ENSG00000089916 |
| ZNF148 | 0.155930317 | 1.114139836 | 0.266581866 | 0.432735676 | 252.3783752 | 281.3998854 | ENSG00000163848 |
| TOMM70 | 0.155805862 | 1.114043728 | 0.014014739 | 0.041864095 | 1635.719447 | 1823.188343 | ENSG00000154174 |
| EZH1 | 0.155696142 | 1.113959006 | 0.24775502 | 0.411311711 | 376.2285937 | 418.8114651 | ENSG00000108799 |
| NA | 0.155595433 | 1.113881247 | 0.860547828 | 0.921699125 | 5.296128817 | 5.921247548 | ENSG00000248375 |
| CYB561 | 0.155592494 | 1.113878978 | 0.069021747 | 0.154287199 | 882.3528681 | 982.9401553 | ENSG00000008283 |
| ROCK2 | 0.155541264 | 1.113839425 | 0.084798234 | 0.181829228 | 630.5202479 | 702.2970551 | ENSG00000134318 |
| CRIP2 | 0.15551033 | 1.113815542 | 0.260779385 | 0.426149522 | 229.5677955 | 255.4797407 | ENSG00000182809 |
| NA | 0.155147813 | 1.113535701 | 0.89902355 | 0.944354114 | 2.961047286 | 3.32549228 | ENSG00000225674 |
| NA | 0.155074392 | 1.113479033 | 0.864808028 | 0.924215371 | 4.961156576 | 5.525728761 | ENSG00000214646 |
| RASGEF1C | 0.154885061 | 1.113332916 | 0.712233724 | 0.826597327 | 22.98581885 | 25.61984523 | ENSG00000146090 |
| NA | 0.154884404 | 1.113332409 | 0.737094405 | 0.843705033 | 19.40717852 | 21.56246536 | ENSG00000280239 |
| CD2AP | 0.154646079 | 1.113148508 | 0.015534135 | 0.045563174 | 1299.695834 | 1446.988795 | ENSG00000198087 |
| NA | 0.154231397 | 1.112828596 | 0.344541409 | 0.517123353 | 188.7519777 | 210.1093809 | ENSG00000250182 |
| TUT7 | 0.154213256 | 1.112814602 | 0.177898112 | 0.321844929 | 399.4904635 | 444.5186783 | ENSG00000083223 |
| NA | 0.154184964 | 1.112792779 | 0.557539488 | 0.713428546 | 79.74209911 | 88.45816718 | ENSG00000187653 |
| GPX2 | 0.154175608 | 1.112785563 | 0.840028511 | 0.908905319 | 7.611674027 | 8.478825526 | ENSG00000176153 |
| TNIK | 0.154106667 | 1.112732388 | 0.021105425 | 0.058717156 | 1586.085138 | 1764.307123 | ENSG00000154310 |
| RP2 | 0.154018847 | 1.112664656 | 0.28287583 | 0.451211977 | 250.9471374 | 279.436031 | ENSG00000102218 |
| ZNF43 | 0.153651243 | 1.112381181 | 0.234963274 | 0.396074495 | 268.6339854 | 298.803082 | ENSG00000198521 |

| | | | | | | | |
|---|---|---|---|---|---|---|---|
| CIDECP1 | 0.153588234 | 1.112332599 | 0.577002697 | 0.728847501 | 58.2590566 | 64.82345211 | ENSG00000186162 |
| TENM3 | 0.153513889 | 1.11227528 | 0.008475182 | 0.027391938 | 1915.444387 | 2131.118786 | ENSG00000218336 |
| ASMTL | 0.153418888 | 1.11220204 | 0.007150688 | 0.023787536 | 2249.148909 | 2500.782239 | ENSG00000169093 |
| SPDYE3 | 0.153390886 | 1.112180452 | 0.668886835 | 0.796659624 | 42.29331273 | 46.89139483 | ENSG00000214300 |
| PAPOLG | 0.153340947 | 1.112141955 | 0.145190841 | 0.275378676 | 423.6055263 | 470.8989321 | ENSG00000115421 |
| MARCHF2 | 0.15312026 | 1.111971845 | 0.256018324 | 0.421151998 | 241.6287751 | 268.8588579 | ENSG00000099785 |
| RSPH1 | 0.152915838 | 1.111814296 | 0.864371439 | 0.923879632 | 5.380619642 | 5.994918514 | ENSG00000160188 |
| HPDL | 0.152825057 | 1.111744338 | 0.141767097 | 0.270534085 | 566.2168989 | 629.8780903 | ENSG00000186603 |
| RAB3IL1 | 0.152618161 | 1.111584915 | 0.254461708 | 0.419306466 | 248.2634414 | 276.1831131 | ENSG00000167994 |
| KCNB1 | 0.152556454 | 1.111537371 | 0.720283058 | 0.832084584 | 26.67359258 | 29.56160873 | ENSG00000158445 |
| UNC5B-AS1 | 0.152553864 | 1.111535375 | 0.819914439 | 0.897162192 | 14.3812676 | 15.85390099 | ENSG00000237512 |
| TCF4 | 0.152340641 | 1.111371108 | 0.046126423 | 0.1118995 | 1190.564432 | 1323.571223 | ENSG00000196628 |
| ATP8B2 | 0.15227904 | 1.111323656 | 0.009645284 | 0.030507817 | 1801.953852 | 2002.049032 | ENSG00000143515 |
| FLAD1 | 0.152236644 | 1.111290998 | 0.025711477 | 0.069264082 | 1946.32505 | 2162.213741 | ENSG00000160688 |
| COG5 | 0.152235283 | 1.111289949 | 0.194921393 | 0.345070074 | 442.4350603 | 491.3441925 | ENSG00000164597 |
| PRUNE2 | 0.152135855 | 1.111213364 | 0.165307042 | 0.304607713 | 439.4455915 | 488.3846319 | ENSG00000106772 |
| DLGAP1 | 0.152084875 | 1.111174098 | 0.237798334 | 0.399631297 | 287.1854924 | 319.1383721 | ENSG00000170579 |
| CCT4 | 0.152075072 | 1.111166548 | 0.000288487 | 0.001403871 | 5357.527227 | 5952.753697 | ENSG00000115484 |
| ERCC6L2 | 0.152053762 | 1.111150134 | 0.230186423 | 0.39047892 | 283.6154249 | 314.9454877 | ENSG00000182150 |
| ELOVL4 | 0.152032768 | 1.111133966 | 0.227480143 | 0.386823515 | 302.515146 | 336.480256 | ENSG00000118402 |
| SPINT1-AS1 | 0.151974466 | 1.111089063 | 0.68128335 | 0.805555447 | 35.0285363 | 38.85889625 | ENSG00000261183 |
| NDST3 | 0.151965228 | 1.111081949 | 0.840864145 | 0.909051234 | 9.054238691 | 10.00794194 | ENSG00000164100 |
| CCN1 | 0.151952041 | 1.111071793 | 0.027645464 | 0.07361058 | 1686.684518 | 1873.852169 | ENSG00000142871 |
| GALT | 0.151927284 | 1.111052727 | 0.529645952 | 0.690716585 | 83.10845479 | 92.25741082 | ENSG00000213930 |
| GGPS1 | 0.151878944 | 1.1110155 | 0.324077098 | 0.495543064 | 185.4624911 | 206.1423557 | ENSG00000152904 |
| ADORA1 | 0.15177837 | 1.110938051 | 0.447798394 | 0.618226496 | 142.4132284 | 158.1461631 | ENSG00000163485 |
| SCIN | 0.151662166 | 1.110848572 | 0.826381106 | 0.901120226 | 9.759789188 | 10.81188296 | ENSG00000006747 |
| PSMD5 | 0.15162414 | 1.110819293 | 0.163665184 | 0.302217388 | 453.7377135 | 503.94202 | ENSG00000095261 |
| NA | 0.151593734 | 1.110795882 | 0.759279042 | 0.857014416 | 17.0965394 | 18.9371351 | ENSG00000275457 |
| CCDC171 | 0.151539976 | 1.110754492 | 0.598923492 | 0.745552887 | 50.93299444 | 56.5387481 | ENSG00000164989 |
| GPC2 | 0.15143465 | 1.110673403 | 0.042006202 | 0.103690548 | 933.5013384 | 1037.113496 | ENSG00000213420 |
| NA | 0.151229412 | 1.110515409 | 0.83795789 | 0.907638723 | 7.655652752 | 8.501948799 | ENSG00000217261 |
| RNASET2 | 0.151185302 | 1.110481456 | 0.306379107 | 0.477411902 | 263.4712833 | 292.6807218 | ENSG00000026297 |
| RFK | 0.151176464 | 1.110474653 | 0.326953952 | 0.498953302 | 254.2822296 | 282.4305338 | ENSG00000135002 |
| NA | 0.151146551 | 1.110451629 | 0.785658828 | 0.874662076 | 14.15223753 | 15.63691667 | ENSG00000253477 |
| NCKAP5-AS2 | 0.151114696 | 1.11042711 | 0.829183211 | 0.90286078 | 10.20580984 | 11.31441582 | ENSG00000226953 |
| TRAP1 | 0.151109184 | 1.110422868 | 0.001756188 | 0.007049234 | 3516.068313 | 3904.174303 | ENSG00000126602 |
| DNAJC15 | 0.150971041 | 1.110316546 | 0.163058072 | 0.301298206 | 439.8440036 | 487.9165572 | ENSG00000120675 |
| TSGA10IP | 0.150953111 | 1.110302747 | 0.610255738 | 0.753967174 | 48.45404939 | 53.77414516 | ENSG00000175513 |
| BTN3A3 | 0.150895519 | 1.110258425 | 0.716024178 | 0.829598944 | 29.73691525 | 33.10140172 | ENSG00000111801 |
| GLB1 | 0.150788333 | 1.11017594 | 0.040750658 | 0.101160848 | 932.564204 | 1034.867588 | ENSG00000170266 |
| MBTD1 | 0.150592874 | 1.110025542 | 0.029961681 | 0.078456739 | 1350.161718 | 1498.150241 | ENSG00000011258 |
| ATP5MC1 | 0.1505762 | 1.110012713 | 0.066574111 | 0.149910256 | 1559.773525 | 1730.388901 | ENSG00000159199 |
| B3GALT5 | 0.150535328 | 1.109981266 | 0.057881115 | 0.134156505 | 827.9412799 | 918.7530805 | ENSG00000183778 |
| LRP1 | 0.150431226 | 1.109901176 | 0.005572193 | 0.019260558 | 6186.863898 | 6865.256497 | ENSG00000123384 |
| BSCL2 | 0.15041398 | 1.109887907 | 0.791442404 | 0.878616299 | 12.67964865 | 14.0615537 | ENSG00000168000 |
| TMEM14B | 0.15038418 | 1.109864982 | 0.183285607 | 0.329183748 | 420.6838398 | 466.5831556 | ENSG00000137210 |
| MUS81 | 0.150233997 | 1.109749452 | 0.13095075 | 0.254167514 | 535.9517364 | 594.3629997 | ENSG00000172732 |
| HDAC1 | 0.150197478 | 1.109721362 | 0.005059788 | 0.017721201 | 3956.778562 | 4390.43309 | ENSG00000116478 |
| DLG4 | 0.150167923 | 1.109698629 | 0.047727716 | 0.115113399 | 995.0920333 | 1103.80519 | ENSG00000132535 |
| WDFY1 | 0.150159878 | 1.10969244 | 0.059764292 | 0.137480677 | 1140.138359 | 1265.951623 | ENSG00000085449 |
| KAT6B | 0.150035254 | 1.109596586 | 0.14066378 | 0.269017681 | 551.9605831 | 612.6908304 | ENSG00000156650 |
| TRAPPC12 | 0.150015903 | 1.109581703 | 0.063991813 | 0.145160428 | 1000.027346 | 1109.660848 | ENSG00000171853 |
| GTF2E1 | 0.149978462 | 1.109552907 | 0.025689194 | 0.069213721 | 1208.027044 | 1339.79629 | ENSG00000153767 |
| INPP1 | 0.149774826 | 1.109396305 | 0.589042538 | 0.738542806 | 55.96516289 | 62.18601191 | ENSG00000151689 |
| B3GALNT1 | 0.149747473 | 1.109375272 | 0.412904126 | 0.585186558 | 133.5629769 | 148.4089699 | ENSG00000169255 |
| TMEM18 | 0.149716311 | 1.10935131 | 0.240824071 | 0.403065864 | 303.7626087 | 336.5663095 | ENSG00000151353 |
| OSBPL2 | 0.149711656 | 1.109347731 | 0.07100914 | 0.157816762 | 701.9103554 | 778.3373595 | ENSG00000130703 |
| PPIEL | 0.149515973 | 1.109197272 | 0.597433061 | 0.744425518 | 51.2979909 | 56.8455419 | ENSG00000243970 |
| WASF2 | 0.149472845 | 1.109164114 | 0.00319401 | 0.011835506 | 3021.794261 | 3352.33687 | ENSG00000158195 |
| HUWE1 | 0.149319687 | 1.10904637 | 0.012754734 | 0.038650892 | 8092.609071 | 8974.517781 | ENSG00000086758 |
| PIM3 | 0.149139597 | 1.108907938 | 0.040890544 | 0.101455882 | 1152.700814 | 1278.901425 | ENSG00000198355 |
| DNM1L | 0.149115204 | 1.108889188 | 0.014575146 | 0.043216899 | 1640.011847 | 1819.370265 | ENSG00000087470 |
| ATP5PO | 0.149104861 | 1.108881239 | 0.533393451 | 0.693924025 | 88.51543186 | 98.04719449 | ENSG00000241837 |
| ZBTB1 | 0.149075936 | 1.108859007 | 0.210478712 | 0.365939568 | 353.259296 | 391.5928611 | ENSG00000126804 |
| ABR | 0.148849614 | 1.108685069 | 0.008852666 | 0.028405985 | 1762.143038 | 1953.306382 | ENSG00000159842 |
| NA | 0.148783052 | 1.108633918 | 0.076194644 | 0.166708731 | 1009.736824 | 1119.118684 | ENSG00000250899 |
| HADHB | 0.148714456 | 1.108581207 | 0.046936485 | 0.113564748 | 996.2924765 | 1104.951061 | ENSG00000138029 |
| ZNF480 | 0.148586726 | 1.108483062 | 0.124856112 | 0.245424828 | 551.1492778 | 611.0127603 | ENSG00000198464 |
| RANBP2 | 0.148511751 | 1.108425457 | 0.015029492 | 0.044340942 | 3659.201855 | 4055.556838 | ENSG00000153201 |
| CSNK1E | 0.14850975 | 1.10842392 | 0.023987197 | 0.065284568 | 1447.648036 | 1604.343388 | ENSG00000213923 |
| ARRDC1 | 0.14846974 | 1.108393181 | 0.320286556 | 0.491635992 | 249.4772514 | 276.7429485 | ENSG00000197070 |
| SLC6A11 | 0.148302624 | 1.108264796 | 0.439053317 | 0.610126606 | 117.5853937 | 130.4497486 | ENSG00000132164 |
| PROM2 | 0.148279394 | 1.108246951 | 0.503699954 | 0.66897596 | 99.98625184 | 110.857111 | ENSG00000155066 |
| FMN2 | 0.148226951 | 1.108206666 | 0.530162226 | 0.691084499 | 81.56521746 | 90.43791169 | ENSG00000155816 |
| LOC105374338 | 0.148131645 | 1.108133459 | 0.879342408 | 0.93354215 | 4.754653887 | 5.237757081 | ENSG00000283183 |
| EXOC6B | 0.148119041 | 1.108123779 | 0.035380099 | 0.089970669 | 1187.804142 | 1316.069318 | ENSG00000144036 |

| | | | | | | | |
|---|---|---|---|---|---|---|---|
| ZNF239 | 0.148020554 | 1.108048134 | 0.357838346 | 0.531045429 | 167.8684555 | 186.1716523 | ENSG00000196793 |
| ACOX1 | 0.147999404 | 1.10803189 | 0.198549643 | 0.349983659 | 369.7937481 | 409.8236429 | ENSG00000161533 |
| THAP9-AS1 | 0.147854923 | 1.10792093 | 0.291054313 | 0.460752401 | 282.4223778 | 312.768938 | ENSG00000251022 |
| AASDH | 0.14778572 | 1.107867787 | 0.323086493 | 0.494655678 | 204.1731676 | 226.4566794 | ENSG00000157426 |
| SSR1 | 0.147585348 | 1.107713929 | 0.015468312 | 0.045406748 | 1648.692041 | 1827.005888 | ENSG00000124783 |
| PTGER4 | 0.147550893 | 1.107687474 | 0.811568592 | 0.891807113 | 10.62997883 | 11.78764625 | ENSG00000171522 |
| RWDD4 | 0.147449209 | 1.107609405 | 0.291410925 | 0.461107051 | 249.9399422 | 277.0739074 | ENSG00000182552 |
| GRAMD4 | 0.147379996 | 1.107556268 | 0.120873851 | 0.239919923 | 640.7741419 | 710.1278902 | ENSG00000075240 |
| IFITM2 | 0.147344539 | 1.107529049 | 0.043977002 | 0.107659944 | 1079.372559 | 1194.98327 | ENSG00000185201 |
| MRRF | 0.147252699 | 1.107458548 | 0.176773005 | 0.320410415 | 400.8292764 | 443.9174916 | ENSG00000148187 |
| GPR107 | 0.147245124 | 1.107452733 | 0.01133528 | 0.035071193 | 1799.8069 | 1993.245559 | ENSG00000148358 |
| NA | 0.147242284 | 1.107450553 | 0.753999217 | 0.854204775 | 20.35695291 | 22.51133727 | ENSG00000273373 |
| KRT8 | 0.147237994 | 1.107447259 | 0.000185303 | 0.000940458 | 8533.608667 | 9449.696013 | ENSG00000170421 |
| ZNF747 | 0.147211011 | 1.107426547 | 0.618355265 | 0.760031311 | 48.29643151 | 53.37270769 | ENSG00000169955 |
| NTNG1 | 0.146938529 | 1.107217407 | 0.759047295 | 0.857004645 | 18.46019504 | 20.4310183 | ENSG00000162631 |
| DNAH11 | 0.146897731 | 1.107186096 | 0.493609567 | 0.660027875 | 113.4242908 | 125.4357002 | ENSG00000105877 |
| TCFL5 | 0.146503946 | 1.10688393 | 0.11526996 | 0.231199084 | 599.5497365 | 663.6414705 | ENSG00000101190 |
| NA | 0.146439677 | 1.106834622 | 0.729717357 | 0.838015093 | 23.76469657 | 26.29741701 | ENSG00000260362 |
| CTNNA1 | 0.146320179 | 1.106742947 | 0.000114754 | 0.000612538 | 8847.349255 | 9791.072211 | ENSG00000044115 |
| NPC1 | 0.146309849 | 1.106735022 | 0.044222476 | 0.108123674 | 976.3499684 | 1080.987265 | ENSG00000141458 |
| LPCAT2 | 0.146130655 | 1.106597566 | 0.536415983 | 0.696038255 | 91.08377382 | 100.9276621 | ENSG00000087253 |
| CEP128 | 0.146045394 | 1.106532169 | 0.304039702 | 0.474763631 | 237.8588628 | 262.9607336 | ENSG00000100629 |
| NA | 0.146044245 | 1.106531288 | 0.76468622 | 0.860801883 | 18.95557606 | 21.00266052 | ENSG00000212664 |
| NA | 0.145636977 | 1.106218962 | 0.345204736 | 0.517592458 | 199.6067563 | 220.7766056 | ENSG00000233476 |
| BMPR2 | 0.145568036 | 1.106166101 | 0.182286287 | 0.327785678 | 432.044087 | 478.0137087 | ENSG00000204217 |
| ATL2 | 0.145556911 | 1.106157571 | 0.020537576 | 0.057493889 | 1372.210516 | 1517.465714 | ENSG00000119787 |
| C14orf28 | 0.14535246 | 1.106000824 | 0.635120344 | 0.77136449 | 58.81592551 | 65.17835204 | ENSG00000179476 |
| PCSK6 | 0.145346167 | 1.105996 | 0.553667696 | 0.710018183 | 79.01334551 | 87.21272129 | ENSG00000140479 |
| LIMA1 | 0.145308698 | 1.105967276 | 0.043767526 | 0.107274683 | 1121.673696 | 1240.573855 | ENSG00000050405 |
| TNFRSF18 | 0.145248415 | 1.105921064 | 0.74432804 | 0.848613954 | 23.20280943 | 25.76503659 | ENSG00000186891 |
| VWC2 | 0.145103666 | 1.105810109 | 0.834544386 | 0.906384177 | 10.44512764 | 11.53355072 | ENSG00000188730 |
| NA | 0.145065688 | 1.105781 | 0.839311224 | 0.908339569 | 8.657102848 | 9.608382474 | ENSG00000266947 |
| TMLHE | 0.14501884 | 1.105745093 | 0.409122194 | 0.581304872 | 141.7296048 | 156.889946 | ENSG00000185973 |
| TRMT10B | 0.145003924 | 1.105733661 | 0.391620422 | 0.564598636 | 145.5295782 | 161.0062153 | ENSG00000165275 |
| PRIM1 | 0.14496732 | 1.105705606 | 0.310678836 | 0.481660807 | 263.3761045 | 291.573743 | ENSG00000198056 |
| ATP5F1A | 0.144947747 | 1.105690605 | 0.000429355 | 0.002005931 | 14425.68029 | 15949.60148 | ENSG00000152234 |
| ITGB3 | 0.144930612 | 1.105677474 | 0.90654431 | 0.94946231 | 2.947092772 | 3.277095161 | ENSG00000259207 |
| FAM189A2 | 0.144870602 | 1.105631483 | 0.288457702 | 0.457731145 | 289.8566974 | 320.6419605 | ENSG00000135063 |
| RNF144A-AS1 | 0.144835825 | 1.105604831 | 0.748563564 | 0.851045481 | 22.98581885 | 25.36979037 | ENSG00000228203 |
| TOM1L1 | 0.144814781 | 1.105588704 | 0.096317403 | 0.201219301 | 725.6862525 | 802.0268871 | ENSG00000141198 |
| NA | 0.144762464 | 1.105548613 | 0.468201804 | 0.637228799 | 126.55359 | 139.9785536 | ENSG00000229150 |
| NA | 0.144588486 | 1.1054153 | 0.872778105 | 0.929283719 | 5.018414094 | 5.527879334 | ENSG00000229759 |
| NA | 0.144542566 | 1.105380116 | 0.737563583 | 0.844011097 | 22.88266759 | 25.21410661 | ENSG00000262801 |
| SPPL2B | 0.144471036 | 1.105325311 | 0.156535962 | 0.292128785 | 759.487015 | 839.4056882 | ENSG00000005206 |
| TIMMDC1 | 0.144382437 | 1.105257433 | 0.01876319 | 0.053221687 | 1715.371523 | 1895.652597 | ENSG00000113845 |
| NA | 0.14435821 | 1.105238873 | 0.85119724 | 0.915330155 | 7.353495625 | 8.091909034 | ENSG00000273382 |
| ACTN4 | 0.144326132 | 1.105214298 | 0.0010266 | 0.004361185 | 9357.509556 | 10341.25355 | ENSG00000130402 |
| NA | 0.144271955 | 1.105172795 | 0.877566141 | 0.932734348 | 4.988389884 | 5.496153767 | ENSG00000235554 |
| GOLGA6L10 | 0.144256132 | 1.105160675 | 0.65198305 | 0.78368777 | 45.87618343 | 50.58121337 | ENSG00000278662 |
| ZNF542P | 0.144254296 | 1.105159268 | 0.623137322 | 0.762604966 | 63.18584646 | 70.03027712 | ENSG00000240225 |
| STXBP5L | 0.144226034 | 1.105137618 | 0.422105756 | 0.593804067 | 134.1346762 | 148.3922983 | ENSG00000145087 |
| NA | 0.144194798 | 1.105113691 | 0.494147681 | 0.660564266 | 96.76096879 | 106.7830596 | ENSG00000215158 |
| PBX2 | 0.14419141 | 1.105111096 | 0.006489143 | 0.021946363 | 3070.717646 | 3392.895949 | ENSG00000204304 |
| DDX17 | 0.144165361 | 1.105091142 | 0.005544202 | 0.01918554 | 7441.009584 | 8222.347551 | ENSG00000100201 |
| UFL1 | 0.144106119 | 1.105045764 | 0.216191674 | 0.373304116 | 342.6087048 | 378.6358631 | ENSG00000014123 |
| FBXL6 | 0.14391964 | 1.104902938 | 0.207655485 | 0.362392072 | 524.4641213 | 579.6674371 | ENSG00000182325 |
| LOC105369147 | 0.143855039 | 1.104853465 | 0.741418809 | 0.846815283 | 21.94876274 | 24.28405682 | ENSG00000247081 |
| IL1RAPL2 | 0.143791791 | 1.104805028 | 0.81042233 | 0.890970374 | 11.28317789 | 12.50286229 | ENSG00000189108 |
| HUS1B | 0.143651445 | 1.104697558 | 0.915967277 | 0.954976206 | 3.021095707 | 3.348615553 | ENSG00000188996 |
| ALCAM | 0.143628744 | 1.104680175 | 0.183935317 | 0.329905502 | 525.8293286 | 581.3333604 | ENSG00000170017 |
| SEPSECS | 0.143618438 | 1.104672284 | 0.57978721 | 0.731024514 | 68.23651209 | 75.44469067 | ENSG00000109618 |
| CDC5L | 0.143504552 | 1.104585085 | 0.024765502 | 0.0670999 | 1370.015707 | 1513.139458 | ENSG00000096401 |
| SYNE2 | 0.143451435 | 1.104544417 | 0.05644457 | 0.131506304 | 1391.407813 | 1536.685096 | ENSG00000054654 |
| RAB39A | 0.14333527 | 1.104455483 | 0.413323896 | 0.585592779 | 141.3865721 | 156.0397584 | ENSG00000179331 |
| CUL5 | 0.143258303 | 1.104396562 | 0.22166472 | 0.38009964 | 321.4819226 | 355.0242834 | ENSG00000166266 |
| C6orf136 | 0.143255234 | 1.104394213 | 0.305822646 | 0.476814407 | 215.7803543 | 238.3056384 | ENSG00000204564 |
| DENND11 | 0.143171646 | 1.104330228 | 0.068697695 | 0.153687492 | 945.214392 | 1043.772326 | ENSG00000257093 |
| NDRG3 | 0.143105124 | 1.104279309 | 0.109215174 | 0.22135637 | 647.5271902 | 715.0790448 | ENSG00000101079 |
| MAD2L1BP | 0.143099592 | 1.104275074 | 0.17782385 | 0.321740752 | 422.7954362 | 466.8378147 | ENSG00000124688 |
| ZMYM3 | 0.143088216 | 1.104266367 | 0.007376907 | 0.024417969 | 2658.438511 | 2935.775341 | ENSG00000147130 |
| ZKSCAN3 | 0.143085593 | 1.10426436 | 0.328843169 | 0.501083221 | 201.7894014 | 222.9268458 | ENSG00000189298 |
| C2orf68 | 0.143022233 | 1.104215864 | 0.085176228 | 0.182489407 | 835.0857332 | 922.2449789 | ENSG00000168887 |
| NA | 0.143001432 | 1.104199943 | 0.819705983 | 0.897041177 | 10.98939348 | 12.14713833 | ENSG00000242858 |
| STYX | 0.142870702 | 1.104099891 | 0.178768285 | 0.323055642 | 545.461771 | 602.6016444 | ENSG00000198252 |
| RITA1 | 0.142744603 | 1.10400339 | 0.133829358 | 0.258661671 | 569.3356251 | 628.5826482 | ENSG00000139405 |
| CDCA7L | 0.142561497 | 1.10386328 | 0.006050416 | 0.020688111 | 2647.57471 | 2922.174163 | ENSG00000164649 |
| N4BP1 | 0.142467453 | 1.103791326 | 0.051969721 | 0.122999644 | 1148.078358 | 1267.113785 | ENSG00000102921 |

| | | | | | | | |
|---|---|---|---|---|---|---|---|
| PCDH18 | 0.142449369 | 1.10377749 | 0.084882621 | 0.181961401 | 858.7268152 | 947.6731136 | ENSG00000189184 |
| CARMIL1 | 0.142444804 | 1.103773997 | 0.053929051 | 0.126716847 | 956.4282729 | 1055.917798 | ENSG00000079691 |
| ZDHHC23 | 0.142203824 | 1.103589644 | 0.212517407 | 0.368510602 | 366.6446584 | 404.8512673 | ENSG00000184307 |
| CLIP1 | 0.142161713 | 1.103557432 | 0.146142115 | 0.276666098 | 483.2783627 | 533.3914653 | ENSG00000130779 |
| MHENCR | 0.14214789 | 1.103546858 | 0.667514046 | 0.795712381 | 47.54112032 | 52.42625893 | ENSG00000232442 |
| CALCOCO1 | 0.14191882 | 1.103371652 | 0.320147564 | 0.491560542 | 296.5931252 | 326.690395 | ENSG00000012822 |
| OSBP | 0.141892068 | 1.103351192 | 0.014520235 | 0.043073944 | 1667.170295 | 1839.239382 | ENSG00000110048 |
| FBXL16 | 0.141813609 | 1.103291189 | 0.28146748 | 0.449886284 | 288.5426151 | 318.1588591 | ENSG00000127585 |
| ASNSD1 | 0.14178459 | 1.103268998 | 0.44222402 | 0.61325164 | 128.410148 | 141.6816307 | ENSG00000138381 |
| SETD1B | 0.141704263 | 1.103207571 | 0.185882267 | 0.33236495 | 470.3693958 | 518.7795573 | ENSG00000139718 |
| PHC1 | 0.141683563 | 1.103191743 | 0.00113128 | 0.004761841 | 14889.28573 | 16424.21708 | ENSG00000111752 |
| SGMS1 | 0.14166386 | 1.103176676 | 0.258643968 | 0.423844784 | 394.9895206 | 435.7070464 | ENSG00000198964 |
| TIPARP | 0.141468649 | 1.103027415 | 0.476300657 | 0.644703619 | 131.9802155 | 145.8554204 | ENSG00000163659 |
| MYL12B | 0.141395201 | 1.102971261 | 0.025051351 | 0.067674781 | 2563.276435 | 2826.087122 | ENSG00000118680 |
| FAM98C | 0.141338329 | 1.102927782 | 0.491629287 | 0.657856494 | 113.9829867 | 125.7976033 | ENSG00000130244 |
| HEXB | 0.141324349 | 1.102917094 | 0.064409997 | 0.145888606 | 990.5008787 | 1092.673815 | ENSG00000049860 |
| AFTPH | 0.141135448 | 1.102772693 | 0.21046441 | 0.365939568 | 354.6715994 | 391.3834251 | ENSG00000119844 |
| SMIM13 | 0.140940972 | 1.102624048 | 0.296866763 | 0.467034501 | 261.0992685 | 287.9783077 | ENSG00000224531 |
| CECR2 | 0.140897304 | 1.102590674 | 0.013168894 | 0.039731175 | 3315.292851 | 3654.645627 | ENSG00000099954 |
| NOB1 | 0.140872652 | 1.102571834 | 0.062344771 | 0.142229539 | 1389.431359 | 1531.206844 | ENSG00000141101 |
| NA | 0.140856942 | 1.102559828 | 0.800935655 | 0.885251398 | 13.62663213 | 15.02709716 | ENSG00000270605 |
| AVL9 | 0.140844508 | 1.102550325 | 0.065155034 | 0.147195896 | 854.5413962 | 942.1390978 | ENSG00000105778 |
| LYPLA1 | 0.140796427 | 1.102513581 | 0.042254134 | 0.104209218 | 1935.163862 | 2134.453074 | ENSG00000120992 |
| LRR1 | 0.140757299 | 1.10248368 | 0.391197972 | 0.56427397 | 163.6825099 | 180.5848955 | ENSG00000165501 |
| NOTCH2 | 0.140672523 | 1.102418897 | 0.026631446 | 0.071314162 | 1729.376692 | 1906.016184 | ENSG00000134250 |
| SLC25A46 | 0.140641093 | 1.10239488 | 0.199313278 | 0.35068884 | 455.5499805 | 502.6336013 | ENSG00000164209 |
| TMPRSS6 | 0.140611622 | 1.102372361 | 0.835336783 | 0.906737673 | 11.18261737 | 12.34906864 | ENSG00000187045 |
| ZRANB1 | 0.14059503 | 1.102359683 | 0.342582347 | 0.515382987 | 216.4902473 | 238.6110751 | ENSG00000019995 |
| ICA1 | 0.140455391 | 1.10225299 | 0.223602154 | 0.382132278 | 377.7106824 | 415.8511353 | ENSG00000003147 |
| COL4A2 | 0.140446402 | 1.102246123 | 0.005040574 | 0.017663526 | 3353.713888 | 3696.44559 | ENSG00000134871 |
| FRRS1 | 0.140355897 | 1.102176978 | 0.596666986 | 0.74400866 | 67.5108617 | 74.47672405 | ENSG00000156869 |
| PGM3 | 0.140316257 | 1.102146694 | 0.108443481 | 0.220139239 | 650.0715653 | 716.5965843 | ENSG00000013375 |
| TAF6 | 0.140243496 | 1.10209111 | 0.036307207 | 0.091976669 | 1556.189492 | 1714.546099 | ENSG00000106290 |
| NA | 0.140233029 | 1.102083114 | 0.86418767 | 0.923808287 | 6.2647638 | 6.951859682 | ENSG00000214832 |
| BMP8A | 0.140145068 | 1.102015922 | 0.724277358 | 0.83450006 | 27.04329495 | 29.77859305 | ENSG00000183682 |
| C8orf88 | 0.140066286 | 1.101955745 | 0.435406826 | 0.606135187 | 132.3014334 | 145.7979001 | ENSG00000253250 |
| SLC35F6 | 0.139881647 | 1.101814724 | 0.394009042 | 0.566676241 | 185.8588632 | 204.8485127 | ENSG00000213699 |
| RPS28 | 0.139809988 | 1.101759998 | 0.246271217 | 0.409700942 | 6773.542607 | 7462.8132 | ENSG00000233927 |
| MDH2 | 0.139801529 | 1.101753538 | 0.000669321 | 0.002964837 | 5730.207859 | 6313.336678 | ENSG00000146701 |
| GRB14 | 0.139737894 | 1.101704942 | 0.388393768 | 0.561741203 | 166.8232269 | 183.7049619 | ENSG00000115290 |
| TRIM59 | 0.139709083 | 1.101682941 | 0.218144546 | 0.375700959 | 398.4022584 | 439.2303856 | ENSG00000213186 |
| ZMAT5 | 0.139703881 | 1.101678968 | 0.528156746 | 0.689681974 | 115.575397 | 127.0578611 | ENSG00000100319 |
| RPL7A | 0.139588139 | 1.101590589 | 0.000360229 | 0.001716696 | 25776.24371 | 28393.24502 | ENSG00000148303 |
| GNAQ | 0.139485413 | 1.101512153 | 0.03385621 | 0.086625289 | 1317.850442 | 1451.702489 | ENSG00000156052 |
| KDM7A | 0.139441085 | 1.101478309 | 0.250617439 | 0.414778742 | 320.3667212 | 352.8356238 | ENSG00000006459 |
| CCSER2 | 0.139408224 | 1.101453221 | 0.263487921 | 0.42919408 | 309.6317513 | 341.1657261 | ENSG00000107771 |
| TUT1 | 0.139335215 | 1.101397482 | 0.359543403 | 0.532601991 | 203.8500347 | 224.7511791 | ENSG00000149016 |
| NA | 0.139308088 | 1.101376773 | 0.842735466 | 0.909954609 | 9.849186098 | 10.78230797 | ENSG00000270933 |
| NA | 0.139293091 | 1.101365323 | 0.907024292 | 0.949810314 | 3.290437721 | 3.651641249 | ENSG00000284698 |
| PLEKHA1 | 0.139289422 | 1.101362523 | 0.115891586 | 0.23218035 | 679.3622598 | 748.4947212 | ENSG00000107679 |
| SCRN1 | 0.139243123 | 1.101327178 | 0.007927617 | 0.025912751 | 3049.590012 | 3358.529667 | ENSG00000136193 |
| RPL39 | 0.139017367 | 1.101154854 | 0.273438659 | 0.440235784 | 4189.942903 | 4613.30069 | ENSG00000198918 |
| TBC1D3 | 0.138923906 | 1.101083521 | 0.707706896 | 0.823921166 | 36.53999965 | 40.13150609 | ENSG00000274512 |
| NA | 0.138881348 | 1.10105104 | 0.848993306 | 0.913958172 | 8.099558222 | 8.89961816 | ENSG00000264769 |
| MIER2 | 0.138804265 | 1.100992213 | 0.065026806 | 0.146957838 | 899.2285996 | 989.6400516 | ENSG00000105556 |
| NA | 0.138796188 | 1.100986049 | 0.906283611 | 0.949343894 | 2.985489691 | 3.300218434 | ENSG00000279415 |
| PCNP | 0.13877671 | 1.100971184 | 0.007247195 | 0.024054653 | 3117.554154 | 3432.836416 | ENSG00000081154 |
| PMPCB | 0.138640072 | 1.100866916 | 0.036434417 | 0.092202065 | 1932.268904 | 2126.954343 | ENSG00000105819 |
| LZIC | 0.138577982 | 1.100819539 | 0.239851944 | 0.40194927 | 340.8453978 | 375.4284287 | ENSG00000162441 |
| PPM1A | 0.138542588 | 1.100792532 | 0.10118923 | 0.208930903 | 902.3983912 | 994.2203187 | ENSG00000100614 |
| KDM6A | 0.138446469 | 1.100719195 | 0.067481432 | 0.151546484 | 1041.151102 | 1145.791864 | ENSG00000147050 |
| KATNA1 | 0.138311891 | 1.100616522 | 0.275795096 | 0.442773236 | 277.6770605 | 305.4132177 | ENSG00000186625 |
| LINC02585 | 0.138279739 | 1.100591993 | 0.872860217 | 0.929319855 | 5.647170753 | 6.22212267 | ENSG00000228350 |
| NA | 0.138278162 | 1.100590791 | 0.856194492 | 0.918981135 | 7.465219758 | 8.161278853 | ENSG00000277010 |
| NA | 0.138277744 | 1.100590472 | 0.878428219 | 0.933290675 | 5.687682854 | 6.264068068 | ENSG00000264304 |
| RAP1GDS1 | 0.138267098 | 1.10058235 | 0.164973582 | 0.304022294 | 473.5394612 | 521.0071876 | ENSG00000138698 |
| BLCAP | 0.138106266 | 1.100459664 | 0.066503962 | 0.149822249 | 1330.974849 | 1464.869342 | ENSG00000166619 |
| GALNT17 | 0.138032178 | 1.100403152 | 0.013536625 | 0.040603943 | 1974.951345 | 2173.741457 | ENSG00000185274 |
| ETFA | 0.137980037 | 1.100363384 | 0.038140101 | 0.095839609 | 1886.166043 | 2074.747826 | ENSG00000140374 |
| CD247 | 0.137931149 | 1.100326096 | 0.7191345 | 0.831454818 | 32.00233635 | 35.15267873 | ENSG00000198821 |
| CHTF18 | 0.137828513 | 1.10024782 | 0.062142016 | 0.141880644 | 2117.222838 | 2329.505802 | ENSG00000127586 |
| MRPL28 | 0.137786367 | 1.100215679 | 0.018283713 | 0.052060679 | 2028.624542 | 2231.660187 | ENSG00000086504 |
| TMEM120A | 0.137666652 | 1.100124386 | 0.27228892 | 0.439080963 | 310.5276596 | 341.4407943 | ENSG00000189077 |
| KCNG1 | 0.137497032 | 1.099995051 | 0.14087759 | 0.269236139 | 650.3440248 | 715.5934329 | ENSG00000026559 |
| TEF | 0.137474838 | 1.099978129 | 0.374204454 | 0.547457767 | 229.9437923 | 252.6508505 | ENSG00000167074 |
| SHISA7 | 0.13745817 | 1.09996542 | 0.697319131 | 0.817153648 | 33.47492522 | 36.80842485 | ENSG00000187902 |
| NA | 0.137452571 | 1.099961151 | 0.903498562 | 0.947455466 | 3.314880125 | 3.674764522 | ENSG00000224680 |

| | | | | | | | |
|---|---|---|---|---|---|---|---|
| NA | 0.137447723 | 1.099957455 | 0.875980424 | 0.931600831 | 5.399480241 | 5.923398121 | ENSG00000233597 |
| CLTC | 0.13740975 | 1.099928503 | 0.001312508 | 0.005447417 | 7507.986582 | 8257.942105 | ENSG00000141367 |
| PPP1R10 | 0.137273027 | 1.099824269 | 0.053992242 | 0.126826582 | 1076.984401 | 1184.561569 | ENSG00000204569 |
| PEX7 | 0.137264916 | 1.099818086 | 0.396933743 | 0.569472215 | 161.8125098 | 177.9117012 | ENSG00000112357 |
| BRAF | 0.137211173 | 1.099777116 | 0.238224768 | 0.400103953 | 449.369056 | 494.6642813 | ENSG00000157764 |
| MSMO1 | 0.137176672 | 1.099750816 | 0.048286539 | 0.116094599 | 1829.044569 | 2012.322996 | ENSG00000052802 |
| ACTN1 | 0.137097428 | 1.099690411 | 0.002512055 | 0.009627999 | 7982.215871 | 8778.755816 | ENSG00000072110 |
| FNBP1 | 0.137032646 | 1.099641032 | 0.085258378 | 0.182604571 | 826.4220847 | 908.9753052 | ENSG00000187239 |
| MICALL1 | 0.136980386 | 1.0996012 | 0.064377096 | 0.145831207 | 1122.965363 | 1235.237141 | ENSG00000100139 |
| NA | 0.136901161 | 1.099540817 | 0.750785654 | 0.852585092 | 26.0837084 | 28.71221467 | ENSG00000287678 |
| WNK1 | 0.136819643 | 1.099478691 | 0.010917899 | 0.03395956 | 3823.71655 | 4204.278018 | ENSG00000060237 |
| ENPP5 | 0.136771851 | 1.099442269 | 0.46849594 | 0.637314377 | 138.4287913 | 152.2119937 | ENSG00000112796 |
| POLR2M | 0.136516464 | 1.099247662 | 0.455163356 | 0.624906149 | 202.4153303 | 223.025227 | ENSG00000255529 |
| COL18A1 | 0.136413541 | 1.099169243 | 0.013204998 | 0.039802665 | 7511.099774 | 8256.397052 | ENSG00000182871 |
| CRKL | 0.136373204 | 1.099138512 | 0.00518168 | 0.018088995 | 2748.35208 | 3021.2581 | ENSG00000099942 |
| FYTTD1 | 0.136345973 | 1.099117766 | 0.156211609 | 0.291692785 | 546.1002887 | 600.1419573 | ENSG00000122068 |
| TRAF6 | 0.136322395 | 1.099099803 | 0.374481503 | 0.54773847 | 194.3053211 | 213.6860247 | ENSG00000175104 |
| RIOK3 | 0.136199235 | 1.099005979 | 0.225500386 | 0.384558695 | 347.6449034 | 382.0928619 | ENSG00000101782 |
| KCNT1 | 0.136071408 | 1.098908608 | 0.848346635 | 0.913505717 | 9.807798116 | 10.84737664 | ENSG00000107147 |
| PRRC2B | 0.135881433 | 1.098763913 | 0.003032002 | 0.011302483 | 6112.633643 | 6715.36166 | ENSG00000130723 |
| SEPTIN11 | 0.135834489 | 1.09872816 | 0.008869296 | 0.028445151 | 2732.182825 | 3002.390807 | ENSG00000138758 |
| NA | 0.135797594 | 1.098700062 | 0.873755483 | 0.929913771 | 5.72328887 | 6.287191341 | ENSG00000256591 |
| OPLAH | 0.135754623 | 1.098667338 | 0.124229275 | 0.244666354 | 853.4745161 | 937.3743596 | ENSG00000178814 |
| ZPLD2P | 0.135730423 | 1.098648908 | 0.725818821 | 0.835365268 | 38.50450293 | 42.16557851 | ENSG00000236155 |
| GGA1 | 0.135544856 | 1.098507604 | 0.05224646 | 0.123409631 | 1032.894364 | 1134.607337 | ENSG00000100083 |
| ARHGAP32 | 0.135495594 | 1.098470095 | 0.118387354 | 0.236028364 | 627.5138576 | 689.3279898 | ENSG00000134909 |
| STK11 | 0.135476097 | 1.09845525 | 0.02229133 | 0.061466844 | 2275.251191 | 2498.799339 | ENSG00000118046 |
| CCNT1 | 0.135463395 | 1.098445579 | 0.125339623 | 0.246099405 | 747.9620151 | 820.9368825 | ENSG00000129315 |
| SNAP23 | 0.135328226 | 1.098342668 | 0.327360268 | 0.499415336 | 346.8798539 | 380.5785453 | ENSG00000092531 |
| NOA1 | 0.135210662 | 1.098253169 | 0.086465917 | 0.184739617 | 833.7884077 | 915.4528742 | ENSG00000084092 |
| MINDY2 | 0.135177556 | 1.098227967 | 0.231463859 | 0.391887775 | 457.8415589 | 503.1487708 | ENSG00000128923 |
| AIFM2 | 0.13509613 | 1.098165985 | 0.316757432 | 0.48786967 | 268.5485066 | 295.1078962 | ENSG00000042286 |
| LOC283335 | 0.135041521 | 1.098124418 | 0.657514573 | 0.788028928 | 49.17307884 | 53.87308997 | ENSG00000257337 |
| SLMAP | 0.135018906 | 1.098107204 | 0.077978365 | 0.169860055 | 978.0887961 | 1074.733919 | ENSG00000163681 |
| NA | 0.134968679 | 1.098068974 | 0.905716018 | 0.949058539 | 3.624734239 | 4.017585042 | ENSG00000237846 |
| MTX2 | 0.134909307 | 1.098023786 | 0.081160036 | 0.175423895 | 822.7291295 | 903.0694085 | ENSG00000128654 |
| NA | 0.134853201 | 1.097981085 | 0.179964204 | 0.324699725 | 584.0416893 | 640.8446031 | ENSG00000111788 |
| PACC1 | 0.134749014 | 1.097901795 | 0.252660917 | 0.417265794 | 339.8401177 | 373.1088078 | ENSG00000065600 |
| SCAF8 | 0.134716611 | 1.097877137 | 0.084016854 | 0.180406181 | 884.2300513 | 970.8109419 | ENSG00000213079 |
| ELMOD2 | 0.134706752 | 1.097869634 | 0.18772485 | 0.334942664 | 453.769965 | 498.1895896 | ENSG00000179387 |
| SAMD3 | 0.134606682 | 1.097793485 | 0.753334137 | 0.853912734 | 23.62015732 | 25.97771976 | ENSG00000164483 |
| DDIT4L | 0.134568239 | 1.097764232 | 0.535068868 | 0.695272365 | 89.60069706 | 98.41204167 | ENSG00000145358 |
| MRPS6 | 0.134563986 | 1.097760996 | 0.174464969 | 0.317330163 | 468.579544 | 514.3984274 | ENSG00000243927 |
| LONP1 | 0.134542246 | 1.097744454 | 0.003178949 | 0.011784226 | 4589.436303 | 5038.186427 | ENSG00000196365 |
| KIF20B | 0.134497663 | 1.097710531 | 0.082957172 | 0.178727586 | 1046.299715 | 1149.11572 | ENSG00000138182 |
| ACOT2 | 0.134469823 | 1.097689349 | 0.474628944 | 0.643027387 | 148.1193715 | 162.6944926 | ENSG00000119673 |
| SLC25A25 | 0.134453262 | 1.097676749 | 0.174838385 | 0.317802136 | 525.1639395 | 576.2069612 | ENSG00000148339 |
| SCRN2 | 0.134415927 | 1.097648343 | 0.181219624 | 0.326415291 | 534.3911389 | 585.9656418 | ENSG00000141295 |
| MYH9 | 0.134255753 | 1.097526484 | 0.000195156 | 0.000986572 | 15512.31803 | 17024.71713 | ENSG00000100345 |
| GPHA2 | 0.13424797 | 1.097520563 | 0.859037743 | 0.920821105 | 7.731770869 | 8.441181275 | ENSG00000149735 |
| NA | 0.134148355 | 1.097444784 | 0.911185846 | 0.951993281 | 3.312089222 | 3.655942396 | ENSG00000234322 |
| RHOBTB2 | 0.134062768 | 1.097379666 | 0.418782078 | 0.590895631 | 199.5653314 | 219.3491481 | ENSG00000008853 |
| METTL5 | 0.134042361 | 1.097364159 | 0.119762422 | 0.238154508 | 794.7001787 | 872.0246331 | ENSG00000138382 |
| ATP2A1 | 0.133930885 | 1.097279369 | 0.61489856 | 0.757375056 | 59.05121311 | 64.89146485 | ENSG00000196296 |
| NA | 0.13388033 | 1.097240919 | 0.62888675 | 0.766771224 | 67.15739228 | 73.88976736 | ENSG00000259959 |
| AHSP | 0.13381495 | 1.097191195 | 0.861901102 | 0.922689647 | 7.740143577 | 8.449250532 | ENSG00000169877 |
| B3GALT6 | 0.133729514 | 1.097126222 | 0.151834715 | 0.285259397 | 547.61538 | 600.9513145 | ENSG00000176022 |
| TXLNA | 0.133674707 | 1.097084543 | 0.06481041 | 0.146594375 | 1777.989197 | 1949.59189 | ENSG00000084652 |
| VASP | 0.13356385 | 1.097000247 | 0.042716701 | 0.105186685 | 2810.743283 | 3084.005693 | ENSG00000125753 |
| KNTC1 | 0.133476195 | 1.096933598 | 0.089856507 | 0.190485921 | 1111.515829 | 1218.779805 | ENSG00000184445 |
| ANXA5 | 0.13347503 | 1.096932712 | 0.011734745 | 0.035995384 | 3869.623643 | 4244.554176 | ENSG00000164111 |
| SLC8B1 | 0.133389147 | 1.096867413 | 0.250721294 | 0.414889339 | 361.9033954 | 396.790575 | ENSG00000089060 |
| RHOV | 0.133287087 | 1.096789821 | 0.622620391 | 0.762504441 | 66.38094205 | 72.82259548 | ENSG00000104140 |
| M6PR | 0.133236448 | 1.096751324 | 0.047240972 | 0.114172587 | 1438.237387 | 1576.76694 | ENSG00000003056 |
| ASAH2 | 0.133184801 | 1.096712062 | 0.753249505 | 0.853907046 | 22.98581885 | 25.21195604 | ENSG00000188611 |
| AGTRAP | 0.133181088 | 1.09670924 | 0.049281338 | 0.117921892 | 1136.660177 | 1246.733811 | ENSG00000177674 |
| KAT7 | 0.13311386 | 1.096658136 | 0.009551491 | 0.030270709 | 4676.040204 | 5128.616244 | ENSG00000136504 |
| NA | 0.132959227 | 1.096540598 | 0.719509662 | 0.831663685 | 39.07042026 | 42.9012221 | ENSG00000288398 |
| DKK1 | 0.132689936 | 1.096335939 | 0.912055052 | 0.952535122 | 2.961047286 | 3.253971888 | ENSG00000107984 |
| TMEM132C | 0.132642987 | 1.096300262 | 0.758131341 | 0.856402715 | 22.78768887 | 24.95947803 | ENSG00000181234 |
| PLOD1 | 0.132414542 | 1.096126681 | 0.01651262 | 0.0479087 | 1923.776517 | 2109.153321 | ENSG00000083444 |
| ASTN2 | 0.132364316 | 1.096088521 | 0.189742137 | 0.337792134 | 457.2753903 | 501.3096623 | ENSG00000148219 |
| PKP2 | 0.132307874 | 1.09604564 | 0.183943993 | 0.329905502 | 457.588799 | 501.642263 | ENSG00000057294 |
| N4BP2 | 0.132209129 | 1.095970624 | 0.134567517 | 0.25972406 | 634.3270055 | 695.5576182 | ENSG00000078177 |
| NA | 0.132172711 | 1.095942958 | 0.804036158 | 0.887065768 | 15.03917259 | 16.50807693 | ENSG00000227885 |
| NETO2 | 0.132060431 | 1.095857668 | 0.247464387 | 0.411006068 | 400.6716217 | 439.2578406 | ENSG00000171208 |
| INO80C | 0.131964572 | 1.095784857 | 0.397965778 | 0.570346957 | 183.4802033 | 200.9132009 | ENSG00000153391 |

| | | | | | | | |
|---|---|---|---|---|---|---|---|
| DCLRE1A | 0.131883229 | 1.095723075 | 0.299462489 | 0.469954021 | 338.5098776 | 370.5676225 | ENSG00000198924 |
| GOSR1 | 0.131795517 | 1.09565646 | 0.136685433 | 0.262770316 | 808.8342456 | 886.7755775 | ENSG00000108587 |
| BLOC1S2 | 0.131766275 | 1.095634253 | 0.116804355 | 0.233741994 | 753.4144118 | 826.089931 | ENSG00000196072 |
| GPR180 | 0.131758477 | 1.095628331 | 0.361140182 | 0.534249291 | 222.8887501 | 244.1809119 | ENSG00000152749 |
| MEDAG | 0.131723072 | 1.095601444 | 0.735183472 | 0.842286754 | 35.29105921 | 38.53758147 | ENSG00000102802 |
| LRP4 | 0.131659329 | 1.095553037 | 0.080283265 | 0.173898997 | 971.9900967 | 1064.251468 | ENSG00000134569 |
| RASD2 | 0.131639725 | 1.095538151 | 0.430220764 | 0.601622962 | 174.5593402 | 190.9595869 | ENSG00000100302 |
| MTUS1 | 0.131626348 | 1.095527993 | 0.5821362 | 0.732591091 | 89.13689185 | 97.5685663 | ENSG00000129422 |
| GNRH1 | 0.131527926 | 1.095453257 | 0.889754661 | 0.939066482 | 5.393898435 | 5.887371406 | ENSG00000147437 |
| SMS | 0.131493965 | 1.095427471 | 0.008398649 | 0.027202541 | 4035.676922 | 4421.55774 | ENSG00000102172 |
| TUSC3 | 0.131433515 | 1.095381572 | 0.052780943 | 0.124428347 | 1191.65424 | 1305.078622 | ENSG00000104723 |
| CHP1 | 0.131241112 | 1.095235498 | 0.05752384 | 0.133456692 | 1734.141057 | 1899.65955 | ENSG00000187446 |
| NA | 0.131198141 | 1.095202877 | 0.696241092 | 0.816684112 | 41.96536176 | 45.86749488 | ENSG00000279483 |
| PPFIA3 | 0.131089803 | 1.095120636 | 0.112522921 | 0.226963579 | 749.8880477 | 821.3071458 | ENSG00000177380 |
| DOT1L | 0.131045503 | 1.095087009 | 0.03287082 | 0.084581963 | 2290.342105 | 2508.300186 | ENSG00000104885 |
| AP3B1 | 0.131030696 | 1.09507577 | 0.062987326 | 0.143390195 | 1395.286156 | 1527.938224 | ENSG00000132842 |
| NA | 0.130972922 | 1.095031918 | 0.89586313 | 0.942392226 | 5.144092742 | 5.597249153 | ENSG00000237840 |
| ADO | 0.130795611 | 1.094897344 | 0.096093853 | 0.200882803 | 825.176574 | 903.7216759 | ENSG00000181915 |
| BDNF-AS | 0.130662255 | 1.094796142 | 0.790289734 | 0.87763484 | 18.56613721 | 20.38262118 | ENSG00000245573 |
| CABP1 | 0.130648467 | 1.094785679 | 0.538775484 | 0.698254475 | 101.0065625 | 110.660548 | ENSG00000157782 |
| GALNT1 | 0.130442328 | 1.094629261 | 0.077551067 | 0.169101123 | 1264.776843 | 1384.702518 | ENSG00000141429 |
| ARHGAP11A | 0.130392834 | 1.094591709 | 0.065867842 | 0.148649571 | 1274.396434 | 1394.842427 | ENSG00000198826 |
| APLP2 | 0.130386393 | 1.094586822 | 0.000514591 | 0.002351807 | 12094.17644 | 13238.34175 | ENSG00000084234 |
| CREB1 | 0.130374175 | 1.094577553 | 0.067864823 | 0.152198198 | 1157.4321 | 1266.523557 | ENSG00000118260 |
| C12orf29 | 0.130355038 | 1.094563033 | 0.379728973 | 0.553016176 | 216.1225479 | 236.6119633 | ENSG00000133641 |
| OTUD4 | 0.130286159 | 1.094510777 | 0.062145329 | 0.141880644 | 1294.634041 | 1417.252638 | ENSG00000164164 |
| CA2 | 0.13016663 | 1.094420099 | 0.504672149 | 0.669852101 | 122.7385349 | 134.4141024 | ENSG00000104267 |
| ACTC1 | 0.129951243 | 1.09425672 | 0.406492903 | 0.578762609 | 255.6602031 | 280.0872629 | ENSG00000159251 |
| MAN2B1 | 0.129889569 | 1.094209942 | 0.071382816 | 0.158498296 | 951.9807944 | 1041.638478 | ENSG00000104774 |
| PINK1 | 0.12977384 | 1.094122171 | 0.4344241 | 0.605525673 | 195.4928616 | 214.0721477 | ENSG00000158828 |
| CCL2 | 0.12970409 | 1.094069275 | 0.710865374 | 0.825872368 | 35.07722295 | 38.36228208 | ENSG00000108691 |
| FBXO21 | 0.129626923 | 1.094010757 | 0.055302597 | 0.129471165 | 1350.088242 | 1476.967968 | ENSG00000135108 |
| TFB1M | 0.129563696 | 1.093962811 | 0.297224547 | 0.467451388 | 344.4533114 | 376.6386719 | ENSG00000029639 |
| ETV5 | 0.129554813 | 1.093956076 | 0.097168197 | 0.202557996 | 1006.017497 | 1100.195942 | ENSG00000244405 |
| AGBL3 | 0.129412686 | 1.093848311 | 0.718919134 | 0.831355294 | 37.2209956 | 40.80020301 | ENSG00000146856 |
| CPLANE1 | 0.129225184 | 1.093706156 | 0.512453287 | 0.676780746 | 123.9168638 | 135.2263973 | ENSG00000197603 |
| TRIM66 | 0.129072477 | 1.093590395 | 0.660268944 | 0.790200893 | 52.07108664 | 56.89367856 | ENSG00000166436 |
| ZNF334 | 0.128976186 | 1.093517407 | 0.376749963 | 0.550235282 | 253.5472042 | 277.0246984 | ENSG00000198185 |
| NA | 0.128938018 | 1.093488477 | 0.912868769 | 0.95298184 | 3.018304804 | 3.302369007 | ENSG00000248569 |
| ANKRD46 | 0.128923866 | 1.093477751 | 0.27335755 | 0.440191783 | 411.5392403 | 450.1293702 | ENSG00000186106 |
| COPB1 | 0.128879878 | 1.093444411 | 0.063848847 | 0.144906836 | 1433.153368 | 1567.48413 | ENSG00000129083 |
| SNORD46 | 0.128826535 | 1.093403982 | 0.826155971 | 0.901078467 | 12.0592647 | 13.19280448 | ENSG00000200913 |
| CTTNBP2NL | 0.12878863 | 1.093375255 | 0.25906616 | 0.424176311 | 564.2394902 | 617.1745695 | ENSG00000143079 |
| FOXD3 | 0.128666957 | 1.093283047 | 0.278941403 | 0.446709166 | 436.0491378 | 477.3404686 | ENSG00000187140 |
| KCNH7 | 0.12864224 | 1.093264316 | 0.851169141 | 0.915330155 | 9.612459038 | 10.54650152 | ENSG00000184611 |
| NA | 0.128385764 | 1.093069978 | 0.699001839 | 0.818280507 | 38.47447872 | 42.04134775 | ENSG00000280063 |
| ARHGEF10L | 0.128373733 | 1.093060862 | 0.039421668 | 0.098578233 | 1357.909397 | 1484.046784 | ENSG00000074964 |
| SS18 | 0.12835004 | 1.093042911 | 0.031448484 | 0.08160676 | 1718.311182 | 1878.274978 | ENSG00000141380 |
| XAB2 | 0.12822314 | 1.092946771 | 0.024028305 | 0.065378002 | 2091.045714 | 2285.20445 | ENSG00000076924 |
| STXBP4 | 0.128195538 | 1.09292586 | 0.407205229 | 0.57939195 | 181.5844846 | 198.4645271 | ENSG00000166263 |
| ZNF470 | 0.1281459 | 1.092888257 | 0.428075524 | 0.599404368 | 183.011343 | 200.2356291 | ENSG00000197016 |
| MMAB | 0.12813859 | 1.09288272 | 0.227490165 | 0.386823515 | 412.6087993 | 450.7102025 | ENSG00000139428 |
| WAC | 0.127832516 | 1.092650885 | 0.039015035 | 0.097758027 | 2328.35612 | 2544.649035 | ENSG00000095787 |
| ZNF318 | 0.127793149 | 1.09262107 | 0.127550476 | 0.249424858 | 747.8208176 | 816.6179014 | ENSG00000171467 |
| GNB4 | 0.127783231 | 1.092613559 | 0.247334911 | 0.410932543 | 513.5252139 | 561.0154985 | ENSG00000114450 |
| ETV1 | 0.127778553 | 1.092610015 | 0.056642977 | 0.131888917 | 1383.145569 | 1511.888299 | ENSG00000006468 |
| NA | 0.127748196 | 1.092587026 | 0.920457976 | 0.95753639 | 3.263204413 | 3.580120856 | ENSG00000287735 |
| DACH1 | 0.127622281 | 1.092491671 | 0.882738483 | 0.935911408 | 7.405171337 | 8.134114897 | ENSG00000276644 |
| DNAJC13 | 0.127607619 | 1.092480568 | 0.124313321 | 0.244806889 | 726.1878173 | 793.7193629 | ENSG00000138246 |
| CCDC17 | 0.12756772 | 1.092450355 | 0.829279296 | 0.90286078 | 12.30137341 | 13.45980346 | ENSG00000159588 |
| GLI4 | 0.127353015 | 1.092287787 | 0.640163979 | 0.774888832 | 56.57525911 | 61.76952042 | ENSG00000250571 |
| ZNF304 | 0.127321987 | 1.092264295 | 0.370936513 | 0.544659482 | 237.2635969 | 258.8538906 | ENSG00000131845 |
| DGCR5 | 0.127159436 | 1.092141235 | 0.885738575 | 0.937037652 | 6.322021318 | 6.918516579 | ENSG00000273032 |
| LDLR | 0.127058232 | 1.092064624 | 0.025139862 | 0.067894873 | 3588.072467 | 3917.004248 | ENSG00000130164 |
| SLC25A37 | 0.12698075 | 1.092005975 | 0.128717195 | 0.251019351 | 909.1268948 | 992.4801546 | ENSG00000147454 |
| AGK | 0.126965329 | 1.091994303 | 0.177596807 | 0.321360098 | 589.4942847 | 643.4534917 | ENSG00000006530 |
| FBXL5 | 0.126949253 | 1.091982135 | 0.076175659 | 0.166686097 | 1104.13902 | 1206.213783 | ENSG00000118564 |
| ZNF688 | 0.126874493 | 1.09192555 | 0.663396884 | 0.792273218 | 55.2435427 | 60.50981402 | ENSG00000229809 |
| STYK1 | 0.126825538 | 1.091888498 | 0.765420688 | 0.861069997 | 25.25036407 | 27.63320541 | ENSG00000060140 |
| NA | 0.126737288 | 1.091821709 | 0.512803467 | 0.677104153 | 119.7281272 | 130.6559801 | ENSG00000242474 |
| NUDT16P1 | 0.126715387 | 1.091805135 | 0.503320254 | 0.668747918 | 118.5765561 | 129.3489732 | ENSG00000246082 |
| AFG3L2 | 0.12664046 | 1.091748433 | 0.029142186 | 0.076706497 | 2156.935793 | 2354.931665 | ENSG00000141385 |
| NA | 0.126622886 | 1.091735134 | 0.913862482 | 0.953761152 | 2.979907885 | 3.253971888 | ENSG00000225912 |
| NA | 0.126576728 | 1.091700205 | 0.906455323 | 0.949420655 | 3.630316045 | 3.996612343 | ENSG00000285877 |
| SEMA6C | 0.126562617 | 1.091689527 | 0.141039465 | 0.269438039 | 847.7587113 | 925.8254397 | ENSG00000143434 |
| DAG1 | 0.126536191 | 1.09166953 | 0.011321873 | 0.03503533 | 2870.255495 | 3133.587365 | ENSG00000173402 |
| GRID2 | 0.126455823 | 1.091608719 | 0.228600062 | 0.388265874 | 497.3721914 | 542.7688692 | ENSG00000152208 |

| | | | | | | | |
|---|---|---|---|---|---|---|---|
| CDK9 | 0.126277178 | 1.091473556 | 0.031567687 | 0.081862111 | 2249.563181 | 2454.784147 | ENSG00000136807 |
| GIPC2 | 0.126260923 | 1.091461259 | 0.784305559 | 0.873764212 | 22.19107161 | 24.12864563 | ENSG00000137960 |
| NA | 0.126258644 | 1.091459535 | 0.84581425 | 0.912000597 | 10.78847259 | 11.71182472 | ENSG00000236782 |
| NSUN6 | 0.126254226 | 1.091456192 | 0.436553946 | 0.6073972 | 173.1384271 | 188.7249412 | ENSG00000241058 |
| SOX11 | 0.126148913 | 1.091376522 | 0.101759191 | 0.209860654 | 1778.365362 | 1941.510691 | ENSG00000176887 |
| NA | 0.126067037 | 1.091314585 | 0.910852407 | 0.951993281 | 3.303716514 | 3.622066255 | ENSG00000227155 |
| NCAPD3 | 0.125995613 | 1.091260559 | 0.057936924 | 0.134242631 | 1648.432359 | 1799.115171 | ENSG00000151503 |
| MRPL41 | 0.125908431 | 1.091194616 | 0.12222355 | 0.241851933 | 1590.682667 | 1735.781325 | ENSG00000182154 |
| RSBN1 | 0.125835326 | 1.091139324 | 0.359051769 | 0.532109371 | 282.2850599 | 308.2272597 | ENSG00000081019 |
| KIF11 | 0.125804037 | 1.09111566 | 0.033637814 | 0.086187684 | 1671.151052 | 1823.835806 | ENSG00000138160 |
| NA | 0.125740937 | 1.091067937 | 0.759188963 | 0.857014416 | 25.84283899 | 28.28712089 | ENSG00000273270 |
| NA | 0.125691032 | 1.091030196 | 0.804681179 | 0.887171265 | 18.72463097 | 20.44984043 | ENSG00000240731 |
| ZNF461 | 0.12566538 | 1.091010798 | 0.651167687 | 0.783195701 | 58.98924966 | 64.4031804 | ENSG00000197808 |
| LOC100996419 | 0.125417439 | 1.090823313 | 0.832265298 | 0.904825849 | 12.02086778 | 13.10676311 | ENSG00000245317 |
| GS1-124K5.4 | 0.125366277 | 1.090784631 | 0.798885152 | 0.883907026 | 19.29353937 | 20.94404356 | ENSG00000237310 |
| NA | 0.125153254 | 1.090623581 | 0.913278828 | 0.953256842 | 3.012722998 | 3.283546882 | ENSG00000237804 |
| GPX7 | 0.125056832 | 1.090550692 | 0.416653723 | 0.588753807 | 183.0536068 | 199.6478668 | ENSG00000116157 |
| GMEB1 | 0.125019223 | 1.090522263 | 0.224194183 | 0.382737167 | 513.2677496 | 560.171539 | ENSG00000162419 |
| LINC01182 | 0.124959284 | 1.090476956 | 0.917081468 | 0.955518152 | 3.037165403 | 3.308820728 | ENSG00000250634 |
| CCDC180 | 0.124934931 | 1.090458549 | 0.830531916 | 0.903705296 | 12.08649801 | 13.1696812 | ENSG00000197816 |
| ZBTB24 | 0.124882201 | 1.090418694 | 0.234028491 | 0.394878301 | 465.3934216 | 507.1655929 | ENSG00000112365 |
| C1orf112 | 0.124854611 | 1.090397841 | 0.253102817 | 0.417707733 | 384.441642 | 419.3427795 | ENSG00000000460 |
| TRAPPC6B | 0.124834755 | 1.090382834 | 0.491644657 | 0.657856494 | 138.3878037 | 150.7837122 | ENSG00000182400 |
| RPL17 | 0.124823375 | 1.090374233 | 0.648790232 | 0.781652034 | 57.9792267 | 63.23356817 | ENSG00000265681 |
| KPNA5 | 0.124762037 | 1.090327875 | 0.505672816 | 0.670442219 | 171.4614138 | 187.2861368 | ENSG00000196911 |
| TPK1 | 0.124583886 | 1.090193245 | 0.808978542 | 0.890279831 | 20.97175505 | 22.95336307 | ENSG00000196511 |
| CDKN1B | 0.124444846 | 1.090088182 | 0.121366017 | 0.24048174 | 846.3154325 | 922.3420578 | ENSG00000111276 |
| MYL6 | 0.124442541 | 1.09008644 | 0.018631587 | 0.05289506 | 2603.627923 | 2837.295917 | ENSG00000092841 |
| PCYOX1 | 0.124309856 | 1.0898619 | 0.107617747 | 0.218900671 | 1274.219957 | 1389.627744 | ENSG00000116005 |
| NNAT | 0.124223126 | 1.089920665 | 0.241415666 | 0.403705757 | 407.4333309 | 444.0581214 | ENSG00000053438 |
| MPZL2 | 0.124181554 | 1.089889259 | 0.737208648 | 0.843705033 | 30.33373267 | 33.05138707 | ENSG00000149573 |
| BCL10 | 0.124153605 | 1.089868145 | 0.431601838 | 0.602593258 | 214.1016261 | 233.3741542 | ENSG00000142867 |
| NA | 0.124095755 | 1.089824444 | 0.859539162 | 0.921084878 | 8.77719969 | 9.539012655 | ENSG00000261211 |
| RAB21 | 0.124083267 | 1.089815011 | 0.214491411 | 0.370965915 | 657.5354848 | 716.9786297 | ENSG00000080371 |
| UBAC2-AS1 | 0.123915362 | 1.089688182 | 0.755668728 | 0.855091825 | 29.46912484 | 32.05680165 | ENSG00000228889 |
| CLUAP1 | 0.12389298 | 1.089671277 | 0.566699737 | 0.7202656 | 95.06600769 | 103.4094307 | ENSG00000103351 |
| NPAS4 | 0.123779393 | 1.089585488 | 0.884634938 | 0.936840831 | 6.989174896 | 7.648253591 | ENSG00000174576 |
| NA | 0.123764819 | 1.089574481 | 0.722604633 | 0.83371806 | 39.08581424 | 42.67508402 | ENSG00000278768 |
| IKBKB | 0.123756385 | 1.089568111 | 0.135507138 | 0.260963182 | 688.266014 | 749.7598438 | ENSG00000104365 |
| DCAF10 | 0.123713578 | 1.089535782 | 0.29830498 | 0.468730257 | 311.532176 | 339.4607772 | ENSG00000122741 |
| USP11 | 0.123702115 | 1.089527126 | 0.005160792 | 0.018025863 | 5078.344663 | 5532.862061 | ENSG00000102226 |
| RPL13A | 0.123611774 | 1.089458902 | 0.006336558 | 0.021501942 | 48992.58845 | 53374.4299 | ENSG00000142541 |
| SNUPN | 0.12358987 | 1.089442361 | 0.328716917 | 0.501000083 | 290.433591 | 316.3006497 | ENSG00000169371 |
| TTC19 | 0.123530865 | 1.089397804 | 0.027897988 | 0.07411928 | 1936.55749 | 2110.0769 | ENSG00000011295 |
| BCKDK | 0.12340315 | 1.08930137 | 0.089798526 | 0.190425785 | 1037.420151 | 1129.952805 | ENSG00000103507 |
| FRMD4A | 0.123361847 | 1.089270184 | 0.237399633 | 0.399135117 | 467.4306005 | 508.8321103 | ENSG00000151474 |
| LOC101928069 | 0.123348913 | 1.089260419 | 0.85447972 | 0.91780447 | 9.307511009 | 10.14506405 | ENSG00000255517 |
| ZNF182 | 0.123215497 | 1.089159692 | 0.500779112 | 0.666686425 | 157.3631162 | 171.3531976 | ENSG00000147118 |
| DENND2B | 0.123129121 | 1.089094485 | 0.279320584 | 0.447167916 | 411.4525222 | 447.900698 | ENSG00000166444 |
| AGA | 0.123082581 | 1.089059352 | 0.521333322 | 0.684299983 | 119.1821096 | 129.8853822 | ENSG00000038002 |
| SCAI | 0.123062556 | 1.089044236 | 0.344960622 | 0.517524352 | 262.0762009 | 285.3556489 | ENSG00000173611 |
| TXN | 0.123052751 | 1.089036835 | 0.04871751 | 0.116964466 | 2666.298543 | 2903.38497 | ENSG00000136810 |
| BORCS8 | 0.123044655 | 1.089030723 | 0.707063287 | 0.823594399 | 42.31967016 | 46.05705479 | ENSG00000254901 |
| FOXJ3 | 0.12296227 | 1.088968536 | 0.134659178 | 0.259797001 | 752.7212131 | 820.0036039 | ENSG00000198815 |
| MARCHF6 | 0.122935865 | 1.088948605 | 0.07356469 | 0.162335441 | 1637.334532 | 1782.365289 | ENSG00000145495 |
| LARP7 | 0.122772253 | 1.088825118 | 0.160672479 | 0.297917416 | 872.5676612 | 950.4979447 | ENSG00000174720 |
| NUDT14 | 0.122503583 | 1.088622367 | 0.302894894 | 0.473589351 | 335.6569629 | 365.2984308 | ENSG00000183828 |
| BOLA3 | 0.122474639 | 1.088600526 | 0.376027023 | 0.549415837 | 221.122577 | 240.7623814 | ENSG00000163170 |
| PRDM14 | 0.122424837 | 1.088562948 | 0.031950622 | 0.082710607 | 2191.301034 | 2386.090942 | ENSG00000147596 |
| PALD1 | 0.122350579 | 1.08850692 | 0.028401265 | 0.07517684 | 1845.200493 | 2008.199591 | ENSG00000107719 |
| ARHGAP42 | 0.12218423 | 1.088381418 | 0.22939491 | 0.389375909 | 518.6179575 | 564.6010476 | ENSG00000165895 |
| PON2 | 0.122111638 | 1.088326655 | 0.359597001 | 0.53263097 | 269.6775503 | 293.7516681 | ENSG00000105854 |
| USP12 | 0.122006739 | 1.088247525 | 0.204774231 | 0.358336328 | 542.1122729 | 589.9842624 | ENSG00000152484 |
| FAM107B | 0.121845102 | 1.088125606 | 0.188527685 | 0.336138447 | 616.0921979 | 670.4927741 | ENSG00000065809 |
| MED9 | 0.121746312 | 1.088051099 | 0.247651422 | 0.411207552 | 405.9404419 | 441.711609 | ENSG00000141026 |
| ASCC1 | 0.12156124 | 1.08791153 | 0.207723389 | 0.362412219 | 572.5529358 | 623.0219099 | ENSG00000138303 |
| PRKAA2 | 0.121501336 | 1.087866358 | 0.441872099 | 0.612973015 | 230.9843662 | 251.5543335 | ENSG00000162409 |
| FOXD1 | 0.121336326 | 1.087741939 | 0.892131177 | 0.940288616 | 5.690473756 | 6.165123255 | ENSG00000251493 |
| MLXIP | 0.121183396 | 1.087626642 | 0.086865968 | 0.185368516 | 1505.72735 | 1637.671527 | ENSG00000175727 |
| PRMT3 | 0.121047482 | 1.087524183 | 0.15718817 | 0.292977487 | 665.8685277 | 724.3053265 | ENSG00000185238 |
| FIBCD1 | 0.120953198 | 1.087453113 | 0.5382812 | 0.697942104 | 132.6426631 | 144.1674278 | ENSG00000130720 |
| GLUD1P3 | 0.120895957 | 1.087409967 | 0.816350768 | 0.894937925 | 18.53419798 | 20.13391129 | ENSG00000250959 |
| IP6K2 | 0.120884975 | 1.08740169 | 0.05561824 | 0.130068064 | 1369.442017 | 1489.014574 | ENSG00000068745 |
| CDR2 | 0.120833001 | 1.087362516 | 0.133094534 | 0.257396155 | 985.2481409 | 1070.889077 | ENSG00000140743 |
| TRAM2 | 0.120769309 | 1.087314512 | 0.184523971 | 0.330699734 | 737.511256 | 801.7206995 | ENSG00000065308 |
| TAX1BP1 | 0.120661883 | 1.087233551 | 0.090858175 | 0.192039559 | 1076.042748 | 1170.052016 | ENSG00000106052 |
| PDCD6IP | 0.120645194 | 1.087220975 | 0.070342969 | 0.156705353 | 1454.453563 | 1581.288154 | ENSG00000170248 |

| | | | | | | | |
|---|---|---|---|---|---|---|---|
| ZNF550 | 0.120509825 | 1.087118965 | 0.454251571 | 0.624131491 | 175.324827 | 190.41377 | ENSG00000251369 |
| ABHD10 | 0.120439071 | 1.087065651 | 0.2330948 | 0.393923066 | 470.7410259 | 511.6510106 | ENSG00000144827 |
| NA | 0.12039862 | 1.087035171 | 0.302557878 | 0.473215828 | 529.0946851 | 574.8002097 | ENSG00000281398 |
| SLC6A17 | 0.120397077 | 1.087034008 | 0.864398086 | 0.923879632 | 8.986493283 | 9.751695833 | ENSG00000197106 |
| LRRC41 | 0.12024269 | 1.086917689 | 0.11694983 | 0.2339603 | 1215.39512 | 1321.689115 | ENSG00000132128 |
| NA | 0.120206458 | 1.086890392 | 0.814933775 | 0.894197569 | 17.99954415 | 19.53028305 | ENSG00000228205 |
| PWWP2B | 0.120044854 | 1.08676865 | 0.25635996 | 0.421525713 | 444.8415682 | 483.7239332 | ENSG00000171813 |
| PTK2 | 0.120029156 | 1.086756825 | 0.016138463 | 0.047042475 | 3239.020168 | 3519.263528 | ENSG00000169398 |
| ZYG11B | 0.119999871 | 1.086734766 | 0.234699543 | 0.395802857 | 458.4020435 | 498.1384968 | ENSG00000162378 |
| NME7 | 0.119997836 | 1.086733233 | 0.348024574 | 0.5206873 | 296.9904996 | 323.0000008 | ENSG00000143156 |
| UQCRB | 0.119936504 | 1.086687034 | 0.040962719 | 0.101595758 | 1706.714444 | 1854.547745 | ENSG00000156467 |
| NA | 0.119755181 | 1.086550464 | 0.895577048 | 0.94232447 | 6.014282385 | 6.522997791 | ENSG00000287686 |
| IGSF3 | 0.119677306 | 1.086491815 | 0.038768479 | 0.097216006 | 1804.875951 | 1960.533025 | ENSG00000143061 |
| FOPNL | 0.119661305 | 1.086479765 | 0.31236008 | 0.483373841 | 488.1662062 | 530.9667204 | ENSG00000133393 |
| TRDMT1 | 0.119598151 | 1.086432205 | 0.414847173 | 0.586974975 | 214.3117956 | 233.0896475 | ENSG00000107614 |
| FECH | 0.119584646 | 1.086422035 | 0.125505764 | 0.246350396 | 818.5091294 | 889.2398689 | ENSG00000066926 |
| AP5B1 | 0.119581674 | 1.086419797 | 0.157181456 | 0.292977487 | 747.0586484 | 811.5282919 | ENSG00000254470 |
| CHST12 | 0.119541487 | 1.086389535 | 0.263874506 | 0.429641531 | 430.4033554 | 467.6135378 | ENSG00000136213 |
| SMC6 | 0.119489976 | 1.086350746 | 0.055906335 | 0.130551884 | 1396.126261 | 1516.402583 | ENSG00000163029 |
| ZNF638 | 0.119417476 | 1.086296155 | 0.043186755 | 0.106115869 | 2353.328917 | 2556.573154 | ENSG00000075292 |
| AP1S1 | 0.119385698 | 1.086272227 | 0.097515136 | 0.203083719 | 1529.481509 | 1661.772033 | ENSG00000106367 |
| FAM110D | 0.119360837 | 1.086253509 | 0.769978164 | 0.864539245 | 31.29658569 | 34.11802592 | ENSG00000197245 |
| TEPSIN | 0.119337122 | 1.086235653 | 0.490981001 | 0.657452395 | 152.0561872 | 165.0270476 | ENSG00000167302 |
| AURKA | 0.119168857 | 1.08610897 | 0.035503918 | 0.090190424 | 2155.630532 | 2340.536172 | ENSG00000087586 |
| SSR4 | 0.119131411 | 1.086080779 | 0.078185615 | 0.170215406 | 1538.367553 | 1670.135964 | ENSG00000180879 |
| MAFK | 0.119003392 | 1.08598441 | 0.187141073 | 0.33417925 | 660.5741137 | 717.1534814 | ENSG00000198517 |
| TRERF1 | 0.118973025 | 1.085961551 | 0.115642492 | 0.231777592 | 891.835543 | 968.4265675 | ENSG00000124496 |
| YPEL3 | 0.118854914 | 1.085872649 | 0.359411943 | 0.532479518 | 286.5834566 | 311.4901369 | ENSG00000090238 |
| PLK5 | 0.118779593 | 1.085815959 | 0.829798893 | 0.903182315 | 14.21420097 | 15.5108079 | ENSG00000185988 |
| EPB41L4A | 0.118767131 | 1.08580658 | 0.725245113 | 0.835021231 | 38.26921533 | 41.56623931 | ENSG00000129595 |
| TAZ | 0.118745784 | 1.085790514 | 0.343370113 | 0.516165577 | 364.8847797 | 395.5650479 | ENSG00000102125 |
| RBBP9 | 0.118668322 | 1.085732216 | 0.507813128 | 0.672586547 | 198.9708293 | 216.2032685 | ENSG00000089050 |
| FAM234A | 0.118664047 | 1.085728999 | 0.023942594 | 0.06519997 | 2398.87147 | 2605.048042 | ENSG00000167930 |
| ANO6 | 0.118652151 | 1.085720046 | 0.091400511 | 0.193016667 | 1273.438199 | 1382.610319 | ENSG00000177119 |
| NA | 0.118578461 | 1.085664591 | 0.039429249 | 0.098578233 | 2642.494311 | 2869.667013 | ENSG00000215030 |
| ARHGAP28 | 0.118480217 | 1.085590662 | 0.467740386 | 0.637008457 | 228.4537085 | 248.48725 | ENSG00000088756 |
| RASSF1 | 0.11840903 | 1.085537098 | 0.344223948 | 0.516879611 | 289.766913 | 314.5101925 | ENSG00000068028 |
| CHAMP1 | 0.118393382 | 1.085525323 | 0.286456274 | 0.455604583 | 594.0389708 | 645.3706276 | ENSG00000198824 |
| TULP3 | 0.118375731 | 1.085512042 | 0.236494779 | 0.397925929 | 470.3623873 | 510.295334 | ENSG00000078246 |
| ZNF473 | 0.118355395 | 1.085496742 | 0.408422025 | 0.580707327 | 210.0600565 | 227.9645627 | ENSG00000142528 |
| FAM124A | 0.118075395 | 1.085286088 | 0.076275402 | 0.166847577 | 1494.006387 | 1621.740175 | ENSG00000150510 |
| NA | 0.117961837 | 1.085200665 | 0.920942946 | 0.95778265 | 2.982698788 | 3.228698041 | ENSG00000259531 |
| HJURP | 0.117911627 | 1.085162898 | 0.066558596 | 0.149892818 | 1452.241306 | 1575.5142 | ENSG00000123485 |
| RWDD2A | 0.117905431 | 1.085158237 | 0.592015663 | 0.740787017 | 113.9571048 | 123.6167525 | ENSG00000013392 |
| NA | 0.117842055 | 1.085110568 | 0.891580621 | 0.939954256 | 5.970979382 | 6.463847803 | ENSG00000226330 |
| MZF1 | 0.117780069 | 1.085063947 | 0.486271793 | 0.653750725 | 167.5221195 | 181.8359996 | ENSG00000099326 |
| RRAGA | 0.117539053 | 1.084882692 | 0.167092015 | 0.307104807 | 762.7263774 | 827.5443346 | ENSG00000155876 |
| MTERF2 | 0.117466882 | 1.084828422 | 0.667042211 | 0.795444847 | 56.69168917 | 61.60146625 | ENSG00000120832 |
| LMO4 | 0.117453251 | 1.084818172 | 0.055843451 | 0.130468211 | 1690.057868 | 1832.948689 | ENSG00000143013 |
| STX18-AS1 | 0.117370086 | 1.084755639 | 0.546570042 | 0.704645318 | 119.9539661 | 130.024667 | ENSG00000247708 |
| SLC16A4 | 0.117310862 | 1.08471111 | 0.881665915 | 0.935376247 | 6.934032559 | 7.549308779 | ENSG00000168679 |
| MIER3 | 0.11726688 | 1.084678041 | 0.320648584 | 0.492055671 | 380.951923 | 413.2136645 | ENSG00000155545 |
| PNPO | 0.117256893 | 1.084670533 | 0.674995341 | 0.800623163 | 57.46239433 | 62.28118861 | ENSG00000108439 |
| NA | 0.117029609 | 1.084499666 | 0.921437711 | 0.958038961 | 3.039956306 | 3.285697455 | ENSG00000279684 |
| WAS | 0.116972715 | 1.084456899 | 0.7793791 | 0.870480335 | 25.36978519 | 27.49446578 | ENSG00000015285 |
| SRFBP1 | 0.116938079 | 1.084430864 | 0.33341274 | 0.506001443 | 324.5049333 | 352.0843505 | ENSG00000151304 |
| CRCP | 0.116932525 | 1.084426689 | 0.30864705 | 0.479700644 | 398.6830763 | 432.7345276 | ENSG00000241258 |
| SH2D7 | 0.116877133 | 1.084385053 | 0.782860778 | 0.872832206 | 24.8862435 | 26.93466093 | ENSG00000183476 |
| APPL1 | 0.116842482 | 1.084359009 | 0.145765724 | 0.276170357 | 899.9149645 | 976.4055075 | ENSG00000157500 |
| NA | 0.116774273 | 1.084307743 | 0.85431098 | 0.91767432 | 10.01238578 | 10.87265049 | ENSG00000234782 |
| UNC5B | 0.116768302 | 1.084303254 | 0.101845963 | 0.209994706 | 1610.969833 | 1747.306031 | ENSG00000107731 |
| MKRN2 | 0.116548403 | 1.084137995 | 0.210178755 | 0.365736818 | 537.4350886 | 582.9175862 | ENSG00000075975 |
| NA | 0.116532015 | 1.08412568 | 0.888160594 | 0.938179871 | 6.055470207 | 6.541819917 | ENSG00000215895 |
| PNISR | 0.116509421 | 1.084108702 | 0.048096482 | 0.115776047 | 2018.951883 | 2188.601323 | ENSG00000132424 |
| CBWD2 | 0.116438899 | 1.08405571 | 0.24107922 | 0.403358803 | 533.8378758 | 578.7709483 | ENSG00000136682 |
| NA | 0.116376586 | 1.084008888 | 0.915883971 | 0.954940963 | 3.905239865 | 4.265761897 | ENSG00000228492 |
| DENND4C | 0.116361601 | 1.083997629 | 0.362696802 | 0.535987744 | 303.0582095 | 328.523579 | ENSG00000137145 |
| SYT16 | 0.116344702 | 1.083984932 | 0.744921054 | 0.848905485 | 34.39188449 | 37.36580655 | ENSG00000139973 |
| MAT2A | 0.116181699 | 1.083862464 | 0.050131555 | 0.119541141 | 4064.770414 | 4404.567159 | ENSG00000168906 |
| NA | 0.116168985 | 1.083852913 | 0.703859525 | 0.821872411 | 46.9325757 | 50.94526705 | ENSG00000239569 |
| NFYA | 0.116166804 | 1.083851274 | 0.142707219 | 0.271870922 | 830.3989 | 900.4480641 | ENSG00000001167 |
| TGFBR3 | 0.116067664 | 1.083776796 | 0.583071574 | 0.733248614 | 110.1825377 | 119.2665668 | ENSG00000069702 |
| ZNF706 | 0.116045862 | 1.083760418 | 0.134651307 | 0.259797001 | 817.978581 | 886.8500846 | ENSG00000120963 |
| AGGF1 | 0.115966195 | 1.083700573 | 0.500181835 | 0.666413964 | 168.0259102 | 181.8937925 | ENSG00000164252 |
| SPTAN1 | 0.115812337 | 1.083585007 | 0.014913969 | 0.04405227 | 6123.111085 | 6633.344751 | ENSG00000197694 |
| PIGBOS1 | 0.115763465 | 1.0835483 | 0.759260495 | 0.857014416 | 30.84817446 | 33.34365989 | ENSG00000225973 |
| NA | 0.115760016 | 1.083545711 | 0.872684042 | 0.929234854 | 8.708778561 | 9.391670723 | ENSG00000179131 |

| | | | | | | | |
|---|---|---|---|---|---|---|---|
| GNG11 | 0.115707856 | 1.083506536 | 0.698947442 | 0.818280507 | 49.33069672 | 53.31570827 | ENSG00000127920 |
| ZNF12 | 0.115631069 | 1.083448868 | 0.325723938 | 0.497509145 | 370.7977889 | 401.8043325 | ENSG00000164631 |
| LIPG | 0.115603411 | 1.083428097 | 0.411350113 | 0.583577271 | 236.5481974 | 256.3772593 | ENSG00000101670 |
| SIPA1 | 0.115588057 | 1.083416567 | 0.141627453 | 0.270294341 | 1403.711643 | 1521.112837 | ENSG00000213445 |
| NA | 0.115556049 | 1.083392531 | 0.892838944 | 0.940687531 | 6.047097499 | 6.528916475 | ENSG00000279833 |
| MRPS36 | 0.115546741 | 1.083385541 | 0.355148882 | 0.52823262 | 339.7965904 | 368.1649348 | ENSG00000134056 |
| NA | 0.115490398 | 1.083343231 | 0.824941395 | 0.900593514 | 16.61491273 | 18.01837122 | ENSG00000213178 |
| CRPPA | 0.115469875 | 1.08332782 | 0.864088017 | 0.923752947 | 9.691368058 | 10.49433629 | ENSG00000214960 |
| GJA1 | 0.115422141 | 1.083291977 | 0.003090505 | 0.011487271 | 12487.51927 | 13528.99311 | ENSG00000152661 |
| EI24 | 0.11525434 | 1.083165985 | 0.014276559 | 0.042507963 | 3257.314174 | 3528.314244 | ENSG00000149547 |
| CNEP1R1 | 0.115252329 | 1.083164475 | 0.452791106 | 0.622979333 | 207.3545968 | 224.6127 | ENSG00000205423 |
| RACGAP1 | 0.115244966 | 1.083158947 | 0.123207399 | 0.24318356 | 1149.705311 | 1244.897919 | ENSG00000161800 |
| NA | 0.115145018 | 1.083083911 | 0.811868717 | 0.892001334 | 20.23773194 | 21.80419049 | ENSG00000225234 |
| IFI6 | 0.115106841 | 1.08305525 | 0.656449824 | 0.7869973 | 65.33762841 | 70.90950664 | ENSG00000126709 |
| LEMD2 | 0.11508342 | 1.083037668 | 0.077270704 | 0.168699542 | 1407.101551 | 1524.356261 | ENSG00000161904 |
| AP1S2 | 0.115007124 | 1.082980393 | 0.104609323 | 0.214308047 | 4654.569099 | 5042.022688 | ENSG00000182287 |
| TACC1 | 0.114910374 | 1.082907769 | 0.327629351 | 0.499746803 | 401.5559276 | 434.5016131 | ENSG00000147526 |
| FBXO2 | 0.114803485 | 1.082827539 | 0.167523543 | 0.307569659 | 966.6640828 | 1046.601985 | ENSG00000116661 |
| NA | 0.114712222 | 1.082759044 | 0.473155003 | 0.641887001 | 179.282018 | 194.3646567 | ENSG00000277072 |
| KRAS | 0.114603985 | 1.082677813 | 0.206846148 | 0.361273792 | 931.7420006 | 1009.727892 | ENSG00000133703 |
| MAT2B | 0.114570894 | 1.08265298 | 0.137335947 | 0.263879259 | 1155.194676 | 1250.631194 | ENSG00000038274 |
| MRE11 | 0.1144256 | 1.082543952 | 0.217829209 | 0.375392467 | 618.7410516 | 670.0961524 | ENSG00000020922 |
| SELENOV | 0.114415722 | 1.08253654 | 0.641341641 | 0.775703377 | 80.29462551 | 86.84220378 | ENSG00000186838 |
| SMAD2 | 0.114412801 | 1.082534348 | 0.14085608 | 0.269221702 | 945.5964462 | 1023.489939 | ENSG00000175387 |
| NA | 0.114363738 | 1.082497533 | 0.895333198 | 0.942250941 | 6.270345606 | 6.781394477 | ENSG00000230897 |
| CD248 | 0.11430362 | 1.082452426 | 0.644156517 | 0.777623489 | 112.1938858 | 121.4848319 | ENSG00000174807 |
| BRCA1 | 0.114231202 | 1.082398092 | 0.232861007 | 0.393631413 | 755.9082994 | 818.2559467 | ENSG00000012048 |
| POLG2 | 0.114219761 | 1.082389509 | 0.25412875 | 0.41892656 | 481.7712069 | 521.3930502 | ENSG00000256525 |
| ACSL3 | 0.114173207 | 1.082354582 | 0.043479074 | 0.106738969 | 2222.263336 | 2406.113459 | ENSG00000123983 |
| DDX11 | 0.11412315 | 1.082317028 | 0.057382322 | 0.133238974 | 2375.828603 | 2570.798384 | ENSG00000013573 |
| RUVBL1 | 0.114081982 | 1.082286144 | 0.027912385 | 0.074128372 | 3784.382542 | 4095.513768 | ENSG00000175792 |
| FUBP1 | 0.113977284 | 1.082207604 | 0.011479879 | 0.035461704 | 4245.266836 | 4593.866626 | ENSG00000162613 |
| ATP6V1C2 | 0.113910991 | 1.082157877 | 0.802054616 | 0.885936266 | 22.09888379 | 23.87024895 | ENSG00000143882 |
| ARRDC1-AS1 | 0.113868047 | 1.082125666 | 0.254792033 | 0.419567948 | 644.3658978 | 696.8354143 | ENSG00000203993 |
| WARS2-AS1 | 0.113855666 | 1.082116379 | 0.727195886 | 0.836082516 | 41.17667188 | 44.61639078 | ENSG00000231365 |
| ARV1 | 0.113843922 | 1.08210757 | 0.526780095 | 0.688723471 | 139.4608292 | 151.0869984 | ENSG00000173409 |
| WSB1 | 0.113641011 | 1.081955386 | 0.082320813 | 0.177535026 | 1621.414632 | 1754.120095 | ENSG00000109046 |
| PCDHA11 | 0.113451934 | 1.081813596 | 0.425134002 | 0.596782098 | 217.6726064 | 235.261666 | ENSG00000249158 |
| GABRA1 | 0.113413063 | 1.081784449 | 0.887762053 | 0.937989799 | 6.379278836 | 6.886791011 | ENSG00000022355 |
| PEX11B | 0.11330128 | 1.081700633 | 0.571207128 | 0.723801675 | 109.9644711 | 118.989881 | ENSG00000131779 |
| LINC02021 | 0.113297101 | 1.0816975 | 0.835339895 | 0.906737673 | 18.47694046 | 20.03952809 | ENSG00000249846 |
| LOC283177 | 0.113286661 | 1.081689672 | 0.824402732 | 0.90032117 | 17.81815959 | 19.18531195 | ENSG00000255545 |
| NA | 0.113256557 | 1.081667101 | 0.92910481 | 0.962113616 | 3.701528078 | 4.013283894 | ENSG00000255893 |
| DSN1 | 0.113231604 | 1.081648392 | 0.246650013 | 0.410164964 | 505.767361 | 546.826831 | ENSG00000149636 |
| LRAT | 0.113102376 | 1.08155151 | 0.297353235 | 0.467577539 | 508.5725564 | 550.5727944 | ENSG00000121207 |
| NA | 0.113052038 | 1.081513773 | 0.807076011 | 0.888912551 | 21.73109644 | 23.44247156 | ENSG00000235587 |
| PLEKHF1 | 0.113008776 | 1.081481343 | 0.295011535 | 0.464956497 | 430.7028097 | 466.1276812 | ENSG00000166289 |
| LAMP2 | 0.112898179 | 1.081398439 | 0.166569816 | 0.306378559 | 779.9326138 | 843.8219297 | ENSG00000005893 |
| NA | 0.112846852 | 1.081359966 | 0.884113032 | 0.936545193 | 7.035268802 | 7.622979745 | ENSG00000279792 |
| PLEKHA8 | 0.112706588 | 1.081254838 | 0.149444889 | 0.281627327 | 971.3171612 | 1049.838734 | ENSG00000106086 |
| RAB43 | 0.112660554 | 1.081220338 | 0.533152042 | 0.693856577 | 162.547572 | 175.9859999 | ENSG00000172780 |
| NA | 0.112622477 | 1.081191801 | 0.794891749 | 0.880588753 | 23.32079109 | 25.21840776 | ENSG00000235238 |
| CRACR2B | 0.112585687 | 1.08116423 | 0.364184356 | 0.537897589 | 301.420469 | 325.8230029 | ENSG00000177685 |
| ALDH6A1 | 0.112563886 | 1.081147893 | 0.287159512 | 0.456459636 | 507.7785587 | 548.8175768 | ENSG00000119711 |
| PAWR | 0.112549648 | 1.081137223 | 0.155805541 | 0.29109699 | 2224.955945 | 2406.387399 | ENSG00000177425 |
| VMP1 | 0.112424105 | 1.081043146 | 0.123953384 | 0.244272626 | 1163.71257 | 1257.665872 | ENSG00000062716 |
| NA | 0.112332535 | 1.080974533 | 0.598186431 | 0.745019978 | 98.25520916 | 106.1025132 | ENSG00000213442 |
| ZNF93 | 0.112269535 | 1.08092733 | 0.22793235 | 0.387370537 | 640.9304836 | 692.9385218 | ENSG00000184635 |
| ZFYVE28 | 0.112254039 | 1.08091572 | 0.580430683 | 0.731378652 | 123.6172974 | 133.7093666 | ENSG00000159733 |
| TIMM17A | 0.112183506 | 1.080862875 | 0.08615203 | 0.184232216 | 1518.332019 | 1640.312966 | ENSG00000134375 |
| NA | 0.112017964 | 1.080738859 | 0.81569617 | 0.894616739 | 18.51466166 | 19.93010298 | ENSG00000270871 |
| NGB | 0.111985843 | 1.080714797 | 0.841934699 | 0.909545964 | 13.88760144 | 15.03946757 | ENSG00000165553 |
| ZBED3-AS1 | 0.111954593 | 1.080691388 | 0.809397435 | 0.890310719 | 21.71223584 | 23.39837558 | ENSG00000250802 |
| STN1 | 0.111854849 | 1.080616675 | 0.48842318 | 0.655318651 | 210.7193498 | 227.4144262 | ENSG00000107960 |
| YEATS2 | 0.111805466 | 1.080579686 | 0.128782667 | 0.251058627 | 2061.3634 | 2226.911254 | ENSG00000163872 |
| CCDC97 | 0.111762727 | 1.080547675 | 0.221705519 | 0.380103196 | 612.524609 | 661.8386429 | ENSG00000142039 |
| BCL9L | 0.111752529 | 1.080540037 | 0.104338342 | 0.213990108 | 1474.126719 | 1593.11047 | ENSG00000186174 |
| NUP37 | 0.111652766 | 1.08046532 | 0.294006625 | 0.463789501 | 513.4049041 | 555.0125476 | ENSG00000075188 |
| RABL2A | 0.111579603 | 1.080410528 | 0.624077563 | 0.763058842 | 92.63120462 | 100.3092525 | ENSG00000144134 |
| RAB5A | 0.111439461 | 1.080305583 | 0.123211659 | 0.24318356 | 1205.246506 | 1302.92866 | ENSG00000144566 |
| SDC2 | 0.111333829 | 1.080226487 | 0.100019854 | 0.207181682 | 1321.445189 | 1426.995774 | ENSG00000169439 |
| TTLL1 | 0.111261071 | 1.080172011 | 0.570990172 | 0.723801675 | 127.6079039 | 137.7820731 | ENSG00000100271 |
| PPP2R2C | 0.111255344 | 1.080167723 | 0.338079007 | 0.510699225 | 346.4060875 | 373.9587349 | ENSG00000074211 |
| RIBC1 | 0.111089919 | 1.080043874 | 0.839192598 | 0.908262141 | 15.12645432 | 16.26179018 | ENSG00000158423 |
| WDR70 | 0.111010155 | 1.079984162 | 0.281064762 | 0.449400772 | 736.2076473 | 794.4270612 | ENSG00000082068 |
| NA | 0.11100083 | 1.079977181 | 0.883256608 | 0.936254916 | 6.999662786 | 7.572432051 | ENSG00000255031 |
| ARHGDIG | 0.110943069 | 1.079933943 | 0.840260994 | 0.908946853 | 14.96305447 | 16.22172278 | ENSG00000242173 |

| | | | | | | | |
|---|---|---|---|---|---|---|---|
| INTS3 | 0.110841272 | 1.079857745 | 0.114292314 | 0.229687076 | 2103.830589 | 2271.734415 | ENSG00000143624 |
| THSD7B | 0.110754155 | 1.079792541 | 0.568019132 | 0.721278112 | 111.0924757 | 119.9196582 | ENSG00000144229 |
| TEFM | 0.11070194 | 1.07975346 | 0.579508247 | 0.730863859 | 115.8050659 | 124.9485123 | ENSG00000172171 |
| NA | 0.110669246 | 1.079728991 | 0.928067355 | 0.961460037 | 3.402161854 | 3.670463374 | ENSG00000103832 |
| MAP3K12 | 0.110660307 | 1.079722302 | 0.356547747 | 0.529904668 | 325.0417023 | 350.6835361 | ENSG00000139625 |
| CRYL1 | 0.110547539 | 1.079637908 | 0.779995033 | 0.870819155 | 30.94103799 | 33.40737149 | ENSG00000165475 |
| FAM171B | 0.110467704 | 1.079578166 | 0.371558528 | 0.545038336 | 306.772616 | 331.3707462 | ENSG00000144369 |
| ADD3 | 0.110386349 | 1.079517289 | 0.174778799 | 0.317751183 | 1470.764116 | 1587.778734 | ENSG00000148700 |
| NA | 0.110335856 | 1.079479507 | 0.920444153 | 0.95753639 | 3.714806871 | 4.019735616 | ENSG00000244009 |
| ADGRL3 | 0.110307521 | 1.079458306 | 0.450998393 | 0.621354807 | 252.1449515 | 272.0528558 | ENSG00000150471 |
| NA | 0.110221924 | 1.079394263 | 0.91944344 | 0.9570718 | 3.336531627 | 3.598942982 | ENSG00000254458 |
| MTRF1L | 0.110194959 | 1.079374088 | 0.368506389 | 0.542384802 | 297.4956928 | 321.3351194 | ENSG00000112031 |
| YDJC | 0.110130693 | 1.079326008 | 0.218124945 | 0.375700742 | 804.2874331 | 868.5816891 | ENSG00000161179 |
| SFT2D1 | 0.10985967 | 1.079123266 | 0.145154063 | 0.275343818 | 950.5172298 | 1025.604177 | ENSG00000198818 |
| WBP4 | 0.109728744 | 1.079025339 | 0.451327514 | 0.621541909 | 208.9822881 | 225.6871476 | ENSG00000120688 |
| EEF2KMT | 0.109627112 | 1.078949329 | 0.357462981 | 0.530760367 | 329.3005749 | 355.5004763 | ENSG00000118894 |
| NACC2 | 0.109595825 | 1.07892593 | 0.213012826 | 0.369136762 | 651.8281776 | 703.5406536 | ENSG00000148411 |
| TEP1 | 0.109513926 | 1.078864684 | 0.401266476 | 0.573764151 | 358.6925793 | 386.6239931 | ENSG00000129566 |
| ARFGEF2 | 0.109497359 | 1.078852294 | 0.181125686 | 0.326307024 | 837.5665422 | 902.9255742 | ENSG00000124198 |
| GALC | 0.109451406 | 1.078817931 | 0.422017961 | 0.593804067 | 240.3384466 | 259.477407 | ENSG00000054983 |
| EGLN3 | 0.109310825 | 1.078712813 | 0.311699989 | 0.48270085 | 430.4695492 | 464.6163145 | ENSG00000129521 |
| TET3 | 0.109264307 | 1.078678032 | 0.264124121 | 0.42986741 | 577.1241027 | 622.210209 | ENSG00000187605 |
| MCF2L | 0.109203957 | 1.07863291 | 0.270250461 | 0.436798968 | 544.9930086 | 587.9119885 | ENSG00000126217 |
| NDUFA8 | 0.109096022 | 1.078552215 | 0.118820228 | 0.236622383 | 1437.470036 | 1550.173748 | ENSG00000119421 |
| HSP90AA1 | 0.109043544 | 1.078512983 | 0.001310525 | 0.005440358 | 43224.63585 | 46619.07444 | ENSG00000080824 |
| HNRNPH1 | 0.108860815 | 1.078376389 | 0.006819032 | 0.022897468 | 10631.15645 | 11464.78252 | ENSG00000169045 |
| GPATCH8 | 0.108608316 | 1.07818767 | 0.092340723 | 0.194448737 | 1596.627822 | 1720.951814 | ENSG00000186566 |
| CCDC138 | 0.10845056 | 1.078069778 | 0.344247019 | 0.516879611 | 377.8122452 | 407.6718341 | ENSG00000163006 |
| FAM200A | 0.108435493 | 1.078058519 | 0.523641049 | 0.686069036 | 163.2909558 | 176.2435725 | ENSG00000221909 |
| COX7A2 | 0.108374889 | 1.078013234 | 0.127932625 | 0.249868206 | 2059.563901 | 2219.198718 | ENSG00000112695 |
| NA | 0.108358603 | 1.078001065 | 0.440357159 | 0.611233988 | 227.9841582 | 245.914085 | ENSG00000224578 |
| RERE | 0.108329572 | 1.077979373 | 0.039338287 | 0.098427354 | 2637.28451 | 2843.366603 | ENSG00000142599 |
| RPS18 | 0.10825066 | 1.077920412 | 0.058628763 | 0.135396094 | 24111.07669 | 25988.18606 | ENSG00000231500 |
| HDGF | 0.108133248 | 1.077832689 | 0.019425567 | 0.05485853 | 10189.55387 | 10982.09078 | ENSG00000143321 |
| LINC00862 | 0.108033654 | 1.077758286 | 0.830776669 | 0.903920627 | 18.05680167 | 19.48859811 | ENSG00000203721 |
| NA | 0.107916149 | 1.077670508 | 0.929133821 | 0.962113616 | 3.067189614 | 3.285697455 | ENSG00000228107 |
| HIPK1 | 0.107915887 | 1.077670312 | 0.17305987 | 0.315488371 | 1127.267098 | 1214.531775 | ENSG00000163349 |
| PTPN12 | 0.107901298 | 1.077659414 | 0.110395035 | 0.223184931 | 2194.743959 | 2366.387073 | ENSG00000127947 |
| TGS1 | 0.107898421 | 1.077657266 | 0.151537942 | 0.284868294 | 1013.903346 | 1093.14189 | ENSG00000137574 |
| NA | 0.107841685 | 1.077614886 | 0.673337075 | 0.79993506 | 65.54010089 | 70.61482278 | ENSG00000278041 |
| KLHL15 | 0.107838998 | 1.077612879 | 0.395715683 | 0.568283425 | 328.0904317 | 353.4749878 | ENSG00000174010 |
| SRSF5 | 0.107822447 | 1.077600516 | 0.018576176 | 0.052768814 | 4485.474358 | 4832.795464 | ENSG00000100650 |
| NA | 0.107814547 | 1.077594616 | 0.863967972 | 0.923726994 | 11.87632854 | 12.86235436 | ENSG00000230658 |
| PTGFRN | 0.10779504 | 1.077580045 | 0.163142747 | 0.301425795 | 980.8249685 | 1057.235055 | ENSG00000134247 |
| ZNF414 | 0.107587501 | 1.077425041 | 0.399216387 | 0.571735956 | 304.6416097 | 328.3738381 | ENSG00000133250 |
| AASDHPPT | 0.107515164 | 1.07737102 | 0.248683665 | 0.412498416 | 536.082121 | 577.4841021 | ENSG00000149313 |
| LAMTOR5 | 0.107475876 | 1.077341681 | 0.451062921 | 0.621387568 | 294.8848485 | 317.8289178 | ENSG00000134248 |
| NPIPB4 | 0.107438666 | 1.077313895 | 0.676016096 | 0.801291952 | 70.74181457 | 76.4142627 | ENSG00000185864 |
| ZNF711 | 0.1074233 | 1.07730242 | 0.321324748 | 0.492740517 | 407.495882 | 439.1758157 | ENSG00000147180 |
| PAPOLA | 0.107341213 | 1.077241126 | 0.020271678 | 0.056856782 | 4481.316573 | 4827.419391 | ENSG00000090060 |
| SNX24 | 0.107307027 | 1.0772156 | 0.491127587 | 0.65748385 | 206.3903796 | 222.380217 | ENSG00000064652 |
| MEGF6 | 0.107277532 | 1.077193576 | 0.192923124 | 0.342475383 | 986.071436 | 1062.223103 | ENSG00000162591 |
| ZNF721 | 0.107272434 | 1.07718977 | 0.231317838 | 0.391674924 | 646.6856364 | 697.0765026 | ENSG00000182903 |
| NINJ1 | 0.107174762 | 1.077116846 | 0.276080736 | 0.443084315 | 534.6850497 | 576.3827694 | ENSG00000131669 |
| NA | 0.107159036 | 1.077105105 | 0.923515923 | 0.95901091 | 3.943636784 | 4.26791247 | ENSG00000213609 |
| USP42 | 0.107126224 | 1.077080608 | 0.265407728 | 0.43144691 | 555.3623446 | 598.2902301 | ENSG00000106346 |
| XPNPEP3 | 0.107108151 | 1.077067115 | 0.461803255 | 0.631594938 | 223.2327708 | 240.4741493 | ENSG00000196236 |
| NA | 0.107096731 | 1.077058589 | 0.880488643 | 0.934399077 | 10.5399062 | 11.36308551 | ENSG00000260948 |
| NA | 0.107096107 | 1.077058123 | 0.881492867 | 0.93532122 | 10.0263403 | 10.78607608 | ENSG00000249042 |
| RAD52 | 0.107036407 | 1.077013555 | 0.50384303 | 0.669027801 | 184.7586924 | 198.7221303 | ENSG00000002016 |
| ITPKC | 0.107007518 | 1.076991989 | 0.350143161 | 0.522800066 | 362.278204 | 390.2689101 | ENSG00000086544 |
| KIAA1109 | 0.106997062 | 1.076984183 | 0.346982865 | 0.519733399 | 644.3205292 | 693.6262281 | ENSG00000138688 |
| SNX8 | 0.106942938 | 1.07694378 | 0.116466094 | 0.233186033 | 1231.78134 | 1326.208215 | ENSG00000106266 |
| CORO6 | 0.106668966 | 1.076739284 | 0.706273288 | 0.8232458 | 57.18447945 | 61.48720696 | ENSG00000167549 |
| NA | 0.106473164 | 1.07659316 | 0.919465641 | 0.9570718 | 4.349145335 | 4.711828377 | ENSG00000253392 |
| ESYT2 | 0.106381626 | 1.076524853 | 0.150029497 | 0.282390381 | 1014.726602 | 1092.323985 | ENSG00000117868 |
| ERI1 | 0.106307758 | 1.076469734 | 0.293163544 | 0.462882097 | 455.803365 | 490.8209843 | ENSG00000104626 |
| HIF1AN | 0.106210635 | 1.076397268 | 0.070716523 | 0.157346302 | 1683.565226 | 1811.875357 | ENSG00000166135 |
| QTRT1 | 0.106160984 | 1.076360224 | 0.089519139 | 0.189895943 | 1384.297941 | 1489.952705 | ENSG00000213339 |
| MCM3AP-AS1 | 0.106155141 | 1.076355865 | 0.664509979 | 0.793258787 | 69.76289186 | 75.07821383 | ENSG00000215424 |
| FITM2 | 0.106046394 | 1.076274735 | 0.596612863 | 0.74400866 | 110.0590864 | 118.3313918 | ENSG00000197296 |
| RAB3C | 0.105969026 | 1.076217019 | 0.461837425 | 0.631596893 | 239.158603 | 257.382325 | ENSG00000152932 |
| FAT1 | 0.105945582 | 1.07619953 | 0.18084996 | 0.325962495 | 2544.531042 | 2738.273218 | ENSG00000083857 |
| NET1 | 0.105902287 | 1.076167234 | 0.083793181 | 0.180086259 | 1873.691547 | 2016.725378 | ENSG00000173848 |
| TMEM106A | 0.105761693 | 1.076062364 | 0.841625481 | 0.909490034 | 14.70978215 | 15.85577899 | ENSG00000184988 |
| IGFBP5 | 0.105756818 | 1.076058728 | 0.826335475 | 0.901120226 | 23.78375732 | 25.54643472 | ENSG00000115461 |
| RMI1 | 0.105718389 | 1.076030065 | 0.393577812 | 0.566478368 | 383.8633729 | 413.1983622 | ENSG00000178966 |

| | | | | | | | |
|---|---|---|---|---|---|---|---|
| USP37 | 0.105455804 | 1.075834235 | 0.403857006 | 0.576115175 | 307.4075549 | 330.4353229 | ENSG00000135913 |
| COX5B | 0.105384671 | 1.075781191 | 0.099779987 | 0.206884752 | 2000.74451 | 2151.631945 | ENSG00000135940 |
| NUTM2A | 0.105350029 | 1.07575536 | 0.869143806 | 0.926984177 | 10.65163033 | 11.46364786 | ENSG00000184923 |
| PRKDC | 0.1052692 | 1.075695091 | 0.369346768 | 0.543069245 | 8158.872115 | 8776.171015 | ENSG00000253729 |
| TNFSF11 | 0.105157417 | 1.075611747 | 0.529891844 | 0.690872098 | 152.7065954 | 164.2795424 | ENSG00000120659 |
| ENO2 | 0.104959052 | 1.075463865 | 0.05302635 | 0.12495357 | 2460.240105 | 2645.73155 | ENSG00000111674 |
| NOP53 | 0.10492452 | 1.075438123 | 0.023197148 | 0.063501659 | 7973.664689 | 8575.777234 | ENSG00000105373 |
| ZNF507 | 0.104750183 | 1.075308174 | 0.202991341 | 0.35579738 | 708.5722551 | 761.8677083 | ENSG00000168813 |
| GTPBP1 | 0.10471129 | 1.075279185 | 0.050171232 | 0.119606183 | 2513.544162 | 2703.061107 | ENSG00000100226 |
| PLAGL1 | 0.104694806 | 1.0752669 | 0.831743726 | 0.904411714 | 17.47481464 | 18.7712315 | ENSG00000118495 |
| CC2D1B | 0.104647014 | 1.07523128 | 0.353457893 | 0.526407489 | 398.5367853 | 428.9576137 | ENSG00000154222 |
| WDR1 | 0.104510049 | 1.075129206 | 0.005480815 | 0.018985527 | 10911.06271 | 11730.52866 | ENSG00000071127 |
| PIAS2 | 0.104500063 | 1.075121764 | 0.233282494 | 0.394121995 | 738.4639973 | 794.3054651 | ENSG00000078043 |
| TANC1 | 0.104475736 | 1.075103635 | 0.378063991 | 0.551598279 | 330.3416612 | 355.1708439 | ENSG00000115183 |
| SH2B3 | 0.104450597 | 1.075084901 | 0.132129829 | 0.255889622 | 1072.535571 | 1153.29094 | ENSG00000111252 |
| SDC4 | 0.10443391 | 1.075072467 | 0.162943876 | 0.301116038 | 984.2655272 | 1058.23691 | ENSG00000124145 |
| NA | 0.104426729 | 1.075067116 | 0.885707618 | 0.937037652 | 8.297688203 | 8.971138553 | ENSG00000278238 |
| IL17REL | 0.104372531 | 1.075026729 | 0.932895961 | 0.964184087 | 3.048329015 | 3.256122461 | ENSG00000188263 |
| ENDOV | 0.104353603 | 1.075012625 | 0.57683579 | 0.728827562 | 141.0957788 | 151.3534764 | ENSG00000173818 |
| AVPR2 | 0.104352643 | 1.07501191 | 0.926860283 | 0.960672598 | 3.312089222 | 3.552696436 | ENSG00000126895 |
| NA | 0.104311013 | 1.07498089 | 0.742742838 | 0.847821391 | 43.46777469 | 46.75370916 | ENSG00000214654 |
| NA | 0.104215069 | 1.074909402 | 0.920240981 | 0.957436234 | 3.690364466 | 3.971338496 | ENSG00000258645 |
| PXDN | 0.104132614 | 1.07484797 | 0.011612922 | 0.035735344 | 6507.983947 | 6994.946827 | ENSG00000130508 |
| KCNK12 | 0.103984354 | 1.074737517 | 0.531548799 | 0.692377304 | 209.5581809 | 225.6392835 | ENSG00000184261 |
| NA | 0.103925546 | 1.074693709 | 0.840605022 | 0.909025813 | 16.41610702 | 17.58252457 | ENSG00000228393 |
| CCDC174 | 0.103904713 | 1.07467819 | 0.408329427 | 0.580691704 | 315.4782033 | 339.2327796 | ENSG00000154781 |
| NA | 0.103853008 | 1.074639676 | 0.930033967 | 0.962618589 | 3.072771419 | 3.277095161 | ENSG00000250938 |
| RBM38 | 0.103707119 | 1.07453101 | 0.169331605 | 0.310210666 | 1437.456783 | 1544.371698 | ENSG00000132819 |
| TMF1 | 0.103644116 | 1.074484087 | 0.454722638 | 0.62439002 | 227.9345977 | 245.0047475 | ENSG00000144747 |
| DNPH1 | 0.103612981 | 1.074460898 | 0.183599578 | 0.329594217 | 896.6840755 | 963.8537271 | ENSG00000112667 |
| TMEM243 | 0.103378913 | 1.074286588 | 0.333227915 | 0.505846687 | 433.7963936 | 465.986791 | ENSG00000135185 |
| LINC00638 | 0.103366007 | 1.074276978 | 0.848466977 | 0.913547795 | 14.61691862 | 15.71918993 | ENSG00000258701 |
| ARRDC4 | 0.103142097 | 1.07411026 | 0.580476839 | 0.731378652 | 135.3031929 | 145.1383385 | ENSG00000140450 |
| KRR1 | 0.103026675 | 1.07402433 | 0.315816145 | 0.486925292 | 486.972445 | 523.2028504 | ENSG00000111615 |
| NA | 0.102957764 | 1.07397303 | 0.386422953 | 0.55977211 | 12364.21871 | 13278.88617 | ENSG00000211459 |
| PIP5K1C | 0.102936452 | 1.073957165 | 0.126242638 | 0.247494587 | 1373.601919 | 1475.395083 | ENSG00000186111 |
| ALDH1B1 | 0.102917104 | 1.073942762 | 0.047306335 | 0.114301918 | 3116.148685 | 3346.036522 | ENSG00000137124 |
| BOD1L1 | 0.102829082 | 1.07387724 | 0.271504802 | 0.438476278 | 958.290011 | 1028.708038 | ENSG00000038219 |
| NA | 0.102541837 | 1.07366345 | 0.789794879 | 0.877393205 | 31.38127668 | 33.78353512 | ENSG00000263272 |
| TRAPPC8 | 0.102526019 | 1.073651678 | 0.173441368 | 0.31609423 | 1061.395567 | 1139.363249 | ENSG00000153339 |
| ENPEP | 0.102518113 | 1.073645794 | 0.851941934 | 0.91580219 | 14.0621649 | 15.07979543 | ENSG00000138792 |
| CSRNP2 | 0.102409113 | 1.07356468 | 0.167356656 | 0.307474024 | 960.4818309 | 1031.040624 | ENSG00000110925 |
| PTOV1-AS1 | 0.102249761 | 1.073446106 | 0.861658459 | 0.922498125 | 14.29378571 | 15.34249326 | ENSG00000268006 |
| TOPORS | 0.102242672 | 1.073440832 | 0.263719357 | 0.429498533 | 666.2068913 | 715.0680009 | ENSG00000197579 |
| NR1D2 | 0.102127434 | 1.073355092 | 0.37345539 | 0.54695686 | 343.5345123 | 368.8408953 | ENSG00000174738 |
| DNAJA2 | 0.102118604 | 1.073348523 | 0.050677597 | 0.120619558 | 2947.889665 | 3164.353424 | ENSG00000069345 |
| SNHG6 | 0.102106142 | 1.073339252 | 0.197516406 | 0.348589084 | 1036.62812 | 1112.037171 | ENSG00000245910 |
| CYFIP2 | 0.102095686 | 1.073331472 | 0.078799281 | 0.17137733 | 2024.933901 | 2173.507335 | ENSG00000055163 |
| MAN1A2 | 0.102006342 | 1.073265004 | 0.337575719 | 0.510333846 | 434.1453204 | 465.7665837 | ENSG00000198162 |
| LOC101927572 | 0.101943097 | 1.073217955 | 0.909484622 | 0.951250697 | 5.846176615 | 6.213520375 | ENSG00000267698 |
| IFIT5 | 0.101891886 | 1.07317986 | 0.723007192 | 0.833883281 | 54.16039268 | 58.26952225 | ENSG00000152778 |
| APOBEC3B-AS1 | 0.101856118 | 1.073153254 | 0.934739899 | 0.965262427 | 3.075562322 | 3.277095161 | ENSG00000249310 |
| MAGED2 | 0.101839674 | 1.073141022 | 0.045608525 | 0.110847594 | 3207.483389 | 3441.110743 | ENSG00000102316 |
| CYP27B1 | 0.101799382 | 1.073111052 | 0.747686225 | 0.850499144 | 45.67438667 | 49.09483585 | ENSG00000111012 |
| AKIRIN2 | 0.101793021 | 1.07310632 | 0.102110628 | 0.210416181 | 1751.079455 | 1878.625025 | ENSG00000135334 |
| CDC42 | 0.101786884 | 1.073101756 | 0.062409693 | 0.142293509 | 3647.655383 | 3913.913501 | ENSG00000070831 |
| GAS2L3 | 0.101721179 | 1.073052884 | 0.572659133 | 0.725196003 | 161.0688746 | 172.8304092 | ENSG00000139354 |
| DCAF12L2 | 0.101525111 | 1.072907062 | 0.750429907 | 0.852363598 | 41.61411966 | 44.72152685 | ENSG00000198354 |
| CEP72 | 0.101444001 | 1.072846743 | 0.299629903 | 0.470122852 | 474.3669115 | 508.9530939 | ENSG00000112877 |
| CTDSP1 | 0.101395111 | 1.072810388 | 0.317184672 | 0.48837793 | 456.755767 | 489.751916 | ENSG00000144579 |
| FBXO34 | 0.101394277 | 1.072809767 | 0.3824176 | 0.555848245 | 366.6940188 | 393.5919729 | ENSG00000178974 |
| POLE2 | 0.101392876 | 1.072808725 | 0.483583316 | 0.651361944 | 266.069235 | 285.1628539 | ENSG00000100479 |
| RBKS | 0.101167792 | 1.072641363 | 0.846188215 | 0.912075406 | 16.23596176 | 17.48599079 | ENSG00000171174 |
| RUNDC3A | 0.10104988 | 1.072553699 | 0.666621467 | 0.795139722 | 92.14594806 | 98.83981907 | ENSG00000108309 |
| UBASH3B | 0.100963253 | 1.072489299 | 0.42083335 | 0.592835997 | 285.9650245 | 306.6968288 | ENSG00000154127 |
| LINC01347 | 0.100907958 | 1.072448194 | 0.906028499 | 0.949270868 | 6.820393404 | 7.290379055 | ENSG00000214837 |
| VSIG10 | 0.100446708 | 1.072105372 | 0.149775847 | 0.28210563 | 1341.984255 | 1439.374467 | ENSG00000176834 |
| NOS1AP | 0.100333619 | 1.072021336 | 0.505228464 | 0.670313776 | 234.6070605 | 251.7498548 | ENSG00000198929 |
| WDR89 | 0.100263307 | 1.07196909 | 0.440098486 | 0.611138777 | 288.4613538 | 309.5644172 | ENSG00000140006 |
| LOC283922 | 0.100256835 | 1.071964281 | 0.749563549 | 0.851881137 | 49.83465061 | 53.22321518 | ENSG00000214331 |
| BTBD7 | 0.100174141 | 1.071902839 | 0.318012842 | 0.489294883 | 602.2240334 | 645.4571651 | ENSG00000011114 |
| BAZ1A | 0.100000851 | 1.071774095 | 0.257833432 | 0.42290964 | 617.7252835 | 662.3954673 | ENSG00000198604 |
| FAM219A | 0.099721591 | 1.071566654 | 0.351026829 | 0.523659992 | 440.0717205 | 471.9556116 | ENSG00000164970 |
| B3GAT3 | 0.099256819 | 1.071221498 | 0.318257747 | 0.489477014 | 567.4872396 | 607.8389232 | ENSG00000149541 |
| SECISBP2L | 0.099119749 | 1.071119727 | 0.335360653 | 0.507965009 | 460.5230133 | 493.100792 | ENSG00000138593 |
| NA | 0.099097698 | 1.071103355 | 0.829803891 | 0.903182315 | 19.63812362 | 21.0827832 | ENSG00000227939 |
| UBQLN1 | 0.099077453 | 1.071088324 | 0.040011547 | 0.099628173 | 3282.005129 | 3515.105144 | ENSG00000135018 |

| | | | | | | | |
|---|---|---|---|---|---|---|---|
| STAG3 | 0.099050676 | 1.071068445 | 0.745574294 | 0.849098159 | 52.33158038 | 56.04669553 | ENSG00000066923 |
| ERO1A | 0.099022963 | 1.071047871 | 0.307185346 | 0.478127522 | 526.5312619 | 563.9750654 | ENSG00000197930 |
| FKBP3 | 0.098935275 | 1.070982773 | 0.126930228 | 0.24843862 | 2447.523124 | 2621.586409 | ENSG00000100442 |
| COL4A1 | 0.098762202 | 1.070854301 | 0.051727408 | 0.122658033 | 2878.812086 | 3082.787853 | ENSG00000187498 |
| NA | 0.098332081 | 1.070535087 | 0.937025154 | 0.966565868 | 3.064398711 | 3.253971888 | ENSG00000274721 |
| WSCD1 | 0.098307476 | 1.070516829 | 0.087284694 | 0.186076809 | 1885.841397 | 2018.277957 | ENSG00000179314 |
| IVD | 0.098274008 | 1.070491995 | 0.108551513 | 0.220289 | 2209.355345 | 2364.917914 | ENSG00000128928 |
| CYP2A13 | 0.098252946 | 1.070476367 | 0.805494549 | 0.887728253 | 27.61287907 | 29.45674523 | ENSG00000197838 |
| CEP41 | 0.098078787 | 1.070347149 | 0.543529494 | 0.702010409 | 165.5683043 | 177.251334 | ENSG00000106477 |
| EIF4E3 | 0.098018971 | 1.070302772 | 0.506818544 | 0.671592013 | 231.4681833 | 247.8664293 | ENSG00000163412 |
| TUBA1C | 0.097928345 | 1.070235541 | 0.040593091 | 0.10086055 | 3589.464285 | 3840.705213 | ENSG00000167553 |
| FAM83H-AS1 | 0.097836196 | 1.070167184 | 0.751174307 | 0.852757227 | 49.33819355 | 52.95243598 | ENSG00000203499 |
| PRSS36 | 0.097817179 | 1.070153078 | 0.860458583 | 0.921699125 | 16.18795283 | 17.23378537 | ENSG00000178226 |
| NDRG4 | 0.097700048 | 1.070066197 | 0.272143324 | 0.439080963 | 581.0510693 | 621.6797003 | ENSG00000103034 |
| SNHG7 | 0.097665729 | 1.070040742 | 0.246275057 | 0.409700942 | 778.1521739 | 833.004177 | ENSG00000233016 |
| ABCG4 | 0.097652294 | 1.070030778 | 0.749675433 | 0.851958101 | 50.79383684 | 54.43451235 | ENSG00000172350 |
| CRBN | 0.09758301 | 1.069979392 | 0.411962446 | 0.584092867 | 314.1291027 | 336.0126838 | ENSG00000113851 |
| NA | 0.097463342 | 1.069890643 | 0.890017496 | 0.939292495 | 8.602636233 | 9.251040975 | ENSG00000236540 |
| ZNF639 | 0.097298499 | 1.069768404 | 0.374527447 | 0.547764139 | 423.7472377 | 453.475308 | ENSG00000121864 |
| TNK2 | 0.097275359 | 1.069751245 | 0.272214997 | 0.439080963 | 865.0148957 | 925.25679 | ENSG00000061938 |
| PPP1CB | 0.097249114 | 1.069731785 | 0.134900707 | 0.260132901 | 1665.333199 | 1781.998012 | ENSG00000213639 |
| EFCAB14 | 0.097213238 | 1.069705184 | 0.234173823 | 0.395088965 | 864.2446917 | 924.3664739 | ENSG00000159658 |
| NA | 0.097207751 | 1.069701116 | 0.795015827 | 0.880657498 | 35.47136773 | 38.00655811 | ENSG00000266066 |
| FAM149B1 | 0.097181673 | 1.06968178 | 0.485893815 | 0.653385722 | 281.5574945 | 300.8777548 | ENSG00000138286 |
| MECP2 | 0.097162876 | 1.069667844 | 0.251053647 | 0.415215698 | 782.4338477 | 836.4606122 | ENSG00000169057 |
| MIR1180 | 0.097142229 | 1.069652535 | 0.937484663 | 0.966854672 | 3.094422921 | 3.281396308 | ENSG00000221540 |
| PAK2 | 0.097108841 | 1.06962778 | 0.15178752 | 0.285198505 | 1417.345157 | 1516.867678 | ENSG00000180370 |
| APPL2 | 0.097105088 | 1.069624998 | 0.406399693 | 0.578761733 | 356.7318692 | 381.7478787 | ENSG00000136044 |
| SLC30A2 | 0.097063939 | 1.069594491 | 0.85399137 | 0.917433175 | 15.95613186 | 17.10284241 | ENSG00000158014 |
| DCAKD | 0.097024205 | 1.069565032 | 0.241399126 | 0.403705757 | 725.9796877 | 776.4807371 | ENSG00000172992 |
| NR2C2 | 0.096901558 | 1.06947411 | 0.275590575 | 0.442481713 | 730.0129826 | 780.4275831 | ENSG00000177463 |
| CAMK2G | 0.096831807 | 1.069422405 | 0.222780905 | 0.381201564 | 808.839525 | 864.7555114 | ENSG00000148660 |
| UBE2G2 | 0.09670811 | 1.069330716 | 0.071494904 | 0.158657333 | 2337.365701 | 2498.68663 | ENSG00000184787 |
| NA | 0.09667288 | 1.069304603 | 0.761020412 | 0.857878106 | 43.46219288 | 46.53672484 | ENSG00000263266 |
| SPATS2 | 0.096616178 | 1.069262578 | 0.250366528 | 0.4145413 | 749.8847301 | 801.2700409 | ENSG00000123352 |
| RNASEH2C | 0.096445424 | 1.06913603 | 0.293165059 | 0.462882097 | 615.5111747 | 657.8355973 | ENSG00000172922 |
| GTF2IRD2 | 0.096375446 | 1.069084173 | 0.791978669 | 0.878982065 | 31.39802209 | 33.60661819 | ENSG00000196275 |
| NA | 0.09617794 | 1.068937824 | 0.631235247 | 0.768662993 | 118.4983228 | 126.5418734 | ENSG00000239883 |
| PROB1 | 0.09613137 | 1.06890332 | 0.560351944 | 0.715498368 | 176.1454561 | 188.3046695 | ENSG00000228672 |
| ZNF317 | 0.096111368 | 1.068888501 | 0.260684401 | 0.426030394 | 896.7359002 | 958.166748 | ENSG00000130803 |
| CFDP1 | 0.096105808 | 1.068884381 | 0.32145068 | 0.492855275 | 722.9308562 | 772.3937517 | ENSG00000153774 |
| NA | 0.096062987 | 1.068852656 | 0.899474618 | 0.944674599 | 7.698955755 | 8.230648672 | ENSG00000257907 |
| ATR | 0.09606094 | 1.068851139 | 0.340359399 | 0.512946626 | 573.2764326 | 612.0963312 | ENSG00000175054 |
| GET1 | 0.095893727 | 1.068727262 | 0.673386668 | 0.799944714 | 90.06171137 | 96.35647562 | ENSG00000182093 |
| ETV6 | 0.095890745 | 1.068725054 | 0.392667272 | 0.565589811 | 423.9960043 | 452.9050349 | ENSG00000139083 |
| HAUS8 | 0.095816729 | 1.068670226 | 0.3902377 | 0.56343653 | 395.6741081 | 423.0253223 | ENSG00000131351 |
| C1orf115 | 0.095789279 | 1.068649892 | 0.39682704 | 0.569416775 | 382.9098054 | 408.9191701 | ENSG00000162817 |
| NA | 0.095773941 | 1.068638531 | 0.927520968 | 0.961048823 | 3.72317958 | 3.971338496 | ENSG00000254701 |
| NR2E3 | 0.09576479 | 1.068631752 | 0.903792298 | 0.947711976 | 6.733111675 | 7.155940565 | ENSG00000278570 |
| MICU1 | 0.095660687 | 1.068554644 | 0.259737251 | 0.425022589 | 821.9556703 | 878.3034641 | ENSG00000107745 |
| SLC30A7 | 0.095651187 | 1.068547608 | 0.499833818 | 0.666088297 | 216.0308357 | 230.9144244 | ENSG00000162695 |
| COPB2 | 0.095630991 | 1.068532649 | 0.057847887 | 0.134095601 | 2988.874064 | 3194.298194 | ENSG00000184432 |
| TMCO3 | 0.095620662 | 1.068525 | 0.253123394 | 0.417707733 | 985.8872349 | 1053.401069 | ENSG00000150403 |
| LRRCC1 | 0.095496394 | 1.068432965 | 0.530735162 | 0.691644398 | 249.0730559 | 266.0649283 | ENSG00000133739 |
| CHST2 | 0.095465682 | 1.06841022 | 0.253093186 | 0.417707733 | 847.9241894 | 906.614284 | ENSG00000175040 |
| NA | 0.095330054 | 1.068309784 | 0.93107358 | 0.963281155 | 3.750412887 | 3.998762916 | ENSG00000226007 |
| STRN | 0.09521137 | 1.068221902 | 0.261014945 | 0.426318784 | 726.5642144 | 776.1037373 | ENSG00000115808 |
| FASTK | 0.095110876 | 1.068147496 | 0.3213742 | 0.492777179 | 863.2762441 | 921.636105 | ENSG00000164896 |
| TAPT1 | 0.095098858 | 1.068138598 | 0.603250076 | 0.748716284 | 131.5828042 | 140.5625541 | ENSG00000169762 |
| CHST14 | 0.095075532 | 1.068121328 | 0.379479735 | 0.552827637 | 430.2390797 | 459.9450929 | ENSG00000169105 |
| EIF3A | 0.095048192 | 1.068101086 | 0.022307834 | 0.061494773 | 11673.35521 | 12467.67016 | ENSG00000107581 |
| PRDX1 | 0.095019306 | 1.068079701 | 0.069181462 | 0.154590477 | 14193.50677 | 15158.80806 | ENSG00000117450 |
| SYNE1 | 0.094933054 | 1.068015847 | 0.752425708 | 0.853444012 | 55.35143679 | 59.0121627 | ENSG00000131018 |
| NBR2 | 0.094847902 | 1.067952812 | 0.758554805 | 0.856649148 | 49.89382315 | 53.14524307 | ENSG00000198496 |
| OSBP2 | 0.094640098 | 1.067798997 | 0.603249901 | 0.748716284 | 150.4823621 | 160.9292972 | ENSG00000184792 |
| CHPF2 | 0.094638359 | 1.067797709 | 0.300578431 | 0.471189543 | 620.5516038 | 662.8523172 | ENSG00000033100 |
| GPBP1 | 0.094609272 | 1.067776181 | 0.110261692 | 0.223008838 | 1754.427486 | 1872.807805 | ENSG00000062194 |
| UGGT2 | 0.094546003 | 1.067729355 | 0.48487467 | 0.652736724 | 351.540307 | 375.5416341 | ENSG00000102595 |
| MYL12A | 0.094542117 | 1.067726479 | 0.117327169 | 0.234538813 | 1734.15057 | 1851.236186 | ENSG00000101608 |
| NA | 0.094495233 | 1.067691781 | 0.922374306 | 0.958444532 | 4.615696446 | 4.960005232 | ENSG00000272562 |
| YTHDF2 | 0.094445708 | 1.06765513 | 0.058721859 | 0.135583887 | 2717.873357 | 2901.967908 | ENSG00000198492 |
| LRP10 | 0.094377437 | 1.067604607 | 0.342652813 | 0.515408609 | 621.6979806 | 663.8275596 | ENSG00000197324 |
| H2BC4 | 0.094354832 | 1.06758788 | 0.876296826 | 0.931744589 | 12.06696169 | 12.8962305 | ENSG00000180596 |
| TGOLN2 | 0.094220772 | 1.06748868 | 0.148527352 | 0.280191781 | 1705.595637 | 1820.933494 | ENSG00000152291 |
| UBE3A | 0.094193634 | 1.067468601 | 0.157582864 | 0.293542973 | 1446.320816 | 1544.513278 | ENSG00000114062 |
| NA | 0.094156336 | 1.067441004 | 0.817510837 | 0.895649793 | 26.72574385 | 28.51432504 | ENSG00000248367 |
| JPH1 | 0.094068999 | 1.067376386 | 0.304594153 | 0.475321614 | 574.9875898 | 613.7744071 | ENSG00000104369 |

| | | | | | | | |
|---|---|---|---|---|---|---|---|
| IL17RB | 0.094041165 | 1.067355793 | 0.673100906 | 0.799703737 | 112.1050894 | 119.8403169 | ENSG00000056736 |
| GGA3 | 0.094018327 | 1.067338897 | 0.13588607 | 0.26155786 | 1448.084897 | 1545.043829 | ENSG00000125447 |
| DGKZ | 0.093959469 | 1.067295353 | 0.088796787 | 0.188766311 | 3291.558068 | 3513.422524 | ENSG00000149091 |
| MICU2 | 0.093918658 | 1.067265162 | 0.358561201 | 0.531627402 | 844.0867588 | 901.5165626 | ENSG00000165487 |
| LATS2 | 0.093881964 | 1.067238017 | 0.30011565 | 0.470731849 | 593.0755669 | 633.3103057 | ENSG00000150457 |
| GEMIN2 | 0.093840091 | 1.067207042 | 0.389355066 | 0.562629833 | 414.9185994 | 442.4612527 | ENSG00000092208 |
| EPHA1 | 0.093719104 | 1.067117548 | 0.050825729 | 0.120912461 | 4786.825481 | 5107.656149 | ENSG00000146904 |
| KMT5C | 0.093696346 | 1.067100714 | 0.299051078 | 0.469481961 | 702.5713778 | 750.2395139 | ENSG00000133247 |
| RBM20 | 0.093528048 | 1.066976238 | 0.886064478 | 0.937222823 | 13.27299945 | 14.09059566 | ENSG00000203867 |
| LRRC3-DT | 0.093516407 | 1.066967629 | 0.929691172 | 0.962375709 | 4.0386155 | 4.312008443 | ENSG00000229356 |
| RIOX1 | 0.093514109 | 1.06696593 | 0.353753405 | 0.526806927 | 594.5694198 | 634.7122352 | ENSG00000170468 |
| ZNF695 | 0.093356952 | 1.066849709 | 0.708037537 | 0.82397288 | 69.23046536 | 73.8160964 | ENSG00000197472 |
| TTC5 | 0.093291001 | 1.06680094 | 0.486913229 | 0.654258193 | 266.3028219 | 284.255667 | ENSG00000136319 |
| MTREX | 0.09326871 | 1.066784457 | 0.079706857 | 0.172922287 | 2745.199723 | 2928.535746 | ENSG00000039123 |
| ABLIM2 | 0.093247123 | 1.066768495 | 0.7321663 | 0.840066719 | 64.62179018 | 68.99509127 | ENSG00000163995 |
| NA | 0.093124175 | 1.066677587 | 0.834235643 | 0.906156773 | 26.38606569 | 28.10024459 | ENSG00000232611 |
| UGGT1 | 0.093097508 | 1.066657871 | 0.138049978 | 0.26488185 | 1926.68261 | 2056.022525 | ENSG00000136731 |
| PTGIS | 0.093040626 | 1.066615816 | 0.230555643 | 0.390830308 | 875.215737 | 933.1910825 | ENSG00000124212 |
| CLDN15 | 0.093024375 | 1.066603801 | 0.72139735 | 0.8327734 | 63.46040686 | 67.71872542 | ENSG00000106404 |
| METAP1 | 0.092976941 | 1.066568733 | 0.12491803 | 0.24549523 | 2339.996835 | 2495.524266 | ENSG00000164024 |
| PPP1CC | 0.092886815 | 1.066502106 | 0.032770137 | 0.084415079 | 7156.882463 | 7633.590228 | ENSG00000186298 |
| CYYR1 | 0.092730596 | 1.066386629 | 0.273914869 | 0.440744994 | 672.7438633 | 717.56615 | ENSG00000166265 |
| NRARP | 0.092678881 | 1.066348404 | 0.387046826 | 0.56046542 | 758.2906871 | 808.2950322 | ENSG00000198435 |
| ZBTB44 | 0.092658769 | 1.066333538 | 0.161301509 | 0.298653172 | 1526.263597 | 1627.696534 | ENSG00000196323 |
| HBS1L | 0.092565145 | 1.066264341 | 0.148314139 | 0.279899047 | 1384.583665 | 1476.408454 | ENSG00000112339 |
| NCKAP5L | 0.092551104 | 1.066253963 | 0.247551125 | 0.411079345 | 1055.380474 | 1125.817466 | ENSG00000167566 |
| SIGLEC10 | 0.092442853 | 1.066173961 | 0.809357572 | 0.890310719 | 31.97039712 | 33.95993112 | ENSG00000142512 |
| DNAJC18 | 0.092391458 | 1.06613598 | 0.404791789 | 0.576936712 | 364.3807889 | 388.5437941 | ENSG00000170464 |
| PARD3B | 0.092369016 | 1.066119396 | 0.721693362 | 0.833065261 | 68.96882033 | 73.58753517 | ENSG00000116117 |
| ZNF14 | 0.092284904 | 1.066057241 | 0.686721778 | 0.809455567 | 81.97977444 | 87.52648775 | ENSG00000105708 |
| ZNF566 | 0.092226192 | 1.066013857 | 0.622987057 | 0.762604966 | 168.3307091 | 179.6497511 | ENSG00000186017 |
| RBM47 | 0.092079061 | 1.065905147 | 0.12251728 | 0.242250832 | 1625.152256 | 1732.448799 | ENSG00000163694 |
| EHBP1 | 0.091878886 | 1.065757262 | 0.274849289 | 0.441594219 | 761.7774037 | 812.1160057 | ENSG00000115504 |
| TMCO1 | 0.091824883 | 1.065717369 | 0.216709561 | 0.373930602 | 937.1657969 | 998.9795329 | ENSG00000143183 |
| NA | 0.091716528 | 1.065637331 | 0.027354377 | 0.072944515 | 508827.2462 | 542224.4661 | N_multimapping |
| STAT1 | 0.09171358 | 1.065635153 | 0.169751178 | 0.310837207 | 1653.91911 | 1763.227532 | ENSG00000115415 |
| CDK5R1 | 0.091711802 | 1.06563384 | 0.462886902 | 0.632628496 | 337.186409 | 359.3481413 | ENSG00000176749 |
| PKNOX2 | 0.091687701 | 1.065616038 | 0.753583213 | 0.853959904 | 53.85332947 | 57.48278582 | ENSG00000165495 |
| FOXN3 | 0.091627919 | 1.065571882 | 0.211512773 | 0.367263087 | 2029.209292 | 2161.996197 | ENSG00000053254 |
| SLC39A11 | 0.091627486 | 1.065571562 | 0.434924698 | 0.605992526 | 406.7940352 | 433.2580452 | ENSG00000133195 |
| HMG20A | 0.091551695 | 1.065515585 | 0.18169771 | 0.327093181 | 1162.274424 | 1237.913001 | ENSG00000140382 |
| MRPS5 | 0.091502496 | 1.065479249 | 0.169552066 | 0.310561042 | 1170.861717 | 1247.511976 | ENSG00000144029 |
| NA | 0.091475583 | 1.065459373 | 0.925271512 | 0.960029077 | 4.310748416 | 4.608582417 | ENSG00000231434 |
| CCDC102A | 0.091437476 | 1.065431231 | 0.602782648 | 0.748328726 | 213.2337149 | 227.5523723 | ENSG00000135736 |
| TNFRSF12A | 0.091417112 | 1.065416192 | 0.260828053 | 0.426173785 | 956.9749052 | 1019.508175 | ENSG00000006327 |
| ATP2A2 | 0.091382842 | 1.065390884 | 0.016366919 | 0.047557498 | 8193.435345 | 8729.680958 | ENSG00000174437 |
| ESYT3 | 0.091317404 | 1.065342561 | 0.59124913 | 0.74009946 | 216.0348148 | 229.7950994 | ENSG00000158220 |
| ZBED5 | 0.09125301 | 1.065295011 | 0.38056996 | 0.553955715 | 654.8953544 | 698.4437319 | ENSG00000236287 |
| ST6GAL2 | 0.091234313 | 1.065281205 | 0.438692126 | 0.609712522 | 360.7609607 | 384.2638023 | ENSG00000144057 |
| PNPLA8 | 0.091162591 | 1.065228247 | 0.541400217 | 0.700278174 | 206.0737924 | 219.6037646 | ENSG00000135241 |
| MRPS30 | 0.09107286 | 1.065161996 | 0.191856544 | 0.340895698 | 1198.483007 | 1276.929471 | ENSG00000112996 |
| FUS | 0.091021637 | 1.065124177 | 0.030171897 | 0.07888941 | 18424.13514 | 19624.16263 | ENSG00000089280 |
| ZNF865 | 0.090958702 | 1.065077715 | 0.473917623 | 0.642289228 | 608.8320314 | 648.7811131 | ENSG00000261221 |
| ESRP2 | 0.090945438 | 1.065067922 | 0.249692251 | 0.413815567 | 1217.269289 | 1296.577024 | ENSG00000103067 |
| TXNDC11 | 0.090841818 | 1.064991428 | 0.536335059 | 0.696038255 | 230.1416852 | 245.0792362 | ENSG00000153066 |
| C2CD5 | 0.090803613 | 1.064963225 | 0.217212784 | 0.374664861 | 1165.006929 | 1240.486421 | ENSG00000111731 |
| GPX4 | 0.090797807 | 1.064958939 | 0.041042322 | 0.101740868 | 4072.664028 | 4337.862464 | ENSG00000167468 |
| SC5D | 0.090788378 | 1.06495198 | 0.140114978 | 0.268177204 | 1741.9715 | 1855.43388 | ENSG00000109929 |
| RPL18 | 0.090786928 | 1.064950909 | 0.032470139 | 0.083774435 | 14958.35005 | 15928.72679 | ENSG00000063177 |
| UFM1 | 0.090751186 | 1.064924526 | 0.360365362 | 0.533400461 | 504.2956111 | 537.4531463 | ENSG00000120686 |
| MFSD6 | 0.09072178 | 1.06490282 | 0.71837655 | 0.831109947 | 77.48928108 | 82.42612534 | ENSG00000151690 |
| NDUFA3 | 0.090647111 | 1.064847706 | 0.338916345 | 0.511503303 | 874.2016966 | 930.3297159 | ENSG00000170906 |
| NA | 0.090591489 | 1.064806652 | 0.557747523 | 0.713430492 | 267.0992457 | 284.2537769 | ENSG00000279865 |
| C14orf93 | 0.090508465 | 1.064745376 | 0.694219045 | 0.815154885 | 83.8842293 | 89.46617697 | ENSG00000100802 |
| NA | 0.090278847 | 1.064575926 | 0.783410779 | 0.873217619 | 42.40416099 | 45.2202916 | ENSG00000264204 |
| FAM110A | 0.090267896 | 1.064567845 | 0.49539708 | 0.661638401 | 253.4090858 | 269.7840735 | ENSG00000125898 |
| TM2D3 | 0.090234607 | 1.064543281 | 0.396698917 | 0.569356358 | 407.9656693 | 434.5131779 | ENSG00000184277 |
| NA | 0.090223904 | 1.064535384 | 0.907346408 | 0.949958569 | 7.056920304 | 7.473487239 | ENSG00000269934 |
| UPRT | 0.090202427 | 1.064519536 | 0.438365635 | 0.609346553 | 352.5624816 | 375.2611802 | ENSG00000094841 |
| MAT1A | 0.090111676 | 1.064452576 | 0.708562333 | 0.824320104 | 94.65646893 | 100.5714173 | ENSG00000151224 |
| MRPL2 | 0.090053007 | 1.06440929 | 0.860590153 | 0.921699125 | 20.36165883 | 21.80445096 | ENSG00000112651 |
| UTF1 | 0.090032736 | 1.064394334 | 0.493881929 | 0.660300526 | 486.9568381 | 518.49774 | ENSG00000171794 |
| ZNF224 | 0.08998087 | 1.064356069 | 0.743838736 | 0.848323562 | 60.36127802 | 64.10016682 | ENSG00000267680 |
| NA | 0.089903619 | 1.064299079 | 0.881700361 | 0.935376247 | 14.02376798 | 14.86685179 | ENSG00000277304 |
| NA | 0.089886061 | 1.064286125 | 0.859074464 | 0.920821105 | 16.90869715 | 18.0328922 | ENSG00000260628 |
| IREB2 | 0.08972397 | 1.064166557 | 0.158479493 | 0.29484308 | 1934.440526 | 2059.256342 | ENSG00000136381 |
| DEF6 | 0.089490695 | 1.063994501 | 0.528290531 | 0.689768008 | 222.3395782 | 236.7214005 | ENSG00000023892 |

| | | | | | | | |
|---|---|---|---|---|---|---|---|
| GOPC | 0.089471678 | 1.063980476 | 0.402265547 | 0.574546188 | 479.8429101 | 510.2681637 | ENSG00000047932 |
| OPTN | 0.089450408 | 1.06396479 | 0.531117729 | 0.69204945 | 252.6201205 | 268.4574325 | ENSG00000123240 |
| ANPEP | 0.08938808 | 1.063918825 | 0.724505185 | 0.834613013 | 72.56891201 | 77.21553222 | ENSG00000166825 |
| DGCR8 | 0.089384926 | 1.063916499 | 0.274332207 | 0.441089633 | 1097.4609 | 1167.035033 | ENSG00000128191 |
| CEP95 | 0.089342891 | 1.063885501 | 0.42759937 | 0.599042806 | 401.8819267 | 427.3338841 | ENSG00000258890 |
| LINC01775 | 0.08932163 | 1.063869823 | 0.926829577 | 0.960672598 | 4.658999449 | 4.978827358 | ENSG00000267201 |
| LIMS1 | 0.08929881 | 1.063852994 | 0.307376138 | 0.478385888 | 767.5145733 | 817.1454234 | ENSG00000169756 |
| ATP5F1D | 0.089253277 | 1.063819419 | 0.328342721 | 0.500557881 | 1565.428988 | 1665.800033 | ENSG00000099624 |
| SND1 | 0.089149357 | 1.063742792 | 0.020032396 | 0.056275658 | 9098.765206 | 9678.862629 | ENSG00000197157 |
| SLC51B | 0.088962847 | 1.063605282 | 0.932666196 | 0.964180982 | 3.72317958 | 3.943914076 | ENSG00000186198 |
| PCDHB4 | 0.088885057 | 1.063547934 | 0.896188824 | 0.942608288 | 9.544037909 | 10.20798214 | ENSG00000081818 |
| PARM1 | 0.088883023 | 1.063546435 | 0.584005062 | 0.733964094 | 208.9403365 | 222.3772424 | ENSG00000169116 |
| SALL3 | 0.088852043 | 1.063523596 | 0.260910885 | 0.426219991 | 816.6971009 | 868.9621233 | ENSG00000256463 |
| ENTPD1 | 0.088817297 | 1.063497983 | 0.565524135 | 0.719578265 | 182.5915165 | 194.1837112 | ENSG00000138185 |
| NA | 0.088649546 | 1.063374331 | 0.832923589 | 0.905233741 | 28.25559024 | 30.02918095 | ENSG00000228623 |
| NSUN4 | 0.088640144 | 1.0633674 | 0.404290903 | 0.576551456 | 466.9081863 | 496.2730115 | ENSG00000117481 |
| POLK | 0.088624943 | 1.063356197 | 0.637631909 | 0.773474008 | 132.6801089 | 140.8615512 | ENSG00000122008 |
| ATP5ME | 0.088560832 | 1.063308944 | 0.836047886 | 0.906924912 | 26.29300199 | 28.02012191 | ENSG00000169020 |
| CCDC106 | 0.088514098 | 1.0632745 | 0.425150401 | 0.596782098 | 452.4613396 | 480.9953507 | ENSG00000173581 |
| ZNF791 | 0.088486244 | 1.063253972 | 0.433033178 | 0.604046794 | 366.3852407 | 389.5104037 | ENSG00000173875 |
| NA | 0.088482993 | 1.063251576 | 0.863388736 | 0.923414764 | 16.0950893 | 17.10122488 | ENSG00000233426 |
| SLX4 | 0.088344941 | 1.063149838 | 0.457588012 | 0.627475856 | 373.9914962 | 397.9217558 | ENSG00000188827 |
| RNASEK | 0.088335934 | 1.0631432 | 0.859512185 | 0.921084878 | 19.26351516 | 20.43263584 | ENSG00000219200 |
| TUBG2 | 0.088224484 | 1.063061074 | 0.533974481 | 0.694458418 | 318.6792811 | 338.4086836 | ENSG00000037042 |
| SASS6 | 0.088208566 | 1.063049345 | 0.567955199 | 0.721244404 | 229.4464225 | 244.1695955 | ENSG00000156876 |
| HMGXB3 | 0.0881162 | 1.062981287 | 0.223177247 | 0.381710459 | 1066.447923 | 1133.298198 | ENSG00000113716 |
| OTULIN | 0.08797106 | 1.062874353 | 0.397652655 | 0.570131354 | 464.428892 | 494.0561035 | ENSG00000154124 |
| PGAP4 | 0.087888573 | 1.062813585 | 0.337712451 | 0.510343519 | 592.2474779 | 629.200779 | ENSG00000165152 |
| FLVCR1 | 0.087882229 | 1.062808911 | 0.179075792 | 0.323459841 | 1495.60657 | 1590.292568 | ENSG00000162769 |
| ANKLE1 | 0.087871818 | 1.062801241 | 0.424487497 | 0.596219927 | 404.8794559 | 429.9812532 | ENSG00000160117 |
| PI4KAP1 | 0.087789335 | 1.06274048 | 0.621994469 | 0.762199274 | 157.2859221 | 167.198763 | ENSG00000274602 |
| PAIP2 | 0.087769649 | 1.062725979 | 0.207744807 | 0.362416809 | 1916.263439 | 2036.103 | ENSG00000120727 |
| UBXN4 | 0.087767751 | 1.06272458 | 0.142751697 | 0.271928803 | 1825.371749 | 1939.841144 | ENSG00000144224 |
| CKS2 | 0.087675957 | 1.062656965 | 0.528572601 | 0.689851434 | 3488.726698 | 3707.035508 | ENSG00000123975 |
| LINGO2 | 0.087664887 | 1.06264881 | 0.757919654 | 0.856332907 | 60.32220538 | 64.17435871 | ENSG00000174482 |
| CINP | 0.087663365 | 1.06264769 | 0.371492303 | 0.545038336 | 517.9299772 | 550.3447967 | ENSG00000100865 |
| RDH10 | 0.087654476 | 1.062641142 | 0.604168619 | 0.749359253 | 149.0734884 | 158.4486374 | ENSG00000121039 |
| CKAP5 | 0.087591298 | 1.062594609 | 0.115393402 | 0.231337265 | 4635.644122 | 4925.763311 | ENSG00000175216 |
| PRRC2A | 0.087482692 | 1.062514619 | 0.011640337 | 0.035782032 | 14840.93878 | 15768.80942 | ENSG00000204469 |
| MEPCE | 0.087366558 | 1.062429093 | 0.211364608 | 0.367071899 | 1528.896956 | 1624.257728 | ENSG00000146834 |
| NKX3-1 | 0.087330603 | 1.062402615 | 0.800778084 | 0.885210591 | 35.48061632 | 37.63254506 | ENSG00000167034 |
| GLRB | 0.08715006 | 1.062269671 | 0.914756621 | 0.954332917 | 6.760344983 | 7.126365571 | ENSG00000109738 |
| NA | 0.087123616 | 1.0622502 | 0.805475546 | 0.887728253 | 39.96572803 | 42.54064553 | ENSG00000259485 |
| JADE2 | 0.087111911 | 1.062241582 | 0.309057134 | 0.480149889 | 881.4055482 | 936.5453684 | ENSG00000043143 |
| DRAXIN | 0.087097855 | 1.062231233 | 0.243529481 | 0.406491998 | 1041.368294 | 1106.235042 | ENSG00000162490 |
| CD151 | 0.087088358 | 1.06222424 | 0.22566855 | 0.3847227 | 1146.276843 | 1217.461574 | ENSG00000177697 |
| KCNIP2 | 0.086989754 | 1.062151643 | 0.778792765 | 0.870181942 | 44.18102217 | 46.87498374 | ENSG00000120049 |
| RYBP | 0.086925757 | 1.062104528 | 0.295583441 | 0.465667633 | 1087.500867 | 1155.760277 | ENSG00000163602 |
| FER | 0.086851465 | 1.062049836 | 0.590626274 | 0.739778362 | 190.5848202 | 202.6797292 | ENSG00000151422 |
| ZNF543 | 0.08682116 | 1.062027576 | 0.725801017 | 0.835365268 | 81.27075732 | 86.39854834 | ENSG00000178229 |
| NPR1 | 0.086814457 | 1.062022592 | 0.356850676 | 0.530314027 | 600.2887026 | 637.1650044 | ENSG00000169418 |
| CARMIL2 | 0.086805543 | 1.062016031 | 0.272221837 | 0.439080963 | 1036.086181 | 1100.165616 | ENSG00000159753 |
| GMIP | 0.086752408 | 1.061976917 | 0.39110167 | 0.564219438 | 564.7073767 | 599.7889596 | ENSG00000089639 |
| NA | 0.08673778 | 1.061966149 | 0.860950248 | 0.921931237 | 17.87175032 | 19.03392934 | ENSG00000231369 |
| SLC41A3 | 0.086731199 | 1.061961305 | 0.382726584 | 0.556045981 | 525.9566837 | 558.516568 | ENSG00000114544 |
| FZD4 | 0.086624879 | 1.061883046 | 0.535515361 | 0.695478386 | 228.7078936 | 242.8693192 | ENSG00000174804 |
| RPL27 | 0.086556149 | 1.061832459 | 0.108661642 | 0.220442921 | 12821.62453 | 13613.2797 | ENSG00000131469 |
| DPF2 | 0.086525383 | 1.061809815 | 0.12247242 | 0.242220162 | 2098.836782 | 2228.366935 | ENSG00000133884 |
| KMT2B | 0.086435013 | 1.061743306 | 0.158006511 | 0.294218495 | 2198.449477 | 2333.239141 | ENSG00000272333 |
| MTG2 | 0.08643057 | 1.061740036 | 0.3372565 | 0.510118057 | 600.9127164 | 638.2356413 | ENSG00000101181 |
| EP300 | 0.086412499 | 1.061726737 | 0.104680075 | 0.214394663 | 2931.956922 | 3112.257943 | ENSG00000100393 |
| AHCYL1 | 0.086346773 | 1.061678368 | 0.061413133 | 0.140536268 | 3650.159998 | 3875.012217 | ENSG00000168710 |
| TRIB3 | 0.086217223 | 1.061583037 | 0.470387011 | 0.639300819 | 353.446701 | 375.2351249 | ENSG00000101255 |
| ULK3 | 0.086105137 | 1.061500563 | 0.303781172 | 0.474475151 | 711.6109721 | 755.0513652 | ENSG00000140474 |
| ZNF554 | 0.085967046 | 1.061398964 | 0.599926284 | 0.746319594 | 161.9652216 | 171.8866746 | ENSG00000172006 |
| NA | 0.085946333 | 1.061383726 | 0.906115482 | 0.949270868 | 8.276036702 | 8.786152369 | ENSG00000267541 |
| TYW1B | 0.085900515 | 1.061350018 | 0.772064423 | 0.865873774 | 54.34456815 | 57.82990749 | ENSG00000277149 |
| SELENOT | 0.085784591 | 1.061264739 | 0.262310194 | 0.427817548 | 910.089887 | 966.1365158 | ENSG00000198843 |
| FAM177A1 | 0.085739645 | 1.061231677 | 0.444398684 | 0.615161812 | 450.8221596 | 478.8571962 | ENSG00000151327 |
| CROCCP3 | 0.085681039 | 1.061188568 | 0.684747243 | 0.807913843 | 97.63654008 | 103.6742791 | ENSG00000080947 |
| NA | 0.085663585 | 1.06117573 | 0.936676773 | 0.966331442 | 3.714806871 | 3.922941376 | ENSG00000251666 |
| TTC23L | 0.085653592 | 1.061168379 | 0.89212437 | 0.940288616 | 11.41164744 | 12.06969925 | ENSG00000205838 |
| CMPK1 | 0.085518617 | 1.061069104 | 0.392832858 | 0.565659377 | 670.0795177 | 711.4695968 | ENSG00000162368 |
| STARD5 | 0.085507928 | 1.061061242 | 0.870184405 | 0.927494743 | 16.92755774 | 17.97212467 | ENSG00000172345 |
| NEMF | 0.085471522 | 1.061034467 | 0.351205832 | 0.523781345 | 624.6762617 | 662.9135809 | ENSG00000165525 |
| RAB9B | 0.085435881 | 1.061008255 | 0.84937868 | 0.914208788 | 20.74639176 | 22.00208012 | ENSG00000123570 |
| CELF1 | 0.08535756 | 1.060950657 | 0.113822358 | 0.22896216 | 2661.156837 | 2822.935257 | ENSG00000149187 |

| | | | | | | | |
|---|---|---|---|---|---|---|---|
| PCDHA2 | 0.085316485 | 1.060920451 | 0.830899418 | 0.903961401 | 27.31821877 | 29.05933634 | ENSG00000204969 |
| COL22A1 | 0.085266047 | 1.06088336 | 0.680481935 | 0.804854507 | 112.4263073 | 119.2143894 | ENSG00000169436 |
| PARP9 | 0.085265588 | 1.060883023 | 0.693182298 | 0.814433261 | 110.1887952 | 116.7966477 | ENSG00000138496 |
| DUBR | 0.085264407 | 1.060882155 | 0.804918926 | 0.887347334 | 38.60506345 | 40.8365023 | ENSG00000243701 |
| NCK1 | 0.0850677 | 1.060737517 | 0.542531505 | 0.701331855 | 271.5831565 | 288.5136201 | ENSG00000158092 |
| DSC3 | 0.084958057 | 1.060656905 | 0.723478972 | 0.834137153 | 81.31752695 | 86.29851903 | ENSG00000134762 |
| TMEM161B | 0.084920876 | 1.06062957 | 0.586947696 | 0.736586521 | 192.9217657 | 204.5640365 | ENSG00000164180 |
| ERAP1 | 0.084750329 | 1.060504196 | 0.53230973 | 0.693135363 | 291.1476263 | 308.7217354 | ENSG00000164307 |
| KATNAL1 | 0.084633432 | 1.06041827 | 0.591432908 | 0.740202594 | 180.2789623 | 191.2499575 | ENSG00000102781 |
| UBE2C | 0.084614457 | 1.060404323 | 0.144036893 | 0.273593981 | 2333.152735 | 2473.609076 | ENSG00000175063 |
| RBM7 | 0.084597961 | 1.060392198 | 0.527265107 | 0.689118264 | 321.3860311 | 341.0512063 | ENSG00000076053 |
| IFI27L1 | 0.084550606 | 1.060357393 | 0.603806354 | 0.749117524 | 188.4957144 | 199.8153695 | ENSG00000165948 |
| TMEM259 | 0.084289455 | 1.060165468 | 0.211018541 | 0.366635924 | 2579.71153 | 2734.967168 | ENSG00000182087 |
| NA | 0.084136426 | 1.060053021 | 0.916019003 | 0.954978523 | 8.278827604 | 8.773781965 | ENSG00000213757 |
| THUMPD3 | 0.084064456 | 1.06000014 | 0.46362215 | 0.633319281 | 388.5409817 | 411.9077605 | ENSG00000134077 |
| LRP5L | 0.084022765 | 1.059969509 | 0.743126517 | 0.847927029 | 76.78643347 | 81.26861094 | ENSG00000100068 |
| EPC2 | 0.083990315 | 1.059945667 | 0.413303407 | 0.585592779 | 435.3132366 | 461.4591553 | ENSG00000135999 |
| NA | 0.083873987 | 1.059860205 | 0.775936707 | 0.868533369 | 51.02306708 | 54.06075977 | ENSG00000233966 |
| LAMB1 | 0.083825688 | 1.059824724 | 0.098310387 | 0.20438686 | 2483.241342 | 2632.045888 | ENSG00000091136 |
| CAPZA1 | 0.083817157 | 1.059818457 | 0.167741908 | 0.307830018 | 1976.41899 | 2094.266005 | ENSG00000116489 |
| GALK1 | 0.083763871 | 1.059779313 | 0.339721213 | 0.512242083 | 613.2552024 | 649.7468928 | ENSG00000108479 |
| ESCO2 | 0.083672477 | 1.059712179 | 0.527776833 | 0.689499096 | 259.5305496 | 275.2993099 | ENSG00000171320 |
| CD164L2 | 0.08364741 | 1.059693766 | 0.782281592 | 0.872362229 | 48.12119233 | 51.03184146 | ENSG00000174950 |
| ANKS6 | 0.083609624 | 1.059666012 | 0.166363587 | 0.306086786 | 1900.49446 | 2013.982366 | ENSG00000165138 |
| UBAP2 | 0.083532665 | 1.059609486 | 0.251773528 | 0.416085251 | 1529.897904 | 1620.643676 | ENSG00000137073 |
| LARP4B | 0.083499299 | 1.059584981 | 0.198038852 | 0.349338652 | 1246.115713 | 1320.28535 | ENSG00000107929 |
| DCAF4 | 0.083474144 | 1.059566506 | 0.531832558 | 0.692691725 | 335.6651355 | 355.8126558 | ENSG00000119599 |
| NA | 0.083337374 | 1.059466061 | 0.731363206 | 0.839305628 | 88.83964081 | 94.06588452 | ENSG00000259429 |
| NA | 0.083250304 | 1.059402122 | 0.913398687 | 0.953328685 | 7.416334949 | 7.814157185 | ENSG00000225774 |
| MED14 | 0.083158066 | 1.059334392 | 0.19277005 | 0.34223514 | 1579.581646 | 1673.244102 | ENSG00000180182 |
| NMNAT3 | 0.083054756 | 1.059258537 | 0.778555523 | 0.870118451 | 49.35793003 | 52.19501409 | ENSG00000163864 |
| CELSR2 | 0.082977733 | 1.059201987 | 0.23861153 | 0.400524808 | 1587.748222 | 1681.076088 | ENSG00000143126 |
| SCO2 | 0.082973017 | 1.059198524 | 0.947864783 | 0.972984731 | 3.390998242 | 3.567217414 | ENSG00000284194 |
| RABL2B | 0.082929914 | 1.059166879 | 0.545822135 | 0.704108719 | 282.6602689 | 298.9883713 | ENSG00000079974 |
| D2HGDH | 0.082906832 | 1.059149933 | 0.588917091 | 0.738474911 | 204.9471392 | 217.0437877 | ENSG00000180902 |
| PRDM15 | 0.082693212 | 1.058993117 | 0.508391708 | 0.673029402 | 352.6113905 | 373.1609851 | ENSG00000141956 |
| TFRC | 0.082657292 | 1.05896675 | 0.146361101 | 0.276917616 | 3200.400349 | 3389.369112 | ENSG00000072274 |
| RAP2B | 0.082650095 | 1.058961468 | 0.224094737 | 0.382635119 | 1273.907237 | 1349.292503 | ENSG00000181467 |
| ALG5 | 0.082632336 | 1.058948433 | 0.570324256 | 0.723300587 | 247.1973491 | 261.5771117 | ENSG00000120697 |
| EPS15 | 0.082574858 | 1.058906244 | 0.294952641 | 0.464901658 | 790.0934058 | 836.5345374 | ENSG00000085832 |
| PDPR | 0.082568041 | 1.05890124 | 0.27641855 | 0.443545848 | 944.4640991 | 1000.273333 | ENSG00000090857 |
| TAF12 | 0.082556489 | 1.058892762 | 0.523988616 | 0.686376342 | 344.1965966 | 364.4078728 | ENSG00000120656 |
| NA | 0.082517802 | 1.058864367 | 0.816546203 | 0.895101307 | 40.67339371 | 43.00716378 | ENSG00000242299 |
| PFN1 | 0.082467715 | 1.058827606 | 0.035624722 | 0.090461565 | 14195.72345 | 15031.01476 | ENSG00000108518 |
| NA | 0.082447969 | 1.058813115 | 0.884412176 | 0.936795019 | 13.68947146 | 14.51999059 | ENSG00000225093 |
| CLPB | 0.082436313 | 1.05880456 | 0.411321742 | 0.583577271 | 475.4984707 | 503.3311039 | ENSG00000162129 |
| PPP2R5C | 0.082212605 | 1.058640391 | 0.260304888 | 0.425662591 | 1215.590323 | 1287.67863 | ENSG00000078304 |
| NA | 0.082060611 | 1.058528865 | 0.865409268 | 0.924704231 | 17.93738055 | 19.01510721 | ENSG00000286827 |
| CASP3 | 0.08202154 | 1.058500199 | 0.366976491 | 0.54082074 | 1551.062263 | 1642.475763 | ENSG00000164305 |
| RECQL5 | 0.081984965 | 1.058473364 | 0.371246479 | 0.544907238 | 581.2568962 | 615.1193492 | ENSG00000108469 |
| DCK | 0.081984444 | 1.058472982 | 0.613893737 | 0.756749457 | 258.7331747 | 274.1928578 | ENSG00000156136 |
| OSBPL1A | 0.081732374 | 1.05828806 | 0.219371767 | 0.377410217 | 1193.840314 | 1263.722915 | ENSG00000141447 |
| SNAI1 | 0.08167965 | 1.058249385 | 0.629992523 | 0.767682759 | 165.5627225 | 175.3527845 | ENSG00000124216 |
| SYNDIG1L | 0.081667144 | 1.058240211 | 0.854763418 | 0.918046799 | 22.97553112 | 24.2023166 | ENSG00000183379 |
| EIF1B | 0.081534711 | 1.058143074 | 0.285922196 | 0.454942683 | 1124.866162 | 1189.866763 | ENSG00000114784 |
| DSE | 0.081517433 | 1.058130401 | 0.235374702 | 0.396594757 | 1088.034644 | 1151.238046 | ENSG00000111817 |
| NA | 0.081502723 | 1.058119613 | 0.884011857 | 0.936493029 | 13.2972417 | 14.0825264 | ENSG00000279798 |
| RPS9 | 0.081492044 | 1.058111781 | 0.105737846 | 0.216010674 | 30092.38548 | 31840.46839 | ENSG00000170889 |
| NA | 0.08146237 | 1.058090017 | 0.812406627 | 0.892246907 | 40.69033929 | 42.95687655 | ENSG00000260007 |
| AKTIP | 0.081419335 | 1.058058455 | 0.544212384 | 0.702704223 | 251.6199097 | 266.168459 | ENSG00000166971 |
| FOXRED2 | 0.08124083 | 1.057927549 | 0.435307729 | 0.606135187 | 695.7028506 | 736.3976707 | ENSG00000100350 |
| ACE | 0.081231926 | 1.05792102 | 0.770426979 | 0.864741191 | 55.84873284 | 59.05007953 | ENSG00000159640 |
| CD9 | 0.081213845 | 1.057907761 | 0.179480866 | 0.324130817 | 2727.106901 | 2884.769215 | ENSG00000010278 |
| CASP6 | 0.081158558 | 1.057867221 | 0.394248533 | 0.566892054 | 569.917737 | 602.6551852 | ENSG00000138794 |
| SPATA7 | 0.081134101 | 1.057849288 | 0.722458959 | 0.833649706 | 95.21545302 | 100.5429083 | ENSG00000042317 |
| FBN1 | 0.081062805 | 1.057797012 | 0.607848894 | 0.751797117 | 193.2958986 | 204.338171 | ENSG00000166147 |
| CGNL1 | 0.081016936 | 1.057763381 | 0.166715513 | 0.306529638 | 1982.921356 | 2096.727128 | ENSG00000128849 |
| PRR16 | 0.080989207 | 1.05774305 | 0.908581573 | 0.950821695 | 8.390551738 | 8.859823336 | ENSG00000184838 |
| RNH1 | 0.080941329 | 1.057707948 | 0.326162832 | 0.497942956 | 1260.722421 | 1333.929128 | ENSG00000023191 |
| OCRL | 0.080894633 | 1.057673714 | 0.271908907 | 0.438871713 | 1271.036748 | 1344.114986 | ENSG00000122126 |
| STON2 | 0.080879778 | 1.057662823 | 0.570137058 | 0.723168543 | 222.3623058 | 235.1207637 | ENSG00000140022 |
| SOX2-OT | 0.080871985 | 1.05765711 | 0.819052093 | 0.896676348 | 35.70597961 | 37.78795625 | ENSG00000242808 |
| METTL8 | 0.080828816 | 1.057625463 | 0.397719544 | 0.570147896 | 646.2059858 | 683.5510975 | ENSG00000123600 |
| ZNF682 | 0.080816303 | 1.05761629 | 0.631845343 | 0.768920547 | 178.2343988 | 188.2035862 | ENSG00000197124 |
| CDH3 | 0.080757074 | 1.057572871 | 0.129312528 | 0.251773111 | 3842.128306 | 4063.051935 | ENSG00000062038 |
| NA | 0.080703912 | 1.057533901 | 0.94733256 | 0.972798489 | 3.097213824 | 3.230848615 | ENSG00000250144 |
| ACSL1 | 0.080694374 | 1.057526909 | 0.521905588 | 0.684810277 | 315.4609824 | 333.9608432 | ENSG00000151726 |

| | | | | | | | |
|---|---|---|---|---|---|---|---|
| MICAL2 | 0.080678797 | 1.057515491 | 0.788113323 | 0.876396218 | 49.98601096 | 52.76450572 | ENSG00000133816 |
| TMOD4 | 0.080634115 | 1.057482739 | 0.911001307 | 0.951993281 | 8.09676732 | 8.533674367 | ENSG00000163157 |
| MAP4K5 | 0.080602372 | 1.057459472 | 0.398022842 | 0.570365372 | 504.3672235 | 533.4422855 | ENSG00000012983 |
| NA | 0.080590861 | 1.057451034 | 0.935207522 | 0.965468934 | 4.656208547 | 4.932580812 | ENSG00000250327 |
| MAPK15 | 0.08055572 | 1.057425278 | 0.62820188 | 0.766129882 | 198.6628391 | 209.8889437 | ENSG00000181085 |
| FCHSD1 | 0.080413942 | 1.057321367 | 0.414433572 | 0.586734041 | 659.9073094 | 696.9741292 | ENSG00000197948 |
| LOC389641 | 0.080214126 | 1.057174936 | 0.846242292 | 0.912075406 | 30.05968474 | 31.62687368 | ENSG00000246582 |
| PRKX | 0.080186317 | 1.057154558 | 0.181716823 | 0.327097064 | 2313.209914 | 2445.640488 | ENSG00000183943 |
| N6AMT1 | 0.080101415 | 1.057092347 | 0.569922265 | 0.722981099 | 228.2275062 | 241.5031318 | ENSG00000156239 |
| PALLD | 0.080077756 | 1.057075011 | 0.482894066 | 0.650797082 | 367.0296667 | 388.1273027 | ENSG00000129116 |
| SERP2 | 0.079923137 | 1.056961727 | 0.852846084 | 0.916417806 | 22.73812834 | 24.05308456 | ENSG00000151778 |
| SHCBP1 | 0.0798563 | 1.056912761 | 0.45349612 | 0.623637985 | 429.2414965 | 454.0587754 | ENSG00000171241 |
| DCBLD1 | 0.079838356 | 1.056899616 | 0.548942943 | 0.706401062 | 266.7433729 | 282.0755976 | ENSG00000164465 |
| GCK | 0.079750399 | 1.056835181 | 0.63969565 | 0.774742503 | 168.9266876 | 178.2873439 | ENSG00000106633 |
| DDIAS | 0.079733301 | 1.056822656 | 0.488864037 | 0.655460281 | 385.689231 | 407.979403 | ENSG00000165490 |
| NA | 0.07960548 | 1.056729028 | 0.605623808 | 0.750292827 | 183.381758 | 193.6706859 | ENSG00000159247 |
| TSPY26P | 0.079521229 | 1.056667318 | 0.546607897 | 0.704645318 | 262.9000965 | 277.6918204 | ENSG00000235217 |
| TDRD12 | 0.079520956 | 1.056667118 | 0.855295895 | 0.918419475 | 23.29076688 | 24.65053365 | ENSG00000173809 |
| ZNF330 | 0.079497711 | 1.056650093 | 0.348115433 | 0.520782847 | 1016.996491 | 1074.594913 | ENSG00000109445 |
| DELE1 | 0.079496464 | 1.05664918 | 0.31250312 | 0.483543379 | 918.2272033 | 969.6510712 | ENSG00000081791 |
| STRN3 | 0.07947163 | 1.056630991 | 0.391708299 | 0.564630112 | 640.1238586 | 676.1117711 | ENSG00000196792 |
| STRA6 | 0.079364687 | 1.056552669 | 0.557536977 | 0.713428546 | 267.2480025 | 282.112733 | ENSG00000137868 |
| SEC24B | 0.079062718 | 1.056331546 | 0.418567581 | 0.590679386 | 583.4968741 | 616.9116539 | ENSG00000138802 |
| WBP1L | 0.078954378 | 1.056252223 | 0.355186033 | 0.528247148 | 691.1631459 | 730.3887095 | ENSG00000166272 |
| NKTR | 0.078849 | 1.056175075 | 0.272279296 | 0.439080963 | 1123.616921 | 1186.602832 | ENSG00000114857 |
| GOLIM4 | 0.078809413 | 1.056146094 | 0.267163776 | 0.43346143 | 1272.097473 | 1343.743905 | ENSG00000173905 |
| NA | 0.078767329 | 1.056115286 | 0.908810646 | 0.950942162 | 8.980911477 | 9.488464962 | ENSG00000226453 |
| NPM3 | 0.078746632 | 1.056100135 | 0.215958307 | 0.373068121 | 2627.49308 | 2774.275741 | ENSG00000107833 |
| RAPH1 | 0.0786573 | 1.056034743 | 0.70932025 | 0.824903361 | 104.6929848 | 110.6011254 | ENSG00000173166 |
| NME6 | 0.078641331 | 1.056023055 | 0.553292418 | 0.709662844 | 260.9402992 | 275.7440498 | ENSG00000172113 |
| GNL3L | 0.078550866 | 1.055956838 | 0.28228096 | 0.450635717 | 1130.831422 | 1194.401148 | ENSG00000130119 |
| BET1L | 0.078520591 | 1.055934679 | 0.438168213 | 0.609159916 | 464.2857411 | 490.0382885 | ENSG00000177951 |
| NA | 0.078322353 | 1.055789595 | 0.886675011 | 0.937442292 | 14.98749687 | 15.87245054 | ENSG00000240914 |
| XBP1 | 0.0783042 | 1.05577631 | 0.125387851 | 0.246169045 | 2851.752209 | 3011.390249 | ENSG00000100219 |
| ANKRD10 | 0.078165686 | 1.05567495 | 0.233225465 | 0.39410936 | 1351.790612 | 1427.036635 | ENSG00000088448 |
| SNHG18 | 0.078008185 | 1.055559706 | 0.705051997 | 0.82256777 | 105.8757315 | 111.8277492 | ENSG00000250786 |
| RARS2 | 0.078006188 | 1.055558245 | 0.365228287 | 0.538931466 | 768.6330397 | 811.0252202 | ENSG00000146282 |
| TOP2A | 0.077985416 | 1.055543047 | 0.114780822 | 0.23040966 | 7878.426098 | 8316.47436 | ENSG00000131747 |
| PCDHB7 | 0.077966707 | 1.055529359 | 0.891423924 | 0.939943124 | 13.80119559 | 14.57053828 | ENSG00000113212 |
| ITGA2 | 0.077869058 | 1.055457918 | 0.675470062 | 0.800890779 | 143.9112108 | 151.7616382 | ENSG00000164171 |
| TMEM165 | 0.077842286 | 1.055438332 | 0.211729085 | 0.367551955 | 1527.417159 | 1612.427482 | ENSG00000134851 |
| ZDHHC20 | 0.077662058 | 1.05530649 | 0.286942374 | 0.456227259 | 1603.515789 | 1693.261493 | ENSG00000180776 |
| RAB7B | 0.077625772 | 1.055279948 | 0.9099793 | 0.951510145 | 9.666925654 | 10.22088559 | ENSG00000276600 |
| ZFP3 | 0.077578612 | 1.055245452 | 0.841389444 | 0.909364321 | 28.02309355 | 29.60193659 | ENSG00000180787 |
| CYP2C8 | 0.077425723 | 1.055133629 | 0.919945931 | 0.957261824 | 7.468010661 | 7.82275948 | ENSG00000138115 |
| FBXL20 | 0.077385556 | 1.055104253 | 0.649020494 | 0.781685316 | 154.1435782 | 162.4866742 | ENSG00000108306 |
| ATXN2 | 0.07730199 | 1.055043139 | 0.333189278 | 0.505846687 | 963.5558289 | 1016.7973 | ENSG00000204842 |
| HSPA12B | 0.077194444 | 1.054964494 | 0.585167319 | 0.735021666 | 246.3797991 | 259.5411612 | ENSG00000132622 |
| FAM168B | 0.077023531 | 1.054839522 | 0.158941763 | 0.295503619 | 2165.443763 | 2284.349034 | ENSG00000152102 |
| FLT1 | 0.077006494 | 1.054827065 | 0.185926381 | 0.332379374 | 2078.4442 | 2192.870339 | ENSG00000102755 |
| NANS | 0.076953761 | 1.05478851 | 0.343720451 | 0.516450754 | 851.9133551 | 898.1986676 | ENSG00000095380 |
| ANTXR2 | 0.076670702 | 1.054581579 | 0.65135304 | 0.783266107 | 163.4933531 | 172.2173852 | ENSG00000163297 |
| SLC35A5 | 0.076411385 | 1.054392041 | 0.436243381 | 0.607096456 | 511.3932437 | 539.5331927 | ENSG00000138459 |
| ELOVL5 | 0.076408663 | 1.054390051 | 0.106254294 | 0.216767301 | 3817.129265 | 4024.831621 | ENSG00000012660 |
| ZNF813 | 0.076342009 | 1.054341339 | 0.475462024 | 0.643857075 | 453.0122633 | 477.5905355 | ENSG00000198346 |
| NFIB | 0.076336121 | 1.054337035 | 0.814679831 | 0.894020628 | 41.41878059 | 43.67424316 | ENSG00000147862 |
| HAUS3 | 0.076230933 | 1.054260166 | 0.564521655 | 0.71882424 | 254.0294697 | 267.8551371 | ENSG00000214367 |
| COL8A2 | 0.076014549 | 1.054102053 | 0.681986559 | 0.805942337 | 137.4118351 | 144.842316 | ENSG00000171812 |
| HIC2 | 0.075996225 | 1.054088665 | 0.252906044 | 0.417599118 | 1615.991525 | 1703.159369 | ENSG00000169635 |
| PDE2A | 0.075936482 | 1.054045016 | 0.528651515 | 0.689907751 | 319.7090916 | 337.1796125 | ENSG00000186642 |
| FN3K | 0.075691249 | 1.053865861 | 0.574710701 | 0.726856628 | 263.9847981 | 278.3365579 | ENSG00000167363 |
| CDPF1 | 0.075659033 | 1.053842329 | 0.612676039 | 0.75594074 | 193.8937551 | 204.3247345 | ENSG00000205643 |
| LUZP2 | 0.075648367 | 1.053834537 | 0.843857575 | 0.910501509 | 30.15637831 | 31.86724174 | ENSG00000187398 |
| FOSL2 | 0.075590693 | 1.05379241 | 0.696742006 | 0.816923967 | 123.5523429 | 130.0182152 | ENSG00000075426 |
| ZFR | 0.075537975 | 1.053753903 | 0.192505787 | 0.341860364 | 2184.967825 | 2302.907643 | ENSG00000056097 |
| LRRC40 | 0.075522409 | 1.053742533 | 0.429001496 | 0.600353127 | 792.8779874 | 835.7017664 | ENSG00000066557 |
| RECK | 0.075463485 | 1.053699497 | 0.57875733 | 0.730346546 | 266.8949334 | 281.2947614 | ENSG00000122707 |
| ARMC9 | 0.075440208 | 1.053682496 | 0.367118736 | 0.540947951 | 768.1183736 | 809.5361285 | ENSG00000135931 |
| NA | 0.075409289 | 1.053659914 | 0.833759658 | 0.905786877 | 33.79873385 | 35.65332148 | ENSG00000287271 |
| CDC40 | 0.075119011 | 1.053447934 | 0.423342313 | 0.595023429 | 604.0339851 | 636.3252482 | ENSG00000168438 |
| PCMTD2 | 0.074997194 | 1.053358987 | 0.353219368 | 0.52613349 | 923.7022381 | 972.6990111 | ENSG00000203880 |
| TBC1D15 | 0.074990787 | 1.053354309 | 0.423890232 | 0.595619962 | 789.0565115 | 830.882403 | ENSG00000121749 |
| NA | 0.074973044 | 1.053341354 | 0.950859418 | 0.974707957 | 3.44825576 | 3.596792408 | ENSG00000287809 |
| USP41 | 0.074961255 | 1.053332747 | 0.907792742 | 0.950253936 | 10.24612178 | 10.82855451 | ENSG00000161133 |
| NA | 0.074953681 | 1.053327218 | 0.93918455 | 0.967771015 | 5.410643853 | 5.715016091 | ENSG00000215630 |
| CLTCL1 | 0.074918744 | 1.05330171 | 0.643627079 | 0.777277371 | 190.1676214 | 199.9976721 | ENSG00000070371 |
| DDX55 | 0.07488335 | 1.053275869 | 0.379067344 | 0.552436638 | 760.8308205 | 800.7242177 | ENSG00000111364 |

| | | | | | | | |
|---|---|---|---|---|---|---|---|
| GSAP | 0.074871621 | 1.053267306 | 0.89038604 | 0.939388982 | 14.35594931 | 15.13411123 | ENSG00000186088 |
| SERF2 | 0.074725331 | 1.05316051 | 0.445431731 | 0.61592886 | 632.5837089 | 665.4550815 | ENSG00000140264 |
| TICRR | 0.074694196 | 1.053137782 | 0.370835432 | 0.544593964 | 874.4776709 | 920.5989927 | ENSG00000140534 |
| TMEM171 | 0.074649214 | 1.053104946 | 0.85859447 | 0.920821105 | 28.45112948 | 29.83505943 | ENSG00000157111 |
| PRCC | 0.074632964 | 1.053093085 | 0.167778157 | 0.307867252 | 2118.320881 | 2231.240945 | ENSG00000143294 |
| NDUFS1 | 0.074556191 | 1.053037046 | 0.198488764 | 0.34990832 | 2051.38094 | 2160.193449 | ENSG00000023228 |
| AGFG1 | 0.07431438 | 1.05286056 | 0.357117332 | 0.530452601 | 1440.571442 | 1516.936799 | ENSG00000173744 |
| MBLAC2 | 0.074174741 | 1.052758659 | 0.746600351 | 0.849865308 | 92.57135635 | 97.56076961 | ENSG00000176055 |
| C11orf87 | 0.074173074 | 1.052757442 | 0.888185111 | 0.938179871 | 15.95055005 | 16.85304802 | ENSG00000185742 |
| ADAM9 | 0.074170853 | 1.052755821 | 0.448028985 | 0.618456326 | 511.1026505 | 538.1602931 | ENSG00000168615 |
| TMEM160 | 0.074060782 | 1.052675504 | 0.473735612 | 0.642182893 | 640.4349904 | 674.4035808 | ENSG00000130748 |
| DIPK1A | 0.073980609 | 1.052617007 | 0.711295004 | 0.826054537 | 117.7449266 | 124.1036799 | ENSG00000154511 |
| SLC2A1 | 0.073963841 | 1.052604773 | 0.160556639 | 0.297874411 | 4953.728891 | 5213.337625 | ENSG00000117394 |
| MEAF6 | 0.073925059 | 1.052576477 | 0.444729526 | 0.615347158 | 686.4673523 | 723.0435185 | ENSG00000163875 |
| TMEM219 | 0.073874289 | 1.052539437 | 0.425697122 | 0.597462581 | 680.6869918 | 716.4642658 | ENSG00000149932 |
| STARD9 | 0.073816662 | 1.052497395 | 0.613720634 | 0.756749457 | 220.8143625 | 232.2494193 | ENSG00000159433 |
| TTC12 | 0.073632511 | 1.052363058 | 0.737135369 | 0.843705033 | 100.0031974 | 105.2622729 | ENSG00000149292 |
| CRLS1 | 0.073627228 | 1.052359205 | 0.614862144 | 0.757375056 | 317.0577494 | 333.9700212 | ENSG00000088766 |
| FSIP1 | 0.073607099 | 1.052344522 | 0.910179005 | 0.951615803 | 9.694158961 | 10.21013272 | ENSG00000150667 |
| OSBPL3 | 0.073590494 | 1.05233241 | 0.517832554 | 0.681564084 | 399.3444479 | 420.3443618 | ENSG00000070882 |
| DESI1 | 0.073517735 | 1.052279339 | 0.372815774 | 0.546296171 | 759.3674065 | 799.323609 | ENSG00000100418 |
| CRIPT | 0.073251144 | 1.05208491 | 0.649400505 | 0.78189896 | 211.3707602 | 222.6910209 | ENSG00000119878 |
| NFIC | 0.073150314 | 1.052011383 | 0.752442888 | 0.853444012 | 84.1668501 | 88.6665371 | ENSG00000141905 |
| NDC80 | 0.073098983 | 1.051973952 | 0.414631063 | 0.586837715 | 628.0253084 | 660.9599101 | ENSG00000080986 |
| CAMKK2 | 0.073095404 | 1.051971342 | 0.291123615 | 0.460824292 | 1132.750007 | 1191.596205 | ENSG00000110931 |
| GLE1 | 0.073085268 | 1.051963952 | 0.546321295 | 0.704470255 | 429.0271705 | 450.8475667 | ENSG00000119392 |
| LYSMD4 | 0.073066044 | 1.051949935 | 0.572479224 | 0.725080212 | 394.6278403 | 414.8611483 | ENSG00000183060 |
| EIF2A | 0.073037385 | 1.051929038 | 0.355468494 | 0.528544992 | 1171.567255 | 1232.412521 | ENSG00000144895 |
| EVC | 0.072992882 | 1.051896589 | 0.5560744 | 0.712252257 | 322.0172521 | 338.680753 | ENSG00000072840 |
| RBMX | 0.072846789 | 1.051790076 | 0.056778512 | 0.132140702 | 11443.92368 | 12035.94155 | ENSG00000147274 |
| NA | 0.072825348 | 1.051774444 | 0.950282438 | 0.974271818 | 4.926226281 | 5.187209388 | ENSG00000234241 |
| LDB2 | 0.072786907 | 1.05174642 | 0.412074334 | 0.584208567 | 830.9548036 | 873.4835677 | ENSG00000169744 |
| U2SURP | 0.072731919 | 1.051706334 | 0.182615076 | 0.328223928 | 2417.752941 | 2542.362515 | ENSG00000163714 |
| CCDC82 | 0.072717553 | 1.051695861 | 0.580950977 | 0.731628608 | 329.2259715 | 346.6086485 | ENSG00000149231 |
| PSMG2 | 0.072663613 | 1.05165654 | 0.416781188 | 0.588807116 | 634.5339838 | 667.6090288 | ENSG00000128789 |
| MTIF2 | 0.072567573 | 1.051586534 | 0.400048521 | 0.572585667 | 835.7521983 | 879.3475496 | ENSG00000085760 |
| RPS4X | 0.072534612 | 1.051562509 | 0.056607374 | 0.131837842 | 29329.48733 | 30841.28655 | ENSG00000198034 |
| ZNF844 | 0.072526563 | 1.051556642 | 0.771635792 | 0.86554402 | 69.92908261 | 73.47919456 | ENSG00000223547 |
| C10orf88 | 0.072409272 | 1.051471154 | 0.685654982 | 0.808494514 | 146.7921978 | 154.4366984 | ENSG00000119965 |
| CENPB | 0.072348996 | 1.051427225 | 0.242933496 | 0.405751522 | 1807.771145 | 1901.210591 | ENSG00000125817 |
| SLC46A3 | 0.072253138 | 1.051357366 | 0.934859605 | 0.965304601 | 5.919303669 | 6.259766921 | ENSG00000139508 |
| TOPBP1 | 0.072159574 | 1.051289184 | 0.31803787 | 0.489294883 | 1143.957706 | 1202.758196 | ENSG00000163781 |
| VANGL1 | 0.072154041 | 1.051285152 | 0.472669677 | 0.64152607 | 485.5937077 | 510.412822 | ENSG00000173218 |
| IMMT | 0.072140989 | 1.051275641 | 0.156886052 | 0.292640576 | 2562.705926 | 2694.227326 | ENSG00000132305 |
| NA | 0.072075354 | 1.051227815 | 0.756154558 | 0.855290388 | 89.09136152 | 93.46441896 | ENSG00000269837 |
| SVIL | 0.072059724 | 1.051216426 | 0.43897977 | 0.610068349 | 641.8030623 | 674.4896348 | ENSG00000197321 |
| NA | 0.071887897 | 1.051091232 | 0.926697062 | 0.960672598 | 8.486206175 | 8.887247756 | ENSG00000231305 |
| EIF2AK1 | 0.07184074 | 1.051056876 | 0.167336764 | 0.307474024 | 2702.919827 | 2841.387857 | ENSG00000086232 |
| LINC01783 | 0.071756101 | 1.050995215 | 0.916560302 | 0.955284715 | 9.323580705 | 9.791490657 | ENSG00000233421 |
| CAPN2 | 0.071730347 | 1.050976453 | 0.275104942 | 0.441922677 | 1218.688249 | 1280.945687 | ENSG00000162909 |
| XRN2 | 0.071704975 | 1.05095797 | 0.118635185 | 0.23632707 | 3539.291621 | 3719.89954 | ENSG00000088930 |
| NA | 0.071693591 | 1.050949678 | 0.950195486 | 0.974234447 | 3.693155369 | 3.851420984 | ENSG00000234789 |
| NA | 0.07165928 | 1.050924684 | 0.93362551 | 0.964473049 | 6.61022393 | 6.995955654 | ENSG00000232150 |
| P2RX5 | 0.071599575 | 1.050881193 | 0.832542382 | 0.904974086 | 37.4346317 | 39.30470227 | ENSG00000083454 |
| SNTG2 | 0.071551163 | 1.050845929 | 0.624204103 | 0.763058842 | 201.8138438 | 212.0544679 | ENSG00000172554 |
| ACTR2 | 0.071520929 | 1.050823907 | 0.213901509 | 0.370310868 | 5595.772393 | 5880.364186 | ENSG00000138071 |
| NA | 0.07142371 | 1.050753098 | 0.948119174 | 0.972984731 | 3.744831081 | 3.899818104 | ENSG00000277767 |
| PSEN1 | 0.071420374 | 1.050750668 | 0.313179298 | 0.484330755 | 1008.677189 | 1059.965261 | ENSG00000080815 |
| WASF1 | 0.071394534 | 1.050731849 | 0.358197444 | 0.531369199 | 879.1482854 | 923.7526506 | ENSG00000112290 |
| UBE2K | 0.071385723 | 1.05072543 | 0.257601659 | 0.422747223 | 3388.118532 | 3561.079694 | ENSG00000078140 |
| CREBL2 | 0.071291004 | 1.050656449 | 0.488894148 | 0.655460281 | 542.8015535 | 570.8389815 | ENSG00000111269 |
| HNRNPU | 0.071204207 | 1.050593241 | 0.132037356 | 0.255787572 | 21036.81687 | 22101.97702 | ENSG00000153187 |
| PUS1 | 0.071145881 | 1.050550767 | 0.335816354 | 0.508575387 | 1963.853396 | 2063.127078 | ENSG00000177192 |
| SREBF2 | 0.071037344 | 1.050471735 | 0.101331712 | 0.209180317 | 8009.363264 | 8414.068687 | ENSG00000198911 |
| SCARB2 | 0.070929281 | 1.050393054 | 0.337724606 | 0.510343519 | 1025.922974 | 1078.06533 | ENSG00000138760 |
| MASTL | 0.070888512 | 1.050363371 | 0.321590215 | 0.492990849 | 1051.466083 | 1104.453647 | ENSG00000120539 |
| PLEKHO2 | 0.070785211 | 1.050288165 | 0.482064352 | 0.650071312 | 602.2298906 | 632.3278913 | ENSG00000241839 |
| SRGAP2C | 0.070783342 | 1.050286804 | 0.74110912 | 0.846561821 | 139.6048929 | 146.5026174 | ENSG00000171943 |
| PGM2L1 | 0.070699752 | 1.050225952 | 0.457445034 | 0.627327439 | 942.4101009 | 990.443889 | ENSG00000165434 |
| CNNM4 | 0.070680387 | 1.050211855 | 0.523129899 | 0.685763719 | 458.5924396 | 482.189407 | ENSG00000158158 |
| UBE3C | 0.070509712 | 1.05008762 | 0.219893733 | 0.37791083 | 1920.84194 | 2017.603596 | ENSG00000009335 |
| ZNF331 | 0.070465235 | 1.050055247 | 0.328339693 | 0.500557881 | 1159.651051 | 1217.138902 | ENSG00000130844 |
| TMEM187 | 0.070353984 | 1.049974277 | 0.698669439 | 0.818040309 | 128.6676507 | 135.1484131 | ENSG00000177854 |
| ATPAF1 | 0.070301926 | 1.049936391 | 0.281295088 | 0.449657237 | 1362.503093 | 1431.066881 | ENSG00000123472 |
| HIP1R | 0.070280012 | 1.049920442 | 0.423463122 | 0.595149868 | 636.5431784 | 668.0863183 | ENSG00000130787 |
| PRR11 | 0.070194048 | 1.049857884 | 0.517877059 | 0.681564084 | 630.1149267 | 661.5801851 | ENSG00000068489 |
| KIF18A | 0.07015572 | 1.049829993 | 0.541068165 | 0.700176533 | 347.2649133 | 364.4033233 | ENSG00000121621 |

| | | | | | | | |
|---|---|---|---|---|---|---|---|
| RIF1 | 0.070107267 | 1.049794735 | 0.396835087 | 0.569416775 | 1372.243559 | 1440.852374 | ENSG00000080345 |
| TIMP1 | 0.070096088 | 1.0497866 | 0.487414559 | 0.654511643 | 513.3759319 | 539.2613595 | ENSG00000102265 |
| FANCE | 0.070050095 | 1.049753134 | 0.482424081 | 0.650390869 | 495.914396 | 520.5011777 | ENSG00000112039 |
| LTN1 | 0.069944864 | 1.049676567 | 0.473692526 | 0.642182893 | 575.7431011 | 604.3644167 | ENSG00000198862 |
| PPP3CB | 0.069882666 | 1.049631314 | 0.444755867 | 0.615347158 | 582.9973877 | 611.7647966 | ENSG00000107758 |
| ZNF510 | 0.069609953 | 1.049432921 | 0.626414115 | 0.765013591 | 273.161449 | 286.5782504 | ENSG00000081386 |
| NA | 0.069582591 | 1.049413017 | 0.938282167 | 0.967367581 | 5.296128817 | 5.571975306 | ENSG00000260855 |
| CMTM4 | 0.069545521 | 1.049386053 | 0.415645143 | 0.58779696 | 1261.82688 | 1323.653811 | ENSG00000183723 |
| ZNF852 | 0.069375643 | 1.049262494 | 0.669667775 | 0.797187392 | 167.5345593 | 175.6125199 | ENSG00000178917 |
| EIF3G | 0.069365838 | 1.049255364 | 0.18669351 | 0.333627096 | 4277.978115 | 4488.083441 | ENSG00000130811 |
| ZXDC | 0.069197443 | 1.049132899 | 0.548203922 | 0.705732523 | 471.2574438 | 493.841585 | ENSG00000070476 |
| KAT5 | 0.069190087 | 1.04912755 | 0.639769932 | 0.774742503 | 213.719721 | 224.1042789 | ENSG00000172977 |
| CHD2 | 0.069132973 | 1.049086017 | 0.322113294 | 0.493596597 | 1349.975754 | 1416.42315 | ENSG00000173575 |
| VGLL4 | 0.069114051 | 1.049072258 | 0.457231427 | 0.627209803 | 601.6355376 | 630.6984424 | ENSG00000144560 |
| CDC37L1-DT | 0.069056079 | 1.049030103 | 0.942697175 | 0.969948861 | 5.021204997 | 5.268949611 | ENSG00000273061 |
| WWP1 | 0.068940508 | 1.048946072 | 0.330356694 | 0.502714442 | 1146.253351 | 1202.894991 | ENSG00000123124 |
| CYP26C1 | 0.068938863 | 1.048944876 | 0.951919737 | 0.975432043 | 4.104245726 | 4.282433449 | ENSG00000187553 |
| PARP8 | 0.068890282 | 1.048909554 | 0.547958822 | 0.705605342 | 401.7207542 | 421.11121 | ENSG00000151883 |
| NA | 0.068862165 | 1.048889112 | 0.893646002 | 0.941175276 | 16.25549808 | 17.09692373 | ENSG00000287016 |
| ZNF335 | 0.068836863 | 1.048870716 | 0.488581226 | 0.655439529 | 593.8058352 | 622.9308165 | ENSG00000198026 |
| FHOD1 | 0.068728733 | 1.048792107 | 0.386741927 | 0.56019211 | 1393.037394 | 1460.944226 | ENSG00000135723 |
| ZNF222 | 0.068711692 | 1.048779719 | 0.843583211 | 0.910409159 | 37.94887332 | 39.892192 | ENSG00000159885 |
| NA | 0.06865945 | 1.048741741 | 0.742864301 | 0.847821391 | 98.04204863 | 102.9362003 | ENSG00000272645 |
| BABAM2 | 0.068641014 | 1.04872834 | 0.685439465 | 0.808289793 | 230.7490658 | 242.145513 | ENSG00000158019 |
| PRDX5 | 0.068610548 | 1.048706194 | 0.176935558 | 0.320554455 | 4516.988476 | 4736.721588 | ENSG00000126432 |
| MRPS16 | 0.068594692 | 1.048694668 | 0.289470327 | 0.4588825 | 1690.238828 | 1772.341294 | ENSG00000182180 |
| UQCC2 | 0.068554195 | 1.048665231 | 0.504314905 | 0.669493983 | 558.2406145 | 584.8597291 | ENSG00000137288 |
| NA | 0.068454481 | 1.048592754 | 0.948092337 | 0.972984731 | 4.321912028 | 4.516089325 | ENSG00000242085 |
| DR1 | 0.068362037 | 1.048525565 | 0.352594142 | 0.525405035 | 1079.164656 | 1132.164539 | ENSG00000117505 |
| NA | 0.068188597 | 1.048399519 | 0.879289066 | 0.933536895 | 22.15335041 | 23.1399789 | ENSG00000253954 |
| ZSCAN29 | 0.068182164 | 1.048394844 | 0.396695478 | 0.569356358 | 810.9841366 | 850.0234133 | ENSG00000140265 |
| B2M | 0.068122071 | 1.048351176 | 0.44302186 | 0.613960827 | 831.5191565 | 871.8587174 | ENSG00000166710 |
| RPS17 | 0.068063262 | 1.048308442 | 0.705006133 | 0.822564008 | 183.6976694 | 192.3534713 | ENSG00000182774 |
| SPTY2D1 | 0.0679 | 1.048189818 | 0.566051109 | 0.72000935 | 368.3921711 | 386.4166651 | ENSG00000179119 |
| SNORD17 | 0.067898731 | 1.048188896 | 0.88784198 | 0.938001384 | 21.23639114 | 22.15723085 | ENSG00000212232 |
| ATP5F1E | 0.067879378 | 1.048174835 | 0.479685402 | 0.647830516 | 889.1957775 | 931.7206562 | ENSG00000124172 |
| RACK1 | 0.067704057 | 1.048047466 | 0.039872035 | 0.099377141 | 25217.83317 | 26428.75334 | ENSG00000204628 |
| CDK14 | 0.067610903 | 1.047979796 | 0.431204889 | 0.602387828 | 813.2318981 | 851.93884 | ENSG00000058091 |
| LOC652276 | 0.067457259 | 1.047868194 | 0.785467757 | 0.874499827 | 79.76287473 | 83.54761303 | ENSG00000215154 |
| METTL15 | 0.067388399 | 1.04781818 | 0.620235841 | 0.76139303 | 298.0754894 | 312.2010334 | ENSG00000169519 |
| WDR63 | 0.067301714 | 1.047755223 | 0.942838946 | 0.96999126 | 5.658334364 | 5.944370821 | ENSG00000162643 |
| TRAPPC11 | 0.06729032 | 1.047746949 | 0.424442871 | 0.596219927 | 720.7356591 | 755.0699027 | ENSG00000168538 |
| TIMM9 | 0.067264174 | 1.047727961 | 0.694392885 | 0.815309381 | 180.5788041 | 189.1763749 | ENSG00000100575 |
| DCTN4 | 0.066944654 | 1.047495941 | 0.381638022 | 0.555092684 | 1057.298217 | 1107.71862 | ENSG00000132912 |
| LRP2BP | 0.066875868 | 1.047445999 | 0.878853541 | 0.933405785 | 24.30828668 | 25.569558 | ENSG00000109771 |
| CASC2 | 0.066869914 | 1.047441676 | 0.939226009 | 0.967771015 | 5.314989415 | 5.576276454 | ENSG00000177640 |
| ZDHHC13 | 0.066861968 | 1.047435907 | 0.553929874 | 0.710113365 | 470.6396137 | 493.2193587 | ENSG00000177054 |
| VPS13A | 0.066818355 | 1.047404243 | 0.55491225 | 0.711094739 | 735.6968993 | 770.5674735 | ENSG00000197969 |
| WDR33 | 0.066690257 | 1.047311248 | 0.205911943 | 0.359912083 | 2247.596625 | 2354.015591 | ENSG00000136709 |
| MKNK2 | 0.066683173 | 1.047306106 | 0.161206717 | 0.29853497 | 3560.669448 | 3729.341002 | ENSG00000099875 |
| HELQ | 0.06663069 | 1.047268007 | 0.747574824 | 0.850422569 | 112.099032 | 117.250232 | ENSG00000163312 |
| HILPDA | 0.066629638 | 1.047267243 | 0.597336994 | 0.744394571 | 304.1602201 | 318.4728982 | ENSG00000135245 |
| PHLDB3 | 0.066458526 | 1.047143038 | 0.688699058 | 0.810814899 | 169.3527716 | 177.3828284 | ENSG00000176531 |
| TMED8 | 0.066431426 | 1.047123369 | 0.561907552 | 0.716583499 | 449.1125172 | 470.5576865 | ENSG00000100580 |
| YBX3 | 0.066427277 | 1.047120358 | 0.262644575 | 0.428218095 | 5399.347005 | 5654.724843 | ENSG00000060138 |
| MAP7D3 | 0.066253418 | 1.046994177 | 0.348639022 | 0.521366113 | 1048.324415 | 1097.343171 | ENSG00000129680 |
| CCDC43 | 0.066202307 | 1.046957085 | 0.622252675 | 0.762199274 | 285.3157677 | 298.9221755 | ENSG00000180329 |
| SNX13 | 0.066128517 | 1.046903537 | 0.55783498 | 0.713430492 | 411.1412797 | 430.4956171 | ENSG00000071189 |
| ZMYM5 | 0.065905932 | 1.046742029 | 0.717569868 | 0.83069133 | 167.750674 | 175.3506582 | ENSG00000132950 |
| TUT4 | 0.065883007 | 1.046725396 | 0.415513266 | 0.587747169 | 921.7160847 | 965.3742597 | ENSG00000134744 |
| IFITM3 | 0.065725941 | 1.046611445 | 0.177303126 | 0.32103567 | 3207.620935 | 3356.765368 | ENSG00000142089 |
| SYPL1 | 0.065597883 | 1.046518549 | 0.377681102 | 0.551123006 | 915.0984889 | 957.8765647 | ENSG00000008282 |
| CBLB | 0.065384718 | 1.046363932 | 0.560511874 | 0.715655209 | 425.2100514 | 444.5915679 | ENSG00000114423 |
| RDH13 | 0.065354479 | 1.046342001 | 0.655157992 | 0.786083644 | 220.4009568 | 230.5804972 | ENSG00000160439 |
| GSTCD | 0.065215503 | 1.046241211 | 0.566198841 | 0.72000935 | 354.4972361 | 371.1169533 | ENSG00000138780 |
| PTPRJ | 0.065174273 | 1.046211311 | 0.500323513 | 0.666503001 | 569.119323 | 595.2628867 | ENSG00000149177 |
| SENP5 | 0.065118466 | 1.046170842 | 0.54127931 | 0.700276251 | 536.9174172 | 561.7302062 | ENSG00000119231 |
| LOC100506282 | 0.065110687 | 1.046165201 | 0.885772941 | 0.937037652 | 22.28026836 | 23.35454008 | ENSG00000247137 |
| RIOK2 | 0.065091814 | 1.046151516 | 0.520754711 | 0.683672188 | 463.8362538 | 485.2998171 | ENSG00000058729 |
| FGFR3 | 0.065087893 | 1.046148672 | 0.440272414 | 0.611204314 | 704.3715912 | 736.6248628 | ENSG00000068078 |
| NA | 0.06490741 | 1.046017806 | 0.818641011 | 0.896378916 | 61.72792472 | 64.40775412 | ENSG00000270872 |
| MTOR | 0.0648761 | 1.045995105 | 0.196986903 | 0.347896581 | 3888.629731 | 4066.760741 | ENSG00000198793 |
| VWA2 | 0.064856164 | 1.045980651 | 0.765203468 | 0.861065676 | 95.20301321 | 99.64432365 | ENSG00000165816 |
| C3orf67 | 0.064838399 | 1.045967771 | 0.750930989 | 0.852639177 | 110.0430167 | 114.9704558 | ENSG00000163689 |
| TKT | 0.064816367 | 1.045951798 | 0.060181248 | 0.138242016 | 23770.9526 | 24863.19355 | ENSG00000163931 |
| SLC22A18 | 0.064792459 | 1.045934465 | 0.786280612 | 0.875152278 | 77.56096868 | 81.11130964 | ENSG00000110628 |
| CTSL | 0.064776484 | 1.045922883 | 0.499445269 | 0.665800476 | 703.0806381 | 735.6745849 | ENSG00000135047 |

| | | | | | | | |
|---|---|---|---|---|---|---|---|
| RHPN2 | 0.0647386 | 1.045895419 | 0.401289466 | 0.573764151 | 843.621375 | 882.5055387 | ENSG00000131941 |
| NA | 0.064667228 | 1.045843678 | 0.939848419 | 0.968257188 | 7.274586605 | 7.654705312 | ENSG00000223343 |
| GNB1 | 0.064545526 | 1.045755457 | 0.10142585 | 0.209337105 | 10616.97956 | 11101.68673 | ENSG00000078369 |
| RPS12 | 0.064519389 | 1.045736511 | 0.147432683 | 0.278575312 | 14233.3073 | 14883.40271 | ENSG00000112306 |
| C20orf96 | 0.064375958 | 1.045632551 | 0.519565607 | 0.682988384 | 561.8419696 | 586.9728397 | ENSG00000196476 |
| QDPR | 0.064335408 | 1.045603161 | 0.421792831 | 0.593754051 | 893.8019372 | 934.5946537 | ENSG00000151552 |
| TRIM47 | 0.064333761 | 1.045601967 | 0.800522299 | 0.88513072 | 79.60302953 | 83.14054787 | ENSG00000132481 |
| SEPTIN7 | 0.064195597 | 1.045501837 | 0.302240333 | 0.472989952 | 2702.680275 | 2826.805396 | ENSG00000122545 |
| HACD3 | 0.064076433 | 1.045415484 | 0.153010858 | 0.287189381 | 4368.802561 | 4568.153923 | ENSG00000074696 |
| SUN2 | 0.064052493 | 1.045398137 | 0.632993341 | 0.769822396 | 251.4369735 | 262.83975 | ENSG00000100242 |
| TMEM50B | 0.064005986 | 1.045364437 | 0.694942933 | 0.815746479 | 208.6750616 | 217.8799873 | ENSG00000142188 |
| NA | 0.063971176 | 1.045339215 | 0.951244021 | 0.974894991 | 4.074221516 | 4.223816498 | ENSG00000277491 |
| ARF6 | 0.063966342 | 1.045335712 | 0.27285562 | 0.43966403 | 2391.934585 | 2501.167296 | ENSG00000165527 |
| ISCU | 0.063918395 | 1.045300972 | 0.33346101 | 0.506001443 | 1415.030799 | 1478.551976 | ENSG00000136003 |
| LLGL2 | 0.063843078 | 1.045246402 | 0.268522997 | 0.434752592 | 1794.252081 | 1875.558978 | ENSG00000073350 |
| NA | 0.063723817 | 1.045160001 | 0.8160422 | 0.894698714 | 59.29910378 | 62.00773791 | ENSG00000255121 |
| NA | 0.063626284 | 1.045089345 | 0.946416654 | 0.972272003 | 4.694605466 | 4.88848484 | ENSG00000287817 |
| LPIN3 | 0.063561663 | 1.045042535 | 0.749999654 | 0.8520254 | 133.0232537 | 138.8049191 | ENSG00000132793 |
| CYBRD1 | 0.063544755 | 1.045030287 | 0.728727132 | 0.837326529 | 129.2356832 | 135.1338921 | ENSG00000071967 |
| CUL3 | 0.063528096 | 1.04501822 | 0.315756459 | 0.48687218 | 1654.221679 | 1728.759241 | ENSG00000036257 |
| SFPQ | 0.06337263 | 1.044905614 | 0.087092942 | 0.18577067 | 10725.98509 | 11207.69114 | ENSG00000116560 |
| NA | 0.063276289 | 1.044835839 | 0.878096721 | 0.933092643 | 27.6043062 | 28.87650073 | ENSG00000228175 |
| SRPRA | 0.0632705 | 1.044831646 | 0.364261855 | 0.537935965 | 1695.143512 | 1770.612435 | ENSG00000182934 |
| CLBA1 | 0.063128497 | 1.04472881 | 0.825118914 | 0.900593514 | 53.43897265 | 55.84235419 | ENSG00000140104 |
| EPCAM | 0.063119946 | 1.044722618 | 0.306821948 | 0.477828426 | 3451.814127 | 3607.198765 | ENSG00000119888 |
| RUNDC3B | 0.063001832 | 1.044637089 | 0.940054867 | 0.968314739 | 7.236189686 | 7.606308193 | ENSG00000105784 |
| ZFX | 0.062960215 | 1.044606955 | 0.561451636 | 0.716333565 | 421.3713318 | 440.4989732 | ENSG00000005889 |
| NA | 0.062922205 | 1.044579434 | 0.884901789 | 0.936917696 | 21.97878695 | 22.98429513 | ENSG00000226396 |
| ZC3H4 | 0.062633551 | 1.044370456 | 0.445043281 | 0.615568217 | 1172.386271 | 1224.610832 | ENSG00000130749 |
| PUM1 | 0.062593381 | 1.044341377 | 0.203444466 | 0.356462052 | 2864.722061 | 2991.423114 | ENSG00000134644 |
| PHLDA1 | 0.062342392 | 1.044159707 | 0.602326178 | 0.748099041 | 465.8546233 | 486.536636 | ENSG00000139289 |
| VPS35 | 0.062250893 | 1.044093485 | 0.28921397 | 0.458667108 | 1607.158419 | 1677.92369 | ENSG00000069329 |
| NA | 0.062215892 | 1.044068155 | 0.944531676 | 0.970904277 | 5.320571221 | 5.555303755 | ENSG00000257556 |
| KANK1 | 0.062162977 | 1.044029861 | 0.386880755 | 0.560279267 | 1133.919663 | 1184.134815 | ENSG00000107104 |
| CENPP | 0.062158395 | 1.044026546 | 0.776081948 | 0.868612024 | 98.12394871 | 102.4753402 | ENSG00000188312 |
| GATAD1 | 0.062117938 | 1.043997269 | 0.458300083 | 0.628184376 | 685.4226361 | 715.8257075 | ENSG00000157259 |
| DNAJB4 | 0.062006179 | 1.043916398 | 0.704479823 | 0.822230239 | 173.3341296 | 181.0032771 | ENSG00000162616 |
| ATRN | 0.062002766 | 1.043913929 | 0.484706812 | 0.652556292 | 861.722066 | 899.0238233 | ENSG00000088812 |
| ZNF155 | 0.061910797 | 1.043847383 | 0.84472275 | 0.911129246 | 43.89194368 | 45.85082333 | ENSG00000204920 |
| TIMELESS | 0.061878925 | 1.043824323 | 0.182232369 | 0.327734379 | 3834.461367 | 4002.392981 | ENSG00000111602 |
| TMEM47 | 0.061786002 | 1.043757093 | 0.535179119 | 0.695283348 | 582.9555752 | 608.8283592 | ENSG00000147027 |
| TMEM26 | 0.061659732 | 1.043665744 | 0.870444731 | 0.927669704 | 31.44479172 | 32.79299038 | ENSG00000196932 |
| STK32C | 0.061567219 | 1.04359882 | 0.534288783 | 0.694679848 | 555.4448083 | 579.8994328 | ENSG00000165752 |
| NANOGP8 | 0.06152611 | 1.043569084 | 0.707284904 | 0.823594399 | 172.0929983 | 179.5599295 | ENSG00000255192 |
| TJAP1 | 0.061516905 | 1.043562426 | 0.454546566 | 0.62428149 | 748.848476 | 781.7093588 | ENSG00000137221 |
| ACADM | 0.061414018 | 1.043488006 | 0.427310546 | 0.598681658 | 1062.340662 | 1108.558376 | ENSG00000117054 |
| NUMA1 | 0.06140192 | 1.043479256 | 0.323877261 | 0.495473136 | 1893.22423 | 1975.608828 | ENSG00000137497 |
| BTF3L4 | 0.061371837 | 1.043457497 | 0.36493591 | 0.538636469 | 1298.685787 | 1355.347668 | ENSG00000134717 |
| CAPZA2 | 0.061359531 | 1.043448597 | 0.356995291 | 0.53036551 | 1340.576894 | 1398.98343 | ENSG00000198898 |
| RN7SL1 | 0.061293982 | 1.043401189 | 0.844296006 | 0.910821734 | 389.7314366 | 406.4798737 | ENSG00000276168 |
| PROC | 0.061262293 | 1.043378271 | 0.900208721 | 0.945238757 | 18.01349867 | 18.84490247 | ENSG00000115718 |
| AIFM1 | 0.061246014 | 1.043366498 | 0.188933095 | 0.336724717 | 3605.401895 | 3761.489467 | ENSG00000156709 |
| PDGFD | 0.06114005 | 1.043289866 | 0.723930818 | 0.834405189 | 173.863565 | 181.6579861 | ENSG00000170962 |
| TFG | 0.061116257 | 1.043272661 | 0.3462656 | 0.518820169 | 1351.827132 | 1410.789675 | ENSG00000114354 |
| C12orf45 | 0.0610943 | 1.043256783 | 0.560641775 | 0.715726323 | 439.8319272 | 458.8722749 | ENSG00000151131 |
| MRPL9 | 0.061060858 | 1.0432326 | 0.450935382 | 0.621349645 | 854.9322618 | 892.4916781 | ENSG00000143436 |
| AGO2 | 0.061027465 | 1.043208453 | 0.307128996 | 0.478078388 | 2054.940608 | 2143.779719 | ENSG00000123908 |
| NPIPB5 | 0.060910051 | 1.043123555 | 0.843430714 | 0.91029706 | 45.88993779 | 47.94377317 | ENSG00000243716 |
| HCN3 | 0.060832811 | 1.043067709 | 0.656447137 | 0.7869973 | 236.2595192 | 246.3469997 | ENSG00000143630 |
| C9orf16 | 0.060696557 | 1.042969202 | 0.532965103 | 0.693660087 | 559.434451 | 583.5758454 | ENSG00000171159 |
| ZSCAN9 | 0.060694282 | 1.042967558 | 0.706428956 | 0.823341598 | 222.5434149 | 231.7511876 | ENSG00000137185 |
| FNBP1L | 0.060691956 | 1.042965876 | 0.284738005 | 0.453619693 | 2935.914476 | 3062.456675 | ENSG00000137942 |
| RAB18 | 0.060642454 | 1.04293009 | 0.454184839 | 0.624130737 | 951.0980144 | 991.8993712 | ENSG00000099246 |
| GLIS3 | 0.060477335 | 1.042810732 | 0.957649572 | 0.978289446 | 3.807670405 | 3.927242524 | ENSG00000107249 |
| PCSK5 | 0.060465338 | 1.04280206 | 0.6817577 | 0.805819974 | 234.3488068 | 244.1215015 | ENSG00000099139 |
| ISM2 | 0.060400416 | 1.042755134 | 0.888028831 | 0.938117528 | 23.93559324 | 25.01862802 | ENSG00000100593 |
| SH3BGRL3 | 0.060387266 | 1.04274563 | 0.493330433 | 0.659791832 | 757.4660179 | 790.3325701 | ENSG00000142669 |
| SELENOM | 0.060352317 | 1.04272037 | 0.726059429 | 0.835411015 | 172.5829607 | 180.2837478 | ENSG00000198832 |
| F3 | 0.060327489 | 1.042702426 | 0.805064119 | 0.887405972 | 80.02017725 | 83.52233919 | ENSG00000117525 |
| KIAA0232 | 0.060192005 | 1.042604509 | 0.502359958 | 0.668228848 | 670.6124424 | 698.9457681 | ENSG00000170871 |
| TSC22D2 | 0.060175915 | 1.042592882 | 0.512155909 | 0.676434317 | 666.5608874 | 695.1959634 | ENSG00000196428 |
| GLO1 | 0.060148361 | 1.04257297 | 0.319881338 | 0.491386438 | 3350.898818 | 3493.626283 | ENSG00000124767 |
| CYB5R3 | 0.060068497 | 1.042515257 | 0.303898943 | 0.474620669 | 2137.58018 | 2228.913049 | ENSG00000100243 |
| BTG3 | 0.059996889 | 1.042463513 | 0.700262905 | 0.819160251 | 198.9724178 | 207.4232831 | ENSG00000154640 |
| RNMT | 0.059981843 | 1.042452641 | 0.364375258 | 0.537974382 | 1281.339903 | 1335.263511 | ENSG00000101654 |
| DUSP3 | 0.059969385 | 1.042443639 | 0.534227394 | 0.694668387 | 626.6686229 | 653.223725 | ENSG00000108861 |
| NDUFS4 | 0.059953388 | 1.042432081 | 0.54609902 | 0.704338851 | 557.9630631 | 581.3280053 | ENSG00000164258 |

| | | | | | | | |
|---|---|---|---|---|---|---|---|
| DSTNP2 | 0.059889357 | 1.042385815 | 0.87983627 | 0.933860879 | 26.79069836 | 27.90235497 | ENSG00000248593 |
| SNX17 | 0.059741867 | 1.042279255 | 0.271659605 | 0.438542715 | 2040.495513 | 2126.63664 | ENSG00000115234 |
| IER3IP1 | 0.059719982 | 1.042263445 | 0.79304106 | 0.879581159 | 96.86515919 | 101.1737005 | ENSG00000134049 |
| TMEM100 | 0.059686326 | 1.04223913 | 0.915404631 | 0.954649348 | 14.11663151 | 14.64609936 | ENSG00000166292 |
| PCBD2 | 0.059608562 | 1.042182953 | 0.718655799 | 0.831214704 | 169.9729923 | 176.9536939 | ENSG00000132570 |
| C20orf194 | 0.059574967 | 1.042158685 | 0.690128778 | 0.811785015 | 232.2249708 | 241.6324877 | ENSG00000088854 |
| TSPAN7 | 0.059520243 | 1.042119154 | 0.847697349 | 0.912959443 | 48.62283088 | 50.78366461 | ENSG00000156298 |
| LRRC73 | 0.059517026 | 1.042116831 | 0.940992733 | 0.968764627 | 6.640248141 | 6.951859682 | ENSG00000204052 |
| WNK3 | 0.059501445 | 1.042105576 | 0.636661408 | 0.772685449 | 356.4863819 | 371.5401507 | ENSG00000196632 |
| ARHGEF10 | 0.05947706 | 1.042087962 | 0.837325449 | 0.907463215 | 57.95306943 | 60.30600571 | ENSG00000104728 |
| NA | 0.059322414 | 1.041976264 | 0.838728106 | 0.908136413 | 55.86691771 | 58.08857675 | ENSG00000183199 |
| TRIM33 | 0.059243255 | 1.041919094 | 0.382722697 | 0.556045981 | 1166.093784 | 1215.088388 | ENSG00000197323 |
| RPS27A | 0.059242628 | 1.041918641 | 0.166257423 | 0.305949818 | 10451.4319 | 10888.69741 | ENSG00000143947 |
| PLEKHG1 | 0.059016773 | 1.04175554 | 0.841083022 | 0.909135014 | 48.82116102 | 50.82319897 | ENSG00000120278 |
| LMAN1 | 0.058850288 | 1.04163533 | 0.389551092 | 0.562782155 | 1954.918201 | 2037.025646 | ENSG00000074695 |
| TMEM170A | 0.058738353 | 1.041554516 | 0.551779594 | 0.708528644 | 572.5945991 | 596.6607691 | ENSG00000166822 |
| NRL | 0.058655408 | 1.041494635 | 0.924032723 | 0.959186142 | 10.99218438 | 11.44482573 | ENSG00000129535 |
| SNCB | 0.058634439 | 1.041479498 | 0.692498566 | 0.813828195 | 227.6858311 | 237.4798885 | ENSG00000074317 |
| SLC25A30 | 0.058618757 | 1.041468177 | 0.754491205 | 0.854310609 | 138.4152371 | 143.9679086 | ENSG00000174032 |
| EIF3M | 0.058553887 | 1.041421349 | 0.20296862 | 0.355789881 | 3826.127561 | 3984.812043 | ENSG00000149100 |
| LAMB2 | 0.058503125 | 1.041384706 | 0.286237451 | 0.455352416 | 2141.268578 | 2230.042606 | ENSG00000172037 |
| SOCS6 | 0.058468583 | 1.041359773 | 0.58433237 | 0.734259754 | 411.2028798 | 428.4136685 | ENSG00000170677 |
| MYOZ3 | 0.058463356 | 1.041356 | 0.613087763 | 0.756206907 | 367.7089478 | 382.7411497 | ENSG00000164591 |
| RAD21 | 0.058390947 | 1.041303736 | 0.234891867 | 0.395991395 | 3784.347676 | 3941.576275 | ENSG00000164754 |
| NA | 0.058385937 | 1.04130012 | 0.943625966 | 0.970490404 | 6.300369816 | 6.581614742 | ENSG00000273674 |
| CEP55 | 0.058221703 | 1.041181587 | 0.648882954 | 0.781685316 | 367.7092232 | 383.2895954 | ENSG00000138180 |
| DENND6B | 0.058177183 | 1.041149458 | 0.889298744 | 0.938893502 | 28.59546857 | 29.65384136 | ENSG00000205593 |
| STEAP1 | 0.058151622 | 1.041131011 | 0.793583001 | 0.880030448 | 95.22486487 | 99.26384677 | ENSG00000164647 |
| GNPDA1 | 0.058113713 | 1.041103654 | 0.308580688 | 0.4796801 | 2264.382157 | 2356.572078 | ENSG00000113552 |
| PPIP5K1 | 0.058000296 | 1.041021811 | 0.742913274 | 0.847821391 | 179.3637548 | 186.672828 | ENSG00000168781 |
| RINT1 | 0.057976462 | 1.041004613 | 0.579957839 | 0.731144078 | 424.4250794 | 441.7328727 | ENSG00000135249 |
| LRRC28 | 0.05796678 | 1.040997627 | 0.744357302 | 0.848613954 | 153.6726396 | 159.7398089 | ENSG00000168904 |
| WDR86 | 0.057935722 | 1.040975217 | 0.67383779 | 0.800111161 | 369.5014031 | 384.2718589 | ENSG00000187260 |
| JTB | 0.05791363 | 1.040959277 | 0.914440797 | 0.954106627 | 15.47470535 | 16.06792913 | ENSG00000143543 |
| SUPT5H | 0.057882476 | 1.040936798 | 0.239531447 | 0.401634171 | 4974.922273 | 5178.450608 | ENSG00000196235 |
| SLC2A13 | 0.057881511 | 1.040936102 | 0.730263285 | 0.83844239 | 163.9239429 | 170.7960885 | ENSG00000151229 |
| FAM86B2 | 0.057724453 | 1.040822787 | 0.921614824 | 0.958171467 | 13.8661501 | 14.35382653 | ENSG00000145002 |
| AAK1 | 0.057675525 | 1.040787489 | 0.648152257 | 0.781171369 | 328.6119331 | 342.4079553 | ENSG00000115977 |
| DCAF15 | 0.057603863 | 1.040735792 | 0.323741491 | 0.495343996 | 1992.76615 | 2073.442273 | ENSG00000132017 |
| MRPS9 | 0.057601692 | 1.040734226 | 0.513421834 | 0.677642351 | 740.6933737 | 770.2771761 | ENSG00000135972 |
| EPHX4 | 0.057522238 | 1.040676911 | 0.926547478 | 0.960672598 | 11.66424404 | 12.14767136 | ENSG00000172031 |
| FANCB | 0.05745897 | 1.040631274 | 0.773887966 | 0.867313818 | 112.0904592 | 116.6027682 | ENSG00000181544 |
| GRK5 | 0.057457208 | 1.040630003 | 0.793945064 | 0.880229553 | 91.14940405 | 94.85074294 | ENSG00000198873 |
| MPP6 | 0.057444597 | 1.040620906 | 0.385745652 | 0.559336995 | 1481.23763 | 1542.071613 | ENSG00000105926 |
| ZNF235 | 0.057435982 | 1.040614693 | 0.851227151 | 0.915330155 | 47.42556614 | 49.24568543 | ENSG00000159917 |
| GGA2 | 0.057418762 | 1.040602272 | 0.299667917 | 0.470144257 | 2669.132167 | 2776.93018 | ENSG00000103365 |
| HSD11B2 | 0.057393729 | 1.040584216 | 0.541426 | 0.700278174 | 668.8844803 | 696.1559339 | ENSG00000176387 |
| RPL13 | 0.057340917 | 1.040546125 | 0.118799788 | 0.236606102 | 18987.16246 | 19756.87587 | ENSG00000167526 |
| CCN2 | 0.057264062 | 1.040490694 | 0.53551206 | 0.695478386 | 885.4224738 | 920.8143653 | ENSG00000118523 |
| ZNF383 | 0.057264049 | 1.040490685 | 0.862872977 | 0.92296549 | 43.81044391 | 45.67227676 | ENSG00000188283 |
| VDAC1 | 0.05722663 | 1.040463698 | 0.190522536 | 0.33886873 | 5918.236438 | 6158.666378 | ENSG00000213585 |
| DTWD1 | 0.057182727 | 1.040432036 | 0.638372739 | 0.773863705 | 353.5396526 | 367.9812935 | ENSG00000104047 |
| ZNF226 | 0.056933413 | 1.040252253 | 0.704654904 | 0.822353161 | 209.400712 | 217.6328949 | ENSG00000167380 |
| SOD1 | 0.056908335 | 1.040234171 | 0.254559457 | 0.419386609 | 3426.697522 | 3564.34731 | ENSG00000142168 |
| GDI1 | 0.056804789 | 1.040159513 | 0.296056558 | 0.466298747 | 2727.656938 | 2836.409133 | ENSG00000203879 |
| NA | 0.056789442 | 1.040148448 | 0.949926371 | 0.974062065 | 5.366665128 | 5.571975306 | ENSG00000284526 |
| ZWILCH | 0.056731656 | 1.040106787 | 0.542310513 | 0.701093158 | 768.5996994 | 799.8019098 | ENSG00000174442 |
| TUBGCP3 | 0.056499871 | 1.039939695 | 0.483482678 | 0.651277749 | 782.5402655 | 813.5432454 | ENSG00000126216 |
| ALDH7A1 | 0.056458123 | 1.039909603 | 0.218736808 | 0.376519298 | 3786.592021 | 3937.306013 | ENSG00000164904 |
| MTHFSD | 0.056354865 | 1.039835175 | 0.741680394 | 0.847013763 | 148.0336414 | 153.8903422 | ENSG00000103248 |
| FSTL1 | 0.056344667 | 1.039827826 | 0.222616885 | 0.381026377 | 4125.920815 | 4291.126995 | ENSG00000163430 |
| NA | 0.056285823 | 1.039785414 | 0.910843553 | 0.951993281 | 16.73712475 | 17.38704598 | ENSG00000230629 |
| ANO5 | 0.056249194 | 1.039759015 | 0.72629658 | 0.83557892 | 238.599707 | 248.1121525 | ENSG00000171714 |
| PRKAR1A | 0.056211356 | 1.039731745 | 0.403998891 | 0.576189754 | 1459.286352 | 1517.94456 | ENSG00000108946 |
| GRK6 | 0.0561637 | 1.039697401 | 0.326408338 | 0.498238903 | 1859.259079 | 1932.99393 | ENSG00000198055 |
| NDUFB4 | 0.055999967 | 1.039579412 | 0.368271827 | 0.542191239 | 1649.737958 | 1715.356425 | ENSG00000065518 |
| WDCP | 0.055979681 | 1.039564794 | 0.516395322 | 0.680447331 | 723.8613086 | 752.4400535 | ENSG00000163026 |
| WDFY3 | 0.055973253 | 1.039560162 | 0.514735414 | 0.678633192 | 784.0288119 | 814.5706466 | ENSG00000163625 |
| NA | 0.055929933 | 1.039528948 | 0.895391097 | 0.942250941 | 26.93033152 | 27.94430037 | ENSG00000232472 |
| SCN3B | 0.05591293 | 1.039516696 | 0.921967787 | 0.958383477 | 12.96226946 | 13.5146523 | ENSG00000166257 |
| NA | 0.055613196 | 1.039300749 | 0.932770763 | 0.964184087 | 10.65163033 | 11.04930695 | ENSG00000213228 |
| LY6L | 0.055564132 | 1.039265404 | 0.947772328 | 0.972984731 | 5.67440406 | 5.893823127 | ENSG00000261667 |
| MED21 | 0.055493255 | 1.039214348 | 0.574274893 | 0.726479147 | 500.7589224 | 520.5511618 | ENSG00000152944 |
| SSR4P1 | 0.055410305 | 1.039154599 | 0.926025647 | 0.960479636 | 11.70264096 | 12.16219234 | ENSG00000235374 |
| TINF2 | 0.055350864 | 1.039111785 | 0.637063369 | 0.773076018 | 385.1002596 | 399.8095645 | ENSG00000092330 |
| NANOG | 0.055345011 | 1.039107569 | 0.430972405 | 0.602237499 | 1649.404222 | 1714.352364 | ENSG00000111704 |
| CREB3L2 | 0.055183834 | 1.038991487 | 0.48966777 | 0.656121481 | 1132.541726 | 1176.97693 | ENSG00000182158 |

| | | | | | | | |
|---|---|---|---|---|---|---|---|
| MAGEL2 | 0.055179109 | 1.038988085 | 0.775944635 | 0.868533369 | 133.5184346 | 138.7142855 | ENSG00000254585 |
| RUNX3 | 0.0551715 | 1.038982605 | 0.884950678 | 0.936918036 | 32.01928192 | 33.1863586 | ENSG00000020633 |
| TRMT10C | 0.055169139 | 1.038980905 | 0.55900944 | 0.714356707 | 740.3742852 | 769.3269592 | ENSG00000174173 |
| THAP3 | 0.055134072 | 1.038955651 | 0.692547851 | 0.813836537 | 224.2581877 | 233.0493196 | ENSG00000041988 |
| NFE2L3 | 0.055129108 | 1.038952076 | 0.243801696 | 0.406708624 | 3850.152626 | 4000.780403 | ENSG00000050344 |
| GALNT11 | 0.055127925 | 1.038951224 | 0.596174918 | 0.743763199 | 565.8892971 | 588.0848768 | ENSG00000178234 |
| NA | 0.055104923 | 1.038934659 | 0.94425915 | 0.970809901 | 6.950777976 | 7.252734804 | ENSG00000265399 |
| OXSR1 | 0.055009031 | 1.038865606 | 0.358490789 | 0.531604721 | 1569.568197 | 1630.572519 | ENSG00000172939 |
| CCT8 | 0.054995424 | 1.038855808 | 0.223111642 | 0.381632088 | 6829.636762 | 7094.305613 | ENSG00000156261 |
| NA | 0.054902848 | 1.038789149 | 0.957090603 | 0.978044405 | 5.162953341 | 5.344771151 | ENSG00000271335 |
| BET1 | 0.054747004 | 1.038676941 | 0.630911433 | 0.768462714 | 398.3141377 | 414.0488044 | ENSG00000105829 |
| COX4I1 | 0.054708001 | 1.038648861 | 0.236945719 | 0.398476146 | 7359.451531 | 7643.12734 | ENSG00000131143 |
| SHLD2 | 0.054671638 | 1.038622682 | 0.58019448 | 0.731346818 | 583.0074383 | 606.1124139 | ENSG00000122376 |
| NA | 0.054624473 | 1.038588728 | 0.869176807 | 0.926984177 | 46.23994058 | 47.87225278 | ENSG00000226562 |
| CKAP2 | 0.054609665 | 1.038578068 | 0.387455095 | 0.560846119 | 1426.424336 | 1481.250946 | ENSG00000136108 |
| SLC12A7 | 0.054601849 | 1.038572442 | 0.440922944 | 0.611931261 | 1447.902474 | 1503.800553 | ENSG00000113504 |
| NA | 0.054553409 | 1.038537571 | 0.747553888 | 0.850422569 | 184.0462087 | 191.1585489 | ENSG00000204959 |
| NA | 0.054520133 | 1.038513617 | 0.948056559 | 0.972984731 | 6.003794495 | 6.238794221 | ENSG00000243964 |
| PNMA6A | 0.054485548 | 1.038488722 | 0.799291091 | 0.884159303 | 103.77351 | 107.8669031 | ENSG00000235961 |
| FYCO1 | 0.054484536 | 1.038487993 | 0.623888271 | 0.763014769 | 363.688317 | 377.7977855 | ENSG00000163820 |
| LYG1 | 0.054426855 | 1.038446474 | 0.946091708 | 0.972141843 | 6.681435963 | 6.958311403 | ENSG00000144214 |
| TRIP4 | 0.054313234 | 1.038364693 | 0.674389447 | 0.800199707 | 304.9160068 | 316.4237838 | ENSG00000103671 |
| FAM72D | 0.05429872 | 1.038354247 | 0.821161333 | 0.898166734 | 94.14027539 | 97.96596308 | ENSG00000215784 |
| TXNDC5 | 0.054033559 | 1.038163419 | 0.773608035 | 0.867201615 | 269.5882144 | 279.5086601 | ENSG00000239264 |
| GCLC | 0.054029637 | 1.038160597 | 0.549842639 | 0.706992892 | 759.7030303 | 789.0277138 | ENSG00000001084 |
| YIPF5 | 0.054023948 | 1.038156503 | 0.589170551 | 0.738633689 | 480.9717537 | 499.4667674 | ENSG00000145817 |
| NA | 0.053986329 | 1.038129433 | 0.822810019 | 0.899004762 | 74.03870998 | 76.94180894 | ENSG00000261505 |
| NA | 0.053963043 | 1.038112677 | 0.911945286 | 0.952533246 | 22.89084014 | 23.86190712 | ENSG00000269867 |
| FAM174B | 0.053949129 | 1.038102665 | 0.643335249 | 0.777118365 | 462.7182246 | 480.5605949 | ENSG00000185442 |
| NA | 0.053930216 | 1.038089056 | 0.856947891 | 0.91963628 | 58.65582903 | 60.74536001 | ENSG00000280287 |
| DSC2 | 0.053794223 | 1.037991207 | 0.528994623 | 0.69021544 | 690.3586745 | 716.6132616 | ENSG00000134755 |
| SRP54 | 0.053647183 | 1.03788542 | 0.439919081 | 0.611021599 | 1151.967392 | 1196.008503 | ENSG00000100883 |
| EPB41L5 | 0.05351425 | 1.037789792 | 0.614540392 | 0.757127264 | 538.8397446 | 559.2126893 | ENSG00000115109 |
| SMG5 | 0.053467896 | 1.037756448 | 0.238620684 | 0.400524808 | 4502.656605 | 4672.469094 | ENSG00000198952 |
| PHLPP2 | 0.053276373 | 1.037618691 | 0.659499423 | 0.789485573 | 329.6197656 | 341.8938761 | ENSG00000040199 |
| BCAT2 | 0.053275531 | 1.037618086 | 0.527985198 | 0.689522636 | 776.5901001 | 806.2491166 | ENSG00000105552 |
| PKD2 | 0.05315789 | 1.037533479 | 0.502977213 | 0.668568415 | 852.8200649 | 884.9816369 | ENSG00000118762 |
| CNOT2 | 0.053123473 | 1.037508728 | 0.442377572 | 0.613376392 | 1204.763839 | 1249.683673 | ENSG00000111596 |
| MANBA | 0.053108938 | 1.037498276 | 0.518472757 | 0.68211539 | 746.0409412 | 774.0541084 | ENSG00000109323 |
| PDE5A | 0.053098451 | 1.037490734 | 0.652411529 | 0.784007401 | 342.9906468 | 355.5795266 | ENSG00000138735 |
| PRKCA | 0.053048256 | 1.037454637 | 0.560340306 | 0.715498368 | 574.776181 | 596.5432932 | ENSG00000154229 |
| PDE8B | 0.053022543 | 1.037436147 | 0.903368759 | 0.947370846 | 24.30058969 | 25.22055833 | ENSG00000113231 |
| RAI14 | 0.05298315 | 1.03740782 | 0.379287987 | 0.55266597 | 2395.044185 | 2484.971053 | ENSG00000039560 |
| DDI2 | 0.052834243 | 1.03730075 | 0.469838402 | 0.638774384 | 1096.653149 | 1138.069927 | ENSG00000197312 |
| CCNYL1 | 0.052792936 | 1.037271051 | 0.557040939 | 0.713111471 | 799.9448703 | 830.4564731 | ENSG00000163249 |
| IDI1 | 0.052752125 | 1.037241709 | 0.456429512 | 0.626243365 | 1196.709025 | 1241.676248 | ENSG00000067064 |
| FXYD5 | 0.052694549 | 1.037200315 | 0.4851259 | 0.652828023 | 879.3465275 | 912.0305876 | ENSG00000089327 |
| NUP214 | 0.052675067 | 1.037186308 | 0.475045474 | 0.643501316 | 1064.479865 | 1103.565603 | ENSG00000126883 |
| CORO1A | 0.05266942 | 1.037182249 | 0.703677246 | 0.821808799 | 248.0777143 | 257.469172 | ENSG00000102879 |
| FAM83D | 0.052413184 | 1.036998052 | 0.373945335 | 0.547214396 | 1794.816559 | 1860.80163 | ENSG00000101447 |
| ABHD13 | 0.052398397 | 1.036987423 | 0.725185477 | 0.835021231 | 209.0051789 | 216.9604178 | ENSG00000139826 |
| ARC | 0.052396506 | 1.036986064 | 0.56843291 | 0.721613534 | 655.7833655 | 680.2178512 | ENSG00000198576 |
| TMCC1 | 0.052324717 | 1.036934465 | 0.643785346 | 0.777369999 | 401.3472729 | 416.0654055 | ENSG00000172765 |
| MRM3 | 0.052261681 | 1.036889159 | 0.647250303 | 0.780406104 | 348.9013776 | 361.6900615 | ENSG00000171861 |
| CDAN1 | 0.052095811 | 1.036769952 | 0.634948293 | 0.771283454 | 411.4233739 | 426.3626825 | ENSG00000140326 |
| NA | 0.052089884 | 1.036765692 | 0.929233454 | 0.962113616 | 12.01807688 | 12.45661574 | ENSG00000281571 |
| SCN2A | 0.051937107 | 1.036655908 | 0.893105402 | 0.940852791 | 28.47816263 | 29.46911048 | ENSG00000136531 |
| AURKB | 0.051909236 | 1.036635881 | 0.377403262 | 0.550884256 | 2312.644558 | 2396.565905 | ENSG00000178999 |
| DNM2 | 0.051833082 | 1.036581163 | 0.258023141 | 0.423078999 | 4384.549886 | 4544.33733 | ENSG00000079805 |
| SERTAD3 | 0.051772123 | 1.036537364 | 0.653826701 | 0.784925689 | 376.7546505 | 390.3218809 | ENSG00000167565 |
| PITPNM2 | 0.051747979 | 1.036520018 | 0.757308307 | 0.855942967 | 163.5709859 | 169.5213402 | ENSG00000090975 |
| GFOD1 | 0.051626952 | 1.036433069 | 0.729095628 | 0.837550393 | 198.6302241 | 205.7844811 | ENSG00000145990 |
| TBC1D31 | 0.051558824 | 1.036384127 | 0.695528268 | 0.816191958 | 436.6942623 | 452.4352695 | ENSG00000156787 |
| CLUHP3 | 0.051533108 | 1.036365653 | 0.75583597 | 0.855130591 | 207.3165122 | 214.3950737 | ENSG00000131797 |
| ARMC1 | 0.051443182 | 1.036301057 | 0.542303612 | 0.701093158 | 769.9728165 | 797.5186124 | ENSG00000104442 |
| NTAN1 | 0.051434193 | 1.0362946 | 0.703155647 | 0.821457229 | 291.9539006 | 302.8475463 | ENSG00000157045 |
| UFSP2 | 0.051291373 | 1.036192016 | 0.659306676 | 0.789485573 | 373.4347153 | 387.1778793 | ENSG00000109775 |
| SLC25A35 | 0.051267851 | 1.036175122 | 0.884836633 | 0.936917696 | 41.33728082 | 42.67618063 | ENSG00000125434 |
| CNTROB | 0.051195524 | 1.036123177 | 0.519596575 | 0.682988384 | 870.0410437 | 901.5107049 | ENSG00000170037 |
| NA | 0.051160807 | 1.036098243 | 0.900924552 | 0.94573301 | 27.50394584 | 28.42129891 | ENSG00000143429 |
| SRRM2 | 0.051087489 | 1.036045591 | 0.333434818 | 0.506001443 | 13142.51587 | 13615.31606 | ENSG00000167978 |
| TTK | 0.050994696 | 1.035978955 | 0.516585912 | 0.68046834 | 1002.209428 | 1038.826011 | ENSG00000112742 |
| FUT10 | 0.050862024 | 1.035883689 | 0.727332209 | 0.836171969 | 229.3427076 | 237.5952565 | ENSG00000172728 |
| MKI67 | 0.050718349 | 1.035780533 | 0.405494965 | 0.577746259 | 3151.408766 | 3263.551111 | ENSG00000148773 |
| SON | 0.050710305 | 1.035774758 | 0.253652305 | 0.418330051 | 4896.283258 | 5071.881048 | ENSG00000159140 |
| HSPA4 | 0.050670609 | 1.035746259 | 0.172417339 | 0.314600408 | 11915.67196 | 12342.50418 | ENSG00000170606 |
| TRPC4AP | 0.050648723 | 1.035730546 | 0.37379811 | 0.547183931 | 2664.576756 | 2758.946181 | ENSG00000100991 |

| | | | | | | | |
|---|---|---|---|---|---|---|---|
| CAPRIN1 | 0.050586525 | 1.035685895 | 0.228572056 | 0.38825249 | 7493.866824 | 7761.7933 | ENSG00000135387 |
| ARSJ | 0.05055275 | 1.035661648 | 0.799368742 | 0.884159303 | 120.2215932 | 124.4042824 | ENSG00000180801 |
| NA | 0.050527565 | 1.035643569 | 0.849487841 | 0.914224265 | 65.46629812 | 67.72626164 | ENSG00000186076 |
| GID8 | 0.050491852 | 1.035617933 | 0.322999643 | 0.494582906 | 2511.806586 | 2601.31093 | ENSG00000101193 |
| GPR61 | 0.050485153 | 1.035613124 | 0.953375647 | 0.97624978 | 5.756103983 | 5.946521394 | ENSG00000156097 |
| SMARCC1 | 0.050369219 | 1.035529906 | 0.187245443 | 0.334303733 | 8348.541709 | 8644.684568 | ENSG00000173473 |
| ZGRF1 | 0.050326939 | 1.035499559 | 0.635170543 | 0.77136449 | 427.5145962 | 442.5171981 | ENSG00000138658 |
| ATIC | 0.050281202 | 1.035466732 | 0.256056769 | 0.421179325 | 4417.698372 | 4573.9244 | ENSG00000138363 |
| L3HYPDH | 0.050186186 | 1.035398538 | 0.80268009 | 0.88621529 | 108.7496972 | 112.5109792 | ENSG00000126790 |
| POR | 0.049887911 | 1.035184493 | 0.336715729 | 0.50947557 | 3068.595277 | 3176.680631 | ENSG00000127948 |
| NA | 0.04983179 | 1.035144225 | 0.952115048 | 0.975457419 | 6.05826111 | 6.264068068 | ENSG00000256356 |
| GTF2H3 | 0.049810467 | 1.035128926 | 0.638330933 | 0.773863705 | 443.3241617 | 458.7418771 | ENSG00000111358 |
| NA | 0.04980531 | 1.035125225 | 0.953690719 | 0.976468747 | 5.756103983 | 5.942220247 | ENSG00000238035 |
| NA | 0.049794114 | 1.035117193 | 0.855487886 | 0.918462615 | 64.90085635 | 67.22237171 | ENSG00000227946 |
| AXL | 0.049689747 | 1.035042313 | 0.564324951 | 0.718716091 | 683.1598057 | 707.4633224 | ENSG00000167601 |
| PTMS | 0.04963813 | 1.035005282 | 0.292615109 | 0.46227535 | 7521.651072 | 7785.397344 | ENSG00000159335 |
| RRBP1 | 0.049527714 | 1.034926071 | 0.3050954 | 0.47594981 | 5853.210965 | 6057.224124 | ENSG00000125844 |
| SLC12A6 | 0.049494914 | 1.034902542 | 0.742700357 | 0.847821391 | 249.9029336 | 258.4519079 | ENSG00000140199 |
| TSC22D3 | 0.049444022 | 1.034866036 | 0.595421383 | 0.74325572 | 634.6876452 | 656.4105082 | ENSG00000157514 |
| SEPTIN9 | 0.049415132 | 1.034845313 | 0.214405015 | 0.370942915 | 10905.12836 | 11285.80124 | ENSG00000184640 |
| BLVRA | 0.049414412 | 1.034844796 | 0.588521087 | 0.73812888 | 619.6015029 | 640.9451228 | ENSG00000106605 |
| FAM161B | 0.049384584 | 1.034823401 | 0.949093239 | 0.973673562 | 7.359077431 | 7.639651296 | ENSG00000156050 |
| ING3 | 0.04931073 | 1.034770428 | 0.711747521 | 0.826281605 | 261.3154201 | 270.4425506 | ENSG00000071243 |
| TBRG4 | 0.049275392 | 1.034745082 | 0.345856116 | 0.518367674 | 3026.067826 | 3130.764497 | ENSG00000136270 |
| TCAP | 0.049253866 | 1.034729644 | 0.911860069 | 0.952533246 | 22.87987669 | 23.61508733 | ENSG00000173991 |
| GPN3 | 0.049227313 | 1.034710599 | 0.539810889 | 0.698985895 | 885.7017402 | 916.4412683 | ENSG00000111231 |
| RRAS2 | 0.049174109 | 1.034672442 | 0.355021646 | 0.528182278 | 3087.326115 | 3194.548297 | ENSG00000133818 |
| PC | 0.04913415 | 1.034643784 | 0.581899007 | 0.732447768 | 752.2107632 | 777.6431467 | ENSG00000173599 |
| MAN1A1 | 0.049114236 | 1.034629503 | 0.777141892 | 0.869299514 | 150.1622202 | 155.4592413 | ENSG00000111885 |
| NA | 0.049110803 | 1.034627041 | 0.957097557 | 0.978044405 | 5.342222723 | 5.504756061 | ENSG00000263050 |
| DNA2 | 0.048980053 | 1.034533278 | 0.582937729 | 0.733248614 | 823.7253995 | 852.9906853 | ENSG00000138346 |
| HPRT1 | 0.048925193 | 1.034493939 | 0.502893257 | 0.668548951 | 1321.016855 | 1367.089558 | ENSG00000165704 |
| TRAF2 | 0.048886279 | 1.034466036 | 0.477690695 | 0.646041166 | 1334.935393 | 1380.778002 | ENSG00000127191 |
| PRKD2 | 0.04884409 | 1.034435785 | 0.508064304 | 0.672809644 | 1212.93695 | 1254.900245 | ENSG00000105287 |
| NA | 0.048788474 | 1.034395909 | 0.95198593 | 0.975448057 | 6.322021318 | 6.539669343 | ENSG00000199476 |
| GOLGA8B | 0.048757732 | 1.034373868 | 0.770694743 | 0.864868483 | 162.3239605 | 167.8384301 | ENSG00000215252 |
| DPY30 | 0.048638804 | 1.034288603 | 0.495902656 | 0.662159152 | 1491.945026 | 1542.976037 | ENSG00000162961 |
| HNRNPA1 | 0.048607743 | 1.034266335 | 0.189632029 | 0.337665359 | 62692.82581 | 64840.75287 | ENSG00000135486 |
| NA | 0.04858982 | 1.034253486 | 0.956614709 | 0.978042701 | 5.707219174 | 5.887371406 | ENSG00000287787 |
| TEX2 | 0.04857818 | 1.034245141 | 0.633017425 | 0.769822396 | 469.084537 | 485.4896375 | ENSG00000136478 |
| ING1 | 0.048389035 | 1.034109555 | 0.604725214 | 0.74971598 | 753.4852732 | 778.7377552 | ENSG00000153487 |
| IST1 | 0.04833871 | 1.034073483 | 0.328736667 | 0.501000083 | 3415.49002 | 3531.125118 | ENSG00000182149 |
| ADGRG6 | 0.048338309 | 1.034073196 | 0.836679571 | 0.907107819 | 91.05654052 | 94.18068896 | ENSG00000112414 |
| IWS1 | 0.048222135 | 1.03398993 | 0.392287227 | 0.565211211 | 2035.918377 | 2105.094645 | ENSG00000163166 |
| CC2D2A | 0.048021815 | 1.033846369 | 0.732638934 | 0.84029936 | 245.0822492 | 253.1224755 | ENSG00000048342 |
| CREBBP | 0.047996375 | 1.033828138 | 0.401974526 | 0.574301425 | 1774.973647 | 1834.927919 | ENSG00000005339 |
| NCAPD2 | 0.047960909 | 1.033802724 | 0.292506997 | 0.462255467 | 5235.209344 | 5412.182157 | ENSG00000010292 |
| NPAT | 0.047880906 | 1.033745397 | 0.731224377 | 0.839239274 | 264.8921085 | 274.2259467 | ENSG00000149308 |
| ATP5F1C | 0.047785813 | 1.033677262 | 0.382457073 | 0.555863738 | 2857.001365 | 2952.882994 | ENSG00000165629 |
| ARHGAP26 | 0.047718155 | 1.033628787 | 0.637908247 | 0.773642659 | 507.0315322 | 523.8790651 | ENSG00000145819 |
| BAZ2A | 0.047496256 | 1.033469818 | 0.405701972 | 0.577977752 | 3653.569227 | 3774.482564 | ENSG00000076108 |
| GPANK1 | 0.047467458 | 1.033449189 | 0.634737993 | 0.771175881 | 502.5510143 | 519.0435816 | ENSG00000204438 |
| NA | 0.047241797 | 1.033287553 | 0.916382088 | 0.955150575 | 23.98447805 | 24.74653439 | ENSG00000231113 |
| SP1 | 0.047165788 | 1.033233115 | 0.52050659 | 0.683532513 | 1401.571457 | 1447.81692 | ENSG00000185591 |
| ZBTB21 | 0.04716229 | 1.03323061 | 0.710653629 | 0.825806828 | 372.0134756 | 384.1605073 | ENSG00000173276 |
| DCP2 | 0.047107878 | 1.033191642 | 0.559177488 | 0.714471647 | 1463.32936 | 1512.482446 | ENSG00000172795 |
| TPI1P2 | 0.047064335 | 1.033160459 | 0.966105277 | 0.982484721 | 4.686232757 | 4.8126633 | ENSG00000230359 |
| NA | 0.047006659 | 1.033119156 | 0.942459777 | 0.969808053 | 11.15279332 | 11.47870188 | ENSG00000267655 |
| UBE2E1 | 0.047005555 | 1.033118365 | 0.432694504 | 0.603681903 | 2019.8032 | 2086.877747 | ENSG00000170142 |
| SPRYD7 | 0.047001565 | 1.033115508 | 0.802077491 | 0.885936266 | 124.3590175 | 128.4850581 | ENSG00000123178 |
| TMSB4X | 0.04696781 | 1.033091337 | 0.301714445 | 0.472509608 | 16679.09787 | 17230.015 | ENSG00000205542 |
| NA | 0.046964803 | 1.033089184 | 0.817862356 | 0.895882273 | 107.5157504 | 111.002575 | ENSG00000272711 |
| WDR59 | 0.046875589 | 1.033025301 | 0.476397008 | 0.644743559 | 1413.209871 | 1459.292199 | ENSG00000103091 |
| SBSPON | 0.046830447 | 1.032992978 | 0.867721265 | 0.926302413 | 55.86113575 | 57.79441381 | ENSG00000164764 |
| XYLT2 | 0.04681347 | 1.032980822 | 0.506355389 | 0.671163122 | 1241.439619 | 1282.40573 | ENSG00000015532 |
| AGPS | 0.046808199 | 1.032977048 | 0.523582593 | 0.686069036 | 1630.756226 | 1685.580993 | ENSG00000018510 |
| NEU3 | 0.046772321 | 1.032951359 | 0.604472474 | 0.749600272 | 626.4496931 | 646.6658092 | ENSG00000162139 |
| TMEM254 | 0.046759686 | 1.032942313 | 0.781905744 | 0.87209434 | 176.8865246 | 182.5953116 | ENSG00000133678 |
| FNIP2 | 0.046680797 | 1.032885831 | 0.752112625 | 0.853347831 | 223.7098917 | 230.9445508 | ENSG00000052795 |
| CCK | 0.046536766 | 1.032782719 | 0.95565592 | 0.977429542 | 8.257376262 | 8.445482423 | ENSG00000187094 |
| RPS6 | 0.046532936 | 1.032779977 | 0.27338851 | 0.440191783 | 22516.79927 | 23253.38942 | ENSG00000137154 |
| TMEM268 | 0.046505207 | 1.032760127 | 0.675593339 | 0.800938489 | 379.4081221 | 391.8034242 | ENSG00000157693 |
| TSKU | 0.046341987 | 1.032643292 | 0.582985044 | 0.733248614 | 893.8800456 | 923.2713751 | ENSG00000182704 |
| EIF2B3 | 0.046317412 | 1.032625702 | 0.592078762 | 0.740787017 | 791.3994533 | 816.8934784 | ENSG00000070785 |
| NA | 0.046177415 | 1.032525502 | 0.924834389 | 0.959760084 | 20.08761089 | 20.74588137 | ENSG00000286651 |
| TMEM248 | 0.046170671 | 1.032520676 | 0.527521121 | 0.689318563 | 1158.335443 | 1196.26905 | ENSG00000106609 |
| CCNI | 0.046114107 | 1.032480194 | 0.440093956 | 0.611138777 | 2906.218654 | 3001.761154 | ENSG00000118816 |

| | | | | | | | |
|---|---|---|---|---|---|---|---|
| SPCS1 | 0.046098262 | 1.032468855 | 0.539069966 | 0.698545187 | 901.4256748 | 930.7577232 | ENSG00000114902 |
| UQCRC1 | 0.045993534 | 1.032393908 | 0.29142176 | 0.461107051 | 5888.224988 | 6078.737852 | ENSG00000010256 |
| BLMH | 0.045959407 | 1.032369487 | 0.441895999 | 0.612973015 | 2025.214546 | 2090.97316 | ENSG00000108578 |
| SEPTIN7P2 | 0.04593968 | 1.032355371 | 0.745447707 | 0.849098159 | 227.1085869 | 234.3388615 | ENSG00000214765 |
| TRIP6 | 0.045939103 | 1.032354958 | 0.454462201 | 0.624210062 | 2935.659219 | 3031.004005 | ENSG00000087077 |
| IRAK2 | 0.045837986 | 1.032282604 | 0.922093413 | 0.958413349 | 22.86380699 | 23.529579 | ENSG00000134070 |
| OLIG2 | 0.045799772 | 1.032255262 | 0.813499604 | 0.893185108 | 146.4021584 | 151.4819841 | ENSG00000205927 |
| VAMP3 | 0.045693335 | 1.032179108 | 0.615855861 | 0.758212125 | 609.8900491 | 629.763007 | ENSG00000049245 |
| XRRA1 | 0.045684316 | 1.032172656 | 0.869708849 | 0.927295274 | 58.98155268 | 60.85667522 | ENSG00000166435 |
| EIF1 | 0.045620285 | 1.032126846 | 0.270262427 | 0.436798968 | 5354.392355 | 5526.563141 | ENSG00000173812 |
| LRRC37A11P | 0.045592016 | 1.032106622 | 0.946354902 | 0.97226034 | 10.13248263 | 10.38893975 | ENSG00000214553 |
| GOLT1B | 0.045541322 | 1.032070356 | 0.706440834 | 0.823341598 | 391.2042655 | 404.1642813 | ENSG00000111711 |
| PAQR3 | 0.045483465 | 1.032028967 | 0.664647801 | 0.793374217 | 545.7615248 | 563.6481045 | ENSG00000163291 |
| C4orf33 | 0.045456705 | 1.032009825 | 0.897161127 | 0.943168136 | 33.73031272 | 34.82033851 | ENSG00000151470 |
| VPS54 | 0.045373665 | 1.031950425 | 0.71471396 | 0.828621912 | 359.0227719 | 370.5947864 | ENSG00000143952 |
| NA | 0.045326315 | 1.031916556 | 0.942428964 | 0.969808053 | 11.16116603 | 11.46364786 | ENSG00000234589 |
| VTI1A | 0.045284854 | 1.031886901 | 0.745141121 | 0.84900581 | 255.0849861 | 263.3845009 | ENSG00000151532 |
| LENG8 | 0.045277951 | 1.031881963 | 0.721799916 | 0.833127234 | 3501.627143 | 3612.860915 | ENSG00000167615 |
| NA | 0.045273544 | 1.031878812 | 0.942517373 | 0.969815588 | 12.12489493 | 12.48404016 | ENSG00000271833 |
| TENT5B | 0.04516844 | 1.031803639 | 0.500870165 | 0.666686425 | 2638.850053 | 2722.957183 | ENSG00000158246 |
| IL20RA | 0.045121778 | 1.031770267 | 0.888590505 | 0.938505307 | 45.86657142 | 47.15703674 | ENSG00000016402 |
| SGTB | 0.045012276 | 1.031691958 | 0.737808951 | 0.844191725 | 248.3697839 | 256.2393132 | ENSG00000197860 |
| DPPA4 | 0.044986161 | 1.031673283 | 0.300555265 | 0.471189543 | 12923.1796 | 13331.84236 | ENSG00000121570 |
| MIEF2 | 0.044907574 | 1.031617086 | 0.751483734 | 0.852957914 | 237.3476874 | 244.7498584 | ENSG00000177427 |
| PTEN | 0.044794565 | 1.031536281 | 0.621409225 | 0.761915061 | 731.8514211 | 755.5240568 | ENSG00000171862 |
| MAP4 | 0.04478003 | 1.031525889 | 0.272242208 | 0.439080963 | 6653.488566 | 6862.888536 | ENSG00000047849 |
| FADS3 | 0.044473482 | 1.03130673 | 0.618931355 | 0.760351829 | 744.1019436 | 767.1138498 | ENSG00000221968 |
| PHKB | 0.044439252 | 1.031282262 | 0.556516503 | 0.712676571 | 1015.839413 | 1047.693861 | ENSG00000102893 |
| BRI3 | 0.044354327 | 1.031221556 | 0.688715615 | 0.810814899 | 664.1121412 | 685.0143944 | ENSG00000164713 |
| NA | 0.044329013 | 1.031203463 | 0.96043216 | 0.979722803 | 5.666031352 | 5.820152161 | ENSG00000272502 |
| SIK3 | 0.044312171 | 1.031191424 | 0.593567381 | 0.741757358 | 847.1708159 | 873.2682436 | ENSG00000160584 |
| TRIM65 | 0.044310418 | 1.031190171 | 0.529736762 | 0.690716585 | 1436.558148 | 1481.408587 | ENSG00000141569 |
| LDHB | 0.044275272 | 1.031165051 | 0.86806859 | 0.926365612 | 69.41184992 | 71.4827785 | ENSG00000111716 |
| CDC20 | 0.044262434 | 1.031155875 | 0.315663469 | 0.486845538 | 4694.306227 | 4840.16527 | ENSG00000117399 |
| TNRC6C | 0.044244872 | 1.031143322 | 0.687285103 | 0.80992162 | 691.5584533 | 712.5754853 | ENSG00000078687 |
| HDAC4 | 0.044214943 | 1.031121931 | 0.707685326 | 0.823921166 | 426.5231457 | 439.7982903 | ENSG00000068024 |
| TBC1D22A | 0.04418986 | 1.031104004 | 0.653270965 | 0.784551466 | 486.7562423 | 501.8571089 | ENSG00000054611 |
| EPS8L1 | 0.044156969 | 1.031080497 | 0.625610558 | 0.764322563 | 774.4632489 | 798.6495632 | ENSG00000131037 |
| RHOT1 | 0.04415264 | 1.031077403 | 0.643175675 | 0.777006705 | 684.7551191 | 706.464381 | ENSG00000126858 |
| CYP4V2 | 0.044103 | 1.031041927 | 0.878875262 | 0.933405785 | 62.55218372 | 64.64947926 | ENSG00000145476 |
| ERN1 | 0.043899558 | 1.030896544 | 0.836494128 | 0.907107819 | 109.2262176 | 112.730102 | ENSG00000178607 |
| PDCD6 | 0.043786604 | 1.030815834 | 0.845968555 | 0.912024708 | 88.34944127 | 91.05821148 | ENSG00000249915 |
| SNHG20 | 0.043714701 | 1.03076446 | 0.875281395 | 0.931024117 | 70.16225503 | 72.2315981 | ENSG00000234912 |
| ATP10D | 0.043642167 | 1.030712638 | 0.855138741 | 0.918307807 | 79.48340825 | 82.0236218 | ENSG00000145246 |
| MTERF4 | 0.043604979 | 1.03068607 | 0.649226912 | 0.781787538 | 516.3797186 | 532.1409368 | ENSG00000122085 |
| NA | 0.043581021 | 1.030668954 | 0.951586522 | 0.975142395 | 8.297688203 | 8.586372634 | ENSG00000217130 |
| NA | 0.043542804 | 1.030641652 | 0.978864591 | 0.988963906 | 9.731004278 | 10.03804998 | ENSG00000280056 |
| P3H1 | 0.043514384 | 1.030621349 | 0.523250924 | 0.68582916 | 1580.835506 | 1629.711349 | ENSG00000117385 |
| MRM1 | 0.043514103 | 1.030621149 | 0.652646238 | 0.784094076 | 538.4487783 | 554.6159685 | ENSG00000278619 |
| NA | 0.043409264 | 1.030546258 | 0.957007252 | 0.978044405 | 6.376487933 | 6.558491469 | ENSG00000259287 |
| LCMT2 | 0.043392264 | 1.030534115 | 0.725861507 | 0.835365268 | 310.1743023 | 319.3860465 | ENSG00000168806 |
| CAP1 | 0.043387742 | 1.030530885 | 0.42290335 | 0.594457999 | 3899.89515 | 4017.944407 | ENSG00000131236 |
| HMGXB4 | 0.04336852 | 1.030517154 | 0.543318105 | 0.701925366 | 1023.79208 | 1055.271703 | ENSG00000100281 |
| DIP2B | 0.043284718 | 1.030457296 | 0.54005918 | 0.699260463 | 1183.832596 | 1219.247197 | ENSG00000066084 |
| KDM5C | 0.043187534 | 1.030387883 | 0.385063516 | 0.558557808 | 6419.93671 | 6614.331869 | ENSG00000126012 |
| ATP6V1B2 | 0.043036203 | 1.030279807 | 0.504332931 | 0.669493983 | 1833.647874 | 1889.303221 | ENSG00000147416 |
| TP53I13 | 0.043028599 | 1.030274377 | 0.661751553 | 0.791282576 | 488.5671579 | 503.4628588 | ENSG00000167543 |
| ZNF680 | 0.042966595 | 1.030230099 | 0.798794878 | 0.883907026 | 158.9655772 | 163.925969 | ENSG00000173041 |
| H2AZ2 | 0.042677571 | 1.030023727 | 0.420889938 | 0.59287242 | 3009.367647 | 3099.522279 | ENSG00000105968 |
| TMOD2 | 0.042643246 | 1.029999221 | 0.842472619 | 0.909770449 | 104.7564999 | 107.7727924 | ENSG00000128872 |
| NUTF2 | 0.042631825 | 1.029991067 | 0.437452891 | 0.60829696 | 3332.609821 | 3432.289266 | ENSG00000102898 |
| WDR27 | 0.042549956 | 1.02993262 | 0.732418737 | 0.84020366 | 388.7855179 | 400.1112452 | ENSG00000184465 |
| ZCCHC9 | 0.04251591 | 1.029908314 | 0.648188798 | 0.781171369 | 574.662742 | 591.9045843 | ENSG00000131732 |
| APPBP2 | 0.042330978 | 1.029776304 | 0.728370201 | 0.83711585 | 347.0823775 | 357.2347397 | ENSG00000062725 |
| SLC30A9 | 0.042266666 | 1.0297304 | 0.485207061 | 0.652865272 | 1635.762148 | 1684.673043 | ENSG00000014824 |
| PPP1R26 | 0.042254771 | 1.02972191 | 0.481105772 | 0.649112564 | 1685.953973 | 1735.903387 | ENSG00000196422 |
| SYMPK | 0.042226061 | 1.029701418 | 0.466273501 | 0.63554597 | 2477.649651 | 2550.367607 | ENSG00000125755 |
| CRLF3 | 0.042157044 | 1.02965216 | 0.752410912 | 0.853444012 | 336.7276731 | 346.933713 | ENSG00000176390 |
| KCNJ4 | 0.042156739 | 1.029651942 | 0.895031818 | 0.942161195 | 42.00355852 | 43.25291749 | ENSG00000168135 |
| EAF1 | 0.041964136 | 1.029514491 | 0.704750388 | 0.822374465 | 401.517694 | 413.3717778 | ENSG00000144597 |
| ATG4B | 0.041916678 | 1.029480625 | 0.496449209 | 0.66272242 | 1834.085207 | 1887.530829 | ENSG00000168397 |
| MAFF | 0.041850078 | 1.029433101 | 0.827611828 | 0.90179958 | 132.3402306 | 136.0628758 | ENSG00000185022 |
| ZNF770 | 0.041799043 | 1.029396686 | 0.527303485 | 0.689118264 | 1589.416214 | 1636.407913 | ENSG00000198146 |
| ZNF746 | 0.041791403 | 1.029391235 | 0.647754489 | 0.780794292 | 637.9144189 | 656.9544229 | ENSG00000181220 |
| TRIM35 | 0.041520971 | 1.029198294 | 0.689678885 | 0.811485516 | 494.5859461 | 509.4997106 | ENSG00000104228 |
| C4orf3 | 0.041430646 | 1.029133859 | 0.595313303 | 0.74321699 | 912.4637288 | 939.4812058 | ENSG00000164096 |
| BNIPL | 0.041427777 | 1.029131813 | 0.92849454 | 0.961846113 | 22.73621332 | 23.28517026 | ENSG00000163141 |

| | | | | | | | |
|---|---|---|---|---|---|---|---|
| ARHGEF16 | 0.041365944 | 1.029087706 | 0.659937313 | 0.78990201 | 718.2869526 | 739.2486061 | ENSG00000130762 |
| EPOR | 0.04132119 | 1.029055783 | 0.763710142 | 0.859922296 | 258.4675364 | 265.8912586 | ENSG00000187266 |
| DHX32 | 0.041305761 | 1.029044778 | 0.743139406 | 0.847927029 | 316.8820957 | 326.2856983 | ENSG00000089876 |
| TOP1MT | 0.041192273 | 1.028963833 | 0.459877574 | 0.629630822 | 1994.418297 | 2052.046983 | ENSG00000184428 |
| RUNDC1 | 0.041112504 | 1.028906941 | 0.682510567 | 0.806342221 | 493.4899176 | 507.6877535 | ENSG00000198863 |
| NPAS2 | 0.041067027 | 1.028874508 | 0.927093262 | 0.960744034 | 22.21810475 | 22.88319975 | ENSG00000170485 |
| PPP1R14A | 0.040947326 | 1.028789145 | 0.736260742 | 0.843250331 | 367.85166 | 378.1551391 | ENSG00000167641 |
| SMDT1 | 0.040903836 | 1.028758133 | 0.762361647 | 0.858936955 | 270.9198698 | 278.7197489 | ENSG00000183172 |
| TWISTNB | 0.040890123 | 1.028748354 | 0.687137351 | 0.809846447 | 648.4207846 | 667.7767982 | ENSG00000105849 |
| NA | 0.040873385 | 1.028736419 | 0.929203611 | 0.962113616 | 21.07020039 | 21.66786189 | ENSG00000249936 |
| WIPF2 | 0.040767311 | 1.028660784 | 0.59923692 | 0.745793794 | 847.1962733 | 871.526153 | ENSG00000171475 |
| MCU | 0.040683718 | 1.028601183 | 0.59064156 | 0.739778362 | 915.7766059 | 942.3133006 | ENSG00000156026 |
| RNF217 | 0.040640876 | 1.028570638 | 0.867137435 | 0.925884113 | 84.38192566 | 86.89192744 | ENSG00000146373 |
| PTPN18 | 0.040635884 | 1.028567079 | 0.725955543 | 0.83538585 | 356.1248521 | 366.5210018 | ENSG00000072135 |
| LOC112267983 | 0.04051526 | 1.028481084 | 0.932839043 | 0.964184087 | 21.92364462 | 22.55758382 | ENSG00000259826 |
| SBF1 | 0.040466602 | 1.028446397 | 0.431421648 | 0.602534436 | 3272.075545 | 3365.34206 | ENSG00000100241 |
| MRPS25 | 0.040397002 | 1.028396782 | 0.640205482 | 0.774890447 | 809.9244651 | 833.3749123 | ENSG00000131368 |
| SLC66A3 | 0.040387009 | 1.02838966 | 0.792641813 | 0.879340572 | 202.6381397 | 208.4063038 | ENSG00000162976 |
| SHFL | 0.040115274 | 1.028195978 | 0.837474355 | 0.907512338 | 150.4400982 | 154.8902948 | ENSG00000130813 |
| DPY19L4 | 0.040066306 | 1.02816108 | 0.788170545 | 0.876396218 | 252.1393328 | 259.6607635 | ENSG00000156162 |
| ABCB8 | 0.040000097 | 1.028113896 | 0.708786869 | 0.824432167 | 474.4848051 | 487.8888602 | ENSG00000197150 |
| N4BP2L2-IT2 | 0.039949508 | 1.028077845 | 0.962171398 | 0.980632996 | 7.911715972 | 8.146757875 | ENSG00000281026 |
| FAM118A | 0.039866623 | 1.028018782 | 0.661671152 | 0.791256982 | 667.1426615 | 686.0812937 | ENSG00000100376 |
| NDUFA1 | 0.039837361 | 1.027997931 | 0.664155714 | 0.793032178 | 602.2215179 | 619.0869196 | ENSG00000125356 |
| HSDL2 | 0.039806164 | 1.027975702 | 0.706959284 | 0.823594399 | 424.8468578 | 436.6507689 | ENSG00000119471 |
| MTIF3 | 0.039803031 | 1.027973469 | 0.756490437 | 0.855569973 | 294.2421004 | 302.5536801 | ENSG00000122033 |
| SDC3 | 0.039499795 | 1.027757425 | 0.512544153 | 0.676808081 | 1766.468142 | 1815.03577 | ENSG00000162512 |
| LOC102723996 | 0.039451314 | 1.027722889 | 0.911035201 | 0.951993281 | 34.78064763 | 35.70790986 | ENSG00000277117 |
| RPS3A | 0.039440526 | 1.027715203 | 0.514928642 | 0.678795165 | 6165.684825 | 6335.474012 | ENSG00000145425 |
| CCNB1IP1 | 0.03940241 | 1.027688052 | 0.581306987 | 0.731840572 | 1770.785234 | 1819.284242 | ENSG00000100814 |
| FAM53C | 0.039311099 | 1.02762301 | 0.55385173 | 0.710086055 | 1630.839501 | 1675.247897 | ENSG00000120709 |
| TNPO1 | 0.039057778 | 1.027442586 | 0.433151192 | 0.604144288 | 4270.586938 | 4388.279063 | ENSG00000083312 |
| SIAH1 | 0.038979296 | 1.027386696 | 0.873656374 | 0.929910898 | 116.4556373 | 119.8129167 | ENSG00000196470 |
| TMEM269 | 0.038700691 | 1.027188311 | 0.96194059 | 0.980632996 | 6.714251076 | 6.882489863 | ENSG00000274386 |
| UQCRFS1 | 0.038699627 | 1.027187554 | 0.51586233 | 0.679933055 | 2054.627037 | 2110.512033 | ENSG00000169021 |
| WWC2-AS2 | 0.038665448 | 1.027163219 | 0.929699534 | 0.962375709 | 22.60754361 | 23.23462256 | ENSG00000251359 |
| EXOSC7 | 0.038645281 | 1.027148861 | 0.563704181 | 0.718162551 | 1222.857762 | 1255.963104 | ENSG00000075914 |
| MCOLN2 | 0.038616421 | 1.027128313 | 0.918638814 | 0.956469201 | 34.94663822 | 35.86521116 | ENSG00000153898 |
| NA | 0.038589587 | 1.027109209 | 0.967811936 | 0.983248479 | 6.069424722 | 6.213520375 | ENSG00000278616 |
| NXN | 0.038444934 | 1.02700623 | 0.432314076 | 0.603412984 | 3499.896819 | 3595.027704 | ENSG00000167693 |
| WDR75 | 0.038263983 | 1.026877426 | 0.534967886 | 0.695187975 | 1777.581299 | 1825.387755 | ENSG00000115368 |
| UGT8 | 0.038085384 | 1.026750311 | 0.72565513 | 0.835289284 | 450.4266123 | 462.734685 | ENSG00000174607 |
| SETX | 0.038047911 | 1.026723642 | 0.609665408 | 0.753415245 | 1692.896051 | 1737.76417 | ENSG00000107290 |
| TAF4 | 0.038041269 | 1.026718915 | 0.630718533 | 0.768276265 | 838.5959766 | 860.7828464 | ENSG00000130699 |
| LOC100506100 | 0.037995006 | 1.026685992 | 0.951182168 | 0.974894991 | 12.93033023 | 13.23474988 | ENSG00000223478 |
| C19orf25 | 0.037985143 | 1.026678973 | 0.654284003 | 0.785425805 | 956.7342984 | 982.5572611 | ENSG00000119559 |
| RBPMS | 0.03790015 | 1.02661849 | 0.50205513 | 0.66794107 | 2251.938824 | 2312.198436 | ENSG00000157110 |
| MMP15 | 0.037784165 | 1.026535959 | 0.532737659 | 0.693457638 | 2054.850313 | 2110.220171 | ENSG00000102996 |
| CNOT6 | 0.037725051 | 1.026493898 | 0.659425983 | 0.789485573 | 857.8622954 | 880.3829896 | ENSG00000113300 |
| BMT2 | 0.037720051 | 1.02649034 | 0.88368423 | 0.936493029 | 67.29164379 | 69.22014485 | ENSG00000164603 |
| ABI2 | 0.037700814 | 1.026476653 | 0.527982057 | 0.689522636 | 1704.553563 | 1749.355339 | ENSG00000138443 |
| MAP1S | 0.037688671 | 1.026468013 | 0.525507878 | 0.687805989 | 1951.108103 | 2002.587052 | ENSG00000130479 |
| ID2 | 0.037535909 | 1.02635933 | 0.778601802 | 0.870119764 | 283.8390733 | 291.0069534 | ENSG00000115738 |
| LINC00476 | 0.037514367 | 1.026344005 | 0.923072506 | 0.958820213 | 31.74415795 | 32.60934916 | ENSG00000175611 |
| GRPEL2 | 0.037487355 | 1.026324789 | 0.732876342 | 0.840491526 | 408.9095496 | 419.8539268 | ENSG00000164284 |
| RAP2C | 0.037400684 | 1.026263133 | 0.740922977 | 0.846431059 | 777.0464227 | 798.0821548 | ENSG00000123728 |
| RHCG | 0.037305481 | 1.026195412 | 0.959712726 | 0.979463454 | 8.414994142 | 8.649290732 | ENSG00000140519 |
| MPV17L | 0.037098981 | 1.026048539 | 0.866819406 | 0.925595769 | 94.39287198 | 96.91977601 | ENSG00000156968 |
| SLC7A10 | 0.037087215 | 1.026040171 | 0.872201651 | 0.928970491 | 84.13857765 | 86.18882134 | ENSG00000130876 |
| ZNF628 | 0.037085443 | 1.02603891 | 0.819505657 | 0.896989371 | 305.8370105 | 314.059837 | ENSG00000197483 |
| MTCL1 | 0.037034176 | 1.02600245 | 0.59274204 | 0.741158069 | 1369.113679 | 1404.537444 | ENSG00000168502 |
| ATP1A2 | 0.03702146 | 1.025993407 | 0.607856539 | 0.751797117 | 2021.91737 | 2073.659555 | ENSG00000018625 |
| LOC100506258 | 0.037010925 | 1.025985915 | 0.952197328 | 0.975457419 | 13.56745959 | 13.88919839 | ENSG00000251364 |
| LOC105370689 | 0.036991047 | 1.025971779 | 0.957223353 | 0.978061213 | 9.911149541 | 10.21228329 | ENSG00000287346 |
| FBXO5 | 0.036967462 | 1.025955006 | 0.658392829 | 0.788836474 | 744.9544367 | 764.512109 | ENSG00000112029 |
| SNX16 | 0.036840653 | 1.025864832 | 0.844153893 | 0.910770288 | 126.1637877 | 129.3559579 | ENSG00000104497 |
| IGSF9 | 0.036668808 | 1.025742644 | 0.649060982 | 0.781685316 | 1077.924275 | 1105.629814 | ENSG00000085552 |
| NUTM2D | 0.036663471 | 1.025738849 | 0.94930855 | 0.973739099 | 13.69437754 | 14.01745773 | ENSG00000214562 |
| ATPAF2 | 0.036612317 | 1.02570248 | 0.809335854 | 0.890310719 | 213.8351872 | 219.466382 | ENSG00000171953 |
| PCGF3 | 0.036596154 | 1.025690989 | 0.681329379 | 0.805560496 | 663.0963107 | 679.6077712 | ENSG00000185619 |
| ELP4 | 0.036514998 | 1.025633292 | 0.814824252 | 0.894128251 | 191.7463157 | 196.5326467 | ENSG00000109911 |
| TADA2B | 0.03642114 | 1.025566569 | 0.709148815 | 0.824803437 | 571.1157527 | 585.7798379 | ENSG00000173011 |
| ZBTB48 | 0.036358726 | 1.025522202 | 0.745446295 | 0.849098159 | 369.3768758 | 378.6781116 | ENSG00000204859 |
| LOC105371050 | 0.036183462 | 1.025397625 | 0.966136289 | 0.982484721 | 7.138620227 | 7.307583645 | ENSG00000260874 |
| PHF21A | 0.036159776 | 1.025380791 | 0.704046338 | 0.82190951 | 607.7245369 | 623.1588141 | ENSG00000135365 |
| RCHY1 | 0.03611764 | 1.025350844 | 0.75675108 | 0.855714249 | 366.5493674 | 376.1764364 | ENSG00000163743 |
| CDKL2 | 0.036103913 | 1.025341088 | 0.953138022 | 0.976058264 | 14.49211585 | 14.84237145 | ENSG00000138769 |

| | | | | | | | |
|---|---|---|---|---|---|---|---|
| FBXO45 | 0.03596853 | 1.025244874 | 0.706039288 | 0.823224556 | 564.4840775 | 578.9777007 | ENSG00000174013 |
| USP40 | 0.035924259 | 1.025213413 | 0.844836944 | 0.91120147 | 140.9184244 | 144.2179755 | ENSG00000085982 |
| KHNYN | 0.035893329 | 1.025191433 | 0.716550262 | 0.829919376 | 655.6500764 | 671.466932 | ENSG00000100441 |
| MYO18A | 0.035869253 | 1.025174325 | 0.541115328 | 0.700176533 | 1806.617408 | 1852.347521 | ENSG00000196535 |
| PLEKHA6 | 0.035850912 | 1.025161292 | 0.746458263 | 0.849803857 | 407.8366362 | 418.0562122 | ENSG00000143850 |
| TMEM87B | 0.03580948 | 1.025131852 | 0.813272189 | 0.893085546 | 201.6249624 | 206.5437931 | ENSG00000153214 |
| HSPBAP1 | 0.035779832 | 1.025110785 | 0.831676559 | 0.904389657 | 166.865654 | 170.9988428 | ENSG00000169087 |
| LIN7A | 0.035762844 | 1.025098714 | 0.806988134 | 0.888866512 | 221.7378547 | 227.5300426 | ENSG00000111052 |
| SEMA5A | 0.035743249 | 1.025084791 | 0.815197191 | 0.89428314 | 299.1241094 | 306.5631717 | ENSG00000112902 |
| PRKCE | 0.035664925 | 1.025029141 | 0.922208629 | 0.95841945 | 33.57249468 | 34.46514758 | ENSG00000171132 |
| FLYWCH1 | 0.03543328 | 1.024864571 | 0.720180318 | 0.832069677 | 595.5300327 | 610.9245442 | ENSG00000059122 |
| RAB6A | 0.035430071 | 1.024862291 | 0.667669654 | 0.795831769 | 1147.02765 | 1176.154184 | ENSG00000175582 |
| DAD1 | 0.035405045 | 1.024844513 | 0.576892218 | 0.728827622 | 1391.359516 | 1425.828319 | ENSG00000129562 |
| NA | 0.035384806 | 1.024830137 | 0.971034104 | 0.984662474 | 5.081253418 | 5.159784967 | ENSG00000271943 |
| SEC24D | 0.035355486 | 1.024809309 | 0.534839535 | 0.695114833 | 2141.55016 | 2194.882949 | ENSG00000150961 |
| ANKRD27 | 0.035346942 | 1.02480324 | 0.567658682 | 0.721028596 | 1677.238111 | 1718.517915 | ENSG00000105186 |
| SNIP1 | 0.035330077 | 1.02479126 | 0.760319232 | 0.857637604 | 379.4817617 | 388.4532033 | ENSG00000163877 |
| NA | 0.035310696 | 1.024777493 | 0.956647542 | 0.978042701 | 10.94329957 | 11.27247042 | ENSG00000233221 |
| SPG7 | 0.035214847 | 1.024709412 | 0.551858052 | 0.708528644 | 2271.165844 | 2326.764468 | ENSG00000197912 |
| AFMID | 0.035030108 | 1.024578205 | 0.74751211 | 0.850422569 | 519.4438788 | 531.8007878 | ENSG00000183077 |
| ARHGEF7 | 0.034994242 | 1.024552734 | 0.612489474 | 0.755807232 | 1141.877886 | 1169.602466 | ENSG00000102606 |
| TRAK1 | 0.034934369 | 1.024510215 | 0.612836294 | 0.756041756 | 1197.780722 | 1226.514173 | ENSG00000182606 |
| SLC2A11 | 0.034795468 | 1.024411581 | 0.864086218 | 0.923752947 | 117.9471237 | 120.8112581 | ENSG00000133460 |
| GLT8D1 | 0.034616047 | 1.024284188 | 0.697283202 | 0.817153648 | 645.581072 | 660.9472913 | ENSG00000016864 |
| RPA3 | 0.034435548 | 1.024156045 | 0.719048633 | 0.831405369 | 522.9710819 | 535.5949305 | ENSG00000106399 |
| PLTP | 0.034382446 | 1.02411835 | 0.583061896 | 0.733248614 | 2263.944024 | 2318.98154 | ENSG00000100979 |
| KCTD9 | 0.034276185 | 1.024042922 | 0.781248049 | 0.871562348 | 323.2269198 | 330.9620756 | ENSG00000104756 |
| TRPC5 | 0.034226131 | 1.024007393 | 0.912751952 | 0.952911457 | 52.02307771 | 53.2105722 | ENSG00000072315 |
| C3orf38 | 0.034222544 | 1.024004847 | 0.752155917 | 0.853347831 | 426.1776983 | 436.4598519 | ENSG00000179021 |
| ESRRA | 0.03400728 | 1.023852067 | 0.649138825 | 0.781730262 | 1257.820008 | 1287.9651 | ENSG00000173153 |
| YWHAZ | 0.033872161 | 1.02375618 | 0.422499148 | 0.594054742 | 12058.50692 | 12346.25692 | ENSG00000164924 |
| PAXBP1 | 0.03384049 | 1.023733707 | 0.657869309 | 0.788356121 | 936.1995014 | 958.2868707 | ENSG00000159086 |
| GSTK1 | 0.033737039 | 1.0236603 | 0.754681534 | 0.854475964 | 420.3803185 | 430.1105786 | ENSG00000197448 |
| ZDHHC4 | 0.03370771 | 1.02363949 | 0.768661333 | 0.86353454 | 370.3834321 | 379.0098882 | ENSG00000136247 |
| SNORD94 | 0.033671176 | 1.023613568 | 0.969552656 | 0.984039579 | 5.75331308 | 5.845426008 | ENSG00000208772 |
| CCDC66 | 0.033541617 | 1.023521649 | 0.813501754 | 0.893185108 | 286.6561081 | 293.7790805 | ENSG00000180376 |
| TSPO | 0.033491706 | 1.02348624 | 0.80340883 | 0.886596456 | 308.1560081 | 315.0899161 | ENSG00000100300 |
| C8orf58 | 0.033444661 | 1.023452865 | 0.860241254 | 0.921581275 | 147.1515244 | 150.5143022 | ENSG00000241852 |
| NIBAN2 | 0.03341274 | 1.023430221 | 0.580395173 | 0.731378652 | 2942.67923 | 3011.559 | ENSG00000136830 |
| ZSWIM7 | 0.03340365 | 1.023423772 | 0.755963145 | 0.855174165 | 427.4094178 | 437.2759334 | ENSG00000214941 |
| PAXIP1-AS1 | 0.033403369 | 1.023423573 | 0.846902414 | 0.912357987 | 186.4009017 | 191.0068874 | ENSG00000273344 |
| ADNP2 | 0.033251351 | 1.02331574 | 0.63013974 | 0.767813651 | 1419.528337 | 1452.280372 | ENSG00000101544 |
| POP5 | 0.033250013 | 1.02331479 | 0.745573595 | 0.849098159 | 513.5763772 | 525.4872687 | ENSG00000167272 |
| MTRF1 | 0.033157963 | 1.023249501 | 0.843367076 | 0.91029706 | 173.0658636 | 177.1233351 | ENSG00000120662 |
| RAB8B | 0.033154419 | 1.023246987 | 0.743647624 | 0.84825605 | 578.118858 | 591.9916797 | ENSG00000166128 |
| KRIT1 | 0.033132294 | 1.023231295 | 0.911922468 | 0.952533246 | 51.43674818 | 52.68034235 | ENSG00000001631 |
| POLR3G | 0.032969831 | 1.023116075 | 0.592494527 | 0.741062844 | 3688.519066 | 3775.029592 | ENSG00000113356 |
| RIC3 | 0.032943389 | 1.023097323 | 0.835014694 | 0.906690783 | 192.2957999 | 196.544193 | ENSG00000166405 |
| NA | 0.032829013 | 1.023016215 | 0.962577849 | 0.980727225 | 8.687127059 | 8.897467587 | ENSG00000231731 |
| TPRN | 0.032783085 | 1.022983648 | 0.721069618 | 0.832444884 | 740.8009811 | 757.9302885 | ENSG00000176058 |
| RPL18A | 0.032706345 | 1.022929235 | 0.351860412 | 0.524554808 | 18540.86313 | 18965.46581 | ENSG00000105640 |
| CCR10 | 0.032689932 | 1.022917598 | 0.948376557 | 0.973145327 | 19.91795352 | 20.43048527 | ENSG00000184451 |
| LINC01816 | 0.032589174 | 1.02284616 | 0.948012434 | 0.972984731 | 20.68287672 | 21.21775473 | ENSG00000231327 |
| PITPNA | 0.032523272 | 1.022799437 | 0.552552043 | 0.70909389 | 2235.118847 | 2285.482153 | ENSG00000174238 |
| PROM1 | 0.032522306 | 1.022798752 | 0.620927622 | 0.761882379 | 1280.017598 | 1309.201101 | ENSG00000007062 |
| SETD7 | 0.032347434 | 1.022674784 | 0.843388653 | 0.91029706 | 186.506093 | 190.6917823 | ENSG00000145391 |
| SCO1 | 0.032317224 | 1.02265337 | 0.60584009 | 0.750413593 | 1644.582689 | 1681.949108 | ENSG00000133028 |
| PKP4 | 0.03227778 | 1.02262541 | 0.546370627 | 0.704470255 | 2354.79187 | 2408.217489 | ENSG00000144283 |
| FEM1B | 0.03217519 | 1.022552694 | 0.631824858 | 0.768920547 | 1388.756808 | 1419.99954 | ENSG00000169018 |
| CENPC | 0.032159765 | 1.022541761 | 0.826263678 | 0.901094048 | 265.7945497 | 272.1321728 | ENSG00000145241 |
| KLF16 | 0.032104328 | 1.02250247 | 0.697212661 | 0.817153648 | 850.4754225 | 869.6389258 | ENSG00000129911 |
| RHNO1 | 0.03207545 | 1.022482003 | 0.720983516 | 0.832444884 | 643.5948292 | 657.7879631 | ENSG00000171792 |
| S1PR3 | 0.032065378 | 1.022474864 | 0.812191822 | 0.892246907 | 271.8422476 | 277.7969259 | ENSG00000213694 |
| MFSD8 | 0.031892051 | 1.022352031 | 0.867943492 | 0.926334601 | 124.2752024 | 127.1930931 | ENSG00000164073 |
| SELENOI | 0.031809451 | 1.022293499 | 0.709200847 | 0.824814224 | 1254.823718 | 1283.584108 | ENSG00000138018 |
| KPNA6 | 0.03166977 | 1.022194526 | 0.625227047 | 0.763999174 | 1759.147024 | 1798.891873 | ENSG00000025800 |
| TEX45 | 0.031432664 | 1.022026542 | 0.965219596 | 0.982221968 | 8.306060912 | 8.489578395 | ENSG00000198723 |
| CXorf40A | 0.031386099 | 1.021993555 | 0.875195676 | 0.930984272 | 112.5963281 | 115.1381874 | ENSG00000197620 |
| ZNF655 | 0.031370908 | 1.021982794 | 0.761561832 | 0.858336603 | 517.0862202 | 528.6600028 | ENSG00000197343 |
| WDSUB1 | 0.031350713 | 1.021968489 | 0.885792267 | 0.937037652 | 103.3120202 | 105.6139562 | ENSG00000196151 |
| BCLAF3 | 0.031301969 | 1.021933961 | 0.846532704 | 0.91226537 | 174.1115678 | 178.0504408 | ENSG00000173681 |
| MMS19 | 0.031297106 | 1.021930516 | 0.545501795 | 0.703850921 | 2501.631313 | 2556.354129 | ENSG00000155229 |
| WDR20 | 0.031234479 | 1.021886154 | 0.755349144 | 0.855081294 | 493.5756477 | 504.6225238 | ENSG00000140153 |
| FLT4 | 0.031230272 | 1.021883175 | 0.738908085 | 0.844948184 | 577.6211986 | 590.1052828 | ENSG00000037280 |
| CNOT11 | 0.031159334 | 1.02183293 | 0.673015437 | 0.799651441 | 1116.80922 | 1141.80658 | ENSG00000158435 |
| NPTN | 0.031124859 | 1.021808512 | 0.626818252 | 0.765264913 | 1375.682189 | 1406.081324 | ENSG00000156642 |
| TTBK2 | 0.031066298 | 1.021767036 | 0.846805208 | 0.912319442 | 188.0902058 | 192.2281439 | ENSG00000128881 |

| | | | | | | | |
|---|---|---|---|---|---|---|---|
| UBXN7 | 0.030904998 | 1.021652804 | 0.685200858 | 0.808079699 | 1348.68749 | 1377.828812 | ENSG00000163960 |
| PTPRA | 0.0308976 | 1.021647566 | 0.619744291 | 0.761011274 | 1625.18885 | 1660.787189 | ENSG00000132670 |
| IGFBP4 | 0.030806589 | 1.021583118 | 0.477025562 | 0.645322589 | 4962.792681 | 5069.234904 | ENSG00000141753 |
| KLHL21 | 0.030801507 | 1.021579519 | 0.687494124 | 0.810099598 | 950.532523 | 971.2164627 | ENSG00000162413 |
| C11orf74 | 0.030724132 | 1.021524731 | 0.874120307 | 0.930045494 | 151.8479696 | 155.0908744 | ENSG00000166352 |
| NA | 0.030713422 | 1.021517147 | 0.930710964 | 0.963034317 | 34.39679058 | 35.14648748 | ENSG00000213430 |
| SNRPA | 0.030679792 | 1.021493336 | 0.626727805 | 0.765264913 | 2098.08186 | 2143.580904 | ENSG00000077312 |
| TMEM131L | 0.03064181 | 1.021466443 | 0.724232126 | 0.83450006 | 659.2866884 | 673.28213 | ENSG00000121210 |
| TPBG | 0.030575389 | 1.021419416 | 0.669164368 | 0.796782229 | 1075.244059 | 1098.331039 | ENSG00000146242 |
| CISD1 | 0.030476862 | 1.021349662 | 0.78543513 | 0.874499827 | 396.639664 | 405.3215537 | ENSG00000122873 |
| ITPA | 0.030473548 | 1.021347316 | 0.703504516 | 0.821706565 | 854.5309453 | 872.3456932 | ENSG00000125877 |
| TRMT9B | 0.030440415 | 1.02132386 | 0.926928097 | 0.960672598 | 42.1787977 | 43.07034234 | ENSG00000250305 |
| SESN3 | 0.030433387 | 1.021318885 | 0.793939264 | 0.880229553 | 480.4488143 | 491.1927978 | ENSG00000149212 |
| BICDL1 | 0.030364371 | 1.021270028 | 0.774327799 | 0.867655528 | 655.6828419 | 669.2490005 | ENSG00000135127 |
| TEAD1 | 0.030240185 | 1.021182121 | 0.656781446 | 0.787297016 | 1385.059822 | 1414.650177 | ENSG00000187079 |
| LOC729732 | 0.03004108 | 1.021041199 | 0.885551876 | 0.937037652 | 104.6417847 | 107.0215376 | ENSG00000283674 |
| ZNF275 | 0.030027218 | 1.021031389 | 0.775720098 | 0.868533369 | 443.8883641 | 453.1771349 | ENSG00000063587 |
| CDH1 | 0.02998565 | 1.02100197 | 0.520287429 | 0.68346886 | 4154.10996 | 4242.122235 | ENSG00000039068 |
| TPT1 | 0.029785571 | 1.020860383 | 0.571393623 | 0.723942885 | 5384.273579 | 5496.155918 | ENSG00000133112 |
| RPS24 | 0.029670911 | 1.020779252 | 0.484994579 | 0.652807035 | 11365.06334 | 11599.62621 | ENSG00000138326 |
| PGAP6 | 0.029653798 | 1.020767144 | 0.563921531 | 0.718392013 | 3524.079464 | 3597.066429 | ENSG00000129925 |
| TANK | 0.029628992 | 1.020749593 | 0.825565216 | 0.900739697 | 267.5456539 | 272.8078972 | ENSG00000136560 |
| LCORL | 0.029586148 | 1.02071928 | 0.857379699 | 0.919997317 | 194.2872242 | 198.6013645 | ENSG00000178177 |
| PTK2B | 0.029545699 | 1.020690662 | 0.887870264 | 0.938001384 | 109.6561686 | 111.88072 | ENSG00000120899 |
| ZNF431 | 0.029490546 | 1.020651643 | 0.766582015 | 0.861879803 | 560.6374224 | 572.4022956 | ENSG00000196705 |
| NA | 0.029356151 | 1.020556568 | 0.962282028 | 0.980632996 | 12.48921566 | 12.66876588 | ENSG00000276488 |
| RAPGEF1 | 0.02932132 | 1.020531929 | 0.581216167 | 0.731773991 | 3296.228604 | 3364.249087 | ENSG00000107263 |
| NSUN5 | 0.029283294 | 1.020505031 | 0.661684601 | 0.791256982 | 1570.307028 | 1603.147678 | ENSG00000130305 |
| NME2 | 0.029225981 | 1.02046449 | 0.948449949 | 0.973168872 | 33.4946617 | 33.99516433 | ENSG00000243678 |
| CAMK1D | 0.029207588 | 1.02045148 | 0.802679888 | 0.88621529 | 372.2411926 | 379.8394062 | ENSG00000183049 |
| SEL1L3 | 0.029187385 | 1.02043719 | 0.611970831 | 0.755360502 | 1770.78909 | 1806.979021 | ENSG00000091490 |
| LYRM4-AS1 | 0.02912986 | 1.020396503 | 0.96789271 | 0.983248479 | 9.957919169 | 10.17894019 | ENSG00000272142 |
| ATG16L1 | 0.029123173 | 1.020391774 | 0.768428249 | 0.863402025 | 845.6633705 | 862.3321299 | ENSG00000085978 |
| PDCD2 | 0.029035486 | 1.020329756 | 0.647066145 | 0.780388481 | 1391.749118 | 1420.279194 | ENSG00000071994 |
| SCAF11 | 0.029027654 | 1.020324217 | 0.660312208 | 0.790203648 | 1484.92325 | 1514.989586 | ENSG00000139218 |
| SRSF10 | 0.028973776 | 1.020286114 | 0.575716664 | 0.727938007 | 3732.498977 | 3808.532728 | ENSG00000188529 |
| LOC554206 | 0.028969696 | 1.020283228 | 0.956771519 | 0.978044405 | 15.46633264 | 15.7423132 | ENSG00000262587 |
| SLC47A1 | 0.028924846 | 1.02025151 | 0.931323751 | 0.963382625 | 40.34053665 | 41.24626949 | ENSG00000142494 |
| SLU7 | 0.028859437 | 1.020205255 | 0.790770284 | 0.87802173 | 513.6414949 | 524.2829257 | ENSG00000164609 |
| NA | 0.028823894 | 1.020180121 | 0.959227006 | 0.979226764 | 15.86905029 | 16.16095526 | ENSG00000161132 |
| WDR91 | 0.028809357 | 1.020169841 | 0.835986446 | 0.906924912 | 258.2415726 | 263.7622515 | ENSG00000105875 |
| RELN | 0.028781466 | 1.020150119 | 0.976904861 | 0.988018857 | 6.572702892 | 6.678408981 | ENSG00000189056 |
| PRDX3 | 0.028547754 | 1.019984871 | 0.54689845 | 0.70479907 | 4308.429026 | 4393.831466 | ENSG00000165672 |
| SPATA3-AS1 | 0.028545177 | 1.019983049 | 0.970335909 | 0.98427465 | 11.18070235 | 11.33781166 | ENSG00000238062 |
| PRKD3 | 0.028532421 | 1.019974031 | 0.697041221 | 0.817125802 | 1097.761818 | 1119.713043 | ENSG00000115825 |
| UBE2A | 0.028501322 | 1.019952044 | 0.718129651 | 0.831037781 | 822.2168781 | 839.0119684 | ENSG00000077721 |
| NA | 0.028461916 | 1.019924186 | 0.966884829 | 0.98288324 | 9.400374543 | 9.596012069 | ENSG00000250421 |
| TSPAN12 | 0.028434957 | 1.019905127 | 0.822377363 | 0.898735651 | 332.4561734 | 339.3852162 | ENSG00000106025 |
| STAM | 0.028216113 | 1.019750428 | 0.794606765 | 0.880471663 | 393.5980809 | 401.2719826 | ENSG00000136738 |
| SEC22C | 0.028142261 | 1.019698228 | 0.758187393 | 0.856402715 | 697.7188521 | 711.3015794 | ENSG00000093183 |
| DENND5A | 0.028115563 | 1.019679358 | 0.660965223 | 0.790739852 | 1721.578564 | 1754.49366 | ENSG00000184014 |
| RASSF3 | 0.028055691 | 1.019637043 | 0.734697734 | 0.841995884 | 771.2301808 | 786.5793004 | ENSG00000153179 |
| TCEAL8 | 0.028049964 | 1.019632995 | 0.622244524 | 0.762199274 | 2133.718192 | 2176.422999 | ENSG00000180964 |
| TMEM140 | 0.027915099 | 1.019537683 | 0.867687132 | 0.926302413 | 174.7632153 | 178.2687822 | ENSG00000146859 |
| UCHL5 | 0.027911687 | 1.019535271 | 0.73531577 | 0.842388296 | 735.2859594 | 749.5676003 | ENSG00000116750 |
| SLC9A6 | 0.027886773 | 1.019517665 | 0.74755533 | 0.850422569 | 679.5505023 | 692.4470023 | ENSG00000198689 |
| SGSH | 0.027781884 | 1.019443545 | 0.819757343 | 0.897041177 | 346.2085963 | 352.9120089 | ENSG00000181523 |
| MSH3 | 0.027717841 | 1.019398292 | 0.792777414 | 0.879440431 | 423.8364345 | 432.2489694 | ENSG00000113318 |
| CDK17 | 0.027631746 | 1.01933746 | 0.818410451 | 0.89619415 | 319.5249161 | 325.7845225 | ENSG00000059758 |
| CRYBG2 | 0.02756567 | 1.019290775 | 0.827167097 | 0.90146774 | 302.4889888 | 308.5475205 | ENSG00000176092 |
| MMP17 | 0.027561752 | 1.019288007 | 0.725147339 | 0.835021231 | 1210.927905 | 1234.132331 | ENSG00000198598 |
| NA | 0.027537188 | 1.019270651 | 0.933459618 | 0.964413047 | 40.54983024 | 41.41270612 | ENSG00000227615 |
| SPACA6 | 0.027526471 | 1.01926308 | 0.918471342 | 0.956469201 | 72.9117814 | 74.21027022 | ENSG00000182310 |
| MAPRE1 | 0.027431499 | 1.019195985 | 0.558678784 | 0.714203884 | 4654.112372 | 4744.411671 | ENSG00000101367 |
| MYO19 | 0.02723018 | 1.019053772 | 0.697629982 | 0.817418609 | 1423.909897 | 1450.32621 | ENSG00000278259 |
| AAAS | 0.02713511 | 1.018986621 | 0.67456184 | 0.80030617 | 1401.632032 | 1427.763186 | ENSG00000094914 |
| DHX36 | 0.027130806 | 1.018983582 | 0.684396156 | 0.807602535 | 1682.625841 | 1715.23541 | ENSG00000174953 |
| LINC00574 | 0.027082296 | 1.018949319 | 0.973636479 | 0.986264509 | 7.005244592 | 7.107543445 | ENSG00000231690 |
| TRAPPC3 | 0.027079926 | 1.018947645 | 0.748157541 | 0.850784433 | 965.7888238 | 984.1798577 | ENSG00000054116 |
| KCNQ3 | 0.027036259 | 1.018916805 | 0.908844572 | 0.950942162 | 78.97841522 | 80.42351802 | ENSG00000184156 |
| AAMDC | 0.027004322 | 1.018894249 | 0.933138762 | 0.964383363 | 51.07924855 | 52.12402673 | ENSG00000087884 |
| MYADM | 0.026965931 | 1.018867136 | 0.747287613 | 0.850346567 | 810.2070859 | 825.8331519 | ENSG00000179820 |
| LRRC8B | 0.026928411 | 1.018840639 | 0.80991572 | 0.890464161 | 410.1836082 | 418.1882213 | ENSG00000197147 |
| PNPLA2 | 0.026908096 | 1.018826292 | 0.792922782 | 0.879500539 | 537.5543975 | 547.5813025 | ENSG00000177666 |
| SPTLC1 | 0.026872869 | 1.018801416 | 0.699280089 | 0.81841198 | 1105.818824 | 1126.77463 | ENSG00000090054 |
| SHROOM4 | 0.026866936 | 1.018797225 | 0.872243145 | 0.928970491 | 166.0443492 | 169.1981537 | ENSG00000158352 |
| SSH2 | 0.026823628 | 1.018766643 | 0.842034427 | 0.909602763 | 262.2655579 | 267.4335389 | ENSG00000141298 |

| | | | | | | | |
|---|---|---|---|---|---|---|---|
| RPL36A | 0.026817493 | 1.018762311 | 0.946953772 | 0.9726684 | 28.32034459 | 28.86951597 | ENSG00000241343 |
| PNMA3 | 0.02680408 | 1.018752839 | 0.829287759 | 0.90286078 | 343.7972334 | 350.3113768 | ENSG00000183837 |
| CDCA3 | 0.026764828 | 1.018725122 | 0.761528961 | 0.858336603 | 622.9744795 | 634.6616996 | ENSG00000111665 |
| EXOSC2 | 0.026681594 | 1.01866635 | 0.707062778 | 0.823594399 | 1160.559126 | 1181.801267 | ENSG00000130713 |
| SYT3 | 0.026597363 | 1.018606877 | 0.840846546 | 0.909051234 | 338.8170672 | 345.1419598 | ENSG00000213023 |
| TDP1 | 0.026557749 | 1.018578908 | 0.715350912 | 0.829067855 | 1147.651649 | 1169.116642 | ENSG00000042088 |
| SP3 | 0.026360651 | 1.018439762 | 0.706041692 | 0.823224556 | 1306.580945 | 1331.153662 | ENSG00000172845 |
| IGSF8 | 0.026312671 | 1.018405892 | 0.67770701 | 0.802753649 | 2695.25101 | 2744.332517 | ENSG00000162729 |
| GORASP2 | 0.026276413 | 1.018380297 | 0.638997863 | 0.774449191 | 2251.588395 | 2293.377178 | ENSG00000115806 |
| ZNF350-AS1 | 0.026274478 | 1.018378931 | 0.967893475 | 0.983248479 | 10.2817278 | 10.49003514 | ENSG00000269235 |
| ST3GAL2 | 0.026255922 | 1.018365833 | 0.700725017 | 0.819551735 | 1283.569918 | 1306.537878 | ENSG00000157350 |
| NFRKB | 0.026044682 | 1.018216735 | 0.74806757 | 0.850782425 | 935.4430261 | 952.7918077 | ENSG00000170322 |
| EDEM2 | 0.026037313 | 1.018211534 | 0.804614491 | 0.887163817 | 434.2567563 | 442.0170763 | ENSG00000088298 |
| ROR1 | 0.025922042 | 1.018130182 | 0.661940315 | 0.791399412 | 1671.577448 | 1702.193201 | ENSG00000185483 |
| PDIK1L | 0.025916218 | 1.018126072 | 0.925403829 | 0.960029077 | 58.20546587 | 59.29851685 | ENSG00000175087 |
| ZNF230 | 0.025790048 | 1.018037036 | 0.931596449 | 0.963467756 | 58.02205414 | 59.09013482 | ENSG00000159882 |
| DNAJC16 | 0.025694346 | 1.017969506 | 0.833216747 | 0.90535002 | 351.4855011 | 357.8792473 | ENSG00000116138 |
| MON2 | 0.025682532 | 1.01796117 | 0.822325733 | 0.898735651 | 502.7077051 | 511.7015156 | ENSG00000061987 |
| DIDO1 | 0.025665867 | 1.017949412 | 0.6791935 | 0.803872602 | 2971.715899 | 3024.701845 | ENSG00000101191 |
| RALA | 0.025546579 | 1.017865247 | 0.719280547 | 0.831573836 | 1857.31578 | 1891.367686 | ENSG00000006451 |
| EXOC3-AS1 | 0.025512164 | 1.017840967 | 0.957781389 | 0.978372294 | 22.5153558 | 22.96977416 | ENSG00000221990 |
| AHDC1 | 0.025445289 | 1.017793787 | 0.777386851 | 0.869419662 | 922.334115 | 938.4837679 | ENSG00000126705 |
| CORO2B | 0.025429527 | 1.017782667 | 0.886970622 | 0.937513028 | 148.1870289 | 150.8022739 | ENSG00000103647 |
| LUC7L3 | 0.025363458 | 1.017736058 | 0.598518999 | 0.74523821 | 3096.023541 | 3151.477428 | ENSG00000108848 |
| SCAMP1 | 0.025202242 | 1.017622336 | 0.767671631 | 0.862752895 | 827.9091548 | 842.800465 | ENSG00000085365 |
| UBE2D3 | 0.025077766 | 1.017534539 | 0.613994695 | 0.75677096 | 6159.08781 | 6266.731654 | ENSG00000109332 |
| ALG8 | 0.025025713 | 1.017497827 | 0.751543953 | 0.852976078 | 955.8154979 | 972.0529591 | ENSG00000159063 |
| NA | 0.024921329 | 1.01742421 | 0.95640719 | 0.977900626 | 22.15805633 | 22.59737864 | ENSG00000226084 |
| NA | 0.024846555 | 1.017371479 | 0.972105706 | 0.985334762 | 8.382179029 | 8.508400521 | ENSG00000224837 |
| STIMATE-MUSTN1 | 0.024772205 | 1.017319049 | 0.971915143 | 0.98524514 | 8.692708865 | 8.830248342 | ENSG00000248592 |
| TPST2 | 0.024759292 | 1.017309944 | 0.714773704 | 0.828621912 | 2688.710325 | 2735.745787 | ENSG00000128294 |
| TMEM86A | 0.024704198 | 1.017271095 | 0.950678995 | 0.974574795 | 28.66022291 | 29.15963822 | ENSG00000151117 |
| PI4KB | 0.024696477 | 1.017265651 | 0.7432872 | 0.847935038 | 962.3160249 | 978.8346083 | ENSG00000143393 |
| PLS3 | 0.024686575 | 1.017258669 | 0.55669088 | 0.712805242 | 6774.382462 | 6892.432732 | ENSG00000102024 |
| CSNK2A1 | 0.024591905 | 1.017191918 | 0.607078494 | 0.751272767 | 4257.609336 | 4331.304705 | ENSG00000101266 |
| DYM | 0.024474147 | 1.017108895 | 0.777606271 | 0.869490355 | 918.9465465 | 935.4385647 | ENSG00000141627 |
| GOLGA2P5 | 0.024427934 | 1.017076315 | 0.896984602 | 0.943140937 | 154.2666661 | 156.5721509 | ENSG00000238105 |
| PPM1F | 0.024387668 | 1.017047929 | 0.775738716 | 0.868533369 | 775.3728234 | 788.4967025 | ENSG00000100034 |
| UBXN1 | 0.024382252 | 1.01704411 | 0.704506953 | 0.822230239 | 1895.199234 | 1927.233558 | ENSG00000162191 |
| ZNF697 | 0.024274567 | 1.016968199 | 0.777146826 | 0.869299514 | 680.8328215 | 692.5058676 | ENSG00000143067 |
| VAMP5 | 0.024231223 | 1.016937647 | 0.950493117 | 0.974436027 | 32.28216625 | 32.87741422 | ENSG00000168899 |
| NA | 0.024170906 | 1.016895131 | 0.978349302 | 0.988805804 | 6.118309531 | 6.171574976 | ENSG00000272311 |
| SCAF1 | 0.024123771 | 1.016861907 | 0.700462048 | 0.819293844 | 3642.244119 | 3703.788801 | ENSG00000126461 |
| DCTN2 | 0.024103094 | 1.016847334 | 0.718531826 | 0.831206371 | 1785.403128 | 1814.418426 | ENSG00000175203 |
| CTDSPL | 0.024074234 | 1.016826993 | 0.74993125 | 0.8520254 | 932.2927468 | 947.6930445 | ENSG00000144677 |
| SLC24A2 | 0.024001318 | 1.016775602 | 0.919347765 | 0.957052384 | 101.7451283 | 103.5054009 | ENSG00000155886 |
| JKAMP | 0.023991986 | 1.016769026 | 0.817207309 | 0.895520689 | 451.8542585 | 459.5358466 | ENSG00000050130 |
| CUL4B | 0.023932396 | 1.016727029 | 0.736447982 | 0.843334799 | 1581.989454 | 1609.055143 | ENSG00000158290 |
| RBSN | 0.023899679 | 1.016703972 | 0.792044138 | 0.878982065 | 671.4137865 | 682.672983 | ENSG00000131381 |
| SLC4A7 | 0.023828582 | 1.01665387 | 0.778663622 | 0.870138442 | 1246.872201 | 1267.462166 | ENSG00000033867 |
| KCTD11 | 0.023821697 | 1.016649017 | 0.880285211 | 0.934285982 | 180.1972624 | 183.2820066 | ENSG00000213859 |
| RPL31 | 0.023652429 | 1.016529744 | 0.626786574 | 0.765264913 | 11242.50131 | 11427.01329 | ENSG00000071082 |
| METTL26 | 0.02354683 | 1.016455341 | 0.754164015 | 0.854240973 | 1162.707906 | 1182.251604 | ENSG00000130731 |
| OST4 | 0.023529695 | 1.016443268 | 0.783809205 | 0.873560771 | 775.5710783 | 788.1594976 | ENSG00000228474 |
| SIRT3 | 0.02352677 | 1.016441208 | 0.845936433 | 0.912024708 | 382.7668688 | 388.8764191 | ENSG00000142082 |
| CNTLN | 0.023311186 | 1.016289331 | 0.883968469 | 0.936493029 | 206.1459314 | 209.7560801 | ENSG00000044459 |
| NA | 0.023293327 | 1.01627675 | 0.947586482 | 0.972984731 | 37.36429555 | 37.94013237 | ENSG00000280018 |
| PLA2G3 | 0.023140056 | 1.016168787 | 0.772260109 | 0.866042888 | 1053.908271 | 1070.448918 | ENSG00000100078 |
| GRASP | 0.023025958 | 1.016088425 | 0.904711678 | 0.948315202 | 130.1763212 | 132.3042205 | ENSG00000161835 |
| DTX2 | 0.022951576 | 1.016036039 | 0.811102149 | 0.891616136 | 645.0171449 | 655.0075369 | ENSG00000091073 |
| IRGQ | 0.022938912 | 1.01602712 | 0.831506248 | 0.904389657 | 426.4079677 | 433.2577848 | ENSG00000167378 |
| AATK | 0.022899847 | 1.015999609 | 0.907981471 | 0.950348357 | 113.6906417 | 115.435034 | ENSG00000181409 |
| SLC26A6 | 0.022788178 | 1.015920971 | 0.767268462 | 0.862450524 | 1401.604987 | 1423.225416 | ENSG00000225697 |
| FAM174A | 0.022741143 | 1.01588785 | 0.925327347 | 0.960029077 | 83.28270594 | 84.79172027 | ENSG00000174132 |
| RAB2B | 0.022736813 | 1.015884801 | 0.892618619 | 0.940596809 | 244.7331592 | 248.5732793 | ENSG00000129472 |
| TCF24 | 0.022679746 | 1.015844618 | 0.945100599 | 0.971385564 | 42.40416099 | 43.11605585 | ENSG00000261787 |
| TCF20 | 0.022552135 | 1.015754767 | 0.686334511 | 0.809048522 | 2277.157511 | 2312.502323 | ENSG00000100207 |
| SAT2 | 0.022536133 | 1.0157435 | 0.826866704 | 0.901242194 | 548.4444556 | 556.747877 | ENSG00000141504 |
| RPL32 | 0.022479688 | 1.01570376 | 0.607101299 | 0.751272767 | 8696.883325 | 8832.206949 | ENSG00000144713 |
| CHURC1 | 0.022434684 | 1.015672077 | 0.922883644 | 0.958789508 | 89.07999775 | 90.60327013 | ENSG00000258289 |
| FBXO6 | 0.022372447 | 1.015628262 | 0.957422552 | 0.97816113 | 25.8672814 | 26.36033511 | ENSG00000116663 |
| FLCN | 0.022349483 | 1.015612096 | 0.836076545 | 0.906924912 | 407.006283 | 413.317753 | ENSG00000154803 |
| PSTK | 0.022343909 | 1.015608172 | 0.921067223 | 0.957860259 | 90.47455349 | 91.88742635 | ENSG00000179988 |
| TIAL1 | 0.022162685 | 1.015480605 | 0.717665039 | 0.830701804 | 1828.72788 | 1856.979444 | ENSG00000151923 |
| NA | 0.022127055 | 1.015455526 | 0.962486812 | 0.980727225 | 20.45472252 | 20.77383882 | ENSG00000235381 |
| LOC100129503 | 0.022126909 | 1.015455423 | 0.98650412 | 0.992213294 | 2.952674577 | 3.035370027 | ENSG00000280351 |
| SPTBN1 | 0.022109899 | 1.01544345 | 0.621746555 | 0.762165785 | 5896.369198 | 5986.401559 | ENSG00000115306 |

| | | | | | | | |
|---|---|---|---|---|---|---|---|
| VGLL3 | 0.022029204 | 1.015386655 | 0.927754736 | 0.961187788 | 76.06047077 | 77.31473749 | ENSG00000206538 |
| TXNDC9 | 0.021995423 | 1.015362879 | 0.869870397 | 0.927365008 | 287.6432402 | 291.8913018 | ENSG00000115514 |
| STX3 | 0.021978343 | 1.015350859 | 0.720513523 | 0.832149671 | 1614.818051 | 1639.609762 | ENSG00000166900 |
| CCDC61 | 0.021967815 | 1.01534345 | 0.854848097 | 0.918046799 | 379.7243218 | 385.4008891 | ENSG00000104983 |
| MAP3K11 | 0.021960156 | 1.015338059 | 0.766031297 | 0.861545881 | 1601.244699 | 1625.837938 | ENSG00000173327 |
| PML | 0.021928398 | 1.015315709 | 0.816694506 | 0.895136415 | 860.7357742 | 873.9017373 | ENSG00000140464 |
| VAPB | 0.02187676 | 1.015279368 | 0.74485777 | 0.848883514 | 1237.033678 | 1255.758193 | ENSG00000124164 |
| RNF111 | 0.021872744 | 1.015276542 | 0.825287353 | 0.900593514 | 555.7595543 | 563.7739712 | ENSG00000157450 |
| SPAG7 | 0.021762623 | 1.015199049 | 0.734638637 | 0.841995884 | 1469.836783 | 1492.516202 | ENSG00000091640 |
| SYNJ2 | 0.021669399 | 1.015133451 | 0.833015882 | 0.905233741 | 573.7468091 | 582.898764 | ENSG00000078269 |
| DIMT1 | 0.021603855 | 1.015087333 | 0.787289092 | 0.875923427 | 1108.355992 | 1125.773351 | ENSG00000086189 |
| CHMP2A | 0.021598312 | 1.015083433 | 0.787312389 | 0.875923427 | 1110.802045 | 1127.248722 | ENSG00000130724 |
| UQCRC2 | 0.021556266 | 1.015053849 | 0.679443104 | 0.803970859 | 3529.640626 | 3582.112851 | ENSG00000140740 |
| VMA21 | 0.021474081 | 1.014996028 | 0.85767615 | 0.920264231 | 580.417496 | 589.6833873 | ENSG00000160131 |
| RTKN | 0.021444916 | 1.014975509 | 0.74456428 | 0.848722265 | 1388.454902 | 1409.08234 | ENSG00000114993 |
| MAPK8IP3 | 0.021400005 | 1.014943913 | 0.766329416 | 0.861746474 | 1295.025107 | 1313.865404 | ENSG00000138834 |
| CEP97 | 0.021390232 | 1.014937038 | 0.897966944 | 0.943860966 | 185.0238921 | 187.9177409 | ENSG00000182504 |
| BUB1 | 0.021362607 | 1.014917604 | 0.672855589 | 0.799564127 | 3676.246834 | 3731.577337 | ENSG00000169679 |
| KLHL2 | 0.021237157 | 1.014829355 | 0.869651272 | 0.927285136 | 277.1509795 | 281.1197346 | ENSG00000109466 |
| EFNA5 | 0.021201856 | 1.014804524 | 0.916660604 | 0.95533764 | 122.6132196 | 124.6863354 | ENSG00000184349 |
| ARHGEF4 | 0.021119425 | 1.014746543 | 0.815362393 | 0.894413506 | 626.5372503 | 635.4048973 | ENSG00000136002 |
| C1orf198 | 0.020956178 | 1.014631727 | 0.790255982 | 0.87763484 | 815.8580115 | 827.8263755 | ENSG00000119280 |
| FERMT3 | 0.020933114 | 1.014615506 | 0.955654908 | 0.977429542 | 29.74895474 | 30.16550955 | ENSG00000149781 |
| CASP8AP2 | 0.020851691 | 1.014558244 | 0.832783593 | 0.905185277 | 515.2314508 | 522.5760868 | ENSG00000118412 |
| C11orf95 | 0.020846585 | 1.014554654 | 0.775241312 | 0.868326084 | 1195.415052 | 1212.222217 | ENSG00000188070 |
| NA | 0.020770914 | 1.014501441 | 0.923992355 | 0.959186142 | 98.92515364 | 100.1551983 | ENSG00000234742 |
| ELF4 | 0.020743435 | 1.014482118 | 0.89896319 | 0.944354114 | 163.0483716 | 165.3728122 | ENSG00000102034 |
| NA | 0.020736318 | 1.014477114 | 0.965790292 | 0.982433028 | 20.69750695 | 20.99835937 | ENSG00000267523 |
| PMM2 | 0.020654681 | 1.014419709 | 0.868678596 | 0.926811503 | 299.1184027 | 303.2718708 | ENSG00000140650 |
| GABPB2 | 0.020508205 | 1.014316721 | 0.949871867 | 0.974057952 | 47.17979065 | 47.92037732 | ENSG00000143458 |
| MFSD14B | 0.020491139 | 1.014304722 | 0.791910566 | 0.878982065 | 1197.25071 | 1214.209691 | ENSG00000148110 |
| GLS | 0.020394073 | 1.014236481 | 0.808454725 | 0.889922973 | 924.4393417 | 938.1203977 | ENSG00000115419 |
| RNF144B | 0.02034145 | 1.014199487 | 0.926811526 | 0.960672598 | 106.706485 | 108.1201746 | ENSG00000137393 |
| GMDS | 0.020320467 | 1.014184737 | 0.802697619 | 0.88621529 | 788.7946677 | 800.3034008 | ENSG00000112699 |
| FKBP5 | 0.020258179 | 1.01414095 | 0.724449951 | 0.834603608 | 2018.760511 | 2046.921062 | ENSG00000096060 |
| HSD17B4 | 0.020201287 | 1.014100959 | 0.639756151 | 0.774742503 | 7549.208738 | 7656.077116 | ENSG00000133835 |
| RAC1 | 0.020154889 | 1.014068345 | 0.695047444 | 0.815746479 | 5618.004757 | 5697.312468 | ENSG00000136238 |
| TTC13 | 0.02013011 | 1.014050928 | 0.908488891 | 0.950776282 | 142.2344346 | 144.159619 | ENSG00000143643 |
| ARSK | 0.020115426 | 1.014040607 | 0.864606849 | 0.924051564 | 380.3005269 | 385.786709 | ENSG00000164291 |
| NA | 0.020079267 | 1.014015192 | 0.964124094 | 0.981782342 | 27.82795463 | 28.06180684 | ENSG00000214106 |
| NA | 0.020072072 | 1.014010135 | 0.970407652 | 0.984285926 | 15.92052584 | 16.21742164 | ENSG00000286018 |
| TLCD3A | 0.019952613 | 1.013926175 | 0.840953167 | 0.90907448 | 505.2393282 | 512.1527621 | ENSG00000167695 |
| NA | 0.019903784 | 1.01389186 | 0.96343314 | 0.981487825 | 27.04608586 | 27.49042509 | ENSG00000243176 |
| NIPBL | 0.019880927 | 1.013875796 | 0.777632942 | 0.869490355 | 1312.102286 | 1330.598407 | ENSG00000164190 |
| ENTPD3-AS1 | 0.019878399 | 1.01387402 | 0.965377016 | 0.982271747 | 20.40304681 | 20.67489401 | ENSG00000223797 |
| LHX8 | 0.019835076 | 1.013843574 | 0.977889846 | 0.988393222 | 9.370350333 | 9.484696852 | ENSG00000162624 |
| LRPPRC | 0.019798165 | 1.013817635 | 0.669464705 | 0.797003432 | 6294.684526 | 6382.258836 | ENSG00000138095 |
| SMAGP | 0.019770447 | 1.013798157 | 0.925377148 | 0.960029077 | 99.88050983 | 101.1693994 | ENSG00000170545 |
| PANK1 | 0.019725688 | 1.013766706 | 0.836973587 | 0.907285764 | 674.5756666 | 684.3060778 | ENSG00000152782 |
| FAM219B | 0.019714726 | 1.013759002 | 0.890441934 | 0.939388982 | 296.4150071 | 299.9802546 | ENSG00000178761 |
| PTGER1 | 0.019626408 | 1.013696944 | 0.974357869 | 0.986590643 | 11.68589554 | 11.86131722 | ENSG00000160951 |
| KIAA1671 | 0.019563043 | 1.013652423 | 0.806738392 | 0.88864217 | 875.2359121 | 887.6569576 | ENSG00000197077 |
| PPP1R9B | 0.019422772 | 1.013553872 | 0.83782481 | 0.907638723 | 638.2446483 | 647.2110994 | ENSG00000108819 |
| LTO1 | 0.019355823 | 1.013506838 | 0.872603048 | 0.9291999 | 340.5433287 | 345.13386 | ENSG00000149716 |
| PLEKHB2 | 0.019302072 | 1.013469078 | 0.707230191 | 0.823594399 | 2633.29528 | 2669.279104 | ENSG00000115762 |
| PPP6R1 | 0.019258069 | 1.013438168 | 0.774843159 | 0.86807286 | 2728.893229 | 2766.456847 | ENSG00000105063 |
| GLB1L2 | 0.019257202 | 1.013437558 | 0.897898178 | 0.943840113 | 211.6846814 | 214.3630755 | ENSG00000149328 |
| ATG14 | 0.019243194 | 1.013427719 | 0.873554297 | 0.929897741 | 368.6459559 | 373.4005476 | ENSG00000126775 |
| TMEM175 | 0.019193643 | 1.013392912 | 0.938282868 | 0.967367581 | 71.75809507 | 72.82553955 | ENSG00000127419 |
| SYNGR2 | 0.019120233 | 1.013341348 | 0.804454675 | 0.887069454 | 1112.751281 | 1127.492585 | ENSG00000108639 |
| NA | 0.019099743 | 1.013326956 | 0.948229413 | 0.973046099 | 53.49343926 | 54.20192255 | ENSG00000249855 |
| RPS25 | 0.019065064 | 1.013302598 | 0.657816476 | 0.78834178 | 10255.73814 | 10390.8343 | ENSG00000118181 |
| NA | 0.018956774 | 1.013226541 | 0.987900708 | 0.993048438 | 2.952674577 | 3.03106888 | ENSG00000229896 |
| ELP3 | 0.018837502 | 1.013142778 | 0.829686768 | 0.903182315 | 672.4177762 | 680.9005176 | ENSG00000134014 |
| CCDC148 | 0.018814935 | 1.013126931 | 0.977645615 | 0.988353504 | 9.661343848 | 9.772668532 | ENSG00000153237 |
| EARS2 | 0.01877315 | 1.013097588 | 0.755546437 | 0.855091825 | 1600.418762 | 1621.675021 | ENSG00000103356 |
| RPL15 | 0.018764854 | 1.013091762 | 0.587824704 | 0.737351347 | 19273.8989 | 19525.59586 | ENSG00000174748 |
| MRPL14 | 0.018763399 | 1.01309074 | 0.781714464 | 0.871981827 | 1680.668833 | 1702.384161 | ENSG00000180992 |
| ACTB | 0.018753697 | 1.013083928 | 0.605558004 | 0.750277421 | 97224.52851 | 98497.25472 | ENSG00000075624 |
| THRAP3 | 0.018509412 | 1.012912401 | 0.668171583 | 0.796028624 | 4789.927314 | 4851.212936 | ENSG00000054118 |
| ZIC5 | 0.018463819 | 1.012880391 | 0.8462714 | 0.912075406 | 884.0122743 | 895.7128699 | ENSG00000139800 |
| DHX15 | 0.018452129 | 1.012872184 | 0.660532828 | 0.790418633 | 6067.948972 | 6146.370408 | ENSG00000109606 |
| LRIG3 | 0.018388895 | 1.01282779 | 0.904989264 | 0.948451561 | 261.9132766 | 265.6489578 | ENSG00000139263 |
| KCNG3 | 0.018386248 | 1.012825932 | 0.912243345 | 0.952535122 | 164.7950084 | 167.0688861 | ENSG00000171126 |
| NA | 0.01835 | 1.012800485 | 0.980688991 | 0.989984072 | 7.789028387 | 7.843732179 | ENSG00000228929 |
| ATAD2B | 0.018324535 | 1.012782608 | 0.876016747 | 0.931600831 | 346.4363871 | 350.9515769 | ENSG00000119778 |
| FBXO30 | 0.018288691 | 1.012757446 | 0.817351698 | 0.895589009 | 990.9338477 | 1003.735724 | ENSG00000118496 |

| | | | | | | | |
|---|---|---|---|---|---|---|---|
| HDLBP | 0.018285999 | 1.012755555 | 0.637971459 | 0.773642659 | 10482.73099 | 10616.11145 | ENSG00000115677 |
| SMARCC2 | 0.01825902 | 1.012736617 | 0.789161226 | 0.877042816 | 2230.73581 | 2258.361314 | ENSG00000139613 |
| PRX | 0.018217994 | 1.012707818 | 0.87807355 | 0.933092643 | 333.8201413 | 338.2596878 | ENSG00000105227 |
| BORA | 0.01820182 | 1.012696465 | 0.917046815 | 0.955518152 | 164.5165299 | 166.6118063 | ENSG00000136122 |
| MYLK | 0.017951857 | 1.012521019 | 0.867485457 | 0.926153187 | 421.4690503 | 426.6366662 | ENSG00000065534 |
| NA | 0.017930027 | 1.012505698 | 0.982333787 | 0.990351054 | 7.451940965 | 7.500911659 | ENSG00000237950 |
| IL3RA | 0.017925621 | 1.012502606 | 0.965825443 | 0.982433028 | 26.3607474 | 26.74563406 | ENSG00000185291 |
| MPHOSPH8 | 0.017922625 | 1.012500503 | 0.814270971 | 0.893724471 | 908.7757919 | 920.4080331 | ENSG00000196199 |
| ERRFI1 | 0.017882192 | 1.012472127 | 0.777466127 | 0.869419662 | 1738.279094 | 1759.532649 | ENSG00000116285 |
| NA | 0.017841693 | 1.012443706 | 0.96126267 | 0.980267388 | 35.4666618 | 35.93484144 | ENSG00000247679 |
| USP4 | 0.017795852 | 1.012411536 | 0.814261591 | 0.893724471 | 963.6143754 | 975.1047039 | ENSG00000114316 |
| SPATA13 | 0.017678505 | 1.012329191 | 0.869615884 | 0.927285136 | 394.1053012 | 398.890528 | ENSG00000182957 |
| PIAS1 | 0.017675794 | 1.012327289 | 0.858745825 | 0.920821105 | 524.5974217 | 531.2264557 | ENSG00000033800 |
| ZC3H13 | 0.017658488 | 1.012315146 | 0.73926516 | 0.84515611 | 2369.885834 | 2398.460977 | ENSG00000123200 |
| NAGS | 0.017633671 | 1.012297732 | 0.912356429 | 0.952601638 | 176.5990858 | 178.6073016 | ENSG00000161653 |
| FURIN | 0.017626895 | 1.012292977 | 0.800264207 | 0.884896069 | 1237.921703 | 1253.095001 | ENSG00000140564 |
| PRSS8 | 0.017609684 | 1.012280901 | 0.751447331 | 0.852957914 | 2067.456219 | 2092.760363 | ENSG00000052344 |
| TMEM150C | 0.017573222 | 1.012255317 | 0.94765658 | 0.972984731 | 63.50582505 | 64.24105703 | ENSG00000249242 |
| BBS7 | 0.017553207 | 1.012241275 | 0.888951579 | 0.938578344 | 311.2918701 | 315.2329263 | ENSG00000138686 |
| RAD23B | 0.017545333 | 1.01223575 | 0.713369719 | 0.827616986 | 6207.871321 | 6284.86153 | ENSG00000119318 |
| C2orf76 | 0.017477624 | 1.01218244 | 0.940498531 | 0.968508435 | 85.88541463 | 86.88279211 | ENSG00000186132 |
| BHMT | 0.017460481 | 1.012176217 | 0.968482188 | 0.983367014 | 25.58533631 | 25.9373919 | ENSG00000145692 |
| MPP1 | 0.017458649 | 1.012174932 | 0.896420466 | 0.942785335 | 281.7272399 | 284.9187178 | ENSG00000130830 |
| COPG1 | 0.017399718 | 1.012133588 | 0.742914976 | 0.847821391 | 2436.090248 | 2465.562365 | ENSG00000181789 |
| LGR4 | 0.017392663 | 1.012128638 | 0.806707483 | 0.88864217 | 1375.094747 | 1392.547298 | ENSG00000205213 |
| NA | 0.01734457 | 1.012094898 | 0.988713151 | 0.993554482 | 3.249249898 | 3.327642854 | ENSG00000188985 |
| TCEANC | 0.017328875 | 1.012083888 | 0.958578621 | 0.97877302 | 40.88500264 | 41.32613171 | ENSG00000176896 |
| GPR89A | 0.017324038 | 1.012080495 | 0.93021627 | 0.962755624 | 130.3177941 | 131.7126901 | ENSG00000117262 |
| GINS1 | 0.017322974 | 1.012079749 | 0.785464966 | 0.874499827 | 1529.580066 | 1547.989062 | ENSG00000101003 |
| RASAL1 | 0.017007402 | 1.011858392 | 0.919817194 | 0.957231132 | 200.3367492 | 202.7837871 | ENSG00000111344 |
| ARAP1 | 0.016932017 | 1.011805521 | 0.750679984 | 0.852585092 | 2540.636466 | 2570.384 | ENSG00000186635 |
| NA | 0.01688749 | 1.011774294 | 0.927435551 | 0.961011936 | 131.7410978 | 133.240189 | ENSG00000248124 |
| AP1G1 | 0.016863429 | 1.01175742 | 0.794711139 | 0.880471663 | 1467.520259 | 1484.821641 | ENSG00000166747 |
| HDHD3 | 0.016703815 | 1.011645489 | 0.907924962 | 0.950340773 | 245.9891209 | 248.6692922 | ENSG00000119431 |
| RBM15B | 0.016634225 | 1.011596692 | 0.784038612 | 0.873614559 | 2775.631979 | 2808.68558 | ENSG00000259956 |
| UPK3B | 0.016594124 | 1.011568575 | 0.946728878 | 0.972540966 | 80.1470952 | 81.21348952 | ENSG00000243566 |
| FAM229A | 0.016578425 | 1.011557567 | 0.97146369 | 0.984994536 | 20.49870125 | 20.70231843 | ENSG00000225828 |
| TMEM116 | 0.016566086 | 1.011548916 | 0.931421354 | 0.963382625 | 131.8118343 | 133.2673408 | ENSG00000198270 |
| RAB33A | 0.016518451 | 1.011515517 | 0.94345598 | 0.970367312 | 103.4237074 | 104.7428081 | ENSG00000134594 |
| LINC00237 | 0.016450196 | 1.011467662 | 0.982941852 | 0.990647551 | 11.8821105 | 12.01887898 | ENSG00000225127 |
| CCDC149 | 0.016392205 | 1.011427006 | 0.9267558 | 0.960672598 | 180.8366702 | 183.3129266 | ENSG00000181982 |
| ESRRB | 0.016312148 | 1.011370882 | 0.973239564 | 0.986017774 | 19.88860503 | 20.21833513 | ENSG00000119715 |
| DOK5 | 0.016218644 | 1.011305335 | 0.903106786 | 0.947302105 | 253.5340887 | 256.2562573 | ENSG00000101134 |
| RMND1 | 0.016201639 | 1.011293415 | 0.884895247 | 0.936917696 | 363.5341658 | 367.5830911 | ENSG00000155906 |
| CMIP | 0.016094418 | 1.011218259 | 0.79895738 | 0.883907026 | 1621.191724 | 1639.494406 | ENSG00000153815 |
| NCBP3 | 0.016051728 | 1.011188336 | 0.802572581 | 0.88621529 | 1630.245236 | 1647.953144 | ENSG00000074356 |
| POMGNT1 | 0.016030558 | 1.011173498 | 0.816056373 | 0.894698714 | 1140.121637 | 1152.575452 | ENSG00000085998 |
| ODF2 | 0.015945275 | 1.011113726 | 0.87359581 | 0.929897741 | 518.040462 | 523.6282046 | ENSG00000136811 |
| FBL | 0.015836162 | 1.011037257 | 0.703407673 | 0.821643198 | 11854.3771 | 11985.1603 | ENSG00000105202 |
| NA | 0.015760009 | 1.01098389 | 0.896736542 | 0.942978721 | 454.7932157 | 459.2597181 | ENSG00000225178 |
| FAM167A | 0.015721211 | 1.010956703 | 0.966032728 | 0.982484721 | 36.19019702 | 36.60570104 | ENSG00000154319 |
| ZYX | 0.015467954 | 1.010779251 | 0.820522846 | 0.897675167 | 3071.669766 | 3105.91219 | ENSG00000159840 |
| TTC30B | 0.01535954 | 1.010703297 | 0.96761834 | 0.9832144 | 31.8550062 | 32.30255536 | ENSG00000196659 |
| NA | 0.015342727 | 1.010691518 | 0.963805142 | 0.981666578 | 37.4562832 | 37.88528353 | ENSG00000179611 |
| NORAD | 0.015336304 | 1.010687018 | 0.758212568 | 0.856402715 | 3988.107566 | 4031.506529 | ENSG00000260032 |
| C1orf122 | 0.015307852 | 1.010667087 | 0.872441904 | 0.92913088 | 553.3178932 | 559.1078685 | ENSG00000197982 |
| MGAT5 | 0.015249175 | 1.010625982 | 0.821402673 | 0.898330756 | 1246.306659 | 1259.39747 | ENSG00000152127 |
| GPR21 | 0.015226487 | 1.010610088 | 0.853574654 | 0.917087644 | 809.3884968 | 818.148308 | ENSG00000011454 |
| PNPT1 | 0.015218026 | 1.010604162 | 0.836624881 | 0.907107819 | 1008.297363 | 1019.383375 | ENSG00000138035 |
| CPEB2 | 0.015102455 | 1.010523208 | 0.972935475 | 0.98595118 | 25.31185195 | 25.59672195 | ENSG00000137449 |
| TMEM14C | 0.015046163 | 1.010483779 | 0.824169087 | 0.900132812 | 1716.658659 | 1734.19006 | ENSG00000111843 |
| NUDT15 | 0.015019203 | 1.010464896 | 0.866632205 | 0.92554835 | 803.9030252 | 813.0369081 | ENSG00000136159 |
| ASXL2 | 0.015013421 | 1.010460846 | 0.863963299 | 0.923726994 | 838.2291531 | 846.7728696 | ENSG00000143970 |
| INPP5F | 0.015001677 | 1.010452621 | 0.797277386 | 0.882655842 | 1888.553916 | 1908.771906 | ENSG00000198825 |
| NA | 0.014987824 | 1.010442919 | 0.983863263 | 0.99111274 | 8.148443032 | 8.184402126 | ENSG00000227388 |
| NA | 0.014930458 | 1.010402741 | 0.984385537 | 0.991219407 | 8.115627918 | 8.154827132 | ENSG00000286895 |
| NECAB3 | 0.01492549 | 1.010399261 | 0.901980532 | 0.946583965 | 334.2549984 | 337.7450571 | ENSG00000125967 |
| SALL2 | 0.014849069 | 1.010345741 | 0.743305364 | 0.847935038 | 8680.529515 | 8770.415319 | ENSG00000165821 |
| NR3C2 | 0.014833786 | 1.010335038 | 0.975800558 | 0.987428655 | 19.23070004 | 19.37083117 | ENSG00000151623 |
| QPRT | 0.014791145 | 1.010305177 | 0.791003349 | 0.878179421 | 4174.061384 | 4216.789309 | ENSG00000103485 |
| NA | 0.014783811 | 1.010300041 | 0.943358547 | 0.970367312 | 111.6933233 | 112.8911714 | ENSG00000254285 |
| TRIP12 | 0.014760126 | 1.010283455 | 0.766173444 | 0.861621307 | 2721.396943 | 2749.652359 | ENSG00000153827 |
| NA | 0.014697913 | 1.01023989 | 0.939043127 | 0.967737644 | 129.6530311 | 130.8278145 | ENSG00000226259 |
| SLC29A3 | 0.014558588 | 1.010142332 | 0.918600108 | 0.956469201 | 334.3834551 | 337.7792363 | ENSG00000198246 |
| FGFRL1 | 0.01445967 | 1.010073075 | 0.866303455 | 0.925300921 | 733.7874887 | 740.9613516 | ENSG00000127418 |
| ATP6V1C1 | 0.014367661 | 1.010008659 | 0.874022354 | 0.929992567 | 588.5881768 | 594.4519609 | ENSG00000155097 |
| AK2 | 0.014337558 | 1.009987584 | 0.790321987 | 0.87763484 | 2531.217481 | 2556.460816 | ENSG00000004455 |

| | | | | | | | |
|---|---|---|---|---|---|---|---|
| GPR176 | 0.014329637 | 1.009982039 | 0.78806698 | 0.876396218 | 3618.619424 | 3655.244263 | ENSG00000166073 |
| PSME4 | 0.014295974 | 1.009958473 | 0.794259794 | 0.880349251 | 2918.445206 | 2948.097185 | ENSG00000068878 |
| KAT6A | 0.014282566 | 1.009949087 | 0.897037474 | 0.943140937 | 635.0607149 | 640.9623395 | ENSG00000083168 |
| BCAS4 | 0.014250991 | 1.009926983 | 0.900758897 | 0.945687078 | 366.2301014 | 369.7714902 | ENSG00000124243 |
| NA | 0.014238933 | 1.009918542 | 0.973737537 | 0.986270798 | 23.62785431 | 23.85734551 | ENSG00000227486 |
| MAPKAPK5-AS1 | 0.014156309 | 1.009860705 | 0.883020569 | 0.936107589 | 556.7606041 | 561.9864766 | ENSG00000234608 |
| MFN1 | 0.014119195 | 1.009834726 | 0.869145 | 0.926984177 | 689.9658812 | 696.7401581 | ENSG00000171109 |
| NA | 0.014107075 | 1.009826243 | 0.935293522 | 0.965468934 | 170.6340885 | 172.0598234 | ENSG00000230202 |
| MTCH2 | 0.014055958 | 1.009790464 | 0.812266534 | 0.892246907 | 1628.252661 | 1644.309814 | ENSG00000109919 |
| PAN3 | 0.014039537 | 1.00977897 | 0.886479293 | 0.937353813 | 557.5126119 | 562.8272315 | ENSG00000152520 |
| HMGB2 | 0.014030257 | 1.009772475 | 0.787336697 | 0.875923427 | 4137.058625 | 4176.982327 | ENSG00000164104 |
| ACOT13 | 0.014023167 | 1.009767512 | 0.917629044 | 0.955922674 | 259.0501991 | 261.4781363 | ENSG00000112304 |
| FAM104A | 0.013880452 | 1.009667629 | 0.909432984 | 0.951248262 | 378.1925476 | 382.2909999 | ENSG00000133193 |
| DHPS | 0.013761392 | 1.009584308 | 0.964721938 | 0.981988895 | 49.27902101 | 49.61782048 | ENSG00000095059 |
| PLK4 | 0.013716531 | 1.009552915 | 0.865997878 | 0.925076974 | 756.1188692 | 763.3010965 | ENSG00000142731 |
| KIRREL2 | 0.013661049 | 1.009514092 | 0.933808043 | 0.964558308 | 174.137762 | 175.6775885 | ENSG00000126259 |
| NA | 0.013566506 | 1.009447938 | 0.943829124 | 0.970647597 | 149.3645941 | 150.7877408 | ENSG00000185986 |
| NA | 0.013552151 | 1.009437894 | 0.981719523 | 0.990314032 | 12.36911882 | 12.48619074 | ENSG00000279342 |
| GNB5 | 0.013468913 | 1.009379655 | 0.889469999 | 0.93892015 | 517.8992772 | 522.6145551 | ENSG00000069966 |
| NNT | 0.013436725 | 1.009357135 | 0.903002823 | 0.947296072 | 400.3209048 | 403.8472982 | ENSG00000112992 |
| IPO7 | 0.013216493 | 1.009203065 | 0.79441811 | 0.880349251 | 5603.62641 | 5656.040917 | ENSG00000205339 |
| ZFP90 | 0.013209693 | 1.009198308 | 0.91816205 | 0.956334113 | 293.2321896 | 296.021547 | ENSG00000184939 |
| TMIE | 0.013188962 | 1.009183807 | 0.982944625 | 0.990647551 | 11.45495045 | 11.54592112 | ENSG00000181585 |
| FBXO42 | 0.013149155 | 1.009155961 | 0.886709321 | 0.937442292 | 567.7899092 | 572.9438235 | ENSG00000037637 |
| WDR41 | 0.013057297 | 1.00909171 | 0.882297796 | 0.935715397 | 818.3238907 | 825.6164217 | ENSG00000164253 |
| TM9SF3 | 0.013027465 | 1.009070843 | 0.818382187 | 0.89619415 | 2971.685165 | 2999.697234 | ENSG00000077147 |
| MYCL | 0.012997171 | 1.009049655 | 0.903352024 | 0.947370846 | 473.189183 | 477.6773399 | ENSG00000116990 |
| PER2 | 0.012968088 | 1.009029314 | 0.947748664 | 0.972984731 | 137.1237205 | 138.4744384 | ENSG00000132326 |
| CROCCP2 | 0.012839959 | 1.008939704 | 0.895771566 | 0.942392226 | 577.1268808 | 581.9571921 | ENSG00000215908 |
| FBXO27 | 0.012838672 | 1.008938804 | 0.928679274 | 0.961846113 | 219.8238759 | 221.8655742 | ENSG00000161243 |
| CDC16 | 0.012782299 | 1.00889938 | 0.841814408 | 0.90951788 | 1456.333738 | 1469.605287 | ENSG00000130177 |
| AFDN | 0.012593541 | 1.008767388 | 0.779840494 | 0.870697012 | 3902.702837 | 3936.987363 | ENSG00000130396 |
| ASAP3 | 0.012565709 | 1.008747927 | 0.84073898 | 0.909051234 | 1834.831708 | 1850.258364 | ENSG00000088280 |
| PYM1 | 0.012526678 | 1.008720637 | 0.855632179 | 0.918479793 | 1247.56953 | 1257.997182 | ENSG00000170473 |
| NEK9 | 0.012446503 | 1.00866458 | 0.869956523 | 0.927405575 | 1172.398897 | 1182.92373 | ENSG00000119638 |
| NELFE | 0.012358357 | 1.008602955 | 0.852331323 | 0.916006883 | 1387.01768 | 1398.654647 | ENSG00000204356 |
| NSMCE3 | 0.012284111 | 1.00855105 | 0.905358641 | 0.948787133 | 487.6009504 | 491.6800283 | ENSG00000185115 |
| TIPIN | 0.012239419 | 1.008519808 | 0.92080742 | 0.957693333 | 303.8571871 | 306.2736251 | ENSG00000075131 |
| SLC10A3 | 0.012122096 | 1.008437796 | 0.947255034 | 0.972798489 | 193.0144802 | 194.9352328 | ENSG00000126903 |
| ZNF341 | 0.012096951 | 1.00842022 | 0.929346593 | 0.962113616 | 247.6692516 | 249.6571654 | ENSG00000131061 |
| NA | 0.011945588 | 1.008314425 | 0.974558235 | 0.986628334 | 38.02623074 | 38.45907632 | ENSG00000233225 |
| SPTSSA | 0.01181394 | 1.008222419 | 0.862009245 | 0.922689647 | 1322.704556 | 1333.531125 | ENSG00000165389 |
| P3H4 | 0.011768417 | 1.008190606 | 0.88446141 | 0.936795019 | 942.2313313 | 950.4552548 | ENSG00000141696 |
| RNF114 | 0.011751597 | 1.008178852 | 0.82443489 | 0.90032117 | 2180.68019 | 2198.295251 | ENSG00000124226 |
| ARID5A | 0.011617754 | 1.008085325 | 0.934345931 | 0.964907246 | 252.7223959 | 254.7024185 | ENSG00000196843 |
| HEATR6 | 0.011559356 | 1.008044519 | 0.898406351 | 0.944070801 | 605.829731 | 610.6567517 | ENSG00000068097 |
| SMAD4 | 0.011497919 | 1.008001593 | 0.871701001 | 0.928708447 | 1038.634962 | 1046.803643 | ENSG00000141646 |
| WNT11 | 0.011453041 | 1.007970237 | 0.992020911 | 0.995323003 | 3.921309561 | 3.990160622 | ENSG00000085741 |
| INTS14 | 0.011428845 | 1.007953333 | 0.895885699 | 0.942392226 | 711.1484814 | 716.7590229 | ENSG00000138614 |
| MAPKAPK2 | 0.011304545 | 1.007866493 | 0.867871634 | 0.926309156 | 1170.250106 | 1179.267303 | ENSG00000162889 |
| CAD | 0.01116265 | 1.00776737 | 0.825471242 | 0.900739054 | 6250.118243 | 6298.108652 | ENSG00000084774 |
| LRP5 | 0.011116903 | 1.007735415 | 0.836186234 | 0.906992895 | 2224.503587 | 2241.744471 | ENSG00000162337 |
| PIGG | 0.011050222 | 1.007688838 | 0.920760154 | 0.957693333 | 601.9480946 | 605.8597423 | ENSG00000174227 |
| ACSF3 | 0.011004633 | 1.007656997 | 0.926242062 | 0.960497534 | 362.1638892 | 364.9849006 | ENSG00000176715 |
| VPS37D | 0.010963061 | 1.00762796 | 0.960356507 | 0.979722803 | 111.205239 | 112.3343106 | ENSG00000176428 |
| ANKRD16 | 0.010848042 | 1.007547631 | 0.939965144 | 0.968314739 | 243.0207032 | 245.0386357 | ENSG00000134461 |
| GUF1 | 0.010754306 | 1.00748217 | 0.90720416 | 0.949947101 | 652.1944374 | 657.537884 | ENSG00000151806 |
| NA | 0.010721707 | 1.007459405 | 0.989382695 | 0.993903094 | 8.752081564 | 8.779700648 | ENSG00000285416 |
| GFM2 | 0.010676906 | 1.00742812 | 0.921683196 | 0.95819091 | 541.9828763 | 546.1564434 | ENSG00000164347 |
| MAK16 | 0.010645634 | 1.007406283 | 0.926089623 | 0.960494351 | 436.2490196 | 439.6848245 | ENSG00000198042 |
| GPC5 | 0.010602476 | 1.007376147 | 0.986746702 | 0.992302071 | 10.72284237 | 10.7715551 | ENSG00000179399 |
| INO80 | 0.010531084 | 1.007326298 | 0.892899766 | 0.940687531 | 875.1748246 | 881.293206 | ENSG00000128908 |
| MAPKAP1 | 0.010494837 | 1.00730099 | 0.864394309 | 0.923879632 | 2219.439078 | 2235.612649 | ENSG00000119487 |
| DCAF1 | 0.010485862 | 1.007294723 | 0.887758388 | 0.937989799 | 1123.790309 | 1131.581115 | ENSG00000145041 |
| VAMP8 | 0.010466727 | 1.007281363 | 0.930415215 | 0.962909866 | 368.2085961 | 370.5751587 | ENSG00000118640 |
| NA | 0.010252966 | 1.007132128 | 0.976759722 | 0.987990029 | 35.8964126 | 36.25507173 | ENSG00000124097 |
| CPSF3 | 0.010236874 | 1.007120894 | 0.862503156 | 0.922764098 | 1887.447987 | 1900.888676 | ENSG00000119203 |
| PIK3C2B | 0.010234127 | 1.007118977 | 0.908994778 | 0.950996164 | 816.1435496 | 822.4789208 | ENSG00000133056 |
| NA | 0.010215187 | 1.007105755 | 0.973077623 | 0.98595118 | 48.34810722 | 48.62511306 | ENSG00000272923 |
| THOC1 | 0.010138067 | 1.007051921 | 0.902140456 | 0.946601282 | 873.2349511 | 879.2599207 | ENSG00000079134 |
| LSM14B | 0.010116996 | 1.007037213 | 0.868864831 | 0.926958935 | 1994.4638 | 2007.766198 | ENSG00000149657 |
| PDXP | 0.01004124 | 1.006984335 | 0.943405853 | 0.970367312 | 288.498987 | 290.557664 | ENSG00000241360 |
| AQP3 | 0.009959778 | 1.006927477 | 0.967872408 | 0.983248479 | 75.8462711 | 76.31452439 | ENSG00000165272 |
| NA | 0.009925578 | 1.006903607 | 0.906400772 | 0.949415068 | 895.8668107 | 901.6529338 | ENSG00000237550 |
| MIOS | 0.009908551 | 1.006891724 | 0.900839827 | 0.945695533 | 854.5859627 | 860.7951741 | ENSG00000164654 |
| WDHD1 | 0.009794957 | 1.006812446 | 0.882704722 | 0.935911408 | 1604.076212 | 1615.273528 | ENSG00000198554 |
| CCDC117 | 0.009775348 | 1.006798762 | 0.902763841 | 0.947148379 | 1094.889441 | 1102.238586 | ENSG00000159873 |

| | | | | | | | |
|---|---|---|---|---|---|---|---|
| TNKS1BP1 | 0.009773914 | 1.006797761 | 0.841654982 | 0.909490034 | 4509.724515 | 4539.65458 | ENSG00000149115 |
| SPNS2 | 0.009715007 | 1.006756654 | 0.974849024 | 0.986718185 | 51.94090223 | 52.12779484 | ENSG00000183018 |
| NA | 0.009661124 | 1.006719053 | 0.989220084 | 0.993856811 | 9.212732452 | 9.225767129 | ENSG00000228060 |
| GABPA | 0.009629685 | 1.006697115 | 0.928745077 | 0.961852218 | 469.1160007 | 472.2841958 | ENSG00000154727 |
| CUTC | 0.009617143 | 1.006688363 | 0.937074449 | 0.966565868 | 409.8648916 | 412.1817443 | ENSG00000119929 |
| LOC100288123 | 0.009611989 | 1.006684767 | 0.980793281 | 0.990031039 | 26.70899843 | 26.87981209 | ENSG00000267244 |
| FOXK2 | 0.009609536 | 1.006683055 | 0.881799182 | 0.935391472 | 1547.325161 | 1557.827914 | ENSG00000141568 |
| NA | 0.00955349 | 1.006643948 | 0.971836133 | 0.985216819 | 63.46407365 | 63.82697658 | ENSG00000272994 |
| CTDSP2 | 0.009542099 | 1.006636 | 0.839999938 | 0.908905319 | 3473.309515 | 3496.10692 | ENSG00000175215 |
| NA | 0.009457838 | 1.006577209 | 0.988606955 | 0.993499519 | 10.47236095 | 10.50455612 | ENSG00000286158 |
| TFPI2 | 0.009379 | 1.006522205 | 0.965230646 | 0.982221968 | 97.81477032 | 98.31498697 | ENSG00000105825 |
| KBTBD6 | 0.009327397 | 1.006486204 | 0.938768615 | 0.967624505 | 372.7328685 | 374.9016881 | ENSG00000165572 |
| TMEM184C | 0.009290483 | 1.006460451 | 0.912036314 | 0.952535122 | 739.0188263 | 744.2098952 | ENSG00000164168 |
| COX17 | 0.009269962 | 1.006446135 | 0.931603549 | 0.963467756 | 478.5234717 | 481.9039011 | ENSG00000138495 |
| CNNM2 | 0.009260263 | 1.006439369 | 0.922443309 | 0.958464606 | 552.2053195 | 555.7312713 | ENSG00000148842 |
| PTGES2 | 0.009214839 | 1.006407681 | 0.892579783 | 0.940596809 | 2116.310536 | 2130.369834 | ENSG00000148334 |
| TC2N | 0.009183286 | 1.006385671 | 0.939147647 | 0.967771015 | 409.7056852 | 412.4650933 | ENSG00000165929 |
| ABHD12 | 0.009099307 | 1.006327091 | 0.888459128 | 0.938417928 | 1521.787372 | 1532.012033 | ENSG00000100997 |
| NA | 0.009074463 | 1.006309761 | 0.955313951 | 0.977404477 | 172.9198849 | 173.922625 | ENSG00000220785 |
| SNAP29 | 0.009072113 | 1.006308123 | 0.954525712 | 0.976960713 | 190.5460599 | 191.8355876 | ENSG00000099940 |
| TBK1 | 0.00906887 | 1.00630586 | 0.941186871 | 0.96886597 | 439.376533 | 442.6505216 | ENSG00000183735 |
| LRRFIP1 | 0.008862844 | 1.006162164 | 0.883678779 | 0.936493029 | 1776.147507 | 1786.761237 | ENSG00000124831 |
| METTL21A | 0.008772103 | 1.006098882 | 0.91538486 | 0.954649348 | 721.7824904 | 726.3601112 | ENSG00000144401 |
| CHFR | 0.008704101 | 1.00605146 | 0.904936919 | 0.948448229 | 1084.834554 | 1091.086682 | ENSG00000072609 |
| NA | 0.008678159 | 1.00603337 | 0.934075542 | 0.964679656 | 445.6333613 | 448.2703731 | ENSG00000249353 |
| NA | 0.008640119 | 1.006006843 | 0.987139345 | 0.992541702 | 17.14051812 | 17.26953951 | ENSG00000269982 |
| CUEDC1 | 0.008386148 | 1.005829762 | 0.952922114 | 0.975888969 | 233.8963285 | 235.3262017 | ENSG00000180891 |
| GPR173 | 0.008370002 | 1.005818505 | 0.955887896 | 0.977576918 | 222.5841144 | 223.8300042 | ENSG00000184194 |
| BPGM | 0.008331656 | 1.005791772 | 0.947202275 | 0.972798489 | 365.8022667 | 368.3184558 | ENSG00000172331 |
| DAGLB | 0.008328169 | 1.005789341 | 0.937464867 | 0.966854672 | 529.3169451 | 532.6106781 | ENSG00000164535 |
| TAF9B | 0.008285508 | 1.005759599 | 0.938736376 | 0.967624505 | 443.2934859 | 445.8528797 | ENSG00000187325 |
| TMEM134 | 0.00821696 | 1.005711813 | 0.946952227 | 0.9726684 | 356.455682 | 358.4809912 | ENSG00000172663 |
| ATAD3C | 0.008175792 | 1.005683115 | 0.985995826 | 0.992041493 | 21.635442 | 21.8297248 | ENSG00000215915 |
| ZNF7 | 0.008081863 | 1.005617641 | 0.949433762 | 0.973815754 | 280.0103903 | 281.6618018 | ENSG00000147789 |
| NA | 0.008003806 | 1.005563233 | 0.984216214 | 0.991219407 | 26.81793166 | 26.94488076 | ENSG00000268573 |
| CNOT3 | 0.007961548 | 1.00553378 | 0.9152894 | 0.954649348 | 1824.598517 | 1834.758363 | ENSG00000088038 |
| CHD8 | 0.007876757 | 1.005474683 | 0.87385191 | 0.929913791 | 3956.026729 | 3977.414396 | ENSG00000100888 |
| RPS3 | 0.007876567 | 1.005474551 | 0.85080057 | 0.915101063 | 39063.02656 | 39275.7142 | ENSG00000149273 |
| TMEM30A | 0.007847159 | 1.005454056 | 0.923870059 | 0.959186142 | 1140.520777 | 1147.477712 | ENSG00000112697 |
| RCL1 | 0.007833827 | 1.005444764 | 0.936274345 | 0.966102155 | 497.2633334 | 500.1607502 | ENSG00000120158 |
| SPIDR | 0.007791984 | 1.005415603 | 0.919915391 | 0.957261824 | 1084.409695 | 1089.969689 | ENSG00000164808 |
| TTC39B | 0.007772033 | 1.0054017 | 0.962002516 | 0.980632996 | 184.0784602 | 184.8915183 | ENSG00000155158 |
| COX2 | 0.007654858 | 1.005320045 | 0.965902432 | 0.982454066 | 28857.76904 | 29011.36201 | ENSG00000198712 |
| NA | 0.007460773 | 1.005184808 | 0.989869934 | 0.994130074 | 13.95323166 | 14.02122584 | ENSG00000260167 |
| RNF146 | 0.007386452 | 1.005133027 | 0.936486764 | 0.966238764 | 586.9611244 | 589.8528169 | ENSG00000118518 |
| NA | 0.007383877 | 1.005131234 | 0.985904942 | 0.992024423 | 25.3914367 | 25.48110559 | ENSG00000277283 |
| PRMT9 | 0.00734377 | 1.005103291 | 0.96046079 | 0.979722803 | 253.0897076 | 254.5077029 | ENSG00000164169 |
| KLF8 | 0.007343634 | 1.005103196 | 0.963770768 | 0.981666578 | 184.0815265 | 185.093176 | ENSG00000102349 |
| ACSS1 | 0.007333297 | 1.005095995 | 0.981493554 | 0.990314032 | 49.53373278 | 49.70386185 | ENSG00000154930 |
| MIS12 | 0.007248527 | 1.005036939 | 0.950962096 | 0.974761413 | 423.2686902 | 425.1677901 | ENSG00000167842 |
| UXS1 | 0.007240338 | 1.005031234 | 0.921389466 | 0.958038961 | 1051.315375 | 1056.845697 | ENSG00000115652 |
| FZR1 | 0.007232876 | 1.005026036 | 0.923984762 | 0.959186142 | 1004.691577 | 1009.65324 | ENSG00000105325 |
| PPP1R26-AS1 | 0.007171227 | 1.00498309 | 0.9837103 | 0.99105679 | 39.85890998 | 40.15220622 | ENSG00000225361 |
| GPR108 | 0.007161817 | 1.004976535 | 0.933406905 | 0.964413047 | 674.3679246 | 677.5449173 | ENSG00000125734 |
| RFXANK | 0.007156223 | 1.004972638 | 0.937997669 | 0.967280325 | 602.3314022 | 605.4335277 | ENSG00000064490 |
| ANXA4 | 0.007113255 | 1.004942708 | 0.960739518 | 0.979808711 | 219.1603891 | 220.2100885 | ENSG00000196975 |
| ADPGK | 0.007071675 | 1.004913744 | 0.932869195 | 0.964184087 | 736.5188671 | 739.7252831 | ENSG00000159322 |
| GOLPH3 | 0.00700755 | 1.004869079 | 0.90682568 | 0.949653884 | 1994.631584 | 2004.537693 | ENSG00000113384 |
| RAPGEF2 | 0.007006554 | 1.004868385 | 0.920559153 | 0.957590004 | 1242.178822 | 1247.885195 | ENSG00000109756 |
| PHF6 | 0.007002653 | 1.004865669 | 0.905784964 | 0.949079232 | 2005.866301 | 2016.367503 | ENSG00000156531 |
| SPATA24 | 0.006964557 | 1.004839134 | 0.970345513 | 0.98427465 | 147.7243509 | 148.4342558 | ENSG00000170469 |
| MYORG | 0.006916465 | 1.004805639 | 0.944391162 | 0.970811569 | 546.7020875 | 549.1884095 | ENSG00000164976 |
| TPM4 | 0.006874212 | 1.004776211 | 0.856349159 | 0.919096007 | 9100.121338 | 9144.0168 | ENSG00000167460 |
| LOC101927989 | 0.006827848 | 1.004743921 | 0.984123156 | 0.991219407 | 42.80476345 | 42.99183719 | ENSG00000265778 |
| AOX1 | 0.006806837 | 1.004729288 | 0.964184853 | 0.981782342 | 245.6034 | 246.9183572 | ENSG00000138356 |
| RNF126 | 0.00680098 | 1.004725209 | 0.922255926 | 0.95841945 | 1686.003636 | 1694.255183 | ENSG00000070423 |
| VDAC2 | 0.006797641 | 1.004722883 | 0.884751532 | 0.936912872 | 4112.939508 | 4132.780786 | ENSG00000165637 |
| DDX19B | 0.006745715 | 1.004686722 | 0.959412777 | 0.979260946 | 262.9498572 | 263.9736202 | ENSG00000157349 |
| PROSER1 | 0.00666272 | 1.004628926 | 0.910101627 | 0.951586476 | 2180.784279 | 2191.354895 | ENSG00000120685 |
| ANAPC11 | 0.006646919 | 1.004617923 | 0.907548541 | 0.950087463 | 2218.953775 | 2229.260691 | ENSG00000141552 |
| CLASP2 | 0.006477317 | 1.004499828 | 0.935669224 | 0.965653488 | 1374.375039 | 1380.093626 | ENSG00000163539 |
| SLC18B1 | 0.006432433 | 1.004468577 | 0.940847245 | 0.968717133 | 694.7656778 | 697.8514322 | ENSG00000146409 |
| HDAC7 | 0.006414521 | 1.004456106 | 0.926204742 | 0.960497534 | 2060.260492 | 2069.42312 | ENSG00000061273 |
| PPP1R3F | 0.006370737 | 1.004425623 | 0.973488612 | 0.986218297 | 124.7360534 | 125.1714032 | ENSG00000049769 |
| PCK2 | 0.006342368 | 1.004405872 | 0.927127206 | 0.960744034 | 1997.675202 | 2005.428058 | ENSG00000100889 |
| NA | 0.006257637 | 1.004346884 | 0.985779954 | 0.991950399 | 35.28161046 | 35.48715742 | ENSG00000214485 |
| CASQ1 | 0.006190077 | 1.004299853 | 0.989955872 | 0.994130074 | 19.8811082 | 19.87095299 | ENSG00000143318 |

| | | | | | | | |
|---|---|---|---|---|---|---|---|
| NA | 0.006065932 | 1.004213435 | 0.993336616 | 0.996228576 | 9.158265837 | 9.141343294 | ENSG00000273893 |
| MIDN | 0.006028759 | 1.004187561 | 0.924417721 | 0.959482527 | 3523.170167 | 3538.974793 | ENSG00000167470 |
| NA | 0.006026148 | 1.004185743 | 0.968008662 | 0.983248479 | 247.8135538 | 249.1035518 | ENSG00000277200 |
| DNAJC30 | 0.006009166 | 1.004173923 | 0.983919967 | 0.99111274 | 58.59231398 | 58.80861486 | ENSG00000176410 |
| ANKFY1 | 0.005966317 | 1.004144099 | 0.934019444 | 0.964679656 | 1390.384698 | 1395.210055 | ENSG00000185722 |
| COL4A6 | 0.005884791 | 1.004087357 | 0.933525147 | 0.964421015 | 1168.469527 | 1173.006991 | ENSG00000197565 |
| DIXDC1 | 0.005846361 | 1.004060611 | 0.961982207 | 0.980632996 | 337.4779661 | 339.2187795 | ENSG00000150764 |
| NA | 0.005832697 | 1.004051101 | 0.991271833 | 0.994933659 | 15.03079988 | 15.08786469 | ENSG00000258741 |
| CD101 | 0.005824 | 1.004045049 | 0.992365207 | 0.99546135 | 12.35025822 | 12.37219191 | ENSG00000134256 |
| NBEA | 0.00543718 | 1.003775877 | 0.955406712 | 0.977429542 | 618.6482618 | 620.9241137 | ENSG00000172915 |
| RAPGEFL1 | 0.00540122 | 1.003750857 | 0.966803374 | 0.982852229 | 368.813261 | 370.2611381 | ENSG00000108352 |
| NA | 0.005335139 | 1.003704882 | 0.989420715 | 0.993903094 | 30.82460793 | 30.80060289 | ENSG00000287769 |
| PLPP5 | 0.005274972 | 1.003663025 | 0.955445425 | 0.977429542 | 576.8853357 | 579.2081215 | ENSG00000147535 |
| KDM3B | 0.005239564 | 1.003638392 | 0.916244333 | 0.955110203 | 3334.977743 | 3346.304151 | ENSG00000120733 |
| ZNF789 | 0.00521555 | 1.003621686 | 0.982100344 | 0.990314032 | 79.19049971 | 79.48271537 | ENSG00000198556 |
| KCTD1 | 0.005206173 | 1.003615163 | 0.964395805 | 0.981853466 | 342.8051951 | 344.0158679 | ENSG00000134504 |
| TMEM25 | 0.005149065 | 1.003575437 | 0.970140632 | 0.98427465 | 281.0921888 | 282.0807349 | ENSG00000149582 |
| TFCP2 | 0.005038839 | 1.003498763 | 0.940544141 | 0.968508435 | 1427.599988 | 1433.077304 | ENSG00000135457 |
| NAA30 | 0.004995346 | 1.003468511 | 0.965245747 | 0.982221968 | 616.7627329 | 619.2538893 | ENSG00000139977 |
| ARHGAP44 | 0.004926993 | 1.003420969 | 0.960062311 | 0.979722803 | 530.1861837 | 531.9287988 | ENSG00000006740 |
| REPS1 | 0.004839333 | 1.003360002 | 0.940512476 | 0.968508435 | 1380.920097 | 1385.953052 | ENSG00000135597 |
| SAMD1 | 0.004838914 | 1.00335971 | 0.941351705 | 0.968977949 | 1755.734725 | 1761.671428 | ENSG00000141858 |
| CCNT2 | 0.004806413 | 1.003337107 | 0.953550057 | 0.976376547 | 844.599896 | 846.8592204 | ENSG00000082258 |
| HEATR1 | 0.004756206 | 1.003302191 | 0.92328349 | 0.958830585 | 2911.437459 | 2920.966052 | ENSG00000119285 |
| YPEL5 | 0.004745311 | 1.003294614 | 0.962033081 | 0.980632996 | 490.6386794 | 492.2113426 | ENSG00000119801 |
| SEC61G | 0.004662181 | 1.003236804 | 0.948996883 | 0.973626489 | 1527.503214 | 1533.004389 | ENSG00000132432 |
| NA | 0.004617322 | 1.003205611 | 0.99477653 | 0.997056698 | 9.422026045 | 9.393821296 | ENSG00000260293 |
| FOXN4 | 0.004588366 | 1.003185476 | 0.99449647 | 0.997056698 | 12.5078761 | 12.63568324 | ENSG00000139445 |
| NAMPT | 0.004500864 | 1.003124632 | 0.944303573 | 0.970809901 | 1672.437602 | 1678.306492 | ENSG00000105835 |
| MN1 | 0.00447768 | 1.003108513 | 0.975959946 | 0.987518554 | 203.9235621 | 204.5035416 | ENSG00000169184 |
| ZNF660 | 0.004437273 | 1.003080418 | 0.97908707 | 0.988983938 | 163.845234 | 164.3723081 | ENSG00000144792 |
| FRAT2 | 0.004311384 | 1.002992893 | 0.958391059 | 0.978684131 | 1399.821669 | 1404.276698 | ENSG00000181274 |
| PPP2CB | 0.004225366 | 1.002933094 | 0.947306475 | 0.972798489 | 1688.109649 | 1693.128522 | ENSG00000104695 |
| ZCCHC3 | 0.004185817 | 1.002905601 | 0.94099381 | 0.968764627 | 2633.475277 | 2641.508062 | ENSG00000247315 |
| IMPA1 | 0.00397673 | 1.002760261 | 0.973870199 | 0.986345882 | 486.558727 | 488.5201428 | ENSG00000133731 |
| PIK3CD | 0.003746145 | 1.002600004 | 0.966553243 | 0.98274283 | 633.2446433 | 635.181437 | ENSG00000171608 |
| NA | 0.003693546 | 1.002563451 | 0.997642262 | 0.998625824 | 2.977116982 | 3.01009618 | ENSG00000284052 |
| STAT5A | 0.003651002 | 1.002533887 | 0.977721216 | 0.988361337 | 264.9309057 | 265.7430926 | ENSG00000126561 |
| FAM71F1 | 0.003445652 | 1.002391199 | 0.993099014 | 0.996145647 | 28.29610234 | 28.31454531 | ENSG00000135248 |
| SOS2 | 0.003304088 | 1.002292844 | 0.983044277 | 0.990681803 | 235.0169484 | 235.3321019 | ENSG00000100485 |
| NPM1 | 0.003263511 | 1.002264654 | 0.925441953 | 0.960029077 | 46349.58036 | 46453.82536 | ENSG00000181163 |
| MFSD10 | 0.003187127 | 1.00221159 | 0.959016987 | 0.979115993 | 2287.540078 | 2293.291761 | ENSG00000109736 |
| RMST | 0.003109547 | 1.002157698 | 0.988156358 | 0.993201912 | 121.0847616 | 121.1250488 | ENSG00000255794 |
| NDUFB10 | 0.003100553 | 1.002151451 | 0.967339395 | 0.983086307 | 1683.842767 | 1687.165432 | ENSG00000140990 |
| HS2ST1 | 0.003085692 | 1.002141128 | 0.982097282 | 0.990314032 | 347.3775517 | 348.2703376 | ENSG00000153936 |
| SLC7A6OS | 0.003049994 | 1.002116331 | 0.9843119 | 0.991219407 | 219.7201241 | 220.3095969 | ENSG00000103061 |
| SLC1A1 | 0.003023903 | 1.002098208 | 0.981318559 | 0.990314032 | 322.2123911 | 322.7706703 | ENSG00000106688 |
| ATP2C1 | 0.003002857 | 1.00208359 | 0.962129069 | 0.980632996 | 1910.518148 | 1915.275779 | ENSG00000017260 |
| ARFIP2 | 0.002966605 | 1.00205841 | 0.969059202 | 0.983849327 | 1168.536886 | 1170.762204 | ENSG00000132254 |
| DSCC1 | 0.00282162 | 1.001957712 | 0.969448897 | 0.984039579 | 936.4327618 | 938.2314709 | ENSG00000136982 |
| MRPL40 | 0.002621274 | 1.00181858 | 0.984338693 | 0.991219407 | 260.8322418 | 261.470346 | ENSG00000185608 |
| WDFY2 | 0.00260386 | 1.001806488 | 0.974067334 | 0.98649375 | 850.3639724 | 851.3546096 | ENSG00000139668 |
| ACTL6B | 0.002540499 | 1.001762491 | 0.995504555 | 0.997339499 | 26.02318442 | 26.00676172 | ENSG00000077080 |
| FLVCR2 | 0.002465114 | 1.001710147 | 0.992214001 | 0.995401491 | 71.26530479 | 71.40775046 | ENSG00000119686 |
| RHOA | 0.002236603 | 1.001551497 | 0.956872319 | 0.978044405 | 9296.322994 | 9311.466339 | ENSG00000067560 |
| GAS8 | 0.002179215 | 1.001511658 | 0.990673699 | 0.994592055 | 134.5249909 | 134.5423738 | ENSG00000141013 |
| KRT18 | 0.002159493 | 1.001497967 | 0.968263496 | 0.983351992 | 3280.696515 | 3285.695362 | ENSG00000111057 |
| NA | 0.002142134 | 1.001485917 | 0.997143588 | 0.998385683 | 13.39568704 | 13.38398192 | ENSG00000286615 |
| SSB | 0.002067226 | 1.001433919 | 0.96744737 | 0.983138775 | 4071.202502 | 4077.19764 | ENSG00000138385 |
| PANX1 | 0.001996339 | 1.001384715 | 0.983346419 | 0.990812403 | 690.5412231 | 691.5074956 | ENSG00000110218 |
| RAB10 | 0.001993834 | 1.001382976 | 0.969552094 | 0.984039579 | 3119.505156 | 3124.802993 | ENSG00000084733 |
| MOCS2 | 0.001955695 | 1.001356503 | 0.986835948 | 0.992312792 | 324.5801756 | 325.0792658 | ENSG00000164172 |
| C5orf15 | 0.001899876 | 1.001317761 | 0.986024757 | 0.992041493 | 629.7483929 | 631.2563267 | ENSG00000113583 |
| PDPK1 | 0.001885038 | 1.001307463 | 0.984879026 | 0.991457564 | 517.3226349 | 518.1640795 | ENSG00000140992 |
| ZC2HC1A | 0.001843771 | 1.001278822 | 0.993817788 | 0.996607525 | 94.97609832 | 95.25943775 | ENSG00000104427 |
| ASTE1 | 0.001806369 | 1.001252864 | 0.994953205 | 0.997124125 | 60.73397145 | 60.86123683 | ENSG00000034533 |
| NA | 0.001756587 | 1.001218315 | 0.993796839 | 0.996607525 | 90.93903442 | 91.2114721 | ENSG00000205746 |
| PUS7 | 0.001748499 | 1.001212702 | 0.980966124 | 0.990125113 | 1108.81668 | 1110.617716 | ENSG00000091127 |
| PRRC1 | 0.001642292 | 1.001138998 | 0.981774041 | 0.990314032 | 1275.476044 | 1277.142917 | ENSG00000164244 |
| NA | 0.001603279 | 1.001111926 | 0.998208662 | 0.99900125 | 10.30146428 | 10.25852984 | ENSG00000287906 |
| VAV2 | 0.001553428 | 1.001077334 | 0.978710133 | 0.988963238 | 1780.452149 | 1782.601132 | ENSG00000160293 |
| RNF216 | 0.001542867 | 1.001070006 | 0.982846437 | 0.990647551 | 1114.497066 | 1115.934288 | ENSG00000011275 |
| DGLUCY | 0.00138422 | 1.000959929 | 0.990249299 | 0.994284865 | 365.8966808 | 366.1432327 | ENSG00000133943 |
| ASB6 | 0.001361593 | 1.00094423 | 0.990749746 | 0.994616639 | 441.099653 | 441.3577936 | ENSG00000148331 |
| CLUH | 0.001346022 | 1.000933427 | 0.981989384 | 0.990314032 | 4599.869592 | 4604.272184 | ENSG00000132361 |
| NA | 0.00122437 | 1.000849029 | 0.999063656 | 0.999445059 | 4.675744867 | 4.711828377 | ENSG00000285676 |
| PRR12 | 0.001062652 | 1.000736846 | 0.988021324 | 0.993117933 | 2234.233317 | 2236.116641 | ENSG00000126464 |

| | | | | | | | |
|---|---|---|---|---|---|---|---|
| PSMB3 | 0.0010297 | 1.000713989 | 0.986744763 | 0.992302071 | 2975.298004 | 2976.361044 | ENSG00000277791 |
| CEP57 | 0.000699638 | 1.00048507 | 0.99325313 | 0.996221629 | 952.1273736 | 952.516418 | ENSG00000166037 |
| PARVA | 0.000648558 | 1.000449647 | 0.995427791 | 0.997339499 | 399.001428 | 398.8588025 | ENSG00000197702 |
| RDH11 | 0.000623337 | 1.000432158 | 0.991623156 | 0.995130952 | 1817.96131 | 1819.024301 | ENSG00000072042 |
| WDR60 | 0.000548914 | 1.000380551 | 0.996090615 | 0.997849304 | 450.162889 | 449.7925047 | ENSG00000126870 |
| POLB | 0.000462499 | 1.000320632 | 0.996348851 | 0.997983846 | 491.0995304 | 491.1557171 | ENSG00000070501 |
| KIF4A | 0.000458553 | 1.000317895 | 0.994532689 | 0.997056698 | 1194.497581 | 1194.622439 | ENSG00000090889 |
| NA | 0.000451697 | 1.000313142 | 0.999432357 | 0.999639632 | 11.47939285 | 11.42600361 | ENSG00000225689 |
| PIGB | 0.000438282 | 1.00030384 | 0.999141827 | 0.999445059 | 29.34153117 | 29.2903086 | ENSG00000069943 |
| PCBD1 | 0.000427684 | 1.000296492 | 0.996280734 | 0.997983846 | 683.9176693 | 683.9474219 | ENSG00000166228 |
| TMEM69 | 0.000426059 | 1.000295365 | 0.996494368 | 0.997994645 | 556.1746379 | 556.302653 | ENSG00000159596 |
| LARGE2 | 0.00038145 | 1.000264436 | 0.995090821 | 0.997210235 | 2136.112233 | 2136.900331 | ENSG00000165905 |
| FARP2 | 0.000293223 | 1.000203267 | 0.997615619 | 0.998625824 | 486.2038039 | 486.2747685 | ENSG00000006607 |
| FNBP4 | 0.000187464 | 1.000129948 | 0.997989151 | 0.998921222 | 1093.275103 | 1093.285513 | ENSG00000109920 |
| PNMA8B | 0.000113733 | 1.000078837 | 0.999608684 | 0.999707619 | 81.86055348 | 81.91633515 | ENSG00000204851 |
| RPRM | -3.09E-05 | 0.999978615 | 0.999801982 | 0.999801982 | 321.2085504 | 321.0458212 | ENSG00000177519 |
| WSB2 | -7.86E-05 | 0.999945547 | 0.999186015 | 0.999445059 | 983.7299705 | 983.4087447 | ENSG00000176871 |
| SH3D19 | -8.03E-05 | 0.999944332 | 0.999060148 | 0.999445059 | 1474.55414 | 1475.167527 | ENSG00000109686 |
| CTU1 | -0.00013402 | 0.999907109 | 0.999156254 | 0.999445059 | 340.5332042 | 340.8239901 | ENSG00000142544 |
| ZNF121 | -0.000221882 | 0.999846215 | 0.997523993 | 0.998625824 | 1509.61394 | 1509.325329 | ENSG00000197961 |
| TP53 | -0.000292932 | 0.999796976 | 0.995530008 | 0.997339499 | 3594.03256 | 3592.888303 | ENSG00000141510 |
| PCNX1 | -0.000299846 | 0.999792184 | 0.996820657 | 0.998114154 | 1044.669457 | 1044.670583 | ENSG00000100731 |
| QKI | -0.000334499 | 0.99976817 | 0.995391044 | 0.997339499 | 2265.111751 | 2264.909341 | ENSG00000112531 |
| KIAA1324L | -0.000382328 | 0.999735026 | 0.996431852 | 0.997983846 | 828.8437722 | 828.5039105 | ENSG00000164659 |
| TNFAIP2 | -0.000394741 | 0.999726424 | 0.996782931 | 0.998114154 | 569.3146862 | 568.7434028 | ENSG00000185215 |
| ATP6AP2 | -0.000448798 | 0.999688966 | 0.994819195 | 0.997056698 | 1261.907132 | 1262.137623 | ENSG00000182220 |
| NA | -0.000514238 | 0.999643621 | 0.999625387 | 0.999707619 | 3.995312496 | 4.021886189 | ENSG00000280106 |
| NA | -0.000520281 | 0.999639434 | 0.999655796 | 0.999707619 | 2.982698788 | 3.005795033 | ENSG00000239636 |
| CYSTM1 | -0.00056294 | 0.999609876 | 0.996637257 | 0.998034122 | 313.0649013 | 312.9469274 | ENSG00000120306 |
| NA | -0.000571348 | 0.99960405 | 0.998971129 | 0.999445059 | 21.71223584 | 21.7221777 | ENSG00000272578 |
| KCMF1 | -0.000636769 | 0.999558723 | 0.991446364 | 0.995005296 | 2151.70313 | 2151.43981 | ENSG00000176407 |
| FARSB | -0.00066049 | 0.999542288 | 0.992122033 | 0.995372693 | 1304.256276 | 1304.147748 | ENSG00000116120 |
| SHROOM1 | -0.000742368 | 0.999485562 | 0.996568622 | 0.998017198 | 155.4881731 | 155.4006365 | ENSG00000164403 |
| C1orf50 | -0.000774099 | 0.99946358 | 0.998224462 | 0.99900125 | 43.5599625 | 43.44194436 | ENSG00000164008 |
| CNIH1 | -0.000807358 | 0.999440539 | 0.991256408 | 0.994933659 | 1041.704142 | 1041.447022 | ENSG00000100528 |
| TRIM44 | -0.00086463 | 0.999400863 | 0.988305991 | 0.993300556 | 1675.063025 | 1674.057227 | ENSG00000166326 |
| PPP2R3B | -0.000939865 | 0.999348747 | 0.990089171 | 0.994212181 | 936.1810042 | 935.4983452 | ENSG00000167393 |
| USF1 | -0.00101192 | 0.999298837 | 0.991056932 | 0.994873252 | 1365.612345 | 1363.973182 | ENSG00000158773 |
| KIF14 | -0.001055645 | 0.99926855 | 0.991247736 | 0.994933659 | 650.8346856 | 650.4203812 | ENSG00000118193 |
| NA | -0.001130565 | 0.999216659 | 0.999152323 | 0.999445059 | 3.965288286 | 3.992311195 | ENSG00000279970 |
| CHST8 | -0.001145347 | 0.999206421 | 0.99139425 | 0.995004759 | 498.7291381 | 497.7736617 | ENSG00000124302 |
| SMAD7 | -0.001193931 | 0.999172772 | 0.989174394 | 0.993856811 | 1073.53388 | 1072.885421 | ENSG00000101665 |
| SH3BP4 | -0.00123532 | 0.999144108 | 0.985175987 | 0.991584323 | 1469.611043 | 1468.76315 | ENSG00000130147 |
| FBXO38 | -0.001296701 | 0.999101599 | 0.988597701 | 0.993499519 | 620.7879674 | 620.4624602 | ENSG00000145868 |
| ZNF529 | -0.001390402 | 0.999036711 | 0.994471384 | 0.997056698 | 144.1149226 | 143.9929035 | ENSG00000186020 |
| CDS2 | -0.001474289 | 0.998978623 | 0.981798373 | 0.990314032 | 1469.373164 | 1467.622574 | ENSG00000101290 |
| DYNLT1 | -0.001528783 | 0.99894089 | 0.986292761 | 0.992155898 | 1066.804597 | 1064.901634 | ENSG00000146425 |
| KIFBP | -0.001546409 | 0.998928686 | 0.983901924 | 0.99111274 | 1044.93537 | 1044.367346 | ENSG00000198954 |
| STIL | -0.001711849 | 0.998814141 | 0.984581917 | 0.991323276 | 639.2272747 | 638.496982 | ENSG00000123473 |
| MADD | -0.00175716 | 0.998782771 | 0.977758933 | 0.988361337 | 1431.419347 | 1429.549893 | ENSG00000110514 |
| TECR | -0.00178241 | 0.99876529 | 0.975273462 | 0.986989841 | 1940.170697 | 1938.290522 | ENSG00000099797 |
| C5AR1 | -0.001903357 | 0.998681563 | 0.998756799 | 0.999378464 | 3.306507417 | 3.323341707 | ENSG00000197405 |
| SELENON | -0.00197489 | 0.998632047 | 0.979164841 | 0.988983938 | 2009.680101 | 2007.037497 | ENSG00000162430 |
| PXMP4 | -0.001979416 | 0.998628914 | 0.994452971 | 0.997056698 | 60.01860879 | 59.92123979 | ENSG00000101417 |
| NA | -0.002040002 | 0.998586977 | 0.998715342 | 0.999378464 | 3.591919126 | 3.605394703 | ENSG00000278948 |
| FAM120A | -0.002069215 | 0.998566757 | 0.976000661 | 0.987518554 | 1485.270986 | 1482.605612 | ENSG00000048828 |
| ARPIN | -0.002093981 | 0.998549616 | 0.990264637 | 0.994284865 | 161.5642557 | 161.1570649 | ENSG00000242498 |
| SPATA41 | -0.00211151 | 0.998537484 | 0.997568878 | 0.998625824 | 10.5399062 | 10.58897995 | ENSG00000189419 |
| NA | -0.0021219 | 0.998530292 | 0.998530865 | 0.999256052 | 3.282065012 | 3.302369007 | ENSG00000272540 |
| PEG13 | -0.002174113 | 0.998494155 | 0.998078287 | 0.998958609 | 5.666031352 | 5.696193966 | ENSG00000282164 |
| NA | -0.002281953 | 0.998419521 | 0.997537376 | 0.998625824 | 8.537006006 | 8.582071487 | ENSG00000259319 |
| CHMP5 | -0.002316278 | 0.998395766 | 0.977607508 | 0.988353504 | 814.7700803 | 813.5413431 | ENSG00000086065 |
| SFSWAP | -0.002353492 | 0.998370013 | 0.970023744 | 0.98427465 | 1584.506496 | 1582.056329 | ENSG00000061936 |
| C2orf49 | -0.00254583 | 0.998236921 | 0.982044827 | 0.990314032 | 393.0381088 | 392.5404065 | ENSG00000135974 |
| IMP4 | -0.002638485 | 0.998172813 | 0.964464515 | 0.9818716 | 2080.828642 | 2077.334531 | ENSG00000136718 |
| ADNP | -0.002655116 | 0.998161306 | 0.96514962 | 0.982221968 | 2520.208124 | 2515.687431 | ENSG00000101126 |
| SMC1A | -0.00281272 | 0.99805227 | 0.952916116 | 0.975888969 | 4637.32811 | 4628.180603 | ENSG00000072501 |
| ZC3H7A | -0.002858264 | 0.998020764 | 0.97012996 | 0.98427465 | 925.6070436 | 923.895382 | ENSG00000122299 |
| RBM5 | -0.002976524 | 0.997938957 | 0.972987818 | 0.98595118 | 625.6311423 | 624.3303044 | ENSG00000003756 |
| BORCS5 | -0.002983659 | 0.997934022 | 0.98521055 | 0.991584323 | 186.5316866 | 186.3668583 | ENSG00000165714 |
| NA | -0.003065178 | 0.997877636 | 0.987703157 | 0.992901595 | 113.8943535 | 113.6889332 | ENSG00000284648 |
| VPS39 | -0.003133041 | 0.997830698 | 0.964016551 | 0.981778233 | 1339.474744 | 1336.061582 | ENSG00000166887 |
| TMEM59 | -0.003277166 | 0.997731019 | 0.962453302 | 0.980727225 | 1501.863938 | 1498.811965 | ENSG00000116209 |
| MAP3K10 | -0.003309259 | 0.997708825 | 0.972291798 | 0.985471606 | 588.0566773 | 586.6886545 | ENSG00000130758 |
| CLP1 | -0.003409082 | 0.997639794 | 0.969998363 | 0.98427465 | 628.8563501 | 627.1828572 | ENSG00000172409 |
| POLR2D | -0.003439223 | 0.997618952 | 0.960208211 | 0.979722803 | 1423.689555 | 1420.417394 | ENSG00000144231 |
| GARNL3 | -0.00352434 | 0.997560095 | 0.984940569 | 0.991467779 | 128.964226 | 128.6219198 | ENSG00000136895 |

| | | | | | | | |
|---|---|---|---|---|---|---|---|
| C6orf118 | -0.003557099 | 0.997537444 | 0.994646073 | 0.997056698 | 16.0734378 | 15.99264063 | ENSG00000112539 |
| KDM2B | -0.003560608 | 0.997535017 | 0.941192695 | 0.96886597 | 2891.905353 | 2884.613678 | ENSG00000089094 |
| SLC25A3 | -0.003571729 | 0.997527328 | 0.928689322 | 0.961846113 | 9681.752465 | 9657.107016 | ENSG00000075415 |
| SNAPC2 | -0.0036206 | 0.997493538 | 0.970840238 | 0.984569395 | 488.8588271 | 487.6304756 | ENSG00000104976 |
| TIAM2 | -0.003668039 | 0.997460739 | 0.981599314 | 0.990314032 | 203.056839 | 202.5528269 | ENSG00000146426 |
| GTPBP4 | -0.003684763 | 0.997449176 | 0.944107923 | 0.97077906 | 2469.711328 | 2462.942718 | ENSG00000107937 |
| C1GALT1C1 | -0.003832066 | 0.997347339 | 0.973102245 | 0.98595118 | 373.6245847 | 372.7167782 | ENSG00000171155 |
| FANCG | -0.003910457 | 0.997293148 | 0.954716032 | 0.977051829 | 1183.731035 | 1180.461389 | ENSG00000221829 |
| DNAJC9-AS1 | -0.004119158 | 0.997148889 | 0.99472117 | 0.997056698 | 12.17098884 | 12.06969925 | ENSG00000227540 |
| MSL2 | -0.004121947 | 0.997146962 | 0.965468591 | 0.982271747 | 779.6546237 | 777.4400524 | ENSG00000174579 |
| KBTBD2 | -0.004219039 | 0.997079857 | 0.964626859 | 0.981985052 | 569.9670605 | 568.5333727 | ENSG00000170852 |
| POC5 | -0.004227351 | 0.997074112 | 0.980140508 | 0.989527451 | 207.2292305 | 206.2160572 | ENSG00000152359 |
| TMCC2 | -0.004308325 | 0.997018151 | 0.973044416 | 0.98595118 | 301.6499746 | 300.9006055 | ENSG00000133069 |
| CAVIN1 | -0.004321135 | 0.997009298 | 0.951416779 | 0.975020244 | 1142.230806 | 1139.203035 | ENSG00000177469 |
| PLCXD1 | -0.004422398 | 0.996939321 | 0.929325058 | 0.962113616 | 3244.796686 | 3234.596271 | ENSG00000182378 |
| CSNK1A1 | -0.004498732 | 0.996886574 | 0.936304281 | 0.966102155 | 3244.70196 | 3234.617982 | ENSG00000113712 |
| MEIS2 | -0.004512892 | 0.996876789 | 0.996414613 | 0.997983846 | 6.077121709 | 6.083110458 | ENSG00000134138 |
| CEP76 | -0.004514906 | 0.996875398 | 0.975019525 | 0.986836393 | 224.9096719 | 224.0125794 | ENSG00000101624 |
| NA | -0.004600447 | 0.996816292 | 0.995461773 | 0.997339499 | 8.290191376 | 8.186552699 | ENSG00000234337 |
| RAB3A | -0.004604253 | 0.996813663 | 0.977807066 | 0.988361337 | 202.476855 | 201.6849138 | ENSG00000105649 |
| GAK | -0.004623436 | 0.996800408 | 0.932293927 | 0.963923366 | 2243.672537 | 2236.131635 | ENSG00000178950 |
| FAM120C | -0.004629268 | 0.996796379 | 0.976968177 | 0.988031093 | 176.2017498 | 175.7195339 | ENSG00000184083 |
| ZZZ3 | -0.00480542 | 0.996674678 | 0.956309037 | 0.977852096 | 742.8859815 | 740.2625282 | ENSG00000036549 |
| HYDIN | -0.004892378 | 0.996614606 | 0.991790811 | 0.995195658 | 19.76101136 | 19.66902269 | ENSG00000157423 |
| DPM3 | -0.004906286 | 0.996604998 | 0.966191914 | 0.982489496 | 355.9213405 | 354.5731223 | ENSG00000179085 |
| ORC4 | -0.00500589 | 0.996536194 | 0.951235829 | 0.974894991 | 903.292109 | 900.3483258 | ENSG00000115947 |
| ST13 | -0.005038814 | 0.996513453 | 0.898804053 | 0.944277801 | 7384.396836 | 7359.348017 | ENSG00000100380 |
| DGKE | -0.005092956 | 0.996476055 | 0.976537397 | 0.987906179 | 181.0852735 | 180.3907435 | ENSG00000153933 |
| ZNF157 | -0.005119429 | 0.99645777 | 0.992253936 | 0.995401491 | 18.28371656 | 18.18668585 | ENSG00000147117 |
| DICER1 | -0.005173463 | 0.99642045 | 0.948871467 | 0.973549595 | 1138.519515 | 1134.305359 | ENSG00000100697 |
| COMTD1 | -0.005185047 | 0.99641245 | 0.964224159 | 0.981782342 | 458.1742158 | 456.5470384 | ENSG00000165644 |
| STAG2 | -0.005188246 | 0.996410241 | 0.922300485 | 0.95841945 | 3604.714129 | 3592.496556 | ENSG00000101972 |
| SNX4 | -0.005387374 | 0.996272721 | 0.957590733 | 0.978281143 | 573.2704647 | 571.6125661 | ENSG00000114520 |
| FANCC | -0.005411999 | 0.996255716 | 0.964966296 | 0.982175108 | 317.8514064 | 316.8055936 | ENSG00000158169 |
| KIFC1 | -0.005557578 | 0.99615519 | 0.919739448 | 0.95720186 | 2250.922155 | 2241.561102 | ENSG00000237649 |
| NA | -0.005631401 | 0.996104219 | 0.978633348 | 0.98893745 | 115.3030639 | 114.6824342 | ENSG00000224631 |
| ADGRL2 | -0.005648518 | 0.996092401 | 0.932578125 | 0.96416553 | 2198.191535 | 2189.775982 | ENSG00000117114 |
| NA | -0.005670463 | 0.996077249 | 0.995399361 | 0.997339499 | 4.604532834 | 4.615034138 | ENSG00000219928 |
| DNAJC28 | -0.005730116 | 0.996036063 | 0.987515151 | 0.992867816 | 33.08548637 | 33.00540099 | ENSG00000177692 |
| NQO1 | -0.005864157 | 0.995943526 | 0.935811565 | 0.965748724 | 1001.190518 | 997.0352821 | ENSG00000181019 |
| UBE2J2 | -0.005899121 | 0.995919389 | 0.926570818 | 0.960672598 | 1784.462043 | 1777.064427 | ENSG00000160087 |
| AP1AR | -0.006072823 | 0.995799487 | 0.947865869 | 0.972984731 | 604.5342722 | 602.1187762 | ENSG00000138660 |
| MIS18BP1 | -0.006285889 | 0.995652432 | 0.932111551 | 0.963838118 | 1091.587165 | 1087.550851 | ENSG00000129534 |
| BEST2 | -0.006286842 | 0.995651774 | 0.981792916 | 0.990314032 | 58.72269855 | 58.36013734 | ENSG00000039987 |
| CCDC71 | -0.006372193 | 0.995592872 | 0.949835735 | 0.974057952 | 591.7487566 | 589.1636988 | ENSG00000177352 |
| REV1 | -0.006442886 | 0.995544089 | 0.946189168 | 0.972141843 | 595.690352 | 592.7034491 | ENSG00000135945 |
| NA | -0.006484986 | 0.995515038 | 0.99483424 | 0.997056698 | 4.658999449 | 4.663431258 | ENSG00000233155 |
| CHCHD4 | -0.006546021 | 0.995472922 | 0.955693122 | 0.977429542 | 379.4078098 | 377.7760193 | ENSG00000163528 |
| LOC101927060 | -0.006706018 | 0.995362529 | 0.964323501 | 0.98183167 | 204.9419577 | 203.8254368 | ENSG00000262879 |
| NDEL1 | -0.006832556 | 0.99527523 | 0.9385554 | 0.967596848 | 641.1313292 | 638.2894241 | ENSG00000166579 |
| NA | -0.006834981 | 0.995273557 | 0.988965915 | 0.99374992 | 22.04075039 | 22.03595626 | ENSG00000229344 |
| RBP7 | -0.006867681 | 0.995250999 | 0.985500098 | 0.991772257 | 32.54959324 | 32.28562334 | ENSG00000162444 |
| BTAF1 | -0.006885318 | 0.995238832 | 0.918811222 | 0.956597079 | 1315.029904 | 1308.155459 | ENSG00000095564 |
| ATP1B3 | -0.007031969 | 0.99513767 | 0.885673912 | 0.937037652 | 3409.928394 | 3393.651358 | ENSG00000069849 |
| TCF12 | -0.007063838 | 0.995115688 | 0.910910672 | 0.951993281 | 1770.981388 | 1762.972438 | ENSG00000140262 |
| FAM83F | -0.007118427 | 0.995078035 | 0.92774948 | 0.961187788 | 885.2118402 | 880.9990125 | ENSG00000133477 |
| STAC | -0.007140659 | 0.995062701 | 0.958168068 | 0.978611827 | 256.97222 | 255.6563972 | ENSG00000144681 |
| DDX20 | -0.00723182 | 0.994999827 | 0.925367756 | 0.960029077 | 873.943981 | 869.518972 | ENSG00000064703 |
| MND1 | -0.007252071 | 0.99498586 | 0.958098725 | 0.978592815 | 314.4907958 | 313.3921577 | ENSG00000121211 |
| ENTPD2 | -0.007287643 | 0.994961328 | 0.974178331 | 0.98650629 | 97.17784111 | 96.50004945 | ENSG00000054179 |
| SYNGR3 | -0.007300888 | 0.994952193 | 0.940738973 | 0.968657354 | 644.218694 | 641.2581016 | ENSG00000127561 |
| ANKRD34B | -0.007331942 | 0.994930777 | 0.976680514 | 0.987990029 | 72.17967331 | 71.91269436 | ENSG00000189127 |
| TXNL1 | -0.007353559 | 0.99491587 | 0.903057264 | 0.947301669 | 1585.756425 | 1577.772793 | ENSG00000091164 |
| ARHGAP10 | -0.007360504 | 0.99491108 | 0.965830821 | 0.982433028 | 172.1982647 | 171.4935548 | ENSG00000071205 |
| HK2 | -0.007375093 | 0.994901019 | 0.910521955 | 0.951922774 | 2468.260639 | 2454.859102 | ENSG00000159399 |
| CDKN2D | -0.007383365 | 0.994895315 | 0.983559594 | 0.99095671 | 35.27044685 | 35.13626765 | ENSG00000129355 |
| C3orf52 | -0.007425125 | 0.994866517 | 0.952691151 | 0.975846222 | 295.2007599 | 293.522586 | ENSG00000114529 |
| HPS5 | -0.007471574 | 0.994834487 | 0.943929184 | 0.970698752 | 453.2896146 | 450.7757921 | ENSG00000110756 |
| GRHPR | -0.007473173 | 0.994833385 | 0.887057026 | 0.937552986 | 2410.576224 | 2397.994998 | ENSG00000137106 |
| MOSPD3 | -0.00752656 | 0.994796571 | 0.958839058 | 0.978986147 | 239.1407816 | 238.2467367 | ENSG00000106330 |
| CD164 | -0.007575269 | 0.994762985 | 0.926016929 | 0.960479636 | 1583.20199 | 1575.575717 | ENSG00000135535 |
| ACTR6 | -0.007584471 | 0.99475664 | 0.944339215 | 0.970809901 | 406.3068778 | 404.292026 | ENSG00000075089 |
| LOC102723582 | -0.007588972 | 0.994753537 | 0.994136102 | 0.996874912 | 4.94508688 | 4.941183107 | ENSG00000272758 |
| SLC4A2 | -0.007723393 | 0.994660856 | 0.898997353 | 0.944354114 | 2462.619367 | 2449.627361 | ENSG00000164889 |
| LPP-AS2 | -0.007790143 | 0.994614836 | 0.994729599 | 0.997056698 | 2.982698788 | 2.984822334 | ENSG00000270959 |
| PPP4R3A | -0.007847419 | 0.99457535 | 0.890449791 | 0.939388982 | 2102.791533 | 2091.450884 | ENSG00000100796 |
| DAPK3 | -0.007946044 | 0.994507362 | 0.907362842 | 0.949958569 | 2078.440785 | 2067.169306 | ENSG00000167657 |

| | | | | | | | |
|---|---|---|---|---|---|---|---|
| PTPRK | -0.008068812 | 0.994422737 | 0.908306007 | 0.950636457 | 1149.457308 | 1143.025686 | ENSG00000152894 |
| WASH2P | -0.008096407 | 0.994403716 | 0.977595016 | 0.988353504 | 55.4828605 | 55.31427491 | ENSG00000146556 |
| ZBTB11-AS1 | -0.008097134 | 0.994403215 | 0.983206206 | 0.990755899 | 29.04407996 | 28.88672056 | ENSG00000256628 |
| MEX3C | -0.008269801 | 0.994284209 | 0.88775784 | 0.937989799 | 1846.787523 | 1836.506759 | ENSG00000176624 |
| TAB3 | -0.008331913 | 0.994241403 | 0.952728617 | 0.975846222 | 290.0430761 | 288.388329 | ENSG00000157625 |
| MTF2 | -0.008413635 | 0.994185085 | 0.890624319 | 0.939403184 | 1989.310574 | 1978.21496 | ENSG00000143033 |
| GDI2 | -0.008422439 | 0.994179018 | 0.837199938 | 0.907378167 | 9169.278639 | 9116.115996 | ENSG00000057608 |
| CKMT2-AS1 | -0.008428225 | 0.994175031 | 0.968353539 | 0.983367014 | 120.5740746 | 119.7731098 | ENSG00000247572 |
| ISCA1 | -0.008486704 | 0.994134734 | 0.920804044 | 0.957693333 | 739.2032388 | 734.9376274 | ENSG00000135070 |
| RSRC1 | -0.008530658 | 0.994104446 | 0.947280054 | 0.972798489 | 316.8966507 | 314.6556564 | ENSG00000174891 |
| ZNF768 | -0.008598994 | 0.99405736 | 0.909365352 | 0.951248262 | 984.2229752 | 978.8060688 | ENSG00000169957 |
| NUP155 | -0.00869129 | 0.993993766 | 0.89236188 | 0.940480388 | 1784.586042 | 1774.265317 | ENSG00000113569 |
| POU5F1P4 | -0.008813209 | 0.99390977 | 0.983365003 | 0.990812403 | 30.35067825 | 30.16738755 | ENSG00000237872 |
| LEMD1 | -0.008832175 | 0.993896704 | 0.986486778 | 0.992213294 | 18.37849512 | 18.34639819 | ENSG00000186007 |
| G3BP1 | -0.008926247 | 0.993831898 | 0.820648755 | 0.897762002 | 7172.481084 | 7128.275782 | ENSG00000145907 |
| NOTCH2NLC | -0.008983097 | 0.993792737 | 0.993278047 | 0.996221629 | 4.30516661 | 4.297487464 | ENSG00000286219 |
| PMS1 | -0.009002596 | 0.993779305 | 0.918530184 | 0.956469201 | 910.9099909 | 906.0385706 | ENSG00000064933 |
| SMG8 | -0.00900653 | 0.993776595 | 0.92308377 | 0.958820213 | 638.5144275 | 634.4683716 | ENSG00000167447 |
| PLCG1-AS1 | -0.009029869 | 0.993760519 | 0.989010664 | 0.99374992 | 11.16587196 | 11.14825176 | ENSG00000226648 |
| NME4 | -0.009066135 | 0.993735538 | 0.85073297 | 0.915101063 | 5236.728407 | 5203.966783 | ENSG00000103202 |
| NRDC | -0.009090368 | 0.993718847 | 0.850871794 | 0.915101063 | 3612.039602 | 3589.271294 | ENSG00000078618 |
| BPTF | -0.009095215 | 0.993715508 | 0.848746055 | 0.913782797 | 3616.085425 | 3593.335561 | ENSG00000171634 |
| TAX1BP3 | -0.00913184 | 0.993690281 | 0.991941611 | 0.995295205 | 5.995421786 | 5.984165645 | ENSG00000213977 |
| GRIK5 | -0.009182229 | 0.993655575 | 0.899692929 | 0.944800016 | 1471.835877 | 1462.061994 | ENSG00000105737 |
| DOCK5 | -0.009314358 | 0.993564576 | 0.943316321 | 0.970367312 | 273.9736174 | 272.0682008 | ENSG00000147459 |
| SNX25 | -0.00937306 | 0.993524149 | 0.938802833 | 0.967624505 | 337.0490912 | 335.1299346 | ENSG00000109762 |
| POGLUT1 | -0.009388444 | 0.993513555 | 0.939762014 | 0.968219879 | 326.5481824 | 324.1287764 | ENSG00000163389 |
| ETNK1 | -0.009429537 | 0.993485257 | 0.918545011 | 0.956469201 | 894.5764341 | 888.6789611 | ENSG00000139163 |
| REEP4 | -0.009570506 | 0.993388186 | 0.907584358 | 0.950087463 | 848.9917581 | 843.6507015 | ENSG00000168476 |
| ZNF738 | -0.009582002 | 0.99338027 | 0.899492315 | 0.944674599 | 930.7409366 | 924.2393112 | ENSG00000172687 |
| ANKRD12 | -0.009639104 | 0.993340952 | 0.93609726 | 0.965991882 | 326.4259334 | 324.3285561 | ENSG00000101745 |
| C6orf47 | -0.009860343 | 0.993188634 | 0.950024242 | 0.974110644 | 229.4131318 | 228.2086867 | ENSG00000204439 |
| MBTPS1 | -0.009915177 | 0.993150886 | 0.882706225 | 0.935911408 | 1576.279205 | 1565.713791 | ENSG00000140943 |
| PDCL | -0.00991742 | 0.993149342 | 0.922233662 | 0.95841945 | 495.2594821 | 491.9470336 | ENSG00000136940 |
| PDE6D | -0.009955994 | 0.993122788 | 0.929340827 | 0.962113616 | 509.1738413 | 505.8588101 | ENSG00000156973 |
| NA | -0.009956222 | 0.993122631 | 0.991739035 | 0.995195471 | 5.758894886 | 5.740289938 | ENSG00000227014 |
| TAFA4 | -0.009979429 | 0.993106656 | 0.952333438 | 0.975545048 | 172.7909398 | 171.5209913 | ENSG00000163377 |
| POLRMT | -0.009996345 | 0.993095011 | 0.885127461 | 0.936934918 | 3894.971209 | 3868.358756 | ENSG00000099821 |
| UBIAD1 | -0.010070243 | 0.993044144 | 0.915879483 | 0.954940963 | 674.1589945 | 669.0932983 | ENSG00000120942 |
| GTDC1 | -0.010186067 | 0.992964422 | 0.942117498 | 0.969611004 | 259.607068 | 257.9103738 | ENSG00000121964 |
| ARMT1 | -0.010209238 | 0.992948475 | 0.908896976 | 0.950945415 | 1011.167 | 1004.909601 | ENSG00000146476 |
| LIPE | -0.010219375 | 0.992941498 | 0.944033907 | 0.970754696 | 242.9489276 | 241.4302848 | ENSG00000079435 |
| LRBA | -0.01022012 | 0.992940985 | 0.878677085 | 0.933298075 | 2141.86215 | 2125.89423 | ENSG00000198589 |
| LONP2 | -0.01023852 | 0.992928321 | 0.909320057 | 0.951248262 | 653.6790658 | 649.2099202 | ENSG00000102910 |
| NA | -0.010330396 | 0.99286509 | 0.981428377 | 0.990314032 | 23.08637938 | 22.8788986 | ENSG00000224094 |
| NA | -0.01033064 | 0.992864922 | 0.984790669 | 0.991420352 | 15.52917196 | 15.34034269 | ENSG00000265095 |
| NUDT2 | -0.010515369 | 0.9927378 | 0.94922736 | 0.973739099 | 199.7497439 | 198.3053541 | ENSG00000164978 |
| PABPC1 | -0.010624746 | 0.992662539 | 0.829850246 | 0.903182315 | 27059.00378 | 26861.94498 | ENSG00000070756 |
| H2AJ | -0.010695105 | 0.992614129 | 0.930746535 | 0.963034317 | 330.3947764 | 327.9471453 | ENSG00000246705 |
| ATP5F1B | -0.010718035 | 0.992598353 | 0.769720367 | 0.864350405 | 20491.65552 | 20339.07542 | ENSG00000110955 |
| STYXL1 | -0.010722609 | 0.992595205 | 0.917861403 | 0.956072588 | 462.9152275 | 459.60578 | ENSG00000127952 |
| COA1 | -0.010739949 | 0.992583275 | 0.887338047 | 0.937747248 | 2008.417643 | 1992.81617 | ENSG00000106603 |
| TMEM120B | -0.010754032 | 0.992573586 | 0.89952453 | 0.944674599 | 718.8144219 | 713.4198152 | ENSG00000188735 |
| RAB9A | -0.010801902 | 0.992540652 | 0.923866014 | 0.959186142 | 422.688418 | 419.9547375 | ENSG00000123595 |
| NUDT22 | -0.010977434 | 0.992419898 | 0.923997425 | 0.959186142 | 367.300248 | 364.7630758 | ENSG00000149761 |
| AGTPBP1 | -0.011010375 | 0.992397238 | 0.895863708 | 0.942392226 | 1098.971072 | 1091.444174 | ENSG00000135049 |
| CDV3 | -0.011023215 | 0.992388406 | 0.817132311 | 0.895489372 | 3223.59694 | 3198.790991 | ENSG00000091527 |
| GTF3C3 | -0.011068925 | 0.992356964 | 0.886391223 | 0.937353813 | 1087.945796 | 1079.420231 | ENSG00000119041 |
| CCAR1 | -0.011085152 | 0.992345802 | 0.837697516 | 0.907563032 | 2508.333463 | 2489.396885 | ENSG00000060339 |
| PHF8 | -0.011225397 | 0.99224934 | 0.888896634 | 0.938578344 | 966.8716488 | 958.9922005 | ENSG00000172943 |
| RNF149 | -0.01122664 | 0.992248485 | 0.92497556 | 0.959854951 | 370.9909007 | 368.2006889 | ENSG00000163162 |
| NA | -0.011242906 | 0.992237298 | 0.989942433 | 0.994130074 | 5.611564737 | 5.592948006 | ENSG00000279392 |
| NA | -0.011244074 | 0.992236495 | 0.986466015 | 0.992213294 | 12.78279992 | 12.7257653 | ENSG00000232600 |
| ERLIN1 | -0.011245809 | 0.992235301 | 0.879093126 | 0.93343161 | 1005.893948 | 998.2003143 | ENSG00000107566 |
| MAD2L1 | -0.01124826 | 0.992233616 | 0.838267448 | 0.907821104 | 2279.777843 | 2262.475783 | ENSG00000164109 |
| NA | -0.011393711 | 0.992133585 | 0.961321846 | 0.980275914 | 78.3927614 | 77.81485931 | ENSG00000264577 |
| SPATA20 | -0.011470787 | 0.992080581 | 0.902851289 | 0.947188613 | 580.3132914 | 575.6541106 | ENSG00000006282 |
| CNP | -0.011518892 | 0.992047502 | 0.852245946 | 0.916006883 | 1621.926386 | 1608.923243 | ENSG00000173786 |
| RFTN1 | -0.011589797 | 0.991998747 | 0.95464167 | 0.977027559 | 111.6374173 | 110.7654115 | ENSG00000131378 |
| NECAB1 | -0.011591701 | 0.991997438 | 0.954911503 | 0.977148196 | 114.9647003 | 114.2135049 | ENSG00000123119 |
| NA | -0.011604555 | 0.991988599 | 0.926419119 | 0.960629501 | 409.3076863 | 405.7923731 | ENSG00000224114 |
| TRAPPC9 | -0.0116292 | 0.991971654 | 0.913995581 | 0.95379686 | 485.1165357 | 481.0700941 | ENSG00000167632 |
| KCNQ2 | -0.011656611 | 0.991952806 | 0.835748758 | 0.906924912 | 2396.557889 | 2376.907187 | ENSG00000075043 |
| RNF185 | -0.011742122 | 0.991894014 | 0.89946469 | 0.944674599 | 569.4107041 | 564.6402973 | ENSG00000138942 |
| CNTNAP3C | -0.011744227 | 0.991892566 | 0.981502257 | 0.990314032 | 20.56433148 | 20.34497693 | ENSG00000283378 |
| NA | -0.011907106 | 0.991780588 | 0.98007686 | 0.989514979 | 24.40518041 | 24.28028871 | ENSG00000234825 |
| CISD2 | -0.012086162 | 0.991657504 | 0.938867566 | 0.967624505 | 285.1620309 | 283.094124 | ENSG00000145354 |

| | | | | | | | |
|---|---|---|---|---|---|---|---|
| NT5DC2 | -0.012186854 | 0.991588295 | 0.83515176 | 0.906737673 | 7119.977624 | 7059.80493 | ENSG00000168268 |
| NA | -0.012204375 | 0.991576252 | 0.990585152 | 0.994554917 | 3.98973069 | 3.97348907 | ENSG00000243960 |
| ZNF570 | -0.012294926 | 0.991514018 | 0.960163884 | 0.979722803 | 81.54723274 | 80.87873826 | ENSG00000171827 |
| UBA2 | -0.012339533 | 0.991483361 | 0.815672182 | 0.894616739 | 4080.384519 | 4045.875076 | ENSG00000126261 |
| AZGP1 | -0.012376059 | 0.99145826 | 0.986231959 | 0.992146471 | 8.978120574 | 8.935644875 | ENSG00000160862 |
| DIP2C | -0.012390414 | 0.991448394 | 0.904422916 | 0.948084184 | 461.5803567 | 457.6655393 | ENSG00000151240 |
| HIVEP3 | -0.012408 | 0.991436309 | 0.978805013 | 0.988963906 | 25.04809175 | 24.79493151 | ENSG00000127124 |
| NA | -0.012478735 | 0.9913877 | 0.989934954 | 0.994130074 | 4.604532834 | 4.58330857 | ENSG00000259488 |
| SEC23IP | -0.012491723 | 0.991378775 | 0.845215922 | 0.911559253 | 1525.230708 | 1512.18494 | ENSG00000107651 |
| PARG | -0.012507712 | 0.991367788 | 0.858207724 | 0.920629773 | 1239.550996 | 1228.893423 | ENSG00000227345 |
| SGK1 | -0.012528702 | 0.991353364 | 0.924272121 | 0.959383023 | 339.1847282 | 335.716946 | ENSG00000118515 |
| TRAF3IP2-AS1 | -0.012545714 | 0.991341675 | 0.965266916 | 0.982221968 | 63.45829168 | 62.98833539 | ENSG00000231889 |
| ZNF764 | -0.012548922 | 0.99133947 | 0.945409445 | 0.97154771 | 142.6598429 | 141.5256865 | ENSG00000169951 |
| ACAD8 | -0.012558574 | 0.991332838 | 0.860032638 | 0.921437017 | 1428.706842 | 1415.71131 | ENSG00000151498 |
| LSM1 | -0.012633276 | 0.991281509 | 0.931409733 | 0.963382625 | 354.3302944 | 351.1796236 | ENSG00000175324 |
| NA | -0.012719242 | 0.991222443 | 0.962950975 | 0.981055572 | 72.6981453 | 72.15925367 | ENSG00000230074 |
| CBLL1 | -0.012785081 | 0.991177208 | 0.891349493 | 0.93992692 | 585.8450712 | 580.8077043 | ENSG00000105879 |
| TEC | -0.012824224 | 0.991150316 | 0.965559728 | 0.982312663 | 49.08492123 | 48.69071477 | ENSG00000135605 |
| NA | -0.012887445 | 0.991106884 | 0.989940648 | 0.994130074 | 3.98973069 | 3.971338496 | ENSG00000234286 |
| NA | -0.012894973 | 0.991101712 | 0.97518636 | 0.986953469 | 25.85208758 | 25.56929753 | ENSG00000267565 |
| RAD18 | -0.012919371 | 0.991084951 | 0.894617783 | 0.941915276 | 502.6254928 | 498.1070438 | ENSG00000070950 |
| SNX5 | -0.012931092 | 0.991076899 | 0.840333707 | 0.908946853 | 1929.497389 | 1912.1985 | ENSG00000089006 |
| UBTD2 | -0.013046951 | 0.990997312 | 0.87544647 | 0.931097025 | 877.7721019 | 869.8803599 | ENSG00000168246 |
| RAB3B | -0.013146772 | 0.990928746 | 0.826842577 | 0.901242194 | 1582.285366 | 1567.972148 | ENSG00000169213 |
| ADD1 | -0.013382043 | 0.990767161 | 0.789578913 | 0.87730485 | 3505.449444 | 3472.421397 | ENSG00000087274 |
| CNGA1 | -0.013388003 | 0.990763069 | 0.982310633 | 0.990351054 | 12.2930007 | 12.22511044 | ENSG00000198515 |
| NA | -0.013404511 | 0.990751732 | 0.969217444 | 0.983955795 | 44.27157036 | 43.87348986 | ENSG00000230715 |
| THY1 | -0.013466596 | 0.990709096 | 0.746934562 | 0.850095258 | 6343.900894 | 6283.92106 | ENSG00000154096 |
| KEAP1 | -0.013469038 | 0.990707419 | 0.826149574 | 0.901078467 | 2187.462854 | 2167.20701 | ENSG00000079999 |
| ZNF579 | -0.013517221 | 0.990674332 | 0.945406339 | 0.97154771 | 785.6480709 | 778.4675211 | ENSG00000218891 |
| ACTR3 | -0.013546899 | 0.990653954 | 0.809760555 | 0.890395056 | 3441.16919 | 3410.034485 | ENSG00000115091 |
| GPATCH11 | -0.013565306 | 0.990641314 | 0.954109698 | 0.976742208 | 87.97731141 | 87.16941882 | ENSG00000152133 |
| RPL37A | -0.013682119 | 0.990561106 | 0.788540813 | 0.876555309 | 13585.2919 | 13455.84128 | ENSG00000197756 |
| RABIF | -0.013694899 | 0.990552332 | 0.935338359 | 0.965468934 | 178.5112376 | 176.8544765 | ENSG00000183155 |
| ZSCAN32 | -0.013716632 | 0.99053741 | 0.92371792 | 0.959169039 | 279.4451855 | 276.7552762 | ENSG00000140987 |
| AGAP1 | -0.013973735 | 0.990360902 | 0.835666599 | 0.906888184 | 1399.930977 | 1386.564744 | ENSG00000157985 |
| TCF19 | -0.014058071 | 0.990303009 | 0.904055105 | 0.947822284 | 409.7509785 | 406.0542653 | ENSG00000137310 |
| TNFAIP8 | -0.014181772 | 0.990218102 | 0.911230933 | 0.951993281 | 277.4117487 | 274.7435457 | ENSG00000145779 |
| POU5F1 | -0.014181812 | 0.990218074 | 0.683134967 | 0.80675584 | 52923.55386 | 52404.97403 | ENSG00000204531 |
| L3MBTL1 | -0.014379739 | 0.990082233 | 0.949811728 | 0.974057952 | 92.24718431 | 91.20314238 | ENSG00000185513 |
| CD320 | -0.014460534 | 0.990026787 | 0.825228859 | 0.900593514 | 1577.097221 | 1562.238837 | ENSG00000167775 |
| ZMYM2 | -0.014461066 | 0.990026422 | 0.789792034 | 0.877393205 | 2574.433974 | 2548.374298 | ENSG00000121741 |
| CDK5RAP2 | -0.01446977 | 0.990020449 | 0.829864005 | 0.903182315 | 1496.404419 | 1480.873257 | ENSG00000136861 |
| YIPF4 | -0.014605355 | 0.989927411 | 0.903350989 | 0.947370846 | 359.6994253 | 356.4307987 | ENSG00000119820 |
| SERTM1 | -0.014627107 | 0.989912486 | 0.9719829 | 0.985262053 | 26.69783482 | 26.51197819 | ENSG00000180440 |
| PTCHD1 | -0.014651389 | 0.989895825 | 0.957944315 | 0.97848691 | 75.50236257 | 74.70177764 | ENSG00000165186 |
| NA | -0.014739666 | 0.989835256 | 0.93811245 | 0.967346979 | 161.3584287 | 159.6610006 | ENSG00000280128 |
| CHAF1A | -0.014745125 | 0.989831511 | 0.755653582 | 0.855091825 | 4711.798053 | 4664.025223 | ENSG00000167670 |
| TMTC1 | -0.014777597 | 0.989809232 | 0.901767023 | 0.946462875 | 379.0977044 | 374.9057593 | ENSG00000133687 |
| POLD1 | -0.014778139 | 0.98980886 | 0.802554562 | 0.88621529 | 4020.875965 | 3980.039399 | ENSG00000062822 |
| RNF181 | -0.014810466 | 0.989786681 | 0.867802363 | 0.926309156 | 679.1678335 | 671.9385207 | ENSG00000168894 |
| NA | -0.014812981 | 0.989784955 | 0.986082053 | 0.992047399 | 6.643039044 | 6.594518184 | ENSG00000231563 |
| ZNF875 | -0.014827402 | 0.989775062 | 0.88398287 | 0.936493029 | 520.1064017 | 514.4452069 | ENSG00000181666 |
| CDKN3 | -0.014835289 | 0.989769651 | 0.90406577 | 0.947822284 | 510.1090095 | 505.1728475 | ENSG00000100526 |
| NAT16 | -0.014845014 | 0.989762979 | 0.987657804 | 0.992901595 | 5.042856499 | 5.006251778 | ENSG00000167011 |
| CDH22 | -0.014855138 | 0.989756033 | 0.9797961 | 0.989307559 | 12.95947856 | 12.86880608 | ENSG00000149654 |
| TCERG1 | -0.014859914 | 0.989752757 | 0.788513051 | 0.876555309 | 2697.374406 | 2669.617344 | ENSG00000113649 |
| C22orf46 | -0.014911774 | 0.98971718 | 0.909828339 | 0.951403864 | 344.3037028 | 340.2773733 | ENSG00000184208 |
| HCG18 | -0.014923705 | 0.989708994 | 0.890658039 | 0.939403184 | 513.5301441 | 507.8313457 | ENSG00000231074 |
| PCCA | -0.01497788 | 0.98967183 | 0.840346847 | 0.908946853 | 972.4319381 | 962.3240898 | ENSG00000175198 |
| TMEM161A | -0.015138061 | 0.989561954 | 0.829238954 | 0.90286078 | 1126.67898 | 1114.843156 | ENSG00000064545 |
| CRTC3-AS1 | -0.015237929 | 0.989493456 | 0.989718469 | 0.994130074 | 3.284855915 | 3.260423609 | ENSG00000259736 |
| SLC37A4 | -0.015272764 | 0.989469564 | 0.863909235 | 0.923726994 | 775.766954 | 766.9939202 | ENSG00000137700 |
| ZKSCAN2 | -0.015286603 | 0.989460072 | 0.912146981 | 0.952535122 | 255.0672015 | 252.274699 | ENSG00000155592 |
| NEK6 | -0.015316895 | 0.989439297 | 0.889677531 | 0.939036453 | 425.1719057 | 420.8826367 | ENSG00000119408 |
| ATP2A1-AS1 | -0.015403811 | 0.98937969 | 0.946013764 | 0.972116951 | 85.90686597 | 84.95359529 | ENSG00000260442 |
| SLC9A3R2 | -0.015507054 | 0.98930889 | 0.901325872 | 0.946051331 | 356.4221784 | 352.8541855 | ENSG00000065054 |
| CYP20A1 | -0.015597067 | 0.989247166 | 0.916938564 | 0.955518152 | 241.3047663 | 238.6866483 | ENSG00000119004 |
| BTNL8 | -0.015653741 | 0.989208307 | 0.984741192 | 0.991420352 | 7.29902901 | 7.239831362 | ENSG00000113303 |
| NA | -0.015676583 | 0.989192645 | 0.989285023 | 0.993870294 | 3.975776176 | 3.941763502 | ENSG00000272848 |
| ST7L | -0.015710104 | 0.98916966 | 0.916155148 | 0.955068844 | 237.8613414 | 235.3589812 | ENSG00000007341 |
| UMPS | -0.015722145 | 0.989161405 | 0.812700728 | 0.892508809 | 1332.88252 | 1318.964343 | ENSG00000114491 |
| CTC-338M12.4 | -0.015801342 | 0.989107107 | 0.962084855 | 0.980632996 | 41.97720109 | 41.49229577 | ENSG00000233937 |
| NA | -0.015808208 | 0.989102399 | 0.982060349 | 0.990314032 | 8.651521043 | 8.58422206 | ENSG00000184100 |
| OFD1 | -0.015813638 | 0.989098676 | 0.895640171 | 0.942339489 | 344.3382328 | 340.2771008 | ENSG00000046651 |
| ANGPTL2 | -0.01581696 | 0.989096399 | 0.886959391 | 0.937513028 | 386.269303 | 382.2598194 | ENSG00000136859 |
| ALDH2 | -0.015895256 | 0.989042721 | 0.877362145 | 0.93256891 | 596.8463537 | 590.0538806 | ENSG00000111275 |

| | | | | | | | |
|---|---|---|---|---|---|---|---|
| NA | -0.015925686 | 0.98902186 | 0.93540205 | 0.965468934 | 130.1023182 | 128.6103734 | ENSG00000279457 |
| ZNF418 | -0.015954572 | 0.989002058 | 0.912981114 | 0.953047547 | 221.3168769 | 218.6299035 | ENSG00000196724 |
| SPSB1 | -0.015969326 | 0.988991944 | 0.94428164 | 0.970809901 | 101.6418649 | 100.3947729 | ENSG00000171621 |
| RBBP8NL | -0.016081062 | 0.98891535 | 0.977199188 | 0.988195006 | 14.5813126 | 14.45922306 | ENSG00000130701 |
| MIR1244-2 | -0.016156229 | 0.988863827 | 0.940445588 | 0.968508435 | 98.10142133 | 97.12895154 | ENSG00000283498 |
| NA | -0.016232218 | 0.988811743 | 0.982635347 | 0.990542848 | 8.738802771 | 8.662194174 | ENSG00000261790 |
| DYNC1LI2 | -0.016258485 | 0.98879374 | 0.799339725 | 0.884159303 | 1481.423943 | 1464.739089 | ENSG00000135720 |
| NA | -0.016339687 | 0.988738087 | 0.977232706 | 0.988195006 | 13.60709581 | 13.49152903 | ENSG00000229994 |
| NCOR2 | -0.016351977 | 0.988729664 | 0.753128441 | 0.853870159 | 6357.768931 | 6286.919665 | ENSG00000196498 |
| TYRO3 | -0.016423082 | 0.988680935 | 0.736986247 | 0.843705033 | 3551.359294 | 3510.635247 | ENSG00000092445 |
| TSC22D4 | -0.016444873 | 0.988666002 | 0.858916034 | 0.920821105 | 731.0109434 | 723.1314437 | ENSG00000166925 |
| PTP4A2 | -0.016454875 | 0.988659148 | 0.764286041 | 0.860451825 | 2786.857573 | 2756.006013 | ENSG00000184007 |
| LRRC34 | -0.01647855 | 0.988642924 | 0.956219753 | 0.97781263 | 52.65115865 | 52.04524901 | ENSG00000171757 |
| CCDC130 | -0.016575276 | 0.988576642 | 0.886412312 | 0.937353813 | 350.7440323 | 346.9089906 | ENSG00000104957 |
| B4GALNT4 | -0.016596814 | 0.988561884 | 0.828438456 | 0.90234353 | 2175.658298 | 2150.723245 | ENSG00000182272 |
| RSBN1L | -0.016706778 | 0.988486537 | 0.877637916 | 0.932759244 | 494.9820059 | 489.6720722 | ENSG00000187257 |
| EDEM1 | -0.016709564 | 0.988484628 | 0.891667767 | 0.939954256 | 384.5716632 | 380.3249523 | ENSG00000134109 |
| NA | -0.016762193 | 0.988448569 | 0.985733157 | 0.991950399 | 5.021204997 | 4.976676784 | ENSG00000271366 |
| GRM7 | -0.016817793 | 0.988410476 | 0.986724205 | 0.992302071 | 4.310748416 | 4.272213618 | ENSG00000196277 |
| CPLANE2 | -0.016820688 | 0.988408493 | 0.945188249 | 0.971423895 | 76.77391842 | 75.78321004 | ENSG00000132881 |
| FAM234B | -0.016864451 | 0.98837851 | 0.854184154 | 0.917589181 | 725.4089764 | 717.4147432 | ENSG00000084444 |
| LSM6 | -0.016967904 | 0.988307639 | 0.832008863 | 0.904649024 | 882.9566827 | 872.9557731 | ENSG00000164167 |
| MPPED1 | -0.016972952 | 0.98830418 | 0.982202505 | 0.990351054 | 8.218779183 | 8.142456727 | ENSG00000186732 |
| NDUFS8 | -0.016990832 | 0.988291932 | 0.795870435 | 0.88140172 | 1485.492472 | 1468.66987 | ENSG00000110717 |
| TUBB4B | -0.017131959 | 0.98819526 | 0.709726245 | 0.825105495 | 7272.19154 | 7186.267058 | ENSG00000188229 |
| TRUB1 | -0.017145214 | 0.988186181 | 0.867998828 | 0.926342409 | 511.4110652 | 505.1885014 | ENSG00000165832 |
| PPP1R13L | -0.017238201 | 0.988122491 | 0.885160898 | 0.936934918 | 321.4169681 | 317.4896292 | ENSG00000104881 |
| EZR | -0.017266307 | 0.988103241 | 0.703774425 | 0.821822786 | 6442.669664 | 6366.530246 | ENSG00000092820 |
| CES3 | -0.017313556 | 0.988070881 | 0.957156203 | 0.978044405 | 46.34154024 | 45.75564663 | ENSG00000172828 |
| NA | -0.017336392 | 0.988055241 | 0.835593779 | 0.906888184 | 851.0487103 | 841.1613541 | ENSG00000232445 |
| RPSA | -0.017364915 | 0.988035706 | 0.673781626 | 0.800111161 | 18690.05286 | 18465.28907 | ENSG00000168028 |
| NFATC4 | -0.017402742 | 0.9880098 | 0.8639241 | 0.923726994 | 598.9924289 | 591.4773889 | ENSG00000100968 |
| POLDIP3 | -0.017452959 | 0.987975411 | 0.733515099 | 0.840974669 | 3485.730433 | 3442.867566 | ENSG00000100227 |
| CAST | -0.017485554 | 0.98795309 | 0.874684826 | 0.930543483 | 442.7652897 | 437.2019835 | ENSG00000153113 |
| STXBP1 | -0.01758718 | 0.987883499 | 0.79526531 | 0.880883273 | 1344.048941 | 1327.955872 | ENSG00000136854 |
| ABCD3 | -0.017673934 | 0.987824096 | 0.837178097 | 0.907378167 | 681.9246565 | 673.5579612 | ENSG00000117528 |
| SPATA6 | -0.017742047 | 0.98777746 | 0.914361966 | 0.954075978 | 189.6182124 | 187.4157716 | ENSG00000132122 |
| TDH | -0.017759252 | 0.98776568 | 0.974412001 | 0.986590643 | 16.26128005 | 15.96306563 | ENSG00000154316 |
| VEGFA | -0.01779802 | 0.987739137 | 0.838822582 | 0.908136413 | 928.3832028 | 917.1793961 | ENSG00000112715 |
| NA | -0.017798531 | 0.987738787 | 0.987609784 | 0.992901595 | 3.614246349 | 3.577970282 | ENSG00000265791 |
| RREB1 | -0.017801429 | 0.987736803 | 0.827758317 | 0.901908257 | 763.2114835 | 753.6709478 | ENSG00000124782 |
| VTA1 | -0.017817947 | 0.987725494 | 0.812415993 | 0.892246907 | 1117.131602 | 1104.077198 | ENSG00000009844 |
| ESF1 | -0.017879369 | 0.987683443 | 0.8320721 | 0.904666791 | 836.3638973 | 826.7365587 | ENSG00000089048 |
| NPW | -0.01796396 | 0.987625533 | 0.94196155 | 0.969525003 | 83.49758134 | 82.49631919 | ENSG00000183971 |
| METTL3 | -0.018052509 | 0.987564917 | 0.770072684 | 0.864544733 | 2016.257148 | 1990.333469 | ENSG00000165819 |
| NA | -0.018072237 | 0.987551413 | 0.984361215 | 0.991219407 | 5.698846465 | 5.639194551 | ENSG00000286129 |
| NTN5 | -0.018098602 | 0.987533365 | 0.982068766 | 0.990314032 | 6.953568879 | 6.882489863 | ENSG00000142233 |
| SEL1L | -0.018105254 | 0.987528812 | 0.846003341 | 0.912024708 | 676.2923161 | 668.0451117 | ENSG00000071537 |
| APBA3 | -0.01821105 | 0.987456397 | 0.89516044 | 0.942194459 | 341.9528781 | 337.4729877 | ENSG00000011132 |
| NA | -0.018212746 | 0.987455237 | 0.980989129 | 0.990125113 | 7.301819913 | 7.227460957 | ENSG00000285752 |
| ATP6V0E2-AS1 | -0.018399058 | 0.987327723 | 0.969266082 | 0.983955795 | 19.73935986 | 19.53054351 | ENSG00000204934 |
| GPR83 | -0.018406842 | 0.987322396 | 0.985332505 | 0.99165533 | 4.909480864 | 4.856579272 | ENSG00000123901 |
| NA | -0.018416779 | 0.987315595 | 0.986860241 | 0.992312792 | 3.714806871 | 3.674764522 | ENSG00000205274 |
| MTMR7 | -0.018428387 | 0.987307652 | 0.933278644 | 0.964413047 | 96.8692263 | 95.54472582 | ENSG00000003987 |
| NA | -0.018501812 | 0.987257404 | 0.984591466 | 0.991323276 | 5.333850014 | 5.275401332 | ENSG00000276141 |
| DSTN | -0.01850826 | 0.987252992 | 0.755614852 | 0.855091825 | 2512.986407 | 2482.108001 | ENSG00000125868 |
| PXMP2 | -0.018597897 | 0.987191654 | 0.977320195 | 0.988231674 | 11.3076203 | 11.18374544 | ENSG00000176894 |
| HNRNPL | -0.018738577 | 0.987095396 | 0.623707834 | 0.763014769 | 10087.83974 | 9957.17224 | ENSG00000104824 |
| PAFAH1B2 | -0.018745528 | 0.98709064 | 0.772850468 | 0.866604183 | 1874.983916 | 1850.896019 | ENSG00000168092 |
| TOLLIP-AS1 | -0.018858478 | 0.987013363 | 0.973744939 | 0.986270798 | 14.39713714 | 14.23632005 | ENSG00000255153 |
| METTL14 | -0.019012966 | 0.986907676 | 0.840116673 | 0.908905319 | 612.4590157 | 604.6402848 | ENSG00000145388 |
| NSMCE2 | -0.019131098 | 0.986826869 | 0.885215998 | 0.936941831 | 273.5239299 | 269.8370443 | ENSG00000156831 |
| SUGT1 | -0.019224894 | 0.986762713 | 0.761021617 | 0.857878106 | 1825.970558 | 1802.778576 | ENSG00000165416 |
| SLC25A51 | -0.019245968 | 0.986748299 | 0.943431182 | 0.970367312 | 72.78785451 | 71.57150348 | ENSG00000122696 |
| NA | -0.019259806 | 0.986738834 | 0.93335308 | 0.964413047 | 91.05442533 | 89.72593073 | ENSG00000228409 |
| SNX10 | -0.019271199 | 0.986731042 | 0.923292569 | 0.958830585 | 122.8691196 | 121.0384744 | ENSG00000086300 |
| PIP4K2A | -0.019292796 | 0.986716271 | 0.836683385 | 0.907107819 | 828.7980048 | 818.5849361 | ENSG00000150867 |
| PDHA1 | -0.019314315 | 0.986701553 | 0.712901283 | 0.827173021 | 2468.057279 | 2434.684075 | ENSG00000131828 |
| NA | -0.019440876 | 0.986614998 | 0.976773927 | 0.987990029 | 11.38094751 | 11.245046 | ENSG00000253570 |
| NA | -0.01948271 | 0.98658639 | 0.931415692 | 0.963382625 | 87.78966932 | 86.61553266 | ENSG00000174977 |
| SPON1 | -0.019538114 | 0.986548502 | 0.91964406 | 0.957154225 | 137.934174 | 135.9045206 | ENSG00000262655 |
| H3P47 | -0.019577471 | 0.98652159 | 0.968457405 | 0.983367014 | 17.67082944 | 17.4650181 | ENSG00000179101 |
| PFDN6 | -0.019642891 | 0.986476856 | 0.784354692 | 0.873764876 | 1193.796185 | 1177.205818 | ENSG00000204220 |
| NA | -0.019834751 | 0.986345676 | 0.971711736 | 0.985194256 | 13.94764986 | 13.78165128 | ENSG00000280294 |
| VEGFB | -0.019837971 | 0.986343474 | 0.883746677 | 0.936493029 | 401.8671346 | 396.6494428 | ENSG00000173511 |
| NA | -0.019871439 | 0.986320594 | 0.985199701 | 0.991584323 | 4.596160126 | 4.539212598 | ENSG00000287742 |
| NA | -0.019872368 | 0.986319958 | 0.932229488 | 0.963908402 | 79.65605668 | 78.54459633 | ENSG00000180867 |

| | | | | | | | |
|---|---|---|---|---|---|---|---|
| NA | -0.01987622 | 0.986317325 | 0.976482968 | 0.987902923 | 11.15558422 | 11.01784184 | ENSG00000286389 |
| REXO2 | -0.019877603 | 0.986316379 | 0.84423776 | 0.910809833 | 591.5238689 | 583.7736861 | ENSG00000076043 |
| SREK1IP1 | -0.019915902 | 0.986290196 | 0.842461921 | 0.909770449 | 545.9597669 | 538.808302 | ENSG00000153006 |
| LINC01829 | -0.019941688 | 0.986272567 | 0.970754532 | 0.984534234 | 14.92465755 | 14.74504417 | ENSG00000236780 |
| SLC39A10 | -0.019959447 | 0.986260427 | 0.731541885 | 0.839460741 | 2746.522294 | 2709.632119 | ENSG00000196950 |
| NA | -0.02003425 | 0.986209291 | 0.973122724 | 0.98595118 | 13.61056244 | 13.44313191 | ENSG00000272853 |
| RN7SL3 | -0.020109183 | 0.98615807 | 0.975809439 | 0.987428655 | 11.33552933 | 11.19449831 | ENSG00000278771 |
| NKX2-4 | -0.020194586 | 0.986099693 | 0.981807923 | 0.990314032 | 5.345013626 | 5.279702479 | ENSG00000125816 |
| DHRS2 | -0.020248071 | 0.986063137 | 0.967919592 | 0.983248479 | 29.97519391 | 29.58015828 | ENSG00000100867 |
| NBPF10 | -0.020261149 | 0.986054198 | 0.970906229 | 0.984584559 | 17.00925767 | 16.7922805 | ENSG00000271425 |
| CHD3 | -0.0202761 | 0.986043979 | 0.758008665 | 0.856383318 | 1552.053521 | 1529.914595 | ENSG00000170004 |
| INO80E | -0.020435316 | 0.985935165 | 0.776855493 | 0.869225644 | 1237.855308 | 1219.697304 | ENSG00000169592 |
| RB1 | -0.020487945 | 0.9858992 | 0.857341227 | 0.919997317 | 360.432246 | 355.4133505 | ENSG00000139687 |
| RANBP6 | -0.020505931 | 0.985886909 | 0.841164467 | 0.909172105 | 507.1950584 | 500.3771831 | ENSG00000137040 |
| SETMAR | -0.020574729 | 0.985839895 | 0.858887022 | 0.920821105 | 408.7585385 | 402.7198797 | ENSG00000170364 |
| MAP1LC3A | -0.020576357 | 0.985838783 | 0.931210162 | 0.963370796 | 81.6420113 | 80.5961338 | ENSG00000101460 |
| ARL6IP6 | -0.020706982 | 0.985749527 | 0.79781888 | 0.883153957 | 885.4374547 | 873.0423597 | ENSG00000177917 |
| CTTNBP2 | -0.020736726 | 0.985729204 | 0.848480423 | 0.913547795 | 455.3421874 | 448.3862804 | ENSG00000077063 |
| TLCD1 | -0.02075936 | 0.985713739 | 0.897101509 | 0.943156859 | 232.1307189 | 228.8239161 | ENSG00000160606 |
| NA | -0.020810952 | 0.98567849 | 0.974534428 | 0.986628334 | 10.32012472 | 10.18700944 | ENSG00000272267 |
| KLHDC2 | -0.020941125 | 0.985589557 | 0.79987983 | 0.884673879 | 737.6695254 | 727.3442042 | ENSG00000165516 |
| WTAP | -0.020946972 | 0.985585563 | 0.750801697 | 0.852585092 | 1419.541994 | 1399.594632 | ENSG00000146457 |
| ZFAND5 | -0.020962243 | 0.985575131 | 0.736286796 | 0.843250331 | 2395.024666 | 2361.103173 | ENSG00000107372 |
| SAT1 | -0.020973114 | 0.985567704 | 0.830197628 | 0.903443473 | 700.211489 | 689.7455353 | ENSG00000130066 |
| FBXL12 | -0.020987939 | 0.985557576 | 0.804199428 | 0.887069454 | 743.335757 | 732.402918 | ENSG00000127452 |
| HAUS7 | -0.021026517 | 0.985531222 | 0.979379391 | 0.989069618 | 6.349254625 | 6.266218642 | ENSG00000213397 |
| WDR25 | -0.021043772 | 0.985519435 | 0.891304586 | 0.93992692 | 206.0434559 | 203.1586178 | ENSG00000176473 |
| NR1D1 | -0.021045917 | 0.98551797 | 0.9157831 | 0.954939016 | 120.5983168 | 118.8527468 | ENSG00000126368 |
| BCAM | -0.021095092 | 0.985484379 | 0.754291538 | 0.854310609 | 1842.924086 | 1816.197343 | ENSG00000187244 |
| PDIA3 | -0.021105331 | 0.985477385 | 0.62294777 | 0.762604966 | 9328.562378 | 9193.580354 | ENSG00000167004 |
| COG1 | -0.021124931 | 0.985463997 | 0.877073862 | 0.932365232 | 281.09314 | 277.1707259 | ENSG00000166685 |
| ZDHHC8 | -0.021183107 | 0.985424259 | 0.802682875 | 0.88621529 | 883.6158511 | 870.7585355 | ENSG00000099904 |
| DNAJC21 | -0.021289943 | 0.985351288 | 0.780217135 | 0.870840926 | 1057.18483 | 1041.928291 | ENSG00000168724 |
| KCNV2 | -0.02133633 | 0.985319606 | 0.982552033 | 0.990510622 | 4.675744867 | 4.612883564 | ENSG00000168263 |
| TMEM223 | -0.021344681 | 0.985313903 | 0.851950967 | 0.91580219 | 351.5098314 | 346.3091305 | ENSG00000168569 |
| FASTKD5 | -0.021358452 | 0.985304497 | 0.824555491 | 0.900350935 | 572.9110742 | 564.6690788 | ENSG00000215251 |
| SPRYD3 | -0.021388782 | 0.985283784 | 0.878603672 | 0.933298075 | 252.8817286 | 249.287193 | ENSG00000167778 |
| TNKS2 | -0.021422526 | 0.985260738 | 0.704084046 | 0.82190951 | 2236.554366 | 2203.506874 | ENSG00000107854 |
| XPA | -0.021440826 | 0.985248241 | 0.926239693 | 0.960497534 | 85.14473367 | 83.98401114 | ENSG00000136936 |
| NR2C2AP | -0.021601995 | 0.985138182 | 0.851430986 | 0.915396263 | 360.7532637 | 355.4773531 | ENSG00000184162 |
| ZNF33A | -0.021692804 | 0.985076175 | 0.812318704 | 0.892246907 | 596.1149597 | 587.3790992 | ENSG00000189180 |
| AP3S2 | -0.021758301 | 0.985031455 | 0.882311082 | 0.935715397 | 220.3647872 | 217.1405698 | ENSG00000157823 |
| ZNF10 | -0.021788647 | 0.985010735 | 0.933763356 | 0.964558308 | 77.20155403 | 75.98782396 | ENSG00000256223 |
| KLHL5 | -0.021799506 | 0.985003321 | 0.86258533 | 0.922764098 | 330.807018 | 326.2432198 | ENSG00000109790 |
| WDR74 | -0.021867406 | 0.984956963 | 0.748676905 | 0.851124178 | 1628.009414 | 1602.903336 | ENSG00000133316 |
| ZBTB14 | -0.021894041 | 0.984938779 | 0.876967141 | 0.932303158 | 230.0205492 | 226.4884171 | ENSG00000198081 |
| LAMTOR2 | -0.021910718 | 0.984927394 | 0.800648712 | 0.885210591 | 717.8035358 | 707.0688696 | ENSG00000116586 |
| MAGED1 | -0.02204321 | 0.984836946 | 0.723005549 | 0.833883281 | 2712.939757 | 2672.027617 | ENSG00000179222 |
| NA | -0.022116726 | 0.984786762 | 0.958579579 | 0.97877302 | 26.36932027 | 25.85135053 | ENSG00000260588 |
| MRPL42 | -0.022140171 | 0.984770759 | 0.742251548 | 0.847565681 | 1650.253439 | 1625.588257 | ENSG00000198015 |
| PACS2 | -0.022179147 | 0.984744154 | 0.766523669 | 0.861864435 | 1738.33669 | 1711.290056 | ENSG00000179364 |
| NA | -0.022260258 | 0.984688792 | 0.935179788 | 0.965468934 | 62.84022307 | 61.78834254 | ENSG00000213300 |
| BLOC1S5 | -0.022288117 | 0.984669777 | 0.88350248 | 0.936464084 | 208.0351044 | 205.0141437 | ENSG00000188428 |
| NA | -0.022298921 | 0.984662404 | 0.954407839 | 0.976891899 | 46.97472742 | 46.19364385 | ENSG00000249264 |
| ZNF777 | -0.022300237 | 0.984661505 | 0.847470916 | 0.9128175 | 409.4655668 | 403.3458498 | ENSG00000196453 |
| RXRB | -0.022406154 | 0.984589218 | 0.805828752 | 0.887979418 | 582.596585 | 573.7899703 | ENSG00000204231 |
| NPIPB3 | -0.022537148 | 0.984499823 | 0.958332418 | 0.978676055 | 24.29779879 | 23.9412363 | ENSG00000169246 |
| NA | -0.022545269 | 0.984494281 | 0.976884508 | 0.988018857 | 7.326262317 | 7.218858663 | ENSG00000197149 |
| NA | -0.022680407 | 0.984402068 | 0.984320528 | 0.991219407 | 3.638688754 | 3.584422004 | ENSG00000198658 |
| PSMA3-AS1 | -0.022715865 | 0.984377874 | 0.885098416 | 0.936934918 | 212.7876432 | 209.7138621 | ENSG00000257621 |
| NA | -0.022798281 | 0.984321641 | 0.983081321 | 0.990681803 | 4.455287663 | 4.387829982 | ENSG00000220884 |
| PREP | -0.022851028 | 0.984285654 | 0.738216765 | 0.844463914 | 1887.442542 | 1857.83961 | ENSG00000085377 |
| LINC00641 | -0.022876194 | 0.984268484 | 0.904094026 | 0.947822284 | 136.7979089 | 134.5985676 | ENSG00000258441 |
| TIFA | -0.022987551 | 0.984192515 | 0.945075753 | 0.971385564 | 44.56188816 | 44.01464053 | ENSG00000145365 |
| ABHD17B | -0.022995478 | 0.984187107 | 0.788401489 | 0.876551964 | 889.0436165 | 875.6114661 | ENSG00000107362 |
| RPL5 | -0.02299578 | 0.984186901 | 0.566946102 | 0.720424369 | 20191.68954 | 19871.75528 | ENSG00000122406 |
| PCCB | -0.02312289 | 0.984100192 | 0.726478764 | 0.835688839 | 2054.094635 | 2020.959457 | ENSG00000114054 |
| SAMHD1 | -0.023124169 | 0.98409932 | 0.756022498 | 0.855191158 | 1094.663515 | 1077.31147 | ENSG00000101347 |
| SMC3 | -0.023181746 | 0.984060046 | 0.675223805 | 0.800795673 | 2655.133858 | 2613.687865 | ENSG00000108055 |
| RPL23 | -0.023226536 | 0.984029495 | 0.642361358 | 0.776574103 | 15124.58467 | 14881.95944 | ENSG00000125691 |
| SLC39A14 | -0.023269104 | 0.98400046 | 0.575811301 | 0.727976591 | 7216.567486 | 7099.981332 | ENSG00000104635 |
| NEO1 | -0.02336869 | 0.983932539 | 0.715496294 | 0.829186532 | 1470.368119 | 1447.071317 | ENSG00000067141 |
| PSMC2 | -0.023479888 | 0.983856705 | 0.641934188 | 0.776203612 | 2770.319653 | 2725.21217 | ENSG00000161057 |
| PLXNA3 | -0.023565212 | 0.983798519 | 0.826248655 | 0.901094048 | 509.1813622 | 500.4742926 | ENSG00000130827 |
| ACO2 | -0.02357368 | 0.983792744 | 0.691391612 | 0.812873938 | 1758.09202 | 1729.361028 | ENSG00000100412 |
| TCEA1 | -0.023607025 | 0.983770006 | 0.688572108 | 0.810794393 | 3516.698962 | 3460.609093 | ENSG00000187735 |
| ASL | -0.023630156 | 0.983754233 | 0.835665741 | 0.906888184 | 421.2210844 | 414.2940372 | ENSG00000126522 |

| | | | | | | | |
|---|---|---|---|---|---|---|---|
| NDUFB6 | -0.023632048 | 0.983752943 | 0.790890403 | 0.878104562 | 775.4171145 | 762.6612179 | ENSG00000165264 |
| NA | -0.023689085 | 0.983714052 | 0.97947044 | 0.989109787 | 4.988389884 | 4.909457539 | ENSG00000279838 |
| SLC1A3 | -0.023703877 | 0.983703966 | 0.811311638 | 0.891693984 | 556.444968 | 546.9903236 | ENSG00000079215 |
| PEX1 | -0.023707349 | 0.983701598 | 0.822015353 | 0.898645482 | 444.9768958 | 437.7593531 | ENSG00000127980 |
| CRYBB3 | -0.023772553 | 0.98365714 | 0.968221954 | 0.983351992 | 12.02924049 | 11.83819395 | ENSG00000100053 |
| NA | -0.023841862 | 0.983609884 | 0.982882146 | 0.990647551 | 3.935264075 | 3.872393683 | ENSG00000272384 |
| CACNG5 | -0.024028209 | 0.983482844 | 0.916873632 | 0.955508035 | 90.92854653 | 89.59201316 | ENSG00000075429 |
| NEU1 | -0.024048825 | 0.98346879 | 0.752856172 | 0.853762178 | 978.493778 | 961.8960456 | ENSG00000204386 |
| SYNE3 | -0.024058303 | 0.983462329 | 0.98197778 | 0.990314032 | 3.660340256 | 3.601093555 | ENSG00000176438 |
| CEP70 | -0.024068228 | 0.983455564 | 0.824056461 | 0.900060762 | 419.11392 | 412.4317807 | ENSG00000114107 |
| TNFSF12 | -0.024085576 | 0.983443738 | 0.954011266 | 0.976693273 | 30.14868133 | 29.6581425 | ENSG00000239697 |
| SESTD1 | -0.024093332 | 0.983438451 | 0.817497108 | 0.895649793 | 502.8345862 | 494.7433805 | ENSG00000187231 |
| ACTR10 | -0.024147401 | 0.983401594 | 0.822333624 | 0.898735651 | 547.6279688 | 538.1640549 | ENSG00000131966 |
| CHPF | -0.024200051 | 0.983365707 | 0.688672016 | 0.810814899 | 2561.178686 | 2519.450509 | ENSG00000123989 |
| NA | -0.02420034 | 0.983365509 | 0.97441872 | 0.986590643 | 9.598504524 | 9.44060088 | ENSG00000271862 |
| NUP43 | -0.024207748 | 0.98336046 | 0.716698108 | 0.829980982 | 1261.689703 | 1241.065029 | ENSG00000120253 |
| NA | -0.0243342 | 0.983274272 | 0.979540349 | 0.989128605 | 5.045647402 | 4.962155806 | ENSG00000286181 |
| CCNY | -0.024391067 | 0.983235515 | 0.728575555 | 0.83720223 | 1300.1798 | 1278.656701 | ENSG00000108100 |
| VARS1 | -0.024424581 | 0.983212675 | 0.614282961 | 0.756906783 | 7084.446293 | 6965.519803 | ENSG00000204394 |
| HRK | -0.024458913 | 0.983189277 | 0.933467441 | 0.964413047 | 55.99259636 | 54.92332984 | ENSG00000135116 |
| NA | -0.024486176 | 0.983170698 | 0.968789462 | 0.983627365 | 11.91193455 | 11.71397529 | ENSG00000231856 |
| BAHD1 | -0.024547278 | 0.983129059 | 0.71828042 | 0.831072807 | 1250.682012 | 1229.629139 | ENSG00000140320 |
| HARBI1 | -0.024575192 | 0.983110037 | 0.918339399 | 0.956467193 | 79.91403493 | 78.72366383 | ENSG00000180423 |
| SPCS3 | -0.024599987 | 0.983093141 | 0.789689274 | 0.877376939 | 1174.947555 | 1155.909727 | ENSG00000129128 |
| RFX8 | -0.024629561 | 0.983072988 | 0.976465784 | 0.987902923 | 7.579534635 | 7.452514539 | ENSG00000196460 |
| DGKA | -0.024744335 | 0.982994783 | 0.879776543 | 0.933848866 | 208.0256926 | 204.5048681 | ENSG00000065357 |
| RFT1 | -0.024770012 | 0.982977288 | 0.804713223 | 0.887171265 | 467.4442427 | 459.3898618 | ENSG00000163933 |
| NA | -0.024796563 | 0.982959198 | 0.977572621 | 0.988353504 | 5.345013626 | 5.254428633 | ENSG00000242951 |
| NAAA | -0.024833199 | 0.982934236 | 0.940124689 | 0.968334955 | 47.50359928 | 46.57223062 | ENSG00000138744 |
| P2RX7 | -0.024840142 | 0.982929506 | 0.981248989 | 0.990314032 | 4.283515108 | 4.210913056 | ENSG00000089041 |
| NCKAP1 | -0.024857233 | 0.982917862 | 0.614866183 | 0.757375056 | 3657.506061 | 3596.016298 | ENSG00000061676 |
| MPZ | -0.024875159 | 0.982905649 | 0.931718709 | 0.9635352 | 69.85750716 | 68.45788877 | ENSG00000158887 |
| NA | -0.024945656 | 0.98285762 | 0.979820135 | 0.989307559 | 4.368005934 | 4.293186317 | ENSG00000273156 |
| IL22RA1 | -0.025109057 | 0.982746307 | 0.978917057 | 0.988965119 | 4.691814563 | 4.610732991 | ENSG00000142677 |
| REEP6 | -0.025163516 | 0.982709211 | 0.811599486 | 0.891807113 | 470.765068 | 462.4948863 | ENSG00000115255 |
| NA | -0.025205351 | 0.982680715 | 0.979267263 | 0.989008155 | 4.672953964 | 4.591910865 | ENSG00000258682 |
| RPS14 | -0.025293069 | 0.982620969 | 0.795982084 | 0.881474764 | 22575.47778 | 22182.54546 | ENSG00000164587 |
| NA | -0.025315804 | 0.982605484 | 0.9461791 | 0.972141843 | 35.13639549 | 34.65443492 | ENSG00000180015 |
| ZNF703 | -0.025437305 | 0.982522735 | 0.936777088 | 0.966383252 | 46.27311911 | 45.59860579 | ENSG00000183779 |
| NA | -0.02547796 | 0.982495047 | 0.96987952 | 0.984267759 | 9.658552945 | 9.488464962 | ENSG00000215210 |
| POLR1E | -0.025542191 | 0.982451306 | 0.763595899 | 0.85991903 | 706.6886511 | 694.5853625 | ENSG00000137054 |
| NA | -0.025579812 | 0.982425687 | 0.967492951 | 0.983138775 | 10.72005146 | 10.52982996 | ENSG00000248590 |
| TRIM26 | -0.02559376 | 0.982416189 | 0.708152282 | 0.824041843 | 1192.684801 | 1171.688177 | ENSG00000234127 |
| IFNAR1 | -0.025697429 | 0.982345597 | 0.859110631 | 0.920821105 | 228.8904294 | 224.9490809 | ENSG00000142166 |
| AJM1 | -0.025784077 | 0.9822866 | 0.932693033 | 0.964180982 | 53.33562122 | 52.38135734 | ENSG00000232434 |
| CWF19L1 | -0.025829018 | 0.982256001 | 0.808341248 | 0.889848837 | 429.1137396 | 421.2773741 | ENSG00000095485 |
| LOC283683 | -0.025861207 | 0.982234085 | 0.969765089 | 0.984203405 | 9.971197962 | 9.791490657 | ENSG00000274253 |
| AK3 | -0.025945207 | 0.982176898 | 0.794331992 | 0.880349251 | 531.5337696 | 522.4297988 | ENSG00000147853 |
| ID3 | -0.025993982 | 0.982143692 | 0.618786603 | 0.760275316 | 3402.062924 | 3340.933791 | ENSG00000117318 |
| METAP2 | -0.02609853 | 0.982072522 | 0.585554716 | 0.735192012 | 3376.780483 | 3316.682701 | ENSG00000111142 |
| TBCA | -0.026119732 | 0.982058089 | 0.649709108 | 0.781978702 | 2673.602149 | 2625.841976 | ENSG00000171530 |
| ZNF586 | -0.02619732 | 0.982005275 | 0.837027254 | 0.907292957 | 282.6227109 | 277.5890954 | ENSG00000083828 |
| NA | -0.026214603 | 0.981993512 | 0.960762095 | 0.979808711 | 22.4937043 | 22.17846401 | ENSG00000262691 |
| MPHOSPH10 | -0.026271635 | 0.981954692 | 0.764537792 | 0.860685022 | 780.3524403 | 766.3813802 | ENSG00000124383 |
| BFAR | -0.026325416 | 0.981918087 | 0.667407751 | 0.795634837 | 1866.253146 | 1831.607825 | ENSG00000103429 |
| WRNIP1 | -0.026362606 | 0.981892776 | 0.720740816 | 0.83226452 | 1151.748174 | 1131.083192 | ENSG00000124535 |
| MACROD2 | -0.026515699 | 0.981788587 | 0.95686686 | 0.978044405 | 23.84938755 | 23.2980737 | ENSG00000172264 |
| BVES | -0.026572854 | 0.981749692 | 0.911016863 | 0.951993281 | 82.90251568 | 81.49580298 | ENSG00000112276 |
| GLYATL1 | -0.026631443 | 0.981709823 | 0.893800524 | 0.941276663 | 117.5785726 | 115.3756114 | ENSG00000166840 |
| DCAF13 | -0.02667853 | 0.981677783 | 0.64098558 | 0.775396803 | 2288.925382 | 2247.101001 | ENSG00000164934 |
| NA | -0.026704776 | 0.981659924 | 0.979156753 | 0.988983938 | 4.019754901 | 3.943914076 | ENSG00000232702 |
| CRHR1 | -0.026767411 | 0.981617306 | 0.973615309 | 0.986264509 | 6.972429478 | 6.840544465 | ENSG00000120088 |
| LARGE1 | -0.02681681 | 0.981583695 | 0.665376294 | 0.794047268 | 1529.748835 | 1501.530594 | ENSG00000133424 |
| GTF2F2 | -0.026859908 | 0.981554373 | 0.730426934 | 0.838530468 | 880.4893257 | 864.461621 | ENSG00000188342 |
| NA | -0.026863411 | 0.981551989 | 0.97484638 | 0.986718185 | 6.661899643 | 6.535368196 | ENSG00000279789 |
| STK39 | -0.026887098 | 0.981535873 | 0.789531588 | 0.877302797 | 760.552379 | 746.4488057 | ENSG00000198648 |
| ARHGEF35 | -0.02690692 | 0.981522388 | 0.860911599 | 0.921931237 | 203.4126749 | 199.6849596 | ENSG00000213214 |
| NA | -0.027135593 | 0.981366825 | 0.963477007 | 0.981487825 | 12.04810109 | 11.81507068 | ENSG00000280303 |
| KARS1 | -0.027217352 | 0.981311211 | 0.511463571 | 0.675751233 | 5286.108024 | 5187.668099 | ENSG00000065427 |
| NA | -0.027248383 | 0.981290105 | 0.948044307 | 0.972984731 | 25.74036345 | 25.24153103 | ENSG00000223509 |
| SKI | -0.027250973 | 0.981288343 | 0.632441387 | 0.769403216 | 2275.024942 | 2232.915307 | ENSG00000157933 |
| NAPG | -0.027254082 | 0.981286228 | 0.806463436 | 0.888475268 | 415.1821226 | 407.4298422 | ENSG00000134265 |
| OSBPL7 | -0.02734313 | 0.981225662 | 0.879042564 | 0.93343161 | 186.0263302 | 182.1409154 | ENSG00000006025 |
| GMPS | -0.027437535 | 0.981161456 | 0.594252399 | 0.742276808 | 3577.625515 | 3511.136573 | ENSG00000163655 |
| GPX8 | -0.027453331 | 0.981150713 | 0.72563535 | 0.835289284 | 967.9997287 | 949.6773869 | ENSG00000164294 |
| NUB1 | -0.027472693 | 0.981137546 | 0.719629518 | 0.831678231 | 945.8414338 | 927.8115991 | ENSG00000013374 |
| MMADHC | -0.02752433 | 0.981102429 | 0.675350843 | 0.800828482 | 1616.029472 | 1586.069024 | ENSG00000168288 |

| | | | | | | | |
|---|---|---|---|---|---|---|---|
| TRMT10A | -0.027552582 | 0.981083217 | 0.930785073 | 0.963034317 | 45.87598327 | 45.11919621 | ENSG00000145331 |
| R3HDM1 | -0.027553577 | 0.98108254 | 0.719497293 | 0.831663685 | 1207.521564 | 1184.999294 | ENSG00000048991 |
| TWF2 | -0.027588215 | 0.981058985 | 0.776477461 | 0.868903457 | 748.8019946 | 734.3800215 | ENSG00000247596 |
| MICB-DT | -0.0276395 | 0.981024111 | 0.982342409 | 0.990351054 | 3.374928546 | 3.308820728 | ENSG00000286940 |
| ZNF782 | -0.027665672 | 0.981006315 | 0.908748481 | 0.950942162 | 89.62533962 | 87.74214553 | ENSG00000196597 |
| INSIG2 | -0.027730855 | 0.980961992 | 0.884015243 | 0.936493029 | 137.9626467 | 135.367015 | ENSG00000125629 |
| HECW2 | -0.027781857 | 0.980927314 | 0.888799567 | 0.938578344 | 120.348311 | 117.9603533 | ENSG00000138411 |
| CERS4 | -0.027787264 | 0.980923638 | 0.674325306 | 0.800199707 | 1220.235517 | 1196.981582 | ENSG00000090661 |
| SRPK1 | -0.027788383 | 0.980922877 | 0.671908647 | 0.79887764 | 1520.621006 | 1491.203264 | ENSG00000096063 |
| NSRP1 | -0.02781009 | 0.980908118 | 0.76284246 | 0.859328111 | 570.4978843 | 559.5789241 | ENSG00000126653 |
| CMTM6 | -0.02782819 | 0.980895812 | 0.753434394 | 0.853915921 | 943.0980301 | 925.3632221 | ENSG00000091317 |
| TSPAN2 | -0.027848432 | 0.980882049 | 0.847040579 | 0.912404925 | 290.8509758 | 284.9219528 | ENSG00000134198 |
| COX6B1 | -0.02786752 | 0.980869071 | 0.648651284 | 0.781605019 | 2739.438538 | 2686.192496 | ENSG00000126267 |
| TRIM45 | -0.027908173 | 0.980841432 | 0.815990953 | 0.894698714 | 367.9299175 | 360.5032447 | ENSG00000134253 |
| NOC3L | -0.027965919 | 0.980802174 | 0.776446127 | 0.868903457 | 549.5156925 | 539.0516567 | ENSG00000173145 |
| TRRAP | -0.027981114 | 0.980791843 | 0.62551589 | 0.764255306 | 2828.007734 | 2772.621103 | ENSG00000196367 |
| NA | -0.027992855 | 0.980783862 | 0.97050615 | 0.984334076 | 8.433854741 | 8.264524813 | ENSG00000230454 |
| DYNC2LI1 | -0.028011828 | 0.980770963 | 0.914296217 | 0.954058977 | 68.69090545 | 67.45496152 | ENSG00000138036 |
| B3GLCT | -0.028030034 | 0.980758586 | 0.885650727 | 0.937037652 | 129.2236437 | 126.8984093 | ENSG00000187676 |
| RAB29 | -0.028037196 | 0.980753717 | 0.803599664 | 0.88672909 | 369.0245945 | 361.8874181 | ENSG00000117280 |
| DLGAP3 | -0.028333601 | 0.98055224 | 0.803056331 | 0.886394088 | 601.8287388 | 590.4879103 | ENSG00000116544 |
| ARL15 | -0.028405082 | 0.980503658 | 0.873179869 | 0.929557577 | 205.7881436 | 202.1962789 | ENSG00000185305 |
| ADNP-AS1 | -0.028465322 | 0.980462718 | 0.966655037 | 0.982753218 | 9.361977624 | 9.168767714 | ENSG00000259456 |
| SMYD4 | -0.028493562 | 0.980443526 | 0.788145579 | 0.876396218 | 586.1247747 | 574.2000038 | ENSG00000186532 |
| NCS1 | -0.028503908 | 0.980436495 | 0.683392265 | 0.806868328 | 1421.740006 | 1394.098218 | ENSG00000107130 |
| DCTN1 | -0.028571037 | 0.980390876 | 0.571066614 | 0.723801675 | 4577.964331 | 4487.96966 | ENSG00000204843 |
| ST6GAL1 | -0.028604067 | 0.98036843 | 0.581045668 | 0.731654829 | 2758.165046 | 2704.015818 | ENSG00000073849 |
| SINHCAF | -0.028700693 | 0.980302772 | 0.568429927 | 0.721613534 | 6689.441763 | 6557.704199 | ENSG00000139146 |
| TMEM86B | -0.028703426 | 0.980300914 | 0.978624087 | 0.98893745 | 5.443458966 | 5.332400746 | ENSG00000180089 |
| SNHG15 | -0.028708058 | 0.980297767 | 0.766813155 | 0.862039194 | 554.065983 | 543.257172 | ENSG00000232956 |
| FKBP1B | -0.028715128 | 0.980292963 | 0.914940957 | 0.954473611 | 67.18011982 | 65.92261003 | ENSG00000119782 |
| DGKQ | -0.028744232 | 0.980273187 | 0.770719718 | 0.864868483 | 626.5748466 | 614.1446825 | ENSG00000145214 |
| NA | -0.028814067 | 0.980225738 | 0.980695434 | 0.989984072 | 3.681991758 | 3.605394703 | ENSG00000213985 |
| TRAIP | -0.029169957 | 0.979983961 | 0.819524009 | 0.896989371 | 371.3440067 | 363.7994287 | ENSG00000183763 |
| ZC3HC1 | -0.029190508 | 0.979970002 | 0.753489988 | 0.85392876 | 609.1289417 | 596.5462799 | ENSG00000091732 |
| KCNK3 | -0.029206339 | 0.979959249 | 0.967271195 | 0.983068785 | 9.022099299 | 8.830248342 | ENSG00000171303 |
| EIF3H | -0.029295007 | 0.979899022 | 0.540930463 | 0.700109467 | 3766.789757 | 3690.513007 | ENSG00000147677 |
| PRC1 | -0.029347507 | 0.979863364 | 0.967178125 | 0.983045516 | 8.390551738 | 8.209675972 | ENSG00000198901 |
| DCHS2 | -0.029409837 | 0.979821031 | 0.928585038 | 0.961846113 | 45.93479239 | 44.95464969 | ENSG00000197410 |
| LOC728673 | -0.029499165 | 0.979760365 | 0.962550164 | 0.980727225 | 11.79550449 | 11.54161997 | ENSG00000170161 |
| MYH10 | -0.029672726 | 0.979642503 | 0.454275583 | 0.624131491 | 10350.18809 | 10139.19473 | ENSG00000133026 |
| SDAD1 | -0.029704836 | 0.9796207 | 0.695061162 | 0.815746479 | 1195.672869 | 1171.96104 | ENSG00000198301 |
| ALOX12-AS1 | -0.029738629 | 0.979597754 | 0.953811176 | 0.97654025 | 16.44613124 | 16.08513372 | ENSG00000215067 |
| MIR124-2HG | -0.029822614 | 0.979540729 | 0.979164915 | 0.988983938 | 4.329609015 | 4.236186903 | ENSG00000254377 |
| NA | -0.029968511 | 0.979441675 | 0.959280305 | 0.979229356 | 16.61423701 | 16.36261299 | ENSG00000280187 |
| NLN | -0.029975625 | 0.979436846 | 0.527332114 | 0.689118264 | 3859.699469 | 3781.166976 | ENSG00000123213 |
| ATG7 | -0.029998218 | 0.979421507 | 0.816717435 | 0.895136415 | 290.1188308 | 284.1553651 | ENSG00000197548 |
| UBXN2B | -0.03018815 | 0.979292574 | 0.749824677 | 0.852024107 | 606.8529959 | 594.7044084 | ENSG00000215114 |
| SECTM1 | -0.03020222 | 0.979283024 | 0.974851583 | 0.986718185 | 4.969529285 | 4.858909846 | ENSG00000141574 |
| NA | -0.030261085 | 0.979243068 | 0.942782845 | 0.969985273 | 28.80388628 | 28.29303957 | ENSG00000224858 |
| NAA35 | -0.030337418 | 0.979191257 | 0.712805797 | 0.827111978 | 778.6052768 | 762.47213 | ENSG00000135040 |
| NAGK | -0.030354311 | 0.979179792 | 0.720514315 | 0.832149671 | 842.4859105 | 824.5280777 | ENSG00000124357 |
| EML1 | -0.030355436 | 0.979179028 | 0.753405881 | 0.853915921 | 622.6507589 | 609.2822587 | ENSG00000066629 |
| FOS | -0.030356388 | 0.979178382 | 0.801547272 | 0.885657066 | 376.5116659 | 368.2026095 | ENSG00000170345 |
| ZFAND2A | -0.030448826 | 0.979115645 | 0.890543572 | 0.939403184 | 93.79780632 | 91.83419504 | ENSG00000178381 |
| USP6 | -0.030457157 | 0.979109991 | 0.954400061 | 0.976891899 | 16.90120032 | 16.52259791 | ENSG00000129204 |
| TMEM144 | -0.030474175 | 0.979098441 | 0.926958672 | 0.960672598 | 41.019054 | 40.09358927 | ENSG00000164124 |
| SLC43A1 | -0.030489137 | 0.979088287 | 0.778310263 | 0.869988497 | 590.047599 | 577.234865 | ENSG00000149150 |
| BISPR | -0.030651111 | 0.978978369 | 0.955939051 | 0.977577409 | 16.16351043 | 15.79931261 | ENSG00000282851 |
| CRACD | -0.030655315 | 0.978975517 | 0.756722428 | 0.855714249 | 602.0722727 | 589.8379686 | ENSG00000109265 |
| NA | -0.030680034 | 0.978958743 | 0.96603693 | 0.982484721 | 8.77719969 | 8.577770339 | ENSG00000226015 |
| RNF19B | -0.030760578 | 0.978904091 | 0.756731316 | 0.855714249 | 609.2672746 | 596.8040704 | ENSG00000116514 |
| SLC35F5 | -0.030879933 | 0.978823109 | 0.777479581 | 0.869419662 | 410.9685802 | 402.4120198 | ENSG00000115084 |
| PKN2 | -0.031038346 | 0.978715636 | 0.746504927 | 0.849806831 | 619.3654274 | 606.1129469 | ENSG00000065243 |
| RNF11 | -0.031170351 | 0.978626089 | 0.746364472 | 0.849747228 | 706.8749048 | 692.2267523 | ENSG00000123091 |
| NA | -0.031190884 | 0.978612161 | 0.967023322 | 0.982972226 | 8.12679153 | 7.938375845 | ENSG00000242083 |
| BAIAP2 | -0.031195588 | 0.97860897 | 0.778858417 | 0.870204896 | 415.9345307 | 407.1308635 | ENSG00000175866 |
| NEXMIF | -0.031225068 | 0.978588974 | 0.955610627 | 0.977429542 | 13.78724108 | 13.46410461 | ENSG00000050030 |
| NA | -0.031310702 | 0.978530889 | 0.970264731 | 0.98427465 | 7.015732482 | 6.852914869 | ENSG00000243014 |
| ARPC3 | -0.031381129 | 0.978483122 | 0.714404592 | 0.828468829 | 859.5206369 | 841.0113954 | ENSG00000111229 |
| NSF | -0.031423565 | 0.978454341 | 0.761683117 | 0.858423147 | 528.8882335 | 518.048972 | ENSG00000073969 |
| ATP6V1E1 | -0.031475451 | 0.978419152 | 0.643202533 | 0.777006705 | 1285.148324 | 1257.524449 | ENSG00000131100 |
| PIKFYVE | -0.03153566 | 0.97837832 | 0.713871074 | 0.828099032 | 1080.487723 | 1056.669846 | ENSG00000115020 |
| CUL4A | -0.031618079 | 0.978322428 | 0.549822641 | 0.706992892 | 2928.363363 | 2864.161723 | ENSG00000139842 |
| RPL4 | -0.031782363 | 0.97821103 | 0.414401684 | 0.586731955 | 33695.41486 | 32960.10843 | ENSG00000174444 |
| DISP3 | -0.031803632 | 0.978196609 | 0.832356213 | 0.904873694 | 213.7772539 | 209.2583999 | ENSG00000204624 |
| GLRX | -0.031855366 | 0.978161532 | 0.805638018 | 0.887835648 | 403.1226705 | 394.4368117 | ENSG00000173221 |

| | | | | | | | |
|---|---|---|---|---|---|---|---|
| NA | -0.031860671 | 0.978157935 | 0.974675027 | 0.986694792 | 4.955574771 | 4.837937146 | ENSG00000235545 |
| PACSIN1 | -0.031878984 | 0.978145519 | 0.777107475 | 0.869299514 | 420.630012 | 411.2095007 | ENSG00000124507 |
| TMEM63B | -0.031919775 | 0.978117863 | 0.647270627 | 0.780406104 | 1123.384 | 1098.586224 | ENSG00000137216 |
| MBD4 | -0.032027803 | 0.978044625 | 0.743821286 | 0.848323562 | 621.843921 | 607.6891645 | ENSG00000129071 |
| PRDM13 | -0.032037797 | 0.978037849 | 0.979192018 | 0.988983938 | 3.037165403 | 2.963849634 | ENSG00000112238 |
| SGCE | -0.032039088 | 0.978036974 | 0.780998089 | 0.871424971 | 371.1198827 | 362.6591126 | ENSG00000127990 |
| MEAK7 | -0.032059248 | 0.978023308 | 0.787713615 | 0.876235135 | 476.0013344 | 465.1175087 | ENSG00000140950 |
| SMPD2 | -0.032064365 | 0.978019839 | 0.864982534 | 0.924350658 | 131.1221533 | 128.1118691 | ENSG00000135587 |
| NADK2 | -0.032123792 | 0.977979553 | 0.79490812 | 0.880588753 | 352.8561411 | 345.1873823 | ENSG00000152620 |
| LINC01409 | -0.032139 | 0.977969244 | 0.96719734 | 0.983045516 | 8.403830531 | 8.203224251 | ENSG00000237491 |
| TMEM205 | -0.032173458 | 0.977945886 | 0.887231091 | 0.937685585 | 89.78766343 | 87.7615007 | ENSG00000105518 |
| MAP2K4 | -0.032224754 | 0.977911115 | 0.697117428 | 0.817153648 | 729.9772275 | 713.7521249 | ENSG00000065559 |
| POLR1D | -0.032231942 | 0.977906243 | 0.561013334 | 0.715963762 | 2307.17442 | 2255.861649 | ENSG00000186184 |
| UBE2T | -0.032236723 | 0.977903002 | 0.73040602 | 0.838530468 | 722.8713713 | 706.9338739 | ENSG00000077152 |
| LPAR4 | -0.032242774 | 0.977898901 | 0.88758852 | 0.937960566 | 103.8056494 | 101.6816006 | ENSG00000147145 |
| GRSF1 | -0.032310243 | 0.977853169 | 0.523904803 | 0.686313158 | 2748.397686 | 2687.694127 | ENSG00000132463 |
| SEC62 | -0.032320166 | 0.977846444 | 0.660954328 | 0.790739852 | 1536.592873 | 1502.883872 | ENSG00000008952 |
| DNAJB9 | -0.032320829 | 0.977845994 | 0.829349174 | 0.902876688 | 225.8535522 | 220.9274551 | ENSG00000128590 |
| ZNF408 | -0.032427992 | 0.977773363 | 0.792045266 | 0.878982065 | 352.8230506 | 345.2965228 | ENSG00000175213 |
| PRKCG | -0.032434782 | 0.977768761 | 0.822362951 | 0.898735651 | 221.6106613 | 216.4861334 | ENSG00000126583 |
| RAP1B | -0.032516359 | 0.977713475 | 0.615768599 | 0.758204637 | 1493.630953 | 1460.441886 | ENSG00000127314 |
| NCOR1 | -0.032540515 | 0.977697105 | 0.523647322 | 0.686069036 | 2639.945477 | 2581.272704 | ENSG00000141027 |
| CRADD | -0.032584767 | 0.977667116 | 0.895378686 | 0.942250941 | 70.79994797 | 69.20321283 | ENSG00000169372 |
| G3BP2 | -0.032630918 | 0.977635841 | 0.467966622 | 0.63706857 | 5605.470816 | 5480.943992 | ENSG00000138757 |
| DLGAP4-AS1 | -0.03268542 | 0.977598909 | 0.958290182 | 0.978676055 | 12.38028243 | 12.07615097 | ENSG00000232907 |
| NA | -0.032708019 | 0.977583595 | 0.959693664 | 0.979463454 | 10.37180043 | 10.11333848 | ENSG00000255618 |
| NA | -0.032711823 | 0.977581018 | 0.922575279 | 0.9585501 | 40.55349703 | 39.55072855 | ENSG00000231007 |
| RPS11 | -0.032716991 | 0.977577516 | 0.577828732 | 0.729699795 | 24323.22237 | 23776.65559 | ENSG00000142534 |
| NA | -0.032814571 | 0.977511398 | 0.962272551 | 0.980632996 | 9.430398754 | 9.196192135 | ENSG00000265511 |
| STT3B | -0.032863895 | 0.977477978 | 0.526028212 | 0.688206865 | 2826.534873 | 2763.603373 | ENSG00000163527 |
| COX7C | -0.032882249 | 0.977465543 | 0.522357885 | 0.685217324 | 5055.854587 | 4940.726492 | ENSG00000127184 |
| SHISA5 | -0.032974136 | 0.977403289 | 0.51762922 | 0.681556816 | 3132.636156 | 3061.774209 | ENSG00000164054 |
| UACA | -0.03317065 | 0.977270163 | 0.755628614 | 0.855091825 | 474.0556179 | 463.4076888 | ENSG00000137831 |
| GFRA3 | -0.033289654 | 0.977189554 | 0.954334027 | 0.976891899 | 14.42995225 | 14.06800542 | ENSG00000146013 |
| NA | -0.033405055 | 0.977111391 | 0.92023903 | 0.957436234 | 47.28429336 | 46.22536942 | ENSG00000231793 |
| ZNF618 | -0.033428119 | 0.977095771 | 0.632114536 | 0.769151098 | 1285.230437 | 1255.297072 | ENSG00000157657 |
| SAFB | -0.033511697 | 0.977039167 | 0.563166144 | 0.717666672 | 5094.406479 | 4977.442304 | ENSG00000160633 |
| TBL2 | -0.033563411 | 0.977004146 | 0.70189683 | 0.820302242 | 657.5117791 | 642.2005281 | ENSG00000106638 |
| RFC3 | -0.033632141 | 0.976957602 | 0.64599352 | 0.779464783 | 1070.651966 | 1045.670306 | ENSG00000133119 |
| NA | -0.033676637 | 0.976927471 | 0.971807338 | 0.985216819 | 5.363874225 | 5.227004212 | ENSG00000269680 |
| TUFM | -0.033677996 | 0.976926551 | 0.414682949 | 0.586837715 | 5725.5526 | 5593.639837 | ENSG00000178952 |
| DOCK9 | -0.033737082 | 0.976886542 | 0.723331831 | 0.834108096 | 565.0449013 | 551.8456952 | ENSG00000088387 |
| RELA | -0.033758831 | 0.976871814 | 0.786785626 | 0.875461817 | 543.8799848 | 531.58759 | ENSG00000173039 |
| SPRY1 | -0.033777494 | 0.976859178 | 0.744527281 | 0.848722265 | 504.2284662 | 492.7883278 | ENSG00000164056 |
| RPLP1 | -0.034041845 | 0.9766802 | 0.445102554 | 0.615606063 | 23513.47323 | 22964.07478 | ENSG00000137818 |
| FDFT1 | -0.03407358 | 0.976658716 | 0.426528067 | 0.598106633 | 12595.91559 | 12301.292 | ENSG00000079459 |
| PAFAH1B1 | -0.034165149 | 0.976596729 | 0.575961011 | 0.728056082 | 1884.874366 | 1841.648225 | ENSG00000007168 |
| AKIRIN1 | -0.034169439 | 0.976593825 | 0.50432482 | 0.669493983 | 4909.46634 | 4795.855751 | ENSG00000174574 |
| CERT1 | -0.034200822 | 0.976572581 | 0.64713465 | 0.780388481 | 1341.154941 | 1310.558346 | ENSG00000113163 |
| FBXL7 | -0.034263257 | 0.976530319 | 0.740899347 | 0.846431059 | 482.3816154 | 471.0462925 | ENSG00000183580 |
| RNF19A | -0.034276416 | 0.976521412 | 0.846773057 | 0.912319442 | 149.9132904 | 146.2756922 | ENSG00000034677 |
| CYP2S1 | -0.034281431 | 0.976518018 | 0.578722476 | 0.730346546 | 7278.506808 | 7108.208438 | ENSG00000167600 |
| MSH6 | -0.034397463 | 0.976439483 | 0.404458255 | 0.576716994 | 6436.726989 | 6285.135744 | ENSG00000116062 |
| NUDCD1 | -0.034519865 | 0.976356642 | 0.62694536 | 0.765371658 | 1343.031235 | 1311.957325 | ENSG00000120526 |
| H2BW2 | -0.034571303 | 0.976321832 | 0.978820187 | 0.988963906 | 3.045538112 | 2.966000208 | ENSG00000101812 |
| CIAPIN1 | -0.034589531 | 0.976309496 | 0.614044794 | 0.75677096 | 2010.636333 | 1962.050886 | ENSG00000005194 |
| ATOX1 | -0.034606357 | 0.97629811 | 0.784217603 | 0.873763531 | 415.0591596 | 404.9341092 | ENSG00000177556 |
| NAA20 | -0.034648697 | 0.976269458 | 0.711643775 | 0.826210885 | 658.3507563 | 642.9429144 | ENSG00000173418 |
| RARG | -0.034735599 | 0.976210653 | 0.641732607 | 0.776057154 | 1247.588704 | 1218.258809 | ENSG00000172819 |
| LEPROTL1 | -0.034742638 | 0.97620589 | 0.688118881 | 0.81055757 | 860.3385262 | 840.4999451 | ENSG00000104660 |
| RANGAP1 | -0.034813827 | 0.976157721 | 0.479021869 | 0.647184796 | 6067.157408 | 5922.855134 | ENSG00000100401 |
| NA | -0.034956234 | 0.97606137 | 0.921344195 | 0.958038961 | 37.59715567 | 36.7299197 | ENSG00000225285 |
| PNPLA3 | -0.03510812 | 0.975958616 | 0.716249078 | 0.829809679 | 595.6116432 | 580.9206371 | ENSG00000100344 |
| FAM210A | -0.035127142 | 0.975945749 | 0.756898243 | 0.855780331 | 369.0546187 | 360.0872741 | ENSG00000177150 |
| EXTL3-AS1 | -0.0351867 | 0.97590546 | 0.969497381 | 0.984039579 | 8.219655064 | 8.007745663 | ENSG00000246339 |
| GPR75 | -0.035280946 | 0.97584171 | 0.952116383 | 0.975457419 | 17.42102375 | 17.06545862 | ENSG00000119737 |
| COX5A | -0.035285121 | 0.975838886 | 0.526766402 | 0.688723471 | 3124.953356 | 3049.635595 | ENSG00000178741 |
| PSMG4 | -0.035302983 | 0.975826804 | 0.682346644 | 0.806269086 | 776.7859885 | 757.54067 | ENSG00000180822 |
| MPLKIP | -0.035392326 | 0.975766375 | 0.689384451 | 0.811401055 | 876.7756956 | 855.557417 | ENSG00000168303 |
| CALCA | -0.03557092 | 0.975645591 | 0.968244468 | 0.983351992 | 6.104355017 | 5.9379191 | ENSG00000110680 |
| SETD3 | -0.035677001 | 0.975573855 | 0.554285498 | 0.710479837 | 1745.141165 | 1702.783666 | ENSG00000183576 |
| ARL13B | -0.035723886 | 0.97554215 | 0.862302592 | 0.922713518 | 119.9227906 | 116.9079445 | ENSG00000169379 |
| SUN1 | -0.035756915 | 0.975519816 | 0.541254146 | 0.700276251 | 2187.728011 | 2133.310245 | ENSG00000164828 |
| STK25 | -0.035832772 | 0.975468525 | 0.435030812 | 0.606064807 | 4349.255836 | 4242.044985 | ENSG00000115694 |
| NA | -0.035880736 | 0.975436095 | 0.886248829 | 0.937353813 | 77.37611749 | 75.37747142 | ENSG00000233328 |
| ELP5 | -0.035905531 | 0.975419331 | 0.651500666 | 0.783266107 | 821.1723252 | 801.0702491 | ENSG00000170291 |
| FBXO9 | -0.035941006 | 0.975395346 | 0.699283845 | 0.81841198 | 612.5761726 | 597.8952201 | ENSG00000112146 |

| | | | | | | | |
|---|---|---|---|---|---|---|---|
| RPS7 | -0.035964217 | 0.975379653 | 0.407100008 | 0.579327696 | 11751.02601 | 11460.34338 | ENSG00000171863 |
| NA | -0.036035273 | 0.975331615 | 0.88181169 | 0.935391472 | 88.88641043 | 86.57169715 | ENSG00000215492 |
| SF3B1 | -0.036050874 | 0.975321068 | 0.348868588 | 0.521545563 | 8616.659411 | 8403.476933 | ENSG00000115524 |
| LARS1 | -0.036151488 | 0.975253051 | 0.383465653 | 0.556806306 | 6356.73223 | 6199.711522 | ENSG00000133706 |
| JMY | -0.036154948 | 0.975250712 | 0.79033085 | 0.87763484 | 290.7945203 | 283.7542002 | ENSG00000152409 |
| DNAH5 | -0.036332694 | 0.975130565 | 0.949291681 | 0.973739099 | 14.43206743 | 14.02552699 | ENSG00000039139 |
| KDM4C | -0.036408444 | 0.975079366 | 0.662946434 | 0.792029457 | 724.2015872 | 706.291529 | ENSG00000107077 |
| CNOT10 | -0.036417865 | 0.975072999 | 0.651052939 | 0.783181022 | 834.4169031 | 813.7866001 | ENSG00000182973 |
| NA | -0.036428121 | 0.975066067 | 0.955512417 | 0.977429542 | 10.76123928 | 10.456159 | ENSG00000284294 |
| KCTD7 | -0.036492685 | 0.975022431 | 0.780240245 | 0.870840926 | 269.8238413 | 263.0319936 | ENSG00000243335 |
| MARK2 | -0.03650664 | 0.975013 | 0.572948963 | 0.725403106 | 1471.542269 | 1434.603773 | ENSG00000072518 |
| PPP1CA | -0.03654098 | 0.974989792 | 0.409609818 | 0.581847666 | 5063.247406 | 4935.587637 | ENSG00000172531 |
| PLK1 | -0.036546267 | 0.974986219 | 0.435332355 | 0.606135187 | 3229.283998 | 3148.639984 | ENSG00000166851 |
| FLNA | -0.036606339 | 0.974945623 | 0.32521358 | 0.49692635 | 42988.08914 | 41910.92823 | ENSG00000196924 |
| NAA25 | -0.036925329 | 0.97473008 | 0.62920224 | 0.76709332 | 1028.3652 | 1003.074527 | ENSG00000111300 |
| TBC1D2B | -0.036939938 | 0.974720209 | 0.68424086 | 0.807468675 | 583.9520681 | 569.1515525 | ENSG00000167202 |
| H2BU1 | -0.037001565 | 0.974678573 | 0.964071346 | 0.981782206 | 6.700296562 | 6.505793202 | ENSG00000196890 |
| CEP126 | -0.037022713 | 0.974664286 | 0.917667928 | 0.955922674 | 36.62102386 | 35.70817032 | ENSG00000110318 |
| LINC01278 | -0.03707662 | 0.974627868 | 0.754425624 | 0.854310609 | 361.6665562 | 352.7511695 | ENSG00000235437 |
| LOC100131496 | -0.037172957 | 0.974562789 | 0.978432455 | 0.988838039 | 3.064398711 | 2.978370613 | ENSG00000267882 |
| PRSS23 | -0.037240923 | 0.974516877 | 0.828713945 | 0.902525353 | 151.9367661 | 148.0933133 | ENSG00000150687 |
| AHCY | -0.03726736 | 0.97449902 | 0.371480186 | 0.545038336 | 7522.688158 | 7330.611171 | ENSG00000101444 |
| NA | -0.037270533 | 0.974496877 | 0.934047381 | 0.964679656 | 23.20280943 | 22.66889903 | ENSG00000230409 |
| ZFP36L1 | -0.037356663 | 0.9744387 | 0.547604506 | 0.705307928 | 2383.668885 | 2323.618466 | ENSG00000185650 |
| TERF2IP | -0.037381395 | 0.974421996 | 0.766061712 | 0.861545881 | 327.6486045 | 319.6003045 | ENSG00000166848 |
| RPL9 | -0.037424548 | 0.97439285 | 0.456091661 | 0.625868846 | 10097.84362 | 9837.785543 | ENSG00000163682 |
| RPL34 | -0.037445764 | 0.974378521 | 0.488805365 | 0.655460281 | 6172.868469 | 6013.372266 | ENSG00000109475 |
| PFN2 | -0.037457388 | 0.97437067 | 0.493738578 | 0.660154624 | 2471.835438 | 2408.086515 | ENSG00000070087 |
| CTAGE4 | -0.037535017 | 0.974318242 | 0.957369995 | 0.978159238 | 9.838698208 | 9.551916097 | ENSG00000225932 |
| KIAA1549 | -0.037549289 | 0.974308604 | 0.682201533 | 0.806147 | 676.985627 | 659.6577189 | ENSG00000122778 |
| NA | -0.03757196 | 0.974293293 | 0.956864477 | 0.978044405 | 9.767486175 | 9.480395705 | ENSG00000287837 |
| NA | -0.0376265 | 0.974256462 | 0.960474711 | 0.979722803 | 8.178467242 | 7.938375845 | ENSG00000279161 |
| STRN4 | -0.037680619 | 0.974219915 | 0.425148213 | 0.596782098 | 3378.739479 | 3292.106693 | ENSG00000090372 |
| AGAP2 | -0.037686748 | 0.974215777 | 0.895510788 | 0.942306148 | 53.19753966 | 51.76561916 | ENSG00000135439 |
| DNAI1 | -0.037727365 | 0.974188349 | 0.957122979 | 0.978044405 | 9.484865369 | 9.20479443 | ENSG00000122735 |
| NA | -0.037798725 | 0.974140164 | 0.940029904 | 0.968314739 | 19.74494166 | 19.16648983 | ENSG00000272070 |
| NA | -0.037810511 | 0.974132206 | 0.968459012 | 0.983367014 | 5.312198513 | 5.15548382 | ENSG00000270820 |
| CITED1 | -0.037929833 | 0.974051642 | 0.971144085 | 0.984722236 | 4.014173095 | 3.893366382 | ENSG00000125931 |
| RAF1 | -0.037951139 | 0.974037257 | 0.470862322 | 0.639766521 | 2730.098888 | 2659.106967 | ENSG00000132155 |
| NA | -0.037963876 | 0.974028657 | 0.974181993 | 0.98650629 | 3.360974032 | 3.260423609 | ENSG00000227141 |
| BECN1 | -0.038005954 | 0.974000249 | 0.680453972 | 0.804854507 | 666.1081578 | 648.6009432 | ENSG00000126581 |
| PDZD7 | -0.03809873 | 0.973937615 | 0.915068412 | 0.954554953 | 42.01260695 | 41.02174894 | ENSG00000186862 |
| LOC285074 | -0.038123508 | 0.973920889 | 0.8868945 | 0.937513028 | 64.22166328 | 62.58557138 | ENSG00000285793 |
| GAR1 | -0.038144817 | 0.973906503 | 0.602101941 | 0.747868685 | 1054.407482 | 1027.07252 | ENSG00000109534 |
| TMEM214 | -0.038184862 | 0.973879471 | 0.618465348 | 0.760118185 | 1115.477214 | 1085.78253 | ENSG00000119777 |
| ETAA1 | -0.038192214 | 0.973874508 | 0.719454205 | 0.831663685 | 468.5147527 | 456.5518299 | ENSG00000143971 |
| TSTD2 | -0.038260807 | 0.973828206 | 0.767619304 | 0.862744348 | 359.2111394 | 349.5318977 | ENSG00000136925 |
| LMBR1 | -0.038315376 | 0.973791373 | 0.643570944 | 0.777277371 | 999.6020247 | 973.2128546 | ENSG00000105983 |
| SOX7 | -0.038478641 | 0.973681178 | 0.955212361 | 0.977404214 | 9.794719483 | 9.497067257 | ENSG00000171056 |
| SH3KBP1 | -0.038501664 | 0.97366564 | 0.699639747 | 0.818679507 | 598.1129907 | 581.8937225 | ENSG00000147010 |
| INCENP | -0.038506774 | 0.973662191 | 0.501647307 | 0.667582657 | 2371.439484 | 2308.205659 | ENSG00000149503 |
| NAP1L1 | -0.038518719 | 0.97365413 | 0.301740133 | 0.472511478 | 14559.69961 | 14176.94931 | ENSG00000187109 |
| CCDC112 | -0.038546824 | 0.973635162 | 0.753650271 | 0.853959904 | 313.0171678 | 304.6651916 | ENSG00000164221 |
| ATF4 | -0.038555317 | 0.973629431 | 0.354056286 | 0.527135897 | 10546.84216 | 10267.52419 | ENSG00000128272 |
| KIF19 | -0.038564982 | 0.973622908 | 0.930730179 | 0.963034317 | 27.60171546 | 26.78463539 | ENSG00000196169 |
| TGIF1 | -0.038683019 | 0.973543253 | 0.453905638 | 0.623889823 | 2710.219883 | 2638.428274 | ENSG00000177426 |
| MTA1 | -0.03875378 | 0.973495503 | 0.471867445 | 0.640771144 | 3577.318579 | 3482.350949 | ENSG00000182979 |
| C1orf54 | -0.0387806 | 0.973477406 | 0.84083036 | 0.909051234 | 127.2201799 | 123.7622164 | ENSG00000118292 |
| HS3ST5 | -0.038859556 | 0.973424131 | 0.839982741 | 0.908905319 | 177.630323 | 173.2646689 | ENSG00000249853 |
| PCDHGA12 | -0.038878851 | 0.973411112 | 0.966593876 | 0.98274283 | 5.377828739 | 5.212483234 | ENSG00000253159 |
| ZNF227 | -0.038913317 | 0.973387857 | 0.784410504 | 0.873776593 | 230.8675358 | 224.7807662 | ENSG00000131115 |
| WASL | -0.038962617 | 0.973354595 | 0.734794765 | 0.842023481 | 683.3606528 | 665.5394869 | ENSG00000106299 |
| CACNB2 | -0.038968563 | 0.973350583 | 0.890348148 | 0.939388982 | 54.70764958 | 53.31811931 | ENSG00000165995 |
| MRPL18 | -0.039060451 | 0.973288591 | 0.590694794 | 0.739797005 | 1428.916573 | 1390.662985 | ENSG00000112110 |
| ZNF316 | -0.039174754 | 0.973211482 | 0.664737795 | 0.793432545 | 806.174816 | 784.266719 | ENSG00000205903 |
| ZSWIM1 | -0.0392387 | 0.973168346 | 0.793783238 | 0.8802019 | 234.08158 | 227.5760408 | ENSG00000168612 |
| C19orf44 | -0.039257215 | 0.973155857 | 0.87170825 | 0.928708447 | 98.09986973 | 95.47724611 | ENSG00000105072 |
| TMEM229B | -0.039312902 | 0.973118295 | 0.923205998 | 0.958830585 | 28.91070433 | 28.15429993 | ENSG00000198133 |
| PPCDC | -0.039444174 | 0.973029754 | 0.773021316 | 0.866695002 | 261.2276628 | 253.9500912 | ENSG00000138621 |
| EPHX3 | -0.039490529 | 0.97299849 | 0.694558741 | 0.815404861 | 485.4288684 | 472.2653921 | ENSG00000105131 |
| PCF11 | -0.039527485 | 0.972973566 | 0.630656738 | 0.768249503 | 854.5047383 | 831.4726496 | ENSG00000165494 |
| COPS7A | -0.039656893 | 0.972886295 | 0.615632869 | 0.758134291 | 848.832174 | 826.0014665 | ENSG00000111652 |
| HNRNPC | -0.03969832 | 0.972858359 | 0.279527914 | 0.447438943 | 22312.41236 | 21705.82257 | ENSG00000092199 |
| SLC20A2 | -0.039780244 | 0.972803116 | 0.649055378 | 0.781685316 | 733.4570222 | 713.4792194 | ENSG00000168575 |
| SLC37A3 | -0.039896991 | 0.972724398 | 0.614476445 | 0.757096833 | 1077.987276 | 1048.425004 | ENSG00000157800 |
| RBM41 | -0.03991757 | 0.972710523 | 0.717772322 | 0.83075518 | 512.656627 | 498.6937216 | ENSG00000089682 |
| SARAF | -0.039924434 | 0.972705895 | 0.543491403 | 0.702008213 | 1580.469834 | 1537.987596 | ENSG00000133872 |

| | | | | | | | |
|---|---|---|---|---|---|---|---|
| ACADS | -0.039964609 | 0.972678808 | 0.847636217 | 0.912944574 | 135.4928622 | 131.6710173 | ENSG00000122971 |
| TLCD3B | -0.04006014 | 0.972614402 | 0.739049193 | 0.84500301 | 428.1117402 | 416.8529722 | ENSG00000149926 |
| POU2F3 | -0.04007947 | 0.972601371 | 0.931982551 | 0.963756387 | 24.75286787 | 24.10309921 | ENSG00000137709 |
| POSTN | -0.040165128 | 0.972543626 | 0.934911125 | 0.965304601 | 20.23494104 | 19.59535172 | ENSG00000133110 |
| TBC1D32 | -0.040168565 | 0.972541309 | 0.870039018 | 0.927442266 | 78.77960951 | 76.55327491 | ENSG00000146350 |
| KMT5B | -0.040341765 | 0.972424559 | 0.600122404 | 0.746371046 | 922.3452786 | 897.1421996 | ENSG00000110066 |
| MIPOL1 | -0.040372393 | 0.972403915 | 0.816086183 | 0.894698714 | 188.7500627 | 183.3266478 | ENSG00000151338 |
| LHFPL6 | -0.040405463 | 0.972381625 | 0.862136794 | 0.922689647 | 100.6065237 | 97.91946818 | ENSG00000183722 |
| TUBG1 | -0.040418465 | 0.972372862 | 0.481755392 | 0.649807249 | 1772.736648 | 1723.654323 | ENSG00000131462 |
| PTPN23 | -0.040453263 | 0.972349409 | 0.520743816 | 0.683672188 | 1763.015331 | 1714.027452 | ENSG00000076201 |
| WARS2 | -0.04064592 | 0.97221957 | 0.836243492 | 0.907004003 | 121.7085 | 118.1983104 | ENSG00000116874 |
| PRMT5 | -0.040684038 | 0.972193883 | 0.607058231 | 0.751272767 | 1410.028481 | 1370.478072 | ENSG00000100462 |
| SOCS5 | -0.04070544 | 0.972179461 | 0.717220145 | 0.830493388 | 373.8109875 | 363.5488224 | ENSG00000171150 |
| MKKS | -0.04072902 | 0.972163571 | 0.776924543 | 0.869252486 | 288.3719938 | 280.1090596 | ENSG00000125863 |
| APEH | -0.040765604 | 0.972138919 | 0.358882348 | 0.53194002 | 5466.90946 | 5313.564006 | ENSG00000164062 |
| NA | -0.040830339 | 0.9720953 | 0.970275395 | 0.98427465 | 4.381960449 | 4.242638624 | ENSG00000261360 |
| TBL3 | -0.040858359 | 0.97207642 | 0.539747731 | 0.698951029 | 1946.478514 | 1892.297319 | ENSG00000183751 |
| NA | -0.040970369 | 0.972000951 | 0.973029031 | 0.98595118 | 3.641479657 | 3.525272016 | ENSG00000279742 |
| ZDHHC2 | -0.040982949 | 0.971992476 | 0.754457664 | 0.854310609 | 391.427038 | 380.6508345 | ENSG00000104219 |
| SETD2 | -0.040995425 | 0.97198407 | 0.52841346 | 0.689817823 | 1359.220538 | 1320.649428 | ENSG00000181555 |
| CAPN10 | -0.041011445 | 0.971973277 | 0.585515402 | 0.735192012 | 1017.329237 | 989.0912601 | ENSG00000142330 |
| SMN1 | -0.041022178 | 0.971966046 | 0.821204959 | 0.898166734 | 160.4735719 | 156.1817147 | ENSG00000172062 |
| TOB1-AS1 | -0.041039831 | 0.971954153 | 0.922905105 | 0.958789508 | 24.32993818 | 23.67853847 | ENSG00000229980 |
| SORT1 | -0.041095828 | 0.971916429 | 0.589268181 | 0.738633689 | 1362.977847 | 1325.211375 | ENSG00000134243 |
| IDH3B | -0.041210834 | 0.971838954 | 0.502976166 | 0.668568415 | 1962.020139 | 1906.432537 | ENSG00000101365 |
| S1PR5 | -0.041217762 | 0.971834287 | 0.762044641 | 0.858680092 | 278.5818681 | 270.8230275 | ENSG00000180739 |
| CNN2 | -0.041272036 | 0.971797727 | 0.312129553 | 0.483126914 | 9394.580109 | 9130.630863 | ENSG00000064666 |
| USP46 | -0.041277684 | 0.971793923 | 0.582453166 | 0.732900921 | 1098.265008 | 1067.522045 | ENSG00000109189 |
| NFX1 | -0.041294805 | 0.971782391 | 0.580949054 | 0.731628608 | 981.6803007 | 953.8939219 | ENSG00000086102 |
| NA | -0.041295483 | 0.971781934 | 0.843678716 | 0.910436526 | 124.0926665 | 120.4590418 | ENSG00000213621 |
| TRIM37 | -0.041494945 | 0.971647588 | 0.514199385 | 0.678170569 | 1683.264783 | 1636.072938 | ENSG00000108395 |
| SYT17 | -0.041518133 | 0.971631971 | 0.894700377 | 0.941915276 | 67.69917951 | 65.84357184 | ENSG00000103528 |
| FLT3 | -0.041587336 | 0.971585365 | 0.940278544 | 0.968441718 | 15.52638106 | 15.01204315 | ENSG00000122025 |
| ARMC8 | -0.041609821 | 0.971570223 | 0.601307191 | 0.747122054 | 784.1461931 | 761.8929821 | ENSG00000114098 |
| CPSF4 | -0.041620221 | 0.971563219 | 0.557604811 | 0.713428546 | 1152.860733 | 1119.835463 | ENSG00000160917 |
| ACTR3B | -0.041674837 | 0.971526439 | 0.612806008 | 0.756041756 | 837.0585581 | 812.833948 | ENSG00000133627 |
| MEA1 | -0.041726577 | 0.971491597 | 0.569235498 | 0.72234719 | 1233.127575 | 1197.500557 | ENSG00000124733 |
| RAD9A | -0.041975979 | 0.971323668 | 0.729348108 | 0.837690781 | 384.2567908 | 372.974945 | ENSG00000172613 |
| MOV10 | -0.042041721 | 0.971279407 | 0.571186088 | 0.723801675 | 1188.494849 | 1153.950266 | ENSG00000155363 |
| LOC105371267 | -0.042102796 | 0.971238289 | 0.893585167 | 0.941175276 | 47.83578062 | 46.36384859 | ENSG00000277639 |
| DPY19L1 | -0.042128914 | 0.971220707 | 0.739018181 | 0.84500301 | 378.0452175 | 367.643283 | ENSG00000173852 |
| RASA3 | -0.042206317 | 0.9711686 | 0.639362262 | 0.77469615 | 757.9513994 | 736.6181143 | ENSG00000185989 |
| NA | -0.042217511 | 0.971161065 | 0.970321105 | 0.98427465 | 3.393789145 | 3.277095161 | ENSG00000261770 |
| RSL24D1 | -0.042411781 | 0.9710303 | 0.528516657 | 0.689851434 | 2660.065806 | 2583.306157 | ENSG00000137876 |
| ZNF517 | -0.042514406 | 0.970961229 | 0.801745754 | 0.885808137 | 195.2066109 | 189.2000617 | ENSG00000197363 |
| NA | -0.042607493 | 0.970898582 | 0.959339505 | 0.97923797 | 6.433745451 | 6.211369801 | ENSG00000175886 |
| DNMT3A | -0.042679618 | 0.970850044 | 0.491326133 | 0.65765837 | 4067.67509 | 3948.715867 | ENSG00000119772 |
| VPS52 | -0.04271751 | 0.970824546 | 0.649640675 | 0.781978702 | 659.4817762 | 640.1471005 | ENSG00000223501 |
| ANKRD24 | -0.042782101 | 0.970781082 | 0.729975538 | 0.838161902 | 445.1882677 | 432.3285896 | ENSG00000089847 |
| MTFR1 | -0.04297418 | 0.970651841 | 0.63461075 | 0.771166991 | 708.1153831 | 687.4388241 | ENSG00000066855 |
| TMEM129 | -0.04298038 | 0.97064767 | 0.571339539 | 0.723921908 | 1122.770097 | 1089.412914 | ENSG00000168936 |
| VWA8 | -0.043041541 | 0.970606522 | 0.663156709 | 0.792133503 | 532.3478899 | 516.3663587 | ENSG00000102763 |
| DDX46 | -0.043082323 | 0.970579085 | 0.38944134 | 0.562665735 | 3535.387595 | 3430.549254 | ENSG00000145833 |
| USP18 | -0.04333663 | 0.970408014 | 0.816945488 | 0.895335497 | 133.5319135 | 129.5802059 | ENSG00000184979 |
| RUBCN | -0.043403974 | 0.970362717 | 0.775857235 | 0.868533369 | 230.5875426 | 223.7944984 | ENSG00000145016 |
| KPNA2 | -0.043411002 | 0.97035799 | 0.246941103 | 0.410490377 | 11531.80459 | 11190.23879 | ENSG00000182481 |
| NA | -0.043481218 | 0.970310764 | 0.960597464 | 0.979764445 | 5.731661579 | 5.527879334 | ENSG00000238186 |
| NA | -0.04349878 | 0.970298952 | 0.954816466 | 0.977102777 | 7.772958691 | 7.498761085 | ENSG00000254331 |
| NPIPB11 | -0.043518476 | 0.970285706 | 0.935440353 | 0.965468934 | 17.37425412 | 16.90306268 | ENSG00000254206 |
| NA | -0.043544643 | 0.970268107 | 0.967979994 | 0.983248479 | 4.011382192 | 3.87024311 | ENSG00000183298 |
| PPP2R3A | -0.04361801 | 0.970218766 | 0.736340828 | 0.84326215 | 300.9564636 | 292.1910866 | ENSG00000073711 |
| AP3M1 | -0.043663511 | 0.970188167 | 0.575909926 | 0.728039212 | 992.6806207 | 963.5958514 | ENSG00000185009 |
| PRKCZ-AS1 | -0.043673721 | 0.970181301 | 0.876232526 | 0.931727575 | 55.61432112 | 53.93654111 | ENSG00000182873 |
| HTRA4 | -0.043754174 | 0.9701272 | 0.928669628 | 0.961846113 | 20.48263155 | 19.90078845 | ENSG00000169495 |
| MAMDC2 | -0.043761095 | 0.970122546 | 0.768680479 | 0.86353454 | 205.3821964 | 199.1119908 | ENSG00000165072 |
| NA | -0.04383933 | 0.970069939 | 0.760654336 | 0.857664551 | 286.1804124 | 277.5127287 | ENSG00000132967 |
| R3HDM4 | -0.043842369 | 0.970067896 | 0.445499654 | 0.615978628 | 1846.235197 | 1791.038842 | ENSG00000198858 |
| TMED5 | -0.043869221 | 0.970049841 | 0.738212631 | 0.844463914 | 361.0879989 | 350.6606428 | ENSG00000117500 |
| SLC36A1 | -0.043884122 | 0.970039822 | 0.6882159 | 0.810622354 | 386.2855729 | 374.7183195 | ENSG00000123643 |
| RPL8 | -0.043888768 | 0.970036698 | 0.390906378 | 0.564148645 | 51303.48005 | 49765.6566 | ENSG00000161016 |
| NCCRP1 | -0.044109228 | 0.969888476 | 0.859149814 | 0.920821105 | 79.9309805 | 77.37283352 | ENSG00000188505 |
| OPRL1 | -0.044134135 | 0.969871732 | 0.961609679 | 0.980465768 | 6.036609608 | 5.824453309 | ENSG00000125510 |
| CDH13 | -0.044149423 | 0.969861454 | 0.936637532 | 0.966331442 | 15.19679047 | 14.79559186 | ENSG00000140945 |
| GPALPP1 | -0.044188922 | 0.969834902 | 0.694522534 | 0.815404861 | 434.3365923 | 421.6817435 | ENSG00000133114 |
| EIF2B4 | -0.044196174 | 0.969830026 | 0.565820373 | 0.719812768 | 1207.649495 | 1171.174104 | ENSG00000115211 |
| DIS3 | -0.04421067 | 0.969820281 | 0.506109917 | 0.670929522 | 1218.510531 | 1181.740185 | ENSG00000083520 |
| SLBP | -0.04421095 | 0.969820094 | 0.536568131 | 0.696188849 | 1083.329626 | 1050.784686 | ENSG00000163950 |

| | | | | | | | |
|---|---|---|---|---|---|---|---|
| CALY | -0.044230187 | 0.969807162 | 0.929825778 | 0.962454745 | 19.99542308 | 19.41922829 | ENSG00000130643 |
| NA | -0.044301214 | 0.969759417 | 0.962183551 | 0.980632996 | 5.369456031 | 5.176456519 | ENSG00000231510 |
| MYNN | -0.044301267 | 0.969759382 | 0.75718446 | 0.855942967 | 226.2881211 | 219.449983 | ENSG00000085274 |
| NA | -0.044307626 | 0.969755108 | 0.826616701 | 0.9012243 | 117.0741431 | 113.4472081 | ENSG00000226085 |
| KDM5A | -0.044343199 | 0.969731196 | 0.582168961 | 0.732591091 | 896.6038008 | 869.3065308 | ENSG00000073614 |
| CSTF1 | -0.044345669 | 0.969729536 | 0.553867859 | 0.710086055 | 1010.634359 | 980.0303305 | ENSG00000101138 |
| BLM | -0.044459675 | 0.969652908 | 0.599757209 | 0.746157379 | 747.3924693 | 725.0917904 | ENSG00000197299 |
| VPS37C | -0.044640773 | 0.969531197 | 0.662397153 | 0.791698422 | 653.0624878 | 633.5695327 | ENSG00000167987 |
| CWC25 | -0.044671984 | 0.969510223 | 0.707912198 | 0.823963032 | 379.5805704 | 368.2095395 | ENSG00000273559 |
| LINC01012 | -0.044744982 | 0.969461169 | 0.881503038 | 0.93532122 | 51.38439674 | 49.80280666 | ENSG00000281706 |
| RASL10A | -0.044755661 | 0.969453993 | 0.90973825 | 0.951403864 | 30.48576875 | 29.66055354 | ENSG00000100276 |
| PSMB2 | -0.044890835 | 0.969363164 | 0.392747393 | 0.565659377 | 2834.697257 | 2748.175747 | ENSG00000126067 |
| LOC101929243 | -0.044900279 | 0.969356818 | 0.938627332 | 0.967619295 | 15.8152594 | 15.25430132 | ENSG00000272288 |
| EEF1AKMT2 | -0.044931187 | 0.969336051 | 0.895079997 | 0.942161195 | 43.77742864 | 42.45972934 | ENSG00000203791 |
| ING5 | -0.044950246 | 0.969323245 | 0.530729924 | 0.691644398 | 1148.533052 | 1112.989242 | ENSG00000168395 |
| MIF4GD | -0.044959925 | 0.969316742 | 0.749965916 | 0.8520254 | 254.1004063 | 246.1270408 | ENSG00000125457 |
| ZNF549 | -0.0449783 | 0.969304396 | 0.782244527 | 0.872362229 | 188.3701478 | 182.3635459 | ENSG00000121406 |
| DHX9 | -0.044997144 | 0.969291736 | 0.283608853 | 0.452194262 | 7658.806147 | 7423.037909 | ENSG00000135829 |
| IQSEC1 | -0.045069703 | 0.969242988 | 0.545847592 | 0.704108719 | 1178.298302 | 1142.557308 | ENSG00000144711 |
| CYB5R2 | -0.045112204 | 0.969214435 | 0.66807834 | 0.796028624 | 526.7023588 | 510.627892 | ENSG00000166394 |
| PRKACA | -0.045272178 | 0.969106968 | 0.611866073 | 0.755316639 | 1244.788024 | 1206.02532 | ENSG00000072062 |
| NA | -0.045301275 | 0.969087423 | 0.963678409 | 0.981641155 | 4.781887194 | 4.606431843 | ENSG00000236154 |
| NA | -0.045316913 | 0.969076919 | 0.895409478 | 0.942250941 | 37.66278589 | 36.5640161 | ENSG00000248103 |
| OCLN | -0.045339293 | 0.969061886 | 0.68790979 | 0.810410245 | 442.8853866 | 429.3601051 | ENSG00000197822 |
| TMEM50A | -0.045357414 | 0.969049714 | 0.470377432 | 0.639300819 | 1624.748373 | 1574.303956 | ENSG00000183726 |
| FAM184B | -0.045375753 | 0.969037396 | 0.910599214 | 0.951951955 | 26.74181355 | 25.8954465 | ENSG00000047662 |
| SLC24A1 | -0.045440465 | 0.968993931 | 0.834831578 | 0.906542977 | 153.7998699 | 149.160461 | ENSG00000074621 |
| MTMR9 | -0.04546727 | 0.968975928 | 0.67385544 | 0.800111161 | 479.1210529 | 464.5746296 | ENSG00000104643 |
| FBXW7 | -0.045507347 | 0.96894901 | 0.69849891 | 0.817989588 | 424.1622462 | 411.1067573 | ENSG00000109670 |
| CA13 | -0.045578685 | 0.9689011 | 0.903343094 | 0.947370846 | 33.62157965 | 32.51738911 | ENSG00000185015 |
| TTC28-AS1 | -0.04559811 | 0.968888054 | 0.811175118 | 0.891639582 | 149.989772 | 145.1655025 | ENSG00000235954 |
| NA | -0.045601512 | 0.968885769 | 0.863002284 | 0.923052621 | 70.48016955 | 68.42103802 | ENSG00000204860 |
| NA | -0.045620622 | 0.968872935 | 0.912691175 | 0.952899576 | 27.28907044 | 26.4194851 | ENSG00000213839 |
| MET | -0.045708488 | 0.968813928 | 0.696013851 | 0.816616178 | 380.5755628 | 368.935248 | ENSG00000105976 |
| SF1 | -0.045738413 | 0.968793834 | 0.328989885 | 0.5012276 | 6165.494602 | 5973.658917 | ENSG00000168066 |
| PGP | -0.045785306 | 0.968762344 | 0.587031543 | 0.736643824 | 1696.748019 | 1644.467419 | ENSG00000184207 |
| PSRC1 | -0.045805117 | 0.968749042 | 0.479147999 | 0.647285997 | 1370.14678 | 1327.064054 | ENSG00000134222 |
| TRIM28 | -0.045817236 | 0.968740904 | 0.37852311 | 0.551934188 | 31216.17494 | 30240.26259 | ENSG00000130726 |
| VPS28 | -0.045895378 | 0.968688435 | 0.614750913 | 0.757338264 | 672.8750485 | 651.6080647 | ENSG00000160948 |
| PAXX | -0.04590655 | 0.968680933 | 0.679318395 | 0.803872602 | 560.3547647 | 542.9692493 | ENSG00000148362 |
| SPECC1 | -0.045979132 | 0.9686322 | 0.455767272 | 0.625485559 | 1697.102461 | 1644.24844 | ENSG00000128487 |
| RTL8B | -0.045989185 | 0.968625451 | 0.830954721 | 0.903961401 | 106.8721633 | 103.5688826 | ENSG00000212747 |
| SPIN1 | -0.045990093 | 0.968624841 | 0.432643212 | 0.603654 | 2786.677863 | 2699.372474 | ENSG00000106723 |
| FLOT2 | -0.046028565 | 0.968599011 | 0.340242398 | 0.512822572 | 4428.976985 | 4289.5907 | ENSG00000132589 |
| LASP1 | -0.046078619 | 0.968565406 | 0.32485974 | 0.496503663 | 5594.715561 | 5418.85428 | ENSG00000002834 |
| CRK | -0.046107283 | 0.968546163 | 0.440000515 | 0.611090709 | 1775.116723 | 1719.392862 | ENSG00000167193 |
| SMIM12 | -0.046133837 | 0.968528336 | 0.545785469 | 0.704108719 | 1026.510179 | 994.0596135 | ENSG00000163866 |
| PAPSS1 | -0.046146539 | 0.968519808 | 0.441753743 | 0.612907901 | 2676.149015 | 2591.737107 | ENSG00000138801 |
| ALKBH2 | -0.0461512 | 0.96851668 | 0.662817677 | 0.791973727 | 461.4594209 | 446.9375351 | ENSG00000189046 |
| EPN2 | -0.046164418 | 0.968507806 | 0.622894371 | 0.762604966 | 609.0159414 | 589.6667462 | ENSG00000072134 |
| SASH1 | -0.046242998 | 0.968455055 | 0.665867378 | 0.794436736 | 574.2386979 | 555.76542 | ENSG00000111961 |
| PIK3C2A | -0.046323377 | 0.9684011 | 0.428440125 | 0.599776083 | 1873.572401 | 1814.285253 | ENSG00000011405 |
| LPCAT1 | -0.046337478 | 0.968391635 | 0.257759211 | 0.422861888 | 7287.584666 | 7057.421022 | ENSG00000153395 |
| ZSCAN25 | -0.0463422 | 0.968388465 | 0.698165867 | 0.817947148 | 391.7837739 | 379.3817813 | ENSG00000197037 |
| UTP14A | -0.04643905 | 0.968323458 | 0.547514035 | 0.705307928 | 1088.779981 | 1054.576697 | ENSG00000156697 |
| MLX | -0.046550985 | 0.968248331 | 0.590780538 | 0.739856359 | 806.1605108 | 780.3039465 | ENSG00000108788 |
| MED10 | -0.046581543 | 0.968227823 | 0.665149322 | 0.793874626 | 466.2070535 | 451.2097608 | ENSG00000133398 |
| CENPA | -0.046646728 | 0.968184077 | 0.71010628 | 0.82536966 | 294.0303282 | 284.5245561 | ENSG00000115163 |
| RPS5 | -0.046649389 | 0.968182291 | 0.298596839 | 0.46906992 | 16085.81546 | 15573.04721 | ENSG00000083845 |
| PGLS | -0.046702913 | 0.968146372 | 0.693314433 | 0.814467988 | 407.8849205 | 395.203381 | ENSG00000130313 |
| NA | -0.046708508 | 0.968142617 | 0.866679042 | 0.92554835 | 58.64291365 | 56.83263846 | ENSG00000242622 |
| HSP90AB1 | -0.046746979 | 0.968116801 | 0.17746494 | 0.321211875 | 83227.70916 | 80573.3565 | ENSG00000096384 |
| ICE1 | -0.04690018 | 0.968014001 | 0.421996111 | 0.593804067 | 2068.766634 | 2002.533172 | ENSG00000164151 |
| NUDT4 | -0.046926634 | 0.967996252 | 0.502001307 | 0.667915526 | 1417.449072 | 1372.608605 | ENSG00000173598 |
| GRK2 | -0.046979349 | 0.967960883 | 0.400756348 | 0.573217966 | 2247.38911 | 2175.777159 | ENSG00000173020 |
| FBH1 | -0.046994563 | 0.967950675 | 0.534538967 | 0.694864637 | 1079.275153 | 1044.321105 | ENSG00000134452 |
| OARD1 | -0.047065532 | 0.967903061 | 0.619554904 | 0.760827147 | 607.5349557 | 588.20537 | ENSG00000124596 |
| FLJ16779 | -0.047083052 | 0.967891307 | 0.939556867 | 0.968060221 | 13.89405912 | 13.37376209 | ENSG00000275620 |
| NA | -0.047286419 | 0.967754879 | 0.963992062 | 0.981778233 | 4.365215031 | 4.196392078 | ENSG00000274099 |
| EIF2AK3 | -0.047306452 | 0.967741441 | 0.707815505 | 0.82394816 | 322.7532783 | 312.2446269 | ENSG00000172071 |
| HPCAL1 | -0.047354158 | 0.967709441 | 0.740138669 | 0.84572929 | 233.295681 | 225.8057201 | ENSG00000115756 |
| ITCH | -0.047425449 | 0.967661623 | 0.577319753 | 0.729200246 | 712.777649 | 689.6393452 | ENSG00000078747 |
| NAIF1 | -0.047449351 | 0.967645591 | 0.742883982 | 0.847821391 | 250.0048967 | 241.7290215 | ENSG00000171169 |
| CCNL1 | -0.047490802 | 0.967617789 | 0.546881557 | 0.70479907 | 865.538711 | 837.1720723 | ENSG00000163660 |
| TRAF4 | -0.047561117 | 0.96757063 | 0.327912029 | 0.500098897 | 3320.920033 | 3212.981028 | ENSG00000076604 |
| YWHAG | -0.047645768 | 0.967513859 | 0.247670963 | 0.411207552 | 7871.550804 | 7615.421899 | ENSG00000170027 |
| NUP62CL | -0.047732699 | 0.967455562 | 0.893525026 | 0.941175276 | 40.38260035 | 39.08636087 | ENSG00000198088 |

| | | | | | | | |
|---|---|---|---|---|---|---|---|
| ZNF707 | -0.04780674 | 0.967405912 | 0.701259198 | 0.819997315 | 314.5293929 | 304.1787973 | ENSG00000181135 |
| SLC35E1 | -0.047835107 | 0.967386891 | 0.639877751 | 0.774742503 | 531.8843601 | 514.5559891 | ENSG00000127526 |
| MVK | -0.047842092 | 0.967382207 | 0.618790251 | 0.760275316 | 524.1149561 | 506.948336 | ENSG00000110921 |
| TFDP2 | -0.047882276 | 0.967355262 | 0.519390382 | 0.682856939 | 1054.837233 | 1020.460276 | ENSG00000114126 |
| KHDRBS1 | -0.047911129 | 0.967335916 | 0.330130777 | 0.502529221 | 7162.845927 | 6929.677005 | ENSG00000121774 |
| ZRANB2 | -0.048045804 | 0.96724562 | 0.454169438 | 0.624130737 | 1830.812907 | 1771.107073 | ENSG00000132485 |
| ERICH1 | -0.048213846 | 0.967132964 | 0.56158206 | 0.716452584 | 842.1530534 | 814.1501513 | ENSG00000104714 |
| PVR | -0.048246881 | 0.967110819 | 0.648284891 | 0.78123837 | 512.1700574 | 495.8178097 | ENSG00000073008 |
| EYA3 | -0.048254577 | 0.96710566 | 0.618869095 | 0.760323762 | 722.2608748 | 698.4690237 | ENSG00000158161 |
| ARL1 | -0.048310935 | 0.967067881 | 0.584899144 | 0.734780502 | 953.2492101 | 922.5885808 | ENSG00000120805 |
| ADM2 | -0.04833006 | 0.967055061 | 0.697307095 | 0.817153648 | 440.7628041 | 426.3110745 | ENSG00000128165 |
| BID | -0.048541051 | 0.966913642 | 0.58302731 | 0.733248614 | 742.5695064 | 718.3475071 | ENSG00000015475 |
| ZEB1-AS1 | -0.04856717 | 0.966896137 | 0.923059627 | 0.958820213 | 17.91014724 | 17.33649829 | ENSG00000237036 |
| EXO5 | -0.048588121 | 0.966882096 | 0.875018537 | 0.930847171 | 59.55046107 | 57.37282768 | ENSG00000164002 |
| OTUD3 | -0.048637123 | 0.966849255 | 0.634468275 | 0.771091003 | 487.9698138 | 471.59426 | ENSG00000169914 |
| TGFB1 | -0.048756201 | 0.966769456 | 0.708458829 | 0.824299111 | 300.0433344 | 290.2471084 | ENSG00000105329 |
| PGAM5 | -0.048787392 | 0.966748555 | 0.373090974 | 0.546534559 | 2384.660822 | 2305.440177 | ENSG00000247077 |
| IMPAD1 | -0.048821034 | 0.966726012 | 0.639900937 | 0.774742503 | 662.2583499 | 640.8340496 | ENSG00000104331 |
| GALNT14 | -0.048856754 | 0.966702077 | 0.681652952 | 0.805794907 | 437.3761852 | 423.1721375 | ENSG00000158089 |
| ZNF720 | -0.048952683 | 0.9666378 | 0.736635343 | 0.843449216 | 293.4398932 | 283.5199992 | ENSG00000197302 |
| UBA52 | -0.04896846 | 0.966627229 | 0.209799992 | 0.365209497 | 9495.950616 | 9179.303567 | ENSG00000221983 |
| PLEKHG6 | -0.048977674 | 0.966621056 | 0.584217237 | 0.734162903 | 637.3091776 | 615.8780977 | ENSG00000008323 |
| PTMA | -0.049090362 | 0.966545556 | 0.219003657 | 0.376844131 | 41127.67638 | 39751.55495 | ENSG00000187514 |
| NA | -0.0491677 | 0.966493745 | 0.843432152 | 0.91029706 | 90.0194844 | 86.83709071 | ENSG00000213612 |
| BCL7B | -0.049228607 | 0.966452942 | 0.593778949 | 0.741925624 | 612.3153154 | 591.7199013 | ENSG00000106635 |
| PTK7 | -0.049372793 | 0.966356358 | 0.269634935 | 0.435974176 | 4974.157308 | 4806.084871 | ENSG00000112655 |
| PIGO | -0.04946009 | 0.966297886 | 0.592724664 | 0.741158069 | 593.6647243 | 573.5692179 | ENSG00000165282 |
| NA | -0.049463095 | 0.966295873 | 0.966351212 | 0.982599685 | 3.399370951 | 3.260423609 | ENSG00000285730 |
| BRF1 | -0.049573967 | 0.966221616 | 0.616135083 | 0.758317079 | 587.173172 | 567.5928732 | ENSG00000185024 |
| MED30 | -0.049578677 | 0.966218461 | 0.747262998 | 0.850346567 | 203.0033363 | 196.1190871 | ENSG00000164758 |
| DMXL1 | -0.049596507 | 0.96620652 | 0.624206591 | 0.763058842 | 667.9840648 | 645.1885365 | ENSG00000172869 |
| UBALD1 | -0.049623248 | 0.966188611 | 0.567161924 | 0.720569064 | 1137.781114 | 1099.661217 | ENSG00000153443 |
| SYNCRIP | -0.049657323 | 0.966165791 | 0.281146649 | 0.449457198 | 3990.011281 | 3855.770942 | ENSG00000135316 |
| PSKH2 | -0.049717496 | 0.966125494 | 0.955307496 | 0.977404477 | 5.786128194 | 5.54670146 | ENSG00000147613 |
| NA | -0.049722953 | 0.96612184 | 0.956922874 | 0.978044405 | 5.402271144 | 5.178607093 | ENSG00000222881 |
| AAR2 | -0.049813447 | 0.966061241 | 0.577593042 | 0.72944991 | 832.433437 | 803.7160311 | ENSG00000131043 |
| RFC4 | -0.04986651 | 0.96602571 | 0.442677098 | 0.613571195 | 1393.208629 | 1346.097947 | ENSG00000163918 |
| DYNLT3 | -0.04994033 | 0.965976281 | 0.858502243 | 0.920821105 | 72.78570243 | 70.44435757 | ENSG00000165169 |
| NA | -0.050013797 | 0.965927092 | 0.952148121 | 0.975457419 | 8.465793974 | 8.234949819 | ENSG00000233672 |
| STRADB | -0.050137695 | 0.965844141 | 0.619218269 | 0.760607426 | 738.2048422 | 713.4522917 | ENSG00000082146 |
| PAQR8 | -0.050164361 | 0.965826289 | 0.715897089 | 0.829501516 | 304.781355 | 294.50451 | ENSG00000170915 |
| FAM199X | -0.050183351 | 0.965813576 | 0.456645392 | 0.626495004 | 1218.516276 | 1176.869189 | ENSG00000123575 |
| ISY1 | -0.050204819 | 0.965799205 | 0.835333064 | 0.906737673 | 88.44865035 | 85.48004492 | ENSG00000240682 |
| C11orf1 | -0.050305972 | 0.965731491 | 0.799303005 | 0.884159303 | 123.325428 | 119.1848144 | ENSG00000137720 |
| SLC4A5 | -0.050306983 | 0.965730814 | 0.964732442 | 0.981988895 | 3.423813356 | 3.279245734 | ENSG00000188687 |
| CASC3 | -0.050309816 | 0.965728918 | 0.314207395 | 0.485375949 | 3760.566871 | 3631.64176 | ENSG00000108349 |
| SPAG16 | -0.050331588 | 0.965714344 | 0.7592451 | 0.857014416 | 209.3231303 | 202.273718 | ENSG00000144451 |
| CLASRP | -0.050372005 | 0.96568729 | 0.468399905 | 0.637287157 | 1459.674037 | 1409.179697 | ENSG00000104859 |
| HTRA2 | -0.050375429 | 0.965684998 | 0.688115127 | 0.81055757 | 326.8924543 | 315.6757762 | ENSG00000115317 |
| ECE1 | -0.050377442 | 0.965683651 | 0.396543216 | 0.569259948 | 2113.51335 | 2041.452654 | ENSG00000117298 |
| NA | -0.050484071 | 0.96561228 | 0.825291179 | 0.900593514 | 89.89380576 | 86.70184661 | ENSG00000139239 |
| METTL1 | -0.050498491 | 0.965602629 | 0.681749176 | 0.805819974 | 317.8200307 | 306.8027705 | ENSG00000037897 |
| ZFP37 | -0.050506118 | 0.965597524 | 0.85549406 | 0.918462615 | 86.2170835 | 83.47151892 | ENSG00000136866 |
| SLC25A19 | -0.050530413 | 0.965581264 | 0.718287124 | 0.831072807 | 269.4752269 | 259.8291329 | ENSG00000125454 |
| TMA16 | -0.050590978 | 0.965540729 | 0.621015057 | 0.761892741 | 728.4893086 | 703.4427932 | ENSG00000198498 |
| HDAC5 | -0.050601141 | 0.965533927 | 0.39419957 | 0.566882815 | 2420.011255 | 2336.199774 | ENSG00000108840 |
| LDB1 | -0.050664449 | 0.965491559 | 0.348914334 | 0.521573531 | 2638.472151 | 2547.284634 | ENSG00000198728 |
| ZRANB3 | -0.050673022 | 0.965485821 | 0.768481121 | 0.863411142 | 166.2916763 | 160.3224644 | ENSG00000121988 |
| STAT5B | -0.050677339 | 0.965482932 | 0.427180403 | 0.598608428 | 1548.488221 | 1494.819933 | ENSG00000173757 |
| CDCA2 | -0.05071904 | 0.965455026 | 0.492585242 | 0.658932245 | 998.6190732 | 964.5148878 | ENSG00000184661 |
| NA | -0.050738379 | 0.965442084 | 0.804482924 | 0.887069454 | 132.1579333 | 127.7018293 | ENSG00000273209 |
| BRINP2 | -0.050747659 | 0.965435874 | 0.902193311 | 0.946601282 | 29.9668212 | 28.8587631 | ENSG00000198797 |
| APBA1 | -0.050806491 | 0.965396505 | 0.920262436 | 0.957436234 | 19.89697774 | 19.20036597 | ENSG00000107282 |
| TMEM258 | -0.050809489 | 0.965394499 | 0.614258421 | 0.756906783 | 780.7681599 | 753.1622421 | ENSG00000134825 |
| SH2D6 | -0.050812133 | 0.96539273 | 0.961397633 | 0.980301377 | 4.071430613 | 3.89766753 | ENSG00000152292 |
| HS3ST3B1 | -0.050856485 | 0.965363051 | 0.755912608 | 0.855167143 | 183.0577492 | 176.8955979 | ENSG00000125430 |
| MVD | -0.051002459 | 0.96526538 | 0.485139348 | 0.652828023 | 2568.328822 | 2479.222367 | ENSG00000167508 |
| GSTO1 | -0.051020972 | 0.965252993 | 0.561844342 | 0.716559985 | 987.1923426 | 952.4857953 | ENSG00000148834 |
| GSN | -0.051078966 | 0.965214192 | 0.45436004 | 0.624142047 | 1259.32531 | 1215.608664 | ENSG00000148180 |
| NA | -0.051155075 | 0.965163274 | 0.873099211 | 0.929523006 | 43.2452023 | 41.69045797 | ENSG00000225177 |
| CDK7 | -0.051388058 | 0.965007421 | 0.509813885 | 0.674379942 | 1146.634216 | 1106.273443 | ENSG00000134058 |
| SEZ6L2 | -0.05147479 | 0.964949408 | 0.389951102 | 0.563191338 | 1722.649275 | 1662.623402 | ENSG00000174938 |
| PSPN | -0.051512795 | 0.964923989 | 0.881988249 | 0.935527317 | 39.44502872 | 38.17970693 | ENSG00000125650 |
| PEX6 | -0.051578108 | 0.964880307 | 0.528298692 | 0.689768008 | 855.7341183 | 826.0455562 | ENSG00000124587 |
| TMEM135 | -0.051583547 | 0.964876669 | 0.634880533 | 0.771274526 | 570.7185928 | 551.1952327 | ENSG00000166575 |
| ZNHIT2 | -0.051588537 | 0.964873332 | 0.80873827 | 0.890133506 | 100.6312784 | 97.02544511 | ENSG00000174276 |
| APOL2 | -0.051609183 | 0.964859523 | 0.658906126 | 0.789304402 | 406.9987861 | 392.5702725 | ENSG00000128335 |

| | | | | | | | |
|---|---|---|---|---|---|---|---|
| ABL1 | -0.05163368 | 0.964843141 | 0.272436289 | 0.439171761 | 3959.22552 | 3820.228966 | ENSG00000097007 |
| CHKB-DT | -0.051679118 | 0.964812753 | 0.862073885 | 0.922689647 | 54.61825267 | 52.69728648 | ENSG00000205559 |
| ETFDH | -0.05175944 | 0.964759038 | 0.755183387 | 0.854964446 | 206.7097705 | 199.1714318 | ENSG00000171503 |
| TBCEL | -0.05192245 | 0.964650036 | 0.80649476 | 0.888475268 | 121.4508852 | 117.3074918 | ENSG00000154114 |
| TYW1 | -0.051963065 | 0.96462288 | 0.605687118 | 0.750292827 | 503.2465524 | 485.2151265 | ENSG00000198874 |
| NA | -0.052158109 | 0.964492477 | 0.913280439 | 0.953256842 | 19.69538113 | 18.98123107 | ENSG00000216285 |
| TMPO-AS1 | -0.052184843 | 0.964474605 | 0.839089805 | 0.908201842 | 67.40894973 | 65.05144976 | ENSG00000257167 |
| SLC2A4RG | -0.052187755 | 0.964472658 | 0.641359642 | 0.775703377 | 458.4932305 | 442.3593638 | ENSG00000125520 |
| MAPK8 | -0.052206916 | 0.964459849 | 0.556951678 | 0.71304452 | 820.5465724 | 791.2284527 | ENSG00000107643 |
| KIF1B | -0.052211372 | 0.96445687 | 0.408883733 | 0.581185335 | 2482.294057 | 2393.951224 | ENSG00000054523 |
| PDXDC1 | -0.052305269 | 0.964394101 | 0.343467159 | 0.516242264 | 2878.516711 | 2775.910358 | ENSG00000179889 |
| SIK2 | -0.052377755 | 0.964345647 | 0.578630566 | 0.730282123 | 554.8524824 | 534.9294566 | ENSG00000170145 |
| YIPF6 | -0.052404633 | 0.964327681 | 0.564250344 | 0.718668519 | 659.955994 | 636.3056083 | ENSG00000181704 |
| ATRAID | -0.052427231 | 0.964312577 | 0.597312893 | 0.744394571 | 562.586756 | 542.3333135 | ENSG00000138085 |
| ESRP1 | -0.052512197 | 0.964255786 | 0.31624765 | 0.487356879 | 2850.914871 | 2749.410343 | ENSG00000104413 |
| USP48 | -0.052633523 | 0.964174699 | 0.345092879 | 0.517566141 | 2101.895237 | 2027.150399 | ENSG00000090686 |
| ARPP19 | -0.052674315 | 0.964147437 | 0.459270704 | 0.629112487 | 2299.735597 | 2218.081652 | ENSG00000128989 |
| GCFC2 | -0.052790921 | 0.964069513 | 0.779551117 | 0.870508482 | 127.3582615 | 122.8937277 | ENSG00000005436 |
| TTYH3 | -0.052823352 | 0.964047842 | 0.244532924 | 0.407611219 | 6920.120795 | 6671.822311 | ENSG00000136295 |
| UBE3B | -0.052901602 | 0.963995554 | 0.570708027 | 0.72361071 | 583.518087 | 562.6016512 | ENSG00000151148 |
| DDX24 | -0.052906127 | 0.963992531 | 0.301851925 | 0.472648173 | 3398.844785 | 3275.825936 | ENSG00000089737 |
| NA | -0.05295697 | 0.963958559 | 0.942377417 | 0.969808053 | 10.58408508 | 10.13646175 | ENSG00000234911 |
| TMEM216 | -0.053080207 | 0.963876219 | 0.66735991 | 0.795634837 | 342.2458988 | 330.1664338 | ENSG00000187049 |
| MSANTD1 | -0.05315578 | 0.96382573 | 0.949561143 | 0.973894626 | 7.086944514 | 6.77709333 | ENSG00000188981 |
| HDAC2 | -0.053156091 | 0.963825522 | 0.184395173 | 0.330530317 | 7453.246048 | 7183.912165 | ENSG00000196591 |
| KCNH2 | -0.053200212 | 0.963796047 | 0.643915975 | 0.777434452 | 458.3511698 | 442.1031177 | ENSG00000055118 |
| QRICH1 | -0.053253511 | 0.96376044 | 0.335655285 | 0.508371368 | 2276.824656 | 2194.932339 | ENSG00000198218 |
| SEC22B | -0.053280038 | 0.96374272 | 0.524321888 | 0.686719638 | 729.4217101 | 702.9501892 | ENSG00000265808 |
| ZNF283 | -0.053287934 | 0.963737445 | 0.83604596 | 0.906924912 | 71.55179254 | 68.79747421 | ENSG00000167637 |
| PCNX4 | -0.053317524 | 0.963717679 | 0.603465393 | 0.748838988 | 571.2674765 | 551.0333819 | ENSG00000126773 |
| CSTF3 | -0.053439931 | 0.963635915 | 0.510507641 | 0.674950511 | 930.9826066 | 896.9247006 | ENSG00000176102 |
| ANKRD36C | -0.053469287 | 0.963616307 | 0.898777455 | 0.944277801 | 27.07611007 | 26.15033554 | ENSG00000174501 |
| NUP54 | -0.053481614 | 0.963608073 | 0.460929843 | 0.630758148 | 1430.566742 | 1378.159625 | ENSG00000138750 |
| MEGF10 | -0.053533618 | 0.963573339 | 0.557264735 | 0.713303297 | 605.5557711 | 583.3238699 | ENSG00000145794 |
| NA | -0.053659404 | 0.96348933 | 0.96546612 | 0.982271747 | 3.804879502 | 3.643038954 | ENSG00000239268 |
| ZNF189 | -0.05370443 | 0.963459261 | 0.691620611 | 0.81298395 | 270.6077373 | 260.6067035 | ENSG00000136870 |
| IL17RC | -0.053741287 | 0.963434648 | 0.758616267 | 0.856668407 | 173.5514325 | 167.3049715 | ENSG00000163702 |
| ELMO1 | -0.053823234 | 0.963379925 | 0.731064166 | 0.839112204 | 191.5146949 | 184.602741 | ENSG00000155849 |
| LIN28A | -0.053868465 | 0.963349721 | 0.159024122 | 0.295628249 | 23448.98002 | 22590.12164 | ENSG00000131914 |
| PSME3IP1 | -0.053904688 | 0.963325534 | 0.447685593 | 0.618115 | 1188.890158 | 1145.161133 | ENSG00000172775 |
| PPP4R3B | -0.053979453 | 0.963275613 | 0.429132429 | 0.600492796 | 1216.715773 | 1172.017512 | ENSG00000275052 |
| GRK3 | -0.053989439 | 0.963268946 | 0.545060646 | 0.703422861 | 827.8415215 | 797.6890471 | ENSG00000100077 |
| FOXQ1 | -0.054022316 | 0.963246994 | 0.960617112 | 0.979764445 | 4.049779111 | 3.868092536 | ENSG00000164379 |
| GOSR2 | -0.054085319 | 0.963204929 | 0.61708309 | 0.759048006 | 444.7369775 | 428.6830969 | ENSG00000108433 |
| ICA1L | -0.054179373 | 0.963142137 | 0.897593659 | 0.943571428 | 32.97567726 | 31.86670871 | ENSG00000163596 |
| RAB1A | -0.054205771 | 0.963124514 | 0.38831068 | 0.561716673 | 3351.764341 | 3228.568044 | ENSG00000138069 |
| DSP | -0.054215709 | 0.96311788 | 0.345993961 | 0.518493707 | 2442.624077 | 2351.917207 | ENSG00000096696 |
| PNMA6F | -0.054230843 | 0.963107777 | 0.938177213 | 0.967362051 | 10.56522449 | 10.09666693 | ENSG00000225110 |
| USPL1 | -0.054239904 | 0.963101728 | 0.611339411 | 0.754893995 | 456.6427667 | 439.7477426 | ENSG00000132952 |
| MFSD14A | -0.054266268 | 0.963084128 | 0.868951327 | 0.926984177 | 43.92966487 | 42.35727688 | ENSG00000156875 |
| NFXL1 | -0.054346829 | 0.96303035 | 0.684529518 | 0.807710498 | 293.5062375 | 282.8163721 | ENSG00000170448 |
| DUOX2 | -0.054437602 | 0.962969759 | 0.95909769 | 0.979146567 | 4.460869468 | 4.261460749 | ENSG00000140279 |
| TEKT3 | -0.054469015 | 0.962948792 | 0.952892671 | 0.975888969 | 6.082703515 | 5.811549866 | ENSG00000125409 |
| DENND1C | -0.054483936 | 0.962938833 | 0.587622365 | 0.73714547 | 497.4940031 | 478.9239129 | ENSG00000205744 |
| STAT3 | -0.054595331 | 0.962864484 | 0.32583622 | 0.497601847 | 3045.222719 | 2931.062314 | ENSG00000168610 |
| EPC1 | -0.054653116 | 0.962825919 | 0.715619635 | 0.829279653 | 254.3853297 | 244.9945276 | ENSG00000120616 |
| ZNF696 | -0.054829308 | 0.962708339 | 0.592025171 | 0.740787017 | 621.6564805 | 599.1199232 | ENSG00000185730 |
| UBAP1 | -0.054883988 | 0.962671852 | 0.495924288 | 0.662159152 | 1007.152174 | 969.7900229 | ENSG00000165006 |
| NA | -0.054887266 | 0.962669664 | 0.598408912 | 0.745200847 | 645.9688329 | 621.2379349 | ENSG00000218283 |
| PELP1 | -0.054896039 | 0.962663811 | 0.243628623 | 0.406491998 | 5871.534285 | 5652.284627 | ENSG00000141456 |
| HSF2 | -0.055087796 | 0.962535866 | 0.698962361 | 0.818280507 | 238.6400189 | 229.7410319 | ENSG00000025156 |
| RPL10 | -0.055098972 | 0.96252841 | 0.1512029 | 0.284321586 | 10258.2951 | 9873.76786 | ENSG00000147403 |
| ECEL1 | -0.05515625 | 0.962490196 | 0.697714944 | 0.817468508 | 262.3332153 | 252.7143016 | ENSG00000171551 |
| FAM189A1 | -0.055269939 | 0.962414351 | 0.717271588 | 0.830495541 | 205.3564026 | 197.6793961 | ENSG00000104059 |
| RMND5A | -0.055270416 | 0.962414033 | 0.372947462 | 0.546447629 | 1729.311138 | 1664.920966 | ENSG00000153561 |
| EDRF1-AS1 | -0.055332403 | 0.962372683 | 0.960418571 | 0.979722803 | 4.161503245 | 3.97348907 | ENSG00000236991 |
| TMEM51 | -0.055403898 | 0.962324992 | 0.652630665 | 0.784094076 | 331.8238621 | 319.493321 | ENSG00000171729 |
| CEBPA-DT | -0.055519679 | 0.962247765 | 0.83424106 | 0.906156773 | 79.69879768 | 76.57478065 | ENSG00000267296 |
| NA | -0.055581309 | 0.962206661 | 0.911860694 | 0.952533246 | 19.21395462 | 18.46175409 | ENSG00000269983 |
| CCDC51 | -0.055597019 | 0.962196182 | 0.57192154 | 0.724529807 | 576.9180386 | 555.2083172 | ENSG00000164051 |
| RTN4R | -0.055661435 | 0.962153222 | 0.652260621 | 0.78392372 | 574.8634131 | 553.410299 | ENSG00000040608 |
| EFCAB7 | -0.055783002 | 0.962072151 | 0.861091366 | 0.922031169 | 56.12577184 | 54.15996504 | ENSG00000203965 |
| TMSB10 | -0.055813683 | 0.962051691 | 0.283768932 | 0.452374719 | 7319.705281 | 7040.507261 | ENSG00000034510 |
| ZNF561 | -0.055899002 | 0.961994798 | 0.543309212 | 0.701925366 | 770.5901342 | 741.1562055 | ENSG00000171469 |
| PPP1R18 | -0.055942971 | 0.96196548 | 0.52041049 | 0.68346886 | 847.0593062 | 814.7693908 | ENSG00000146112 |
| GART | -0.05604925 | 0.961894618 | 0.231777042 | 0.392349151 | 4363.078693 | 4196.364213 | ENSG00000159131 |
| GPR135 | -0.056110305 | 0.961853911 | 0.921262009 | 0.958011181 | 14.74538817 | 14.19222408 | ENSG00000181619 |

| | | | | | | | |
|---|---|---|---|---|---|---|---|
| NA | -0.056141952 | 0.961832812 | 0.701730044 | 0.820229901 | 212.2074959 | 204.0340793 | ENSG00000214182 |
| N4BP3 | -0.056175343 | 0.961810551 | 0.276774341 | 0.443981238 | 2613.952581 | 2513.977491 | ENSG00000145911 |
| NA | -0.056185234 | 0.961803956 | 0.924519043 | 0.959536066 | 14.16551632 | 13.63887096 | ENSG00000287707 |
| ARID4A | -0.056219791 | 0.961780918 | 0.696353453 | 0.816716589 | 249.6358701 | 240.2326725 | ENSG00000032219 |
| PLD2 | -0.056231855 | 0.961772876 | 0.466227702 | 0.635528448 | 1133.792469 | 1090.236501 | ENSG00000129219 |
| SNRK | -0.056233516 | 0.961771769 | 0.760557157 | 0.857655235 | 180.5016227 | 173.6472842 | ENSG00000163788 |
| NFKB2 | -0.056319113 | 0.961714707 | 0.527019242 | 0.68889607 | 688.743098 | 662.4756206 | ENSG00000077150 |
| PAIP1 | -0.056351973 | 0.961692802 | 0.400281048 | 0.57275046 | 2095.300095 | 2016.077636 | ENSG00000172239 |
| PCBP3 | -0.056416849 | 0.961649558 | 0.764115184 | 0.860309677 | 142.7062491 | 137.2150046 | ENSG00000183570 |
| DNAJA3 | -0.056454018 | 0.961624783 | 0.342044469 | 0.514835246 | 1889.503652 | 1816.597393 | ENSG00000103423 |
| ADGRV1 | -0.056528111 | 0.961575397 | 0.524093993 | 0.686467763 | 887.6842139 | 853.1469142 | ENSG00000164199 |
| KRT19 | -0.056622061 | 0.96151278 | 0.38054404 | 0.553955715 | 2321.523638 | 2231.920547 | ENSG00000171345 |
| SMTNL2 | -0.056663298 | 0.961485298 | 0.824700535 | 0.900458344 | 86.59431961 | 83.18947803 | ENSG00000188176 |
| ATMIN | -0.056694975 | 0.961464187 | 0.449094679 | 0.619306988 | 1324.809655 | 1273.923682 | ENSG00000166454 |
| OXR1 | -0.056745324 | 0.961430633 | 0.683439349 | 0.806868328 | 257.569513 | 247.8019 | ENSG00000164830 |
| NUP93 | -0.056917471 | 0.961315919 | 0.237661674 | 0.399436431 | 3772.823189 | 3626.365788 | ENSG00000102900 |
| MAN1B1 | -0.056972186 | 0.961279461 | 0.420114185 | 0.59212554 | 1112.663184 | 1069.380891 | ENSG00000177239 |
| SLCO2A1 | -0.057193237 | 0.961132184 | 0.871155589 | 0.928357582 | 36.97677172 | 35.62212895 | ENSG00000174640 |
| C14orf119 | -0.057207585 | 0.961122625 | 0.49323359 | 0.659708048 | 733.8695889 | 705.0447687 | ENSG00000179933 |
| VAT1 | -0.057282745 | 0.961072555 | 0.158936738 | 0.295503619 | 8339.341395 | 8014.601018 | ENSG00000108828 |
| ANKRD11 | -0.057303317 | 0.961058851 | 0.22181311 | 0.380200981 | 3945.359293 | 3791.414905 | ENSG00000167522 |
| BUD13 | -0.057309679 | 0.961054613 | 0.511123432 | 0.675579454 | 673.1974929 | 646.9217828 | ENSG00000137656 |
| FTH1 | -0.057508365 | 0.960922267 | 0.64892762 | 0.781685316 | 3342.26673 | 3211.780678 | ENSG00000167996 |
| KCTD6 | -0.057601289 | 0.960860376 | 0.701614349 | 0.820229901 | 205.9409803 | 197.9154751 | ENSG00000168301 |
| FGD1 | -0.057634835 | 0.960838034 | 0.518732708 | 0.682248095 | 701.3589193 | 674.2414517 | ENSG00000102302 |
| PLEKHO1 | -0.057705025 | 0.960791289 | 0.472381229 | 0.641288268 | 832.0778283 | 799.3832983 | ENSG00000023902 |
| TMC5 | -0.057749358 | 0.960761764 | 0.898312621 | 0.944069992 | 26.08024178 | 24.96566929 | ENSG00000103534 |
| SRP68 | -0.057797655 | 0.960729601 | 0.237568895 | 0.399315288 | 3205.607459 | 3079.794609 | ENSG00000167881 |
| ZNF134 | -0.057840312 | 0.960701196 | 0.414971162 | 0.587107348 | 1226.661691 | 1178.897349 | ENSG00000213762 |
| ANKRD40 | -0.057900801 | 0.960660917 | 0.425994333 | 0.597705773 | 1166.298884 | 1120.287957 | ENSG00000154945 |
| PRIMA1 | -0.057942744 | 0.960632988 | 0.767245132 | 0.862450524 | 139.2727868 | 133.9785283 | ENSG00000175785 |
| PACS1 | -0.057949001 | 0.960628821 | 0.472469567 | 0.641363058 | 856.381084 | 822.6918765 | ENSG00000175115 |
| DPYSL3 | -0.057997019 | 0.960596849 | 0.095683197 | 0.200132771 | 20110.78936 | 19318.01599 | ENSG00000113657 |
| AGAP6 | -0.058001944 | 0.960593569 | 0.90217477 | 0.946601282 | 21.51622104 | 20.60552419 | ENSG00000204149 |
| B3GALT1 | -0.058041434 | 0.960567276 | 0.643628051 | 0.777277371 | 318.3777755 | 305.8909977 | ENSG00000172318 |
| SEC16A | -0.058143307 | 0.96049945 | 0.331448553 | 0.504137352 | 2107.222027 | 2023.229439 | ENSG00000148396 |
| ST6GALNAC4 | -0.058158513 | 0.960489327 | 0.548547208 | 0.706033106 | 509.993982 | 489.8987249 | ENSG00000136840 |
| TAF9 | -0.058202879 | 0.96045979 | 0.645234167 | 0.778635808 | 522.6883731 | 502.1303118 | ENSG00000273841 |
| ZNF436 | -0.058229982 | 0.960441747 | 0.869087637 | 0.926984177 | 41.26885969 | 39.53459003 | ENSG00000125945 |
| ZBTB45 | -0.058240089 | 0.960435018 | 0.572278989 | 0.724874194 | 552.3800093 | 530.8750513 | ENSG00000119574 |
| NA | -0.058267712 | 0.960416629 | 0.934930857 | 0.965304601 | 10.40182464 | 10.02702453 | ENSG00000214776 |
| ZNF527 | -0.058384748 | 0.96033872 | 0.840451119 | 0.90896129 | 53.60005715 | 51.52389402 | ENSG00000189164 |
| KHDRBS3 | -0.058493884 | 0.960266076 | 0.596138709 | 0.743763199 | 661.8626265 | 635.9697605 | ENSG00000131773 |
| TUBE1 | -0.058502282 | 0.960260486 | 0.67052465 | 0.797822181 | 272.9961341 | 261.9435764 | ENSG00000074935 |
| IFNGR1 | -0.058512886 | 0.960253428 | 0.57071883 | 0.72361071 | 509.9273127 | 489.908369 | ENSG00000027697 |
| GHET1 | -0.058594773 | 0.960198926 | 0.930710526 | 0.963034317 | 10.3005884 | 9.932380869 | ENSG00000281189 |
| CRAMP1 | -0.058632165 | 0.96017404 | 0.623635137 | 0.763014769 | 370.2649621 | 355.4577311 | ENSG00000007545 |
| CLGN | -0.058688257 | 0.960136709 | 0.737173287 | 0.843705033 | 170.4500763 | 163.8130241 | ENSG00000153132 |
| ROCK1 | -0.058765434 | 0.960085347 | 0.430164501 | 0.601587893 | 1317.593428 | 1264.938204 | ENSG00000067900 |
| APCDD1L | -0.0588295 | 0.960042714 | 0.744224263 | 0.848593706 | 144.0660378 | 138.4572338 | ENSG00000198768 |
| MRTFA | -0.058922986 | 0.959980505 | 0.431024446 | 0.602266594 | 1056.663367 | 1014.13223 | ENSG00000196588 |
| EEFSEC | -0.058970162 | 0.959949114 | 0.627780767 | 0.765906719 | 366.8700969 | 352.4653789 | ENSG00000132394 |
| DYNC1LI1 | -0.058979205 | 0.959943098 | 0.361336671 | 0.534305207 | 1514.685998 | 1454.551298 | ENSG00000144635 |
| HS3ST2 | -0.058999557 | 0.959929556 | 0.850228105 | 0.914765777 | 46.18459808 | 44.30288479 | ENSG00000122254 |
| PTCD1 | -0.059002891 | 0.959927337 | 0.878359545 | 0.933269111 | 29.40070371 | 28.21560049 | ENSG00000106246 |
| KDM8 | -0.059061625 | 0.959888258 | 0.727479514 | 0.836291478 | 169.0813144 | 162.3261562 | ENSG00000155666 |
| TMBIM1 | -0.059180344 | 0.959809273 | 0.526340977 | 0.688475982 | 603.9884036 | 579.6980214 | ENSG00000135926 |
| RNF103 | -0.059249685 | 0.959763142 | 0.806419526 | 0.888475268 | 75.57346246 | 72.48675971 | ENSG00000239305 |
| TBC1D17 | -0.0593885 | 0.959670799 | 0.534035126 | 0.694490469 | 636.5951295 | 611.1665482 | ENSG00000104946 |
| NA | -0.059402591 | 0.959661425 | 0.804207606 | 0.887069454 | 77.71791084 | 74.66413339 | ENSG00000262539 |
| DCAF7 | -0.05948406 | 0.959607235 | 0.281916218 | 0.450277013 | 3069.581616 | 2945.717354 | ENSG00000136485 |
| FGD4 | -0.059540041 | 0.95957 | 0.597564818 | 0.744461977 | 481.5367583 | 462.6575185 | ENSG00000139132 |
| ZNRF2 | -0.059549059 | 0.959564002 | 0.682777712 | 0.806531434 | 226.0430213 | 217.0418975 | ENSG00000180233 |
| GTPBP2 | -0.05955451 | 0.959560376 | 0.399088229 | 0.571594849 | 1317.87556 | 1264.497057 | ENSG00000172432 |
| TUBGCP6 | -0.059562822 | 0.959554848 | 0.448099482 | 0.618471411 | 1079.55621 | 1035.586627 | ENSG00000128159 |
| NDUFAF2 | -0.059566282 | 0.959552547 | 0.587265649 | 0.736860691 | 490.8159585 | 470.6913741 | ENSG00000164182 |
| MGRN1 | -0.059584298 | 0.959540564 | 0.356061464 | 0.5292635 | 1898.949573 | 1822.509784 | ENSG00000102858 |
| EMD | -0.059590444 | 0.959536476 | 0.37308941 | 0.546534559 | 1249.598299 | 1198.878545 | ENSG00000102119 |
| PYCR3 | -0.059602801 | 0.959528258 | 0.633714671 | 0.770452621 | 309.0965339 | 296.467353 | ENSG00000104524 |
| CEP83 | -0.059620426 | 0.959516535 | 0.605251939 | 0.7500773 | 372.6960984 | 357.5657413 | ENSG00000173588 |
| NA | -0.059624582 | 0.959513772 | 0.778236755 | 0.869988497 | 114.6236707 | 109.9668377 | ENSG00000234797 |
| ARHGEF9 | -0.05963976 | 0.959503677 | 0.446964854 | 0.617340804 | 856.6805751 | 821.6655655 | ENSG00000131089 |
| LSM7 | -0.059671568 | 0.959482522 | 0.379674177 | 0.552985621 | 3468.625564 | 3327.441777 | ENSG00000130332 |
| CBARP | -0.059780691 | 0.959409952 | 0.631606088 | 0.768901632 | 496.611561 | 476.4206565 | ENSG00000099625 |
| ABCE1 | -0.059782609 | 0.959408676 | 0.297288208 | 0.467513395 | 2032.629529 | 1950.780214 | ENSG00000164163 |
| PRKAB1 | -0.0597874 | 0.95940549 | 0.62714115 | 0.765475153 | 398.566483 | 382.4679168 | ENSG00000111725 |
| DAPK1 | -0.059922505 | 0.959315648 | 0.230922724 | 0.391211932 | 3285.858005 | 3152.627455 | ENSG00000196730 |

| | | | | | | | |
|---|---|---|---|---|---|---|---|
| GASK1A | -0.060062083 | 0.959222841 | 0.777718929 | 0.869536099 | 112.261468 | 107.5531245 | ENSG00000144649 |
| RNF145 | -0.060072825 | 0.959215699 | 0.293649094 | 0.463339169 | 1969.105956 | 1888.495585 | ENSG00000145860 |
| DDX10 | -0.060191442 | 0.959136836 | 0.406710685 | 0.578944497 | 1116.766479 | 1071.214578 | ENSG00000178105 |
| PHF21B | -0.060208574 | 0.959125446 | 0.529954065 | 0.690906526 | 561.113216 | 538.0175248 | ENSG00000056487 |
| NA | -0.060227407 | 0.959112926 | 0.960434345 | 0.979722803 | 3.069980516 | 2.911151367 | ENSG00000227201 |
| C14orf132 | -0.060274177 | 0.959081833 | 0.365499786 | 0.539179963 | 1304.991287 | 1251.912164 | ENSG00000227051 |
| NA | -0.060543419 | 0.958902862 | 0.937104423 | 0.966565868 | 9.500734905 | 9.143493868 | ENSG00000278840 |
| TASOR2 | -0.060545057 | 0.958901773 | 0.342270448 | 0.514994089 | 1565.415718 | 1500.719432 | ENSG00000108021 |
| WFS1 | -0.060631044 | 0.958844623 | 0.47782182 | 0.646081043 | 1185.891918 | 1137.758094 | ENSG00000109501 |
| FAM49B | -0.060651255 | 0.95883119 | 0.31984228 | 0.491386438 | 1827.788018 | 1752.619779 | ENSG00000153310 |
| TLN1 | -0.06074231 | 0.958770676 | 0.172763792 | 0.315067717 | 4266.58942 | 4090.244333 | ENSG00000137076 |
| KCP | -0.060826122 | 0.958714979 | 0.726173413 | 0.835487047 | 167.8120739 | 160.7171897 | ENSG00000135253 |
| CCDC9 | -0.060892828 | 0.958670651 | 0.424502654 | 0.596219927 | 1058.855936 | 1015.026241 | ENSG00000105321 |
| TOX4 | -0.0610626 | 0.958557845 | 0.36344931 | 0.536988841 | 1243.668032 | 1192.284778 | ENSG00000092203 |
| CALU | -0.061079093 | 0.958546887 | 0.222001246 | 0.380373572 | 4695.345234 | 4500.722762 | ENSG00000128595 |
| ARGLU1 | -0.061080679 | 0.958545833 | 0.265006355 | 0.430987613 | 3047.49381 | 2920.285536 | ENSG00000134884 |
| LRSAM1 | -0.061194811 | 0.958470005 | 0.568634511 | 0.721774468 | 435.4677002 | 417.4958438 | ENSG00000148356 |
| NA | -0.061251297 | 0.958432479 | 0.916992784 | 0.955518152 | 13.11797232 | 12.57868383 | ENSG00000274114 |
| BACE1 | -0.061258015 | 0.958428015 | 0.760466838 | 0.857655235 | 132.8763991 | 127.3705552 | ENSG00000186318 |
| COG8 | -0.061502449 | 0.958265644 | 0.786257873 | 0.875152278 | 97.61142195 | 93.49561149 | ENSG00000213380 |
| F2R | -0.061661933 | 0.958159717 | 0.48943735 | 0.655949418 | 648.2773582 | 621.4570577 | ENSG00000181104 |
| RPAIN | -0.061699305 | 0.958134898 | 0.551102667 | 0.70799957 | 545.4324083 | 522.1996811 | ENSG00000129197 |
| STK35 | -0.061717327 | 0.958122929 | 0.388570361 | 0.561912357 | 1429.204001 | 1369.645889 | ENSG00000125834 |
| USP10 | -0.061719665 | 0.958121376 | 0.1680746 | 0.308295477 | 4936.605173 | 4729.971113 | ENSG00000103194 |
| TMEM74 | -0.061756779 | 0.958096728 | 0.720519732 | 0.832149671 | 161.7749887 | 154.9814493 | ENSG00000164841 |
| C15orf39 | -0.061777784 | 0.958082779 | 0.395960691 | 0.568508312 | 1515.775556 | 1452.792318 | ENSG00000167173 |
| PDIA6 | -0.061786461 | 0.958077016 | 0.119316839 | 0.237439771 | 10240.84875 | 9811.438928 | ENSG00000143870 |
| KCNQ1OT1 | -0.062034823 | 0.957912097 | 0.873813758 | 0.929913791 | 40.2268975 | 38.6324856 | ENSG00000269821 |
| MRPL1 | -0.062082634 | 0.957880351 | 0.618672886 | 0.760227954 | 408.1428363 | 390.6746729 | ENSG00000169288 |
| GSDME | -0.062128004 | 0.957850229 | 0.719530709 | 0.831663685 | 156.9403498 | 150.4780029 | ENSG00000105928 |
| TAF15 | -0.062145787 | 0.957838422 | 0.196158257 | 0.346623517 | 10102.06059 | 9675.882702 | ENSG00000270647 |
| PHF12 | -0.062154524 | 0.957832621 | 0.339621257 | 0.512166484 | 1315.21886 | 1259.786549 | ENSG00000109118 |
| CSKMT | -0.062290463 | 0.957742373 | 0.794304323 | 0.880349251 | 85.20111531 | 81.66574726 | ENSG00000214756 |
| MOB3B | -0.06233559 | 0.957712416 | 0.550930724 | 0.707889978 | 508.3147272 | 486.507352 | ENSG00000120162 |
| CMAS | -0.062336298 | 0.957711946 | 0.378554988 | 0.551938952 | 1590.901545 | 1523.528974 | ENSG00000111726 |
| SLC2A4 | -0.062349813 | 0.957702974 | 0.747791215 | 0.850568416 | 120.1173659 | 115.0145018 | ENSG00000181856 |
| BAG4 | -0.062381725 | 0.95768179 | 0.535515834 | 0.695478386 | 601.1929977 | 576.0490779 | ENSG00000156735 |
| FAM221A | -0.06249506 | 0.95760656 | 0.671814169 | 0.798838096 | 233.5809295 | 223.4468558 | ENSG00000188732 |
| NIPSNAP3A | -0.062576946 | 0.957552208 | 0.725894355 | 0.835365268 | 181.5577637 | 173.76075 | ENSG00000136783 |
| SWAP70 | -0.062770379 | 0.957423831 | 0.344659028 | 0.51721964 | 1365.854416 | 1307.468449 | ENSG00000133789 |
| BDH2 | -0.062836654 | 0.957379849 | 0.701313802 | 0.819997315 | 219.8442 | 210.3753259 | ENSG00000164039 |
| ADPRH | -0.062867805 | 0.957359177 | 0.745082628 | 0.848999786 | 122.4440747 | 117.1550553 | ENSG00000144843 |
| POLR3E | -0.062877914 | 0.957352469 | 0.345555052 | 0.51801591 | 2410.234444 | 2306.768867 | ENSG00000058600 |
| LXN | -0.062905607 | 0.957334093 | 0.802314813 | 0.886147662 | 72.32209738 | 69.14836399 | ENSG00000079257 |
| LOC283194 | -0.062920549 | 0.957324178 | 0.93326398 | 0.964413047 | 8.877560052 | 8.537975515 | ENSG00000255240 |
| LOC100288175 | -0.062972948 | 0.957289408 | 0.865933266 | 0.925059178 | 36.32935462 | 34.67299658 | ENSG00000217801 |
| MST1 | -0.062983049 | 0.957282706 | 0.834655438 | 0.906453755 | 61.34618286 | 58.64568587 | ENSG00000173531 |
| PDZD8 | -0.063078668 | 0.957219261 | 0.477100516 | 0.645339499 | 878.9294551 | 841.9026375 | ENSG00000165650 |
| TAF1A | -0.063116134 | 0.957194403 | 0.691993519 | 0.813284207 | 206.6057433 | 197.7562837 | ENSG00000143498 |
| SRRD | -0.063117716 | 0.957193353 | 0.669114301 | 0.796782229 | 231.1720083 | 221.4079492 | ENSG00000100104 |
| ADAMTSL1 | -0.063201017 | 0.957138087 | 0.860737884 | 0.921806169 | 34.56278117 | 33.12505803 | ENSG00000178031 |
| IL17D | -0.063246091 | 0.957108183 | 0.909390521 | 0.951248262 | 18.16621046 | 17.45049712 | ENSG00000172458 |
| PUS10 | -0.063430845 | 0.956985622 | 0.858587176 | 0.920821105 | 39.28549582 | 37.58817652 | ENSG00000162927 |
| PARP14 | -0.06350598 | 0.956935784 | 0.586336948 | 0.735915815 | 384.4480997 | 367.9925673 | ENSG00000173193 |
| PHETA1 | -0.063547925 | 0.956907962 | 0.551882792 | 0.708528644 | 489.8419802 | 469.1197672 | ENSG00000198324 |
| LRRC47 | -0.063570978 | 0.956892672 | 0.233923902 | 0.39477762 | 2353.674292 | 2252.436957 | ENSG00000130764 |
| COMMD1 | -0.063579146 | 0.956887255 | 0.607062176 | 0.751272767 | 348.3865355 | 333.3056255 | ENSG00000173163 |
| NR2F6 | -0.063640844 | 0.956846333 | 0.296833184 | 0.467034501 | 2632.176849 | 2519.155001 | ENSG00000160113 |
| GNA12 | -0.063650832 | 0.956839709 | 0.166232616 | 0.305933351 | 3815.584287 | 3651.051476 | ENSG00000146535 |
| EIF2S3 | -0.06375565 | 0.956770193 | 0.133529724 | 0.258134273 | 11749.24693 | 11241.50953 | ENSG00000130741 |
| STEAP3 | -0.063761449 | 0.956766347 | 0.514207104 | 0.678170569 | 609.8360069 | 583.2268458 | ENSG00000115107 |
| TIMM29 | -0.063799639 | 0.956741021 | 0.563512463 | 0.718013139 | 498.8152827 | 477.2746185 | ENSG00000142444 |
| NA | -0.063861029 | 0.95670031 | 0.924805001 | 0.959760084 | 9.655762042 | 9.274164248 | ENSG00000226468 |
| GAPVD1 | -0.063906331 | 0.956670269 | 0.406462286 | 0.578761733 | 898.8913121 | 860.1436818 | ENSG00000165219 |
| PALB2 | -0.063909082 | 0.956668445 | 0.567673128 | 0.721028596 | 433.0640342 | 414.030267 | ENSG00000083093 |
| SDHAF1 | -0.063950658 | 0.956640876 | 0.599702012 | 0.746136828 | 343.9927357 | 329.022659 | ENSG00000205138 |
| SNX1 | -0.063953612 | 0.956638917 | 0.312829801 | 0.48386784 | 1455.692228 | 1392.612646 | ENSG00000028528 |
| PEX2 | -0.063975358 | 0.956624498 | 0.558962812 | 0.714356707 | 463.4952242 | 443.5302904 | ENSG00000164751 |
| SLC2A12 | -0.064054733 | 0.956571867 | 0.765460026 | 0.861069997 | 103.9114283 | 99.3646817 | ENSG00000146411 |
| FAM3A | -0.064107506 | 0.956536877 | 0.567796524 | 0.721137711 | 420.5301272 | 402.055254 | ENSG00000071889 |
| GAS7 | -0.064158265 | 0.956503223 | 0.617374033 | 0.759357464 | 329.9046137 | 315.7550627 | ENSG00000007237 |
| EXOC2 | -0.064168588 | 0.956496379 | 0.338192268 | 0.510770143 | 1275.621008 | 1220.01883 | ENSG00000112685 |
| STT3A | -0.064267865 | 0.956430561 | 0.197384435 | 0.34843916 | 3607.466634 | 3451.099983 | ENSG00000134910 |
| NA | -0.064279567 | 0.956422804 | 0.783695847 | 0.873484897 | 96.80542307 | 92.61636986 | ENSG00000227232 |
| SCRN3 | -0.064438129 | 0.956317692 | 0.74462829 | 0.848722265 | 120.4977563 | 115.1346919 | ENSG00000144306 |
| MED4 | -0.064446498 | 0.956312144 | 0.554912664 | 0.711094739 | 410.3680207 | 392.4382266 | ENSG00000136146 |
| NTNG2 | -0.064477857 | 0.956291358 | 0.855520954 | 0.918462615 | 39.90308887 | 38.13184285 | ENSG00000196358 |

| | | | | | | | |
|---|---|---|---|---|---|---|---|
| ZNF200 | -0.064554792 | 0.956240363 | 0.738265715 | 0.844463914 | 149.6327479 | 143.1803786 | ENSG00000010539 |
| NA | -0.064573565 | 0.95622792 | 0.960237351 | 0.979722803 | 3.124447132 | 2.957397913 | ENSG00000275481 |
| BAIAP2L2 | -0.064588701 | 0.956217888 | 0.7414627 | 0.846815283 | 121.5290305 | 116.2336078 | ENSG00000128298 |
| L2HGDH | -0.064622785 | 0.956195297 | 0.422137081 | 0.593804067 | 809.8792599 | 774.3525419 | ENSG00000087299 |
| RSF1 | -0.064819731 | 0.956064773 | 0.359740557 | 0.532802694 | 1423.562274 | 1361.21778 | ENSG00000048649 |
| PANK2 | -0.064826728 | 0.956060136 | 0.463143843 | 0.632822974 | 720.9301975 | 689.5213057 | ENSG00000125779 |
| NCOA1 | -0.064843995 | 0.956048694 | 0.52149892 | 0.684416332 | 471.1625162 | 450.4163001 | ENSG00000084676 |
| BTN2A3P | -0.064955974 | 0.95597449 | 0.844613127 | 0.911061945 | 61.14526197 | 58.28109915 | ENSG00000124549 |
| OTUD6B-AS1 | -0.065016509 | 0.955934379 | 0.647239356 | 0.780406104 | 285.0647738 | 272.6772084 | ENSG00000253738 |
| PRKAG2 | -0.065034426 | 0.955922507 | 0.683782487 | 0.80712525 | 253.4500862 | 242.1234437 | ENSG00000106617 |
| AHNAK | -0.06509857 | 0.955880006 | 0.831021969 | 0.90398358 | 87.94134197 | 84.19425906 | ENSG00000124942 |
| CHMP1B | -0.065142682 | 0.95585078 | 0.392829477 | 0.565659377 | 895.8779133 | 856.1535638 | ENSG00000255112 |
| RALGAPB | -0.065169725 | 0.955832862 | 0.38126827 | 0.554721035 | 1290.61416 | 1233.156222 | ENSG00000170471 |
| PUSL1 | -0.06530629 | 0.955742388 | 0.604755086 | 0.74971598 | 338.4551739 | 323.3694949 | ENSG00000169972 |
| AIMP2 | -0.065316019 | 0.955735943 | 0.485823605 | 0.653375848 | 672.6107758 | 642.5898982 | ENSG00000106305 |
| HADHA | -0.065444469 | 0.955650853 | 0.189910129 | 0.337997629 | 4390.524642 | 4195.723263 | ENSG00000084754 |
| DCBLD2 | -0.065487306 | 0.955622478 | 0.367303112 | 0.541114382 | 1161.191213 | 1109.865637 | ENSG00000057019 |
| MBP | -0.065498765 | 0.955614887 | 0.485661893 | 0.653203903 | 630.1069302 | 601.9034031 | ENSG00000197971 |
| C17orf80 | -0.065671908 | 0.955500208 | 0.540681546 | 0.699925359 | 540.2207334 | 516.5044985 | ENSG00000141219 |
| TMED2 | -0.065791124 | 0.955421254 | 0.240501712 | 0.402701027 | 3093.248274 | 2956.498069 | ENSG00000086598 |
| CAPZB | -0.065864181 | 0.955372873 | 0.129112531 | 0.251536037 | 7927.588541 | 7573.575903 | ENSG00000077549 |
| PPTC7 | -0.065881085 | 0.955361679 | 0.492514446 | 0.65888323 | 600.9510012 | 574.5618948 | ENSG00000196850 |
| TIMM17B | -0.065900705 | 0.955348687 | 0.396859081 | 0.569416775 | 1020.876627 | 975.3652512 | ENSG00000126768 |
| ITPRIP | -0.065930148 | 0.95532919 | 0.474875407 | 0.643316115 | 653.4625636 | 623.8651858 | ENSG00000148841 |
| PTGES3 | -0.066048038 | 0.955251129 | 0.145924958 | 0.276396483 | 9115.278697 | 8708.681161 | ENSG00000110958 |
| INIP | -0.066068315 | 0.955237703 | 0.612350786 | 0.755684431 | 282.463178 | 269.7324597 | ENSG00000148153 |
| ERP44 | -0.066107381 | 0.955211837 | 0.518731403 | 0.682248095 | 708.5627935 | 677.3399697 | ENSG00000023318 |
| RELCH | -0.066262145 | 0.955109372 | 0.490587904 | 0.657126181 | 607.9792487 | 580.6520448 | ENSG00000134444 |
| LEMD3 | -0.066298682 | 0.955085184 | 0.379419756 | 0.552781989 | 969.4776608 | 925.757378 | ENSG00000174106 |
| TRPV6 | -0.066304327 | 0.955081447 | 0.871819029 | 0.928775176 | 29.28985545 | 27.91472537 | ENSG00000165125 |
| ZSCAN16-AS1 | -0.066326314 | 0.955066891 | 0.849022011 | 0.913958172 | 41.75096192 | 39.80589016 | ENSG00000269293 |
| FBXO10 | -0.066334809 | 0.955061268 | 0.722129691 | 0.833369458 | 149.2017578 | 142.4210666 | ENSG00000147912 |
| ATP6V1E2 | -0.066340419 | 0.955057554 | 0.779450459 | 0.870480335 | 79.38583879 | 75.86279969 | ENSG00000250565 |
| TMBIM6 | -0.066387681 | 0.955026267 | 0.138458025 | 0.265453564 | 11631.26564 | 11108.0215 | ENSG00000139644 |
| ND5 | -0.066388836 | 0.955025503 | 0.748774921 | 0.851185445 | 17434.7001 | 16650.63927 | ENSG00000198786 |
| EIF2B5 | -0.066391962 | 0.955023433 | 0.332799242 | 0.505553994 | 1259.062901 | 1202.207521 | ENSG00000145191 |
| PPP1R8 | -0.066410384 | 0.955011238 | 0.359278585 | 0.532322826 | 1058.673964 | 1010.991712 | ENSG00000117751 |
| CXXC5 | -0.066458227 | 0.954979569 | 0.235069304 | 0.39618399 | 2967.314493 | 2834.541164 | ENSG00000171604 |
| NA | -0.066468006 | 0.954973095 | 0.917539349 | 0.955891969 | 13.53675966 | 12.91290205 | ENSG00000225356 |
| TDRD3 | -0.066555556 | 0.954915145 | 0.628028932 | 0.766071576 | 254.4845388 | 242.9335759 | ENSG00000083544 |
| SLC16A1-AS1 | -0.066794337 | 0.95475711 | 0.733261919 | 0.840784336 | 126.1651391 | 120.495862 | ENSG00000226419 |
| BAG6 | -0.066871331 | 0.954706157 | 0.225590558 | 0.384644463 | 6973.197314 | 6658.052564 | ENSG00000204463 |
| RAB3GAP1 | -0.067065879 | 0.954577423 | 0.395435738 | 0.568050549 | 919.3658463 | 877.6178963 | ENSG00000115839 |
| TRIO | -0.067203205 | 0.954486564 | 0.311131083 | 0.48209393 | 1502.427413 | 1434.019978 | ENSG00000038382 |
| NA | -0.067273058 | 0.954440351 | 0.030636027 | 0.079933526 | 2300495.314 | 2195685.29 | N_ambiguous |
| CCDC127 | -0.067435367 | 0.954332978 | 0.505569216 | 0.670442219 | 465.7405085 | 444.4259553 | ENSG00000164366 |
| VPS41 | -0.06756132 | 0.954249665 | 0.416186481 | 0.588305208 | 897.546605 | 856.7961202 | ENSG00000006715 |
| RBM27 | -0.06757405 | 0.954241245 | 0.681449661 | 0.80560396 | 346.5051858 | 331.0989251 | ENSG00000091009 |
| RTTN | -0.067651752 | 0.954189852 | 0.551670398 | 0.708474594 | 387.8009012 | 369.9465171 | ENSG00000176225 |
| TRMT13 | -0.067714577 | 0.954148301 | 0.703736175 | 0.821822786 | 146.4143981 | 139.7319942 | ENSG00000122435 |
| PRKRIP1 | -0.067744531 | 0.954128491 | 0.468442895 | 0.637287157 | 675.9209741 | 644.4018733 | ENSG00000128563 |
| TRIT1 | -0.067756423 | 0.954120626 | 0.47336758 | 0.641990664 | 566.4071175 | 540.3801573 | ENSG00000043514 |
| CEP192 | -0.0678448 | 0.95406218 | 0.42883692 | 0.600166354 | 708.0147104 | 675.261323 | ENSG00000101639 |
| LCA5 | -0.067941511 | 0.953998226 | 0.793459022 | 0.879943547 | 67.87518242 | 64.76294505 | ENSG00000135338 |
| DMKN | -0.067942125 | 0.95399782 | 0.215630356 | 0.372601684 | 2674.836883 | 2550.87011 | ENSG00000161249 |
| GNB2 | -0.067947327 | 0.95399438 | 0.240660394 | 0.402826868 | 3085.468545 | 2944.532501 | ENSG00000172354 |
| RPS13 | -0.067949217 | 0.953993131 | 0.096455434 | 0.201376815 | 9037.917769 | 8621.212457 | ENSG00000110700 |
| VEZT | -0.067996987 | 0.953961543 | 0.39239082 | 0.565318246 | 837.4801747 | 799.2396939 | ENSG00000028203 |
| DACT2 | -0.06801251 | 0.953951279 | 0.726784854 | 0.835941249 | 132.9995751 | 126.7913952 | ENSG00000164488 |
| CBWD3 | -0.068013925 | 0.953950343 | 0.912200325 | 0.952535122 | 13.56466869 | 12.91935378 | ENSG00000196873 |
| LINC01126 | -0.068040282 | 0.953932915 | 0.922095887 | 0.958413349 | 9.636901443 | 9.21501426 | ENSG00000279873 |
| GPR137B | -0.068105289 | 0.953889932 | 0.742979697 | 0.847845096 | 121.7999241 | 116.2252659 | ENSG00000077585 |
| PPARA | -0.068146301 | 0.953862817 | 0.695567663 | 0.816191958 | 151.0275038 | 144.1679488 | ENSG00000186951 |
| GATD1 | -0.068199932 | 0.953827358 | 0.357378377 | 0.530689421 | 1044.568384 | 996.2122522 | ENSG00000177225 |
| MLYCD | -0.068201694 | 0.953826193 | 0.525973244 | 0.688181623 | 508.3293575 | 485.003231 | ENSG00000103150 |
| C16orf86 | -0.068231547 | 0.953806456 | 0.880437675 | 0.93439639 | 22.08213838 | 21.0704128 | ENSG00000159761 |
| UNC50 | -0.068317501 | 0.953749631 | 0.528853218 | 0.690124294 | 489.1732239 | 466.3100264 | ENSG00000115446 |
| NA | -0.068320957 | 0.953747346 | 0.867301527 | 0.926008067 | 27.49278223 | 26.15814434 | ENSG00000259495 |
| AQR | -0.068340822 | 0.953734214 | 0.243578567 | 0.406491998 | 2037.549522 | 1943.771627 | ENSG00000021776 |
| LIG4 | -0.068343538 | 0.953732419 | 0.74839442 | 0.85095348 | 116.4372892 | 111.0100928 | ENSG00000174405 |
| GPATCH1 | -0.068506572 | 0.953624646 | 0.61049431 | 0.754165326 | 283.0480681 | 269.7905252 | ENSG00000076650 |
| XRCC4 | -0.068521493 | 0.953614784 | 0.67907649 | 0.803872602 | 167.8630739 | 159.9436172 | ENSG00000152422 |
| PCDHGC3 | -0.068591491 | 0.953568516 | 0.729915348 | 0.838142677 | 130.3252909 | 124.2053267 | ENSG00000240184 |
| MTFR1L | -0.068881469 | 0.953376871 | 0.673767611 | 0.800111161 | 188.6876989 | 179.7761019 | ENSG00000117640 |
| UTP11 | -0.068908815 | 0.9533588 | 0.358223606 | 0.531369199 | 986.694899 | 940.9684074 | ENSG00000183520 |
| C9orf129 | -0.068959613 | 0.953325232 | 0.678170049 | 0.80312234 | 167.9788282 | 160.1269858 | ENSG00000204352 |
| EIF5B | -0.069000838 | 0.953297991 | 0.19050755 | 0.33886873 | 4076.20669 | 3886.581571 | ENSG00000158417 |

| | | | | | | | |
|---|---|---|---|---|---|---|---|
| FXN | -0.069011064 | 0.953291235 | 0.701708943 | 0.820229901 | 156.1853509 | 148.8846234 | ENSG00000165060 |
| LOC107984875 | -0.06908125 | 0.953244859 | 0.882908261 | 0.936039968 | 22.22089566 | 21.17931698 | ENSG00000235560 |
| KIF23 | -0.069191471 | 0.953172034 | 0.32546422 | 0.497151814 | 1372.629155 | 1308.371843 | ENSG00000137807 |
| ZNF277 | -0.069214313 | 0.953156943 | 0.604195419 | 0.749359253 | 308.1288259 | 294.058171 | ENSG00000198839 |
| ATF2 | -0.069245799 | 0.953136141 | 0.557621462 | 0.713428546 | 451.6870798 | 430.5310923 | ENSG00000115966 |
| NA | -0.069301289 | 0.953099481 | 0.54515667 | 0.703452664 | 499.8681814 | 476.1412935 | ENSG00000272888 |
| TOR1B | -0.069415589 | 0.953023974 | 0.495724145 | 0.661983558 | 468.1287052 | 446.081974 | ENSG00000136816 |
| ZNF532 | -0.069448627 | 0.95300215 | 0.154988384 | 0.289970025 | 4419.578282 | 4211.414204 | ENSG00000074657 |
| TLK1 | -0.069550822 | 0.952934645 | 0.36511284 | 0.538815161 | 1166.214095 | 1111.833023 | ENSG00000198586 |
| PCBP2 | -0.069680177 | 0.952849206 | 0.23230784 | 0.392902905 | 2093.332698 | 1994.927663 | ENSG00000197111 |
| LINC02387 | -0.069709662 | 0.952829733 | 0.898281557 | 0.944069992 | 19.94394753 | 18.91884601 | ENSG00000256232 |
| ZBTB22 | -0.069714266 | 0.952826692 | 0.579201214 | 0.730715494 | 332.48288 | 316.4001396 | ENSG00000236104 |
| NUP42 | -0.069725235 | 0.952819448 | 0.567505621 | 0.720910769 | 323.0737324 | 307.6290413 | ENSG00000136243 |
| LRRC23 | -0.06973471 | 0.95281319 | 0.892531526 | 0.940596809 | 20.71445253 | 19.63783016 | ENSG00000010626 |
| RSPRY1 | -0.069754236 | 0.952800295 | 0.458403122 | 0.628280968 | 580.6027333 | 553.4608409 | ENSG00000159579 |
| SNTB1 | -0.06975855 | 0.952797445 | 0.583365369 | 0.733474635 | 385.9291493 | 367.2956951 | ENSG00000172164 |
| SHOC2 | -0.069781772 | 0.952782109 | 0.540941608 | 0.700109467 | 395.1666876 | 376.6227755 | ENSG00000108061 |
| DDX39B | -0.069803788 | 0.952767569 | 0.386889229 | 0.560279267 | 1461.352963 | 1391.68182 | ENSG00000198563 |
| NA | -0.069829042 | 0.952750892 | 0.848337783 | 0.913505717 | 38.84525713 | 36.94716449 | ENSG00000284946 |
| SYNJ1 | -0.069996656 | 0.952640206 | 0.813889409 | 0.893509025 | 50.42692537 | 48.09756682 | ENSG00000159082 |
| ERC2 | -0.070231393 | 0.952485217 | 0.717541493 | 0.83069133 | 121.8174332 | 115.9983344 | ENSG00000187672 |
| PEX11G | -0.070302118 | 0.952438525 | 0.85870307 | 0.920821105 | 31.59981886 | 29.97407164 | ENSG00000104883 |
| GPAT4 | -0.070394254 | 0.9523777 | 0.303524522 | 0.474227874 | 1209.182445 | 1151.293719 | ENSG00000158669 |
| ZNF131 | -0.070403156 | 0.952371824 | 0.453761036 | 0.623824412 | 608.6094448 | 579.6889123 | ENSG00000172262 |
| PGAM1 | -0.070458229 | 0.952335469 | 0.227218243 | 0.38649741 | 13553.79134 | 12908.29981 | ENSG00000171314 |
| MCCC1 | -0.070590813 | 0.952247953 | 0.487522229 | 0.654565097 | 532.6904839 | 506.9824726 | ENSG00000078070 |
| LINC02609 | -0.070736601 | 0.952151731 | 0.938883183 | 0.967624505 | 6.496584775 | 6.116726136 | ENSG00000233593 |
| CALM2 | -0.07075871 | 0.952137139 | 0.145388486 | 0.275618051 | 4473.037824 | 4258.980438 | ENSG00000143933 |
| MTRR | -0.070827632 | 0.952091654 | 0.381135051 | 0.554569036 | 796.1833307 | 757.8762027 | ENSG00000124275 |
| SMCR8 | -0.070869071 | 0.952064307 | 0.337816005 | 0.51044165 | 1026.950469 | 977.8230909 | ENSG00000176994 |
| NT5M | -0.070944342 | 0.952014636 | 0.664342876 | 0.793206562 | 194.2164877 | 185.1168324 | ENSG00000205309 |
| ARF4 | -0.071080774 | 0.951924611 | 0.229853563 | 0.390050571 | 2131.876876 | 2029.834891 | ENSG00000168374 |
| ZNF192P1 | -0.071118673 | 0.951899604 | 0.906757397 | 0.949633928 | 13.0418542 | 12.38294478 | ENSG00000226314 |
| UBE2G1 | -0.071159727 | 0.951872517 | 0.190241371 | 0.33846226 | 2876.697576 | 2739.137595 | ENSG00000132388 |
| TMEM179 | -0.071187787 | 0.951854003 | 0.640134834 | 0.774888832 | 235.2072324 | 224.0876074 | ENSG00000258986 |
| SLC11A2 | -0.071218178 | 0.951833952 | 0.300484789 | 0.471119316 | 1207.233047 | 1149.025148 | ENSG00000110911 |
| STC2 | -0.071248312 | 0.951814072 | 0.276437288 | 0.443545848 | 1959.265942 | 1864.449133 | ENSG00000113739 |
| MLLT10 | -0.071282238 | 0.951791689 | 0.387122894 | 0.560491896 | 947.5859923 | 901.3324615 | ENSG00000078403 |
| POLR2L | -0.0712953 | 0.951783072 | 0.334751273 | 0.50740055 | 1455.351986 | 1384.292551 | ENSG00000177700 |
| EXOSC6 | -0.07138982 | 0.951720716 | 0.473984122 | 0.642334225 | 546.5146086 | 520.2696724 | ENSG00000223496 |
| NRXN2 | -0.071415836 | 0.951703554 | 0.500317543 | 0.666503001 | 504.6863773 | 480.5538521 | ENSG00000110076 |
| EEA1 | -0.071441761 | 0.951686452 | 0.567434501 | 0.720867894 | 317.5329042 | 302.0635056 | ENSG00000102189 |
| NDUFB9 | -0.071528812 | 0.95162903 | 0.392237203 | 0.565211211 | 924.7910451 | 879.4336636 | ENSG00000147684 |
| PIP4P1 | -0.071547399 | 0.95161677 | 0.42729415 | 0.598681658 | 651.2506452 | 619.5227784 | ENSG00000165782 |
| RALBP1 | -0.071616186 | 0.951571399 | 0.181550779 | 0.326889684 | 2329.956389 | 2217.635126 | ENSG00000017797 |
| ANK2 | -0.071712511 | 0.951507867 | 0.46664247 | 0.635824261 | 566.6179627 | 539.0475855 | ENSG00000145362 |
| IRF3 | -0.071757011 | 0.951478518 | 0.232039883 | 0.392656262 | 1801.40597 | 1714.220192 | ENSG00000126456 |
| PTPRD | -0.071788935 | 0.951457464 | 0.467742719 | 0.637008457 | 789.8132621 | 750.9509434 | ENSG00000153707 |
| DUSP18 | -0.071822473 | 0.951435345 | 0.904442136 | 0.948084184 | 12.0809162 | 11.47009958 | ENSG00000167065 |
| TCEANC2 | -0.071878798 | 0.951398201 | 0.623863938 | 0.763014769 | 241.4596302 | 229.5985242 | ENSG00000116205 |
| MIEN1 | -0.07191906 | 0.95137165 | 0.739759001 | 0.845477132 | 108.8796062 | 103.6683483 | ENSG00000141741 |
| SHISAL2B | -0.072022043 | 0.951303741 | 0.711012666 | 0.825872368 | 141.6962261 | 134.850773 | ENSG00000145642 |
| AIP | -0.072038821 | 0.951292678 | 0.422118867 | 0.593804067 | 684.9115856 | 651.1329805 | ENSG00000110711 |
| ODF2L | -0.07207073 | 0.951271638 | 0.562977464 | 0.717521026 | 303.2573786 | 288.464702 | ENSG00000122417 |
| VAPA | -0.072085789 | 0.951261709 | 0.233310118 | 0.394121995 | 1814.359955 | 1726.284457 | ENSG00000101558 |
| GDPD3 | -0.072095395 | 0.951255375 | 0.915667542 | 0.954870131 | 11.98314659 | 11.3813746 | ENSG00000102886 |
| RPP25 | -0.072223397 | 0.951170979 | 0.40496313 | 0.577138278 | 775.1110521 | 736.6861634 | ENSG00000178718 |
| NKAPD1 | -0.072242002 | 0.951158712 | 0.491240092 | 0.657588829 | 504.087845 | 479.6302357 | ENSG00000150776 |
| TSC1 | -0.072411086 | 0.951047243 | 0.375219228 | 0.548567953 | 832.8584578 | 791.9487819 | ENSG00000165699 |
| LAMTOR4 | -0.072434233 | 0.951031985 | 0.312150425 | 0.483126914 | 1415.43975 | 1345.845221 | ENSG00000188186 |
| NA | -0.072521842 | 0.950974234 | 0.449017563 | 0.61924491 | 646.3307133 | 614.0731437 | ENSG00000227063 |
| MOAP1 | -0.072652122 | 0.950888362 | 0.434808148 | 0.605929638 | 625.771376 | 594.8517808 | ENSG00000165943 |
| DLC1 | -0.072692826 | 0.950861534 | 0.769361704 | 0.864048241 | 74.4500005 | 70.76216471 | ENSG00000164741 |
| EIF3L | -0.072777745 | 0.950805567 | 0.171444548 | 0.313283772 | 3486.608808 | 3315.153894 | ENSG00000100129 |
| MRPL45 | -0.072779054 | 0.950804704 | 0.296212284 | 0.46646785 | 1258.128259 | 1196.693308 | ENSG00000278845 |
| NA | -0.072800639 | 0.950790479 | 0.912217451 | 0.952535122 | 10.00680398 | 9.518039956 | ENSG00000260296 |
| ZNF621 | -0.072856837 | 0.950753443 | 0.54980618 | 0.706992892 | 359.1687875 | 341.3754531 | ENSG00000172888 |
| RPUSD2 | -0.072962077 | 0.950684091 | 0.417562921 | 0.589649828 | 821.7997063 | 781.327828 | ENSG00000166133 |
| PPP2R3C | -0.073071986 | 0.950611668 | 0.674016831 | 0.800111161 | 194.3463087 | 185.064122 | ENSG00000092020 |
| ATP5MC2 | -0.073099524 | 0.950593522 | 0.139195048 | 0.266707557 | 3038.093586 | 2887.664368 | ENSG00000135390 |
| FOCAD | -0.07318203 | 0.950539161 | 0.271736123 | 0.438629533 | 1285.710775 | 1222.002101 | ENSG00000188352 |
| ECHDC2 | -0.073199524 | 0.950527635 | 0.631187383 | 0.76865323 | 220.7578928 | 209.631601 | ENSG00000121310 |
| ZSWIM3 | -0.073201905 | 0.950526066 | 0.70824308 | 0.82409779 | 124.6463442 | 118.5817192 | ENSG00000132801 |
| RPL10A | -0.073313364 | 0.950452633 | 0.054554327 | 0.12799897 | 17033.29203 | 16188.00518 | ENSG00000198755 |
| BRIP1 | -0.073367569 | 0.950416924 | 0.528329899 | 0.689768008 | 398.2821984 | 378.7565925 | ENSG00000136492 |
| AKAP8 | -0.073518851 | 0.950317268 | 0.334608821 | 0.50730421 | 1027.080715 | 976.0116489 | ENSG00000105127 |
| NDNF | -0.073538554 | 0.950304289 | 0.837511905 | 0.907512338 | 40.55628793 | 38.54296711 | ENSG00000173376 |

| | | | | | | | |
|---|---|---|---|---|---|---|---|
| NA | -0.073586596 | 0.950272644 | 0.611702709 | 0.755174514 | 289.6056652 | 274.7422313 | ENSG00000244716 |
| MAPK14 | -0.073653124 | 0.950228825 | 0.294734642 | 0.464709929 | 1224.163409 | 1162.818116 | ENSG00000112062 |
| TTI2 | -0.07369921 | 0.950198471 | 0.550098407 | 0.707133229 | 359.2535537 | 341.178127 | ENSG00000129696 |
| PIP5KL1 | -0.073700063 | 0.950197909 | 0.672851939 | 0.799564127 | 166.9653386 | 158.8296352 | ENSG00000167103 |
| RIMKLB | -0.073731645 | 0.950177109 | 0.314318147 | 0.485469286 | 1190.156444 | 1130.680609 | ENSG00000166532 |
| IRF7 | -0.073744471 | 0.950168661 | 0.651905419 | 0.783643285 | 184.7166287 | 175.603112 | ENSG00000185507 |
| PIAS4 | -0.073805751 | 0.950128303 | 0.287843876 | 0.457208416 | 1519.921314 | 1444.088203 | ENSG00000105229 |
| CACNB3 | -0.073815289 | 0.950122021 | 0.553313972 | 0.709662844 | 315.4470278 | 299.9353225 | ENSG00000167535 |
| WDR3 | -0.073866676 | 0.95008818 | 0.223419585 | 0.38195562 | 1557.42692 | 1479.810585 | ENSG00000065183 |
| KIF5A | -0.073872489 | 0.950084351 | 0.414689323 | 0.586837715 | 796.7682577 | 756.2470996 | ENSG00000155980 |
| LDLRAP1 | -0.073878338 | 0.9500805 | 0.434139744 | 0.605304265 | 586.2942462 | 557.222483 | ENSG00000157978 |
| QSOX2 | -0.074099353 | 0.949934962 | 0.272313293 | 0.439083562 | 1476.688249 | 1402.359011 | ENSG00000165661 |
| RAD51D | -0.074131688 | 0.949913671 | 0.669838497 | 0.797264277 | 168.1774707 | 159.6620851 | ENSG00000185379 |
| NA | -0.074164954 | 0.949891769 | 0.84187883 | 0.909536543 | 36.92250526 | 34.98409153 | ENSG00000226415 |
| ZDHHC5 | -0.074165367 | 0.949891497 | 0.217070156 | 0.374502887 | 1942.03131 | 1844.610009 | ENSG00000156599 |
| EXTL3 | -0.074268115 | 0.949823848 | 0.105231299 | 0.21524939 | 3947.547443 | 3749.036576 | ENSG00000012232 |
| FAM43B | -0.074371646 | 0.949755689 | 0.749509382 | 0.851869763 | 83.24865153 | 79.00975751 | ENSG00000183114 |
| ZBED6CL | -0.074459948 | 0.94969756 | 0.915406337 | 0.954649348 | 8.975329671 | 8.537975515 | ENSG00000188707 |
| NFYB | -0.074489264 | 0.949678262 | 0.365763176 | 0.539475109 | 908.3291707 | 863.1596782 | ENSG00000120837 |
| TRIM36 | -0.074530895 | 0.949650858 | 0.566344581 | 0.720099744 | 306.0145508 | 290.5173119 | ENSG00000152503 |
| HBEGF | -0.074604034 | 0.949602716 | 0.762668175 | 0.85923213 | 74.92564504 | 71.2101213 | ENSG00000113070 |
| DEPDC1B | -0.074619182 | 0.949592745 | 0.306286077 | 0.477305494 | 1068.021787 | 1014.645721 | ENSG00000035499 |
| NA | -0.074634398 | 0.94958273 | 0.903881559 | 0.947754057 | 12.06205561 | 11.42170246 | ENSG00000269906 |
| RCC1L | -0.074648603 | 0.94957338 | 0.305322055 | 0.476264879 | 2311.365175 | 2195.012632 | ENSG00000274523 |
| SLC25A42 | -0.074756971 | 0.949502056 | 0.71483369 | 0.828621912 | 137.8949012 | 130.8202782 | ENSG00000181035 |
| PDE9A | -0.074797343 | 0.949475486 | 0.313934919 | 0.485110424 | 964.3909008 | 915.4571874 | ENSG00000160191 |
| ZNF282 | -0.07479804 | 0.949475027 | 0.288372109 | 0.457670617 | 1624.298839 | 1542.299417 | ENSG00000170265 |
| STK24 | -0.074804673 | 0.949470661 | 0.313308315 | 0.484452605 | 2144.039414 | 2035.612014 | ENSG00000102572 |
| ANGEL1 | -0.074844801 | 0.949444253 | 0.354701338 | 0.527892571 | 776.7455275 | 737.6253364 | ENSG00000013523 |
| ZNF496 | -0.074909013 | 0.949401996 | 0.154313132 | 0.289183469 | 2494.450554 | 2368.226092 | ENSG00000162714 |
| PAXBP1-AS1 | -0.074956002 | 0.949371074 | 0.862120157 | 0.922689647 | 25.38373971 | 24.16009863 | ENSG00000238197 |
| LONRF1 | -0.074965566 | 0.94936478 | 0.299436683 | 0.469954021 | 1142.334069 | 1084.354218 | ENSG00000154359 |
| GGN | -0.075019988 | 0.949328968 | 0.85219886 | 0.916006883 | 34.48108124 | 32.78492112 | ENSG00000179168 |
| NOL4L | -0.075049227 | 0.949309729 | 0.414556663 | 0.586779117 | 642.0055732 | 609.5269343 | ENSG00000197183 |
| GKAP1 | -0.075056042 | 0.949305244 | 0.744217308 | 0.848593706 | 95.31302248 | 90.39407618 | ENSG00000165113 |
| STXBP2 | -0.075062101 | 0.949301257 | 0.508377048 | 0.673029402 | 459.872317 | 436.1520468 | ENSG00000076944 |
| CNBP | -0.075249862 | 0.949177718 | 0.052061254 | 0.123107827 | 8141.603173 | 7727.606348 | ENSG00000169714 |
| RAB11FIP4 | -0.075282211 | 0.949156435 | 0.178094194 | 0.32204866 | 2205.809632 | 2094.287783 | ENSG00000131242 |
| SCNN1D | -0.075364954 | 0.949101999 | 0.81332914 | 0.893097247 | 44.30670082 | 41.97197793 | ENSG00000162572 |
| TMEM164 | -0.075400451 | 0.949078647 | 0.286681079 | 0.455908547 | 1189.293151 | 1128.746615 | ENSG00000157600 |
| ANAPC5 | -0.075438865 | 0.949053377 | 0.198949918 | 0.350401065 | 1981.129452 | 1880.185607 | ENSG00000089053 |
| CDC23 | -0.07544377 | 0.94905015 | 0.351072017 | 0.523662744 | 909.0546195 | 862.2025377 | ENSG00000094880 |
| ABLIM1 | -0.075451595 | 0.949045003 | 0.21968327 | 0.377710335 | 1532.110637 | 1454.116809 | ENSG00000099204 |
| TPI1 | -0.075525236 | 0.948996561 | 0.066839986 | 0.15038607 | 13509.93532 | 12819.59007 | ENSG00000111669 |
| VIRMA | -0.075610443 | 0.948940514 | 0.288437997 | 0.457731145 | 1095.368152 | 1039.131733 | ENSG00000164944 |
| WDR43 | -0.075626345 | 0.948930054 | 0.16018056 | 0.297376882 | 2559.585886 | 2428.419195 | ENSG00000163811 |
| HIKESHI | -0.07563804 | 0.948922362 | 0.342750869 | 0.515435532 | 833.6593135 | 791.205602 | ENSG00000149196 |
| SCAND1 | -0.075640687 | 0.948920621 | 0.324586025 | 0.496240945 | 1021.377665 | 969.36336 | ENSG00000171222 |
| USP27X | -0.07571021 | 0.948874894 | 0.639564937 | 0.774741394 | 188.1786388 | 178.6317514 | ENSG00000273820 |
| NBPF11 | -0.075806984 | 0.948811247 | 0.583331216 | 0.733474635 | 281.8941064 | 267.1052515 | ENSG00000263956 |
| NFKBID | -0.07584933 | 0.948783397 | 0.771338202 | 0.865310843 | 88.78864081 | 84.02328504 | ENSG00000167604 |
| UQCRH | -0.07587756 | 0.948764832 | 0.129152664 | 0.251563414 | 4448.441155 | 4219.410501 | ENSG00000173660 |
| COX6C | -0.075905308 | 0.948746585 | 0.396535542 | 0.569259948 | 661.7490513 | 628.1212367 | ENSG00000164919 |
| RPS8 | -0.076076078 | 0.948634289 | 0.079253476 | 0.172209823 | 18117.21203 | 17185.70515 | ENSG00000142937 |
| ANKRD6 | -0.076102309 | 0.948617041 | 0.696325506 | 0.816716589 | 169.7622592 | 160.6991732 | ENSG00000135299 |
| C2CD2 | -0.07617145 | 0.94857158 | 0.510422431 | 0.674884107 | 380.1745969 | 360.6911628 | ENSG00000157617 |
| CNOT4 | -0.076191794 | 0.948558204 | 0.563580425 | 0.718052307 | 338.8596945 | 321.4846361 | ENSG00000080802 |
| OSTM1 | -0.076292922 | 0.948491716 | 0.454361835 | 0.624142047 | 490.9974936 | 465.9209288 | ENSG00000081087 |
| CABIN1 | -0.076296525 | 0.948489347 | 0.108212192 | 0.219762227 | 3358.383276 | 3185.218007 | ENSG00000099991 |
| NA | -0.076305077 | 0.948483724 | 0.818744855 | 0.896404784 | 41.56822592 | 39.37837323 | ENSG00000233762 |
| PLXNB1 | -0.076310568 | 0.948480115 | 0.221233704 | 0.379597917 | 3736.390574 | 3543.427123 | ENSG00000164050 |
| SCPEP1 | -0.076334295 | 0.948464516 | 0.370180658 | 0.54400511 | 882.6935087 | 836.9359506 | ENSG00000121064 |
| ZNF485 | -0.076388791 | 0.948428689 | 0.699823716 | 0.818845109 | 131.5747069 | 124.9425815 | ENSG00000198298 |
| PDK2 | -0.076665574 | 0.948246749 | 0.647989602 | 0.781028891 | 191.4733069 | 181.5426607 | ENSG00000005882 |
| PPP2R5E | -0.076685846 | 0.948233425 | 0.309276116 | 0.480412718 | 1234.085448 | 1170.199376 | ENSG00000154001 |
| C9orf40 | -0.076724319 | 0.948208139 | 0.526246538 | 0.68839913 | 331.9216317 | 314.8686001 | ENSG00000135045 |
| PIK3CD-AS2 | -0.076743964 | 0.948195227 | 0.757637341 | 0.856114219 | 76.28758582 | 72.19448689 | ENSG00000231789 |
| COA7 | -0.076874354 | 0.948109534 | 0.233766837 | 0.394643919 | 1452.510323 | 1377.021485 | ENSG00000162377 |
| CAMKMT | -0.076904512 | 0.948089714 | 0.548418166 | 0.70596122 | 321.8743525 | 305.1835599 | ENSG00000143919 |
| NR1H2 | -0.076930504 | 0.948072634 | 0.312519768 | 0.483543379 | 1228.401507 | 1164.607028 | ENSG00000131408 |
| TMEM231 | -0.076961876 | 0.948052018 | 0.683038702 | 0.806740929 | 132.1927003 | 125.1867298 | ENSG00000205084 |
| NIFK | -0.076982897 | 0.948038204 | 0.376903768 | 0.550363378 | 695.4211795 | 659.0968907 | ENSG00000155438 |
| YTHDF1 | -0.077065491 | 0.94798393 | 0.255585345 | 0.420583203 | 1786.016255 | 1692.669879 | ENSG00000149658 |
| IRF1 | -0.07709737 | 0.947962983 | 0.760630749 | 0.857664551 | 71.24297756 | 67.59827487 | ENSG00000125347 |
| ARNTL | -0.077113788 | 0.947952195 | 0.663281055 | 0.792185928 | 149.985066 | 142.0521482 | ENSG00000133794 |
| MORF4L1 | -0.077171022 | 0.947914589 | 0.093342333 | 0.196108359 | 10533.52024 | 9984.703277 | ENSG00000185787 |
| ING2 | -0.077173326 | 0.947913076 | 0.549621794 | 0.706883924 | 294.0717162 | 278.6019514 | ENSG00000168556 |

| | | | | | | | |
|---|---|---|---|---|---|---|---|
| ITGA9 | -0.077214979 | 0.947885708 | 0.616617002 | 0.75876498 | 196.3675939 | 186.3023226 | ENSG00000144668 |
| ALAS1 | -0.077263212 | 0.947854018 | 0.262296042 | 0.427817548 | 1164.082712 | 1103.424137 | ENSG00000023330 |
| PDCD10 | -0.077312939 | 0.947821349 | 0.313507529 | 0.484682942 | 1193.733495 | 1130.89116 | ENSG00000114209 |
| NA | -0.077315714 | 0.947819525 | 0.826600327 | 0.9012243 | 47.52082026 | 45.0307438 | ENSG00000175741 |
| PPIP5K2 | -0.077523375 | 0.947683106 | 0.425839144 | 0.597618429 | 577.2732598 | 546.8972364 | ENSG00000145725 |
| PSME1 | -0.077585545 | 0.947642268 | 0.37363923 | 0.547087239 | 795.8583197 | 753.5892076 | ENSG00000092010 |
| PCDHGA10 | -0.077645848 | 0.947602659 | 0.603886731 | 0.749169062 | 255.4849128 | 242.3877285 | ENSG00000253846 |
| NF1 | -0.077684466 | 0.947577294 | 0.401988411 | 0.574301425 | 663.40151 | 628.7280632 | ENSG00000196712 |
| CDCA4 | -0.077688104 | 0.947574904 | 0.304743667 | 0.475478007 | 1508.235808 | 1429.178316 | ENSG00000170779 |
| ATP1B2 | -0.077827082 | 0.947483627 | 0.550980665 | 0.707889978 | 313.8141423 | 297.496596 | ENSG00000129244 |
| NDC1 | -0.077852534 | 0.947466912 | 0.230060892 | 0.390300296 | 1826.653106 | 1731.557405 | ENSG00000058804 |
| PPHLN1 | -0.077880131 | 0.947448788 | 0.23489345 | 0.395991395 | 1423.462064 | 1348.739434 | ENSG00000134283 |
| DPYSL2 | -0.077980744 | 0.947382715 | 0.122917402 | 0.242726952 | 2771.245311 | 2625.537981 | ENSG00000092964 |
| NA | -0.078065182 | 0.947327268 | 0.90609907 | 0.949270868 | 10.35784591 | 9.793641231 | ENSG00000203706 |
| SOWAHD | -0.07806556 | 0.947327021 | 0.754144141 | 0.854240973 | 85.95172057 | 81.27614716 | ENSG00000187808 |
| NA | -0.078146298 | 0.947274006 | 0.898020507 | 0.943865839 | 12.13538282 | 11.44912688 | ENSG00000229944 |
| DEPTOR | -0.07829902 | 0.947173734 | 0.842306356 | 0.909743682 | 36.45570899 | 34.42858783 | ENSG00000155792 |
| LOC101927151 | -0.078324175 | 0.947157219 | 0.733710801 | 0.84114905 | 91.15989194 | 86.22378198 | ENSG00000267575 |
| VDAC3 | -0.07852235 | 0.947027123 | 0.138172407 | 0.265037675 | 2791.328993 | 2644.01497 | ENSG00000078668 |
| BTF3 | -0.078680828 | 0.946923099 | 0.091337967 | 0.192926828 | 11182.23352 | 10588.4493 | ENSG00000145741 |
| EHMT2 | -0.078715573 | 0.946900294 | 0.098628978 | 0.204938777 | 5173.217999 | 4897.865919 | ENSG00000204371 |
| SS18L1 | -0.078749265 | 0.946878181 | 0.469134976 | 0.638003724 | 489.8139079 | 463.407471 | ENSG00000184402 |
| ZNHIT1 | -0.078759074 | 0.946871743 | 0.292286874 | 0.461985166 | 1216.60431 | 1151.477639 | ENSG00000106400 |
| MRPS7 | -0.078798869 | 0.946845625 | 0.270705526 | 0.437405167 | 1577.473654 | 1493.134067 | ENSG00000125445 |
| RNF34 | -0.078844939 | 0.94681539 | 0.250775339 | 0.414897691 | 1244.954781 | 1178.600787 | ENSG00000170633 |
| FAM193A | -0.078854193 | 0.946809316 | 0.379527107 | 0.552847309 | 754.5139807 | 714.2299538 | ENSG00000125386 |
| TPST1 | -0.078869581 | 0.946799218 | 0.534617989 | 0.694920532 | 332.6287338 | 315.2471626 | ENSG00000169902 |
| ZNF585A | -0.078886398 | 0.946788181 | 0.760058054 | 0.857543567 | 66.94426864 | 63.33251298 | ENSG00000196967 |
| PRR14 | -0.078890154 | 0.946785716 | 0.596438336 | 0.743899395 | 229.7327101 | 217.4745397 | ENSG00000156858 |
| SERP1 | -0.078896254 | 0.946781713 | 0.117538634 | 0.234821644 | 4095.136071 | 3878.256139 | ENSG00000120742 |
| NA | -0.07894876 | 0.946747256 | 0.800701713 | 0.885210591 | 48.22781022 | 45.7585907 | ENSG00000235944 |
| DNAJB12 | -0.078997656 | 0.94671517 | 0.409995311 | 0.582116696 | 554.9081514 | 525.5966754 | ENSG00000148719 |
| PDP2 | -0.079031735 | 0.946692807 | 0.371614632 | 0.545038336 | 653.2039848 | 618.4160416 | ENSG00000172840 |
| NR2C1 | -0.079125092 | 0.946631548 | 0.439667882 | 0.610760757 | 496.4095258 | 469.8527329 | ENSG00000120798 |
| FMNL2 | -0.079227699 | 0.946564224 | 0.227819277 | 0.3872466 | 1340.867012 | 1269.126618 | ENSG00000157827 |
| ASIC3 | -0.079244476 | 0.946553217 | 0.886860593 | 0.937513028 | 16.68353402 | 15.68961493 | ENSG00000213199 |
| DDX41 | -0.07925021 | 0.946549455 | 0.188178595 | 0.335597049 | 1963.643249 | 1858.214471 | ENSG00000183258 |
| RRP8 | -0.079268528 | 0.946537436 | 0.53602998 | 0.695818206 | 315.9981517 | 299.1364946 | ENSG00000132275 |
| NPRL3 | -0.079390183 | 0.946457623 | 0.407628378 | 0.579908483 | 650.5771092 | 615.9413004 | ENSG00000103148 |
| ARHGAP23 | -0.079580093 | 0.946333044 | 0.311584744 | 0.482599863 | 2077.495452 | 1965.782784 | ENSG00000275832 |
| PHACTR1 | -0.079655314 | 0.946283704 | 0.667378048 | 0.795634837 | 144.4584677 | 136.8332012 | ENSG00000112137 |
| VPS36 | -0.07973783 | 0.946229582 | 0.500799097 | 0.666686425 | 349.2680889 | 330.5022817 | ENSG00000136100 |
| NA | -0.07980184 | 0.946187601 | 0.948096503 | 0.972984731 | 3.774855292 | 3.525272016 | ENSG00000260923 |
| TRAPPC4 | -0.079875545 | 0.946139263 | 0.379311464 | 0.55266597 | 682.23371 | 645.6967765 | ENSG00000196655 |
| TECTA | -0.079880367 | 0.9461361 | 0.837926571 | 0.907638723 | 30.1789057 | 28.49980406 | ENSG00000109927 |
| BCL2L12 | -0.079899238 | 0.946123724 | 0.18037205 | 0.325313877 | 2158.730529 | 2043.012672 | ENSG00000126453 |
| NADSYN1 | -0.080027467 | 0.946039635 | 0.362203965 | 0.535423501 | 703.6283564 | 665.3703424 | ENSG00000172890 |
| HNRNPH3 | -0.080040896 | 0.94603083 | 0.145158189 | 0.275343818 | 4829.934661 | 4570.141452 | ENSG00000096746 |
| HGS | -0.080075504 | 0.946008136 | 0.232179812 | 0.392824132 | 1744.417632 | 1650.62609 | ENSG00000185359 |
| ZNF32-AS2 | -0.080080522 | 0.946004846 | 0.878531838 | 0.933298075 | 16.41889793 | 15.55920502 | ENSG00000230565 |
| HSP90B1 | -0.08014749 | 0.945960934 | 0.027074715 | 0.072350508 | 14965.82207 | 14157.20112 | ENSG00000166598 |
| FLII | -0.080194841 | 0.945929887 | 0.052195726 | 0.123350208 | 6031.605777 | 5705.91341 | ENSG00000177731 |
| CLSTN3 | -0.080251324 | 0.945892854 | 0.355346288 | 0.528404011 | 964.2338849 | 912.5931008 | ENSG00000139182 |
| CSE1L | -0.080352184 | 0.945826727 | 0.120118339 | 0.238690037 | 7151.49777 | 6765.480225 | ENSG00000124207 |
| KLK13 | -0.08036851 | 0.945816025 | 0.778954359 | 0.870236815 | 58.49366848 | 55.24087652 | ENSG00000167759 |
| RBFOX2 | -0.080396993 | 0.945797352 | 0.102948599 | 0.211770252 | 3760.009303 | 3556.83628 | ENSG00000100320 |
| ICK | -0.080444816 | 0.945766 | 0.613890709 | 0.756749457 | 237.499123 | 224.7366703 | ENSG00000112144 |
| MAPK1IP1L | -0.080559118 | 0.945691072 | 0.26001474 | 0.425329563 | 1361.795051 | 1288.361921 | ENSG00000168175 |
| PLRG1 | -0.080613085 | 0.945655697 | 0.201315373 | 0.353469998 | 1660.814507 | 1570.24528 | ENSG00000171566 |
| NACC1 | -0.080641567 | 0.945637028 | 0.188429176 | 0.336012871 | 4980.271444 | 4710.030743 | ENSG00000160877 |
| NA | -0.080686689 | 0.945607453 | 0.91456018 | 0.95417958 | 8.28440941 | 7.841581606 | ENSG00000227470 |
| GAB2 | -0.080804615 | 0.945530162 | 0.530001518 | 0.690921698 | 293.2514505 | 277.1061902 | ENSG00000033327 |
| BRAP | -0.080996276 | 0.945404557 | 0.403908639 | 0.576146225 | 568.0046213 | 536.8169805 | ENSG00000089234 |
| ZNF76 | -0.081147857 | 0.945305231 | 0.488174285 | 0.655030266 | 394.2919042 | 372.3777379 | ENSG00000065029 |
| NA | -0.0811975 | 0.945272703 | 0.801777817 | 0.885808137 | 44.7441486 | 42.27877173 | ENSG00000244398 |
| MIR924HG | -0.08125268 | 0.94523655 | 0.770268948 | 0.864714751 | 59.42669745 | 56.2023793 | ENSG00000267374 |
| PPRC1 | -0.081260073 | 0.945231706 | 0.217774858 | 0.375355778 | 2792.95015 | 2639.514185 | ENSG00000148840 |
| LRRN2 | -0.08129538 | 0.945208573 | 0.51705536 | 0.680947225 | 303.7173537 | 287.1152046 | ENSG00000170382 |
| PRELID3B | -0.081321938 | 0.945191173 | 0.426084405 | 0.597708465 | 758.7588618 | 717.6182663 | ENSG00000101166 |
| ATG12 | -0.081342041 | 0.945178003 | 0.392274997 | 0.565211211 | 606.8559869 | 573.884856 | ENSG00000145782 |
| TEAD4 | -0.081433398 | 0.945118152 | 0.228335853 | 0.387953755 | 1972.34626 | 1864.055608 | ENSG00000197905 |
| ZNF513 | -0.081455808 | 0.945103472 | 0.3768956 | 0.550363378 | 615.8601895 | 581.8439868 | ENSG00000163795 |
| FGD5-AS1 | -0.08155501 | 0.945038487 | 0.235057135 | 0.39618399 | 1587.305318 | 1499.453541 | ENSG00000225733 |
| RHOC | -0.081570383 | 0.945028417 | 0.494268273 | 0.660679688 | 435.1390224 | 411.0497337 | ENSG00000155366 |
| KCNK5 | -0.081645849 | 0.944978985 | 0.17353537 | 0.316235672 | 1831.443513 | 1730.091001 | ENSG00000164626 |
| SRD5A1 | -0.081689201 | 0.944950589 | 0.408906264 | 0.581185335 | 581.7397991 | 549.5155819 | ENSG00000145545 |
| PPIA | -0.081782299 | 0.944889613 | 0.034419243 | 0.087863123 | 24694.78632 | 23332.84593 | ENSG00000196262 |

| | | | | | | | |
|---|---|---|---|---|---|---|---|
| MIPEP | -0.081809062 | 0.944872085 | 0.427196217 | 0.598608428 | 553.5070869 | 523.4431942 | ENSG00000027001 |
| TP53RK | -0.081811232 | 0.944870664 | 0.47822637 | 0.646448384 | 472.0943569 | 445.9058932 | ENSG00000172315 |
| SELENOF | -0.081837964 | 0.944853156 | 0.220962679 | 0.379375663 | 2179.779938 | 2059.80803 | ENSG00000183291 |
| NA | -0.081840286 | 0.944851635 | 0.315317375 | 0.486545151 | 941.2737463 | 889.5232363 | ENSG00000179899 |
| MRPL51 | -0.08189988 | 0.944812607 | 0.164578607 | 0.303555397 | 2326.274209 | 2197.226771 | ENSG00000111639 |
| BUB3 | -0.081913876 | 0.944803441 | 0.082007293 | 0.177017197 | 5290.377535 | 4998.268245 | ENSG00000154473 |
| MARF1 | -0.081958209 | 0.944774408 | 0.418067221 | 0.590016442 | 470.2294872 | 444.2162162 | ENSG00000166783 |
| PIN1 | -0.081975185 | 0.944763291 | 0.221837128 | 0.380200981 | 1974.50878 | 1865.544342 | ENSG00000127445 |
| UBE2V2 | -0.08198566 | 0.944756432 | 0.280137348 | 0.448252288 | 1270.420823 | 1201.098603 | ENSG00000169139 |
| SHPRH | -0.082059166 | 0.944708297 | 0.439447295 | 0.610586126 | 556.4967686 | 525.2154171 | ENSG00000146414 |
| PTCD3 | -0.082156104 | 0.944644822 | 0.15142433 | 0.284682463 | 2349.38976 | 2219.186529 | ENSG00000132300 |
| PPP1R12C | -0.082203376 | 0.94461387 | 0.27468606 | 0.441470364 | 1813.186846 | 1712.846578 | ENSG00000125503 |
| AP2A1 | -0.082219344 | 0.944603415 | 0.113849448 | 0.228992774 | 3621.986048 | 3421.361374 | ENSG00000196961 |
| C1orf159 | -0.082304916 | 0.944547388 | 0.573034987 | 0.725403106 | 258.904696 | 244.4140348 | ENSG00000131591 |
| TPPP3 | -0.082321868 | 0.94453629 | 0.910871182 | 0.951993281 | 7.976670478 | 7.545007631 | ENSG00000159713 |
| SLC25A28 | -0.082427887 | 0.944466882 | 0.580939724 | 0.731628608 | 232.9577177 | 219.9936678 | ENSG00000155287 |
| FAM102A | -0.08248045 | 0.944432471 | 0.329251466 | 0.501507307 | 958.6601904 | 905.0373704 | ENSG00000167106 |
| SNRPB | -0.082488409 | 0.944427261 | 0.11290125 | 0.227370082 | 3300.318017 | 3116.176699 | ENSG00000125835 |
| UQCRHL | -0.082569981 | 0.944373863 | 0.397297159 | 0.569875789 | 772.084738 | 728.7255067 | ENSG00000233954 |
| BMS1P1 | -0.082578385 | 0.944368362 | 0.836762409 | 0.907107819 | 30.98481656 | 29.3112813 | ENSG00000204177 |
| ZNF624 | -0.082668631 | 0.94430929 | 0.8427765 | 0.909954609 | 31.14175871 | 29.56617034 | ENSG00000197566 |
| ZNF449 | -0.08271442 | 0.94427932 | 0.707260501 | 0.823594399 | 133.516883 | 126.0073729 | ENSG00000173275 |
| CD81 | -0.082722981 | 0.944273717 | 0.17624442 | 0.319712752 | 3424.774528 | 3233.570256 | ENSG00000110651 |
| EML4 | -0.082756317 | 0.944251897 | 0.13751745 | 0.264122772 | 2268.414955 | 2142.540755 | ENSG00000143924 |
| NA | -0.082932454 | 0.944136622 | 0.882581346 | 0.935899117 | 15.70632616 | 14.8504407 | ENSG00000236969 |
| LMTK3 | -0.08298915 | 0.94409952 | 0.516469062 | 0.680447331 | 357.7008676 | 337.6749059 | ENSG00000142235 |
| NA | -0.083016288 | 0.94408176 | 0.878623863 | 0.933298075 | 16.70430964 | 15.77403877 | ENSG00000225200 |
| SDHD | -0.083020184 | 0.944079211 | 0.543020893 | 0.701682367 | 407.1582452 | 384.9061224 | ENSG00000204370 |
| RPS6KA1 | -0.083079417 | 0.944040451 | 0.180784306 | 0.325874767 | 1975.519539 | 1865.010834 | ENSG00000117676 |
| THAP7 | -0.083173038 | 0.943979191 | 0.558892439 | 0.714356707 | 273.9927534 | 258.4841908 | ENSG00000184436 |
| GLIS2 | -0.083220785 | 0.94394795 | 0.497542775 | 0.663906597 | 367.022221 | 346.7331582 | ENSG00000126603 |
| STK26 | -0.083243751 | 0.943932923 | 0.109860303 | 0.222336876 | 4238.449049 | 4001.729777 | ENSG00000134602 |
| STAR | -0.083276144 | 0.943911729 | 0.754115371 | 0.854240973 | 69.04493845 | 65.01301201 | ENSG00000147465 |
| DNAJC10 | -0.083325046 | 0.943879735 | 0.134398889 | 0.259502449 | 2247.684708 | 2122.11219 | ENSG00000077232 |
| MRPL49 | -0.083387721 | 0.94383873 | 0.293289335 | 0.462998736 | 1153.802572 | 1088.230137 | ENSG00000149792 |
| ACIN1 | -0.083392568 | 0.94383556 | 0.105430043 | 0.215595018 | 6106.148015 | 5762.712962 | ENSG00000100813 |
| SMARCAL1 | -0.083417108 | 0.943819505 | 0.411460303 | 0.583638288 | 541.7370128 | 511.1689174 | ENSG00000138375 |
| NA | -0.083426441 | 0.9438134 | 0.935385838 | 0.965468934 | 4.958365673 | 4.730650503 | ENSG00000214018 |
| PHAX | -0.083448251 | 0.943799132 | 0.238738997 | 0.400653675 | 1136.632393 | 1072.461411 | ENSG00000164902 |
| ITPK1 | -0.083454031 | 0.94379535 | 0.081439791 | 0.175949716 | 4066.388326 | 3838.568925 | ENSG00000100605 |
| SETD5 | -0.083488796 | 0.943772608 | 0.135352778 | 0.260744003 | 3805.268166 | 3591.766546 | ENSG00000168137 |
| CLNS1A | -0.083674218 | 0.943651317 | 0.120874073 | 0.239919923 | 2424.080132 | 2287.175326 | ENSG00000074201 |
| RBM4 | -0.083838011 | 0.943544188 | 0.707787749 | 0.82394816 | 112.1797681 | 105.6900624 | ENSG00000173933 |
| MFF | -0.083934196 | 0.943481284 | 0.332948242 | 0.505700696 | 717.2204969 | 676.585032 | ENSG00000168958 |
| ACRV1 | -0.084150881 | 0.943339589 | 0.911156746 | 0.951993281 | 8.602636233 | 8.108580586 | ENSG00000134940 |
| CASC4 | -0.084164538 | 0.943330659 | 0.411356784 | 0.583577271 | 537.7553425 | 507.6737292 | ENSG00000166734 |
| CCDC78 | -0.084185736 | 0.943316798 | 0.669701875 | 0.797187392 | 136.9209599 | 129.3360697 | ENSG00000162004 |
| RANBP9 | -0.084188361 | 0.943315082 | 0.397733177 | 0.570147896 | 709.7110471 | 669.4761378 | ENSG00000010017 |
| TMEM18-DT | -0.084310554 | 0.943235189 | 0.92315121 | 0.958830585 | 6.691923853 | 6.337739034 | ENSG00000233296 |
| CRY1 | -0.084375072 | 0.943193008 | 0.357910447 | 0.531111572 | 610.5798422 | 575.8154526 | ENSG00000008405 |
| DHX8 | -0.084526655 | 0.943093912 | 0.197176393 | 0.348135622 | 1456.328408 | 1373.930975 | ENSG00000067596 |
| ZFP36L2 | -0.084726284 | 0.942963423 | 0.317491609 | 0.488688314 | 1530.553122 | 1443.981704 | ENSG00000152518 |
| HNRNPK | -0.084744455 | 0.942951546 | 0.016209388 | 0.04719929 | 18501.5724 | 17446.59899 | ENSG00000165119 |
| TOMM40L | -0.084808158 | 0.942909911 | 0.544539936 | 0.702938966 | 357.5391443 | 336.8443708 | ENSG00000158882 |
| SUMO3 | -0.084809697 | 0.942908905 | 0.112817072 | 0.227271734 | 3198.924297 | 3016.548398 | ENSG00000184900 |
| SKA2 | -0.084939077 | 0.942824349 | 0.283358383 | 0.451832251 | 848.3431386 | 799.8309639 | ENSG00000182628 |
| KLC1 | -0.084959319 | 0.942811121 | 0.381744629 | 0.555170669 | 548.8886876 | 517.3929302 | ENSG00000126214 |
| NA | -0.084981477 | 0.942796641 | 0.868212736 | 0.926463417 | 17.8914868 | 16.8465963 | ENSG00000229546 |
| ASTN1 | -0.08498782 | 0.942792495 | 0.535185446 | 0.695283348 | 267.7778383 | 252.2542593 | ENSG00000152092 |
| ZHX2 | -0.085039899 | 0.942758463 | 0.346673112 | 0.519309753 | 752.2558052 | 709.1567127 | ENSG00000178764 |
| MFAP2 | -0.085043459 | 0.942756137 | 0.174636 | 0.317551426 | 1879.075811 | 1771.969959 | ENSG00000117122 |
| MTX3 | -0.085110693 | 0.942712202 | 0.432740632 | 0.603702599 | 467.8112294 | 441.342176 | ENSG00000177034 |
| TFAM | -0.085246459 | 0.942623492 | 0.161856809 | 0.29942268 | 2053.767488 | 1936.458615 | ENSG00000108064 |
| KLF10 | -0.085280916 | 0.942600978 | 0.371344692 | 0.54495085 | 541.6365403 | 510.5244161 | ENSG00000155090 |
| NELFB | -0.085291124 | 0.942594309 | 0.129338393 | 0.251798057 | 3305.693648 | 3116.122965 | ENSG00000188986 |
| USP47 | -0.085538133 | 0.942432938 | 0.365783755 | 0.539475109 | 603.3745157 | 568.9711279 | ENSG00000170242 |
| IPPK | -0.085542666 | 0.942429977 | 0.622099544 | 0.762199274 | 161.725228 | 152.5755144 | ENSG00000127080 |
| PPP1R7 | -0.085601049 | 0.942391839 | 0.247231143 | 0.410795519 | 1029.945645 | 970.3420553 | ENSG00000115685 |
| EFNB2 | -0.085645041 | 0.942363103 | 0.60255488 | 0.748190409 | 170.0272587 | 160.0952603 | ENSG00000125266 |
| EGR2 | -0.08579545 | 0.942264862 | 0.83907753 | 0.908201842 | 25.52269715 | 24.02996129 | ENSG00000122877 |
| COMMD2 | -0.085923686 | 0.942181111 | 0.334406649 | 0.507077399 | 749.5480572 | 706.2348327 | ENSG00000114744 |
| PAIP2B | -0.086064742 | 0.942088996 | 0.28993051 | 0.459388047 | 927.43543 | 873.2663351 | ENSG00000124374 |
| TBCK | -0.086088932 | 0.9420732 | 0.572656029 | 0.725196003 | 274.8585125 | 258.9238055 | ENSG00000145348 |
| NA | -0.086271612 | 0.941953918 | 0.704104327 | 0.82190951 | 99.67648575 | 93.96452867 | ENSG00000214223 |
| PLEKHA7 | -0.086287553 | 0.94194351 | 0.465422524 | 0.634879501 | 458.0946168 | 430.9443977 | ENSG00000166689 |
| TPD52L1 | -0.086391853 | 0.941875415 | 0.698244579 | 0.817989588 | 105.2368872 | 98.86618952 | ENSG00000111907 |
| BRD3 | -0.086414454 | 0.94186066 | 0.151637328 | 0.285027349 | 2050.07621 | 1931.460494 | ENSG00000169925 |

| | | | | | | | |
|---|---|---|---|---|---|---|---|
| PRKD1 | -0.086423699 | 0.941854624 | 0.589479046 | 0.738801992 | 187.7435064 | 176.6348023 | ENSG00000184304 |
| GSR | -0.086424648 | 0.941854004 | 0.138028317 | 0.264866632 | 2099.790237 | 1977.461119 | ENSG00000104687 |
| CNTN1 | -0.086461918 | 0.941829673 | 0.442599684 | 0.613507976 | 406.0930785 | 382.4087789 | ENSG00000018236 |
| RASA1 | -0.086509431 | 0.941798656 | 0.231203676 | 0.391515986 | 1043.96283 | 983.5348477 | ENSG00000145715 |
| NRF1 | -0.086517699 | 0.941793259 | 0.430554129 | 0.60191461 | 435.2448155 | 409.9976463 | ENSG00000106459 |
| LAGE3 | -0.086520085 | 0.941791701 | 0.24202671 | 0.404447095 | 1391.315039 | 1310.399543 | ENSG00000196976 |
| B3GNTL1 | -0.086538857 | 0.941779447 | 0.629618862 | 0.767469826 | 142.7645457 | 134.4732645 | ENSG00000175711 |
| NDFIP1 | -0.086656674 | 0.94170254 | 0.256613351 | 0.421771164 | 1291.540872 | 1216.48524 | ENSG00000131507 |
| FN3KRP | -0.086732468 | 0.941653067 | 0.33840522 | 0.51101175 | 663.555498 | 624.7314693 | ENSG00000141560 |
| YEATS4 | -0.086793772 | 0.941613055 | 0.347561481 | 0.52027691 | 596.6147712 | 561.9426411 | ENSG00000127337 |
| HSD17B14 | -0.086804128 | 0.941606296 | 0.617464836 | 0.759372316 | 187.5644371 | 176.570515 | ENSG00000087076 |
| RNFT1 | -0.086806365 | 0.941604836 | 0.826130816 | 0.901078467 | 28.91840132 | 27.28393317 | ENSG00000189050 |
| CDKN2AIP | -0.086864185 | 0.941567099 | 0.443259294 | 0.614069302 | 508.0785268 | 478.585412 | ENSG00000168564 |
| ZNF280A | -0.086870314 | 0.941563099 | 0.910710797 | 0.951993281 | 7.29902901 | 6.884640437 | ENSG00000169548 |
| NA | -0.086943245 | 0.941515503 | 0.653566188 | 0.78472259 | 124.365275 | 117.0001771 | ENSG00000232533 |
| RPS6KA4 | -0.08696438 | 0.94150171 | 0.378613441 | 0.551982458 | 657.2232273 | 618.7003062 | ENSG00000162302 |
| SPINDOC | -0.086987388 | 0.941486695 | 0.123848765 | 0.244094265 | 2678.524336 | 2521.908646 | ENSG00000168005 |
| TAF3 | -0.087098722 | 0.941414042 | 0.463590203 | 0.633319281 | 412.1057724 | 388.1749063 | ENSG00000165632 |
| NA | -0.087201079 | 0.941347253 | 0.626624336 | 0.765221882 | 144.1472621 | 135.6684231 | ENSG00000182484 |
| TRMT61A | -0.087260241 | 0.941308651 | 0.418060375 | 0.590016442 | 652.2346884 | 614.2780365 | ENSG00000166166 |
| PDIA4 | -0.087277037 | 0.941297692 | 0.036281231 | 0.091922936 | 10136.23048 | 9541.675915 | ENSG00000155660 |
| LOC150776 | -0.087404019 | 0.941214846 | 0.455767774 | 0.625485559 | 361.5495626 | 340.2047748 | ENSG00000152117 |
| PACSIN2 | -0.08749053 | 0.941158407 | 0.114765248 | 0.230402373 | 2408.600922 | 2267.553579 | ENSG00000100266 |
| DLGAP5 | -0.087535014 | 0.941129388 | 0.118142193 | 0.235661363 | 2861.221728 | 2692.793883 | ENSG00000126787 |
| SGPL1 | -0.087576554 | 0.94110229 | 0.207019183 | 0.361412403 | 1509.70763 | 1420.189742 | ENSG00000166224 |
| ZNF605 | -0.087602089 | 0.941085633 | 0.569031057 | 0.722135263 | 226.050555 | 212.5352161 | ENSG00000196458 |
| NA | -0.087617334 | 0.941075689 | 0.78889827 | 0.876902133 | 48.25332866 | 45.27487997 | ENSG00000275964 |
| ZNF143 | -0.087634111 | 0.941064745 | 0.2665575 | 0.432732536 | 837.1955267 | 788.0691245 | ENSG00000166478 |
| LTA4H | -0.087647499 | 0.941056012 | 0.109437298 | 0.221783267 | 2871.076022 | 2702.497945 | ENSG00000111144 |
| NA | -0.087886001 | 0.940900453 | 0.852625462 | 0.916271951 | 21.19500316 | 19.98065068 | ENSG00000276672 |
| NA | -0.087925515 | 0.940874683 | 0.88209804 | 0.93559233 | 12.94061796 | 12.22726101 | ENSG00000274422 |
| ZNF407 | -0.0879387 | 0.940866084 | 0.61189614 | 0.755316639 | 208.8502398 | 196.7219219 | ENSG00000215421 |
| ZFP64 | -0.087945328 | 0.940861761 | 0.332445951 | 0.505176435 | 630.9389983 | 593.8125664 | ENSG00000020256 |
| ARMC10 | -0.088011997 | 0.940818284 | 0.255193039 | 0.420152677 | 936.6208553 | 881.2243571 | ENSG00000170632 |
| MVB12A | -0.088111927 | 0.940753119 | 0.39970076 | 0.572292601 | 502.7000593 | 473.1211953 | ENSG00000141971 |
| MED11 | -0.08827869 | 0.940644383 | 0.725124002 | 0.835021231 | 76.51957006 | 72.13774794 | ENSG00000161920 |
| TET1 | -0.088329968 | 0.94061095 | 0.236873904 | 0.398390103 | 1374.635107 | 1292.988706 | ENSG00000138336 |
| NOM1 | -0.088364941 | 0.940588148 | 0.249149581 | 0.413164666 | 1001.84806 | 941.8339457 | ENSG00000146909 |
| ZNF251 | -0.088453545 | 0.940530383 | 0.547607057 | 0.705307928 | 227.8487554 | 214.3937045 | ENSG00000198169 |
| USP13 | -0.088552632 | 0.940465788 | 0.349968552 | 0.522609746 | 676.754557 | 636.7452352 | ENSG00000058056 |
| LSM4 | -0.088598401 | 0.940435952 | 0.049777855 | 0.11888877 | 5156.868411 | 4849.305523 | ENSG00000130520 |
| GJC1 | -0.088688376 | 0.940377303 | 0.149813165 | 0.28214837 | 1793.93816 | 1687.558927 | ENSG00000182963 |
| RAB2A | -0.088688964 | 0.94037692 | 0.201046977 | 0.353159464 | 1578.961502 | 1485.149395 | ENSG00000104388 |
| FIZ1 | -0.088762264 | 0.940329143 | 0.480802502 | 0.648892574 | 453.4625526 | 426.7407362 | ENSG00000179943 |
| BCAS3 | -0.088781639 | 0.940316515 | 0.471377712 | 0.640286399 | 319.539271 | 300.304774 | ENSG00000141376 |
| MAU2 | -0.088995117 | 0.940177385 | 0.266650392 | 0.432781462 | 943.7495385 | 887.4292873 | ENSG00000129933 |
| SERTAD2 | -0.089095191 | 0.94011217 | 0.595360318 | 0.743227587 | 183.0214674 | 172.2208807 | ENSG00000179833 |
| ZNF250 | -0.089135753 | 0.940085739 | 0.565109163 | 0.719287468 | 203.3876689 | 191.0069179 | ENSG00000196150 |
| NUP107 | -0.089143148 | 0.94008092 | 0.140923813 | 0.269297798 | 1948.429313 | 1832.338004 | ENSG00000111581 |
| SLC12A8 | -0.089147745 | 0.940077925 | 0.585444933 | 0.735192012 | 234.7203362 | 220.5951576 | ENSG00000221955 |
| RPS6KL1 | -0.089190049 | 0.94005036 | 0.591007438 | 0.739910841 | 171.6253261 | 161.2963376 | ENSG00000198208 |
| OPA3 | -0.089386276 | 0.939922508 | 0.286791873 | 0.456025554 | 793.8381489 | 746.0907254 | ENSG00000125741 |
| CSNK2A2 | -0.089475786 | 0.939864194 | 0.249778976 | 0.413888183 | 932.3826193 | 876.1325664 | ENSG00000070770 |
| SDHAF2 | -0.089533353 | 0.939826692 | 0.766700638 | 0.861962935 | 53.74930232 | 50.64062382 | ENSG00000167985 |
| RPA2 | -0.089538475 | 0.939823355 | 0.215021387 | 0.37178252 | 1187.502223 | 1115.811165 | ENSG00000117748 |
| INTS9 | -0.089604813 | 0.939780141 | 0.41594063 | 0.588047687 | 443.870767 | 417.1441606 | ENSG00000104299 |
| AP3S1 | -0.089629587 | 0.939764003 | 0.231563384 | 0.392021873 | 1035.736717 | 973.8170095 | ENSG00000177879 |
| USP44 | -0.089644498 | 0.939754291 | 0.072303898 | 0.160121054 | 4227.13283 | 3972.793096 | ENSG00000136014 |
| RNF168 | -0.089657332 | 0.93974593 | 0.186686618 | 0.333627096 | 1211.266081 | 1138.231021 | ENSG00000163961 |
| ZFHX2 | -0.089723328 | 0.939702943 | 0.347290241 | 0.520032293 | 672.2424632 | 632.1732125 | ENSG00000136367 |
| PAGR1 | -0.089754924 | 0.939682363 | 0.553296994 | 0.709662844 | 199.8919678 | 187.8021366 | ENSG00000280789 |
| SLC35E3 | -0.089803247 | 0.939650889 | 0.455098809 | 0.624862002 | 353.7958279 | 332.7721363 | ENSG00000175782 |
| SDHAF3 | -0.089954851 | 0.939552152 | 0.384505058 | 0.557957506 | 454.0590435 | 426.5135136 | ENSG00000196636 |
| PIK3R1 | -0.090055803 | 0.939486409 | 0.536361268 | 0.696038255 | 250.3681783 | 235.1110769 | ENSG00000145675 |
| ZZEF1 | -0.090074281 | 0.939474377 | 0.360962154 | 0.534119883 | 616.1288558 | 578.9169268 | ENSG00000074755 |
| CACUL1 | -0.090111773 | 0.939449962 | 0.306147358 | 0.477156663 | 726.7775751 | 682.4807026 | ENSG00000151893 |
| GTF2A2 | -0.090129975 | 0.93943811 | 0.213552875 | 0.369840068 | 1352.981581 | 1270.705725 | ENSG00000140307 |
| SNHG17 | -0.090202189 | 0.939391087 | 0.410965216 | 0.583279133 | 451.2355895 | 423.9250341 | ENSG00000196756 |
| THUMPD3-AS1 | -0.090215126 | 0.939382664 | 0.578490377 | 0.730200737 | 190.5242082 | 178.9452879 | ENSG00000206573 |
| RNF175 | -0.09027107 | 0.939346237 | 0.6016325 | 0.747381838 | 156.8501139 | 147.2412356 | ENSG00000145428 |
| NA | -0.090281544 | 0.939339418 | 0.858996982 | 0.920821105 | 18.60752519 | 17.41877155 | ENSG00000227376 |
| ANAPC10 | -0.090293802 | 0.939331437 | 0.86349608 | 0.923478372 | 21.71415086 | 20.50899042 | ENSG00000164162 |
| SEH1L | -0.090316803 | 0.939316461 | 0.261896172 | 0.427395234 | 1239.496468 | 1164.261754 | ENSG00000085415 |
| ZNF114 | -0.090331827 | 0.939306679 | 0.550974916 | 0.707889978 | 243.9638566 | 228.9981431 | ENSG00000178150 |
| ZUP1 | -0.090508707 | 0.939191524 | 0.539364966 | 0.698681435 | 272.3934978 | 255.9013574 | ENSG00000153975 |
| NAP1L5 | -0.090515873 | 0.939186859 | 0.774299289 | 0.867655528 | 47.04150893 | 44.14263941 | ENSG00000177432 |
| TOP2B | -0.090532091 | 0.9391763 | 0.124847154 | 0.245424828 | 4342.847287 | 4079.869125 | ENSG00000077097 |

| | | | | | | | |
|---|---|---|---|---|---|---|---|
| YIPF3 | -0.090762797 | 0.939026126 | 0.203565204 | 0.356608822 | 1189.935275 | 1117.13001 | ENSG00000137207 |
| GRIPAP1 | -0.090983476 | 0.938882501 | 0.179075046 | 0.323459841 | 1394.593861 | 1308.775498 | ENSG00000068400 |
| PEX14 | -0.090986049 | 0.938880826 | 0.258912269 | 0.424104321 | 832.6957848 | 782.0142504 | ENSG00000142655 |
| SMARCAD1 | -0.091029325 | 0.938852663 | 0.176997762 | 0.320606932 | 1407.053054 | 1321.753123 | ENSG00000163104 |
| KLC4 | -0.091130405 | 0.938786887 | 0.359143593 | 0.532204567 | 508.5221058 | 477.1369328 | ENSG00000137171 |
| FBXO33 | -0.091341566 | 0.938649491 | 0.502479274 | 0.668228848 | 285.0886526 | 267.6725925 | ENSG00000165355 |
| CBFA2T3 | -0.091374725 | 0.938627917 | 0.807883588 | 0.889446541 | 33.89717919 | 31.74087251 | ENSG00000129993 |
| FAM98B | -0.09141449 | 0.938602046 | 0.182771415 | 0.328448319 | 1285.808759 | 1207.361629 | ENSG00000171262 |
| GID4 | -0.09142684 | 0.938594011 | 0.435326963 | 0.606135187 | 369.4681877 | 346.6845006 | ENSG00000141034 |
| OCIAD1 | -0.091458747 | 0.938573253 | 0.107912705 | 0.21924634 | 1894.089963 | 1777.668818 | ENSG00000109180 |
| NA | -0.091541969 | 0.938519113 | 0.907276082 | 0.949958569 | 7.005244592 | 6.583765315 | ENSG00000214653 |
| QTRT2 | -0.091687982 | 0.938424132 | 0.202765212 | 0.355497928 | 1253.205412 | 1175.718169 | ENSG00000151576 |
| MIIP | -0.091920257 | 0.938273057 | 0.155313305 | 0.290329339 | 1410.447355 | 1323.361072 | ENSG00000116691 |
| KISS1R | -0.091930844 | 0.938266171 | 0.80902622 | 0.890279831 | 32.82384134 | 30.73338365 | ENSG00000116014 |
| DNTTIP2 | -0.091970253 | 0.938240542 | 0.174193377 | 0.316985609 | 1645.30033 | 1544.034208 | ENSG00000067334 |
| RAB7A | -0.092050547 | 0.938188325 | 0.037833577 | 0.09516854 | 4367.055484 | 4097.112483 | ENSG00000075785 |
| DTX3 | -0.092063559 | 0.938179863 | 0.302146466 | 0.472955816 | 814.9157708 | 763.9158049 | ENSG00000178498 |
| FEZ1 | -0.092087682 | 0.938164176 | 0.216310198 | 0.373375092 | 975.1737916 | 914.9578654 | ENSG00000149557 |
| BOP1 | -0.092089782 | 0.938162811 | 0.111068106 | 0.224381069 | 4378.404722 | 4107.582607 | ENSG00000261236 |
| RPL38 | -0.092114932 | 0.938146456 | 0.161042668 | 0.298341927 | 7486.583883 | 7022.76734 | ENSG00000172809 |
| NA | -0.092202144 | 0.938089746 | 0.860341906 | 0.921637923 | 16.80275498 | 15.7213405 | ENSG00000254851 |
| PGGT1B | -0.092214834 | 0.938081495 | 0.478743893 | 0.64701194 | 316.9665865 | 297.0806618 | ENSG00000164219 |
| TTC7B | -0.092230201 | 0.938071503 | 0.452420868 | 0.622561732 | 325.7689043 | 305.4164344 | ENSG00000165914 |
| PAICS | -0.092249345 | 0.938059055 | 0.014061675 | 0.041984796 | 10517.52284 | 9866.188964 | ENSG00000128050 |
| ADHFE1 | -0.092369482 | 0.937980943 | 0.937940599 | 0.96727318 | 4.217884882 | 3.998762916 | ENSG00000147576 |
| BNIP2 | -0.092375593 | 0.937976971 | 0.347128224 | 0.519910759 | 582.2919124 | 546.3417448 | ENSG00000140299 |
| EHD3 | -0.092441811 | 0.937933919 | 0.328440575 | 0.500667494 | 635.882809 | 596.6344719 | ENSG00000013016 |
| ZCWPW1 | -0.092484415 | 0.937906222 | 0.842263401 | 0.909743682 | 25.34342777 | 23.72101691 | ENSG00000078487 |
| FBXO8 | -0.092490123 | 0.937902511 | 0.650069428 | 0.782358329 | 114.3509742 | 107.1723872 | ENSG00000164117 |
| IRX2 | -0.092603084 | 0.937829078 | 0.542140898 | 0.700967832 | 285.6081637 | 268.1909545 | ENSG00000170561 |
| RHOT2 | -0.092604484 | 0.937828168 | 0.133895004 | 0.25871079 | 4438.75699 | 4162.783992 | ENSG00000140983 |
| LINC01003 | -0.092638432 | 0.9378061 | 0.853349148 | 0.916896425 | 23.01795825 | 21.68049276 | ENSG00000261455 |
| NCKIPSD | -0.092713273 | 0.937757452 | 0.230991729 | 0.391294471 | 976.8630829 | 916.1930915 | ENSG00000213672 |
| SLC25A20 | -0.092761519 | 0.937726092 | 0.685793741 | 0.808559288 | 98.55736629 | 92.49267213 | ENSG00000178537 |
| KATNBL1 | -0.092819746 | 0.937688247 | 0.421451554 | 0.593403542 | 361.2121629 | 338.9004394 | ENSG00000134152 |
| NA | -0.092861025 | 0.937661417 | 0.866287387 | 0.925300921 | 16.76156716 | 15.67724453 | ENSG00000278864 |
| CDYL | -0.092894925 | 0.937639385 | 0.135030967 | 0.260228011 | 1785.526907 | 1674.303216 | ENSG00000153046 |
| ZCCHC18 | -0.092958789 | 0.937597879 | 0.674172835 | 0.800127035 | 104.2956488 | 97.73233143 | ENSG00000166707 |
| MRPL39 | -0.092980644 | 0.937583676 | 0.250338495 | 0.414530464 | 1098.507693 | 1029.958367 | ENSG00000154719 |
| ALG2 | -0.092993588 | 0.937575263 | 0.291778411 | 0.461500395 | 727.9265697 | 682.7990612 | ENSG00000119523 |
| PECR | -0.093031925 | 0.937550349 | 0.527998983 | 0.689522636 | 269.4883935 | 252.7694109 | ENSG00000115425 |
| HEBP2 | -0.093066301 | 0.93752801 | 0.267369374 | 0.433722048 | 848.0183419 | 795.05649 | ENSG00000051620 |
| GAPDH | -0.093201866 | 0.937439918 | 0.016213304 | 0.047203569 | 38751.94289 | 36327.79832 | ENSG00000111640 |
| MACC1 | -0.093240743 | 0.937414657 | 0.498066807 | 0.66440963 | 250.0993119 | 234.3829574 | ENSG00000183742 |
| NA | -0.093249372 | 0.93740905 | 0.840311276 | 0.908946853 | 21.45896352 | 20.12369146 | ENSG00000260276 |
| AKR1A1 | -0.093320614 | 0.937362761 | 0.049308701 | 0.11794418 | 3458.846999 | 3242.187984 | ENSG00000117448 |
| SRSF3 | -0.093342883 | 0.937348292 | 0.03841554 | 0.096431254 | 12981.09534 | 12166.90413 | ENSG00000112081 |
| GTSE1 | -0.093449215 | 0.937279208 | 0.223941842 | 0.382441755 | 1278.104957 | 1197.609648 | ENSG00000075218 |
| NA | -0.093471629 | 0.937264647 | 0.537395141 | 0.69697269 | 226.6236938 | 212.2448213 | ENSG00000283041 |
| C16orf91 | -0.093539428 | 0.937220601 | 0.50993014 | 0.674379942 | 267.6385174 | 250.6033537 | ENSG00000174109 |
| ARID2 | -0.093543823 | 0.937217746 | 0.2802832 | 0.448374094 | 767.7528136 | 719.7467466 | ENSG00000189079 |
| ACYP2 | -0.093597608 | 0.937182807 | 0.745471318 | 0.849098159 | 58.67034712 | 54.96983685 | ENSG00000170634 |
| ND3 | -0.093642368 | 0.937153731 | 0.098186072 | 0.204216443 | 5089.19813 | 4769.245852 | ENSG00000198840 |
| MARK4 | -0.093645958 | 0.937151398 | 0.148061632 | 0.279647516 | 1600.976545 | 1501.028328 | ENSG00000007047 |
| EIF4EBP2 | -0.093705838 | 0.937112502 | 0.036330165 | 0.091991812 | 6971.160105 | 6533.624247 | ENSG00000148730 |
| NA | -0.093758813 | 0.937078093 | 0.750354317 | 0.852327924 | 55.15905188 | 51.72582433 | ENSG00000272686 |
| ND6 | -0.0937871 | 0.93705972 | 0.668028395 | 0.796028624 | 3929.959798 | 3682.714745 | ENSG00000198695 |
| CPOX | -0.093820423 | 0.937038076 | 0.263015532 | 0.428597094 | 877.2182 | 821.8449177 | ENSG00000080819 |
| RAB31 | -0.093883687 | 0.936996986 | 0.27317343 | 0.440029112 | 748.6613833 | 701.2535517 | ENSG00000168461 |
| NA | -0.093946369 | 0.936956277 | 0.925662344 | 0.960206069 | 4.936714172 | 4.665581831 | ENSG00000232187 |
| IGBP1 | -0.094004694 | 0.936918398 | 0.313062012 | 0.484188189 | 562.7815074 | 527.2134507 | ENSG00000089289 |
| MYC | -0.094011698 | 0.93691385 | 0.19593852 | 0.346393877 | 1338.976399 | 1254.766334 | ENSG00000136997 |
| MRPL22 | -0.09406906 | 0.936876599 | 0.255583432 | 0.420583203 | 854.1790274 | 800.3592915 | ENSG00000082515 |
| SH3GL3 | -0.094233557 | 0.936769782 | 0.523588845 | 0.686069036 | 241.0403673 | 225.4693819 | ENSG00000140600 |
| CEP85 | -0.094313104 | 0.936718132 | 0.219745291 | 0.377715976 | 1236.136404 | 1157.275213 | ENSG00000130695 |
| ISG20 | -0.094323941 | 0.936711095 | 0.836738678 | 0.907107819 | 21.06182768 | 19.73677497 | ENSG00000172183 |
| MRPL52 | -0.094725646 | 0.936450313 | 0.219744743 | 0.377715976 | 996.69134 | 933.1886899 | ENSG00000172590 |
| ZNF484 | -0.094770857 | 0.936420967 | 0.635174402 | 0.77136449 | 121.7698999 | 114.0070129 | ENSG00000127081 |
| RALGAPA2 | -0.094819063 | 0.936389678 | 0.293464775 | 0.463173493 | 633.7669823 | 593.3936455 | ENSG00000188559 |
| OXA1L | -0.094833201 | 0.936380502 | 0.079845634 | 0.173146303 | 2290.303106 | 2145.09004 | ENSG00000155463 |
| TACC3 | -0.094833532 | 0.936380287 | 0.036177874 | 0.091680414 | 5040.034561 | 4719.184893 | ENSG00000013810 |
| SUB1 | -0.09489182 | 0.936342456 | 0.109667723 | 0.222076015 | 2319.803646 | 2172.323722 | ENSG00000113387 |
| CDK11B | -0.094942317 | 0.936309683 | 0.134634406 | 0.259797001 | 1628.753024 | 1525.625358 | ENSG00000248333 |
| RHPN1 | -0.094943259 | 0.936309072 | 0.34170271 | 0.514541174 | 927.0382076 | 867.8396548 | ENSG00000158106 |
| RAMAC | -0.094952745 | 0.936302915 | 0.700133751 | 0.819108508 | 96.92752296 | 90.88694067 | ENSG00000169612 |
| AKR1C2 | -0.095027005 | 0.936254722 | 0.919535122 | 0.957092481 | 5.559889024 | 5.23130536 | ENSG00000151632 |
| LRGUK | -0.095160228 | 0.936168269 | 0.788480667 | 0.876555309 | 34.76457793 | 32.55718393 | ENSG00000155530 |

| | | | | | | | |
|---|---|---|---|---|---|---|---|
| DEK | -0.095177303 | 0.93615719 | 0.051970872 | 0.122999644 | 4760.373414 | 4457.40405 | ENSG00000124795 |
| BIRC5 | -0.095182988 | 0.9361535 | 0.055916893 | 0.130560734 | 3901.039738 | 3651.298871 | ENSG00000089685 |
| ARL5A | -0.095187769 | 0.936150398 | 0.452042593 | 0.622260144 | 569.5899508 | 533.6219103 | ENSG00000162980 |
| NA | -0.095387351 | 0.936020901 | 0.941983678 | 0.969525003 | 3.481070874 | 3.207725342 | ENSG00000253390 |
| CCDC6 | -0.095390641 | 0.936018766 | 0.207289963 | 0.361852383 | 1311.532928 | 1228.074489 | ENSG00000108091 |
| METRNL | -0.095528242 | 0.935929495 | 0.419794827 | 0.591805174 | 366.1321316 | 342.8965002 | ENSG00000176845 |
| CSNK1D | -0.095535593 | 0.935924726 | 0.077674897 | 0.169294592 | 2938.338732 | 2749.69551 | ENSG00000141551 |
| LPAR3 | -0.095637013 | 0.935858933 | 0.373465127 | 0.54695686 | 420.0794007 | 393.2814294 | ENSG00000171517 |
| RNGTT | -0.09568737 | 0.935826268 | 0.34997886 | 0.522609746 | 696.4188748 | 651.4268646 | ENSG00000111880 |
| NKILA | -0.095766279 | 0.935775084 | 0.836433123 | 0.907107678 | 23.95935992 | 22.49224257 | ENSG00000278709 |
| RB1CC1 | -0.095777477 | 0.935767821 | 0.257833809 | 0.42290964 | 909.437639 | 851.6661707 | ENSG00000023287 |
| MRAP2 | -0.095798837 | 0.935753966 | 0.678184797 | 0.80312234 | 97.50567994 | 91.25395054 | ENSG00000135324 |
| KIF7 | -0.095886063 | 0.935697391 | 0.283806419 | 0.452397094 | 726.2461508 | 679.4225309 | ENSG00000166813 |
| USP6NL | -0.095937317 | 0.93566415 | 0.258422044 | 0.42358906 | 726.4548808 | 679.8252643 | ENSG00000148429 |
| CYTB | -0.095987553 | 0.93563157 | 0.651578411 | 0.783299004 | 22087.47321 | 20665.79549 | ENSG00000198727 |
| PPP6C | -0.096117046 | 0.935547594 | 0.225968408 | 0.384982474 | 962.2356154 | 900.8866128 | ENSG00000119414 |
| PMPCA | -0.096179672 | 0.935506983 | 0.061051011 | 0.139890137 | 2794.838648 | 2614.666942 | ENSG00000165688 |
| DMAC2 | -0.096192911 | 0.935498399 | 0.128241689 | 0.250319783 | 1447.054988 | 1353.89198 | ENSG00000105341 |
| TGFBI | -0.09619455 | 0.935497336 | 0.50408201 | 0.669299061 | 265.0454208 | 248.176434 | ENSG00000120708 |
| TOMM22 | -0.096245856 | 0.935464068 | 0.102311249 | 0.21075134 | 2090.29412 | 1955.628971 | ENSG00000100216 |
| PRMT6 | -0.096257305 | 0.935456644 | 0.209486519 | 0.364762563 | 1277.381613 | 1195.045146 | ENSG00000198890 |
| SRBD1 | -0.096275726 | 0.9354447 | 0.345151725 | 0.517592458 | 572.5110722 | 535.9363697 | ENSG00000068784 |
| TLE1 | -0.096280953 | 0.93544131 | 0.04873577 | 0.116979189 | 3623.818676 | 3390.217362 | ENSG00000196781 |
| PPFIA2 | -0.096308396 | 0.935423517 | 0.83156167 | 0.904389657 | 22.1421868 | 20.68511384 | ENSG00000139220 |
| SLC38A10 | -0.09630963 | 0.935422716 | 0.256814391 | 0.42195779 | 1211.602843 | 1132.904097 | ENSG00000157637 |
| NA | -0.096425389 | 0.935347663 | 0.860058832 | 0.921437017 | 17.93758071 | 16.71000724 | ENSG00000273340 |
| SYT12 | -0.09645883 | 0.935325982 | 0.580421285 | 0.731378652 | 177.5269332 | 165.8629746 | ENSG00000173227 |
| STRBP | -0.096475667 | 0.935315067 | 0.066914587 | 0.150536361 | 2465.560476 | 2306.499844 | ENSG00000165209 |
| RHBDD3 | -0.096665542 | 0.935191977 | 0.181610819 | 0.326967272 | 1656.298647 | 1549.301825 | ENSG00000100263 |
| SLITRK2 | -0.096666955 | 0.935191061 | 0.862443557 | 0.922764098 | 15.44747204 | 14.41459405 | ENSG00000185985 |
| RNF6 | -0.096678481 | 0.93518359 | 0.507043986 | 0.67183892 | 292.4498452 | 273.8516427 | ENSG00000127870 |
| HLA-L | -0.096766432 | 0.935126579 | 0.740058221 | 0.84572929 | 57.09739788 | 53.36221528 | ENSG00000243753 |
| RETREG3 | -0.096857077 | 0.935067827 | 0.297934536 | 0.468224452 | 597.6086123 | 558.8241858 | ENSG00000141699 |
| TFIP11 | -0.096887749 | 0.935047948 | 0.271612236 | 0.438522505 | 654.2831047 | 611.6559108 | ENSG00000100109 |
| PTENP1 | -0.096997455 | 0.934976847 | 0.916302583 | 0.955119313 | 5.314989415 | 5.004101204 | ENSG00000237984 |
| CTBP1 | -0.097001287 | 0.934974363 | 0.150410115 | 0.282996346 | 1890.240551 | 1766.708818 | ENSG00000159692 |
| SLAIN2 | -0.097125887 | 0.934893617 | 0.271418061 | 0.438372891 | 653.1898302 | 610.7575866 | ENSG00000109171 |
| SPIN2B | -0.097139338 | 0.934884901 | 0.621570928 | 0.762041612 | 131.8834098 | 123.2604771 | ENSG00000186787 |
| NPLOC4 | -0.09721074 | 0.934838632 | 0.057255308 | 0.133073752 | 3130.460705 | 2926.435122 | ENSG00000182446 |
| GIT1 | -0.097216453 | 0.934834931 | 0.159847767 | 0.296901914 | 2491.208338 | 2328.86075 | ENSG00000108262 |
| NA | -0.097227147 | 0.934828001 | 0.871186424 | 0.928357582 | 12.3174431 | 11.54592112 | ENSG00000251689 |
| CYTH3 | -0.097255192 | 0.934809829 | 0.325410232 | 0.497108709 | 541.4821889 | 506.1822997 | ENSG00000008256 |
| UNK | -0.097295881 | 0.934783464 | 0.334248419 | 0.506917159 | 513.4433891 | 479.9429603 | ENSG00000132478 |
| TOGARAM1 | -0.097296522 | 0.934783049 | 0.6011719 | 0.747033702 | 145.4026603 | 136.0244075 | ENSG00000198718 |
| ZIC2 | -0.097443496 | 0.934687823 | 0.08383371 | 0.180139871 | 3981.040761 | 3722.033866 | ENSG00000043355 |
| NECTIN2 | -0.09753049 | 0.934631463 | 0.050506329 | 0.120271273 | 4369.296204 | 4084.305329 | ENSG00000130202 |
| ADAM15 | -0.097580079 | 0.934599338 | 0.234454904 | 0.395459435 | 946.1355079 | 884.4342451 | ENSG00000143537 |
| PLA2G12A | -0.097598571 | 0.934587359 | 0.491035929 | 0.657452395 | 298.2176764 | 278.7135513 | ENSG00000123739 |
| PFDN5 | -0.097639672 | 0.934560734 | 0.137184993 | 0.26361547 | 1623.704984 | 1516.606955 | ENSG00000123349 |
| AP2B1 | -0.097711995 | 0.934513885 | 0.031999752 | 0.082826676 | 5526.255532 | 5163.879069 | ENSG00000006125 |
| STX1A | -0.09783115 | 0.934436705 | 0.599390467 | 0.745845417 | 141.2760873 | 131.8987545 | ENSG00000106089 |
| SNPH | -0.097837644 | 0.934432499 | 0.535134527 | 0.695283348 | 182.4087805 | 170.471557 | ENSG00000101298 |
| ZFP42 | -0.097893327 | 0.934396433 | 0.111204693 | 0.224562937 | 4549.657717 | 4250.479781 | ENSG00000179059 |
| NA | -0.097914322 | 0.934382836 | 0.876895644 | 0.932278526 | 13.34122042 | 12.46951919 | ENSG00000267248 |
| SEPTIN4 | -0.098092891 | 0.93426719 | 0.611627231 | 0.755129658 | 123.9241604 | 115.7310749 | ENSG00000108387 |
| P3H3 | -0.098124764 | 0.93424655 | 0.185058136 | 0.331387575 | 1172.232493 | 1095.232763 | ENSG00000110811 |
| CHUK | -0.098130857 | 0.934242604 | 0.342789928 | 0.515454089 | 507.0984017 | 473.2954833 | ENSG00000213341 |
| MSI2 | -0.098205673 | 0.934194157 | 0.245844781 | 0.409196866 | 1019.767595 | 953.1670863 | ENSG00000153944 |
| PPP2R1B | -0.098259511 | 0.934159296 | 0.101563724 | 0.209536973 | 2022.53752 | 1889.687224 | ENSG00000137713 |
| SLC25A16 | -0.098302723 | 0.934131316 | 0.668411744 | 0.796192093 | 87.93420857 | 82.1397712 | ENSG00000122912 |
| LOC389906 | -0.098353398 | 0.934098505 | 0.392138996 | 0.565166482 | 446.7320927 | 417.5832239 | ENSG00000285756 |
| TRIM52-AS1 | -0.098470837 | 0.93402247 | 0.619455277 | 0.760753231 | 136.252567 | 127.4746131 | ENSG00000248275 |
| KANSL2 | -0.098516571 | 0.933992862 | 0.320192734 | 0.49159077 | 602.0330609 | 561.845047 | ENSG00000139620 |
| ZNF35 | -0.098677143 | 0.933888914 | 0.496527607 | 0.662747811 | 267.1575296 | 249.5095446 | ENSG00000169981 |
| CYP4X1 | -0.098763686 | 0.933832895 | 0.731261882 | 0.839239274 | 55.35335181 | 51.66048309 | ENSG00000186377 |
| PLXDC1 | -0.098985812 | 0.933689127 | 0.862024435 | 0.922689647 | 14.52772187 | 13.54422729 | ENSG00000161381 |
| C2orf42 | -0.099028896 | 0.933661245 | 0.620266904 | 0.76139303 | 118.8591769 | 110.8678639 | ENSG00000115998 |
| YJEFN3 | -0.099045287 | 0.933650636 | 0.63827104 | 0.773863705 | 114.1331077 | 106.4230345 | ENSG00000250067 |
| RGS19 | -0.099073411 | 0.933632436 | 0.485541435 | 0.653129864 | 243.7418478 | 227.7631654 | ENSG00000171700 |
| SLIT2 | -0.099135115 | 0.933592505 | 0.5249387 | 0.687247537 | 199.3893654 | 186.3109249 | ENSG00000145147 |
| PLP2 | -0.09921733 | 0.933539304 | 0.668145835 | 0.796028624 | 90.71454701 | 84.60647362 | ENSG00000102007 |
| NA | -0.099229151 | 0.933531655 | 0.888915768 | 0.938578344 | 9.917607228 | 9.198342708 | ENSG00000286099 |
| NA | -0.099238953 | 0.933525313 | 0.7595221 | 0.857189541 | 45.05276342 | 41.95530637 | ENSG00000255310 |
| PDCD2L | -0.099283556 | 0.933496452 | 0.659314718 | 0.789485573 | 96.12942123 | 89.68316129 | ENSG00000126249 |
| AFAP1 | -0.099326425 | 0.933468714 | 0.616733695 | 0.758837926 | 123.2199614 | 115.0924739 | ENSG00000196526 |
| GTF3C2 | -0.099492286 | 0.933361403 | 0.091660861 | 0.193386117 | 2071.60975 | 1933.172792 | ENSG00000115207 |
| TSPAN9 | -0.099530714 | 0.933336542 | 0.184917535 | 0.331220443 | 1252.9917 | 1169.067463 | ENSG00000011105 |

| | | | | | | | |
|---|---|---|---|---|---|---|---|
| ISOC2 | -0.099680579 | 0.933239594 | 0.143680879 | 0.273267065 | 1832.644235 | 1709.971919 | ENSG00000063241 |
| WDR6 | -0.099710834 | 0.933220022 | 0.075280045 | 0.165310311 | 4949.411624 | 4617.888495 | ENSG00000178252 |
| NA | -0.099711612 | 0.933219519 | 0.915222011 | 0.954649348 | 5.710010077 | 5.349072298 | ENSG00000259834 |
| CIB2 | -0.099724031 | 0.933211486 | 0.174573371 | 0.317467469 | 1106.517553 | 1032.951792 | ENSG00000136425 |
| MPP5 | -0.099734501 | 0.933204713 | 0.301371994 | 0.472097506 | 538.1131303 | 502.1558213 | ENSG00000072415 |
| PKIG | -0.099746263 | 0.933197105 | 0.360583353 | 0.533641232 | 425.2738177 | 396.8473203 | ENSG00000168734 |
| ARL6IP1 | -0.099893409 | 0.93310193 | 0.161074358 | 0.298347137 | 1552.512007 | 1449.240615 | ENSG00000170540 |
| AP4B1 | -0.099912023 | 0.93308989 | 0.535645559 | 0.695553209 | 178.8692128 | 166.8973549 | ENSG00000134262 |
| C2orf27A | -0.099921561 | 0.933083722 | 0.752310138 | 0.85344398 | 48.82654267 | 45.63652262 | ENSG00000287151 |
| CD99 | -0.100148059 | 0.932937243 | 0.105536247 | 0.215758768 | 1668.057905 | 1556.40955 | ENSG00000002586 |
| MRPS10 | -0.100265666 | 0.932861193 | 0.232605748 | 0.393337074 | 757.5259158 | 706.7886946 | ENSG00000048544 |
| DHRS4L2 | -0.100328437 | 0.932820606 | 0.707274978 | 0.823594399 | 74.05880988 | 69.16288497 | ENSG00000187630 |
| FIS1 | -0.100384946 | 0.932784069 | 0.186714818 | 0.333634267 | 1072.27469 | 1000.163981 | ENSG00000214253 |
| RPS20 | -0.100407566 | 0.932769444 | 0.008926696 | 0.028601876 | 14234.37056 | 13276.10305 | ENSG00000008988 |
| TIMM21 | -0.100454421 | 0.932739151 | 0.229974303 | 0.390187711 | 856.6805751 | 799.2399728 | ENSG00000075336 |
| PCDH8 | -0.100464344 | 0.932732735 | 0.850854712 | 0.915101063 | 17.15850284 | 16.05125758 | ENSG00000136099 |
| NA | -0.100478677 | 0.932723469 | 0.826728055 | 0.901230101 | 21.08347918 | 19.63137843 | ENSG00000233913 |
| FERMT1 | -0.100489503 | 0.932716469 | 0.274854688 | 0.441594219 | 608.4516637 | 567.5608629 | ENSG00000101311 |
| ARAP3 | -0.100581785 | 0.93265681 | 0.116249863 | 0.232849768 | 2631.16113 | 2453.093526 | ENSG00000120318 |
| TMEM11 | -0.100696362 | 0.932582743 | 0.23227727 | 0.392885648 | 715.9716444 | 667.6558268 | ENSG00000178307 |
| LYSMD2 | -0.100803489 | 0.932513497 | 0.763328009 | 0.859724464 | 43.47682312 | 40.59450459 | ENSG00000140280 |
| TTC28 | -0.100967432 | 0.932407534 | 0.386221608 | 0.559606508 | 383.2426624 | 357.2704938 | ENSG00000100154 |
| DRG1 | -0.10101543 | 0.932376514 | 0.107616291 | 0.218900671 | 1850.435562 | 1725.641852 | ENSG00000185721 |
| CYTH1 | -0.101095308 | 0.932324893 | 0.344543386 | 0.517123353 | 455.9811197 | 425.0059214 | ENSG00000108669 |
| CBR1 | -0.101120747 | 0.932308453 | 0.243733204 | 0.406629528 | 848.3243901 | 790.4895862 | ENSG00000159228 |
| CHID1 | -0.101161885 | 0.932281869 | 0.093729541 | 0.196674495 | 1756.234349 | 1637.768788 | ENSG00000177830 |
| CMSS1 | -0.101257891 | 0.932219831 | 0.216559624 | 0.373738746 | 897.8526291 | 837.1085664 | ENSG00000184220 |
| ILRUN | -0.101397903 | 0.932129365 | 0.067255269 | 0.151126546 | 3018.801561 | 2814.233832 | ENSG00000196821 |
| TXLNG | -0.101430575 | 0.932108256 | 0.071206416 | 0.158199122 | 2726.815694 | 2541.446101 | ENSG00000086712 |
| IGF2BP2 | -0.101487251 | 0.932071639 | 0.04923474 | 0.117883502 | 3018.289197 | 2813.87377 | ENSG00000073792 |
| RPS15 | -0.101523543 | 0.932048192 | 0.017066107 | 0.049262646 | 14374.25384 | 13396.71284 | ENSG00000115268 |
| CCNB3 | -0.101537158 | 0.932039396 | 0.926653225 | 0.960672598 | 4.550066219 | 4.272213618 | ENSG00000147082 |
| POLR1C | -0.101648924 | 0.931967193 | 0.426151698 | 0.597752666 | 339.3702182 | 316.341529 | ENSG00000171453 |
| FMNL3 | -0.101745144 | 0.931905038 | 0.233348283 | 0.394142816 | 832.7535179 | 775.7236183 | ENSG00000161791 |
| NEIL2 | -0.101789234 | 0.931876559 | 0.337140952 | 0.510020867 | 494.7299586 | 460.97705 | ENSG00000154328 |
| ANKDD1A | -0.10180294 | 0.931867706 | 0.718868807 | 0.831346931 | 57.99741158 | 54.04167718 | ENSG00000166839 |
| HYAL2 | -0.101856684 | 0.931832992 | 0.222226372 | 0.380657871 | 880.1290351 | 819.9584598 | ENSG00000068001 |
| COA8 | -0.101951687 | 0.931771632 | 0.634061278 | 0.770742252 | 117.2873789 | 109.38417 | ENSG00000256053 |
| NA | -0.102098438 | 0.931676857 | 0.879638497 | 0.933753715 | 10.84206333 | 10.13861232 | ENSG00000227543 |
| QSER1 | -0.102106018 | 0.931671962 | 0.119833076 | 0.23824589 | 1870.067127 | 1743.081517 | ENSG00000060749 |
| MROH1 | -0.102125443 | 0.931659417 | 0.297516614 | 0.467741477 | 520.0213984 | 484.2313124 | ENSG00000179832 |
| NRAS | -0.102238103 | 0.931586667 | 0.09449998 | 0.198044444 | 1768.154242 | 1647.627716 | ENSG00000213281 |
| PAMR1 | -0.102299826 | 0.931546812 | 0.640648253 | 0.775170981 | 119.6751 | 111.5314356 | ENSG00000149090 |
| DNMT3B | -0.102395634 | 0.931484951 | 0.011584203 | 0.035666062 | 40429.30632 | 37658.28278 | ENSG00000088305 |
| SEC23B | -0.102415623 | 0.931472045 | 0.063195148 | 0.143795424 | 2266.614704 | 2111.442567 | ENSG00000101310 |
| RARS1 | -0.102497611 | 0.931419111 | 0.079676939 | 0.172915719 | 2260.069673 | 2105.925956 | ENSG00000113643 |
| KIF2A | -0.10260212 | 0.931351642 | 0.160126402 | 0.297304949 | 1045.841715 | 973.9708458 | ENSG00000068796 |
| CIP2A | -0.102623793 | 0.93133765 | 0.218816864 | 0.376623494 | 789.4592306 | 735.6388554 | ENSG00000163507 |
| APMAP | -0.102642294 | 0.931325706 | 0.08350937 | 0.179616374 | 2026.198687 | 1887.848321 | ENSG00000101474 |
| NA | -0.102670459 | 0.931307525 | 0.81506983 | 0.894245129 | 29.4705643 | 27.49204263 | ENSG00000246596 |
| CARD19 | -0.102746542 | 0.931258412 | 0.374313248 | 0.547575405 | 395.0650879 | 368.0546798 | ENSG00000165233 |
| CCDC40 | -0.102757687 | 0.931251218 | 0.522946853 | 0.685570353 | 196.8266932 | 183.5702388 | ENSG00000141519 |
| SNHG3 | -0.102848443 | 0.931192638 | 0.291262984 | 0.461007074 | 1215.014664 | 1130.837451 | ENSG00000242125 |
| PET100 | -0.102895278 | 0.931162408 | 0.862522964 | 0.922764098 | 12.99787547 | 12.0949731 | ENSG00000229833 |
| FAM229B | -0.102953873 | 0.93112459 | 0.615766842 | 0.758204637 | 109.9304167 | 102.3153674 | ENSG00000203778 |
| SLF1 | -0.103003026 | 0.931092867 | 0.367474092 | 0.541239183 | 411.2779218 | 383.1896088 | ENSG00000133302 |
| LOC100130111 | -0.10316604 | 0.930987666 | 0.7989517 | 0.883907026 | 30.03783307 | 28.00129978 | ENSG00000256802 |
| KIN | -0.103216422 | 0.930955154 | 0.338783803 | 0.511352842 | 459.8594896 | 428.4072046 | ENSG00000151657 |
| ZNF286B | -0.1032519 | 0.930932262 | 0.798001786 | 0.883255061 | 31.35212835 | 29.06767817 | ENSG00000249459 |
| NA | -0.103270664 | 0.930920153 | 0.840390652 | 0.908946853 | 16.95479105 | 15.83265572 | ENSG00000227398 |
| SF3B3 | -0.103452063 | 0.930803111 | 0.007618523 | 0.025062914 | 7945.839179 | 7396.020029 | ENSG00000189091 |
| PHLPP1 | -0.103478257 | 0.930786211 | 0.112280155 | 0.226564806 | 1572.357696 | 1463.293646 | ENSG00000081913 |
| ZBED3 | -0.103591165 | 0.930713368 | 0.62080488 | 0.761828685 | 105.0968538 | 97.86272923 | ENSG00000132846 |
| ADD2 | -0.103597124 | 0.930709524 | 0.0087417 | 0.028124772 | 7161.513193 | 6665.187107 | ENSG00000075340 |
| GOLGA7 | -0.103704766 | 0.930640085 | 0.309743789 | 0.480906836 | 621.8265879 | 578.4727568 | ENSG00000147533 |
| MIER1 | -0.103736803 | 0.930619419 | 0.329966305 | 0.50235814 | 551.5070287 | 513.8598251 | ENSG00000198160 |
| ADH5 | -0.103796376 | 0.930580992 | 0.022609317 | 0.06223692 | 6138.697917 | 5711.975251 | ENSG00000197894 |
| PEX11A | -0.103838778 | 0.930553642 | 0.711555498 | 0.826158117 | 58.11818414 | 54.04382775 | ENSG00000166821 |
| UTP4 | -0.103869313 | 0.930533946 | 0.040639988 | 0.100963572 | 2771.784269 | 2578.862523 | ENSG00000141076 |
| NA | -0.103965215 | 0.930472092 | 0.818243167 | 0.89619415 | 24.32802316 | 22.52370768 | ENSG00000224831 |
| CXCL2 | -0.104118725 | 0.930373091 | 0.902051907 | 0.946601282 | 6.964056769 | 6.46169723 | ENSG00000081041 |
| CHMP2B | -0.104125226 | 0.930368898 | 0.20561478 | 0.359513706 | 779.6940966 | 725.3776236 | ENSG00000083937 |
| WDR36 | -0.10425417 | 0.930285748 | 0.225077162 | 0.384040645 | 794.2290272 | 738.877476 | ENSG00000134987 |
| DDX3X | -0.104262298 | 0.930280507 | 0.009288561 | 0.029563706 | 7580.233607 | 7052.536916 | ENSG00000215301 |
| ANKIB1 | -0.104263659 | 0.93027963 | 0.192389281 | 0.341684921 | 896.4743304 | 834.3996058 | ENSG00000001629 |
| CKMT1B | -0.10430298 | 0.930254275 | 0.570283588 | 0.723296561 | 164.7061239 | 153.0802099 | ENSG00000237289 |
| NA | -0.104385424 | 0.930201116 | 0.904896461 | 0.948448229 | 6.436536354 | 5.986316219 | ENSG00000255142 |

| | | | | | | | |
|---|---|---|---|---|---|---|---|
| FAM89A | -0.104412717 | 0.930183518 | 0.797621123 | 0.882985716 | 30.60095951 | 28.47399718 | ENSG00000182118 |
| ZNF362 | -0.104603886 | 0.93006027 | 0.298618407 | 0.46906992 | 600.8217679 | 558.8365868 | ENSG00000160094 |
| SURF2 | -0.10467176 | 0.930016515 | 0.453414351 | 0.623576157 | 306.0574166 | 284.7020366 | ENSG00000148291 |
| NA | -0.104717562 | 0.929986989 | 0.765659498 | 0.861244162 | 46.67228212 | 43.45940941 | ENSG00000274471 |
| ZBTB9 | -0.104754932 | 0.9299629 | 0.177283056 | 0.32103567 | 864.8099574 | 804.2491929 | ENSG00000213588 |
| FASTKD2 | -0.104757043 | 0.92996154 | 0.191846491 | 0.340895698 | 1004.371698 | 934.141051 | ENSG00000118246 |
| SEMA6B | -0.104772462 | 0.929951601 | 0.114452332 | 0.229965622 | 3102.874464 | 2886.339109 | ENSG00000167680 |
| NA | -0.104788067 | 0.929941542 | 0.918591513 | 0.956469201 | 6.008700579 | 5.590797432 | ENSG00000276744 |
| SORD | -0.104895277 | 0.929872438 | 0.137670126 | 0.264310748 | 1482.020562 | 1377.990021 | ENSG00000140263 |
| PPM1J | -0.105025094 | 0.92978877 | 0.465661124 | 0.635040226 | 256.3785057 | 238.2096682 | ENSG00000155367 |
| MFSD12 | -0.105029229 | 0.929786105 | 0.213738183 | 0.370127763 | 968.9072023 | 901.5921542 | ENSG00000161091 |
| RNF44 | -0.105031836 | 0.929784425 | 0.028783425 | 0.075969496 | 4008.863807 | 3727.439484 | ENSG00000146083 |
| ATG4C | -0.105083898 | 0.929750872 | 0.365706167 | 0.539443161 | 432.609604 | 402.4585084 | ENSG00000125703 |
| INSIG1 | -0.105092998 | 0.929745008 | 0.119829401 | 0.23824589 | 1350.010559 | 1255.483906 | ENSG00000186480 |
| PSMB8 | -0.105107151 | 0.929735887 | 0.425920552 | 0.597689195 | 308.1282993 | 286.5322459 | ENSG00000204264 |
| ZNF345 | -0.105115083 | 0.929730775 | 0.701710635 | 0.820229901 | 59.8161363 | 55.6678483 | ENSG00000251247 |
| STX7 | -0.105176001 | 0.929691518 | 0.281808131 | 0.450277013 | 624.772792 | 581.1838253 | ENSG00000079950 |
| NPR2 | -0.105409422 | 0.929541111 | 0.444602098 | 0.615289998 | 278.2792596 | 258.7979451 | ENSG00000159899 |
| NPRL2 | -0.105447656 | 0.929516477 | 0.407073044 | 0.579327696 | 323.0006054 | 300.3348699 | ENSG00000114388 |
| LOC105372401 | -0.105559388 | 0.929444491 | 0.918931035 | 0.956670189 | 4.577299527 | 4.286734596 | ENSG00000286177 |
| NUDT3 | -0.105618843 | 0.929406189 | 0.484606755 | 0.652511755 | 269.890674 | 251.2295537 | ENSG00000272325 |
| LOC728554 | -0.105676268 | 0.929369196 | 0.299473577 | 0.469954021 | 559.639527 | 520.2215236 | ENSG00000170089 |
| KCNB2 | -0.105759185 | 0.929315783 | 0.79571285 | 0.881277793 | 28.61153826 | 26.49934732 | ENSG00000182674 |
| CLTA | -0.105885163 | 0.929234637 | 0.027912938 | 0.074128372 | 4152.11367 | 3858.113814 | ENSG00000122705 |
| NA | -0.105941196 | 0.929198548 | 0.906278028 | 0.949343894 | 5.611564737 | 5.224853639 | ENSG00000230457 |
| LSM14A | -0.106025223 | 0.92914443 | 0.022655818 | 0.06234713 | 4766.439446 | 4429.279628 | ENSG00000257103 |
| HNRNPA2B1 | -0.106149142 | 0.929064625 | 0.00338999 | 0.012480211 | 43761.9562 | 40657.65677 | ENSG00000122566 |
| BICD1 | -0.106202937 | 0.929029983 | 0.252634461 | 0.417265794 | 695.0933917 | 645.9787927 | ENSG00000151746 |
| REEP1 | -0.106218363 | 0.929020049 | 0.789107519 | 0.877033655 | 29.40907642 | 27.35115242 | ENSG00000068615 |
| CASTOR3 | -0.106282337 | 0.928978854 | 0.573589863 | 0.72591498 | 136.6619425 | 126.770162 | ENSG00000239521 |
| RASD1 | -0.106344664 | 0.928938722 | 0.736844957 | 0.843639151 | 45.71836539 | 42.54656421 | ENSG00000108551 |
| NDUFC1 | -0.106424838 | 0.928887099 | 0.268321822 | 0.43470554 | 616.5142276 | 573.203795 | ENSG00000109390 |
| LINC01664 | -0.106444933 | 0.928874161 | 0.917402318 | 0.955800827 | 5.380619642 | 5.012703499 | ENSG00000235478 |
| RPL7 | -0.106471381 | 0.928857133 | 0.004448778 | 0.015857608 | 27885.77339 | 25901.20388 | ENSG00000147604 |
| DDOST | -0.10658569 | 0.92878354 | 0.008569926 | 0.02765977 | 6754.759604 | 6273.896071 | ENSG00000244038 |
| CYB5R4 | -0.106597082 | 0.928776206 | 0.469950931 | 0.638843169 | 254.2210668 | 236.037377 | ENSG00000065615 |
| CSDE1 | -0.106597754 | 0.928775773 | 0.002794156 | 0.010552478 | 13560.26956 | 12595.24767 | ENSG00000009307 |
| SMIM8 | -0.106642831 | 0.928746754 | 0.613695341 | 0.756749457 | 121.5088184 | 112.7234019 | ENSG00000111850 |
| SCMH1 | -0.106665499 | 0.928732162 | 0.330994732 | 0.503526487 | 432.114223 | 401.5714885 | ENSG00000010803 |
| GLOD4 | -0.106794657 | 0.928649021 | 0.088963042 | 0.189028093 | 1683.923014 | 1563.859362 | ENSG00000167699 |
| MFSD4B | -0.106826808 | 0.928628325 | 0.598669736 | 0.74530235 | 113.8789595 | 105.782816 | ENSG00000173214 |
| NOL9 | -0.106920814 | 0.928567818 | 0.285513636 | 0.45451754 | 570.9612139 | 529.8092302 | ENSG00000162408 |
| PSKH1 | -0.106924366 | 0.928565532 | 0.21538786 | 0.372316059 | 823.3029823 | 764.5705202 | ENSG00000159792 |
| SEZ6 | -0.106965914 | 0.92853879 | 0.676930467 | 0.802129339 | 74.4351701 | 69.21369313 | ENSG00000063015 |
| PSMB1 | -0.107072184 | 0.928470396 | 0.023527836 | 0.064315603 | 3759.656695 | 3490.688671 | ENSG00000008018 |
| DIAPH1 | -0.107075768 | 0.928468089 | 0.019675462 | 0.055450598 | 4217.141305 | 3915.352149 | ENSG00000131504 |
| NDUFA12 | -0.107213087 | 0.92837972 | 0.152440152 | 0.286257468 | 1024.517655 | 951.4191381 | ENSG00000184752 |
| ORMDL1 | -0.107236913 | 0.928364388 | 0.388100006 | 0.561526829 | 366.724043 | 340.8301571 | ENSG00000128699 |
| PKD1P6-NPIPP1 | -0.107434376 | 0.92823733 | 0.775884023 | 0.868533369 | 34.08203038 | 31.68628413 | ENSG00000270580 |
| NBN | -0.107440986 | 0.928233077 | 0.304241207 | 0.474962946 | 701.8340883 | 651.5496956 | ENSG00000104320 |
| RABEPK | -0.107470263 | 0.928214241 | 0.154602909 | 0.289501526 | 982.8372294 | 912.0290006 | ENSG00000136933 |
| CEP57L1 | -0.107489578 | 0.928201814 | 0.642674906 | 0.776758449 | 110.0260711 | 102.32045 | ENSG00000183137 |
| NA | -0.107539263 | 0.928169848 | 0.921754539 | 0.95821344 | 4.612905543 | 4.307707295 | ENSG00000230910 |
| ZC3H14 | -0.107552642 | 0.928161241 | 0.101129274 | 0.208874179 | 1655.696469 | 1536.296623 | ENSG00000100722 |
| EPB41L3 | -0.10770285 | 0.928064609 | 0.29425224 | 0.464138999 | 475.0260926 | 440.8571082 | ENSG00000082397 |
| A4GALT | -0.107710356 | 0.92805978 | 0.467840446 | 0.63706857 | 214.9004405 | 199.4717434 | ENSG00000128274 |
| TRIM74 | -0.107710906 | 0.928059427 | 0.690875372 | 0.812514589 | 64.67673236 | 60.05164971 | ENSG00000174353 |
| SLC12A5 | -0.107760781 | 0.928027343 | 0.800770058 | 0.885210591 | 26.92128309 | 24.92130074 | ENSG00000124140 |
| RPS19 | -0.10782756 | 0.927984388 | 0.009422587 | 0.029924038 | 17561.33636 | 16295.91186 | ENSG00000105372 |
| TSEN2 | -0.107893123 | 0.927942217 | 0.469389046 | 0.638304249 | 265.3240114 | 245.8503976 | ENSG00000154743 |
| FAM81A | -0.108020817 | 0.927860088 | 0.807227097 | 0.888970029 | 26.41453829 | 24.45426156 | ENSG00000157470 |
| PIH1D1 | -0.108037885 | 0.927849111 | 0.107797483 | 0.219150727 | 2048.749724 | 1900.613366 | ENSG00000104872 |
| SUPT7L | -0.108075838 | 0.927824702 | 0.334925127 | 0.507544432 | 458.5796732 | 425.7487917 | ENSG00000119760 |
| LOC93622 | -0.108087419 | 0.927817254 | 0.315241316 | 0.486505618 | 446.6557872 | 414.5795978 | ENSG00000170846 |
| ERCC6L | -0.108283289 | 0.927691297 | 0.192253352 | 0.341506393 | 780.4537134 | 724.2943011 | ENSG00000186871 |
| PRMT7 | -0.108343496 | 0.927652582 | 0.116854449 | 0.233793734 | 1168.439429 | 1084.111384 | ENSG00000132600 |
| HSPA9 | -0.108376534 | 0.927631339 | 0.005305508 | 0.018464503 | 12422.62295 | 11524.7955 | ENSG00000113013 |
| PPM1K | -0.108405603 | 0.927612648 | 0.571931984 | 0.724529807 | 127.995591 | 118.6556628 | ENSG00000163644 |
| LRWD1 | -0.108411309 | 0.92760898 | 0.170514339 | 0.311879408 | 925.1017013 | 858.6124269 | ENSG00000161036 |
| NA | -0.108545719 | 0.927522562 | 0.557821144 | 0.713430492 | 133.2269655 | 123.5164506 | ENSG00000228709 |
| ZFYVE21 | -0.108547887 | 0.927521168 | 0.197010975 | 0.347907242 | 835.7030892 | 775.5874954 | ENSG00000100711 |
| LNX2 | -0.108552443 | 0.927518239 | 0.310330182 | 0.481279598 | 424.6341728 | 393.9224535 | ENSG00000139517 |
| LAMA1 | -0.108574588 | 0.927504002 | 0.060153562 | 0.138222741 | 2769.813494 | 2568.516236 | ENSG00000101680 |
| RAB32 | -0.108847813 | 0.927328363 | 0.564406958 | 0.718750324 | 130.652366 | 121.1500501 | ENSG00000118508 |
| HPS1 | -0.108878106 | 0.927308892 | 0.23616359 | 0.397542042 | 816.8538045 | 757.4151069 | ENSG00000107521 |
| NA | -0.108941148 | 0.927268372 | 0.80972513 | 0.890395056 | 24.99150995 | 23.14159643 | ENSG00000235023 |
| SLC16A13 | -0.108988687 | 0.927237818 | 0.628427382 | 0.766356446 | 98.14731507 | 91.04558062 | ENSG00000174327 |

| | | | | | | | |
|---|---|---|---|---|---|---|---|
| NA | -0.109041602 | 0.927203809 | 0.868256292 | 0.926463417 | 11.26922338 | 10.45830957 | ENSG00000257511 |
| PARL | -0.109268449 | 0.927058028 | 0.731952297 | 0.839881737 | 65.55693433 | 60.85908626 | ENSG00000175193 |
| PNN | -0.109329454 | 0.927018828 | 0.042538278 | 0.1048028 | 4578.743053 | 4243.667957 | ENSG00000100941 |
| NA | -0.109395341 | 0.926976493 | 0.751699491 | 0.853002052 | 38.95802041 | 36.16634674 | ENSG00000224043 |
| INVS | -0.10942858 | 0.926955136 | 0.539144642 | 0.698545187 | 168.2919857 | 155.699361 | ENSG00000119509 |
| WDPCP | -0.109528139 | 0.92689117 | 0.471704774 | 0.64060882 | 214.8591646 | 199.0743466 | ENSG00000143951 |
| NA | -0.109674279 | 0.926797283 | 0.791887874 | 0.878982065 | 27.54983959 | 25.57144811 | ENSG00000255753 |
| CCT5 | -0.109776874 | 0.926731378 | 0.007020246 | 0.023438484 | 11785.57742 | 10920.82422 | ENSG00000150753 |
| NA | -0.109777987 | 0.926730663 | 0.80944741 | 0.890310719 | 23.21763983 | 21.52884968 | ENSG00000268034 |
| AGER | -0.109867955 | 0.926672873 | 0.622203316 | 0.762199274 | 113.0244393 | 104.9194524 | ENSG00000204305 |
| CBX3 | -0.109989777 | 0.926594628 | 0.033295901 | 0.085470556 | 3899.71148 | 3613.977621 | ENSG00000122565 |
| ZNRF3 | -0.110023646 | 0.926572875 | 0.240559734 | 0.402763221 | 689.6384426 | 638.4940806 | ENSG00000183579 |
| PBDC1 | -0.110157927 | 0.926486637 | 0.255528551 | 0.420561494 | 744.9965146 | 690.3000276 | ENSG00000102390 |
| DDX42 | -0.110204145 | 0.926456957 | 0.020113497 | 0.056478816 | 4102.751659 | 3800.505737 | ENSG00000198231 |
| FADS6 | -0.110209838 | 0.926453301 | 0.82144806 | 0.898330756 | 21.08627009 | 19.5896935 | ENSG00000172782 |
| YWHAH | -0.110243417 | 0.926431738 | 0.034092271 | 0.087132221 | 3125.621137 | 2896.357488 | ENSG00000128245 |
| IRS1 | -0.110253332 | 0.92642537 | 0.119928348 | 0.238390934 | 1288.571564 | 1193.998662 | ENSG00000169047 |
| CASP2 | -0.110381431 | 0.926343116 | 0.24706367 | 0.410587979 | 580.836306 | 537.8521848 | ENSG00000106144 |
| CCT6P3 | -0.110389499 | 0.926337935 | 0.820376511 | 0.897565975 | 25.83255126 | 23.88530296 | ENSG00000234585 |
| AP4S1 | -0.110440338 | 0.926305293 | 0.531187873 | 0.692094101 | 176.4448224 | 163.5231929 | ENSG00000100478 |
| PHTF2 | -0.110681208 | 0.926150651 | 0.354988644 | 0.528182278 | 393.6190566 | 364.4570876 | ENSG00000006576 |
| CDC14B | -0.110708241 | 0.926133297 | 0.097068452 | 0.202393807 | 1886.447301 | 1747.805728 | ENSG00000081377 |
| NA | -0.110714298 | 0.926129409 | 0.890166566 | 0.93936831 | 7.086944514 | 6.531067048 | ENSG00000230530 |
| LINC01668 | -0.110807349 | 0.926069678 | 0.911117321 | 0.951993281 | 4.909480864 | 4.564486445 | ENSG00000283051 |
| AP3M2 | -0.110857052 | 0.926037773 | 0.332111651 | 0.50474796 | 459.6465903 | 425.5377382 | ENSG00000070718 |
| PALM3 | -0.110865082 | 0.926032619 | 0.118249714 | 0.235813816 | 1741.232986 | 1612.220027 | ENSG00000187867 |
| FKBP4 | -0.11088951 | 0.92601694 | 0.003782651 | 0.013770735 | 9955.182109 | 9218.59759 | ENSG00000004478 |
| NA | -0.110896648 | 0.926012358 | 0.791618134 | 0.878760814 | 27.41877929 | 25.42866779 | ENSG00000238072 |
| KIDINS220 | -0.110951791 | 0.925976964 | 0.099571138 | 0.206562729 | 1325.289556 | 1227.297638 | ENSG00000134313 |
| MRPL16 | -0.111244791 | 0.925788925 | 0.130266564 | 0.253247231 | 1550.146474 | 1434.414425 | ENSG00000166902 |
| GNAI1 | -0.111273973 | 0.925770199 | 0.31074931 | 0.481688017 | 617.2584631 | 571.9924736 | ENSG00000127955 |
| GIN1 | -0.111294869 | 0.92575679 | 0.680030879 | 0.804468981 | 67.1244139 | 62.05964267 | ENSG00000145723 |
| PAK3 | -0.111299139 | 0.92575405 | 0.482824956 | 0.650794873 | 218.7474958 | 202.2968534 | ENSG00000077264 |
| SPRED2 | -0.111307725 | 0.92574854 | 0.250466134 | 0.414635046 | 541.6131371 | 501.6513984 | ENSG00000198369 |
| SIN3A | -0.111320326 | 0.925740455 | 0.041331516 | 0.102339402 | 2561.07378 | 2370.297064 | ENSG00000169375 |
| NCK2 | -0.111339945 | 0.925727866 | 0.109464097 | 0.221814274 | 1603.17128 | 1484.853143 | ENSG00000071051 |
| NA | -0.111360686 | 0.925714557 | 0.849993691 | 0.91461559 | 15.54436578 | 14.42104577 | ENSG00000287001 |
| SLC38A4 | -0.111429372 | 0.925670485 | 0.414797846 | 0.586948232 | 333.4026302 | 308.6510085 | ENSG00000139209 |
| RNF157 | -0.111480723 | 0.925637538 | 0.383888601 | 0.557230624 | 330.7509487 | 306.1012698 | ENSG00000141576 |
| ZFR2 | -0.111497387 | 0.925626846 | 0.389702668 | 0.562916823 | 327.1621086 | 302.7399929 | ENSG00000105278 |
| ARFGAP1 | -0.111673492 | 0.925513865 | 0.060940788 | 0.139654162 | 2950.279637 | 2729.608991 | ENSG00000101199 |
| LGALS3 | -0.111681807 | 0.925508531 | 0.708050305 | 0.82397288 | 63.06952855 | 58.37035717 | ENSG00000131981 |
| SLCO4C1 | -0.11171647 | 0.925486294 | 0.523796183 | 0.686217465 | 164.8145447 | 152.6967889 | ENSG00000173930 |
| ERI2 | -0.111774055 | 0.925449354 | 0.529735446 | 0.690716585 | 145.4047754 | 134.5071406 | ENSG00000196678 |
| NCBP2 | -0.111784194 | 0.92544285 | 0.047249717 | 0.114179417 | 2240.938493 | 2073.882603 | ENSG00000114503 |
| PWP1 | -0.111793883 | 0.925436635 | 0.071313044 | 0.158381295 | 1618.414914 | 1497.974972 | ENSG00000136045 |
| NA | -0.111871293 | 0.925386981 | 0.846281624 | 0.912075406 | 16.08113478 | 14.90690708 | ENSG00000271870 |
| TIMP3 | -0.112099154 | 0.925240836 | 0.598683594 | 0.74530235 | 100.0232973 | 92.61260175 | ENSG00000100234 |
| RAP1A | -0.112152736 | 0.925206473 | 0.217928771 | 0.375504184 | 605.2918107 | 560.0271412 | ENSG00000116473 |
| ZNF684 | -0.112168075 | 0.925196636 | 0.61214829 | 0.755482865 | 101.2001868 | 93.67896802 | ENSG00000117010 |
| CRTAP | -0.112279612 | 0.92512511 | 0.034453162 | 0.087926437 | 2643.422844 | 2446.13906 | ENSG00000170275 |
| NA | -0.112311521 | 0.925104649 | 0.728834621 | 0.837400159 | 49.92105646 | 46.21756063 | ENSG00000214389 |
| ZCCHC8 | -0.112356352 | 0.925075902 | 0.258768026 | 0.423927095 | 628.9490376 | 581.5944713 | ENSG00000033030 |
| RPL35A | -0.112363778 | 0.925071141 | 0.006122795 | 0.020886796 | 9996.362392 | 9246.575081 | ENSG00000182899 |
| MGST1 | -0.112370026 | 0.925067134 | 0.042160354 | 0.104017828 | 3634.592144 | 3363.441765 | ENSG00000008394 |
| CENPH | -0.112389702 | 0.925054518 | 0.226021291 | 0.385038567 | 685.112745 | 634.3390277 | ENSG00000153044 |
| PPP2R5B | -0.11240724 | 0.925043272 | 0.551918508 | 0.708528644 | 132.8555114 | 122.8386184 | ENSG00000068971 |
| INTS1 | -0.112412242 | 0.925040065 | 0.035866466 | 0.090991582 | 5905.863892 | 5463.224741 | ENSG00000164880 |
| NA | -0.112424694 | 0.925032081 | 0.576107347 | 0.728193345 | 116.862334 | 108.1675056 | ENSG00000124224 |
| DNAJC19 | -0.112437839 | 0.925023653 | 0.174155914 | 0.316978587 | 841.3796454 | 778.0821979 | ENSG00000205981 |
| VBP1 | -0.112439389 | 0.925022659 | 0.126497049 | 0.247917766 | 1183.847228 | 1095.71992 | ENSG00000155959 |
| COPZ1 | -0.11254761 | 0.924953273 | 0.100796647 | 0.20834331 | 1975.735001 | 1826.890823 | ENSG00000111481 |
| ZNF85 | -0.112548715 | 0.924952564 | 0.482872202 | 0.650797082 | 191.0747316 | 176.9146926 | ENSG00000105750 |
| GCAT | -0.11266353 | 0.924878957 | 0.248962912 | 0.412890606 | 698.9463682 | 646.4848822 | ENSG00000100116 |
| CMTM3 | -0.112686278 | 0.924864373 | 0.320599159 | 0.492018963 | 459.8733192 | 425.7049989 | ENSG00000140931 |
| GHDC | -0.112780855 | 0.924803745 | 0.468716203 | 0.637569051 | 230.9012268 | 213.5666218 | ENSG00000167925 |
| ADGRA1 | -0.112936046 | 0.924704269 | 0.876087704 | 0.931624933 | 9.884591955 | 9.168767714 | ENSG00000197177 |
| TCP1 | -0.113021447 | 0.924649532 | 0.005855203 | 0.020123415 | 7442.302862 | 6882.203949 | ENSG00000120438 |
| LGMN | -0.113050917 | 0.924630644 | 0.074521834 | 0.164053485 | 1598.095957 | 1477.026912 | ENSG00000100600 |
| SLC41A2 | -0.113175417 | 0.924550856 | 0.759039732 | 0.857004645 | 37.12880779 | 34.37265449 | ENSG00000136052 |
| ITGA4 | -0.113227591 | 0.92451742 | 0.627008242 | 0.765399987 | 90.62782886 | 83.9081896 | ENSG00000115232 |
| ESPN | -0.113344561 | 0.924442466 | 0.272148989 | 0.439080963 | 518.6002482 | 479.8426463 | ENSG00000187017 |
| MCMBP | -0.113385182 | 0.924416437 | 0.041480011 | 0.102641213 | 2232.976677 | 2064.194085 | ENSG00000197771 |
| OSER1 | -0.113405358 | 0.924403509 | 0.226279196 | 0.385443502 | 614.0168591 | 567.4877036 | ENSG00000132823 |
| NA | -0.113446943 | 0.924376864 | 0.78020556 | 0.870840926 | 27.97699964 | 25.86157036 | ENSG00000273253 |
| UQCR11 | -0.113536223 | 0.924319662 | 0.44492875 | 0.615542205 | 273.5121659 | 252.8229574 | ENSG00000127540 |
| CPSF1 | -0.113644639 | 0.924250203 | 0.020029369 | 0.056275352 | 4772.778863 | 4411.317931 | ENSG00000071894 |

| | | | | | | | |
|---|---|---|---|---|---|---|---|
| CUL2 | -0.1136554 | 0.92424331 | 0.208530522 | 0.363425991 | 813.7756387 | 752.5120216 | ENSG00000108094 |
| LEMD1-DT | -0.113696671 | 0.92421687 | 0.745232166 | 0.849009255 | 41.435526 | 38.37974713 | ENSG00000224717 |
| RAB35 | -0.113747865 | 0.924184075 | 0.107613572 | 0.218900671 | 1644.81565 | 1520.2723 | ENSG00000111737 |
| NA | -0.113890106 | 0.924092961 | 0.699056284 | 0.818294586 | 60.32990236 | 55.65468439 | ENSG00000259295 |
| SUOX | -0.113966163 | 0.924044245 | 0.355282442 | 0.528349799 | 449.9633707 | 415.2967403 | ENSG00000139531 |
| RTL8A | -0.114014012 | 0.924013598 | 0.239708205 | 0.401825771 | 582.7128149 | 538.3786587 | ENSG00000203950 |
| TOB2 | -0.114207416 | 0.923889735 | 0.062544593 | 0.142550536 | 2064.287182 | 1907.532992 | ENSG00000183864 |
| EXOC3L1 | -0.11424347 | 0.923866647 | 0.677272205 | 0.802386422 | 69.11104424 | 63.92807196 | ENSG00000179044 |
| SNRPA1 | -0.11424958 | 0.923862734 | 0.071870555 | 0.159307695 | 2391.61911 | 2209.332491 | ENSG00000131876 |
| CTTN | -0.114256893 | 0.923858051 | 0.01538519 | 0.04520879 | 5277.31066 | 4875.613961 | ENSG00000085733 |
| UBE2Z | -0.11431978 | 0.923817781 | 0.034308472 | 0.087626734 | 2469.05525 | 2280.963126 | ENSG00000159202 |
| ACACA | -0.114355885 | 0.923794662 | 0.007626918 | 0.025073458 | 6039.200075 | 5578.940191 | ENSG00000278540 |
| ZNF202 | -0.114409759 | 0.923760166 | 0.135385238 | 0.260780492 | 989.7900345 | 914.0200616 | ENSG00000166261 |
| TTLL5 | -0.114451739 | 0.923733286 | 0.210314959 | 0.365925206 | 704.5759418 | 651.4028872 | ENSG00000119685 |
| VEZF1 | -0.114459005 | 0.923728634 | 0.145340625 | 0.275581482 | 943.2098153 | 871.6339421 | ENSG00000136451 |
| GFM1 | -0.114595506 | 0.923641239 | 0.127518798 | 0.249388192 | 944.8581317 | 872.7027621 | ENSG00000168827 |
| TVP23C | -0.114653848 | 0.923603888 | 0.760778197 | 0.857754074 | 32.04284845 | 29.62936101 | ENSG00000175106 |
| CBSL | -0.114694678 | 0.92357775 | 0.178083662 | 0.32204866 | 897.4312751 | 828.3335126 | ENSG00000274276 |
| NA | -0.114732029 | 0.923553839 | 0.886568496 | 0.937396155 | 7.119759628 | 6.535368196 | ENSG00000234645 |
| GRN | -0.114884023 | 0.923456544 | 0.051148683 | 0.121542059 | 2675.208853 | 2470.905804 | ENSG00000030582 |
| RHOBTB3 | -0.115023363 | 0.923367358 | 0.386170414 | 0.559606508 | 281.6622485 | 260.3539529 | ENSG00000164292 |
| TMEM150A | -0.115086119 | 0.923327193 | 0.308045497 | 0.479079787 | 380.8698228 | 351.8071623 | ENSG00000168890 |
| NA | -0.115100134 | 0.923318223 | 0.595673436 | 0.743377944 | 111.4382992 | 102.9805688 | ENSG00000224861 |
| AK9 | -0.115106263 | 0.923314301 | 0.786704305 | 0.875421825 | 29.27290988 | 27.11372843 | ENSG00000155085 |
| NTM | -0.115107368 | 0.923313593 | 0.424646551 | 0.596335198 | 305.6821693 | 282.2237267 | ENSG00000182667 |
| PHOX2A | -0.115174748 | 0.923270472 | 0.898700448 | 0.944271805 | 5.720497967 | 5.277551906 | ENSG00000165462 |
| NA | -0.115264729 | 0.923212889 | 0.863571504 | 0.923507838 | 9.955128266 | 9.219315408 | ENSG00000287225 |
| KRI1 | -0.115266192 | 0.923211953 | 0.137376042 | 0.26393001 | 1572.140895 | 1451.326751 | ENSG00000129347 |
| GAB1 | -0.115315284 | 0.923180538 | 0.430094663 | 0.601577447 | 283.216374 | 261.1208069 | ENSG00000109458 |
| PDIA5 | -0.115427875 | 0.923108494 | 0.18765044 | 0.334840869 | 726.8338446 | 670.7390609 | ENSG00000065485 |
| PCMTD1 | -0.115428037 | 0.923108391 | 0.358413524 | 0.531531004 | 337.1843818 | 311.598738 | ENSG00000168300 |
| NA | -0.115428774 | 0.923107919 | 0.536245014 | 0.696003671 | 239.3446808 | 221.0963149 | ENSG00000237973 |
| SLITRK5 | -0.115461961 | 0.923086685 | 0.528561521 | 0.689851434 | 141.7686774 | 130.8893755 | ENSG00000165300 |
| ZNF416 | -0.115467265 | 0.923083291 | 0.511009334 | 0.675474926 | 155.26301 | 143.1497191 | ENSG00000083817 |
| SLC26A11 | -0.11547844 | 0.923076141 | 0.291323861 | 0.461065601 | 414.266915 | 382.5114918 | ENSG00000181045 |
| MTA3 | -0.115540857 | 0.923036206 | 0.02267377 | 0.062367057 | 3093.962619 | 2855.857028 | ENSG00000057935 |
| FAM160A2 | -0.115555788 | 0.923026653 | 0.22850978 | 0.388180888 | 536.9751134 | 495.674248 | ENSG00000051009 |
| NA | -0.115595815 | 0.923001044 | 0.871547252 | 0.928639494 | 10.27403081 | 9.497067257 | ENSG00000205940 |
| TEX9 | -0.115662711 | 0.922958246 | 0.755787403 | 0.855125795 | 44.9185119 | 41.64204874 | ENSG00000151575 |
| POU5F1P3 | -0.115662714 | 0.922958245 | 0.63521312 | 0.77136449 | 89.64120916 | 82.69580793 | ENSG00000235602 |
| BCAT1 | -0.115680114 | 0.922947113 | 0.024506375 | 0.066491206 | 5010.949231 | 4625.865087 | ENSG00000060982 |
| NOP58 | -0.115700771 | 0.922933898 | 0.051887937 | 0.122893825 | 2309.541815 | 2132.16475 | ENSG00000055044 |
| FLYWCH2 | -0.115764976 | 0.922892826 | 0.508902308 | 0.673520474 | 227.8096459 | 210.4239956 | ENSG00000162076 |
| ASB8 | -0.115777848 | 0.922884591 | 0.632341545 | 0.769378791 | 84.68284349 | 78.09529477 | ENSG00000177981 |
| NAA50 | -0.115821888 | 0.92285642 | 0.031185975 | 0.081079503 | 4532.213668 | 4183.391344 | ENSG00000121579 |
| TMED7 | -0.115845179 | 0.922841521 | 0.358790595 | 0.531844887 | 500.8233275 | 462.7691063 | ENSG00000134970 |
| DACH2 | -0.11594123 | 0.922780082 | 0.683215643 | 0.806801725 | 58.42891413 | 53.82737646 | ENSG00000126733 |
| SHMT1 | -0.115951993 | 0.922773198 | 0.169078623 | 0.309870389 | 1001.612109 | 924.1271846 | ENSG00000176974 |
| IDE | -0.115967128 | 0.922763518 | 0.10086751 | 0.208422787 | 1073.225452 | 990.4474577 | ENSG00000119912 |
| RPS29 | -0.115982281 | 0.922753826 | 0.089939399 | 0.190588295 | 4013.167197 | 3702.918741 | ENSG00000213741 |
| FAIM | -0.115986885 | 0.922750881 | 0.584803966 | 0.73470878 | 112.3210039 | 103.821621 | ENSG00000158234 |
| NPHP4 | -0.115993749 | 0.922746491 | 0.302701658 | 0.473402325 | 451.4492014 | 416.4897363 | ENSG00000131697 |
| FAM83G | -0.116049425 | 0.922710881 | 0.195181952 | 0.345436241 | 1045.034631 | 964.5437425 | ENSG00000188522 |
| CLDN10 | -0.116073846 | 0.922695262 | 0.245961105 | 0.409319847 | 542.230855 | 500.197843 | ENSG00000134873 |
| TSPAN4 | -0.116169543 | 0.92263406 | 0.101146272 | 0.208886921 | 1634.839058 | 1508.140797 | ENSG00000214063 |
| MTERF3 | -0.116226723 | 0.922597493 | 0.241103642 | 0.403358803 | 497.8629417 | 459.4406699 | ENSG00000156469 |
| MT1F | -0.116227347 | 0.922597094 | 0.3611887 | 0.534250207 | 391.5129541 | 360.9909719 | ENSG00000198417 |
| TACO1 | -0.116267111 | 0.922571666 | 0.232627907 | 0.393337074 | 736.6357358 | 679.3047697 | ENSG00000136463 |
| EPHB4 | -0.116398374 | 0.92248773 | 0.022235926 | 0.061340376 | 4752.937327 | 4384.55053 | ENSG00000196411 |
| NA | -0.116426455 | 0.922469774 | 0.835624145 | 0.906888184 | 15.0691968 | 13.93759551 | ENSG00000276449 |
| SNRPD2 | -0.116431338 | 0.922466652 | 0.093466372 | 0.196283452 | 5827.923757 | 5375.199076 | ENSG00000125743 |
| CCNB1 | -0.116654656 | 0.922323872 | 0.017005572 | 0.049102603 | 5762.992012 | 5314.458701 | ENSG00000134057 |
| TRNP1 | -0.116793519 | 0.922235101 | 0.160639311 | 0.297913186 | 994.5128997 | 917.4785863 | ENSG00000253368 |
| RANBP17 | -0.116909011 | 0.922161276 | 0.502461127 | 0.668228848 | 170.6541004 | 157.3279432 | ENSG00000204764 |
| LINC02693 | -0.117039637 | 0.922077784 | 0.355032896 | 0.528182278 | 322.4368785 | 297.4893308 | ENSG00000212719 |
| CIAO1 | -0.117056299 | 0.922067135 | 0.066660369 | 0.150086972 | 1730.776029 | 1595.858941 | ENSG00000144021 |
| BRD1 | -0.117059177 | 0.922065296 | 0.088901917 | 0.188960653 | 1269.429595 | 1170.355866 | ENSG00000100425 |
| NDUFAB1 | -0.117113858 | 0.922030348 | 0.064068432 | 0.145251394 | 1528.39827 | 1408.774722 | ENSG00000004779 |
| PGBD1 | -0.117156569 | 0.922003052 | 0.489090451 | 0.655621145 | 201.7268744 | 185.9777912 | ENSG00000137338 |
| DNAJC24 | -0.117202225 | 0.921973874 | 0.417620156 | 0.589687486 | 245.399488 | 226.4550619 | ENSG00000170946 |
| DNAAF2 | -0.117248222 | 0.92194448 | 0.143482172 | 0.273023731 | 1008.653762 | 929.9145509 | ENSG00000165506 |
| NA | -0.117306245 | 0.921907401 | 0.870091762 | 0.927447241 | 10.10804022 | 9.33923292 | ENSG00000275888 |
| NKD2 | -0.117311805 | 0.921903848 | 0.359817148 | 0.532875218 | 329.828771 | 304.0451707 | ENSG00000145506 |
| DPP7 | -0.117330519 | 0.92189189 | 0.347486184 | 0.52027691 | 311.7961363 | 287.3418573 | ENSG00000176978 |
| MIR4500HG | -0.117337957 | 0.921887137 | 0.900366 | 0.945352447 | 5.636007141 | 5.187209388 | ENSG00000228824 |
| SH3BP5L | -0.117529557 | 0.921764712 | 0.092239173 | 0.194292233 | 1260.310578 | 1161.730238 | ENSG00000175137 |
| HMGN3-AS1 | -0.117533502 | 0.921762191 | 0.860554953 | 0.921699125 | 10.61670004 | 9.78718951 | ENSG00000270362 |

| | | | | | | | |
|---|---|---|---|---|---|---|---|
| NA | -0.117537137 | 0.921759869 | 0.06009888 | 0.138146165 | 1968.163855 | 1814.277258 | ENSG00000198618 |
| PPP1R1B | -0.117576635 | 0.921734634 | 0.344009199 | 0.516676027 | 314.3027903 | 289.7999453 | ENSG00000131771 |
| GALNT7 | -0.117672891 | 0.921673138 | 0.094362639 | 0.197778104 | 1273.945234 | 1174.739643 | ENSG00000109586 |
| CEBPZ | -0.11771437 | 0.921646639 | 0.058270093 | 0.134766618 | 2583.126599 | 2381.301359 | ENSG00000115816 |
| EXT1 | -0.117831415 | 0.921571869 | 0.164386896 | 0.303288793 | 843.6856779 | 777.6030183 | ENSG00000182197 |
| SMARCD2 | -0.117906298 | 0.921524037 | 0.019717829 | 0.055553768 | 2913.053061 | 2684.976558 | ENSG00000108604 |
| HS6ST2 | -0.117919648 | 0.921515509 | 0.259931887 | 0.425271432 | 477.4556646 | 439.9760007 | ENSG00000171004 |
| DUSP8 | -0.117931807 | 0.921507743 | 0.725222277 | 0.835021231 | 45.00579363 | 41.55197879 | ENSG00000184545 |
| SIRT7 | -0.117939626 | 0.921502749 | 0.25741682 | 0.422551725 | 453.9364312 | 418.4716192 | ENSG00000187531 |
| TLN2 | -0.118004087 | 0.921461576 | 0.107826918 | 0.219167044 | 1085.794704 | 1000.760855 | ENSG00000171914 |
| MED15 | -0.118013128 | 0.921455802 | 0.069153257 | 0.154545351 | 1451.147455 | 1336.998865 | ENSG00000099917 |
| SSBP3 | -0.118186972 | 0.921344773 | 0.031186041 | 0.081079503 | 3435.680548 | 3165.681339 | ENSG00000157216 |
| FOXN3-AS1 | -0.118332473 | 0.921251857 | 0.798393592 | 0.883638026 | 23.70185724 | 21.7746155 | ENSG00000258920 |
| PUM2 | -0.118417015 | 0.921197873 | 0.065020726 | 0.146957838 | 1909.930016 | 1760.020618 | ENSG00000055917 |
| TAOK2 | -0.1184389 | 0.921183899 | 0.078222564 | 0.170276629 | 1612.645307 | 1485.377775 | ENSG00000149930 |
| NBPF20 | -0.118477182 | 0.921159455 | 0.859233153 | 0.920859264 | 10.35784591 | 9.549765524 | ENSG00000162825 |
| DUSP11 | -0.118538827 | 0.921120096 | 0.354334515 | 0.527428019 | 322.7988839 | 297.4944575 | ENSG00000144048 |
| GPKOW | -0.118558607 | 0.921107467 | 0.253766701 | 0.418437253 | 497.463103 | 458.0347303 | ENSG00000068394 |
| ABRACL | -0.118581163 | 0.921093066 | 0.070798216 | 0.157473583 | 1497.053965 | 1379.370117 | ENSG00000146386 |
| RFWD3 | -0.118642318 | 0.921054022 | 0.026455142 | 0.070940525 | 2715.697533 | 2501.451778 | ENSG00000168411 |
| PPP1R1A | -0.118674496 | 0.921033479 | 0.363478315 | 0.536988841 | 310.2387812 | 285.6000577 | ENSG00000135447 |
| VPS26C | -0.118834764 | 0.920931168 | 0.092376652 | 0.1944949 | 1350.526478 | 1243.950089 | ENSG00000157538 |
| NA | -0.118952325 | 0.920856127 | 0.909763731 | 0.951403864 | 4.901108155 | 4.518239899 | ENSG00000218582 |
| GPC3 | -0.119022812 | 0.920811137 | 0.017235676 | 0.049662945 | 3073.412006 | 2830.54537 | ENSG00000147257 |
| TBKBP1 | -0.119195951 | 0.920700636 | 0.241813635 | 0.404161049 | 672.1236057 | 619.2116897 | ENSG00000198933 |
| GCSH | -0.119210666 | 0.920691245 | 0.794702193 | 0.880471663 | 22.04999898 | 20.27507408 | ENSG00000140905 |
| CCNB2 | -0.119365675 | 0.920592328 | 0.051915375 | 0.122913538 | 1677.521632 | 1543.811014 | ENSG00000157456 |
| XPO7 | -0.119427692 | 0.920552756 | 0.014838261 | 0.043882399 | 3394.996692 | 3125.955697 | ENSG00000130227 |
| RNF4 | -0.119444311 | 0.920542151 | 0.04864976 | 0.116816346 | 2031.428908 | 1869.593956 | ENSG00000063978 |
| WDR61 | -0.119512357 | 0.920498734 | 0.228006339 | 0.387462147 | 610.2998491 | 561.9509402 | ENSG00000140395 |
| CHD1 | -0.119600228 | 0.920442671 | 0.09536318 | 0.199571611 | 1300.030494 | 1197.273819 | ENSG00000153922 |
| RING1 | -0.119640289 | 0.920417112 | 0.138565986 | 0.265607754 | 955.649283 | 879.7576136 | ENSG00000204227 |
| BTBD3 | -0.119646728 | 0.920413004 | 0.130406141 | 0.253441956 | 943.9829238 | 869.0516602 | ENSG00000132640 |
| TMEM81 | -0.119649352 | 0.92041133 | 0.665503673 | 0.794116512 | 60.77727446 | 55.86977861 | ENSG00000174529 |
| MRPL15 | -0.119699594 | 0.920379277 | 0.079428688 | 0.172493395 | 1272.419366 | 1170.904409 | ENSG00000137547 |
| KLC2 | -0.119715516 | 0.92036912 | 0.099371381 | 0.206279864 | 1111.856285 | 1023.167613 | ENSG00000174996 |
| DOCK11 | -0.119759599 | 0.920340997 | 0.302811104 | 0.4735351 | 461.8935653 | 425.2035137 | ENSG00000147251 |
| ZMYM6 | -0.119790975 | 0.920320982 | 0.672543071 | 0.799434862 | 66.22312401 | 60.98923571 | ENSG00000163867 |
| RADIL | -0.119821297 | 0.920301639 | 0.391572786 | 0.564598636 | 243.901693 | 224.5385081 | ENSG00000157927 |
| EIPR1 | -0.119822414 | 0.920300927 | 0.21927397 | 0.377275612 | 578.099634 | 531.9059176 | ENSG00000032389 |
| COPS8 | -0.119868987 | 0.920271218 | 0.124020546 | 0.244355056 | 1201.564089 | 1106.053447 | ENSG00000198612 |
| PAFAH2 | -0.119884633 | 0.920261238 | 0.417705593 | 0.589692058 | 232.4405219 | 213.7731321 | ENSG00000158006 |
| SLC38A9 | -0.119896861 | 0.920253437 | 0.274472362 | 0.441200428 | 437.2709188 | 402.1165546 | ENSG00000177058 |
| EIF4G3 | -0.119968895 | 0.92020749 | 0.040110073 | 0.09982756 | 2819.863508 | 2594.774919 | ENSG00000075151 |
| IFFO1 | -0.120125871 | 0.920107371 | 0.284100737 | 0.452754012 | 413.3537858 | 380.2015819 | ENSG00000010295 |
| VSTM4 | -0.120211054 | 0.920053045 | 0.707331781 | 0.823594399 | 48.15871337 | 44.40989886 | ENSG00000165633 |
| GRAMD2A | -0.120226362 | 0.920043282 | 0.775434492 | 0.868441613 | 25.02153416 | 23.05581553 | ENSG00000175318 |
| GRB2 | -0.120243152 | 0.920032575 | 0.024800526 | 0.067157068 | 2755.094202 | 2534.032679 | ENSG00000177885 |
| SNX29 | -0.120278765 | 0.920009865 | 0.406555571 | 0.578809116 | 229.5997347 | 211.2545555 | ENSG00000048471 |
| NA | -0.120344826 | 0.919967738 | 0.752986511 | 0.853786015 | 31.55093405 | 29.02761077 | ENSG00000267787 |
| RAE1 | -0.120372659 | 0.91994999 | 0.129859372 | 0.25265932 | 846.7011677 | 778.9124789 | ENSG00000101146 |
| AGPAT1 | -0.120433582 | 0.919911143 | 0.201931308 | 0.354357943 | 560.65811 | 515.8377343 | ENSG00000204310 |
| CACNB1 | -0.120444914 | 0.919903917 | 0.453712282 | 0.623824412 | 235.4737082 | 216.4934091 | ENSG00000067191 |
| DDX21 | -0.120517451 | 0.919857667 | 0.005314801 | 0.018483473 | 7121.547431 | 6551.825061 | ENSG00000165732 |
| PLXNA1 | -0.120528467 | 0.919850643 | 0.012638584 | 0.038342781 | 3811.176259 | 3505.666427 | ENSG00000114554 |
| PPARD | -0.120623605 | 0.919789986 | 0.155143029 | 0.290152278 | 716.9463368 | 659.3700688 | ENSG00000112033 |
| KAT2A | -0.120635928 | 0.919782129 | 0.11673654 | 0.233630522 | 1176.978574 | 1082.394889 | ENSG00000108773 |
| TSTA3 | -0.120891435 | 0.919619246 | 0.052096879 | 0.123176969 | 2121.07022 | 1950.213522 | ENSG00000104522 |
| UBR2 | -0.120918467 | 0.919602016 | 0.196970366 | 0.347896581 | 707.0573881 | 649.7670414 | ENSG00000024048 |
| ATP13A2 | -0.120936957 | 0.919590229 | 0.062554556 | 0.142556401 | 2303.561701 | 2118.433222 | ENSG00000159363 |
| ZNF287 | -0.12107669 | 0.919501167 | 0.584020986 | 0.733964094 | 102.5888484 | 94.36273106 | ENSG00000141040 |
| ARHGEF40 | -0.121142295 | 0.919459354 | 0.052472973 | 0.123868836 | 2311.840592 | 2126.104321 | ENSG00000165801 |
| FAM149A | -0.121145289 | 0.919457446 | 0.365896263 | 0.539558539 | 274.970725 | 252.8350247 | ENSG00000109794 |
| GINS4 | -0.121163326 | 0.919445951 | 0.183022421 | 0.328802898 | 780.2305022 | 717.0929685 | ENSG00000147536 |
| CRNKL1 | -0.121192222 | 0.919427535 | 0.153950945 | 0.288588834 | 803.5591905 | 738.5430036 | ENSG00000101343 |
| APOO | -0.121276911 | 0.919373565 | 0.207680858 | 0.362403567 | 635.3087049 | 583.8546391 | ENSG00000184831 |
| B9D2 | -0.12150769 | 0.91922651 | 0.706642912 | 0.823527362 | 54.04261118 | 49.61835352 | ENSG00000123810 |
| WAPL | -0.121528059 | 0.919213531 | 0.043846956 | 0.10740973 | 1806.737378 | 1660.951736 | ENSG00000062650 |
| LRRC37A2 | -0.121613889 | 0.919158847 | 0.569493598 | 0.72257965 | 106.1557615 | 97.50161963 | ENSG00000238083 |
| HLA-DPB2 | -0.121664648 | 0.919126508 | 0.302948208 | 0.473613024 | 410.8941386 | 377.2979362 | ENSG00000224557 |
| HSD17B12 | -0.121718289 | 0.919092335 | 0.258750575 | 0.423927095 | 527.214136 | 485.0177457 | ENSG00000149084 |
| NUS1 | -0.121744522 | 0.919075622 | 0.032076699 | 0.082959058 | 1997.548676 | 1836.425807 | ENSG00000153989 |
| PITPNB | -0.121748948 | 0.919072803 | 0.092934717 | 0.195358356 | 1858.494109 | 1708.583917 | ENSG00000180957 |
| POMP | -0.121752141 | 0.919070769 | 0.022422451 | 0.061766599 | 2896.03055 | 2662.133371 | ENSG00000132963 |
| MAP3K6 | -0.121832635 | 0.919019492 | 0.373120485 | 0.5465351 | 271.9487534 | 250.0905889 | ENSG00000142733 |
| KCNK1 | -0.121841783 | 0.919013664 | 0.394498599 | 0.567042134 | 305.0032644 | 280.088862 | ENSG00000135750 |
| ZFYVE1 | -0.121876024 | 0.918991852 | 0.307070703 | 0.478026221 | 392.1244159 | 360.5645516 | ENSG00000165861 |

| | | | | | | | |
|---|---|---|---|---|---|---|---|
| FAM13A | -0.121912535 | 0.918968595 | 0.300415309 | 0.471086955 | 343.2372613 | 315.325159 | ENSG00000138640 |
| DPH1 | -0.121919127 | 0.918964396 | 0.373629566 | 0.547087239 | 278.6718527 | 256.188214 | ENSG00000108963 |
| PPP4R2 | -0.122029517 | 0.918894083 | 0.143810175 | 0.273351275 | 921.1821947 | 846.9852986 | ENSG00000163605 |
| CLSTN1 | -0.122037729 | 0.918888852 | 0.020622517 | 0.057673876 | 3662.937952 | 3365.832186 | ENSG00000171603 |
| NARS1 | -0.122141859 | 0.918822532 | 0.003894066 | 0.014107124 | 5740.821218 | 5275.024995 | ENSG00000134440 |
| GABBR1 | -0.122145174 | 0.918820421 | 0.114301795 | 0.229687076 | 1566.424311 | 1438.586434 | ENSG00000204681 |
| MPST | -0.122145651 | 0.918820117 | 0.059451822 | 0.136859796 | 2201.77411 | 2023.23208 | ENSG00000128309 |
| NA | -0.122179014 | 0.918798869 | 0.859692831 | 0.921147212 | 9.595713621 | 8.832398915 | ENSG00000215006 |
| LDHC | -0.122202276 | 0.918784055 | 0.854842216 | 0.918046799 | 11.82465282 | 10.94632145 | ENSG00000166796 |
| B4GALNT3 | -0.122259211 | 0.918747796 | 0.133876806 | 0.25870154 | 1086.380494 | 997.6241652 | ENSG00000139044 |
| CDK6 | -0.122342836 | 0.918694543 | 0.34182953 | 0.514691966 | 359.4345138 | 330.6100771 | ENSG00000105810 |
| AGPAT4 | -0.122430093 | 0.91863898 | 0.160628423 | 0.297913186 | 704.983239 | 647.3256434 | ENSG00000026652 |
| ABCD4 | -0.122494903 | 0.918597713 | 0.174315721 | 0.317178323 | 1164.623008 | 1069.029263 | ENSG00000119688 |
| NEDD1 | -0.122496657 | 0.918596596 | 0.319331687 | 0.49073748 | 410.880874 | 377.8870314 | ENSG00000139350 |
| SPAG5 | -0.122509807 | 0.918588224 | 0.059658231 | 0.137269435 | 1760.635455 | 1617.076992 | ENSG00000076382 |
| STK19 | -0.122563993 | 0.918553723 | 0.41342622 | 0.585608724 | 216.6533348 | 198.9452938 | ENSG00000204344 |
| PIP4K2B | -0.122892787 | 0.918344406 | 0.017955846 | 0.051368467 | 2513.558529 | 2308.508109 | ENSG00000276293 |
| DRD1 | -0.122986629 | 0.918284673 | 0.9010509 | 0.945814175 | 4.607323737 | 4.242638624 | ENSG00000184845 |
| NA | -0.122986673 | 0.918284645 | 0.872504873 | 0.929146647 | 8.218779183 | 7.597705898 | ENSG00000228960 |
| PCDHA5 | -0.123001845 | 0.918274988 | 0.865477566 | 0.924725988 | 9.585225731 | 8.81357679 | ENSG00000204965 |
| AP5M1 | -0.123090758 | 0.918218396 | 0.14550104 | 0.275777222 | 746.9651091 | 686.0078647 | ENSG00000053770 |
| LINC00174 | -0.12312377 | 0.918197386 | 0.694095199 | 0.815108692 | 52.63249821 | 48.16667618 | ENSG00000179406 |
| KXD1 | -0.123147536 | 0.918182261 | 0.131001699 | 0.254240822 | 807.8069747 | 741.6273344 | ENSG00000105700 |
| NA | -0.123191822 | 0.918154076 | 0.849757854 | 0.914412828 | 10.69002725 | 9.797942378 | ENSG00000228038 |
| PCID2 | -0.123243728 | 0.918121042 | 0.129735632 | 0.252494964 | 1129.819558 | 1036.942807 | ENSG00000126226 |
| SENP6 | -0.123271504 | 0.918103366 | 0.069866177 | 0.15579568 | 1499.784731 | 1377.592254 | ENSG00000112701 |
| FUT4 | -0.123298753 | 0.918086026 | 0.494521272 | 0.660880488 | 151.223043 | 138.8742583 | ENSG00000196371 |
| PROSER2 | -0.123374073 | 0.918038095 | 0.344661162 | 0.51721964 | 281.2839733 | 258.2943583 | ENSG00000148426 |
| NA | -0.123489229 | 0.91796482 | 0.913931226 | 0.953781298 | 3.970870091 | 3.672613948 | ENSG00000254528 |
| TAC3 | -0.123529867 | 0.917938964 | 0.546863312 | 0.70479907 | 115.9265142 | 106.4302797 | ENSG00000166863 |
| FKBP8 | -0.12373132 | 0.917810795 | 0.005209409 | 0.018169355 | 8303.174231 | 7621.234194 | ENSG00000105701 |
| ZNF606 | -0.123754941 | 0.917795768 | 0.446535483 | 0.616880263 | 180.2461472 | 165.488156 | ENSG00000166704 |
| SMU1 | -0.123848064 | 0.917736527 | 0.039208258 | 0.098191161 | 1839.723017 | 1688.051004 | ENSG00000122692 |
| FPGS | -0.12388002 | 0.9177162 | 0.055884156 | 0.130531694 | 1597.851723 | 1466.490067 | ENSG00000136877 |
| NA | -0.124037277 | 0.917616172 | 0.894074193 | 0.94151347 | 5.377828739 | 4.926129091 | ENSG00000259005 |
| CRISPLD1 | -0.124232056 | 0.917492293 | 0.29521582 | 0.465202449 | 435.7704961 | 400.0394586 | ENSG00000121005 |
| TRIB2 | -0.124308037 | 0.917443973 | 0.031108815 | 0.080911441 | 1942.035577 | 1781.436899 | ENSG00000071575 |
| CWC22 | -0.124338116 | 0.917424846 | 0.13051169 | 0.253570451 | 918.5438288 | 843.1975892 | ENSG00000163510 |
| ADGRB3 | -0.124379302 | 0.917398655 | 0.768907542 | 0.863739323 | 26.21082651 | 24.01920842 | ENSG00000135298 |
| NA | -0.124404658 | 0.917382532 | 0.822357093 | 0.898735651 | 14.64694283 | 13.44743305 | ENSG00000232888 |
| EDC3 | -0.124438004 | 0.917361328 | 0.083688965 | 0.179902365 | 1031.189479 | 945.8246152 | ENSG00000179151 |
| TRNT1 | -0.124499145 | 0.917322451 | 0.206889762 | 0.361317255 | 524.9979489 | 481.6599948 | ENSG00000072756 |
| ADGRL1 | -0.124529353 | 0.917303244 | 0.032308823 | 0.083425179 | 4912.859078 | 4507.643098 | ENSG00000072071 |
| NA | -0.124603693 | 0.917255978 | 0.823474303 | 0.899475838 | 15.64069594 | 14.32640211 | ENSG00000257740 |
| RPS6KA3 | -0.124622563 | 0.917243981 | 0.292531798 | 0.462256792 | 374.7265953 | 343.8684954 | ENSG00000177189 |
| PLPPR2 | -0.12467287 | 0.917211997 | 0.307634089 | 0.478710108 | 344.8518739 | 316.3116751 | ENSG00000105520 |
| MITD1 | -0.124760349 | 0.917156383 | 0.280922285 | 0.449247434 | 376.5734292 | 345.4989861 | ENSG00000158411 |
| RABEP1 | -0.124767271 | 0.917151982 | 0.091601981 | 0.193315154 | 1109.388304 | 1017.380525 | ENSG00000029725 |
| AZI2 | -0.124772491 | 0.917148664 | 0.498326184 | 0.664584198 | 165.8799102 | 152.2582282 | ENSG00000163512 |
| PSMD12 | -0.124852312 | 0.917097921 | 0.044179419 | 0.108045787 | 2032.424278 | 1864.649434 | ENSG00000197170 |
| RNF138 | -0.124948334 | 0.917036884 | 0.272390227 | 0.439134203 | 741.7682788 | 680.7251998 | ENSG00000134758 |
| LINC01097 | -0.124960359 | 0.91702924 | 0.80953451 | 0.890310719 | 18.55026767 | 16.91785623 | ENSG00000281202 |
| XKR6 | -0.125272596 | 0.916830792 | 0.390034885 | 0.563270173 | 240.4567917 | 220.6575121 | ENSG00000171044 |
| PHF23 | -0.125276382 | 0.916828386 | 0.058582447 | 0.135342993 | 1524.458339 | 1397.928957 | ENSG00000040633 |
| DNAJC14 | -0.125355217 | 0.916778288 | 0.29894871 | 0.46935945 | 342.0162299 | 313.4755397 | ENSG00000135392 |
| CCDC84 | -0.125373334 | 0.916766776 | 0.43139738 | 0.602534436 | 204.2980073 | 187.1877434 | ENSG00000186166 |
| PKN1 | -0.125384092 | 0.916759939 | 0.006709164 | 0.022599352 | 5194.580897 | 4762.379071 | ENSG00000123143 |
| EHD4 | -0.125391678 | 0.916755119 | 0.050302123 | 0.119814577 | 1678.098415 | 1538.610368 | ENSG00000103966 |
| DIAPH3 | -0.125429187 | 0.916731284 | 0.174156656 | 0.316978587 | 673.0984599 | 617.3638995 | ENSG00000139734 |
| PPME1 | -0.125433223 | 0.91672872 | 0.113631712 | 0.228697898 | 875.0321492 | 801.8534899 | ENSG00000214517 |
| HS6ST1 | -0.125457177 | 0.916713499 | 0.031269736 | 0.081242353 | 2481.479349 | 2274.970934 | ENSG00000136720 |
| GLRX3 | -0.125463718 | 0.916709343 | 0.037830302 | 0.09516854 | 2503.661813 | 2295.564511 | ENSG00000108010 |
| NCR1 | -0.125520051 | 0.916673549 | 0.884005509 | 0.936493029 | 6.850417614 | 6.234493074 | ENSG00000189430 |
| MED8 | -0.125630733 | 0.916603225 | 0.427976461 | 0.599309953 | 214.4585622 | 196.6713558 | ENSG00000159479 |
| SIX1 | -0.125670826 | 0.916577753 | 0.879234744 | 0.933530598 | 7.571161926 | 6.998106228 | ENSG00000126778 |
| NDUFB11 | -0.125707204 | 0.916554641 | 0.048439438 | 0.116412756 | 1639.366433 | 1503.050012 | ENSG00000147123 |
| LRRTM4 | -0.125783685 | 0.916506054 | 0.604815625 | 0.74974285 | 102.9354967 | 94.07826703 | ENSG00000176204 |
| LYRM4 | -0.125858065 | 0.916458803 | 0.183318601 | 0.329212356 | 663.6922522 | 608.3989401 | ENSG00000214113 |
| MRPL47 | -0.125881139 | 0.916444146 | 0.118596349 | 0.236274106 | 933.8996255 | 855.9000197 | ENSG00000136522 |
| SYNGAP1 | -0.125896907 | 0.916434129 | 0.054645108 | 0.12811847 | 2351.598401 | 2154.564948 | ENSG00000197283 |
| POM121C | -0.125897335 | 0.916433858 | 0.098433887 | 0.204599517 | 1372.695323 | 1258.281665 | ENSG00000272391 |
| PMS2P3 | -0.125921541 | 0.916418481 | 0.564706703 | 0.718964956 | 98.13020624 | 89.81250513 | ENSG00000127957 |
| UNKL | -0.125931582 | 0.916412103 | 0.194793945 | 0.344907755 | 622.2033868 | 570.1262497 | ENSG00000059145 |
| CAMKV | -0.125982407 | 0.916379819 | 0.057502815 | 0.13344001 | 2281.793333 | 2090.434407 | ENSG00000164076 |
| TRMT61B | -0.125983209 | 0.91637931 | 0.233927892 | 0.39477762 | 469.2764463 | 429.8250364 | ENSG00000171103 |
| STK33 | -0.126002518 | 0.916367045 | 0.386408051 | 0.55977211 | 280.8629075 | 257.0838852 | ENSG00000130413 |
| RBM10 | -0.126003458 | 0.916366448 | 0.030656431 | 0.079972035 | 2301.832874 | 2109.389473 | ENSG00000182872 |

| | | | | | | | |
|---|---|---|---|---|---|---|---|
| GABPB1 | -0.126029328 | 0.916350016 | 0.230696135 | 0.390999749 | 528.4563803 | 484.3694885 | ENSG00000104064 |
| GUCD1 | -0.126118087 | 0.916293641 | 0.058557725 | 0.13530208 | 1418.414665 | 1299.832044 | ENSG00000138867 |
| ZFP82 | -0.126118139 | 0.916293608 | 0.249427827 | 0.413448377 | 404.7532648 | 370.8725567 | ENSG00000181007 |
| ADK | -0.126126049 | 0.916288585 | 0.199181298 | 0.350616517 | 511.8493888 | 468.7987371 | ENSG00000156110 |
| RAB5B | -0.126252417 | 0.916208329 | 0.085396048 | 0.182858826 | 1355.782194 | 1242.398413 | ENSG00000111540 |
| IER2 | -0.126259649 | 0.916203736 | 0.09968908 | 0.206762933 | 1553.090115 | 1422.473064 | ENSG00000160888 |
| RASGEF1B | -0.126370232 | 0.916133512 | 0.640876926 | 0.775313983 | 62.04507551 | 56.90254132 | ENSG00000138670 |
| ZNF446 | -0.126497842 | 0.916052481 | 0.753631131 | 0.853959904 | 32.48742964 | 29.81649777 | ENSG00000083838 |
| NA | -0.126565182 | 0.916009723 | 0.781397876 | 0.871679086 | 22.553077 | 20.71468884 | ENSG00000269929 |
| LINC01529 | -0.12658681 | 0.915995992 | 0.753941414 | 0.854189454 | 31.15379821 | 28.49496988 | ENSG00000225872 |
| NA | -0.126775184 | 0.915876397 | 0.894738839 | 0.941915276 | 5.012832289 | 4.587609718 | ENSG00000234026 |
| SRP72 | -0.126809297 | 0.915854741 | 0.022931605 | 0.062911078 | 2399.402017 | 2196.85217 | ENSG00000174780 |
| SF3B6 | -0.126911354 | 0.915789955 | 0.105419662 | 0.215595018 | 1035.357491 | 948.5531487 | ENSG00000115128 |
| SCOC | -0.126921752 | 0.915783355 | 0.27466479 | 0.441470364 | 479.1321044 | 439.1580596 | ENSG00000153130 |
| QRSL1 | -0.12693815 | 0.915772946 | 0.185533721 | 0.331954277 | 665.3196071 | 609.3916717 | ENSG00000130348 |
| KCTD2 | -0.1269778 | 0.915747778 | 0.084793422 | 0.181829228 | 1157.300339 | 1059.889972 | ENSG00000180901 |
| CDK2 | -0.126978624 | 0.915747255 | 0.260846938 | 0.426173785 | 397.228509 | 363.502588 | ENSG00000123374 |
| NA | -0.127089568 | 0.915676836 | 0.53669483 | 0.696306408 | 112.2641467 | 102.8060629 | ENSG00000288066 |
| SBDS | -0.127108791 | 0.915664635 | 0.310394408 | 0.481304972 | 321.3979073 | 294.3383644 | ENSG00000126524 |
| ZMYND8 | -0.127272541 | 0.91556071 | 0.01194251 | 0.03648764 | 3294.895096 | 3015.924725 | ENSG00000101040 |
| NA | -0.127289279 | 0.915550089 | 0.643919324 | 0.777434452 | 62.66845052 | 57.25962235 | ENSG00000237638 |
| HES6 | -0.127320681 | 0.91553016 | 0.089105675 | 0.189247778 | 1896.707406 | 1736.648765 | ENSG00000144485 |
| NA | -0.127470016 | 0.915435398 | 0.727164318 | 0.836082516 | 36.02420643 | 33.06616851 | ENSG00000271888 |
| CCDC152 | -0.127474782 | 0.915432374 | 0.3906007 | 0.563876233 | 220.0469238 | 201.5439931 | ENSG00000198865 |
| VRK2 | -0.127510574 | 0.915409663 | 0.308969858 | 0.48007593 | 349.9592974 | 320.385987 | ENSG00000028116 |
| NAT2 | -0.127597554 | 0.915354475 | 0.782869059 | 0.872832206 | 26.20292936 | 24.05255152 | ENSG00000156006 |
| ECT2 | -0.127694582 | 0.915292915 | 0.037159762 | 0.09375477 | 1613.410543 | 1477.09464 | ENSG00000114346 |
| LINC01090 | -0.127772252 | 0.91524364 | 0.515054118 | 0.678914178 | 140.339904 | 128.2387714 | ENSG00000231689 |
| KRTDAP | -0.127821668 | 0.915212291 | 0.881461018 | 0.93532122 | 6.814811598 | 6.186095954 | ENSG00000188508 |
| WDYHV1 | -0.127852908 | 0.915192473 | 0.372666393 | 0.546160251 | 255.4521346 | 233.6193443 | ENSG00000156795 |
| SCN5A | -0.127857681 | 0.915189445 | 0.261328317 | 0.426793478 | 391.177069 | 357.805328 | ENSG00000183873 |
| DNAJC5 | -0.127924818 | 0.915146857 | 0.036348799 | 0.092009537 | 1819.456263 | 1664.742232 | ENSG00000101152 |
| RBBP8 | -0.127938144 | 0.915138404 | 0.136789546 | 0.262907954 | 797.0517927 | 729.1785395 | ENSG00000101773 |
| ERCC4 | -0.128110972 | 0.915028781 | 0.526568712 | 0.688570564 | 112.235474 | 102.673242 | ENSG00000175595 |
| POLR1B | -0.128119321 | 0.915023486 | 0.042864421 | 0.105471626 | 1678.744341 | 1536.339944 | ENSG00000125630 |
| PITHD1 | -0.128122785 | 0.915021289 | 0.050571721 | 0.12041213 | 1630.656303 | 1492.095143 | ENSG00000057757 |
| B4GALT2 | -0.12816523 | 0.914994369 | 0.023810143 | 0.064920685 | 1934.164061 | 1769.773993 | ENSG00000117411 |
| RNF41 | -0.128202355 | 0.914970824 | 0.060821555 | 0.139527878 | 1223.11067 | 1118.876601 | ENSG00000181852 |
| LOC100996756 | -0.128211722 | 0.914964883 | 0.855337945 | 0.918419475 | 8.953678169 | 8.21397712 | ENSG00000287978 |
| MCTS1 | -0.128224888 | 0.914956533 | 0.167418403 | 0.30749961 | 642.3746878 | 587.9964365 | ENSG00000232119 |
| PTBP1 | -0.128248107 | 0.914941808 | 0.005685767 | 0.019603955 | 19781.5562 | 18099.66992 | ENSG00000011304 |
| INTS6 | -0.12830227 | 0.914907459 | 0.174344295 | 0.317200395 | 618.5203431 | 565.7913721 | ENSG00000102786 |
| PDPN | -0.128401384 | 0.914844606 | 0.024013933 | 0.065348113 | 2136.498004 | 1954.216537 | ENSG00000162493 |
| TRAF7 | -0.128430212 | 0.914826326 | 0.009822065 | 0.030965427 | 3641.034335 | 3330.427411 | ENSG00000131653 |
| FAM189B | -0.12847266 | 0.914799409 | 0.103944077 | 0.213408386 | 1349.095992 | 1234.750729 | ENSG00000160767 |
| TRAPPC10 | -0.128484786 | 0.914791721 | 0.157425809 | 0.293335389 | 732.966384 | 670.4554088 | ENSG00000160218 |
| RPL24 | -0.128616924 | 0.914707938 | 0.011056861 | 0.034314336 | 15968.0839 | 14604.85959 | ENSG00000114391 |
| NDUFAF4 | -0.128753851 | 0.914621126 | 0.182039209 | 0.327554669 | 620.6159578 | 568.0550476 | ENSG00000123545 |
| EXOC3 | -0.128965664 | 0.914486854 | 0.082201577 | 0.177339172 | 1022.874333 | 935.6593903 | ENSG00000180104 |
| FARSA | -0.128981848 | 0.914476595 | 0.017895654 | 0.051240516 | 3851.611665 | 3522.281524 | ENSG00000179115 |
| DEPDC7 | -0.129185816 | 0.914347316 | 0.498318955 | 0.664584198 | 130.1554334 | 119.1331822 | ENSG00000121690 |
| GNA11 | -0.129199624 | 0.914338565 | 0.208736251 | 0.363631034 | 642.7336269 | 587.762024 | ENSG00000088256 |
| NA | -0.129207937 | 0.914333296 | 0.73880341 | 0.844878569 | 32.61522347 | 29.71514192 | ENSG00000259570 |
| GM2A | -0.129251948 | 0.914305404 | 0.12209012 | 0.241642301 | 846.7286253 | 774.3517363 | ENSG00000196743 |
| MBD1 | -0.129308556 | 0.914269529 | 0.093527439 | 0.19636894 | 1080.2954 | 987.1808975 | ENSG00000141644 |
| HCCS | -0.129502239 | 0.914146796 | 0.132542445 | 0.256534194 | 722.7432283 | 661.0478416 | ENSG00000004961 |
| PYCR1 | -0.129522236 | 0.914134125 | 0.012720368 | 0.038564925 | 3917.588925 | 3581.504486 | ENSG00000183010 |
| ANAPC1 | -0.129543546 | 0.914120623 | 0.022879505 | 0.06281932 | 2175.306853 | 1987.898159 | ENSG00000153107 |
| SPAG9 | -0.12960227 | 0.914083415 | 0.0647227 | 0.14644213 | 1245.907147 | 1139.213515 | ENSG00000008294 |
| NA | -0.12963622 | 0.914061905 | 0.597439007 | 0.744425518 | 80.05877433 | 73.07613955 | ENSG00000240489 |
| NELFA | -0.129647453 | 0.914054788 | 0.050988552 | 0.121225062 | 1676.70441 | 1533.450783 | ENSG00000185049 |
| BEX3 | -0.129672188 | 0.914039117 | 0.001687048 | 0.006800009 | 8256.645959 | 7546.915403 | ENSG00000166681 |
| HNRNPA3 | -0.129704921 | 0.914018378 | 0.000639794 | 0.002851068 | 15583.16831 | 14244.25822 | ENSG00000170144 |
| NA | -0.129715331 | 0.914011783 | 0.74251107 | 0.84771149 | 29.9263091 | 27.32157742 | ENSG00000287038 |
| DOK4 | -0.129773621 | 0.913974854 | 0.103970433 | 0.213439781 | 868.8338177 | 794.4227416 | ENSG00000125170 |
| NA | -0.12978824 | 0.913965593 | 0.894748713 | 0.941915276 | 4.661790352 | 4.263611323 | ENSG00000188451 |
| CFAP410 | -0.129834636 | 0.913936201 | 0.712495468 | 0.826851355 | 36.66970851 | 33.56037165 | ENSG00000160226 |
| SLC16A1 | -0.129881227 | 0.913906686 | 0.00060844 | 0.002726459 | 9802.27204 | 8959.173443 | ENSG00000155380 |
| TPMT | -0.129915431 | 0.913885019 | 0.174503368 | 0.317370083 | 635.8216462 | 580.8036878 | ENSG00000137364 |
| EMG1 | -0.129928813 | 0.913876543 | 0.058861443 | 0.135857394 | 1340.723097 | 1224.965175 | ENSG00000126749 |
| NIF3L1 | -0.129953552 | 0.913860872 | 0.126617718 | 0.248053458 | 851.1048038 | 777.6275291 | ENSG00000196290 |
| ZNF573 | -0.130023815 | 0.913816366 | 0.698480425 | 0.817989588 | 39.87306466 | 36.38118049 | ENSG00000189144 |
| LOC102724450 | -0.130061399 | 0.91379256 | 0.890290355 | 0.939388982 | 5.081253418 | 4.636006837 | ENSG00000226944 |
| HSDL1 | -0.130075563 | 0.913783588 | 0.098550668 | 0.204809834 | 1235.816038 | 1129.209274 | ENSG00000103160 |
| PIGA | -0.130107245 | 0.913763522 | 0.476020489 | 0.644414824 | 150.2554472 | 137.4771694 | ENSG00000165195 |
| CBX2 | -0.130151451 | 0.913735523 | 0.032086668 | 0.082973714 | 4130.339925 | 3774.358698 | ENSG00000173894 |
| NA | -0.130186631 | 0.913713242 | 0.891462309 | 0.939943124 | 5.056811014 | 4.612883564 | ENSG00000274423 |

| | | | | | | | |
|---|---|---|---|---|---|---|---|
| APOB | -0.130220897 | 0.91369154 | 0.761992745 | 0.858671771 | 24.87228899 | 22.69632346 | ENSG00000084674 |
| RABGGTB | -0.13028497 | 0.913650962 | 0.116997024 | 0.234006179 | 969.1346296 | 885.4995574 | ENSG00000137955 |
| CYBC1 | -0.130385927 | 0.913587029 | 0.090608219 | 0.19161622 | 987.9750518 | 902.9150575 | ENSG00000178927 |
| KIF16B | -0.130424916 | 0.91356234 | 0.407043549 | 0.579327696 | 247.0637861 | 225.6209944 | ENSG00000089177 |
| MAK | -0.130478392 | 0.913528478 | 0.869532798 | 0.927261316 | 8.515354505 | 7.797485633 | ENSG00000111837 |
| NBDY | -0.130486515 | 0.913523334 | 0.142429533 | 0.27150278 | 668.4619878 | 610.5535057 | ENSG00000204272 |
| XRCC1 | -0.130504641 | 0.913511856 | 0.182541662 | 0.328153127 | 658.2141511 | 600.9389625 | ENSG00000073050 |
| RPL19 | -0.130652895 | 0.913417987 | 0.000843024 | 0.003647999 | 15851.25198 | 14478.36315 | ENSG00000108298 |
| PLCG1 | -0.130692165 | 0.913393124 | 0.012924144 | 0.039072193 | 4583.483527 | 4185.652785 | ENSG00000124181 |
| TBC1D20 | -0.130786865 | 0.91333317 | 0.173761167 | 0.316527543 | 588.3594221 | 537.4292416 | ENSG00000125875 |
| GPR62 | -0.130851249 | 0.913292412 | 0.858023281 | 0.92058549 | 9.410862433 | 8.598743039 | ENSG00000180929 |
| IGF2R | -0.130890418 | 0.913267616 | 0.009048942 | 0.028920333 | 4253.58764 | 3884.092781 | ENSG00000197081 |
| SCOC-AS1 | -0.131089593 | 0.913141541 | 0.891657625 | 0.939954256 | 5.081253418 | 4.63170569 | ENSG00000196951 |
| SAMD14 | -0.131109768 | 0.913128771 | 0.505524863 | 0.670442219 | 122.8778557 | 112.3241029 | ENSG00000167100 |
| NBPF8 | -0.131120932 | 0.913121705 | 0.664469539 | 0.793258787 | 55.99028102 | 51.13697753 | ENSG00000270231 |
| PRKAR2A | -0.131151483 | 0.913102369 | 0.134508748 | 0.259636608 | 806.5516645 | 736.7933831 | ENSG00000114302 |
| ZNF665 | -0.131185101 | 0.913081092 | 0.62860603 | 0.766477398 | 60.80921369 | 55.53932849 | ENSG00000197497 |
| HIF1A | -0.131347056 | 0.912978597 | 0.022861456 | 0.062778697 | 2741.687979 | 2503.4896 | ENSG00000100644 |
| LRIG1 | -0.131364049 | 0.912967843 | 0.00844422 | 0.027308875 | 2967.757794 | 2709.431916 | ENSG00000144749 |
| XKR8 | -0.131374863 | 0.912961 | 0.28513291 | 0.454098817 | 418.614672 | 382.3491264 | ENSG00000158156 |
| BSN | -0.131391894 | 0.912950222 | 0.46451334 | 0.634087662 | 160.0961726 | 146.1426107 | ENSG00000164061 |
| PCDHB2 | -0.131446964 | 0.912915374 | 0.458812149 | 0.628707569 | 171.8648072 | 157.1693153 | ENSG00000112852 |
| DBR1 | -0.131474691 | 0.912897829 | 0.147468973 | 0.278603855 | 648.0325836 | 591.3803037 | ENSG00000138231 |
| HIBCH | -0.131483342 | 0.912892355 | 0.134488196 | 0.259622915 | 688.4598384 | 628.2156015 | ENSG00000198130 |
| ZNF212 | -0.13148929 | 0.912888591 | 0.201560702 | 0.353836336 | 563.8446455 | 514.3137367 | ENSG00000170260 |
| MANBAL | -0.131539297 | 0.912856949 | 0.103472672 | 0.212644222 | 1153.317292 | 1052.522694 | ENSG00000101363 |
| SRSF11 | -0.13155032 | 0.912849974 | 0.007416686 | 0.024528593 | 3717.040374 | 3392.921119 | ENSG00000116754 |
| NA | -0.131555548 | 0.912846666 | 0.782679964 | 0.872756022 | 20.49311944 | 18.65561514 | ENSG00000220472 |
| UBL7-AS1 | -0.13157007 | 0.912837478 | 0.639820762 | 0.774742503 | 63.27555567 | 57.84092082 | ENSG00000247240 |
| PDE4DIP | -0.131691153 | 0.912760868 | 0.143931377 | 0.273446936 | 840.9766907 | 767.7247417 | ENSG00000178104 |
| GRAMD2B | -0.131700603 | 0.91275489 | 0.45561901 | 0.625398202 | 158.3248179 | 144.2878784 | ENSG00000155324 |
| NDUFV1 | -0.131795539 | 0.912694828 | 0.033809201 | 0.086534868 | 2672.204669 | 2439.417464 | ENSG00000167792 |
| NUP35 | -0.13189982 | 0.912628859 | 0.10630362 | 0.216838282 | 823.22595 | 751.2190269 | ENSG00000163002 |
| MARCHF8 | -0.131925198 | 0.912612805 | 0.316235509 | 0.487356879 | 396.072469 | 361.020559 | ENSG00000165406 |
| NEFM | -0.131951047 | 0.912596454 | 0.212244359 | 0.368182145 | 551.9888925 | 503.6545877 | ENSG00000104722 |
| PRKCD | -0.132036162 | 0.912542615 | 0.157018395 | 0.292801402 | 718.3332083 | 655.2809512 | ENSG00000163932 |
| SPINK1 | -0.132050319 | 0.91253366 | 0.849051081 | 0.913958172 | 9.331953413 | 8.514852242 | ENSG00000164266 |
| NA | -0.132090112 | 0.91250849 | 0.893655462 | 0.941175276 | 4.670163061 | 4.261460749 | ENSG00000256968 |
| WDR86-AS1 | -0.132150411 | 0.912470352 | 0.75818005 | 0.856402715 | 26.96717684 | 24.6338621 | ENSG00000243836 |
| FBLN1 | -0.132183156 | 0.912449642 | 0.002185071 | 0.008525933 | 5669.836654 | 5173.603353 | ENSG00000077942 |
| POLR3K | -0.132202835 | 0.912437196 | 0.097990222 | 0.203928756 | 1028.034457 | 937.5727518 | ENSG00000161980 |
| NA | -0.132294029 | 0.912379522 | 0.872205642 | 0.928970491 | 7.902667542 | 7.244132509 | ENSG00000229413 |
| ZFPL1 | -0.132338515 | 0.912351389 | 0.610552383 | 0.75418877 | 95.54520687 | 87.32104979 | ENSG00000162300 |
| UPF1 | -0.132437422 | 0.912288842 | 0.004174794 | 0.01499617 | 4472.927046 | 4080.21351 | ENSG00000005007 |
| ZNRF1 | -0.132438557 | 0.912288125 | 0.23655808 | 0.397997725 | 579.1200441 | 528.4578305 | ENSG00000186187 |
| MAP1LC3B2 | -0.132467387 | 0.912269894 | 0.852291502 | 0.916006883 | 9.400374543 | 8.575619766 | ENSG00000258102 |
| RNF220 | -0.132549385 | 0.912218045 | 0.058049139 | 0.134449027 | 1533.245363 | 1398.553309 | ENSG00000187147 |
| PPIL4 | -0.132597442 | 0.912187659 | 0.165938685 | 0.305567313 | 583.0639818 | 531.7225552 | ENSG00000131013 |
| ZNF90 | -0.132653439 | 0.912152255 | 0.124332029 | 0.24481618 | 697.2514312 | 636.1238693 | ENSG00000213988 |
| GDF11 | -0.132756484 | 0.912087106 | 0.182355424 | 0.327879438 | 557.2638453 | 508.3983652 | ENSG00000135414 |
| ZNF567 | -0.132773186 | 0.912076547 | 0.444305131 | 0.615130776 | 168.9378512 | 154.1820699 | ENSG00000189042 |
| GPR150 | -0.132777113 | 0.912074064 | 0.888814566 | 0.938578344 | 5.383410545 | 4.88848484 | ENSG00000178015 |
| ORAI2 | -0.132868817 | 0.91201609 | 0.140246426 | 0.268375539 | 869.3195257 | 793.1754483 | ENSG00000160991 |
| WDR11 | -0.132873039 | 0.912013421 | 0.069913964 | 0.155884221 | 1218.030093 | 1110.966673 | ENSG00000120008 |
| PHF20 | -0.132893835 | 0.912000275 | 0.051334986 | 0.12188586 | 1576.2746 | 1437.331452 | ENSG00000025293 |
| TLE3 | -0.132969795 | 0.911952259 | 0.058369429 | 0.134978411 | 1316.288807 | 1200.391802 | ENSG00000140332 |
| EEF1D | -0.132979053 | 0.911946407 | 0.051153367 | 0.121542059 | 1400.465706 | 1276.678914 | ENSG00000104529 |
| AIF1L | -0.133092036 | 0.911874991 | 0.006281189 | 0.021351616 | 5698.671064 | 5195.79677 | ENSG00000126878 |
| LINC00888 | -0.133107413 | 0.911865272 | 0.394752458 | 0.567322481 | 195.752992 | 178.323631 | ENSG00000240024 |
| TPP1 | -0.133147474 | 0.911839951 | 0.277877574 | 0.445375232 | 358.3633153 | 326.823749 | ENSG00000166340 |
| SNRPD1 | -0.133182241 | 0.911817977 | 0.009252626 | 0.029468195 | 3003.793852 | 2738.182629 | ENSG00000167088 |
| IQCE | -0.133188139 | 0.91181425 | 0.393827275 | 0.566676241 | 197.2916866 | 179.8083789 | ENSG00000106012 |
| RPRD1B | -0.133207832 | 0.911801803 | 0.213204495 | 0.369402543 | 450.2199094 | 410.4794791 | ENSG00000101413 |
| ZRSR2 | -0.133263797 | 0.911766433 | 0.22155051 | 0.379938741 | 506.7191016 | 462.1211521 | ENSG00000169249 |
| HAUS6 | -0.133337361 | 0.911719943 | 0.085465774 | 0.18294721 | 1297.500624 | 1183.238084 | ENSG00000147874 |
| NA | -0.133345054 | 0.911715082 | 0.668514193 | 0.796264961 | 50.01324427 | 45.48703011 | ENSG00000228232 |
| TTYH1 | -0.13338583 | 0.911689314 | 0.212605433 | 0.368629463 | 446.4358056 | 406.9090387 | ENSG00000167614 |
| NAPSB | -0.133444319 | 0.911652353 | 0.889350338 | 0.938896589 | 4.996762593 | 4.541363172 | ENSG00000131401 |
| AIFM3 | -0.133469487 | 0.911636449 | 0.853974907 | 0.917433175 | 9.671831738 | 8.802823921 | ENSG00000183773 |
| MAN2C1 | -0.133474982 | 0.911632977 | 0.154943941 | 0.289942699 | 769.8730439 | 701.3458269 | ENSG00000140400 |
| KCNMB3 | -0.133498351 | 0.911618211 | 0.81601643 | 0.894698714 | 16.76503378 | 15.33200086 | ENSG00000171121 |
| ZNF326 | -0.133660244 | 0.911515918 | 0.10956313 | 0.22194501 | 893.9386545 | 815.0877189 | ENSG00000162664 |
| RPL21P28 | -0.133729176 | 0.911472367 | 0.792263462 | 0.879072498 | 17.64638703 | 16.10825699 | ENSG00000220749 |
| THEMIS2 | -0.133744963 | 0.911462393 | 0.268568359 | 0.434766942 | 338.3881923 | 308.4808464 | ENSG00000130775 |
| IPO8 | -0.133868244 | 0.91138451 | 0.050064458 | 0.119437574 | 1364.675999 | 1243.702996 | ENSG00000133704 |
| IFT140 | -0.133951504 | 0.911331915 | 0.176073616 | 0.319472925 | 635.5644927 | 578.64029 | ENSG00000187535 |
| EIF4A1 | -0.133961 | 0.911325916 | 0.551520774 | 0.708395183 | 148.8791885 | 135.5482757 | ENSG00000161960 |

| | | | | | | | |
|---|---|---|---|---|---|---|---|
| NPIPB12 | -0.134242137 | 0.911148344 | 0.85060653 | 0.915070855 | 9.636901443 | 8.763029096 | ENSG00000169203 |
| DHX16 | -0.134253789 | 0.911140985 | 0.06598679 | 0.148876296 | 1539.620111 | 1402.583853 | ENSG00000204560 |
| NRIP3 | -0.134284803 | 0.911121398 | 0.430807357 | 0.602050472 | 186.3400143 | 169.9606702 | ENSG00000175352 |
| ZDHHC12 | -0.134306228 | 0.911107868 | 0.263935827 | 0.429641531 | 394.1087309 | 359.262633 | ENSG00000160446 |
| UCHL1 | -0.134360824 | 0.911073389 | 0.00102639 | 0.004361185 | 11506.46772 | 10482.44574 | ENSG00000154277 |
| RGPD8 | -0.134398109 | 0.911049843 | 0.792888792 | 0.879500539 | 17.34702081 | 15.83695686 | ENSG00000169629 |
| PRTFDC1 | -0.134401983 | 0.911047397 | 0.402926236 | 0.57525533 | 212.5358103 | 193.9395508 | ENSG00000099256 |
| NA | -0.13442716 | 0.911031498 | 0.713993517 | 0.828174365 | 38.78836304 | 35.39251376 | ENSG00000284610 |
| TES | -0.134500799 | 0.910984998 | 0.171863894 | 0.313823289 | 539.6305738 | 491.5144036 | ENSG00000135269 |
| KRBA1 | -0.13451938 | 0.910973265 | 0.299567139 | 0.470062606 | 323.106911 | 294.4964408 | ENSG00000133619 |
| PKIA | -0.134535329 | 0.910963194 | 0.282485964 | 0.450766644 | 353.9142979 | 322.2474436 | ENSG00000171033 |
| MRAS | -0.134562327 | 0.910946147 | 0.294613082 | 0.464556234 | 328.2527186 | 299.2623429 | ENSG00000158186 |
| RBBP7 | -0.134587438 | 0.910930291 | 0.001552265 | 0.006305486 | 6979.272042 | 6358.380119 | ENSG00000102054 |
| TFAP4 | -0.134620194 | 0.91090961 | 0.178440615 | 0.322614611 | 621.6079208 | 566.2318172 | ENSG00000090447 |
| SNU13 | -0.13485399 | 0.910762004 | 0.011558366 | 0.035624292 | 2781.696948 | 2533.565888 | ENSG00000100138 |
| NA | -0.134855244 | 0.910761212 | 0.899849578 | 0.944913081 | 4.280724206 | 3.904119251 | ENSG00000236058 |
| LOC101927974 | -0.134961191 | 0.910694331 | 0.67100814 | 0.798249756 | 47.81133822 | 43.46936878 | ENSG00000241764 |
| MPRIP | -0.135105704 | 0.910603113 | 0.003158859 | 0.011716507 | 3667.156552 | 3339.345754 | ENSG00000133030 |
| BICRA | -0.135149989 | 0.910575161 | 0.221421115 | 0.379808687 | 602.8372343 | 549.0660625 | ENSG00000063169 |
| NBL1 | -0.135201732 | 0.910542503 | 0.452548514 | 0.62268997 | 166.1637703 | 151.1045061 | ENSG00000158747 |
| DDX31 | -0.135273723 | 0.910497069 | 0.163437253 | 0.30188319 | 633.8559277 | 576.7766949 | ENSG00000125485 |
| ZNF692 | -0.135315181 | 0.910470904 | 0.171775596 | 0.313769816 | 677.155997 | 616.2040046 | ENSG00000171163 |
| NBPF12 | -0.135320254 | 0.910467702 | 0.478184765 | 0.646437442 | 171.6424718 | 156.1225831 | ENSG00000268043 |
| NKIRAS2 | -0.135428449 | 0.910399425 | 0.091665681 | 0.193386117 | 885.8295581 | 806.0047321 | ENSG00000168256 |
| NEPRO | -0.135543522 | 0.910326812 | 0.17211799 | 0.314246464 | 708.3074572 | 645.0522811 | ENSG00000163608 |
| RAB38 | -0.135572254 | 0.910308683 | 0.445337947 | 0.615843321 | 171.1409085 | 155.9537049 | ENSG00000123892 |
| NA | -0.135578688 | 0.910304623 | 0.719919512 | 0.83186387 | 33.22723471 | 30.20368684 | ENSG00000127589 |
| HSD17B7 | -0.135672153 | 0.910245651 | 0.121243526 | 0.240359497 | 772.7076005 | 703.4115702 | ENSG00000132196 |
| ZNF3 | -0.135700192 | 0.91022796 | 0.145942384 | 0.276396483 | 865.3844334 | 787.2912693 | ENSG00000166526 |
| CIAO2A | -0.135765475 | 0.910186772 | 0.090227481 | 0.190999488 | 1036.900504 | 943.9351826 | ENSG00000166797 |
| NA | -0.135831909 | 0.910144861 | 0.885987467 | 0.937192752 | 5.081253418 | 4.608582417 | ENSG00000254577 |
| ECD | -0.135936195 | 0.910079073 | 0.05532074 | 0.129482212 | 1164.225023 | 1059.702291 | ENSG00000122882 |
| RNF5 | -0.135946186 | 0.91007277 | 0.020643532 | 0.057706762 | 2252.266562 | 2049.613588 | ENSG00000204308 |
| NA | -0.135948906 | 0.910071054 | 0.879633996 | 0.933753715 | 5.758894886 | 5.201730366 | ENSG00000251050 |
| WLS | -0.136071274 | 0.909993866 | 0.682533935 | 0.806342221 | 44.42891284 | 40.322944 | ENSG00000116729 |
| ID4 | -0.136089807 | 0.909982176 | 0.549315295 | 0.706644528 | 111.9714384 | 101.709316 | ENSG00000172201 |
| MRPL27 | -0.136191447 | 0.909918069 | 0.119035815 | 0.236978299 | 869.9248777 | 791.2860463 | ENSG00000108826 |
| FGF22 | -0.136221591 | 0.909899057 | 0.711532758 | 0.826158117 | 34.63610838 | 31.55776433 | ENSG00000070388 |
| NA | -0.136453093 | 0.909753062 | 0.811392463 | 0.891703555 | 13.67830785 | 12.43564304 | ENSG00000258674 |
| DEAF1 | -0.136503371 | 0.909721358 | 0.038412443 | 0.096431254 | 1284.446492 | 1168.632076 | ENSG00000177030 |
| NA | -0.136515155 | 0.909713927 | 0.241891596 | 0.404256326 | 394.1867257 | 358.5492828 | ENSG00000264538 |
| WDR77 | -0.136527251 | 0.9097063 | 0.074535692 | 0.164056702 | 933.1839122 | 848.6529126 | ENSG00000116455 |
| IRF2 | -0.13657473 | 0.909676362 | 0.372543036 | 0.546020947 | 230.6634606 | 209.6805496 | ENSG00000168310 |
| RHEB | -0.136626313 | 0.909643837 | 0.014662821 | 0.043450151 | 2661.961874 | 2422.163935 | ENSG00000106615 |
| INHBE | -0.136660835 | 0.909622071 | 0.303155039 | 0.473842477 | 320.8222658 | 291.5326338 | ENSG00000139269 |
| APEX2 | -0.136694812 | 0.909600649 | 0.161900129 | 0.299474101 | 651.5986961 | 592.45397 | ENSG00000169188 |
| GUSBP3 | -0.136731907 | 0.909577261 | 0.769173654 | 0.863937639 | 22.98302795 | 20.88327604 | ENSG00000253203 |
| ZNF749 | -0.136797994 | 0.909535596 | 0.325391044 | 0.497108709 | 249.2921248 | 226.934211 | ENSG00000186230 |
| MEGF8 | -0.136812738 | 0.909526301 | 0.007528433 | 0.024847049 | 3427.389792 | 3117.071007 | ENSG00000105429 |
| RSPO4 | -0.136845933 | 0.909505374 | 0.318971674 | 0.49045772 | 301.051042 | 273.9998392 | ENSG00000101282 |
| TAF13 | -0.136873883 | 0.909487754 | 0.254575563 | 0.419386609 | 436.1298866 | 396.987126 | ENSG00000197780 |
| DMRTA1 | -0.136908194 | 0.909466124 | 0.451287764 | 0.621531539 | 145.4429722 | 132.3440153 | ENSG00000176399 |
| FNDC5 | -0.137010778 | 0.909401458 | 0.722774195 | 0.833863823 | 31.47414021 | 28.62133911 | ENSG00000160097 |
| MED27 | -0.137023289 | 0.909393572 | 0.189487603 | 0.337494724 | 573.0154633 | 520.6565767 | ENSG00000160563 |
| C16orf72 | -0.137058722 | 0.909371237 | 0.078651827 | 0.171095218 | 1011.80523 | 920.6586331 | ENSG00000182831 |
| MTO1 | -0.137214493 | 0.909273055 | 0.221841389 | 0.380200981 | 521.1434209 | 473.6125154 | ENSG00000135297 |
| PWWP3A | -0.137220147 | 0.909269492 | 0.090822729 | 0.191985675 | 861.8723886 | 783.4546661 | ENSG00000160953 |
| CHEK1 | -0.137345358 | 0.90919058 | 0.027437758 | 0.073138287 | 1804.891233 | 1640.558137 | ENSG00000149554 |
| ATP11C | -0.137419009 | 0.909144166 | 0.018910464 | 0.053592148 | 1910.525206 | 1737.2149 | ENSG00000101974 |
| PTRH2 | -0.137479994 | 0.909105736 | 0.140712213 | 0.269017681 | 647.8112746 | 588.6931214 | ENSG00000141378 |
| NA | -0.137507913 | 0.909088143 | 0.890571759 | 0.939403184 | 5.059601916 | 4.58330857 | ENSG00000279668 |
| SMC2 | -0.137618073 | 0.90901873 | 0.056105645 | 0.130848767 | 1362.977773 | 1239.512523 | ENSG00000136824 |
| FDPS | -0.137621992 | 0.909016261 | 0.058988757 | 0.136053576 | 1833.620002 | 1665.960829 | ENSG00000160752 |
| COA4 | -0.137760346 | 0.908929091 | 0.04250284 | 0.104728864 | 1316.065459 | 1196.117698 | ENSG00000181924 |
| SNRNP70 | -0.137763091 | 0.908927362 | 0.005244421 | 0.018284859 | 9752.600896 | 8864.094014 | ENSG00000104852 |
| NA | -0.138031357 | 0.908758365 | 0.824797482 | 0.900513228 | 11.99363448 | 10.94417088 | ENSG00000286277 |
| ZNF254 | -0.138063669 | 0.908738011 | 0.344059435 | 0.516678387 | 279.9941204 | 254.7351796 | ENSG00000213096 |
| SCN4B | -0.13822438 | 0.908636787 | 0.464896168 | 0.634475556 | 172.0291198 | 156.1543087 | ENSG00000177098 |
| C6orf226 | -0.138319191 | 0.908577075 | 0.737999963 | 0.844360197 | 33.48732813 | 30.46476713 | ENSG00000221821 |
| WDR82 | -0.138444752 | 0.908498003 | 0.008930849 | 0.028604516 | 3659.559718 | 3324.526228 | ENSG00000164091 |
| PAFAH1B3 | -0.138452087 | 0.908493384 | 0.165609708 | 0.305051208 | 564.8282614 | 513.2567726 | ENSG00000079462 |
| NA | -0.138478733 | 0.908476604 | 0.846791308 | 0.912319442 | 8.651521043 | 7.864704878 | ENSG00000232346 |
| LRRC4B | -0.138484828 | 0.908472767 | 0.20976382 | 0.365209497 | 636.8016464 | 578.9755743 | ENSG00000131409 |
| HNRNPUL2 | -0.138538156 | 0.908439186 | 0.099591712 | 0.206583195 | 830.3099034 | 754.5329301 | ENSG00000214753 |
| RPL35 | -0.138588682 | 0.908407371 | 0.011297646 | 0.034965971 | 18382.21144 | 16697.66447 | ENSG00000136942 |
| RPGR | -0.13859620 1 | 0.908402637 | 0.404639568 | 0.576913252 | 166.5830332 | 151.3596556 | ENSG00000156313 |
| CHD4 | -0.13875748 | 0.908301092 | 0.309096843 | 0.48017291 | 271.3672419 | 246.533052 | ENSG00000111642 |

| | | | | | | | |
|---|---|---|---|---|---|---|---|
| EXTL2 | -0.1387646 | 0.90829661 | 0.124509901 | 0.245068922 | 740.6202481 | 673.3054711 | ENSG00000162694 |
| MTF1 | -0.138766864 | 0.908295184 | 0.25750147 | 0.422654714 | 329.2150449 | 298.952314 | ENSG00000188786 |
| TMEM107 | -0.138780628 | 0.908286518 | 0.606467314 | 0.751018163 | 63.22320424 | 57.37362118 | ENSG00000179029 |
| COG6 | -0.138875551 | 0.90822676 | 0.273712068 | 0.440492159 | 324.0406157 | 294.3402361 | ENSG00000133103 |
| NFE2L1 | -0.138949714 | 0.908180072 | 0.001712792 | 0.006890817 | 6699.044436 | 6083.383248 | ENSG00000082641 |
| C11orf58 | -0.138952807 | 0.908178125 | 0.019379197 | 0.054743606 | 5262.534202 | 4779.523476 | ENSG00000110696 |
| HERPUD2 | -0.138991599 | 0.908153706 | 0.254794387 | 0.419567948 | 362.0467465 | 329.1210281 | ENSG00000122557 |
| TMEM138 | -0.138995127 | 0.908151485 | 0.29681219 | 0.467034501 | 299.273542 | 271.5508865 | ENSG00000149483 |
| PSMA2 | -0.139058767 | 0.908111426 | 0.621362084 | 0.761915061 | 67.71023098 | 61.67351968 | ENSG00000106588 |
| SMARCB1 | -0.139146705 | 0.908056075 | 0.004245646 | 0.015215076 | 3762.295789 | 3416.318804 | ENSG00000099956 |
| NA | -0.139185315 | 0.908031773 | 0.893468728 | 0.941175276 | 4.691814563 | 4.251240919 | ENSG00000269696 |
| MTHFD1L | -0.139208342 | 0.90801728 | 0.007129767 | 0.023734313 | 2909.286601 | 2641.34721 | ENSG00000120254 |
| NLGN3 | -0.139286175 | 0.907968294 | 0.275501848 | 0.442376074 | 377.8755601 | 343.1584351 | ENSG00000196338 |
| MAST2 | -0.139327054 | 0.907942567 | 0.047590782 | 0.11482764 | 1509.999453 | 1371.235766 | ENSG00000086015 |
| PI4KA | -0.139392761 | 0.907901216 | 0.028926472 | 0.076232317 | 1634.503033 | 1484.411923 | ENSG00000241973 |
| KIAA1191 | -0.139441347 | 0.907870641 | 0.023964789 | 0.065235435 | 1839.338923 | 1669.956483 | ENSG00000122203 |
| YARS2 | -0.139470523 | 0.907852281 | 0.092525047 | 0.194709358 | 757.7372366 | 688.0792967 | ENSG00000139131 |
| NA | -0.139471141 | 0.907851892 | 0.849276481 | 0.914149793 | 8.986493283 | 8.148908448 | ENSG00000260793 |
| FLJ45513 | -0.13947779 | 0.907847708 | 0.870294408 | 0.927560742 | 7.282959314 | 6.653135134 | ENSG00000204584 |
| NA | -0.139568086 | 0.907790889 | 0.892706648 | 0.940638188 | 4.395239242 | 3.992311195 | ENSG00000251775 |
| ADIPOR2 | -0.139580193 | 0.907783271 | 0.035136923 | 0.089458075 | 1616.361991 | 1467.76072 | ENSG00000006831 |
| C17orf107 | -0.139616992 | 0.907760116 | 0.78396965 | 0.873614559 | 20.76736754 | 18.88684787 | ENSG00000205710 |
| CORO1C | -0.139700065 | 0.907707847 | 0.006104585 | 0.020846797 | 2946.086795 | 2674.013534 | ENSG00000110880 |
| EIF4E2 | -0.139702727 | 0.907706172 | 0.028077877 | 0.074474127 | 2266.48298 | 2057.693647 | ENSG00000135930 |
| SFXN1 | -0.139797275 | 0.907646687 | 0.013845179 | 0.041428161 | 2162.381392 | 1962.578334 | ENSG00000164466 |
| SNHG26 | -0.139950776 | 0.90755012 | 0.453282674 | 0.623522252 | 141.5111748 | 128.5060308 | ENSG00000228649 |
| ANO8 | -0.140057625 | 0.907482907 | 0.086911236 | 0.185444602 | 1298.69471 | 1178.434957 | ENSG00000074855 |
| TSPAN13 | -0.140082952 | 0.907466976 | 0.183809973 | 0.329796678 | 682.6210975 | 619.9539966 | ENSG00000106537 |
| RPL11 | -0.1400845 | 0.907466003 | 0.000109207 | 0.000585847 | 15554.31186 | 14114.37667 | ENSG00000142676 |
| TBC1D30 | -0.140143206 | 0.907429077 | 0.517872492 | 0.681564084 | 108.0853345 | 97.9775521 | ENSG00000111490 |
| MRPL57 | -0.140270621 | 0.907348939 | 0.097031688 | 0.202364295 | 985.3493161 | 893.9786555 | ENSG00000173141 |
| TMEM14A | -0.140368084 | 0.907287644 | 0.45180358 | 0.621975512 | 152.4981776 | 138.1133593 | ENSG00000096092 |
| JOSD1 | -0.140377818 | 0.907281522 | 0.108677371 | 0.220451647 | 762.1809972 | 691.3053417 | ENSG00000100221 |
| PPP2R2D | -0.140468395 | 0.907224562 | 0.108290338 | 0.219874636 | 751.1332702 | 681.723808 | ENSG00000175470 |
| ZNF609 | -0.140608548 | 0.907136432 | 0.016868389 | 0.048764888 | 2417.921721 | 2192.637283 | ENSG00000180357 |
| BCR | -0.140704103 | 0.907076352 | 0.006952333 | 0.023264085 | 4373.334417 | 3967.633452 | ENSG00000186716 |
| SUPT3H | -0.140751468 | 0.907046572 | 0.30237696 | 0.473086289 | 256.8976535 | 232.9796893 | ENSG00000196284 |
| WASH5P | -0.14077693 | 0.907030563 | 0.709749208 | 0.825105495 | 31.45527961 | 28.49120177 | ENSG00000282458 |
| SLC25A29 | -0.140847187 | 0.906986393 | 0.021638257 | 0.059991896 | 1924.132414 | 1744.652477 | ENSG00000197119 |
| TRIM24 | -0.140848421 | 0.906985618 | 0.00898851 | 0.028760547 | 3307.410722 | 3000.490918 | ENSG00000122779 |
| UQCRQ | -0.14086758 | 0.906973573 | 0.329665655 | 0.502019273 | 254.5008086 | 230.4931108 | ENSG00000164405 |
| ACP6 | -0.140957912 | 0.906916786 | 0.258247427 | 0.423410769 | 342.2084146 | 310.0078122 | ENSG00000162836 |
| KIAA1143 | -0.141006995 | 0.906885932 | 0.063069164 | 0.143559557 | 919.9692094 | 834.3837704 | ENSG00000163807 |
| ZNF319 | -0.141110565 | 0.906820829 | 0.37871718 | 0.552091976 | 218.6825924 | 198.3344082 | ENSG00000166188 |
| TMEM39A | -0.141136542 | 0.906804502 | 0.088124522 | 0.187597677 | 799.5266835 | 725.1574042 | ENSG00000176142 |
| DLGAP4 | -0.14122406 | 0.906749494 | 0.06835899 | 0.153054003 | 1013.373798 | 919.0284397 | ENSG00000080845 |
| GCDH | -0.141246583 | 0.906735338 | 0.043759417 | 0.107274683 | 1208.002452 | 1095.148884 | ENSG00000105607 |
| SLC9A3R1 | -0.141344994 | 0.906673489 | 0.006744681 | 0.022691254 | 2885.834027 | 2616.643767 | ENSG00000109062 |
| NA | -0.141375092 | 0.906654573 | 0.714559298 | 0.828587115 | 30.89773499 | 27.99699863 | ENSG00000243679 |
| EXOSC8 | -0.141411209 | 0.906631877 | 0.048159454 | 0.115884249 | 1384.282561 | 1255.762761 | ENSG00000120699 |
| CSRP2 | -0.141413761 | 0.906630272 | 0.008188024 | 0.026641114 | 2501.556171 | 2267.52833 | ENSG00000175183 |
| NRBP1 | -0.141414303 | 0.906629932 | 0.004737854 | 0.016751823 | 3069.250376 | 2782.375253 | ENSG00000115216 |
| HAUS1 | -0.141625975 | 0.906496921 | 0.064013258 | 0.145160428 | 977.8507787 | 886.4309095 | ENSG00000152240 |
| LAPTM4A | -0.1416424 | 0.906486601 | 0.002847798 | 0.010721972 | 4987.967265 | 4521.509857 | ENSG00000068697 |
| AMMECR1L | -0.141712607 | 0.906442489 | 0.09409842 | 0.19734279 | 777.2877151 | 704.3997344 | ENSG00000144233 |
| METTL9 | -0.141829739 | 0.906368898 | 0.020622798 | 0.057673876 | 2487.556874 | 2255.410954 | ENSG00000197006 |
| TBP | -0.141918112 | 0.906313379 | 0.148423785 | 0.280056291 | 517.7527861 | 469.4281722 | ENSG00000112592 |
| RBFA | -0.141964939 | 0.906283963 | 0.408917555 | 0.581185335 | 162.3381152 | 147.0164546 | ENSG00000101546 |
| GPM6B | -0.142048968 | 0.906231178 | 0.014820049 | 0.043848707 | 2024.888346 | 1835.722416 | ENSG00000046653 |
| DPYS | -0.14210098 | 0.906198508 | 0.846025719 | 0.912024708 | 8.705987658 | 7.885677578 | ENSG00000147647 |
| TRIM41 | -0.142113267 | 0.90619079 | 0.104389345 | 0.214026449 | 727.6544609 | 659.1832046 | ENSG00000146063 |
| RBM14 | -0.142175576 | 0.906151653 | 0.030590466 | 0.079832344 | 1952.811248 | 1769.874616 | ENSG00000239306 |
| CDC42BPB | -0.142181049 | 0.906148215 | 0.004858882 | 0.017120127 | 3728.1794 | 3377.39192 | ENSG00000198752 |
| PKD1 | -0.142188229 | 0.906143706 | 0.06402674 | 0.145173935 | 1188.637525 | 1076.672651 | ENSG00000008710 |
| USP54 | -0.1422514 | 0.90610403 | 0.137751415 | 0.264440496 | 611.9544982 | 554.1846835 | ENSG00000166348 |
| PLD3 | -0.142278874 | 0.906086774 | 0.006207751 | 0.021135495 | 4864.926603 | 4408.716413 | ENSG00000105223 |
| NSMF | -0.142330155 | 0.906054567 | 0.024612659 | 0.066723271 | 2291.478819 | 2076.398841 | ENSG00000165802 |
| LUC7L2 | -0.142510958 | 0.905941025 | 0.171869942 | 0.313823289 | 622.4301526 | 563.974042 | ENSG00000146963 |
| GCC1 | -0.142515496 | 0.905938176 | 0.107007626 | 0.218004447 | 740.5382586 | 670.6981694 | ENSG00000179562 |
| H4-16 | -0.142635394 | 0.905862889 | 0.797091204 | 0.882500369 | 17.34981171 | 15.70681952 | ENSG00000197837 |
| HNRNPUL1 | -0.142709813 | 0.905816163 | 0.000961626 | 0.004114157 | 7011.180215 | 6351.569332 | ENSG00000105323 |
| COMMD7 | -0.142800989 | 0.905758918 | 0.267415711 | 0.433724271 | 300.241189 | 272.0230387 | ENSG00000149600 |
| ATP5PF | -0.142873829 | 0.905713189 | 0.15954557 | 0.296454785 | 517.7816079 | 468.6973691 | ENSG00000154723 |
| ZNF16 | -0.142929038 | 0.90567853 | 0.447966206 | 0.618413917 | 155.8519303 | 141.0876893 | ENSG00000170631 |
| NA | -0.142971631 | 0.905651791 | 0.486980517 | 0.654262257 | 109.3994296 | 98.97535417 | ENSG00000197744 |
| DKKL1 | -0.142972564 | 0.905651206 | 0.677093929 | 0.802273754 | 38.08224896 | 34.49849069 | ENSG00000104901 |
| FMR1 | -0.142998754 | 0.905634765 | 0.107162381 | 0.218204506 | 678.8098214 | 614.9644284 | ENSG00000102081 |

| | | | | | | | |
|---|---|---|---|---|---|---|---|
| OGDH | -0.143012601 | 0.905626073 | 0.002841757 | 0.01070919 | 3923.256392 | 3552.591704 | ENSG00000105953 |
| OSMR | -0.143064402 | 0.905593556 | 0.550433426 | 0.707422466 | 84.4952014 | 76.40593298 | ENSG00000145623 |
| MBD6 | -0.143101608 | 0.905570202 | 0.199821226 | 0.351358243 | 503.4437696 | 456.0186865 | ENSG00000166987 |
| SNX3 | -0.143112677 | 0.905563254 | 0.025386128 | 0.068492838 | 1617.656137 | 1465.59654 | ENSG00000112335 |
| ADAMTSL3 | -0.143121359 | 0.905557804 | 0.714112798 | 0.828230024 | 29.45237942 | 26.68648408 | ENSG00000156218 |
| GFER | -0.143171903 | 0.905526079 | 0.183068773 | 0.328855546 | 535.3955943 | 484.78117 | ENSG00000127554 |
| ATXN7L3 | -0.143282629 | 0.905456583 | 0.007032566 | 0.023471492 | 3020.540688 | 2735.064138 | ENSG00000087152 |
| LYN | -0.143296668 | 0.905447772 | 0.057382481 | 0.133238974 | 1031.581784 | 934.3934927 | ENSG00000254087 |
| AUNIP | -0.143297471 | 0.905447268 | 0.273038391 | 0.43988505 | 339.603003 | 307.2840886 | ENSG00000127423 |
| SERINC2 | -0.143309164 | 0.90543993 | 0.073170764 | 0.161669591 | 981.9560025 | 889.4254307 | ENSG00000168528 |
| NA | -0.143335499 | 0.905423402 | 0.790183649 | 0.87763484 | 18.68276743 | 16.86300739 | ENSG00000198237 |
| NA | -0.143336123 | 0.90542301 | 0.879092611 | 0.93343161 | 5.43787716 | 4.886334266 | ENSG00000232545 |
| NA | -0.143392431 | 0.905387673 | 0.852856196 | 0.916417806 | 8.945305461 | 8.081156166 | ENSG00000229582 |
| PCBP1 | -0.143452137 | 0.905350204 | 0.433308345 | 0.604319786 | 193.7882516 | 175.6251629 | ENSG00000169564 |
| HMGCR | -0.143455151 | 0.905348313 | 0.000701106 | 0.003086497 | 5126.675948 | 4641.096182 | ENSG00000113161 |
| URGCP | -0.143458357 | 0.905346301 | 0.168175597 | 0.308449843 | 467.6900182 | 423.2751472 | ENSG00000106608 |
| ARL3 | -0.143520824 | 0.905307101 | 0.085775961 | 0.183570453 | 831.0083574 | 752.0227074 | ENSG00000138175 |
| BHLHE40 | -0.143536835 | 0.905297054 | 0.630342869 | 0.76801265 | 71.09356913 | 64.21389307 | ENSG00000134107 |
| NA | -0.143562954 | 0.905280665 | 0.68986832 | 0.811588653 | 36.9804385 | 33.37968661 | ENSG00000250994 |
| NA | -0.143596142 | 0.90525984 | 0.909823884 | 0.951403864 | 3.290437721 | 3.005795033 | ENSG00000204805 |
| RPF2 | -0.143609017 | 0.905251761 | 0.149968369 | 0.28233042 | 959.4235521 | 869.1468001 | ENSG00000197498 |
| CEBPG | -0.143678521 | 0.90520815 | 0.044614818 | 0.108939846 | 1217.694659 | 1102.590088 | ENSG00000153879 |
| SMIM26 | -0.143710968 | 0.905187792 | 0.651074286 | 0.783181022 | 54.02059626 | 48.82945441 | ENSG00000232388 |
| ABTB1 | -0.143794322 | 0.905135495 | 0.525637721 | 0.687835929 | 105.9153677 | 96.10075044 | ENSG00000114626 |
| UNC13B | -0.143855121 | 0.905097351 | 0.015977331 | 0.046671517 | 2508.350819 | 2270.167097 | ENSG00000198722 |
| THRA | -0.143993782 | 0.905010363 | 0.191231518 | 0.339973017 | 505.2904162 | 457.4267767 | ENSG00000126351 |
| ZNF672 | -0.144050356 | 0.904974875 | 0.069304687 | 0.154806963 | 1003.456266 | 908.1848554 | ENSG00000171161 |
| TMEM70 | -0.144106732 | 0.904939512 | 0.136612891 | 0.26272548 | 729.0988796 | 660.1100256 | ENSG00000175606 |
| RTN2 | -0.144111865 | 0.904936292 | 0.277009684 | 0.444235147 | 290.5063177 | 262.8817075 | ENSG00000125744 |
| KLF3 | -0.144133387 | 0.904922793 | 0.283115963 | 0.451520341 | 309.8879778 | 280.8019153 | ENSG00000109787 |
| PPP3R1 | -0.144180872 | 0.904893008 | 0.298465456 | 0.468944218 | 455.5703189 | 412.5729566 | ENSG00000221823 |
| SLC35C2 | -0.144234515 | 0.904859363 | 0.061473328 | 0.140657333 | 992.9623273 | 898.0258034 | ENSG00000080189 |
| SLC25A21 | -0.144313484 | 0.904809835 | 0.567018867 | 0.72043476 | 79.74716846 | 72.2810613 | ENSG00000183032 |
| NA | -0.144370225 | 0.90477425 | 0.898411268 | 0.944070801 | 3.635897851 | 3.308820728 | ENSG00000241511 |
| PTS | -0.14444693 | 0.904726146 | 0.26257378 | 0.428175045 | 343.0814832 | 310.2532934 | ENSG00000150787 |
| POTEF | -0.144499945 | 0.904692901 | 0.896007922 | 0.942469401 | 3.630316045 | 3.304519581 | ENSG00000196604 |
| EIF4H | -0.144559255 | 0.904655709 | 0.000361786 | 0.001722839 | 10673.84297 | 9656.211364 | ENSG00000106682 |
| ZFP62 | -0.144638842 | 0.904605804 | 0.104470377 | 0.214147067 | 629.9022545 | 569.7011375 | ENSG00000196670 |
| PDZD11 | -0.144713416 | 0.904559046 | 0.198888036 | 0.35035605 | 397.1042699 | 359.1497065 | ENSG00000120509 |
| MAGT1 | -0.14474186 | 0.904541212 | 0.043975522 | 0.107659944 | 1098.399898 | 993.8347714 | ENSG00000102158 |
| TANGO2 | -0.14474833 | 0.904537155 | 0.453744224 | 0.623824412 | 136.4849515 | 123.4952174 | ENSG00000183597 |
| CSGALNACT2 | -0.144837235 | 0.904481416 | 0.382193191 | 0.555605791 | 171.0872057 | 154.778174 | ENSG00000169826 |
| RAD51AP1 | -0.144868307 | 0.904461936 | 0.107688104 | 0.218974513 | 720.2684383 | 651.3738511 | ENSG00000111247 |
| RUSC2 | -0.144892241 | 0.90444693 | 0.310072609 | 0.481262427 | 310.1072326 | 280.1830032 | ENSG00000198853 |
| CATSPERG | -0.144898727 | 0.904442864 | 0.815733163 | 0.894616739 | 12.40751574 | 11.23052502 | ENSG00000099338 |
| LPAL2 | -0.144929678 | 0.904423461 | 0.756997218 | 0.855820859 | 21.69337524 | 19.59158361 | ENSG00000213071 |
| NA | -0.144935192 | 0.904420004 | 0.027958613 | 0.074229232 | 1390.675054 | 1257.520178 | ENSG00000188856 |
| RRAGC | -0.144964703 | 0.904401504 | 0.495031039 | 0.661429731 | 123.0354736 | 111.5712305 | ENSG00000116954 |
| NA | -0.145170851 | 0.904272283 | 0.787920807 | 0.876337918 | 20.11416848 | 18.21357723 | ENSG00000214077 |
| BASP1 | -0.145446892 | 0.904099279 | 0.017090464 | 0.049323696 | 4922.517785 | 4451.190452 | ENSG00000176788 |
| MED22 | -0.145550069 | 0.904034623 | 0.039427305 | 0.098578233 | 1909.938015 | 1726.638437 | ENSG00000148297 |
| SELENOS | -0.145559422 | 0.904022495 | 0.09521315 | 0.199344136 | 688.2487561 | 622.415889 | ENSG00000131871 |
| IL17RD | -0.145807165 | 0.903873533 | 0.013465298 | 0.040442016 | 2223.604048 | 2009.867108 | ENSG00000144730 |
| MYO1B | -0.145838365 | 0.903853986 | 0.011816138 | 0.03618751 | 2073.950338 | 1874.006272 | ENSG00000128641 |
| IQCB1 | -0.145948953 | 0.903784705 | 0.088176979 | 0.187667928 | 836.5162968 | 756.580197 | ENSG00000173226 |
| ZNF827 | -0.145978856 | 0.903765972 | 0.131595543 | 0.255136645 | 603.2400626 | 544.773385 | ENSG00000151612 |
| AUP1 | -0.146141427 | 0.903664137 | 0.070286096 | 0.156623436 | 1018.988178 | 920.3957296 | ENSG00000115307 |
| SLC38A1 | -0.146214286 | 0.903618501 | 0.000485573 | 0.002232942 | 6893.31185 | 6229.916578 | ENSG00000111371 |
| LMAN2 | -0.146376756 | 0.903516745 | 0.006223088 | 0.021176503 | 3257.748233 | 2943.689619 | ENSG00000169223 |
| NA | -0.146423954 | 0.903487187 | 0.838725113 | 0.908136413 | 10.19532195 | 9.158547884 | ENSG00000214433 |
| DOLK | -0.146541256 | 0.903413729 | 0.1703393 | 0.311677458 | 528.2284135 | 477.2187036 | ENSG00000175283 |
| IGF2BP1 | -0.14658612 | 0.903385636 | 8.75E-05 | 0.000479422 | 9129.069451 | 8247.432007 | ENSG00000159217 |
| GPR19 | -0.146610226 | 0.903370541 | 0.300460879 | 0.471119316 | 247.9787054 | 224.1529365 | ENSG00000183150 |
| DNMT1 | -0.146690858 | 0.903320054 | 0.000646185 | 0.002876451 | 7106.584572 | 6419.349207 | ENSG00000130816 |
| RSPH3 | -0.146693805 | 0.903318208 | 0.65488914 | 0.785858821 | 45.82729862 | 41.38689923 | ENSG00000130363 |
| NA | -0.146940932 | 0.903163488 | 0.820223696 | 0.897449681 | 11.33764451 | 10.27735196 | ENSG00000270878 |
| AGPAT2 | -0.146955624 | 0.90315429 | 0.124801933 | 0.245424828 | 1004.362952 | 907.3518845 | ENSG00000169692 |
| BRF2 | -0.146965864 | 0.903147879 | 0.334959934 | 0.507557307 | 217.4401097 | 196.4339502 | ENSG00000104221 |
| AGO3 | -0.146968454 | 0.903146258 | 0.232012088 | 0.392643669 | 326.6584301 | 295.130468 | ENSG00000126070 |
| UPF2 | -0.147001688 | 0.903125453 | 0.049067246 | 0.117511638 | 1223.181721 | 1105.246823 | ENSG00000151461 |
| GUCY1A2 | -0.147004511 | 0.903123686 | 0.723914039 | 0.834405189 | 31.77609718 | 28.72002346 | ENSG00000152402 |
| TMEM17 | -0.147100809 | 0.903063406 | 0.739765098 | 0.845477132 | 24.6509559 | 22.25885927 | ENSG00000186889 |
| ARL8B | -0.147132447 | 0.903043602 | 0.080803985 | 0.174771799 | 766.4955388 | 692.3717017 | ENSG00000134108 |
| TWSG1 | -0.147139587 | 0.903039133 | 0.158521816 | 0.294864946 | 598.8743862 | 540.8394119 | ENSG00000128791 |
| DVL2 | -0.147178427 | 0.903014822 | 0.015254635 | 0.044893542 | 1695.546621 | 1531.13284 | ENSG00000004975 |
| NA | -0.147193522 | 0.903005373 | 0.442139427 | 0.61325164 | 160.905675 | 144.9850963 | ENSG00000267383 |
| SMAP1 | -0.147218055 | 0.902990018 | 0.310746772 | 0.481688017 | 263.5915803 | 238.1198466 | ENSG00000112305 |

| | | | | | | | |
|---|---|---|---|---|---|---|---|
| RESF1 | -0.147272939 | 0.902955667 | 0.048879563 | 0.117236808 | 1301.164016 | 1174.96443 | ENSG00000174718 |
| CLDND2 | -0.147357739 | 0.902902594 | 0.804214539 | 0.887069454 | 13.37335982 | 12.03582311 | ENSG00000160318 |
| NLE1 | -0.14738988 | 0.902882479 | 0.037496147 | 0.094467568 | 1516.17542 | 1368.960532 | ENSG00000073536 |
| KIAA1958 | -0.147392408 | 0.902880896 | 0.115291688 | 0.231218625 | 648.5397159 | 585.4931743 | ENSG00000165185 |
| TMEM19 | -0.147442061 | 0.902849822 | 0.443099686 | 0.61402457 | 130.8628989 | 118.2958981 | ENSG00000139291 |
| NRIP2 | -0.147449423 | 0.902845216 | 0.676155705 | 0.80140819 | 43.43352011 | 39.26946905 | ENSG00000053702 |
| COL9A3 | -0.147473309 | 0.902830268 | 0.039405749 | 0.098570579 | 1853.690331 | 1673.588916 | ENSG00000092758 |
| GPR160 | -0.147473515 | 0.902830139 | 0.181147479 | 0.326315812 | 577.639859 | 521.4602205 | ENSG00000173890 |
| NA | -0.147481622 | 0.902825065 | 0.885403797 | 0.937037652 | 4.68902366 | 4.21306363 | ENSG00000232004 |
| SOGA1 | -0.147658735 | 0.902714237 | 0.073224554 | 0.161769912 | 944.5230331 | 852.3039903 | ENSG00000149639 |
| RTL6 | -0.147692798 | 0.902692923 | 0.046398321 | 0.112460109 | 1106.830999 | 998.6585212 | ENSG00000188636 |
| NDUFA5 | -0.147779298 | 0.902638802 | 0.052217939 | 0.123372476 | 1126.064668 | 1016.970722 | ENSG00000128609 |
| FICD | -0.147830768 | 0.9026066 | 0.430093438 | 0.601577447 | 138.1078616 | 124.5554046 | ENSG00000198855 |
| FANCI | -0.147873113 | 0.902580107 | 0.02174089 | 0.060207223 | 1805.24579 | 1629.235101 | ENSG00000140525 |
| PFKFB1 | -0.14787934 | 0.902576211 | 0.889456764 | 0.93892015 | 4.458078566 | 4.017585042 | ENSG00000158571 |
| NOSIP | -0.147957822 | 0.902527113 | 0.035974334 | 0.091229246 | 1787.053439 | 1613.256721 | ENSG00000142546 |
| USP22 | -0.147961648 | 0.902524719 | 0.000996593 | 0.004250557 | 6308.074954 | 5692.215505 | ENSG00000124422 |
| RNF115 | -0.147986832 | 0.902508965 | 0.121305497 | 0.240429914 | 547.6951649 | 494.4468002 | ENSG00000265491 |
| SIN3B | -0.14799896 | 0.902501378 | 0.028579666 | 0.075552662 | 2228.68186 | 2010.940381 | ENSG00000127511 |
| MRPL50 | -0.148111195 | 0.90243117 | 0.04942926 | 0.118173237 | 950.3700118 | 857.4694456 | ENSG00000136897 |
| TRAPPC1 | -0.14823455 | 0.902354013 | 0.080891619 | 0.174882912 | 883.2406166 | 796.4917989 | ENSG00000170043 |
| RPL36 | -0.148236146 | 0.902353014 | 0.014351479 | 0.042678339 | 5777.050602 | 5212.228056 | ENSG00000130255 |
| TMEM41A | -0.148323606 | 0.902298314 | 0.277583761 | 0.445015236 | 274.3246224 | 247.5940631 | ENSG00000163900 |
| LINC00843 | -0.14838378 | 0.90226068 | 0.889556896 | 0.938960499 | 4.011382192 | 3.622066255 | ENSG00000178440 |
| LARP1B | -0.148430016 | 0.902231764 | 0.07588081 | 0.166229469 | 738.3887551 | 666.1992968 | ENSG00000138709 |
| ATP5MPL | -0.148484656 | 0.902197594 | 0.07224582 | 0.160010805 | 811.6249328 | 732.2463981 | ENSG00000156411 |
| ZNHIT3 | -0.148491818 | 0.902193115 | 0.110379235 | 0.223176378 | 588.3650919 | 531.0258703 | ENSG00000273611 |
| MEST | -0.148520255 | 0.902175332 | 0.006815399 | 0.022889253 | 2195.967379 | 1981.606642 | ENSG00000106484 |
| ITGB1BP1 | -0.148555311 | 0.90215341 | 0.208742423 | 0.363631034 | 371.0252674 | 334.5787562 | ENSG00000119185 |
| NA | -0.148559979 | 0.902150492 | 0.886479859 | 0.937353813 | 4.014173095 | 3.624216828 | ENSG00000197670 |
| OPA1 | -0.148573738 | 0.902141888 | 0.01143643 | 0.035355796 | 1976.599098 | 1783.377358 | ENSG00000198836 |
| HEBP1 | -0.148662186 | 0.902086582 | 0.109549305 | 0.22194031 | 737.2285358 | 665.2697558 | ENSG00000013583 |
| NA | -0.148666236 | 0.902084049 | 0.900782751 | 0.945687078 | 4.299584804 | 3.87024311 | ENSG00000260532 |
| NA | -0.148696518 | 0.902065115 | 0.905541022 | 0.948926713 | 3.728761385 | 3.376039973 | ENSG00000287972 |
| SRGAP2B | -0.148862376 | 0.901961416 | 0.327436356 | 0.499491914 | 220.0823666 | 198.374445 | ENSG00000196369 |
| NA | -0.14895246 | 0.901905098 | 0.834307558 | 0.90617798 | 9.794719483 | 8.779700648 | ENSG00000267108 |
| LRRC8A | -0.148953451 | 0.901904478 | 0.007025489 | 0.023451932 | 3993.814659 | 3602.10251 | ENSG00000136802 |
| DHODH | -0.148994632 | 0.901878734 | 0.198932479 | 0.350401065 | 410.0996298 | 370.1000566 | ENSG00000102967 |
| MORN4 | -0.149013135 | 0.901867167 | 0.223228487 | 0.381764249 | 334.4311134 | 301.4475069 | ENSG00000171160 |
| RBL1 | -0.149107098 | 0.901808431 | 0.375502049 | 0.548898237 | 181.2062093 | 163.3070263 | ENSG00000080839 |
| KHDRBS2 | -0.149219529 | 0.901738154 | 0.35737699 | 0.530689421 | 191.8878639 | 173.2541765 | ENSG00000112232 |
| PPIL3 | -0.149339737 | 0.901663023 | 0.174948074 | 0.317910521 | 487.6368077 | 439.6057679 | ENSG00000240344 |
| PPP5C | -0.149344427 | 0.901660092 | 0.014504007 | 0.043052286 | 3497.152584 | 3152.38931 | ENSG00000011485 |
| POFUT2 | -0.149424033 | 0.901610341 | 0.069310442 | 0.154806963 | 889.7576135 | 802.0750238 | ENSG00000186866 |
| CTH | -0.149473854 | 0.901579205 | 0.250220977 | 0.414430782 | 349.2599533 | 314.5688094 | ENSG00000116761 |
| NA | -0.149563197 | 0.901523374 | 0.737483207 | 0.843969185 | 31.59491277 | 28.4062449 | ENSG00000238045 |
| ASAP1 | -0.14963668 | 0.901477457 | 0.076643939 | 0.167549876 | 807.9858806 | 728.0763765 | ENSG00000153317 |
| TRIB1 | -0.149636838 | 0.901477358 | 0.046204833 | 0.112075623 | 1075.561861 | 969.9903294 | ENSG00000173334 |
| ERGIC2 | -0.149765967 | 0.901396675 | 0.234771293 | 0.39588925 | 386.7504542 | 348.9664226 | ENSG00000087502 |
| GIGYF1 | -0.149782627 | 0.901386266 | 0.024563849 | 0.066609673 | 2934.976842 | 2645.081332 | ENSG00000146830 |
| NA | -0.149801388 | 0.901374544 | 0.86167092 | 0.922498125 | 7.22011999 | 6.528916475 | ENSG00000229525 |
| GATD3B | -0.149809958 | 0.90136919 | 0.416328935 | 0.58842417 | 151.5157514 | 136.6035981 | ENSG00000280071 |
| NA | -0.149851068 | 0.901343505 | 0.85155875 | 0.915482604 | 7.277377508 | 6.585915889 | ENSG00000226179 |
| NA | -0.149872026 | 0.901330411 | 0.901977314 | 0.946583965 | 3.651967547 | 3.302369007 | ENSG00000251229 |
| DHX30 | -0.149890046 | 0.901319154 | 0.002713952 | 0.010291892 | 3432.426479 | 3093.469908 | ENSG00000132153 |
| NA | -0.149922202 | 0.901299064 | 0.656829892 | 0.787306166 | 42.60998796 | 38.45907632 | ENSG00000236565 |
| SLC15A3 | -0.150227919 | 0.901108093 | 0.27370175 | 0.440492159 | 435.2337512 | 392.3040791 | ENSG00000110446 |
| DBF4B | -0.150233537 | 0.901104584 | 0.075775201 | 0.166092422 | 910.6996439 | 820.1348437 | ENSG00000161692 |
| POLG | -0.150260794 | 0.90108756 | 0.018115606 | 0.051681183 | 1889.182035 | 1702.658696 | ENSG00000140521 |
| CPNE3 | -0.150274465 | 0.901079021 | 0.064892258 | 0.146739719 | 905.8071966 | 816.1298348 | ENSG00000085719 |
| KYAT3 | -0.150289453 | 0.901069659 | 0.072130372 | 0.159813228 | 756.4798865 | 681.8437497 | ENSG00000137944 |
| LMNB1 | -0.150402765 | 0.900998891 | 0.001904968 | 0.007563947 | 4821.964634 | 4345.481451 | ENSG00000113368 |
| NA | -0.150485987 | 0.900946918 | 0.835987432 | 0.906924912 | 8.741593674 | 7.8501839 | ENSG00000254332 |
| NCAPG2 | -0.150515331 | 0.900928594 | 0.006663934 | 0.022458761 | 2420.600249 | 2180.472594 | ENSG00000146918 |
| NA | -0.15068074 | 0.900825305 | 0.894596333 | 0.941915276 | 3.65475845 | 3.304519581 | ENSG00000260361 |
| SLC51A | -0.150724771 | 0.900797813 | 0.547624382 | 0.705307928 | 80.94271833 | 72.86964763 | ENSG00000163959 |
| SPATA2L | -0.150753676 | 0.900779765 | 0.31910336 | 0.490581998 | 326.7279656 | 294.6247185 | ENSG00000158792 |
| STX1B | -0.150911878 | 0.900680993 | 0.260860507 | 0.426173785 | 298.8285605 | 269.1788461 | ENSG00000099365 |
| MGAT1 | -0.150960471 | 0.900650657 | 0.005476743 | 0.01897483 | 2967.534448 | 2673.06825 | ENSG00000131446 |
| MLF2 | -0.150998127 | 0.900627149 | 0.001759208 | 0.007058422 | 5953.350685 | 5361.399438 | ENSG00000089693 |
| TM9SF4 | -0.151096251 | 0.900565896 | 0.028091019 | 0.074498741 | 1870.001105 | 1683.567155 | ENSG00000101337 |
| EIF4E | -0.151110468 | 0.900557021 | 0.009099068 | 0.029046851 | 2149.397661 | 1936.132515 | ENSG00000151247 |
| POLDIP2 | -0.151112337 | 0.900555854 | 0.004743734 | 0.016763394 | 2936.590268 | 2644.900743 | ENSG00000004142 |
| DMWD | -0.151181951 | 0.900512401 | 0.124173726 | 0.244606897 | 712.6702944 | 641.687527 | ENSG00000185800 |
| VGF | -0.151223091 | 0.900486722 | 0.365077128 | 0.538803677 | 241.2108267 | 217.3984271 | ENSG00000128564 |
| GMFB | -0.151233174 | 0.900480429 | 0.050619655 | 0.120511386 | 1334.992164 | 1202.899855 | ENSG00000197045 |
| AP2S1 | -0.151292883 | 0.900443161 | 0.011477467 | 0.035461704 | 2869.82105 | 2583.889711 | ENSG00000042753 |

| | | | | | | | |
|---|---|---|---|---|---|---|---|
| NRSN2 | -0.151310666 | 0.900432063 | 0.16203277 | 0.299559412 | 646.8490746 | 582.7493142 | ENSG00000125841 |
| RBX1 | -0.151411748 | 0.900368976 | 0.020526054 | 0.057478314 | 2009.252768 | 1808.523494 | ENSG00000100387 |
| SUCLG1 | -0.151457477 | 0.900340438 | 0.080524683 | 0.174324055 | 718.7258256 | 647.0981361 | ENSG00000163541 |
| PXK | -0.151471827 | 0.900331483 | 0.350975459 | 0.523640184 | 187.1810557 | 168.5509504 | ENSG00000168297 |
| LRRC69 | -0.151499089 | 0.900314469 | 0.8337367 | 0.905786877 | 8.714360366 | 7.816307759 | ENSG00000214954 |
| NUP85 | -0.151542528 | 0.900287361 | 0.039554067 | 0.098826255 | 1446.898432 | 1302.048438 | ENSG00000125450 |
| NA | -0.151568183 | 0.900271352 | 0.891015746 | 0.939677714 | 3.981357982 | 3.58227143 | ENSG00000260418 |
| TAP2 | -0.151570826 | 0.900269703 | 0.4078715 | 0.580211571 | 141.638493 | 127.4410095 | ENSG00000204267 |
| MSL1 | -0.151574614 | 0.900267339 | 0.011999562 | 0.036623198 | 1878.292927 | 1691.007905 | ENSG00000188895 |
| CYB5A | -0.151577749 | 0.900265383 | 0.021699343 | 0.060118057 | 1721.492097 | 1549.672282 | ENSG00000166347 |
| MSH4 | -0.151586771 | 0.900259753 | 0.883965576 | 0.936493029 | 4.716256968 | 4.221665925 | ENSG00000057468 |
| ACTL6A | -0.151637292 | 0.900228228 | 0.017746726 | 0.050884674 | 1762.524117 | 1586.387414 | ENSG00000136518 |
| KCNS3 | -0.151753529 | 0.9001557 | 0.189478072 | 0.337494724 | 524.8255759 | 472.9412916 | ENSG00000170745 |
| FAF1 | -0.151777197 | 0.900140933 | 0.048937065 | 0.117340421 | 1085.498302 | 976.9535055 | ENSG00000185104 |
| ANP32AP1 | -0.15182594 | 0.900110521 | 0.800225521 | 0.884896069 | 13.88568642 | 12.43779362 | ENSG00000259516 |
| CDC14A | -0.151927086 | 0.900047417 | 0.393939988 | 0.566676241 | 164.7778996 | 148.3963269 | ENSG00000079335 |
| UBE3D | -0.152158602 | 0.899902994 | 0.257655913 | 0.42280029 | 278.4121596 | 250.4073236 | ENSG00000118420 |
| ACSL6 | -0.152229817 | 0.899858574 | 0.845546813 | 0.911814164 | 7.957809879 | 7.155940565 | ENSG00000164398 |
| UBN1 | -0.15224422 | 0.89984959 | 0.021938578 | 0.060632823 | 1694.452057 | 1525.094904 | ENSG00000118900 |
| JMJD8 | -0.152271009 | 0.899832881 | 0.105643329 | 0.215931921 | 731.8403455 | 658.8879635 | ENSG00000161999 |
| NA | -0.152292104 | 0.899819724 | 0.246973242 | 0.410493875 | 273.2397575 | 245.8587152 | ENSG00000184319 |
| AMER1 | -0.152363868 | 0.899774965 | 0.080882661 | 0.174882912 | 777.9640434 | 700.4195026 | ENSG00000184675 |
| TBC1D10B | -0.152436506 | 0.899729664 | 0.057817971 | 0.134042359 | 1032.582121 | 929.4329786 | ENSG00000169221 |
| ZC3H8 | -0.152583634 | 0.899637913 | 0.444389362 | 0.615161812 | 130.2250938 | 117.2717377 | ENSG00000144161 |
| NA | -0.1525955 | 0.899630514 | 0.739743632 | 0.845477132 | 22.55798308 | 20.32777234 | ENSG00000229212 |
| COTL1 | -0.152745786 | 0.899536803 | 0.003959078 | 0.014318441 | 2858.654051 | 2571.639865 | ENSG00000103187 |
| INTS10 | -0.152758521 | 0.899528863 | 0.035141428 | 0.089458075 | 1095.833909 | 985.6006456 | ENSG00000104613 |
| CTNNB1 | -0.152780835 | 0.89951495 | 2.72E-05 | 0.00016558 | 10890.5762 | 9796.252731 | ENSG00000168036 |
| CFL2 | -0.152834392 | 0.899481558 | 0.359224457 | 0.532283508 | 217.8998478 | 196.2390168 | ENSG00000165410 |
| RNF14 | -0.152963077 | 0.89940133 | 0.201147811 | 0.353294068 | 338.9406675 | 304.7138492 | ENSG00000013561 |
| TTC37 | -0.152963133 | 0.899401295 | 0.023530758 | 0.064315603 | 1447.006803 | 1301.487239 | ENSG00000198677 |
| HSPA14 | -0.15297793 | 0.899392071 | 0.217788462 | 0.375355778 | 430.1360022 | 386.815952 | ENSG00000284024 |
| UBLCP1 | -0.153074804 | 0.899331681 | 0.123620943 | 0.243741988 | 616.4249187 | 554.4699715 | ENSG00000164332 |
| IL4R | -0.153116788 | 0.899305509 | 0.06116931 | 0.140096005 | 895.693613 | 805.845256 | ENSG00000077238 |
| EFNA3 | -0.153244063 | 0.899226176 | 0.561770322 | 0.7165506 | 69.40925918 | 62.32932526 | ENSG00000143590 |
| INAFM1 | -0.153289586 | 0.899197802 | 0.460444595 | 0.630228231 | 121.9197086 | 109.6044015 | ENSG00000257704 |
| LRIF1 | -0.153298886 | 0.899192006 | 0.453139165 | 0.623386113 | 163.7171279 | 147.4864379 | ENSG00000121931 |
| BCOR | -0.153448286 | 0.899098893 | 0.021372314 | 0.059365486 | 1683.540309 | 1514.091589 | ENSG00000183337 |
| CHIC2 | -0.153694081 | 0.898945725 | 0.428147514 | 0.599418949 | 135.2120442 | 121.7321726 | ENSG00000109220 |
| NA | -0.153745743 | 0.898913535 | 0.80004779 | 0.884808916 | 12.37190972 | 11.09555349 | ENSG00000214391 |
| ABHD5 | -0.153767288 | 0.898900111 | 0.246056213 | 0.409442802 | 292.7701872 | 263.4455289 | ENSG00000011198 |
| SMIM4 | -0.153879 | 0.898830509 | 0.259037066 | 0.424164675 | 261.2128324 | 234.6851896 | ENSG00000168273 |
| PPIG | -0.153882334 | 0.898828432 | 0.007220726 | 0.023987432 | 2046.94688 | 1839.985234 | ENSG00000138398 |
| GANAB | -0.153924086 | 0.89880242 | 0.000123286 | 0.000652129 | 12354.41311 | 11103.87258 | ENSG00000089597 |
| SEC24C | -0.153987186 | 0.89876311 | 0.017222972 | 0.049636428 | 1411.371172 | 1268.268361 | ENSG00000176986 |
| SPTLC2 | -0.154023957 | 0.898740203 | 0.020400967 | 0.057169531 | 1462.320349 | 1314.571636 | ENSG00000100596 |
| GTF2IRD1 | -0.154252754 | 0.898597682 | 0.045968423 | 0.111592629 | 967.5779502 | 869.8832982 | ENSG00000006704 |
| SLC35E2B | -0.154324017 | 0.898553296 | 0.026034256 | 0.069996771 | 1472.338755 | 1323.073398 | ENSG00000189339 |
| RPGRIP1L | -0.154380222 | 0.898518291 | 0.296139108 | 0.466390688 | 235.4367749 | 211.3621147 | ENSG00000103494 |
| TASOR | -0.154425215 | 0.898490269 | 0.089450351 | 0.18981264 | 843.4724933 | 758.2326896 | ENSG00000163946 |
| PPOX | -0.154468584 | 0.89846326 | 0.11249851 | 0.22693807 | 579.3709386 | 520.2462581 | ENSG00000143224 |
| TEDC1 | -0.154469228 | 0.898462859 | 0.257292061 | 0.422420822 | 379.8233179 | 341.1682103 | ENSG00000185347 |
| NA | -0.154473169 | 0.898460405 | 0.670496727 | 0.797822181 | 36.53075106 | 32.90215503 | ENSG00000204685 |
| TONSL | -0.154534325 | 0.89842232 | 0.240224704 | 0.402411841 | 2847.847684 | 2558.580665 | ENSG00000160949 |
| GYS1 | -0.154706969 | 0.898314814 | 0.018773256 | 0.053242413 | 1485.632781 | 1335.089011 | ENSG00000104812 |
| TRIM23 | -0.154756895 | 0.898283727 | 0.342036791 | 0.514835246 | 199.5687228 | 179.3416002 | ENSG00000113595 |
| SPIRE1 | -0.154851397 | 0.898224889 | 0.25319228 | 0.417775294 | 314.1730062 | 281.8067327 | ENSG00000134278 |
| LCN12 | -0.154890759 | 0.898200382 | 0.794355406 | 0.880349251 | 13.52348087 | 12.08206966 | ENSG00000184925 |
| NOP14 | -0.154948852 | 0.898164215 | 0.047330447 | 0.114331537 | 1153.979303 | 1036.701912 | ENSG00000087269 |
| MZF1-AS1 | -0.154962083 | 0.898155978 | 0.719915923 | 0.83186387 | 27.75654244 | 24.90946338 | ENSG00000267858 |
| DDX1 | -0.155057724 | 0.898096438 | 0.002606084 | 0.009939497 | 4929.106052 | 4427.611227 | ENSG00000079785 |
| TMED10 | -0.155164182 | 0.898030169 | 0.002511136 | 0.009626854 | 3147.687028 | 2827.367704 | ENSG00000170348 |
| ABCD1 | -0.155237855 | 0.897984311 | 0.239439493 | 0.401523107 | 314.1690143 | 282.3299657 | ENSG00000101986 |
| MDFIC | -0.155257963 | 0.897971795 | 0.27210534 | 0.439080963 | 263.8299342 | 237.2518908 | ENSG00000135272 |
| SPATA5 | -0.155267072 | 0.897966126 | 0.315750507 | 0.48687218 | 252.0510999 | 226.5504991 | ENSG00000145375 |
| GNS | -0.155299863 | 0.897945716 | 0.115805539 | 0.232056161 | 893.6873824 | 802.4939505 | ENSG00000135677 |
| ATP6V0A1 | -0.155303098 | 0.897943703 | 0.053995476 | 0.126826582 | 813.6136031 | 730.3451771 | ENSG00000033627 |
| SPINT1 | -0.15532243 | 0.89793167 | 0.004138253 | 0.01487997 | 3046.651444 | 2735.688266 | ENSG00000166145 |
| NA | -0.155341322 | 0.897919912 | 0.145593688 | 0.275925713 | 459.2324848 | 412.2887399 | ENSG00000184809 |
| PGS1 | -0.155342368 | 0.897919261 | 0.02906422 | 0.076532606 | 1144.587576 | 1027.447363 | ENSG00000087157 |
| SRD5A3-AS1 | -0.155404133 | 0.89788082 | 0.813582553 | 0.893222983 | 11.57975322 | 10.39109033 | ENSG00000249700 |
| C19orf48 | -0.15545614 | 0.897848454 | 0.01451883 | 0.043073944 | 1977.194953 | 1775.266203 | ENSG00000167747 |
| MPP7 | -0.155670092 | 0.897715312 | 0.314654241 | 0.485871685 | 240.4927981 | 215.8526392 | ENSG00000150054 |
| ABCC5 | -0.155692545 | 0.897701341 | 0.071478529 | 0.158657333 | 1262.523871 | 1132.430279 | ENSG00000114770 |
| UBE2B | -0.155723677 | 0.897681969 | 0.18987865 | 0.337972785 | 351.7105521 | 315.9688425 | ENSG00000119048 |
| HINFP | -0.155841098 | 0.89760891 | 0.097130477 | 0.202501247 | 646.8639036 | 580.4850324 | ENSG00000172273 |
| NA | -0.156099667 | 0.897448049 | 0.230833686 | 0.391144963 | 408.2351235 | 366.0956902 | ENSG00000214826 |

| | | | | | | | |
|---|---|---|---|---|---|---|---|
| NAA15 | -0.156268824 | 0.897342829 | 0.004323954 | 0.015464109 | 2918.553004 | 2619.846294 | ENSG00000164134 |
| HS3ST3A1 | -0.156276433 | 0.897338096 | 0.413419322 | 0.585608724 | 162.1825756 | 145.810531 | ENSG00000153976 |
| USP30 | -0.156277422 | 0.897337481 | 0.468040096 | 0.63706857 | 116.7162801 | 104.4693574 | ENSG00000135093 |
| DAZAP1 | -0.156311034 | 0.897316575 | 0.00028775 | 0.001400701 | 9025.424692 | 8098.862272 | ENSG00000071626 |
| MIR4482 | -0.156361414 | 0.897285241 | 0.859145356 | 0.920821105 | 6.261972897 | 5.668769546 | ENSG00000266852 |
| CNOT8 | -0.156368381 | 0.897280907 | 0.040979661 | 0.101624712 | 999.684751 | 896.7824592 | ENSG00000155508 |
| CCDC58 | -0.156370556 | 0.897279555 | 0.08858523 | 0.18839132 | 685.4941363 | 615.2690717 | ENSG00000160124 |
| FAM53A | -0.156403523 | 0.897259051 | 0.61214585 | 0.755482865 | 50.73677949 | 45.48864765 | ENSG00000174137 |
| LSM8 | -0.156454803 | 0.897227159 | 0.139881476 | 0.267756852 | 656.0636468 | 588.8323878 | ENSG00000128534 |
| NA | -0.156513033 | 0.897190946 | 0.666521709 | 0.795139722 | 44.15725549 | 39.67897578 | ENSG00000246465 |
| GAREM2 | -0.156593066 | 0.897141176 | 0.060650743 | 0.139193735 | 867.2514198 | 778.5177415 | ENSG00000157833 |
| SLC18A2 | -0.156609519 | 0.897130945 | 0.56515568 | 0.719299216 | 72.48972759 | 64.93099921 | ENSG00000165646 |
| LOC101929331 | -0.156915555 | 0.896940659 | 0.894453035 | 0.941860999 | 3.333740724 | 3.01009618 | ENSG00000224307 |
| MNS1 | -0.157009652 | 0.896882159 | 0.600418038 | 0.746526209 | 58.18401453 | 52.0396029 | ENSG00000138587 |
| NELL2 | -0.157022964 | 0.896873884 | 0.025008828 | 0.067617934 | 1296.689718 | 1162.515914 | ENSG00000184613 |
| SREK1 | -0.157078654 | 0.896839264 | 0.069585078 | 0.155330448 | 714.8420002 | 641.3226251 | ENSG00000153914 |
| FUNDC1 | -0.157092348 | 0.896830751 | 0.297647313 | 0.467887411 | 214.1338777 | 192.1625422 | ENSG00000069509 |
| TRA2B | -0.157115175 | 0.896816561 | 0.001961158 | 0.00775419 | 7028.429571 | 6303.84894 | ENSG00000136527 |
| PHLDB1 | -0.157227072 | 0.896747006 | 0.030453532 | 0.079522997 | 1578.698041 | 1415.125982 | ENSG00000019144 |
| TWNK | -0.157272368 | 0.896718851 | 0.022855218 | 0.062778697 | 1716.205343 | 1538.377845 | ENSG00000107815 |
| TMEM255A | -0.157294038 | 0.896705382 | 0.638385286 | 0.773863705 | 47.22636012 | 42.41966194 | ENSG00000125355 |
| COX1 | -0.15745193 | 0.89660725 | 0.481839999 | 0.649875929 | 94440.68809 | 84676.24828 | ENSG00000198804 |
| MAP4K4 | -0.157506619 | 0.896573262 | 0.000410949 | 0.00192792 | 4786.874581 | 4292.066272 | ENSG00000071054 |
| DDX18 | -0.15751509 | 0.896567998 | 0.002332472 | 0.009018985 | 3254.223808 | 2918.16194 | ENSG00000088205 |
| ANKMY1 | -0.157584426 | 0.89652491 | 0.453830061 | 0.623840007 | 113.4972177 | 101.7743847 | ENSG00000144504 |
| AARS2 | -0.157620308 | 0.896502612 | 0.04792294 | 0.115502428 | 881.6239512 | 790.5269642 | ENSG00000124608 |
| NEK8 | -0.157632499 | 0.896495037 | 0.231878457 | 0.392474763 | 286.9800288 | 257.4855526 | ENSG00000160602 |
| DCUN1D4 | -0.157641793 | 0.896489262 | 0.112699573 | 0.227201115 | 718.518298 | 644.123812 | ENSG00000109184 |
| CAMSAP1 | -0.157658182 | 0.896479078 | 0.0078742 | 0.025785298 | 1813.618622 | 1626.402988 | ENSG00000130559 |
| CDIP1 | -0.157763151 | 0.896413853 | 0.019591557 | 0.055262572 | 1765.81268 | 1583.434844 | ENSG00000089486 |
| TRAPPC13 | -0.157764702 | 0.896412889 | 0.371097148 | 0.544770952 | 178.5492711 | 159.9763967 | ENSG00000113597 |
| KLHL23 | -0.157775274 | 0.89640632 | 0.039866747 | 0.099377141 | 1281.744425 | 1149.864558 | ENSG00000213160 |
| UBFD1 | -0.157995837 | 0.896269286 | 0.001936994 | 0.007669653 | 3220.52681 | 2886.270048 | ENSG00000103353 |
| PEX13 | -0.158127637 | 0.896187409 | 0.199717103 | 0.351207168 | 473.5784345 | 424.2887968 | ENSG00000162928 |
| EPN1 | -0.158147571 | 0.896175026 | 0.01170648 | 0.035925819 | 4474.577538 | 4010.73775 | ENSG00000063245 |
| SORCS1 | -0.158193594 | 0.896146438 | 0.379980159 | 0.553233522 | 156.9941407 | 140.8892361 | ENSG00000108018 |
| DTD1 | -0.158209244 | 0.896136717 | 0.157377504 | 0.293302042 | 407.7076911 | 365.1650584 | ENSG00000125821 |
| FBXO36 | -0.158353753 | 0.896046959 | 0.608486633 | 0.752263243 | 56.4274534 | 50.45940575 | ENSG00000153832 |
| PABPC4 | -0.158383673 | 0.896028376 | 0.002408785 | 0.009289858 | 4669.222867 | 4183.550488 | ENSG00000090621 |
| DCTN6 | -0.158493805 | 0.895959978 | 0.518652347 | 0.682248095 | 80.06348026 | 71.85192683 | ENSG00000104671 |
| NA | -0.158511807 | 0.895948799 | 0.838432002 | 0.907948341 | 7.652861849 | 6.850764296 | ENSG00000226049 |
| SUPT6H | -0.158546667 | 0.89592715 | 0.003881797 | 0.014078538 | 3449.0565 | 3089.968607 | ENSG00000109111 |
| BTN2A1 | -0.158619401 | 0.895881983 | 0.206989364 | 0.361393051 | 320.7337448 | 287.0926264 | ENSG00000112763 |
| TTC38 | -0.158720253 | 0.895819358 | 0.044895737 | 0.109492246 | 929.3421874 | 832.3548231 | ENSG00000075234 |
| TYK2 | -0.158735031 | 0.895810182 | 0.118208141 | 0.235768533 | 690.7223948 | 618.7142942 | ENSG00000105397 |
| NA | -0.158768226 | 0.89578957 | 0.838142199 | 0.907736423 | 7.982252283 | 7.130666718 | ENSG00000178162 |
| THUMPD1 | -0.158867636 | 0.895727847 | 0.045007864 | 0.109714628 | 1108.439229 | 992.589374 | ENSG00000066654 |
| FAM200B | -0.15887117 | 0.895725653 | 0.303142988 | 0.473842477 | 196.9551628 | 176.5315442 | ENSG00000237765 |
| ZNF775 | -0.158879395 | 0.895720546 | 0.108835948 | 0.220680499 | 624.0661286 | 559.2963502 | ENSG00000196456 |
| ATG2B | -0.159018206 | 0.895634368 | 0.198408019 | 0.349797944 | 357.2467993 | 319.8998531 | ENSG00000066739 |
| SRA1 | -0.159050777 | 0.895614147 | 0.059427558 | 0.136820268 | 1003.968806 | 899.49517 | ENSG00000213523 |
| SUZ12 | -0.159137576 | 0.895560265 | 0.050689865 | 0.120633875 | 858.4971478 | 769.2919622 | ENSG00000178691 |
| SRF | -0.159180639 | 0.895533534 | 0.002682458 | 0.010186477 | 3146.500254 | 2818.093364 | ENSG00000112658 |
| NA | -0.159196973 | 0.895523395 | 0.877720087 | 0.932795186 | 4.376378643 | 3.895516956 | ENSG00000273026 |
| COX11 | -0.159314878 | 0.89545021 | 0.083836803 | 0.180139871 | 770.7511449 | 690.0225299 | ENSG00000166260 |
| EBAG9 | -0.159377832 | 0.895411137 | 0.262270763 | 0.427817548 | 268.3842308 | 240.5980954 | ENSG00000147654 |
| YIF1B | -0.159398091 | 0.895398564 | 0.015446406 | 0.045354128 | 1503.64106 | 1345.986941 | ENSG00000167645 |
| DVL1 | -0.159464643 | 0.89535726 | 0.013480852 | 0.040476128 | 2290.057433 | 2050.651312 | ENSG00000107404 |
| CHRDL1 | -0.159477419 | 0.895349331 | 0.076784985 | 0.167791021 | 792.7070268 | 709.3153648 | ENSG00000101938 |
| EMSY | -0.159487707 | 0.895342946 | 0.117379325 | 0.234555102 | 577.2157141 | 516.5467712 | ENSG00000158636 |
| LY6G5C | -0.15962383 | 0.895258471 | 0.781034563 | 0.871424971 | 14.60084892 | 13.08148926 | ENSG00000204428 |
| SH3BP1 | -0.159668342 | 0.89523085 | 0.481373539 | 0.649337595 | 95.1813617 | 85.27812672 | ENSG00000100092 |
| SNX12 | -0.159698071 | 0.895212403 | 0.011111023 | 0.034463952 | 1968.81027 | 1762.6412 | ENSG00000147164 |
| INSR | -0.159725419 | 0.895195433 | 0.060570775 | 0.139054006 | 724.2958022 | 648.2422383 | ENSG00000171105 |
| TAC1 | -0.159906342 | 0.895083177 | 0.652891864 | 0.784340326 | 37.97822181 | 33.96423226 | ENSG00000006128 |
| PHF13 | -0.159957597 | 0.895051377 | 0.017889943 | 0.051234395 | 1319.233158 | 1180.258956 | ENSG00000116273 |
| ESCO1 | -0.160016501 | 0.895014834 | 0.105995846 | 0.21640024 | 729.3866549 | 652.9594339 | ENSG00000141446 |
| NA | -0.160070164 | 0.894981543 | 0.328722133 | 0.501000083 | 180.1269263 | 161.2105567 | ENSG00000078319 |
| MED28 | -0.160126352 | 0.894946687 | 0.007198848 | 0.023922994 | 2248.474598 | 2012.054435 | ENSG00000118579 |
| SLC35B4 | -0.160151833 | 0.894930881 | 0.045971003 | 0.111592629 | 969.4620667 | 867.5120266 | ENSG00000205060 |
| PPAT | -0.16016391 | 0.89492339 | 0.026781365 | 0.071655938 | 1782.806806 | 1595.795955 | ENSG00000128059 |
| MRPL37 | -0.160202841 | 0.89489924 | 0.003437473 | 0.012640545 | 3067.407299 | 2745.378098 | ENSG00000116221 |
| NAB1 | -0.160301504 | 0.894838042 | 0.143506442 | 0.273042979 | 471.6469337 | 421.9151326 | ENSG00000138386 |
| ZBTB49 | -0.16031357 | 0.894830558 | 0.465807893 | 0.635135713 | 108.2417131 | 96.74339212 | ENSG00000168826 |
| COMMD10 | -0.160343542 | 0.894811968 | 0.228206393 | 0.38776795 | 301.967926 | 270.365372 | ENSG00000145781 |
| ABI3BP | -0.160407559 | 0.894772264 | 0.884505983 | 0.936795019 | 3.687573563 | 3.302369007 | ENSG00000154175 |
| WNT5A | -0.160409556 | 0.894771025 | 0.559523797 | 0.714772106 | 63.54402181 | 56.98858269 | ENSG00000114251 |

| | | | | | | | |
|---|---|---|---|---|---|---|---|
| HSPA13 | -0.160447185 | 0.894747688 | 0.05793915 | 0.134242631 | 773.4090839 | 692.1926221 | ENSG00000155304 |
| MKRN1 | -0.160468217 | 0.894734644 | 0.008949626 | 0.028650387 | 2655.488866 | 2375.96324 | ENSG00000133606 |
| NA | -0.16047693 | 0.89472924 | 0.855605167 | 0.918479793 | 6.29478801 | 5.675221267 | ENSG00000286360 |
| RSL1D1 | -0.160516328 | 0.894704806 | 0.000440045 | 0.002049964 | 4612.560991 | 4126.815358 | ENSG00000171490 |
| NPHP1 | -0.160520688 | 0.894702103 | 0.658709834 | 0.789118264 | 36.59849647 | 32.76288235 | ENSG00000144061 |
| PTPRF | -0.160535917 | 0.894692658 | 1.33E-05 | 8.52E-05 | 10224.2215 | 9147.952222 | ENSG00000142949 |
| CCDC181 | -0.160577975 | 0.894666576 | 0.52560987 | 0.687835929 | 75.64814112 | 67.64452142 | ENSG00000117477 |
| GCH1 | -0.160627227 | 0.894636034 | 0.319912398 | 0.491395021 | 201.0655539 | 179.9955157 | ENSG00000131979 |
| NA | -0.160635978 | 0.894630607 | 0.897035581 | 0.943140937 | 3.290437721 | 2.957397913 | ENSG00000243024 |
| PUM3 | -0.160723941 | 0.894576062 | 0.008546461 | 0.027597888 | 1746.24643 | 1562.415536 | ENSG00000080608 |
| NAF1 | -0.160724208 | 0.894575897 | 0.106352946 | 0.216899743 | 609.4725011 | 545.2987928 | ENSG00000145414 |
| LSM10 | -0.160725656 | 0.894574999 | 0.077028139 | 0.168265182 | 836.3076789 | 748.1033163 | ENSG00000181817 |
| PCYT1B | -0.160752511 | 0.894558347 | 0.021891681 | 0.060520552 | 1164.047581 | 1041.600822 | ENSG00000102230 |
| ACAA2 | -0.16077283 | 0.894545748 | 0.016344067 | 0.047499196 | 1824.952698 | 1632.553355 | ENSG00000167315 |
| TMEM41B | -0.160861258 | 0.89449092 | 0.167470153 | 0.307523935 | 523.775329 | 468.583073 | ENSG00000166471 |
| HHIP | -0.160892491 | 0.894471555 | 0.892879404 | 0.940687531 | 3.314880125 | 2.980521186 | ENSG00000164161 |
| GSPT2 | -0.160949486 | 0.894436219 | 0.049064403 | 0.117511638 | 793.4178979 | 709.7305176 | ENSG00000189369 |
| ARHGAP33 | -0.160999431 | 0.894405254 | 0.13223308 | 0.256063877 | 575.1560831 | 514.307043 | ENSG00000004777 |
| PSMA7 | -0.161014857 | 0.894395691 | 0.000998836 | 0.004259182 | 7244.169799 | 6478.968754 | ENSG00000101182 |
| SLC25A17 | -0.161042392 | 0.894378621 | 0.060627871 | 0.139168521 | 706.0210219 | 631.698383 | ENSG00000100372 |
| CPPED1 | -0.161127685 | 0.894325746 | 0.265771964 | 0.431904761 | 252.8654956 | 226.280556 | ENSG00000103381 |
| ATXN7L1 | -0.161159102 | 0.894306271 | 0.039330591 | 0.098420863 | 914.2652936 | 817.3002952 | ENSG00000146776 |
| SMCHD1 | -0.161174135 | 0.894296952 | 0.059086274 | 0.136247482 | 1008.524263 | 901.7069465 | ENSG00000101596 |
| NA | -0.161251978 | 0.8942487 | 0.883151342 | 0.936194777 | 3.663131159 | 3.277095161 | ENSG00000255966 |
| SLC12A9-AS1 | -0.161257424 | 0.894245325 | 0.778387191 | 0.869988497 | 16.10904381 | 14.36404636 | ENSG00000236305 |
| SYNPO2 | -0.161317804 | 0.894207899 | 0.383523134 | 0.556806306 | 142.1207954 | 127.0651064 | ENSG00000172403 |
| ZNF32 | -0.161393363 | 0.894161068 | 0.368166726 | 0.542077874 | 157.6831459 | 140.913456 | ENSG00000169740 |
| ZFAND2B | -0.16144025 | 0.894132009 | 0.322371007 | 0.493873817 | 202.3279733 | 181.0600039 | ENSG00000158552 |
| RNASEH2B | -0.161462768 | 0.894118053 | 0.074386295 | 0.163848597 | 722.199526 | 646.2683761 | ENSG00000136104 |
| NA | -0.161465076 | 0.894116622 | 0.834140549 | 0.906149642 | 8.006694688 | 7.132817292 | ENSG00000251259 |
| DLG3 | -0.161483442 | 0.89410524 | 0.024784278 | 0.06712885 | 1242.430543 | 1110.55564 | ENSG00000082458 |
| CLIC1 | -0.161508105 | 0.894089955 | 0.001118846 | 0.004713618 | 4650.244699 | 4156.7685 | ENSG00000213719 |
| ZNF207 | -0.161523177 | 0.894080615 | 0.025999237 | 0.06991724 | 2710.172374 | 2423.826969 | ENSG00000010244 |
| TMEM167B | -0.161574441 | 0.894048846 | 0.244927963 | 0.408086441 | 256.964523 | 229.802072 | ENSG00000215717 |
| DGCR2 | -0.161663779 | 0.893993484 | 0.001710793 | 0.006885649 | 3059.055394 | 2734.727266 | ENSG00000070413 |
| FXR2 | -0.161722115 | 0.893957336 | 0.015004212 | 0.044284923 | 1317.942029 | 1177.918896 | ENSG00000129245 |
| AKNA | -0.161808467 | 0.893903829 | 0.027971642 | 0.074253604 | 1166.041108 | 1042.522052 | ENSG00000106948 |
| CARD6 | -0.161829095 | 0.893891048 | 0.85050039 | 0.915007697 | 7.015732482 | 6.285040767 | ENSG00000132357 |
| EPS15L1 | -0.161948394 | 0.893817134 | 0.045949996 | 0.111569713 | 932.874846 | 833.386253 | ENSG00000127527 |
| EHBP1L1 | -0.161955147 | 0.89381295 | 0.381974102 | 0.555370998 | 151.0395433 | 135.0335902 | ENSG00000173442 |
| PCAT1 | -0.162035638 | 0.893763084 | 0.724244943 | 0.83450006 | 21.65218742 | 19.3686806 | ENSG00000253438 |
| RPS21 | -0.16207006 | 0.89374176 | 0.004286735 | 0.015350919 | 6777.530579 | 6056.956839 | ENSG00000171858 |
| EPM2AIP1 | -0.162090121 | 0.893729332 | 0.304791126 | 0.475513595 | 235.9460352 | 210.46353 | ENSG00000178567 |
| DHRS7 | -0.162102156 | 0.893721877 | 0.086660585 | 0.184991629 | 803.5993392 | 717.9544299 | ENSG00000100612 |
| YY1 | -0.162186936 | 0.893669358 | 0.003335891 | 0.012299821 | 2912.843958 | 2603.591301 | ENSG00000100811 |
| MLST8 | -0.162215926 | 0.893651401 | 0.016615487 | 0.04813476 | 1347.473769 | 1204.418498 | ENSG00000167965 |
| FAM20C | -0.162253691 | 0.893628008 | 0.043616976 | 0.106995814 | 1206.663753 | 1078.40795 | ENSG00000177706 |
| AKT1 | -0.162281425 | 0.89361083 | 0.028162342 | 0.074636591 | 2147.511211 | 1918.150571 | ENSG00000142208 |
| FAM131A | -0.162375094 | 0.893552812 | 0.189144123 | 0.337069679 | 387.3423286 | 346.3830803 | ENSG00000175182 |
| SLCO5A1 | -0.162441649 | 0.893511592 | 0.465145645 | 0.634636441 | 106.5701183 | 95.22556161 | ENSG00000137571 |
| FREM1 | -0.162563802 | 0.893435941 | 0.549294448 | 0.706644528 | 63.67057634 | 56.84769248 | ENSG00000164946 |
| GNAO1 | -0.162607339 | 0.89340898 | 0.049786622 | 0.118894988 | 1094.240236 | 977.927415 | ENSG00000087258 |
| CDCA8 | -0.16280161 | 0.893288683 | 0.022402448 | 0.061720312 | 1282.084014 | 1145.37112 | ENSG00000134690 |
| MCRIP1 | -0.162846806 | 0.893260699 | 0.009980567 | 0.031372841 | 1907.709601 | 1704.126475 | ENSG00000225663 |
| ATP5MF | -0.16301611 | 0.893155879 | 0.238989619 | 0.400969624 | 329.4393691 | 294.1947784 | ENSG00000241468 |
| SHQ1 | -0.163019779 | 0.893153608 | 0.096938714 | 0.202238844 | 702.1411884 | 627.0607523 | ENSG00000144736 |
| SLC31A1 | -0.16305485 | 0.893131896 | 0.005435814 | 0.018856732 | 1978.603499 | 1766.499363 | ENSG00000136868 |
| NUP50 | -0.163057178 | 0.893130455 | 0.002695772 | 0.010228982 | 2559.422524 | 2286.631562 | ENSG00000093000 |
| SMARCD1 | -0.16310192 | 0.893102757 | 0.000413339 | 0.001937248 | 4194.216724 | 3746.352375 | ENSG00000066117 |
| TMEM98 | -0.163141429 | 0.893078299 | 0.012095534 | 0.036855937 | 1730.905099 | 1545.264327 | ENSG00000006042 |
| EP400P1 | -0.163143658 | 0.893076919 | 0.185167301 | 0.331433852 | 321.9414974 | 287.5179807 | ENSG00000185684 |
| PRPF6 | -0.163295595 | 0.892982869 | 0.002171015 | 0.008479663 | 3415.288178 | 3050.03201 | ENSG00000101161 |
| COX7A1 | -0.163426062 | 0.892902118 | 0.891524233 | 0.939954256 | 3.300925611 | 2.959548487 | ENSG00000161281 |
| TMEM132B | -0.163441958 | 0.89289228 | 0.003540102 | 0.012987443 | 2101.667445 | 1876.427709 | ENSG00000139364 |
| CRYBG1 | -0.163488982 | 0.892863177 | 0.079781479 | 0.173045256 | 618.2715013 | 552.153543 | ENSG00000112297 |
| WDR83 | -0.163495931 | 0.892858877 | 0.676204697 | 0.801417018 | 38.3198519 | 34.06909576 | ENSG00000123154 |
| TOM1L2 | -0.163503029 | 0.892854484 | 0.480967371 | 0.649016617 | 86.08613535 | 76.92298682 | ENSG00000175662 |
| NA | -0.163561346 | 0.892818393 | 0.580485124 | 0.731378652 | 53.40125145 | 47.73324056 | ENSG00000268222 |
| UTP15 | -0.16356349 | 0.892817066 | 0.086849173 | 0.185366016 | 607.816137 | 542.6941627 | ENSG00000164338 |
| RPS15A | -0.163584275 | 0.892804204 | 0.071731344 | 0.159072232 | 668.6077536 | 597.2585092 | ENSG00000134419 |
| NA | -0.163609326 | 0.892788701 | 0.774124877 | 0.86752893 | 17.15100601 | 15.23064501 | ENSG00000273723 |
| FBXO11 | -0.163668765 | 0.892751919 | 0.076208778 | 0.166720745 | 605.298832 | 540.2825695 | ENSG00000138081 |
| MLEC | -0.163674277 | 0.892748508 | 3.74E-05 | 0.000220662 | 6711.39328 | 5991.512674 | ENSG00000110917 |
| UBA1 | -0.163742506 | 0.892706289 | 4.54E-06 | 3.18E-05 | 19372.83528 | 17293.55755 | ENSG00000130985 |
| DTD2 | -0.163768871 | 0.892689975 | 0.572894966 | 0.725403106 | 57.62559402 | 51.48167605 | ENSG00000129480 |
| CCDC57 | -0.163815696 | 0.892661001 | 0.498890281 | 0.665244516 | 95.31050698 | 85.04689399 | ENSG00000176155 |
| SLC7A6 | -0.163839395 | 0.892646338 | 0.18677847 | 0.333717094 | 346.8555747 | 309.2549519 | ENSG00000103064 |

| | | | | | | | |
|---|---|---|---|---|---|---|---|
| CEP170B | -0.163892847 | 0.892613266 | 0.028098786 | 0.074509096 | 1993.534854 | 1779.105854 | ENSG00000099814 |
| DOCK7 | -0.16402023 | 0.892534456 | 0.070868816 | 0.15757611 | 761.8269642 | 680.2745232 | ENSG00000116641 |
| COMMD8 | -0.164152049 | 0.892452909 | 0.623129206 | 0.762604966 | 44.45786101 | 39.76959087 | ENSG00000169019 |
| MED31 | -0.164226095 | 0.892407105 | 0.621878707 | 0.762199274 | 45.64802924 | 40.66844813 | ENSG00000108590 |
| TRMT112 | -0.16426516 | 0.892382941 | 0.002293802 | 0.008885886 | 4077.167647 | 3637.683626 | ENSG00000173113 |
| RGL2 | -0.164304656 | 0.892358511 | 0.025949548 | 0.06980794 | 1030.853343 | 920.0423619 | ENSG00000237441 |
| ATG5 | -0.164314177 | 0.892352622 | 0.056214978 | 0.131066369 | 739.7792806 | 659.8575476 | ENSG00000057663 |
| NA | -0.16434035 | 0.892336433 | 0.628173147 | 0.766129882 | 40.07398554 | 35.75200583 | ENSG00000218226 |
| ZBTB11 | -0.164361517 | 0.892323341 | 0.225962961 | 0.384982474 | 264.8882784 | 236.3680997 | ENSG00000066422 |
| ZNF800 | -0.164366595 | 0.8923202 | 0.572112349 | 0.724710705 | 56.70285278 | 50.64922612 | ENSG00000048405 |
| ARFGAP2 | -0.164378862 | 0.892312613 | 0.019976612 | 0.056184404 | 1683.314781 | 1501.17982 | ENSG00000149182 |
| NA | -0.164383752 | 0.892309589 | 0.605318719 | 0.7500773 | 52.22053197 | 46.54990085 | ENSG00000229132 |
| UBXN2A | -0.164395153 | 0.892302537 | 0.127776702 | 0.249690069 | 520.316185 | 464.4934529 | ENSG00000173960 |
| CAB39 | -0.164435102 | 0.892277829 | 0.038323409 | 0.096250971 | 1257.543293 | 1122.621517 | ENSG00000135932 |
| RBMXL1 | -0.164487817 | 0.892245227 | 0.094348724 | 0.197770427 | 569.5943159 | 508.1416045 | ENSG00000213516 |
| GVINP1 | -0.164504201 | 0.892235094 | 0.872196132 | 0.928970491 | 4.400821048 | 3.895516956 | ENSG00000254838 |
| FKBP14 | -0.164513795 | 0.89222916 | 0.383482012 | 0.556806306 | 176.8349 | 158.0595703 | ENSG00000106080 |
| APBB2 | -0.164641351 | 0.892150277 | 0.039775262 | 0.099224696 | 884.4409476 | 789.4113525 | ENSG00000163697 |
| ARPC1A | -0.164689715 | 0.89212037 | 0.425414262 | 0.597109031 | 113.1112454 | 101.0163992 | ENSG00000241685 |
| ABCA17P | -0.164718 | 0.892102879 | 0.828367387 | 0.902331876 | 7.734561772 | 6.88033929 | ENSG00000238098 |
| NAE1 | -0.164743243 | 0.89208727 | 0.005959085 | 0.0204186 | 2186.731244 | 1950.695082 | ENSG00000159593 |
| FAF2 | -0.164784938 | 0.892061489 | 0.014360804 | 0.042686329 | 1278.674206 | 1140.948051 | ENSG00000113194 |
| MYO1E | -0.164805138 | 0.892048999 | 0.012805833 | 0.038775284 | 1337.348169 | 1192.869063 | ENSG00000157483 |
| DBNDD1 | -0.164829883 | 0.892033698 | 0.055825269 | 0.13044153 | 1365.670081 | 1218.835013 | ENSG00000003249 |
| SNX22 | -0.164858699 | 0.892015881 | 0.507855992 | 0.67259714 | 89.40400654 | 79.6712092 | ENSG00000157734 |
| PITPNC1 | -0.164875354 | 0.892005584 | 0.02034403 | 0.057034834 | 1163.17544 | 1037.467124 | ENSG00000154217 |
| CDK16 | -0.164907443 | 0.891985743 | 0.000653409 | 0.002903018 | 4537.828269 | 4048.569595 | ENSG00000102225 |
| TNS3 | -0.164909299 | 0.891984596 | 0.035515295 | 0.090207446 | 896.8449087 | 799.9826375 | ENSG00000136205 |
| SAP130 | -0.165008594 | 0.891923206 | 0.011932946 | 0.036473734 | 1464.989388 | 1306.373059 | ENSG00000136715 |
| DNAH9 | -0.165085638 | 0.891875576 | 0.763031984 | 0.859491417 | 15.12645432 | 13.46625518 | ENSG00000007174 |
| MCUR1 | -0.165104191 | 0.891864107 | 0.069683657 | 0.155514511 | 746.4054124 | 665.9753398 | ENSG00000050393 |
| ZSCAN21 | -0.165109993 | 0.89186052 | 0.300114684 | 0.470731849 | 226.6802756 | 202.1188398 | ENSG00000166529 |
| NDUFS5 | -0.165155101 | 0.891832635 | 0.001326872 | 0.005491672 | 5315.301535 | 4739.536508 | ENSG00000168653 |
| NA | -0.16515857 | 0.891830491 | 0.22096651 | 0.379375663 | 303.4151966 | 270.3382628 | ENSG00000185596 |
| MFSD13A | -0.165303281 | 0.891741039 | 0.228993059 | 0.388858804 | 264.990754 | 236.3632655 | ENSG00000138111 |
| FAAH | -0.165324207 | 0.891728105 | 0.209943878 | 0.365426992 | 304.121623 | 270.9058643 | ENSG00000117480 |
| LIN7B | -0.165334624 | 0.891721666 | 0.622675737 | 0.762523814 | 48.18191647 | 43.05178068 | ENSG00000104863 |
| ULBP3 | -0.165423274 | 0.891666874 | 0.516577001 | 0.68046834 | 78.31664328 | 70.00042955 | ENSG00000131019 |
| WDR12 | -0.165461972 | 0.891642956 | 0.00605061 | 0.020688111 | 1919.023665 | 1710.934997 | ENSG00000138442 |
| NA | -0.165599905 | 0.891557713 | 0.71779728 | 0.83075518 | 23.29355779 | 20.84105807 | ENSG00000280649 |
| ANKS3 | -0.165662201 | 0.891519216 | 0.18719512 | 0.334244822 | 321.6648588 | 286.6078191 | ENSG00000168096 |
| TULP2 | -0.165667206 | 0.891516123 | 0.716299283 | 0.829818008 | 22.46599543 | 19.99517165 | ENSG00000104804 |
| NA | -0.165730709 | 0.891476882 | 0.890186723 | 0.93936831 | 3.303716514 | 2.95524734 | ENSG00000121089 |
| NA | -0.165789911 | 0.8914403 | 0.828074046 | 0.90215036 | 7.740143577 | 6.878188716 | ENSG00000272140 |
| DNASE2 | -0.165858885 | 0.891397682 | 0.128098426 | 0.250090753 | 444.42676 | 396.0732084 | ENSG00000105612 |
| PHF10 | -0.165893157 | 0.891376507 | 0.085838295 | 0.183663105 | 1054.611079 | 940.2039338 | ENSG00000130024 |
| MAPKAPK5 | -0.165899653 | 0.891372493 | 0.053070059 | 0.124987731 | 865.2076952 | 771.766213 | ENSG00000089022 |
| CHAC2 | -0.165920092 | 0.891359865 | 0.339031828 | 0.511557527 | 202.6575639 | 180.5857195 | ENSG00000143942 |
| RFC1 | -0.165927881 | 0.891355053 | 0.003117863 | 0.011580034 | 1976.385286 | 1761.581 | ENSG00000035928 |
| NA | -0.165992044 | 0.891315411 | 0.690697918 | 0.812355416 | 27.92464821 | 24.91323149 | ENSG00000232931 |
| KHSRP | -0.166034948 | 0.891288905 | 0.00051343 | 0.002347612 | 13177.22142 | 11744.54926 | ENSG00000088247 |
| SH3BGRL | -0.166075175 | 0.891264054 | 0.082331868 | 0.177539019 | 656.8906456 | 585.9365572 | ENSG00000131171 |
| MRPL20-AS1 | -0.166187137 | 0.891194889 | 0.137511725 | 0.264122772 | 504.5791078 | 449.3752318 | ENSG00000224870 |
| NA | -0.166240765 | 0.891161761 | 0.885771618 | 0.937037652 | 3.306507417 | 2.957397913 | ENSG00000278743 |
| ELOVL6 | -0.166245967 | 0.891158548 | 0.002262812 | 0.008788424 | 2475.841203 | 2207.095076 | ENSG00000170522 |
| NA | -0.166287256 | 0.891133044 | 0.885053735 | 0.936934918 | 3.665922062 | 3.262574183 | ENSG00000256282 |
| ENOX1 | -0.166347259 | 0.891095982 | 0.701318527 | 0.819997315 | 26.22948695 | 23.46208718 | ENSG00000120658 |
| FIBP | -0.166431018 | 0.891044249 | 0.015818317 | 0.046291134 | 1332.087321 | 1187.025266 | ENSG00000172500 |
| FKBPL | -0.166443284 | 0.891036673 | 0.358233203 | 0.531369199 | 160.2863302 | 142.593955 | ENSG00000204315 |
| GABARAPL2 | -0.166551848 | 0.890969625 | 0.041832742 | 0.103354948 | 837.1337265 | 746.1813284 | ENSG00000034713 |
| NA | -0.166555163 | 0.890967577 | 0.473031182 | 0.641764156 | 85.11538519 | 75.86279969 | ENSG00000234771 |
| TRAM1 | -0.166666742 | 0.890898671 | 0.010890817 | 0.033893014 | 1723.943022 | 1536.460879 | ENSG00000067167 |
| MYT1 | -0.166964577 | 0.890714771 | 0.497829411 | 0.66419719 | 80.27990726 | 71.43679242 | ENSG00000196132 |
| MIR647 | -0.166988093 | 0.890700252 | 0.890838799 | 0.939542468 | 4.335190821 | 3.838517542 | ENSG00000207554 |
| ZNF256 | -0.166999463 | 0.890693232 | 0.333163667 | 0.505846687 | 228.1699222 | 202.8214555 | ENSG00000152454 |
| TSC2 | -0.167003413 | 0.890690793 | 0.007782099 | 0.025541165 | 2195.09077 | 1955.421159 | ENSG00000103197 |
| DMPK | -0.167007901 | 0.890688023 | 0.143755588 | 0.273351275 | 420.1279732 | 374.1929359 | ENSG00000104936 |
| DENND2C | -0.167021878 | 0.890679393 | 0.52533021 | 0.687620103 | 74.11781916 | 66.0629672 | ENSG00000175984 |
| CLPP | -0.167059515 | 0.890656158 | 0.023531156 | 0.064315603 | 1152.551681 | 1026.938627 | ENSG00000125656 |
| BIN1 | -0.167094092 | 0.890634812 | 0.017416499 | 0.050101654 | 1421.451867 | 1265.735022 | ENSG00000136717 |
| ANKRD28 | -0.16713701 | 0.890608317 | 0.023783699 | 0.064858826 | 1040.017278 | 926.2706574 | ENSG00000206560 |
| ZC2HC1C | -0.167154216 | 0.890597696 | 0.674903305 | 0.800563221 | 31.94228793 | 28.44442219 | ENSG00000119703 |
| TLNRD1 | -0.167216724 | 0.890559109 | 0.034341105 | 0.087688404 | 1321.750153 | 1177.479039 | ENSG00000140406 |
| MICALL2 | -0.167224521 | 0.890554297 | 0.321952626 | 0.493428784 | 186.4777325 | 165.9285763 | ENSG00000164877 |
| LAG3 | -0.167282771 | 0.89051834 | 0.553275203 | 0.709662844 | 61.4838641 | 54.83916647 | ENSG00000089692 |
| ZC3H3 | -0.167307054 | 0.890503351 | 0.082587686 | 0.178011067 | 796.1272387 | 709.401152 | ENSG00000014164 |
| LMTK2 | -0.167328007 | 0.890490419 | 0.038048635 | 0.095672082 | 913.8228276 | 813.8438358 | ENSG00000164715 |

| | | | | | | | |
|---|---|---|---|---|---|---|---|
| TRAK2 | -0.167452197 | 0.890413766 | 0.189676101 | 0.337705737 | 325.6889193 | 290.2949419 | ENSG00000115993 |
| NA | -0.167501863 | 0.890383114 | 0.742312956 | 0.847585631 | 18.44691625 | 16.40052982 | ENSG00000261094 |
| MAGED4B | -0.167621753 | 0.890309125 | 0.596286228 | 0.743851064 | 46.05333763 | 41.00266635 | ENSG00000187243 |
| PPP1R16A | -0.167623758 | 0.890307887 | 0.242833919 | 0.405655449 | 253.2462495 | 225.4021626 | ENSG00000160972 |
| TIMM8A | -0.167651844 | 0.890290555 | 0.149914362 | 0.282283841 | 440.5639841 | 392.0572472 | ENSG00000126953 |
| SERGEF | -0.167699914 | 0.890260892 | 0.289296046 | 0.458748272 | 218.9713075 | 195.0096973 | ENSG00000129158 |
| IFNLR1 | -0.167753293 | 0.890227953 | 0.198659364 | 0.350049122 | 296.4575975 | 264.0077447 | ENSG00000185436 |
| UTP6 | -0.167824768 | 0.89018385 | 0.009364692 | 0.029766729 | 1740.008561 | 1548.681198 | ENSG00000108651 |
| KLHDC10 | -0.167937351 | 0.890114385 | 0.151352792 | 0.284575703 | 453.6096315 | 404.15809 | ENSG00000128607 |
| DDX54 | -0.167957098 | 0.890102202 | 0.003761615 | 0.013699323 | 3490.530185 | 3106.460334 | ENSG00000123064 |
| STAT2 | -0.168021894 | 0.890062226 | 0.078993271 | 0.171702443 | 613.8980257 | 546.1653793 | ENSG00000170581 |
| GLIPR1L1 | -0.168060188 | 0.890038601 | 0.760234897 | 0.857637004 | 15.12366341 | 13.42215921 | ENSG00000173401 |
| ZNF419 | -0.168202274 | 0.889950949 | 0.340461835 | 0.512992991 | 173.9747625 | 154.7295043 | ENSG00000105136 |
| HECTD3 | -0.168209804 | 0.889946303 | 0.04906214 | 0.117511638 | 1049.414513 | 934.0491278 | ENSG00000126107 |
| CADPS2 | -0.168278112 | 0.889904167 | 0.115717871 | 0.231904577 | 459.2360266 | 408.9344846 | ENSG00000081803 |
| NA | -0.168289182 | 0.889897339 | 0.888948169 | 0.938578344 | 4.362424128 | 3.857872705 | ENSG00000278993 |
| POC1A | -0.168320036 | 0.889878308 | 0.030042402 | 0.078636089 | 957.2940689 | 851.3726382 | ENSG00000164087 |
| APEX1 | -0.168351532 | 0.889858881 | 0.000109853 | 0.000588333 | 9201.501034 | 8187.533801 | ENSG00000100823 |
| NA | -0.16835536 | 0.889856519 | 0.875364247 | 0.931060908 | 4.016963998 | 3.552696436 | ENSG00000231977 |
| VPS51 | -0.168493081 | 0.889771577 | 0.010685903 | 0.033323918 | 2139.585483 | 1903.65821 | ENSG00000149823 |
| NUP98 | -0.168521871 | 0.889753821 | 0.000878066 | 0.0037852 | 3155.460625 | 2807.529634 | ENSG00000110713 |
| BCLAF1 | -0.168563262 | 0.889728295 | 0.000683198 | 0.003020151 | 3038.387535 | 2703.714198 | ENSG00000029363 |
| SURF4 | -0.168750996 | 0.889612524 | 0.000310752 | 0.001503945 | 6462.147827 | 5749.258053 | ENSG00000148248 |
| SOBP | -0.168821357 | 0.889569138 | 0.339128143 | 0.511622821 | 175.8710079 | 156.3175224 | ENSG00000112320 |
| CNOT1 | -0.168849572 | 0.889551741 | 0.000489816 | 0.002248701 | 6495.885029 | 5778.431162 | ENSG00000125107 |
| ZFP41 | -0.168855304 | 0.889548207 | 0.250963828 | 0.41513833 | 274.4056466 | 244.0776844 | ENSG00000181638 |
| RPL21 | -0.168912903 | 0.889512692 | 4.50E-05 | 0.000260933 | 12117.41096 | 10777.67673 | ENSG00000122026 |
| NA | -0.168994097 | 0.889462633 | 0.558708766 | 0.714203884 | 63.33927088 | 56.2357224 | ENSG00000223546 |
| PRKACB | -0.169124538 | 0.889382216 | 0.107188095 | 0.21823383 | 523.204781 | 465.5154628 | ENSG00000142875 |
| PGRMC2 | -0.169268517 | 0.889293461 | 0.011269625 | 0.034896041 | 1491.732243 | 1326.977989 | ENSG00000164040 |
| NA | -0.169297421 | 0.889275645 | 0.801213952 | 0.885438723 | 11.33764451 | 10.04826981 | ENSG00000218537 |
| ZBTB46 | -0.169332655 | 0.889253927 | 0.124729297 | 0.245375574 | 538.8312328 | 479.0016003 | ENSG00000130584 |
| RXRA | -0.169333408 | 0.889253463 | 0.008718731 | 0.028064881 | 1634.030942 | 1452.451388 | ENSG00000186350 |
| NUP153 | -0.169421865 | 0.889198941 | 0.006126764 | 0.020896643 | 2363.402429 | 2102.056221 | ENSG00000124789 |
| ZNF862 | -0.169502733 | 0.889149099 | 0.532481739 | 0.693264847 | 66.4321422 | 58.944683 | ENSG00000106479 |
| ZNF420 | -0.169530213 | 0.889132163 | 0.502117152 | 0.667977518 | 82.73931599 | 73.63378172 | ENSG00000197050 |
| EPAS1 | -0.169610122 | 0.889082917 | 0.524828116 | 0.687196035 | 71.02427212 | 62.99829475 | ENSG00000116016 |
| NOTO | -0.169641401 | 0.889063641 | 0.815165411 | 0.89428314 | 9.296347397 | 8.312921933 | ENSG00000214513 |
| CENPT | -0.169725397 | 0.88901188 | 0.021305538 | 0.05920558 | 1074.445445 | 954.8786027 | ENSG00000102901 |
| TALDO1 | -0.169826088 | 0.888949835 | 0.000345591 | 0.001653472 | 5249.746866 | 4666.514866 | ENSG00000177156 |
| NA | -0.169929057 | 0.88888639 | 2.86E-05 | 0.000173089 | 2461511.144 | 2188003.061 | N_unmapped |
| CMTM8 | -0.169933455 | 0.888883681 | 0.413735118 | 0.585917205 | 145.671602 | 129.7038915 | ENSG00000170293 |
| RPAP3 | -0.169938459 | 0.888880597 | 0.052188861 | 0.123350208 | 780.7618158 | 694.5261883 | ENSG00000005175 |
| MBOAT7 | -0.170035351 | 0.888820902 | 0.009325932 | 0.029663075 | 1954.804587 | 1737.709189 | ENSG00000125505 |
| MAPK6 | -0.170134483 | 0.88875983 | 0.004797864 | 0.016929871 | 2832.894428 | 2518.591041 | ENSG00000069956 |
| AKAP12 | -0.170166085 | 0.888740362 | 0.000612093 | 0.002740921 | 3692.104309 | 3281.047414 | ENSG00000131016 |
| ZNF367 | -0.170217173 | 0.888708891 | 0.292279006 | 0.461985166 | 184.6836134 | 164.2687895 | ENSG00000165244 |
| STAU1 | -0.170219948 | 0.888707182 | 0.000480792 | 0.002213067 | 3314.18442 | 2945.874698 | ENSG00000124214 |
| CPD | -0.170349162 | 0.888627589 | 0.046373155 | 0.112413236 | 1518.176967 | 1349.178152 | ENSG00000108582 |
| C5 | -0.170379556 | 0.888608868 | 0.825173279 | 0.900593514 | 8.175676339 | 7.22315981 | ENSG00000106804 |
| SETD9 | -0.170479002 | 0.888547618 | 0.416502648 | 0.588583443 | 123.0550468 | 109.0843915 | ENSG00000155542 |
| BRPF1 | -0.170490251 | 0.88854069 | 0.041721455 | 0.103119615 | 834.5994759 | 741.3966652 | ENSG00000156983 |
| NA | -0.170516415 | 0.888524575 | 0.786201709 | 0.875152278 | 12.70130015 | 11.33753909 | ENSG00000231566 |
| HABP4 | -0.17053481 | 0.888513247 | 0.156457273 | 0.29203511 | 405.1201762 | 360.3636752 | ENSG00000130956 |
| CHST10 | -0.170584733 | 0.888482501 | 0.04503815 | 0.109742446 | 821.9561601 | 730.5621615 | ENSG00000115526 |
| ZNF607 | -0.170587958 | 0.888480515 | 0.206382645 | 0.360594838 | 292.2487242 | 259.7541049 | ENSG00000198182 |
| LOC100128568 | -0.170595681 | 0.888475759 | 0.804478855 | 0.887069454 | 10.61949094 | 9.42339629 | ENSG00000266983 |
| SPEG | -0.170611346 | 0.888466111 | 0.029412082 | 0.077248261 | 1192.920615 | 1059.852129 | ENSG00000072195 |
| LDLRAD3 | -0.170865881 | 0.888309373 | 0.118413431 | 0.236055957 | 416.2024333 | 369.8723131 | ENSG00000179241 |
| HINT3 | -0.17090754 | 0.888283723 | 0.241372418 | 0.403703426 | 231.9882068 | 206.2176442 | ENSG00000111911 |
| SPDL1 | -0.170941646 | 0.888262723 | 0.029391157 | 0.077203813 | 947.786327 | 842.3178451 | ENSG00000040275 |
| PXDC1 | -0.170954021 | 0.888255104 | 0.634899148 | 0.771274526 | 40.81147527 | 36.40241366 | ENSG00000168994 |
| WBP11 | -0.171105569 | 0.888161802 | 0.000286688 | 0.001396661 | 3493.407841 | 3102.764401 | ENSG00000084463 |
| ZNF174 | -0.171109114 | 0.88815962 | 0.310310767 | 0.481279598 | 222.9245194 | 198.1149944 | ENSG00000103343 |
| AJAP1 | -0.171226138 | 0.88808758 | 0.185577719 | 0.332002206 | 321.9404214 | 286.0154952 | ENSG00000196581 |
| EIF4G1 | -0.171230208 | 0.888085075 | 0.000187276 | 0.000949723 | 20928.39032 | 18586.00323 | ENSG00000114867 |
| STIP1 | -0.171270913 | 0.888060018 | 6.18E-05 | 0.000349161 | 12266.81295 | 10893.43086 | ENSG00000168439 |
| ZNF451 | -0.171274115 | 0.888058047 | 0.120903332 | 0.239937179 | 496.4907132 | 441.3214332 | ENSG00000112200 |
| RUVBL2 | -0.171499111 | 0.88791956 | 0.002554568 | 0.009770058 | 4136.469583 | 3672.469102 | ENSG00000183207 |
| PCGF1 | -0.171514878 | 0.887909857 | 0.333561358 | 0.506034301 | 179.532224 | 159.1835117 | ENSG00000115289 |
| PFDN4 | -0.171608556 | 0.887852204 | 0.092072497 | 0.194000353 | 578.7626083 | 513.9781799 | ENSG00000101132 |
| TNFRSF25 | -0.171718421 | 0.887784594 | 0.372180374 | 0.54561377 | 146.7585068 | 130.2136938 | ENSG00000215788 |
| TRAM1L1 | -0.171813875 | 0.887725857 | 0.719581243 | 0.831672264 | 26.02298426 | 23.03726598 | ENSG00000174599 |
| NA | -0.171818833 | 0.887722806 | 0.84050346 | 0.908966938 | 6.687017769 | 5.946521394 | ENSG00000268521 |
| CWC27 | -0.17185057 | 0.887703278 | 0.01730861 | 0.049843319 | 1172.794132 | 1041.310724 | ENSG00000153015 |
| KIF22 | -0.171857493 | 0.887699018 | 0.001207275 | 0.005059643 | 3080.217981 | 2733.7023 | ENSG00000079616 |
| ZIK1 | -0.171903444 | 0.887670745 | 0.072411095 | 0.160284846 | 717.4304662 | 636.6511856 | ENSG00000171649 |

| | | | | | | | |
|---|---|---|---|---|---|---|---|
| CACNA1E | -0.172020592 | 0.887598668 | 0.694147399 | 0.815120372 | 24.68935282 | 21.88646376 | ENSG00000198216 |
| PER1 | -0.172024464 | 0.887596286 | 0.084247029 | 0.180840039 | 893.4461155 | 793.2663543 | ENSG00000179094 |
| TMED4 | -0.172034243 | 0.88759027 | 0.023724645 | 0.064743545 | 985.6685678 | 874.9422298 | ENSG00000158604 |
| ULBP1 | -0.172036953 | 0.887588602 | 0.801549257 | 0.885657066 | 9.63411054 | 8.582071487 | ENSG00000111981 |
| SNN | -0.17214837 | 0.887520058 | 0.001101998 | 0.004647714 | 2575.135857 | 2285.366283 | ENSG00000184602 |
| ERLIN2 | -0.172245155 | 0.887460519 | 0.024337574 | 0.066107594 | 1264.767091 | 1122.131336 | ENSG00000147475 |
| TMEM53 | -0.172255538 | 0.887454132 | 0.328281996 | 0.500544419 | 155.0090619 | 137.4306624 | ENSG00000126106 |
| CTAGE15 | -0.172279525 | 0.887439377 | 0.826762174 | 0.901230101 | 7.383519835 | 6.531067048 | ENSG00000271079 |
| PLEKHM1 | -0.172293219 | 0.887430954 | 0.237343879 | 0.399076159 | 298.2826678 | 264.3508805 | ENSG00000225190 |
| SNRPN | -0.17235977 | 0.887390018 | 0.465593592 | 0.635023048 | 86.18824747 | 76.34463242 | ENSG00000128739 |
| NA | -0.172415724 | 0.887355602 | 0.745187644 | 0.849008672 | 16.66937934 | 14.81441399 | ENSG00000268869 |
| NA | -0.172491181 | 0.887309192 | 0.885704176 | 0.937037652 | 3.360974032 | 2.986972907 | ENSG00000232385 |
| TMEM87A | -0.172527923 | 0.887286595 | 0.157487417 | 0.293417985 | 378.5320625 | 335.9817518 | ENSG00000103978 |
| SPPL2A | -0.172564848 | 0.887263885 | 0.281850321 | 0.450277013 | 288.7546258 | 256.4662569 | ENSG00000138600 |
| ATXN10 | -0.172587187 | 0.887250147 | 0.005080109 | 0.01777301 | 2033.973146 | 1804.925556 | ENSG00000130638 |
| YBX1 | -0.172592467 | 0.887246899 | 0.00086229 | 0.003723568 | 27825.67389 | 24688.7091 | ENSG00000065978 |
| TOP3A | -0.172600886 | 0.887241722 | 0.014867736 | 0.043949356 | 1159.241727 | 1028.490073 | ENSG00000177302 |
| STIM2 | -0.17265271 | 0.887209851 | 0.10373078 | 0.213106547 | 441.6556701 | 391.930587 | ENSG00000109689 |
| ZSCAN16 | -0.172788083 | 0.887126605 | 0.621009117 | 0.761892741 | 42.42234586 | 37.58253041 | ENSG00000196812 |
| UBAC2 | -0.172794019 | 0.887122955 | 0.020727367 | 0.057865651 | 1176.25191 | 1043.026977 | ENSG00000134882 |
| RFLNA | -0.172880308 | 0.887069897 | 0.31404677 | 0.485226851 | 219.5010183 | 194.8878834 | ENSG00000178882 |
| RPL37 | -0.172985515 | 0.887005211 | 0.001298576 | 0.005397723 | 9042.986425 | 8020.315805 | ENSG00000145592 |
| MFF-DT | -0.173122114 | 0.88692123 | 0.802758652 | 0.886231954 | 9.955128266 | 8.838850636 | ENSG00000236432 |
| ELOB | -0.173177072 | 0.886887444 | 0.009445103 | 0.029977867 | 2846.702796 | 2524.41733 | ENSG00000103363 |
| NECTIN3 | -0.173182685 | 0.886883994 | 0.076082278 | 0.166519539 | 528.2757836 | 468.3596553 | ENSG00000177707 |
| RPL29 | -0.17319101 | 0.886878876 | 0.000139334 | 0.000726458 | 17539.98572 | 15554.9731 | ENSG00000162244 |
| NMT1 | -0.173195905 | 0.886875867 | 0.000179165 | 0.000911704 | 3669.270462 | 3253.932092 | ENSG00000136448 |
| PCDHB18P | -0.173310623 | 0.886805349 | 0.878664594 | 0.933298075 | 3.714806871 | 3.283546882 | ENSG00000146001 |
| TM6SF1 | -0.173397774 | 0.88675178 | 0.877350206 | 0.93256891 | 3.660340256 | 3.232999189 | ENSG00000136404 |
| SCFD1 | -0.173478592 | 0.886702106 | 0.056693845 | 0.131959575 | 794.2485777 | 704.6159068 | ENSG00000092108 |
| PKD1P5-LOC10537675 | -0.173479334 | 0.886701651 | 0.19614583 | 0.346623517 | 299.0914448 | 265.3656024 | ENSG00000254681 |
| GMEB2 | -0.173604308 | 0.886624843 | 0.046886084 | 0.113513078 | 790.3830479 | 700.3929264 | ENSG00000101216 |
| THAP12 | -0.173618187 | 0.886616313 | 0.030002293 | 0.078552421 | 905.9058805 | 803.3847264 | ENSG00000137492 |
| ABHD2 | -0.173653988 | 0.886594312 | 0.060525822 | 0.138967345 | 660.644562 | 586.0207269 | ENSG00000140526 |
| HEATR5B | -0.17379139 | 0.886509877 | 0.093686794 | 0.196660712 | 478.8908204 | 424.6560855 | ENSG00000008869 |
| CAMK1G | -0.173812562 | 0.886496867 | 0.709913737 | 0.825195583 | 21.80230847 | 19.27188636 | ENSG00000008118 |
| NA | -0.173838577 | 0.886480882 | 0.81973528 | 0.897041177 | 9.016517493 | 8.041621805 | ENSG00000279039 |
| AKT2 | -0.174046549 | 0.8863531 | 0.001797541 | 0.007192774 | 3078.766355 | 2728.162657 | ENSG00000105221 |
| AVPI1 | -0.174048996 | 0.886351597 | 0.299292726 | 0.469823092 | 178.5457676 | 158.4459538 | ENSG00000119986 |
| TMEM184B | -0.174176476 | 0.88627328 | 0.021657571 | 0.060028189 | 1784.957149 | 1582.723371 | ENSG00000198792 |
| C12orf49 | -0.174197299 | 0.886260488 | 0.00431572 | 0.015440384 | 1591.332108 | 1410.185768 | ENSG00000111412 |
| ARAF | -0.174231655 | 0.886239383 | 0.044493051 | 0.1086888 | 945.1577733 | 837.26273 | ENSG00000078061 |
| TMPO | -0.174264675 | 0.8862191 | 0.002377127 | 0.009175103 | 3187.833555 | 2825.896913 | ENSG00000120802 |
| DPP3 | -0.174483674 | 0.886084583 | 0.002276801 | 0.008830273 | 2397.803714 | 2125.230683 | ENSG00000254986 |
| HEMK1 | -0.174509933 | 0.886068455 | 0.306596202 | 0.477595877 | 172.6219581 | 152.9922784 | ENSG00000114735 |
| CHDH | -0.174527812 | 0.886057475 | 0.154180138 | 0.288962309 | 362.0676343 | 320.8331501 | ENSG00000016391 |
| NA | -0.174542722 | 0.886048318 | 0.67363426 | 0.800111161 | 29.01214073 | 25.64511907 | ENSG00000286067 |
| SYAP1 | -0.174711104 | 0.88594491 | 0.004084744 | 0.014723244 | 1810.048806 | 1604.062952 | ENSG00000169895 |
| CSPP1 | -0.174820272 | 0.885877873 | 0.208783673 | 0.363637219 | 270.8673182 | 240.1896731 | ENSG00000104218 |
| ZNF879 | -0.174845742 | 0.885862233 | 0.594088807 | 0.742168577 | 47.21519651 | 41.9636361 | ENSG00000234284 |
| TMEM106C | -0.174850497 | 0.885859314 | 0.128491195 | 0.25065463 | 420.4657732 | 372.5110861 | ENSG00000134291 |
| MTMR12 | -0.174861592 | 0.885852501 | 0.009931661 | 0.031244564 | 1293.531667 | 1145.869315 | ENSG00000150712 |
| TRIM15 | -0.174952935 | 0.885796416 | 0.822951488 | 0.89905749 | 9.433189656 | 8.386592899 | ENSG00000204610 |
| IRF8 | -0.174974953 | 0.885782897 | 0.639640568 | 0.774741394 | 35.83165825 | 31.80620165 | ENSG00000140968 |
| CCT7 | -0.174975118 | 0.885782796 | 8.01E-06 | 5.38E-05 | 15113.86389 | 13387.40445 | ENSG00000135624 |
| EMC1 | -0.174994461 | 0.88577092 | 0.004133896 | 0.014871043 | 1634.020605 | 1447.41232 | ENSG00000127463 |
| NA | -0.175095955 | 0.885708608 | 0.856046351 | 0.918873256 | 5.548725413 | 4.955704085 | ENSG00000224669 |
| SCRT1 | -0.175107063 | 0.885701788 | 0.833004677 | 0.905233741 | 7.078571806 | 6.259766921 | ENSG00000261678 |
| BMS1 | -0.175516454 | 0.88545049 | 0.000400501 | 0.001886709 | 3428.553942 | 3036.020689 | ENSG00000165733 |
| MSH2 | -0.175618976 | 0.88538757 | 6.77E-05 | 0.000379768 | 6804.563866 | 6024.514774 | ENSG00000095002 |
| EFEMP2 | -0.17582598 | 0.88526054 | 0.499720876 | 0.666029806 | 89.86697277 | 79.40097515 | ENSG00000172638 |
| FASN | -0.175851363 | 0.885244964 | 0.000899952 | 0.003866585 | 35295.16818 | 31244.84679 | ENSG00000169710 |
| CACTIN | -0.175936435 | 0.885192765 | 0.064331259 | 0.145744771 | 921.0721101 | 815.4313819 | ENSG00000105298 |
| API5 | -0.175960865 | 0.885177776 | 0.001178692 | 0.00494738 | 3258.030854 | 2884.689197 | ENSG00000166181 |
| ELP6 | -0.176086396 | 0.885100759 | 0.044023034 | 0.107731619 | 836.2020987 | 739.8083683 | ENSG00000163832 |
| TNIP1 | -0.176112082 | 0.885085001 | 0.051658591 | 0.122531154 | 825.5976014 | 730.7073223 | ENSG00000145901 |
| SLC25A33 | -0.176133305 | 0.88507198 | 0.04510597 | 0.109879943 | 724.004972 | 640.827659 | ENSG00000171612 |
| FNDC3A | -0.17615637 | 0.88505783 | 0.092567343 | 0.194755874 | 558.9673424 | 495.0496165 | ENSG00000102531 |
| SUV39H1 | -0.176170012 | 0.885049461 | 0.14507431 | 0.275292987 | 332.6394218 | 294.2633484 | ENSG00000101945 |
| PARD3 | -0.176447108 | 0.884879488 | 0.007124906 | 0.023724672 | 1638.615037 | 1450.015351 | ENSG00000148498 |
| RAD1 | -0.176473677 | 0.884863191 | 0.047525339 | 0.114743784 | 747.8963863 | 662.2983821 | ENSG00000113456 |
| PSMC4 | -0.176486378 | 0.884855401 | 0.003269215 | 0.012074753 | 3346.312399 | 2960.476091 | ENSG00000013275 |
| TRMT2A | -0.176576786 | 0.884799953 | 0.051362669 | 0.121934194 | 678.3278824 | 600.223195 | ENSG00000099899 |
| RAB14 | -0.176587444 | 0.884793417 | 0.069763156 | 0.155619918 | 944.4661035 | 835.9034783 | ENSG00000119396 |
| KIF3C | -0.176606381 | 0.884781803 | 0.001998479 | 0.007882368 | 2136.815218 | 1890.805773 | ENSG00000084731 |
| LINC00242 | -0.176625134 | 0.884770302 | 0.821558296 | 0.898400379 | 7.8218435 | 6.878188716 | ENSG00000229214 |
| UBC | -0.176641713 | 0.884760134 | 7.23E-05 | 0.000403076 | 10416.09991 | 9215.315412 | ENSG00000150991 |

| | | | | | | | |
|---|---|---|---|---|---|---|---|
| BMP2 | -0.176646137 | 0.884757421 | 0.562598539 | 0.717227624 | 64.26899648 | 57.03913038 | ENSG00000125845 |
| GREB1 | -0.176825645 | 0.884647342 | 0.0358732 | 0.0909967 | 958.2566834 | 847.1482765 | ENSG00000196208 |
| NTN4 | -0.176826182 | 0.884647012 | 0.652614059 | 0.784094076 | 30.39099019 | 26.8712098 | ENSG00000074527 |
| ANKRD17 | -0.176870529 | 0.88461982 | 0.001494358 | 0.006104968 | 2842.887076 | 2514.742122 | ENSG00000132466 |
| TTYH2 | -0.176908074 | 0.884596798 | 0.401767172 | 0.574195474 | 127.3410405 | 112.5144868 | ENSG00000141540 |
| CAND1 | -0.17696005 | 0.88456493 | 0.000191327 | 0.000968485 | 4960.892757 | 4388.333943 | ENSG00000111530 |
| NA | -0.176977315 | 0.884554344 | 0.691248278 | 0.812804495 | 25.67772429 | 22.63502289 | ENSG00000231167 |
| HMGN1 | -0.176980736 | 0.884552246 | 9.26E-05 | 0.000504533 | 7867.325902 | 6959.546904 | ENSG00000205581 |
| ZC3H7B | -0.177156955 | 0.884444209 | 0.002923101 | 0.010957936 | 3821.034097 | 3380.168354 | ENSG00000100403 |
| SCNN1G | -0.177160612 | 0.884441967 | 0.84383821 | 0.910501509 | 7.987834089 | 7.126626035 | ENSG00000166828 |
| SZRD1 | -0.177173497 | 0.884434068 | 0.001934286 | 0.007663651 | 2504.518658 | 2214.468377 | ENSG00000055070 |
| BEND4 | -0.17723478 | 0.8843965 | 0.016567164 | 0.048021978 | 1271.248059 | 1125.022575 | ENSG00000188848 |
| CDH6 | -0.177270435 | 0.884374643 | 0.622023153 | 0.762199274 | 35.92364591 | 31.79572135 | ENSG00000113361 |
| DNALI1 | -0.177394354 | 0.884298683 | 0.301989181 | 0.472786341 | 250.9207657 | 221.518192 | ENSG00000163879 |
| IBTK | -0.177479563 | 0.884246456 | 0.060844609 | 0.139533208 | 685.4429233 | 606.51977 | ENSG00000005700 |
| ASS1 | -0.177639186 | 0.884148627 | 0.002949468 | 0.011037475 | 2281.090246 | 2015.922061 | ENSG00000130707 |
| MED20 | -0.177658225 | 0.884136959 | 0.06236677 | 0.142262901 | 999.4411644 | 882.79279 | ENSG00000124641 |
| SLIT1 | -0.177659053 | 0.884136451 | 0.674562144 | 0.80030617 | 33.75176406 | 29.9200163 | ENSG00000187122 |
| SLC44A1 | -0.177706604 | 0.884107311 | 0.001926779 | 0.007635475 | 2007.326562 | 1774.942925 | ENSG00000070214 |
| ARPC2 | -0.177743641 | 0.884084614 | 0.000344209 | 0.00164731 | 4220.042329 | 3731.000357 | ENSG00000163466 |
| MTRNR2L2 | -0.177764731 | 0.88407169 | 0.656632957 | 0.787167933 | 31.18852834 | 27.64987697 | ENSG00000269028 |
| WDR45B | -0.177781822 | 0.884061217 | 0.001959259 | 0.00774986 | 2631.816908 | 2325.929389 | ENSG00000141580 |
| NA | -0.177813097 | 0.884042052 | 0.834813787 | 0.906542977 | 6.29478801 | 5.578427028 | ENSG00000278434 |
| TROAP | -0.177825188 | 0.884034644 | 0.045031882 | 0.109742446 | 693.9426965 | 613.727597 | ENSG00000135451 |
| BAG5 | -0.177845906 | 0.884021948 | 0.024142444 | 0.065642268 | 929.205984 | 821.7496926 | ENSG00000166170 |
| ACER3 | -0.177956078 | 0.883954442 | 0.2235467 | 0.382071356 | 261.7446584 | 231.7234785 | ENSG00000078124 |
| CHORDC1 | -0.177971401 | 0.883945053 | 0.004112529 | 0.014812321 | 1907.104198 | 1686.020185 | ENSG00000110172 |
| ZBTB12 | -0.178067014 | 0.883886473 | 0.028931745 | 0.076235801 | 945.557737 | 835.3066176 | ENSG00000204366 |
| NA | -0.178119558 | 0.883854282 | 0.843255496 | 0.91029706 | 6.243112298 | 5.527879334 | ENSG00000234791 |
| YAE1 | -0.178132714 | 0.883846222 | 0.18997144 | 0.338075559 | 280.3652991 | 247.9011053 | ENSG00000241127 |
| PTH2R | -0.178158835 | 0.88383022 | 0.838810395 | 0.908136413 | 5.938164268 | 5.277551906 | ENSG00000144407 |
| MTPN | -0.178220681 | 0.883792332 | 0.007537702 | 0.024860628 | 2097.234675 | 1853.872499 | ENSG00000105887 |
| AURKAIP1 | -0.178234697 | 0.883783746 | 0.005804578 | 0.01997077 | 2830.238507 | 2501.980155 | ENSG00000175756 |
| SNAPC1 | -0.178295579 | 0.883746451 | 0.372788282 | 0.546296171 | 121.5826212 | 107.6249054 | ENSG00000023608 |
| HAUS5 | -0.178306976 | 0.883739469 | 0.059330338 | 0.13669432 | 767.7787707 | 678.0749293 | ENSG00000249115 |
| HMCES | -0.178490744 | 0.883626907 | 0.013519327 | 0.040560083 | 1453.120796 | 1284.095614 | ENSG00000183624 |
| LOC100507373 | -0.178604446 | 0.88355727 | 0.553962782 | 0.710113365 | 67.16301098 | 59.12295699 | ENSG00000267169 |
| BTBD10 | -0.178626566 | 0.883543723 | 0.079318124 | 0.172272678 | 527.3276119 | 466.0930115 | ENSG00000148925 |
| PPP2CA | -0.178655716 | 0.883525871 | 0.001071171 | 0.004533558 | 2308.219487 | 2039.823739 | ENSG00000113575 |
| NA | -0.178744201 | 0.883471683 | 0.884543117 | 0.936795019 | 3.330949821 | 2.940726362 | ENSG00000275719 |
| ZCCHC24 | -0.178792381 | 0.883442179 | 0.1819229 | 0.327437458 | 289.8119308 | 256.0411631 | ENSG00000165424 |
| TRPM7 | -0.178879432 | 0.883388874 | 0.033132551 | 0.085107861 | 831.8303011 | 734.9647545 | ENSG00000092439 |
| CMC2 | -0.179124299 | 0.88323895 | 0.397980426 | 0.570346957 | 111.7233475 | 98.55939572 | ENSG00000103121 |
| NA | -0.179250253 | 0.883161843 | 0.405390422 | 0.577661887 | 109.9445713 | 96.93026841 | ENSG00000235552 |
| LSM12 | -0.179404536 | 0.883067402 | 0.052289078 | 0.123449835 | 981.4389458 | 867.200405 | ENSG00000161654 |
| NA | -0.179415823 | 0.883060493 | 0.830328916 | 0.903535375 | 6.324812221 | 5.597249153 | ENSG00000275155 |
| HAUS4 | -0.179496276 | 0.88301125 | 0.375918486 | 0.549340468 | 116.9379766 | 103.1994311 | ENSG00000092036 |
| USP14 | -0.179597545 | 0.88294927 | 0.002670239 | 0.010154067 | 2242.725368 | 1980.634883 | ENSG00000101557 |
| TSNARE1 | -0.179619805 | 0.882935647 | 0.187099636 | 0.334136185 | 339.2448277 | 299.3040399 | ENSG00000171045 |
| MID1 | -0.179642581 | 0.882921708 | 0.010546439 | 0.032931589 | 1265.847977 | 1117.534863 | ENSG00000101871 |
| TMEM9B | -0.179647861 | 0.882918477 | 0.230494947 | 0.390774064 | 245.8504389 | 216.9870487 | ENSG00000175348 |
| SHROOM2 | -0.179867816 | 0.882783876 | 0.02754384 | 0.073400777 | 900.4499434 | 794.7104712 | ENSG00000146950 |
| RTN4 | -0.17988183 | 0.882775301 | 0.003203726 | 0.011861422 | 6377.174887 | 5629.99475 | ENSG00000115310 |
| GSTZ1 | -0.179923092 | 0.882750053 | 0.137975999 | 0.264792577 | 462.109954 | 407.7565 | ENSG00000100577 |
| ARID3B | -0.179969746 | 0.882721507 | 0.001035394 | 0.00439467 | 5394.188867 | 4761.592287 | ENSG00000179361 |
| DENND10 | -0.18014866 | 0.882612045 | 0.03321451 | 0.085284321 | 792.2634848 | 699.1224245 | ENSG00000119979 |
| RTCA | -0.180148909 | 0.882611892 | 0.031887281 | 0.082579881 | 776.3697678 | 685.0090087 | ENSG00000137996 |
| APBB3 | -0.180246407 | 0.882552247 | 0.288208473 | 0.457486189 | 161.0943562 | 142.15056 | ENSG00000113108 |
| NEFL | -0.180410637 | 0.882451787 | 0.080671874 | 0.174564344 | 462.1026957 | 407.7823374 | ENSG00000277586 |
| MRPS12 | -0.180550695 | 0.882366122 | 0.013342807 | 0.040124097 | 1168.036963 | 1031.070211 | ENSG00000128626 |
| STK4 | -0.180679175 | 0.882287546 | 0.027837836 | 0.073990038 | 927.6761376 | 818.8046282 | ENSG00000101109 |
| LOC101927420 | -0.18077972 | 0.882226059 | 0.68946587 | 0.811401055 | 23.93280234 | 21.20700186 | ENSG00000224046 |
| NA | -0.180785134 | 0.882222748 | 0.443149738 | 0.614029679 | 93.85351224 | 82.832397 | ENSG00000213707 |
| GOT1 | -0.180793101 | 0.882217876 | 0.011908674 | 0.036418871 | 1492.492195 | 1316.676132 | ENSG00000120053 |
| WASHC2A | -0.180837448 | 0.882190758 | 0.019253689 | 0.054452854 | 941.4048066 | 830.1505702 | ENSG00000099290 |
| TMEM126A | -0.180871396 | 0.882169999 | 0.131387163 | 0.254809467 | 381.6883736 | 336.7730498 | ENSG00000171202 |
| URAHP | -0.180882636 | 0.882163126 | 0.792167768 | 0.879067442 | 10.12410992 | 8.922741433 | ENSG00000222019 |
| MARK3 | -0.180900157 | 0.882152413 | 0.004554238 | 0.016190165 | 1865.645994 | 1646.112351 | ENSG00000075413 |
| TBC1D25 | -0.180949138 | 0.882122463 | 0.297823078 | 0.468117187 | 179.235961 | 157.9057766 | ENSG00000068354 |
| CYCS | -0.180962808 | 0.882114105 | 0.000742125 | 0.003249492 | 5939.57625 | 5240.107158 | ENSG00000172115 |
| FAM111A-DT | -0.180998008 | 0.882092583 | 0.593116634 | 0.74153033 | 40.19408238 | 35.40918531 | ENSG00000245571 |
| MRFAP1L1 | -0.181089081 | 0.882036901 | 0.007316568 | 0.0242557 | 1494.690373 | 1318.489483 | ENSG00000178988 |
| DGUOK-AS1 | -0.181196622 | 0.881971155 | 0.838841052 | 0.908136413 | 6.36320914 | 5.620372426 | ENSG00000237883 |
| ABHD17A | -0.181198576 | 0.88196996 | 0.029179544 | 0.076782119 | 1611.564725 | 1421.895764 | ENSG00000129968 |
| CBFB | -0.181300749 | 0.8819075 | 0.038831827 | 0.097349549 | 674.4596 | 595.0332715 | ENSG00000067955 |
| FUBP3 | -0.181312349 | 0.88190041 | 0.002285537 | 0.00885703 | 2798.96924 | 2468.262172 | ENSG00000107164 |
| KIAA1841 | -0.181321154 | 0.881895027 | 0.19501641 | 0.345174931 | 263.4008592 | 232.5481317 | ENSG00000162929 |

| | | | | | | | |
|---|---|---|---|---|---|---|---|
| FAM50A | -0.181335751 | 0.881886104 | 0.015158383 | 0.044672326 | 1336.078704 | 1178.094511 | ENSG00000071859 |
| C5orf30 | -0.181393192 | 0.881850993 | 0.122375699 | 0.242128164 | 404.3697711 | 356.7513015 | ENSG00000181751 |
| VKORC1 | -0.181443466 | 0.881820263 | 0.757536948 | 0.856114219 | 13.0851572 | 11.5394694 | ENSG00000167397 |
| NA | -0.181498216 | 0.881786799 | 0.828380982 | 0.902331876 | 7.149783838 | 6.285040767 | ENSG00000272787 |
| ZNF730 | -0.181505816 | 0.881782154 | 0.489545361 | 0.656003026 | 79.17598162 | 69.7923201 | ENSG00000183850 |
| FAM131B | -0.181661168 | 0.881687207 | 0.018007561 | 0.051475324 | 1159.628425 | 1022.246917 | ENSG00000159784 |
| FIP1L1 | -0.181715034 | 0.881654288 | 0.246855473 | 0.41040025 | 211.3266182 | 186.1792006 | ENSG00000145216 |
| CD40 | -0.181945111 | 0.881513695 | 0.127811295 | 0.249698206 | 386.5270428 | 340.3997203 | ENSG00000101017 |
| TP53I3 | -0.182040645 | 0.881455325 | 0.080124162 | 0.17365186 | 522.5993524 | 460.3876759 | ENSG00000115129 |
| ADORA2B | -0.182056304 | 0.881445757 | 0.467463245 | 0.636762707 | 88.80631323 | 78.32626704 | ENSG00000170425 |
| ACOT8 | -0.182086353 | 0.881427398 | 0.10572683 | 0.216010674 | 395.148815 | 348.1942313 | ENSG00000101473 |
| RPL27A | -0.182087184 | 0.88142689 | 4.90E-05 | 0.000282319 | 14201.11726 | 12515.80738 | ENSG00000166441 |
| GTF3C4 | -0.182110368 | 0.881412726 | 0.005007666 | 0.017557776 | 1817.938707 | 1602.241103 | ENSG00000125484 |
| SOD2 | -0.182121248 | 0.881406079 | 0.043599006 | 0.106979562 | 761.2184707 | 670.7982167 | ENSG00000112096 |
| CCT6A | -0.182125992 | 0.881403181 | 0.000112582 | 0.000601447 | 10212.93225 | 9001.233991 | ENSG00000146731 |
| IKZF2 | -0.182128202 | 0.881401831 | 0.668191981 | 0.796028624 | 28.67437759 | 25.17216121 | ENSG00000030419 |
| NA | -0.182142705 | 0.88139297 | 0.883929404 | 0.936493029 | 3.36934674 | 2.966000208 | ENSG00000213026 |
| NFATC3 | -0.182257198 | 0.881323025 | 0.10320236 | 0.212241373 | 411.7045552 | 362.7282099 | ENSG00000072736 |
| BTBD2 | -0.182273155 | 0.881313277 | 0.001826943 | 0.007293715 | 2848.472721 | 2510.862265 | ENSG00000133243 |
| PIN4 | -0.182384686 | 0.881245148 | 0.049277709 | 0.117921892 | 646.3049195 | 569.4131537 | ENSG00000102309 |
| PAM16 | -0.182584648 | 0.881123013 | 0.503303216 | 0.668747918 | 63.84079729 | 56.29057124 | ENSG00000217930 |
| HMGA1 | -0.182606646 | 0.881109578 | 4.87E-05 | 0.000280611 | 54557.43527 | 48071.82692 | ENSG00000137309 |
| CPLX2 | -0.182646236 | 0.881085399 | 0.615116129 | 0.757594665 | 34.6717144 | 30.58630218 | ENSG00000145920 |
| EMC10 | -0.182735965 | 0.881030602 | 0.018189669 | 0.05187713 | 1593.284948 | 1403.199603 | ENSG00000161671 |
| NA | -0.182809313 | 0.88098581 | 0.876858408 | 0.932278526 | 3.312089222 | 2.911151367 | ENSG00000275552 |
| NA | -0.182841 | 0.88096646 | 0.568569188 | 0.721739041 | 44.82584852 | 39.51119419 | ENSG00000278635 |
| FAM117A | -0.182873312 | 0.88094673 | 0.084766237 | 0.181812927 | 650.4259248 | 572.7854924 | ENSG00000121104 |
| DERL1 | -0.183051685 | 0.880837817 | 0.033470589 | 0.085804803 | 841.1971336 | 740.9045884 | ENSG00000136986 |
| GPR4 | -0.183060066 | 0.880832701 | 0.828502581 | 0.902362426 | 7.038059705 | 6.17372555 | ENSG00000177464 |
| CWF19L2 | -0.183089764 | 0.880814569 | 0.094526812 | 0.198079158 | 432.1883139 | 380.6616242 | ENSG00000152404 |
| ZMIZ2 | -0.183102443 | 0.880806827 | 0.028200773 | 0.074707651 | 1700.501057 | 1497.808354 | ENSG00000122515 |
| MZT2B | -0.183161761 | 0.880770613 | 0.511306914 | 0.675720602 | 83.5330241 | 73.30255021 | ENSG00000152082 |
| KBTBD8 | -0.183247307 | 0.880718388 | 0.147355195 | 0.278470715 | 352.8860901 | 310.9601491 | ENSG00000163376 |
| RUFY2 | -0.183274223 | 0.880701957 | 0.348438039 | 0.521225047 | 135.257099 | 119.1337153 | ENSG00000204130 |
| IK | -0.183308041 | 0.880681313 | 0.007410526 | 0.024512423 | 1542.419597 | 1357.674285 | ENSG00000113141 |
| ZDHHC22 | -0.183319641 | 0.880674232 | 0.007361998 | 0.024376984 | 1533.20639 | 1350.18795 | ENSG00000177108 |
| LLPH | -0.183360344 | 0.880649386 | 0.238107498 | 0.4000133 | 224.957644 | 198.3152951 | ENSG00000139233 |
| HDAC6 | -0.183389465 | 0.88063161 | 0.025022304 | 0.067624722 | 844.1482197 | 743.277186 | ENSG00000094631 |
| GSKIP | -0.183410542 | 0.880618744 | 0.053785916 | 0.126422789 | 715.4387935 | 630.0372633 | ENSG00000100744 |
| APTR | -0.183528969 | 0.88054646 | 0.253348877 | 0.41793682 | 184.1374326 | 162.0930334 | ENSG00000214293 |
| CENPN | -0.183629471 | 0.88048512 | 0.076304503 | 0.166892309 | 488.7817579 | 430.3479905 | ENSG00000166451 |
| AOC2 | -0.183639189 | 0.88047919 | 0.521339322 | 0.684299983 | 63.70897326 | 55.99320377 | ENSG00000131480 |
| CTSO | -0.183649887 | 0.880472661 | 0.71762235 | 0.830701804 | 20.79248567 | 18.24772595 | ENSG00000256043 |
| LINC00367 | -0.183668135 | 0.880461524 | 0.873710241 | 0.929913771 | 3.393789145 | 2.98267176 | ENSG00000233780 |
| ACP1 | -0.183682227 | 0.880452924 | 0.000892234 | 0.003841352 | 2691.762004 | 2370.151738 | ENSG00000143727 |
| USP19 | -0.183739611 | 0.880417904 | 0.007545831 | 0.024874681 | 2226.706831 | 1959.758502 | ENSG00000172046 |
| DMAC1 | -0.183756923 | 0.880407339 | 0.019700998 | 0.055514455 | 1074.012715 | 945.1425123 | ENSG00000137038 |
| CYP46A1 | -0.183802845 | 0.880379316 | 0.770695178 | 0.864868483 | 15.11808161 | 13.23233884 | ENSG00000036530 |
| POLQ | -0.183834298 | 0.880360122 | 0.03331915 | 0.08550748 | 744.5822188 | 655.7399091 | ENSG00000051341 |
| IQCC | -0.183841733 | 0.880355586 | 0.507391219 | 0.67212003 | 66.37939045 | 58.59137007 | ENSG00000160051 |
| GSE1 | -0.183848946 | 0.880351184 | 0.00072258 | 0.003171624 | 2695.508587 | 2372.49269 | ENSG00000131149 |
| PAK4 | -0.183850231 | 0.8803504 | 0.010263578 | 0.032136776 | 2980.511515 | 2624.289918 | ENSG00000130669 |
| PRADC1 | -0.183868249 | 0.880339405 | 0.323163408 | 0.494734172 | 140.7015971 | 123.8590107 | ENSG00000135617 |
| GNAI2 | -0.183894678 | 0.880323278 | 6.18E-05 | 0.000349161 | 7105.709806 | 6255.124155 | ENSG00000114353 |
| MYLK3 | -0.184029642 | 0.880240928 | 0.768115439 | 0.8631511 | 11.99921628 | 10.60135036 | ENSG00000140795 |
| CENPK | -0.184053257 | 0.88022652 | 0.071548512 | 0.158721521 | 546.1784851 | 480.8856409 | ENSG00000123219 |
| ZNF71 | -0.184054055 | 0.880226032 | 0.182105003 | 0.327581836 | 283.4468067 | 249.6800161 | ENSG00000197951 |
| SLC7A8 | -0.184119409 | 0.880186159 | 1.86E-05 | 0.000116622 | 6340.497557 | 5580.791005 | ENSG00000092068 |
| FSD1 | -0.184153901 | 0.880165116 | 0.001368626 | 0.005646312 | 2043.08806 | 1798.189155 | ENSG00000105255 |
| MEI4 | -0.184235005 | 0.880115637 | 0.809202564 | 0.890310719 | 8.910375166 | 7.864704878 | ENSG00000269964 |
| ERGIC3 | -0.18437112 | 0.880032604 | 0.000602059 | 0.002700377 | 2874.426521 | 2528.884143 | ENSG00000125991 |
| NA | -0.18437859 | 0.880028048 | 0.81875757 | 0.896404784 | 7.879101018 | 6.88033929 | ENSG00000253106 |
| ALDH18A1 | -0.18445079 | 0.879984007 | 5.98E-05 | 0.000339527 | 4538.464369 | 3993.22262 | ENSG00000059573 |
| MIR1915HG | -0.184644422 | 0.879865908 | 0.213496749 | 0.369776063 | 225.9662275 | 198.9541565 | ENSG00000204682 |
| GSPT1 | -0.184719605 | 0.879820057 | 0.000109707 | 0.000587878 | 4886.8837 | 4300.306352 | ENSG00000103342 |
| RECQL4 | -0.184733255 | 0.879811733 | 0.010386896 | 0.032470202 | 1979.549697 | 1742.088181 | ENSG00000160957 |
| CALCOCO2 | -0.184817325 | 0.879760465 | 0.126977446 | 0.248505824 | 387.9579555 | 341.2695235 | ENSG00000136436 |
| NA | -0.184842486 | 0.879745121 | 0.837378485 | 0.907469713 | 5.95981577 | 5.25872978 | ENSG00000204620 |
| CDC42EP1 | -0.184852545 | 0.879738987 | 0.166689361 | 0.306510767 | 397.6690743 | 349.8793346 | ENSG00000128283 |
| STK11IP | -0.184927057 | 0.879693552 | 0.0610853 | 0.13995208 | 626.6337437 | 550.9293972 | ENSG00000144589 |
| NA | -0.184988217 | 0.87965626 | 0.506355714 | 0.671163122 | 61.75061537 | 54.41003201 | ENSG00000253636 |
| PLPP6 | -0.185048666 | 0.879619403 | 0.252049014 | 0.416504842 | 203.2469597 | 179.0334493 | ENSG00000205808 |
| FAM98A | -0.185097937 | 0.879589363 | 0.00067091 | 0.002970511 | 2466.412093 | 2169.32842 | ENSG00000119812 |
| SAR1A | -0.18510083 | 0.8795876 | 0.005005442 | 0.017557731 | 2036.611412 | 1791.94538 | ENSG00000079332 |
| CTCF | -0.185193846 | 0.879530891 | 0.00023871 | 0.001185315 | 2745.112643 | 2414.398236 | ENSG00000102974 |
| NA | -0.185223051 | 0.879513086 | 0.551893992 | 0.708528644 | 49.16171507 | 43.16902669 | ENSG00000259985 |
| RRAGD | -0.185250956 | 0.879496075 | 0.11492379 | 0.230648651 | 390.7399847 | 343.9230901 | ENSG00000025039 |

| | | | | | | | |
|---|---|---|---|---|---|---|---|
| PCGF6 | -0.185255054 | 0.879493577 | 0.086433569 | 0.184731883 | 424.3538673 | 373.2978347 | ENSG00000156374 |
| MLLT11 | -0.185262856 | 0.87948882 | 0.132001404 | 0.255759156 | 427.063069 | 375.6166558 | ENSG00000213190 |
| TMUB1 | -0.185371075 | 0.879422851 | 0.073091225 | 0.161530855 | 932.1690116 | 820.3356837 | ENSG00000164897 |
| PPP1R37 | -0.185509331 | 0.879338578 | 0.022119984 | 0.061072936 | 893.4345374 | 785.7568099 | ENSG00000104866 |
| DYNLL1 | -0.185887184 | 0.879108303 | 0.000102333 | 0.0005522 | 4658.230949 | 4095.137107 | ENSG00000088986 |
| UGT3A2 | -0.185896271 | 0.879102766 | 0.056021751 | 0.130710644 | 588.2888858 | 517.3020425 | ENSG00000168671 |
| NA | -0.185905752 | 0.879096988 | 0.624984792 | 0.763774412 | 34.9515443 | 30.71026038 | ENSG00000226221 |
| UBE2N | -0.185928187 | 0.879083318 | 0.000708058 | 0.003113552 | 3873.043957 | 3405.115012 | ENSG00000177889 |
| ENC1 | -0.185987741 | 0.87904703 | 0.096706004 | 0.201834419 | 573.6889497 | 503.8898732 | ENSG00000171617 |
| HDHD2 | -0.186052674 | 0.879007467 | 0.781024599 | 0.871424971 | 10.42905795 | 9.124671742 | ENSG00000167220 |
| IARS1 | -0.186075503 | 0.878993558 | 9.15E-06 | 6.08E-05 | 8112.283851 | 7130.548575 | ENSG00000196305 |
| CXorf40B | -0.186270352 | 0.87887485 | 0.044848026 | 0.109431226 | 691.5783034 | 607.4033555 | ENSG00000197021 |
| CHCHD1 | -0.186316487 | 0.878846745 | 0.020730635 | 0.057866399 | 892.3259938 | 784.1376235 | ENSG00000172586 |
| DPM2 | -0.186387799 | 0.878803305 | 0.03129232 | 0.081279136 | 1015.440702 | 891.9314192 | ENSG00000136908 |
| NXT1 | -0.186388314 | 0.878802991 | 0.139652319 | 0.267397805 | 385.163187 | 338.0930333 | ENSG00000132661 |
| ELP1 | -0.186439759 | 0.878771655 | 0.00434992 | 0.015545445 | 1531.507487 | 1345.972693 | ENSG00000070061 |
| FBXW2 | -0.186442994 | 0.878769684 | 0.012892447 | 0.039000813 | 1095.435748 | 963.0562073 | ENSG00000119402 |
| COMT | -0.186467261 | 0.878754903 | 0.286250518 | 0.455352416 | 187.2251224 | 164.3905971 | ENSG00000093010 |
| GIPC1 | -0.1865451 | 0.878707492 | 0.005556045 | 0.019215072 | 2459.821796 | 2161.846343 | ENSG00000123159 |
| NA | -0.186550013 | 0.8787045 | 0.751663598 | 0.853002052 | 13.0795754 | 11.47225015 | ENSG00000184844 |
| RALY | -0.186622681 | 0.878660241 | 0.000129656 | 0.000680849 | 5143.952439 | 4519.932798 | ENSG00000125970 |
| SCAF4 | -0.186748373 | 0.878583693 | 0.009147772 | 0.029187837 | 1585.215576 | 1393.063589 | ENSG00000156304 |
| UPF3A | -0.186852674 | 0.878520177 | 0.003636364 | 0.01328833 | 1662.891485 | 1460.451907 | ENSG00000169062 |
| LOC145783 | -0.186864138 | 0.878513196 | 0.804268603 | 0.887069454 | 10.17299473 | 8.918440286 | ENSG00000285331 |
| NA | -0.186938989 | 0.878467618 | 0.886337232 | 0.937353813 | 3.328158918 | 2.915452515 | ENSG00000200090 |
| HCFC1 | -0.186974442 | 0.87844603 | 0.000398759 | 0.001879878 | 4958.221895 | 4356.085898 | ENSG00000172534 |
| NKAIN4 | -0.187083244 | 0.878379784 | 0.34136437 | 0.514071824 | 122.677135 | 107.761234 | ENSG00000101198 |
| NA | -0.187119965 | 0.878357427 | 0.818425812 | 0.89619415 | 6.733111675 | 5.893823127 | ENSG00000287878 |
| GNPTAB | -0.187144065 | 0.878342754 | 0.008140712 | 0.026509535 | 1176.597633 | 1033.485802 | ENSG00000111670 |
| RNF135 | -0.187157714 | 0.878334444 | 0.255632332 | 0.420624643 | 188.5176413 | 165.6865907 | ENSG00000181481 |
| HSPB11 | -0.187160269 | 0.878332889 | 0.109888549 | 0.222370712 | 402.4157927 | 353.6164295 | ENSG00000081870 |
| RTCB | -0.187160843 | 0.878332539 | 0.000274601 | 0.001342874 | 3421.627829 | 3004.708703 | ENSG00000100220 |
| TTLL10 | -0.187229325 | 0.878290848 | 0.755761166 | 0.855125795 | 13.00345728 | 11.39642862 | ENSG00000162571 |
| AGPAT5 | -0.187376789 | 0.878201079 | 0.011567779 | 0.035638269 | 1811.217287 | 1591.28988 | ENSG00000155189 |
| MBD2 | -0.187390767 | 0.87819257 | 0.122656471 | 0.242433689 | 428.3677523 | 376.0189173 | ENSG00000134046 |
| NEGR1 | -0.187454558 | 0.87815374 | 0.562672061 | 0.717247589 | 47.66121716 | 41.96578667 | ENSG00000172260 |
| BCL2L2 | -0.187673379 | 0.878020556 | 0.100525096 | 0.207924966 | 419.8438377 | 368.6744587 | ENSG00000129473 |
| STRIP2 | -0.187678338 | 0.878017538 | 0.215020696 | 0.37178252 | 232.794518 | 204.2188171 | ENSG00000128578 |
| DCUN1D2 | -0.187702639 | 0.878002749 | 0.076432823 | 0.167135071 | 449.7588951 | 394.7220934 | ENSG00000150401 |
| PISD | -0.187754653 | 0.877971094 | 0.011628911 | 0.035767432 | 993.5078325 | 872.0555893 | ENSG00000241878 |
| NA | -0.187883237 | 0.877892846 | 0.590883122 | 0.739910841 | 38.63508766 | 33.88006889 | ENSG00000213235 |
| TGFBRAP1 | -0.18795469 | 0.877849367 | 0.02724556 | 0.072726455 | 844.6254528 | 741.3942299 | ENSG00000135966 |
| CENPVL3 | -0.187979168 | 0.877834473 | 0.315929207 | 0.487060683 | 131.6936525 | 115.5947463 | ENSG00000224109 |
| THOP1 | -0.188021549 | 0.877808686 | 0.000736851 | 0.00322839 | 3647.399836 | 3201.722704 | ENSG00000172009 |
| CADM1 | -0.18803033 | 0.877803343 | 0.127649359 | 0.249491771 | 402.5581287 | 353.5521543 | ENSG00000182985 |
| NA | -0.188202504 | 0.877698591 | 0.842784666 | 0.909954609 | 5.638798044 | 4.97452621 | ENSG00000272189 |
| GMPPA | -0.188214823 | 0.877691096 | 0.014863702 | 0.043944166 | 966.7718009 | 848.3111766 | ENSG00000144591 |
| LARP1 | -0.188267583 | 0.877658999 | 1.11E-06 | 8.68E-06 | 9653.952834 | 8473.120864 | ENSG00000155506 |
| ANGPTL4 | -0.188427101 | 0.877561962 | 0.625878617 | 0.764553217 | 40.92080882 | 35.81330639 | ENSG00000167772 |
| ALKBH4 | -0.188662121 | 0.877419016 | 0.14192711 | 0.270759085 | 372.8453947 | 327.3451224 | ENSG00000160993 |
| PHF20L1 | -0.188773162 | 0.877351485 | 0.090576228 | 0.191569566 | 477.6421185 | 419.5333934 | ENSG00000129292 |
| GEMIN6 | -0.188832778 | 0.877315232 | 0.15830794 | 0.294694439 | 344.3763543 | 301.7045771 | ENSG00000152147 |
| UIMC1 | -0.188887061 | 0.877282222 | 0.029708926 | 0.077911213 | 846.3957313 | 743.0198375 | ENSG00000087206 |
| ATN1 | -0.188973962 | 0.877229381 | 0.008453472 | 0.027334215 | 2467.995958 | 2165.281624 | ENSG00000111676 |
| C8orf37 | -0.188976504 | 0.877227835 | 0.560562806 | 0.715672872 | 46.98983322 | 41.19599437 | ENSG00000156172 |
| AASS | -0.188978739 | 0.877226477 | 0.000321345 | 0.001546286 | 4286.95562 | 3760.439328 | ENSG00000008311 |
| NA | -0.189009344 | 0.877207868 | 0.726942673 | 0.83605886 | 15.77965338 | 13.83650012 | ENSG00000253931 |
| TP73-AS1 | -0.189065377 | 0.877173798 | 0.01624067 | 0.047261845 | 1153.589399 | 1012.132518 | ENSG00000227372 |
| RAB28 | -0.189088191 | 0.877159927 | 0.093281389 | 0.196023015 | 392.5885335 | 344.4457958 | ENSG00000157869 |
| GSTP1 | -0.18930231 | 0.877029752 | 3.67E-07 | 3.11E-06 | 15232.45744 | 13358.72038 | ENSG00000084207 |
| DROSHA | -0.189305444 | 0.877027847 | 0.000362375 | 0.001724789 | 2625.37972 | 2301.949642 | ENSG00000113360 |
| BMP7 | -0.189306549 | 0.877027175 | 0.148905486 | 0.280795282 | 394.0371171 | 345.3876525 | ENSG00000101144 |
| HERPUD1 | -0.189317076 | 0.877020775 | 0.065519047 | 0.147948955 | 495.6550904 | 434.649246 | ENSG00000051108 |
| TRIM29 | -0.189466242 | 0.876930102 | 0.83307217 | 0.905243916 | 5.97935209 | 5.252278059 | ENSG00000137699 |
| PLIN2 | -0.189543644 | 0.876883055 | 0.02770075 | 0.073696754 | 737.7482712 | 646.9787701 | ENSG00000147872 |
| MAL2 | -0.189581442 | 0.876860081 | 0.014226635 | 0.042378939 | 1847.702633 | 1620.620492 | ENSG00000147676 |
| NA | -0.189600118 | 0.87684873 | 0.869241174 | 0.927001575 | 3.363764935 | 2.936425214 | ENSG00000237417 |
| PPWD1 | -0.189639423 | 0.876824841 | 0.020380959 | 0.057121762 | 821.2906703 | 720.2138221 | ENSG00000113593 |
| ADRA2B | -0.18968507 | 0.876797099 | 0.189635918 | 0.337665359 | 258.3154634 | 226.7962649 | ENSG00000274286 |
| NA | -0.18976182 | 0.876750455 | 0.732461808 | 0.84020366 | 18.23973784 | 15.89154524 | ENSG00000236698 |
| ANKRD39 | -0.189770272 | 0.876745319 | 0.50312801 | 0.668630645 | 63.75977309 | 55.81251873 | ENSG00000213337 |
| CDC42EP3 | -0.18983192 | 0.876707856 | 0.050201857 | 0.119634838 | 562.0484354 | 492.8203565 | ENSG00000163171 |
| CHCHD2 | -0.189856861 | 0.876692699 | 0.002843483 | 0.010713599 | 3454.145441 | 3028.304495 | ENSG00000106153 |
| UBP1 | -0.189951717 | 0.87663506 | 0.003743523 | 0.013638585 | 1911.566667 | 1676.327039 | ENSG00000153560 |
| CUTA | -0.19005408 | 0.876572862 | 0.000632473 | 0.00282497 | 3255.304665 | 2852.533881 | ENSG00000112514 |
| BANF1 | -0.190096583 | 0.876547038 | 0.000572227 | 0.002582723 | 4746.340943 | 4160.052949 | ENSG00000175334 |
| NA | -0.190342537 | 0.876397615 | 0.867853192 | 0.926309156 | 3.366555838 | 2.936425214 | ENSG00000240399 |

| | | | | | | | |
|---|---|---|---|---|---|---|---|
| MED19 | -0.190368583 | 0.876381793 | 0.139331047 | 0.266888614 | 366.3905854 | 321.1762432 | ENSG00000156603 |
| DAXX | -0.190442137 | 0.876337112 | 0.00380823 | 0.013842956 | 2353.492845 | 2062.626622 | ENSG00000204209 |
| MAPK1 | -0.190582832 | 0.876251655 | 0.001543275 | 0.006279544 | 1877.822762 | 1645.326699 | ENSG00000100030 |
| UBL4A | -0.190686078 | 0.876188948 | 0.018261193 | 0.052049988 | 1000.228517 | 876.0538313 | ENSG00000102178 |
| DCLRE1B | -0.190727963 | 0.87616351 | 0.168063252 | 0.308295477 | 271.0666364 | 237.6474096 | ENSG00000118655 |
| CHAC1 | -0.190730216 | 0.876162142 | 0.05555103 | 0.129958151 | 772.3060214 | 676.3941698 | ENSG00000128965 |
| SEMA4D | -0.190749114 | 0.876150665 | 0.01318778 | 0.03976328 | 1253.83662 | 1098.831372 | ENSG00000187764 |
| PRDM4 | -0.190768427 | 0.876138936 | 0.077638442 | 0.169242473 | 440.7722897 | 386.0432277 | ENSG00000110851 |
| CSTF2 | -0.190781886 | 0.876130763 | 0.003472649 | 0.012760165 | 1551.077953 | 1358.892332 | ENSG00000101811 |
| PMS2CL | -0.190811304 | 0.876112898 | 0.258782099 | 0.423927095 | 172.8840787 | 151.4271474 | ENSG00000187953 |
| FCF1 | -0.190869913 | 0.876077307 | 0.005305518 | 0.018464503 | 1247.690666 | 1093.047029 | ENSG00000119616 |
| HHEX | -0.191031237 | 0.875979348 | 0.786494787 | 0.875289658 | 10.44233674 | 9.097247322 | ENSG00000152804 |
| SF3A2 | -0.191140721 | 0.875912874 | 0.002519477 | 0.009651159 | 2061.080133 | 1805.819399 | ENSG00000104897 |
| CD82 | -0.191232451 | 0.875857183 | 0.334922071 | 0.507544432 | 120.9686949 | 105.8911871 | ENSG00000085117 |
| RPS16 | -0.191302282 | 0.87581479 | 0.000129684 | 0.000680849 | 20595.40439 | 18036.94426 | ENSG00000105193 |
| BLOC1S6 | -0.191403972 | 0.87575306 | 0.185184478 | 0.331433852 | 444.8037349 | 389.4891763 | ENSG00000104164 |
| NA | -0.191636146 | 0.875612135 | 0.725052234 | 0.835021231 | 15.41465693 | 13.49583017 | ENSG00000235508 |
| ZNF792 | -0.1917499 | 0.875543097 | 0.305389286 | 0.476331235 | 140.9655575 | 123.5651082 | ENSG00000180884 |
| SUMO2 | -0.191754422 | 0.875540353 | 4.25E-05 | 0.000247784 | 4782.752039 | 4187.719855 | ENSG00000188612 |
| NHSL1 | -0.191789964 | 0.875518784 | 0.089743573 | 0.190330175 | 410.2102027 | 359.3059112 | ENSG00000135540 |
| GPM6A | -0.191851242 | 0.875481597 | 0.617713466 | 0.759443988 | 33.10223178 | 29.01093922 | ENSG00000150625 |
| NA | -0.191898055 | 0.87545319 | 0.865896679 | 0.925059178 | 3.780437098 | 3.281396308 | ENSG00000188460 |
| KDM1A | -0.191994117 | 0.875394899 | 7.50E-06 | 5.06E-05 | 8329.262431 | 7292.361313 | ENSG00000004487 |
| ALKBH1 | -0.192116315 | 0.875320755 | 0.102090336 | 0.210408619 | 354.9008296 | 310.6350968 | ENSG00000100601 |
| SENP3 | -0.192133541 | 0.875310304 | 0.009931642 | 0.031244564 | 1066.032817 | 933.3050813 | ENSG00000161956 |
| ASPSCR1 | -0.192182948 | 0.875280329 | 0.014764138 | 0.043696685 | 1012.714128 | 886.4906167 | ENSG00000169696 |
| TMUB2 | -0.192193776 | 0.875273759 | 0.104286479 | 0.213906482 | 357.2402536 | 312.5731869 | ENSG00000168591 |
| MRPS18A | -0.192253295 | 0.87523765 | 0.071752026 | 0.159099808 | 533.0464319 | 466.1080777 | ENSG00000096080 |
| ZSCAN5A | -0.192269516 | 0.87522781 | 0.451682405 | 0.621853074 | 74.41786111 | 65.0950127 | ENSG00000131848 |
| PSMC6 | -0.192335763 | 0.875187621 | 0.001921319 | 0.007617875 | 1574.081581 | 1377.765452 | ENSG00000100519 |
| UBL3 | -0.192489844 | 0.875094155 | 0.075307608 | 0.165349314 | 561.4521816 | 491.6915625 | ENSG00000122042 |
| RPA1 | -0.192520579 | 0.875075512 | 0.000143058 | 0.000743461 | 4674.302941 | 4089.908703 | ENSG00000132383 |
| LRTOMT | -0.192536985 | 0.875065561 | 0.679623996 | 0.804135581 | 21.73946915 | 19.00435434 | ENSG00000184154 |
| FAM111A | -0.192546129 | 0.875060015 | 0.089101104 | 0.189247778 | 452.3551461 | 395.6118338 | ENSG00000166801 |
| H4C5 | -0.192549848 | 0.875057759 | 0.700880384 | 0.819683752 | 19.63812362 | 17.26308779 | ENSG00000276966 |
| ADGRA3 | -0.19258507 | 0.875036396 | 0.001758378 | 0.007056555 | 1805.402583 | 1579.706872 | ENSG00000152990 |
| BRD4 | -0.192632093 | 0.875007876 | 0.00220447 | 0.008587729 | 2572.169444 | 2250.862213 | ENSG00000141867 |
| NIP7 | -0.192750744 | 0.874935915 | 0.007759057 | 0.025477441 | 1257.772074 | 1100.953679 | ENSG00000132603 |
| PLCXD2 | -0.192830038 | 0.874887828 | 0.533350368 | 0.693924025 | 65.91913988 | 57.71401855 | ENSG00000240891 |
| PDF | -0.192855913 | 0.874872137 | 0.226975578 | 0.386228563 | 331.4395806 | 290.2024794 | ENSG00000258429 |
| SCRIB | -0.192952227 | 0.874813733 | 0.000534025 | 0.002431408 | 4367.401125 | 3821.204566 | ENSG00000180900 |
| ARID1A | -0.193109954 | 0.874718097 | 2.48E-05 | 0.000151609 | 4158.722067 | 3637.820537 | ENSG00000117713 |
| DYNLL2 | -0.193143357 | 0.874697844 | 0.00058031 | 0.002612546 | 3131.036162 | 2739.160458 | ENSG00000264364 |
| SAP18 | -0.193161525 | 0.874686829 | 0.000410666 | 0.001927064 | 4149.455516 | 3628.800189 | ENSG00000150459 |
| THOC3 | -0.193178863 | 0.874676317 | 0.001031339 | 0.004380102 | 2322.408568 | 2030.932251 | ENSG00000051596 |
| CTR9 | -0.19317976 | 0.874675773 | 0.002883581 | 0.010835053 | 1622.328327 | 1419.630906 | ENSG00000198730 |
| ATG9A | -0.193222561 | 0.874649825 | 0.028121429 | 0.07455889 | 675.6085804 | 590.9952468 | ENSG00000198925 |
| MAP2K7 | -0.193348911 | 0.874573227 | 0.004068112 | 0.014677004 | 1444.762497 | 1264.051778 | ENSG00000076984 |
| MAD2L2 | -0.193374882 | 0.874557483 | 8.46E-05 | 0.000464955 | 4475.451072 | 3913.549732 | ENSG00000116670 |
| KCNJ2-AS1 | -0.193379072 | 0.874554943 | 0.882429594 | 0.935789637 | 3.347695239 | 2.915452515 | ENSG00000267365 |
| LINC01252 | -0.193386834 | 0.874550238 | 0.830925297 | 0.903961401 | 7.128132336 | 6.198999397 | ENSG00000247157 |
| CNOT6L | -0.193464315 | 0.874503271 | 0.169958729 | 0.311128662 | 261.1135112 | 228.4579418 | ENSG00000138767 |
| YWHAE | -0.193467619 | 0.874501268 | 4.49E-05 | 0.000260665 | 22260.54271 | 19466.50718 | ENSG00000108953 |
| POLE | -0.193477549 | 0.874495249 | 0.004098627 | 0.014770523 | 2309.499776 | 2019.15248 | ENSG00000177084 |
| PARP6 | -0.193479749 | 0.874493915 | 0.114683714 | 0.230262649 | 363.5365932 | 317.9760114 | ENSG00000137817 |
| RPS10 | -0.193544017 | 0.87445496 | 0.722915701 | 0.833877469 | 16.57930671 | 14.56838771 | ENSG00000124614 |
| CYP51A1 | -0.193762387 | 0.87432261 | 0.176023881 | 0.319442774 | 266.0320405 | 232.7301738 | ENSG00000001630 |
| NA | -0.193848134 | 0.874270646 | 0.538355591 | 0.697944739 | 53.53462709 | 46.71473837 | ENSG00000249565 |
| NECTIN1 | -0.193915879 | 0.874229594 | 0.001791193 | 0.007170346 | 3610.54087 | 3156.042697 | ENSG00000110400 |
| NA | -0.193932919 | 0.874219268 | 0.831660881 | 0.904389657 | 6.75755408 | 5.883070259 | ENSG00000243230 |
| NA | -0.194046569 | 0.874150403 | 0.483487048 | 0.651277749 | 62.72080195 | 54.72946879 | ENSG00000218426 |
| COLGALT1 | -0.194068253 | 0.874137265 | 0.000199369 | 0.001005234 | 2624.624723 | 2294.443884 | ENSG00000130309 |
| TCTA | -0.194083109 | 0.874128263 | 0.34271855 | 0.515435532 | 112.0115501 | 97.90897578 | ENSG00000145022 |
| NA | -0.194105156 | 0.874114905 | 0.746758162 | 0.849969325 | 15.64627774 | 13.6509809 | ENSG00000273066 |
| ALAD | -0.194191561 | 0.874062554 | 0.087217793 | 0.186009133 | 398.7168553 | 348.3674107 | ENSG00000148218 |
| OAZ2 | -0.194237468 | 0.874034742 | 1.74E-05 | 0.000109668 | 6244.112077 | 5457.574326 | ENSG00000180304 |
| ALOX15 | -0.194286142 | 0.874005255 | 0.723441673 | 0.834137153 | 16.52504026 | 14.38340152 | ENSG00000161905 |
| SEMA4B | -0.194462078 | 0.873898676 | 0.002788833 | 0.01054063 | 1450.114042 | 1267.224822 | ENSG00000185033 |
| IRAK1BP1 | -0.194477567 | 0.873889294 | 0.307997305 | 0.479079787 | 139.5200018 | 122.1411278 | ENSG00000146243 |
| CEBPD | -0.194496008 | 0.873878123 | 0.54452897 | 0.702938966 | 47.60395965 | 41.61840454 | ENSG00000221869 |
| ELOA | -0.194547108 | 0.873847172 | 0.011091211 | 0.034409867 | 1258.533667 | 1100.100554 | ENSG00000011007 |
| NA | -0.194729382 | 0.873736775 | 0.874340445 | 0.930228409 | 3.344904336 | 2.90685022 | ENSG00000239486 |
| HNRNPF | -0.194888514 | 0.873640405 | 4.78E-06 | 3.33E-05 | 9361.685077 | 8178.20812 | ENSG00000169813 |
| NA | -0.195035733 | 0.873551259 | 0.75437132 | 0.854310609 | 13.18850863 | 11.4765513 | ENSG00000225616 |
| CD59 | -0.195156813 | 0.873477949 | 0.058243482 | 0.134737379 | 496.6302711 | 433.8124829 | ENSG00000085063 |
| PMS2P4 | -0.195185669 | 0.873460478 | 0.758268487 | 0.856402715 | 13.02510878 | 11.32920937 | ENSG00000067601 |
| RAVER1 | -0.195227182 | 0.873435344 | 0.016865402 | 0.048763561 | 1058.88486 | 924.7448492 | ENSG00000161847 |

| | | | | | | | |
|---|---|---|---|---|---|---|---|
| BMF | -0.195269794 | 0.873409547 | 0.051885871 | 0.122893825 | 533.7613063 | 465.8803405 | ENSG00000104081 |
| CADPS | -0.195277118 | 0.873405113 | 0.208394357 | 0.363221498 | 239.4504724 | 209.2056895 | ENSG00000163618 |
| CD200 | -0.195524548 | 0.873255332 | 0.007444762 | 0.024596148 | 1106.318197 | 965.9719508 | ENSG00000091972 |
| PCYT1A | -0.195554007 | 0.873237501 | 0.054635699 | 0.128111982 | 521.8847407 | 455.8030287 | ENSG00000161217 |
| DEDD2 | -0.1956831 | 0.873159367 | 0.005757301 | 0.019832873 | 1332.394085 | 1163.91376 | ENSG00000160570 |
| AMFR | -0.195843989 | 0.873061998 | 0.073662654 | 0.162495856 | 549.7535566 | 479.8996336 | ENSG00000159461 |
| RBM42 | -0.195871486 | 0.873045358 | 0.001520333 | 0.006196649 | 2224.993489 | 1942.713937 | ENSG00000126254 |
| CCDC12 | -0.195970417 | 0.872985492 | 0.004706668 | 0.016662936 | 1211.559241 | 1057.884682 | ENSG00000160799 |
| PSMA4 | -0.195971217 | 0.872985008 | 0.000138756 | 0.000723834 | 5346.853747 | 4667.583196 | ENSG00000041357 |
| TMED9 | -0.195995123 | 0.872970542 | 0.000120843 | 0.000640258 | 4121.981748 | 3598.320472 | ENSG00000184840 |
| C15orf62 | -0.196034298 | 0.872946838 | 0.696941963 | 0.817090264 | 18.11194401 | 15.80738187 | ENSG00000188277 |
| MOB3C | -0.196037944 | 0.872944632 | 0.380984248 | 0.554433247 | 93.81666692 | 81.8577182 | ENSG00000142961 |
| ATXN7L2 | -0.196055312 | 0.872934123 | 0.236698995 | 0.39820008 | 223.9337786 | 195.6593237 | ENSG00000162650 |
| NA | -0.196089081 | 0.87291369 | 0.771202994 | 0.865209477 | 11.2454567 | 9.842038351 | ENSG00000224664 |
| NA | -0.196212991 | 0.872838721 | 0.694093927 | 0.815108692 | 19.49725115 | 16.94528065 | ENSG00000264016 |
| FAM126B | -0.196243044 | 0.872820539 | 0.458558047 | 0.628404012 | 70.16560951 | 61.31941325 | ENSG00000155744 |
| TCF7L1 | -0.196304208 | 0.872783536 | 0.000947383 | 0.004058619 | 3794.485604 | 3312.216736 | ENSG00000152284 |
| TAMM41 | -0.196359839 | 0.872749881 | 0.226992913 | 0.386228563 | 230.6227483 | 201.4802573 | ENSG00000144559 |
| ROGDI | -0.196375564 | 0.872740369 | 0.205496123 | 0.359371381 | 260.1435247 | 226.8979359 | ENSG00000067836 |
| H4C4 | -0.19645248 | 0.87269384 | 0.851416222 | 0.915396263 | 5.200474379 | 4.560185297 | ENSG00000277157 |
| NA | -0.19650131 | 0.872664304 | 0.822626574 | 0.898906153 | 5.968188479 | 5.20388094 | ENSG00000226002 |
| RNF166 | -0.196557831 | 0.872630115 | 0.122663987 | 0.242433689 | 446.978595 | 390.1471572 | ENSG00000158717 |
| RNF165 | -0.196577474 | 0.872618235 | 0.188632196 | 0.336281646 | 292.3571307 | 254.7502763 | ENSG00000141622 |
| KDM4A | -0.196639697 | 0.872580599 | 0.012214122 | 0.037182046 | 1035.464897 | 903.5862081 | ENSG00000066135 |
| POU5F1B | -0.196742774 | 0.872518258 | 0.240435192 | 0.402654967 | 175.1895364 | 152.7664313 | ENSG00000212993 |
| PDLIM7 | -0.196765693 | 0.872504397 | 0.005063161 | 0.017726577 | 1568.708584 | 1368.844879 | ENSG00000196923 |
| CHMP7 | -0.196888305 | 0.872430248 | 0.002889207 | 0.010854079 | 1399.392456 | 1220.660115 | ENSG00000147457 |
| C6orf132 | -0.196921115 | 0.872410407 | 0.007447499 | 0.024600977 | 1707.1157 | 1489.204486 | ENSG00000188112 |
| ITGAL | -0.196952676 | 0.872391322 | 0.8691004 | 0.926984177 | 3.363764935 | 2.915452515 | ENSG00000005844 |
| MRPL30 | -0.197038533 | 0.872339406 | 0.037678028 | 0.094826723 | 683.1816062 | 596.1646943 | ENSG00000185414 |
| NA | -0.197239105 | 0.872218137 | 0.759018433 | 0.857004645 | 12.06975259 | 10.52122767 | ENSG00000236603 |
| TMEM39B | -0.197273434 | 0.872197383 | 0.056108082 | 0.130848767 | 492.695846 | 429.5886906 | ENSG00000121775 |
| ERCC5 | -0.197371868 | 0.872137875 | 0.555125864 | 0.711320693 | 44.84729987 | 39.22484004 | ENSG00000134899 |
| ICOSLG | -0.197381189 | 0.872132241 | 0.54450245 | 0.702938966 | 44.54322772 | 38.85512814 | ENSG00000160223 |
| AMPD2 | -0.197486892 | 0.872068343 | 0.017824965 | 0.051078638 | 950.5603199 | 828.8473435 | ENSG00000116337 |
| AOC3 | -0.197513072 | 0.872052519 | 0.626078145 | 0.764748528 | 34.89882945 | 30.37146843 | ENSG00000131471 |
| SMOX | -0.197517187 | 0.872050032 | 0.120602842 | 0.239480127 | 348.7970382 | 304.44307 | ENSG00000088826 |
| MSL3 | -0.197518 | 0.87204954 | 0.125645295 | 0.246574098 | 297.0625875 | 259.0582319 | ENSG00000005302 |
| UBE4B | -0.19753057 | 0.872041942 | 0.00054021 | 0.002455514 | 3391.848351 | 2957.644982 | ENSG00000130939 |
| FAHD2A | -0.197585185 | 0.87200893 | 0.027654234 | 0.073623769 | 713.0515337 | 621.56973 | ENSG00000115042 |
| ZNF785 | -0.197621442 | 0.871987016 | 0.343476341 | 0.516242264 | 111.5524509 | 97.19644336 | ENSG00000197162 |
| ZNHIT6 | -0.197629412 | 0.871982199 | 0.038084409 | 0.095737077 | 608.0560426 | 530.2724891 | ENSG00000117174 |
| AP5S1 | -0.197776142 | 0.871893518 | 0.503685317 | 0.66897596 | 56.2855049 | 49.05854867 | ENSG00000125843 |
| KIF15 | -0.197816057 | 0.871869395 | 0.03497049 | 0.089128779 | 697.8877585 | 608.2822698 | ENSG00000163808 |
| NA | -0.19786202 | 0.871841619 | 0.629516196 | 0.767393171 | 28.58988676 | 24.83740995 | ENSG00000285679 |
| SIGIRR | -0.197878212 | 0.871831834 | 0.021226297 | 0.059027892 | 857.524846 | 747.4883901 | ENSG00000185187 |
| FANCD2 | -0.197901854 | 0.871817547 | 0.002126268 | 0.008328494 | 1864.340231 | 1625.204582 | ENSG00000144554 |
| MED29 | -0.197921851 | 0.871805463 | 0.016977046 | 0.049034913 | 1127.814995 | 983.4063889 | ENSG00000063322 |
| PLCB3 | -0.197941638 | 0.871793506 | 0.001173668 | 0.004928436 | 4934.471715 | 4301.632759 | ENSG00000149782 |
| FAM168A | -0.197952017 | 0.871787234 | 0.024559137 | 0.066606258 | 793.9593729 | 691.8802064 | ENSG00000054965 |
| NA | -0.19800208 | 0.871756983 | 0.593412545 | 0.741659955 | 54.47478946 | 47.29363791 | ENSG00000243302 |
| ILF3 | -0.198028293 | 0.871741144 | 1.18E-06 | 9.23E-06 | 23035.85376 | 20080.18425 | ENSG00000129351 |
| PSMD11 | -0.19807513 | 0.871712843 | 5.96E-05 | 0.000338725 | 3147.966144 | 2743.570127 | ENSG00000108671 |
| SLC15A1 | -0.198150584 | 0.871667253 | 0.560090185 | 0.715353533 | 45.28438423 | 39.44182437 | ENSG00000088386 |
| NA | -0.198163231 | 0.871659612 | 0.835200691 | 0.906737673 | 5.595495041 | 4.886334266 | ENSG00000229237 |
| TYMS | -0.198327287 | 0.871560497 | 0.13399234 | 0.258872932 | 307.2675215 | 267.5384266 | ENSG00000176890 |
| ZNF140 | -0.198497794 | 0.871457496 | 0.253099217 | 0.417707733 | 176.6675069 | 153.7174659 | ENSG00000196387 |
| IKZF5 | -0.198598097 | 0.87139691 | 0.247364324 | 0.41094602 | 172.8226789 | 150.777521 | ENSG00000095574 |
| FOXC1 | -0.198722419 | 0.871321822 | 0.879039664 | 0.93343161 | 3.36934674 | 2.919753662 | ENSG00000054598 |
| CCDC153 | -0.198771437 | 0.871292218 | 0.6026174 | 0.748219879 | 34.04662452 | 29.60408717 | ENSG00000248712 |
| GPR89B | -0.19882275 | 0.871261229 | 0.332718888 | 0.505471733 | 122.0944722 | 106.4638833 | ENSG00000188092 |
| KLHL3 | -0.198887851 | 0.871221914 | 0.112947258 | 0.227435664 | 370.6824349 | 322.9118273 | ENSG00000146021 |
| TUBGCP2 | -0.199081977 | 0.871104693 | 0.002442592 | 0.009405199 | 1884.951197 | 1641.486122 | ENSG00000130640 |
| UBB | -0.199145398 | 0.8710664 | 0.000405843 | 0.001908144 | 6752.401909 | 5880.941531 | ENSG00000170315 |
| YWHAQ | -0.199154356 | 0.871060991 | 1.22E-06 | 9.53E-06 | 10838.81305 | 9442.004522 | ENSG00000134308 |
| CEP104 | -0.199336123 | 0.870951252 | 0.029217659 | 0.076855982 | 654.184259 | 569.7664787 | ENSG00000116198 |
| ATXN2L | -0.19939766 | 0.870914103 | 0.000187999 | 0.000952889 | 3987.683722 | 3473.16059 | ENSG00000168488 |
| CDCA7 | -0.19951527 | 0.870843108 | 0.000570792 | 0.002576915 | 2193.809875 | 1909.857694 | ENSG00000144354 |
| KLF13 | -0.199673873 | 0.870747377 | 0.003141094 | 0.011659084 | 1361.384322 | 1185.557445 | ENSG00000169926 |
| GPI | -0.199696257 | 0.870733867 | 9.86E-06 | 6.50E-05 | 6551.449611 | 5704.88739 | ENSG00000105220 |
| NA | -0.199728093 | 0.870714653 | 0.740950769 | 0.846431059 | 13.71112296 | 11.98768645 | ENSG00000271780 |
| AFG3L1P | -0.199751452 | 0.870700555 | 0.212804006 | 0.36880802 | 186.6421715 | 162.4154143 | ENSG00000223959 |
| OLA1 | -0.199915822 | 0.870601359 | 0.001231042 | 0.005149185 | 2157.144389 | 1878.426039 | ENSG00000138430 |
| PPP1R11 | -0.199944689 | 0.870583939 | 0.050009364 | 0.119338279 | 487.7353652 | 424.6614833 | ENSG00000204619 |
| ZNF219 | -0.200014089 | 0.870542062 | 0.00280742 | 0.010598422 | 2526.056796 | 2199.665733 | ENSG00000165804 |
| PPP1R3E | -0.200033135 | 0.87053057 | 0.535891857 | 0.69573254 | 50.03278059 | 43.41882108 | ENSG00000235194 |
| NA | -0.200144417 | 0.870463423 | 0.543414067 | 0.70198947 | 46.78245465 | 40.84268144 | ENSG00000260719 |

| | | | | | | | |
|---|---|---|---|---|---|---|---|
| POLR3F | -0.200157689 | 0.870455416 | 0.132057315 | 0.255800549 | 296.741733 | 258.4615884 | ENSG00000132664 |
| ESS2 | -0.200233964 | 0.870409396 | 0.047595346 | 0.11482764 | 551.0646749 | 479.8717003 | ENSG00000100056 |
| KNDC1 | -0.200287989 | 0.870376803 | 0.06113935 | 0.140059281 | 471.4378403 | 409.9465593 | ENSG00000171798 |
| PPP1R12A | -0.200369927 | 0.870327371 | 0.003089082 | 0.011484194 | 1317.1523 | 1146.183378 | ENSG00000058272 |
| NA | -0.200376983 | 0.870323114 | 0.468051052 | 0.63706857 | 67.20822901 | 58.37519136 | ENSG00000273247 |
| ANKRD9 | -0.200382104 | 0.870320025 | 0.005569056 | 0.019253165 | 1087.715853 | 946.4035331 | ENSG00000156381 |
| C19orf12 | -0.200382384 | 0.870319856 | 0.130448792 | 0.253489474 | 360.2657799 | 313.2211774 | ENSG00000131943 |
| GTF2I | -0.200394372 | 0.870312624 | 0.002705065 | 0.010260206 | 1632.363779 | 1420.854387 | ENSG00000263001 |
| NA | -0.20050283 | 0.870247199 | 0.463874152 | 0.633573794 | 65.92029116 | 57.40964789 | ENSG00000227827 |
| MLLT1 | -0.200618927 | 0.870177171 | 0.004926578 | 0.017323845 | 1799.270884 | 1565.967209 | ENSG00000130382 |
| CAMK4 | -0.200697341 | 0.870129876 | 0.239650064 | 0.401786355 | 161.2200348 | 140.2474305 | ENSG00000152495 |
| LHFPL2 | -0.200720103 | 0.870116147 | 0.0012984 | 0.005397723 | 1530.529352 | 1331.573147 | ENSG00000145685 |
| CSF1 | -0.200741215 | 0.870103415 | 0.280544092 | 0.448754236 | 153.2555671 | 133.3722164 | ENSG00000184371 |
| MYCBP | -0.200754755 | 0.870095248 | 0.241805647 | 0.404161049 | 245.9685838 | 214.1329152 | ENSG00000214114 |
| POLR2H | -0.20077464 | 0.870083256 | 0.00576331 | 0.019846484 | 1456.375064 | 1266.746465 | ENSG00000163882 |
| MACO1 | -0.200775691 | 0.870082622 | 0.106602446 | 0.217316685 | 322.3789452 | 280.6707119 | ENSG00000204178 |
| KCTD10 | -0.200812451 | 0.870060453 | 0.004049108 | 0.014619378 | 1363.050148 | 1185.66421 | ENSG00000110906 |
| E2F5 | -0.200893599 | 0.870011515 | 0.106218907 | 0.216718026 | 526.4535184 | 458.5554912 | ENSG00000133740 |
| ZNF33B | -0.200935632 | 0.869986167 | 0.080148424 | 0.173684929 | 396.9666639 | 345.3341607 | ENSG00000196693 |
| DDX5 | -0.200989058 | 0.869953951 | 3.38E-07 | 2.87E-06 | 9709.525416 | 8446.853876 | ENSG00000108654 |
| TMEM218 | -0.201008728 | 0.86994209 | 0.09643921 | 0.201364736 | 347.252986 | 301.9320417 | ENSG00000150433 |
| H3C13 | -0.201116083 | 0.869877357 | 0.836650506 | 0.907107819 | 5.32894393 | 4.65913011 | ENSG00000183598 |
| GNAI3 | -0.201291804 | 0.869771413 | 0.000810915 | 0.003519316 | 2165.766822 | 1884.560703 | ENSG00000065135 |
| GJC3 | -0.201381777 | 0.869717171 | 0.704236236 | 0.822013749 | 16.99876978 | 14.80419416 | ENSG00000176402 |
| SRSF1 | -0.201407189 | 0.869701852 | 0.02151692 | 0.059706977 | 1470.707813 | 1280.040335 | ENSG00000136450 |
| RABGGTA | -0.201471865 | 0.869662864 | 0.158372754 | 0.294729771 | 255.5699672 | 222.4829299 | ENSG00000100949 |
| TPPP | -0.201505009 | 0.869642885 | 0.456850467 | 0.626731785 | 65.20512865 | 56.63528187 | ENSG00000171368 |
| RLN1 | -0.201524184 | 0.869631326 | 0.650627021 | 0.782789439 | 23.410188 | 20.34874504 | ENSG00000107014 |
| NA | -0.201548113 | 0.869616903 | 0.862666832 | 0.922796155 | 3.780437098 | 3.251821314 | ENSG00000234106 |
| RTKN2 | -0.201839196 | 0.869441463 | 0.061342806 | 0.140425309 | 524.1988223 | 455.9266959 | ENSG00000182010 |
| GPR26 | -0.201889376 | 0.869411223 | 0.702027969 | 0.820379304 | 16.60654002 | 14.47374404 | ENSG00000154478 |
| THOC6 | -0.202032878 | 0.869324748 | 0.004643202 | 0.016474528 | 1179.635287 | 1025.053266 | ENSG00000131652 |
| PCP4 | -0.202033177 | 0.869324568 | 0.742017291 | 0.847348344 | 15.27869055 | 13.26432487 | ENSG00000183036 |
| NA | -0.202136449 | 0.869262342 | 0.819445608 | 0.896989371 | 6.022655094 | 5.222703065 | ENSG00000261614 |
| NOL11 | -0.202235894 | 0.869202426 | 0.000617635 | 0.002764454 | 2086.318219 | 1813.548593 | ENSG00000130935 |
| NA | -0.202528432 | 0.869026194 | 0.824519722 | 0.900350935 | 5.728870676 | 4.985279079 | ENSG00000286162 |
| NA | -0.202643288 | 0.868957012 | 0.837868327 | 0.907638723 | 5.227707687 | 4.560185297 | ENSG00000276259 |
| SPCS2 | -0.202654636 | 0.868950176 | 0.001012975 | 0.004311848 | 1992.203638 | 1731.795126 | ENSG00000118363 |
| NA | -0.20271612 | 0.868913145 | 0.881661136 | 0.935376247 | 3.456628469 | 2.989123481 | ENSG00000254016 |
| PRKAR2B | -0.202766086 | 0.868883052 | 0.016739193 | 0.048442209 | 856.0013693 | 743.8482831 | ENSG00000005249 |
| USP15 | -0.202777454 | 0.868876205 | 0.053782437 | 0.126422789 | 463.4614452 | 402.6104546 | ENSG00000135655 |
| PAXIP1 | -0.202798194 | 0.868863714 | 0.012969437 | 0.039202978 | 906.7010649 | 787.7260314 | ENSG00000157212 |
| TPRA1 | -0.202830296 | 0.868844381 | 0.047987667 | 0.115577979 | 515.4824589 | 448.2415979 | ENSG00000163870 |
| CEMIP2 | -0.202836308 | 0.86884076 | 0.004375261 | 0.01562732 | 1532.874997 | 1332.438322 | ENSG00000135048 |
| CCDC134 | -0.202891328 | 0.868807626 | 0.354231873 | 0.527356618 | 108.2967433 | 94.15166542 | ENSG00000100147 |
| LOC100287049 | -0.202919342 | 0.868790756 | 0.869817254 | 0.927359602 | 3.429395161 | 2.959548487 | ENSG00000228606 |
| DNAJB2 | -0.203095414 | 0.868684732 | 0.014529645 | 0.043095231 | 1092.661955 | 949.108211 | ENSG00000135924 |
| ANO10 | -0.203150587 | 0.868651512 | 0.024898397 | 0.067403168 | 981.1010549 | 852.5844258 | ENSG00000160746 |
| AMDHD2 | -0.203245142 | 0.868594581 | 0.100728149 | 0.208224039 | 327.5543015 | 284.6299526 | ENSG00000162066 |
| NA | -0.20329589 | 0.868564028 | 0.565661358 | 0.719705399 | 37.80174333 | 32.81343004 | ENSG00000213058 |
| ATG3 | -0.203308439 | 0.868556473 | 0.020354107 | 0.057054792 | 744.5206188 | 646.7714846 | ENSG00000144848 |
| PRR19 | -0.20330962 | 0.868555762 | 0.516349096 | 0.680435167 | 64.6536414 | 56.20184626 | ENSG00000188368 |
| FBXL14 | -0.203347215 | 0.868533129 | 0.077801874 | 0.169533034 | 373.3796851 | 324.4455174 | ENSG00000171823 |
| ZNF133 | -0.203349746 | 0.868531605 | 0.131116073 | 0.254386016 | 282.1449512 | 245.0676656 | ENSG00000125846 |
| UNC5CL | -0.203402565 | 0.868499808 | 0.607758859 | 0.751797117 | 33.91025782 | 29.46534753 | ENSG00000124602 |
| RPS6KB1 | -0.203443157 | 0.868475372 | 0.068860749 | 0.154034404 | 447.3079471 | 388.1684482 | ENSG00000108443 |
| ABHD17C | -0.203624009 | 0.868366509 | 0.03600147 | 0.091286061 | 554.706667 | 481.6296384 | ENSG00000136379 |
| NUDT5 | -0.203679635 | 0.868333028 | 0.004353016 | 0.015553626 | 1082.506806 | 939.9869858 | ENSG00000165609 |
| NA | -0.20379534 | 0.86826339 | 0.043776515 | 0.107278042 | 671.9856732 | 583.2169169 | ENSG00000250251 |
| STAMBP | -0.203859397 | 0.868224839 | 0.006480583 | 0.021927997 | 992.9431544 | 861.8312082 | ENSG00000124356 |
| NA | -0.203882716 | 0.868210806 | 0.69696855 | 0.817090264 | 18.05189559 | 15.62831437 | ENSG00000275807 |
| CCDC28B | -0.203978181 | 0.868153357 | 0.163557559 | 0.302047565 | 275.0495219 | 238.7866655 | ENSG00000160050 |
| KIAA2013 | -0.203982434 | 0.868150798 | 0.0059509 | 0.0203978 | 1561.612891 | 1355.943542 | ENSG00000116685 |
| NA | -0.204017821 | 0.868129504 | 0.573774303 | 0.726100766 | 39.46496536 | 34.17879344 | ENSG00000264350 |
| GTF2E2 | -0.204021563 | 0.868127252 | 0.016590863 | 0.048070643 | 844.8307531 | 732.9963143 | ENSG00000197265 |
| HCG11 | -0.204036559 | 0.868118229 | 0.1499478 | 0.282319247 | 306.3518895 | 265.8348043 | ENSG00000228223 |
| NPC2 | -0.204054851 | 0.868107222 | 0.008813857 | 0.028309709 | 1016.571246 | 881.9548997 | ENSG00000119655 |
| RASSF4 | -0.204084654 | 0.868089289 | 0.159651737 | 0.296623486 | 240.3543531 | 208.4259499 | ENSG00000107551 |
| RNF214 | -0.204126243 | 0.868064265 | 0.062407056 | 0.142293509 | 479.0980357 | 415.4612626 | ENSG00000167257 |
| PARD6G | -0.204136149 | 0.868058304 | 0.073530818 | 0.162303221 | 380.0911453 | 330.11482 | ENSG00000178184 |
| COX14 | -0.204307108 | 0.867955446 | 0.51393387 | 0.678086197 | 70.94966871 | 61.56921976 | ENSG00000178449 |
| C1QBP | -0.204316987 | 0.867949502 | 9.62E-06 | 6.36E-05 | 6643.378043 | 5766.469841 | ENSG00000108561 |
| RPL41 | -0.20443965 | 0.867875709 | 1.91E-05 | 0.000119424 | 9781.628145 | 8488.042724 | ENSG00000229117 |
| NA | -0.204458734 | 0.867864229 | 0.311824061 | 0.482815472 | 116.142829 | 100.8416329 | ENSG00000229320 |
| CYHR1 | -0.204500734 | 0.867838964 | 0.015575667 | 0.045657223 | 1044.140423 | 906.1779591 | ENSG00000187954 |
| CLDN12 | -0.204543178 | 0.867813432 | 0.018034013 | 0.051519335 | 692.961183 | 601.4153791 | ENSG00000157224 |
| C16orf87 | -0.204601534 | 0.867778331 | 0.084585759 | 0.181506548 | 430.0715744 | 373.4854739 | ENSG00000155330 |

| | | | | | | | |
|---|---|---|---|---|---|---|---|
| GRIP1 | -0.204628688 | 0.867761998 | 0.100550674 | 0.207946296 | 334.6781282 | 290.2912043 | ENSG00000155974 |
| HAUS2 | -0.204639868 | 0.867755273 | 0.053057527 | 0.124973473 | 464.1804378 | 402.9500585 | ENSG00000137814 |
| SUGT1P4-STRA6LP | -0.204701409 | 0.867718258 | 0.795352275 | 0.88092902 | 8.374482042 | 7.278008651 | ENSG00000254876 |
| ANO2 | -0.204746911 | 0.867690891 | 0.40346183 | 0.575849523 | 78.01747722 | 67.65312372 | ENSG00000047617 |
| USP1 | -0.204782177 | 0.867669681 | 0.004480753 | 0.01595684 | 1322.723643 | 1148.368276 | ENSG00000162607 |
| BTBD6 | -0.204914123 | 0.867590329 | 0.169117811 | 0.309912767 | 245.1098459 | 212.5140014 | ENSG00000184887 |
| BUD23 | -0.205072458 | 0.867495117 | 0.000318817 | 0.001535656 | 2315.517877 | 2008.798131 | ENSG00000071462 |
| ELAC2 | -0.20522935 | 0.867400783 | 7.05E-05 | 0.000393862 | 3404.999253 | 2953.778762 | ENSG00000006744 |
| ID1 | -0.205270088 | 0.867376289 | 0.003037834 | 0.011322034 | 1645.424331 | 1427.158541 | ENSG00000125968 |
| NA | -0.205302947 | 0.867356534 | 0.631647676 | 0.768901632 | 29.94863632 | 25.92287092 | ENSG00000223703 |
| ATF1 | -0.205304842 | 0.867355395 | 0.062796554 | 0.143057194 | 454.4986065 | 394.1190166 | ENSG00000123268 |
| HNRNPD | -0.205310507 | 0.867351989 | 3.69E-05 | 0.000218271 | 19636.40168 | 17032.4568 | ENSG00000138668 |
| SELENOW | -0.205331828 | 0.867339172 | 0.000280329 | 0.001369072 | 3274.729651 | 2839.22864 | ENSG00000178980 |
| FBXO25 | -0.205337157 | 0.867335968 | 0.195724551 | 0.346159807 | 192.1964787 | 166.6161075 | ENSG00000147364 |
| GLCCI1 | -0.205447823 | 0.867269438 | 0.135213285 | 0.260527315 | 337.6984233 | 293.1738163 | ENSG00000106415 |
| ELMO2 | -0.205497489 | 0.867239582 | 0.058375661 | 0.134978411 | 510.8181899 | 443.3815914 | ENSG00000062598 |
| PNRC2 | -0.205511878 | 0.867230933 | 0.001074143 | 0.00454514 | 2466.156769 | 2138.719236 | ENSG00000189266 |
| BNC2 | -0.205532385 | 0.867218606 | 0.064265124 | 0.145645971 | 436.7605441 | 378.6167805 | ENSG00000173068 |
| BRPF3 | -0.205694981 | 0.867120873 | 0.003283456 | 0.012125027 | 1276.461248 | 1106.746177 | ENSG00000096070 |
| ATAT1 | -0.205750771 | 0.867087342 | 0.081269631 | 0.175621423 | 369.4080512 | 320.4225467 | ENSG00000137343 |
| ZNF500 | -0.205797018 | 0.867059547 | 0.213955292 | 0.370370741 | 226.1817786 | 196.1027187 | ENSG00000103199 |
| ZNF234 | -0.205821167 | 0.867045034 | 0.272590306 | 0.439346611 | 146.5765586 | 127.1575994 | ENSG00000263002 |
| MUL1 | -0.205848006 | 0.867028904 | 0.022076356 | 0.060969934 | 749.6104451 | 649.6883063 | ENSG00000090432 |
| HMOX2 | -0.205848382 | 0.867028678 | 0.005542739 | 0.01918554 | 1106.86359 | 959.6076114 | ENSG00000103415 |
| UBE2M | -0.205864458 | 0.867019017 | 0.000187091 | 0.000949034 | 2866.679181 | 2485.212614 | ENSG00000130725 |
| SFMBT1 | -0.205885709 | 0.867006246 | 0.069782365 | 0.155644768 | 409.6997031 | 355.4101338 | ENSG00000163935 |
| PRPS2 | -0.205915015 | 0.866988634 | 0.018264808 | 0.052049988 | 785.1057427 | 681.2115883 | ENSG00000101911 |
| ZNF787 | -0.205932119 | 0.866978355 | 0.003794766 | 0.01380442 | 1679.811986 | 1456.822013 | ENSG00000142409 |
| HIGD1A | -0.206009727 | 0.866931789 | 0.267335789 | 0.433704037 | 210.2377373 | 182.5251178 | ENSG00000181061 |
| RIPK2 | -0.206052257 | 0.866906162 | 0.108716106 | 0.22050306 | 342.5870533 | 297.1658733 | ENSG00000104312 |
| COPE | -0.206083691 | 0.866887274 | 0.011705589 | 0.035925819 | 811.5576388 | 703.5785704 | ENSG00000105669 |
| RAD54B | -0.206145383 | 0.866850205 | 0.398511227 | 0.570915742 | 90.94222564 | 78.84950002 | ENSG00000197275 |
| TMX1 | -0.20618326 | 0.866827447 | 0.007040427 | 0.023493664 | 1642.900545 | 1424.735285 | ENSG00000139921 |
| RARA | -0.206185884 | 0.86682587 | 0.104013039 | 0.213504527 | 361.2727749 | 313.04858 | ENSG00000131759 |
| GAD2 | -0.206231898 | 0.866798223 | 0.803668623 | 0.88672909 | 8.075115818 | 7.023380075 | ENSG00000136750 |
| MAPK9 | -0.20626393 | 0.866778978 | 0.009034552 | 0.028888704 | 1037.538609 | 899.8659479 | ENSG00000050748 |
| TBCB | -0.206331347 | 0.866738475 | 0.00094762 | 0.004058734 | 2135.001251 | 1850.956853 | ENSG00000105254 |
| FAM86DP | -0.206347186 | 0.866728959 | 0.211980987 | 0.367844312 | 193.5610102 | 167.7448588 | ENSG00000244026 |
| HTR1B | -0.206363554 | 0.866719126 | 0.682425041 | 0.806312334 | 19.75822046 | 17.13079987 | ENSG00000135312 |
| EIF2S1 | -0.206435703 | 0.866675783 | 0.001524895 | 0.006213932 | 2115.316232 | 1834.201369 | ENSG00000134001 |
| MZT2A | -0.20644719 | 0.866668882 | 0.004819243 | 0.016995983 | 1804.198701 | 1564.013241 | ENSG00000173272 |
| MTAP | -0.20649154 | 0.86664224 | 0.022352636 | 0.061609474 | 715.4625474 | 620.4213509 | ENSG00000099810 |
| PPIL1 | -0.206609458 | 0.866571408 | 0.002346157 | 0.009068266 | 1371.737188 | 1188.892109 | ENSG00000137168 |
| PCMT1 | -0.206614997 | 0.866568081 | 0.002153964 | 0.008421588 | 1459.305537 | 1264.695486 | ENSG00000120265 |
| ZDHHC7 | -0.20670894 | 0.866511655 | 0.002143719 | 0.008388334 | 1515.061292 | 1312.825832 | ENSG00000153786 |
| MTHFD2L | -0.206765713 | 0.866477557 | 0.399752553 | 0.572292601 | 98.82754729 | 85.69865889 | ENSG00000163738 |
| NA | -0.206911627 | 0.866389926 | 0.674168465 | 0.800127035 | 20.36811652 | 17.61209956 | ENSG00000284428 |
| COG3 | -0.206911776 | 0.866389836 | 0.075872821 | 0.166229469 | 388.6698146 | 337.0075297 | ENSG00000136152 |
| ABCB1 | -0.206930688 | 0.866378479 | 0.715280054 | 0.82903554 | 21.12957309 | 18.3614522 | ENSG00000085563 |
| RBM22 | -0.206964144 | 0.866358388 | 0.014464622 | 0.042948595 | 759.4201951 | 657.8366207 | ENSG00000086589 |
| BRCC3 | -0.207013322 | 0.866328857 | 0.046029755 | 0.111693083 | 465.2721607 | 403.0659474 | ENSG00000185515 |
| DAAM2 | -0.207039998 | 0.866312838 | 0.199216135 | 0.350627549 | 187.1417829 | 162.066675 | ENSG00000146122 |
| SLC49A3 | -0.207043035 | 0.866311014 | 0.360539449 | 0.533617194 | 126.1815722 | 109.2037881 | ENSG00000169026 |
| ZNF708 | -0.207130405 | 0.866258552 | 0.174123677 | 0.316978377 | 239.3437296 | 207.5633798 | ENSG00000182141 |
| CBX6 | -0.207250384 | 0.866186514 | 0.012975379 | 0.039208311 | 1023.070072 | 886.1456704 | ENSG00000183741 |
| NA | -0.207260834 | 0.86618024 | 0.543567136 | 0.702012025 | 41.69669547 | 36.01308613 | ENSG00000181524 |
| DGUOK | -0.207398936 | 0.866097329 | 0.007831713 | 0.025671381 | 954.5387365 | 826.5212282 | ENSG00000114956 |
| CCDC115 | -0.207416366 | 0.866086865 | 0.104358206 | 0.214008095 | 354.1268069 | 306.3222948 | ENSG00000136710 |
| PSMB5 | -0.20745961 | 0.866060905 | 2.24E-05 | 0.000138354 | 3994.522287 | 3459.931159 | ENSG00000100804 |
| CCDC30 | -0.207676605 | 0.865930651 | 0.770566569 | 0.864847549 | 9.87063744 | 8.607345333 | ENSG00000186409 |
| UBE2O | -0.207685898 | 0.865925073 | 0.000156584 | 0.000806585 | 2714.072772 | 2349.878143 | ENSG00000175931 |
| CLASP1 | -0.207695619 | 0.865919238 | 0.00197207 | 0.007794142 | 1433.372323 | 1241.173952 | ENSG00000074054 |
| TPM2 | -0.207709456 | 0.865910934 | 0.000281417 | 0.001374037 | 2927.532889 | 2535.284559 | ENSG00000198467 |
| SMN1 | -0.207723468 | 0.865902524 | 0.521782406 | 0.684695217 | 47.6730565 | 41.27611706 | ENSG00000205571 |
| HSPH1 | -0.207792683 | 0.865860982 | 1.05E-05 | 6.90E-05 | 4830.812556 | 4183.269138 | ENSG00000120694 |
| MPP3 | -0.207883314 | 0.86580659 | 0.259018626 | 0.424164675 | 145.5034209 | 125.893071 | ENSG00000161647 |
| FOXO3B | -0.207885191 | 0.865805463 | 0.143887316 | 0.273390153 | 316.554457 | 274.2603438 | ENSG00000240445 |
| TADA3 | -0.207886502 | 0.865804677 | 0.003124557 | 0.01160043 | 1285.007052 | 1112.505791 | ENSG00000171148 |
| ERCC2 | -0.207929389 | 0.865778939 | 0.001422048 | 0.005846704 | 2651.997618 | 2295.353337 | ENSG00000104884 |
| RNF167 | -0.207937669 | 0.86577397 | 0.000170981 | 0.000874442 | 2247.907606 | 1946.059305 | ENSG00000108523 |
| POU4F1 | -0.207965388 | 0.865757336 | 0.822572828 | 0.898898341 | 5.679985866 | 4.913758686 | ENSG00000152192 |
| ARHGEF18 | -0.208045262 | 0.865709405 | 0.025971399 | 0.069856979 | 617.5634353 | 534.8635823 | ENSG00000104880 |
| DNAJC2 | -0.208082434 | 0.865687099 | 0.017586307 | 0.050514809 | 850.5712402 | 736.586134 | ENSG00000105821 |
| GLB1L3 | -0.208109847 | 0.865670651 | 0.006093242 | 0.020819118 | 1536.153609 | 1329.851067 | ENSG00000166105 |
| QRFP | -0.208237888 | 0.865593825 | 0.818415419 | 0.89619415 | 5.758894886 | 4.983128505 | ENSG00000188710 |
| SAP30L-AS1 | -0.208728333 | 0.865299616 | 0.681040586 | 0.805367127 | 19.13783651 | 16.59196772 | ENSG00000245275 |
| MTMR2 | -0.208739866 | 0.865292698 | 0.000941959 | 0.004038072 | 1518.415472 | 1314.157871 | ENSG00000087053 |

| | | | | | | | |
|---|---|---|---|---|---|---|---|
| NUDCD2 | -0.208811269 | 0.865249874 | 0.007423146 | 0.02453734 | 992.0241311 | 858.6441525 | ENSG00000170584 |
| DNAJC17 | -0.208826979 | 0.865240452 | 0.217695203 | 0.3752621 | 256.7921116 | 221.7892622 | ENSG00000104129 |
| DPY19L2 | -0.208855364 | 0.865223429 | 0.53534997 | 0.695403425 | 45.02206348 | 38.95784106 | ENSG00000177990 |
| PIK3C3 | -0.208868475 | 0.865215565 | 0.030130742 | 0.078792483 | 638.3717153 | 552.5391024 | ENSG00000078142 |
| POM121 | -0.208913379 | 0.865188636 | 0.000425067 | 0.001987871 | 1980.644097 | 1713.834081 | ENSG00000196313 |
| LOC220729 | -0.208925181 | 0.865181558 | 0.149438869 | 0.281627327 | 312.7093777 | 270.2161826 | ENSG00000214135 |
| ASB3 | -0.208959759 | 0.865160822 | 0.074447578 | 0.163946146 | 415.2408195 | 359.6312362 | ENSG00000115239 |
| PTPN5 | -0.208968336 | 0.865155679 | 0.137640324 | 0.264279833 | 267.3387509 | 231.3542996 | ENSG00000110786 |
| RAB1B | -0.208996905 | 0.865138547 | 0.009598983 | 0.030395539 | 904.2051133 | 782.2275093 | ENSG00000174903 |
| RNF122 | -0.209028329 | 0.865119702 | 0.094019084 | 0.197250914 | 311.5257183 | 269.5480066 | ENSG00000133874 |
| GTPBP3 | -0.209037837 | 0.865114001 | 0.00706582 | 0.023562096 | 1209.583537 | 1047.12494 | ENSG00000130299 |
| SKP1 | -0.209063664 | 0.865098514 | 0.001251965 | 0.00522763 | 1589.556837 | 1375.497445 | ENSG00000113558 |
| KCNAB1 | -0.209170356 | 0.86503454 | 0.706218305 | 0.823231454 | 18.78400367 | 16.16284537 | ENSG00000169282 |
| CHD9 | -0.209448595 | 0.864867725 | 0.043739793 | 0.107269813 | 496.0163449 | 428.7919827 | ENSG00000177200 |
| RAB23 | -0.209468796 | 0.864855615 | 0.109141405 | 0.221230099 | 292.4155154 | 252.9291354 | ENSG00000112210 |
| GOLGA2 | -0.20967343 | 0.864732951 | 0.002820331 | 0.010636756 | 1189.553621 | 1028.596796 | ENSG00000167110 |
| NA | -0.209703663 | 0.86471483 | 0.282458763 | 0.450766644 | 177.8451473 | 153.7784939 | ENSG00000260404 |
| NYAP1 | -0.209795784 | 0.864659617 | 0.186091641 | 0.332613161 | 207.0250062 | 178.8119155 | ENSG00000166924 |
| GHRL | -0.209887537 | 0.864604628 | 0.868664524 | 0.926811503 | 3.363764935 | 2.881576373 | ENSG00000157017 |
| ANAPC13 | -0.210088615 | 0.86448413 | 0.040575637 | 0.100830169 | 495.1726759 | 428.2996575 | ENSG00000129055 |
| LAP3 | -0.210120365 | 0.864465105 | 0.000627076 | 0.00280346 | 1906.794869 | 1648.423673 | ENSG00000002549 |
| SWSAP1 | -0.210180538 | 0.86442905 | 0.620215664 | 0.76139303 | 31.73087915 | 27.37401523 | ENSG00000173928 |
| NR1I3 | -0.210222442 | 0.864403943 | 0.661141017 | 0.790816567 | 24.0696446 | 20.84966036 | ENSG00000143257 |
| RPF1 | -0.210303446 | 0.86435541 | 0.007506261 | 0.024786595 | 973.9355904 | 841.909677 | ENSG00000117133 |
| CCDC85B | -0.210388427 | 0.864304497 | 0.005312372 | 0.01848169 | 1638.116906 | 1416.523253 | ENSG00000175602 |
| SAMD12 | -0.210429269 | 0.86428003 | 0.336040608 | 0.508875059 | 120.5348018 | 104.2555776 | ENSG00000177570 |
| EMC2 | -0.210492627 | 0.864242074 | 0.021754575 | 0.060227827 | 641.800598 | 554.8181472 | ENSG00000104412 |
| MDH1 | -0.210523348 | 0.864223671 | 2.65E-06 | 1.93E-05 | 4259.812654 | 3681.290414 | ENSG00000014641 |
| FAU | -0.210601477 | 0.86417687 | 0.00056985 | 0.002573866 | 2019.806304 | 1745.087971 | ENSG00000149806 |
| TXNDC17 | -0.210615002 | 0.864168769 | 0.015796863 | 0.046235363 | 1024.814394 | 885.229366 | ENSG00000129235 |
| VHL | -0.210724214 | 0.864103354 | 0.015316408 | 0.045052988 | 794.0067061 | 686.0718979 | ENSG00000134086 |
| CYP2D7 | -0.210817862 | 0.864047265 | 0.601586071 | 0.747372288 | 34.4528088 | 29.73852566 | ENSG00000205702 |
| PLPPR3 | -0.210819942 | 0.864046019 | 0.006733109 | 0.022664177 | 1632.076817 | 1410.852345 | ENSG00000129951 |
| ZNF767P | -0.210839971 | 0.864034024 | 0.17128385 | 0.313019775 | 198.4745213 | 171.527443 | ENSG00000133624 |
| RPL14 | -0.210938397 | 0.863975078 | 1.81E-07 | 1.60E-06 | 18513.36974 | 15994.65222 | ENSG00000188846 |
| MRFAP1 | -0.211084698 | 0.863887468 | 1.43E-07 | 1.29E-06 | 10379.19312 | 8967.115945 | ENSG00000179010 |
| KLHL17 | -0.21109865 | 0.863879114 | 0.052521814 | 0.123953794 | 699.1057251 | 603.7805078 | ENSG00000187961 |
| PSMF1 | -0.211173148 | 0.863834506 | 0.000399654 | 0.001883179 | 1814.136768 | 1567.178433 | ENSG00000125818 |
| FAAP100 | -0.211194408 | 0.863821777 | 0.01292279 | 0.039072193 | 1244.747941 | 1075.207259 | ENSG00000185504 |
| ANKRD26 | -0.211278407 | 0.863771483 | 0.256351179 | 0.421525713 | 168.5945062 | 145.6271439 | ENSG00000107890 |
| NA | -0.211314316 | 0.863749984 | 0.580332762 | 0.731378652 | 35.05345627 | 30.31877016 | ENSG00000264575 |
| GPN2 | -0.211353496 | 0.863726527 | 0.036337028 | 0.091991812 | 539.3986279 | 465.9104485 | ENSG00000142751 |
| SMIM30 | -0.211397311 | 0.863700296 | 0.089905128 | 0.19056805 | 423.065417 | 365.568919 | ENSG00000214194 |
| SFRP5 | -0.211417035 | 0.863688488 | 0.420824535 | 0.592835997 | 75.6369775 | 65.41927156 | ENSG00000120057 |
| MALSU1 | -0.211419789 | 0.863686839 | 0.009139957 | 0.029170712 | 934.5366186 | 807.3119995 | ENSG00000156928 |
| NA | -0.211436734 | 0.863676695 | 0.820766706 | 0.89784012 | 5.636007141 | 4.856759272 | ENSG00000237082 |
| NA | -0.211490688 | 0.863644395 | 0.651474844 | 0.783266107 | 22.16172312 | 19.14362702 | ENSG00000257489 |
| GATAD2A | -0.211501444 | 0.863637957 | 1.26E-05 | 8.12E-05 | 4035.191084 | 3484.746294 | ENSG00000167491 |
| DRAP1 | -0.211509853 | 0.863632923 | 0.001093912 | 0.004618658 | 3050.045936 | 2633.946891 | ENSG00000175550 |
| ARPC4 | -0.211619273 | 0.863567424 | 0.002671158 | 0.010155558 | 1368.338231 | 1181.881887 | ENSG00000241553 |
| KCTD5 | -0.21170214 | 0.863517822 | 0.008680299 | 0.027964538 | 938.3572797 | 810.344165 | ENSG00000167977 |
| MYDGF | -0.211784036 | 0.863468806 | 0.005940143 | 0.020364547 | 1989.31653 | 1717.145065 | ENSG00000074842 |
| E4F1 | -0.211838261 | 0.863436352 | 0.014511926 | 0.043069164 | 896.992527 | 774.1232604 | ENSG00000167967 |
| LRCH2 | -0.211849808 | 0.863429441 | 0.193613704 | 0.343353739 | 215.6665519 | 185.9995695 | ENSG00000130224 |
| LDLRAD4 | -0.211887649 | 0.863406794 | 0.516798421 | 0.680701785 | 44.62213674 | 38.50800647 | ENSG00000168675 |
| GNPDA2 | -0.211904566 | 0.86339667 | 0.264362252 | 0.430146103 | 140.8184275 | 121.564924 | ENSG00000163281 |
| LRRC8D | -0.212018764 | 0.86332833 | 0.014949871 | 0.044144798 | 797.9419943 | 688.9555943 | ENSG00000171492 |
| STMP1 | -0.212075106 | 0.863294615 | 0.008833197 | 0.028357663 | 912.3954951 | 788.158934 | ENSG00000243317 |
| TWIST2 | -0.212100572 | 0.863279376 | 0.45956297 | 0.629447053 | 75.48530485 | 65.31334199 | ENSG00000233608 |
| SLC25A21-AS1 | -0.212154413 | 0.863247159 | 0.779366428 | 0.870480335 | 7.982252283 | 6.903462563 | ENSG00000258708 |
| COPS4 | -0.212187715 | 0.863227233 | 0.019338768 | 0.054645403 | 821.6331152 | 709.6632678 | ENSG00000138663 |
| NLGN4X | -0.212204354 | 0.863217277 | 1.28E-06 | 9.92E-06 | 4943.685895 | 4267.488027 | ENSG00000146938 |
| CHCHD5 | -0.212230838 | 0.863201431 | 0.161952887 | 0.299530539 | 284.3313512 | 245.1881588 | ENSG00000125611 |
| PIK3R4 | -0.212297154 | 0.863161753 | 0.006119728 | 0.020883928 | 1017.166974 | 877.9752072 | ENSG00000196455 |
| FANCM | -0.212319613 | 0.863148316 | 0.146339057 | 0.276903065 | 230.972639 | 199.3795108 | ENSG00000187790 |
| ATP5PB | -0.212519328 | 0.863028838 | 0.001062359 | 0.004500212 | 3123.008638 | 2694.631187 | ENSG00000116459 |
| LOC100506098 | -0.212622055 | 0.862967388 | 0.527943736 | 0.689522636 | 41.07332046 | 35.44521203 | ENSG00000233834 |
| MICAL1 | -0.212678727 | 0.862933489 | 0.005662827 | 0.019538831 | 1586.47086 | 1368.404538 | ENSG00000135596 |
| ARHGAP18 | -0.212711829 | 0.86291369 | 0.268273958 | 0.434677719 | 129.9714581 | 112.1135703 | ENSG00000146376 |
| C11orf68 | -0.21291029 | 0.862794993 | 0.027301326 | 0.072835001 | 710.1281466 | 613.0448879 | ENSG00000175573 |
| NPTX1 | -0.213023313 | 0.862727403 | 0.378373595 | 0.551850761 | 112.9709734 | 97.18327945 | ENSG00000171246 |
| ARIH2 | -0.213053943 | 0.862709087 | 0.001209742 | 0.005066681 | 2053.670993 | 1772.022675 | ENSG00000177479 |
| MYLPF | -0.213067987 | 0.862700689 | 0.816683108 | 0.895136415 | 5.668822255 | 4.877731971 | ENSG00000180209 |
| ENTPD6 | -0.213078292 | 0.862694526 | 0.005683821 | 0.019600748 | 1176.677945 | 1015.158275 | ENSG00000197586 |
| RPL30 | -0.213174295 | 0.862637121 | 0.000715635 | 0.003145435 | 2962.752533 | 2555.063855 | ENSG00000156482 |
| KLHL22 | -0.213188217 | 0.862628797 | 0.221832884 | 0.380200981 | 157.2789888 | 135.7377929 | ENSG00000099910 |
| TRIM16L | -0.213204809 | 0.862618876 | 0.293009785 | 0.462746767 | 118.2017474 | 101.9475335 | ENSG00000108448 |

| | | | | | | | |
|---|---|---|---|---|---|---|---|
| C16orf70 | -0.213290031 | 0.862567922 | 0.087810501 | 0.187011744 | 375.8407207 | 324.1879569 | ENSG00000125149 |
| FRMD3 | -0.213303804 | 0.862559687 | 0.560277517 | 0.715498368 | 36.82665066 | 31.8209952 | ENSG00000172159 |
| MAFG | -0.213376886 | 0.862515994 | 0.008677979 | 0.027961734 | 1036.556381 | 893.5363329 | ENSG00000197063 |
| EIF3K | -0.213423084 | 0.862488375 | 7.57E-05 | 0.000420898 | 3403.889855 | 2935.142248 | ENSG00000178982 |
| POLA1 | -0.213467263 | 0.862461963 | 8.41E-05 | 0.000462053 | 2335.156001 | 2013.918648 | ENSG00000101868 |
| GNPNAT1 | -0.21352297 | 0.862428661 | 0.013444019 | 0.040395207 | 755.5786335 | 651.5252705 | ENSG00000100522 |
| ATP6V1FNB | -0.213694448 | 0.86232616 | 0.739425122 | 0.845288891 | 11.72429246 | 10.06494136 | ENSG00000272899 |
| HLA-C | -0.213697916 | 0.862324087 | 1.46E-05 | 9.30E-05 | 5958.237627 | 5138.249568 | ENSG00000204525 |
| DGKD | -0.213705461 | 0.862319577 | 0.010164144 | 0.031903107 | 858.9432806 | 741.0285161 | ENSG00000077044 |
| MLLT3 | -0.213815052 | 0.862254076 | 0.223687781 | 0.382177042 | 152.6521287 | 131.6164289 | ENSG00000171843 |
| MIGA1 | -0.213887956 | 0.862210504 | 0.364323458 | 0.537974382 | 95.19396478 | 82.19246947 | ENSG00000180488 |
| NA | -0.213959849 | 0.862167539 | 0.631746602 | 0.768920547 | 23.5817604 | 20.37401889 | ENSG00000253771 |
| TARS3 | -0.214022059 | 0.862130363 | 0.108339044 | 0.21995038 | 323.1559079 | 278.892322 | ENSG00000185418 |
| AACS | -0.214043446 | 0.862117582 | 0.001924482 | 0.007627942 | 1532.955195 | 1321.601226 | ENSG00000081760 |
| TAF1D | -0.214089076 | 0.862090316 | 0.022960178 | 0.062969122 | 1114.542657 | 960.3236631 | ENSG00000166012 |
| SUV39H2 | -0.214202959 | 0.862022267 | 0.022826456 | 0.062718297 | 679.8428984 | 586.0987048 | ENSG00000152455 |
| FHAD1 | -0.214274641 | 0.861979437 | 0.546744509 | 0.704747316 | 37.32223185 | 32.20576112 | ENSG00000142621 |
| DLG5 | -0.214326663 | 0.861948356 | 0.000207809 | 0.00104397 | 2697.429386 | 2324.808441 | ENSG00000151208 |
| SLC6A2 | -0.214472941 | 0.861860965 | 0.819309019 | 0.896906724 | 5.682776769 | 4.884183692 | ENSG00000103546 |
| FGFR2 | -0.214701427 | 0.861724479 | 0.000587298 | 0.002642158 | 1643.385373 | 1416.064258 | ENSG00000066468 |
| H2BC6 | -0.214767632 | 0.861684936 | 0.648670176 | 0.781605019 | 22.07723229 | 19.00435434 | ENSG00000274290 |
| PSMC1 | -0.214787847 | 0.861672862 | 0.009895674 | 0.031161844 | 1654.184172 | 1425.350502 | ENSG00000100764 |
| NA | -0.214839877 | 0.861641787 | 0.310123444 | 0.481279598 | 152.1237324 | 131.0509964 | ENSG00000237517 |
| KIFC2 | -0.214936028 | 0.861584363 | 0.120267249 | 0.238862916 | 267.0243541 | 230.1067153 | ENSG00000167702 |
| MTURN | -0.214956763 | 0.86157198 | 0.21274571 | 0.368744168 | 165.2676988 | 142.4097502 | ENSG00000180354 |
| TMEM170B | -0.214988113 | 0.861553258 | 0.182973739 | 0.328746055 | 193.8607399 | 167.2119269 | ENSG00000205269 |
| STARD3NL | -0.214991033 | 0.861551515 | 0.033406934 | 0.085709957 | 664.3899056 | 572.3272428 | ENSG00000010270 |
| CPTP | -0.215028168 | 0.861529338 | 0.172414408 | 0.314600408 | 332.1867945 | 286.2760546 | ENSG00000224051 |
| BCAR3 | -0.215038608 | 0.861523104 | 0.127317771 | 0.249045541 | 248.8793196 | 214.3025685 | ENSG00000137936 |
| UBE2D1 | -0.215039538 | 0.861522549 | 0.03887488 | 0.097432158 | 689.7292805 | 594.5463378 | ENSG00000072401 |
| ENOPH1 | -0.215212312 | 0.861419381 | 0.000539393 | 0.002452377 | 1954.926586 | 1683.92938 | ENSG00000145293 |
| FAM155B | -0.215256653 | 0.861392906 | 0.015796673 | 0.046235363 | 770.1703219 | 663.4306837 | ENSG00000130054 |
| SLC38A2 | -0.215319618 | 0.861355312 | 1.77E-07 | 1.57E-06 | 6625.62522 | 5707.410927 | ENSG00000134294 |
| POLE3 | -0.215401096 | 0.861306667 | 0.006534311 | 0.022087507 | 1002.390323 | 863.2368874 | ENSG00000148229 |
| MFN2 | -0.215444255 | 0.861280901 | 0.000392444 | 0.001855546 | 2677.305079 | 2305.140539 | ENSG00000116688 |
| MOGS | -0.215455474 | 0.861274203 | 0.00186801 | 0.007437726 | 1961.761562 | 1690.221969 | ENSG00000115275 |
| IFT122 | -0.215618578 | 0.861176838 | 0.033437236 | 0.085764887 | 636.5093241 | 547.8222405 | ENSG00000163913 |
| DENR | -0.215639732 | 0.86116421 | 0.000240643 | 0.001193991 | 2600.636581 | 2239.95746 | ENSG00000139726 |
| PGAP3 | -0.215791335 | 0.861073721 | 0.073147911 | 0.161637612 | 487.2671806 | 419.5399183 | ENSG00000161395 |
| PAQR9 | -0.215912792 | 0.861001233 | 0.653476441 | 0.784700531 | 29.46095229 | 25.29746437 | ENSG00000188582 |
| IDH1 | -0.215930205 | 0.860990841 | 5.46E-08 | 5.22E-07 | 10387.44093 | 8943.918324 | ENSG00000138413 |
| GTPBP6 | -0.216026065 | 0.860933634 | 0.000210064 | 0.001054198 | 2053.106191 | 1767.808521 | ENSG00000178605 |
| DDA1 | -0.21602663 | 0.860933297 | 0.002936141 | 0.011002545 | 1226.243066 | 1055.864324 | ENSG00000130311 |
| TIGD3 | -0.216027631 | 0.860932699 | 0.623549627 | 0.763008988 | 24.40394111 | 21.00266052 | ENSG00000173825 |
| PREX2 | -0.216178761 | 0.860842517 | 0.003413422 | 0.01256168 | 1199.746501 | 1032.588271 | ENSG00000046889 |
| MAPKBP1 | -0.216202071 | 0.860828608 | 0.089133645 | 0.189265509 | 317.9354217 | 273.4287421 | ENSG00000137802 |
| NA | -0.216242259 | 0.860804629 | 0.529436654 | 0.690558654 | 41.31283842 | 35.52748529 | ENSG00000224905 |
| POP1 | -0.216336542 | 0.860748375 | 0.050092901 | 0.119478506 | 450.601339 | 387.7551918 | ENSG00000104356 |
| ANK3 | -0.216341776 | 0.860745253 | 0.010865399 | 0.03382353 | 836.8304792 | 720.3847962 | ENSG00000151150 |
| LRRC3 | -0.216345141 | 0.860743245 | 0.106203325 | 0.216718026 | 303.1880816 | 261.1036087 | ENSG00000160233 |
| JADE3 | -0.21637353 | 0.860726308 | 0.018254354 | 0.052042071 | 619.3544512 | 533.0857681 | ENSG00000102221 |
| NUDT21 | -0.216390082 | 0.860716433 | 1.35E-06 | 1.04E-05 | 4690.295935 | 4037.778442 | ENSG00000167005 |
| SNX33 | -0.216407317 | 0.86070615 | 0.005066904 | 0.017736463 | 991.1087603 | 852.8380188 | ENSG00000173548 |
| RALGPS1 | -0.216550544 | 0.860620706 | 0.083052793 | 0.178833574 | 412.1695486 | 354.549708 | ENSG00000136828 |
| NACA | -0.216551646 | 0.860620049 | 1.13E-05 | 7.31E-05 | 4938.240332 | 4249.477023 | ENSG00000196531 |
| EID2 | -0.216782084 | 0.860482595 | 0.015983658 | 0.046681081 | 658.6091349 | 566.6845896 | ENSG00000176396 |
| CRB3 | -0.216822571 | 0.860458447 | 0.077382861 | 0.168853295 | 340.0945171 | 292.7968776 | ENSG00000130545 |
| PPM1M | -0.216879237 | 0.860424651 | 0.29068807 | 0.460474918 | 120.2936442 | 103.6514163 | ENSG00000164088 |
| IRF2BPL | -0.217078533 | 0.860305799 | 0.00020627 | 0.001037319 | 2066.457611 | 1777.896065 | ENSG00000119669 |
| SNHG11 | -0.217079018 | 0.86030551 | 0.314175663 | 0.485365797 | 102.193064 | 87.89432165 | ENSG00000174365 |
| NA | -0.217105352 | 0.860289806 | 0.308650736 | 0.479700644 | 101.5531437 | 87.38344696 | ENSG00000287086 |
| RBM45 | -0.217228744 | 0.86021623 | 0.246450814 | 0.409922629 | 170.5795339 | 146.5996905 | ENSG00000155636 |
| PNCK | -0.217286203 | 0.86018197 | 0.360640724 | 0.533685195 | 84.01848081 | 72.22567942 | ENSG00000130822 |
| CCDC91 | -0.217399224 | 0.860114586 | 0.166688046 | 0.306510767 | 194.1929212 | 167.0766949 | ENSG00000123106 |
| QARS1 | -0.217459088 | 0.860078897 | 2.28E-06 | 1.68E-05 | 6215.103283 | 5344.252506 | ENSG00000172053 |
| SPPL3 | -0.217674571 | 0.859950444 | 0.037011499 | 0.093454036 | 660.1051895 | 567.7896783 | ENSG00000157837 |
| CEP78 | -0.217677313 | 0.859948809 | 0.006035241 | 0.02064969 | 861.434153 | 740.7626074 | ENSG00000148019 |
| NA | -0.217747771 | 0.859906812 | 0.711797032 | 0.826289358 | 13.68388965 | 11.74570086 | ENSG00000259891 |
| ULK1 | -0.217781363 | 0.85988679 | 0.00051812 | 0.002366812 | 1901.66956 | 1635.451311 | ENSG00000177169 |
| STAP2 | -0.217848442 | 0.85984681 | 0.092064705 | 0.194000353 | 308.5981618 | 265.5513821 | ENSG00000178078 |
| RHEBL1 | -0.218060867 | 0.859720214 | 0.327297939 | 0.499399236 | 118.6814591 | 102.0300672 | ENSG00000167550 |
| LEO1 | -0.218095767 | 0.859699417 | 0.006922333 | 0.023184485 | 1012.912672 | 871.0561455 | ENSG00000166477 |
| ALKBH7 | -0.218453017 | 0.859486559 | 0.006369251 | 0.021597684 | 1124.614531 | 966.523487 | ENSG00000125652 |
| AFG1L | -0.218491873 | 0.859463411 | 0.339415153 | 0.511895686 | 95.49017668 | 82.11637535 | ENSG00000135537 |
| BRD2 | -0.218600581 | 0.859398652 | 0.000347765 | 0.001662225 | 3645.629326 | 3132.104641 | ENSG00000204256 |
| AFP | -0.218623202 | 0.859385177 | 0.654821599 | 0.785826654 | 26.08283252 | 22.41616057 | ENSG00000081051 |
| CNOT7 | -0.218628198 | 0.859382201 | 0.001553917 | 0.006309538 | 1872.021406 | 1609.541574 | ENSG00000198791 |

| | | | | | | | |
|---|---|---|---|---|---|---|---|
| TRAF3 | -0.218668895 | 0.859357959 | 0.029115785 | 0.076647462 | 543.2274743 | 466.6646659 | ENSG00000131323 |
| PPIF | -0.218720806 | 0.859327038 | 0.000544709 | 0.002474795 | 2606.771335 | 2240.476218 | ENSG00000108179 |
| MAP7 | -0.218740004 | 0.859315603 | 6.80E-05 | 0.000380877 | 2378.552036 | 2044.083812 | ENSG00000135525 |
| LAMTOR3 | -0.218784392 | 0.859289165 | 0.068004867 | 0.152473088 | 370.4605894 | 318.4194064 | ENSG00000109270 |
| TIMM50 | -0.218784846 | 0.859288894 | 5.49E-05 | 0.000313675 | 3408.758751 | 2928.580485 | ENSG00000105197 |
| ELOVL1 | -0.218869616 | 0.859238405 | 0.023947447 | 0.06520398 | 558.2621411 | 479.4428384 | ENSG00000066322 |
| PKMYT1 | -0.218870133 | 0.859238097 | 0.014200021 | 0.042312729 | 1459.387639 | 1253.881017 | ENSG00000127564 |
| ADSL | -0.218905109 | 0.859217267 | 0.400595207 | 0.573114961 | 85.46555124 | 73.38267289 | ENSG00000239900 |
| SLC29A1 | -0.218906318 | 0.859216547 | 2.08E-06 | 1.55E-05 | 4709.16155 | 4045.957872 | ENSG00000112759 |
| TRIP13 | -0.2189578 | 0.859185887 | 0.001887831 | 0.007508898 | 1190.655143 | 1022.844699 | ENSG00000071539 |
| ITGA6 | -0.218969299 | 0.859179038 | 0.000758534 | 0.003312104 | 2461.112357 | 2114.583757 | ENSG00000091409 |
| WASHC3 | -0.218988085 | 0.859167851 | 0.418961167 | 0.590975423 | 64.07520901 | 55.02389219 | ENSG00000120860 |
| MRPS26 | -0.219065229 | 0.859121911 | 0.000811869 | 0.003522664 | 2124.707908 | 1825.440606 | ENSG00000125901 |
| SART1 | -0.219107527 | 0.859096722 | 0.001036438 | 0.004397715 | 2848.069358 | 2447.233766 | ENSG00000175467 |
| NA | -0.219131651 | 0.859082357 | 0.59272826 | 0.741158069 | 34.16883655 | 29.27578762 | ENSG00000286797 |
| DLAT | -0.219214566 | 0.859032985 | 5.10E-05 | 0.000292525 | 2373.458619 | 2039.22065 | ENSG00000150768 |
| SLC43A2 | -0.219305791 | 0.858978668 | 0.047615726 | 0.114862444 | 547.6097981 | 470.3985988 | ENSG00000167703 |
| STARD7 | -0.219332201 | 0.858962944 | 1.01E-05 | 6.65E-05 | 3983.512981 | 3421.442339 | ENSG00000084090 |
| CLIC4 | -0.219374874 | 0.858937537 | 1.08E-05 | 7.03E-05 | 6648.480959 | 5711.370591 | ENSG00000169504 |
| NREP | -0.219526404 | 0.858847325 | 0.116362545 | 0.233002891 | 300.5303043 | 257.8023057 | ENSG00000134986 |
| FRMPD3 | -0.219574436 | 0.858818732 | 0.308039072 | 0.479079787 | 101.4883894 | 87.13311953 | ENSG00000147234 |
| PSMA3 | -0.219579369 | 0.858815795 | 0.000192723 | 0.000975291 | 2485.403714 | 2135.445837 | ENSG00000100567 |
| ZNF384 | -0.219622494 | 0.858790125 | 0.017934799 | 0.05132476 | 856.7322636 | 735.7122354 | ENSG00000126746 |
| APBA2 | -0.219706662 | 0.858740023 | 0.006960125 | 0.023281386 | 847.090131 | 727.3090136 | ENSG00000034053 |
| HSPA12A | -0.219732309 | 0.858724757 | 0.175770353 | 0.319162828 | 210.705083 | 180.6370729 | ENSG00000165868 |
| CHCHD7 | -0.219754197 | 0.858711729 | 0.009356991 | 0.029747152 | 728.8376959 | 625.8467778 | ENSG00000170791 |
| ZNF649-AS1 | -0.219797512 | 0.858685948 | 0.858153262 | 0.920622544 | 3.426604259 | 2.90685022 | ENSG00000268095 |
| RBM28 | -0.219871352 | 0.858642 | 0.002141346 | 0.008380749 | 1441.749997 | 1237.263448 | ENSG00000106344 |
| CD83 | -0.219876483 | 0.858638946 | 0.16201768 | 0.299559412 | 240.2313404 | 206.4865212 | ENSG00000112149 |
| CRAT | -0.219969077 | 0.858583839 | 0.088955464 | 0.189028093 | 323.3423107 | 277.5993152 | ENSG00000095321 |
| IL21R | -0.220048232 | 0.858536733 | 0.620054715 | 0.761295539 | 29.29388566 | 25.13130031 | ENSG00000103522 |
| DNAJC11 | -0.22005927 | 0.858530165 | 0.001034635 | 0.004392416 | 1900.817181 | 1631.604029 | ENSG00000007923 |
| SF3B2 | -0.220112148 | 0.858498698 | 3.67E-07 | 3.11E-06 | 7501.712536 | 6440.050714 | ENSG00000087365 |
| LOC101928445 | -0.220252936 | 0.858414924 | 0.780177724 | 0.870840926 | 7.946646267 | 6.813120045 | ENSG00000244945 |
| DNAJC4 | -0.220380397 | 0.858339087 | 0.231107624 | 0.39138769 | 169.9294891 | 145.6959928 | ENSG00000110011 |
| ZC3H12B | -0.220437988 | 0.858304824 | 0.519873702 | 0.683212997 | 45.24252069 | 38.82178504 | ENSG00000102053 |
| NDUFB2 | -0.220477408 | 0.858281372 | 0.074311667 | 0.163702907 | 381.2904499 | 326.9652028 | ENSG00000090266 |
| FAM184A | -0.220528113 | 0.858251208 | 0.142519822 | 0.27162121 | 245.8969331 | 210.8246091 | ENSG00000111879 |
| APTX | -0.22058726 | 0.858216022 | 0.01351565 | 0.040555359 | 828.8410182 | 710.7742995 | ENSG00000137074 |
| LAPTM4B | -0.220725541 | 0.858133766 | 4.35E-08 | 4.22E-07 | 11211.00926 | 9621.558975 | ENSG00000104341 |
| RER1 | -0.220776707 | 0.858103333 | 0.002810841 | 0.01060926 | 1165.389748 | 1000.035722 | ENSG00000157916 |
| NA | -0.22082829 | 0.858072653 | 0.653449532 | 0.784700531 | 19.95914134 | 17.18511567 | ENSG00000257851 |
| NDUFAF6 | -0.220845214 | 0.858062586 | 0.204286079 | 0.357546975 | 195.1397045 | 167.6206217 | ENSG00000156170 |
| MYLK2 | -0.220917976 | 0.858019312 | 0.811215919 | 0.891639582 | 5.347804529 | 4.58330857 | ENSG00000101306 |
| ZBTB7C | -0.220954584 | 0.85799754 | 0.773405139 | 0.867024555 | 10.32869759 | 8.775399501 | ENSG00000184828 |
| IFT88 | -0.221044465 | 0.857944088 | 0.326052036 | 0.497852606 | 94.30646614 | 80.94272243 | ENSG00000032742 |
| FBXO44 | -0.221059338 | 0.857935243 | 0.01442612 | 0.042847464 | 701.3888044 | 601.696366 | ENSG00000132879 |
| C7orf26 | -0.221092697 | 0.857915405 | 0.014981867 | 0.044225738 | 779.7158745 | 668.9929844 | ENSG00000146576 |
| ATP23 | -0.221185346 | 0.857860312 | 0.27443543 | 0.441177824 | 124.298969 | 106.5891986 | ENSG00000166896 |
| NA | -0.221190327 | 0.857857351 | 0.653142916 | 0.784504704 | 20.42258313 | 17.55966176 | ENSG00000180211 |
| EPHX2 | -0.221264662 | 0.85781315 | 0.136167025 | 0.261998612 | 257.6720254 | 220.7000026 | ENSG00000120915 |
| CLK4 | -0.221273673 | 0.857807793 | 0.161055195 | 0.298341927 | 220.4623567 | 189.2564855 | ENSG00000113240 |
| CIC | -0.22132595 | 0.85777671 | 0.002644403 | 0.010065743 | 1766.806481 | 1515.688445 | ENSG00000079432 |
| FKBP15 | -0.22141848 | 0.857721696 | 0.033056922 | 0.084967003 | 474.0619634 | 406.5266838 | ENSG00000119321 |
| PSIP1 | -0.221423275 | 0.857718846 | 1.50E-06 | 1.15E-05 | 8220.532571 | 7051.923509 | ENSG00000164985 |
| CNPY3 | -0.221515003 | 0.857664313 | 0.000168254 | 0.000862551 | 2237.312207 | 1918.896337 | ENSG00000137161 |
| MAP2K2 | -0.221633328 | 0.857593973 | 0.000193938 | 0.000980722 | 3255.908028 | 2792.022903 | ENSG00000126934 |
| MTMR4 | -0.221658388 | 0.857579077 | 0.024206644 | 0.065789006 | 811.9411935 | 695.8628555 | ENSG00000108389 |
| NA | -0.221736582 | 0.857532597 | 0.684906261 | 0.808006768 | 17.25503316 | 14.73052319 | ENSG00000234769 |
| SLC52A2 | -0.221738589 | 0.857531404 | 0.005438026 | 0.018861012 | 979.6379034 | 839.7673066 | ENSG00000185803 |
| LYAR | -0.221874105 | 0.857450858 | 0.000449102 | 0.002085608 | 1669.360875 | 1431.920583 | ENSG00000145220 |
| NA | -0.221883267 | 0.857445412 | 0.60076576 | 0.746737905 | 27.44389742 | 23.46182672 | ENSG00000273142 |
| ARHGAP35 | -0.221944037 | 0.857409296 | 4.32E-06 | 3.04E-05 | 3602.566389 | 3088.88738 | ENSG00000160007 |
| HSF1 | -0.222172842 | 0.857273325 | 0.00016868 | 0.000864042 | 2228.949477 | 1910.953436 | ENSG00000185122 |
| LOC105378083 | -0.222172958 | 0.857273256 | 0.815820847 | 0.894662041 | 5.356177238 | 4.585459144 | ENSG00000271913 |
| RAB11FIP5 | -0.222215467 | 0.857247997 | 0.015184006 | 0.044733455 | 713.481611 | 611.5962399 | ENSG00000135631 |
| NA | -0.222229788 | 0.857239487 | 0.371543797 | 0.545038336 | 81.26807856 | 69.66380029 | ENSG00000204745 |
| LOC100506405 | -0.222239 | 0.857234013 | 0.589217232 | 0.738633689 | 31.49579171 | 27.00026264 | ENSG00000228496 |
| COL7A1 | -0.222279971 | 0.857209669 | 0.135710896 | 0.261264278 | 285.0479772 | 244.0403127 | ENSG00000114270 |
| NA | -0.222281376 | 0.857208835 | 0.52931611 | 0.690494798 | 38.28125482 | 32.72953925 | ENSG00000178715 |
| FOXL2 | -0.222305936 | 0.857194242 | 0.60081071 | 0.746745662 | 30.10490276 | 25.73788474 | ENSG00000183770 |
| CCNF | -0.222338016 | 0.857175181 | 0.003052899 | 0.011365008 | 1204.491232 | 1032.706099 | ENSG00000162063 |
| ACOT1 | -0.222490746 | 0.857084442 | 0.553853357 | 0.710086055 | 34.96358379 | 29.92755252 | ENSG00000184227 |
| AP3D1 | -0.222572108 | 0.857036107 | 1.40E-06 | 1.07E-05 | 7335.694409 | 6286.772948 | ENSG00000065000 |
| MRGBP | -0.222723852 | 0.856945968 | 0.02897232 | 0.076311446 | 757.8206273 | 649.1728822 | ENSG00000101189 |
| DAPK2 | -0.222728642 | 0.856943123 | 0.477814082 | 0.646081043 | 48.94596379 | 41.96605924 | ENSG00000035664 |
| PKHD1 | -0.222799747 | 0.856900888 | 0.769814623 | 0.864405931 | 8.09676732 | 6.928736409 | ENSG00000170927 |

| | | | | | | | |
|---|---|---|---|---|---|---|---|
| 2-Mar | -0.222832603 | 0.856881374 | 0.163958397 | 0.302613991 | 215.2850854 | 184.4403451 | ENSG00000117791 |
| TTF2 | -0.222941454 | 0.856816724 | 0.000268659 | 0.001317076 | 1939.004194 | 1661.517205 | ENSG00000116830 |
| NA | -0.22302441 | 0.856767458 | 0.576666856 | 0.728661841 | 36.61910883 | 31.28484666 | ENSG00000288075 |
| ANKRD13D | -0.223142632 | 0.856697253 | 0.045369895 | 0.110425264 | 473.9847693 | 406.0013187 | ENSG00000172932 |
| ND4 | -0.223147551 | 0.856694332 | 0.483129227 | 0.65104741 | 25995.33064 | 22270.09227 | ENSG00000198886 |
| LARS2 | -0.223188941 | 0.856669755 | 3.50E-05 | 0.00020853 | 2298.76731 | 1969.366478 | ENSG00000011376 |
| PORCN | -0.223295897 | 0.856606247 | 0.089061741 | 0.189196128 | 309.0216551 | 264.7151946 | ENSG00000102312 |
| RSRP1 | -0.2233073 | 0.856599476 | 0.026103519 | 0.070134121 | 707.6619025 | 605.4948404 | ENSG00000117616 |
| PDHX | -0.22331104 | 0.856597256 | 0.008098626 | 0.026390304 | 859.1543628 | 735.6001388 | ENSG00000110435 |
| GNAS | -0.223363791 | 0.856565935 | 1.28E-10 | 1.74E-09 | 17725.79451 | 15183.82446 | ENSG00000087460 |
| DNAL4 | -0.223401978 | 0.856543263 | 0.250584612 | 0.414759996 | 187.2025439 | 160.150921 | ENSG00000100246 |
| PIMREG | -0.223513156 | 0.856477258 | 0.00063647 | 0.002836909 | 1996.617249 | 1709.392542 | ENSG00000129195 |
| RBM11 | -0.223572052 | 0.856442294 | 0.506742925 | 0.671592013 | 47.64669907 | 40.77923031 | ENSG00000185272 |
| COX7B | -0.223648859 | 0.8563967 | 0.012075631 | 0.03680692 | 1052.526519 | 900.7178378 | ENSG00000131174 |
| ENO1 | -0.223706127 | 0.856362705 | 1.90E-11 | 2.83E-10 | 56560.5248 | 48436.30218 | ENSG00000074800 |
| LCMT1 | -0.223750863 | 0.856336151 | 0.001805149 | 0.007218211 | 1091.014827 | 934.2424069 | ENSG00000205629 |
| PSMB6 | -0.223785065 | 0.85631585 | 3.96E-05 | 0.000232724 | 3029.835324 | 2594.415306 | ENSG00000142507 |
| FABP5 | -0.22380205 | 0.856305769 | 0.00231863 | 0.008972655 | 1079.209398 | 924.3702484 | ENSG00000164687 |
| NOP56 | -0.22394441 | 0.856221276 | 0.00012926 | 0.00067926 | 5123.600997 | 4386.211704 | ENSG00000101361 |
| CHERP | -0.223965148 | 0.856208968 | 0.002363256 | 0.009130697 | 1747.836627 | 1496.398604 | ENSG00000085872 |
| KNOP1 | -0.224018014 | 0.856177594 | 0.000771033 | 0.003359839 | 1284.938243 | 1100.002396 | ENSG00000103550 |
| BRK1 | -0.224143158 | 0.856103329 | 0.000266396 | 0.001306981 | 3977.957225 | 3405.422976 | ENSG00000254999 |
| HMGA2 | -0.224168392 | 0.856088356 | 0.000202429 | 0.001019597 | 3473.228422 | 2973.840813 | ENSG00000149948 |
| GRINA | -0.224168689 | 0.85608818 | 0.002629433 | 0.010018642 | 1059.807038 | 907.247251 | ENSG00000178719 |
| CC2D1A | -0.224192668 | 0.856073951 | 0.007852299 | 0.025722312 | 1050.789407 | 899.8953172 | ENSG00000132024 |
| GOLGA6L4 | -0.224318417 | 0.855999336 | 0.728962363 | 0.837497049 | 11.90635275 | 10.24185829 | ENSG00000184206 |
| GRPEL1 | -0.224434814 | 0.855930277 | 0.002103766 | 0.008248728 | 1086.502282 | 930.2261547 | ENSG00000109519 |
| MFHAS1 | -0.224450978 | 0.855920687 | 0.000412648 | 0.001934953 | 1675.506692 | 1433.579303 | ENSG00000147324 |
| COPS5 | -0.224518743 | 0.855880485 | 0.005875787 | 0.020186966 | 998.0922782 | 853.8895852 | ENSG00000121022 |
| NA | -0.224584256 | 0.855841619 | 0.447110762 | 0.617477482 | 58.23269918 | 49.7699966 | ENSG00000261971 |
| SP2 | -0.224589212 | 0.85583868 | 0.094598113 | 0.198185512 | 335.8251198 | 287.5698855 | ENSG00000167182 |
| NA | -0.224599786 | 0.855832407 | 0.431137748 | 0.602337652 | 64.18884816 | 54.93516721 | ENSG00000279041 |
| SCAMP2 | -0.22462764 | 0.855815883 | 0.005790086 | 0.019928019 | 984.0191002 | 841.770738 | ENSG00000140497 |
| LIX1L | -0.22472356 | 0.855758985 | 0.517450542 | 0.681374635 | 48.67036425 | 41.66141601 | ENSG00000271601 |
| PRPF18 | -0.224737366 | 0.855750796 | 0.303408099 | 0.474136735 | 116.0768354 | 99.11866752 | ENSG00000165630 |
| GRIA3 | -0.224752535 | 0.855741798 | 0.196062149 | 0.346540081 | 207.3981 | 177.4032375 | ENSG00000125675 |
| VTI1B | -0.224830563 | 0.855695517 | 0.027282763 | 0.072805615 | 489.0906481 | 418.6135755 | ENSG00000100568 |
| TLK2 | -0.224879738 | 0.855666351 | 0.002175886 | 0.008494174 | 1074.306287 | 919.2481198 | ENSG00000146872 |
| USP36 | -0.224900596 | 0.85565398 | 0.001855543 | 0.007395718 | 1359.066676 | 1162.784022 | ENSG00000055483 |
| ZNF571-AS1 | -0.224944822 | 0.85562775 | 0.814457353 | 0.893878181 | 5.380619642 | 4.594061439 | ENSG00000267470 |
| ABCC8 | -0.224951253 | 0.855623935 | 0.504751834 | 0.669911774 | 45.11684204 | 38.68223979 | ENSG00000006071 |
| LSAMP | -0.224998086 | 0.855596161 | 0.290339243 | 0.459997893 | 113.8512507 | 97.27147139 | ENSG00000185565 |
| PNPLA5 | -0.225037283 | 0.855572915 | 0.13936299 | 0.266923296 | 211.268685 | 180.6330322 | ENSG00000100341 |
| KAT14 | -0.225041602 | 0.855570354 | 0.028736515 | 0.075866446 | 562.8551469 | 481.1674519 | ENSG00000149474 |
| ZNF713 | -0.225045377 | 0.855568115 | 0.685055121 | 0.808079699 | 15.01193928 | 12.87310723 | ENSG00000178665 |
| RASSF7 | -0.225061856 | 0.855558343 | 0.039654171 | 0.098973814 | 552.4298069 | 472.3259736 | ENSG00000099849 |
| EXOSC10 | -0.225151132 | 0.855505401 | 0.003385477 | 0.012465975 | 1249.460717 | 1068.817851 | ENSG00000171824 |
| ZNF589 | -0.225154551 | 0.855503374 | 0.00016705 | 0.000856609 | 2659.028533 | 2273.926649 | ENSG00000164048 |
| NA | -0.225221706 | 0.855463553 | 0.768258701 | 0.863261801 | 8.121209724 | 6.933037557 | ENSG00000256633 |
| NUP188 | -0.225302064 | 0.855415904 | 2.22E-05 | 0.000137324 | 3243.903634 | 2775.165833 | ENSG00000095319 |
| EEF1AKMT4 | -0.225402531 | 0.855356337 | 0.580806121 | 0.731628608 | 50.40144383 | 42.94262815 | ENSG00000284753 |
| CHCHD6 | -0.225519 | 0.855287286 | 0.009702201 | 0.030650571 | 849.8040883 | 726.909963 | ENSG00000159685 |
| DNAJC12 | -0.22564336 | 0.855213564 | 0.598504143 | 0.74523821 | 25.32648219 | 21.6802323 | ENSG00000108176 |
| GPS1 | -0.225645956 | 0.855212025 | 2.31E-06 | 1.70E-05 | 3607.635715 | 3085.22309 | ENSG00000169727 |
| CD63 | -0.225707881 | 0.855175318 | 3.47E-06 | 2.48E-05 | 6943.328697 | 5937.44032 | ENSG00000135404 |
| C22orf39 | -0.225736449 | 0.855158384 | 0.070450055 | 0.156898178 | 347.0331676 | 296.9938027 | ENSG00000242259 |
| LOXL1-AS1 | -0.225845064 | 0.855094004 | 0.393879983 | 0.566676241 | 83.95177454 | 71.90759971 | ENSG00000261801 |
| VCPKMT | -0.225957723 | 0.855027233 | 0.252775763 | 0.417419727 | 132.932105 | 113.6677001 | ENSG00000100483 |
| HMGN3 | -0.226108698 | 0.854937761 | 0.014358764 | 0.042686329 | 870.4263785 | 744.3298553 | ENSG00000118418 |
| RAB11A | -0.22611441 | 0.854934376 | 4.03E-05 | 0.000236582 | 2673.364323 | 2286.38874 | ENSG00000103769 |
| TIGD5 | -0.226114438 | 0.85493436 | 0.156798839 | 0.29250618 | 218.9595434 | 187.2501406 | ENSG00000179886 |
| CXCL6 | -0.226134686 | 0.854922361 | 0.302845044 | 0.47354979 | 110.6429884 | 94.44395036 | ENSG00000124875 |
| TMEM35A | -0.226387604 | 0.854772498 | 0.265282449 | 0.431279599 | 117.8095177 | 100.7321957 | ENSG00000126950 |
| PRCP | -0.226415553 | 0.854755939 | 0.001875409 | 0.007462566 | 1532.690307 | 1310.194256 | ENSG00000137509 |
| ANP32A | -0.226428421 | 0.854748315 | 4.69E-08 | 4.52E-07 | 7724.107359 | 6602.319348 | ENSG00000140350 |
| TMEM234 | -0.226540296 | 0.854682036 | 0.418644661 | 0.590744946 | 61.89303943 | 52.88442324 | ENSG00000160055 |
| NA | -0.226546703 | 0.85467824 | 0.827163988 | 0.90146774 | 4.92555056 | 4.21306363 | ENSG00000271737 |
| MXRA7 | -0.226691198 | 0.854592643 | 0.014045844 | 0.041946004 | 901.2736246 | 769.7017657 | ENSG00000182534 |
| TRIM27 | -0.226696391 | 0.854589567 | 5.37E-05 | 0.000307255 | 2219.113156 | 1896.471871 | ENSG00000204713 |
| SLC25A26 | -0.226809593 | 0.854522513 | 0.024077523 | 0.065493442 | 567.5222068 | 485.1718182 | ENSG00000144741 |
| TBC1D14 | -0.226930444 | 0.854450935 | 0.00827991 | 0.026876617 | 858.3685519 | 733.3942504 | ENSG00000132405 |
| GTF2IP4 | -0.226950923 | 0.854438806 | 0.066228263 | 0.149323215 | 587.3084725 | 501.8430967 | ENSG00000233369 |
| SPINT2 | -0.226960331 | 0.854433234 | 0.000427401 | 0.00199782 | 2714.309651 | 2319.080212 | ENSG00000167642 |
| E2F1 | -0.226975657 | 0.854424158 | 0.008208934 | 0.026695642 | 1102.98935 | 941.964132 | ENSG00000101412 |
| C1D | -0.227274202 | 0.854247365 | 0.079879184 | 0.173179291 | 307.0101068 | 262.2632736 | ENSG00000197223 |
| ACLY | -0.227376488 | 0.854186802 | 4.80E-09 | 5.34E-08 | 14386.28613 | 12287.94508 | ENSG00000131473 |
| NUP210 | -0.227447404 | 0.854144815 | 2.67E-07 | 2.30E-06 | 4465.463267 | 3814.324626 | ENSG00000132182 |

| | | | | | | | |
|---|---|---|---|---|---|---|---|
| TRMT6 | -0.227638687 | 0.854031574 | 0.029503329 | 0.077445738 | 586.6080184 | 501.0964703 | ENSG00000089195 |
| MYO10 | -0.227900889 | 0.853876373 | 2.35E-06 | 1.73E-05 | 4200.900636 | 3586.040203 | ENSG00000145555 |
| NA | -0.228250655 | 0.853669384 | 0.842470614 | 0.909770449 | 4.231839396 | 3.647340101 | ENSG00000223571 |
| FBXW11 | -0.228317426 | 0.853629875 | 0.000307223 | 0.00148761 | 1538.366916 | 1313.376793 | ENSG00000072803 |
| MON1B | -0.22846752 | 0.853541071 | 0.002373749 | 0.009165836 | 965.0008849 | 823.5912547 | ENSG00000103111 |
| PSMD10 | -0.22848833 | 0.853528759 | 0.00099916 | 0.004259622 | 2308.71992 | 1970.076265 | ENSG00000101843 |
| ND4L | -0.228589788 | 0.853468736 | 0.322179981 | 0.493659572 | 3567.953201 | 3045.159312 | ENSG00000212907 |
| CDC7 | -0.228649224 | 0.853433576 | 0.002367009 | 0.009143368 | 1086.703115 | 927.7303919 | ENSG00000097046 |
| NA | -0.228661472 | 0.853426331 | 0.513726996 | 0.677859608 | 42.86346043 | 36.6059615 | ENSG00000196656 |
| NA | -0.228778929 | 0.853356852 | 0.46524297 | 0.634724337 | 52.51798317 | 44.7505688 | ENSG00000276291 |
| TM9SF2 | -0.228856297 | 0.85331109 | 0.000215218 | 0.001076705 | 2862.175797 | 2443.279201 | ENSG00000125304 |
| METTL2A | -0.228918748 | 0.853274153 | 0.009506823 | 0.030148959 | 770.452328 | 657.2843336 | ENSG00000087995 |
| PRPF4 | -0.228938732 | 0.853262334 | 0.000315262 | 0.001521188 | 1597.555297 | 1363.089778 | ENSG00000136875 |
| HSPD1 | -0.228941186 | 0.853260882 | 8.14E-11 | 1.13E-09 | 28720.67688 | 24506.23428 | ENSG00000144381 |
| ATP9B | -0.229100535 | 0.853166643 | 0.051267839 | 0.121784003 | 419.2129289 | 358.0367906 | ENSG00000166377 |
| SH3GL1 | -0.229232437 | 0.853088643 | 5.50E-05 | 0.000313871 | 3765.006833 | 3212.134966 | ENSG00000141985 |
| NA | -0.229358397 | 0.853014164 | 0.701919502 | 0.820302242 | 12.69571835 | 10.82640394 | ENSG00000273284 |
| SOCS3 | -0.229467928 | 0.852949405 | 0.027988604 | 0.074288411 | 546.7924483 | 466.2543535 | ENSG00000184557 |
| TREML2 | -0.229705126 | 0.852809181 | 0.771111586 | 0.865157241 | 9.514013698 | 8.127935749 | ENSG00000112195 |
| PODXL | -0.229823659 | 0.852739116 | 2.40E-09 | 2.78E-08 | 25017.97765 | 21333.95613 | ENSG00000128567 |
| TGFBR1 | -0.229843184 | 0.852727575 | 0.007161148 | 0.023818225 | 861.0953138 | 734.565728 | ENSG00000106799 |
| HMGCS1 | -0.229845799 | 0.85272603 | 8.45E-07 | 6.73E-06 | 5907.830546 | 5037.997261 | ENSG00000112972 |
| FNTB | -0.229939323 | 0.852670753 | 0.403607583 | 0.576014935 | 66.52009965 | 56.70061101 | ENSG00000257365 |
| LZTS3 | -0.230060626 | 0.852599063 | 0.001088953 | 0.00460175 | 1293.281625 | 1102.456759 | ENSG00000088899 |
| NA | -0.230194317 | 0.852520058 | 0.835865458 | 0.906924912 | 4.26465451 | 3.672613948 | ENSG00000279058 |
| TBC1D1 | -0.230435742 | 0.852377407 | 0.000432737 | 0.002019825 | 2250.608858 | 1918.663904 | ENSG00000065882 |
| TOP1 | -0.230499974 | 0.852339457 | 1.28E-06 | 9.92E-06 | 3923.602793 | 3344.515771 | ENSG00000198900 |
| MRPS23 | -0.230571623 | 0.852297129 | 0.000445284 | 0.002071876 | 1538.013608 | 1310.815967 | ENSG00000181610 |
| SRSF7 | -0.230723887 | 0.852207181 | 1.93E-06 | 1.45E-05 | 7110.4897 | 6059.437488 | ENSG00000115875 |
| EXTL1 | -0.230751409 | 0.852190924 | 0.596704652 | 0.74400866 | 25.49825474 | 21.71787655 | ENSG00000158008 |
| PSMD14 | -0.230933535 | 0.85208335 | 5.10E-05 | 0.000292557 | 2434.73652 | 2075.204391 | ENSG00000115233 |
| LRRC37A | -0.231035477 | 0.852023143 | 0.663692409 | 0.792577086 | 18.15313183 | 15.44358865 | ENSG00000176681 |
| PSPH | -0.231107473 | 0.851980625 | 0.002293905 | 0.008885886 | 1102.129973 | 938.6194719 | ENSG00000146733 |
| C5orf51 | -0.231155974 | 0.851951983 | 0.03988668 | 0.099386972 | 469.3729398 | 400.2407948 | ENSG00000205765 |
| NKRF | -0.231159673 | 0.851949799 | 0.041691831 | 0.103072807 | 390.0361859 | 332.2731111 | ENSG00000186416 |
| NDUFB8 | -0.231163664 | 0.851947442 | 0.595589835 | 0.743369792 | 24.68656192 | 21.05535878 | ENSG00000166136 |
| DDX19A | -0.231197684 | 0.851927353 | 0.056929079 | 0.132427212 | 375.0021196 | 319.3478386 | ENSG00000168872 |
| EDC4 | -0.23121516 | 0.851917033 | 0.002472671 | 0.00950963 | 1707.179364 | 1454.047282 | ENSG00000038358 |
| SORBS3 | -0.231235117 | 0.851905248 | 0.004684192 | 0.016592499 | 980.682807 | 835.9650336 | ENSG00000120896 |
| ACTR8 | -0.231298258 | 0.851867965 | 0.013908181 | 0.041597321 | 678.2544799 | 577.6371023 | ENSG00000113812 |
| SET | -0.231361418 | 0.851830671 | 4.04E-08 | 3.94E-07 | 23183.11088 | 19749.12943 | ENSG00000119335 |
| LOC100287042 | -0.231382733 | 0.851818086 | 0.569339417 | 0.722431539 | 28.9437196 | 24.58708252 | ENSG00000263843 |
| RPUSD4 | -0.231401157 | 0.851807208 | 0.003221518 | 0.011921408 | 1128.53455 | 961.340282 | ENSG00000165526 |
| RNASEH1 | -0.231408782 | 0.851802706 | 0.015594088 | 0.045704277 | 609.7061859 | 519.4525432 | ENSG00000171865 |
| BMP4 | -0.231427478 | 0.851791667 | 0.326095173 | 0.497879064 | 89.81960266 | 76.64953611 | ENSG00000125378 |
| NA | -0.231441576 | 0.851783344 | 0.179818139 | 0.324496888 | 184.0212027 | 156.4691654 | ENSG00000254682 |
| TMEM33 | -0.231464968 | 0.851769533 | 0.008212539 | 0.026702864 | 1109.456451 | 945.4589255 | ENSG00000109133 |
| NA | -0.231582652 | 0.851700055 | 0.803089309 | 0.886394088 | 5.40785295 | 4.585459144 | ENSG00000218418 |
| NA | -0.231637523 | 0.851667662 | 0.765385767 | 0.861069997 | 7.767376885 | 6.606888588 | ENSG00000224765 |
| ERAL1 | -0.231769064 | 0.851590013 | 0.00115814 | 0.004868527 | 1331.239295 | 1133.535343 | ENSG00000132591 |
| DOCK10 | -0.231831848 | 0.851552954 | 0.436777198 | 0.607576358 | 54.97140978 | 46.79969524 | ENSG00000135905 |
| RAD17 | -0.23183492 | 0.85155114 | 0.050942281 | 0.121129982 | 469.3466576 | 399.9807927 | ENSG00000152942 |
| CCDC136 | -0.231861356 | 0.851535537 | 0.01431743 | 0.042596783 | 658.5667447 | 560.4092356 | ENSG00000128596 |
| ZC3H15 | -0.231874209 | 0.851527951 | 2.69E-05 | 0.00016372 | 2272.108799 | 1934.947316 | ENSG00000065548 |
| ARMCX4 | -0.23190542 | 0.851509529 | 0.047466185 | 0.11461637 | 404.4153384 | 344.0758419 | ENSG00000196440 |
| NA | -0.232075196 | 0.851409329 | 0.750167596 | 0.852166005 | 9.274695895 | 7.934074697 | ENSG00000228519 |
| LSR | -0.232105359 | 0.851391529 | 0.000130077 | 0.000682251 | 3817.313308 | 3250.356927 | ENSG00000105699 |
| NA | -0.232159785 | 0.851359411 | 0.808627625 | 0.890062508 | 4.988389884 | 4.244789198 | ENSG00000280486 |
| HTT | -0.232230971 | 0.851317403 | 1.10E-05 | 7.17E-05 | 3200.296074 | 2724.287396 | ENSG00000197386 |
| TAF8 | -0.232359768 | 0.851241406 | 0.031199329 | 0.08110312 | 469.2207404 | 399.1150486 | ENSG00000137413 |
| SRSF9 | -0.232423819 | 0.851203614 | 0.00017499 | 0.000893257 | 3869.51343 | 3294.325757 | ENSG00000111786 |
| TMEM67 | -0.232463431 | 0.851180243 | 0.198971193 | 0.350403968 | 171.6628471 | 146.135868 | ENSG00000164953 |
| TOMM5 | -0.232612578 | 0.851092251 | 0.60896946 | 0.752715438 | 23.86246618 | 20.23312868 | ENSG00000175768 |
| GLI2 | -0.23273492 | 0.851020081 | 0.011655934 | 0.035810577 | 1146.333525 | 976.0132785 | ENSG00000074047 |
| RAB15 | -0.232980583 | 0.850875181 | 0.000466292 | 0.002155066 | 2607.220358 | 2218.593819 | ENSG00000139998 |
| NA | -0.233006805 | 0.850859716 | 0.483147309 | 0.65104741 | 44.55092471 | 37.93368065 | ENSG00000213713 |
| TPCN2 | -0.233051895 | 0.850833124 | 0.100834016 | 0.208398221 | 290.0943884 | 246.5368506 | ENSG00000162341 |
| MAP3K4 | -0.233057691 | 0.850829706 | 0.000469392 | 0.002166279 | 1357.661982 | 1154.964194 | ENSG00000085511 |
| DKK3 | -0.233148681 | 0.850776046 | 0.015231052 | 0.044830976 | 567.5562612 | 482.8600182 | ENSG00000050165 |
| GK5 | -0.233218732 | 0.850734737 | 0.193631713 | 0.343354066 | 162.0840791 | 137.9001126 | ENSG00000175066 |
| ALDH3A2 | -0.233329221 | 0.850669586 | 0.000100684 | 0.000544686 | 2010.323328 | 1710.490239 | ENSG00000072210 |
| LOC645513 | -0.233560677 | 0.850533121 | 0.236409235 | 0.397816692 | 134.1877033 | 114.2592184 | ENSG00000245958 |
| RNPS1 | -0.233587582 | 0.850517259 | 3.71E-09 | 4.19E-08 | 7371.664609 | 6269.733653 | ENSG00000205937 |
| MGC12916 | -0.233749137 | 0.850422023 | 0.275819804 | 0.442776054 | 129.2953313 | 110.2521136 | ENSG00000266709 |
| XPOT | -0.233755374 | 0.850418346 | 1.45E-07 | 1.30E-06 | 5475.664872 | 4656.72309 | ENSG00000184575 |
| CNRIP1 | -0.23382802 | 0.850375525 | 0.430460894 | 0.601827882 | 58.68083501 | 49.82216183 | ENSG00000119865 |
| PAOX | -0.233829992 | 0.850374363 | 0.760003629 | 0.857538689 | 10.47148507 | 8.977590274 | ENSG00000148832 |

| | | | | | | | |
|---|---|---|---|---|---|---|---|
| ARHGAP17 | -0.233840972 | 0.850367891 | 0.000510892 | 0.002337116 | 1377.667981 | 1171.355589 | ENSG00000140750 |
| CARM1 | -0.233922531 | 0.850319819 | 2.47E-06 | 1.81E-05 | 5050.912766 | 4295.17936 | ENSG00000142453 |
| MARCHF9 | -0.233949272 | 0.850304058 | 0.310170876 | 0.481279598 | 102.0358095 | 86.96506537 | ENSG00000139266 |
| TRABD | -0.234110498 | 0.850209039 | 0.000181267 | 0.000921673 | 1885.945877 | 1603.776629 | ENSG00000170638 |
| NA | -0.234177145 | 0.850169763 | 0.825931437 | 0.901037344 | 5.361083322 | 4.539212598 | ENSG00000257195 |
| TMEM230 | -0.23430605 | 0.850093804 | 0.001434283 | 0.005891984 | 1303.126005 | 1107.690153 | ENSG00000089063 |
| TRMT44 | -0.234317784 | 0.85008689 | 0.067238635 | 0.151106768 | 324.7275057 | 275.8690862 | ENSG00000155275 |
| SMG1P7 | -0.234362032 | 0.850060818 | 0.325262987 | 0.49696248 | 85.33026058 | 72.59807493 | ENSG00000261556 |
| LRRC29 | -0.234365255 | 0.850058919 | 0.775827197 | 0.868533369 | 7.277377508 | 6.188246528 | ENSG00000125122 |
| METRN | -0.234423356 | 0.850024685 | 0.053519235 | 0.125953343 | 396.3460798 | 336.6324748 | ENSG00000103260 |
| ERICH2 | -0.234503847 | 0.849977262 | 0.681974043 | 0.805942337 | 14.59526712 | 12.4103692 | ENSG00000204334 |
| NA | -0.234634041 | 0.849900561 | 0.81504878 | 0.894245129 | 5.002344398 | 4.246939771 | ENSG00000226054 |
| C9orf163 | -0.234673507 | 0.849877311 | 0.784500198 | 0.873826048 | 7.410753143 | 6.303862893 | ENSG00000196366 |
| OAZ1 | -0.234680312 | 0.849873302 | 1.89E-08 | 1.93E-07 | 5227.930891 | 4442.993854 | ENSG00000104904 |
| UBE2L3 | -0.234695519 | 0.849864344 | 5.97E-07 | 4.86E-06 | 3818.049593 | 3244.880571 | ENSG00000185651 |
| SSH3 | -0.23480591 | 0.849799317 | 0.024401601 | 0.06623488 | 542.9638021 | 461.3024176 | ENSG00000172830 |
| GRIN3B | -0.234833658 | 0.849782973 | 0.529181253 | 0.690412252 | 47.16698742 | 40.15757344 | ENSG00000116032 |
| XPO5 | -0.234876828 | 0.849757546 | 4.28E-06 | 3.01E-05 | 3229.707802 | 2744.343863 | ENSG00000124571 |
| TCEAL9 | -0.234952534 | 0.849712955 | 0.000422413 | 0.001975938 | 1386.067181 | 1178.104409 | ENSG00000185222 |
| ZNF263 | -0.23498126 | 0.849696036 | 0.005119375 | 0.017890916 | 920.9092597 | 782.1713339 | ENSG00000006194 |
| SERBP1 | -0.234995975 | 0.84968737 | 4.88E-09 | 5.43E-08 | 22421.61709 | 19051.64816 | ENSG00000142864 |
| METTL2B | -0.235022356 | 0.849671833 | 0.018274917 | 0.052054399 | 523.0838452 | 444.6913246 | ENSG00000165055 |
| SCARB1 | -0.235175772 | 0.849581483 | 0.000313049 | 0.001513163 | 1391.076383 | 1181.590977 | ENSG00000073060 |
| ST3GAL4 | -0.235209865 | 0.849561407 | 0.062027815 | 0.141690774 | 418.0917695 | 355.2533898 | ENSG00000110080 |
| DCAF12 | -0.235250414 | 0.849537529 | 0.004673555 | 0.016560903 | 861.4395346 | 732.2251591 | ENSG00000198876 |
| C1S | -0.235349333 | 0.849479282 | 0.149448911 | 0.281627327 | 193.8352583 | 164.593321 | ENSG00000182326 |
| NDUFA4 | -0.23540093 | 0.849448902 | 4.21E-06 | 2.97E-05 | 3316.187983 | 2817.070524 | ENSG00000189043 |
| NA | -0.235410041 | 0.849443537 | 0.124364184 | 0.24481618 | 240.9644124 | 204.5525023 | ENSG00000183458 |
| TBC1D24 | -0.235475649 | 0.849404909 | 0.067150894 | 0.150958383 | 317.938488 | 270.0266227 | ENSG00000162065 |
| ADCY6 | -0.235526332 | 0.849375069 | 0.027929835 | 0.074163035 | 461.1754487 | 391.7308072 | ENSG00000174233 |
| GYG2 | -0.235558212 | 0.8493563 | 4.38E-06 | 3.08E-05 | 3274.409999 | 2780.707658 | ENSG00000056998 |
| CD276 | -0.235562359 | 0.849353858 | 2.32E-06 | 1.71E-05 | 4411.135524 | 3746.947716 | ENSG00000103855 |
| PREB | -0.235623595 | 0.849317808 | 0.000216768 | 0.001083334 | 1609.441349 | 1366.767256 | ENSG00000138073 |
| STK16 | -0.235705149 | 0.849269798 | 0.06196767 | 0.141570147 | 370.7371769 | 315.0549007 | ENSG00000115661 |
| PCDHGC4 | -0.235787341 | 0.849221416 | 0.375320861 | 0.548674957 | 72.74156044 | 61.71037043 | ENSG00000242419 |
| SLC25A1 | -0.235883419 | 0.849164863 | 1.13E-05 | 7.32E-05 | 5185.388642 | 4403.798718 | ENSG00000100075 |
| KNSTRN | -0.235950727 | 0.849125246 | 0.000977684 | 0.004179148 | 1183.287981 | 1004.81257 | ENSG00000128944 |
| ZNF821 | -0.236025635 | 0.849081159 | 0.370808781 | 0.544593964 | 76.66518535 | 64.96702593 | ENSG00000102984 |
| SCP2 | -0.23605323 | 0.849064919 | 0.009601908 | 0.030395539 | 685.4429361 | 581.8235471 | ENSG00000116171 |
| TMEM203 | -0.23611033 | 0.849031314 | 0.001604625 | 0.006500381 | 1000.953003 | 849.8714671 | ENSG00000187713 |
| TM2D2 | -0.236182408 | 0.848988897 | 0.001547891 | 0.006292137 | 1179.228614 | 1001.481414 | ENSG00000169490 |
| HAP1 | -0.236265373 | 0.848940076 | 0.179701761 | 0.324405749 | 167.8489192 | 142.5600667 | ENSG00000173805 |
| ZNF280B | -0.236315107 | 0.848910811 | 0.120912076 | 0.239937179 | 229.5848163 | 195.0696529 | ENSG00000275004 |
| NA | -0.236358014 | 0.848885564 | 0.685160062 | 0.808079699 | 15.49424167 | 13.11913351 | ENSG00000226239 |
| CARS1 | -0.236365339 | 0.848881254 | 0.000315062 | 0.001520608 | 1667.527055 | 1415.186944 | ENSG00000110619 |
| NA | -0.236467453 | 0.848821172 | 0.278875467 | 0.446699032 | 102.4651969 | 86.99599743 | ENSG00000196295 |
| ECHDC1 | -0.236593037 | 0.848747287 | 0.002478613 | 0.009528682 | 1024.821015 | 870.022874 | ENSG00000093144 |
| WRN | -0.236641921 | 0.848718529 | 0.00691068 | 0.023152818 | 758.2767965 | 643.7820396 | ENSG00000165392 |
| R3HCC1 | -0.236682731 | 0.848694521 | 0.02295238 | 0.062956686 | 535.3558701 | 454.4831063 | ENSG00000104679 |
| RPL36AL | -0.236704591 | 0.848681662 | 2.81E-05 | 0.000170753 | 3718.087218 | 3154.576159 | ENSG00000165502 |
| ZC3H18 | -0.236721402 | 0.848671773 | 2.88E-06 | 2.08E-05 | 3710.274114 | 3149.293124 | ENSG00000158545 |
| NATD1 | -0.236762407 | 0.848647651 | 0.043967264 | 0.107659944 | 385.3639077 | 326.7379987 | ENSG00000274180 |
| RNFT2 | -0.236787216 | 0.848633058 | 0.120034151 | 0.238571895 | 231.15729 | 196.383663 | ENSG00000135119 |
| SMNDC1 | -0.236838658 | 0.848602799 | 0.01640243 | 0.047631985 | 680.2501459 | 577.403961 | ENSG00000119953 |
| LSM2 | -0.236838989 | 0.848602604 | 0.001858764 | 0.007407025 | 1188.443425 | 1008.431741 | ENSG00000204392 |
| CIZ1 | -0.236893622 | 0.848570469 | 0.000398177 | 0.00187851 | 2407.392821 | 2042.737144 | ENSG00000148337 |
| C3orf70 | -0.236981597 | 0.848518725 | 0.395900501 | 0.568468083 | 62.94528936 | 53.43132464 | ENSG00000187068 |
| SLC35A1 | -0.23707722 | 0.848462487 | 0.552358224 | 0.708958311 | 34.07174265 | 28.80874844 | ENSG00000164414 |
| NDUFS3 | -0.237096001 | 0.848451442 | 0.000887181 | 0.003822715 | 1664.428455 | 1412.2645 | ENSG00000213619 |
| NA | -0.237143481 | 0.848423519 | 0.485923934 | 0.653385722 | 44.25694013 | 37.51477813 | ENSG00000230555 |
| RNF20 | -0.237165459 | 0.848410594 | 0.000170969 | 0.000874442 | 1555.297689 | 1319.621215 | ENSG00000155827 |
| ALG1L2 | -0.237386017 | 0.8482809 | 0.81784147 | 0.895882273 | 4.71904787 | 4.017585042 | ENSG00000251287 |
| GLT8D2 | -0.237441233 | 0.848248434 | 0.350829314 | 0.523517225 | 74.3906278 | 63.04669187 | ENSG00000120820 |
| MOCOS | -0.237444436 | 0.848246551 | 0.012072038 | 0.036801783 | 634.9608173 | 538.1622074 | ENSG00000075643 |
| IPO11 | -0.237465989 | 0.848233879 | 0.003141578 | 0.011659084 | 904.654876 | 767.8032042 | ENSG00000086200 |
| AKR7A2 | -0.237488438 | 0.84822068 | 0.00861632 | 0.027795555 | 724.4813802 | 614.9243916 | ENSG00000053371 |
| COQ7 | -0.23760214 | 0.848153833 | 0.105985448 | 0.21640024 | 226.7397235 | 192.4631447 | ENSG00000167186 |
| PLS1 | -0.237614586 | 0.848146516 | 0.041255433 | 0.102177242 | 559.4933481 | 474.7301555 | ENSG00000120756 |
| ZNF780B | -0.237724928 | 0.848081649 | 0.252984018 | 0.417622232 | 130.8560778 | 111.1386005 | ENSG00000128000 |
| NA | -0.237795101 | 0.848040399 | 0.821076459 | 0.898128032 | 4.585672235 | 3.89766753 | ENSG00000242797 |
| HMGN4 | -0.237832904 | 0.848018178 | 0.002791718 | 0.010549469 | 999.7560141 | 848.2041015 | ENSG00000182952 |
| ASCC2 | -0.237850023 | 0.848008116 | 0.005005833 | 0.017557731 | 946.1369842 | 802.164888 | ENSG00000100325 |
| NA | -0.237858273 | 0.848003267 | 0.678409946 | 0.803241041 | 14.32871601 | 12.16649349 | ENSG00000258199 |
| ZNF557 | -0.237860897 | 0.848001724 | 0.302143612 | 0.472955816 | 97.63327361 | 82.78776799 | ENSG00000130544 |
| SMIM15 | -0.23797198 | 0.847936433 | 0.051566262 | 0.12234224 | 413.2924612 | 350.60474 | ENSG00000188725 |
| NA | -0.237998107 | 0.847921077 | 0.40098907 | 0.573508315 | 62.52949308 | 52.92018949 | ENSG00000280206 |
| NA | -0.237998576 | 0.847920802 | 0.826122272 | 0.901078467 | 4.931132366 | 4.171118231 | ENSG00000259146 |

| | | | | | | | |
|---|---|---|---|---|---|---|---|
| GRTP1 | -0.238064216 | 0.847882224 | 0.000671979 | 0.002973883 | 1140.611036 | 967.0794811 | ENSG00000139835 |
| PFKFB4 | -0.23812278 | 0.847847806 | 0.063637022 | 0.144562151 | 368.9471619 | 312.4032547 | ENSG00000114268 |
| MAGI1 | -0.238190725 | 0.847807877 | 0.000111574 | 0.000596555 | 2172.319801 | 1842.029011 | ENSG00000151276 |
| ARHGEF17 | -0.238241587 | 0.847777988 | 0.051335992 | 0.12188586 | 346.547911 | 293.880721 | ENSG00000110237 |
| BMP2K | -0.238294053 | 0.847747158 | 0.127636972 | 0.249491771 | 205.9502289 | 174.6547731 | ENSG00000138756 |
| NA | -0.23836423 | 0.847705921 | 0.739087439 | 0.84500301 | 9.285859507 | 7.894279872 | ENSG00000219507 |
| NA | -0.238386473 | 0.847692852 | 0.495176487 | 0.661481172 | 43.400705 | 36.80842485 | ENSG00000276791 |
| C10orf82 | -0.238521763 | 0.847613363 | 0.239860833 | 0.40194927 | 151.1676125 | 128.3215656 | ENSG00000165863 |
| BTRC | -0.238575768 | 0.847581634 | 0.00366281 | 0.01336472 | 792.4939174 | 671.8003141 | ENSG00000166167 |
| UBR1 | -0.238613535 | 0.847559447 | 0.017729589 | 0.050850654 | 517.747041 | 438.843215 | ENSG00000159459 |
| PTGS1 | -0.238768033 | 0.847468686 | 0.443596089 | 0.614403515 | 52.60438902 | 44.6438152 | ENSG00000095303 |
| NA | -0.23891153 | 0.847384397 | 0.708524207 | 0.824320104 | 12.0592647 | 10.22680427 | ENSG00000283078 |
| CTNNBL1 | -0.238944602 | 0.847364972 | 0.000380048 | 0.001802681 | 1530.57701 | 1296.815975 | ENSG00000132792 |
| PBRM1 | -0.238977332 | 0.847345749 | 0.000606234 | 0.00271847 | 1227.407115 | 1040.454594 | ENSG00000163939 |
| GCHFR | -0.239004766 | 0.847329636 | 0.785944458 | 0.874929571 | 7.751307189 | 6.549889174 | ENSG00000137880 |
| NA | -0.239040301 | 0.847308766 | 0.77138654 | 0.86531475 | 7.026896093 | 5.971795241 | ENSG00000267191 |
| TM2D1 | -0.239095865 | 0.847276133 | 0.196234129 | 0.346694073 | 156.8523923 | 133.0275058 | ENSG00000162604 |
| DAAM1 | -0.23918232 | 0.84722536 | 0.041851989 | 0.103389258 | 452.3939065 | 383.5625315 | ENSG00000100592 |
| PRDX2 | -0.23929705 | 0.847157988 | 3.38E-05 | 0.000201786 | 2428.225577 | 2056.860997 | ENSG00000167815 |
| PSMA1 | -0.239300771 | 0.847155802 | 0.253782378 | 0.418437253 | 122.8457163 | 104.0652363 | ENSG00000129084 |
| ARL10 | -0.239394529 | 0.847100749 | 0.077109103 | 0.168422974 | 281.9796364 | 238.7038408 | ENSG00000175414 |
| UTP18 | -0.239450413 | 0.847067937 | 2.65E-05 | 0.000161818 | 2060.782205 | 1745.983042 | ENSG00000011260 |
| CYP3A5 | -0.239460436 | 0.847062052 | 0.745687921 | 0.849177432 | 8.953678169 | 7.625130318 | ENSG00000106258 |
| JPH3 | -0.23946978 | 0.847056566 | 0.00125976 | 0.005254493 | 1323.740099 | 1121.697428 | ENSG00000154118 |
| UBBP4 | -0.239495834 | 0.847041269 | 0.154886653 | 0.289892153 | 173.5320963 | 146.9193878 | ENSG00000263563 |
| NXPE3 | -0.239549575 | 0.847009717 | 0.012769448 | 0.038689403 | 614.157168 | 519.7768263 | ENSG00000144815 |
| PSMG3 | -0.239570831 | 0.846997237 | 0.004712205 | 0.016676416 | 773.3861931 | 654.9381428 | ENSG00000157778 |
| PUF60 | -0.239665589 | 0.846941607 | 2.08E-06 | 1.55E-05 | 4079.874688 | 3455.161047 | ENSG00000179950 |
| SYNRG | -0.23992478 | 0.846789461 | 0.003204078 | 0.011861422 | 835.6117262 | 707.5012208 | ENSG00000275066 |
| PABPC1L | -0.240112288 | 0.846679411 | 0.0172162 | 0.049629066 | 889.2352235 | 752.1169033 | ENSG00000101104 |
| TUBB | -0.240367725 | 0.846529515 | 1.38E-13 | 2.56E-12 | 33110.26979 | 28028.39165 | ENSG00000196230 |
| ZKSCAN7 | -0.240404951 | 0.846507672 | 0.181294207 | 0.326458186 | 166.8333513 | 141.126127 | ENSG00000196345 |
| NA | -0.240419721 | 0.846499006 | 0.637384621 | 0.77331992 | 18.25560737 | 15.5108079 | ENSG00000269119 |
| ATL1 | -0.240594519 | 0.84639645 | 0.26707726 | 0.433388957 | 102.157458 | 86.55072445 | ENSG00000198513 |
| LOC728752 | -0.240640651 | 0.846369386 | 0.627481645 | 0.765687167 | 19.8971779 | 16.81702131 | ENSG00000267309 |
| LRRC8C | -0.240712212 | 0.846327405 | 0.342058128 | 0.514835246 | 81.72283534 | 69.26881454 | ENSG00000171488 |
| MRGPRF | -0.240713351 | 0.846326737 | 0.289044244 | 0.458473319 | 113.6743719 | 96.27984847 | ENSG00000172935 |
| LOC100505715 | -0.240762919 | 0.846297659 | 0.726064024 | 0.835411015 | 10.8590089 | 9.12897289 | ENSG00000267058 |
| POMT1 | -0.240798074 | 0.846277037 | 0.013487998 | 0.040491279 | 576.427876 | 487.5266966 | ENSG00000130714 |
| SDHAP1 | -0.240911378 | 0.846210576 | 0.127460029 | 0.249298531 | 230.6846365 | 194.9019018 | ENSG00000185485 |
| HMGB1 | -0.241174965 | 0.846055984 | 4.51E-06 | 3.16E-05 | 15713.69774 | 13294.93878 | ENSG00000189403 |
| GABPB1-IT1 | -0.241195181 | 0.846044128 | 0.142678341 | 0.271870922 | 181.8021509 | 153.8290416 | ENSG00000285410 |
| RAP2A | -0.241334547 | 0.845962404 | 0.007733005 | 0.025400544 | 887.3167162 | 751.1009447 | ENSG00000125249 |
| TMEM260 | -0.241439439 | 0.845900899 | 0.077869006 | 0.169660153 | 277.0149763 | 234.4880935 | ENSG00000070269 |
| PHKA1 | -0.241527154 | 0.84584947 | 0.060653286 | 0.139193735 | 379.2573623 | 320.7691233 | ENSG00000067177 |
| DCTD | -0.241604949 | 0.845803861 | 0.000175055 | 0.000893257 | 1531.280721 | 1294.89269 | ENSG00000129187 |
| PRKRA | -0.241626462 | 0.845791249 | 0.009762042 | 0.030806403 | 840.9140132 | 711.3332624 | ENSG00000180228 |
| LZTS2 | -0.241683987 | 0.845757525 | 8.90E-06 | 5.92E-05 | 3282.63096 | 2776.672359 | ENSG00000107816 |
| HSPA8 | -0.241706127 | 0.845744546 | 1.94E-13 | 3.56E-12 | 46536.79572 | 39358.88048 | ENSG00000109971 |
| MICA | -0.241762188 | 0.845711682 | 0.057554769 | 0.133512393 | 344.7302624 | 291.4772277 | ENSG00000204520 |
| POLR2G | -0.241781746 | 0.845700217 | 0.000640795 | 0.00285421 | 1301.152026 | 1099.87255 | ENSG00000168002 |
| HDGFL2 | -0.24182352 | 0.84567573 | 3.16E-06 | 2.27E-05 | 4431.719943 | 3748.263913 | ENSG00000167674 |
| SLC27A6 | -0.241877785 | 0.845643921 | 0.25771287 | 0.422821821 | 116.7627374 | 98.64490405 | ENSG00000113396 |
| NA | -0.242005372 | 0.845569139 | 0.559078809 | 0.714392879 | 31.34008885 | 26.47999216 | ENSG00000273619 |
| BAG2 | -0.242228011 | 0.845438659 | 0.025049209 | 0.067674781 | 460.1014351 | 388.9887761 | ENSG00000112208 |
| PCDHB16 | -0.242252769 | 0.845424151 | 0.304530701 | 0.47526984 | 97.36285556 | 82.43849575 | ENSG00000272674 |
| SMARCA4 | -0.242357987 | 0.845362495 | 1.82E-08 | 1.87E-07 | 10684.92069 | 9032.581464 | ENSG00000127616 |
| RNF144A | -0.242362425 | 0.845359894 | 0.01933039 | 0.054635819 | 485.0194177 | 410.1866427 | ENSG00000151692 |
| MBNL3 | -0.242721155 | 0.84514972 | 0.319261805 | 0.490669175 | 91.40391567 | 77.19349344 | ENSG00000076770 |
| NSMAF | -0.242730846 | 0.845144042 | 0.00863933 | 0.027860469 | 611.3945261 | 516.706744 | ENSG00000035681 |
| TRMT11 | -0.242798425 | 0.845104455 | 0.039570106 | 0.098853524 | 411.581155 | 348.1070934 | ENSG00000066651 |
| RSPH9 | -0.242806149 | 0.84509993 | 0.679974248 | 0.804451319 | 16.70239462 | 14.01530716 | ENSG00000172426 |
| HUS1 | -0.242963037 | 0.845008034 | 0.105058446 | 0.215032624 | 282.028233 | 238.4217757 | ENSG00000136273 |
| BAZ1B | -0.243011836 | 0.844979452 | 1.66E-07 | 1.48E-06 | 3615.833443 | 3055.460853 | ENSG00000009954 |
| RASL10B | -0.243156074 | 0.844894977 | 0.001544946 | 0.006284483 | 1049.733816 | 886.9668097 | ENSG00000270885 |
| MAPRE2 | -0.243199816 | 0.84486936 | 0.000337476 | 0.001618256 | 1267.723246 | 1070.81669 | ENSG00000166974 |
| NA | -0.243214327 | 0.844860863 | 0.570434313 | 0.723392606 | 27.31072194 | 23.02624053 | ENSG00000260290 |
| CREB3 | -0.243229972 | 0.844851701 | 0.010320647 | 0.032284027 | 601.0608103 | 508.1257265 | ENSG00000107175 |
| ELOC | -0.243238891 | 0.844846477 | 0.000564165 | 0.002551174 | 1475.67719 | 1246.603456 | ENSG00000154582 |
| PLEKHJ1 | -0.243296747 | 0.844812597 | 2.67E-05 | 0.000162654 | 2094.0588 | 1768.815247 | ENSG00000104886 |
| EIF2S2 | -0.2433249 | 0.844796112 | 2.67E-06 | 1.94E-05 | 5397.32205 | 4560.821968 | ENSG00000125977 |
| SIMC1 | -0.243332629 | 0.844791586 | 0.045345688 | 0.110380274 | 381.3381707 | 321.9659358 | ENSG00000170085 |
| SELENOP | -0.243341881 | 0.844786169 | 0.520422688 | 0.68346886 | 38.1064912 | 32.29825421 | ENSG00000250722 |
| ADCY3 | -0.243369895 | 0.844769765 | 0.004554664 | 0.016190165 | 815.8700141 | 689.2683189 | ENSG00000138031 |
| RAB40C | -0.243374019 | 0.84476735 | 0.004044707 | 0.014606222 | 787.3418523 | 664.7788362 | ENSG00000197562 |
| CTBP2 | -0.243391653 | 0.844757025 | 2.02E-07 | 1.78E-06 | 3766.143445 | 3181.302861 | ENSG00000175029 |
| ZNF497 | -0.243417584 | 0.844741841 | 0.603315939 | 0.748749857 | 24.03682948 | 20.28797752 | ENSG00000174586 |

| | | | | | | | |
|---|---|---|---|---|---|---|---|
| GTF2H1 | -0.243427065 | 0.844736289 | 0.05220304 | 0.123352382 | 445.2044126 | 376.4074208 | ENSG00000110768 |
| ERVK3-1 | -0.243469012 | 0.844711728 | 0.632752013 | 0.769635504 | 18.68344315 | 15.80308072 | ENSG00000142396 |
| SBK1 | -0.243610791 | 0.84462872 | 0.000109815 | 0.000588295 | 4612.59565 | 3896.521301 | ENSG00000188322 |
| PTPN2 | -0.24377997 | 0.84452968 | 0.008409486 | 0.0272285 | 752.1435302 | 634.9838079 | ENSG00000175354 |
| BACE2 | -0.243987409 | 0.844408257 | 0.077642211 | 0.169242473 | 333.7773139 | 282.191456 | ENSG00000182240 |
| ARMCX1 | -0.244006105 | 0.844397314 | 0.019489793 | 0.055017989 | 681.8764602 | 575.4766791 | ENSG00000126947 |
| ARR3 | -0.244025575 | 0.844385919 | 0.809134472 | 0.890310719 | 4.672953964 | 3.950365797 | ENSG00000120500 |
| EDA | -0.24412646 | 0.844326874 | 0.109656839 | 0.222076015 | 225.7326916 | 190.5541271 | ENSG00000158813 |
| SDC1 | -0.244197244 | 0.844285449 | 9.18E-05 | 0.000501087 | 2086.113971 | 1761.673778 | ENSG00000115884 |
| SETD4 | -0.244287762 | 0.844232479 | 0.019603719 | 0.055278132 | 593.0226647 | 500.3419925 | ENSG00000185917 |
| ENTR1 | -0.244319824 | 0.844213717 | 0.000195451 | 0.000987546 | 1996.750435 | 1685.237265 | ENSG00000165689 |
| BICD2 | -0.244336789 | 0.84420379 | 0.004867051 | 0.017145777 | 1029.815374 | 869.486223 | ENSG00000185963 |
| TERF1 | -0.244349338 | 0.844196447 | 4.14E-09 | 4.66E-08 | 11008.47857 | 9294.249885 | ENSG00000147601 |
| IMP3 | -0.244357562 | 0.844191634 | 0.047327198 | 0.114331537 | 349.8981218 | 295.0691796 | ENSG00000177971 |
| TBRG1 | -0.244399548 | 0.844167067 | 0.038690841 | 0.097059169 | 396.9104824 | 334.8062393 | ENSG00000154144 |
| SYBU | -0.244463662 | 0.844129553 | 0.70617149 | 0.823231454 | 12.46477325 | 10.49380325 | ENSG00000147642 |
| NRM | -0.24449312 | 0.844112317 | 0.039745271 | 0.099162704 | 505.8007028 | 427.0171615 | ENSG00000137404 |
| CCDC126 | -0.244676948 | 0.844004766 | 0.450463259 | 0.62088359 | 48.11261946 | 40.67436681 | ENSG00000169193 |
| GADD45GIP1 | -0.244693927 | 0.843994834 | 0.000796769 | 0.003461816 | 2001.633413 | 1689.439855 | ENSG00000179271 |
| ATG4A | -0.244731515 | 0.843972845 | 0.282541552 | 0.450766644 | 95.48160381 | 80.67734099 | ENSG00000101844 |
| BEND6 | -0.244753118 | 0.843960207 | 0.465058756 | 0.634608349 | 45.00320288 | 38.00089989 | ENSG00000151917 |
| NA | -0.244761503 | 0.843955302 | 0.821900661 | 0.898581801 | 4.607323737 | 3.889065235 | ENSG00000223723 |
| PPP2R1A | -0.245015651 | 0.843806642 | 9.70E-10 | 1.18E-08 | 8957.49996 | 7558.569646 | ENSG00000105568 |
| MLPH | -0.245141307 | 0.843733151 | 0.180102103 | 0.32491814 | 155.6260914 | 131.234074 | ENSG00000115648 |
| RABL3 | -0.245398308 | 0.843582862 | 0.044577403 | 0.108867285 | 411.04172 | 346.5417203 | ENSG00000144840 |
| ZFP69 | -0.245678467 | 0.843419062 | 0.304324671 | 0.475054802 | 108.1654828 | 91.3475097 | ENSG00000187815 |
| MAFG-DT | -0.245706547 | 0.843402646 | 0.32321863 | 0.494779447 | 86.63789801 | 73.18907231 | ENSG00000265688 |
| CDCA5 | -0.245710142 | 0.843400544 | 1.52E-05 | 9.67E-05 | 2693.582967 | 2271.3029 | ENSG00000146670 |
| ZNF117 | -0.245751749 | 0.843376221 | 0.202395213 | 0.354978275 | 151.6614788 | 127.78383 | ENSG00000152926 |
| NA | -0.245774885 | 0.843362696 | 0.594293218 | 0.742279732 | 24.43029854 | 20.66629172 | ENSG00000280273 |
| NA | -0.245787123 | 0.843355542 | 0.298780963 | 0.46917245 | 88.14138698 | 74.25975184 | ENSG00000213976 |
| CASKIN2 | -0.245860354 | 0.843312735 | 0.005007447 | 0.017557776 | 883.7384265 | 745.3946162 | ENSG00000177303 |
| CROCC | -0.245945463 | 0.843262987 | 7.88E-05 | 0.000435845 | 2290.267039 | 1931.416744 | ENSG00000058453 |
| SEMA6A-AS2 | -0.246055483 | 0.843198682 | 0.76910878 | 0.863915073 | 7.682886059 | 6.44072453 | ENSG00000249167 |
| ATP8A1 | -0.246204725 | 0.84311146 | 0.050069549 | 0.119437574 | 327.8535044 | 276.3914951 | ENSG00000124406 |
| ACTR5 | -0.246296891 | 0.8430576 | 0.017370619 | 0.049999494 | 557.8694358 | 469.8559922 | ENSG00000101442 |
| SUDS3 | -0.246311165 | 0.843049259 | 2.15E-05 | 0.000133489 | 2255.053536 | 1901.426727 | ENSG00000111707 |
| STAB2 | -0.246353685 | 0.843024413 | 0.675376008 | 0.800828482 | 15.23239648 | 12.90268222 | ENSG00000136011 |
| DHRS4-AS1 | -0.246377585 | 0.843010447 | 0.12143382 | 0.240585377 | 192.3240724 | 162.139813 | ENSG00000215256 |
| ATOH7 | -0.24638024 | 0.843008896 | 0.771769747 | 0.865643944 | 6.588572428 | 5.576276454 | ENSG00000179774 |
| MRPL13 | -0.246413408 | 0.842989515 | 0.00699292 | 0.023363427 | 772.2854218 | 650.8976887 | ENSG00000172172 |
| PPIH | -0.246456119 | 0.842964559 | 0.089152746 | 0.189285233 | 239.9623987 | 202.2608267 | ENSG00000171960 |
| DDX39A | -0.246693819 | 0.842825682 | 1.58E-06 | 1.20E-05 | 3394.00961 | 2860.438574 | ENSG00000123136 |
| IKZF4 | -0.246754457 | 0.842790258 | 0.080695495 | 0.174595872 | 259.500162 | 218.7820675 | ENSG00000123411 |
| SNRNP40 | -0.246950972 | 0.842675466 | 3.42E-05 | 0.000203868 | 2139.247016 | 1802.088992 | ENSG00000060688 |
| GPR157 | -0.246954987 | 0.842673121 | 0.247457101 | 0.411006068 | 118.6042649 | 99.8602419 | ENSG00000180758 |
| P4HB | -0.247007009 | 0.842642736 | 5.48E-09 | 6.04E-08 | 7702.669469 | 6490.124027 | ENSG00000185624 |
| TOR1A | -0.247078184 | 0.842601165 | 0.000561033 | 0.002538687 | 1188.14685 | 1001.440305 | ENSG00000136827 |
| RNASEH2A | -0.247080677 | 0.842599709 | 0.727211059 | 0.836082516 | 11.26643248 | 9.509437661 | ENSG00000104889 |
| UFD1 | -0.247097186 | 0.842590067 | 0.000956763 | 0.004094257 | 1207.565241 | 1017.25976 | ENSG00000070010 |
| RNF10 | -0.247194184 | 0.842533419 | 1.34E-06 | 1.03E-05 | 4005.22793 | 3374.703859 | ENSG00000022840 |
| NUTM2A-AS1 | -0.247208229 | 0.842525216 | 0.034506693 | 0.088051404 | 391.7131254 | 330.323202 | ENSG00000223482 |
| SGPP1 | -0.247230428 | 0.842512252 | 0.107653355 | 0.218950012 | 226.1720544 | 190.8243306 | ENSG00000126821 |
| ADRA1A | -0.247242926 | 0.842504953 | 0.723208772 | 0.834065907 | 9.337535219 | 7.866855452 | ENSG00000120907 |
| MAGED4 | -0.247314137 | 0.842463369 | 0.503529128 | 0.668887233 | 34.61445688 | 29.18114396 | ENSG00000154545 |
| NA | -0.247433057 | 0.842393928 | 0.750787741 | 0.852585092 | 10.76614537 | 9.009055378 | ENSG00000248971 |
| NA | -0.247458551 | 0.842379043 | 0.556821628 | 0.712925336 | 30.54880823 | 25.7381452 | ENSG00000215859 |
| TSPAN3 | -0.247512244 | 0.842347692 | 1.25E-06 | 9.71E-06 | 2935.331552 | 2472.623638 | ENSG00000140391 |
| NA | -0.247519642 | 0.842343373 | 0.825180635 | 0.900593514 | 4.370796837 | 3.702188942 | ENSG00000232946 |
| GABRB1 | -0.247667535 | 0.842257027 | 0.827521385 | 0.901751966 | 3.95970648 | 3.376039973 | ENSG00000163288 |
| PSMG3-AS1 | -0.247825482 | 0.842164822 | 0.217979355 | 0.375517033 | 125.9778604 | 106.2213647 | ENSG00000230487 |
| ZNF382 | -0.248134864 | 0.841984241 | 0.347791765 | 0.520501609 | 80.47034025 | 67.68297128 | ENSG00000161298 |
| CHMP4A | -0.248137513 | 0.841982695 | 0.606567607 | 0.751094147 | 20.69124942 | 17.43974425 | ENSG00000254505 |
| UBTD1 | -0.248261429 | 0.841910378 | 0.017688451 | 0.050762855 | 626.9942486 | 528.3274021 | ENSG00000165886 |
| SYP | -0.24829531 | 0.841890607 | 0.131025396 | 0.254261232 | 209.9128896 | 177.0238572 | ENSG00000102003 |
| TECPR1 | -0.248306336 | 0.841884172 | 0.034251901 | 0.087505419 | 387.1274901 | 325.6689367 | ENSG00000205356 |
| TCEAL4 | -0.248426405 | 0.841814109 | 0.000557132 | 0.00252529 | 1291.884328 | 1087.387401 | ENSG00000133142 |
| CCT2 | -0.248491696 | 0.841776012 | 6.75E-10 | 8.36E-09 | 8861.274772 | 7459.906343 | ENSG00000166226 |
| NA | -0.248778629 | 0.841608611 | 0.733407058 | 0.840900776 | 9.427607851 | 7.934074697 | ENSG00000254244 |
| NSFL1C | -0.248830356 | 0.841578436 | 9.23E-06 | 6.12E-05 | 2699.570205 | 2271.272791 | ENSG00000088833 |
| IL17RA | -0.248849676 | 0.841567166 | 0.048027196 | 0.115652558 | 334.7140096 | 281.5416238 | ENSG00000177663 |
| NA | -0.248944228 | 0.841512013 | 0.304536338 | 0.47526984 | 88.63226224 | 74.46032508 | ENSG00000260727 |
| CACNA2D3 | -0.248983669 | 0.841489008 | 0.306729517 | 0.477764967 | 85.99905378 | 72.37947307 | ENSG00000157445 |
| RDH14 | -0.249112281 | 0.841413995 | 0.083280439 | 0.179243886 | 244.8773492 | 206.0928925 | ENSG00000240857 |
| USP3 | -0.249244721 | 0.841336756 | 0.082198942 | 0.177339172 | 252.6152144 | 212.4841233 | ENSG00000140455 |
| LINC01224 | -0.249250918 | 0.841333142 | 0.014948853 | 0.044144798 | 547.0924022 | 460.6008558 | ENSG00000269416 |
| NA | -0.249432824 | 0.841227067 | 0.703623 | 0.821795199 | 12.0348223 | 10.09666693 | ENSG00000225259 |

| | | | | | | | |
|---|---|---|---|---|---|---|---|
| SPAST | -0.249454396 | 0.841214489 | 0.020118915 | 0.05648581 | 579.4214261 | 487.7575957 | ENSG00000021574 |
| MRPL36 | -0.249481411 | 0.841198737 | 0.003695835 | 0.013469934 | 747.7245385 | 629.0295019 | ENSG00000171421 |
| RAB39B | -0.249503319 | 0.841185963 | 0.063442134 | 0.144204336 | 286.4399053 | 240.9981879 | ENSG00000155961 |
| FBXO17 | -0.249680582 | 0.841082613 | 0.112957394 | 0.227435664 | 237.0736395 | 199.1845773 | ENSG00000269190 |
| TBC1D7 | -0.249707153 | 0.841067123 | 0.021635887 | 0.059991896 | 545.7797835 | 458.9822025 | ENSG00000145979 |
| NA | -0.249793596 | 0.841016729 | 0.512911487 | 0.677200431 | 35.19644391 | 29.69873083 | ENSG00000264070 |
| SAAL1 | -0.249807757 | 0.841008474 | 0.002037963 | 0.008024975 | 952.5959599 | 801.2267447 | ENSG00000166788 |
| GHRHR | -0.249823107 | 0.840999526 | 0.549509084 | 0.706799555 | 31.78487021 | 26.66766195 | ENSG00000106128 |
| TJP2 | -0.249968368 | 0.840914852 | 0.342102192 | 0.514861397 | 85.31439105 | 71.77073807 | ENSG00000119139 |
| LINGO3 | -0.249977657 | 0.840909438 | 0.643014108 | 0.776973702 | 16.37559492 | 13.75852801 | ENSG00000220008 |
| CD44 | -0.250000877 | 0.840895904 | 0.27765694 | 0.445095565 | 100.3678816 | 84.57930966 | ENSG00000026508 |
| RBP1 | -0.250109173 | 0.840832785 | 0.042071086 | 0.103824141 | 365.3329908 | 306.8627445 | ENSG00000114115 |
| NA | -0.250330197 | 0.840703977 | 0.626166932 | 0.764808554 | 18.29121339 | 15.38228809 | ENSG00000251495 |
| G6PD | -0.25033564 | 0.840700805 | 0.000441285 | 0.002054751 | 1358.488431 | 1142.258057 | ENSG00000160211 |
| PSMD7 | -0.250371535 | 0.840679888 | 0.000212945 | 0.001066991 | 2017.542647 | 1695.77912 | ENSG00000103035 |
| FBXO7 | -0.250488624 | 0.840611661 | 0.000116415 | 0.000620207 | 1409.59152 | 1184.905153 | ENSG00000100225 |
| MAPK3 | -0.250489228 | 0.84061131 | 0.001342863 | 0.005550712 | 912.5560288 | 767.0651496 | ENSG00000102882 |
| KCTD21 | -0.250549831 | 0.840575999 | 0.443690143 | 0.614489665 | 51.45301803 | 43.10557556 | ENSG00000188997 |
| GINM1 | -0.250589869 | 0.840552671 | 0.082474002 | 0.177785896 | 261.5381557 | 220.1420637 | ENSG00000055211 |
| PHACTR2 | -0.250665781 | 0.840508445 | 0.014070514 | 0.041998187 | 550.9538266 | 463.5429393 | ENSG00000112419 |
| SLC47A2 | -0.250697425 | 0.840490009 | 0.812154731 | 0.892246907 | 4.648511559 | 3.901968677 | ENSG00000180638 |
| NCAPH2 | -0.250751034 | 0.840458778 | 9.37E-05 | 0.000509767 | 1545.213391 | 1298.595734 | ENSG00000025770 |
| HHAT | -0.250885997 | 0.840380157 | 0.746194563 | 0.849654073 | 9.133823432 | 7.692349563 | ENSG00000054392 |
| CAPN1 | -0.250953576 | 0.840340793 | 1.28E-07 | 1.16E-06 | 4604.290624 | 3869.128433 | ENSG00000014216 |
| TIMM22 | -0.251019367 | 0.840302472 | 0.002520205 | 0.009652031 | 783.5024428 | 658.1996204 | ENSG00000177370 |
| FAM8A1 | -0.251079067 | 0.8402677 | 0.086537741 | 0.184811199 | 267.9675587 | 225.4204212 | ENSG00000137414 |
| VRTN | -0.251126758 | 0.840239924 | 1.24E-06 | 9.60E-06 | 5643.399093 | 4741.755471 | ENSG00000133980 |
| NA | -0.251185677 | 0.84020561 | 0.092062988 | 0.194000353 | 221.8493034 | 186.3821849 | ENSG00000263146 |
| PSMG1 | -0.251206882 | 0.84019326 | 0.003335811 | 0.012299821 | 977.9355335 | 821.3423912 | ENSG00000183527 |
| STUB1 | -0.251239161 | 0.840174462 | 0.003852469 | 0.013985317 | 1052.342508 | 884.6971913 | ENSG00000103266 |
| ZBTB2 | -0.251336553 | 0.840117746 | 0.034004972 | 0.086955186 | 476.8966436 | 400.960082 | ENSG00000181472 |
| TMEM242 | -0.25135227 | 0.840108594 | 0.064010225 | 0.145160428 | 305.7220057 | 256.5240619 | ENSG00000215712 |
| NA | -0.251397782 | 0.840082092 | 0.76138215 | 0.858234373 | 6.99408098 | 5.866398707 | ENSG00000260095 |
| NSD3 | -0.251417056 | 0.840070868 | 0.00139717 | 0.005755457 | 1256.373612 | 1055.086184 | ENSG00000147548 |
| KLF12 | -0.251537999 | 0.840000447 | 0.089201681 | 0.189368288 | 250.5479233 | 210.6334195 | ENSG00000118922 |
| RRM1 | -0.251615348 | 0.839955413 | 1.42E-05 | 9.07E-05 | 2128.826769 | 1788.302179 | ENSG00000167325 |
| CAMSAP3 | -0.251632935 | 0.839945173 | 0.003610635 | 0.013209322 | 1310.710835 | 1100.964796 | ENSG00000076826 |
| FKBP1A | -0.251696109 | 0.839908394 | 0.000509678 | 0.002332669 | 2970.480568 | 2495.538526 | ENSG00000088832 |
| TMEM167A | -0.251977726 | 0.839744458 | 3.70E-05 | 0.000218692 | 3339.861284 | 2805.592785 | ENSG00000174695 |
| OLFM2 | -0.25197944 | 0.83974346 | 3.54E-05 | 0.000210165 | 2072.549688 | 1741.042901 | ENSG00000105088 |
| LRRC55 | -0.252008173 | 0.839726736 | 0.283028526 | 0.451418214 | 98.02577877 | 82.44441443 | ENSG00000183908 |
| PARN | -0.252359076 | 0.839522516 | 0.00107775 | 0.004558405 | 1260.582437 | 1058.332917 | ENSG00000140694 |
| HCK | -0.25238327 | 0.839508438 | 0.615849356 | 0.758212125 | 18.72674615 | 15.69391608 | ENSG00000101336 |
| GPRIN2 | -0.2525554 | 0.83940828 | 0.124368802 | 0.24481618 | 299.5641111 | 251.2772 | ENSG00000204175 |
| DOP1B | -0.252726594 | 0.83930868 | 0.067797101 | 0.152096044 | 253.2715678 | 212.6290542 | ENSG00000142197 |
| NA | -0.25282941 | 0.839248867 | 0.617705091 | 0.759443988 | 22.05857185 | 18.45960351 | ENSG00000247373 |
| NA | -0.252969908 | 0.83916714 | 0.561762907 | 0.7165506 | 26.74113783 | 22.48794142 | ENSG00000268297 |
| EPG5 | -0.253047451 | 0.839122037 | 0.03544629 | 0.09008556 | 440.5336845 | 369.5004385 | ENSG00000152223 |
| CD8B | -0.253072082 | 0.839107711 | 0.79604007 | 0.881488375 | 6.428163645 | 5.422743264 | ENSG00000172116 |
| SRP19 | -0.253077121 | 0.83910478 | 0.226550913 | 0.38577047 | 123.434161 | 103.6944278 | ENSG00000153037 |
| SEC11A | -0.253121529 | 0.839078952 | 5.45E-07 | 4.46E-06 | 3582.437653 | 3006.028267 | ENSG00000140612 |
| CACNG7 | -0.253123197 | 0.839077982 | 0.000111678 | 0.000596948 | 1374.6899 | 1153.626485 | ENSG00000105605 |
| NA | -0.253200303 | 0.839033138 | 0.446079163 | 0.616426443 | 43.86558625 | 36.78261797 | ENSG00000261377 |
| ZCCHC17 | -0.253218437 | 0.839022592 | 0.006010683 | 0.020573473 | 668.6903421 | 560.8859432 | ENSG00000121766 |
| DPY19L3 | -0.253255011 | 0.839001322 | 0.006107425 | 0.020849112 | 931.5570103 | 781.6730348 | ENSG00000178904 |
| FRG1 | -0.253324176 | 0.8389611 | 0.052025729 | 0.123038904 | 368.5123559 | 309.3753783 | ENSG00000109536 |
| BRD8 | -0.253364736 | 0.838937514 | 0.000163712 | 0.000840608 | 1565.436666 | 1312.688752 | ENSG00000112983 |
| GIGYF2 | -0.25337388 | 0.838932196 | 0.000318432 | 0.001534953 | 1670.374913 | 1400.601593 | ENSG00000204120 |
| IFT20 | -0.253381311 | 0.838927875 | 0.125862886 | 0.246925754 | 203.5704049 | 170.7670649 | ENSG00000109083 |
| ZNF354A | -0.253600719 | 0.838800299 | 0.126394269 | 0.2477415 | 218.4004472 | 183.5202241 | ENSG00000169131 |
| SMARCA5 | -0.25361492 | 0.838792042 | 2.72E-08 | 2.71E-07 | 4648.090854 | 3899.288628 | ENSG00000153147 |
| CCNJ | -0.25361562 | 0.838791635 | 0.002176051 | 0.008494174 | 926.8085146 | 777.8600642 | ENSG00000107443 |
| CLCNKA | -0.253616832 | 0.838790931 | 0.593556355 | 0.741757358 | 21.94126591 | 18.32946617 | ENSG00000186510 |
| NR4A1 | -0.253640354 | 0.838777255 | 0.032936007 | 0.084727097 | 506.1109204 | 424.587564 | ENSG00000123358 |
| AOPEP | -0.253741606 | 0.83871839 | 0.101569656 | 0.209536973 | 210.7067978 | 176.7953081 | ENSG00000148120 |
| ATP5MC3 | -0.253751817 | 0.838712454 | 5.63E-08 | 5.36E-07 | 5143.923525 | 4313.886129 | ENSG00000154518 |
| CBX3P2 | -0.253846453 | 0.838657439 | 0.812326432 | 0.892246907 | 4.628975239 | 3.872393683 | ENSG00000266405 |
| COQ3 | -0.253888095 | 0.838633232 | 0.057394732 | 0.133238974 | 283.0896194 | 237.3772061 | ENSG00000132423 |
| SLC25A39 | -0.253974485 | 0.838583015 | 2.44E-08 | 2.44E-07 | 5791.300182 | 4856.184306 | ENSG00000013306 |
| SPATA2 | -0.254222006 | 0.838439153 | 0.017984598 | 0.051436751 | 509.1629531 | 426.6697609 | ENSG00000158480 |
| RASA2 | -0.254269536 | 0.838411531 | 0.05595576 | 0.130619866 | 324.3843609 | 272.1719677 | ENSG00000155903 |
| FCHO1 | -0.254286902 | 0.838401439 | 0.011705897 | 0.035925819 | 722.3605736 | 605.7844296 | ENSG00000130475 |
| HIP1 | -0.254319377 | 0.838382567 | 4.28E-07 | 3.57E-06 | 3543.930545 | 2971.226254 | ENSG00000127946 |
| CTIF | -0.254412929 | 0.838328203 | 0.025225856 | 0.068117579 | 497.5020876 | 416.7510891 | ENSG00000134030 |
| PPP4C | -0.254437937 | 0.838313671 | 6.52E-07 | 5.28E-06 | 4685.670702 | 3928.294752 | ENSG00000149923 |
| ZNF385C | -0.254499009 | 0.838278185 | 0.202291455 | 0.354863743 | 122.2450689 | 102.5046548 | ENSG00000187595 |
| TBC1D16 | -0.254546235 | 0.838250745 | 1.12E-07 | 1.02E-06 | 6533.053365 | 5475.988121 | ENSG00000167291 |

| | | | | | | | |
|---|---|---|---|---|---|---|---|
| C3orf14 | -0.25474073 | 0.838137745 | 0.233419904 | 0.394179858 | 144.0219469 | 120.8271362 | ENSG00000114405 |
| MRPS22 | -0.254758341 | 0.838127514 | 0.009188101 | 0.029297134 | 608.6186182 | 509.877516 | ENSG00000175110 |
| YME1L1 | -0.254809684 | 0.838097687 | 2.14E-07 | 1.87E-06 | 3532.514069 | 2961.089774 | ENSG00000136758 |
| FTO | -0.254830565 | 0.838085557 | 0.002405416 | 0.009280575 | 813.4038339 | 681.3922976 | ENSG00000140718 |
| NA | -0.254922767 | 0.838031997 | 0.731040939 | 0.839112204 | 8.597054427 | 7.229611531 | ENSG00000223612 |
| ARVCF | -0.254973907 | 0.838002291 | 0.000398443 | 0.001879211 | 1696.835748 | 1421.764051 | ENSG00000099889 |
| NUBP2 | -0.255064767 | 0.837949516 | 0.000385229 | 0.001824863 | 1939.931168 | 1625.277024 | ENSG00000095906 |
| NA | -0.255108199 | 0.83792429 | 0.565210218 | 0.719310343 | 24.33551998 | 20.42886773 | ENSG00000229036 |
| JUP | -0.255137343 | 0.837907363 | 3.12E-05 | 0.000187842 | 3218.987191 | 2697.720146 | ENSG00000173801 |
| NA | -0.255226674 | 0.837855482 | 0.761899884 | 0.858617284 | 8.349363916 | 7.034132943 | ENSG00000272121 |
| DENND1A | -0.255247029 | 0.837843661 | 0.002569795 | 0.009818559 | 978.1154147 | 819.1931986 | ENSG00000119522 |
| SOCS7 | -0.255264897 | 0.837833284 | 8.33E-05 | 0.000458688 | 1565.195297 | 1311.198304 | ENSG00000274211 |
| NA | -0.255307147 | 0.837808748 | 0.498926873 | 0.665247326 | 35.87775216 | 30.09640019 | ENSG00000259917 |
| CNNM3 | -0.25531326 | 0.837805198 | 0.003779927 | 0.013763415 | 987.741242 | 827.8677816 | ENSG00000168763 |
| ZNRD1 | -0.255347753 | 0.837785167 | 0.260561308 | 0.425937479 | 116.9351857 | 97.79633402 | ENSG00000066379 |
| NA | -0.255481108 | 0.837707731 | 0.254029179 | 0.418808382 | 104.5167817 | 87.63539193 | ENSG00000180385 |
| NA | -0.255563547 | 0.837659863 | 0.804309371 | 0.887069454 | 6.583666344 | 5.517126466 | ENSG00000225972 |
| MGAT2 | -0.25559033 | 0.837644313 | 0.794412807 | 0.880349251 | 4.694605466 | 3.920790803 | ENSG00000168282 |
| EVI5L | -0.255638172 | 0.837616536 | 0.016154153 | 0.047059765 | 536.9216859 | 449.7234259 | ENSG00000142459 |
| PTOV1 | -0.25565381 | 0.837607456 | 0.002595806 | 0.009902253 | 1000.198467 | 837.7714115 | ENSG00000104960 |
| EPS8 | -0.255686544 | 0.837588452 | 0.25264041 | 0.417265794 | 107.4658264 | 89.99398367 | ENSG00000151491 |
| CHMP4B | -0.255775761 | 0.837536656 | 7.73E-06 | 5.20E-05 | 2317.078236 | 1940.794862 | ENSG00000101421 |
| CNOT9 | -0.255912756 | 0.83745713 | 1.33E-05 | 8.52E-05 | 1793.724472 | 1502.020787 | ENSG00000144580 |
| ENOSF1 | -0.255935535 | 0.837443907 | 0.016685814 | 0.048305602 | 450.5040819 | 377.1092003 | ENSG00000132199 |
| CCDC3 | -0.256150839 | 0.837318938 | 0.467275171 | 0.636551467 | 41.88222238 | 34.99914555 | ENSG00000151468 |
| NXT2 | -0.25620448 | 0.837287806 | 0.058201764 | 0.13472165 | 390.9060264 | 327.7422282 | ENSG00000101888 |
| PTPA | -0.256204674 | 0.837287694 | 1.12E-08 | 1.19E-07 | 5873.929138 | 4918.413778 | ENSG00000119383 |
| RARB | -0.25621225 | 0.837283297 | 0.804396998 | 0.887069454 | 4.642929753 | 3.874544257 | ENSG00000077092 |
| NA | -0.256237103 | 0.837268873 | 0.802627596 | 0.88621529 | 4.316330222 | 3.626367402 | ENSG00000287733 |
| SPA17 | -0.256270683 | 0.837249386 | 0.729421122 | 0.837724764 | 8.689917962 | 7.301131924 | ENSG00000064199 |
| DBF4 | -0.256313349 | 0.837224625 | 0.004485414 | 0.015970492 | 852.668919 | 714.4113955 | ENSG00000006634 |
| ZNF700 | -0.256317798 | 0.837222043 | 0.285455231 | 0.454462067 | 101.7395465 | 85.05604775 | ENSG00000196757 |
| AP2M1 | -0.256338065 | 0.837210282 | 9.44E-09 | 1.01E-07 | 12824.34292 | 10736.56621 | ENSG00000161203 |
| MED16 | -0.256466021 | 0.837136031 | 2.41E-05 | 0.000147769 | 2308.827031 | 1932.935901 | ENSG00000175221 |
| NUBP1 | -0.256751072 | 0.836970645 | 0.073275605 | 0.161808573 | 300.7223401 | 251.5433628 | ENSG00000103274 |
| CCDC47 | -0.25675799 | 0.836966631 | 3.84E-05 | 0.000225908 | 2009.7881 | 1682.740382 | ENSG00000108588 |
| GUCY1B1 | -0.25679004 | 0.836948037 | 0.3495783 | 0.522201873 | 72.00407072 | 60.17586837 | ENSG00000061918 |
| NA | -0.256912514 | 0.83687699 | 0.517659554 | 0.681556816 | 31.27022826 | 26.17104778 | ENSG00000234964 |
| PNP | -0.257030996 | 0.836808264 | 0.000172825 | 0.000882843 | 1470.315856 | 1229.93649 | ENSG00000198805 |
| C2CD2L | -0.25708152 | 0.836778959 | 0.394460992 | 0.567042134 | 51.5156572 | 43.13110987 | ENSG00000172375 |
| GNAL | -0.257383658 | 0.836603734 | 0.018878727 | 0.053517932 | 560.194593 | 468.9769563 | ENSG00000141404 |
| FAM214B | -0.257387045 | 0.83660177 | 0.179863314 | 0.324548049 | 138.4077034 | 115.9776221 | ENSG00000005238 |
| GMPR2 | -0.257474262 | 0.836551195 | 8.72E-05 | 0.000478023 | 1492.802263 | 1248.906369 | ENSG00000100938 |
| TMEM97 | -0.257506355 | 0.836532586 | 1.39E-05 | 8.86E-05 | 2340.069963 | 1957.945327 | ENSG00000109084 |
| RRP9 | -0.257550864 | 0.836506778 | 0.00012761 | 0.0006715 | 2122.544562 | 1775.408463 | ENSG00000114767 |
| ADAL | -0.257621016 | 0.836466103 | 0.178613049 | 0.322849397 | 195.3330406 | 163.3538059 | ENSG00000168803 |
| MAML1 | -0.257631887 | 0.836459801 | 0.000104215 | 0.000561726 | 1772.510722 | 1482.758612 | ENSG00000161021 |
| TMEM237 | -0.257690414 | 0.836425868 | 0.005827487 | 0.020035299 | 655.4485934 | 548.5376864 | ENSG00000155755 |
| MED6 | -0.257694096 | 0.836423733 | 0.055192948 | 0.129261522 | 318.06453 | 266.3918829 | ENSG00000133997 |
| ASF1B | -0.257730301 | 0.836402743 | 0.000766471 | 0.003343734 | 978.520611 | 818.6922653 | ENSG00000105011 |
| BRD7 | -0.257739919 | 0.836397167 | 6.64E-06 | 4.53E-05 | 2363.716199 | 1977.371516 | ENSG00000166164 |
| ZNF346 | -0.257753017 | 0.836389574 | 0.012321695 | 0.037480501 | 532.2542385 | 445.132036 | ENSG00000113761 |
| ARF5 | -0.257826556 | 0.836346942 | 0.00012078 | 0.000640157 | 2162.59155 | 1808.306026 | ENSG00000004059 |
| HMOX1 | -0.257838999 | 0.836339728 | 0.006625884 | 0.022350193 | 857.2427888 | 716.7577206 | ENSG00000100292 |
| BST2 | -0.25792953 | 0.836287248 | 0.001462477 | 0.005993762 | 1007.075329 | 842.0360341 | ENSG00000130303 |
| GUSBP1 | -0.258003034 | 0.836244642 | 0.280888067 | 0.449229953 | 84.41071057 | 70.50135699 | ENSG00000183666 |
| NA | -0.258067676 | 0.836207173 | 0.72845709 | 0.837165836 | 8.645939237 | 7.248433657 | ENSG00000236814 |
| EED | -0.258089671 | 0.836194425 | 0.003238086 | 0.011978021 | 736.761139 | 615.8248906 | ENSG00000074266 |
| GABRA5 | -0.258138912 | 0.836165884 | 0.022858797 | 0.062778697 | 479.625356 | 401.0119747 | ENSG00000186297 |
| RHOG | -0.258199858 | 0.836130562 | 0.022641787 | 0.062317409 | 452.7425962 | 378.7878093 | ENSG00000177105 |
| H3-5 | -0.258333047 | 0.836053374 | 0.668954255 | 0.796690735 | 14.40762503 | 11.98527542 | ENSG00000188375 |
| EXOSC3 | -0.258420917 | 0.836002455 | 0.019784206 | 0.055700106 | 441.8098214 | 369.4372783 | ENSG00000107371 |
| NA | -0.258454032 | 0.835983265 | 0.642282152 | 0.776527012 | 15.67072015 | 13.09601024 | ENSG00000267265 |
| SARS1 | -0.258494409 | 0.835959869 | 6.07E-06 | 4.16E-05 | 3879.745861 | 3242.406593 | ENSG00000031698 |
| NA | -0.258510069 | 0.835950795 | 0.773353601 | 0.867017162 | 7.111386919 | 5.929849843 | ENSG00000262692 |
| TESK2 | -0.25858361 | 0.835908184 | 0.217583999 | 0.375200219 | 124.8929956 | 104.3066888 | ENSG00000070759 |
| C11orf49 | -0.258655593 | 0.835866477 | 0.028812736 | 0.076015449 | 403.7137301 | 337.14361 | ENSG00000149179 |
| RPN2 | -0.258664296 | 0.835861435 | 3.80E-07 | 3.21E-06 | 5154.190345 | 4308.707485 | ENSG00000118705 |
| BANP | -0.258669526 | 0.835858405 | 0.008553935 | 0.027617398 | 691.0303466 | 577.797596 | ENSG00000172530 |
| NA | -0.258682389 | 0.835850952 | 0.73486638 | 0.842023481 | 9.054238691 | 7.563829757 | ENSG00000223529 |
| NKX1-2 | -0.258726033 | 0.835825667 | 0.01226318 | 0.037319608 | 511.3520559 | 427.2626179 | ENSG00000229544 |
| NA | -0.258785635 | 0.835791137 | 0.751173853 | 0.852757227 | 7.054129401 | 5.872850428 | ENSG00000226945 |
| FAM156A | -0.258850074 | 0.835753807 | 0.745960931 | 0.849438187 | 8.77719969 | 7.353830191 | ENSG00000268350 |
| PQBP1 | -0.258960791 | 0.835689671 | 0.001324271 | 0.005483263 | 1240.365968 | 1036.027829 | ENSG00000102103 |
| NA | -0.25896715 | 0.835685987 | 0.62378117 | 0.763014769 | 17.3197875 | 14.50762018 | ENSG00000228532 |
| FDX1 | -0.259091412 | 0.835614011 | 0.0189855 | 0.053781099 | 539.5888366 | 451.337717 | ENSG00000137714 |
| C17orf100 | -0.259107392 | 0.835604755 | 0.513098969 | 0.677401602 | 39.86680713 | 33.21997427 | ENSG00000256806 |

| | | | | | | | |
|---|---|---|---|---|---|---|---|
| COL17A1 | -0.259107785 | 0.835604528 | 0.387642625 | 0.560949207 | 53.8095509 | 44.94604739 | ENSG00000065618 |
| TCTN1 | -0.25916 | 0.835574285 | 0.274083379 | 0.440918439 | 92.91836809 | 77.54706683 | ENSG00000204852 |
| MRPL11 | -0.25920086 | 0.835550621 | 4.61E-05 | 0.000266859 | 1898.745133 | 1586.038735 | ENSG00000174547 |
| LOC100128398 | -0.259245423 | 0.835524812 | 0.208260335 | 0.363053507 | 135.4217382 | 113.0610972 | ENSG00000176593 |
| FNDC7 | -0.259299157 | 0.835493693 | 0.578074228 | 0.729895171 | 23.1142884 | 19.26113349 | ENSG00000143107 |
| COX6A1 | -0.259353041 | 0.835462489 | 0.210919165 | 0.366521726 | 118.93721 | 99.46362651 | ENSG00000111775 |
| CCNA2 | -0.25936733 | 0.835454214 | 6.87E-06 | 4.66E-05 | 2670.297109 | 2231.550373 | ENSG00000145386 |
| WDR24 | -0.259414851 | 0.835426695 | 0.014374748 | 0.042714613 | 487.190699 | 407.0015318 | ENSG00000127580 |
| MARVELD3 | -0.259687629 | 0.835268751 | 0.003823669 | 0.01389384 | 712.3691253 | 594.8870262 | ENSG00000140832 |
| ZBTB33 | -0.25997618 | 0.835101708 | 0.00201724 | 0.007949861 | 838.6319091 | 700.851569 | ENSG00000177485 |
| ZNRD2 | -0.260006078 | 0.835084401 | 0.002785209 | 0.01053106 | 736.1547096 | 614.6082141 | ENSG00000173465 |
| LOC101929798 | -0.260042553 | 0.835063289 | 0.719869806 | 0.83186387 | 9.098217416 | 7.591254177 | ENSG00000275585 |
| NA | -0.260065251 | 0.83505015 | 0.389371116 | 0.562629833 | 53.45360288 | 44.62850072 | ENSG00000227086 |
| PROS1 | -0.260223362 | 0.834958639 | 0.641198419 | 0.775605638 | 17.33027539 | 14.4930992 | ENSG00000184500 |
| HSD17B10 | -0.260400716 | 0.834856002 | 7.29E-05 | 0.000406215 | 2410.741314 | 2011.973883 | ENSG00000072506 |
| ILVBL | -0.26041565 | 0.83484736 | 0.000596943 | 0.002681796 | 1413.624555 | 1180.709863 | ENSG00000105135 |
| SDF2 | -0.260485124 | 0.834807158 | 0.010252548 | 0.03211788 | 645.0921486 | 537.9301812 | ENSG00000132581 |
| TANGO6 | -0.260595918 | 0.83474305 | 0.02147444 | 0.059597672 | 438.4883956 | 365.9133571 | ENSG00000103047 |
| OTUD5 | -0.260705574 | 0.834679606 | 5.29E-05 | 0.000302734 | 1606.840155 | 1341.069724 | ENSG00000068308 |
| SNF8 | -0.26071407 | 0.83467469 | 0.038841252 | 0.097360524 | 401.3605886 | 334.5830758 | ENSG00000159210 |
| CDK13 | -0.260745722 | 0.834656378 | 0.000517825 | 0.002366026 | 1275.405658 | 1065.272909 | ENSG00000065883 |
| SEC13 | -0.260758281 | 0.834649112 | 5.65E-07 | 4.61E-06 | 2827.907138 | 2359.903538 | ENSG00000157020 |
| DUSP1 | -0.260816887 | 0.834615207 | 0.115567999 | 0.231652355 | 197.4097563 | 164.6377079 | ENSG00000120129 |
| CCZ1 | -0.260834092 | 0.834605254 | 0.036325084 | 0.091991812 | 347.3323706 | 289.9432586 | ENSG00000122674 |
| MCF2 | -0.260905956 | 0.834563681 | 0.573118628 | 0.725447081 | 30.6952625 | 25.73787263 | ENSG00000101977 |
| HEPH | -0.260915764 | 0.834558008 | 5.33E-05 | 0.000304957 | 1686.507925 | 1407.296364 | ENSG00000089472 |
| ZNF518B | -0.260939297 | 0.834544395 | 0.033610795 | 0.086141338 | 417.6407292 | 348.185096 | ENSG00000178163 |
| TRUB2 | -0.261065172 | 0.834471584 | 0.003571508 | 0.013082084 | 684.6163249 | 571.3348569 | ENSG00000167112 |
| ABHD6 | -0.261077556 | 0.834464421 | 0.147112755 | 0.278094282 | 159.326268 | 132.8728881 | ENSG00000163686 |
| DPAGT1 | -0.261112984 | 0.83444393 | 0.000549092 | 0.002491774 | 1056.654118 | 882.0003285 | ENSG00000172269 |
| BCL7C | -0.26118082 | 0.834404694 | 0.223997588 | 0.382503096 | 119.013128 | 99.30903935 | ENSG00000099385 |
| RTEL1-TNFRSF6B | -0.261219411 | 0.834382375 | 0.468335971 | 0.637276519 | 50.67114926 | 42.1714972 | ENSG00000026036 |
| LRRC1 | -0.261234067 | 0.834373899 | 0.085420596 | 0.182880845 | 292.5086912 | 244.132218 | ENSG00000137269 |
| NFYC | -0.2612919 | 0.834340452 | 0.005723636 | 0.019720426 | 796.336193 | 664.5975876 | ENSG00000066136 |
| MTG1 | -0.261390071 | 0.83428368 | 0.305527605 | 0.476469929 | 78.54288245 | 65.60399728 | ENSG00000148824 |
| USP39 | -0.261598732 | 0.834163024 | 2.33E-06 | 1.71E-05 | 2671.689311 | 2228.730177 | ENSG00000168883 |
| PPP1R14B | -0.261635927 | 0.834141518 | 2.32E-06 | 1.71E-05 | 6414.686068 | 5350.0108 | ENSG00000173457 |
| NA | -0.261661574 | 0.83412669 | 0.76954175 | 0.864200134 | 6.61022393 | 5.506906635 | ENSG00000231579 |
| DHCR7 | -0.261676227 | 0.834118218 | 2.68E-09 | 3.09E-08 | 5251.792108 | 4380.758689 | ENSG00000172893 |
| HNRNPAB | -0.261789328 | 0.834052829 | 6.59E-09 | 7.19E-08 | 17460.459 | 14563.55907 | ENSG00000197451 |
| RRN3 | -0.261826047 | 0.834031601 | 0.001445814 | 0.005933036 | 807.4742809 | 673.7252097 | ENSG00000085721 |
| EIF2AK4 | -0.262061006 | 0.833895781 | 1.27E-06 | 9.83E-06 | 3747.358537 | 3125.176951 | ENSG00000128829 |
| NA | -0.262089991 | 0.833879027 | 0.178752942 | 0.323055642 | 147.1887331 | 122.9609469 | ENSG00000286456 |
| PRPF39 | -0.262119503 | 0.83386197 | 0.006813463 | 0.022886735 | 642.3412708 | 535.3819748 | ENSG00000185246 |
| RPP30 | -0.262223729 | 0.83380173 | 0.001606654 | 0.006506138 | 826.3738387 | 689.2155901 | ENSG00000148688 |
| NA | -0.262450245 | 0.833670826 | 0.809591552 | 0.890310719 | 3.95970648 | 3.329793428 | ENSG00000228903 |
| LOC101927837 | -0.262459079 | 0.833665721 | 0.80363907 | 0.88672909 | 4.289096914 | 3.577970282 | ENSG00000270049 |
| HPF1 | -0.262582931 | 0.833594156 | 0.019897145 | 0.055977224 | 475.9653791 | 397.020166 | ENSG00000056050 |
| NUTM2G | -0.262604393 | 0.833581755 | 0.498124404 | 0.66440963 | 33.04497427 | 27.54017928 | ENSG00000188152 |
| TSPAN15 | -0.2626315 | 0.833566094 | 0.028734246 | 0.075866446 | 355.1731627 | 296.035244 | ENSG00000099282 |
| RPL6 | -0.262645705 | 0.833557886 | 7.93E-12 | 1.23E-10 | 19339.78562 | 16120.31763 | ENSG00000089009 |
| MSANTD3 | -0.262732798 | 0.833507567 | 0.021290273 | 0.059181756 | 442.3063906 | 368.3566637 | ENSG00000066697 |
| HSPBP1 | -0.262932052 | 0.833392457 | 0.001997148 | 0.007879689 | 1672.929581 | 1394.63281 | ENSG00000133265 |
| C9orf135 | -0.26318305 | 0.833247477 | 0.133237367 | 0.257620733 | 179.4543541 | 149.7797189 | ENSG00000204711 |
| SREBF1 | -0.263222174 | 0.833224881 | 0.000178368 | 0.000908129 | 2253.844044 | 1877.956674 | ENSG00000072310 |
| FAM209B | -0.263262439 | 0.833201626 | 0.809547434 | 0.890310719 | 3.98973069 | 3.352916701 | ENSG00000213714 |
| SRC | -0.263323748 | 0.833166219 | 0.000146276 | 0.000757735 | 1369.962367 | 1141.553327 | ENSG00000197122 |
| MALT1 | -0.263513164 | 0.833056838 | 0.049277842 | 0.117921892 | 315.0545979 | 262.7045059 | ENSG00000172175 |
| NA | -0.263526217 | 0.833049301 | 0.773813794 | 0.867313818 | 5.905349155 | 4.955704085 | ENSG00000228057 |
| DPY19L2P3 | -0.263526971 | 0.833048865 | 0.472689418 | 0.64152607 | 40.54233342 | 33.85720608 | ENSG00000227855 |
| ZNF142 | -0.263566573 | 0.833025998 | 0.00753513 | 0.024856394 | 676.9035779 | 563.9888355 | ENSG00000115568 |
| ZC4H2 | -0.263586172 | 0.833014681 | 0.006972127 | 0.023306066 | 586.0286079 | 487.8227192 | ENSG00000126970 |
| ZNF830 | -0.263592815 | 0.833010846 | 0.030100086 | 0.07874434 | 430.0621995 | 357.9456546 | ENSG00000198783 |
| ENGASE | -0.263604399 | 0.833004157 | 0.002628163 | 0.010015783 | 892.4789668 | 742.9179849 | ENSG00000167280 |
| WHAMM | -0.263626433 | 0.832991435 | 0.166306297 | 0.306010566 | 139.3190809 | 116.2387024 | ENSG00000156232 |
| TOMM34 | -0.263644825 | 0.832980816 | 0.000395448 | 0.001867005 | 1176.825862 | 980.266379 | ENSG00000025772 |
| ZNF385D | -0.263741137 | 0.832925209 | 0.263460351 | 0.429186276 | 90.16015671 | 75.18737848 | ENSG00000151789 |
| SLC25A41 | -0.263871057 | 0.832850205 | 0.693373538 | 0.814467988 | 10.97053288 | 9.12897289 | ENSG00000181240 |
| PTPN1 | -0.263946699 | 0.832806539 | 0.000419641 | 0.0019644 | 1162.143853 | 968.4496545 | ENSG00000196396 |
| SIGMAR1 | -0.264110199 | 0.832712162 | 7.91E-06 | 5.31E-05 | 2714.319625 | 2260.479452 | ENSG00000147955 |
| MFSD2A | -0.264171573 | 0.832676738 | 0.009377375 | 0.029797224 | 510.1222514 | 424.8424409 | ENSG00000168389 |
| ECM1 | -0.264249666 | 0.832631667 | 0.655920984 | 0.786579245 | 17.18940293 | 14.22100557 | ENSG00000143369 |
| CCDC25 | -0.264260745 | 0.832625273 | 0.001996017 | 0.007877487 | 842.7995946 | 702.0563961 | ENSG00000147419 |
| NA | -0.264301149 | 0.832601955 | 0.678341079 | 0.803241041 | 13.58073839 | 11.29989484 | ENSG00000270938 |
| CZIB | -0.264324139 | 0.832588687 | 0.002591976 | 0.009891555 | 807.7378537 | 672.4700892 | ENSG00000162384 |
| DHRS3 | -0.264391723 | 0.832549685 | 0.077124945 | 0.168438505 | 252.103051 | 209.9287264 | ENSG00000162496 |
| AGAP4 | -0.264450633 | 0.83251569 | 0.310305069 | 0.481279598 | 105.8614135 | 87.93977469 | ENSG00000188234 |

| | | | | | | | |
|---|---|---|---|---|---|---|---|
| HINT1 | -0.264497405 | 0.8324887 | 3.09E-06 | 2.22E-05 | 3268.686848 | 2720.210511 | ENSG00000169567 |
| PLP1 | -0.26455381 | 0.832456153 | 4.26E-05 | 0.000248476 | 1466.140273 | 1220.744284 | ENSG00000123560 |
| PLBD2 | -0.264593841 | 0.832433055 | 0.00026032 | 0.001280101 | 1121.314617 | 933.399725 | ENSG00000151176 |
| CDK5RAP1 | -0.26461332 | 0.832421815 | 0.00152919 | 0.006229854 | 885.2110139 | 736.5113849 | ENSG00000101391 |
| RTN3 | -0.265037014 | 0.832177384 | 2.26E-06 | 1.67E-05 | 3734.599282 | 3108.646544 | ENSG00000133318 |
| WDR19 | -0.265042749 | 0.832174076 | 0.035633584 | 0.09047216 | 326.3629308 | 271.5530249 | ENSG00000157796 |
| AP5Z1 | -0.265282472 | 0.832035811 | 0.008277341 | 0.026872802 | 512.6018722 | 426.5748142 | ENSG00000242802 |
| PSMD3 | -0.265311577 | 0.832019025 | 8.32E-08 | 7.74E-07 | 7480.424324 | 6223.558995 | ENSG00000108344 |
| EMP2 | -0.265330141 | 0.832008319 | 0.014158983 | 0.042203484 | 468.5330497 | 390.026664 | ENSG00000213853 |
| DIS3L | -0.265610065 | 0.831846902 | 0.000858741 | 0.003711507 | 1104.824669 | 918.376699 | ENSG00000166938 |
| INTS6L | -0.265683863 | 0.831804351 | 0.046434857 | 0.112534526 | 299.8216379 | 249.5375083 | ENSG00000165359 |
| YJU2 | -0.265702316 | 0.831793712 | 0.020425086 | 0.057228807 | 532.3584289 | 442.6621411 | ENSG00000105248 |
| SF3A1 | -0.265771108 | 0.831754051 | 6.22E-09 | 6.80E-08 | 4446.761815 | 3698.789085 | ENSG00000099995 |
| LETM1 | -0.265938599 | 0.831657493 | 3.14E-05 | 0.000188883 | 2277.236046 | 1893.477448 | ENSG00000168924 |
| ZNF274 | -0.265975193 | 0.831636398 | 0.01181515 | 0.03618751 | 562.9433798 | 467.9749079 | ENSG00000171606 |
| EEPD1 | -0.266015099 | 0.831613395 | 0.035179883 | 0.089508722 | 351.9999429 | 292.6366138 | ENSG00000122547 |
| SLC9A1 | -0.266129501 | 0.831547453 | 0.008949242 | 0.028650387 | 871.5874254 | 724.7926117 | ENSG00000090020 |
| NA | -0.266143196 | 0.831539559 | 0.526068901 | 0.688213425 | 26.81793166 | 22.25025697 | ENSG00000203279 |
| SSR3 | -0.266201156 | 0.831506153 | 4.53E-06 | 3.18E-05 | 2477.669152 | 2060.686266 | ENSG00000114850 |
| DNPEP | -0.266206018 | 0.83150335 | 0.00010218 | 0.000551785 | 1501.509927 | 1248.232819 | ENSG00000123992 |
| NA | -0.266508272 | 0.831329164 | 0.809480292 | 0.890310719 | 4.743490275 | 3.92509195 | ENSG00000273819 |
| MYBL1 | -0.26663154 | 0.831258135 | 0.322440763 | 0.493902237 | 77.99303481 | 64.84845338 | ENSG00000185697 |
| GYPC | -0.266706703 | 0.831214829 | 0.010220434 | 0.032033609 | 513.1740471 | 426.5396115 | ENSG00000136732 |
| AKAP5 | -0.266819159 | 0.831150039 | 0.295434228 | 0.465508592 | 82.83967635 | 68.95582948 | ENSG00000179841 |
| JMJD6 | -0.266893692 | 0.831107101 | 0.009969652 | 0.031348745 | 555.0408755 | 461.2281951 | ENSG00000070495 |
| SLC3A2 | -0.266916177 | 0.831094148 | 2.75E-12 | 4.48E-11 | 9416.304118 | 7825.771152 | ENSG00000168003 |
| DHX34 | -0.267082913 | 0.830998102 | 0.00028296 | 0.00138087 | 1596.904401 | 1327.286473 | ENSG00000134815 |
| RSRC2 | -0.267199294 | 0.830931069 | 6.99E-06 | 4.74E-05 | 2105.955791 | 1749.751657 | ENSG00000111011 |
| AP1S3 | -0.267408063 | 0.830810836 | 0.24233464 | 0.404891524 | 93.39931903 | 77.58120344 | ENSG00000152056 |
| EIF3B | -0.267449867 | 0.830786762 | 1.28E-09 | 1.54E-08 | 11647.22964 | 9675.823897 | ENSG00000106263 |
| NA | -0.267459447 | 0.830781245 | 0.759682648 | 0.857270428 | 6.340881917 | 5.277551906 | ENSG00000268947 |
| ZNF774 | -0.267468582 | 0.830775985 | 0.539103222 | 0.698545187 | 29.08738297 | 24.10283875 | ENSG00000196391 |
| HDAC8 | -0.267480054 | 0.830769379 | 0.343988473 | 0.516676027 | 65.19763183 | 54.02877374 | ENSG00000147099 |
| NA | -0.267555545 | 0.830725909 | 0.710921941 | 0.825872368 | 11.29713241 | 9.349725324 | ENSG00000286548 |
| NAXD | -0.267637686 | 0.830678612 | 0.00187108 | 0.007448412 | 793.9785089 | 659.6117939 | ENSG00000213995 |
| ANKH | -0.267716825 | 0.830633046 | 0.042726808 | 0.105186685 | 341.0707611 | 283.2616267 | ENSG00000154122 |
| FBLL1 | -0.267754629 | 0.830611281 | 0.114621671 | 0.23023393 | 186.9386347 | 155.3073378 | ENSG00000188573 |
| NA | -0.26786065 | 0.830550243 | 0.782929267 | 0.872832206 | 6.079912612 | 5.079922744 | ENSG00000261461 |
| NRDE2 | -0.267866409 | 0.830546928 | 0.090481728 | 0.191432663 | 199.3993777 | 165.5701688 | ENSG00000119720 |
| HNRNPA0 | -0.267964447 | 0.83049049 | 1.89E-09 | 2.23E-08 | 13289.92411 | 11037.27573 | ENSG00000177733 |
| ARHGAP1 | -0.268127725 | 0.830396504 | 0.001742447 | 0.007001363 | 941.3323056 | 781.9371023 | ENSG00000175220 |
| CANT1 | -0.268202419 | 0.830353512 | 0.000267758 | 0.00131333 | 1090.624312 | 905.7318499 | ENSG00000171302 |
| RGS12 | -0.26830339 | 0.830295399 | 0.004162752 | 0.014962482 | 619.108 | 514.2537569 | ENSG00000159788 |
| UBA5 | -0.268348762 | 0.830269288 | 0.001831543 | 0.00730536 | 813.1255698 | 675.4462971 | ENSG00000081307 |
| MAGOH | -0.268443889 | 0.830214544 | 0.000416294 | 0.001950628 | 1563.279016 | 1297.333356 | ENSG00000162385 |
| DOLPP1 | -0.268569845 | 0.830142064 | 0.001224773 | 0.005125183 | 805.8413201 | 668.8418501 | ENSG00000167130 |
| TBCCD1 | -0.26860587 | 0.830121336 | 0.068975864 | 0.154202502 | 291.3712748 | 241.824707 | ENSG00000113838 |
| PCDHGB1 | -0.268631723 | 0.83010646 | 0.556019414 | 0.712229118 | 26.68667121 | 22.09216217 | ENSG00000254221 |
| NA | -0.268901379 | 0.829951318 | 0.049129278 | 0.117645592 | 309.9817172 | 257.2898256 | ENSG00000210196 |
| NA | -0.268960467 | 0.829917327 | 0.675909333 | 0.801214633 | 12.79416369 | 10.6352265 | ENSG00000233270 |
| SLC39A6 | -0.269009275 | 0.82988925 | 0.000393617 | 0.001860638 | 1105.828148 | 918.0042609 | ENSG00000141424 |
| TNIP2 | -0.269018469 | 0.829883962 | 0.000699626 | 0.00308139 | 1443.082968 | 1198.023802 | ENSG00000168884 |
| ZNF598 | -0.269022394 | 0.829881704 | 0.001051492 | 0.004456137 | 1399.315627 | 1161.68699 | ENSG00000167962 |
| DERA | -0.269029586 | 0.829877567 | 0.042128229 | 0.103951862 | 288.9737173 | 239.6408937 | ENSG00000023697 |
| STK38 | -0.269078418 | 0.829849478 | 0.006892513 | 0.02310399 | 557.6981019 | 463.1573798 | ENSG00000112079 |
| NPIPB15 | -0.269140501 | 0.829813768 | 0.608208609 | 0.752015913 | 16.99318797 | 14.10995082 | ENSG00000196436 |
| CALHM2 | -0.269212499 | 0.829772357 | 0.403812905 | 0.576115175 | 48.24408007 | 39.96990365 | ENSG00000138172 |
| IFT52 | -0.269244654 | 0.829753863 | 0.009837639 | 0.03100946 | 559.0074158 | 463.8050977 | ENSG00000101052 |
| NOC2L | -0.269396794 | 0.829666366 | 2.84E-06 | 2.06E-05 | 5488.904322 | 4553.865464 | ENSG00000188976 |
| THAP11 | -0.26961553 | 0.829540584 | 0.001827304 | 0.007293715 | 957.6320095 | 794.6195894 | ENSG00000168286 |
| PIK3AP1 | -0.269641863 | 0.829525443 | 0.198842922 | 0.350308566 | 114.7603128 | 95.24546823 | ENSG00000155629 |
| CHM | -0.269679047 | 0.829504063 | 0.004673357 | 0.016560903 | 653.9925626 | 542.3069857 | ENSG00000188419 |
| SMARCA5-AS1 | -0.269697266 | 0.829493588 | 0.311814452 | 0.482815472 | 71.24173826 | 59.06298297 | ENSG00000245112 |
| CTDSPL2 | -0.269732327 | 0.82947343 | 0.002583387 | 0.009862681 | 730.8031631 | 606.2258797 | ENSG00000137770 |
| XKRX | -0.269770504 | 0.82945148 | 0.431999292 | 0.603017247 | 48.9964002 | 40.54746454 | ENSG00000182489 |
| MCM3AP | -0.269808899 | 0.829429406 | 3.70E-06 | 2.63E-05 | 1885.404463 | 1563.662593 | ENSG00000160294 |
| NA | -0.269811364 | 0.829427988 | 0.592923649 | 0.741337099 | 23.0737763 | 19.16890087 | ENSG00000235316 |
| SDF4 | -0.269915052 | 0.829368379 | 9.32E-06 | 6.17E-05 | 2904.163252 | 2408.882528 | ENSG00000078808 |
| MTMR14 | -0.269968578 | 0.829337609 | 3.36E-05 | 0.000201113 | 1643.6011 | 1363.098411 | ENSG00000163719 |
| ADCY2 | -0.27002471 | 0.829305342 | 6.22E-05 | 0.000351157 | 2001.619616 | 1659.373974 | ENSG00000078295 |
| CCNDBP1 | -0.270073649 | 0.829277211 | 0.013112633 | 0.039567621 | 478.1601149 | 396.7580438 | ENSG00000166946 |
| SRI | -0.270209859 | 0.829198919 | 0.000701893 | 0.003089258 | 906.934889 | 751.7551754 | ENSG00000075142 |
| CCDC144B | -0.270214161 | 0.829196447 | 0.802049313 | 0.885936266 | 4.406402853 | 3.651641249 | ENSG00000154874 |
| EXOSC5 | -0.27022706 | 0.829189033 | 0.0019052 | 0.007563947 | 1206.114104 | 999.4485475 | ENSG00000077348 |
| ISL2 | -0.270228037 | 0.829188471 | 0.778394145 | 0.869988497 | 5.892070362 | 4.907306965 | ENSG00000159556 |
| NA | -0.270257133 | 0.829171748 | 0.655457216 | 0.786295949 | 12.99508457 | 10.75703412 | ENSG00000214535 |
| VPS16 | -0.270285398 | 0.829155504 | 0.013433954 | 0.04037931 | 428.3605436 | 355.0001062 | ENSG00000215305 |

| | | | | | | | |
|---|---|---|---|---|---|---|---|
| NLK | -0.270465969 | 0.829051731 | 0.003650935 | 0.013328953 | 667.2441348 | 552.9354695 | ENSG00000087095 |
| NA | -0.270491252 | 0.829037202 | 0.76021198 | 0.857637004 | 6.820393404 | 5.641345125 | ENSG00000276007 |
| ZNF582-AS1 | -0.27070542 | 0.828914141 | 0.549144586 | 0.706566278 | 26.33650516 | 21.81090268 | ENSG00000267454 |
| GUSBP1 | -0.270843392 | 0.828834872 | 0.5473699 | 0.705176488 | 26.46900491 | 21.88646376 | ENSG00000215190 |
| TENT4A | -0.270888098 | 0.828809188 | 0.001642328 | 0.006639177 | 741.3603655 | 614.7022942 | ENSG00000112941 |
| MCM10 | -0.270964939 | 0.828765046 | 0.000446612 | 0.002077046 | 1136.910696 | 941.8059946 | ENSG00000065328 |
| PRPF8 | -0.270968682 | 0.828762895 | 2.14E-11 | 3.16E-10 | 11554.12777 | 9575.380745 | ENSG00000174231 |
| NA | -0.271002997 | 0.828743183 | 0.652532734 | 0.784094076 | 13.85566221 | 11.42170246 | ENSG00000282381 |
| NCL | -0.271132211 | 0.82866896 | 7.32E-13 | 1.27E-11 | 36994.37791 | 30656.43884 | ENSG00000115053 |
| NDE1 | -0.271168049 | 0.828648376 | 0.021094051 | 0.058693976 | 438.690868 | 363.77493 | ENSG00000072864 |
| RILPL2 | -0.271225085 | 0.828615616 | 0.198065581 | 0.349353856 | 113.3739665 | 93.80104822 | ENSG00000150977 |
| NA | -0.271443789 | 0.828490012 | 0.528439572 | 0.689817823 | 26.23594463 | 21.70335557 | ENSG00000206228 |
| C11orf80 | -0.27153171 | 0.828439524 | 0.082115758 | 0.177231493 | 212.6799861 | 176.1048329 | ENSG00000173715 |
| VSTM2B | -0.27156916 | 0.828418019 | 0.529693237 | 0.690716585 | 26.88547691 | 22.30887392 | ENSG00000187135 |
| TMEM176A | -0.271655243 | 0.82836859 | 0.720385054 | 0.832149671 | 8.270454896 | 6.859366591 | ENSG00000002933 |
| OXNAD1 | -0.271699549 | 0.828343152 | 0.012628169 | 0.038321536 | 436.74878 | 361.8739816 | ENSG00000154814 |
| LIN28B | -0.27176419 | 0.828306038 | 3.94E-05 | 0.000231702 | 2355.05651 | 1951.506685 | ENSG00000187772 |
| LINGO1 | -0.271795775 | 0.828287904 | 0.000313786 | 0.001515965 | 2180.418235 | 1806.300456 | ENSG00000169783 |
| RBM39 | -0.271803589 | 0.828283418 | 3.12E-10 | 4.07E-09 | 4941.94757 | 4093.058672 | ENSG00000131051 |
| ZNF577 | -0.27188443 | 0.828237006 | 0.136637368 | 0.262746359 | 152.5663986 | 126.2609049 | ENSG00000161551 |
| FRMD5 | -0.271930521 | 0.828210546 | 0.038455124 | 0.09651806 | 339.2140398 | 281.2458191 | ENSG00000171877 |
| CPSF7 | -0.271955001 | 0.828196493 | 1.25E-05 | 8.05E-05 | 2175.939165 | 1801.883063 | ENSG00000149532 |
| LRIG2-DT | -0.272242253 | 0.828031609 | 0.807266444 | 0.888970029 | 4.746281178 | 3.906269825 | ENSG00000238198 |
| PSMC3IP | -0.272342176 | 0.82797426 | 0.042931162 | 0.105581979 | 329.50669 | 272.7831563 | ENSG00000131470 |
| IKBKG | -0.272572475 | 0.827842101 | 0.195475517 | 0.345860608 | 109.3666145 | 90.45028209 | ENSG00000269335 |
| PELO | -0.272596798 | 0.827828144 | 0.001267394 | 0.005281765 | 890.3931544 | 737.0679731 | ENSG00000152684 |
| POU3F1 | -0.272610202 | 0.827820452 | 0.00102484 | 0.004356381 | 1127.880589 | 933.9961017 | ENSG00000185668 |
| CDIPT | -0.272673477 | 0.827784146 | 0.002742198 | 0.010382678 | 955.6250664 | 791.5371551 | ENSG00000103502 |
| WASHC2C | -0.272908351 | 0.827649392 | 0.000594681 | 0.002672253 | 930.0149228 | 769.3132501 | ENSG00000172661 |
| PSMC5 | -0.272938089 | 0.827632332 | 7.33E-05 | 0.000408237 | 1808.324847 | 1496.588963 | ENSG00000087191 |
| PPM1E | -0.272938739 | 0.827631959 | 0.020949612 | 0.058367845 | 369.9766473 | 306.3204047 | ENSG00000175175 |
| SHMT2 | -0.273018994 | 0.827585921 | 2.70E-09 | 3.10E-08 | 7971.385789 | 6595.930264 | ENSG00000182199 |
| FAM204A | -0.273072364 | 0.827555306 | 0.11288158 | 0.227354204 | 181.3813364 | 150.2421965 | ENSG00000165669 |
| SFTPD | -0.273085631 | 0.827547696 | 0.765367827 | 0.861069997 | 6.330394027 | 5.237757081 | ENSG00000133661 |
| EIF1AD | -0.273108632 | 0.827534502 | 0.003453488 | 0.012692177 | 623.2803034 | 515.7774877 | ENSG00000175376 |
| DHX37 | -0.27318834 | 0.827488783 | 1.58E-05 | 1.00E-04 | 2703.368415 | 2237.262598 | ENSG00000150990 |
| THAP4 | -0.273226797 | 0.827466725 | 2.84E-05 | 0.000172027 | 1761.438514 | 1457.374058 | ENSG00000176946 |
| NA | -0.273250669 | 0.827453033 | 0.689098052 | 0.811166128 | 10.31733381 | 8.552496493 | ENSG00000271992 |
| OGFOD1 | -0.273269124 | 0.827442449 | 0.000418713 | 0.001960534 | 1169.006422 | 966.8697904 | ENSG00000087263 |
| HNRNPA1L2 | -0.273329179 | 0.827408005 | 0.199130586 | 0.350591233 | 109.6136293 | 90.71915907 | ENSG00000139675 |
| C1orf174 | -0.273361512 | 0.827389462 | 0.004544469 | 0.016159879 | 784.3067779 | 649.1115147 | ENSG00000198912 |
| ILF3-DT | -0.273600253 | 0.827252555 | 0.016531421 | 0.047948804 | 400.4905622 | 331.1516234 | ENSG00000267100 |
| CUL7 | -0.273646577 | 0.827225993 | 0.01158486 | 0.035666062 | 596.8194327 | 493.1379153 | ENSG00000044090 |
| IZUMO4 | -0.273710159 | 0.827189536 | 0.73562445 | 0.842640381 | 9.403165446 | 7.728115815 | ENSG00000099840 |
| MAZ | -0.273720839 | 0.827183412 | 0.02870812 | 0.07582261 | 528.0909196 | 436.6123553 | ENSG00000103495 |
| ABHD14A | -0.273729862 | 0.827178239 | 0.753009966 | 0.853786015 | 7.470801564 | 6.118876709 | ENSG00000248487 |
| SAC3D1 | -0.273734341 | 0.827175671 | 0.013822529 | 0.041373222 | 491.5518597 | 406.7915201 | ENSG00000168061 |
| SNRPF | -0.2737924 | 0.827142383 | 5.96E-06 | 4.09E-05 | 2374.300561 | 1963.606845 | ENSG00000139343 |
| C1orf109 | -0.273800425 | 0.827137782 | 0.003509407 | 0.012887869 | 758.0922718 | 627.0035414 | ENSG00000116922 |
| AMN1 | -0.273822242 | 0.827125274 | 0.371034917 | 0.544721049 | 69.60914092 | 57.53144341 | ENSG00000151743 |
| KPNA3 | -0.273842246 | 0.827113806 | 0.000240977 | 0.001195345 | 1556.040262 | 1287.753391 | ENSG00000102753 |
| NA | -0.273846693 | 0.827111256 | 0.376355434 | 0.549729134 | 56.70775887 | 47.05216113 | ENSG00000243199 |
| C17orf58 | -0.273856302 | 0.827105747 | 0.025615202 | 0.069043295 | 398.2658164 | 329.2713689 | ENSG00000186665 |
| ANKRD52 | -0.273865691 | 0.827100364 | 2.48E-05 | 0.000151708 | 1645.110083 | 1360.721245 | ENSG00000139645 |
| GNL2 | -0.273876527 | 0.827094152 | 3.63E-05 | 0.000214922 | 2787.652682 | 2305.461169 | ENSG00000134697 |
| SMPDL3A | -0.273888236 | 0.82708744 | 0.480808172 | 0.648892574 | 33.79526723 | 27.95290266 | ENSG00000172594 |
| TMEM178B | -0.273905858 | 0.827077337 | 0.017390455 | 0.050042541 | 435.0716035 | 359.6382757 | ENSG00000261115 |
| ZEB1 | -0.273909119 | 0.827075468 | 0.724065256 | 0.834405189 | 8.009485591 | 6.657436282 | ENSG00000148516 |
| VCP | -0.273950364 | 0.827051823 | 1.14E-14 | 2.39E-13 | 15157.17726 | 12535.70268 | ENSG00000165280 |
| NA | -0.273952727 | 0.827050468 | 0.336074035 | 0.50888573 | 61.6230217 | 50.94580009 | ENSG00000286132 |
| MAST1 | -0.27397282 | 0.827038949 | 0.019332544 | 0.054635819 | 379.7238335 | 314.2757126 | ENSG00000105613 |
| GPD1L | -0.27430952 | 0.826845956 | 0.104121342 | 0.213681363 | 180.8182099 | 149.6495816 | ENSG00000152642 |
| SLC7A5 | -0.27434038 | 0.826828269 | 7.37E-07 | 5.93E-06 | 9999.356882 | 8267.624035 | ENSG00000103257 |
| TRO | -0.274348931 | 0.826823368 | 0.000280188 | 0.001368729 | 1224.771052 | 1012.66602 | ENSG00000067445 |
| PACRGL | -0.274458907 | 0.826760342 | 0.067733466 | 0.151970958 | 223.9618126 | 185.1491094 | ENSG00000163138 |
| DDX27 | -0.274459112 | 0.826760225 | 2.03E-05 | 0.000126069 | 1875.864582 | 1551.417589 | ENSG00000124228 |
| TMEM145 | -0.274485094 | 0.826745336 | 0.176864685 | 0.320456151 | 141.17186 | 116.652795 | ENSG00000167619 |
| SDK2 | -0.274535193 | 0.826716626 | 0.045159431 | 0.1099824 | 303.4635178 | 251.0246673 | ENSG00000069188 |
| DM1-AS | -0.274673418 | 0.826637423 | 0.633557484 | 0.770323801 | 15.08247559 | 12.44586288 | ENSG00000267395 |
| FOXI2 | -0.274723953 | 0.826608468 | 0.623586113 | 0.763008988 | 14.67975794 | 12.13476792 | ENSG00000186766 |
| NA | -0.274860436 | 0.826530272 | 0.789381024 | 0.877236554 | 4.343563529 | 3.577970282 | ENSG00000260364 |
| SLC35G1 | -0.274970908 | 0.826466984 | 0.101103909 | 0.208844149 | 175.5215175 | 145.1485462 | ENSG00000176273 |
| PRDM11 | -0.275125882 | 0.82637821 | 0.033845199 | 0.086615512 | 303.0341305 | 250.3651178 | ENSG00000019485 |
| AEBP1 | -0.275211268 | 0.826329302 | 1.81E-05 | 0.000113586 | 1942.651745 | 1604.895735 | ENSG00000106624 |
| ORC6 | -0.27532394 | 0.82626477 | 8.27E-05 | 0.000455793 | 1281.694099 | 1059.00242 | ENSG00000091651 |
| DUS1L | -0.275324998 | 0.826264163 | 1.51E-05 | 9.61E-05 | 1935.883764 | 1599.534523 | ENSG00000169718 |
| B3GAT1 | -0.275346138 | 0.826252056 | 0.000575727 | 0.002593727 | 1302.124869 | 1075.69501 | ENSG00000109956 |

| | | | | | | | |
|---|---|---|---|---|---|---|---|
| ARHGEF26 | -0.275366089 | 0.82624063 | 0.018292724 | 0.052078651 | 363.7331716 | 300.4916503 | ENSG00000114790 |
| AVEN | -0.27539797 | 0.826222372 | 0.020013642 | 0.05624562 | 420.3309212 | 347.4263355 | ENSG00000169857 |
| CFL1 | -0.275416735 | 0.826211625 | 6.25E-11 | 8.79E-10 | 9486.835791 | 7838.493738 | ENSG00000172757 |
| LOC101928053 | -0.27549285 | 0.826168037 | 0.823065203 | 0.899130802 | 3.573058527 | 2.98267176 | ENSG00000246308 |
| PPP6R2 | -0.275512932 | 0.826156536 | 0.000360471 | 0.001717425 | 1078.245306 | 890.9956748 | ENSG00000100239 |
| PCDHB13 | -0.275548524 | 0.826136155 | 0.413705789 | 0.585917205 | 44.35558563 | 36.64843994 | ENSG00000187372 |
| LIN54 | -0.275671159 | 0.826065933 | 0.106311569 | 0.216838282 | 193.4644799 | 159.9070453 | ENSG00000189308 |
| ATG101 | -0.275675096 | 0.826063679 | 0.005360793 | 0.018619924 | 867.4206655 | 716.458705 | ENSG00000123395 |
| RIC8B | -0.275721519 | 0.826037098 | 0.027748387 | 0.073772621 | 334.6419217 | 276.5595493 | ENSG00000111785 |
| FAM83H | -0.275772929 | 0.826007663 | 8.14E-05 | 0.000449015 | 2295.443533 | 1896.066435 | ENSG00000180921 |
| RUFY1 | -0.275811233 | 0.825985733 | 0.013219455 | 0.039827661 | 569.8588399 | 470.1466596 | ENSG00000176783 |
| KCNJ8 | -0.275845803 | 0.82596594 | 0.661967805 | 0.791399412 | 14.98123935 | 12.34906864 | ENSG00000121361 |
| ZNF512 | -0.275917162 | 0.825925087 | 0.013320317 | 0.040072148 | 461.5313598 | 381.4690608 | ENSG00000243943 |
| NCOA6 | -0.275958612 | 0.825901358 | 0.030580951 | 0.079818309 | 320.1276788 | 264.5199765 | ENSG00000198646 |
| TTC25 | -0.276188538 | 0.825769743 | 0.525587303 | 0.687835929 | 26.56445919 | 21.96390283 | ENSG00000204815 |
| BCCIP | -0.276194964 | 0.825766064 | 8.57E-07 | 6.82E-06 | 2233.485988 | 1844.078937 | ENSG00000107949 |
| PPID | -0.276211633 | 0.825756524 | 0.000137963 | 0.000720672 | 1190.912045 | 983.4668591 | ENSG00000171497 |
| LRRC32 | -0.276296576 | 0.825707906 | 0.3907356 | 0.563986567 | 59.40851257 | 48.99886565 | ENSG00000137507 |
| SLC45A4 | -0.276306898 | 0.825701999 | 0.000696581 | 0.003069379 | 900.6035822 | 743.3823221 | ENSG00000022567 |
| TNC | -0.276562225 | 0.825555879 | 0.148622256 | 0.28031294 | 137.934174 | 113.8975757 | ENSG00000041982 |
| STARD3 | -0.276605068 | 0.825531363 | 0.000161233 | 0.000828651 | 1468.627642 | 1212.567188 | ENSG00000131748 |
| SNTA1 | -0.276818781 | 0.825409083 | 0.06211353 | 0.141852978 | 248.656547 | 205.366893 | ENSG00000101400 |
| HSPB8 | -0.276869072 | 0.82538031 | 0.089275163 | 0.189503431 | 190.9880503 | 157.4575596 | ENSG00000152137 |
| TARDBP | -0.276872879 | 0.825378132 | 2.75E-07 | 2.37E-06 | 2897.186778 | 2392.070182 | ENSG00000120948 |
| GTF3A | -0.27694808 | 0.82533511 | 0.000185932 | 0.000943404 | 1242.187733 | 1025.411643 | ENSG00000122034 |
| ME1 | -0.276970156 | 0.825322481 | 0.092246258 | 0.194292233 | 210.8493852 | 174.1691844 | ENSG00000065833 |
| VPS26A | -0.276978617 | 0.825317641 | 0.002607931 | 0.009944574 | 895.5571852 | 739.5978172 | ENSG00000122958 |
| FBXW5 | -0.276992186 | 0.825309879 | 5.23E-05 | 0.000299731 | 1839.340339 | 1518.060244 | ENSG00000159069 |
| SERPING1 | -0.277093551 | 0.825251894 | 0.005095718 | 0.017821157 | 722.486001 | 596.4801331 | ENSG00000149131 |
| MERTK | -0.277134484 | 0.825228479 | 0.075160903 | 0.165121253 | 231.9074197 | 191.1744085 | ENSG00000153208 |
| RETREG1 | -0.277145366 | 0.825222255 | 0.490903411 | 0.65741185 | 34.11416977 | 28.24328538 | ENSG00000154153 |
| COPS3 | -0.277364921 | 0.825096679 | 2.24E-05 | 0.000138212 | 1689.015706 | 1392.985653 | ENSG00000141030 |
| TUBA1B | -0.27738546 | 0.825084933 | 4.72E-09 | 5.26E-08 | 4436.408225 | 3660.060568 | ENSG00000123416 |
| GPATCH3 | -0.277418665 | 0.825065942 | 0.03230882 | 0.083425179 | 374.0415323 | 308.6913974 | ENSG00000198746 |
| MDC1 | -0.277691172 | 0.824910113 | 4.28E-05 | 0.000249208 | 1556.224009 | 1283.401128 | ENSG00000137337 |
| MZT1 | -0.277716992 | 0.824895349 | 0.00856386 | 0.027644816 | 605.8271545 | 499.9725411 | ENSG00000204899 |
| ELAVL1 | -0.277799153 | 0.824848373 | 3.23E-07 | 2.75E-06 | 3228.989362 | 2663.21899 | ENSG00000066044 |
| SH2B1 | -0.277801777 | 0.824846873 | 0.000670839 | 0.002970511 | 1240.311163 | 1022.710043 | ENSG00000178188 |
| ATP6V0B | -0.27800651 | 0.824729827 | 1.47E-05 | 9.33E-05 | 1831.402502 | 1510.101398 | ENSG00000117410 |
| ATP6V0A2 | -0.278017963 | 0.82472328 | 0.000633916 | 0.002828264 | 808.6910067 | 666.9201711 | ENSG00000185344 |
| NKAP | -0.278047889 | 0.824706173 | 0.114601139 | 0.230216659 | 163.3414803 | 134.7598791 | ENSG00000101882 |
| NA | -0.278053095 | 0.824703197 | 0.807241979 | 0.888970029 | 4.381960449 | 3.603244129 | ENSG00000275092 |
| ZNF184 | -0.278087139 | 0.824683736 | 0.107154901 | 0.218204506 | 169.947674 | 140.2057456 | ENSG00000096654 |
| SCGB3A2 | -0.278232522 | 0.824600635 | 0.000283211 | 0.001381748 | 1087.958413 | 896.8930114 | ENSG00000164265 |
| PYDC1 | -0.278234898 | 0.824599277 | 0.683668415 | 0.807039982 | 9.985152477 | 8.262374239 | ENSG00000169900 |
| RFFL | -0.278295252 | 0.824564782 | 0.078395955 | 0.170634816 | 249.2256556 | 205.5502617 | ENSG00000092871 |
| DNAAF5 | -0.278368519 | 0.824522908 | 8.91E-05 | 0.000486788 | 1491.841039 | 1230.668323 | ENSG00000164818 |
| CCDC22 | -0.278409016 | 0.824499763 | 0.06803652 | 0.152509006 | 292.7423903 | 241.2270401 | ENSG00000101997 |
| RMDN3 | -0.278532658 | 0.824429105 | 0.002550215 | 0.009755344 | 870.9671153 | 718.3257593 | ENSG00000137824 |
| PTGR1 | -0.278680496 | 0.824344627 | 0.012176212 | 0.037078343 | 610.9534243 | 503.8333215 | ENSG00000106853 |
| WAKMAR2 | -0.278691642 | 0.824338258 | 0.809839517 | 0.890431127 | 4.354727141 | 3.577970282 | ENSG00000237499 |
| SAMD4B | -0.27878209 | 0.824286579 | 6.74E-08 | 6.35E-07 | 3305.897909 | 2725.097911 | ENSG00000179134 |
| ERCC3 | -0.278889935 | 0.824224964 | 4.05E-06 | 2.86E-05 | 1687.883071 | 1391.422061 | ENSG00000163161 |
| GMNN | -0.278904834 | 0.824216452 | 1.23E-06 | 9.60E-06 | 2005.173754 | 1652.590471 | ENSG00000112312 |
| ATG13 | -0.278945844 | 0.824193023 | 0.000204827 | 0.0010306 | 1110.244124 | 915.0347715 | ENSG00000175224 |
| LINC01572 | -0.278962126 | 0.824183721 | 0.458993978 | 0.628912056 | 38.00700672 | 31.30151821 | ENSG00000261008 |
| SH3GL2 | -0.278973244 | 0.82417737 | 0.205708391 | 0.359644785 | 115.0120335 | 94.92577775 | ENSG00000107295 |
| CHCHD3 | -0.278997577 | 0.824163469 | 2.98E-06 | 2.15E-05 | 3031.069959 | 2498.16722 | ENSG00000106554 |
| CSNK1G2 | -0.279012555 | 0.824154913 | 2.32E-06 | 1.71E-05 | 4437.690082 | 3657.554738 | ENSG00000133275 |
| SARS2 | -0.279031234 | 0.824144242 | 0.673963117 | 0.800111161 | 12.58957602 | 10.35291304 | ENSG00000104835 |
| NAV2 | -0.279097034 | 0.824106655 | 7.86E-05 | 0.000434851 | 1320.008457 | 1087.864902 | ENSG00000166833 |
| SNRPG | -0.279098575 | 0.824105774 | 2.17E-05 | 0.000134642 | 1960.157322 | 1615.438899 | ENSG00000143977 |
| TTF1 | -0.279135335 | 0.824084776 | 0.001940886 | 0.007683489 | 653.5217361 | 538.6160584 | ENSG00000125482 |
| OGFR | -0.279245395 | 0.824021911 | 0.000135901 | 0.000710673 | 1334.102287 | 1099.686746 | ENSG00000060491 |
| PKNOX1 | -0.27926318 | 0.824011753 | 0.004189016 | 0.015037273 | 616.7719914 | 508.170095 | ENSG00000160199 |
| FANCA | -0.279276626 | 0.824004073 | 0.000689812 | 0.003045116 | 1262.644645 | 1039.797819 | ENSG00000187741 |
| ZFP14 | -0.279310459 | 0.823984749 | 0.140679409 | 0.269017681 | 167.5626316 | 138.3883849 | ENSG00000142065 |
| FAM161A | -0.279314103 | 0.823982668 | 0.040915246 | 0.101491065 | 303.8979746 | 250.2599938 | ENSG00000170264 |
| HDDC3 | -0.279396881 | 0.823935392 | 0.163874659 | 0.302488379 | 124.6348172 | 102.5259 | ENSG00000184508 |
| NA | -0.279598942 | 0.823820001 | 0.194388939 | 0.344411923 | 120.5159412 | 99.2609027 | ENSG00000213866 |
| PBX3 | -0.279663753 | 0.823782993 | 0.007574842 | 0.02494399 | 569.2026129 | 468.8675675 | ENSG00000167081 |
| DDX52 | -0.279748259 | 0.823734741 | 0.003051421 | 0.011363894 | 927.6677152 | 764.3707221 | ENSG00000278053 |
| TIMM10B | -0.279768427 | 0.823723226 | 0.013688339 | 0.041009745 | 415.3305287 | 342.2767398 | ENSG00000132286 |
| GBA2 | -0.279784814 | 0.823713869 | 2.99E-08 | 2.96E-07 | 3091.820561 | 2546.471491 | ENSG00000070610 |
| TMEM229A | -0.279844348 | 0.823679879 | 0.734711354 | 0.841995884 | 9.475616779 | 7.831361775 | ENSG00000234224 |
| NA | -0.279931014 | 0.8236304 | 0.807977875 | 0.889499583 | 3.954124674 | 3.262574183 | ENSG00000213087 |
| TBPL1 | -0.280038684 | 0.823568934 | 0.042464871 | 0.104670386 | 307.0960243 | 253.1229964 | ENSG00000028839 |

| | | | | | | | |
|---|---|---|---|---|---|---|---|
| CEBPB | -0.28019069 | 0.823482166 | 0.021325295 | 0.059251946 | 423.1936112 | 348.613679 | ENSG00000172216 |
| TPRG1L | -0.280489187 | 0.823311803 | 0.00132171 | 0.005475006 | 735.311353 | 605.4671128 | ENSG00000158109 |
| ZFYVE27 | -0.280491943 | 0.82331023 | 0.011774956 | 0.036084301 | 593.2665252 | 487.9445389 | ENSG00000155256 |
| DHX33 | -0.280573147 | 0.82326389 | 6.87E-07 | 5.55E-06 | 2307.272764 | 1899.763984 | ENSG00000005100 |
| GTPBP8 | -0.280717885 | 0.823181301 | 0.113708757 | 0.228805219 | 173.689514 | 143.196214 | ENSG00000163607 |
| RGS8 | -0.280775984 | 0.82314815 | 0.6962199 | 0.816684112 | 9.565689411 | 7.915252572 | ENSG00000135824 |
| SART3 | -0.280796767 | 0.823136293 | 7.42E-07 | 5.97E-06 | 2734.836984 | 2250.44188 | ENSG00000075856 |
| TPRKB | -0.280812069 | 0.823127562 | 0.012188611 | 0.037110243 | 436.9513405 | 359.6310062 | ENSG00000144034 |
| PDCL3 | -0.280948445 | 0.823049757 | 0.000770999 | 0.003359839 | 779.5112357 | 641.798818 | ENSG00000115539 |
| NEIL3 | -0.281023145 | 0.823007142 | 0.061694976 | 0.14105476 | 222.704975 | 183.1338712 | ENSG00000109674 |
| ETF1 | -0.281092421 | 0.822967623 | 1.23E-08 | 1.30E-07 | 4892.70113 | 4027.388939 | ENSG00000120705 |
| KIAA0100 | -0.281094905 | 0.822966206 | 6.70E-08 | 6.31E-07 | 2672.028013 | 2199.35436 | ENSG00000007202 |
| NA | -0.281115835 | 0.822954267 | 0.495062121 | 0.661429731 | 31.28629796 | 25.69970745 | ENSG00000274021 |
| NA | -0.281185033 | 0.822914796 | 0.607308681 | 0.751432991 | 18.82922169 | 15.58447886 | ENSG00000236166 |
| EWSR1 | -0.281236075 | 0.822885682 | 9.32E-10 | 1.13E-08 | 10072.43898 | 8287.746787 | ENSG00000182944 |
| LOC101928725 | -0.281270308 | 0.822866156 | 0.79029834 | 0.87763484 | 4.365215031 | 3.573669135 | ENSG00000245975 |
| SLCO4A1 | -0.281373156 | 0.822807497 | 0.005475849 | 0.01897483 | 584.6795214 | 481.1515738 | ENSG00000101187 |
| KCNK17 | -0.281459756 | 0.822758108 | 0.393906162 | 0.566676241 | 54.42522893 | 44.61425231 | ENSG00000124780 |
| DHRS11 | -0.281549329 | 0.822707027 | 0.001694085 | 0.006825521 | 726.6580418 | 597.5368247 | ENSG00000278535 |
| PRORP | -0.281696764 | 0.822622955 | 0.249349981 | 0.413425912 | 90.47108686 | 74.43316112 | ENSG00000100890 |
| SSNA1 | -0.281736279 | 0.822600424 | 8.03E-06 | 5.38E-05 | 2104.994792 | 1731.257148 | ENSG00000176101 |
| CDCP1 | -0.281759452 | 0.822587211 | 0.008287343 | 0.026896219 | 457.9424317 | 376.6190379 | ENSG00000163814 |
| PPP1R13B | -0.281795569 | 0.822566619 | 0.039714411 | 0.099098526 | 276.798298 | 227.6954131 | ENSG00000088808 |
| ACSM3 | -0.281840785 | 0.822540839 | 0.600601212 | 0.746663049 | 19.26909696 | 15.81813474 | ENSG00000005187 |
| CMTM7 | -0.281841764 | 0.822540281 | 0.026113656 | 0.070151585 | 363.7229591 | 299.4551073 | ENSG00000153551 |
| ARHGAP45 | -0.281886968 | 0.822514508 | 7.92E-05 | 0.00043778 | 1886.28667 | 1551.527899 | ENSG00000180448 |
| ATP6V1A | -0.281914315 | 0.822498917 | 0.000609905 | 0.002732391 | 913.6850726 | 751.581512 | ENSG00000114573 |
| ZNF630 | -0.281972728 | 0.822465616 | 0.564606996 | 0.718885458 | 30.30805097 | 24.96619022 | ENSG00000221994 |
| RPS19BP1 | -0.282007959 | 0.822445531 | 0.001186655 | 0.004978635 | 1383.059341 | 1137.732602 | ENSG00000187051 |
| ST7 | -0.282020127 | 0.822438595 | 0.078608218 | 0.171038927 | 212.6602496 | 174.7994435 | ENSG00000004866 |
| RNF130 | -0.282060439 | 0.822415615 | 0.000370515 | 0.001760975 | 1104.902979 | 909.0038383 | ENSG00000113269 |
| BABAM1 | -0.282156198 | 0.822361028 | 0.059111743 | 0.136288352 | 234.904912 | 193.3654975 | ENSG00000105393 |
| PRPS1 | -0.282157003 | 0.82236057 | 1.26E-09 | 1.51E-08 | 7002.825503 | 5758.485915 | ENSG00000147224 |
| RCE1 | -0.282562855 | 0.82212926 | 0.000630324 | 0.002816022 | 869.8608743 | 714.9163941 | ENSG00000173653 |
| HSBP1 | -0.282594606 | 0.822111166 | 2.18E-05 | 0.000134831 | 3538.223617 | 2908.843042 | ENSG00000230989 |
| MED12 | -0.282714512 | 0.822042841 | 6.60E-05 | 0.000371026 | 1257.177797 | 1033.738843 | ENSG00000184634 |
| LSM3 | -0.282722329 | 0.822038387 | 6.61E-05 | 0.000371582 | 1633.647961 | 1343.109587 | ENSG00000170860 |
| SNRNP25 | -0.282722596 | 0.822038235 | 2.45E-05 | 0.0001499 | 1466.017874 | 1205.228642 | ENSG00000161981 |
| LINC00309 | -0.282753319 | 0.82202073 | 0.673864735 | 0.800111161 | 10.33898532 | 8.485277248 | ENSG00000230923 |
| CLTB | -0.282753488 | 0.822020633 | 0.003641274 | 0.013298716 | 748.6651892 | 615.4089263 | ENSG00000175416 |
| RRP1 | -0.282863082 | 0.821958191 | 5.99E-05 | 0.000339723 | 2153.148061 | 1769.481878 | ENSG00000160214 |
| SEMA3B | -0.282889822 | 0.821942956 | 0.238259091 | 0.400126763 | 89.93798464 | 73.80802714 | ENSG00000012171 |
| F13A1 | -0.282958602 | 0.821903771 | 0.520642421 | 0.683664349 | 25.27779754 | 20.73996268 | ENSG00000124491 |
| PER3 | -0.282983081 | 0.821889826 | 0.116632458 | 0.233470661 | 160.2531147 | 131.8815378 | ENSG00000049246 |
| NA | -0.283000708 | 0.821879784 | 0.513476989 | 0.677668782 | 27.61479409 | 22.75332287 | ENSG00000248121 |
| TXNDC12 | -0.283113851 | 0.821815331 | 0.12092008 | 0.239937179 | 162.9681111 | 133.9360498 | ENSG00000117862 |
| SAMD13 | -0.283201809 | 0.821765228 | 0.712707964 | 0.827048204 | 8.338876025 | 6.836243318 | ENSG00000203943 |
| MEMO1 | -0.283272404 | 0.821725018 | 0.201039866 | 0.353159464 | 120.8547435 | 99.35743647 | ENSG00000162959 |
| RFESD | -0.283346779 | 0.821682657 | 0.679043975 | 0.803872602 | 10.03961909 | 8.264524813 | ENSG00000175449 |
| NA | -0.283351829 | 0.821679781 | 0.135437933 | 0.260855948 | 149.8886479 | 123.2513296 | ENSG00000254615 |
| LINC02728 | -0.283365217 | 0.821672156 | 0.753238346 | 0.853907046 | 5.940955171 | 4.886334266 | ENSG00000251323 |
| PATZ1 | -0.283454502 | 0.821621306 | 5.34E-06 | 3.69E-05 | 2336.380863 | 1920.384296 | ENSG00000100105 |
| BRAT1 | -0.28345487 | 0.821621096 | 5.78E-05 | 0.000329098 | 1717.383885 | 1411.331234 | ENSG00000106009 |
| C12orf65 | -0.2838458 | 0.82139849 | 0.038576787 | 0.096785642 | 334.7007308 | 275.2078707 | ENSG00000130921 |
| GLA | -0.283960539 | 0.821333166 | 0.004125203 | 0.014845198 | 672.8303813 | 552.4415758 | ENSG00000102393 |
| NA | -0.283971259 | 0.821327063 | 0.760910748 | 0.857853384 | 5.638798044 | 4.663431258 | ENSG00000280374 |
| RRP7BP | -0.284100517 | 0.82125348 | 0.045301549 | 0.110286746 | 260.5609848 | 213.8435985 | ENSG00000182841 |
| ANAPC7 | -0.284134072 | 0.821234378 | 0.000203009 | 0.001022253 | 1032.906879 | 848.1958207 | ENSG00000196510 |
| ZNF670-ZNF695 | -0.284270351 | 0.821156807 | 0.688346548 | 0.810627756 | 9.652971139 | 7.963649691 | ENSG00000135747 |
| NA | -0.28431577 | 0.821130956 | 0.793871507 | 0.880229553 | 4.460869468 | 3.647340101 | ENSG00000270035 |
| NA | -0.284326255 | 0.821124988 | 0.456381857 | 0.62622252 | 33.94797902 | 27.90612308 | ENSG00000227896 |
| CCSER1 | -0.284492469 | 0.821030392 | 0.685186765 | 0.808079699 | 10.10524932 | 8.310771359 | ENSG00000184305 |
| VPS25 | -0.284568533 | 0.820987105 | 3.71E-06 | 2.63E-05 | 1946.004996 | 1597.171351 | ENSG00000131475 |
| ASB7 | -0.284573775 | 0.820984122 | 0.076577623 | 0.167432723 | 209.6231723 | 172.2133445 | ENSG00000183475 |
| ECI2 | -0.284643443 | 0.820944477 | 0.041274017 | 0.102210149 | 279.6675578 | 229.4044027 | ENSG00000198721 |
| NA | -0.284683766 | 0.820921532 | 0.832532029 | 0.904974086 | 3.975776176 | 3.264724756 | ENSG00000272097 |
| TTC26 | -0.284696784 | 0.820914125 | 0.110942107 | 0.224173477 | 156.1969148 | 128.125033 | ENSG00000105948 |
| CEP152 | -0.28469838 | 0.820913217 | 0.049589124 | 0.118482016 | 271.8605078 | 223.1250201 | ENSG00000103995 |
| ECPAS | -0.284705486 | 0.820909173 | 4.02E-10 | 5.14E-09 | 4665.5829 | 3830.03734 | ENSG00000136813 |
| RTF2 | -0.284739094 | 0.82089005 | 0.000989751 | 0.004225449 | 988.421071 | 811.0477799 | ENSG00000022277 |
| RASGEF1A | -0.285073325 | 0.820699896 | 0.008654757 | 0.027896228 | 470.9813829 | 386.6931025 | ENSG00000198915 |
| RAB5C | -0.285155178 | 0.820653334 | 0.021439489 | 0.059534934 | 389.8051287 | 319.7310176 | ENSG00000108774 |
| GOLGA2P7 | -0.285235878 | 0.82060743 | 0.106137644 | 0.216620957 | 204.9570394 | 168.2578657 | ENSG00000225151 |
| METTL17 | -0.285325749 | 0.820556313 | 0.000781902 | 0.003403354 | 780.4224499 | 640.0941539 | ENSG00000165792 |
| SNORA14B | -0.285533964 | 0.820437895 | 0.809055814 | 0.890279831 | 3.940845881 | 3.232999189 | ENSG00000207181 |
| PHB | -0.285536336 | 0.820436547 | 1.36E-06 | 1.05E-05 | 5969.50905 | 4897.050608 | ENSG00000167085 |
| PPA1 | -0.285600832 | 0.82039987 | 1.58E-07 | 1.41E-06 | 2778.111424 | 2279.460047 | ENSG00000180817 |

| | | | | | | | |
|---|---|---|---|---|---|---|---|
| MSRB1 | -0.285644061 | 0.820375287 | 0.000254063 | 0.001252845 | 969.1269454 | 795.3369133 | ENSG00000198736 |
| DPP9-AS1 | -0.285703597 | 0.820341434 | 0.757278054 | 0.855942967 | 5.671613158 | 4.684403957 | ENSG00000205790 |
| GTF3C5 | -0.285729346 | 0.820326792 | 9.10E-08 | 8.44E-07 | 2968.547148 | 2435.004851 | ENSG00000148308 |
| GPR68 | -0.285732486 | 0.820325007 | 0.627324543 | 0.765592317 | 14.89271832 | 12.18316504 | ENSG00000119714 |
| MTMR8 | -0.285779549 | 0.820298247 | 0.187475787 | 0.334591119 | 111.2144876 | 91.21928089 | ENSG00000102043 |
| CLMP | -0.285979644 | 0.820184484 | 0.010090308 | 0.031686819 | 486.3063547 | 399.3018822 | ENSG00000166250 |
| SRRM1 | -0.286084843 | 0.820124679 | 5.56E-07 | 4.55E-06 | 3233.842887 | 2652.216662 | ENSG00000133226 |
| NA | -0.286095324 | 0.820118721 | 0.593768985 | 0.741925624 | 17.56680229 | 14.43609979 | ENSG00000227542 |
| LGALSL | -0.286132632 | 0.820097513 | 0.007552487 | 0.024888117 | 442.3079933 | 362.7053592 | ENSG00000119862 |
| EMC3 | -0.286175094 | 0.820073376 | 0.003008136 | 0.011224361 | 592.9604131 | 486.0755339 | ENSG00000125037 |
| GLYR1 | -0.286228138 | 0.820043225 | 7.16E-09 | 7.78E-08 | 3323.377303 | 2725.95014 | ENSG00000140632 |
| ADSS1 | -0.286233126 | 0.820040389 | 0.053056872 | 0.124973473 | 235.4300418 | 192.8667449 | ENSG00000185100 |
| DCAF5 | -0.286332168 | 0.819984095 | 0.000624303 | 0.002791706 | 858.5240546 | 703.5465843 | ENSG00000139990 |
| NA | -0.286360749 | 0.819967851 | 0.720598219 | 0.832149671 | 8.705987658 | 7.096790577 | ENSG00000231747 |
| GNL3 | -0.286459227 | 0.819911882 | 2.09E-08 | 2.11E-07 | 3350.247662 | 2746.763362 | ENSG00000163938 |
| ACO1 | -0.286469882 | 0.819905826 | 4.90E-06 | 3.41E-05 | 2185.259666 | 1791.975544 | ENSG00000122729 |
| CAMTA1 | -0.286471004 | 0.819905189 | 6.07E-05 | 0.000343545 | 1378.372817 | 1129.854629 | ENSG00000171735 |
| SLC39A7 | -0.286541012 | 0.819865403 | 4.92E-07 | 4.05E-06 | 3656.729198 | 2997.375403 | ENSG00000112473 |
| SERPINH1 | -0.286597467 | 0.819833321 | 1.59E-11 | 2.38E-10 | 16439.76424 | 13477.54956 | ENSG00000149257 |
| NA | -0.28664551 | 0.81980602 | 0.671692655 | 0.798817768 | 12.76413948 | 10.42711704 | ENSG00000226243 |
| SAPCD2 | -0.286653101 | 0.819801707 | 0.000222271 | 0.001109973 | 1139.905499 | 934.96568 | ENSG00000186193 |
| PLA2G7 | -0.286719191 | 0.819764152 | 0.246660295 | 0.410164964 | 80.55068873 | 66.03743289 | ENSG00000146070 |
| NBAS | -0.286749712 | 0.81974681 | 0.00113718 | 0.004785633 | 865.3637089 | 709.2962459 | ENSG00000151779 |
| TYMSOS | -0.286836866 | 0.819697291 | 0.4678872 | 0.63706857 | 31.21787683 | 25.64726965 | ENSG00000176912 |
| DIP2A | -0.286856466 | 0.819686154 | 0.000686171 | 0.003031127 | 811.5684759 | 664.9974139 | ENSG00000160305 |
| ZMIZ1 | -0.286986757 | 0.819612131 | 6.64E-06 | 4.53E-05 | 1508.438529 | 1236.072269 | ENSG00000108175 |
| PRPF40A | -0.287007208 | 0.819600513 | 4.65E-09 | 5.19E-08 | 3741.412992 | 3067.106991 | ENSG00000196504 |
| MRPL4 | -0.287096536 | 0.819549767 | 0.00049474 | 0.002268606 | 892.5071783 | 731.8944301 | ENSG00000105364 |
| PCNA | -0.287121248 | 0.819535739 | 1.82E-09 | 2.15E-08 | 5768.356739 | 4726.986036 | ENSG00000132646 |
| CLCN7 | -0.28713682 | 0.819526883 | 7.31E-05 | 0.000407443 | 1400.135291 | 1147.538589 | ENSG00000103249 |
| PDLIM4 | -0.287208183 | 0.819486346 | 0.042845749 | 0.10543913 | 258.6687327 | 211.999086 | ENSG00000131435 |
| CDK5RAP3 | -0.287345604 | 0.819408291 | 0.221330714 | 0.379730595 | 89.25802783 | 73.07775709 | ENSG00000108465 |
| TSEN54 | -0.287353694 | 0.819403697 | 0.000105979 | 0.000570434 | 1179.915243 | 966.7372363 | ENSG00000182173 |
| PDE7A | -0.287438725 | 0.819355403 | 2.42E-05 | 0.000148711 | 1396.367606 | 1144.158974 | ENSG00000205268 |
| HYAL3 | -0.287518733 | 0.819309965 | 0.434695437 | 0.605816333 | 37.83455845 | 31.0339862 | ENSG00000186792 |
| PSAT1 | -0.287641636 | 0.819240171 | 2.66E-09 | 3.06E-08 | 8848.356145 | 7249.364731 | ENSG00000135069 |
| NA | -0.287694473 | 0.819210168 | 0.763726541 | 0.859922296 | 5.578749623 | 4.591910865 | ENSG00000281332 |
| BRI3BP | -0.288016162 | 0.819027523 | 0.030782545 | 0.080203413 | 374.21925 | 306.8576256 | ENSG00000184992 |
| COQ8B | -0.288040641 | 0.819013626 | 0.000274276 | 0.001341886 | 969.742348 | 794.0697134 | ENSG00000123815 |
| TK1 | -0.288057456 | 0.81900408 | 1.04E-05 | 6.84E-05 | 2105.747025 | 1724.450553 | ENSG00000167900 |
| RPLP2 | -0.288061299 | 0.819001898 | 7.15E-08 | 6.70E-07 | 10931.54383 | 8952.498483 | ENSG00000177600 |
| YAP1 | -0.288164768 | 0.818943162 | 1.28E-06 | 9.93E-06 | 2252.261633 | 1844.607531 | ENSG00000137693 |
| SUMO1 | -0.28821525 | 0.818914507 | 4.13E-07 | 3.45E-06 | 4375.191469 | 3583.704643 | ENSG00000116030 |
| TSPYL1 | -0.288283519 | 0.818875756 | 8.63E-05 | 0.000473623 | 1064.465398 | 871.4175335 | ENSG00000189241 |
| FCHSD2 | -0.288439712 | 0.818787106 | 0.138257978 | 0.265175448 | 156.0052425 | 127.590502 | ENSG00000137478 |
| KANK2 | -0.288468139 | 0.818770972 | 3.67E-05 | 0.000216926 | 1196.917992 | 980.0335351 | ENSG00000197256 |
| MRPS2 | -0.288484674 | 0.818761589 | 0.00362453 | 0.013250105 | 809.375871 | 663.1244045 | ENSG00000122140 |
| MED26 | -0.288567528 | 0.818714568 | 0.128789392 | 0.251058627 | 145.1017424 | 118.843884 | ENSG00000105085 |
| KIAA0930 | -0.28867875 | 0.818651453 | 4.78E-06 | 3.33E-05 | 2406.456173 | 1969.880085 | ENSG00000100364 |
| SPRY2 | -0.288703609 | 0.818637347 | 0.006586433 | 0.022228637 | 931.1551032 | 762.0959664 | ENSG00000136158 |
| DTYMK | -0.288733701 | 0.818620272 | 5.67E-06 | 3.90E-05 | 1649.479253 | 1349.899148 | ENSG00000168393 |
| DNTTIP1 | -0.288767383 | 0.81860116 | 0.001536257 | 0.006256268 | 821.377289 | 672.4480383 | ENSG00000101457 |
| NA | -0.288770095 | 0.818599622 | 0.481332552 | 0.649327711 | 33.08760155 | 27.01022201 | ENSG00000237854 |
| ZNF30 | -0.288809558 | 0.81857723 | 0.146934477 | 0.277800993 | 123.9871999 | 101.3734803 | ENSG00000168661 |
| AKAP6 | -0.288970131 | 0.818486127 | 0.716564164 | 0.829919376 | 7.592813428 | 6.236643648 | ENSG00000151320 |
| NSG1 | -0.28899394 | 0.818472619 | 0.000210334 | 0.001055006 | 1089.902716 | 892.1771302 | ENSG00000168824 |
| NLGN2 | -0.28906982 | 0.818429572 | 5.67E-08 | 5.39E-07 | 2768.408167 | 2266.173398 | ENSG00000169992 |
| GRHL2 | -0.289114855 | 0.818404025 | 0.002860059 | 0.010765543 | 694.0270993 | 568.2309169 | ENSG00000083307 |
| C5orf49 | -0.289237283 | 0.818334578 | 0.621311016 | 0.761915061 | 13.68109875 | 11.21977215 | ENSG00000215217 |
| HADH | -0.28929901 | 0.818299565 | 0.000128132 | 0.000673697 | 1266.347449 | 1035.774775 | ENSG00000138796 |
| AHCYL2 | -0.28941693 | 0.818232683 | 0.004792094 | 0.016920004 | 604.5408789 | 494.5968137 | ENSG00000158467 |
| MAPK7 | -0.289471237 | 0.818201883 | 0.001800266 | 0.007202184 | 754.9000793 | 618.0428589 | ENSG00000166484 |
| ZNF321P | -0.289491973 | 0.818190123 | 0.57405381 | 0.72631154 | 18.8957278 | 15.40756194 | ENSG00000213801 |
| BSDC1 | -0.28949716 | 0.818187181 | 0.00015853 | 0.000815955 | 1062.474952 | 869.6085026 | ENSG00000160058 |
| NRXN3 | -0.289559972 | 0.81815156 | 0.34350141 | 0.516242264 | 59.20815527 | 48.59579853 | ENSG00000021645 |
| SLC35F2 | -0.289596869 | 0.818130636 | 0.001738643 | 0.006987532 | 637.3059864 | 521.5161601 | ENSG00000110660 |
| NA | -0.28963617 | 0.81810835 | 0.716377585 | 0.829858884 | 8.779990593 | 7.137118439 | ENSG00000205976 |
| GPHN | -0.289813838 | 0.818007606 | 0.028602727 | 0.075575291 | 346.0447961 | 283.2490021 | ENSG00000171723 |
| DCUN1D5 | -0.289957952 | 0.817925897 | 1.86E-06 | 1.40E-05 | 2133.825461 | 1745.586258 | ENSG00000137692 |
| EDF1 | -0.290168154 | 0.817806733 | 3.41E-06 | 2.44E-05 | 4432.275621 | 3624.156272 | ENSG00000107223 |
| RAET1K | -0.290368341 | 0.817693263 | 0.738234561 | 0.844463914 | 6.403721241 | 5.206031513 | ENSG00000218358 |
| XPO4 | -0.290402048 | 0.817674159 | 0.000480438 | 0.002212214 | 1058.769833 | 866.4682448 | ENSG00000132953 |
| MEN1 | -0.290416635 | 0.817665891 | 5.58E-06 | 3.85E-05 | 1746.318369 | 1428.615772 | ENSG00000133895 |
| ADCY1 | -0.290459979 | 0.817641326 | 9.50E-05 | 0.000516609 | 1247.763067 | 1020.71193 | ENSG00000164742 |
| ZNF530 | -0.290513686 | 0.817610888 | 0.094233104 | 0.197549533 | 159.4997555 | 130.386037 | ENSG00000183647 |
| EEF1E1 | -0.29061065 | 0.817555938 | 0.282026426 | 0.45037846 | 93.69461816 | 76.82698608 | ENSG00000124802 |
| SPRED3 | -0.290653897 | 0.817531431 | 0.134122818 | 0.259073123 | 130.7082721 | 106.9032376 | ENSG00000188766 |

| | | | | | | | |
|---|---|---|---|---|---|---|---|
| DDR1 | -0.290698393 | 0.817506217 | 1.80E-09 | 2.13E-08 | 6561.752577 | 5364.598023 | ENSG00000204580 |
| NA | -0.290929643 | 0.817375189 | 0.524504081 | 0.686869156 | 23.84081468 | 19.50096852 | ENSG00000233325 |
| ZBTB26 | -0.290950491 | 0.817363377 | 0.114104148 | 0.229385485 | 152.1336568 | 124.4188034 | ENSG00000171448 |
| KIAA0895L | -0.291031226 | 0.817317638 | 0.004871995 | 0.017160061 | 620.6647162 | 506.9462159 | ENSG00000196123 |
| PEX5 | -0.291105075 | 0.817275802 | 4.38E-07 | 3.65E-06 | 1941.962138 | 1586.925161 | ENSG00000139197 |
| TAF4B | -0.291127144 | 0.8172633 | 0.011570697 | 0.035639519 | 434.6299253 | 355.0573781 | ENSG00000141384 |
| INPP4A | -0.291131 | 0.817261116 | 0.00021095 | 0.001057545 | 908.281599 | 742.1750293 | ENSG00000040933 |
| ZNF271P | -0.291141383 | 0.817255234 | 0.008226528 | 0.026739335 | 480.836226 | 393.1690602 | ENSG00000257267 |
| FYN | -0.291171929 | 0.817237931 | 4.07E-06 | 2.88E-05 | 1739.538585 | 1421.478969 | ENSG00000010810 |
| CEP89 | -0.291196174 | 0.817224197 | 0.006021406 | 0.020606518 | 475.9804849 | 388.8613529 | ENSG00000121289 |
| USP31 | -0.291213491 | 0.817214388 | 0.014293463 | 0.042538598 | 401.5930484 | 328.3275674 | ENSG00000103404 |
| DKC1 | -0.291403061 | 0.817107013 | 1.92E-08 | 1.96E-07 | 4577.254472 | 3739.863247 | ENSG00000130826 |
| PARK7 | -0.291460211 | 0.817074645 | 3.02E-10 | 3.94E-09 | 6342.610432 | 5182.146193 | ENSG00000116288 |
| CIR1 | -0.291641984 | 0.816971704 | 0.02990003 | 0.078327196 | 295.7139255 | 241.8002388 | ENSG00000138433 |
| TMEM126B | -0.291668697 | 0.816956577 | 0.000295365 | 0.001434512 | 899.7475472 | 735.1642616 | ENSG00000171204 |
| LGALS7 | -0.291814272 | 0.816874146 | 0.785445484 | 0.874499827 | 4.016963998 | 3.277095161 | ENSG00000205076 |
| FUCA2 | -0.291918325 | 0.816815232 | 0.000395435 | 0.001867005 | 914.5081533 | 747.1143223 | ENSG00000001036 |
| NA | -0.291935133 | 0.816805716 | 0.311401743 | 0.482458358 | 60.10433892 | 49.02682311 | ENSG00000287222 |
| ZNF45 | -0.291976143 | 0.816782498 | 0.018375302 | 0.05225976 | 373.4640126 | 305.0324683 | ENSG00000124459 |
| PRICKLE1 | -0.292018491 | 0.816758523 | 0.036712609 | 0.092808667 | 267.2693417 | 218.3814541 | ENSG00000139174 |
| COX8A | -0.292137622 | 0.816691081 | 0.03045544 | 0.079522997 | 281.8933938 | 230.114524 | ENSG00000176340 |
| CRYBB2P1 | -0.292159962 | 0.816678435 | 0.042726051 | 0.105186685 | 287.3096194 | 234.4671513 | ENSG00000100058 |
| NDUFA6 | -0.292306008 | 0.816595766 | 3.63E-05 | 0.000215197 | 1952.057013 | 1593.624264 | ENSG00000184983 |
| CALR | -0.292476991 | 0.816498992 | 3.19E-12 | 5.15E-11 | 29130.16747 | 23783.9793 | ENSG00000179218 |
| NFKBIL1 | -0.292522969 | 0.81647297 | 0.117829848 | 0.235282498 | 142.3084375 | 116.0550733 | ENSG00000204498 |
| C5orf22 | -0.292531996 | 0.816467862 | 0.002188316 | 0.008534319 | 596.2892108 | 487.0445729 | ENSG00000082213 |
| C2orf88 | -0.292565113 | 0.81644912 | 0.683132683 | 0.80675584 | 10.38787013 | 8.445482423 | ENSG00000187699 |
| RABGEF1 | -0.292575894 | 0.816443019 | 0.448901024 | 0.61917272 | 35.04161693 | 28.64231181 | ENSG00000154710 |
| CELSR3 | -0.29298274 | 0.816212811 | 5.59E-05 | 0.00031877 | 1515.525986 | 1236.256207 | ENSG00000008300 |
| LDAH | -0.293138204 | 0.816124861 | 0.003340068 | 0.012312871 | 513.5520469 | 419.0631497 | ENSG00000118961 |
| MRPL43 | -0.29319481 | 0.81609284 | 0.000683217 | 0.003020151 | 899.0921577 | 733.7067024 | ENSG00000055950 |
| PEMT | -0.293258772 | 0.816056659 | 0.144511116 | 0.274413224 | 124.0845692 | 101.3947134 | ENSG00000133027 |
| CEP131 | -0.293293917 | 0.81603678 | 0.002246211 | 0.008732701 | 901.80464 | 735.9717835 | ENSG00000141577 |
| KPNA4 | -0.293381509 | 0.815987236 | 2.09E-06 | 1.56E-05 | 2076.41747 | 1695.008994 | ENSG00000186432 |
| NFKB1 | -0.293509071 | 0.81591509 | 0.021364378 | 0.059351989 | 331.9388896 | 270.9386317 | ENSG00000109320 |
| ZNF432 | -0.293528203 | 0.815904271 | 0.14719612 | 0.278224609 | 123.7547032 | 101.0051133 | ENSG00000256087 |
| NA | -0.293733178 | 0.815788357 | 0.724453757 | 0.834603608 | 7.263422993 | 5.963192946 | ENSG00000269038 |
| HECTD2 | -0.293790266 | 0.815756077 | 0.06848594 | 0.153302654 | 216.762268 | 177.0542257 | ENSG00000165338 |
| SEPTIN6 | -0.293858891 | 0.815717274 | 0.000170404 | 0.000871954 | 929.7098257 | 758.6312014 | ENSG00000125354 |
| RRP1B | -0.293884797 | 0.815702627 | 4.68E-07 | 3.87E-06 | 3453.690044 | 2817.581485 | ENSG00000160208 |
| RPL7L1 | -0.293932298 | 0.81567577 | 1.43E-08 | 1.49E-07 | 2930.037201 | 2390.567176 | ENSG00000146223 |
| FBXO31 | -0.293948092 | 0.815666841 | 0.007087984 | 0.023627837 | 535.0988557 | 436.3235844 | ENSG00000103264 |
| UNC119 | -0.293984428 | 0.815646297 | 0.000540939 | 0.002458245 | 964.6057762 | 786.2886815 | ENSG00000109103 |
| ACAP3 | -0.294013105 | 0.815630084 | 0.000575999 | 0.002594348 | 864.1776092 | 704.8931072 | ENSG00000131584 |
| EXOSC4 | -0.294041731 | 0.815613901 | 0.040201279 | 0.100015845 | 241.5116693 | 196.9795066 | ENSG00000178896 |
| ZNF773 | -0.294134585 | 0.815561408 | 0.246865611 | 0.41040025 | 86.307944 | 70.3706866 | ENSG00000152439 |
| TIMM23 | -0.294198584 | 0.81552523 | 1.36E-07 | 1.23E-06 | 2452.12196 | 1999.534689 | ENSG00000265354 |
| NDN | -0.294420087 | 0.815400029 | 0.00095117 | 0.004072039 | 737.1002167 | 601.3180518 | ENSG00000182636 |
| NA | -0.294479499 | 0.81536645 | 0.488786229 | 0.655460281 | 28.48932624 | 23.15880102 | ENSG00000275437 |
| PRELID3A | -0.29454313 | 0.815330489 | 0.107086985 | 0.218108623 | 175.7320504 | 143.1618291 | ENSG00000141391 |
| TUBB1 | -0.294644269 | 0.815273333 | 0.750937723 | 0.852639177 | 5.64437985 | 4.617184712 | ENSG00000101162 |
| ZNF22 | -0.294796627 | 0.815187239 | 0.018046816 | 0.051531214 | 332.3209949 | 271.1029178 | ENSG00000165512 |
| COPS7B | -0.294961999 | 0.815093802 | 4.81E-06 | 3.35E-05 | 1613.358102 | 1314.655291 | ENSG00000144524 |
| NA | -0.29511159 | 0.81500929 | 0.016073454 | 0.046881314 | 399.4538326 | 325.9291808 | ENSG00000259781 |
| PHRF1 | -0.295216349 | 0.814950112 | 2.25E-06 | 1.66E-05 | 2741.304399 | 2234.109994 | ENSG00000070047 |
| CDK12 | -0.295324121 | 0.814889236 | 2.14E-06 | 1.59E-05 | 1951.278122 | 1589.817769 | ENSG00000167258 |
| IDH3A | -0.295415425 | 0.814837665 | 0.008270118 | 0.026858393 | 439.5961881 | 358.3919815 | ENSG00000166411 |
| WWC1 | -0.295651463 | 0.814704362 | 3.01E-05 | 0.000181291 | 1605.145181 | 1307.831206 | ENSG00000113645 |
| HCFC2 | -0.295657506 | 0.814700949 | 0.48776535 | 0.654754809 | 31.61289749 | 25.85107796 | ENSG00000111727 |
| NUDT16L1 | -0.295783895 | 0.81462958 | 0.000662783 | 0.002939253 | 955.6986065 | 778.5664722 | ENSG00000168101 |
| POLR2J | -0.295858236 | 0.814587603 | 0.005199212 | 0.018140349 | 469.7355456 | 382.4733267 | ENSG00000005075 |
| EDRF1 | -0.29590866 | 0.814559133 | 0.005276654 | 0.01838063 | 471.5001672 | 384.137142 | ENSG00000107938 |
| ZNF429 | -0.296046471 | 0.814481327 | 0.264892462 | 0.430899779 | 67.84448249 | 55.29975393 | ENSG00000197013 |
| TMEM159 | -0.296082252 | 0.814461127 | 0.006246361 | 0.021244454 | 473.7420217 | 385.5151605 | ENSG00000011638 |
| VPS53 | -0.296148186 | 0.814423905 | 0.00056114 | 0.002538687 | 806.8784514 | 657.434105 | ENSG00000141252 |
| B4GALT1 | -0.296186764 | 0.814402128 | 0.003050028 | 0.011360898 | 557.639642 | 454.2252121 | ENSG00000086062 |
| LINC00449 | -0.296230338 | 0.814377531 | 0.821910522 | 0.898581801 | 3.282065012 | 2.709221058 | ENSG00000203441 |
| MPV17L2 | -0.296326831 | 0.814323064 | 0.000875176 | 0.003773583 | 862.9384568 | 702.6684209 | ENSG00000254858 |
| MYD88 | -0.296368236 | 0.814299693 | 0.004450116 | 0.015859447 | 683.9462555 | 557.3582664 | ENSG00000172936 |
| ARL2 | -0.296468558 | 0.81424307 | 0.005645602 | 0.019485221 | 484.4431246 | 394.2969996 | ENSG00000213465 |
| KAT8 | -0.296583271 | 0.81417833 | 0.000145457 | 0.000754305 | 996.1908896 | 810.6678723 | ENSG00000103510 |
| KLHL7 | -0.29661952 | 0.814157873 | 5.70E-07 | 4.65E-06 | 1799.06055 | 1464.610321 | ENSG00000122550 |
| NA | -0.296650805 | 0.814140219 | 0.782907499 | 0.872832206 | 4.044197305 | 3.283546882 | ENSG00000270953 |
| TMEM127 | -0.296684471 | 0.81412122 | 0.000524949 | 0.002393473 | 822.6863532 | 670.0993754 | ENSG00000135956 |
| KLHL9 | -0.296825664 | 0.814041549 | 0.002753589 | 0.010419672 | 705.0013486 | 573.6243209 | ENSG00000198642 |
| ZNF569 | -0.29718876 | 0.813836697 | 0.136698711 | 0.262770316 | 151.7718146 | 123.8199972 | ENSG00000196437 |
| XRCC6 | -0.297316587 | 0.813764592 | 1.29E-14 | 2.68E-13 | 16554.06083 | 13470.85087 | ENSG00000196419 |

| | | | | | | | |
|---|---|---|---|---|---|---|---|
| CCDC142 | -0.297322167 | 0.813761444 | 0.173286878 | 0.315842513 | 139.1760932 | 113.2837461 | ENSG00000135637 |
| ADAMTS2 | -0.297416674 | 0.813708139 | 0.022439415 | 0.061804505 | 294.4079277 | 239.515318 | ENSG00000087116 |
| PPM1B | -0.29751997 | 0.81364988 | 0.013966787 | 0.041740246 | 486.7957295 | 396.5380849 | ENSG00000138032 |
| MRPL46 | -0.297746799 | 0.813521963 | 0.391478166 | 0.564598636 | 40.87730566 | 33.25787899 | ENSG00000259494 |
| DDN | -0.297755612 | 0.813516994 | 0.679112639 | 0.803872602 | 10.15971593 | 8.245702688 | ENSG00000181418 |
| NA | -0.297755913 | 0.813516824 | 0.621229899 | 0.761915061 | 14.15502843 | 11.48462056 | ENSG00000230882 |
| ANKRD61 | -0.297849007 | 0.813464331 | 0.696431543 | 0.816741277 | 9.19020507 | 7.494459938 | ENSG00000157999 |
| NTF3 | -0.298014556 | 0.813370992 | 0.609964926 | 0.753704418 | 17.22221804 | 13.9120612 | ENSG00000185652 |
| AMBRA1 | -0.298120567 | 0.813311226 | 5.44E-05 | 0.000310838 | 1122.959943 | 912.8523089 | ENSG00000110497 |
| NAT14 | -0.298235001 | 0.813246717 | 0.0002642 | 0.001297195 | 1175.148801 | 955.886885 | ENSG00000090971 |
| NA | -0.298317814 | 0.813200037 | 0.388651349 | 0.561987346 | 44.4683489 | 36.278195 | ENSG00000228065 |
| NSD2 | -0.298368659 | 0.813171378 | 7.71E-09 | 8.35E-08 | 5184.197877 | 4215.349857 | ENSG00000109685 |
| ZNF211 | -0.298374672 | 0.813167989 | 0.020548194 | 0.057515266 | 337.6830421 | 274.5564089 | ENSG00000121417 |
| COPG2 | -0.298435974 | 0.813133437 | 9.36E-05 | 0.000509539 | 1050.403621 | 854.1842691 | ENSG00000158623 |
| CREBZF | -0.298509601 | 0.81309194 | 1.38E-05 | 8.81E-05 | 1316.971316 | 1070.453158 | ENSG00000137504 |
| CTPS1 | -0.2985765 | 0.813054237 | 1.68E-07 | 1.50E-06 | 2926.835572 | 2379.405906 | ENSG00000171793 |
| DIPK1B | -0.29864269 | 0.813016936 | 0.016107526 | 0.046973587 | 434.2971804 | 352.8396766 | ENSG00000165716 |
| SNORD116-17 | -0.298700314 | 0.812984463 | 0.828745742 | 0.902525353 | 3.621943337 | 2.953096766 | ENSG00000206656 |
| NA | -0.2987423 | 0.812960803 | 0.575668731 | 0.72792511 | 17.48597825 | 14.22341661 | ENSG00000232499 |
| COPS2 | -0.298750619 | 0.812956115 | 0.000181997 | 0.000924897 | 1146.415075 | 932.4849713 | ENSG00000166200 |
| MFSD5 | -0.298844939 | 0.812902968 | 0.013283728 | 0.039976743 | 445.1983041 | 362.0468884 | ENSG00000182544 |
| RCN2 | -0.298867368 | 0.81289033 | 7.39E-08 | 6.92E-07 | 2767.713254 | 2250.547252 | ENSG00000117906 |
| KCNH4 | -0.299070482 | 0.812775893 | 0.580031738 | 0.731189456 | 18.20393166 | 14.86066053 | ENSG00000089558 |
| CHST11 | -0.299168124 | 0.812720886 | 0.004900815 | 0.017245826 | 508.2761301 | 413.2338316 | ENSG00000171310 |
| NA | -0.29918834 | 0.812709498 | 0.670801783 | 0.798102701 | 10.50238516 | 8.485277248 | ENSG00000225022 |
| GPN1 | -0.299229995 | 0.812686033 | 0.002691467 | 0.010216666 | 682.0381196 | 554.1838779 | ENSG00000198522 |
| C11orf45 | -0.299361393 | 0.812612018 | 0.622038855 | 0.762199274 | 12.99787547 | 10.60350093 | ENSG00000174370 |
| NA | -0.299378741 | 0.812602247 | 0.796290395 | 0.881714959 | 3.657549353 | 2.986972907 | ENSG00000231050 |
| ZCCHC2 | -0.299527113 | 0.812518681 | 0.020788861 | 0.058002912 | 371.0603609 | 301.4803291 | ENSG00000141664 |
| MAN1C1 | -0.299570914 | 0.812494012 | 0.013613794 | 0.040799084 | 347.4654339 | 282.3942224 | ENSG00000117643 |
| FLRT1 | -0.299632278 | 0.812459454 | 0.34326134 | 0.516042277 | 48.12956504 | 39.11137425 | ENSG00000126500 |
| AGAP9 | -0.299738709 | 0.812399519 | 0.31338109 | 0.484526298 | 56.20535658 | 45.64459187 | ENSG00000204172 |
| CENPI | -0.299758305 | 0.812388485 | 0.016864535 | 0.048763561 | 343.8292606 | 279.303198 | ENSG00000102384 |
| BYSL | -0.299764567 | 0.812384959 | 1.94E-05 | 0.000121179 | 1223.1342 | 993.5562687 | ENSG00000112578 |
| ZC3H12C | -0.299818215 | 0.81235475 | 0.356386824 | 0.529706312 | 50.70464009 | 41.13307627 | ENSG00000149289 |
| OTUB2 | -0.299931539 | 0.812290942 | 0.179037094 | 0.323459841 | 109.0010545 | 88.50440161 | ENSG00000089723 |
| UBR5 | -0.30004736 | 0.812225732 | 1.76E-05 | 0.000110489 | 4257.105879 | 3457.210245 | ENSG00000104517 |
| CTDP1 | -0.300132098 | 0.812178027 | 1.75E-05 | 0.000109919 | 1764.87283 | 1433.48283 | ENSG00000060069 |
| NOL7 | -0.300212811 | 0.81213259 | 7.61E-05 | 0.00042262 | 2136.334584 | 1735.646571 | ENSG00000225921 |
| STAG3L5P | -0.300387273 | 0.812034387 | 0.166955718 | 0.306883541 | 118.0432537 | 95.67272469 | ENSG00000242294 |
| LYPLA2 | -0.300565853 | 0.811933877 | 4.27E-08 | 4.15E-07 | 3699.427255 | 3003.083955 | ENSG00000011009 |
| HIRIP3 | -0.300626862 | 0.811899543 | 0.007139252 | 0.0237591 | 552.9578909 | 448.7264294 | ENSG00000149929 |
| PDCD7 | -0.300628754 | 0.811898478 | 0.000450223 | 0.002089308 | 744.7810245 | 604.7489406 | ENSG00000090470 |
| UBL7 | -0.300667515 | 0.811876665 | 1.92E-05 | 0.000119675 | 1236.50557 | 1003.701321 | ENSG00000138629 |
| STOML2 | -0.300789154 | 0.811808216 | 5.71E-09 | 6.27E-08 | 4333.656876 | 3517.122024 | ENSG00000165283 |
| ANAPC15 | -0.300805677 | 0.811798918 | 0.118768561 | 0.236568333 | 150.2209172 | 121.9394701 | ENSG00000110200 |
| CCDC32 | -0.300853031 | 0.811772273 | 0.344414702 | 0.517076312 | 56.43909257 | 45.75886328 | ENSG00000128891 |
| NA | -0.300856871 | 0.811770112 | 0.622562231 | 0.76248162 | 12.95947856 | 10.55295324 | ENSG00000236683 |
| RFC5 | -0.301028865 | 0.811673341 | 0.002067543 | 0.008124868 | 586.4509129 | 475.890966 | ENSG00000111445 |
| AHSA1 | -0.301181255 | 0.811587609 | 3.98E-10 | 5.09E-09 | 3836.820556 | 3114.393705 | ENSG00000100591 |
| CLCN6 | -0.301261324 | 0.811542568 | 0.000934044 | 0.004005031 | 748.3461106 | 606.8556421 | ENSG00000011021 |
| NIBAN3 | -0.301408931 | 0.811459541 | 0.743301828 | 0.847935038 | 5.698846465 | 4.63170569 | ENSG00000167483 |
| SF3B5 | -0.301589349 | 0.811358069 | 4.43E-06 | 3.11E-05 | 1721.788184 | 1396.565235 | ENSG00000169976 |
| MBIP | -0.301850104 | 0.811211436 | 0.008415414 | 0.027238549 | 432.133447 | 350.9281811 | ENSG00000151332 |
| NA | -0.30189485 | 0.811186276 | 0.578906841 | 0.730487433 | 17.30304209 | 14.0196083 | ENSG00000237214 |
| ZNF580 | -0.301933227 | 0.811164698 | 0.04925949 | 0.117921892 | 237.1048661 | 192.3894859 | ENSG00000213015 |
| TMEM38B | -0.301964785 | 0.811146955 | 0.016918673 | 0.048888276 | 325.2738867 | 263.9746742 | ENSG00000095209 |
| CXorf56 | -0.301968002 | 0.811145146 | 0.003356045 | 0.012362319 | 558.3390102 | 453.1408356 | ENSG00000018610 |
| MCRS1 | -0.301988756 | 0.811133478 | 2.38E-06 | 1.75E-05 | 1950.433038 | 1581.863031 | ENSG00000187778 |
| TTC14 | -0.302061425 | 0.811092621 | 0.0258766 | 0.069650549 | 333.0388361 | 270.2750416 | ENSG00000163728 |
| ZNF276 | -0.302090618 | 0.811076209 | 0.128339147 | 0.250433996 | 152.5142104 | 123.7681351 | ENSG00000158805 |
| KCNC3 | -0.302121649 | 0.811058764 | 0.089617186 | 0.190083028 | 188.1669116 | 152.4408275 | ENSG00000131398 |
| NAA10 | -0.302210575 | 0.811008773 | 0.01282892 | 0.038826906 | 388.7742052 | 315.5246661 | ENSG00000102030 |
| SNAP91 | -0.30222176 | 0.811002485 | 0.560334885 | 0.715498368 | 21.07211541 | 17.18726625 | ENSG00000065609 |
| NA | -0.302379054 | 0.810914068 | 0.61786781 | 0.759577359 | 15.87096531 | 12.94247705 | ENSG00000269737 |
| NDST1 | -0.302386232 | 0.810910033 | 1.69E-06 | 1.28E-05 | 2737.900082 | 2220.371088 | ENSG00000070614 |
| EIF3J | -0.302451451 | 0.810873376 | 1.06E-05 | 6.92E-05 | 1694.007727 | 1374.146615 | ENSG00000104131 |
| SAP30BP | -0.302484566 | 0.810854764 | 9.02E-07 | 7.15E-06 | 1621.040238 | 1314.355512 | ENSG00000161526 |
| PMP22 | -0.30268407 | 0.810742642 | 0.478857226 | 0.647119772 | 26.72506813 | 21.66571132 | ENSG00000109099 |
| WASH8P | -0.302752579 | 0.810704143 | 0.046252095 | 0.112176158 | 303.1728508 | 245.8651791 | ENSG00000226210 |
| INPP5E | -0.302826101 | 0.810662829 | 0.003052212 | 0.011364645 | 597.1090762 | 484.0753497 | ENSG00000148384 |
| POFUT1 | -0.302848217 | 0.810650402 | 7.68E-05 | 0.000426321 | 1081.604165 | 876.6918872 | ENSG00000101346 |
| COX3 | -0.302877912 | 0.810633717 | 0.083859923 | 0.180149418 | 49580.98252 | 40192.03171 | ENSG00000198938 |
| GBF1 | -0.302878904 | 0.810633159 | 9.32E-05 | 0.000507726 | 1286.07108 | 1042.471807 | ENSG00000107862 |
| NA | -0.302932508 | 0.81060304 | 0.744699451 | 0.848753226 | 7.31856533 | 5.948671968 | ENSG00000279179 |
| VILL | -0.302957495 | 0.810589001 | 0.184741841 | 0.330997943 | 101.028214 | 81.86982814 | ENSG00000136059 |
| RRS1 | -0.302985662 | 0.810573176 | 0.000598146 | 0.002686433 | 1321.459009 | 1071.040206 | ENSG00000179041 |

| | | | | | | | |
|---|---|---|---|---|---|---|---|
| NGDN | -0.303016409 | 0.810555901 | 0.000260098 | 0.001279334 | 930.618337 | 754.3783614 | ENSG00000129460 |
| KLHDC3 | -0.303311173 | 0.810390309 | 5.64E-09 | 6.21E-08 | 5352.810333 | 4337.004359 | ENSG00000124702 |
| NOC4L | -0.30336341 | 0.810360967 | 0.000762504 | 0.003327932 | 1241.473198 | 1005.648092 | ENSG00000184967 |
| IFT172 | -0.303433985 | 0.810321326 | 0.014224289 | 0.042378497 | 369.0568092 | 299.2991631 | ENSG00000138002 |
| MMP14 | -0.303434652 | 0.810320952 | 4.51E-07 | 3.74E-06 | 2155.494567 | 1746.641611 | ENSG00000157227 |
| EIF4G2 | -0.303451973 | 0.810311223 | 7.38E-15 | 1.57E-13 | 17927.7941 | 14527.34106 | ENSG00000110321 |
| STX4 | -0.303535753 | 0.810264168 | 0.001649601 | 0.006665783 | 801.8318671 | 649.7837556 | ENSG00000103496 |
| MMP24 | -0.303656943 | 0.810196107 | 0.017883679 | 0.05122406 | 340.3633836 | 275.9010601 | ENSG00000125966 |
| ZNF668 | -0.30367044 | 0.810188527 | 0.102703559 | 0.21145301 | 168.3813457 | 136.5799297 | ENSG00000167394 |
| IRS2 | -0.30379547 | 0.810118316 | 0.011805054 | 0.036165046 | 468.4214889 | 379.9318567 | ENSG00000185950 |
| ZNF180 | -0.303817786 | 0.810105784 | 0.082612719 | 0.17804513 | 181.8606477 | 147.3100845 | ENSG00000167384 |
| CLPTM1L | -0.303868525 | 0.810077294 | 5.68E-07 | 4.63E-06 | 2737.207359 | 2217.6137 | ENSG00000049656 |
| CALM1 | -0.303884839 | 0.810068134 | 8.47E-10 | 1.04E-08 | 6315.770778 | 5116.594422 | ENSG00000198668 |
| NA | -0.304066552 | 0.809966109 | 0.784876582 | 0.874043418 | 4.011382192 | 3.230848615 | ENSG00000267219 |
| THAP7-AS1 | -0.304128316 | 0.809931433 | 0.646005025 | 0.779464783 | 10.92655415 | 8.878645461 | ENSG00000230513 |
| GAD1 | -0.304199395 | 0.809891531 | 0.709793463 | 0.825105495 | 7.290656301 | 5.923398121 | ENSG00000128683 |
| COQ10A | -0.304323797 | 0.809821698 | 0.056615418 | 0.131840661 | 212.8760009 | 172.419576 | ENSG00000135469 |
| RABAC1 | -0.304350435 | 0.809806745 | 0.50381697 | 0.669027801 | 28.61835937 | 23.1523493 | ENSG00000105404 |
| PYURF | -0.304369887 | 0.809795826 | 0.001473016 | 0.006031831 | 770.881379 | 624.4106938 | ENSG00000145337 |
| MAP3K3 | -0.30454711 | 0.809696356 | 0.000228741 | 0.001140808 | 884.6248349 | 716.3177906 | ENSG00000198909 |
| ADRA2C | -0.304571595 | 0.809682614 | 0.000733101 | 0.003214148 | 716.2261561 | 579.8255198 | ENSG00000184160 |
| NA | -0.304734363 | 0.809591269 | 0.775966648 | 0.868533369 | 5.661125267 | 4.58330857 | ENSG00000284624 |
| NAP1L4 | -0.304859865 | 0.809520845 | 1.57E-11 | 2.35E-10 | 5155.703787 | 4173.724491 | ENSG00000205531 |
| FAM241B | -0.304967953 | 0.809460197 | 0.008545152 | 0.027597888 | 473.5575709 | 383.1221354 | ENSG00000171224 |
| RCBTB1 | -0.30506671 | 0.809404789 | 0.000184219 | 0.000935204 | 973.2817028 | 788.3804678 | ENSG00000136144 |
| SIRT6 | -0.305131485 | 0.809368449 | 0.000944822 | 0.004048549 | 651.2140511 | 527.145971 | ENSG00000077463 |
| PDSS1 | -0.305249857 | 0.809302043 | 0.03182029 | 0.082428524 | 274.8116307 | 222.5627921 | ENSG00000148459 |
| BRIX1 | -0.305254386 | 0.809299502 | 0.0048509 | 0.01709825 | 591.6155442 | 479.2753052 | ENSG00000113460 |
| NA | -0.305334116 | 0.809254778 | 0.73964747 | 0.845477132 | 6.871193235 | 5.592948006 | ENSG00000224225 |
| SYF2 | -0.305343183 | 0.809249692 | 0.002011969 | 0.007930711 | 689.1947516 | 558.22375 | ENSG00000117614 |
| ADGRB1 | -0.305347152 | 0.809247466 | 0.002033629 | 0.008011177 | 596.4856259 | 482.5121151 | ENSG00000181790 |
| H2BC10 | -0.305427846 | 0.809202204 | 0.767478692 | 0.862636569 | 5.363874225 | 4.36470671 | ENSG00000278588 |
| C13orf42 | -0.305431173 | 0.809200337 | 7.79E-05 | 0.00043133 | 1342.778462 | 1086.696671 | ENSG00000226792 |
| ACSS2 | -0.305456888 | 0.809185914 | 0.000134451 | 0.000703469 | 963.6944868 | 779.7892668 | ENSG00000131069 |
| NA | -0.305480953 | 0.809172417 | 0.597210927 | 0.744333634 | 14.97633326 | 12.12401505 | ENSG00000257576 |
| TCOF1 | -0.30557356 | 0.809120477 | 3.73E-10 | 4.79E-09 | 8577.488012 | 6940.004283 | ENSG00000070814 |
| MINCR | -0.305768071 | 0.809011395 | 0.317824929 | 0.489084293 | 59.56286399 | 48.26534841 | ENSG00000253716 |
| LOC101929709 | -0.305800641 | 0.808993131 | 0.243566932 | 0.406491998 | 82.75952803 | 67.00700493 | ENSG00000251136 |
| ZNF268 | -0.305865472 | 0.808956778 | 0.025317892 | 0.068339533 | 294.3621972 | 238.3965018 | ENSG00000090612 |
| SNORA5C | -0.305987797 | 0.80888819 | 0.765713934 | 0.861255175 | 6.077121709 | 4.896554097 | ENSG00000201772 |
| EIF5 | -0.306080361 | 0.808836294 | 5.11E-10 | 6.45E-09 | 4440.119804 | 3592.005486 | ENSG00000100664 |
| NA | -0.306091928 | 0.808829808 | 0.269639101 | 0.435974176 | 70.78599346 | 57.17519852 | ENSG00000237004 |
| IFT43 | -0.30629647 | 0.808715142 | 0.043997029 | 0.107695304 | 212.5827801 | 171.9821118 | ENSG00000119650 |
| H1-5 | -0.306380993 | 0.808667764 | 0.803321602 | 0.886596456 | 3.681991758 | 2.984822334 | ENSG00000184357 |
| SAE1 | -0.306441376 | 0.808633918 | 4.43E-15 | 9.59E-14 | 8205.466158 | 6634.936585 | ENSG00000142230 |
| FAM13B | -0.306766311 | 0.808451812 | 0.049893012 | 0.119119565 | 221.9339575 | 179.5204072 | ENSG00000031003 |
| CNN3 | -0.306778819 | 0.808444803 | 3.82E-12 | 6.09E-11 | 4745.855576 | 3837.190615 | ENSG00000117519 |
| NA | -0.30690661 | 0.808373195 | 0.667937805 | 0.79602077 | 10.57359719 | 8.483126674 | ENSG00000233045 |
| NA | -0.30696658 | 0.808339594 | 0.646582639 | 0.780015364 | 11.0494419 | 8.956617575 | ENSG00000281849 |
| MAGOHB | -0.307043149 | 0.808296694 | 0.00612116 | 0.020884912 | 463.8944384 | 374.9667689 | ENSG00000111196 |
| ARL16 | -0.307051235 | 0.808292163 | 0.006965417 | 0.023291707 | 437.8316546 | 353.7603064 | ENSG00000214087 |
| CCP110 | -0.307055247 | 0.808289916 | 0.014417807 | 0.042829368 | 369.2687304 | 298.491987 | ENSG00000103540 |
| ACOX3 | -0.30710804 | 0.808260338 | 0.030297474 | 0.079174826 | 253.2310557 | 204.5769521 | ENSG00000087008 |
| WIZ | -0.307233215 | 0.808190213 | 0.000753839 | 0.003294587 | 1303.627032 | 1053.793734 | ENSG00000011451 |
| CDK2AP1 | -0.307358874 | 0.808119822 | 5.75E-05 | 0.000327304 | 3099.951453 | 2505.202334 | ENSG00000111328 |
| CALM3 | -0.307442698 | 0.80807287 | 3.46E-12 | 5.56E-11 | 9748.110701 | 7877.496806 | ENSG00000160014 |
| GPR162 | -0.307452806 | 0.808067208 | 0.085329459 | 0.182736523 | 217.5466154 | 175.9988728 | ENSG00000250510 |
| RPL13AP6 | -0.307500721 | 0.808040371 | 0.78948236 | 0.877298629 | 3.630316045 | 2.938575748 | ENSG00000234118 |
| OSTC | -0.307567059 | 0.808003216 | 2.13E-06 | 1.58E-05 | 2015.561699 | 1628.466375 | ENSG00000198856 |
| ALG12 | -0.307722886 | 0.807915948 | 0.023083105 | 0.06321638 | 292.6809905 | 236.4388266 | ENSG00000182858 |
| TTL | -0.30776462 | 0.807892577 | 1.87E-05 | 0.000117364 | 1132.235953 | 914.8024663 | ENSG00000114999 |
| ZHX3 | -0.307951074 | 0.807788172 | 0.105074327 | 0.215042313 | 137.9872012 | 111.4214775 | ENSG00000174306 |
| ZMYM4 | -0.307954428 | 0.807786294 | 2.25E-07 | 1.97E-06 | 2044.726425 | 1651.793993 | ENSG00000146463 |
| CLEC16A | -0.308004886 | 0.807758042 | 0.007097916 | 0.023649854 | 447.2080623 | 361.1111925 | ENSG00000038532 |
| SLC25A22 | -0.30823147 | 0.807631189 | 0.00095136 | 0.004072039 | 1215.96627 | 981.9956272 | ENSG00000177542 |
| DACT3 | -0.308420308 | 0.807525482 | 0.293472065 | 0.463173493 | 61.09318594 | 49.38872621 | ENSG00000197380 |
| XRCC2 | -0.308468438 | 0.807498543 | 0.008817258 | 0.02831592 | 573.5344123 | 463.4127956 | ENSG00000196584 |
| NA | -0.308489218 | 0.807486912 | 0.160105895 | 0.297295487 | 124.9825557 | 101.0118376 | ENSG00000279207 |
| EMC8 | -0.308525115 | 0.807466821 | 0.003326534 | 0.012270013 | 690.4169584 | 557.649721 | ENSG00000131148 |
| TRIM8 | -0.308635662 | 0.807404951 | 0.002500929 | 0.00959917 | 583.2076324 | 470.8968121 | ENSG00000171206 |
| MOXD1 | -0.308853742 | 0.807282911 | 0.221200976 | 0.379575607 | 83.93937163 | 67.75019053 | ENSG00000079931 |
| ATP6 | -0.309003414 | 0.807199164 | 0.055176985 | 0.129239828 | 24782.17933 | 20004.16869 | ENSG00000198899 |
| SKAP2 | -0.309158347 | 0.807112483 | 0.214435914 | 0.370942915 | 80.34119498 | 64.82183457 | ENSG00000005020 |
| NA | -0.309205266 | 0.807086235 | 0.749814542 | 0.852024107 | 5.377828739 | 4.36470671 | ENSG00000240616 |
| PSMA5 | -0.309219223 | 0.807078427 | 7.87E-10 | 9.70E-09 | 3523.327842 | 2843.329819 | ENSG00000143106 |
| CYP2J2 | -0.30928367 | 0.807042374 | 0.482201705 | 0.650181945 | 30.18815429 | 24.26093355 | ENSG00000134716 |
| ZMYND11 | -0.309351145 | 0.807004629 | 0.00087248 | 0.003762801 | 625.0409827 | 504.393169 | ENSG00000015171 |

| | | | | | | | |
|---|---|---|---|---|---|---|---|
| THG1L | -0.3094777 | 0.806933841 | 0.023154508 | 0.063402926 | 311.7407938 | 251.8296743 | ENSG00000113272 |
| CCDC93 | -0.309483002 | 0.806930875 | 0.002676831 | 0.010169112 | 609.1671513 | 491.2377483 | ENSG00000125633 |
| CYB561D1 | -0.309672073 | 0.806825131 | 0.101717632 | 0.209797375 | 152.4768896 | 123.1835773 | ENSG00000174151 |
| NA | -0.309887776 | 0.806704509 | 0.698656619 | 0.818040309 | 7.29902901 | 5.895973701 | ENSG00000267742 |
| PWWP2A | -0.309907949 | 0.806693228 | 3.11E-06 | 2.24E-05 | 1377.390542 | 1111.49425 | ENSG00000170234 |
| ZNF891 | -0.31016168 | 0.806551365 | 0.243595873 | 0.406491998 | 78.64710976 | 63.31637446 | ENSG00000214029 |
| IL1R1 | -0.310170407 | 0.806546487 | 0.671833978 | 0.798838096 | 8.959259975 | 7.25058423 | ENSG00000115594 |
| RCOR2 | -0.310576982 | 0.806319221 | 2.35E-06 | 1.73E-05 | 2787.248485 | 2247.111026 | ENSG00000167771 |
| MAP1LC3B | -0.310585594 | 0.806314407 | 0.000108416 | 0.000582252 | 884.3919748 | 713.1925805 | ENSG00000140941 |
| TMTC2 | -0.310612812 | 0.806299195 | 0.12715135 | 0.248764517 | 118.9756069 | 95.94239518 | ENSG00000179104 |
| MOCS1 | -0.310656619 | 0.806274713 | 0.00048219 | 0.002218971 | 749.207452 | 604.1885906 | ENSG00000124615 |
| LINC01980 | -0.310691376 | 0.806255289 | 0.230921954 | 0.391211932 | 74.35434606 | 59.97770617 | ENSG00000225548 |
| NONO | -0.310731728 | 0.806232738 | 2.77E-15 | 6.15E-14 | 15923.78309 | 12837.95772 | ENSG00000147140 |
| NA | -0.310738703 | 0.80622884 | 0.570889259 | 0.723744466 | 19.43229665 | 15.61379339 | ENSG00000285531 |
| NA | -0.310762687 | 0.806215437 | 0.784867681 | 0.874043418 | 3.668712964 | 2.961699061 | ENSG00000258515 |
| VPS33A | -0.310796963 | 0.806196283 | 0.032059341 | 0.082927766 | 257.1994983 | 207.507186 | ENSG00000139719 |
| GS1-124K5.11 | -0.310819249 | 0.806183829 | 0.018300085 | 0.052082889 | 333.6017993 | 268.7685275 | ENSG00000229180 |
| TAOK3 | -0.310895763 | 0.806141075 | 0.033308797 | 0.085492284 | 285.8284946 | 230.4107949 | ENSG00000135090 |
| ULK4P3 | -0.310984857 | 0.806091292 | 0.743322245 | 0.847935038 | 6.121100434 | 4.905156391 | ENSG00000178081 |
| TOX | -0.310996979 | 0.80608452 | 0.043820302 | 0.10735807 | 248.993271 | 200.7951493 | ENSG00000198846 |
| PRKCZ | -0.311034725 | 0.80606343 | 0.000108239 | 0.000581462 | 911.0754675 | 734.23775 | ENSG00000067606 |
| NCOA2 | -0.311050078 | 0.806054852 | 0.000418616 | 0.001960534 | 745.6748176 | 601.1177269 | ENSG00000140396 |
| NA | -0.311210611 | 0.805965165 | 0.798611849 | 0.883828878 | 3.614246349 | 2.913301941 | ENSG00000224282 |
| CRY2 | -0.311327517 | 0.805899857 | 0.002813727 | 0.010615999 | 514.9329235 | 414.7952493 | ENSG00000121671 |
| ZFAND3 | -0.311402741 | 0.805857838 | 2.35E-06 | 1.73E-05 | 1610.828245 | 1298.038558 | ENSG00000156639 |
| RASSF2 | -0.311439811 | 0.805837131 | 0.023865196 | 0.06504429 | 306.9908828 | 247.4348596 | ENSG00000101265 |
| PIGW | -0.311710207 | 0.805686112 | 0.024710509 | 0.066960308 | 326.6965147 | 263.5925798 | ENSG00000277161 |
| SSTR2 | -0.311810565 | 0.805630069 | 0.070338945 | 0.156705353 | 194.2377758 | 156.5267042 | ENSG00000180616 |
| ZNF135 | -0.311960206 | 0.80554651 | 0.394940989 | 0.567550976 | 40.75230273 | 32.82686652 | ENSG00000176293 |
| GPR153 | -0.312024423 | 0.805510654 | 0.010976732 | 0.034098573 | 360.3922095 | 290.3492698 | ENSG00000158292 |
| DUSP28 | -0.312171574 | 0.805428499 | 0.487057846 | 0.654305913 | 30.56084772 | 24.48006845 | ENSG00000188542 |
| UNC5A | -0.31217943 | 0.805424113 | 0.031502385 | 0.081714738 | 315.975136 | 254.4905409 | ENSG00000113763 |
| DHRS4 | -0.312180222 | 0.805423671 | 0.080572137 | 0.174387646 | 190.8304325 | 153.6429772 | ENSG00000157326 |
| CS | -0.312315081 | 0.805348386 | 1.05E-09 | 1.27E-08 | 3054.039744 | 2459.845986 | ENSG00000062485 |
| GPR27 | -0.312353562 | 0.805326905 | 0.001915286 | 0.007596171 | 518.8794392 | 417.8007717 | ENSG00000170837 |
| NA | -0.312459159 | 0.805267962 | 0.736590078 | 0.843447451 | 5.731661579 | 4.606431843 | ENSG00000231120 |
| SPOCK2 | -0.312536309 | 0.8052249 | 0.278604148 | 0.44635434 | 68.62846645 | 55.16316487 | ENSG00000107742 |
| ENPP4 | -0.31254823 | 0.805218246 | 0.181278901 | 0.326458186 | 99.82919754 | 80.48051744 | ENSG00000001561 |
| HNRNPH2 | -0.312632487 | 0.805171221 | 2.08E-06 | 1.55E-05 | 1434.301649 | 1155.23759 | ENSG00000126945 |
| MAEA | -0.312649855 | 0.805161528 | 1.36E-07 | 1.23E-06 | 2479.832773 | 1996.533814 | ENSG00000090316 |
| CCKBR | -0.312763516 | 0.805098097 | 0.001914426 | 0.007594322 | 526.72744 | 424.14658 | ENSG00000110148 |
| PLPPR5 | -0.312908803 | 0.805017023 | 0.262960625 | 0.428588493 | 61.94383926 | 49.91680549 | ENSG00000117598 |
| WDR5 | -0.313019422 | 0.804955301 | 4.04E-08 | 3.94E-07 | 2568.651826 | 2067.478002 | ENSG00000196363 |
| REPIN1 | -0.313056857 | 0.804934414 | 5.54E-08 | 5.29E-07 | 2642.29663 | 2127.049679 | ENSG00000214022 |
| CNPPD1 | -0.313169651 | 0.804871485 | 0.000529906 | 0.002414362 | 725.4529921 | 583.564529 | ENSG00000115649 |
| CELSR1 | -0.313211018 | 0.804848406 | 6.20E-05 | 0.000350497 | 982.2891335 | 790.5721141 | ENSG00000075275 |
| NUDT16 | -0.313289826 | 0.804804442 | 0.000289212 | 0.001406399 | 1038.889259 | 835.46822 | ENSG00000198585 |
| RAB11FIP2 | -0.313576945 | 0.80464429 | 0.07506599 | 0.165006609 | 207.6693443 | 167.2073653 | ENSG00000107560 |
| IFRD2 | -0.313598168 | 0.804632453 | 6.83E-05 | 0.000382327 | 1699.591983 | 1367.442763 | ENSG00000214706 |
| AGBL5 | -0.313716352 | 0.804566541 | 0.000109511 | 0.000586991 | 1142.491272 | 919.0106589 | ENSG00000084693 |
| CLK3 | -0.313849353 | 0.804492372 | 0.000178923 | 0.000910711 | 838.9149671 | 674.545865 | ENSG00000179335 |
| KDM4D | -0.313862635 | 0.804484965 | 0.442459265 | 0.613445571 | 38.41634532 | 31.0033267 | ENSG00000186280 |
| MAGIX | -0.313924169 | 0.804450654 | 0.553519175 | 0.709874911 | 17.15379691 | 13.76067858 | ENSG00000269313 |
| CERCAM | -0.313954616 | 0.804433676 | 0.083846856 | 0.180141408 | 154.4275505 | 124.2564074 | ENSG00000167123 |
| PPAN | -0.313993778 | 0.80441184 | 0.08870019 | 0.18859422 | 143.7312657 | 115.6141015 | ENSG00000130810 |
| VPS26B | -0.314015774 | 0.804399576 | 0.000767291 | 0.003346556 | 832.6330207 | 670.4483508 | ENSG00000151502 |
| VPS37B | -0.314060889 | 0.804374422 | 0.046277632 | 0.112223984 | 228.1078849 | 183.677798 | ENSG00000139722 |
| ZNF343 | -0.314082258 | 0.804362507 | 0.134979347 | 0.260196387 | 117.2671669 | 94.36756525 | ENSG00000088876 |
| PRMT1 | -0.314167199 | 0.80431515 | 4.48E-11 | 6.38E-10 | 15887.62002 | 12778.65433 | ENSG00000126457 |
| SEMA4A | -0.31429727 | 0.804242638 | 0.012411834 | 0.037724387 | 332.7991549 | 267.505344 | ENSG00000196189 |
| NA | -0.314438829 | 0.804163729 | 0.538322808 | 0.697944739 | 23.85967528 | 19.08206599 | ENSG00000225159 |
| RAMP3 | -0.314461272 | 0.804151219 | 0.55845904 | 0.7140266 | 17.89898363 | 14.41512709 | ENSG00000122679 |
| LHFPL4 | -0.314477443 | 0.804142206 | 0.000717387 | 0.003152417 | 793.4630279 | 637.8396321 | ENSG00000156959 |
| PDCD11 | -0.314563061 | 0.804094484 | 5.08E-10 | 6.42E-09 | 4506.16038 | 3623.261934 | ENSG00000148843 |
| WNK2 | -0.314711305 | 0.804011864 | 4.78E-07 | 3.95E-06 | 2473.188096 | 1989.149784 | ENSG00000165238 |
| TRA2A | -0.314743742 | 0.803993787 | 1.96E-07 | 1.73E-06 | 1765.298912 | 1419.476331 | ENSG00000164548 |
| TAF1C | -0.314818883 | 0.803951913 | 7.87E-05 | 0.000435673 | 1568.968763 | 1261.142844 | ENSG00000103168 |
| DDAH2 | -0.314844693 | 0.803937531 | 5.77E-06 | 3.96E-05 | 1384.17086 | 1112.730579 | ENSG00000213722 |
| ZNF107 | -0.315062091 | 0.803816395 | 0.011053799 | 0.034310351 | 432.6903911 | 348.1044219 | ENSG00000196247 |
| GPRC5C | -0.315067393 | 0.803813441 | 0.021451548 | 0.059559846 | 311.1520368 | 250.3207492 | ENSG00000170412 |
| CTNS | -0.315210813 | 0.803733537 | 0.028829939 | 0.076030569 | 293.10372 | 235.6620374 | ENSG00000040531 |
| DHDDS | -0.315269154 | 0.803701036 | 0.000171413 | 0.000876418 | 875.0234642 | 703.0507821 | ENSG00000117682 |
| ZMYND19 | -0.315818908 | 0.803394835 | 6.95E-05 | 0.00038843 | 1139.341707 | 914.9829214 | ENSG00000165724 |
| NECAB2 | -0.315873698 | 0.803364325 | 0.04849622 | 0.116520186 | 225.64178 | 181.0549093 | ENSG00000103154 |
| ATP13A1 | -0.316000747 | 0.803293581 | 2.14E-05 | 0.000132937 | 2167.584184 | 1741.42622 | ENSG00000105726 |
| APBB1 | -0.316021976 | 0.803281761 | 0.000138368 | 0.000722395 | 952.1009054 | 765.1375582 | ENSG00000166313 |
| MRPL3 | -0.316152679 | 0.803208989 | 2.89E-07 | 2.48E-06 | 2575.009631 | 2069.203039 | ENSG00000114686 |

| | | | | | | | |
|---|---|---|---|---|---|---|---|
| ZNF300 | -0.316297016 | 0.803128635 | 0.067652795 | 0.151860608 | 198.8665509 | 159.5800965 | ENSG00000145908 |
| STK40 | -0.316308203 | 0.803122407 | 5.00E-05 | 0.00028761 | 968.5809888 | 777.8383101 | ENSG00000196182 |
| TMEM189 | -0.316310605 | 0.80312107 | 0.011989308 | 0.036607428 | 317.481704 | 254.9164282 | ENSG00000240849 |
| LOC100289495 | -0.316342473 | 0.80310333 | 0.667782008 | 0.795884265 | 9.770277078 | 7.803937354 | ENSG00000286760 |
| ATP6V1F | -0.316360455 | 0.80309332 | 0.015162281 | 0.044676274 | 364.4262823 | 292.78103 | ENSG00000128524 |
| CDC123 | -0.316371379 | 0.803087239 | 1.29E-07 | 1.17E-06 | 3213.701926 | 2580.844169 | ENSG00000151465 |
| HECTD4 | -0.316644676 | 0.802935121 | 0.000558183 | 0.002528866 | 770.4302521 | 618.2851229 | ENSG00000173064 |
| NA | -0.316657144 | 0.802928181 | 0.673865559 | 0.800111161 | 10.13248263 | 8.073086909 | ENSG00000262227 |
| ARSA | -0.316727895 | 0.802888806 | 0.022055158 | 0.060928834 | 266.4521793 | 213.9113508 | ENSG00000100299 |
| DLEU2 | -0.316778177 | 0.802860824 | 0.65588898 | 0.786579245 | 10.22726118 | 8.255922518 | ENSG00000231607 |
| MORC4 | -0.316860297 | 0.802815125 | 0.043764842 | 0.107274683 | 200.0665313 | 160.5434773 | ENSG00000133131 |
| TUB | -0.316889026 | 0.802799139 | 1.00E-05 | 6.60E-05 | 1374.526086 | 1103.473861 | ENSG00000166402 |
| GRAMD1B | -0.317012651 | 0.80273035 | 0.003325407 | 0.012268201 | 460.6360889 | 369.4932054 | ENSG00000023171 |
| GSS | -0.317047406 | 0.802711012 | 0.000169681 | 0.000868484 | 909.4560979 | 729.5976841 | ENSG00000100983 |
| MRPS21 | -0.317259835 | 0.802592826 | 1.49E-05 | 9.48E-05 | 1132.534205 | 908.529275 | ENSG00000266472 |
| NUP160 | -0.317408048 | 0.802510377 | 7.96E-10 | 9.79E-09 | 3569.917259 | 2864.932224 | ENSG00000030066 |
| NA | -0.317457568 | 0.802482831 | 0.702948307 | 0.821305541 | 7.465219758 | 5.990617367 | ENSG00000275325 |
| ST6GALNAC3 | -0.317512213 | 0.802452436 | 0.008765127 | 0.02818859 | 399.5964057 | 320.8073006 | ENSG00000184005 |
| SLC9A7 | -0.317590247 | 0.802409033 | 0.003615833 | 0.013220819 | 484.9346514 | 389.1565577 | ENSG00000065923 |
| RAD54B | -0.31771258 | 0.802340996 | 0.097313551 | 0.202817169 | 140.0290619 | 112.3622802 | ENSG00000265817 |
| WDR55 | -0.317785208 | 0.802300606 | 0.001921548 | 0.007617875 | 507.1474498 | 406.9143938 | ENSG00000120314 |
| NA | -0.317842046 | 0.802268998 | 0.776164525 | 0.868654049 | 3.663131159 | 2.93427464 | ENSG00000226709 |
| PNKD | -0.317963119 | 0.802201674 | 0.000349156 | 0.001668458 | 703.4474985 | 564.199356 | ENSG00000127838 |
| GHITM | -0.317972082 | 0.80219669 | 3.63E-09 | 4.11E-08 | 2800.580671 | 2246.653843 | ENSG00000165678 |
| TRIR | -0.317990978 | 0.802186183 | 1.06E-07 | 9.77E-07 | 3761.62143 | 3017.707107 | ENSG00000123144 |
| ACVR2B-AS1 | -0.318069452 | 0.80214255 | 0.369874942 | 0.54372152 | 41.81380125 | 33.5474682 | ENSG00000229589 |
| CALN1 | -0.318220421 | 0.802058615 | 0.002165116 | 0.008461761 | 559.7870445 | 449.421963 | ENSG00000183166 |
| WDR81 | -0.318241247 | 0.802047037 | 0.006961124 | 0.023281386 | 615.373908 | 493.3355871 | ENSG00000167716 |
| PHF2 | -0.318306577 | 0.802010719 | 0.009004408 | 0.028801864 | 442.5069112 | 354.6110512 | ENSG00000197724 |
| NA | -0.318350668 | 0.801986208 | 0.022684793 | 0.062373482 | 289.1551771 | 231.9458606 | ENSG00000284691 |
| ARHGAP22 | -0.31841158 | 0.801952348 | 0.014145352 | 0.04216937 | 426.4571279 | 341.6860513 | ENSG00000128805 |
| HLA-DPB1 | -0.318507575 | 0.801898989 | 0.021047432 | 0.058606525 | 302.515146 | 242.937865 | ENSG00000223865 |
| NA | -0.318511106 | 0.801897027 | 0.648780365 | 0.781652034 | 12.10814951 | 9.762176127 | ENSG00000146677 |
| SNRPC | -0.318562602 | 0.801868404 | 2.90E-08 | 2.88E-07 | 2447.210527 | 1961.951941 | ENSG00000124562 |
| LAMTOR1 | -0.318691657 | 0.801796677 | 1.61E-05 | 0.000101699 | 1057.271911 | 847.725286 | ENSG00000149357 |
| TAP1 | -0.318709351 | 0.801786843 | 0.00490814 | 0.017265303 | 404.9135103 | 324.5528041 | ENSG00000168394 |
| RAB8A | -0.318730084 | 0.801775321 | 0.012977299 | 0.039208311 | 410.7873348 | 329.5200666 | ENSG00000167461 |
| POLR2B | -0.3188631 | 0.801701401 | 2.22E-09 | 2.58E-08 | 3023.557605 | 2424.178634 | ENSG00000047315 |
| INSYN2A | -0.318897948 | 0.801682036 | 0.638863233 | 0.774334671 | 13.5903504 | 10.94417088 | ENSG00000188916 |
| GUSB | -0.318954352 | 0.801650694 | 0.000141798 | 0.000737509 | 951.1976253 | 762.9012829 | ENSG00000169919 |
| GARS1 | -0.31896286 | 0.801645966 | 2.26E-10 | 3.00E-09 | 4864.759515 | 3899.171801 | ENSG00000106105 |
| MICAL3 | -0.318983747 | 0.801634361 | 0.000711502 | 0.00312798 | 882.5986564 | 706.9947141 | ENSG00000243156 |
| GNA14 | -0.319134798 | 0.801550433 | 0.019986596 | 0.05620429 | 305.7186512 | 244.9321426 | ENSG00000156049 |
| FBN3 | -0.319174537 | 0.801528355 | 1.04E-06 | 8.15E-06 | 1942.714845 | 1556.603229 | ENSG00000142449 |
| NA | -0.319198417 | 0.801515088 | 0.569922428 | 0.722981099 | 15.566693 | 12.52598556 | ENSG00000269069 |
| GPT2 | -0.31931993 | 0.801447582 | 3.83E-07 | 3.23E-06 | 2802.437658 | 2245.657829 | ENSG00000166123 |
| E2F3 | -0.31940804 | 0.801398636 | 3.04E-06 | 2.19E-05 | 1325.576631 | 1062.321169 | ENSG00000112242 |
| OTUB1 | -0.319422694 | 0.801390496 | 6.50E-08 | 6.15E-07 | 2089.987094 | 1674.403288 | ENSG00000167770 |
| PICK1 | -0.319650186 | 0.801264139 | 0.001065626 | 0.00451207 | 633.1984742 | 506.9228138 | ENSG00000100151 |
| VKORC1L1 | -0.319693268 | 0.801240211 | 4.76E-06 | 3.32E-05 | 1556.194162 | 1247.174274 | ENSG00000196715 |
| CENPO | -0.319887428 | 0.801132386 | 0.001727375 | 0.006946588 | 617.6686776 | 495.0195269 | ENSG00000138092 |
| MXD4 | -0.319978191 | 0.801081987 | 0.000127596 | 0.0006715 | 1062.572933 | 851.491738 | ENSG00000123933 |
| ERCC8 | -0.320148608 | 0.800987366 | 0.103716336 | 0.213099566 | 152.5520438 | 122.2083592 | ENSG00000049167 |
| CABLES1 | -0.320203146 | 0.800957087 | 0.004134229 | 0.014871043 | 444.2260024 | 355.9928015 | ENSG00000134508 |
| SNORA55 | -0.320235569 | 0.800939086 | 0.780908754 | 0.871424971 | 3.668712964 | 2.932124067 | ENSG00000201457 |
| NMRK2 | -0.32029023 | 0.800908741 | 9.24E-05 | 0.000504081 | 837.430414 | 670.801167 | ENSG00000077009 |
| PYGB | -0.320328236 | 0.800887642 | 5.86E-09 | 6.43E-08 | 2711.105128 | 2170.6968 | ENSG00000100994 |
| CCDC150 | -0.320492799 | 0.800796293 | 0.022977086 | 0.06299758 | 268.2499282 | 214.8583326 | ENSG00000144395 |
| MFAP1 | -0.320601291 | 0.800736075 | 3.37E-07 | 2.87E-06 | 1637.04518 | 1310.892322 | ENSG00000140259 |
| ANKRD45 | -0.320601382 | 0.800736024 | 0.735043952 | 0.842176925 | 5.375037836 | 4.312008443 | ENSG00000183831 |
| ZNF836 | -0.320620622 | 0.800725345 | 0.238862853 | 0.400791798 | 70.84604188 | 56.84150122 | ENSG00000196267 |
| NHLRC3 | -0.320645493 | 0.800711542 | 0.038825071 | 0.097345263 | 227.4268138 | 182.0120925 | ENSG00000188811 |
| AKR1C3 | -0.32066383 | 0.800701365 | 0.417249497 | 0.589333206 | 44.39378239 | 35.68960866 | ENSG00000196139 |
| EIF3I | -0.320876227 | 0.800583492 | 3.77E-13 | 6.74E-12 | 5672.715436 | 4541.246208 | ENSG00000084623 |
| LOC441242 | -0.321032781 | 0.800496622 | 0.499963321 | 0.666214853 | 21.3772636 | 17.13456798 | ENSG00000272693 |
| RBPMS2 | -0.321034935 | 0.800495426 | 3.59E-08 | 3.52E-07 | 3657.608133 | 2928.106405 | ENSG00000166831 |
| NA | -0.321162895 | 0.800424429 | 0.797078476 | 0.882500369 | 3.330949821 | 2.688248359 | ENSG00000229251 |
| GALM | -0.321186557 | 0.800411302 | 0.007098266 | 0.023649854 | 392.0835404 | 313.5973594 | ENSG00000143891 |
| ASTL | -0.3212265 | 0.800389141 | 0.695011505 | 0.815746479 | 6.975220381 | 5.599399727 | ENSG00000188886 |
| OS9 | -0.321242576 | 0.800380223 | 9.13E-07 | 7.23E-06 | 2034.853835 | 1627.934328 | ENSG00000135506 |
| FAM43A | -0.321359472 | 0.800315374 | 0.126554249 | 0.248004675 | 123.7239153 | 98.92103836 | ENSG00000185112 |
| C19orf53 | -0.321369029 | 0.800310072 | 2.40E-06 | 1.76E-05 | 2078.54899 | 1662.873555 | ENSG00000104979 |
| SRM | -0.321372565 | 0.800308111 | 7.20E-07 | 5.80E-06 | 3381.665396 | 2706.939437 | ENSG00000116649 |
| ANO1 | -0.321395102 | 0.800295609 | 0.104443211 | 0.214114131 | 166.6252602 | 133.3144051 | ENSG00000131620 |
| SNX7 | -0.322060698 | 0.799926473 | 0.004647032 | 0.016485086 | 445.1115349 | 355.8500333 | ENSG00000162627 |
| POLR3A | -0.322292431 | 0.799797995 | 2.61E-06 | 1.90E-05 | 1493.13815 | 1194.835638 | ENSG00000148606 |
| PEX10 | -0.322304895 | 0.799791085 | 0.006178325 | 0.021046455 | 468.3125684 | 374.584705 | ENSG00000157911 |

| | | | | | | | |
|---|---|---|---|---|---|---|---|
| LUC7L | -0.322331432 | 0.799776374 | 1.28E-08 | 1.34E-07 | 2069.390148 | 1655.094533 | ENSG00000007392 |
| NA | -0.322423492 | 0.799725341 | 0.727125667 | 0.836082516 | 6.139961033 | 4.861060419 | ENSG00000245322 |
| KDELR2 | -0.322477562 | 0.799695369 | 2.27E-05 | 0.000139674 | 1249.022132 | 998.9198741 | ENSG00000136240 |
| PLCE1 | -0.322698523 | 0.799572898 | 0.422488719 | 0.594054742 | 30.30929027 | 24.2356597 | ENSG00000138193 |
| PCOLCE2 | -0.322773614 | 0.799531282 | 0.034995702 | 0.089169475 | 275.4647546 | 220.5235824 | ENSG00000163710 |
| SLC12A2 | -0.322948255 | 0.799434503 | 0.014233808 | 0.042393762 | 319.2619949 | 255.5179059 | ENSG00000064651 |
| RPTOR | -0.323014232 | 0.799397944 | 4.47E-05 | 0.000259645 | 1016.944102 | 812.4787813 | ENSG00000141564 |
| NA | -0.32301786 | 0.799395934 | 0.577495651 | 0.729374663 | 15.24846618 | 12.25038429 | ENSG00000271966 |
| METTL6 | -0.323029618 | 0.799389419 | 0.029333905 | 0.077095418 | 265.3826714 | 212.4066842 | ENSG00000206562 |
| SELENOK | -0.323083492 | 0.799359568 | 0.000664379 | 0.002945655 | 629.8483004 | 503.2490911 | ENSG00000113811 |
| CAB39L | -0.323096221 | 0.799352516 | 0.218850801 | 0.376635584 | 76.7166609 | 61.37264456 | ENSG00000102547 |
| AGO1 | -0.323096995 | 0.799352087 | 1.26E-08 | 1.33E-07 | 2536.461309 | 2027.646499 | ENSG00000092847 |
| SOX12 | -0.323268961 | 0.799256811 | 1.23E-07 | 1.12E-06 | 4163.580541 | 3328.380926 | ENSG00000177732 |
| DDB1 | -0.323304013 | 0.799237393 | 4.39E-16 | 1.06E-14 | 10085.51147 | 8059.838247 | ENSG00000167986 |
| REXO4 | -0.323479344 | 0.799140267 | 3.88E-07 | 3.26E-06 | 1612.979641 | 1288.85238 | ENSG00000148300 |
| EIF4A3 | -0.323484063 | 0.799137653 | 1.34E-10 | 1.83E-09 | 5240.572085 | 4188.011904 | ENSG00000141543 |
| LINC01122 | -0.323503224 | 0.79912704 | 0.488719947 | 0.655460281 | 24.31195346 | 19.37083117 | ENSG00000233723 |
| ZBTB8A | -0.323556674 | 0.799097433 | 0.123729921 | 0.243907001 | 127.827173 | 102.2951882 | ENSG00000160062 |
| ARHGEF19 | -0.323559461 | 0.79909589 | 6.81E-05 | 0.000381501 | 1481.652211 | 1183.684824 | ENSG00000142632 |
| ZNF213 | -0.323810974 | 0.798956591 | 0.033607663 | 0.086141338 | 246.031311 | 196.7716577 | ENSG00000085644 |
| REEP2 | -0.323998359 | 0.798852825 | 5.54E-05 | 0.000316162 | 944.1236714 | 754.5757117 | ENSG00000132563 |
| NA | -0.324039969 | 0.798829785 | 0.057665222 | 0.133736452 | 174.7386096 | 139.4881185 | ENSG00000247092 |
| MRPL33 | -0.324077396 | 0.798809062 | 0.010685005 | 0.033323918 | 396.3387831 | 316.790509 | ENSG00000243147 |
| PRKY | -0.324111323 | 0.798790277 | 0.639981693 | 0.774765411 | 10.3005884 | 8.258073092 | ENSG00000099725 |
| RAB11B | -0.324269444 | 0.798702733 | 4.37E-06 | 3.07E-05 | 1749.878387 | 1398.276314 | ENSG00000185236 |
| SMARCA1 | -0.324385872 | 0.798638279 | 2.57E-10 | 3.39E-09 | 3330.25225 | 2659.786065 | ENSG00000102038 |
| SLC39A13 | -0.324394235 | 0.798633649 | 0.001097323 | 0.004632049 | 543.7942675 | 434.1711935 | ENSG00000165915 |
| MESD | -0.324396889 | 0.79863218 | 1.52E-06 | 1.16E-05 | 1373.357032 | 1096.757317 | ENSG00000117899 |
| PCDHA4 | -0.324440648 | 0.798607957 | 0.149468127 | 0.281636025 | 107.5103687 | 85.69865889 | ENSG00000204967 |
| PRSS12 | -0.324486319 | 0.798582677 | 0.04477767 | 0.109287201 | 207.4640056 | 165.6599598 | ENSG00000164099 |
| CLDN7 | -0.324601563 | 0.798518887 | 8.59E-07 | 6.84E-06 | 1477.898018 | 1179.765838 | ENSG00000181885 |
| CLDN3 | -0.324605686 | 0.798516605 | 0.021557337 | 0.059810527 | 342.2862235 | 273.2144414 | ENSG00000165215 |
| GOT2 | -0.324611232 | 0.798513536 | 3.69E-12 | 5.92E-11 | 4814.722382 | 3844.250931 | ENSG00000125166 |
| NA | -0.324625721 | 0.798505516 | 0.728297187 | 0.837081806 | 5.756103983 | 4.568787592 | ENSG00000217325 |
| NA | -0.324713638 | 0.798456857 | 0.632488738 | 0.7694123 | 10.75286658 | 8.588523208 | ENSG00000242588 |
| FAM122B | -0.32476715 | 0.798427241 | 3.38E-05 | 0.000201826 | 1076.400899 | 859.2913495 | ENSG00000156504 |
| NA | -0.324892701 | 0.798357761 | 0.784308802 | 0.873764212 | 3.309298319 | 2.662974513 | ENSG00000231173 |
| TXNDC15 | -0.324943781 | 0.798329495 | 0.000598392 | 0.002686433 | 622.4951312 | 497.0616502 | ENSG00000113621 |
| NA | -0.325015242 | 0.798289953 | 0.2863009 | 0.455395008 | 56.59411971 | 45.07215616 | ENSG00000254510 |
| CASKIN1 | -0.325075448 | 0.798256639 | 0.043880107 | 0.107477291 | 204.6944305 | 163.4852882 | ENSG00000167971 |
| UMAD1 | -0.32524844 | 0.798160927 | 0.233367853 | 0.394142816 | 76.69413352 | 61.17717808 | ENSG00000219545 |
| NUP62 | -0.325290144 | 0.798137855 | 2.14E-08 | 2.16E-07 | 2392.611687 | 1910.169347 | ENSG00000213024 |
| RNF25 | -0.32537833 | 0.798089069 | 0.00166133 | 0.006703351 | 500.7491102 | 399.6355916 | ENSG00000163481 |
| LRFN3 | -0.325676733 | 0.797924012 | 0.007000307 | 0.023380006 | 427.5041836 | 341.3106754 | ENSG00000126243 |
| CAPN15 | -0.325698204 | 0.797912137 | 5.09E-05 | 0.000292502 | 2259.572754 | 1802.919388 | ENSG00000103326 |
| FPGT | -0.32580009 | 0.797855789 | 0.370025755 | 0.543818897 | 38.17423661 | 30.50402892 | ENSG00000254685 |
| MBLAC1 | -0.325875508 | 0.797814081 | 0.415514794 | 0.587747169 | 31.30603444 | 24.90462919 | ENSG00000214309 |
| NA | -0.325877155 | 0.79781317 | 0.66115233 | 0.790816567 | 10.25728539 | 8.201606715 | ENSG00000234630 |
| ACOX2 | -0.325913584 | 0.797793025 | 0.529572599 | 0.69068927 | 18.16920152 | 14.47374404 | ENSG00000168306 |
| NA | -0.325961431 | 0.797766567 | 0.662583009 | 0.791840467 | 8.989284186 | 7.155940565 | ENSG00000282787 |
| DEF8 | -0.32615818 | 0.797657778 | 1.71E-06 | 1.29E-05 | 1435.562792 | 1145.040688 | ENSG00000140995 |
| PRLR | -0.326169506 | 0.797651516 | 0.394211444 | 0.566882815 | 37.09898374 | 29.6057047 | ENSG00000113494 |
| USP5 | -0.326225994 | 0.797620285 | 5.88E-11 | 8.28E-10 | 4350.783039 | 3469.507059 | ENSG00000111667 |
| PRPF19 | -0.326313287 | 0.797572025 | 4.62E-10 | 5.88E-09 | 4348.481664 | 3467.542187 | ENSG00000110107 |
| GUCY2D | -0.326370658 | 0.797540309 | 0.779465933 | 0.870480335 | 3.668712964 | 2.913301941 | ENSG00000132518 |
| NCBP1 | -0.326550838 | 0.797440709 | 4.34E-07 | 3.62E-06 | 1919.856434 | 1531.728084 | ENSG00000136937 |
| COPS9 | -0.326566264 | 0.797432183 | 3.78E-05 | 0.000222692 | 1006.856274 | 802.4998997 | ENSG00000172428 |
| ZPR1 | -0.326642663 | 0.797389955 | 1.45E-07 | 1.30E-06 | 1601.178503 | 1276.755214 | ENSG00000109917 |
| PROZ | -0.326687722 | 0.797365051 | 0.683487934 | 0.806876307 | 8.218779183 | 6.590217036 | ENSG00000126231 |
| IRX6 | -0.326755629 | 0.79732752 | 0.639349753 | 0.77469615 | 11.0305813 | 8.760878523 | ENSG00000159387 |
| IFT46 | -0.32676466 | 0.79732253 | 0.034676804 | 0.08845038 | 235.0203029 | 187.4829603 | ENSG00000118096 |
| AKAP8L | -0.326841031 | 0.797280323 | 7.96E-06 | 5.35E-05 | 1241.215881 | 988.9815746 | ENSG00000011243 |
| CASD1 | -0.326914734 | 0.797239594 | 0.008370521 | 0.027120545 | 383.4990011 | 305.8124741 | ENSG00000127995 |
| APLP1 | -0.326991237 | 0.797197319 | 0.000114961 | 0.000613175 | 830.8122546 | 661.9045235 | ENSG00000105290 |
| NOP10 | -0.327089887 | 0.797142809 | 5.94E-07 | 4.83E-06 | 1655.127175 | 1318.89879 | ENSG00000182117 |
| TSEN34 | -0.327187112 | 0.79708909 | 0.002269802 | 0.008808439 | 502.8337983 | 400.9815878 | ENSG00000170892 |
| C4orf19 | -0.327265858 | 0.797045585 | 0.615918509 | 0.758212125 | 13.52607161 | 10.80758181 | ENSG00000154274 |
| LMO2 | -0.327324424 | 0.797013229 | 0.370869256 | 0.544602178 | 40.42099727 | 32.12993958 | ENSG00000135363 |
| HNRNPDL | -0.327426858 | 0.796956642 | 1.51E-16 | 3.82E-15 | 7864.648082 | 6268.272736 | ENSG00000152795 |
| PRKCH | -0.327569366 | 0.796877923 | 0.028342609 | 0.075062777 | 292.6946696 | 233.4249197 | ENSG00000027075 |
| ATP8A2 | -0.32761621 | 0.796852049 | 0.01270248 | 0.0385228 | 343.341852 | 273.3897713 | ENSG00000132932 |
| NTMT1 | -0.328093372 | 0.796588539 | 0.000800172 | 0.003475822 | 789.1562104 | 628.3861094 | ENSG00000148335 |
| LSS | -0.328095251 | 0.796587501 | 1.18E-08 | 1.25E-07 | 5528.553115 | 4404.203185 | ENSG00000160285 |
| PCSK4 | -0.328119089 | 0.796574339 | 0.098375954 | 0.204501134 | 137.2851173 | 109.4196637 | ENSG00000115257 |
| PDE3B | -0.328126816 | 0.796570073 | 0.019362941 | 0.054705695 | 380.631432 | 303.5818386 | ENSG00000152270 |
| OTUD6B | -0.328317001 | 0.796465071 | 0.004905817 | 0.017260281 | 486.1818032 | 387.6390182 | ENSG00000155100 |
| PCM1 | -0.32834476 | 0.796449746 | 4.40E-08 | 4.26E-07 | 2086.150476 | 1661.931806 | ENSG00000078674 |

| | | | | | | | |
|---|---|---|---|---|---|---|---|
| NA | -0.328384513 | 0.796427801 | 0.699509739 | 0.818577032 | 8.20570055 | 6.470299524 | ENSG00000244459 |
| SLC16A2 | -0.328404056 | 0.796417012 | 5.00E-09 | 5.54E-08 | 2430.636804 | 1936.093575 | ENSG00000147100 |
| NA | -0.328436091 | 0.796399328 | 0.531481285 | 0.692377304 | 18.59915248 | 14.75364646 | ENSG00000272449 |
| LOX | -0.328457755 | 0.796387369 | 0.400658482 | 0.573162979 | 32.60039307 | 26.02396631 | ENSG00000113083 |
| MROH6 | -0.32859538 | 0.796311402 | 0.230258949 | 0.390533268 | 72.04226747 | 57.25962235 | ENSG00000204839 |
| NA | -0.328790915 | 0.796203482 | 0.420128016 | 0.59212554 | 33.55941605 | 26.72896251 | ENSG00000260565 |
| PUS3 | -0.32889223 | 0.796147569 | 0.007883263 | 0.025797461 | 371.6047758 | 295.7730792 | ENSG00000110060 |
| CELF2 | -0.329043414 | 0.796064143 | 0.001016708 | 0.004325829 | 555.8775743 | 442.589512 | ENSG00000048740 |
| ZNF592 | -0.329117695 | 0.796023156 | 1.61E-06 | 1.22E-05 | 1499.929107 | 1194.087133 | ENSG00000166716 |
| CNTN4 | -0.329160745 | 0.795999403 | 0.487999401 | 0.65488671 | 22.23138355 | 17.73201708 | ENSG00000144619 |
| ALG3 | -0.329181677 | 0.795987855 | 1.89E-05 | 0.000118048 | 1050.973657 | 836.432097 | ENSG00000214160 |
| ZNF74 | -0.329279362 | 0.79593396 | 1.38E-07 | 1.24E-06 | 1920.010459 | 1527.87003 | ENSG00000185252 |
| NA | -0.329318606 | 0.795912309 | 0.440232842 | 0.611204314 | 26.34400198 | 20.96071512 | ENSG00000237126 |
| HDAC10 | -0.32939153 | 0.795872079 | 0.732994849 | 0.840528065 | 7.048547595 | 5.611770131 | ENSG00000100429 |
| ADPRHL2 | -0.329392042 | 0.795871797 | 0.000334987 | 0.001607521 | 682.2055255 | 543.0321369 | ENSG00000116863 |
| TMSB15B | -0.329519447 | 0.795801516 | 0.320976323 | 0.492323626 | 51.23371211 | 40.76255876 | ENSG00000158427 |
| GATB | -0.329533819 | 0.795793588 | 0.003164008 | 0.011731093 | 437.3404159 | 347.8578625 | ENSG00000059691 |
| TSR2 | -0.329562277 | 0.795777892 | 2.44E-05 | 0.000149864 | 1356.060475 | 1079.230447 | ENSG00000158526 |
| ZNF286A | -0.329593269 | 0.795760796 | 0.004175063 | 0.01499617 | 482.7211446 | 384.5786348 | ENSG00000187607 |
| RAB40B | -0.329668125 | 0.795719509 | 0.358343217 | 0.531467595 | 41.80004689 | 33.18797613 | ENSG00000141542 |
| HDAC3 | -0.329711977 | 0.795695323 | 5.29E-07 | 4.34E-06 | 1635.406475 | 1300.979642 | ENSG00000171720 |
| ZKSCAN4 | -0.329747237 | 0.795675875 | 0.054626839 | 0.128106777 | 182.8659647 | 145.3332841 | ENSG00000187626 |
| DIRAS1 | -0.329751988 | 0.795673255 | 0.001010664 | 0.004304918 | 939.1883598 | 747.2308295 | ENSG00000176490 |
| NA | -0.329767555 | 0.79566467 | 0.667375513 | 0.795634837 | 8.670381641 | 6.905613136 | ENSG00000270587 |
| DBP | -0.3297892 | 0.795652732 | 0.104624216 | 0.214308047 | 164.1824847 | 130.7769942 | ENSG00000105516 |
| LMNTD2-AS1 | -0.32994568 | 0.795566437 | 0.619276269 | 0.760630237 | 16.9466185 | 13.43883076 | ENSG00000254815 |
| SIPA1L3 | -0.329992599 | 0.795540565 | 3.54E-05 | 0.00021021 | 1029.698193 | 819.1004329 | ENSG00000105738 |
| NINL | -0.330187658 | 0.795433012 | 1.59E-07 | 1.42E-06 | 1885.961708 | 1499.802304 | ENSG00000101004 |
| MPHOSPH6 | -0.330299608 | 0.79537129 | 0.0022219 | 0.008645153 | 643.5644316 | 512.3046414 | ENSG00000135698 |
| OSCP1 | -0.330632489 | 0.795187791 | 0.512011838 | 0.676336645 | 22.18596536 | 17.66264726 | ENSG00000116885 |
| CCNJL | -0.330652814 | 0.795176588 | 1.42E-05 | 9.08E-05 | 1134.691895 | 902.2127876 | ENSG00000135083 |
| TFDP1 | -0.330745184 | 0.795125678 | 3.23E-08 | 3.18E-07 | 2661.982673 | 2116.178967 | ENSG00000198176 |
| NDUFAF5 | -0.331105501 | 0.794927118 | 0.030244069 | 0.079056686 | 264.3659764 | 209.9572169 | ENSG00000101247 |
| H2AZ1 | -0.331213731 | 0.794867485 | 2.69E-13 | 4.89E-12 | 5037.470698 | 4003.834245 | ENSG00000164032 |
| EOGT | -0.331283341 | 0.794829134 | 0.014895554 | 0.044011354 | 314.9444254 | 250.6388168 | ENSG00000163378 |
| KPNB1 | -0.331325138 | 0.794806106 | 3.23E-17 | 8.68E-16 | 11124.81543 | 8842.084821 | ENSG00000108424 |
| NDUFS6 | -0.331412643 | 0.7947579 | 3.39E-08 | 3.33E-07 | 3213.38137 | 2553.298144 | ENSG00000145494 |
| C4orf46 | -0.331428068 | 0.794749403 | 0.001165486 | 0.004895144 | 508.6207144 | 404.3522421 | ENSG00000205208 |
| MIS18A | -0.331554619 | 0.794679692 | 8.75E-06 | 5.83E-05 | 1865.336154 | 1483.070161 | ENSG00000159055 |
| RNF8 | -0.331695759 | 0.794601951 | 2.01E-05 | 0.000125054 | 992.1125641 | 788.6708748 | ENSG00000112130 |
| AADAT | -0.331765192 | 0.79456371 | 0.02459871 | 0.066694831 | 251.208419 | 199.5508121 | ENSG00000109576 |
| GCA | -0.33180074 | 0.794544132 | 0.003300042 | 0.012179284 | 495.6915113 | 394.0985464 | ENSG00000115271 |
| FAM122A | -0.331805279 | 0.794541632 | 0.426055748 | 0.597708465 | 28.12712069 | 22.33629835 | ENSG00000187866 |
| NA | -0.331810852 | 0.794538563 | 0.674883997 | 0.800563221 | 8.779990593 | 6.98358525 | ENSG00000284602 |
| BAG3 | -0.331883382 | 0.794498619 | 0.00160672 | 0.006506138 | 644.777502 | 512.6550102 | ENSG00000151929 |
| PPT2 | -0.331904007 | 0.794487261 | 0.112765809 | 0.227271734 | 118.7867255 | 94.26970494 | ENSG00000221988 |
| PCED1A | -0.331940205 | 0.794467328 | 0.001865738 | 0.007431748 | 503.8255853 | 400.1682204 | ENSG00000132635 |
| PXYLP1 | -0.33217849 | 0.794336119 | 0.006086881 | 0.020801067 | 447.9753009 | 356.0460086 | ENSG00000155893 |
| MOB2 | -0.332206603 | 0.79432064 | 0.00564628 | 0.019485221 | 441.3134523 | 350.5840461 | ENSG00000182208 |
| AP1M1 | -0.332225583 | 0.79431019 | 7.47E-05 | 0.000415419 | 800.2081039 | 635.4140389 | ENSG00000072958 |
| AVP | -0.332296903 | 0.794270924 | 0.696711702 | 0.816923967 | 6.613014833 | 5.277551906 | ENSG00000101200 |
| TBL1X | -0.332383762 | 0.794223106 | 4.24E-08 | 4.13E-07 | 1949.29183 | 1548.54997 | ENSG00000101849 |
| CDC34 | -0.332419838 | 0.794203246 | 5.51E-09 | 6.08E-08 | 2798.756754 | 2222.590685 | ENSG00000099804 |
| PSMD9 | -0.332423881 | 0.79420102 | 0.600703403 | 0.746708509 | 14.48565816 | 11.56904439 | ENSG00000110801 |
| UCKL1-AS1 | -0.332458559 | 0.79418193 | 0.605661875 | 0.750292827 | 12.70409106 | 10.04396866 | ENSG00000280213 |
| NA | -0.332701369 | 0.794048278 | 0.672466489 | 0.799434862 | 9.19108095 | 7.280159224 | ENSG00000242615 |
| PTPRZ1 | -0.332786868 | 0.794001222 | 2.57E-12 | 4.21E-11 | 5874.543296 | 4664.350638 | ENSG00000106278 |
| GMCL1 | -0.332892726 | 0.793942964 | 0.000438314 | 0.002043381 | 715.5096052 | 568.4788028 | ENSG00000087338 |
| DOK2 | -0.332969939 | 0.793900473 | 0.780123504 | 0.870840926 | 3.33932253 | 2.665125086 | ENSG00000147443 |
| NA | -0.333001585 | 0.793883059 | 0.533725002 | 0.694180758 | 18.53053119 | 14.73859245 | ENSG00000272476 |
| CEP162 | -0.333130393 | 0.793812182 | 0.062839175 | 0.143096761 | 171.7715802 | 136.3365991 | ENSG00000135315 |
| GTF3C6 | -0.33314783 | 0.793802588 | 5.53E-06 | 3.81E-05 | 1301.005162 | 1033.154267 | ENSG00000155115 |
| LOC100288748 | -0.333183167 | 0.793783145 | 0.466480529 | 0.635722431 | 25.99104502 | 20.52325093 | ENSG00000261437 |
| HLA-A | -0.333461387 | 0.79363008 | 1.73E-11 | 2.57E-10 | 6245.539557 | 4956.020001 | ENSG00000206503 |
| CCN3 | -0.33353186 | 0.793591314 | 0.760482602 | 0.857655235 | 4.582881332 | 3.676915095 | ENSG00000136999 |
| SRP14 | -0.333681621 | 0.793508938 | 1.09E-12 | 1.84E-11 | 5930.865938 | 4705.904635 | ENSG00000140319 |
| SMOC1 | -0.333751991 | 0.793470235 | 0.389581994 | 0.562784652 | 33.52572505 | 26.68003236 | ENSG00000198732 |
| EIF3E | -0.333755543 | 0.793468281 | 2.29E-10 | 3.04E-09 | 6230.583037 | 4944.881331 | ENSG00000104408 |
| NDUFA11 | -0.333765841 | 0.793462617 | 0.174759827 | 0.317746637 | 89.87406928 | 71.2921341 | ENSG00000174886 |
| CYP27C1 | -0.333861684 | 0.793409906 | 0.111173284 | 0.224523014 | 121.4209731 | 96.19838082 | ENSG00000186684 |
| PTTG1IP | -0.334047212 | 0.793307882 | 9.49E-11 | 1.31E-09 | 3762.095016 | 2984.737022 | ENSG00000183255 |
| CCDC59 | -0.334099611 | 0.793279069 | 0.004198092 | 0.015064248 | 594.242383 | 471.485671 | ENSG00000133773 |
| ICMT | -0.334120966 | 0.793267327 | 2.53E-13 | 4.59E-12 | 4052.785371 | 3215.05976 | ENSG00000116237 |
| PSME2 | -0.334126011 | 0.793264553 | 1.64E-05 | 0.000103485 | 1280.814823 | 1015.36476 | ENSG00000100911 |
| CPS1 | -0.334221853 | 0.793211856 | 0.00378463 | 0.01377274 | 396.2913377 | 314.317658 | ENSG00000021826 |
| NOP16 | -0.334259303 | 0.793191266 | 0.000168912 | 0.00086478 | 714.4415099 | 566.727068 | ENSG00000048162 |
| SHKBP1 | -0.334278856 | 0.793180516 | 9.59E-07 | 7.56E-06 | 1589.962121 | 1261.047637 | ENSG00000160410 |

| | | | | | | | |
|---|---|---|---|---|---|---|---|
| FTSJ3 | -0.334729283 | 0.792932914 | 1.59E-08 | 1.65E-07 | 2724.689384 | 2160.17372 | ENSG00000108592 |
| LOC105377623 | -0.334731441 | 0.792931728 | 0.667696707 | 0.795831769 | 8.354945721 | 6.653135134 | ENSG00000250326 |
| ND2 | -0.334847821 | 0.792867766 | 0.148287015 | 0.279875237 | 13332.14062 | 10570.69754 | ENSG00000198763 |
| ASIC1 | -0.334873681 | 0.792853554 | 0.053308563 | 0.125518796 | 172.1030858 | 136.4817905 | ENSG00000110881 |
| CPXM2 | -0.334960738 | 0.792805712 | 0.051838673 | 0.122822383 | 185.6716215 | 147.2162222 | ENSG00000121898 |
| MARS1 | -0.334966898 | 0.792802327 | 7.60E-16 | 1.79E-14 | 9515.903005 | 7543.289895 | ENSG00000166986 |
| SEC61B | -0.335087859 | 0.792735858 | 5.24E-07 | 4.30E-06 | 1686.412397 | 1336.724391 | ENSG00000106803 |
| PLEKHM2 | -0.335139983 | 0.792707218 | 1.02E-07 | 9.40E-07 | 1880.617782 | 1490.372729 | ENSG00000116786 |
| TRMT1 | -0.335186783 | 0.792681503 | 1.91E-08 | 1.95E-07 | 2859.101062 | 2266.161974 | ENSG00000104907 |
| MTCP1 | -0.335271489 | 0.792634963 | 0.760009277 | 0.857538689 | 4.618487349 | 3.700038368 | ENSG00000214827 |
| LSM5 | -0.335366512 | 0.792582758 | 0.000547904 | 0.00248697 | 686.9666882 | 544.728465 | ENSG00000106355 |
| MEX3B | -0.335399773 | 0.792564485 | 0.00108577 | 0.004590312 | 618.0749371 | 490.3716706 | ENSG00000183496 |
| NUP205 | -0.335451815 | 0.792535896 | 1.57E-11 | 2.35E-10 | 5671.230818 | 4494.179429 | ENSG00000155561 |
| EDN1 | -0.335649294 | 0.792427419 | 0.713818654 | 0.828088019 | 6.760344983 | 5.378647292 | ENSG00000078401 |
| NA | -0.335699101 | 0.792400063 | 0.729771946 | 0.838027895 | 5.75331308 | 4.518239899 | ENSG00000213695 |
| NA | -0.335829272 | 0.792328569 | 0.726427427 | 0.835679616 | 5.369456031 | 4.238337476 | ENSG00000144158 |
| RNF224 | -0.335862648 | 0.792310239 | 0.724014122 | 0.834405189 | 5.429504452 | 4.28888517 | ENSG00000233198 |
| ANAPC4 | -0.335919773 | 0.792278867 | 0.000437438 | 0.00203979 | 620.3641987 | 491.2485012 | ENSG00000053900 |
| HTR3A | -0.336027298 | 0.792219821 | 0.032498148 | 0.083835487 | 207.442154 | 164.2701466 | ENSG00000166736 |
| C11orf96 | -0.336051807 | 0.792206363 | 0.63592368 | 0.772030427 | 10.00959488 | 7.959348544 | ENSG00000187479 |
| TMEM30B | -0.336083754 | 0.79218882 | 0.006274351 | 0.021332128 | 521.2006174 | 412.9921307 | ENSG00000182107 |
| ZNF84 | -0.336178686 | 0.792136694 | 0.0013177 | 0.005461914 | 523.6804624 | 415.1643855 | ENSG00000198040 |
| SLC7A1 | -0.336198358 | 0.792125893 | 2.71E-12 | 4.41E-11 | 3724.749268 | 2949.953402 | ENSG00000139514 |
| UBXN11 | -0.336229854 | 0.7921086 | 0.044362875 | 0.108411986 | 211.7103631 | 167.5572011 | ENSG00000158062 |
| NA | -0.336252938 | 0.792095926 | 0.246443829 | 0.409922629 | 67.82985226 | 53.81258291 | ENSG00000254612 |
| RBFADN | -0.336287275 | 0.792077074 | 0.75160975 | 0.85300057 | 5.018414094 | 3.988010048 | ENSG00000261126 |
| TUG1 | -0.336314501 | 0.792062126 | 3.08E-08 | 3.04E-07 | 1900.88913 | 1505.179836 | ENSG00000253352 |
| PCGF2 | -0.336461203 | 0.791981589 | 0.000262565 | 0.001290813 | 707.7478343 | 560.7689697 | ENSG00000277258 |
| DNAJC25 | -0.336512819 | 0.791953254 | 0.092696807 | 0.195006991 | 125.0890104 | 99.13532697 | ENSG00000059769 |
| LOC101927751 | -0.336571002 | 0.791921315 | 0.742822931 | 0.847821391 | 5.878115847 | 4.711828377 | ENSG00000259583 |
| NA | -0.336631679 | 0.791888009 | 0.589721814 | 0.738962228 | 14.54937337 | 11.48031941 | ENSG00000279259 |
| PPM1L | -0.336634817 | 0.791886287 | 0.026264553 | 0.070473258 | 280.9174252 | 222.5692254 | ENSG00000163590 |
| PGAP2 | -0.336672263 | 0.791865734 | 0.002262822 | 0.008788424 | 682.4839671 | 540.3296828 | ENSG00000148985 |
| AP2A2 | -0.33671147 | 0.791844214 | 1.22E-07 | 1.11E-06 | 1651.327038 | 1307.114348 | ENSG00000183020 |
| ZNF571 | -0.336751737 | 0.791822113 | 0.383646821 | 0.556921576 | 37.3981498 | 29.74497738 | ENSG00000180479 |
| LTBP3 | -0.336962867 | 0.791706243 | 0.009140892 | 0.029170712 | 303.0732031 | 239.8890706 | ENSG00000168056 |
| ANXA11 | -0.337080847 | 0.791641502 | 1.02E-08 | 1.09E-07 | 1896.263272 | 1501.075404 | ENSG00000122359 |
| IPO5 | -0.337244801 | 0.791551542 | 3.84E-17 | 1.02E-15 | 11764.69534 | 9311.930791 | ENSG00000065150 |
| CLEC18A | -0.337319317 | 0.791510658 | 0.722343177 | 0.833565964 | 5.377828739 | 4.238337476 | ENSG00000157322 |
| ZNF614 | -0.337370471 | 0.791482594 | 0.003094707 | 0.01149968 | 505.3117042 | 400.1789612 | ENSG00000142556 |
| FBRSL1 | -0.337383698 | 0.791475338 | 0.023421664 | 0.06405264 | 3137.017395 | 2482.991751 | ENSG00000112787 |
| PSMC3 | -0.337395362 | 0.791468939 | 2.56E-11 | 3.73E-10 | 4472.241486 | 3539.415547 | ENSG00000165916 |
| FAT2 | -0.337706856 | 0.79129807 | 0.267712847 | 0.434023745 | 52.37256805 | 41.50950036 | ENSG00000086570 |
| ORC2 | -0.337782885 | 0.791256371 | 1.66E-05 | 0.000105096 | 1174.77936 | 930.1422091 | ENSG00000115942 |
| MCUB | -0.337981989 | 0.791147178 | 0.015439117 | 0.045339625 | 408.996431 | 323.490509 | ENSG00000005059 |
| STXBP6 | -0.337992413 | 0.791141462 | 0.068090765 | 0.152595137 | 154.9681495 | 122.786441 | ENSG00000168952 |
| TRIM74 | -0.338099307 | 0.791082846 | 0.726688541 | 0.835880307 | 5.761685789 | 4.513938752 | ENSG00000155428 |
| RRP12 | -0.338221849 | 0.791015654 | 3.28E-06 | 2.34E-05 | 2081.542516 | 1646.699423 | ENSG00000052749 |
| C1QL3 | -0.338420827 | 0.790906564 | 0.337420316 | 0.510173133 | 49.8624475 | 39.42730339 | ENSG00000165985 |
| GDE1 | -0.338487484 | 0.790870023 | 1.02E-05 | 6.70E-05 | 1118.313146 | 884.755784 | ENSG00000006007 |
| NA | -0.338495533 | 0.79086561 | 0.699193858 | 0.818405966 | 7.02410519 | 5.540249739 | ENSG00000272277 |
| B4GALT7 | -0.338531482 | 0.790845903 | 0.003411769 | 0.012557991 | 413.0020682 | 326.5188638 | ENSG00000027847 |
| CCDC85C | -0.338767354 | 0.790716616 | 1.73E-05 | 0.000109118 | 934.4347677 | 738.5919764 | ENSG00000205476 |
| PHF14 | -0.338919871 | 0.790633028 | 2.93E-06 | 2.11E-05 | 1356.394194 | 1072.064305 | ENSG00000106443 |
| RIC8A | -0.339146713 | 0.790508723 | 6.68E-07 | 5.40E-06 | 2021.879215 | 1597.731241 | ENSG00000177963 |
| LIME1 | -0.339150857 | 0.790506452 | 0.629943961 | 0.767672076 | 11.18194165 | 8.880796035 | ENSG00000203896 |
| GTF2IP1 | -0.339166197 | 0.790498047 | 0.060158527 | 0.138222741 | 205.380757 | 162.4627332 | ENSG00000277053 |
| MOSMO | -0.339174154 | 0.790493687 | 0.108052479 | 0.219484087 | 127.566516 | 100.6348805 | ENSG00000185716 |
| SYNJ2BP | -0.339259279 | 0.790447046 | 0.0028827 | 0.010835053 | 421.7431863 | 333.5188175 | ENSG00000213463 |
| CITED4 | -0.339291826 | 0.790429214 | 0.047768629 | 0.115188078 | 249.0166005 | 196.8235745 | ENSG00000179862 |
| WIPI2 | -0.33932212 | 0.790412616 | 1.35E-08 | 1.41E-07 | 1924.043632 | 1521.078082 | ENSG00000157954 |
| PRDM10 | -0.339433102 | 0.790351815 | 0.001235871 | 0.005167141 | 483.3452322 | 381.8465751 | ENSG00000170325 |
| LINC00662 | -0.339467901 | 0.790332751 | 0.024395882 | 0.066228674 | 270.826004 | 213.7502814 | ENSG00000261824 |
| LRRC59 | -0.339492495 | 0.790319278 | 9.78E-11 | 1.35E-09 | 3242.537261 | 2562.505053 | ENSG00000108829 |
| MNAT1 | -0.339547754 | 0.790289007 | 0.004013131 | 0.014503489 | 404.2636657 | 319.2408551 | ENSG00000020426 |
| MCOLN1 | -0.339645712 | 0.790235349 | 0.008765919 | 0.02818859 | 350.3118029 | 276.7001975 | ENSG00000090674 |
| KLHDC4 | -0.339646979 | 0.790234655 | 6.27E-06 | 4.29E-05 | 1297.640521 | 1025.775969 | ENSG00000104731 |
| CACNA1C-AS2 | -0.339801928 | 0.790149787 | 0.662423111 | 0.791698422 | 8.349363916 | 6.613340309 | ENSG00000256271 |
| CBX8 | -0.33999096 | 0.790046262 | 0.349081049 | 0.521661052 | 40.83679356 | 32.27781455 | ENSG00000141570 |
| RCC1 | -0.340127374 | 0.789971563 | 1.07E-08 | 1.14E-07 | 3482.101861 | 2750.119878 | ENSG00000180198 |
| SQLE | -0.340144492 | 0.78996219 | 7.16E-14 | 1.37E-12 | 4344.424963 | 3431.903538 | ENSG00000104549 |
| ZNF385B | -0.340180603 | 0.789942417 | 0.318127149 | 0.489370877 | 44.67169727 | 35.28522712 | ENSG00000144331 |
| NA | -0.340240584 | 0.789909576 | 0.775547253 | 0.868517477 | 3.336531627 | 2.642001813 | ENSG00000225536 |
| RPL31P11 | -0.340782209 | 0.789613079 | 0.727776652 | 0.836583208 | 5.342222723 | 4.194241504 | ENSG00000213075 |
| GPT | -0.340806279 | 0.789599905 | 0.687575287 | 0.810099598 | 7.032477899 | 5.532180482 | ENSG00000167701 |
| NA | -0.340888366 | 0.789554979 | 0.72917724 | 0.837555591 | 4.958365673 | 3.920790803 | ENSG00000261634 |
| MANEAL | -0.341035312 | 0.789474563 | 0.159757371 | 0.296762585 | 90.16469937 | 71.2119993 | ENSG00000185090 |

| | | | | | | | |
|---|---|---|---|---|---|---|---|
| DVL3 | -0.341182868 | 0.789393821 | 2.45E-08 | 2.45E-07 | 2376.323537 | 1876.323718 | ENSG00000161202 |
| CD34 | -0.341348741 | 0.789303067 | 0.510213578 | 0.6746542 | 19.89630202 | 15.72510861 | ENSG00000174059 |
| PRELID1 | -0.341401835 | 0.789274019 | 1.21E-07 | 1.11E-06 | 3693.143849 | 2915.281465 | ENSG00000169230 |
| ARID1B | -0.341477094 | 0.789232847 | 4.55E-08 | 4.40E-07 | 2005.413096 | 1582.136754 | ENSG00000049618 |
| EHMT1 | -0.341556164 | 0.789189593 | 1.82E-10 | 2.44E-09 | 3445.406733 | 2718.677136 | ENSG00000181090 |
| EFEMP1 | -0.341940729 | 0.788979254 | 0.049539087 | 0.118377125 | 232.9026875 | 184.02787 | ENSG00000115380 |
| SDR39U1 | -0.342064843 | 0.788911382 | 0.152171779 | 0.28580915 | 92.22378104 | 72.79570409 | ENSG00000100445 |
| NA | -0.342293229 | 0.788786503 | 0.031050709 | 0.080782094 | 277.3229522 | 218.330125 | ENSG00000255769 |
| RABL6 | -0.342294364 | 0.788785883 | 6.03E-08 | 5.71E-07 | 3268.742246 | 2578.289372 | ENSG00000196642 |
| PRKCB | -0.342308121 | 0.788778361 | 0.010175106 | 0.031927127 | 325.6468925 | 256.8800526 | ENSG00000166501 |
| RAB3D | -0.342318985 | 0.788772421 | 0.031238746 | 0.081194645 | 223.07245 | 176.1083284 | ENSG00000105514 |
| HTATSF1 | -0.34243885 | 0.788706889 | 9.50E-09 | 1.02E-07 | 2092.039079 | 1649.76352 | ENSG00000102241 |
| PPIB | -0.342589164 | 0.788624719 | 2.84E-08 | 2.82E-07 | 2171.839965 | 1712.841417 | ENSG00000166794 |
| SLC66A1 | -0.342723089 | 0.788551514 | 0.008012848 | 0.026150541 | 390.630688 | 307.7151132 | ENSG00000040487 |
| BEX2 | -0.342751554 | 0.788535956 | 3.49E-05 | 0.000208101 | 1048.510193 | 826.4166799 | ENSG00000133134 |
| ATP5MD | -0.342871095 | 0.788470621 | 8.42E-06 | 5.63E-05 | 1271.132881 | 1001.943679 | ENSG00000173915 |
| EBNA1BP2 | -0.342989699 | 0.788405803 | 6.00E-07 | 4.88E-06 | 1438.066165 | 1133.370003 | ENSG00000117395 |
| TMEM271 | -0.343038263 | 0.788379265 | 0.805860539 | 0.887979418 | 4.273027218 | 3.403464394 | ENSG00000273238 |
| SUMF1 | -0.343051881 | 0.788371823 | 0.013500799 | 0.040523403 | 273.8254114 | 215.8841106 | ENSG00000144455 |
| SS18L2 | -0.343314759 | 0.788228184 | 0.000572347 | 0.002582723 | 883.146491 | 696.2914201 | ENSG00000008324 |
| RGMA | -0.343425041 | 0.788167933 | 5.03E-05 | 0.000288877 | 918.921028 | 724.5581076 | ENSG00000182175 |
| LAS1L | -0.343458918 | 0.788149425 | 5.46E-09 | 6.02E-08 | 2590.516362 | 2042.044452 | ENSG00000001497 |
| SMG9 | -0.343485996 | 0.788134633 | 0.001079981 | 0.004566837 | 529.0716821 | 416.8726121 | ENSG00000105771 |
| CDC26 | -0.343634262 | 0.78805364 | 0.004218414 | 0.015131541 | 353.984546 | 278.9942536 | ENSG00000176386 |
| DYNLRB1 | -0.343692319 | 0.788021928 | 0.610698484 | 0.75432094 | 11.62652284 | 9.168767714 | ENSG00000125971 |
| NA | -0.343749542 | 0.787990673 | 0.698496414 | 0.817989588 | 6.61022393 | 5.210332661 | ENSG00000235510 |
| ZNF622 | -0.343857344 | 0.787931794 | 2.94E-06 | 2.12E-05 | 1293.469643 | 1019.219665 | ENSG00000173545 |
| CMTR1 | -0.343900518 | 0.787908215 | 1.27E-06 | 9.86E-06 | 1316.144429 | 1036.983637 | ENSG00000137200 |
| ARMC5 | -0.343944444 | 0.787884226 | 0.01579051 | 0.046230797 | 300.0939709 | 236.3000685 | ENSG00000140691 |
| SORD2P | -0.344067819 | 0.787816851 | 0.211755349 | 0.367551955 | 65.70753094 | 51.74464646 | ENSG00000259479 |
| PAQR4 | -0.344085651 | 0.787807113 | 0.010931984 | 0.033980369 | 367.4594686 | 289.727904 | ENSG00000162073 |
| CDO1 | -0.34413532 | 0.787779992 | 0.004688703 | 0.01660543 | 361.6849413 | 284.9816359 | ENSG00000129596 |
| UQCC3 | -0.344137847 | 0.787778612 | 0.176051346 | 0.319462563 | 96.23919344 | 75.84209956 | ENSG00000204922 |
| H2BC20P | -0.344153273 | 0.787770188 | 0.329871266 | 0.502292729 | 44.78936663 | 35.29787009 | ENSG00000261716 |
| MIR4664 | -0.344161243 | 0.787765836 | 0.184413754 | 0.330532911 | 77.02631485 | 60.69534535 | ENSG00000265660 |
| DBI | -0.344201667 | 0.787743763 | 1.80E-09 | 2.13E-08 | 2014.27595 | 1586.615944 | ENSG00000155368 |
| NA | -0.344212093 | 0.787738071 | 0.773840466 | 0.867313818 | 3.720388677 | 2.904699646 | ENSG00000287934 |
| SEPHS1 | -0.344247541 | 0.787718716 | 3.46E-20 | 1.20E-18 | 15894.42845 | 12520.53749 | ENSG00000086475 |
| HARS1 | -0.344297916 | 0.787691211 | 2.00E-07 | 1.76E-06 | 1625.481334 | 1280.261167 | ENSG00000170445 |
| LRFN1 | -0.34439723 | 0.787636989 | 0.013180488 | 0.039751438 | 275.2117576 | 216.7719545 | ENSG00000128011 |
| SUPT16H | -0.344432184 | 0.787617906 | 7.67E-19 | 2.33E-17 | 9863.590473 | 7768.419843 | ENSG00000092201 |
| TSSC4 | -0.344587323 | 0.787533215 | 0.001413789 | 0.005816466 | 726.8064253 | 572.4622938 | ENSG00000184281 |
| KBTBD4 | -0.344684762 | 0.787480027 | 0.330212865 | 0.502574863 | 50.05826214 | 39.40123604 | ENSG00000123444 |
| NA | -0.344692485 | 0.787475812 | 0.583927201 | 0.733964094 | 14.48354298 | 11.43030476 | ENSG00000261499 |
| ADH6 | -0.344776437 | 0.787429989 | 0.737625607 | 0.844032006 | 5.423922646 | 4.249090345 | ENSG00000172955 |
| BCAP31 | -0.344786752 | 0.787424359 | 1.31E-11 | 1.98E-10 | 4002.590217 | 3151.628587 | ENSG00000185825 |
| MCM7 | -0.34482466 | 0.787403669 | 3.92E-14 | 7.73E-13 | 12332.41041 | 9709.596068 | ENSG00000166508 |
| NA | -0.344844744 | 0.787392707 | 0.340164763 | 0.51274562 | 46.93169982 | 37.02864425 | ENSG00000260257 |
| KIT | -0.344861891 | 0.787383349 | 0.000563264 | 0.002547694 | 559.5792273 | 440.5841905 | ENSG00000157404 |
| TMED3 | -0.34562417 | 0.786967428 | 7.75E-06 | 5.21E-05 | 1140.347964 | 896.9172008 | ENSG00000166557 |
| LOC100129534 | -0.345626947 | 0.786965913 | 0.480075152 | 0.648175374 | 21.70453885 | 17.08832143 | ENSG00000269896 |
| ANKRD54 | -0.345653709 | 0.786951315 | 0.022998726 | 0.06304795 | 222.4176113 | 175.185536 | ENSG00000100124 |
| TFEB | -0.345691741 | 0.78693057 | 0.219679898 | 0.377710335 | 71.01673839 | 55.91146354 | ENSG00000112561 |
| GRK4 | -0.345705552 | 0.786923037 | 0.388181097 | 0.561602036 | 31.20180713 | 24.58708252 | ENSG00000125388 |
| GTF2F1 | -0.345867158 | 0.786834893 | 1.16E-07 | 1.06E-06 | 2751.745272 | 2165.266806 | ENSG00000125651 |
| ECHS1 | -0.346169759 | 0.786669874 | 6.01E-11 | 8.45E-10 | 3020.672123 | 2375.732916 | ENSG00000127884 |
| FAM117B | -0.346364583 | 0.786563648 | 0.000467521 | 0.002158678 | 546.557385 | 429.8957327 | ENSG00000138439 |
| NELFCD | -0.346577507 | 0.786447569 | 4.61E-07 | 3.82E-06 | 1426.71642 | 1122.445151 | ENSG00000101158 |
| RANGRF | -0.346658412 | 0.786403468 | 0.613919911 | 0.756749457 | 10.90211175 | 8.63261918 | ENSG00000108961 |
| CAAP1 | -0.346935726 | 0.78625232 | 0.000140597 | 0.000732052 | 672.5782361 | 528.8315647 | ENSG00000120159 |
| PSMD8 | -0.347143247 | 0.786139231 | 8.60E-12 | 1.33E-10 | 3154.604541 | 2479.589231 | ENSG00000099341 |
| FAM131C | -0.34715763 | 0.786131394 | 0.344530386 | 0.517123353 | 41.22623241 | 32.3259512 | ENSG00000185519 |
| EFNA2 | -0.347192267 | 0.786112521 | 0.015204988 | 0.044781591 | 323.5423557 | 254.6883879 | ENSG00000099617 |
| ZNF396 | -0.347287098 | 0.78606085 | 0.639586924 | 0.774741394 | 10.5371153 | 8.357017905 | ENSG00000186496 |
| NA | -0.347548676 | 0.78591834 | 0.784033848 | 0.873614559 | 3.363764935 | 2.644152387 | ENSG00000213169 |
| CTPS2 | -0.347637995 | 0.785869685 | 3.06E-09 | 3.50E-08 | 2174.232741 | 1708.877504 | ENSG00000047230 |
| CHMP6 | -0.347713065 | 0.785828793 | 0.00726485 | 0.024104959 | 322.5063756 | 253.2741186 | ENSG00000176108 |
| INTS11 | -0.34782221 | 0.785769344 | 3.38E-08 | 3.32E-07 | 3643.862373 | 2862.997594 | ENSG00000127054 |
| MCFD2 | -0.347868123 | 0.785744339 | 9.86E-09 | 1.05E-07 | 1769.380016 | 1390.208061 | ENSG00000180398 |
| WASHC1 | -0.347895599 | 0.785729374 | 0.105695957 | 0.215993718 | 113.3582602 | 89.02791928 | ENSG00000181404 |
| P4HTM | -0.347935141 | 0.785707839 | 0.037723603 | 0.094916658 | 255.042884 | 200.0025184 | ENSG00000178467 |
| DBN1 | -0.347970781 | 0.785688429 | 2.38E-11 | 3.49E-10 | 14310.47981 | 11243.77316 | ENSG00000113758 |
| FAM171A2 | -0.348003081 | 0.785670839 | 0.000195967 | 0.000989635 | 971.5851532 | 763.295251 | ENSG00000161682 |
| ZNF544 | -0.348197128 | 0.785565171 | 0.000263468 | 0.001294922 | 616.3451948 | 484.0396019 | ENSG00000198131 |
| USP7 | -0.348254201 | 0.785534094 | 2.05E-16 | 5.14E-15 | 5441.352673 | 4274.564985 | ENSG00000187555 |
| CBWD5 | -0.348356112 | 0.785478607 | 0.093725692 | 0.196674495 | 115.1499518 | 90.44839198 | ENSG00000147996 |
| SLC19A1 | -0.348467742 | 0.785417832 | 6.33E-06 | 4.32E-05 | 1125.183063 | 883.614717 | ENSG00000173638 |

| | | | | | | | |
|---|---|---|---|---|---|---|---|
| ATP1A1 | -0.348496654 | 0.785402092 | 1.09E-17 | 3.03E-16 | 7042.560901 | 5531.094128 | ENSG00000163399 |
| NA | -0.348578688 | 0.785357434 | 0.686101778 | 0.808823601 | 7.054129401 | 5.506906635 | ENSG00000228169 |
| LINC01948 | -0.348712582 | 0.78528455 | 0.763634487 | 0.85991903 | 3.747621984 | 2.911151367 | ENSG00000248727 |
| ADRM1 | -0.348744785 | 0.785267021 | 1.03E-06 | 8.14E-06 | 1585.688068 | 1245.376554 | ENSG00000130706 |
| NHP2 | -0.348891239 | 0.78518731 | 6.91E-06 | 4.68E-05 | 1864.140678 | 1463.347235 | ENSG00000145912 |
| PPP2R5D | -0.349087715 | 0.785080385 | 9.25E-07 | 7.31E-06 | 1446.158716 | 1135.081428 | ENSG00000112640 |
| CDC25C | -0.349149402 | 0.785046817 | 0.002822961 | 0.010644594 | 383.5870465 | 301.1877716 | ENSG00000158402 |
| NA | -0.349156387 | 0.785043016 | 0.238116267 | 0.4000133 | 57.34421251 | 45.07671777 | ENSG00000235363 |
| CKB | -0.349567705 | 0.784819229 | 1.67E-10 | 2.25E-09 | 7453.302211 | 5849.671501 | ENSG00000166165 |
| PFDN1 | -0.349723879 | 0.784734276 | 0.000272543 | 0.001334421 | 617.3551056 | 484.4407668 | ENSG00000113068 |
| PLAUR | -0.349772006 | 0.784708098 | 0.004793446 | 0.016920469 | 343.0670403 | 269.2554612 | ENSG00000011422 |
| PSMD13 | -0.349918436 | 0.784628456 | 4.17E-08 | 4.06E-07 | 3161.198629 | 2479.392735 | ENSG00000185627 |
| ATP5MG | -0.349997217 | 0.784585611 | 0.107579727 | 0.218892577 | 120.9422254 | 94.8865092 | ENSG00000167283 |
| PAF1 | -0.350081337 | 0.784539865 | 8.67E-06 | 5.78E-05 | 1133.736464 | 889.1627634 | ENSG00000006712 |
| SNX11 | -0.350151539 | 0.78450169 | 0.001257135 | 0.005245813 | 528.5106836 | 414.4572508 | ENSG00000002919 |
| SLC33A1 | -0.350231128 | 0.784458413 | 0.193965516 | 0.343788016 | 96.46946281 | 75.88886704 | ENSG00000169359 |
| POLE4 | -0.350244305 | 0.784451248 | 0.000757476 | 0.003308232 | 492.9241124 | 386.7312798 | ENSG00000115350 |
| NA | -0.350252389 | 0.784446853 | 0.539377532 | 0.698681435 | 16.8467337 | 13.20947603 | ENSG00000239218 |
| CHPT1 | -0.350283068 | 0.784430172 | 0.011494476 | 0.035495428 | 318.6917351 | 250.0494612 | ENSG00000111666 |
| MIR4292 | -0.35031002 | 0.784415517 | 0.374455839 | 0.54773847 | 33.23839832 | 26.08473383 | ENSG00000265806 |
| CFL1P1 | -0.35033997 | 0.784399233 | 0.686090264 | 0.808823601 | 7.573952829 | 5.986316219 | ENSG00000223820 |
| NSDHL | -0.350580535 | 0.784268448 | 2.64E-06 | 1.92E-05 | 1345.915324 | 1055.625271 | ENSG00000147383 |
| SERHL2 | -0.350636846 | 0.784237837 | 0.268403799 | 0.43470554 | 53.73901459 | 42.06044244 | ENSG00000183569 |
| SHLD1 | -0.350651397 | 0.784229927 | 0.193167629 | 0.342846327 | 84.94612813 | 66.49237424 | ENSG00000171984 |
| ISOC1 | -0.350673941 | 0.784217673 | 2.84E-05 | 0.000171781 | 1059.778629 | 831.034113 | ENSG00000066583 |
| TEX13C | -0.350771847 | 0.784164455 | 0.634708443 | 0.771175881 | 10.83436634 | 8.531523793 | ENSG00000282815 |
| BSG | -0.350856117 | 0.784118652 | 5.30E-20 | 1.79E-18 | 21153.97737 | 16587.63547 | ENSG00000172270 |
| WDR48 | -0.35108626 | 0.783993577 | 6.87E-06 | 4.67E-05 | 1146.93581 | 898.9146587 | ENSG00000114742 |
| NPY1R | -0.351136152 | 0.783966465 | 0.123010099 | 0.24288514 | 120.6218834 | 94.43264603 | ENSG00000164128 |
| BIN3 | -0.351174216 | 0.783945781 | 0.008441146 | 0.027303512 | 336.1299415 | 263.2957575 | ENSG00000147439 |
| NDUFB7 | -0.351232685 | 0.783914011 | 1.03E-05 | 6.77E-05 | 1467.060012 | 1149.89583 | ENSG00000099795 |
| CCDC144NL | -0.351279574 | 0.783888533 | 0.566973343 | 0.720424369 | 13.73835627 | 10.72315798 | ENSG00000205212 |
| SIPA1L1 | -0.351282146 | 0.783887135 | 5.15E-08 | 4.95E-07 | 1658.114855 | 1300.308716 | ENSG00000197555 |
| UBE2D2 | -0.351310619 | 0.783871665 | 8.11E-09 | 8.77E-08 | 3421.634595 | 2682.9016 | ENSG00000131508 |
| NF2 | -0.351436362 | 0.783803347 | 0.000101053 | 0.000546204 | 789.5726312 | 619.0532613 | ENSG00000186575 |
| PPIL2 | -0.351561751 | 0.783735227 | 0.000123556 | 0.0006532 | 856.8752257 | 671.5763755 | ENSG00000100023 |
| NA | -0.35161261 | 0.783707598 | 0.777984662 | 0.869782794 | 3.426604259 | 2.686097786 | ENSG00000251634 |
| ITGA7 | -0.35169287 | 0.783664001 | 5.79E-05 | 0.00032936 | 1664.607807 | 1303.969602 | ENSG00000135424 |
| CLK1 | -0.351847326 | 0.783580105 | 0.011151906 | 0.034575928 | 312.9844648 | 245.2964688 | ENSG00000013441 |
| MRPL20 | -0.351956519 | 0.783520801 | 1.72E-06 | 1.29E-05 | 1312.952101 | 1028.340853 | ENSG00000242485 |
| CCZ1B | -0.351985646 | 0.783504982 | 0.006119789 | 0.020883928 | 380.0567785 | 298.0053017 | ENSG00000146574 |
| UBE2I | -0.351999629 | 0.783497389 | 3.85E-11 | 5.52E-10 | 3204.814052 | 2511.047184 | ENSG00000103275 |
| SAMM50 | -0.352162855 | 0.783408749 | 3.38E-07 | 2.87E-06 | 1479.479602 | 1159.183425 | ENSG00000100347 |
| STX10 | -0.352172939 | 0.783403273 | 4.20E-05 | 0.00024519 | 953.6408379 | 746.580379 | ENSG00000104915 |
| ZNF214 | -0.352236102 | 0.783368975 | 0.357313113 | 0.530674206 | 33.50215853 | 26.1984722 | ENSG00000149050 |
| LRRC17 | -0.352304802 | 0.783331673 | 0.526591754 | 0.688570564 | 16.32324349 | 12.79298454 | ENSG00000128606 |
| NA | -0.352499853 | 0.783225774 | 0.780851459 | 0.871422278 | 3.396580048 | 2.658673365 | ENSG00000273291 |
| NA | -0.352525577 | 0.783211809 | 0.684041917 | 0.807283288 | 6.711460173 | 5.247976912 | ENSG00000268670 |
| CCNL2 | -0.35256574 | 0.783190006 | 3.93E-07 | 3.30E-06 | 2158.26364 | 1689.667016 | ENSG00000221978 |
| NA | -0.352642758 | 0.783148196 | 0.169863691 | 0.311013711 | 82.42667098 | 64.5679995 | ENSG00000283567 |
| NA | -0.352681035 | 0.783127418 | 0.635446552 | 0.771599386 | 10.02566458 | 7.837280484 | ENSG00000275202 |
| SNX32 | -0.352753066 | 0.78308832 | 0.467620589 | 0.636932062 | 23.80030258 | 18.5797936 | ENSG00000172803 |
| VAC14 | -0.352768488 | 0.783079948 | 2.92E-07 | 2.51E-06 | 1427.711712 | 1118.207371 | ENSG00000103043 |
| KLHDC8B | -0.352771864 | 0.783078116 | 0.00405162 | 0.014625713 | 371.3715665 | 291.0053359 | ENSG00000185909 |
| NA | -0.352830915 | 0.783046064 | 0.659179841 | 0.789485573 | 7.913831154 | 6.211369801 | ENSG00000272438 |
| NUDCD3 | -0.352895188 | 0.78301118 | 2.03E-08 | 2.06E-07 | 1669.315931 | 1307.020461 | ENSG00000015676 |
| TIMM8B | -0.352945981 | 0.782983613 | 1.05E-05 | 6.87E-05 | 1403.025139 | 1098.226793 | ENSG00000150779 |
| SPECC1L | -0.352981048 | 0.782964582 | 7.79E-05 | 0.00043133 | 744.1574493 | 582.7272449 | ENSG00000100014 |
| CFAP221 | -0.353012775 | 0.782947363 | 0.71899748 | 0.831396055 | 4.99397169 | 3.899818104 | ENSG00000163075 |
| RANBP3 | -0.353015562 | 0.782945851 | 2.47E-09 | 2.86E-08 | 2509.978339 | 1965.348687 | ENSG00000031823 |
| NA | -0.353029467 | 0.782938304 | 0.19467656 | 0.344826509 | 76.78575775 | 60.19441792 | ENSG00000244313 |
| SPOP | -0.353053059 | 0.782925502 | 0.0003335 | 0.001601582 | 611.3966044 | 479.062622 | ENSG00000121067 |
| NUTM2B-AS1 | -0.353073117 | 0.782914616 | 0.070634168 | 0.157217461 | 164.9743899 | 129.0994514 | ENSG00000225484 |
| EML5 | -0.353209472 | 0.782840624 | 0.545136844 | 0.703452664 | 15.61067173 | 12.28426043 | ENSG00000165521 |
| NSA2 | -0.353239284 | 0.782824447 | 5.83E-08 | 5.54E-07 | 1509.951518 | 1182.008541 | ENSG00000164346 |
| ERH | -0.353239857 | 0.782824136 | 1.60E-09 | 1.90E-08 | 3318.076232 | 2597.105123 | ENSG00000100632 |
| KIF21A | -0.353357949 | 0.78276006 | 9.31E-07 | 7.35E-06 | 1225.583396 | 959.8027869 | ENSG00000139116 |
| KLHL4 | -0.353371621 | 0.782752643 | 2.48E-05 | 0.000151936 | 813.4419938 | 636.6274988 | ENSG00000102271 |
| KLF15 | -0.353393668 | 0.782740681 | 0.010816928 | 0.03368895 | 386.0310996 | 302.4598484 | ENSG00000163884 |
| ARRB2 | -0.353460129 | 0.782704623 | 4.03E-07 | 3.37E-06 | 1795.929235 | 1405.814101 | ENSG00000141480 |
| PID1 | -0.353916258 | 0.782457198 | 0.183901638 | 0.329890878 | 70.87307503 | 55.53717792 | ENSG00000153823 |
| SOX8 | -0.353966777 | 0.7824298 | 0.001785296 | 0.007151193 | 436.6432013 | 341.6698886 | ENSG00000005513 |
| NA | -0.353978135 | 0.78242364 | 0.729042633 | 0.837539389 | 4.664581255 | 3.674764522 | ENSG00000229431 |
| DNAJC7 | -0.354094835 | 0.782360352 | 2.06E-06 | 1.54E-05 | 1397.5783 | 1093.589641 | ENSG00000168259 |
| SIAH2 | -0.354389411 | 0.782200622 | 0.000469228 | 0.00216604 | 522.4452763 | 408.7223466 | ENSG00000181788 |
| NA | -0.354430089 | 0.782178568 | 0.677605962 | 0.802732536 | 6.75755408 | 5.275401332 | ENSG00000268049 |
| SNRNP27 | -0.354486928 | 0.782147752 | 0.046913731 | 0.113523931 | 184.7169921 | 144.5339047 | ENSG00000124380 |

| | | | | | | | |
|---|---|---|---|---|---|---|---|
| ANKS1A | -0.35450099 | 0.782140129 | 4.97E-09 | 5.51E-08 | 1882.284008 | 1471.765759 | ENSG00000064999 |
| ENY2 | -0.354507199 | 0.782136763 | 6.68E-10 | 8.28E-09 | 2572.434077 | 2011.319409 | ENSG00000120533 |
| GANC | -0.35451873 | 0.782130511 | 0.083346804 | 0.179346704 | 148.8978858 | 116.2731237 | ENSG00000214013 |
| MELK | -0.354622327 | 0.78207435 | 7.20E-06 | 4.87E-05 | 1140.553429 | 892.4567111 | ENSG00000165304 |
| KIAA0513 | -0.354654784 | 0.782056756 | 0.007950832 | 0.025974514 | 324.4538581 | 253.5526823 | ENSG00000135709 |
| HSPA5 | -0.354677958 | 0.782044194 | 1.93E-19 | 6.19E-18 | 21243.11824 | 16612.84678 | ENSG00000044574 |
| UNC119B | -0.35470974 | 0.782026966 | 1.66E-09 | 1.97E-08 | 2141.045329 | 1673.709584 | ENSG00000175970 |
| ATP2B1 | -0.354786107 | 0.781985571 | 2.84E-05 | 0.000171781 | 884.5479289 | 692.2224454 | ENSG00000070961 |
| ERGIC1 | -0.354814631 | 0.78197011 | 3.95E-11 | 5.64E-10 | 3046.562785 | 2381.762329 | ENSG00000113719 |
| NA | -0.354838839 | 0.78195699 | 0.004618708 | 0.016402706 | 65203.56423 | 50986.49163 | ENSG00000210082 |
| ITIH5 | -0.354899459 | 0.781924133 | 0.026558255 | 0.0711478 | 217.631694 | 170.1236419 | ENSG00000123243 |
| DTNBP1 | -0.355062965 | 0.78183552 | 0.018867037 | 0.053492652 | 228.4679001 | 178.6938639 | ENSG00000047579 |
| HYOU1 | -0.355151141 | 0.781787736 | 8.06E-15 | 1.71E-13 | 9674.575828 | 7562.607623 | ENSG00000149428 |
| PITPNM1 | -0.355376171 | 0.781665804 | 3.80E-07 | 3.21E-06 | 2210.781153 | 1728.66169 | ENSG00000110697 |
| NA | -0.355519639 | 0.781588075 | 0.379901711 | 0.553191719 | 34.86561401 | 27.31135759 | ENSG00000165121 |
| RBM23 | -0.355684394 | 0.781498823 | 5.47E-06 | 3.77E-05 | 1008.220956 | 788.11296 | ENSG00000100461 |
| PRPF4B | -0.355700378 | 0.781490165 | 3.11E-08 | 3.07E-07 | 1907.813701 | 1490.73004 | ENSG00000112739 |
| ZNF23 | -0.355827919 | 0.78142108 | 0.383538551 | 0.556806306 | 45.34096603 | 35.27610389 | ENSG00000167377 |
| LY6G5B | -0.355865738 | 0.781400596 | 0.366436536 | 0.540190067 | 32.40226309 | 25.3399428 | ENSG00000240053 |
| C17orf75 | -0.355871307 | 0.78139758 | 8.89E-05 | 0.000486215 | 745.6733412 | 582.699802 | ENSG00000108666 |
| RDH5 | -0.355943313 | 0.781358581 | 0.743765354 | 0.848323562 | 5.062392819 | 3.948215223 | ENSG00000135437 |
| TMEM43 | -0.355955409 | 0.78135203 | 2.72E-05 | 0.0001655 | 789.8401335 | 617.1114641 | ENSG00000170876 |
| UNC79 | -0.355995233 | 0.781330462 | 0.189566851 | 0.337604702 | 73.09037506 | 57.1189926 | ENSG00000133958 |
| CPE | -0.35600092 | 0.781327382 | 0.000429617 | 0.002006229 | 536.1447601 | 419.045121 | ENSG00000109472 |
| DPYSL4 | -0.356012368 | 0.781321182 | 3.50E-05 | 0.00020853 | 890.205863 | 695.6840116 | ENSG00000151640 |
| COL20A1 | -0.35606499 | 0.781292684 | 0.698475895 | 0.817989588 | 8.38285475 | 6.53751877 | ENSG00000101203 |
| DCLRE1C | -0.356107248 | 0.7812698 | 0.044110585 | 0.107904805 | 156.7975623 | 122.4772362 | ENSG00000152457 |
| NA | -0.356201568 | 0.781218723 | 0.243265015 | 0.406164567 | 55.56176952 | 43.51425825 | ENSG00000261578 |
| ITSN1 | -0.356217846 | 0.781209909 | 7.52E-12 | 1.17E-10 | 3229.819116 | 2523.308649 | ENSG00000205726 |
| VAV3 | -0.356266247 | 0.781183701 | 0.008846954 | 0.028392379 | 276.4495714 | 215.9104505 | ENSG00000134215 |
| NKAIN3-IT1 | -0.356310545 | 0.781159715 | 0.681082335 | 0.805367128 | 6.352045528 | 4.980977931 | ENSG00000274956 |
| ABT1 | -0.356369777 | 0.781127644 | 2.81E-06 | 2.04E-05 | 1376.109274 | 1074.584803 | ENSG00000146109 |
| ASNS | -0.356401952 | 0.781110223 | 0.375624231 | 0.549022281 | 31.41197661 | 24.48813771 | ENSG00000070669 |
| ARHGAP21 | -0.356418586 | 0.781101217 | 0.000116786 | 0.000621838 | 783.9688629 | 612.2361369 | ENSG00000107863 |
| SCARF2 | -0.356488119 | 0.781063572 | 0.006284537 | 0.021359231 | 397.4375656 | 310.3882891 | ENSG00000244486 |
| HNRNPR | -0.356526682 | 0.781042695 | 3.46E-18 | 9.95E-17 | 6410.854176 | 5007.611792 | ENSG00000125944 |
| NA | -0.356597343 | 0.781004441 | 0.357492021 | 0.530760367 | 37.4899742 | 29.18544511 | ENSG00000224086 |
| SORL1 | -0.356692604 | 0.780952873 | 7.79E-07 | 6.24E-06 | 2393.640479 | 1868.704942 | ENSG00000137642 |
| GPR158 | -0.356777842 | 0.780906734 | 0.466851663 | 0.636064371 | 22.9371342 | 17.92775613 | ENSG00000151025 |
| ELMOD3 | -0.35686364 | 0.780860294 | 0.048773053 | 0.117024997 | 177.3304301 | 138.3585616 | ENSG00000115459 |
| LRATD2 | -0.356887805 | 0.780847215 | 1.76E-05 | 0.00011085 | 844.2425227 | 659.4141342 | ENSG00000168672 |
| NA | -0.357022257 | 0.780774447 | 0.027333396 | 0.072900393 | 200.5905851 | 156.5678377 | ENSG00000248641 |
| CSTF2T | -0.357367012 | 0.780587891 | 2.29E-07 | 2.00E-06 | 1345.315003 | 1050.181307 | ENSG00000177613 |
| EXOC7 | -0.357520123 | 0.780505053 | 6.62E-08 | 6.24E-07 | 1810.533446 | 1412.382297 | ENSG00000182473 |
| CERS5 | -0.357654888 | 0.780432148 | 3.15E-05 | 0.00018917 | 824.9563538 | 643.8135232 | ENSG00000139624 |
| NECAP2 | -0.357705458 | 0.780404792 | 3.49E-05 | 0.000207763 | 1043.298578 | 813.7059323 | ENSG00000157191 |
| MRPS28 | -0.357834868 | 0.780334793 | 0.145124743 | 0.275334521 | 104.2607185 | 81.28448899 | ENSG00000147586 |
| NUP88 | -0.357857942 | 0.780322312 | 2.63E-08 | 2.62E-07 | 1760.276054 | 1373.3529 | ENSG00000108559 |
| HEY1 | -0.357915111 | 0.780291391 | 0.22780428 | 0.3872466 | 75.30634777 | 58.64757598 | ENSG00000164683 |
| NA | -0.35795871 | 0.780267811 | 0.066343584 | 0.149530795 | 141.0428269 | 110.1658118 | ENSG00000260708 |
| NA | -0.35806602 | 0.780209776 | 0.764843145 | 0.860878062 | 3.774855292 | 2.909000794 | ENSG00000181201 |
| CFAP46 | -0.358069942 | 0.780207655 | 0.081753112 | 0.176508034 | 120.4857168 | 94.09358151 | ENSG00000171811 |
| NA | -0.358077451 | 0.780203594 | 0.766384467 | 0.861758146 | 3.390998242 | 2.637700666 | ENSG00000228835 |
| NA | -0.358109458 | 0.780186285 | 0.661666477 | 0.791256982 | 7.922203862 | 6.183945381 | ENSG00000267868 |
| CHRNA7 | -0.358149846 | 0.780164444 | 0.117166485 | 0.234272249 | 95.40201907 | 74.4259159 | ENSG00000175344 |
| CHST6 | -0.358153706 | 0.780162357 | 0.000461564 | 0.002134748 | 527.0616485 | 411.383431 | ENSG00000183196 |
| LOC105371899 | -0.358322913 | 0.78007086 | 0.73705562 | 0.843705033 | 5.162953341 | 4.021886189 | ENSG00000267568 |
| CDKN1C | -0.358339008 | 0.780062158 | 0.012992692 | 0.039248673 | 310.1231533 | 242.1874278 | ENSG00000129757 |
| PRKCSH | -0.358361747 | 0.780049863 | 1.33E-15 | 3.06E-14 | 7533.572065 | 5876.574383 | ENSG00000130175 |
| MYO9B | -0.35841901 | 0.780018902 | 1.01E-08 | 1.07E-07 | 2370.258506 | 1849.344682 | ENSG00000099331 |
| KLF9 | -0.358526262 | 0.779960916 | 0.422144668 | 0.593804067 | 24.86391628 | 19.35362659 | ENSG00000119138 |
| MARVELD1 | -0.35863124 | 0.779904164 | 5.11E-06 | 3.54E-05 | 1351.290973 | 1053.349261 | ENSG00000155254 |
| CCDC63 | -0.358746063 | 0.779842095 | 0.51140536 | 0.675720602 | 16.01618028 | 12.50501286 | ENSG00000173093 |
| HNRNPLL | -0.358777657 | 0.779825017 | 3.40E-07 | 2.89E-06 | 1917.425413 | 1496.016691 | ENSG00000143889 |
| RP9 | -0.359069792 | 0.779667124 | 0.037181321 | 0.093784632 | 183.4765365 | 142.9827495 | ENSG00000164610 |
| COPRS | -0.359103256 | 0.77964904 | 0.000142195 | 0.000739373 | 825.7025044 | 643.6159245 | ENSG00000172301 |
| SCLY | -0.359290932 | 0.779547624 | 0.137845786 | 0.264595328 | 114.0191563 | 89.13409721 | ENSG00000132330 |
| ZNF445 | -0.359402086 | 0.779487565 | 0.000658988 | 0.002925113 | 636.9929907 | 496.3950796 | ENSG00000185219 |
| DUS3L | -0.359412691 | 0.779481836 | 4.20E-06 | 2.96E-05 | 1133.527772 | 884.0448812 | ENSG00000141994 |
| ZNF583 | -0.359478631 | 0.779446209 | 0.362193157 | 0.535423501 | 33.23560742 | 25.83467898 | ENSG00000198440 |
| ARPC1B | -0.359744209 | 0.779302738 | 0.379221888 | 0.552618933 | 31.06372557 | 24.16628988 | ENSG00000130429 |
| SPTBN2 | -0.360107387 | 0.779106584 | 1.04E-08 | 1.11E-07 | 2183.75066 | 1701.091693 | ENSG00000173898 |
| PFAS | -0.360328524 | 0.778987172 | 7.42E-12 | 1.16E-10 | 5802.527027 | 4519.391435 | ENSG00000178921 |
| SLC12A9 | -0.360495764 | 0.778896876 | 4.51E-05 | 0.000261594 | 1248.382412 | 972.1621912 | ENSG00000146828 |
| LPAR2 | -0.360631843 | 0.778823411 | 0.00015236 | 0.000786507 | 851.8073263 | 663.4419817 | ENSG00000064547 |
| BAP1 | -0.360635298 | 0.778821546 | 1.11E-09 | 1.34E-08 | 3154.981177 | 2456.366301 | ENSG00000163930 |
| ADRA2A | -0.360637017 | 0.778820618 | 0.40374767 | 0.576115175 | 29.23750402 | 22.77161196 | ENSG00000150594 |

| | | | | | | | |
|---|---|---|---|---|---|---|---|
| ZBTB8OS | -0.36067459 | 0.778800335 | 0.001320062 | 0.00547053 | 454.9674668 | 354.1240812 | ENSG00000176261 |
| NA | -0.360684564 | 0.778794951 | 0.757684955 | 0.856117874 | 4.229048493 | 3.329793428 | ENSG00000265566 |
| NGEF | -0.360735221 | 0.778767606 | 0.120165407 | 0.2387098 | 90.35445664 | 70.36449535 | ENSG00000066248 |
| SLC29A4 | -0.360809487 | 0.778727518 | 0.009193575 | 0.0293049 | 333.524317 | 259.8250922 | ENSG00000164638 |
| NEDD8 | -0.36086972 | 0.778695006 | 0.028509759 | 0.075391606 | 214.2799685 | 166.7099576 | ENSG00000129559 |
| AVIL | -0.360921683 | 0.77866696 | 0.464047512 | 0.633683583 | 20.26967118 | 15.83480629 | ENSG00000135407 |
| FAM207A | -0.361094247 | 0.778573828 | 0.000126813 | 0.000668217 | 1324.423425 | 1031.644555 | ENSG00000160256 |
| DOCK8 | -0.361166535 | 0.778534817 | 0.333284978 | 0.505893501 | 43.93488326 | 34.09652018 | ENSG00000107099 |
| MMS22L | -0.361187077 | 0.778523732 | 0.001830813 | 0.007305264 | 483.6759372 | 377.0311856 | ENSG00000146263 |
| LOC101928837 | -0.361334051 | 0.778444424 | 0.6680948 | 0.796028624 | 9.364768527 | 7.311351754 | ENSG00000255100 |
| ZNF322 | -0.361487803 | 0.778361467 | 0.013231689 | 0.039858293 | 265.94795 | 207.1100376 | ENSG00000181315 |
| TARS1 | -0.361506004 | 0.778351648 | 3.82E-20 | 1.32E-18 | 8690.760877 | 6764.643209 | ENSG00000113407 |
| PBX4 | -0.36156587 | 0.77831935 | 0.382225514 | 0.555610911 | 35.91355834 | 27.83890383 | ENSG00000105717 |
| CLSPN | -0.361641565 | 0.778278514 | 5.05E-06 | 3.51E-05 | 1021.464093 | 794.5190149 | ENSG00000092853 |
| ALKBH5 | -0.361649121 | 0.778274438 | 1.12E-07 | 1.03E-06 | 1365.787585 | 1063.399627 | ENSG00000091542 |
| TSPOAP1 | -0.361838025 | 0.778172539 | 0.014245046 | 0.042420682 | 277.3381461 | 215.5326999 | ENSG00000005379 |
| RTL8C | -0.361993721 | 0.778088563 | 1.06E-05 | 6.92E-05 | 984.7997438 | 766.5177151 | ENSG00000134590 |
| SMIM3 | -0.362028926 | 0.778069576 | 0.000119894 | 0.000636106 | 688.0210391 | 534.9929382 | ENSG00000256235 |
| CYP2T1P | -0.362067164 | 0.778048954 | 0.550081364 | 0.707133229 | 14.33987962 | 11.16277274 | ENSG00000233622 |
| NA | -0.36216537 | 0.777995993 | 0.526586909 | 0.688570564 | 17.17803916 | 13.37806323 | ENSG00000283103 |
| USP16 | -0.362205962 | 0.777974104 | 0.002568266 | 0.009814664 | 377.9450573 | 294.1821292 | ENSG00000156256 |
| EEF1B2 | -0.362254613 | 0.777947869 | 2.59E-15 | 5.77E-14 | 7202.814301 | 5602.947133 | ENSG00000114942 |
| ITFG2 | -0.362438554 | 0.777848688 | 0.006538201 | 0.022090699 | 314.5061017 | 244.8079239 | ENSG00000111203 |
| C19orf73 | -0.362497536 | 0.777816888 | 0.402019928 | 0.574301425 | 29.30524943 | 22.7845154 | ENSG00000221916 |
| ARNTL2 | -0.362545176 | 0.777791204 | 0.006499625 | 0.021977962 | 312.3971472 | 243.1744955 | ENSG00000029153 |
| MFSD9 | -0.362578315 | 0.777773338 | 0.004572592 | 0.016247904 | 387.6341469 | 301.5797827 | ENSG00000135953 |
| TIMM44 | -0.362590281 | 0.777766887 | 6.30E-10 | 7.84E-09 | 2946.776681 | 2292.004649 | ENSG00000104980 |
| NA | -0.362637389 | 0.777741491 | 0.663797551 | 0.792653578 | 7.668931545 | 5.994918514 | ENSG00000232811 |
| NA | -0.362764619 | 0.777672906 | 0.672046703 | 0.798992504 | 6.790369193 | 5.256579206 | ENSG00000273306 |
| CNR1 | -0.36285609 | 0.7776236 | 0.428464787 | 0.599776083 | 29.40137943 | 22.8159805 | ENSG00000118432 |
| DOHH | -0.362895061 | 0.777602595 | 0.001529448 | 0.006229854 | 709.5168494 | 551.6494779 | ENSG00000129932 |
| PPP4R1 | -0.363012828 | 0.777539122 | 1.91E-09 | 2.25E-08 | 1853.402164 | 1441.000328 | ENSG00000154845 |
| FAM110B | -0.363027831 | 0.777531036 | 0.099544034 | 0.206528711 | 110.126995 | 85.73656361 | ENSG00000169122 |
| RTRAF | -0.363156348 | 0.777461776 | 3.50E-11 | 5.05E-10 | 3281.047035 | 2551.800892 | ENSG00000087302 |
| EFHC1 | -0.363328903 | 0.777368793 | 0.029178315 | 0.076782119 | 229.4124049 | 178.0106581 | ENSG00000096093 |
| LRRFIP2 | -0.363436196 | 0.777310982 | 0.000286703 | 0.001396661 | 611.6110922 | 475.5970636 | ENSG00000093167 |
| RASGRP1 | -0.363761419 | 0.777135775 | 0.203865517 | 0.357005237 | 70.26752147 | 54.62164911 | ENSG00000172575 |
| LINC02449 | -0.364046808 | 0.776982059 | 0.622099053 | 0.762199274 | 9.642483249 | 7.477788386 | ENSG00000215241 |
| POLR2E | -0.364081265 | 0.776963502 | 1.25E-12 | 2.11E-11 | 5802.500276 | 4508.774309 | ENSG00000099817 |
| NA | -0.364098176 | 0.776954395 | 0.616794292 | 0.758837926 | 10.09687661 | 7.803937354 | ENSG00000241556 |
| RAC3 | -0.364344559 | 0.776821718 | 5.43E-07 | 4.45E-06 | 1784.012293 | 1386.082197 | ENSG00000169750 |
| MSRA | -0.36440172 | 0.77679094 | 0.210408951 | 0.365939568 | 71.20401706 | 55.22744004 | ENSG00000175806 |
| HACD2 | -0.364419635 | 0.776781294 | 3.27E-05 | 0.000195738 | 886.8483044 | 689.2693486 | ENSG00000206527 |
| PALM2AKAP2 | -0.364432254 | 0.7767745 | 9.74E-10 | 1.18E-08 | 2122.732153 | 1648.836953 | ENSG00000157654 |
| SLC23A2 | -0.364477302 | 0.776750246 | 0.000217801 | 0.001088217 | 665.3984281 | 516.8455142 | ENSG00000089057 |
| NA | -0.364563868 | 0.77670364 | 0.714605916 | 0.828587115 | 5.10290492 | 3.943914076 | ENSG00000270108 |
| THAP1 | -0.364664699 | 0.776649357 | 0.156946927 | 0.29272582 | 95.73958205 | 74.40600927 | ENSG00000131931 |
| SCAP | -0.364867209 | 0.776540347 | 2.92E-11 | 4.25E-10 | 3707.062447 | 2879.02428 | ENSG00000114650 |
| ZAN | -0.364932167 | 0.776505384 | 0.709776384 | 0.825105495 | 5.108486726 | 3.946064649 | ENSG00000146839 |
| GSK3A | -0.365143643 | 0.776391569 | 1.07E-06 | 8.40E-06 | 1153.292474 | 895.7690579 | ENSG00000105723 |
| ING4 | -0.36527432 | 0.776321248 | 0.004791412 | 0.016920004 | 343.6524187 | 266.6554048 | ENSG00000111653 |
| DHX29 | -0.365343836 | 0.776283842 | 3.06E-06 | 2.20E-05 | 1110.96821 | 862.6558373 | ENSG00000067248 |
| NA | -0.365346525 | 0.776282395 | 0.260665328 | 0.426030394 | 48.35839495 | 37.62420323 | ENSG00000238123 |
| CTNND2 | -0.365348108 | 0.776281543 | 0.001783856 | 0.007146908 | 420.8422087 | 326.482837 | ENSG00000169862 |
| SH3BGRL2 | -0.365372698 | 0.776268312 | 0.000646236 | 0.002876451 | 500.0155627 | 388.2926548 | ENSG00000198478 |
| NA | -0.365582404 | 0.776155484 | 0.77042134 | 0.864741191 | 4.566811636 | 3.546244715 | ENSG00000237609 |
| CHRNB1 | -0.365747392 | 0.776066727 | 0.007395977 | 0.024468566 | 330.3513853 | 256.5834787 | ENSG00000170175 |
| NA | -0.366007438 | 0.775926854 | 0.79420329 | 0.880349251 | 3.374928546 | 2.608125672 | ENSG00000256944 |
| AXIN1 | -0.366057948 | 0.775899689 | 3.82E-10 | 4.89E-09 | 2085.064097 | 1617.736287 | ENSG00000103126 |
| HS3ST1 | -0.366194236 | 0.775826394 | 0.369793762 | 0.543643612 | 31.88590629 | 24.74517732 | ENSG00000002587 |
| CCNI2 | -0.366225408 | 0.775809631 | 0.476645493 | 0.644944112 | 21.35648798 | 16.54572118 | ENSG00000205089 |
| USP35 | -0.366575893 | 0.775621181 | 0.007665012 | 0.025190076 | 266.2177675 | 206.4781914 | ENSG00000118369 |
| SVIP | -0.366651938 | 0.775580299 | 0.067848537 | 0.152193736 | 127.0309863 | 98.61827313 | ENSG00000198168 |
| SKA1 | -0.366752194 | 0.775526404 | 0.004056406 | 0.01463751 | 333.9713015 | 259.2668744 | ENSG00000154839 |
| RPN1 | -0.366953685 | 0.775418099 | 1.42E-16 | 3.61E-15 | 5181.040357 | 4017.584685 | ENSG00000163902 |
| EPB41 | -0.367102028 | 0.775338372 | 3.10E-11 | 4.51E-10 | 2203.595836 | 1708.557856 | ENSG00000159023 |
| PIGU | -0.367108841 | 0.775334711 | 0.000231428 | 0.001152121 | 710.9464461 | 551.2022664 | ENSG00000101464 |
| UBL5 | -0.367111883 | 0.775333076 | 1.11E-09 | 1.34E-08 | 2496.509358 | 1934.771248 | ENSG00000198258 |
| DDX51 | -0.367160051 | 0.77530719 | 1.24E-05 | 7.98E-05 | 963.2463141 | 746.6188289 | ENSG00000185163 |
| TBC1D22B | -0.367254189 | 0.775256601 | 0.000480492 | 0.002212214 | 486.2507737 | 376.8822567 | ENSG00000065491 |
| NA | -0.367357469 | 0.775201104 | 0.631670367 | 0.768901632 | 8.063952206 | 6.240944795 | ENSG00000269935 |
| SHANK3 | -0.367491401 | 0.775129142 | 0.000648839 | 0.002884708 | 596.5247227 | 462.371449 | ENSG00000251322 |
| MTUS2 | -0.367740044 | 0.774995563 | 0.537992823 | 0.697661975 | 15.24376025 | 11.74570086 | ENSG00000132938 |
| SLC25A38 | -0.367883541 | 0.774918482 | 1.02E-05 | 6.73E-05 | 1099.700113 | 852.0474532 | ENSG00000144659 |
| SNRPB2 | -0.367888515 | 0.774915811 | 6.67E-08 | 6.29E-07 | 1536.043286 | 1190.227879 | ENSG00000125870 |
| NA | -0.367951248 | 0.774882115 | 0.484639906 | 0.652511755 | 21.15170015 | 16.44247522 | ENSG00000280433 |
| ANKRD13A | -0.36809509 | 0.774804861 | 0.00432258 | 0.015462061 | 319.7927066 | 247.9599948 | ENSG00000076513 |

| | | | | | | | |
|---|---|---|---|---|---|---|---|
| SEC61A1 | -0.368301544 | 0.774693992 | 7.09E-14 | 1.36E-12 | 5196.362773 | 4025.977984 | ENSG00000058262 |
| ATP9A | -0.368346786 | 0.774669698 | 0.008768218 | 0.028191283 | 259.2744864 | 200.9083667 | ENSG00000054793 |
| LINC01089 | -0.368543303 | 0.774564184 | 0.039606076 | 0.098879359 | 191.7106217 | 148.4100665 | ENSG00000212694 |
| TSPAN5 | -0.368561544 | 0.77455439 | 0.000103456 | 0.00055779 | 699.3987585 | 542.2093311 | ENSG00000168785 |
| LOC107986257 | -0.368575473 | 0.774546912 | 0.723592834 | 0.834209608 | 4.730211482 | 3.672613948 | ENSG00000286176 |
| PARP11 | -0.368634717 | 0.774515106 | 0.06460608 | 0.146229717 | 127.4287978 | 98.6911506 | ENSG00000111224 |
| NA | -0.368641401 | 0.774511518 | 0.440328162 | 0.611233988 | 21.67662983 | 16.8256236 | ENSG00000220842 |
| NA | -0.368694498 | 0.774483013 | 0.022661421 | 0.062353654 | 199.7409709 | 154.8410921 | ENSG00000273015 |
| NA | -0.368721507 | 0.774468514 | 0.550674355 | 0.707658325 | 13.5393504 | 10.52767939 | ENSG00000233406 |
| SNORD3B-2 | -0.368864505 | 0.774391754 | 0.304710231 | 0.475464294 | 43.38886567 | 33.51386464 | ENSG00000262074 |
| PSORS1C1 | -0.368904026 | 0.774370541 | 0.724012145 | 0.834405189 | 4.612905543 | 3.573669135 | ENSG00000204540 |
| TOE1 | -0.369094571 | 0.774268272 | 0.030733331 | 0.080107647 | 241.520793 | 187.0630038 | ENSG00000132773 |
| ENDOD1 | -0.369129311 | 0.774249627 | 0.083027881 | 0.178833574 | 118.1989565 | 91.54460583 | ENSG00000149218 |
| ACOT7 | -0.36934992 | 0.774131243 | 1.27E-12 | 2.14E-11 | 2771.686588 | 2145.30734 | ENSG00000097021 |
| CD24 | -0.369367282 | 0.774121926 | 6.09E-13 | 1.06E-11 | 32428.82105 | 25104.8915 | ENSG00000272398 |
| DAB2IP | -0.369559587 | 0.774018746 | 1.94E-07 | 1.71E-06 | 1995.254345 | 1544.399389 | ENSG00000136848 |
| TTC9C | -0.36963084 | 0.773980519 | 0.001874421 | 0.007460174 | 447.3229777 | 345.8197737 | ENSG00000162222 |
| PREX1 | -0.369684645 | 0.773951654 | 2.82E-05 | 0.0001709 | 845.4517389 | 654.4296976 | ENSG00000124126 |
| TGIF2 | -0.369704744 | 0.773940872 | 3.95E-06 | 2.80E-05 | 987.3141558 | 764.2112765 | ENSG00000118707 |
| ZIC3 | -0.369724956 | 0.77393003 | 2.96E-07 | 2.54E-06 | 1676.741731 | 1297.825954 | ENSG00000156925 |
| XRCC5 | -0.369780602 | 0.773900179 | 1.24E-23 | 5.46E-22 | 14303.29792 | 11069.65179 | ENSG00000079246 |
| NA | -0.36984752 | 0.773864283 | 0.624970437 | 0.763774412 | 9.628528734 | 7.422939545 | ENSG00000259577 |
| IFT81 | -0.369878751 | 0.773847531 | 0.004405923 | 0.015725192 | 315.9033362 | 244.5363633 | ENSG00000122970 |
| SALL1 | -0.369979854 | 0.773793302 | 5.74E-06 | 3.94E-05 | 1112.310176 | 860.5053181 | ENSG00000103449 |
| HAS3 | -0.370071816 | 0.773743979 | 2.49E-08 | 2.49E-07 | 2622.127745 | 2028.691475 | ENSG00000103044 |
| VOPP1 | -0.370137348 | 0.773708834 | 1.50E-07 | 1.34E-06 | 1645.60086 | 1273.498056 | ENSG00000154978 |
| NIPBL-DT | -0.370167327 | 0.773692757 | 0.091946783 | 0.193851956 | 121.2827794 | 93.94166586 | ENSG00000285967 |
| NAA60 | -0.370310802 | 0.773615818 | 9.56E-05 | 0.00051953 | 773.7331679 | 598.5723015 | ENSG00000122390 |
| BOLA2B | -0.37034654 | 0.773596654 | 0.491593462 | 0.657856494 | 19.45797835 | 15.1029187 | ENSG00000169627 |
| SSBP2 | -0.370375884 | 0.77358092 | 0.001843902 | 0.00735084 | 382.9298173 | 296.1842034 | ENSG00000145687 |
| NR3C1 | -0.370466107 | 0.773532543 | 0.013567616 | 0.040673332 | 252.8442076 | 195.7918537 | ENSG00000113580 |
| ZNF136 | -0.370548885 | 0.773488161 | 0.075675879 | 0.165931279 | 118.0258325 | 91.26606048 | ENSG00000196646 |
| USP2 | -0.370707408 | 0.773403175 | 0.040671937 | 0.101004421 | 161.6749038 | 125.0186756 | ENSG00000036672 |
| NDUFS7 | -0.370753154 | 0.773378652 | 0.175740667 | 0.319138964 | 67.08725628 | 51.90865994 | ENSG00000115286 |
| ANKLE2 | -0.370800702 | 0.773353163 | 2.08E-09 | 2.44E-08 | 1687.558535 | 1304.825102 | ENSG00000176915 |
| DXO | -0.370807777 | 0.773349371 | 0.037655773 | 0.094783079 | 162.066658 | 125.3300311 | ENSG00000204348 |
| PLXNC1 | -0.371234465 | 0.773120681 | 0.197704748 | 0.348844991 | 64.0263242 | 49.49869647 | ENSG00000136040 |
| UBE2E3 | -0.37137643 | 0.773044607 | 1.25E-06 | 9.69E-06 | 1272.598311 | 984.4799393 | ENSG00000170035 |
| FBXW9 | -0.37186856 | 0.772780952 | 0.032780533 | 0.084428473 | 182.3886806 | 140.7429907 | ENSG00000132004 |
| VWA1 | -0.371915267 | 0.772755934 | 0.000394759 | 0.001865124 | 1230.277379 | 950.8016498 | ENSG00000179403 |
| LHFPL5 | -0.371928491 | 0.772748851 | 0.446303308 | 0.616603675 | 24.22746263 | 18.6760548 | ENSG00000197753 |
| TMEM208 | -0.371948475 | 0.772738147 | 0.000110996 | 0.000594079 | 576.8932328 | 445.7076883 | ENSG00000168701 |
| DUSP9 | -0.371958209 | 0.772732933 | 0.157140296 | 0.292944864 | 74.6460153 | 57.5793196 | ENSG00000130829 |
| NA | -0.372071029 | 0.772672507 | 0.622352973 | 0.762273727 | 9.290765591 | 7.176913264 | ENSG00000184441 |
| NA | -0.372137031 | 0.772637159 | 0.60393072 | 0.749175456 | 9.751416479 | 7.503062233 | ENSG00000257303 |
| TMEM132A | -0.372138555 | 0.772636343 | 7.01E-10 | 8.68E-09 | 3022.235654 | 2335.434514 | ENSG00000006118 |
| TIMM10 | -0.372163779 | 0.772622834 | 1.84E-06 | 1.38E-05 | 1092.972322 | 844.122587 | ENSG00000134809 |
| EXOSC9 | -0.372332021 | 0.772532739 | 2.49E-06 | 1.82E-05 | 1013.762912 | 783.440666 | ENSG00000123737 |
| CNDP2 | -0.372347125 | 0.772524651 | 1.37E-18 | 4.10E-17 | 6543.111779 | 5054.362685 | ENSG00000133313 |
| NA | -0.372470417 | 0.772458634 | 0.371618332 | 0.545038336 | 29.26665235 | 22.62695364 | ENSG00000174403 |
| LRRC14 | -0.372533164 | 0.772425039 | 0.000862882 | 0.003724739 | 573.7952695 | 443.4354105 | ENSG00000160959 |
| GON7 | -0.372646671 | 0.772364269 | 0.054512054 | 0.127915342 | 140.5377217 | 108.5148999 | ENSG00000170270 |
| PRSS16 | -0.372746717 | 0.77231071 | 0.015512602 | 0.045513857 | 247.2224431 | 190.861987 | ENSG00000112812 |
| PABPN1 | -0.372932574 | 0.772211222 | 6.92E-08 | 6.51E-07 | 2110.442306 | 1629.370467 | ENSG00000100836 |
| AARSD1 | -0.372957224 | 0.772198029 | 0.578220016 | 0.73000277 | 11.89307395 | 9.221465981 | ENSG00000266967 |
| FGF12 | -0.372969337 | 0.772191545 | 0.015877224 | 0.046421269 | 235.6584714 | 182.0647908 | ENSG00000114279 |
| RBM8A | -0.373041034 | 0.772153171 | 1.17E-05 | 7.55E-05 | 862.2835301 | 665.7836415 | ENSG00000265241 |
| NA | -0.373202518 | 0.772066747 | 0.557380987 | 0.713357435 | 14.13058603 | 10.91459588 | ENSG00000260526 |
| FAM3B | -0.373228429 | 0.77205288 | 0.426224589 | 0.59781144 | 29.56063693 | 22.84528292 | ENSG00000183844 |
| PSMD6-AS2 | -0.373264381 | 0.772033641 | 0.67155145 | 0.798748321 | 6.760344983 | 5.176456519 | ENSG00000239653 |
| ABCC6P1 | -0.373280942 | 0.772024779 | 0.744014511 | 0.848473867 | 4.310748416 | 3.357217848 | ENSG00000256340 |
| PIGN | -0.373287392 | 0.772021327 | 0.002139837 | 0.008378243 | 437.01438 | 337.4008795 | ENSG00000197563 |
| GLRX5 | -0.373376548 | 0.771973619 | 2.13E-12 | 3.51E-11 | 3046.890048 | 2352.727563 | ENSG00000182512 |
| NOL10 | -0.373382408 | 0.771970483 | 3.78E-05 | 0.000222544 | 787.6333966 | 608.2032437 | ENSG00000115761 |
| PIGT | -0.373447147 | 0.771935843 | 6.05E-06 | 4.15E-05 | 1042.227744 | 804.2656282 | ENSG00000124155 |
| COL9A1 | -0.373447544 | 0.771935631 | 0.000862418 | 0.003723568 | 500.7868683 | 386.1177348 | ENSG00000112280 |
| XKR4 | -0.373491876 | 0.77191191 | 0.467080657 | 0.636331423 | 20.16093811 | 15.58878001 | ENSG00000206579 |
| POLR2K | -0.373493217 | 0.771911193 | 0.001023249 | 0.004350783 | 702.3280809 | 542.617747 | ENSG00000147669 |
| NSUN2 | -0.373560835 | 0.771875015 | 4.36E-12 | 6.91E-11 | 3752.194408 | 2896.381763 | ENSG00000037474 |
| FAM210B | -0.373586438 | 0.771861317 | 0.000523204 | 0.002386648 | 473.2647376 | 365.2672262 | ENSG00000124098 |
| TAF1B | -0.373628414 | 0.771838859 | 0.007575908 | 0.02494399 | 367.6305143 | 284.1308726 | ENSG00000115750 |
| TNFRSF8 | -0.374250542 | 0.771506094 | 8.09E-06 | 5.42E-05 | 1123.460632 | 866.8348069 | ENSG00000120949 |
| GALR3 | -0.374316203 | 0.771470981 | 0.561132357 | 0.716063622 | 13.28695397 | 10.20583157 | ENSG00000128310 |
| FRMD4B | -0.374490329 | 0.771377875 | 0.039641615 | 0.098955278 | 153.8965634 | 118.7328293 | ENSG00000114541 |
| CBS | -0.374494226 | 0.771375791 | 5.43E-06 | 3.75E-05 | 1008.616829 | 778.1558752 | ENSG00000160200 |
| PIGF | -0.374662206 | 0.771285981 | 0.126020959 | 0.247160465 | 89.31181873 | 68.81682938 | ENSG00000151665 |
| CD3EAP | -0.37466729 | 0.771283263 | 0.031784246 | 0.082357273 | 196.7080358 | 151.6979145 | ENSG00000117877 |

| | | | | | | | |
|---|---|---|---|---|---|---|---|
| PSMB7 | -0.374675922 | 0.771278648 | 7.69E-13 | 1.33E-11 | 3957.797798 | 3052.035417 | ENSG00000136930 |
| IMPA2 | -0.374798304 | 0.771213225 | 1.15E-10 | 1.57E-09 | 2493.566671 | 1922.567358 | ENSG00000141401 |
| RNF121 | -0.374924707 | 0.771145657 | 0.00010101 | 0.00054613 | 679.08467 | 523.354524 | ENSG00000137522 |
| POMT2 | -0.37496604 | 0.771123564 | 2.67E-05 | 0.000162851 | 724.3318838 | 558.6988706 | ENSG00000009830 |
| NA | -0.375012425 | 0.771098772 | 0.053029645 | 0.12495357 | 149.8461086 | 115.6272654 | ENSG00000188002 |
| ACVR1B | -0.375104826 | 0.771049386 | 8.67E-07 | 6.89E-06 | 1035.123667 | 798.4720401 | ENSG00000135503 |
| CASZ1 | -0.375106922 | 0.771048266 | 0.073261336 | 0.161797458 | 142.6972759 | 110.1970043 | ENSG00000130940 |
| TPX2 | -0.375141948 | 0.771029547 | 3.20E-15 | 7.04E-14 | 6223.764764 | 4797.83775 | ENSG00000088325 |
| RNF26 | -0.375311864 | 0.770938743 | 8.15E-08 | 7.60E-07 | 1843.538787 | 1421.941422 | ENSG00000173456 |
| SULF2 | -0.375329864 | 0.770929124 | 2.22E-05 | 0.000137169 | 739.7664405 | 570.597832 | ENSG00000196562 |
| EIF5A | -0.375341502 | 0.770922905 | 1.05E-11 | 1.61E-10 | 19866.34097 | 15315.55516 | ENSG00000132507 |
| SENP1 | -0.37540998 | 0.770886314 | 0.00114502 | 0.004816524 | 411.1592133 | 316.782755 | ENSG00000079387 |
| PINX1 | -0.37563371 | 0.770766776 | 0.018559667 | 0.052737448 | 308.1005535 | 237.2597423 | ENSG00000254093 |
| MED25 | -0.37567418 | 0.770745155 | 2.99E-07 | 2.56E-06 | 1504.924857 | 1159.828774 | ENSG00000104973 |
| CDK2AP2 | -0.375802388 | 0.770676664 | 2.43E-06 | 1.78E-05 | 932.5053949 | 718.3335739 | ENSG00000167797 |
| TRMU | -0.375976844 | 0.770583476 | 2.30E-07 | 2.00E-06 | 1294.0993 | 996.9202293 | ENSG00000100416 |
| TXNL4A | -0.376014397 | 0.770563418 | 4.36E-09 | 4.88E-08 | 2099.309101 | 1618.266735 | ENSG00000141759 |
| NA | -0.376258543 | 0.770433028 | 0.54462292 | 0.702999047 | 14.14107392 | 10.88932204 | ENSG00000283692 |
| NA | -0.376337234 | 0.770391006 | 0.532666275 | 0.693411506 | 17.42293877 | 13.47700805 | ENSG00000251188 |
| MOK | -0.376741927 | 0.770174933 | 0.417163606 | 0.589258485 | 23.96840835 | 18.52520522 | ENSG00000080823 |
| DHX57 | -0.376831281 | 0.770127233 | 4.93E-06 | 3.43E-05 | 877.5784025 | 676.0037093 | ENSG00000163214 |
| NPDC1 | -0.376839017 | 0.770123104 | 0.00240068 | 0.009264157 | 534.0154928 | 411.1538584 | ENSG00000107281 |
| NA | -0.376844263 | 0.770120303 | 0.641856006 | 0.776157727 | 7.658443655 | 5.900274849 | ENSG00000250299 |
| TIMM13 | -0.3768895 | 0.770096156 | 1.74E-10 | 2.34E-09 | 4350.745075 | 3350.729861 | ENSG00000099800 |
| SLC35C1 | -0.376947774 | 0.770065051 | 0.000851725 | 0.003683997 | 487.189623 | 375.5612497 | ENSG00000181830 |
| CNMD | -0.377279916 | 0.769887784 | 8.54E-13 | 1.47E-11 | 3338.680262 | 2570.3154 | ENSG00000136110 |
| PCBP4 | -0.377318128 | 0.769867392 | 0.003531461 | 0.012961455 | 339.2620487 | 261.4496337 | ENSG00000090097 |
| DUSP6 | -0.377333128 | 0.769859388 | 2.91E-09 | 3.33E-08 | 2309.911904 | 1778.575975 | ENSG00000139318 |
| FAM92A | -0.377387903 | 0.769830159 | 0.016296912 | 0.04738977 | 239.8381595 | 184.5489704 | ENSG00000188343 |
| POP7 | -0.377440044 | 0.769802337 | 1.02E-08 | 1.09E-07 | 1552.947753 | 1195.381249 | ENSG00000172336 |
| H2AC4 | -0.377448087 | 0.769798046 | 0.759108643 | 0.857014416 | 3.453837566 | 2.637700666 | ENSG00000278463 |
| UBA6-AS1 | -0.377578049 | 0.769728703 | 0.134612988 | 0.259785829 | 79.81020794 | 61.46057604 | ENSG00000248049 |
| PIF1 | -0.377582233 | 0.769726471 | 1.21E-06 | 9.46E-06 | 1860.84065 | 1431.692695 | ENSG00000140451 |
| PNPLA6 | -0.377741287 | 0.769641615 | 3.75E-12 | 6.01E-11 | 2813.261348 | 2165.408181 | ENSG00000032444 |
| POLD2 | -0.377880773 | 0.769567206 | 3.52E-14 | 7.00E-13 | 3518.761132 | 2708.377605 | ENSG00000106628 |
| HGH1 | -0.37800539 | 0.769500735 | 7.96E-05 | 0.000439798 | 962.1669941 | 740.1509831 | ENSG00000235173 |
| KIAA1549L | -0.378163046 | 0.76941665 | 0.080221724 | 0.173804726 | 114.8184462 | 88.43959341 | ENSG00000110427 |
| ESYT1 | -0.378180013 | 0.769407601 | 1.70E-10 | 2.28E-09 | 2490.137752 | 1915.882237 | ENSG00000139641 |
| NA | -0.378222333 | 0.769385031 | 0.488820492 | 0.655460281 | 20.71828257 | 15.96521621 | ENSG00000262519 |
| NOMO2 | -0.378244871 | 0.769373012 | 8.02E-09 | 8.67E-08 | 1511.00567 | 1162.850102 | ENSG00000185164 |
| COL14A1 | -0.378290097 | 0.769348894 | 2.30E-06 | 1.70E-05 | 996.8817219 | 767.025058 | ENSG00000187955 |
| RNF216P1 | -0.378342656 | 0.769320866 | 0.001011499 | 0.004306518 | 420.2073947 | 323.1826065 | ENSG00000196204 |
| IP6K1 | -0.378437431 | 0.769270329 | 2.09E-07 | 1.83E-06 | 1256.666583 | 966.8627757 | ENSG00000176095 |
| NA | -0.378577651 | 0.769195565 | 0.274064048 | 0.440918439 | 41.01817812 | 31.59379104 | ENSG00000272655 |
| PLA2G15 | -0.378747088 | 0.769105232 | 0.029068772 | 0.076534145 | 188.2671598 | 144.9366687 | ENSG00000103066 |
| NDOR1 | -0.37877429 | 0.769090731 | 8.47E-05 | 0.000465246 | 797.5313795 | 613.402563 | ENSG00000188566 |
| TRIM17 | -0.378834167 | 0.769058811 | 0.345567713 | 0.51801591 | 35.57260397 | 27.30894656 | ENSG00000162931 |
| SPATA5L1 | -0.378899882 | 0.769023781 | 0.002423745 | 0.009338219 | 360.5061368 | 277.4282864 | ENSG00000171763 |
| PIP4K2C | -0.378903999 | 0.769021587 | 8.77E-05 | 0.000480405 | 647.8306988 | 497.9172539 | ENSG00000166908 |
| SMIM20 | -0.379025245 | 0.76895696 | 1.02E-05 | 6.73E-05 | 814.1946021 | 626.2232019 | ENSG00000250317 |
| PDCD5 | -0.379051092 | 0.768943183 | 6.27E-06 | 4.29E-05 | 1176.45974 | 904.1774718 | ENSG00000105185 |
| USP43 | -0.379156886 | 0.768886798 | 0.013713222 | 0.041058786 | 235.1652056 | 180.929061 | ENSG00000154914 |
| DMAP1 | -0.379235473 | 0.768844916 | 5.44E-07 | 4.46E-06 | 1019.788595 | 784.2860073 | ENSG00000178028 |
| RPP40 | -0.37927922 | 0.768821603 | 0.098028076 | 0.203974131 | 107.0313328 | 82.29168685 | ENSG00000124787 |
| LRPAP1 | -0.37933813 | 0.76879021 | 4.87E-11 | 6.91E-10 | 2217.073661 | 1704.658329 | ENSG00000163956 |
| NUDT11 | -0.379351846 | 0.768782901 | 1.50E-05 | 9.51E-05 | 822.0868939 | 631.5895336 | ENSG00000196368 |
| NA | -0.379487617 | 0.768710555 | 0.490249199 | 0.656722903 | 18.7754308 | 14.50331903 | ENSG00000264964 |
| HSPE1 | -0.379597798 | 0.76865185 | 0.003587621 | 0.013130109 | 547.9841823 | 421.1187705 | ENSG00000115541 |
| FBXL15 | -0.379601309 | 0.768649979 | 0.003704919 | 0.013500488 | 322.5813056 | 248.1188952 | ENSG00000107872 |
| ZNF17 | -0.379880264 | 0.76850137 | 0.21244958 | 0.368458586 | 53.0201853 | 40.79858547 | ENSG00000186272 |
| RNLS | -0.380299907 | 0.768277865 | 0.062398024 | 0.142293509 | 141.8529681 | 109.2365555 | ENSG00000184719 |
| PLOD3 | -0.380589459 | 0.768123685 | 1.36E-09 | 1.62E-08 | 1692.493949 | 1300.086661 | ENSG00000106397 |
| LRRC20 | -0.380667857 | 0.768081946 | 3.09E-05 | 0.00018584 | 716.8758374 | 550.2746455 | ENSG00000172731 |
| PTDSS1 | -0.380875446 | 0.767971434 | 2.20E-09 | 2.56E-08 | 1961.080038 | 1505.5722 | ENSG00000156471 |
| RPP21 | -0.380942685 | 0.767935643 | 0.454380328 | 0.624142047 | 20.47425884 | 15.67456092 | ENSG00000241370 |
| MAP7D2 | -0.380942809 | 0.767935577 | 0.104626675 | 0.214308047 | 99.64888902 | 76.40486691 | ENSG00000184368 |
| ZDHHC14 | -0.38103009 | 0.767889119 | 0.015743231 | 0.046113372 | 212.1605261 | 162.9378353 | ENSG00000175048 |
| GABRA3 | -0.381213356 | 0.76779158 | 0.014738789 | 0.043635049 | 211.9072538 | 162.5877696 | ENSG00000011677 |
| NA | -0.38126994 | 0.767761467 | 0.055774121 | 0.130362672 | 142.2414558 | 109.221241 | ENSG00000230330 |
| RPL22L1 | -0.381317526 | 0.767736144 | 1.02E-06 | 8.00E-06 | 1215.757665 | 933.5094716 | ENSG00000163584 |
| NOL6 | -0.381502135 | 0.76763791 | 1.90E-10 | 2.54E-09 | 2370.436898 | 1819.56801 | ENSG00000165271 |
| MID1IP1 | -0.381762661 | 0.767499299 | 5.88E-16 | 1.41E-14 | 4535.384855 | 3480.661993 | ENSG00000165175 |
| NA | -0.381763253 | 0.767498985 | 0.786332006 | 0.875158987 | 3.483861777 | 2.658673365 | ENSG00000287923 |
| CENPBD1P1 | -0.38186813 | 0.767443193 | 0.001224575 | 0.005125183 | 428.5867459 | 328.9352789 | ENSG00000213753 |
| ZNF841 | -0.38200546 | 0.767370144 | 0.032770957 | 0.084415079 | 170.4388247 | 130.8256518 | ENSG00000197608 |
| NA | -0.382051182 | 0.767345824 | 0.597529182 | 0.744461977 | 9.413653336 | 7.202187111 | ENSG00000231233 |
| CHRAC1 | -0.382081464 | 0.767329718 | 5.71E-05 | 0.000324941 | 685.0817697 | 526.0182799 | ENSG00000104472 |

| | | | | | | | |
|---|---|---|---|---|---|---|---|
| CREM | -0.382097335 | 0.767321277 | 0.002672531 | 0.010156775 | 330.5937822 | 253.810255 | ENSG00000095794 |
| C3orf18 | -0.382315783 | 0.7672051 | 0.029272906 | 0.076966557 | 203.4118359 | 155.8558567 | ENSG00000088543 |
| MXD3 | -0.382364942 | 0.767178959 | 0.03675188 | 0.092860277 | 161.8865127 | 124.0206131 | ENSG00000213347 |
| KRBOX4 | -0.382823166 | 0.766935329 | 0.053018619 | 0.12495357 | 136.2518913 | 104.6594382 | ENSG00000147121 |
| TDP2 | -0.383121709 | 0.76677664 | 2.35E-07 | 2.04E-06 | 1205.172628 | 924.2307152 | ENSG00000111802 |
| CERK | -0.383255754 | 0.7667054 | 1.12E-11 | 1.71E-10 | 2479.6104 | 1901.145449 | ENSG00000100422 |
| TRABD2B | -0.383263165 | 0.766701462 | 0.043769583 | 0.107274683 | 156.4887473 | 120.0463 | ENSG00000269113 |
| RPL26 | -0.383361106 | 0.766649414 | 7.23E-05 | 0.000403063 | 653.4683455 | 500.8262426 | ENSG00000161970 |
| TNFAIP3 | -0.383383038 | 0.766637759 | 0.267452731 | 0.433747847 | 51.91625966 | 39.79648226 | ENSG00000118503 |
| ERP29 | -0.383497069 | 0.766577166 | 3.91E-14 | 7.70E-13 | 4069.617114 | 3119.440915 | ENSG00000089248 |
| UQCR10 | -0.383798665 | 0.76641693 | 0.058507553 | 0.135234748 | 146.0884743 | 111.8939144 | ENSG00000184076 |
| FADS2 | -0.383963211 | 0.766329522 | 3.59E-13 | 6.43E-12 | 11181.44198 | 8568.865207 | ENSG00000134824 |
| ABCF1 | -0.384002883 | 0.766308449 | 1.15E-14 | 2.41E-13 | 5784.535895 | 4433.444399 | ENSG00000204574 |
| STX5 | -0.384130939 | 0.766240433 | 0.000685507 | 0.003028887 | 485.6499772 | 372.2774239 | ENSG00000162236 |
| RETREG2 | -0.384206841 | 0.766200121 | 2.46E-07 | 2.13E-06 | 1277.067691 | 978.6025578 | ENSG00000144567 |
| ZSWIM8 | -0.384318254 | 0.766140953 | 7.48E-05 | 0.000416154 | 640.5448364 | 490.5803558 | ENSG00000214655 |
| MYPOP | -0.384527911 | 0.766029623 | 0.015230922 | 0.044830976 | 220.5466473 | 168.9610023 | ENSG00000176182 |
| TDG | -0.38456625 | 0.766009267 | 6.83E-07 | 5.52E-06 | 1278.13173 | 979.041397 | ENSG00000139372 |
| ZNF48 | -0.384596807 | 0.765993042 | 3.21E-05 | 0.000192106 | 1046.881064 | 802.1705341 | ENSG00000180035 |
| MAP6D1 | -0.384645253 | 0.76596732 | 0.319812259 | 0.491386438 | 40.77654498 | 31.33271074 | ENSG00000180834 |
| NA | -0.384649146 | 0.765965254 | 0.465072659 | 0.634608349 | 23.400576 | 17.91323515 | ENSG00000269918 |
| ABCC6 | -0.384704945 | 0.765935629 | 0.469766659 | 0.638727701 | 18.33451639 | 14.01745773 | ENSG00000091262 |
| SPATC1L | -0.384733966 | 0.765920222 | 1.41E-05 | 9.02E-05 | 945.8595449 | 724.2658774 | ENSG00000160284 |
| SYNDIG1 | -0.384924702 | 0.765818968 | 0.059236644 | 0.136527372 | 125.9184877 | 96.49789887 | ENSG00000101463 |
| PSMD2 | -0.385024743 | 0.765765865 | 4.00E-17 | 1.06E-15 | 11140.53502 | 8530.423294 | ENSG00000175166 |
| SPATA33 | -0.385123591 | 0.7657134 | 0.019595953 | 0.055266891 | 209.695899 | 160.6459298 | ENSG00000167523 |
| UTP3 | -0.385167446 | 0.765690124 | 8.98E-07 | 7.13E-06 | 1450.714303 | 1110.704085 | ENSG00000132467 |
| ZNF512B | -0.385214666 | 0.765665063 | 9.25E-05 | 0.00050414 | 1205.910506 | 923.0184373 | ENSG00000196700 |
| NA | -0.385380725 | 0.765576937 | 0.705126639 | 0.822605104 | 4.697396369 | 3.580120856 | ENSG00000228201 |
| EMC7 | -0.385534842 | 0.765495159 | 6.26E-06 | 4.29E-05 | 776.2156165 | 594.2984277 | ENSG00000134153 |
| NA | -0.385702419 | 0.765406247 | 0.760425205 | 0.857655235 | 3.481070874 | 2.63985124 | ENSG00000285969 |
| RABEP2 | -0.385826048 | 0.76534066 | 0.000399194 | 0.00188147 | 533.046395 | 407.7240236 | ENSG00000177548 |
| MRPS11 | -0.385924422 | 0.765288475 | 0.000229157 | 0.001142587 | 556.3948197 | 425.6990802 | ENSG00000181991 |
| METTL25 | -0.386029073 | 0.765232964 | 0.161961695 | 0.299530539 | 65.30848008 | 49.95660032 | ENSG00000127720 |
| CRMP1 | -0.386033011 | 0.765230875 | 2.22E-18 | 6.51E-17 | 6206.631401 | 4749.693998 | ENSG00000072832 |
| ZNF112 | -0.386091607 | 0.765199796 | 0.271531469 | 0.438482635 | 40.72227852 | 31.15632686 | ENSG00000062370 |
| NA | -0.386271229 | 0.765104531 | 0.48766935 | 0.654717059 | 17.50483885 | 13.31891325 | ENSG00000237398 |
| MRPS17 | -0.386346337 | 0.765064699 | 0.071435244 | 0.158579666 | 111.7652111 | 85.51957928 | ENSG00000239789 |
| CLINT1 | -0.386426654 | 0.765022108 | 7.60E-12 | 1.18E-10 | 2248.009443 | 1719.97602 | ENSG00000113282 |
| NA | -0.386562564 | 0.764950042 | 0.411190312 | 0.583469827 | 25.41675498 | 19.3686806 | ENSG00000177337 |
| TRPC1 | -0.386629976 | 0.7649143 | 0.028585761 | 0.075552662 | 166.2442309 | 127.0331203 | ENSG00000144935 |
| PNO1 | -0.386724466 | 0.764864203 | 4.20E-05 | 0.000245291 | 679.6300502 | 519.8593416 | ENSG00000115946 |
| NA | -0.386901186 | 0.764770518 | 0.227340653 | 0.386637444 | 47.7352201 | 36.55379627 | ENSG00000185275 |
| URB1 | -0.387002817 | 0.764716646 | 3.50E-09 | 3.97E-08 | 2011.465751 | 1537.820991 | ENSG00000142207 |
| C12orf73 | -0.387096337 | 0.764667076 | 0.056292637 | 0.131199862 | 135.103111 | 103.4131988 | ENSG00000204954 |
| PCDHB14 | -0.38712224 | 0.764653347 | 0.213976536 | 0.370374281 | 51.12900924 | 39.103305 | ENSG00000120327 |
| IGHMBP2 | -0.387160365 | 0.76463314 | 0.000246711 | 0.001220959 | 540.1903458 | 412.6966412 | ENSG00000132740 |
| SNRNP200 | -0.387238745 | 0.7645916 | 2.90E-23 | 1.24E-21 | 12509.21151 | 9563.742034 | ENSG00000144028 |
| MTCH1 | -0.387382137 | 0.764515609 | 5.25E-18 | 1.50E-16 | 5800.519246 | 4434.139527 | ENSG00000137409 |
| PA2G4 | -0.387443672 | 0.764483001 | 2.41E-14 | 4.90E-13 | 3095.878863 | 2366.910192 | ENSG00000170515 |
| LDOC1 | -0.387517274 | 0.764444001 | 2.09E-05 | 0.000129878 | 887.8214211 | 678.9277826 | ENSG00000182195 |
| SLC4A1AP | -0.387749351 | 0.76432104 | 4.12E-05 | 0.000241125 | 621.3112574 | 475.0057936 | ENSG00000163798 |
| FAAP20 | -0.387756081 | 0.764317474 | 2.08E-07 | 1.82E-06 | 1023.459423 | 782.2624394 | ENSG00000162585 |
| PTPRU | -0.3878735 | 0.76425527 | 8.82E-10 | 1.08E-08 | 1854.538267 | 1417.43021 | ENSG00000060656 |
| THSD1 | -0.387944277 | 0.764217777 | 0.618548912 | 0.760172457 | 11.76847135 | 8.950165854 | ENSG00000136114 |
| SSRP1 | -0.38795331 | 0.764212992 | 1.16E-19 | 3.82E-18 | 8314.558303 | 6354.105217 | ENSG00000149136 |
| AATF | -0.388013083 | 0.76418133 | 4.44E-09 | 4.97E-08 | 2084.504489 | 1592.774168 | ENSG00000275700 |
| MRPL34 | -0.388098427 | 0.764136126 | 4.83E-08 | 4.65E-07 | 1233.819134 | 942.8056441 | ENSG00000130312 |
| RAB36 | -0.388159557 | 0.764103748 | 0.290925217 | 0.460661443 | 42.48095483 | 32.42489601 | ENSG00000100228 |
| PEPD | -0.388357299 | 0.763999024 | 4.45E-09 | 4.98E-08 | 2319.974745 | 1773.073157 | ENSG00000124299 |
| KIF1C | -0.388416772 | 0.76396753 | 2.44E-11 | 3.58E-10 | 4766.567153 | 3641.717545 | ENSG00000129250 |
| AKIP1 | -0.388503209 | 0.76392176 | 0.002092592 | 0.008211592 | 401.6227972 | 306.3973229 | ENSG00000166452 |
| MAIP1 | -0.388638344 | 0.763850208 | 0.008777339 | 0.028211203 | 237.0787089 | 181.0543578 | ENSG00000162972 |
| LRFN4 | -0.388770156 | 0.763780421 | 0.00011877 | 0.000630836 | 889.3295663 | 679.5806072 | ENSG00000173621 |
| CCNQ | -0.388852654 | 0.763736747 | 0.002833603 | 0.010680547 | 403.0553637 | 308.1065608 | ENSG00000262919 |
| NA | -0.388873124 | 0.763725911 | 0.555659163 | 0.711814923 | 11.67819856 | 8.952316427 | ENSG00000278126 |
| MAPKAPK3 | -0.388993804 | 0.763662028 | 6.15E-12 | 9.62E-11 | 2151.57854 | 1642.929942 | ENSG00000114738 |
| SLC35A3 | -0.389324949 | 0.763486764 | 0.063282615 | 0.143943512 | 134.9278718 | 103.1703892 | ENSG00000117620 |
| SMG6 | -0.389347518 | 0.76347482 | 2.90E-05 | 0.000175151 | 802.4429499 | 612.1689361 | ENSG00000070366 |
| MCM8 | -0.389429534 | 0.763431418 | 0.021074643 | 0.058648434 | 240.5796795 | 183.8450281 | ENSG00000125885 |
| MIR4442 | -0.389466986 | 0.7634116 | 0.408497658 | 0.580759696 | 30.49434162 | 23.36340283 | ENSG00000283971 |
| KIAA1586 | -0.389508183 | 0.763389801 | 0.023886016 | 0.065091841 | 250.6483347 | 191.4578186 | ENSG00000168116 |
| PDZRN3 | -0.389519065 | 0.763384043 | 0.001261567 | 0.005260892 | 373.7133059 | 285.3043077 | ENSG00000121440 |
| SRPRB | -0.389619376 | 0.763330966 | 3.78E-05 | 0.000222692 | 634.4463755 | 484.2885602 | ENSG00000144867 |
| NFATC2IP | -0.389892675 | 0.763186377 | 1.10E-06 | 8.59E-06 | 1149.844768 | 877.4683917 | ENSG00000176953 |
| LOC100505501 | -0.389925874 | 0.763168815 | 0.090109985 | 0.19085548 | 102.1988459 | 77.82911983 | ENSG00000167912 |
| NA | -0.389963434 | 0.763148946 | 0.654637399 | 0.785703358 | 6.917962863 | 5.323798452 | ENSG00000225335 |

| | | | | | | | |
|---|---|---|---|---|---|---|---|
| SLITRK4 | -0.389972391 | 0.763144209 | 0.028147088 | 0.074616666 | 200.4465214 | 153.1638097 | ENSG00000179542 |
| BAIAP2-DT | -0.390462582 | 0.762884956 | 0.003078991 | 0.011453302 | 368.0114683 | 280.6231509 | ENSG00000226137 |
| H3-3B | -0.390544704 | 0.762841532 | 6.34E-18 | 1.81E-16 | 14268.7416 | 10885.13317 | ENSG00000132475 |
| GABRB3 | -0.390818252 | 0.762696903 | 1.36E-16 | 3.45E-15 | 4032.529816 | 3074.925571 | ENSG00000166206 |
| CBX1 | -0.390841581 | 0.762684571 | 2.13E-22 | 8.59E-21 | 7540.959124 | 5751.411714 | ENSG00000108468 |
| TACR3 | -0.390927723 | 0.762639033 | 0.181120084 | 0.326307024 | 59.73376066 | 45.53327666 | ENSG00000169836 |
| EIF2S3B | -0.39100385 | 0.762598791 | 0.369417993 | 0.543132575 | 25.42704271 | 19.38750273 | ENSG00000180574 |
| CST6 | -0.391265823 | 0.762460327 | 0.659061711 | 0.789441755 | 6.052679304 | 4.610732991 | ENSG00000175315 |
| NA | -0.391319325 | 0.762432052 | 0.386154717 | 0.559606508 | 27.18695832 | 20.8093325 | ENSG00000260285 |
| MIR22HG | -0.391432362 | 0.762372317 | 0.337585932 | 0.510333846 | 34.47657548 | 26.1774995 | ENSG00000186594 |
| GET3 | -0.391450407 | 0.762362781 | 4.14E-10 | 5.29E-09 | 2389.938765 | 1821.210543 | ENSG00000198356 |
| NA | -0.391477728 | 0.762348344 | 0.329220817 | 0.501507307 | 31.08749226 | 23.72478502 | ENSG00000240429 |
| CENPU | -0.391810878 | 0.762172321 | 1.93E-07 | 1.70E-06 | 1236.716341 | 942.4529006 | ENSG00000151725 |
| DENND2A | -0.391859299 | 0.762146741 | 0.001633537 | 0.006606407 | 382.9666257 | 292.1107035 | ENSG00000146966 |
| ARL6 | -0.391926152 | 0.762111425 | 0.130072029 | 0.252945515 | 89.47693343 | 68.38903988 | ENSG00000113966 |
| TLL2 | -0.391978508 | 0.762083768 | 0.020551512 | 0.057516207 | 175.9710048 | 134.0968283 | ENSG00000095587 |
| DCAF11 | -0.391987731 | 0.762078896 | 9.02E-07 | 7.15E-06 | 1582.744352 | 1205.623416 | ENSG00000100897 |
| NA | -0.392025331 | 0.762059035 | 0.621920555 | 0.762199274 | 8.738802771 | 6.687011276 | ENSG00000234297 |
| C8orf33 | -0.392044179 | 0.762049079 | 3.11E-08 | 3.07E-07 | 1219.510505 | 929.6528949 | ENSG00000182307 |
| NA | -0.392246725 | 0.761942099 | 0.662741804 | 0.791973727 | 6.491002969 | 4.909457539 | ENSG00000260160 |
| NA | -0.392346145 | 0.761889594 | 0.317080616 | 0.488289469 | 34.46867833 | 26.2404176 | ENSG00000285728 |
| AGFG2 | -0.392394133 | 0.761864251 | 0.075342573 | 0.165407258 | 111.4361841 | 84.97886914 | ENSG00000106351 |
| WDR4 | -0.392490038 | 0.761813607 | 1.21E-08 | 1.27E-07 | 1455.878043 | 1108.628867 | ENSG00000160193 |
| L3MBTL2 | -0.392568273 | 0.761772296 | 5.80E-05 | 0.000329654 | 559.3039031 | 426.0419007 | ENSG00000100395 |
| CGRRF1 | -0.392711376 | 0.761696739 | 0.012572808 | 0.038171552 | 208.2810801 | 158.6290619 | ENSG00000100532 |
| TIMP2 | -0.39281985 | 0.76163947 | 0.000381477 | 0.001807682 | 548.1150666 | 417.8037158 | ENSG00000035862 |
| NA | -0.392852754 | 0.761622099 | 0.70620315 | 0.823231454 | 4.730211482 | 3.580120856 | ENSG00000233747 |
| LIMK1 | -0.392904051 | 0.761595019 | 7.37E-14 | 1.41E-12 | 3237.404394 | 2465.896584 | ENSG00000106683 |
| CLSTN2 | -0.393128478 | 0.761476554 | 0.232646105 | 0.393337074 | 54.75777369 | 41.79317089 | ENSG00000158258 |
| NA | -0.39325499 | 0.761409782 | 0.561202939 | 0.716063622 | 11.65096525 | 8.887247756 | ENSG00000272525 |
| TRIM31 | -0.393513961 | 0.761273117 | 0.655782687 | 0.786539656 | 7.370241042 | 5.618221852 | ENSG00000204616 |
| LOC102724474 | -0.393526907 | 0.761266286 | 0.501445372 | 0.667359964 | 15.92889855 | 12.14767136 | ENSG00000283828 |
| BCL9 | -0.393540949 | 0.761258877 | 1.09E-08 | 1.15E-07 | 1589.477001 | 1210.041839 | ENSG00000116128 |
| MTNR1A | -0.393544212 | 0.761257155 | 0.581417617 | 0.731932083 | 10.45908216 | 7.970101412 | ENSG00000168412 |
| WRAP73 | -0.393552279 | 0.761252898 | 3.69E-05 | 0.000218037 | 646.8434914 | 492.0771951 | ENSG00000116213 |
| CCAR2 | -0.393552631 | 0.761252712 | 3.88E-14 | 7.65E-13 | 3712.506831 | 2825.87424 | ENSG00000158941 |
| NA | -0.393610202 | 0.761222335 | 0.160709173 | 0.297956815 | 68.11573953 | 51.95328895 | ENSG00000261188 |
| CCDC24 | -0.393664179 | 0.761193856 | 0.037207456 | 0.093838283 | 159.2269468 | 121.0153511 | ENSG00000159214 |
| SLC2A6 | -0.393847741 | 0.761097011 | 0.001151705 | 0.00484359 | 642.888329 | 489.1287147 | ENSG00000160326 |
| TMEM250 | -0.393848486 | 0.761096617 | 5.24E-08 | 5.02E-07 | 1457.088713 | 1109.26287 | ENSG00000238227 |
| CCDC137 | -0.393899001 | 0.761069969 | 9.63E-08 | 8.90E-07 | 1671.411872 | 1272.037819 | ENSG00000185298 |
| COASY | -0.394067054 | 0.76098132 | 1.21E-09 | 1.45E-08 | 2132.686817 | 1623.113063 | ENSG00000068120 |
| NA | -0.394082867 | 0.76097298 | 0.607924205 | 0.751808824 | 9.569356194 | 7.246283083 | ENSG00000226976 |
| AXIN2 | -0.39436603 | 0.760823635 | 0.007531714 | 0.024849375 | 274.6006223 | 209.1067447 | ENSG00000168646 |
| PNPLA4 | -0.394461627 | 0.760773223 | 0.000968854 | 0.004144161 | 511.8648197 | 389.4203095 | ENSG00000006757 |
| TCF3 | -0.394477842 | 0.760764672 | 1.53E-13 | 2.84E-12 | 7351.136676 | 5592.363733 | ENSG00000071564 |
| CHAD | -0.394575011 | 0.760713434 | 0.755477305 | 0.855091825 | 3.453837566 | 2.593604694 | ENSG00000136457 |
| PTPN13 | -0.394635817 | 0.760681373 | 4.27E-14 | 8.39E-13 | 2658.376775 | 2022.490184 | ENSG00000163629 |
| PLEKHG3 | -0.394738877 | 0.760627035 | 3.87E-07 | 3.26E-06 | 1340.612063 | 1019.730843 | ENSG00000126822 |
| CDADC1 | -0.394769472 | 0.760610905 | 0.168434711 | 0.308836993 | 65.00823797 | 49.42287493 | ENSG00000102543 |
| SLAIN1 | -0.394811286 | 0.76058886 | 0.000118996 | 0.000631515 | 569.0822775 | 432.8442558 | ENSG00000139737 |
| VPS50 | -0.394815627 | 0.760586571 | 0.04686712 | 0.113496562 | 129.8681067 | 98.66587675 | ENSG00000004766 |
| DPCD | -0.395064336 | 0.760455464 | 0.003261044 | 0.012049187 | 362.9753818 | 275.7491871 | ENSG00000166171 |
| STMN3 | -0.395081415 | 0.760446461 | 7.92E-07 | 6.34E-06 | 2124.483548 | 1615.919521 | ENSG00000197457 |
| MYBBP1A | -0.395143194 | 0.760413898 | 4.53E-11 | 6.45E-10 | 4928.588368 | 3747.649133 | ENSG00000132382 |
| MAP7D1 | -0.395311092 | 0.760325408 | 1.93E-09 | 2.26E-08 | 3257.832627 | 2477.463241 | ENSG00000116871 |
| NA | -0.395339712 | 0.760310325 | 0.588903905 | 0.738474911 | 10.46957005 | 7.963649691 | ENSG00000229932 |
| CENPW | -0.395397903 | 0.760279658 | 1.29E-05 | 8.26E-05 | 698.4019632 | 531.0097502 | ENSG00000203760 |
| YRDC | -0.395402323 | 0.760277329 | 0.001289553 | 0.005365999 | 357.5914588 | 272.0380927 | ENSG00000196449 |
| VIPR2 | -0.395591844 | 0.760177461 | 0.224657328 | 0.38339212 | 53.409424 | 40.65769526 | ENSG00000106018 |
| KLHL25 | -0.3958434 | 0.760044924 | 0.000150784 | 0.000779622 | 808.5186095 | 614.6773297 | ENSG00000183655 |
| ELOVL7 | -0.396152291 | 0.759882211 | 0.114207408 | 0.229545229 | 78.69300351 | 59.7545427 | ENSG00000164181 |
| GNG2 | -0.396185035 | 0.759864964 | 0.002283904 | 0.00885471 | 330.6864456 | 251.4341677 | ENSG00000186469 |
| NA | -0.396229812 | 0.759841381 | 0.521449239 | 0.684397691 | 17.21875142 | 13.02233927 | ENSG00000284617 |
| SMCO4 | -0.396243017 | 0.759834426 | 1.64E-05 | 0.00010375 | 748.8123192 | 568.9657422 | ENSG00000166002 |
| PTPN11 | -0.396261666 | 0.759824604 | 1.67E-15 | 3.79E-14 | 3605.230598 | 2739.593578 | ENSG00000179295 |
| SLC35F3 | -0.396435685 | 0.759732959 | 0.20892575 | 0.363851824 | 53.92944759 | 40.99783217 | ENSG00000183780 |
| DPP9 | -0.396491858 | 0.759703378 | 1.26E-07 | 1.14E-06 | 1371.18251 | 1041.970849 | ENSG00000142002 |
| UROD | -0.396525836 | 0.759685486 | 9.82E-10 | 1.19E-08 | 1675.608767 | 1272.669113 | ENSG00000126088 |
| SPSB3 | -0.396559173 | 0.759667932 | 0.475524002 | 0.643878257 | 17.20248156 | 13.08955852 | ENSG00000162032 |
| DNAJC1 | -0.396576658 | 0.759658725 | 0.001361399 | 0.005617699 | 399.2659392 | 303.1282785 | ENSG00000136770 |
| NA | -0.396596726 | 0.759648158 | 0.5463466 | 0.704470255 | 11.69705915 | 8.895317013 | ENSG00000275582 |
| NA | -0.396927109 | 0.759474216 | 0.642525379 | 0.776675047 | 6.939614365 | 5.304976326 | ENSG00000267199 |
| RPL23A | -0.397022831 | 0.759423827 | 5.24E-08 | 5.02E-07 | 1630.944581 | 1238.408858 | ENSG00000198242 |
| PHB2 | -0.397080162 | 0.759393649 | 3.92E-20 | 1.34E-18 | 8034.565805 | 6100.731378 | ENSG00000215021 |
| RBM44 | -0.397203699 | 0.759328625 | 0.559400869 | 0.714709727 | 10.39903374 | 7.889978725 | ENSG00000177483 |
| NPY | -0.397281867 | 0.759287484 | 0.659367518 | 0.789485573 | 8.167303631 | 6.152752851 | ENSG00000122585 |

| | | | | | | | |
|---|---|---|---|---|---|---|---|
| AUTS2 | -0.397405353 | 0.759222497 | 2.63E-10 | 3.46E-09 | 1939.885212 | 1473.226027 | ENSG00000158321 |
| FBXW8 | -0.397516606 | 0.759163952 | 0.001579148 | 0.006406591 | 480.5332299 | 364.4439663 | ENSG00000174989 |
| NA | -0.397588672 | 0.759126031 | 0.706773374 | 0.823594399 | 4.37358774 | 3.329793428 | ENSG00000277173 |
| ARHGDIA | -0.398043575 | 0.758886705 | 8.04E-10 | 9.88E-09 | 3804.960298 | 2887.666102 | ENSG00000141522 |
| APIP | -0.398178453 | 0.75881576 | 0.003783905 | 0.013772701 | 322.5969749 | 245.154773 | ENSG00000149089 |
| FAM171A1 | -0.398303046 | 0.75875023 | 3.00E-07 | 2.57E-06 | 1553.218895 | 1178.450495 | ENSG00000148468 |
| NAPB | -0.398330323 | 0.758735885 | 0.006901118 | 0.023128818 | 249.4547609 | 189.3734589 | ENSG00000125814 |
| DBNL | -0.398518087 | 0.758637143 | 4.35E-06 | 3.06E-05 | 1366.114524 | 1036.483607 | ENSG00000136279 |
| ZBED5-AS1 | -0.398832477 | 0.75847184 | 0.514652235 | 0.678586366 | 13.42012944 | 10.15958502 | ENSG00000247271 |
| MCC | -0.398839807 | 0.758467986 | 0.000150736 | 0.000779586 | 600.2778655 | 455.3292774 | ENSG00000171444 |
| MCAT | -0.398842648 | 0.758466493 | 0.000574423 | 0.002589066 | 446.4963295 | 338.979738 | ENSG00000100294 |
| GDF1 | -0.398846965 | 0.758464223 | 0.021172491 | 0.058886753 | 183.5152969 | 139.3453382 | ENSG00000223802 |
| BNIP1 | -0.398906536 | 0.758432905 | 0.100522397 | 0.207924966 | 110.0528288 | 83.25752131 | ENSG00000113734 |
| UBXN6 | -0.398918968 | 0.75842637 | 4.59E-07 | 3.80E-06 | 965.9624617 | 732.8729075 | ENSG00000167671 |
| ZNF615 | -0.398957334 | 0.758406201 | 0.027219746 | 0.072667605 | 160.3491695 | 121.5670746 | ENSG00000197619 |
| CSNK2B | -0.398991654 | 0.75838816 | 0.231025752 | 0.391317744 | 49.00110612 | 37.2243833 | ENSG00000204435 |
| NA | -0.399041094 | 0.758362171 | 0.503517547 | 0.668887233 | 15.0691968 | 11.38567575 | ENSG00000275180 |
| NA | -0.399213844 | 0.758271369 | 0.18523696 | 0.331484805 | 59.09539199 | 44.79466478 | ENSG00000255857 |
| FST | -0.399256761 | 0.758248813 | 0.035172874 | 0.089502693 | 152.6577106 | 115.6864459 | ENSG00000134363 |
| SLC44A3 | -0.399264521 | 0.758244734 | 0.155739553 | 0.291008496 | 68.56052088 | 52.01458951 | ENSG00000143036 |
| NA | -0.399268965 | 0.758242399 | 0.539394672 | 0.698681435 | 12.19822214 | 9.221465981 | ENSG00000250539 |
| DPM1 | -0.399364471 | 0.758192205 | 1.59E-06 | 1.21E-05 | 1092.715821 | 829.1233989 | ENSG00000000419 |
| TMEM115 | -0.399460912 | 0.758141523 | 0.000285207 | 0.001390428 | 471.091192 | 357.1054264 | ENSG00000126062 |
| NA | -0.399645633 | 0.758044458 | 0.538600298 | 0.698168146 | 12.58678512 | 9.585259201 | ENSG00000279520 |
| NA | -0.399691983 | 0.758020104 | 0.337691743 | 0.510343519 | 33.9500942 | 25.68088532 | ENSG00000253341 |
| TIGD1 | -0.39969246 | 0.758019854 | 0.06629474 | 0.149438166 | 118.0749927 | 89.59443631 | ENSG00000221944 |
| ARHGAP39 | -0.399781775 | 0.757972927 | 0.000318574 | 0.001535252 | 575.2363947 | 436.2144013 | ENSG00000147799 |
| FAM104B | -0.399932137 | 0.757893933 | 0.034164871 | 0.087306204 | 147.1569941 | 111.6973392 | ENSG00000182518 |
| NA | -0.400163761 | 0.757772263 | 0.58997311 | 0.739181092 | 9.119868918 | 6.884640437 | ENSG00000227008 |
| EFTUD2 | -0.400203669 | 0.757751302 | 5.68E-14 | 1.10E-12 | 4551.942305 | 3448.740211 | ENSG00000108883 |
| TRIM3 | -0.400364241 | 0.757666969 | 0.001610466 | 0.006518568 | 326.2616946 | 247.1716408 | ENSG00000110171 |
| AP4M1 | -0.400400282 | 0.757648042 | 0.011617009 | 0.035742219 | 236.7084429 | 179.095053 | ENSG00000221838 |
| SFRP2 | -0.40062681 | 0.757529087 | 1.64E-16 | 4.13E-15 | 7063.442985 | 5351.957188 | ENSG00000145423 |
| NA | -0.400647443 | 0.757518253 | 0.390810767 | 0.564052858 | 31.13406173 | 23.62530717 | ENSG00000279413 |
| RELT | -0.400773577 | 0.757452026 | 0.046292719 | 0.112232355 | 143.6883261 | 108.936777 | ENSG00000054967 |
| NA | -0.400787615 | 0.757444656 | 0.395903204 | 0.568468083 | 22.06885958 | 16.69280265 | ENSG00000225630 |
| MREG | -0.401158546 | 0.757249935 | 0.00045384 | 0.002103564 | 456.8975906 | 346.1225147 | ENSG00000118242 |
| MSL3P1 | -0.40122494 | 0.757215087 | 0.640840027 | 0.775313983 | 7.429613742 | 5.622523 | ENSG00000224287 |
| ARF3 | -0.401318044 | 0.757166221 | 8.88E-19 | 2.69E-17 | 4420.603829 | 3346.901012 | ENSG00000134287 |
| AAGAB | -0.401462926 | 0.757090187 | 7.53E-07 | 6.05E-06 | 964.6198429 | 730.1408358 | ENSG00000103591 |
| NA | -0.401540533 | 0.757049462 | 0.113679593 | 0.228770397 | 82.81100359 | 62.69526906 | ENSG00000237238 |
| FNDC11 | -0.401627032 | 0.757004073 | 0.160221688 | 0.297424613 | 68.06302467 | 51.59352431 | ENSG00000125531 |
| LOC101928000 | -0.401958187 | 0.756830331 | 0.338028033 | 0.510682025 | 32.08702733 | 24.17704275 | ENSG00000234327 |
| ALG14 | -0.402457467 | 0.756568457 | 0.186078254 | 0.332613161 | 56.0938326 | 42.38255073 | ENSG00000172339 |
| MBD3 | -0.4027402 | 0.756420202 | 1.98E-05 | 0.000123598 | 1034.056313 | 782.3157317 | ENSG00000071655 |
| NOL4 | -0.403078993 | 0.75624259 | 0.433953472 | 0.605088285 | 22.6585436 | 17.17651338 | ENSG00000101746 |
| TIGD7 | -0.403080354 | 0.756241877 | 0.238831943 | 0.400774793 | 44.2101705 | 33.47648085 | ENSG00000140993 |
| OSGEP | -0.403183052 | 0.756188046 | 6.79E-05 | 0.000380866 | 546.6397606 | 413.1351899 | ENSG00000092094 |
| XPO6 | -0.40336549 | 0.756092427 | 3.73E-17 | 9.94E-16 | 4271.17383 | 3229.84973 | ENSG00000169180 |
| ACBD7 | -0.403507473 | 0.75601802 | 0.002052178 | 0.00807271 | 342.075966 | 258.8549445 | ENSG00000176244 |
| FBLN5 | -0.403532645 | 0.756004829 | 0.240342465 | 0.402539198 | 45.85799855 | 34.57620234 | ENSG00000140092 |
| ARMC2 | -0.403533822 | 0.756004212 | 0.20526813 | 0.359037764 | 58.34721421 | 43.9598038 | ENSG00000118690 |
| TUT1 | -0.403564536 | 0.755988118 | 0.488075561 | 0.654943354 | 17.76090207 | 13.3985029 | ENSG00000255508 |
| RANBP10 | -0.403624865 | 0.755956505 | 9.51E-05 | 0.000516932 | 586.3952823 | 443.2687134 | ENSG00000141084 |
| SLC6A3 | -0.403713729 | 0.755909943 | 0.373729285 | 0.547176938 | 27.1284615 | 20.5568545 | ENSG00000142319 |
| LRRC37B | -0.403977418 | 0.755771794 | 0.023570935 | 0.064396957 | 195.0269412 | 147.534835 | ENSG00000185158 |
| NA | -0.404116688 | 0.755698839 | 0.582159233 | 0.732591091 | 9.927894958 | 7.50736338 | ENSG00000235558 |
| COA3 | -0.404202807 | 0.755653731 | 1.43E-06 | 1.09E-05 | 1069.188928 | 808.3476636 | ENSG00000183978 |
| HMGN5 | -0.404294541 | 0.755605684 | 3.37E-05 | 0.000201515 | 738.6708265 | 558.4372694 | ENSG00000198157 |
| KIAA1328 | -0.404655797 | 0.755416501 | 0.176450656 | 0.319976461 | 61.86493024 | 46.86664191 | ENSG00000150477 |
| YARS1 | -0.40485933 | 0.755309936 | 1.58E-21 | 5.96E-20 | 6909.491464 | 5218.088365 | ENSG00000134684 |
| INPP5A | -0.405252406 | 0.755104172 | 0.000101818 | 0.000550034 | 560.7109739 | 423.2584572 | ENSG00000068383 |
| S100A1 | -0.405287811 | 0.755085642 | 0.517950776 | 0.6816146 | 13.42012944 | 10.09236578 | ENSG00000160678 |
| DNAJA1 | -0.405346381 | 0.755054988 | 4.85E-21 | 1.77E-19 | 7027.110253 | 5306.719548 | ENSG00000086061 |
| DIABLO | -0.405382654 | 0.755036004 | 0.459050416 | 0.628944714 | 24.37092584 | 18.47627507 | ENSG00000184047 |
| SLC39A3 | -0.405446889 | 0.755002387 | 0.001193438 | 0.005003829 | 483.7142972 | 365.5850454 | ENSG00000141873 |
| MAST3 | -0.405586482 | 0.754929338 | 0.006347968 | 0.021533084 | 320.3332545 | 241.9817357 | ENSG00000099308 |
| TELO2 | -0.405726009 | 0.75485633 | 1.29E-07 | 1.17E-06 | 2164.952452 | 1634.117547 | ENSG00000100726 |
| UBE2L6 | -0.405772059 | 0.754832236 | 9.60E-09 | 1.03E-07 | 1287.671975 | 972.06075 | ENSG00000156587 |
| RAB6B | -0.405864196 | 0.75478403 | 0.000248171 | 0.001227868 | 442.5040081 | 334.0762113 | ENSG00000154917 |
| B3GNT5 | -0.405902604 | 0.754763937 | 0.063885393 | 0.144972724 | 106.7894243 | 80.61065478 | ENSG00000176597 |
| JDP2 | -0.405940277 | 0.754744228 | 6.72E-06 | 4.57E-05 | 700.7446296 | 528.9479929 | ENSG00000140044 |
| NA | -0.406099569 | 0.754660899 | 0.701788867 | 0.820248957 | 4.346354432 | 3.277095161 | ENSG00000240695 |
| NCDN | -0.406215859 | 0.754600071 | 7.59E-06 | 5.12E-05 | 1213.443434 | 915.6349894 | ENSG00000020129 |
| TNPO3 | -0.40621746 | 0.754599234 | 1.56E-17 | 4.27E-16 | 3750.978743 | 2830.830621 | ENSG00000064419 |
| NA | -0.406323895 | 0.754543565 | 0.037176696 | 0.093784632 | 164.4669694 | 124.0200801 | ENSG00000204791 |
| GABRA2 | -0.406423286 | 0.754491584 | 0.430272897 | 0.601652251 | 23.38103968 | 17.65430543 | ENSG00000151834 |

| | | | | | | | |
|---|---|---|---|---|---|---|---|
| TATDN2 | -0.406959086 | 0.754211427 | 0.000607247 | 0.002721747 | 517.81918 | 390.3399522 | ENSG00000157014 |
| NA | -0.407450205 | 0.753954724 | 0.727074742 | 0.836082516 | 4.381960449 | 3.302369007 | ENSG00000269473 |
| SEPTIN1 | -0.407919308 | 0.75370961 | 0.524788465 | 0.687190761 | 12.72016075 | 9.612683621 | ENSG00000180096 |
| GULP1 | -0.408135589 | 0.753596626 | 2.25E-10 | 2.99E-09 | 1949.066767 | 1468.729516 | ENSG00000144366 |
| PERP | -0.408164517 | 0.753581516 | 5.45E-06 | 3.76E-05 | 981.7754569 | 740.1159492 | ENSG00000112378 |
| NABP2 | -0.408200808 | 0.75356256 | 1.94E-08 | 1.97E-07 | 1160.341622 | 874.3757249 | ENSG00000139579 |
| POLR3C | -0.408201559 | 0.753562167 | 4.19E-05 | 0.00024455 | 638.0928365 | 480.6460969 | ENSG00000186141 |
| TTC33 | -0.408330662 | 0.753494736 | 0.020135269 | 0.056523498 | 197.7848403 | 149.232266 | ENSG00000113638 |
| ARMH3 | -0.40833384 | 0.753493076 | 3.36E-05 | 0.000200747 | 553.8784289 | 417.363035 | ENSG00000120029 |
| MINPP1 | -0.40841864 | 0.753448788 | 0.000168724 | 0.000864042 | 713.7407164 | 537.7928049 | ENSG00000107789 |
| DTX1 | -0.408453434 | 0.753430617 | 0.002526023 | 0.009670474 | 346.3149388 | 260.7032678 | ENSG00000135144 |
| CXXC1 | -0.408845342 | 0.753225975 | 2.51E-11 | 3.68E-10 | 2101.48899 | 1582.480908 | ENSG00000154832 |
| CA11 | -0.408893073 | 0.753201056 | 1.42E-07 | 1.28E-06 | 1059.788065 | 798.2109961 | ENSG00000063180 |
| EXOC6 | -0.408983076 | 0.753154068 | 0.036593675 | 0.09255652 | 143.5086933 | 108.2140126 | ENSG00000138190 |
| OGFOD3 | -0.409008968 | 0.753140552 | 1.89E-05 | 0.000118048 | 696.4606758 | 524.1358748 | ENSG00000181396 |
| NA | -0.409012032 | 0.753138952 | 0.683416973 | 0.806868328 | 5.657658643 | 4.263611323 | ENSG00000243824 |
| SETD1A | -0.409132812 | 0.753075903 | 1.73E-07 | 1.54E-06 | 1791.928291 | 1349.799428 | ENSG00000099381 |
| USP25 | -0.409139219 | 0.753072559 | 8.86E-06 | 5.90E-05 | 1009.472036 | 760.4835106 | ENSG00000155313 |
| WDR7 | -0.409490607 | 0.75288916 | 0.00182584 | 0.007290889 | 319.4245558 | 240.2923982 | ENSG00000091157 |
| USB1 | -0.409644924 | 0.752808632 | 7.25E-09 | 7.87E-08 | 1718.664315 | 1293.794938 | ENSG00000103005 |
| PCDH19 | -0.409692724 | 0.75278369 | 0.170165065 | 0.311447275 | 63.00150774 | 47.34040539 | ENSG00000165194 |
| RIOK1 | -0.409730347 | 0.752764059 | 1.76E-05 | 0.000110489 | 753.7755399 | 567.5083946 | ENSG00000124784 |
| NA | -0.40976276 | 0.752747147 | 0.287533709 | 0.45693058 | 33.13158027 | 24.91108091 | ENSG00000226360 |
| C11orf24 | -0.409779716 | 0.7527383 | 4.49E-06 | 3.15E-05 | 783.5868598 | 590.1300052 | ENSG00000171067 |
| UBE2V1 | -0.40985519 | 0.752698922 | 0.274180295 | 0.440988166 | 38.68656321 | 29.0902684 | ENSG00000244687 |
| RRM2 | -0.409923878 | 0.752663086 | 3.13E-16 | 7.70E-15 | 3890.651871 | 2928.340426 | ENSG00000171848 |
| FRG2C | -0.410200743 | 0.752518658 | 0.782978078 | 0.87283618 | 3.617037252 | 2.759768752 | ENSG00000172969 |
| CABLES2 | -0.410210073 | 0.752513791 | 0.000296677 | 0.001439796 | 450.4429574 | 339.0824815 | ENSG00000149679 |
| SLC25A23 | -0.410255907 | 0.752489884 | 6.43E-07 | 5.22E-06 | 1529.779099 | 1151.663412 | ENSG00000125648 |
| BRMS1 | -0.410477546 | 0.752374289 | 2.16E-06 | 1.61E-05 | 1040.382675 | 782.1651669 | ENSG00000174744 |
| LINC00863 | -0.410539616 | 0.75234192 | 0.14364331 | 0.27322255 | 93.96049354 | 70.79686488 | ENSG00000224914 |
| GOLGA8H | -0.410692655 | 0.752262117 | 0.484934562 | 0.652771798 | 15.85018969 | 11.92853647 | ENSG00000261794 |
| HRAT92 | -0.41071499 | 0.752250471 | 0.464251793 | 0.633820335 | 18.69114013 | 14.04004796 | ENSG00000223855 |
| RAB5IF | -0.410855787 | 0.75217706 | 0.006878931 | 0.023068446 | 258.3943724 | 194.6270819 | ENSG00000101084 |
| NOP9 | -0.411030419 | 0.752086018 | 6.81E-06 | 4.63E-05 | 991.0579605 | 745.29486 | ENSG00000196943 |
| PNKP | -0.411073261 | 0.752063684 | 4.76E-09 | 5.30E-08 | 1324.286378 | 995.5771588 | ENSG00000039650 |
| NA | -0.411116875 | 0.752040949 | 0.628538923 | 0.766444017 | 7.347913819 | 5.49830434 | ENSG00000267394 |
| CAPN5 | -0.411177381 | 0.75200941 | 6.86E-08 | 6.45E-07 | 1171.254222 | 880.5142783 | ENSG00000149260 |
| DDX56 | -0.411400736 | 0.751892994 | 1.60E-08 | 1.66E-07 | 1583.447374 | 1190.124427 | ENSG00000136271 |
| GPR3 | -0.41145747 | 0.751863426 | 0.006784969 | 0.022814368 | 244.1776929 | 183.5223868 | ENSG00000181773 |
| TMEM201 | -0.411490555 | 0.751846185 | 1.85E-05 | 0.000116149 | 797.9948482 | 600.0992489 | ENSG00000188807 |
| NTHL1 | -0.411591634 | 0.75179351 | 5.20E-09 | 5.75E-08 | 1272.198597 | 956.7055954 | ENSG00000065057 |
| EIF3C | -0.411610389 | 0.751783737 | 2.81E-07 | 2.42E-06 | 1217.729575 | 915.4136371 | ENSG00000184110 |
| KDELR1 | -0.411638402 | 0.751769139 | 2.50E-16 | 6.17E-15 | 3588.605763 | 2698.283113 | ENSG00000105438 |
| SNORD12C | -0.411658162 | 0.751758843 | 0.706925074 | 0.823594399 | 4.776305388 | 3.552696436 | ENSG00000209042 |
| NA | -0.411852075 | 0.751657805 | 0.247780025 | 0.411317827 | 49.05066666 | 36.92350818 | ENSG00000236397 |
| SYN3 | -0.411933363 | 0.751615455 | 0.000356208 | 0.001697952 | 450.2025252 | 338.053208 | ENSG00000185666 |
| PET117 | -0.412046971 | 0.751556269 | 0.705606658 | 0.822866524 | 5.747731275 | 4.322761311 | ENSG00000232838 |
| RWDD1 | -0.412054395 | 0.751552402 | 7.30E-06 | 4.94E-05 | 716.2015135 | 538.5523347 | ENSG00000111832 |
| SBDSP1 | -0.412206686 | 0.751473073 | 0.01657009 | 0.048021978 | 191.8123093 | 144.3171808 | ENSG00000225648 |
| UBTF | -0.412266139 | 0.751442105 | 1.47E-13 | 2.73E-12 | 3503.949651 | 2633.201349 | ENSG00000108312 |
| ANKRD35 | -0.412432434 | 0.751355494 | 0.020700938 | 0.057808597 | 179.8982965 | 135.0311792 | ENSG00000198483 |
| ARPC5L | -0.412502284 | 0.751319117 | 3.63E-10 | 4.68E-09 | 1619.147874 | 1216.36536 | ENSG00000136950 |
| DHRS1 | -0.41261682 | 0.751259472 | 0.001902902 | 0.007559491 | 296.5427641 | 222.6896944 | ENSG00000157379 |
| ZDHHC18 | -0.412717171 | 0.751207217 | 1.81E-07 | 1.60E-06 | 1081.325798 | 812.2873376 | ENSG00000204160 |
| ILK | -0.412838307 | 0.751144145 | 0.662061689 | 0.791462605 | 6.139961033 | 4.581157997 | ENSG00000166333 |
| BZW1 | -0.412868506 | 0.751128421 | 4.58E-14 | 8.95E-13 | 3570.734347 | 2681.809123 | ENSG00000082153 |
| TPGS2 | -0.412998256 | 0.751060871 | 9.23E-14 | 1.75E-12 | 2446.671193 | 1837.917824 | ENSG00000134779 |
| NA | -0.4131617 | 0.750975788 | 0.518750457 | 0.682248095 | 13.84920452 | 10.40346073 | ENSG00000286511 |
| CCDC124 | -0.413390717 | 0.750856586 | 2.64E-09 | 3.04E-08 | 2760.823052 | 2072.864707 | ENSG00000007080 |
| SLC8A1 | -0.413480545 | 0.750809836 | 0.047881775 | 0.115417634 | 125.2721467 | 93.92903499 | ENSG00000183023 |
| WDR46 | -0.413726562 | 0.750681814 | 2.35E-10 | 3.10E-09 | 2325.780405 | 1745.412631 | ENSG00000227057 |
| CASTOR2 | -0.41377695 | 0.750655596 | 0.000279214 | 0.001364318 | 408.9447922 | 307.1219226 | ENSG00000274070 |
| ST3GAL1 | -0.413892794 | 0.750595323 | 0.007907595 | 0.025855155 | 225.4074435 | 169.265906 | ENSG00000008513 |
| NA | -0.414143828 | 0.750464728 | 0.63728296 | 0.773245209 | 7.843294843 | 5.921247548 | ENSG00000270761 |
| OTULINL | -0.414175743 | 0.750448127 | 0.013538156 | 0.040603943 | 217.6005937 | 163.4358066 | ENSG00000145569 |
| CHSY3 | -0.414276203 | 0.750395872 | 0.146238211 | 0.276739389 | 63.05422259 | 47.36299562 | ENSG00000198108 |
| NA | -0.414643414 | 0.750204898 | 0.579163984 | 0.730715494 | 8.744384577 | 6.535368196 | ENSG00000280300 |
| TM9SF1 | -0.415148179 | 0.749942464 | 0.172295314 | 0.314480925 | 60.41678377 | 45.42007133 | ENSG00000100926 |
| JPT1 | -0.415156626 | 0.749938073 | 1.30E-17 | 3.58E-16 | 4538.997586 | 3403.681201 | ENSG00000189159 |
| RAMP1 | -0.415274212 | 0.749876952 | 0.506808047 | 0.671592013 | 14.76424877 | 11.0116627 | ENSG00000132329 |
| KIF13A | -0.415302936 | 0.749862022 | 1.86E-07 | 1.65E-06 | 1007.829928 | 755.6969452 | ENSG00000137177 |
| ZNF771 | -0.415364395 | 0.749830079 | 0.00350348 | 0.012868553 | 267.9469832 | 200.9965587 | ENSG00000179965 |
| NA | -0.415477311 | 0.749771394 | 0.444152721 | 0.614997856 | 17.53207216 | 13.09816081 | ENSG00000217094 |
| ZFYVE9 | -0.415767929 | 0.749620375 | 2.17E-05 | 0.00013469 | 755.0630036 | 566.6227012 | ENSG00000157077 |
| BBOF1 | -0.41578424 | 0.749611899 | 0.658626395 | 0.789067307 | 6.547384606 | 4.936881959 | ENSG00000119636 |
| ATAD2 | -0.415841031 | 0.749582392 | 1.31E-05 | 8.39E-05 | 648.9357757 | 486.6027955 | ENSG00000156802 |

| | | | | | | | |
|---|---|---|---|---|---|---|---|
| POLR2I | -0.415865025 | 0.749569925 | 0.211172953 | 0.3668051 | 48.63351893 | 36.39838508 | ENSG00000105258 |
| RCAN2 | -0.416203883 | 0.749393888 | 0.004566227 | 0.016228276 | 249.3120614 | 186.7053898 | ENSG00000172348 |
| ELMSAN1 | -0.416219609 | 0.74938572 | 0.002304891 | 0.008923069 | 308.737044 | 231.1642188 | ENSG00000156030 |
| DNAJC8 | -0.416225471 | 0.749382675 | 3.42E-13 | 6.16E-12 | 3868.974273 | 2898.840256 | ENSG00000126698 |
| APC2 | -0.416443747 | 0.749269304 | 0.20424445 | 0.357512429 | 52.67963126 | 39.53269992 | ENSG00000115266 |
| LIN52 | -0.416488236 | 0.749246198 | 0.006266869 | 0.021310447 | 222.6775415 | 166.8048618 | ENSG00000205659 |
| COL11A2 | -0.416840528 | 0.749063262 | 0.102721631 | 0.21145301 | 80.90240639 | 60.51144366 | ENSG00000204248 |
| CTDNEP1 | -0.416955938 | 0.749003342 | 0.000143444 | 0.000744894 | 607.0266593 | 455.0479752 | ENSG00000175826 |
| SSBP1 | -0.417046372 | 0.748956393 | 1.97E-12 | 3.26E-11 | 3494.905639 | 2617.615489 | ENSG00000106028 |
| ARRDC3-AS1 | -0.417103374 | 0.748926802 | 0.654471189 | 0.785579064 | 5.707219174 | 4.26791247 | ENSG00000281357 |
| NEURL4 | -0.41716575 | 0.748894422 | 7.32E-07 | 5.89E-06 | 1220.613955 | 913.7488041 | ENSG00000215041 |
| NA | -0.417231794 | 0.74886014 | 0.468324696 | 0.637276519 | 15.42861144 | 11.56689382 | ENSG00000273716 |
| NA | -0.417393669 | 0.74877612 | 0.710256966 | 0.825445335 | 3.954124674 | 2.98267176 | ENSG00000226390 |
| RTN4RL1 | -0.417532634 | 0.748703999 | 0.582752346 | 0.733133919 | 8.354945721 | 6.264068068 | ENSG00000185924 |
| PSMD6 | -0.417549126 | 0.748695441 | 9.61E-15 | 2.03E-13 | 2577.312696 | 1929.602024 | ENSG00000163636 |
| SEC14L2 | -0.417559092 | 0.748690269 | 0.188534446 | 0.336138447 | 51.63766906 | 38.66369024 | ENSG00000100003 |
| SLC44A2 | -0.417567476 | 0.748685918 | 3.15E-14 | 6.31E-13 | 3157.471535 | 2364.043731 | ENSG00000129353 |
| ULK2 | -0.417658217 | 0.748638829 | 0.008927059 | 0.028601876 | 206.4616797 | 154.5216674 | ENSG00000083290 |
| TMEM266 | -0.417745754 | 0.748593407 | 0.027712459 | 0.073717739 | 143.0566153 | 106.9852504 | ENSG00000169758 |
| NA | -0.417870979 | 0.748528432 | 0.163470928 | 0.301916484 | 63.46407365 | 47.36541877 | ENSG00000228222 |
| MTDH | -0.41790588 | 0.748510324 | 2.74E-10 | 3.59E-09 | 1648.386353 | 1233.829554 | ENSG00000147649 |
| UGDH | -0.418392351 | 0.748257972 | 1.84E-08 | 1.89E-07 | 1857.176823 | 1390.249753 | ENSG00000109814 |
| KCND2 | -0.418527657 | 0.748187798 | 0.002480253 | 0.009533088 | 286.5775113 | 214.5942777 | ENSG00000184408 |
| ABCC1 | -0.418544306 | 0.748179164 | 1.20E-11 | 1.82E-10 | 2633.183753 | 1969.900525 | ENSG00000103222 |
| LIMK2 | -0.418719245 | 0.748088446 | 6.24E-07 | 5.07E-06 | 1194.754809 | 893.8399222 | ENSG00000182541 |
| POLR1A | -0.418918989 | 0.747984879 | 7.43E-14 | 1.42E-12 | 2867.695512 | 2144.561253 | ENSG00000068654 |
| HAT1 | -0.41901403 | 0.747935605 | 5.50E-08 | 5.25E-07 | 1849.886191 | 1384.406199 | ENSG00000128708 |
| RBM17 | -0.419087014 | 0.74789777 | 3.39E-14 | 6.77E-13 | 2695.323172 | 2015.525222 | ENSG00000134453 |
| NIPA2 | -0.41908707 | 0.747897741 | 5.78E-11 | 8.16E-10 | 1955.664678 | 1462.801605 | ENSG00000140157 |
| LINC01838 | -0.419271426 | 0.747802176 | 0.708666686 | 0.824391789 | 3.98973069 | 3.005795033 | ENSG00000269037 |
| YKT6 | -0.419361495 | 0.747755491 | 8.77E-13 | 1.50E-11 | 3551.806009 | 2656.075681 | ENSG00000106636 |
| NRCAM | -0.41955292 | 0.747656281 | 0.045446128 | 0.110568956 | 124.3528721 | 92.87583262 | ENSG00000091129 |
| LRRC6 | -0.419553041 | 0.747656218 | 0.175310267 | 0.318477292 | 56.85038309 | 42.45837227 | ENSG00000129295 |
| TRIM25 | -0.419595108 | 0.747634418 | 2.22E-08 | 2.24E-07 | 1065.302172 | 796.5232398 | ENSG00000121060 |
| GSTM1 | -0.419595275 | 0.747634332 | 0.105615443 | 0.215897798 | 80.22197402 | 59.90969342 | ENSG00000134184 |
| LOC339192 | -0.419720515 | 0.747569433 | 0.672506766 | 0.799434862 | 5.33943182 | 4.019735616 | ENSG00000267121 |
| CASP9 | -0.419772212 | 0.747542645 | 0.000255284 | 0.001258223 | 476.6103176 | 355.9519343 | ENSG00000132906 |
| MKS1 | -0.41980289 | 0.747526749 | 0.002075264 | 0.008151888 | 350.2717663 | 261.6085768 | ENSG00000011143 |
| NEK4 | -0.420023255 | 0.747412577 | 3.20E-06 | 2.29E-05 | 762.8672868 | 570.1267827 | ENSG00000114904 |
| JPT2 | -0.420024276 | 0.747412048 | 4.79E-21 | 1.76E-19 | 4767.523218 | 3563.315037 | ENSG00000206053 |
| SNW1 | -0.420116236 | 0.747364408 | 1.72E-08 | 1.77E-07 | 1120.58077 | 837.6471381 | ENSG00000100603 |
| NA | -0.420157246 | 0.747343164 | 0.369045736 | 0.542833495 | 23.35293049 | 17.49029194 | ENSG00000261189 |
| ATPSCKMT | -0.420165282 | 0.747339001 | 0.013068754 | 0.039447557 | 182.1036323 | 136.1034763 | ENSG00000150756 |
| NA | -0.420198124 | 0.747321988 | 0.662140289 | 0.791507517 | 5.780546388 | 4.314159016 | ENSG00000279641 |
| ALPL | -0.420278683 | 0.747280259 | 8.41E-22 | 3.25E-20 | 5738.737583 | 4288.757506 | ENSG00000162551 |
| EPO | -0.420414918 | 0.747209696 | 0.612633198 | 0.75593623 | 8.248803394 | 6.158671534 | ENSG00000130427 |
| RPS26 | -0.420453834 | 0.747189541 | 6.04E-12 | 9.46E-11 | 3549.035342 | 2651.170331 | ENSG00000197728 |
| WRAP53 | -0.420475347 | 0.747178399 | 2.10E-06 | 1.56E-05 | 763.7912673 | 570.4529438 | ENSG00000141499 |
| RFC2 | -0.420553273 | 0.747138042 | 1.46E-11 | 2.20E-10 | 1776.070188 | 1326.619636 | ENSG00000049541 |
| NA | -0.420571639 | 0.747128531 | 0.675657864 | 0.800965762 | 5.687682854 | 4.238337476 | ENSG00000235513 |
| LMNB2 | -0.420575188 | 0.747126693 | 4.49E-20 | 1.53E-18 | 13108.70624 | 9793.9373 | ENSG00000176619 |
| NUDT17 | -0.420581337 | 0.747123509 | 0.343852372 | 0.516608731 | 29.18774333 | 21.77891665 | ENSG00000186364 |
| FAM118B | -0.420612552 | 0.747107344 | 1.72E-07 | 1.53E-06 | 909.9245721 | 679.6687992 | ENSG00000197798 |
| SLC29A2 | -0.420748367 | 0.747037015 | 8.83E-06 | 5.88E-05 | 814.7651614 | 608.3613627 | ENSG00000174669 |
| ZNF175 | -0.420829211 | 0.746995154 | 0.039192074 | 0.098167386 | 145.3970785 | 108.5638484 | ENSG00000105497 |
| SPRYD4 | -0.420888924 | 0.746964237 | 0.011080469 | 0.034382071 | 191.5951555 | 143.101843 | ENSG00000176422 |
| SCFD2 | -0.420929177 | 0.746943396 | 0.000897909 | 0.003860469 | 331.3132872 | 247.4518158 | ENSG00000184178 |
| NA | -0.42101888 | 0.746896954 | 0.462567071 | 0.632280973 | 16.72105505 | 12.48995885 | ENSG00000223803 |
| BBOX1-AS1 | -0.421115866 | 0.746846745 | 0.174368086 | 0.317213763 | 55.55773932 | 41.52402134 | ENSG00000254560 |
| ABHD4 | -0.421269916 | 0.746767002 | 1.97E-07 | 1.73E-06 | 881.5730393 | 658.4429631 | ENSG00000100439 |
| C8orf48 | -0.421306108 | 0.746748268 | 0.646879871 | 0.780276345 | 5.731661579 | 4.26791247 | ENSG00000164743 |
| TMX2 | -0.421469946 | 0.746663469 | 2.73E-07 | 2.35E-06 | 1136.782377 | 849.0887287 | ENSG00000213593 |
| NA | -0.421496651 | 0.746649649 | 0.586081703 | 0.73573906 | 8.393342641 | 6.268369215 | ENSG00000248578 |
| CHN2 | -0.421715043 | 0.746536631 | 0.650965252 | 0.783147532 | 5.671613158 | 4.217364777 | ENSG00000106069 |
| SMYD5 | -0.421853515 | 0.746464981 | 7.59E-09 | 8.22E-08 | 1585.071851 | 1183.586841 | ENSG00000135632 |
| RAD54L | -0.421896699 | 0.746442637 | 4.39E-07 | 3.65E-06 | 993.0451033 | 740.736298 | ENSG00000085999 |
| TVP23A | -0.421932508 | 0.74642411 | 0.103448126 | 0.212616429 | 82.98333972 | 62.100231 | ENSG00000166676 |
| KMT5A | -0.422375543 | 0.746194927 | 0.001165428 | 0.004895144 | 338.6983587 | 252.9170254 | ENSG00000183955 |
| C20orf204 | -0.422674732 | 0.746040196 | 0.552642441 | 0.709127666 | 11.35650511 | 8.443331849 | ENSG00000196421 |
| NA | -0.422914906 | 0.745916008 | 0.608647897 | 0.752414392 | 8.371015417 | 6.240944795 | ENSG00000232439 |
| NA | -0.422924736 | 0.745910926 | 0.464114174 | 0.63372215 | 14.91907575 | 11.14825176 | ENSG00000258559 |
| KCNN1 | -0.423243308 | 0.745746234 | 0.006481439 | 0.021927997 | 237.8113805 | 177.3242114 | ENSG00000105642 |
| LINC02043 | -0.423257435 | 0.745738931 | 0.395292994 | 0.567972378 | 20.6123404 | 15.37206826 | ENSG00000232233 |
| NA | -0.4235696 | 0.745577589 | 0.421074867 | 0.593046306 | 20.66210109 | 15.31291827 | ENSG00000238000 |
| PHTF1 | -0.423811488 | 0.745452593 | 2.84E-05 | 0.000171834 | 620.779995 | 462.9457506 | ENSG00000116793 |
| CCDC28A | -0.423867743 | 0.745423526 | 0.019796516 | 0.055718499 | 171.9355805 | 127.9647876 | ENSG00000024862 |
| CDC42EP4 | -0.423974841 | 0.745368191 | 0.000331113 | 0.001591306 | 460.3029948 | 343.0030239 | ENSG00000179604 |

| | | | | | | | |
|---|---|---|---|---|---|---|---|
| LINC01134 | -0.424101129 | 0.745302948 | 0.5678338 | 0.721137711 | 9.313092814 | 6.979284103 | ENSG00000236423 |
| EFHD1 | -0.424130691 | 0.745287676 | 0.026522769 | 0.071072473 | 149.5961027 | 111.6032286 | ENSG00000115468 |
| RARRES2 | -0.424182982 | 0.745260663 | 2.17E-18 | 6.36E-17 | 4687.236668 | 3493.001932 | ENSG00000106538 |
| CDS1 | -0.424197975 | 0.745252918 | 0.000272348 | 0.001333809 | 416.4248055 | 310.6095504 | ENSG00000163624 |
| LOC102724064 | -0.424209622 | 0.745246902 | 0.232215487 | 0.392850036 | 47.77505648 | 35.66407435 | ENSG00000255468 |
| SRCAP | -0.424247738 | 0.745227213 | 0.214133486 | 0.370601966 | 50.65104936 | 37.68686087 | ENSG00000080603 |
| ANKMY2 | -0.42433987 | 0.745179623 | 0.001667113 | 0.006725276 | 320.7400023 | 238.7554851 | ENSG00000106524 |
| MON1A | -0.424442105 | 0.745126819 | 0.005674284 | 0.01957136 | 312.4705497 | 232.9364174 | ENSG00000164077 |
| R3HDM2 | -0.424544905 | 0.745073726 | 0.008114818 | 0.026429674 | 217.8013513 | 162.321031 | ENSG00000179912 |
| TRIM9 | -0.424621093 | 0.745034381 | 0.049469136 | 0.118239264 | 116.5776861 | 86.94355963 | ENSG00000100505 |
| NA | -0.424742005 | 0.744971942 | 0.710804603 | 0.825872368 | 3.926891366 | 2.936425214 | ENSG00000215283 |
| LYPD6 | -0.42474401 | 0.744970906 | 0.012059357 | 0.036774748 | 186.5102353 | 138.9444522 | ENSG00000187123 |
| ASGR2 | -0.424790126 | 0.744947094 | 0.70131161 | 0.819997315 | 5.268895509 | 3.937462355 | ENSG00000161944 |
| RBBP4 | -0.424799481 | 0.744942263 | 4.53E-14 | 8.88E-13 | 4063.910501 | 3027.002165 | ENSG00000162521 |
| OAF | -0.424852007 | 0.744915142 | 0.219476292 | 0.377522706 | 48.81557922 | 36.46156365 | ENSG00000184232 |
| CD99P1 | -0.424867085 | 0.744907356 | 0.090769213 | 0.191917027 | 91.56117013 | 68.21318902 | ENSG00000223773 |
| LINC02762 | -0.424919628 | 0.744880227 | 0.578581825 | 0.730268384 | 8.801642095 | 6.512244923 | ENSG00000250303 |
| NA | -0.425021251 | 0.74482776 | 0.527124218 | 0.688986604 | 12.63566993 | 9.377149745 | ENSG00000268912 |
| FAM76B | -0.425074564 | 0.744800236 | 0.00131402 | 0.005450176 | 311.9533539 | 232.4075141 | ENSG00000077458 |
| KCNC4 | -0.425319657 | 0.744673716 | 0.000725412 | 0.003181882 | 405.8465904 | 302.1955451 | ENSG00000116396 |
| NA | -0.425453176 | 0.744604801 | 0.711943588 | 0.826368397 | 3.932473172 | 2.938575788 | ENSG00000285677 |
| NA | -0.425622721 | 0.744517301 | 0.579844983 | 0.731049577 | 9.606877233 | 7.143037123 | ENSG00000184616 |
| SRSF2 | -0.425721936 | 0.744466102 | 2.13E-11 | 3.15E-10 | 2644.504444 | 1968.944153 | ENSG00000161547 |
| MRPL58 | -0.425773141 | 0.744439679 | 1.83E-05 | 0.000114791 | 597.5006302 | 444.8400357 | ENSG00000167862 |
| CCNH | -0.425985352 | 0.744330185 | 0.002575526 | 0.009838508 | 277.9978781 | 207.0992847 | ENSG00000134480 |
| C9orf85 | -0.426005215 | 0.744319937 | 0.004853342 | 0.017103732 | 253.9414001 | 189.0427483 | ENSG00000155621 |
| PHLDB2 | -0.42602898 | 0.744307676 | 1.98E-06 | 1.48E-05 | 781.8588323 | 582.3139458 | ENSG00000144824 |
| LRRN1 | -0.426265529 | 0.744185647 | 8.25E-16 | 1.94E-14 | 2733.768089 | 2034.515248 | ENSG00000175928 |
| ZNF266 | -0.426334682 | 0.744149977 | 0.001447292 | 0.005937839 | 331.2732876 | 246.2577417 | ENSG00000174652 |
| RMC1 | -0.426592804 | 0.744016848 | 0.009255616 | 0.029468575 | 220.9924678 | 164.6661863 | ENSG00000141452 |
| LBP | -0.426738776 | 0.743941572 | 0.666896046 | 0.79531971 | 5.846176615 | 4.339432863 | ENSG00000129988 |
| ADCY4 | -0.426957599 | 0.743828742 | 0.6159712 | 0.758212125 | 7.116968725 | 5.281853053 | ENSG00000129467 |
| RIPK4 | -0.427115432 | 0.74374737 | 0.034932565 | 0.089043884 | 127.5480188 | 94.97900229 | ENSG00000183421 |
| SLC35F1 | -0.427244739 | 0.743680712 | 0.002986384 | 0.011158307 | 264.5121184 | 196.9144259 | ENSG00000196376 |
| DNAJB5 | -0.427244786 | 0.743680688 | 9.99E-09 | 1.07E-07 | 1401.044164 | 1041.621546 | ENSG00000137094 |
| SRRT | -0.427353149 | 0.743624831 | 1.05E-15 | 2.46E-14 | 6774.555813 | 5037.13958 | ENSG00000087087 |
| CHMP1A | -0.427550447 | 0.743523143 | 1.37E-11 | 2.06E-10 | 2092.114936 | 1555.399717 | ENSG00000131165 |
| NA | -0.427701501 | 0.743445298 | 0.098037849 | 0.203974131 | 82.96303966 | 61.7211233 | ENSG00000206567 |
| NDUFA2 | -0.427749436 | 0.743420597 | 0.006044816 | 0.020675628 | 247.9767152 | 184.1295227 | ENSG00000131495 |
| NA | -0.428007418 | 0.74328767 | 0.440172087 | 0.611196986 | 18.90689141 | 13.96906061 | ENSG00000276570 |
| KIF1A | -0.428285926 | 0.743144195 | 5.40E-24 | 2.47E-22 | 14268.26422 | 10603.14761 | ENSG00000130294 |
| ZNF423 | -0.428347759 | 0.743112345 | 4.82E-10 | 6.10E-09 | 1522.170078 | 1131.350452 | ENSG00000102935 |
| LINC01918 | -0.428418511 | 0.743075902 | 0.159206234 | 0.295881257 | 57.15657042 | 42.52129037 | ENSG00000226508 |
| ZNF350 | -0.428486752 | 0.743040755 | 0.069701305 | 0.155519844 | 90.90641946 | 67.48480908 | ENSG00000256683 |
| NA | -0.428617397 | 0.742973471 | 0.118504264 | 0.236180365 | 72.50926391 | 53.77897934 | ENSG00000274317 |
| PTPN9 | -0.429110649 | 0.742719495 | 3.27E-10 | 4.24E-09 | 1304.879199 | 969.0909633 | ENSG00000169410 |
| AMIGO1 | -0.429137902 | 0.742705465 | 0.038074264 | 0.09572405 | 125.7066034 | 93.39235342 | ENSG00000181754 |
| BMP6 | -0.429319108 | 0.742612184 | 0.073497745 | 0.162280531 | 98.33371786 | 73.14792041 | ENSG00000153162 |
| LINC01950 | -0.429443993 | 0.742547904 | 0.090135622 | 0.190859232 | 90.86187716 | 67.45281094 | ENSG00000251027 |
| FAM228B | -0.429492816 | 0.742522776 | 0.507184071 | 0.671891767 | 12.49200656 | 9.276314822 | ENSG00000219626 |
| EEF1AKMT1 | -0.429606685 | 0.742464172 | 0.104610706 | 0.214308047 | 76.77930006 | 57.12570479 | ENSG00000150456 |
| RNPEPL1 | -0.429821606 | 0.742353574 | 0.002476665 | 0.00952309 | 361.404623 | 268.2754273 | ENSG00000142327 |
| TMEM94 | -0.42993369 | 0.742295902 | 2.85E-10 | 3.72E-09 | 1465.158271 | 1087.554395 | ENSG00000177728 |
| FASTKD1 | -0.430008424 | 0.742257451 | 7.75E-06 | 5.21E-05 | 714.8831 | 530.973693 | ENSG00000138399 |
| C12orf75 | -0.430010894 | 0.74225618 | 0.000179988 | 0.000915651 | 460.0209376 | 341.7024261 | ENSG00000235162 |
| SLTM | -0.430606287 | 0.741949918 | 2.05E-16 | 5.14E-15 | 3795.941286 | 2816.915417 | ENSG00000137776 |
| CSK | -0.430734366 | 0.741884053 | 2.14E-08 | 2.16E-07 | 1697.984542 | 1259.928046 | ENSG00000103653 |
| PANK4 | -0.430737721 | 0.741882327 | 4.46E-08 | 4.32E-07 | 973.5459002 | 722.084402 | ENSG00000157881 |
| ADAMTS15 | -0.431015321 | 0.74173959 | 0.00408007 | 0.014711894 | 255.809285 | 189.9375649 | ENSG00000166106 |
| SCD | -0.431027767 | 0.741733191 | 4.30E-37 | 4.06E-35 | 30264.81029 | 22449.33652 | ENSG00000099194 |
| SNURF | -0.431107998 | 0.741691943 | 0.693380735 | 0.814467988 | 4.395239242 | 3.230848615 | ENSG00000273173 |
| KBTBD11 | -0.431351115 | 0.741566966 | 0.000374161 | 0.001776069 | 359.9895798 | 267.0108683 | ENSG00000176595 |
| MGAT4A | -0.431994078 | 0.741236548 | 0.079287672 | 0.172225931 | 90.90729534 | 67.30868566 | ENSG00000071073 |
| TASP1 | -0.432261694 | 0.741099063 | 0.013477152 | 0.040471317 | 176.6842523 | 130.982129 | ENSG00000089123 |
| TMEM151B | -0.432442094 | 0.741006399 | 0.00092148 | 0.0039582 | 414.7873773 | 307.4964992 | ENSG00000178233 |
| AP1B1 | -0.432527572 | 0.740962496 | 8.75E-13 | 1.50E-11 | 2943.980359 | 2181.366473 | ENSG00000100280 |
| LINC00632 | -0.432851657 | 0.740796066 | 0.053544139 | 0.125996583 | 106.6260244 | 79.06486681 | ENSG00000203930 |
| EPDR1 | -0.432917395 | 0.740762312 | 0.011223648 | 0.034781591 | 197.9468887 | 146.6123335 | ENSG00000086289 |
| DYRK1B | -0.432917507 | 0.740762255 | 0.006312702 | 0.021436073 | 268.3378856 | 198.5645685 | ENSG00000105204 |
| GEMIN7-AS1 | -0.433281662 | 0.7405753 | 0.612912185 | 0.75608703 | 7.086944514 | 5.222703065 | ENSG00000267348 |
| IFT22 | -0.43331948 | 0.740555887 | 0.004248003 | 0.015220695 | 244.7195312 | 181.3619451 | ENSG00000128581 |
| CA8 | -0.433347637 | 0.740541434 | 0.032198585 | 0.083207355 | 131.6595981 | 97.54329245 | ENSG00000178538 |
| FBXO15 | -0.433352825 | 0.740538771 | 0.465521405 | 0.634969485 | 16.29947681 | 12.05464524 | ENSG00000141665 |
| CDC25A | -0.433450859 | 0.740488452 | 1.28E-13 | 2.38E-12 | 2161.225292 | 1600.573193 | ENSG00000164045 |
| NA | -0.433551078 | 0.740437014 | 0.330270119 | 0.502622347 | 25.71592105 | 19.04253163 | ENSG00000282022 |
| SNAPC4 | -0.433708321 | 0.740356317 | 1.52E-06 | 1.16E-05 | 1071.021394 | 792.6248151 | ENSG00000165684 |
| TRAPPC2 | -0.433784968 | 0.740316984 | 0.014334924 | 0.042642253 | 169.7795682 | 125.700276 | ENSG00000196459 |

| | | | | | | | |
|---|---|---|---|---|---|---|---|
| NCAPG | -0.43381097 | 0.740303642 | 9.72E-08 | 8.98E-07 | 1219.297846 | 902.7494265 | ENSG00000109805 |
| ITGAE | -0.433869288 | 0.740273717 | 0.000462851 | 0.002140187 | 401.1776524 | 296.6399688 | ENSG00000083457 |
| RRAGB | -0.433934071 | 0.740240476 | 0.022964071 | 0.062970846 | 147.0356579 | 108.9117636 | ENSG00000083750 |
| CLN6 | -0.4340444 | 0.740183869 | 8.09E-06 | 5.42E-05 | 956.2870512 | 707.8191074 | ENSG00000128973 |
| TBC1D4 | -0.434394431 | 0.740004305 | 1.66E-08 | 1.71E-07 | 1250.933808 | 925.5955514 | ENSG00000136111 |
| C2CD3 | -0.434520887 | 0.739939445 | 4.11E-05 | 0.000240452 | 550.1052004 | 407.20345 | ENSG00000168014 |
| LIPT2 | -0.434655319 | 0.7398705 | 0.278969489 | 0.44671706 | 42.60719705 | 31.43973692 | ENSG00000175536 |
| PES1 | -0.434799033 | 0.739796801 | 1.95E-11 | 2.89E-10 | 4539.117927 | 3358.086509 | ENSG00000100029 |
| MBNL2 | -0.435072423 | 0.739656623 | 0.374957771 | 0.548268806 | 22.6341012 | 16.76432304 | ENSG00000139793 |
| ANKS1B | -0.435323801 | 0.739527755 | 0.19470696 | 0.344848692 | 45.22366009 | 33.37753604 | ENSG00000185046 |
| TCF7 | -0.435407918 | 0.739484638 | 0.00169866 | 0.006841096 | 278.7383588 | 206.0493295 | ENSG00000081059 |
| TXNRD1 | -0.435416333 | 0.739480324 | 2.95E-18 | 8.52E-17 | 3618.667748 | 2675.944973 | ENSG00000198431 |
| PLAC1 | -0.435861978 | 0.739251936 | 0.570899359 | 0.723744466 | 8.447809256 | 6.215670948 | ENSG00000170965 |
| METTL16 | -0.435886095 | 0.739239579 | 3.63E-11 | 5.21E-10 | 1716.442583 | 1268.634256 | ENSG00000127804 |
| NA | -0.436178764 | 0.73908963 | 0.7031633 | 0.821457229 | 4.865502139 | 3.624216828 | ENSG00000213018 |
| WASF3 | -0.43625285 | 0.739051676 | 5.56E-05 | 0.000317443 | 530.7077957 | 392.1870936 | ENSG00000132970 |
| MEX3D | -0.436592486 | 0.738877711 | 2.62E-11 | 3.82E-10 | 2264.832537 | 1673.995635 | ENSG00000181588 |
| CLDN5 | -0.436688555 | 0.738828511 | 0.362549311 | 0.535810829 | 21.75832975 | 16.05340815 | ENSG00000184113 |
| SLC27A5 | -0.43680298 | 0.738769914 | 0.009410408 | 0.029892341 | 235.6992717 | 174.0030203 | ENSG00000083807 |
| NA | -0.436848142 | 0.738746788 | 0.707956417 | 0.823963032 | 3.965288286 | 2.929973493 | ENSG00000270890 |
| CACFD1 | -0.436854618 | 0.738743472 | 9.01E-05 | 0.000491964 | 671.8659525 | 496.574717 | ENSG00000160325 |
| NUFIP1 | -0.43705231 | 0.738642249 | 4.71E-05 | 0.000271993 | 477.7950673 | 352.7805145 | ENSG00000083635 |
| PPP4R4 | -0.437153656 | 0.738590363 | 0.027319597 | 0.072873665 | 136.8279331 | 100.9672086 | ENSG00000119698 |
| TOMM40 | -0.437512862 | 0.73840649 | 1.37E-11 | 2.06E-10 | 5732.092309 | 4232.833346 | ENSG00000130204 |
| ANKRD13B | -0.437820566 | 0.738249016 | 1.35E-05 | 8.67E-05 | 1020.200936 | 753.248835 | ENSG00000198720 |
| SHISAL1 | -0.437841076 | 0.738238521 | 0.001682943 | 0.006785611 | 286.6120413 | 211.6987439 | ENSG00000138944 |
| MAP2K5 | -0.437997604 | 0.738158429 | 0.001548001 | 0.006292137 | 279.586897 | 206.4843706 | ENSG00000137764 |
| PRR7 | -0.438167742 | 0.738071383 | 0.00870151 | 0.028023509 | 202.9810843 | 149.838348 | ENSG00000131188 |
| RCC2 | -0.438249371 | 0.738029623 | 5.30E-33 | 3.98E-31 | 15095.75578 | 11140.711 | ENSG00000179051 |
| PPP6R3 | -0.438268681 | 0.738019744 | 3.33E-13 | 6.01E-12 | 2057.284176 | 1518.63237 | ENSG00000110075 |
| NA | -0.438422855 | 0.73794088 | 0.693318525 | 0.814467988 | 4.45249676 | 3.253971888 | ENSG00000259790 |
| PPT1 | -0.438641199 | 0.737829205 | 1.27E-13 | 2.38E-12 | 3664.36542 | 2703.215106 | ENSG00000131238 |
| TXN2 | -0.438680319 | 0.737809199 | 5.49E-08 | 5.25E-07 | 1119.500574 | 826.2198263 | ENSG00000100348 |
| CROT | -0.43875264 | 0.737772214 | 0.11230173 | 0.226564806 | 78.86477606 | 58.26333099 | ENSG00000005469 |
| ENTPD7 | -0.438932186 | 0.737680402 | 0.001866144 | 0.007431831 | 321.7280984 | 237.2486558 | ENSG00000198018 |
| NA | -0.439331296 | 0.737476357 | 0.501750162 | 0.667627423 | 12.00970417 | 8.857672762 | ENSG00000225808 |
| CWC15 | -0.439470221 | 0.737405345 | 7.68E-06 | 5.18E-05 | 834.8426365 | 615.7477062 | ENSG00000150316 |
| LMF2 | -0.439579636 | 0.737349422 | 2.52E-07 | 2.18E-06 | 908.4680771 | 669.5237472 | ENSG00000100258 |
| TNNI3 | -0.439581958 | 0.737348235 | 0.229657026 | 0.389786528 | 42.46979121 | 31.3606682 | ENSG00000129991 |
| PPP2R2B | -0.439623114 | 0.737327201 | 1.48E-08 | 1.54E-07 | 1495.079538 | 1103.171332 | ENSG00000156475 |
| TSPYL2 | -0.439996727 | 0.737136281 | 7.63E-06 | 5.14E-05 | 644.0220772 | 474.6726599 | ENSG00000184205 |
| PRPF31 | -0.440009863 | 0.737129569 | 3.26E-07 | 2.78E-06 | 1128.780815 | 832.2639781 | ENSG00000105618 |
| L3MBTL3 | -0.440106816 | 0.737080034 | 0.000120791 | 0.000640157 | 427.846302 | 315.1100647 | ENSG00000198945 |
| LOC100507557 | -0.440110445 | 0.73707818 | 0.587280759 | 0.736860691 | 7.941064461 | 5.847576582 | ENSG00000235652 |
| STAC2 | -0.440204665 | 0.737030044 | 0.003890443 | 0.014097729 | 241.0963486 | 177.9138396 | ENSG00000141750 |
| INPP5K | -0.440312897 | 0.736974754 | 0.000500126 | 0.002292214 | 386.1088943 | 284.388258 | ENSG00000132376 |
| ZNF233 | -0.440314034 | 0.736974173 | 0.325775752 | 0.49754889 | 25.42146091 | 18.73520479 | ENSG00000159915 |
| HIGD2A | -0.440354922 | 0.736953286 | 0.022034722 | 0.060881099 | 149.4082605 | 110.0789769 | ENSG00000146066 |
| SF3A3 | -0.440463768 | 0.736897688 | 1.31E-19 | 4.28E-18 | 5117.918676 | 3770.641933 | ENSG00000183431 |
| DDX23 | -0.44048498 | 0.736886853 | 5.58E-14 | 1.08E-12 | 4897.864207 | 3608.488211 | ENSG00000174243 |
| AGAP3 | -0.440626221 | 0.736814715 | 3.26E-12 | 5.24E-11 | 1849.226381 | 1362.274296 | ENSG00000133612 |
| NA | -0.440662575 | 0.736796148 | 0.424069235 | 0.595741306 | 18.46019504 | 13.64317211 | ENSG00000255389 |
| KCNA2 | -0.440738424 | 0.736757413 | 0.394006084 | 0.566676241 | 22.77951632 | 16.69710379 | ENSG00000177301 |
| NIPAL4 | -0.440927044 | 0.736661095 | 0.711437052 | 0.826133147 | 4.041406403 | 2.978370613 | ENSG00000172548 |
| DDRGK1 | -0.440949679 | 0.736649537 | 1.57E-09 | 1.87E-08 | 1306.687087 | 962.6685584 | ENSG00000198171 |
| VPS9D1 | -0.440990485 | 0.736628701 | 0.095788858 | 0.200332054 | 85.5996026 | 62.98566388 | ENSG00000075399 |
| RPS23 | -0.441140393 | 0.736552163 | 7.16E-11 | 9.98E-10 | 2310.444384 | 1701.472109 | ENSG00000186468 |
| PODNL1 | -0.441238096 | 0.736502284 | 0.361729219 | 0.534844666 | 24.21263224 | 17.9299067 | ENSG00000132000 |
| TTC1 | -0.441519145 | 0.736358821 | 9.16E-10 | 1.12E-08 | 1510.040364 | 1111.615276 | ENSG00000113312 |
| PPM1G | -0.441925458 | 0.736151466 | 9.60E-19 | 2.89E-17 | 8682.94069 | 6392.024759 | ENSG00000115241 |
| RNF7 | -0.441950968 | 0.736138449 | 2.38E-06 | 1.75E-05 | 670.8685568 | 494.146446 | ENSG00000114125 |
| ERBB3 | -0.442074853 | 0.73607524 | 1.89E-09 | 2.22E-08 | 1265.942406 | 931.6326942 | ENSG00000065361 |
| CTSC | -0.442148329 | 0.736037753 | 2.19E-30 | 1.41E-28 | 7961.12887 | 5859.692946 | ENSG00000109861 |
| MTM1 | -0.44219729 | 0.736012774 | 0.00050212 | 0.002300808 | 349.1846741 | 257.159131 | ENSG00000171100 |
| CIB1 | -0.442365675 | 0.735926875 | 0.003640891 | 0.013298716 | 230.1071553 | 169.219399 | ENSG00000185043 |
| HLA-B | -0.442409923 | 0.735904304 | 1.43E-05 | 9.10E-05 | 560.4281288 | 412.5307255 | ENSG00000234745 |
| RIPPLY3 | -0.442414392 | 0.735902024 | 0.261935192 | 0.427415395 | 40.99508716 | 30.17626241 | ENSG00000183145 |
| MAF1 | -0.442454033 | 0.735881804 | 3.54E-16 | 8.67E-15 | 2871.51471 | 2113.386848 | ENSG00000179632 |
| SEPTIN2 | -0.442539297 | 0.735838314 | 8.10E-30 | 5.12E-28 | 9333.411339 | 6868.564931 | ENSG00000168385 |
| CCNE1 | -0.442760444 | 0.735725528 | 2.48E-09 | 2.87E-08 | 1229.997445 | 905.115003 | ENSG00000105173 |
| NA | -0.442791674 | 0.735709602 | 0.023965763 | 0.065235435 | 135.734183 | 99.95998022 | ENSG00000211772 |
| DHX40 | -0.442925487 | 0.735641367 | 9.29E-08 | 8.61E-07 | 940.4616901 | 691.7637481 | ENSG00000108406 |
| YY2 | -0.442926023 | 0.735641093 | 0.538188902 | 0.697869336 | 11.50575028 | 8.4809761 | ENSG00000230797 |
| NA | -0.442954608 | 0.735626518 | 0.299925412 | 0.470509972 | 33.46567663 | 24.6844098 | ENSG00000241506 |
| RAN | -0.443175004 | 0.735514147 | 8.65E-35 | 7.01E-33 | 20207.31212 | 14862.46889 | ENSG00000132341 |
| ACP2 | -0.443476965 | 0.735360218 | 0.001010685 | 0.004304918 | 326.696827 | 240.0573973 | ENSG00000134575 |
| TEAD2 | -0.443537305 | 0.735329462 | 2.79E-14 | 5.61E-13 | 2788.584987 | 2050.620889 | ENSG00000074219 |

| | | | | | | | |
|---|---|---|---|---|---|---|---|
| TSR3 | -0.443683664 | 0.735254868 | 1.81E-07 | 1.60E-06 | 1163.080275 | 855.5964426 | ENSG00000007520 |
| ACOXL | -0.443721831 | 0.735235417 | 7.75E-05 | 0.000429861 | 433.7477842 | 318.8525815 | ENSG00000153093 |
| BCAS2 | -0.443730602 | 0.735230947 | 3.43E-09 | 3.90E-08 | 1206.682449 | 887.1681522 | ENSG00000116752 |
| ELK1 | -0.443966806 | 0.735110582 | 5.80E-08 | 5.51E-07 | 1066.119208 | 783.2336716 | ENSG00000126767 |
| TMEM65 | -0.444177497 | 0.735003234 | 0.001519904 | 0.006196211 | 394.7322692 | 290.3188829 | ENSG00000164983 |
| IGDCC3 | -0.444326083 | 0.734927538 | 1.31E-12 | 2.21E-11 | 2277.554497 | 1673.71955 | ENSG00000174498 |
| LINC00539 | -0.444446478 | 0.73486621 | 0.604547907 | 0.749603656 | 8.376597223 | 6.114575562 | ENSG00000224429 |
| C12orf43 | -0.444502405 | 0.734837723 | 2.67E-05 | 0.000162654 | 541.74401 | 397.9607508 | ENSG00000157895 |
| ITGA1 | -0.444942137 | 0.73461378 | 0.01893142 | 0.053643658 | 149.4077849 | 109.7794283 | ENSG00000213949 |
| CASP7 | -0.444980754 | 0.734594116 | 0.038394284 | 0.096402985 | 126.0078847 | 92.47736976 | ENSG00000165806 |
| CAPNS1 | -0.445222419 | 0.734471075 | 1.00E-16 | 2.57E-15 | 2980.013627 | 2189.04421 | ENSG00000126247 |
| ACAD9 | -0.445348638 | 0.73440682 | 4.15E-06 | 2.93E-05 | 612.2244038 | 449.6862905 | ENSG00000177646 |
| TSN | -0.445383409 | 0.73438912 | 1.50E-14 | 3.08E-13 | 2538.849194 | 1864.353218 | ENSG00000211460 |
| ASGR1 | -0.445413019 | 0.734374048 | 0.006989124 | 0.023354789 | 203.8422625 | 149.632377 | ENSG00000141505 |
| DOP1A | -0.445843657 | 0.734154873 | 0.005822466 | 0.020021604 | 224.2851456 | 164.519117 | ENSG00000083097 |
| NFKBIB | -0.445869468 | 0.734141738 | 3.92E-07 | 3.29E-06 | 1041.932836 | 764.8388579 | ENSG00000104825 |
| NA | -0.445889222 | 0.734131686 | 0.646948015 | 0.78030975 | 5.323362124 | 3.89766753 | ENSG00000231006 |
| SFXN2 | -0.44623841 | 0.73395402 | 0.000334647 | 0.001606688 | 371.2793276 | 272.3682824 | ENSG00000156398 |
| DNAJC9 | -0.446545338 | 0.73379789 | 1.64E-05 | 0.000103588 | 558.6236709 | 409.6526873 | ENSG00000213551 |
| UHRF1BP1 | -0.446570519 | 0.733785083 | 1.69E-08 | 1.75E-07 | 991.1825631 | 727.2234747 | ENSG00000065060 |
| RFXAP | -0.446636334 | 0.733751608 | 0.015053797 | 0.044404096 | 163.5551164 | 119.8715336 | ENSG00000133111 |
| EXOC3L4 | -0.44671732 | 0.73371042 | 0.684995583 | 0.808062728 | 4.041406403 | 2.959548487 | ENSG00000205436 |
| EGFL8 | -0.446734293 | 0.733701789 | 0.473431091 | 0.641990664 | 17.66891442 | 12.95753106 | ENSG00000241404 |
| PIGZ | -0.446813731 | 0.73366139 | 0.583725616 | 0.733831923 | 7.598395234 | 5.599399727 | ENSG00000119227 |
| VPS4A | -0.446858224 | 0.733638764 | 1.07E-06 | 8.40E-06 | 678.3011616 | 497.7440625 | ENSG00000132612 |
| MIR25 | -0.447076838 | 0.733527603 | 0.377561495 | 0.551044564 | 24.48352586 | 17.8957701 | ENSG00000207547 |
| KCNV1 | -0.447150925 | 0.733489935 | 0.707089063 | 0.823594399 | 5.10290492 | 3.769408187 | ENSG00000164794 |
| CACTIN-AS1 | -0.447404355 | 0.733361099 | 0.591166936 | 0.740052002 | 8.006694688 | 5.856178876 | ENSG00000226800 |
| SFXN4 | -0.447414664 | 0.733355858 | 0.00033951 | 0.001627204 | 335.5627479 | 246.1243572 | ENSG00000183605 |
| CTNNBIP1 | -0.447454788 | 0.733335462 | 1.49E-07 | 1.33E-06 | 1006.402395 | 737.9697986 | ENSG00000178585 |
| ZKSCAN5 | -0.447487694 | 0.733318737 | 1.50E-06 | 1.14E-05 | 676.4684823 | 495.9595177 | ENSG00000196652 |
| DOCK4 | -0.447730793 | 0.73319518 | 5.17E-07 | 4.26E-06 | 699.8101612 | 513.1357585 | ENSG00000128512 |
| APOBR | -0.448260617 | 0.732925966 | 0.714342575 | 0.828446713 | 4.06863971 | 2.978370613 | ENSG00000184730 |
| GATC | -0.448314265 | 0.732898713 | 0.003638427 | 0.013293351 | 244.2000201 | 178.7484644 | ENSG00000257218 |
| LINC00852 | -0.448686253 | 0.732709764 | 0.460990531 | 0.630796449 | 13.96651046 | 10.28380368 | ENSG00000231177 |
| EBP | -0.448762037 | 0.732671276 | 0.000392419 | 0.001855546 | 353.0874865 | 258.6283161 | ENSG00000147155 |
| ZNF8-ERVK3-1 | -0.448962394 | 0.732569532 | 0.621140968 | 0.761915061 | 6.289206204 | 4.63170569 | ENSG00000267216 |
| FEM1A | -0.449071596 | 0.732514084 | 0.146655004 | 0.277364871 | 54.3323654 | 39.76797334 | ENSG00000141965 |
| TBC1D13 | -0.449323573 | 0.732386157 | 9.29E-07 | 7.35E-06 | 958.9130255 | 702.478328 | ENSG00000107021 |
| COQ2 | -0.449366408 | 0.732364412 | 0.001141439 | 0.004802509 | 395.2299656 | 289.5039286 | ENSG00000173085 |
| NA | -0.449415193 | 0.732339647 | 0.698466175 | 0.817989588 | 4.906689961 | 3.60969585 | ENSG00000213261 |
| HDDC2 | -0.449521097 | 0.73228589 | 5.32E-09 | 5.88E-08 | 1297.679404 | 950.4987935 | ENSG00000111906 |
| CPNE5 | -0.44954594 | 0.73227328 | 0.409053884 | 0.581250624 | 17.05044549 | 12.48619074 | ENSG00000124772 |
| LOC101927811 | -0.449763705 | 0.732162757 | 0.534495253 | 0.694855702 | 10.67953936 | 7.793184486 | ENSG00000271133 |
| DNAJB1 | -0.449811744 | 0.732138378 | 3.70E-18 | 1.06E-16 | 3464.720784 | 2536.258916 | ENSG00000132002 |
| ESPL1 | -0.449882719 | 0.73210236 | 7.00E-10 | 8.67E-09 | 1953.993383 | 1430.303384 | ENSG00000135476 |
| EIF1AX | -0.449900913 | 0.732093128 | 6.61E-14 | 1.28E-12 | 3018.418659 | 2210.230202 | ENSG00000173674 |
| PLEKHD1 | -0.450036004 | 0.732024579 | 0.31638792 | 0.487378373 | 28.13828431 | 20.53562134 | ENSG00000175985 |
| RELB | -0.450383614 | 0.731848223 | 0.001911296 | 0.007583464 | 291.9794702 | 213.9629525 | ENSG00000104856 |
| LAT | -0.450478881 | 0.731799897 | 0.691373575 | 0.812873938 | 4.049779111 | 2.95524734 | ENSG00000213658 |
| TNFRSF1A | -0.450681544 | 0.731697105 | 3.01E-09 | 3.44E-08 | 1133.266915 | 829.6824223 | ENSG00000067182 |
| BEGAIN | -0.450741965 | 0.731666462 | 0.083484589 | 0.179583096 | 78.65113996 | 57.54623696 | ENSG00000183092 |
| NUDC | -0.450896639 | 0.731588022 | 1.38E-19 | 4.48E-18 | 6548.801734 | 4791.092783 | ENSG00000090273 |
| NDUFAF8 | -0.451040097 | 0.731515279 | 4.28E-06 | 3.02E-05 | 615.6249772 | 450.5601465 | ENSG00000224877 |
| GRM2 | -0.451223013 | 0.731422538 | 0.02500167 | 0.067617934 | 137.0121085 | 100.0756087 | ENSG00000164082 |
| NA | -0.451362999 | 0.731351571 | 0.443478745 | 0.614285091 | 14.8571123 | 10.83070509 | ENSG00000270157 |
| NA | -0.451577031 | 0.731243078 | 0.323425749 | 0.495057219 | 29.08670725 | 21.23201524 | ENSG00000231503 |
| NA | -0.451604493 | 0.731229159 | 0.642844535 | 0.776866124 | 7.998997701 | 5.833055603 | ENSG00000280233 |
| NME1 | -0.451798163 | 0.731131004 | 5.18E-05 | 0.000297056 | 558.0179172 | 407.5323252 | ENSG00000239672 |
| OTX2 | -0.451919734 | 0.731069397 | 4.70E-06 | 3.28E-05 | 731.6277216 | 535.0109547 | ENSG00000165588 |
| TSPAN6 | -0.451997216 | 0.731030135 | 4.41E-12 | 6.98E-11 | 1742.407582 | 1273.494772 | ENSG00000000003 |
| ATP1A3 | -0.45213154 | 0.730962075 | 0.225842544 | 0.384910463 | 37.19443802 | 27.22478318 | ENSG00000105409 |
| ZNF26 | -0.452146044 | 0.730954726 | 0.009885676 | 0.031135441 | 184.5300257 | 135.0069898 | ENSG00000198393 |
| RIT2 | -0.452174127 | 0.730940497 | 0.453610177 | 0.623750369 | 13.92041655 | 10.21013272 | ENSG00000152214 |
| EXD2 | -0.452502492 | 0.730774151 | 1.99E-06 | 1.49E-05 | 684.3433899 | 499.9399978 | ENSG00000081177 |
| ZNF324 | -0.452609675 | 0.730719861 | 0.041788665 | 0.103272501 | 113.2276755 | 82.83940018 | ENSG00000083812 |
| MCM3 | -0.452660128 | 0.730694307 | 3.50E-23 | 1.49E-21 | 11421.24142 | 8344.529405 | ENSG00000112118 |
| GFRA1 | -0.452758027 | 0.730644725 | 0.272272409 | 0.439080963 | 35.74505225 | 26.19443151 | ENSG00000151892 |
| IMPDH1 | -0.453310488 | 0.730364988 | 3.44E-10 | 4.46E-09 | 1464.603256 | 1070.27864 | ENSG00000106348 |
| MIR1244-3 | -0.453535019 | 0.730251327 | 0.308031721 | 0.479079787 | 30.71220808 | 22.39114719 | ENSG00000283429 |
| TM6SF2 | -0.453551736 | 0.730242866 | 0.430594334 | 0.601927196 | 15.38463272 | 11.26816927 | ENSG00000213996 |
| NA | -0.453589229 | 0.730223889 | 0.690296007 | 0.811932215 | 4.044197305 | 2.940726362 | ENSG00000260593 |
| ADAM23 | -0.453861091 | 0.730086298 | 3.66E-09 | 4.14E-08 | 1030.934091 | 752.3305978 | ENSG00000114948 |
| MORF4L2 | -0.454057582 | 0.729986869 | 5.92E-25 | 2.92E-23 | 6967.529498 | 5086.052064 | ENSG00000123562 |
| ABCA3 | -0.454069553 | 0.729980812 | 3.12E-05 | 0.000187539 | 480.9458719 | 350.9077657 | ENSG00000167972 |
| PNMA8A | -0.454145952 | 0.729942156 | 4.73E-09 | 5.28E-08 | 1076.867645 | 785.7105633 | ENSG00000182013 |
| BNC2-AS1 | -0.454411789 | 0.729807666 | 0.38476632 | 0.558210655 | 20.66469184 | 15.08409658 | ENSG00000234779 |

| | | | | | | | |
|---|---|---|---|---|---|---|---|
| THYN1 | -0.45446209 | 0.729782221 | 5.90E-10 | 7.39E-09 | 1183.577684 | 863.644504 | ENSG00000151500 |
| EHD1 | -0.454487214 | 0.729769512 | 1.73E-09 | 2.04E-08 | 1510.330358 | 1102.531223 | ENSG00000110047 |
| TNPO2 | -0.454576846 | 0.729724175 | 3.47E-16 | 8.51E-15 | 2830.491412 | 2065.557371 | ENSG00000105576 |
| C20orf27 | -0.454709527 | 0.729657067 | 1.43E-12 | 2.40E-11 | 2413.193955 | 1761.045446 | ENSG00000101220 |
| STRAP | -0.454760113 | 0.729631483 | 1.91E-20 | 6.73E-19 | 4322.222242 | 3154.402258 | ENSG00000023734 |
| MRPS34 | -0.454789544 | 0.729616599 | 9.56E-11 | 1.32E-09 | 2044.673552 | 1491.80748 | ENSG00000074071 |
| NA | -0.454886361 | 0.729567637 | 0.259576278 | 0.424831253 | 31.10981948 | 22.70654329 | ENSG00000128692 |
| STX2 | -0.455044087 | 0.72948788 | 5.99E-05 | 0.000339817 | 491.0309092 | 358.3951982 | ENSG00000111450 |
| NA | -0.455096874 | 0.729461189 | 0.683922063 | 0.807191222 | 4.016963998 | 2.913301941 | ENSG00000213305 |
| HERC2 | -0.455290577 | 0.729363254 | 1.89E-13 | 3.49E-12 | 4380.145706 | 3193.599256 | ENSG00000128731 |
| NA | -0.455303509 | 0.729356717 | 0.239866413 | 0.40194927 | 37.34523479 | 27.25220761 | ENSG00000218175 |
| NA | -0.455350337 | 0.729333043 | 0.068909684 | 0.154090265 | 93.71503036 | 68.50707939 | ENSG00000182057 |
| KLHL13 | -0.45551513 | 0.729249739 | 2.56E-06 | 1.87E-05 | 612.6255698 | 446.6315774 | ENSG00000003096 |
| POLD3 | -0.455592972 | 0.729210393 | 7.34E-08 | 6.87E-07 | 787.4552912 | 574.345474 | ENSG00000077514 |
| SRRM4 | -0.455596926 | 0.729208394 | 0.085132541 | 0.182416068 | 77.82193798 | 56.72724193 | ENSG00000139767 |
| CANX | -0.455605493 | 0.729204064 | 2.63E-35 | 2.22E-33 | 11809.45577 | 8611.961227 | ENSG00000127022 |
| PEBP1 | -0.456034805 | 0.728987102 | 2.03E-24 | 9.55E-23 | 8378.853745 | 6107.785702 | ENSG00000089220 |
| CETN2 | -0.456122564 | 0.728942759 | 2.54E-05 | 0.000155505 | 533.609158 | 389.0119178 | ENSG00000147400 |
| NA | -0.456208949 | 0.728899114 | 0.442198304 | 0.61325164 | 15.34065399 | 11.19879945 | ENSG00000229644 |
| SCLT1 | -0.456429465 | 0.72878771 | 0.00618614 | 0.021069355 | 193.6018857 | 141.2653997 | ENSG00000151466 |
| PRR5 | -0.456756097 | 0.728622728 | 0.003789312 | 0.013787179 | 232.7789239 | 169.6520105 | ENSG00000186654 |
| NA | -0.456787749 | 0.728606743 | 0.67477392 | 0.800508191 | 5.79171 | 4.177569952 | ENSG00000273891 |
| GINS3 | -0.456972045 | 0.728513673 | 0.002494292 | 0.009577505 | 245.7665116 | 178.9657092 | ENSG00000181938 |
| NA | -0.456981619 | 0.728508838 | 0.124910972 | 0.24549523 | 71.02722628 | 51.80002834 | ENSG00000213888 |
| CIAO3 | -0.457005538 | 0.72849676 | 0.000144626 | 0.000750602 | 384.1385338 | 279.7102752 | ENSG00000103245 |
| NOMO1 | -0.457059771 | 0.728469376 | 4.80E-13 | 8.45E-12 | 2183.826588 | 1591.282574 | ENSG00000103512 |
| MARCHF5 | -0.457067801 | 0.728465321 | 3.92E-05 | 0.000230399 | 570.0328909 | 415.1888595 | ENSG00000198060 |
| NA | -0.457111647 | 0.728443182 | 0.522455238 | 0.685277029 | 10.41789434 | 7.589103603 | ENSG00000287009 |
| NA | -0.457129215 | 0.728434312 | 0.150515978 | 0.283112688 | 58.26503873 | 42.35512631 | ENSG00000264164 |
| NA | -0.457142322 | 0.728427694 | 0.402198894 | 0.574514541 | 16.70989144 | 12.18316504 | ENSG00000198406 |
| NA | -0.457193692 | 0.728401757 | 0.419051513 | 0.591059643 | 17.50954477 | 12.78491529 | ENSG00000235245 |
| NXPH3 | -0.457241011 | 0.728377867 | 0.018075848 | 0.051605917 | 170.7159517 | 124.3061616 | ENSG00000182575 |
| MAP9 | -0.457250644 | 0.728373003 | 0.01795806 | 0.051368467 | 138.426564 | 100.9104697 | ENSG00000164114 |
| CBY1 | -0.457307917 | 0.728344089 | 0.024325414 | 0.06608387 | 159.2766195 | 116.0432238 | ENSG00000100211 |
| IRF4 | -0.457390592 | 0.728302351 | 0.177933475 | 0.321875691 | 47.57114453 | 34.62675003 | ENSG00000137265 |
| CNKSR2 | -0.457532558 | 0.728230688 | 0.000292661 | 0.001422813 | 366.2371979 | 266.9928396 | ENSG00000149970 |
| LOC100129215 | -0.457697705 | 0.728147331 | 0.292004278 | 0.46172578 | 27.87085731 | 20.32132062 | ENSG00000226180 |
| NA | -0.457743194 | 0.728124372 | 0.202316228 | 0.354872009 | 41.63856207 | 30.41179629 | ENSG00000240253 |
| FAM218A | -0.457771242 | 0.728110217 | 0.483895903 | 0.651646482 | 13.58632019 | 9.850640645 | ENSG00000250486 |
| FOXP1 | -0.457781262 | 0.72810516 | 0.063540296 | 0.144359422 | 86.87059487 | 63.35804729 | ENSG00000114861 |
| TIA1 | -0.458274238 | 0.727856405 | 8.22E-10 | 1.01E-08 | 1657.724001 | 1206.857981 | ENSG00000116001 |
| DHCR24 | -0.458487344 | 0.727748898 | 8.83E-24 | 3.98E-22 | 14228.78088 | 10354.74295 | ENSG00000116133 |
| TRPC3 | -0.458673027 | 0.727655239 | 0.468257049 | 0.637259029 | 13.47180516 | 9.745244111 | ENSG00000138741 |
| TESK1 | -0.458776103 | 0.727603252 | 0.000473383 | 0.002182609 | 327.1745868 | 238.2150233 | ENSG00000107140 |
| POLA2 | -0.458963056 | 0.727508971 | 0.031425513 | 0.081559205 | 126.7149868 | 92.12405683 | ENSG00000014138 |
| TMEM104 | -0.458998686 | 0.727491004 | 1.59E-05 | 0.000100437 | 686.2939259 | 499.0941419 | ENSG00000109066 |
| SHISA9 | -0.459197501 | 0.727390757 | 3.56E-05 | 0.000211513 | 473.491977 | 344.4977006 | ENSG00000237515 |
| NA | -0.459473997 | 0.727251364 | 0.065009195 | 0.146957838 | 85.01973075 | 61.93836809 | ENSG00000269958 |
| ORC1 | -0.459563471 | 0.727206263 | 1.98E-11 | 2.93E-10 | 1468.498482 | 1067.588556 | ENSG00000085840 |
| IPO13 | -0.459676469 | 0.727149307 | 1.64E-06 | 1.24E-05 | 666.0535421 | 484.3781213 | ENSG00000117408 |
| RAD51 | -0.459810137 | 0.727081939 | 2.57E-07 | 2.22E-06 | 854.5919207 | 621.3796371 | ENSG00000051180 |
| RFX2 | -0.459825186 | 0.727074354 | 0.027832734 | 0.073986672 | 120.8213648 | 87.97685538 | ENSG00000087903 |
| NA | -0.460004699 | 0.726983891 | 0.459842826 | 0.629627933 | 13.59939883 | 9.936682016 | ENSG00000260747 |
| GGCT | -0.460051651 | 0.726960232 | 0.000117624 | 0.000625952 | 412.4646234 | 299.9985316 | ENSG00000006625 |
| DNAAF3 | -0.460303514 | 0.726833332 | 0.202027686 | 0.354494824 | 43.42917761 | 31.68198299 | ENSG00000167646 |
| STARD10 | -0.460386866 | 0.72679134 | 0.000285058 | 0.001390052 | 359.4329111 | 261.2428629 | ENSG00000214530 |
| FRRS1L | -0.460445694 | 0.726761705 | 0.156023248 | 0.291397471 | 51.92028987 | 37.77826946 | ENSG00000260230 |
| VWA7 | -0.460508199 | 0.726730218 | 0.495157603 | 0.661481172 | 12.54280639 | 9.120370595 | ENSG00000204396 |
| OAT | -0.460529642 | 0.726719416 | 9.15E-18 | 2.57E-16 | 2549.181382 | 1852.409064 | ENSG00000065154 |
| NA | -0.460795801 | 0.726585358 | 0.6897059 | 0.811485516 | 4.071430613 | 2.940726362 | ENSG00000272108 |
| ADAMTS9 | -0.460811155 | 0.726577626 | 0.315465719 | 0.486735118 | 26.26720814 | 19.14147645 | ENSG00000163638 |
| GMPPB | -0.460931349 | 0.726517096 | 4.96E-05 | 0.000285371 | 609.8106276 | 443.0993327 | ENSG00000173540 |
| BANCR | -0.461171232 | 0.726396305 | 0.54266368 | 0.701455713 | 9.103799222 | 6.611189736 | ENSG00000278910 |
| TSC22D1-AS1 | -0.461188947 | 0.726387385 | 0.366517346 | 0.540226629 | 24.19723826 | 17.5594013 | ENSG00000278156 |
| NA | -0.461476545 | 0.726242596 | 0.551589378 | 0.708436131 | 9.44646845 | 6.817421192 | ENSG00000287234 |
| NA | -0.461508619 | 0.72622645 | 0.655281043 | 0.786182387 | 4.939505075 | 3.596792408 | ENSG00000287780 |
| TCEAL1 | -0.461641731 | 0.726159447 | 0.091591658 | 0.193314516 | 67.77885227 | 49.21234233 | ENSG00000172465 |
| NA | -0.461718858 | 0.726120627 | 0.602521224 | 0.748190409 | 6.34367282 | 4.612883564 | ENSG00000232546 |
| AK6 | -0.461757981 | 0.726100937 | 0.002880986 | 0.010831632 | 239.675999 | 174.0223755 | ENSG00000085231 |
| PPP1R32 | -0.461763521 | 0.726098149 | 0.398215393 | 0.570556532 | 18.16920152 | 13.18420218 | ENSG00000162148 |
| NPY5R | -0.461769896 | 0.72609494 | 0.31487388 | 0.486150732 | 26.85074678 | 19.52032368 | ENSG00000164129 |
| MAP3K5 | -0.461915233 | 0.726021797 | 0.060920609 | 0.139641096 | 87.99425699 | 63.94259294 | ENSG00000197442 |
| NA | -0.46195467 | 0.726001951 | 0.002081402 | 0.008174332 | 254.7270478 | 184.8944808 | ENSG00000159712 |
| DERL2 | -0.462122625 | 0.725917436 | 1.42E-05 | 9.08E-05 | 529.9140507 | 384.5329455 | ENSG00000072849 |
| HNRNPM | -0.462329918 | 0.725813141 | 2.58E-18 | 7.49E-17 | 15206.81585 | 11037.52487 | ENSG00000099783 |
| C4orf48 | -0.46242746 | 0.72576407 | 0.007230624 | 0.024007911 | 233.669777 | 169.7778467 | ENSG00000243449 |
| MGAT4B | -0.462436228 | 0.725759659 | 8.09E-12 | 1.25E-10 | 1859.70122 | 1349.642939 | ENSG00000161013 |

| | | | | | | | |
|---|---|---|---|---|---|---|---|
| DERL3 | -0.462536529 | 0.725709203 | 0.025257197 | 0.068192665 | 120.2966353 | 87.26998117 | ENSG00000099958 |
| TTC27 | -0.462769199 | 0.725592174 | 8.28E-08 | 7.71E-07 | 837.5484837 | 608.0545326 | ENSG00000018699 |
| COMMD5 | -0.462904326 | 0.725524216 | 2.00E-06 | 1.49E-05 | 778.4074861 | 564.5309027 | ENSG00000170619 |
| NA | -0.46295592 | 0.725498271 | 0.575541263 | 0.727811636 | 8.170094534 | 5.891672554 | ENSG00000267799 |
| LOC728975 | -0.463160177 | 0.725395562 | 0.351370933 | 0.523906065 | 24.15413542 | 17.54890889 | ENSG00000254614 |
| WWP2 | -0.46340492 | 0.725272514 | 6.44E-11 | 9.02E-10 | 1460.499784 | 1059.113226 | ENSG00000198373 |
| NA | -0.463518487 | 0.725215424 | 0.199455206 | 0.350906556 | 43.13714495 | 31.22139553 | ENSG00000277531 |
| DUT | -0.463737227 | 0.725105475 | 2.25E-12 | 3.70E-11 | 2062.074061 | 1494.789098 | ENSG00000128951 |
| ACTR1A | -0.464000455 | 0.724973188 | 3.73E-14 | 7.38E-13 | 2515.917965 | 1823.844258 | ENSG00000138107 |
| GALNS | -0.46400689 | 0.724969954 | 6.00E-05 | 0.000340065 | 466.3816681 | 338.0217734 | ENSG00000141012 |
| ARMC6 | -0.464242779 | 0.724851427 | 1.09E-10 | 1.50E-09 | 1744.893001 | 1264.841804 | ENSG00000105676 |
| BRD3OS | -0.464261126 | 0.724842209 | 0.000402412 | 0.001894321 | 345.8266543 | 250.895887 | ENSG00000235106 |
| STAG3L4 | -0.464413822 | 0.724765495 | 0.038737335 | 0.097163168 | 134.1780161 | 97.1284185 | ENSG00000106610 |
| MIF | -0.464434292 | 0.724755212 | 0.600440654 | 0.746526209 | 6.357627334 | 4.610732991 | ENSG00000240972 |
| NA | -0.464535615 | 0.724704313 | 0.243629697 | 0.406491998 | 37.51979825 | 27.13659124 | ENSG00000271976 |
| NA | -0.464557644 | 0.724693247 | 0.646744407 | 0.780161729 | 4.985598981 | 3.622066255 | ENSG00000215414 |
| H2BC8 | -0.464639901 | 0.724651929 | 0.241062753 | 0.403358803 | 31.74348222 | 23.01171955 | ENSG00000273802 |
| NA | -0.464745794 | 0.724598742 | 0.411093529 | 0.583375405 | 16.30159199 | 11.82959165 | ENSG00000236924 |
| GPRC5B | -0.464772448 | 0.724585355 | 1.77E-12 | 2.93E-11 | 3129.151669 | 2267.315484 | ENSG00000167191 |
| NA | -0.465040659 | 0.72445066 | 0.400371166 | 0.572836917 | 16.3127556 | 11.83604337 | ENSG00000277159 |
| DPH2 | -0.465071642 | 0.724435102 | 1.89E-10 | 2.52E-09 | 1223.262069 | 886.4787552 | ENSG00000132768 |
| SMARCE1 | -0.465370843 | 0.724284877 | 0.200021324 | 0.351678031 | 40.31956087 | 29.22523993 | ENSG00000073584 |
| CACHD1 | -0.465386985 | 0.724276773 | 1.55E-12 | 2.59E-11 | 1737.733717 | 1259.131295 | ENSG00000158966 |
| SHISAL2A | -0.465439201 | 0.724250559 | 0.43304998 | 0.604046794 | 14.88626063 | 10.79897952 | ENSG00000182183 |
| METTL22 | -0.465496998 | 0.724221545 | 2.34E-05 | 0.000143828 | 670.2983948 | 485.1108024 | ENSG00000067365 |
| SERPINE3 | -0.465742866 | 0.724098132 | 0.617656436 | 0.759443988 | 6.270345606 | 4.534911451 | ENSG00000253309 |
| XXYLT1 | -0.465823695 | 0.724057564 | 3.84E-07 | 3.24E-06 | 740.2945245 | 536.0576991 | ENSG00000173950 |
| BCL7A | -0.465869983 | 0.724034334 | 2.25E-05 | 0.000138764 | 537.2744796 | 389.4246107 | ENSG00000110987 |
| SRSF8 | -0.466107359 | 0.723915214 | 6.08E-05 | 0.000343835 | 525.22479 | 380.6129361 | ENSG00000263465 |
| RIMS2 | -0.46652385 | 0.723706257 | 3.58E-05 | 0.000212414 | 433.6148089 | 313.6739744 | ENSG00000176406 |
| PHYHIP | -0.466589093 | 0.72367353 | 0.002736911 | 0.010368763 | 257.8505822 | 186.5432664 | ENSG00000168490 |
| NAA40 | -0.466599732 | 0.723668193 | 3.01E-07 | 2.58E-06 | 957.286521 | 692.2921304 | ENSG00000110583 |
| P2RY8 | -0.466684197 | 0.723625826 | 0.647515996 | 0.780555588 | 4.972320188 | 3.603244129 | ENSG00000182162 |
| TYMP | -0.466726046 | 0.723604836 | 0.740204931 | 0.84572929 | 3.564685818 | 2.585002399 | ENSG00000025708 |
| TRMT12 | -0.466778856 | 0.723578348 | 6.39E-05 | 0.000360133 | 440.9546623 | 318.7778445 | ENSG00000183665 |
| GOLM1 | -0.466868074 | 0.723533603 | 3.75E-21 | 1.38E-19 | 4101.873599 | 2967.510938 | ENSG00000135052 |
| HRC | -0.466903566 | 0.723515803 | 0.100380217 | 0.207732713 | 77.30645706 | 55.867095 | ENSG00000130528 |
| NUDT10 | -0.467015168 | 0.723459836 | 9.36E-06 | 6.20E-05 | 585.4924904 | 423.3343093 | ENSG00000122824 |
| FAM174C | -0.467058695 | 0.72343801 | 0.000107504 | 0.000577678 | 610.3022297 | 441.8982127 | ENSG00000228300 |
| PEX3 | -0.467259009 | 0.72333757 | 0.00688704 | 0.023089658 | 200.6326119 | 145.333805 | ENSG00000034693 |
| NA | -0.467262999 | 0.723335569 | 0.672716983 | 0.799543055 | 4.607323737 | 3.376039973 | ENSG00000235649 |
| NA | -0.467275744 | 0.723329179 | 0.56268856 | 0.717247589 | 8.964841781 | 6.46169723 | ENSG00000286064 |
| BCORL1 | -0.467284736 | 0.723324671 | 8.43E-05 | 0.000463052 | 396.6089641 | 286.9103118 | ENSG00000085185 |
| GTF3C1 | -0.467348073 | 0.723292916 | 2.13E-21 | 7.95E-20 | 5394.266659 | 3901.122093 | ENSG00000077235 |
| YTHDC2 | -0.467440293 | 0.723246683 | 2.29E-07 | 1.99E-06 | 784.8575256 | 567.8407532 | ENSG00000047188 |
| VPS11 | -0.467512316 | 0.723210578 | 6.97E-05 | 0.000389889 | 480.4734058 | 347.560483 | ENSG00000160695 |
| NA | -0.467600028 | 0.72316661 | 0.280185569 | 0.448292262 | 30.01917263 | 21.64635616 | ENSG00000269352 |
| NA | -0.467636342 | 0.723148407 | 0.314363913 | 0.4855011 | 26.38818087 | 19.00865549 | ENSG00000234743 |
| ARIH1 | -0.467732069 | 0.723100426 | 2.79E-10 | 3.66E-09 | 1302.331921 | 941.9825594 | ENSG00000166233 |
| NA | -0.467762988 | 0.723084929 | 0.519769911 | 0.683169677 | 9.973988865 | 7.221009236 | ENSG00000257718 |
| ZBTB17 | -0.467787958 | 0.723072414 | 1.91E-05 | 0.000119598 | 577.8788645 | 417.6426891 | ENSG00000116809 |
| SHBG | -0.467911527 | 0.723010485 | 0.461336588 | 0.631135673 | 14.90599711 | 10.69788413 | ENSG00000129214 |
| DPH3 | -0.467993738 | 0.722969285 | 1.05E-06 | 8.25E-06 | 658.9067494 | 476.6306198 | ENSG00000154813 |
| ST14 | -0.468117843 | 0.722907096 | 3.78E-14 | 7.48E-13 | 1963.902342 | 1419.984813 | ENSG00000149418 |
| ATP6V1H | -0.468411511 | 0.722759959 | 1.92E-06 | 1.44E-05 | 627.8759511 | 454.1545036 | ENSG00000047249 |
| DYNC1I2 | -0.468463078 | 0.722734126 | 9.78E-14 | 1.86E-12 | 1858.676641 | 1343.439032 | ENSG00000077380 |
| TEX10 | -0.468631219 | 0.722649899 | 3.18E-13 | 5.73E-12 | 2089.529935 | 1510.279666 | ENSG00000136891 |
| PARPBP | -0.468782965 | 0.722573893 | 7.56E-05 | 0.000420408 | 416.8617777 | 301.314686 | ENSG00000185480 |
| RRP7A | -0.468904311 | 0.722513119 | 3.19E-12 | 5.15E-11 | 2033.285932 | 1468.99419 | ENSG00000189306 |
| WDTC1 | -0.469072286 | 0.722429001 | 8.69E-08 | 8.07E-07 | 858.4866088 | 620.1048888 | ENSG00000142784 |
| TAF6L | -0.469091674 | 0.722419293 | 0.000390508 | 0.001847299 | 360.39664 | 260.2889148 | ENSG00000162227 |
| RECQL | -0.469441462 | 0.72224416 | 1.04E-05 | 6.84E-05 | 645.0632005 | 465.7695157 | ENSG00000004700 |
| LEXM | -0.469518836 | 0.722205426 | 0.653134977 | 0.784504704 | 5.375037836 | 3.847119837 | ENSG00000162398 |
| SYT13 | -0.469561153 | 0.722184242 | 6.55E-05 | 0.000368589 | 486.321073 | 351.5863793 | ENSG00000019505 |
| SLIRP | -0.469668578 | 0.72213047 | 1.05E-11 | 1.61E-10 | 1487.064932 | 1073.343912 | ENSG00000119705 |
| SEC63 | -0.46970009 | 0.722114697 | 2.56E-14 | 5.20E-13 | 2221.046246 | 1604.635364 | ENSG00000025796 |
| CENPM | -0.469752645 | 0.722088391 | 1.38E-07 | 1.24E-06 | 729.2150952 | 526.4382727 | ENSG00000100162 |
| TSPAN17 | -0.469859825 | 0.722034748 | 1.94E-14 | 3.96E-13 | 2009.38746 | 1450.844027 | ENSG00000048140 |
| ADAMTS7 | -0.469869671 | 0.722029821 | 2.77E-16 | 6.83E-15 | 2419.625056 | 1746.921768 | ENSG00000136378 |
| NA | -0.47002356 | 0.721952808 | 0.585137885 | 0.735021666 | 8.504190893 | 6.141999982 | ENSG00000235586 |
| GPRASP1 | -0.470420028 | 0.721754435 | 0.15815986 | 0.294447198 | 54.80521904 | 39.56094838 | ENSG00000198932 |
| GTSE1-DT | -0.470455156 | 0.721736861 | 0.414531806 | 0.586779117 | 15.90936223 | 11.51634613 | ENSG00000277232 |
| ZNF354B | -0.47065376 | 0.721637512 | 0.005717175 | 0.019705203 | 193.5824616 | 139.5674477 | ENSG00000178338 |
| FGF21 | -0.470675852 | 0.721626462 | 0.494348605 | 0.660741283 | 11.8227378 | 8.491728969 | ENSG00000105550 |
| GEMIN5 | -0.470692009 | 0.72161838 | 8.89E-12 | 1.37E-10 | 1605.707557 | 1158.849225 | ENSG00000082516 |
| NA | -0.470727956 | 0.7216004 | 0.679259583 | 0.803872602 | 4.071430613 | 2.909000794 | ENSG00000269910 |
| NA | -0.470840036 | 0.721544343 | 0.383213411 | 0.556669422 | 17.70364455 | 12.75749087 | ENSG00000260806 |

| | | | | | | | |
|---|---|---|---|---|---|---|---|
| MICOS10 | -0.470965432 | 0.72148163 | 0.074592865 | 0.164117622 | 73.28794146 | 52.84247784 | ENSG00000173436 |
| ZNF511 | -0.471047919 | 0.72144038 | 0.033621859 | 0.086158247 | 107.1730812 | 77.24887532 | ENSG00000198546 |
| NA | -0.471274856 | 0.721326906 | 0.443322173 | 0.614112311 | 13.67830785 | 9.881833175 | ENSG00000223969 |
| NCBP2AS2 | -0.471350404 | 0.721289134 | 5.65E-08 | 5.38E-07 | 877.5623087 | 632.9823093 | ENSG00000270170 |
| THOC5 | -0.471569131 | 0.721179788 | 2.87E-11 | 4.18E-10 | 1470.768396 | 1060.566242 | ENSG00000100296 |
| SYTL4 | -0.471709538 | 0.721109604 | 0.617640978 | 0.759443988 | 5.908140058 | 4.28888517 | ENSG00000102362 |
| TTLL12 | -0.471714588 | 0.72110708 | 2.44E-18 | 7.11E-17 | 6322.240756 | 4559.177358 | ENSG00000100304 |
| LZTFL1 | -0.471889239 | 0.721019789 | 0.036608279 | 0.092581321 | 133.2465018 | 96.21044812 | ENSG00000163818 |
| SUPV3L1 | -0.471925355 | 0.72100174 | 1.33E-07 | 1.21E-06 | 818.5236347 | 590.0160185 | ENSG00000156502 |
| GRWD1 | -0.471998127 | 0.720965372 | 2.66E-16 | 6.58E-15 | 2659.723136 | 1917.507366 | ENSG00000105447 |
| NMT2 | -0.472024407 | 0.720952239 | 0.000263678 | 0.001295301 | 354.8869872 | 255.9295632 | ENSG00000152465 |
| FBXL17 | -0.472112322 | 0.720908307 | 0.057171775 | 0.132911631 | 85.45334849 | 61.63104125 | ENSG00000145743 |
| TMEM132E | -0.472293111 | 0.720817973 | 0.188671364 | 0.336289317 | 40.25692171 | 29.03191192 | ENSG00000181291 |
| ATCAY | -0.472431277 | 0.720748944 | 2.55E-11 | 3.73E-10 | 1368.709821 | 986.6390545 | ENSG00000167654 |
| FAM136A | -0.472508606 | 0.720710313 | 3.94E-13 | 7.02E-12 | 2330.763166 | 1679.901357 | ENSG00000035141 |
| KIF24 | -0.472514449 | 0.720707394 | 0.010689516 | 0.033329797 | 168.2885943 | 121.3726805 | ENSG00000186638 |
| LRRC3B | -0.472700981 | 0.720614216 | 0.157758219 | 0.293841248 | 52.11909557 | 37.55295541 | ENSG00000179796 |
| NA | -0.47281002 | 0.720559754 | 0.634277225 | 0.770955955 | 4.996762593 | 3.598942982 | ENSG00000270558 |
| FOXRED1 | -0.472863728 | 0.72053293 | 1.58E-10 | 2.13E-09 | 1337.539854 | 963.2732648 | ENSG00000110074 |
| HSPA14 | -0.472889353 | 0.720520132 | 0.00062781 | 0.002806091 | 412.1321539 | 297.4360768 | ENSG00000187522 |
| SNORA11 | -0.473125089 | 0.720402409 | 0.735666855 | 0.842640381 | 3.647061463 | 2.633399519 | ENSG00000221716 |
| CDC45 | -0.4734246 | 0.720252866 | 1.27E-05 | 8.19E-05 | 598.7317478 | 431.2793547 | ENSG00000093009 |
| STRIP1 | -0.473501746 | 0.720214352 | 1.26E-10 | 1.73E-09 | 1218.14218 | 876.9451708 | ENSG00000143093 |
| NAP1L2 | -0.473790066 | 0.720070433 | 4.22E-05 | 0.000245931 | 533.0396492 | 384.2712958 | ENSG00000186462 |
| TSR1 | -0.474080573 | 0.719925451 | 2.63E-12 | 4.29E-11 | 3592.347652 | 2586.049796 | ENSG00000167721 |
| TSC22D1 | -0.474224134 | 0.719853815 | 2.66E-14 | 5.38E-13 | 2047.982096 | 1473.644978 | ENSG00000102804 |
| LINS1 | -0.47423738 | 0.719847207 | 0.065999175 | 0.148876296 | 86.08681107 | 62.11313444 | ENSG00000140471 |
| NA | -0.474252708 | 0.719839558 | 0.616174643 | 0.758317387 | 7.236189686 | 5.216784382 | ENSG00000272599 |
| MANF | -0.474385148 | 0.71977348 | 1.11E-16 | 2.85E-15 | 2400.323206 | 1727.329621 | ENSG00000145050 |
| NA | -0.474543879 | 0.719694292 | 0.499824209 | 0.666088297 | 12.27992207 | 8.760878523 | ENSG00000182165 |
| PRPF38B | -0.474626526 | 0.719653064 | 4.50E-15 | 9.71E-14 | 2469.927145 | 1777.648954 | ENSG00000134186 |
| NA | -0.47464961 | 0.71964155 | 0.383348307 | 0.556767998 | 18.43017083 | 13.31272199 | ENSG00000276248 |
| HLA-F-AS1 | -0.474736549 | 0.719598184 | 0.259286824 | 0.424465557 | 30.59278696 | 21.96443587 | ENSG00000214922 |
| IQCK | -0.474750973 | 0.71959099 | 0.210679028 | 0.366243952 | 36.9600263 | 26.63378581 | ENSG00000174628 |
| COQ4 | -0.474784048 | 0.719574493 | 1.01E-05 | 6.64E-05 | 565.7581119 | 407.2040014 | ENSG00000167113 |
| GABRQ | -0.474822614 | 0.719555257 | 0.000103204 | 0.000556742 | 379.2142978 | 273.0487983 | ENSG00000268089 |
| MRPL21 | -0.47490306 | 0.719515135 | 1.63E-09 | 1.94E-08 | 1100.120191 | 791.232524 | ENSG00000197345 |
| SRSF4 | -0.47495867 | 0.719487401 | 7.75E-18 | 2.19E-16 | 3993.85726 | 2873.481819 | ENSG00000116350 |
| ZNF165 | -0.475115867 | 0.71940901 | 0.071561629 | 0.158732367 | 80.12843476 | 57.58792189 | ENSG00000197279 |
| ST8SIA6 | -0.475486732 | 0.719224099 | 0.198294869 | 0.349656157 | 43.33160815 | 31.17783259 | ENSG00000148488 |
| CLPTM1 | -0.475863051 | 0.719036518 | 1.22E-13 | 2.29E-12 | 2592.207882 | 1863.986777 | ENSG00000104853 |
| TCTN2 | -0.475995241 | 0.718970638 | 0.01252025 | 0.038023952 | 154.6459805 | 111.1348629 | ENSG00000168778 |
| AKR1B1 | -0.476264657 | 0.718836387 | 4.27E-23 | 1.79E-21 | 4307.026085 | 3095.62142 | ENSG00000085662 |
| CNKSR3 | -0.47642483 | 0.718756583 | 0.088659955 | 0.188529453 | 80.91951523 | 58.32060298 | ENSG00000153721 |
| GEMIN7 | -0.476458225 | 0.718739946 | 0.631780574 | 0.768920547 | 5.078462515 | 3.649490675 | ENSG00000142252 |
| OLFM1 | -0.476617816 | 0.718660443 | 5.33E-12 | 8.38E-11 | 1464.670587 | 1052.940542 | ENSG00000130558 |
| SETD6 | -0.476831249 | 0.718554132 | 2.23E-06 | 1.65E-05 | 702.1210516 | 504.3808413 | ENSG00000103037 |
| STIM1 | -0.476935997 | 0.718501963 | 2.76E-06 | 2.01E-05 | 553.1111151 | 397.4729873 | ENSG00000167323 |
| PGRMC1 | -0.477167086 | 0.718386883 | 2.09E-18 | 6.13E-17 | 6577.533411 | 4725.80054 | ENSG00000101856 |
| NA | -0.477198245 | 0.718371368 | 0.691108535 | 0.812689713 | 4.591254041 | 3.323341707 | ENSG00000284642 |
| FBXO3 | -0.477329661 | 0.718305934 | 4.81E-05 | 0.000277319 | 422.2675155 | 303.6181563 | ENSG00000110429 |
| LZTS1 | -0.477386537 | 0.718277616 | 1.46E-05 | 9.28E-05 | 488.7071545 | 351.0787823 | ENSG00000061337 |
| NRIP1 | -0.47753438 | 0.718204014 | 0.01525929 | 0.044900391 | 200.2483162 | 143.7783302 | ENSG00000180530 |
| LAYN | -0.477684628 | 0.718129221 | 0.024226912 | 0.065825543 | 116.0555473 | 83.37661479 | ENSG00000204381 |
| NA | -0.477741419 | 0.718100952 | 0.608308727 | 0.752091498 | 6.316439512 | 4.513938752 | ENSG00000279799 |
| CFAP20 | -0.477881422 | 0.718031269 | 2.42E-06 | 1.77E-05 | 564.829901 | 405.5579237 | ENSG00000070761 |
| NA | -0.478269536 | 0.717838131 | 0.427926608 | 0.599283643 | 15.18092093 | 10.88555393 | ENSG00000276728 |
| FEN1 | -0.478394121 | 0.717776144 | 3.03E-15 | 6.68E-14 | 2387.025692 | 1713.816604 | ENSG00000168496 |
| KIF18B | -0.478483376 | 0.717731738 | 9.34E-13 | 1.59E-11 | 1619.598274 | 1162.097563 | ENSG00000186185 |
| MBOAT2 | -0.478672143 | 0.717637834 | 0.000225352 | 0.001124486 | 399.9880861 | 287.2913096 | ENSG00000143797 |
| TMEM255B | -0.478816105 | 0.717566227 | 0.068524091 | 0.153352464 | 76.30903717 | 54.67246938 | ENSG00000184497 |
| GLCE | -0.478859851 | 0.717544469 | 0.000293667 | 0.001427343 | 401.7927058 | 288.1923358 | ENSG00000138604 |
| ALYREF | -0.479198241 | 0.717376186 | 1.02E-15 | 2.38E-14 | 2968.604916 | 2129.185156 | ENSG00000183684 |
| TMEM52 | -0.479248617 | 0.717351137 | 0.008367478 | 0.027115239 | 184.4248714 | 132.2848653 | ENSG00000178821 |
| ADCK2 | -0.47943022 | 0.717260844 | 0.00925397 | 0.029468195 | 155.6397336 | 111.6449136 | ENSG00000133597 |
| DHFR | -0.479738553 | 0.717107567 | 1.87E-10 | 2.49E-09 | 1432.83179 | 1027.412948 | ENSG00000228716 |
| FUNDC2 | -0.480237594 | 0.716859556 | 1.33E-06 | 1.02E-05 | 801.1512573 | 574.434962 | ENSG00000165775 |
| RASA4B | -0.480301059 | 0.716828022 | 0.647115404 | 0.780388481 | 5.07288071 | 3.630668549 | ENSG00000170667 |
| PCDHB11 | -0.480620052 | 0.716669543 | 0.58072886 | 0.731628608 | 7.734561772 | 5.511207782 | ENSG00000197479 |
| TPD52L2 | -0.480862302 | 0.716549213 | 1.78E-20 | 6.29E-19 | 2939.573066 | 2106.775895 | ENSG00000101150 |
| ARHGEF25 | -0.481346785 | 0.716308623 | 0.112815715 | 0.227271734 | 64.9155746 | 46.41492932 | ENSG00000240771 |
| TUSC2 | -0.481489562 | 0.716237737 | 9.86E-08 | 9.09E-07 | 992.2751392 | 711.1397044 | ENSG00000114383 |
| SLC1A5 | -0.481506015 | 0.716229569 | 2.50E-23 | 1.08E-21 | 7459.24958 | 5342.022471 | ENSG00000105281 |
| PMEL | -0.481623392 | 0.716171299 | 7.25E-13 | 1.25E-11 | 2062.822774 | 1477.075533 | ENSG00000185664 |
| PDXDC2P-NPIPB14P | -0.481682194 | 0.71614211 | 0.285539336 | 0.454519093 | 29.16696771 | 20.80315335 | ENSG00000196696 |
| ZDHHC9 | -0.481709337 | 0.716128636 | 2.30E-12 | 3.78E-11 | 1737.659812 | 1244.836594 | ENSG00000188706 |
| NA | -0.481808344 | 0.716079492 | 0.652321242 | 0.783947737 | 4.640138851 | 3.355067274 | ENSG00000253838 |

| | | | | | | | |
|---|---|---|---|---|---|---|---|
| PURG | -0.481998022 | 0.715985352 | 0.013593481 | 0.040744536 | 144.6323554 | 103.5288152 | ENSG00000172733 |
| MPC1 | -0.48210737 | 0.715931087 | 0.002579217 | 0.009850658 | 220.5885989 | 158.1501733 | ENSG00000060762 |
| SP110 | -0.4821166 | 0.715926506 | 0.047453665 | 0.11461463 | 86.31305024 | 61.79102616 | ENSG00000135899 |
| POP4 | -0.482189621 | 0.715890271 | 3.53E-05 | 0.000209986 | 443.4170252 | 317.2333952 | ENSG00000105171 |
| UBE2J1 | -0.482258193 | 0.715856245 | 1.46E-05 | 9.30E-05 | 581.3423013 | 416.533275 | ENSG00000198833 |
| ZNF674-AS1 | -0.482315749 | 0.715827687 | 0.078810215 | 0.17138179 | 74.48349133 | 53.28907736 | ENSG00000230844 |
| PAK1IP1 | -0.48257411 | 0.715699506 | 2.66E-10 | 3.50E-09 | 1191.87172 | 853.0050006 | ENSG00000111845 |
| MRPL48 | -0.482670563 | 0.715651659 | 0.000122815 | 0.000650173 | 376.3798419 | 269.2436238 | ENSG00000175581 |
| SLC15A2 | -0.482995678 | 0.715490404 | 0.12288916 | 0.242720875 | 59.66581509 | 42.81786434 | ENSG00000163406 |
| STBD1 | -0.483101346 | 0.715438 | 0.395525119 | 0.56813664 | 17.86058671 | 12.81395724 | ENSG00000118804 |
| ZGPAT | -0.483362984 | 0.715308265 | 0.006310486 | 0.021432323 | 174.4389311 | 124.8646094 | ENSG00000197114 |
| KCTD15 | -0.483452411 | 0.715263927 | 4.33E-10 | 5.52E-09 | 1232.608316 | 881.7731486 | ENSG00000153885 |
| CENPQ | -0.483705883 | 0.715138271 | 0.003446782 | 0.012672359 | 196.4087818 | 140.4316232 | ENSG00000031691 |
| UHRF2 | -0.484115961 | 0.714935026 | 2.66E-09 | 3.06E-08 | 1080.541651 | 772.6150435 | ENSG00000147854 |
| NA | -0.484118707 | 0.714933665 | 0.522530843 | 0.685293349 | 9.726974075 | 6.972832381 | ENSG00000273456 |
| KAZALD1 | -0.484181278 | 0.714902659 | 0.31031667 | 0.481279598 | 23.37458199 | 16.69495322 | ENSG00000107821 |
| NFKBIE | -0.484238195 | 0.714874455 | 0.000215807 | 0.001079371 | 389.0208566 | 278.2145203 | ENSG00000146232 |
| UPP1 | -0.484246944 | 0.714870119 | 1.67E-07 | 1.49E-06 | 752.4549502 | 538.1807143 | ENSG00000183696 |
| ATXN7L3B | -0.484310758 | 0.714838499 | 7.07E-15 | 1.51E-13 | 1776.027561 | 1269.859257 | ENSG00000253719 |
| NA | -0.484340068 | 0.714823977 | 0.234850768 | 0.395988651 | 33.35973446 | 23.79819552 | ENSG00000235494 |
| NA | -0.484625643 | 0.714682495 | 0.674024492 | 0.800111161 | 3.663131159 | 2.616727967 | ENSG00000277749 |
| NAT10 | -0.484760067 | 0.714615907 | 5.39E-21 | 1.97E-19 | 3463.742739 | 2474.952195 | ENSG00000135372 |
| GRB10 | -0.485154329 | 0.714420642 | 1.16E-15 | 2.69E-14 | 3029.240829 | 2163.574179 | ENSG00000106070 |
| KIF20A | -0.485159011 | 0.714418324 | 1.26E-11 | 1.91E-10 | 1733.979446 | 1238.649717 | ENSG00000112984 |
| LMF1 | -0.485393302 | 0.714302313 | 0.007530585 | 0.024849375 | 160.6972204 | 114.8900106 | ENSG00000260807 |
| GNL1 | -0.485461304 | 0.714268645 | 5.92E-15 | 1.27E-13 | 2122.046025 | 1515.398608 | ENSG00000204590 |
| PCDHA3 | -0.48566574 | 0.714167437 | 0.257854086 | 0.42290964 | 29.07689508 | 20.80503135 | ENSG00000255408 |
| TOX2 | -0.485730954 | 0.714135155 | 0.066351434 | 0.149531017 | 77.03257238 | 54.97549507 | ENSG00000124191 |
| CSNK2A3 | -0.485732069 | 0.714134603 | 0.659341065 | 0.789485573 | 4.601741931 | 3.308820728 | ENSG00000254598 |
| LINC01841 | -0.485764873 | 0.714118366 | 0.596725005 | 0.74400866 | 6.019864191 | 4.314159016 | ENSG00000266913 |
| NA | -0.485950612 | 0.714026433 | 0.366382977 | 0.540190067 | 18.3834012 | 13.11913351 | ENSG00000248334 |
| HERC2P9 | -0.485952551 | 0.714025473 | 3.30E-05 | 0.000197239 | 444.7330722 | 317.2976582 | ENSG00000206149 |
| POMGNT2 | -0.486324446 | 0.713841437 | 1.91E-06 | 1.43E-05 | 612.7394844 | 437.4305511 | ENSG00000144647 |
| CHRNA5 | -0.486364333 | 0.713821701 | 0.002457191 | 0.009455751 | 218.8761286 | 156.3597161 | ENSG00000169684 |
| SVBP | -0.486421545 | 0.713793394 | 0.065388834 | 0.147672207 | 83.86779618 | 59.82176194 | ENSG00000177868 |
| FABP6 | -0.486466643 | 0.713771082 | 0.157871972 | 0.293996352 | 42.7573181 | 30.51048064 | ENSG00000170231 |
| COL2A1 | -0.48651765 | 0.713745846 | 2.62E-05 | 0.00016002 | 422.8433571 | 301.7911515 | ENSG00000139219 |
| NA | -0.486529517 | 0.713739975 | 0.42364774 | 0.595322593 | 16.10258613 | 11.56259267 | ENSG00000214222 |
| C7orf31 | -0.486549912 | 0.713729886 | 0.210043277 | 0.365534041 | 36.44454538 | 25.94169305 | ENSG00000153790 |
| MFSD11 | -0.486584418 | 0.713712815 | 0.004116885 | 0.014825242 | 183.381758 | 130.8955546 | ENSG00000092931 |
| TCTEX1D1 | -0.486645471 | 0.713682612 | 0.320923316 | 0.492320611 | 21.17076091 | 15.07119314 | ENSG00000152760 |
| NA | -0.48677895 | 0.713616585 | 0.395574364 | 0.568165069 | 16.18516193 | 11.53516825 | ENSG00000279656 |
| LAMP1 | -0.486821004 | 0.713595783 | 5.06E-23 | 2.12E-21 | 4653.41523 | 3320.650957 | ENSG00000185896 |
| PARP12 | -0.486835794 | 0.713588468 | 0.015835687 | 0.046320885 | 181.3341153 | 129.4723862 | ENSG00000059378 |
| HSPA1L | -0.486937016 | 0.713538403 | 0.122856823 | 0.242681852 | 61.10000705 | 43.65757161 | ENSG00000204390 |
| SSU72 | -0.486957866 | 0.713528091 | 1.31E-19 | 4.28E-18 | 3744.836353 | 2672.101064 | ENSG00000160075 |
| TRNAU1AP | -0.486966925 | 0.713523611 | 1.24E-05 | 8.02E-05 | 495.7812574 | 353.4726073 | ENSG00000180098 |
| DOK1 | -0.487088019 | 0.713463723 | 0.000466995 | 0.002157282 | 284.3026784 | 202.8703736 | ENSG00000115325 |
| CELF4 | -0.487148349 | 0.713433888 | 0.168967061 | 0.309695351 | 45.92506824 | 32.68759385 | ENSG00000101489 |
| ZNF740 | -0.48742809 | 0.713295566 | 7.02E-08 | 6.60E-07 | 719.7789145 | 513.7095391 | ENSG00000139651 |
| SYVN1 | -0.487664768 | 0.713178557 | 3.76E-06 | 2.67E-05 | 735.1575523 | 524.485147 | ENSG00000162298 |
| RBM6 | -0.487927392 | 0.713048744 | 4.00E-17 | 1.06E-15 | 2631.011772 | 1875.260332 | ENSG00000004534 |
| NA | -0.488059496 | 0.712983455 | 0.624102729 | 0.763058842 | 6.395348532 | 4.532760877 | ENSG00000275956 |
| NA | -0.488416042 | 0.712807271 | 0.309437219 | 0.480585571 | 24.60294698 | 17.54299021 | ENSG00000234062 |
| UTP20 | -0.488570623 | 0.7127309 | 5.37E-13 | 9.41E-12 | 1775.534257 | 1264.942888 | ENSG00000120800 |
| PI4K2A | -0.488583651 | 0.712724463 | 6.40E-07 | 5.20E-06 | 574.3244153 | 409.2399334 | ENSG00000155252 |
| MAX | -0.489068498 | 0.712484978 | 2.15E-06 | 1.59E-05 | 579.1455384 | 412.9119838 | ENSG00000125952 |
| NA | -0.489181184 | 0.71242933 | 0.417934838 | 0.589959093 | 16.20960434 | 11.52871653 | ENSG00000214439 |
| NA | -0.489228503 | 0.712405963 | 0.467943976 | 0.63706857 | 12.98459668 | 9.288685226 | ENSG00000255224 |
| SGTA | -0.489497707 | 0.712273042 | 9.65E-19 | 2.90E-17 | 3300.218311 | 2350.519149 | ENSG00000104969 |
| NA | -0.489709691 | 0.712168391 | 0.57918419 | 0.730715494 | 7.451940965 | 5.304976326 | ENSG00000256897 |
| TCF15 | -0.489752685 | 0.712147168 | 0.000306587 | 0.001484902 | 339.8616201 | 242.2054565 | ENSG00000125878 |
| DMC1 | -0.490210604 | 0.711921164 | 0.042915797 | 0.105571109 | 91.10243426 | 64.98180737 | ENSG00000100206 |
| ABCB9 | -0.490519249 | 0.711768875 | 0.040150688 | 0.099906796 | 113.4062691 | 80.68112121 | ENSG00000150967 |
| ABCF3 | -0.490588241 | 0.711734838 | 5.91E-12 | 9.28E-11 | 1337.968841 | 951.9540027 | ENSG00000161204 |
| PIGS | -0.490592473 | 0.71173275 | 3.16E-12 | 5.11E-11 | 1284.165423 | 913.8592526 | ENSG00000087111 |
| TLCD2 | -0.490627551 | 0.711715445 | 0.270528788 | 0.437156211 | 28.60942308 | 20.30464907 | ENSG00000185561 |
| MRPL54 | -0.490638711 | 0.711709939 | 2.86E-05 | 0.000172857 | 474.7145237 | 337.5549947 | ENSG00000183617 |
| HDGFL3 | -0.490660644 | 0.71169912 | 1.10E-17 | 3.04E-16 | 2365.413815 | 1683.600807 | ENSG00000166503 |
| NEK10 | -0.490695737 | 0.711681808 | 0.614066347 | 0.75677096 | 6.109936822 | 4.360405562 | ENSG00000163491 |
| SLC35A4 | -0.491004455 | 0.711529533 | 2.41E-16 | 5.98E-15 | 2299.13023 | 1635.893416 | ENSG00000176087 |
| TMEM240 | -0.491015872 | 0.711523903 | 0.069570956 | 0.155330448 | 79.85909275 | 56.82510224 | ENSG00000205090 |
| FSIP2 | -0.491029195 | 0.711517332 | 0.44179044 | 0.612914734 | 13.78165927 | 9.75976509 | ENSG00000188738 |
| NA | -0.491109673 | 0.711477642 | 0.680548798 | 0.804884242 | 3.744831081 | 2.662974513 | ENSG00000236875 |
| ERG28 | -0.491508779 | 0.711280847 | 3.54E-10 | 4.58E-09 | 1244.982002 | 885.6751236 | ENSG00000133935 |
| SLC10A4 | -0.49211308 | 0.710982975 | 0.060110876 | 0.138146165 | 84.66781293 | 60.23422485 | ENSG00000145248 |
| SMARCA2 | -0.492178188 | 0.71095089 | 0.000646827 | 0.002878417 | 341.7193791 | 242.5552923 | ENSG00000080503 |

| | | | | | | | |
|---|---|---|---|---|---|---|---|
| SLC4A10 | -0.492369816 | 0.710856463 | 0.595778537 | 0.743461011 | 6.460978759 | 4.558034724 | ENSG00000144290 |
| DLST | -0.492416546 | 0.710833438 | 7.81E-16 | 1.84E-14 | 2908.923318 | 2067.437353 | ENSG00000119689 |
| GALNT18 | -0.492568677 | 0.710758486 | 0.001591578 | 0.006452948 | 253.9466937 | 180.4631 | ENSG00000110328 |
| NOLC1 | -0.492709475 | 0.710689123 | 1.33E-20 | 4.75E-19 | 6777.021717 | 4816.115822 | ENSG00000166197 |
| NBPF9 | -0.492765309 | 0.71066162 | 0.002646768 | 0.010072755 | 213.4774121 | 151.5575573 | ENSG00000269713 |
| KIAA0753 | -0.492874181 | 0.710607992 | 1.23E-06 | 9.56E-06 | 547.137457 | 388.7252848 | ENSG00000198920 |
| DUSP22 | -0.493271619 | 0.710412258 | 1.15E-06 | 8.95E-06 | 758.5080667 | 538.2692336 | ENSG00000112679 |
| E2F4 | -0.493617506 | 0.710241957 | 1.29E-14 | 2.67E-13 | 2327.013402 | 1652.666905 | ENSG00000205250 |
| BTG3-AS1 | -0.493725481 | 0.710188802 | 0.511371421 | 0.675720602 | 9.800301289 | 6.960461977 | ENSG00000280594 |
| MAP2K6 | -0.493753103 | 0.710175205 | 5.83E-09 | 6.40E-08 | 973.5247371 | 691.3407864 | ENSG00000108984 |
| LOC100128108 | -0.493815667 | 0.710144409 | 0.349163168 | 0.521743352 | 17.98905626 | 12.77416242 | ENSG00000259658 |
| TCN2 | -0.493827827 | 0.710138423 | 8.46E-13 | 1.45E-11 | 1352.97894 | 960.4718111 | ENSG00000185339 |
| DDX11L1 | -0.49384432 | 0.710130305 | 0.669196293 | 0.796782229 | 3.72317958 | 2.637700666 | ENSG00000223972 |
| RCBTB2 | -0.494082338 | 0.710013156 | 0.00033485 | 0.001607264 | 316.0340331 | 224.598179 | ENSG00000136161 |
| NPEPL1 | -0.49409263 | 0.710008091 | 0.014968992 | 0.044194493 | 154.1025025 | 109.3468168 | ENSG00000215440 |
| MAP2K3 | -0.494128745 | 0.709990318 | 1.89E-08 | 1.93E-07 | 936.3139284 | 664.7393082 | ENSG00000034152 |
| NA | -0.494772555 | 0.709673551 | 0.476532385 | 0.644836296 | 11.15837513 | 7.89858102 | ENSG00000232450 |
| EFL1 | -0.494954312 | 0.709584149 | 8.89E-11 | 1.23E-09 | 1104.014943 | 783.2258323 | ENSG00000140598 |
| SWI5 | -0.495130151 | 0.709497668 | 0.000150838 | 0.000779694 | 336.1378017 | 238.5277358 | ENSG00000175854 |
| RTP1 | -0.495756813 | 0.709189552 | 0.070347227 | 0.156705353 | 78.4926462 | 55.59821802 | ENSG00000175077 |
| FBXL19-AS1 | -0.495845532 | 0.709145941 | 0.263963062 | 0.429641531 | 28.31005686 | 20.0290478 | ENSG00000260852 |
| MMD | -0.495883652 | 0.709127204 | 2.63E-06 | 1.92E-05 | 620.8513334 | 440.7403952 | ENSG00000108960 |
| SP6 | -0.495997257 | 0.709071366 | 0.090036327 | 0.19072041 | 62.76968676 | 44.51529539 | ENSG00000189120 |
| RFLNB | -0.496154725 | 0.708993976 | 3.63E-10 | 4.68E-09 | 1176.516772 | 834.6706333 | ENSG00000183688 |
| EYA1 | -0.49619861 | 0.70897241 | 0.369158499 | 0.54295796 | 19.60598422 | 13.97739033 | ENSG00000104313 |
| PRPSAP1 | -0.49630961 | 0.708917864 | 1.18E-08 | 1.25E-07 | 839.0025754 | 594.912009 | ENSG00000161542 |
| YIF1A | -0.496324394 | 0.708910599 | 8.31E-13 | 1.43E-11 | 1364.825333 | 967.3244534 | ENSG00000174851 |
| SNHG21 | -0.496333909 | 0.708905924 | 0.500851649 | 0.666686425 | 9.655762042 | 6.827641023 | ENSG00000250988 |
| NA | -0.496419423 | 0.708863906 | 0.233430651 | 0.394179858 | 35.85697654 | 25.36790026 | ENSG00000224945 |
| NA | -0.496564294 | 0.708792727 | 0.581941161 | 0.732447768 | 6.022655094 | 4.265761897 | ENSG00000252061 |
| AMPH | -0.49667065 | 0.708740477 | 0.000123757 | 0.000654079 | 329.1748962 | 233.4555913 | ENSG00000078053 |
| USH1G | -0.496720017 | 0.708716225 | 0.065947934 | 0.148812913 | 71.99270694 | 51.04206129 | ENSG00000182040 |
| ZNF514 | -0.496733671 | 0.708709518 | 0.002921844 | 0.010955353 | 221.0804253 | 156.8119738 | ENSG00000144026 |
| SCYL1 | -0.497052702 | 0.708552814 | 3.98E-15 | 8.68E-14 | 1771.930724 | 1255.532346 | ENSG00000142186 |
| CAMK1 | -0.497127713 | 0.708515975 | 0.31824543 | 0.489477014 | 25.73689683 | 18.16760326 | ENSG00000134072 |
| ZDHHC3 | -0.49719795 | 0.708481482 | 1.20E-14 | 2.50E-13 | 1632.300689 | 1156.789018 | ENSG00000163812 |
| ASMT | -0.497245048 | 0.708458353 | 0.465389338 | 0.634879126 | 13.32938109 | 9.416944569 | ENSG00000196433 |
| NA | -0.497261981 | 0.708450038 | 0.509916406 | 0.674379942 | 11.20426887 | 7.997525833 | ENSG00000256542 |
| LGI2 | -0.497570917 | 0.708298348 | 0.002992706 | 0.011176763 | 237.0653179 | 168.1944024 | ENSG00000153012 |
| GAL3ST3 | -0.497696912 | 0.708236493 | 0.000744842 | 0.003258959 | 294.4647849 | 208.7553218 | ENSG00000175229 |
| CKAP4 | -0.497756536 | 0.708207223 | 2.77E-16 | 6.83E-15 | 2709.325804 | 1919.07088 | ENSG00000136026 |
| RND2 | -0.498006506 | 0.708084526 | 2.87E-18 | 8.30E-17 | 3361.599387 | 2380.558961 | ENSG00000108830 |
| BRSK2 | -0.498324213 | 0.70792861 | 1.54E-08 | 1.60E-07 | 916.8016978 | 649.1905956 | ENSG00000174672 |
| LINC01456 | -0.498325167 | 0.707928142 | 0.482597063 | 0.650578613 | 11.18002663 | 7.887828151 | ENSG00000225882 |
| LINC00665 | -0.498331367 | 0.7079251 | 4.84E-14 | 9.42E-13 | 1936.741001 | 1370.950218 | ENSG00000232677 |
| SNRPD3 | -0.498418728 | 0.707882233 | 6.06E-08 | 5.74E-07 | 764.4826263 | 541.3317433 | ENSG00000100028 |
| TMCC3 | -0.498462987 | 0.707860518 | 0.067419622 | 0.151460572 | 75.35176595 | 53.26542105 | ENSG00000057704 |
| EIF4EBP1 | -0.498576737 | 0.707804708 | 5.17E-16 | 1.24E-14 | 3070.960445 | 2173.545295 | ENSG00000187840 |
| TMEM147 | -0.498604651 | 0.707791013 | 4.05E-12 | 6.44E-11 | 1292.850285 | 915.2256821 | ENSG00000105677 |
| NA | -0.498633336 | 0.70777694 | 0.640373341 | 0.775044991 | 4.615696446 | 3.277095161 | ENSG00000213755 |
| YIPF1 | -0.498673677 | 0.70775715 | 3.20E-06 | 2.29E-05 | 502.5652442 | 355.712366 | ENSG00000058799 |
| NOMO3 | -0.499081337 | 0.707557188 | 0.02129149 | 0.059181756 | 126.6398325 | 89.64847954 | ENSG00000103226 |
| ARTN | -0.499135872 | 0.707530442 | 0.001474478 | 0.006035255 | 226.0076155 | 159.9551635 | ENSG00000117407 |
| TAF11 | -0.499163268 | 0.707517007 | 3.62E-06 | 2.58E-05 | 483.7723922 | 342.2466439 | ENSG00000064995 |
| NA | -0.499234528 | 0.707482061 | 0.68294937 | 0.806684809 | 3.687573563 | 2.595755268 | ENSG00000241478 |
| ANTXR1 | -0.499329037 | 0.707435716 | 8.94E-12 | 1.38E-10 | 1185.662886 | 839.1509806 | ENSG00000169604 |
| NOX1 | -0.499336002 | 0.707432301 | 0.566949689 | 0.720424369 | 6.91517196 | 4.905156391 | ENSG00000007952 |
| ZNF358 | -0.499884242 | 0.70716352 | 0.003800329 | 0.013819442 | 197.3288954 | 139.7467757 | ENSG00000198816 |
| NA | -0.500227809 | 0.706995134 | 0.665288606 | 0.793991741 | 3.720388677 | 2.616727967 | ENSG00000235272 |
| FBXL19 | -0.500483384 | 0.7068699 | 1.71E-15 | 3.87E-14 | 3407.723802 | 2409.280718 | ENSG00000099364 |
| TARBP2 | -0.500988908 | 0.706622255 | 2.23E-11 | 3.28E-10 | 1405.587374 | 993.2236801 | ENSG00000139546 |
| WNT4 | -0.501190582 | 0.706523483 | 0.343590093 | 0.516335318 | 19.44836634 | 13.69345933 | ENSG00000162552 |
| MARCKS | -0.50149908 | 0.70637242 | 7.23E-17 | 1.88E-15 | 3024.582535 | 2137.328811 | ENSG00000277443 |
| LMAN2L | -0.501582357 | 0.706331647 | 8.66E-08 | 8.04E-07 | 698.9552903 | 493.577518 | ENSG00000114988 |
| NA | -0.501926146 | 0.706163351 | 0.40843072 | 0.580707327 | 14.85065462 | 10.52982996 | ENSG00000273796 |
| SNORD3A | -0.501997517 | 0.706128418 | 0.354929525 | 0.528150684 | 17.59961741 | 12.43349247 | ENSG00000263934 |
| RNF40 | -0.502283269 | 0.70598857 | 5.23E-10 | 6.59E-09 | 1863.283942 | 1315.078101 | ENSG00000103549 |
| CACNG4 | -0.50246564 | 0.705899331 | 0.009384391 | 0.029814607 | 137.3303354 | 96.97945903 | ENSG00000075461 |
| NA | -0.502650203 | 0.705809032 | 0.244944943 | 0.408086441 | 31.28091631 | 22.05907953 | ENSG00000277103 |
| JRK | -0.502783967 | 0.705743594 | 2.35E-09 | 2.72E-08 | 983.2593953 | 693.4598889 | ENSG00000234616 |
| H1-3 | -0.502828459 | 0.70572183 | 0.671124406 | 0.798289611 | 4.70855998 | 3.331944001 | ENSG00000124575 |
| TKFC | -0.503000156 | 0.705637845 | 1.28E-07 | 1.16E-06 | 651.6483815 | 459.7367109 | ENSG00000149476 |
| PRPF38A | -0.503625251 | 0.705332171 | 4.09E-15 | 8.91E-14 | 1647.818734 | 1162.237617 | ENSG00000134748 |
| KPTN | -0.503826341 | 0.705233865 | 0.000408892 | 0.001920609 | 293.0383268 | 206.6021375 | ENSG00000118162 |
| KALRN | -0.503897376 | 0.705199142 | 0.00278161 | 0.010520208 | 234.1390377 | 165.2219505 | ENSG00000160145 |
| PHF5A | -0.50415223 | 0.705074578 | 3.89E-15 | 8.50E-14 | 1605.762512 | 1132.084471 | ENSG00000100410 |
| ATP6V0C | -0.504212328 | 0.705045208 | 0.633084102 | 0.769822396 | 4.670163061 | 3.300218434 | ENSG00000185883 |

| | | | | | | | |
|---|---|---|---|---|---|---|---|
| CYGB | -0.504274185 | 0.705014979 | 0.096161485 | 0.20095456 | 59.90620893 | 42.14056513 | ENSG00000161544 |
| RAVER2 | -0.504421393 | 0.704943045 | 1.07E-08 | 1.14E-07 | 830.8490119 | 585.7779173 | ENSG00000162437 |
| MRPS15 | -0.504472177 | 0.704918231 | 1.29E-15 | 2.97E-14 | 1978.817298 | 1394.195866 | ENSG00000116898 |
| LIG1 | -0.504695423 | 0.704809159 | 3.58E-14 | 7.11E-13 | 2232.348948 | 1573.169354 | ENSG00000105486 |
| SDCBP2 | -0.504710558 | 0.704801765 | 0.037444194 | 0.094364628 | 87.9704903 | 61.93595705 | ENSG00000125775 |
| TMEM222 | -0.505108709 | 0.704607283 | 1.20E-05 | 7.75E-05 | 491.8796347 | 346.6471168 | ENSG00000186501 |
| DCLK3 | -0.505490681 | 0.704420753 | 0.067429192 | 0.151464432 | 71.29341397 | 50.1730516 | ENSG00000163673 |
| SCAMP5 | -0.505512947 | 0.704409882 | 2.83E-15 | 6.28E-14 | 2022.30212 | 1424.457631 | ENSG00000198794 |
| MAGEH1 | -0.505514487 | 0.70440913 | 0.000405285 | 0.001906041 | 268.7385761 | 189.3535707 | ENSG00000187601 |
| CDC6 | -0.505544245 | 0.7043946 | 2.06E-20 | 7.24E-19 | 3098.457595 | 2182.660001 | ENSG00000094804 |
| TTC7A | -0.505800238 | 0.704269623 | 3.07E-06 | 2.21E-05 | 570.8667988 | 401.8592424 | ENSG00000068724 |
| NA | -0.505985883 | 0.704179004 | 0.409031431 | 0.581250624 | 17.55930547 | 12.27754824 | ENSG00000255201 |
| NA | -0.506216496 | 0.704066451 | 0.531861999 | 0.692691725 | 7.941064461 | 5.620372426 | ENSG00000225345 |
| YPEL1 | -0.506338824 | 0.704006754 | 0.012891479 | 0.039000813 | 145.7189721 | 102.7028048 | ENSG00000100027 |
| LINC01128 | -0.506446406 | 0.703954259 | 0.023502139 | 0.06426361 | 111.3366996 | 78.46741771 | ENSG00000228794 |
| UBOX5 | -0.506477231 | 0.703939218 | 0.018989624 | 0.053784882 | 140.6474939 | 99.16894264 | ENSG00000185019 |
| SEC11C | -0.506530227 | 0.70391336 | 1.23E-07 | 1.12E-06 | 726.3216812 | 511.5318745 | ENSG00000166562 |
| NA | -0.506607628 | 0.703875596 | 0.422197739 | 0.593804067 | 12.99508457 | 9.131123463 | ENSG00000271119 |
| NA | -0.506668829 | 0.703845737 | 0.522659213 | 0.685379572 | 8.417785045 | 5.912645253 | ENSG00000249492 |
| IGSF22 | -0.506739339 | 0.703811339 | 0.548070506 | 0.705654951 | 6.996871883 | 4.930430238 | ENSG00000179057 |
| ZER1 | -0.507319439 | 0.703528396 | 1.79E-07 | 1.59E-06 | 694.361596 | 488.247292 | ENSG00000160445 |
| CARMIL3 | -0.507396855 | 0.703490646 | 0.000193948 | 0.000980722 | 343.5369157 | 241.7152819 | ENSG00000186648 |
| ADAT3 | -0.507537383 | 0.703422124 | 0.009527038 | 0.030208101 | 146.073844 | 102.8135991 | ENSG00000213638 |
| OVOS2 | -0.507549103 | 0.70341641 | 0.0683452 | 0.153040888 | 80.09246533 | 56.13651712 | ENSG00000177359 |
| LOC145694 | -0.507572424 | 0.703405039 | 0.546967336 | 0.70479907 | 8.015067397 | 5.666618972 | ENSG00000259426 |
| RNF182 | -0.507899693 | 0.703245493 | 0.041440598 | 0.102569998 | 106.6258243 | 75.09406135 | ENSG00000180537 |
| ADAMTS3 | -0.507985813 | 0.703203515 | 0.030018664 | 0.078584618 | 105.3329051 | 74.13658715 | ENSG00000156140 |
| HDAC11 | -0.508064931 | 0.703164952 | 0.000181493 | 0.000922577 | 365.948957 | 257.2234062 | ENSG00000163517 |
| NA | -0.508080164 | 0.703157527 | 0.655781061 | 0.786539656 | 4.672953964 | 3.287848029 | ENSG00000241889 |
| KIF5C | -0.508222541 | 0.703088138 | 2.34E-14 | 4.75E-13 | 2220.486599 | 1561.549112 | ENSG00000168280 |
| NAB2 | -0.50834131 | 0.703030259 | 3.44E-06 | 2.46E-05 | 695.3674525 | 488.8799685 | ENSG00000166886 |
| NA | -0.508398171 | 0.703002551 | 0.254538879 | 0.419386609 | 35.41374679 | 24.80621742 | ENSG00000273162 |
| UBE2R2 | -0.508538021 | 0.702934407 | 4.99E-17 | 1.31E-15 | 2280.506247 | 1603.792954 | ENSG00000107341 |
| ITPKA | -0.508756216 | 0.702828102 | 0.420008626 | 0.592063311 | 13.17243893 | 9.248890402 | ENSG00000137825 |
| TEKT2 | -0.508782035 | 0.702815525 | 0.304215221 | 0.474960815 | 20.97175505 | 14.74289359 | ENSG00000092850 |
| DUSP7 | -0.509255304 | 0.702585007 | 5.59E-08 | 5.33E-07 | 679.0577873 | 477.3711097 | ENSG00000164086 |
| KANSL3 | -0.50936781 | 0.70253022 | 1.36E-11 | 2.05E-10 | 1222.843719 | 858.655772 | ENSG00000114982 |
| ATP6AP1 | -0.509567565 | 0.702432954 | 1.27E-15 | 2.94E-14 | 1663.260951 | 1168.526147 | ENSG00000071553 |
| WDR35 | -0.509637464 | 0.702398922 | 1.01E-08 | 1.07E-07 | 770.0896469 | 540.6883082 | ENSG00000118965 |
| NRP1 | -0.509767653 | 0.70233554 | 0.1613362 | 0.298672048 | 43.46075342 | 30.51048064 | ENSG00000099250 |
| NA | -0.509932814 | 0.702255141 | 0.666807232 | 0.795262956 | 4.763026595 | 3.350766127 | ENSG00000235314 |
| MYH6 | -0.509936285 | 0.702253451 | 0.566501599 | 0.720251917 | 6.98359309 | 4.903005818 | ENSG00000197616 |
| NA | -0.50997628 | 0.702233983 | 0.663089619 | 0.792102411 | 3.720388677 | 2.589303546 | ENSG00000287967 |
| PRR5L | -0.509979497 | 0.702232418 | 0.016424382 | 0.047685994 | 134.3933671 | 94.24873224 | ENSG00000135362 |
| L1TD1 | -0.510075121 | 0.702185874 | 3.35E-36 | 2.97E-34 | 15127.38983 | 10623.18943 | ENSG00000240563 |
| NA | -0.510095104 | 0.702176148 | 0.622158477 | 0.762199274 | 4.672953964 | 3.281396308 | ENSG00000260669 |
| IGSF11 | -0.510373352 | 0.702040735 | 0.291964405 | 0.461700578 | 23.79404505 | 16.66161012 | ENSG00000144847 |
| PRRG2 | -0.510389613 | 0.702032822 | 0.132598284 | 0.256616523 | 49.94203224 | 34.98032342 | ENSG00000126460 |
| MAPK10 | -0.510419063 | 0.702018491 | 0.176823225 | 0.320441224 | 42.27118567 | 29.60193659 | ENSG00000109339 |
| CCDC96 | -0.51053492 | 0.701962117 | 0.615280344 | 0.757748539 | 6.014282385 | 4.200693225 | ENSG00000173013 |
| PLA2R1 | -0.510792721 | 0.701836692 | 0.074970672 | 0.164834651 | 70.93611451 | 49.69741013 | ENSG00000153246 |
| EIF3CL | -0.511321806 | 0.701579352 | 0.000508744 | 0.002328947 | 361.0551327 | 253.1625613 | ENSG00000205609 |
| CHAF1B | -0.511386885 | 0.701547705 | 2.59E-08 | 2.58E-07 | 821.8359738 | 576.1870241 | ENSG00000159259 |
| NA | -0.5114933 | 0.70149596 | 0.615970318 | 0.758212125 | 5.660449546 | 3.990160622 | ENSG00000258702 |
| CRYM | -0.511542043 | 0.701472259 | 0.002151583 | 0.008415691 | 272.9358856 | 191.3440618 | ENSG00000103316 |
| NA | -0.51166077 | 0.701414534 | 0.685219414 | 0.808079699 | 3.695946272 | 2.570481421 | ENSG00000288095 |
| CSAD | -0.511774659 | 0.701359165 | 0.088407771 | 0.188055389 | 62.98735306 | 44.0522969 | ENSG00000139631 |
| ZNF329 | -0.512015165 | 0.701242254 | 0.000577504 | 0.002600522 | 287.480553 | 201.5939956 | ENSG00000181894 |
| THBS4 | -0.512032089 | 0.701234028 | 1.02E-05 | 6.69E-05 | 423.8560077 | 297.0064456 | ENSG00000113296 |
| VPS33B | -0.512304745 | 0.701101514 | 0.268581788 | 0.434766942 | 25.28617025 | 17.65834611 | ENSG00000184056 |
| BCL2 | -0.512567549 | 0.700973812 | 0.043258689 | 0.106252021 | 83.97988373 | 58.98043714 | ENSG00000171791 |
| ZW10 | -0.51257192 | 0.700971688 | 1.72E-07 | 1.53E-06 | 695.0344435 | 487.1427242 | ENSG00000086827 |
| TSPEAR-AS1 | -0.512669023 | 0.700924509 | 0.072488527 | 0.160437836 | 71.3618351 | 50.01951841 | ENSG00000235890 |
| NDUFAF7 | -0.512678403 | 0.700919952 | 3.27E-10 | 4.24E-09 | 997.8884656 | 699.1740688 | ENSG00000003509 |
| CELF5 | -0.512928894 | 0.700798264 | 0.000177751 | 0.000905465 | 293.6218286 | 205.7269487 | ENSG00000161082 |
| ZNF66 | -0.513124317 | 0.700703342 | 0.013464433 | 0.040442016 | 121.9570664 | 85.39374309 | ENSG00000160229 |
| CDC37 | -0.513264514 | 0.700635253 | 2.98E-17 | 8.03E-16 | 5244.236132 | 3674.218317 | ENSG00000105401 |
| BMERB1 | -0.513377314 | 0.700580475 | 6.53E-05 | 0.000367714 | 336.7202643 | 235.9112804 | ENSG00000166780 |
| BARX1 | -0.514264617 | 0.700149728 | 0.125323393 | 0.246092586 | 45.47625669 | 31.8209952 | ENSG00000131668 |
| THADA | -0.514405728 | 0.700081249 | 1.11E-07 | 1.02E-06 | 730.1270732 | 511.3753851 | ENSG00000115970 |
| BOD1 | -0.514533987 | 0.700019013 | 1.53E-21 | 5.78E-20 | 3421.766124 | 2394.395292 | ENSG00000145919 |
| DNAJC3-DT | -0.514796285 | 0.699891753 | 0.105730948 | 0.216010674 | 54.94070985 | 38.52252745 | ENSG00000247400 |
| ASPHD2 | -0.514929405 | 0.699827176 | 0.000929503 | 0.003990678 | 234.5470489 | 164.1932285 | ENSG00000128203 |
| PSENEN | -0.515049493 | 0.699768926 | 0.191136876 | 0.339836081 | 34.14698488 | 23.83207166 | ENSG00000205155 |
| ZNF610 | -0.515227085 | 0.699682792 | 0.100388485 | 0.207732713 | 69.72102832 | 48.74798676 | ENSG00000167554 |
| CDK20 | -0.515695511 | 0.69945565 | 0.031757854 | 0.08229994 | 92.63997764 | 64.82854676 | ENSG00000156345 |
| TACC2 | -0.515731723 | 0.699438093 | 1.07E-10 | 1.47E-09 | 1132.767979 | 792.1593508 | ENSG00000138162 |

| | | | | | | | |
|---|---|---|---|---|---|---|---|
| RPL13AP20 | -0.515776725 | 0.699416276 | 0.386879051 | 0.560279267 | 15.59480219 | 10.87910221 | ENSG00000234498 |
| CCDC92 | -0.516444375 | 0.699092676 | 0.258601166 | 0.423810644 | 27.15848571 | 18.93068338 | ENSG00000119242 |
| GNB3 | -0.516504809 | 0.699063392 | 0.014283464 | 0.04252196 | 135.4009625 | 94.43373053 | ENSG00000111664 |
| JAM3 | -0.516798059 | 0.698921311 | 1.16E-13 | 2.18E-12 | 1403.604434 | 981.1585304 | ENSG00000166086 |
| PASK | -0.517148426 | 0.698751594 | 1.38E-15 | 3.16E-14 | 1729.433749 | 1207.869311 | ENSG00000115687 |
| SHE | -0.517501001 | 0.69858085 | 0.215553042 | 0.372534826 | 35.87966718 | 25.16113577 | ENSG00000169291 |
| INSYN1 | -0.517541368 | 0.698561303 | 0.001313386 | 0.005449806 | 222.4534175 | 155.4541467 | ENSG00000205363 |
| CRNDE | -0.517555692 | 0.698554367 | 0.035718615 | 0.090640331 | 96.6385566 | 67.50954989 | ENSG00000245694 |
| ZMAT2 | -0.517579534 | 0.698542823 | 2.44E-12 | 4.00E-11 | 1678.120465 | 1171.675558 | ENSG00000146007 |
| AAMP | -0.517751857 | 0.698459391 | 4.53E-17 | 1.20E-15 | 3955.169461 | 2762.10568 | ENSG00000127837 |
| NA | -0.517861504 | 0.698406309 | 0.40550953 | 0.577746259 | 13.99374377 | 9.743093538 | ENSG00000276564 |
| ALKBH6 | -0.518156109 | 0.698263705 | 0.244447655 | 0.407504462 | 36.33003034 | 25.2305177 | ENSG00000239382 |
| CHRM4 | -0.518260175 | 0.698213339 | 0.329530795 | 0.501853522 | 18.78400367 | 13.08148926 | ENSG00000180720 |
| RPIA | -0.51886759 | 0.697919434 | 2.51E-07 | 2.17E-06 | 598.4119325 | 417.768501 | ENSG00000153574 |
| STS | -0.518974985 | 0.697867482 | 0.0210715 | 0.058648147 | 119.2117335 | 83.28143809 | ENSG00000101846 |
| SMIM7 | -0.519208074 | 0.697754741 | 0.00099127 | 0.00423066 | 230.9036543 | 161.0293386 | ENSG00000214046 |
| CCDC86 | -0.519598131 | 0.697566116 | 3.88E-12 | 6.19E-11 | 1535.355195 | 1070.985048 | ENSG00000110104 |
| SMAP2 | -0.519880813 | 0.697429448 | 0.004304013 | 0.015407072 | 173.8762802 | 121.2965985 | ENSG00000084070 |
| LOC102724159 | -0.520026015 | 0.697359258 | 4.42E-09 | 4.95E-08 | 941.8628426 | 656.7016965 | ENSG00000275464 |
| ZNF584 | -0.520512144 | 0.697124316 | 0.035705431 | 0.090618795 | 99.84335222 | 69.75923746 | ENSG00000171574 |
| ELOF1 | -0.52066512 | 0.697050401 | 3.61E-10 | 4.66E-09 | 1477.866719 | 1030.01816 | ENSG00000130165 |
| PSME3 | -0.520768403 | 0.6970005 | 4.05E-26 | 2.10E-24 | 6313.428082 | 4400.000577 | ENSG00000131467 |
| LOC101927124 | -0.521103415 | 0.696838667 | 0.589561775 | 0.738857675 | 6.197218551 | 4.293186317 | ENSG00000250365 |
| CCND3 | -0.521143439 | 0.696819335 | 1.06E-08 | 1.12E-07 | 886.0162633 | 617.3768577 | ENSG00000112576 |
| ZNF324B | -0.521392746 | 0.696698931 | 0.086455998 | 0.184739617 | 70.66341802 | 49.10909637 | ENSG00000249471 |
| NA | -0.52144678 | 0.696672837 | 0.650331833 | 0.782531898 | 4.730211482 | 3.287848029 | ENSG00000260518 |
| GLDC | -0.521534587 | 0.696630437 | 2.36E-25 | 1.18E-23 | 4392.912576 | 3060.043623 | ENSG00000178445 |
| TEDC2 | -0.522034193 | 0.696389235 | 2.05E-09 | 2.40E-08 | 1055.781264 | 735.3796895 | ENSG00000162062 |
| TTC34 | -0.522219457 | 0.696299814 | 0.611513702 | 0.755037815 | 4.697396369 | 3.256122461 | ENSG00000215912 |
| LRRC42 | -0.522235948 | 0.696291855 | 1.58E-12 | 2.64E-11 | 1166.071145 | 812.0789371 | ENSG00000116212 |
| NPEPPS | -0.522434402 | 0.696196081 | 4.39E-15 | 9.50E-14 | 1713.328523 | 1193.195206 | ENSG00000141279 |
| CHRNA3 | -0.522490133 | 0.696169188 | 0.019585315 | 0.055253045 | 113.0183819 | 78.681991 | ENSG00000080644 |
| ARMH1 | -0.522554779 | 0.696137994 | 0.623583797 | 0.763008988 | 4.670163061 | 3.235149762 | ENSG00000198520 |
| NA | -0.522572655 | 0.696129368 | 0.522851193 | 0.685570353 | 8.423366851 | 5.820152161 | ENSG00000260144 |
| MMEL1 | -0.523163064 | 0.695844542 | 0.533685496 | 0.694176178 | 8.486206175 | 5.868549281 | ENSG00000142606 |
| UBAC1 | -0.523204603 | 0.695824507 | 5.01E-17 | 1.32E-15 | 1767.532594 | 1229.916548 | ENSG00000130560 |
| NA | -0.523535252 | 0.69566505 | 0.459712791 | 0.629539253 | 10.45629125 | 7.27155693 | ENSG00000273188 |
| PNMT | -0.523565662 | 0.695650387 | 0.04086142 | 0.101396661 | 91.42365215 | 63.64978708 | ENSG00000141744 |
| WARS1 | -0.52392252 | 0.695478336 | 1.11E-17 | 3.06E-16 | 2903.72849 | 2018.467305 | ENSG00000140105 |
| RPE | -0.524177797 | 0.695355285 | 8.04E-12 | 1.25E-10 | 1099.608139 | 764.7382408 | ENSG00000197713 |
| LTV1 | -0.524311139 | 0.69529102 | 0.000254886 | 0.001256583 | 287.299495 | 199.9909478 | ENSG00000135521 |
| CDH4 | -0.524538951 | 0.695181237 | 1.24E-05 | 7.98E-05 | 480.7656017 | 334.350486 | ENSG00000179242 |
| ZNF790 | -0.525003079 | 0.694957627 | 0.113159745 | 0.227795538 | 47.20769968 | 32.75911424 | ENSG00000197863 |
| CARHSP1 | -0.525109123 | 0.694906546 | 1.28E-14 | 2.67E-13 | 1592.487522 | 1106.242268 | ENSG00000153048 |
| EVA1A | -0.525113162 | 0.694904601 | 0.153249242 | 0.28752491 | 44.24230989 | 30.73553422 | ENSG00000115363 |
| ANKRD44 | -0.525180023 | 0.694872397 | 0.323572159 | 0.495163455 | 19.05055478 | 13.29174929 | ENSG00000065413 |
| CDH23 | -0.525365768 | 0.694782938 | 0.000545843 | 0.002479364 | 246.9680436 | 171.4806634 | ENSG00000107736 |
| DCTN5 | -0.525469426 | 0.69473302 | 8.94E-17 | 2.30E-15 | 1855.97859 | 1289.52453 | ENSG00000166847 |
| FTSJ1 | -0.525477834 | 0.694728971 | 8.27E-10 | 1.02E-08 | 1018.590642 | 707.3861801 | ENSG00000068438 |
| C1QTNF4 | -0.525575515 | 0.694681935 | 0.010742477 | 0.033484105 | 145.8383563 | 101.4444676 | ENSG00000172247 |
| NA | -0.525827461 | 0.694560629 | 0.639922609 | 0.774742503 | 4.259072704 | 2.984822334 | ENSG00000271275 |
| UBR7 | -0.526058202 | 0.694449551 | 2.04E-08 | 2.07E-07 | 722.8259162 | 502.0273015 | ENSG00000012963 |
| NOL8 | -0.526341409 | 0.694313242 | 3.66E-06 | 2.60E-05 | 584.7604817 | 406.1037101 | ENSG00000198000 |
| FBRS | -0.526819335 | 0.694083272 | 3.86E-06 | 2.74E-05 | 516.2811483 | 358.5186539 | ENSG00000156860 |
| MATR3 | -0.527090294 | 0.693952926 | 0.313901419 | 0.485097514 | 22.62572849 | 15.69606665 | ENSG00000015479 |
| H1-2 | -0.527715545 | 0.693652238 | 0.006382546 | 0.021638961 | 155.3832701 | 107.9908492 | ENSG00000187837 |
| LINC02724 | -0.52786663 | 0.6935796 | 0.607172976 | 0.751313271 | 5.587122332 | 3.868092536 | ENSG00000181908 |
| HSD17B6 | -0.528115283 | 0.693460069 | 0.099182424 | 0.205933499 | 54.76020117 | 37.99874932 | ENSG00000025423 |
| RAB34 | -0.528118269 | 0.693458634 | 8.65E-23 | 3.60E-21 | 3443.106345 | 2387.42707 | ENSG00000109113 |
| NA | -0.528176036 | 0.693430868 | 0.448947446 | 0.619192478 | 11.32715662 | 7.8501839 | ENSG00000236675 |
| SYT11 | -0.528577561 | 0.693237902 | 0.000439482 | 0.002048334 | 287.7900579 | 199.6440987 | ENSG00000132718 |
| NA | -0.528948848 | 0.693059516 | 0.645237262 | 0.778635808 | 4.368005934 | 3.056342726 | ENSG00000229222 |
| COX16 | -0.529096295 | 0.692988687 | 0.22904995 | 0.388858804 | 30.72384725 | 21.27019253 | ENSG00000133983 |
| TOR2A | -0.529315531 | 0.692883386 | 2.06E-05 | 0.000128409 | 369.0219669 | 255.9365479 | ENSG00000160404 |
| SMPD4 | -0.529945158 | 0.692581061 | 6.51E-16 | 1.55E-14 | 2875.418474 | 1991.23209 | ENSG00000136699 |
| PAK6 | -0.529949062 | 0.692579187 | 0.000971169 | 0.004153139 | 211.1194398 | 146.179431 | ENSG00000137843 |
| NA | -0.530179955 | 0.692468353 | 0.409710342 | 0.581926242 | 17.76648388 | 12.22915112 | ENSG00000271032 |
| CYFIP1 | -0.530328805 | 0.692396912 | 4.59E-21 | 1.69E-19 | 2374.102079 | 1643.683565 | ENSG00000273749 |
| GNG5 | -0.530426926 | 0.692349822 | 0.015276762 | 0.044944946 | 123.152216 | 85.24640115 | ENSG00000174021 |
| PSMD1 | -0.530627903 | 0.69225338 | 9.36E-32 | 6.56E-30 | 5668.888369 | 3924.209276 | ENSG00000173692 |
| NA | -0.530779513 | 0.692180636 | 0.458498798 | 0.628367456 | 10.47515185 | 7.229611531 | ENSG00000226525 |
| MAP3K13 | -0.531106274 | 0.692023879 | 0.017692745 | 0.050767626 | 124.9543954 | 86.29610799 | ENSG00000073803 |
| CCDC34 | -0.531446745 | 0.691860583 | 8.32E-07 | 6.63E-06 | 592.8368496 | 410.3275144 | ENSG00000109881 |
| NCLN | -0.531671058 | 0.69175302 | 4.61E-13 | 8.14E-12 | 4440.003292 | 3071.406388 | ENSG00000125912 |
| ERI3 | -0.531794982 | 0.691693603 | 5.87E-16 | 1.41E-14 | 1619.659037 | 1120.623514 | ENSG00000117419 |
| NT5C3B | -0.531844289 | 0.691669963 | 4.81E-14 | 9.38E-13 | 1519.767013 | 1051.381318 | ENSG00000141698 |
| STAMBPL1 | -0.531848961 | 0.691667723 | 0.007227485 | 0.02400162 | 135.2869231 | 93.57599464 | ENSG00000138134 |

| | | | | | | | |
|---|---|---|---|---|---|---|---|
| ORMDL3 | -0.532195284 | 0.691501706 | 5.12E-13 | 8.98E-12 | 1606.525795 | 1110.624247 | ENSG00000172057 |
| NA | -0.53220355 | 0.691497744 | 0.000381276 | 0.001807245 | 239.727987 | 165.6903588 | ENSG00000259203 |
| NA | -0.532612675 | 0.691301674 | 0.611358088 | 0.754893995 | 5.786128194 | 4.00091349 | ENSG00000275329 |
| NA | -0.532890373 | 0.691168622 | 0.36875499 | 0.542529936 | 16.28273139 | 11.32301811 | ENSG00000229692 |
| NA | -0.532924341 | 0.691152348 | 0.459580205 | 0.629447053 | 10.00401308 | 6.930886983 | ENSG00000213763 |
| NA | -0.533318127 | 0.690963723 | 0.373936982 | 0.547214396 | 19.30681816 | 13.23879056 | ENSG00000228944 |
| CDKN2AIPNL | -0.533333196 | 0.690956506 | 5.47E-13 | 9.58E-12 | 1303.937461 | 901.0127458 | ENSG00000237190 |
| NA | -0.533361186 | 0.690943101 | 0.533265335 | 0.693924025 | 7.698955755 | 5.349072298 | ENSG00000225137 |
| SCNN1A | -0.533373137 | 0.690937377 | 2.27E-14 | 4.63E-13 | 1690.939974 | 1168.767872 | ENSG00000111319 |
| PHF3 | -0.533473271 | 0.690889422 | 2.57E-15 | 5.73E-14 | 1693.827854 | 1170.26118 | ENSG00000118482 |
| BAK1 | -0.53355747 | 0.690849101 | 4.04E-12 | 6.43E-11 | 1180.53838 | 815.2345094 | ENSG00000030110 |
| ATXN7 | -0.533561168 | 0.690847331 | 0.516459637 | 0.680447331 | 8.545378715 | 5.9379191 | ENSG00000163635 |
| GRM8 | -0.533826929 | 0.69072008 | 0.102221596 | 0.210611643 | 51.49420585 | 35.51458184 | ENSG00000179603 |
| GPAT3 | -0.534253779 | 0.690515747 | 0.194578755 | 0.344716573 | 34.95222002 | 24.24667304 | ENSG00000138678 |
| ZNF793 | -0.534385713 | 0.690452603 | 0.006326882 | 0.021476665 | 166.0303946 | 114.5122415 | ENSG00000188227 |
| GK | -0.534451226 | 0.69042125 | 0.001123 | 0.004730085 | 213.0015674 | 147.1543886 | ENSG00000198814 |
| NA | -0.534924239 | 0.690194921 | 0.560759509 | 0.715829254 | 8.026231008 | 5.521427613 | ENSG00000181227 |
| STOX1 | -0.535222111 | 0.690052431 | 0.044992317 | 0.109700049 | 76.27813707 | 52.75669693 | ENSG00000165730 |
| MYB | -0.535320654 | 0.690005299 | 0.033858 | 0.086625289 | 84.99807925 | 58.63788919 | ENSG00000118513 |
| FRG1-DT | -0.535763315 | 0.689793617 | 0.345196057 | 0.517592458 | 18.71346736 | 12.95538049 | ENSG00000245685 |
| CNTN3 | -0.536190295 | 0.689589496 | 0.585569709 | 0.735192012 | 5.64437985 | 3.876694831 | ENSG00000113805 |
| KCNJ11 | -0.536247975 | 0.689561927 | 0.000573473 | 0.002586595 | 242.4572133 | 167.0812687 | ENSG00000187486 |
| SAP30L | -0.536332834 | 0.689521368 | 2.78E-07 | 2.39E-06 | 551.9571406 | 380.5874018 | ENSG00000164576 |
| NA | -0.536450254 | 0.689465251 | 0.251389675 | 0.415633636 | 24.32224119 | 16.80250033 | ENSG00000243547 |
| NA | -0.536765879 | 0.689314429 | 0.460562206 | 0.630344485 | 11.0305813 | 7.627280892 | ENSG00000286868 |
| NA | -0.536918875 | 0.689241333 | 0.32025933 | 0.491635992 | 20.79160979 | 14.33930555 | ENSG00000268262 |
| PRRT3 | -0.536951764 | 0.68922562 | 0.001629724 | 0.006593751 | 220.0773114 | 151.9256396 | ENSG00000163704 |
| NA | -0.537307086 | 0.689055891 | 0.347207777 | 0.519949172 | 16.62540062 | 11.45934671 | ENSG00000260401 |
| NA | -0.537654741 | 0.688889865 | 0.688271906 | 0.810627756 | 3.780437098 | 2.572631995 | ENSG00000253476 |
| PTPRO | -0.537739804 | 0.688849249 | 0.249583411 | 0.413670726 | 28.33920519 | 19.45446151 | ENSG00000151490 |
| ANKRD42 | -0.537881877 | 0.688781416 | 0.015833475 | 0.046320885 | 110.0978467 | 75.86871837 | ENSG00000137494 |
| KIF26A | -0.537942896 | 0.688752284 | 7.92E-05 | 0.000437989 | 4104.889243 | 2827.253483 | ENSG00000066735 |
| ENOX2 | -0.538022778 | 0.688714149 | 2.12E-07 | 1.86E-06 | 541.9789342 | 373.1069298 | ENSG00000165675 |
| NA | -0.538358335 | 0.68855398 | 0.401719952 | 0.574170526 | 12.34746731 | 8.485277248 | ENSG00000272716 |
| PAQR5 | -0.538781856 | 0.688351876 | 0.000693966 | 0.003059254 | 221.120662 | 152.0498703 | ENSG00000137819 |
| ACTR3C | -0.53893555 | 0.688278548 | 0.485944869 | 0.653385722 | 9.084262902 | 6.240944795 | ENSG00000106526 |
| FAM53B | -0.539077444 | 0.688210857 | 3.72E-08 | 3.64E-07 | 647.716347 | 445.6555536 | ENSG00000189319 |
| B3GAT2 | -0.539094218 | 0.688202855 | 0.248247433 | 0.411917319 | 27.06195539 | 18.66583497 | ENSG00000112309 |
| SIX5 | -0.539283313 | 0.688112658 | 0.002562764 | 0.009797518 | 173.1293787 | 118.9850468 | ENSG00000177045 |
| ZNF613 | -0.539297589 | 0.688105848 | 0.004340118 | 0.015516164 | 162.3353243 | 111.8172568 | ENSG00000176024 |
| CA12 | -0.53937098 | 0.688070845 | 0.015916006 | 0.046513509 | 121.6962604 | 83.72939468 | ENSG00000074410 |
| PSMA6 | -0.539441646 | 0.688037143 | 0.202883623 | 0.355673207 | 31.51512787 | 21.75175269 | ENSG00000100902 |
| NCOA7 | -0.539878698 | 0.68782874 | 0.001935559 | 0.007665543 | 188.3825508 | 129.5223946 | ENSG00000111912 |
| SLC35B1 | -0.539954783 | 0.687792465 | 2.39E-10 | 3.16E-09 | 1063.705908 | 731.3578154 | ENSG00000121073 |
| NDRG2 | -0.540024672 | 0.687759148 | 8.81E-06 | 5.87E-05 | 404.9548983 | 278.4338973 | ENSG00000165795 |
| NA | -0.540425629 | 0.68756803 | 0.557615061 | 0.713428546 | 8.0583704 | 5.51550893 | ENSG00000272087 |
| DONSON | -0.54043794 | 0.687562163 | 4.60E-07 | 3.82E-06 | 513.1055138 | 353.0464169 | ENSG00000159147 |
| CELF3 | -0.540807859 | 0.687385889 | 0.283865518 | 0.452453912 | 21.56211479 | 14.89023553 | ENSG00000159409 |
| STAG1 | -0.540916689 | 0.687334038 | 1.69E-07 | 1.50E-06 | 574.4551633 | 394.9603293 | ENSG00000118007 |
| SETBP1 | -0.540917405 | 0.687333697 | 0.027680666 | 0.073663641 | 93.59533383 | 64.30289063 | ENSG00000152217 |
| NA | -0.541004523 | 0.687292193 | 0.244774613 | 0.40790835 | 24.87420401 | 17.1619924 | ENSG00000287737 |
| WDFY3-AS2 | -0.541173071 | 0.687211902 | 0.086274679 | 0.184419219 | 56.03589936 | 38.48703377 | ENSG00000180769 |
| NA | -0.541185434 | 0.687206013 | 0.026641265 | 0.071330553 | 103.3015323 | 70.92053208 | ENSG00000235501 |
| RUFY4 | -0.541348675 | 0.68712826 | 0.533419696 | 0.693924025 | 6.681435963 | 4.587609718 | ENSG00000188282 |
| PGD | -0.541701808 | 0.68696009 | 5.55E-37 | 5.22E-35 | 10743.77204 | 7380.363062 | ENSG00000142657 |
| CCDC69 | -0.541777881 | 0.686923868 | 4.06E-05 | 0.000238004 | 316.6770325 | 217.5342106 | ENSG00000198624 |
| C9orf116 | -0.541873541 | 0.686878322 | 0.017918681 | 0.051293853 | 101.6451314 | 69.82593578 | ENSG00000160345 |
| PRRT4 | -0.541927371 | 0.686852694 | 0.167717454 | 0.307814424 | 42.17936128 | 29.0588033 | ENSG00000224940 |
| NA | -0.542271376 | 0.686688936 | 0.483356437 | 0.651192752 | 8.651521043 | 5.969644667 | ENSG00000234692 |
| UNC45A | -0.542345869 | 0.68665348 | 1.02E-24 | 4.92E-23 | 3277.123946 | 2250.270827 | ENSG00000140553 |
| ASF1A | -0.542488215 | 0.686585734 | 1.84E-06 | 1.38E-05 | 467.9343173 | 321.3060895 | ENSG00000111875 |
| NIPSNAP3B | -0.542556463 | 0.686553255 | 0.432358985 | 0.603432006 | 10.95936927 | 7.528336079 | ENSG00000165028 |
| PCDH7 | -0.542588555 | 0.686537983 | 0.014519116 | 0.043073944 | 117.6117511 | 80.81582016 | ENSG00000169851 |
| NA | -0.542883066 | 0.686397848 | 0.290894647 | 0.460650848 | 19.60809941 | 13.48937845 | ENSG00000240376 |
| CFAP43 | -0.543087799 | 0.686300448 | 0.510739925 | 0.675211341 | 7.650070946 | 5.25872978 | ENSG00000197748 |
| POLR2F | -0.543148466 | 0.686271588 | 2.36E-18 | 6.89E-17 | 2080.422695 | 1427.638737 | ENSG00000100142 |
| ZNF205 | -0.5434411 | 0.686132401 | 6.29E-05 | 0.00035511 | 359.0027231 | 246.5341365 | ENSG00000122386 |
| BEND3 | -0.544102222 | 0.685818049 | 3.62E-11 | 5.20E-10 | 1165.048981 | 799.2760238 | ENSG00000178409 |
| ATP6V1G1 | -0.544440521 | 0.68565725 | 4.92E-15 | 1.06E-13 | 2029.837891 | 1391.799345 | ENSG00000136888 |
| VPS29 | -0.544644234 | 0.685560439 | 3.47E-06 | 2.48E-05 | 407.8935814 | 279.8444411 | ENSG00000111237 |
| WFDC2 | -0.544681811 | 0.685542584 | 2.85E-06 | 2.06E-05 | 449.3638745 | 308.2243156 | ENSG00000101443 |
| RAB27A | -0.544794991 | 0.685488805 | 0.006431906 | 0.021775692 | 143.0047394 | 97.94742564 | ENSG00000069974 |
| C1orf52 | -0.544807792 | 0.685482722 | 7.61E-08 | 7.12E-07 | 619.0314063 | 424.0218405 | ENSG00000162642 |
| MINDY3 | -0.54490428 | 0.685436878 | 0.0004748 | 0.002188093 | 249.0064377 | 170.6898863 | ENSG00000148481 |
| SOCS2 | -0.545044881 | 0.685370081 | 6.32E-05 | 0.000356498 | 343.7571727 | 235.5362012 | ENSG00000120833 |
| AUH | -0.54531844 | 0.685240135 | 2.87E-06 | 2.08E-05 | 426.7310126 | 292.5511117 | ENSG00000148090 |
| AKAP17A | -0.545353331 | 0.685223563 | 1.43E-16 | 3.62E-15 | 2313.886157 | 1585.556109 | ENSG00000197976 |

| | | | | | | | |
|---|---|---|---|---|---|---|---|
| NA | -0.545358132 | 0.685221283 | 0.49067419 | 0.657196125 | 9.729764978 | 6.703682827 | ENSG00000272836 |
| NAALAD2 | -0.545421206 | 0.685191326 | 0.100510305 | 0.207924966 | 51.81502342 | 35.51619938 | ENSG00000077616 |
| AMD1 | -0.54553855 | 0.685135597 | 6.58E-15 | 1.41E-13 | 4129.266992 | 2829.920435 | ENSG00000123505 |
| BRINP1 | -0.545567641 | 0.685121782 | 2.16E-08 | 2.18E-07 | 929.1720559 | 636.8584774 | ENSG00000078725 |
| USF2 | -0.545820515 | 0.685001705 | 1.27E-24 | 6.06E-23 | 2745.338369 | 1880.785461 | ENSG00000105698 |
| TNFRSF19 | -0.546479259 | 0.684689001 | 0.475906258 | 0.644305399 | 9.620156026 | 6.615490883 | ENSG00000127863 |
| TUBD1 | -0.546617122 | 0.684623575 | 0.000313517 | 0.001515044 | 284.94758 | 195.114294 | ENSG00000108423 |
| PRAF2 | -0.546737761 | 0.684566329 | 0.193874269 | 0.343689444 | 31.18506172 | 21.41323332 | ENSG00000243279 |
| HCN4 | -0.546815226 | 0.684529573 | 1.18E-08 | 1.25E-07 | 646.1327977 | 442.4677044 | ENSG00000138622 |
| TACSTD2 | -0.546828073 | 0.684523477 | 0.244244879 | 0.40730705 | 30.38732341 | 20.78809934 | ENSG00000184292 |
| ALDH4A1 | -0.547227595 | 0.68433394 | 1.45E-15 | 3.31E-14 | 1510.796426 | 1033.824612 | ENSG00000159423 |
| CENPX | -0.547389546 | 0.684257124 | 1.70E-15 | 3.86E-14 | 1793.244937 | 1226.950045 | ENSG00000169689 |
| SLC6A9 | -0.547525646 | 0.684192576 | 4.37E-07 | 3.64E-06 | 581.4231622 | 397.5953522 | ENSG00000196517 |
| NEXN | -0.547698203 | 0.684110746 | 0.017204225 | 0.049601958 | 108.4597428 | 74.31647868 | ENSG00000162614 |
| SETDB2 | -0.547811132 | 0.684057199 | 0.003628743 | 0.013262994 | 168.2535888 | 114.9537343 | ENSG00000136169 |
| CDR2L | -0.547900082 | 0.684015024 | 2.05E-11 | 3.04E-10 | 1433.380449 | 980.6562885 | ENSG00000109089 |
| EPHB3 | -0.548199117 | 0.683873259 | 0.037529604 | 0.094527168 | 80.12843476 | 54.79238689 | ENSG00000182580 |
| LAT2 | -0.54865718 | 0.68365616 | 0.348545024 | 0.521344659 | 15.92889855 | 10.9506226 | ENSG00000086730 |
| U2AF2 | -0.548784698 | 0.683595736 | 4.83E-19 | 1.49E-17 | 9082.902399 | 6208.741572 | ENSG00000063244 |
| B3GNT7 | -0.548816925 | 0.683580465 | 3.42E-20 | 1.19E-18 | 2338.283061 | 1597.752383 | ENSG00000156966 |
| MPDU1 | -0.549101876 | 0.683445463 | 2.53E-08 | 2.53E-07 | 697.1346008 | 476.3577448 | ENSG00000129255 |
| SERF1A | -0.549347691 | 0.683329023 | 0.362442241 | 0.53569363 | 14.42437044 | 9.842038351 | ENSG00000205572 |
| IQCD | -0.549380864 | 0.683313311 | 0.251729352 | 0.416047886 | 24.89114959 | 16.9286091 | ENSG00000166578 |
| NRG2 | -0.549601811 | 0.683208671 | 0.417656132 | 0.589692058 | 12.10256771 | 8.279045791 | ENSG00000158458 |
| NAPA | -0.549739382 | 0.683143525 | 4.29E-06 | 3.02E-05 | 467.8979603 | 319.5955009 | ENSG00000105402 |
| ARHGAP29 | -0.549885297 | 0.683074435 | 0.028012755 | 0.074337974 | 84.44823161 | 57.64707188 | ENSG00000137962 |
| KIF3A | -0.550220495 | 0.682915747 | 6.07E-05 | 0.000343362 | 306.8965429 | 209.7560801 | ENSG00000131437 |
| EEF1AKMT3 | -0.550349405 | 0.682854728 | 0.167570849 | 0.307603887 | 37.99496723 | 25.96965051 | ENSG00000123427 |
| HTR7P1 | -0.550357327 | 0.682850979 | 0.030373823 | 0.079363594 | 84.14954111 | 57.47418353 | ENSG00000183935 |
| TGFB3 | -0.550467373 | 0.682798894 | 0.420200244 | 0.592160342 | 12.0997768 | 8.26882596 | ENSG00000119699 |
| TMEM102 | -0.550501185 | 0.682782892 | 0.060915644 | 0.139641096 | 86.70967362 | 59.38885937 | ENSG00000181284 |
| GNRHR2 | -0.550953521 | 0.682568849 | 0.387583675 | 0.56093888 | 14.32871601 | 9.755463942 | ENSG00000211451 |
| NA | -0.551271457 | 0.682418443 | 0.229867676 | 0.390050571 | 26.03972967 | 17.79708575 | ENSG00000249129 |
| KEL | -0.551677019 | 0.682226633 | 0.175926346 | 0.319295808 | 33.17488328 | 22.59092692 | ENSG00000197993 |
| SLC25A13 | -0.551841735 | 0.682148746 | 1.48E-21 | 5.60E-20 | 2238.652834 | 1527.487348 | ENSG00000004864 |
| ALG1 | -0.551853928 | 0.682142981 | 2.60E-07 | 2.24E-06 | 551.7500502 | 376.222453 | ENSG00000033011 |
| OGG1 | -0.552349704 | 0.681908605 | 6.11E-08 | 5.78E-07 | 614.9455087 | 419.3820534 | ENSG00000114026 |
| PPM1H | -0.552852267 | 0.681671104 | 5.08E-19 | 1.56E-17 | 2030.013265 | 1384.024092 | ENSG00000111110 |
| WASHC4 | -0.553582126 | 0.681326334 | 9.18E-11 | 1.27E-09 | 886.7153291 | 604.04475 | ENSG00000136051 |
| MLLT6 | -0.55381116 | 0.681218179 | 6.18E-06 | 4.23E-05 | 389.9290797 | 265.6100112 | ENSG00000275023 |
| MRPL12 | -0.554155837 | 0.681055447 | 0.024348222 | 0.066127206 | 99.1606045 | 67.58213636 | ENSG00000262814 |
| DAP | -0.554328193 | 0.680974088 | 7.28E-13 | 1.26E-11 | 1040.180166 | 708.4888214 | ENSG00000112977 |
| ATP8 | -0.554351452 | 0.680963109 | 0.06673787 | 0.150226399 | 60.16418718 | 41.01718733 | ENSG00000228253 |
| RFNG | -0.554394318 | 0.680942876 | 2.33E-06 | 1.71E-05 | 393.9852044 | 268.2627659 | ENSG00000169733 |
| LOC339260 | -0.554931295 | 0.680689474 | 4.67E-07 | 3.87E-06 | 477.0834595 | 324.7324046 | ENSG00000233098 |
| GAS6-DT | -0.555074216 | 0.680622044 | 0.377598781 | 0.551044564 | 14.07544369 | 9.598162643 | ENSG00000272695 |
| ACPP | -0.5550807 | 0.680618985 | 0.319992069 | 0.491478265 | 18.92575201 | 12.86235436 | ENSG00000014257 |
| CYB5B | -0.555103615 | 0.680608175 | 1.43E-19 | 4.66E-18 | 2781.646975 | 1893.147215 | ENSG00000103018 |
| ZBTB40 | -0.555281147 | 0.680524427 | 9.24E-10 | 1.13E-08 | 734.8903752 | 499.8268167 | ENSG00000184677 |
| LOC100287015 | -0.555485149 | 0.680428206 | 0.002534918 | 0.009700677 | 197.8813706 | 134.9007572 | ENSG00000246089 |
| ZDHHC16 | -0.555619865 | 0.680364672 | 4.69E-11 | 6.67E-10 | 1230.507595 | 837.0610665 | ENSG00000171307 |
| ZNF616 | -0.555634059 | 0.680357978 | 3.89E-08 | 3.81E-07 | 573.6774369 | 390.4143861 | ENSG00000204611 |
| LOC653160 | -0.556623584 | 0.679891489 | 0.57112273 | 0.723801675 | 5.350595432 | 3.651641249 | ENSG00000241014 |
| CYB561D2 | -0.556685371 | 0.679862372 | 0.023706193 | 0.0647115 | 86.81333735 | 58.98877897 | ENSG00000114395 |
| GPC1 | -0.556897544 | 0.679762394 | 8.20E-13 | 1.41E-11 | 1790.127788 | 1217.37257 | ENSG00000063660 |
| LINC00960 | -0.556960111 | 0.679732914 | 0.12507417 | 0.24570324 | 47.79718354 | 32.55450032 | ENSG00000242516 |
| MROH8 | -0.557039364 | 0.679695575 | 0.130661633 | 0.253810649 | 38.75151772 | 26.38130781 | ENSG00000101353 |
| SCAMP4 | -0.557265039 | 0.679589261 | 8.33E-10 | 1.02E-08 | 887.5152579 | 603.1047714 | ENSG00000227500 |
| NA | -0.557632791 | 0.679416052 | 0.306482174 | 0.477495365 | 17.3358572 | 11.76452298 | ENSG00000267493 |
| PTPRT | -0.557671163 | 0.679397981 | 0.011638665 | 0.035782032 | 116.5335072 | 79.30014022 | ENSG00000196090 |
| WFIKKN1 | -0.557694029 | 0.679387214 | 0.001349641 | 0.005576337 | 204.30344 | 138.9557381 | ENSG00000127578 |
| SGSM2 | -0.55782413 | 0.67932595 | 6.44E-09 | 7.03E-08 | 802.3179342 | 544.6260736 | ENSG00000141258 |
| RANBP1 | -0.557860917 | 0.679308628 | 3.95E-35 | 3.26E-33 | 8067.364831 | 5479.584276 | ENSG00000099901 |
| TOM1 | -0.558242613 | 0.679128926 | 3.53E-09 | 4.00E-08 | 660.428135 | 448.6113219 | ENSG00000100284 |
| NA | -0.558319955 | 0.679092519 | 0.471714701 | 0.64060882 | 10.11294631 | 6.83194217 | ENSG00000273014 |
| ILKAP | -0.558446159 | 0.679033116 | 1.57E-07 | 1.41E-06 | 610.1484518 | 414.1988663 | ENSG00000132323 |
| KRT10 | -0.558518351 | 0.678999138 | 5.45E-07 | 4.46E-06 | 579.4387351 | 393.7492805 | ENSG00000186395 |
| TSPAN14 | -0.559151721 | 0.67870111 | 6.48E-19 | 1.98E-17 | 1869.621106 | 1269.441166 | ENSG00000108219 |
| EIF6 | -0.559158756 | 0.678697801 | 4.51E-13 | 7.98E-12 | 1526.911978 | 1035.790708 | ENSG00000242372 |
| SH3BP5 | -0.559197455 | 0.678679596 | 0.020679648 | 0.057774236 | 100.3601846 | 68.15969726 | ENSG00000131370 |
| CNOT2-DT | -0.559374013 | 0.678596544 | 0.383360495 | 0.556767998 | 13.55908688 | 9.149945589 | ENSG00000257815 |
| RNF24 | -0.559477643 | 0.678547801 | 7.06E-06 | 4.79E-05 | 362.2104586 | 245.7578803 | ENSG00000101236 |
| C19orf54 | -0.559889181 | 0.678354269 | 8.07E-08 | 7.52E-07 | 532.3376022 | 361.0275437 | ENSG00000188493 |
| NA | -0.560015914 | 0.678294682 | 0.576952724 | 0.7288321 | 5.372246934 | 3.653791822 | ENSG00000278291 |
| OIP5 | -0.560114087 | 0.678248526 | 0.021055205 | 0.058619709 | 89.07509167 | 60.45308718 | ENSG00000104147 |
| TRAPPC2L | -0.56011932 | 0.678246066 | 2.86E-12 | 4.63E-11 | 1031.169667 | 699.0536183 | ENSG00000167515 |
| UBXN8 | -0.560630587 | 0.67800575 | 0.006621161 | 0.022341929 | 130.6501387 | 88.62889285 | ENSG00000104691 |

| | | | | | | | |
|---|---|---|---|---|---|---|---|
| CCDC88A | -0.560752817 | 0.677948309 | 1.21E-11 | 1.84E-10 | 1167.189449 | 791.7250243 | ENSG00000115355 |
| PTX3 | -0.5608413 | 0.677906731 | 0.248227518 | 0.411917319 | 24.78433154 | 16.81272016 | ENSG00000163661 |
| FARS2 | -0.56088159 | 0.677887799 | 1.59E-05 | 0.000100578 | 362.6594335 | 245.7654287 | ENSG00000145982 |
| NA | -0.560934429 | 0.677862972 | 0.645112659 | 0.778582852 | 4.2751424 | 2.888028095 | ENSG00000267676 |
| ATP6V1D | -0.561163275 | 0.677755455 | 1.91E-11 | 2.84E-10 | 879.2944643 | 595.7813097 | ENSG00000100554 |
| NA | -0.561269758 | 0.677705433 | 0.087343501 | 0.186181599 | 51.01469437 | 34.55764068 | ENSG00000179967 |
| SAP30 | -0.56129359 | 0.677694238 | 3.67E-05 | 0.000217093 | 333.8099288 | 226.2104048 | ENSG00000164105 |
| NA | -0.561457675 | 0.677617165 | 0.194949373 | 0.345087938 | 36.17644266 | 24.44780984 | ENSG00000233757 |
| ARL2BP | -0.561904849 | 0.677407165 | 0.40713663 | 0.579337076 | 13.46622335 | 9.063904218 | ENSG00000102931 |
| SAP30-DT | -0.562010295 | 0.677357655 | 0.425003579 | 0.596709217 | 10.75565748 | 7.301131924 | ENSG00000272870 |
| VDR | -0.562145246 | 0.677294298 | 0.475707644 | 0.644081706 | 8.711569464 | 5.887371406 | ENSG00000111424 |
| CYB561A3 | -0.562659211 | 0.677053052 | 0.00020625 | 0.001037319 | 258.8177024 | 175.1212913 | ENSG00000162144 |
| MAF | -0.5627 | 0.67703391 | 0.005464327 | 0.018942019 | 134.9750418 | 91.43490815 | ENSG00000178573 |
| ZWINT | -0.562767682 | 0.677002149 | 2.69E-22 | 1.08E-20 | 2178.70122 | 1475.143628 | ENSG00000122952 |
| HIRA | -0.56287034 | 0.676953977 | 9.41E-05 | 0.000511862 | 328.7037574 | 222.5531238 | ENSG00000100084 |
| ARHGAP8 | -0.563399233 | 0.676705851 | 0.14863517 | 0.28031294 | 35.49457083 | 24.02350956 | ENSG00000241484 |
| PIP5K1B | -0.563569693 | 0.6766259 | 0.017455764 | 0.050203051 | 103.9643434 | 70.20237197 | ENSG00000107242 |
| DLL4 | -0.56366427 | 0.676581545 | 0.490252788 | 0.656722903 | 8.681545253 | 5.856178876 | ENSG00000128917 |
| PHKG2 | -0.563980293 | 0.676433355 | 3.57E-06 | 2.55E-05 | 410.1877874 | 277.24682 | ENSG00000156873 |
| NA | -0.564294841 | 0.67628589 | 0.100632672 | 0.208093566 | 46.16294658 | 31.21709438 | ENSG00000232640 |
| WIPF3 | -0.564354397 | 0.676257973 | 3.59E-07 | 3.04E-06 | 477.3991708 | 323.0486889 | ENSG00000122574 |
| C1QL2 | -0.564366928 | 0.676252099 | 0.278888764 | 0.446699032 | 19.39322401 | 13.09816081 | ENSG00000144119 |
| NA | -0.564477923 | 0.676200073 | 0.445911274 | 0.616326889 | 9.770277078 | 6.604738015 | ENSG00000226900 |
| PCDH9 | -0.564551958 | 0.676165373 | 0.370019339 | 0.543818897 | 13.66714423 | 9.276314822 | ENSG00000184226 |
| NA | -0.564709264 | 0.676091651 | 0.535841408 | 0.695713864 | 6.733111675 | 4.513938752 | ENSG00000263345 |
| PCSK7 | -0.56480656 | 0.676046056 | 1.10E-11 | 1.68E-10 | 927.6157769 | 627.1304315 | ENSG00000160613 |
| SYT10 | -0.564895948 | 0.676004171 | 0.305998228 | 0.477011056 | 26.36585364 | 17.80191993 | ENSG00000110975 |
| SLC6A8 | -0.565152559 | 0.675883941 | 3.36E-18 | 9.69E-17 | 3127.07818 | 2113.376441 | ENSG00000130821 |
| GJB3 | -0.565407837 | 0.675764357 | 0.381638809 | 0.555092684 | 13.05234209 | 8.788302943 | ENSG00000188910 |
| TMA7 | -0.565449932 | 0.67574464 | 0.004223671 | 0.01514758 | 139.7247896 | 94.49555202 | ENSG00000232112 |
| CCDC120 | -0.565485348 | 0.675728051 | 5.43E-06 | 3.75E-05 | 402.4680192 | 272.0302961 | ENSG00000147144 |
| CALCRL | -0.565888076 | 0.675539448 | 0.398202947 | 0.570556532 | 11.70543186 | 7.919553719 | ENSG00000064989 |
| DDX58 | -0.566097245 | 0.675441513 | 0.010300893 | 0.032237919 | 117.8938083 | 79.55880948 | ENSG00000107201 |
| DZANK1 | -0.56618105 | 0.675402278 | 0.000220228 | 0.001100058 | 234.7063064 | 158.513706 | ENSG00000089091 |
| POLR3H | -0.566285211 | 0.675353516 | 2.16E-10 | 2.88E-09 | 955.545043 | 645.4854504 | ENSG00000100413 |
| CYLD | -0.566562263 | 0.675223835 | 0.000106762 | 0.000574011 | 317.235328 | 214.2912826 | ENSG00000083799 |
| TUBB8 | -0.566580143 | 0.675215467 | 0.552579361 | 0.70909389 | 7.807888986 | 5.254428633 | ENSG00000261456 |
| TMPRSS3 | -0.566787868 | 0.675118254 | 0.636525608 | 0.77256924 | 3.929682269 | 2.688248359 | ENSG00000160183 |
| CCDC184 | -0.566984251 | 0.675026361 | 0.130713298 | 0.253868927 | 43.27925671 | 29.24674567 | ENSG00000177875 |
| PLEKHA3 | -0.567105991 | 0.674969402 | 0.000774907 | 0.003375193 | 237.3856599 | 160.1896435 | ENSG00000116095 |
| CA5BP1 | -0.567249419 | 0.674902303 | 0.001935234 | 0.007665543 | 174.2968193 | 117.7782991 | ENSG00000186312 |
| CDC25B | -0.567455032 | 0.674806123 | 1.05E-08 | 1.12E-07 | 905.7879741 | 611.268164 | ENSG00000101224 |
| C9orf78 | -0.567542973 | 0.67476499 | 4.20E-15 | 9.12E-14 | 1224.843879 | 826.3988996 | ENSG00000136819 |
| NA | -0.567657625 | 0.674711368 | 0.436899259 | 0.607641853 | 12.00970417 | 8.060183466 | ENSG00000261143 |
| CEND1 | -0.567709928 | 0.674686908 | 0.009821708 | 0.030965427 | 152.0022842 | 102.4143243 | ENSG00000184524 |
| LLGL1 | -0.568377893 | 0.674374601 | 2.77E-17 | 7.48E-16 | 3339.755366 | 2252.30882 | ENSG00000131899 |
| ZNF488 | -0.568464089 | 0.674334311 | 0.00015435 | 0.000795929 | 249.2142918 | 167.9320077 | ENSG00000265763 |
| NA | -0.568825389 | 0.674165456 | 0.16474954 | 0.303783539 | 31.69180651 | 21.39226062 | ENSG00000260947 |
| NA | -0.569758365 | 0.673729621 | 0.458168246 | 0.628137563 | 10.15413413 | 6.781394477 | ENSG00000220804 |
| NA | -0.569796252 | 0.673711928 | 0.562364192 | 0.71705443 | 5.353386335 | 3.601093555 | ENSG00000253919 |
| SPX | -0.570095604 | 0.673572151 | 0.591543037 | 0.740282629 | 5.869743138 | 3.990160622 | ENSG00000134548 |
| NA | -0.570441297 | 0.673410772 | 0.133185841 | 0.257546919 | 48.32191305 | 32.51497807 | ENSG00000260920 |
| MCM6 | -0.571252681 | 0.673032146 | 7.33E-37 | 6.83E-35 | 5392.290391 | 3628.952172 | ENSG00000076003 |
| SUSD5 | -0.572164366 | 0.67260697 | 0.018534712 | 0.052682057 | 97.068032 | 65.38162731 | ENSG00000173705 |
| FAM155A | -0.572442115 | 0.672477491 | 0.000210159 | 0.001054401 | 291.9787064 | 196.7439486 | ENSG00000204442 |
| FOXP4 | -0.572477319 | 0.672461082 | 5.52E-07 | 4.52E-06 | 858.9954702 | 577.5040451 | ENSG00000137166 |
| IFIH1 | -0.572586928 | 0.672409994 | 0.226793078 | 0.386012463 | 24.87998598 | 16.67613109 | ENSG00000115267 |
| ASIC2 | -0.572618941 | 0.672395073 | 0.192355037 | 0.341655558 | 28.65464111 | 19.2315585 | ENSG00000108684 |
| RHPN1-AS1 | -0.572674548 | 0.672369157 | 0.230399798 | 0.390703454 | 24.29991397 | 16.32040713 | ENSG00000254389 |
| NA | -0.57274124 | 0.672338076 | 0.426089203 | 0.597708465 | 10.63835154 | 7.128516145 | ENSG00000253893 |
| MDM1 | -0.572781513 | 0.672319308 | 0.000892287 | 0.003841352 | 203.2804506 | 136.8318562 | ENSG00000111554 |
| NA | -0.572877079 | 0.672274774 | 0.399753398 | 0.572292601 | 11.74315306 | 7.887828151 | ENSG00000255921 |
| IRX3 | -0.572918388 | 0.672255525 | 0.091468209 | 0.193117349 | 54.64636186 | 36.67398636 | ENSG00000177508 |
| FXYD7 | -0.572984042 | 0.672224933 | 0.140290178 | 0.268406012 | 39.02556566 | 26.31838971 | ENSG00000221946 |
| PRKAR1B | -0.573347944 | 0.672055394 | 5.37E-14 | 1.04E-12 | 2045.945108 | 1375.016497 | ENSG00000188191 |
| NR6A1 | -0.57351232 | 0.671978826 | 4.13E-32 | 2.96E-30 | 5759.745803 | 3871.125466 | ENSG00000148200 |
| MAP3K14 | -0.573574118 | 0.671950043 | 0.00402426 | 0.014537466 | 155.6117366 | 104.6492184 | ENSG00000006062 |
| SLC35E4 | -0.573578685 | 0.671947915 | 0.000574654 | 0.002486418 | 208.7302551 | 140.2197457 | ENSG00000100036 |
| NA | -0.573962753 | 0.671769056 | 0.261372777 | 0.426829953 | 20.04363217 | 13.46625518 | ENSG00000231993 |
| NA | -0.57399426 | 0.671754386 | 0.18383203 | 0.329796678 | 29.89070308 | 20.0623909 | ENSG00000277459 |
| VPS18 | -0.574159956 | 0.671677238 | 1.94E-13 | 3.56E-12 | 1346.997412 | 904.6902429 | ENSG00000104142 |
| BAG1 | -0.574179604 | 0.67166809 | 1.83E-10 | 2.45E-09 | 1235.632779 | 829.8717644 | ENSG00000107262 |
| NA | -0.574539928 | 0.671500357 | 0.294371331 | 0.464283454 | 20.57828599 | 13.78380185 | ENSG00000267904 |
| NA | -0.574730655 | 0.671411589 | 0.59801115 | 0.744849761 | 4.428054355 | 2.957397913 | ENSG00000204434 |
| FLJ37453 | -0.574844206 | 0.671358746 | 0.008386173 | 0.027166693 | 112.9611244 | 75.91764853 | ENSG00000179743 |
| TXNL4B | -0.574928814 | 0.671319375 | 0.005107001 | 0.017854141 | 155.4454337 | 104.5924794 | ENSG00000140830 |
| NA | -0.574934297 | 0.671316824 | 0.048544338 | 0.116592235 | 70.54089369 | 47.30196764 | ENSG00000273117 |

| | | | | | | | |
|---|---|---|---|---|---|---|---|
| STK36 | -0.575322345 | 0.671136281 | 1.14E-07 | 1.04E-06 | 708.9339354 | 475.588225 | ENSG00000163482 |
| ASAP2 | -0.575354962 | 0.671121108 | 2.91E-07 | 2.50E-06 | 500.5976746 | 336.0987131 | ENSG00000151693 |
| G0S2 | -0.575900607 | 0.670867329 | 0.256229731 | 0.421427892 | 21.88034161 | 14.67352378 | ENSG00000123689 |
| PANX2 | -0.575989012 | 0.670826222 | 0.001036567 | 0.004397715 | 221.5003398 | 148.8647231 | ENSG00000073150 |
| FZD1 | -0.576036023 | 0.670804363 | 0.087640867 | 0.18671231 | 59.28044334 | 39.67871532 | ENSG00000157240 |
| NA | -0.57612711 | 0.670762012 | 0.640584756 | 0.775170981 | 3.902448962 | 2.63985124 | ENSG00000223349 |
| SLC10A7 | -0.57636527 | 0.670651291 | 0.143830098 | 0.273362208 | 37.30145623 | 24.99067056 | ENSG00000120519 |
| ALKAL1 | -0.576922506 | 0.670392305 | 0.499110177 | 0.665399753 | 7.419125852 | 4.980977931 | ENSG00000196711 |
| FZD8 | -0.577008586 | 0.670352306 | 1.29E-06 | 9.95E-06 | 425.1873749 | 285.0644727 | ENSG00000177283 |
| NA | -0.577167463 | 0.670278488 | 0.301453379 | 0.472177423 | 17.21942714 | 11.51634613 | ENSG00000230383 |
| NA | -0.577221882 | 0.670253205 | 0.117666976 | 0.235029368 | 59.13274977 | 39.54507033 | ENSG00000238178 |
| WDR45 | -0.577412719 | 0.670164551 | 0.005931487 | 0.020350659 | 123.7965668 | 83.03352169 | ENSG00000196998 |
| NA | -0.577720938 | 0.670021392 | 0.016872598 | 0.048769745 | 93.96244547 | 62.88429592 | ENSG00000263823 |
| NA | -0.577776656 | 0.669995515 | 0.522293704 | 0.685179724 | 7.274586605 | 4.86751214 | ENSG00000278867 |
| NA | -0.577837104 | 0.669967444 | 0.655938018 | 0.786579245 | 4.324702931 | 2.8794258 | ENSG00000275630 |
| NA | -0.577919991 | 0.669928953 | 0.000302024 | 0.001465008 | 219.7046549 | 147.1723988 | ENSG00000234664 |
| TOLLIP | -0.577961863 | 0.66990951 | 1.67E-09 | 1.99E-08 | 871.7913743 | 583.8455401 | ENSG00000078902 |
| NA | -0.578412925 | 0.669700094 | 0.49794625 | 0.664307131 | 7.296238107 | 4.879882545 | ENSG00000271918 |
| LSG1 | -0.57842127 | 0.66969622 | 3.47E-20 | 1.20E-18 | 1763.962127 | 1181.537982 | ENSG00000041802 |
| NA | -0.57879636 | 0.669522127 | 0.527673992 | 0.689424922 | 6.324812221 | 4.221665925 | ENSG00000259408 |
| HDX | -0.578977635 | 0.669438006 | 0.010980103 | 0.034103553 | 106.7898998 | 71.47281913 | ENSG00000165259 |
| NA | -0.57905809 | 0.669400675 | 0.522274639 | 0.685179724 | 7.280168411 | 4.865361567 | ENSG00000235489 |
| TMEM200A | -0.579178821 | 0.669344659 | 3.06E-06 | 2.20E-05 | 439.911475 | 294.712359 | ENSG00000164484 |
| NA | -0.579382691 | 0.669250079 | 0.0059606 | 0.020420163 | 129.2461711 | 86.58245002 | ENSG00000240356 |
| NA | -0.579602996 | 0.66914789 | 0.522203892 | 0.68515509 | 6.352045528 | 4.238337476 | ENSG00000260992 |
| PCTP | -0.580008373 | 0.668959895 | 8.25E-08 | 7.69E-07 | 495.0635425 | 331.211046 | ENSG00000141179 |
| LTBP1 | -0.580127384 | 0.668904713 | 6.64E-16 | 1.58E-14 | 1316.697319 | 880.8350053 | ENSG00000049323 |
| PIGQ | -0.580734821 | 0.668623135 | 3.23E-07 | 2.76E-06 | 469.4873668 | 313.8901532 | ENSG00000007541 |
| KCNN2 | -0.580838631 | 0.668575025 | 0.000125827 | 0.000663931 | 325.4002042 | 217.6234807 | ENSG00000080709 |
| TEX261 | -0.580846994 | 0.66857115 | 8.60E-07 | 6.84E-06 | 420.5674481 | 281.2533796 | ENSG00000144043 |
| DHRS13 | -0.580885979 | 0.668553083 | 5.35E-07 | 4.39E-06 | 556.0877949 | 371.9564186 | ENSG00000167536 |
| THAP10 | -0.581411232 | 0.668309722 | 0.018298427 | 0.052082889 | 100.0219459 | 66.82821 | ENSG00000129028 |
| RHOF | -0.581540436 | 0.668249873 | 0.02976522 | 0.078027022 | 84.28953769 | 56.39355674 | ENSG00000139725 |
| TCTN3 | -0.581673842 | 0.668188083 | 7.91E-10 | 9.73E-09 | 840.6056851 | 561.8082026 | ENSG00000119977 |
| RCN1 | -0.58180677 | 0.66812652 | 1.21E-09 | 1.46E-08 | 910.7986301 | 609.0133639 | ENSG00000049449 |
| SLIT3 | -0.581806936 | 0.668126443 | 0.049049598 | 0.117511638 | 68.31454523 | 45.77096111 | ENSG00000184347 |
| ASXL3 | -0.581924021 | 0.668072222 | 0.394489826 | 0.567042134 | 12.82331202 | 8.63261918 | ENSG00000141431 |
| CXCL12 | -0.582074156 | 0.668002702 | 2.18E-11 | 3.22E-10 | 823.3493133 | 550.191282 | ENSG00000107562 |
| NA | -0.582103994 | 0.667988887 | 0.381094979 | 0.554552557 | 13.42503553 | 9.002864121 | ENSG00000250838 |
| KHDC1 | -0.582105591 | 0.667988147 | 0.002125949 | 0.008328494 | 157.6420702 | 105.3472298 | ENSG00000135314 |
| NA | -0.582912569 | 0.66761461 | 0.617027789 | 0.759048006 | 4.368005934 | 2.890178668 | ENSG00000260306 |
| C5orf60 | -0.583039475 | 0.667555886 | 0.613869576 | 0.756749457 | 3.986939787 | 2.688248359 | ENSG00000204661 |
| HEG1 | -0.583139457 | 0.667509624 | 2.13E-05 | 0.000132204 | 290.9114629 | 194.1855892 | ENSG00000173706 |
| PDGFA | -0.583248188 | 0.667459318 | 6.30E-11 | 8.84E-10 | 777.753272 | 519.1836357 | ENSG00000197461 |
| EDNRA | -0.583280696 | 0.667444279 | 0.014365565 | 0.042693901 | 94.39777807 | 63.02733671 | ENSG00000151617 |
| CCDC107 | -0.58332343 | 0.667424509 | 0.128371343 | 0.250471485 | 41.94737704 | 27.9125748 | ENSG00000159884 |
| RBM15-AS1 | -0.583367401 | 0.667404167 | 0.551692777 | 0.708474594 | 5.973770284 | 4.021886189 | ENSG00000227963 |
| NA | -0.583371891 | 0.66740209 | 0.393547761 | 0.56647738 | 13.39501132 | 8.966837405 | ENSG00000272205 |
| PRXL2A | -0.583494767 | 0.667345249 | 2.57E-15 | 5.73E-14 | 1419.741238 | 948.0519914 | ENSG00000122378 |
| ANO4 | -0.583618451 | 0.667288039 | 0.059346869 | 0.136715599 | 65.37602533 | 43.77936712 | ENSG00000151572 |
| H2AC12 | -0.583937775 | 0.667140359 | 0.602480984 | 0.748190409 | 4.398030145 | 2.90685022 | ENSG00000274997 |
| FRMD6-AS1 | -0.584037399 | 0.667094291 | 0.455779923 | 0.625485559 | 8.330503317 | 5.548852033 | ENSG00000273888 |
| AMN | -0.584207473 | 0.667015655 | 0.144944504 | 0.275144369 | 43.03291765 | 28.63881628 | ENSG00000166126 |
| NA | -0.584211188 | 0.667013937 | 0.610196241 | 0.753941951 | 3.956915577 | 2.662974513 | ENSG00000227698 |
| MED24 | -0.584215665 | 0.667011867 | 5.03E-28 | 2.88E-26 | 2818.792526 | 1880.151464 | ENSG00000008838 |
| QPCTL | -0.584268021 | 0.666987662 | 3.59E-08 | 3.53E-07 | 561.9488757 | 374.4967735 | ENSG00000011478 |
| NA | -0.584296197 | 0.666974635 | 0.277028087 | 0.444235147 | 18.83567938 | 12.57438268 | ENSG00000277182 |
| ZNF302 | -0.584668262 | 0.666802648 | 5.18E-06 | 3.58E-05 | 461.5243385 | 307.8626666 | ENSG00000089335 |
| NA | -0.585232802 | 0.666541773 | 0.563229461 | 0.717699949 | 5.982142993 | 4.017585042 | ENSG00000277351 |
| NA | -0.585238248 | 0.666539256 | 0.072206707 | 0.159942541 | 53.53462709 | 35.60304636 | ENSG00000279673 |
| HMGN2 | -0.585390183 | 0.666469065 | 1.53E-26 | 8.19E-25 | 6622.39424 | 4414.017082 | ENSG00000198830 |
| MLH1 | -0.585497723 | 0.666419387 | 8.66E-14 | 1.65E-12 | 1217.909445 | 811.4634778 | ENSG00000076242 |
| SEMA4G | -0.585541983 | 0.666398943 | 0.00576764 | 0.019854308 | 123.4427339 | 82.13090844 | ENSG00000095539 |
| CIPC | -0.585786565 | 0.666285977 | 6.98E-07 | 5.62E-06 | 439.851976 | 292.943426 | ENSG00000198894 |
| NA | -0.58586553 | 0.666249509 | 0.146910917 | 0.277794404 | 34.500142 | 22.96117186 | ENSG00000215835 |
| PLAGL2 | -0.586029884 | 0.666173613 | 2.35E-13 | 4.28E-12 | 1141.99707 | 761.1127037 | ENSG00000126003 |
| ALKBH8 | -0.58606397 | 0.666157874 | 1.22E-05 | 7.85E-05 | 315.8103094 | 210.2113124 | ENSG00000137760 |
| NA | -0.586147741 | 0.666119194 | 0.51894901 | 0.682462701 | 9.015841772 | 6.04116506 | ENSG00000265800 |
| CNPY4 | -0.58632164 | 0.666038907 | 0.000592999 | 0.002665317 | 185.0816251 | 123.3521645 | ENSG00000166997 |
| NACAD | -0.586327402 | 0.666036247 | 0.003890742 | 0.014097729 | 147.7150143 | 98.47523235 | ENSG00000136274 |
| DBX1 | -0.586816653 | 0.665810417 | 0.489251416 | 0.655745784 | 7.916622057 | 5.3071269 | ENSG00000109851 |
| ITPRIPL1 | -0.587720081 | 0.665393611 | 1.97E-06 | 1.48E-05 | 403.8164938 | 268.6072039 | ENSG00000198885 |
| PEF1 | -0.587989989 | 0.665269137 | 3.23E-08 | 3.18E-07 | 546.6976938 | 363.4563414 | ENSG00000162517 |
| ETNPPL | -0.588510978 | 0.665028937 | 0.131424626 | 0.2548565 | 39.30348054 | 26.10329549 | ENSG00000164089 |
| SERAC1 | -0.588629516 | 0.664974297 | 4.15E-05 | 0.000242508 | 276.2753713 | 183.8977448 | ENSG00000122335 |
| NA | -0.588809792 | 0.664891209 | 0.491526622 | 0.657856494 | 8.371015417 | 5.553153181 | ENSG00000235997 |
| KCTD17 | -0.588835451 | 0.664879384 | 3.78E-11 | 5.42E-10 | 905.9376834 | 602.4731552 | ENSG00000100379 |

| | | | | | | | |
|---|---|---|---|---|---|---|---|
| CTSD | -0.588869167 | 0.664863845 | 4.76E-08 | 4.59E-07 | 697.8870331 | 463.8532649 | ENSG00000117984 |
| UNQ6494 | -0.589298441 | 0.664666045 | 0.468049197 | 0.63706857 | 8.466669855 | 5.620372426 | ENSG00000237372 |
| SLC43A3 | -0.589359884 | 0.664637738 | 0.000252314 | 0.001245493 | 303.3411057 | 201.39023 | ENSG00000134802 |
| LARP6 | -0.589962902 | 0.66435999 | 8.29E-08 | 7.72E-07 | 486.8480298 | 323.5985465 | ENSG00000166173 |
| IFRD1 | -0.59031315 | 0.664198721 | 4.01E-14 | 7.89E-13 | 1043.975909 | 693.3079427 | ENSG00000006652 |
| RIMS3 | -0.590502457 | 0.664111572 | 1.99E-13 | 3.64E-12 | 1060.662671 | 704.4370755 | ENSG00000117016 |
| GDAP2 | -0.590749501 | 0.663997861 | 0.000132426 | 0.000693814 | 233.8238771 | 155.4194465 | ENSG00000196505 |
| KATNB1 | -0.591005459 | 0.663880067 | 3.94E-17 | 1.05E-15 | 1613.075646 | 1070.968091 | ENSG00000140854 |
| CAPN10-DT | -0.591124587 | 0.663825251 | 0.033648905 | 0.086204653 | 68.8137932 | 45.70589244 | ENSG00000260942 |
| TMED1 | -0.59116677 | 0.663805842 | 0.000457575 | 0.002117823 | 231.9275196 | 153.9185785 | ENSG00000099203 |
| LOC100130264 | -0.591578418 | 0.663616463 | 0.643534142 | 0.777277371 | 3.935264075 | 2.621029114 | ENSG00000179447 |
| LOC105371067 | -0.591853463 | 0.663489959 | 0.324004996 | 0.495543064 | 16.66937934 | 11.01758138 | ENSG00000285567 |
| CISH | -0.592291129 | 0.663288709 | 0.047731692 | 0.115113399 | 62.18315707 | 41.25676189 | ENSG00000114737 |
| BCL2L1 | -0.592727019 | 0.663088335 | 1.82E-09 | 2.15E-08 | 696.0182723 | 461.644123 | ENSG00000171552 |
| NA | -0.592830393 | 0.663040824 | 0.262911965 | 0.428545392 | 20.71569183 | 13.70582974 | ENSG00000239881 |
| SCD5 | -0.592899341 | 0.663009138 | 0.111083518 | 0.224388706 | 43.93054076 | 29.19807597 | ENSG00000145284 |
| SLITRK3 | -0.592950425 | 0.662985662 | 0.045489558 | 0.110660663 | 61.87541813 | 41.08655715 | ENSG00000121871 |
| BBS1 | -0.593125591 | 0.66290517 | 0.511226932 | 0.675669961 | 7.829340328 | 5.20388094 | ENSG00000174483 |
| MIGA2 | -0.593597513 | 0.662688361 | 2.54E-09 | 2.94E-08 | 734.1820579 | 486.2148371 | ENSG00000148343 |
| GDAP1 | -0.594220154 | 0.662402419 | 9.10E-10 | 1.11E-08 | 836.4888249 | 553.8738979 | ENSG00000104381 |
| C18orf65 | -0.594310474 | 0.66236095 | 0.419630102 | 0.591664088 | 10.38228832 | 6.859366591 | ENSG00000275763 |
| EMC9 | -0.594412179 | 0.662314258 | 1.52E-06 | 1.16E-05 | 452.7289298 | 299.9305005 | ENSG00000100908 |
| PTRHD1 | -0.594916105 | 0.662082955 | 0.005574829 | 0.019266218 | 133.9260583 | 88.53130511 | ENSG00000184924 |
| PCDHGB7 | -0.595393994 | 0.661863678 | 0.00028908 | 0.001406112 | 208.2690406 | 137.8675935 | ENSG00000254122 |
| SIVA1 | -0.59547697 | 0.661825612 | 8.97E-12 | 1.38E-10 | 1655.015665 | 1094.865292 | ENSG00000184990 |
| GPRASP2 | -0.595584638 | 0.661776223 | 0.000216608 | 0.001082816 | 259.9474972 | 172.2190027 | ENSG00000158301 |
| EFCAB11 | -0.5960658 | 0.661555547 | 0.025933337 | 0.069793527 | 75.54343825 | 50.00284686 | ENSG00000140025 |
| NA | -0.59618467 | 0.66150104 | 0.296595375 | 0.46691867 | 16.47615545 | 10.90437605 | ENSG00000226074 |
| C19orf47 | -0.596429605 | 0.661388743 | 9.59E-06 | 6.34E-05 | 381.6929305 | 252.7906562 | ENSG00000160392 |
| ATAD3A | -0.596787915 | 0.661224499 | 8.85E-14 | 1.68E-12 | 1912.790129 | 1264.582384 | ENSG00000197785 |
| CHD7 | -0.596862882 | 0.661190141 | 2.88E-23 | 1.24E-21 | 3022.178052 | 1997.763587 | ENSG00000171316 |
| ARL6IP5 | -0.596965357 | 0.661143178 | 5.50E-10 | 6.91E-09 | 622.3422562 | 411.3116501 | ENSG00000144746 |
| DHRSX | -0.597008092 | 0.661123594 | 3.91E-06 | 2.77E-05 | 336.3714497 | 222.4458187 | ENSG00000169084 |
| ATP8B1 | -0.597050254 | 0.661104274 | 0.050193765 | 0.119630333 | 63.27344049 | 41.86307375 | ENSG00000081923 |
| WDR76 | -0.597060185 | 0.661099723 | 1.25E-06 | 9.67E-06 | 401.7405787 | 265.2819842 | ENSG00000092470 |
| SLC27A4 | -0.597124245 | 0.661070369 | 3.20E-10 | 4.16E-09 | 896.6836752 | 592.6889708 | ENSG00000167114 |
| TMEM117 | -0.597138306 | 0.661063926 | 0.195738572 | 0.346159807 | 28.9177256 | 19.18557242 | ENSG00000139173 |
| IL12RB2 | -0.597408046 | 0.660940339 | 0.07815196 | 0.170161338 | 46.62974286 | 30.83447903 | ENSG00000081985 |
| WNT5B | -0.597438827 | 0.660926237 | 0.055683671 | 0.130205296 | 69.30610792 | 45.77364472 | ENSG00000111186 |
| GSTM2 | -0.597714973 | 0.660799741 | 0.098980123 | 0.205557708 | 41.87038304 | 27.68537064 | ENSG00000213366 |
| CACNA2D2 | -0.597809491 | 0.660756451 | 1.44E-12 | 2.41E-11 | 894.2818362 | 591.0079082 | ENSG00000007402 |
| SRCIN1 | -0.598128257 | 0.660610471 | 0.01300566 | 0.039275545 | 120.6279407 | 79.57575361 | ENSG00000277363 |
| NA | -0.598189806 | 0.660582289 | 0.29753023 | 0.467741477 | 19.33684237 | 12.69188916 | ENSG00000279069 |
| LRP3 | -0.598284498 | 0.660538933 | 1.72E-20 | 6.10E-19 | 1833.772302 | 1211.505886 | ENSG00000130881 |
| NA | -0.598585987 | 0.66040091 | 0.302470914 | 0.473186844 | 16.38117673 | 10.80543124 | ENSG00000279382 |
| C5orf38 | -0.598931259 | 0.660242879 | 0.160955042 | 0.298269329 | 32.92699261 | 21.81036964 | ENSG00000186493 |
| NA | -0.598981081 | 0.660220078 | 0.522474447 | 0.685277029 | 6.409303046 | 4.194241504 | ENSG00000268279 |
| TAFA5 | -0.599412535 | 0.660022662 | 1.33E-15 | 3.05E-14 | 1124.599161 | 742.4060015 | ENSG00000219438 |
| PEX12 | -0.599878623 | 0.659809464 | 0.04797453 | 0.115577979 | 70.14029122 | 46.34797054 | ENSG00000108733 |
| CCDC189 | -0.5999644 | 0.659770235 | 0.435511903 | 0.606209693 | 8.950887267 | 5.923398121 | ENSG00000196118 |
| YTHDF3-AS1 | -0.600348387 | 0.659594655 | 0.27488432 | 0.441605048 | 18.02187138 | 11.93068704 | ENSG00000270673 |
| ZNF138 | -0.60052021 | 0.659516103 | 9.53E-06 | 6.30E-05 | 310.9379493 | 205.1026082 | ENSG00000197008 |
| NA | -0.600753971 | 0.65940925 | 0.592512224 | 0.741062844 | 3.986939787 | 2.637700666 | ENSG00000286430 |
| ASRGL1 | -0.600986394 | 0.659303025 | 0.000198019 | 0.000999475 | 252.3104665 | 166.2047291 | ENSG00000162174 |
| TBC1D10A | -0.601049828 | 0.659274037 | 0.012312368 | 0.03746339 | 93.96782711 | 61.95908032 | ENSG00000099992 |
| XYLB | -0.601093921 | 0.659253888 | 0.023627567 | 0.064524264 | 80.32791619 | 53.05703902 | ENSG00000093217 |
| HEXIM2 | -0.601188216 | 0.6592108 | 0.149509821 | 0.281659566 | 30.50550523 | 20.12584203 | ENSG00000168517 |
| NA | -0.601539491 | 0.659050312 | 0.596322467 | 0.743851064 | 3.95970648 | 2.614577393 | ENSG00000284731 |
| KLHL18 | -0.60158743 | 0.659028412 | 8.03E-12 | 1.25E-10 | 864.1245068 | 569.6317371 | ENSG00000114648 |
| OSBPL11 | -0.601904 | 0.658883818 | 3.45E-12 | 5.55E-11 | 859.1375564 | 566.1686081 | ENSG00000144909 |
| NCKAP5 | -0.601952392 | 0.658861718 | 0.253745828 | 0.418437253 | 22.1072565 | 14.60871557 | ENSG00000176771 |
| NUCB1 | -0.602045836 | 0.658819044 | 1.46E-25 | 7.38E-24 | 2452.336086 | 1615.281967 | ENSG00000104805 |
| NA | -0.602615801 | 0.658558816 | 0.366470893 | 0.540199434 | 16.02387727 | 10.5723084 | ENSG00000227252 |
| TBXAS1 | -0.602913986 | 0.658422715 | 0.191065305 | 0.33977146 | 25.3998094 | 16.70785666 | ENSG00000059377 |
| NA | -0.603018268 | 0.658375124 | 0.637542886 | 0.773414653 | 4.552857122 | 3.035370027 | ENSG00000231460 |
| NA | -0.603106935 | 0.658334662 | 0.052263698 | 0.123435235 | 59.12521604 | 38.84626538 | ENSG00000272849 |
| RBCK1 | -0.603514286 | 0.658148805 | 2.55E-11 | 3.73E-10 | 1310.083505 | 862.2665645 | ENSG00000125826 |
| RPUSD1 | -0.603564674 | 0.658125819 | 9.19E-10 | 1.12E-08 | 923.004237 | 607.8335318 | ENSG00000007376 |
| NA | -0.60363277 | 0.658094755 | 0.606886977 | 0.751272767 | 3.962497383 | 2.610276246 | ENSG00000188512 |
| CTSA | -0.603662039 | 0.658081404 | 5.17E-10 | 6.53E-09 | 652.0035289 | 429.2176095 | ENSG00000064601 |
| TMEM158 | -0.604008486 | 0.657923393 | 0.001291235 | 0.005370678 | 334.8801507 | 220.4757425 | ENSG00000249992 |
| TMEM200C | -0.604156051 | 0.657856101 | 0.000155752 | 0.000802727 | 231.3021927 | 152.3733236 | ENSG00000206432 |
| NA | -0.604493367 | 0.657702306 | 0.518695815 | 0.682248095 | 5.995421786 | 3.950365797 | ENSG00000249014 |
| AK1 | -0.604638489 | 0.65763615 | 0.120142337 | 0.238693456 | 36.00622171 | 23.6258402 | ENSG00000106992 |
| NA | -0.604661836 | 0.657625508 | 0.047837695 | 0.115340205 | 65.23303768 | 42.82190503 | ENSG00000279495 |
| ZNF438 | -0.604921708 | 0.657507061 | 0.145308113 | 0.275573536 | 34.69683252 | 22.90443291 | ENSG00000183621 |
| GNB1L | -0.604922658 | 0.657506628 | 1.50E-06 | 1.14E-05 | 394.8619397 | 259.6572985 | ENSG00000185838 |

| | | | | | | | |
|---|---|---|---|---|---|---|---|
| PWP2 | -0.605862089 | 0.657078622 | 0.089834225 | 0.190460691 | 45.31353256 | 29.75116863 | ENSG00000241945 |
| MIR4713HG | -0.605908351 | 0.657057553 | 0.536086879 | 0.695845242 | 6.958474964 | 4.57308874 | ENSG00000259240 |
| NA | -0.605939224 | 0.657043492 | 0.562162919 | 0.716861771 | 4.955574771 | 3.256122461 | ENSG00000272416 |
| GNG7 | -0.606575508 | 0.656753774 | 1.01E-07 | 9.31E-07 | 464.0080264 | 304.7587387 | ENSG00000176533 |
| NA | -0.606647551 | 0.656720979 | 0.0042442 | 0.015212721 | 131.7521493 | 86.66203967 | ENSG00000274712 |
| SEMA4F | -0.607046258 | 0.656539512 | 0.000189216 | 0.000958302 | 242.3251139 | 159.0003914 | ENSG00000135622 |
| GNAZ | -0.60726166 | 0.656441494 | 4.17E-05 | 0.000243468 | 283.2979249 | 186.1824051 | ENSG00000128266 |
| NA | -0.607399722 | 0.656378677 | 0.376984174 | 0.550439124 | 12.40751574 | 8.098360755 | ENSG00000258748 |
| ECEL1P2 | -0.607643111 | 0.656267953 | 0.19618166 | 0.34663312 | 25.65607278 | 16.81487073 | ENSG00000244280 |
| STAM-AS1 | -0.607870265 | 0.656164631 | 0.486790477 | 0.654233584 | 7.051338498 | 4.633856264 | ENSG00000260589 |
| GNG13 | -0.608327672 | 0.655956626 | 0.411609605 | 0.583807138 | 10.0263403 | 6.581614742 | ENSG00000127588 |
| UBE2Q2P1 | -0.608659937 | 0.655805571 | 0.410357906 | 0.582545763 | 9.973988865 | 6.539669343 | ENSG00000189136 |
| ICAM5 | -0.608729762 | 0.655773832 | 0.008651472 | 0.0278903 | 108.140677 | 70.76405482 | ENSG00000105376 |
| PDAP1 | -0.609128011 | 0.655592833 | 3.47E-29 | 2.12E-27 | 6533.853133 | 4283.397221 | ENSG00000106244 |
| NA | -0.609453322 | 0.655445022 | 0.115348913 | 0.231285301 | 40.3864304 | 26.49934732 | ENSG00000272667 |
| CASR | -0.609576872 | 0.655388893 | 0.349606742 | 0.522203922 | 13.54704739 | 8.924892007 | ENSG00000036828 |
| MCM4 | -0.609639021 | 0.65536066 | 9.23E-35 | 7.45E-33 | 8852.130384 | 5801.404537 | ENSG00000104738 |
| HMG20B | -0.610014148 | 0.655190276 | 2.12E-22 | 8.54E-21 | 1804.490581 | 1182.673439 | ENSG00000064961 |
| MISP3 | -0.610059353 | 0.655169748 | 0.06772004 | 0.151958508 | 49.21059988 | 32.24823955 | ENSG00000141854 |
| H2BC13 | -0.61020171 | 0.655105102 | 0.33527948 | 0.507881936 | 13.02231788 | 8.52937322 | ENSG00000185130 |
| SLCO4A1-AS1 | -0.610382282 | 0.655023113 | 0.571172636 | 0.723801675 | 6.857914442 | 4.482213184 | ENSG00000232803 |
| SLC22A31 | -0.610599978 | 0.65492428 | 0.001313649 | 0.005449806 | 174.744667 | 114.3705457 | ENSG00000259803 |
| HEY2 | -0.610784578 | 0.654840485 | 0.000364576 | 0.001734255 | 184.5993228 | 120.9545715 | ENSG00000135547 |
| ARHGAP6 | -0.610836147 | 0.654817078 | 0.173104239 | 0.315539439 | 27.96025422 | 18.29343946 | ENSG00000047648 |
| FUCA1 | -0.610836188 | 0.654817059 | 0.00018745 | 0.000950355 | 225.3861924 | 147.3654664 | ENSG00000179163 |
| ZNF689 | -0.611149753 | 0.654674752 | 3.80E-09 | 4.29E-08 | 600.6740743 | 393.1187851 | ENSG00000156853 |
| DPH7 | -0.611186675 | 0.654657998 | 1.18E-11 | 1.79E-10 | 880.020214 | 576.3709321 | ENSG00000148399 |
| CFAP73 | -0.611431743 | 0.654546802 | 0.445587615 | 0.616056094 | 8.033927996 | 5.256579206 | ENSG00000186710 |
| NA | -0.611449512 | 0.65453874 | 0.487280038 | 0.654467675 | 7.528059082 | 4.882033118 | ENSG00000263551 |
| MDK | -0.611501955 | 0.654514947 | 1.54E-40 | 1.85E-38 | 6900.051246 | 4517.068051 | ENSG00000110492 |
| NA | -0.611625024 | 0.654459116 | 0.421916191 | 0.593797713 | 9.040959898 | 5.914795827 | ENSG00000233859 |
| NA | -0.611664371 | 0.654441268 | 0.626249222 | 0.764860636 | 3.929682269 | 2.5640297 | ENSG00000230956 |
| NA | -0.611833621 | 0.654364496 | 0.19955075 | 0.351010625 | 23.66346032 | 15.5317806 | ENSG00000246889 |
| MTHFS | -0.611894632 | 0.654336824 | 0.322826576 | 0.494414694 | 18.36645562 | 12.03152196 | ENSG00000136371 |
| NA | -0.611911965 | 0.654328963 | 0.546145265 | 0.704351405 | 5.021204997 | 3.283546882 | ENSG00000253973 |
| EMILIN3 | -0.61229771 | 0.654154032 | 4.28E-05 | 0.000249467 | 269.5691296 | 176.1102307 | ENSG00000183798 |
| NA | -0.612532016 | 0.654047801 | 0.605025515 | 0.749906657 | 3.965288286 | 2.587152973 | ENSG00000276071 |
| TTN-AS1 | -0.612618898 | 0.654008414 | 0.125965581 | 0.247102098 | 35.87496126 | 23.50833373 | ENSG00000237298 |
| ARG2 | -0.612963974 | 0.653852002 | 4.35E-08 | 4.22E-07 | 536.690102 | 350.7746905 | ENSG00000081181 |
| NA | -0.613294428 | 0.653702252 | 0.475232475 | 0.643709429 | 8.172885437 | 5.349072298 | ENSG00000279744 |
| NPEPPSP1 | -0.613760332 | 0.653491179 | 0.014360153 | 0.042686329 | 91.83474251 | 59.89517244 | ENSG00000274487 |
| LOC102724250 | -0.613767278 | 0.653488033 | 0.003933056 | 0.014232337 | 143.8911109 | 94.03604906 | ENSG00000271254 |
| SLCO3A1 | -0.614035788 | 0.653366419 | 1.02E-07 | 9.36E-07 | 434.0443595 | 283.6025755 | ENSG00000176463 |
| EIF2B1 | -0.614273924 | 0.653258581 | 9.57E-16 | 2.24E-14 | 1364.095503 | 891.0263585 | ENSG00000111361 |
| LURAP1 | -0.614333469 | 0.65323162 | 0.311418354 | 0.482458358 | 14.62808223 | 9.596012069 | ENSG00000171357 |
| EIF2B2 | -0.615273017 | 0.652806344 | 2.08E-11 | 3.08E-10 | 727.1606826 | 474.836134 | ENSG00000119718 |
| NA | -0.6152781 | 0.652804044 | 0.488890967 | 0.655460281 | 8.622372713 | 5.592948006 | ENSG00000248690 |
| SPHK1 | -0.615534488 | 0.652688042 | 0.083061989 | 0.178833574 | 44.79929094 | 29.24029395 | ENSG00000176170 |
| C1QL4 | -0.615605448 | 0.652655939 | 0.174418691 | 0.317275904 | 25.72361803 | 16.81487073 | ENSG00000186897 |
| FAHD1 | -0.616040343 | 0.652459229 | 8.13E-07 | 6.50E-06 | 377.0226042 | 246.060373 | ENSG00000180185 |
| FLRT2 | -0.616395702 | 0.652298537 | 0.01211551 | 0.036905147 | 110.6365307 | 72.33832117 | ENSG00000185070 |
| ADGRB2 | -0.617008473 | 0.652021538 | 4.31E-17 | 1.14E-15 | 1238.152085 | 807.238068 | ENSG00000121753 |
| NA | -0.617091293 | 0.651984109 | 0.338702397 | 0.511352842 | 13.59102612 | 8.887247756 | ENSG00000259394 |
| TRAF3IP1 | -0.617120915 | 0.651970722 | 7.65E-09 | 8.29E-08 | 539.0379868 | 351.2858073 | ENSG00000204104 |
| NA | -0.617140941 | 0.651961672 | 0.142797068 | 0.271988373 | 31.26464646 | 20.43693699 | ENSG00000261342 |
| NA | -0.617374902 | 0.651855953 | 0.296666264 | 0.466954055 | 15.69516255 | 10.26283098 | ENSG00000287630 |
| SKP2 | -0.617990474 | 0.651577877 | 3.18E-19 | 9.96E-18 | 1738.052881 | 1132.174857 | ENSG00000145604 |
| OXCT1 | -0.618088257 | 0.651533716 | 3.14E-13 | 5.66E-12 | 885.2606128 | 577.1071208 | ENSG00000083720 |
| STX18 | -0.618361031 | 0.65141054 | 4.37E-09 | 4.90E-08 | 599.3925189 | 390.0982086 | ENSG00000168818 |
| SPSB4 | -0.618383485 | 0.651400402 | 1.73E-10 | 2.32E-09 | 717.5394122 | 467.1376237 | ENSG00000175093 |
| NA | -0.618865871 | 0.651182633 | 0.557245985 | 0.713303297 | 6.069424722 | 3.946064649 | ENSG00000233542 |
| NA | -0.61908507 | 0.651083702 | 0.594549448 | 0.742536915 | 4.598951028 | 3.03106888 | ENSG00000272338 |
| LETM2 | -0.619150841 | 0.65105402 | 0.183793584 | 0.329796678 | 24.79337997 | 16.15235296 | ENSG00000165046 |
| SCG3 | -0.619201625 | 0.651031103 | 6.54E-13 | 1.14E-11 | 1098.789851 | 715.6727194 | ENSG00000104112 |
| FAM27E3 | -0.619397854 | 0.650942559 | 0.480891281 | 0.648959336 | 7.005244592 | 4.541363172 | ENSG00000274026 |
| NA | -0.619415654 | 0.650934527 | 0.508286487 | 0.672982473 | 6.077121709 | 3.946064649 | ENSG00000240207 |
| NA | -0.619779431 | 0.650770414 | 0.55069031 | 0.707658325 | 5.084044321 | 3.302369007 | ENSG00000272696 |
| TBC1D19 | -0.620417183 | 0.650482801 | 0.099902525 | 0.207049808 | 40.40616688 | 26.29688397 | ENSG00000109680 |
| DKFZP434A062 | -0.620563882 | 0.650416661 | 0.561663303 | 0.716508847 | 5.086835224 | 3.302369007 | ENSG00000262075 |
| SPHK2 | -0.620609728 | 0.650395992 | 1.93E-07 | 1.70E-06 | 853.3484002 | 554.6362025 | ENSG00000063176 |
| MGAT5B | -0.620829104 | 0.6502971 | 6.78E-06 | 4.61E-05 | 306.8448672 | 199.6209754 | ENSG00000167889 |
| EME2 | -0.62113955 | 0.650157181 | 1.30E-06 | 1.01E-05 | 571.1884795 | 371.1358302 | ENSG00000197774 |
| NA | -0.621188414 | 0.650135161 | 0.508724919 | 0.673424306 | 7.015732482 | 4.543513746 | ENSG00000259448 |
| ZNF699 | -0.621233537 | 0.650114827 | 0.149372484 | 0.281565818 | 37.19898068 | 24.1832219 | ENSG00000196110 |
| GLIDR | -0.621347221 | 0.6500636 | 0.113816625 | 0.22896216 | 36.8573506 | 24.01490727 | ENSG00000278175 |
| HLF | -0.621605289 | 0.649947328 | 0.516499418 | 0.680447331 | 7.204250453 | 4.680102809 | ENSG00000108924 |
| NA | -0.621641304 | 0.649931103 | 0.471027248 | 0.639945534 | 8.015067397 | 5.187209388 | ENSG00000240409 |

| | | | | | | | |
|---|---|---|---|---|---|---|---|
| PLA2G4B | -0.621809858 | 0.649855174 | 0.593324781 | 0.741598312 | 4.0386155 | 2.614577393 | ENSG00000243708 |
| FAM32A | -0.622276691 | 0.649644925 | 2.17E-21 | 8.10E-20 | 1816.533925 | 1179.972609 | ENSG00000105058 |
| TFPT | -0.622298646 | 0.649635039 | 2.88E-05 | 0.000174309 | 336.667037 | 218.7331679 | ENSG00000105619 |
| RNF208 | -0.622522226 | 0.649534371 | 0.011848155 | 0.03626829 | 115.3307359 | 75.11019987 | ENSG00000212864 |
| EFR3B | -0.622646929 | 0.649478229 | 7.03E-08 | 6.60E-07 | 454.2549703 | 294.8919901 | ENSG00000084710 |
| SLC22A5 | -0.622710694 | 0.649449524 | 0.005719858 | 0.019710931 | 115.0820062 | 74.89321554 | ENSG00000197375 |
| H3C12 | -0.622774004 | 0.649421024 | 0.257567744 | 0.422727528 | 17.74973846 | 11.53516825 | ENSG00000197153 |
| NA | -0.623039247 | 0.649301638 | 0.041522699 | 0.10272049 | 69.86113704 | 45.29936032 | ENSG00000272106 |
| TMEM191A | -0.62307312 | 0.649286393 | 0.02561942 | 0.069045016 | 76.58240938 | 49.69202448 | ENSG00000226287 |
| PCDH1 | -0.623353793 | 0.649160088 | 5.55E-32 | 3.93E-30 | 5583.427717 | 3624.468953 | ENSG00000156453 |
| ZNF100 | -0.623483424 | 0.649101761 | 0.002040835 | 0.008033002 | 147.0560701 | 95.47561646 | ENSG00000197020 |
| NA | -0.623488075 | 0.649099669 | 0.062828129 | 0.143096761 | 61.63506119 | 40.02286238 | ENSG00000275234 |
| CHRNB4 | -0.623501264 | 0.649093735 | 0.012639145 | 0.038342781 | 105.7178013 | 68.72944936 | ENSG00000117971 |
| ST3GAL6 | -0.623584015 | 0.649056505 | 0.05490548 | 0.128666378 | 57.41139434 | 37.27493099 | ENSG00000064225 |
| DUSP26 | -0.62366615 | 0.649019554 | 0.047170998 | 0.114046338 | 54.74816168 | 35.58610224 | ENSG00000133878 |
| HCRTR1 | -0.623748793 | 0.648982377 | 0.574272971 | 0.726479147 | 5.682776769 | 3.718860494 | ENSG00000121764 |
| RNU6-8 | -0.623830522 | 0.648945613 | 0.542974096 | 0.701682367 | 5.045647402 | 3.258273035 | ENSG00000202337 |
| TRIM39 | -0.623896024 | 0.648916149 | 1.21E-05 | 7.80E-05 | 283.4695342 | 183.8533884 | ENSG00000204599 |
| TMEM198 | -0.623915745 | 0.648907279 | 2.63E-05 | 0.000160489 | 261.9574555 | 169.8356702 | ENSG00000188760 |
| URM1 | -0.623935796 | 0.648898261 | 2.26E-15 | 5.08E-14 | 1307.657337 | 848.8709509 | ENSG00000167118 |
| EPN3 | -0.624704754 | 0.648552489 | 0.011545107 | 0.035606181 | 88.96077679 | 57.61319574 | ENSG00000049283 |
| MT1E | -0.624786228 | 0.648515864 | 2.24E-05 | 0.000138625 | 289.0356808 | 187.6464408 | ENSG00000169715 |
| UNG | -0.625321657 | 0.648275224 | 9.64E-36 | 8.26E-34 | 4695.992278 | 3044.10886 | ENSG00000076248 |
| NA | -0.625584785 | 0.648156999 | 0.009925548 | 0.031235521 | 93.64680939 | 60.73514017 | ENSG00000272086 |
| RPS6KA2 | -0.625591311 | 0.648154067 | 6.29E-12 | 9.83E-11 | 779.3624278 | 504.9298506 | ENSG00000071242 |
| RPL26L1 | -0.625617268 | 0.648142405 | 4.08E-10 | 5.22E-09 | 595.4288703 | 385.9566412 | ENSG00000037241 |
| ABCA1 | -0.625925701 | 0.648003854 | 0.000123008 | 0.000651013 | 219.3467919 | 142.1556547 | ENSG00000165029 |
| NA | -0.626180904 | 0.647889237 | 0.244447754 | 0.407504462 | 19.61089031 | 12.67736818 | ENSG00000228218 |
| C7orf50 | -0.626283213 | 0.647843293 | 1.16E-21 | 4.45E-20 | 2085.157776 | 1350.716829 | ENSG00000146540 |
| PGAM1P5 | -0.626304505 | 0.647833732 | 0.078597494 | 0.171034885 | 48.06201979 | 31.03802688 | ENSG00000257150 |
| NA | -0.627238269 | 0.647414566 | 0.385040776 | 0.558557808 | 10.69281815 | 6.937338704 | ENSG00000250616 |
| KLLN | -0.627243575 | 0.647412186 | 0.409896275 | 0.582018919 | 9.582434828 | 6.213520375 | ENSG00000227268 |
| NA | -0.62766545 | 0.647222896 | 0.596964282 | 0.744192035 | 4.019754901 | 2.585002399 | ENSG00000258982 |
| ABITRAM | -0.627682642 | 0.647215183 | 0.002233802 | 0.00868796 | 164.2022212 | 106.529485 | ENSG00000119328 |
| NA | -0.627980884 | 0.647081401 | 0.097450606 | 0.20301388 | 38.2184155 | 24.75324658 | ENSG00000123965 |
| MCTP1 | -0.628144003 | 0.647008243 | 0.013702923 | 0.041047064 | 83.6937963 | 54.18041682 | ENSG00000175471 |
| NXF1 | -0.628989029 | 0.646629383 | 7.34E-13 | 1.27E-11 | 1077.228649 | 696.3659335 | ENSG00000162231 |
| NA | -0.629186137 | 0.646541044 | 0.421598098 | 0.593523236 | 8.629869541 | 5.595098579 | ENSG00000283213 |
| CYP2U1 | -0.629507506 | 0.646397039 | 0.001513679 | 0.006176053 | 182.0767624 | 117.9388171 | ENSG00000155016 |
| NA | -0.629825964 | 0.64625437 | 0.101643663 | 0.209667226 | 36.07376696 | 23.31474525 | ENSG00000257354 |
| NA | -0.630333242 | 0.646027175 | 0.33875435 | 0.511352842 | 12.65520625 | 8.167730574 | ENSG00000262454 |
| MGAT4C | -0.63078702 | 0.645824009 | 0.048974893 | 0.117392478 | 63.12140442 | 40.71899582 | ENSG00000182050 |
| MYCN | -0.630788921 | 0.645823158 | 2.03E-18 | 5.96E-17 | 1827.058602 | 1180.167015 | ENSG00000134323 |
| HTRA3 | -0.63093361 | 0.645758391 | 9.25E-05 | 0.000504168 | 236.1626255 | 152.3426762 | ENSG00000170801 |
| MAOA | -0.63125467 | 0.645614699 | 0.000208695 | 0.001047876 | 198.7398331 | 128.3691692 | ENSG00000189221 |
| NUDT8 | -0.631373979 | 0.64556131 | 0.019000467 | 0.053799795 | 103.9115916 | 67.1091785 | ENSG00000167799 |
| MTHFD1 | -0.631556471 | 0.645479655 | 2.07E-42 | 2.61E-40 | 5425.486862 | 3501.800667 | ENSG00000100714 |
| MARCKSL1 | -0.63170713 | 0.645412252 | 1.97E-09 | 2.31E-08 | 12246.62068 | 7904.226673 | ENSG00000175130 |
| NA | -0.631916686 | 0.645318511 | 0.500635644 | 0.666686425 | 7.040850608 | 4.511788178 | ENSG00000238287 |
| CDC27 | -0.632445188 | 0.645082155 | 2.10E-09 | 2.45E-08 | 795.5153524 | 513.2855051 | ENSG00000004897 |
| NA | -0.632558387 | 0.645031541 | 0.581936255 | 0.732447768 | 4.999553496 | 3.201273621 | ENSG00000244265 |
| ATP6V0E1 | -0.632575669 | 0.645023814 | 1.67E-18 | 4.96E-17 | 1448.810323 | 934.7357802 | ENSG00000113732 |
| ZSCAN18 | -0.632862857 | 0.644895426 | 2.15E-06 | 1.59E-05 | 341.9885962 | 220.5220075 | ENSG00000121413 |
| NA | -0.63315327 | 0.644765623 | 0.607658808 | 0.751721564 | 4.109827532 | 2.637700666 | ENSG00000287760 |
| DNAL1 | -0.633541889 | 0.644591966 | 0.000538741 | 0.002449989 | 180.576777 | 116.5196893 | ENSG00000119661 |
| FJX1 | -0.634272233 | 0.644265732 | 4.70E-08 | 4.53E-07 | 532.0408778 | 342.6559022 | ENSG00000179431 |
| TUBA1A | -0.634452673 | 0.644185158 | 1.68E-41 | 2.08E-39 | 5994.689562 | 3861.585386 | ENSG00000167552 |
| DUSP16 | -0.634790457 | 0.64403435 | 1.80E-22 | 7.30E-21 | 2595.337207 | 1671.580498 | ENSG00000111266 |
| CFTR | -0.63542404 | 0.643751574 | 0.350168454 | 0.522800066 | 11.74594396 | 7.551459352 | ENSG00000001626 |
| FAM47E | -0.635479037 | 0.643727034 | 0.556326568 | 0.712480636 | 5.160162438 | 3.304519581 | ENSG00000189157 |
| ERICH6-AS1 | -0.635678266 | 0.643638145 | 0.404312275 | 0.576551456 | 9.622946929 | 6.186095954 | ENSG00000240137 |
| RWDD3 | -0.636051407 | 0.643471695 | 0.216069847 | 0.373193967 | 23.50372726 | 15.10668681 | ENSG00000122481 |
| NA | -0.636057279 | 0.643469076 | 0.449964931 | 0.620273841 | 7.658443655 | 4.936881959 | ENSG00000268230 |
| DNAJB11 | -0.636129125 | 0.643437032 | 2.27E-16 | 5.63E-15 | 1337.843964 | 861.4732668 | ENSG00000090520 |
| SLC4A8 | -0.63634341 | 0.643341468 | 1.40E-06 | 1.07E-05 | 339.6806728 | 218.3900564 | ENSG00000050438 |
| NA | -0.636502053 | 0.643270729 | 0.597032365 | 0.744207247 | 4.098663921 | 2.61887854 | ENSG00000287523 |
| LRRC8E | -0.636782427 | 0.643145727 | 0.291743075 | 0.461500395 | 16.79908819 | 10.84522606 | ENSG00000171017 |
| LRP4-AS1 | -0.636928145 | 0.64308077 | 0.514893606 | 0.678795165 | 5.614355639 | 3.628517976 | ENSG00000247675 |
| UPF3B | -0.637066439 | 0.643019129 | 1.63E-15 | 3.71E-14 | 1104.815071 | 710.7670058 | ENSG00000125351 |
| NISCH | -0.637297927 | 0.642915961 | 2.62E-23 | 1.13E-21 | 3441.781011 | 2212.181924 | ENSG00000010322 |
| NA | -0.637346751 | 0.642894204 | 0.372127292 | 0.54561377 | 12.17098884 | 7.763609492 | ENSG00000230373 |
| FMC1 | -0.637404202 | 0.642868603 | 0.621298183 | 0.761915061 | 3.638688754 | 2.368551112 | ENSG00000164898 |
| NA | -0.637492221 | 0.642829383 | 0.259955458 | 0.425271432 | 17.36309051 | 11.16277274 | ENSG00000259343 |
| NA | -0.637841522 | 0.642673762 | 0.605178604 | 0.750048214 | 4.095873018 | 2.614577393 | ENSG00000257043 |
| P2RY11 | -0.637924438 | 0.642636826 | 0.047551657 | 0.114762986 | 52.62237374 | 33.86124677 | ENSG00000244165 |
| TAF7 | -0.638102802 | 0.642557381 | 4.63E-33 | 3.50E-31 | 3145.762336 | 2020.998718 | ENSG00000178913 |
| SLC20A1 | -0.638208278 | 0.642510404 | 5.00E-31 | 3.34E-29 | 2571.88985 | 1652.575666 | ENSG00000144136 |

| | | | | | | | |
|---|---|---|---|---|---|---|---|
| LOC112694756 | -0.63831604 | 0.642462414 | 0.39013535 | 0.563373085 | 9.694158961 | 6.219972096 | ENSG00000285043 |
| SLC35D1 | -0.638372386 | 0.642437322 | 0.007544649 | 0.024874681 | 104.8308662 | 67.46409685 | ENSG00000116704 |
| SEPT5-GP1BB | -0.638790412 | 0.642251201 | 0.518065585 | 0.681701302 | 6.760344983 | 4.358254988 | ENSG00000284874 |
| NA | -0.638806082 | 0.642244225 | 0.184554845 | 0.330724339 | 24.61901667 | 15.77780688 | ENSG00000244932 |
| AACSP1 | -0.639046765 | 0.642137089 | 0.52336887 | 0.685937148 | 5.592704138 | 3.603244129 | ENSG00000250420 |
| UBE2F | -0.639076598 | 0.642123811 | 0.001999105 | 0.007883224 | 136.9473542 | 88.00159618 | ENSG00000184182 |
| NAGPA | -0.639205721 | 0.642066342 | 6.43E-05 | 0.000362567 | 234.467264 | 150.4712907 | ENSG00000103174 |
| MYL6B | -0.639343208 | 0.642005157 | 4.55E-14 | 8.91E-13 | 1221.076607 | 783.7566015 | ENSG00000196465 |
| NA | -0.639506085 | 0.64193268 | 0.255685408 | 0.420676094 | 20.48107995 | 13.20355735 | ENSG00000286606 |
| GET4 | -0.63960173 | 0.641890125 | 0.000124426 | 0.000657437 | 212.2582957 | 136.2051047 | ENSG00000239857 |
| PTPRG-AS1 | -0.640199627 | 0.64162416 | 0.208602831 | 0.36351917 | 29.58296415 | 18.92529773 | ENSG00000241472 |
| SFXN5 | -0.64021073 | 0.641619223 | 1.32E-10 | 1.80E-09 | 956.1828608 | 613.2108952 | ENSG00000144040 |
| SLC25A27 | -0.640264324 | 0.641595388 | 0.037539916 | 0.094540798 | 74.97069981 | 47.94297967 | ENSG00000153291 |
| VPS8 | -0.640303186 | 0.641578105 | 1.11E-11 | 1.70E-10 | 763.3444077 | 489.8392902 | ENSG00000156931 |
| CYP7B1 | -0.640400835 | 0.641534682 | 0.426385848 | 0.59799414 | 10.25537037 | 6.535368196 | ENSG00000172817 |
| COMMD4 | -0.640449751 | 0.64151293 | 8.43E-19 | 2.56E-17 | 1695.669998 | 1087.720862 | ENSG00000140365 |
| SCN4A | -0.640657704 | 0.641420468 | 8.89E-06 | 5.91E-05 | 282.2402053 | 180.9355248 | ENSG00000007314 |
| DRG2 | -0.640693451 | 0.641404575 | 3.98E-11 | 5.69E-10 | 664.5173871 | 426.4712835 | ENSG00000108591 |
| ZDHHC6 | -0.64070409 | 0.641399845 | 1.62E-11 | 2.42E-10 | 695.171625 | 445.8776326 | ENSG00000023041 |
| C17orf49 | -0.64098958 | 0.641272933 | 0.343638074 | 0.516367197 | 12.77530309 | 8.163429427 | ENSG00000258315 |
| NBPF1 | -0.641191155 | 0.64118334 | 2.83E-08 | 2.81E-07 | 439.7180879 | 282.1204992 | ENSG00000219481 |
| NA | -0.641197839 | 0.641180369 | 0.086069594 | 0.184076334 | 40.55628793 | 26.00461115 | ENSG00000256663 |
| NA | -0.641219047 | 0.641170944 | 0.573020963 | 0.725403106 | 4.585672235 | 2.95524734 | ENSG00000236762 |
| NA | -0.641302939 | 0.641133661 | 0.470720301 | 0.639618604 | 6.681435963 | 4.293186317 | ENSG00000229816 |
| CUL1 | -0.642012092 | 0.640818591 | 1.23E-21 | 4.68E-20 | 2714.503149 | 1739.207192 | ENSG00000055130 |
| NA | -0.642165412 | 0.640750493 | 0.47324612 | 0.641965466 | 6.684226866 | 4.291035743 | ENSG00000260751 |
| RIPOR3 | -0.64228618 | 0.640696858 | 0.045750108 | 0.111126333 | 70.01875492 | 44.89871635 | ENSG00000042062 |
| LPCAT3 | -0.642616565 | 0.640550152 | 1.41E-10 | 1.92E-09 | 602.3476721 | 385.7566131 | ENSG00000111684 |
| NA | -0.643185261 | 0.640297703 | 0.564426347 | 0.718750324 | 5.054020111 | 3.203424194 | ENSG00000270926 |
| CFAP36 | -0.643687348 | 0.640074905 | 4.01E-05 | 0.000235646 | 260.9499112 | 167.1175558 | ENSG00000163001 |
| CNTF | -0.643785979 | 0.640031147 | 0.434423786 | 0.605525673 | 7.71291027 | 4.934731385 | ENSG00000242689 |
| NA | -0.643964911 | 0.639951772 | 0.347794622 | 0.520501609 | 12.77183646 | 8.136005006 | ENSG00000286753 |
| GPLD1 | -0.644018707 | 0.639927909 | 0.003931267 | 0.014228533 | 137.5769867 | 88.07633323 | ENSG00000112293 |
| ADAMTS8 | -0.644105909 | 0.639889231 | 1.57E-10 | 2.12E-09 | 616.5237884 | 394.3903168 | ENSG00000134917 |
| BUD31 | -0.644233674 | 0.639832564 | 8.96E-13 | 1.53E-11 | 814.4704018 | 521.1445702 | ENSG00000106245 |
| NA | -0.644267034 | 0.63981777 | 0.400069211 | 0.572585667 | 11.23728415 | 7.128516145 | ENSG00000226471 |
| ZFP69B | -0.644620572 | 0.639660999 | 0.003488566 | 0.012816211 | 140.4296644 | 90.0259697 | ENSG00000187801 |
| GPR45 | -0.645032844 | 0.639478232 | 0.517820368 | 0.681564084 | 5.587122332 | 3.575819709 | ENSG00000135973 |
| NA | -0.645195069 | 0.63940633 | 0.384220357 | 0.557628266 | 10.23284299 | 6.562792616 | ENSG00000279694 |
| DCTN3 | -0.645618738 | 0.639218586 | 6.54E-07 | 5.29E-06 | 355.2450504 | 226.8955128 | ENSG00000137100 |
| NRG3 | -0.645767755 | 0.639152564 | 0.062471207 | 0.142416929 | 50.52728573 | 32.41440361 | ENSG00000185737 |
| RIDA | -0.646682119 | 0.638747605 | 9.51E-10 | 1.15E-08 | 574.4400575 | 366.957945 | ENSG00000132541 |
| CCM2L | -0.647434482 | 0.638414586 | 0.003084728 | 0.01147243 | 127.9700726 | 81.82169149 | ENSG00000101331 |
| MIR1915 | -0.647466831 | 0.638400271 | 0.563142859 | 0.717666672 | 4.58009043 | 2.932124067 | ENSG00000222071 |
| MRTO4 | -0.647500551 | 0.63838535 | 7.59E-22 | 2.96E-20 | 1833.382588 | 1170.336553 | ENSG00000053372 |
| NA | -0.647865375 | 0.638223937 | 0.412946173 | 0.585186558 | 10.95099656 | 7.044352774 | ENSG00000218690 |
| PIGH | -0.647936998 | 0.638192254 | 0.008997308 | 0.028783926 | 96.70283539 | 61.75231583 | ENSG00000100564 |
| PCDHA12 | -0.648179889 | 0.638084817 | 0.000214233 | 0.001072331 | 176.9848578 | 112.9403741 | ENSG00000251664 |
| BEX1 | -0.648305571 | 0.638029232 | 2.51E-17 | 6.81E-16 | 1325.416586 | 845.3984382 | ENSG00000133169 |
| FBN2 | -0.648574689 | 0.637910226 | 1.03E-10 | 1.42E-09 | 939.8937825 | 599.5848118 | ENSG00000138829 |
| TMEM199 | -0.648734586 | 0.637839529 | 0.01589125 | 0.046455236 | 89.55412759 | 57.03833688 | ENSG00000244045 |
| KSR2 | -0.649302856 | 0.637588337 | 0.012673279 | 0.038440288 | 85.83720554 | 54.59072915 | ENSG00000171435 |
| GABRA4 | -0.649861706 | 0.637341405 | 0.45744726 | 0.627327439 | 7.753422371 | 4.932580812 | ENSG00000109158 |
| DHX38 | -0.649964485 | 0.637296002 | 1.42E-27 | 8.00E-26 | 2382.01058 | 1517.707216 | ENSG00000140829 |
| ARRDC3 | -0.650079313 | 0.63724528 | 0.011749903 | 0.036024695 | 81.76489904 | 52.06649428 | ENSG00000113369 |
| COL9A2 | -0.650569177 | 0.637028941 | 0.001017264 | 0.004327243 | 156.6762262 | 99.89033783 | ENSG00000049089 |
| LOC730101 | -0.650763657 | 0.636943074 | 1.51E-05 | 9.57E-05 | 275.9380838 | 175.9561339 | ENSG00000216775 |
| TNFSF13 | -0.650798187 | 0.636927829 | 0.044878431 | 0.109472922 | 54.73017696 | 34.8165704 | ENSG00000161955 |
| GORASP1 | -0.651087707 | 0.636800023 | 2.89E-14 | 5.79E-13 | 1061.354606 | 675.8646787 | ENSG00000114745 |
| OBI1 | -0.651532332 | 0.636603797 | 0.000988266 | 0.00422119 | 177.8369506 | 113.2256622 | ENSG00000152193 |
| SH2B2 | -0.651588157 | 0.636579165 | 1.71E-07 | 1.52E-06 | 450.8323593 | 287.1509708 | ENSG00000160999 |
| NA | -0.651617726 | 0.636566118 | 0.256474875 | 0.421579483 | 17.56122049 | 11.2240733 | ENSG00000249572 |
| NA | -0.651769463 | 0.636499169 | 0.267401715 | 0.433724271 | 16.41610702 | 10.40346073 | ENSG00000260005 |
| LOC100507437 | -0.652595921 | 0.636134651 | 0.069583717 | 0.155330448 | 54.10505019 | 34.52591511 | ENSG00000251602 |
| NA | -0.653184229 | 0.635875298 | 0.509937505 | 0.674379942 | 5.726079773 | 3.645189528 | ENSG00000242262 |
| CPNE1 | -0.653495859 | 0.63573796 | 4.93E-20 | 1.67E-18 | 1374.955173 | 873.7648393 | ENSG00000214078 |
| SLITRK1 | -0.65366977 | 0.635661329 | 0.517394525 | 0.681347381 | 5.677194963 | 3.607545277 | ENSG00000178235 |
| ZMPSTE24 | -0.654282912 | 0.635391232 | 2.86E-13 | 5.19E-12 | 1095.089538 | 696.037585 | ENSG00000084073 |
| SPICE1 | -0.654422802 | 0.635329625 | 0.457710259 | 0.627598878 | 6.730320772 | 4.265761897 | ENSG00000163611 |
| IQGAP1 | -0.654668816 | 0.635221296 | 2.31E-31 | 1.57E-29 | 3401.01211 | 2160.714774 | ENSG00000140575 |
| SCT | -0.654780399 | 0.635172167 | 0.221454985 | 0.379808687 | 18.66390683 | 11.88874164 | ENSG00000070031 |
| PCNX3 | -0.654835051 | 0.635148106 | 1.15E-19 | 3.80E-18 | 1884.900235 | 1197.066885 | ENSG00000197136 |
| CNIH2 | -0.654905209 | 0.63511722 | 0.00028178 | 0.001375461 | 220.7763531 | 140.1009126 | ENSG00000174871 |
| NA | -0.655938771 | 0.634662378 | 0.391093064 | 0.564219438 | 9.808673997 | 6.186095954 | ENSG00000260236 |
| KCNK10 | -0.655960818 | 0.634652679 | 0.105200192 | 0.215220596 | 33.6325431 | 21.3772066 | ENSG00000100433 |
| POU6F1 | -0.656171597 | 0.634559963 | 0.005329699 | 0.018525264 | 102.9948325 | 65.42599585 | ENSG00000184271 |
| ANKRD18B | -0.656795816 | 0.634285464 | 0.003571787 | 0.013082084 | 115.2025034 | 73.01079831 | ENSG00000230453 |

| | | | | | | | |
|---|---|---|---|---|---|---|---|
| RAPGEF4 | -0.65685058 | 0.634261387 | 0.039951626 | 0.099523535 | 55.07264603 | 35.01607756 | ENSG00000091428 |
| SLC25A53 | -0.657325083 | 0.634052812 | 1.51E-05 | 9.61E-05 | 253.4199371 | 160.8672031 | ENSG00000269743 |
| FBXW4P1 | -0.657767271 | 0.633858504 | 0.302302139 | 0.473007588 | 15.40137814 | 9.71136797 | ENSG00000230701 |
| PTDSS2 | -0.658071555 | 0.633724829 | 1.31E-20 | 4.69E-19 | 1498.947244 | 950.2812819 | ENSG00000174915 |
| CENPV | -0.658111548 | 0.633707261 | 3.21E-24 | 1.49E-22 | 2294.728659 | 1454.713676 | ENSG00000166582 |
| NA | -0.65812556 | 0.633701107 | 0.585305378 | 0.735147214 | 3.665922062 | 2.341126692 | ENSG00000199804 |
| PJA1 | -0.658147841 | 0.63369132 | 1.97E-25 | 9.84E-24 | 1824.895579 | 1156.25293 | ENSG00000181191 |
| PRRT2 | -0.658150801 | 0.633690019 | 0.001399803 | 0.005765074 | 172.5138128 | 109.0970345 | ENSG00000167371 |
| MX1 | -0.65822519 | 0.633657346 | 0.031599994 | 0.081923866 | 62.97810447 | 39.86612465 | ENSG00000157601 |
| NA | -0.658331841 | 0.633610505 | 0.208719858 | 0.363631034 | 19.61926302 | 12.42919132 | ENSG00000269883 |
| TNFRSF11A | -0.65868291 | 0.633456339 | 2.75E-05 | 0.000167045 | 224.0424365 | 141.9991774 | ENSG00000141655 |
| CTBP1-AS | -0.658906615 | 0.633358123 | 0.236110008 | 0.397486531 | 17.70364455 | 11.24289543 | ENSG00000280927 |
| NRTN | -0.659132739 | 0.63325886 | 0.015342368 | 0.045103987 | 92.43375036 | 58.51797168 | ENSG00000171119 |
| SCHIP1 | -0.659143147 | 0.633254291 | 0.005557944 | 0.019218192 | 108.3321491 | 68.64662464 | ENSG00000283154 |
| NA | -0.659690686 | 0.633014001 | 0.160619491 | 0.297913186 | 27.54637296 | 17.4983612 | ENSG00000228158 |
| H4C12 | -0.659944477 | 0.632902654 | 0.373983143 | 0.547214396 | 10.33061261 | 6.528916475 | ENSG00000273542 |
| HOMER2 | -0.660190801 | 0.632794602 | 2.19E-09 | 2.55E-08 | 502.4192655 | 317.8117496 | ENSG00000103942 |
| FCN3 | -0.660198106 | 0.632791398 | 0.532783038 | 0.693469913 | 4.672953964 | 2.959548487 | ENSG00000142748 |
| NA | -0.660355681 | 0.632722287 | 0.429801079 | 0.601297601 | 7.764585982 | 4.882033118 | ENSG00000249631 |
| FBXL13 | -0.660546595 | 0.632638563 | 0.332662993 | 0.505426621 | 12.90010586 | 8.167730574 | ENSG00000161040 |
| PDIA2 | -0.66061069 | 0.632610458 | 0.265245077 | 0.431255186 | 16.08113478 | 10.15743445 | ENSG00000185615 |
| INPPL1 | -0.66083239 | 0.632513251 | 5.37E-25 | 2.66E-23 | 2426.393825 | 1534.272329 | ENSG00000165458 |
| NA | -0.661239115 | 0.632334958 | 0.54885284 | 0.70633223 | 6.218870053 | 3.893366382 | ENSG00000219797 |
| POMC | -0.661266569 | 0.632322925 | 0.014805317 | 0.043811839 | 80.77306096 | 51.09664967 | ENSG00000115138 |
| NUBPL | -0.6613265 | 0.632296658 | 0.000172937 | 0.000883038 | 200.1572796 | 126.7074923 | ENSG00000151413 |
| EXOSC1 | -0.661440526 | 0.632246686 | 3.55E-11 | 5.12E-10 | 710.557771 | 448.895265 | ENSG00000171311 |
| CRELD2 | -0.661574677 | 0.632187898 | 3.52E-12 | 5.65E-11 | 1057.497801 | 668.4014845 | ENSG00000184164 |
| WDR90 | -0.661904273 | 0.632043486 | 3.00E-11 | 4.36E-10 | 802.2373998 | 507.0919403 | ENSG00000161996 |
| FCRLB | -0.662277784 | 0.631879872 | 0.63678219 | 0.772783419 | 4.569602539 | 2.881576373 | ENSG00000162746 |
| NA | -0.662303185 | 0.631868747 | 0.495594684 | 0.661856494 | 6.754763177 | 4.251240919 | ENSG00000233514 |
| NA | -0.662385597 | 0.631832653 | 0.578370346 | 0.730097 | 4.582881332 | 2.888028095 | ENSG00000280107 |
| MGC16275 | -0.662681183 | 0.631703214 | 0.389859436 | 0.563101107 | 9.343117025 | 5.900274849 | ENSG00000246731 |
| NA | -0.663198929 | 0.631476552 | 0.590287927 | 0.739479472 | 3.720388677 | 2.366400538 | ENSG00000233668 |
| MAGEF1 | -0.663662693 | 0.631273593 | 1.75E-19 | 5.64E-18 | 1795.30969 | 1133.560096 | ENSG00000177383 |
| GRID1 | -0.663783011 | 0.631220948 | 0.342262787 | 0.514994089 | 11.41164744 | 7.176913264 | ENSG00000182771 |
| EFS | -0.663801099 | 0.631213034 | 2.33E-14 | 4.75E-13 | 1105.345696 | 697.732363 | ENSG00000100842 |
| WNT2B | -0.664669197 | 0.630833335 | 0.212051236 | 0.367900026 | 21.76092049 | 13.76282915 | ENSG00000134245 |
| NA | -0.664821736 | 0.630766639 | 0.427082978 | 0.59858019 | 8.801642095 | 5.509057209 | ENSG00000255639 |
| NA | -0.664836557 | 0.630760159 | 0.470174115 | 0.639101526 | 7.900552361 | 5.033676198 | ENSG00000267890 |
| USP49 | -0.665411273 | 0.630508938 | 1.50E-11 | 2.25E-10 | 704.7008183 | 443.9097255 | ENSG00000164663 |
| NA | -0.665643698 | 0.630407368 | 0.403224446 | 0.575553295 | 8.390551738 | 5.300675179 | ENSG00000279759 |
| NA | -0.665758009 | 0.63035742 | 0.503280357 | 0.668747918 | 7.234074504 | 4.566637019 | ENSG00000225883 |
| CHST1 | -0.665767686 | 0.630353192 | 0.0405159 | 0.100694695 | 59.5536154 | 37.64785954 | ENSG00000175264 |
| ATP7B | -0.665851434 | 0.630316601 | 5.70E-06 | 3.92E-05 | 317.9219427 | 200.2482905 | ENSG00000123191 |
| MRPS18C | -0.665990323 | 0.630255923 | 0.002792903 | 0.010549814 | 119.7506177 | 75.57644554 | ENSG00000163319 |
| ZFYVE26 | -0.666117152 | 0.630200519 | 3.61E-16 | 8.82E-15 | 988.0085554 | 622.5129742 | ENSG00000072121 |
| NA | -0.666482879 | 0.630040782 | 0.449423543 | 0.619627613 | 7.304610816 | 4.617184712 | ENSG00000243485 |
| DTX4 | -0.666528923 | 0.630020674 | 3.11E-13 | 5.61E-12 | 899.6355349 | 566.4337533 | ENSG00000110042 |
| NA | -0.666568146 | 0.630003546 | 0.003557907 | 0.013043631 | 103.6922104 | 65.28027146 | ENSG00000277013 |
| FGFR1 | -0.666700255 | 0.629945859 | 7.24E-65 | 1.86E-62 | 9960.326261 | 6274.20026 | ENSG00000077782 |
| INSRR | -0.666938306 | 0.629841923 | 0.536995436 | 0.696555874 | 4.637347948 | 2.911151367 | ENSG00000027644 |
| MAMDC4 | -0.667182913 | 0.629735143 | 2.87E-06 | 2.08E-05 | 345.0557489 | 217.3887588 | ENSG00000177943 |
| NA | -0.66748059 | 0.629605221 | 0.53570979 | 0.695589794 | 4.702978174 | 2.957397913 | ENSG00000213856 |
| NA | -0.667575854 | 0.629563649 | 0.360353727 | 0.533400461 | 11.08225701 | 7.006708523 | ENSG00000268087 |
| ST3GAL5 | -0.667593875 | 0.629555785 | 0.031285256 | 0.08127173 | 64.86333531 | 40.80665473 | ENSG00000115525 |
| CARD9 | -0.667874569 | 0.629433309 | 0.16048964 | 0.297809112 | 25.44590331 | 15.96091506 | ENSG00000187796 |
| TSPAN33 | -0.667960921 | 0.629395635 | 1.16E-13 | 2.19E-12 | 975.4719569 | 613.920695 | ENSG00000158457 |
| THEM6 | -0.66822289 | 0.629281358 | 0.000389841 | 0.001845048 | 197.5425315 | 124.2220104 | ENSG00000130193 |
| GDF6 | -0.668276563 | 0.629257947 | 0.360000843 | 0.533095927 | 12.39914303 | 7.757690809 | ENSG00000156466 |
| NA | -0.668328472 | 0.629235307 | 0.000329756 | 0.001585177 | 161.8001069 | 101.7033973 | ENSG00000226067 |
| NA | -0.668371925 | 0.629216355 | 0.502426492 | 0.668228848 | 6.248694104 | 3.950365797 | ENSG00000230105 |
| RASGRF1 | -0.66863111 | 0.629103324 | 0.193280963 | 0.342984368 | 20.38139531 | 12.83116183 | ENSG00000058335 |
| KREMEN1 | -0.669050498 | 0.628920472 | 0.000371859 | 0.001766445 | 220.1619513 | 138.3166346 | ENSG00000183762 |
| ZSCAN1 | -0.669102858 | 0.628897647 | 0.313882423 | 0.485097514 | 14.03704677 | 8.807125069 | ENSG00000152467 |
| NA | -0.669144448 | 0.628879517 | 0.245709538 | 0.409112956 | 249.1536174 | 156.711151 | ENSG00000198744 |
| EYA2 | -0.669222267 | 0.628845596 | 0.124850963 | 0.245424828 | 31.82864877 | 19.98656936 | ENSG00000064655 |
| TOP3B | -0.669224665 | 0.628844551 | 0.296425218 | 0.466750186 | 13.61825942 | 8.600893612 | ENSG00000100038 |
| NA | -0.669549579 | 0.628702943 | 0.121160463 | 0.240265881 | 30.23596305 | 19.03392934 | ENSG00000255836 |
| CDK4 | -0.670132 | 0.628449185 | 5.76E-30 | 3.68E-28 | 2437.900701 | 1531.668207 | ENSG00000135446 |
| AREL1 | -0.670171677 | 0.628431901 | 1.38E-17 | 3.78E-16 | 1113.497441 | 699.448313 | ENSG00000119682 |
| FBXW4 | -0.670307053 | 0.628372934 | 3.23E-12 | 5.21E-11 | 724.5900381 | 455.4833194 | ENSG00000107829 |
| NA | -0.670590389 | 0.628249538 | 0.322100855 | 0.493596597 | 12.48921566 | 7.793184486 | ENSG00000267390 |
| NA | -0.670645182 | 0.628225678 | 0.441411655 | 0.61252138 | 8.2543852 | 5.174305945 | ENSG00000286311 |
| NA | -0.671067911 | 0.628041626 | 0.185902734 | 0.3323679 | 21.40449691 | 13.42430978 | ENSG00000217648 |
| REC8 | -0.671125671 | 0.628016483 | 1.56E-10 | 2.11E-09 | 596.8417982 | 374.6481561 | ENSG00000100918 |
| NA | -0.671353624 | 0.627917261 | 0.416482639 | 0.588583443 | 8.469460758 | 5.323798452 | ENSG00000259172 |
| NA | -0.671414649 | 0.627890701 | 0.487505813 | 0.654565097 | 6.229157783 | 3.92509195 | ENSG00000227440 |

| | | | | | | | |
|---|---|---|---|---|---|---|---|
| TMEM88 | -0.671769112 | 0.62773645 | 0.145466586 | 0.275739011 | 28.03704806 | 17.53627802 | ENSG00000167874 |
| NA | -0.672879162 | 0.627253638 | 0.481043346 | 0.649073734 | 6.291997107 | 3.964886775 | ENSG00000217835 |
| UCK1 | -0.672896836 | 0.627245954 | 1.01E-16 | 2.59E-15 | 1263.555298 | 792.3131508 | ENSG00000130717 |
| FHOD3 | -0.672965246 | 0.627216212 | 3.07E-07 | 2.62E-06 | 334.4523646 | 209.9238617 | ENSG00000134775 |
| SEMA7A | -0.67351167 | 0.626978697 | 1.72E-13 | 3.17E-12 | 793.8206156 | 497.8164069 | ENSG00000138623 |
| PRXL2B | -0.673606442 | 0.626937512 | 2.01E-22 | 8.13E-21 | 2055.725296 | 1288.66176 | ENSG00000157870 |
| JPH4 | -0.673985038 | 0.62677301 | 1.17E-28 | 6.95E-27 | 3328.475031 | 2086.466429 | ENSG00000092051 |
| CRB2 | -0.673999969 | 0.626766524 | 1.45E-05 | 9.24E-05 | 327.0492715 | 204.9942492 | ENSG00000148204 |
| AQP4-AS1 | -0.674063221 | 0.626739045 | 0.52320798 | 0.68581947 | 5.747731275 | 3.58227143 | ENSG00000260372 |
| NA | -0.674097498 | 0.626724154 | 0.428571103 | 0.599881378 | 7.347913819 | 4.610732991 | ENSG00000183889 |
| NA | -0.674593496 | 0.626508724 | 0.269844251 | 0.436232754 | 14.76703967 | 9.267712527 | ENSG00000273137 |
| PNMA1 | -0.674646247 | 0.626485817 | 3.33E-29 | 2.05E-27 | 2166.232704 | 1357.309681 | ENSG00000176903 |
| HSD17B1 | -0.674827582 | 0.626407077 | 0.154957993 | 0.289942699 | 25.20937641 | 15.76758705 | ENSG00000108786 |
| TTLL4 | -0.674883127 | 0.626382961 | 3.61E-26 | 1.89E-24 | 2066.273311 | 1294.754763 | ENSG00000135912 |
| PARD6A | -0.675157753 | 0.626263736 | 8.83E-05 | 0.000483271 | 215.3061492 | 134.6725112 | ENSG00000102981 |
| NA | -0.67591804 | 0.625933788 | 0.496843935 | 0.663111689 | 5.79171 | 3.603244129 | ENSG00000230185 |
| ARL9 | -0.676078371 | 0.62586423 | 0.267849588 | 0.434172457 | 15.81737458 | 9.896887191 | ENSG00000196503 |
| TRH | -0.676329386 | 0.625755345 | 0.549312676 | 0.706644528 | 6.177482071 | 3.865941962 | ENSG00000170893 |
| BEND5 | -0.676964228 | 0.625480049 | 0.003142169 | 0.011659084 | 115.1031822 | 71.99524019 | ENSG00000162373 |
| DLEU7 | -0.677104278 | 0.625419333 | 0.006553528 | 0.0221292 | 112.5561794 | 70.58335767 | ENSG00000186047 |
| MVB12B | -0.677479437 | 0.62525672 | 4.17E-07 | 3.48E-06 | 311.430264 | 194.7115179 | ENSG00000196814 |
| LKAAEAR1 | -0.677609185 | 0.62520049 | 0.503052337 | 0.668622201 | 6.953568879 | 4.39213113 | ENSG00000171695 |
| NA | -0.677832555 | 0.625103699 | 0.361050828 | 0.534210119 | 9.957919169 | 6.234493074 | ENSG00000241627 |
| FMNL1 | -0.678291863 | 0.624904718 | 0.000107219 | 0.000576304 | 208.6095947 | 130.2465039 | ENSG00000184922 |
| DPP4 | -0.678318737 | 0.624893077 | 0.011974568 | 0.03657363 | 86.2102624 | 53.9537457 | ENSG00000197635 |
| NA | -0.678368935 | 0.624871335 | 0.596093613 | 0.743757964 | 3.774855292 | 2.366400538 | ENSG00000241651 |
| BIVM | -0.67882142 | 0.624675382 | 0.005152739 | 0.018000995 | 93.10900124 | 58.12433089 | ENSG00000134897 |
| NA | -0.679318917 | 0.624460007 | 0.117372665 | 0.234555102 | 35.32682848 | 22.1746959 | ENSG00000261754 |
| NA | -0.679821145 | 0.624242659 | 0.258009935 | 0.423078999 | 14.74259727 | 9.196192135 | ENSG00000278376 |
| NA | -0.679888932 | 0.624213328 | 0.461690564 | 0.63153036 | 7.39747435 | 4.617184712 | ENSG00000285763 |
| NA | -0.680216469 | 0.624071629 | 0.147439424 | 0.278575312 | 25.3998094 | 15.90202554 | ENSG00000253683 |
| MPP2 | -0.680321035 | 0.624026398 | 4.53E-28 | 2.61E-26 | 2453.331315 | 1530.554256 | ENSG00000108852 |
| KIF9 | -0.680686885 | 0.623868173 | 0.007846071 | 0.025706278 | 90.30258077 | 56.31935273 | ENSG00000088727 |
| WDR62 | -0.680984094 | 0.623739663 | 6.30E-21 | 2.29E-19 | 1912.659042 | 1192.690359 | ENSG00000075702 |
| PTBP2 | -0.681290703 | 0.623607117 | 5.34E-14 | 1.04E-12 | 824.5569026 | 514.4298498 | ENSG00000117569 |
| OTX1 | -0.681386407 | 0.62356575 | 0.172429103 | 0.314600408 | 24.11506278 | 14.9867693 | ENSG00000115507 |
| PARP3 | -0.681986117 | 0.623306595 | 0.067956607 | 0.152382995 | 40.52347282 | 25.21787472 | ENSG00000041880 |
| NA | -0.682328937 | 0.6231585 | 0.000633833 | 0.002828264 | 159.0757866 | 98.95169785 | ENSG00000276710 |
| NA | -0.682664474 | 0.623013585 | 0.301341653 | 0.472097506 | 13.83612589 | 8.630468606 | ENSG00000256238 |
| EXOSC10-AS1 | -0.682672862 | 0.623009962 | 0.379068155 | 0.552436638 | 8.953678169 | 5.595098579 | ENSG00000230337 |
| DPP10-AS1 | -0.683024931 | 0.622857944 | 0.030718213 | 0.080089883 | 56.24220189 | 35.02872054 | ENSG00000235026 |
| MAP3K8 | -0.683390931 | 0.62269995 | 0.01899539 | 0.053793315 | 81.13056058 | 50.65378773 | ENSG00000107968 |
| EID1 | -0.683752968 | 0.622543707 | 4.61E-29 | 2.79E-27 | 2915.413815 | 1815.030117 | ENSG00000255302 |
| NA | -0.683865663 | 0.622495079 | 0.570580164 | 0.723530004 | 3.72317958 | 2.315852845 | ENSG00000259416 |
| NA | -0.684259385 | 0.622325218 | 0.475411119 | 0.643857075 | 6.814811598 | 4.196392078 | ENSG00000284720 |
| NA | -0.684428898 | 0.622252101 | 0.543257437 | 0.701925366 | 5.247244007 | 3.277095161 | ENSG00000232748 |
| ARSB | -0.684537041 | 0.622205459 | 1.45E-07 | 1.30E-06 | 351.2683984 | 218.4586327 | ENSG00000113273 |
| C12orf10 | -0.684727685 | 0.622123244 | 5.25E-11 | 7.43E-10 | 742.0299096 | 461.8278011 | ENSG00000139637 |
| SIRT2 | -0.6848621 | 0.622065284 | 0.000239667 | 0.001189764 | 206.3115955 | 128.3737308 | ENSG00000068903 |
| AMER3 | -0.685091341 | 0.621966447 | 0.422182462 | 0.593804067 | 7.955018976 | 4.976676784 | ENSG00000178171 |
| ME3 | -0.685123687 | 0.621952502 | 0.057165026 | 0.132911631 | 47.04796662 | 29.21663763 | ENSG00000151376 |
| HTRA1 | -0.685167785 | 0.621933492 | 5.93E-11 | 8.34E-10 | 649.3634993 | 404.2874218 | ENSG00000166033 |
| NA | -0.685288095 | 0.621881629 | 0.156978816 | 0.292756994 | 25.18684903 | 15.66003994 | ENSG00000213740 |
| LOC642361 | -0.685296102 | 0.621878178 | 0.044874587 | 0.109472922 | 55.34198804 | 34.41083178 | ENSG00000272447 |
| NA | -0.685534086 | 0.621775603 | 0.133004855 | 0.257248511 | 27.54513366 | 17.11789643 | ENSG00000215102 |
| ANAPC2 | -0.685897601 | 0.621618954 | 3.03E-14 | 6.07E-13 | 1381.211905 | 858.2444115 | ENSG00000176248 |
| OTUD7A | -0.686281988 | 0.621453354 | 0.000862042 | 0.003723568 | 150.5766651 | 93.59428373 | ENSG00000169918 |
| SLC25A10 | -0.686673205 | 0.621284857 | 0.003201466 | 0.011856301 | 110.0517528 | 68.46245038 | ENSG00000183048 |
| CPEB3 | -0.686751696 | 0.621251056 | 0.164478869 | 0.303429459 | 24.73823763 | 15.33012286 | ENSG00000107864 |
| EPHB6 | -0.687336891 | 0.620999111 | 8.55E-13 | 1.47E-11 | 776.1096375 | 481.7775439 | ENSG00000106123 |
| NA | -0.687702925 | 0.620841574 | 0.431868285 | 0.60292163 | 7.350704722 | 4.543513746 | ENSG00000250308 |
| CLDN6 | -0.687783518 | 0.620806893 | 7.27E-59 | 1.67E-56 | 14600.46626 | 9064.501592 | ENSG00000184697 |
| LRP2 | -0.688130678 | 0.620657525 | 0.131170622 | 0.254466258 | 28.00634813 | 17.38489541 | ENSG00000081479 |
| ATG2A | -0.688583086 | 0.620462926 | 1.92E-13 | 3.53E-12 | 911.8909535 | 565.6838555 | ENSG00000110046 |
| DGCR6L | -0.688742943 | 0.62039418 | 2.68E-14 | 5.40E-13 | 1342.406334 | 832.6640827 | ENSG00000128185 |
| SNHG19 | -0.688893022 | 0.620329646 | 1.18E-07 | 1.08E-06 | 425.50903 | 263.6490888 | ENSG00000260260 |
| NA | -0.689189167 | 0.620202322 | 0.143269577 | 0.272699892 | 28.20582955 | 17.58278503 | ENSG00000256667 |
| DGCR11 | -0.68954627 | 0.620048826 | 0.118125402 | 0.235652866 | 37.97869738 | 23.58981349 | ENSG00000273311 |
| SUFU | -0.690378572 | 0.619691218 | 3.48E-10 | 4.50E-09 | 519.3575993 | 321.8215258 | ENSG00000107882 |
| ZNF664 | -0.69105435 | 0.619401014 | 2.14E-22 | 8.59E-21 | 1731.370856 | 1072.561125 | ENSG00000179195 |
| MED14OS | -0.691508995 | 0.619205849 | 0.568784171 | 0.721916932 | 3.750412887 | 2.315852845 | ENSG00000234636 |
| AKT1S1 | -0.691955455 | 0.619014258 | 1.72E-16 | 4.33E-15 | 1497.604778 | 927.2029188 | ENSG00000204673 |
| ELMOD1 | -0.692333455 | 0.618852092 | 0.110614183 | 0.223557696 | 32.06238477 | 19.91961058 | ENSG00000110675 |
| NRGN | -0.692994017 | 0.618568805 | 0.40024553 | 0.57274212 | 7.97946138 | 4.953553511 | ENSG00000154146 |
| ZNF433-AS1 | -0.693210223 | 0.618476111 | 0.016481386 | 0.047825272 | 75.96445292 | 46.89380587 | ENSG00000219665 |
| ATF5 | -0.69323323 | 0.618466248 | 1.78E-05 | 0.000111589 | 220.1446934 | 136.1766142 | ENSG00000169136 |
| KCNQ1 | -0.693274986 | 0.618448348 | 0.01469263 | 0.043525111 | 75.68989252 | 46.94084592 | ENSG00000053918 |

| | | | | | | | |
|---|---|---|---|---|---|---|---|
| TMX4 | -0.693329806 | 0.618424849 | 5.30E-06 | 3.66E-05 | 288.9913017 | 178.981563 | ENSG00000125827 |
| CHKB | -0.694206426 | 0.618049191 | 0.47414839 | 0.642434437 | 6.458187856 | 3.992311195 | ENSG00000100288 |
| LZTR1 | -0.694215746 | 0.618045199 | 0.000654324 | 0.002905745 | 170.0422524 | 105.0237766 | ENSG00000099949 |
| NA | -0.694569395 | 0.617893715 | 0.351354934 | 0.523906065 | 10.0668524 | 6.196848823 | ENSG00000215007 |
| NA | -0.694618042 | 0.617872881 | 0.001380099 | 0.00568878 | 120.7250346 | 74.68080494 | ENSG00000276529 |
| NA | -0.694738262 | 0.617821395 | 0.307767432 | 0.478878974 | 13.19389027 | 8.161278853 | ENSG00000267980 |
| LINC00511 | -0.694821738 | 0.617785649 | 0.269179661 | 0.435576741 | 17.14733923 | 10.6771719 | ENSG00000227036 |
| COL21A1 | -0.695129892 | 0.617653706 | 0.008988149 | 0.028760547 | 77.18201771 | 47.70634918 | ENSG00000124749 |
| NA | -0.695152117 | 0.617644191 | 0.394639787 | 0.567202811 | 8.033927996 | 4.980977931 | ENSG00000278987 |
| DNLZ | -0.695389945 | 0.617542381 | 0.474157554 | 0.642434437 | 7.315098706 | 4.488664905 | ENSG00000213221 |
| PCYT2 | -0.695679495 | 0.617418452 | 4.36E-15 | 9.45E-14 | 1253.770341 | 773.9934019 | ENSG00000185813 |
| EXT2 | -0.695899996 | 0.617324093 | 4.35E-28 | 2.51E-26 | 2127.819975 | 1313.705328 | ENSG00000151348 |
| UBA7 | -0.695958102 | 0.61729923 | 0.480617333 | 0.648725789 | 6.322021318 | 3.889065235 | ENSG00000182179 |
| FANK1 | -0.696614728 | 0.617018338 | 0.450962437 | 0.621349645 | 6.978011284 | 4.335131715 | ENSG00000203780 |
| SMIM10L1 | -0.696867144 | 0.616910393 | 2.70E-08 | 2.69E-07 | 365.3646177 | 225.5123812 | ENSG00000256537 |
| APOM | -0.697666249 | 0.616568782 | 0.267096641 | 0.433388957 | 16.93892152 | 10.36581648 | ENSG00000204444 |
| DDX49 | -0.697756409 | 0.616530252 | 5.12E-14 | 9.96E-13 | 1046.832404 | 645.4902967 | ENSG00000105671 |
| NA | -0.697757311 | 0.616529866 | 0.153204745 | 0.287469385 | 22.42269242 | 13.83434955 | ENSG00000250132 |
| SPC25 | -0.698040171 | 0.616408999 | 2.04E-09 | 2.40E-08 | 596.0492301 | 367.3392154 | ENSG00000152253 |
| PGAM4 | -0.698264136 | 0.616313315 | 0.451139698 | 0.621416345 | 8.10725521 | 5.010552925 | ENSG00000226784 |
| KCNK15 | -0.698499538 | 0.61621276 | 0.39753473 | 0.570131354 | 8.025555287 | 4.955704085 | ENSG00000124249 |
| NA | -0.698803191 | 0.616083076 | 0.000629768 | 0.002814189 | 148.0729142 | 91.27223962 | ENSG00000280079 |
| MFSD1 | -0.698848792 | 0.616063603 | 0.000445285 | 0.002071876 | 154.9798767 | 95.60172523 | ENSG00000118855 |
| HES1 | -0.699594661 | 0.615745182 | 8.30E-05 | 0.000456923 | 222.7232351 | 137.3204195 | ENSG00000114315 |
| ARMCX5 | -0.699770681 | 0.615670061 | 7.21E-05 | 0.000402441 | 188.0797179 | 115.7316079 | ENSG00000125962 |
| SNORA11F | -0.699864788 | 0.615629902 | 0.20645253 | 0.360684274 | 21.46263031 | 13.16322948 | ENSG00000221164 |
| PIM2 | -0.700071644 | 0.615541638 | 1.35E-45 | 1.94E-43 | 5394.586272 | 3319.983865 | ENSG00000102096 |
| RTN1 | -0.700111329 | 0.615524706 | 0.006879517 | 0.023068446 | 89.8194394 | 55.20295969 | ENSG00000139970 |
| CTAGE8 | -0.70052139 | 0.615349779 | 0.461501636 | 0.631316697 | 6.360418237 | 3.895516956 | ENSG00000244693 |
| CPXM1 | -0.700819934 | 0.615222455 | 3.43E-28 | 2.00E-26 | 1878.514685 | 1155.535285 | ENSG00000088882 |
| SLC26A8 | -0.700914672 | 0.615182056 | 0.411661251 | 0.583837465 | 7.938273558 | 4.879882545 | ENSG00000112053 |
| ANKRD36B | -0.702340186 | 0.614574501 | 0.016024725 | 0.046746253 | 65.12342873 | 40.0075479 | ENSG00000196912 |
| NRSN1 | -0.702945078 | 0.614316876 | 0.500790157 | 0.666686425 | 5.306616707 | 3.256122461 | ENSG00000152954 |
| DEXI | -0.703412278 | 0.614117969 | 0.016436922 | 0.047710603 | 73.95297985 | 45.44104403 | ENSG00000182108 |
| EAPP | -0.703483308 | 0.614087734 | 1.73E-08 | 1.78E-07 | 386.014516 | 236.8348784 | ENSG00000129518 |
| KANK4 | -0.703825398 | 0.61394214 | 0.333549248 | 0.506034301 | 11.29992331 | 6.974982955 | ENSG00000132854 |
| NAT8L | -0.703893029 | 0.61391336 | 6.57E-15 | 1.41E-13 | 1015.821593 | 624.0727196 | ENSG00000185818 |
| PJVK | -0.705300151 | 0.613314876 | 0.268390983 | 0.43470554 | 15.55640527 | 9.507820125 | ENSG00000204311 |
| RORB | -0.705621237 | 0.613178392 | 0.000719373 | 0.003159704 | 156.2264266 | 95.69584797 | ENSG00000198963 |
| NA | -0.706144035 | 0.612956231 | 0.087246404 | 0.186015736 | 38.97987207 | 23.80894839 | ENSG00000215105 |
| NA | -0.706187735 | 0.612937664 | 0.125043611 | 0.245691639 | 26.69225302 | 16.33546115 | ENSG00000249637 |
| NA | -0.707868179 | 0.612224134 | 0.573886598 | 0.726147603 | 4.776305388 | 2.894479816 | ENSG00000270988 |
| NA | -0.708137795 | 0.61210973 | 0.499574929 | 0.665927305 | 5.32894393 | 3.253971888 | ENSG00000212440 |
| NA | -0.708255255 | 0.612059895 | 0.511996067 | 0.676336645 | 5.331734833 | 3.256122461 | ENSG00000259151 |
| NA | -0.708429099 | 0.611986147 | 0.296796393 | 0.467034501 | 12.42358543 | 7.616528024 | ENSG00000280087 |
| SEC61A2 | -0.708803293 | 0.611827435 | 2.95E-10 | 3.86E-09 | 459.4568841 | 281.1046806 | ENSG00000065665 |
| TTC4 | -0.708939421 | 0.611769708 | 0.382111828 | 0.555529374 | 9.068193206 | 5.517126466 | ENSG00000243725 |
| VPS9D1-AS1 | -0.709151127 | 0.611679942 | 0.006129468 | 0.02090217 | 89.48319096 | 54.77356476 | ENSG00000261373 |
| ARMH4 | -0.709212041 | 0.611654116 | 1.72E-06 | 1.30E-05 | 307.7452812 | 188.1753378 | ENSG00000139971 |
| DCHS1 | -0.709694528 | 0.611449592 | 3.04E-15 | 6.70E-14 | 848.926389 | 518.9158738 | ENSG00000166341 |
| CLCF1 | -0.709699015 | 0.61144769 | 0.238026368 | 0.399944836 | 15.63232323 | 9.598162643 | ENSG00000175505 |
| ZNF253 | -0.710403894 | 0.611149019 | 5.27E-08 | 5.05E-07 | 358.7801876 | 219.4090916 | ENSG00000256771 |
| TTLL3 | -0.71161841 | 0.610634747 | 0.098764091 | 0.205153234 | 38.08033394 | 23.30855399 | ENSG00000214021 |
| MEGF11 | -0.711772738 | 0.610569429 | 0.008364411 | 0.027109854 | 78.45906735 | 47.89752662 | ENSG00000157890 |
| NA | -0.712049715 | 0.61045222 | 0.539705739 | 0.698943569 | 5.290547011 | 3.216327637 | ENSG00000213453 |
| ABLIM3 | -0.712430829 | 0.610290979 | 0.189307002 | 0.33732878 | 19.89159609 | 12.08852138 | ENSG00000173210 |
| CH17-340M24.3 | -0.71270809 | 0.610173703 | 0.302959137 | 0.473613024 | 11.2887597 | 6.903462563 | ENSG00000197180 |
| FAXC | -0.712882134 | 0.610100097 | 1.03E-05 | 6.74E-05 | 220.4817808 | 134.5404837 | ENSG00000146267 |
| ZNF772 | -0.713201737 | 0.609964956 | 0.013273147 | 0.039964456 | 76.40680678 | 46.70934061 | ENSG00000197128 |
| CYB5D2 | -0.713290782 | 0.609927309 | 0.001865463 | 0.007431748 | 113.6241356 | 69.38443091 | ENSG00000167740 |
| ANOS1 | -0.713436324 | 0.609865781 | 3.85E-36 | 3.40E-34 | 4541.34789 | 2770.347092 | ENSG00000011201 |
| ATP4A | -0.713786735 | 0.609717671 | 0.199481494 | 0.350920801 | 19.26263927 | 11.73064684 | ENSG00000105675 |
| NT5C3A | -0.713907645 | 0.609666574 | 5.90E-09 | 6.47E-08 | 412.5945324 | 251.6312638 | ENSG00000122643 |
| PCDHA6 | -0.713997908 | 0.609628431 | 0.00111401 | 0.004696321 | 116.0236081 | 70.78743856 | ENSG00000081842 |
| CCDC74B | -0.714013532 | 0.609621829 | 2.78E-05 | 0.000169137 | 201.0229266 | 122.6960985 | ENSG00000152076 |
| NA | -0.71463178 | 0.60936064 | 0.14573009 | 0.276129964 | 23.57338769 | 14.35382653 | ENSG00000231982 |
| NA | -0.714900107 | 0.609247315 | 0.516302509 | 0.680420256 | 4.313539319 | 2.637700666 | ENSG00000235254 |
| TMEM123 | -0.715255066 | 0.609097435 | 1.66E-22 | 6.76E-21 | 1940.942144 | 1182.890944 | ENSG00000152558 |
| TMEM62 | -0.715512559 | 0.608988733 | 0.054754198 | 0.128358638 | 41.03703872 | 25.02669728 | ENSG00000137842 |
| P2RX2 | -0.715610125 | 0.60894755 | 0.042988651 | 0.105709887 | 49.52256917 | 30.11066071 | ENSG00000187848 |
| SH3RF3 | -0.71598407 | 0.608789731 | 0.048855112 | 0.117207308 | 42.01962822 | 25.58220098 | ENSG00000172985 |
| RNF150 | -0.716453868 | 0.608591518 | 2.44E-05 | 0.000149864 | 203.9300197 | 123.9875305 | ENSG00000170153 |
| NBPF15 | -0.717020219 | 0.608352653 | 2.03E-06 | 1.51E-05 | 306.2849689 | 186.2367209 | ENSG00000266338 |
| FLJ30679 | -0.717039644 | 0.608344462 | 0.583363984 | 0.733474635 | 3.8020886 | 2.292729572 | ENSG00000280278 |
| FGFR1OP | -0.717306724 | 0.608231852 | 4.35E-05 | 0.000253094 | 209.0393454 | 127.2718708 | ENSG00000213066 |
| GCGR | -0.717323288 | 0.608224869 | 0.418865861 | 0.590927405 | 8.110046113 | 4.91590926 | ENSG00000215644 |
| NASP | -0.717343942 | 0.608216162 | 4.27E-96 | 3.17E-93 | 18181.00777 | 11058.0248 | ENSG00000132780 |

| | | | | | | | |
|---|---|---|---|---|---|---|---|
| PMM1 | -0.717371331 | 0.608204615 | 5.59E-05 | 0.000318501 | 182.6058713 | 110.9711099 | ENSG00000100417 |
| ABCG1 | -0.717458705 | 0.608167782 | 0.001539928 | 0.006269893 | 128.9942502 | 78.42789546 | ENSG00000160179 |
| AJUBA | -0.718870611 | 0.607572884 | 3.93E-11 | 5.62E-10 | 857.5538581 | 521.3749789 | ENSG00000129474 |
| KTN1 | -0.71888902 | 0.607565131 | 2.85E-35 | 2.40E-33 | 2706.515449 | 1644.716855 | ENSG00000126777 |
| REM2 | -0.719200786 | 0.607433851 | 0.009811903 | 0.030943504 | 74.85829995 | 45.47223656 | ENSG00000139890 |
| TRAM2-AS1 | -0.719541871 | 0.607290257 | 0.018047011 | 0.051531214 | 66.45706016 | 40.34875088 | ENSG00000225791 |
| NOL12 | -0.719647032 | 0.607245992 | 0.011661965 | 0.035823403 | 67.12074712 | 40.79428432 | ENSG00000273899 |
| MCM2 | -0.719825603 | 0.607170834 | 2.88E-39 | 3.18E-37 | 4933.799769 | 2995.175663 | ENSG00000073111 |
| MACROH2A1 | -0.719898729 | 0.60714006 | 8.05E-31 | 5.28E-29 | 2767.669226 | 1680.280174 | ENSG00000113648 |
| NA | -0.719950467 | 0.607118287 | 0.254845749 | 0.419616707 | 19.21933627 | 11.63438564 | ENSG00000284636 |
| NA | -0.721627672 | 0.606412891 | 0.248149095 | 0.411859605 | 14.02376798 | 8.485277248 | ENSG00000261324 |
| MIR4453HG | -0.721661838 | 0.60639853 | 0.072787243 | 0.161006616 | 38.75921471 | 23.43010115 | ENSG00000268471 |
| PHKA2 | -0.721853956 | 0.606317784 | 2.19E-15 | 4.93E-14 | 765.1804045 | 463.9747879 | ENSG00000044446 |
| DHX35 | -0.72191884 | 0.606290516 | 1.03E-11 | 1.58E-10 | 554.1973683 | 335.7620959 | ENSG00000101452 |
| SEPTIN8 | -0.722394778 | 0.606090537 | 1.99E-23 | 8.68E-22 | 1618.077289 | 980.2889998 | ENSG00000164402 |
| RNF38 | -0.722874221 | 0.605889151 | 3.50E-13 | 6.28E-12 | 665.6578699 | 403.3044375 | ENSG00000137075 |
| SLC49A4 | -0.723731485 | 0.605529233 | 9.20E-07 | 7.28E-06 | 292.5538724 | 177.1636935 | ENSG00000138463 |
| SLC13A4 | -0.72376164 | 0.605516576 | 0.023673504 | 0.064631413 | 56.56621068 | 34.28418997 | ENSG00000164707 |
| NA | -0.723826238 | 0.605489464 | 0.462019075 | 0.631715787 | 6.031027803 | 3.670463374 | ENSG00000227038 |
| NA | -0.724132464 | 0.605360957 | 0.273037957 | 0.43988505 | 13.49345666 | 8.112881733 | ENSG00000250397 |
| NA | -0.725724815 | 0.604693168 | 0.46638254 | 0.635649681 | 5.913721863 | 3.580120856 | ENSG00000272008 |
| KCNK13 | -0.725798349 | 0.604662348 | 0.216393827 | 0.373486027 | 16.79159137 | 10.11548905 | ENSG00000152315 |
| GFOD2 | -0.725935146 | 0.604605016 | 1.29E-15 | 2.97E-14 | 932.7116733 | 563.6632196 | ENSG00000141098 |
| NOS1 | -0.726607622 | 0.604323261 | 6.70E-05 | 0.000376234 | 180.8283344 | 109.3228694 | ENSG00000089250 |
| PITRM1-AS1 | -0.726768924 | 0.604255697 | 0.538711206 | 0.698233482 | 4.37358774 | 2.644152387 | ENSG00000237399 |
| APCDD1 | -0.726797495 | 0.604243731 | 0.000647432 | 0.002879781 | 120.6281409 | 72.87797735 | ENSG00000154856 |
| NA | -0.727069206 | 0.604129941 | 9.36E-07 | 7.39E-06 | 269.5481538 | 162.7620029 | ENSG00000267321 |
| ALDH16A1 | -0.727101767 | 0.604116306 | 2.74E-17 | 7.41E-16 | 1123.367292 | 678.5355047 | ENSG00000161618 |
| ASPRV1 | -0.728072832 | 0.603709818 | 0.246735544 | 0.410254731 | 14.66301252 | 8.853371614 | ENSG00000244617 |
| CHN1 | -0.728206343 | 0.603653951 | 5.76E-19 | 1.77E-17 | 938.8499067 | 566.7189988 | ENSG00000128656 |
| ENKD1 | -0.728743972 | 0.603429038 | 1.92E-05 | 0.000119679 | 225.3373076 | 136.0502328 | ENSG00000124074 |
| ARHGEF28 | -0.72913596 | 0.603265105 | 0.140721446 | 0.269017681 | 22.39266821 | 13.535625 | ENSG00000214944 |
| SLC27A1 | -0.729349811 | 0.60317569 | 8.02E-06 | 5.38E-05 | 243.0123674 | 146.5459262 | ENSG00000130304 |
| POLR2C | -0.729411006 | 0.603150105 | 6.77E-28 | 3.86E-26 | 1541.246412 | 929.5434698 | ENSG00000102978 |
| TMEM45A | -0.729953673 | 0.602923274 | 3.29E-07 | 2.80E-06 | 285.16218 | 172.0832072 | ENSG00000181458 |
| TFAP2E | -0.730562733 | 0.602668793 | 0.333005707 | 0.505711815 | 10.95187244 | 6.558491469 | ENSG00000116819 |
| MCM5 | -0.730590593 | 0.602657155 | 2.29E-38 | 2.34E-36 | 8900.855631 | 5363.42806 | ENSG00000100297 |
| ARNT2 | -0.730761549 | 0.602585746 | 2.99E-08 | 2.96E-07 | 348.8380627 | 210.064231 | ENSG00000172379 |
| GSTM4 | -0.731028969 | 0.60247406 | 3.86E-07 | 3.25E-06 | 302.4394282 | 182.2495165 | ENSG00000168765 |
| KCTD21-AS1 | -0.731089368 | 0.602448837 | 0.010220666 | 0.032033609 | 79.22331482 | 47.77625204 | ENSG00000246174 |
| EFNB3 | -0.731165485 | 0.602417053 | 3.74E-12 | 5.99E-11 | 592.9592249 | 357.2629882 | ENSG00000108947 |
| SLC4A3 | -0.731830759 | 0.602139323 | 1.14E-08 | 1.21E-07 | 368.5292135 | 221.7045353 | ENSG00000114923 |
| LBHD1 | -0.732000518 | 0.602068475 | 0.388339218 | 0.561716673 | 7.704537561 | 4.65913011 | ENSG00000162194 |
| SAFB2 | -0.732392542 | 0.601904896 | 1.37E-24 | 6.51E-23 | 2510.981118 | 1511.801816 | ENSG00000130254 |
| POLR3B | -0.733475441 | 0.601453271 | 8.23E-12 | 1.27E-10 | 542.2439337 | 326.0152343 | ENSG00000013503 |
| NA | -0.733480091 | 0.601451332 | 0.385727403 | 0.559336995 | 9.260741381 | 5.582728175 | ENSG00000255129 |
| NA | -0.733768169 | 0.601331246 | 0.400206934 | 0.572729374 | 7.584440719 | 4.564486445 | ENSG00000236264 |
| ULK4 | -0.733953433 | 0.601254031 | 0.000171811 | 0.000878222 | 175.4900539 | 105.5475426 | ENSG00000168038 |
| KLHL14 | -0.734163793 | 0.601166369 | 0.04889482 | 0.117258824 | 42.40348527 | 25.42840732 | ENSG00000197705 |
| MYO5C | -0.734624134 | 0.600974576 | 3.64E-10 | 4.69E-09 | 523.7626748 | 314.8927957 | ENSG00000128833 |
| LSMEM1 | -0.735498669 | 0.600610387 | 0.436918778 | 0.607641853 | 6.031027803 | 3.624216828 | ENSG00000181016 |
| CCDC88B | -0.735505156 | 0.600607686 | 5.62E-09 | 6.20E-08 | 498.597165 | 299.1873575 | ENSG00000168071 |
| ZNF599 | -0.735673151 | 0.600537753 | 0.030080452 | 0.078714324 | 62.7085623 | 37.8349963 | ENSG00000153896 |
| SLC38A3 | -0.736039239 | 0.600385384 | 0.275259817 | 0.442061033 | 12.0781253 | 7.254885378 | ENSG00000188338 |
| NA | -0.73613957 | 0.600343632 | 0.509684178 | 0.674323947 | 4.40361195 | 2.637700666 | ENSG00000243554 |
| NA | -0.736702707 | 0.600109342 | 0.403986176 | 0.576189754 | 7.114177822 | 4.242638624 | ENSG00000272501 |
| NA | -0.736856712 | 0.600045285 | 0.040497987 | 0.100694695 | 47.87051076 | 28.68668036 | ENSG00000259642 |
| LINC01431 | -0.737378374 | 0.599828354 | 0.487753081 | 0.654754809 | 5.435086257 | 3.226547467 | ENSG00000232645 |
| NA | -0.737501904 | 0.599776996 | 0.18348538 | 0.329419866 | 17.99742897 | 10.78015739 | ENSG00000206120 |
| TCF25 | -0.738514211 | 0.599356294 | 1.80E-24 | 8.51E-23 | 1351.17437 | 809.5195117 | ENSG00000141002 |
| NA | -0.738574974 | 0.599331051 | 0.391681609 | 0.564630112 | 8.330503317 | 5.033676198 | ENSG00000213938 |
| NA | -0.738739082 | 0.59926288 | 0.180695184 | 0.325776668 | 19.20346673 | 11.4765513 | ENSG00000279738 |
| FAM78A | -0.738838762 | 0.599221477 | 1.40E-05 | 8.96E-05 | 204.042783 | 122.1658808 | ENSG00000126882 |
| NA | -0.739472781 | 0.598958196 | 0.382717373 | 0.556045981 | 8.324921511 | 5.027224477 | ENSG00000278341 |
| NR0B1 | -0.739602185 | 0.598904474 | 0.352408787 | 0.525250553 | 9.837822327 | 5.923398121 | ENSG00000169297 |
| C12orf57 | -0.740427011 | 0.598562163 | 0.002677569 | 0.010169911 | 101.7884313 | 60.79564723 | ENSG00000111678 |
| KIF3B | -0.741264344 | 0.598214861 | 8.19E-24 | 3.71E-22 | 1238.930799 | 741.3828951 | ENSG00000101350 |
| RBM25 | -0.741488301 | 0.598122004 | 3.07E-38 | 3.10E-36 | 3446.956928 | 2060.820269 | ENSG00000119707 |
| LANCL2 | -0.742722212 | 0.59761066 | 4.71E-09 | 5.26E-08 | 524.5595004 | 313.2335783 | ENSG00000132434 |
| GJA3 | -0.743028082 | 0.597483972 | 0.250428398 | 0.414608155 | 14.99798477 | 9.019535672 | ENSG00000121743 |
| IGSF5 | -0.744615909 | 0.596826744 | 0.050740299 | 0.120724112 | 38.99362642 | 23.27226681 | ENSG00000183067 |
| NA | -0.745461469 | 0.596477048 | 0.153344454 | 0.287675567 | 23.61756658 | 14.02122584 | ENSG00000259556 |
| NA | -0.745503475 | 0.596459681 | 0.526923124 | 0.688817103 | 4.340772627 | 2.568330847 | ENSG00000270607 |
| SNHG9 | -0.745848012 | 0.596317255 | 0.226859486 | 0.386091429 | 15.3462358 | 9.131123463 | ENSG00000255198 |
| SPEF2 | -0.745990491 | 0.596258366 | 0.379987775 | 0.553233522 | 8.379388126 | 5.029375051 | ENSG00000152582 |
| NA | -0.746341131 | 0.596113466 | 0.297013876 | 0.467196239 | 11.01383588 | 6.567093764 | ENSG00000261526 |
| BCL6B | -0.746448547 | 0.596069084 | 5.05E-09 | 5.59E-08 | 383.1433412 | 228.5076839 | ENSG00000161940 |

| | | | | | | | |
|---|---|---|---|---|---|---|---|
| FOXP4-AS1 | -0.746521814 | 0.596038814 | 0.375643828 | 0.549022281 | 7.740143577 | 4.608582417 | ENSG00000234753 |
| BEND7 | -0.746532439 | 0.596034424 | 0.000504101 | 0.002309334 | 131.3647376 | 78.4437614 | ENSG00000165626 |
| EML3 | -0.746547341 | 0.596028267 | 3.72E-13 | 6.66E-12 | 651.9330295 | 388.254508 | ENSG00000149499 |
| NA | -0.747261129 | 0.595733449 | 0.448773967 | 0.61904173 | 6.055470207 | 3.594641834 | ENSG00000275720 |
| TSPAN18 | -0.747606877 | 0.595590696 | 6.88E-13 | 1.19E-11 | 591.0980219 | 352.0082685 | ENSG00000157570 |
| MYO5B | -0.748036606 | 0.595413317 | 2.09E-09 | 2.45E-08 | 406.8559617 | 242.2167424 | ENSG00000167306 |
| ALOXE3 | -0.748138308 | 0.595371345 | 0.445037146 | 0.615568217 | 6.142751936 | 3.651641249 | ENSG00000179148 |
| PCDHB10 | -0.748756647 | 0.595116223 | 0.01854473 | 0.052702767 | 65.22542871 | 38.83415544 | ENSG00000120324 |
| DGKG | -0.74883874 | 0.595082361 | 0.273463807 | 0.440239531 | 12.10814951 | 7.172612117 | ENSG00000058866 |
| LINC01426 | -0.749246358 | 0.594914251 | 0.279017357 | 0.446756627 | 12.67127594 | 7.536405336 | ENSG00000234380 |
| LINC01534 | -0.749626047 | 0.594757701 | 0.273356752 | 0.440191783 | 15.12924522 | 9.040508372 | ENSG00000225975 |
| ITFG1 | -0.749877748 | 0.594653945 | 2.10E-08 | 2.12E-07 | 355.7101318 | 211.5271821 | ENSG00000129636 |
| MKX | -0.749928614 | 0.59463298 | 0.063793436 | 0.144832197 | 38.69617522 | 23.10851379 | ENSG00000150051 |
| SMO | -0.749991198 | 0.594607185 | 6.43E-36 | 5.57E-34 | 2762.064457 | 1642.244325 | ENSG00000128602 |
| NA | -0.750639465 | 0.594340062 | 0.001729812 | 0.006954941 | 108.2764432 | 64.34268545 | ENSG00000261061 |
| MMP25 | -0.750885136 | 0.594238862 | 7.90E-12 | 1.23E-10 | 504.9914375 | 300.1767872 | ENSG00000008516 |
| MSANTD2 | -0.75089046 | 0.59423667 | 7.06E-07 | 5.69E-06 | 279.2816367 | 166.0073725 | ENSG00000120458 |
| PLPBP | -0.750959349 | 0.594208295 | 9.44E-18 | 2.64E-16 | 849.174531 | 504.5480593 | ENSG00000147471 |
| NA | -0.750966189 | 0.594205478 | 0.43429984 | 0.605483719 | 6.760344983 | 4.040708315 | ENSG00000141028 |
| CCNC | -0.751309612 | 0.594064049 | 1.66E-12 | 2.76E-11 | 707.355092 | 420.3475784 | ENSG00000112237 |
| UBXN10 | -0.751328121 | 0.594056427 | 0.071390373 | 0.158498296 | 35.38304686 | 20.95211282 | ENSG00000162543 |
| SETSIP | -0.7532588 | 0.593261966 | 0.204886954 | 0.35846855 | 16.55977039 | 9.844188924 | ENSG00000230667 |
| TMPRSS5 | -0.753283172 | 0.593251944 | 0.377206371 | 0.550691067 | 8.916157131 | 5.25872978 | ENSG00000166682 |
| CYP27A1 | -0.753451402 | 0.59318277 | 0.000312786 | 0.001512268 | 130.7627387 | 77.57528475 | ENSG00000135929 |
| HSPA1B | -0.75424087 | 0.592858259 | 1.37E-22 | 5.63E-21 | 1305.939522 | 774.2409847 | ENSG00000204388 |
| NA | -0.75447752 | 0.592761018 | 0.340112792 | 0.51274562 | 11.49334737 | 6.751819483 | ENSG00000248873 |
| TENM4 | -0.754828034 | 0.59261702 | 4.36E-07 | 3.63E-06 | 287.7839494 | 170.4365964 | ENSG00000149256 |
| SHF | -0.755001306 | 0.592545849 | 1.42E-13 | 2.64E-12 | 720.3012407 | 427.0345776 | ENSG00000138606 |
| TLR2 | -0.755512152 | 0.592336071 | 0.166835328 | 0.306720701 | 23.01883413 | 13.65769308 | ENSG00000137462 |
| BORCS7 | -0.755519073 | 0.592333229 | 0.092077509 | 0.194000353 | 32.81036239 | 19.51979064 | ENSG00000166275 |
| EGFL6 | -0.756040856 | 0.592119037 | 0.064533704 | 0.146117333 | 33.32480417 | 19.77441922 | ENSG00000198759 |
| H4C6 | -0.756143068 | 0.592077088 | 0.480413005 | 0.648495366 | 5.677194963 | 3.40131382 | ENSG00000274618 |
| ADI1 | -0.756416768 | 0.591964774 | 2.78E-18 | 8.04E-17 | 867.318214 | 513.4858547 | ENSG00000182551 |
| PLEKHB1 | -0.756483254 | 0.591937494 | 9.80E-06 | 6.47E-05 | 211.392812 | 124.984539 | ENSG00000021300 |
| C16orf54 | -0.760741218 | 0.590193028 | 0.145797096 | 0.276202669 | 21.83512359 | 12.9048328 | ENSG00000185905 |
| NA | -0.760762207 | 0.590184441 | 0.472851095 | 0.64165521 | 5.018414094 | 2.963849634 | ENSG00000228818 |
| NAA16 | -0.760969581 | 0.590099614 | 6.39E-10 | 7.96E-09 | 487.8068143 | 287.645416 | ENSG00000172766 |
| C1orf216 | -0.761672852 | 0.589812028 | 1.50E-08 | 1.56E-07 | 345.759109 | 203.846149 | ENSG00000142686 |
| RIN3 | -0.761699123 | 0.589801288 | 0.016197234 | 0.04717102 | 74.83210579 | 44.11253138 | ENSG00000100599 |
| NA | -0.761750256 | 0.589780384 | 0.50083302 | 0.666686425 | 4.952783868 | 2.915452515 | ENSG00000256312 |
| NA | -0.761966241 | 0.589692095 | 0.308233684 | 0.47929518 | 10.68165454 | 6.335588461 | ENSG00000213467 |
| H1-10 | -0.762238503 | 0.58958082 | 1.88E-31 | 1.28E-29 | 2351.701779 | 1386.415536 | ENSG00000184897 |
| ZNF252P-AS1 | -0.762880815 | 0.589318387 | 0.513661333 | 0.677859608 | 4.977901994 | 2.929973493 | ENSG00000255559 |
| LOXL1 | -0.762977321 | 0.589278967 | 1.08E-06 | 8.45E-06 | 327.1750382 | 192.8764133 | ENSG00000129038 |
| ATP6V0D1 | -0.763086685 | 0.589234299 | 6.67E-19 | 2.03E-17 | 1346.271523 | 792.8767542 | ENSG00000159720 |
| NA | -0.763483901 | 0.589072087 | 0.531898874 | 0.692692984 | 3.962497383 | 2.366400538 | ENSG00000124593 |
| LRRC10B | -0.764362626 | 0.588713401 | 0.06342741 | 0.144187859 | 33.37647988 | 19.62492671 | ENSG00000204950 |
| NA | -0.764533388 | 0.588643723 | 0.236797234 | 0.39833061 | 15.58343842 | 9.18758984 | ENSG00000238207 |
| GPR182 | -0.764538541 | 0.588641621 | 0.311133141 | 0.48209393 | 11.59024111 | 6.806668324 | ENSG00000166856 |
| SLC35A2 | -0.764762341 | 0.588550314 | 1.99E-13 | 3.64E-12 | 686.5228069 | 403.9645626 | ENSG00000102100 |
| SHISA6 | -0.764814073 | 0.58852921 | 0.000303735 | 0.001472248 | 166.7093876 | 98.36363245 | ENSG00000188803 |
| ST8SIA4 | -0.764952281 | 0.588472833 | 0.40131666 | 0.573764151 | 6.670272351 | 3.918640229 | ENSG00000113532 |
| ACKR1 | -0.765077201 | 0.58842188 | 0.533322459 | 0.693924025 | 3.935264075 | 2.343277265 | ENSG00000213088 |
| NA | -0.765287968 | 0.588335922 | 0.280762664 | 0.449066624 | 13.53675966 | 7.970101412 | ENSG00000274943 |
| BTN1A1 | -0.765624053 | 0.588198882 | 0.293207504 | 0.462907436 | 10.64114244 | 6.285040767 | ENSG00000124557 |
| ZFP30 | -0.766037978 | 0.588030145 | 0.016372945 | 0.047567844 | 57.86491182 | 33.93680784 | ENSG00000120784 |
| LINC00648 | -0.766708552 | 0.587756889 | 0.100677961 | 0.208164903 | 31.73423363 | 18.68519013 | ENSG00000259129 |
| DNAJC3 | -0.766827063 | 0.587708609 | 1.20E-14 | 2.51E-13 | 764.2313811 | 449.1800016 | ENSG00000102580 |
| P2RY2 | -0.7671735 | 0.587567498 | 0.011976342 | 0.03657363 | 63.37121011 | 37.25610887 | ENSG00000175591 |
| LYPD3 | -0.767272423 | 0.587527211 | 0.004444636 | 0.015845772 | 82.1964016 | 48.22367559 | ENSG00000124466 |
| NA | -0.767340527 | 0.587499477 | 0.214439655 | 0.370942915 | 15.69795346 | 9.240288107 | ENSG00000235459 |
| MARCHF4 | -0.767868711 | 0.587284428 | 0.004125504 | 0.014845198 | 86.29821985 | 50.61508951 | ENSG00000144583 |
| GCLM | -0.767986444 | 0.587236504 | 2.07E-10 | 2.76E-09 | 465.6862052 | 273.6403287 | ENSG00000023909 |
| PLEK2 | -0.767988339 | 0.587235732 | 0.30069838 | 0.471320485 | 10.12690082 | 5.942220247 | ENSG00000100558 |
| RCAN1 | -0.768272237 | 0.587120185 | 1.40E-05 | 8.94E-05 | 181.3432518 | 106.4512524 | ENSG00000159200 |
| NCAPH | -0.768717069 | 0.586939184 | 1.71E-19 | 5.52E-18 | 1369.485368 | 803.5592628 | ENSG00000121152 |
| LINC00652 | -0.768839752 | 0.586889275 | 0.442902571 | 0.613839608 | 6.145542839 | 3.573669135 | ENSG00000179935 |
| NA | -0.769404524 | 0.58665957 | 0.335208561 | 0.507826995 | 8.932026668 | 5.254428633 | ENSG00000272604 |
| SLC66A2 | -0.769749502 | 0.586519304 | 4.02E-09 | 4.53E-08 | 467.6335755 | 274.2977518 | ENSG00000122490 |
| NA | -0.769851913 | 0.586477671 | 0.473522662 | 0.642069703 | 4.985598981 | 2.911151367 | ENSG00000286459 |
| PRSS53 | -0.769959631 | 0.586433884 | 0.171852361 | 0.313823289 | 17.91014724 | 10.53413111 | ENSG00000151006 |
| YPEL2 | -0.769980805 | 0.586425277 | 0.008089431 | 0.02636925 | 79.5522297 | 46.5875451 | ENSG00000175155 |
| B4GAT1 | -0.770355877 | 0.586272838 | 5.03E-17 | 1.32E-15 | 770.2577158 | 451.7539665 | ENSG00000174684 |
| ANKHD1 | -0.770443975 | 0.586237038 | 0.142850718 | 0.272063698 | 24.11872956 | 14.06800542 | ENSG00000131503 |
| NA | -0.771034866 | 0.58599698 | 0.205602692 | 0.359513706 | 15.82853819 | 9.292986374 | ENSG00000264107 |
| ZNF460-AS1 | -0.771228374 | 0.585918386 | 0.468958263 | 0.637853335 | 5.638798044 | 3.323341707 | ENSG00000267871 |
| ZNF576 | -0.771644268 | 0.585749504 | 0.000378742 | 0.001796931 | 127.2807919 | 74.64477822 | ENSG00000124444 |

| | | | | | | | |
|---|---|---|---|---|---|---|---|
| CCBE1 | -0.77178384 | 0.585692839 | 0.000244759 | 0.001211923 | 129.7132797 | 75.93862123 | ENSG00000183287 |
| GFAP | -0.771939138 | 0.585629795 | 0.00956402 | 0.030305439 | 63.87919421 | 37.37629895 | ENSG00000131095 |
| DFFB | -0.772792573 | 0.585283465 | 8.65E-07 | 6.87E-06 | 235.9496651 | 137.9899342 | ENSG00000169598 |
| NDUFC2 | -0.772969611 | 0.585211648 | 0.016023626 | 0.046746253 | 67.0430774 | 39.39368771 | ENSG00000151366 |
| NA | -0.773164219 | 0.585132712 | 0.390465439 | 0.563723154 | 6.763135886 | 3.946064649 | ENSG00000285600 |
| LOC101929295 | -0.773403116 | 0.585035828 | 0.344431632 | 0.517076312 | 8.382179029 | 4.882033118 | ENSG00000245552 |
| PHC2 | -0.773539857 | 0.58498038 | 8.54E-09 | 9.20E-08 | 339.5976966 | 198.6968443 | ENSG00000134686 |
| NA | -0.773741748 | 0.584898523 | 0.020005239 | 0.056240031 | 50.5519283 | 29.54923832 | ENSG00000227473 |
| PTENP1-AS | -0.773918849 | 0.584826727 | 0.416783017 | 0.588807116 | 6.654202655 | 3.87024311 | ENSG00000281128 |
| GLYCTK | -0.773948677 | 0.584814636 | 0.005461051 | 0.018934066 | 73.99473126 | 43.27389019 | ENSG00000168237 |
| PDS5A | -0.774019819 | 0.584785798 | 1.49E-45 | 2.12E-43 | 2616.855356 | 1530.353058 | ENSG00000121892 |
| TIAF1 | -0.774036893 | 0.584778878 | 0.172581306 | 0.314764676 | 19.06383357 | 11.13373078 | ENSG00000221995 |
| MYBL2 | -0.774370024 | 0.584643863 | 1.84E-35 | 1.57E-33 | 6574.235971 | 3843.109229 | ENSG00000101057 |
| NMNAT1 | -0.774533255 | 0.584577718 | 0.011557988 | 0.035624292 | 60.57739272 | 35.4387603 | ENSG00000173614 |
| C1QL1 | -0.774800505 | 0.584469439 | 2.41E-07 | 2.09E-06 | 288.9490748 | 169.0421789 | ENSG00000131094 |
| CHRD | -0.774962323 | 0.584403886 | 0.131752076 | 0.255414461 | 20.76525236 | 12.11164465 | ENSG00000090539 |
| GINS2 | -0.77508957 | 0.584352344 | 7.61E-24 | 3.46E-22 | 1602.76672 | 935.8721283 | ENSG00000131153 |
| CHST7 | -0.775421643 | 0.584217855 | 1.46E-12 | 2.45E-11 | 689.875059 | 403.3568632 | ENSG00000147119 |
| CPT1B | -0.776184484 | 0.583909025 | 0.230842642 | 0.391144963 | 17.78813538 | 10.32118747 | ENSG00000205560 |
| NA | -0.777038015 | 0.583563674 | 0.459980675 | 0.629716856 | 5.021204997 | 2.911151367 | ENSG00000235078 |
| NA | -0.777068702 | 0.583551261 | 0.46414819 | 0.63372374 | 5.056811014 | 2.936425214 | ENSG00000273226 |
| PRR18 | -0.777716578 | 0.583289262 | 0.336214349 | 0.508978342 | 8.966956963 | 5.229154786 | ENSG00000176381 |
| CAVIN4 | -0.7781816 | 0.583101282 | 0.00011051 | 0.00059169 | 159.9368398 | 93.3114067 | ENSG00000170681 |
| NA | -0.778238742 | 0.583078187 | 0.039582075 | 0.098870621 | 41.00701451 | 23.89498976 | ENSG00000260924 |
| ADGRG2 | -0.778322524 | 0.583044327 | 1.12E-06 | 8.73E-06 | 260.7987879 | 151.9890907 | ENSG00000173698 |
| NA | -0.778732683 | 0.582878591 | 0.270759031 | 0.437418346 | 11.8632499 | 6.884640437 | ENSG00000274104 |
| NA | -0.778792034 | 0.582854612 | 0.532452512 | 0.693264847 | 3.956915577 | 2.322304566 | ENSG00000260367 |
| PRRT1 | -0.779143348 | 0.582712697 | 0.063334935 | 0.144045535 | 32.67160511 | 19.05113393 | ENSG00000204314 |
| COL26A1 | -0.779866468 | 0.582420698 | 1.34E-39 | 1.52E-37 | 2889.785824 | 1683.3954 | ENSG00000160963 |
| PEX26 | -0.77992277 | 0.582397969 | 9.50E-30 | 5.97E-28 | 1552.149139 | 904.0782423 | ENSG00000215193 |
| HRAS | -0.780133409 | 0.582312943 | 5.76E-17 | 1.51E-15 | 799.5941918 | 465.6832444 | ENSG00000174775 |
| SRRM2-AS1 | -0.781148713 | 0.581903282 | 0.180697145 | 0.325776668 | 16.3288253 | 9.49921783 | ENSG00000205913 |
| AQP5 | -0.781190567 | 0.5818864 | 0.024132636 | 0.065624851 | 50.88219461 | 29.53848546 | ENSG00000161798 |
| ARFRP1 | -0.781563304 | 0.581736082 | 4.91E-17 | 1.29E-15 | 737.0287662 | 428.5583389 | ENSG00000101246 |
| FAM222A | -0.782609277 | 0.581314469 | 0.010933397 | 0.033980369 | 64.7535262 | 37.58844909 | ENSG00000139438 |
| NA | -0.782670568 | 0.581289773 | 0.003252471 | 0.012021167 | 84.92841882 | 49.41830121 | ENSG00000278133 |
| RPGRIP1 | -0.782696296 | 0.581279407 | 0.229010052 | 0.388858804 | 13.00066638 | 7.568130904 | ENSG00000092200 |
| ND1 | -0.783008481 | 0.581153637 | 0.000776026 | 0.003379304 | 18308.1132 | 10639.81274 | ENSG00000198888 |
| C10orf143 | -0.783092748 | 0.581119694 | 0.032927693 | 0.084717007 | 48.57067961 | 28.19085969 | ENSG00000237489 |
| CCDC160 | -0.784358336 | 0.580610136 | 0.0021824 | 0.008517233 | 95.98871202 | 55.80632747 | ENSG00000203952 |
| TM7SF2 | -0.784443862 | 0.580575718 | 1.12E-19 | 3.72E-18 | 1094.188197 | 635.2984347 | ENSG00000149809 |
| SPATA31C2 | -0.785306462 | 0.58022869 | 0.479651017 | 0.647829431 | 5.018414094 | 2.885877521 | ENSG00000177910 |
| SLC38A7 | -0.785382228 | 0.580198219 | 5.47E-16 | 1.32E-14 | 730.2082479 | 423.6935288 | ENSG00000103042 |
| LOC101928415 | -0.785710208 | 0.580066332 | 0.018717425 | 0.053099684 | 53.72996616 | 31.16439611 | ENSG00000286724 |
| GRIA1 | -0.785727173 | 0.580059512 | 0.101527073 | 0.209493932 | 27.26674322 | 15.80093015 | ENSG00000155511 |
| SRPK3 | -0.785945067 | 0.57997191 | 0.0527549 | 0.124390498 | 37.16999561 | 21.54956192 | ENSG00000184343 |
| DSCAML1 | -0.786278142 | 0.579838028 | 0.262831305 | 0.428450118 | 12.39144604 | 7.155940565 | ENSG00000177103 |
| SYS1 | -0.786787715 | 0.57963326 | 3.43E-07 | 2.91E-06 | 255.0071531 | 147.8155614 | ENSG00000204070 |
| NA | -0.786944333 | 0.579570338 | 0.250324882 | 0.414530464 | 14.62317615 | 8.460003401 | ENSG00000228302 |
| C1QTNF3 | -0.78782843 | 0.579215281 | 0.352563971 | 0.525400662 | 8.033927996 | 4.684403957 | ENSG00000082196 |
| KPNA1 | -0.787883854 | 0.57919303 | 6.52E-14 | 1.26E-12 | 644.6012365 | 373.68257 | ENSG00000114030 |
| NA | -0.787967621 | 0.579159401 | 0.17685945 | 0.320456151 | 17.08605151 | 9.911408169 | ENSG00000263731 |
| NYNRIN | -0.788292576 | 0.579028965 | 3.78E-29 | 2.30E-27 | 1364.156852 | 789.7574324 | ENSG00000205978 |
| MAPK8IP2 | -0.788656762 | 0.578882817 | 2.88E-05 | 0.000174004 | 255.851424 | 148.0718197 | ENSG00000008735 |
| GGT7 | -0.78869298 | 0.578868284 | 3.49E-22 | 1.39E-20 | 1585.049396 | 916.7995007 | ENSG00000131067 |
| NA | -0.78929129 | 0.578628268 | 0.502802092 | 0.668473822 | 4.019754901 | 2.341126692 | ENSG00000273521 |
| NA | -0.789378526 | 0.57859328 | 0.509213464 | 0.673839823 | 3.956915577 | 2.297030719 | ENSG00000250859 |
| GFRA2 | -0.790557562 | 0.578120621 | 0.000196518 | 0.000992156 | 141.7491411 | 82.09917076 | ENSG00000168546 |
| NA | -0.791085522 | 0.577909094 | 0.104588945 | 0.214308047 | 24.35438058 | 14.05510198 | ENSG00000260051 |
| NA | -0.791295952 | 0.577824807 | 6.94E-07 | 5.60E-06 | 257.0085386 | 148.322668 | ENSG00000262001 |
| SPACA9 | -0.791359967 | 0.577799168 | 0.2163017 | 0.373375092 | 14.26866758 | 8.28979866 | ENSG00000165698 |
| EPHA4 | -0.791424544 | 0.577773306 | 8.35E-05 | 0.000459339 | 178.2436473 | 103.0773509 | ENSG00000116106 |
| DNAAF1 | -0.791455514 | 0.577760903 | 0.392556464 | 0.565514655 | 7.287865398 | 4.192090931 | ENSG00000154099 |
| RBM12B-AS1 | -0.792149243 | 0.57748315 | 0.335216887 | 0.507826995 | 9.767486175 | 5.654248567 | ENSG00000279331 |
| NA | -0.792448508 | 0.577363372 | 0.368515764 | 0.542384802 | 7.375147127 | 4.244789198 | ENSG00000287620 |
| NA | -0.793669202 | 0.57687506 | 0.155465306 | 0.290580489 | 18.78400367 | 10.81780164 | ENSG00000269044 |
| UBQLN2 | -0.794176266 | 0.576672341 | 1.82E-20 | 6.45E-19 | 1087.469916 | 627.1312371 | ENSG00000188021 |
| NA | -0.794527107 | 0.576532121 | 0.418046111 | 0.590016442 | 5.728870676 | 3.302369007 | ENSG00000287806 |
| U2AF1L4 | -0.794941668 | 0.576366477 | 0.100895971 | 0.208459267 | 28.48162925 | 16.39837925 | ENSG00000161265 |
| HAPLN3 | -0.795017919 | 0.576336015 | 5.19E-39 | 5.60E-37 | 2002.041685 | 1153.916347 | ENSG00000140511 |
| RGPD6 | -0.795284455 | 0.576229548 | 0.44998872 | 0.620273841 | 5.619937445 | 3.226547467 | ENSG00000183054 |
| ADAMTS19-AS1 | -0.795368908 | 0.576195817 | 0.016891876 | 0.048818155 | 56.52223196 | 32.58675893 | ENSG00000249421 |
| ZMYND15 | -0.795411914 | 0.576178641 | 0.453418835 | 0.623576157 | 5.630425335 | 3.232999189 | ENSG00000141497 |
| ESR2 | -0.795558976 | 0.576119911 | 0.368585778 | 0.542405084 | 9.546828812 | 5.48808451 | ENSG00000140009 |
| NA | -0.795975122 | 0.575953753 | 0.420771682 | 0.592835708 | 6.226366881 | 3.598942982 | ENSG00000270504 |
| ACAT2 | -0.796665183 | 0.575678332 | 6.30E-29 | 3.77E-27 | 2588.908921 | 1490.282975 | ENSG00000120437 |
| TAF5 | -0.797152667 | 0.575483844 | 5.78E-13 | 1.01E-11 | 615.1066301 | 354.064677 | ENSG00000148835 |

| | | | | | | | |
|---|---|---|---|---|---|---|---|
| ERG | -0.797590907 | 0.575309059 | 0.094122397 | 0.19734279 | 28.12644497 | 16.21742164 | ENSG00000157554 |
| NA | -0.797607698 | 0.575302363 | 0.269268848 | 0.435631109 | 12.08370711 | 6.970681808 | ENSG00000286835 |
| MORN2 | -0.797758385 | 0.575242277 | 0.002883218 | 0.010835053 | 80.00131665 | 46.09711008 | ENSG00000188010 |
| KLRG1 | -0.798504696 | 0.574944779 | 0.023051827 | 0.063157618 | 53.53271206 | 30.64895981 | ENSG00000139187 |
| MMP2 | -0.798711334 | 0.574862436 | 1.37E-28 | 8.03E-27 | 2059.175227 | 1184.447491 | ENSG00000087245 |
| C1R | -0.799066897 | 0.574720774 | 0.000662556 | 0.002938923 | 123.5642702 | 70.83961589 | ENSG00000159403 |
| PTPRD-AS1 | -0.79990033 | 0.574388858 | 0.349339282 | 0.521966083 | 8.0178583 | 4.610732991 | ENSG00000225706 |
| NA | -0.801599267 | 0.573712849 | 0.347696282 | 0.520438313 | 9.076565914 | 5.172155372 | ENSG00000269086 |
| VASH1 | -0.802181899 | 0.573481201 | 4.10E-11 | 5.85E-10 | 429.1902212 | 246.1883598 | ENSG00000071246 |
| MDH1B | -0.803218675 | 0.573069224 | 0.061914978 | 0.141466521 | 35.97320644 | 20.60069001 | ENSG00000138400 |
| ANG | -0.803249119 | 0.573057131 | 0.513197551 | 0.677439028 | 4.046988208 | 2.322304566 | ENSG00000214274 |
| ABCF2 | -0.803618647 | 0.572910369 | 4.06E-11 | 5.80E-10 | 404.9709311 | 231.9931732 | ENSG00000033050 |
| CCDC15 | -0.804034986 | 0.57274506 | 0.007573899 | 0.02494399 | 67.21448653 | 38.45100706 | ENSG00000149548 |
| SUSD4 | -0.804076743 | 0.572728482 | 0.004456604 | 0.015879636 | 80.41069215 | 45.9788101 | ENSG00000143502 |
| NA | -0.804353579 | 0.572618593 | 0.417715099 | 0.589692058 | 6.842044905 | 3.874544257 | ENSG00000228477 |
| MTA2 | -0.804588456 | 0.572525376 | 8.83E-56 | 1.79E-53 | 4418.582343 | 2529.381625 | ENSG00000149480 |
| TP53BP1 | -0.804667735 | 0.572493916 | 2.35E-37 | 2.27E-35 | 2709.455172 | 1550.675821 | ENSG00000067369 |
| NA | -0.805066722 | 0.57233561 | 0.420144855 | 0.59212554 | 5.786128194 | 3.302369007 | ENSG00000287601 |
| TMEM99 | -0.805070178 | 0.572334239 | 0.000651564 | 0.002896157 | 125.6296463 | 71.86644781 | ENSG00000167920 |
| BRD9 | -0.805185077 | 0.572288659 | 1.61E-21 | 6.04E-20 | 1119.104012 | 640.0301513 | ENSG00000028310 |
| NA | -0.805314147 | 0.572237462 | 0.29718606 | 0.467428967 | 9.726974075 | 5.54670146 | ENSG00000225416 |
| SOAT1 | -0.805361155 | 0.572218817 | 0.000198873 | 0.001003523 | 125.5054824 | 71.78606466 | ENSG00000057252 |
| RTL5 | -0.805366749 | 0.572216598 | 0.000532418 | 0.002424663 | 127.1149135 | 72.629007 | ENSG00000242732 |
| SLC22A4 | -0.805803004 | 0.572043592 | 0.330940109 | 0.503483095 | 8.621496832 | 4.951402937 | ENSG00000197208 |
| NA | -0.806464644 | 0.571781305 | 0.218562742 | 0.376253244 | 13.68109875 | 7.791033912 | ENSG00000234618 |
| LTBP2 | -0.806699454 | 0.571688251 | 0.043036375 | 0.105813753 | 44.66803049 | 25.67280396 | ENSG00000119681 |
| DNAJB6 | -0.806742908 | 0.571671032 | 1.63E-63 | 4.08E-61 | 8716.27997 | 4982.263466 | ENSG00000105993 |
| NA | -0.807722339 | 0.571283062 | 0.3801514 | 0.55342998 | 7.933367474 | 4.516089325 | ENSG00000262902 |
| WWOX | -0.808145579 | 0.57111549 | 1.54E-06 | 1.17E-05 | 214.8419437 | 122.7450287 | ENSG00000186153 |
| NA | -0.808317772 | 0.571047329 | 0.247545896 | 0.411079345 | 13.78425001 | 7.915252572 | ENSG00000279089 |
| SLC24A3 | -0.80840361 | 0.571013354 | 0.00013285 | 0.000695659 | 128.1336726 | 73.16809961 | ENSG00000185052 |
| KCNQ4 | -0.808831011 | 0.570844215 | 0.000129824 | 0.000681108 | 133.7984646 | 76.45835867 | ENSG00000117013 |
| UVRAG | -0.808920405 | 0.570808845 | 8.70E-12 | 1.34E-10 | 430.6212219 | 245.8000983 | ENSG00000198382 |
| HIPK2 | -0.809883702 | 0.570427839 | 8.05E-22 | 3.13E-20 | 905.8026171 | 516.6645381 | ENSG00000064393 |
| NA | -0.810046975 | 0.570363286 | 0.252185289 | 0.41665865 | 15.73076857 | 9.011466415 | ENSG00000236801 |
| NA | -0.810252405 | 0.570282076 | 0.500729258 | 0.666686425 | 4.104245726 | 2.343277265 | ENSG00000224713 |
| NA | -0.811316332 | 0.569861672 | 0.112661131 | 0.227147354 | 21.38563631 | 12.21058946 | ENSG00000262049 |
| ACAA1 | -0.811334607 | 0.569854454 | 2.32E-12 | 3.80E-11 | 593.3636574 | 337.97795 | ENSG00000060971 |
| SPANXA2-OT1 | -0.811832907 | 0.569657663 | 0.464665058 | 0.634249885 | 4.642929753 | 2.665125086 | ENSG00000277215 |
| NA | -0.812140479 | 0.569536229 | 0.576911399 | 0.728827622 | 3.970870091 | 2.252934747 | ENSG00000255122 |
| ZNF594 | -0.812186117 | 0.569518212 | 1.90E-05 | 0.000119061 | 210.4571554 | 120.1699735 | ENSG00000180626 |
| DLD | -0.812574311 | 0.569364989 | 1.83E-32 | 1.34E-30 | 1859.892505 | 1059.623743 | ENSG00000091140 |
| RND1 | -0.812602762 | 0.569353761 | 2.17E-06 | 1.61E-05 | 226.8433504 | 129.1742068 | ENSG00000172602 |
| NA | -0.812655212 | 0.569333063 | 0.457317079 | 0.627282691 | 5.668822255 | 3.201273621 | ENSG00000185065 |
| SLC4A4 | -0.813117149 | 0.569150797 | 0.025360893 | 0.068434325 | 42.08804935 | 23.94553745 | ENSG00000080493 |
| FOXM1 | -0.813406887 | 0.569036505 | 1.34E-38 | 1.40E-36 | 3102.913947 | 1765.751622 | ENSG00000111206 |
| LBH | -0.814163059 | 0.568738329 | 1.13E-22 | 4.68E-21 | 1182.675694 | 672.4080135 | ENSG00000213626 |
| ATAD3B | -0.81422116 | 0.568715425 | 3.01E-23 | 1.29E-21 | 2689.424273 | 1529.436312 | ENSG00000160072 |
| EPHB2 | -0.814297388 | 0.568685377 | 7.19E-20 | 2.39E-18 | 900.0553614 | 512.2368891 | ENSG00000133216 |
| RET | -0.816937639 | 0.567645587 | 1.34E-09 | 1.60E-08 | 420.7700839 | 238.8662672 | ENSG00000165731 |
| NA | -0.817380683 | 0.567471293 | 0.076038032 | 0.166479363 | 28.54379286 | 16.2281745 | ENSG00000274964 |
| FRY | -0.817400647 | 0.56746344 | 4.06E-05 | 0.000238005 | 180.8782584 | 102.764911 | ENSG00000073910 |
| CNPY2 | -0.817893153 | 0.567269753 | 0.057803663 | 0.134025296 | 30.81324416 | 17.50911407 | ENSG00000257727 |
| PDE4D | -0.819280037 | 0.566724691 | 0.082992752 | 0.178784274 | 26.01461155 | 14.73267376 | ENSG00000113448 |
| DUSP4 | -0.820655936 | 0.566184463 | 1.10E-10 | 1.51E-09 | 401.3336818 | 227.4544815 | ENSG00000120875 |
| CFAP298 | -0.820800535 | 0.566127718 | 0.001467169 | 0.006010438 | 91.36830965 | 51.67312607 | ENSG00000159079 |
| NA | -0.821220253 | 0.56596304 | 0.087020568 | 0.185645431 | 24.42001081 | 13.83219897 | ENSG00000234518 |
| SDF2L1 | -0.822653862 | 0.565400921 | 3.71E-19 | 1.15E-17 | 1683.48562 | 951.9637137 | ENSG00000128228 |
| SKIV2L | -0.822696928 | 0.565384043 | 6.13E-27 | 3.32E-25 | 1344.477928 | 760.4267717 | ENSG00000204351 |
| TEX14 | -0.822983201 | 0.565271865 | 0.084688498 | 0.181698271 | 26.9135861 | 15.23520662 | ENSG00000121101 |
| BRICD5 | -0.823043412 | 0.565248275 | 0.001384797 | 0.005706927 | 89.85241777 | 50.80652742 | ENSG00000182685 |
| SCYL2 | -0.823135032 | 0.565212379 | 6.88E-16 | 1.63E-14 | 707.2513772 | 400.034315 | ENSG00000136021 |
| PM20D2 | -0.824338465 | 0.5647411 | 1.70E-07 | 1.52E-06 | 280.6856042 | 158.8083899 | ENSG00000146281 |
| RUBCNL | -0.825576984 | 0.564256492 | 0.002422047 | 0.00933354 | 83.12787897 | 46.88950472 | ENSG00000102445 |
| LRRC37A3 | -0.825989033 | 0.564095357 | 0.085027203 | 0.182251086 | 28.64694412 | 16.21365353 | ENSG00000176809 |
| KHK | -0.826111362 | 0.564047528 | 2.24E-07 | 1.95E-06 | 277.5013088 | 156.4578795 | ENSG00000138030 |
| LANCL3 | -0.826706719 | 0.56381481 | 0.263690263 | 0.429487408 | 11.00055709 | 6.186095954 | ENSG00000147036 |
| NA | -0.82675871 | 0.563794493 | 0.201519489 | 0.353796183 | 19.08037883 | 10.76993756 | ENSG00000235043 |
| SEPTIN3 | -0.826961588 | 0.563715215 | 5.35E-35 | 4.37E-33 | 1714.337982 | 966.3104823 | ENSG00000100167 |
| PIANP | -0.827083739 | 0.563667488 | 7.71E-07 | 6.18E-06 | 237.8516925 | 134.0836644 | ENSG00000139200 |
| GPR1 | -0.827446312 | 0.563525847 | 0.275909381 | 0.442873166 | 11.68378036 | 6.594518184 | ENSG00000183671 |
| ZNF181 | -0.827477519 | 0.563513657 | 0.000558388 | 0.002529201 | 131.4861859 | 74.06695686 | ENSG00000197841 |
| HSBP1L1 | -0.827524124 | 0.563495454 | 0.167342405 | 0.307474024 | 15.68678984 | 8.825947194 | ENSG00000226742 |
| TCEAL2 | -0.827564067 | 0.563479853 | 1.25E-10 | 1.71E-09 | 395.8398233 | 222.9674462 | ENSG00000184905 |
| NA | -0.827938596 | 0.56333359 | 0.37625202 | 0.549619697 | 6.923544669 | 3.89766753 | ENSG00000273554 |
| ARMC7 | -0.828215603 | 0.563225437 | 3.70E-09 | 4.19E-08 | 329.5074779 | 185.3515848 | ENSG00000125449 |
| COMMD9 | -0.828282367 | 0.563199373 | 9.32E-10 | 1.13E-08 | 367.9708682 | 207.1974482 | ENSG00000110442 |

| | | | | | | | |
|---|---|---|---|---|---|---|---|
| NA | -0.828650649 | 0.563055621 | 0.383367856 | 0.556767998 | 6.991290077 | 3.939612928 | ENSG00000259863 |
| NA | -0.829482612 | 0.562731016 | 0.04142458 | 0.10254351 | 35.17845919 | 19.83141863 | ENSG00000257122 |
| MITF | -0.829549604 | 0.562704886 | 0.134929114 | 0.260161673 | 19.84741721 | 11.21547101 | ENSG00000187098 |
| KCNJ10 | -0.82956614 | 0.562698437 | 0.102767431 | 0.211487786 | 26.62450761 | 14.93890522 | ENSG00000177807 |
| FZD2 | -0.829698022 | 0.562647001 | 8.66E-18 | 2.44E-16 | 1468.615104 | 826.5750231 | ENSG00000180340 |
| EPB41L1 | -0.829921349 | 0.56255991 | 6.44E-16 | 1.54E-14 | 927.6571279 | 521.5710758 | ENSG00000088367 |
| FOXI3 | -0.830397634 | 0.56237422 | 2.22E-05 | 0.000137324 | 172.9182453 | 97.29915628 | ENSG00000214336 |
| NA | -0.830738839 | 0.562241231 | 2.92E-05 | 0.000176627 | 199.6355412 | 111.9506534 | ENSG00000282936 |
| C1QTNF1 | -0.830844408 | 0.562200091 | 0.001358901 | 0.005609791 | 92.21752352 | 51.8957565 | ENSG00000173918 |
| EFHD2 | -0.831302031 | 0.562021789 | 4.51E-23 | 1.89E-21 | 1352.550643 | 759.9038476 | ENSG00000142634 |
| MRO | -0.83133619 | 0.562008482 | 0.056874341 | 0.132321016 | 30.9466198 | 17.33434771 | ENSG00000134042 |
| ZAR1 | -0.831372883 | 0.561994189 | 0.097033318 | 0.202364295 | 23.40807282 | 13.16322948 | ENSG00000182223 |
| ETHE1 | -0.83154625 | 0.561926658 | 0.002744702 | 0.010388081 | 76.72503361 | 43.12250757 | ENSG00000105755 |
| NA | -0.831946138 | 0.561770924 | 0.448654014 | 0.618920522 | 5.345013626 | 3.028918306 | ENSG00000276116 |
| ERBB2 | -0.832013255 | 0.56174479 | 1.64E-12 | 2.74E-11 | 13599.06045 | 7639.18773 | ENSG00000141736 |
| ZNF528 | -0.832063595 | 0.56172519 | 0.382601488 | 0.556031739 | 6.403721241 | 3.573669135 | ENSG00000167555 |
| NA | -0.832090093 | 0.561714873 | 0.306823015 | 0.477828426 | 8.744384577 | 4.88848484 | ENSG00000275120 |
| ULK4P2 | -0.834851958 | 0.560640566 | 0.324754319 | 0.496421201 | 8.755548189 | 4.884183692 | ENSG00000260128 |
| BCO2 | -0.836014472 | 0.560188988 | 0.486746838 | 0.654233584 | 4.607323737 | 2.570481421 | ENSG00000197580 |
| SLC30A3 | -0.836920156 | 0.559837427 | 2.33E-09 | 2.71E-08 | 312.9601104 | 175.25974 | ENSG00000115194 |
| DZIP3 | -0.837022376 | 0.559797762 | 5.09E-16 | 1.23E-14 | 599.190522 | 335.3687277 | ENSG00000198919 |
| NA | -0.837358583 | 0.559667321 | 0.476363399 | 0.644743305 | 4.670163061 | 2.608125672 | ENSG00000260773 |
| GATA5 | -0.837843136 | 0.559479379 | 0.268856829 | 0.435110493 | 9.955128266 | 5.578427028 | ENSG00000130700 |
| TMSB15B | -0.838127959 | 0.559368935 | 0.272743649 | 0.439557038 | 9.979570671 | 5.592948006 | ENSG00000269226 |
| DLX4 | -0.838302701 | 0.559301188 | 0.278288886 | 0.445997416 | 10.62439703 | 5.973945815 | ENSG00000108813 |
| RADX | -0.838488926 | 0.559228997 | 4.47E-07 | 3.72E-06 | 232.0257648 | 129.8609018 | ENSG00000147231 |
| SLC34A3 | -0.838754009 | 0.559126253 | 0.366068879 | 0.539771823 | 7.056920304 | 3.939612928 | ENSG00000198569 |
| SCN9A | -0.839142256 | 0.558975805 | 0.003530193 | 0.012959267 | 77.9979409 | 43.52475066 | ENSG00000169432 |
| CLCN2 | -0.839387183 | 0.558880916 | 1.04E-07 | 9.52E-07 | 263.2801988 | 147.2261816 | ENSG00000114859 |
| FKBP7 | -0.839772895 | 0.558731516 | 0.013983291 | 0.041776624 | 50.67202514 | 28.30379244 | ENSG00000079150 |
| PCSK1N | -0.839818266 | 0.558713945 | 0.000405297 | 0.001906041 | 151.3269453 | 84.74815733 | ENSG00000102109 |
| RGS7BP | -0.840027457 | 0.558632937 | 0.019490588 | 0.055017989 | 53.42270279 | 29.89367638 | ENSG00000186479 |
| SLC25A48 | -0.840153523 | 0.558584125 | 0.022060309 | 0.060934339 | 43.57949882 | 24.33460452 | ENSG00000145832 |
| PAK5 | -0.840263408 | 0.558541581 | 0.171228046 | 0.312977093 | 18.04352288 | 10.02514653 | ENSG00000101349 |
| HYKK | -0.840557266 | 0.558427825 | 0.12124505 | 0.240359497 | 21.3367515 | 11.97908416 | ENSG00000188266 |
| FOXO6 | -0.840587747 | 0.558416027 | 8.88E-05 | 0.000485866 | 132.3277908 | 73.97150759 | ENSG00000204060 |
| HPCA | -0.840686951 | 0.55837763 | 0.028179091 | 0.074660464 | 47.91381376 | 26.86555158 | ENSG00000121905 |
| ZNF69 | -0.840771932 | 0.55834474 | 0.000283779 | 0.001384169 | 110.3547858 | 61.51731499 | ENSG00000198429 |
| SOWAHB | -0.841443478 | 0.558084902 | 0.200908005 | 0.353076098 | 14.69861854 | 8.199456142 | ENSG00000186212 |
| MAN1B1-DT | -0.841839587 | 0.557931694 | 0.284353819 | 0.453045055 | 10.02566458 | 5.599399727 | ENSG00000268996 |
| NA | -0.842356478 | 0.557731833 | 1.08E-08 | 1.15E-07 | 518.3034358 | 289.0205088 | ENSG00000284976 |
| MAPK4 | -0.84404572 | 0.557079171 | 4.44E-06 | 3.12E-05 | 209.5859266 | 116.5662206 | ENSG00000141639 |
| PPP1R14C | -0.845582103 | 0.556486231 | 0.287830268 | 0.457208416 | 9.479283563 | 5.25872978 | ENSG00000198729 |
| SEPTIN5 | -0.845678141 | 0.556449188 | 0.361266671 | 0.534290242 | 13.17012359 | 7.37480289 | ENSG00000184702 |
| PCDHGA2 | -0.847411969 | 0.55578085 | 0.166540436 | 0.306353728 | 16.05948328 | 8.956617575 | ENSG00000081853 |
| BAD | -0.847870143 | 0.555604372 | 0.057303668 | 0.133151072 | 32.97934404 | 18.30796043 | ENSG00000002330 |
| ATP10A | -0.848228709 | 0.5554663 | 1.88E-10 | 2.51E-09 | 357.8720014 | 198.7011455 | ENSG00000206190 |
| XKR7 | -0.848233765 | 0.555464353 | 1.59E-07 | 1.42E-06 | 284.6198661 | 158.0031223 | ENSG00000260903 |
| SLC2A8 | -0.848840301 | 0.555230875 | 2.47E-19 | 7.80E-18 | 842.7686576 | 467.8789192 | ENSG00000136856 |
| TAGLN3 | -0.849276037 | 0.555063204 | 0.000413146 | 0.001936818 | 108.9105063 | 60.32671795 | ENSG00000144834 |
| EVL | -0.850378061 | 0.554639373 | 1.70E-23 | 7.42E-22 | 1232.938468 | 683.6436453 | ENSG00000196405 |
| NA | -0.850568006 | 0.554566354 | 0.193480924 | 0.34321292 | 17.63310824 | 9.758147554 | ENSG00000275734 |
| HTR7 | -0.851229272 | 0.554312224 | 0.006404526 | 0.021694419 | 62.90689244 | 34.96122872 | ENSG00000148680 |
| POGLUT2 | -0.851235963 | 0.554309653 | 1.64E-15 | 3.72E-14 | 635.7275817 | 352.7100481 | ENSG00000134901 |
| NA | -0.852533703 | 0.553811262 | 0.151978035 | 0.285500854 | 15.98615607 | 8.857672762 | ENSG00000226361 |
| ZDHHC15 | -0.852593469 | 0.55378832 | 0.000750781 | 0.003283456 | 115.4053393 | 63.82778219 | ENSG00000102383 |
| MT2A | -0.853404326 | 0.553477154 | 2.10E-17 | 5.72E-16 | 676.8841153 | 374.7704726 | ENSG00000125148 |
| NOVA1 | -0.853441557 | 0.553462871 | 1.06E-05 | 6.92E-05 | 205.0857844 | 113.7155336 | ENSG00000139910 |
| TPR | -0.854039885 | 0.553233382 | 4.48E-50 | 7.51E-48 | 3544.78047 | 1960.445644 | ENSG00000047410 |
| EFNB1 | -0.855428088 | 0.5527013 | 5.80E-11 | 8.17E-10 | 449.2117646 | 248.5827177 | ENSG00000090776 |
| NA | -0.855608939 | 0.55263202 | 0.168798073 | 0.309415016 | 15.35460851 | 8.455702254 | ENSG00000282886 |
| GLT1D1 | -0.857534353 | 0.551894972 | 2.71E-10 | 3.56E-09 | 341.5204485 | 188.6579703 | ENSG00000151948 |
| NA | -0.858278924 | 0.551610214 | 0.242078433 | 0.404498488 | 10.77172717 | 5.942220247 | ENSG00000267500 |
| C2orf81 | -0.858443249 | 0.551547388 | 0.444649451 | 0.615289998 | 4.809120502 | 2.637700666 | ENSG00000284308 |
| TP53TG5 | -0.858458812 | 0.551541439 | 0.314316075 | 0.485469286 | 8.426157754 | 4.661280684 | ENSG00000124251 |
| TRIL | -0.858941552 | 0.551356918 | 4.79E-07 | 3.95E-06 | 202.8942029 | 111.7747904 | ENSG00000255690 |
| SHLD3 | -0.860487135 | 0.550766557 | 0.072823657 | 0.161066484 | 25.70754834 | 14.18792293 | ENSG00000253251 |
| NA | -0.860928835 | 0.550597958 | 0.378167541 | 0.551707633 | 5.938164268 | 3.279245734 | ENSG00000242457 |
| NA | -0.862882546 | 0.549852838 | 0.21670145 | 0.373930602 | 11.96919207 | 6.604738015 | ENSG00000226608 |
| CCNE2 | -0.863497425 | 0.54961854 | 0.010597793 | 0.033065672 | 61.15514938 | 33.56441233 | ENSG00000175305 |
| CEP290 | -0.863864094 | 0.549478869 | 1.40E-06 | 1.07E-05 | 210.3393228 | 115.6439369 | ENSG00000198707 |
| TFE3 | -0.864821767 | 0.549114241 | 4.44E-53 | 8.13E-51 | 3074.174175 | 1687.666838 | ENSG00000068323 |
| SLC16A14 | -0.865080362 | 0.549015825 | 0.001751427 | 0.007033052 | 97.88375503 | 53.68701929 | ENSG00000163053 |
| NA | -0.86532563 | 0.548922499 | 0.061807776 | 0.141271779 | 27.00557376 | 14.86926283 | ENSG00000288559 |
| SHD | -0.866318646 | 0.548544799 | 0.004740708 | 0.016758843 | 68.65163265 | 37.70406546 | ENSG00000105251 |
| DCTPP1 | -0.866377079 | 0.548522582 | 9.84E-24 | 4.41E-22 | 1065.071953 | 584.3520468 | ENSG00000179958 |
| E2F8 | -0.866380502 | 0.548521281 | 4.61E-08 | 4.45E-07 | 255.8452914 | 140.4211308 | ENSG00000129173 |

| | | | | | | | |
|---|---|---|---|---|---|---|---|
| CDH24 | -0.866494925 | 0.548477778 | 1.85E-16 | 4.64E-15 | 669.5313707 | 367.2722751 | ENSG00000139880 |
| TCEA2 | -0.866559138 | 0.548453366 | 6.73E-22 | 2.63E-20 | 867.9733281 | 476.0108835 | ENSG00000171703 |
| TXNDC16 | -0.867243684 | 0.548193192 | 3.14E-09 | 3.58E-08 | 373.0445879 | 204.8275279 | ENSG00000087301 |
| EZH2 | -0.867586761 | 0.548062845 | 1.54E-31 | 1.06E-29 | 1614.973167 | 884.8937786 | ENSG00000106462 |
| NA | -0.867723972 | 0.548010723 | 0.417316967 | 0.589345652 | 6.011491482 | 3.300218434 | ENSG00000259051 |
| LOC107985164 | -0.868367649 | 0.547766275 | 0.011230305 | 0.0347922 | 52.23919241 | 28.69124197 | ENSG00000267327 |
| NA | -0.868823545 | 0.547593207 | 0.208206419 | 0.363025129 | 12.0809162 | 6.634313009 | ENSG00000286808 |
| SOX18 | -0.868830946 | 0.547590398 | 0.390181145 | 0.563397041 | 5.984258175 | 3.279245734 | ENSG00000203883 |
| NA | -0.86930462 | 0.547410639 | 0.1276447 | 0.249491771 | 18.01070776 | 9.871613345 | ENSG00000263968 |
| BOLA3-AS1 | -0.869337512 | 0.547398159 | 2.82E-07 | 2.42E-06 | 215.2950608 | 117.950394 | ENSG00000225439 |
| GRHL1 | -0.869449209 | 0.54735578 | 0.001683124 | 0.006785611 | 86.6094254 | 47.34391303 | ENSG00000134317 |
| MYL5 | -0.869967177 | 0.547159299 | 0.001639003 | 0.006627125 | 86.07285656 | 47.01561349 | ENSG00000215375 |
| ZNF578 | -0.870069204 | 0.547120605 | 0.04199494 | 0.103676015 | 38.9917114 | 21.28524655 | ENSG00000258405 |
| CUX2 | -0.870372744 | 0.547005505 | 1.20E-09 | 1.44E-08 | 297.6381168 | 162.7797405 | ENSG00000111249 |
| HKDC1 | -0.870444531 | 0.546978287 | 0.396956761 | 0.569472215 | 6.074330806 | 3.331944001 | ENSG00000156510 |
| NA | -0.870544759 | 0.546940288 | 0.411049192 | 0.5833554 | 5.353386335 | 2.909000794 | ENSG00000248476 |
| NA | -0.870921113 | 0.546797627 | 0.142879385 | 0.272091432 | 16.83557009 | 9.181671157 | ENSG00000232000 |
| NA | -0.87143644 | 0.546602347 | 0.137628169 | 0.264279833 | 24.91068591 | 13.52299413 | ENSG00000256092 |
| TMEM121B | -0.871509999 | 0.546574478 | 1.76E-06 | 1.32E-05 | 244.5772193 | 133.4668722 | ENSG00000183307 |
| NA | -0.87164848 | 0.546522016 | 0.209425183 | 0.364688681 | 14.285413 | 7.784582191 | ENSG00000253333 |
| NA | -0.871934522 | 0.546413668 | 0.486991434 | 0.654262257 | 4.310748416 | 2.391674385 | ENSG00000223834 |
| MAATS1 | -0.872656393 | 0.546140332 | 0.403039072 | 0.575331266 | 5.372246934 | 2.913301941 | ENSG00000183833 |
| UBE2QL1 | -0.87329974 | 0.545896843 | 0.030816144 | 0.080258436 | 37.28191991 | 20.3024985 | ENSG00000215218 |
| TUBB3 | -0.873323791 | 0.545887743 | 0.191331261 | 0.340088865 | 13.25884478 | 7.27155693 | ENSG00000258947 |
| KIF6 | -0.873343998 | 0.545880097 | 0.053725851 | 0.126362521 | 33.42448881 | 18.23052136 | ENSG00000164627 |
| NA | -0.873596726 | 0.545784479 | 0.434906206 | 0.605992526 | 5.908140058 | 3.214177063 | ENSG00000235772 |
| GAS1 | -0.873736498 | 0.545731605 | 0.000213607 | 0.001069754 | 138.4344612 | 75.53988579 | ENSG00000180447 |
| NA | -0.87421385 | 0.545551066 | 0.008714089 | 0.028054655 | 53.52326331 | 29.24621263 | ENSG00000277534 |
| LINC02523 | -0.874534348 | 0.545429884 | 0.159967401 | 0.297066916 | 20.85253409 | 11.30340249 | ENSG00000224506 |
| NA | -0.874876607 | 0.545300504 | 0.472280566 | 0.641196734 | 4.277933303 | 2.366400538 | ENSG00000273711 |
| FIG4 | -0.875072589 | 0.545226433 | 8.88E-18 | 2.50E-16 | 704.6502187 | 384.1589082 | ENSG00000112367 |
| NA | -0.875338355 | 0.545126003 | 0.162586095 | 0.300512442 | 2517.119494 | 1372.149992 | ENSG00000248527 |
| SPINK2 | -0.87565789 | 0.545005279 | 0.196075112 | 0.346540081 | 12.69013654 | 6.930886983 | ENSG00000128040 |
| PDRG1 | -0.875816679 | 0.544945297 | 8.39E-22 | 3.25E-20 | 996.3220891 | 542.9869564 | ENSG00000088356 |
| VIPR1 | -0.876326817 | 0.544752638 | 0.001276324 | 0.005314388 | 81.41174193 | 44.38677558 | ENSG00000114812 |
| NA | -0.876680742 | 0.544619015 | 0.342947207 | 0.515650395 | 6.673063254 | 3.651641249 | ENSG00000270175 |
| HINT2 | -0.877436403 | 0.544333827 | 0.381845949 | 0.555226517 | 6.112727725 | 3.331944001 | ENSG00000137133 |
| PIK3R3 | -0.877736813 | 0.544220492 | 5.10E-08 | 4.90E-07 | 278.4512834 | 151.7059716 | ENSG00000117461 |
| NA | -0.87792186 | 0.544150693 | 0.372386599 | 0.545833134 | 6.003794495 | 3.258273035 | ENSG00000260577 |
| SNORD62A | -0.878152549 | 0.544063689 | 0.465979801 | 0.635311034 | 4.289096914 | 2.366400538 | ENSG00000235284 |
| MACROH2A2 | -0.878747477 | 0.543839378 | 1.10E-34 | 8.79E-33 | 1862.763278 | 1012.41408 | ENSG00000099284 |
| NA | -0.87899117 | 0.543747523 | 0.005531268 | 0.019149982 | 58.00666017 | 31.50129795 | ENSG00000234009 |
| NA | -0.879752065 | 0.54346082 | 0.171697289 | 0.313656478 | 15.16485123 | 8.232799245 | ENSG00000256338 |
| SIRPA | -0.88007407 | 0.543339535 | 1.86E-08 | 1.91E-07 | 269.8699352 | 146.6787472 | ENSG00000198053 |
| NA | -0.880431231 | 0.54320504 | 0.000489052 | 0.002246628 | 106.0502949 | 57.48037479 | ENSG00000186301 |
| GNA15 | -0.88053725 | 0.543165122 | 0.331702198 | 0.504364081 | 8.330503317 | 4.490815479 | ENSG00000060558 |
| NA | -0.880543735 | 0.543162681 | 0.100696205 | 0.208180312 | 22.60004678 | 12.30308255 | ENSG00000219891 |
| ST8SIA1 | -0.8807146 | 0.543098356 | 0.003159633 | 0.011717125 | 74.2960125 | 40.43479225 | ENSG00000111728 |
| ACADSB | -0.880943175 | 0.543012316 | 1.22E-14 | 2.54E-13 | 499.7733787 | 271.5231774 | ENSG00000196177 |
| NA | -0.880972891 | 0.543001131 | 0.406708726 | 0.578944497 | 5.402271144 | 2.90685022 | ENSG00000284728 |
| PROKR1 | -0.881536152 | 0.542789173 | 0.005793008 | 0.019934521 | 61.81536971 | 33.65716588 | ENSG00000169618 |
| LAMA4 | -0.882424629 | 0.542455001 | 0.062837809 | 0.143096761 | 27.93090573 | 15.2077822 | ENSG00000112769 |
| SSTR3 | -0.883124025 | 0.542192092 | 0.213132056 | 0.369310203 | 15.19140882 | 8.224196951 | ENSG00000278195 |
| HM13 | -0.884056922 | 0.541841604 | 9.64E-39 | 1.03E-36 | 2447.71646 | 1326.227092 | ENSG00000101294 |
| H2AW | -0.884142311 | 0.541809536 | 3.21E-11 | 4.65E-10 | 371.4354819 | 201.2377507 | ENSG00000181218 |
| SYPL2 | -0.884335888 | 0.541736842 | 0.001265507 | 0.005275043 | 81.63989612 | 44.21200923 | ENSG00000143028 |
| PPP1R14B-AS1 | -0.884454907 | 0.541692151 | 0.338152395 | 0.51074991 | 6.643039044 | 3.598942982 | ENSG00000256940 |
| SH2D5 | -0.885662019 | 0.541239104 | 4.38E-20 | 1.49E-18 | 734.6748241 | 397.3210959 | ENSG00000189410 |
| ABHD8 | -0.886276779 | 0.54100852 | 2.91E-07 | 2.50E-06 | 237.7087048 | 128.465182 | ENSG00000127220 |
| CDK5 | -0.886864916 | 0.540788015 | 4.94E-12 | 7.81E-11 | 420.6590724 | 227.3321651 | ENSG00000164885 |
| HDAC9 | -0.88715241 | 0.54068026 | 2.02E-07 | 1.78E-06 | 215.11444 | 116.4277293 | ENSG00000048052 |
| NA | -0.887888626 | 0.540404418 | 0.03810143 | 0.095767386 | 34.62302975 | 18.66153382 | ENSG00000284727 |
| TLL1 | -0.887924815 | 0.540390863 | 0.07694712 | 0.168107236 | 24.85554357 | 13.38398192 | ENSG00000038295 |
| CD2BP2 | -0.88805298 | 0.540342858 | 2.65E-47 | 4.06E-45 | 2389.012872 | 1290.677277 | ENSG00000169217 |
| TMEM109 | -0.888222752 | 0.540279275 | 2.43E-26 | 1.29E-24 | 1046.896895 | 565.7276484 | ENSG00000110108 |
| NA | -0.889147855 | 0.539932942 | 0.376206502 | 0.549602004 | 6.031027803 | 3.235149762 | ENSG00000268157 |
| TTPA | -0.891733027 | 0.5389663 | 0.42065065 | 0.592708473 | 5.584331429 | 3.056342726 | ENSG00000137561 |
| MAPK8IP1 | -0.892498905 | 0.538680257 | 8.54E-28 | 4.85E-26 | 1268.804771 | 683.5513943 | ENSG00000121653 |
| HEXA | -0.892938345 | 0.538516202 | 0.003219617 | 0.011916662 | 75.68374713 | 40.74426967 | ENSG00000213614 |
| NA | -0.893547101 | 0.538289019 | 0.350471386 | 0.523090406 | 6.664690545 | 3.575819709 | ENSG00000279722 |
| LOC729603 | -0.895639798 | 0.537508771 | 0.164785035 | 0.303801379 | 16.0573681 | 8.678865726 | ENSG00000213073 |
| NA | -0.895922974 | 0.537403278 | 0.118117168 | 0.235652866 | 18.92575201 | 10.11763963 | ENSG00000266777 |
| RPL13P5 | -0.896017153 | 0.537368197 | 0.239321722 | 0.401387179 | 10.3061702 | 5.521427613 | ENSG00000240370 |
| NA | -0.896584365 | 0.537156966 | 0.283284896 | 0.451752412 | 9.269114089 | 5.027224477 | ENSG00000277342 |
| NA | -0.896908506 | 0.537036293 | 0.104164948 | 0.213748112 | 18.93324884 | 10.18700944 | ENSG00000215302 |
| JAM2 | -0.897250325 | 0.536909068 | 1.14E-05 | 7.41E-05 | 146.7343767 | 78.67715682 | ENSG00000154721 |
| PRR36 | -0.897531931 | 0.536804276 | 5.43E-15 | 1.17E-13 | 746.922408 | 401.0770491 | ENSG00000183248 |

| | | | | | | | |
|---|---|---|---|---|---|---|---|
| NKAIN3 | -0.898004413 | 0.536628502 | 0.1555133 | 0.290642033 | 14.69092155 | 7.887828151 | ENSG00000185942 |
| WTIP | -0.898615089 | 0.536401401 | 1.19E-12 | 2.01E-11 | 444.9307139 | 238.4180502 | ENSG00000142279 |
| TRAPPC5 | -0.898651683 | 0.536387796 | 0.007394123 | 0.024468566 | 51.46677239 | 27.6181514 | ENSG00000181029 |
| HMX3 | -0.899084577 | 0.536226871 | 0.301258468 | 0.47202543 | 7.356286528 | 3.946064649 | ENSG00000188620 |
| NPHP3 | -0.899691523 | 0.536001327 | 0.050932369 | 0.121121342 | 28.14598129 | 15.05452158 | ENSG00000113971 |
| GREM1 | -0.900516503 | 0.535694912 | 0.34788717 | 0.520562473 | 6.714251076 | 3.580120856 | ENSG00000166923 |
| NLRP2 | -0.901376036 | 0.535375849 | 2.12E-66 | 5.76E-64 | 3921.792349 | 2099.188997 | ENSG00000022556 |
| WNT3A | -0.901796652 | 0.535219783 | 0.404686284 | 0.576913252 | 5.554307219 | 3.007945607 | ENSG00000154342 |
| NA | -0.90195671 | 0.535160407 | 0.291655441 | 0.461401109 | 7.995531076 | 4.291035743 | ENSG00000281468 |
| EBLN2 | -0.902414996 | 0.534990435 | 0.111491569 | 0.225042783 | 19.50195708 | 10.43303573 | ENSG00000255423 |
| SEMA5B | -0.902506645 | 0.534956451 | 1.18E-16 | 3.01E-15 | 806.0041195 | 430.7118384 | ENSG00000082684 |
| TMEM191B | -0.903412551 | 0.534620643 | 0.30224183 | 0.472989952 | 8.556542327 | 4.587609718 | ENSG00000278558 |
| NA | -0.903449865 | 0.534606816 | 0.017602547 | 0.050553927 | 43.56833521 | 23.24914354 | ENSG00000279348 |
| DZIP1 | -0.903588705 | 0.534555369 | 7.58E-25 | 3.71E-23 | 1076.872826 | 575.9527925 | ENSG00000134874 |
| NA | -0.903997264 | 0.534404009 | 0.371305685 | 0.54495085 | 6.10714592 | 3.230848615 | ENSG00000250012 |
| NA | -0.904749989 | 0.534125257 | 0.32128648 | 0.492721002 | 7.903343264 | 4.217364777 | ENSG00000251400 |
| LOC101929240 | -0.905892022 | 0.533702612 | 0.447166865 | 0.617487186 | 4.291887817 | 2.297030719 | ENSG00000265298 |
| STEAP3-AS1 | -0.906143283 | 0.53360967 | 0.24517906 | 0.408370682 | 10.52192148 | 5.599399727 | ENSG00000229867 |
| CLEC14A | -0.90648121 | 0.533484696 | 0.324060218 | 0.495543064 | 7.293447204 | 3.872393683 | ENSG00000176435 |
| COL12A1 | -0.906513035 | 0.533472928 | 0.009990201 | 0.031392891 | 50.62784626 | 27.09222269 | ENSG00000111799 |
| VARS2 | -0.907623734 | 0.533062377 | 0.221440306 | 0.379808687 | 11.0738843 | 5.887371406 | ENSG00000137411 |
| NA | -0.907708467 | 0.53303107 | 0.390955254 | 0.564176976 | 5.617146542 | 3.03106888 | ENSG00000254783 |
| PSORS1C2 | -0.908271658 | 0.532823029 | 0.36845506 | 0.542378219 | 6.681435963 | 3.531723737 | ENSG00000204538 |
| CAMK2N2 | -0.908777476 | 0.53263625 | 5.53E-05 | 0.000315963 | 146.6486097 | 78.20339334 | ENSG00000163888 |
| ATAD1 | -0.909196208 | 0.532481679 | 3.32E-19 | 1.04E-17 | 809.4016776 | 431.2140561 | ENSG00000138138 |
| TP53INP2 | -0.909219959 | 0.532472913 | 2.69E-05 | 0.00016362 | 146.3878405 | 77.88880285 | ENSG00000078804 |
| HTR1A | -0.909543115 | 0.532353655 | 0.023539771 | 0.064330037 | 35.81770374 | 19.06565491 | ENSG00000178394 |
| RNF139-AS1 | -0.909881599 | 0.532228769 | 0.170323604 | 0.311677458 | 14.0893982 | 7.452514539 | ENSG00000245149 |
| ZNF585B | -0.909979092 | 0.532192804 | 0.002284479 | 0.00885471 | 66.73497505 | 35.49145857 | ENSG00000245680 |
| GPR137C | -0.910235126 | 0.532098364 | 0.00123013 | 0.005146484 | 77.79191377 | 41.41647423 | ENSG00000180998 |
| MPPED2 | -0.9109333 | 0.531840925 | 0.003965898 | 0.01434042 | 85.91974444 | 45.66692165 | ENSG00000066382 |
| LINC00115 | -0.911520081 | 0.531624655 | 0.139622522 | 0.267378594 | 17.18382112 | 9.071973475 | ENSG00000225880 |
| FAAP24 | -0.911607403 | 0.531592479 | 0.000246505 | 0.001220251 | 97.11268644 | 51.61853769 | ENSG00000131944 |
| NUCB2 | -0.912543882 | 0.531247524 | 3.12E-09 | 3.56E-08 | 316.5724928 | 168.1764164 | ENSG00000070081 |
| NA | -0.914638841 | 0.530476652 | 0.312823643 | 0.48386784 | 7.470801564 | 3.950365797 | ENSG00000276571 |
| NA | -0.914645243 | 0.530474298 | 0.380651851 | 0.554033112 | 6.21587899 | 3.329793428 | ENSG00000267898 |
| LYPD6B | -0.915018933 | 0.530336911 | 0.009052224 | 0.028926031 | 59.88946352 | 31.70161072 | ENSG00000150556 |
| NA | -0.915850861 | 0.530031181 | 0.435309545 | 0.606135187 | 4.286306011 | 2.267455725 | ENSG00000258477 |
| NA | -0.916997248 | 0.529610178 | 0.143171945 | 0.272621656 | 16.02734389 | 8.460003401 | ENSG00000269688 |
| BBS9 | -0.917066697 | 0.529584684 | 6.75E-11 | 9.43E-10 | 455.6435808 | 241.4966679 | ENSG00000122507 |
| LOC105374618 | -0.918112606 | 0.52920089 | 0.165443142 | 0.304800272 | 16.6114461 | 8.76894778 | ENSG00000248994 |
| RBP4 | -0.918201254 | 0.529168374 | 0.160490567 | 0.297809112 | 15.37414483 | 8.091909034 | ENSG00000138207 |
| SLC16A9 | -0.919567126 | 0.528667621 | 3.39E-10 | 4.40E-09 | 295.9518919 | 156.4355498 | ENSG00000165449 |
| FAM133B | -0.919597088 | 0.528656641 | 4.26E-20 | 1.46E-18 | 798.0155486 | 422.0517401 | ENSG00000234545 |
| NA | -0.920381946 | 0.528369119 | 0.290792578 | 0.46052702 | 9.2397656 | 4.879882545 | ENSG00000273729 |
| CTB-178M22.2 | -0.920483643 | 0.528331875 | 0.083438668 | 0.179504332 | 22.84965232 | 12.06754868 | ENSG00000253978 |
| ZNF443 | -0.920814837 | 0.528210602 | 0.003661012 | 0.01336076 | 82.02240172 | 43.19995876 | ENSG00000180855 |
| VSX2 | -0.920924863 | 0.52817032 | 0.418832532 | 0.590923599 | 4.346354432 | 2.292729572 | ENSG00000119614 |
| NXPH2 | -0.920986276 | 0.528147837 | 0.002733316 | 0.010357179 | 62.71155336 | 33.11430516 | ENSG00000144227 |
| GPAM | -0.922454439 | 0.527610639 | 1.64E-21 | 6.14E-20 | 759.7849914 | 400.8740222 | ENSG00000119927 |
| LINC02570 | -0.922759489 | 0.52749909 | 2.49E-07 | 2.16E-06 | 199.37262 | 105.0415326 | ENSG00000237923 |
| NDUFA9 | -0.923122708 | 0.527366302 | 0.001954287 | 0.007731778 | 70.36250019 | 37.06305343 | ENSG00000139180 |
| PDSS2 | -0.923556516 | 0.52720775 | 1.27E-08 | 1.33E-07 | 289.7348488 | 152.9239931 | ENSG00000164494 |
| SOX21-AS1 | -0.923676708 | 0.52716383 | 0.014754793 | 0.043675725 | 44.84662415 | 23.69305945 | ENSG00000227640 |
| IGFLR1 | -0.924097619 | 0.527010051 | 0.365376463 | 0.539080485 | 7.372356224 | 3.855722131 | ENSG00000126246 |
| NA | -0.92413171 | 0.526997598 | 0.153158028 | 0.28740968 | 16.09979522 | 8.506249947 | ENSG00000269388 |
| TMX3 | -0.924208091 | 0.526969697 | 2.38E-10 | 3.15E-09 | 387.0248896 | 204.0356784 | ENSG00000166479 |
| LOC105376114 | -0.924606006 | 0.526824372 | 0.157795306 | 0.293881951 | 14.42157954 | 7.61437745 | ENSG00000254473 |
| NA | -0.925524668 | 0.526489014 | 0.377588952 | 0.551044564 | 6.267554703 | 3.323341707 | ENSG00000233461 |
| H2BC21 | -0.92579799 | 0.526389278 | 0.001304691 | 0.005419639 | 81.20926944 | 42.84045458 | ENSG00000184678 |
| RIMS4 | -0.926197305 | 0.526243603 | 1.90E-49 | 3.07E-47 | 2079.892408 | 1094.579932 | ENSG00000101098 |
| COBLL1 | -0.926357224 | 0.526185273 | 0.001953957 | 0.007731778 | 85.20928786 | 45.0221294 | ENSG00000082438 |
| PRC1-AS1 | -0.926423507 | 0.526161099 | 0.361155472 | 0.534249291 | 5.67440406 | 3.007945607 | ENSG00000258725 |
| NA | -0.926538614 | 0.52611912 | 0.081654111 | 0.176353499 | 23.96282655 | 12.69645077 | ENSG00000271524 |
| BICC1 | -0.926823415 | 0.52601527 | 0.021619013 | 0.059964397 | 37.61689215 | 19.79754249 | ENSG00000122870 |
| NA | -0.92730916 | 0.525838194 | 0.091436414 | 0.19307135 | 19.94865345 | 10.50025497 | ENSG00000272931 |
| NBPF25P | -0.92807003 | 0.525560943 | 0.219430205 | 0.377477089 | 11.93446194 | 6.245245943 | ENSG00000272150 |
| NA | -0.92810675 | 0.525547566 | 0.400076834 | 0.572585667 | 5.657658643 | 2.986972907 | ENSG00000229670 |
| BLOC1S1 | -0.928143434 | 0.525534203 | 0.123483874 | 0.243546408 | 17.47269946 | 9.154246736 | ENSG00000135441 |
| PIGL | -0.928259132 | 0.525492059 | 1.14E-08 | 1.20E-07 | 267.1823723 | 140.6305668 | ENSG00000108474 |
| NA | -0.929078819 | 0.525193578 | 0.498126405 | 0.66440963 | 4.299584804 | 2.246483026 | ENSG00000274425 |
| NA | -0.929709682 | 0.524963971 | 0.181115633 | 0.326307024 | 13.80889258 | 7.241981936 | ENSG00000280046 |
| MSC | -0.931238025 | 0.524408137 | 0.150469483 | 0.283080437 | 18.0441986 | 9.433616121 | ENSG00000178860 |
| LCA5L | -0.931622198 | 0.524268511 | 0.450656229 | 0.621079016 | 4.980017175 | 2.608125672 | ENSG00000157578 |
| TCEAL3 | -0.931805533 | 0.524201892 | 3.81E-12 | 6.09E-11 | 367.052082 | 192.2313911 | ENSG00000196507 |
| GPX3 | -0.931808887 | 0.524200674 | 3.73E-21 | 1.38E-19 | 722.188838 | 378.5737811 | ENSG00000211445 |
| QRFPR | -0.931983276 | 0.524137314 | 0.052013757 | 0.123036053 | 29.33439776 | 15.35486367 | ENSG00000186867 |

| | | | | | | | |
|---|---|---|---|---|---|---|---|
| ACAN | -0.932511072 | 0.523945598 | 0.051831107 | 0.122819541 | 30.44458093 | 15.92918949 | ENSG00000157766 |
| KCNMA1 | -0.932705441 | 0.523875014 | 3.02E-05 | 0.00018201 | 132.4470487 | 69.28333552 | ENSG00000156113 |
| OLIG1 | -0.932755769 | 0.523856739 | 0.02452967 | 0.066545052 | 36.82473564 | 19.22348924 | ENSG00000184221 |
| NA | -0.933330164 | 0.523648212 | 0.255932837 | 0.421047275 | 8.801642095 | 4.589760291 | ENSG00000255139 |
| RGS11 | -0.934369496 | 0.523271106 | 0.066771762 | 0.150262111 | 24.00612955 | 12.62493038 | ENSG00000076344 |
| NTRK3 | -0.93611986 | 0.522636627 | 2.67E-06 | 1.94E-05 | 179.7217811 | 94.16913048 | ENSG00000140538 |
| ZNF501 | -0.93657721 | 0.522470972 | 2.71E-05 | 0.000165147 | 143.4846143 | 75.10428118 | ENSG00000186446 |
| VAT1L | -0.936869569 | 0.522365105 | 7.51E-50 | 1.24E-47 | 2706.424152 | 1414.006833 | ENSG00000171724 |
| SMPDL3B | -0.937348136 | 0.522191856 | 9.16E-20 | 3.04E-18 | 607.8998641 | 317.4810269 | ENSG00000130768 |
| LINC01963 | -0.937819175 | 0.522021389 | 0.008155553 | 0.026553378 | 52.18193489 | 27.28555071 | ENSG00000260804 |
| HMSD | -0.939141774 | 0.521543042 | 0.092566683 | 0.194755874 | 24.78971319 | 13.02690088 | ENSG00000221887 |
| NA | -0.939240449 | 0.521507372 | 0.221399022 | 0.379808687 | 10.0479918 | 5.224853639 | ENSG00000287811 |
| LOC101929748 | -0.939618906 | 0.521370584 | 0.118039904 | 0.235554752 | 19.51332085 | 10.14668158 | ENSG00000288062 |
| NA | -0.940304761 | 0.521122784 | 0.386312392 | 0.559696008 | 4.996762593 | 2.59145412 | ENSG00000273271 |
| FUT8 | -0.941129369 | 0.520825009 | 4.43E-16 | 1.07E-14 | 484.9581299 | 252.7056993 | ENSG00000033170 |
| NTN3 | -0.94131145 | 0.52075928 | 0.363502766 | 0.536988841 | 5.72328887 | 2.986972907 | ENSG00000162068 |
| FIGNL2 | -0.941783667 | 0.520588855 | 2.85E-06 | 2.07E-05 | 160.6870959 | 83.73825743 | ENSG00000261308 |
| SNAP25 | -0.94196628 | 0.520522965 | 0.045594173 | 0.110845266 | 54.76499511 | 28.49120177 | ENSG00000132639 |
| SLFN11 | -0.942437966 | 0.520352809 | 0.093566872 | 0.196430355 | 19.71212655 | 10.30692696 | ENSG00000172716 |
| MEGF9 | -0.94252898 | 0.520319983 | 0.00597342 | 0.020459615 | 58.51061406 | 30.50428938 | ENSG00000106780 |
| PCDHGB4 | -0.942739373 | 0.520244109 | 5.67E-05 | 0.000322977 | 124.0599266 | 64.56290486 | ENSG00000253953 |
| LINC01273 | -0.943385485 | 0.520011169 | 0.143441874 | 0.272973975 | 15.16696642 | 7.862554305 | ENSG00000231742 |
| NA | -0.943751257 | 0.519879345 | 0.268507309 | 0.434752592 | 9.508431892 | 4.985279079 | ENSG00000225695 |
| NA | -0.944433478 | 0.519633563 | 0.166661543 | 0.306510767 | 16.8291124 | 8.72055066 | ENSG00000288060 |
| NA | -0.944613223 | 0.519568826 | 0.114155996 | 0.229465804 | 16.41331612 | 8.510551094 | ENSG00000286340 |
| KCND3 | -0.944775607 | 0.519510349 | 0.252097427 | 0.416549162 | 10.23284299 | 5.372195571 | ENSG00000171385 |
| GLIPR2 | -0.945474692 | 0.519258671 | 1.38E-07 | 1.24E-06 | 219.7867551 | 114.0532595 | ENSG00000122694 |
| LINC01806 | -0.945916718 | 0.5190996 | 0.3163162 | 0.487378373 | 6.349254625 | 3.304519581 | ENSG00000227403 |
| RAX | -0.947134911 | 0.518661464 | 0.425963922 | 0.597705773 | 4.45249676 | 2.292729572 | ENSG00000134438 |
| LOC105371414 | -0.947199921 | 0.518638093 | 0.083899143 | 0.180193539 | 22.78980405 | 11.80485084 | ENSG00000260279 |
| NA | -0.947699636 | 0.51845848 | 0.396773968 | 0.569416775 | 5.138510936 | 2.660823939 | ENSG00000280114 |
| NUP210L | -0.94830655 | 0.518240421 | 0.28703264 | 0.456295564 | 7.587231622 | 3.948215223 | ENSG00000143552 |
| NA | -0.94839767 | 0.51820769 | 0.263957306 | 0.429641531 | 8.270454896 | 4.31630959 | ENSG00000266490 |
| LIPE-AS1 | -0.948660999 | 0.518113113 | 0.16929257 | 0.310174088 | 12.66848504 | 6.567093764 | ENSG00000213904 |
| TMEM221 | -0.94870306 | 0.518098007 | 0.011255779 | 0.03486436 | 46.98935766 | 24.32868583 | ENSG00000188051 |
| ANKRD36 | -0.948796943 | 0.518064293 | 0.001372296 | 0.005660246 | 72.68641811 | 37.55080484 | ENSG00000135976 |
| NA | -0.949919109 | 0.517661486 | 0.027539264 | 0.073398722 | 33.6053098 | 17.42092212 | ENSG00000261762 |
| TRPC4 | -0.950712442 | 0.517376904 | 2.11E-09 | 2.46E-08 | 256.2208879 | 132.4394646 | ENSG00000133107 |
| CBLN1 | -0.950984604 | 0.517279311 | 3.90E-08 | 3.82E-07 | 248.2554321 | 128.587238 | ENSG00000102924 |
| NA | -0.951051172 | 0.517255444 | 0.409832802 | 0.58197163 | 5.160162438 | 2.665125086 | ENSG00000283538 |
| SPTB | -0.951161178 | 0.517216004 | 0.001706267 | 0.006868865 | 66.15769394 | 34.2610667 | ENSG00000070182 |
| ATF7IP2 | -0.95148435 | 0.517100158 | 6.08E-10 | 7.58E-09 | 265.0356086 | 137.1047739 | ENSG00000166669 |
| NA | -0.951551228 | 0.517076188 | 0.290776041 | 0.46052702 | 7.677304254 | 3.990160622 | ENSG00000279822 |
| NA | -0.951902282 | 0.516950382 | 0.274119327 | 0.440926869 | 7.647280043 | 3.971338496 | ENSG00000254893 |
| NPPC | -0.952318876 | 0.516801128 | 0.058164537 | 0.134687264 | 24.6293044 | 12.69834088 | ENSG00000163273 |
| RRP36 | -0.952945603 | 0.516576671 | 4.10E-36 | 3.60E-34 | 1648.251751 | 850.9037515 | ENSG00000124541 |
| UTS2R | -0.953081092 | 0.51652816 | 0.259652658 | 0.424920209 | 8.270454896 | 4.293186317 | ENSG00000181408 |
| ZEB2 | -0.95341302 | 0.516409333 | 0.216109892 | 0.373229715 | 12.05579808 | 6.213520375 | ENSG00000169554 |
| NA | -0.954429249 | 0.516045704 | 0.661660087 | 0.791256982 | 7.457522771 | 3.838517542 | ENSG00000224195 |
| PTPMT1 | -0.955336116 | 0.515721424 | 0.315710545 | 0.48687218 | 7.09252632 | 3.670463374 | ENSG00000110536 |
| PINLYP | -0.955510648 | 0.515659038 | 0.003196041 | 0.011840163 | 69.05765366 | 35.59040338 | ENSG00000234465 |
| NA | -0.956039442 | 0.515470067 | 0.041012515 | 0.101680046 | 27.33987027 | 14.08682756 | ENSG00000258568 |
| NA | -0.956085335 | 0.51545367 | 0.374934226 | 0.548268806 | 5.108486726 | 2.614577393 | ENSG00000256393 |
| SLC1A2 | -0.956293008 | 0.515379476 | 0.014837752 | 0.043882399 | 40.11728854 | 20.66199057 | ENSG00000110436 |
| LNP1 | -0.956315059 | 0.515371599 | 1.30E-07 | 1.18E-06 | 190.0717668 | 97.88585251 | ENSG00000206535 |
| DRD2 | -0.957138196 | 0.515077635 | 0.144696647 | 0.274711446 | 13.40896583 | 6.903462563 | ENSG00000149295 |
| NA | -0.957431927 | 0.514972777 | 0.339790443 | 0.512301479 | 5.728870676 | 2.93427464 | ENSG00000262410 |
| DANCR | -0.957576277 | 0.514921253 | 6.83E-40 | 7.94E-38 | 1503.515956 | 774.4283635 | ENSG00000226950 |
| LINC01315 | -0.959050037 | 0.514395513 | 1.08E-15 | 2.52E-14 | 450.8623452 | 231.8057638 | ENSG00000229891 |
| WDR73 | -0.959640415 | 0.514185056 | 2.00E-05 | 0.000124627 | 121.0984038 | 62.22392873 | ENSG00000177082 |
| NA | -0.960437597 | 0.513901014 | 1.01E-13 | 1.90E-12 | 382.0793272 | 196.3670035 | ENSG00000166492 |
| ZNF704 | -0.960547704 | 0.513861794 | 8.03E-06 | 5.38E-05 | 151.6127573 | 77.84579138 | ENSG00000164684 |
| NCAM1 | -0.961056288 | 0.513680677 | 5.99E-09 | 6.57E-08 | 251.811582 | 129.3500392 | ENSG00000149294 |
| H2BC17 | -0.961079439 | 0.513672434 | 0.251641101 | 0.415973306 | 10.69281815 | 5.447756647 | ENSG00000274641 |
| H4C3 | -0.962003721 | 0.513343449 | 0.062526178 | 0.142525405 | 22.31096829 | 11.44267516 | ENSG00000197061 |
| FAM163A | -0.962976299 | 0.5129975 | 0.046903304 | 0.113513708 | 29.07968598 | 14.99563206 | ENSG00000143340 |
| CAMK2N1 | -0.963146153 | 0.512937107 | 0.000127888 | 0.000672738 | 108.3314734 | 55.46754764 | ENSG00000162545 |
| GDF10 | -0.963424612 | 0.512838113 | 0.277022793 | 0.444235147 | 8.243221588 | 4.223816498 | ENSG00000266524 |
| ATP2B1-AS1 | -0.963575453 | 0.512784496 | 0.243604856 | 0.406491998 | 8.923653959 | 4.58330857 | ENSG00000271614 |
| VEGFC | -0.963803751 | 0.512703357 | 0.042231183 | 0.104165932 | 29.04966177 | 14.96820764 | ENSG00000150630 |
| CAMK2A | -0.963955766 | 0.512649337 | 0.130914085 | 0.254121917 | 24.4898714 | 12.57008153 | ENSG00000070808 |
| TBC1D9 | -0.96413452 | 0.512585822 | 0.000679196 | 0.003004442 | 76.95856945 | 39.52356459 | ENSG00000109436 |
| NA | -0.965186527 | 0.512212183 | 0.002914669 | 0.010934514 | 63.2236798 | 32.35929431 | ENSG00000271109 |
| SLC24A4 | -0.965286309 | 0.512176758 | 0.071635182 | 0.158877248 | 21.91873853 | 11.29344312 | ENSG00000140090 |
| NA | -0.965955345 | 0.511939296 | 0.025322222 | 0.068339533 | 34.58231749 | 17.71964667 | ENSG00000255397 |
| ZNF823 | -0.966798768 | 0.511640095 | 0.003661032 | 0.01336076 | 57.75386341 | 29.55569005 | ENSG00000197933 |
| CCDC81 | -0.967483888 | 0.511397181 | 0.193432413 | 0.343158423 | 12.28550387 | 6.259766921 | ENSG00000149201 |

| | | | | | | | |
|---|---|---|---|---|---|---|---|
| GLDN | -0.967687782 | 0.511324911 | 0.133518466 | 0.258134273 | 15.49635685 | 7.923854867 | ENSG00000186417 |
| ZNF649 | -0.968205941 | 0.511141296 | 5.71E-16 | 1.37E-14 | 524.1057075 | 268.0374213 | ENSG00000198093 |
| DHDH | -0.968686226 | 0.510971161 | 0.037514798 | 0.094502215 | 28.31775385 | 14.42749749 | ENSG00000104808 |
| PARP16 | -0.97027585 | 0.510408461 | 9.45E-06 | 6.26E-05 | 136.0181553 | 69.36696585 | ENSG00000138617 |
| NAA38 | -0.970311518 | 0.510395842 | 2.52E-13 | 4.59E-12 | 386.7735451 | 197.3954651 | ENSG00000183011 |
| GATD3A | -0.970915859 | 0.510182084 | 0.032709669 | 0.084290973 | 29.53878527 | 15.09001526 | ENSG00000160221 |
| NA | -0.971060687 | 0.510130871 | 0.329897851 | 0.502293563 | 6.499375678 | 3.304519581 | ENSG00000249485 |
| C3orf80 | -0.971294673 | 0.510048141 | 0.0095678 | 0.030307458 | 43.08112673 | 21.95798415 | ENSG00000180044 |
| EXOG | -0.971668201 | 0.509916101 | 1.46E-08 | 1.52E-07 | 221.9981852 | 113.1823597 | ENSG00000157036 |
| CTXN1 | -0.97355953 | 0.509248055 | 1.42E-14 | 2.93E-13 | 814.2818611 | 414.5855528 | ENSG00000178531 |
| NA | -0.974745576 | 0.508829572 | 0.332506109 | 0.505228051 | 7.622837638 | 3.857872705 | ENSG00000213683 |
| MSI1 | -0.976504541 | 0.508209574 | 9.52E-37 | 8.74E-35 | 1512.235423 | 768.2694925 | ENSG00000135097 |
| TRABD2A | -0.977910608 | 0.507714508 | 5.70E-06 | 3.92E-05 | 171.9347799 | 87.33613434 | ENSG00000186854 |
| DOK3 | -0.978264267 | 0.507590064 | 5.27E-05 | 0.000301773 | 123.6022668 | 62.63370803 | ENSG00000146094 |
| SOX21 | -0.97872586 | 0.507427685 | 0.002312587 | 0.008951065 | 60.97731946 | 31.02376637 | ENSG00000125285 |
| NA | -0.979648236 | 0.507103369 | 0.264858747 | 0.43088127 | 7.767376885 | 3.918640229 | ENSG00000235370 |
| LOC105375304 | -0.980631694 | 0.506757804 | 0.377213818 | 0.550691067 | 5.79171 | 2.904699646 | ENSG00000225792 |
| ANKHD1-EIF4EBP3 | -0.980787648 | 0.506703027 | 0.345948288 | 0.51846554 | 6.536896716 | 3.352916701 | ENSG00000254996 |
| RNF39 | -0.981962333 | 0.506290622 | 0.002975294 | 0.011121179 | 60.44401708 | 30.71052084 | ENSG00000204618 |
| CAMK2B | -0.982870606 | 0.505971979 | 0.165327058 | 0.3046155 | 15.77897766 | 8.020649106 | ENSG00000058404 |
| NA | -0.983124063 | 0.505883096 | 0.070097483 | 0.156275343 | 20.66959792 | 10.43088515 | ENSG00000285693 |
| LRRC45 | -0.983370486 | 0.505796695 | 1.28E-19 | 4.20E-18 | 919.099873 | 464.8126477 | ENSG00000169683 |
| TSSC2 | -0.983445081 | 0.505770543 | 3.80E-48 | 5.92E-46 | 1907.763826 | 964.7778767 | ENSG00000223756 |
| MAN2A1 | -0.983914488 | 0.505606008 | 1.38E-43 | 1.82E-41 | 1637.025443 | 828.0742372 | ENSG00000112893 |
| TAF2 | -0.986401134 | 0.50473529 | 1.10E-24 | 5.31E-23 | 763.5762798 | 385.6759147 | ENSG00000064313 |
| H2AC11 | -0.989399708 | 0.503687311 | 0.002098341 | 0.008230804 | 61.78881213 | 31.14772456 | ENSG00000196787 |
| NA | -0.990373988 | 0.503347276 | 0.283907086 | 0.452482782 | 7.938273558 | 4.042858888 | ENSG00000228395 |
| RAB11FIP3 | -0.990921751 | 0.503156201 | 8.32E-38 | 8.19E-36 | 1427.650312 | 718.3870841 | ENSG00000090565 |
| NA | -0.991403859 | 0.502988089 | 0.129922666 | 0.252756973 | 14.36990383 | 7.229611531 | ENSG00000261485 |
| MAP2 | -0.99333164 | 0.502316427 | 1.34E-07 | 1.21E-06 | 185.147819 | 93.10061363 | ENSG00000078018 |
| ZNF487 | -0.993405835 | 0.502290594 | 0.074825709 | 0.164609733 | 18.96327305 | 9.545464376 | ENSG00000243660 |
| NA | -0.993628792 | 0.502212975 | 0.092280262 | 0.194342634 | 17.72250515 | 8.92059086 | ENSG00000207425 |
| ADGRG5 | -0.993719136 | 0.502181527 | 1.43E-11 | 2.16E-10 | 377.2642615 | 189.542325 | ENSG00000159618 |
| RHBDL3 | -0.993955275 | 0.502099337 | 9.96E-18 | 2.78E-16 | 551.583259 | 276.8362472 | ENSG00000141314 |
| SNORC | -0.994263138 | 0.501992203 | 0.083658988 | 0.179857968 | 21.73476322 | 10.89147261 | ENSG00000182600 |
| ZSWIM5 | -0.994454454 | 0.501925639 | 3.76E-07 | 3.18E-06 | 206.9084882 | 103.986428 | ENSG00000162415 |
| KCNIP3 | -0.994541268 | 0.501895436 | 0.000590203 | 0.002653985 | 84.63694974 | 42.46805906 | ENSG00000115041 |
| NA | -0.995176839 | 0.501674377 | 0.099870904 | 0.207027015 | 22.50526823 | 11.28484082 | ENSG00000250770 |
| SPNS1 | -0.995186128 | 0.501671147 | 0.142693318 | 0.271870922 | 13.02231788 | 6.510094349 | ENSG00000169682 |
| CNTNAP3B | -0.997301194 | 0.50093621 | 0.008982097 | 0.028749564 | 46.13850418 | 23.18218476 | ENSG00000154529 |
| ZHX1 | -0.998033129 | 0.50068213 | 0.000138716 | 0.000723823 | 96.79011712 | 48.39549787 | ENSG00000165156 |
| ANKRD23 | -0.999506014 | 0.500171232 | 0.37975249 | 0.553016176 | 4.637347948 | 2.343277265 | ENSG00000163126 |
| IFT57 | -0.999656285 | 0.500119137 | 5.09E-18 | 1.46E-16 | 590.237971 | 294.9742754 | ENSG00000114446 |
| NA | -0.999870617 | 0.500044843 | 0.09781945 | 0.203629546 | 19.69538113 | 9.850107608 | ENSG00000271851 |
| ZNF778 | -1.000064462 | 0.49997766 | 7.61E-16 | 1.79E-14 | 525.7937594 | 262.6757487 | ENSG00000170100 |
| NA | -1.001536392 | 0.499467811 | 0.058168655 | 0.134687264 | 24.49333802 | 12.26328773 | ENSG00000230562 |
| FAM111B | -1.003014545 | 0.498956329 | 8.67E-17 | 2.23E-15 | 648.4790812 | 323.8166153 | ENSG00000189057 |
| NA | -1.003022034 | 0.498953739 | 0.078113724 | 0.170116488 | 18.49301015 | 9.223616555 | ENSG00000269993 |
| PHACTR3 | -1.004158641 | 0.4985608 | 0.172495367 | 0.314637682 | 13.81918031 | 6.920134115 | ENSG00000087495 |
| ZNF558 | -1.004549432 | 0.49842577 | 1.06E-08 | 1.13E-07 | 205.7388202 | 102.5509013 | ENSG00000167785 |
| CXXC5-AS1 | -1.006855045 | 0.497629858 | 0.342267004 | 0.514994089 | 5.312198513 | 2.662974513 | ENSG00000250635 |
| TREX2 | -1.007124934 | 0.497536774 | 0.197523575 | 0.348589084 | 10.58946673 | 5.298524605 | ENSG00000183479 |
| NA | -1.007781052 | 0.497310552 | 0.120144806 | 0.238693456 | 15.26262085 | 7.568130904 | ENSG00000228501 |
| YWHAB | -1.009007662 | 0.496887908 | 7.33E-115 | 8.84E-112 | 6564.644323 | 3261.835308 | ENSG00000166913 |
| KIAA0895 | -1.009175791 | 0.496830005 | 0.317209742 | 0.48837793 | 5.935373365 | 2.961699061 | ENSG00000164542 |
| WWTR1-AS1 | -1.009519658 | 0.496711599 | 0.426604279 | 0.598161981 | 3.998103399 | 1.998306171 | ENSG00000241313 |
| PLXNA4 | -1.010408779 | 0.496405574 | 0.004102576 | 0.014781993 | 50.84571272 | 25.19690202 | ENSG00000221866 |
| NA | -1.010415202 | 0.496403364 | 0.123981483 | 0.244303043 | 15.91215313 | 7.931924124 | ENSG00000262877 |
| ZNF846 | -1.010701955 | 0.496304708 | 0.000470538 | 0.002171047 | 78.10946487 | 38.79005947 | ENSG00000196605 |
| NA | -1.010732568 | 0.496294177 | 0.337358408 | 0.510173133 | 5.290547011 | 2.637700666 | ENSG00000197320 |
| LOC101927278 | -1.010779473 | 0.496278041 | 0.12607397 | 0.247214165 | 17.80632026 | 8.807125069 | ENSG00000287261 |
| IL18R1 | -1.010903622 | 0.496235337 | 0.375915085 | 0.549340468 | 6.575493795 | 3.224396894 | ENSG00000115604 |
| NA | -1.011409409 | 0.496061395 | 0.312824542 | 0.48386784 | 5.943746074 | 2.959548487 | ENSG00000228110 |
| BNC1 | -1.011660584 | 0.495975037 | 0.404763945 | 0.576936712 | 4.730211482 | 2.364249965 | ENSG00000169594 |
| NA | -1.012065427 | 0.495835878 | 0.295896822 | 0.466123273 | 6.643039044 | 3.308820728 | ENSG00000287966 |
| E2F2 | -1.012828048 | 0.495573845 | 9.18E-09 | 9.85E-08 | 232.6948334 | 115.3223801 | ENSG00000007968 |
| NA | -1.013092368 | 0.495483057 | 0.409814482 | 0.58197163 | 4.046988208 | 2.021429444 | ENSG00000270039 |
| NA | -1.013342465 | 0.495397171 | 0.364421821 | 0.537974382 | 5.317780318 | 2.644152387 | ENSG00000275601 |
| PCDHB15 | -1.014811783 | 0.494892889 | 8.65E-05 | 0.000474408 | 109.1061208 | 53.91907605 | ENSG00000113248 |
| NA | -1.015120479 | 0.494787007 | 0.261616794 | 0.427011555 | 7.924994765 | 3.927242524 | ENSG00000285646 |
| LINC01018 | -1.015685639 | 0.494593217 | 0.076909001 | 0.168042988 | 22.78402209 | 11.36496351 | ENSG00000250056 |
| NA | -1.016492756 | 0.494316594 | 0.359549828 | 0.532601991 | 5.32894393 | 2.642001813 | ENSG00000284959 |
| PVALB | -1.016938668 | 0.494163833 | 0.038122825 | 0.095808678 | 28.00443311 | 13.81552742 | ENSG00000100362 |
| NA | -1.017443756 | 0.493990857 | 0.003003033 | 0.011209657 | 56.89069503 | 28.00103932 | ENSG00000235859 |
| HID1 | -1.018481213 | 0.49363575 | 2.76E-11 | 4.02E-10 | 268.5998189 | 132.4728077 | ENSG00000167861 |
| NA | -1.019328926 | 0.49334578 | 0.412841465 | 0.585167134 | 4.071430613 | 2.021429444 | ENSG00000261226 |
| IGFBP6 | -1.02030128 | 0.493013385 | 5.82E-09 | 6.39E-08 | 229.2695806 | 112.8723613 | ENSG00000167779 |

| | | | | | | | |
|---|---|---|---|---|---|---|---|
| ERICH5 | -1.020671944 | 0.492886734 | 0.013930447 | 0.041651001 | 39.43541671 | 19.4735441 | ENSG00000177459 |
| CALML4 | -1.021317899 | 0.492666097 | 0.251166868 | 0.415331738 | 8.031137093 | 3.964886775 | ENSG00000129007 |
| NA | -1.021494658 | 0.492605739 | 0.402663284 | 0.574965019 | 4.041406403 | 1.998306171 | ENSG00000242952 |
| NA | -1.023206859 | 0.492021458 | 0.365247524 | 0.538931466 | 4.667372158 | 2.297030719 | ENSG00000240522 |
| NA | -1.023370633 | 0.491965607 | 0.303707506 | 0.474434827 | 5.970979382 | 2.93427464 | ENSG00000233588 |
| PPP2R5A | -1.023526165 | 0.491912573 | 3.32E-31 | 2.24E-29 | 890.0444519 | 437.7596441 | ENSG00000066027 |
| NA | -1.023534426 | 0.491909756 | 0.278472471 | 0.446217496 | 6.673063254 | 3.283546882 | ENSG00000214784 |
| EMC6 | -1.024550667 | 0.491563375 | 0.061603442 | 0.140918953 | 23.51353943 | 11.57334554 | ENSG00000127774 |
| NA | -1.024817957 | 0.49147231 | 0.339724494 | 0.512242083 | 5.287756108 | 2.59145412 | ENSG00000197332 |
| DDX47 | -1.025228487 | 0.491332478 | 0.301932915 | 0.472736618 | 7.280168411 | 3.569367988 | ENSG00000213782 |
| NA | -1.02543417 | 0.491262435 | 0.356920938 | 0.530336733 | 4.670163061 | 2.292729572 | ENSG00000262766 |
| PLPPR1 | -1.026152758 | 0.491017804 | 0.005638243 | 0.019464451 | 71.19485649 | 35.04403502 | ENSG00000148123 |
| NA | -1.026462111 | 0.490912528 | 0.356951085 | 0.530340679 | 5.405062047 | 2.658673365 | ENSG00000260060 |
| ANKRD53 | -1.026685375 | 0.490836562 | 0.229873254 | 0.390050571 | 8.003903785 | 3.922941376 | ENSG00000144031 |
| ARX | -1.026710194 | 0.490828119 | 0.393933751 | 0.566676241 | 4.016963998 | 1.970881751 | ENSG00000004848 |
| STIMATE | -1.027218142 | 0.490655337 | 0.170335521 | 0.311677458 | 11.34322632 | 5.557454328 | ENSG00000213533 |
| NA | -1.028652289 | 0.490167831 | 0.322424145 | 0.493902237 | 5.345013626 | 2.614577393 | ENSG00000251654 |
| NA | -1.028673635 | 0.490160579 | 0.387259925 | 0.560605674 | 4.694605466 | 2.299181293 | ENSG00000237991 |
| TRIM7 | -1.02991027 | 0.489740608 | 0.365864481 | 0.539552917 | 5.467901371 | 2.686097786 | ENSG00000146054 |
| MXRA5 | -1.030009256 | 0.489707007 | 3.76E-07 | 3.18E-06 | 152.2112143 | 74.49528572 | ENSG00000101825 |
| KCTD8 | -1.030297264 | 0.489609256 | 6.24E-07 | 5.07E-06 | 144.7402126 | 70.87752061 | ENSG00000183783 |
| CLIC2 | -1.031113464 | 0.489332339 | 0.268354762 | 0.43470554 | 7.29902901 | 3.55484701 | ENSG00000155962 |
| NA | -1.031355623 | 0.489250211 | 0.281485016 | 0.449886284 | 7.462428855 | 3.653791822 | ENSG00000244471 |
| LOC102724532 | -1.032832507 | 0.488749622 | 0.035044936 | 0.089271341 | 28.35739007 | 13.90802051 | ENSG00000264920 |
| ADAMTS17 | -1.034606562 | 0.488148986 | 0.008350778 | 0.027079316 | 53.97342631 | 26.25091 | ENSG00000140470 |
| SLC25A18 | -1.035736963 | 0.487766654 | 0.141053069 | 0.269438039 | 13.57986251 | 6.680559554 | ENSG00000182902 |
| NA | -1.036377237 | 0.487550229 | 0.406246565 | 0.578703471 | 4.022545804 | 1.952059625 | ENSG00000286443 |
| MFGE8 | -1.036818745 | 0.487401047 | 6.89E-190 | 4.43E-186 | 19998.02296 | 9747.035294 | ENSG00000140545 |
| ROM1 | -1.037140859 | 0.487292236 | 0.134972369 | 0.260196387 | 13.45505974 | 6.531067048 | ENSG00000149489 |
| EPHA10 | -1.037187757 | 0.487276396 | 0.009423489 | 0.029924038 | 38.53106051 | 18.7922042 | ENSG00000183317 |
| UHRF1 | -1.03725233 | 0.487254586 | 4.18E-40 | 4.95E-38 | 2537.673382 | 1236.097374 | ENSG00000276043 |
| GPR155 | -1.038023491 | 0.486994205 | 0.007295661 | 0.024198865 | 43.24173568 | 21.05750936 | ENSG00000163328 |
| NA | -1.038645249 | 0.48678437 | 0.085803379 | 0.183608761 | 17.47481464 | 8.464304548 | ENSG00000266865 |
| EMX1 | -1.039135668 | 0.486618924 | 8.85E-07 | 7.03E-06 | 163.4120166 | 79.35876928 | ENSG00000135638 |
| DDX11-AS1 | -1.040226796 | 0.486251028 | 0.017685799 | 0.050762795 | 34.46588743 | 16.75625378 | ENSG00000245614 |
| C9orf24 | -1.040265557 | 0.486237964 | 0.022679224 | 0.062367057 | 32.27379354 | 15.67456092 | ENSG00000164972 |
| NA | -1.040504239 | 0.486157526 | 0.320809725 | 0.492224641 | 6.033818705 | 2.915452515 | ENSG00000286529 |
| PINK1-AS | -1.042448428 | 0.485502817 | 0.048747645 | 0.116993134 | 28.00510883 | 13.62381694 | ENSG00000117242 |
| HUNK | -1.042525522 | 0.485476874 | 7.77E-15 | 1.65E-13 | 403.411536 | 196.0615547 | ENSG00000142149 |
| NA | -1.042528389 | 0.485475909 | 0.394419729 | 0.567042134 | 4.077012419 | 1.970881751 | ENSG00000259155 |
| NA | -1.042607003 | 0.485449456 | 0.288504554 | 0.457767836 | 6.711460173 | 3.235149762 | ENSG00000259969 |
| NA | -1.042971288 | 0.485326893 | 0.257256989 | 0.422420822 | 9.560107605 | 4.677952236 | ENSG00000277383 |
| NGRN | -1.043660481 | 0.485095102 | 1.55E-19 | 5.00E-18 | 566.6746709 | 274.9360802 | ENSG00000182768 |
| TSNAXIP1 | -1.043885178 | 0.485019556 | 0.120387007 | 0.239076153 | 16.22914066 | 7.835129884 | ENSG00000102904 |
| ECRG4 | -1.04394317 | 0.48500006 | 0.020569382 | 0.05754952 | 31.24598602 | 15.15131582 | ENSG00000119147 |
| NA | -1.045243658 | 0.484563063 | 0.426629552 | 0.598161981 | 4.079803322 | 1.968731177 | ENSG00000260368 |
| PCBP1-AS1 | -1.048354934 | 0.483519194 | 2.13E-09 | 2.49E-08 | 232.70213 | 112.5308674 | ENSG00000179818 |
| LINC01465 | -1.049389585 | 0.483172555 | 0.187862594 | 0.335137973 | 10.23284299 | 4.980977931 | ENSG00000221949 |
| DLX3 | -1.049543839 | 0.483120897 | 0.33349706 | 0.506016343 | 6.022655094 | 2.881576373 | ENSG00000064195 |
| EPHB1 | -1.051080182 | 0.482606689 | 5.00E-12 | 7.89E-11 | 285.3030524 | 137.6073188 | ENSG00000154928 |
| C2orf78 | -1.051868557 | 0.482343036 | 0.290957752 | 0.460675147 | 6.877650922 | 3.302369007 | ENSG00000187833 |
| LINC00545 | -1.05229955 | 0.482198962 | 0.069758719 | 0.155619918 | 18.41900722 | 8.882946609 | ENSG00000236094 |
| VN1R1 | -1.054760623 | 0.481377087 | 0.311947361 | 0.48296762 | 6.902093326 | 3.308820728 | ENSG00000178201 |
| TMEM121 | -1.05524221 | 0.481216425 | 0.000707027 | 0.003110437 | 69.18668679 | 33.30627611 | ENSG00000184986 |
| TRPM3 | -1.055936272 | 0.480984973 | 9.57E-08 | 8.85E-07 | 194.2886125 | 93.37138072 | ENSG00000083067 |
| FAM151B | -1.056268833 | 0.480874112 | 0.269570095 | 0.435935678 | 7.64448914 | 3.721011067 | ENSG00000152380 |
| RTN4RL2 | -1.057393534 | 0.480499377 | 7.02E-11 | 9.80E-10 | 352.129727 | 169.0395195 | ENSG00000186907 |
| NA | -1.057913881 | 0.480326103 | 0.098257237 | 0.204328614 | 15.0475453 | 7.227460957 | ENSG00000255182 |
| AMBP | -1.058358443 | 0.480178115 | 0.010215042 | 0.032031596 | 39.27433221 | 18.91427229 | ENSG00000106927 |
| ITGB1BP2 | -1.06065491 | 0.47941438 | 0.023900368 | 0.065112554 | 28.00423295 | 13.41140634 | ENSG00000147166 |
| LIMCH1 | -1.063202754 | 0.478568467 | 0.024643255 | 0.066796829 | 28.1920752 | 13.4850773 | ENSG00000064042 |
| SLC25A12 | -1.063981259 | 0.478310292 | 3.30E-11 | 4.78E-10 | 273.9485745 | 131.0974849 | ENSG00000115840 |
| ANGPT1 | -1.064183668 | 0.47824319 | 0.001736932 | 0.00698211 | 56.16608378 | 26.87551094 | ENSG00000154188 |
| ALKBH3 | -1.064186693 | 0.478242188 | 5.30E-09 | 5.86E-08 | 194.3629789 | 92.85002573 | ENSG00000166199 |
| NA | -1.065719878 | 0.477734219 | 0.292963963 | 0.462717133 | 7.498034872 | 3.544094141 | ENSG00000274292 |
| NA | -1.06586157 | 0.477687301 | 0.048874644 | 0.117236808 | 23.91135099 | 11.34588092 | ENSG00000204121 |
| MAN2A2 | -1.06600175 | 0.477640889 | 1.23E-16 | 3.13E-15 | 418.8960904 | 199.8350157 | ENSG00000196547 |
| ABAT | -1.06655774 | 0.47745685 | 3.03E-12 | 4.90E-11 | 292.9608573 | 139.6938169 | ENSG00000183044 |
| NAP1L3 | -1.066743078 | 0.477395516 | 8.64E-46 | 1.24E-43 | 1522.714105 | 726.5604666 | ENSG00000186310 |
| N4BP2L1 | -1.066794256 | 0.477378581 | 0.035172365 | 0.089502693 | 28.45930203 | 13.64532268 | ENSG00000139597 |
| PRSS27 | -1.068314523 | 0.4768758 | 0.028844525 | 0.076047524 | 30.77185618 | 14.66492148 | ENSG00000172382 |
| NA | -1.069127982 | 0.476606991 | 0.394276641 | 0.566892054 | 4.776305388 | 2.2486336 | ENSG00000267940 |
| SCUBE2 | -1.070872816 | 0.476030918 | 0.100846466 | 0.208401625 | 14.50327946 | 6.88033929 | ENSG00000175356 |
| NA | -1.071480946 | 0.475830302 | 0.139759232 | 0.267564015 | 11.73545607 | 5.576276454 | ENSG00000225506 |
| EPOP | -1.071851232 | 0.47570819 | 2.19E-06 | 1.62E-05 | 151.7718146 | 72.08882989 | ENSG00000273604 |
| OR2A20P | -1.074047103 | 0.474984683 | 0.220392849 | 0.378660354 | 8.278827604 | 3.941763502 | ENSG00000170356 |
| CABP7 | -1.075017835 | 0.474665193 | 0.238357377 | 0.400256978 | 8.346573013 | 3.975639643 | ENSG00000100314 |

| | | | | | | | |
|---|---|---|---|---|---|---|---|
| C2orf15 | -1.075730383 | 0.474430813 | 0.025045483 | 0.067674781 | 28.26029617 | 13.39688536 | ENSG00000273045 |
| CP | -1.076306011 | 0.474241555 | 0.081499597 | 0.176059208 | 18.69480692 | 8.811426216 | ENSG00000047457 |
| C19orf57 | -1.076745479 | 0.474097116 | 6.93E-05 | 0.000387762 | 90.97117381 | 43.24672623 | ENSG00000132016 |
| NA | -1.076975126 | 0.474021655 | 0.004991429 | 0.017516764 | 43.50617161 | 20.56304576 | ENSG00000261663 |
| C1orf229 | -1.077694385 | 0.473785389 | 0.217672789 | 0.375256994 | 8.349363916 | 3.967037349 | ENSG00000221953 |
| NA | -1.078016046 | 0.473679767 | 0.445621407 | 0.616058661 | 4.29467872 | 2.069826564 | ENSG00000280149 |
| BTBD17 | -1.078033144 | 0.473674153 | 2.11E-16 | 5.26E-15 | 496.418313 | 235.0210375 | ENSG00000204347 |
| ENHO | -1.078984473 | 0.47336191 | 1.18E-07 | 1.08E-06 | 187.7623301 | 88.96579469 | ENSG00000168913 |
| CES4A | -1.079021592 | 0.473349731 | 0.287427941 | 0.456811041 | 6.261972897 | 2.986972907 | ENSG00000172824 |
| NOG | -1.079614292 | 0.473155306 | 0.125759331 | 0.246747687 | 13.20525405 | 6.219972096 | ENSG00000183691 |
| SLC39A1 | -1.079712886 | 0.473122971 | 0.000803913 | 0.003490497 | 59.67717886 | 28.25754589 | ENSG00000143570 |
| TIMM23B | -1.07992502 | 0.473053408 | 0.002340557 | 0.009048432 | 50.70184919 | 24.01060612 | ENSG00000204152 |
| KIAA0319 | -1.080238945 | 0.472950485 | 0.043071887 | 0.105887571 | 23.49256365 | 11.05575867 | ENSG00000137261 |
| NA | -1.080626398 | 0.472823485 | 0.430337647 | 0.601699177 | 4.164294147 | 1.945607904 | ENSG00000235052 |
| ADCYAP1R1 | -1.081470566 | 0.472546902 | 5.37E-31 | 3.57E-29 | 858.8173776 | 406.1120582 | ENSG00000078549 |
| NA | -1.081893909 | 0.472408259 | 0.078627654 | 0.171061923 | 18.86829433 | 8.971138553 | ENSG00000213144 |
| NA | -1.082094094 | 0.472342713 | 0.267615974 | 0.433939628 | 6.88514775 | 3.256122461 | ENSG00000073905 |
| NA | -1.082988561 | 0.472049953 | 0.250823546 | 0.41494186 | 7.008035494 | 3.327642854 | ENSG00000260269 |
| GABRG3 | -1.083149383 | 0.471997335 | 0.309769894 | 0.480908662 | 5.587122332 | 2.662974513 | ENSG00000182256 |
| RBM19 | -1.083183873 | 0.471986051 | 2.84E-49 | 4.53E-47 | 1815.013429 | 856.4878126 | ENSG00000122965 |
| NA | -1.085528338 | 0.471219668 | 0.14094502 | 0.269311646 | 13.96371956 | 6.596668758 | ENSG00000276386 |
| NA | -1.08680765 | 0.470801999 | 0.323554745 | 0.495163455 | 5.693264659 | 2.711371632 | ENSG00000230626 |
| H2AC8 | -1.087066838 | 0.470717424 | 0.029480084 | 0.077395252 | 27.12479472 | 12.79298454 | ENSG00000277075 |
| HESX1 | -1.087365892 | 0.47061986 | 6.98E-07 | 5.62E-06 | 138.7908847 | 65.33350276 | ENSG00000163666 |
| NA | -1.089420836 | 0.469949996 | 0.000703042 | 0.003093608 | 67.14259878 | 31.51420139 | ENSG00000206791 |
| ALOX12B | -1.089629267 | 0.469882106 | 0.345318135 | 0.517722246 | 5.710010077 | 2.709221058 | ENSG00000179477 |
| GALR2 | -1.090820571 | 0.469494261 | 0.213891625 | 0.370310868 | 8.366109333 | 3.916489655 | ENSG00000182687 |
| NA | -1.090983913 | 0.469441108 | 0.000607182 | 0.002721747 | 64.87976842 | 30.45348123 | ENSG00000274372 |
| RTEL1 | -1.091893096 | 0.469145361 | 0.030261065 | 0.079090369 | 28.10479347 | 13.20785849 | ENSG00000258366 |
| MTRNR2L8 | -1.0926262 | 0.468907025 | 0.355103421 | 0.528213834 | 4.898317252 | 2.315852845 | ENSG00000255823 |
| NA | -1.093222334 | 0.468713309 | 0.285638935 | 0.454566961 | 6.338091014 | 2.98267176 | ENSG00000269752 |
| NA | -1.093408552 | 0.468652813 | 0.263926125 | 0.429641531 | 6.297578913 | 2.959548487 | ENSG00000226445 |
| NA | -1.093628426 | 0.468581393 | 0.017945586 | 0.051348012 | 33.21048929 | 15.61620443 | ENSG00000231806 |
| METTL7A | -1.09385443 | 0.468507994 | 7.31E-33 | 5.45E-31 | 944.8186445 | 442.6637039 | ENSG00000185432 |
| PCLAF | -1.093868743 | 0.468503346 | 4.66E-06 | 3.26E-05 | 118.0340051 | 55.42827374 | ENSG00000166803 |
| NA | -1.093951641 | 0.468476426 | 0.274389534 | 0.441140803 | 6.340881917 | 2.98267176 | ENSG00000285454 |
| MYCNOS | -1.094613148 | 0.468261669 | 0.000826543 | 0.003581501 | 65.73611569 | 30.74360348 | ENSG00000233718 |
| LOC101927551 | -1.095310112 | 0.468035507 | 0.145322149 | 0.275573536 | 12.72295166 | 5.976096388 | ENSG00000230922 |
| NA | -1.096295977 | 0.467715785 | 0.167440784 | 0.30751144 | 11.8227378 | 5.479482215 | ENSG00000286747 |
| PRSS35 | -1.096968149 | 0.46749792 | 0.000143449 | 0.000744894 | 85.13010344 | 39.84756299 | ENSG00000146250 |
| ARL4C | -1.097197883 | 0.467423482 | 3.33E-23 | 1.42E-21 | 620.9694641 | 290.5159548 | ENSG00000188042 |
| NA | -1.097425327 | 0.467349797 | 0.040506205 | 0.100694695 | 21.76391155 | 10.15528388 | ENSG00000280604 |
| NA | -1.098624872 | 0.466961375 | 0.270472277 | 0.437101508 | 6.322021318 | 2.95524734 | ENSG00000232186 |
| NA | -1.098822852 | 0.466897299 | 0.263104855 | 0.428678696 | 6.291997107 | 2.936425214 | ENSG00000226666 |
| FXYD6 | -1.09886168 | 0.466884734 | 1.44E-06 | 1.10E-05 | 128.3403754 | 60.01373288 | ENSG00000137726 |
| MAP1A | -1.099968005 | 0.466526842 | 1.55E-06 | 1.18E-05 | 154.3607321 | 72.03209094 | ENSG00000166963 |
| TSFM | -1.100329371 | 0.466410001 | 0.291003849 | 0.46071032 | 5.671613158 | 2.660823939 | ENSG00000123297 |
| CDK8 | -1.100698849 | 0.466290567 | 3.81E-22 | 1.51E-20 | 714.593722 | 333.6225843 | ENSG00000132964 |
| KLK4 | -1.101782558 | 0.465940436 | 0.105666691 | 0.215956789 | 13.3188932 | 6.190397102 | ENSG00000167749 |
| NA | -1.102076521 | 0.465845505 | 0.039797005 | 0.099266101 | 25.86795712 | 12.02345271 | ENSG00000272079 |
| NA | -1.102779876 | 0.465618448 | 0.01148249 | 0.035464092 | 34.37513908 | 15.95446334 | ENSG00000254252 |
| NA | -1.103092762 | 0.465517477 | 0.292615434 | 0.46227535 | 5.64437985 | 2.635550092 | ENSG00000217275 |
| ENTPD3 | -1.104425973 | 0.465087486 | 0.25691992 | 0.422059285 | 6.322021318 | 2.93427464 | ENSG00000168032 |
| RBM3 | -1.105297554 | 0.464806596 | 3.75E-50 | 6.41E-48 | 1708.113395 | 794.1079091 | ENSG00000102317 |
| GPR137 | -1.105568701 | 0.464719246 | 3.84E-21 | 1.41E-19 | 672.5157474 | 312.4387605 | ENSG00000173264 |
| NA | -1.10626429 | 0.464495238 | 0.319207591 | 0.490669175 | 4.958365673 | 2.318003419 | ENSG00000268189 |
| HSP90B2P | -1.107407984 | 0.464127156 | 0.242737154 | 0.40552892 | 8.485330294 | 3.967037349 | ENSG00000259706 |
| FAM186B | -1.108045851 | 0.463921994 | 0.377797801 | 0.551251598 | 4.26465451 | 1.998306171 | ENSG00000135436 |
| PDGFRL | -1.108274919 | 0.463848339 | 8.94E-05 | 0.000488372 | 80.8975003 | 37.57795669 | ENSG00000104213 |
| FOXD2-AS1 | -1.108452669 | 0.463791193 | 0.262598212 | 0.428178691 | 6.387651545 | 2.959548487 | ENSG00000237424 |
| CCDC177 | -1.109707635 | 0.463387928 | 1.45E-10 | 1.97E-09 | 253.3846064 | 117.5150804 | ENSG00000267909 |
| BRWD1-AS2 | -1.111730513 | 0.462738643 | 0.326215696 | 0.497984252 | 4.942295977 | 2.292729572 | ENSG00000255568 |
| NA | -1.11213104 | 0.462610193 | 0.336101408 | 0.508887236 | 5.0239959 | 2.338976118 | ENSG00000260209 |
| SLC17A6 | -1.112181005 | 0.462594172 | 0.19133194 | 0.340088865 | 12.0376132 | 5.561222438 | ENSG00000091664 |
| PANO1 | -1.112344405 | 0.462541781 | 0.259174845 | 0.42431825 | 6.376487933 | 2.938575788 | ENSG00000274897 |
| EGR1 | -1.113188326 | 0.462271291 | 1.13E-15 | 2.62E-14 | 651.655001 | 300.7782891 | ENSG00000120738 |
| NA | -1.114324866 | 0.461907262 | 0.252984988 | 0.417622232 | 7.059711207 | 3.237300336 | ENSG00000251593 |
| NA | -1.114834457 | 0.461744136 | 0.34906375 | 0.521661052 | 4.969529285 | 2.299181293 | ENSG00000272829 |
| CFAP69 | -1.115615002 | 0.461494385 | 0.004637705 | 0.016458052 | 44.94574521 | 20.80341382 | ENSG00000105792 |
| NA | -1.115639795 | 0.461486454 | 0.33628225 | 0.509041187 | 5.714916161 | 2.637700666 | ENSG00000286408 |
| CEP135 | -1.116194071 | 0.461309187 | 1.13E-13 | 2.13E-12 | 302.3812068 | 139.5612565 | ENSG00000174799 |
| ADGRF3 | -1.116387267 | 0.461247416 | 5.31E-05 | 0.000304061 | 85.98345965 | 39.64967336 | ENSG00000173567 |
| NA | -1.116942704 | 0.46106987 | 0.006676452 | 0.022497021 | 40.78232694 | 18.8551223 | ENSG00000213222 |
| MATN1-AS1 | -1.117009203 | 0.461048618 | 0.429891207 | 0.601380078 | 4.229048493 | 1.954210199 | ENSG00000186056 |
| DQX1 | -1.117823467 | 0.460788473 | 0.092024407 | 0.193973211 | 14.24422518 | 6.585915889 | ENSG00000144045 |
| ADAM33 | -1.118188839 | 0.46067179 | 1.24E-14 | 2.59E-13 | 339.8284416 | 156.5984972 | ENSG00000149451 |
| SLC5A2 | -1.118612531 | 0.46053652 | 0.122506897 | 0.242250832 | 12.13259192 | 5.569824733 | ENSG00000140675 |

| | | | | | | | |
|---|---|---|---|---|---|---|---|
| GBX1 | -1.118648775 | 0.46052495 | 0.381804605 | 0.555208249 | 4.267445412 | 1.975182898 | ENSG00000164900 |
| ZNF236-DT | -1.119641649 | 0.460208122 | 0.073603064 | 0.162401546 | 20.42449815 | 9.374999171 | ENSG00000264278 |
| NA | -1.121026054 | 0.45976672 | 0.082704233 | 0.178222448 | 16.32045259 | 7.47993896 | ENSG00000234373 |
| NCAM2 | -1.121626673 | 0.459575351 | 0.017568894 | 0.050487344 | 30.92496829 | 14.23847063 | ENSG00000154654 |
| NA | -1.122077695 | 0.459431699 | 0.361160386 | 0.534249291 | 4.2751424 | 1.973032325 | ENSG00000286585 |
| CLIP3 | -1.122488398 | 0.459300928 | 1.42E-15 | 3.24E-14 | 347.3884399 | 159.6623577 | ENSG00000105270 |
| ADPRHL1 | -1.123041784 | 0.459124783 | 0.009703128 | 0.030650571 | 33.6514037 | 15.4812329 | ENSG00000153531 |
| NA | -1.123810238 | 0.458880295 | 0.326656132 | 0.498538247 | 4.955574771 | 2.267455725 | ENSG00000232470 |
| NA | -1.124283276 | 0.45872986 | 0.349986797 | 0.522609746 | 4.286306011 | 1.975182898 | ENSG00000280129 |
| NA | -1.124409417 | 0.458689753 | 0.063353564 | 0.144068559 | 17.97298657 | 8.281196365 | ENSG00000198225 |
| LOC644656 | -1.124951978 | 0.458517284 | 0.172993931 | 0.31542778 | 10.66000304 | 4.869662714 | ENSG00000268403 |
| TOMM20L | -1.127040265 | 0.457854064 | 0.369019828 | 0.542833495 | 4.379169546 | 2.021429444 | ENSG00000196860 |
| NA | -1.128782142 | 0.457301596 | 0.20229312 | 0.354863743 | 8.736011868 | 4.021886189 | ENSG00000280219 |
| IZUMO1 | -1.128994861 | 0.457234173 | 0.087567495 | 0.186576602 | 14.99240296 | 6.834092744 | ENSG00000182264 |
| ABCA5 | -1.129509552 | 0.457071081 | 0.002005751 | 0.007907817 | 50.96860046 | 23.24967658 | ENSG00000154265 |
| MAGEE1 | -1.129525236 | 0.457066112 | 2.58E-14 | 5.22E-13 | 337.2383729 | 154.0003124 | ENSG00000198934 |
| PTN | -1.130338729 | 0.456808459 | 9.44E-32 | 6.60E-30 | 824.227163 | 376.7636657 | ENSG00000105894 |
| FNDC4 | -1.131432723 | 0.456462193 | 4.78E-10 | 6.06E-09 | 211.4025362 | 96.48231182 | ENSG00000115226 |
| RNF31 | -1.131509186 | 0.456438001 | 0.213482972 | 0.369776063 | 7.97946138 | 3.670463374 | ENSG00000092098 |
| LRRN3 | -1.131774144 | 0.456354182 | 0.285841865 | 0.454852381 | 5.726079773 | 2.593604694 | ENSG00000173114 |
| NA | -1.13331691 | 0.455866434 | 0.282320538 | 0.450660759 | 6.433745451 | 2.904699646 | ENSG00000286681 |
| DLL1 | -1.133465188 | 0.455819583 | 4.60E-05 | 0.000266504 | 96.91388074 | 44.21925445 | ENSG00000198719 |
| NA | -1.133976203 | 0.455658156 | 0.212252015 | 0.368182145 | 9.386420029 | 4.276514765 | ENSG00000235790 |
| D21S2088E | -1.134147779 | 0.455603969 | 7.37E-31 | 4.85E-29 | 757.7552213 | 345.494685 | ENSG00000228592 |
| NA | -1.134252499 | 0.4555709 | 0.344841754 | 0.517450394 | 4.319121125 | 1.970881751 | ENSG00000278952 |
| KRT7 | -1.135940293 | 0.455038244 | 2.12E-11 | 3.14E-10 | 281.7153637 | 128.4613897 | ENSG00000135480 |
| OLIG3 | -1.138139047 | 0.454345266 | 0.38428669 | 0.55768258 | 4.250699995 | 1.922484631 | ENSG00000177468 |
| CRELD1 | -1.138356492 | 0.454276792 | 0.022766508 | 0.062584409 | 25.30415497 | 11.51419555 | ENSG00000163703 |
| NA | -1.138410128 | 0.454259903 | 0.030671787 | 0.079985083 | 23.79472078 | 10.77585625 | ENSG00000212769 |
| NA | -1.13846498 | 0.454242632 | 0.273384646 | 0.440191783 | 6.469351467 | 2.90685022 | ENSG00000263603 |
| ZNF493 | -1.138577652 | 0.454207158 | 4.07E-08 | 3.97E-07 | 156.9124407 | 71.3596138 | ENSG00000196268 |
| LRP11 | -1.139980309 | 0.453765771 | 0.036752279 | 0.092860277 | 21.03459437 | 9.564286502 | ENSG00000120256 |
| CDH15 | -1.14037456 | 0.453641785 | 0.068012592 | 0.152473088 | 17.22134216 | 7.793184486 | ENSG00000129910 |
| ZNF25 | -1.140388755 | 0.453637322 | 0.148271878 | 0.279874051 | 10.15692503 | 4.587609718 | ENSG00000175395 |
| NA | -1.140850932 | 0.45349202 | 0.173906384 | 0.316700858 | 8.657102848 | 3.920790803 | ENSG00000273391 |
| TMEM163 | -1.141547743 | 0.453273039 | 0.006656776 | 0.02244248 | 36.79750234 | 16.62128225 | ENSG00000152128 |
| NA | -1.1422094 | 0.453065204 | 0.32008842 | 0.491508852 | 5.086835224 | 2.294880146 | ENSG00000249593 |
| NA | -1.142372185 | 0.453014086 | 0.357657002 | 0.530817142 | 4.351936238 | 1.970881751 | ENSG00000246575 |
| NA | -1.142399532 | 0.453005499 | 0.183438238 | 0.329365883 | 8.673172544 | 3.922941376 | ENSG00000280254 |
| NA | -1.142483111 | 0.452979256 | 0.381615233 | 0.555092684 | 4.2751424 | 1.926785779 | ENSG00000254717 |
| LTB4R | -1.143006284 | 0.452815019 | 3.34E-06 | 2.38E-05 | 112.6879523 | 51.03560957 | ENSG00000213903 |
| GLYATL2 | -1.14435191 | 0.452392868 | 0.126073033 | 0.247214165 | 11.62373194 | 5.275401332 | ENSG00000156689 |
| NSMCE4A | -1.145401763 | 0.45206378 | 1.27E-36 | 1.16E-34 | 992.1474433 | 448.240023 | ENSG00000107672 |
| UBE2E1-AS1 | -1.147387263 | 0.451442058 | 0.222619268 | 0.381026377 | 7.287865398 | 3.306670155 | ENSG00000223791 |
| NA | -1.14739616 | 0.451439274 | 0.015325832 | 0.045068692 | 29.1369435 | 13.15300965 | ENSG00000267480 |
| NA | -1.148323492 | 0.451149192 | 0.334720737 | 0.50740055 | 4.337981724 | 1.947758478 | ENSG00000272744 |
| OVOL2 | -1.149107528 | 0.450904081 | 1.24E-10 | 1.70E-09 | 223.7282398 | 100.7886742 | ENSG00000125850 |
| H3C6 | -1.149467739 | 0.450791514 | 0.124851795 | 0.245424828 | 12.93503615 | 5.797028888 | ENSG00000274750 |
| NA | -1.149717519 | 0.450713473 | 0.378499004 | 0.551934188 | 4.29467872 | 1.922484631 | ENSG00000276434 |
| NA | -1.149848898 | 0.45067243 | 0.176486019 | 0.319992786 | 8.687127059 | 3.895516956 | ENSG00000228150 |
| CRTC3 | -1.150079175 | 0.450600502 | 6.26E-10 | 7.80E-09 | 262.6730936 | 118.6042973 | ENSG00000140577 |
| NA | -1.151449893 | 0.450172585 | 0.221042035 | 0.379404031 | 7.307401718 | 3.302369007 | ENSG00000235027 |
| GNG8 | -1.152613059 | 0.449809782 | 0.225696273 | 0.3847227 | 7.941064461 | 3.559148157 | ENSG00000167414 |
| ARSI | -1.153294526 | 0.449597361 | 7.33E-08 | 6.87E-07 | 181.0812944 | 81.27480219 | ENSG00000183876 |
| SHISA2 | -1.153638563 | 0.44949016 | 3.86E-20 | 1.32E-18 | 440.4465292 | 198.047763 | ENSG00000180730 |
| ZNF428 | -1.153710012 | 0.449467899 | 3.67E-09 | 4.16E-08 | 227.1599134 | 102.2510923 | ENSG00000131116 |
| SEM1 | -1.155305989 | 0.448970952 | 1.74E-54 | 3.32E-52 | 2252.45642 | 1011.127737 | ENSG00000127922 |
| RTF1 | -1.156809793 | 0.448503207 | 1.69E-74 | 6.40E-72 | 2332.565919 | 1046.573749 | ENSG00000137815 |
| PARTICL | -1.156852489 | 0.448489934 | 0.001299546 | 0.00540059 | 48.99205769 | 21.9746557 | ENSG00000286532 |
| NA | -1.157234007 | 0.448371348 | 0.180713085 | 0.325776668 | 9.512098676 | 4.236186903 | ENSG00000260774 |
| LOC399900 | -1.157550594 | 0.448272967 | 0.208187504 | 0.363024961 | 8.039509801 | 3.594641834 | ENSG00000279491 |
| NA | -1.158965115 | 0.447833664 | 0.075929569 | 0.166317397 | 15.29456009 | 6.829791597 | ENSG00000272927 |
| MTSS1 | -1.159790575 | 0.447577502 | 2.00E-06 | 1.50E-05 | 109.8459258 | 49.11124694 | ENSG00000170873 |
| C18orf32 | -1.16012481 | 0.447473822 | 0.022119775 | 0.061072936 | 30.85308054 | 13.79670529 | ENSG00000177576 |
| TSPYL4 | -1.162442992 | 0.44675538 | 1.25E-26 | 6.73E-25 | 884.3885962 | 394.6234638 | ENSG00000187189 |
| SMKR1 | -1.162849779 | 0.446629429 | 0.030826183 | 0.080273745 | 24.78779816 | 11.00090983 | ENSG00000240204 |
| FRMPD1 | -1.162916977 | 0.446608626 | 0.130456871 | 0.253489474 | 12.50508519 | 5.616071279 | ENSG00000070601 |
| CCDC74A | -1.164248415 | 0.446196649 | 9.11E-15 | 1.93E-13 | 305.3047331 | 136.2594205 | ENSG00000163040 |
| KIAA1324 | -1.164402055 | 0.446149134 | 0.00393013 | 0.014227086 | 41.86759214 | 18.6341094 | ENSG00000116299 |
| AMOTL1 | -1.164785947 | 0.446030433 | 3.74E-63 | 9.14E-61 | 1742.021149 | 776.9198188 | ENSG00000166025 |
| CCT6B | -1.165470902 | 0.445818719 | 0.071147233 | 0.15808584 | 15.39579633 | 6.834092744 | ENSG00000132141 |
| NA | -1.166308424 | 0.445559984 | 0.205343045 | 0.359136235 | 7.326262317 | 3.253971888 | ENSG00000248126 |
| SCML1 | -1.166653021 | 0.445453572 | 0.001051324 | 0.004456137 | 53.71544807 | 23.85089378 | ENSG00000047634 |
| MAPRE3 | -1.167351558 | 0.445237941 | 2.73E-08 | 2.71E-07 | 156.08643 | 69.40137503 | ENSG00000084764 |
| NA | -1.167481517 | 0.445197835 | 0.081304335 | 0.175676739 | 15.49057489 | 6.933037557 | ENSG00000250046 |
| NA | -1.167714344 | 0.445125994 | 1.15E-07 | 1.05E-06 | 158.3350688 | 70.55782336 | ENSG00000225138 |
| CCDC154 | -1.17135471 | 0.444004219 | 0.185074312 | 0.331387575 | 9.551734896 | 4.244789198 | ENSG00000197599 |

| | | | | | | | |
|---|---|---|---|---|---|---|---|
| NA | -1.171925795 | 0.443828496 | 0.266040783 | 0.432257454 | 6.754763177 | 3.012246754 | ENSG00000188801 |
| KCNJ12 | -1.172929175 | 0.443519925 | 7.58E-07 | 6.08E-06 | 112.4880706 | 49.89368222 | ENSG00000184185 |
| LOC100291105 | -1.173054421 | 0.443481423 | 0.244629845 | 0.407737545 | 6.754763177 | 3.01009618 | ENSG00000218018 |
| ARAP2 | -1.173244157 | 0.443423103 | 0.024811226 | 0.067176611 | 26.57024115 | 11.72849627 | ENSG00000047365 |
| RBM24 | -1.173534072 | 0.443334004 | 0.198252984 | 0.349620469 | 8.085603708 | 3.552696436 | ENSG00000112183 |
| SFRP1 | -1.173885523 | 0.443226018 | 1.90E-176 | 7.32E-173 | 14611.66082 | 6477.0009 | ENSG00000104332 |
| SLC5A6 | -1.174444284 | 0.443054388 | 7.19E-88 | 4.08E-85 | 4016.68699 | 1779.39431 | ENSG00000138074 |
| KCNAB3 | -1.174654699 | 0.442989774 | 0.000653576 | 0.002903095 | 57.07064014 | 25.28992815 | ENSG00000170049 |
| H2BC7 | -1.174659012 | 0.44298845 | 0.042768467 | 0.105275807 | 22.24832912 | 9.827517372 | ENSG00000277224 |
| NA | -1.177081449 | 0.442245249 | 0.129936442 | 0.252758285 | 11.817156 | 5.187209388 | ENSG00000231043 |
| PCGF5 | -1.177824808 | 0.442017438 | 2.88E-06 | 2.08E-05 | 131.2660921 | 58.22998789 | ENSG00000180628 |
| SEMA6D | -1.178057711 | 0.441946086 | 0.000793269 | 0.003448942 | 53.42290295 | 23.59196406 | ENSG00000137872 |
| H3C8 | -1.178346107 | 0.44185775 | 0.000633944 | 0.002828264 | 52.75084329 | 23.3104441 | ENSG00000273983 |
| CNTFR | -1.179044314 | 0.44164396 | 1.07E-05 | 6.98E-05 | 113.9394835 | 50.22979055 | ENSG00000122756 |
| ST8SIA2 | -1.181279424 | 0.440960268 | 1.75E-05 | 0.000110294 | 90.28862626 | 39.9131647 | ENSG00000140557 |
| SMAD9 | -1.181387815 | 0.44092714 | 0.01280211 | 0.038770095 | 30.55506576 | 13.4517342 | ENSG00000120693 |
| TMEM233 | -1.182296697 | 0.440649448 | 0.230502207 | 0.390774064 | 6.730320772 | 2.963849634 | ENSG00000224982 |
| NA | -1.182821714 | 0.440489119 | 0.370268955 | 0.544051977 | 5.220210859 | 2.27605802 | ENSG00000286299 |
| ACP7 | -1.182876945 | 0.440472256 | 0.103842355 | 0.213222233 | 15.54524166 | 6.798066029 | ENSG00000183760 |
| ATP6V0E2 | -1.183519901 | 0.440275997 | 5.03E-29 | 3.04E-27 | 650.6473701 | 286.4104809 | ENSG00000171130 |
| NOX4 | -1.18378086 | 0.440196366 | 0.017535485 | 0.050398843 | 27.02443435 | 11.95596089 | ENSG00000086991 |
| SLC16A11 | -1.184427843 | 0.439999002 | 0.106701919 | 0.217496483 | 14.29398587 | 6.28073962 | ENSG00000174326 |
| NA | -1.184564613 | 0.439957291 | 0.127063027 | 0.248622868 | 11.92588907 | 5.25872978 | ENSG00000279452 |
| NA | -1.185350602 | 0.439717665 | 0.315493805 | 0.486739523 | 5.274477315 | 2.343277265 | ENSG00000279072 |
| ABCB7 | -1.185879825 | 0.439556393 | 2.54E-38 | 2.58E-36 | 861.409235 | 378.7840291 | ENSG00000131269 |
| NA | -1.186907156 | 0.439243501 | 0.144229642 | 0.27393266 | 9.691368058 | 4.246939771 | ENSG00000213790 |
| LCMT1-AS2 | -1.187453057 | 0.439077327 | 0.265778934 | 0.431904761 | 5.910930961 | 2.59145412 | ENSG00000260034 |
| H2AX | -1.187971917 | 0.438919443 | 3.20E-57 | 6.71E-55 | 3581.892341 | 1572.475067 | ENSG00000188486 |
| NA | -1.188574431 | 0.438736174 | 0.23299807 | 0.393794093 | 6.724738966 | 2.938575788 | ENSG00000265451 |
| WNT7B | -1.190711885 | 0.438086637 | 0.141404406 | 0.269948778 | 10.56801539 | 4.615034138 | ENSG00000188064 |
| LOC106699570 | -1.192491713 | 0.437546511 | 0.264943321 | 0.430946172 | 5.987049078 | 2.61887854 | ENSG00000246777 |
| NRN1 | -1.193837905 | 0.437138422 | 0.125307662 | 0.246086746 | 11.42771714 | 5.031525625 | ENSG00000124785 |
| CD53 | -1.19436515 | 0.436978696 | 0.185091817 | 0.331387575 | 8.969747865 | 3.914339082 | ENSG00000143119 |
| SLC2A9 | -1.19490036 | 0.436816616 | 0.256398461 | 0.421525713 | 6.675854157 | 2.892329242 | ENSG00000109667 |
| PARP2 | -1.195481511 | 0.436640691 | 9.46E-36 | 8.15E-34 | 763.8033949 | 333.5180239 | ENSG00000129484 |
| SYT9 | -1.201667286 | 0.434772536 | 1.20E-05 | 7.77E-05 | 88.49175319 | 38.52037688 | ENSG00000170743 |
| GPD1 | -1.203678849 | 0.434166751 | 0.095558777 | 0.199945829 | 12.0809162 | 5.227004212 | ENSG00000167588 |
| ZNF442 | -1.203717449 | 0.434155134 | 0.00775637 | 0.025472953 | 33.32288915 | 14.43609979 | ENSG00000198342 |
| THBD | -1.203738421 | 0.434148823 | 0.037980686 | 0.095513677 | 19.65486903 | 8.540126088 | ENSG00000178726 |
| HSF2BP | -1.204925616 | 0.433791709 | 0.025948212 | 0.06980794 | 25.48621525 | 11.0240331 | ENSG00000160207 |
| NA | -1.207116812 | 0.433133357 | 0.094822095 | 0.198568501 | 13.65877153 | 5.925548695 | ENSG00000235082 |
| NA | -1.20734063 | 0.433066166 | 0.385798009 | 0.559340709 | 4.623393433 | 2.023580018 | ENSG00000226009 |
| CMPK2 | -1.207576809 | 0.432995276 | 0.050997691 | 0.121231849 | 19.52081768 | 8.428277833 | ENSG00000134326 |
| WDR34 | -1.207738083 | 0.432946876 | 3.79E-45 | 5.34E-43 | 1372.096542 | 594.1941278 | ENSG00000119333 |
| CYP26A1 | -1.208489339 | 0.432721486 | 3.74E-05 | 0.000220496 | 76.06413756 | 32.92957945 | ENSG00000095596 |
| GRPR | -1.209572722 | 0.432396658 | 2.79E-12 | 4.52E-11 | 232.7315154 | 100.5848537 | ENSG00000126010 |
| FBXO43 | -1.210268718 | 0.432188108 | 0.011653985 | 0.035810295 | 28.86481058 | 12.44801345 | ENSG00000156509 |
| HERC2P2 | -1.210377676 | 0.432155469 | 1.77E-53 | 3.29E-51 | 1862.202828 | 804.2685907 | ENSG00000276550 |
| NA | -1.21057572 | 0.432096149 | 0.10873593 | 0.22050306 | 13.55071418 | 5.794878315 | ENSG00000213642 |
| MC5R | -1.211349971 | 0.431864319 | 0.323817756 | 0.495421392 | 4.618487349 | 2.021429444 | ENSG00000176136 |
| NA | -1.212217932 | 0.431604577 | 0.19430686 | 0.34432929 | 7.710119367 | 3.355067274 | ENSG00000229020 |
| ITGB8 | -1.212726079 | 0.431452583 | 0.002974726 | 0.011121179 | 43.73241077 | 18.91857344 | ENSG00000105855 |
| RSPO2 | -1.213125157 | 0.431333251 | 0.083074963 | 0.178841547 | 14.45997646 | 6.219972096 | ENSG00000147655 |
| NA | -1.214683903 | 0.430867473 | 0.025586667 | 0.068976019 | 21.41566052 | 9.246739828 | ENSG00000224825 |
| NA | -1.215131853 | 0.430733712 | 0.154003319 | 0.288658961 | 10.00189789 | 4.337282289 | ENSG00000279605 |
| SCUBE3 | -1.215222172 | 0.430706747 | 1.18E-05 | 7.62E-05 | 85.43448789 | 36.76594641 | ENSG00000146197 |
| ADAM11 | -1.215966173 | 0.430484688 | 7.32E-14 | 1.40E-12 | 257.9956339 | 111.0410127 | ENSG00000073670 |
| DPH5 | -1.220074291 | 0.429260613 | 2.14E-34 | 1.69E-32 | 734.6060395 | 315.1963666 | ENSG00000117543 |
| CDHR1 | -1.221340673 | 0.428883978 | 9.39E-24 | 4.21E-22 | 528.5570402 | 226.8438685 | ENSG00000148600 |
| NA | -1.223548452 | 0.428228152 | 0.300273357 | 0.470902636 | 5.279383399 | 2.246483026 | ENSG00000235084 |
| VWA5B2 | -1.224268188 | 0.42801457 | 0.098713585 | 0.205070404 | 12.32302491 | 5.296374031 | ENSG00000145198 |
| NA | -1.22431345 | 0.428001142 | 0.309362929 | 0.480508877 | 5.331734833 | 2.273907447 | ENSG00000236972 |
| NA | -1.224468982 | 0.427955003 | 0.113912981 | 0.229072795 | 11.59861381 | 5.006251778 | ENSG00000253549 |
| NA | -1.225135367 | 0.427757375 | 0.075628172 | 0.165864377 | 13.8284289 | 5.893823127 | ENSG00000287431 |
| DLG1 | -1.225440543 | 0.4276669 | 2.25E-54 | 4.26E-52 | 1224.700416 | 523.922628 | ENSG00000075711 |
| TNKS2-AS1 | -1.22655726 | 0.427335993 | 0.283645238 | 0.452214899 | 6.145542839 | 2.608125672 | ENSG00000228701 |
| SLC1A4 | -1.227477421 | 0.427063522 | 4.40E-05 | 0.00025547 | 76.80844839 | 32.83923693 | ENSG00000115902 |
| NA | -1.22840513 | 0.426788992 | 0.292558507 | 0.462261131 | 4.640138851 | 1.996155598 | ENSG00000226855 |
| NA | -1.228898015 | 0.426643208 | 0.260345002 | 0.425692103 | 5.347804529 | 2.267455725 | ENSG00000260176 |
| SLC27A2 | -1.229071148 | 0.426592011 | 3.62E-05 | 0.000214331 | 81.43530845 | 34.76118852 | ENSG00000140284 |
| PRR22 | -1.229194576 | 0.426555516 | 0.003843047 | 0.013956367 | 35.4450103 | 15.1238914 | ENSG00000212123 |
| MYOM2 | -1.231242948 | 0.425950312 | 0.008475583 | 0.027391938 | 31.54726727 | 13.4517342 | ENSG00000036448 |
| TM4SF18 | -1.232172176 | 0.425676049 | 3.73E-05 | 0.000219929 | 80.38258296 | 34.2277236 | ENSG00000163762 |
| NA | -1.233818663 | 0.42519052 | 0.087224022 | 0.186009133 | 12.43753995 | 5.300675179 | ENSG00000287710 |
| TCHH | -1.234099243 | 0.425107836 | 0.185531054 | 0.331954277 | 10.16442186 | 4.369007857 | ENSG00000159450 |
| CCDC157 | -1.234839608 | 0.424889734 | 3.17E-05 | 0.000190297 | 76.35704609 | 32.42059487 | ENSG00000187860 |
| NA | -1.236037317 | 0.424537142 | 0.254714333 | 0.41951355 | 5.432295355 | 2.294880146 | ENSG00000229539 |

| | | | | | | | |
|---|---|---|---|---|---|---|---|
| NA | -1.236361866 | 0.424441649 | 0.294842102 | 0.464765403 | 5.347804529 | 2.250784174 | ENSG00000269846 |
| JAKMIP2 | -1.238842283 | 0.423712536 | 1.22E-15 | 2.83E-14 | 286.4938964 | 121.5184049 | ENSG00000176049 |
| SMIM27 | -1.239963626 | 0.423383331 | 0.060029926 | 0.13800945 | 14.79706388 | 6.266218642 | ENSG00000235453 |
| H2AC16 | -1.24050555 | 0.423224324 | 0.214203608 | 0.370667545 | 7.018523385 | 2.980521186 | ENSG00000276903 |
| BMI1 | -1.242253302 | 0.422711919 | 0.003565307 | 0.013068276 | 35.66546751 | 15.06904256 | ENSG00000168283 |
| NA | -1.2422662 | 0.42270814 | 0.018689342 | 0.053027813 | 24.90721928 | 10.51100784 | ENSG00000220848 |
| GRM3 | -1.243650413 | 0.422302762 | 0.001960747 | 0.007754154 | 41.90232228 | 17.70244208 | ENSG00000198822 |
| OR7E14P | -1.245212739 | 0.421845688 | 0.041164612 | 0.102004691 | 19.30103619 | 8.108580586 | ENSG00000184669 |
| NA | -1.246138462 | 0.421575093 | 0.06318812 | 0.143795424 | 15.5827627 | 6.590217036 | ENSG00000229097 |
| NA | -1.246821342 | 0.421375593 | 0.296886843 | 0.467034501 | 4.642929753 | 1.952059625 | ENSG00000275882 |
| C1QTNF2 | -1.25154481 | 0.419998241 | 0.011994297 | 0.036616866 | 26.67060151 | 11.24074485 | ENSG00000145861 |
| PCDHB12 | -1.252817626 | 0.419627861 | 5.04E-06 | 3.50E-05 | 90.54979572 | 37.92481789 | ENSG00000120328 |
| NA | -1.253773166 | 0.419350021 | 0.321515117 | 0.492914895 | 4.760235692 | 2.000456745 | ENSG00000274767 |
| BBS5 | -1.255048201 | 0.418979569 | 0.055852432 | 0.13047339 | 15.61278691 | 6.53751877 | ENSG00000163093 |
| CDH7 | -1.255942396 | 0.418719962 | 0.010370496 | 0.032424186 | 28.37692639 | 11.93068704 | ENSG00000081138 |
| GNG3 | -1.256622781 | 0.418522538 | 0.287939634 | 0.457322862 | 4.71904787 | 1.970881751 | ENSG00000162188 |
| MAP3K2-DT | -1.256756375 | 0.418483784 | 0.106854873 | 0.217739238 | 11.89048321 | 4.978827358 | ENSG00000236682 |
| CHKB-CPT1B | -1.25709375 | 0.418385933 | 0.245205531 | 0.408379513 | 6.975220381 | 2.902549073 | ENSG00000254413 |
| L3MBTL4 | -1.257467115 | 0.41827767 | 0.003805345 | 0.013835077 | 36.23226072 | 15.1658368 | ENSG00000154655 |
| FLJ38576 | -1.260224924 | 0.417478867 | 0.063950702 | 0.145086793 | 17.5779659 | 7.421049436 | ENSG00000279943 |
| PLAC9 | -1.260499579 | 0.417399397 | 7.47E-05 | 0.000415661 | 65.48774946 | 27.39524839 | ENSG00000189129 |
| NA | -1.26217623 | 0.416914591 | 0.066773957 | 0.150262111 | 13.38452343 | 5.57412588 | ENSG00000259529 |
| NA | -1.262606645 | 0.416790227 | 0.068220742 | 0.15285091 | 13.34891741 | 5.551002607 | ENSG00000279329 |
| NA | -1.263175898 | 0.416625804 | 0.114653391 | 0.230249379 | 10.33619441 | 4.339432863 | ENSG00000271781 |
| ASXL1 | -1.263587191 | 0.416507047 | 1.17E-39 | 1.35E-37 | 1011.641691 | 421.2857159 | ENSG00000171456 |
| PYGO1 | -1.263940168 | 0.416405154 | 0.018269165 | 0.052049988 | 27.48488508 | 11.44912688 | ENSG00000171016 |
| ADAMTS19 | -1.265848679 | 0.415854665 | 5.13E-29 | 3.08E-27 | 575.6324161 | 239.4239094 | ENSG00000145808 |
| SFXN3 | -1.266541045 | 0.415655139 | 0.002455126 | 0.009449687 | 40.54791522 | 16.9095144 | ENSG00000107819 |
| TMEFF2 | -1.267603947 | 0.415349019 | 0.00253362 | 0.00969763 | 38.08503986 | 15.86223071 | ENSG00000144339 |
| STK32B | -1.271558232 | 0.414212147 | 5.67E-07 | 4.63E-06 | 129.0008712 | 53.64452874 | ENSG00000152953 |
| FLJ13224 | -1.273015541 | 0.41379395 | 0.282791563 | 0.451114863 | 4.749072081 | 1.947758478 | ENSG00000177340 |
| ISM1 | -1.274289833 | 0.413428619 | 0.005670142 | 0.019560572 | 31.01137414 | 12.84353224 | ENSG00000101230 |
| FGFBP3 | -1.27629252 | 0.412855114 | 2.61E-72 | 8.98E-70 | 2172.34029 | 897.3538167 | ENSG00000174721 |
| PKD1L1 | -1.277159128 | 0.412607192 | 0.20101134 | 0.353159464 | 7.903343264 | 3.262574183 | ENSG00000158683 |
| TTI1 | -1.27827127 | 0.412289244 | 4.19E-45 | 5.86E-43 | 1377.243346 | 567.4143203 | ENSG00000101407 |
| NA | -1.281024763 | 0.411503109 | 0.017530573 | 0.050392234 | 24.07734158 | 9.945284311 | ENSG00000203497 |
| GDPD2 | -1.281849599 | 0.411267907 | 2.54E-19 | 8.01E-18 | 372.1009958 | 152.8825929 | ENSG00000130055 |
| GRIA2 | -1.283245802 | 0.410870085 | 0.028684702 | 0.075781511 | 25.74180291 | 10.60995265 | ENSG00000120251 |
| KCND1 | -1.285371387 | 0.410265178 | 3.60E-15 | 7.88E-14 | 305.7524686 | 125.3655368 | ENSG00000102057 |
| ZNF732 | -1.285723188 | 0.410165147 | 8.04E-11 | 1.12E-09 | 180.3235046 | 74.03119061 | ENSG00000186777 |
| NKAIN1 | -1.288443503 | 0.409392477 | 5.20E-61 | 1.22E-58 | 2113.480034 | 865.6952359 | ENSG00000084628 |
| NA | -1.28853415 | 0.409366755 | 0.16385125 | 0.30247766 | 8.022764384 | 3.283546882 | ENSG00000272275 |
| HACL1 | -1.288913778 | 0.409259049 | 3.39E-35 | 2.82E-33 | 722.8876923 | 295.5445606 | ENSG00000131373 |
| MAGI2-AS3 | -1.290303243 | 0.40886508 | 2.41E-08 | 2.41E-07 | 129.2307771 | 52.82418875 | ENSG00000234456 |
| NA | -1.292867474 | 0.408139013 | 0.635630856 | 0.7717746 | 4.743490275 | 1.931086926 | ENSG00000267681 |
| MIOX | -1.292917534 | 0.408124851 | 0.221036177 | 0.379404031 | 6.398139435 | 2.59145412 | ENSG00000100253 |
| UCP1 | -1.294174434 | 0.407769441 | 0.184849129 | 0.33112866 | 8.860814635 | 3.632819123 | ENSG00000109424 |
| KIAA1755 | -1.294810735 | 0.407589634 | 0.274262996 | 0.441074338 | 5.56547083 | 2.265305152 | ENSG00000149633 |
| GRIK1 | -1.294890718 | 0.407567038 | 0.112574687 | 0.227020519 | 9.71301956 | 3.969187922 | ENSG00000171189 |
| HSPA6 | -1.295292925 | 0.407453428 | 0.284306643 | 0.453007306 | 6.330394027 | 2.557577979 | ENSG00000173110 |
| NA | -1.299783114 | 0.406187257 | 0.000448365 | 0.002083192 | 48.5406554 | 19.70935055 | ENSG00000272841 |
| SLC6A12 | -1.300616809 | 0.4059526 | 0.116531292 | 0.233292356 | 9.726974075 | 3.948215223 | ENSG00000111181 |
| H4C1 | -1.301452185 | 0.405717606 | 0.225964094 | 0.384982474 | 5.649961656 | 2.292729572 | ENSG00000278637 |
| KCNC1 | -1.303399519 | 0.405170342 | 0.034908974 | 0.089007272 | 18.622719 | 7.553609926 | ENSG00000129159 |
| S1PR1 | -1.304859847 | 0.404760428 | 0.005761533 | 0.019843908 | 32.45202378 | 13.07503754 | ENSG00000170989 |
| GALNT16 | -1.304983145 | 0.404725837 | 3.81E-13 | 6.79E-12 | 254.1793906 | 102.7396677 | ENSG00000100626 |
| NA | -1.30511248 | 0.404689555 | 0.001453334 | 0.005960092 | 42.21843392 | 17.12649872 | ENSG00000286707 |
| NA | -1.305607209 | 0.404550803 | 0.095314648 | 0.199513333 | 11.38652932 | 4.608582417 | ENSG00000233967 |
| TEK | -1.308436588 | 0.403758185 | 0.000584438 | 0.002630517 | 48.65217937 | 19.62062557 | ENSG00000120156 |
| LTB4R2 | -1.30923759 | 0.403534076 | 0.026061436 | 0.070050321 | 19.47001785 | 7.820608906 | ENSG00000213906 |
| ALDH1L1 | -1.309264952 | 0.403526423 | 0.001831736 | 0.00730536 | 38.21004279 | 15.38443866 | ENSG00000144908 |
| H2AC13 | -1.30953382 | 0.403451226 | 0.051013438 | 0.121254341 | 14.69092155 | 5.942220247 | ENSG00000196747 |
| ADM | -1.309697025 | 0.403405588 | 9.22E-58 | 2.00E-55 | 1154.018813 | 465.7154909 | ENSG00000148926 |
| TTLL7 | -1.312161171 | 0.402717153 | 2.06E-06 | 1.54E-05 | 97.54993407 | 39.25253704 | ENSG00000137941 |
| H2BC15 | -1.313387198 | 0.402375063 | 0.026650866 | 0.071336459 | 18.73511886 | 7.521884358 | ENSG00000233822 |
| ANAPC1P1 | -1.317649379 | 0.401188073 | 0.089425128 | 0.189779994 | 12.23574318 | 4.91590926 | ENSG00000233673 |
| PDE1B | -1.319586604 | 0.400649726 | 0.00024209 | 0.001200557 | 66.08301528 | 26.41439045 | ENSG00000123360 |
| SOX9 | -1.322782857 | 0.39976308 | 9.46E-08 | 8.76E-07 | 121.3690972 | 48.49874383 | ENSG00000125398 |
| NA | -1.32501012 | 0.399146393 | 0.013216984 | 0.039826436 | 26.46688973 | 10.59113053 | ENSG00000283544 |
| ZNF626 | -1.326565804 | 0.398716219 | 6.05E-09 | 6.63E-08 | 134.2155002 | 53.57489846 | ENSG00000188171 |
| SNORD62B | -1.328633708 | 0.398145123 | 0.120990112 | 0.240051449 | 9.073775011 | 3.603244129 | ENSG00000231587 |
| CXCL16 | -1.32962311 | 0.397872169 | 0.002590717 | 0.009888705 | 39.59168315 | 15.76920458 | ENSG00000161921 |
| ZNF20 | -1.33125585 | 0.39742214 | 0.187039202 | 0.334059183 | 7.389101641 | 2.917603089 | ENSG00000132010 |
| DRC3 | -1.331458082 | 0.397366434 | 0.044618409 | 0.108939846 | 16.53062206 | 6.543970491 | ENSG00000171962 |
| NA | -1.331721859 | 0.397293788 | 0.26441772 | 0.430200071 | 6.539687619 | 2.597905841 | ENSG00000286004 |
| NUPR1 | -1.335638163 | 0.396216767 | 0.239891129 | 0.40194927 | 4.963947479 | 1.973032325 | ENSG00000176046 |
| UGT3A1 | -1.337811165 | 0.39562043 | 0.000111966 | 0.000598322 | 68.13443687 | 26.89003192 | ENSG00000145626 |

| | | | | | | | |
|---|---|---|---|---|---|---|---|
| NA | -1.337960312 | 0.395579533 | 0.23916408 | 0.401192546 | 5.021204997 | 1.998306171 | ENSG00000230832 |
| NA | -1.339925452 | 0.395041068 | 0.02936417 | 0.077153936 | 18.3212376 | 7.257035951 | ENSG00000215014 |
| NA | -1.342385089 | 0.39436814 | 0.263460882 | 0.429186276 | 4.988389884 | 1.968731177 | ENSG00000269925 |
| ODC1 | -1.34420041 | 0.393872225 | 1.95E-135 | 3.76E-132 | 4470.751577 | 1760.72819 | ENSG00000115758 |
| GPR6 | -1.346239652 | 0.393315882 | 0.194729354 | 0.344851433 | 6.607433027 | 2.589303546 | ENSG00000146360 |
| NA | -1.346691198 | 0.393192798 | 0.177320339 | 0.32103567 | 9.90903436 | 3.885297125 | ENSG00000248489 |
| LOC730183 | -1.347020347 | 0.393103102 | 0.039855761 | 0.099377141 | 17.45786906 | 6.876038142 | ENSG00000261840 |
| ELP2 | -1.348747329 | 0.392632818 | 1.50E-91 | 9.62E-89 | 2018.188311 | 792.4878866 | ENSG00000134759 |
| CYP39A1 | -1.350285319 | 0.392214474 | 0.000429449 | 0.002005931 | 51.0237428 | 19.96344609 | ENSG00000146233 |
| NA | -1.350655946 | 0.392113727 | 0.240276399 | 0.40246349 | 5.081253418 | 1.998306171 | ENSG00000213113 |
| NA | -1.35908983 | 0.389828148 | 0.309533426 | 0.480618909 | 4.291887817 | 1.701732197 | ENSG00000259081 |
| NA | -1.359410693 | 0.389741457 | 3.61E-11 | 5.20E-10 | 161.8039369 | 63.11176054 | ENSG00000278922 |
| NA | -1.361553304 | 0.389163065 | 0.244441964 | 0.407504462 | 5.045647402 | 1.952059625 | ENSG00000286223 |
| NA | -1.361669381 | 0.389131754 | 0.128483671 | 0.25065463 | 8.41220324 | 3.253971888 | ENSG00000279601 |
| H3C1 | -1.36469321 | 0.388317004 | 0.260225593 | 0.42560508 | 5.105695823 | 1.977333472 | ENSG00000275714 |
| SCN7A | -1.365088138 | 0.38821072 | 0.238525404 | 0.400479522 | 5.105695823 | 1.975182898 | ENSG00000136546 |
| ZAR1L | -1.368976935 | 0.387165704 | 0.251078653 | 0.415221458 | 5.165744244 | 1.998306171 | ENSG00000189167 |
| NA | -1.369964065 | 0.386900885 | 0.232260488 | 0.392885648 | 6.014282385 | 2.345427839 | ENSG00000270532 |
| SYCE3 | -1.37172782 | 0.386428172 | 0.241680999 | 0.404044384 | 5.135720033 | 1.975182898 | ENSG00000217442 |
| HGSNAT | -1.371781143 | 0.386413889 | 2.15E-08 | 2.17E-07 | 124.3743234 | 48.13090992 | ENSG00000165102 |
| NA | -1.374477889 | 0.385692263 | 0.150514166 | 0.283112688 | 8.463878952 | 3.235149762 | ENSG00000276408 |
| NA | -1.375322098 | 0.385466637 | 0.195752285 | 0.346159807 | 5.935373365 | 2.292729572 | ENSG00000285669 |
| CPNE9 | -1.383992787 | 0.383156906 | 9.81E-08 | 9.06E-07 | 109.1421271 | 41.81844474 | ENSG00000144550 |
| HERC2P3 | -1.388146979 | 0.382055205 | 1.98E-07 | 1.75E-06 | 109.3900178 | 41.7426232 | ENSG00000180229 |
| NA | -1.389436621 | 0.381713834 | 0.195867018 | 0.346299207 | 5.973770284 | 2.271756873 | ENSG00000189014 |
| NA | -1.389618597 | 0.381665689 | 0.173059309 | 0.315488371 | 9.291441313 | 3.518820294 | ENSG00000272764 |
| NA | -1.389741 | 0.381633309 | 0.014620799 | 0.043338941 | 21.56578157 | 8.216127694 | ENSG00000288586 |
| NA | -1.389945546 | 0.381579204 | 0.142163643 | 0.271076291 | 7.677304254 | 2.909000794 | ENSG00000270184 |
| NA | -1.3916142 | 0.381138116 | 0.014462303 | 0.042948319 | 20.68634334 | 7.892129299 | ENSG00000270354 |
| NA | -1.393340915 | 0.380682218 | 0.094118903 | 0.19734279 | 10.27614599 | 3.901968677 | ENSG00000286478 |
| NA | -1.394363689 | 0.380412435 | 0.136241083 | 0.262114963 | 7.761795079 | 2.936425214 | ENSG00000256167 |
| ST8SIA5 | -1.396849702 | 0.379757483 | 2.68E-06 | 1.95E-05 | 83.8332293 | 31.87746157 | ENSG00000101638 |
| ACSM1 | -1.397468662 | 0.379594591 | 0.170647969 | 0.312035068 | 6.920753766 | 2.642001813 | ENSG00000166743 |
| DYNC2H1 | -1.398415764 | 0.379345475 | 1.19E-21 | 4.51E-20 | 396.6718162 | 150.4279883 | ENSG00000187240 |
| PCDHGB6 | -1.400030613 | 0.378921101 | 2.07E-09 | 2.42E-08 | 141.6353018 | 53.58539087 | ENSG00000253305 |
| PLCL2 | -1.400206049 | 0.378875026 | 3.11E-11 | 4.51E-10 | 211.9608445 | 80.42217937 | ENSG00000154822 |
| RNF103-CHMP3 | -1.400496166 | 0.378798844 | 0.19417773 | 0.34413253 | 6.074330806 | 2.292729572 | ENSG00000249884 |
| STKLD1 | -1.401276105 | 0.378594116 | 0.077368397 | 0.168853295 | 12.86382412 | 4.854608698 | ENSG00000198870 |
| LOXL3 | -1.402659762 | 0.378231189 | 1.15E-25 | 5.85E-24 | 441.1027193 | 166.8823008 | ENSG00000115318 |
| ACD | -1.406416582 | 0.377247545 | 8.23E-45 | 1.13E-42 | 808.874982 | 305.0128463 | ENSG00000102977 |
| NA | -1.406987774 | 0.377098215 | 0.084339595 | 0.180998457 | 11.21754767 | 4.221665925 | ENSG00000235912 |
| DGKI | -1.407603898 | 0.376937204 | 0.011732268 | 0.035993507 | 23.55472725 | 8.861973909 | ENSG00000157680 |
| SLC7A11 | -1.4108399 | 0.376092672 | 1.14E-22 | 4.70E-21 | 471.0520824 | 177.02791 | ENSG00000151012 |
| TEX13B | -1.412710135 | 0.37560544 | 0.007575216 | 0.02494399 | 28.90619856 | 10.80543124 | ENSG00000170925 |
| INCA1 | -1.414188622 | 0.375220714 | 0.007090919 | 0.023633539 | 29.75433639 | 11.2031006 | ENSG00000196388 |
| LRRC49 | -1.414651294 | 0.3751004 | 9.88E-14 | 1.87E-12 | 204.6080246 | 76.70574203 | ENSG00000137821 |
| NA | -1.415015852 | 0.375005627 | 0.176954493 | 0.320558656 | 6.986383993 | 2.61887854 | ENSG00000267342 |
| MAP3K7 | -1.416239721 | 0.374687636 | 8.00E-71 | 2.62E-68 | 1310.456561 | 490.9484317 | ENSG00000135341 |
| TTLL11 | -1.416898516 | 0.374516577 | 0.013207032 | 0.039802665 | 23.50285138 | 8.786152369 | ENSG00000175764 |
| NA | -1.4195576 | 0.373826928 | 0.086217439 | 0.184351654 | 12.15125236 | 4.532760877 | ENSG00000240668 |
| LHFPL3 | -1.419801538 | 0.373763725 | 0.170777788 | 0.312213258 | 6.958474964 | 2.589303546 | ENSG00000187416 |
| PODN | -1.421244841 | 0.373389991 | 0.2238125 | 0.382322406 | 5.314989415 | 2.000456745 | ENSG00000174348 |
| NA | -1.421518291 | 0.373319225 | 0.2202439 | 0.378438148 | 5.266104606 | 1.975182898 | ENSG00000242571 |
| SLC38A5 | -1.422932604 | 0.372953429 | 1.49E-42 | 1.89E-40 | 1166.717523 | 434.7657291 | ENSG00000017483 |
| CLIP4 | -1.423355576 | 0.372844102 | 0.088334489 | 0.187940953 | 9.666925654 | 3.601093555 | ENSG00000115295 |
| RAI2 | -1.424429414 | 0.372566687 | 0.000973424 | 0.004161861 | 39.47784384 | 14.69664705 | ENSG00000131831 |
| COL15A1 | -1.427957567 | 0.371656677 | 0.135910839 | 0.26155786 | 8.839163133 | 3.308820728 | ENSG00000204291 |
| LEAP2 | -1.43038004 | 0.371033141 | 0.132979483 | 0.25722523 | 11.39557775 | 4.204994373 | ENSG00000164406 |
| ZNF521 | -1.430630601 | 0.370968707 | 6.72E-17 | 1.76E-15 | 274.8299278 | 102.1179987 | ENSG00000198795 |
| NA | -1.435985879 | 0.369594225 | 0.22569572 | 0.3847227 | 5.323362124 | 1.968731177 | ENSG00000237233 |
| NA | -1.440812939 | 0.36835968 | 0.208769642 | 0.363637219 | 5.314989415 | 1.952059625 | ENSG00000280007 |
| NA | -1.441474464 | 0.368190813 | 0.105470306 | 0.215646811 | 8.980911477 | 3.331944001 | ENSG00000280828 |
| ADAM20P1 | -1.441936674 | 0.368072871 | 0.223650163 | 0.382177042 | 6.230033664 | 2.265305152 | ENSG00000259158 |
| OGFR-AS1 | -1.443978031 | 0.367552431 | 0.0792823 | 0.172225931 | 12.50440947 | 4.617184712 | ENSG00000229873 |
| LINC02048 | -1.44683102 | 0.366826299 | 0.201715053 | 0.354042862 | 5.380619642 | 1.970881751 | ENSG00000228271 |
| GFI1 | -1.448237751 | 0.366468792 | 0.207805203 | 0.36248939 | 6.39323335 | 2.366400538 | ENSG00000162676 |
| NA | -1.457061655 | 0.364234212 | 0.018208757 | 0.051923893 | 18.01070776 | 6.558491469 | ENSG00000214975 |
| NCR3 | -1.458793533 | 0.363797231 | 0.215452323 | 0.37239412 | 5.350595432 | 1.926785779 | ENSG00000204475 |
| TAS1R3 | -1.459660883 | 0.363578581 | 0.097504492 | 0.203083477 | 9.125450724 | 3.329793428 | ENSG00000169962 |
| LOC107984813 | -1.462187567 | 0.36294238 | 0.225366909 | 0.384433022 | 5.350595432 | 1.920334058 | ENSG00000259804 |
| NA | -1.462958718 | 0.362748432 | 0.004813486 | 0.016981887 | 29.13714366 | 10.55940496 | ENSG00000279233 |
| NA | -1.463602532 | 0.362586588 | 0.214146499 | 0.370601966 | 5.369456031 | 1.924635205 | ENSG00000239345 |
| PDYN | -1.468996357 | 0.361233512 | 0.002039955 | 0.008031177 | 38.91595671 | 14.07392411 | ENSG00000101327 |
| NA | -1.470188486 | 0.36093514 | 0.261514717 | 0.426881063 | 4.58009043 | 1.674307777 | ENSG00000250794 |
| NPY2R | -1.470411799 | 0.360879276 | 0.257415901 | 0.422551725 | 4.631766142 | 1.699581623 | ENSG00000185149 |
| NA | -1.47226188 | 0.360416789 | 0.271089117 | 0.437914936 | 4.591254041 | 1.674307777 | ENSG00000274718 |
| SARM1 | -1.47341295 | 0.360129341 | 1.62E-07 | 1.44E-06 | 95.82618806 | 34.58883321 | ENSG00000004139 |

| | | | | | | | |
|---|---|---|---|---|---|---|---|
| NA | -1.47413713 | 0.359948614 | 0.150929472 | 0.2838351 | 7.347913819 | 2.658673365 | ENSG00000236643 |
| NA | -1.476215511 | 0.359430437 | 0.162747849 | 0.300782598 | 6.357627334 | 2.271756873 | ENSG00000238082 |
| NA | -1.478379478 | 0.358891714 | 0.004682563 | 0.016589775 | 27.58197898 | 9.957654715 | ENSG00000268798 |
| LOC105373383 | -1.47903018 | 0.358729879 | 0.075399521 | 0.165494614 | 10.99776618 | 3.95251637 | ENSG00000260081 |
| FEZF1 | -1.47931698 | 0.358658573 | 1.15E-14 | 2.41E-13 | 226.3028393 | 81.2280105 | ENSG00000128610 |
| NOXRED1 | -1.483220087 | 0.357689559 | 0.252978214 | 0.417622232 | 4.68902366 | 1.701732197 | ENSG00000165555 |
| ARHGAP36 | -1.483384185 | 0.357648877 | 0.004465584 | 0.015908696 | 25.70196653 | 9.194041561 | ENSG00000147256 |
| NA | -1.48521041 | 0.357196436 | 0.11513441 | 0.230999262 | 8.267663993 | 2.963849634 | ENSG00000212789 |
| NA | -1.486459107 | 0.356887405 | 0.155315887 | 0.290329339 | 6.463769662 | 2.294880146 | ENSG00000274818 |
| NA | -1.4883539 | 0.356418987 | 0.089127077 | 0.189265509 | 10.18136744 | 3.619915681 | ENSG00000261338 |
| CDYL2 | -1.488955057 | 0.356270502 | 1.44E-09 | 1.72E-08 | 116.2685077 | 41.46325381 | ENSG00000166446 |
| CTNNAL1 | -1.489000575 | 0.356259261 | 1.78E-81 | 7.82E-79 | 1426.73209 | 508.5527104 | ENSG00000119326 |
| NA | -1.490547165 | 0.355877551 | 0.028830212 | 0.076030569 | 14.73701546 | 5.229154786 | ENSG00000248015 |
| NKX6-1 | -1.493834217 | 0.355067639 | 0.254599155 | 0.41938966 | 4.686232757 | 1.678608924 | ENSG00000163623 |
| SPEF1 | -1.494111208 | 0.354999474 | 0.036572832 | 0.092515933 | 15.92331674 | 5.717166665 | ENSG00000101222 |
| NA | -1.495543529 | 0.354647202 | 0.236386806 | 0.39781365 | 4.642929753 | 1.655485651 | ENSG00000217289 |
| DBNDD2 | -1.497039427 | 0.354279667 | 0.002504954 | 0.009612703 | 29.68120933 | 10.52982996 | ENSG00000244274 |
| LUARIS | -1.497083699 | 0.354268795 | 0.083655425 | 0.179857968 | 9.331953413 | 3.329793428 | ENSG00000231638 |
| LONRF2 | -1.497533262 | 0.354158418 | 0.037108045 | 0.093636534 | 13.88760144 | 4.934731385 | ENSG00000170500 |
| NA | -1.499334881 | 0.353716425 | 0.249404236 | 0.413448377 | 4.612905543 | 1.632362378 | ENSG00000272370 |
| GPR50 | -1.499922863 | 0.353572295 | 3.50E-14 | 6.97E-13 | 197.074696 | 69.6874566 | ENSG00000102195 |
| PAPLN | -1.501110327 | 0.353281393 | 1.46E-11 | 2.20E-10 | 174.5190283 | 61.51033023 | ENSG00000100767 |
| ZNF878 | -1.502684797 | 0.352896054 | 2.07E-05 | 0.000129037 | 58.57237734 | 20.68941499 | ENSG00000257446 |
| NA | -1.502716639 | 0.352888265 | 0.110090311 | 0.222708913 | 8.330503317 | 2.929973493 | ENSG00000232320 |
| CDK5R2 | -1.503746741 | 0.352636388 | 0.046879311 | 0.113511836 | 14.94930012 | 5.254428633 | ENSG00000171450 |
| NA | -1.504034137 | 0.352566148 | 0.006832611 | 0.02293907 | 23.26285786 | 8.21397712 | ENSG00000214189 |
| C11orf94 | -1.50408991 | 0.352552518 | 0.288069188 | 0.457410478 | 4.582881332 | 1.609239105 | ENSG00000234776 |
| ZNF528-AS1 | -1.506519254 | 0.351959357 | 0.173644133 | 0.3163741 | 6.637457238 | 2.368551112 | ENSG00000269834 |
| SPAG1 | -1.510047162 | 0.351099741 | 0.000303745 | 0.001472248 | 42.03569791 | 14.75149589 | ENSG00000104450 |
| CACNA1G | -1.514860419 | 0.34993032 | 1.00E-22 | 4.15E-21 | 343.7950813 | 120.2304927 | ENSG00000006283 |
| NPAS3 | -1.515640658 | 0.349741122 | 0.195717078 | 0.346159807 | 5.617146542 | 1.968731177 | ENSG00000151322 |
| NA | -1.518012867 | 0.349166519 | 0.155128964 | 0.290152278 | 7.715701173 | 2.734494905 | ENSG00000281641 |
| B4GALNT1 | -1.52263443 | 0.34804978 | 0.000172853 | 0.000882843 | 43.40561108 | 15.11313854 | ENSG00000135454 |
| LEF1 | -1.523159741 | 0.347923072 | 0.189458327 | 0.337494724 | 5.758894886 | 2.019278871 | ENSG00000138795 |
| TPP2 | -1.524759584 | 0.347537465 | 1.61E-107 | 1.55E-104 | 1881.721269 | 653.8349137 | ENSG00000134900 |
| LIG3 | -1.525617942 | 0.347330753 | 2.17E-123 | 2.98E-120 | 2315.702901 | 804.1623396 | ENSG00000005156 |
| GRIN2B | -1.526161754 | 0.347199854 | 3.16E-06 | 2.27E-05 | 75.16862963 | 26.21540421 | ENSG00000273079 |
| MID2 | -1.527402032 | 0.346901496 | 2.62E-06 | 1.91E-05 | 73.12474177 | 25.39102354 | ENSG00000080561 |
| NME5 | -1.528382676 | 0.346665777 | 0.589240423 | 0.738633689 | 6.504081603 | 2.252934747 | ENSG00000112981 |
| NA | -1.530168728 | 0.346236871 | 0.175110438 | 0.318144232 | 5.64437985 | 1.945607904 | ENSG00000279812 |
| NA | -1.531047359 | 0.34602607 | 0.162344779 | 0.300095163 | 5.704428271 | 1.973032325 | ENSG00000248559 |
| NA | -1.533751606 | 0.345378072 | 0.238577336 | 0.400521746 | 4.68902366 | 1.60063681 | ENSG00000235748 |
| MYZAP | -1.536658148 | 0.344682953 | 0.122085642 | 0.241642301 | 7.63400125 | 2.642001813 | ENSG00000263155 |
| CRHR2 | -1.537713701 | 0.344430856 | 0.002429411 | 0.009358182 | 29.51924895 | 10.18700944 | ENSG00000106113 |
| LOC101928521 | -1.540516666 | 0.343762322 | 0.027216103 | 0.072667605 | 14.3259251 | 4.930430238 | ENSG00000242686 |
| PRTG | -1.541394883 | 0.343553126 | 1.22E-08 | 1.28E-07 | 105.3028809 | 36.21259329 | ENSG00000166450 |
| MMUT | -1.541429712 | 0.343544833 | 7.20E-32 | 5.09E-30 | 502.0760085 | 172.7712593 | ENSG00000146085 |
| SRSF12 | -1.543479958 | 0.34305696 | 1.71E-11 | 2.54E-10 | 155.8316061 | 53.43132464 | ENSG00000154548 |
| NA | -1.545874287 | 0.342488087 | 0.167871896 | 0.308009963 | 5.786128194 | 1.975182898 | ENSG00000224550 |
| INHBB | -1.546824747 | 0.342262527 | 0.198309471 | 0.349656157 | 5.840594809 | 2.000456745 | ENSG00000163083 |
| ADAP2 | -1.548562712 | 0.341850464 | 0.01771367 | 0.050812552 | 17.3819511 | 5.967494094 | ENSG00000184060 |
| YAF2 | -1.549917972 | 0.341529482 | 0.010484322 | 0.032758836 | 20.99898836 | 7.134967866 | ENSG00000015153 |
| HS6ST3 | -1.554170882 | 0.340524172 | 0.000231417 | 0.001152121 | 48.66892479 | 16.69091254 | ENSG00000185352 |
| LRRC4 | -1.554784006 | 0.340379486 | 5.42E-20 | 1.83E-18 | 299.3601111 | 101.8080004 | ENSG00000128594 |
| DRD5 | -1.556985092 | 0.339860572 | 0.128754626 | 0.251041591 | 7.61446493 | 2.566180274 | ENSG00000169676 |
| NA | -1.560346636 | 0.339069604 | 0.179794859 | 0.324485231 | 5.75331308 | 1.924635205 | ENSG00000254554 |
| NA | -1.562209121 | 0.338632156 | 0.003885444 | 0.014089118 | 25.20638535 | 8.561098787 | ENSG00000278493 |
| HRH3 | -1.565106387 | 0.337952786 | 5.86E-13 | 1.02E-11 | 177.45636 | 59.95270489 | ENSG00000101180 |
| LINC02249 | -1.565122232 | 0.337949074 | 0.138479564 | 0.265468476 | 6.820393404 | 2.290578998 | ENSG00000225930 |
| LRRC46 | -1.56579971 | 0.337790414 | 0.13531742 | 0.260701923 | 6.78757829 | 2.271756873 | ENSG00000141294 |
| NA | -1.565934983 | 0.337758742 | 0.197324816 | 0.348365793 | 5.813361501 | 1.943457331 | ENSG00000268295 |
| FAIM2 | -1.566658177 | 0.337589473 | 0.014290213 | 0.042535487 | 18.41342541 | 6.190397102 | ENSG00000135472 |
| NA | -1.569578953 | 0.336906705 | 0.239663864 | 0.401786355 | 4.841935615 | 1.604937958 | ENSG00000261079 |
| NA | -1.569645497 | 0.336891166 | 0.040000281 | 0.099618566 | 12.61960023 | 4.246939771 | ENSG00000242375 |
| NA | -1.572580069 | 0.336206595 | 0.099835017 | 0.206976604 | 9.973988865 | 3.403464394 | ENSG00000267419 |
| ETV2 | -1.573446775 | 0.336004678 | 0.021270986 | 0.059143642 | 16.6234856 | 5.553153181 | ENSG00000105672 |
| LINC01801 | -1.573700624 | 0.335945562 | 0.009841009 | 0.031015016 | 21.75832975 | 7.370501742 | ENSG00000267767 |
| ZNF763 | -1.57463671 | 0.335727656 | 0.146548123 | 0.277189905 | 7.908925069 | 2.683947212 | ENSG00000197054 |
| P2RX5-TAX1BP3 | -1.575542372 | 0.335516967 | 0.000294403 | 0.001430563 | 38.28125482 | 12.84783338 | ENSG00000257950 |
| TMSB15A | -1.579830304 | 0.334521234 | 3.80E-43 | 4.95E-41 | 649.3598694 | 217.0007762 | ENSG00000158164 |
| PCDHB5 | -1.580405174 | 0.334387964 | 1.45E-18 | 4.33E-17 | 262.38414 | 87.6870057 | ENSG00000113209 |
| DPF1 | -1.583232176 | 0.333733363 | 1.91E-25 | 9.57E-24 | 369.2792694 | 123.1728366 | ENSG00000011332 |
| NA | -1.58563286 | 0.333178483 | 0.058048939 | 0.134449027 | 10.8590089 | 3.598942982 | ENSG00000285602 |
| NA | -1.586509341 | 0.332976129 | 0.23268625 | 0.393370471 | 4.876665751 | 1.625910657 | ENSG00000287574 |
| NA | -1.592370755 | 0.33162605 | 0.20501855 | 0.35866626 | 4.909480864 | 1.630211804 | ENSG00000266648 |
| SLC9A3 | -1.595231479 | 0.33096912 | 0.000456033 | 0.002112204 | 35.643816 | 11.78979683 | ENSG00000066230 |
| NA | -1.595984916 | 0.330796319 | 0.201789267 | 0.354140898 | 4.92555056 | 1.630211804 | ENSG00000233971 |

| | | | | | | | |
|---|---|---|---|---|---|---|---|
| LOC101928718 | -1.597960839 | 0.330343568 | 0.143199971 | 0.272648109 | 6.001003592 | 1.996155598 | ENSG00000284830 |
| SPACA4 | -1.598948621 | 0.330117466 | 0.145941972 | 0.276396483 | 5.95981577 | 1.973032325 | ENSG00000177202 |
| NA | -1.60330804 | 0.32912145 | 0.193595176 | 0.343352444 | 4.996762593 | 1.64903393 | ENSG00000276223 |
| CALCB | -1.604095071 | 0.328941954 | 3.42E-10 | 4.43E-09 | 127.5559159 | 41.90852679 | ENSG00000175868 |
| NA | -1.604945071 | 0.328748207 | 0.212431273 | 0.368458586 | 5.070089807 | 1.678608924 | ENSG00000275017 |
| NA | -1.605677736 | 0.328581296 | 0.196373139 | 0.346907896 | 5.88090675 | 1.920334058 | ENSG00000237080 |
| NA | -1.605923241 | 0.328525386 | 0.185883855 | 0.33236495 | 4.969529285 | 1.630211804 | ENSG00000259137 |
| NA | -1.607880985 | 0.328079877 | 0.100202381 | 0.207448393 | 7.924994765 | 2.589303546 | ENSG00000224080 |
| NA | -1.60788753 | 0.328078389 | 0.202117902 | 0.354620869 | 4.963947479 | 1.623760083 | ENSG00000286281 |
| NA | -1.608807398 | 0.327869272 | 0.066077787 | 0.149036196 | 9.999106991 | 3.277095161 | ENSG00000286416 |
| OR7E12P | -1.61144551 | 0.327270278 | 3.72E-10 | 4.78E-09 | 111.5140539 | 36.49679686 | ENSG00000189398 |
| LRRTM3 | -1.613174721 | 0.326878247 | 0.009081324 | 0.029004605 | 22.31029257 | 7.351679617 | ENSG00000198739 |
| GLTPD2 | -1.614082505 | 0.326672631 | 0.067026069 | 0.150734425 | 11.16866286 | 3.695737221 | ENSG00000182327 |
| PRR29 | -1.615268926 | 0.326404098 | 0.077576851 | 0.169138227 | 8.972538768 | 2.911151367 | ENSG00000224383 |
| BOC | -1.616202274 | 0.326193 | 2.19E-16 | 5.45E-15 | 191.5048827 | 62.52506432 | ENSG00000144857 |
| LOC728392 | -1.617067042 | 0.325997534 | 9.17E-19 | 2.77E-17 | 226.6179118 | 73.80372599 | ENSG00000286190 |
| ABHD16A | -1.617878113 | 0.325814313 | 0.044803173 | 0.109335612 | 11.07946611 | 3.598942982 | ENSG00000204427 |
| ANGPTL8 | -1.617947115 | 0.32579873 | 0.179717186 | 0.324405749 | 5.015623191 | 1.625910657 | ENSG00000130173 |
| NA | -1.620119496 | 0.325308518 | 0.111493853 | 0.225042783 | 7.026896093 | 2.271756873 | ENSG00000280020 |
| NA | -1.620257265 | 0.325277454 | 0.208285812 | 0.363065113 | 5.086835224 | 1.655485651 | ENSG00000270076 |
| NA | -1.622502928 | 0.324771529 | 0.0578953 | 0.134173262 | 11.06060551 | 3.571518561 | ENSG00000274825 |
| NA | -1.623175208 | 0.324620225 | 0.22257003 | 0.381026377 | 4.996762593 | 1.609239105 | ENSG00000280010 |
| NA | -1.624476045 | 0.324327656 | 0.043714351 | 0.107221048 | 12.21130078 | 3.996612343 | ENSG00000271933 |
| NA | -1.62491789 | 0.324228341 | 0.007336604 | 0.024309589 | 21.29277278 | 6.922284688 | ENSG00000286482 |
| NA | -1.62631587 | 0.323914314 | 0.176435482 | 0.319976461 | 5.054020111 | 1.625910657 | ENSG00000224888 |
| SCN2B | -1.628687025 | 0.323382379 | 0.016005154 | 0.04671742 | 18.47414955 | 6.011590066 | ENSG00000149575 |
| ZNF799 | -1.629455509 | 0.323210168 | 0.000168407 | 0.000863109 | 40.64077876 | 13.10461253 | ENSG00000196466 |
| NA | -1.629796777 | 0.323133722 | 0.171561418 | 0.313467637 | 5.075671612 | 1.628061231 | ENSG00000272710 |
| NA | -1.630878831 | 0.322891455 | 0.06377068 | 0.144814619 | 10.16250684 | 3.260423609 | ENSG00000269318 |
| NA | -1.631001057 | 0.322864101 | 0.175736864 | 0.319138964 | 6.063842916 | 1.943457331 | ENSG00000248996 |
| LHB | -1.635028025 | 0.321964153 | 0.170406801 | 0.311771396 | 6.231948686 | 2.044552717 | ENSG00000104826 |
| PAX9 | -1.637543364 | 0.321403297 | 0.19135649 | 0.340101164 | 5.160162438 | 1.653335077 | ENSG00000198807 |
| DCLK2 | -1.639300542 | 0.321012072 | 0.042205113 | 0.104114941 | 12.41867935 | 4.017585042 | ENSG00000170390 |
| NA | -1.644010225 | 0.319965836 | 0.19863457 | 0.350037407 | 5.07288071 | 1.60063681 | ENSG00000286136 |
| CTU2 | -1.64451489 | 0.319853929 | 2.07E-36 | 1.85E-34 | 532.461292 | 170.3656275 | ENSG00000174177 |
| ZGLP1 | -1.645228567 | 0.319695742 | 0.034258133 | 0.08750975 | 15.53734451 | 5.010552925 | ENSG00000220201 |
| NMU | -1.650284826 | 0.318577255 | 7.83E-17 | 2.03E-15 | 217.1967617 | 69.29219828 | ENSG00000109255 |
| NA | -1.65055364 | 0.318517901 | 0.001262945 | 0.005265502 | 28.98124064 | 9.276314822 | ENSG00000285825 |
| H3C10 | -1.650986151 | 0.318422426 | 4.14E-05 | 0.000241781 | 50.44578597 | 16.04050471 | ENSG00000278828 |
| ZNF433 | -1.655665998 | 0.317391193 | 0.017110947 | 0.049362536 | 16.45729485 | 5.185058814 | ENSG00000197647 |
| ABALON | -1.657864239 | 0.316907951 | 0.005603555 | 0.01935509 | 24.74872552 | 7.812006611 | ENSG00000281376 |
| C21orf59-TCP10L | -1.658700415 | 0.316724327 | 0.075199578 | 0.165168513 | 9.228601989 | 2.911151367 | ENSG00000265590 |
| NA | -1.659792856 | 0.316484586 | 0.074593505 | 0.164117622 | 9.247462588 | 2.915452515 | ENSG00000272944 |
| DUSP15 | -1.660929563 | 0.316235325 | 5.04E-06 | 3.50E-05 | 67.12477732 | 21.17769944 | ENSG00000149599 |
| NA | -1.6622311 | 0.31595016 | 0.074577309 | 0.164117622 | 8.324921511 | 2.637700666 | ENSG00000272092 |
| NA | -1.662576014 | 0.315874632 | 0.104899794 | 0.214778536 | 7.247353297 | 2.294880146 | ENSG00000272619 |
| LOC728485 | -1.663363755 | 0.315702206 | 0.096418816 | 0.201343946 | 8.439436547 | 2.683947212 | ENSG00000267260 |
| HMX2 | -1.667757526 | 0.314742188 | 2.39E-13 | 4.35E-12 | 160.5807903 | 50.48951378 | ENSG00000188816 |
| PHKG1 | -1.670117026 | 0.314227854 | 0.093129439 | 0.195725026 | 7.296238107 | 2.294880146 | ENSG00000164776 |
| MIR762HG | -1.674995469 | 0.313167093 | 0.090796861 | 0.191952027 | 7.32905322 | 2.294880146 | ENSG00000260083 |
| MIR3176 | -1.676875243 | 0.312759315 | 0.008886575 | 0.028495832 | 20.8392553 | 6.463847803 | ENSG00000266235 |
| NA | -1.678947209 | 0.312310459 | 0.121562318 | 0.240765779 | 6.297578913 | 1.970881751 | ENSG00000260018 |
| ZNF880 | -1.680274188 | 0.312023331 | 2.69E-15 | 5.99E-14 | 174.82837 | 54.61411289 | ENSG00000221923 |
| NA | -1.683382001 | 0.311351902 | 0.086612659 | 0.184924588 | 9.380838223 | 2.892329242 | ENSG00000258317 |
| LY6H | -1.688616017 | 0.310224382 | 0.143755033 | 0.273351275 | 6.291997107 | 1.943457331 | ENSG00000176956 |
| C8orf76 | -1.68890815 | 0.31016157 | 0.017390762 | 0.050042541 | 15.85278043 | 4.932580812 | ENSG00000189376 |
| NA | -1.689611854 | 0.31001032 | 0.0209087 | 0.058262276 | 13.78724108 | 4.265761897 | ENSG00000231713 |
| JAML | -1.692187934 | 0.309457258 | 0.110180585 | 0.222868161 | 6.379278836 | 1.973032325 | ENSG00000160593 |
| TMEM202-AS1 | -1.695992074 | 0.308642347 | 0.091766953 | 0.1935786 | 7.416334949 | 2.269606299 | ENSG00000261423 |
| NA | -1.699992562 | 0.30778769 | 0.041813275 | 0.103320082 | 11.60698652 | 3.548395288 | ENSG00000259404 |
| SAMD15 | -1.701158233 | 0.307539104 | 0.028586363 | 0.075552662 | 14.99798477 | 4.627404542 | ENSG00000100583 |
| NA | -1.703186663 | 0.307107008 | 0.102781879 | 0.211494958 | 7.468010661 | 2.273907447 | ENSG00000286084 |
| DNAJC9-AS1 | -1.705928243 | 0.306523961 | 0.040021583 | 0.09963303 | 12.91705143 | 3.950365797 | ENSG00000236756 |
| KCNA5 | -1.706901305 | 0.306317288 | 0.033738726 | 0.08640034 | 11.75431667 | 3.580120856 | ENSG00000130037 |
| SHISA3 | -1.708814981 | 0.305911239 | 7.15E-09 | 7.78E-08 | 102.8410447 | 31.4846264 | ENSG00000178343 |
| LOC101928058 | -1.712244503 | 0.305184902 | 0.111241223 | 0.224613193 | 7.465219758 | 2.2486336 | ENSG00000253764 |
| CCDC163 | -1.714976474 | 0.304607533 | 4.97E-05 | 0.00028587 | 43.30525072 | 13.21592775 | ENSG00000280670 |
| NA | -1.718079911 | 0.303952984 | 0.06569239 | 0.14827096 | 8.602636233 | 2.614577393 | ENSG00000248774 |
| SMPD1 | -1.7187171 | 0.303818768 | 8.69E-06 | 5.79E-05 | 53.17867906 | 16.20935238 | ENSG00000166311 |
| H3C2 | -1.718911226 | 0.30377789 | 0.161342668 | 0.298672048 | 5.345013626 | 1.607088532 | ENSG00000286522 |
| ZNF559 | -1.722949279 | 0.302928815 | 3.83E-16 | 9.32E-15 | 190.1529543 | 57.74574412 | ENSG00000188321 |
| ZNF536 | -1.724473501 | 0.302608937 | 0.209782084 | 0.365209497 | 5.363874225 | 1.609239105 | ENSG00000198597 |
| LONRF3 | -1.729564249 | 0.301543021 | 0.200999107 | 0.353159464 | 5.600401125 | 1.72485547 | ENSG00000175556 |
| NA | -1.730943282 | 0.301254922 | 0.037701653 | 0.094873805 | 10.96216017 | 3.329793428 | ENSG00000253200 |
| NA | -1.733773684 | 0.300664474 | 0.077434873 | 0.168900489 | 7.674513351 | 2.318003419 | ENSG00000237410 |
| ZNF876P | -1.734996719 | 0.300409696 | 0.007058399 | 0.023541421 | 18.62830081 | 5.620372426 | ENSG00000198155 |
| H2BC9 | -1.737118078 | 0.299968294 | 1.97E-08 | 2.00E-07 | 89.62271198 | 26.91153766 | ENSG00000275713 |

| | | | | | | | |
|---|---|---|---|---|---|---|---|
| IQUB | -1.738258727 | 0.299731221 | 0.126923234 | 0.24843862 | 6.59973604 | 1.994005024 | ENSG00000164675 |
| CREG2 | -1.741756839 | 0.299005341 | 0.028705745 | 0.07582261 | 13.07120269 | 3.874544257 | ENSG00000175874 |
| SLC12A1 | -1.742056447 | 0.298943253 | 0.102719821 | 0.21145301 | 7.571161926 | 2.246483026 | ENSG00000074803 |
| H2BC5 | -1.743004855 | 0.298746796 | 3.16E-10 | 4.11E-09 | 110.8423577 | 33.11430516 | ENSG00000158373 |
| H2BC11 | -1.74977209 | 0.297348749 | 1.82E-08 | 1.87E-07 | 88.97664633 | 26.36222522 | ENSG00000124635 |
| GRIN1 | -1.751049688 | 0.297085544 | 6.92E-14 | 1.33E-12 | 158.4438387 | 47.1535291 | ENSG00000176884 |
| DPYSL5 | -1.761041031 | 0.295035195 | 7.08E-86 | 3.80E-83 | 1112.929223 | 328.3805688 | ENSG00000157851 |
| NA | -1.766867156 | 0.293846138 | 0.007725952 | 0.025381698 | 18.8957278 | 5.502605488 | ENSG00000286342 |
| NA | -1.768357757 | 0.293542692 | 0.067124658 | 0.150938545 | 8.912490347 | 2.61887854 | ENSG00000286873 |
| NA | -1.776146609 | 0.291962177 | 0.054303838 | 0.127473271 | 9.019308396 | 2.63985124 | ENSG00000283355 |
| NFIX | -1.776385692 | 0.291913798 | 0.022143205 | 0.061119554 | 12.40472483 | 3.624216828 | ENSG00000008441 |
| TNFAIP8L3 | -1.776943211 | 0.291801011 | 0.027638514 | 0.073602233 | 15.67542607 | 4.587609718 | ENSG00000183578 |
| ZNF677 | -1.778031989 | 0.291580877 | 1.10E-29 | 6.89E-28 | 404.3751144 | 118.0479817 | ENSG00000197928 |
| PCDHGA1 | -1.778927585 | 0.291399926 | 0.00872163 | 0.028064881 | 20.32501367 | 5.904575996 | ENSG00000204956 |
| CCDC173 | -1.77920322 | 0.291344258 | 0.020042183 | 0.056294956 | 17.18015434 | 5.075621597 | ENSG00000154479 |
| THSD7A | -1.786261415 | 0.289922375 | 4.36E-05 | 0.000253639 | 43.26876882 | 12.62493038 | ENSG00000005108 |
| PDE10A | -1.787139715 | 0.289745927 | 2.72E-05 | 0.000165555 | 47.49610246 | 13.69130876 | ENSG00000112541 |
| LINC02696 | -1.789660795 | 0.289240044 | 0.082343119 | 0.177543434 | 8.091185514 | 2.368551112 | ENSG00000254427 |
| NA | -1.789801578 | 0.28921182 | 0.070828799 | 0.157523445 | 7.913831154 | 2.292729572 | ENSG00000261572 |
| CCDC183-AS1 | -1.789873639 | 0.289197375 | 0.123266905 | 0.243217946 | 5.641588947 | 1.625910657 | ENSG00000228544 |
| SLC1A6 | -1.79250447 | 0.288670489 | 2.18E-24 | 1.02E-22 | 296.6230245 | 85.75941431 | ENSG00000105143 |
| ZBTB16 | -1.793121311 | 0.28854709 | 0.053790796 | 0.126422789 | 9.158265837 | 2.63985124 | ENSG00000109906 |
| CLEC4G | -1.794677643 | 0.288235983 | 0.002739929 | 0.010376123 | 24.83022529 | 7.124214997 | ENSG00000182566 |
| NA | -1.79511782 | 0.288148054 | 0.121043362 | 0.240083025 | 5.668822255 | 1.625910657 | ENSG00000280327 |
| NA | -1.795565018 | 0.288058749 | 0.079861029 | 0.173159392 | 7.897761458 | 2.267455725 | ENSG00000287430 |
| SHC4 | -1.795814402 | 0.28800896 | 0.064440792 | 0.14594122 | 9.002562979 | 2.5640297 | ENSG00000185634 |
| NA | -1.803004238 | 0.286577205 | 0.163302006 | 0.301662262 | 5.824525113 | 1.67645835 | ENSG00000240882 |
| LOC101928424 | -1.803725117 | 0.286434045 | 0.136851633 | 0.26300108 | 5.704428271 | 1.623760083 | ENSG00000255337 |
| FOXP2 | -1.804332306 | 0.286313518 | 0.173732418 | 0.316505059 | 4.661790352 | 1.356761103 | ENSG00000128573 |
| FEZF1-AS1 | -1.812705066 | 0.284656695 | 3.06E-26 | 1.61E-24 | 290.9657293 | 82.89101395 | ENSG00000230316 |
| RDX | -1.813992801 | 0.284402726 | 9.63E-131 | 1.55E-127 | 1712.504403 | 487.00101 | ENSG00000137710 |
| NA | -1.81495257 | 0.284213587 | 0.045037342 | 0.109742446 | 12.98200594 | 3.718860494 | ENSG00000237436 |
| LINC02805 | -1.821402075 | 0.282945858 | 0.160627693 | 0.297913186 | 4.667372158 | 1.329336683 | ENSG00000232265 |
| NA | -1.826136159 | 0.282018917 | 0.120704448 | 0.239632552 | 7.100899029 | 2.019278871 | ENSG00000260949 |
| NA | -1.82700948 | 0.281848251 | 0.148240628 | 0.279866931 | 4.694605466 | 1.331487256 | ENSG00000285090 |
| POPDC2 | -1.827669769 | 0.281719285 | 0.148079383 | 0.279647516 | 4.697396369 | 1.331487256 | ENSG00000121577 |
| HOGA1 | -1.827669769 | 0.281719285 | 0.148079383 | 0.279647516 | 4.697396369 | 1.331487256 | ENSG00000241935 |
| ADM5 | -1.827817246 | 0.281690488 | 2.86E-08 | 2.84E-07 | 76.73041525 | 21.57053462 | ENSG00000224420 |
| NA | -1.844466485 | 0.278458361 | 0.007541058 | 0.024867443 | 16.47894635 | 4.564486445 | ENSG00000279692 |
| LHX2 | -1.846287389 | 0.278107125 | 0.030199492 | 0.078950861 | 10.66837575 | 2.98267176 | ENSG00000106689 |
| AMH | -1.84871305 | 0.277639925 | 1.14E-25 | 5.79E-24 | 305.3995116 | 84.63847176 | ENSG00000104899 |
| NA | -1.849499571 | 0.277488604 | 0.161835723 | 0.299412385 | 4.727420579 | 1.304062836 | ENSG00000234773 |
| H4C8 | -1.853134744 | 0.276790293 | 8.57E-05 | 0.000470499 | 40.30002455 | 11.15847159 | ENSG00000158406 |
| NA | -1.862607065 | 0.274978921 | 0.197869061 | 0.349071061 | 4.733002385 | 1.287391284 | ENSG00000280231 |
| ZNF440 | -1.862705264 | 0.274960205 | 1.69E-08 | 1.75E-07 | 116.1945416 | 31.9296268 | ENSG00000171295 |
| FOSB | -1.880070851 | 0.271670374 | 0.000125177 | 0.000660682 | 34.87821709 | 9.450820711 | ENSG00000125740 |
| CBLN3 | -1.883114754 | 0.271097788 | 0.031269491 | 0.081242353 | 11.98805267 | 3.224396894 | ENSG00000139899 |
| NA | -1.884516862 | 0.270834445 | 0.122069487 | 0.241642301 | 6.164403438 | 1.678608924 | ENSG00000271855 |
| EFCAB6 | -1.893631213 | 0.269128818 | 0.011836222 | 0.03623751 | 14.57919742 | 3.920790803 | ENSG00000186976 |
| LOC105376748 | -1.893973581 | 0.269064958 | 0.044246536 | 0.10816879 | 9.704646851 | 2.593604694 | ENSG00000233654 |
| WASHC5 | -1.895381518 | 0.268802504 | 5.74E-69 | 1.76E-66 | 944.7733527 | 254.2506817 | ENSG00000164961 |
| NA | -1.901348806 | 0.267692977 | 0.105929134 | 0.216332726 | 6.079912612 | 1.604937958 | ENSG00000236060 |
| SEC14L1P1 | -1.903496536 | 0.267294761 | 0.023008838 | 0.063066708 | 11.0766752 | 2.959548487 | ENSG00000213693 |
| NA | -1.907444121 | 0.266564373 | 3.95E-13 | 7.02E-12 | 119.6204333 | 31.91778944 | ENSG00000206417 |
| BAALC | -1.908429197 | 0.266382424 | 0.000124809 | 0.000659098 | 32.08403627 | 8.554647066 | ENSG00000164929 |
| LOC100128164 | -1.91062953 | 0.26597646 | 0.114001818 | 0.229203656 | 6.118309531 | 1.602787384 | ENSG00000239219 |
| NA | -1.910930715 | 0.265920939 | 0.016533899 | 0.047948804 | 13.6163444 | 3.601093555 | ENSG00000274849 |
| NA | -1.919310351 | 0.264380862 | 0.146489457 | 0.27710611 | 4.933923269 | 1.304062836 | ENSG00000254094 |
| NA | -1.921907128 | 0.263905418 | 0.156623178 | 0.292263274 | 4.950668686 | 1.30621341 | ENSG00000285803 |
| NA | -1.924349684 | 0.26345899 | 0.039244626 | 0.098256727 | 11.14980226 | 2.940726362 | ENSG00000230042 |
| NA | -1.926648883 | 0.263039454 | 0.18247226 | 0.328058934 | 4.922759657 | 1.287391284 | ENSG00000263235 |
| CHRDL2 | -1.926774105 | 0.263016624 | 0.00783649 | 0.025679246 | 14.97912417 | 3.946064649 | ENSG00000054938 |
| NA | -1.936310506 | 0.261283784 | 0.059891439 | 0.13770748 | 9.883916233 | 2.559728552 | ENSG00000223461 |
| NA | -1.943140538 | 0.260049733 | 0.128877649 | 0.251179908 | 4.991180787 | 1.283090137 | ENSG00000261351 |
| NA | -1.951647687 | 0.258520808 | 0.006316309 | 0.021444548 | 18.8159429 | 4.831485425 | ENSG00000237087 |
| NA | -1.952825287 | 0.258309877 | 0.101436057 | 0.209337105 | 6.313648609 | 1.623760083 | ENSG00000236184 |
| PTGER2 | -1.955429189 | 0.257844077 | 4.50E-07 | 3.73E-06 | 53.43406656 | 13.76282915 | ENSG00000125384 |
| ZNF596 | -1.95630218 | 0.2576881 | 0.021850894 | 0.060433778 | 15.07669363 | 3.865941962 | ENSG00000172748 |
| TAS2R14 | -1.956610316 | 0.257633068 | 0.097784283 | 0.203578308 | 6.29478801 | 1.607088532 | ENSG00000212127 |
| TTBK1 | -1.973276827 | 0.254673927 | 7.17E-09 | 7.78E-08 | 79.54345667 | 20.19065024 | ENSG00000146216 |
| NA | -1.982920408 | 0.252977256 | 0.092872788 | 0.195292334 | 6.588572428 | 1.699581623 | ENSG00000279696 |
| NA | -2.002287552 | 0.249603911 | 0.121011101 | 0.240068403 | 5.227707687 | 1.308363983 | ENSG00000272155 |
| KBTBD11-OT1 | -2.009730458 | 0.248319514 | 0.101778522 | 0.209878082 | 5.314989415 | 1.331487256 | ENSG00000253696 |
| PHGDH | -2.011864276 | 0.247952508 | 0 | 0 | 15479.36997 | 3837.192421 | ENSG00000092621 |
| PP2D1 | -2.014375943 | 0.24752121 | 0.067142626 | 0.150958383 | 6.645829947 | 1.653335077 | ENSG00000183977 |
| NA | -2.016722293 | 0.247118977 | 0.107404284 | 0.218581711 | 5.345013626 | 1.329336683 | ENSG00000272918 |
| NA | -2.017681619 | 0.246954709 | 0.015787679 | 0.046229525 | 11.95523756 | 2.957397913 | ENSG00000273008 |

| | | | | | | | |
|---|---|---|---|---|---|---|---|
| NA | -2.020807857 | 0.246420151 | 0.077418853 | 0.168889189 | 6.738693481 | 1.674307777 | ENSG00000228559 |
| NA | -2.02217954 | 0.246185972 | 0.130737338 | 0.253877157 | 5.249359189 | 1.280939563 | ENSG00000261305 |
| ATOH8 | -2.027799159 | 0.245228887 | 0.001975082 | 0.007804447 | 25.18984009 | 6.118876709 | ENSG00000168874 |
| NA | -2.029632042 | 0.244917532 | 0.000831554 | 0.003601595 | 22.80654947 | 5.578427028 | ENSG00000271147 |
| LOC100379224 | -2.029809621 | 0.244887388 | 0.05596335 | 0.130621776 | 7.943855364 | 1.926785779 | ENSG00000186019 |
| NA | -2.035213264 | 0.243971873 | 0.054278388 | 0.127429035 | 7.971088672 | 1.922484631 | ENSG00000260643 |
| IRX5 | -2.0405424 | 0.243072333 | 0.072410811 | 0.160284846 | 6.67864506 | 1.602787384 | ENSG00000176842 |
| NKX2-5 | -2.043504096 | 0.242573844 | 0.047180847 | 0.114055855 | 8.091185514 | 1.945607904 | ENSG00000183072 |
| NA | -2.049948597 | 0.241492686 | 0.001517379 | 0.006187225 | 21.82184479 | 5.275401332 | ENSG00000254461 |
| TMEM215 | -2.050239531 | 0.241443992 | 0.006878041 | 0.023068446 | 14.96516965 | 3.622066255 | ENSG00000188133 |
| GSC2 | -2.053679782 | 0.24086893 | 0.09245826 | 0.194632506 | 6.730320772 | 1.598486237 | ENSG00000063515 |
| ZNF790-AS1 | -2.05733589 | 0.240259288 | 0.090522177 | 0.191487859 | 7.021314288 | 1.72485547 | ENSG00000267254 |
| ALDH3A1 | -2.061612621 | 0.239548117 | 0.009719317 | 0.030691658 | 13.65386544 | 3.260423609 | ENSG00000108602 |
| NA | -2.064362849 | 0.239091898 | 0.123148722 | 0.243114464 | 5.453946856 | 1.28524071 | ENSG00000230747 |
| NA | -2.068690874 | 0.238375708 | 0.023838954 | 0.064981951 | 10.97611468 | 2.610276246 | ENSG00000284634 |
| NA | -2.073916628 | 0.237513821 | 0.048445581 | 0.116413018 | 8.234848879 | 1.952059625 | ENSG00000269425 |
| FABP7 | -2.083014704 | 0.236020699 | 1.26E-18 | 3.78E-17 | 181.4238756 | 42.66381022 | ENSG00000164434 |
| LOC101928963 | -2.088347997 | 0.235149799 | 0.095334615 | 0.199533478 | 5.611564737 | 1.33363783 | ENSG00000272677 |
| MYO16 | -2.098677969 | 0.233472095 | 0.093329629 | 0.196103024 | 5.592704138 | 1.30621341 | ENSG00000041515 |
| NA | -2.100988302 | 0.233098512 | 0.017839397 | 0.051112402 | 11.22033857 | 2.616727967 | ENSG00000277511 |
| NA | -2.101921822 | 0.23294773 | 0.08724104 | 0.186015736 | 5.608773834 | 1.30621341 | ENSG00000236015 |
| BEX5 | -2.102996595 | 0.232774254 | 0.000138088 | 0.000721131 | 28.12153889 | 6.512244923 | ENSG00000184515 |
| NA | -2.103897345 | 0.232628967 | 0.047556649 | 0.114762986 | 8.390551738 | 1.928936353 | ENSG00000241168 |
| SLITRK6 | -2.105917018 | 0.23230353 | 0.030662496 | 0.079977043 | 11.13584774 | 2.568330847 | ENSG00000184564 |
| NA | -2.109097359 | 0.231791994 | 0.062236413 | 0.142049533 | 7.125341433 | 1.651184504 | ENSG00000260498 |
| ZNF737 | -2.123946876 | 0.229418418 | 2.00E-19 | 6.39E-18 | 167.2739534 | 38.33485766 | ENSG00000237440 |
| ADRA1D | -2.128651433 | 0.228671515 | 0.013011242 | 0.039286253 | 12.98950277 | 2.989123481 | ENSG00000171873 |
| SHLD2P1 | -2.139008815 | 0.227035717 | 0.096133285 | 0.200941536 | 5.737243384 | 1.28524071 | ENSG00000165874 |
| SLC9A2 | -2.144052944 | 0.226243313 | 0.113449774 | 0.22835555 | 5.954233964 | 1.379884376 | ENSG00000115616 |
| BMP3 | -2.152014681 | 0.224998193 | 0.058615078 | 0.135385952 | 7.3095169 | 1.651184504 | ENSG00000152785 |
| H4C9 | -2.153463353 | 0.224772376 | 1.00E-13 | 1.89E-12 | 112.3559342 | 25.27272356 | ENSG00000276180 |
| ZNF781 | -2.158551924 | 0.223980971 | 0.088296644 | 0.187881156 | 5.875324944 | 1.331487256 | ENSG00000196381 |
| CDH9 | -2.163548203 | 0.223206632 | 0.065991714 | 0.148876296 | 7.255726006 | 1.604937958 | ENSG00000113100 |
| EDN3 | -2.173058692 | 0.221740055 | 0.033095605 | 0.085046933 | 8.858023732 | 1.973032325 | ENSG00000124205 |
| NA | -2.186517622 | 0.219681057 | 8.99E-05 | 0.000490913 | 30.04361504 | 6.606888588 | ENSG00000206341 |
| LRRC61 | -2.19119721 | 0.218969645 | 0.059258112 | 0.136560536 | 7.424707657 | 1.602787384 | ENSG00000127399 |
| SV2C | -2.193783565 | 0.218577444 | 0.068234431 | 0.152854452 | 5.98983998 | 1.30621341 | ENSG00000122012 |
| LILRB3 | -2.198896808 | 0.217804126 | 0.084802113 | 0.181829228 | 5.952118783 | 1.283090137 | ENSG00000204577 |
| NA | -2.239037893 | 0.211827545 | 0.045216093 | 0.110092597 | 7.655652752 | 1.60063681 | ENSG00000244586 |
| LAMP5 | -2.247141394 | 0.210641062 | 6.77E-06 | 4.60E-05 | 35.94059149 | 7.572432051 | ENSG00000125869 |
| DGKK | -2.247417989 | 0.210600682 | 7.81E-76 | 3.07E-73 | 686.9909915 | 144.4914262 | ENSG00000274588 |
| DAZL | -2.268480988 | 0.207548299 | 0.037027987 | 0.093483431 | 7.971088672 | 1.674307777 | ENSG00000092345 |
| NA | -2.274282162 | 0.206715409 | 0.054106426 | 0.127071715 | 6.403721241 | 1.331487256 | ENSG00000286650 |
| H4C11 | -2.290091266 | 0.20446258 | 0.005373773 | 0.018661649 | 14.36711293 | 2.915452515 | ENSG00000197238 |
| NA | -2.301856336 | 0.202801983 | 0.09001448 | 0.190695073 | 6.635542216 | 1.356761103 | ENSG00000263011 |
| NA | -2.30635972 | 0.202169922 | 0.018099966 | 0.051646019 | 9.707437754 | 1.949909052 | ENSG00000123009 |
| SLC30A10 | -2.307600084 | 0.20199618 | 0.031344573 | 0.081403898 | 8.115627918 | 1.625910657 | ENSG00000196660 |
| NA | -2.307975968 | 0.201943558 | 0.011155828 | 0.034582531 | 11.3048294 | 2.267455725 | ENSG00000278012 |
| NA | -2.321328935 | 0.200083078 | 0.101961 | 0.210174105 | 5.045647402 | 1.034913282 | ENSG00000225330 |
| FAM135B | -2.330336032 | 0.198837802 | 0.009166301 | 0.029237287 | 11.63980164 | 2.341126692 | ENSG00000147724 |
| RSPH4A | -2.336367624 | 0.19800824 | 0.007548788 | 0.024880175 | 13.33563862 | 2.658673365 | ENSG00000111834 |
| LOC102723313 | -2.336870465 | 0.197939238 | 0.025017564 | 0.067621386 | 8.303270009 | 1.651184504 | ENSG00000248896 |
| FAM133A | -2.340078312 | 0.197499607 | 0.000324651 | 0.001561415 | 21.70944494 | 4.314159016 | ENSG00000179083 |
| PCDHGA3 | -2.353599774 | 0.195657216 | 0.01709271 | 0.049323696 | 9.955128266 | 1.924635205 | ENSG00000254245 |
| TEX26-AS1 | -2.354430281 | 0.195544616 | 0.049926115 | 0.119169102 | 6.648620849 | 1.28524071 | ENSG00000224743 |
| FOXB2 | -2.357413323 | 0.195140709 | 0.001323308 | 0.00548045 | 18.3645406 | 3.548395288 | ENSG00000204612 |
| FRZB | -2.374626534 | 0.192826263 | 3.38E-23 | 1.44E-21 | 171.8173987 | 33.04278477 | ENSG00000162998 |
| MT1L | -2.387505368 | 0.191112577 | 0.092883063 | 0.195292334 | 5.2605228 | 1.034913282 | ENSG00000260549 |
| ANKS4B | -2.387977377 | 0.191050061 | 0.048570449 | 0.116640425 | 6.961265867 | 1.354610529 | ENSG00000175311 |
| NA | -2.405586267 | 0.188732363 | 0.079097124 | 0.171908812 | 5.257731897 | 1.009639435 | ENSG00000287754 |
| NA | -2.423328257 | 0.186425581 | 0.04025506 | 0.100123823 | 7.026896093 | 1.304062836 | ENSG00000276436 |
| DLK2 | -2.43896646 | 0.184415719 | 0.011741436 | 0.036010181 | 10.8317756 | 2.021429444 | ENSG00000171462 |
| FBP1 | -2.440643761 | 0.184201439 | 1.70E-06 | 1.28E-05 | 37.34734997 | 6.861517164 | ENSG00000165140 |
| LOC100129484 | -2.44346744 | 0.183841268 | 0.06767541 | 0.151893699 | 5.366665128 | 0.986516162 | ENSG00000234432 |
| NA | -2.470843096 | 0.180385703 | 0.068320303 | 0.153002898 | 5.429504452 | 0.963392889 | ENSG00000228434 |
| NA | -2.476430492 | 0.179688441 | 1.92E-27 | 1.06E-25 | 214.9775977 | 38.52011641 | ENSG00000278630 |
| UNC45B | -2.478887251 | 0.17938271 | 0.034765894 | 0.088665899 | 7.443568256 | 1.356761103 | ENSG00000141161 |
| NA | -2.481586346 | 0.179047422 | 0.008495361 | 0.027442063 | 10.93492686 | 1.947758478 | ENSG00000235100 |
| LTK | -2.51905651 | 0.174457012 | 0.070391465 | 0.156785793 | 5.573167817 | 0.963392889 | ENSG00000062524 |
| ZNF439 | -2.531089133 | 0.173008025 | 0.020226134 | 0.056745541 | 9.715810463 | 1.72485547 | ENSG00000171291 |
| SLC7A4 | -2.536569203 | 0.172352102 | 0.056945912 | 0.132443741 | 5.72328887 | 0.984365589 | ENSG00000099960 |
| POU3F4 | -2.537650793 | 0.172222938 | 8.01E-05 | 0.000442114 | 24.64604982 | 4.215214203 | ENSG00000196767 |
| HAS1 | -2.551964067 | 0.170522727 | 5.66E-07 | 4.62E-06 | 38.57416336 | 6.609039162 | ENSG00000105509 |
| NA | -2.587900568 | 0.166327593 | 0.057309281 | 0.133151072 | 6.061052013 | 1.034913282 | ENSG00000279637 |
| TMEM114 | -2.59262 | 0.165784381 | 0.000393735 | 0.00186074 | 21.77575089 | 3.603244129 | ENSG00000232258 |
| NA | -2.608365423 | 0.163984866 | 0.015158626 | 0.044672326 | 10.11641293 | 1.674307777 | ENSG00000270344 |
| H2AC6 | -2.623929908 | 0.162225227 | 1.86E-65 | 4.84E-63 | 538.4221469 | 87.44474752 | ENSG00000180573 |

| | | | | | | | |
|---|---|---|---|---|---|---|---|
| ZNF229 | -2.652417923 | 0.159053285 | 1.13E-11 | 1.72E-10 | 76.1245494 | 12.07561793 | ENSG00000278318 |
| WDR97 | -2.667434499 | 0.157406334 | 0.002944397 | 0.011022777 | 16.49414016 | 2.572631995 | ENSG00000179698 |
| H2BC12 | -2.694606297 | 0.154469478 | 3.87E-49 | 6.12E-47 | 359.5234224 | 55.659246 | ENSG00000197903 |
| SHPK | -2.720038553 | 0.151770305 | 2.22E-19 | 7.07E-18 | 123.7391091 | 18.88684787 | ENSG00000197417 |
| ZNF568 | -2.723909565 | 0.151363623 | 5.40E-07 | 4.43E-06 | 43.37578703 | 6.5477386 | ENSG00000198453 |
| NA | -2.752136061 | 0.148430959 | 0.01534251 | 0.045103987 | 8.790478483 | 1.283090137 | ENSG00000231831 |
| NA | -2.922657066 | 0.131884135 | 0.007573036 | 0.02494399 | 9.999106991 | 1.329336683 | ENSG00000272468 |
| ZNF44 | -2.924483703 | 0.131717259 | 2.72E-14 | 5.48E-13 | 79.8295441 | 10.55080266 | ENSG00000197857 |
| TMED6 | -2.958869802 | 0.128614945 | 0.006789623 | 0.022818399 | 10.3250308 | 1.354610529 | ENSG00000157315 |
| NA | -3.031315479 | 0.122315956 | 0.039592693 | 0.098879359 | 5.402271144 | 0.664668341 | ENSG00000259869 |
| GPR85 | -3.053427429 | 0.120455533 | 0.044156322 | 0.108002993 | 5.432295355 | 0.643695642 | ENSG00000164604 |
| MT1M | -3.077049882 | 0.118499273 | 3.00E-05 | 0.00018097 | 22.23784123 | 2.637700666 | ENSG00000205364 |
| SIAH3 | -3.083499196 | 0.117970726 | 0.011230679 | 0.0347922 | 8.276036702 | 0.963392889 | ENSG00000215475 |
| CFAP65 | -3.085457994 | 0.117810661 | 0.010952828 | 0.034029795 | 8.450600159 | 1.009639435 | ENSG00000181378 |
| STX16-NPEPL1 | -3.095685652 | 0.116978423 | 0.003576684 | 0.013097533 | 11.25123866 | 1.30621341 | ENSG00000254995 |
| ZNF491 | -3.132989181 | 0.113992501 | 0.031585209 | 0.08189654 | 5.731661579 | 0.641545068 | ENSG00000177599 |
| ZNF829 | -3.185737216 | 0.109899961 | 1.05E-08 | 1.11E-07 | 41.89741619 | 4.63170569 | ENSG00000185869 |
| TBXT | -3.214318303 | 0.107744169 | 0.028413567 | 0.075188768 | 6.052679304 | 0.639394495 | ENSG00000164458 |
| NA | -3.327957128 | 0.099582971 | 0.005247583 | 0.018290067 | 9.879010149 | 0.988666736 | ENSG00000286964 |
| MAP6 | -3.371370854 | 0.096630949 | 7.58E-17 | 1.97E-15 | 104.9437785 | 10.05472153 | ENSG00000171533 |
| NUTM2B | -3.434925975 | 0.092466464 | 0.018846012 | 0.053440897 | 7.163062631 | 0.666818915 | ENSG00000188199 |
| ORM2 | -3.444997815 | 0.091823179 | 2.34E-05 | 0.000144165 | 21.50764817 | 1.998306171 | ENSG00000228278 |
| NA | -3.545154785 | 0.085664734 | 0.009688005 | 0.030617841 | 7.701746658 | 0.666818915 | ENSG00000279412 |
| MTTP | -3.560234162 | 0.08477401 | 0.009095818 | 0.029041282 | 7.707328464 | 0.641545068 | ENSG00000138823 |
| NA | -3.701121742 | 0.076886721 | 0.000127915 | 0.000672738 | 16.97153647 | 1.283090137 | ENSG00000283236 |
| ZNF441 | -3.730096107 | 0.075357969 | 9.79E-05 | 0.000530975 | 17.41544194 | 1.30621341 | ENSG00000197044 |
| PITX3 | -3.77006356 | 0.073298955 | 0.000489132 | 0.002246628 | 13.46622335 | 0.986516162 | ENSG00000107859 |
| ORM1 | -3.833869398 | 0.070127816 | 0.000537511 | 0.002445721 | 13.93927715 | 0.959091742 | ENSG00000229314 |
| NA | -3.928470313 | 0.065676893 | 0.002505988 | 0.009614758 | 10.03682819 | 0.666818915 | ENSG00000240093 |
| ZFP28 | -4.009342624 | 0.062096569 | 0.000175669 | 0.000895571 | 15.93659554 | 1.009639435 | ENSG00000196867 |
| ZNF563 | -4.052224135 | 0.060278021 | 3.74E-10 | 4.80E-09 | 43.66745627 | 2.637700666 | ENSG00000188868 |
| NA | -4.086138571 | 0.058877549 | 0.009768627 | 0.030822143 | 5.995421786 | 0.321847821 | ENSG00000229271 |
| PKIB | -4.663384566 | 0.039462206 | 1.54E-73 | 5.50E-71 | 383.1093379 | 15.1868095 | ENSG00000135549 |
| GDF7 | -4.753744957 | 0.03706638 | 1.37E-06 | 1.05E-05 | 26.61701078 | 0.984365589 | ENSG00000143869 |
| NKX3-2 | -5.47425816 | 0.022494901 | 3.41E-09 | 3.87E-08 | 43.93333166 | 1.009639435 | ENSG00000109705 |
| ZNF354C | -5.52761245 | 0.021678182 | 1.24E-06 | 9.60E-06 | 30.25223291 | 0.639394495 | ENSG00000177932 |
| ZNF502 | -5.649082319 | 0.019927682 | 2.35E-05 | 0.00014454 | 17.72808696 | 0.344971094 | ENSG00000196653 |
| TCEAL5 | -5.707550587 | 0.019136216 | 1.70E-15 | 3.86E-14 | 85.69602077 | 1.623760083 | ENSG00000204065 |
| DMRT2 | -5.741603223 | 0.018689824 | 1.81E-05 | 0.000113774 | 18.93691562 | 0.319697247 | ENSG00000173253 |
| LINC00664 | -5.762739762 | 0.018418 | 2.32E-10 | 3.08E-09 | 53.5108604 | 0.986516162 | ENSG00000268658 |
| ZNF595 | -6.122191821 | 0.014356105 | 1.15E-11 | 1.75E-10 | 68.80418119 | 1.009639435 | ENSG00000272602 |
| LOC105372321 | -6.151800586 | 0.014064474 | 4.63E-43 | 5.96E-41 | 280.1362691 | 3.916489655 | ENSG00000268119 |
| NA | -6.250204183 | 0.013137147 | 7.15E-06 | 4.84E-05 | 13.825638 | 0 | ENSG00000270325 |
| ZNF736 | -6.303595468 | 0.012659854 | 5.13E-20 | 1.74E-18 | 129.6285887 | 1.623760083 | ENSG00000234444 |
| ZNF572 | -6.761092111 | 0.009219523 | 3.97E-07 | 3.32E-06 | 19.66045084 | 0 | ENSG00000180938 |

**Table S3. Total gene expression using RNA-seq data in 3 biological replicates of SF3B1$^{K700E}$ versus SF3B1$^{WT}$ ES cells (K700E ES line 2).** Average baseMean, log2fold change, fold change (FC) and p-values are shown.

| Gene Symbol | log2 FC | FC | p-value | padj | WT baseMean | MT baseMean | Ensembl ID |
|---|---|---|---|---|---|---|---|
| CRYZ | 8.242771432 | 302.9154665 | 4.02E-29 | 4.57E-27 | 1.279874314 | 400.6412587 | ENSG00000116791 |
| ZXDA | 8.190694607 | 292.1761499 | 3.54E-11 | 8.44E-10 | 0 | 53.38454532 | ENSG00000198205 |
| HORMAD1 | 8.146699087 | 283.4006165 | 1.66E-11 | 4.11E-10 | 0.319968578 | 100.866641 | ENSG00000143452 |
| ZNF717 | 7.745476067 | 214.5955081 | 7.43E-10 | 1.45E-08 | 0 | 39.22583136 | ENSG00000227124 |
| LOC105373085 | 7.69043025 | 206.5618834 | 3.06E-25 | 2.81E-23 | 1.348673346 | 273.5252812 | ENSG00000286381 |
| NA | 7.676194491 | 204.5336621 | 1.43E-09 | 2.69E-08 | 0 | 37.35389021 | ENSG00000224848 |
| NA | 7.434273983 | 172.9575262 | 1.70E-09 | 3.17E-08 | 0.319968578 | 61.58966005 | ENSG00000265206 |
| CPA4 | 7.287506785 | 156.2277348 | 0.002121987 | 0.010709174 | 0 | 28.55344195 | ENSG00000128510 |
| KHDC3L | 7.092427378 | 136.4688032 | 5.14E-08 | 7.50E-07 | 0 | 24.94765513 | ENSG00000203908 |
| KRT80 | 7.048836913 | 132.4071219 | 0.071453941 | 0.185792051 | 0 | 24.19907051 | ENSG00000167767 |
| NNMT | 6.809423845 | 112.1607308 | 0.081672409 | 0.205049646 | 0 | 20.49890839 | ENSG00000166741 |
| TUBA3C | 6.737173608 | 106.6820474 | 0.08498025 | 0.211387252 | 0 | 19.49768404 | ENSG00000198033 |
| TYW3 | 6.463206412 | 88.23055235 | 5.68E-25 | 5.06E-23 | 2.050290591 | 175.4681275 | ENSG00000162623 |
| ZNF248 | 6.129825244 | 70.02631396 | 4.86E-15 | 1.84E-13 | 1.283433786 | 92.51505542 | ENSG00000198105 |
| NA | 6.12129724 | 69.61359808 | 4.82E-05 | 0.000389079 | 0 | 12.7353949 | ENSG00000233820 |
| PRAME | 6.044934624 | 66.0247315 | 0.000327191 | 0.002116823 | 0.701617244 | 44.91995124 | ENSG00000185686 |
| LTF | 5.942188469 | 61.48610311 | 3.92E-05 | 0.000323919 | 0 | 11.22692653 | ENSG00000012223 |
| NA | 5.865337566 | 58.29650812 | 7.67E-05 | 0.000586339 | 0 | 10.65888513 | ENSG00000286168 |
| CRYGD | 5.757325535 | 54.09133258 | 0.000122119 | 0.000888878 | 0 | 9.872160761 | ENSG00000118231 |
| NA | 5.686933935 | 51.51547372 | 0.131933201 | 0.292081713 | 0 | 9.415825726 | ENSG00000260604 |
| REG1A | 5.66857426 | 50.86404385 | 0.00016282 | 0.001147337 | 0 | 9.295417497 | ENSG00000115386 |
| PRAM1 | 5.664084035 | 50.70598138 | 0.000143714 | 0.001028828 | 0 | 9.260893626 | ENSG00000133246 |
| DUXAP10 | 5.634629287 | 49.68124032 | 0.134285059 | 0.295705847 | 0 | 9.080633968 | ENSG00000244306 |
| NA | 5.568766571 | 47.46415754 | 0.000251821 | 0.001686398 | 0 | 8.6819749 | ENSG00000275160 |
| DUXAP8 | 5.523224398 | 45.9892383 | 0.139263744 | 0.302994908 | 0 | 8.405899524 | ENSG00000206195 |
| NAP1L6P | 5.507566398 | 45.49280222 | 0.00028954 | 0.001905877 | 0 | 8.316610197 | ENSG00000204118 |
| LOC101928254 | 5.500549355 | 45.27206959 | 1.29E-06 | 1.46E-05 | 0.647056102 | 29.90364589 | ENSG00000219445 |
| NA | 5.454670015 | 43.85501773 | 0.00055917 | 0.003380259 | 0 | 8.022185154 | ENSG00000233309 |
| IL7R | 5.401695098 | 42.27389378 | 0.149169807 | 0.317559668 | 0 | 7.726814153 | ENSG00000168685 |
| MOV10L1 | 5.394077911 | 42.05128292 | 0.000620368 | 0.003693469 | 0 | 7.693519787 | ENSG00000073146 |
| MAGEA6 | 5.393743784 | 42.04154502 | 0.000634001 | 0.003766314 | 0 | 7.691344323 | ENSG00000197172 |
| NA | 5.383920711 | 41.75626338 | 0.000491899 | 0.00303678 | 0 | 7.630052477 | ENSG00000261435 |
| ZNF454 | 5.352430224 | 40.85470201 | 3.84E-06 | 3.98E-05 | 0.670777201 | 27.05318098 | ENSG00000178187 |
| NA | 5.309148702 | 39.64724464 | 1.36E-15 | 5.53E-14 | 2.019450547 | 78.83270717 | ENSG00000226539 |
| PRND | 5.295056549 | 39.26185818 | 0.163542245 | 0.338762235 | 0 | 7.173222985 | ENSG00000171864 |
| A2M | 5.271600942 | 38.62869251 | 0.151397283 | 0.321032941 | 0 | 7.060781563 | ENSG00000175899 |
| LINC00452 | 5.271600942 | 38.62869251 | 0.151397283 | 0.321032941 | 0 | 7.060781563 | ENSG00000229373 |
| NA | 5.259345478 | 38.30193777 | 0.000983458 | 0.005532499 | 0 | 7.004786602 | ENSG00000251598 |
| COL3A1 | 5.202709248 | 36.827441 | 0.038058374 | 0.114284049 | 0.319968578 | 13.11163692 | ENSG00000168542 |
| C6orf15 | 5.202333681 | 36.81785521 | 0.156854242 | 0.328933604 | 0 | 6.729940732 | ENSG00000204542 |
| NA | 5.190482354 | 36.51664593 | 0.001465445 | 0.007802347 | 0 | 6.679242657 | ENSG00000256540 |
| LINC00668 | 5.184308662 | 36.36071496 | 0.001150903 | 0.00634593 | 0 | 6.645948291 | ENSG00000265933 |
| NA | 5.152477829 | 35.56725745 | 0.000261945 | 0.001747998 | 0.323528051 | 12.66880617 | ENSG00000272405 |
| KRT81 | 5.13476368 | 35.13321458 | 9.76E-05 | 0.000725882 | 1.988610503 | 69.83792788 | ENSG00000205426 |
| TDRD1 | 5.132118094 | 35.06884703 | 1.08E-07 | 1.50E-06 | 0.970584153 | 34.68635488 | ENSG00000095627 |
| DPEP3 | 5.10978467 | 34.53014938 | 4.41E-100 | 1.19E-96 | 15.88160346 | 548.4444865 | ENSG00000141096 |
| F7 | 5.093349157 | 34.13900622 | 9.48E-08 | 1.33E-06 | 0.997864724 | 33.92282557 | ENSG00000057593 |
| MAGEA4 | 5.073056547 | 33.66217632 | 1.10E-35 | 1.73E-33 | 5.601980206 | 189.2650926 | ENSG00000147381 |
| GAB4 | 5.070949133 | 33.61304025 | 0.000324424 | 0.002103962 | 0.319968578 | 11.96730375 | ENSG00000215568 |
| CYTOR | 5.06928834 | 33.57436805 | 1.36E-05 | 0.000125081 | 0.670777201 | 22.25741921 | ENSG00000222041 |
| LINC00839 | 4.961274525 | 31.15246729 | 0.044304652 | 0.128211997 | 0.323528051 | 11.09178452 | ENSG00000185904 |
| ANKRD1 | 4.960190706 | 31.12907292 | 0.04831301 | 0.137449713 | 0.323528051 | 11.08308266 | ENSG00000148677 |
| NPAP1 | 4.852613072 | 28.89229842 | 0.000756587 | 0.004393647 | 0.350808622 | 10.28264311 | ENSG00000185823 |
| TBX3 | 4.820399335 | 28.25431553 | 5.67E-05 | 0.000450659 | 0.643496629 | 18.63377717 | ENSG00000135111 |
| DMRTB1 | 4.763140466 | 27.15489662 | 0.001160465 | 0.006389326 | 0.350808622 | 9.673551435 | ENSG00000143006 |
| NA | 4.750003248 | 26.90874587 | 0.00123433 | 0.006727367 | 0.350808622 | 9.568087905 | ENSG00000233854 |
| NA | 4.733605216 | 26.60462596 | 9.53E-05 | 0.000711629 | 0.643496629 | 17.5205629 | ENSG00000286186 |
| NA | 4.718506775 | 26.32764868 | 0.001785231 | 0.009221576 | 0.350808622 | 9.384706824 | ENSG00000270127 |
| MIR205HG | 4.716647294 | 26.29373699 | 1.08E-38 | 1.89E-36 | 9.51040218 | 251.7921965 | ENSG00000230937 |
| LOC101927314 | 4.612675894 | 24.46548593 | 5.07E-14 | 1.68E-12 | 2.690227748 | 64.91213974 | ENSG00000271860 |
| NA | 4.568536082 | 23.72828756 | 9.21E-09 | 1.52E-07 | 1.699481969 | 39.45527604 | ENSG00000267279 |
| NA | 4.546530855 | 23.3691095 | 0.002622351 | 0.012809907 | 0.319968578 | 8.321907083 | ENSG00000287106 |
| ICAM2 | 4.420757578 | 21.41808485 | 0.004317354 | 0.01956641 | 0.350808622 | 7.621350622 | ENSG00000108622 |
| NA | 4.417758917 | 21.37361329 | 0.003782899 | 0.017509818 | 0.350808622 | 7.602054997 | ENSG00000267769 |
| LOC101927793 | 4.360790884 | 20.54607461 | 0.004835744 | 0.021461568 | 0.350808622 | 7.315385849 | ENSG00000259881 |
| OR2W3 | 4.358664362 | 20.51581215 | 0.004630393 | 0.020701935 | 0.319968578 | 7.302333068 | ENSG00000238243 |
| SIX3 | 4.328593149 | 20.09261103 | 2.35E-05 | 0.000206341 | 0.990745779 | 19.9421958 | ENSG00000138083 |
| GIP | 4.327452113 | 20.07672595 | 0.000702779 | 0.004124293 | 0.643496629 | 13.2016613 | ENSG00000159224 |
| NA | 4.310975993 | 19.84874647 | 1.92E-11 | 4.73E-10 | 2.666506649 | 52.59498308 | ENSG00000287839 |
| MEG3 | 4.303574019 | 19.74717015 | 0.008683832 | 0.034951868 | 0.323528051 | 7.038080968 | ENSG00000214548 |
| NA | 4.296871869 | 19.65564591 | 0.00565328 | 0.024481515 | 0.350808622 | 7.00166518 | ENSG00000287477 |

| | | | | | | | |
|---|---|---|---|---|---|---|---|
| IZUMO2 | 4.246100974 | 18.97596007 | 1.69E-08 | 2.70E-07 | 1.968448877 | 37.6242172 | ENSG00000161652 |
| NA | 4.231746025 | 18.78808376 | 0.00835292 | 0.033858101 | 0.350808622 | 6.696646366 | ENSG00000287862 |
| SCG2 | 4.218302549 | 18.61382377 | 2.27E-15 | 8.98E-14 | 4.042460567 | 74.24162961 | ENSG00000171951 |
| TEX101 | 4.155477169 | 17.82063906 | 0.009102016 | 0.036235188 | 0.350808622 | 6.350577289 | ENSG00000131126 |
| C11orf16 | 4.154057553 | 17.80311213 | 0.009102464 | 0.036235188 | 0.319968578 | 6.34310494 | ENSG00000176029 |
| LGALS12 | 4.149642016 | 17.74870692 | 7.99E-05 | 0.000609071 | 0.997864724 | 17.64844348 | ENSG00000133317 |
| LINC01679 | 4.119904818 | 17.38661068 | 8.46E-05 | 0.000640384 | 0.990745779 | 17.24978441 | ENSG00000237989 |
| ANXA1 | 4.100254174 | 17.15139687 | 2.80E-10 | 5.83E-09 | 2.601267089 | 45.25304017 | ENSG00000135046 |
| PITX2 | 4.074062853 | 16.84283224 | 1.51E-14 | 5.29E-13 | 4.284146904 | 72.38434961 | ENSG00000164093 |
| NA | 4.051738025 | 16.58420586 | 7.65E-85 | 9.63E-82 | 26.78812118 | 444.3737236 | ENSG00000287064 |
| NPIPA5 | 4.041305562 | 16.46471417 | 0.000142705 | 0.001023153 | 0.963465208 | 16.23673679 | ENSG00000183793 |
| KRT17 | 4.038527269 | 16.4330375 | 1.06E-13 | 3.39E-12 | 5.278452155 | 86.97004662 | ENSG00000128422 |
| MLKL | 4.006410435 | 16.07125219 | 0.000581388 | 0.003498891 | 3.550494407 | 57.42425013 | ENSG00000168404 |
| DNAJC22 | 3.966491219 | 15.63265827 | 0.000247507 | 0.001662812 | 0.997864724 | 15.52530301 | ENSG00000178401 |
| MPO | 3.888265851 | 14.80759925 | 0.003515681 | 0.016458512 | 0.701617244 | 9.939978998 | ENSG00000005381 |
| LINC01139 | 3.854475941 | 14.46481474 | 2.49E-70 | 1.81E-67 | 29.10529375 | 417.9068087 | ENSG00000215808 |
| CYP2B6 | 3.851905541 | 14.43906622 | 4.67E-156 | 2.94E-152 | 60.50673429 | 874.1315679 | ENSG00000197408 |
| TCL1B | 3.845138652 | 14.37149909 | 1.08E-06 | 1.24E-05 | 2.012331602 | 28.64876321 | ENSG00000213231 |
| CNTN5 | 3.827175716 | 14.19366952 | 0.004733627 | 0.021098548 | 0.639937157 | 9.320010009 | ENSG00000149972 |
| ASB2 | 3.814918256 | 14.07358777 | 1.11E-09 | 2.13E-08 | 2.955635184 | 41.84295216 | ENSG00000100628 |
| SLC22A1 | 3.753544777 | 13.48744131 | 0.004925043 | 0.021800214 | 0.674336673 | 8.967698088 | ENSG00000175003 |
| TEX11 | 3.739480999 | 13.35660088 | 0.006328547 | 0.026923769 | 0.701617244 | 8.986047756 | ENSG00000120498 |
| LINC01414 | 3.721474826 | 13.1909341 | 0.001279372 | 0.006936817 | 0.970584153 | 13.0235771 | ENSG00000253554 |
| LOC101926892 | 3.692943781 | 12.93262993 | 0.006770288 | 0.028463243 | 0.674336673 | 8.594861037 | ENSG00000253967 |
| DPPA5 | 3.684131461 | 12.85387516 | 4.43E-25 | 3.98E-23 | 10.16338202 | 131.5190824 | ENSG00000203909 |
| NA | 3.653622261 | 12.5849036 | 0.007222724 | 0.030050925 | 0.643496629 | 8.264966164 | ENSG00000280355 |
| EPHA3 | 3.61663793 | 12.26638242 | 2.14E-08 | 3.34E-07 | 2.982915755 | 36.55299915 | ENSG00000044524 |
| EBF2 | 3.600286359 | 12.12813959 | 0.009731002 | 0.03816851 | 0.639937157 | 7.951245495 | ENSG00000221818 |
| NA | 3.5871186 | 12.01794734 | 0.009064213 | 0.036151564 | 0.674336673 | 7.999768106 | ENSG00000230650 |
| CER1 | 3.572584253 | 11.89748106 | 3.20E-15 | 1.25E-13 | 6.81661496 | 82.14234499 | ENSG00000147869 |
| GDF15 | 3.552840496 | 11.73576917 | 6.94E-85 | 9.36E-82 | 43.18274214 | 508.061081 | ENSG00000130513 |
| LINC00917 | 3.539800485 | 11.63017165 | 0.01205043 | 0.045696503 | 0.639937157 | 7.622580127 | ENSG00000168367 |
| GPRC5D-AS1 | 3.475584747 | 11.12385352 | 0.00018632 | 0.001294097 | 1.617640255 | 18.27806028 | ENSG00000247498 |
| PRSS3 | 3.453886554 | 10.95780214 | 0.014465117 | 0.052899637 | 0.674336673 | 7.289563833 | ENSG00000010438 |
| NA | 3.417667109 | 10.6861266 | 0.018054481 | 0.063298316 | 0.639937157 | 7.006962066 | ENSG00000286717 |
| NA | 3.417375899 | 10.68396981 | 0.019145481 | 0.066244487 | 0.639937157 | 7.002611139 | ENSG00000277349 |
| MAS1 | 3.397113581 | 10.53496474 | 0.017182365 | 0.060840479 | 0.670777201 | 7.006016107 | ENSG00000130368 |
| NA | 3.393062768 | 10.50542606 | 0.000285736 | 0.001885436 | 1.637801881 | 17.33566877 | ENSG00000272141 |
| S100A2 | 3.391054704 | 10.49081389 | 0.044920699 | 0.129674014 | 2.775628934 | 28.90074053 | ENSG00000196754 |
| OR52N2 | 3.385145892 | 10.44793478 | 0.016272122 | 0.058222626 | 0.670777201 | 6.933846943 | ENSG00000180988 |
| PKP1 | 3.380436192 | 10.41388295 | 5.36E-20 | 3.20E-18 | 9.943052517 | 103.3975939 | ENSG00000081277 |
| TRDN | 3.376301009 | 10.38407648 | 9.34E-07 | 1.09E-05 | 2.955635184 | 30.84764577 | ENSG00000186439 |
| NA | 3.375061303 | 10.37515729 | 0.000213896 | 0.001458805 | 1.692363024 | 17.29083468 | ENSG00000256424 |
| SCARA5 | 3.370952592 | 10.34565148 | 1.54E-05 | 0.000140147 | 2.670066121 | 27.4665012 | ENSG00000168079 |
| AGBL4 | 3.368642321 | 10.32909765 | 0.0037857 | 0.017518485 | 0.994305252 | 10.2515242 | ENSG00000186094 |
| TRIM58 | 3.343907492 | 10.153516 | 1.35E-23 | 1.09E-21 | 12.29315006 | 124.4638813 | ENSG00000162722 |
| F5 | 3.337622578 | 10.10937974 | 0.023182461 | 0.077541758 | 0.639937157 | 6.615775347 | ENSG00000198734 |
| GPAT2 | 3.337487023 | 10.10842991 | 2.84E-09 | 5.06E-08 | 4.297189585 | 43.46603741 | ENSG00000186281 |
| GSTT2 | 3.298223689 | 9.837036047 | 0.006264469 | 0.026699275 | 0.96702468 | 9.696252029 | ENSG00000099984 |
| NA | 3.292485101 | 9.79798512 | 0.001114666 | 0.00616412 | 1.31427383 | 12.94053062 | ENSG00000280202 |
| COL8A1 | 3.287789748 | 9.766148723 | 1.46E-06 | 1.63E-05 | 2.986475227 | 29.08797808 | ENSG00000144810 |
| PCDHGB5 | 3.282925969 | 9.733279396 | 6.31E-11 | 1.46E-09 | 6.013273708 | 58.17616708 | ENSG00000276547 |
| PAX7 | 3.28245932 | 9.730131619 | 1.40E-05 | 0.000128527 | 2.751907835 | 26.09802023 | ENSG00000009709 |
| NA | 3.272723729 | 9.664691817 | 0.001518095 | 0.008037351 | 1.372394445 | 12.94176012 | ENSG00000256417 |
| RASSF6 | 3.269151569 | 9.640791327 | 0.025795842 | 0.084384979 | 0.639937157 | 6.31510746 | ENSG00000169435 |
| NA | 3.268947851 | 9.639430077 | 0.030051611 | 0.095094617 | 0.639937157 | 6.30763511 | ENSG00000201699 |
| STMN2 | 3.267997783 | 9.633084248 | 2.90E-47 | 7.60E-45 | 29.76183306 | 284.7198593 | ENSG00000104435 |
| C1orf87 | 3.267568508 | 9.630218343 | 0.006125344 | 0.026195002 | 1.025145295 | 9.667025044 | ENSG00000162598 |
| NA | 3.266801866 | 9.625102245 | 0.005628556 | 0.024396439 | 1.025145295 | 9.657377231 | ENSG00000257869 |
| HSD17B7P2 | 3.251704375 | 9.524902833 | 0.000125905 | 0.000913102 | 1.995729448 | 18.9484438 | ENSG00000099251 |
| NECTIN4 | 3.249851878 | 9.5126802 | 0.000136659 | 0.000983162 | 1.992169976 | 18.90521808 | ENSG00000143217 |
| GPRC5D | 3.246688378 | 9.491843923 | 0.023775139 | 0.079117796 | 0.674336673 | 6.320404345 | ENSG00000111291 |
| LOC157273 | 3.244720767 | 9.478907351 | 2.02E-09 | 3.72E-08 | 5.191855761 | 49.84678765 | ENSG00000248538 |
| NA | 3.242876992 | 9.466800973 | 0.023089501 | 0.077271883 | 0.674336673 | 6.298933256 | ENSG00000224892 |
| NA | 3.234495158 | 9.411959794 | 0.000497539 | 0.003062703 | 1.672201397 | 15.63199605 | ENSG00000248213 |
| MYL7 | 3.206346012 | 9.230098297 | 2.03E-05 | 0.000180305 | 2.608386034 | 24.33014515 | ENSG00000106631 |
| NA | 3.204169961 | 9.216186814 | 0.042449062 | 0.124306178 | 0.639937157 | 6.03033023 | ENSG00000271267 |
| VAV1 | 3.188480854 | 9.116505065 | 7.47E-39 | 1.33E-36 | 27.81118154 | 251.6480035 | ENSG00000141968 |
| NA | 3.182389306 | 9.07809324 | 0.011115272 | 0.042656351 | 0.997864724 | 9.042042716 | ENSG00000254739 |
| LOC283299 | 3.182363526 | 9.077931019 | 0.002226196 | 0.011142783 | 1.317833303 | 12.01024593 | ENSG00000254951 |
| NCF1 | 3.16888659 | 8.993524381 | 0.028325172 | 0.090787655 | 0.674336673 | 5.989563514 | ENSG00000158517 |
| TMEM82 | 3.142154648 | 8.828416223 | 1.75E-06 | 1.93E-05 | 3.302884333 | 29.19221211 | ENSG00000162460 |
| TBX1 | 3.137191388 | 8.798096273 | 0.000452565 | 0.00282069 | 1.937608833 | 17.32725046 | ENSG00000184058 |
| EVPL | 3.135624518 | 8.788546097 | 1.11E-21 | 7.59E-20 | 15.40862987 | 136.2147931 | ENSG00000167880 |

| | | | | | | | |
|---|---|---|---|---|---|---|---|
| FMOD | 3.126126086 | 8.730874029 | 0.010814284 | 0.04167063 | 0.994305252 | 8.680745395 | ENSG00000122176 |
| NA | 3.107348409 | 8.617971989 | 0.000774855 | 0.004485943 | 1.665082452 | 14.28999951 | ENSG00000236212 |
| ZNF648 | 3.103317351 | 8.593925982 | 0.00079831 | 0.004607605 | 1.665082452 | 14.24053094 | ENSG00000179930 |
| LEFTY2 | 3.089445072 | 8.511686838 | 1.91E-120 | 6.01E-117 | 134.4144698 | 1139.740463 | ENSG00000143768 |
| MCCC2 | 3.08255541 | 8.471135758 | 9.39E-06 | 9.03E-05 | 2.986475227 | 25.25702487 | ENSG00000131844 |
| SRRM5 | 3.073941731 | 8.420709129 | 0.000161906 | 0.001142175 | 2.346538071 | 19.59830219 | ENSG00000226763 |
| NA | 3.07262975 | 8.413054851 | 0.000344789 | 0.002215496 | 1.961329932 | 16.64721913 | ENSG00000259345 |
| TTC22 | 3.067111079 | 8.380934272 | 1.37E-18 | 7.21E-17 | 12.94020616 | 108.2753835 | ENSG00000006555 |
| BTBD16 | 3.065053299 | 8.368988698 | 0.015334173 | 0.055454969 | 0.997864724 | 8.329379432 | ENSG00000138152 |
| LOC102724770 | 3.04682613 | 8.263919093 | 0.011831122 | 0.044958168 | 1.025145295 | 8.290788181 | ENSG00000278817 |
| PRPH2 | 3.026024805 | 8.145621647 | 0.000456931 | 0.002845085 | 2.015891075 | 16.25291099 | ENSG00000112619 |
| GDA | 3.010693712 | 8.059518846 | 0.004205384 | 0.019187889 | 1.31427383 | 10.64800781 | ENSG00000119125 |
| NA | 2.993388002 | 7.963419186 | 0.005323356 | 0.023298163 | 1.345113874 | 10.59513427 | ENSG00000228157 |
| SYT4 | 2.989073274 | 7.939638236 | 4.92E-35 | 7.50E-33 | 27.69702525 | 220.3241896 | ENSG00000132872 |
| CTCFL | 2.939237674 | 7.67005899 | 3.09E-11 | 7.45E-10 | 9.532038347 | 71.92250678 | ENSG00000124092 |
| LINC00707 | 2.937314346 | 7.65984047 | 0.0001116 | 0.000820387 | 2.659387704 | 20.32373471 | ENSG00000238266 |
| NA | 2.926903381 | 7.604763491 | 0.008954023 | 0.035810405 | 1.403234489 | 10.30628966 | ENSG00000241439 |
| KIF25 | 2.90523159 | 7.491380444 | 0.000999751 | 0.005614124 | 2.019450547 | 14.97003084 | ENSG00000125337 |
| LINC01508 | 2.892349044 | 7.424783937 | 0.010239107 | 0.039803528 | 1.372394445 | 9.972043859 | ENSG00000231107 |
| ACTA1 | 2.892002181 | 7.422999034 | 1.41E-163 | 1.33E-159 | 199.4105523 | 1480.560833 | ENSG00000143632 |
| ZIC1 | 2.891364041 | 7.419716382 | 7.17E-07 | 8.50E-06 | 5.080648543 | 37.1673877 | ENSG00000152977 |
| SATB2-AS1 | 2.88062935 | 7.364713231 | 8.16E-05 | 0.000621236 | 2.97579681 | 21.9296998 | ENSG00000225953 |
| NA | 2.880059903 | 7.36180687 | 0.02225638 | 0.075069074 | 0.990745779 | 7.302333068 | ENSG00000287150 |
| LOC400499 | 2.876477905 | 7.343551271 | 1.76E-17 | 8.31E-16 | 16.25613318 | 119.2643745 | ENSG00000188897 |
| CTSLP2 | 2.87643089 | 7.34331196 | 0.000423315 | 0.002656798 | 2.724627264 | 19.69194244 | ENSG00000280913 |
| NINJ2 | 2.870919562 | 7.315312838 | 0.003776498 | 0.017488762 | 1.61052131 | 11.99312577 | ENSG00000171840 |
| NA | 2.851565139 | 7.217829873 | 0.003403335 | 0.016039924 | 1.668641925 | 11.98565342 | ENSG00000235072 |
| NA | 2.829142993 | 7.106518694 | 0.000152049 | 0.001080713 | 2.9900347 | 21.18714712 | ENSG00000234273 |
| NA | 2.820398641 | 7.063575487 | 0.016347576 | 0.05844831 | 1.403234489 | 9.587383531 | ENSG00000157306 |
| FRMD1 | 2.820234895 | 7.062773817 | 0.029863138 | 0.094577454 | 0.963465208 | 6.92419913 | ENSG00000153303 |
| PLA2G4C | 2.818707936 | 7.055302479 | 1.47E-21 | 9.99E-20 | 19.23312519 | 135.5407211 | ENSG00000105499 |
| DUSP27 | 2.810987537 | 7.017647766 | 1.54E-06 | 1.72E-05 | 4.620717636 | 32.46789315 | ENSG00000198842 |
| NA | 2.795802933 | 6.944173205 | 0.028002158 | 0.090012043 | 1.025145295 | 6.978964586 | ENSG00000267733 |
| TCEA3 | 2.792552031 | 6.928543146 | 0.026962523 | 0.087459163 | 1.025145295 | 6.957493496 | ENSG00000204219 |
| SERPINA1 | 2.786127424 | 6.897757568 | 0.059799971 | 0.161932333 | 1.052425867 | 7.067307954 | ENSG00000197249 |
| UNC5C | 2.78576822 | 6.896040368 | 7.68E-16 | 3.21E-14 | 14.33988118 | 98.38929668 | ENSG00000182168 |
| PPP1R36 | 2.771587809 | 6.828590448 | 0.025573034 | 0.083874045 | 1.279874314 | 8.889002534 | ENSG00000165807 |
| RGN | 2.766233541 | 6.803294499 | 3.24E-06 | 3.39E-05 | 5.404176594 | 36.37368545 | ENSG00000130988 |
| LINC-ROR | 2.759764686 | 6.772857708 | 2.49E-16 | 1.07E-14 | 14.58393178 | 98.69140498 | ENSG00000258609 |
| UNC93A | 2.751908731 | 6.736077472 | 1.50E-05 | 0.000137018 | 4.379031299 | 29.18038883 | ENSG00000112494 |
| NA | 2.747095655 | 6.713642216 | 0.001358036 | 0.007312947 | 2.332300181 | 15.58999983 | ENSG00000258636 |
| HLA-DQA1 | 2.746997971 | 6.713187656 | 2.41E-09 | 4.39E-08 | 8.260172702 | 55.51326622 | ENSG00000196735 |
| NA | 2.746419513 | 6.7104965 | 0.012427101 | 0.046810804 | 1.345113874 | 8.927877332 | ENSG00000265055 |
| POT1-AS1 | 2.735323386 | 6.65908237 | 0.025664974 | 0.084082043 | 1.279874314 | 8.690676754 | ENSG00000224897 |
| NA | 2.732622906 | 6.646629361 | 0.0357136 | 0.10893097 | 1.021585823 | 6.680472162 | ENSG00000277504 |
| ZNF385D-AS1 | 2.726027246 | 6.616311897 | 0.016889086 | 0.059986909 | 1.283433786 | 8.658328346 | ENSG00000225542 |
| NA | 2.723621363 | 6.605287528 | 0.005563676 | 0.02414293 | 1.668641925 | 10.9648499 | ENSG00000233058 |
| KIF25-AS1 | 2.721369532 | 6.594985705 | 2.38E-08 | 3.66E-07 | 7.205382571 | 47.82163836 | ENSG00000229921 |
| LINC00504 | 2.702813388 | 6.510703268 | 0.001003542 | 0.005630392 | 3.129997029 | 19.84543412 | ENSG00000248360 |
| MFAP5 | 2.69836558 | 6.490661777 | 0.081269004 | 0.204199725 | 0.970584153 | 6.377345264 | ENSG00000197614 |
| LAIR1 | 2.689104159 | 6.449128245 | 0.017949969 | 0.063000047 | 1.372394445 | 8.667976159 | ENSG00000167613 |
| NA | 2.689062136 | 6.448940399 | 0.040107899 | 0.11896304 | 0.963465208 | 6.334403086 | ENSG00000254683 |
| NTSR2 | 2.687206168 | 6.440649433 | 0.047674389 | 0.135971453 | 0.970584153 | 6.340929476 | ENSG00000169006 |
| ANKDD1B | 2.6865147 | 6.43756324 | 0.007352575 | 0.030471696 | 1.634242409 | 10.59636377 | ENSG00000189045 |
| NA | 2.672685589 | 6.376150085 | 1.30E-10 | 2.84E-09 | 9.561403851 | 61.10946756 | ENSG00000286699 |
| NA | 2.667328928 | 6.352519627 | 0.000308347 | 0.002014856 | 2.982915755 | 18.9322696 | ENSG00000250131 |
| RGS4 | 2.664732755 | 6.341098356 | 0.005034848 | 0.022226064 | 1.95777046 | 12.49675391 | ENSG00000117152 |
| PRSS21 | 2.658266998 | 6.312742912 | 0.009867977 | 0.038607156 | 1.723203067 | 10.62218579 | ENSG00000007038 |
| CACNA1C-AS1 | 2.647570503 | 6.266111766 | 0.040850057 | 0.120696533 | 1.025145295 | 6.296757792 | ENSG00000246627 |
| LOC343052 | 2.644385676 | 6.252294244 | 0.000271412 | 0.001802256 | 3.395404465 | 20.90700436 | ENSG00000231827 |
| CFAP300 | 2.643451691 | 6.248247887 | 3.10E-07 | 3.96E-06 | 6.687610381 | 41.53452838 | ENSG00000137691 |
| FAM90A1 | 2.642486171 | 6.244067665 | 0.001030353 | 0.005755168 | 2.666506649 | 16.57287451 | ENSG00000171847 |
| LINC00304 | 2.625190668 | 6.169658637 | 0.003452658 | 0.016231875 | 2.339419126 | 14.36841152 | ENSG00000180422 |
| NA | 2.616964755 | 6.134580769 | 0.031366984 | 0.098421802 | 1.290552731 | 8.042710285 | ENSG00000235192 |
| NA | 2.608002013 | 6.096587826 | 0.011258076 | 0.043099284 | 2.07401169 | 12.38090752 | ENSG00000248587 |
| MIR4435-2HG | 2.60191228 | 6.07090787 | 5.10E-07 | 6.25E-06 | 8.432170283 | 50.75108236 | ENSG00000172965 |
| NA | 2.598974362 | 6.058557595 | 0.021558063 | 0.073118539 | 1.31427383 | 8.005064992 | ENSG00000213514 |
| NA | 2.588228991 | 6.013600343 | 0.0212247 | 0.07216933 | 1.317833303 | 7.941597682 | ENSG00000229953 |
| NA | 2.585990177 | 6.004275508 | 0.010086313 | 0.039322879 | 1.668641925 | 9.968922436 | ENSG00000266217 |
| SOX15 | 2.563814105 | 5.912687812 | 2.77E-27 | 2.86E-25 | 31.08531076 | 183.8609742 | ENSG00000129194 |
| LINC00989 | 2.562008844 | 5.905293824 | 0.009076939 | 0.036187039 | 1.950651515 | 11.63740888 | ENSG00000250334 |
| PLAAT4 | 2.557108771 | 5.885270684 | 0.007896026 | 0.032332276 | 1.964889405 | 11.63863838 | ENSG00000133321 |
| SP140L | 2.552631183 | 5.867033312 | 3.78E-09 | 6.62E-08 | 9.530563807 | 56.15319325 | ENSG00000185404 |
| NA | 2.549112992 | 5.852743253 | 0.006711036 | 0.028270783 | 1.985051031 | 11.61905921 | ENSG00000260455 |

| | | | | | | | |
|---|---|---|---|---|---|---|---|
| KRT42P | 2.547335399 | 5.845536331 | 0.061668653 | 0.165733956 | 1.052425867 | 5.949742757 | ENSG00000214514 |
| CHRM1 | 2.537980872 | 5.807756116 | 0.071397559 | 0.185671014 | 0.963465208 | 5.699489398 | ENSG00000168539 |
| KCNJ9 | 2.5353758 | 5.797278524 | 0.00622948 | 0.026580148 | 2.274179566 | 13.32547449 | ENSG00000162728 |
| SLC22A2 | 2.533241911 | 5.788710111 | 0.000838668 | 0.004808213 | 2.982915755 | 17.24760895 | ENSG00000112499 |
| NA | 2.530496493 | 5.777704797 | 0.073847516 | 0.190624705 | 0.963465208 | 5.66307361 | ENSG00000286859 |
| LOC105369332 | 2.527048651 | 5.763913366 | 0.030258091 | 0.095620896 | 1.345113874 | 7.691344323 | ENSG00000256690 |
| AADACL3 | 2.523797682 | 5.750939585 | 0.015715562 | 0.056595802 | 1.692363024 | 9.600436312 | ENSG00000188984 |
| NA | 2.523786902 | 5.750896613 | 0.029037478 | 0.092539073 | 1.341554401 | 7.658049957 | ENSG00000229334 |
| LINC02588 | 2.523349182 | 5.74915203 | 0.0291215 | 0.092722899 | 1.31427383 | 7.57623298 | ENSG00000257842 |
| NA | 2.514821428 | 5.715269097 | 1.45E-07 | 1.95E-06 | 7.92360197 | 45.31506706 | ENSG00000287238 |
| NA | 2.513232027 | 5.708976118 | 7.34E-10 | 1.43E-08 | 11.40440762 | 64.58129891 | ENSG00000280623 |
| VSIG1 | 2.512826304 | 5.707370832 | 1.18E-20 | 7.40E-19 | 25.37776775 | 145.1254347 | ENSG00000101842 |
| CGB2 | 2.510910978 | 5.699798737 | 0.003468917 | 0.016296142 | 2.62854766 | 15.06272513 | ENSG00000104818 |
| NA | 2.508170526 | 5.688982038 | 0.060469074 | 0.16332358 | 0.997864724 | 5.664019569 | ENSG00000254186 |
| PLD6 | 2.506477712 | 5.682310676 | 7.29E-06 | 7.19E-05 | 5.911270367 | 33.70972304 | ENSG00000179598 |
| SERPINB9 | 2.50271778 | 5.667520796 | 2.47E-126 | 9.32E-123 | 228.8358408 | 1296.265401 | ENSG00000170542 |
| NA | 2.494815093 | 5.636560548 | 0.000745807 | 0.004343066 | 4.193991036 | 23.93649942 | ENSG00000180066 |
| LINC01162 | 2.493580025 | 5.631737246 | 0.016592629 | 0.059156466 | 1.634242409 | 9.265244554 | ENSG00000232790 |
| SEMA3C | 2.493416157 | 5.631097603 | 7.45E-05 | 0.000571592 | 4.967966786 | 27.97715019 | ENSG00000075223 |
| PIWIL2 | 2.481779171 | 5.585859054 | 5.83E-07 | 7.02E-06 | 8.552860709 | 47.94299254 | ENSG00000197181 |
| LEFTY1 | 2.476284883 | 5.564626602 | 4.82E-89 | 7.00E-86 | 134.7943384 | 748.6697517 | ENSG00000243709 |
| NA | 2.471516051 | 5.546263084 | 9.03E-07 | 1.05E-05 | 7.191144681 | 40.17134429 | ENSG00000277883 |
| NA | 2.471315378 | 5.545491672 | 0.000114264 | 0.000836708 | 4.71679724 | 25.89628948 | ENSG00000270977 |
| MUC4 | 2.468176593 | 5.533439795 | 4.73E-13 | 1.41E-11 | 18.44491156 | 102.3188309 | ENSG00000145113 |
| NEAT1 | 2.466539594 | 5.527164665 | 1.00E-05 | 9.54E-05 | 29.3090211 | 161.8779197 | ENSG00000245532 |
| C14orf39 | 2.463351171 | 5.514962862 | 7.85E-07 | 9.23E-06 | 7.314504856 | 40.24322991 | ENSG00000179008 |
| GJC2 | 2.45494283 | 5.482913945 | 1.84E-19 | 1.06E-17 | 28.17711778 | 154.2784058 | ENSG00000198835 |
| PTPN7 | 2.44252535 | 5.435924241 | 0.044579854 | 0.128827932 | 1.372394445 | 7.317561313 | ENSG00000143851 |
| NA | 2.438685311 | 5.421474613 | 0.007506795 | 0.031014181 | 2.339419126 | 12.61309476 | ENSG00000253854 |
| ATP1A4 | 2.434260738 | 5.404873073 | 0.000582873 | 0.003504476 | 3.919100392 | 21.35607796 | ENSG00000132681 |
| CDKN2C | 2.431142484 | 5.393203547 | 0.050380776 | 0.141921383 | 1.699481969 | 9.071932113 | ENSG00000123080 |
| AGMO | 2.419654573 | 5.350429003 | 0.015644849 | 0.056416453 | 1.995729448 | 10.6545342 | ENSG00000187546 |
| NA | 2.412597573 | 5.324321074 | 0.051212107 | 0.14374861 | 1.317833303 | 7.046782823 | ENSG00000243155 |
| VSIR | 2.408002877 | 5.307391148 | 0.000815704 | 0.004690798 | 3.626412384 | 19.29640479 | ENSG00000107738 |
| NA | 2.406225104 | 5.300855096 | 0.044705357 | 0.129111586 | 1.31427383 | 7.000435675 | ENSG00000199691 |
| EZHIP | 2.39924442 | 5.275268113 | 0.051229571 | 0.143776258 | 1.341554401 | 7.03713501 | ENSG00000187690 |
| MEIOB | 2.393883022 | 5.255700344 | 8.99E-25 | 7.79E-23 | 36.66740868 | 191.955045 | ENSG00000162039 |
| NA | 2.392158688 | 5.249422394 | 0.002408455 | 0.011934837 | 2.986475227 | 15.66529041 | ENSG00000267505 |
| NA | 2.39136084 | 5.246520126 | 0.045270154 | 0.130423624 | 1.348673346 | 7.00913753 | ENSG00000125514 |
| CCDC80 | 2.386078927 | 5.227346984 | 5.76E-33 | 8.00E-31 | 51.72611964 | 270.9814435 | ENSG00000091986 |
| IFIT3 | 2.382905685 | 5.215861946 | 0.001046451 | 0.005827848 | 3.592012868 | 18.86568087 | ENSG00000119917 |
| TREH | 2.382337974 | 5.213809873 | 0.041595447 | 0.12229955 | 1.345113874 | 6.941319293 | ENSG00000118094 |
| NA | 2.378516001 | 5.200015779 | 0.045935835 | 0.131958967 | 1.348673346 | 6.93167148 | ENSG00000229282 |
| NHLRC1 | 2.366367284 | 5.156411085 | 2.03E-08 | 3.18E-07 | 10.55214963 | 54.53975581 | ENSG00000187566 |
| TSKS | 2.364431712 | 5.149497693 | 0.012225867 | 0.046251102 | 2.683108803 | 13.74778012 | ENSG00000126467 |
| LINC02078 | 2.362437882 | 5.142385913 | 0.005761473 | 0.024864126 | 2.662947177 | 13.65820724 | ENSG00000267719 |
| COLEC11 | 2.360083957 | 5.134002353 | 1.69E-05 | 0.000153198 | 6.833217114 | 35.37073714 | ENSG00000118004 |
| CHADL | 2.348131329 | 5.091643216 | 0.008228608 | 0.033447538 | 2.597707617 | 13.37806449 | ENSG00000100399 |
| MS4A15 | 2.34449367 | 5.078821149 | 0.000240311 | 0.00162023 | 5.087767488 | 25.4879826 | ENSG00000166961 |
| H2BW1 | 2.342227577 | 5.070849922 | 0.058393164 | 0.159043973 | 1.283433786 | 6.614829389 | ENSG00000123569 |
| LINC00607 | 2.340993599 | 5.066514536 | 0.006497879 | 0.02756344 | 2.604826561 | 13.32736641 | ENSG00000235770 |
| NA | 2.337230991 | 5.053318073 | 0.034031493 | 0.10497954 | 1.634242409 | 8.317556156 | ENSG00000196796 |
| SLC25A3P1 | 2.333650389 | 5.040791873 | 0.076938832 | 0.196190904 | 1.294112204 | 6.589953331 | ENSG00000236253 |
| TNN | 2.330604741 | 5.030161571 | 4.28E-13 | 1.29E-11 | 20.25827049 | 101.7398168 | ENSG00000120332 |
| NA | 2.329822745 | 5.027435771 | 0.014680755 | 0.053605194 | 2.274179566 | 11.52882393 | ENSG00000249199 |
| NA | 2.325195624 | 5.011337227 | 0.009452727 | 0.037299363 | 2.312138554 | 11.5944667 | ENSG00000229646 |
| SDR16C5 | 2.309281409 | 4.956361474 | 0.050978611 | 0.143306202 | 1.345113874 | 6.596479721 | ENSG00000170786 |
| HR | 2.30487987 | 4.941263085 | 0.031525268 | 0.098846867 | 1.937608833 | 9.705899842 | ENSG00000168453 |
| USH2A | 2.297441709 | 4.915852775 | 0.043820602 | 0.127315704 | 1.603402365 | 8.023414659 | ENSG00000042781 |
| NLRP10 | 2.29560836 | 4.909609772 | 0.027346772 | 0.088371628 | 1.964889405 | 9.715547655 | ENSG00000182261 |
| CSRP1 | 2.289339564 | 4.888322825 | 2.38E-05 | 0.000208006 | 7.318064329 | 35.71077428 | ENSG00000159176 |
| FAT4 | 2.289142662 | 4.887655703 | 5.98E-27 | 6.10E-25 | 48.32507078 | 236.0571827 | ENSG00000196159 |
| LOC101927447 | 2.281709429 | 4.862537677 | 0.078324258 | 0.19870414 | 1.279874314 | 6.336578549 | ENSG00000203527 |
| NA | 2.280324024 | 4.857870473 | 0.045214864 | 0.13030409 | 1.630682936 | 7.984539861 | ENSG00000280164 |
| GPER1 | 2.27806816 | 4.850280423 | 0.048010493 | 0.13669202 | 1.644920826 | 8.010361878 | ENSG00000164850 |
| RERG | 2.277956012 | 4.8499034 | 0.008061241 | 0.032866262 | 2.69378722 | 12.93835516 | ENSG00000134533 |
| CFHR3 | 2.276988012 | 4.84665037 | 0.013857093 | 0.051061699 | 2.697346693 | 12.95547532 | ENSG00000116785 |
| NA | 2.275748016 | 4.842486464 | 0.031843729 | 0.099663378 | 1.637801881 | 7.981418439 | ENSG00000268401 |
| LINC01814 | 2.274761647 | 4.839176793 | 0.053148389 | 0.147777874 | 1.66152298 | 8.034008431 | ENSG00000236008 |
| MIR2052HG | 2.263837172 | 4.802671631 | 0.025052157 | 0.082610439 | 2.046731118 | 9.682253289 | ENSG00000254349 |
| ADGRD1 | 2.260727234 | 4.792329927 | 0.000954335 | 0.005390555 | 5.219136332 | 25.16962746 | ENSG00000111452 |
| MRPS9-AS2 | 2.260023733 | 4.789993617 | 0.063866109 | 0.170083982 | 1.317833303 | 6.333457127 | ENSG00000224509 |
| CHRNG | 2.259245548 | 4.787410604 | 0.058648177 | 0.159546359 | 1.31427383 | 6.320404345 | ENSG00000196811 |
| NA | 2.257404078 | 4.781303804 | 0.041986654 | 0.123219575 | 1.692363024 | 8.000714065 | ENSG00000256654 |

| | | | | | | | |
|---|---|---|---|---|---|---|---|
| NPIPA2 | 2.254439637 | 4.7714893 | 0.073013731 | 0.189092747 | 1.341554401 | 6.366751493 | ENSG00000254852 |
| NA | 2.248934814 | 4.753317643 | 0.074898484 | 0.192383452 | 1.348673346 | 6.358049639 | ENSG00000233695 |
| CCDC87 | 2.24860945 | 4.752245773 | 0.000174643 | 0.001220146 | 5.322334881 | 25.21502865 | ENSG00000182791 |
| HAND2-AS1 | 2.245913783 | 4.743374521 | 0.028070736 | 0.090171091 | 1.934049361 | 9.308186733 | ENSG00000237125 |
| NA | 2.242408516 | 4.731863693 | 0.04850075 | 0.137859222 | 1.72676254 | 8.00194357 | ENSG00000258131 |
| CALHM3 | 2.235517461 | 4.709315768 | 0.027338219 | 0.088359107 | 1.985051031 | 9.362006229 | ENSG00000183128 |
| NA | 2.23343687 | 4.702529096 | 0.079062397 | 0.200012508 | 1.372394445 | 6.341875435 | ENSG00000249577 |
| ARID5B | 2.233408059 | 4.702435185 | 2.35E-05 | 0.000206292 | 7.392787098 | 34.46627446 | ENSG00000150347 |
| DENND2D | 2.231034553 | 4.694705153 | 8.13E-07 | 9.55E-06 | 9.905093528 | 46.57791654 | ENSG00000162777 |
| NA | 2.230067543 | 4.691559437 | 0.017432112 | 0.061556736 | 2.370259169 | 10.99190142 | ENSG00000244349 |
| CCDC146 | 2.229280089 | 4.688999381 | 0.008540568 | 0.034485488 | 3.251882663 | 15.39886285 | ENSG00000135205 |
| NA | 2.227681455 | 4.683806431 | 0.005627909 | 0.024396439 | 2.986475227 | 13.97315742 | ENSG00000259723 |
| DCC | 2.224090075 | 4.672161278 | 1.11E-05 | 0.000104662 | 8.670297148 | 40.32143101 | ENSG00000187323 |
| EID3 | 2.218297671 | 4.65344021 | 1.14E-05 | 0.000107398 | 7.916483025 | 36.91590484 | ENSG00000255150 |
| C2orf80 | 2.215395359 | 4.644088159 | 0.039955306 | 0.118593823 | 2.046731118 | 9.381301855 | ENSG00000188674 |
| TSPYL5 | 2.214899656 | 4.642492746 | 9.53E-33 | 1.30E-30 | 58.57860866 | 271.9482596 | ENSG00000180543 |
| IQSEC3 | 2.207068205 | 4.617359954 | 1.17E-17 | 5.64E-16 | 31.47763785 | 145.2867252 | ENSG00000120645 |
| BCAS1 | 2.206151747 | 4.614427753 | 0.001802791 | 0.009281818 | 3.949940435 | 18.28676214 | ENSG00000064787 |
| NA | 2.204917902 | 4.610483016 | 0.010341755 | 0.040111736 | 2.686668275 | 12.28632131 | ENSG00000286496 |
| RCSD1 | 2.200358822 | 4.595936363 | 4.94E-05 | 0.00039707 | 7.052656893 | 32.14792963 | ENSG00000198771 |
| NA | 2.199512093 | 4.593239763 | 0.057153013 | 0.156470665 | 1.644920826 | 7.566585167 | ENSG00000262089 |
| PTH1R | 2.195925489 | 4.581834946 | 2.64E-06 | 2.82E-05 | 10.2393 | 46.99123676 | ENSG00000160801 |
| NR1I2 | 2.193049496 | 4.572710219 | 6.18E-12 | 1.60E-10 | 19.85882446 | 90.75896066 | ENSG00000144852 |
| LINC02714 | 2.183477879 | 4.542472809 | 0.013609481 | 0.050296488 | 2.635666605 | 12.0136509 | ENSG00000251226 |
| TLE6 | 2.183425356 | 4.54230744 | 1.96E-09 | 3.62E-08 | 17.24095522 | 78.28757726 | ENSG00000104953 |
| FLJ22447 | 2.183287433 | 4.541873211 | 0.039837391 | 0.118383812 | 1.992169976 | 9.04828556 | ENSG00000232774 |
| NA | 2.182454867 | 4.539252893 | 0.005839559 | 0.025126983 | 3.244763718 | 14.87326916 | ENSG00000272129 |
| NA | 2.180969908 | 4.534583067 | 0.081620517 | 0.204946612 | 1.321392775 | 6.003562254 | ENSG00000276216 |
| ANGPT4 | 2.17876949 | 4.527672131 | 0.000229847 | 0.001556352 | 5.642303459 | 25.48892856 | ENSG00000101280 |
| PF4 | 2.176211243 | 4.519650589 | 0.005173379 | 0.022755147 | 3.964178325 | 17.93416667 | ENSG00000163737 |
| NA | 2.169414869 | 4.498409095 | 0.035192446 | 0.107716263 | 2.019450547 | 9.014045236 | ENSG00000276975 |
| DPPA3 | 2.158507305 | 4.464526907 | 1.07E-16 | 4.71E-15 | 34.91781476 | 156.2724791 | ENSG00000187569 |
| ODF3L2 | 2.155104009 | 4.454007565 | 0.089437011 | 0.219415763 | 1.348673346 | 5.943216367 | ENSG00000181781 |
| GRAMD1C | 2.151527826 | 4.442980551 | 0.000143254 | 0.001026696 | 6.705407743 | 29.60080254 | ENSG00000178075 |
| CAV1 | 2.145609117 | 4.424790399 | 1.39E-39 | 2.65E-37 | 78.33779405 | 346.8233624 | ENSG00000105974 |
| OTOF | 2.143141341 | 4.417228124 | 0.002148414 | 0.010802132 | 4.277027959 | 18.95932112 | ENSG00000115155 |
| TGFB2 | 2.13689539 | 4.398145677 | 0.000975389 | 0.005492019 | 5.818750236 | 25.87018392 | ENSG00000092969 |
| LINC01618 | 2.134621978 | 4.391220494 | 0.002068122 | 0.010473717 | 4.307868003 | 18.89897523 | ENSG00000250302 |
| PRDM1 | 2.134544502 | 4.390984683 | 0.011831723 | 0.044958168 | 3.071876414 | 13.27572238 | ENSG00000057657 |
| NA | 2.134309842 | 4.39027053 | 0.077951987 | 0.19807902 | 1.699481969 | 7.381974581 | ENSG00000275216 |
| SPAG17 | 2.13089513 | 4.379891496 | 0.005254479 | 0.023044781 | 3.629971857 | 15.91961115 | ENSG00000155761 |
| NA | 2.129125319 | 4.374521806 | 0.066634137 | 0.175973386 | 1.695922496 | 7.333735516 | ENSG00000225083 |
| RORC | 2.128186513 | 4.371676095 | 2.03E-05 | 0.000179947 | 8.52913961 | 37.44346308 | ENSG00000143365 |
| LCN15 | 2.126618439 | 4.366927076 | 0.01683397 | 0.0598249 | 2.941397294 | 12.91688407 | ENSG00000177984 |
| MIR1281 | 2.124572601 | 4.360738868 | 0.036732384 | 0.111221914 | 1.964889405 | 8.614156663 | ENSG00000284015 |
| IL18 | 2.123688224 | 4.35806654 | 0.009095933 | 0.036232167 | 3.571851242 | 15.73093319 | ENSG00000150782 |
| EMP1 | 2.122921725 | 4.355751729 | 0.000929047 | 0.005260884 | 5.257095321 | 23.00091785 | ENSG00000134531 |
| SERPINA11 | 2.121971085 | 4.352882524 | 0.034589601 | 0.106249937 | 1.985051031 | 8.644329606 | ENSG00000186910 |
| NA | 2.11645936 | 4.336284339 | 0.034212613 | 0.105366143 | 2.424820312 | 10.28699403 | ENSG00000251161 |
| IL1A | 2.114055141 | 4.329064037 | 0.011345632 | 0.043399282 | 3.264925345 | 14.21783034 | ENSG00000115008 |
| PANCR | 2.112842826 | 4.325427796 | 0.083242258 | 0.207995508 | 1.723203067 | 7.31945323 | ENSG00000250103 |
| NA | 2.110224609 | 4.317585085 | 4.11E-06 | 4.23E-05 | 10.05929375 | 43.12242735 | ENSG00000285689 |
| NA | 2.108527799 | 4.312509991 | 0.067705879 | 0.178011494 | 1.72676254 | 7.287388369 | ENSG00000266573 |
| HAO2 | 2.108091108 | 4.31120483 | 0.052721251 | 0.146909055 | 2.346538071 | 10.07873689 | ENSG00000116882 |
| MECOM | 2.107373181 | 4.309059982 | 0.000270772 | 0.001799272 | 7.300266966 | 31.39109467 | ENSG00000085276 |
| RGS7 | 2.102618574 | 4.294882237 | 1.25E-14 | 4.47E-13 | 28.58368275 | 122.4788363 | ENSG00000182901 |
| APOA5 | 2.101127825 | 4.290446592 | 0.085967817 | 0.213141948 | 1.31427383 | 5.666195032 | ENSG00000110243 |
| NA | 2.099509011 | 4.285635082 | 0.114132698 | 0.263431434 | 1.286993259 | 5.582202592 | ENSG00000232807 |
| ANGPTL6 | 2.096425854 | 4.27648611 | 0.043687887 | 0.127059095 | 2.050290591 | 8.623804476 | ENSG00000130812 |
| ADGRE5 | 2.094905842 | 4.27198281 | 8.90E-16 | 3.70E-14 | 33.58248958 | 143.1337907 | ENSG00000123146 |
| SMC1B | 2.094414199 | 4.270527248 | 0.097710353 | 0.234042999 | 1.321392775 | 5.650020829 | ENSG00000077935 |
| SGCD | 2.091093565 | 4.260709129 | 0.06511628 | 0.172616641 | 1.61052131 | 6.95966896 | ENSG00000170624 |
| CDKN1A | 2.088734977 | 4.253749207 | 1.28E-81 | 1.51E-78 | 236.2146949 | 1006.171369 | ENSG00000124762 |
| NA | 2.087912709 | 4.251325462 | 0.053255477 | 0.147923177 | 2.043171646 | 8.552864816 | ENSG00000229835 |
| ARSL | 2.085370671 | 4.243841195 | 1.54E-38 | 2.68E-36 | 88.83301827 | 377.441112 | ENSG00000157399 |
| SNORA65 | 2.082701006 | 4.235995352 | 0.030284398 | 0.095686942 | 2.397539741 | 9.982921177 | ENSG00000201302 |
| LMX1B | 2.080108259 | 4.228389446 | 0.008270573 | 0.033596425 | 3.302884333 | 13.99027758 | ENSG00000136944 |
| ESPNL | 2.077405769 | 4.220476143 | 1.12E-12 | 3.19E-11 | 25.38488669 | 107.2944734 | ENSG00000144488 |
| PAX8-AS1 | 2.069779631 | 4.198225413 | 0.062615362 | 0.167623329 | 1.665082452 | 6.964965846 | ENSG00000189223 |
| PENK | 2.069672156 | 4.197912675 | 0.043522319 | 0.126722015 | 1.988610503 | 8.352080027 | ENSG00000181195 |
| TRIM4 | 2.067367409 | 4.191211743 | 1.62E-18 | 8.48E-17 | 37.63651829 | 157.9205131 | ENSG00000146833 |
| RBM43 | 2.066531087 | 4.188782826 | 2.72E-08 | 4.16E-07 | 18.05881369 | 75.26099271 | ENSG00000184898 |
| NA | 2.062064952 | 4.175835708 | 0.47550785 | 0.67798911 | 1.599842892 | 6.70534822 | ENSG00000163364 |
| NA | 2.057985836 | 4.164045509 | 0.073247685 | 0.189490758 | 1.692363024 | 6.964965846 | ENSG00000253966 |

| | | | | | | | |
|---|---|---|---|---|---|---|---|
| NA | 2.056654257 | 4.160203952 | 0.055375161 | 0.152665682 | 2.012331602 | 8.31973162 | ENSG00000234997 |
| LINC00958 | 2.053214135 | 4.150295717 | 0.027556513 | 0.088942874 | 2.666506649 | 11.04449141 | ENSG00000251381 |
| BEX5 | 2.050392851 | 4.142187475 | 8.38E-14 | 2.71E-12 | 27.84410652 | 115.2430919 | ENSG00000184515 |
| HJV | 2.049417897 | 4.139389186 | 0.034646925 | 0.106339539 | 2.748348363 | 11.16563469 | ENSG00000168509 |
| FCGBP | 2.049155144 | 4.138635361 | 2.45E-06 | 2.63E-05 | 12.04790425 | 49.60958707 | ENSG00000275395 |
| ITGA11 | 2.039404556 | 4.110758322 | 0.018421236 | 0.064304073 | 2.9900347 | 12.26362071 | ENSG00000137809 |
| TNS4 | 2.037611362 | 4.105652041 | 2.67E-06 | 2.85E-05 | 12.46039296 | 50.70613268 | ENSG00000131746 |
| SLC7A14 | 2.033263819 | 4.093298333 | 0.000109506 | 0.000805932 | 7.97460364 | 32.58328804 | ENSG00000013293 |
| NA | 2.029155705 | 4.081659133 | 0.023388282 | 0.078091887 | 3.048155315 | 12.31119736 | ENSG00000236975 |
| TOGARAM2 | 2.028036904 | 4.078495059 | 5.64E-05 | 0.000448256 | 8.305250636 | 33.78718909 | ENSG00000189350 |
| LGR5 | 2.027574066 | 4.077186825 | 0.043504833 | 0.126690636 | 2.254017939 | 9.321239514 | ENSG00000139292 |
| NA | 2.025475417 | 4.071260166 | 0.000310616 | 0.002026949 | 7.423627142 | 29.94876353 | ENSG00000252051 |
| ENPP6 | 2.022244134 | 4.062151745 | 0.017985265 | 0.063097109 | 3.04103637 | 12.20667979 | ENSG00000164303 |
| CFAP44 | 2.020261681 | 4.056573649 | 0.003476885 | 0.016325452 | 4.871887182 | 19.94966815 | ENSG00000206530 |
| NA | 2.018289603 | 4.051032343 | 0.049989527 | 0.141035432 | 1.95777046 | 7.997592643 | ENSG00000227117 |
| COL13A1 | 2.016506031 | 4.046027236 | 9.27E-45 | 2.22E-42 | 119.6194118 | 484.8510327 | ENSG00000197467 |
| NA | 2.014876461 | 4.041459699 | 0.078776703 | 0.199450066 | 1.641361354 | 6.66741938 | ENSG00000226853 |
| SLC39A4 | 2.013133264 | 4.036579382 | 2.35E-12 | 6.44E-11 | 26.2952653 | 105.8023902 | ENSG00000147804 |
| WEE2-AS1 | 2.012860516 | 4.035816322 | 0.05773273 | 0.157669212 | 1.954210987 | 7.946894568 | ENSG00000228775 |
| LOXL4 | 2.012825126 | 4.035717322 | 0.012469136 | 0.046933004 | 4.508035878 | 18.41899364 | ENSG00000138131 |
| ABCC3 | 2.012709723 | 4.035394512 | 0.008575914 | 0.03460601 | 3.622852912 | 14.66936295 | ENSG00000108846 |
| PRDM16 | 2.011234125 | 4.031269197 | 7.58E-08 | 1.08E-06 | 15.55095646 | 62.68185474 | ENSG00000142611 |
| HCLS1 | 2.010098986 | 4.028098566 | 1.91E-07 | 2.51E-06 | 15.61027228 | 62.77454903 | ENSG00000180353 |
| ADGRG3 | 2.009954151 | 4.027694198 | 0.030939326 | 0.097299657 | 3.102716458 | 12.27355207 | ENSG00000182885 |
| PKDREJ | 2.007018056 | 4.019505586 | 0.024016315 | 0.079751734 | 2.944956766 | 11.89078366 | ENSG00000130943 |
| SKAP1 | 2.006668373 | 4.01853145 | 0.150280091 | 0.319275456 | 1.341554401 | 5.367702608 | ENSG00000141293 |
| C8orf31 | 2.006267454 | 4.017414873 | 0.000129031 | 0.000931834 | 8.305250636 | 33.32584072 | ENSG00000177335 |
| LOC105373958 | 2.006213179 | 4.017263738 | 0.017104598 | 0.060638321 | 2.979356283 | 11.97042517 | ENSG00000234949 |
| LOC100130691 | 2.005952861 | 4.016538934 | 0.000267113 | 0.001777457 | 7.59059071 | 30.52768226 | ENSG00000213963 |
| CPB2-AS1 | 2.004643943 | 4.012896488 | 0.010399889 | 0.040320653 | 4.177388883 | 16.99953106 | ENSG00000235903 |
| NA | 2.002490515 | 4.006911136 | 0.00870056 | 0.034989367 | 3.653692956 | 14.61242203 | ENSG00000230982 |
| PTP4A1 | 1.993234185 | 3.981285095 | 0.050380412 | 0.141921383 | 2.019450547 | 7.967419699 | ENSG00000112245 |
| PRELP | 1.993122262 | 3.980976243 | 0.020586938 | 0.070444705 | 3.044595843 | 11.99530123 | ENSG00000188783 |
| GALR1 | 1.991711383 | 3.977084962 | 0.024599008 | 0.081400432 | 2.690227748 | 10.60383612 | ENSG00000166573 |
| NA | 1.986423636 | 3.962534885 | 0.001350508 | 0.00727794 | 5.61858236 | 22.28853811 | ENSG00000285847 |
| FLI1 | 1.984359542 | 3.956869655 | 0.000243416 | 0.001638241 | 7.318064329 | 28.87754548 | ENSG00000151702 |
| KCNE3 | 1.981421539 | 3.948819814 | 9.42E-11 | 2.12E-09 | 26.49334824 | 103.9794661 | ENSG00000175538 |
| NA | 1.981307185 | 3.948506827 | 0.09193463 | 0.223879494 | 1.692363024 | 6.600830648 | ENSG00000245146 |
| NA | 1.980819138 | 3.94717132 | 0.036063109 | 0.109688646 | 2.91055725 | 11.60411451 | ENSG00000189229 |
| NA | 1.976130419 | 3.934363945 | 0.000643524 | 0.003815686 | 6.544088579 | 25.89723544 | ENSG00000204054 |
| TRPM2 | 1.976071443 | 3.934203114 | 0.005223601 | 0.022941326 | 4.886125072 | 19.39127455 | ENSG00000142185 |
| TUSC8 | 1.974589062 | 3.930162765 | 0.015531042 | 0.056068661 | 3.769934186 | 14.57042581 | ENSG00000237361 |
| NA | 1.973954075 | 3.928433326 | 0.090472209 | 0.221284168 | 1.723203067 | 6.634125014 | ENSG00000237531 |
| NLRP1 | 1.972547427 | 3.924604914 | 0.000274027 | 0.001817064 | 8.001884211 | 31.30788023 | ENSG00000091592 |
| HSPA2 | 1.971958662 | 3.923003606 | 3.04E-129 | 1.44E-125 | 652.5103022 | 2558.942635 | ENSG00000126803 |
| DNM3 | 1.961173492 | 3.893785717 | 2.89E-17 | 1.34E-15 | 43.06294144 | 168.2096826 | ENSG00000197959 |
| BTG2 | 1.960814907 | 3.892818028 | 1.45E-23 | 1.16E-21 | 62.66703689 | 244.3399744 | ENSG00000159388 |
| NA | 1.959964578 | 3.890524266 | 0.099230662 | 0.236881691 | 1.606961837 | 6.352752753 | ENSG00000259436 |
| NA | 1.959928647 | 3.890427372 | 0.023879425 | 0.079408864 | 2.955635184 | 11.524473 | ENSG00000245025 |
| NA | 1.955369974 | 3.87815368 | 0.001435549 | 0.007671289 | 6.994536278 | 27.01307668 | ENSG00000277270 |
| MYRFL | 1.955067028 | 3.877339405 | 0.077023846 | 0.196328158 | 1.988610503 | 7.720571308 | ENSG00000166268 |
| PCDHGA11 | 1.949896276 | 3.863467539 | 5.53E-15 | 2.08E-13 | 36.27836173 | 140.3958828 | ENSG00000253873 |
| SH3TC2 | 1.949651906 | 3.862813183 | 0.023293241 | 0.077843366 | 3.368123893 | 12.89446702 | ENSG00000169247 |
| PADI1 | 1.949126924 | 3.861407799 | 0.000417298 | 0.00262689 | 7.947323068 | 30.70871992 | ENSG00000142623 |
| GZMM | 1.946318199 | 3.853897492 | 0.075108521 | 0.192773092 | 1.995729448 | 7.68169651 | ENSG00000197540 |
| NA | 1.94516768 | 3.850825317 | 1.35E-06 | 1.53E-05 | 13.81471071 | 53.34160314 | ENSG00000287535 |
| COL4A4 | 1.944107823 | 3.847997396 | 1.16E-05 | 0.000108927 | 11.55357383 | 44.52108126 | ENSG00000081052 |
| TRADD | 1.935112654 | 3.824079864 | 0.009530458 | 0.037551158 | 4.031782149 | 15.3064521 | ENSG00000102871 |
| BMPER | 1.928352 | 3.806201658 | 0.032718199 | 0.101742661 | 2.917676195 | 11.19892905 | ENSG00000164619 |
| PSTPIP1 | 1.926350058 | 3.800923683 | 0.000432797 | 0.002710904 | 7.221984725 | 27.56912685 | ENSG00000140368 |
| NA | 1.923823174 | 3.794272182 | 0.044204258 | 0.128036413 | 2.611945506 | 9.979799754 | ENSG00000280399 |
| PAPPA2 | 1.9236261 | 3.793753914 | 1.95E-05 | 0.000174529 | 12.65552682 | 48.43508095 | ENSG00000116183 |
| CHODL | 1.92327492 | 3.792830553 | 7.49E-09 | 1.26E-07 | 23.49383033 | 88.52366231 | ENSG00000154645 |
| PDCD1 | 1.921951557 | 3.789353042 | 2.57E-09 | 4.63E-08 | 21.45510788 | 81.42544637 | ENSG00000188389 |
| LINC01152 | 1.919360609 | 3.782553817 | 0.077510369 | 0.197248796 | 2.050290591 | 7.637524826 | ENSG00000256124 |
| KCNH3 | 1.917701205 | 3.778205582 | 0.000343889 | 0.002210461 | 9.58987963 | 36.2573446 | ENSG00000135519 |
| TDRD10 | 1.917027261 | 3.776441034 | 0.025916314 | 0.084676401 | 3.006636854 | 11.30221712 | ENSG00000163239 |
| INMT | 1.916369106 | 3.774718623 | 0.028305729 | 0.090740739 | 3.017315271 | 11.32368821 | ENSG00000241644 |
| SIGLEC15 | 1.914226255 | 3.769116153 | 0.009740645 | 0.038188188 | 3.946380963 | 14.9183868 | ENSG00000197046 |
| TNFAIP6 | 1.913698405 | 3.76773737 | 0.001894672 | 0.009688838 | 5.645862931 | 21.22819738 | ENSG00000123610 |
| SOST | 1.907340738 | 3.751170239 | 0.139492816 | 0.303318576 | 1.665082452 | 6.213994857 | ENSG00000167941 |
| APOBEC3B-AS1 | 1.906836369 | 3.749859051 | 0.034974017 | 0.107186677 | 3.048155315 | 11.31186493 | ENSG00000249310 |
| ASCL5 | 1.906562899 | 3.749148313 | 0.023583865 | 0.078661478 | 4.155142324 | 15.29775025 | ENSG00000232237 |
| LINC01686 | 1.905767105 | 3.747080846 | 0.088628978 | 0.217823823 | 1.930489888 | 7.337802897 | ENSG00000261504 |

| | | | | | | | |
|---|---|---|---|---|---|---|---|
| DDX60L | 1.900335115 | 3.732998981 | 0.000350126 | 0.002245967 | 8.577777016 | 32.03688567 | ENSG00000181381 |
| FOXS1 | 1.893307338 | 3.714858702 | 0.04999159 | 0.141035432 | 2.312138554 | 8.588334646 | ENSG00000179772 |
| SNTG1 | 1.890816932 | 3.708451579 | 0.022071538 | 0.074552135 | 3.33728385 | 12.33484392 | ENSG00000147481 |
| LGALS1 | 1.890747691 | 3.7082736 | 2.14E-41 | 4.55E-39 | 148.0521489 | 548.4196534 | ENSG00000100097 |
| TPRG1 | 1.890213025 | 3.706899559 | 0.03423455 | 0.105382147 | 2.955635184 | 11.00779208 | ENSG00000188001 |
| NA | 1.887177349 | 3.699107812 | 0.051389172 | 0.144138498 | 2.342978598 | 8.611981199 | ENSG00000253252 |
| TBX10 | 1.884805995 | 3.693032592 | 2.92E-05 | 0.000249924 | 11.3700081 | 41.7558383 | ENSG00000167800 |
| PTGS2 | 1.881378418 | 3.684269044 | 0.080045961 | 0.2019057 | 1.961329932 | 7.262512311 | ENSG00000073756 |
| NA | 1.877460435 | 3.674277106 | 0.031116255 | 0.097742926 | 3.010196326 | 11.01337251 | ENSG00000230333 |
| PCDHGC5 | 1.876214551 | 3.671105438 | 0.01768573 | 0.062254428 | 3.622852912 | 13.34571608 | ENSG00000240764 |
| LINC02473 | 1.875890101 | 3.67027993 | 0.069472889 | 0.181623319 | 2.401099213 | 8.677623972 | ENSG00000261121 |
| NA | 1.875436924 | 3.66912721 | 0.060580695 | 0.163534048 | 2.397539741 | 8.656152883 | ENSG00000256139 |
| SLC25A45 | 1.874888073 | 3.667731612 | 0.016077459 | 0.057708694 | 4.249747387 | 15.68988292 | ENSG00000162241 |
| TNFRSF10C | 1.874608587 | 3.667021151 | 1.23E-07 | 1.67E-06 | 18.50659164 | 67.87192465 | ENSG00000173535 |
| KLHDC8A | 1.874218763 | 3.666030434 | 1.14E-18 | 6.14E-17 | 51.38242997 | 188.7865085 | ENSG00000162873 |
| NA | 1.868128468 | 3.650587014 | 0.10982506 | 0.256147669 | 1.637801881 | 6.019736458 | ENSG00000229728 |
| LINC01063 | 1.866668617 | 3.646894883 | 0.122549867 | 0.276709919 | 1.630682936 | 5.991738978 | ENSG00000232065 |
| NA | 1.863549421 | 3.639018589 | 0.024946341 | 0.082347779 | 3.279163235 | 11.99842265 | ENSG00000272654 |
| KCNAB2 | 1.861075133 | 3.632782856 | 6.98E-53 | 2.40E-50 | 188.0508915 | 682.0030005 | ENSG00000069424 |
| NA | 1.858643433 | 3.626664863 | 0.048029228 | 0.13670414 | 2.686668275 | 9.685374711 | ENSG00000273576 |
| NA | 1.857737723 | 3.624388793 | 0.105088027 | 0.247516512 | 1.637801881 | 5.965916961 | ENSG00000230863 |
| ZP3 | 1.857380217 | 3.623490765 | 0.00363049 | 0.016920485 | 6.767087831 | 24.2987427 | ENSG00000188372 |
| ETV3L | 1.854716857 | 3.616807606 | 0.000579861 | 0.003492007 | 8.046962145 | 28.92606809 | ENSG00000253831 |
| C4orf50 | 1.852462135 | 3.611159478 | 0.000197861 | 0.001366714 | 10.90415346 | 39.38339042 | ENSG00000181215 |
| NA | 1.850764382 | 3.606912391 | 0.124774152 | 0.280519032 | 1.672201397 | 6.007913181 | ENSG00000231464 |
| ZCCHC12 | 1.85075994 | 3.606901287 | 2.49E-12 | 6.80E-11 | 35.993072 | 129.3797677 | ENSG00000174460 |
| NA | 1.849226995 | 3.603070785 | 0.085391234 | 0.212158066 | 2.319257499 | 8.363903303 | ENSG00000227242 |
| ADGRE2 | 1.848384392 | 3.600967034 | 2.84E-10 | 5.92E-09 | 28.0691907 | 101.163525 | ENSG00000127507 |
| EMP3 | 1.8455264 | 3.593840546 | 2.22E-12 | 6.11E-11 | 45.84271467 | 164.1180931 | ENSG00000142227 |
| LRRC4C | 1.843700397 | 3.589294739 | 0.000235577 | 0.001592293 | 9.224833118 | 33.18150239 | ENSG00000148948 |
| MIR302C | 1.838986496 | 3.57758612 | 0.063006799 | 0.168467139 | 2.339419126 | 8.335905823 | ENSG00000199102 |
| NA | 1.831594253 | 3.559301765 | 0.032815753 | 0.101895017 | 3.241204246 | 11.66105543 | ENSG00000276627 |
| DLGAP2 | 1.83122699 | 3.558395798 | 2.38E-05 | 0.000208006 | 12.57515965 | 44.7293384 | ENSG00000198010 |
| NA | 1.830570789 | 3.556777653 | 0.042206621 | 0.123692124 | 2.986475227 | 10.60506563 | ENSG00000266904 |
| LHX5 | 1.827395809 | 3.548958759 | 0.133927974 | 0.295207783 | 1.937608833 | 6.942548798 | ENSG00000089116 |
| NA | 1.823547454 | 3.539504609 | 0.073190248 | 0.189420012 | 2.373818642 | 8.306962384 | ENSG00000233332 |
| IP6K3 | 1.821405209 | 3.53425273 | 0.031926185 | 0.09987179 | 3.251882663 | 11.58576485 | ENSG00000161896 |
| KCNMB1 | 1.821316 | 3.534034197 | 0.000147533 | 0.001052971 | 10.50114796 | 37.28155309 | ENSG00000145936 |
| NTN1 | 1.820386654 | 3.531758398 | 2.46E-26 | 2.41E-24 | 87.79983815 | 309.8810685 | ENSG00000065320 |
| DHRS2 | 1.820045525 | 3.530923405 | 0.063728446 | 0.169795668 | 29.68355082 | 104.7869694 | ENSG00000100867 |
| RBPJL | 1.818200645 | 3.526411039 | 0.009071601 | 0.036173392 | 4.589877592 | 16.26255881 | ENSG00000124232 |
| HLA-DQB1 | 1.818053557 | 3.526051527 | 2.54E-49 | 7.74E-47 | 221.1106011 | 778.7121781 | ENSG00000179344 |
| SPATA31E1 | 1.817785941 | 3.525397514 | 0.092333474 | 0.224590115 | 1.964889405 | 6.961844424 | ENSG00000177992 |
| TRIM50 | 1.817601051 | 3.524945743 | 0.06553353 | 0.173503534 | 2.904633514 | 10.3591632 | ENSG00000146755 |
| NA | 1.816822316 | 3.523043569 | 0.000830836 | 0.004767653 | 7.658194534 | 26.89751383 | ENSG00000282221 |
| PDE4B | 1.816713077 | 3.522776816 | 9.44E-42 | 2.03E-39 | 152.4023989 | 536.4951106 | ENSG00000184588 |
| FAM20A | 1.813493858 | 3.514924884 | 0.000363858 | 0.002323795 | 8.597938643 | 30.23883765 | ENSG00000108950 |
| NA | 1.81329012 | 3.51442854 | 0.054857303 | 0.151569885 | 2.92479514 | 10.37940478 | ENSG00000268201 |
| NA | 1.810073872 | 3.506602435 | 0.005363034 | 0.023433803 | 5.322334881 | 18.62072439 | ENSG00000249626 |
| CAVIN3 | 1.809808845 | 3.505958321 | 1.53E-34 | 2.29E-32 | 169.7169073 | 596.1848547 | ENSG00000170955 |
| CD79B | 1.809004331 | 3.50400378 | 0.00017459 | 0.001220146 | 10.38994074 | 36.18422948 | ENSG00000007312 |
| S100B | 1.808842508 | 3.503610767 | 0.140497831 | 0.3047809 | 1.72676254 | 5.923920741 | ENSG00000160307 |
| TNFSF9 | 1.807999747 | 3.501564705 | 1.62E-13 | 5.13E-12 | 40.27810863 | 141.2211984 | ENSG00000125657 |
| NMI | 1.807306059 | 3.499881459 | 5.83E-12 | 1.52E-10 | 36.71902074 | 129.0513859 | ENSG00000123609 |
| NA | 1.806784561 | 3.498616569 | 0.00929485 | 0.036799422 | 5.591301789 | 19.62128633 | ENSG00000273314 |
| IER3 | 1.803241059 | 3.490033922 | 1.70E-53 | 5.93E-51 | 246.7232673 | 861.6564757 | ENSG00000137331 |
| NA | 1.796033225 | 3.472640884 | 0.090799962 | 0.221803996 | 2.019450547 | 6.947845683 | ENSG00000287527 |
| C20orf144 | 1.793493563 | 3.466533165 | 0.077508326 | 0.197248796 | 2.308579082 | 8.028711545 | ENSG00000149609 |
| ISLR | 1.793103981 | 3.465597197 | 0.000649322 | 0.003845231 | 11.37949131 | 39.2787049 | ENSG00000129009 |
| ABCC2 | 1.789407324 | 3.456728569 | 0.001765343 | 0.009136338 | 8.601498115 | 29.77625977 | ENSG00000023839 |
| C6orf223 | 1.789043028 | 3.455855819 | 3.36E-08 | 5.09E-07 | 21.78931435 | 75.40041302 | ENSG00000181577 |
| NA | 1.785631879 | 3.447694348 | 0.105139066 | 0.247605844 | 2.046731118 | 6.952196611 | ENSG00000278000 |
| NA | 1.785325919 | 3.446963254 | 0.140355561 | 0.304598342 | 1.606961837 | 5.612375535 | ENSG00000286667 |
| DEGS2 | 1.784374835 | 3.444691623 | 4.79E-05 | 0.000386928 | 12.87733087 | 44.38355287 | ENSG00000168350 |
| RASSF9 | 1.784349726 | 3.444631671 | 0.004385522 | 0.019808796 | 7.090615882 | 24.21162884 | ENSG00000198774 |
| HIPK4 | 1.782693989 | 3.440680639 | 0.132236206 | 0.292563645 | 1.641361354 | 5.678018309 | ENSG00000160396 |
| NA | 1.782674802 | 3.440634881 | 0.010020072 | 0.039126903 | 5.927872521 | 20.40694914 | ENSG00000232057 |
| NA | 1.781891981 | 3.438768465 | 0.000693334 | 0.004079008 | 8.301691163 | 28.48323734 | ENSG00000230438 |
| NA | 1.78152701 | 3.437898639 | 0.013384922 | 0.049573299 | 4.658676625 | 15.98430797 | ENSG00000227712 |
| HAPLN1 | 1.779866217 | 3.433943298 | 3.86E-09 | 6.75E-08 | 27.02505284 | 92.98756466 | ENSG00000145681 |
| EBF1 | 1.775491315 | 3.423545804 | 0.129476892 | 0.288107825 | 1.634242409 | 5.626374276 | ENSG00000164330 |
| PLAT | 1.773315567 | 3.418386601 | 5.24E-34 | 7.67E-32 | 129.2943616 | 441.0563299 | ENSG00000104368 |
| BIRC7 | 1.773190931 | 3.418091295 | 0.16197546 | 0.336745581 | 1.941168306 | 6.704118715 | ENSG00000101197 |
| LINC00622 | 1.773018597 | 3.417683019 | 0.061863182 | 0.16602049 | 2.601267089 | 8.968644047 | ENSG00000260941 |

| | | | | | | | |
|---|---|---|---|---|---|---|---|
| TMCC1-AS1 | 1.772272512 | 3.415916031 | 0.00016775 | 0.001176805 | 10.81755707 | 37.10609586 | ENSG00000271270 |
| TFF3 | 1.772105455 | 3.415520508 | 0.000207055 | 0.001418804 | 11.60101602 | 39.61455907 | ENSG00000160180 |
| PTPRB | 1.772076947 | 3.415453018 | 0.001642163 | 0.008588272 | 7.726993566 | 26.19601141 | ENSG00000127329 |
| NA | 1.770456718 | 3.411619423 | 0.001267965 | 0.006888813 | 7.579912292 | 25.91993604 | ENSG00000175772 |
| DPPA2 | 1.770192266 | 3.410994115 | 1.12E-22 | 8.33E-21 | 86.93957149 | 296.291952 | ENSG00000163530 |
| HTR2C | 1.768856825 | 3.407838165 | 6.89E-11 | 1.57E-09 | 33.46861262 | 114.0106263 | ENSG00000147246 |
| PTPRH | 1.767682473 | 3.405065318 | 0.076166949 | 0.19482769 | 2.370259169 | 7.983593902 | ENSG00000080031 |
| NA | 1.767274089 | 3.404101582 | 0.146571956 | 0.313833491 | 1.66152298 | 5.644723943 | ENSG00000228852 |
| LINC02188 | 1.761524055 | 3.390561129 | 3.78E-14 | 1.28E-12 | 46.30206076 | 157.0346836 | ENSG00000261175 |
| MX2 | 1.761473059 | 3.390441281 | 6.55E-15 | 2.44E-13 | 56.91975543 | 192.4488144 | ENSG00000183486 |
| NA | 1.759140168 | 3.384963243 | 0.058331264 | 0.158975805 | 2.632107133 | 8.927877332 | ENSG00000280057 |
| NA | 1.75848743 | 3.383432085 | 0.088360487 | 0.217333655 | 2.683108803 | 9.039867252 | ENSG00000280138 |
| ADAMTSL2 | 1.758137607 | 3.382611774 | 2.25E-11 | 5.47E-10 | 37.79191372 | 127.7190372 | ENSG00000197859 |
| SORBS2 | 1.755580592 | 3.376621784 | 1.02E-72 | 8.34E-70 | 399.728839 | 1351.561604 | ENSG00000154556 |
| ANKRD31 | 1.754052684 | 3.373047616 | 0.091157948 | 0.22236087 | 2.401099213 | 7.967419699 | ENSG00000145700 |
| NA | 1.753645148 | 3.372094922 | 0.020314256 | 0.069645209 | 4.362429145 | 14.61648941 | ENSG00000261068 |
| BANK1 | 1.751299747 | 3.366617334 | 0.002509204 | 0.0123434 | 9.230756855 | 31.10631744 | ENSG00000153064 |
| GPA33 | 1.749985936 | 3.363552872 | 0.021572143 | 0.073140022 | 4.351750728 | 14.54677926 | ENSG00000143167 |
| NDUFV2-AS1 | 1.747885521 | 3.358659448 | 0.013967961 | 0.051390016 | 4.620717636 | 15.52218159 | ENSG00000266053 |
| NME9 | 1.74598451 | 3.354236723 | 0.09548826 | 0.230188382 | 2.397539741 | 7.906127853 | ENSG00000181322 |
| DCST2 | 1.744963456 | 3.351863633 | 0.032064019 | 0.100170223 | 4.000942105 | 13.38553684 | ENSG00000163354 |
| FAM124B | 1.742535605 | 3.346227667 | 1.39E-05 | 0.000127409 | 16.47765789 | 55.27237713 | ENSG00000124019 |
| NA | 1.741271643 | 3.343297281 | 0.009740667 | 0.038188188 | 6.300037978 | 21.07375974 | ENSG00000269842 |
| MSLN | 1.740438673 | 3.341367515 | 0.135111262 | 0.297005724 | 2.305019609 | 7.73116508 | ENSG00000102854 |
| LINC01270 | 1.739449184 | 3.339076585 | 0.060485682 | 0.163345073 | 3.73553467 | 12.36255785 | ENSG00000203999 |
| NA | 1.739241026 | 3.338594845 | 0.108043719 | 0.252816969 | 1.992169976 | 6.640651405 | ENSG00000257953 |
| NA | 1.739171967 | 3.338435037 | 0.157181358 | 0.329379709 | 1.72676254 | 5.656547219 | ENSG00000260047 |
| SGMS2 | 1.738552102 | 3.337000961 | 0.086862493 | 0.214655611 | 2.277739038 | 7.672048697 | ENSG00000164023 |
| PADI3 | 1.737388514 | 3.334310628 | 1.62E-09 | 3.02E-08 | 29.31822497 | 97.98243022 | ENSG00000142619 |
| GABRG2 | 1.737320551 | 3.334153558 | 0.002354591 | 0.011683253 | 9.852896649 | 32.95707105 | ENSG00000113327 |
| NA | 1.736175977 | 3.331509429 | 0.050618713 | 0.142485376 | 3.602691286 | 12.07711821 | ENSG00000260495 |
| HLA-DMB | 1.736072783 | 3.33127114 | 0.0031084 | 0.014850022 | 7.882083508 | 26.37023914 | ENSG00000242574 |
| ALPK1 | 1.735617747 | 3.330220601 | 0.021117103 | 0.071932988 | 4.23906897 | 14.20288565 | ENSG00000073331 |
| SYTL2 | 1.735394863 | 3.32970615 | 3.51E-14 | 1.19E-12 | 49.45221941 | 164.4677351 | ENSG00000137501 |
| LINC00346 | 1.728991167 | 3.314959321 | 0.093639964 | 0.227182774 | 2.308579082 | 7.672994656 | ENSG00000255874 |
| RIN1 | 1.72876524 | 3.314440239 | 1.95E-11 | 4.78E-10 | 38.83483022 | 128.5521087 | ENSG00000174791 |
| TEX19 | 1.728474956 | 3.313773407 | 0.015688441 | 0.056530471 | 5.171694134 | 17.33160139 | ENSG00000182459 |
| SLC8A3 | 1.727197221 | 3.310839835 | 0.040021787 | 0.118744924 | 3.739094142 | 12.21850307 | ENSG00000100678 |
| THPO | 1.726324688 | 3.308838065 | 0.009390802 | 0.037070505 | 5.625701305 | 18.6219539 | ENSG00000090534 |
| MYOM1 | 1.724500356 | 3.304656582 | 0.001916343 | 0.009778478 | 8.577777016 | 28.34760087 | ENSG00000101605 |
| NA | 1.723476353 | 3.302311821 | 0.111139063 | 0.258544124 | 2.019450547 | 6.606127534 | ENSG00000286570 |
| ACOXL-AS1 | 1.722830733 | 3.300834336 | 0.013835986 | 0.050999131 | 5.060486917 | 16.53958014 | ENSG00000204581 |
| NA | 1.722511472 | 3.300103958 | 0.075685261 | 0.194016038 | 2.886836151 | 9.663903622 | ENSG00000287595 |
| SERPINE1 | 1.720213905 | 3.294852553 | 7.01E-12 | 1.80E-10 | 44.18564088 | 145.399094 | ENSG00000106366 |
| MSRB3 | 1.716720728 | 3.286884423 | 1.09E-05 | 0.00010315 | 19.28265232 | 63.69887442 | ENSG00000174099 |
| GABRB2 | 1.715699735 | 3.284559124 | 0.000974524 | 0.005488781 | 8.863346079 | 29.23014095 | ENSG00000145864 |
| LOC101928335 | 1.713544276 | 3.279655492 | 0.001674542 | 0.008743081 | 8.772300488 | 28.51000532 | ENSG00000236064 |
| APOBEC3H | 1.710038673 | 3.271695934 | 0.079384477 | 0.200719754 | 2.655828232 | 8.687271785 | ENSG00000100298 |
| LINC00672 | 1.709042567 | 3.269437778 | 4.81E-05 | 0.000388242 | 17.43636841 | 57.14448624 | ENSG00000263874 |
| LOC101927531 | 1.708572847 | 3.268373472 | 0.07634062 | 0.195139769 | 2.62854766 | 8.617278085 | ENSG00000255595 |
| HCN1 | 1.706084619 | 3.262741339 | 6.52E-11 | 1.50E-09 | 41.97019232 | 137.404201 | ENSG00000164588 |
| IFI35 | 1.704590646 | 3.259364379 | 6.07E-07 | 7.30E-06 | 22.86665652 | 74.51430002 | ENSG00000068079 |
| NA | 1.704361893 | 3.258847616 | 0.098323269 | 0.235221046 | 2.366699697 | 7.635349363 | ENSG00000228528 |
| GDNF | 1.698460854 | 3.245545213 | 0.039044686 | 0.116578481 | 3.708254098 | 11.92530753 | ENSG00000168621 |
| ZNF37A | 1.69768527 | 3.243800896 | 3.39E-17 | 1.56E-15 | 61.34920358 | 199.2624641 | ENSG00000075407 |
| DNAH3 | 1.69754156 | 3.24347779 | 0.000147452 | 0.00105279 | 12.29670953 | 39.79510227 | ENSG00000158486 |
| NXPH1 | 1.696467219 | 3.241063347 | 0.074483769 | 0.19176905 | 2.69378722 | 8.665800696 | ENSG00000122584 |
| LINC01140 | 1.694947687 | 3.237651464 | 0.003975659 | 0.01827661 | 7.610752336 | 24.66070243 | ENSG00000267272 |
| PPP1R17 | 1.69483038 | 3.237388218 | 4.65E-07 | 5.74E-06 | 21.77507646 | 70.56461958 | ENSG00000106341 |
| SATB2 | 1.694046079 | 3.235628736 | 4.46E-12 | 1.18E-10 | 47.84022356 | 154.9054683 | ENSG00000119042 |
| MXRA8 | 1.693286071 | 3.233924664 | 3.59E-42 | 7.80E-40 | 168.7899271 | 545.8466614 | ENSG00000162576 |
| PLK2 | 1.692290251 | 3.231693218 | 1.57E-47 | 4.19E-45 | 203.9867768 | 658.4850753 | ENSG00000145632 |
| OAZ3 | 1.690068029 | 3.226719185 | 0.061733101 | 0.165817029 | 3.541011198 | 11.54499813 | ENSG00000143450 |
| LNCTAM34A | 1.689537077 | 3.225531882 | 0.037579786 | 0.113080667 | 3.884700875 | 12.66786021 | ENSG00000234546 |
| JSRP1 | 1.688764389 | 3.223804793 | 0.00935107 | 0.03697987 | 5.700424074 | 18.25971062 | ENSG00000167476 |
| PLA2G2A | 1.688561664 | 3.223351821 | 3.84E-10 | 7.85E-09 | 33.80637856 | 108.8910016 | ENSG00000188257 |
| NA | 1.688163339 | 3.222461984 | 0.018594539 | 0.064738103 | 4.655117152 | 14.97438176 | ENSG00000287935 |
| PI15 | 1.68698254 | 3.219825582 | 0.089011584 | 0.218593466 | 2.713948847 | 8.644329606 | ENSG00000137558 |
| NA | 1.686038162 | 3.217718598 | 0.117272836 | 0.268578404 | 2.246898994 | 7.336856939 | ENSG00000283696 |
| SVIL-AS1 | 1.684954288 | 3.215302084 | 0.162228726 | 0.337107182 | 1.668641925 | 5.352474363 | ENSG00000224597 |
| OVGP1 | 1.684142869 | 3.213494202 | 0.003220413 | 0.01529615 | 8.914347749 | 28.69917774 | ENSG00000085465 |
| NA | 1.682112317 | 3.208974481 | 0.012252257 | 0.046323098 | 5.260654793 | 16.93606375 | ENSG00000271474 |
| RIMBP3B | 1.679030779 | 3.202127558 | 0.18325017 | 0.366977969 | 1.672201397 | 5.334124696 | ENSG00000274600 |
| NA | 1.678461573 | 3.200864427 | 0.057385027 | 0.156901149 | 3.90605771 | 12.58074635 | ENSG00000251396 |

| | | | | | | | |
|---|---|---|---|---|---|---|---|
| DNAH2 | 1.678146958 | 3.200166478 | 0.005947587 | 0.02554473 | 7.647516116 | 24.3918885 | ENSG00000183914 |
| ST7-AS1 | 1.676081287 | 3.195587713 | 0.170760195 | 0.34929273 | 1.672201397 | 5.319179997 | ENSG00000227199 |
| NA | 1.675508772 | 3.194319835 | 0.047823924 | 0.136304679 | 3.664371373 | 11.68375603 | ENSG00000231720 |
| KLK8 | 1.673728478 | 3.190380458 | 0.003916722 | 0.018049608 | 7.318064329 | 23.3049907 | ENSG00000129455 |
| MAP1LC3C | 1.670287331 | 3.182779762 | 0.000815984 | 0.004690979 | 11.44948555 | 36.1724062 | ENSG00000197769 |
| ADAD2 | 1.66783796 | 3.177380706 | 0.022475228 | 0.075577758 | 4.589877592 | 14.6606611 | ENSG00000140955 |
| LINC00698 | 1.667520429 | 3.176681455 | 5.10E-16 | 2.16E-14 | 77.82805668 | 247.6594912 | ENSG00000244342 |
| ALOX15B | 1.667457485 | 3.176542859 | 0.10080253 | 0.239333137 | 2.284857983 | 7.308859458 | ENSG00000179593 |
| LAMP3 | 1.664023338 | 3.168990508 | 0.038778673 | 0.116031797 | 4.011620523 | 12.66445524 | ENSG00000078081 |
| UBE2D3-AS1 | 1.659281317 | 3.158591394 | 0.005231422 | 0.022964995 | 6.939975135 | 21.94275258 | ENSG00000246560 |
| LINC02482 | 1.658568629 | 3.157031441 | 0.118064858 | 0.269945942 | 2.335859653 | 7.357382069 | ENSG00000251580 |
| NA | 1.656454616 | 3.152408761 | 0.05936012 | 0.161065498 | 3.241204246 | 10.32463933 | ENSG00000268366 |
| MYLK4 | 1.655998551 | 3.151412379 | 0.095223428 | 0.229843359 | 2.611945506 | 8.278964904 | ENSG00000145949 |
| COL6A3 | 1.653688337 | 3.146370004 | 9.44E-23 | 7.02E-21 | 105.4464686 | 331.1029504 | ENSG00000163359 |
| NA | 1.653610127 | 3.146199439 | 0.140472746 | 0.3047809 | 2.02301002 | 6.310756533 | ENSG00000287229 |
| NA | 1.649776363 | 3.137849946 | 0.15326019 | 0.324039262 | 2.050290591 | 6.352752753 | ENSG00000234353 |
| GCNT4 | 1.649575508 | 3.137413117 | 1.22E-09 | 2.31E-08 | 34.63344091 | 108.8091846 | ENSG00000176928 |
| NA | 1.64851405 | 3.135105625 | 0.006537874 | 0.027689561 | 6.698288798 | 20.87030503 | ENSG00000279278 |
| NA | 1.646997781 | 3.131812369 | 0.016160421 | 0.057954721 | 5.329453825 | 16.65157006 | ENSG00000285130 |
| IL32 | 1.644724079 | 3.126880488 | 0.010084461 | 0.039322879 | 7.866955895 | 24.30214767 | ENSG00000008517 |
| VIL1 | 1.644276513 | 3.125910588 | 4.97E-34 | 7.40E-32 | 143.1562351 | 447.229601 | ENSG00000127831 |
| SOX1 | 1.641678604 | 3.120286723 | 0.107238537 | 0.251555734 | 2.370259169 | 7.318507271 | ENSG00000182968 |
| LINC00858 | 1.638719913 | 3.113894169 | 0.007052294 | 0.029452137 | 7.283664813 | 22.70119592 | ENSG00000229404 |
| LAMA2 | 1.631559543 | 3.098477609 | 3.35E-14 | 1.14E-12 | 70.81422292 | 218.6300921 | ENSG00000196569 |
| LOC100129434 | 1.630819533 | 3.096888698 | 0.071677822 | 0.186271593 | 3.017315271 | 9.284540179 | ENSG00000233251 |
| SPSB2 | 1.628010855 | 3.090865455 | 6.95E-23 | 5.26E-21 | 113.8288319 | 352.6557458 | ENSG00000111671 |
| MANEA-DT | 1.624228443 | 3.082772536 | 0.227874732 | 0.427222706 | 1.614080782 | 5.037807736 | ENSG00000261366 |
| ADAMTSL5 | 1.624107314 | 3.082513717 | 0.056521962 | 0.155188303 | 3.361004949 | 10.28046764 | ENSG00000185761 |
| DLK1 | 1.622467761 | 3.079012578 | 6.85E-09 | 1.16E-07 | 33.92706898 | 104.7058446 | ENSG00000185559 |
| NA | 1.619347408 | 3.072360289 | 0.077914035 | 0.19807902 | 3.384726047 | 10.30193873 | ENSG00000267413 |
| NA | 1.618649844 | 3.070875119 | 6.44E-09 | 1.10E-07 | 36.35783918 | 111.6722079 | ENSG00000188013 |
| SLC52A1 | 1.616021655 | 3.06528593 | 0.033311265 | 0.103123018 | 4.358869673 | 13.27912735 | ENSG00000132517 |
| LAPTM5 | 1.613725187 | 3.06041052 | 0.13753296 | 0.30037122 | 2.281298511 | 7.0436614 | ENSG00000162511 |
| OXCT2 | 1.612599292 | 3.058023074 | 0.000106141 | 0.000783917 | 17.58344969 | 53.70110387 | ENSG00000198754 |
| LINC01091 | 1.611074166 | 3.054792032 | 0.141785198 | 0.306573559 | 2.281298511 | 7.026257692 | ENSG00000249464 |
| LOC102724057 | 1.61053475 | 3.053650076 | 0.036388564 | 0.110500747 | 4.729839921 | 14.32452338 | ENSG00000230303 |
| KCNG2 | 1.609237612 | 3.050905749 | 0.146633142 | 0.313928906 | 2.261136884 | 6.971492236 | ENSG00000178342 |
| NA | 1.606504146 | 3.045130689 | 0.153725463 | 0.324695826 | 1.954210987 | 6.001386791 | ENSG00000226699 |
| CCDC9B | 1.605672156 | 3.043375094 | 0.000156376 | 0.001107299 | 15.14558669 | 46.20886333 | ENSG00000188549 |
| NA | 1.605082737 | 3.042131965 | 0.188064819 | 0.373971159 | 1.634242409 | 5.007634792 | ENSG00000184414 |
| PTGDR2 | 1.599506225 | 3.030395776 | 0.204439676 | 0.396206468 | 1.637801881 | 4.988339166 | ENSG00000183134 |
| EBI3 | 1.597658102 | 3.026516261 | 0.005076724 | 0.022374314 | 7.541953304 | 22.93215365 | ENSG00000105246 |
| FOXB1 | 1.596609207 | 3.024316665 | 8.30E-14 | 2.69E-12 | 67.62905378 | 204.768641 | ENSG00000171956 |
| TAFA2 | 1.594130812 | 3.019125675 | 0.008412078 | 0.034053174 | 6.578488096 | 19.92072471 | ENSG00000198673 |
| NA | 1.59293425 | 3.01662267 | 2.73E-05 | 0.000235269 | 18.08342452 | 54.69419345 | ENSG00000283662 |
| NA | 1.591773274 | 3.014196088 | 0.149957693 | 0.318877494 | 1.992169976 | 6.001386791 | ENSG00000261757 |
| CFAP100 | 1.591039777 | 3.012663996 | 0.19374087 | 0.381800723 | 1.668641925 | 5.015107141 | ENSG00000163885 |
| LOC91370 | 1.589903698 | 3.010292547 | 0.127638176 | 0.285125024 | 2.305019609 | 6.950021147 | ENSG00000268812 |
| NA | 1.58953241 | 3.009517927 | 0.124881033 | 0.280659086 | 2.305019609 | 6.947845683 | ENSG00000285820 |
| IL6R | 1.589139741 | 3.008698914 | 2.90E-22 | 2.09E-20 | 103.4285182 | 311.7549742 | ENSG00000160712 |
| ADAMTS4 | 1.588677423 | 3.007734919 | 8.87E-40 | 1.71E-37 | 186.043925 | 559.434664 | ENSG00000158859 |
| PCDHGB3 | 1.588048894 | 3.006424844 | 0.041872368 | 0.122941492 | 4.297189585 | 12.92000549 | ENSG00000262209 |
| SCO2 | 1.58762294 | 3.005537333 | 0.088486508 | 0.217558614 | 3.357445476 | 10.04638849 | ENSG00000284194 |
| PRTN3 | 1.587465936 | 3.005210266 | 0.049988312 | 0.141035432 | 3.657252428 | 10.96920083 | ENSG00000196415 |
| NA | 1.587445916 | 3.005168564 | 0.168047722 | 0.345442607 | 1.992169976 | 5.978686196 | ENSG00000287464 |
| LINC02348 | 1.586946757 | 3.004128983 | 0.066799984 | 0.176288107 | 3.330164905 | 9.978570249 | ENSG00000286733 |
| C1orf74 | 1.586555057 | 3.003313456 | 3.73E-24 | 3.14E-22 | 111.6691141 | 335.242443 | ENSG00000162757 |
| NA | 1.584701717 | 2.999457765 | 0.15907443 | 0.332461334 | 1.988610503 | 5.956269148 | ENSG00000259659 |
| NPFFR2 | 1.5844375 | 2.99890849 | 1.38E-18 | 7.26E-17 | 89.19506397 | 266.8363907 | ENSG00000056291 |
| NA | 1.58414868 | 2.998308184 | 0.117349086 | 0.268687862 | 2.339419126 | 6.983315513 | ENSG00000249916 |
| NA | 1.582909993 | 2.995734965 | 0.086009838 | 0.213190155 | 2.9900347 | 8.935349681 | ENSG00000285407 |
| HLA-DRA | 1.582489802 | 2.994862572 | 3.56E-10 | 7.32E-09 | 43.84670589 | 131.0992358 | ENSG00000204287 |
| POLN | 1.582458635 | 2.994797872 | 0.067212199 | 0.177128436 | 3.711813571 | 11.03919453 | ENSG00000130997 |
| GAU1 | 1.581155731 | 2.99209448 | 0.112270436 | 0.2601825 | 2.972237338 | 8.885881111 | ENSG00000255474 |
| LCP1 | 1.580921497 | 2.991608727 | 5.11E-08 | 7.48E-07 | 29.97476428 | 89.84172875 | ENSG00000136167 |
| NA | 1.580726143 | 2.991203663 | 0.084429086 | 0.210345758 | 3.010196326 | 8.958050275 | ENSG00000268942 |
| ALPK2 | 1.580372468 | 2.990470462 | 0.102034451 | 0.241589846 | 3.037476898 | 9.018112617 | ENSG00000198796 |
| SLC14A2 | 1.580125159 | 2.989957876 | 0.207266226 | 0.399768992 | 1.672201397 | 4.975286385 | ENSG00000132874 |
| NA | 1.579968706 | 2.989633647 | 0.195629099 | 0.384638785 | 1.665082452 | 4.958166222 | ENSG00000279735 |
| FAM163B | 1.579289796 | 2.988227102 | 1.27E-11 | 3.18E-10 | 51.2839861 | 153.6840909 | ENSG00000196990 |
| NA | 1.578931813 | 2.987485711 | 0.00099717 | 0.005602963 | 13.13681456 | 39.40876094 | ENSG00000257474 |
| TM4SF1 | 1.578284617 | 2.98614582 | 0.00695926 | 0.029128021 | 7.552631721 | 22.65078139 | ENSG00000169908 |
| B3GNT3 | 1.577702718 | 2.984941625 | 3.19E-05 | 0.000270024 | 17.74329431 | 53.14422332 | ENSG00000179913 |
| NA | 1.576499802 | 2.982453826 | 1.32E-10 | 2.88E-09 | 53.29690252 | 158.6013224 | ENSG00000235152 |

| | | | | | | | |
|---|---|---|---|---|---|---|---|
| SLFN13 | 1.575605963 | 2.980606583 | 9.66E-30 | 1.13E-27 | 187.2484662 | 557.6235335 | ENSG00000154760 |
| RNF212 | 1.574079279 | 2.977454122 | 1.34E-13 | 4.29E-12 | 62.26403138 | 185.7867349 | ENSG00000178222 |
| SH2D2A | 1.573718104 | 2.976708818 | 1.45E-20 | 8.99E-19 | 94.43764207 | 280.97602 | ENSG00000027869 |
| ATP1A1-AS1 | 1.570396909 | 2.969864087 | 0.044248595 | 0.128125531 | 4.531756977 | 13.58291666 | ENSG00000203865 |
| TGM2 | 1.570136138 | 2.969327324 | 6.71E-12 | 1.73E-10 | 50.71905104 | 150.5107519 | ENSG00000198959 |
| SCX | 1.56997474 | 2.968995157 | 2.79E-09 | 4.98E-08 | 39.51008277 | 117.5797864 | ENSG00000260428 |
| LPL | 1.56982112 | 2.968679031 | 7.68E-24 | 6.33E-22 | 112.7761011 | 334.4148363 | ENSG00000175445 |
| TMEM249 | 1.569166653 | 2.96733262 | 0.096401811 | 0.231857896 | 3.017315271 | 8.885881111 | ENSG00000261587 |
| ADAMTS14 | 1.567287147 | 2.963469372 | 1.90E-10 | 4.06E-09 | 60.56632944 | 179.2383263 | ENSG00000138316 |
| ALOX5 | 1.566774248 | 2.962416002 | 1.38E-07 | 1.87E-06 | 31.6377618 | 93.51438785 | ENSG00000012779 |
| LOC105370854 | 1.566719124 | 2.962302814 | 0.203841945 | 0.395475152 | 2.050290591 | 6.002332749 | ENSG00000259370 |
| LINC02204 | 1.565562705 | 2.959929277 | 0.122097639 | 0.27599105 | 2.713948847 | 7.941597682 | ENSG00000280639 |
| SLC9A7P1 | 1.563852671 | 2.956422934 | 0.001291308 | 0.006989498 | 11.0298779 | 32.44112517 | ENSG00000227825 |
| NA | 1.563516156 | 2.955733416 | 0.048020644 | 0.13670031 | 4.562597021 | 13.56456699 | ENSG00000286214 |
| EHBP1L1 | 1.562772636 | 2.954210517 | 2.95E-26 | 2.86E-24 | 149.5523527 | 441.5914464 | ENSG00000173442 |
| LOC105378909 | 1.561454647 | 2.9515129 | 0.158989217 | 0.332393597 | 2.363140224 | 6.903674 | ENSG00000231246 |
| EYS | 1.560894687 | 2.950367537 | 0.218794468 | 0.414977757 | 1.699481969 | 4.949464368 | ENSG00000188107 |
| RAC2 | 1.559509686 | 2.947536515 | 7.46E-08 | 1.07E-06 | 43.78919566 | 129.5967267 | ENSG00000128340 |
| NRK | 1.55229647 | 2.932836142 | 1.63E-20 | 1.00E-18 | 101.6738647 | 298.0099366 | ENSG00000123572 |
| C16orf89 | 1.54868928 | 2.925512292 | 6.54E-10 | 1.29E-08 | 45.1503013 | 131.9382666 | ENSG00000153446 |
| PDE6G | 1.54821088 | 2.924542349 | 2.43E-12 | 6.66E-11 | 53.30015651 | 155.80144 | ENSG00000185527 |
| NA | 1.545378167 | 2.918805681 | 0.004586022 | 0.020547335 | 8.983146781 | 26.13907049 | ENSG00000228192 |
| TMEM232 | 1.544103647 | 2.91622826 | 0.008495815 | 0.034312119 | 7.252824769 | 21.17220242 | ENSG00000186952 |
| FAM66B | 1.540615951 | 2.909186831 | 0.000792893 | 0.004579141 | 13.04073496 | 38.18913718 | ENSG00000215374 |
| TRPC6 | 1.538592158 | 2.905108724 | 0.056389772 | 0.154875051 | 5.098445906 | 14.6768353 | ENSG00000137672 |
| AMY2B | 1.537402654 | 2.902714446 | 0.072894399 | 0.188809591 | 4.771358382 | 13.72385002 | ENSG00000240038 |
| CMKLR1 | 1.53568115 | 2.89925283 | 0.000113111 | 0.000829873 | 21.2703731 | 61.60394234 | ENSG00000174600 |
| RNVU1-15 | 1.534841198 | 2.897565348 | 0.22114993 | 0.418269047 | 2.104851733 | 5.965916961 | ENSG00000207205 |
| FOXA2 | 1.534093661 | 2.896064354 | 0.022277961 | 0.075128446 | 5.622141833 | 16.29367771 | ENSG00000125798 |
| LGR6 | 1.533966351 | 2.895808803 | 1.35E-05 | 0.000125017 | 24.19516824 | 70.01776572 | ENSG00000133067 |
| APCDD1L-DT | 1.533885135 | 2.895645789 | 0.100931696 | 0.239519362 | 3.013755799 | 8.693798176 | ENSG00000231290 |
| TLX2 | 1.533418023 | 2.894708396 | 0.228189976 | 0.427559053 | 1.937608833 | 5.667140991 | ENSG00000115297 |
| TNRC6C-AS1 | 1.532899169 | 2.893667525 | 0.002698317 | 0.013131581 | 9.830370759 | 28.58434994 | ENSG00000204282 |
| SGCG | 1.532884321 | 2.893637743 | 0.180638347 | 0.363606393 | 2.93903303 | 8.527988759 | ENSG00000102683 |
| C12orf56 | 1.527728361 | 2.883314804 | 0.0155578 | 0.056139443 | 7.420067669 | 21.22261695 | ENSG00000185306 |
| FAM183A | 1.526664389 | 2.881189175 | 0.026195117 | 0.085470207 | 6.238357891 | 18.05797987 | ENSG00000186973 |
| PLD5 | 1.526142424 | 2.880146954 | 1.45E-10 | 3.17E-09 | 54.55750493 | 157.0772993 | ENSG00000180287 |
| GJA5 | 1.525891824 | 2.879646707 | 1.45E-07 | 1.94E-06 | 32.54786108 | 93.57083431 | ENSG00000265107 |
| NA | 1.525539404 | 2.878943357 | 0.027065542 | 0.087748113 | 5.673143503 | 16.27343612 | ENSG00000271013 |
| F10 | 1.52515838 | 2.878183112 | 7.50E-09 | 1.26E-07 | 37.0475828 | 106.6771744 | ENSG00000126218 |
| CFD | 1.524615471 | 2.87710021 | 1.42E-06 | 1.60E-05 | 25.86497922 | 74.34101825 | ENSG00000197766 |
| SNORD14E | 1.524529247 | 2.876928263 | 0.118446423 | 0.270315491 | 3.044595843 | 8.690676754 | ENSG00000200879 |
| FRK | 1.524187347 | 2.876246547 | 0.138547614 | 0.301830117 | 2.955635184 | 8.504342205 | ENSG00000111816 |
| HPGD | 1.524014624 | 2.875902216 | 3.38E-05 | 0.000283557 | 18.8063986 | 54.17656657 | ENSG00000164120 |
| NA | 1.52308921 | 2.874058066 | 0.136674096 | 0.299152794 | 2.308579082 | 6.648123755 | ENSG00000227355 |
| VIM | 1.521404914 | 2.870704662 | 1.08E-32 | 1.45E-30 | 210.887955 | 606.0254654 | ENSG00000026025 |
| MYOM3 | 1.520851856 | 2.869604386 | 0.149068189 | 0.317379116 | 2.305019609 | 6.626652665 | ENSG00000142661 |
| INPP5J | 1.520500166 | 2.868904941 | 1.46E-22 | 1.08E-20 | 128.5017505 | 369.324436 | ENSG00000185133 |
| NA | 1.519863547 | 2.867639256 | 0.156863968 | 0.328933604 | 2.335859653 | 6.682647625 | ENSG00000234807 |
| RAB41 | 1.518232267 | 2.8643986 | 0.205100209 | 0.397059708 | 1.95777046 | 5.641318974 | ENSG00000147127 |
| NODAL | 1.516031604 | 2.860032625 | 1.73E-10 | 3.74E-09 | 46.13036866 | 131.7546746 | ENSG00000156574 |
| OR2A7 | 1.515672303 | 2.859320427 | 0.017408994 | 0.0614981 | 6.667448754 | 19.02373439 | ENSG00000243896 |
| CYSRT1 | 1.515603657 | 2.859184378 | 0.003672231 | 0.017078935 | 10.61027024 | 30.33654528 | ENSG00000197191 |
| SLC4A9 | 1.513933057 | 2.85587544 | 0.182775647 | 0.36646333 | 1.988610503 | 5.6845447 | ENSG00000113073 |
| NA | 1.512149218 | 2.852346438 | 0.000932504 | 0.005277295 | 12.2041894 | 34.867109 | ENSG00000241634 |
| ABI3 | 1.511892236 | 2.851838404 | 2.20E-08 | 3.41E-07 | 37.11490729 | 106.1914014 | ENSG00000108798 |
| NA | 1.511419733 | 2.850904541 | 0.02912535 | 0.092722899 | 5.976509927 | 17.03623039 | ENSG00000253348 |
| FAM89B | 1.51133326 | 2.850733668 | 0.083306596 | 0.20812873 | 3.619293439 | 10.35698773 | ENSG00000176973 |
| HS3ST6 | 1.510996384 | 2.850068085 | 0.169438208 | 0.347337414 | 2.346538071 | 6.654650145 | ENSG00000162040 |
| CDKN2A | 1.510586857 | 2.849259172 | 0.151437275 | 0.321047702 | 2.342978598 | 6.641597364 | ENSG00000147889 |
| NA | 1.510308542 | 2.848709565 | 0.05740629 | 0.156936564 | 3.977221007 | 11.33239006 | ENSG00000259349 |
| LOC728755 | 1.510121401 | 2.848340065 | 0.091203123 | 0.222413609 | 4.282951695 | 12.20573383 | ENSG00000258932 |
| NA | 1.510105817 | 2.848309298 | 0.19621359 | 0.385257474 | 1.981491558 | 5.650020829 | ENSG00000284188 |
| NA | 1.509832223 | 2.847769193 | 0.17957211 | 0.362271051 | 1.985051031 | 5.656547219 | ENSG00000236330 |
| LOC105378721 | 1.508744991 | 2.84562389 | 0.197503283 | 0.387118741 | 1.954210987 | 5.588728982 | ENSG00000266993 |
| CABYR | 1.508235967 | 2.844620049 | 2.07E-15 | 8.27E-14 | 81.45830787 | 231.7266163 | ENSG00000154040 |
| CD19 | 1.507901054 | 2.843959763 | 0.019859346 | 0.06836379 | 5.962272037 | 16.95441341 | ENSG00000177455 |
| TAGLN | 1.506999555 | 2.842183209 | 9.46E-73 | 8.12E-70 | 536.0808659 | 1522.956167 | ENSG00000149591 |
| KCTD19 | 1.505850647 | 2.839920704 | 0.021677594 | 0.0734053 | 6.351039648 | 17.96528557 | ENSG00000168676 |
| NA | 1.501264471 | 2.830907228 | 0.15113503 | 0.320658919 | 2.366699697 | 6.631949551 | ENSG00000273812 |
| SNORD116-5 | 1.501057898 | 2.830501911 | 0.134107481 | 0.295424407 | 2.921235667 | 8.343661719 | ENSG00000207191 |
| CNIH3-AS2 | 1.500627322 | 2.829657268 | 0.002914258 | 0.014036155 | 11.13752564 | 31.68194679 | ENSG00000233384 |
| LOC101927021 | 1.500443414 | 2.829296578 | 0.000719457 | 0.004210398 | 13.48406371 | 38.28060197 | ENSG00000229043 |
| MYOCD | 1.499373834 | 2.827199782 | 0.000307534 | 0.002010309 | 16.6837495 | 47.37909115 | ENSG00000141052 |

| | | | | | | | |
|---|---|---|---|---|---|---|---|
| MN1 | 1.499250315 | 2.826957738 | 2.26E-36 | 3.69E-34 | 201.9053668 | 570.6705759 | ENSG00000169184 |
| LHX4 | 1.498914467 | 2.826299721 | 2.86E-08 | 4.36E-07 | 41.07107695 | 116.0051082 | ENSG00000121454 |
| LTB | 1.498708087 | 2.825895442 | 0.176362641 | 0.357398131 | 2.346538071 | 6.584656445 | ENSG00000227507 |
| LRFN5 | 1.49870454 | 2.825888494 | 0.007790178 | 0.031989023 | 8.546936973 | 24.22222261 | ENSG00000165379 |
| COL5A2 | 1.498125333 | 2.824754197 | 1.27E-65 | 7.51E-63 | 465.3195026 | 1314.873304 | ENSG00000204262 |
| NA | 1.496557799 | 2.821686679 | 0.074706899 | 0.192128819 | 4.004501578 | 11.25151905 | ENSG00000270112 |
| PHOX2A | 1.494900475 | 2.818447071 | 0.03087097 | 0.097129148 | 5.662465085 | 15.90684192 | ENSG00000165462 |
| LINC00847 | 1.493492846 | 2.815698471 | 0.029604996 | 0.093885868 | 5.842471335 | 16.58810275 | ENSG00000245060 |
| GREM2 | 1.491804099 | 2.812404482 | 0.063513784 | 0.169462815 | 4.038901094 | 11.26674729 | ENSG00000180875 |
| NA | 1.490809706 | 2.81046667 | 0.181870096 | 0.365120512 | 2.019450547 | 5.626374276 | ENSG00000225087 |
| NRROS | 1.488367241 | 2.80571261 | 6.02E-15 | 2.25E-13 | 80.38007597 | 225.2049327 | ENSG00000174004 |
| PCDH11Y | 1.488269344 | 2.805522229 | 0.04273457 | 0.12489049 | 4.960847841 | 13.91498699 | ENSG00000099715 |
| SLC9A9 | 1.484166405 | 2.797554826 | 0.087775575 | 0.21624636 | 3.33728385 | 9.300714383 | ENSG00000181804 |
| NA | 1.483764184 | 2.79677498 | 0.09729908 | 0.233391734 | 3.704694626 | 10.28481857 | ENSG00000246203 |
| PTAFR | 1.483628892 | 2.796512719 | 2.06E-07 | 2.69E-06 | 43.0193642 | 120.2711363 | ENSG00000169403 |
| NA | 1.483502714 | 2.796268147 | 0.205213822 | 0.397157515 | 2.043171646 | 5.642548479 | ENSG00000267301 |
| GSTT2B | 1.482557831 | 2.794437351 | 0.000115304 | 0.000843342 | 21.0108894 | 58.55865194 | ENSG00000133433 |
| NA | 1.482539658 | 2.794402149 | 0.019325439 | 0.066781504 | 6.285800088 | 17.55915416 | ENSG00000225465 |
| GHR | 1.481593364 | 2.792569845 | 0.066815904 | 0.176305484 | 4.231950025 | 11.8898377 | ENSG00000112964 |
| KCNJ2 | 1.481301941 | 2.792005804 | 0.004126291 | 0.018868041 | 9.626643411 | 26.82439871 | ENSG00000123700 |
| ART3 | 1.480313238 | 2.790093051 | 0.111655165 | 0.259329454 | 2.982915755 | 8.321907083 | ENSG00000156219 |
| MYOZ1 | 1.477389736 | 2.784444884 | 0.000265411 | 0.001768627 | 16.56186386 | 46.08221225 | ENSG00000177791 |
| NA | 1.477051801 | 2.783792735 | 0.117409725 | 0.268744421 | 2.9900347 | 8.31225927 | ENSG00000258623 |
| SLC44A4 | 1.47375146 | 2.777431748 | 3.32E-07 | 4.23E-06 | 32.94463736 | 91.69938762 | ENSG00000204385 |
| NA | 1.470082303 | 2.770376977 | 0.164572547 | 0.340038572 | 2.666506649 | 7.381028622 | ENSG00000230552 |
| NA | 1.46996742 | 2.770156378 | 0.019143164 | 0.066244487 | 9.153949154 | 25.1298067 | ENSG00000227799 |
| DMBX1 | 1.469725037 | 2.769691011 | 0.010333701 | 0.04008873 | 8.870465024 | 24.6628779 | ENSG00000197587 |
| NA | 1.469585101 | 2.769422375 | 0.061803874 | 0.165932067 | 4.331589101 | 11.96200687 | ENSG00000272084 |
| NA | 1.469351427 | 2.768973845 | 0.135998083 | 0.298230426 | 2.62854766 | 7.302333068 | ENSG00000225603 |
| OXCT2P1 | 1.468532482 | 2.767402486 | 0.002905197 | 0.014010378 | 12.06806588 | 33.22321506 | ENSG00000237624 |
| LPO | 1.468477621 | 2.767297253 | 0.03546851 | 0.10830117 | 5.291494837 | 14.65971514 | ENSG00000167419 |
| FLRT3 | 1.46835546 | 2.767062939 | 0.021391233 | 0.072662131 | 6.547648052 | 18.19094642 | ENSG00000125848 |
| GDPGP1 | 1.468110652 | 2.766593442 | 0.078853568 | 0.19955029 | 5.312851672 | 14.69613093 | ENSG00000183208 |
| TBC1D10C | 1.467706515 | 2.765818555 | 0.010963614 | 0.042168603 | 8.921466694 | 24.72104832 | ENSG00000175463 |
| NA | 1.466602525 | 2.763702884 | 0.13422694 | 0.295612334 | 3.353886004 | 9.204898666 | ENSG00000285257 |
| NA | 1.465759221 | 2.762087878 | 0.003872937 | 0.017874 | 10.00117313 | 27.52277971 | ENSG00000213080 |
| AMDHD1 | 1.465486909 | 2.761566576 | 0.033041111 | 0.102426359 | 5.642303459 | 15.54894956 | ENSG00000139344 |
| NR2F1-AS1 | 1.462928887 | 2.756674422 | 0.00986659 | 0.038607156 | 9.350557557 | 25.70950344 | ENSG00000237187 |
| NA | 1.461839338 | 2.754593318 | 0.067656235 | 0.177975761 | 4.579199175 | 12.6893313 | ENSG00000253553 |
| C1orf94 | 1.459286847 | 2.749724056 | 9.13E-18 | 4.44E-16 | 109.1170686 | 300.5032872 | ENSG00000142698 |
| MMP9 | 1.459261107 | 2.749674997 | 1.69E-11 | 4.18E-10 | 66.45593749 | 182.8435031 | ENSG00000100985 |
| ARHGAP24 | 1.455985894 | 2.743439753 | 0.053694152 | 0.148900744 | 4.937126742 | 13.567026 | ENSG00000138639 |
| CATSPER2 | 1.452043691 | 2.735953464 | 0.015037122 | 0.054589782 | 8.454416841 | 23.31057113 | ENSG00000166762 |
| LINC01138 | 1.451932898 | 2.735743361 | 1.86E-06 | 2.04E-05 | 36.04023487 | 98.70495222 | ENSG00000274020 |
| RGS16 | 1.451368078 | 2.734672518 | 1.46E-42 | 3.20E-40 | 324.5106691 | 888.3832394 | ENSG00000143333 |
| NA | 1.449735702 | 2.73158005 | 0.148518627 | 0.316458791 | 3.044595843 | 8.230442293 | ENSG00000237227 |
| EOMES | 1.449019157 | 2.730223688 | 0.071728341 | 0.186359326 | 4.573275438 | 12.53875013 | ENSG00000163508 |
| PNLIPRP2 | 1.447717179 | 2.727760876 | 0.0251 | 0.082730718 | 6.756409413 | 18.26935843 | ENSG00000266200 |
| GJB5 | 1.446500286 | 2.725461019 | 0.008275866 | 0.033601916 | 8.879948232 | 24.27944708 | ENSG00000189280 |
| AIPL1 | 1.445002531 | 2.72263301 | 0.001601695 | 0.008409371 | 13.46746156 | 36.76872863 | ENSG00000129221 |
| LINC01291 | 1.443027639 | 2.718908572 | 0.078765169 | 0.199450066 | 4.062622193 | 10.92938007 | ENSG00000204792 |
| NA | 1.439801469 | 2.712835312 | 0.002499564 | 0.012312014 | 11.87710188 | 32.28356611 | ENSG00000246263 |
| TAC4 | 1.438811285 | 2.710974015 | 0.193475468 | 0.3813971 | 2.332300181 | 6.303284183 | ENSG00000176358 |
| NA | 1.437939749 | 2.709336804 | 0.008018475 | 0.032741385 | 8.979587309 | 24.28597347 | ENSG00000254275 |
| SLC30A1 | 1.437409832 | 2.708341818 | 1.63E-36 | 2.67E-34 | 502.8794913 | 1363.251526 | ENSG00000170385 |
| DANT2 | 1.436811005 | 2.707217887 | 0.048742962 | 0.138360263 | 6.302402242 | 17.07075426 | ENSG00000235244 |
| ATXN1 | 1.436540648 | 2.706710608 | 3.35E-05 | 0.000281292 | 23.27439055 | 62.90271315 | ENSG00000124788 |
| NA | 1.43385709 | 2.701680533 | 0.099557718 | 0.237418485 | 3.935702545 | 10.68347764 | ENSG00000257951 |
| NA | 1.43375268 | 2.701485017 | 0.061296207 | 0.165085406 | 4.297189585 | 11.61376233 | ENSG00000213070 |
| LOC349160 | 1.433179587 | 2.700412097 | 0.023502913 | 0.078433029 | 7.211306308 | 19.54230723 | ENSG00000234352 |
| NA | 1.432546583 | 2.699227511 | 0.044487047 | 0.128599093 | 6.254960045 | 16.9148762 | ENSG00000281852 |
| C22orf15 | 1.430780212 | 2.69592472 | 1.46E-05 | 0.000133476 | 30.9631458 | 83.67987238 | ENSG00000169314 |
| ATP8B3 | 1.428743248 | 2.692120988 | 7.07E-09 | 1.20E-07 | 54.00800297 | 145.0480116 | ENSG00000130270 |
| NA | 1.427369055 | 2.689557914 | 0.107028105 | 0.251239995 | 3.361004949 | 8.979521365 | ENSG00000283491 |
| FAM107A | 1.427245741 | 2.689328035 | 0.045203076 | 0.13029 | 6.739807259 | 18.023456 | ENSG00000168309 |
| GABRP | 1.425452442 | 2.685987223 | 2.77E-09 | 4.95E-08 | 49.05368926 | 132.0335879 | ENSG00000094755 |
| CPA2 | 1.421392809 | 2.678439686 | 0.211752698 | 0.405572993 | 1.985051031 | 5.319179997 | ENSG00000158516 |
| PPP1R3C | 1.420975069 | 2.677664242 | 6.03E-05 | 0.000474853 | 22.03692442 | 59.18637683 | ENSG00000119938 |
| DAND5 | 1.420940925 | 2.677600869 | 0.209538223 | 0.402720267 | 1.985051031 | 5.317004534 | ENSG00000179284 |
| NA | 1.419981637 | 2.675821051 | 0.254813574 | 0.461420495 | 2.012331602 | 5.366473103 | ENSG00000225511 |
| NA | 1.419341395 | 2.674633833 | 0.260847927 | 0.467817009 | 2.346538071 | 6.238587369 | ENSG00000259347 |
| LOC105372273 | 1.419313605 | 2.674582313 | 0.077948663 | 0.19807902 | 4.985764148 | 13.32084002 | ENSG00000267082 |
| OLIG3 | 1.417236333 | 2.670734074 | 0.097615866 | 0.233907913 | 4.204669454 | 11.34326738 | ENSG00000177468 |
| ATOH1 | 1.416610865 | 2.66957645 | 0.039387101 | 0.117341126 | 7.130939134 | 18.84175077 | ENSG00000172238 |

| | | | | | | | |
|---|---|---|---|---|---|---|---|
| NA | 1.416446997 | 2.669273244 | 0.231963447 | 0.43275295 | 1.995729448 | 5.317004534 | ENSG00000215482 |
| S100A4 | 1.415595026 | 2.667697395 | 1.29E-15 | 5.29E-14 | 93.4955337 | 249.2895543 | ENSG00000196154 |
| LINC01184 | 1.41556973 | 2.66765062 | 3.48E-11 | 8.33E-10 | 58.57504919 | 156.2283801 | ENSG00000245937 |
| HAPLN2 | 1.414900584 | 2.666413606 | 0.03245042 | 0.101059726 | 6.567809678 | 17.56255913 | ENSG00000132702 |
| NA | 1.414205781 | 2.665129768 | 0.0340308 | 0.10497954 | 8.463010327 | 22.41140535 | ENSG00000258417 |
| FKBP2 | 1.41417934 | 2.665080923 | 0.071217058 | 0.185354761 | 4.610039219 | 12.31961567 | ENSG00000173486 |
| NA | 1.412587316 | 2.662141611 | 0.129932885 | 0.288816212 | 2.982915755 | 7.925423479 | ENSG00000236372 |
| PTGER3 | 1.411438788 | 2.660023126 | 0.007598726 | 0.03135969 | 10.19541727 | 27.18541249 | ENSG00000050628 |
| ZFP57 | 1.410389131 | 2.658088484 | 0.030537891 | 0.09632672 | 9.413432853 | 24.89818656 | ENSG00000204644 |
| NA | 1.409088259 | 2.655692777 | 5.19E-05 | 0.000416026 | 22.96510039 | 60.86138966 | ENSG00000263826 |
| NA | 1.407609964 | 2.652972946 | 0.213555608 | 0.40799167 | 2.019450547 | 5.317004534 | ENSG00000232485 |
| FMO5 | 1.407425011 | 2.652632859 | 0.00363577 | 0.016936733 | 11.85574504 | 31.50866502 | ENSG00000131781 |
| CSPG5 | 1.407202089 | 2.652223012 | 3.00E-17 | 1.39E-15 | 109.7285562 | 291.5431568 | ENSG00000114646 |
| NA | 1.406378632 | 2.650709615 | 0.003478834 | 0.016326486 | 13.08728744 | 34.48897506 | ENSG00000285184 |
| NOS3 | 1.406317713 | 2.650597689 | 3.57E-09 | 6.27E-08 | 52.85919202 | 140.2315138 | ENSG00000164867 |
| KLHDC7A | 1.403658065 | 2.645715742 | 6.14E-60 | 2.70E-57 | 531.308364 | 1406.13398 | ENSG00000179023 |
| NA | 1.403639798 | 2.645682241 | 0.006232632 | 0.026581583 | 9.881372429 | 26.19166049 | ENSG00000227012 |
| TLR3 | 1.403296965 | 2.645053614 | 0.003221275 | 0.015296401 | 12.58939754 | 33.30843701 | ENSG00000164342 |
| MARCHF10 | 1.402419429 | 2.643445218 | 0.101139088 | 0.239860816 | 3.622852912 | 9.588613036 | ENSG00000173838 |
| NA | 1.402291119 | 2.643210126 | 0.088115807 | 0.216788302 | 4.297189585 | 11.3840341 | ENSG00000271387 |
| IL15 | 1.400571722 | 2.640061838 | 0.013384631 | 0.049573299 | 10.84987165 | 28.36377507 | ENSG00000164136 |
| NA | 1.400174542 | 2.639335118 | 2.51E-05 | 0.000218834 | 35.77838691 | 94.47723186 | ENSG00000223745 |
| NA | 1.399711894 | 2.638488863 | 0.167482126 | 0.344542477 | 2.662947177 | 7.018785343 | ENSG00000272945 |
| INTS6-AS1 | 1.398778474 | 2.636782319 | 0.058639018 | 0.159544425 | 5.50234113 | 14.64883782 | ENSG00000236778 |
| KCNK3 | 1.394836247 | 2.629587033 | 0.015310411 | 0.05538473 | 8.935704584 | 23.47872397 | ENSG00000171303 |
| NA | 1.394424966 | 2.628837501 | 0.060309166 | 0.162984918 | 5.50234113 | 14.59284286 | ENSG00000248923 |
| PCSK2 | 1.394154715 | 2.628345104 | 0.005655837 | 0.024486585 | 10.65178871 | 27.93543752 | ENSG00000125851 |
| GPR20 | 1.393507949 | 2.62716707 | 0.000509644 | 0.003127926 | 19.75710045 | 51.63736332 | ENSG00000204882 |
| NA | 1.393013946 | 2.626267639 | 0.163221 | 0.338180807 | 2.666506649 | 6.983315513 | ENSG00000225742 |
| OCA2 | 1.391970527 | 2.624368894 | 0.008919215 | 0.035701437 | 9.623083938 | 25.22278455 | ENSG00000104044 |
| FOXJ1 | 1.389820937 | 2.620461543 | 8.65E-18 | 4.24E-16 | 120.5259516 | 316.4618685 | ENSG00000129654 |
| NFASC | 1.38862247 | 2.618285593 | 3.59E-43 | 8.07E-41 | 325.9918809 | 853.4478607 | ENSG00000163531 |
| SPARCL1 | 1.388333654 | 2.617761486 | 0.012763853 | 0.047798011 | 10.04625107 | 26.16460896 | ENSG00000152583 |
| LIPA | 1.38818088 | 2.617484293 | 9.51E-49 | 2.81E-46 | 449.6030303 | 1177.887509 | ENSG00000107798 |
| RAB17 | 1.386564423 | 2.614553195 | 2.44E-48 | 7.08E-46 | 408.5835666 | 1067.8712 | ENSG00000124839 |
| TRPM6 | 1.385827718 | 2.613218427 | 0.00188059 | 0.009629864 | 15.04861737 | 39.1536622 | ENSG00000119121 |
| NA | 1.385767319 | 2.613109025 | 0.260446919 | 0.467408541 | 2.077571162 | 5.338475623 | ENSG00000235660 |
| NA | 1.385625384 | 2.612851956 | 0.166719374 | 0.343197664 | 2.893955096 | 7.658049957 | ENSG00000272195 |
| LMCD1 | 1.385623491 | 2.612848528 | 5.35E-25 | 4.79E-23 | 170.3360986 | 445.5076739 | ENSG00000071282 |
| LYNX1 | 1.385354834 | 2.612362011 | 1.57E-13 | 4.98E-12 | 89.66035995 | 234.2180117 | ENSG00000180155 |
| DBH-AS1 | 1.384280333 | 2.610417082 | 0.011693211 | 0.044494516 | 8.897745595 | 23.28134415 | ENSG00000225756 |
| BHLHE41 | 1.383609123 | 2.609202875 | 0.001712222 | 0.008908605 | 16.80858422 | 43.6203071 | ENSG00000123095 |
| CEACAM1 | 1.382874309 | 2.607874256 | 0.003776137 | 0.017488762 | 12.36787283 | 32.08975921 | ENSG00000079385 |
| KCNN4 | 1.381750263 | 2.605843176 | 0.014618811 | 0.053389335 | 9.990494714 | 25.98747073 | ENSG00000104783 |
| NA | 1.380897684 | 2.604303675 | 0.020503256 | 0.070209244 | 9.646805037 | 25.11346455 | ENSG00000286909 |
| ARHGEF33 | 1.380545421 | 2.60366786 | 0.123369986 | 0.278057307 | 3.306443806 | 8.606684313 | ENSG00000214694 |
| FGF16 | 1.379349526 | 2.601510492 | 4.36E-06 | 4.47E-05 | 30.6562199 | 79.85757807 | ENSG00000196468 |
| WDR66 | 1.377876062 | 2.598854855 | 0.004123804 | 0.018861234 | 15.49520011 | 40.35209841 | ENSG00000158023 |
| NA | 1.377340767 | 2.597890759 | 1.21E-07 | 1.65E-06 | 42.08315341 | 109.2781209 | ENSG00000253320 |
| GGT1 | 1.377046348 | 2.597360647 | 0.000323418 | 0.002098164 | 20.94445464 | 54.26318598 | ENSG00000100031 |
| SLC6A17 | 1.376883493 | 2.597067468 | 0.016992662 | 0.060300007 | 8.897745595 | 23.12546609 | ENSG00000197106 |
| ALG1L | 1.376828894 | 2.596969182 | 3.20E-05 | 0.000270226 | 27.57158014 | 71.52188316 | ENSG00000189366 |
| SMOC2 | 1.376626809 | 2.596605438 | 1.79E-14 | 6.19E-13 | 100.0321978 | 259.9906036 | ENSG00000112562 |
| XCL1 | 1.376018325 | 2.595510501 | 0.233886031 | 0.435080545 | 2.74478889 | 7.048958286 | ENSG00000143184 |
| NA | 1.37343166 | 2.590861077 | 0.270746654 | 0.479166306 | 2.077571162 | 5.284656126 | ENSG00000234558 |
| NPNT | 1.372773098 | 2.589678669 | 0.041505778 | 0.12213098 | 6.905575619 | 17.92858624 | ENSG00000168743 |
| ZNF354C | 1.372731015 | 2.589603132 | 1.51E-05 | 0.000137778 | 29.95132252 | 77.39255149 | ENSG00000177932 |
| NA | 1.370946779 | 2.58640245 | 0.040058025 | 0.118833772 | 7.737671984 | 19.88960581 | ENSG00000253706 |
| LPAR5 | 1.370684957 | 2.585933109 | 0.154009327 | 0.325040918 | 2.948516239 | 7.665522306 | ENSG00000184574 |
| NR5A1 | 1.370554899 | 2.585699999 | 0.055284685 | 0.1525053 | 5.233374222 | 13.60126632 | ENSG00000136931 |
| ERVMER34-1 | 1.369469153 | 2.58375478 | 1.36E-28 | 1.49E-26 | 217.6045998 | 562.1451845 | ENSG00000226887 |
| LINC01694 | 1.369132688 | 2.583152268 | 9.01E-06 | 8.72E-05 | 29.83508129 | 77.00883712 | ENSG00000233922 |
| CFAP45 | 1.367443707 | 2.580129909 | 0.155759377 | 0.327558713 | 2.928354612 | 7.606405924 | ENSG00000213085 |
| GBGT1 | 1.36740704 | 2.580064335 | 4.63E-15 | 1.76E-13 | 109.0704906 | 280.9091477 | ENSG00000148288 |
| NA | 1.367375883 | 2.580008615 | 0.072324821 | 0.187617341 | 5.053367972 | 12.93835516 | ENSG00000242861 |
| LINC02572 | 1.36666949 | 2.578745664 | 0.257756512 | 0.464655567 | 2.342978598 | 6.025979302 | ENSG00000229536 |
| CD79A | 1.366582857 | 2.578590816 | 0.070804179 | 0.184534491 | 4.624277109 | 11.92965846 | ENSG00000105369 |
| POLR3GL | 1.365708774 | 2.577029004 | 0.001816799 | 0.009341206 | 14.54597279 | 37.49822853 | ENSG00000121851 |
| GUCY1B2 | 1.365687282 | 2.576990613 | 0.198968617 | 0.389060538 | 2.721067792 | 6.926374594 | ENSG00000123201 |
| NA | 1.363565136 | 2.57320275 | 1.18E-11 | 2.98E-10 | 94.69887965 | 243.1421482 | ENSG00000251095 |
| NAALADL2 | 1.363105248 | 2.572382622 | 0.098144545 | 0.234852983 | 4.215347871 | 10.933731 | ENSG00000177694 |
| KLHL31 | 1.361495421 | 2.569513836 | 0.06013264 | 0.162600933 | 6.078513268 | 15.47053756 | ENSG00000124743 |
| CPA6 | 1.36105865 | 2.568736044 | 0.304106088 | 0.514194957 | 1.919811471 | 4.995811515 | ENSG00000165078 |
| C3P1 | 1.360823568 | 2.56831751 | 0.24111517 | 0.444668216 | 2.370259169 | 6.047733938 | ENSG00000167798 |

| | | | | | | | |
|---|---|---|---|---|---|---|---|
| TRIM29 | 1.36051595 | 2.567769941 | 0.061258598 | 0.165031188 | 5.921948785 | 15.2352289 | ENSG00000137699 |
| NA | 1.360375633 | 2.56752021 | 6.34E-07 | 7.60E-06 | 36.8803399 | 94.88700884 | ENSG00000250519 |
| WDR72 | 1.360060017 | 2.56695858 | 8.24E-06 | 8.04E-05 | 30.07320815 | 77.21151383 | ENSG00000166415 |
| TNFRSF14 | 1.357664238 | 2.562699356 | 0.119802067 | 0.272404316 | 3.592012868 | 9.24907035 | ENSG00000157873 |
| MINAR1 | 1.354488721 | 2.557064799 | 0.002572778 | 0.012593808 | 16.1520449 | 41.01111015 | ENSG00000169330 |
| LEPR | 1.354017916 | 2.55623047 | 7.24E-06 | 7.16E-05 | 31.87441413 | 81.73727512 | ENSG00000116678 |
| TXNIP | 1.353760278 | 2.555774017 | 2.51E-15 | 9.87E-14 | 112.1862759 | 286.0640313 | ENSG00000265972 |
| PPP1R3B | 1.351685767 | 2.552101605 | 1.38E-47 | 3.72E-45 | 401.1848035 | 1023.329224 | ENSG00000173281 |
| KLK7 | 1.351403148 | 2.551601706 | 0.002310762 | 0.011499009 | 15.03082001 | 38.21250019 | ENSG00000169035 |
| ALPG | 1.348979785 | 2.547319257 | 0.000288 | 0.001897066 | 21.04737385 | 53.75113953 | ENSG00000163286 |
| MCHR1 | 1.34884779 | 2.547086208 | 2.53E-08 | 3.87E-07 | 47.03927273 | 119.7379023 | ENSG00000128285 |
| LGSN | 1.348199342 | 2.545941628 | 0.203812293 | 0.395475152 | 2.57754599 | 6.636300478 | ENSG00000146166 |
| NA | 1.346600813 | 2.543122246 | 0.20906084 | 0.402212098 | 2.370259169 | 5.963741498 | ENSG00000198857 |
| NA | 1.346112611 | 2.54226181 | 0.031757295 | 0.099475289 | 7.589395501 | 19.33310413 | ENSG00000187952 |
| TNNC1 | 1.345674389 | 2.541489709 | 1.77E-05 | 0.000159077 | 28.73640843 | 73.11224114 | ENSG00000114854 |
| POU3F2 | 1.345417252 | 2.541036771 | 0.199156476 | 0.389203606 | 3.048155315 | 7.689168859 | ENSG00000184486 |
| TDRD6 | 1.34513074 | 2.540532183 | 0.10660945 | 0.250386381 | 4.273468486 | 10.86903418 | ENSG00000180113 |
| NA | 1.34395433 | 2.538461413 | 0.194844236 | 0.383534854 | 2.608386034 | 6.671770308 | ENSG00000241170 |
| NA | 1.343396427 | 2.537479957 | 0.137289793 | 0.300013625 | 4.541240186 | 11.59758812 | ENSG00000234694 |
| NA | 1.342868588 | 2.53655174 | 0.127090661 | 0.284110628 | 3.939262018 | 10.0410916 | ENSG00000274922 |
| SEMA3G | 1.342835948 | 2.536494352 | 5.42E-13 | 1.61E-11 | 86.90013797 | 220.594444 | ENSG00000010319 |
| CREB5 | 1.342744063 | 2.536332809 | 0.008108697 | 0.033016978 | 10.81043812 | 27.53677845 | ENSG00000146592 |
| NA | 1.342705181 | 2.536264452 | 0.088356588 | 0.217333655 | 5.189491497 | 13.26172364 | ENSG00000261326 |
| GALNT5 | 1.342453466 | 2.535821975 | 0.330434623 | 0.541830151 | 1.919811471 | 4.913994538 | ENSG00000136542 |
| PRKAR2A-AS1 | 1.340115818 | 2.531716423 | 0.122862805 | 0.277211956 | 3.711813571 | 9.324360936 | ENSG00000224424 |
| CNTNAP5 | 1.339297841 | 2.530281402 | 0.119236954 | 0.271627247 | 3.908421974 | 9.954923696 | ENSG00000155052 |
| LOC100506274 | 1.338688081 | 2.529212196 | 0.203307857 | 0.394885209 | 2.611945506 | 6.645948291 | ENSG00000229727 |
| LOC100507642 | 1.338549758 | 2.528969711 | 0.057382678 | 0.156901149 | 6.820174433 | 17.39194728 | ENSG00000240859 |
| PYY | 1.337808936 | 2.527671421 | 0.0014528 | 0.007745943 | 16.37949335 | 41.2435083 | ENSG00000131096 |
| ENPP2 | 1.335273308 | 2.52323278 | 9.66E-05 | 0.000719704 | 34.24083449 | 86.77241293 | ENSG00000136960 |
| DUSP10 | 1.335212198 | 2.523125903 | 0.00424308 | 0.019308572 | 15.13045908 | 37.97217819 | ENSG00000143507 |
| NA | 1.334875576 | 2.522537254 | 0.043346046 | 0.126286653 | 6.585607041 | 16.66216383 | ENSG00000277287 |
| NA | 1.334578479 | 2.522017836 | 1.08E-11 | 2.71E-10 | 78.3781173 | 197.7776852 | ENSG00000234261 |
| NOD2 | 1.334416742 | 2.521735116 | 0.007973522 | 0.032571913 | 11.56661651 | 29.1760379 | ENSG00000167207 |
| APOBEC3B | 1.332474782 | 2.518342981 | 6.28E-12 | 1.62E-10 | 85.82994089 | 216.4149627 | ENSG00000179750 |
| ARL11 | 1.332129788 | 2.517740839 | 0.004105333 | 0.018794959 | 13.55998169 | 34.13855505 | ENSG00000152213 |
| MVP | 1.33190881 | 2.517355224 | 9.29E-05 | 0.000695526 | 28.35147962 | 71.18451593 | ENSG00000013364 |
| FALEC | 1.331198381 | 2.516115905 | 0.142666452 | 0.30795016 | 3.27204429 | 8.267141628 | ENSG00000228126 |
| GTSF1 | 1.330528622 | 2.51494809 | 0.061734685 | 0.165817029 | 5.291494837 | 13.32206953 | ENSG00000170627 |
| NA | 1.330371852 | 2.514674819 | 0.143609314 | 0.309454803 | 3.739094142 | 9.303835805 | ENSG00000286216 |
| NA | 1.329534366 | 2.513215471 | 0.007634761 | 0.031474016 | 12.37855125 | 31.00068595 | ENSG00000266389 |
| CHRNE | 1.327860502 | 2.510301243 | 0.139226745 | 0.302949312 | 3.302884333 | 8.29826053 | ENSG00000108556 |
| HNMT | 1.327658592 | 2.509949942 | 0.000238767 | 0.001610974 | 20.9385309 | 52.45549014 | ENSG00000150540 |
| HHIP-AS1 | 1.326866108 | 2.508571585 | 0.305697648 | 0.516031107 | 2.250458467 | 5.711312675 | ENSG00000248890 |
| FGD2 | 1.326502616 | 2.507939622 | 0.164426544 | 0.339885648 | 3.766374713 | 9.336184213 | ENSG00000146192 |
| NA | 1.326094038 | 2.507229462 | 0.234373283 | 0.435643647 | 2.621428715 | 6.586831908 | ENSG00000267055 |
| NPL | 1.32578425 | 2.506691146 | 9.22E-08 | 1.30E-06 | 59.81579546 | 150.0712535 | ENSG00000135838 |
| CACNA1S | 1.325526069 | 2.506242595 | 0.274939709 | 0.483718689 | 1.985051031 | 4.974340426 | ENSG00000081248 |
| PLCB2 | 1.325368289 | 2.505968516 | 6.08E-29 | 6.83E-27 | 255.1325545 | 638.8708527 | ENSG00000137841 |
| NA | 1.324435451 | 2.504348694 | 0.006648974 | 0.02806996 | 11.95538412 | 29.90024092 | ENSG00000251611 |
| ARHGDIB | 1.324280628 | 2.504079954 | 0.251011132 | 0.456548706 | 1.992169976 | 4.981812775 | ENSG00000111348 |
| MOGAT3 | 1.323570784 | 2.502848184 | 0.154955034 | 0.326525978 | 3.313562751 | 8.286437254 | ENSG00000106384 |
| LOC100507336 | 1.322938375 | 2.501751296 | 0.145865547 | 0.312959652 | 3.33728385 | 8.324082547 | ENSG00000228793 |
| NA | 1.321626269 | 2.499477031 | 0.219319369 | 0.415703519 | 2.683108803 | 6.678296698 | ENSG00000253807 |
| NUPR1 | 1.3210464 | 2.498472606 | 0.087931833 | 0.216448472 | 4.913405643 | 12.34108676 | ENSG00000176046 |
| LINC02617 | 1.318084407 | 2.493348262 | 0.000513137 | 0.003147318 | 20.32943379 | 50.56162639 | ENSG00000256288 |
| NA | 1.317709227 | 2.492699938 | 0.099738461 | 0.237659577 | 4.685957196 | 11.62870702 | ENSG00000231680 |
| ELF3 | 1.317039752 | 2.491543483 | 2.58E-09 | 4.64E-08 | 58.33900726 | 145.569495 | ENSG00000163435 |
| SCARF1 | 1.314993185 | 2.488011553 | 1.59E-10 | 3.45E-09 | 87.16704609 | 216.7587407 | ENSG00000074660 |
| PRODH | 1.314899761 | 2.487850443 | 2.43E-05 | 0.000212228 | 29.12754031 | 72.4860293 | ENSG00000100033 |
| HGC6.3 | 1.314688417 | 2.48748602 | 0.191963409 | 0.379193768 | 2.662947177 | 6.596479721 | ENSG00000235994 |
| TMPRSS9 | 1.314610922 | 2.487352405 | 0.006022464 | 0.02580763 | 13.48406371 | 33.64769615 | ENSG00000178297 |
| NA | 1.31360095 | 2.485611722 | 0.279404794 | 0.488473282 | 1.992169976 | 4.935465628 | ENSG00000237713 |
| ADARB2 | 1.313547212 | 2.485519138 | 1.95E-07 | 2.55E-06 | 62.92143484 | 155.854808 | ENSG00000185736 |
| CALHM5 | 1.311486567 | 2.481971531 | 0.255451662 | 0.462309979 | 2.015891075 | 4.965638572 | ENSG00000178033 |
| TENM2 | 1.311317572 | 2.481680813 | 1.09E-05 | 0.000103143 | 42.21126827 | 104.6737071 | ENSG00000145934 |
| NA | 1.310672195 | 2.480570903 | 0.279906112 | 0.488994296 | 2.046731118 | 5.023808996 | ENSG00000249193 |
| FGF17 | 1.309061512 | 2.477803039 | 0.294818158 | 0.504909849 | 2.043171646 | 5.012931678 | ENSG00000158815 |
| MYO18B | 1.308740579 | 2.477251903 | 0.00070225 | 0.004123215 | 19.58393382 | 48.4503092 | ENSG00000133454 |
| SLC22A20P | 1.307763985 | 2.475575561 | 4.98E-13 | 1.48E-11 | 113.9675996 | 281.5320702 | ENSG00000197847 |
| NA | 1.306703576 | 2.473756634 | 0.18783405 | 0.373590945 | 3.340843322 | 8.219848522 | ENSG00000287778 |
| PTCH2 | 1.306683253 | 2.473721786 | 0.043258568 | 0.126051235 | 7.238586879 | 17.9534623 | ENSG00000117425 |
| HPSE2 | 1.305518138 | 2.471724825 | 0.015897811 | 0.057169991 | 10.82259108 | 26.51502897 | ENSG00000172987 |
| C5orf58 | 1.305273581 | 2.471305869 | 0.094848234 | 0.229289414 | 4.736958866 | 11.59011577 | ENSG00000234511 |

| | | | | | | | |
|---|---|---|---|---|---|---|---|
| NA | 1.304354848 | 2.469732599 | 0.070445253 | 0.183700441 | 5.703983547 | 14.02480145 | ENSG00000267364 |
| WFDC1 | 1.304287378 | 2.469617101 | 0.009627524 | 0.0378704 | 11.35340595 | 27.94508533 | ENSG00000103175 |
| NA | 1.304269107 | 2.469585824 | 0.043576617 | 0.12685018 | 7.052656893 | 17.32412904 | ENSG00000242798 |
| TPCN2 | 1.304154949 | 2.469390418 | 0.04286725 | 0.125181385 | 7.452992645 | 18.5519602 | ENSG00000287725 |
| MICAL2 | 1.303161344 | 2.467690298 | 4.59E-07 | 5.67E-06 | 49.4901784 | 121.9852348 | ENSG00000133816 |
| CR2 | 1.300814535 | 2.463679409 | 2.30E-18 | 1.20E-16 | 134.1149677 | 330.2412916 | ENSG00000117322 |
| NA | 1.300619668 | 2.46334666 | 0.072238093 | 0.187469615 | 6.183796748 | 15.34627286 | ENSG00000213225 |
| TNNI1 | 1.299760905 | 2.461880789 | 0.000120618 | 0.000878664 | 27.8533104 | 68.86645465 | ENSG00000159173 |
| NA | 1.298322027 | 2.459426647 | 0.108047962 | 0.252816969 | 4.351750728 | 10.65765562 | ENSG00000228878 |
| PWWP3B | 1.29817678 | 2.459179051 | 0.002577324 | 0.012612793 | 17.42688521 | 42.91490525 | ENSG00000157502 |
| NEU4 | 1.29619188 | 2.455797969 | 0.057145041 | 0.156470665 | 6.343920703 | 15.49853504 | ENSG00000204099 |
| CTHRC1 | 1.293055828 | 2.450465487 | 0.000613775 | 0.00366576 | 19.92998776 | 48.74662616 | ENSG00000164932 |
| ACTG2 | 1.291423649 | 2.447694745 | 1.28E-05 | 0.000118937 | 31.73236687 | 77.72121686 | ENSG00000163017 |
| RNASEL | 1.290887449 | 2.446785191 | 0.006767437 | 0.028463243 | 11.96250306 | 29.21708817 | ENSG00000135828 |
| HSD52 | 1.290872754 | 2.446760268 | 0.230985332 | 0.431331045 | 2.312138554 | 5.668370496 | ENSG00000224609 |
| CD27-AS1 | 1.290769903 | 2.446585843 | 0.000222515 | 0.001511162 | 24.1963373 | 59.17795852 | ENSG00000215039 |
| RGS2 | 1.290375039 | 2.445916307 | 2.68E-21 | 1.77E-19 | 188.9722535 | 462.0744509 | ENSG00000116741 |
| NABP1 | 1.289742363 | 2.444843915 | 3.04E-05 | 0.000258958 | 31.1303887 | 76.10503686 | ENSG00000173559 |
| LINC00886 | 1.288407707 | 2.442583204 | 0.153885022 | 0.324923819 | 4.124302281 | 9.926926216 | ENSG00000240875 |
| TMPRSS4 | 1.284513251 | 2.43599851 | 0.327611221 | 0.538978822 | 2.046731118 | 4.921466888 | ENSG00000137648 |
| ARIH2OS | 1.284385664 | 2.435783088 | 0.127422465 | 0.284710513 | 3.922659864 | 9.600436312 | ENSG00000221883 |
| NA | 1.284319969 | 2.435672175 | 0.179183746 | 0.361714629 | 3.275603762 | 8.024360618 | ENSG00000226070 |
| CD7 | 1.283010277 | 2.433462051 | 0.007077958 | 0.02953972 | 13.68779106 | 33.5092218 | ENSG00000173762 |
| NA | 1.282671498 | 2.432890685 | 0.108530766 | 0.253663801 | 5.111488587 | 12.30032005 | ENSG00000283538 |
| NA | 1.282176473 | 2.432056042 | 0.237702893 | 0.439841186 | 2.308579082 | 5.616726463 | ENSG00000270571 |
| AHNAK | 1.281635131 | 2.431143633 | 0.054918763 | 0.151695311 | 87.08192424 | 211.7029957 | ENSG00000124942 |
| LOC729867 | 1.279478594 | 2.427512281 | 0.082636635 | 0.206810608 | 5.360293869 | 12.94488155 | ENSG00000227741 |
| SOX3 | 1.278587364 | 2.426013139 | 1.02E-35 | 1.62E-33 | 382.1607968 | 927.3548454 | ENSG00000134595 |
| DDR2 | 1.276489872 | 2.422488593 | 0.003889805 | 0.017943081 | 14.95253776 | 36.15434008 | ENSG00000162733 |
| AFDN-DT | 1.274930823 | 2.419872143 | 0.000560624 | 0.00338784 | 27.67953337 | 66.82702308 | ENSG00000198221 |
| ZNF503 | 1.27488734 | 2.41979921 | 0.220599478 | 0.417562507 | 3.064757469 | 7.344329288 | ENSG00000165655 |
| ANXA2 | 1.274738966 | 2.419550358 | 1.45E-69 | 1.01E-66 | 850.7786567 | 2058.097923 | ENSG00000182718 |
| PLA2G4D | 1.273549159 | 2.417555749 | 0.361487773 | 0.573611502 | 2.050290591 | 4.892523449 | ENSG00000159337 |
| NA | 1.273235915 | 2.417030897 | 0.073650495 | 0.190220131 | 5.652981876 | 13.64326254 | ENSG00000287608 |
| OPTC | 1.272153059 | 2.415217405 | 0.004773088 | 0.021229338 | 18.85473052 | 45.80613688 | ENSG00000188770 |
| OTOGL | 1.271502615 | 2.414128743 | 0.12392808 | 0.278982214 | 4.706118822 | 11.29379881 | ENSG00000165899 |
| ITGB2 | 1.270969833 | 2.413237378 | 0.025109465 | 0.082741625 | 8.932145111 | 21.55374132 | ENSG00000160255 |
| ADAMTS5 | 1.270600575 | 2.412619788 | 8.55E-06 | 8.34E-05 | 34.37751668 | 82.97787551 | ENSG00000154736 |
| NA | 1.270375063 | 2.412242695 | 0.007676604 | 0.031625799 | 13.26965795 | 32.00749073 | ENSG00000284620 |
| RFPL2 | 1.269745765 | 2.411190713 | 0.19882802 | 0.389030323 | 3.32304596 | 8.007240455 | ENSG00000128253 |
| HHLA1 | 1.268080409 | 2.408408992 | 2.47E-11 | 5.98E-10 | 87.62222232 | 210.6988772 | ENSG00000132297 |
| CCDC110 | 1.267587051 | 2.40758553 | 0.00776909 | 0.031932511 | 13.22813949 | 31.85255863 | ENSG00000168491 |
| NA | 1.267323393 | 2.407145574 | 0.23166755 | 0.432316371 | 2.624988188 | 6.350577289 | ENSG00000253649 |
| NA | 1.266953077 | 2.406527779 | 0.003107739 | 0.014850022 | 16.34717877 | 39.54607842 | ENSG00000249602 |
| NA | 1.266076118 | 2.405065387 | 0.132649384 | 0.293143565 | 5.026087401 | 12.05318811 | ENSG00000284630 |
| LYST | 1.265521531 | 2.404141033 | 0.162053479 | 0.336860754 | 3.333724377 | 7.995417179 | ENSG00000143669 |
| SPIB | 1.263949315 | 2.401522482 | 6.71E-11 | 1.54E-09 | 97.26015454 | 233.8437545 | ENSG00000269404 |
| GDF3 | 1.262939525 | 2.399842166 | 1.08E-13 | 3.45E-12 | 114.2914587 | 274.4618087 | ENSG00000184344 |
| PHF24 | 1.261362428 | 2.397220187 | 0.024804022 | 0.081989855 | 9.285317997 | 22.23812358 | ENSG00000122733 |
| PNMA6F | 1.260818258 | 2.396316151 | 0.022138578 | 0.074725119 | 10.46466351 | 24.87766143 | ENSG00000225110 |
| NA | 1.259997595 | 2.394953417 | 0.006001497 | 0.025752845 | 13.62402604 | 32.58300449 | ENSG00000279393 |
| NA | 1.259957244 | 2.394886433 | 0.001022784 | 0.005717964 | 20.30571269 | 48.52531624 | ENSG00000256969 |
| NA | 1.259707197 | 2.394471388 | 0.250783732 | 0.456317828 | 2.370259169 | 5.612375535 | ENSG00000273355 |
| NA | 1.258560386 | 2.392568759 | 0.229362185 | 0.429159305 | 2.608386034 | 6.273111239 | ENSG00000267293 |
| LOC105373159 | 1.25586064 | 2.38809568 | 1.62E-06 | 1.80E-05 | 62.4517414 | 149.0720367 | ENSG00000177788 |
| LRRC15 | 1.255719454 | 2.387861985 | 0.03735787 | 0.112675145 | 7.906999816 | 18.91609539 | ENSG00000172061 |
| TUBA3E | 1.25484338 | 2.386412402 | 0.216451181 | 0.411898484 | 2.635666605 | 6.30110872 | ENSG00000152086 |
| APOBEC3D | 1.253022302 | 2.383401992 | 0.085149168 | 0.211695899 | 5.564021218 | 13.33606827 | ENSG00000243811 |
| NA | 1.252848696 | 2.383115205 | 0.000771217 | 0.004467619 | 22.17452249 | 52.85916254 | ENSG00000268970 |
| SYNPO | 1.252401162 | 2.382376061 | 4.86E-11 | 1.14E-09 | 93.1509543 | 221.50392 | ENSG00000171992 |
| MAG | 1.251680363 | 2.381186075 | 0.092844652 | 0.225630068 | 5.311656463 | 12.63551181 | ENSG00000105695 |
| IL2RB | 1.251266158 | 2.380502523 | 7.81E-06 | 7.67E-05 | 38.26844678 | 90.87218008 | ENSG00000100385 |
| NA | 1.250847269 | 2.379811441 | 0.281844532 | 0.490709708 | 2.401099213 | 5.633846625 | ENSG00000226191 |
| LINC01780 | 1.248319809 | 2.375645895 | 0.01306311 | 0.048687419 | 11.92810355 | 28.35244625 | ENSG00000232650 |
| SSTR1 | 1.248268991 | 2.375562215 | 0.006905718 | 0.028942407 | 13.81115124 | 32.90665652 | ENSG00000139874 |
| PEX5L | 1.247719187 | 2.374657073 | 4.23E-05 | 0.000346352 | 34.85969415 | 82.97598359 | ENSG00000114757 |
| POLD4 | 1.247155248 | 2.373729017 | 8.44E-05 | 0.000639386 | 28.12255664 | 66.77348712 | ENSG00000175482 |
| TAPBPL | 1.247108952 | 2.373652847 | 0.021813125 | 0.073837761 | 10.19778154 | 24.27386664 | ENSG00000139192 |
| NA | 1.246526683 | 2.372695038 | 0.297642112 | 0.507812023 | 1.95777046 | 4.682090846 | ENSG00000213433 |
| NA | 1.246508868 | 2.372665739 | 0.074367563 | 0.191591463 | 6.03107107 | 14.22435673 | ENSG00000280310 |
| NA | 1.245553077 | 2.371094359 | 0.298960449 | 0.50928202 | 1.961329932 | 4.68426631 | ENSG00000235313 |
| OAS3 | 1.245146609 | 2.370426417 | 0.001791236 | 0.009236679 | 22.33701071 | 52.81309894 | ENSG00000111331 |
| MUC19 | 1.244775922 | 2.369817436 | 0.028386325 | 0.090891106 | 9.772250144 | 23.32116491 | ENSG00000205592 |
| CD1D | 1.244626539 | 2.369572067 | 1.40E-05 | 0.00012824 | 43.01814284 | 101.9390885 | ENSG00000158473 |

| | | | | | | | |
|---|---|---|---|---|---|---|---|
| SULT2B1 | 1.243317961 | 2.367423751 | 0.043098272 | 0.125661705 | 7.552631721 | 17.94504399 | ENSG00000088002 |
| NEURL1 | 1.243040527 | 2.366968534 | 0.001602968 | 0.008413577 | 19.13348612 | 45.31071613 | ENSG00000107954 |
| BAHCC1 | 1.240150983 | 2.362232525 | 7.62E-05 | 0.000583803 | 29.08009811 | 68.74387096 | ENSG00000266074 |
| NA | 1.239123604 | 2.36055092 | 0.044104735 | 0.127876062 | 7.572793347 | 17.91174962 | ENSG00000287406 |
| NA | 1.238783094 | 2.35999384 | 0.174062421 | 0.354297136 | 3.691651944 | 8.672327087 | ENSG00000214252 |
| SBK3 | 1.238768898 | 2.359970618 | 0.012327025 | 0.046512661 | 11.25732635 | 26.56232207 | ENSG00000231274 |
| NA | 1.23853021 | 2.359580204 | 0.186122109 | 0.371006547 | 3.932143073 | 9.314713123 | ENSG00000272848 |
| F8 | 1.23822829 | 2.359086453 | 0.000114143 | 0.000836148 | 28.08935233 | 66.2799583 | ENSG00000185010 |
| NA | 1.237881082 | 2.358518769 | 0.197478355 | 0.387118741 | 4.155142324 | 9.659552695 | ENSG00000237976 |
| KCNJ1 | 1.237841815 | 2.358454577 | 0.233164365 | 0.434097722 | 2.69378722 | 6.306405605 | ENSG00000151704 |
| NOS2 | 1.237669598 | 2.358173061 | 4.86E-09 | 8.43E-08 | 67.25127007 | 158.4742722 | ENSG00000007171 |
| NA | 1.236477337 | 2.356225042 | 0.012229872 | 0.046256987 | 11.5571333 | 27.26940493 | ENSG00000262468 |
| INPP4B | 1.236126131 | 2.355651518 | 0.090132377 | 0.220657239 | 5.680262448 | 13.34259466 | ENSG00000109452 |
| NA | 1.235925113 | 2.355323316 | 0.003672627 | 0.017078935 | 19.13348612 | 45.11059376 | ENSG00000287315 |
| NA | 1.234176217 | 2.352469823 | 0.320077372 | 0.530559219 | 1.992169976 | 4.689563196 | ENSG00000259868 |
| MYO1F | 1.233656577 | 2.351622647 | 1.06E-09 | 2.03E-08 | 87.70167362 | 205.9173978 | ENSG00000142347 |
| IRX1 | 1.232198177 | 2.349246626 | 0.012428349 | 0.046810804 | 11.80830284 | 27.83432491 | ENSG00000170549 |
| NA | 1.232004539 | 2.348931332 | 0.146026981 | 0.31312814 | 4.235509498 | 10.02709286 | ENSG00000230593 |
| GABRR1 | 1.231433639 | 2.348002003 | 0.218051526 | 0.414192624 | 2.959194656 | 6.966195351 | ENSG00000146276 |
| FZD4-DT | 1.231010128 | 2.347312835 | 0.304502335 | 0.514680623 | 1.985051031 | 4.663741179 | ENSG00000246523 |
| TRIM10 | 1.23022327 | 2.34603294 | 0.299753608 | 0.509848409 | 1.985051031 | 4.660619757 | ENSG00000204613 |
| PTCHD1 | 1.229709399 | 2.34519746 | 1.58E-07 | 2.11E-06 | 74.77812192 | 175.114019 | ENSG00000165186 |
| OPN3 | 1.229168816 | 2.34431887 | 0.002394164 | 0.011870253 | 17.53008375 | 41.0885762 | ENSG00000054277 |
| UCMA | 1.229136617 | 2.344266548 | 0.002963699 | 0.014245216 | 17.11403557 | 40.22705571 | ENSG00000165623 |
| MIR3936HG | 1.228225723 | 2.342786884 | 0.181735443 | 0.364958937 | 3.742653615 | 8.684150363 | ENSG00000233006 |
| BGN | 1.227151539 | 2.341043171 | 2.13E-09 | 3.91E-08 | 75.90616028 | 177.58635 | ENSG00000182492 |
| RHBDL1 | 1.226611726 | 2.340167386 | 1.04E-08 | 1.72E-07 | 63.02999847 | 147.3359726 | ENSG00000103269 |
| NA | 1.225362066 | 2.338141215 | 0.073669759 | 0.190243867 | 6.207517847 | 14.58972144 | ENSG00000259033 |
| SULT1C2 | 1.225066816 | 2.337662759 | 0.008432453 | 0.034107276 | 13.93095194 | 32.53665734 | ENSG00000198203 |
| RHBG | 1.224824802 | 2.337270646 | 0.001218545 | 0.006660554 | 20.17523357 | 47.17518494 | ENSG00000132677 |
| RHBDL2 | 1.223218677 | 2.334670055 | 0.15945563 | 0.333012059 | 4.695440405 | 10.89920713 | ENSG00000158315 |
| NA | 1.222447939 | 2.333423126 | 0.064969245 | 0.172379782 | 7.403465515 | 17.139024 | ENSG00000203362 |
| SIT1 | 1.22196952 | 2.332649457 | 1.85E-10 | 3.98E-09 | 86.86665433 | 203.0646847 | ENSG00000137078 |
| NA | 1.221727675 | 2.332258458 | 0.002534739 | 0.012436619 | 17.92121563 | 41.72254393 | ENSG00000258053 |
| TCAM1P | 1.221138184 | 2.331305682 | 0.05768566 | 0.157567462 | 8.336090679 | 19.41057018 | ENSG00000240280 |
| NA | 1.220604723 | 2.330443801 | 0.290936373 | 0.500577023 | 2.254017939 | 5.312653607 | ENSG00000197927 |
| NEK7 | 1.220454307 | 2.330200841 | 1.74E-14 | 6.01E-13 | 128.0996091 | 298.1067936 | ENSG00000151414 |
| NA | 1.218842105 | 2.327598312 | 0.190107042 | 0.376488975 | 3.275603762 | 7.665522306 | ENSG00000282390 |
| FGF4 | 1.218619909 | 2.327239855 | 0.000474045 | 0.002940977 | 25.28524761 | 59.01186556 | ENSG00000075388 |
| SPOCD1 | 1.217004394 | 2.324635295 | 0.007151118 | 0.029831865 | 16.96486937 | 39.40876094 | ENSG00000134668 |
| CDA | 1.216674919 | 2.324104467 | 6.14E-11 | 1.42E-09 | 90.21161578 | 209.4780669 | ENSG00000158825 |
| NOSTRIN | 1.215720533 | 2.322567512 | 0.09875874 | 0.235963917 | 5.564021218 | 12.99217465 | ENSG00000163072 |
| NAV3 | 1.21570375 | 2.322540494 | 0.000126007 | 0.00091349 | 31.32580189 | 72.81079525 | ENSG00000067798 |
| SLC5A4-AS1 | 1.215695252 | 2.322526812 | 0.07881112 | 0.199510455 | 8.766376751 | 20.24409319 | ENSG00000242082 |
| IRAK1 | 1.215029984 | 2.321456075 | 6.83E-73 | 6.15E-70 | 1072.231524 | 2490.168321 | ENSG00000184216 |
| NUTM2D | 1.214089269 | 2.319942855 | 0.00971544 | 0.038128862 | 13.55522701 | 31.43649586 | ENSG00000214562 |
| LRRC38 | 1.213922182 | 2.319674185 | 0.071037073 | 0.185039328 | 6.871176103 | 16.03283058 | ENSG00000162494 |
| NA | 1.213888316 | 2.319619732 | 0.132416652 | 0.29278638 | 5.491662713 | 12.82541928 | ENSG00000278997 |
| NA | 1.21293606 | 2.318089164 | 0.012772006 | 0.047800106 | 13.46626635 | 31.33493175 | ENSG00000271643 |
| ADGRG1 | 1.212813192 | 2.317891751 | 0.0003328 | 0.002147961 | 31.71693377 | 73.60018953 | ENSG00000205336 |
| NA | 1.212568393 | 2.317498481 | 0.300664999 | 0.510754566 | 2.019450547 | 4.644445553 | ENSG00000224520 |
| GLP1R | 1.212122065 | 2.316781625 | 6.59E-10 | 1.30E-08 | 86.03066743 | 199.0795774 | ENSG00000112164 |
| CACNA2D4 | 1.211544558 | 2.315854408 | 0.11121496 | 0.25865696 | 5.342496507 | 12.3401408 | ENSG00000151062 |
| BASP1-AS1 | 1.211432308 | 2.315674229 | 0.097636247 | 0.233907913 | 5.573504427 | 12.95888029 | ENSG00000215196 |
| S100A11 | 1.211319923 | 2.315493846 | 3.97E-71 | 3.00E-68 | 1182.932471 | 2739.218025 | ENSG00000163191 |
| NA | 1.21102298 | 2.315017309 | 0.211917359 | 0.405682649 | 3.013755799 | 6.93167148 | ENSG00000272115 |
| RASA4 | 1.211013999 | 2.315002897 | 0.193927167 | 0.382048252 | 3.299324861 | 7.651523566 | ENSG00000105808 |
| SYTL3 | 1.210871872 | 2.314774846 | 0.193308176 | 0.381226493 | 3.306443806 | 7.663346843 | ENSG00000164674 |
| CAPN13 | 1.210247304 | 2.313772957 | 1.46E-06 | 1.63E-05 | 47.17211611 | 108.9989241 | ENSG00000162949 |
| METTL27 | 1.209337029 | 2.312313532 | 1.77E-08 | 2.81E-07 | 76.99295951 | 177.9089834 | ENSG00000165171 |
| NA | 1.209245834 | 2.312167372 | 0.30345784 | 0.513550538 | 2.019450547 | 4.630446813 | ENSG00000228043 |
| MYOF | 1.207705979 | 2.309700805 | 3.64E-05 | 0.00030332 | 33.75537689 | 77.95066154 | ENSG00000138119 |
| LOC105370802 | 1.206805118 | 2.308259009 | 0.315103566 | 0.525450848 | 2.291976928 | 5.312653607 | ENSG00000259200 |
| OXSM | 1.204863596 | 2.305154736 | 0.0032261 | 0.015303919 | 19.38020647 | 44.51606792 | ENSG00000151093 |
| NA | 1.20345737 | 2.302908947 | 0.124334835 | 0.27973116 | 4.589877592 | 10.61036251 | ENSG00000272720 |
| SYT5 | 1.203138779 | 2.302400449 | 7.48E-05 | 0.000573815 | 36.25225022 | 83.58015724 | ENSG00000129990 |
| AMIGO2 | 1.202441043 | 2.3012872 | 1.94E-12 | 5.37E-11 | 131.6171785 | 302.3219127 | ENSG00000139211 |
| NA | 1.202136644 | 2.300801695 | 0.269006553 | 0.477013029 | 2.312138554 | 5.32882781 | ENSG00000255621 |
| PHLDA3 | 1.201636555 | 2.300004295 | 2.03E-40 | 4.03E-38 | 904.9827338 | 2082.95405 | ENSG00000174307 |
| NA | 1.20161465 | 2.299969373 | 0.158002515 | 0.330696831 | 4.362429145 | 9.984150681 | ENSG00000232389 |
| CA3 | 1.201339036 | 2.299530026 | 0.049355112 | 0.139678 | 8.911983485 | 20.49746797 | ENSG00000164879 |
| CRABP2 | 1.201125907 | 2.299190342 | 3.68E-76 | 3.66E-73 | 1483.429603 | 3409.046588 | ENSG00000143320 |
| NA | 1.200096711 | 2.297550721 | 0.272815361 | 0.481506365 | 2.57754599 | 5.989563514 | ENSG00000237301 |
| NA | 1.199813702 | 2.297100062 | 0.111817132 | 0.259514137 | 5.407736067 | 12.32614206 | ENSG00000228242 |

| | | | | | | | |
|---|---|---|---|---|---|---|---|
| NA | 1.199396275 | 2.296435519 | 0.201154861 | 0.391993556 | 3.326605432 | 7.6182292 | ENSG00000224728 |
| FAM50B | 1.198091538 | 2.29435962 | 4.37E-08 | 6.46E-07 | 68.85942712 | 158.0404698 | ENSG00000145945 |
| TSHZ2 | 1.198081039 | 2.294342923 | 0.177810204 | 0.3597542 | 3.599131813 | 8.29826053 | ENSG00000182463 |
| NA | 1.1967857 | 2.292283847 | 0.004162463 | 0.019015025 | 18.00425255 | 41.14551712 | ENSG00000284606 |
| LOC105370295 | 1.196643483 | 2.292057891 | 0.016868181 | 0.059935203 | 11.82610021 | 27.18106156 | ENSG00000287376 |
| GNGT2 | 1.195618937 | 2.290430738 | 0.132498679 | 0.292899163 | 4.937126742 | 11.3451593 | ENSG00000167083 |
| NA | 1.194921598 | 2.289323906 | 0.156782651 | 0.328887732 | 4.3553102 | 9.910752013 | ENSG00000261441 |
| NA | 1.194080935 | 2.287990299 | 0.26274301 | 0.469966067 | 3.102716458 | 7.00166518 | ENSG00000228863 |
| LOC100506207 | 1.192700228 | 2.285801663 | 0.127093809 | 0.284110628 | 5.491662713 | 12.67968349 | ENSG00000285219 |
| ILDR1 | 1.191489066 | 2.283883507 | 9.10E-05 | 0.000683934 | 31.33055657 | 71.64766091 | ENSG00000145103 |
| EGFLAM | 1.191438192 | 2.283802972 | 0.203889098 | 0.395485376 | 3.664371373 | 8.360781881 | ENSG00000164318 |
| KLHL32 | 1.19044432 | 2.282230203 | 0.126997056 | 0.283991871 | 4.675278779 | 10.62965814 | ENSG00000186231 |
| B3GALT4 | 1.1896617 | 2.280992495 | 0.000610331 | 0.003652125 | 24.1726162 | 55.12832235 | ENSG00000235863 |
| WNT3 | 1.188641473 | 2.279380022 | 4.47E-10 | 9.06E-09 | 87.1848173 | 198.5218033 | ENSG00000108379 |
| ELN | 1.188186697 | 2.278661613 | 4.12E-18 | 2.09E-16 | 166.6399974 | 379.7855195 | ENSG00000049540 |
| DHRS12 | 1.186097517 | 2.275364251 | 0.000193529 | 0.001339242 | 28.75301058 | 65.48095916 | ENSG00000102796 |
| ACKR1 | 1.185899017 | 2.275051206 | 0.180306244 | 0.363247645 | 3.895379293 | 8.923526404 | ENSG00000213088 |
| NA | 1.185679632 | 2.274705274 | 0.102932425 | 0.243228083 | 6.051232696 | 13.6992575 | ENSG00000262703 |
| MIR210HG | 1.184815313 | 2.273342906 | 0.001926752 | 0.009823629 | 22.76729678 | 51.548852 | ENSG00000247095 |
| LRRC37A6P | 1.184025943 | 2.272099387 | 0.002420823 | 0.011974133 | 22.87138505 | 51.96992812 | ENSG00000230445 |
| CAPN11 | 1.183801014 | 2.271745173 | 0.000348938 | 0.00223911 | 28.61096332 | 64.89674354 | ENSG00000137225 |
| LINC00954 | 1.183443637 | 2.271182498 | 0.112383281 | 0.260261077 | 5.264214265 | 11.99624719 | ENSG00000228784 |
| CD93 | 1.182346279 | 2.269455625 | 0.318557571 | 0.528518441 | 2.346538071 | 5.296479403 | ENSG00000125810 |
| NA | 1.179885131 | 2.265587376 | 0.002257403 | 0.011272094 | 29.26010436 | 66.48542052 | ENSG00000262152 |
| CAV2 | 1.179320066 | 2.264700178 | 0.000993709 | 0.005586842 | 26.3201816 | 59.46869505 | ENSG00000105971 |
| GABRA1 | 1.179222953 | 2.264547739 | 0.077108647 | 0.19649127 | 6.316640132 | 14.27600077 | ENSG00000022355 |
| CYP26B1 | 1.178767118 | 2.263832343 | 0.087528071 | 0.215904863 | 8.183365001 | 18.36299868 | ENSG00000003137 |
| APOBEC3G | 1.177858335 | 2.262406758 | 0.002221774 | 0.011126547 | 20.28079638 | 45.78996267 | ENSG00000239713 |
| FZD10 | 1.177314222 | 2.261553652 | 0.039167947 | 0.116854134 | 10.81191266 | 24.27850112 | ENSG00000111432 |
| ZNF124 | 1.176433133 | 2.260172887 | 2.08E-14 | 7.17E-13 | 159.8545274 | 362.0125277 | ENSG00000196418 |
| PTPRR | 1.17642421 | 2.260158908 | 0.124756752 | 0.280513307 | 5.311656463 | 12.00059812 | ENSG00000153233 |
| SLC12A5-AS1 | 1.176115467 | 2.259675277 | 0.184673526 | 0.368858755 | 3.984339952 | 9.000046496 | ENSG00000204044 |
| STARD13 | 1.176081237 | 2.259621664 | 3.19E-07 | 4.06E-06 | 56.90996674 | 128.6457489 | ENSG00000133121 |
| MESP2 | 1.174103576 | 2.256526272 | 0.02396868 | 0.079635561 | 11.35696542 | 25.52222293 | ENSG00000188095 |
| NA | 1.173949252 | 2.256284907 | 0.079404558 | 0.200729645 | 6.605768667 | 14.9099685 | ENSG00000223374 |
| NA | 1.173602153 | 2.255742131 | 0.273417028 | 0.482117701 | 2.639226078 | 5.957215107 | ENSG00000253541 |
| WNT9A | 1.173020222 | 2.25483243 | 1.88E-07 | 2.47E-06 | 62.16706206 | 140.2566734 | ENSG00000143816 |
| NA | 1.172431837 | 2.253913012 | 0.154240905 | 0.325420567 | 4.531756977 | 10.31064059 | ENSG00000250794 |
| NA | 1.171671285 | 2.25272512 | 0.007907784 | 0.032361516 | 15.13490828 | 34.17307893 | ENSG00000279756 |
| RASGRF2-AS1 | 1.171276863 | 2.252109326 | 0.162214212 | 0.337107182 | 3.977221007 | 8.947172957 | ENSG00000251450 |
| LCN2 | 1.169997928 | 2.250113738 | 0.056218986 | 0.154495861 | 8.205611559 | 18.50561305 | ENSG00000148346 |
| EXOC8 | 1.168718534 | 2.248119203 | 2.12E-11 | 5.18E-10 | 103.1954253 | 231.9526131 | ENSG00000116903 |
| MAP10 | 1.168257035 | 2.247400175 | 0.0001322 | 0.000953262 | 34.69540033 | 77.74486341 | ENSG00000212916 |
| RALYL | 1.167845121 | 2.246758594 | 0.252643604 | 0.458591633 | 3.446406135 | 7.63222794 | ENSG00000184672 |
| NA | 1.167517277 | 2.24624809 | 0.152033017 | 0.321987844 | 4.559037549 | 10.31376201 | ENSG00000223723 |
| CHRNA9 | 1.166261488 | 2.244293702 | 0.002139038 | 0.010775066 | 21.10429926 | 47.42174978 | ENSG00000174343 |
| SNCG | 1.166171146 | 2.244153167 | 9.91E-10 | 1.91E-08 | 92.38882602 | 207.055699 | ENSG00000173267 |
| NA | 1.164976832 | 2.242296147 | 0.024893991 | 0.082218085 | 11.4610537 | 25.78429956 | ENSG00000251271 |
| NA | 1.16477211 | 2.241977983 | 0.250460489 | 0.455942388 | 3.234085301 | 7.316331808 | ENSG00000272750 |
| PSMB9 | 1.16409142 | 2.240920425 | 1.49E-05 | 0.000135681 | 41.66830044 | 93.43909727 | ENSG00000240065 |
| NA | 1.163599792 | 2.240156916 | 0.24443113 | 0.448729647 | 2.979356283 | 6.683593584 | ENSG00000204778 |
| GCNA | 1.163063244 | 2.239323941 | 0.049114893 | 0.139144126 | 9.758012254 | 22.01151677 | ENSG00000147174 |
| RBPMS-AS1 | 1.161972852 | 2.237632095 | 0.159259407 | 0.332663853 | 4.610039219 | 10.34828588 | ENSG00000254109 |
| NA | 1.161791047 | 2.237350131 | 0.108006377 | 0.252813637 | 5.611463415 | 12.55369483 | ENSG00000280322 |
| SPTBN5 | 1.159463044 | 2.233742745 | 3.31E-07 | 4.20E-06 | 73.15069297 | 163.4778826 | ENSG00000137877 |
| LINC01748 | 1.159278576 | 2.23345715 | 0.153552971 | 0.324518483 | 4.300749058 | 9.600436312 | ENSG00000226476 |
| NA | 1.158682449 | 2.232534468 | 0.164291534 | 0.339718123 | 4.320910684 | 9.629379751 | ENSG00000274099 |
| NA | 1.155252785 | 2.227233452 | 0.218350455 | 0.414542705 | 3.564732297 | 8.016888268 | ENSG00000237438 |
| CDK10 | 1.155179836 | 2.227120836 | 1.29E-15 | 5.28E-14 | 263.3236488 | 586.520134 | ENSG00000185324 |
| BVES-AS1 | 1.154832833 | 2.226585224 | 0.298429248 | 0.508695401 | 2.659387704 | 5.893747797 | ENSG00000203808 |
| RASGRP4 | 1.154819503 | 2.226564652 | 1.60E-06 | 1.78E-05 | 52.0496477 | 116.0872087 | ENSG00000171777 |
| NA | 1.154215882 | 2.225633256 | 0.172242752 | 0.351708788 | 4.3553102 | 9.65085084 | ENSG00000224543 |
| NA | 1.152960019 | 2.22369669 | 0.09374292 | 0.227374138 | 6.278681143 | 13.9784543 | ENSG00000246100 |
| IGIP | 1.151998788 | 2.22221559 | 0.11788324 | 0.269584038 | 6.560690733 | 14.62859623 | ENSG00000182700 |
| NA | 1.151090961 | 2.220817684 | 0.228323367 | 0.427681689 | 4.413430815 | 9.741369672 | ENSG00000206549 |
| TMC1 | 1.150782997 | 2.220343668 | 0.263452409 | 0.470744512 | 3.526773308 | 7.903952389 | ENSG00000165091 |
| KLK5 | 1.14819012 | 2.21635675 | 0.09728006 | 0.233375754 | 6.823733905 | 15.2202842 | ENSG00000167754 |
| LINC00857 | 1.148040943 | 2.216127587 | 0.2488499 | 0.453807341 | 3.622852912 | 8.060113994 | ENSG00000237523 |
| GSG1L | 1.147044812 | 2.214597957 | 0.029575748 | 0.09384039 | 10.64466976 | 23.50795096 | ENSG00000169181 |
| SPOCK3 | 1.147004167 | 2.214535565 | 0.165081918 | 0.340755495 | 4.358869673 | 9.591734458 | ENSG00000196104 |
| NA | 1.146977423 | 2.214494514 | 0.010481694 | 0.040579499 | 18.22221779 | 40.30090588 | ENSG00000261799 |
| STAT6 | 1.146578207 | 2.213881813 | 1.25E-52 | 4.20E-50 | 1001.400291 | 2216.433807 | ENSG00000166888 |
| TIGD6 | 1.145437792 | 2.212132486 | 0.000513781 | 0.003150245 | 30.19656833 | 66.72997785 | ENSG00000164296 |
| NA | 1.14517832 | 2.211734664 | 0.014116845 | 0.051858567 | 14.52581116 | 32.14547062 | ENSG00000154608 |

| | | | | | | | |
|---|---|---|---|---|---|---|---|
| HCP5 | 1.144103553 | 2.210087598 | 2.54E-05 | 0.000220494 | 43.0807388 | 95.40463572 | ENSG00000206337 |
| NA | 1.143673392 | 2.209428726 | 0.017918233 | 0.062920384 | 13.46034262 | 29.82277487 | ENSG00000179066 |
| FAXDC2 | 1.142618693 | 2.207814088 | 4.19E-05 | 0.000343143 | 38.29425281 | 84.62895824 | ENSG00000170271 |
| DAGLA | 1.141804835 | 2.206568959 | 3.35E-22 | 2.38E-20 | 257.6997269 | 569.0404402 | ENSG00000134780 |
| DNAH1 | 1.141325462 | 2.205835892 | 0.019396673 | 0.066966393 | 16.68997872 | 36.75522435 | ENSG00000114841 |
| KIFAP3 | 1.140525174 | 2.204612615 | 3.52E-06 | 3.67E-05 | 56.6739248 | 125.0584797 | ENSG00000075945 |
| AZU1 | 1.140435434 | 2.204475487 | 0.070111191 | 0.183051125 | 8.343209624 | 18.35458038 | ENSG00000172232 |
| CPZ | 1.140112097 | 2.203981475 | 0.039117874 | 0.116760082 | 10.85551605 | 24.00147978 | ENSG00000109625 |
| NUDT7 | 1.140058722 | 2.203899936 | 0.025748219 | 0.084307987 | 11.6282966 | 25.59344613 | ENSG00000140876 |
| SNORA66 | 1.139197453 | 2.202584631 | 0.216564923 | 0.412073408 | 3.629971857 | 8.01594231 | ENSG00000207523 |
| ANXA3 | 1.139122059 | 2.202469528 | 2.22E-19 | 1.27E-17 | 196.2239092 | 432.1892929 | ENSG00000138772 |
| WNT9B | 1.137939998 | 2.20066569 | 0.075469093 | 0.193619592 | 7.51111326 | 16.62451854 | ENSG00000158955 |
| BTBD19 | 1.137036518 | 2.199287968 | 0.141412611 | 0.305978134 | 5.315215936 | 11.68687745 | ENSG00000222009 |
| ARHGAP31-AS1 | 1.135295406 | 2.196635363 | 0.057541036 | 0.157236648 | 8.863346079 | 19.51676876 | ENSG00000241155 |
| PCDHB17P | 1.134727406 | 2.195770703 | 0.137932835 | 0.300966099 | 5.274892683 | 11.59541266 | ENSG00000255622 |
| PHYHD1 | 1.134698908 | 2.195727329 | 1.35E-09 | 2.55E-08 | 94.33711327 | 206.8516772 | ENSG00000175287 |
| MEF2C | 1.132892712 | 2.192980087 | 0.019389275 | 0.066953092 | 14.50209007 | 31.84631579 | ENSG00000081189 |
| NA | 1.132309211 | 2.192093311 | 0.205140111 | 0.397096248 | 3.626412384 | 7.960893308 | ENSG00000250049 |
| NA | 1.131375508 | 2.190675061 | 0.192181264 | 0.379479629 | 4.211788399 | 9.307240774 | ENSG00000237118 |
| VCX3A | 1.131346323 | 2.190630745 | 0.357083354 | 0.569121372 | 2.274179566 | 5.024754954 | ENSG00000169059 |
| C11orf86 | 1.131227567 | 2.19045043 | 0.028807496 | 0.091927943 | 11.50613163 | 25.28502235 | ENSG00000173237 |
| NEK11 | 1.130655383 | 2.189581852 | 0.025040405 | 0.082586108 | 12.32635437 | 26.92551131 | ENSG00000114670 |
| TUSC1 | 1.129105881 | 2.187231432 | 4.55E-07 | 5.63E-06 | 63.88286685 | 139.9751855 | ENSG00000198680 |
| RNF227 | 1.128757508 | 2.186703337 | 0.008018406 | 0.032741385 | 18.61334966 | 40.67206192 | ENSG00000179859 |
| NA | 1.127676289 | 2.18506514 | 0.230007671 | 0.429903311 | 3.677414055 | 8.013766846 | ENSG00000280401 |
| NA | 1.126974958 | 2.184003182 | 0.000172985 | 0.001209936 | 36.07107491 | 78.71759582 | ENSG00000281490 |
| GBP1P1 | 1.126440626 | 2.18319444 | 0.236635857 | 0.438640378 | 3.691651944 | 8.030887009 | ENSG00000225492 |
| NA | 1.125896457 | 2.182371119 | 0.180834931 | 0.363839553 | 4.555478076 | 10.02056647 | ENSG00000259488 |
| MASP1 | 1.125498779 | 2.181769632 | 5.78E-05 | 0.000457845 | 42.50721026 | 92.87462878 | ENSG00000127241 |
| INHBA | 1.125108242 | 2.181179107 | 0.042165 | 0.1236277 | 10.78671702 | 23.65730262 | ENSG00000122641 |
| TIRAP | 1.1247658 | 2.180661438 | 0.008413668 | 0.034053174 | 17.3737986 | 37.71395802 | ENSG00000150455 |
| NTSR1 | 1.124465608 | 2.180207741 | 0.004461675 | 0.020104682 | 18.88824031 | 41.15544848 | ENSG00000101188 |
| NA | 1.124363132 | 2.180052883 | 0.201582286 | 0.392587026 | 3.657252428 | 7.953420959 | ENSG00000285813 |
| FMN1 | 1.12363662 | 2.17895533 | 0.000192424 | 0.001333065 | 36.69468925 | 79.72612457 | ENSG00000248905 |
| FHIT | 1.123350806 | 2.178523697 | 0.007311823 | 0.030341413 | 18.09054346 | 39.50597412 | ENSG00000189283 |
| NA | 1.121317652 | 2.17545572 | 0.05944781 | 0.161210784 | 11.48477479 | 25.08300806 | ENSG00000234537 |
| NA | 1.121244148 | 2.175344885 | 0.17262164 | 0.352187449 | 4.269909014 | 9.3265364 | ENSG00000242611 |
| TEX37 | 1.120678281 | 2.174491818 | 0.324036967 | 0.53482242 | 2.284857983 | 5.003283865 | ENSG00000172073 |
| ZBTB44-DT | 1.120400954 | 2.17407386 | 0.241458225 | 0.445045733 | 3.395404465 | 7.306683995 | ENSG00000175773 |
| HPCAL4 | 1.120235928 | 2.173825188 | 5.40E-05 | 0.000431089 | 38.5999835 | 84.02856843 | ENSG00000116983 |
| NA | 1.118581836 | 2.171334262 | 0.134696542 | 0.296301012 | 5.656541349 | 12.24867601 | ENSG00000259834 |
| LAMB3 | 1.116599682 | 2.168353061 | 0.003265046 | 0.01546536 | 23.79808648 | 51.67020619 | ENSG00000196878 |
| ARSD | 1.115774344 | 2.167112942 | 5.58E-09 | 9.61E-08 | 87.5546185 | 189.5918661 | ENSG00000006756 |
| PXDNL | 1.113849648 | 2.16422373 | 0.333505807 | 0.545539816 | 2.288417455 | 4.977461848 | ENSG00000147485 |
| FAM13C | 1.112934783 | 2.162851753 | 0.057637221 | 0.157476695 | 8.625219214 | 18.65089734 | ENSG00000148541 |
| LINC00639 | 1.112740334 | 2.16256026 | 0.205599996 | 0.397732391 | 4.00806105 | 8.636857257 | ENSG00000259070 |
| PCSK1 | 1.112371494 | 2.16200745 | 0.00126447 | 0.006871801 | 26.61731881 | 57.69256961 | ENSG00000175426 |
| NA | 1.111276293 | 2.160366815 | 0.37839659 | 0.590824274 | 2.270620093 | 4.930168742 | ENSG00000233750 |
| CALB1 | 1.109732405 | 2.158056154 | 2.72E-24 | 2.30E-22 | 352.8556146 | 761.4100943 | ENSG00000104327 |
| LINC00471 | 1.108591902 | 2.156350806 | 0.016005844 | 0.057498644 | 14.66933297 | 31.53760846 | ENSG00000181798 |
| LAMC2 | 1.108439526 | 2.156123066 | 2.68E-23 | 2.07E-21 | 274.417571 | 591.3006543 | ENSG00000058085 |
| KIAA1257 | 1.10625338 | 2.152858321 | 0.000631749 | 0.003754115 | 34.90477208 | 75.33526471 | ENSG00000114656 |
| SFR1 | 1.10619342 | 2.152768848 | 1.20E-05 | 0.000112152 | 47.30112069 | 101.8295576 | ENSG00000156384 |
| FGF8 | 1.106065866 | 2.152578524 | 0.003621742 | 0.016887055 | 23.60028287 | 50.69771437 | ENSG00000107831 |
| PARP15 | 1.105553731 | 2.151814525 | 0.315476508 | 0.525747781 | 2.312138554 | 4.979637312 | ENSG00000173200 |
| NA | 1.105141549 | 2.151199835 | 0.383724108 | 0.596037982 | 2.652268759 | 5.713488138 | ENSG00000231536 |
| CCL25 | 1.105065963 | 2.15108713 | 0.080037485 | 0.2019057 | 7.514672732 | 16.2463846 | ENSG00000131142 |
| NA | 1.10497087 | 2.150945349 | 0.136025036 | 0.298247299 | 5.731264118 | 12.222854 | ENSG00000227288 |
| SLC44A5 | 1.104394423 | 2.150086084 | 0.000254678 | 0.001704322 | 32.7159937 | 70.38653539 | ENSG00000137968 |
| LOC285095 | 1.104160327 | 2.149737233 | 0.18136337 | 0.364444269 | 4.596996537 | 9.900158241 | ENSG00000216921 |
| S100A16 | 1.102862549 | 2.147804304 | 1.29E-20 | 8.02E-19 | 249.1157213 | 534.6318417 | ENSG00000188643 |
| NA | 1.102732081 | 2.14761008 | 0.004327762 | 0.019604171 | 20.93497143 | 44.90697109 | ENSG00000205959 |
| SLCO6A1 | 1.102417809 | 2.147142302 | 0.326128581 | 0.537335392 | 2.339419126 | 5.009810256 | ENSG00000205359 |
| NA | 1.102228117 | 2.146860004 | 0.27926767 | 0.488375771 | 3.237644773 | 7.017839384 | ENSG00000215452 |
| NA | 1.102093647 | 2.14665991 | 0.180629387 | 0.363606393 | 4.3553102 | 9.302889847 | ENSG00000271727 |
| SNORA71B | 1.102086843 | 2.146649787 | 0.309292769 | 0.51980012 | 2.608386034 | 5.630725203 | ENSG00000235408 |
| C2orf91 | 1.101549345 | 2.145850169 | 0.269369596 | 0.477414112 | 3.241204246 | 7.02313627 | ENSG00000205086 |
| MYBPC2 | 1.100935479 | 2.144937304 | 4.36E-10 | 8.84E-09 | 101.1228882 | 216.7325398 | ENSG00000086967 |
| XK | 1.100548515 | 2.14436206 | 0.001793122 | 0.009242114 | 28.64625256 | 61.48682349 | ENSG00000047597 |
| SERPINB1 | 1.100378953 | 2.144110045 | 1.10E-16 | 4.84E-15 | 192.0927673 | 411.7755339 | ENSG00000021355 |
| NA | 1.095981853 | 2.137585094 | 9.04E-13 | 2.60E-11 | 131.64151 | 281.4413404 | ENSG00000243004 |
| NA | 1.095423797 | 2.136758404 | 0.009268191 | 0.036719201 | 18.23526047 | 38.96498423 | ENSG00000242759 |
| HECW1 | 1.094853909 | 2.135914517 | 2.30E-08 | 3.56E-07 | 123.3558362 | 263.7994757 | ENSG00000002746 |
| HACD4 | 1.094205866 | 2.134955303 | 0.001714391 | 0.008914173 | 25.22504207 | 53.7928522 | ENSG00000188921 |

| | | | | | | | |
|---|---|---|---|---|---|---|---|
| ANO9 | 1.093159076 | 2.133406784 | 0.000753302 | 0.004377263 | 275.6215797 | 588.0464482 | ENSG00000185101 |
| NMUR1 | 1.093034058 | 2.133221921 | 0.02730636 | 0.088301475 | 12.90817091 | 27.52807659 | ENSG00000171596 |
| NA | 1.092668028 | 2.132680764 | 0.339495981 | 0.551295017 | 2.342978598 | 4.972164962 | ENSG00000272368 |
| PEG13 | 1.09244852 | 2.132356298 | 0.13460997 | 0.296214085 | 5.607903943 | 11.98130249 | ENSG00000282164 |
| NA | 1.091792315 | 2.131386624 | 0.334303615 | 0.546292653 | 2.339419126 | 4.961287645 | ENSG00000231198 |
| CCDC144A | 1.091507188 | 2.130965429 | 0.307963324 | 0.518350034 | 3.422685036 | 7.193748116 | ENSG00000170160 |
| NDRG1 | 1.091150677 | 2.130438901 | 4.61E-05 | 0.000374593 | 45.90647969 | 97.61082267 | ENSG00000104419 |
| MEIS1 | 1.091123549 | 2.130398841 | 0.359421779 | 0.571533011 | 2.363140224 | 5.001108401 | ENSG00000143995 |
| MGARP | 1.090683327 | 2.129748874 | 0.181490581 | 0.364583515 | 4.937126742 | 10.51237133 | ENSG00000137463 |
| SLC6A16 | 1.090404497 | 2.129337296 | 0.108414274 | 0.253454267 | 7.399906043 | 15.67805965 | ENSG00000063127 |
| ZBTB7B | 1.090184973 | 2.129013316 | 7.35E-49 | 2.20E-46 | 743.0087863 | 1582.236844 | ENSG00000160685 |
| NA | 1.089107545 | 2.127423928 | 0.00162439 | 0.008509463 | 26.08294446 | 55.57332857 | ENSG00000224411 |
| METTL11B | 1.088624174 | 2.12671126 | 0.142565372 | 0.307767153 | 5.322334881 | 11.29692023 | ENSG00000203740 |
| LINC02615 | 1.088553972 | 2.126607775 | 0.051316654 | 0.143956479 | 10.37570285 | 21.9847488 | ENSG00000251432 |
| MAFB | 1.08847447 | 2.126490588 | 0.000584359 | 0.003511171 | 32.05945439 | 68.15793138 | ENSG00000204103 |
| TRPV3 | 1.088264759 | 2.126181503 | 0.022767264 | 0.076345309 | 13.87283133 | 29.51122966 | ENSG00000167723 |
| IRF2BP2 | 1.087515874 | 2.125078116 | 9.98E-29 | 1.10E-26 | 457.7908968 | 973.6573989 | ENSG00000168264 |
| TMEM38A | 1.087178824 | 2.124581701 | 3.18E-15 | 1.24E-13 | 187.3323928 | 397.661611 | ENSG00000072954 |
| NUTM2F | 1.086766959 | 2.123975255 | 0.311640896 | 0.522197973 | 2.935473557 | 6.259112499 | ENSG00000130950 |
| CPEB4 | 1.086321901 | 2.12332013 | 3.47E-13 | 1.05E-11 | 151.089041 | 320.8179935 | ENSG00000113742 |
| NA | 1.086066752 | 2.122944643 | 0.203673944 | 0.395311657 | 4.413430815 | 9.270541439 | ENSG00000237821 |
| NA | 1.085711031 | 2.122421258 | 0.014901252 | 0.054242511 | 15.49995479 | 32.96889432 | ENSG00000277011 |
| DYRK3 | 1.084935338 | 2.121280405 | 2.22E-17 | 1.04E-15 | 211.5412141 | 448.9740405 | ENSG00000143479 |
| NA | 1.083172149 | 2.118689466 | 0.253758556 | 0.459951083 | 3.279163235 | 6.975843164 | ENSG00000234322 |
| LINC01976 | 1.082884139 | 2.118266548 | 0.363206239 | 0.575190761 | 2.366699697 | 4.970935457 | ENSG00000261514 |
| PHYHIPL | 1.082874518 | 2.118252421 | 0.169336069 | 0.347297096 | 5.342496507 | 11.27544915 | ENSG00000165443 |
| MEF2D | 1.082406904 | 2.117565954 | 1.06E-32 | 1.43E-30 | 487.6602998 | 1032.599103 | ENSG00000116604 |
| TBCE | 1.08130258 | 2.115945663 | 0.160830253 | 0.335018677 | 5.373336551 | 11.30874351 | ENSG00000284770 |
| SRMS | 1.080788691 | 2.115192096 | 0.07464493 | 0.192000751 | 8.301691163 | 17.51621198 | ENSG00000125508 |
| LOC105372233 | 1.080132938 | 2.114230889 | 0.14733734 | 0.314867834 | 5.622141833 | 11.87148803 | ENSG00000267751 |
| PRDM13 | 1.079926873 | 2.113928929 | 0.286143421 | 0.495345072 | 3.006636854 | 6.338754013 | ENSG00000112238 |
| GLUD1P2 | 1.079745022 | 2.113662485 | 0.390102955 | 0.601690159 | 2.366699697 | 4.962517149 | ENSG00000265366 |
| MYO15B | 1.079483364 | 2.11327917 | 1.59E-06 | 1.77E-05 | 124.9826809 | 264.1621314 | ENSG00000266714 |
| STK32A | 1.078439995 | 2.111751382 | 0.054907526 | 0.151686459 | 9.854091858 | 20.85971125 | ENSG00000169302 |
| DOC2A | 1.078220799 | 2.111430558 | 1.27E-06 | 1.45E-05 | 129.363771 | 273.3086917 | ENSG00000149927 |
| CD247 | 1.078125 | 2.111290357 | 0.000528528 | 0.0032208 | 31.68492467 | 66.94286947 | ENSG00000198821 |
| MYH3 | 1.077461602 | 2.11031974 | 0.001583153 | 0.008344361 | 28.28596073 | 59.85704389 | ENSG00000109063 |
| GNGT1 | 1.076778827 | 2.109321239 | 0.008962923 | 0.03581605 | 19.89558824 | 41.96336039 | ENSG00000127928 |
| NA | 1.076693551 | 2.109196564 | 0.111515763 | 0.259135468 | 6.640168183 | 14.0011549 | ENSG00000227214 |
| C16orf74 | 1.076134837 | 2.108379891 | 0.00257829 | 0.012614252 | 26.52863748 | 55.92157311 | ENSG00000154102 |
| NA | 1.075343217 | 2.107223321 | 0.109686626 | 0.255858688 | 6.582047568 | 13.88916498 | ENSG00000265975 |
| RIMBP3 | 1.074402717 | 2.105850059 | 0.048613292 | 0.138053813 | 10.20845995 | 21.53009477 | ENSG00000275793 |
| RNF152 | 1.074203792 | 2.105559715 | 0.008896173 | 0.035654547 | 23.0027539 | 48.56296153 | ENSG00000176641 |
| GJB7 | 1.073537 | 2.104586782 | 0.049415728 | 0.139786702 | 11.77537787 | 24.65228413 | ENSG00000164411 |
| GRIN2C | 1.073505035 | 2.104540152 | 0.261983466 | 0.468977732 | 3.310003278 | 6.964019887 | ENSG00000161509 |
| NYX | 1.072557802 | 2.103158824 | 0.211824392 | 0.40563523 | 4.273468486 | 9.026814471 | ENSG00000188937 |
| SUCLG2 | 1.07190914 | 2.102213418 | 9.74E-34 | 1.40E-31 | 669.1324506 | 1405.528366 | ENSG00000172340 |
| ALS2CL | 1.07135674 | 2.101408645 | 1.78E-07 | 2.35E-06 | 163.4263793 | 343.4677696 | ENSG00000178038 |
| PTPRVP | 1.071129836 | 2.101078167 | 0.231353195 | 0.431857772 | 4.930007797 | 10.40115942 | ENSG00000243323 |
| CCDC102B | 1.07112622 | 2.101072901 | 0.053224087 | 0.147857736 | 9.653923982 | 20.22791899 | ENSG00000150636 |
| NAALADL1 | 1.07062164 | 2.100338183 | 0.0003299 | 0.002131428 | 34.14058502 | 71.6464314 | ENSG00000168060 |
| NA | 1.069355776 | 2.098496091 | 0.03034569 | 0.095864563 | 14.56496536 | 30.56873252 | ENSG00000254978 |
| CBX7 | 1.067800527 | 2.096235096 | 4.14E-07 | 5.15E-06 | 86.4123161 | 180.9474639 | ENSG00000100307 |
| PPT2-EGFL8 | 1.067732809 | 2.096136705 | 0.206701354 | 0.39921815 | 4.262790069 | 8.970819511 | ENSG00000258388 |
| ANKRD33 | 1.067540764 | 2.095857695 | 0.136439674 | 0.298778204 | 6.722009897 | 14.02574741 | ENSG00000167612 |
| NA | 1.067372866 | 2.095613798 | 0.234039602 | 0.435237606 | 3.919100392 | 8.243495075 | ENSG00000279148 |
| NA | 1.066202836 | 2.093914938 | 0.286109243 | 0.495345072 | 3.04103637 | 6.31510746 | ENSG00000287392 |
| TMEM88B | 1.066099354 | 2.093764751 | 0.014936421 | 0.054328642 | 17.63417203 | 37.08651668 | ENSG00000205116 |
| NA | 1.065371718 | 2.092709007 | 0.096118051 | 0.231293245 | 7.283664813 | 15.22993201 | ENSG00000261235 |
| OXCT1-AS1 | 1.065165865 | 2.092410427 | 0.334693666 | 0.546630623 | 3.003077381 | 6.254761572 | ENSG00000248668 |
| LGALS3BP | 1.064638153 | 2.091645202 | 2.95E-56 | 1.12E-53 | 1246.661339 | 2606.71907 | ENSG00000108679 |
| COX6B2 | 1.064550166 | 2.09151764 | 0.005307865 | 0.02324653 | 27.85953962 | 58.25921356 | ENSG00000160471 |
| PLAC8L1 | 1.064254873 | 2.09108959 | 0.210511644 | 0.404015487 | 4.579199175 | 9.615381011 | ENSG00000173261 |
| ALOX12P2 | 1.064144246 | 2.090929249 | 2.68E-05 | 0.000231661 | 56.33290488 | 117.79043 | ENSG00000262943 |
| LINC01356 | 1.063877273 | 2.090542354 | 6.44E-12 | 1.66E-10 | 164.0366454 | 342.888597 | ENSG00000215866 |
| NA | 1.063772162 | 2.090390049 | 0.331535237 | 0.542983463 | 3.664371373 | 7.647456185 | ENSG00000272263 |
| GAS5 | 1.060886762 | 2.086213432 | 7.46E-58 | 2.94E-55 | 3842.104162 | 8014.312912 | ENSG00000234741 |
| LINC00548 | 1.0606382 | 2.085854029 | 0.199736811 | 0.389994797 | 4.293630113 | 8.963347161 | ENSG00000215483 |
| GCSAM | 1.060085545 | 2.085055152 | 0.018491564 | 0.064430476 | 16.02009124 | 33.57051365 | ENSG00000174500 |
| LOC103611081 | 1.059966744 | 2.084883461 | 0.029075975 | 0.092596929 | 13.72366512 | 28.46706314 | ENSG00000255455 |
| LOC101927468 | 1.059271836 | 2.083879471 | 3.22E-06 | 3.37E-05 | 60.43465511 | 125.7520583 | ENSG00000227733 |
| NA | 1.05917562 | 2.083740497 | 0.23948449 | 0.442444198 | 4.293630113 | 8.950577926 | ENSG00000283312 |
| PTGES3L | 1.059175385 | 2.083740158 | 0.07927999 | 0.200509255 | 8.59437917 | 17.91392509 | ENSG00000267060 |
| NA | 1.057049951 | 2.080672572 | 0.081899004 | 0.205481957 | 8.961789946 | 18.63472313 | ENSG00000287808 |

| | | | | | | | |
|---|---|---|---|---|---|---|---|
| KIRREL1 | 1.056045548 | 2.079224514 | 1.30E-60 | 5.86E-58 | 1546.945058 | 3215.959807 | ENSG00000183853 |
| COLQ | 1.055325142 | 2.078186518 | 0.126767278 | 0.283649294 | 6.516808008 | 13.63361473 | ENSG00000206561 |
| FZD5 | 1.055013281 | 2.077737333 | 3.24E-64 | 1.75E-61 | 1634.875148 | 3396.381371 | ENSG00000163251 |
| FBXO48 | 1.054875476 | 2.077538879 | 0.018117971 | 0.063397791 | 18.2162679 | 37.83001532 | ENSG00000204923 |
| NA | 1.05473565 | 2.077337534 | 0.338189662 | 0.550246554 | 2.748348363 | 5.624198812 | ENSG00000260136 |
| LMOD1 | 1.054499255 | 2.076997177 | 0.027094133 | 0.087810658 | 14.28176056 | 29.64497422 | ENSG00000163431 |
| GSG1 | 1.052014008 | 2.073422334 | 0.182523609 | 0.366113473 | 5.615022888 | 11.65916351 | ENSG00000111305 |
| NA | 1.05081674 | 2.07170235 | 0.006502506 | 0.027570682 | 21.17665776 | 43.89203155 | ENSG00000232325 |
| LRRTM1 | 1.050729343 | 2.071576852 | 0.002299999 | 0.011451486 | 27.06301182 | 56.12888429 | ENSG00000162951 |
| LY6L | 1.048715863 | 2.068687697 | 0.153458054 | 0.324384974 | 5.61858236 | 11.63305795 | ENSG00000261667 |
| LOC112268389 | 1.046151072 | 2.065013297 | 0.012742847 | 0.047757226 | 17.45416578 | 36.09276469 | ENSG00000255052 |
| SERHL | 1.046106054 | 2.064948863 | 0.242349647 | 0.445992309 | 4.386150244 | 9.001938413 | ENSG00000172250 |
| EFHB | 1.045343519 | 2.063857724 | 0.175122892 | 0.355608499 | 5.325894353 | 10.97014678 | ENSG00000163576 |
| SPTLC3 | 1.045310504 | 2.063810494 | 0.247398326 | 0.452330079 | 4.647998207 | 9.571492873 | ENSG00000172296 |
| EPSTI1 | 1.044471965 | 2.062611293 | 0.028112401 | 0.090258874 | 15.35879725 | 31.81368383 | ENSG00000133106 |
| NA | 1.044229124 | 2.062264133 | 0.052219841 | 0.14596976 | 12.97905487 | 26.96098114 | ENSG00000236200 |
| HES4 | 1.043988215 | 2.061919794 | 0.015753025 | 0.056698281 | 19.86118873 | 40.97641833 | ENSG00000188290 |
| NAXE | 1.043800064 | 2.061650904 | 2.59E-59 | 1.06E-56 | 1413.178788 | 2913.268657 | ENSG00000163382 |
| RYR2 | 1.043446111 | 2.061145157 | 1.89E-08 | 2.98E-07 | 108.316982 | 222.7071586 | ENSG00000198626 |
| PRORSD1P | 1.043209194 | 2.060806707 | 0.101214767 | 0.239997826 | 8.874024496 | 18.35212137 | ENSG00000162997 |
| MUC12 | 1.04162462 | 2.058544477 | 0.020672965 | 0.070687842 | 16.0085231 | 32.85255348 | ENSG00000205277 |
| MT1H | 1.037782095 | 2.053068971 | 5.21E-05 | 0.000417289 | 48.45199042 | 99.39848784 | ENSG00000205358 |
| NA | 1.037499199 | 2.052666428 | 0.403692356 | 0.614998476 | 2.257577412 | 4.686441774 | ENSG00000280502 |
| MYEOV | 1.037417445 | 2.052550112 | 0.05377211 | 0.149007527 | 12.72935987 | 26.04904612 | ENSG00000172927 |
| UCP3 | 1.03740126 | 2.052527085 | 0.120861549 | 0.273874532 | 7.259943714 | 14.91214396 | ENSG00000175564 |
| NA | 1.036983369 | 2.051932637 | 0.089450991 | 0.219415763 | 7.927161442 | 16.2840299 | ENSG00000279602 |
| NA | 1.036931599 | 2.051859006 | 0.080065798 | 0.201928766 | 8.901305067 | 18.3038823 | ENSG00000231551 |
| MIRLET7BHG | 1.03612912 | 2.050718005 | 0.316654885 | 0.526735445 | 3.766374713 | 7.640929795 | ENSG00000197182 |
| ALDH1A1 | 1.035502808 | 2.049827927 | 0.007037583 | 0.029403707 | 20.61500285 | 42.18031938 | ENSG00000165092 |
| SH3RF2 | 1.035179041 | 2.049367961 | 0.036186558 | 0.110010165 | 14.50000513 | 29.50716228 | ENSG00000156463 |
| NA | 1.03465114 | 2.048618207 | 0.146157685 | 0.313230582 | 6.705407743 | 13.68743423 | ENSG00000243926 |
| NA | 1.033105541 | 2.046424641 | 0.013365472 | 0.049540126 | 19.21532783 | 39.34529363 | ENSG00000244041 |
| C1RL-AS1 | 1.033036416 | 2.046326592 | 0.002995448 | 0.014390493 | 25.17048093 | 51.50250485 | ENSG00000205885 |
| NEDD9 | 1.032371458 | 2.045383628 | 5.88E-17 | 2.65E-15 | 227.5158964 | 464.9600294 | ENSG00000111859 |
| UPK1B | 1.030789605 | 2.043142182 | 0.335324582 | 0.547377018 | 3.234085301 | 6.665243917 | ENSG00000114638 |
| OR52A1 | 1.030106116 | 2.042174456 | 0.074518406 | 0.191800015 | 8.921466694 | 18.22206532 | ENSG00000182070 |
| KCNN3 | 1.029958735 | 2.041965845 | 1.12E-06 | 1.29E-05 | 74.23156846 | 151.4377043 | ENSG00000143603 |
| NA | 1.029536468 | 2.041368263 | 0.339739636 | 0.551394601 | 3.259001608 | 6.699767788 | ENSG00000279711 |
| LOC101928063 | 1.029312505 | 2.041051387 | 0.39936168 | 0.610828948 | 2.254017939 | 4.64879648 | ENSG00000266921 |
| TUBB6 | 1.029096307 | 2.040745544 | 4.92E-63 | 2.58E-60 | 3053.364317 | 6231.726987 | ENSG00000176014 |
| C16orf92 | 1.02849537 | 2.039895673 | 0.274482843 | 0.483275078 | 3.898938765 | 8.019063732 | ENSG00000167194 |
| CDH11 | 1.028171957 | 2.039438436 | 1.44E-14 | 5.07E-13 | 195.9499083 | 399.8023434 | ENSG00000140937 |
| UHMK1 | 1.027399718 | 2.038347066 | 7.13E-15 | 2.63E-13 | 418.4636654 | 852.5295313 | ENSG00000152332 |
| NA | 1.027210213 | 2.038079338 | 0.265025304 | 0.472628608 | 3.881141403 | 7.983593902 | ENSG00000231991 |
| NA | 1.027176738 | 2.038032049 | 0.425727018 | 0.635611886 | 2.270620093 | 4.661565716 | ENSG00000261386 |
| NA | 1.027011506 | 2.037798645 | 0.005836131 | 0.025126983 | 23.35979174 | 47.440383 | ENSG00000247735 |
| NA | 1.026813202 | 2.037518561 | 0.293109347 | 0.502804708 | 3.557613352 | 7.313210386 | ENSG00000275223 |
| NA | 1.02676127 | 2.03744522 | 0.388366985 | 0.600408663 | 2.594148144 | 5.325706388 | ENSG00000227107 |
| GRIN2D | 1.026660433 | 2.037302817 | 4.30E-27 | 4.42E-25 | 563.2429326 | 1146.728153 | ENSG00000105464 |
| NA | 1.026379946 | 2.036906766 | 0.01760359 | 0.062011527 | 18.44491156 | 37.67056434 | ENSG00000267325 |
| CTSF | 1.02580267 | 2.036091886 | 0.343713783 | 0.555691704 | 2.914116723 | 5.983037123 | ENSG00000174080 |
| PTPN14 | 1.023670267 | 2.033084624 | 4.00E-39 | 7.33E-37 | 1135.397543 | 2307.082461 | ENSG00000152104 |
| CBLC | 1.023093608 | 2.032272144 | 4.49E-14 | 1.50E-12 | 200.5036069 | 407.3494877 | ENSG00000142273 |
| C1orf116 | 1.021847229 | 2.030517173 | 9.59E-08 | 1.34E-06 | 103.7042991 | 210.2167202 | ENSG00000182795 |
| SNORA33 | 1.021148563 | 2.029534076 | 0.076161872 | 0.19482769 | 11.48268986 | 23.12206112 | ENSG00000200534 |
| CASC8 | 1.020828363 | 2.029083679 | 0.424918364 | 0.634856275 | 2.315698027 | 4.71226379 | ENSG00000246228 |
| LOC107984525 | 1.020598988 | 2.0287611 | 0.109411443 | 0.255279849 | 8.077802188 | 16.28620536 | ENSG00000286069 |
| FAM78B | 1.020196329 | 2.028194948 | 6.69E-06 | 6.65E-05 | 72.78770523 | 147.8224806 | ENSG00000188859 |
| RCN3 | 1.019865038 | 2.027729259 | 5.04E-18 | 2.54E-16 | 289.9768678 | 587.6938489 | ENSG00000142552 |
| NA | 1.019349542 | 2.027004851 | 0.178481801 | 0.360688825 | 6.885413993 | 14.01269463 | ENSG00000261172 |
| PSEN2 | 1.018686068 | 2.026072877 | 9.31E-25 | 8.02E-23 | 427.9877735 | 866.3271721 | ENSG00000143801 |
| TOR4A | 1.018026028 | 2.02514615 | 1.88E-13 | 5.86E-12 | 220.23368 | 445.4159982 | ENSG00000198113 |
| POU6F2 | 1.017987734 | 2.025092397 | 0.046574038 | 0.133487738 | 15.51922669 | 31.24346696 | ENSG00000106536 |
| IER5 | 1.017845902 | 2.02489332 | 4.46E-30 | 5.27E-28 | 518.6925762 | 1050.274001 | ENSG00000162783 |
| LIF | 1.017543655 | 2.024469145 | 9.89E-07 | 1.15E-05 | 84.21230063 | 170.6931018 | ENSG00000128342 |
| TGM1 | 1.016000503 | 2.022304866 | 0.021018475 | 0.07168754 | 19.68565783 | 39.95003437 | ENSG00000092295 |
| ACTN3 | 1.015952161 | 2.022237105 | 5.73E-19 | 3.18E-17 | 355.5784362 | 718.92557 | ENSG00000248746 |
| CD8A | 1.015260427 | 2.021267728 | 0.310807625 | 0.521464285 | 3.302884333 | 6.693241397 | ENSG00000153563 |
| NA | 1.01517776 | 2.021151912 | 0.31517531 | 0.525477685 | 2.948516239 | 5.987388051 | ENSG00000227953 |
| BMP8B | 1.015124891 | 2.021077846 | 0.001022143 | 0.005716072 | 33.59553226 | 67.77565743 | ENSG00000116985 |
| SUGP1 | 1.015037792 | 2.020955832 | 1.73E-43 | 3.98E-41 | 876.5422429 | 1771.359919 | ENSG00000105705 |
| DCST1 | 1.014880457 | 2.020735446 | 0.256553249 | 0.463427453 | 3.926219337 | 7.971770626 | ENSG00000163357 |
| LOC101927572 | 1.014850241 | 2.020693123 | 0.195582076 | 0.384625666 | 5.792944205 | 11.57394157 | ENSG00000267698 |
| NA | 1.014477117 | 2.02017058 | 0.18387739 | 0.367930124 | 5.229814749 | 10.61783486 | ENSG00000260015 |

| | | | | | | | |
|---|---|---|---|---|---|---|---|
| NA | 1.013690894 | 2.019069951 | 0.377611848 | 0.590038039 | 2.308579082 | 4.674618497 | ENSG00000245904 |
| NA | 1.013462457 | 2.018750276 | 0.107835272 | 0.252572711 | 8.158169362 | 16.548282 | ENSG00000205267 |
| NA | 1.011595635 | 2.016139739 | 0.216433706 | 0.411898484 | 5.518943284 | 11.17963343 | ENSG00000281332 |
| MGAT3 | 1.011348263 | 2.01579407 | 7.99E-28 | 8.47E-26 | 442.0489247 | 891.210414 | ENSG00000128268 |
| C3orf35 | 1.011191748 | 2.015575392 | 0.196870448 | 0.38627516 | 5.601980206 | 11.31621586 | ENSG00000198590 |
| LOC285097 | 1.010264508 | 2.014280371 | 0.274568817 | 0.483381385 | 4.317351211 | 8.699095062 | ENSG00000280119 |
| BRINP3 | 1.00968826 | 2.013475978 | 0.402207859 | 0.613595236 | 2.301460137 | 4.644445553 | ENSG00000162670 |
| NA | 1.00934793 | 2.013001057 | 0.16063328 | 0.334756068 | 5.911270367 | 11.93277988 | ENSG00000226330 |
| ZBTB41 | 1.009316928 | 2.0129578 | 5.05E-06 | 5.13E-05 | 64.60967973 | 129.9527272 | ENSG00000177888 |
| TSPOAP1-AS1 | 1.009183993 | 2.012772327 | 0.345726214 | 0.557608639 | 2.632107133 | 5.310478143 | ENSG00000265148 |
| APOLD1 | 1.008173079 | 2.011362448 | 0.270552793 | 0.478990297 | 4.29007064 | 8.638749174 | ENSG00000178878 |
| PCLO | 1.00808954 | 2.011245984 | 1.21E-07 | 1.65E-06 | 92.80934955 | 186.5499807 | ENSG00000186472 |
| PRCD | 1.008004813 | 2.01112787 | 0.159678014 | 0.333353891 | 6.620006557 | 13.32301548 | ENSG00000214140 |
| CGAS | 1.007206441 | 2.010015241 | 0.00023295 | 0.001576229 | 51.11285267 | 102.3720606 | ENSG00000164430 |
| FRMPD4 | 1.006627146 | 2.009208308 | 0.015899221 | 0.057169991 | 20.95276879 | 41.99114696 | ENSG00000169933 |
| TTC39C | 1.006451494 | 2.008963696 | 1.62E-21 | 1.10E-19 | 335.9764257 | 674.8354103 | ENSG00000168234 |
| LINC01852 | 1.006402051 | 2.008894848 | 0.302392178 | 0.512490019 | 3.964178325 | 7.969595162 | ENSG00000236914 |
| NA | 1.006166917 | 2.00856746 | 0.39851223 | 0.610274365 | 2.312138554 | 4.647566976 | ENSG00000280122 |
| DARS2 | 1.005499372 | 2.007638297 | 1.08E-30 | 1.31E-28 | 757.0154547 | 1520.633215 | ENSG00000117593 |
| WASIR2 | 1.005339225 | 2.007415451 | 0.127823954 | 0.285371245 | 6.956577289 | 13.96445556 | ENSG00000231439 |
| NA | 1.005155534 | 2.007159874 | 0.350167406 | 0.562322403 | 2.9900347 | 5.998265368 | ENSG00000243746 |
| UNC80 | 1.004801344 | 2.006667164 | 0.087504865 | 0.215875817 | 8.917907221 | 17.90399373 | ENSG00000144406 |
| ZFHX3 | 1.0047545 | 2.006602009 | 0.000212623 | 0.001451694 | 45.35163823 | 91.00129017 | ENSG00000140836 |
| NA | 1.004741743 | 2.006584266 | 0.305319303 | 0.51555392 | 3.313562751 | 6.648123755 | ENSG00000213399 |
| HLA-G | 1.004518498 | 2.006273787 | 0.000127479 | 0.000922388 | 47.87965709 | 95.86893756 | ENSG00000204632 |
| NA | 1.004508403 | 2.006259749 | 0.226590951 | 0.425491709 | 4.906286698 | 9.86563437 | ENSG00000214018 |
| AIF1 | 1.004393174 | 2.006099514 | 0.000950435 | 0.005370727 | 35.46286752 | 71.35323586 | ENSG00000204472 |
| MIR647 | 1.003594352 | 2.004989042 | 0.243556753 | 0.447820979 | 4.29007064 | 8.60450885 | ENSG00000207554 |
| NA | 1.00333002 | 2.00462172 | 0.353265377 | 0.565470134 | 2.982915755 | 5.976510733 | ENSG00000288271 |
| PIWIL3 | 1.003085119 | 2.00428146 | 0.403420646 | 0.614750173 | 2.305019609 | 4.622974464 | ENSG00000184571 |
| ZNF467 | 1.002474702 | 2.00343361 | 0.123968322 | 0.279039542 | 7.318064329 | 14.63919001 | ENSG00000181444 |
| ANKRD65 | 1.00224006 | 2.003107794 | 0.047314698 | 0.13515929 | 13.32065962 | 26.63761266 | ENSG00000235098 |
| RUNX1 | 1.001674272 | 2.002322381 | 0.129285653 | 0.287851869 | 7.338225955 | 14.6606611 | ENSG00000159216 |
| FGD3 | 1.001648332 | 2.00228638 | 7.15E-05 | 0.000551451 | 59.50027608 | 119.4666723 | ENSG00000127084 |
| C3orf62 | 1.001227908 | 2.001702966 | 0.005522751 | 0.024003946 | 24.63819151 | 49.46118136 | ENSG00000188315 |
| MKLN1-AS | 1.000951422 | 2.001319386 | 0.007248908 | 0.030113424 | 23.95437163 | 48.08466095 | ENSG00000236753 |
| NMNAT2 | 1.000487087 | 2.00067536 | 2.01E-22 | 1.46E-20 | 366.6356208 | 732.9801523 | ENSG00000157064 |
| NA | 1.000475252 | 2.000658948 | 0.103915735 | 0.245152862 | 8.298131691 | 16.57627948 | ENSG00000286207 |
| PYCR2 | 1.000288961 | 2.000400626 | 1.32E-24 | 1.13E-22 | 439.9805835 | 880.6435084 | ENSG00000143811 |
| TLR4 | 1.000235725 | 2.000326812 | 0.358806627 | 0.570841373 | 2.666506649 | 5.321355461 | ENSG00000136869 |
| NIT1 | 0.999521329 | 1.999336532 | 3.51E-25 | 3.20E-23 | 496.3816253 | 991.7845124 | ENSG00000158793 |
| SYTL5 | 0.998993791 | 1.998605585 | 0.121955114 | 0.275790005 | 8.701137191 | 17.33812778 | ENSG00000147041 |
| ERN2 | 0.998390903 | 1.997770562 | 0.0126522 | 0.047445751 | 19.60053597 | 39.07307472 | ENSG00000134398 |
| GCC2-AS1 | 0.99777851 | 1.99692273 | 0.071145715 | 0.185220131 | 11.65201769 | 23.21229641 | ENSG00000214184 |
| ERVH48-1 | 0.997748084 | 1.996880617 | 5.92E-16 | 2.51E-14 | 529.1435355 | 1055.462769 | ENSG00000233056 |
| LOC101929563 | 0.997590564 | 1.9966626 | 0.47415273 | 0.677209204 | 2.373818642 | 4.718790181 | ENSG00000283982 |
| TENM1 | 0.997159163 | 1.996065638 | 0.00304042 | 0.014580912 | 38.63140836 | 76.78658125 | ENSG00000009694 |
| NA | 0.99681061 | 1.99558345 | 0.40560976 | 0.616792038 | 2.366699697 | 4.696089587 | ENSG00000258365 |
| LIX1L | 0.996596688 | 1.995287568 | 0.00024811 | 0.00166627 | 48.17323482 | 96.20725074 | ENSG00000271601 |
| ZNF575 | 0.996394173 | 1.995007504 | 0.255975421 | 0.462814344 | 4.004501578 | 7.959947349 | ENSG00000176472 |
| CGN | 0.995887522 | 1.994307013 | 3.46E-68 | 2.18E-65 | 1888.079628 | 3765.971964 | ENSG00000143375 |
| TRIM15 | 0.995880619 | 1.994297471 | 0.096625795 | 0.232270166 | 9.344633821 | 18.57343129 | ENSG00000204610 |
| RD3 | 0.995590649 | 1.993896673 | 0.285988612 | 0.49531512 | 3.650133483 | 7.259390889 | ENSG00000198570 |
| NA | 0.99412823 | 1.991876541 | 0.414761798 | 0.625609309 | 2.363140224 | 4.679915383 | ENSG00000273002 |
| NA | 0.994001542 | 1.991701635 | 0.296954299 | 0.507188382 | 4.059062721 | 8.037413399 | ENSG00000260725 |
| OLFML3 | 0.993418298 | 1.990896605 | 3.86E-39 | 7.15E-37 | 803.3274539 | 1598.691035 | ENSG00000116774 |
| SYCP1 | 0.992892178 | 1.990170699 | 0.366077951 | 0.57862254 | 2.690227748 | 5.323530924 | ENSG00000198765 |
| NPIPB13 | 0.992874348 | 1.990146103 | 0.290395541 | 0.499919762 | 3.653692956 | 7.247567612 | ENSG00000198064 |
| TSLP | 0.992573307 | 1.989730872 | 0.191048218 | 0.377817504 | 5.707543019 | 11.29569073 | ENSG00000145777 |
| ARHGAP9 | 0.992496823 | 1.98962539 | 0.001509783 | 0.008002316 | 31.97997694 | 63.72100792 | ENSG00000123329 |
| GPRIN3 | 0.992428693 | 1.989531433 | 9.31E-05 | 0.000696793 | 49.67374412 | 98.76213373 | ENSG00000185477 |
| CYTL1 | 0.991999196 | 1.988939228 | 1.50E-08 | 2.42E-07 | 114.608121 | 227.8197625 | ENSG00000170891 |
| HMCN1 | 0.991325531 | 1.988010713 | 7.19E-10 | 1.40E-08 | 241.104276 | 478.4212654 | ENSG00000143341 |
| MAP4K3-DT | 0.990302092 | 1.98660093 | 1.92E-05 | 0.000171905 | 62.356857 | 123.7592574 | ENSG00000231312 |
| BRCA2 | 0.989999779 | 1.986184686 | 8.40E-12 | 2.15E-10 | 217.7472051 | 432.7310905 | ENSG00000139618 |
| NA | 0.989327312 | 1.985259105 | 0.129645408 | 0.288376048 | 8.05408109 | 15.89936957 | ENSG00000280099 |
| NA | 0.989064171 | 1.984897035 | 0.107265922 | 0.251579352 | 9.192797866 | 18.31135465 | ENSG00000285852 |
| RAB26 | 0.988986971 | 1.984790824 | 4.08E-05 | 0.000335429 | 74.18532147 | 147.3111395 | ENSG00000167964 |
| ARID3C | 0.987622884 | 1.982915066 | 0.435509442 | 0.644005602 | 2.335859653 | 4.611151187 | ENSG00000205143 |
| STEAP2 | 0.987622332 | 1.982914307 | 8.51E-12 | 2.17E-10 | 196.4125345 | 390.14875 | ENSG00000157214 |
| NA | 0.987203457 | 1.982338667 | 0.188744021 | 0.374651129 | 6.767087831 | 13.32424499 | ENSG00000242242 |
| TAGLN2 | 0.986894678 | 1.981914434 | 5.06E-56 | 1.87E-53 | 1640.236533 | 3250.556078 | ENSG00000158710 |
| NA | 0.985828208 | 1.980449904 | 0.052210286 | 0.14596976 | 12.28247164 | 24.3149169 | ENSG00000280435 |
| KLF7 | 0.985280396 | 1.97969804 | 1.35E-15 | 5.50E-14 | 272.739777 | 540.1525939 | ENSG00000118263 |

| | | | | | | | |
|---|---|---|---|---|---|---|---|
| NA | 0.984828206 | 1.979077633 | 0.241646635 | 0.44519938 | 5.094886433 | 9.99909538 | ENSG00000287516 |
| PADI2 | 0.984503867 | 1.978632758 | 6.97E-09 | 1.18E-07 | 144.8385046 | 285.9428451 | ENSG00000117115 |
| PMFBP1 | 0.983926695 | 1.977841333 | 0.116295965 | 0.26705366 | 8.934509375 | 17.69261517 | ENSG00000118557 |
| NA | 0.983391274 | 1.977107443 | 0.173774894 | 0.353826293 | 7.239782087 | 14.3587637 | ENSG00000260193 |
| LINC01873 | 0.982108307 | 1.975350012 | 0.400916296 | 0.612310747 | 2.370259169 | 4.637919163 | ENSG00000232164 |
| PDLIM3 | 0.981691882 | 1.974779921 | 3.07E-06 | 3.23E-05 | 71.37409781 | 141.0177437 | ENSG00000154553 |
| GADD45G | 0.981006098 | 1.973841435 | 1.06E-11 | 2.67E-10 | 161.3022295 | 318.4846314 | ENSG00000130222 |
| LOC653653 | 0.980098499 | 1.972600082 | 0.062165775 | 0.166737772 | 11.25732635 | 22.18865501 | ENSG00000280852 |
| TRPV1 | 0.97999637 | 1.972460446 | 0.000840162 | 0.004813854 | 48.50182304 | 95.6182327 | ENSG00000196689 |
| A1BG-AS1 | 0.979367797 | 1.971601244 | 0.18808806 | 0.373977997 | 6.186161012 | 12.26702568 | ENSG00000268895 |
| FAHD2CP | 0.979137538 | 1.971286595 | 0.000612535 | 0.003659512 | 42.74948142 | 83.98997717 | ENSG00000231584 |
| SNAI2 | 0.978937109 | 1.971012751 | 0.103820775 | 0.244959438 | 8.894186122 | 17.55262777 | ENSG00000019549 |
| DBH | 0.978657917 | 1.970631355 | 0.091424674 | 0.222833443 | 10.0047326 | 19.68011916 | ENSG00000123454 |
| NA | 0.978234208 | 1.970052679 | 0.10072987 | 0.239221889 | 8.938068848 | 17.60985223 | ENSG00000261559 |
| IRX6 | 0.977660638 | 1.969269604 | 0.07801607 | 0.198161867 | 10.92431509 | 21.49121997 | ENSG00000159387 |
| HFE | 0.976821538 | 1.968124571 | 0.022879672 | 0.076664764 | 17.58108542 | 34.58450723 | ENSG00000010704 |
| RBM15 | 0.976675032 | 1.967924718 | 4.87E-20 | 2.91E-18 | 376.55668 | 740.2666377 | ENSG00000162775 |
| LOC105375431 | 0.976212562 | 1.967293981 | 0.187489776 | 0.37298477 | 5.897032477 | 11.65575855 | ENSG00000274993 |
| NA | 0.975329675 | 1.966090423 | 0.141010061 | 0.30538704 | 7.968679903 | 15.66216899 | ENSG00000285722 |
| HLA-DRB1 | 0.974991662 | 1.965629837 | 3.75E-15 | 1.44E-13 | 278.4499119 | 546.6335537 | ENSG00000196126 |
| LOC101927825 | 0.974099724 | 1.964414972 | 0.229788691 | 0.429616814 | 5.226255277 | 10.32340982 | ENSG00000282556 |
| CDH13 | 0.973937538 | 1.964194149 | 0.072553601 | 0.188055826 | 15.04358335 | 29.67458007 | ENSG00000140945 |
| EFCAB2 | 0.973280875 | 1.963300322 | 9.81E-09 | 1.62E-07 | 116.6628869 | 229.0958327 | ENSG00000203666 |
| IFIT1 | 0.972394959 | 1.962095087 | 0.006047363 | 0.025908451 | 28.6868813 | 56.13607309 | ENSG00000185745 |
| NA | 0.971403828 | 1.960747591 | 0.027208271 | 0.088089866 | 16.54526171 | 32.41312769 | ENSG00000228648 |
| NA | 0.97085468 | 1.960001393 | 0.141005062 | 0.30538704 | 6.929296718 | 13.59379397 | ENSG00000255031 |
| SNHG8 | 0.970426232 | 1.959419404 | 3.14E-22 | 2.25E-20 | 562.6475397 | 1103.487977 | ENSG00000269893 |
| KCNH5 | 0.970127262 | 1.959013394 | 0.00250135 | 0.012317601 | 33.62845724 | 65.94448299 | ENSG00000140015 |
| NA | 0.969325633 | 1.957925177 | 0.285692386 | 0.494933661 | 4.228390553 | 8.34025675 | ENSG00000264281 |
| CYS1 | 0.968736603 | 1.95712595 | 0.164912851 | 0.340518172 | 6.602209194 | 12.92993685 | ENSG00000205795 |
| EXPH5 | 0.9679234 | 1.95602309 | 0.001008872 | 0.005651901 | 37.83223697 | 73.88853969 | ENSG00000110723 |
| RYR1 | 0.967913386 | 1.956009512 | 1.53E-32 | 2.02E-30 | 1045.408952 | 2045.240304 | ENSG00000196218 |
| RRNAD1 | 0.967668728 | 1.955677832 | 5.03E-14 | 1.67E-12 | 243.0158773 | 475.0523136 | ENSG00000143303 |
| MEIS3 | 0.967558759 | 1.955528767 | 2.94E-30 | 3.52E-28 | 633.8202494 | 1238.80523 | ENSG00000105419 |
| ACTRT3 | 0.967127372 | 1.954944122 | 0.000250983 | 0.00168198 | 54.83831931 | 107.1971447 | ENSG00000184378 |
| ASH1L | 0.966664211 | 1.95431661 | 2.79E-21 | 1.84E-19 | 444.0627314 | 867.0190267 | ENSG00000116539 |
| LINC01376 | 0.96596981 | 1.953376181 | 0.192521727 | 0.37987387 | 7.39634657 | 14.39858446 | ENSG00000236204 |
| NA | 0.96558271 | 1.952852127 | 0.08466444 | 0.210879445 | 12.41503569 | 24.37743826 | ENSG00000287625 |
| JPX | 0.96540862 | 1.952616491 | 0.162714528 | 0.337677347 | 6.643727656 | 12.95575886 | ENSG00000225470 |
| GABRG1 | 0.965062997 | 1.952148762 | 0.269846575 | 0.477964037 | 4.246187915 | 8.342432214 | ENSG00000163285 |
| ZFHX4 | 0.964710756 | 1.951672194 | 0.003714013 | 0.017245913 | 32.38178723 | 63.18548288 | ENSG00000091656 |
| NA | 0.964540083 | 1.951441322 | 0.094743591 | 0.229196202 | 10.08301485 | 19.55318454 | ENSG00000234232 |
| SH3BP5-AS1 | 0.964170497 | 1.950941471 | 0.114655166 | 0.264249794 | 9.572082268 | 18.72089104 | ENSG00000224660 |
| PYROXD2 | 0.963556147 | 1.950110868 | 0.002668917 | 0.013003741 | 32.24093518 | 62.77767045 | ENSG00000119943 |
| CADM3 | 0.963383606 | 1.949877657 | 0.000103195 | 0.000763648 | 54.33094621 | 106.1158273 | ENSG00000162706 |
| ITPRID1 | 0.962739362 | 1.949007121 | 0.235849258 | 0.437749338 | 5.280816419 | 10.3053437 | ENSG00000180347 |
| LOC101928053 | 0.962224833 | 1.948312143 | 0.352239905 | 0.564306973 | 3.533892253 | 6.950967106 | ENSG00000246308 |
| NA | 0.961803338 | 1.947743011 | 0.380184065 | 0.592537433 | 3.068316941 | 5.898098724 | ENSG00000250144 |
| ZMAT4 | 0.961032112 | 1.946702078 | 4.09E-11 | 9.68E-10 | 196.4273572 | 382.5867791 | ENSG00000165061 |
| LMO1 | 0.960993659 | 1.946650193 | 0.409286794 | 0.62043515 | 2.751907835 | 5.279359241 | ENSG00000166407 |
| RNF128 | 0.960708336 | 1.946265239 | 0.003075971 | 0.014709982 | 32.8832366 | 63.86674371 | ENSG00000133135 |
| HLA-F | 0.960554173 | 1.946057278 | 0.000774037 | 0.004482578 | 43.42709823 | 84.50034262 | ENSG00000204642 |
| RGS20 | 0.959061125 | 1.94404434 | 0.284477777 | 0.493726967 | 3.919100392 | 7.66769777 | ENSG00000147509 |
| TMEM119 | 0.958858784 | 1.943771703 | 0.356616806 | 0.568713764 | 3.217483147 | 6.322579809 | ENSG00000183160 |
| NA | 0.958749956 | 1.943625083 | 0.059085874 | 0.16052895 | 13.29693852 | 25.77370579 | ENSG00000224616 |
| NFAM1 | 0.958007181 | 1.942624661 | 0.013314133 | 0.049417727 | 24.00804305 | 46.52504301 | ENSG00000235568 |
| FIBIN | 0.957865106 | 1.942433362 | 0.255664619 | 0.4625624 | 5.278452155 | 10.25776705 | ENSG00000176971 |
| POLR2J3 | 0.957374574 | 1.941773026 | 0.199166728 | 0.389203606 | 5.676702975 | 10.99284738 | ENSG00000285437 |
| RIN2 | 0.957336695 | 1.941722044 | 0.020368656 | 0.069798962 | 20.15031726 | 39.14618985 | ENSG00000132669 |
| ACER2 | 0.956552426 | 1.940666784 | 2.52E-05 | 0.000219067 | 82.66345941 | 160.347967 | ENSG00000177076 |
| DPP6 | 0.955958684 | 1.939868266 | 2.02E-10 | 4.30E-09 | 146.5608435 | 284.2975537 | ENSG00000130226 |
| NA | 0.955581335 | 1.939360943 | 0.210624555 | 0.40411068 | 5.700424074 | 11.01337251 | ENSG00000238035 |
| NA | 0.955435472 | 1.939164875 | 0.238135925 | 0.440512973 | 4.620717636 | 8.972994974 | ENSG00000277475 |
| NA | 0.955283969 | 1.938961247 | 0.00047786 | 0.002962698 | 46.65434392 | 90.21144438 | ENSG00000239213 |
| PARVG | 0.954845191 | 1.938371625 | 0.272156077 | 0.480747126 | 4.300749058 | 8.349904563 | ENSG00000138964 |
| PPARG | 0.954767702 | 1.938267515 | 0.32143266 | 0.531913727 | 3.592012868 | 7.006016107 | ENSG00000132170 |
| NA | 0.954538984 | 1.937960256 | 0.259877975 | 0.466786718 | 4.273468486 | 8.306962384 | ENSG00000205871 |
| KCNA1 | 0.954280994 | 1.937613731 | 0.24115922 | 0.444668216 | 5.308096991 | 10.27081983 | ENSG00000111262 |
| NA | 0.953690048 | 1.936820222 | 0.049389268 | 0.139732782 | 14.28176056 | 27.6453634 | ENSG00000279249 |
| NA | 0.95365504 | 1.936773225 | 0.271818912 | 0.480477361 | 4.242628443 | 8.250967424 | ENSG00000183022 |
| LOC101928222 | 0.952713916 | 1.935510207 | 0.264026081 | 0.471457313 | 4.640879262 | 8.980467323 | ENSG00000227210 |
| LOC101929494 | 0.952208419 | 1.934832155 | 1.02E-05 | 9.71E-05 | 76.678356 | 148.6419322 | ENSG00000276170 |
| NA | 0.95218446 | 1.934800023 | 0.315094412 | 0.525450848 | 3.588453396 | 6.985490977 | ENSG00000224358 |
| ZBTB7C | 0.952103138 | 1.934690965 | 0.146693836 | 0.313983685 | 10.23365559 | 19.62942109 | ENSG00000184828 |

| | | | | | | | |
|---|---|---|---|---|---|---|---|
| SNORD15B | 0.951922082 | 1.934448179 | 0.26307708 | 0.470251837 | 4.273468486 | 8.286437254 | ENSG00000207445 |
| EVA1C | 0.951175223 | 1.933447007 | 0.002405913 | 0.011925374 | 33.17356034 | 64.08296766 | ENSG00000166979 |
| YOD1 | 0.951160302 | 1.933427011 | 1.50E-10 | 3.25E-09 | 152.6177206 | 295.2281633 | ENSG00000180667 |
| SYNPR-AS1 | 0.951036371 | 1.933260932 | 0.12350182 | 0.278221624 | 8.930949903 | 17.27030955 | ENSG00000241359 |
| LOC101928059 | 0.950782675 | 1.932921 | 0.265623837 | 0.473175215 | 5.797393401 | 11.30221712 | ENSG00000284738 |
| KBTBD3 | 0.950740817 | 1.93286492 | 0.018425436 | 0.064304723 | 25.67429456 | 49.51783873 | ENSG00000182359 |
| PROS1 | 0.950221924 | 1.932169853 | 0.027301071 | 0.088299491 | 17.15435882 | 33.18367785 | ENSG00000184500 |
| CRYGB | 0.949843998 | 1.931663771 | 0.211342925 | 0.40503462 | 6.085632213 | 11.67722964 | ENSG00000182187 |
| NA | 0.949419871 | 1.931095979 | 0.002651319 | 0.012944713 | 32.87375339 | 63.39792299 | ENSG00000234614 |
| CREB3L3 | 0.949034643 | 1.930580408 | 0.349138596 | 0.561195198 | 3.279163235 | 6.366751493 | ENSG00000060566 |
| CD70 | 0.948802858 | 1.930270264 | 0.007965363 | 0.032545624 | 26.03906174 | 50.30435219 | ENSG00000125726 |
| LINC01123 | 0.948030114 | 1.929236638 | 0.139510049 | 0.303321124 | 7.549072248 | 14.61554345 | ENSG00000204588 |
| CRTC3-AS1 | 0.947891833 | 1.929051732 | 0.357989992 | 0.569904458 | 3.251882663 | 6.316053418 | ENSG00000259736 |
| NA | 0.947767256 | 1.928885165 | 6.85E-08 | 9.83E-07 | 178.6357054 | 344.0507854 | ENSG00000255666 |
| EFCAB13 | 0.9474877 | 1.928511434 | 0.137769576 | 0.300679352 | 7.92360197 | 15.30304713 | ENSG00000178852 |
| USP46-AS1 | 0.947070754 | 1.927954165 | 0.119608861 | 0.27216434 | 9.275834788 | 17.86228105 | ENSG00000248866 |
| FMO4 | 0.946286597 | 1.926906538 | 0.060876684 | 0.164236598 | 14.11095819 | 27.28747105 | ENSG00000076258 |
| APOL6 | 0.946139156 | 1.926709621 | 0.30840896 | 0.51877645 | 3.626412384 | 7.006962066 | ENSG00000221963 |
| ULK4P3 | 0.945748738 | 1.926188291 | 0.232451095 | 0.433361033 | 6.065470586 | 11.6106409 | ENSG00000178081 |
| FAM167A | 0.94522559 | 1.925489946 | 0.003320309 | 0.0156917 | 35.82435456 | 69.08263564 | ENSG00000154319 |
| DNAH10 | 0.945110703 | 1.925336618 | 0.169525057 | 0.347458292 | 6.891337729 | 13.32206953 | ENSG00000197653 |
| NA | 0.944868396 | 1.925013276 | 0.042844293 | 0.125137318 | 14.82917759 | 28.609226 | ENSG00000258744 |
| SLC45A1 | 0.944676365 | 1.924757063 | 1.85E-05 | 0.000166564 | 74.35940399 | 143.1369847 | ENSG00000162426 |
| C4orf51 | 0.944532816 | 1.924565559 | 0.357906478 | 0.569904458 | 3.904862502 | 7.536412223 | ENSG00000237136 |
| C5orf60 | 0.944355004 | 1.924328371 | 0.283431206 | 0.492464654 | 3.946380963 | 7.606405924 | ENSG00000204661 |
| CEBPZOS | 0.943626967 | 1.923357529 | 0.080275305 | 0.20224108 | 11.24072419 | 21.64520611 | ENSG00000218739 |
| NA | 0.943491982 | 1.923177579 | 0.060030283 | 0.162393911 | 13.3491354 | 25.60091848 | ENSG00000181097 |
| KIAA1614 | 0.943315883 | 1.922942845 | 0.001494427 | 0.007938749 | 45.70244685 | 87.83530799 | ENSG00000135835 |
| NA | 0.943306622 | 1.922930502 | 0.354428426 | 0.566515517 | 3.643014538 | 7.011312993 | ENSG00000235888 |
| ZMAT3 | 0.943187027 | 1.922771103 | 3.92E-12 | 1.04E-10 | 199.7082743 | 383.7513311 | ENSG00000172667 |
| NA | 0.943057337 | 1.922598265 | 0.20004248 | 0.390461886 | 6.814530028 | 12.97505449 | ENSG00000256967 |
| MSH5 | 0.943010079 | 1.922535287 | 0.162666957 | 0.337677347 | 7.272986395 | 14.00210086 | ENSG00000204410 |
| NPBWR1 | 0.942994523 | 1.922514558 | 0.273550696 | 0.482218324 | 6.795258125 | 13.11643935 | ENSG00000183729 |
| TOP3B | 0.94253781 | 1.921906045 | 0.074004979 | 0.190874628 | 13.48050424 | 25.98529526 | ENSG00000100038 |
| LRRN4 | 0.942447362 | 1.921785557 | 0.085263563 | 0.211924504 | 11.50493642 | 22.14230786 | ENSG00000125872 |
| PRDX4 | 0.942125684 | 1.921357104 | 1.43E-62 | 6.91E-60 | 2196.625528 | 4219.848584 | ENSG00000123131 |
| NEB | 0.941759268 | 1.92086918 | 0.008443237 | 0.034128968 | 27.60597965 | 52.87824726 | ENSG00000183091 |
| SELENBP1 | 0.94160231 | 1.92066021 | 1.25E-18 | 6.64E-17 | 402.1082509 | 772.0979791 | ENSG00000143416 |
| LINC00624 | 0.941346764 | 1.920320031 | 0.1635824 | 0.338773684 | 7.256384241 | 13.95603725 | ENSG00000278811 |
| NA | 0.940672399 | 1.919422618 | 0.2592098 | 0.466162945 | 6.1473123 | 11.70210569 | ENSG00000224846 |
| LMCD1-AS1 | 0.94050519 | 1.919200169 | 0.117025674 | 0.26824006 | 9.489045346 | 18.29735591 | ENSG00000227110 |
| ZFPM2 | 0.939662867 | 1.918079964 | 0.32854875 | 0.540048678 | 3.275603762 | 6.310756533 | ENSG00000169946 |
| LINC01460 | 0.939432944 | 1.917774304 | 0.287212964 | 0.496341218 | 4.348191255 | 8.306962384 | ENSG00000205334 |
| NA | 0.938658602 | 1.916745247 | 0.049811098 | 0.140694375 | 19.54717004 | 37.44469258 | ENSG00000233290 |
| PGA5 | 0.938596078 | 1.91666218 | 0.393115794 | 0.604796215 | 3.317122223 | 6.364576029 | ENSG00000256713 |
| NA | 0.938102728 | 1.916006862 | 0.287954871 | 0.497213837 | 5.989552609 | 11.42648182 | ENSG00000267141 |
| NA | 0.937819016 | 1.915630109 | 0.1637168 | 0.338975088 | 7.276545868 | 13.94203851 | ENSG00000279145 |
| SYCP2 | 0.937810104 | 1.915618274 | 0.34147053 | 0.553106695 | 3.268484817 | 6.287109979 | ENSG00000196074 |
| LPXN | 0.937578269 | 1.915310468 | 0.056951116 | 0.156098958 | 13.16409514 | 25.28284689 | ENSG00000110031 |
| KCNS1 | 0.937527925 | 1.915243632 | 0.000143633 | 0.001028635 | 59.39972112 | 113.8827457 | ENSG00000124134 |
| PLA2G4A | 0.937362598 | 1.915024167 | 0.005886785 | 0.025318135 | 28.67116887 | 54.96062102 | ENSG00000116711 |
| NA | 0.937068319 | 1.914633581 | 6.53E-05 | 0.000509144 | 67.59912961 | 129.8101128 | ENSG00000179277 |
| RALY-AS1 | 0.936704986 | 1.914151455 | 0.078046037 | 0.198211324 | 13.3965776 | 25.5328167 | ENSG00000285230 |
| MYMX | 0.93567544 | 1.912785952 | 0.0133066 | 0.049399479 | 28.29068926 | 54.24220934 | ENSG00000262179 |
| NA | 0.935082207 | 1.911999581 | 0.287085237 | 0.496231181 | 4.00806105 | 7.634403404 | ENSG00000225579 |
| NA | 0.934933188 | 1.911802096 | 0.133650968 | 0.294756339 | 9.797445783 | 18.60265827 | ENSG00000243829 |
| RGMB-AS1 | 0.93485048 | 1.911692499 | 0.303388186 | 0.513486816 | 4.000942105 | 7.623526086 | ENSG00000246763 |
| NA | 0.934698714 | 1.911491406 | 0.330070353 | 0.541514953 | 3.302884333 | 6.322579809 | ENSG00000225536 |
| RNF112 | 0.934494206 | 1.911220463 | 0.001901796 | 0.009717377 | 45.12983419 | 86.39486877 | ENSG00000128482 |
| PLIN4 | 0.933063959 | 1.909326673 | 0.009268755 | 0.036719201 | 27.29757922 | 52.27305501 | ENSG00000167676 |
| ZDHHC21 | 0.932731117 | 1.908886225 | 5.94E-05 | 0.000468704 | 102.1092365 | 195.2001313 | ENSG00000175893 |
| PPP5D1 | 0.932721778 | 1.908873869 | 0.099120329 | 0.236678141 | 11.28460692 | 21.50181374 | ENSG00000230510 |
| DCAF4L1 | 0.932304307 | 1.908321579 | 0.367501857 | 0.579660284 | 2.959194656 | 5.668370496 | ENSG00000182308 |
| TP73 | 0.932072805 | 1.908015386 | 1.05E-06 | 1.21E-05 | 92.59939298 | 176.7658797 | ENSG00000078900 |
| CILP2 | 0.93202803 | 1.90795617 | 1.11E-18 | 5.99E-17 | 358.5003084 | 683.9853326 | ENSG00000160161 |
| GJD2 | 0.931526072 | 1.907292449 | 0.278338241 | 0.48756309 | 4.848166083 | 9.334008749 | ENSG00000159248 |
| NA | 0.930947998 | 1.906528368 | 0.000519575 | 0.003180022 | 47.11015669 | 89.97518976 | ENSG00000273599 |
| GH2 | 0.930345019 | 1.905731695 | 0.183687427 | 0.367627921 | 7.476713744 | 14.35564228 | ENSG00000136487 |
| FAM71E1 | 0.930219165 | 1.905565456 | 5.58E-07 | 6.75E-06 | 97.67528685 | 186.04775 | ENSG00000142530 |
| DUSP2 | 0.928904105 | 1.903829267 | 2.02E-09 | 3.72E-08 | 141.5584772 | 269.5679605 | ENSG00000158050 |
| FAM178B | 0.928886991 | 1.903806682 | 0.341345855 | 0.552952176 | 3.33728385 | 6.340929476 | ENSG00000168754 |
| NA | 0.928359214 | 1.903110346 | 0.362121376 | 0.574002198 | 4.447830331 | 8.402778101 | ENSG00000270069 |
| NA | 0.928167733 | 1.902857772 | 0.10897623 | 0.254484494 | 11.54645488 | 22.01558416 | ENSG00000233577 |
| NA | 0.928010738 | 1.902650714 | 0.27995739 | 0.488994296 | 5.22033154 | 9.986042599 | ENSG00000286540 |

| | | | | | | | |
|---|---|---|---|---|---|---|---|
| PSMB10 | 0.927497969 | 1.901974585 | 0.18094746 | 0.363879543 | 6.987417333 | 13.25737271 | ENSG00000205220 |
| LOC101059948 | 0.927221205 | 1.901609749 | 0.373656231 | 0.585698698 | 2.97579681 | 5.666195032 | ENSG00000268655 |
| CALB2 | 0.926902512 | 1.901189728 | 0.002157433 | 0.010835936 | 34.77340324 | 66.10310361 | ENSG00000172137 |
| GAL3ST1 | 0.925799661 | 1.899736942 | 3.57E-13 | 1.08E-11 | 218.0313002 | 414.0294654 | ENSG00000128242 |
| NA | 0.925517137 | 1.899364951 | 0.30889887 | 0.519369227 | 4.004501578 | 7.564409703 | ENSG00000280407 |
| MEF2C-AS1 | 0.925285563 | 1.8990601 | 0.152128222 | 0.322114752 | 9.886406442 | 18.60237472 | ENSG00000248309 |
| NA | 0.925266151 | 1.899034547 | 0.360660206 | 0.572695535 | 3.673854582 | 6.950021147 | ENSG00000249863 |
| NA | 0.925018905 | 1.898709122 | 0.285496308 | 0.494732775 | 4.871887182 | 9.315659082 | ENSG00000271737 |
| UBE2D4 | 0.923574075 | 1.896808555 | 0.006753978 | 0.028419982 | 27.11164923 | 51.44215897 | ENSG00000078967 |
| BBC3 | 0.923461396 | 1.896660414 | 7.57E-16 | 3.17E-14 | 279.5643489 | 529.9581262 | ENSG00000105327 |
| NA | 0.923193658 | 1.896308462 | 0.04763128 | 0.135898899 | 17.61548494 | 33.39772634 | ENSG00000234287 |
| BDNF | 0.923128088 | 1.896222276 | 2.77E-05 | 0.000238243 | 66.81150079 | 126.6876399 | ENSG00000176697 |
| PON1 | 0.92257821 | 1.895499676 | 0.146076036 | 0.313151159 | 8.525580138 | 16.20003746 | ENSG00000005421 |
| NA | 0.922435095 | 1.895311652 | 0.060347623 | 0.163065512 | 13.45322367 | 25.59127067 | ENSG00000276805 |
| OR2H2 | 0.922129569 | 1.894910316 | 0.338544313 | 0.550538996 | 4.177388883 | 8.000714065 | ENSG00000204657 |
| COL1A1 | 0.920830004 | 1.893204169 | 2.30E-29 | 2.64E-27 | 1721.290925 | 3260.386711 | ENSG00000108821 |
| MAGEA8 | 0.920367998 | 1.892597989 | 0.414053586 | 0.625152061 | 2.632107133 | 4.996757474 | ENSG00000156009 |
| PLEKHN1 | 0.920266729 | 1.892465145 | 0.005827885 | 0.025099114 | 34.18062895 | 64.86628705 | ENSG00000187583 |
| LAMC3 | 0.920050497 | 1.892181522 | 5.81E-39 | 1.05E-36 | 1030.315893 | 1950.232191 | ENSG00000050555 |
| FKBP6 | 0.919873556 | 1.891949467 | 0.420823811 | 0.630884946 | 2.611945506 | 4.963463108 | ENSG00000077800 |
| NA | 0.919784427 | 1.891832588 | 0.223411413 | 0.421267916 | 5.638743987 | 10.6694789 | ENSG00000267506 |
| RPL39L | 0.919702138 | 1.891724683 | 8.23E-22 | 5.73E-20 | 506.0411675 | 956.8095762 | ENSG00000163923 |
| NA | 0.919644428 | 1.891649013 | 0.074898681 | 0.192383452 | 11.5808544 | 21.90822871 | ENSG00000213513 |
| CDH6 | 0.919495296 | 1.891453483 | 0.002277774 | 0.011353867 | 35.5625066 | 67.40187442 | ENSG00000113361 |
| PURA | 0.918697164 | 1.890407376 | 1.46E-05 | 0.000133438 | 91.49210049 | 173.1764351 | ENSG00000185129 |
| NDUFS2 | 0.918080177 | 1.889599093 | 4.51E-62 | 2.13E-59 | 2160.102531 | 4081.00057 | ENSG00000158864 |
| NA | 0.918070677 | 1.88958665 | 0.124372697 | 0.279783011 | 9.761571726 | 18.5519602 | ENSG00000279453 |
| WIPF1 | 0.917603736 | 1.888975167 | 1.64E-08 | 2.63E-07 | 153.5097432 | 290.3931479 | ENSG00000115935 |
| PKNOX2 | 0.917342838 | 1.888633595 | 0.000374557 | 0.002385668 | 53.31767454 | 100.895868 | ENSG00000165495 |
| NA | 0.917203898 | 1.888451717 | 0.396972585 | 0.608843902 | 2.948516239 | 5.572554779 | ENSG00000231628 |
| NA | 0.917118934 | 1.888340504 | 0.018525255 | 0.064535961 | 21.85099444 | 41.28550452 | ENSG00000225630 |
| RORA-AS1 | 0.91706903 | 1.888275186 | 0.406267877 | 0.617196433 | 2.639226078 | 4.993636052 | ENSG00000245534 |
| NA | 0.915188936 | 1.885816023 | 0.406314181 | 0.617217125 | 2.659387704 | 5.015107141 | ENSG00000248268 |
| TEKT3 | 0.914871705 | 1.8854014 | 0.209837006 | 0.403130403 | 6.023952125 | 11.3140404 | ENSG00000125409 |
| NA | 0.914748481 | 1.88524037 | 0.0063393 | 0.026951303 | 30.73241721 | 57.72926895 | ENSG00000250903 |
| NA | 0.914540892 | 1.884969124 | 0.201464978 | 0.392476389 | 6.340361231 | 11.89948551 | ENSG00000176700 |
| NCF1B | 0.913912442 | 1.884148195 | 0.250514414 | 0.455996598 | 5.318775408 | 10.01839101 | ENSG00000182487 |
| AKT3 | 0.913880346 | 1.884106278 | 1.48E-15 | 6.02E-14 | 281.595673 | 530.2382689 | ENSG00000117020 |
| LINC02614 | 0.913638448 | 1.883790394 | 0.185167593 | 0.369493871 | 7.245705824 | 13.69273111 | ENSG00000241288 |
| NA | 0.913212894 | 1.883234811 | 0.086410559 | 0.213874626 | 13.38798411 | 25.35407009 | ENSG00000203644 |
| LOC101929551 | 0.912399025 | 1.88217272 | 0.273247015 | 0.481952917 | 5.346055979 | 10.04761799 | ENSG00000226506 |
| NA | 0.912364961 | 1.882128279 | 0.121478631 | 0.27487706 | 9.867134539 | 18.62072439 | ENSG00000285761 |
| NA | 0.912321381 | 1.882071427 | 0.023244392 | 0.077693865 | 22.96746466 | 43.13396708 | ENSG00000228956 |
| RASSF10 | 0.91212128 | 1.881810403 | 7.51E-05 | 0.000576164 | 62.91731671 | 118.326858 | ENSG00000189431 |
| TRG-AS1 | 0.911919421 | 1.881547122 | 0.138880983 | 0.302371151 | 9.05786955 | 16.96406123 | ENSG00000281103 |
| CMTM1 | 0.911772952 | 1.881356109 | 0.1234613 | 0.278183408 | 9.892050846 | 18.64966783 | ENSG00000089505 |
| SLC22A3 | 0.911377266 | 1.880840182 | 0.007717943 | 0.031754543 | 39.81550797 | 75.21268101 | ENSG00000146477 |
| SNORA73B | 0.91136961 | 1.880830201 | 0.022555326 | 0.075779635 | 20.8673676 | 39.2560043 | ENSG00000200087 |
| ACAP1 | 0.910626662 | 1.879861874 | 4.36E-24 | 3.64E-22 | 552.0212009 | 1036.97215 | ENSG00000072818 |
| ACOT11 | 0.910591837 | 1.879816498 | 1.46E-06 | 1.64E-05 | 97.66371871 | 183.362643 | ENSG00000162390 |
| NA | 0.910560961 | 1.879776267 | 0.258537619 | 0.465397286 | 5.291494837 | 9.946221842 | ENSG00000223784 |
| AHRR | 0.909618933 | 1.878549241 | 0.121133398 | 0.274260186 | 12.95413857 | 24.47370547 | ENSG00000286169 |
| TAFA3 | 0.908969791 | 1.877704177 | 0.229415335 | 0.429202162 | 7.227908461 | 13.60032037 | ENSG00000184599 |
| RPH3AL | 0.90836901 | 1.876922408 | 0.010241885 | 0.039806134 | 27.60005592 | 51.6892909 | ENSG00000181031 |
| NCMAP | 0.908108758 | 1.876583855 | 0.209796478 | 0.403130403 | 6.787249457 | 12.62368853 | ENSG00000184454 |
| SMS | 0.907969992 | 1.876403364 | 1.10E-60 | 5.06E-58 | 3995.328304 | 7497.70089 | ENSG00000102172 |
| NPAS1 | 0.907951666 | 1.876379528 | 1.89E-07 | 2.48E-06 | 120.2821803 | 225.8365368 | ENSG00000130751 |
| LOC100507291 | 0.907553741 | 1.875862056 | 0.010600618 | 0.040955415 | 27.88798924 | 52.29235064 | ENSG00000248932 |
| ITPR2 | 0.907442739 | 1.875717731 | 8.70E-21 | 5.46E-19 | 842.1622324 | 1578.511519 | ENSG00000123104 |
| NA | 0.907310745 | 1.875546126 | 0.457324789 | 0.663026151 | 2.659387704 | 4.98058327 | ENSG00000276538 |
| TRPV4 | 0.906949509 | 1.875076569 | 0.002172345 | 0.010896345 | 43.07300947 | 80.47877656 | ENSG00000111199 |
| NA | 0.906665679 | 1.874707709 | 0.00626121 | 0.026691409 | 32.87967713 | 61.50857813 | ENSG00000231806 |
| NA | 0.906442423 | 1.874417622 | 0.099046231 | 0.236531117 | 11.71133352 | 21.85128779 | ENSG00000281344 |
| PIPSL | 0.906385967 | 1.874344274 | 0.082113455 | 0.205855913 | 11.86998293 | 22.29411854 | ENSG00000180764 |
| PYGM | 0.906282135 | 1.874209381 | 3.30E-11 | 7.92E-10 | 207.7056832 | 388.914109 | ENSG00000068976 |
| LAMC1 | 0.905763544 | 1.873535798 | 9.61E-46 | 2.39E-43 | 4151.019431 | 7776.220394 | ENSG00000135862 |
| NA | 0.905447878 | 1.873125908 | 0.219335981 | 0.415703519 | 7.207746835 | 13.52909716 | ENSG00000261812 |
| TMEM47 | 0.905292342 | 1.872923978 | 3.12E-18 | 1.60E-16 | 577.0920416 | 1081.149305 | ENSG00000147027 |
| AMER2 | 0.905064366 | 1.87262804 | 0.115726952 | 0.26600578 | 9.592243894 | 17.98051382 | ENSG00000165566 |
| PAQR7 | 0.905008337 | 1.872555316 | 1.65E-07 | 2.19E-06 | 126.5848881 | 236.9976526 | ENSG00000182749 |
| NA | 0.90493308 | 1.872457638 | 0.214996867 | 0.410039975 | 6.967255707 | 13.05904693 | ENSG00000279792 |
| NA | 0.904818487 | 1.872308915 | 0.451321701 | 0.657517808 | 2.64278555 | 4.944167482 | ENSG00000271755 |
| NA | 0.904719884 | 1.872180954 | 0.480061888 | 0.682164692 | 2.335859653 | 4.372721105 | ENSG00000248968 |
| MST1L | 0.903777422 | 1.870958323 | 0.000715741 | 0.004191248 | 49.81014698 | 93.11449928 | ENSG00000186715 |

| | | | | | | | |
|---|---|---|---|---|---|---|---|
| ESRRG | 0.903767971 | 1.870946067 | 0.00594586 | 0.025543121 | 31.57252224 | 58.9409259 | ENSG00000196482 |
| NA | 0.903130433 | 1.870119464 | 0.047720612 | 0.136030754 | 17.72549695 | 33.25584702 | ENSG00000276855 |
| TTC41P | 0.90311339 | 1.870097371 | 0.423897979 | 0.634084225 | 2.659387704 | 4.960341686 | ENSG00000214198 |
| NA | 0.902756388 | 1.869634664 | 0.080990748 | 0.203690276 | 12.25519107 | 22.90510213 | ENSG00000287562 |
| THBS3 | 0.90243407 | 1.869217009 | 0.000136639 | 0.000983162 | 60.31813455 | 112.9384623 | ENSG00000169231 |
| CAPN6 | 0.902407451 | 1.869182521 | 3.59E-09 | 6.30E-08 | 171.9875541 | 321.2586488 | ENSG00000077274 |
| ZNF502 | 0.90238785 | 1.869157126 | 0.046542758 | 0.133433528 | 17.55380485 | 32.78407283 | ENSG00000196653 |
| NA | 0.902250517 | 1.868979205 | 0.012866161 | 0.048114346 | 26.62324254 | 49.909309 | ENSG00000205918 |
| PRSS30P | 0.902170816 | 1.868875956 | 0.287108668 | 0.496231181 | 5.022527928 | 9.362006229 | ENSG00000172460 |
| TMEM151A | 0.902112589 | 1.86880053 | 0.00707743 | 0.02953972 | 31.65052515 | 59.17405909 | ENSG00000179292 |
| NA | 0.902034809 | 1.868699781 | 0.017347288 | 0.061303057 | 25.24492436 | 47.2938692 | ENSG00000255495 |
| NA | 0.901989801 | 1.868641484 | 0.262825782 | 0.469982842 | 5.880430324 | 11.05849015 | ENSG00000232024 |
| HMGB3 | 0.90097227 | 1.867323998 | 5.60E-90 | 9.61E-87 | 6510.825532 | 12158.04375 | ENSG00000029993 |
| ZNF790-AS1 | 0.900818373 | 1.867124815 | 0.191441736 | 0.378415029 | 6.949458344 | 12.9847023 | ENSG00000267254 |
| NLRC5 | 0.900709192 | 1.866983519 | 0.013633524 | 0.050355781 | 29.66308371 | 55.45727126 | ENSG00000140853 |
| NA | 0.90030974 | 1.866466662 | 0.287281704 | 0.496369165 | 4.627836581 | 8.645275565 | ENSG00000261821 |
| TMC8 | 0.899970002 | 1.866027182 | 0.012658188 | 0.04745878 | 25.45249051 | 47.53061828 | ENSG00000167895 |
| PLXDC2 | 0.899480467 | 1.865394109 | 2.33E-19 | 1.32E-17 | 390.1543669 | 727.8455829 | ENSG00000120594 |
| NA | 0.898773064 | 1.864479667 | 0.432350231 | 0.641695544 | 2.683108803 | 4.977461848 | ENSG00000231443 |
| NA | 0.898322132 | 1.863896992 | 0.301197853 | 0.511337778 | 4.671719306 | 8.69597364 | ENSG00000285907 |
| BMPR1B | 0.897773949 | 1.863188899 | 0.079337353 | 0.200627466 | 12.34059226 | 22.92780272 | ENSG00000138696 |
| NA | 0.89767962 | 1.86306708 | 0.124096656 | 0.279261839 | 9.272275316 | 17.27343097 | ENSG00000254198 |
| OPCML | 0.896922519 | 1.862089632 | 0.009083518 | 0.036197981 | 37.55943126 | 69.95815488 | ENSG00000183715 |
| TMEM184A | 0.896775024 | 1.86189927 | 0.023944588 | 0.079577355 | 20.85312971 | 38.81875398 | ENSG00000164855 |
| NA | 0.896735415 | 1.861848152 | 0.224333233 | 0.422596818 | 6.540529107 | 12.21415214 | ENSG00000235510 |
| CYP2A13 | 0.89673163 | 1.861843268 | 0.010111008 | 0.039399727 | 27.34413169 | 50.75247982 | ENSG00000197838 |
| NA | 0.896521525 | 1.86157214 | 0.414385039 | 0.625352522 | 3.550494407 | 6.664297958 | ENSG00000278948 |
| RAB24 | 0.896470101 | 1.861505787 | 0.000183455 | 0.001277015 | 63.27732921 | 117.9996331 | ENSG00000169228 |
| ABHD12B | 0.896143441 | 1.861084346 | 0.000445047 | 0.002780258 | 51.83671647 | 96.34071175 | ENSG00000131969 |
| JMJD4 | 0.895019587 | 1.859635133 | 0.006322275 | 0.026915276 | 34.27907281 | 63.8656298 | ENSG00000081692 |
| NA | 0.893916093 | 1.858213272 | 0.410627786 | 0.621669532 | 3.584893923 | 6.705064674 | ENSG00000237846 |
| CSF2RA | 0.893414082 | 1.857566787 | 0.016079595 | 0.057708694 | 24.33749484 | 45.06046277 | ENSG00000198223 |
| TPO | 0.893287294 | 1.857403546 | 0.053747757 | 0.148961901 | 18.60742593 | 34.51574303 | ENSG00000115705 |
| NA | 0.892962663 | 1.856985645 | 0.016445689 | 0.058743493 | 29.25209569 | 54.27796272 | ENSG00000267432 |
| LOC574538 | 0.892587571 | 1.856502902 | 0.254696633 | 0.461252965 | 8.874303828 | 16.36584687 | ENSG00000249695 |
| OPRK1 | 0.892109816 | 1.855888215 | 0.130878405 | 0.290439328 | 8.948747265 | 16.59775057 | ENSG00000082556 |
| FAM71F1 | 0.891719811 | 1.855386579 | 0.010657324 | 0.041120411 | 28.02202784 | 51.85169535 | ENSG00000135248 |
| NA | 0.891462397 | 1.855055559 | 0.399382319 | 0.610828948 | 3.068316941 | 5.624198812 | ENSG00000235092 |
| NA | 0.891418919 | 1.854999656 | 0.195211721 | 0.38402279 | 7.916483025 | 14.72753337 | ENSG00000287200 |
| SLC17A9 | 0.891418061 | 1.854998553 | 4.12E-18 | 2.09E-16 | 741.9475137 | 1377.003245 | ENSG00000101194 |
| TCIRG1 | 0.890322302 | 1.853590174 | 1.97E-12 | 5.45E-11 | 284.8117078 | 528.1023498 | ENSG00000110719 |
| CCER2 | 0.889990594 | 1.853164042 | 0.483298477 | 0.684395001 | 2.363140224 | 4.358722365 | ENSG00000262484 |
| BOLA1 | 0.889916311 | 1.853068627 | 1.05E-07 | 1.46E-06 | 125.5327415 | 232.3097704 | ENSG00000178096 |
| TMEM220 | 0.889813909 | 1.852937101 | 0.002317857 | 0.011524826 | 40.7321158 | 75.48629738 | ENSG00000187824 |
| METTL23 | 0.889482821 | 1.852511913 | 1.62E-08 | 2.60E-07 | 151.2948533 | 280.6154577 | ENSG00000181038 |
| GAP43 | 0.88938459 | 1.852385784 | 1.06E-17 | 5.15E-16 | 399.1564545 | 738.8489943 | ENSG00000172020 |
| AATBC | 0.889187854 | 1.852133196 | 0.212622475 | 0.406702668 | 6.667448754 | 12.3251961 | ENSG00000215458 |
| NA | 0.888999955 | 1.851891987 | 0.293932913 | 0.50375924 | 4.300749058 | 7.965244235 | ENSG00000230224 |
| MRPL24 | 0.888800681 | 1.85163621 | 2.01E-21 | 1.35E-19 | 432.3751184 | 800.7085596 | ENSG00000143314 |
| PTPN6 | 0.888239936 | 1.850916658 | 9.35E-17 | 4.14E-15 | 341.7886418 | 632.311145 | ENSG00000111679 |
| HEATR4 | 0.888230061 | 1.85090399 | 0.144134027 | 0.3100548 | 8.625219214 | 15.95413502 | ENSG00000187105 |
| BAIAP3 | 0.887900628 | 1.850481392 | 1.79E-10 | 3.85E-09 | 219.817989 | 407.0530028 | ENSG00000007516 |
| KCNIP1 | 0.887397106 | 1.849835659 | 0.026329394 | 0.085803385 | 22.04404337 | 40.87473863 | ENSG00000182132 |
| CACNG6 | 0.887332502 | 1.849752825 | 0.032966843 | 0.102278984 | 20.89820765 | 38.65873591 | ENSG00000130433 |
| SNORD104 | 0.887270313 | 1.84967309 | 0.010572282 | 0.040871423 | 40.61203577 | 75.39059725 | ENSG00000199753 |
| NA | 0.887182483 | 1.849560488 | 0.443733322 | 0.651211433 | 2.686668275 | 4.932344206 | ENSG00000259182 |
| HTR1D | 0.886157468 | 1.848246868 | 0.009781011 | 0.038298651 | 28.80045278 | 53.25933466 | ENSG00000179546 |
| NA | 0.885926348 | 1.847950802 | 0.434735496 | 0.643487883 | 3.075435886 | 5.618901926 | ENSG00000186543 |
| LOC107986257 | 0.885816895 | 1.847810609 | 0.300023013 | 0.510168792 | 4.685957196 | 8.619453548 | ENSG00000286176 |
| CATSPER3 | 0.885721495 | 1.847688423 | 0.394108732 | 0.605790608 | 3.609810231 | 6.701943251 | ENSG00000152705 |
| SVOPL | 0.88523238 | 1.84706211 | 0.006355572 | 0.027014401 | 32.81919225 | 60.52882487 | ENSG00000157703 |
| G2E3-AS1 | 0.88500545 | 1.846771598 | 0.295368957 | 0.505462499 | 5.0498085 | 9.260893626 | ENSG00000257636 |
| ALDH1A3 | 0.884181923 | 1.845717715 | 0.005433125 | 0.023690735 | 38.88466284 | 71.66109255 | ENSG00000184254 |
| NA | 0.884116971 | 1.84563462 | 0.074579133 | 0.191861243 | 15.06610924 | 27.91273692 | ENSG00000231770 |
| SSPO | 0.883827538 | 1.845264386 | 0.086465731 | 0.213879248 | 13.85147449 | 25.59845947 | ENSG00000197558 |
| CYP19A1 | 0.883191943 | 1.844451614 | 0.473860494 | 0.676969168 | 2.686668275 | 4.920520929 | ENSG00000137869 |
| CERS3 | 0.882710931 | 1.843836754 | 0.281414029 | 0.490403904 | 5.438576111 | 9.925980258 | ENSG00000154227 |
| NA | 0.882581435 | 1.843671259 | 0.026531456 | 0.086296062 | 21.83675655 | 40.29298203 | ENSG00000285517 |
| FCRL5 | 0.882013127 | 1.842945141 | 0.0279163 | 0.089800619 | 22.55617115 | 41.5478647 | ENSG00000143297 |
| EPHA8 | 0.881961169 | 1.84287877 | 1.51E-14 | 5.30E-13 | 267.1926372 | 492.5176166 | ENSG00000070886 |
| NA | 0.881672504 | 1.84251007 | 0.302025305 | 0.512282009 | 5.244052639 | 9.712426233 | ENSG00000248375 |
| NA | 0.881664902 | 1.842500361 | 0.182097608 | 0.365414231 | 7.521791677 | 13.91716246 | ENSG00000253540 |
| POTEF | 0.881433397 | 1.842204724 | 0.356894332 | 0.569012197 | 3.592012868 | 6.655596104 | ENSG00000196604 |
| CXCL1 | 0.881100804 | 1.841780079 | 0.350930401 | 0.563021036 | 3.929778809 | 7.255039962 | ENSG00000163739 |

| | | | | | | | |
|---|---|---|---|---|---|---|---|
| GPR78 | 0.879883198 | 1.840226309 | 0.114565389 | 0.26417184 | 11.97674095 | 22.02551551 | ENSG00000155269 |
| GABRD | 0.879066629 | 1.839185031 | 0.004465654 | 0.020117813 | 37.35214444 | 68.73176414 | ENSG00000187730 |
| IFI27L2 | 0.878925108 | 1.839004625 | 1.80E-07 | 2.38E-06 | 128.1974682 | 235.9472432 | ENSG00000119632 |
| NA | 0.878092016 | 1.837942988 | 0.217442185 | 0.413406909 | 8.214205045 | 14.93891194 | ENSG00000234337 |
| VPS13B-DT | 0.877949083 | 1.837760906 | 0.105746005 | 0.248775909 | 11.56661651 | 21.27454453 | ENSG00000253948 |
| LINC02861 | 0.877815371 | 1.837590585 | 0.471697312 | 0.674951033 | 2.363140224 | 4.309253795 | ENSG00000262185 |
| NA | 0.876993088 | 1.836543526 | 0.337837638 | 0.549815908 | 4.004501578 | 7.339978361 | ENSG00000223486 |
| LACTB2 | 0.87693033 | 1.836463637 | 0.001271902 | 0.006907637 | 46.80679026 | 86.12058999 | ENSG00000147592 |
| AGAP1-IT1 | 0.876734153 | 1.836213932 | 0.477978995 | 0.680484985 | 2.346538071 | 4.281256315 | ENSG00000235529 |
| CSGALNACT1 | 0.876659859 | 1.836119376 | 0.296676591 | 0.506875709 | 5.792944205 | 10.51672226 | ENSG00000147408 |
| TNNT2 | 0.876386852 | 1.835771953 | 0.215949245 | 0.411233386 | 6.725569369 | 12.26797164 | ENSG00000118194 |
| CDK18 | 0.876289348 | 1.835647886 | 8.45E-41 | 1.75E-38 | 1252.850806 | 2300.184437 | ENSG00000117266 |
| NA | 0.876244915 | 1.835591353 | 0.040594219 | 0.120122504 | 21.40677596 | 39.15271624 | ENSG00000266651 |
| NA | 0.876118037 | 1.835429928 | 0.230837494 | 0.431150834 | 5.996671554 | 10.98537503 | ENSG00000271714 |
| SH3TC1 | 0.876048795 | 1.835341839 | 0.356886974 | 0.569012197 | 4.508035878 | 8.358606418 | ENSG00000125089 |
| NA | 0.875412727 | 1.834532835 | 0.352221577 | 0.564306973 | 3.602691286 | 6.636300478 | ENSG00000288546 |
| ADCY7 | 0.8745255 | 1.833404984 | 2.67E-05 | 0.000231069 | 91.29343331 | 167.4325334 | ENSG00000121281 |
| PDP1 | 0.8744559 | 1.833316536 | 1.80E-12 | 5.03E-11 | 279.8935213 | 513.0928139 | ENSG00000164951 |
| NDUFV2 | 0.873918133 | 1.832633292 | 0.278566743 | 0.487748658 | 6.425762417 | 11.71487493 | ENSG00000178127 |
| KCNA6 | 0.873655397 | 1.832299573 | 0.000916952 | 0.005203314 | 50.66301536 | 93.0566124 | ENSG00000130035 |
| KCNJ6 | 0.873508331 | 1.8321128 | 0.041160569 | 0.121323138 | 19.75682112 | 36.28562562 | ENSG00000157542 |
| ITGB3 | 0.872697201 | 1.831083017 | 0.48194635 | 0.68352167 | 2.914116723 | 5.382647307 | ENSG00000259207 |
| SEC14L5 | 0.87264633 | 1.831018452 | 5.81E-12 | 1.52E-10 | 241.1559659 | 441.5668836 | ENSG00000103184 |
| CNTNAP2 | 0.872622536 | 1.830988254 | 5.79E-25 | 5.13E-23 | 1492.365306 | 2732.225839 | ENSG00000174469 |
| SSUH2 | 0.871992354 | 1.830188637 | 0.240856857 | 0.444457861 | 6.000231026 | 10.9499052 | ENSG00000125046 |
| DOK7 | 0.87165094 | 1.829755574 | 0.162948103 | 0.338087774 | 9.629007675 | 17.59897491 | ENSG00000175920 |
| NA | 0.871230068 | 1.829221864 | 0.108479942 | 0.253576397 | 13.71626684 | 25.1913821 | ENSG00000236809 |
| FAM27C | 0.870964301 | 1.828884923 | 0.349552642 | 0.561717294 | 3.622852912 | 6.638475942 | ENSG00000231527 |
| GLRX2 | 0.870859519 | 1.828752097 | 6.14E-07 | 7.37E-06 | 175.0069537 | 320.5573144 | ENSG00000023572 |
| NA | 0.870635472 | 1.828468119 | 0.141330026 | 0.305869533 | 9.588684422 | 17.5205629 | ENSG00000286504 |
| LOC101929054 | 0.87043858 | 1.828218595 | 0.091056408 | 0.222228 | 12.52415798 | 22.94303097 | ENSG00000243224 |
| DYDC2 | 0.870358413 | 1.828117009 | 0.000124942 | 0.000906817 | 67.20619214 | 122.8479848 | ENSG00000133665 |
| HES2 | 0.869907201 | 1.827545342 | 0.086121297 | 0.213410404 | 13.76370904 | 25.24955252 | ENSG00000069812 |
| NOTUM | 0.869603339 | 1.827160464 | 0.175069086 | 0.355579662 | 7.634473435 | 13.92898573 | ENSG00000185269 |
| PPIL6 | 0.868801048 | 1.826144652 | 0.209239175 | 0.402391224 | 7.318064329 | 13.35753936 | ENSG00000185250 |
| CA7 | 0.868447626 | 1.82569735 | 0.011363671 | 0.043433097 | 28.23523839 | 51.48103376 | ENSG00000168748 |
| HEYL | 0.868422038 | 1.825664969 | 1.12E-07 | 1.54E-06 | 135.8802997 | 248.2991643 | ENSG00000163909 |
| EDEM3 | 0.868195856 | 1.825378768 | 1.70E-11 | 4.19E-10 | 298.4834031 | 544.5537842 | ENSG00000116406 |
| MSTO1 | 0.867875808 | 1.824973871 | 7.72E-18 | 3.81E-16 | 400.8817425 | 731.185931 | ENSG00000125459 |
| BHLHA15 | 0.867200319 | 1.824119593 | 0.0042172 | 0.019213951 | 35.85310967 | 65.29585411 | ENSG00000180535 |
| NKX3-1 | 0.867027779 | 1.82390145 | 0.004308312 | 0.019534806 | 35.1313308 | 64.02943171 | ENSG00000167034 |
| LOC101926913 | 0.866847873 | 1.823674021 | 0.356614787 | 0.568713764 | 3.622852912 | 6.612653925 | ENSG00000234350 |
| NA | 0.866415853 | 1.823127997 | 0.0047482 | 0.021135401 | 40.27071035 | 73.6877549 | ENSG00000260912 |
| NA | 0.865781293 | 1.822326282 | 0.292074898 | 0.501713156 | 4.889684544 | 8.963347161 | ENSG00000226421 |
| NA | 0.865677016 | 1.822194571 | 0.293431122 | 0.503173226 | 5.274892683 | 9.625028824 | ENSG00000232093 |
| NA | 0.865623255 | 1.82212667 | 0.038721998 | 0.115907766 | 26.93639766 | 49.06055774 | ENSG00000259692 |
| STEAP1B | 0.864824416 | 1.821118014 | 8.38E-05 | 0.000635786 | 75.83942002 | 138.2488122 | ENSG00000105889 |
| CALCA | 0.863807256 | 1.819834502 | 0.260075414 | 0.467011057 | 6.044113751 | 10.93685242 | ENSG00000110680 |
| MAOB | 0.863638755 | 1.819621965 | 0.001465039 | 0.007802347 | 46.79995065 | 85.09013866 | ENSG00000069535 |
| USH1C | 0.863392208 | 1.81931103 | 0.009925103 | 0.038790473 | 32.27741963 | 58.84342917 | ENSG00000006611 |
| SYN1 | 0.862844806 | 1.81862086 | 1.00E-05 | 9.56E-05 | 88.81166144 | 161.65955 | ENSG00000008056 |
| EMC1-AS1 | 0.862431231 | 1.818099593 | 0.411553781 | 0.622721989 | 4.475110903 | 8.067586343 | ENSG00000230424 |
| CFAP74 | 0.862340577 | 1.817985355 | 0.202706349 | 0.394000556 | 7.341785428 | 13.31336767 | ENSG00000142609 |
| SPATA32 | 0.861895669 | 1.817424798 | 0.395716217 | 0.607618567 | 3.306443806 | 6.02285788 | ENSG00000184361 |
| TEX41 | 0.861120682 | 1.816448775 | 0.26578973 | 0.473308727 | 5.318775408 | 9.649904882 | ENSG00000226674 |
| NA | 0.860974474 | 1.816264699 | 0.192500383 | 0.379871445 | 7.314504856 | 13.25425129 | ENSG00000250896 |
| NA | 0.860758001 | 1.815992194 | 0.002108466 | 0.010658024 | 44.14264788 | 80.28886909 | ENSG00000213025 |
| NKAIN2 | 0.860580592 | 1.815768894 | 0.369917013 | 0.582108391 | 3.650133483 | 6.614829389 | ENSG00000188580 |
| ITGA10 | 0.860053468 | 1.81510558 | 0.406174247 | 0.617103833 | 3.306443806 | 6.012264109 | ENSG00000143127 |
| HBQ1 | 0.859598794 | 1.814533629 | 0.010318769 | 0.040047255 | 30.36945563 | 55.17561546 | ENSG00000086506 |
| BFSP1 | 0.859411187 | 1.814297683 | 0.00031856 | 0.002073057 | 61.22523302 | 110.8520212 | ENSG00000125864 |
| SARDH | 0.859191607 | 1.814021565 | 4.59E-14 | 1.52E-12 | 308.4113541 | 559.0180341 | ENSG00000123453 |
| NA | 0.859054854 | 1.813849623 | 0.298677385 | 0.50902648 | 5.346055979 | 9.67478094 | ENSG00000233597 |
| CSDC2 | 0.857992634 | 1.812514622 | 0.007595946 | 0.031355069 | 33.21032412 | 60.12441741 | ENSG00000172346 |
| TUBB8B | 0.857980015 | 1.812498769 | 0.354699577 | 0.566806892 | 3.684532999 | 6.648123755 | ENSG00000173213 |
| KANK4 | 0.857692855 | 1.812138037 | 0.133595272 | 0.294702337 | 11.18972252 | 20.33773345 | ENSG00000132854 |
| PRRG1 | 0.856964429 | 1.811223308 | 7.73E-06 | 7.60E-05 | 96.05973152 | 174.1056058 | ENSG00000130962 |
| LMNA | 0.856612405 | 1.810781415 | 4.57E-30 | 5.36E-28 | 1236.936075 | 2240.968886 | ENSG00000160789 |
| RAG1 | 0.856554593 | 1.810708855 | 0.097590831 | 0.233883608 | 15.34130537 | 27.66756954 | ENSG00000166349 |
| TMCO4 | 0.856386005 | 1.810497274 | 5.63E-06 | 5.68E-05 | 108.1272132 | 195.3851637 | ENSG00000162542 |
| KANK3 | 0.856222898 | 1.810292597 | 0.001508372 | 0.008000103 | 46.56746819 | 84.38088035 | ENSG00000186994 |
| GPR146 | 0.856125415 | 1.810170278 | 0.480489046 | 0.682566243 | 2.93903303 | 5.349352941 | ENSG00000164849 |
| LENG9 | 0.856007633 | 1.810022502 | 6.14E-05 | 0.000482241 | 74.10940349 | 134.1685516 | ENSG00000275183 |
| LOC100507551 | 0.855886555 | 1.809870603 | 0.29919298 | 0.509445445 | 5.765663634 | 10.36663555 | ENSG00000269439 |

| | | | | | | | |
|---|---|---|---|---|---|---|---|
| RDH16 | 0.855673133 | 1.809602884 | 0.288490914 | 0.497704503 | 5.673143503 | 10.22012175 | ENSG00000139547 |
| PKDCC | 0.855661857 | 1.80958874 | 2.14E-22 | 1.55E-20 | 916.8593299 | 1660.323301 | ENSG00000162878 |
| MSS51 | 0.85511117 | 1.808898139 | 0.148355634 | 0.316254222 | 10.2393 | 18.51838228 | ENSG00000166343 |
| ADAMTSL4 | 0.855066178 | 1.808841727 | 1.95E-05 | 0.000174556 | 135.8200685 | 245.2677477 | ENSG00000143382 |
| CCSAP | 0.85458692 | 1.808240936 | 6.65E-12 | 1.72E-10 | 412.5897692 | 745.9248026 | ENSG00000154429 |
| DPEP1 | 0.854474074 | 1.808099502 | 0.001451315 | 0.007742394 | 61.96480925 | 111.8823046 | ENSG00000015413 |
| WDR31 | 0.854215648 | 1.807775652 | 0.233243589 | 0.434160909 | 7.001655223 | 12.62680995 | ENSG00000148225 |
| NKPD1 | 0.854004328 | 1.807510876 | 0.136938465 | 0.299627263 | 11.73149514 | 21.07856216 | ENSG00000179846 |
| HLX | 0.853816239 | 1.80727524 | 0.436245066 | 0.644810651 | 2.931914085 | 5.335354201 | ENSG00000136630 |
| TRHDE | 0.853624976 | 1.80703566 | 0.011549025 | 0.044061281 | 31.13839737 | 56.47466982 | ENSG00000072657 |
| FES | 0.85359276 | 1.806995308 | 1.06E-05 | 0.000100152 | 104.9586723 | 189.725663 | ENSG00000182511 |
| NA | 0.853571057 | 1.806968125 | 0.039674649 | 0.117993031 | 23.43451451 | 42.19403457 | ENSG00000213062 |
| NA | 0.853420755 | 1.806779882 | 0.099313289 | 0.236959123 | 13.26729368 | 23.96790187 | ENSG00000286615 |
| GAL3ST2 | 0.853420058 | 1.806779009 | 0.312392694 | 0.522611975 | 4.58631812 | 8.32625801 | ENSG00000154252 |
| SMG1P6 | 0.853131045 | 1.806417096 | 0.374611741 | 0.58656427 | 3.691651944 | 6.631949551 | ENSG00000254634 |
| MISP | 0.85278373 | 1.805982271 | 0.203868378 | 0.395485376 | 7.518232205 | 13.6519644 | ENSG00000099812 |
| CLVS2 | 0.852620454 | 1.805777892 | 0.077081024 | 0.196447388 | 15.73599672 | 28.32350281 | ENSG00000146352 |
| NA | 0.851702717 | 1.804629553 | 0.499382452 | 0.697805224 | 2.57754599 | 4.702615977 | ENSG00000276931 |
| LOC107984660 | 0.851697991 | 1.804623642 | 0.303023628 | 0.513237777 | 5.005925775 | 9.019342122 | ENSG00000258457 |
| LOC105370941 | 0.851568086 | 1.804461154 | 0.31212223 | 0.52250938 | 4.596996537 | 8.328433474 | ENSG00000261136 |
| TTLL10-AS1 | 0.851515377 | 1.804395229 | 0.383011907 | 0.595421458 | 3.306443806 | 5.965916961 | ENSG00000205231 |
| LINC02525 | 0.851228337 | 1.804036261 | 0.16561358 | 0.341442409 | 9.064988495 | 16.26473427 | ENSG00000230269 |
| OLFM3 | 0.851076811 | 1.803846794 | 0.319067847 | 0.529256214 | 4.559037549 | 8.267141628 | ENSG00000118733 |
| SCNN1G | 0.851028908 | 1.803786899 | 0.204366306 | 0.396166875 | 7.910559288 | 14.29529639 | ENSG00000166828 |
| NA | 0.850536732 | 1.803171642 | 0.116249805 | 0.266980125 | 12.50636062 | 22.60102927 | ENSG00000268912 |
| LTBR | 0.850491225 | 1.803114765 | 0.004268916 | 0.019408726 | 50.54290975 | 90.8040783 | ENSG00000111321 |
| AKR7L | 0.850242697 | 1.802804176 | 0.320632872 | 0.531150323 | 4.569715966 | 8.276789441 | ENSG00000211454 |
| NA | 0.849167787 | 1.801461459 | 0.035038583 | 0.107262708 | 20.87329134 | 37.61021846 | ENSG00000233913 |
| TFAP2C | 0.84829248 | 1.800368814 | 1.00E-05 | 9.54E-05 | 101.5816239 | 182.7177979 | ENSG00000087510 |
| GLIPR1 | 0.848264187 | 1.800333508 | 0.356289308 | 0.568409019 | 4.242628443 | 7.689168859 | ENSG00000139278 |
| C2orf74 | 0.847661694 | 1.799581816 | 0.419664816 | 0.630097794 | 3.333724377 | 5.986442092 | ENSG00000237651 |
| SDCBP2-AS1 | 0.847619998 | 1.799529805 | 0.128010303 | 0.285564312 | 10.83771869 | 19.55100908 | ENSG00000234684 |
| VAMP4 | 0.847516716 | 1.799400982 | 1.15E-06 | 1.32E-05 | 133.6707493 | 240.2717776 | ENSG00000117533 |
| LINC01806 | 0.847041043 | 1.798807797 | 0.234200748 | 0.4354944 | 6.285800088 | 11.31621586 | ENSG00000227403 |
| TMEM169 | 0.847030791 | 1.798795015 | 0.002262453 | 0.011294325 | 50.53075679 | 90.70391166 | ENSG00000163449 |
| CASP8 | 0.846644315 | 1.79831321 | 0.0015695 | 0.008281644 | 49.14648872 | 88.26007262 | ENSG00000064012 |
| PLD1 | 0.846467996 | 1.798093442 | 0.025588867 | 0.0839114 | 22.52177164 | 40.47853857 | ENSG00000075651 |
| ACTC1 | 0.846320295 | 1.797909366 | 4.61E-09 | 8.01E-08 | 253.0671874 | 455.7263553 | ENSG00000159251 |
| THEM4 | 0.846272699 | 1.797850052 | 2.58E-09 | 4.65E-08 | 186.6096981 | 335.0621404 | ENSG00000159445 |
| LOC101929705 | 0.846236379 | 1.797804791 | 0.138153353 | 0.301308012 | 10.84839711 | 19.54013176 | ENSG00000237686 |
| KLKB1 | 0.846108072 | 1.797644909 | 5.12E-07 | 6.27E-06 | 147.7155776 | 265.9811857 | ENSG00000164344 |
| HTR6 | 0.845918849 | 1.797409146 | 0.001544029 | 0.008154078 | 48.83127483 | 87.65221046 | ENSG00000158748 |
| MST1R | 0.84515586 | 1.796458814 | 0.001530032 | 0.008086943 | 56.0167751 | 100.4199538 | ENSG00000164078 |
| LINC02582 | 0.845090813 | 1.796377818 | 0.039051178 | 0.116579432 | 19.62781654 | 35.18735605 | ENSG00000261780 |
| INPP5D | 0.844702861 | 1.795894823 | 8.87E-11 | 2.00E-09 | 253.6721152 | 455.3259723 | ENSG00000168918 |
| CEACAM19 | 0.844516249 | 1.79566254 | 0.081084183 | 0.203898109 | 14.72597904 | 26.55267426 | ENSG00000186567 |
| NA | 0.844495495 | 1.795636708 | 0.147292297 | 0.314867834 | 10.14677986 | 18.30700372 | ENSG00000273295 |
| ATP1B1 | 0.844490014 | 1.795629886 | 2.30E-26 | 2.26E-24 | 793.3592569 | 1425.351469 | ENSG00000143153 |
| TNFRSF11B | 0.84371887 | 1.79467035 | 0.110415486 | 0.257177658 | 12.55736229 | 22.54002097 | ENSG00000164761 |
| NA | 0.843025232 | 1.793807691 | 0.476516669 | 0.679017376 | 2.92479514 | 5.268481923 | ENSG00000229337 |
| NA | 0.842937327 | 1.793698397 | 0.29718832 | 0.5074504 | 5.384014968 | 9.589558994 | ENSG00000240087 |
| RN7SL832P | 0.842531822 | 1.793194305 | 0.522919023 | 0.717284954 | 2.428379784 | 4.285607242 | ENSG00000243819 |
| NA | 0.842190299 | 1.792769859 | 0.385033175 | 0.597138171 | 4.575639702 | 8.218619017 | ENSG00000287460 |
| AMT | 0.841875132 | 1.792378259 | 0.289566201 | 0.498946871 | 5.529621701 | 9.979799754 | ENSG00000145020 |
| FER1L5 | 0.841869892 | 1.792371749 | 0.223311512 | 0.421260403 | 8.559979654 | 15.39233646 | ENSG00000249715 |
| ARMC12 | 0.84135418 | 1.791731154 | 0.163025388 | 0.338120875 | 10.07945537 | 17.97398743 | ENSG00000157343 |
| NA | 0.840513082 | 1.790686871 | 0.010616994 | 0.040994055 | 217.17017 | 388.8650528 | ENSG00000227671 |
| DGKH | 0.839551579 | 1.789493842 | 3.71E-05 | 0.000308473 | 86.96092833 | 155.3803639 | ENSG00000102780 |
| LINC00467 | 0.839183748 | 1.789037649 | 0.001660866 | 0.008676477 | 47.45621063 | 85.05466883 | ENSG00000153363 |
| TSTD3 | 0.839153907 | 1.789000645 | 0.100596551 | 0.239082818 | 12.2516316 | 21.9147551 | ENSG00000279170 |
| ATP2B3 | 0.839042441 | 1.788862427 | 0.089946629 | 0.22037391 | 14.79241381 | 26.51720443 | ENSG00000067842 |
| NA | 0.838812983 | 1.788577934 | 0.043778153 | 0.127231521 | 21.25494001 | 38.00547255 | ENSG00000228106 |
| SAMSN1 | 0.838529104 | 1.788226031 | 0.241674674 | 0.44519938 | 6.725569369 | 11.95642643 | ENSG00000155307 |
| CST3 | 0.83843108 | 1.788104533 | 9.86E-15 | 3.58E-13 | 355.0983954 | 635.5482348 | ENSG00000101439 |
| NA | 0.838276116 | 1.787912478 | 0.002566477 | 0.012566224 | 47.59261349 | 85.17441465 | ENSG00000227081 |
| FN1 | 0.838237297 | 1.787864371 | 2.59E-25 | 2.40E-23 | 2666.960164 | 4766.577664 | ENSG00000115414 |
| RAD51-AS1 | 0.838138086 | 1.787741428 | 0.004800626 | 0.02132169 | 43.55757735 | 77.7663345 | ENSG00000245849 |
| PAH | 0.837760452 | 1.787273538 | 0.025003796 | 0.082494181 | 24.55546008 | 43.85192725 | ENSG00000171759 |
| TSEN15 | 0.837672601 | 1.787164707 | 2.21E-21 | 1.47E-19 | 659.4710755 | 1178.399723 | ENSG00000198860 |
| POLA2 | 0.837416323 | 1.786847267 | 0.2050881 | 0.397059708 | 7.661754006 | 13.67126002 | ENSG00000285816 |
| SP140 | 0.836661062 | 1.785912084 | 0.418513995 | 0.629270445 | 3.38828552 | 6.007913181 | ENSG00000079263 |
| REXO5 | 0.83574391 | 1.784777103 | 0.00021048 | 0.001438622 | 82.87283116 | 148.2371459 | ENSG00000005189 |
| FAM241A | 0.835469807 | 1.784438038 | 0.012269927 | 0.046368735 | 34.68652751 | 62.14739124 | ENSG00000174749 |
| CHCHD10 | 0.835327312 | 1.784261797 | 1.41E-10 | 3.08E-09 | 551.3870628 | 984.8397749 | ENSG00000250479 |

| | | | | | | | |
|---|---|---|---|---|---|---|---|
| NA | 0.835114334 | 1.783998416 | 0.078844199 | 0.19955029 | 15.4382747 | 27.63912056 | ENSG00000267424 |
| NA | 0.834245237 | 1.782924037 | 0.162491448 | 0.337548234 | 9.357676502 | 16.62886947 | ENSG00000286138 |
| PCED1B | 0.834185891 | 1.782850697 | 0.00012824 | 0.000926835 | 77.39777061 | 138.2623595 | ENSG00000179715 |
| AHNAK2 | 0.833913162 | 1.782513696 | 0.000957722 | 0.005405436 | 60.29469278 | 107.4046667 | ENSG00000185567 |
| NA | 0.833661438 | 1.782202707 | 0.006501119 | 0.027570682 | 36.99093673 | 65.76356092 | ENSG00000273230 |
| KCNJ3 | 0.833158915 | 1.781582035 | 0.317141505 | 0.527263189 | 5.087767488 | 8.989169178 | ENSG00000162989 |
| LINC02601 | 0.832532329 | 1.780808432 | 0.458632511 | 0.663997598 | 3.003077381 | 5.338475623 | ENSG00000223714 |
| KLHL12 | 0.832159227 | 1.780347948 | 3.45E-26 | 3.31E-24 | 924.329913 | 1644.558357 | ENSG00000117153 |
| KDELR3 | 0.831340373 | 1.779337734 | 1.07E-06 | 1.23E-05 | 121.0291548 | 215.2248494 | ENSG00000100196 |
| HPX | 0.831278729 | 1.779261708 | 0.457443901 | 0.663147928 | 2.993594172 | 5.319179997 | ENSG00000110169 |
| FCSK | 0.830956319 | 1.778864126 | 1.14E-07 | 1.56E-06 | 149.6745177 | 265.9140299 | ENSG00000157353 |
| NA | 0.830837402 | 1.778717507 | 0.403966509 | 0.615184866 | 3.769934186 | 6.620126274 | ENSG00000239268 |
| PABPC1L2B | 0.829035294 | 1.776497052 | 0.211528667 | 0.405267209 | 8.077802188 | 14.25888061 | ENSG00000184388 |
| KLK10 | 0.82873196 | 1.776123573 | 0.000377867 | 0.00240432 | 60.91802779 | 108.1680281 | ENSG00000129451 |
| MIR573 | 0.82713463 | 1.774158165 | 0.353222703 | 0.565470134 | 4.30430853 | 7.635349363 | ENSG00000207697 |
| GABRE | 0.826508346 | 1.773388156 | 0.003049496 | 0.014612976 | 73.63818378 | 130.4780803 | ENSG00000102287 |
| MIR17HG | 0.826485683 | 1.7733603 | 4.48E-05 | 0.000365336 | 113.3774944 | 201.3915115 | ENSG00000215417 |
| ARL4D | 0.826306066 | 1.773139528 | 1.30E-08 | 2.11E-07 | 228.6163231 | 404.7707473 | ENSG00000175906 |
| NA | 0.826239757 | 1.773058033 | 0.475292732 | 0.677887112 | 3.020874744 | 5.337529665 | ENSG00000232936 |
| QPCT | 0.826076549 | 1.772857464 | 0.000200517 | 0.001383545 | 69.6867688 | 123.3957846 | ENSG00000115828 |
| ANXA9 | 0.825927465 | 1.772674271 | 0.000405841 | 0.002559892 | 65.49514203 | 115.9250152 | ENSG00000143412 |
| GEMIN8 | 0.825696145 | 1.772390065 | 2.75E-08 | 4.20E-07 | 170.2207215 | 301.4227899 | ENSG00000046647 |
| PKP3 | 0.825495886 | 1.772144058 | 6.91E-13 | 2.03E-11 | 349.7295081 | 620.3454793 | ENSG00000184363 |
| TNFRSF18 | 0.825165325 | 1.771738059 | 0.036580969 | 0.1109057 | 22.96243064 | 40.80720394 | ENSG00000186891 |
| ACKR3 | 0.82461452 | 1.771061757 | 0.038616192 | 0.115664836 | 21.72763426 | 38.57285155 | ENSG00000144476 |
| NA | 0.824612674 | 1.77105949 | 0.373732155 | 0.585708628 | 3.911981447 | 6.967141309 | ENSG00000226390 |
| FZD9 | 0.824379054 | 1.770772722 | 2.06E-06 | 2.25E-05 | 130.2083252 | 230.9122734 | ENSG00000188763 |
| NA | 0.824000312 | 1.770307912 | 0.275477733 | 0.484439611 | 5.61858236 | 9.949626811 | ENSG00000278200 |
| NA | 0.823847455 | 1.770120353 | 1.38E-09 | 2.60E-08 | 294.2671916 | 520.7002517 | ENSG00000281406 |
| TMEM37 | 0.82370258 | 1.769942607 | 3.34E-11 | 8.01E-10 | 231.6701746 | 410.2318792 | ENSG00000171227 |
| KCNK7 | 0.823560892 | 1.769768789 | 0.339166927 | 0.550934752 | 4.300749058 | 7.60423046 | ENSG00000173338 |
| NA | 0.823504898 | 1.769700102 | 0.124800029 | 0.280537317 | 14.92970639 | 26.58407671 | ENSG00000267284 |
| DDX28 | 0.82347744 | 1.769666421 | 2.74E-13 | 8.46E-12 | 341.8017362 | 605.3652241 | ENSG00000182810 |
| NA | 0.823451723 | 1.769634875 | 0.000296055 | 0.001942374 | 77.0184862 | 136.4777001 | ENSG00000267640 |
| NA | 0.823396984 | 1.769567732 | 0.013468361 | 0.049852998 | 32.17569562 | 56.92809434 | ENSG00000255026 |
| XPR1 | 0.82275464 | 1.768780028 | 1.66E-33 | 2.36E-31 | 1428.774595 | 2526.723258 | ENSG00000143324 |
| RASIP1 | 0.82209979 | 1.767977347 | 8.06E-08 | 1.14E-06 | 170.6557361 | 301.388266 | ENSG00000105538 |
| CAPN14 | 0.821989856 | 1.767842632 | 0.51543949 | 0.711815339 | 2.581105463 | 4.588450593 | ENSG00000214711 |
| SMIM38 | 0.821966194 | 1.767813638 | 0.453578601 | 0.659622125 | 2.632107133 | 4.67026757 | ENSG00000284713 |
| RNF213-AS1 | 0.821903043 | 1.767736256 | 0.274006559 | 0.482661493 | 6.047673224 | 10.6318336 | ENSG00000263069 |
| UFSP1 | 0.821881779 | 1.767710202 | 0.005897 | 0.025356293 | 41.07196667 | 72.81564063 | ENSG00000176125 |
| ZNF132 | 0.820963715 | 1.766585671 | 0.028592728 | 0.09128887 | 25.5616128 | 45.09281118 | ENSG00000131849 |
| IFITM10 | 0.820499573 | 1.766017419 | 0.470674548 | 0.674100473 | 3.446406135 | 6.02285788 | ENSG00000244242 |
| LINC00605 | 0.819999618 | 1.765405525 | 0.017418057 | 0.061518611 | 31.67808506 | 55.7803562 | ENSG00000251533 |
| NA | 0.819991525 | 1.765395622 | 0.289756629 | 0.499138369 | 6.061911114 | 10.63930595 | ENSG00000272311 |
| HDAC4-AS1 | 0.819507705 | 1.764803681 | 0.362934652 | 0.574867178 | 4.348191255 | 7.64622668 | ENSG00000222020 |
| ZNF79 | 0.819010477 | 1.764195542 | 9.25E-06 | 8.91E-05 | 106.2990315 | 187.740661 | ENSG00000196152 |
| CEP350 | 0.818812842 | 1.763953881 | 3.94E-12 | 1.05E-10 | 552.3550496 | 973.7026712 | ENSG00000135837 |
| NA | 0.818259884 | 1.76327792 | 0.234247306 | 0.435538089 | 7.338225955 | 12.88671112 | ENSG00000269378 |
| HSPA6 | 0.818100154 | 1.763082708 | 0.340656201 | 0.552213936 | 6.269197935 | 11.08648763 | ENSG00000173110 |
| NA | 0.81801324 | 1.762976495 | 0.447673289 | 0.654431651 | 3.426244509 | 5.971213847 | ENSG00000254016 |
| RPS27 | 0.81770328 | 1.762597764 | 5.09E-34 | 7.52E-32 | 10141.68507 | 17876.1448 | ENSG00000177954 |
| CPNE7 | 0.817023177 | 1.761767051 | 0.037158749 | 0.112144817 | 363.0801442 | 639.756897 | ENSG00000178773 |
| TSHZ1 | 0.816170837 | 1.760726511 | 0.251455972 | 0.45709366 | 6.589166513 | 11.62341014 | ENSG00000179981 |
| DMTN | 0.816098361 | 1.760638061 | 0.039202353 | 0.116901381 | 23.5492812 | 41.40853971 | ENSG00000158856 |
| LOC107985177 | 0.816095749 | 1.760634873 | 0.459171662 | 0.664323996 | 3.068316941 | 5.35682529 | ENSG00000287281 |
| SNORD101 | 0.816031165 | 1.760556058 | 0.468285094 | 0.671564029 | 2.639226078 | 4.654093366 | ENSG00000206754 |
| NA | 0.815670478 | 1.760115958 | 0.429890936 | 0.639654139 | 3.537451726 | 6.282759052 | ENSG00000286918 |
| CDHR2 | 0.815339041 | 1.759711645 | 0.160173021 | 0.334091839 | 10.16694149 | 17.95998869 | ENSG00000074276 |
| STPG1 | 0.815310278 | 1.759676562 | 1.14E-07 | 1.56E-06 | 156.6806221 | 275.6437872 | ENSG00000001460 |
| GEM | 0.815281779 | 1.759641801 | 5.82E-05 | 0.000460271 | 84.63902723 | 148.9699352 | ENSG00000164949 |
| AMZ1 | 0.814758849 | 1.759004105 | 0.004500499 | 0.020253665 | 41.25344746 | 72.62306324 | ENSG00000174945 |
| CCDC183 | 0.814497909 | 1.758685982 | 0.168743123 | 0.346419598 | 9.677645081 | 16.97805997 | ENSG00000213213 |
| NA | 0.814306665 | 1.758452866 | 0.046242396 | 0.132658005 | 20.63280021 | 36.21440242 | ENSG00000260528 |
| NA | 0.814299238 | 1.758443814 | 0.250457895 | 0.455942388 | 7.018257377 | 12.30126601 | ENSG00000225518 |
| AP3B2 | 0.814298784 | 1.758443259 | 4.76E-11 | 1.12E-09 | 261.4812298 | 459.3020418 | ENSG00000103723 |
| MUC5B | 0.813728729 | 1.75774858 | 0.423264957 | 0.633337985 | 3.575410714 | 6.330052158 | ENSG00000117983 |
| PKD1L2 | 0.813563349 | 1.757547096 | 0.017782706 | 0.062520019 | 31.010588 | 54.6013312 | ENSG00000166473 |
| NA | 0.813306219 | 1.757233878 | 0.382828086 | 0.59533172 | 3.939262018 | 6.933846943 | ENSG00000260418 |
| NA | 0.813287842 | 1.757211495 | 0.305656997 | 0.516031107 | 5.842471335 | 10.35046134 | ENSG00000279406 |
| NA | 0.813235595 | 1.757147859 | 0.439637217 | 0.647624715 | 3.044595843 | 5.303005794 | ENSG00000250938 |
| CLHC1 | 0.812731054 | 1.756533454 | 0.009153403 | 0.036353738 | 40.14051057 | 70.62714093 | ENSG00000162994 |
| EIF1B-AS1 | 0.812509153 | 1.756263302 | 0.214740158 | 0.409718068 | 7.965120431 | 13.98498069 | ENSG00000280739 |
| OSBPL5 | 0.811969755 | 1.75560679 | 2.14E-11 | 5.21E-10 | 314.3780754 | 551.8464625 | ENSG00000021762 |

| | | | | | | | |
|---|---|---|---|---|---|---|---|
| TMEM204 | 0.811624881 | 1.755187165 | 0.379769777 | 0.592380627 | 3.949940435 | 6.939143829 | ENSG00000131634 |
| POMK | 0.811217992 | 1.754692212 | 0.007200729 | 0.029979225 | 37.94344419 | 66.63227022 | ENSG00000185900 |
| SPATA18 | 0.811029498 | 1.75446297 | 0.047717377 | 0.136030754 | 20.64584289 | 36.12416713 | ENSG00000163071 |
| NA | 0.810688863 | 1.754048772 | 0.023371738 | 0.078050447 | 27.22077152 | 47.65611248 | ENSG00000259495 |
| RALGAPA1P1 | 0.809893523 | 1.753082053 | 0.292439568 | 0.502202531 | 5.700424074 | 9.937803534 | ENSG00000229419 |
| NA | 0.809761559 | 1.752921705 | 0.264224099 | 0.471721696 | 7.896321398 | 13.84811471 | ENSG00000272989 |
| CDX4 | 0.80961788 | 1.752747139 | 0.141693978 | 0.306440122 | 11.02275895 | 19.25034119 | ENSG00000131264 |
| NA | 0.809580094 | 1.752701233 | 0.375937754 | 0.587957307 | 4.000942105 | 6.999489717 | ENSG00000287077 |
| SYT14 | 0.809464149 | 1.752560379 | 4.89E-07 | 6.02E-06 | 205.3734086 | 360.1150052 | ENSG00000143469 |
| ZNF793-AS1 | 0.809437743 | 1.752528302 | 0.157309022 | 0.329574093 | 10.61382972 | 18.58402506 | ENSG00000266916 |
| VLDLR-AS1 | 0.809034671 | 1.752038734 | 0.010258753 | 0.039842869 | 37.84883912 | 66.29133007 | ENSG00000236404 |
| NA | 0.807965476 | 1.750740764 | 0.14239527 | 0.307505394 | 12.7020793 | 22.14107836 | ENSG00000280173 |
| ELL3 | 0.80796268 | 1.750737371 | 0.467563397 | 0.671155346 | 2.666506649 | 4.658444293 | ENSG00000128886 |
| SLC46A1 | 0.807709389 | 1.750430024 | 0.00498188 | 0.022012821 | 41.36968869 | 72.43003434 | ENSG00000076351 |
| TP53I11 | 0.807579399 | 1.750272313 | 4.25E-28 | 4.56E-26 | 1172.7128 | 2053.292636 | ENSG00000175274 |
| NA | 0.807502801 | 1.750179387 | 0.008599435 | 0.034678696 | 36.99302166 | 64.81085919 | ENSG00000267278 |
| GRIK2 | 0.807419388 | 1.750078199 | 0.022523022 | 0.075698038 | 29.04094391 | 50.89993957 | ENSG00000164418 |
| RNF180 | 0.807078065 | 1.749664202 | 0.035481933 | 0.108303911 | 23.99616942 | 41.89242073 | ENSG00000164197 |
| PPP1R15B | 0.806951417 | 1.749510614 | 2.10E-27 | 2.19E-25 | 1010.357047 | 1766.65491 | ENSG00000158615 |
| MYCBPAP | 0.8068225 | 1.749354287 | 0.129704324 | 0.288376048 | 12.98408889 | 22.69372357 | ENSG00000136449 |
| MGMT | 0.806820964 | 1.749352425 | 1.65E-10 | 3.57E-09 | 271.791998 | 475.7200701 | ENSG00000170430 |
| FOSL1 | 0.8060357 | 1.748400505 | 2.63E-07 | 3.40E-06 | 160.4318692 | 280.1525963 | ENSG00000175592 |
| HLA-DPA1 | 0.805696099 | 1.747988991 | 0.013085583 | 0.04873225 | 32.94847616 | 57.48677148 | ENSG00000231389 |
| NA | 0.80447545 | 1.746510662 | 0.470554215 | 0.673979245 | 2.662947177 | 4.637919163 | ENSG00000286786 |
| NIT2 | 0.804432069 | 1.746458147 | 2.13E-11 | 5.19E-10 | 295.714335 | 516.0319259 | ENSG00000114021 |
| ADRA1A | 0.803885087 | 1.745796121 | 0.211457791 | 0.405213637 | 9.244994745 | 16.12039594 | ENSG00000120907 |
| OBSCN | 0.803631593 | 1.745489396 | 1.81E-31 | 2.26E-29 | 1240.427989 | 2164.634164 | ENSG00000154358 |
| MTMR11 | 0.803367842 | 1.745170319 | 0.004499456 | 0.020253665 | 45.62327489 | 79.47558766 | ENSG00000014914 |
| VEPH1 | 0.80308003 | 1.744822198 | 0.006112156 | 0.026144521 | 57.86633915 | 100.9398287 | ENSG00000197415 |
| BTN3A2 | 0.803006686 | 1.744733498 | 0.000229674 | 0.001555739 | 74.05275742 | 129.0103356 | ENSG00000186470 |
| NT5E | 0.802171051 | 1.743723209 | 0.127626571 | 0.285125024 | 12.48619899 | 21.80494064 | ENSG00000135318 |
| VEGFD | 0.801771742 | 1.743240648 | 0.400222056 | 0.611652791 | 3.622852912 | 6.334403086 | ENSG00000165197 |
| LOC102724301 | 0.801702379 | 1.743156837 | 0.155103482 | 0.326765869 | 13.59674547 | 23.71481063 | ENSG00000254854 |
| KIRREL3 | 0.801166108 | 1.742509001 | 0.100170306 | 0.238311443 | 14.31260061 | 24.88201236 | ENSG00000149571 |
| DYRK4 | 0.801115541 | 1.742447926 | 0.017551103 | 0.06186125 | 32.30617474 | 56.1858252 | ENSG00000010219 |
| NA | 0.800119387 | 1.741245213 | 0.308844132 | 0.51932343 | 5.353174924 | 9.27706783 | ENSG00000278918 |
| NA | 0.800036174 | 1.741144783 | 0.336787482 | 0.548704693 | 4.960847841 | 8.642154143 | ENSG00000260412 |
| SMYD3 | 0.800007423 | 1.741110085 | 6.16E-12 | 1.60E-10 | 308.8191137 | 538.0817504 | ENSG00000185420 |
| MPPED1 | 0.799980297 | 1.741077349 | 0.227915854 | 0.427257385 | 8.134448263 | 14.2427064 | ENSG00000186732 |
| LYSMD1 | 0.799665748 | 1.740697785 | 1.34E-05 | 0.000123877 | 105.0550313 | 182.7322482 | ENSG00000163155 |
| TSNAX | 0.799510881 | 1.740510938 | 6.51E-07 | 7.77E-06 | 219.9543913 | 383.3109594 | ENSG00000116918 |
| ANP32C | 0.799493664 | 1.740490167 | 0.460305964 | 0.664999857 | 3.640650274 | 6.344050899 | ENSG00000248546 |
| SNHG5 | 0.799458064 | 1.740447219 | 2.73E-41 | 5.73E-39 | 3749.169288 | 6524.201423 | ENSG00000203875 |
| ATP2B2 | 0.799443166 | 1.740429248 | 0.016555417 | 0.0590684 | 29.71291632 | 51.75370417 | ENSG00000157087 |
| LINC01600 | 0.79893683 | 1.739818524 | 0.246356824 | 0.451212164 | 7.259943714 | 12.65480743 | ENSG00000164385 |
| SPTY2D1OS | 0.798638204 | 1.739458433 | 0.290726873 | 0.500353322 | 6.745730996 | 11.66975729 | ENSG00000247595 |
| GPR39 | 0.798503784 | 1.739296371 | 0.279760269 | 0.488880785 | 6.292919033 | 10.93808192 | ENSG00000183840 |
| CTD-2297D10.2 | 0.798029159 | 1.738724262 | 0.00029499 | 0.001937024 | 137.1453257 | 238.558375 | ENSG00000250579 |
| CLC | 0.797959735 | 1.738640594 | 0.198864915 | 0.389030323 | 8.605057588 | 14.96908488 | ENSG00000105205 |
| NA | 0.796970812 | 1.737449218 | 0.329955413 | 0.541420451 | 5.917194104 | 10.30628966 | ENSG00000274248 |
| CD81-AS1 | 0.796141908 | 1.73645125 | 0.374512521 | 0.58653427 | 5.026087401 | 8.708742875 | ENSG00000238184 |
| NA | 0.795120453 | 1.735222244 | 0.085741852 | 0.212721344 | 16.73950585 | 29.10944917 | ENSG00000260628 |
| CD101 | 0.79477552 | 1.734807421 | 0.126232556 | 0.282854985 | 12.2279105 | 21.21732006 | ENSG00000134256 |
| SLC5A11 | 0.794629143 | 1.734631415 | 0.490165979 | 0.69013064 | 2.690227748 | 4.636973204 | ENSG00000158865 |
| PWRN1 | 0.793706834 | 1.733522827 | 0.269597215 | 0.477611952 | 7.74359572 | 13.36718717 | ENSG00000259905 |
| NA | 0.79277327 | 1.732401432 | 0.450086132 | 0.656361128 | 3.248323191 | 5.672721423 | ENSG00000286415 |
| TMEM196 | 0.792594005 | 1.732186182 | 0.421272354 | 0.631256718 | 3.637090802 | 6.30110872 | ENSG00000173452 |
| NA | 0.792435271 | 1.731995608 | 0.009355315 | 0.036984533 | 42.27767688 | 73.09389147 | ENSG00000237978 |
| ARL8A | 0.792148734 | 1.731651645 | 1.98E-23 | 1.57E-21 | 767.4722587 | 1328.870035 | ENSG00000143862 |
| LAD1 | 0.792063542 | 1.731549394 | 7.49E-16 | 3.14E-14 | 502.3474562 | 869.8010337 | ENSG00000159166 |
| PEAR1 | 0.791769913 | 1.731197011 | 0.054778036 | 0.151417332 | 20.94920932 | 36.21099745 | ENSG00000187800 |
| SLFN5 | 0.791751815 | 1.731175294 | 7.08E-05 | 0.000546543 | 92.60679126 | 160.0938572 | ENSG00000166750 |
| NA | 0.79174414 | 1.731166084 | 0.347574186 | 0.559556672 | 4.593437065 | 7.983593902 | ENSG00000227620 |
| CCN3 | 0.791729538 | 1.731148562 | 0.389470976 | 0.601255569 | 4.53531645 | 7.892129113 | ENSG00000136999 |
| CDC42BPA | 0.791402152 | 1.730755763 | 1.93E-21 | 1.30E-19 | 816.2979153 | 1412.265852 | ENSG00000143776 |
| S100A10 | 0.791110677 | 1.730406125 | 3.58E-21 | 2.34E-19 | 614.9313981 | 1064.006793 | ENSG00000197747 |
| LOC112267895 | 0.790941849 | 1.73020364 | 0.184966588 | 0.36917079 | 9.57564174 | 16.57722543 | ENSG00000261241 |
| NA | 0.790798485 | 1.730031714 | 0.225181276 | 0.423518414 | 8.615736005 | 14.8872679 | ENSG00000248740 |
| COQ8A | 0.790494407 | 1.729667112 | 1.53E-25 | 1.44E-23 | 962.4736279 | 1665.731634 | ENSG00000163050 |
| TNRC18 | 0.790463725 | 1.729630328 | 3.25E-06 | 3.39E-05 | 157.2562094 | 272.1885089 | ENSG00000182095 |
| RASAL2 | 0.790430124 | 1.729590045 | 8.99E-18 | 4.39E-16 | 597.0386066 | 1031.809709 | ENSG00000075391 |
| ZNF469 | 0.789964218 | 1.729031578 | 0.003107719 | 0.014850022 | 55.72586654 | 96.42922307 | ENSG00000225614 |
| MC1R | 0.789654264 | 1.728660146 | 0.116833314 | 0.26789669 | 14.03295528 | 24.15468792 | ENSG00000258839 |
| NA | 0.789449381 | 1.72841467 | 0.453901527 | 0.659968547 | 3.643014538 | 6.291460907 | ENSG00000213985 |

| | | | | | | | |
|---|---|---|---|---|---|---|---|
| ADGRD2 | 0.789418435 | 1.728377595 | 0.046141944 | 0.132389944 | 24.99019534 | 43.32276063 | ENSG00000180264 |
| PSMD4 | 0.789213918 | 1.728132597 | 1.76E-52 | 5.84E-50 | 4329.793616 | 7481.70548 | ENSG00000159352 |
| DPYS | 0.788872036 | 1.727723122 | 0.2133319 | 0.407657787 | 8.618100269 | 14.86144589 | ENSG00000147647 |
| TMEM238 | 0.788707151 | 1.727525673 | 0.002848815 | 0.0137666 | 52.83430186 | 91.36079089 | ENSG00000233493 |
| PTGR2 | 0.78841726 | 1.727178584 | 0.179289338 | 0.361816536 | 9.599362839 | 16.57192855 | ENSG00000140043 |
| IDI2-AS1 | 0.788082827 | 1.72677825 | 0.136410277 | 0.298778204 | 11.95538412 | 20.62875352 | ENSG00000232656 |
| NA | 0.787953817 | 1.726623844 | 0.485661248 | 0.686556829 | 2.690227748 | 4.60680026 | ENSG00000237276 |
| DNER | 0.787698349 | 1.726318126 | 0.001099269 | 0.006089672 | 60.0375995 | 103.5528095 | ENSG00000187957 |
| CACTIN-AS1 | 0.787543457 | 1.726132793 | 0.250025437 | 0.455369888 | 7.927161442 | 13.71760717 | ENSG00000226800 |
| ASB16 | 0.787420291 | 1.725985436 | 0.115189986 | 0.26515878 | 13.30761694 | 22.92780272 | ENSG00000161664 |
| LINC02018 | 0.787370206 | 1.725925517 | 0.131066963 | 0.290755267 | 12.91173038 | 22.29411854 | ENSG00000272690 |
| LINC01775 | 0.787276483 | 1.725813398 | 0.365019678 | 0.577191377 | 4.610039219 | 7.971770626 | ENSG00000267201 |
| CARD14 | 0.787218705 | 1.725744283 | 0.362184802 | 0.574006376 | 4.644438735 | 8.021239196 | ENSG00000141527 |
| TOMM20 | 0.787008005 | 1.725492263 | 3.05E-51 | 9.59E-49 | 4442.025632 | 7663.766886 | ENSG00000173726 |
| TTC39A | 0.786265113 | 1.724603978 | 3.90E-05 | 0.000322546 | 110.2142675 | 189.648975 | ENSG00000085831 |
| NA | 0.785949845 | 1.724227146 | 0.520061125 | 0.715550501 | 2.74478889 | 4.686441774 | ENSG00000280089 |
| LOC148709 | 0.785882673 | 1.724146868 | 0.003967602 | 0.018244012 | 51.36402221 | 88.32940391 | ENSG00000234996 |
| LINC02751 | 0.785771852 | 1.724014432 | 0.143408956 | 0.309128882 | 11.5571333 | 19.95089765 | ENSG00000254946 |
| NA | 0.785537774 | 1.723734734 | 0.004995681 | 0.022068642 | 42.37347716 | 73.03259962 | ENSG00000259820 |
| CYP2A6 | 0.785158439 | 1.723281563 | 0.441337054 | 0.64904972 | 3.248323191 | 5.636022089 | ENSG00000255974 |
| ABHD1 | 0.784559128 | 1.722565841 | 0.350678235 | 0.562807723 | 4.617158164 | 7.963068772 | ENSG00000143994 |
| SNORD46 | 0.784472255 | 1.722462119 | 0.140060345 | 0.304132415 | 11.94234144 | 20.54163966 | ENSG00000200913 |
| SPATS2L | 0.78443814 | 1.722421389 | 1.37E-18 | 7.21E-17 | 934.718841 | 1609.617039 | ENSG00000196141 |
| SLC8A2 | 0.783985149 | 1.721880651 | 5.07E-07 | 6.22E-06 | 206.5933567 | 355.6983233 | ENSG00000118160 |
| GGACT | 0.783982277 | 1.721877224 | 0.209510228 | 0.402719054 | 10.38134726 | 18.0085113 | ENSG00000134864 |
| NA | 0.78374811 | 1.721597765 | 0.125266933 | 0.281225158 | 19.08276378 | 32.79639056 | ENSG00000230805 |
| STING1 | 0.78368766 | 1.72152563 | 0.289133378 | 0.498473968 | 5.96583151 | 10.25464563 | ENSG00000184584 |
| ASPM | 0.783567781 | 1.721382588 | 5.61E-19 | 3.12E-17 | 1063.463797 | 1829.671363 | ENSG00000066279 |
| ZBTB18 | 0.783334951 | 1.721104804 | 8.68E-10 | 1.68E-08 | 268.3167851 | 461.4325593 | ENSG00000179456 |
| ELK4 | 0.783280502 | 1.721039849 | 3.85E-12 | 1.03E-10 | 315.5138436 | 542.5197878 | ENSG00000158711 |
| HSPB7 | 0.783123185 | 1.72085219 | 0.391839068 | 0.603234759 | 4.668159834 | 8.025590123 | ENSG00000173641 |
| DECR2 | 0.782970143 | 1.720669651 | 6.27E-05 | 0.000491282 | 98.8439801 | 169.9425823 | ENSG00000242612 |
| ADAMTS10 | 0.782914146 | 1.720602866 | 1.02E-05 | 9.72E-05 | 178.1318912 | 306.7456287 | ENSG00000142303 |
| NA | 0.782717805 | 1.72036872 | 0.432592697 | 0.641954509 | 4.100581182 | 6.985490977 | ENSG00000261542 |
| LNCPRESS1 | 0.782478188 | 1.720083007 | 8.75E-09 | 1.46E-07 | 248.8672215 | 427.6619662 | ENSG00000232301 |
| CDKN2D | 0.781878708 | 1.719368414 | 0.013904392 | 0.051187815 | 34.9154505 | 60.1530773 | ENSG00000129355 |
| RAB42 | 0.781062753 | 1.718396254 | 4.24E-08 | 6.28E-07 | 175.7334617 | 301.9462808 | ENSG00000188060 |
| H2AC14 | 0.780969945 | 1.718285713 | 0.395201331 | 0.607074763 | 4.208228926 | 7.285212905 | ENSG00000276368 |
| C1orf56 | 0.779992579 | 1.717122041 | 0.000617646 | 0.003683059 | 67.03894924 | 115.2097976 | ENSG00000143443 |
| NA | 0.779769948 | 1.716857081 | 0.016210981 | 0.058091913 | 32.1412961 | 55.17372354 | ENSG00000265519 |
| SCG5 | 0.779454599 | 1.716481846 | 1.35E-12 | 3.80E-11 | 340.6985624 | 584.3818676 | ENSG00000166922 |
| DNAJC6 | 0.779238847 | 1.716225169 | 5.92E-05 | 0.000467583 | 94.66272626 | 162.3885554 | ENSG00000116675 |
| ZYG11A | 0.77890066 | 1.71582291 | 0.021516899 | 0.073018264 | 29.39650721 | 50.4838768 | ENSG00000203995 |
| LINC00910 | 0.778352274 | 1.715170829 | 0.018058218 | 0.063298316 | 36.62650061 | 63.06679861 | ENSG00000188825 |
| SOD3 | 0.778104124 | 1.714875837 | 0.000115569 | 0.000844626 | 84.74339484 | 145.2803668 | ENSG00000109610 |
| CREB3L4 | 0.77762369 | 1.714304859 | 3.20E-10 | 6.63E-09 | 294.8602714 | 504.7971279 | ENSG00000143578 |
| SPATA17 | 0.776927514 | 1.713477816 | 0.373498992 | 0.585549432 | 4.631396053 | 7.927598942 | ENSG00000162814 |
| CHGB | 0.776162795 | 1.712569806 | 7.28E-05 | 0.000559918 | 95.82071494 | 163.9921771 | ENSG00000089199 |
| NA | 0.776053697 | 1.712440305 | 0.202082639 | 0.393314505 | 9.763046267 | 16.62139712 | ENSG00000235493 |
| ABHD14A | 0.775936173 | 1.712300812 | 0.258965899 | 0.465982147 | 7.399906043 | 12.5882187 | ENSG00000248487 |
| SBSN | 0.77526576 | 1.7115053 | 0.800252802 | 0.898320559 | 3.1572776 | 5.391349161 | ENSG00000189001 |
| WDR78 | 0.774994874 | 1.711183971 | 0.39056996 | 0.602017115 | 5.067605862 | 8.61103524 | ENSG00000152763 |
| NA | 0.774853591 | 1.711016404 | 0.409346741 | 0.620469288 | 4.610039219 | 7.8824813 | ENSG00000233155 |
| PROX1 | 0.774739063 | 1.710880581 | 0.147080176 | 0.314569607 | 11.66981506 | 19.90455051 | ENSG00000117707 |
| IFITM1 | 0.7745795 | 1.710691366 | 6.71E-05 | 0.000521984 | 823.7637687 | 1409.023651 | ENSG00000185885 |
| DUSP23 | 0.773058992 | 1.708889357 | 8.27E-18 | 4.07E-16 | 598.4297654 | 1023.443927 | ENSG00000158716 |
| NA | 0.772371164 | 1.708074811 | 0.31628793 | 0.526264103 | 6.653210865 | 11.36038754 | ENSG00000254419 |
| CD74 | 0.772184211 | 1.707853483 | 7.02E-18 | 3.49E-16 | 607.0848832 | 1036.892763 | ENSG00000019582 |
| URB1-AS1 | 0.771678207 | 1.707254583 | 0.319264447 | 0.529397584 | 5.635184514 | 9.610084125 | ENSG00000256073 |
| NA | 0.771468464 | 1.707006396 | 0.275670767 | 0.484643683 | 7.587031237 | 12.97164952 | ENSG00000257027 |
| DEGS1 | 0.770754926 | 1.706162341 | 3.33E-23 | 2.54E-21 | 766.1547826 | 1307.521051 | ENSG00000143753 |
| C21orf91 | 0.770302973 | 1.705627936 | 2.02E-08 | 3.17E-07 | 335.5043157 | 572.1476286 | ENSG00000154642 |
| MRPL55 | 0.770216723 | 1.70552597 | 1.63E-13 | 5.15E-12 | 425.5176215 | 726.1115193 | ENSG00000162910 |
| COL1A2 | 0.770042962 | 1.705320566 | 4.83E-35 | 7.41E-33 | 2327.271441 | 3968.274787 | ENSG00000164692 |
| HS3ST2 | 0.76992581 | 1.705182093 | 0.00532181 | 0.023296799 | 45.73211784 | 77.92077215 | ENSG00000122254 |
| PPL | 0.769441798 | 1.704610115 | 4.17E-25 | 3.78E-23 | 1005.65782 | 1713.734349 | ENSG00000118898 |
| SNORD20 | 0.769370795 | 1.704526223 | 0.436533293 | 0.644964109 | 3.306443806 | 5.636022089 | ENSG00000207280 |
| DHX58 | 0.768428576 | 1.703413367 | 0.001219966 | 0.006666395 | 65.10637442 | 110.7197897 | ENSG00000108771 |
| RNF213 | 0.767420839 | 1.702223931 | 1.84E-15 | 7.39E-14 | 1042.9555 | 1774.239086 | ENSG00000173821 |
| CRYAB | 0.767253749 | 1.702026795 | 0.126285229 | 0.282905877 | 16.42546101 | 28.05802121 | ENSG00000109846 |
| GPR37L1 | 0.766884109 | 1.701590765 | 0.339949139 | 0.551541193 | 5.625701305 | 9.556264628 | ENSG00000170075 |
| MYL9 | 0.76618255 | 1.700763511 | 5.49E-32 | 7.00E-30 | 1430.345403 | 2433.443397 | ENSG00000101335 |
| ILDR2 | 0.766146886 | 1.700721468 | 0.049853455 | 0.140771904 | 24.87423345 | 42.3060245 | ENSG00000143195 |
| SUSD2 | 0.766133078 | 1.700705191 | 1.08E-07 | 1.49E-06 | 215.164067 | 365.896028 | ENSG00000099994 |

| | | | | | | | |
|---|---|---|---|---|---|---|---|
| CTNNA3 | 0.766063309 | 1.700622946 | 0.364410272 | 0.576517382 | 6.038190015 | 10.21047394 | ENSG00000183230 |
| ABL2 | 0.765707371 | 1.700203424 | 1.49E-14 | 5.22E-13 | 537.6116304 | 913.6811333 | ENSG00000143322 |
| TNFRSF19 | 0.765528835 | 1.699993035 | 0.207741394 | 0.400326145 | 9.519885389 | 16.22056259 | ENSG00000127863 |
| NA | 0.765413622 | 1.699857279 | 0.447909337 | 0.654652722 | 3.746213087 | 6.305459647 | ENSG00000254013 |
| RAB4A | 0.765400589 | 1.699841923 | 1.26E-17 | 6.05E-16 | 647.7635808 | 1100.961362 | ENSG00000168118 |
| MIR4479 | 0.765310436 | 1.699735705 | 0.136435845 | 0.298778204 | 15.91956244 | 27.08044341 | ENSG00000266507 |
| PARVB | 0.764968487 | 1.69933288 | 2.48E-09 | 4.50E-08 | 244.8822706 | 415.8729873 | ENSG00000188677 |
| MME | 0.764964244 | 1.699327881 | 1.01E-08 | 1.67E-07 | 236.2407285 | 401.0679152 | ENSG00000196549 |
| H3C13 | 0.764874443 | 1.69922211 | 0.361274177 | 0.573333011 | 5.278452155 | 8.984818251 | ENSG00000183598 |
| SNHG12 | 0.764743774 | 1.699068213 | 2.28E-06 | 2.46E-05 | 320.1046894 | 543.9176973 | ENSG00000197989 |
| GDPD5 | 0.764664977 | 1.698975416 | 8.11E-05 | 0.000617066 | 113.9258018 | 193.3782914 | ENSG00000158555 |
| ATP2C2 | 0.764427369 | 1.698695623 | 0.00137521 | 0.0073991 | 67.22607443 | 114.3139212 | ENSG00000064270 |
| NA | 0.763897638 | 1.698072007 | 0.00010422 | 0.000770631 | 102.5104102 | 174.0692854 | ENSG00000177822 |
| NR1I3 | 0.763403344 | 1.697490316 | 0.055750893 | 0.153455125 | 23.83724068 | 40.44713612 | ENSG00000143257 |
| ZNF562 | 0.763267066 | 1.697329978 | 1.61E-09 | 3.00E-08 | 308.7067119 | 523.3529016 | ENSG00000171466 |
| F8A1 | 0.762939992 | 1.696945218 | 0.000763148 | 0.004427666 | 79.98479981 | 136.0174133 | ENSG00000277203 |
| NA | 0.762676027 | 1.696634763 | 0.241517629 | 0.445088566 | 8.178330988 | 13.93333666 | ENSG00000228589 |
| NA | 0.762674829 | 1.696633354 | 0.39693335 | 0.608843902 | 4.331589101 | 7.339978361 | ENSG00000224272 |
| TMEM220-AS1 | 0.762532556 | 1.696466046 | 0.206275883 | 0.398641021 | 10.03557265 | 16.96964166 | ENSG00000263400 |
| NA | 0.762382282 | 1.696289348 | 0.293100629 | 0.502804708 | 6.26207899 | 10.64583234 | ENSG00000259209 |
| ADAMTS1 | 0.762198362 | 1.696073113 | 2.33E-15 | 9.19E-14 | 496.736857 | 841.9055824 | ENSG00000154734 |
| PCDHGC4 | 0.762046571 | 1.695894673 | 0.002253715 | 0.011259634 | 72.03094261 | 121.9437627 | ENSG00000242419 |
| CEP112 | 0.761164526 | 1.694858142 | 0.007420632 | 0.030711952 | 45.40947951 | 77.15194595 | ENSG00000154240 |
| BIK | 0.761068406 | 1.694745225 | 0.150796425 | 0.320156323 | 13.13325509 | 22.34453307 | ENSG00000100290 |
| ADGRA2 | 0.760926891 | 1.694578994 | 0.000414792 | 0.002611986 | 99.67015215 | 169.36338 | ENSG00000020181 |
| C6orf141 | 0.760906048 | 1.694554512 | 0.001117853 | 0.006178326 | 86.63829 | 146.9448288 | ENSG00000197261 |
| MIR4258 | 0.760887546 | 1.69453278 | 0.37266941 | 0.584685704 | 4.889684544 | 8.342432214 | ENSG00000264349 |
| NA | 0.760872829 | 1.694515495 | 0.175445311 | 0.355933754 | 10.58654915 | 17.94721945 | ENSG00000223551 |
| SP2-AS1 | 0.76055081 | 1.69413731 | 0.447305396 | 0.654123697 | 3.361004949 | 5.666195032 | ENSG00000234494 |
| NA | 0.760418656 | 1.69398213 | 0.0338308 | 0.104462827 | 30.31280956 | 51.26937166 | ENSG00000185495 |
| SNAPC5 | 0.759376391 | 1.692758767 | 2.56E-08 | 3.92E-07 | 203.3260932 | 344.2191788 | ENSG00000174446 |
| CDHR4 | 0.75908049 | 1.692411613 | 0.28781922 | 0.497120576 | 8.714179873 | 14.71325109 | ENSG00000187492 |
| PLCH2 | 0.75858527 | 1.691830774 | 0.21698599 | 0.412749841 | 9.234316327 | 15.65658856 | ENSG00000149527 |
| CES1P1 | 0.75856542 | 1.691807497 | 0.507980699 | 0.704869733 | 2.897514569 | 4.939816555 | ENSG00000228695 |
| ZNF687 | 0.758544144 | 1.691782547 | 1.15E-32 | 1.52E-30 | 1825.088199 | 3088.096168 | ENSG00000143373 |
| NA | 0.758099 | 1.691260627 | 0.271141987 | 0.479538904 | 7.221984725 | 12.24120366 | ENSG00000267249 |
| NA | 0.757989135 | 1.691131839 | 0.369351128 | 0.581613339 | 4.906286698 | 8.344607677 | ENSG00000215861 |
| PCDH11X | 0.757619927 | 1.690699107 | 5.79E-05 | 0.000458592 | 111.7832704 | 188.601687 | ENSG00000102290 |
| NA | 0.757161699 | 1.690162193 | 0.450228299 | 0.656517754 | 3.933338281 | 6.678296698 | ENSG00000254528 |
| TDRKH | 0.756589279 | 1.689491719 | 1.16E-16 | 5.09E-15 | 494.7322809 | 835.6340069 | ENSG00000182134 |
| GPR139 | 0.756276423 | 1.689125383 | 0.243776656 | 0.448050818 | 9.115990165 | 15.29992571 | ENSG00000180269 |
| ARHGAP25 | 0.755731653 | 1.68848768 | 0.099279288 | 0.236959123 | 16.46697947 | 27.84520223 | ENSG00000163219 |
| RP9P | 0.755431933 | 1.688136933 | 0.587617507 | 0.76572348 | 2.5633081 | 4.374896568 | ENSG00000205763 |
| NA | 0.755430677 | 1.688135464 | 0.458720428 | 0.664029993 | 3.364564421 | 5.645669901 | ENSG00000286156 |
| POMZP3 | 0.755426899 | 1.688131043 | 0.143394545 | 0.309128882 | 13.74591168 | 23.31586802 | ENSG00000146707 |
| EPCAM-DT | 0.755130405 | 1.687784144 | 0.087085576 | 0.21509431 | 19.47598059 | 32.90977794 | ENSG00000234690 |
| NA | 0.754679108 | 1.687256262 | 0.404366626 | 0.615476848 | 4.331589101 | 7.286158864 | ENSG00000286592 |
| NA | 0.754099713 | 1.686578786 | 0.345083487 | 0.557034745 | 5.315215936 | 8.947172957 | ENSG00000144158 |
| LOC102724330 | 0.753877915 | 1.686319513 | 0.006752428 | 0.02841979 | 50.26415415 | 84.57920613 | ENSG00000228801 |
| ZNF57 | 0.753526047 | 1.685908276 | 0.38986836 | 0.601542768 | 4.328029629 | 7.271214165 | ENSG00000171970 |
| ZNF286B | 0.753200168 | 1.685527502 | 0.034340921 | 0.105671153 | 31.04526684 | 52.19312995 | ENSG00000249459 |
| NA | 0.753158783 | 1.685479153 | 0.481710202 | 0.683478375 | 2.959194656 | 5.007634792 | ENSG00000233719 |
| FIGNL1 | 0.753039318 | 1.685339589 | 2.08E-06 | 2.27E-05 | 169.8699646 | 286.7412248 | ENSG00000132436 |
| SCAPER | 0.752877449 | 1.685150506 | 6.16E-05 | 0.000483287 | 106.5958638 | 179.466047 | ENSG00000140386 |
| ELFN2 | 0.752548802 | 1.684766671 | 0.000601574 | 0.003606578 | 78.48990933 | 131.9775929 | ENSG00000166897 |
| SLC45A3 | 0.75213381 | 1.684282117 | 5.44E-13 | 1.61E-11 | 466.6189798 | 786.4645081 | ENSG00000158715 |
| NA | 0.752027084 | 1.684157524 | 0.303380714 | 0.513486816 | 6.309521187 | 10.60818705 | ENSG00000261098 |
| GSTM3 | 0.751588024 | 1.683645057 | 0.010326339 | 0.040068403 | 45.63367397 | 76.80632837 | ENSG00000134202 |
| PTER | 0.751414627 | 1.683442712 | 5.16E-05 | 0.000413403 | 118.2179573 | 199.0123263 | ENSG00000165983 |
| ALKAL2 | 0.751185458 | 1.683175322 | 0.030809711 | 0.096957406 | 35.51414853 | 59.71393507 | ENSG00000189292 |
| KRBA2 | 0.751113406 | 1.683091262 | 0.188662851 | 0.374568749 | 10.84839711 | 18.318827 | ENSG00000184619 |
| DISP1 | 0.751087849 | 1.683061447 | 0.00035752 | 0.00228796 | 88.55576336 | 149.1453924 | ENSG00000154309 |
| NA | 0.749897464 | 1.681673306 | 0.367356884 | 0.579625262 | 4.947805159 | 8.342432214 | ENSG00000217716 |
| CD55 | 0.749662432 | 1.681399363 | 8.55E-11 | 1.93E-09 | 312.6916878 | 526.2664979 | ENSG00000196352 |
| DCDC2C | 0.749569742 | 1.681291341 | 0.427081468 | 0.636929126 | 3.919100392 | 6.615775347 | ENSG00000214866 |
| IGSF21 | 0.74930823 | 1.680986607 | 5.64E-10 | 1.12E-08 | 403.6832815 | 678.7589423 | ENSG00000117154 |
| RGL1 | 0.748967279 | 1.680589387 | 0.000325808 | 0.002109318 | 91.01498319 | 152.9362413 | ENSG00000143344 |
| CGREF1 | 0.748908628 | 1.680521066 | 0.002112464 | 0.010667673 | 58.73636835 | 98.61393893 | ENSG00000138028 |
| NA | 0.748532803 | 1.680083344 | 0.184934735 | 0.369146229 | 11.32612538 | 19.00443876 | ENSG00000188459 |
| ROCK1P1 | 0.747584356 | 1.678979198 | 0.439500222 | 0.647473413 | 3.95705938 | 6.656825609 | ENSG00000263006 |
| SSC5D | 0.747527118 | 1.678912587 | 0.061358346 | 0.165182091 | 27.0621221 | 45.34327545 | ENSG00000179954 |
| LINC01836 | 0.747525856 | 1.678911118 | 0.109405722 | 0.255279849 | 15.79620227 | 26.58379316 | ENSG00000267530 |
| NA | 0.747415108 | 1.678782242 | 0.020292724 | 0.069589268 | 37.901036 | 63.56779979 | ENSG00000266869 |
| ZFP92 | 0.747079358 | 1.678391594 | 0.098142205 | 0.234852983 | 18.56234799 | 31.11870781 | ENSG00000189420 |

| | | | | | | | |
|---|---|---|---|---|---|---|---|
| TULP1 | 0.747064148 | 1.6783739 | 0.000148396 | 0.001058329 | 94.04739992 | 158.0073434 | ENSG00000112041 |
| PFKFB2 | 0.746533312 | 1.67775646 | 4.39E-13 | 1.32E-11 | 473.7581297 | 794.1415701 | ENSG00000123836 |
| PARD6G-AS1 | 0.746486327 | 1.67770182 | 0.136966846 | 0.299654644 | 15.15626511 | 25.43823049 | ENSG00000267270 |
| NLRC3 | 0.746315697 | 1.677503407 | 0.429751157 | 0.639615333 | 3.926219337 | 6.606127534 | ENSG00000167984 |
| TMEM79 | 0.746047141 | 1.677191171 | 0.034345223 | 0.105671153 | 32.14721984 | 53.92290824 | ENSG00000163472 |
| NA | 0.745811529 | 1.676917285 | 0.316120183 | 0.526170426 | 6.523926953 | 11.01119705 | ENSG00000236409 |
| NA | 0.745447939 | 1.676494719 | 0.115620856 | 0.265826622 | 155.7471072 | 261.1165051 | ENSG00000235688 |
| OBSCN-AS1 | 0.745413578 | 1.676454791 | 0.486285294 | 0.687181815 | 2.948516239 | 4.955990759 | ENSG00000162913 |
| ERO1B | 0.745336353 | 1.676365055 | 0.001713534 | 0.00891217 | 65.17517345 | 109.0273731 | ENSG00000086619 |
| HMCN2 | 0.745088693 | 1.676077306 | 3.79E-12 | 1.01E-10 | 399.3717762 | 669.2188424 | ENSG00000148357 |
| NA | 0.744689172 | 1.67561322 | 0.481846447 | 0.683503506 | 2.982915755 | 5.001108401 | ENSG00000223668 |
| CREG1 | 0.744484718 | 1.675375775 | 1.51E-13 | 4.80E-12 | 513.0914852 | 860.1071024 | ENSG00000143162 |
| NA | 0.744449993 | 1.675335449 | 0.418929511 | 0.629503762 | 3.980780479 | 6.671770308 | ENSG00000221930 |
| GLRA4 | 0.74415553 | 1.674993539 | 0.227353636 | 0.426330395 | 10.42998466 | 17.54950634 | ENSG00000188828 |
| POGK | 0.744127704 | 1.674961232 | 2.22E-31 | 2.76E-29 | 1346.634221 | 2255.079223 | ENSG00000143157 |
| SP8 | 0.74250192 | 1.673074769 | 8.78E-05 | 0.000661411 | 111.9193939 | 186.9821021 | ENSG00000164651 |
| TMEM86B | 0.742491865 | 1.673063109 | 0.404477523 | 0.615476848 | 5.394693385 | 8.977345901 | ENSG00000180089 |
| NA | 0.742383672 | 1.672937644 | 0.248904101 | 0.453852848 | 8.986706254 | 15.02507984 | ENSG00000262903 |
| NA | 0.742293704 | 1.672833321 | 0.11196778 | 0.259717416 | 18.56679719 | 31.22388779 | ENSG00000239415 |
| TUBA3D | 0.742245734 | 1.6727777 | 0.362827977 | 0.574832747 | 5.411295539 | 8.986993714 | ENSG00000075886 |
| NKX2-5 | 0.741974939 | 1.672463747 | 0.287886228 | 0.49718623 | 8.012562628 | 13.38553684 | ENSG00000183072 |
| NPIPA1 | 0.741017343 | 1.671354009 | 9.21E-05 | 0.000691128 | 113.9895668 | 190.5027825 | ENSG00000183426 |
| CAPN9 | 0.740636531 | 1.670912899 | 0.427228693 | 0.637098378 | 3.94282149 | 6.594304258 | ENSG00000135773 |
| DUOX1 | 0.740551412 | 1.670814319 | 0.073089733 | 0.189263624 | 21.01177913 | 35.21535354 | ENSG00000137857 |
| PCOLCE-AS1 | 0.740539392 | 1.670800397 | 0.180940747 | 0.363879543 | 13.16409514 | 22.0696872 | ENSG00000224729 |
| SPDYC | 0.740235682 | 1.670448705 | 0.493418191 | 0.692854676 | 2.97579681 | 4.969989499 | ENSG00000204710 |
| ACRV1 | 0.739771469 | 1.669911295 | 0.243426535 | 0.447625133 | 8.516096929 | 14.27817623 | ENSG00000134940 |
| NA | 0.739757824 | 1.669895501 | 0.212958556 | 0.407212272 | 10.58298967 | 17.70226298 | ENSG00000228038 |
| SRP14-AS1 | 0.739578508 | 1.669687959 | 0.020009743 | 0.068818794 | 37.02889572 | 61.62709444 | ENSG00000248508 |
| NA | 0.739531896 | 1.669634013 | 0.430334998 | 0.640113141 | 4.351750728 | 7.217394669 | ENSG00000251775 |
| ZFP3 | 0.739316693 | 1.669384977 | 0.042380261 | 0.124143202 | 27.74090797 | 46.30323863 | ENSG00000180787 |
| FAM86EP | 0.739264998 | 1.66932516 | 0.018079641 | 0.063314513 | 44.80925522 | 74.5355602 | ENSG00000251669 |
| CHRNB2 | 0.739216859 | 1.66926946 | 1.32E-05 | 0.000121854 | 132.6337303 | 221.2046495 | ENSG00000160716 |
| NA | 0.739121939 | 1.669159636 | 0.052515097 | 0.146578128 | 28.8404967 | 48.32821996 | ENSG00000247627 |
| PDGFRB | 0.73890072 | 1.668903712 | 5.17E-10 | 1.04E-08 | 297.5140141 | 496.2785761 | ENSG00000113721 |
| CAHM | 0.738393574 | 1.66831715 | 0.327775047 | 0.539201373 | 8.19612835 | 13.73349783 | ENSG00000270419 |
| KCNH7 | 0.738209849 | 1.668104706 | 0.224965669 | 0.423271978 | 9.517521125 | 15.92301612 | ENSG00000184611 |
| EIF4E1B | 0.737989776 | 1.667850268 | 0.015509214 | 0.0560022 | 38.39745136 | 64.02725624 | ENSG00000175766 |
| NA | 0.737819492 | 1.66765342 | 0.325366597 | 0.536548042 | 5.996671554 | 9.980745713 | ENSG00000272462 |
| SORCS2 | 0.737659801 | 1.667468839 | 0.003833493 | 0.017717945 | 64.4614294 | 107.5200616 | ENSG00000184985 |
| RBMS2 | 0.737552628 | 1.667344973 | 1.87E-06 | 2.05E-05 | 163.829969 | 273.3780324 | ENSG00000076067 |
| BLZF1 | 0.73720699 | 1.666945561 | 8.62E-07 | 1.01E-05 | 182.4077239 | 304.15519 | ENSG00000117475 |
| ZBTB37 | 0.736931921 | 1.666627767 | 3.38E-05 | 0.000283557 | 164.1205203 | 272.9203296 | ENSG00000185278 |
| HRH1 | 0.736761882 | 1.666431345 | 0.550951333 | 0.738132515 | 2.57754599 | 4.339426739 | ENSG00000196639 |
| TRAF1 | 0.736658354 | 1.666311767 | 0.055110392 | 0.152157858 | 29.67050814 | 49.3820343 | ENSG00000056558 |
| NA | 0.736607135 | 1.66625261 | 0.372927952 | 0.584945545 | 4.99880683 | 8.310083807 | ENSG00000229970 |
| C2CD4B | 0.736457221 | 1.666079475 | 0.458823802 | 0.664077828 | 4.81880058 | 7.935071291 | ENSG00000205502 |
| SLC50A1 | 0.736330502 | 1.665933141 | 5.47E-20 | 3.25E-18 | 714.6709086 | 1189.954244 | ENSG00000169241 |
| XG | 0.735956836 | 1.66550171 | 0.139734521 | 0.303669325 | 14.00211524 | 23.26299448 | ENSG00000124343 |
| TMEM154 | 0.735881144 | 1.665414332 | 0.174263814 | 0.354554204 | 15.41483294 | 25.58133931 | ENSG00000170006 |
| FGF2 | 0.734729943 | 1.664085941 | 2.90E-22 | 2.09E-20 | 911.7183456 | 1517.791613 | ENSG00000138685 |
| LINC01484 | 0.734502642 | 1.66382378 | 0.000579875 | 0.003492007 | 82.66462847 | 137.6252703 | ENSG00000253686 |
| C1QTNF6 | 0.734449857 | 1.663762905 | 0.002420484 | 0.011974133 | 61.34711865 | 102.019676 | ENSG00000133466 |
| KCNT2 | 0.734438059 | 1.6637493 | 2.00E-05 | 0.000177862 | 146.1239971 | 242.9016153 | ENSG00000162687 |
| CCDC121 | 0.733869481 | 1.663093732 | 0.119522094 | 0.272079887 | 18.27202425 | 30.38941882 | ENSG00000176714 |
| ETV3 | 0.733618087 | 1.662803958 | 1.40E-15 | 5.67E-14 | 721.5856352 | 1199.881316 | ENSG00000117036 |
| GIPC3 | 0.733556744 | 1.662733258 | 0.00216639 | 0.010872249 | 72.93600787 | 121.3506773 | ENSG00000179855 |
| ITGAM | 0.733478024 | 1.662642533 | 0.340403964 | 0.551995651 | 5.993112081 | 9.936574029 | ENSG00000169896 |
| SDE2 | 0.732981823 | 1.662070782 | 3.13E-12 | 8.43E-11 | 400.5264847 | 665.0652855 | ENSG00000143751 |
| MC5R | 0.732880799 | 1.661954401 | 0.413974304 | 0.625132315 | 4.573275438 | 7.636578868 | ENSG00000176136 |
| LOC102723566 | 0.731797253 | 1.660706646 | 0.374924477 | 0.586858579 | 5.601980206 | 9.324360936 | ENSG00000225032 |
| DEPP1 | 0.731331371 | 1.66017045 | 0.462569767 | 0.667504398 | 3.575410714 | 5.975564774 | ENSG00000165507 |
| CDH2 | 0.731322034 | 1.660159706 | 8.71E-15 | 3.20E-13 | 574.568778 | 953.7110928 | ENSG00000170558 |
| CCDC177 | 0.731230251 | 1.660054091 | 4.00E-07 | 5.00E-06 | 250.8419507 | 416.9213165 | ENSG00000267909 |
| NA | 0.73074317 | 1.65949372 | 0.038408114 | 0.115169457 | 31.1597542 | 51.89057015 | ENSG00000268903 |
| LOC102606465 | 0.73068898 | 1.659431388 | 0.078769415 | 0.199450066 | 20.8424513 | 34.61250471 | ENSG00000233184 |
| GAS2L1 | 0.729356178 | 1.657899067 | 7.30E-09 | 1.23E-07 | 291.9820799 | 484.6221254 | ENSG00000185340 |
| CNTN2 | 0.729107652 | 1.657613494 | 0.000209967 | 0.001435861 | 131.2960403 | 217.8659527 | ENSG00000184144 |
| DIAPH2 | 0.72879405 | 1.657253213 | 2.01E-23 | 1.59E-21 | 955.514278 | 1582.875258 | ENSG00000147202 |
| C4A | 0.728792599 | 1.657251546 | 0.318399846 | 0.528431801 | 6.629489766 | 10.98849645 | ENSG00000244731 |
| NEFH | 0.728209008 | 1.656581299 | 4.58E-05 | 0.000372651 | 118.4350067 | 195.9839451 | ENSG00000100285 |
| RPL36A | 0.72770972 | 1.65600809 | 0.042506945 | 0.124432407 | 28.04191013 | 46.49704553 | ENSG00000241343 |
| PCDHGA5 | 0.727708907 | 1.656007156 | 0.323706558 | 0.534510655 | 7.184025736 | 11.94990004 | ENSG00000253485 |
| NA | 0.727657695 | 1.655948373 | 0.471060395 | 0.674448488 | 4.638514998 | 7.689168859 | ENSG00000274356 |

| | | | | | | | |
|---|---|---|---|---|---|---|---|
| GP1BA | 0.727591959 | 1.655872922 | 0.367219242 | 0.57955335 | 5.600784998 | 9.286715643 | ENSG00000185245 |
| CHP2 | 0.727014154 | 1.655209871 | 0.01987938 | 0.06839839 | 37.88323864 | 62.57326978 | ENSG00000166869 |
| NA | 0.726972191 | 1.655161728 | 0.449346816 | 0.655789383 | 3.984339952 | 6.570657705 | ENSG00000251661 |
| SAMD11 | 0.726935906 | 1.6551201 | 0.445603803 | 0.653102663 | 4.447830331 | 7.276511051 | ENSG00000187634 |
| HYAL1 | 0.726641095 | 1.654781914 | 0.000963059 | 0.005429063 | 73.72183109 | 122.0024276 | ENSG00000114378 |
| KCTD13 | 0.726149561 | 1.654218217 | 2.74E-06 | 2.92E-05 | 171.0187494 | 283.2015026 | ENSG00000174943 |
| UAP1L1 | 0.726056851 | 1.654111918 | 1.31E-06 | 1.49E-05 | 202.9924716 | 335.6184968 | ENSG00000197355 |
| VDR | 0.725940151 | 1.653978122 | 0.245368414 | 0.4497508 | 8.625219214 | 14.24488187 | ENSG00000111424 |
| MAMLD1 | 0.725927561 | 1.653963688 | 0.257555186 | 0.464425636 | 9.231952063 | 15.29557478 | ENSG00000013619 |
| EPM2A | 0.725399331 | 1.653358215 | 0.000608447 | 0.003644315 | 79.33921824 | 131.2650452 | ENSG00000112425 |
| P3H2 | 0.725146756 | 1.653068783 | 4.98E-12 | 1.31E-10 | 411.4549691 | 680.5356551 | ENSG00000090530 |
| CALHM1 | 0.724825917 | 1.652701201 | 0.309123014 | 0.519653562 | 7.807360739 | 12.98564826 | ENSG00000185933 |
| SSTR5 | 0.724575853 | 1.652414761 | 0.057857791 | 0.157901735 | 25.36975908 | 41.78383578 | ENSG00000162009 |
| NA | 0.724355854 | 1.6521628 | 0.362156734 | 0.574006376 | 6.258519517 | 10.3753374 | ENSG00000279880 |
| DET1 | 0.724063813 | 1.651828392 | 0.31562409 | 0.525761748 | 6.643727656 | 10.96049897 | ENSG00000140543 |
| PRNP | 0.72358464 | 1.651279849 | 1.17E-12 | 3.33E-11 | 468.1582077 | 773.034516 | ENSG00000171867 |
| FOXD1 | 0.72337912 | 1.651044632 | 0.370475663 | 0.582696169 | 5.631625042 | 9.294187992 | ENSG00000251493 |
| GCK | 0.723345612 | 1.651006284 | 9.56E-07 | 1.11E-05 | 167.2458144 | 276.007995 | ENSG00000106633 |
| LOC105371814 | 0.722686187 | 1.650251818 | 0.529710295 | 0.721915199 | 2.604826561 | 4.328549421 | ENSG00000248278 |
| NA | 0.722467299 | 1.650001457 | 0.529208003 | 0.721629123 | 2.608386034 | 4.332900348 | ENSG00000273568 |
| DCAF6 | 0.72235996 | 1.649878699 | 2.09E-09 | 3.84E-08 | 282.7554673 | 466.5888833 | ENSG00000143164 |
| ICAM3 | 0.722254742 | 1.649758375 | 0.002487977 | 0.012277356 | 65.19386054 | 107.5551526 | ENSG00000076662 |
| NA | 0.722113966 | 1.649597403 | 0.1210795 | 0.274260186 | 18.77911802 | 31.06538277 | ENSG00000282012 |
| NA | 0.721889796 | 1.649341103 | 0.19060203 | 0.377149972 | 11.8972635 | 19.64777076 | ENSG00000281571 |
| LINC00909 | 0.721680677 | 1.649102048 | 0.000846035 | 0.004843096 | 80.87293185 | 133.3546078 | ENSG00000264247 |
| SH3PXD2A | 0.721380932 | 1.648759454 | 6.64E-16 | 2.80E-14 | 567.5680131 | 935.5448677 | ENSG00000107957 |
| BTN3A1 | 0.721228108 | 1.648584811 | 0.002702288 | 0.013145984 | 76.66589814 | 126.3192694 | ENSG00000026950 |
| CD163L1 | 0.721195822 | 1.648547918 | 0.53757235 | 0.727589323 | 2.624988188 | 4.349074552 | ENSG00000177675 |
| NTRK1 | 0.720996577 | 1.648320259 | 0.456501138 | 0.662631902 | 3.595572341 | 5.947567294 | ENSG00000198400 |
| CDH22 | 0.72046862 | 1.647717163 | 0.185629266 | 0.370258628 | 12.8263292 | 21.15914964 | ENSG00000149654 |
| STC1 | 0.720337349 | 1.647567244 | 5.01E-08 | 7.36E-07 | 327.1524876 | 538.1460778 | ENSG00000159167 |
| LDHD | 0.720249001 | 1.647466354 | 0.000355209 | 0.002275482 | 105.3904329 | 173.6212007 | ENSG00000166816 |
| BSND | 0.720042195 | 1.647230211 | 0.337058853 | 0.548974251 | 6.200398902 | 10.25899655 | ENSG00000162399 |
| LURAP1L | 0.719653625 | 1.646786612 | 0.442522036 | 0.650095617 | 4.066181665 | 6.638475942 | ENSG00000153714 |
| NA | 0.718848735 | 1.645868114 | 0.315595846 | 0.525761748 | 6.640168183 | 10.90138259 | ENSG00000215156 |
| IL23A | 0.718570607 | 1.645550848 | 0.101234984 | 0.239997826 | 20.08774745 | 32.94430181 | ENSG00000110944 |
| ELAC1 | 0.71817262 | 1.645096963 | 0.06240203 | 0.167181493 | 23.1617088 | 38.08149819 | ENSG00000141642 |
| COL12A1 | 0.717660471 | 1.644513067 | 0.043011615 | 0.125505926 | 50.11915781 | 82.56469356 | ENSG00000111799 |
| CCDC33 | 0.717520867 | 1.64435394 | 0.133197126 | 0.294030125 | 17.6415703 | 28.9745907 | ENSG00000140481 |
| EBF4 | 0.717336588 | 1.644143917 | 0.033314987 | 0.103123018 | 37.54338777 | 61.50037073 | ENSG00000088881 |
| OMA1 | 0.71729371 | 1.644095053 | 9.67E-05 | 0.000720122 | 112.8638665 | 185.405196 | ENSG00000162600 |
| CHRFAM7A | 0.717084017 | 1.643856104 | 0.144157533 | 0.310070047 | 14.47480949 | 23.82914397 | ENSG00000166664 |
| ZNF34 | 0.716820775 | 1.643556184 | 0.000653466 | 0.00386856 | 87.53534659 | 144.0287211 | ENSG00000196378 |
| PTAR1 | 0.716647313 | 1.643358583 | 1.33E-16 | 5.81E-15 | 702.5289573 | 1154.0591 | ENSG00000188647 |
| MB | 0.716505391 | 1.643196929 | 0.247213123 | 0.45212278 | 9.850532385 | 16.21961663 | ENSG00000198125 |
| GPATCH2 | 0.716439359 | 1.643121722 | 6.13E-09 | 1.05E-07 | 276.1719191 | 453.9286634 | ENSG00000092978 |
| TOR1AIP2 | 0.716045161 | 1.642672822 | 8.36E-13 | 2.43E-11 | 521.1882021 | 856.028543 | ENSG00000169905 |
| LYRM9 | 0.715716588 | 1.642298746 | 0.256261955 | 0.463107876 | 8.859786606 | 14.59284286 | ENSG00000232859 |
| TMEM161B-AS1 | 0.715643236 | 1.642215248 | 0.000616425 | 0.003679261 | 107.865339 | 176.8946109 | ENSG00000247828 |
| PAQR6 | 0.715548113 | 1.642106974 | 0.004975451 | 0.021989562 | 89.13785924 | 146.4375122 | ENSG00000160781 |
| PMVK | 0.715539084 | 1.642096696 | 2.98E-13 | 9.12E-12 | 542.3328763 | 891.1851294 | ENSG00000163344 |
| NA | 0.715402302 | 1.641941016 | 0.453958911 | 0.659970287 | 3.657252428 | 6.001386791 | ENSG00000279368 |
| IER5L | 0.715332239 | 1.641861279 | 0.111424138 | 0.25901612 | 17.76109167 | 29.23203287 | ENSG00000188483 |
| NA | 0.715309564 | 1.641835474 | 0.306512185 | 0.516741226 | 7.22910367 | 11.88331131 | ENSG00000285752 |
| LINC01237 | 0.714759136 | 1.641209187 | 0.385368205 | 0.597559615 | 5.860268697 | 9.683199248 | ENSG00000233806 |
| RNF32 | 0.714583704 | 1.641009629 | 0.161582315 | 0.336214414 | 15.170503 | 24.93242688 | ENSG00000105982 |
| FTCDNL1 | 0.714568817 | 1.640992696 | 0.205616168 | 0.397732391 | 11.5808544 | 19.03243624 | ENSG00000226124 |
| SCYL3 | 0.714368669 | 1.640765053 | 9.82E-05 | 0.000730429 | 108.2215127 | 177.5536227 | ENSG00000000457 |
| GUCY2C | 0.714091259 | 1.640449587 | 0.027656077 | 0.089218484 | 34.52075915 | 56.76474393 | ENSG00000070019 |
| TTC32 | 0.713970591 | 1.640312384 | 0.00350926 | 0.016436199 | 65.12297657 | 106.7138737 | ENSG00000183891 |
| PIFO | 0.713514253 | 1.63979362 | 0.006896103 | 0.028914945 | 51.85807331 | 84.87268521 | ENSG00000173947 |
| FBXW12 | 0.713436155 | 1.639704856 | 0.36211329 | 0.574002198 | 6.718450425 | 10.96579586 | ENSG00000164049 |
| TNFRSF1B | 0.713289017 | 1.639537633 | 0.000295347 | 0.001938698 | 96.39157373 | 158.1250817 | ENSG00000028137 |
| NA | 0.713201972 | 1.639438715 | 0.529269146 | 0.721629123 | 3.075435886 | 4.991460588 | ENSG00000282024 |
| ENO3 | 0.713124368 | 1.63935053 | 9.16E-08 | 1.29E-06 | 269.6524419 | 441.9220037 | ENSG00000108515 |
| NA | 0.712114854 | 1.638203809 | 0.039009638 | 0.116510679 | 29.65271077 | 48.45343062 | ENSG00000123870 |
| NA | 0.71192684 | 1.63799033 | 0.142686151 | 0.307957482 | 14.61121235 | 23.9022591 | ENSG00000277142 |
| NA | 0.711739563 | 1.637777715 | 0.318430177 | 0.528431801 | 7.950882541 | 13.0372923 | ENSG00000284707 |
| RENBP | 0.711452137 | 1.637451456 | 8.15E-08 | 1.15E-06 | 231.4768201 | 379.3620273 | ENSG00000102032 |
| PPFIBP2 | 0.711326581 | 1.637308957 | 5.89E-06 | 5.92E-05 | 168.5636739 | 275.9541029 | ENSG00000166387 |
| NA | 0.711059447 | 1.637005815 | 0.28081543 | 0.489649698 | 9.824726355 | 15.97588966 | ENSG00000286099 |
| CYP24A1 | 0.710947686 | 1.636879007 | 0.558452849 | 0.744063237 | 2.64278555 | 4.332900348 | ENSG00000019186 |
| RNF2 | 0.710940452 | 1.6368708 | 3.32E-11 | 7.96E-10 | 634.406287 | 1039.058766 | ENSG00000121481 |
| ZNF843 | 0.710891357 | 1.636815097 | 0.452003202 | 0.658191107 | 4.262790069 | 7.008191571 | ENSG00000176723 |

| | | | | | | | |
|---|---|---|---|---|---|---|---|
| TRIM72 | 0.710683083 | 1.636578816 | 0.033165618 | 0.102761722 | 36.93401132 | 60.32028418 | ENSG00000177238 |
| TICAM1 | 0.710607759 | 1.636493372 | 0.046946789 | 0.134281976 | 35.43942576 | 57.9543627 | ENSG00000127666 |
| GAB3 | 0.710402151 | 1.63626016 | 0.33230554 | 0.544094861 | 6.31308066 | 10.32246386 | ENSG00000160219 |
| CLCN1 | 0.710221485 | 1.636055267 | 0.148474773 | 0.316436754 | 15.35643299 | 25.24085067 | ENSG00000188037 |
| ZEB1 | 0.709484322 | 1.635219519 | 0.27976355 | 0.488880785 | 7.930720914 | 12.98687777 | ENSG00000148516 |
| NA | 0.709145653 | 1.6348357 | 0.525498177 | 0.718994605 | 2.62854766 | 4.307078331 | ENSG00000236539 |
| TGFB1I1 | 0.70900627 | 1.634677762 | 0.000249509 | 0.001673464 | 100.953281 | 165.0292502 | ENSG00000140682 |
| ADGRL4 | 0.708967311 | 1.634633619 | 0.259445256 | 0.466408725 | 9.864770275 | 16.1290978 | ENSG00000162618 |
| SWT1 | 0.708764554 | 1.634403902 | 0.000184855 | 0.001286067 | 106.3135749 | 173.6444254 | ENSG00000116668 |
| NA | 0.708614798 | 1.634234256 | 0.078414386 | 0.198852649 | 24.40985334 | 39.73163496 | ENSG00000240086 |
| CIITA | 0.70859874 | 1.634216065 | 0.210489349 | 0.404013758 | 11.0061568 | 17.9385176 | ENSG00000179583 |
| SNORD12C | 0.708582527 | 1.6341977 | 0.447060365 | 0.653867333 | 4.729839921 | 7.690398364 | ENSG00000209042 |
| NA | 0.708579425 | 1.634194186 | 0.135942867 | 0.298230426 | 15.88516293 | 25.98245739 | ENSG00000235726 |
| NBPF10 | 0.708548188 | 1.634158804 | 0.113491523 | 0.262079655 | 16.83794972 | 27.53677845 | ENSG00000271425 |
| COLEC12 | 0.707950048 | 1.633481424 | 0.012750059 | 0.047765297 | 50.05630866 | 81.87213359 | ENSG00000158270 |
| NA | 0.707508006 | 1.632981002 | 0.268417649 | 0.476326564 | 8.536258555 | 13.99462851 | ENSG00000267053 |
| PLCD4 | 0.70725961 | 1.632699867 | 0.009629949 | 0.037872049 | 52.64954093 | 85.93482254 | ENSG00000115556 |
| RGPD4 | 0.706834296 | 1.632218609 | 0.460149443 | 0.664999857 | 3.653692956 | 5.947567294 | ENSG00000196862 |
| NA | 0.706833331 | 1.632217517 | 0.218296603 | 0.414533066 | 11.22292683 | 18.35987726 | ENSG00000218537 |
| RGS9 | 0.706443677 | 1.631776735 | 1.85E-06 | 2.03E-05 | 169.6703554 | 276.8405721 | ENSG00000108370 |
| LINC00242 | 0.706121064 | 1.631411881 | 0.321720734 | 0.532255459 | 7.747155192 | 12.5493439 | ENSG00000229214 |
| GNG12 | 0.706026194 | 1.631304605 | 9.64E-13 | 2.77E-11 | 519.6670248 | 847.9071169 | ENSG00000172380 |
| NA | 0.705966773 | 1.631237417 | 0.193596782 | 0.381569482 | 13.402222 | 21.96545317 | ENSG00000233406 |
| MIR635 | 0.705769441 | 1.631014312 | 0.433564029 | 0.642497135 | 5.267773738 | 8.598928418 | ENSG00000207561 |
| NDUFAF1 | 0.705674697 | 1.630907204 | 8.53E-11 | 1.93E-09 | 336.2409434 | 548.2374589 | ENSG00000137806 |
| PLIN5 | 0.70561172 | 1.630836013 | 0.160655246 | 0.334764904 | 13.80759177 | 22.5742613 | ENSG00000214456 |
| PSD4 | 0.70560706 | 1.630830744 | 0.036367271 | 0.11047063 | 39.95905593 | 65.35809192 | ENSG00000125637 |
| P2RX4 | 0.705408373 | 1.630606163 | 0.000249535 | 0.001673464 | 102.9054071 | 167.6183735 | ENSG00000135124 |
| NA | 0.705285165 | 1.630466913 | 0.535420654 | 0.726186373 | 2.62854766 | 4.290904128 | ENSG00000271971 |
| EFR3A | 0.704053899 | 1.629075988 | 9.02E-15 | 3.30E-13 | 682.1429292 | 1111.323034 | ENSG00000132294 |
| SURF1 | 0.704048104 | 1.629069443 | 1.38E-10 | 3.01E-09 | 452.1161492 | 736.2928818 | ENSG00000148290 |
| NTPCR | 0.703928088 | 1.628933929 | 4.26E-21 | 2.76E-19 | 1007.547019 | 1640.362409 | ENSG00000135778 |
| STAC3 | 0.703696026 | 1.62867193 | 0.052669583 | 0.146835707 | 28.20439835 | 45.91907276 | ENSG00000185482 |
| NA | 0.703177442 | 1.628086602 | 0.575975275 | 0.758153886 | 2.69378722 | 4.374896568 | ENSG00000245711 |
| ANKRD29 | 0.703131839 | 1.628035139 | 0.008276065 | 0.033601916 | 59.26481896 | 96.52922176 | ENSG00000154065 |
| PRKG2 | 0.7030193 | 1.627908148 | 0.095479775 | 0.230188382 | 29.84962466 | 48.42325768 | ENSG00000138669 |
| JOSD2 | 0.702748405 | 1.627602504 | 0.027836315 | 0.089631512 | 46.34774908 | 75.74090167 | ENSG00000161677 |
| SLC9B2 | 0.702576928 | 1.62740906 | 0.004608914 | 0.020634908 | 58.58216813 | 95.35537806 | ENSG00000164038 |
| NFE2 | 0.7024817 | 1.627301643 | 0.07646619 | 0.195350252 | 22.88918241 | 37.22773359 | ENSG00000123405 |
| NR0B1 | 0.702207947 | 1.62699289 | 0.266309707 | 0.473743295 | 9.734291155 | 15.94666267 | ENSG00000169297 |
| SYT11 | 0.702024443 | 1.626785958 | 7.41E-08 | 1.06E-06 | 284.8772 | 463.864786 | ENSG00000132718 |
| TEX22 | 0.701901212 | 1.626647008 | 0.205875334 | 0.398085353 | 15.90740949 | 26.0381688 | ENSG00000226174 |
| TRIM56 | 0.701883189 | 1.626626688 | 6.37E-18 | 3.18E-16 | 738.3854256 | 1200.959525 | ENSG00000169871 |
| PAPPA | 0.701753864 | 1.626480881 | 0.421700147 | 0.631747364 | 4.871887182 | 7.979242975 | ENSG00000182752 |
| C1orf21 | 0.701660817 | 1.626375984 | 3.89E-18 | 1.98E-16 | 733.421297 | 1192.952314 | ENSG00000116667 |
| LIN28B-AS1 | 0.700983488 | 1.625612598 | 0.255969922 | 0.462814344 | 9.010427352 | 14.59936925 | ENSG00000203809 |
| LOC100507053 | 0.700706013 | 1.625299973 | 0.043580316 | 0.12685018 | 29.27818105 | 47.50044534 | ENSG00000246090 |
| MIR124-2HG | 0.700546851 | 1.625120676 | 0.479107712 | 0.68145793 | 4.282951695 | 6.969316773 | ENSG00000254377 |
| IRF6 | 0.699287968 | 1.623703227 | 1.29E-10 | 2.83E-09 | 403.1518301 | 654.4091226 | ENSG00000117595 |
| RTL4 | 0.699188105 | 1.623590839 | 0.510487766 | 0.70709707 | 3.264925345 | 5.331003274 | ENSG00000187823 |
| RINL | 0.698973956 | 1.623349856 | 2.51E-06 | 2.69E-05 | 189.8162235 | 308.1295785 | ENSG00000187994 |
| NCAN | 0.698908033 | 1.62327568 | 8.70E-07 | 1.02E-05 | 216.6331253 | 351.9190141 | ENSG00000130287 |
| LOC107985911 | 0.698709636 | 1.623052465 | 0.536310661 | 0.726767138 | 3.302884333 | 5.37829638 | ENSG00000283196 |
| NA | 0.698318813 | 1.622612844 | 0.178309916 | 0.360533486 | 13.54218433 | 22.0158677 | ENSG00000260066 |
| GPR156 | 0.698059863 | 1.622321626 | 0.010574412 | 0.040871423 | 51.33882657 | 83.24591144 | ENSG00000175697 |
| RPL3L | 0.69789755 | 1.622139114 | 0.233716704 | 0.434894073 | 13.78031119 | 22.44687518 | ENSG00000140986 |
| TFR2 | 0.697852703 | 1.62208869 | 6.23E-05 | 0.000487848 | 121.330157 | 196.6388169 | ENSG00000106327 |
| NA | 0.697780002 | 1.622006951 | 0.332693126 | 0.544635028 | 6.980298388 | 11.30968947 | ENSG00000287750 |
| NRP2 | 0.697163185 | 1.621313618 | 5.88E-14 | 1.93E-12 | 689.5079294 | 1118.047048 | ENSG00000118257 |
| CEP170 | 0.696949044 | 1.621072983 | 6.43E-17 | 2.89E-15 | 1001.126699 | 1622.552639 | ENSG00000143702 |
| ANKRD50 | 0.696674122 | 1.620764098 | 3.39E-13 | 1.03E-11 | 632.8958599 | 1025.241866 | ENSG00000151458 |
| IBA57 | 0.696420269 | 1.620478937 | 7.48E-08 | 1.07E-06 | 242.7815887 | 393.55806 | ENSG00000181873 |
| LHX1 | 0.695936877 | 1.619936067 | 0.321028296 | 0.531456561 | 6.956577289 | 11.25709948 | ENSG00000273706 |
| NPTXR | 0.695876611 | 1.6198684 | 4.71E-11 | 1.11E-09 | 444.011476 | 719.30965 | ENSG00000221890 |
| RGS5 | 0.695297701 | 1.619218526 | 2.61E-07 | 3.38E-06 | 219.7011112 | 355.6086254 | ENSG00000143248 |
| LMO7 | 0.695258122 | 1.619174104 | 7.91E-08 | 1.12E-06 | 233.691379 | 378.2062497 | ENSG00000136153 |
| COL11A1 | 0.695018933 | 1.618905679 | 8.93E-09 | 1.48E-07 | 315.8827028 | 510.8320926 | ENSG00000060718 |
| ADORA2A | 0.694846528 | 1.618712227 | 0.431928096 | 0.641510501 | 4.902727226 | 7.970541121 | ENSG00000128271 |
| NUP133 | 0.694753837 | 1.618608231 | 7.63E-24 | 6.32E-22 | 1837.771949 | 2974.735282 | ENSG00000069248 |
| LOC441204 | 0.694334592 | 1.618137934 | 0.506037832 | 0.70340972 | 3.251882663 | 5.291182517 | ENSG00000214870 |
| NA | 0.693706106 | 1.617433173 | 0.468114864 | 0.671437878 | 4.348191255 | 7.018785343 | ENSG00000273828 |
| PTPRD-AS1 | 0.693688434 | 1.617413361 | 0.33906504 | 0.550859767 | 7.941399332 | 12.81794693 | ENSG00000225706 |
| SDCCAG8 | 0.693683894 | 1.61740827 | 0.000950005 | 0.005369901 | 92.83215478 | 149.9352382 | ENSG00000054282 |
| GALNT12 | 0.693175722 | 1.616838659 | 7.35E-08 | 1.05E-06 | 329.6936787 | 533.3334497 | ENSG00000119514 |

| | | | | | | | |
|---|---|---|---|---|---|---|---|
| CCDC162P | 0.692595668 | 1.616188718 | 0.120390158 | 0.273166902 | 19.93354723 | 32.16694171 | ENSG00000203799 |
| SUCLG2-AS1 | 0.691768493 | 1.615262335 | 0.094510147 | 0.228823606 | 23.94605748 | 38.7745823 | ENSG00000241316 |
| NA | 0.691637775 | 1.615115989 | 0.41185134 | 0.622972571 | 4.93356727 | 7.993241715 | ENSG00000226790 |
| PDE4DIP | 0.691434056 | 1.614887939 | 9.70E-17 | 4.29E-15 | 832.5484552 | 1344.897599 | ENSG00000178104 |
| NA | 0.69143073 | 1.614884216 | 0.500560544 | 0.698882588 | 3.302884333 | 5.347177478 | ENSG00000251485 |
| COL25A1 | 0.691208458 | 1.614635435 | 0.000123501 | 0.000897389 | 116.4472859 | 188.0942024 | ENSG00000188517 |
| WDR26 | 0.690846643 | 1.614230549 | 2.31E-23 | 1.79E-21 | 1545.773774 | 2494.912034 | ENSG00000162923 |
| RFTN2 | 0.690051639 | 1.613341265 | 0.132386323 | 0.292753595 | 22.55586567 | 36.51507031 | ENSG00000162944 |
| CACNA1B | 0.689970082 | 1.613250063 | 2.04E-11 | 4.98E-10 | 376.9149386 | 607.8249109 | ENSG00000148408 |
| TUBB4A | 0.689607985 | 1.61284521 | 1.26E-16 | 5.53E-15 | 675.6594039 | 1089.899016 | ENSG00000104833 |
| SPIRE2 | 0.689301578 | 1.612502702 | 9.67E-08 | 1.35E-06 | 230.6453086 | 371.9698378 | ENSG00000204991 |
| RO60 | 0.689154658 | 1.612338498 | 2.09E-06 | 2.27E-05 | 316.0900669 | 509.9630865 | ENSG00000116747 |
| PLAAT5 | 0.688959916 | 1.612120871 | 0.00184562 | 0.009461057 | 72.34615651 | 116.5095144 | ENSG00000168004 |
| STX6 | 0.688905112 | 1.612059633 | 2.82E-20 | 1.71E-18 | 967.2532743 | 1559.95025 | ENSG00000135823 |
| PLPPR4 | 0.687988674 | 1.611035934 | 0.017907881 | 0.06289573 | 41.88062127 | 67.35458132 | ENSG00000117600 |
| LAMA3 | 0.687774898 | 1.610797232 | 0.05898779 | 0.160308594 | 28.62044653 | 45.97145184 | ENSG00000053747 |
| TNFSF4 | 0.687694511 | 1.610707481 | 0.333481914 | 0.545539816 | 7.194704154 | 11.64488123 | ENSG00000117586 |
| HLA-DRB5 | 0.687428439 | 1.61041045 | 0.009320805 | 0.036886714 | 52.24417117 | 84.10771548 | ENSG00000198502 |
| TRANK1 | 0.687386378 | 1.6103635 | 6.84E-07 | 8.15E-06 | 257.0410771 | 413.2697496 | ENSG00000168016 |
| NA | 0.687029846 | 1.609965581 | 0.296808985 | 0.506986043 | 9.364795447 | 15.05090186 | ENSG00000276900 |
| BCAN | 0.687007923 | 1.609941117 | 0.000452945 | 0.002822126 | 108.4095538 | 174.7296422 | ENSG00000132692 |
| JPH2 | 0.686937287 | 1.609862294 | 0.025915914 | 0.084676401 | 38.13440819 | 61.3249135 | ENSG00000149596 |
| NELL1 | 0.686266809 | 1.609114301 | 0.348183827 | 0.560232732 | 6.602209194 | 10.63713049 | ENSG00000165973 |
| NA | 0.686206125 | 1.609046617 | 0.051288708 | 0.143899457 | 34.92968839 | 56.32486665 | ENSG00000214485 |
| MIR34AHG | 0.686170983 | 1.609007425 | 0.006648751 | 0.02806996 | 78.30014054 | 125.9936528 | ENSG00000228526 |
| ELAVL2 | 0.686003908 | 1.6088211 | 2.57E-07 | 3.33E-06 | 271.985989 | 437.754616 | ENSG00000107105 |
| VASH2 | 0.685418139 | 1.608168012 | 8.01E-21 | 5.07E-19 | 1448.944009 | 2330.267905 | ENSG00000143494 |
| ST8SIA3 | 0.684778228 | 1.607454864 | 0.000447616 | 0.002793529 | 134.7777619 | 216.9455041 | ENSG00000177511 |
| NA | 0.684512323 | 1.607158619 | 0.314492761 | 0.524936499 | 7.689034577 | 12.32179114 | ENSG00000249631 |
| DGAT2 | 0.684496023 | 1.607140461 | 0.001087001 | 0.006030555 | 88.37247697 | 141.8707733 | ENSG00000062282 |
| CHRNA4 | 0.684121072 | 1.606722825 | 0.001768217 | 0.009148704 | 84.82554204 | 136.053997 | ENSG00000101204 |
| RASGRF2 | 0.683802717 | 1.606368313 | 2.70E-10 | 5.65E-09 | 357.4338979 | 574.0347683 | ENSG00000113319 |
| HNF4G | 0.683595607 | 1.606137724 | 0.073323927 | 0.189584112 | 27.72194155 | 44.59107496 | ENSG00000164749 |
| NA | 0.68349925 | 1.606030454 | 0.472465324 | 0.675813429 | 3.919100392 | 6.331281663 | ENSG00000228903 |
| MRC1 | 0.683058811 | 1.605540225 | 0.55228202 | 0.739107051 | 2.686668275 | 4.281256315 | ENSG00000260314 |
| NALCN | 0.683036421 | 1.605515308 | 0.195681055 | 0.384660843 | 12.66767978 | 20.29573723 | ENSG00000102452 |
| GPHA2 | 0.682906904 | 1.605371181 | 0.323576135 | 0.534435484 | 7.654635061 | 12.25737787 | ENSG00000149735 |
| SCCPDH | 0.682645712 | 1.605080564 | 7.28E-14 | 2.37E-12 | 570.8030137 | 916.461931 | ENSG00000143653 |
| SH2D3A | 0.681972528 | 1.604331782 | 3.19E-14 | 1.09E-12 | 534.4116653 | 857.3049195 | ENSG00000125731 |
| HYLS1 | 0.681735991 | 1.604068766 | 1.88E-05 | 0.000168953 | 169.0980482 | 271.2654857 | ENSG00000198331 |
| NA | 0.6817228 | 1.6040541 | 0.426414137 | 0.636390955 | 5.597225525 | 9.000992454 | ENSG00000243824 |
| ADA2 | 0.680947777 | 1.603192625 | 4.30E-16 | 1.83E-14 | 939.4856462 | 1505.309636 | ENSG00000093072 |
| NA | 0.680603533 | 1.60281013 | 0.143378184 | 0.309128882 | 17.88089238 | 28.68763801 | ENSG00000282718 |
| TEN1-CDK3 | 0.680363029 | 1.602542956 | 0.291951698 | 0.501592778 | 8.301691163 | 13.28972112 | ENSG00000261408 |
| ZNF91 | 0.680203333 | 1.602365576 | 9.89E-09 | 1.63E-07 | 280.2381269 | 449.1975985 | ENSG00000167232 |
| NA | 0.680066958 | 1.602214115 | 0.587670547 | 0.76572348 | 2.69378722 | 4.286553201 | ENSG00000265174 |
| ATRNL1 | 0.679870888 | 1.601996381 | 1.99E-05 | 0.000177222 | 155.2189615 | 248.7914637 | ENSG00000107518 |
| NUDT6 | 0.67965016 | 1.601751299 | 0.014715638 | 0.053701417 | 45.9939658 | 73.66968878 | ENSG00000170917 |
| NA | 0.679578263 | 1.601671477 | 0.177139381 | 0.358665938 | 15.90296029 | 25.43416311 | ENSG00000273145 |
| NA | 0.678564946 | 1.600546893 | 0.277112154 | 0.486332123 | 9.364795447 | 14.93021008 | ENSG00000259424 |
| TMEM63A | 0.678498453 | 1.600473127 | 0.00039935 | 0.002524007 | 1656.658209 | 2651.347259 | ENSG00000196187 |
| BTNL2 | 0.678070228 | 1.59999814 | 0.437203058 | 0.645246932 | 4.988128412 | 7.967419699 | ENSG00000204290 |
| HS1BP3 | 0.677686278 | 1.599572383 | 1.47E-10 | 3.19E-09 | 386.3918571 | 617.9692476 | ENSG00000118960 |
| EWSAT1 | 0.677587857 | 1.599463264 | 0.466662423 | 0.670694054 | 4.582758647 | 7.372326768 | ENSG00000212766 |
| COLCA2 | 0.677558743 | 1.599430986 | 0.499872764 | 0.698232038 | 3.333724377 | 5.319179997 | ENSG00000214290 |
| PLEK2 | 0.677084062 | 1.598904822 | 0.259405377 | 0.466401 | 10.0284537 | 15.99083436 | ENSG00000100558 |
| NA | 0.67704265 | 1.598858926 | 0.475211169 | 0.677873172 | 4.393269189 | 6.976789122 | ENSG00000212195 |
| APELA | 0.676096862 | 1.597811106 | 2.93E-06 | 3.10E-05 | 228.56334 | 365.669843 | ENSG00000248329 |
| PPARGC1A | 0.675997537 | 1.597701105 | 0.096855127 | 0.232563137 | 22.64869129 | 36.07999545 | ENSG00000109819 |
| RCOR3 | 0.675976165 | 1.597677437 | 4.55E-08 | 6.70E-07 | 245.750572 | 392.7100868 | ENSG00000117625 |
| PTGES | 0.675513787 | 1.59716547 | 0.003881987 | 0.017911392 | 75.46844977 | 120.7526739 | ENSG00000148344 |
| C6orf163 | 0.67549071 | 1.597139923 | 0.236450871 | 0.43845703 | 12.07162535 | 19.17722607 | ENSG00000203872 |
| ADAM19 | 0.675338559 | 1.596971492 | 1.22E-17 | 5.85E-16 | 818.3011209 | 1306.726527 | ENSG00000135074 |
| NA | 0.675312154 | 1.596942263 | 0.008665731 | 0.034893885 | 56.03398765 | 89.60575768 | ENSG00000280385 |
| MAML2 | 0.675204279 | 1.59682286 | 0.000626503 | 0.003727635 | 97.83660601 | 156.0367486 | ENSG00000184384 |
| NA | 0.675191111 | 1.596808285 | 0.567748044 | 0.751047019 | 2.901074041 | 4.679915383 | ENSG00000267422 |
| NA | 0.675141287 | 1.59675314 | 0.540979504 | 0.730264327 | 3.313562751 | 5.282480663 | ENSG00000251550 |
| NA | 0.674902104 | 1.596488438 | 0.575724608 | 0.757933721 | 2.686668275 | 4.253258835 | ENSG00000278982 |
| ASB12 | 0.674723734 | 1.596291065 | 0.276934726 | 0.48615011 | 8.945187792 | 14.27287935 | ENSG00000198881 |
| ISG15 | 0.674704823 | 1.59627014 | 1.85E-09 | 3.43E-08 | 329.3505216 | 525.7640267 | ENSG00000187608 |
| BATF2 | 0.674452788 | 1.5959913 | 0.068176565 | 0.17897059 | 30.34453932 | 48.48483307 | ENSG00000168062 |
| SH2D3C | 0.674397464 | 1.5959301 | 0.001333289 | 0.007194049 | 90.7065566 | 144.8761273 | ENSG00000095370 |
| TCEAL2 | 0.674291734 | 1.595813144 | 2.38E-09 | 4.34E-08 | 391.8935356 | 625.1775116 | ENSG00000184905 |
| SERF1A | 0.674288289 | 1.595809333 | 0.375860906 | 0.587885805 | 6.26207899 | 10.0249174 | ENSG00000172058 |

| | | | | | | | |
|---|---|---|---|---|---|---|---|
| IAH1 | 0.67408739 | 1.595587128 | 7.28E-06 | 7.19E-05 | 188.743636 | 301.2000825 | ENSG00000134330 |
| LINC01869 | 0.674052337 | 1.595548361 | 0.560910303 | 0.746076283 | 2.897514569 | 4.672443033 | ENSG00000180279 |
| SLC5A5 | 0.673936937 | 1.595420739 | 0.039881378 | 0.118457542 | 38.97123308 | 62.03975225 | ENSG00000105641 |
| ZFP36 | 0.673735599 | 1.595198103 | 0.001482885 | 0.007884334 | 95.15649802 | 152.0349 | ENSG00000128016 |
| EHHADH | 0.67367358 | 1.59512953 | 0.042237209 | 0.123762558 | 32.63771146 | 52.14082351 | ENSG00000113790 |
| LGALS8 | 0.673302011 | 1.594718754 | 6.63E-05 | 0.000516349 | 150.4840882 | 239.6190814 | ENSG00000116977 |
| CLU | 0.673104617 | 1.594500574 | 1.39E-40 | 2.80E-38 | 3877.674218 | 6184.062362 | ENSG00000120885 |
| NDST3 | 0.67296465 | 1.594345886 | 0.300885551 | 0.511018736 | 8.961789946 | 14.27287935 | ENSG00000164100 |
| CD47 | 0.672656392 | 1.594005262 | 8.73E-08 | 1.23E-06 | 286.5651405 | 456.5072359 | ENSG00000196776 |
| TNNC2 | 0.672651685 | 1.594000062 | 0.012972086 | 0.048424165 | 53.79212267 | 85.65042417 | ENSG00000101470 |
| LOC105369306 | 0.672625791 | 1.593971452 | 0.044998118 | 0.12983796 | 30.85786232 | 49.1421638 | ENSG00000233393 |
| PCDH9 | 0.672240412 | 1.59354572 | 0.186693581 | 0.371910156 | 13.52794644 | 21.58268476 | ENSG00000184226 |
| GPR89A | 0.672118368 | 1.593410921 | 0.000337731 | 0.002174584 | 129.0360731 | 205.3532558 | ENSG00000117262 |
| DES | 0.671976394 | 1.593254123 | 0.01525625 | 0.055247226 | 46.8601562 | 74.75367606 | ENSG00000175084 |
| NFIA | 0.671898663 | 1.593168283 | 0.189384837 | 0.375552859 | 14.03295528 | 22.29411854 | ENSG00000162599 |
| TTPAL | 0.671826579 | 1.593088681 | 1.18E-08 | 1.92E-07 | 301.7024386 | 480.947815 | ENSG00000124120 |
| MRPS21 | 0.671742664 | 1.592996021 | 5.06E-14 | 1.68E-12 | 1121.302933 | 1786.354424 | ENSG00000266472 |
| SDHC | 0.671740494 | 1.592993626 | 2.27E-30 | 2.73E-28 | 1906.04882 | 3035.850307 | ENSG00000143252 |
| NA | 0.670914501 | 1.592081842 | 0.360998588 | 0.573030729 | 8.72338375 | 13.99368255 | ENSG00000198580 |
| TMEM130 | 0.67089188 | 1.59205688 | 0.004211977 | 0.019204054 | 65.30417803 | 103.9060674 | ENSG00000166448 |
| FAM86B3P | 0.670870874 | 1.592033699 | 0.003453679 | 0.016232637 | 75.76263847 | 120.6696274 | ENSG00000173295 |
| ATF6 | 0.670764008 | 1.591915776 | 5.16E-13 | 1.53E-11 | 475.3410905 | 756.71057 | ENSG00000118217 |
| PLOD2 | 0.670450506 | 1.591569885 | 1.04E-07 | 1.45E-06 | 418.187658 | 665.1145525 | ENSG00000152952 |
| GLMP | 0.66999082 | 1.591062843 | 5.53E-10 | 1.11E-08 | 363.950732 | 579.1624754 | ENSG00000198715 |
| HEPHL1 | 0.669936906 | 1.591003385 | 0.55716298 | 0.74339571 | 2.92479514 | 4.693914123 | ENSG00000181333 |
| SPART | 0.669866109 | 1.590925314 | 1.01E-18 | 5.50E-17 | 922.089015 | 1466.92566 | ENSG00000133104 |
| RAB13 | 0.669765606 | 1.590814487 | 2.40E-06 | 2.58E-05 | 261.0452988 | 415.5357786 | ENSG00000143545 |
| MAPK11 | 0.669453835 | 1.590470745 | 0.014915045 | 0.05427325 | 50.60253048 | 80.68506915 | ENSG00000185386 |
| CNIH3 | 0.668632648 | 1.5895657 | 0.001449729 | 0.007736121 | 99.96164552 | 158.6582204 | ENSG00000143786 |
| NA | 0.668593877 | 1.589522984 | 0.269459095 | 0.477415889 | 10.09013379 | 15.9510136 | ENSG00000215086 |
| UEVLD | 0.667983618 | 1.588850758 | 0.002737676 | 0.013280505 | 79.30954726 | 126.0496048 | ENSG00000151116 |
| ZNF284 | 0.667958187 | 1.588822751 | 0.197356963 | 0.386948322 | 15.11978066 | 23.90566406 | ENSG00000186026 |
| CRYBA4 | 0.667829814 | 1.588681382 | 0.100148773 | 0.238290233 | 25.23691569 | 40.01858766 | ENSG00000196431 |
| SMIM10 | 0.667789644 | 1.588637148 | 0.076297578 | 0.195060888 | 32.51018142 | 51.45984622 | ENSG00000184785 |
| NKAIN3-IT1 | 0.667654959 | 1.588488845 | 0.368175026 | 0.580237451 | 6.289359561 | 10.00127084 | ENSG00000274956 |
| DAP3 | 0.66762379 | 1.588454526 | 3.54E-17 | 1.62E-15 | 1473.107572 | 2339.393845 | ENSG00000132676 |
| C1orf167 | 0.667589468 | 1.588416737 | 0.224610693 | 0.422914571 | 12.60599969 | 20.03583605 | ENSG00000215910 |
| NA | 0.66754087 | 1.588363232 | 0.379316256 | 0.591819852 | 6.292919033 | 10.00439227 | ENSG00000249406 |
| TUBB1 | 0.667401456 | 1.588209749 | 0.415141898 | 0.625944491 | 5.587742316 | 8.874057835 | ENSG00000101162 |
| C8orf34-AS1 | 0.667266088 | 1.588060734 | 0.206209947 | 0.398554379 | 12.54787908 | 19.95307312 | ENSG00000248801 |
| CX3CL1 | 0.667106619 | 1.587885207 | 0.017479292 | 0.061688737 | 47.98257631 | 76.1434172 | ENSG00000006210 |
| NMB | 0.666805451 | 1.587553764 | 8.91E-06 | 8.65E-05 | 178.1766636 | 283.1372573 | ENSG00000197696 |
| SCN3A | 0.666790096 | 1.587536867 | 0.551561105 | 0.738582061 | 2.917676195 | 4.672443033 | ENSG00000153253 |
| SAMD4A | 0.6665174 | 1.587236822 | 0.001699777 | 0.008857698 | 86.17599483 | 136.9275517 | ENSG00000020577 |
| TAF10 | 0.666394907 | 1.587102062 | 0.000539414 | 0.003277614 | 98.6695921 | 156.5321694 | ENSG00000166337 |
| ISLR2 | 0.66606728 | 1.586741683 | 0.064990418 | 0.172379782 | 29.01129908 | 46.08032034 | ENSG00000167178 |
| ANP32E | 0.666012704 | 1.586681658 | 6.96E-23 | 5.26E-21 | 3996.489267 | 6341.507058 | ENSG00000143401 |
| NA | 0.665955186 | 1.586618401 | 0.43369678 | 0.64253265 | 5.029646873 | 7.934125333 | ENSG00000286757 |
| LCTL | 0.665101506 | 1.585679835 | 0.00013029 | 0.00094021 | 124.6769241 | 197.5801898 | ENSG00000188501 |
| NA | 0.664848811 | 1.58540212 | 0.044675723 | 0.129045737 | 39.63845081 | 62.71108172 | ENSG00000260265 |
| GALNT6 | 0.664847213 | 1.585400365 | 1.38E-06 | 1.55E-05 | 215.0652921 | 340.5712278 | ENSG00000139629 |
| CLDN11 | 0.664637539 | 1.585169967 | 0.001593501 | 0.008381523 | 85.9117826 | 136.3090958 | ENSG00000013297 |
| SNORD17 | 0.664312543 | 1.584812916 | 0.114463898 | 0.263974752 | 21.03105103 | 33.23967281 | ENSG00000212232 |
| GALE | 0.664072486 | 1.584549233 | 1.89E-10 | 4.05E-09 | 516.7949073 | 818.46067 | ENSG00000117308 |
| NA | 0.664009399 | 1.584479945 | 0.163951106 | 0.339274231 | 16.54170224 | 26.20906419 | ENSG00000198406 |
| POU2F3 | 0.664003045 | 1.584472966 | 0.100786435 | 0.239325013 | 24.51513682 | 38.82934776 | ENSG00000137709 |
| NA | 0.663944131 | 1.584408264 | 0.490583017 | 0.690413613 | 4.566156493 | 7.258161384 | ENSG00000230910 |
| ZDHHC1 | 0.663762992 | 1.584209345 | 0.01011437 | 0.039399727 | 53.07270806 | 84.01456969 | ENSG00000159714 |
| RORA | 0.663728668 | 1.584171653 | 0.032982313 | 0.102278984 | 47.10568134 | 74.54114063 | ENSG00000069667 |
| NA | 0.663704545 | 1.584145165 | 0.381637243 | 0.594066876 | 6.251400572 | 9.911981517 | ENSG00000260908 |
| ORMDL2 | 0.663371938 | 1.58377999 | 7.89E-06 | 7.74E-05 | 184.9241485 | 292.7329638 | ENSG00000123353 |
| GLUD2 | 0.663181378 | 1.583570809 | 0.072857495 | 0.188765783 | 26.73711951 | 42.34222937 | ENSG00000182890 |
| DISC1 | 0.662483609 | 1.582805089 | 0.026199649 | 0.085470207 | 41.21904795 | 65.30361001 | ENSG00000162946 |
| INAVA | 0.662151163 | 1.582440399 | 7.33E-17 | 3.27E-15 | 880.9281917 | 1394.631253 | ENSG00000163362 |
| RAD51C | 0.66204245 | 1.58232116 | 4.75E-06 | 4.85E-05 | 189.74923 | 300.5642955 | ENSG00000108384 |
| CCDC172 | 0.661960558 | 1.582231346 | 0.077879634 | 0.198028394 | 25.36116559 | 40.20898959 | ENSG00000182645 |
| RBFOX1 | 0.66161899 | 1.581856785 | 0.041351729 | 0.121780719 | 35.03405599 | 55.44639395 | ENSG00000078328 |
| LOC100506691 | 0.661338062 | 1.581548789 | 0.003928494 | 0.018090613 | 76.11133601 | 120.2130814 | ENSG00000256546 |
| MPPE1 | 0.661226575 | 1.581426576 | 0.001815021 | 0.009334604 | 101.0641572 | 159.4890438 | ENSG00000154889 |
| TMEM177 | 0.660762789 | 1.580918274 | 6.74E-16 | 2.83E-14 | 954.9826501 | 1510.294507 | ENSG00000144120 |
| DCXR | 0.660228343 | 1.580332731 | 9.04E-19 | 4.95E-17 | 953.5648467 | 1507.429655 | ENSG00000169738 |
| SNAP47 | 0.660003201 | 1.580086129 | 1.07E-18 | 5.77E-17 | 992.7220621 | 1568.975407 | ENSG00000143740 |
| LOC101927322 | 0.659872146 | 1.579942601 | 0.479167586 | 0.68145793 | 3.980780479 | 6.275286703 | ENSG00000267226 |
| LOC100287049 | 0.659629059 | 1.57967641 | 0.546897178 | 0.734786085 | 3.398963937 | 5.326652347 | ENSG00000228606 |

| | | | | | | | |
|---|---|---|---|---|---|---|---|
| SCN1B | 0.659562518 | 1.579603553 | 0.020875134 | 0.071262997 | 46.25965257 | 72.89573365 | ENSG00000105711 |
| NA | 0.65955127 | 1.579591237 | 0.214806708 | 0.409801463 | 14.10531379 | 22.15195568 | ENSG00000285979 |
| PCDHGA7 | 0.659448368 | 1.579478575 | 0.526637724 | 0.719933477 | 3.38828552 | 5.305181257 | ENSG00000253537 |
| ADORA2A-AS1 | 0.659402109 | 1.579427932 | 0.474421364 | 0.67731814 | 4.606479746 | 7.285212905 | ENSG00000178803 |
| SPINK5 | 0.659025023 | 1.579015161 | 0.012632771 | 0.047403909 | 63.21119992 | 99.87177508 | ENSG00000133710 |
| APOBEC3F | 0.658159574 | 1.578068219 | 0.004597701 | 0.020589894 | 77.43928907 | 122.4317541 | ENSG00000128394 |
| KRTCAP2 | 0.657917778 | 1.577803757 | 0.018596993 | 0.064738103 | 52.11399753 | 82.19192915 | ENSG00000163463 |
| DDIT4L | 0.657760906 | 1.577632204 | 0.00116425 | 0.006406426 | 88.71469211 | 139.9001785 | ENSG00000145358 |
| SPINK1 | 0.657615195 | 1.577472872 | 0.283162133 | 0.492186485 | 9.2378758 | 14.58319505 | ENSG00000164266 |
| NUP50-DT | 0.657360278 | 1.577194164 | 0.026785776 | 0.086990436 | 53.93770325 | 85.09231412 | ENSG00000226328 |
| NA | 0.657278903 | 1.577105206 | 0.00180463 | 0.00928622 | 93.01099197 | 146.5485132 | ENSG00000203761 |
| C5AR2 | 0.656649813 | 1.576417656 | 0.42142258 | 0.631431722 | 5.229814749 | 8.276789441 | ENSG00000134830 |
| ZNFX1 | 0.655715767 | 1.575397364 | 8.34E-09 | 1.39E-07 | 349.1705229 | 550.4017746 | ENSG00000124201 |
| EMILIN1 | 0.655639849 | 1.575314465 | 7.94E-05 | 0.00060565 | 149.0491233 | 234.9879721 | ENSG00000138080 |
| TARBP1 | 0.655530779 | 1.575195373 | 3.59E-13 | 1.08E-11 | 536.7670762 | 845.2187635 | ENSG00000059588 |
| NA | 0.655483135 | 1.575143354 | 0.442210768 | 0.649844397 | 4.65155768 | 7.320682735 | ENSG00000259562 |
| PIWIL4 | 0.655408699 | 1.575062087 | 0.385646946 | 0.597653274 | 6.977934124 | 10.98319957 | ENSG00000134627 |
| ENDOG | 0.655300033 | 1.574943455 | 4.13E-05 | 0.000339305 | 176.5231755 | 278.4764695 | ENSG00000167136 |
| CENPL | 0.655223058 | 1.574859426 | 2.84E-05 | 0.00024281 | 192.0005265 | 302.067592 | ENSG00000120334 |
| CAPG | 0.654647256 | 1.574231002 | 1.10E-14 | 3.97E-13 | 686.2672059 | 1079.986583 | ENSG00000042493 |
| ANKRD33B | 0.654347619 | 1.573904079 | 3.05E-10 | 6.33E-09 | 384.1698483 | 604.5369121 | ENSG00000164236 |
| HHIPL1 | 0.65412249 | 1.573658494 | 0.000253291 | 0.00169564 | 121.7456204 | 191.7586838 | ENSG00000182218 |
| NA | 0.65405567 | 1.573585611 | 0.148757529 | 0.316798697 | 18.76963481 | 29.60080254 | ENSG00000212664 |
| ARHGAP4 | 0.65381224 | 1.573320117 | 5.99E-14 | 1.97E-12 | 723.6760487 | 1138.506024 | ENSG00000089820 |
| C1orf53 | 0.653684393 | 1.5731807 | 0.41145153 | 0.622617159 | 5.700424074 | 8.911703128 | ENSG00000203724 |
| PLEKHG4 | 0.653487268 | 1.572965762 | 1.68E-09 | 3.12E-08 | 403.7080676 | 634.831715 | ENSG00000196155 |
| EFCAB10 | 0.653405412 | 1.572876516 | 0.278573313 | 0.487748658 | 10.34486281 | 16.20968527 | ENSG00000185055 |
| LIN9 | 0.652865529 | 1.572288027 | 3.77E-07 | 4.74E-06 | 296.5758224 | 466.3755496 | ENSG00000183814 |
| WDR88 | 0.652561666 | 1.571956905 | 0.305912306 | 0.516092607 | 9.312598569 | 14.61554345 | ENSG00000166359 |
| NUDT19 | 0.652499098 | 1.571888731 | 0.028223637 | 0.090533227 | 45.1559457 | 71.08775426 | ENSG00000213965 |
| MMRN2 | 0.652427626 | 1.571810861 | 0.030702946 | 0.096718122 | 38.6237046 | 60.77210033 | ENSG00000173269 |
| NA | 0.652423193 | 1.571806032 | 0.354824943 | 0.56695736 | 6.987417333 | 10.96267444 | ENSG00000269934 |
| NA | 0.652161613 | 1.571521068 | 0.147928347 | 0.315735401 | 19.88966451 | 31.25122286 | ENSG00000261490 |
| C3 | 0.652113571 | 1.571468736 | 2.77E-07 | 3.58E-06 | 267.9627225 | 420.5921563 | ENSG00000125730 |
| C2CD4D | 0.652082575 | 1.571434974 | 0.071613762 | 0.186156353 | 31.93133953 | 50.30049572 | ENSG00000225556 |
| NA | 0.651974401 | 1.571317152 | 0.087457065 | 0.215786082 | 26.63183603 | 41.74306906 | ENSG00000233396 |
| THSD4 | 0.651355814 | 1.57064356 | 0.001161076 | 0.006390823 | 96.09877538 | 150.7368643 | ENSG00000187720 |
| NA | 0.651286052 | 1.570567612 | 0.433900158 | 0.642632302 | 5.240493167 | 8.24879196 | ENSG00000213598 |
| GRIP2 | 0.651240026 | 1.570517508 | 0.000700936 | 0.004117689 | 198.7326823 | 312.318678 | ENSG00000144596 |
| CHST13 | 0.651090048 | 1.57035425 | 0.00450909 | 0.020279629 | 78.58868426 | 123.6233374 | ENSG00000180767 |
| NA | 0.650645947 | 1.569870927 | 0.049464364 | 0.139882378 | 34.06586226 | 53.50741256 | ENSG00000229852 |
| GAL | 0.650454799 | 1.569662942 | 2.97E-12 | 8.04E-11 | 821.3350372 | 1290.243566 | ENSG00000069482 |
| CXCL5 | 0.650207592 | 1.569394002 | 0.015999781 | 0.057487796 | 72.44637982 | 113.765248 | ENSG00000163735 |
| LINC00205 | 0.650037212 | 1.56920867 | 9.89E-07 | 1.15E-05 | 256.5752742 | 402.8344358 | ENSG00000223768 |
| PRKAB2 | 0.649922461 | 1.569083862 | 2.21E-12 | 6.10E-11 | 721.8848329 | 1131.656535 | ENSG00000131791 |
| LINC01842 | 0.649722107 | 1.568865971 | 0.51271443 | 0.709022818 | 4.664600361 | 7.303562573 | ENSG00000267147 |
| IDH1-AS1 | 0.649265011 | 1.568368978 | 0.260551493 | 0.46750455 | 10.60671077 | 16.63539586 | ENSG00000231908 |
| CLDN19 | 0.6492561 | 1.56835929 | 0.000107831 | 0.000795777 | 152.4441967 | 238.7706136 | ENSG00000164007 |
| NA | 0.648705034 | 1.56776034 | 0.55191875 | 0.738889411 | 2.959194656 | 4.651917903 | ENSG00000269887 |
| GOLGA6A | 0.64846084 | 1.567494999 | 0.401444061 | 0.612708101 | 5.90059195 | 9.27706783 | ENSG00000159289 |
| TAS2R5 | 0.647772948 | 1.566747779 | 0.519662628 | 0.715241508 | 3.602691286 | 5.673667382 | ENSG00000127366 |
| PIAS3 | 0.647613358 | 1.566574476 | 6.40E-15 | 2.39E-13 | 783.1875619 | 1226.667044 | ENSG00000131788 |
| LACTB | 0.647311316 | 1.566246533 | 4.39E-08 | 6.48E-07 | 290.4145516 | 455.0095293 | ENSG00000103642 |
| IPCEF1 | 0.647168511 | 1.566091506 | 0.403527296 | 0.614813472 | 7.474628812 | 11.61376233 | ENSG00000074706 |
| SLC52A3 | 0.646981421 | 1.565888427 | 6.30E-06 | 6.27E-05 | 185.3660289 | 290.2801393 | ENSG00000101276 |
| ZBED6CL | 0.646845795 | 1.565741227 | 0.301917867 | 0.512145775 | 8.883507705 | 13.93863355 | ENSG00000188707 |
| VGLL3 | 0.646601102 | 1.565475687 | 0.003935909 | 0.018115923 | 75.30031715 | 117.8961341 | ENSG00000206538 |
| KLC3 | 0.646532634 | 1.565401393 | 0.003941032 | 0.018130667 | 85.66623131 | 134.2399428 | ENSG00000104892 |
| PLK3 | 0.646469267 | 1.565332638 | 0.00049799 | 0.003064369 | 120.6487014 | 188.8243647 | ENSG00000173846 |
| SP100 | 0.646311453 | 1.565161418 | 0.624636871 | 0.7923564 | 3.003077381 | 4.699211009 | ENSG00000067066 |
| SLC34A2 | 0.64626918 | 1.565115557 | 0.022789603 | 0.076390071 | 46.86519021 | 73.19763104 | ENSG00000157765 |
| SV2B | 0.64617105 | 1.565009105 | 0.182773559 | 0.36646333 | 15.99428521 | 24.99617774 | ENSG00000185518 |
| SPRTN | 0.646092658 | 1.564924069 | 1.83E-05 | 0.000164104 | 207.0035073 | 323.8635472 | ENSG00000010072 |
| NCALD | 0.646071488 | 1.564901105 | 0.064985779 | 0.172379782 | 30.36709137 | 47.59843652 | ENSG00000104490 |
| PBXIP1 | 0.645921691 | 1.564738628 | 3.90E-15 | 1.50E-13 | 661.1744485 | 1034.449332 | ENSG00000163346 |
| CPNE4 | 0.645878273 | 1.564691537 | 0.056670492 | 0.155487759 | 34.0519037 | 53.12681961 | ENSG00000196353 |
| NPIPB12 | 0.645782368 | 1.564587526 | 0.289366351 | 0.498742179 | 9.541242224 | 14.96908488 | ENSG00000169203 |
| LRRC9 | 0.645424223 | 1.56419917 | 0.196482947 | 0.385675134 | 16.61286554 | 25.96940461 | ENSG00000131951 |
| THAP7-AS1 | 0.645319408 | 1.564085532 | 0.27603446 | 0.485029968 | 10.81399759 | 16.98553232 | ENSG00000230513 |
| RRP15 | 0.645124802 | 1.563874565 | 2.92E-10 | 6.07E-09 | 904.284676 | 1414.801133 | ENSG00000067533 |
| ARL4A | 0.644878967 | 1.563608104 | 3.79E-08 | 5.66E-07 | 323.080765 | 505.4158674 | ENSG00000122644 |
| NA | 0.644091528 | 1.562754902 | 2.14E-05 | 0.000188798 | 179.5069554 | 280.5426691 | ENSG00000249673 |
| PSMB4 | 0.643893061 | 1.562539933 | 8.56E-39 | 1.51E-36 | 5700.126335 | 8906.72536 | ENSG00000159377 |
| RRM2B | 0.643774434 | 1.562411458 | 1.84E-18 | 9.59E-17 | 1206.047191 | 1884.324592 | ENSG00000048392 |

| | | | | | | | |
|---|---|---|---|---|---|---|---|
| METAP1D | 0.643749096 | 1.562384017 | 0.00731928 | 0.03035235 | 66.380325 | 103.6209113 | ENSG00000172878 |
| RNF157-AS1 | 0.643525529 | 1.562141921 | 0.399855882 | 0.611241569 | 6.516808008 | 10.21700033 | ENSG00000267128 |
| PTK6 | 0.643478641 | 1.562091152 | 0.414095759 | 0.625165753 | 5.955153093 | 9.306011269 | ENSG00000101213 |
| CAND2 | 0.643470952 | 1.562082827 | 7.10E-07 | 8.43E-06 | 294.7123254 | 460.7178885 | ENSG00000144712 |
| MELTF | 0.643432137 | 1.5620408 | 0.007193374 | 0.029966164 | 85.70744486 | 133.634467 | ENSG00000163975 |
| NA | 0.643148414 | 1.561733637 | 0.525707553 | 0.719210891 | 3.629971857 | 5.685490658 | ENSG00000287917 |
| C22orf42 | 0.642911906 | 1.561477635 | 0.508724026 | 0.705481952 | 3.595572341 | 5.640373016 | ENSG00000205856 |
| NES | 0.642382982 | 1.560905268 | 1.08E-43 | 2.51E-41 | 6337.300145 | 9892.377601 | ENSG00000132688 |
| VWF | 0.64166203 | 1.560125438 | 0.094750518 | 0.229196202 | 24.28293369 | 37.80324735 | ENSG00000110799 |
| IPP | 0.641423968 | 1.55986802 | 0.005605867 | 0.024314837 | 83.79533658 | 130.7349326 | ENSG00000197429 |
| ARHGAP30 | 0.640916688 | 1.559319636 | 0.17208171 | 0.351515748 | 16.79050752 | 26.23271075 | ENSG00000186517 |
| CTC1 | 0.640691836 | 1.559076626 | 6.21E-12 | 1.61E-10 | 498.4580007 | 777.10536 | ENSG00000178971 |
| NA | 0.640504816 | 1.558874532 | 0.108256258 | 0.2532416 | 43.64033494 | 68.09450703 | ENSG00000261786 |
| NA | 0.640444867 | 1.558809758 | 0.010280344 | 0.039910176 | 57.7420631 | 89.90519606 | ENSG00000274652 |
| MCMDC2 | 0.639955345 | 1.558280926 | 0.275551976 | 0.484515046 | 11.37356758 | 17.67984593 | ENSG00000178460 |
| AOC1 | 0.639859607 | 1.558177521 | 0.538611051 | 0.728316604 | 4.042460567 | 6.253815614 | ENSG00000002726 |
| HEIH | 0.639619457 | 1.55791817 | 0.375385313 | 0.587433845 | 6.582047568 | 10.27081983 | ENSG00000278970 |
| TMEM256 | 0.639507726 | 1.557797519 | 0.145132173 | 0.311704888 | 23.75776323 | 37.09164562 | ENSG00000205544 |
| MRVI1 | 0.639333374 | 1.557609269 | 0.005526591 | 0.024015112 | 71.71186375 | 111.7173255 | ENSG00000072952 |
| ACBD3 | 0.639132898 | 1.557392839 | 2.65E-15 | 1.04E-13 | 740.6322979 | 1153.73528 | ENSG00000182827 |
| C7orf25 | 0.639114293 | 1.557372755 | 0.436933864 | 0.645089622 | 5.522502756 | 8.665800696 | ENSG00000136197 |
| KLHL11 | 0.638624163 | 1.556843755 | 0.011311332 | 0.043285613 | 68.04332252 | 105.6182311 | ENSG00000178502 |
| SCRT2 | 0.637894387 | 1.556056438 | 0.227290106 | 0.426253589 | 13.03865003 | 20.20209697 | ENSG00000215397 |
| LY75 | 0.637840944 | 1.555998796 | 0.102552615 | 0.242543035 | 28.98312878 | 44.97240295 | ENSG00000054219 |
| RALGPS2 | 0.637617166 | 1.555757462 | 4.81E-08 | 7.07E-07 | 583.0842907 | 906.823972 | ENSG00000116191 |
| CD2BP2-DT | 0.637254684 | 1.555366622 | 0.13855257 | 0.301830117 | 20.24995634 | 31.43649586 | ENSG00000260219 |
| NA | 0.637046899 | 1.555142625 | 0.005743191 | 0.024790892 | 67.70527724 | 105.165301 | ENSG00000250320 |
| DENND6B | 0.637038466 | 1.555133535 | 0.092718874 | 0.22538241 | 28.31827532 | 43.98103733 | ENSG00000205593 |
| TMEM267 | 0.63698 | 1.555070514 | 0.003255957 | 0.01543005 | 75.75076485 | 117.8095147 | ENSG00000151881 |
| HUS1B | 0.636645539 | 1.554710043 | 0.583526365 | 0.762833684 | 2.993594172 | 4.646621017 | ENSG00000188996 |
| ABCA11P | 0.636229338 | 1.554261591 | 0.47753133 | 0.680052723 | 4.269909014 | 6.662122495 | ENSG00000251595 |
| FRAT1 | 0.636208597 | 1.554239246 | 0.002337169 | 0.011605953 | 91.4651254 | 142.3232518 | ENSG00000165879 |
| NA | 0.636166482 | 1.554193876 | 0.370259458 | 0.582445426 | 8.58014128 | 13.3650117 | ENSG00000227598 |
| PLEKHA4 | 0.635787454 | 1.553785609 | 3.52E-07 | 4.43E-06 | 355.4586361 | 551.7300263 | ENSG00000105559 |
| ABRAXAS1 | 0.635480312 | 1.553454851 | 3.62E-07 | 4.55E-06 | 296.4363926 | 460.4842085 | ENSG00000163322 |
| STAB1 | 0.635230408 | 1.553185784 | 0.160445526 | 0.334475523 | 21.02868677 | 32.58858492 | ENSG00000010327 |
| FBXL6 | 0.635036617 | 1.552977165 | 1.12E-08 | 1.83E-07 | 519.2323674 | 807.1325785 | ENSG00000182325 |
| ARHGEF11 | 0.634104735 | 1.551974373 | 2.27E-15 | 8.98E-14 | 983.7080503 | 1525.893143 | ENSG00000132694 |
| NRAV | 0.633972824 | 1.551832476 | 0.000545059 | 0.003305535 | 123.2751641 | 191.1096987 | ENSG00000248008 |
| LOC102546294 | 0.633702211 | 1.551541419 | 0.572582658 | 0.755060645 | 4.774917855 | 7.3689218 | ENSG00000247199 |
| PIGC | 0.633359903 | 1.551173329 | 1.48E-10 | 3.22E-09 | 468.8210547 | 727.0872778 | ENSG00000135845 |
| WIF1 | 0.633156538 | 1.550954688 | 0.013297193 | 0.049374792 | 60.36201728 | 93.75783127 | ENSG00000156076 |
| DPH6 | 0.632953507 | 1.550736437 | 0.000202237 | 0.001393377 | 128.1254151 | 198.6004989 | ENSG00000134146 |
| NA | 0.632561979 | 1.550315645 | 0.400040682 | 0.611474567 | 6.656770337 | 10.30193873 | ENSG00000255145 |
| SQSTM1 | 0.632443836 | 1.550188695 | 4.89E-22 | 3.44E-20 | 1428.728399 | 2215.492566 | ENSG00000161011 |
| CLPSL2 | 0.632357543 | 1.550095974 | 0.195770157 | 0.384795942 | 15.69803773 | 24.26232691 | ENSG00000196748 |
| TM6SF1 | 0.632278414 | 1.550010957 | 0.528174155 | 0.721043079 | 3.622852912 | 5.618901926 | ENSG00000136404 |
| NA | 0.632273941 | 1.550006151 | 0.419060605 | 0.629503762 | 6.051232696 | 9.338359676 | ENSG00000229666 |
| ETNPPL | 0.632124755 | 1.549845877 | 0.039034594 | 0.11656678 | 38.90571419 | 60.36568537 | ENSG00000164089 |
| NA | 0.632053023 | 1.549768819 | 0.109333105 | 0.255191655 | 25.5826903 | 39.7456337 | ENSG00000260400 |
| TRIM11 | 0.631570089 | 1.54925013 | 1.39E-11 | 3.46E-10 | 535.8692078 | 830.3365089 | ENSG00000154370 |
| H19 | 0.631542049 | 1.549220019 | 0.02921351 | 0.092956551 | 42.42447883 | 65.6742716 | ENSG00000130600 |
| GORAB | 0.631328378 | 1.548990588 | 1.26E-05 | 0.000117149 | 205.6127568 | 318.8982352 | ENSG00000120370 |
| TNS1 | 0.631222438 | 1.548876846 | 2.67E-05 | 0.00023084 | 165.0511123 | 255.4803541 | ENSG00000079308 |
| ZNF669 | 0.631144067 | 1.548792709 | 4.94E-06 | 5.03E-05 | 222.8527961 | 345.3680512 | ENSG00000188295 |
| NA | 0.63109497 | 1.548740003 | 0.522278565 | 0.716804556 | 3.657252428 | 5.658722683 | ENSG00000215003 |
| NANOS1 | 0.630675041 | 1.548289273 | 0.009787341 | 0.038315493 | 74.76952843 | 115.7585434 | ENSG00000188613 |
| CERS2 | 0.630538872 | 1.548143145 | 1.34E-25 | 1.26E-23 | 2184.760526 | 3382.177691 | ENSG00000143418 |
| MPI | 0.630439532 | 1.548036548 | 1.97E-11 | 4.83E-10 | 539.5987402 | 835.4357968 | ENSG00000178802 |
| ACTN2 | 0.630311854 | 1.547899552 | 0.025084002 | 0.08270101 | 47.62820821 | 73.75434363 | ENSG00000077522 |
| SV2A | 0.630141442 | 1.547716724 | 1.42E-22 | 1.05E-20 | 2621.080884 | 4055.78361 | ENSG00000159164 |
| TATDN3 | 0.629905617 | 1.547463754 | 2.66E-06 | 2.83E-05 | 214.9671532 | 332.8919177 | ENSG00000203705 |
| COL28A1 | 0.629833436 | 1.547386332 | 0.366200898 | 0.578768429 | 7.318064329 | 11.30874351 | ENSG00000215018 |
| NA | 0.62972556 | 1.547270633 | 0.409871703 | 0.62092308 | 6.681686644 | 10.31064059 | ENSG00000258433 |
| SRD5A3 | 0.629622481 | 1.547160086 | 1.90E-08 | 3.00E-07 | 358.1690766 | 554.4377909 | ENSG00000128039 |
| LIFR | 0.629443209 | 1.546967845 | 1.95E-05 | 0.000174556 | 205.2260224 | 317.3111439 | ENSG00000113594 |
| NA | 0.629277905 | 1.546790603 | 0.35355053 | 0.565638915 | 8.618100269 | 13.33729777 | ENSG00000231747 |
| NA | 0.629179543 | 1.546685147 | 0.515821965 | 0.71213523 | 4.331589101 | 6.698821829 | ENSG00000261474 |
| PYROXD1 | 0.629012477 | 1.546506049 | 0.016253487 | 0.058189314 | 57.46241778 | 88.69181515 | ENSG00000121350 |
| ZNF365 | 0.62894925 | 1.546438275 | 0.447041247 | 0.653867333 | 5.591301789 | 8.676678014 | ENSG00000138311 |
| NA | 0.628937072 | 1.546425221 | 0.534257944 | 0.725234422 | 4.066181665 | 6.221467206 | ENSG00000258704 |
| SERPINF2 | 0.628708741 | 1.546180492 | 0.028020794 | 0.090056618 | 46.7403555 | 72.41575206 | ENSG00000167711 |
| PRKN | 0.628580946 | 1.546043537 | 0.409785828 | 0.620855246 | 7.083496937 | 10.8637373 | ENSG00000185345 |
| RPS27L | 0.628564634 | 1.546026056 | 1.33E-09 | 2.50E-08 | 683.9723316 | 1058.002813 | ENSG00000185088 |

| | | | | | | | |
|---|---|---|---|---|---|---|---|
| NA | 0.62855885 | 1.546019857 | 0.401729515 | 0.613063587 | 7.754274137 | 11.9125383 | ENSG00000280195 |
| NA | 0.628397295 | 1.545846741 | 0.118257514 | 0.270243606 | 21.4657863 | 33.21479676 | ENSG00000278600 |
| NA | 0.628332886 | 1.545777729 | 0.349066824 | 0.561170342 | 10.02341969 | 15.6005936 | ENSG00000273654 |
| PDGFC | 0.628312971 | 1.545756392 | 0.119033082 | 0.271252028 | 26.50995039 | 40.7944347 | ENSG00000145431 |
| IARS2 | 0.628099279 | 1.545527451 | 1.95E-32 | 2.55E-30 | 2563.617958 | 3962.282442 | ENSG00000067704 |
| RNASEH2A | 0.627965607 | 1.545384258 | 0.296255303 | 0.506438845 | 11.14700885 | 17.28430829 | ENSG00000104889 |
| NA | 0.627950502 | 1.545368078 | 0.003718866 | 0.017259961 | 79.24077438 | 122.5818408 | ENSG00000282508 |
| NA | 0.627481836 | 1.544866139 | 0.567415216 | 0.751025817 | 3.695211417 | 5.694192512 | ENSG00000284669 |
| NUMBL | 0.627279169 | 1.544649135 | 1.98E-11 | 4.85E-10 | 508.8411541 | 785.749406 | ENSG00000105245 |
| SUMF2 | 0.62716371 | 1.544525521 | 6.93E-15 | 2.57E-13 | 1101.599401 | 1700.422528 | ENSG00000129103 |
| NA | 0.626637847 | 1.543962644 | 0.445187695 | 0.65276507 | 5.60434447 | 8.675448509 | ENSG00000250999 |
| C5orf24 | 0.626627895 | 1.543951993 | 0.000327595 | 0.002117984 | 170.4728325 | 263.6758937 | ENSG00000181904 |
| ATP2A3 | 0.626503126 | 1.543818473 | 9.23E-16 | 3.82E-14 | 1028.883371 | 1588.525572 | ENSG00000074370 |
| NA | 0.626483954 | 1.543797957 | 0.517235148 | 0.713304058 | 3.653692956 | 5.628549739 | ENSG00000279989 |
| LINC02035 | 0.626299462 | 1.543600549 | 0.180143142 | 0.362957777 | 17.27060005 | 26.64848998 | ENSG00000273033 |
| C15orf56 | 0.626198657 | 1.543492697 | 0.553506217 | 0.739980526 | 3.602691286 | 5.560731502 | ENSG00000176753 |
| ELFN1 | 0.626126834 | 1.543415857 | 0.001919544 | 0.009792163 | 111.3194745 | 171.7900262 | ENSG00000225968 |
| NA | 0.625840874 | 1.543109964 | 0.342452414 | 0.554269269 | 7.981722585 | 12.29814458 | ENSG00000235499 |
| ACOT4 | 0.625826128 | 1.543094192 | 0.139582423 | 0.303408616 | 21.49899061 | 33.17592196 | ENSG00000177465 |
| NA | 0.625777028 | 1.543041675 | 0.427058938 | 0.636929126 | 5.580623371 | 8.639978679 | ENSG00000214578 |
| LRRN2 | 0.625717048 | 1.542977525 | 3.79E-08 | 5.66E-07 | 300.6920899 | 464.0194345 | ENSG00000170382 |
| DNAH14 | 0.625245838 | 1.542473643 | 4.44E-07 | 5.50E-06 | 268.1457034 | 413.5508813 | ENSG00000185842 |
| SPATA12 | 0.625067997 | 1.542283514 | 0.542082054 | 0.731177781 | 3.688092472 | 5.669316454 | ENSG00000186451 |
| CR1 | 0.624464889 | 1.541638909 | 0.340896783 | 0.552461663 | 9.313793777 | 14.35346682 | ENSG00000203710 |
| AARD | 0.624454243 | 1.541627533 | 0.266003494 | 0.473433051 | 14.94274907 | 23.13823533 | ENSG00000205002 |
| NAGLU | 0.624438527 | 1.541610739 | 5.77E-07 | 6.96E-06 | 298.7539485 | 460.9332095 | ENSG00000108784 |
| NA | 0.624351329 | 1.541517566 | 0.144112217 | 0.3100432 | 22.54668795 | 34.70035362 | ENSG00000265749 |
| ZNF552 | 0.624171668 | 1.541325609 | 0.000856599 | 0.004892435 | 114.3735536 | 176.1587526 | ENSG00000178935 |
| LTC4S | 0.624149947 | 1.541302404 | 0.307437754 | 0.517738517 | 11.92810355 | 18.41029179 | ENSG00000213316 |
| SPI1 | 0.623918901 | 1.541055586 | 0.456901485 | 0.662717707 | 4.947805159 | 7.634403404 | ENSG00000066336 |
| QRICH2 | 0.623857993 | 1.540990527 | 0.046670878 | 0.133663866 | 38.36661131 | 59.15185296 | ENSG00000129646 |
| LOC100129129 | 0.623819602 | 1.540949521 | 0.415868416 | 0.626739688 | 6.254960045 | 9.667025044 | ENSG00000255020 |
| NUDT18 | 0.62351006 | 1.540618933 | 0.072013972 | 0.186965061 | 32.38534671 | 49.95783161 | ENSG00000275074 |
| ZNF18 | 0.623310803 | 1.540406166 | 3.49E-05 | 0.00029164 | 165.0840373 | 254.3449337 | ENSG00000154957 |
| ZBTB42 | 0.623284544 | 1.540378128 | 0.00250843 | 0.0123434 | 89.17342781 | 137.2304677 | ENSG00000179627 |
| SKOR2 | 0.622866481 | 1.539931824 | 0.426554402 | 0.63654523 | 5.587742316 | 8.625979939 | ENSG00000215474 |
| C5orf47 | 0.622860614 | 1.539925561 | 0.488400994 | 0.688779985 | 4.300749058 | 6.617950811 | ENSG00000185056 |
| PLB1 | 0.622761248 | 1.539819503 | 0.587350493 | 0.765666704 | 3.602691286 | 5.544557299 | ENSG00000163803 |
| NBPF26 | 0.622752549 | 1.539810218 | 0.007821082 | 0.032081044 | 70.03521316 | 107.6822552 | ENSG00000273136 |
| PIGM | 0.622366886 | 1.539398649 | 8.79E-06 | 8.54E-05 | 236.0856641 | 363.62729 | ENSG00000143315 |
| FAM157A | 0.622307459 | 1.53933524 | 0.014243262 | 0.052270502 | 77.7601218 | 119.4900353 | ENSG00000236438 |
| THEM5 | 0.622223458 | 1.539245615 | 0.019019355 | 0.065904683 | 50.23895851 | 77.37514779 | ENSG00000196407 |
| NA | 0.622065431 | 1.539077021 | 1.14E-05 | 0.000107194 | 188.0870967 | 289.4078142 | ENSG00000274265 |
| MAGEB17 | 0.621908602 | 1.538909724 | 0.431521443 | 0.641322299 | 6.265638462 | 9.662674117 | ENSG00000182798 |
| NA | 0.621524369 | 1.538499921 | 0.601213248 | 0.776335877 | 4.07330061 | 6.214940815 | ENSG00000287760 |
| NA | 0.621349132 | 1.538313058 | 0.487359233 | 0.688133017 | 4.300749058 | 6.608302998 | ENSG00000279800 |
| RNF187 | 0.621217091 | 1.538172272 | 2.30E-26 | 2.26E-24 | 2402.666389 | 3695.854524 | ENSG00000168159 |
| CFAP99 | 0.62121567 | 1.538170756 | 0.554786713 | 0.741242673 | 3.868098721 | 6.005737718 | ENSG00000206113 |
| RDM1 | 0.621213771 | 1.538168733 | 0.016181794 | 0.058006721 | 72.80725647 | 111.8420847 | ENSG00000278023 |
| EPRS1 | 0.621021815 | 1.537964086 | 1.89E-33 | 2.67E-31 | 4491.209372 | 6906.850064 | ENSG00000136628 |
| NA | 0.620985156 | 1.537925007 | 0.452464331 | 0.658557911 | 4.971526258 | 7.644051217 | ENSG00000271366 |
| INKA1 | 0.62083061 | 1.537760269 | 0.016537708 | 0.059016366 | 58.8926535 | 90.5547709 | ENSG00000185614 |
| GPSM1 | 0.620492796 | 1.537400237 | 5.26E-11 | 1.23E-09 | 593.2372992 | 912.2451107 | ENSG00000160360 |
| CSMD1 | 0.620260808 | 1.53715304 | 0.433062287 | 0.642298067 | 5.607903943 | 8.630330866 | ENSG00000183117 |
| ZMYM1 | 0.620207656 | 1.537096409 | 9.90E-06 | 9.47E-05 | 218.0164775 | 335.2202665 | ENSG00000197056 |
| MYH11 | 0.619924095 | 1.536794324 | 0.143894703 | 0.309786988 | 20.57348439 | 31.60448074 | ENSG00000133392 |
| COCH | 0.61984655 | 1.536711722 | 1.26E-05 | 0.000117149 | 229.9949724 | 353.3149684 | ENSG00000100473 |
| ZNF808 | 0.619260077 | 1.536087157 | 0.035885713 | 0.109324356 | 57.35001535 | 87.98193738 | ENSG00000198482 |
| LYL1 | 0.619202855 | 1.536026233 | 0.258001802 | 0.464808695 | 13.15461193 | 20.24939008 | ENSG00000104903 |
| PRKAG1 | 0.619169507 | 1.535990728 | 3.39E-09 | 5.98E-08 | 418.4186659 | 642.5345764 | ENSG00000181929 |
| MUC16 | 0.618955291 | 1.535762675 | 0.607613847 | 0.781288725 | 3.020874744 | 4.612097146 | ENSG00000181143 |
| CRABP1 | 0.618528494 | 1.535308413 | 2.00E-13 | 6.23E-12 | 790.1332076 | 1212.907606 | ENSG00000166426 |
| MYO15A | 0.618164596 | 1.534921204 | 0.28642047 | 0.495567068 | 13.93451141 | 21.40554653 | ENSG00000091536 |
| SRR | 0.617111825 | 1.533801541 | 0.06502687 | 0.172443379 | 33.87250784 | 52.05125064 | ENSG00000167720 |
| TMEM26 | 0.617049745 | 1.533735543 | 0.086819601 | 0.214577695 | 31.13750764 | 47.72516023 | ENSG00000196932 |
| TP53BP2 | 0.616509552 | 1.533161369 | 3.04E-18 | 1.56E-16 | 1176.526156 | 1802.971074 | ENSG00000143514 |
| BPHL | 0.616371333 | 1.53301449 | 7.88E-08 | 1.12E-06 | 363.1446693 | 556.2041006 | ENSG00000137274 |
| LGALS3 | 0.616075415 | 1.532700078 | 0.030732227 | 0.096773804 | 62.45621675 | 95.80312683 | ENSG00000131981 |
| APOBEC3C | 0.615619157 | 1.532215433 | 2.14E-08 | 3.34E-07 | 371.9810659 | 570.037976 | ENSG00000244509 |
| TMEFF1 | 0.615521853 | 1.532112094 | 2.43E-05 | 0.00021208 | 251.7902889 | 385.5862894 | ENSG00000241697 |
| YY1AP1 | 0.615304605 | 1.531881399 | 2.05E-11 | 5.00E-10 | 544.1886963 | 833.6962551 | ENSG00000163374 |
| HPDL | 0.615227581 | 1.531799615 | 2.56E-09 | 4.62E-08 | 560.5025917 | 859.2022311 | ENSG00000186603 |
| GBAP1 | 0.615141911 | 1.531708657 | 0.044801626 | 0.129350054 | 49.50177269 | 75.93565452 | ENSG00000160766 |
| ADORA1 | 0.615106 | 1.53167053 | 0.000960285 | 0.005418281 | 140.9911517 | 215.7369685 | ENSG00000163485 |

| | | | | | | | |
|---|---|---|---|---|---|---|---|
| GNMT | 0.614623685 | 1.531158555 | 0.439070448 | 0.647129674 | 6.340361231 | 9.698427493 | ENSG00000124713 |
| PLK5 | 0.614537226 | 1.531066798 | 0.237939855 | 0.44023652 | 14.06943973 | 21.64303065 | ENSG00000185988 |
| NA | 0.614273469 | 1.530786909 | 0.106908095 | 0.25098948 | 27.02416311 | 41.28550452 | ENSG00000213839 |
| PROCR | 0.614118233 | 1.530622204 | 0.000855316 | 0.004890061 | 132.6061966 | 202.9828678 | ENSG00000101000 |
| MYH6 | 0.614058535 | 1.530558868 | 0.415301489 | 0.626135128 | 6.916254036 | 10.59730973 | ENSG00000197616 |
| EDA2R | 0.613723047 | 1.530202989 | 0.001798002 | 0.009264738 | 120.5903014 | 184.7160113 | ENSG00000131080 |
| PGBD2 | 0.612774335 | 1.529197063 | 0.000379028 | 0.002410081 | 135.074568 | 206.6618424 | ENSG00000185220 |
| OCEL1 | 0.612595168 | 1.529007166 | 1.10E-05 | 0.000103649 | 229.0516949 | 350.00862 | ENSG00000099330 |
| CEP19 | 0.612546835 | 1.528955942 | 0.132655807 | 0.293143565 | 28.67737194 | 43.7355864 | ENSG00000174007 |
| FAM227B | 0.612054319 | 1.528434067 | 0.033643376 | 0.103969098 | 48.33841894 | 73.70828003 | ENSG00000166262 |
| NUAK2 | 0.611675543 | 1.528032833 | 5.25E-06 | 5.32E-05 | 219.9030847 | 336.2786427 | ENSG00000163545 |
| AFF3 | 0.611528561 | 1.527877165 | 0.191976257 | 0.379193768 | 23.01401713 | 34.97692345 | ENSG00000144218 |
| ACP5 | 0.611138335 | 1.527463955 | 0.003298587 | 0.015608571 | 95.517654 | 145.6093157 | ENSG00000102575 |
| JAG2 | 0.610818328 | 1.527125181 | 5.20E-10 | 1.04E-08 | 473.9292625 | 723.9601704 | ENSG00000184916 |
| NA | 0.610737741 | 1.527039881 | 0.627208761 | 0.793729923 | 2.632107133 | 4.03970481 | ENSG00000272843 |
| ABCB10 | 0.61059249 | 1.526886146 | 2.26E-06 | 2.44E-05 | 268.5866934 | 410.6121886 | ENSG00000135776 |
| GRHL3 | 0.61057387 | 1.526866439 | 0.300641468 | 0.510754566 | 10.86619447 | 16.62886947 | ENSG00000158055 |
| DDHD2 | 0.610332722 | 1.526611244 | 1.54E-06 | 1.72E-05 | 284.8469453 | 434.3454013 | ENSG00000085788 |
| NA | 0.609783893 | 1.526030602 | 0.368365895 | 0.580392952 | 7.610752336 | 11.61376233 | ENSG00000253930 |
| INKA2-AS1 | 0.609753034 | 1.52599796 | 0.556171355 | 0.742544206 | 3.904862502 | 5.9948604 | ENSG00000227811 |
| ADCK5 | 0.609544342 | 1.525777235 | 0.000168012 | 0.001178205 | 151.6969429 | 231.6193562 | ENSG00000173137 |
| AGMAT | 0.609369734 | 1.525592583 | 3.07E-05 | 0.000261196 | 211.5525029 | 323.1116732 | ENSG00000116771 |
| SLC35D3 | 0.609233793 | 1.525448837 | 0.273903913 | 0.48261573 | 13.34438072 | 20.31938378 | ENSG00000182747 |
| MATN4 | 0.609076951 | 1.525283007 | 0.186871474 | 0.372146766 | 20.56547572 | 31.18841796 | ENSG00000124159 |
| NA | 0.608995755 | 1.525197166 | 0.486169641 | 0.687121215 | 5.009485247 | 7.610756851 | ENSG00000213236 |
| RAB37 | 0.608523359 | 1.524697837 | 0.012790369 | 0.047840389 | 68.92672546 | 105.2029463 | ENSG00000172794 |
| NA | 0.608424249 | 1.524593097 | 0.44370985 | 0.651211433 | 6.37832022 | 9.695022524 | ENSG00000261349 |
| FAM89A | 0.6076619 | 1.523787684 | 0.100848163 | 0.239381291 | 30.29948754 | 46.27261418 | ENSG00000182118 |
| CPT1C | 0.607523043 | 1.52364103 | 2.00E-15 | 8.03E-14 | 1253.174562 | 1909.384306 | ENSG00000169169 |
| ZSCAN31 | 0.607475963 | 1.523591308 | 0.009383233 | 0.037052928 | 71.72106763 | 109.4755008 | ENSG00000235109 |
| NA | 0.607328517 | 1.523435602 | 0.310548926 | 0.521122843 | 11.09008345 | 16.99205871 | ENSG00000256542 |
| LINC01844 | 0.607204692 | 1.523304853 | 0.01955578 | 0.067454046 | 56.40287296 | 85.79861897 | ENSG00000236714 |
| BHLHE23 | 0.607090642 | 1.523184436 | 0.389662922 | 0.601355258 | 7.579912292 | 11.54376863 | ENSG00000125533 |
| TPCN1 | 0.607074305 | 1.523167188 | 1.47E-07 | 1.97E-06 | 519.215383 | 790.4169537 | ENSG00000186815 |
| AGPAT3 | 0.606978229 | 1.523065756 | 0.000409949 | 0.002583212 | 158.5802981 | 241.3807136 | ENSG00000160216 |
| ARID4B | 0.606951478 | 1.523037514 | 1.84E-10 | 3.97E-09 | 529.0418627 | 805.2273953 | ENSG00000054267 |
| ETS1 | 0.606889793 | 1.522972396 | 1.70E-18 | 8.88E-17 | 1541.090382 | 2346.323469 | ENSG00000134954 |
| ASPG | 0.606810313 | 1.522888495 | 0.090936657 | 0.221993119 | 35.07112526 | 53.30473585 | ENSG00000166183 |
| MAN2B2 | 0.606599309 | 1.522665779 | 9.64E-06 | 9.24E-05 | 205.8517478 | 313.4681268 | ENSG00000013288 |
| TLE2 | 0.606478367 | 1.522538138 | 6.97E-08 | 9.99E-07 | 376.0179599 | 572.8060865 | ENSG00000065717 |
| NA | 0.606372815 | 1.522426749 | 0.538470423 | 0.728178586 | 3.94282149 | 6.031559734 | ENSG00000258603 |
| SYNPR | 0.606351505 | 1.522404262 | 0.292550751 | 0.502308753 | 12.43311239 | 18.80968591 | ENSG00000163630 |
| CEBPA | 0.605585426 | 1.521596071 | 0.111691178 | 0.259381197 | 26.6353955 | 40.42661099 | ENSG00000245848 |
| INHA | 0.605390783 | 1.521390797 | 0.535815185 | 0.726437264 | 3.946380963 | 6.031559734 | ENSG00000123999 |
| ZNF490 | 0.605203209 | 1.521193004 | 0.114893952 | 0.264622779 | 24.39414091 | 37.15556443 | ENSG00000188033 |
| SOX6 | 0.604972655 | 1.520949924 | 0.476831592 | 0.679293925 | 5.056927445 | 7.641875753 | ENSG00000110693 |
| FZD7 | 0.604817924 | 1.520786809 | 1.58E-24 | 1.35E-22 | 6608.104681 | 10050.82822 | ENSG00000155760 |
| CACNA1F | 0.604782774 | 1.520749757 | 0.212395187 | 0.406432578 | 17.81000841 | 26.97280442 | ENSG00000102001 |
| ZNF814 | 0.604222869 | 1.520159674 | 5.10E-07 | 6.25E-06 | 270.3389571 | 411.1058854 | ENSG00000204514 |
| CSNK2A3 | 0.604199114 | 1.520134644 | 0.505577911 | 0.702948074 | 4.551918604 | 6.95966896 | ENSG00000254598 |
| NA | 0.603976678 | 1.519900287 | 0.261636283 | 0.468785693 | 14.2639632 | 21.68909425 | ENSG00000286989 |
| NA | 0.603836309 | 1.519752413 | 0.536645845 | 0.726856257 | 3.939262018 | 6.013210067 | ENSG00000267745 |
| MTX1 | 0.603793847 | 1.519707684 | 3.50E-09 | 6.15E-08 | 603.8717714 | 917.2627924 | ENSG00000173171 |
| WWTR1-AS1 | 0.603724057 | 1.51963417 | 0.571088846 | 0.753841189 | 3.960618853 | 6.03685662 | ENSG00000241313 |
| MX1 | 0.603304537 | 1.519192341 | 0.013297336 | 0.049374792 | 62.34706831 | 94.78684219 | ENSG00000157601 |
| KCNF1 | 0.603243526 | 1.519128096 | 0.030193515 | 0.095495651 | 46.80114586 | 71.01492268 | ENSG00000162975 |
| SSPN | 0.603126458 | 1.519004831 | 0.297862886 | 0.508112549 | 11.7300206 | 17.9073987 | ENSG00000123096 |
| ARNT | 0.602961308 | 1.518830956 | 6.59E-14 | 2.16E-12 | 799.2465018 | 1213.596371 | ENSG00000143437 |
| TSTD1 | 0.602685944 | 1.518541087 | 0.001287227 | 0.006973401 | 143.6579638 | 217.9712912 | ENSG00000215845 |
| PLEKHF2 | 0.602570337 | 1.518419408 | 0.006645581 | 0.02806996 | 111.4650812 | 169.1309389 | ENSG00000175895 |
| NA | 0.602303624 | 1.518138721 | 0.028914058 | 0.092174543 | 52.35420933 | 79.59985236 | ENSG00000260077 |
| CKS1B | 0.602279736 | 1.518113584 | 1.09E-21 | 7.54E-20 | 3357.33 | 5095.685991 | ENSG00000173207 |
| PCCA-DT | 0.602137793 | 1.517964229 | 6.59E-05 | 0.00051368 | 207.7858233 | 315.4549684 | ENSG00000274605 |
| TRIM6 | 0.602033912 | 1.517854931 | 1.20E-05 | 0.000112154 | 220.8446088 | 335.0283515 | ENSG00000121236 |
| SHANK1 | 0.601926316 | 1.517741734 | 0.007865846 | 0.032222667 | 74.11532723 | 112.4835247 | ENSG00000161681 |
| LOC101927483 | 0.601834903 | 1.517645569 | 0.474263554 | 0.67728881 | 5.714661964 | 8.628155403 | ENSG00000286595 |
| ACVRL1 | 0.601742754 | 1.517548636 | 0.349426357 | 0.561562143 | 8.990265726 | 13.61526506 | ENSG00000139567 |
| AGRN | 0.601714437 | 1.51751885 | 1.13E-34 | 1.70E-32 | 8341.292866 | 12658.95489 | ENSG00000188157 |
| NA | 0.6016784 | 1.517480945 | 0.154081128 | 0.325133498 | 22.65817449 | 34.34200977 | ENSG00000275557 |
| LGALS9 | 0.600562211 | 1.516307349 | 0.448664207 | 0.655281496 | 6.409160263 | 9.667025044 | ENSG00000168961 |
| NBEAL2 | 0.600513679 | 1.516256341 | 1.29E-18 | 6.86E-17 | 1909.830601 | 2895.999937 | ENSG00000160796 |
| LOC100505797 | 0.60049856 | 1.516240451 | 0.42879595 | 0.638613353 | 7.685475105 | 11.61934276 | ENSG00000263958 |
| H3C11 | 0.600366382 | 1.516101542 | 0.611261501 | 0.783844194 | 3.302884333 | 5.01851211 | ENSG00000275379 |
| PLAAT1 | 0.600298138 | 1.516029827 | 0.004226013 | 0.019244819 | 91.29549209 | 138.5429537 | ENSG00000127252 |

| | | | | | | | |
|---|---|---|---|---|---|---|---|
| ONECUT2 | 0.600229132 | 1.515957315 | 0.111772877 | 0.259475204 | 28.14243893 | 42.82270542 | ENSG00000119547 |
| NA | 0.599473182 | 1.515163185 | 0.610010647 | 0.782665336 | 3.095597513 | 4.630446813 | ENSG00000275481 |
| PRRC2C | 0.59891644 | 1.51457859 | 2.50E-05 | 0.000217863 | 4530.777469 | 6861.688364 | ENSG00000117523 |
| NA | 0.598878784 | 1.514539058 | 0.533856757 | 0.72489825 | 3.946380963 | 5.996089905 | ENSG00000274220 |
| COQ6 | 0.598670637 | 1.514320562 | 0.334897676 | 0.546869227 | 12.82899894 | 19.33963052 | ENSG00000119723 |
| NDUFB1 | 0.598491122 | 1.514132146 | 1.69E-07 | 2.24E-06 | 442.986837 | 670.8257737 | ENSG00000183648 |
| TUBB8 | 0.598023237 | 1.513641173 | 0.464895999 | 0.669294471 | 7.72935783 | 11.66010947 | ENSG00000261456 |
| RGS6 | 0.597805971 | 1.513413239 | 0.15637222 | 0.328376446 | 23.5647143 | 35.66442713 | ENSG00000182732 |
| NA | 0.597786703 | 1.513393027 | 0.369360515 | 0.581613339 | 8.370490196 | 12.61621618 | ENSG00000239407 |
| CCDC114 | 0.597356165 | 1.512941459 | 0.094682568 | 0.229182298 | 30.90502518 | 46.87953039 | ENSG00000105479 |
| LEMD1-DT | 0.596698171 | 1.512251583 | 0.049911486 | 0.140893633 | 41.02955849 | 62.04410317 | ENSG00000224717 |
| SIAE | 0.596681927 | 1.512234556 | 0.00465048 | 0.020781902 | 97.02881556 | 146.3904596 | ENSG00000110013 |
| SSBP4 | 0.596625034 | 1.512174922 | 2.02E-10 | 4.31E-09 | 798.1004647 | 1207.133659 | ENSG00000130511 |
| B3GNT8 | 0.596564743 | 1.512111729 | 0.269264983 | 0.477336825 | 13.87163612 | 21.01246789 | ENSG00000177191 |
| TBX19 | 0.596494364 | 1.512037965 | 0.257106486 | 0.463974588 | 12.96392726 | 19.56500782 | ENSG00000143178 |
| CCNA1 | 0.596097526 | 1.51162211 | 0.112317673 | 0.260228509 | 30.31400476 | 45.70691619 | ENSG00000133101 |
| TMEM139 | 0.595907481 | 1.511422999 | 0.612727736 | 0.784552396 | 2.659387704 | 4.023530606 | ENSG00000178826 |
| KLK6 | 0.595875799 | 1.511389807 | 0.297038328 | 0.507286021 | 11.02275895 | 16.60522291 | ENSG00000167755 |
| DCAF8 | 0.595863071 | 1.511376473 | 5.91E-06 | 5.93E-05 | 300.5034907 | 453.858073 | ENSG00000132716 |
| SEZ6L | 0.595801302 | 1.511311766 | 0.430953684 | 0.640680179 | 6.582047568 | 9.970151941 | ENSG00000100095 |
| PRR35 | 0.595639856 | 1.511142651 | 0.133684731 | 0.294796374 | 24.99255961 | 37.84931095 | ENSG00000161992 |
| RHOB | 0.595241856 | 1.510725825 | 1.08E-07 | 1.50E-06 | 362.2008326 | 547.427991 | ENSG00000143878 |
| PLXNA2 | 0.594578476 | 1.510031322 | 0.024830182 | 0.082050391 | 64.57975556 | 97.66029124 | ENSG00000076356 |
| NA | 0.594334679 | 1.509776168 | 0.015943849 | 0.05730864 | 67.40043628 | 101.7848189 | ENSG00000229335 |
| NA | 0.594315653 | 1.509756258 | 0.232804286 | 0.433708657 | 16.56898281 | 25.03694445 | ENSG00000230629 |
| KCNA3 | 0.594048718 | 1.509476941 | 0.278649091 | 0.487770759 | 12.56092176 | 18.98391363 | ENSG00000177272 |
| VPS72 | 0.594020825 | 1.509447756 | 4.06E-17 | 1.85E-15 | 1737.858325 | 2622.123431 | ENSG00000163159 |
| KRCC1 | 0.593954768 | 1.509378645 | 0.000538551 | 0.00327448 | 136.6889543 | 206.2592112 | ENSG00000172086 |
| CXCL3 | 0.593355487 | 1.508751794 | 0.038111949 | 0.114426733 | 43.30966179 | 65.35837546 | ENSG00000163734 |
| TMOD1 | 0.593055394 | 1.508437994 | 0.241084802 | 0.444661245 | 15.1942241 | 22.94492289 | ENSG00000136842 |
| NA | 0.593042531 | 1.508424545 | 0.539665105 | 0.729167597 | 3.984339952 | 6.013210067 | ENSG00000279529 |
| GSTO2 | 0.592885418 | 1.508260283 | 0.000543051 | 0.003295476 | 140.6249623 | 212.0029879 | ENSG00000065621 |
| CCS | 0.592576869 | 1.507937746 | 6.43E-09 | 1.10E-07 | 653.8646978 | 986.5312291 | ENSG00000173992 |
| GADD45B | 0.591697268 | 1.507018647 | 0.001895441 | 0.009690147 | 110.3094568 | 166.3628284 | ENSG00000099860 |
| DCAF12L1 | 0.591693684 | 1.507014903 | 0.000619946 | 0.003693276 | 135.4906423 | 204.2230893 | ENSG00000198889 |
| NA | 0.591477587 | 1.506789189 | 0.406078895 | 0.61705825 | 7.75783361 | 11.61376233 | ENSG00000198416 |
| TADA2A | 0.591321691 | 1.506626375 | 1.16E-05 | 0.000108859 | 289.0554274 | 435.1001037 | ENSG00000276234 |
| C15orf40 | 0.591231118 | 1.506531792 | 0.000200875 | 0.001385255 | 162.0133299 | 244.2003861 | ENSG00000169609 |
| RTN4IP1 | 0.59118447 | 1.50648308 | 7.73E-07 | 9.11E-06 | 300.3922829 | 452.4311677 | ENSG00000130347 |
| STIMATE-MUSTN1 | 0.591026569 | 1.506318207 | 0.385510313 | 0.597632758 | 8.60861706 | 12.97164952 | ENSG00000248592 |
| JUN | 0.591004212 | 1.506294864 | 5.91E-06 | 5.93E-05 | 363.9734849 | 548.2357779 | ENSG00000177606 |
| LOC107985206 | 0.590849915 | 1.506133774 | 0.44162067 | 0.649302802 | 6.596285458 | 9.94310042 | ENSG00000286391 |
| JAKMIP3 | 0.590707254 | 1.505984847 | 0.024030859 | 0.079786002 | 56.2095447 | 84.68807462 | ENSG00000188385 |
| LINC01456 | 0.590545644 | 1.505816157 | 0.334313221 | 0.546292653 | 11.07020115 | 16.59652106 | ENSG00000225882 |
| COQ5 | 0.590446557 | 1.505712738 | 2.53E-09 | 4.57E-08 | 508.7304794 | 765.3445894 | ENSG00000110871 |
| PARS2 | 0.590332584 | 1.505593791 | 2.43E-06 | 2.62E-05 | 278.248295 | 419.0730215 | ENSG00000162396 |
| NA | 0.590017469 | 1.505264973 | 0.543725919 | 0.732348903 | 4.00806105 | 6.027208807 | ENSG00000286001 |
| ZNF425 | 0.589958295 | 1.505203234 | 9.67E-05 | 0.000719809 | 184.6285114 | 278.0468595 | ENSG00000204947 |
| SCN8A | 0.58917908 | 1.504390478 | 1.04E-09 | 2.00E-08 | 502.8444303 | 756.2886862 | ENSG00000196876 |
| NA | 0.588678461 | 1.503868541 | 0.643832051 | 0.805803452 | 2.686668275 | 4.033178419 | ENSG00000279196 |
| FOXG1 | 0.588667427 | 1.503857039 | 0.483169885 | 0.68431553 | 5.315215936 | 7.98794483 | ENSG00000176165 |
| MBTPS2 | 0.58831047 | 1.503484995 | 0.000824461 | 0.004736824 | 154.5256323 | 232.4178609 | ENSG00000012174 |
| PDE12 | 0.588079696 | 1.503244516 | 3.75E-07 | 4.71E-06 | 377.9128028 | 568.3906115 | ENSG00000174840 |
| NA | 0.587987819 | 1.503148786 | 0.038905967 | 0.116274599 | 57.49651181 | 86.41612895 | ENSG00000223459 |
| C1orf35 | 0.5877365 | 1.502886959 | 1.20E-12 | 3.40E-11 | 868.6115214 | 1306.042297 | ENSG00000143793 |
| TRPV6 | 0.587680659 | 1.50282879 | 0.121970013 | 0.275790673 | 29.01040935 | 43.49621036 | ENSG00000165125 |
| NFKBIZ | 0.587483394 | 1.502623317 | 0.001102252 | 0.00610083 | 124.2217478 | 186.615907 | ENSG00000144802 |
| MYO7A | 0.586982996 | 1.502102222 | 0.002921948 | 0.014069606 | 106.416468 | 159.9959089 | ENSG00000137474 |
| CST6 | 0.586971652 | 1.502090412 | 0.4442432 | 0.651664815 | 5.993112081 | 8.989169178 | ENSG00000175315 |
| NA | 0.586770519 | 1.501881012 | 0.323305745 | 0.53426925 | 10.56994699 | 15.86919662 | ENSG00000225205 |
| NA | 0.586617516 | 1.501721741 | 0.577255769 | 0.759282257 | 4.184507827 | 6.341875435 | ENSG00000265566 |
| RPS6KC1 | 0.586199065 | 1.501286233 | 6.56E-08 | 9.46E-07 | 470.6195653 | 705.9819774 | ENSG00000136643 |
| NA | 0.586164211 | 1.501249963 | 0.175475538 | 0.355945814 | 29.83271702 | 44.82782403 | ENSG00000237886 |
| IMPDH2 | 0.585699974 | 1.500766963 | 2.53E-27 | 2.63E-25 | 4142.803297 | 6217.597914 | ENSG00000178035 |
| GAMT | 0.585434873 | 1.500491215 | 1.13E-08 | 1.85E-07 | 429.8696254 | 645.1330437 | ENSG00000130005 |
| F12 | 0.585084755 | 1.500127116 | 0.040660023 | 0.120279533 | 47.81085806 | 71.62278485 | ENSG00000131187 |
| TARS2 | 0.58485982 | 1.499893244 | 2.19E-16 | 9.48E-15 | 1210.661454 | 1815.522419 | ENSG00000143374 |
| RNF115 | 0.58483558 | 1.499868044 | 2.93E-10 | 6.09E-09 | 542.2412726 | 813.3092936 | ENSG00000265491 |
| RTL10 | 0.584074504 | 1.499077015 | 2.21E-10 | 4.69E-09 | 532.13985 | 797.6489642 | ENSG00000215012 |
| CRYL1 | 0.583552448 | 1.498534655 | 0.094889779 | 0.229328138 | 30.63633761 | 45.80831234 | ENSG00000165475 |
| TRPV2 | 0.583459383 | 1.498437991 | 0.118836346 | 0.270904935 | 28.2127125 | 42.25873139 | ENSG00000187688 |
| PPFIA4 | 0.582991194 | 1.497951792 | 0.150251764 | 0.319262984 | 139.5126102 | 208.951425 | ENSG00000143847 |
| ZNF670 | 0.582923042 | 1.49788103 | 0.000117915 | 0.00086044 | 214.377659 | 321.4429759 | ENSG00000277462 |
| DDX25 | 0.58274272 | 1.497693823 | 3.91E-05 | 0.000323092 | 202.0023362 | 302.710799 | ENSG00000109832 |

| | | | | | | | |
|---|---|---|---|---|---|---|---|
| CXCL14 | 0.582504667 | 1.497446715 | 0.400924087 | 0.612310747 | 7.968679903 | 11.90913333 | ENSG00000145824 |
| ASPDH | 0.582493697 | 1.497435329 | 0.581060463 | 0.761150628 | 3.364564421 | 5.019458068 | ENSG00000204653 |
| COL6A2 | 0.582473287 | 1.497414144 | 1.18E-18 | 6.33E-17 | 3292.694526 | 4931.503538 | ENSG00000142173 |
| DESI2 | 0.582370268 | 1.497307222 | 4.83E-15 | 1.83E-13 | 2740.870312 | 4103.451163 | ENSG00000121644 |
| NA | 0.582156867 | 1.497085759 | 0.261773366 | 0.468843082 | 13.92739247 | 20.80901318 | ENSG00000271936 |
| GALNTL6 | 0.581798195 | 1.496713611 | 0.36961414 | 0.581680252 | 10.74519856 | 16.13127326 | ENSG00000174473 |
| SNAPIN | 0.581725484 | 1.496638179 | 1.43E-07 | 1.92E-06 | 350.721956 | 524.5762569 | ENSG00000143553 |
| TNXB | 0.581534562 | 1.496440131 | 0.000244253 | 0.001642703 | 160.7242779 | 240.4318386 | ENSG00000168477 |
| ARPC5 | 0.581168914 | 1.49606091 | 3.63E-21 | 2.37E-19 | 2335.101449 | 3493.533399 | ENSG00000162704 |
| JUND | 0.581094703 | 1.495983956 | 4.09E-14 | 1.38E-12 | 1052.198364 | 1574.271434 | ENSG00000130522 |
| SLC7A2 | 0.581087902 | 1.495976904 | 1.91E-09 | 3.54E-08 | 518.264044 | 774.8521205 | ENSG00000003989 |
| MYEF2 | 0.580943245 | 1.495826912 | 8.87E-13 | 2.56E-11 | 992.5381664 | 1483.663514 | ENSG00000104177 |
| FAM86B1 | 0.580605398 | 1.495476663 | 0.062923929 | 0.168340898 | 40.72263259 | 60.93885571 | ENSG00000186523 |
| IDS | 0.580008614 | 1.494858174 | 0.000393808 | 0.00249624 | 151.6263644 | 226.3931111 | ENSG00000010404 |
| TYMP | 0.579949422 | 1.494796843 | 0.614999161 | 0.785862315 | 3.523213836 | 5.317004534 | ENSG00000025708 |
| NA | 0.579120895 | 1.493938641 | 0.524424332 | 0.718312522 | 4.893244017 | 7.358328028 | ENSG00000268496 |
| NA | 0.578723885 | 1.493527586 | 0.464999035 | 0.669294471 | 6.623566029 | 9.869985297 | ENSG00000268521 |
| ZMIZ1-AS1 | 0.578515333 | 1.493311701 | 0.641946869 | 0.804214244 | 2.683108803 | 3.991182199 | ENSG00000224596 |
| NA | 0.57849684 | 1.49329256 | 0.313380448 | 0.523687264 | 10.8863561 | 16.27967897 | ENSG00000251615 |
| RNPEP | 0.578027056 | 1.492806379 | 5.73E-17 | 2.58E-15 | 1074.616606 | 1604.47088 | ENSG00000176393 |
| CACNA1E | 0.577997899 | 1.492776209 | 0.133480631 | 0.294552667 | 24.44158311 | 36.51100293 | ENSG00000198216 |
| THNSL1 | 0.577837242 | 1.492609984 | 0.000374002 | 0.002382936 | 150.160865 | 224.3223927 | ENSG00000185875 |
| EFCAB12 | 0.577596094 | 1.492360514 | 0.607929428 | 0.781467763 | 3.371683366 | 5.009810256 | ENSG00000172771 |
| NPTX2 | 0.577542064 | 1.492304625 | 3.98E-10 | 8.12E-09 | 770.8677666 | 1150.921283 | ENSG00000106236 |
| CITED1 | 0.577497484 | 1.492258513 | 0.553263005 | 0.739863889 | 3.973661534 | 5.912097464 | ENSG00000125931 |
| NA | 0.577442649 | 1.492201796 | 0.417495464 | 0.628430894 | 9.370719184 | 13.93740404 | ENSG00000214992 |
| HLA-E | 0.577360406 | 1.492116732 | 1.73E-14 | 6.00E-13 | 994.6050848 | 1483.358778 | ENSG00000204592 |
| KIF12 | 0.576830337 | 1.491568606 | 0.085426543 | 0.212189959 | 35.68942625 | 53.30662777 | ENSG00000136883 |
| IL17D | 0.576664634 | 1.491397299 | 0.250698085 | 0.456242973 | 17.98617586 | 26.92863274 | ENSG00000172458 |
| RABGAP1L | 0.576531732 | 1.491259917 | 4.34E-15 | 1.65E-13 | 1229.742385 | 1833.805928 | ENSG00000152061 |
| KTN1-AS1 | 0.576155346 | 1.490870911 | 0.021042543 | 0.071730761 | 66.02150771 | 98.17016222 | ENSG00000186615 |
| NAT9 | 0.575950705 | 1.490659452 | 1.57E-09 | 2.95E-08 | 748.8624947 | 1116.521413 | ENSG00000109065 |
| ABCC10 | 0.57588978 | 1.490596503 | 2.96E-08 | 4.51E-07 | 461.5887226 | 688.3933986 | ENSG00000124574 |
| DDTL | 0.575787659 | 1.490490995 | 0.106515948 | 0.250224367 | 30.26389282 | 45.22097531 | ENSG00000099974 |
| PYGO2 | 0.575671678 | 1.490371176 | 5.64E-15 | 2.11E-13 | 1037.22625 | 1545.758358 | ENSG00000163348 |
| CCDC169 | 0.57551307 | 1.490207336 | 0.028894784 | 0.09215977 | 63.53709224 | 94.61507348 | ENSG00000242715 |
| LOC105372493 | 0.574912229 | 1.489586836 | 0.665363659 | 0.82085806 | 2.879717206 | 4.330724884 | ENSG00000286787 |
| B3GALNT1 | 0.574154211 | 1.488804386 | 0.001044556 | 0.005820724 | 132.2266067 | 197.0691619 | ENSG00000169255 |
| CSPG4 | 0.574121384 | 1.488770511 | 0.00030385 | 0.001988291 | 160.1374012 | 238.4523741 | ENSG00000173546 |
| NA | 0.57390844 | 1.488550782 | 0.571886983 | 0.754459355 | 4.197550509 | 6.292406865 | ENSG00000278383 |
| ACSBG1 | 0.573844188 | 1.488484489 | 0.007935985 | 0.032446652 | 94.8453761 | 141.0881163 | ENSG00000103740 |
| GTPBP10 | 0.573521784 | 1.48815189 | 6.12E-05 | 0.000481348 | 207.5613234 | 309.1017212 | ENSG00000105793 |
| NA | 0.572887542 | 1.487497807 | 0.464268704 | 0.668593683 | 6.224120001 | 9.289837065 | ENSG00000259032 |
| C1orf43 | 0.572641418 | 1.487244062 | 9.32E-24 | 7.62E-22 | 2132.752755 | 3171.269799 | ENSG00000143612 |
| MGST2 | 0.572393012 | 1.486988007 | 0.01408606 | 0.051774085 | 69.27783957 | 102.9757827 | ENSG00000085871 |
| NA | 0.572388097 | 1.486982942 | 0.632801124 | 0.797821895 | 2.890395624 | 4.346899088 | ENSG00000230773 |
| BEST4 | 0.572352932 | 1.486946698 | 0.078431931 | 0.198870437 | 34.1678656 | 50.75247982 | ENSG00000142959 |
| MIXL1 | 0.572270633 | 1.486861876 | 0.258248433 | 0.465051236 | 21.80683238 | 32.5615334 | ENSG00000185155 |
| UROS | 0.572199911 | 1.486788991 | 2.62E-12 | 7.14E-11 | 701.7924342 | 1043.517764 | ENSG00000188690 |
| LINC02076 | 0.572198251 | 1.486787281 | 0.344398578 | 0.556370157 | 10.44658682 | 15.60711999 | ENSG00000220161 |
| NA | 0.571923944 | 1.486504617 | 0.218389482 | 0.414542705 | 18.46151371 | 27.45401551 | ENSG00000217165 |
| NA | 0.571863423 | 1.48644226 | 0.436320767 | 0.644810651 | 6.88185452 | 10.25682109 | ENSG00000275494 |
| ADCY10 | 0.571695163 | 1.486268907 | 0.423406088 | 0.633448781 | 6.93285619 | 10.32246386 | ENSG00000143199 |
| PDCD4 | 0.571623463 | 1.486195043 | 9.58E-06 | 9.20E-05 | 253.3642735 | 376.3633676 | ENSG00000150593 |
| LYRM4-AS1 | 0.571330585 | 1.485893365 | 0.369589525 | 0.581680252 | 9.85765133 | 14.70700824 | ENSG00000272142 |
| ALG10 | 0.571248427 | 1.48580875 | 0.165675174 | 0.341494833 | 23.99944956 | 35.70063201 | ENSG00000139133 |
| LGI4 | 0.571227891 | 1.4857876 | 0.046292966 | 0.132782907 | 63.73161571 | 94.69608277 | ENSG00000153902 |
| ELOA-AS1 | 0.571027162 | 1.485580889 | 0.364368652 | 0.576499831 | 9.850532385 | 14.69518497 | ENSG00000236810 |
| HIVEP1 | 0.571015015 | 1.485568382 | 1.02E-06 | 1.18E-05 | 329.600827 | 490.0384757 | ENSG00000095951 |
| MMP19 | 0.570898437 | 1.485448344 | 0.238470813 | 0.441046059 | 17.69969092 | 26.21218562 | ENSG00000123342 |
| NA | 0.57018211 | 1.484710973 | 0.144431329 | 0.310446807 | 29.52459592 | 43.79905371 | ENSG00000236333 |
| DCP1B | 0.570004077 | 1.484527766 | 6.77E-07 | 8.07E-06 | 320.3169318 | 475.1866776 | ENSG00000151065 |
| NA | 0.569886036 | 1.484406307 | 0.479155115 | 0.68145793 | 5.5533428 | 8.276789441 | ENSG00000230457 |
| SYCE1L | 0.569419701 | 1.483926567 | 0.019545592 | 0.067431221 | 67.93628516 | 100.8263687 | ENSG00000205078 |
| LOC101927741 | 0.569287903 | 1.483791008 | 0.353759921 | 0.565878037 | 9.657483454 | 14.30617371 | ENSG00000287299 |
| NSUN7 | 0.569177823 | 1.483677797 | 0.000342777 | 0.002204066 | 164.2463227 | 243.641498 | ENSG00000179299 |
| PCYOX1L | 0.569024763 | 1.483520397 | 1.07E-08 | 1.75E-07 | 478.0238944 | 708.6763962 | ENSG00000145882 |
| FCHO2 | 0.568849029 | 1.483339702 | 0.002855564 | 0.013792156 | 128.2386811 | 190.403918 | ENSG00000157107 |
| NA | 0.568747324 | 1.483235135 | 1.98E-08 | 3.11E-07 | 408.2786995 | 605.4628888 | ENSG00000224032 |
| PCDHGA4 | 0.568550988 | 1.483033295 | 0.371318195 | 0.58329264 | 8.925026166 | 13.22842927 | ENSG00000262576 |
| EFNA4 | 0.568381512 | 1.482859091 | 1.22E-05 | 0.000114106 | 232.6351143 | 344.8901294 | ENSG00000243364 |
| NA | 0.568354472 | 1.482831299 | 0.464941602 | 0.669294471 | 6.241917363 | 9.274892367 | ENSG00000235008 |
| CALCRL | 0.568353971 | 1.482830783 | 0.326773991 | 0.5382579 | 11.59153282 | 17.15831962 | ENSG00000064989 |
| FAM86C2P | 0.568161212 | 1.482632675 | 0.018665752 | 0.064929588 | 70.36171587 | 104.5344547 | ENSG00000160172 |

| | | | | | | | |
|---|---|---|---|---|---|---|---|
| SLC5A3 | 0.5678108 | 1.482272606 | 1.66E-08 | 2.65E-07 | 528.4436971 | 782.7668479 | ENSG00000198743 |
| PARP1 | 0.567772769 | 1.482233533 | 4.22E-43 | 9.38E-41 | 18644.37223 | 27634.61701 | ENSG00000143799 |
| GOLT1A | 0.567665761 | 1.482123596 | 0.017985049 | 0.063097109 | 68.80367077 | 102.0184465 | ENSG00000174567 |
| AIM2 | 0.567591734 | 1.482047548 | 0.209516267 | 0.402719054 | 17.93901299 | 26.54832333 | ENSG00000163568 |
| WDR92 | 0.567314731 | 1.481763017 | 0.0333844 | 0.103304015 | 60.30626092 | 89.53096155 | ENSG00000243667 |
| CADM2 | 0.567238101 | 1.481684314 | 0.004433903 | 0.019989078 | 106.5220308 | 157.8036051 | ENSG00000175161 |
| NMRAL1 | 0.567061823 | 1.481503283 | 2.86E-13 | 8.76E-12 | 860.1078568 | 1274.466506 | ENSG00000153406 |
| LUZP2 | 0.567033062 | 1.481473748 | 0.128921023 | 0.287209334 | 29.84784464 | 44.30391135 | ENSG00000187398 |
| HMGCL | 0.566688183 | 1.481119642 | 0.0013468 | 0.007260726 | 213.8980478 | 316.1988459 | ENSG00000117305 |
| COL16A1 | 0.566478869 | 1.480904768 | 0.000178947 | 0.001246557 | 302.3992756 | 447.8435084 | ENSG00000084636 |
| COL20A1 | 0.566265854 | 1.480686128 | 0.492962365 | 0.692446816 | 8.306445844 | 12.28944273 | ENSG00000101203 |
| C2orf88 | 0.566129926 | 1.480546627 | 0.353962374 | 0.566010113 | 10.28318272 | 15.19228672 | ENSG00000187699 |
| NA | 0.56548486 | 1.479884784 | 0.43219509 | 0.641566127 | 10.19333234 | 14.9949069 | ENSG00000225913 |
| PTPRN2 | 0.565191276 | 1.479583663 | 0.000197953 | 0.001366855 | 207.3750362 | 306.5727555 | ENSG00000155093 |
| NUFIP2 | 0.565007962 | 1.479395674 | 4.13E-17 | 1.88E-15 | 2129.229691 | 3148.817341 | ENSG00000108256 |
| SERTAD1 | 0.564639665 | 1.479018056 | 0.000266702 | 0.001775353 | 185.3704519 | 274.3204239 | ENSG00000197019 |
| NA | 0.564535478 | 1.47891125 | 0.583918898 | 0.763005183 | 4.252111651 | 6.310756533 | ENSG00000260532 |
| LOC102724788 | 0.564273097 | 1.478642307 | 8.50E-05 | 0.000643319 | 204.6415622 | 302.7773877 | ENSG00000277196 |
| PHOSPHO1 | 0.564082768 | 1.478447248 | 0.302694165 | 0.512817736 | 12.778887 | 18.9711444 | ENSG00000173868 |
| SERPINB6 | 0.563485374 | 1.477835176 | 1.01E-07 | 1.41E-06 | 445.5193056 | 657.7468735 | ENSG00000124570 |
| ABHD11 | 0.563389175 | 1.477736637 | 1.73E-05 | 0.000156145 | 286.0782339 | 423.0530858 | ENSG00000106077 |
| NA | 0.563257982 | 1.477602264 | 0.644631225 | 0.806145123 | 3.613369703 | 5.357771249 | ENSG00000273062 |
| NA | 0.562989626 | 1.47732744 | 0.396256344 | 0.608200671 | 8.359811778 | 12.31214332 | ENSG00000286403 |
| LOC101926933 | 0.562980574 | 1.477318171 | 0.309192977 | 0.519692741 | 16.70065714 | 24.62457019 | ENSG00000257285 |
| STRADA | 0.562752692 | 1.477084838 | 0.137828779 | 0.300773803 | 27.97814511 | 41.26403343 | ENSG00000266173 |
| MCAM | 0.562702527 | 1.477033478 | 1.23E-09 | 2.33E-08 | 524.19873 | 774.1207943 | ENSG00000076706 |
| TRIM5 | 0.562702042 | 1.477032982 | 4.04E-05 | 0.000332213 | 250.3543565 | 369.5125671 | ENSG00000132256 |
| CCDC113 | 0.562637951 | 1.476967367 | 0.000679387 | 0.004004441 | 146.8384039 | 216.7636587 | ENSG00000103021 |
| EPS8L2 | 0.56259775 | 1.476926212 | 2.13E-07 | 2.78E-06 | 585.6511321 | 865.1400686 | ENSG00000177106 |
| GNPAT | 0.562359411 | 1.476682237 | 1.28E-14 | 4.56E-13 | 1087.286929 | 1605.447426 | ENSG00000116906 |
| FZD6 | 0.562259046 | 1.476579512 | 0.001117889 | 0.006178326 | 146.7518075 | 216.6811067 | ENSG00000164930 |
| NA | 0.56202984 | 1.476344941 | 0.148762025 | 0.316798697 | 29.18713546 | 42.93904627 | ENSG00000260776 |
| A2M-AS1 | 0.56196649 | 1.476280115 | 0.628379636 | 0.794839211 | 2.928354612 | 4.358722365 | ENSG00000245105 |
| LOC100130357 | 0.561909904 | 1.476222213 | 0.524585231 | 0.718404067 | 5.673143503 | 8.369200189 | ENSG00000215022 |
| NA | 0.561875473 | 1.476186982 | 0.525961914 | 0.719293069 | 4.954924104 | 7.325033662 | ENSG00000226054 |
| CALHM2 | 0.561758711 | 1.476067515 | 0.050349823 | 0.141897688 | 47.76341586 | 70.47270329 | ENSG00000138172 |
| NA | 0.56175638 | 1.476065129 | 0.389426839 | 0.601255569 | 9.169076767 | 13.57326884 | ENSG00000225173 |
| PILRB | 0.561754908 | 1.476063623 | 0.09082395 | 0.221803996 | 36.53692956 | 53.8939648 | ENSG00000121716 |
| SERPINA5 | 0.561617053 | 1.475922586 | 0.055511108 | 0.152951164 | 45.91003916 | 67.61542845 | ENSG00000188488 |
| DENND3 | 0.561600287 | 1.475905435 | 0.014375888 | 0.052654894 | 117.6984574 | 173.9058826 | ENSG00000105339 |
| PCDH8 | 0.561552519 | 1.475856568 | 0.284847276 | 0.494087503 | 16.97999698 | 25.13784615 | ENSG00000136099 |
| CYB5R1 | 0.561456864 | 1.475758717 | 3.20E-05 | 0.00027006 | 240.1527878 | 354.7307198 | ENSG00000159348 |
| LLPH-DT | 0.56137291 | 1.475672842 | 0.317951208 | 0.528054575 | 14.77817592 | 21.84286948 | ENSG00000239335 |
| SPIN2A | 0.561009072 | 1.475300734 | 0.040468698 | 0.119826176 | 49.41754056 | 73.02172231 | ENSG00000147059 |
| VSTM2A | 0.560685358 | 1.474969741 | 0.384415536 | 0.596719332 | 9.261596898 | 13.68090784 | ENSG00000170419 |
| EEF1A2 | 0.560646353 | 1.474929864 | 1.31E-14 | 4.67E-13 | 1278.703528 | 1886.383682 | ENSG00000101210 |
| TST | 0.56049562 | 1.474775771 | 1.24E-08 | 2.01E-07 | 438.4462334 | 646.7330495 | ENSG00000128311 |
| NA | 0.560481297 | 1.47476113 | 0.000268992 | 0.001789334 | 187.2917379 | 276.227624 | ENSG00000258649 |
| EDIL3 | 0.560475 | 1.474754693 | 8.03E-09 | 1.35E-07 | 765.0005299 | 1127.145696 | ENSG00000164176 |
| FNDC1 | 0.560317213 | 1.474593409 | 0.337533431 | 0.549415519 | 16.4589708 | 24.17426709 | ENSG00000164694 |
| COLGALT2 | 0.560120676 | 1.47439254 | 0.002452493 | 0.012108585 | 108.4975986 | 159.9939014 | ENSG00000198756 |
| MARCHF1 | 0.559935222 | 1.474203023 | 0.134076569 | 0.295418971 | 28.94753406 | 42.63402745 | ENSG00000145416 |
| LACC1 | 0.559774973 | 1.474039283 | 0.059105337 | 0.16055873 | 45.08506174 | 66.42930997 | ENSG00000179630 |
| CMYA5 | 0.559076038 | 1.473325336 | 0.203436124 | 0.395053079 | 18.96059881 | 27.88379348 | ENSG00000164309 |
| ZDHHC19 | 0.558871735 | 1.473116711 | 0.024413375 | 0.080842017 | 60.91802779 | 89.76258169 | ENSG00000163958 |
| EVC2 | 0.558772828 | 1.473015721 | 8.21E-05 | 0.000623922 | 197.7134607 | 291.2908006 | ENSG00000173040 |
| MTHFS | 0.558699907 | 1.472941269 | 0.279949734 | 0.488994296 | 18.17091063 | 26.91680946 | ENSG00000136371 |
| TP53INP1 | 0.558662498 | 1.472903077 | 0.002593075 | 0.012680017 | 109.7789474 | 161.5398042 | ENSG00000164938 |
| LRRC75B | 0.558361436 | 1.472595743 | 0.000259368 | 0.001732319 | 204.8188987 | 301.9437491 | ENSG00000178026 |
| LOC283683 | 0.558270366 | 1.472502789 | 0.382436638 | 0.594918939 | 9.867134539 | 14.54337429 | ENSG00000274253 |
| SLC30A2 | 0.558190542 | 1.472421318 | 0.257029907 | 0.463948031 | 15.79620227 | 23.3276913 | ENSG00000158014 |
| FOLR1 | 0.558120653 | 1.47234999 | 0.374612214 | 0.58656427 | 10.11238035 | 14.96785537 | ENSG00000110195 |
| NA | 0.558100942 | 1.472329874 | 0.488396804 | 0.688779985 | 9.620414189 | 14.25982656 | ENSG00000280056 |
| RDH12 | 0.557969705 | 1.472195948 | 0.414177618 | 0.625189379 | 11.964588 | 17.73896231 | ENSG00000139988 |
| SBK2 | 0.557694962 | 1.471915613 | 0.007783069 | 0.031966779 | 92.78651818 | 136.646631 | ENSG00000187550 |
| IZUMO4 | 0.557657959 | 1.471877861 | 0.429804305 | 0.639626028 | 9.313793777 | 13.70985128 | ENSG00000099840 |
| RAP2C-AS1 | 0.557509407 | 1.471726311 | 0.43649851 | 0.644963176 | 8.607421852 | 12.68062945 | ENSG00000232160 |
| PUS7L | 0.557336038 | 1.471549465 | 0.00904033 | 0.036079169 | 105.8779755 | 156.020858 | ENSG00000129317 |
| INTS12 | 0.557064095 | 1.471272109 | 0.003040489 | 0.014580912 | 105.7694119 | 155.6846476 | ENSG00000138785 |
| MRTFB | 0.55670539 | 1.470906344 | 2.51E-05 | 0.000218912 | 287.6124801 | 422.7503448 | ENSG00000186260 |
| ANKRD34A | 0.556515168 | 1.470712415 | 0.647791334 | 0.808876289 | 2.904633514 | 4.302727404 | ENSG00000272031 |
| ADCK1 | 0.55603377 | 1.47022175 | 8.99E-06 | 8.71E-05 | 247.7878199 | 364.3028751 | ENSG00000063761 |
| UNC93B1 | 0.556032974 | 1.470220939 | 2.47E-06 | 2.65E-05 | 433.4678669 | 637.7628198 | ENSG00000110057 |
| SNORA79B | 0.55602393 | 1.470211723 | 0.622458712 | 0.791034769 | 2.928354612 | 4.337251275 | ENSG00000222489 |

| | | | | | | | |
|---|---|---|---|---|---|---|---|
| CHRM3 | 0.555851941 | 1.470036464 | 0.000648549 | 0.00384186 | 153.9408672 | 226.0277464 | ENSG00000133019 |
| NA | 0.555668685 | 1.469849746 | 0.452629951 | 0.658748198 | 6.987417333 | 10.23752546 | ENSG00000270728 |
| M1AP | 0.555618774 | 1.469798897 | 0.013361648 | 0.049535674 | 90.3601716 | 132.5367645 | ENSG00000159374 |
| NA | 0.555613497 | 1.469793521 | 0.55044953 | 0.737721905 | 4.273468486 | 6.292406865 | ENSG00000275665 |
| WSCD2 | 0.555438155 | 1.469614897 | 0.050116443 | 0.141303229 | 50.11826808 | 73.54928055 | ENSG00000075035 |
| CDH15 | 0.555325488 | 1.469500131 | 0.269222241 | 0.477305859 | 17.04523654 | 25.091499 | ENSG00000129910 |
| BLVRB | 0.555247047 | 1.469420235 | 2.55E-07 | 3.30E-06 | 383.720622 | 563.8165385 | ENSG00000090013 |
| RTL9 | 0.555035778 | 1.469205068 | 0.549764365 | 0.73701285 | 4.297189585 | 6.319458387 | ENSG00000243978 |
| NA | 0.555004129 | 1.469172838 | 0.12542575 | 0.281481318 | 32.37230402 | 47.62397499 | ENSG00000287763 |
| TCAF1 | 0.55485734 | 1.469023363 | 7.07E-06 | 7.00E-05 | 362.1055916 | 531.3001953 | ENSG00000198420 |
| NA | 0.554172913 | 1.468326611 | 0.677671913 | 0.82756234 | 2.966313601 | 4.370545641 | ENSG00000254477 |
| CR1L | 0.55410101 | 1.468253432 | 0.325873708 | 0.537056024 | 12.16978988 | 17.8996428 | ENSG00000197721 |
| PPDPF | 0.554016606 | 1.468167535 | 1.19E-10 | 2.63E-09 | 824.4745697 | 1211.306706 | ENSG00000125534 |
| TOR3A | 0.553748737 | 1.467894962 | 8.04E-12 | 2.06E-10 | 764.6658682 | 1122.311154 | ENSG00000186283 |
| TOB1 | 0.5533623 | 1.467501828 | 1.24E-06 | 1.41E-05 | 343.5284726 | 504.4654567 | ENSG00000141232 |
| NA | 0.553325583 | 1.46746448 | 0.641283133 | 0.803884545 | 2.931914085 | 4.327603462 | ENSG00000225674 |
| GALNT10 | 0.553212622 | 1.467349584 | 6.03E-10 | 1.20E-08 | 617.0474869 | 905.0529912 | ENSG00000164574 |
| GRID2IP | 0.553034939 | 1.467168875 | 0.144001843 | 0.309946981 | 30.82465801 | 45.22220481 | ENSG00000215045 |
| ANKAR | 0.552785462 | 1.466915188 | 0.359886228 | 0.572043718 | 12.86309298 | 18.87750414 | ENSG00000151687 |
| LIMD2 | 0.552379627 | 1.466502598 | 1.67E-13 | 5.25E-12 | 1473.474909 | 2161.998743 | ENSG00000136490 |
| ABHD14B | 0.552272483 | 1.46639369 | 0.052726167 | 0.146909055 | 74.60256486 | 109.4415869 | ENSG00000114779 |
| PIGX | 0.552121976 | 1.466240718 | 7.22E-05 | 0.000556342 | 212.9892211 | 312.4601651 | ENSG00000163964 |
| SULT1A2 | 0.551983087 | 1.466099569 | 0.518705166 | 0.714579245 | 6.416279208 | 9.36853262 | ENSG00000197165 |
| SNORD116-17 | 0.551967435 | 1.466083664 | 0.629159511 | 0.795425152 | 3.581334451 | 5.265360501 | ENSG00000206656 |
| NA | 0.551844026 | 1.46595826 | 0.184005721 | 0.368057883 | 24.09433396 | 35.29979748 | ENSG00000269973 |
| NA | 0.551510918 | 1.465619819 | 0.440474736 | 0.648201153 | 7.878524036 | 11.56429376 | ENSG00000219102 |
| DPYD | 0.551292997 | 1.465398452 | 0.380852712 | 0.59343263 | 9.28887747 | 13.59596944 | ENSG00000188641 |
| FADD | 0.551160851 | 1.465264233 | 3.51E-07 | 4.43E-06 | 415.7053274 | 609.0959475 | ENSG00000168040 |
| ZFAS1 | 0.550983082 | 1.465083694 | 0.001877378 | 0.009616025 | 130.5060734 | 191.2346985 | ENSG00000177410 |
| NUF2 | 0.550881025 | 1.464980057 | 1.14E-07 | 1.57E-06 | 416.9463269 | 610.663226 | ENSG00000143228 |
| NBPF14 | 0.550831907 | 1.464930181 | 0.218523523 | 0.41475545 | 18.10122188 | 26.57632081 | ENSG00000270629 |
| PILRA | 0.550510444 | 1.4646038 | 0.467557426 | 0.671155346 | 7.079937464 | 10.31158654 | ENSG00000085514 |
| LINC00173 | 0.550418118 | 1.464510075 | 0.280097211 | 0.489029772 | 16.42101181 | 23.97102329 | ENSG00000196668 |
| LOC100507071 | 0.550316821 | 1.46440725 | 0.503858979 | 0.70156874 | 5.853149752 | 8.619453548 | ENSG00000254064 |
| RASEF | 0.550183551 | 1.46427198 | 0.011788476 | 0.044820893 | 85.84593265 | 125.4827929 | ENSG00000165105 |
| SLC13A3 | 0.550067552 | 1.464154251 | 6.89E-05 | 0.000534666 | 232.9805834 | 341.3647621 | ENSG00000158296 |
| IFI6 | 0.549958662 | 1.464043746 | 0.028061035 | 0.090155264 | 64.68292796 | 94.73709008 | ENSG00000126709 |
| ZBTB4 | 0.549585581 | 1.463665193 | 1.06E-07 | 1.48E-06 | 468.9322351 | 686.7430079 | ENSG00000174282 |
| MICOS13 | 0.549327392 | 1.463403274 | 1.48E-05 | 0.000135609 | 291.6537454 | 426.5568438 | ENSG00000174917 |
| CYB5RL | 0.549069154 | 1.463141354 | 0.000172873 | 0.001209597 | 211.3288933 | 309.3281397 | ENSG00000215883 |
| NA | 0.549007565 | 1.463078892 | 0.347152371 | 0.55928788 | 14.63612866 | 21.41888285 | ENSG00000189212 |
| TMEM171 | 0.548927933 | 1.462998138 | 0.169681585 | 0.347741372 | 28.17503285 | 41.02510889 | ENSG00000157111 |
| NFATC1 | 0.548697497 | 1.462764477 | 0.002037125 | 0.010327813 | 156.6026709 | 229.4570574 | ENSG00000131196 |
| CCDC125 | 0.548686303 | 1.462753128 | 0.019362695 | 0.066898005 | 69.95426117 | 102.3804789 | ENSG00000183323 |
| NA | 0.54856248 | 1.462627589 | 0.627774936 | 0.794139912 | 2.921235667 | 4.297430518 | ENSG00000269889 |
| DDX59 | 0.548251736 | 1.462312586 | 1.91E-06 | 2.09E-05 | 330.8913018 | 483.6688863 | ENSG00000118197 |
| NA | 0.548105655 | 1.462164525 | 0.1123505 | 0.260240303 | 32.46362895 | 47.44445038 | ENSG00000257261 |
| NA | 0.548000595 | 1.462058051 | 0.492899643 | 0.69243545 | 5.700424074 | 8.292963644 | ENSG00000263004 |
| WAKMAR2 | 0.547776424 | 1.461830889 | 0.608959174 | 0.782305701 | 4.314986947 | 6.302054678 | ENSG00000237499 |
| NA | 0.547554866 | 1.46160641 | 0.314467847 | 0.524936499 | 15.46791954 | 22.66355062 | ENSG00000283341 |
| CACNA1C | 0.547487114 | 1.461537772 | 0.025421753 | 0.083493878 | 62.78119318 | 91.6436762 | ENSG00000151067 |
| NA | 0.547289631 | 1.461337723 | 0.624370951 | 0.7923564 | 3.568291769 | 5.227715207 | ENSG00000279400 |
| CFP | 0.547203065 | 1.461250041 | 0.625301265 | 0.792803919 | 3.664371373 | 5.353703868 | ENSG00000126759 |
| NA | 0.547074169 | 1.461119493 | 0.467904035 | 0.671349859 | 7.51111326 | 11.04014049 | ENSG00000272316 |
| NA | 0.546667887 | 1.46070808 | 0.442831443 | 0.650402224 | 8.677416093 | 12.65168601 | ENSG00000227001 |
| CLDN23 | 0.546613554 | 1.46065307 | 0.001388856 | 0.007464014 | 134.4180287 | 196.4597867 | ENSG00000253958 |
| TIMP4 | 0.546367803 | 1.460404281 | 0.023529818 | 0.078495074 | 72.92176998 | 106.6086937 | ENSG00000157150 |
| MCEE | 0.546331396 | 1.460367428 | 0.016104788 | 0.057777152 | 96.31504536 | 140.5443838 | ENSG00000124370 |
| PDLIM2 | 0.546094645 | 1.460127797 | 0.006167751 | 0.026340566 | 170.1705573 | 248.4161676 | ENSG00000120913 |
| TCEAL7 | 0.545818445 | 1.459848286 | 0.011386573 | 0.043503022 | 83.57353253 | 122.0550176 | ENSG00000182916 |
| POLH | 0.545748441 | 1.459777451 | 0.00211193 | 0.010667673 | 132.7544469 | 193.7452418 | ENSG00000170734 |
| SOCS1 | 0.545457519 | 1.459483114 | 8.29E-05 | 0.000629232 | 391.8166043 | 572.4530395 | ENSG00000185338 |
| RAB3C | 0.545426925 | 1.459452164 | 2.45E-05 | 0.000213858 | 236.7866972 | 345.4789069 | ENSG00000152932 |
| GNAS-AS1 | 0.545374228 | 1.459398856 | 0.371070482 | 0.583194578 | 10.21913837 | 14.93361505 | ENSG00000235590 |
| NA | 0.54506923 | 1.459090359 | 0.432175541 | 0.641566127 | 8.192568878 | 12.00807047 | ENSG00000267541 |
| INA | 0.545068299 | 1.459089418 | 1.15E-06 | 1.32E-05 | 404.9292883 | 591.3621344 | ENSG00000148798 |
| MDFI | 0.544729029 | 1.458746332 | 4.80E-18 | 2.42E-16 | 2107.787221 | 3075.555058 | ENSG00000112559 |
| ABCB6 | 0.544512135 | 1.458527042 | 0.404589293 | 0.615551577 | 10.11741436 | 14.68988808 | ENSG00000115657 |
| NA | 0.544260179 | 1.458272343 | 0.007216752 | 0.030032693 | 91.68573424 | 133.6656585 | ENSG00000279088 |
| NA | 0.544226433 | 1.458238233 | 0.615607218 | 0.786373262 | 2.952075711 | 4.321077071 | ENSG00000278876 |
| NBEAL1 | 0.544223734 | 1.458235505 | 0.052159851 | 0.145888445 | 53.26486727 | 77.47983331 | ENSG00000144426 |
| NA | 0.54414494 | 1.458155865 | 0.566728375 | 0.750531324 | 4.324470156 | 6.290231402 | ENSG00000273156 |
| CDC42SE1 | 0.54393915 | 1.457947884 | 4.97E-15 | 1.88E-13 | 1420.364009 | 2070.12317 | ENSG00000197622 |
| TSACC | 0.543831623 | 1.457839224 | 0.198281673 | 0.388277622 | 22.01083906 | 32.15852341 | ENSG00000163467 |

| | | | | | | | |
|---|---|---|---|---|---|---|---|
| RIOX2 | 0.543283219 | 1.457285169 | 2.33E-05 | 0.000204819 | 280.4795083 | 408.4108041 | ENSG00000170854 |
| CNST | 0.543009798 | 1.457009009 | 7.99E-07 | 9.39E-06 | 425.6667616 | 620.3486733 | ENSG00000162852 |
| LINC00939 | 0.542809046 | 1.456806279 | 0.651975527 | 0.811317727 | 2.966313601 | 4.330724884 | ENSG00000249267 |
| NA | 0.542675019 | 1.456670948 | 0.668221425 | 0.822110685 | 2.74478889 | 3.946064556 | ENSG00000269907 |
| MSX2 | 0.542594551 | 1.456589703 | 0.508128667 | 0.704897972 | 5.921948785 | 8.647451029 | ENSG00000120149 |
| MTHFR | 0.542462579 | 1.456456466 | 1.64E-05 | 0.000148962 | 260.6559469 | 379.4708528 | ENSG00000177000 |
| NA | 0.542061818 | 1.456051938 | 0.064207916 | 0.170746959 | 49.51745897 | 71.99533835 | ENSG00000228232 |
| NA | 0.54193392 | 1.455922862 | 0.463976073 | 0.668542004 | 6.585607041 | 9.598260848 | ENSG00000244363 |
| DBIL5P | 0.541540318 | 1.455525704 | 0.50420719 | 0.701743148 | 6.860497685 | 10.04203756 | ENSG00000231784 |
| CD37 | 0.541519265 | 1.455504465 | 0.25084095 | 0.456327073 | 18.75923573 | 27.19070937 | ENSG00000104894 |
| RAMP3 | 0.541100435 | 1.455081977 | 0.251247674 | 0.456802959 | 17.71601374 | 25.84124048 | ENSG00000122679 |
| NA | 0.541048328 | 1.455029424 | 0.488169048 | 0.688710031 | 5.938550939 | 8.658328346 | ENSG00000259553 |
| MMP24OS | 0.541040223 | 1.45502125 | 0.000174714 | 0.001220146 | 239.6296762 | 349.1680964 | ENSG00000126005 |
| ZNF135 | 0.540863721 | 1.454843251 | 0.07810982 | 0.198293314 | 40.35166234 | 58.70495482 | ENSG00000176293 |
| MYT1L | 0.540635714 | 1.454613341 | 0.576475628 | 0.758517416 | 4.317351211 | 6.261287963 | ENSG00000186487 |
| NA | 0.540609658 | 1.45458707 | 0.267504301 | 0.47528636 | 16.01444684 | 23.22534919 | ENSG00000279347 |
| MMP23B | 0.540535257 | 1.454512058 | 0.168411898 | 0.346040551 | 27.99474727 | 40.67017001 | ENSG00000189409 |
| EML2 | 0.540488168 | 1.454464584 | 3.02E-06 | 3.18E-05 | 371.6359022 | 540.2437657 | ENSG00000125746 |
| ADSS2 | 0.540432222 | 1.454408183 | 1.85E-17 | 8.72E-16 | 1511.027903 | 2198.015367 | ENSG00000035687 |
| TACR2 | 0.540329913 | 1.454305047 | 0.61973441 | 0.788817438 | 2.955635184 | 4.309253795 | ENSG00000075073 |
| FHDC1 | 0.540031476 | 1.454004239 | 8.31E-13 | 2.41E-11 | 874.818735 | 1271.590123 | ENSG00000137460 |
| CES4A | 0.539731732 | 1.453702177 | 0.539395394 | 0.729040189 | 6.19683943 | 9.066635228 | ENSG00000172824 |
| RPP25L | 0.539563334 | 1.453532504 | 1.21E-06 | 1.38E-05 | 394.0499994 | 572.8503012 | ENSG00000164967 |
| DACT2 | 0.53937031 | 1.453338043 | 0.002932863 | 0.014111367 | 131.6714342 | 191.2407734 | ENSG00000164488 |
| LOC100268168 | 0.539300688 | 1.45326791 | 0.620408932 | 0.789409963 | 2.955635184 | 4.304902868 | ENSG00000204758 |
| LINC01137 | 0.539186573 | 1.453152963 | 0.419620428 | 0.630081242 | 9.435958744 | 13.64014112 | ENSG00000233621 |
| HAX1 | 0.539086554 | 1.453052223 | 8.82E-13 | 2.55E-11 | 1279.700251 | 1859.103585 | ENSG00000143575 |
| SDR42E2 | 0.538929054 | 1.4528936 | 0.600969407 | 0.776307564 | 3.633531329 | 5.27283285 | ENSG00000183921 |
| CIAO2B | 0.538811347 | 1.452775066 | 1.11E-11 | 2.78E-10 | 1116.464172 | 1621.851469 | ENSG00000166595 |
| ALPK3 | 0.538684022 | 1.452646857 | 2.32E-16 | 1.00E-14 | 1728.998031 | 2510.773655 | ENSG00000136383 |
| NA | 0.538379587 | 1.452340355 | 0.128432448 | 0.286357353 | 33.80162388 | 49.05192852 | ENSG00000261167 |
| CRIM1 | 0.538348275 | 1.452308834 | 2.97E-06 | 3.14E-05 | 368.2784562 | 534.8184692 | ENSG00000150938 |
| TMEM54 | 0.538205834 | 1.45216545 | 0.086003041 | 0.213190155 | 51.20862679 | 74.24491899 | ENSG00000121900 |
| B4GALT3 | 0.538188642 | 1.452148146 | 1.24E-10 | 2.72E-09 | 693.7638542 | 1007.139685 | ENSG00000158850 |
| SESN1 | 0.537709042 | 1.451665483 | 4.05E-06 | 4.18E-05 | 319.6049672 | 464.1703946 | ENSG00000080546 |
| IVNS1ABP | 0.537607225 | 1.451563037 | 4.19E-25 | 3.78E-23 | 5712.223238 | 8290.311788 | ENSG00000116679 |
| ENAH | 0.537515608 | 1.45147086 | 1.38E-17 | 6.56E-16 | 6488.888168 | 9418.02392 | ENSG00000154380 |
| KCTD18 | 0.537451137 | 1.451405998 | 0.018665382 | 0.064929588 | 80.63272006 | 116.7588218 | ENSG00000155729 |
| NA | 0.537153092 | 1.451106184 | 0.622775844 | 0.79114079 | 2.986475227 | 4.335075811 | ENSG00000287836 |
| SLC25A44 | 0.537041392 | 1.450993837 | 3.48E-10 | 7.18E-09 | 752.518578 | 1091.728516 | ENSG00000160785 |
| NA | 0.537018025 | 1.450970336 | 0.562248299 | 0.747224428 | 4.593437065 | 6.699767788 | ENSG00000226855 |
| NA | 0.536578005 | 1.450527859 | 0.558489002 | 0.744063237 | 4.3553102 | 6.282759052 | ENSG00000270441 |
| CYP2F1 | 0.536479067 | 1.450428387 | 0.569919145 | 0.752843159 | 4.365988618 | 6.298933256 | ENSG00000197446 |
| TACR1 | 0.535875308 | 1.449821519 | 0.524026072 | 0.718166487 | 5.27133321 | 7.662400884 | ENSG00000115353 |
| NA | 0.535516299 | 1.449460782 | 0.630825019 | 0.796625039 | 2.9900347 | 4.332900348 | ENSG00000248569 |
| ATP10B | 0.535391042 | 1.449334943 | 0.523708729 | 0.717880479 | 5.765663634 | 8.286437254 | ENSG00000118322 |
| PIP5K1A | 0.535246934 | 1.449190179 | 3.82E-19 | 2.15E-17 | 1913.398918 | 2772.989004 | ENSG00000143398 |
| AFP | 0.535185279 | 1.449128248 | 0.207699022 | 0.400285339 | 25.82081717 | 37.56510081 | ENSG00000081051 |
| BBS1 | 0.535048645 | 1.448991011 | 0.482464129 | 0.683726181 | 7.745680652 | 11.3140404 | ENSG00000174483 |
| TRIM54 | 0.534970042 | 1.448912067 | 0.007942145 | 0.032457782 | 101.0985823 | 146.7169496 | ENSG00000138100 |
| INTS7 | 0.53492466 | 1.448866491 | 6.38E-09 | 1.09E-07 | 556.7786519 | 806.8991485 | ENSG00000143493 |
| CA4 | 0.534838491 | 1.448779955 | 0.057152103 | 0.156470665 | 50.39760792 | 73.16265567 | ENSG00000167434 |
| AK7 | 0.534614401 | 1.448554938 | 0.037824283 | 0.113671482 | 67.05138153 | 96.88321469 | ENSG00000140057 |
| TBXA2R | 0.534588709 | 1.448529141 | 0.318069907 | 0.528112378 | 14.13704355 | 20.50276486 | ENSG00000006638 |
| ABCG2 | 0.534161892 | 1.448100662 | 0.100687929 | 0.239192486 | 40.02546454 | 58.12887397 | ENSG00000118777 |
| NA | 0.534159786 | 1.448098548 | 0.356528681 | 0.568669268 | 16.59361979 | 24.10834077 | ENSG00000253878 |
| FGR | 0.533648625 | 1.447585564 | 0.528994261 | 0.72146 | 5.754985217 | 8.260615237 | ENSG00000000938 |
| NBPF20 | 0.533643646 | 1.447580567 | 0.406677373 | 0.617595253 | 10.25234268 | 14.81509874 | ENSG00000162825 |
| LOC101927752 | 0.533490001 | 1.44742641 | 0.021383015 | 0.072655316 | 93.5705358 | 135.2373106 | ENSG00000253982 |
| NA | 0.533355649 | 1.447291624 | 0.668524517 | 0.822213318 | 3.020874744 | 4.365248755 | ENSG00000206762 |
| NA | 0.533189594 | 1.44712505 | 0.536420335 | 0.726836969 | 5.236933694 | 7.597704069 | ENSG00000251292 |
| NEK2 | 0.532792697 | 1.446726988 | 3.72E-08 | 5.57E-07 | 584.1378339 | 844.411682 | ENSG00000117650 |
| IGSF9B | 0.53264437 | 1.446578255 | 2.31E-10 | 4.88E-09 | 1490.983047 | 2155.682643 | ENSG00000080854 |
| CRYBB1 | 0.532607328 | 1.446541114 | 0.122565579 | 0.276709919 | 32.14485558 | 46.4789794 | ENSG00000100122 |
| NA | 0.532541632 | 1.446475244 | 0.119700263 | 0.272255747 | 35.21909625 | 50.90618242 | ENSG00000249898 |
| INSRR | 0.532456766 | 1.446390158 | 0.55707088 | 0.743325342 | 4.589877592 | 6.668648885 | ENSG00000027644 |
| THRB | 0.532368919 | 1.446302089 | 0.042524503 | 0.124441017 | 56.573396 | 81.77631787 | ENSG00000151090 |
| PCDH7 | 0.532300505 | 1.446233505 | 0.004169202 | 0.019031999 | 116.4292092 | 168.5530091 | ENSG00000169851 |
| STX17 | 0.532151173 | 1.446083815 | 2.54E-05 | 0.000220746 | 294.638188 | 425.6341041 | ENSG00000136874 |
| USP21 | 0.53213716 | 1.446069769 | 3.00E-08 | 4.57E-07 | 537.0020008 | 776.7955387 | ENSG00000143258 |
| LOC105369779 | 0.531949943 | 1.445882126 | 0.018633212 | 0.064840282 | 74.01123896 | 106.9103075 | ENSG00000258056 |
| PHF11 | 0.531921731 | 1.445853852 | 0.02349285 | 0.078413304 | 76.58314054 | 110.5056686 | ENSG00000136147 |
| C1orf131 | 0.531293342 | 1.445224224 | 6.98E-05 | 0.000539845 | 220.3526427 | 318.4236231 | ENSG00000143633 |
| LINC01503 | 0.531239305 | 1.445170093 | 0.07995623 | 0.201760209 | 41.6137393 | 60.21682816 | ENSG00000233901 |

| | | | | | | | |
|---|---|---|---|---|---|---|---|
| PALLD | 0.530915821 | 1.444846091 | 1.54E-06 | 1.72E-05 | 363.3947477 | 525.1054235 | ENSG00000129116 |
| PDZK1 | 0.530864617 | 1.444794811 | 0.497927597 | 0.696960538 | 7.85983695 | 11.24622216 | ENSG00000174827 |
| MRPS14 | 0.530741168 | 1.444671188 | 1.01E-07 | 1.41E-06 | 470.4205403 | 679.1158363 | ENSG00000120333 |
| SETDB1 | 0.53068689 | 1.444616837 | 8.64E-10 | 1.68E-08 | 641.745657 | 927.0413332 | ENSG00000143379 |
| APRT | 0.530513802 | 1.444443528 | 3.98E-19 | 2.23E-17 | 2029.702098 | 2932.041307 | ENSG00000198931 |
| LIN7B | 0.53047334 | 1.444403018 | 0.071076189 | 0.185115685 | 47.70649045 | 68.93862382 | ENSG00000104863 |
| H6PD | 0.530345376 | 1.444274908 | 1.51E-05 | 0.000137878 | 290.9234254 | 420.0416866 | ENSG00000049239 |
| FAM160B2 | 0.529956723 | 1.443885882 | 5.66E-06 | 5.70E-05 | 343.5729663 | 496.1820683 | ENSG00000158863 |
| ZNF185 | 0.529907626 | 1.443836746 | 1.97E-05 | 0.000176255 | 293.9531729 | 424.0709132 | ENSG00000147394 |
| TFB2M | 0.529821671 | 1.443750725 | 1.74E-08 | 2.77E-07 | 523.6240841 | 756.2394583 | ENSG00000162851 |
| EPPK1 | 0.529808351 | 1.443737395 | 0.000176868 | 0.001233437 | 280.4522277 | 404.5412167 | ENSG00000261150 |
| NA | 0.529805906 | 1.443734949 | 7.73E-22 | 5.41E-20 | 2347.376654 | 3389.117801 | ENSG00000249790 |
| BDKRB2 | 0.529803889 | 1.44373293 | 0.37077289 | 0.582969447 | 13.94518983 | 20.08168874 | ENSG00000168398 |
| NA | 0.529764157 | 1.44369317 | 0.593278495 | 0.770515396 | 3.884700875 | 5.654371756 | ENSG00000215283 |
| KITLG | 0.529744169 | 1.443673168 | 0.011219839 | 0.042961609 | 87.06977128 | 125.6400684 | ENSG00000049130 |
| TRIM22 | 0.529513818 | 1.44344268 | 3.44E-05 | 0.000288265 | 452.8566942 | 653.3379381 | ENSG00000132274 |
| OTUD1 | 0.529436676 | 1.4433655 | 0.005253028 | 0.023043772 | 115.736491 | 166.9318158 | ENSG00000165312 |
| FDXR | 0.529353367 | 1.443282154 | 2.18E-11 | 5.31E-10 | 845.5844366 | 1220.067851 | ENSG00000161513 |
| 1-Mar | 0.529170396 | 1.44309912 | 5.07E-06 | 5.15E-05 | 330.4830353 | 477.1550015 | ENSG00000186205 |
| BCL2L13 | 0.529117061 | 1.443045772 | 4.82E-10 | 9.72E-09 | 777.9380919 | 1121.931244 | ENSG00000099968 |
| NIPAL1 | 0.529092743 | 1.443021448 | 0.0109743 | 0.042192521 | 107.2847694 | 155.0796234 | ENSG00000163293 |
| SOX2 | 0.528910747 | 1.442839422 | 3.60E-19 | 2.03E-17 | 5549.403377 | 8007.326154 | ENSG00000181449 |
| SLFN12 | 0.528859083 | 1.442787754 | 0.010505323 | 0.040645979 | 98.55485156 | 142.0737335 | ENSG00000172123 |
| FRMPD2 | 0.528822478 | 1.442751147 | 0.622481743 | 0.791034769 | 3.695211417 | 5.305181257 | ENSG00000170324 |
| NA | 0.528173699 | 1.442102489 | 0.596137146 | 0.772634 | 3.711813571 | 5.317004534 | ENSG00000281741 |
| DHTKD1 | 0.528012721 | 1.441941586 | 1.55E-15 | 6.26E-14 | 1488.444096 | 2145.388916 | ENSG00000181192 |
| SLC19A2 | 0.527470948 | 1.441400197 | 8.27E-09 | 1.38E-07 | 618.9539503 | 891.8539342 | ENSG00000117479 |
| NA | 0.5272627 | 1.441192152 | 0.39939392 | 0.610828948 | 10.84839711 | 15.68770746 | ENSG00000257681 |
| DNAJC5G | 0.527215093 | 1.441144594 | 0.598992798 | 0.77503937 | 3.711813571 | 5.312653607 | ENSG00000163793 |
| GLRB | 0.52715633 | 1.441085896 | 0.482755732 | 0.683859246 | 6.694729326 | 9.602611776 | ENSG00000109738 |
| TRIM62 | 0.527144283 | 1.441073862 | 3.76E-05 | 0.000312395 | 318.7462268 | 459.7541415 | ENSG00000116525 |
| CLEC2L | 0.526942559 | 1.440872379 | 0.30225955 | 0.512445547 | 16.00140416 | 23.03326625 | ENSG00000236279 |
| BAMBI | 0.526711917 | 1.440642047 | 6.81E-10 | 1.34E-08 | 614.7736646 | 885.649231 | ENSG00000095739 |
| NA | 0.526349835 | 1.440280525 | 0.470772046 | 0.674161267 | 7.787199113 | 11.28821838 | ENSG00000235734 |
| TRPS1 | 0.526147698 | 1.440078741 | 0.016252744 | 0.058189314 | 77.3921262 | 111.4545865 | ENSG00000104447 |
| UCKL1 | 0.526108277 | 1.440039392 | 3.84E-08 | 5.73E-07 | 641.5615064 | 924.2073191 | ENSG00000198276 |
| GRB7 | 0.52607924 | 1.440010409 | 1.85E-08 | 2.92E-07 | 552.2158017 | 795.555146 | ENSG00000141738 |
| PTH2 | 0.526048346 | 1.439979573 | 0.605337465 | 0.779667429 | 3.926219337 | 5.685490658 | ENSG00000142538 |
| COP1 | 0.52598613 | 1.439917476 | 7.74E-13 | 2.26E-11 | 1036.478866 | 1491.96128 | ENSG00000143207 |
| TCERG1L | 0.525772328 | 1.439704101 | 9.84E-05 | 0.00073149 | 226.0755926 | 325.5104376 | ENSG00000176769 |
| LY6E | 0.525690064 | 1.439622009 | 2.49E-16 | 1.07E-14 | 1637.998 | 2358.47683 | ENSG00000160932 |
| PELI3 | 0.525533944 | 1.43946623 | 0.005779714 | 0.024937148 | 140.4150319 | 201.9917427 | ENSG00000174516 |
| GPRIN1 | 0.525456613 | 1.439389075 | 3.40E-07 | 4.31E-06 | 515.9976717 | 743.0905752 | ENSG00000169258 |
| RPAP2 | 0.525324609 | 1.439257379 | 2.90E-05 | 0.000248457 | 277.0656955 | 398.7515349 | ENSG00000122484 |
| PHLDA2 | 0.52508516 | 1.43901852 | 0.012641595 | 0.047424817 | 98.53229952 | 141.7639123 | ENSG00000181649 |
| ELF1 | 0.525012143 | 1.438945692 | 0.001229124 | 0.006702858 | 196.8377348 | 283.1352271 | ENSG00000120690 |
| NAV1 | 0.524877626 | 1.43881153 | 3.67E-10 | 7.53E-09 | 1239.999456 | 1783.572198 | ENSG00000134369 |
| SULT1A1 | 0.524711086 | 1.438645448 | 0.023653566 | 0.07883617 | 77.50864676 | 111.3604947 | ENSG00000196502 |
| NENF | 0.524629024 | 1.438563619 | 1.67E-05 | 0.000151632 | 306.6807789 | 441.1574855 | ENSG00000117691 |
| NA | 0.524598375 | 1.438533058 | 0.253876344 | 0.460076265 | 18.41051204 | 26.53990503 | ENSG00000278974 |
| ORC5 | 0.524562441 | 1.438497228 | 1.09E-06 | 1.25E-05 | 373.6449792 | 537.5007289 | ENSG00000164815 |
| IL17RE | 0.52430218 | 1.438237748 | 0.033651978 | 0.103978667 | 68.78114488 | 98.95849496 | ENSG00000163701 |
| ZMYND10 | 0.524156235 | 1.438092261 | 0.634673044 | 0.799639765 | 2.986475227 | 4.281256315 | ENSG00000004838 |
| VWA5A | 0.523874653 | 1.437811605 | 1.89E-08 | 2.98E-07 | 535.4335566 | 769.6225361 | ENSG00000110002 |
| ZNF8-ERVK3-1 | 0.523607624 | 1.437545505 | 0.495826596 | 0.695207823 | 6.224120001 | 8.986993714 | ENSG00000267216 |
| ZC3HAV1L | 0.523592466 | 1.437530401 | 0.024417414 | 0.080842017 | 69.543247 | 99.95934044 | ENSG00000146858 |
| PSPC1 | 0.523263055 | 1.437202206 | 4.50E-20 | 2.70E-18 | 2550.417632 | 3664.934553 | ENSG00000121390 |
| AK1 | 0.523170489 | 1.437109996 | 0.112331629 | 0.260228509 | 35.65650127 | 51.12096595 | ENSG00000106992 |
| LOXL2 | 0.523130281 | 1.437069944 | 5.65E-12 | 1.48E-10 | 942.8926182 | 1354.476135 | ENSG00000134013 |
| NR4A2 | 0.523072381 | 1.437012271 | 0.545560075 | 0.733772667 | 6.268002726 | 9.028989934 | ENSG00000153234 |
| ARHGAP31 | 0.522818988 | 1.436759898 | 0.001221002 | 0.006670123 | 193.696449 | 278.0387974 | ENSG00000031081 |
| IRAK3 | 0.52263442 | 1.4365761 | 0.072882512 | 0.188804696 | 48.46978779 | 69.5117512 | ENSG00000090376 |
| STUM | 0.522526233 | 1.436468376 | 0.123879115 | 0.278938486 | 41.48000619 | 59.57255021 | ENSG00000203685 |
| NA | 0.522430281 | 1.436372842 | 0.433552983 | 0.642497135 | 8.563539126 | 12.33484392 | ENSG00000232346 |
| ZNF805 | 0.522385833 | 1.43632859 | 0.011378044 | 0.043479231 | 95.98055956 | 137.780443 | ENSG00000204524 |
| NA | 0.522172005 | 1.436115721 | 0.671047293 | 0.823813829 | 3.04103637 | 4.347845047 | ENSG00000286608 |
| NA | 0.521750674 | 1.435696373 | 0.593183371 | 0.77044484 | 3.926219337 | 5.666195032 | ENSG00000276071 |
| NCSTN | 0.521546621 | 1.435493324 | 4.19E-14 | 1.40E-12 | 1468.276721 | 2107.768255 | ENSG00000162736 |
| TRAF5 | 0.521501772 | 1.4354487 | 1.26E-05 | 0.000117183 | 335.6944161 | 481.9543066 | ENSG00000082512 |
| UBAP2L | 0.521488917 | 1.435435909 | 1.18E-21 | 8.11E-20 | 5547.72971 | 7963.584505 | ENSG00000143569 |
| TENT5C | 0.521292091 | 1.435240087 | 0.19936818 | 0.389556967 | 23.68687926 | 33.91847464 | ENSG00000183508 |
| C8orf34 | 0.521289128 | 1.435237139 | 0.291330771 | 0.500936139 | 15.25470898 | 21.86623249 | ENSG00000165084 |
| TAB1 | 0.521202458 | 1.435150919 | 7.11E-13 | 2.09E-11 | 955.8479258 | 1371.388951 | ENSG00000100324 |
| RPL12 | 0.521006806 | 1.434956304 | 1.86E-23 | 1.48E-21 | 14154.26616 | 20309.9292 | ENSG00000197958 |

| | | | | | | | |
|---|---|---|---|---|---|---|---|
| PGF | 0.520922539 | 1.434872491 | 0.01008566 | 0.039322879 | 102.5540136 | 147.2999357 | ENSG00000119630 |
| NSL1 | 0.520765323 | 1.434716137 | 1.91E-10 | 4.08E-09 | 942.3624393 | 1351.955973 | ENSG00000117697 |
| NA | 0.520089649 | 1.434044357 | 0.28010776 | 0.489029772 | 16.41241833 | 23.61653591 | ENSG00000278784 |
| ALX1 | 0.519817306 | 1.433773672 | 0.215127614 | 0.410165068 | 24.888192 | 35.78557041 | ENSG00000180318 |
| ARRDC2 | 0.519746807 | 1.433703611 | 0.015533557 | 0.056068661 | 95.86014846 | 137.2250982 | ENSG00000105643 |
| ILK | 0.519672459 | 1.433629728 | 0.518838111 | 0.71467963 | 6.08207274 | 8.667976159 | ENSG00000166333 |
| NA | 0.519509263 | 1.433467567 | 0.366359473 | 0.578876192 | 10.87923715 | 15.62329419 | ENSG00000242858 |
| HELZ2 | 0.519114787 | 1.433075667 | 3.00E-08 | 4.57E-07 | 671.3008392 | 962.4304951 | ENSG00000130589 |
| INE1 | 0.518995389 | 1.432957071 | 0.245206187 | 0.44954071 | 19.94778512 | 28.53828634 | ENSG00000224975 |
| NA | 0.518958581 | 1.432920511 | 0.305711201 | 0.516031107 | 15.42759628 | 22.14230786 | ENSG00000258011 |
| SAYSD1 | 0.518116973 | 1.43208485 | 0.00183188 | 0.009408498 | 143.3667759 | 205.4262553 | ENSG00000112167 |
| ZNF658 | 0.518116499 | 1.432084379 | 0.131738038 | 0.291866829 | 30.99990958 | 44.43954783 | ENSG00000274349 |
| NA | 0.517849679 | 1.431819545 | 0.473435234 | 0.676566356 | 6.936415663 | 9.931277143 | ENSG00000214653 |
| PLCD1 | 0.517603541 | 1.431575284 | 1.52E-06 | 1.71E-05 | 383.3398113 | 548.3693639 | ENSG00000187091 |
| NA | 0.51753235 | 1.431504644 | 0.410127281 | 0.621140669 | 9.241435272 | 13.22842927 | ENSG00000250138 |
| LHFPL1 | 0.517438738 | 1.431411761 | 0.095621477 | 0.230391885 | 46.33793481 | 66.12285074 | ENSG00000182508 |
| SERP2 | 0.517435449 | 1.431408497 | 0.199653322 | 0.38987209 | 22.51109322 | 32.22634164 | ENSG00000151778 |
| APOA1 | 0.517207044 | 1.431181897 | 0.636904923 | 0.800725077 | 3.04103637 | 4.321077071 | ENSG00000118137 |
| NA | 0.516913388 | 1.430890614 | 0.577963821 | 0.759450456 | 4.879006127 | 7.034959546 | ENSG00000259146 |
| SYTL1 | 0.51668894 | 1.43066802 | 0.011786102 | 0.044820893 | 117.3929538 | 168.073522 | ENSG00000142765 |
| NA | 0.515878809 | 1.429864868 | 0.435023575 | 0.643690371 | 8.621659742 | 12.33266845 | ENSG00000179131 |
| APC2 | 0.515874776 | 1.429860871 | 0.058396129 | 0.159043973 | 52.14809156 | 74.61368866 | ENSG00000115266 |
| DENND4B | 0.515834325 | 1.42982078 | 1.05E-14 | 3.79E-13 | 1745.77278 | 2496.928665 | ENSG00000198837 |
| PIPOX | 0.515619034 | 1.429607427 | 5.49E-09 | 9.47E-08 | 818.3113642 | 1169.255896 | ENSG00000179761 |
| NTNG1 | 0.515510936 | 1.429500313 | 0.279426893 | 0.488473282 | 18.2791432 | 26.05290259 | ENSG00000162631 |
| OGFRL1 | 0.515356013 | 1.429346816 | 0.003239061 | 0.015357687 | 138.6384106 | 198.0705542 | ENSG00000119900 |
| GATM | 0.515340525 | 1.429331471 | 0.000128595 | 0.000929041 | 330.8405801 | 472.3142174 | ENSG00000171766 |
| NA | 0.514423808 | 1.428423534 | 0.50745814 | 0.704540578 | 7.147261956 | 10.26211797 | ENSG00000229525 |
| NA | 0.514263147 | 1.428264471 | 0.64033929 | 0.803013745 | 3.04103637 | 4.309253795 | ENSG00000222335 |
| ADH6 | 0.514002649 | 1.428006602 | 0.556659162 | 0.742985954 | 5.369777078 | 7.63222794 | ENSG00000172955 |
| NPHS1 | 0.513758517 | 1.427764976 | 0.179814154 | 0.362604409 | 27.30973217 | 38.93386533 | ENSG00000161270 |
| PBX1 | 0.513640418 | 1.427648103 | 1.77E-13 | 5.55E-12 | 3370.517874 | 4811.339836 | ENSG00000185630 |
| NA | 0.513585322 | 1.427593583 | 0.653984956 | 0.81254148 | 3.068316941 | 4.344723624 | ENSG00000287445 |
| CHKA | 0.513481125 | 1.427490481 | 4.35E-08 | 6.43E-07 | 671.4277856 | 958.4424546 | ENSG00000110721 |
| KIFC3 | 0.513454246 | 1.427463885 | 0.001246516 | 0.006784375 | 176.0024025 | 251.3516865 | ENSG00000140859 |
| EXD3 | 0.5132685 | 1.427280112 | 0.006605775 | 0.027952064 | 125.1662466 | 178.6289081 | ENSG00000187609 |
| NAPEPLD | 0.512969261 | 1.426984102 | 0.004294234 | 0.019491144 | 134.4094096 | 191.4714476 | ENSG00000161048 |
| NA | 0.512935834 | 1.426951038 | 0.578966878 | 0.759924207 | 4.668159834 | 6.645948291 | ENSG00000254428 |
| ELMO3 | 0.512915703 | 1.426931128 | 8.58E-05 | 0.000648038 | 257.1982781 | 366.6591488 | ENSG00000102890 |
| IKZF3 | 0.512785083 | 1.42680194 | 0.407596592 | 0.61857302 | 10.68618822 | 15.18046344 | ENSG00000161405 |
| HARS2 | 0.512383353 | 1.426404691 | 2.16E-06 | 2.34E-05 | 427.494586 | 609.5030881 | ENSG00000112855 |
| LINC01238 | 0.512176225 | 1.426199917 | 0.366360038 | 0.578876192 | 11.676934 | 16.60957384 | ENSG00000237940 |
| DUSP12 | 0.511981608 | 1.426007538 | 4.12E-07 | 5.14E-06 | 416.6225195 | 594.2969706 | ENSG00000081721 |
| CD99L2 | 0.511959179 | 1.425985369 | 4.24E-07 | 5.28E-06 | 502.6214316 | 716.0912137 | ENSG00000102181 |
| CENPF | 0.511908711 | 1.425935486 | 5.88E-13 | 1.74E-11 | 2926.816502 | 4172.210847 | ENSG00000117724 |
| NA | 0.511760305 | 1.425788812 | 0.089246846 | 0.219114234 | 44.50413492 | 63.57962306 | ENSG00000185834 |
| C8orf82 | 0.511647257 | 1.425677093 | 1.12E-07 | 1.54E-06 | 620.3513383 | 885.1759867 | ENSG00000213563 |
| CITED2 | 0.511502297 | 1.42553385 | 2.49E-09 | 4.51E-08 | 853.7564599 | 1216.795816 | ENSG00000164442 |
| ETNK2 | 0.51120096 | 1.425236129 | 2.78E-07 | 3.58E-06 | 430.9523071 | 614.0098896 | ENSG00000143845 |
| HES3 | 0.510779334 | 1.424819666 | 4.43E-05 | 0.000361677 | 307.654922 | 438.8746105 | ENSG00000173673 |
| LAMA5 | 0.510724137 | 1.424765153 | 2.84E-12 | 7.70E-11 | 3581.356205 | 5103.150807 | ENSG00000130702 |
| LOC339874 | 0.510710002 | 1.424751194 | 0.660901334 | 0.817732696 | 3.071876414 | 4.339426739 | ENSG00000250608 |
| GATAD2B | 0.510678544 | 1.424720128 | 2.83E-06 | 3.00E-05 | 432.7760645 | 616.8078896 | ENSG00000143614 |
| ENTPD1-AS1 | 0.509990478 | 1.424040797 | 0.452993115 | 0.659225936 | 8.643016576 | 12.2928477 | ENSG00000226688 |
| LYPD5 | 0.509923214 | 1.423974404 | 0.503598422 | 0.701361077 | 6.309521187 | 8.977345901 | ENSG00000159871 |
| B3GNT9 | 0.509829837 | 1.423882242 | 0.001999066 | 0.010153944 | 149.1962046 | 212.5671299 | ENSG00000237172 |
| ASH1L-AS1 | 0.509681826 | 1.423736168 | 0.090898264 | 0.221956778 | 52.1457273 | 74.37386112 | ENSG00000235919 |
| NA | 0.509630346 | 1.423685366 | 0.599565507 | 0.77551184 | 3.95705938 | 5.640373016 | ENSG00000280106 |
| ZBTB6 | 0.509365372 | 1.423423907 | 0.000654299 | 0.003871348 | 189.3550973 | 269.3917252 | ENSG00000186130 |
| USP53 | 0.509207792 | 1.42326844 | 8.07E-06 | 7.90E-05 | 327.4867196 | 465.8128481 | ENSG00000145390 |
| MDM4 | 0.509168676 | 1.423229852 | 6.48E-06 | 6.44E-05 | 368.2464215 | 523.8001692 | ENSG00000198625 |
| PHF7 | 0.509152906 | 1.423214295 | 0.014505253 | 0.053025881 | 116.543671 | 165.8232789 | ENSG00000010318 |
| UROC1 | 0.509141987 | 1.423203523 | 0.606796302 | 0.780598101 | 3.953499908 | 5.632900666 | ENSG00000159650 |
| EEF2KMT | 0.50867962 | 1.422747476 | 1.03E-05 | 9.80E-05 | 326.0319248 | 463.860603 | ENSG00000118894 |
| MYH2 | 0.508146737 | 1.422222058 | 0.116369967 | 0.267158623 | 39.94657191 | 56.68794029 | ENSG00000125414 |
| HPN | 0.507874798 | 1.421954003 | 0.001016709 | 0.005689054 | 222.2872768 | 316.0038524 | ENSG00000105707 |
| LMO3 | 0.507765706 | 1.421846484 | 0.00725096 | 0.030115324 | 139.5209761 | 198.5742253 | ENSG00000048540 |
| SIX4 | 0.507608924 | 1.421691976 | 5.26E-07 | 6.41E-06 | 480.9739369 | 683.7321461 | ENSG00000100625 |
| TRIM32 | 0.507280509 | 1.421368378 | 3.38E-07 | 4.28E-06 | 433.3118878 | 615.6817811 | ENSG00000119401 |
| PHPT1 | 0.506901519 | 1.42099504 | 1.62E-13 | 5.13E-12 | 1333.07218 | 1894.849716 | ENSG00000054148 |
| KCNAB1 | 0.506839218 | 1.420933677 | 0.297298863 | 0.507547372 | 18.59911177 | 26.35262452 | ENSG00000169282 |
| TRIM46 | 0.506688464 | 1.420785205 | 3.43E-05 | 0.000286852 | 475.9564947 | 676.7516091 | ENSG00000163462 |
| SCARA3 | 0.50663522 | 1.420732771 | 6.44E-06 | 6.41E-05 | 392.7776274 | 558.4644859 | ENSG00000168077 |
| TMEM168 | 0.506316534 | 1.42041897 | 0.000500491 | 0.003078753 | 226.9705643 | 322.3611241 | ENSG00000146802 |

| | | | | | | | |
|---|---|---|---|---|---|---|---|
| SYT1 | 0.506204063 | 1.420308241 | 2.51E-08 | 3.86E-07 | 605.4948546 | 860.0964133 | ENSG00000067715 |
| LTBP2 | 0.505669486 | 1.419782057 | 0.191356076 | 0.378285302 | 44.22212533 | 62.89545172 | ENSG00000119681 |
| NA | 0.505604241 | 1.41971785 | 0.685978435 | 0.831762616 | 3.095597513 | 4.356546901 | ENSG00000260329 |
| KIZ | 0.50515773 | 1.419278517 | 1.35E-06 | 1.53E-05 | 465.8852756 | 661.3371915 | ENSG00000088970 |
| OSGEPL1 | 0.504982605 | 1.419106246 | 0.018600432 | 0.064738142 | 82.00572489 | 116.3707565 | ENSG00000128694 |
| ZNF496 | 0.50494592 | 1.419070161 | 6.85E-21 | 4.37E-19 | 2469.699843 | 3504.340536 | ENSG00000162714 |
| MKNK1 | 0.504653009 | 1.418782075 | 0.00047258 | 0.002933816 | 202.1523665 | 286.5563307 | ENSG00000079277 |
| SLC6A15 | 0.504491179 | 1.418622937 | 8.91E-07 | 1.04E-05 | 467.4254972 | 663.1540906 | ENSG00000072041 |
| FUT9 | 0.504344671 | 1.418478881 | 0.032131143 | 0.100263819 | 75.28488405 | 106.859893 | ENSG00000172461 |
| OXER1 | 0.504290446 | 1.418425567 | 0.398451676 | 0.610274365 | 12.42807838 | 17.72061265 | ENSG00000162881 |
| SYNPO2L | 0.504135251 | 1.418272991 | 0.673407207 | 0.825137853 | 3.099156985 | 4.351250015 | ENSG00000166317 |
| LINC02367 | 0.503836009 | 1.417978845 | 0.507139254 | 0.704350311 | 6.554766997 | 9.338359676 | ENSG00000260423 |
| NA | 0.503777839 | 1.417921673 | 0.630879735 | 0.796625039 | 3.275603762 | 4.668092106 | ENSG00000266473 |
| NA | 0.503732861 | 1.417877468 | 0.679681358 | 0.828703108 | 3.213923675 | 4.580978244 | ENSG00000188985 |
| SLC23A1 | 0.503624528 | 1.417771002 | 0.532926703 | 0.724314769 | 5.645862931 | 8.005064992 | ENSG00000170482 |
| NA | 0.503587077 | 1.417734199 | 0.402556752 | 0.61393481 | 10.33182013 | 14.61336799 | ENSG00000269947 |
| THBS1 | 0.50357819 | 1.417725466 | 2.63E-06 | 2.81E-05 | 427.6758653 | 606.3988063 | ENSG00000137801 |
| IL13RA1 | 0.50357767 | 1.417724955 | 7.85E-08 | 1.12E-06 | 550.4753344 | 779.950653 | ENSG00000131724 |
| STK17A | 0.503442143 | 1.41759178 | 8.29E-05 | 0.000629228 | 279.9854828 | 397.0388768 | ENSG00000164543 |
| SLC16A12 | 0.503434934 | 1.417584696 | 0.039453664 | 0.117483828 | 75.37592964 | 106.9679835 | ENSG00000152779 |
| RICTOR | 0.503200201 | 1.417354067 | 0.009632936 | 0.037875911 | 189.0200012 | 267.6304062 | ENSG00000164327 |
| TMEM176B | 0.503170576 | 1.417324963 | 0.304857368 | 0.51500415 | 15.21438573 | 21.5655646 | ENSG00000106565 |
| NA | 0.502941135 | 1.417099575 | 0.484788971 | 0.68583712 | 8.232892131 | 11.70522712 | ENSG00000254910 |
| NA | 0.502935057 | 1.417093605 | 0.379237027 | 0.591794021 | 13.1427383 | 18.68977213 | ENSG00000237350 |
| DRAM1 | 0.502254156 | 1.416424945 | 0.005295644 | 0.023198386 | 127.7565298 | 181.059032 | ENSG00000136048 |
| NA | 0.502177466 | 1.416349653 | 0.514418236 | 0.710768831 | 6.371201275 | 8.989169178 | ENSG00000226329 |
| SF3B4 | 0.502116846 | 1.416290141 | 2.42E-13 | 7.47E-12 | 2130.078271 | 3017.580345 | ENSG00000143368 |
| NA | 0.501878835 | 1.416056506 | 0.253387555 | 0.459455015 | 20.12659616 | 28.5230581 | ENSG00000253981 |
| GBA | 0.501738705 | 1.41591897 | 0.001002087 | 0.005625568 | 187.7852303 | 266.2705248 | ENSG00000177628 |
| CAMK1G | 0.500370352 | 1.41457665 | 0.247962098 | 0.452966166 | 21.58914647 | 30.49098292 | ENSG00000008118 |
| MRGPRG-AS1 | 0.500285822 | 1.414493769 | 0.6361324 | 0.80066251 | 3.991458897 | 5.636022089 | ENSG00000236301 |
| ARHGEF2 | 0.500219922 | 1.414429159 | 1.65E-15 | 6.66E-14 | 1465.226734 | 2072.412311 | ENSG00000116584 |
| CRACR2A | 0.500194926 | 1.414404653 | 0.04151971 | 0.12215294 | 66.32576385 | 93.79358465 | ENSG00000130038 |
| NA | 0.500149649 | 1.414360265 | 0.041101829 | 0.121244616 | 69.07263768 | 97.78353734 | ENSG00000234160 |
| KANSL1-AS1 | 0.500079888 | 1.414291876 | 0.462814307 | 0.667551225 | 10.41722131 | 14.68742907 | ENSG00000214401 |
| MRPS31 | 0.499971973 | 1.414186089 | 8.64E-07 | 1.01E-05 | 478.251369 | 676.1247645 | ENSG00000102738 |
| NAPSB | 0.499848893 | 1.414065446 | 0.605642813 | 0.779795526 | 4.947805159 | 7.012542498 | ENSG00000131401 |
| TET2 | 0.499739542 | 1.413958269 | 0.134835576 | 0.296468722 | 34.51867422 | 48.74539665 | ENSG00000168769 |
| PGPEP1 | 0.499693585 | 1.413913228 | 1.26E-08 | 2.04E-07 | 609.4898474 | 861.5777051 | ENSG00000130517 |
| MDGA1 | 0.49969049 | 1.413910195 | 0.216077458 | 0.411436055 | 30.70989132 | 43.36352735 | ENSG00000112139 |
| OXTR | 0.499512127 | 1.413735402 | 0.072365598 | 0.187697339 | 58.97093575 | 83.25534834 | ENSG00000180914 |
| RUSC1 | 0.49945539 | 1.413679805 | 2.37E-11 | 5.74E-10 | 1311.285665 | 1854.440489 | ENSG00000160753 |
| NA | 0.499452837 | 1.413677303 | 0.613390599 | 0.784815818 | 4.231950025 | 6.027208807 | ENSG00000249119 |
| PNMA3 | 0.499378424 | 1.413604388 | 1.34E-05 | 0.000123951 | 340.4091546 | 480.8794297 | ENSG00000183837 |
| CD177 | 0.499360028 | 1.413586364 | 0.139706628 | 0.30364365 | 35.23452935 | 49.71578565 | ENSG00000204936 |
| NA | 0.499259323 | 1.413487694 | 0.24200699 | 0.445535388 | 24.48310157 | 34.63492176 | ENSG00000249456 |
| MDM2 | 0.499203555 | 1.413433056 | 3.98E-05 | 0.000328061 | 400.1605479 | 565.3663703 | ENSG00000135679 |
| NFIB | 0.499063704 | 1.413296049 | 0.103524782 | 0.244383188 | 41.00820166 | 57.92787827 | ENSG00000147862 |
| SMARCD3 | 0.498835741 | 1.413072749 | 0.004638499 | 0.020733267 | 132.6491634 | 187.3513662 | ENSG00000082014 |
| RCAN3 | 0.498593037 | 1.412835048 | 0.00018224 | 0.00126903 | 254.2396939 | 359.1679495 | ENSG00000117602 |
| PHYH | 0.498275156 | 1.412523781 | 0.000141726 | 0.001017293 | 265.2013576 | 374.2341522 | ENSG00000107537 |
| NA | 0.498196406 | 1.41244668 | 0.427832944 | 0.637657464 | 9.633762356 | 13.5819707 | ENSG00000273456 |
| PLGRKT | 0.4981925 | 1.412442855 | 0.001137998 | 0.006282108 | 179.5273963 | 253.2710458 | ENSG00000107020 |
| RAB9B | 0.498186468 | 1.41243695 | 0.28173191 | 0.490709708 | 20.54383955 | 29.04430086 | ENSG00000123570 |
| NA | 0.49815417 | 1.412405329 | 0.229644371 | 0.429517079 | 22.79310281 | 32.21763979 | ENSG00000224063 |
| SGIP1 | 0.498010206 | 1.412264395 | 0.400381168 | 0.611796938 | 10.61382972 | 14.99273143 | ENSG00000118473 |
| ATP6AP1L | 0.497876354 | 1.412133373 | 0.392599385 | 0.604159012 | 12.78036154 | 17.96774458 | ENSG00000205464 |
| MTMR10 | 0.497854109 | 1.412111599 | 0.000420343 | 0.002641433 | 203.4811832 | 287.5310706 | ENSG00000166912 |
| NA | 0.497809573 | 1.412068008 | 0.155591111 | 0.327409423 | 31.82132753 | 44.94367043 | ENSG00000272977 |
| SNCA | 0.49734369 | 1.411612089 | 0.000402935 | 0.002542409 | 200.3814158 | 282.7712301 | ENSG00000145335 |
| FOXH1 | 0.497256625 | 1.411526902 | 6.99E-10 | 1.37E-08 | 1143.188735 | 1614.701711 | ENSG00000160973 |
| NA | 0.497225612 | 1.41149656 | 0.446304965 | 0.653521693 | 9.125194042 | 12.94923247 | ENSG00000231073 |
| FBLN2 | 0.497171767 | 1.41144388 | 8.98E-10 | 1.73E-08 | 1167.298374 | 1646.988912 | ENSG00000163520 |
| MALAT1 | 0.496753627 | 1.411034856 | 6.20E-09 | 1.06E-07 | 1063.080802 | 1501.000425 | ENSG00000251562 |
| VWA7 | 0.496688855 | 1.410971508 | 0.389911836 | 0.601542768 | 12.41028101 | 17.58280071 | ENSG00000204396 |
| FEZF1 | 0.49664036 | 1.41092408 | 0.000264393 | 0.001762466 | 224.0333107 | 316.2191601 | ENSG00000128610 |
| PLEKHH1 | 0.49658463 | 1.410869578 | 8.92E-08 | 1.26E-06 | 674.699368 | 951.5234367 | ENSG00000054690 |
| NA | 0.496394875 | 1.410684021 | 0.24844125 | 0.453402881 | 21.45866735 | 30.32718101 | ENSG00000220842 |
| RFX5 | 0.495821949 | 1.410123919 | 2.31E-10 | 4.88E-09 | 1059.444804 | 1493.542425 | ENSG00000143390 |
| NA | 0.495778642 | 1.41008159 | 0.602190644 | 0.77717285 | 4.00806105 | 5.628549739 | ENSG00000255216 |
| MINDY1 | 0.495668026 | 1.409973479 | 0.00144699 | 0.007723688 | 170.4505604 | 240.122258 | ENSG00000143409 |
| HASPIN | 0.495645932 | 1.409951886 | 1.97E-05 | 0.000176255 | 312.988189 | 441.001113 | ENSG00000177602 |
| HOMEZ | 0.49552206 | 1.409830831 | 0.002514663 | 0.012367035 | 183.2051676 | 258.5280309 | ENSG00000215271 |
| IGFBPL1 | 0.495373659 | 1.409685818 | 3.19E-12 | 8.57E-11 | 1329.811198 | 1874.379926 | ENSG00000137142 |

| | | | | | | | |
|---|---|---|---|---|---|---|---|
| SYNM | 0.495246966 | 1.409562029 | 0.035026874 | 0.107253389 | 87.11540788 | 122.5031452 | ENSG00000182253 |
| NA | 0.495115499 | 1.409433586 | 0.636742974 | 0.800725077 | 3.299324861 | 4.658444293 | ENSG00000260163 |
| C6orf99 | 0.495066205 | 1.409385431 | 0.263145838 | 0.470286743 | 21.32462876 | 30.11625395 | ENSG00000203711 |
| NA | 0.494948575 | 1.40927052 | 0.538823539 | 0.728551767 | 5.676702975 | 7.977067512 | ENSG00000231120 |
| SLC16A8 | 0.494916725 | 1.409239409 | 0.32550756 | 0.536686771 | 16.72170849 | 23.65512716 | ENSG00000100156 |
| AKAP7 | 0.494915142 | 1.409237863 | 0.000129384 | 0.000934027 | 258.7328815 | 364.3175362 | ENSG00000118507 |
| RBIS | 0.49487526 | 1.409198906 | 0.047540394 | 0.135701383 | 61.29019324 | 86.35031822 | ENSG00000176731 |
| NA | 0.494701681 | 1.409029367 | 0.609526734 | 0.782469742 | 4.218907344 | 5.987388051 | ENSG00000265460 |
| TUBB3 | 0.494640802 | 1.40896991 | 0.361705627 | 0.573767965 | 13.12257668 | 18.5292596 | ENSG00000258947 |
| RBFOX3 | 0.494076086 | 1.408418503 | 5.58E-06 | 5.63E-05 | 456.5314112 | 643.4127453 | ENSG00000167281 |
| TDH | 0.493948974 | 1.408294416 | 0.327312612 | 0.538722214 | 16.1034075 | 22.57861222 | ENSG00000154316 |
| RBP5 | 0.493823316 | 1.40817176 | 0.261807614 | 0.468843082 | 21.30001794 | 29.96087035 | ENSG00000139194 |
| PET117 | 0.493386285 | 1.407745252 | 0.569944329 | 0.752843159 | 5.689745657 | 7.985769366 | ENSG00000232838 |
| LINC01234 | 0.493355882 | 1.407715585 | 0.12794601 | 0.285475 | 37.07249911 | 52.2095877 | ENSG00000249550 |
| NA | 0.49325272 | 1.407614928 | 0.571788768 | 0.754382485 | 4.93356727 | 6.955318033 | ENSG00000286459 |
| PMEPA1 | 0.49315789 | 1.407522407 | 1.44E-06 | 1.62E-05 | 519.5825656 | 731.7563159 | ENSG00000124225 |
| MORN1 | 0.493143765 | 1.407508627 | 0.027766662 | 0.089483518 | 94.65471759 | 133.178699 | ENSG00000116151 |
| IFIT5 | 0.493101252 | 1.407467152 | 0.074862512 | 0.192383452 | 53.61628628 | 75.59060404 | ENSG00000152778 |
| ZNF285 | 0.49306649 | 1.407433238 | 0.13104047 | 0.290730648 | 38.72481821 | 54.31918094 | ENSG00000267508 |
| COL5A3 | 0.492809284 | 1.407182342 | 0.057183248 | 0.156530748 | 69.06404419 | 97.39112112 | ENSG00000080573 |
| DUSP19 | 0.492592665 | 1.406971071 | 0.430857256 | 0.640589378 | 10.52842853 | 14.80980185 | ENSG00000162999 |
| NA | 0.492580996 | 1.406959691 | 0.348511531 | 0.560425772 | 14.10027977 | 19.89584865 | ENSG00000287670 |
| ELOVL2 | 0.492549602 | 1.406929075 | 0.121207574 | 0.274362344 | 42.8393318 | 60.24482564 | ENSG00000197977 |
| PROSER3 | 0.492368218 | 1.406752198 | 0.003446672 | 0.016211806 | 143.6301246 | 202.0186263 | ENSG00000167595 |
| PLA2G6 | 0.492318778 | 1.406703991 | 0.011020654 | 0.042344873 | 126.8149801 | 178.317195 | ENSG00000184381 |
| UBR4 | 0.492278446 | 1.406664666 | 0.000430577 | 0.002697895 | 4397.878707 | 6185.830616 | ENSG00000127481 |
| HCG23 | 0.492271634 | 1.406658024 | 0.15161446 | 0.321279071 | 33.0229196 | 46.49146509 | ENSG00000225914 |
| ZDHHC24 | 0.491868067 | 1.406264593 | 0.002150628 | 0.010807888 | 165.4049217 | 232.8085961 | ENSG00000174165 |
| CHTOP | 0.491560749 | 1.405965067 | 4.44E-14 | 1.48E-12 | 1744.604185 | 2452.953041 | ENSG00000160679 |
| PTCD2 | 0.491530507 | 1.405935595 | 0.001908869 | 0.009750874 | 156.3570941 | 219.7570215 | ENSG00000049883 |
| BTN2A2 | 0.49092391 | 1.405344578 | 0.002128498 | 0.010733429 | 211.7591276 | 297.3737638 | ENSG00000124508 |
| MGST3 | 0.490846732 | 1.405269401 | 1.14E-10 | 2.53E-09 | 1193.518279 | 1677.577343 | ENSG00000143198 |
| TMEM269 | 0.490778786 | 1.405203219 | 0.537712542 | 0.727674766 | 6.650846601 | 9.343656562 | ENSG00000274386 |
| SHARPIN | 0.490549348 | 1.404979761 | 1.47E-08 | 2.38E-07 | 717.2914987 | 1008.293069 | ENSG00000179526 |
| TCAIM | 0.490416673 | 1.40485056 | 0.002231196 | 0.011163236 | 191.4353638 | 269.1905616 | ENSG00000179152 |
| NA | 0.490180422 | 1.404620526 | 0.352898102 | 0.565169662 | 17.38803649 | 24.38585656 | ENSG00000263412 |
| PRR16 | 0.490084238 | 1.404526882 | 0.465581546 | 0.669803268 | 8.308810108 | 11.66540636 | ENSG00000184838 |
| NA | 0.490069201 | 1.404512244 | 0.606640279 | 0.780515171 | 4.246187915 | 5.997035864 | ENSG00000226986 |
| CADM4 | 0.489969277 | 1.404414967 | 5.65E-06 | 5.70E-05 | 584.1443675 | 820.4406814 | ENSG00000105767 |
| ZMAT1 | 0.489877493 | 1.404325621 | 0.299344839 | 0.509612123 | 18.48998949 | 26.01669771 | ENSG00000166432 |
| ZSWIM9 | 0.489534821 | 1.403992103 | 7.46E-06 | 7.36E-05 | 383.0696753 | 537.406973 | ENSG00000185453 |
| NA | 0.489399648 | 1.403860563 | 0.308252517 | 0.518716402 | 17.68873317 | 24.92277907 | ENSG00000231369 |
| RIMS1 | 0.489115351 | 1.403583946 | 0.130619935 | 0.290036153 | 44.66634381 | 62.76440676 | ENSG00000079841 |
| UNC13D | 0.488442342 | 1.402929334 | 0.002907814 | 0.014015841 | 159.6015523 | 223.999403 | ENSG00000092929 |
| NSMCE1 | 0.488363551 | 1.402852717 | 0.002413621 | 0.011951032 | 201.8095666 | 283.1382462 | ENSG00000169189 |
| TMEM241 | 0.488296454 | 1.402787475 | 0.015715125 | 0.056595802 | 111.8144415 | 157.0079156 | ENSG00000134490 |
| ZNF839 | 0.488269405 | 1.402761174 | 0.006767283 | 0.028463243 | 120.6923048 | 169.3733114 | ENSG00000022976 |
| CTBS | 0.487964964 | 1.402465192 | 0.504090724 | 0.701693513 | 7.399906043 | 10.33211168 | ENSG00000117151 |
| CLYBL | 0.487961915 | 1.402462227 | 0.008985568 | 0.035897046 | 114.0417375 | 159.7926222 | ENSG00000125246 |
| ACADVL | 0.487723878 | 1.402230848 | 0.011991612 | 0.045501628 | 2570.398928 | 3604.35215 | ENSG00000072778 |
| PLCD3 | 0.487540432 | 1.402052559 | 1.50E-07 | 2.01E-06 | 674.9652367 | 946.7100603 | ENSG00000161714 |
| JAK1 | 0.487407761 | 1.40192363 | 2.62E-08 | 4.01E-07 | 657.7372469 | 921.6298378 | ENSG00000162434 |
| KDR | 0.48692989 | 1.401459341 | 2.98E-12 | 8.07E-11 | 3297.609962 | 4619.994334 | ENSG00000128052 |
| STEAP1 | 0.486730983 | 1.401266133 | 0.017784355 | 0.062520019 | 94.26920396 | 132.1200393 | ENSG00000164647 |
| NA | 0.486593419 | 1.401132526 | 0.659594508 | 0.816601696 | 3.361004949 | 4.69826505 | ENSG00000259376 |
| GRIK4 | 0.486180607 | 1.400731663 | 0.020764048 | 0.0709479 | 112.6210112 | 157.4805928 | ENSG00000149403 |
| LYRM1 | 0.485875651 | 1.40043561 | 0.000336772 | 0.002169887 | 234.395666 | 327.9169149 | ENSG00000102897 |
| MFAP3L | 0.485665773 | 1.400231893 | 0.021979847 | 0.074322185 | 89.0159736 | 124.4962297 | ENSG00000198948 |
| COLCA1 | 0.485472748 | 1.400044563 | 0.581983739 | 0.761962097 | 5.009485247 | 6.999489717 | ENSG00000196167 |
| SBNO2 | 0.485448837 | 1.400021359 | 1.84E-12 | 5.13E-11 | 1387.882406 | 1943.915643 | ENSG00000064932 |
| GBX2 | 0.485401826 | 1.399975739 | 0.10021295 | 0.238382865 | 46.35929165 | 64.73746052 | ENSG00000168505 |
| MTFP1 | 0.485254924 | 1.399833194 | 0.247818695 | 0.452799082 | 23.58220617 | 33.09750995 | ENSG00000242114 |
| NA | 0.485193136 | 1.399773243 | 0.653873755 | 0.81254148 | 3.313562751 | 4.632622277 | ENSG00000238230 |
| HPS6 | 0.485065082 | 1.399649004 | 6.49E-07 | 7.76E-06 | 527.4906304 | 738.157453 | ENSG00000166189 |
| ACAD10 | 0.485007035 | 1.399592691 | 0.000472802 | 0.002934229 | 300.7197283 | 420.7961781 | ENSG00000111271 |
| MKRN2OS | 0.484882495 | 1.399471876 | 0.581026663 | 0.761150628 | 5.005925775 | 6.990787862 | ENSG00000225526 |
| NA | 0.484657442 | 1.399253583 | 0.547451886 | 0.735112974 | 5.945669884 | 8.334959865 | ENSG00000243964 |
| ANGEL2 | 0.484608957 | 1.399206559 | 1.63E-06 | 1.81E-05 | 619.577668 | 867.2716008 | ENSG00000174606 |
| KCTD12 | 0.48452869 | 1.399128714 | 0.001304706 | 0.007057968 | 263.0193665 | 367.6100836 | ENSG00000178695 |
| SRXN1 | 0.48435984 | 1.398964972 | 0.051465488 | 0.144266847 | 76.13950631 | 106.4917858 | ENSG00000271303 |
| OSCAR | 0.483795425 | 1.398417772 | 0.090334919 | 0.221038475 | 50.96757699 | 71.44034973 | ENSG00000170909 |
| BROX | 0.483765557 | 1.398388821 | 0.000135343 | 0.000975185 | 256.4996356 | 358.6273158 | ENSG00000162819 |
| NPIPB6 | 0.48375974 | 1.398383183 | 0.339821033 | 0.551428098 | 15.94920728 | 22.28447073 | ENSG00000198156 |
| SNORD94 | 0.483612516 | 1.398240488 | 0.571065306 | 0.753841189 | 5.696864602 | 7.922302056 | ENSG00000208772 |

| | | | | | | | |
|---|---|---|---|---|---|---|---|
| FTCD | 0.483312763 | 1.397950002 | 0.13182209 | 0.291931389 | 42.90101189 | 59.92363262 | ENSG00000160282 |
| PLXNB3 | 0.482808835 | 1.397461788 | 0.131281911 | 0.291029168 | 47.66021731 | 66.65641123 | ENSG00000198753 |
| MEGF10 | 0.482716438 | 1.397372291 | 2.34E-05 | 0.000205592 | 599.5455996 | 837.3338396 | ENSG00000145794 |
| CCDC71L | 0.482226055 | 1.396897394 | 3.90E-06 | 4.04E-05 | 424.3121645 | 593.0830226 | ENSG00000253276 |
| MTFMT | 0.482046485 | 1.396723536 | 0.000400774 | 0.002530465 | 346.0743667 | 483.7518898 | ENSG00000103707 |
| ANTKMT | 0.481855416 | 1.396538568 | 4.45E-05 | 0.000363015 | 383.2784617 | 535.8482284 | ENSG00000103254 |
| UAP1 | 0.481661287 | 1.396350662 | 2.13E-10 | 4.52E-09 | 876.8841526 | 1224.604467 | ENSG00000117143 |
| LINC01535 | 0.481374903 | 1.396073505 | 0.51638351 | 0.712545856 | 8.425051338 | 11.72357678 | ENSG00000226686 |
| ZNF678 | 0.48124163 | 1.395944544 | 0.001469486 | 0.00782166 | 188.1825914 | 262.8877719 | ENSG00000181450 |
| SYDE1 | 0.481191597 | 1.395896134 | 9.24E-06 | 8.91E-05 | 440.4315893 | 615.2648676 | ENSG00000105137 |
| SLC44A3-AS1 | 0.481157525 | 1.395863167 | 0.210085544 | 0.403443716 | 29.11597216 | 40.50095562 | ENSG00000224081 |
| MACROD1 | 0.480479858 | 1.395207652 | 3.87E-05 | 0.000320082 | 369.1944274 | 514.8455239 | ENSG00000133315 |
| MATN3 | 0.480330852 | 1.395063558 | 0.009129992 | 0.036283649 | 129.3371008 | 180.6912082 | ENSG00000132031 |
| CDHR3 | 0.480202423 | 1.394939375 | 0.011202445 | 0.042912408 | 132.5314477 | 185.0487441 | ENSG00000128536 |
| NA | 0.480073155 | 1.394814392 | 0.648771934 | 0.809250856 | 4.055503248 | 5.614550999 | ENSG00000257043 |
| AK5 | 0.480023059 | 1.394765959 | 0.031368855 | 0.098421802 | 78.61059976 | 109.5638441 | ENSG00000154027 |
| CPNE8 | 0.479939204 | 1.394684892 | 0.011786634 | 0.044820893 | 113.1126718 | 157.6254256 | ENSG00000139117 |
| GPR179 | 0.47881531 | 1.393598823 | 0.66111068 | 0.817849134 | 3.326605432 | 4.620799 | ENSG00000277399 |
| NA | 0.478768901 | 1.393553994 | 0.296651589 | 0.506875709 | 23.52347517 | 32.67352332 | ENSG00000259048 |
| PROC | 0.478712248 | 1.393499272 | 0.306837228 | 0.517146011 | 17.83225497 | 24.8571363 | ENSG00000115718 |
| UGDH-AS1 | 0.478507213 | 1.393301242 | 0.521408238 | 0.716145653 | 7.776520696 | 10.89173478 | ENSG00000249348 |
| NT5DC1 | 0.478096143 | 1.392904302 | 1.78E-08 | 2.82E-07 | 666.4354344 | 928.4250948 | ENSG00000178425 |
| SPSB1 | 0.47807405 | 1.392882972 | 0.021971808 | 0.074308309 | 100.6190484 | 140.1624661 | ENSG00000171621 |
| ZNF486 | 0.4779763 | 1.3927886 | 2.71E-05 | 0.00023364 | 362.3968829 | 504.7738376 | ENSG00000256229 |
| NA | 0.477760622 | 1.392580399 | 0.315729166 | 0.525792898 | 20.75852465 | 28.79232354 | ENSG00000272419 |
| NA | 0.477729389 | 1.392550251 | 0.466478226 | 0.670531382 | 8.597938643 | 11.98659938 | ENSG00000133519 |
| NA | 0.477446354 | 1.39227708 | 0.684770624 | 0.831472593 | 3.310003278 | 4.597152447 | ENSG00000186676 |
| LINC00869 | 0.477294364 | 1.39213041 | 0.074526265 | 0.191800015 | 57.8132264 | 80.39594099 | ENSG00000277147 |
| LOC101927837 | 0.477151591 | 1.391992648 | 0.619928406 | 0.788958013 | 4.246187915 | 5.923920741 | ENSG00000270049 |
| RAB3GAP2 | 0.476895312 | 1.391745398 | 7.08E-10 | 1.38E-08 | 1624.381242 | 2260.0012 | ENSG00000118873 |
| SLC27A3 | 0.476864296 | 1.391715477 | 6.92E-07 | 8.23E-06 | 626.7133095 | 872.7224155 | ENSG00000143554 |
| NA | 0.476740411 | 1.391595975 | 0.579979646 | 0.760671964 | 5.033206346 | 6.969316773 | ENSG00000271943 |
| LRCH4 | 0.476599145 | 1.391459719 | 0.000154812 | 0.001098279 | 268.9087214 | 374.1775902 | ENSG00000077454 |
| STOM | 0.476509419 | 1.391373182 | 7.19E-07 | 8.52E-06 | 642.5856836 | 893.8681632 | ENSG00000148175 |
| TPM1 | 0.476508299 | 1.391372102 | 3.80E-22 | 2.69E-20 | 4123.191569 | 5736.916177 | ENSG00000140416 |
| NT5C2 | 0.476335268 | 1.391205236 | 1.41E-07 | 1.90E-06 | 641.5492756 | 892.355169 | ENSG00000076685 |
| ADAMTS16 | 0.476316522 | 1.391187159 | 0.001328655 | 0.007171093 | 213.5577156 | 297.3732264 | ENSG00000145536 |
| LINC01841 | 0.47624211 | 1.391115407 | 0.551638865 | 0.738582061 | 5.958712565 | 8.28426179 | ENSG00000266913 |
| NA | 0.47614939 | 1.391026004 | 0.544247146 | 0.73258071 | 5.996671554 | 8.330608937 | ENSG00000215895 |
| OR2A42 | 0.475898586 | 1.390784204 | 0.666941287 | 0.82154232 | 4.127861753 | 5.692017049 | ENSG00000212807 |
| NA | 0.47582122 | 1.390709623 | 0.628487605 | 0.794869303 | 4.284146904 | 5.963741498 | ENSG00000275630 |
| B3GALNT2 | 0.475793325 | 1.390682733 | 9.00E-07 | 1.05E-05 | 541.222331 | 752.1001539 | ENSG00000162885 |
| SLC66A1L | 0.475544617 | 1.390443012 | 0.447434742 | 0.654212225 | 10.22269784 | 14.21688439 | ENSG00000174899 |
| RPRD2 | 0.475407077 | 1.390310461 | 4.27E-08 | 6.31E-07 | 794.5964448 | 1104.283799 | ENSG00000163125 |
| GFI1B | 0.474957727 | 1.389877494 | 0.674895265 | 0.825790913 | 3.353886004 | 4.640094626 | ENSG00000165702 |
| NA | 0.474891412 | 1.389813609 | 0.058345846 | 0.158975805 | 71.74598394 | 99.86948402 | ENSG00000246695 |
| ZNF888 | 0.474781365 | 1.389707599 | 0.009892196 | 0.038677872 | 118.7087283 | 164.825133 | ENSG00000213793 |
| ESD | 0.474772817 | 1.389699365 | 1.21E-09 | 2.31E-08 | 806.4610621 | 1120.784327 | ENSG00000139684 |
| POGZ | 0.474520811 | 1.389456638 | 8.60E-13 | 2.49E-11 | 1328.256075 | 1845.459765 | ENSG00000143442 |
| AHCTF1 | 0.474212873 | 1.389160094 | 1.05E-10 | 2.35E-09 | 1524.121421 | 2116.718836 | ENSG00000153207 |
| FNIP1 | 0.473837334 | 1.388798538 | 0.007769934 | 0.031932511 | 164.7684917 | 228.8612591 | ENSG00000217128 |
| FAM181B | 0.473383439 | 1.388361668 | 0.202045059 | 0.393281894 | 30.57109805 | 42.35499861 | ENSG00000182103 |
| PITRM1-AS1 | 0.473382882 | 1.388361133 | 0.617271643 | 0.787224317 | 4.331589101 | 6.00885914 | ENSG00000237399 |
| NA | 0.473158286 | 1.388145012 | 0.622564082 | 0.791044071 | 4.317351211 | 5.989563514 | ENSG00000278993 |
| LINC01122 | 0.473030679 | 1.388022235 | 0.277414018 | 0.486558495 | 24.07920635 | 33.35573012 | ENSG00000233723 |
| NA | 0.473012742 | 1.388004978 | 0.438023323 | 0.646255505 | 12.28603111 | 17.06734929 | ENSG00000270012 |
| CLIC6 | 0.472806131 | 1.387806214 | 0.051672931 | 0.144744712 | 68.78734795 | 95.3556616 | ENSG00000159212 |
| SLITRK2 | 0.472524553 | 1.387535374 | 0.344855813 | 0.556903268 | 15.29622744 | 21.16785149 | ENSG00000185985 |
| VANGL2 | 0.47220941 | 1.387232314 | 4.40E-14 | 1.47E-12 | 2027.755487 | 2812.340005 | ENSG00000162738 |
| CEP68 | 0.472202267 | 1.387225446 | 2.95E-07 | 3.77E-06 | 579.781432 | 804.5148773 | ENSG00000011523 |
| SLC4A5 | 0.472154328 | 1.38717935 | 0.660416053 | 0.817373256 | 3.391844992 | 4.672443033 | ENSG00000188687 |
| IRX2 | 0.47209541 | 1.387122701 | 0.000519767 | 0.003180022 | 282.7234576 | 392.7405433 | ENSG00000170561 |
| SNX21 | 0.471989358 | 1.387020738 | 0.000669168 | 0.003947904 | 223.7643438 | 310.5242325 | ENSG00000124104 |
| RAB20 | 0.471959275 | 1.386991816 | 4.81E-05 | 0.000388305 | 323.5966793 | 448.7681994 | ENSG00000139832 |
| IGFBP3 | 0.471880141 | 1.386915739 | 0.08307522 | 0.207770565 | 55.18083993 | 76.53488747 | ENSG00000146674 |
| GPATCH4 | 0.471450329 | 1.386502607 | 1.04E-11 | 2.63E-10 | 1825.195716 | 2531.336084 | ENSG00000160818 |
| NEK5 | 0.470964637 | 1.386035911 | 0.546565853 | 0.734549967 | 6.929296718 | 9.60885462 | ENSG00000197168 |
| INTS5 | 0.470366128 | 1.385461027 | 1.23E-07 | 1.68E-06 | 622.0742365 | 862.3302845 | ENSG00000185085 |
| EFNA1 | 0.470037525 | 1.385145496 | 0.086448687 | 0.213879248 | 56.11374443 | 77.63353591 | ENSG00000169242 |
| ACBD6 | 0.469869628 | 1.384984306 | 6.98E-10 | 1.37E-08 | 1127.990186 | 1562.222828 | ENSG00000230124 |
| TLR5 | 0.469868362 | 1.38498309 | 0.061978979 | 0.166307617 | 64.55539792 | 89.23169112 | ENSG00000187554 |
| GIPR | 0.46975899 | 1.384878097 | 0.447553277 | 0.654334879 | 10.34130334 | 14.28564858 | ENSG00000010310 |
| SMIM29 | 0.469714872 | 1.384835748 | 0.033390661 | 0.103306459 | 81.64215292 | 112.9857554 | ENSG00000186577 |
| PITPNM3 | 0.469633331 | 1.38475748 | 0.041703222 | 0.12254012 | 105.2193251 | 145.8418092 | ENSG00000091622 |

| | | | | | | | |
|---|---|---|---|---|---|---|---|
| GPRC5A | 0.469020406 | 1.384169294 | 0.402179871 | 0.613595236 | 13.03509056 | 18.00416037 | ENSG00000013588 |
| NA | 0.469019257 | 1.384168191 | 0.550757558 | 0.738029978 | 6.020392653 | 8.302611457 | ENSG00000261461 |
| NA | 0.468764129 | 1.383923436 | 0.661487747 | 0.818194662 | 3.602691286 | 5.010756214 | ENSG00000254595 |
| PNMA5 | 0.468762159 | 1.383921546 | 0.548575251 | 0.736074416 | 7.204187363 | 10.01526958 | ENSG00000198883 |
| PPP1R15A | 0.468716047 | 1.383877312 | 6.50E-05 | 0.000507384 | 369.0717038 | 510.855215 | ENSG00000087074 |
| ITLN2 | 0.46869781 | 1.383859819 | 0.039631625 | 0.117902211 | 90.32429754 | 124.7957407 | ENSG00000158764 |
| KCNK17 | 0.468621789 | 1.383786901 | 0.089374066 | 0.21936954 | 53.89056653 | 74.47760068 | ENSG00000124780 |
| PDGFRA | 0.4682203 | 1.383401859 | 0.387526815 | 0.599823285 | 14.05520184 | 19.50154051 | ENSG00000134853 |
| KDF1 | 0.468086874 | 1.383273922 | 0.000101177 | 0.000749599 | 348.9533957 | 482.6430921 | ENSG00000175707 |
| PLEKHH3 | 0.467921386 | 1.383115259 | 3.87E-07 | 4.85E-06 | 553.5403706 | 765.6181331 | ENSG00000068137 |
| LPAR1 | 0.467899923 | 1.383094683 | 0.055556577 | 0.153054115 | 68.40925875 | 94.47624294 | ENSG00000198121 |
| PLSCR1 | 0.467892184 | 1.383087264 | 5.59E-07 | 6.76E-06 | 569.4296778 | 787.1901812 | ENSG00000188313 |
| NME3 | 0.467773865 | 1.382973838 | 1.22E-08 | 1.99E-07 | 1241.514533 | 1717.72892 | ENSG00000103024 |
| ARHGAP5 | 0.467761154 | 1.382961653 | 3.97E-05 | 0.000327459 | 404.201229 | 558.5633574 | ENSG00000100852 |
| C1orf127 | 0.467371505 | 1.382588188 | 0.656750992 | 0.814576546 | 3.561172824 | 4.953815295 | ENSG00000175262 |
| POU2F2 | 0.467329732 | 1.382548156 | 0.049632727 | 0.140274475 | 99.02396019 | 136.7585779 | ENSG00000028277 |
| CASK | 0.466729792 | 1.381973347 | 7.91E-09 | 1.33E-07 | 1062.89846 | 1468.438436 | ENSG00000147044 |
| GPC4 | 0.466630395 | 1.381878137 | 1.26E-18 | 6.68E-17 | 16236.80445 | 22437.99254 | ENSG00000076716 |
| NA | 0.466627213 | 1.381875089 | 0.186047088 | 0.370974475 | 31.24543472 | 43.09660533 | ENSG00000259612 |
| SACS | 0.466206102 | 1.38147179 | 1.53E-06 | 1.71E-05 | 1110.405489 | 1533.320505 | ENSG00000151835 |
| SOSTDC1 | 0.466115422 | 1.381384961 | 0.67659768 | 0.827022925 | 3.333724377 | 4.576627316 | ENSG00000171243 |
| SHC2 | 0.465623585 | 1.380914105 | 0.015582724 | 0.056213894 | 111.9202836 | 154.5815756 | ENSG00000129946 |
| C1RL | 0.465581079 | 1.38087342 | 0.004015045 | 0.018444205 | 188.2312033 | 259.4859866 | ENSG00000139178 |
| ATRIP | 0.465493199 | 1.380789308 | 3.12E-05 | 0.000264703 | 345.8139154 | 477.5574444 | ENSG00000164053 |
| NA | 0.465457933 | 1.380755556 | 0.259199797 | 0.466162945 | 27.2631797 | 37.69220339 | ENSG00000228158 |
| USHBP1 | 0.465265876 | 1.380571757 | 0.020806239 | 0.071057749 | 115.2685252 | 159.046991 | ENSG00000130307 |
| NA | 0.464867912 | 1.380190982 | 0.531207631 | 0.72312089 | 8.961789946 | 12.37873206 | ENSG00000223529 |
| GPR89B | 0.464813577 | 1.380139002 | 0.013682635 | 0.050477939 | 120.8737594 | 166.8346026 | ENSG00000188092 |
| NA | 0.46469228 | 1.38002297 | 0.624654598 | 0.7923564 | 4.30430853 | 5.926096204 | ENSG00000240695 |
| SMTN | 0.464639274 | 1.379972267 | 0.00035232 | 0.00225774 | 271.3726959 | 374.5992763 | ENSG00000183963 |
| PRUNE1 | 0.464488133 | 1.379827705 | 4.20E-07 | 5.23E-06 | 689.3713707 | 950.6497867 | ENSG00000143363 |
| HHLA3 | 0.464443489 | 1.379785007 | 0.038781716 | 0.116031797 | 76.82691181 | 106.0154927 | ENSG00000197568 |
| PCDHGA12 | 0.464413134 | 1.379755976 | 0.60056614 | 0.776296479 | 5.325894353 | 7.345558793 | ENSG00000253159 |
| NA | 0.464353543 | 1.379698986 | 0.562782964 | 0.747671915 | 6.03107107 | 8.281140368 | ENSG00000226564 |
| LPP | 0.4642068 | 1.379558658 | 0.000400643 | 0.002530465 | 403.0461894 | 555.4451925 | ENSG00000145012 |
| SH3BP5L | 0.463970638 | 1.379332849 | 9.08E-12 | 2.31E-10 | 1247.793801 | 1720.857906 | ENSG00000175137 |
| PTPN3 | 0.463940429 | 1.379303967 | 0.004314835 | 0.019559689 | 154.0977628 | 212.7457085 | ENSG00000070159 |
| OXLD1 | 0.46356838 | 1.378948312 | 6.03E-06 | 6.04E-05 | 482.3270333 | 665.3986509 | ENSG00000204237 |
| ZCCHC4 | 0.463269403 | 1.378662574 | 0.010222048 | 0.039745395 | 129.9529858 | 179.1091006 | ENSG00000168228 |
| CRPPA | 0.463248266 | 1.378642376 | 0.465185588 | 0.669437616 | 9.595803367 | 13.21225507 | ENSG00000214960 |
| GAREM1 | 0.462912299 | 1.378321363 | 0.046907722 | 0.134260509 | 82.05525202 | 113.1589942 | ENSG00000141441 |
| PEX11B | 0.46287021 | 1.378281153 | 0.018261352 | 0.063818491 | 108.8599754 | 150.1361909 | ENSG00000131779 |
| FBF1 | 0.462764022 | 1.378179709 | 0.107364181 | 0.251716022 | 51.44794886 | 70.88072663 | ENSG00000188878 |
| NADK | 0.462406866 | 1.377838567 | 4.07E-12 | 1.08E-10 | 1823.132638 | 2511.391033 | ENSG00000008130 |
| MAD2L1BP | 0.462296793 | 1.377733446 | 8.12E-06 | 7.94E-05 | 418.5672211 | 576.8160175 | ENSG00000124688 |
| FOXD3-AS1 | 0.462288957 | 1.377725963 | 1.60E-09 | 3.00E-08 | 1563.847558 | 2154.591096 | ENSG00000230798 |
| OTUD7B | 0.462287422 | 1.377724497 | 3.49E-07 | 4.42E-06 | 601.3275076 | 828.3510421 | ENSG00000264522 |
| NA | 0.462129781 | 1.377573964 | 0.352010635 | 0.564131097 | 16.68878351 | 22.93338316 | ENSG00000233830 |
| LINC02175 | 0.461844368 | 1.377301461 | 0.403114545 | 0.614382873 | 13.0659306 | 17.92016793 | ENSG00000262155 |
| NIBAN1 | 0.461451893 | 1.376926826 | 2.71E-06 | 2.88E-05 | 565.3742007 | 778.0714075 | ENSG00000135842 |
| KCNH6 | 0.461437551 | 1.376913139 | 0.120736354 | 0.273698333 | 54.12986245 | 74.48630254 | ENSG00000173826 |
| NA | 0.461425042 | 1.3769012 | 0.643074941 | 0.805122663 | 4.606479746 | 6.366751493 | ENSG00000250327 |
| RGS14 | 0.461199784 | 1.376686232 | 1.68E-05 | 0.000152477 | 380.2424338 | 523.3744157 | ENSG00000169220 |
| FH | 0.461103527 | 1.376594382 | 3.53E-11 | 8.42E-10 | 1490.406192 | 2051.406916 | ENSG00000091483 |
| NA | 0.46099305 | 1.37648897 | 0.084931771 | 0.211350177 | 58.75060624 | 80.83687983 | ENSG00000275131 |
| PLAAT3 | 0.460472677 | 1.375992568 | 2.29E-09 | 4.19E-08 | 1078.581518 | 1483.6331 | ENSG00000176485 |
| SELENOH | 0.460380445 | 1.375904602 | 0.005050835 | 0.022281012 | 179.111322 | 246.2769685 | ENSG00000211450 |
| NA | 0.460362146 | 1.375887151 | 0.371352623 | 0.583298203 | 15.692114 | 21.51392057 | ENSG00000267102 |
| NA | 0.460256198 | 1.375786112 | 0.617227909 | 0.787224317 | 4.3553102 | 5.961566034 | ENSG00000265458 |
| SYCE3 | 0.459986869 | 1.375529299 | 0.607100109 | 0.78089399 | 5.087767488 | 6.93167148 | ENSG00000217442 |
| DACT1 | 0.45994941 | 1.375493584 | 2.17E-05 | 0.00019103 | 397.3395878 | 546.6250198 | ENSG00000165617 |
| ALG1L9P | 0.459799362 | 1.375350532 | 0.223418866 | 0.421267916 | 29.76747746 | 40.98672855 | ENSG00000248671 |
| TUBB2A | 0.459713201 | 1.375268396 | 6.31E-16 | 2.66E-14 | 2796.060226 | 3844.630044 | ENSG00000137267 |
| CACNB4 | 0.459678016 | 1.375234856 | 0.209864762 | 0.403142715 | 32.58195511 | 44.90630868 | ENSG00000182389 |
| KLK1 | 0.459114769 | 1.374698051 | 0.24195893 | 0.445490337 | 25.35760612 | 34.94145362 | ENSG00000167748 |
| LINC02693 | 0.458872751 | 1.374467459 | 8.55E-05 | 0.000645753 | 319.2351921 | 438.9143883 | ENSG00000212719 |
| NA | 0.458702657 | 1.374305419 | 0.582677435 | 0.762472699 | 5.295054309 | 7.266863238 | ENSG00000260176 |
| C1QTNF12 | 0.458693509 | 1.374296705 | 0.302320093 | 0.512445547 | 23.71149009 | 32.66047054 | ENSG00000184163 |
| RPL17 | 0.458676316 | 1.374280326 | 0.081415937 | 0.204487295 | 57.40401783 | 78.96276321 | ENSG00000265681 |
| FEM1C | 0.458296974 | 1.373919021 | 0.000814016 | 0.004685368 | 339.9331279 | 466.6028391 | ENSG00000145780 |
| CPLX2 | 0.457900511 | 1.37354151 | 0.181447053 | 0.36453485 | 34.32295554 | 47.20535788 | ENSG00000145920 |
| COX20 | 0.457883129 | 1.373524961 | 0.418042179 | 0.628811354 | 12.37499177 | 16.93946871 | ENSG00000203667 |
| PITRM1 | 0.457587028 | 1.373243086 | 6.43E-08 | 9.29E-07 | 686.1246 | 941.8813604 | ENSG00000107959 |
| TMEM64 | 0.457521296 | 1.373180519 | 7.55E-05 | 0.000579042 | 509.9849566 | 700.264118 | ENSG00000180694 |

| | | | | | | | |
|---|---|---|---|---|---|---|---|
| COG2 | 0.457485111 | 1.373146078 | 8.63E-07 | 1.01E-05 | 696.9895992 | 956.6748497 | ENSG00000135775 |
| GTF2H2C | 0.457407755 | 1.373072453 | 0.001038073 | 0.005788013 | 256.6230213 | 352.7016711 | ENSG00000183474 |
| CTSZ | 0.457323918 | 1.372992665 | 1.88E-06 | 2.06E-05 | 729.3435984 | 1002.187122 | ENSG00000101160 |
| IMMP1L | 0.457054671 | 1.372736451 | 0.044148609 | 0.127953728 | 83.91722221 | 115.2788453 | ENSG00000148950 |
| MSH2-OT1 | 0.456746517 | 1.372443271 | 0.553499532 | 0.739980526 | 6.26207899 | 8.597982459 | ENSG00000235760 |
| CLBA1 | 0.456647507 | 1.372349085 | 0.099428181 | 0.237203285 | 52.90071048 | 72.59383625 | ENSG00000140104 |
| HAUS7 | 0.456643225 | 1.372345012 | 0.546998533 | 0.734791389 | 6.285800088 | 8.625979939 | ENSG00000213397 |
| SFT2D2 | 0.456620906 | 1.372323781 | 0.004168105 | 0.019031999 | 185.152539 | 254.1295604 | ENSG00000213064 |
| DYNC1I1 | 0.456206312 | 1.371929467 | 0.323189183 | 0.534123367 | 17.87377343 | 24.50387842 | ENSG00000158560 |
| DOCK3 | 0.45602461 | 1.371756688 | 0.000779262 | 0.004508688 | 260.4987459 | 357.0335528 | ENSG00000088538 |
| NA | 0.455820763 | 1.371562879 | 0.642008681 | 0.804214244 | 5.48810324 | 7.561288281 | ENSG00000224669 |
| NA | 0.455750735 | 1.371496304 | 0.622292529 | 0.791007229 | 5.369777078 | 7.343383329 | ENSG00000219470 |
| FOXO4 | 0.45568994 | 1.371438512 | 1.85E-07 | 2.43E-06 | 619.7048414 | 849.8058463 | ENSG00000184481 |
| ETS2 | 0.455410204 | 1.371172617 | 1.14E-07 | 1.56E-06 | 771.6574292 | 1058.05006 | ENSG00000157557 |
| ASB16-AS1 | 0.455388913 | 1.371152382 | 0.053728272 | 0.14895124 | 77.96565474 | 106.9865441 | ENSG00000267080 |
| DNASE1L1 | 0.455151351 | 1.370926619 | 0.167211595 | 0.344023419 | 35.12065239 | 48.12665717 | ENSG00000013563 |
| RASSF1 | 0.455108668 | 1.37088606 | 0.00096487 | 0.005437652 | 286.8809387 | 393.4368871 | ENSG00000068028 |
| ARHGAP28 | 0.454514393 | 1.370321481 | 0.007242398 | 0.030106249 | 226.137019 | 310.1965428 | ENSG00000088756 |
| POLM | 0.454472874 | 1.370282045 | 0.001076569 | 0.00598158 | 237.355471 | 325.2866686 | ENSG00000122678 |
| APOE | 0.454317505 | 1.370134483 | 4.15E-14 | 1.39E-12 | 16757.84133 | 22961.64676 | ENSG00000130203 |
| CORO2A | 0.454277983 | 1.370096949 | 8.71E-11 | 1.96E-09 | 1383.428829 | 1894.660237 | ENSG00000106789 |
| DTNA | 0.454111378 | 1.369938737 | 0.003739283 | 0.017348247 | 179.4310112 | 245.7243233 | ENSG00000134769 |
| HLA-DOA | 0.454108114 | 1.369935638 | 6.11E-06 | 6.12E-05 | 456.3229297 | 625.0064255 | ENSG00000204252 |
| CDC73 | 0.453664926 | 1.369514866 | 6.55E-09 | 1.11E-07 | 1149.218508 | 1573.537505 | ENSG00000134371 |
| DMD | 0.453413787 | 1.369276486 | 0.00808687 | 0.032949415 | 136.4105041 | 186.7099987 | ENSG00000198947 |
| NA | 0.453394069 | 1.369257772 | 0.00919932 | 0.036497724 | 213.7368065 | 292.6190523 | ENSG00000244879 |
| GDPD1 | 0.453019661 | 1.368902469 | 0.045359959 | 0.130582748 | 77.6779746 | 106.2237498 | ENSG00000153982 |
| COPS6 | 0.45301221 | 1.368895399 | 1.57E-13 | 4.98E-12 | 2214.30742 | 3031.60161 | ENSG00000168090 |
| PLEKHM3 | 0.452837389 | 1.36872953 | 0.011664433 | 0.044393957 | 129.2226135 | 176.9545577 | ENSG00000178385 |
| IPMK | 0.452791151 | 1.368685663 | 0.00672968 | 0.028336679 | 185.6569368 | 254.1955391 | ENSG00000151151 |
| NA | 0.452642988 | 1.368545108 | 0.431733658 | 0.6413842 | 11.93522249 | 16.34532174 | ENSG00000231711 |
| FRS2 | 0.452493252 | 1.368403076 | 0.001310317 | 0.00708426 | 333.6601434 | 456.1821732 | ENSG00000166225 |
| RAD51B | 0.451860133 | 1.367802691 | 0.103279963 | 0.243896724 | 51.39458293 | 70.20890269 | ENSG00000182185 |
| GRM1 | 0.451720066 | 1.367669901 | 0.42949679 | 0.639369926 | 13.29337905 | 18.12530365 | ENSG00000152822 |
| MAP3K15 | 0.451571611 | 1.367529174 | 0.111529388 | 0.259135468 | 48.41967584 | 66.29830797 | ENSG00000180815 |
| PLEKHA8P1 | 0.45140832 | 1.3673744 | 0.030944228 | 0.097299657 | 111.869282 | 153.0353893 | ENSG00000134297 |
| KLHL20 | 0.45135565 | 1.36732448 | 0.001436939 | 0.007676548 | 252.2323701 | 344.6175317 | ENSG00000076321 |
| MRPL38 | 0.451313748 | 1.367284768 | 0.249198929 | 0.454215016 | 31.06069994 | 42.32293375 | ENSG00000204316 |
| ECH1 | 0.451298375 | 1.367270198 | 2.54E-06 | 2.72E-05 | 596.8095077 | 816.6452909 | ENSG00000104823 |
| SLC25A5-AS1 | 0.451040088 | 1.367025437 | 0.372042625 | 0.583930264 | 17.37023913 | 23.71235162 | ENSG00000224281 |
| SYT3 | 0.451026548 | 1.367012607 | 0.000200911 | 0.001385255 | 335.4124321 | 458.9098365 | ENSG00000213023 |
| NA | 0.45092092 | 1.366912523 | 0.639174102 | 0.802206613 | 4.362429145 | 5.921745277 | ENSG00000272800 |
| NA | 0.450784576 | 1.366783348 | 0.731207475 | 0.858455957 | 2.935473557 | 4.03970481 | ENSG00000224635 |
| NA | 0.450571388 | 1.366581392 | 0.22233732 | 0.41980007 | 28.44372042 | 38.89499053 | ENSG00000267519 |
| NA | 0.450245154 | 1.366272405 | 0.508659787 | 0.70544467 | 11.10432134 | 15.21404135 | ENSG00000272153 |
| PID1 | 0.450133561 | 1.366166727 | 0.061594197 | 0.165580984 | 70.16154799 | 96.02509916 | ENSG00000153823 |
| SMG1P5 | 0.449996353 | 1.366036804 | 0.239032939 | 0.44178284 | 27.3984135 | 37.44223357 | ENSG00000183604 |
| CCHCR1 | 0.449789044 | 1.365840524 | 1.11E-05 | 0.000104682 | 860.8008038 | 1175.297707 | ENSG00000204536 |
| DHRS7B | 0.449717368 | 1.365772668 | 0.000547809 | 0.003317953 | 241.9073896 | 330.4343635 | ENSG00000109016 |
| KIAA1217 | 0.449653484 | 1.365712192 | 8.19E-05 | 0.000622954 | 364.7348014 | 498.2132792 | ENSG00000120549 |
| NA | 0.449501154 | 1.365567998 | 0.490060332 | 0.69013064 | 9.759486795 | 13.24677894 | ENSG00000226121 |
| NA | 0.449275018 | 1.365353967 | 0.10257088 | 0.242555855 | 88.71736128 | 121.3221854 | ENSG00000248131 |
| KMT2A | 0.449097854 | 1.365186312 | 3.15E-05 | 0.00026678 | 750.7454906 | 1024.336562 | ENSG00000118058 |
| XRCC3 | 0.449090056 | 1.365178932 | 1.74E-05 | 0.000156664 | 439.6155114 | 599.9330653 | ENSG00000126215 |
| TIGD2 | 0.449071847 | 1.365161702 | 0.013485952 | 0.049888775 | 120.9885523 | 165.2054424 | ENSG00000180346 |
| SMOC1 | 0.448872758 | 1.364973325 | 0.196001544 | 0.385010315 | 33.18512849 | 45.38165579 | ENSG00000198732 |
| LRRC75A | 0.448734825 | 1.364842829 | 0.000140277 | 0.001007312 | 330.710432 | 451.0469344 | ENSG00000181350 |
| NLRP6 | 0.448650011 | 1.364762595 | 0.567592492 | 0.751047019 | 6.32731855 | 8.611981199 | ENSG00000174885 |
| AZIN1 | 0.448362159 | 1.364490319 | 9.14E-14 | 2.95E-12 | 4749.821309 | 6480.612359 | ENSG00000155096 |
| NA | 0.448296323 | 1.364428054 | 0.396800239 | 0.608788083 | 13.91671405 | 19.0022633 | ENSG00000230699 |
| SUCO | 0.448158956 | 1.364298144 | 2.35E-08 | 3.61E-07 | 859.4207574 | 1171.974255 | ENSG00000094975 |
| TENT5A | 0.44801308 | 1.364160203 | 0.07159561 | 0.186134786 | 65.80087272 | 89.64311942 | ENSG00000112773 |
| HSPG2 | 0.448012239 | 1.364159408 | 3.55E-08 | 5.33E-07 | 2803.494331 | 3823.640576 | ENSG00000142798 |
| DDX60 | 0.447782801 | 1.363942477 | 0.11013389 | 0.256665781 | 48.70079571 | 66.40471745 | ENSG00000137628 |
| NOL3 | 0.447740259 | 1.363902258 | 0.089999309 | 0.220447418 | 56.70860365 | 77.27063021 | ENSG00000140939 |
| GADD45A | 0.447701782 | 1.363865883 | 3.41E-05 | 0.000285834 | 404.3192497 | 551.3358861 | ENSG00000116717 |
| NA | 0.447565426 | 1.363736984 | 0.700285653 | 0.84040536 | 3.929778809 | 5.389173698 | ENSG00000287830 |
| BORCS8 | 0.447548605 | 1.363721083 | 0.153555582 | 0.324518483 | 41.90050357 | 57.23434266 | ENSG00000254901 |
| SLC19A3 | 0.447487581 | 1.3636634 | 0.021052349 | 0.071738292 | 127.8404309 | 173.9762122 | ENSG00000135917 |
| FGFR3 | 0.447463063 | 1.363640226 | 2.15E-07 | 2.80E-06 | 697.3828422 | 950.8462433 | ENSG00000068078 |
| NA | 0.4473387 | 1.363522682 | 0.278850784 | 0.488008252 | 22.10216398 | 30.15702067 | ENSG00000185641 |
| NUCKS1 | 0.447236196 | 1.363425808 | 1.18E-15 | 4.84E-14 | 12044.68187 | 16421.6933 | ENSG00000069275 |
| ARF1 | 0.447123853 | 1.363319641 | 5.51E-21 | 3.55E-19 | 6523.443353 | 8893.397172 | ENSG00000143761 |
| FLAD1 | 0.446856088 | 1.363066631 | 2.00E-13 | 6.23E-12 | 1926.998866 | 2626.006437 | ENSG00000160688 |

| | | | | | | | |
|---|---|---|---|---|---|---|---|
| NA | 0.446793692 | 1.363007681 | 0.371572737 | 0.583546871 | 15.61975549 | 21.26461317 | ENSG00000268218 |
| GAN | 0.446618285 | 1.362841972 | 0.018077608 | 0.063314513 | 122.762757 | 167.2139231 | ENSG00000261609 |
| NA | 0.44659482 | 1.362819806 | 0.657305698 | 0.814996921 | 3.650133483 | 4.965638572 | ENSG00000231203 |
| CSRNP1 | 0.446134708 | 1.362385238 | 0.009283312 | 0.036761449 | 140.7726278 | 191.8834429 | ENSG00000144655 |
| NA | 0.4461337 | 1.362384286 | 0.618701328 | 0.788459044 | 4.617158164 | 6.294582329 | ENSG00000272572 |
| CSRNP3 | 0.446131529 | 1.362382236 | 0.039542211 | 0.117694551 | 98.14086216 | 133.6639346 | ENSG00000178662 |
| NA | 0.445757595 | 1.362029164 | 0.466096167 | 0.670237274 | 10.27014004 | 13.99897943 | ENSG00000255618 |
| MYRF | 0.445505624 | 1.361791302 | 0.002412724 | 0.011951032 | 265.4916813 | 361.2654359 | ENSG00000124920 |
| LINC01409 | 0.445089924 | 1.361398971 | 0.533029594 | 0.724314769 | 8.318293317 | 11.31091898 | ENSG00000237491 |
| GTF2A1 | 0.445019218 | 1.361332251 | 7.98E-05 | 0.000608369 | 679.9405489 | 926.0734532 | ENSG00000165417 |
| EFHC2 | 0.445013238 | 1.361326608 | 0.153047758 | 0.323662587 | 39.67257099 | 53.99581246 | ENSG00000183690 |
| SLC26A1 | 0.444947404 | 1.361264488 | 0.431610928 | 0.641354345 | 15.46466555 | 20.9748226 | ENSG00000145217 |
| SLC25A4 | 0.444742368 | 1.361071039 | 1.00E-11 | 2.54E-10 | 1455.082851 | 1980.45644 | ENSG00000151729 |
| ZFP2 | 0.444486318 | 1.360829497 | 0.44659258 | 0.653638757 | 11.18852731 | 15.25045714 | ENSG00000198939 |
| TRIM13 | 0.444279095 | 1.360634047 | 0.001061739 | 0.005909504 | 245.0466167 | 333.4627017 | ENSG00000204977 |
| ATP6V0C | 0.4441383 | 1.360501267 | 0.62508984 | 0.792695516 | 4.624277109 | 6.291460907 | ENSG00000185883 |
| LOC101928673 | 0.444052167 | 1.360420044 | 0.112016425 | 0.259785114 | 49.05396859 | 66.68949468 | ENSG00000203739 |
| RAET1K | 0.443965566 | 1.360338383 | 0.568781925 | 0.752014213 | 6.340361231 | 8.588334646 | ENSG00000218358 |
| HS3ST4 | 0.443915734 | 1.360291397 | 0.037122834 | 0.112095724 | 89.63071512 | 121.9932743 | ENSG00000182601 |
| CLDN4 | 0.443864232 | 1.360242838 | 0.000167618 | 0.001176319 | 303.9098785 | 413.3721347 | ENSG00000189143 |
| DOC2B | 0.443862016 | 1.360240748 | 0.010592772 | 0.04093401 | 141.9113707 | 193.2062392 | ENSG00000272636 |
| CLEC11A | 0.443701721 | 1.360089623 | 0.008054614 | 0.032846336 | 147.9412466 | 201.3088009 | ENSG00000105472 |
| UCK2 | 0.443672634 | 1.360062202 | 1.25E-10 | 2.76E-09 | 1346.031022 | 1830.167532 | ENSG00000143179 |
| ARHGEF39 | 0.443417378 | 1.359821587 | 0.001852129 | 0.009491849 | 213.621455 | 290.4099845 | ENSG00000137135 |
| NA | 0.443383357 | 1.359789521 | 0.417031705 | 0.627991687 | 15.69923294 | 21.30471747 | ENSG00000188383 |
| NEMP2 | 0.443380745 | 1.359787059 | 0.05664461 | 0.155439327 | 75.8302423 | 103.0455655 | ENSG00000189362 |
| SLC38A3 | 0.443282891 | 1.359694832 | 0.428563355 | 0.638382944 | 11.95894359 | 16.22803494 | ENSG00000188338 |
| SUGCT | 0.443165624 | 1.359584316 | 0.127915237 | 0.285473796 | 52.70646634 | 71.59790879 | ENSG00000175600 |
| HPSE | 0.44306748 | 1.359491829 | 0.027177755 | 0.088036348 | 97.19519431 | 132.2093286 | ENSG00000173083 |
| ZBTB47 | 0.443038305 | 1.359464336 | 0.025480772 | 0.083673166 | 100.0609791 | 135.9314134 | ENSG00000114853 |
| LOC440173 | 0.442867181 | 1.359303095 | 0.184412397 | 0.368649107 | 38.40101083 | 52.22046502 | ENSG00000269994 |
| NA | 0.442829885 | 1.359267955 | 0.329127108 | 0.540813141 | 23.83159627 | 32.28668753 | ENSG00000285796 |
| ALDH3B1 | 0.442616767 | 1.359067175 | 0.327018059 | 0.538425113 | 23.942498 | 32.64429634 | ENSG00000006534 |
| PXMP2 | 0.442316633 | 1.358784468 | 0.472634945 | 0.675985379 | 11.19208679 | 15.22123016 | ENSG00000176894 |
| PRELID2 | 0.442070065 | 1.358552261 | 0.119598744 | 0.27216434 | 49.77427292 | 67.70812274 | ENSG00000186314 |
| NLRX1 | 0.442006854 | 1.358492738 | 5.71E-06 | 5.75E-05 | 560.0925463 | 760.8843553 | ENSG00000160703 |
| ITGAV | 0.441982033 | 1.358469367 | 6.02E-06 | 6.03E-05 | 751.7003624 | 1020.653751 | ENSG00000138448 |
| NA | 0.441941554 | 1.358431251 | 0.45477915 | 0.660692258 | 12.45891842 | 16.98799133 | ENSG00000279520 |
| TRIB3 | 0.441771072 | 1.358270736 | 0.000130905 | 0.000944289 | 349.9148277 | 475.6130506 | ENSG00000101255 |
| C16orf58 | 0.441677823 | 1.358182947 | 3.89E-05 | 0.000321696 | 368.5500928 | 500.6225646 | ENSG00000140688 |
| LINC02585 | 0.441477209 | 1.357994097 | 0.596089672 | 0.772625495 | 5.591301789 | 7.599879533 | ENSG00000228350 |
| CBWD6 | 0.441339486 | 1.357864467 | 0.232554041 | 0.433456678 | 33.71268937 | 45.79213814 | ENSG00000215126 |
| STON1 | 0.441117444 | 1.357655497 | 0.006644892 | 0.02806996 | 163.8670121 | 222.3621081 | ENSG00000243244 |
| NA | 0.441046153 | 1.357588409 | 0.582933744 | 0.762575961 | 6.591530777 | 8.958996234 | ENSG00000279789 |
| MAP11 | 0.440676319 | 1.357240438 | 0.001593754 | 0.008381523 | 258.2338481 | 350.8878707 | ENSG00000146826 |
| CPLX1 | 0.440531745 | 1.357104434 | 0.325750152 | 0.536984748 | 26.44532123 | 36.05372193 | ENSG00000168993 |
| GGCX | 0.440487666 | 1.35706297 | 1.76E-06 | 1.94E-05 | 598.7506243 | 812.8817044 | ENSG00000115486 |
| LOC101929140 | 0.440201066 | 1.356793408 | 0.596653831 | 0.773144476 | 5.642303459 | 7.652753071 | ENSG00000278390 |
| FHL2 | 0.439668802 | 1.356292929 | 1.20E-06 | 1.37E-05 | 658.6235466 | 892.845817 | ENSG00000115641 |
| RFX8 | 0.439616519 | 1.356243778 | 0.579067891 | 0.760003969 | 7.505189523 | 10.20082613 | ENSG00000196460 |
| LOC283177 | 0.439515889 | 1.356149182 | 0.348253304 | 0.560296784 | 17.64632498 | 23.87426162 | ENSG00000255545 |
| NA | 0.439426416 | 1.356065078 | 0.578935256 | 0.759924207 | 6.381879692 | 8.60450885 | ENSG00000214719 |
| EME1 | 0.43940449 | 1.356044469 | 0.013354826 | 0.049520102 | 155.8239149 | 211.3113442 | ENSG00000154920 |
| LRRC27 | 0.439314026 | 1.355959442 | 0.134125966 | 0.295424407 | 47.54428157 | 64.36150204 | ENSG00000148814 |
| ASIC4 | 0.438978588 | 1.355644206 | 0.576087555 | 0.758153886 | 6.371201275 | 8.586159182 | ENSG00000072182 |
| PDZD2 | 0.438954121 | 1.355621216 | 0.002345487 | 0.011644196 | 275.5782295 | 373.1097341 | ENSG00000133401 |
| CHMP3 | 0.438877076 | 1.355548823 | 0.046548058 | 0.133433528 | 94.50232297 | 128.3429056 | ENSG00000115561 |
| MPG | 0.438773637 | 1.355451636 | 0.000114877 | 0.000840544 | 379.7739617 | 514.974802 | ENSG00000103152 |
| BCL2L14 | 0.438578232 | 1.35526806 | 0.570784391 | 0.75364383 | 6.843895532 | 9.331833286 | ENSG00000121380 |
| CD46 | 0.438422174 | 1.355121467 | 5.14E-07 | 6.29E-06 | 1543.118096 | 2090.495781 | ENSG00000117335 |
| NA | 0.438126434 | 1.354843707 | 0.707736846 | 0.845910689 | 2.917676195 | 3.979358922 | ENSG00000214192 |
| TTC17 | 0.43798226 | 1.354708319 | 4.13E-08 | 6.14E-07 | 976.171131 | 1321.948454 | ENSG00000052841 |
| CTF1 | 0.437920859 | 1.354650664 | 0.406786644 | 0.617595253 | 13.96179198 | 18.87097775 | ENSG00000150281 |
| SENP8 | 0.437899148 | 1.354630279 | 0.12889877 | 0.287209334 | 46.06156963 | 62.35347291 | ENSG00000166192 |
| TMEM106B | 0.437742453 | 1.354483157 | 0.001523514 | 0.008057003 | 361.0316069 | 489.2617123 | ENSG00000106460 |
| SGSM1 | 0.437730413 | 1.354471852 | 0.019654428 | 0.067720099 | 119.821639 | 162.1947915 | ENSG00000167037 |
| LOXHD1 | 0.437627393 | 1.354375136 | 0.447020765 | 0.653867333 | 12.02418315 | 16.22368401 | ENSG00000167210 |
| NA | 0.437396364 | 1.354158267 | 0.726182143 | 0.855720453 | 2.897514569 | 3.94824002 | ENSG00000277369 |
| ZNF595 | 0.43713819 | 1.353915959 | 0.095583517 | 0.230359203 | 68.140318 | 92.1009572 | ENSG00000272602 |
| MRI1 | 0.43700675 | 1.353792613 | 0.003869007 | 0.017860227 | 195.7274677 | 264.9220019 | ENSG00000037757 |
| ACSF2 | 0.436950206 | 1.353739554 | 0.001508312 | 0.008000103 | 253.8692823 | 343.4549145 | ENSG00000167107 |
| FAM72A | 0.436800546 | 1.35359913 | 0.037125092 | 0.112095724 | 101.4475853 | 137.1884715 | ENSG00000196550 |
| LMLN | 0.436727701 | 1.353530785 | 0.096837803 | 0.232555036 | 63.69810592 | 86.28401304 | ENSG00000185621 |
| KLF11 | 0.436417269 | 1.35323957 | 0.00294386 | 0.014157066 | 213.2130845 | 288.1623684 | ENSG00000172059 |

| | | | | | | | |
|---|---|---|---|---|---|---|---|
| BCL2L11 | 0.436406049 | 1.353229046 | 0.003304666 | 0.015633422 | 241.2853016 | 326.963646 | ENSG00000153094 |
| FAM184B | 0.435966128 | 1.352816469 | 0.25183809 | 0.457549413 | 26.47883102 | 35.808271 | ENSG00000047662 |
| ZNF600 | 0.435946053 | 1.352797644 | 0.043615566 | 0.126915249 | 86.07310177 | 116.3975244 | ENSG00000189190 |
| NR2E3 | 0.435751949 | 1.352615648 | 0.572952698 | 0.755337641 | 6.667448754 | 9.008464804 | ENSG00000278570 |
| TSPAN11 | 0.435408787 | 1.352293951 | 0.005531319 | 0.024030125 | 168.2973506 | 227.3571846 | ENSG00000110900 |
| SCAMP3 | 0.435182534 | 1.352081891 | 2.69E-09 | 4.81E-08 | 966.071539 | 1306.544841 | ENSG00000116521 |
| NA | 0.434605144 | 1.351540875 | 0.436698136 | 0.645089622 | 16.31900847 | 22.04821611 | ENSG00000268649 |
| SLC51B | 0.434384845 | 1.351334511 | 0.667982422 | 0.822028414 | 3.688092472 | 4.953815295 | ENSG00000186198 |
| UTP25 | 0.434226364 | 1.351186073 | 1.04E-05 | 9.86E-05 | 509.9991684 | 689.3168506 | ENSG00000117597 |
| CAMSAP2 | 0.434182075 | 1.351144595 | 1.16E-05 | 0.00010886 | 678.6025028 | 916.5342252 | ENSG00000118200 |
| BAZ2B | 0.433916409 | 1.350895811 | 0.357680365 | 0.569801407 | 17.39248569 | 23.56271641 | ENSG00000226266 |
| PABPC5 | 0.433712465 | 1.350704857 | 0.190198202 | 0.376548176 | 45.10878284 | 60.84103248 | ENSG00000174740 |
| MTR | 0.433528394 | 1.350532533 | 1.71E-06 | 1.89E-05 | 677.1103082 | 913.8292985 | ENSG00000116984 |
| NA | 0.433447756 | 1.350457049 | 0.702828718 | 0.842126015 | 2.952075711 | 4.005180939 | ENSG00000259531 |
| NA | 0.433354868 | 1.350370103 | 0.342062814 | 0.553733606 | 19.7642194 | 26.59249502 | ENSG00000234338 |
| NA | 0.433251987 | 1.350273809 | 0.700445663 | 0.840454689 | 3.722491988 | 4.998932938 | ENSG00000233799 |
| NA | 0.433160288 | 1.350187987 | 0.630703372 | 0.796625039 | 4.937126742 | 6.692295438 | ENSG00000273381 |
| NA | 0.433019571 | 1.350056299 | 0.068436554 | 0.179518185 | 70.44650666 | 94.9902969 | ENSG00000242071 |
| NA | 0.432971192 | 1.350011028 | 0.628772647 | 0.795176551 | 4.675278779 | 6.284934516 | ENSG00000270060 |
| IL21R | 0.432934873 | 1.349977043 | 0.246413045 | 0.451227611 | 28.99706119 | 39.21277858 | ENSG00000103522 |
| DCXR-DT | 0.432843131 | 1.349891199 | 0.72317044 | 0.854154341 | 2.979356283 | 4.033178419 | ENSG00000264569 |
| CTNNA2 | 0.432387392 | 1.349464843 | 0.00255564 | 0.012516406 | 200.2986843 | 270.0853038 | ENSG00000066032 |
| NA | 0.432349333 | 1.349429244 | 0.118410414 | 0.270315491 | 56.02358856 | 75.51815133 | ENSG00000238278 |
| PBLD | 0.432298659 | 1.349381848 | 0.207853358 | 0.40048679 | 35.73923271 | 48.25822626 | ENSG00000108187 |
| CFAP157 | 0.432160144 | 1.349252298 | 0.230867029 | 0.431163355 | 36.45124904 | 49.33814617 | ENSG00000160401 |
| ZNF672 | 0.432034733 | 1.349135015 | 6.69E-09 | 1.14E-07 | 993.4432823 | 1340.809269 | ENSG00000171161 |
| C14orf93 | 0.431922805 | 1.349030349 | 0.060709011 | 0.163807641 | 83.03887886 | 112.1611022 | ENSG00000100802 |
| RIBC2 | 0.431868816 | 1.348979866 | 0.3143955 | 0.524872614 | 21.21105728 | 28.61887381 | ENSG00000128408 |
| EEF1AKNMT | 0.431780337 | 1.348897138 | 5.28E-11 | 1.23E-09 | 2595.31773 | 3501.261912 | ENSG00000010165 |
| NA | 0.431675923 | 1.348799516 | 0.337253991 | 0.549215845 | 19.43921681 | 26.27224796 | ENSG00000227939 |
| FRMD6 | 0.431025494 | 1.348191555 | 0.001712307 | 0.008908605 | 259.2948151 | 349.6417195 | ENSG00000139926 |
| TMIE | 0.43095805 | 1.348128531 | 0.459922789 | 0.664802287 | 11.34272753 | 15.24080933 | ENSG00000181585 |
| CNN1 | 0.430934133 | 1.348106182 | 0.000285646 | 0.001885436 | 333.854387 | 450.4211217 | ENSG00000130176 |
| RPRD1A | 0.430679509 | 1.347868273 | 1.02E-10 | 2.27E-09 | 2411.243677 | 3249.855493 | ENSG00000141425 |
| CENPP | 0.43064542 | 1.347836425 | 0.04039858 | 0.119638925 | 97.14803145 | 130.8863442 | ENSG00000188312 |
| USP51 | 0.430410233 | 1.347616719 | 0.053646886 | 0.148797718 | 90.64042732 | 122.0644545 | ENSG00000247746 |
| NA | 0.430185516 | 1.347406829 | 0.590610746 | 0.768530555 | 5.90059195 | 7.981418439 | ENSG00000235280 |
| NAT1 | 0.430131695 | 1.347356563 | 0.111943236 | 0.259717416 | 56.46099358 | 75.97431841 | ENSG00000171428 |
| RGS10 | 0.430121054 | 1.347346626 | 0.038010881 | 0.114159587 | 98.46586475 | 132.6400526 | ENSG00000148908 |
| KC6 | 0.430025865 | 1.347257731 | 0.139106891 | 0.302828084 | 54.84543826 | 73.99572718 | ENSG00000267313 |
| RIT1 | 0.430024191 | 1.347256167 | 0.000109139 | 0.000803543 | 449.7550945 | 605.2881666 | ENSG00000143622 |
| PEA15 | 0.42993905 | 1.347176661 | 3.10E-12 | 8.36E-11 | 1742.826219 | 2347.925443 | ENSG00000162734 |
| ZNF784 | 0.429770765 | 1.347019527 | 0.001719495 | 0.008933333 | 285.6371916 | 384.5481251 | ENSG00000179922 |
| ND3 | 0.429707378 | 1.346960345 | 1.36E-09 | 2.56E-08 | 5038.519503 | 6787.713542 | ENSG00000198840 |
| NA | 0.42969479 | 1.346948593 | 0.598067475 | 0.774094349 | 5.897032477 | 7.971770626 | ENSG00000204620 |
| RIPOR2 | 0.429572023 | 1.346833978 | 9.62E-07 | 1.12E-05 | 616.9351624 | 830.8705506 | ENSG00000111913 |
| DEDD | 0.429469938 | 1.34673868 | 0.000138128 | 0.000993352 | 351.633887 | 473.607764 | ENSG00000158796 |
| PCP4L1 | 0.429442943 | 1.34671348 | 0.34605358 | 0.558088971 | 23.82063852 | 32.05211392 | ENSG00000248485 |
| NA | 0.429290259 | 1.346570962 | 0.713283935 | 0.848794399 | 2.944956766 | 3.979358922 | ENSG00000284052 |
| TMC7 | 0.429245055 | 1.34652877 | 0.087786721 | 0.21624636 | 68.94602351 | 92.74457274 | ENSG00000170537 |
| CA2 | 0.429219481 | 1.346504901 | 0.020240437 | 0.069473039 | 121.5315201 | 163.5982478 | ENSG00000104267 |
| SRGAP2 | 0.429197799 | 1.346484665 | 7.41E-07 | 8.75E-06 | 878.6236535 | 1182.327511 | ENSG00000266028 |
| CCR10 | 0.429173179 | 1.346461687 | 0.447339558 | 0.654123697 | 19.71054798 | 26.58663104 | ENSG00000184451 |
| THBS2 | 0.429143034 | 1.346433553 | 3.87E-05 | 0.000320082 | 548.2936999 | 738.603952 | ENSG00000186340 |
| CATSPER4 | 0.429085394 | 1.34637976 | 0.561139507 | 0.746223477 | 7.67835616 | 10.31499151 | ENSG00000188782 |
| RAMP2 | 0.429056543 | 1.346352835 | 0.073876433 | 0.190647232 | 78.48459599 | 105.8492318 | ENSG00000131477 |
| CEMIP | 0.428729224 | 1.346047409 | 0.239589126 | 0.442594214 | 30.11236235 | 40.54853227 | ENSG00000103888 |
| NA | 0.428653317 | 1.345976589 | 0.463268381 | 0.667951089 | 11.63897501 | 15.65441309 | ENSG00000213885 |
| CMBL | 0.428244529 | 1.34559526 | 5.76E-05 | 0.000456567 | 524.0644115 | 705.4712557 | ENSG00000164237 |
| CYTB | 0.428215064 | 1.345567778 | 0.067037616 | 0.176742341 | 21867.96207 | 29425.04589 | ENSG00000198727 |
| AKAP3 | 0.428100116 | 1.345460574 | 0.508955204 | 0.705543494 | 9.854091858 | 13.295018 | ENSG00000111254 |
| NA | 0.42796938 | 1.345338655 | 0.63185817 | 0.797156893 | 4.93356727 | 6.655596104 | ENSG00000240950 |
| ZNF837 | 0.427753952 | 1.345137779 | 0.043157102 | 0.12581381 | 97.449644 | 131.2459175 | ENSG00000152475 |
| DGAT1 | 0.427641052 | 1.345032517 | 3.78E-05 | 0.000313226 | 509.2779737 | 685.5264033 | ENSG00000185000 |
| KIAA0040 | 0.427302455 | 1.344716879 | 0.000266049 | 0.001772253 | 388.9263846 | 522.5675224 | ENSG00000235750 |
| N4BP2L2-IT2 | 0.427272813 | 1.34468925 | 0.593096501 | 0.770384994 | 7.835836519 | 10.56401537 | ENSG00000281026 |
| NUDT17 | 0.426559751 | 1.344024793 | 0.273132611 | 0.481840226 | 28.89149837 | 38.95505287 | ENSG00000186364 |
| ZNF8 | 0.426316506 | 1.343798203 | 0.008901848 | 0.035669589 | 170.2613497 | 228.9846208 | ENSG00000278129 |
| NA | 0.426297106 | 1.343780132 | 0.712578116 | 0.848221729 | 2.986475227 | 4.019179679 | ENSG00000257605 |
| CKMT2 | 0.426294068 | 1.343777303 | 0.2601164 | 0.467028585 | 27.25517103 | 36.5915904 | ENSG00000131730 |
| RPRML | 0.426127706 | 1.343622357 | 0.304280841 | 0.514398324 | 31.25580765 | 42.03285963 | ENSG00000179673 |
| NMRK1 | 0.42610628 | 1.343602402 | 0.147990977 | 0.315762016 | 52.6967038 | 70.96471907 | ENSG00000106733 |
| MAN1B1-DT | 0.425864006 | 1.343376788 | 0.548303646 | 0.73594274 | 9.921695682 | 13.34448657 | ENSG00000268996 |
| S100A14 | 0.42545737 | 1.342998199 | 0.339091596 | 0.550859767 | 23.76605123 | 31.96237309 | ENSG00000189334 |

| | | | | | | | |
|---|---|---|---|---|---|---|---|
| MAP2 | 0.425235913 | 1.342792062 | 0.007750458 | 0.031874432 | 183.2833975 | 246.2490664 | ENSG00000078018 |
| NA | 0.425044921 | 1.342614307 | 0.665543755 | 0.82085806 | 3.977221007 | 5.347177478 | ENSG00000231066 |
| RIMBP2 | 0.424948064 | 1.342524172 | 0.30148192 | 0.511636058 | 24.24733897 | 32.50676795 | ENSG00000060709 |
| NA | 0.424835551 | 1.342419476 | 0.132459509 | 0.292846854 | 52.36842107 | 70.42077571 | ENSG00000225077 |
| HVCN1 | 0.424618066 | 1.342217122 | 0.03691451 | 0.111630186 | 93.78822171 | 125.8295244 | ENSG00000122986 |
| LOC105369147 | 0.424538387 | 1.342142995 | 0.31572716 | 0.525792898 | 21.72763426 | 29.21614221 | ENSG00000247081 |
| TEX30 | 0.424382584 | 1.341998058 | 0.053828474 | 0.149119955 | 84.01449702 | 112.7343452 | ENSG00000151287 |
| CALD1 | 0.424330344 | 1.341949465 | 2.42E-17 | 1.12E-15 | 3745.235057 | 5025.752637 | ENSG00000122786 |
| LINC02762 | 0.42427311 | 1.341896229 | 0.536439167 | 0.726836969 | 8.717739345 | 11.65887997 | ENSG00000250303 |
| FBN1 | 0.424205409 | 1.34183326 | 0.013104567 | 0.048764483 | 191.3831414 | 256.6015876 | ENSG00000166147 |
| WFIKKN2 | 0.424173256 | 1.341803355 | 0.726260708 | 0.855720453 | 3.217483147 | 4.363073292 | ENSG00000173714 |
| MDN1 | 0.424086221 | 1.341722409 | 0.006876574 | 0.028852275 | 5156.219333 | 6917.844967 | ENSG00000112159 |
| NA | 0.42377956 | 1.341437241 | 0.424308286 | 0.634226465 | 13.88231453 | 18.64560045 | ENSG00000277304 |
| APOOL | 0.423529908 | 1.341205131 | 0.00873457 | 0.035096246 | 153.6467047 | 205.9936343 | ENSG00000155008 |
| NA | 0.423366017 | 1.341052778 | 0.673017412 | 0.824962394 | 3.984339952 | 5.347177478 | ENSG00000232906 |
| FBXO28 | 0.423340429 | 1.341028993 | 1.46E-06 | 1.63E-05 | 725.3803104 | 973.3031224 | ENSG00000143756 |
| KMT2C | 0.423309684 | 1.341000415 | 0.002789036 | 0.013501907 | 486.4560129 | 651.6582643 | ENSG00000055609 |
| NA | 0.423223129 | 1.340919964 | 0.363603789 | 0.575675515 | 18.39538443 | 24.58446589 | ENSG00000263571 |
| ACAT1 | 0.423203943 | 1.340902131 | 2.55E-05 | 0.000221519 | 453.2193497 | 607.4511279 | ENSG00000075239 |
| POLR2A | 0.422910419 | 1.340629345 | 5.12E-10 | 1.03E-08 | 5815.875876 | 7796.521544 | ENSG00000181222 |
| NA | 0.422803595 | 1.340530083 | 0.252513562 | 0.458442474 | 28.39508302 | 38.06532398 | ENSG00000260025 |
| ZDHHC11B | 0.422557317 | 1.340301264 | 0.562858284 | 0.747719379 | 6.93285619 | 9.302889847 | ENSG00000206077 |
| DECR1 | 0.422539694 | 1.340284892 | 4.41E-05 | 0.000360413 | 403.5734965 | 540.9492925 | ENSG00000104325 |
| NA | 0.422266715 | 1.340031315 | 0.581962838 | 0.761962097 | 7.160304638 | 9.653026304 | ENSG00000231940 |
| NA | 0.422259722 | 1.34002482 | 0.703572382 | 0.842524014 | 2.982915755 | 3.999884053 | ENSG00000272583 |
| SPDEF | 0.422040801 | 1.339821493 | 0.723486258 | 0.854420486 | 3.766374713 | 5.008580751 | ENSG00000124664 |
| NA | 0.421896975 | 1.33968793 | 0.433585484 | 0.642497135 | 14.67289244 | 19.61759781 | ENSG00000177855 |
| VPS13D | 0.421885545 | 1.339677316 | 7.00E-05 | 0.000541101 | 751.8716972 | 1006.686764 | ENSG00000048707 |
| LTBP4 | 0.421722519 | 1.339525939 | 1.83E-08 | 2.89E-07 | 1781.441941 | 2386.826494 | ENSG00000090006 |
| ZNF232 | 0.421675823 | 1.339482584 | 5.98E-05 | 0.000471278 | 452.7381137 | 606.7950493 | ENSG00000167840 |
| TAF5L | 0.421671122 | 1.339478219 | 7.44E-07 | 8.78E-06 | 694.4002313 | 930.2921906 | ENSG00000135801 |
| CES2 | 0.421191537 | 1.33903302 | 8.94E-05 | 0.000672629 | 377.1969482 | 505.0020957 | ENSG00000172831 |
| ALS2 | 0.420975424 | 1.33883245 | 2.20E-07 | 2.86E-06 | 1182.391128 | 1582.069792 | ENSG00000003393 |
| DIPK2A | 0.420956454 | 1.338814846 | 0.001341015 | 0.007231603 | 415.0526264 | 555.7772628 | ENSG00000181744 |
| ALDH9A1 | 0.42083053 | 1.338697994 | 1.14E-10 | 2.53E-09 | 1871.330867 | 2504.777783 | ENSG00000143149 |
| PTGDR | 0.4207454 | 1.338619003 | 0.726545163 | 0.855892047 | 3.20680473 | 4.337251275 | ENSG00000168229 |
| NA | 0.420705137 | 1.338581645 | 0.651008018 | 0.810534186 | 7.270622131 | 9.757543875 | ENSG00000233483 |
| IFT80 | 0.420613146 | 1.338496296 | 0.055388218 | 0.152679389 | 90.13038446 | 120.879114 | ENSG00000068885 |
| POU2F1 | 0.420520189 | 1.338410055 | 1.12E-05 | 0.000104885 | 1018.168297 | 1362.645796 | ENSG00000143190 |
| FBXL3 | 0.420469618 | 1.338363141 | 0.053449294 | 0.148330626 | 98.23429817 | 131.578555 | ENSG00000005812 |
| NRBP2 | 0.420043027 | 1.337967458 | 0.017315911 | 0.061238012 | 204.7960418 | 274.0774023 | ENSG00000185189 |
| CDKL3 | 0.419831128 | 1.337770955 | 0.602491067 | 0.777454308 | 5.955153093 | 7.967419699 | ENSG00000006837 |
| KCTD3 | 0.419812864 | 1.337754019 | 3.51E-10 | 7.24E-09 | 1875.140871 | 2508.126368 | ENSG00000136636 |
| NT5DC3 | 0.41952225 | 1.337484572 | 0.00170568 | 0.008881261 | 318.1122139 | 425.4266249 | ENSG00000111696 |
| PRR13 | 0.419495884 | 1.337460129 | 0.000144385 | 0.001032064 | 438.183548 | 586.0122143 | ENSG00000205352 |
| EMBP1 | 0.419419836 | 1.33738963 | 0.204697518 | 0.396605379 | 40.26982063 | 53.94031195 | ENSG00000231752 |
| NIPSNAP1 | 0.41927529 | 1.337255642 | 1.19E-14 | 4.28E-13 | 3673.220108 | 4911.053616 | ENSG00000184117 |
| PGGHG | 0.419247428 | 1.337229816 | 0.206749044 | 0.399262489 | 1476.067481 | 1973.88811 | ENSG00000142102 |
| NGF | 0.41917135 | 1.337159302 | 0.458523486 | 0.663948476 | 13.79691335 | 18.4592659 | ENSG00000134259 |
| GALNT3 | 0.419060319 | 1.337056396 | 0.00157334 | 0.008299588 | 360.9484921 | 482.5512781 | ENSG00000115339 |
| SMTNL2 | 0.418903197 | 1.336910788 | 0.081300484 | 0.204251647 | 85.71751231 | 114.6131916 | ENSG00000188176 |
| BTBD1 | 0.418745317 | 1.336764492 | 9.24E-06 | 8.91E-05 | 720.4568105 | 963.6812877 | ENSG00000064726 |
| RILP | 0.418645883 | 1.336672362 | 0.017114588 | 0.06066235 | 153.5239293 | 205.0169298 | ENSG00000167705 |
| B4GALT4 | 0.418392883 | 1.336437975 | 1.94E-05 | 0.000173518 | 492.8293509 | 658.2994234 | ENSG00000121578 |
| GON4L | 0.418373079 | 1.33641963 | 1.07E-07 | 1.48E-06 | 1007.570155 | 1345.836302 | ENSG00000116580 |
| BORCS6 | 0.418333383 | 1.336382859 | 0.170327029 | 0.348638065 | 41.94319108 | 56.08877998 | ENSG00000196544 |
| LRRC8C-DT | 0.41768354 | 1.335781038 | 0.756066865 | 0.873131704 | 3.508086222 | 4.623920423 | ENSG00000231999 |
| FKRP | 0.417670392 | 1.335768865 | 1.98E-06 | 2.16E-05 | 647.1349847 | 864.5828709 | ENSG00000181027 |
| RPUSD3 | 0.417626064 | 1.335727823 | 5.63E-10 | 1.12E-08 | 1298.228757 | 1733.725154 | ENSG00000156990 |
| NA | 0.417611472 | 1.335714312 | 0.678493083 | 0.828057969 | 3.94282149 | 5.267535964 | ENSG00000270177 |
| JHY | 0.417546917 | 1.335654546 | 0.436986143 | 0.645089622 | 16.4103334 | 21.82518223 | ENSG00000109944 |
| NA | 0.417254992 | 1.335384307 | 0.371742436 | 0.583764835 | 17.8773329 | 23.85931692 | ENSG00000259953 |
| PCAT6 | 0.416962337 | 1.335113448 | 0.183200604 | 0.366977969 | 65.26563423 | 87.37104911 | ENSG00000228288 |
| CCDC184 | 0.416878201 | 1.335035589 | 0.205048467 | 0.397040942 | 42.83785726 | 57.21209357 | ENSG00000177875 |
| SLC22A17 | 0.416260071 | 1.334463708 | 2.12E-07 | 2.77E-06 | 1148.429989 | 1532.864378 | ENSG00000092096 |
| RAB19 | 0.41614349 | 1.334355877 | 0.299605526 | 0.509826127 | 29.77579162 | 39.70770486 | ENSG00000146955 |
| NA | 0.416124157 | 1.334337996 | 0.603829379 | 0.77848488 | 6.193279957 | 8.309137848 | ENSG00000228981 |
| SHB | 0.415974684 | 1.334199758 | 0.065862936 | 0.174253508 | 83.52133565 | 111.5158783 | ENSG00000107338 |
| NA | 0.415545404 | 1.333802819 | 0.357997172 | 0.569904458 | 19.21532783 | 25.6429147 | ENSG00000280239 |
| NA | 0.415440662 | 1.333705987 | 0.28120265 | 0.490170948 | 30.52868986 | 40.5678279 | ENSG00000287769 |
| CASC9 | 0.415418394 | 1.333685401 | 0.239294243 | 0.442179234 | 35.40769599 | 47.07255928 | ENSG00000249395 |
| NA | 0.415081584 | 1.333374077 | 0.711382848 | 0.847591497 | 2.979356283 | 3.967535646 | ENSG00000157021 |
| SOX13 | 0.414995687 | 1.333294691 | 1.76E-12 | 4.93E-11 | 2523.313588 | 3364.997024 | ENSG00000143842 |
| RHBDF2 | 0.414681793 | 1.333004632 | 0.000163445 | 0.001150879 | 504.4877013 | 672.893024 | ENSG00000129667 |

| | | | | | | | |
|---|---|---|---|---|---|---|---|
| LOC105371083 | 0.414627098 | 1.332954096 | 0.584313087 | 0.763175369 | 6.925737245 | 9.223248333 | ENSG00000263080 |
| PRKG1 | 0.414603691 | 1.33293247 | 0.018749335 | 0.065149887 | 153.9886405 | 205.3734544 | ENSG00000185532 |
| CNTNAP3 | 0.414560727 | 1.332892775 | 0.083919435 | 0.209355156 | 67.29634801 | 89.65399674 | ENSG00000106714 |
| GMPR | 0.414492844 | 1.33283006 | 0.408004482 | 0.61908763 | 16.36734039 | 21.88770358 | ENSG00000137198 |
| PLEKHO1 | 0.414437883 | 1.332779285 | 1.12E-07 | 1.55E-06 | 823.8137832 | 1097.824555 | ENSG00000023902 |
| NPIPB2 | 0.41429107 | 1.332643665 | 0.610651035 | 0.783380531 | 5.996671554 | 7.976121553 | ENSG00000234719 |
| TBILA | 0.414216195 | 1.332574503 | 0.612671935 | 0.784534139 | 6.942339399 | 9.241598001 | ENSG00000261488 |
| NA | 0.414161893 | 1.332524347 | 0.061319986 | 0.165125901 | 94.622403 | 126.1847898 | ENSG00000229638 |
| PPA2 | 0.414116577 | 1.332482492 | 3.05E-05 | 0.000259439 | 532.9473617 | 709.8078446 | ENSG00000138777 |
| MYH14 | 0.414013981 | 1.332387737 | 1.28E-09 | 2.42E-08 | 1462.377856 | 1948.681693 | ENSG00000105357 |
| TEX45 | 0.413515583 | 1.331927526 | 0.560129067 | 0.74535213 | 8.223408922 | 10.98197006 | ENSG00000198723 |
| NA | 0.413411828 | 1.33183174 | 0.607302646 | 0.781101305 | 5.985993136 | 7.955596422 | ENSG00000261251 |
| FLNB-AS1 | 0.413329644 | 1.331755874 | 0.669652652 | 0.82299482 | 3.977221007 | 5.28683159 | ENSG00000244161 |
| DGCR5 | 0.413027696 | 1.331477174 | 0.609026128 | 0.782305701 | 6.258519517 | 8.362957345 | ENSG00000273032 |
| MIAT | 0.412872904 | 1.331334323 | 0.059217122 | 0.160769888 | 1060.305682 | 1411.584853 | ENSG00000225783 |
| NA | 0.4127967 | 1.331264003 | 0.236589507 | 0.43859751 | 41.56512804 | 55.36987385 | ENSG00000205534 |
| MIR4482 | 0.41275704 | 1.331227406 | 0.600963399 | 0.776307564 | 6.19683943 | 8.288612717 | ENSG00000266852 |
| ZNF674 | 0.412726706 | 1.331199417 | 0.218148075 | 0.414334346 | 36.99093673 | 49.13223245 | ENSG00000251192 |
| NA | 0.412620082 | 1.331101036 | 0.281413358 | 0.490403904 | 31.29170786 | 41.6076435 | ENSG00000238045 |
| SPDYA | 0.412572769 | 1.331057384 | 0.684489472 | 0.831291555 | 4.031782149 | 5.3502989 | ENSG00000163806 |
| PAK3 | 0.412421206 | 1.330917556 | 0.008821402 | 0.035392432 | 216.6144388 | 287.9035811 | ENSG00000077264 |
| SMAD1 | 0.412413552 | 1.330910495 | 0.00183534 | 0.009421148 | 244.8633041 | 325.6986211 | ENSG00000170365 |
| ZNF702P | 0.412386648 | 1.330885676 | 0.075854794 | 0.194324331 | 85.86937442 | 114.1334506 | ENSG00000242779 |
| NA | 0.412280772 | 1.330788009 | 0.69321746 | 0.836043703 | 4.273468486 | 5.714717643 | ENSG00000287733 |
| FGD6 | 0.41207109 | 1.330594605 | 1.45E-05 | 0.000132886 | 556.4011475 | 740.21824 | ENSG00000180263 |
| PAQR9-AS1 | 0.410977145 | 1.329586045 | 0.653594057 | 0.812465428 | 4.960847841 | 6.582480981 | ENSG00000241570 |
| PC | 0.410949121 | 1.329560218 | 1.37E-06 | 1.55E-05 | 744.7559642 | 989.6438823 | ENSG00000173599 |
| RHBDF1 | 0.410895807 | 1.329511086 | 9.90E-05 | 0.000735048 | 570.8617453 | 759.2474955 | ENSG00000007384 |
| THUMPD2 | 0.410820058 | 1.329441281 | 0.000454689 | 0.002832056 | 325.4756088 | 432.9898051 | ENSG00000138050 |
| TPRN | 0.410693258 | 1.32932444 | 6.16E-06 | 6.16E-05 | 733.4347231 | 975.3557021 | ENSG00000176058 |
| RNU6-8 | 0.410687103 | 1.32931877 | 0.637679739 | 0.801236319 | 4.995247357 | 6.620126274 | ENSG00000202337 |
| CCDC3 | 0.410649129 | 1.32928378 | 0.211095262 | 0.404769977 | 41.47644672 | 55.08727209 | ENSG00000151468 |
| VPS45 | 0.410595447 | 1.329234319 | 6.73E-05 | 0.000522742 | 475.5970409 | 632.3614642 | ENSG00000136631 |
| RBM46 | 0.410320626 | 1.328981135 | 0.649243437 | 0.809530306 | 5.751425744 | 7.582759371 | ENSG00000151962 |
| TTN | 0.409942263 | 1.328632641 | 0.056141468 | 0.154327747 | 89.7816352 | 119.0781125 | ENSG00000155657 |
| TAF12 | 0.409923689 | 1.328615536 | 0.00103195 | 0.00576238 | 340.8290677 | 452.4434854 | ENSG00000120656 |
| SFN | 0.409856349 | 1.328553522 | 0.020316413 | 0.069645209 | 143.0221965 | 190.0273198 | ENSG00000175793 |
| TVP23B | 0.409594687 | 1.328312584 | 0.004735572 | 0.021102237 | 240.8128343 | 320.1014738 | ENSG00000171928 |
| NA | 0.409404731 | 1.328137699 | 0.251190586 | 0.456743136 | 44.0975961 | 58.70204431 | ENSG00000269981 |
| NA | 0.40923726 | 1.327983535 | 0.505000659 | 0.70232988 | 10.2155789 | 13.57761977 | ENSG00000225507 |
| TADA1 | 0.409086554 | 1.327844819 | 2.34E-05 | 0.00020516 | 527.8393803 | 700.5992574 | ENSG00000152382 |
| LOC101929240 | 0.409056044 | 1.327816739 | 0.678535984 | 0.828057969 | 4.249747387 | 5.673667382 | ENSG00000265298 |
| GTF2H2 | 0.408982675 | 1.327749214 | 0.001513883 | 0.008020479 | 289.7573757 | 384.5414605 | ENSG00000145736 |
| CCDC159 | 0.408941544 | 1.32771136 | 0.248129643 | 0.453053109 | 32.31921742 | 42.8118281 | ENSG00000183401 |
| PNLDC1 | 0.408851325 | 1.327628334 | 0.718311767 | 0.851388915 | 3.251882663 | 4.351250015 | ENSG00000146453 |
| BMP1 | 0.408801702 | 1.32758267 | 0.004675518 | 0.020878967 | 222.7469284 | 295.6236283 | ENSG00000168487 |
| ARMCX3 | 0.408684806 | 1.327475106 | 0.000289949 | 0.001907239 | 376.3574797 | 499.5494117 | ENSG00000102401 |
| PLEKHG5 | 0.408621924 | 1.327417247 | 6.18E-06 | 6.17E-05 | 800.6402787 | 1063.128008 | ENSG00000171680 |
| CA10 | 0.408620708 | 1.327416128 | 0.55813571 | 0.743916196 | 7.985282057 | 10.56401537 | ENSG00000154975 |
| NA | 0.408386763 | 1.327200894 | 0.566296579 | 0.750233591 | 8.31236958 | 11.03701906 | ENSG00000248578 |
| CRTC1 | 0.408357923 | 1.327174363 | 0.000319817 | 0.002077653 | 364.9438159 | 484.29699 | ENSG00000105662 |
| NA | 0.408281083 | 1.327103677 | 0.497589722 | 0.696660074 | 10.2630211 | 13.62179145 | ENSG00000220323 |
| LOC105376199 | 0.408066373 | 1.326906186 | 0.72810289 | 0.856731016 | 3.237644773 | 4.327603462 | ENSG00000228317 |
| RELA-DT | 0.408031429 | 1.326874047 | 0.443104455 | 0.650550076 | 15.59959387 | 20.71274596 | ENSG00000285533 |
| TOR1AIP1 | 0.407544908 | 1.326426659 | 2.82E-05 | 0.00024146 | 525.0115806 | 696.1378141 | ENSG00000143337 |
| CC2D1B | 0.407521542 | 1.326405176 | 0.000325413 | 0.002107485 | 394.5562813 | 523.6952298 | ENSG00000154222 |
| SLC38A6 | 0.407503026 | 1.326388153 | 0.168848196 | 0.346597629 | 49.50769643 | 65.72184825 | ENSG00000139974 |
| LINC00667 | 0.407491077 | 1.326377167 | 0.038213436 | 0.114658522 | 129.5485581 | 171.8849686 | ENSG00000263753 |
| GPAA1 | 0.407486902 | 1.326373329 | 6.54E-09 | 1.11E-07 | 1190.99306 | 1580.254522 | ENSG00000197858 |
| LOC100288637 | 0.407433646 | 1.326324367 | 0.009573103 | 0.037711316 | 174.1137365 | 230.693663 | ENSG00000187951 |
| XPC | 0.407409093 | 1.326301795 | 9.78E-06 | 9.35E-05 | 689.0419451 | 913.5203935 | ENSG00000154767 |
| NA | 0.407314375 | 1.326214722 | 0.002192415 | 0.010988258 | 236.375683 | 313.2914401 | ENSG00000214796 |
| IQSEC2 | 0.407305824 | 1.326206861 | 4.77E-09 | 8.27E-08 | 1507.864422 | 2000.415292 | ENSG00000124313 |
| NA | 0.407290385 | 1.326192669 | 0.619476875 | 0.788725553 | 6.227679473 | 8.279910863 | ENSG00000226666 |
| NA | 0.407245158 | 1.326151095 | 0.095846068 | 0.230844662 | 71.63266564 | 94.93089697 | ENSG00000216866 |
| LOC105376805 | 0.407069831 | 1.325989941 | 0.612069252 | 0.784199347 | 5.993112081 | 7.915775666 | ENSG00000238142 |
| UBE2Q1 | 0.406925577 | 1.325857363 | 9.40E-09 | 1.55E-07 | 1348.50372 | 1787.723834 | ENSG00000160714 |
| NA | 0.406913864 | 1.325846598 | 0.481801967 | 0.683503506 | 13.23525843 | 17.5342781 | ENSG00000237742 |
| H3-3A | 0.406722542 | 1.325670784 | 1.82E-18 | 9.51E-17 | 8257.454662 | 10947.19017 | ENSG00000163041 |
| NA | 0.406679136 | 1.3256309 | 0.520688781 | 0.715714945 | 10.45370576 | 13.90751464 | ENSG00000219201 |
| STX12 | 0.406642314 | 1.325597065 | 0.008277263 | 0.033601916 | 174.8316504 | 231.7212039 | ENSG00000117758 |
| NA | 0.40661964 | 1.325576232 | 0.221697354 | 0.418934663 | 78.97239228 | 104.6635124 | ENSG00000187653 |
| NRXN1 | 0.406533831 | 1.325497391 | 0.004666819 | 0.020845051 | 208.9105554 | 276.8855218 | ENSG00000179915 |
| TSSK6 | 0.406383984 | 1.325359725 | 0.215570479 | 0.410719179 | 41.59950141 | 55.19585704 | ENSG00000178093 |

| | | | | | | | |
|---|---|---|---|---|---|---|---|
| ILF2 | 0.406338499 | 1.32531794 | 8.98E-16 | 3.73E-14 | 9093.957672 | 12051.23799 | ENSG00000143621 |
| GPM6A | 0.406268895 | 1.325254 | 0.253821198 | 0.460020472 | 32.778869 | 43.41656884 | ENSG00000150625 |
| GEMIN2 | 0.406252604 | 1.325239036 | 0.000192816 | 0.001335288 | 410.8274648 | 544.0238099 | ENSG00000092208 |
| PGBD5 | 0.406161035 | 1.325154924 | 8.64E-07 | 1.01E-05 | 822.633652 | 1090.26975 | ENSG00000177614 |
| MYLIP | 0.40586443 | 1.324882512 | 9.17E-06 | 8.86E-05 | 566.4654753 | 750.4813899 | ENSG00000007944 |
| NA | 0.405842866 | 1.324862709 | 0.15990779 | 0.333686036 | 54.46854427 | 72.27531316 | ENSG00000214176 |
| SDHAF4 | 0.405578516 | 1.324619972 | 0.007503023 | 0.031005381 | 176.2016806 | 233.3318034 | ENSG00000154079 |
| C10orf105 | 0.405446577 | 1.324498837 | 0.56105829 | 0.746220565 | 8.043402672 | 10.60166066 | ENSG00000214688 |
| NA | 0.405210899 | 1.324282485 | 0.422019959 | 0.632126184 | 16.71814902 | 22.1951814 | ENSG00000234176 |
| ATP7A | 0.405191205 | 1.324264408 | 0.16557916 | 0.341442003 | 57.52201293 | 75.93667312 | ENSG00000165240 |
| APH1B | 0.405176999 | 1.324251368 | 0.000411975 | 0.002595114 | 343.1076703 | 454.1392343 | ENSG00000138613 |
| DENND6A | 0.404803239 | 1.323908338 | 0.003826309 | 0.017693401 | 228.7872034 | 302.7857631 | ENSG00000174839 |
| RC3H1 | 0.404669338 | 1.323785467 | 0.007229205 | 0.030064643 | 210.2099809 | 277.949719 | ENSG00000135870 |
| ECI1 | 0.404645284 | 1.323763396 | 1.18E-09 | 2.26E-08 | 1747.816661 | 2313.195341 | ENSG00000167969 |
| LINC01772 | 0.404532582 | 1.323659989 | 0.669519335 | 0.82295975 | 5.522502756 | 7.359273987 | ENSG00000226029 |
| ASAH2 | 0.404532314 | 1.323659743 | 0.330659509 | 0.542057711 | 22.75633903 | 30.138671 | ENSG00000188611 |
| NA | 0.40448524 | 1.323616554 | 0.651209812 | 0.810631949 | 4.995247357 | 6.582480981 | ENSG00000276603 |
| BTNL8 | 0.40446489 | 1.323597884 | 0.583382598 | 0.762833684 | 7.225544198 | 9.574614295 | ENSG00000113303 |
| SLC35B2 | 0.404157016 | 1.323315456 | 1.63E-05 | 0.000147918 | 663.6377604 | 877.9038266 | ENSG00000157593 |
| IQCG | 0.404137793 | 1.323297823 | 0.137182366 | 0.299917687 | 70.66477739 | 93.35292936 | ENSG00000114473 |
| PFDN2 | 0.404074732 | 1.323239983 | 3.45E-07 | 4.36E-06 | 1235.563726 | 1635.095084 | ENSG00000143256 |
| CCDC7 | 0.404049387 | 1.323216736 | 0.578324438 | 0.759450456 | 7.252824769 | 9.603557734 | ENSG00000216937 |
| LRRC29 | 0.403899104 | 1.323078906 | 0.600565223 | 0.776296479 | 7.205382571 | 9.544441352 | ENSG00000125122 |
| NA | 0.403865214 | 1.323047826 | 0.687970715 | 0.83320004 | 4.222466816 | 5.612375535 | ENSG00000279058 |
| C5AR1 | 0.403595263 | 1.322800287 | 0.719779006 | 0.852321847 | 3.27204429 | 4.354371437 | ENSG00000197405 |
| ZMAT5 | 0.403565827 | 1.322773297 | 0.062257285 | 0.166841042 | 114.4417422 | 151.0740635 | ENSG00000100319 |
| LINC01876 | 0.403479919 | 1.322694533 | 0.375713265 | 0.587736266 | 25.02250993 | 33.07029047 | ENSG00000226383 |
| LIAS | 0.403258988 | 1.322491993 | 0.013057481 | 0.048687419 | 215.9018894 | 285.8819954 | ENSG00000121897 |
| NA | 0.403070839 | 1.322319532 | 0.715797155 | 0.850071015 | 3.268484817 | 4.349074552 | ENSG00000121089 |
| PUSL1 | 0.402834538 | 1.322102965 | 0.000781564 | 0.004520621 | 335.0883198 | 443.3163793 | ENSG00000169972 |
| HMBS | 0.402675619 | 1.321957338 | 4.13E-07 | 5.15E-06 | 1059.518943 | 1400.053197 | ENSG00000256269 |
| HMMR | 0.402637497 | 1.321922407 | 2.50E-06 | 2.69E-05 | 839.7785015 | 1110.08977 | ENSG00000072571 |
| PPP1R1A | 0.402601362 | 1.321889297 | 0.001327894 | 0.007169032 | 307.1599294 | 405.7089558 | ENSG00000135447 |
| NA | 0.402578689 | 1.321868522 | 0.037031149 | 0.111893319 | 125.2700033 | 165.817204 | ENSG00000229150 |
| RILPL1 | 0.402569616 | 1.32186021 | 0.017752121 | 0.062429946 | 172.0717339 | 227.4205793 | ENSG00000188026 |
| ENTPD8 | 0.402397727 | 1.321702727 | 0.548089167 | 0.735759449 | 9.579201213 | 12.68715584 | ENSG00000188833 |
| MIA3 | 0.402251695 | 1.321568949 | 1.65E-06 | 1.83E-05 | 1001.078341 | 1322.927373 | ENSG00000154305 |
| ADAM8 | 0.402244038 | 1.321561935 | 0.061696899 | 0.165762689 | 88.95874271 | 117.6224451 | ENSG00000151651 |
| NA | 0.40223362 | 1.321552391 | 0.736038043 | 0.861271552 | 3.020874744 | 3.971886573 | ENSG00000272328 |
| SLC17A5 | 0.402005424 | 1.321343373 | 0.001764109 | 0.009132457 | 263.8746253 | 348.6504695 | ENSG00000119899 |
| SMPD3 | 0.40174215 | 1.321102266 | 0.286928867 | 0.496139433 | 36.02274299 | 47.49826987 | ENSG00000103056 |
| NA | 0.401298191 | 1.320695787 | 0.181559808 | 0.364683789 | 44.4460143 | 58.82006617 | ENSG00000232573 |
| ZFPM1 | 0.401257752 | 1.320658769 | 0.036275941 | 0.110228663 | 135.1967591 | 178.7031801 | ENSG00000179588 |
| NA | 0.401017607 | 1.320438956 | 0.01732799 | 0.061253636 | 193.5292061 | 255.5836852 | ENSG00000213553 |
| PRKAA1 | 0.401001226 | 1.320423963 | 0.00116574 | 0.006410892 | 357.9469154 | 472.5207077 | ENSG00000132356 |
| ZNF506 | 0.400810381 | 1.320249304 | 0.656484041 | 0.814352412 | 5.278452155 | 6.978018627 | ENSG00000081665 |
| SLC5A12 | 0.400776694 | 1.320218476 | 0.162589557 | 0.337640618 | 49.12276762 | 64.8115216 | ENSG00000148942 |
| ENG | 0.40065308 | 1.320105361 | 0.08069381 | 0.203105771 | 82.13948415 | 108.5430406 | ENSG00000106991 |
| COL24A1 | 0.400631813 | 1.320085902 | 0.349685696 | 0.561787696 | 21.94707404 | 28.94318825 | ENSG00000171502 |
| H2BC13 | 0.400508218 | 1.319972816 | 0.500048316 | 0.698373943 | 12.89156876 | 17.05145863 | ENSG00000185130 |
| C18orf25 | 0.400506884 | 1.319971595 | 0.00031893 | 0.00207442 | 375.5238832 | 495.4269869 | ENSG00000152242 |
| PLPP3 | 0.400136301 | 1.31963258 | 0.000104869 | 0.000775125 | 422.0296452 | 556.7903237 | ENSG00000162407 |
| KIF13B | 0.39982235 | 1.31934544 | 0.002051885 | 0.010397062 | 280.703345 | 370.1169814 | ENSG00000197892 |
| NA | 0.399624988 | 1.319164965 | 0.234334782 | 0.435614957 | 38.92142662 | 51.30824645 | ENSG00000286388 |
| TMEM9 | 0.399576112 | 1.319120274 | 3.44E-08 | 5.19E-07 | 2337.779855 | 3084.354705 | ENSG00000116857 |
| MFNG | 0.39933118 | 1.318896341 | 0.114647889 | 0.264249794 | 69.85490143 | 92.17456678 | ENSG00000100060 |
| COX2 | 0.399167771 | 1.318746963 | 0.062558993 | 0.167506959 | 28572.04172 | 37679.49408 | ENSG00000198712 |
| PBK | 0.39900074 | 1.318594291 | 0.024006583 | 0.079741572 | 160.2246085 | 210.9910542 | ENSG00000168078 |
| GLUL | 0.398900526 | 1.3185027 | 1.20E-14 | 4.28E-13 | 5726.755755 | 7550.154036 | ENSG00000135821 |
| FSTL5 | 0.39862349 | 1.318249537 | 0.698445802 | 0.839388378 | 4.286511168 | 5.660898146 | ENSG00000168843 |
| MARCHF3 | 0.3986106 | 1.318237759 | 0.018079263 | 0.063314513 | 163.1912009 | 215.1783867 | ENSG00000173926 |
| S100A13 | 0.39860414 | 1.318231856 | 0.266373643 | 0.473812399 | 33.87754185 | 44.65047489 | ENSG00000189171 |
| PHF19 | 0.398214898 | 1.317876242 | 0.000600338 | 0.003600309 | 343.1734941 | 452.3119593 | ENSG00000119403 |
| ERFE | 0.398211381 | 1.31787303 | 0.262987432 | 0.470180598 | 33.03003854 | 43.58266181 | ENSG00000178752 |
| PARD3B | 0.398148396 | 1.317815495 | 0.096789437 | 0.232494087 | 68.28709379 | 89.93224758 | ENSG00000116117 |
| NA | 0.398135733 | 1.317803929 | 0.710567259 | 0.847218763 | 3.27204429 | 4.335075811 | ENSG00000278743 |
| HSD11B1L | 0.398116932 | 1.317786755 | 0.160027404 | 0.333861857 | 78.66073786 | 103.5530204 | ENSG00000167733 |
| NAA80 | 0.398113944 | 1.317784027 | 0.091750781 | 0.223518254 | 78.42350072 | 103.2089159 | ENSG00000243477 |
| MEX3A | 0.398048546 | 1.317724292 | 1.22E-09 | 2.31E-08 | 2084.895832 | 2747.317218 | ENSG00000254726 |
| NA | 0.397980053 | 1.317661733 | 0.17467372 | 0.355005726 | 47.04372192 | 62.12469064 | ENSG00000178458 |
| ATG10 | 0.397822433 | 1.317517781 | 0.016414296 | 0.058653545 | 171.8591076 | 226.5504296 | ENSG00000152348 |
| FGF1 | 0.397788608 | 1.317486892 | 0.618452147 | 0.788301079 | 6.316640132 | 8.313488775 | ENSG00000113578 |
| NA | 0.397520587 | 1.317242155 | 0.209318715 | 0.4025032 | 42.43396203 | 55.89792655 | ENSG00000196656 |
| PI4K2B | 0.397491044 | 1.317215181 | 0.001687629 | 0.008806539 | 417.8466369 | 550.7208648 | ENSG00000038210 |

| | | | | | | | |
|---|---|---|---|---|---|---|---|
| TECPR2 | 0.397171269 | 1.316923251 | 2.51E-06 | 2.69E-05 | 734.570946 | 966.932966 | ENSG00000196663 |
| NA | 0.397071784 | 1.316832442 | 0.68315012 | 0.830790283 | 4.280587431 | 5.647845365 | ENSG00000242085 |
| LRRC32 | 0.39688507 | 1.316662028 | 0.153004441 | 0.32360722 | 58.82504968 | 77.53130939 | ENSG00000137507 |
| C17orf113 | 0.396836278 | 1.316617499 | 0.202341601 | 0.393732085 | 46.99210986 | 61.74069273 | ENSG00000267221 |
| SMG1P3 | 0.396781992 | 1.316567959 | 0.189673931 | 0.375943765 | 60.95654544 | 80.02215798 | ENSG00000180747 |
| TMCO6 | 0.396579177 | 1.316382887 | 0.002668854 | 0.013003741 | 312.7939972 | 411.5837345 | ENSG00000113119 |
| KLHL28 | 0.396354525 | 1.31617792 | 0.049593823 | 0.140185503 | 109.5049465 | 144.0622561 | ENSG00000179454 |
| SMIM10L2A | 0.39588966 | 1.31575389 | 0.011315407 | 0.043292434 | 168.7768844 | 222.0722449 | ENSG00000178947 |
| DPP8 | 0.395337758 | 1.315250645 | 0.000207508 | 0.001421393 | 423.1790921 | 556.8947983 | ENSG00000074603 |
| ZNF691 | 0.395251422 | 1.315171938 | 0.000471969 | 0.002931954 | 368.8789604 | 484.9255882 | ENSG00000164011 |
| SIL1 | 0.39486943 | 1.314823757 | 0.094650082 | 0.229133035 | 73.11006423 | 96.29029722 | ENSG00000120725 |
| EBF3 | 0.394866712 | 1.314821281 | 0.695193066 | 0.837250559 | 4.25330686 | 5.602727723 | ENSG00000108001 |
| IL1RAP | 0.394387735 | 1.314384831 | 0.036687514 | 0.11113951 | 113.4741589 | 149.0128773 | ENSG00000196083 |
| NBPF12 | 0.393969511 | 1.314003858 | 0.017530681 | 0.061800805 | 169.9651028 | 222.9922194 | ENSG00000268043 |
| B3GALT6 | 0.393953761 | 1.313989512 | 2.91E-05 | 0.000249048 | 542.1508374 | 712.7364809 | ENSG00000176022 |
| NA | 0.393932177 | 1.313969855 | 0.479084626 | 0.68145793 | 13.41673922 | 17.53768307 | ENSG00000261884 |
| CLDN1 | 0.393749864 | 1.313803818 | 0.122435443 | 0.276545072 | 65.8432809 | 86.46711057 | ENSG00000163347 |
| LINC02021 | 0.393741988 | 1.313796646 | 0.436993991 | 0.645089622 | 18.30050003 | 24.03789557 | ENSG00000249846 |
| NA | 0.393662848 | 1.313724579 | 0.559710623 | 0.745115652 | 8.832506035 | 11.66010947 | ENSG00000230207 |
| MYO1D | 0.393649148 | 1.313712104 | 8.39E-05 | 0.000636044 | 637.6910958 | 837.7220135 | ENSG00000176658 |
| TNMD | 0.393617519 | 1.313683303 | 0.170888216 | 0.349455762 | 50.5838434 | 66.50299218 | ENSG00000000005 |
| NA | 0.3932964 | 1.313390933 | 0.640041382 | 0.802921583 | 5.291494837 | 6.943494756 | ENSG00000242951 |
| GPS2 | 0.393260456 | 1.313358211 | 0.350926222 | 0.563021036 | 24.786468 | 32.58016662 | ENSG00000132522 |
| FOXD3 | 0.393248923 | 1.313347711 | 0.00091768 | 0.005205882 | 431.650416 | 567.5501105 | ENSG00000187140 |
| PEAK1 | 0.393197821 | 1.313301192 | 0.000113824 | 0.000834457 | 868.6523005 | 1139.911767 | ENSG00000173517 |
| TATDN1 | 0.393194429 | 1.313298104 | 0.000523915 | 0.003201985 | 354.6322097 | 465.7147616 | ENSG00000147687 |
| ZSWIM6 | 0.393084531 | 1.313198067 | 0.00288732 | 0.013934844 | 278.1278839 | 365.1033873 | ENSG00000130449 |
| TMIGD2 | 0.39302355 | 1.31314256 | 0.490897135 | 0.69054665 | 13.35714407 | 17.63662021 | ENSG00000167664 |
| GS1-124K5.4 | 0.392940257 | 1.31306675 | 0.434750432 | 0.643487883 | 19.11123956 | 25.03382303 | ENSG00000237310 |
| IDH3G | 0.39288754 | 1.31301877 | 3.86E-05 | 0.000319586 | 524.2488675 | 688.6197218 | ENSG00000067829 |
| NA | 0.392740909 | 1.312885326 | 0.532961532 | 0.724314769 | 11.36644863 | 14.85832446 | ENSG00000230793 |
| ZNF391 | 0.392540198 | 1.312702687 | 0.030423089 | 0.09607693 | 132.6906819 | 173.98586 | ENSG00000124613 |
| LANCL3 | 0.392374882 | 1.312552275 | 0.539339888 | 0.729040189 | 10.89347504 | 14.32140196 | ENSG00000147036 |
| CLEC18A | 0.392044324 | 1.31225157 | 0.649269067 | 0.809530306 | 5.325894353 | 6.976789122 | ENSG00000157322 |
| NA | 0.391705427 | 1.311943352 | 0.72223036 | 0.853577843 | 3.295765389 | 4.335075811 | ENSG00000206147 |
| ZSWIM4 | 0.391705318 | 1.311943252 | 0.000299911 | 0.001963874 | 417.7894333 | 547.9217307 | ENSG00000132003 |
| RABAC1 | 0.39164673 | 1.311889975 | 0.33149346 | 0.542983463 | 28.3215293 | 37.21685627 | ENSG00000105404 |
| CENPBD1 | 0.391532351 | 1.31178597 | 0.018473358 | 0.064390803 | 151.6382119 | 198.7180692 | ENSG00000177946 |
| LYG1 | 0.390935058 | 1.311242988 | 0.619405565 | 0.788725553 | 6.616447084 | 8.693798176 | ENSG00000144214 |
| ZNF37BP | 0.390398999 | 1.310755863 | 0.026503957 | 0.086223466 | 139.4177514 | 183.0038047 | ENSG00000234420 |
| ZNF619 | 0.390204875 | 1.310579504 | 0.080178496 | 0.202095424 | 93.09845194 | 121.9159762 | ENSG00000177873 |
| PLEKHG4B | 0.390077595 | 1.310463885 | 3.43E-08 | 5.19E-07 | 1282.847434 | 1680.91666 | ENSG00000153404 |
| DSCAM | 0.38987437 | 1.310279299 | 0.642976435 | 0.805106085 | 6.360522857 | 8.306962384 | ENSG00000171587 |
| NA | 0.389095257 | 1.309571888 | 0.437780587 | 0.645947838 | 16.22292887 | 21.24219612 | ENSG00000279382 |
| TULP4 | 0.389042187 | 1.309523715 | 0.0156692 | 0.056471916 | 238.0671301 | 311.3497458 | ENSG00000130338 |
| AKAP13 | 0.388979507 | 1.309466822 | 0.00031045 | 0.002026564 | 722.0078603 | 945.2820411 | ENSG00000170776 |
| LOC100506606 | 0.388908081 | 1.309401994 | 0.772488065 | 0.882863539 | 3.102716458 | 4.03223246 | ENSG00000257176 |
| C18orf21 | 0.38859947 | 1.309121926 | 0.067953331 | 0.178508646 | 106.3405761 | 139.3584536 | ENSG00000141428 |
| KIF17 | 0.388590701 | 1.309113969 | 0.001615398 | 0.008469402 | 294.7437764 | 385.9246026 | ENSG00000117245 |
| TMC5 | 0.388557317 | 1.309083676 | 0.317684493 | 0.527777749 | 25.8210965 | 33.77536582 | ENSG00000103534 |
| NANP | 0.388274233 | 1.308826835 | 0.020060501 | 0.068968243 | 177.945935 | 232.8187681 | ENSG00000170191 |
| ATP10D | 0.388185336 | 1.308746189 | 0.092812234 | 0.225580315 | 78.68000918 | 103.1165052 | ENSG00000145246 |
| NA | 0.388179343 | 1.308740752 | 0.692626404 | 0.835855442 | 4.280587431 | 5.600552259 | ENSG00000277879 |
| NA | 0.388112123 | 1.308679775 | 0.310460689 | 0.521021071 | 30.59245488 | 40.024379 | ENSG00000259802 |
| USP18 | 0.388022105 | 1.308598122 | 0.036637375 | 0.111058881 | 132.2165131 | 172.8521206 | ENSG00000184979 |
| NKIRAS1 | 0.387974996 | 1.308555392 | 0.092481114 | 0.224833398 | 80.33560843 | 105.3190036 | ENSG00000197885 |
| CRLS1 | 0.387959827 | 1.308541633 | 0.007911165 | 0.032361516 | 313.8262341 | 411.1376667 | ENSG00000088766 |
| DOK6 | 0.38783997 | 1.308432926 | 0.453649729 | 0.659673145 | 15.72740324 | 20.66639881 | ENSG00000206052 |
| NA | 0.387733564 | 1.308336426 | 0.166861137 | 0.343414621 | 61.13182316 | 79.7999021 | ENSG00000270872 |
| SNORD83A | 0.387634495 | 1.308246586 | 0.554444098 | 0.740972103 | 9.868329748 | 12.93211231 | ENSG00000209482 |
| ERMAP | 0.387538281 | 1.308159342 | 0.227788812 | 0.427104022 | 37.05470175 | 48.48454952 | ENSG00000164010 |
| NCOA5 | 0.3873346 | 1.307974668 | 2.78E-07 | 3.58E-06 | 934.7039131 | 1222.911599 | ENSG00000124160 |
| NA | 0.387303715 | 1.307946667 | 0.207622896 | 0.400179464 | 42.62108723 | 55.7803562 | ENSG00000261655 |
| IRAK2 | 0.387263374 | 1.307910094 | 0.369417633 | 0.581613339 | 22.64157234 | 29.56003582 | ENSG00000134070 |
| KDM5B | 0.387225683 | 1.307875925 | 3.37E-10 | 6.96E-09 | 5008.751938 | 6549.487132 | ENSG00000117139 |
| BAX | 0.386534473 | 1.307249459 | 1.47E-09 | 2.76E-08 | 2858.332013 | 3736.373881 | ENSG00000087088 |
| ACOT1 | 0.386149884 | 1.306901023 | 0.2557312 | 0.462594226 | 34.62067756 | 45.16062942 | ENSG00000184227 |
| TOX3 | 0.385912167 | 1.306685699 | 0.003471442 | 0.016303948 | 279.4991616 | 365.1583636 | ENSG00000103460 |
| MMP11 | 0.385887664 | 1.306663506 | 0.007480288 | 0.030938511 | 197.1758063 | 257.5076789 | ENSG00000099953 |
| ZNF281 | 0.38584948 | 1.306628923 | 6.39E-09 | 1.09E-07 | 1715.071379 | 2240.713057 | ENSG00000162702 |
| INSM1 | 0.385657234 | 1.30645482 | 0.390424895 | 0.601865789 | 19.81374653 | 25.90905872 | ENSG00000173404 |
| NA | 0.385361661 | 1.306187187 | 0.428827037 | 0.638613353 | 17.79785545 | 23.29193792 | ENSG00000275765 |
| SLC9A8 | 0.385338081 | 1.306165838 | 0.011413125 | 0.043586827 | 204.2996526 | 266.7273543 | ENSG00000197818 |
| ZNF629 | 0.385207281 | 1.306047422 | 2.03E-06 | 2.21E-05 | 868.8811516 | 1134.947349 | ENSG00000102870 |

| | | | | | | | |
|---|---|---|---|---|---|---|---|
| VIPAS39 | 0.385031077 | 1.305887917 | 0.000214511 | 0.00146247 | 401.0208407 | 523.6932653 | ENSG00000151445 |
| EDRF1-AS1 | 0.385008023 | 1.30586705 | 0.72216977 | 0.853559651 | 4.124302281 | 5.334124696 | ENSG00000236991 |
| NA | 0.384926405 | 1.305793174 | 0.535310268 | 0.72613054 | 10.95515513 | 14.29747186 | ENSG00000261087 |
| ZNF526 | 0.384906791 | 1.305775422 | 0.000172567 | 0.001207907 | 472.55453 | 617.2702471 | ENSG00000167625 |
| KIF19 | 0.384837387 | 1.305712606 | 0.338278057 | 0.550263392 | 27.3298938 | 35.63642965 | ENSG00000196169 |
| SLC25A11 | 0.384818667 | 1.305695663 | 2.64E-06 | 2.82E-05 | 1422.242199 | 1856.012368 | ENSG00000108528 |
| SH3PXD2B | 0.384392398 | 1.305309931 | 1.11E-10 | 2.48E-09 | 2526.135692 | 3297.440869 | ENSG00000174705 |
| GJA4 | 0.384377366 | 1.30529633 | 0.146009826 | 0.31312691 | 68.43209013 | 89.0497075 | ENSG00000187513 |
| FAM193B | 0.384322153 | 1.305246376 | 0.001064255 | 0.005918275 | 947.1502251 | 1236.634509 | ENSG00000146067 |
| ZNF548 | 0.384218689 | 1.305152772 | 0.067716949 | 0.178011494 | 87.93715689 | 114.7990317 | ENSG00000188785 |
| RBM14-RBM4 | 0.384118636 | 1.305062261 | 0.74920461 | 0.869654826 | 3.075435886 | 3.971886573 | ENSG00000248643 |
| ITPR3 | 0.383876464 | 1.304843211 | 2.35E-10 | 4.95E-09 | 2184.870722 | 2850.741355 | ENSG00000096433 |
| ADCY10P1 | 0.383869176 | 1.304836619 | 0.189206043 | 0.375292104 | 65.46549719 | 85.4217575 | ENSG00000161912 |
| CDKL5 | 0.383685077 | 1.304670123 | 0.004364042 | 0.019735378 | 290.1734239 | 378.335673 | ENSG00000008086 |
| NA | 0.383621754 | 1.304612859 | 0.745884261 | 0.867900448 | 3.302884333 | 4.310199754 | ENSG00000254458 |
| NA | 0.383580169 | 1.304575255 | 0.099654103 | 0.237532214 | 87.10953587 | 113.8656256 | ENSG00000176654 |
| OSGIN1 | 0.383296829 | 1.304319066 | 0.447688774 | 0.654431651 | 17.89037558 | 23.35540523 | ENSG00000140961 |
| PABPC4L | 0.383195675 | 1.304227618 | 0.154799871 | 0.326313671 | 57.17893365 | 74.57698932 | ENSG00000254535 |
| TK2 | 0.382996431 | 1.304047509 | 0.258184587 | 0.4649834 | 39.89201077 | 52.00445199 | ENSG00000166548 |
| IKBKE | 0.38288727 | 1.303948843 | 0.005534974 | 0.024040473 | 227.7997116 | 297.1436137 | ENSG00000263528 |
| CCKBR | 0.382806605 | 1.303875938 | 5.08E-05 | 0.000408028 | 521.4673149 | 680.1370913 | ENSG00000110148 |
| NEBL | 0.382645598 | 1.303730431 | 0.09952525 | 0.237374887 | 78.11006627 | 101.992908 | ENSG00000078114 |
| ELAVL3 | 0.382637565 | 1.303723171 | 0.404976964 | 0.615978573 | 20.60551964 | 26.80917047 | ENSG00000196361 |
| CCM2 | 0.381798454 | 1.302965111 | 9.43E-06 | 9.07E-05 | 628.2280573 | 818.5697791 | ENSG00000136280 |
| SULT4A1 | 0.381736853 | 1.302909478 | 0.016517054 | 0.058976096 | 214.0179003 | 278.5665892 | ENSG00000130540 |
| FAM86JP | 0.381718062 | 1.302892507 | 0.369970905 | 0.582144687 | 26.92599858 | 34.93492723 | ENSG00000171084 |
| ZCCHC14 | 0.381654345 | 1.302834966 | 1.16E-06 | 1.33E-05 | 847.2833599 | 1103.711733 | ENSG00000140948 |
| NA | 0.381391371 | 1.302597508 | 0.526960816 | 0.72003871 | 12.07518482 | 15.68458604 | ENSG00000258072 |
| NME7 | 0.381371009 | 1.302579123 | 0.002097282 | 0.010604332 | 294.0400742 | 383.0825036 | ENSG00000143156 |
| RHBDD1 | 0.381305285 | 1.302519783 | 0.001063673 | 0.005916782 | 345.8159748 | 450.0496821 | ENSG00000144468 |
| LRRC7 | 0.381280348 | 1.30249727 | 0.561771469 | 0.746766189 | 8.94162832 | 11.65358308 | ENSG00000033122 |
| C17orf50 | 0.381236861 | 1.302458009 | 0.733049547 | 0.859729886 | 3.575410714 | 4.693914123 | ENSG00000270806 |
| UBR3 | 0.380789477 | 1.302054176 | 0.000574687 | 0.003464079 | 425.7728836 | 553.882707 | ENSG00000144357 |
| RAPGEF5 | 0.380653992 | 1.301931904 | 0.007246247 | 0.030108994 | 239.1985004 | 311.7503397 | ENSG00000136237 |
| AR | 0.380416544 | 1.301717642 | 0.008692763 | 0.034972874 | 264.1865852 | 343.5673856 | ENSG00000169083 |
| ZNF296 | 0.380399572 | 1.301702328 | 0.036814346 | 0.111398641 | 125.9918083 | 164.1398048 | ENSG00000170684 |
| NA | 0.380361703 | 1.30166816 | 0.594235504 | 0.771358959 | 7.685475105 | 9.975448827 | ENSG00000260123 |
| NA | 0.380179081 | 1.3015034 | 0.614550333 | 0.785501391 | 7.39634657 | 9.560615555 | ENSG00000286084 |
| FOXO3 | 0.380045657 | 1.30138304 | 0.001598344 | 0.00839631 | 364.3317703 | 473.7298103 | ENSG00000118689 |
| VSNL1 | 0.379998123 | 1.301340162 | 1.44E-06 | 1.62E-05 | 1301.236361 | 1693.426819 | ENSG00000163032 |
| CAMKK1 | 0.379922935 | 1.301272343 | 0.012556084 | 0.047197779 | 188.9530077 | 245.925487 | ENSG00000004660 |
| SNHG29 | 0.379882402 | 1.301235784 | 2.47E-10 | 5.19E-09 | 3404.719628 | 4429.361295 | ENSG00000175061 |
| TESMIN | 0.379752672 | 1.30111878 | 0.0001085 | 0.000800088 | 544.2610286 | 708.0656033 | ENSG00000132749 |
| NA | 0.379650297 | 1.301026454 | 0.726461003 | 0.855887253 | 3.310003278 | 4.302727404 | ENSG00000269191 |
| TRAPPC12 | 0.379295217 | 1.300706281 | 2.77E-06 | 2.95E-05 | 990.0487937 | 1288.428545 | ENSG00000171853 |
| MACF1 | 0.379033844 | 1.300470654 | 1.73E-06 | 1.91E-05 | 4702.889856 | 6114.783003 | ENSG00000127603 |
| PACSIN1 | 0.378987752 | 1.300429107 | 0.000335639 | 0.002164803 | 416.4341991 | 541.6966906 | ENSG00000124507 |
| ABCG4 | 0.378755526 | 1.300219797 | 0.205201221 | 0.397157515 | 50.2825619 | 65.55624974 | ENSG00000172350 |
| RASSF10-DT | 0.378680317 | 1.300152017 | 0.714616842 | 0.84957753 | 4.375471827 | 5.666195032 | ENSG00000254670 |
| FHL3 | 0.37857006 | 1.300052658 | 0.000733763 | 0.00428482 | 391.6832997 | 509.1842993 | ENSG00000183386 |
| KLHL8 | 0.378557267 | 1.300041129 | 0.017806103 | 0.062584824 | 217.7732911 | 282.7556754 | ENSG00000145332 |
| CRTC2 | 0.378303504 | 1.299812479 | 0.000144524 | 0.001032667 | 537.2911293 | 698.6640504 | ENSG00000160741 |
| LOC107987373 | 0.378294573 | 1.299804432 | 0.023945441 | 0.079577355 | 154.6519677 | 200.8467901 | ENSG00000276345 |
| SGPP2 | 0.378070332 | 1.299602417 | 0.728653883 | 0.857148832 | 3.340843322 | 4.332900348 | ENSG00000163082 |
| NA | 0.377906816 | 1.299455128 | 0.453458428 | 0.6596002 | 16.34509384 | 21.16350056 | ENSG00000259366 |
| ZPLD2P | 0.37783442 | 1.299389921 | 0.247846657 | 0.452799082 | 38.12848445 | 49.53429648 | ENSG00000236155 |
| NA | 0.377660347 | 1.299233148 | 0.440009071 | 0.647767794 | 16.63778184 | 21.57615837 | ENSG00000254851 |
| SZT2 | 0.377580953 | 1.299161652 | 0.005372246 | 0.023468626 | 265.8833713 | 345.417826 | ENSG00000198198 |
| MTUS2 | 0.377495805 | 1.299084977 | 0.481325703 | 0.683198091 | 15.09961904 | 19.51336379 | ENSG00000132938 |
| SLA | 0.377448573 | 1.299042448 | 0.411366476 | 0.622588228 | 18.93924198 | 24.57169665 | ENSG00000155926 |
| NXPH4 | 0.377291232 | 1.298900781 | 0.148885568 | 0.317026041 | 64.41604598 | 83.80581809 | ENSG00000182379 |
| PHOSPHO2 | 0.377191473 | 1.298810968 | 0.391061235 | 0.602430167 | 23.64388626 | 30.75088409 | ENSG00000144362 |
| C12orf4 | 0.377161211 | 1.298783725 | 0.027895576 | 0.089776406 | 134.6386637 | 174.8749265 | ENSG00000047621 |
| SMUG1 | 0.377130772 | 1.298756322 | 0.000815546 | 0.004690798 | 345.9254281 | 449.2771674 | ENSG00000123415 |
| CFAP73 | 0.377004732 | 1.298642862 | 0.600917549 | 0.776307564 | 7.954442013 | 10.33740856 | ENSG00000186710 |
| RPL23P8 | 0.376824122 | 1.298480296 | 0.728641741 | 0.857148832 | 3.330164905 | 4.314550681 | ENSG00000219451 |
| AMZ2P1 | 0.376550265 | 1.298233838 | 0.066796059 | 0.176288107 | 123.1212432 | 159.5946753 | ENSG00000214174 |
| PTPN21 | 0.376291673 | 1.29800116 | 0.005309537 | 0.023248459 | 233.4968555 | 302.9826914 | ENSG00000070778 |
| LIN37 | 0.376112418 | 1.297839893 | 0.731676567 | 0.858723011 | 3.588453396 | 4.688617237 | ENSG00000267796 |
| NA | 0.376070842 | 1.297802492 | 0.685327825 | 0.831721353 | 5.862632961 | 7.642821712 | ENSG00000235001 |
| LINC02593 | 0.375947627 | 1.297691657 | 0.592677474 | 0.769999594 | 9.272275316 | 12.05564712 | ENSG00000223764 |
| COPA | 0.375945107 | 1.29768939 | 6.59E-13 | 1.94E-11 | 4364.690232 | 5662.97009 | ENSG00000122218 |
| NA | 0.375785341 | 1.297545691 | 0.256657932 | 0.463515831 | 39.14028158 | 50.91828924 | ENSG00000227097 |
| KLF5 | 0.375693174 | 1.297462799 | 0.361805894 | 0.573839281 | 22.78006013 | 29.59645161 | ENSG00000102554 |

| | | | | | | | |
|---|---|---|---|---|---|---|---|
| ITGB5 | 0.375543309 | 1.297328027 | 2.53E-14 | 8.70E-13 | 7090.346723 | 9198.033675 | ENSG00000082781 |
| MYSM1 | 0.375508726 | 1.297296929 | 1.14E-05 | 0.00010718 | 1032.212849 | 1338.398563 | ENSG00000162601 |
| NOVA2 | 0.375222717 | 1.297039771 | 0.000150679 | 0.001072186 | 576.6615285 | 748.3685893 | ENSG00000104967 |
| TBC1D8-AS1 | 0.375203293 | 1.297022308 | 0.723220005 | 0.854159462 | 3.330164905 | 4.307078331 | ENSG00000272902 |
| QSOX1 | 0.375125456 | 1.296952332 | 3.29E-13 | 1.00E-11 | 3297.911727 | 4277.563044 | ENSG00000116260 |
| TMEM185B | 0.374972207 | 1.296814572 | 0.000222411 | 0.001511162 | 425.870184 | 552.2788447 | ENSG00000226479 |
| USP9X | 0.374963137 | 1.296806419 | 2.19E-07 | 2.85E-06 | 4853.164846 | 6293.060177 | ENSG00000124486 |
| SPPL2B | 0.374955574 | 1.296799621 | 0.002603421 | 0.012727312 | 751.9351117 | 975.6646242 | ENSG00000005206 |
| SMIM14 | 0.374853126 | 1.296707536 | 0.011570344 | 0.044106973 | 202.4898531 | 262.4253619 | ENSG00000163683 |
| SHTN1 | 0.374806818 | 1.296665915 | 0.004288168 | 0.019477624 | 320.2893463 | 414.7643348 | ENSG00000187164 |
| NR2F2 | 0.374685011 | 1.296556442 | 0.679250424 | 0.828368191 | 4.617158164 | 6.001386791 | ENSG00000185551 |
| ZNF337-AS1 | 0.374637229 | 1.2965135 | 0.367453633 | 0.579660284 | 24.08841022 | 31.28205821 | ENSG00000213742 |
| INO80D | 0.374455281 | 1.296349999 | 0.017895064 | 0.062862404 | 226.9029343 | 293.9169592 | ENSG00000114933 |
| SPATA3-AS1 | 0.374403303 | 1.296303294 | 0.592351453 | 0.76989382 | 11.07851531 | 14.31894295 | ENSG00000238062 |
| NDUFB5 | 0.37394733 | 1.295893654 | 0.000243772 | 0.001640057 | 730.1773702 | 946.6425256 | ENSG00000136521 |
| FBXO4 | 0.373779665 | 1.295743058 | 0.512922518 | 0.70906535 | 13.67858719 | 17.70878937 | ENSG00000151876 |
| NA | 0.373765418 | 1.295730263 | 0.624697508 | 0.79235761 | 6.691169853 | 8.639978679 | ENSG00000268049 |
| TTC13 | 0.373221431 | 1.295241783 | 0.033993209 | 0.104912855 | 140.8227398 | 182.2501638 | ENSG00000143643 |
| MAVS | 0.373221176 | 1.295241554 | 1.30E-06 | 1.47E-05 | 1077.876777 | 1395.72582 | ENSG00000088888 |
| LINC01023 | 0.373205096 | 1.295227117 | 0.617428689 | 0.787368546 | 9.278199052 | 12.02954156 | ENSG00000272523 |
| POLR2J3 | 0.373071437 | 1.295107126 | 0.639591247 | 0.802516823 | 6.94709408 | 9.010640267 | ENSG00000168255 |
| NA | 0.372622413 | 1.2947041 | 0.678899402 | 0.828284464 | 4.617158164 | 5.989563514 | ENSG00000229431 |
| NA | 0.372589347 | 1.294674426 | 0.432580911 | 0.641954509 | 18.20086095 | 23.56366237 | ENSG00000254297 |
| ASH2L | 0.372506739 | 1.294600296 | 1.44E-07 | 1.94E-06 | 1354.277821 | 1752.568826 | ENSG00000129691 |
| TNFSF11 | 0.372225424 | 1.294347883 | 0.024316988 | 0.080567937 | 151.1874849 | 195.6636981 | ENSG00000120659 |
| LCA10 | 0.371989153 | 1.294135924 | 0.784568443 | 0.890365666 | 3.102716458 | 3.975007995 | ENSG00000273769 |
| VWA2 | 0.371818874 | 1.293983189 | 0.076037658 | 0.194707955 | 94.24520353 | 122.1110126 | ENSG00000165816 |
| EMID1 | 0.371782144 | 1.293950245 | 0.041155256 | 0.121323138 | 126.8802197 | 163.9863131 | ENSG00000186998 |
| VWA5B1 | 0.371636046 | 1.293819217 | 0.337359935 | 0.549275142 | 32.24833346 | 41.56470132 | ENSG00000158816 |
| HIF3A | 0.371620218 | 1.293805022 | 0.004399593 | 0.019867604 | 267.9814357 | 346.4757803 | ENSG00000124440 |
| CUL9 | 0.371605442 | 1.293791771 | 0.034629624 | 0.106320998 | 184.6087076 | 238.8353105 | ENSG00000112659 |
| ZNF786 | 0.371518559 | 1.293713858 | 0.000643385 | 0.003815686 | 392.4166728 | 507.7149892 | ENSG00000197362 |
| CTSH | 0.371510858 | 1.293706952 | 0.00162123 | 0.008497621 | 357.3683795 | 461.9763144 | ENSG00000103811 |
| ITPKB | 0.371500826 | 1.293697956 | 0.004177059 | 0.019063257 | 294.5127946 | 381.0663195 | ENSG00000143772 |
| NAAA | 0.371415765 | 1.293621682 | 0.221102402 | 0.418221041 | 47.0273991 | 60.84710737 | ENSG00000138744 |
| CYC1 | 0.371359071 | 1.293570848 | 3.43E-12 | 9.17E-11 | 4752.852068 | 6148.286422 | ENSG00000179091 |
| SSR4P1 | 0.371355599 | 1.293567735 | 0.539535235 | 0.729048576 | 11.58797334 | 15.00550067 | ENSG00000235374 |
| ZNF860 | 0.371329326 | 1.293544178 | 0.137249081 | 0.299959367 | 69.1375979 | 89.56470742 | ENSG00000197385 |
| MAMSTR | 0.371288345 | 1.293507434 | 0.305776755 | 0.516058292 | 31.52004603 | 40.8943178 | ENSG00000176909 |
| KCNH8 | 0.371118728 | 1.293355366 | 0.248824487 | 0.453807341 | 40.68614814 | 52.50808013 | ENSG00000183960 |
| GALNT2 | 0.370957805 | 1.293211108 | 5.70E-10 | 1.14E-08 | 3274.397983 | 4233.429629 | ENSG00000143641 |
| COMMD6 | 0.370951213 | 1.2932052 | 0.00050288 | 0.003092444 | 437.0924999 | 565.5731277 | ENSG00000188243 |
| NA | 0.370916649 | 1.293174217 | 0.001076966 | 0.005981914 | 683.1668008 | 883.4585355 | ENSG00000197582 |
| ARFGEF3 | 0.370726173 | 1.293003493 | 0.000370024 | 0.002360775 | 858.1087699 | 1108.796989 | ENSG00000112379 |
| PTPDC1 | 0.370690626 | 1.292971635 | 0.035883984 | 0.109324356 | 133.0023363 | 171.807219 | ENSG00000158079 |
| ITSN2 | 0.3706874 | 1.292968744 | 0.006942129 | 0.029075652 | 233.2729665 | 301.4209706 | ENSG00000198399 |
| FOXC1 | 0.370653679 | 1.292938522 | 0.731496262 | 0.858577685 | 3.33728385 | 4.297430518 | ENSG00000054598 |
| BCL11A | 0.370406782 | 1.292717273 | 0.001609143 | 0.008438954 | 370.5369233 | 479.3431224 | ENSG00000119866 |
| ETFB | 0.370366528 | 1.292681205 | 0.148180087 | 0.316058386 | 69.43529377 | 89.83039993 | ENSG00000105379 |
| TNFRSF10B | 0.370264312 | 1.29258962 | 1.00E-06 | 1.16E-05 | 1904.339298 | 2460.831718 | ENSG00000120889 |
| SCN3B | 0.370256023 | 1.292582194 | 0.50311895 | 0.701106949 | 12.82988867 | 16.62574805 | ENSG00000166257 |
| NA | 0.370234996 | 1.292563355 | 0.481964055 | 0.68352167 | 14.63849292 | 18.87532868 | ENSG00000232648 |
| TMEM132C | 0.370213467 | 1.292544067 | 0.388334437 | 0.600408663 | 22.56684957 | 29.11815103 | ENSG00000181234 |
| EGFL7 | 0.370197001 | 1.292529314 | 2.62E-06 | 2.80E-05 | 931.643631 | 1204.529284 | ENSG00000172889 |
| NA | 0.37009554 | 1.292438417 | 0.739315853 | 0.863587169 | 3.326605432 | 4.283431778 | ENSG00000227141 |
| ZNF337 | 0.370045275 | 1.292393388 | 0.390251283 | 0.601716195 | 20.78789015 | 26.89656787 | ENSG00000130684 |
| LOC283335 | 0.369758117 | 1.292136172 | 0.232067208 | 0.432848318 | 48.6951513 | 62.79043969 | ENSG00000257337 |
| NA | 0.36954605 | 1.291946251 | 0.035256633 | 0.107853095 | 147.6553465 | 190.4689937 | ENSG00000267809 |
| RGS8 | 0.369529291 | 1.291931243 | 0.580386449 | 0.760941272 | 9.465324247 | 12.28319988 | ENSG00000135824 |
| NA | 0.369508457 | 1.291912586 | 0.738908435 | 0.863507784 | 3.578970187 | 4.64879648 | ENSG00000265791 |
| SLC16A5 | 0.369471475 | 1.291879469 | 0.008694606 | 0.034972874 | 276.2025059 | 357.1377142 | ENSG00000170190 |
| NA | 0.369366061 | 1.291785078 | 0.499474687 | 0.69783084 | 13.6169071 | 17.56038366 | ENSG00000260267 |
| CHRNA10 | 0.369308292 | 1.291733354 | 0.656411199 | 0.814315542 | 5.676702975 | 7.313210386 | ENSG00000129749 |
| NA | 0.369082546 | 1.291531245 | 0.613046526 | 0.78462019 | 7.972239376 | 10.28264311 | ENSG00000229976 |
| A2ML1 | 0.368942591 | 1.291405961 | 0.030940796 | 0.097299657 | 158.2641683 | 204.2581076 | ENSG00000166535 |
| OPLAH | 0.368930348 | 1.291395001 | 7.61E-06 | 7.49E-05 | 844.9675835 | 1091.255054 | ENSG00000178814 |
| GPR161 | 0.368917014 | 1.291383066 | 5.77E-05 | 0.000457468 | 585.722932 | 756.3478323 | ENSG00000143147 |
| SLC4A11 | 0.368914765 | 1.291381053 | 0.000595125 | 0.003570185 | 586.0494614 | 756.9162861 | ENSG00000088836 |
| FGF19 | 0.368814306 | 1.291291133 | 0.001195045 | 0.006547265 | 353.9429986 | 456.8838568 | ENSG00000162344 |
| TMEM192 | 0.368768887 | 1.291250481 | 0.007657818 | 0.031562178 | 249.4146123 | 321.6570244 | ENSG00000170088 |
| BTG1 | 0.368478204 | 1.290990338 | 0.018058604 | 0.063298316 | 231.8410287 | 299.7738826 | ENSG00000133639 |
| NDUFA13 | 0.367816164 | 1.29039805 | 0.299122385 | 0.509389038 | 32.96271405 | 42.43709913 | ENSG00000186010 |
| TECTA | 0.367750327 | 1.290339165 | 0.313466198 | 0.523784211 | 29.88015922 | 38.51685659 | ENSG00000109927 |
| NA | 0.367738965 | 1.290329002 | 0.654058325 | 0.81257098 | 5.66958403 | 7.292685255 | ENSG00000273759 |

| | | | | | | | |
|---|---|---|---|---|---|---|---|
| PLLP | 0.367599148 | 1.290203958 | 0.132384403 | 0.292753595 | 88.10175619 | 113.8182598 | ENSG00000102934 |
| EXO5 | 0.36750293 | 1.290117913 | 0.176380756 | 0.357398131 | 58.97241029 | 75.89127193 | ENSG00000164002 |
| HM13-AS1 | 0.367371138 | 1.290000065 | 0.687459297 | 0.832702766 | 4.655117152 | 6.003562254 | ENSG00000230613 |
| TIMM17A | 0.367252323 | 1.28989383 | 1.33E-07 | 1.80E-06 | 1503.261094 | 1938.737133 | ENSG00000134375 |
| CYTH2 | 0.367213505 | 1.289859123 | 0.028515489 | 0.091165568 | 1518.99806 | 1959.229658 | ENSG00000105443 |
| BRMS1L | 0.366931575 | 1.289607085 | 0.050802743 | 0.142896904 | 148.5746752 | 191.8764221 | ENSG00000100916 |
| NOTCH3 | 0.366925007 | 1.289601214 | 8.92E-10 | 1.72E-08 | 3599.499554 | 4642.603142 | ENSG00000074181 |
| HDAC10 | 0.366870911 | 1.289552859 | 0.634650757 | 0.799639765 | 6.976738915 | 8.993520105 | ENSG00000100429 |
| RAB29 | 0.3668233 | 1.289510303 | 0.000814926 | 0.004689178 | 365.3732384 | 471.0133438 | ENSG00000117280 |
| TAC1 | 0.366821578 | 1.289508764 | 0.288049915 | 0.497332477 | 37.59499983 | 48.55208421 | ENSG00000006128 |
| PLEKHA5 | 0.366812436 | 1.289500592 | 1.86E-07 | 2.44E-06 | 1244.183432 | 1603.850111 | ENSG00000052126 |
| TXNRD2 | 0.366690764 | 1.289391845 | 0.054313644 | 0.150265626 | 130.5609139 | 168.3778784 | ENSG00000184470 |
| PLPP7 | 0.366631629 | 1.289338994 | 0.159012398 | 0.332405262 | 75.67929607 | 97.67199892 | ENSG00000160539 |
| KCNV2 | 0.366577393 | 1.289290525 | 0.695904141 | 0.837581566 | 4.631396053 | 5.971213847 | ENSG00000168263 |
| TDRG1 | 0.366532609 | 1.289250503 | 0.504433776 | 0.70195504 | 13.98551308 | 18.02033458 | ENSG00000204091 |
| GPBAR1 | 0.36649305 | 1.289215152 | 0.743484818 | 0.866496593 | 3.38828552 | 4.342548161 | ENSG00000179921 |
| PSD3 | 0.366352093 | 1.289089197 | 0.00385162 | 0.017793017 | 357.3076148 | 460.6305637 | ENSG00000156011 |
| NA | 0.366276251 | 1.289021432 | 0.452206261 | 0.658283762 | 16.47409841 | 21.27766595 | ENSG00000237356 |
| ZNF397 | 0.366065315 | 1.288832978 | 0.000495881 | 0.003056912 | 724.3263077 | 933.021031 | ENSG00000186812 |
| ST20 | 0.365933348 | 1.288715091 | 0.580242416 | 0.76091091 | 9.279394261 | 11.95425097 | ENSG00000180953 |
| TIGD4 | 0.36575916 | 1.288559504 | 0.420455337 | 0.630683006 | 25.01152603 | 32.29633534 | ENSG00000169989 |
| UBQLN4 | 0.365726531 | 1.288530361 | 7.38E-09 | 1.24E-07 | 1794.147274 | 2312.560309 | ENSG00000160803 |
| PDGFB | 0.365633687 | 1.288447441 | 0.052995351 | 0.147460911 | 126.6168972 | 163.0878098 | ENSG00000100311 |
| PPIC | 0.365583781 | 1.288402872 | 0.134815337 | 0.296458738 | 71.42037096 | 92.07875106 | ENSG00000168938 |
| ART5 | 0.365536101 | 1.288360291 | 0.063098535 | 0.168616929 | 121.2598834 | 156.387874 | ENSG00000167311 |
| ADCY8 | 0.365175149 | 1.288037993 | 0.305781958 | 0.516058292 | 34.22598621 | 43.98829877 | ENSG00000155897 |
| NA | 0.365149788 | 1.288015351 | 0.277661034 | 0.486828394 | 35.02576799 | 45.13575336 | ENSG00000246067 |
| NA | 0.365122029 | 1.287990568 | 0.174462684 | 0.354785108 | 58.73962234 | 75.73900975 | ENSG00000244627 |
| CEP85L | 0.365045419 | 1.287922175 | 0.144368824 | 0.31038311 | 83.85732215 | 107.7508085 | ENSG00000111860 |
| ATP5F1E | 0.365031444 | 1.287909699 | 0.000338664 | 0.002179107 | 880.2868769 | 1134.000188 | ENSG00000124172 |
| NA | 0.365001297 | 1.287882788 | 0.618956191 | 0.788624184 | 8.070683243 | 10.36010916 | ENSG00000227388 |
| ADAR | 0.364746597 | 1.287655439 | 1.03E-10 | 2.31E-09 | 7868.492836 | 10130.92088 | ENSG00000160710 |
| ALCAM | 0.364473822 | 1.287412 | 0.001452015 | 0.007743943 | 520.6050673 | 670.4961289 | ENSG00000170017 |
| EPHA2 | 0.364465396 | 1.287404482 | 6.47E-08 | 9.33E-07 | 1227.402102 | 1580.26438 | ENSG00000142627 |
| TDRD7 | 0.364435182 | 1.28737752 | 0.175139775 | 0.355608499 | 57.8900341 | 74.48268666 | ENSG00000196116 |
| OR2A20P | 0.364400462 | 1.287346538 | 0.605728368 | 0.779795526 | 8.19612835 | 10.57054176 | ENSG00000170356 |
| ALG9 | 0.364307745 | 1.287263808 | 0.2957921 | 0.505935495 | 33.47929103 | 43.04373179 | ENSG00000086848 |
| ZNF608 | 0.36426 | 1.287221207 | 7.17E-06 | 7.09E-05 | 892.3496567 | 1148.173857 | ENSG00000168916 |
| NA | 0.364036678 | 1.287021968 | 0.745584896 | 0.867742215 | 3.857420304 | 5.012931678 | ENSG00000285589 |
| ERCC1 | 0.363938839 | 1.286934688 | 7.10E-05 | 0.00054745 | 558.7230486 | 719.0108217 | ENSG00000012061 |
| NA | 0.363926342 | 1.286923541 | 0.016534973 | 0.059016366 | 194.8393618 | 250.8913364 | ENSG00000196810 |
| NA | 0.363848121 | 1.286853768 | 0.435478748 | 0.644005602 | 20.42759832 | 26.3562404 | ENSG00000259605 |
| CNKSR1 | 0.363748352 | 1.286764779 | 0.005557373 | 0.024121432 | 265.9439084 | 342.2074088 | ENSG00000142675 |
| LOC400622 | 0.363691414 | 1.286713996 | 0.724078867 | 0.854559399 | 3.595572341 | 4.646621017 | ENSG00000267665 |
| ATP2B4 | 0.36365012 | 1.286677167 | 0.000313949 | 0.002045867 | 521.3207667 | 670.3825962 | ENSG00000058668 |
| TMEM147-AS1 | 0.363636227 | 1.286664776 | 0.015732672 | 0.056635818 | 271.0272267 | 348.6786053 | ENSG00000236144 |
| MORN3 | 0.36332406 | 1.286386401 | 0.581873141 | 0.761938307 | 10.8863561 | 14.04002969 | ENSG00000139714 |
| JAKMIP2-AS1 | 0.363143083 | 1.286225042 | 0.005382879 | 0.023504202 | 256.4631511 | 329.6754257 | ENSG00000280780 |
| LOC112268124 | 0.363128097 | 1.286211681 | 0.611945639 | 0.784174879 | 8.005443683 | 10.26211797 | ENSG00000258738 |
| UBE2T | 0.36303227 | 1.286126251 | 8.01E-05 | 0.000609985 | 715.7530316 | 920.2706687 | ENSG00000077152 |
| PLAU | 0.362419839 | 1.2855804 | 0.014976068 | 0.054441394 | 207.0536454 | 266.0692456 | ENSG00000122861 |
| ZNF587B | 0.362383086 | 1.28554765 | 0.004530236 | 0.020350502 | 320.8033047 | 412.9199397 | ENSG00000269343 |
| NA | 0.362359141 | 1.285526313 | 0.744600454 | 0.867208114 | 3.864539249 | 5.015107141 | ENSG00000228492 |
| MOCS3 | 0.3623491 | 1.285517367 | 0.004019351 | 0.018459497 | 318.23936 | 409.3998692 | ENSG00000124217 |
| FAM131C | 0.362346608 | 1.285515146 | 0.255586586 | 0.462495201 | 40.82583114 | 52.42749266 | ENSG00000185519 |
| ARHGEF4 | 0.362203803 | 1.285387906 | 3.50E-05 | 0.000292913 | 620.3335159 | 797.1550792 | ENSG00000136002 |
| DSEL | 0.362022437 | 1.285226326 | 0.021024495 | 0.071695124 | 181.173233 | 232.7166369 | ENSG00000171451 |
| RGS17 | 0.361720248 | 1.284957149 | 0.081921608 | 0.205511367 | 92.8778431 | 119.3551339 | ENSG00000091844 |
| C4orf19 | 0.361705869 | 1.284944341 | 0.547001756 | 0.734791389 | 13.39273879 | 17.27683594 | ENSG00000154274 |
| NA | 0.361635843 | 1.284881974 | 0.694233727 | 0.836829255 | 4.671719306 | 5.989563514 | ENSG00000237493 |
| CCDC191 | 0.36146399 | 1.284728928 | 0.587436848 | 0.765666704 | 9.551920642 | 12.29379366 | ENSG00000163617 |
| FKBP11 | 0.361308601 | 1.284590561 | 0.000617055 | 0.003680693 | 450.5015621 | 578.8032809 | ENSG00000134285 |
| PYY2 | 0.361246824 | 1.284535555 | 0.634575813 | 0.799623956 | 7.025376322 | 9.003167918 | ENSG00000237575 |
| NA | 0.361111235 | 1.284414836 | 0.046026128 | 0.13215804 | 150.9384002 | 194.0102719 | ENSG00000263072 |
| RGS3 | 0.361082346 | 1.284389117 | 0.002440197 | 0.012057331 | 428.279214 | 549.7906458 | ENSG00000138835 |
| ARHGEF3 | 0.360772564 | 1.284113357 | 0.06354184 | 0.169513723 | 105.3806442 | 135.3923154 | ENSG00000163947 |
| PEG10 | 0.360641004 | 1.283996263 | 9.74E-07 | 1.13E-05 | 2740.634346 | 3519.237053 | ENSG00000242265 |
| GABARAPL1 | 0.36048337 | 1.283855977 | 1.10E-06 | 1.26E-05 | 1195.35208 | 1533.861951 | ENSG00000139112 |
| NA | 0.360470819 | 1.283844808 | 0.746899478 | 0.86838024 | 3.361004949 | 4.285607242 | ENSG00000223901 |
| NA | 0.360143646 | 1.283553692 | 0.632004069 | 0.797206785 | 7.805275808 | 9.93562807 | ENSG00000253106 |
| EMC4 | 0.360135977 | 1.283546869 | 4.89E-06 | 4.98E-05 | 1012.869021 | 1299.921026 | ENSG00000128463 |
| C11orf54 | 0.359942188 | 1.283374468 | 0.20717848 | 0.399708096 | 54.8762783 | 70.47610826 | ENSG00000182919 |
| NA | 0.359916274 | 1.283351417 | 0.079506822 | 0.200921495 | 100.6866784 | 129.203859 | ENSG00000232098 |
| CYBA | 0.359882263 | 1.283321162 | 4.60E-09 | 8.01E-08 | 3033.509203 | 3892.396783 | ENSG00000051523 |

| | | | | | | | |
|---|---|---|---|---|---|---|---|
| TBX6 | 0.359876883 | 1.283316377 | 0.333805655 | 0.545793922 | 39.17704536 | 50.39725739 | ENSG00000149922 |
| ZNF292 | 0.359667797 | 1.283130402 | 0.006102859 | 0.026110668 | 368.4779875 | 472.573275 | ENSG00000188994 |
| ZKSCAN1 | 0.359299547 | 1.282802924 | 8.95E-06 | 8.68E-05 | 772.1292082 | 990.1733755 | ENSG00000106261 |
| FAM49A | 0.35923587 | 1.282746305 | 0.035935168 | 0.109404426 | 149.1271007 | 191.0392106 | ENSG00000197872 |
| NA | 0.359099667 | 1.282625208 | 0.034623131 | 0.106318346 | 342.9205201 | 439.1588503 | ENSG00000274090 |
| PPP1R14C | 0.359043284 | 1.282575082 | 0.617896115 | 0.787824768 | 9.388516546 | 12.01119189 | ENSG00000198729 |
| MRM2 | 0.358841179 | 1.282395421 | 0.583355889 | 0.762833684 | 10.84959232 | 13.94080901 | ENSG00000122687 |
| NA | 0.358769865 | 1.282332032 | 0.729756654 | 0.857642318 | 3.602691286 | 4.632622277 | ENSG00000198658 |
| CLK2 | 0.358703466 | 1.282273015 | 1.34E-07 | 1.82E-06 | 1248.469079 | 1601.066541 | ENSG00000176444 |
| EPHA5 | 0.358651725 | 1.282227028 | 0.730636016 | 0.858105398 | 3.629971857 | 4.663741179 | ENSG00000145242 |
| CLCN4 | 0.358611144 | 1.282190962 | 0.009521496 | 0.037523674 | 278.6242737 | 356.8168774 | ENSG00000073464 |
| NA | 0.358353374 | 1.28196189 | 0.736457091 | 0.861442686 | 3.657252428 | 4.693914123 | ENSG00000177173 |
| ASB8 | 0.358338749 | 1.281948894 | 0.102791721 | 0.243017232 | 83.84516919 | 107.3921811 | ENSG00000177981 |
| ISG20L2 | 0.358295448 | 1.281910418 | 5.03E-08 | 7.38E-07 | 1427.383286 | 1829.969249 | ENSG00000143319 |
| VSTM2L | 0.358031975 | 1.281676331 | 0.193436036 | 0.3813971 | 62.27142966 | 79.72752203 | ENSG00000132821 |
| ANKRD49 | 0.35792738 | 1.281583412 | 0.137054905 | 0.299790131 | 71.83967312 | 92.16537047 | ENSG00000168876 |
| GRIN2A | 0.357717239 | 1.281396752 | 0.164219765 | 0.339644099 | 75.13986214 | 96.07645965 | ENSG00000183454 |
| INKA2 | 0.35751487 | 1.281217021 | 0.01152216 | 0.043985445 | 305.1499627 | 390.388337 | ENSG00000197852 |
| NA | 0.357415276 | 1.281128577 | 0.740051078 | 0.863848501 | 3.660811901 | 4.693914123 | ENSG00000240036 |
| RPP38 | 0.357327995 | 1.281051074 | 0.065596661 | 0.173646332 | 118.8279442 | 152.3408648 | ENSG00000152464 |
| COL6A1 | 0.356983926 | 1.280745591 | 4.20E-08 | 6.22E-07 | 6700.992897 | 8582.563882 | ENSG00000142156 |
| ACTG1 | 0.356941551 | 1.280707974 | 2.17E-17 | 1.02E-15 | 55180.1601 | 70670.0242 | ENSG00000184009 |
| SLC9A3R2 | 0.356731993 | 1.280521959 | 0.004277231 | 0.019440557 | 352.8690144 | 452.2078932 | ENSG00000065054 |
| OTX2 | 0.356559535 | 1.280368896 | 0.000108545 | 0.00080011 | 724.3655647 | 927.7598605 | ENSG00000165588 |
| ZNF701 | 0.356505557 | 1.280320992 | 0.133639657 | 0.294756339 | 75.14642242 | 96.18265823 | ENSG00000167562 |
| JTB | 0.356157733 | 1.280012353 | 0.487896516 | 0.688634208 | 15.32350801 | 19.56623733 | ENSG00000143543 |
| SEMA3D | 0.356099486 | 1.279960675 | 0.112229651 | 0.260119902 | 91.89213134 | 117.4393046 | ENSG00000153993 |
| MFAP3 | 0.35604408 | 1.27991152 | 0.007173936 | 0.029900057 | 240.0753436 | 307.1587677 | ENSG00000037749 |
| ETFRF1 | 0.355514496 | 1.279441777 | 0.208750911 | 0.401738843 | 49.05752806 | 62.72943139 | ENSG00000205707 |
| PNMA6A | 0.35534102 | 1.27928794 | 0.096012286 | 0.231120626 | 102.7352151 | 131.5724074 | ENSG00000235961 |
| TMEM183A | 0.355279376 | 1.279233279 | 3.10E-09 | 5.50E-08 | 2402.357728 | 3072.646673 | ENSG00000163444 |
| CARF | 0.355225701 | 1.279185687 | 0.383420482 | 0.595734937 | 25.73569531 | 32.98383902 | ENSG00000138380 |
| DENND5B | 0.355087999 | 1.279063597 | 0.014386945 | 0.052685177 | 231.6220958 | 296.0729425 | ENSG00000170456 |
| TPM3P9 | 0.35508564 | 1.279061506 | 0.018073244 | 0.063314513 | 203.515862 | 260.2312818 | ENSG00000241015 |
| BCL11B | 0.3550781 | 1.279054821 | 0.146087462 | 0.313151159 | 74.00384068 | 94.62878867 | ENSG00000127152 |
| LOC105374338 | 0.355076258 | 1.279053187 | 0.701296357 | 0.84099452 | 4.709678295 | 5.997035864 | ENSG00000283183 |
| MUC1 | 0.355015544 | 1.278999362 | 0.424151087 | 0.634226465 | 22.30589133 | 28.62757567 | ENSG00000185499 |
| MFSD14C | 0.354501839 | 1.278544025 | 0.000296113 | 0.001942374 | 529.2870823 | 676.3843821 | ENSG00000196312 |
| IL27RA | 0.354475309 | 1.278520514 | 0.00072124 | 0.004218215 | 454.763488 | 581.8105269 | ENSG00000104998 |
| HMBOX1 | 0.354431577 | 1.278481759 | 0.057218065 | 0.156580663 | 124.1330665 | 158.5184439 | ENSG00000147421 |
| NA | 0.354285552 | 1.278352362 | 0.618177697 | 0.788004442 | 8.52913961 | 10.92625865 | ENSG00000250731 |
| FAM151A | 0.35408312 | 1.278173002 | 0.633693091 | 0.798671518 | 7.314504856 | 9.350182953 | ENSG00000162391 |
| KCNQ5 | 0.353364295 | 1.277536309 | 0.357721233 | 0.569801407 | 26.52271374 | 33.86465514 | ENSG00000185760 |
| HBP1 | 0.353338533 | 1.277513497 | 0.002085353 | 0.010552487 | 373.7466777 | 477.0011834 | ENSG00000105856 |
| NA | 0.353262293 | 1.277445987 | 2.79E-05 | 0.000240041 | 1197.192972 | 1529.244066 | ENSG00000196205 |
| RGL3 | 0.353134266 | 1.27733263 | 0.234495899 | 0.435785775 | 607.7462551 | 776.4068376 | ENSG00000205517 |
| CAMK1 | 0.352895028 | 1.277120831 | 0.384641929 | 0.59697262 | 25.47621161 | 32.54129182 | ENSG00000134072 |
| ANKRD13C | 0.352767362 | 1.277007822 | 0.00178647 | 0.009222932 | 390.0387105 | 498.0476283 | ENSG00000118454 |
| DDX50 | 0.352754903 | 1.276996794 | 2.86E-06 | 3.03E-05 | 900.5179714 | 1149.942918 | ENSG00000107625 |
| DUS2 | 0.352753453 | 1.27699551 | 0.00493696 | 0.021839891 | 286.3596331 | 365.5947406 | ENSG00000167264 |
| ATP6V0E2-AS1 | 0.352544883 | 1.276810909 | 0.43732232 | 0.645322083 | 19.54597483 | 24.96818026 | ENSG00000204934 |
| TMEM45B | 0.352515334 | 1.276784758 | 0.780715986 | 0.888276674 | 3.695211417 | 4.714439254 | ENSG00000151715 |
| PGBD4 | 0.352398164 | 1.276681066 | 0.449064207 | 0.655579587 | 20.63635968 | 26.32606746 | ENSG00000182405 |
| ZBTB5 | 0.352373783 | 1.276659491 | 0.003060045 | 0.014652372 | 336.5873028 | 429.5374803 | ENSG00000168795 |
| C17orf107 | 0.352125907 | 1.276440162 | 0.451584674 | 0.657835279 | 20.55568703 | 26.22277939 | ENSG00000205710 |
| APLF | 0.351889472 | 1.27623099 | 0.155817367 | 0.327558713 | 64.47327688 | 82.30531653 | ENSG00000169621 |
| MRPL41 | 0.351857031 | 1.276202293 | 4.66E-05 | 0.000377549 | 1574.686698 | 2009.877342 | ENSG00000182154 |
| NA | 0.351825881 | 1.276174738 | 0.700635671 | 0.840545199 | 5.171694134 | 6.652474682 | ENSG00000276259 |
| CLUAP1 | 0.351814537 | 1.276164703 | 0.105754461 | 0.248775909 | 94.13338592 | 119.9811778 | ENSG00000103351 |
| ARV1 | 0.351749776 | 1.276107419 | 0.050569061 | 0.142366834 | 138.0705526 | 176.3065912 | ENSG00000173409 |
| TEX15 | 0.351605806 | 1.275980079 | 0.012988961 | 0.048467996 | 282.0001787 | 359.8265394 | ENSG00000133863 |
| CRISPLD2 | 0.351557632 | 1.275937473 | 0.077786974 | 0.197846061 | 105.7278934 | 134.92773 | ENSG00000103196 |
| SLCO4A1-AS1 | 0.351349547 | 1.275753454 | 0.705883078 | 0.844296784 | 6.785774916 | 8.722741615 | ENSG00000232803 |
| SRGAP3 | 0.351309327 | 1.275717888 | 0.050369986 | 0.141921383 | 133.0314225 | 169.8110132 | ENSG00000196220 |
| BCKDHB | 0.351306253 | 1.27571517 | 0.004109693 | 0.018801246 | 433.3133362 | 552.7539942 | ENSG00000083123 |
| NA | 0.351276662 | 1.275689004 | 0.767361658 | 0.879694476 | 3.371683366 | 4.267257575 | ENSG00000103832 |
| PTPRE | 0.351120271 | 1.275550724 | 0.407877986 | 0.618945422 | 25.95482961 | 33.17062507 | ENSG00000132334 |
| NA | 0.351102134 | 1.275534688 | 0.688268653 | 0.833414711 | 4.940686215 | 6.31510746 | ENSG00000261351 |
| ZNF169 | 0.351091819 | 1.275525569 | 0.169949618 | 0.348064043 | 65.57403466 | 83.79231381 | ENSG00000175787 |
| NA | 0.351069262 | 1.275505626 | 0.672274938 | 0.82439555 | 6.278681143 | 8.027765586 | ENSG00000270083 |
| EIF4ENIF1 | 0.351019878 | 1.275461965 | 0.00355681 | 0.016622211 | 313.0448613 | 399.1099217 | ENSG00000184708 |
| NA | 0.350709634 | 1.275187714 | 0.338823897 | 0.550644635 | 39.59281421 | 50.68298058 | ENSG00000259865 |
| C4orf47 | 0.35061632 | 1.275105237 | 0.534043678 | 0.725099924 | 13.31710015 | 16.95441341 | ENSG00000205129 |
| ND5 | 0.350571683 | 1.275065786 | 0.128016281 | 0.285564312 | 17261.42967 | 22009.6713 | ENSG00000198786 |

| | | | | | | | |
|---|---|---|---|---|---|---|---|
| OPRD1 | 0.350244987 | 1.274777081 | 0.181256171 | 0.364306384 | 66.47965859 | 84.79788908 | ENSG00000116329 |
| PRAG1 | 0.350105821 | 1.274654119 | 2.06E-06 | 2.25E-05 | 1103.579572 | 1407.263455 | ENSG00000275342 |
| F11R | 0.350005635 | 1.274565605 | 3.98E-08 | 5.93E-07 | 1672.861943 | 2132.162474 | ENSG00000158769 |
| PIM3 | 0.349873821 | 1.274449159 | 2.24E-05 | 0.000196892 | 1141.164504 | 1454.814324 | ENSG00000198355 |
| TRMT5 | 0.349767207 | 1.274354981 | 0.00054424 | 0.003301628 | 537.207151 | 684.113685 | ENSG00000126814 |
| OLFML2A | 0.349638375 | 1.274241187 | 0.01002408 | 0.039128719 | 247.006141 | 314.5400799 | ENSG00000185585 |
| NA | 0.349583866 | 1.274193043 | 0.093311452 | 0.226586904 | 97.43599093 | 124.0785585 | ENSG00000197258 |
| ITPKC | 0.34952744 | 1.274143209 | 0.001965129 | 0.010003092 | 358.6963064 | 456.871918 | ENSG00000086544 |
| RGPD2 | 0.349382617 | 1.274015312 | 0.369394085 | 0.581613339 | 33.55904782 | 42.62409609 | ENSG00000185304 |
| GOLGA8A | 0.349310577 | 1.273951696 | 0.378106741 | 0.590532551 | 324.8048322 | 413.8223987 | ENSG00000175265 |
| MCM9 | 0.349310334 | 1.273951482 | 0.040138622 | 0.119035471 | 149.5885317 | 190.7564409 | ENSG00000111877 |
| NR1H3 | 0.34912212 | 1.273785293 | 0.006281635 | 0.026760362 | 305.7585005 | 389.6084655 | ENSG00000025434 |
| MAT1A | 0.348916449 | 1.273603715 | 0.135859126 | 0.298129165 | 93.7449238 | 119.1322156 | ENSG00000151224 |
| TRIM47 | 0.348733667 | 1.273442366 | 0.171149583 | 0.349952347 | 78.79889401 | 100.2648108 | ENSG00000132481 |
| CORO6 | 0.348718798 | 1.273429242 | 0.307182593 | 0.517497169 | 56.61727873 | 72.16911458 | ENSG00000167549 |
| DYRK2 | 0.3486031 | 1.273327122 | 0.000851157 | 0.004869937 | 741.3928214 | 943.7997737 | ENSG00000127334 |
| CD248 | 0.348341794 | 1.273096513 | 0.121901589 | 0.275701977 | 111.0710264 | 141.6272142 | ENSG00000174807 |
| PROK2 | 0.348257883 | 1.273022469 | 0.398844455 | 0.610475104 | 25.42076075 | 32.27268879 | ENSG00000163421 |
| AIFM2 | 0.347887182 | 1.272695407 | 0.013072537 | 0.048702874 | 265.8543368 | 338.6118596 | ENSG00000042286 |
| NA | 0.347802681 | 1.272620866 | 0.665155118 | 0.820741875 | 6.251400572 | 7.972716585 | ENSG00000279799 |
| NA | 0.347698853 | 1.272529281 | 0.508143699 | 0.704897972 | 14.92169772 | 18.99696641 | ENSG00000276449 |
| REPS2 | 0.347553127 | 1.272400749 | 0.094243339 | 0.228323984 | 103.1165327 | 130.9623698 | ENSG00000169891 |
| RBM4B | 0.347509895 | 1.272362621 | 0.00086712 | 0.004939839 | 423.4586857 | 538.6136166 | ENSG00000173914 |
| RAB30 | 0.347503734 | 1.272357187 | 0.120246031 | 0.272933542 | 84.10673782 | 107.100215 | ENSG00000137502 |
| SUSD3 | 0.347318484 | 1.272193821 | 0.13473345 | 0.296347683 | 99.30833405 | 126.379427 | ENSG00000157303 |
| NA | 0.346699318 | 1.271647946 | 0.556210238 | 0.742544206 | 15.49403105 | 19.75881472 | ENSG00000273066 |
| NDUFAF3 | 0.346665324 | 1.271617983 | 0.053181753 | 0.147783608 | 139.7901962 | 177.9179688 | ENSG00000178057 |
| NA | 0.346499245 | 1.271471607 | 0.488043411 | 0.688687126 | 18.37758706 | 23.32021895 | ENSG00000205041 |
| CCT3 | 0.346486542 | 1.271460411 | 2.64E-17 | 1.23E-15 | 13132.27823 | 16697.24071 | ENSG00000163468 |
| PALM | 0.346427259 | 1.271408165 | 0.001321572 | 0.007136942 | 463.6137914 | 589.7541844 | ENSG00000099864 |
| NA | 0.346396216 | 1.271380808 | 0.717877604 | 0.851146028 | 4.744077811 | 5.991738978 | ENSG00000273001 |
| TRMT1L | 0.346249576 | 1.271251588 | 0.000938147 | 0.00530764 | 605.4720493 | 769.7275348 | ENSG00000121486 |
| ADAP1 | 0.34621575 | 1.271221782 | 0.278996373 | 0.488082123 | 43.17325894 | 54.9036801 | ENSG00000105963 |
| RBM34 | 0.34619105 | 1.271200018 | 0.744268048 | 0.867034858 | 3.94282149 | 5.037807736 | ENSG00000188739 |
| BEX4 | 0.346173248 | 1.271184332 | 3.03E-05 | 0.000258392 | 983.2608042 | 1249.807307 | ENSG00000102409 |
| C19orf81 | 0.346132723 | 1.271148626 | 0.114349222 | 0.263770009 | 92.79956086 | 118.1041506 | ENSG00000235034 |
| CCDC106 | 0.346118184 | 1.271135815 | 0.001166464 | 0.006412588 | 447.9233789 | 569.5211568 | ENSG00000173581 |
| ORC3 | 0.346071722 | 1.271094879 | 8.69E-06 | 8.45E-05 | 864.8866919 | 1099.341303 | ENSG00000135336 |
| NA | 0.345688357 | 1.270757157 | 0.341259498 | 0.552859708 | 30.78550381 | 39.09766724 | ENSG00000280142 |
| MTRES1 | 0.345606476 | 1.270685037 | 0.029553774 | 0.093786426 | 179.6812911 | 228.249089 | ENSG00000130349 |
| CYP2R1 | 0.345549777 | 1.270635099 | 0.034194889 | 0.105345919 | 184.6137154 | 234.5175657 | ENSG00000186104 |
| IQCN | 0.345453647 | 1.270550437 | 0.740177795 | 0.863921982 | 3.629971857 | 4.604624797 | ENSG00000130518 |
| DLL4 | 0.345368354 | 1.270475323 | 0.609677893 | 0.782504214 | 8.59437917 | 10.9110304 | ENSG00000128917 |
| ALOX12 | 0.345249855 | 1.270370974 | 0.179237622 | 0.361750828 | 61.01855659 | 77.57687853 | ENSG00000108839 |
| TNFRSF10A | 0.344931673 | 1.270090829 | 0.022875786 | 0.076664764 | 200.0187335 | 253.903784 | ENSG00000104689 |
| NA | 0.344861218 | 1.270028804 | 0.735185975 | 0.860969065 | 3.895379293 | 4.979637312 | ENSG00000272384 |
| NID2 | 0.344761155 | 1.26994072 | 0.013214359 | 0.049124639 | 214.5860893 | 272.4316461 | ENSG00000087303 |
| SIDT2 | 0.344560052 | 1.269763711 | 0.003578488 | 0.016705342 | 401.5780987 | 509.950825 | ENSG00000149577 |
| LNPEP | 0.344530035 | 1.269737292 | 0.274810529 | 0.483588343 | 58.38227959 | 73.90298993 | ENSG00000113441 |
| FBXO10 | 0.344261847 | 1.269501277 | 0.045120017 | 0.130090308 | 147.7221123 | 187.6420074 | ENSG00000147912 |
| PABPC1L2A | 0.344162151 | 1.269413553 | 0.725550102 | 0.855466921 | 5.013044719 | 6.358049639 | ENSG00000186288 |
| CPQ | 0.344102497 | 1.269361066 | 0.490330593 | 0.690212675 | 16.50493846 | 20.9897673 | ENSG00000104324 |
| ND4L | 0.344003027 | 1.269273549 | 0.098611726 | 0.235761797 | 3532.255012 | 4483.700419 | ENSG00000212907 |
| RASSF5 | 0.343667565 | 1.268978446 | 0.265908632 | 0.473407364 | 44.39381743 | 56.45632015 | ENSG00000266094 |
| NA | 0.343588434 | 1.268908845 | 0.676974898 | 0.827165564 | 6.234798418 | 7.921072551 | ENSG00000226445 |
| YBEY | 0.343524493 | 1.268852608 | 0.00306654 | 0.014672312 | 331.3791498 | 420.5547219 | ENSG00000182362 |
| ALOX15 | 0.343220102 | 1.268584923 | 0.543980099 | 0.73244858 | 16.36169599 | 20.7416894 | ENSG00000161905 |
| MFF-DT | 0.343093162 | 1.268473308 | 0.612202858 | 0.784199347 | 9.854091858 | 12.49987533 | ENSG00000236432 |
| FLNC | 0.343079943 | 1.268461686 | 2.73E-05 | 0.000235269 | 755.7690123 | 958.8621332 | ENSG00000128591 |
| HKDC1 | 0.342876474 | 1.268282802 | 0.684182658 | 0.831079275 | 6.013273708 | 7.592407184 | ENSG00000156510 |
| MYO1C | 0.342856083 | 1.268264877 | 2.97E-09 | 5.28E-08 | 2053.834417 | 2604.890127 | ENSG00000197879 |
| NA | 0.342828024 | 1.268240211 | 0.703345574 | 0.842499163 | 4.975085731 | 6.304230142 | ENSG00000287252 |
| PLCB1 | 0.342652111 | 1.268085578 | 0.076485028 | 0.195350252 | 137.4077323 | 174.0136895 | ENSG00000182621 |
| NA | 0.34245 | 1.267907941 | 0.384161089 | 0.596568265 | 25.38844616 | 32.24469131 | ENSG00000280120 |
| SNHG7 | 0.342419845 | 1.26788144 | 0.000654347 | 0.003871348 | 770.3674607 | 977.2886851 | ENSG00000233016 |
| TMEM229A | 0.342181578 | 1.267672062 | 0.632974455 | 0.79797881 | 9.372804116 | 11.96512829 | ENSG00000234224 |
| RMDN2 | 0.341962954 | 1.267479975 | 0.361571202 | 0.573611572 | 32.14249131 | 40.69086309 | ENSG00000115841 |
| NA | 0.341957811 | 1.267475457 | 0.751909475 | 0.870846753 | 3.691651944 | 4.662795221 | ENSG00000226823 |
| SLC37A2 | 0.341878736 | 1.267405987 | 0.235853827 | 0.437749338 | 47.94817679 | 60.80321924 | ENSG00000134955 |
| LINC00551 | 0.341844108 | 1.267375567 | 0.582303044 | 0.762184154 | 11.8441769 | 14.90344211 | ENSG00000272274 |
| GPSM3 | 0.341721258 | 1.26726765 | 0.125131237 | 0.281020741 | 83.60467806 | 105.8552637 | ENSG00000213654 |
| KLF4 | 0.341604847 | 1.267165399 | 0.238792416 | 0.441511139 | 54.88815193 | 69.52735831 | ENSG00000136826 |
| MYCBP2 | 0.341370211 | 1.266959327 | 0.002439589 | 0.012057331 | 697.6335766 | 883.1876914 | ENSG00000005810 |
| FMN2 | 0.341051863 | 1.266679788 | 0.153992008 | 0.325040918 | 80.73978298 | 102.343496 | ENSG00000155816 |

| | | | | | | | |
|---|---|---|---|---|---|---|---|
| RNF215 | 0.340975901 | 1.266613096 | 0.017866385 | 0.062785019 | 242.5526663 | 307.085412 | ENSG00000099999 |
| SMG1 | 0.340911188 | 1.266556282 | 0.0003939 | 0.00249624 | 1759.967452 | 2228.003094 | ENSG00000157106 |
| TXNDC5 | 0.340867541 | 1.266517965 | 0.042547413 | 0.124478374 | 266.9827335 | 337.8775869 | ENSG00000239264 |
| NT5C | 0.340739042 | 1.266405163 | 0.019102209 | 0.066155367 | 219.0799134 | 277.4305791 | ENSG00000125458 |
| PAQR9 | 0.340729618 | 1.26639689 | 0.390439914 | 0.601865789 | 29.17645704 | 36.79019972 | ENSG00000188582 |
| IDO1 | 0.340665309 | 1.266340441 | 0.003146001 | 0.014987949 | 400.0965814 | 507.1538961 | ENSG00000131203 |
| SEMG1 | 0.340400351 | 1.266107893 | 0.396741881 | 0.608754018 | 24.11805506 | 30.56003066 | ENSG00000124233 |
| MORC2 | 0.340314529 | 1.266032578 | 5.49E-08 | 7.97E-07 | 2260.804825 | 2862.048121 | ENSG00000133422 |
| TNFRSF10D | 0.340236044 | 1.265963705 | 0.003053283 | 0.01462741 | 334.0263846 | 423.0345979 | ENSG00000173530 |
| URB2 | 0.340168523 | 1.265904457 | 6.41E-06 | 6.37E-05 | 1177.481664 | 1490.428954 | ENSG00000135763 |
| FAM160A1 | 0.340072832 | 1.265820495 | 0.000263634 | 0.001758025 | 624.9839046 | 790.8610437 | ENSG00000164142 |
| IL20RA | 0.33989475 | 1.265664256 | 0.286306426 | 0.495500079 | 45.42310701 | 57.30573382 | ENSG00000016402 |
| PIGV | 0.339643122 | 1.265443524 | 0.035243844 | 0.107838626 | 166.989637 | 211.3458251 | ENSG00000060642 |
| CMC1 | 0.339585202 | 1.265392721 | 0.007486976 | 0.030959391 | 299.601374 | 378.7690669 | ENSG00000187118 |
| ACHE | 0.339460458 | 1.265283312 | 0.173754879 | 0.353826293 | 80.51947963 | 101.9764503 | ENSG00000087085 |
| C12orf60 | 0.339420551 | 1.265248313 | 0.387937358 | 0.600150584 | 25.1503193 | 31.80403602 | ENSG00000182993 |
| NA | 0.339300696 | 1.265143204 | 0.509342344 | 0.706028345 | 14.94897829 | 18.87532868 | ENSG00000282386 |
| ANKEF1 | 0.339198607 | 1.265053682 | 0.026452667 | 0.086086268 | 186.3220958 | 235.7799934 | ENSG00000132623 |
| ZNF624 | 0.339136103 | 1.264998876 | 0.367870094 | 0.579853662 | 30.82554773 | 39.08395204 | ENSG00000197566 |
| RP1L1 | 0.33910845 | 1.264974629 | 0.638797042 | 0.802053197 | 7.57635282 | 9.588613036 | ENSG00000183638 |
| INF2 | 0.33907235 | 1.264942976 | 6.87E-08 | 9.85E-07 | 1812.225691 | 2292.935924 | ENSG00000203485 |
| NA | 0.338845633 | 1.264744209 | 0.746986338 | 0.86838024 | 3.677414055 | 4.630446813 | ENSG00000244009 |
| CCDC50 | 0.338794476 | 1.264699362 | 0.002141369 | 0.010778187 | 403.2108404 | 509.8662952 | ENSG00000152492 |
| FAM114A1 | 0.338726951 | 1.26464017 | 0.035733486 | 0.10893097 | 181.1737917 | 228.7556276 | ENSG00000197712 |
| HTATIP2 | 0.338473846 | 1.264418322 | 1.38E-05 | 0.000127349 | 846.6692027 | 1070.646331 | ENSG00000109854 |
| PLEKHD1 | 0.338459802 | 1.264406013 | 0.380991104 | 0.593545089 | 27.86546335 | 35.21412403 | ENSG00000175985 |
| NA | 0.338377094 | 1.264333529 | 0.577485044 | 0.759450456 | 13.65962077 | 17.23361021 | ENSG00000232022 |
| SLC48A1 | 0.338343022 | 1.264303669 | 0.016738202 | 0.05956302 | 251.2143438 | 317.529398 | ENSG00000211584 |
| FAS | 0.338258445 | 1.264229553 | 0.427519145 | 0.637380521 | 23.35119825 | 29.59304664 | ENSG00000026103 |
| LFNG | 0.33825017 | 1.264222302 | 0.00070309 | 0.004124839 | 534.9771852 | 676.3917819 | ENSG00000106003 |
| PALM3 | 0.338240327 | 1.264213677 | 4.08E-07 | 5.09E-06 | 1723.820184 | 2179.828551 | ENSG00000187867 |
| PRR14L | 0.338106564 | 1.264096467 | 0.000324974 | 0.002106086 | 904.4509269 | 1142.563485 | ENSG00000183530 |
| NEK3 | 0.338041624 | 1.264039567 | 0.003429337 | 0.016142332 | 537.7110431 | 679.4300514 | ENSG00000136098 |
| CBARP | 0.338001013 | 1.264003986 | 0.002656817 | 0.012961505 | 491.6827289 | 621.9264433 | ENSG00000099625 |
| DHFR2 | 0.337946877 | 1.263956556 | 0.223109316 | 0.420989977 | 56.92153488 | 72.09172116 | ENSG00000178700 |
| NOD1 | 0.337875288 | 1.263893838 | 0.134678303 | 0.296295404 | 86.11106076 | 108.7877135 | ENSG00000106100 |
| NA | 0.3376624 | 1.263707349 | 0.292857599 | 0.502691852 | 43.18866588 | 54.59743178 | ENSG00000272335 |
| TMEM141 | 0.337582429 | 1.263637301 | 0.201123523 | 0.391972933 | 60.52778564 | 76.61765041 | ENSG00000244187 |
| FBXL18 | 0.337455618 | 1.263526233 | 0.001415811 | 0.007585132 | 437.7309631 | 553.1018169 | ENSG00000155034 |
| CBWD3 | 0.337190181 | 1.263293782 | 0.56792772 | 0.751177685 | 13.43809606 | 16.89406753 | ENSG00000196873 |
| NA | 0.337149684 | 1.263258322 | 0.733011242 | 0.859729886 | 3.949940435 | 5.007634792 | ENSG00000228887 |
| IQCA1 | 0.337140957 | 1.26325068 | 0.061225881 | 0.16496658 | 139.4717021 | 176.1997302 | ENSG00000132321 |
| TGDS | 0.336981054 | 1.263110674 | 0.016478844 | 0.058850788 | 228.8473828 | 289.0133904 | ENSG00000088451 |
| CXCR2 | 0.336779408 | 1.262934141 | 0.550693825 | 0.73799694 | 12.8512455 | 16.24950603 | ENSG00000180871 |
| NA | 0.336713689 | 1.262876612 | 0.495613379 | 0.695207823 | 18.41882619 | 23.3276913 | ENSG00000233871 |
| NAPRT | 0.336674268 | 1.262842105 | 0.002698632 | 0.013131581 | 500.3152418 | 632.1997448 | ENSG00000147813 |
| SLC9A5 | 0.336356386 | 1.262563882 | 0.026797376 | 0.087013144 | 259.1359131 | 327.0643071 | ENSG00000135740 |
| STXBP5L | 0.336265739 | 1.262484556 | 0.05950068 | 0.161307829 | 132.7956599 | 167.7950602 | ENSG00000145087 |
| LRRC69 | 0.336022803 | 1.262271983 | 0.64782917 | 0.808876289 | 8.628778686 | 10.86061587 | ENSG00000214954 |
| PLCH1 | 0.336004431 | 1.262255909 | 0.000333678 | 0.002152892 | 578.1824271 | 729.6739194 | ENSG00000114805 |
| ZNF41 | 0.33590301 | 1.262167176 | 0.107858444 | 0.252572711 | 99.05035104 | 124.8603649 | ENSG00000147124 |
| TM7SF3 | 0.335769891 | 1.26205072 | 6.07E-05 | 0.000477452 | 1048.058375 | 1322.99701 | ENSG00000064115 |
| SCOC-AS1 | 0.33574479 | 1.262028762 | 0.7138284 | 0.84906134 | 5.033206346 | 6.330052158 | ENSG00000196951 |
| SKIDA1 | 0.33537807 | 1.261708006 | 0.204315269 | 0.396149266 | 58.80961658 | 74.19913893 | ENSG00000180592 |
| IRAK4 | 0.335246702 | 1.261593124 | 0.325404754 | 0.536564112 | 35.21881692 | 44.52448623 | ENSG00000198001 |
| NA | 0.335200118 | 1.261552388 | 0.545233335 | 0.733646706 | 14.04007423 | 17.67861643 | ENSG00000268575 |
| MAPK13 | 0.33512805 | 1.261489371 | 0.009250857 | 0.036671377 | 303.5385772 | 382.4963758 | ENSG00000156711 |
| ABCA7 | 0.335058573 | 1.261428621 | 0.003210913 | 0.015262547 | 520.8961506 | 657.4402596 | ENSG00000064687 |
| IGSF9 | 0.335022231 | 1.261396846 | 2.00E-05 | 0.000177941 | 1067.225956 | 1346.541485 | ENSG00000085552 |
| AEN | 0.334831333 | 1.261229948 | 5.87E-10 | 1.17E-08 | 3055.655738 | 3853.308718 | ENSG00000181026 |
| PDE2A | 0.334824618 | 1.261224079 | 0.00582502 | 0.025092494 | 316.5262512 | 399.3692988 | ENSG00000186642 |
| INHBC | 0.334736551 | 1.261147091 | 0.682046516 | 0.830032766 | 6.371201275 | 8.013766846 | ENSG00000175189 |
| LITAF | 0.334713822 | 1.261127222 | 2.75E-05 | 0.000236764 | 1973.108746 | 2489.41289 | ENSG00000189067 |
| TTC12 | 0.334442896 | 1.260890415 | 0.118848154 | 0.270904935 | 99.02398635 | 124.8404068 | ENSG00000149292 |
| NA | 0.334373156 | 1.260829466 | 0.750118992 | 0.869986929 | 3.922659864 | 4.960341686 | ENSG00000255062 |
| NA | 0.334321732 | 1.260784525 | 0.23271763 | 0.433590415 | 63.73901399 | 80.22996362 | ENSG00000137970 |
| RUFY4 | 0.334195978 | 1.260674632 | 0.673530964 | 0.825146432 | 6.616447084 | 8.358606418 | ENSG00000188282 |
| ATP5F1D | 0.333942997 | 1.260453589 | 0.000221639 | 0.001506717 | 1549.590868 | 1954.21219 | ENSG00000099624 |
| FRMD8 | 0.333783518 | 1.260314262 | 0.00053488 | 0.003255698 | 564.2716885 | 711.4762584 | ENSG00000126391 |
| NA | 0.333676805 | 1.260221043 | 0.673436576 | 0.825137853 | 6.847455004 | 8.684150363 | ENSG00000225335 |
| SPX | 0.333674331 | 1.260218882 | 0.729331627 | 0.85751636 | 5.804512346 | 7.378853159 | ENSG00000134548 |
| MIEF1 | 0.333598463 | 1.260152612 | 1.17E-05 | 0.000109392 | 1216.327504 | 1532.487164 | ENSG00000100335 |
| SH3BP2 | 0.33351561 | 1.260080244 | 0.000238114 | 0.001607144 | 834.6470794 | 1051.339942 | ENSG00000087266 |
| ADCY9 | 0.333512443 | 1.260077478 | 0.002718278 | 0.013206763 | 435.9081471 | 549.0624051 | ENSG00000162104 |

| | | | | | | | |
|---|---|---|---|---|---|---|---|
| PFKFB3 | 0.333204116 | 1.259808208 | 2.18E-05 | 0.000191786 | 1073.634629 | 1352.086544 | ENSG00000170525 |
| VASN | 0.333184586 | 1.259791154 | 0.014362652 | 0.05261662 | 338.0771849 | 426.3747875 | ENSG00000168140 |
| PRR15 | 0.333169247 | 1.25977776 | 0.058608736 | 0.159485011 | 161.9635235 | 204.0942854 | ENSG00000176532 |
| MRPS18B | 0.333115484 | 1.259730814 | 2.81E-05 | 0.000241266 | 954.9283683 | 1203.517246 | ENSG00000204568 |
| LINC02323 | 0.332836714 | 1.259487422 | 0.650572684 | 0.810373005 | 7.954442013 | 10.03361925 | ENSG00000259230 |
| NA | 0.332695223 | 1.259363904 | 4.08E-05 | 0.000335429 | 999.6560873 | 1258.921231 | ENSG00000250899 |
| NA | 0.332491664 | 1.259186226 | 0.022926401 | 0.076780474 | 329.3519961 | 414.8095384 | ENSG00000229191 |
| NA | 0.332447318 | 1.259147521 | 0.624596222 | 0.7923564 | 8.979587309 | 11.28944789 | ENSG00000269427 |
| KIF21B | 0.332446853 | 1.259147115 | 3.89E-07 | 4.88E-06 | 1982.961846 | 2497.106891 | ENSG00000116852 |
| BCS1L | 0.332030497 | 1.258783782 | 0.000957516 | 0.005405436 | 535.7923739 | 674.6073537 | ENSG00000074582 |
| RHOU | 0.33193472 | 1.258700218 | 0.043586488 | 0.12685018 | 197.2594013 | 248.4150537 | ENSG00000116574 |
| SSR2 | 0.331920643 | 1.258687937 | 2.52E-05 | 0.000219435 | 1212.645388 | 1526.130588 | ENSG00000163479 |
| GALT | 0.331880502 | 1.258652916 | 0.163945959 | 0.339274231 | 82.30227786 | 103.4893422 | ENSG00000213930 |
| ITGA9-AS1 | 0.331745923 | 1.25853551 | 0.474494182 | 0.67731814 | 21.10785873 | 26.60460184 | ENSG00000235257 |
| NA | 0.331736032 | 1.258526882 | 0.750319149 | 0.870058771 | 4.222466816 | 5.354649827 | ENSG00000230847 |
| NA | 0.331711106 | 1.258505138 | 0.700129686 | 0.840395926 | 6.663889282 | 8.391900784 | ENSG00000272669 |
| SMG7 | 0.331675363 | 1.258473959 | 3.12E-07 | 3.98E-06 | 1732.829801 | 2180.491195 | ENSG00000116698 |
| ZNF197 | 0.331550644 | 1.25836517 | 0.007455866 | 0.030851016 | 318.0270654 | 400.5156691 | ENSG00000186448 |
| SEC14L6 | 0.331511126 | 1.258330701 | 0.376745315 | 0.588878934 | 30.63842254 | 38.64473717 | ENSG00000214491 |
| JAK2 | 0.331404265 | 1.258237499 | 0.318230944 | 0.528240431 | 41.66357191 | 52.4337355 | ENSG00000096968 |
| CRYZL1 | 0.331360432 | 1.258199271 | 0.028360102 | 0.09086877 | 220.0071735 | 276.6406606 | ENSG00000205758 |
| TJP3 | 0.331257134 | 1.258109187 | 0.013837413 | 0.050999131 | 351.1649793 | 441.6070629 | ENSG00000105289 |
| LOC100379224 | 0.33108451 | 1.257958658 | 0.672926276 | 0.82494078 | 7.861921882 | 9.906684632 | ENSG00000186019 |
| ITPA | 0.33091124 | 1.257807585 | 6.31E-05 | 0.00049369 | 846.0541039 | 1063.894372 | ENSG00000125877 |
| STXBP3 | 0.330802427 | 1.25771272 | 0.000164131 | 0.00115528 | 630.5719518 | 793.1494038 | ENSG00000116266 |
| PRIMA1 | 0.33076494 | 1.25768004 | 0.086528208 | 0.21399754 | 137.8793093 | 173.5990242 | ENSG00000175785 |
| ELOVL4 | 0.330631201 | 1.257563457 | 0.020727281 | 0.070838688 | 299.4753958 | 376.7084048 | ENSG00000118402 |
| ACRBP | 0.330231941 | 1.25721548 | 0.577974356 | 0.759450456 | 11.83558342 | 14.9022126 | ENSG00000111644 |
| LINC00641 | 0.329959507 | 1.256978094 | 0.085419343 | 0.212189959 | 135.4517936 | 170.2117524 | ENSG00000258441 |
| ATG4D | 0.329870706 | 1.256900726 | 0.003222903 | 0.015296439 | 599.5394221 | 753.758827 | ENSG00000130734 |
| MIR100HG | 0.329847197 | 1.256880245 | 0.711673441 | 0.847786093 | 6.071394323 | 7.57623298 | ENSG00000255248 |
| FEZ1 | 0.329541615 | 1.256614049 | 1.08E-05 | 0.000101865 | 965.5048506 | 1213.044409 | ENSG00000149557 |
| ZNF81 | 0.329526236 | 1.256600654 | 0.109846341 | 0.256147669 | 113.5900946 | 142.7017419 | ENSG00000197779 |
| SPRY4-AS1 | 0.329442945 | 1.256528109 | 0.522466355 | 0.71692967 | 15.38162863 | 19.26433993 | ENSG00000231185 |
| NA | 0.329429706 | 1.256516578 | 0.66204109 | 0.818450247 | 7.940204123 | 9.984150681 | ENSG00000272275 |
| KLHDC9 | 0.329181929 | 1.256300796 | 0.136000023 | 0.298230426 | 96.89299694 | 121.7297572 | ENSG00000162755 |
| VSTM4 | 0.329054343 | 1.256189698 | 0.286054592 | 0.495345072 | 47.66586172 | 59.97014772 | ENSG00000165633 |
| MCOLN2 | 0.328986021 | 1.25613021 | 0.344293723 | 0.556339122 | 34.59548192 | 43.53168019 | ENSG00000153898 |
| NA | 0.328936344 | 1.256086958 | 0.345723425 | 0.557608639 | 36.25108116 | 45.63457907 | ENSG00000214820 |
| USP20 | 0.328790408 | 1.255959904 | 0.002018962 | 0.010246734 | 444.1490223 | 557.4381746 | ENSG00000136878 |
| ZSCAN22 | 0.328400701 | 1.255620685 | 0.244319055 | 0.448705182 | 58.99793698 | 74.21363213 | ENSG00000182318 |
| SLC22A7 | 0.328363042 | 1.25558791 | 0.718654578 | 0.851423561 | 5.587742316 | 7.048958286 | ENSG00000137204 |
| FBLN7 | 0.328257611 | 1.255496156 | 0.737036686 | 0.861960479 | 3.973661534 | 4.991460588 | ENSG00000144152 |
| ADPRM | 0.328148214 | 1.255400957 | 0.160161093 | 0.334091839 | 84.53966749 | 106.0284299 | ENSG00000170222 |
| SREBF2-AS1 | 0.327985626 | 1.255259485 | 0.282705925 | 0.491655782 | 53.65185485 | 67.46467932 | ENSG00000184068 |
| C11orf65 | 0.327898296 | 1.255183503 | 0.742520141 | 0.865919752 | 4.23906897 | 5.359000754 | ENSG00000166323 |
| DAPK2 | 0.327830113 | 1.255124183 | 0.279235473 | 0.488364689 | 48.45407535 | 60.93980167 | ENSG00000035664 |
| MPZL1 | 0.327695568 | 1.255007136 | 8.75E-07 | 1.02E-05 | 1850.389386 | 2322.804443 | ENSG00000197965 |
| KRTCAP3 | 0.327517369 | 1.25485213 | 0.001012035 | 0.005666257 | 469.2178048 | 588.8082926 | ENSG00000157992 |
| NA | 0.327360487 | 1.254715682 | 0.755363256 | 0.872799503 | 4.246187915 | 5.359946713 | ENSG00000234134 |
| CDH8 | 0.327294859 | 1.254658606 | 0.201520893 | 0.392544823 | 67.1196219 | 84.21072 | ENSG00000150394 |
| JUNB | 0.327241476 | 1.254612182 | 0.117416479 | 0.268744421 | 94.28967108 | 118.1727039 | ENSG00000171223 |
| HROB | 0.327041028 | 1.254437878 | 0.00532738 | 0.023310375 | 338.2028837 | 424.4367724 | ENSG00000125319 |
| ROMO1 | 0.327039088 | 1.254436192 | 0.00139654 | 0.007501043 | 1002.629059 | 1257.49425 | ENSG00000125995 |
| SYNGR1 | 0.326784333 | 1.2542147 | 0.002277988 | 0.011353867 | 602.7999239 | 756.7259795 | ENSG00000100321 |
| AGA | 0.326717891 | 1.254156939 | 0.084271125 | 0.210010582 | 117.9854487 | 148.0334076 | ENSG00000038002 |
| GSDMD | 0.326623561 | 1.254074939 | 0.000176209 | 0.001229754 | 704.092673 | 883.0513332 | ENSG00000104518 |
| FSTL4 | 0.326443719 | 1.253918619 | 0.16446444 | 0.339926775 | 96.89330243 | 121.3915394 | ENSG00000053108 |
| RPS28 | 0.326441212 | 1.253916441 | 2.02E-05 | 0.000179245 | 6705.372282 | 8408.498842 | ENSG00000233927 |
| AIG1 | 0.326317101 | 1.253808574 | 0.001721185 | 0.008937194 | 444.2314738 | 557.1855349 | ENSG00000146416 |
| UFC1 | 0.326156834 | 1.253669298 | 0.001953184 | 0.009947648 | 512.6504957 | 642.4079253 | ENSG00000143222 |
| LBHD1 | 0.326103086 | 1.253622594 | 0.664740829 | 0.820498729 | 7.62735449 | 9.530442612 | ENSG00000162194 |
| SLC36A4 | 0.325907402 | 1.253452566 | 0.006309443 | 0.026866705 | 350.2315161 | 439.1085514 | ENSG00000180773 |
| GLI1 | 0.325809268 | 1.253367308 | 0.010422174 | 0.040382182 | 285.9827392 | 358.403168 | ENSG00000111087 |
| UGP2 | 0.325778585 | 1.253340651 | 7.79E-13 | 2.27E-11 | 7946.271695 | 9958.93635 | ENSG00000169764 |
| BATF3 | 0.325690518 | 1.253264146 | 0.27254376 | 0.481206251 | 48.15070893 | 60.41231606 | ENSG00000123685 |
| ITGB1 | 0.325612827 | 1.253196657 | 6.42E-10 | 1.27E-08 | 5920.374461 | 7419.040181 | ENSG00000150093 |
| KLHL24 | 0.325588393 | 1.253175433 | 0.00936859 | 0.037016314 | 332.1839401 | 415.7725371 | ENSG00000114796 |
| PCIF1 | 0.325416692 | 1.253026296 | 0.001033007 | 0.005766578 | 633.1263098 | 793.0063606 | ENSG00000100982 |
| RHOV | 0.325165334 | 1.252808003 | 0.219521919 | 0.415897673 | 65.72706583 | 82.4157934 | ENSG00000104140 |
| GPR45 | 0.325138553 | 1.252784748 | 0.714838415 | 0.849668508 | 5.529621701 | 6.962790382 | ENSG00000135973 |
| ZNF398 | 0.325105053 | 1.252755658 | 3.71E-06 | 3.85E-05 | 1529.271583 | 1915.55673 | ENSG00000197024 |
| NA | 0.325058483 | 1.25271522 | 0.347311688 | 0.559353678 | 36.70597806 | 46.11550662 | ENSG00000275764 |
| NA | 0.3249349 | 1.252607914 | 0.605964928 | 0.779911925 | 10.37214338 | 12.94488155 | ENSG00000226525 |

| | | | | | | | |
|---|---|---|---|---|---|---|---|
| CLIP4 | 0.324892026 | 1.25257069 | 0.622816528 | 0.79114079 | 9.572082268 | 12.01894779 | ENSG00000115295 |
| FAM20B | 0.324848174 | 1.252532618 | 9.36E-05 | 0.000700482 | 936.5780891 | 1172.67725 | ENSG00000116199 |
| GVQW2 | 0.324847674 | 1.252532184 | 0.678622542 | 0.828107179 | 6.660329809 | 8.334959865 | ENSG00000279968 |
| RBAK | 0.324797316 | 1.252488465 | 0.025763438 | 0.084308183 | 258.5102133 | 323.8062274 | ENSG00000146587 |
| SMURF1 | 0.324788446 | 1.252480764 | 0.000126587 | 0.000916988 | 694.7130816 | 869.9707198 | ENSG00000198742 |
| PHC3 | 0.324778947 | 1.252472518 | 0.020900456 | 0.071328419 | 238.5902674 | 298.7590157 | ENSG00000173889 |
| TMEM106A | 0.324671221 | 1.252378999 | 0.59255975 | 0.769952585 | 14.56377015 | 18.23360505 | ENSG00000184988 |
| ADAM12 | 0.324588899 | 1.252307539 | 0.69634099 | 0.837900181 | 7.123540857 | 8.979521365 | ENSG00000148848 |
| GPR150 | 0.32446806 | 1.25220265 | 0.71591787 | 0.850107455 | 5.333013298 | 6.671770308 | ENSG00000178015 |
| ATP6 | 0.324389802 | 1.252134728 | 0.075856975 | 0.194324331 | 24536.33328 | 30723.04307 | ENSG00000198899 |
| NA | 0.324236462 | 1.252001648 | 0.745868112 | 0.867900448 | 4.000942105 | 5.003283865 | ENSG00000273091 |
| NOP2 | 0.32418366 | 1.251955826 | 0.700265546 | 0.84040536 | 8.063564298 | 10.08308782 | ENSG00000111641 |
| SCHIP1 | 0.32416241 | 1.251937386 | 0.614217958 | 0.785289156 | 9.295996415 | 11.62776107 | ENSG00000151967 |
| ZNF236 | 0.324057204 | 1.251846095 | 0.01836587 | 0.064134514 | 228.5579749 | 286.2411696 | ENSG00000130856 |
| SYDE2 | 0.323856488 | 1.251671942 | 0.126821118 | 0.283736147 | 114.3883763 | 142.8331954 | ENSG00000097096 |
| FAH | 0.323559466 | 1.251414275 | 0.004717174 | 0.021050054 | 332.4140838 | 415.93449 | ENSG00000103876 |
| NA | 0.32343765 | 1.251308614 | 0.725322612 | 0.855358906 | 5.309292199 | 6.638475942 | ENSG00000237938 |
| FAM72C | 0.32338985 | 1.251267156 | 0.232580141 | 0.433462544 | 61.06986374 | 76.34933093 | ENSG00000263513 |
| NA | 0.323130775 | 1.251042478 | 0.66058596 | 0.817400463 | 7.712755676 | 9.600436312 | ENSG00000228929 |
| PDS5B | 0.323112975 | 1.251027043 | 0.000209205 | 0.00143163 | 834.4843107 | 1043.988463 | ENSG00000083642 |
| GTF2E1 | 0.323032198 | 1.250956998 | 3.28E-05 | 0.000275993 | 1196.057587 | 1495.55934 | ENSG00000153767 |
| ZCRB1 | 0.322615757 | 1.250595955 | 0.00602145 | 0.02580763 | 431.7372656 | 540.3800943 | ENSG00000139168 |
| RFXANK | 0.322429371 | 1.250434398 | 0.000435228 | 0.002725226 | 596.3231342 | 745.9615652 | ENSG00000064490 |
| SUSD6 | 0.322370371 | 1.250383262 | 0.116435732 | 0.267244627 | 105.8740849 | 132.2073211 | ENSG00000100647 |
| NA | 0.322300195 | 1.250322441 | 0.475123766 | 0.677841249 | 20.4880832 | 25.65473798 | ENSG00000241860 |
| APOB | 0.322216934 | 1.250250285 | 0.451176545 | 0.657452882 | 24.63018284 | 30.69734814 | ENSG00000084674 |
| PNRC1 | 0.322086074 | 1.250136886 | 0.036808609 | 0.111398641 | 189.6537091 | 237.1088645 | ENSG00000146278 |
| USP40 | 0.321884107 | 1.249961888 | 0.069082539 | 0.18089625 | 139.5348568 | 174.1806129 | ENSG00000085982 |
| PSTPIP2 | 0.321730956 | 1.249829204 | 0.245743787 | 0.450220354 | 60.73593661 | 75.73806379 | ENSG00000152229 |
| PTPRN | 0.321720683 | 1.249820304 | 0.087572109 | 0.21597407 | 116.6014861 | 145.6715535 | ENSG00000054356 |
| ASPH | 0.321677477 | 1.249782875 | 1.57E-05 | 0.000143069 | 1457.061472 | 1820.465373 | ENSG00000198363 |
| GUSBP11 | 0.321157283 | 1.249332321 | 0.446795313 | 0.653682173 | 24.78290852 | 31.01373873 | ENSG00000228315 |
| NA | 0.321088143 | 1.249272449 | 0.690510963 | 0.834528492 | 7.5125878 | 9.306011269 | ENSG00000267466 |
| ZFPL1 | 0.320773398 | 1.248999931 | 0.167946212 | 0.345271524 | 94.57729892 | 118.1431934 | ENSG00000162300 |
| MGLL | 0.320737623 | 1.248968961 | 0.586146152 | 0.764723322 | 14.47984351 | 18.02156408 | ENSG00000074416 |
| ZNF644 | 0.32068638 | 1.248924599 | 0.000659513 | 0.003898249 | 946.6875193 | 1182.695806 | ENSG00000122482 |
| CAP2 | 0.320624081 | 1.248870668 | 0.009602656 | 0.037789848 | 306.5055012 | 382.5169736 | ENSG00000112186 |
| MYO6 | 0.320538371 | 1.248796476 | 0.000164457 | 0.00115714 | 805.5894805 | 1005.565742 | ENSG00000196586 |
| METTL26 | 0.320423536 | 1.248697079 | 0.000149123 | 0.001062713 | 1151.043739 | 1438.04443 | ENSG00000130731 |
| NA | 0.320160025 | 1.248469022 | 0.659950763 | 0.816938918 | 8.043402672 | 10.01744505 | ENSG00000256633 |
| TRPC4 | 0.320091514 | 1.248409736 | 0.019291201 | 0.066675389 | 253.6922769 | 316.5336158 | ENSG00000133107 |
| PMS2 | 0.32008728 | 1.248406073 | 0.012052649 | 0.045696503 | 285.2099325 | 355.9450037 | ENSG00000122512 |
| SMIM10L2B | 0.320013149 | 1.248341927 | 0.074715067 | 0.192128819 | 127.4439595 | 159.1121394 | ENSG00000196972 |
| NA | 0.319953596 | 1.248290398 | 0.729574902 | 0.857588823 | 5.597225525 | 7.004786602 | ENSG00000229670 |
| LINC01405 | 0.319889079 | 1.248234576 | 0.071308827 | 0.185487155 | 167.6529381 | 209.0441489 | ENSG00000185847 |
| RSAD1 | 0.319880878 | 1.24822748 | 2.47E-05 | 0.00021563 | 890.9233371 | 1112.133072 | ENSG00000136444 |
| LINC00574 | 0.319637442 | 1.248016875 | 0.679218572 | 0.828368191 | 6.936415663 | 8.67450255 | ENSG00000231690 |
| ITPR1 | 0.31952601 | 1.247920484 | 0.087887856 | 0.216396631 | 176.2096637 | 219.4656749 | ENSG00000150995 |
| SELENOO | 0.319491816 | 1.247890906 | 0.001739338 | 0.009021532 | 503.0283538 | 627.4615208 | ENSG00000073169 |
| KNL1 | 0.319473696 | 1.247875234 | 0.007034246 | 0.029396912 | 365.9087557 | 456.4729753 | ENSG00000137812 |
| SEMA6D | 0.319328094 | 1.247749299 | 0.26017634 | 0.467028585 | 52.8876678 | 66.00104505 | ENSG00000137872 |
| TLCD1 | 0.319232804 | 1.247666889 | 0.03297242 | 0.102278984 | 229.784431 | 286.8971029 | ENSG00000160606 |
| ALMS1 | 0.319076406 | 1.24753164 | 3.03E-05 | 0.000257927 | 1032.75907 | 1287.898208 | ENSG00000116127 |
| MARCHF2 | 0.318954024 | 1.247425818 | 0.021203088 | 0.072147806 | 239.2192468 | 298.5963707 | ENSG00000099785 |
| PTPN4 | 0.318928319 | 1.247403593 | 0.009603044 | 0.037789848 | 481.5785325 | 600.4499439 | ENSG00000088179 |
| SNPH | 0.318855327 | 1.247340483 | 0.040682029 | 0.120325781 | 180.5899158 | 225.2658254 | ENSG00000101298 |
| SCNM1 | 0.318675219 | 1.247184773 | 0.000394711 | 0.002500541 | 660.237959 | 823.0477958 | ENSG00000163156 |
| C6orf47 | 0.31853384 | 1.247062559 | 0.024088342 | 0.079948742 | 227.107272 | 283.4204261 | ENSG00000204439 |
| MTSS2 | 0.318443491 | 1.246984464 | 0.017943523 | 0.062997476 | 349.5575372 | 435.9909465 | ENSG00000132613 |
| NA | 0.317907385 | 1.246521171 | 0.792760975 | 0.89406764 | 3.797214757 | 4.693914123 | ENSG00000286799 |
| TGFBI | 0.317903508 | 1.246517821 | 0.013643845 | 0.050374195 | 262.4132702 | 327.1661548 | ENSG00000120708 |
| SPR | 0.317859691 | 1.246479963 | 0.004362779 | 0.019734391 | 396.9502616 | 495.180801 | ENSG00000116096 |
| ND6 | 0.317803122 | 1.246431088 | 0.182289102 | 0.365720779 | 3890.736614 | 4849.771754 | ENSG00000198695 |
| GZF1 | 0.31777626 | 1.246407881 | 0.002419053 | 0.011971651 | 423.4417781 | 527.7514877 | ENSG00000125812 |
| LOC100291105 | 0.317571924 | 1.246231358 | 0.688538804 | 0.833582621 | 6.687610381 | 8.309137848 | ENSG00000218018 |
| LBR | 0.317555946 | 1.246217556 | 6.75E-06 | 6.70E-05 | 1921.203663 | 2393.664041 | ENSG00000143815 |
| POLI | 0.317534455 | 1.246198992 | 0.007049909 | 0.029448691 | 320.7190731 | 399.7277153 | ENSG00000101751 |
| LENG1 | 0.317532462 | 1.246197271 | 0.080249759 | 0.202203696 | 145.8016643 | 181.4944857 | ENSG00000105617 |
| TRIM2 | 0.317485983 | 1.246157122 | 0.000878696 | 0.004998246 | 756.7184667 | 942.7632677 | ENSG00000109654 |
| ZNF724 | 0.317469286 | 1.2461427 | 0.184637785 | 0.368858755 | 78.68860267 | 98.02951241 | ENSG00000196081 |
| CENPB | 0.317396077 | 1.246079466 | 1.72E-05 | 0.000155244 | 1789.692298 | 2230.247568 | ENSG00000125817 |
| IGSF10 | 0.317361116 | 1.246049271 | 0.444154088 | 0.651584708 | 23.6652431 | 29.56098178 | ENSG00000152580 |
| CIRBP | 0.317339639 | 1.246030721 | 6.63E-05 | 0.000516364 | 2747.941248 | 3423.883052 | ENSG00000099622 |
| SPSB3 | 0.317310695 | 1.246005723 | 0.526655069 | 0.719933477 | 17.02863439 | 21.30566343 | ENSG00000162032 |

| | | | | | | | |
|---|---|---|---|---|---|---|---|
| NA | 0.316957423 | 1.245700651 | 0.479218392 | 0.681478865 | 20.98213429 | 26.21530704 | ENSG00000276672 |
| RPL13AP5 | 0.316918569 | 1.245667103 | 0.029038207 | 0.092539073 | 206.8887406 | 257.7341177 | ENSG00000236552 |
| SDC2 | 0.316892434 | 1.245644538 | 5.58E-06 | 5.63E-05 | 1308.373147 | 1629.123448 | ENSG00000169439 |
| FA2H | 0.316857013 | 1.245613955 | 0.465998376 | 0.670198716 | 25.84008907 | 32.17252215 | ENSG00000103089 |
| ARHGDIG | 0.316703855 | 1.245481727 | 0.57831591 | 0.759450456 | 14.81613491 | 18.44744263 | ENSG00000242173 |
| PDZRN3 | 0.316589255 | 1.245382796 | 0.005557445 | 0.024121432 | 369.9767429 | 460.8285763 | ENSG00000121440 |
| FSD1L | 0.316498131 | 1.245304137 | 0.330503301 | 0.54184867 | 41.30917766 | 51.46921049 | ENSG00000106701 |
| LYRM7 | 0.316495513 | 1.245301877 | 0.06546465 | 0.173345472 | 173.3127595 | 215.5105297 | ENSG00000186687 |
| ZBTB7A | 0.31642267 | 1.245239003 | 0.002946843 | 0.014167801 | 460.792779 | 574.2198733 | ENSG00000178951 |
| FLNB | 0.316350281 | 1.245176523 | 1.51E-07 | 2.02E-06 | 4166.166538 | 5186.454932 | ENSG00000136068 |
| NA | 0.316275416 | 1.245111909 | 0.708555773 | 0.846314323 | 5.560461745 | 6.943494756 | ENSG00000254783 |
| NA | 0.316004628 | 1.244878229 | 0.213060451 | 0.407334139 | 78.86652399 | 98.20886907 | ENSG00000180867 |
| MRS2 | 0.315881969 | 1.244772393 | 4.29E-06 | 4.40E-05 | 1375.935207 | 1712.183928 | ENSG00000124532 |
| DUSP13 | 0.315827558 | 1.244725447 | 0.454548309 | 0.66052207 | 26.83111418 | 33.29443827 | ENSG00000079393 |
| SOX10 | 0.315742583 | 1.244652135 | 0.08679241 | 0.214538567 | 124.1894071 | 154.6650736 | ENSG00000100146 |
| TRPM4 | 0.315730303 | 1.244641541 | 0.021041168 | 0.071730761 | 251.9705477 | 313.8971268 | ENSG00000130529 |
| MCM3AP-AS1 | 0.31542858 | 1.244381266 | 0.189414837 | 0.375552859 | 69.06551873 | 85.99242587 | ENSG00000215424 |
| CD164L2 | 0.315314691 | 1.244283036 | 0.323936056 | 0.534702597 | 47.63649621 | 59.3381875 | ENSG00000174950 |
| WIPI1 | 0.315254191 | 1.244230857 | 0.014554375 | 0.053184865 | 345.3646635 | 429.3208572 | ENSG00000070540 |
| RPSAP52 | 0.315065567 | 1.244068192 | 0.753670985 | 0.871590725 | 4.038901094 | 5.003283865 | ENSG00000241749 |
| AFF4 | 0.314979226 | 1.24399374 | 0.00085124 | 0.004869937 | 998.6474919 | 1242.203975 | ENSG00000072364 |
| GASK1A | 0.31495936 | 1.243976611 | 0.13190283 | 0.29206061 | 111.1679702 | 138.2157288 | ENSG00000144649 |
| SCUBE1 | 0.314829394 | 1.243864551 | 0.023867803 | 0.079384195 | 230.8863845 | 286.9333077 | ENSG00000159307 |
| MYRIP | 0.314778263 | 1.243820468 | 0.359354614 | 0.571533011 | 37.11162715 | 46.14227459 | ENSG00000170011 |
| RPS17 | 0.314578647 | 1.24364838 | 0.068636844 | 0.179878823 | 181.8840542 | 226.2064204 | ENSG00000182774 |
| ATP5IF1 | 0.314465943 | 1.243551231 | 3.59E-05 | 0.000299154 | 2119.164471 | 2635.667338 | ENSG00000130770 |
| ABO | 0.314414232 | 1.243506658 | 0.555216085 | 0.741583972 | 15.55451593 | 19.37292489 | ENSG00000175164 |
| FOXO1 | 0.314385647 | 1.243482019 | 0.003489866 | 0.016361997 | 654.890759 | 814.3808249 | ENSG00000150907 |
| RAP1GAP | 0.314264729 | 1.243377803 | 0.011556825 | 0.044080861 | 418.3169152 | 520.3962514 | ENSG00000076864 |
| HLCS | 0.314196163 | 1.243318711 | 0.008704849 | 0.034999166 | 329.1037751 | 409.1303726 | ENSG00000159267 |
| PLPP2 | 0.31402225 | 1.243168841 | 8.99E-06 | 8.71E-05 | 1599.262705 | 1988.022333 | ENSG00000141934 |
| NA | 0.313912209 | 1.243074023 | 0.644403556 | 0.805989478 | 9.310234305 | 11.53412081 | ENSG00000250421 |
| RRAS | 0.313578108 | 1.242786184 | 0.01733006 | 0.061253636 | 274.5949075 | 341.3112489 | ENSG00000126458 |
| NA | 0.313566953 | 1.242776574 | 0.663015388 | 0.819386533 | 8.043402672 | 9.953977738 | ENSG00000279583 |
| HEXIM1 | 0.313354364 | 1.242593458 | 3.91E-06 | 4.04E-05 | 1285.478956 | 1597.573848 | ENSG00000186834 |
| ALG10B | 0.313348192 | 1.242588142 | 0.164158029 | 0.339590796 | 89.45780166 | 111.0926267 | ENSG00000175548 |
| NA | 0.313337128 | 1.242578613 | 0.739948241 | 0.863848501 | 5.287935364 | 6.552308037 | ENSG00000263050 |
| SFI1 | 0.313182025 | 1.242445031 | 0.014767101 | 0.0538372 | 296.3267122 | 368.0766468 | ENSG00000198089 |
| NA | 0.312815714 | 1.242129605 | 0.081781478 | 0.205241623 | 141.0576126 | 174.9936108 | ENSG00000258634 |
| EGLN3 | 0.312781444 | 1.2421001 | 0.003923127 | 0.018070306 | 426.156058 | 529.5787627 | ENSG00000129521 |
| NA | 0.312685298 | 1.242017325 | 0.15428895 | 0.32548557 | 97.28562951 | 120.7847387 | ENSG00000213442 |
| NECAB1 | 0.312432725 | 1.241799904 | 0.131262423 | 0.291029168 | 113.8196541 | 141.3838434 | ENSG00000123119 |
| NA | 0.312322478 | 1.241705013 | 0.615474557 | 0.786256984 | 11.7537417 | 14.67030891 | ENSG00000230658 |
| YDJC | 0.312156058 | 1.241561785 | 0.000523356 | 0.003199889 | 796.2443915 | 989.3163215 | ENSG00000161179 |
| SLC25A15 | 0.312097145 | 1.241511087 | 0.001518714 | 0.008038372 | 659.4843975 | 818.500026 | ENSG00000102743 |
| PCNX2 | 0.312064453 | 1.241482954 | 2.68E-05 | 0.000231485 | 1351.665421 | 1677.974678 | ENSG00000135749 |
| DPP10 | 0.312051532 | 1.241471836 | 0.045893287 | 0.131876861 | 196.3499647 | 244.0082602 | ENSG00000175497 |
| MRRF | 0.311924817 | 1.241362799 | 0.006326368 | 0.026920563 | 396.873149 | 492.6550496 | ENSG00000148187 |
| PRXL2C | 0.311698792 | 1.241168331 | 0.151524616 | 0.32116076 | 92.85175774 | 115.3417455 | ENSG00000158122 |
| NA | 0.311451016 | 1.240955185 | 0.353528636 | 0.565638915 | 38.19461373 | 47.47708233 | ENSG00000248092 |
| SLC25A40 | 0.311255677 | 1.240787173 | 0.076919452 | 0.196190904 | 141.2227962 | 175.3182087 | ENSG00000075303 |
| HOOK2 | 0.311188621 | 1.240729503 | 5.69E-05 | 0.000452226 | 1654.852831 | 2053.57908 | ENSG00000095066 |
| AFF1 | 0.311107376 | 1.240659634 | 0.000160826 | 0.001134978 | 934.3219078 | 1158.618775 | ENSG00000172493 |
| CPM | 0.311039128 | 1.240600944 | 0.563749738 | 0.748513341 | 15.31046533 | 18.97521178 | ENSG00000135678 |
| RAB12 | 0.311030866 | 1.240593839 | 0.009694697 | 0.038063273 | 326.0686886 | 404.7027611 | ENSG00000206418 |
| JAK3 | 0.310974706 | 1.240545548 | 0.189051315 | 0.375142805 | 76.62882887 | 95.068498 | ENSG00000105639 |
| GLIS3 | 0.310852109 | 1.240440134 | 0.778619541 | 0.886993808 | 3.773493658 | 4.626095886 | ENSG00000107249 |
| CDT1 | 0.310843778 | 1.240432971 | 4.59E-06 | 4.69E-05 | 2317.886028 | 2875.932039 | ENSG00000167513 |
| SH2D4A | 0.310816393 | 1.240409425 | 0.053316625 | 0.148027707 | 165.3402408 | 204.9644125 | ENSG00000104611 |
| LINC01829 | 0.310776433 | 1.240375069 | 0.592517943 | 0.769952585 | 14.77461645 | 18.39846851 | ENSG00000236780 |
| ZDHHC17 | 0.310725699 | 1.240331449 | 0.012757457 | 0.047783532 | 409.4168314 | 507.8599673 | ENSG00000186908 |
| TUFT1 | 0.310649692 | 1.240266106 | 0.01449472 | 0.052997637 | 339.45957 | 420.5217111 | ENSG00000143367 |
| SNX27 | 0.310526673 | 1.240160352 | 5.25E-05 | 0.00042023 | 1083.651898 | 1343.579931 | ENSG00000143376 |
| OLIG2 | 0.310454375 | 1.240098206 | 0.112474542 | 0.260399869 | 144.9257369 | 179.9826123 | ENSG00000205927 |
| COQ9 | 0.310376526 | 1.240031291 | 0.000782369 | 0.00452251 | 646.6925201 | 801.9370008 | ENSG00000088682 |
| TBCC | 0.310220878 | 1.239897515 | 0.002726543 | 0.013242451 | 452.5227659 | 561.2564742 | ENSG00000124659 |
| IFT74 | 0.309915719 | 1.239635279 | 0.039383544 | 0.117341126 | 185.5368312 | 229.8697878 | ENSG00000096872 |
| DMXL2 | 0.309692637 | 1.239443611 | 0.068273462 | 0.179200046 | 179.9099086 | 222.8771081 | ENSG00000104093 |
| CDC42SE2 | 0.309582447 | 1.239348948 | 0.001823675 | 0.009368906 | 580.5574148 | 719.2840888 | ENSG00000158985 |
| RPL13P5 | 0.309467614 | 1.239250305 | 0.630589426 | 0.796625039 | 10.20134101 | 12.68403442 | ENSG00000240370 |
| NA | 0.309423405 | 1.239212331 | 0.805726086 | 0.900901829 | 2.979356283 | 3.704513051 | ENSG00000239246 |
| NA | 0.309261693 | 1.239073435 | 0.679383489 | 0.828446904 | 6.98385786 | 8.642154143 | ENSG00000226945 |
| CCL26 | 0.309115511 | 1.238947891 | 0.265175249 | 0.47279405 | 56.38863507 | 69.84932933 | ENSG00000006606 |
| PYCR3 | 0.308834387 | 1.238706493 | 0.011169274 | 0.042802707 | 306.0346126 | 379.0255631 | ENSG00000104524 |

| | | | | | | | |
|---|---|---|---|---|---|---|---|
| TDGF1P3 | 0.308754951 | 1.238638291 | 0.001040023 | 0.005797177 | 523.4977493 | 648.5517382 | ENSG00000225366 |
| FAM83E | 0.308736634 | 1.238622564 | 0.218002199 | 0.414158635 | 77.45708643 | 96.08705342 | ENSG00000105523 |
| PLIN3 | 0.308718453 | 1.238606955 | 6.46E-07 | 7.73E-06 | 1692.236121 | 2096.005223 | ENSG00000105355 |
| CEP164 | 0.308284325 | 1.238234297 | 0.017587707 | 0.061978698 | 343.5964075 | 426.0053031 | ENSG00000110274 |
| MOB3A | 0.308012119 | 1.238000691 | 0.081573112 | 0.204854817 | 132.9498601 | 164.6236858 | ENSG00000172081 |
| FAM110C | 0.307947591 | 1.237945319 | 0.057687129 | 0.157567462 | 161.1459443 | 199.4992131 | ENSG00000184731 |
| YPEL4 | 0.307917658 | 1.237919635 | 0.536622271 | 0.726856257 | 20.44539568 | 25.36589337 | ENSG00000166793 |
| ARID3A | 0.307861559 | 1.237871499 | 5.27E-07 | 6.42E-06 | 2127.923885 | 2634.159034 | ENSG00000116017 |
| C2CD4C | 0.307732976 | 1.237761177 | 0.020469477 | 0.070106289 | 378.5942584 | 469.0310117 | ENSG00000183186 |
| LIPG | 0.307550115 | 1.237604301 | 0.037127813 | 0.112095724 | 234.1943546 | 290.0481403 | ENSG00000101670 |
| TIPRL | 0.307502893 | 1.237563792 | 0.004342562 | 0.019647648 | 538.8289088 | 666.4786302 | ENSG00000143155 |
| CUEDC1 | 0.307427293 | 1.237498943 | 0.031798083 | 0.099570021 | 231.5562976 | 286.7809729 | ENSG00000180891 |
| NA | 0.307297724 | 1.237387807 | 0.640369976 | 0.803013745 | 9.905093528 | 12.25955333 | ENSG00000213326 |
| RLIM | 0.307216698 | 1.237318314 | 0.000269885 | 0.001794008 | 765.8184656 | 947.172164 | ENSG00000131263 |
| SLC29A3 | 0.30718115 | 1.237287827 | 0.024653141 | 0.081550984 | 331.082396 | 409.8696359 | ENSG00000198246 |
| LYSMD2 | 0.307065464 | 1.237188617 | 0.35246177 | 0.564614516 | 43.05256851 | 53.30010138 | ENSG00000140280 |
| NA | 0.306909827 | 1.237055156 | 0.668310471 | 0.822110685 | 9.481036677 | 11.6644604 | ENSG00000226976 |
| PCDH12 | 0.306827215 | 1.236984322 | 0.537226439 | 0.72743395 | 17.95205567 | 22.15290164 | ENSG00000113555 |
| BBS12 | 0.306820085 | 1.236978208 | 0.28257181 | 0.491604664 | 53.64028671 | 66.32384644 | ENSG00000181004 |
| CUEDC2 | 0.306504462 | 1.23670762 | 0.00076892 | 0.004455679 | 915.242625 | 1131.650598 | ENSG00000107874 |
| IL17RB | 0.306391661 | 1.236610929 | 0.119689502 | 0.272255747 | 110.9983369 | 137.2536628 | ENSG00000056736 |
| CCDC171 | 0.306313321 | 1.236543781 | 0.288767593 | 0.497979727 | 50.42636304 | 62.26730501 | ENSG00000164989 |
| RYK | 0.306041162 | 1.236310534 | 6.11E-05 | 0.000480501 | 915.6472353 | 1131.994868 | ENSG00000163785 |
| CIART | 0.306024754 | 1.236296474 | 0.119808856 | 0.272404316 | 121.5487326 | 150.5100895 | ENSG00000159208 |
| REXO1 | 0.306012742 | 1.236286179 | 5.71E-05 | 0.000453313 | 1006.647421 | 1245.029977 | ENSG00000079313 |
| SLCO1A2 | 0.306008759 | 1.236282767 | 0.338262892 | 0.550263392 | 43.65246175 | 53.88119557 | ENSG00000084453 |
| NA | 0.305962807 | 1.23624339 | 0.195036049 | 0.383792413 | 72.78089177 | 89.99789035 | ENSG00000204055 |
| GAS2 | 0.305743943 | 1.23605586 | 0.681580052 | 0.829785926 | 6.987417333 | 8.616332126 | ENSG00000148935 |
| SPATA25 | 0.305570826 | 1.235907548 | 0.611763035 | 0.784141186 | 11.79881964 | 14.63606858 | ENSG00000149634 |
| DDIT3 | 0.305523522 | 1.235867024 | 0.439343576 | 0.647293137 | 25.98686486 | 32.17469761 | ENSG00000175197 |
| NA | 0.305444337 | 1.235799193 | 0.684753954 | 0.831472593 | 7.018257377 | 8.649626492 | ENSG00000230530 |
| ZNF853 | 0.305365559 | 1.235731715 | 0.021004758 | 0.071653693 | 251.9506916 | 311.5109637 | ENSG00000236609 |
| JAZF1 | 0.305192402 | 1.235583407 | 0.118082351 | 0.269945942 | 130.4787667 | 161.1947537 | ENSG00000153814 |
| GPX1 | 0.305014641 | 1.235431174 | 6.22E-06 | 6.20E-05 | 4239.592085 | 5237.545004 | ENSG00000233276 |
| LMBRD1 | 0.304962023 | 1.235386116 | 0.03163891 | 0.099170261 | 280.9930839 | 347.0633579 | ENSG00000168216 |
| MTHFD2 | 0.304818542 | 1.235263259 | 1.02E-05 | 9.72E-05 | 2449.45479 | 3024.95664 | ENSG00000065911 |
| ZNF627 | 0.304641207 | 1.235111432 | 0.016774439 | 0.059680724 | 271.1120431 | 334.7938209 | ENSG00000198551 |
| LINC02249 | 0.304601823 | 1.235077714 | 0.700654621 | 0.840545199 | 6.756409413 | 8.286437254 | ENSG00000225930 |
| NA | 0.304568463 | 1.235049156 | 0.610716718 | 0.783411576 | 13.64390834 | 16.94259014 | ENSG00000226450 |
| DLG2 | 0.304544774 | 1.235028877 | 0.11675075 | 0.267758973 | 118.5880117 | 146.2601201 | ENSG00000150672 |
| NA | 0.304190048 | 1.234725248 | 0.608554819 | 0.782019444 | 11.28460692 | 13.91063607 | ENSG00000246877 |
| MCRIP2 | 0.304185135 | 1.234721043 | 0.000294577 | 0.00193499 | 821.1876771 | 1014.543341 | ENSG00000172366 |
| PGP | 0.30412509 | 1.234669656 | 0.000811522 | 0.004674626 | 1679.640342 | 2074.87562 | ENSG00000184207 |
| ZBTB24 | 0.304108066 | 1.234655086 | 0.004836666 | 0.021461568 | 460.755327 | 568.6850592 | ENSG00000112365 |
| DAB1 | 0.304060077 | 1.234614018 | 0.004038919 | 0.018531337 | 440.30734 | 543.4112703 | ENSG00000173406 |
| ERMP1 | 0.303904023 | 1.234480479 | 0.00035693 | 0.002284957 | 844.2559243 | 1042.11645 | ENSG00000099219 |
| FOSL2 | 0.303801769 | 1.234392986 | 0.107355237 | 0.251716022 | 122.3387002 | 150.8927423 | ENSG00000075426 |
| NA | 0.303316499 | 1.233977851 | 0.719652686 | 0.852291545 | 7.076377992 | 8.706850957 | ENSG00000224418 |
| PYCARD | 0.303299272 | 1.233963116 | 0.140791959 | 0.305089632 | 106.4131878 | 131.2846244 | ENSG00000103490 |
| PPP1R10 | 0.303240705 | 1.233913024 | 2.70E-05 | 0.000233256 | 1066.254124 | 1316.067811 | ENSG00000204569 |
| PINK1 | 0.303092698 | 1.233786442 | 0.052574078 | 0.146612739 | 193.5307324 | 238.9446305 | ENSG00000158828 |
| LHFPL6 | 0.303064999 | 1.233762755 | 0.174622122 | 0.354968674 | 99.57791135 | 123.0745916 | ENSG00000183722 |
| NA | 0.302701264 | 1.233451735 | 0.653582226 | 0.812465428 | 9.984570978 | 12.29909054 | ENSG00000271646 |
| NME2 | 0.302562834 | 1.233333388 | 0.434555738 | 0.643382278 | 33.17503489 | 40.75962729 | ENSG00000243678 |
| DZIP1L | 0.302508661 | 1.233287077 | 0.085219016 | 0.211841663 | 153.5888896 | 189.318345 | ENSG00000158163 |
| DSTYK | 0.302440861 | 1.23322912 | 0.00794021 | 0.032456896 | 482.2588191 | 594.4678754 | ENSG00000133059 |
| LIMCH1 | 0.302279467 | 1.233091166 | 0.490396717 | 0.690222811 | 27.91171034 | 34.43347455 | ENSG00000064042 |
| PLBD1 | 0.302157801 | 1.232987181 | 0.063557206 | 0.169519847 | 178.119154 | 219.5470404 | ENSG00000121316 |
| NA | 0.3021499 | 1.232980429 | 0.626042368 | 0.793253855 | 11.27156424 | 13.87204481 | ENSG00000236364 |
| ZNF468 | 0.301930332 | 1.232792792 | 0.034526079 | 0.106135066 | 204.6418416 | 252.2177259 | ENSG00000204604 |
| COX19 | 0.301851386 | 1.232725334 | 0.017265896 | 0.061072569 | 288.1033027 | 355.4269253 | ENSG00000240230 |
| NA | 0.301851021 | 1.232725022 | 0.757645751 | 0.874053636 | 4.307868003 | 5.307356721 | ENSG00000253392 |
| GFPT1 | 0.301835264 | 1.232711559 | 1.37E-05 | 0.000126138 | 1883.629662 | 2321.410694 | ENSG00000198380 |
| ADIPOR1 | 0.301714311 | 1.232608215 | 8.04E-06 | 7.87E-05 | 1478.085937 | 1821.547756 | ENSG00000159346 |
| ACVR1 | 0.301568728 | 1.232483838 | 0.008077565 | 0.032918605 | 395.8672233 | 487.6246713 | ENSG00000115170 |
| PTPRS | 0.301512273 | 1.23243561 | 2.18E-08 | 3.39E-07 | 3809.974359 | 4696.035939 | ENSG00000105426 |
| UPK3B | 0.301428952 | 1.232364434 | 0.212611624 | 0.406702668 | 79.33298902 | 97.91962532 | ENSG00000243566 |
| RPL32P3 | 0.301418221 | 1.232355268 | 0.265687366 | 0.473175215 | 102.9552135 | 126.9107167 | ENSG00000251474 |
| NA | 0.301403267 | 1.232342494 | 0.568378446 | 0.751668554 | 14.83629653 | 18.32535339 | ENSG00000282572 |
| HAGHL | 0.301307246 | 1.232260476 | 0.025386899 | 0.083411847 | 568.1507973 | 700.5197151 | ENSG00000103253 |
| NA | 0.301285477 | 1.232241883 | 0.796689074 | 0.896204025 | 2.97579681 | 3.674340108 | ENSG00000271959 |
| PRLR | 0.301140912 | 1.232118412 | 0.424172671 | 0.634226465 | 36.72285954 | 45.18690294 | ENSG00000113494 |
| PLPP1 | 0.301137107 | 1.232115162 | 0.002604284 | 0.012728237 | 468.4933562 | 577.4523716 | ENSG00000067113 |
| PPFIA3 | 0.300817724 | 1.231842428 | 0.002070423 | 0.010482558 | 742.4044177 | 914.7519358 | ENSG00000177380 |

| | | | | | | | |
|---|---|---|---|---|---|---|---|
| PLSCR4 | 0.300603741 | 1.231659732 | 0.512887026 | 0.70906535 | 19.0990866 | 23.56177045 | ENSG00000114698 |
| AIDA | 0.300594223 | 1.231651607 | 0.000189348 | 0.001314648 | 1776.595728 | 2188.512487 | ENSG00000186063 |
| DST | 0.300580477 | 1.231639871 | 0.003330822 | 0.015733511 | 1314.630673 | 1618.215515 | ENSG00000151914 |
| DBT | 0.300449039 | 1.231527667 | 0.003212473 | 0.015266117 | 582.0273889 | 716.7660731 | ENSG00000137992 |
| NA | 0.300350296 | 1.23144338 | 0.650225147 | 0.810100403 | 9.704925652 | 11.89295912 | ENSG00000279861 |
| GGNBP2 | 0.300238745 | 1.231348167 | 0.000337668 | 0.002174584 | 778.0573862 | 957.6899705 | ENSG00000278311 |
| APOM | 0.30023762 | 1.231347207 | 0.558145303 | 0.743916196 | 16.7741847 | 20.56311075 | ENSG00000204444 |
| NA | 0.300050518 | 1.231187524 | 0.640316701 | 0.803013745 | 11.053599 | 13.5430959 | ENSG00000234589 |
| C5orf34 | 0.300045272 | 1.231183047 | 0.179916959 | 0.362695534 | 89.02873695 | 109.6341214 | ENSG00000172244 |
| CCNK | 0.299773486 | 1.230951129 | 0.015028953 | 0.05458111 | 532.0976694 | 654.3593275 | ENSG00000090061 |
| ICAM1 | 0.299623462 | 1.230823131 | 0.257013984 | 0.463948031 | 65.99867633 | 81.24301124 | ENSG00000090339 |
| TENT2 | 0.299503749 | 1.230721003 | 0.003333624 | 0.015738061 | 453.2833941 | 557.5648983 | ENSG00000164329 |
| PRICKLE2 | 0.299458486 | 1.230682392 | 0.290890934 | 0.500544446 | 58.18508637 | 71.60594823 | ENSG00000163637 |
| TTC8 | 0.299351279 | 1.230590943 | 0.100556363 | 0.23901912 | 151.3511421 | 186.2107215 | ENSG00000165533 |
| CSNK1G1 | 0.299192766 | 1.230455742 | 0.05277308 | 0.146978624 | 216.2120431 | 266.4053833 | ENSG00000169118 |
| NA | 0.299181156 | 1.230445839 | 0.715607879 | 0.849899679 | 5.962272037 | 7.346504752 | ENSG00000164616 |
| MTCP1 | 0.299167887 | 1.230434522 | 0.754794195 | 0.872411882 | 4.573275438 | 5.654371756 | ENSG00000214827 |
| LINC00664 | 0.298831745 | 1.23014787 | 0.296690711 | 0.506875709 | 52.98494261 | 65.20488378 | ENSG00000268658 |
| ARL14EP | 0.298794427 | 1.23011605 | 0.007802191 | 0.032031386 | 462.3347027 | 568.6645341 | ENSG00000152219 |
| DOK5 | 0.298739042 | 1.230068827 | 0.022310527 | 0.075197985 | 251.024575 | 308.7148127 | ENSG00000101134 |
| NA | 0.298594768 | 1.229945823 | 0.000224766 | 0.001524131 | 873.7271282 | 1074.653434 | ENSG00000235655 |
| UBE2S | 0.298563709 | 1.229919344 | 6.65E-08 | 9.56E-07 | 7052.246475 | 8674.372675 | ENSG00000108106 |
| BCHE | 0.298529732 | 1.229890379 | 0.336290514 | 0.548195628 | 50.88273447 | 62.5208907 | ENSG00000114200 |
| SCAT2 | 0.298364388 | 1.229749431 | 0.80070962 | 0.898320559 | 3.217483147 | 3.993357662 | ENSG00000257596 |
| TRIM68 | 0.298239634 | 1.229643096 | 0.093458872 | 0.226859999 | 167.134023 | 205.2114718 | ENSG00000167333 |
| FOXP3 | 0.298109132 | 1.229531871 | 0.755345769 | 0.872799503 | 4.300749058 | 5.279359241 | ENSG00000049768 |
| SETSIP | 0.298076395 | 1.229503972 | 0.563207794 | 0.74797322 | 16.38750202 | 20.19557058 | ENSG00000230667 |
| PDLIM5 | 0.297907985 | 1.229360456 | 0.00051853 | 0.003177299 | 815.0987659 | 1001.655147 | ENSG00000163110 |
| CCDC14 | 0.297844603 | 1.229306448 | 0.005203981 | 0.022860475 | 731.418196 | 899.3003783 | ENSG00000175455 |
| MLXIPL | 0.297826194 | 1.229290762 | 0.21159065 | 0.40534484 | 111.618749 | 137.4263774 | ENSG00000009950 |
| BICRAL | 0.297765459 | 1.229239012 | 0.043171297 | 0.125816349 | 263.0986163 | 323.6413116 | ENSG00000112624 |
| CLTCL1 | 0.297704382 | 1.229186972 | 0.06791042 | 0.17842074 | 188.3047308 | 231.1565674 | ENSG00000070371 |
| NA | 0.297646954 | 1.229138045 | 0.566842905 | 0.750531324 | 16.80858422 | 20.55658436 | ENSG00000273702 |
| DDX47 | 0.29760838 | 1.229105181 | 0.717027025 | 0.850889492 | 7.208942044 | 8.862234558 | ENSG00000213782 |
| AKAP1 | 0.297597485 | 1.229095899 | 1.65E-07 | 2.19E-06 | 2306.734032 | 2834.873985 | ENSG00000121057 |
| SHISA4 | 0.297473921 | 1.228990634 | 0.16385071 | 0.339215157 | 107.3116928 | 131.7276231 | ENSG00000198892 |
| SCRT1 | 0.297409363 | 1.22893564 | 0.705498809 | 0.844050998 | 7.007578959 | 8.574335906 | ENSG00000261678 |
| MYO1G | 0.297265419 | 1.22881303 | 0.802504773 | 0.899050996 | 3.234085301 | 4.007356402 | ENSG00000136286 |
| ERC1 | 0.29720322 | 1.228760053 | 0.000219128 | 0.001490715 | 1076.714567 | 1322.252209 | ENSG00000082805 |
| NA | 0.297190386 | 1.228749122 | 0.800758713 | 0.898320559 | 2.9900347 | 3.676515571 | ENSG00000241429 |
| SIPA1L2 | 0.297052319 | 1.228631536 | 0.006330316 | 0.026925229 | 409.840915 | 503.3328312 | ENSG00000116991 |
| IRF2BP1 | 0.296939488 | 1.22853545 | 0.00100392 | 0.00563084 | 904.373916 | 1111.809324 | ENSG00000170604 |
| DDAH1 | 0.296885378 | 1.228489373 | 0.000519601 | 0.003180022 | 777.8178104 | 955.4734304 | ENSG00000153904 |
| PLCG2 | 0.296725303 | 1.228353073 | 0.022418385 | 0.075478452 | 277.9938715 | 341.3366491 | ENSG00000197943 |
| SDHB | 0.296612419 | 1.228256964 | 0.001172037 | 0.006434277 | 916.2637568 | 1125.547166 | ENSG00000117118 |
| NA | 0.296455489 | 1.228123367 | 0.757931875 | 0.874279969 | 4.351750728 | 5.32882781 | ENSG00000213862 |
| MRPS35 | 0.296353899 | 1.228036889 | 0.000668064 | 0.003942619 | 826.6449142 | 1015.726622 | ENSG00000061794 |
| NA | 0.296346675 | 1.22803074 | 0.39353778 | 0.605164003 | 45.1292238 | 55.2379259 | ENSG00000279369 |
| NA | 0.296144236 | 1.227858435 | 0.795614844 | 0.895554092 | 2.982915755 | 3.664692295 | ENSG00000237804 |
| BLOC1S3 | 0.296140456 | 1.227855218 | 0.135336813 | 0.297270462 | 121.4386945 | 149.2921897 | ENSG00000189114 |
| SMC6 | 0.296001612 | 1.227737056 | 2.52E-05 | 0.000219435 | 1382.24239 | 1696.760733 | ENSG00000163029 |
| NA | 0.295915015 | 1.227663363 | 0.592250547 | 0.769870087 | 14.56969389 | 17.8685239 | ENSG00000267939 |
| TRIM38 | 0.295809268 | 1.227573382 | 0.331407283 | 0.542962899 | 46.50105958 | 57.16652442 | ENSG00000112343 |
| DEPDC4 | 0.295599331 | 1.227394761 | 0.440657962 | 0.64836974 | 27.88918445 | 34.19672548 | ENSG00000166153 |
| NA | 0.295512555 | 1.227320938 | 0.379282442 | 0.591815989 | 43.52671115 | 53.33602271 | ENSG00000269001 |
| NA | 0.295384927 | 1.227212368 | 0.406778547 | 0.617595253 | 39.63817147 | 48.73000045 | ENSG00000236017 |
| NA | 0.295353516 | 1.227185649 | 0.801978309 | 0.898712919 | 2.9900347 | 3.66998918 | ENSG00000213891 |
| KAT2B | 0.295302639 | 1.227142372 | 0.60531883 | 0.779667429 | 12.9461299 | 15.84772553 | ENSG00000114166 |
| HMGN3-AS1 | 0.2952771 | 1.227120649 | 0.638585452 | 0.802000816 | 10.51182638 | 12.92558592 | ENSG00000270362 |
| TRPA1 | 0.295237426 | 1.227086904 | 0.612375534 | 0.784314137 | 12.18283257 | 14.9883805 | ENSG00000104321 |
| ZNF641 | 0.295179074 | 1.227037273 | 0.098692533 | 0.235925126 | 133.995167 | 164.3543048 | ENSG00000167528 |
| CACYBP | 0.29504737 | 1.226925262 | 5.10E-08 | 7.47E-07 | 4544.184954 | 5574.994792 | ENSG00000116161 |
| NA | 0.294986838 | 1.226873784 | 0.622336294 | 0.791009618 | 12.21011314 | 15.01325656 | ENSG00000287858 |
| LY6H | 0.294958396 | 1.226849597 | 0.717182171 | 0.851020125 | 6.227679473 | 7.676399624 | ENSG00000176956 |
| CHIC1 | 0.29487766 | 1.226780943 | 0.121774306 | 0.275447093 | 120.1647439 | 147.4645882 | ENSG00000204116 |
| MRPL10 | 0.294872386 | 1.226776458 | 1.26E-05 | 0.000116926 | 1634.456353 | 2004.481388 | ENSG00000159111 |
| USO1 | 0.294871626 | 1.226775812 | 3.02E-06 | 3.18E-05 | 3419.873662 | 4195.655036 | ENSG00000138768 |
| TRMO | 0.294774842 | 1.226693515 | 0.283864738 | 0.493051563 | 61.01619232 | 74.89345255 | ENSG00000136932 |
| CARD11 | 0.294630801 | 1.226571046 | 2.33E-05 | 0.000204428 | 1204.508827 | 1477.646091 | ENSG00000198286 |
| LOC105371267 | 0.294593813 | 1.2265396 | 0.322274426 | 0.532891388 | 47.3580461 | 58.06691971 | ENSG00000277639 |
| PFKP | 0.294591224 | 1.226537399 | 1.18E-07 | 1.61E-06 | 3685.642183 | 4520.090596 | ENSG00000067057 |
| GPX7 | 0.29441751 | 1.22638972 | 0.055606865 | 0.15312564 | 181.2262935 | 222.2696248 | ENSG00000116157 |
| ARHGEF1 | 0.294388779 | 1.226365298 | 0.000117041 | 0.000854727 | 1343.036844 | 1646.526506 | ENSG00000076928 |
| BCL3 | 0.29436233 | 1.226342815 | 0.316795055 | 0.526875791 | 50.39552299 | 61.7752166 | ENSG00000069399 |

| | | | | | | | |
|---|---|---|---|---|---|---|---|
| RIPK1 | 0.294338103 | 1.226322221 | 0.005922371 | 0.025448003 | 445.7280926 | 546.4575294 | ENSG00000137275 |
| CNIH4 | 0.293856017 | 1.225912506 | 9.08E-05 | 0.000682339 | 1505.473541 | 1845.489981 | ENSG00000143771 |
| NA | 0.293599447 | 1.225694508 | 0.695126651 | 0.837250559 | 7.3453449 | 8.982642787 | ENSG00000225774 |
| RPS6KB2 | 0.293453292 | 1.225570343 | 0.001735137 | 0.009004687 | 564.1180475 | 691.2172635 | ENSG00000175634 |
| SLC12A4 | 0.293445166 | 1.22556344 | 7.44E-05 | 0.000571107 | 1189.471727 | 1457.559527 | ENSG00000124067 |
| KLHL29 | 0.2934249 | 1.225546225 | 0.030211054 | 0.09551913 | 262.5360455 | 322.0395522 | ENSG00000119771 |
| CYB561 | 0.293237443 | 1.225386993 | 0.000526196 | 0.003211767 | 873.5878808 | 1070.80595 | ENSG00000008283 |
| SLC39A11 | 0.293221576 | 1.225373516 | 0.013334496 | 0.049464145 | 402.8057244 | 493.2420305 | ENSG00000133195 |
| AKR1C1 | 0.293125696 | 1.225292082 | 0.584045656 | 0.763089955 | 15.95632622 | 19.50494548 | ENSG00000187134 |
| SPRY3 | 0.293016964 | 1.225199738 | 0.248245259 | 0.453204315 | 64.1378752 | 78.58368331 | ENSG00000168939 |
| TMEM8B | 0.292990451 | 1.225177223 | 0.033639087 | 0.103969098 | 231.3540709 | 283.4297904 | ENSG00000137103 |
| WDR17 | 0.292960907 | 1.225152133 | 0.33673794 | 0.548688178 | 46.47049886 | 56.92279745 | ENSG00000150627 |
| VWDE | 0.292927979 | 1.225124171 | 0.152427614 | 0.32256785 | 207.13841 | 253.7273941 | ENSG00000146530 |
| CCDC90B | 0.29288113 | 1.225084388 | 0.002850554 | 0.013771479 | 493.4853315 | 604.4671293 | ENSG00000137500 |
| HCFC1R1 | 0.292844153 | 1.225052989 | 0.01296682 | 0.048414075 | 388.0068015 | 475.6208494 | ENSG00000103145 |
| CORO1A | 0.292806891 | 1.225021348 | 0.052775327 | 0.146978624 | 245.5913378 | 300.8675474 | ENSG00000102879 |
| TDGF1 | 0.292510957 | 1.22477009 | 3.19E-09 | 5.64E-08 | 5658.849 | 6930.747779 | ENSG00000241186 |
| RAB3IL1 | 0.292429322 | 1.224700788 | 0.029927535 | 0.094749624 | 245.783497 | 301.2126408 | ENSG00000167994 |
| GOLGA6L10 | 0.292426669 | 1.224698536 | 0.370078068 | 0.582264793 | 45.41687779 | 55.64426822 | ENSG00000278662 |
| H2BW2 | 0.292278221 | 1.224572526 | 0.808391052 | 0.902753534 | 3.017315271 | 3.690514311 | ENSG00000101812 |
| WBP1 | 0.292272889 | 1.224568001 | 0.281788135 | 0.490709708 | 81.6397625 | 99.98471096 | ENSG00000239779 |
| RELL2 | 0.292225209 | 1.22452753 | 0.025533221 | 0.083811693 | 347.864206 | 426.1324486 | ENSG00000164620 |
| CDK11A | 0.292134886 | 1.224450868 | 0.644273433 | 0.805989478 | 10.30809903 | 12.59786651 | ENSG00000008128 |
| GLI3 | 0.292069174 | 1.224395098 | 0.004375071 | 0.019771051 | 457.5195139 | 560.1201602 | ENSG00000106571 |
| SYP | 0.29206836 | 1.224394407 | 0.063780388 | 0.169886126 | 207.7920258 | 254.7051695 | ENSG00000102003 |
| MOSPD3 | 0.292066473 | 1.224392806 | 0.031193342 | 0.097936202 | 236.7493486 | 290.0682663 | ENSG00000106330 |
| BDH1 | 0.291884451 | 1.224238336 | 0.003392557 | 0.015997115 | 496.1927201 | 607.2197186 | ENSG00000161267 |
| CELSR2 | 0.291789281 | 1.22415758 | 0.000361188 | 0.002307521 | 1572.093118 | 1923.900853 | ENSG00000143126 |
| NA | 0.291557567 | 1.223960981 | 0.758204141 | 0.874316737 | 4.624277109 | 5.671491918 | ENSG00000279019 |
| D2HGDH | 0.29153699 | 1.223943524 | 0.065205551 | 0.17282903 | 202.9207496 | 248.506802 | ENSG00000180902 |
| NFKBIA | 0.291373204 | 1.22380458 | 0.025206135 | 0.082987771 | 289.9815963 | 354.611709 | ENSG00000100906 |
| RAD50 | 0.29128247 | 1.223727615 | 0.193460288 | 0.3813971 | 114.7688559 | 140.1165704 | ENSG00000113522 |
| OLFML2B | 0.291233417 | 1.223686008 | 0.081193872 | 0.204065248 | 184.2762283 | 225.6384516 | ENSG00000162745 |
| MATN1 | 0.291064887 | 1.22354307 | 0.805668094 | 0.900901829 | 3.013755799 | 3.681812457 | ENSG00000162510 |
| TSSK4 | 0.291023227 | 1.223507739 | 0.522187917 | 0.716804556 | 24.81166363 | 30.26494321 | ENSG00000139908 |
| TUBGCP5 | 0.290940231 | 1.223437355 | 0.012329865 | 0.046514082 | 388.5168967 | 475.2375436 | ENSG00000275835 |
| NA | 0.290848911 | 1.223359916 | 0.71585926 | 0.850091312 | 6.234798418 | 7.655874493 | ENSG00000262619 |
| B9D1 | 0.2908375 | 1.223350239 | 0.036694174 | 0.111141856 | 314.1850264 | 384.3494432 | ENSG00000108641 |
| E2F6 | 0.290674308 | 1.223211867 | 0.010402897 | 0.040324039 | 449.1925493 | 549.1187992 | ENSG00000169016 |
| PPP1R3D | 0.290674285 | 1.223211847 | 0.22847926 | 0.427931253 | 78.08987849 | 95.66162638 | ENSG00000132825 |
| RUNX1T1 | 0.29063062 | 1.223174826 | 0.075520716 | 0.193725717 | 186.6061647 | 227.9474322 | ENSG00000079102 |
| NA | 0.290486978 | 1.223053047 | 0.798021942 | 0.897070233 | 2.982915755 | 3.644167164 | ENSG00000287486 |
| SYT2 | 0.290360828 | 1.222946106 | 0.00028984 | 0.001907186 | 835.4661058 | 1021.750594 | ENSG00000143858 |
| POM121L9P | 0.290109902 | 1.22273342 | 0.723703111 | 0.854472967 | 6.258519517 | 7.675170119 | ENSG00000128262 |
| H1-0 | 0.289999889 | 1.222640183 | 0.000100436 | 0.00074469 | 3372.830794 | 4122.927701 | ENSG00000189060 |
| SLC6A11 | 0.28995437 | 1.222601608 | 0.12088944 | 0.273904869 | 116.4140816 | 142.3432825 | ENSG00000132164 |
| RPL28 | 0.289935773 | 1.222585849 | 4.46E-07 | 5.53E-06 | 13592.96287 | 16618.89484 | ENSG00000108107 |
| FKBP9 | 0.289810638 | 1.222479809 | 0.037750896 | 0.113505121 | 271.8554581 | 332.2525805 | ENSG00000122642 |
| DEPDC5 | 0.289542787 | 1.222252865 | 0.079656411 | 0.201191831 | 192.2731308 | 235.3224288 | ENSG00000100150 |
| CFLAR | 0.289457508 | 1.222180619 | 0.202530378 | 0.393861208 | 96.11279255 | 117.201369 | ENSG00000003402 |
| PSD | 0.289411273 | 1.222141451 | 0.156133988 | 0.328095066 | 143.6926683 | 175.7863374 | ENSG00000059915 |
| PDE3A | 0.289304628 | 1.222051113 | 0.091310358 | 0.222617622 | 195.7230446 | 238.8941003 | ENSG00000172572 |
| ZNF628 | 0.289296684 | 1.222044384 | 0.07429367 | 0.191436206 | 302.7154304 | 370.3343919 | ENSG00000197483 |
| NETO1 | 0.289233308 | 1.221990702 | 0.041956172 | 0.123149256 | 268.2292989 | 327.3970696 | ENSG00000166342 |
| SLC41A2 | 0.289162871 | 1.221931042 | 0.458492766 | 0.663948476 | 36.74986078 | 44.96398464 | ENSG00000136052 |
| PACSIN3 | 0.289103406 | 1.221880677 | 4.04E-05 | 0.000332315 | 1154.409753 | 1410.26699 | ENSG00000165912 |
| TRIOBP | 0.288981181 | 1.221777165 | 7.76E-05 | 0.000592541 | 1348.917734 | 1648.664947 | ENSG00000100106 |
| NDUFB10 | 0.288667157 | 1.221511256 | 2.72E-05 | 0.000234291 | 1666.981355 | 2036.630399 | ENSG00000140990 |
| NA | 0.288650922 | 1.22149751 | 0.759269969 | 0.874958102 | 4.906286698 | 6.029384271 | ENSG00000285872 |
| RAPGEF6 | 0.288378244 | 1.221266661 | 0.30831752 | 0.518758988 | 52.65518534 | 64.4324417 | ENSG00000158987 |
| RUSC1-AS1 | 0.288313976 | 1.221212258 | 0.310142193 | 0.520717933 | 60.71638537 | 74.26555971 | ENSG00000225855 |
| FLVCR1-DT | 0.288163172 | 1.221084612 | 0.059125228 | 0.160589664 | 184.4322085 | 225.2121215 | ENSG00000198468 |
| LETMD1 | 0.288108202 | 1.221038087 | 0.010738456 | 0.041403787 | 367.5480582 | 448.6487801 | ENSG00000050426 |
| ZNF225 | 0.288053776 | 1.220992023 | 0.161490015 | 0.336059387 | 100.816852 | 123.1405179 | ENSG00000256294 |
| FGFR4 | 0.288003217 | 1.220949235 | 1.44E-06 | 1.62E-05 | 2575.578532 | 3145.312909 | ENSG00000160867 |
| PIDD1 | 0.28788346 | 1.220847889 | 0.009731606 | 0.03816851 | 557.2695013 | 680.6554665 | ENSG00000177595 |
| MUTYH | 0.287856039 | 1.220824685 | 0.526034237 | 0.719293069 | 19.58985755 | 23.88513893 | ENSG00000132781 |
| CEBPB-AS1 | 0.287849799 | 1.220819405 | 0.698678374 | 0.839454411 | 8.134448263 | 9.990393526 | ENSG00000277449 |
| RPS6KA6 | 0.287789168 | 1.220768099 | 0.063766304 | 0.169872571 | 253.1235286 | 309.1225822 | ENSG00000072133 |
| LINC00339 | 0.287782628 | 1.220762565 | 0.537387923 | 0.727443977 | 20.4349966 | 24.83784067 | ENSG00000218510 |
| GPCPD1 | 0.287736987 | 1.220723946 | 0.054301452 | 0.150253909 | 188.6511158 | 230.2553941 | ENSG00000125772 |
| ZNF217 | 0.287638332 | 1.220640473 | 0.000526377 | 0.003211835 | 1246.615878 | 1520.980922 | ENSG00000171940 |
| PTP4A3 | 0.287564431 | 1.220577948 | 0.0027097 | 0.013175254 | 841.4550492 | 1026.568028 | ENSG00000184489 |
| NA | 0.287403901 | 1.220442141 | 0.689778614 | 0.83399945 | 9.231952063 | 11.27762461 | ENSG00000228906 |

| | | | | | | | |
|---|---|---|---|---|---|---|---|
| PVT1 | 0.287271778 | 1.220330377 | 0.059773188 | 0.161907151 | 193.6552877 | 236.5070343 | ENSG00000249859 |
| FUZ | 0.287151536 | 1.220228672 | 0.015158478 | 0.054924762 | 334.1808641 | 407.8614388 | ENSG00000010361 |
| NETO2 | 0.287135054 | 1.220214732 | 0.017444476 | 0.061588881 | 396.6836851 | 484.2164752 | ENSG00000171208 |
| NA | 0.286950956 | 1.220059034 | 0.579760306 | 0.760542746 | 16.84626387 | 20.56093528 | ENSG00000279716 |
| SELENOM | 0.286856354 | 1.219979034 | 0.089862658 | 0.220196742 | 170.8452773 | 208.7252993 | ENSG00000198832 |
| ZBTB34 | 0.286811643 | 1.219941226 | 0.027219821 | 0.088112155 | 366.8935521 | 447.0297919 | ENSG00000177125 |
| NA | 0.286716185 | 1.219860509 | 0.568680828 | 0.752014213 | 19.90863092 | 24.27850112 | ENSG00000214077 |
| KCTD20 | 0.286700611 | 1.219847341 | 0.000467399 | 0.002905477 | 900.6204817 | 1098.137613 | ENSG00000112078 |
| NA | 0.286601425 | 1.219763478 | 0.718401438 | 0.851388915 | 6.231238946 | 7.620404664 | ENSG00000272702 |
| CCDC138 | 0.286484887 | 1.219664953 | 0.01158342 | 0.044139 | 374.0364422 | 456.4482875 | ENSG00000163006 |
| PCNT | 0.286380719 | 1.219576891 | 0.000481095 | 0.002980802 | 1114.406231 | 1358.392616 | ENSG00000160299 |
| PACC1 | 0.286281091 | 1.219492674 | 0.018691677 | 0.064995824 | 336.4897749 | 410.160824 | ENSG00000065600 |
| MINK1 | 0.286258017 | 1.21947317 | 8.62E-07 | 1.01E-05 | 2488.462384 | 3035.129531 | ENSG00000141503 |
| RBM18 | 0.286118707 | 1.21935542 | 0.089371799 | 0.21936954 | 189.8675823 | 231.6799427 | ENSG00000119446 |
| SFMBT2 | 0.286011069 | 1.219264449 | 0.147719515 | 0.315539309 | 143.1911933 | 174.2476531 | ENSG00000198879 |
| CDK5R1 | 0.285893998 | 1.219165513 | 0.027613986 | 0.089105527 | 333.8571091 | 407.0268113 | ENSG00000176749 |
| GRASP | 0.285709945 | 1.219009986 | 0.148555356 | 0.316501343 | 128.8931878 | 157.2329368 | ENSG00000161835 |
| JPH1 | 0.285679572 | 1.218984323 | 0.002170229 | 0.010888622 | 569.2437734 | 694.0228513 | ENSG00000104369 |
| GVQW3 | 0.285643357 | 1.218953724 | 0.135139217 | 0.297032602 | 132.592238 | 161.6221156 | ENSG00000179240 |
| NA | 0.285621402 | 1.218935174 | 0.816330151 | 0.906477444 | 3.020874744 | 3.672164644 | ENSG00000243289 |
| HIBADH | 0.285527764 | 1.218856062 | 0.003177156 | 0.015124929 | 533.4594889 | 650.3909319 | ENSG00000106049 |
| CTSV | 0.285404553 | 1.218751972 | 5.54E-06 | 5.60E-05 | 1547.721703 | 1886.068666 | ENSG00000136943 |
| CRYBG3 | 0.285354626 | 1.218709796 | 0.180467684 | 0.363417808 | 152.321142 | 185.2998575 | ENSG00000080200 |
| RDH10 | 0.285197006 | 1.218576654 | 0.087097815 | 0.215096408 | 147.5919125 | 179.8340387 | ENSG00000121039 |
| RASL11A | 0.285152619 | 1.218539163 | 0.41048406 | 0.621501761 | 44.14264788 | 53.78365589 | ENSG00000122035 |
| LOC112268411 | 0.285135874 | 1.21852502 | 0.6708384 | 0.823718022 | 9.030588979 | 10.96920083 | ENSG00000236106 |
| SEMA4C | 0.285010432 | 1.218419074 | 1.01E-06 | 1.17E-05 | 2406.974458 | 2932.623588 | ENSG00000168758 |
| NSUN6 | 0.284998965 | 1.218409389 | 0.07876928 | 0.199450066 | 171.4341872 | 208.7410947 | ENSG00000241058 |
| TMEM176A | 0.284869312 | 1.218299898 | 0.699058144 | 0.839692979 | 8.185449933 | 10.02179597 | ENSG00000002933 |
| NA | 0.284822508 | 1.218260374 | 0.56543411 | 0.749559397 | 18.06117796 | 21.93934761 | ENSG00000236698 |
| TMPPE | 0.28450935 | 1.217995962 | 0.517527908 | 0.713551474 | 22.11996135 | 26.99115409 | ENSG00000188167 |
| SEC22A | 0.284387203 | 1.217892843 | 0.336225173 | 0.548193198 | 56.44263755 | 68.86160927 | ENSG00000121542 |
| XKR4 | 0.284353231 | 1.217864166 | 0.533789063 | 0.724879869 | 19.9608278 | 24.28162254 | ENSG00000206579 |
| KMT2D | 0.284275521 | 1.217798568 | 0.001480199 | 0.007874242 | 1994.081061 | 2428.068877 | ENSG00000167548 |
| UVSSA | 0.284151346 | 1.217693754 | 0.327519824 | 0.538922356 | 336.3017861 | 409.5319984 | ENSG00000163945 |
| ORAI1 | 0.28392687 | 1.217504302 | 0.006011295 | 0.025777318 | 541.5394022 | 659.5873396 | ENSG00000276045 |
| NA | 0.28391239 | 1.217492082 | 0.701017478 | 0.840766866 | 8.212730504 | 10.04326706 | ENSG00000278238 |
| SFTPD | 0.283774585 | 1.217375794 | 0.736380955 | 0.861418484 | 6.269197935 | 7.640929795 | ENSG00000133661 |
| TTC9B | 0.283695347 | 1.217308933 | 0.817747887 | 0.907141028 | 2.993594172 | 3.634519351 | ENSG00000174521 |
| NA | 0.283684222 | 1.217299546 | 0.632769278 | 0.797821895 | 14.01635313 | 17.05145863 | ENSG00000253477 |
| C8orf88 | 0.283570344 | 1.217203464 | 0.129057505 | 0.287411673 | 130.9932849 | 159.4023814 | ENSG00000253250 |
| RNF141 | 0.283565584 | 1.217199448 | 0.016184136 | 0.058006721 | 450.8900236 | 549.0480275 | ENSG00000110315 |
| BICDL2 | 0.283455791 | 1.217106818 | 0.050157282 | 0.141397263 | 288.1222436 | 350.6819046 | ENSG00000162069 |
| TNK1 | 0.283374828 | 1.217038517 | 0.064655564 | 0.171756975 | 198.1218052 | 241.4261577 | ENSG00000174292 |
| NCEH1 | 0.283321648 | 1.216993657 | 0.257445919 | 0.464361621 | 74.15832023 | 90.17646901 | ENSG00000144959 |
| KLHL36 | 0.2831417 | 1.21684187 | 0.001143778 | 0.006310324 | 919.4518227 | 1118.162749 | ENSG00000135686 |
| P4HA2 | 0.283095927 | 1.216803263 | 0.109113045 | 0.254709501 | 144.71695 | 176.1029686 | ENSG00000072682 |
| NA | 0.283035883 | 1.216752621 | 0.809072149 | 0.902949338 | 3.299324861 | 4.030056997 | ENSG00000230216 |
| CPVL | 0.28284731 | 1.216593592 | 0.003911322 | 0.018033522 | 551.8480082 | 670.8694604 | ENSG00000106066 |
| EPHA6 | 0.282721304 | 1.216487338 | 0.51919686 | 0.714901185 | 23.57775698 | 28.68886751 | ENSG00000080224 |
| PPP1R12B | 0.282679861 | 1.216452394 | 0.031915593 | 0.099855199 | 259.0712578 | 315.1109592 | ENSG00000077157 |
| COL5A1 | 0.282638007 | 1.216417104 | 0.002532998 | 0.012434537 | 714.7718207 | 869.5852518 | ENSG00000130635 |
| NA | 0.282458193 | 1.216265502 | 0.676419583 | 0.826926971 | 10.47534193 | 12.68185895 | ENSG00000233045 |
| NCKAP5L | 0.282427958 | 1.216240013 | 0.001027731 | 0.005742217 | 1044.791871 | 1271.055102 | ENSG00000167566 |
| SPATA2L | 0.28229312 | 1.216126345 | 0.055471147 | 0.152885671 | 323.4695843 | 393.7892287 | ENSG00000158792 |
| SMTNL1 | 0.282272752 | 1.216109176 | 0.715296764 | 0.849695403 | 9.644440773 | 11.73975099 | ENSG00000214872 |
| TRIML2 | 0.28221063 | 1.216056811 | 0.015002116 | 0.05450461 | 515.2788163 | 626.5982669 | ENSG00000179046 |
| CNFN | 0.282133299 | 1.215991631 | 0.192396004 | 0.379705144 | 91.94641315 | 111.8153167 | ENSG00000105427 |
| ZNF347 | 0.281943394 | 1.215831578 | 0.082363875 | 0.206374126 | 196.3013534 | 238.969055 | ENSG00000197937 |
| C2orf92 | 0.281907857 | 1.215801629 | 0.597788708 | 0.77397777 | 15.33655069 | 18.60795516 | ENSG00000228486 |
| ZMYND12 | 0.2818828 | 1.215780513 | 0.727957602 | 0.856731016 | 6.023952125 | 7.292685255 | ENSG00000066185 |
| PLEKHA6 | 0.281678088 | 1.215608011 | 0.009023373 | 0.036019106 | 403.8048099 | 490.8233081 | ENSG00000143850 |
| CYREN | 0.281645301 | 1.215580385 | 0.079693029 | 0.201230492 | 180.7939486 | 219.7318619 | ENSG00000122783 |
| SAXO2 | 0.281589127 | 1.215533056 | 0.517163018 | 0.713270264 | 21.91267453 | 26.62796485 | ENSG00000188659 |
| VPS37D | 0.281496602 | 1.215455102 | 0.157920965 | 0.330608653 | 110.0852885 | 133.9277352 | ENSG00000176428 |
| CENPE | 0.281419261 | 1.215389944 | 0.000639466 | 0.003795195 | 770.5174917 | 936.1039534 | ENSG00000138778 |
| NA | 0.281377838 | 1.215355049 | 0.620552924 | 0.789501351 | 14.0199126 | 17.01788072 | ENSG00000230882 |
| AVPR1B | 0.281115812 | 1.215134333 | 0.665788825 | 0.821041006 | 14.356204 | 17.4804586 | ENSG00000198049 |
| LOC107986163 | 0.281083021 | 1.215106714 | 0.723885238 | 0.85451761 | 6.26207899 | 7.613878273 | ENSG00000272970 |
| NA | 0.281068832 | 1.215094764 | 0.571742278 | 0.75437385 | 22.29579773 | 26.97062896 | ENSG00000250770 |
| UQCC1 | 0.280829368 | 1.214893094 | 0.173759247 | 0.353826293 | 910.3349946 | 1106.041045 | ENSG00000101019 |
| CISD3 | 0.280801502 | 1.214869629 | 0.038370795 | 0.115075823 | 268.9437051 | 327.0003454 | ENSG00000277972 |
| FRAS1 | 0.28079647 | 1.214865391 | 0.134122021 | 0.295424407 | 2784.707703 | 3382.792213 | ENSG00000138759 |
| FBXO41 | 0.280658031 | 1.21474882 | 0.00835869 | 0.03387422 | 506.9100538 | 615.4927469 | ENSG00000163013 |

| | | | | | | | |
|---|---|---|---|---|---|---|---|
| NA | 0.280641964 | 1.214735292 | 0.459343534 | 0.664371804 | 46.49419381 | 56.43702452 | ENSG00000249264 |
| C6orf62 | 0.280587938 | 1.214689803 | 0.000235397 | 0.001591651 | 961.426847 | 1168.170112 | ENSG00000112308 |
| TNFAIP8L1 | 0.28043201 | 1.214558525 | 0.060559167 | 0.163520142 | 213.6564126 | 259.6849224 | ENSG00000185361 |
| NA | 0.280391246 | 1.214524208 | 0.373786624 | 0.585708628 | 44.2968481 | 53.83418601 | ENSG00000244398 |
| NA | 0.280368911 | 1.214505405 | 0.810848756 | 0.90389462 | 3.541011198 | 4.340372697 | ENSG00000268047 |
| MYLK | 0.280245294 | 1.214401345 | 0.011568353 | 0.044106973 | 417.3069504 | 506.5237418 | ENSG00000065534 |
| RCCD1 | 0.280079358 | 1.214261676 | 0.004933299 | 0.021828808 | 491.9047082 | 597.1409817 | ENSG00000166965 |
| REEP3 | 0.279816448 | 1.214040414 | 0.042182905 | 0.123661001 | 366.5893727 | 445.4431224 | ENSG00000165476 |
| OBSL1 | 0.279699942 | 1.213942377 | 9.22E-05 | 0.000691128 | 3300.023061 | 4006.564632 | ENSG00000124006 |
| FBXL5 | 0.279534222 | 1.213802942 | 0.000420717 | 0.002642251 | 1093.129452 | 1327.038607 | ENSG00000118564 |
| FRG2C | 0.279353449 | 1.213650859 | 0.82636677 | 0.912467781 | 3.582529659 | 4.377072032 | ENSG00000172969 |
| IGFBP7 | 0.279319783 | 1.213622538 | 0.727881441 | 0.856731016 | 6.296478506 | 7.63970029 | ENSG00000163453 |
| NA | 0.279306988 | 1.213611775 | 0.822786548 | 0.910164243 | 3.310003278 | 4.026652028 | ENSG00000276545 |
| DDIAS | 0.279145062 | 1.213475568 | 0.012018734 | 0.045595377 | 381.8464727 | 463.5847385 | ENSG00000165490 |
| KIFC2 | 0.27910507 | 1.213441931 | 0.097319115 | 0.233410145 | 264.375516 | 321.0527484 | ENSG00000167702 |
| PRKCQ | 0.279100835 | 1.213438369 | 0.004729905 | 0.021089193 | 507.0334396 | 615.0931716 | ENSG00000065675 |
| ZNF776 | 0.279035675 | 1.213383565 | 0.047504565 | 0.135619607 | 269.5904819 | 327.5137369 | ENSG00000152443 |
| MLF1 | 0.279028542 | 1.213377565 | 0.009644216 | 0.03789659 | 485.6894872 | 588.8311385 | ENSG00000178053 |
| PLEKHG2 | 0.278937242 | 1.21330078 | 0.000250551 | 0.001679681 | 1182.296678 | 1434.828768 | ENSG00000090924 |
| ITPRIPL2 | 0.278928088 | 1.213293082 | 0.166602136 | 0.343021641 | 106.3349056 | 128.9210463 | ENSG00000205730 |
| CALHM6 | 0.278789502 | 1.213176537 | 0.72304716 | 0.854154341 | 6.316640132 | 7.658049957 | ENSG00000188820 |
| OTOG | 0.278784721 | 1.213172517 | 0.373576904 | 0.585622963 | 43.31469581 | 52.42966812 | ENSG00000188162 |
| FAM120AOS | 0.278730673 | 1.213127068 | 0.001785781 | 0.009221894 | 767.4174967 | 930.845927 | ENSG00000188938 |
| GCNT1 | 0.278649937 | 1.213059182 | 0.066713064 | 0.176157188 | 188.0381799 | 228.143131 | ENSG00000187210 |
| NRL | 0.278617107 | 1.213031577 | 0.652776112 | 0.811801095 | 10.88279663 | 13.19295944 | ENSG00000129535 |
| NIPSNAP2 | 0.278558086 | 1.212981953 | 0.003144433 | 0.014984261 | 571.9381716 | 693.819934 | ENSG00000146729 |
| PPM1N | 0.278554869 | 1.212979248 | 0.096034771 | 0.231122295 | 179.2370726 | 217.2783751 | ENSG00000213889 |
| HEATR5A | 0.278518731 | 1.212948865 | 0.092068399 | 0.224116094 | 208.9760743 | 253.183721 | ENSG00000129493 |
| FANCF | 0.278361997 | 1.212817098 | 0.027061842 | 0.087748113 | 332.449678 | 403.5387331 | ENSG00000183161 |
| ZFP36L2 | 0.278170479 | 1.212656107 | 0.001289089 | 0.006981488 | 1515.219789 | 1838.40487 | ENSG00000152518 |
| SLC41A1 | 0.278090769 | 1.212589108 | 0.000396208 | 0.002507502 | 940.1843655 | 1140.18814 | ENSG00000133065 |
| UBA3 | 0.277956149 | 1.212475965 | 0.000109618 | 0.000806443 | 1109.714959 | 1345.59122 | ENSG00000144744 |
| ABLIM2 | 0.277950546 | 1.212471256 | 0.314668764 | 0.524986448 | 63.9744711 | 77.59042577 | ENSG00000163995 |
| PCDHGA10 | 0.27774656 | 1.212299834 | 0.059039611 | 0.160426339 | 252.9473873 | 306.6706084 | ENSG00000253846 |
| TMEM144 | 0.277501088 | 1.212093581 | 0.403746418 | 0.614998476 | 40.61734911 | 49.14774424 | ENSG00000164124 |
| RPS12 | 0.277434329 | 1.212037494 | 6.17E-07 | 7.40E-06 | 14091.44966 | 17079.79809 | ENSG00000112306 |
| IQCE | 0.27737141 | 1.211984636 | 0.071247666 | 0.185383326 | 195.3446761 | 236.7413243 | ENSG00000106012 |
| IL34 | 0.277102454 | 1.211758711 | 0.187348521 | 0.372782301 | 100.7175184 | 122.0685649 | ENSG00000157368 |
| FOXA3 | 0.277073846 | 1.211734683 | 0.005733177 | 0.024753324 | 515.5486201 | 625.0462463 | ENSG00000170608 |
| LOC100134368 | 0.277066209 | 1.211728268 | 0.520207037 | 0.715550501 | 24.83243565 | 30.20431377 | ENSG00000236829 |
| LOC283194 | 0.276998168 | 1.211671121 | 0.726180382 | 0.855720453 | 8.78862331 | 10.71270462 | ENSG00000255240 |
| CHST12 | 0.27675222 | 1.211464576 | 0.009187032 | 0.036464291 | 426.1124807 | 516.3945051 | ENSG00000136213 |
| PPARGC1B | 0.27668068 | 1.211404504 | 0.034049187 | 0.105016969 | 308.1409127 | 372.9990893 | ENSG00000155846 |
| PPIAP46 | 0.276612117 | 1.211346934 | 0.77210967 | 0.882599617 | 4.975085731 | 6.039032084 | ENSG00000260266 |
| CACNB3 | 0.276467894 | 1.211225844 | 0.02768428 | 0.089263724 | 312.303505 | 378.5226403 | ENSG00000167535 |
| ABCB4 | 0.276314555 | 1.211097114 | 0.577803021 | 0.759450456 | 18.94872519 | 22.92250584 | ENSG00000005471 |
| KREMEN2 | 0.276281376 | 1.211069262 | 0.058343515 | 0.158975805 | 202.8996721 | 245.7438298 | ENSG00000131650 |
| CDON | 0.276192333 | 1.210994517 | 0.033588207 | 0.103848379 | 350.2154988 | 423.8956966 | ENSG00000064309 |
| CPT2 | 0.27612502 | 1.210938016 | 0.030486057 | 0.096222038 | 265.7478843 | 321.9121231 | ENSG00000157184 |
| ARMC4 | 0.276108244 | 1.210923935 | 0.115204013 | 0.26515878 | 135.84262 | 164.534113 | ENSG00000169126 |
| RALGDS | 0.276034434 | 1.210861985 | 0.382385042 | 0.594896975 | 47.66736241 | 57.70250097 | ENSG00000160271 |
| NA | 0.275946274 | 1.210787993 | 0.728836702 | 0.857302489 | 6.540529107 | 7.943773146 | ENSG00000231579 |
| NOCT | 0.275878919 | 1.210731467 | 0.019180127 | 0.066327908 | 399.5402399 | 483.8940003 | ENSG00000151014 |
| MPC2 | 0.27570942 | 1.210589229 | 0.143095814 | 0.308594775 | 122.1560242 | 147.9228581 | ENSG00000143158 |
| SYNE4 | 0.275527814 | 1.21043685 | 0.039878538 | 0.118457542 | 269.134111 | 325.7808372 | ENSG00000181392 |
| AKAP11 | 0.275471637 | 1.210389718 | 0.013175364 | 0.048989319 | 482.8759249 | 584.3955172 | ENSG00000023516 |
| PCDHAC1 | 0.275471283 | 1.210389421 | 0.288614231 | 0.49785164 | 65.52659247 | 79.4734122 | ENSG00000248383 |
| COX15 | 0.275420894 | 1.210347146 | 0.026148672 | 0.085361741 | 293.9549524 | 355.7322277 | ENSG00000014919 |
| NA | 0.275396332 | 1.21032654 | 0.765691897 | 0.878910348 | 4.967966786 | 6.025033344 | ENSG00000205578 |
| MAP1B | 0.275370468 | 1.210304842 | 0.005241485 | 0.023003821 | 1369.466654 | 1656.779787 | ENSG00000131711 |
| SNX4 | 0.275344329 | 1.210282914 | 0.008833238 | 0.035432392 | 567.5116714 | 687.0142379 | ENSG00000114520 |
| SETD1B | 0.275344308 | 1.210282896 | 0.010495452 | 0.040616108 | 465.7216183 | 563.6711815 | ENSG00000139718 |
| CHURC1 | 0.27531975 | 1.210262295 | 0.20447065 | 0.396206468 | 88.18832643 | 106.7671987 | ENSG00000258289 |
| ATP5ME | 0.275296331 | 1.210242648 | 0.512215012 | 0.708615611 | 26.03669747 | 31.55595813 | ENSG00000169020 |
| FAM174B | 0.275182453 | 1.210147123 | 0.010845264 | 0.041779655 | 458.0696001 | 554.6530689 | ENSG00000185442 |
| CXCR4 | 0.275062643 | 1.210046629 | 0.53372619 | 0.724879869 | 22.5870112 | 27.32975081 | ENSG00000121966 |
| RNF175 | 0.275040053 | 1.210027683 | 0.101400211 | 0.240279246 | 155.3073379 | 187.7249383 | ENSG00000145428 |
| ESYT3 | 0.275033806 | 1.210022442 | 0.094419033 | 0.228690962 | 213.9360847 | 258.6504037 | ENSG00000158220 |
| EGLN2 | 0.274972547 | 1.209971065 | 0.240896184 | 0.444487 | 91.58195145 | 110.6493445 | ENSG00000269858 |
| TLE5 | 0.274777928 | 1.209807851 | 5.25E-06 | 5.32E-05 | 3181.12528 | 3848.508884 | ENSG00000104964 |
| NA | 0.274735299 | 1.209772104 | 0.809169684 | 0.902949338 | 3.279163235 | 3.977183459 | ENSG00000259594 |
| FGFR1OP2 | 0.274664739 | 1.209712937 | 0.008608523 | 0.034700523 | 439.9283605 | 532.5137145 | ENSG00000111790 |
| SSH1 | 0.274601578 | 1.209659977 | 0.001780911 | 0.009201779 | 679.6784227 | 822.1113206 | ENSG00000084112 |
| NA | 0.274594517 | 1.209654057 | 0.630134257 | 0.796418514 | 12.67123925 | 15.29775025 | ENSG00000233270 |

| | | | | | | | |
|---|---|---|---|---|---|---|---|
| CCND1 | 0.274561787 | 1.209626614 | 0.001280431 | 0.006940569 | 2021.242086 | 2445.419499 | ENSG00000110092 |
| NA | 0.274538958 | 1.209607473 | 0.830301302 | 0.914737804 | 3.071876414 | 3.694865238 | ENSG00000226803 |
| APOC1 | 0.274274284 | 1.209385581 | 0.195860617 | 0.38481354 | 100.0076132 | 120.960931 | ENSG00000130208 |
| TRAPPC6A | 0.273968908 | 1.209129617 | 0.002847106 | 0.013761864 | 680.866335 | 823.5840785 | ENSG00000007255 |
| NA | 0.273914354 | 1.209083896 | 0.314936025 | 0.52531062 | 58.81909979 | 71.19255538 | ENSG00000260469 |
| PI4KB | 0.27383818 | 1.209020059 | 0.000474742 | 0.002944338 | 952.7897548 | 1151.973028 | ENSG00000143393 |
| NSD1 | 0.273781518 | 1.208972575 | 8.92E-06 | 8.66E-05 | 3809.02297 | 4604.584884 | ENSG00000165671 |
| C1orf122 | 0.273666312 | 1.208876037 | 0.005033531 | 0.022225444 | 547.8088533 | 662.2790456 | ENSG00000197982 |
| CCNG2 | 0.27359337 | 1.208814918 | 0.080132575 | 0.202023809 | 181.6091374 | 219.497193 | ENSG00000138764 |
| FBXO46 | 0.273506691 | 1.208742293 | 0.00717538 | 0.029900057 | 531.5939331 | 642.7952126 | ENSG00000177051 |
| TFPI | 0.273418248 | 1.208668194 | 0.249178382 | 0.454215016 | 77.23792598 | 93.36860911 | ENSG00000003436 |
| NA | 0.273224058 | 1.208505516 | 0.645573848 | 0.807127582 | 13.52794644 | 16.39384435 | ENSG00000234129 |
| SLC12A9-AS1 | 0.27322125 | 1.208503164 | 0.59631214 | 0.772807766 | 15.95396196 | 19.25469212 | ENSG00000236305 |
| RWDD2A | 0.273131739 | 1.208428186 | 0.244346523 | 0.448705182 | 112.8470112 | 136.2820013 | ENSG00000013392 |
| NHLH2 | 0.273004739 | 1.208321813 | 0.214919957 | 0.409934692 | 95.59624114 | 115.6887606 | ENSG00000177551 |
| TMBIM4 | 0.272941594 | 1.208268927 | 0.413252348 | 0.624641475 | 49.28586623 | 59.63334759 | ENSG00000155957 |
| GHRL | 0.272930527 | 1.208259658 | 0.826045539 | 0.91237981 | 3.330164905 | 4.026652028 | ENSG00000157017 |
| NA | 0.27267711 | 1.20804744 | 0.497931304 | 0.696960538 | 26.07938499 | 31.5289066 | ENSG00000237126 |
| PNPLA2 | 0.272669007 | 1.208040654 | 0.009753266 | 0.038200904 | 532.1653244 | 643.0801078 | ENSG00000177666 |
| NA | 0.272630092 | 1.208008069 | 0.464556081 | 0.668939504 | 28.8348523 | 34.79929076 | ENSG00000260086 |
| TIMP1 | 0.272610124 | 1.20799135 | 0.016913145 | 0.060061063 | 508.2524974 | 614.2717686 | ENSG00000102265 |
| LPCAT4 | 0.272604532 | 1.207986668 | 0.001233935 | 0.006727152 | 861.4331668 | 1040.864649 | ENSG00000176454 |
| ZNF783 | 0.272367349 | 1.207788088 | 0.053304841 | 0.148016751 | 248.2866257 | 300.0329534 | ENSG00000204946 |
| NA | 0.272331141 | 1.207757776 | 0.708721174 | 0.846326173 | 8.563539126 | 10.38063429 | ENSG00000184100 |
| GOLGA6D | 0.272329213 | 1.207756162 | 0.749936663 | 0.869935742 | 6.112912784 | 7.332506011 | ENSG00000140478 |
| NA | 0.272286384 | 1.207720308 | 0.58706364 | 0.765497261 | 16.50493846 | 19.96366689 | ENSG00000268869 |
| PLA2G3 | 0.272265924 | 1.20770318 | 0.000310759 | 0.002027177 | 1043.441367 | 1260.054721 | ENSG00000100078 |
| ZBTB48 | 0.272037134 | 1.207511672 | 0.015117326 | 0.0548072 | 365.7035799 | 441.511551 | ENSG00000204859 |
| COX10 | 0.271823434 | 1.207332822 | 0.012513586 | 0.047075513 | 545.57655 | 658.7717874 | ENSG00000006695 |
| EFNB2 | 0.271821168 | 1.207330925 | 0.087590437 | 0.21597407 | 168.3480729 | 203.1704318 | ENSG00000125266 |
| LOC106660606 | 0.271735928 | 1.207259593 | 0.726263565 | 0.855720453 | 6.574928623 | 7.951245495 | ENSG00000260778 |
| RBM33 | 0.271631312 | 1.207172054 | 0.000117311 | 0.000856369 | 1373.946013 | 1657.885628 | ENSG00000184863 |
| SALL4 | 0.271622433 | 1.207164623 | 6.15E-06 | 6.15E-05 | 3164.498248 | 3819.223831 | ENSG00000101115 |
| P2RY8 | 0.271597994 | 1.207144175 | 0.771357952 | 0.882479852 | 4.924084061 | 5.954093685 | ENSG00000182162 |
| NA | 0.271500619 | 1.207062701 | 2.02E-05 | 0.000179573 | 8802.200536 | 10624.35582 | ENSG00000240342 |
| GRAMD4 | 0.271350373 | 1.206937001 | 0.004323273 | 0.019588536 | 634.4023709 | 765.8701307 | ENSG00000075240 |
| BSPRY | 0.271299991 | 1.206894853 | 0.091278709 | 0.222569195 | 166.5083231 | 201.0351845 | ENSG00000119411 |
| CSKMT | 0.271285665 | 1.206882869 | 0.241310437 | 0.444860173 | 84.3389148 | 101.9340752 | ENSG00000214756 |
| MRPS27 | 0.271262623 | 1.206863593 | 6.37E-05 | 0.000498179 | 1785.701959 | 2154.167158 | ENSG00000113048 |
| LOC101929657 | 0.271198313 | 1.206809797 | 0.818888074 | 0.907979173 | 3.068316941 | 3.674340108 | ENSG00000273523 |
| PHIP | 0.271170356 | 1.206786411 | 0.002871254 | 0.013860847 | 1064.416304 | 1284.161776 | ENSG00000146247 |
| ZNF106 | 0.270997922 | 1.206642182 | 0.000259412 | 0.001732319 | 1600.339158 | 1930.104129 | ENSG00000103994 |
| TACR3 | 0.270981456 | 1.20662841 | 0.30775773 | 0.518096339 | 59.13194942 | 71.40175847 | ENSG00000169836 |
| NA | 0.270891439 | 1.206553124 | 0.451326165 | 0.657517808 | 31.97196827 | 38.49198053 | ENSG00000136149 |
| FAN1 | 0.270759484 | 1.206442773 | 0.009490398 | 0.037432806 | 678.7080668 | 818.2106048 | ENSG00000198690 |
| DNAJC28 | 0.270598308 | 1.206307999 | 0.436967761 | 0.645089622 | 32.75751216 | 39.52555329 | ENSG00000177692 |
| TTC30B | 0.270503357 | 1.206228608 | 0.464218144 | 0.668593683 | 31.53428392 | 38.11167113 | ENSG00000196659 |
| SNX18 | 0.270475428 | 1.206205257 | 0.012294308 | 0.046402492 | 486.5530334 | 586.8485323 | ENSG00000178996 |
| NA | 0.270453259 | 1.206186722 | 0.699488547 | 0.839970842 | 8.5433775 | 10.34393495 | ENSG00000283213 |
| CNGA4 | 0.270441494 | 1.206176886 | 0.831954493 | 0.915381416 | 3.048155315 | 3.653814977 | ENSG00000132259 |
| ALG5 | 0.269988942 | 1.205798585 | 0.068442199 | 0.179518185 | 244.7696411 | 294.9877257 | ENSG00000120697 |
| ERF | 0.269964057 | 1.205777787 | 0.000370939 | 0.002365815 | 1417.327446 | 1709.4965 | ENSG00000105722 |
| NA | 0.269874717 | 1.205703121 | 0.714083665 | 0.849104018 | 7.747155192 | 9.284540179 | ENSG00000226478 |
| CDAN1 | 0.269847602 | 1.205680459 | 0.015999244 | 0.057487796 | 407.355863 | 491.0362427 | ENSG00000140326 |
| NA | 0.269798586 | 1.205639497 | 0.431714655 | 0.6413842 | 40.2733801 | 48.63323878 | ENSG00000242299 |
| RASSF8 | 0.269797637 | 1.205638704 | 0.028123721 | 0.090279868 | 385.4209675 | 464.585709 | ENSG00000123094 |
| STAC | 0.269703439 | 1.205559987 | 0.040827997 | 0.120663023 | 254.4051568 | 306.7915111 | ENSG00000144681 |
| LRMDA | 0.269601773 | 1.205475035 | 0.824023144 | 0.910998541 | 3.595572341 | 4.357776406 | ENSG00000148655 |
| AVPR2 | 0.269519267 | 1.205406097 | 0.806071169 | 0.901122005 | 3.279163235 | 3.957887833 | ENSG00000126895 |
| C12orf29 | 0.26948611 | 1.205378394 | 0.064981748 | 0.172379782 | 213.9662876 | 257.82017 | ENSG00000133641 |
| LHPP | 0.269395566 | 1.205302746 | 0.00452128 | 0.020324771 | 826.5368575 | 996.2499076 | ENSG00000107902 |
| NA | 0.269247323 | 1.205178902 | 0.344994506 | 0.556999337 | 53.14862604 | 63.94666877 | ENSG00000239665 |
| GPR180 | 0.269167343 | 1.205112092 | 0.058913719 | 0.160153388 | 220.6687724 | 266.0318838 | ENSG00000152749 |
| BTD | 0.269008028 | 1.20497902 | 0.043077835 | 0.125650946 | 280.2885443 | 337.6585681 | ENSG00000169814 |
| NA | 0.26876591 | 1.204776814 | 0.655896071 | 0.814050821 | 12.0843887 | 14.62424531 | ENSG00000238058 |
| PRR3 | 0.268595171 | 1.20463424 | 0.097719303 | 0.234042999 | 167.6177005 | 202.0415378 | ENSG00000204576 |
| NA | 0.26847044 | 1.204530095 | 0.019468981 | 0.067191468 | 542.1445047 | 652.8522605 | ENSG00000230623 |
| HARBI1 | 0.268374271 | 1.204449805 | 0.27346815 | 0.482117812 | 79.11385473 | 95.3327501 | ENSG00000180423 |
| LYSMD3 | 0.268373755 | 1.204449374 | 0.1266623 | 0.283501308 | 193.9153557 | 233.8337076 | ENSG00000176018 |
| RASSF4 | 0.268212056 | 1.204314386 | 0.061125406 | 0.164766384 | 237.9841194 | 286.4460647 | ENSG00000107551 |
| EGFR | 0.268033094 | 1.204165003 | 0.098865158 | 0.236188301 | 159.2747703 | 191.71735 | ENSG00000146648 |
| NA | 0.268013837 | 1.20414893 | 0.774648384 | 0.884208808 | 4.675278779 | 5.600552259 | ENSG00000217950 |
| PABPC3 | 0.268008553 | 1.20414452 | 0.688812091 | 0.83375313 | 10.26658057 | 12.38648795 | ENSG00000151846 |
| RIOX1 | 0.267774715 | 1.203949363 | 0.006866266 | 0.028815431 | 588.6073254 | 708.7260757 | ENSG00000170468 |

| | | | | | | | |
|---|---|---|---|---|---|---|---|
| PANK3 | 0.267750833 | 1.203929434 | 0.000383616 | 0.002437612 | 1526.325657 | 1837.689428 | ENSG00000120137 |
| GSTA4 | 0.26765739 | 1.203851458 | 0.079608014 | 0.201112894 | 238.1418785 | 286.9209603 | ENSG00000170899 |
| KCNQ3 | 0.26765547 | 1.203849855 | 0.255972898 | 0.462814344 | 78.19574679 | 94.03804667 | ENSG00000184156 |
| NA | 0.267404896 | 1.203640784 | 0.717919252 | 0.851146028 | 8.023241046 | 9.627204287 | ENSG00000264769 |
| NBPF11 | 0.267392362 | 1.203630326 | 0.053115956 | 0.147709443 | 279.1278342 | 335.5275561 | ENSG00000263956 |
| BAIAP2L1 | 0.267375957 | 1.20361664 | 0.006368835 | 0.027058593 | 738.2208786 | 888.5661596 | ENSG00000006453 |
| HPAT5 | 0.2672071 | 1.203475773 | 0.000700999 | 0.004117689 | 965.2367996 | 1161.808912 | ENSG00000280707 |
| LEMD2 | 0.267158661 | 1.203435367 | 0.001145359 | 0.006317201 | 1393.033556 | 1676.777809 | ENSG00000161904 |
| BAIAP2 | 0.267152594 | 1.203430306 | 0.015667049 | 0.056471916 | 411.7740479 | 495.6751602 | ENSG00000175866 |
| SNRPE | 0.267047887 | 1.203342968 | 0.000149485 | 0.001064888 | 1941.315202 | 2335.590925 | ENSG00000182004 |
| RN7SL3 | 0.267019706 | 1.203319462 | 0.694515153 | 0.837010823 | 11.22768151 | 13.49268137 | ENSG00000278771 |
| ODR4 | 0.267004352 | 1.203306656 | 0.034906732 | 0.107049957 | 289.3078439 | 348.115396 | ENSG00000157181 |
| LCAT | 0.266833668 | 1.203164302 | 0.191228865 | 0.3780734 | 108.7879223 | 130.9725121 | ENSG00000213398 |
| MFSD8 | 0.266718205 | 1.203068013 | 0.146708607 | 0.313983685 | 123.0412071 | 148.0592296 | ENSG00000164073 |
| LOC101930085 | 0.266622285 | 1.202988028 | 0.452101228 | 0.658283091 | 40.30185588 | 48.52124886 | ENSG00000271122 |
| SYNGR2 | 0.266547238 | 1.202925452 | 0.00066583 | 0.003931893 | 1101.612825 | 1325.518918 | ENSG00000108639 |
| AMPD3 | 0.266464959 | 1.202856849 | 0.199062541 | 0.389201376 | 114.2359555 | 137.3769815 | ENSG00000133805 |
| NA | 0.266464755 | 1.202856679 | 0.665589024 | 0.82085806 | 11.6045755 | 13.95357824 | ENSG00000258815 |
| BTBD9 | 0.266398114 | 1.202801118 | 0.120059758 | 0.272712115 | 141.0134244 | 169.5902704 | ENSG00000183826 |
| LOC339803 | 0.266365865 | 1.202774232 | 0.358267109 | 0.570123925 | 60.39611131 | 72.83616576 | ENSG00000212978 |
| LRRK1 | 0.26635945 | 1.202768884 | 0.272101188 | 0.480723209 | 106.5010056 | 128.0032473 | ENSG00000154237 |
| SHFL | 0.266135842 | 1.202582477 | 0.15559966 | 0.327409423 | 148.9148054 | 179.4424005 | ENSG00000130813 |
| BCAR1 | 0.266130426 | 1.202577962 | 3.09E-05 | 0.000262618 | 2843.828095 | 3420.936949 | ENSG00000050820 |
| PTMS | 0.266047491 | 1.202508832 | 1.34E-07 | 1.81E-06 | 7446.438878 | 8955.329838 | ENSG00000159335 |
| PXN | 0.2658841 | 1.202372651 | 0.000983388 | 0.005532499 | 1014.712715 | 1219.686523 | ENSG00000089159 |
| SNHG6 | 0.265599789 | 1.202135724 | 0.000884882 | 0.005030401 | 1026.317543 | 1233.853551 | ENSG00000245910 |
| NA | 0.265466353 | 1.202024542 | 0.771741825 | 0.882491333 | 5.005925775 | 6.011034604 | ENSG00000270926 |
| TMEM215 | 0.265446285 | 1.202007823 | 0.637560751 | 0.801140114 | 14.81138023 | 17.80070566 | ENSG00000188133 |
| GEMIN4 | 0.265403919 | 1.201972525 | 7.02E-05 | 0.000542839 | 1640.1445 | 1971.566805 | ENSG00000179409 |
| TMEM44 | 0.265320283 | 1.201902846 | 0.082913994 | 0.207394805 | 206.6094007 | 248.2478468 | ENSG00000145014 |
| EDN3 | 0.265258558 | 1.201851425 | 0.749470422 | 0.869822328 | 8.763707002 | 10.5695958 | ENSG00000124205 |
| NA | 0.26512461 | 1.201739843 | 0.745061201 | 0.867396409 | 6.694729326 | 8.035237936 | ENSG00000141028 |
| LOC100996419 | 0.265063749 | 1.201689148 | 0.647787582 | 0.808876289 | 11.90082298 | 14.32017245 | ENSG00000245317 |
| ABCA2 | 0.264986121 | 1.20162449 | 0.015271613 | 0.055292257 | 763.7546777 | 918.1754031 | ENSG00000107331 |
| SLC10A3 | 0.264957028 | 1.201600258 | 0.127904163 | 0.285473796 | 191.0546049 | 229.9754623 | ENSG00000126903 |
| SLC4A2 | 0.264783826 | 1.20145601 | 2.01E-05 | 0.000179122 | 2438.037852 | 2929.723453 | ENSG00000164889 |
| NA | 0.264602672 | 1.201305157 | 0.686965731 | 0.83231846 | 10.42878945 | 12.57734138 | ENSG00000260948 |
| UPK1A | 0.264551809 | 1.201262804 | 0.751337685 | 0.870629467 | 7.543427844 | 8.982642787 | ENSG00000105668 |
| NECAB3 | 0.264350208 | 1.201094953 | 0.037361491 | 0.112675145 | 330.9218625 | 397.6067206 | ENSG00000125967 |
| SMAD3 | 0.26415629 | 1.20093352 | 0.017187395 | 0.060840479 | 452.3351088 | 543.0853702 | ENSG00000166949 |
| NA | 0.264130641 | 1.20091217 | 0.498657917 | 0.697463691 | 31.05803019 | 37.3611946 | ENSG00000235174 |
| CRIP3 | 0.264104364 | 1.200890296 | 0.251156601 | 0.456725317 | 171.773707 | 206.2352717 | ENSG00000146215 |
| SLC4A10 | 0.263951811 | 1.200763319 | 0.745835394 | 0.867900448 | 6.398481846 | 7.641875753 | ENSG00000144290 |
| IKZF1 | 0.263437316 | 1.200335178 | 0.426897853 | 0.636856462 | 44.33391736 | 53.10724044 | ENSG00000185811 |
| FUT2 | 0.263371611 | 1.200280512 | 0.171239271 | 0.350097827 | 112.1750126 | 134.6002941 | ENSG00000176920 |
| ZNF311 | 0.263220334 | 1.200154661 | 0.314153713 | 0.524608089 | 70.66091243 | 84.67291901 | ENSG00000197935 |
| SUSD1 | 0.262738436 | 1.199753845 | 0.183268912 | 0.366977969 | 108.8899257 | 130.5820604 | ENSG00000106868 |
| SPIN3 | 0.262721992 | 1.19974017 | 0.046469524 | 0.133248855 | 268.0819645 | 321.570237 | ENSG00000204271 |
| LMF1 | 0.262716431 | 1.199735545 | 0.206346131 | 0.398735975 | 114.8246378 | 137.8088193 | ENSG00000103227 |
| GNPTG | 0.26258317 | 1.199624732 | 0.015358648 | 0.055532848 | 397.4691511 | 476.860293 | ENSG00000090581 |
| C18orf54 | 0.262507072 | 1.199561456 | 0.056321899 | 0.154733644 | 376.9368274 | 452.1313505 | ENSG00000166845 |
| EBPL | 0.262385294 | 1.199460205 | 9.14E-05 | 0.00068653 | 1520.173285 | 1823.63118 | ENSG00000123179 |
| PPM1D | 0.262342722 | 1.199424811 | 0.016815654 | 0.059793566 | 411.1741291 | 492.9376311 | ENSG00000170836 |
| NA | 0.262309584 | 1.199397262 | 0.613148293 | 0.784656293 | 17.23264107 | 20.68692394 | ENSG00000172974 |
| SCIN | 0.262202518 | 1.199308255 | 0.706031249 | 0.844411002 | 9.668161872 | 11.55464594 | ENSG00000006747 |
| GXYLT1 | 0.262140746 | 1.199256905 | 0.043642484 | 0.126954453 | 356.8403117 | 428.1485992 | ENSG00000151233 |
| NA | 0.262111322 | 1.199232446 | 0.708382751 | 0.846207797 | 8.856227134 | 10.66077704 | ENSG00000259994 |
| SLC2A14 | 0.262081727 | 1.199207845 | 0.190197543 | 0.376548176 | 107.8318554 | 129.2398959 | ENSG00000173262 |
| USP32 | 0.261976617 | 1.199120479 | 0.000111904 | 0.000821657 | 1645.479774 | 1972.691147 | ENSG00000170832 |
| NA | 0.261954887 | 1.199102417 | 0.79306519 | 0.894207674 | 4.991687885 | 5.976510733 | ENSG00000227775 |
| ADARB1 | 0.261846471 | 1.199012311 | 0.025238158 | 0.083049767 | 336.535437 | 403.6423047 | ENSG00000197381 |
| LOC105377623 | 0.26173462 | 1.198919355 | 0.713867936 | 0.84906134 | 8.270851119 | 9.9011042 | ENSG00000250326 |
| LINC00472 | 0.261633816 | 1.198835588 | 0.746120454 | 0.867969455 | 7.212501516 | 8.684150363 | ENSG00000233237 |
| HSCB | 0.261603178 | 1.198810129 | 0.360338751 | 0.572516706 | 57.21717197 | 68.52502005 | ENSG00000100209 |
| NA | 0.261522605 | 1.198743178 | 0.519861006 | 0.715410267 | 27.32961447 | 32.79438305 | ENSG00000228175 |
| NA | 0.261500692 | 1.198724971 | 0.522483614 | 0.71692967 | 27.11640391 | 32.5077139 | ENSG00000203635 |
| TRIP6 | 0.261454347 | 1.198686464 | 4.73E-05 | 0.000382645 | 2906.220591 | 3484.574114 | ENSG00000087077 |
| NA | 0.261423889 | 1.198661157 | 0.438102957 | 0.646322502 | 41.5452196 | 49.71767756 | ENSG00000279483 |
| CDH26 | 0.26132147 | 1.198576066 | 0.518016666 | 0.713860533 | 25.50112792 | 30.5656111 | ENSG00000124215 |
| NA | 0.261227698 | 1.198498163 | 0.769765923 | 0.881687559 | 4.995247357 | 5.973389311 | ENSG00000286223 |
| MOB4 | 0.261174071 | 1.198453615 | 0.076668198 | 0.195738668 | 340.9780057 | 409.0571622 | ENSG00000115540 |
| ZKSCAN3 | 0.261110836 | 1.198401085 | 0.076995734 | 0.196282997 | 199.7862772 | 239.3918121 | ENSG00000189298 |
| NA | 0.261098224 | 1.19839061 | 0.80966792 | 0.90305216 | 3.330164905 | 3.981534386 | ENSG00000213169 |
| BOLA3 | 0.260334784 | 1.197756617 | 0.068875094 | 0.180398579 | 218.9221276 | 262.1731011 | ENSG00000163170 |

| | | | | | | | |
|---|---|---|---|---|---|---|---|
| TTPA | 0.260277835 | 1.197709338 | 0.765874318 | 0.878910348 | 5.526062229 | 6.664297958 | ENSG00000137561 |
| CFAP97 | 0.260243187 | 1.197680574 | 0.004329261 | 0.019606257 | 602.8156107 | 721.9954397 | ENSG00000164323 |
| FBXL22 | 0.260233974 | 1.197672925 | 0.746268492 | 0.867969455 | 6.660329809 | 7.958717845 | ENSG00000197361 |
| CLEC2D | 0.260216615 | 1.197658515 | 0.558835277 | 0.744312307 | 20.27131317 | 24.24927413 | ENSG00000069493 |
| ZNF524 | 0.260184085 | 1.19763151 | 0.41858911 | 0.629283182 | 45.19746416 | 54.20579356 | ENSG00000171443 |
| NA | 0.26015501 | 1.197607374 | 0.524126723 | 0.718239903 | 29.06941969 | 34.82482923 | ENSG00000256943 |
| KCNC3 | 0.260149248 | 1.197602591 | 0.102305294 | 0.242200761 | 186.3152823 | 223.1599207 | ENSG00000131398 |
| CTSL | 0.260067939 | 1.197535097 | 0.004332294 | 0.019615289 | 696.0270232 | 833.889116 | ENSG00000135047 |
| SPA17 | 0.259996486 | 1.197475788 | 0.715315324 | 0.849695403 | 8.605057588 | 10.30941108 | ENSG00000064199 |
| EMILIN2 | 0.259966675 | 1.197451044 | 0.078483889 | 0.198936301 | 207.828764 | 248.7885531 | ENSG00000132205 |
| PPP1R35 | 0.259948515 | 1.197435972 | 0.094189595 | 0.228252342 | 198.0221661 | 237.4477448 | ENSG00000160813 |
| PLCB4 | 0.259945672 | 1.197433611 | 0.091932817 | 0.223879494 | 228.5095912 | 273.299525 | ENSG00000101333 |
| TMC6 | 0.259828588 | 1.197336436 | 0.000986248 | 0.005546546 | 1405.254172 | 1683.443704 | ENSG00000141524 |
| USP24 | 0.259817416 | 1.197327164 | 0.002791261 | 0.013505755 | 1555.775628 | 1862.153445 | ENSG00000162402 |
| LINC01275 | 0.25964634 | 1.197185192 | 0.819146578 | 0.908070158 | 3.364564421 | 4.016058257 | ENSG00000237595 |
| ALKBH2 | 0.259644006 | 1.197183255 | 0.017708694 | 0.062323648 | 456.9062731 | 546.968126 | ENSG00000189046 |
| NA | 0.259582739 | 1.197132416 | 0.105691666 | 0.248690145 | 186.8641216 | 223.5945011 | ENSG00000250182 |
| MRAP2 | 0.259337399 | 1.196928853 | 0.237588403 | 0.439801704 | 96.53062018 | 115.4404718 | ENSG00000135324 |
| NA | 0.25928905 | 1.196888741 | 0.637920316 | 0.80145796 | 13.63470446 | 16.30455503 | ENSG00000272145 |
| SOX4 | 0.259100257 | 1.196732124 | 5.82E-07 | 7.02E-06 | 5506.989022 | 6589.554711 | ENSG00000124766 |
| LINC02453 | 0.259044118 | 1.196685558 | 0.817642803 | 0.907131026 | 3.564732297 | 4.285607242 | ENSG00000245017 |
| GFPT2 | 0.2590174 | 1.196663396 | 1.39E-05 | 0.000127465 | 2828.374897 | 3384.445024 | ENSG00000131459 |
| FBXO22 | 0.259013437 | 1.196660109 | 0.000269256 | 0.00179046 | 1526.109342 | 1826.202069 | ENSG00000167196 |
| PPP3CB-AS1 | 0.25895776 | 1.196613928 | 0.769778367 | 0.881687559 | 5.253535848 | 6.30110872 | ENSG00000221817 |
| HAS2 | 0.258786383 | 1.196471791 | 0.23552102 | 0.437389563 | 94.44476102 | 112.9876473 | ENSG00000170961 |
| CLCN5 | 0.258621128 | 1.196334748 | 0.033456801 | 0.103494134 | 336.492113 | 402.3194655 | ENSG00000171365 |
| SNORD19 | 0.258610492 | 1.196325928 | 0.811255005 | 0.904187369 | 3.629971857 | 4.358722365 | ENSG00000212493 |
| ANXA7 | 0.258244133 | 1.196022171 | 0.000497557 | 0.003062703 | 992.9862487 | 1187.470469 | ENSG00000138279 |
| NCOA3 | 0.258204262 | 1.195989117 | 0.017950927 | 0.063000047 | 512.048212 | 611.9925259 | ENSG00000124151 |
| FECH | 0.25819393 | 1.195980552 | 0.003898485 | 0.017978728 | 810.3890353 | 968.8737237 | ENSG00000066926 |
| EPHA7 | 0.258125553 | 1.195923869 | 0.115484865 | 0.265643322 | 162.0765107 | 193.6983276 | ENSG00000135333 |
| B4GALNT4 | 0.258012803 | 1.195830409 | 0.002021346 | 0.010256076 | 2153.995217 | 2576.620461 | ENSG00000182272 |
| DGCR11 | 0.257999551 | 1.195819424 | 0.500627212 | 0.698923996 | 37.59947518 | 45.08846025 | ENSG00000273311 |
| RPP14 | 0.257963936 | 1.195789905 | 0.138508143 | 0.301802964 | 204.448234 | 244.4211586 | ENSG00000163684 |
| PHF1 | 0.257934915 | 1.195765851 | 0.00645654 | 0.027406556 | 793.4838128 | 949.1684261 | ENSG00000112511 |
| TMEM179B | 0.257872761 | 1.195714336 | 0.436409922 | 0.644882738 | 44.63789418 | 53.51706037 | ENSG00000185475 |
| PSMD6-AS2 | 0.257835984 | 1.195683855 | 0.740330304 | 0.864046585 | 6.694729326 | 7.977067512 | ENSG00000239653 |
| MIR1180 | 0.257800612 | 1.195654539 | 0.838962047 | 0.919227098 | 3.064757469 | 3.630168424 | ENSG00000221540 |
| NA | 0.257572935 | 1.195465864 | 0.110362393 | 0.257100296 | 192.4181271 | 230.2542802 | ENSG00000189343 |
| CD109 | 0.257482051 | 1.195390556 | 0.207189226 | 0.399708096 | 118.7766115 | 141.7864019 | ENSG00000156535 |
| SDK1 | 0.257357619 | 1.195287459 | 0.030489419 | 0.096222038 | 413.7917196 | 494.0987782 | ENSG00000146555 |
| MICALL1 | 0.257310373 | 1.195248316 | 0.001259427 | 0.006848341 | 1111.786432 | 1329.232258 | ENSG00000100139 |
| MBNL1 | 0.257291451 | 1.195232639 | 0.184091845 | 0.368146783 | 132.3487455 | 158.2955256 | ENSG00000152601 |
| MARCHF7 | 0.256997739 | 1.194989332 | 0.000424653 | 0.002663424 | 1511.055768 | 1805.834454 | ENSG00000136536 |
| NA | 0.256855913 | 1.194871863 | 0.814882456 | 0.905897873 | 3.361004949 | 3.999884053 | ENSG00000261770 |
| SPOPL | 0.256651357 | 1.194702457 | 0.188870437 | 0.374823266 | 129.8907471 | 155.2447274 | ENSG00000144228 |
| LRRC4B | 0.256643415 | 1.19469588 | 0.024009032 | 0.079741572 | 630.3750891 | 753.6782168 | ENSG00000131409 |
| CAMKK2 | 0.256527441 | 1.194599846 | 0.000749607 | 0.004360439 | 1121.459497 | 1339.536234 | ENSG00000110931 |
| RFX1 | 0.256519667 | 1.194593409 | 0.020678207 | 0.070692965 | 523.0161555 | 625.274913 | ENSG00000132005 |
| PTENP1 | 0.256493768 | 1.194571964 | 0.768346024 | 0.880474191 | 5.260654793 | 6.294582329 | ENSG00000237984 |
| TMEM59L | 0.256451268 | 1.194536774 | 0.020171144 | 0.06927297 | 372.1779797 | 444.5244204 | ENSG00000105696 |
| B3GAT3 | 0.256331379 | 1.194437511 | 0.009117992 | 0.036258849 | 561.8175027 | 671.4533628 | ENSG00000149541 |
| PRPF40B | 0.256091112 | 1.194238606 | 0.094173172 | 0.228252342 | 190.9641441 | 228.2210915 | ENSG00000110844 |
| TSPAN12 | 0.255870556 | 1.194056047 | 0.039846763 | 0.118393034 | 329.1079188 | 393.2115395 | ENSG00000106025 |
| PIGB | 0.255830309 | 1.194022737 | 0.535008274 | 0.725887726 | 29.06141102 | 34.63832673 | ENSG00000069943 |
| ABR | 0.255741305 | 1.193949076 | 3.28E-05 | 0.000276232 | 1744.631517 | 2082.948226 | ENSG00000159842 |
| GGPS1 | 0.255727045 | 1.193937275 | 0.096647019 | 0.232270166 | 183.6140707 | 219.356187 | ENSG00000152904 |
| LRRC28 | 0.255537793 | 1.193780666 | 0.141404637 | 0.305978134 | 152.1441104 | 181.5290095 | ENSG00000168904 |
| AKR1E2 | 0.25553772 | 1.193780605 | 0.422858671 | 0.63308118 | 45.93940466 | 54.89518915 | ENSG00000165568 |
| BCAS4 | 0.255368295 | 1.19364042 | 0.022421717 | 0.075478452 | 362.5795066 | 432.8338544 | ENSG00000124243 |
| AKR1C2 | 0.255264135 | 1.193554244 | 0.782844848 | 0.889742909 | 5.50234113 | 6.600830648 | ENSG00000151632 |
| MTFR2 | 0.25522672 | 1.193523291 | 0.161980241 | 0.336745581 | 125.1935011 | 149.3878898 | ENSG00000146410 |
| KIRREL2 | 0.255070095 | 1.193393724 | 0.129974257 | 0.288865836 | 172.4296877 | 205.7252422 | ENSG00000126259 |
| WBP2 | 0.255030844 | 1.193361256 | 6.93E-05 | 0.000537272 | 1565.237523 | 1867.59167 | ENSG00000132471 |
| NA | 0.255018678 | 1.193351193 | 0.514238648 | 0.710572684 | 30.66333885 | 36.67246142 | ENSG00000257270 |
| NA | 0.254890026 | 1.19324478 | 0.789431563 | 0.892025074 | 4.99880683 | 5.944445872 | ENSG00000237017 |
| ACSS3 | 0.254697492 | 1.193085547 | 0.011558826 | 0.044080861 | 555.4183849 | 662.2390139 | ENSG00000111058 |
| ABHD3 | 0.254654372 | 1.193049888 | 0.204763075 | 0.396651008 | 117.0302977 | 139.7493567 | ENSG00000158201 |
| ESRP2 | 0.254630076 | 1.193029797 | 0.000707815 | 0.004149978 | 1205.097204 | 1438.461425 | ENSG00000103067 |
| KBTBD7 | 0.254495588 | 1.192918588 | 0.032432792 | 0.101038152 | 393.3861141 | 469.3336738 | ENSG00000120696 |
| CBWD1 | 0.254313783 | 1.192768269 | 0.051062711 | 0.143429756 | 263.1175572 | 313.7224475 | ENSG00000172785 |
| DNAJC9-AS1 | 0.254299076 | 1.192756109 | 0.675095177 | 0.825884072 | 12.05502319 | 14.34599447 | ENSG00000227540 |
| HAAO | 0.254286755 | 1.192745923 | 0.506299792 | 0.703618483 | 29.81611487 | 35.56426049 | ENSG00000162882 |
| MSRB2 | 0.254211937 | 1.192684069 | 0.070171743 | 0.183113634 | 231.0269834 | 275.5417287 | ENSG00000148450 |

| | | | | | | | |
|---|---|---|---|---|---|---|---|
| ROGDI | 0.254167018 | 1.192646935 | 0.080036259 | 0.2019057 | 257.523254 | 307.2647257 | ENSG00000067836 |
| PATJ | 0.253837676 | 1.192374705 | 0.003295762 | 0.015599115 | 1060.916227 | 1264.204941 | ENSG00000132849 |
| USP34 | 0.253517991 | 1.192110518 | 0.000896864 | 0.005092386 | 1339.352259 | 1595.8171 | ENSG00000115464 |
| RPS2 | 0.253500331 | 1.192095925 | 9.01E-09 | 1.50E-07 | 57349.44606 | 68366.9414 | ENSG00000140988 |
| CPT1A | 0.253371229 | 1.191989254 | 0.195192764 | 0.38402279 | 137.0107029 | 163.1450342 | ENSG00000110090 |
| IMP4 | 0.253191844 | 1.19184105 | 3.89E-05 | 0.000321737 | 2060.03601 | 2455.944142 | ENSG00000136718 |
| GTF2IRD2 | 0.253059518 | 1.191731738 | 0.544204021 | 0.73258071 | 31.08531076 | 37.10971174 | ENSG00000196275 |
| SNTG2 | 0.252962544 | 1.191651636 | 0.095922762 | 0.230999917 | 199.8099983 | 238.1779071 | ENSG00000172554 |
| RMND5B | 0.252912995 | 1.191610709 | 0.000222908 | 0.001513159 | 1630.532803 | 1942.430917 | ENSG00000145916 |
| KLRG2 | 0.252905682 | 1.19160467 | 0.018891849 | 0.065535004 | 476.1367803 | 567.7126103 | ENSG00000188883 |
| CAMK2G | 0.252752688 | 1.19147831 | 0.002728605 | 0.013243306 | 800.8469295 | 953.983325 | ENSG00000148660 |
| LANCL1 | 0.252650547 | 1.191393957 | 0.001592588 | 0.008380055 | 938.5002387 | 1117.714737 | ENSG00000115365 |
| SLC24A2 | 0.252610182 | 1.191360624 | 0.245190631 | 0.44954071 | 100.7507489 | 119.9234588 | ENSG00000155886 |
| AGTRAP | 0.252309256 | 1.191112149 | 0.000496696 | 0.003059398 | 1125.386074 | 1340.293847 | ENSG00000177674 |
| ASIC1 | 0.252297137 | 1.191102143 | 0.139478088 | 0.303318576 | 170.3930501 | 203.162508 | ENSG00000110881 |
| ANKRD35 | 0.25224026 | 1.191055185 | 0.127895335 | 0.285473796 | 178.1244412 | 211.9488849 | ENSG00000198483 |
| GTF2H5 | 0.252142024 | 1.190974088 | 0.021615458 | 0.07324743 | 411.9341719 | 490.7824687 | ENSG00000272047 |
| CLN5 | 0.251881648 | 1.190759161 | 0.644343946 | 0.805989478 | 14.23075889 | 16.95223795 | ENSG00000102805 |
| GTF2IRD2B | 0.25179093 | 1.190684288 | 0.313877473 | 0.524285873 | 69.75765277 | 83.1243893 | ENSG00000174428 |
| BBS4 | 0.251678962 | 1.190591881 | 0.010658428 | 0.041120411 | 631.9510824 | 751.9947763 | ENSG00000140463 |
| TMSB15B-AS1 | 0.25164018 | 1.190559877 | 0.80912892 | 0.902949338 | 3.629971857 | 4.330724884 | ENSG00000231728 |
| TDRD5 | 0.251469445 | 1.190418989 | 0.695148338 | 0.837250559 | 10.37214338 | 12.31214332 | ENSG00000162782 |
| CHD5 | 0.251300311 | 1.190279438 | 0.298713706 | 0.509042441 | 73.16254044 | 87.11528795 | ENSG00000116254 |
| KLF16 | 0.251252227 | 1.190239768 | 0.002305538 | 0.011476035 | 841.9915852 | 1002.587766 | ENSG00000129911 |
| PROCA1 | 0.25109161 | 1.190107265 | 0.094755343 | 0.229196202 | 237.2810276 | 282.4233121 | ENSG00000167525 |
| NA | 0.251023464 | 1.190051051 | 0.548926143 | 0.736359583 | 24.51274641 | 29.14397304 | ENSG00000259363 |
| ARHGEF16 | 0.250992052 | 1.190025141 | 0.005815406 | 0.025056795 | 711.0940744 | 846.6948608 | ENSG00000130762 |
| MRPL23 | 0.250814458 | 1.189878659 | 0.08263368 | 0.206810608 | 218.4562468 | 260.0888784 | ENSG00000214026 |
| ZFAND6 | 0.25064021 | 1.189734954 | 0.000144251 | 0.001031496 | 2204.887961 | 2622.352127 | ENSG00000086666 |
| NEXMIF | 0.250463423 | 1.189589174 | 0.657452739 | 0.81507221 | 13.65130661 | 16.18726822 | ENSG00000050030 |
| SH3GLB1 | 0.25046048 | 1.189586747 | 0.005717966 | 0.024704595 | 820.4425088 | 976.238839 | ENSG00000097033 |
| INTS4 | 0.250458812 | 1.189585372 | 0.010021542 | 0.039126903 | 565.0255032 | 672.1930776 | ENSG00000149262 |
| NNAT | 0.250377846 | 1.189518612 | 0.021392716 | 0.072662131 | 403.3685478 | 479.7812757 | ENSG00000053438 |
| SGTB | 0.250309926 | 1.189462613 | 0.069330195 | 0.181440614 | 245.8928986 | 292.3811034 | ENSG00000197860 |
| NA | 0.250244585 | 1.189408743 | 0.566046067 | 0.750054727 | 22.99238096 | 27.31452257 | ENSG00000268034 |
| FAM135A | 0.250223619 | 1.189391457 | 0.055331363 | 0.15261177 | 529.0822894 | 629.3819915 | ENSG00000082269 |
| SPARC | 0.250107486 | 1.189295719 | 2.06E-06 | 2.25E-05 | 4060.589058 | 4828.404374 | ENSG00000113140 |
| NTAN1 | 0.249984367 | 1.189194229 | 0.054400131 | 0.150460818 | 289.0638195 | 343.7917872 | ENSG00000157045 |
| FAM53A | 0.249839794 | 1.189075066 | 0.384487038 | 0.59678127 | 50.22828009 | 59.75687725 | ENSG00000174137 |
| CBFA2T3 | 0.249765903 | 1.189014165 | 0.495711653 | 0.695207823 | 33.5694469 | 39.87351428 | ENSG00000129993 |
| TUBB2B | 0.249762521 | 1.189011378 | 3.81E-06 | 3.95E-05 | 10569.24829 | 12566.75496 | ENSG00000137285 |
| SKA3 | 0.249621983 | 1.188895558 | 0.001419977 | 0.007605294 | 961.3853024 | 1143.234353 | ENSG00000165480 |
| SGCB | 0.249533676 | 1.188822788 | 0.025354977 | 0.083370606 | 653.6185873 | 777.6449095 | ENSG00000163069 |
| MNT | 0.249533666 | 1.18882278 | 0.089472255 | 0.21943942 | 262.8889652 | 312.8940534 | ENSG00000070444 |
| FAM86C1 | 0.249509156 | 1.188802583 | 0.211187521 | 0.404860048 | 106.5730324 | 126.6481027 | ENSG00000158483 |
| LINC00894 | 0.249372301 | 1.188689818 | 0.461040473 | 0.665806327 | 63.2423193 | 75.17089571 | ENSG00000235703 |
| TEX264 | 0.249292263 | 1.188623873 | 0.096569316 | 0.232231189 | 213.2801296 | 253.5210585 | ENSG00000164081 |
| LINC01529 | 0.249277626 | 1.188611815 | 0.546399283 | 0.734378368 | 30.85193858 | 36.68712257 | ENSG00000225872 |
| NIPA1 | 0.249235738 | 1.188577304 | 0.023363806 | 0.078050447 | 443.570206 | 527.4999322 | ENSG00000170113 |
| LINC01607 | 0.249022646 | 1.18840176 | 0.842011014 | 0.921498493 | 3.095597513 | 3.634519351 | ENSG00000272138 |
| CHST15 | 0.248955388 | 1.188346357 | 0.035463987 | 0.10830117 | 354.2194417 | 421.0853063 | ENSG00000182022 |
| NA | 0.248831798 | 1.188244561 | 0.398876612 | 0.610475104 | 46.92183629 | 55.79028756 | ENSG00000245248 |
| SNRNP35 | 0.248791367 | 1.188211262 | 0.069950589 | 0.182662732 | 260.2998511 | 309.482525 | ENSG00000184209 |
| KIAA0232 | 0.248752514 | 1.188179262 | 0.009102526 | 0.036235188 | 664.00764 | 788.4877225 | ENSG00000170871 |
| VCPIP1 | 0.24871546 | 1.188148746 | 0.042999996 | 0.125491412 | 318.6376114 | 378.2690117 | ENSG00000175073 |
| NLGN3 | 0.248705751 | 1.18814075 | 0.036883165 | 0.111571133 | 374.1061571 | 444.7807057 | ENSG00000196338 |
| FSTL3 | 0.248693714 | 1.188130836 | 0.161480329 | 0.336059387 | 214.8286392 | 255.4615959 | ENSG00000070404 |
| PCDHA13 | 0.248576467 | 1.188034282 | 0.809894699 | 0.90313851 | 4.822360053 | 5.675842845 | ENSG00000239389 |
| SLC13A5 | 0.248569607 | 1.188028633 | 0.520228137 | 0.715550501 | 28.59672543 | 33.94647212 | ENSG00000141485 |
| C9orf72 | 0.248497312 | 1.187969101 | 0.111576967 | 0.259211591 | 253.5522878 | 301.5061926 | ENSG00000147894 |
| SNHG32 | 0.248490931 | 1.187963847 | 0.0073529 | 0.030471696 | 608.6793872 | 722.9240863 | ENSG00000204387 |
| TPSG1 | 0.248140114 | 1.187675007 | 0.464992563 | 0.669294471 | 40.06014339 | 47.60808433 | ENSG00000116176 |
| BNC2-AS1 | 0.24809628 | 1.187638922 | 0.602236964 | 0.777179518 | 20.46199784 | 24.3525622 | ENSG00000234779 |
| ZFAND1 | 0.248080325 | 1.187625788 | 0.042511935 | 0.124432407 | 388.6894 | 461.6785571 | ENSG00000104231 |
| TMTC4 | 0.248000121 | 1.187559766 | 0.003062543 | 0.014656898 | 824.0279102 | 978.1270926 | ENSG00000125247 |
| PCDH18 | 0.24790919 | 1.187484917 | 0.00805415 | 0.032846336 | 850.2332728 | 1008.985056 | ENSG00000189184 |
| CHST2 | 0.247748593 | 1.187352737 | 0.006945168 | 0.029081929 | 839.4487698 | 997.0668642 | ENSG00000175040 |
| ZNF267 | 0.247728643 | 1.187336319 | 0.06684313 | 0.176352684 | 242.4073644 | 287.7967201 | ENSG00000185947 |
| AXL | 0.247602592 | 1.187232583 | 0.004726694 | 0.021082574 | 676.3426129 | 803.2341 | ENSG00000167601 |
| RASL12 | 0.247571057 | 1.187206632 | 0.050309419 | 0.141804981 | 296.5025481 | 352.0030065 | ENSG00000103710 |
| ZBTB3 | 0.247438007 | 1.18709715 | 0.014915444 | 0.05427325 | 472.1805589 | 560.6267847 | ENSG00000185670 |
| CTNND1 | 0.247430744 | 1.187091173 | 0.225242536 | 0.423586423 | 107.5581338 | 127.5938265 | ENSG00000198561 |
| PLXNB2 | 0.247398579 | 1.187064707 | 6.96E-05 | 0.000538531 | 3763.757191 | 4468.632744 | ENSG00000196576 |
| FTX | 0.247390402 | 1.187057979 | 0.150502118 | 0.319675233 | 152.7111304 | 181.4253426 | ENSG00000230590 |

| | | | | | | | |
|---|---|---|---|---|---|---|---|
| ZNF75A | 0.2473283 | 1.187006882 | 0.107863326 | 0.252572711 | 184.6558181 | 219.280234 | ENSG00000162086 |
| LOC653160 | 0.247306481 | 1.186988931 | 0.779544254 | 0.887571291 | 5.298613782 | 6.280583589 | ENSG00000241014 |
| LIN7C | 0.247279652 | 1.186966857 | 0.062380972 | 0.167148788 | 303.7022862 | 360.3377555 | ENSG00000148943 |
| AARS1 | 0.247248429 | 1.186941169 | 2.92E-07 | 3.74E-06 | 11083.40333 | 13154.25943 | ENSG00000090861 |
| STK17B | 0.24711157 | 1.186828577 | 0.167014677 | 0.343693166 | 153.8362203 | 182.5010795 | ENSG00000081320 |
| SSR4 | 0.247007277 | 1.186742784 | 0.001193756 | 0.006543226 | 1523.092079 | 1807.551228 | ENSG00000180879 |
| TEFM | 0.246939666 | 1.186687169 | 0.202996042 | 0.394401261 | 114.6458006 | 136.1308007 | ENSG00000172171 |
| STK11 | 0.246880716 | 1.18663868 | 0.000347491 | 0.002230578 | 2252.728802 | 2673.285631 | ENSG00000118046 |
| LUZP1 | 0.246802691 | 1.186574505 | 0.000191471 | 0.001327435 | 1734.074971 | 2057.142683 | ENSG00000169641 |
| MTMR3 | 0.246441722 | 1.186277656 | 0.330180287 | 0.541601209 | 68.10917247 | 80.75674386 | ENSG00000100330 |
| NA | 0.246425447 | 1.186264274 | 0.827529637 | 0.913263802 | 3.364564421 | 3.965360182 | ENSG00000273291 |
| NA | 0.246422865 | 1.186262151 | 0.727557186 | 0.856710154 | 9.255673162 | 10.99407688 | ENSG00000284655 |
| SLITRK1 | 0.246379968 | 1.186226879 | 0.780268508 | 0.88783339 | 5.622141833 | 6.682647625 | ENSG00000178235 |
| MPND | 0.246329228 | 1.18618516 | 0.086608275 | 0.21413947 | 294.6864676 | 349.3942314 | ENSG00000008382 |
| ALKAL1 | 0.246236432 | 1.186108866 | 0.762495277 | 0.877069272 | 7.348904373 | 8.714323307 | ENSG00000196711 |
| KANSL1L | 0.246195788 | 1.18607545 | 0.143776088 | 0.309708159 | 165.7711373 | 196.8225245 | ENSG00000144445 |
| SCN11A | 0.24611973 | 1.186012923 | 0.785582339 | 0.890476216 | 5.904151422 | 7.036189051 | ENSG00000168356 |
| WSCD1 | 0.24609003 | 1.185988507 | 0.000231195 | 0.001564915 | 1867.140968 | 2213.694243 | ENSG00000179314 |
| ZNF623 | 0.246084978 | 1.185984355 | 0.007771344 | 0.031932511 | 647.5142174 | 767.5756322 | ENSG00000183309 |
| SNX19 | 0.246032186 | 1.185940957 | 0.004559443 | 0.020452514 | 689.2175539 | 817.1392283 | ENSG00000120451 |
| SLC8B1 | 0.246030273 | 1.185939385 | 0.047151766 | 0.134775404 | 358.3336503 | 424.9805275 | ENSG00000089060 |
| BTK | 0.245880817 | 1.185816534 | 0.780999534 | 0.888429637 | 5.546223855 | 6.594304258 | ENSG00000010671 |
| CACNA1H | 0.245759518 | 1.185716836 | 0.00060655 | 0.003634106 | 1922.298828 | 2279.333797 | ENSG00000196557 |
| FUOM | 0.24570055 | 1.185668373 | 0.062064873 | 0.166514438 | 267.8888894 | 317.3844699 | ENSG00000148803 |
| NA | 0.245582671 | 1.185571499 | 0.715062988 | 0.849670014 | 12.44735028 | 14.72318245 | ENSG00000262580 |
| ATAD5 | 0.245428791 | 1.185445051 | 0.056264889 | 0.154599509 | 330.8720043 | 391.8082511 | ENSG00000176208 |
| NA | 0.245282006 | 1.185324446 | 0.150140533 | 0.319162772 | 302.234728 | 358.5374726 | ENSG00000234072 |
| ALG6 | 0.24521961 | 1.185273182 | 0.05463467 | 0.151087384 | 399.947896 | 473.6516818 | ENSG00000088035 |
| PLEC | 0.245097265 | 1.185172672 | 0.000374743 | 0.00238605 | 4785.509843 | 5671.717464 | ENSG00000178209 |
| THAP8 | 0.244972215 | 1.185069947 | 0.19183432 | 0.379032342 | 130.2035967 | 154.5311611 | ENSG00000161277 |
| NA | 0.244967328 | 1.185065933 | 0.394268441 | 0.60593746 | 49.24109379 | 58.36541214 | ENSG00000265415 |
| MTUS1 | 0.244913776 | 1.185021945 | 0.320888638 | 0.53143751 | 88.27550764 | 104.3984394 | ENSG00000129422 |
| MATK | 0.244575981 | 1.184744515 | 0.00715867 | 0.029856776 | 670.2590394 | 794.5118528 | ENSG00000007264 |
| ASB13 | 0.244501553 | 1.184683396 | 0.007318117 | 0.03035235 | 637.5386227 | 755.50154 | ENSG00000196372 |
| MOB1B | 0.244474941 | 1.184661543 | 0.10662469 | 0.250386381 | 300.1091037 | 355.72405 | ENSG00000173542 |
| CLMN | 0.244416701 | 1.18461372 | 0.24462649 | 0.448950732 | 113.7647619 | 134.5199902 | ENSG00000165959 |
| MMP16 | 0.244355007 | 1.184563064 | 0.046033223 | 0.132158319 | 438.9123412 | 519.6975736 | ENSG00000156103 |
| POLE2 | 0.244307296 | 1.18452389 | 0.094779691 | 0.229196202 | 263.4692299 | 311.6022176 | ENSG00000100479 |
| NA | 0.244300475 | 1.18451829 | 0.605726535 | 0.779795526 | 19.06112761 | 22.65078139 | ENSG00000198134 |
| IGFBP5 | 0.244227784 | 1.184458609 | 0.681705826 | 0.829832056 | 23.55431521 | 27.86622182 | ENSG00000115461 |
| KLHL42 | 0.244227304 | 1.184458215 | 0.041004325 | 0.121013701 | 353.8125974 | 419.3319768 | ENSG00000087448 |
| PMM2 | 0.244191316 | 1.184428669 | 0.044347964 | 0.128275587 | 296.1437308 | 350.7510773 | ENSG00000140650 |
| LOC100506688 | 0.24402927 | 1.184295639 | 0.832530896 | 0.91561881 | 3.629971857 | 4.298376477 | ENSG00000215246 |
| STOX2 | 0.243948874 | 1.184229645 | 0.015666632 | 0.056471916 | 818.2305157 | 968.8722176 | ENSG00000173320 |
| NA | 0.243873035 | 1.184167394 | 0.599022097 | 0.77503937 | 27.33553821 | 32.45873979 | ENSG00000248121 |
| TMEM259 | 0.243819755 | 1.184123663 | 0.131829012 | 0.291931389 | 2554.109648 | 3024.601259 | ENSG00000182087 |
| KATNAL2 | 0.243785648 | 1.184095669 | 0.198239202 | 0.388274977 | 134.5502878 | 159.1936728 | ENSG00000167216 |
| ACOX2 | 0.243616236 | 1.183956632 | 0.618549174 | 0.788371543 | 17.99357413 | 21.26272125 | ENSG00000168306 |
| KCNG1 | 0.243399879 | 1.18377909 | 0.012197578 | 0.046153328 | 643.8214475 | 762.6027321 | ENSG00000026559 |
| HDHD3 | 0.243141729 | 1.183567289 | 0.100692627 | 0.239192486 | 243.5780909 | 288.0420984 | ENSG00000119431 |
| MECR | 0.243127249 | 1.183555409 | 0.007814887 | 0.032068576 | 568.0472415 | 672.336427 | ENSG00000116353 |
| NA | 0.243100801 | 1.183533712 | 0.814933325 | 0.905897873 | 3.919100392 | 4.662795221 | ENSG00000267778 |
| LRRC58 | 0.243056785 | 1.183497604 | 0.066474208 | 0.175673862 | 454.7251712 | 538.2119611 | ENSG00000163428 |
| HSD11B2 | 0.242968071 | 1.18342483 | 0.00811339 | 0.033028965 | 662.1955013 | 784.1802356 | ENSG00000176387 |
| KIAA0556 | 0.242906955 | 1.183374699 | 0.058711422 | 0.159695407 | 317.0063437 | 374.9725946 | ENSG00000047578 |
| PEX16 | 0.242880446 | 1.183352955 | 0.019526053 | 0.067376122 | 461.8051615 | 546.4554695 | ENSG00000121680 |
| ADAMTS20 | 0.242866451 | 1.183341476 | 0.184857427 | 0.369108957 | 140.358639 | 165.8092372 | ENSG00000173157 |
| BRWD1 | 0.242847092 | 1.183325597 | 0.032235368 | 0.100522609 | 639.0349104 | 755.6867271 | ENSG00000185658 |
| ALDH1B1 | 0.242811212 | 1.183296168 | 0.000126387 | 0.000915896 | 3085.18529 | 3649.894891 | ENSG00000137124 |
| PUS1 | 0.242639543 | 1.183155374 | 0.000375371 | 0.002389245 | 1944.166065 | 2301.112072 | ENSG00000177192 |
| SNRNP48 | 0.242602782 | 1.183125227 | 0.060256096 | 0.162911435 | 511.4483176 | 605.4306857 | ENSG00000168566 |
| TERF2 | 0.2422842 | 1.182863993 | 0.006019597 | 0.02580706 | 673.2084721 | 796.5535755 | ENSG00000132604 |
| AEBP2 | 0.242234732 | 1.182823434 | 0.005812882 | 0.025056795 | 816.4681323 | 966.1009036 | ENSG00000139154 |
| NFKBID | 0.242170811 | 1.182771029 | 0.3104527 | 0.521021071 | 87.93181797 | 103.7974825 | ENSG00000167604 |
| ABI3BP | 0.241895105 | 1.182545017 | 0.819735651 | 0.908474441 | 3.650133483 | 4.311429258 | ENSG00000154175 |
| ICOSLG | 0.241842557 | 1.182501946 | 0.425153346 | 0.635106861 | 44.10379917 | 52.11906887 | ENSG00000160223 |
| FRMD3 | 0.241837212 | 1.182497564 | 0.487680061 | 0.6884316 | 36.45005383 | 43.22571541 | ENSG00000172159 |
| NA | 0.241587853 | 1.182293196 | 0.087143881 | 0.215153903 | 225.7236149 | 266.8606044 | ENSG00000224578 |
| ABHD18 | 0.241562338 | 1.182272287 | 0.087277773 | 0.215456307 | 231.024593 | 273.1047722 | ENSG00000164074 |
| FAM72D | 0.241515195 | 1.182233654 | 0.263858022 | 0.471246336 | 93.20968531 | 110.2538068 | ENSG00000215784 |
| FAM162A | 0.241358208 | 1.182105016 | 0.005717366 | 0.024704595 | 752.5081795 | 889.1300907 | ENSG00000114023 |
| PPCS | 0.241311653 | 1.182066871 | 0.045685181 | 0.131298835 | 354.0058739 | 418.1945488 | ENSG00000127125 |
| C1orf112 | 0.241200551 | 1.181975844 | 0.029809238 | 0.094430262 | 380.6309482 | 449.9601092 | ENSG00000000460 |
| LEXM | 0.241124218 | 1.181913307 | 0.783943204 | 0.890065126 | 5.322334881 | 6.268760312 | ENSG00000162398 |

| | | | | | | | |
|---|---|---|---|---|---|---|---|
| FANCE | 0.241093138 | 1.181887845 | 0.014809674 | 0.053981992 | 490.9605405 | 580.3814891 | ENSG00000112039 |
| LRR1 | 0.24108837 | 1.181883939 | 0.158158611 | 0.330950179 | 162.0534 | 191.6208719 | ENSG00000165501 |
| DAPK3 | 0.24108113 | 1.181878008 | 0.000493175 | 0.003042666 | 2057.593645 | 2432.548256 | ENSG00000167657 |
| AZIN2 | 0.241051497 | 1.181853733 | 0.604108715 | 0.778603606 | 18.56471226 | 21.93622619 | ENSG00000142920 |
| BACH1 | 0.24092098 | 1.181746818 | 0.024456319 | 0.080956632 | 607.8390028 | 717.9520963 | ENSG00000156273 |
| SOWAHC | 0.240880026 | 1.181713272 | 0.018944094 | 0.065692107 | 468.142853 | 553.1289934 | ENSG00000198142 |
| NA | 0.240872755 | 1.181707316 | 0.474096158 | 0.677209204 | 38.69130842 | 45.70407832 | ENSG00000278768 |
| LPIN2 | 0.240696065 | 1.181562599 | 0.04698575 | 0.134361884 | 365.253717 | 431.2750442 | ENSG00000101577 |
| RELL1 | 0.240585136 | 1.181471752 | 0.312228831 | 0.522574273 | 74.31313084 | 87.814404 | ENSG00000181826 |
| ASMT | 0.240489909 | 1.18139377 | 0.698437478 | 0.839388378 | 13.1901805 | 15.59624267 | ENSG00000196433 |
| SCML2 | 0.240465351 | 1.18137366 | 0.010105768 | 0.039390595 | 635.6709302 | 750.7201332 | ENSG00000102098 |
| NA | 0.240440219 | 1.181353081 | 0.813362374 | 0.905307269 | 3.919100392 | 4.651917903 | ENSG00000284731 |
| SIPA1 | 0.240334937 | 1.181266873 | 0.003660208 | 0.017029569 | 1389.588928 | 1642.175708 | ENSG00000213445 |
| TBX22 | 0.240321386 | 1.181255778 | 0.71537696 | 0.849695403 | 9.279394261 | 10.95737755 | ENSG00000122145 |
| CA9 | 0.240172954 | 1.18113425 | 0.761926879 | 0.87673587 | 7.338225955 | 8.653031461 | ENSG00000107159 |
| SPP1 | 0.240058063 | 1.181040193 | 0.070662432 | 0.184215907 | 288.1831118 | 340.4619078 | ENSG00000118785 |
| RAD23A | 0.240040125 | 1.181025509 | 0.000621466 | 0.003698827 | 1283.70038 | 1516.257745 | ENSG00000179262 |
| ENSA | 0.240021988 | 1.181010661 | 3.34E-07 | 4.24E-06 | 5170.546086 | 6106.090869 | ENSG00000143420 |
| RTEL1-TNFRSF6B | 0.240021418 | 1.181010194 | 0.453266766 | 0.659471716 | 50.15948106 | 59.34376794 | ENSG00000026036 |
| NA | 0.239944094 | 1.180946897 | 0.172418319 | 0.351936874 | 173.0049957 | 204.4241803 | ENSG00000254635 |
| HELZ | 0.239893557 | 1.18090553 | 0.051950295 | 0.145410007 | 416.5685432 | 491.6242288 | ENSG00000198265 |
| DNAH17 | 0.239709415 | 1.180754812 | 0.398484685 | 0.610274365 | 52.23112849 | 61.67627946 | ENSG00000187775 |
| NA | 0.239380103 | 1.180485322 | 0.66947695 | 0.82295975 | 12.98052941 | 15.30427664 | ENSG00000267750 |
| EAF2 | 0.239346762 | 1.180458041 | 0.329409767 | 0.541082909 | 82.12969546 | 97.11634788 | ENSG00000145088 |
| BACH2 | 0.239279883 | 1.18040332 | 0.386491017 | 0.598661652 | 55.31813251 | 65.4100195 | ENSG00000112182 |
| APH1A | 0.239222464 | 1.18035634 | 0.00010558 | 0.000780081 | 1873.049595 | 2210.334749 | ENSG00000117362 |
| TSPAN32 | 0.239159868 | 1.180305128 | 0.796425831 | 0.896014644 | 5.387574441 | 6.333457127 | ENSG00000064201 |
| SCTR | 0.239133716 | 1.180283733 | 0.728951078 | 0.857335811 | 9.606481784 | 11.3529152 | ENSG00000080293 |
| HSPB1 | 0.2389228 | 1.180111193 | 5.27E-05 | 0.000421309 | 2687.274082 | 3171.625338 | ENSG00000106211 |
| CLPX | 0.238610858 | 1.179856055 | 0.001278475 | 0.006933947 | 1158.216224 | 1366.553682 | ENSG00000166855 |
| TMPRSS2 | 0.238392202 | 1.179677249 | 0.086175028 | 0.213459524 | 234.2219145 | 276.3962713 | ENSG00000184012 |
| NA | 0.23822021 | 1.179536621 | 0.578065669 | 0.759450456 | 25.42668448 | 29.90269993 | ENSG00000231167 |
| FAT2 | 0.23821194 | 1.179529859 | 0.43514219 | 0.643815439 | 51.84947982 | 61.1251473 | ENSG00000086570 |
| NA | 0.238134936 | 1.179466903 | 0.409011558 | 0.620117475 | 66.47198098 | 78.59399354 | ENSG00000259959 |
| ZNF574 | 0.238039881 | 1.179389194 | 0.001556136 | 0.008213423 | 980.3022717 | 1156.333979 | ENSG00000105732 |
| MEIOC | 0.238020398 | 1.179373267 | 0.273111605 | 0.481840226 | 103.3451502 | 121.714529 | ENSG00000180336 |
| HOMER3 | 0.238010748 | 1.179365378 | 0.054819102 | 0.15150867 | 378.2502638 | 446.2623764 | ENSG00000051128 |
| REV3L | 0.237739427 | 1.179143601 | 0.317767786 | 0.527796362 | 93.07119752 | 109.6778939 | ENSG00000009413 |
| NA | 0.237720392 | 1.179128044 | 0.142538703 | 0.307767153 | 197.627423 | 232.9718512 | ENSG00000233476 |
| DSN1 | 0.23769714 | 1.17910904 | 0.018286042 | 0.063879375 | 500.7621567 | 590.279093 | ENSG00000149636 |
| TBC1D9B | 0.237678027 | 1.179093419 | 0.000190587 | 0.00132199 | 1672.235301 | 1971.688234 | ENSG00000197226 |
| AP4E1 | 0.237657045 | 1.179076271 | 0.153838532 | 0.32486198 | 219.9929351 | 259.4304431 | ENSG00000081014 |
| ELL | 0.237614048 | 1.179041131 | 0.020020166 | 0.068842103 | 489.7654826 | 577.7397758 | ENSG00000105656 |
| SLC7A7 | 0.237346415 | 1.178822429 | 0.269182465 | 0.477280147 | 100.3649559 | 118.5175435 | ENSG00000155465 |
| STEAP3 | 0.237309583 | 1.178792334 | 0.014117283 | 0.051858567 | 603.8031287 | 711.670705 | ENSG00000115107 |
| SUPT5H | 0.237275837 | 1.178764761 | 1.73E-06 | 1.91E-05 | 4925.305379 | 5806.218372 | ENSG00000196235 |
| OSGIN2 | 0.237166494 | 1.178675425 | 0.097002417 | 0.232887191 | 242.3706268 | 285.731961 | ENSG00000164823 |
| TPGS1 | 0.237156175 | 1.178666994 | 0.403018997 | 0.61430727 | 53.59284451 | 63.21643382 | ENSG00000141933 |
| POT1 | 0.237097302 | 1.178618897 | 0.07413943 | 0.191143576 | 306.9387102 | 361.3068423 | ENSG00000128513 |
| NA | 0.237055022 | 1.178584356 | 0.706930884 | 0.845211019 | 15.91836724 | 18.78993878 | ENSG00000271870 |
| NA | 0.23700549 | 1.178543893 | 0.81598001 | 0.906477444 | 3.926219337 | 4.644445553 | ENSG00000270890 |
| TFRC | 0.236921387 | 1.17847519 | 0.000351169 | 0.002251128 | 3168.711563 | 3733.453396 | ENSG00000072274 |
| ANO7 | 0.236757034 | 1.178340945 | 0.360230037 | 0.572445662 | 65.63126555 | 77.28179108 | ENSG00000146205 |
| AKR7A3 | 0.236541084 | 1.178164578 | 0.775895496 | 0.884990423 | 5.934991466 | 7.013488457 | ENSG00000162482 |
| NA | 0.236523567 | 1.178150273 | 0.780752159 | 0.888276674 | 5.673143503 | 6.676121235 | ENSG00000286162 |
| NA | 0.236475492 | 1.178111015 | 0.316692578 | 0.526751746 | 88.02670236 | 103.4635202 | ENSG00000215492 |
| NID1 | 0.236470136 | 1.178106641 | 0.004790668 | 0.021289151 | 1295.38563 | 1525.51172 | ENSG00000116962 |
| MAFK | 0.236434956 | 1.178077913 | 0.006841694 | 0.028731463 | 653.984804 | 770.6113903 | ENSG00000198517 |
| NA | 0.236260767 | 1.177935682 | 0.723665225 | 0.854472967 | 9.870694012 | 11.65887997 | ENSG00000257718 |
| THAP5 | 0.236199168 | 1.177885388 | 0.208371871 | 0.40125449 | 141.4573374 | 166.7320496 | ENSG00000177683 |
| COA6 | 0.236156338 | 1.177850421 | 0.043732423 | 0.12711818 | 371.6679369 | 437.8824442 | ENSG00000168275 |
| NA | 0.236084358 | 1.177791656 | 0.666132677 | 0.821155848 | 13.80759177 | 16.31202738 | ENSG00000280294 |
| MFSD3 | 0.235937764 | 1.177671985 | 0.00228276 | 0.011371649 | 1262.619749 | 1487.632911 | ENSG00000167700 |
| IQCH-AS1 | 0.235931842 | 1.177667151 | 0.74613357 | 0.867969455 | 8.185449933 | 9.689725638 | ENSG00000259673 |
| INTS4P2 | 0.235924962 | 1.177661535 | 0.814664708 | 0.905846657 | 3.949940435 | 4.665916643 | ENSG00000273024 |
| NA | 0.235792842 | 1.177553692 | 0.826155389 | 0.912403332 | 3.926219337 | 4.636973204 | ENSG00000279970 |
| SH3RF1 | 0.235746767 | 1.177516085 | 0.129351718 | 0.287965011 | 258.5531801 | 304.3116648 | ENSG00000154447 |
| ALDOC | 0.235721131 | 1.177495161 | 0.005277021 | 0.023127532 | 866.6634618 | 1020.431436 | ENSG00000109107 |
| MRPL28 | 0.235708455 | 1.177484815 | 0.000133603 | 0.00096301 | 2008.392798 | 2364.797525 | ENSG00000086504 |
| KIAA1211L | 0.235565023 | 1.177367756 | 0.764569082 | 0.87816748 | 7.338225955 | 8.615102621 | ENSG00000196872 |
| S1PR2 | 0.235564168 | 1.177367058 | 0.075840449 | 0.194324331 | 253.9538193 | 299.0729473 | ENSG00000267534 |
| ETFBKMT | 0.235487407 | 1.177304416 | 0.687424759 | 0.832702766 | 15.60699214 | 18.25658919 | ENSG00000139160 |
| METTL18 | 0.235444755 | 1.177269611 | 0.320646736 | 0.531150323 | 76.83492048 | 90.53452932 | ENSG00000171806 |
| NA | 0.235369376 | 1.177208102 | 0.423170828 | 0.633337985 | 54.61946435 | 64.21860413 | ENSG00000279407 |

| | | | | | | | |
|---|---|---|---|---|---|---|---|
| BHMT | 0.23535816 | 1.17719895 | 0.60009591 | 0.776007344 | 25.33508023 | 29.87158103 | ENSG00000145692 |
| GEN1 | 0.235319285 | 1.177167229 | 0.129838879 | 0.288641227 | 220.365406 | 259.4115466 | ENSG00000178295 |
| LOC100507557 | 0.235275208 | 1.177131265 | 0.761897478 | 0.87673587 | 7.85836241 | 9.273662862 | ENSG00000235652 |
| GCLC | 0.235194749 | 1.177065619 | 0.010085828 | 0.039322879 | 752.0690195 | 885.6491356 | ENSG00000001084 |
| TMCO1 | 0.235025147 | 1.176927251 | 0.002528852 | 0.012417415 | 927.8164142 | 1092.063163 | ENSG00000143183 |
| NIFK-AS1 | 0.234995581 | 1.176903132 | 0.397336398 | 0.609116478 | 70.47823643 | 83.03218946 | ENSG00000236859 |
| NPC1 | 0.234940903 | 1.176858529 | 0.002143248 | 0.010781898 | 966.6406183 | 1137.607013 | ENSG00000141458 |
| HDHD5 | 0.234803894 | 1.176746771 | 0.000204822 | 0.001405547 | 2104.22755 | 2476.061062 | ENSG00000069998 |
| NA | 0.234755008 | 1.176706897 | 0.752589093 | 0.871085164 | 7.569233875 | 8.907352201 | ENSG00000254893 |
| SNX24 | 0.234748911 | 1.176701924 | 0.136392898 | 0.298778204 | 204.3595788 | 240.4854472 | ENSG00000064652 |
| CXorf38 | 0.234738676 | 1.176693576 | 0.114726514 | 0.264349708 | 194.2955042 | 228.7517711 | ENSG00000185753 |
| SOX5 | 0.23471965 | 1.176678058 | 0.650013112 | 0.810064757 | 16.99214994 | 19.96678831 | ENSG00000134532 |
| ADRB2 | 0.234662055 | 1.176631084 | 0.619417431 | 0.788725553 | 18.88112136 | 22.19707332 | ENSG00000169252 |
| POU5F1P4 | 0.234571313 | 1.176557079 | 0.549150333 | 0.736475695 | 30.05452107 | 35.29376554 | ENSG00000237872 |
| EDEM2 | 0.234296707 | 1.176333152 | 0.029271859 | 0.093110839 | 429.9589177 | 505.7198449 | ENSG00000088298 |
| OVCH2 | 0.234253363 | 1.176297811 | 0.819601084 | 0.908449728 | 4.231950025 | 5.011985719 | ENSG00000183378 |
| MPHOSPH9 | 0.23422195 | 1.176272199 | 0.029263407 | 0.093099638 | 455.130847 | 535.0776353 | ENSG00000051825 |
| VRK3 | 0.233980184 | 1.176075096 | 0.034212135 | 0.105366143 | 524.9238413 | 617.3997588 | ENSG00000105053 |
| CACNA1I | 0.233470674 | 1.175659821 | 0.350443363 | 0.562526384 | 87.89152029 | 103.5687002 | ENSG00000100346 |
| DHRS4L2 | 0.233387442 | 1.175591997 | 0.358248464 | 0.570123925 | 73.30191795 | 86.35561511 | ENSG00000187630 |
| MARVELD2 | 0.233371702 | 1.17557917 | 0.06306366 | 0.168547583 | 302.2367613 | 355.0467839 | ENSG00000152939 |
| BRWD3 | 0.233151043 | 1.175399381 | 0.005723037 | 0.024715195 | 848.2310604 | 996.8377031 | ENSG00000165288 |
| NAGA | 0.233150014 | 1.175398542 | 0.01716594 | 0.060803084 | 549.7838107 | 646.161036 | ENSG00000198951 |
| NA | 0.232977493 | 1.175257993 | 0.805897689 | 0.901034744 | 5.11504806 | 5.951918221 | ENSG00000271335 |
| COL8A2 | 0.232977265 | 1.175257807 | 0.222300165 | 0.419771884 | 136.0451783 | 159.8814171 | ENSG00000171812 |
| KCNE5 | 0.232885237 | 1.175182842 | 0.454942768 | 0.660704703 | 52.18251723 | 61.34449267 | ENSG00000176076 |
| ANXA6 | 0.232839861 | 1.17514588 | 1.50E-05 | 0.000137024 | 2705.996048 | 3179.664158 | ENSG00000197043 |
| CPNE5 | 0.232741172 | 1.175065496 | 0.648814874 | 0.809250968 | 16.88302765 | 19.80778883 | ENSG00000124772 |
| SLC6A6 | 0.232643452 | 1.174985906 | 0.00040221 | 0.002538682 | 3384.105034 | 3975.183082 | ENSG00000131389 |
| CCDC8 | 0.232629421 | 1.174974479 | 0.006948855 | 0.029090917 | 708.4927556 | 832.6018849 | ENSG00000169515 |
| NA | 0.232614773 | 1.174962549 | 0.115328385 | 0.265348021 | 224.3826192 | 263.4827922 | ENSG00000283041 |
| IPO9 | 0.232477496 | 1.174850753 | 1.25E-05 | 0.000116474 | 3682.65958 | 4326.342239 | ENSG00000198700 |
| PRICKLE3 | 0.232463903 | 1.174839684 | 0.146686403 | 0.313983685 | 193.4299242 | 227.4633832 | ENSG00000012211 |
| CYB5R3 | 0.232328594 | 1.174729502 | 0.000279551 | 0.001849794 | 2116.220359 | 2486.728722 | ENSG00000100243 |
| ZNF654 | 0.232224001 | 1.174644339 | 0.278289003 | 0.48756309 | 92.62667356 | 108.795753 | ENSG00000175105 |
| NA | 0.232220369 | 1.174641382 | 0.432036614 | 0.641532588 | 49.12841203 | 57.7840344 | ENSG00000212802 |
| DCLK1 | 0.232216522 | 1.174638249 | 0.009342872 | 0.036958552 | 718.7074971 | 844.0275458 | ENSG00000133083 |
| USP8 | 0.232041548 | 1.174495795 | 0.013153482 | 0.048917589 | 838.6334514 | 984.6434863 | ENSG00000138592 |
| LOC102723996 | 0.232001965 | 1.174463571 | 0.506416427 | 0.703694948 | 34.43207782 | 40.40854487 | ENSG00000277117 |
| GRAMD1A | 0.231846857 | 1.174337308 | 0.000160479 | 0.001133375 | 3720.040024 | 4369.057546 | ENSG00000089351 |
| GSTCD | 0.231797382 | 1.174297037 | 0.080185386 | 0.202095424 | 350.9568544 | 412.1019581 | ENSG00000138780 |
| PRPF3 | 0.231669358 | 1.174192835 | 0.001392946 | 0.007483863 | 1224.040234 | 1437.031128 | ENSG00000117360 |
| DCAF12L2 | 0.231468901 | 1.174029697 | 0.466960216 | 0.670914738 | 41.19413164 | 48.41673129 | ENSG00000198354 |
| TCFL5 | 0.231442315 | 1.174008061 | 0.014459029 | 0.052887616 | 593.5976434 | 697.049225 | ENSG00000101190 |
| CNTNAP1 | 0.231424065 | 1.173993211 | 0.083491722 | 0.208542121 | 353.403617 | 414.454398 | ENSG00000108797 |
| C2orf27A | 0.231208629 | 1.173817913 | 0.457045589 | 0.662777555 | 48.33902933 | 56.87862577 | ENSG00000287151 |
| ABCB1 | 0.230739199 | 1.173436033 | 0.616320657 | 0.786912014 | 20.91717407 | 24.50387842 | ENSG00000085563 |
| MTERF1 | 0.230737509 | 1.173434659 | 0.156290375 | 0.328277581 | 185.9893639 | 218.3003128 | ENSG00000127989 |
| LINC00888 | 0.230729621 | 1.173428243 | 0.121561625 | 0.275031901 | 193.8100466 | 227.3313626 | ENSG00000240024 |
| SLIT1 | 0.230685799 | 1.1733926 | 0.567038004 | 0.750684381 | 33.41021267 | 39.34557718 | ENSG00000187122 |
| NA | 0.230674137 | 1.173383115 | 0.751467053 | 0.870641429 | 8.830421103 | 10.28046764 | ENSG00000286811 |
| PDHB | 0.23057304 | 1.173300893 | 0.001068398 | 0.005939565 | 2205.463801 | 2587.186855 | ENSG00000168291 |
| QTRT1 | 0.230544451 | 1.173277643 | 0.001196776 | 0.006554846 | 1370.523938 | 1608.228732 | ENSG00000213339 |
| FCGRT | 0.230306574 | 1.173084204 | 0.052734788 | 0.146909055 | 437.9264541 | 514.1799858 | ENSG00000104870 |
| JCAD | 0.230235157 | 1.173026135 | 0.119311985 | 0.271732614 | 220.6304824 | 258.5233535 | ENSG00000165757 |
| ANKRD24 | 0.230076898 | 1.172897465 | 0.049689357 | 0.140392506 | 440.7052847 | 517.3152803 | ENSG00000089847 |
| HDGF | 0.229759584 | 1.17263952 | 1.02E-05 | 9.71E-05 | 10088.69977 | 11830.0854 | ENSG00000143321 |
| DTL | 0.229583251 | 1.172496203 | 0.000943557 | 0.005336651 | 1242.287978 | 1456.592266 | ENSG00000143476 |
| NA | 0.229560416 | 1.172477645 | 0.786493071 | 0.891157476 | 5.642303459 | 6.596479721 | ENSG00000217241 |
| HPS4 | 0.229423774 | 1.172366601 | 0.023234283 | 0.077693865 | 762.8979194 | 894.2029167 | ENSG00000100099 |
| POPDC3 | 0.229368474 | 1.172321664 | 0.418957689 | 0.629503762 | 52.89122727 | 62.01080881 | ENSG00000132429 |
| TAF1A | 0.22935039 | 1.17230697 | 0.136100399 | 0.298311982 | 204.5484056 | 239.6783133 | ENSG00000143498 |
| ZDHHC4 | 0.229219682 | 1.172200764 | 0.045622107 | 0.131157481 | 366.6969693 | 429.6824381 | ENSG00000136247 |
| STYX | 0.229171073 | 1.172161269 | 0.050692516 | 0.142629345 | 539.9771343 | 633.0282615 | ENSG00000198252 |
| GPR162 | 0.22912832 | 1.172126534 | 0.169160334 | 0.346974358 | 215.3511399 | 252.5131922 | ENSG00000250510 |
| SLC35B3 | 0.229072186 | 1.172080928 | 0.162820233 | 0.337859587 | 182.0106422 | 213.3371856 | ENSG00000124786 |
| PNMT | 0.228955182 | 1.171985875 | 0.328157643 | 0.539669465 | 90.51498221 | 106.0709206 | ENSG00000141744 |
| EDARADD | 0.228860439 | 1.171908912 | 0.291984347 | 0.501603237 | 90.95447216 | 106.6339486 | ENSG00000186197 |
| NSUN5 | 0.228724701 | 1.171798656 | 0.002273406 | 0.011337015 | 1554.571853 | 1822.219612 | ENSG00000130305 |
| LRP12 | 0.22857179 | 1.171674464 | 0.130711993 | 0.290138222 | 202.8169407 | 237.4957729 | ENSG00000147650 |
| NA | 0.228571074 | 1.171673883 | 0.77152051 | 0.882491333 | 8.544852041 | 9.919453867 | ENSG00000248690 |
| YPEL3 | 0.228523627 | 1.17163535 | 0.081369969 | 0.204399025 | 283.7029146 | 332.7268863 | ENSG00000090238 |
| RASAL2-AS1 | 0.228458927 | 1.171582807 | 0.590518014 | 0.768530555 | 24.69778667 | 28.85295297 | ENSG00000224687 |
| UNC5B | 0.228355987 | 1.171499215 | 0.005020766 | 0.022174265 | 1594.895501 | 1868.775507 | ENSG00000107731 |

| | | | | | | | |
|---|---|---|---|---|---|---|---|
| ZNF517 | 0.228354458 | 1.171497973 | 0.127800367 | 0.28535232 | 193.2768675 | 226.2676169 | ENSG00000197363 |
| ARNTL | 0.228206939 | 1.17137819 | 0.183618234 | 0.367528388 | 148.493977 | 173.827355 | ENSG00000133794 |
| NBEA | 0.22811784 | 1.17130585 | 0.038179099 | 0.114573698 | 612.5338328 | 716.9045045 | ENSG00000172915 |
| DLG4 | 0.227943798 | 1.171164556 | 0.00595193 | 0.025557573 | 985.1895402 | 1153.940924 | ENSG00000132535 |
| DENND2B | 0.227856185 | 1.171093435 | 0.031939432 | 0.099896685 | 407.3745501 | 476.9534387 | ENSG00000166444 |
| SYT7 | 0.227548654 | 1.170843826 | 0.022430033 | 0.07549299 | 546.7908014 | 639.8059695 | ENSG00000011347 |
| SEPHS2 | 0.227441125 | 1.170756563 | 0.000358666 | 0.00229296 | 1725.298422 | 2019.845724 | ENSG00000179918 |
| ACTR3C | 0.227434762 | 1.170751399 | 0.772893934 | 0.883060194 | 8.99262999 | 10.48125243 | ENSG00000106526 |
| C11orf96 | 0.22722707 | 1.170582869 | 0.719833726 | 0.852321847 | 9.908653 | 11.60193905 | ENSG00000187479 |
| FAM76A | 0.227169793 | 1.170536395 | 0.145775494 | 0.312801977 | 193.1205823 | 225.8219483 | ENSG00000009780 |
| CAPS | 0.22698544 | 1.17038683 | 0.747080479 | 0.868435054 | 8.223408922 | 9.652080345 | ENSG00000105519 |
| LINC00240 | 0.226978919 | 1.170381539 | 0.720621728 | 0.852637136 | 10.23574052 | 12.00494904 | ENSG00000224843 |
| TET3 | 0.226953848 | 1.170361201 | 0.048525019 | 0.137907448 | 571.3986854 | 668.2327901 | ENSG00000187605 |
| TMEM185A | 0.226895562 | 1.170313918 | 0.072248553 | 0.187470998 | 336.8334122 | 393.9615215 | ENSG00000269556 |
| FAM120B | 0.226682024 | 1.170140709 | 0.030258431 | 0.095620896 | 508.1347822 | 594.2603962 | ENSG00000112584 |
| ACVR1C | 0.226421777 | 1.169929647 | 0.663611476 | 0.819640466 | 15.65415501 | 18.28458668 | ENSG00000123612 |
| LOC100289495 | 0.226393324 | 1.169906574 | 0.72807092 | 0.856731016 | 9.674085608 | 11.28821838 | ENSG00000286760 |
| NA | 0.226337198 | 1.169861061 | 0.837245601 | 0.918172233 | 4.522273768 | 5.326652347 | ENSG00000237609 |
| AMOTL2 | 0.226244907 | 1.169786226 | 0.003985781 | 0.018318683 | 1096.862772 | 1283.043468 | ENSG00000114019 |
| DUS4L | 0.226228802 | 1.169773168 | 0.504535916 | 0.701993721 | 35.47146101 | 41.46046729 | ENSG00000105865 |
| CNOT3 | 0.226109088 | 1.169676105 | 0.001775825 | 0.009183038 | 1806.41063 | 2113.54402 | ENSG00000088038 |
| KANK1 | 0.225771534 | 1.169402462 | 0.006607851 | 0.027954586 | 1122.668923 | 1312.80477 | ENSG00000107104 |
| ENDOV | 0.225608168 | 1.169270051 | 0.230036883 | 0.429903311 | 139.7163376 | 163.1788957 | ENSG00000173818 |
| NA | 0.225513717 | 1.169193502 | 0.730650506 | 0.858105398 | 11.04292058 | 12.84376894 | ENSG00000267655 |
| CA14 | 0.225414642 | 1.169113213 | 0.006325848 | 0.026920563 | 899.404704 | 1050.870637 | ENSG00000118298 |
| TERT | 0.225272584 | 1.168998099 | 0.031288781 | 0.098219518 | 524.8120493 | 613.7565578 | ENSG00000164362 |
| DENND1B | 0.225134245 | 1.16888601 | 0.471578636 | 0.674832351 | 53.51275667 | 62.36747165 | ENSG00000213047 |
| ZYX | 0.225038714 | 1.168808613 | 0.002270676 | 0.011329381 | 3040.89018 | 3555.436187 | ENSG00000159840 |
| YIPF2 | 0.225007375 | 1.168783223 | 0.009940689 | 0.038835315 | 861.4947679 | 1007.189816 | ENSG00000130733 |
| ZXDB | 0.224978367 | 1.168759723 | 0.137416816 | 0.300186995 | 196.384949 | 229.5601345 | ENSG00000198455 |
| ACOT2 | 0.224849635 | 1.168655439 | 0.222122949 | 0.419521142 | 146.6281418 | 171.5132885 | ENSG00000119673 |
| ARMCX6 | 0.224830873 | 1.168640241 | 0.093450162 | 0.226859999 | 276.6540971 | 323.1389357 | ENSG00000198960 |
| NA | 0.224768709 | 1.168589887 | 0.620976525 | 0.78986608 | 22.96243064 | 26.90744519 | ENSG00000230409 |
| FKBP3 | 0.2247183 | 1.168549056 | 0.00074332 | 0.004330626 | 2423.079663 | 2831.361083 | ENSG00000100442 |
| C1orf159 | 0.224686842 | 1.168523576 | 0.115545128 | 0.265717212 | 256.3525804 | 299.6523605 | ENSG00000131591 |
| CCDC149 | 0.224581655 | 1.168438382 | 0.18085051 | 0.363839553 | 179.0093704 | 209.4992545 | ENSG00000181982 |
| NDFIP2 | 0.224463519 | 1.168342707 | 0.072234499 | 0.187469615 | 409.6617456 | 478.7864621 | ENSG00000102471 |
| RIOK2 | 0.224263858 | 1.168181026 | 0.028384643 | 0.090891106 | 459.2382678 | 536.2875159 | ENSG00000058729 |
| FAR1 | 0.224236627 | 1.168158977 | 0.008560256 | 0.034550211 | 944.4390161 | 1103.135798 | ENSG00000197601 |
| SPNS2 | 0.224070191 | 1.168024221 | 0.474892197 | 0.677776537 | 51.43638072 | 59.8808584 | ENSG00000183018 |
| PDXP | 0.223960487 | 1.167935407 | 0.13750655 | 0.300348277 | 285.6251427 | 333.8884349 | ENSG00000241360 |
| TRPT1 | 0.223921364 | 1.167903735 | 0.124573818 | 0.280202067 | 220.8635752 | 257.900474 | ENSG00000149743 |
| MTMR6 | 0.223638631 | 1.167674877 | 0.142826179 | 0.308189258 | 263.0876324 | 307.4370912 | ENSG00000139505 |
| BBS7 | 0.223619476 | 1.167659373 | 0.080918773 | 0.203536365 | 308.1649387 | 360.0855805 | ENSG00000138686 |
| SH3BGRL3 | 0.223562918 | 1.167613599 | 0.01915779 | 0.066262796 | 749.915635 | 876.1784195 | ENSG00000142669 |
| NA | 0.223553174 | 1.167605712 | 0.695414943 | 0.837344504 | 13.84555075 | 16.15709528 | ENSG00000235454 |
| GDAP1L1 | 0.223444443 | 1.167517717 | 0.226433958 | 0.425281481 | 142.8332913 | 166.8954 | ENSG00000124194 |
| NA | 0.223403235 | 1.16748437 | 0.547577003 | 0.735228703 | 30.95958633 | 36.20910553 | ENSG00000234964 |
| LRATD1 | 0.22321808 | 1.167334544 | 0.244416673 | 0.448729647 | 114.9245562 | 134.1311899 | ENSG00000162981 |
| LOC283731 | 0.223191846 | 1.167313318 | 0.801603302 | 0.898520796 | 6.282240616 | 7.345275247 | ENSG00000248540 |
| NBPF3 | 0.222890831 | 1.167069786 | 0.025333105 | 0.083318651 | 533.628765 | 622.5167635 | ENSG00000142794 |
| LOC286059 | 0.222837408 | 1.16702657 | 0.701390404 | 0.841053894 | 11.95538412 | 13.93333666 | ENSG00000246130 |
| COL23A1 | 0.222819869 | 1.167012382 | 0.100434139 | 0.238808487 | 330.2125683 | 385.3254126 | ENSG00000050767 |
| ZSCAN10 | 0.222655748 | 1.166879631 | 0.001076589 | 0.00598158 | 5131.597286 | 5989.011781 | ENSG00000130182 |
| SEPSECS | 0.222640378 | 1.1668672 | 0.388711526 | 0.600524064 | 67.55791664 | 78.90894371 | ENSG00000109618 |
| STKLD1 | 0.222254982 | 1.166555529 | 0.714917581 | 0.849668508 | 12.72669012 | 14.89691571 | ENSG00000198870 |
| COL4A6 | 0.22225209 | 1.16655319 | 0.003029967 | 0.014537837 | 1156.920891 | 1349.261726 | ENSG00000197565 |
| GABPB2 | 0.222245471 | 1.166547838 | 0.462995349 | 0.667710361 | 46.70743052 | 54.48158539 | ENSG00000143458 |
| AKAP9 | 0.222210884 | 1.166519872 | 0.051462009 | 0.144266847 | 505.9308239 | 589.7539306 | ENSG00000127914 |
| TYSND1 | 0.222190083 | 1.166503052 | 0.014230717 | 0.052244769 | 767.3619156 | 894.868076 | ENSG00000156521 |
| PPIE | 0.222156184 | 1.166475644 | 0.011404297 | 0.043561924 | 681.4392276 | 794.5438224 | ENSG00000084072 |
| DNAJC15 | 0.221966927 | 1.166322632 | 0.05417311 | 0.149920748 | 435.4955805 | 507.6595317 | ENSG00000120675 |
| PIP4P1 | 0.221960589 | 1.166317509 | 0.023760663 | 0.079091803 | 644.7911425 | 752.2269862 | ENSG00000165782 |
| WDR18 | 0.22195749 | 1.166315003 | 0.000619369 | 0.003691003 | 1767.527844 | 2062.007644 | ENSG00000065268 |
| USF3 | 0.221939122 | 1.166300154 | 0.168467996 | 0.346080507 | 219.4837569 | 255.6204142 | ENSG00000176542 |
| COX5B | 0.221846075 | 1.166224936 | 0.001316854 | 0.007113498 | 1980.85198 | 2309.904851 | ENSG00000135940 |
| LARP7 | 0.221814074 | 1.166199067 | 0.012873983 | 0.048134067 | 863.8238141 | 1007.808546 | ENSG00000174720 |
| NA | 0.221749625 | 1.166146971 | 0.821464964 | 0.909334444 | 4.273468486 | 5.001108401 | ENSG00000255130 |
| IKBIP | 0.221616219 | 1.166039142 | 0.132956284 | 0.293601429 | 228.4025795 | 266.4116987 | ENSG00000166130 |
| TMEM182 | 0.221284201 | 1.165770824 | 0.435369935 | 0.643950597 | 59.79207436 | 69.75228411 | ENSG00000170417 |
| DNM1 | 0.22127694 | 1.165764957 | 0.066185853 | 0.175034287 | 785.5740916 | 915.9674102 | ENSG00000106976 |
| NEDD4L | 0.221185921 | 1.165691412 | 0.001844653 | 0.009458665 | 1317.443806 | 1535.299815 | ENSG00000049759 |
| FAM160B1 | 0.221061661 | 1.165591015 | 0.198843325 | 0.389030323 | 209.772069 | 244.7621417 | ENSG00000151553 |
| ERV3-1 | 0.221060963 | 1.165590451 | 0.482032175 | 0.683524375 | 46.18581953 | 53.92413775 | ENSG00000213462 |

| | | | | | | | |
|---|---|---|---|---|---|---|---|
| PRDM6 | 0.221003575 | 1.165544086 | 0.821838654 | 0.9095419 | 4.246187915 | 4.967814035 | ENSG00000061455 |
| NA | 0.220912883 | 1.165470819 | 0.670338079 | 0.823292434 | 16.24189529 | 18.91950036 | ENSG00000213189 |
| TEAD1 | 0.220804319 | 1.16538312 | 0.009060778 | 0.0361455 | 1371.342712 | 1597.586024 | ENSG00000187079 |
| C21orf58 | 0.220756325 | 1.165344352 | 0.171939792 | 0.351301847 | 326.0559258 | 379.8569239 | ENSG00000160298 |
| ANKRA2 | 0.220695252 | 1.165295021 | 0.341123489 | 0.552686778 | 83.2286738 | 97.09196628 | ENSG00000164331 |
| EXOC3L4 | 0.220647759 | 1.16525666 | 0.828500747 | 0.91386231 | 4.000942105 | 4.651917903 | ENSG00000205436 |
| TXNRD3 | 0.220542902 | 1.165171971 | 0.198180783 | 0.388241088 | 150.6777213 | 175.6691232 | ENSG00000197763 |
| LRP6 | 0.220472387 | 1.165115022 | 0.043849626 | 0.127380431 | 1161.34398 | 1352.5044 | ENSG00000070018 |
| FNDC10 | 0.220337638 | 1.165006204 | 0.078489509 | 0.198936301 | 540.3779325 | 629.7595514 | ENSG00000228594 |
| NA | 0.220312706 | 1.164986072 | 0.600960025 | 0.776307564 | 24.16669246 | 28.12657449 | ENSG00000278730 |
| MOXD1 | 0.220147169 | 1.164852407 | 0.344375392 | 0.556370157 | 83.10175415 | 96.9000513 | ENSG00000079931 |
| FOXP4-AS1 | 0.220120615 | 1.164830967 | 0.775439176 | 0.884700308 | 7.665313479 | 8.891177997 | ENSG00000234753 |
| NA | 0.220045989 | 1.164770716 | 0.709419427 | 0.846496645 | 12.81328651 | 14.96596346 | ENSG00000260233 |
| MMP24 | 0.220001053 | 1.164734437 | 0.068683844 | 0.179977023 | 336.9526281 | 392.5069156 | ENSG00000125966 |
| ATXN7L2 | 0.21985286 | 1.164614802 | 0.157112625 | 0.32930876 | 221.700478 | 258.4843107 | ENSG00000162650 |
| CCDC102A | 0.219848495 | 1.164611278 | 0.217629814 | 0.413599581 | 211.0528069 | 246.1951515 | ENSG00000135736 |
| NDUFS4 | 0.219804646 | 1.164575882 | 0.033970959 | 0.104861319 | 552.4081362 | 643.1617342 | ENSG00000164258 |
| SNX9 | 0.219675126 | 1.164471335 | 0.009110822 | 0.036245602 | 810.1755193 | 943.1397433 | ENSG00000130340 |
| MCTP2 | 0.219643005 | 1.164445409 | 0.484367759 | 0.685446618 | 49.77457841 | 57.86100599 | ENSG00000140563 |
| FRS3 | 0.219581968 | 1.164396145 | 0.415814144 | 0.626739688 | 73.83186925 | 86.08634967 | ENSG00000137218 |
| IL11RA | 0.219480006 | 1.164313854 | 0.586213339 | 0.764758145 | 34.99996196 | 40.72900284 | ENSG00000137070 |
| PAX5 | 0.219370848 | 1.164225763 | 0.703838425 | 0.842524014 | 12.27891217 | 14.28564858 | ENSG00000196092 |
| XAB2 | 0.219195946 | 1.164084629 | 0.000224743 | 0.001524131 | 2070.293693 | 2409.789203 | ENSG00000076924 |
| SMIM8 | 0.219130542 | 1.164031857 | 0.253190049 | 0.459273276 | 120.2869089 | 140.0818786 | ENSG00000111850 |
| SOX11 | 0.21911533 | 1.164019583 | 0.009174702 | 0.036430666 | 1760.449048 | 2049.601215 | ENSG00000176887 |
| ZNF316 | 0.218850715 | 1.163806102 | 0.008770169 | 0.035216806 | 798.1671264 | 928.9529366 | ENSG00000205903 |
| TRAFD1 | 0.218831983 | 1.163790991 | 0.01245923 | 0.046908403 | 742.5456537 | 864.1836221 | ENSG00000135148 |
| SEPTIN10 | 0.218741893 | 1.163718319 | 0.001004474 | 0.005632269 | 1457.882253 | 1696.220428 | ENSG00000186522 |
| NFS1 | 0.218543169 | 1.163558034 | 0.120126864 | 0.27279892 | 240.3443367 | 279.6763032 | ENSG00000244005 |
| PANO1 | 0.218432709 | 1.16346895 | 0.788253471 | 0.891600791 | 6.31308066 | 7.348680215 | ENSG00000274897 |
| TRIM65 | 0.218307399 | 1.163367897 | 0.002639779 | 0.012891707 | 1422.212481 | 1655.31737 | ENSG00000141569 |
| MIPOL1 | 0.218272615 | 1.163339848 | 0.197114111 | 0.386672898 | 186.8727151 | 217.1744972 | ENSG00000151338 |
| IL20RB | 0.218265063 | 1.163333758 | 0.59058344 | 0.768530555 | 27.09860655 | 31.54072988 | ENSG00000174564 |
| PLAA | 0.217959701 | 1.163087552 | 0.003773093 | 0.017481566 | 1362.910262 | 1585.018396 | ENSG00000137055 |
| NOTCH4 | 0.21782113 | 1.162975843 | 0.538245553 | 0.728027383 | 36.4005267 | 42.27876206 | ENSG00000204301 |
| KIAA0355 | 0.217731405 | 1.162903517 | 0.015447132 | 0.055810042 | 644.820508 | 749.8816922 | ENSG00000166398 |
| NA | 0.217616337 | 1.162810768 | 0.684923613 | 0.831604889 | 14.60172914 | 16.97805997 | ENSG00000178464 |
| NA | 0.217528383 | 1.16273988 | 0.576529467 | 0.758517416 | 31.86640546 | 36.96922988 | ENSG00000254469 |
| MAPK8IP2 | 0.217430163 | 1.162660722 | 0.181923284 | 0.365180797 | 253.2961627 | 294.819194 | ENSG00000008735 |
| SIK1B | 0.217416932 | 1.162650059 | 0.395800303 | 0.607686774 | 78.82500553 | 91.53024587 | ENSG00000275993 |
| NA | 0.217374191 | 1.162615615 | 0.287123014 | 0.496231181 | 113.4738534 | 131.9933156 | ENSG00000234797 |
| METTL21A | 0.217262232 | 1.162525396 | 0.009745481 | 0.03818981 | 714.603051 | 830.848628 | ENSG00000144401 |
| LBP | 0.217231345 | 1.162500507 | 0.816530129 | 0.906477444 | 5.792944205 | 6.694470902 | ENSG00000129988 |
| LOC145783 | 0.217227064 | 1.162497057 | 0.76354737 | 0.877522527 | 10.07233643 | 11.67845914 | ENSG00000285331 |
| SLC7A3 | 0.217180452 | 1.162459498 | 0.000492114 | 0.003037116 | 2860.826322 | 3324.915686 | ENSG00000165349 |
| NUDT1 | 0.21710081 | 1.162395329 | 0.009754025 | 0.038200904 | 895.3654707 | 1040.824901 | ENSG00000106268 |
| LAYN | 0.216941436 | 1.162266926 | 0.29787207 | 0.508112549 | 114.8946059 | 133.6851651 | ENSG00000204381 |
| VPS13B | 0.21665141 | 1.162033298 | 0.085610614 | 0.212479491 | 335.941442 | 390.0147946 | ENSG00000132549 |
| MED17 | 0.216560367 | 1.161959969 | 0.577965128 | 0.759450456 | 30.54709762 | 35.59443343 | ENSG00000042429 |
| PNPLA7 | 0.216464222 | 1.161882536 | 0.671458159 | 0.823943296 | 18.90959714 | 21.97197956 | ENSG00000130653 |
| ANKRD23 | 0.216371841 | 1.161808138 | 0.853016787 | 0.927897127 | 4.589877592 | 5.351244858 | ENSG00000163126 |
| POP5 | 0.216265703 | 1.161722668 | 0.031969086 | 0.099963474 | 508.441071 | 590.6983726 | ENSG00000167272 |
| NQO2 | 0.216116105 | 1.161602211 | 0.002997412 | 0.014396266 | 1203.94946 | 1398.169432 | ENSG00000124588 |
| FUT4 | 0.215930214 | 1.161452549 | 0.274078714 | 0.482743566 | 149.7157307 | 173.8303811 | ENSG00000196371 |
| TCF7L2 | 0.215869268 | 1.161403484 | 0.042825898 | 0.12511867 | 436.6862404 | 507.3670712 | ENSG00000148737 |
| ABRAXAS2 | 0.215767071 | 1.161321216 | 0.057912445 | 0.157954438 | 356.0808015 | 413.3523446 | ENSG00000165660 |
| MFSD4A | 0.21566594 | 1.161239812 | 0.805092898 | 0.90066817 | 6.020392653 | 6.968370814 | ENSG00000174514 |
| ZNF235 | 0.21564144 | 1.161220092 | 0.485379372 | 0.686363866 | 46.96126982 | 54.4212395 | ENSG00000159917 |
| C1QL4 | 0.215440237 | 1.161058156 | 0.591664881 | 0.769356281 | 25.4667284 | 29.58774975 | ENSG00000186897 |
| NA | 0.215422953 | 1.161044246 | 0.567731613 | 0.751047019 | 37.63028907 | 43.81891643 | ENSG00000233225 |
| CNTRL | 0.215205773 | 1.160869478 | 0.106849522 | 0.250883155 | 252.2276416 | 292.6325136 | ENSG00000119397 |
| SLC22A18 | 0.215186992 | 1.160854366 | 0.402869221 | 0.614207252 | 76.79726698 | 89.20267505 | ENSG00000110628 |
| NARS2 | 0.215087114 | 1.160774003 | 0.004412629 | 0.019912187 | 1048.206905 | 1217.161412 | ENSG00000137513 |
| LAG3 | 0.21474652 | 1.160499998 | 0.426868793 | 0.636856462 | 60.86555158 | 70.78614041 | ENSG00000089692 |
| NUDT22 | 0.214637515 | 1.160412317 | 0.062961915 | 0.168409513 | 363.6432475 | 422.2147901 | ENSG00000149761 |
| NA | 0.21457347 | 1.160360805 | 0.411656945 | 0.622828186 | 74.16577024 | 86.24362519 | ENSG00000261963 |
| ERICH2 | 0.21453128 | 1.160326872 | 0.415027711 | 0.625822286 | 59.51123383 | 69.08871053 | ENSG00000239467 |
| TULP3 | 0.214505121 | 1.160305833 | 0.041147069 | 0.121321218 | 465.7010733 | 540.0483075 | ENSG00000078246 |
| POLR3D | 0.214432312 | 1.160247277 | 0.001141461 | 0.006299382 | 1379.484305 | 1600.549481 | ENSG00000168495 |
| FKTN | 0.214389688 | 1.160212999 | 0.147134448 | 0.314606907 | 222.7324112 | 258.3632401 | ENSG00000106692 |
| TPM3 | 0.214244984 | 1.160096634 | 6.33E-05 | 0.000495209 | 13843.29753 | 16059.24794 | ENSG00000143549 |
| SLC30A5 | 0.214234346 | 1.16008808 | 0.033485888 | 0.103567147 | 542.6428036 | 629.566931 | ENSG00000145740 |
| DTNB | 0.214188114 | 1.160050904 | 0.016616227 | 0.059229415 | 648.7985409 | 752.4858031 | ENSG00000138101 |
| NEDD4 | 0.21412349 | 1.159998942 | 0.159696891 | 0.333356448 | 191.4869503 | 222.0869787 | ENSG00000069869 |

| | | | | | | | |
|---|---|---|---|---|---|---|---|
| NOS1AP | 0.214075661 | 1.159960486 | 0.178027564 | 0.360039682 | 232.2621109 | 269.5782277 | ENSG00000198929 |
| DIS3L2 | 0.213959296 | 1.15986693 | 0.077971439 | 0.198101792 | 324.6197658 | 376.3058166 | ENSG00000144535 |
| RNF139 | 0.213871806 | 1.159796594 | 0.041211481 | 0.121435297 | 441.886131 | 512.7865858 | ENSG00000170881 |
| SORCS3 | 0.213843968 | 1.159774215 | 0.521206109 | 0.716053207 | 40.46078463 | 46.92928251 | ENSG00000156395 |
| SNORD3B-2 | 0.2137507 | 1.15969924 | 0.490933509 | 0.69054665 | 42.9552937 | 49.86201589 | ENSG00000262074 |
| NA | 0.213743918 | 1.159693787 | 0.710569128 | 0.847218763 | 12.8263292 | 14.89161883 | ENSG00000236683 |
| TOMM40L | 0.213538108 | 1.159528361 | 0.07610937 | 0.194760087 | 353.9951955 | 410.2003612 | ENSG00000158882 |
| CLUH | 0.21352766 | 1.159519964 | 7.08E-05 | 0.000546318 | 4553.956345 | 5281.057101 | ENSG00000132361 |
| MCL1 | 0.213393446 | 1.159412099 | 1.23E-05 | 0.000114505 | 6990.929081 | 8105.445854 | ENSG00000143384 |
| LDHC | 0.213187012 | 1.159246212 | 0.736751057 | 0.861679799 | 11.70274003 | 13.64761347 | ENSG00000166796 |
| ATP13A3 | 0.21310296 | 1.159178676 | 0.02116819 | 0.072055023 | 1297.910926 | 1504.033895 | ENSG00000133657 |
| SKIL | 0.213011654 | 1.159105315 | 0.025357732 | 0.083370606 | 1592.584653 | 1846.317844 | ENSG00000136603 |
| CEP120 | 0.212930776 | 1.159040337 | 0.101687921 | 0.240920385 | 348.3979694 | 403.8498565 | ENSG00000168944 |
| MYORG | 0.212883809 | 1.159002605 | 0.038168035 | 0.114558701 | 541.2612053 | 627.4000407 | ENSG00000164976 |
| NA | 0.21287238 | 1.158993424 | 0.387065783 | 0.599276623 | 88.99550649 | 103.2261517 | ENSG00000139239 |
| PIM1 | 0.212858131 | 1.158981977 | 0.002446873 | 0.012083997 | 1220.925975 | 1414.840495 | ENSG00000137193 |
| PCSK6 | 0.212727457 | 1.158877005 | 0.405730006 | 0.616875547 | 78.22539162 | 90.61379196 | ENSG00000140479 |
| EP400 | 0.212453 | 1.158656562 | 0.001694959 | 0.008837468 | 2371.424175 | 2746.936353 | ENSG00000183495 |
| RPL3 | 0.212315529 | 1.158546162 | 4.13E-07 | 5.15E-06 | 54880.28678 | 63580.64353 | ENSG00000100316 |
| ANGPTL2 | 0.212267463 | 1.158507563 | 0.071746843 | 0.186374021 | 382.4262043 | 443.0088156 | ENSG00000136859 |
| C11orf45 | 0.21213575 | 1.1584018 | 0.713068177 | 0.848591123 | 12.86784766 | 14.91403588 | ENSG00000174370 |
| RGP1 | 0.21207363 | 1.158351923 | 0.028153865 | 0.090345919 | 534.8425362 | 619.287357 | ENSG00000107185 |
| VRK1 | 0.211916235 | 1.158225556 | 0.003193562 | 0.015195367 | 1083.794861 | 1255.177646 | ENSG00000100749 |
| TLCD5 | 0.211862377 | 1.158182319 | 0.123039558 | 0.277511292 | 249.4870226 | 289.0664749 | ENSG00000181264 |
| PICALM | 0.211750879 | 1.158092813 | 0.008583336 | 0.034628563 | 1157.146131 | 1340.479615 | ENSG00000073921 |
| MPP6 | 0.211720904 | 1.158068751 | 0.006009518 | 0.025775549 | 1466.422986 | 1698.366372 | ENSG00000105926 |
| NA | 0.211687403 | 1.158041859 | 0.828183969 | 0.91367299 | 4.579199175 | 5.326652347 | ENSG00000228952 |
| NA | 0.211676513 | 1.158033118 | 0.533049415 | 0.724314769 | 38.25776836 | 44.2636391 | ENSG00000213235 |
| B4GALT5 | 0.21160984 | 1.157979602 | 0.00196866 | 0.010018364 | 2434.950747 | 2818.761549 | ENSG00000158470 |
| RPAP1 | 0.211269137 | 1.157706168 | 0.00076612 | 0.004440818 | 1790.300305 | 2072.931589 | ENSG00000103932 |
| SLC25A21 | 0.211247539 | 1.157688837 | 0.378471163 | 0.590891859 | 78.93829767 | 91.53603721 | ENSG00000183032 |
| CYP46A1 | 0.211107082 | 1.157576134 | 0.692143635 | 0.835483064 | 14.96913992 | 17.31448123 | ENSG00000036530 |
| ZNF746 | 0.211072569 | 1.157548441 | 0.027984461 | 0.08997047 | 631.5475177 | 730.8948251 | ENSG00000181220 |
| LYRM2 | 0.211070348 | 1.157546659 | 0.142085723 | 0.307047588 | 253.2916618 | 293.1802408 | ENSG00000083099 |
| MAP1S | 0.211066451 | 1.157543533 | 0.001742291 | 0.009034368 | 1931.645441 | 2236.019504 | ENSG00000130479 |
| NA | 0.210994912 | 1.157486135 | 0.756506776 | 0.873426084 | 8.601498115 | 9.95709916 | ENSG00000231731 |
| IQGAP3 | 0.210747928 | 1.157287995 | 0.022465772 | 0.075559414 | 1065.623366 | 1233.568177 | ENSG00000183856 |
| ARHGAP27 | 0.210663806 | 1.157220517 | 0.111937971 | 0.259717416 | 313.690469 | 362.9935105 | ENSG00000159314 |
| ADAM22 | 0.210636498 | 1.157198612 | 0.069752927 | 0.182247375 | 426.1942446 | 492.8713986 | ENSG00000008277 |
| NBPF25P | 0.210573019 | 1.157147696 | 0.747553179 | 0.868611444 | 11.82401528 | 13.60467129 | ENSG00000272150 |
| PAMR1 | 0.210554651 | 1.157132964 | 0.321052342 | 0.531456561 | 118.4800585 | 136.967824 | ENSG00000149090 |
| BCKDK | 0.210300089 | 1.156928807 | 0.005839679 | 0.025126983 | 1027.09752 | 1188.62215 | ENSG00000103507 |
| RNF207 | 0.210287953 | 1.156919075 | 0.485795862 | 0.686695723 | 61.68488459 | 71.40754982 | ENSG00000158286 |
| SERINC3 | 0.210240602 | 1.156881104 | 0.016234055 | 0.058160398 | 774.9646855 | 896.0944667 | ENSG00000132824 |
| TMEM186 | 0.210090337 | 1.156760615 | 0.183281508 | 0.366977969 | 231.0159739 | 267.0555682 | ENSG00000184857 |
| CDH20 | 0.210080305 | 1.156752571 | 0.785896598 | 0.890641938 | 6.871176103 | 7.983593902 | ENSG00000101542 |
| NA | 0.209824854 | 1.156547768 | 0.760030147 | 0.875513562 | 8.605057588 | 9.949626811 | ENSG00000280211 |
| FASTKD3 | 0.20969502 | 1.15644369 | 0.248090779 | 0.453053109 | 135.3675615 | 156.4916862 | ENSG00000124279 |
| HLTF | 0.209599347 | 1.156367003 | 0.021323052 | 0.07246461 | 881.6827915 | 1019.352522 | ENSG00000071794 |
| ITGA3 | 0.209599136 | 1.156366834 | 0.028393868 | 0.090899847 | 813.7209797 | 941.0867052 | ENSG00000005884 |
| SATL1 | 0.209419938 | 1.156223209 | 0.733737136 | 0.860173437 | 10.61738919 | 12.25832383 | ENSG00000184788 |
| REEP5 | 0.209390272 | 1.156199435 | 0.00655132 | 0.027734067 | 1120.237263 | 1295.224188 | ENSG00000129625 |
| STMN1 | 0.209296569 | 1.156124342 | 0.000111815 | 0.000821327 | 5914.89732 | 6837.515083 | ENSG00000117632 |
| ZEB1-AS1 | 0.209295277 | 1.156123306 | 0.685828798 | 0.831762616 | 17.73025163 | 20.53388376 | ENSG00000237036 |
| CFAP58 | 0.209177165 | 1.15602866 | 0.730446105 | 0.858078794 | 12.38803445 | 14.26947438 | ENSG00000120051 |
| CYP11A1 | 0.209036888 | 1.155916262 | 0.573187494 | 0.755541692 | 34.19634138 | 39.48885395 | ENSG00000140459 |
| TREML2 | 0.208979429 | 1.155870225 | 0.770283933 | 0.881997379 | 9.414322577 | 10.93808192 | ENSG00000112195 |
| PPFIBP1 | 0.20897138 | 1.155863777 | 0.034838182 | 0.106857084 | 542.9698394 | 627.3571914 | ENSG00000110841 |
| NA | 0.208909125 | 1.1558139 | 0.573448267 | 0.755779923 | 30.97499327 | 35.85121318 | ENSG00000254602 |
| FLOT1 | 0.208803051 | 1.155728922 | 9.61E-05 | 0.000716427 | 3025.433136 | 3496.476227 | ENSG00000137312 |
| RBM8A | 0.208667163 | 1.155620068 | 0.014725621 | 0.053717087 | 853.6768072 | 986.8424548 | ENSG00000265241 |
| H2AC20 | 0.208658489 | 1.15561312 | 0.595154556 | 0.772037516 | 28.66285472 | 33.1600313 | ENSG00000184260 |
| SDC4 | 0.208570172 | 1.15554238 | 0.007854152 | 0.032181743 | 974.432773 | 1126.02108 | ENSG00000124145 |
| LIPT1 | 0.208465369 | 1.15545844 | 0.507260782 | 0.704350311 | 44.95249769 | 51.86474813 | ENSG00000144182 |
| TAF9B | 0.20830041 | 1.155326331 | 0.061537465 | 0.165475583 | 438.9207338 | 506.9432096 | ENSG00000187325 |
| NA | 0.208248859 | 1.15528505 | 0.793937738 | 0.894547711 | 6.320199605 | 7.283037442 | ENSG00000273343 |
| UNC13A | 0.208198622 | 1.155244821 | 0.0007653 | 0.004437422 | 1824.12313 | 2107.146539 | ENSG00000130477 |
| NA | 0.208048304 | 1.155124459 | 0.836406677 | 0.917869805 | 4.86832771 | 5.657493178 | ENSG00000271978 |
| ZBTB39 | 0.208016619 | 1.155099091 | 0.00469299 | 0.020952035 | 1041.66246 | 1203.027574 | ENSG00000166860 |
| MAP4K2 | 0.207933654 | 1.155032666 | 0.042987165 | 0.125473356 | 540.1035221 | 624.0793052 | ENSG00000168067 |
| EIF3J-DT | 0.207899322 | 1.15500518 | 0.271768007 | 0.480466094 | 178.6831209 | 206.4423517 | ENSG00000179523 |
| CENPVL3 | 0.207789566 | 1.154917314 | 0.25711234 | 0.463974588 | 130.3817973 | 150.5392009 | ENSG00000224109 |
| TCTE3 | 0.207758758 | 1.154892651 | 0.674922364 | 0.825790913 | 18.15578302 | 21.01123839 | ENSG00000184786 |
| REEP4 | 0.207742696 | 1.154879794 | 0.017082748 | 0.060572231 | 840.4862951 | 971.1454284 | ENSG00000168476 |

| | | | | | | | |
|---|---|---|---|---|---|---|---|
| RBM38 | 0.207690426 | 1.154837953 | 0.004792173 | 0.021289151 | 1423.20567 | 1643.91759 | ENSG00000132819 |
| ZSCAN16 | 0.207686619 | 1.154834905 | 0.529614633 | 0.721836919 | 41.99419275 | 48.52465383 | ENSG00000196812 |
| ELP4 | 0.20764146 | 1.154798757 | 0.169970389 | 0.348068836 | 189.8349105 | 219.2121322 | ENSG00000109911 |
| RBKS | 0.207579078 | 1.154748825 | 0.689737845 | 0.83399945 | 16.06753344 | 18.61542751 | ENSG00000171174 |
| GGH | 0.207370807 | 1.154582135 | 0.025882503 | 0.084624491 | 616.5899465 | 711.6982737 | ENSG00000137563 |
| NA | 0.207187806 | 1.154435689 | 0.520553946 | 0.715685882 | 39.67257099 | 45.81796015 | ENSG00000218226 |
| ADAMTS12 | 0.207056776 | 1.154330844 | 0.087586962 | 0.21597407 | 415.6942924 | 479.3205897 | ENSG00000151388 |
| MOSPD2 | 0.206951122 | 1.154246311 | 0.196687136 | 0.38603581 | 174.8883227 | 201.9703146 | ENSG00000130150 |
| HEXB | 0.206935769 | 1.154234028 | 0.008962039 | 0.03581605 | 980.7035762 | 1131.968858 | ENSG00000049860 |
| EML5 | 0.206907256 | 1.154211216 | 0.69790122 | 0.838947456 | 15.45607206 | 17.85670062 | ENSG00000165521 |
| BRI3 | 0.206902424 | 1.154207351 | 0.06341286 | 0.169217444 | 657.5452434 | 759.3074922 | ENSG00000164713 |
| NA | 0.206690332 | 1.154037682 | 0.675836905 | 0.826595731 | 17.7575322 | 20.5167636 | ENSG00000286827 |
| CENPJ | 0.206675885 | 1.154026125 | 0.036194293 | 0.110015969 | 599.9800038 | 691.9341394 | ENSG00000151849 |
| PCDHGB2 | 0.206578476 | 1.153948209 | 0.637001499 | 0.80075706 | 28.00306142 | 32.27051333 | ENSG00000253910 |
| MFAP4 | 0.206539499 | 1.153917035 | 0.38151887 | 0.593980532 | 77.96090006 | 90.03553565 | ENSG00000166482 |
| TRAPPC2B | 0.206522131 | 1.153903143 | 0.619033156 | 0.788669038 | 26.86047968 | 31.01373873 | ENSG00000256060 |
| ARC | 0.206486522 | 1.153874662 | 0.0354753 | 0.10830117 | 649.2591862 | 749.362246 | ENSG00000198576 |
| XBP1 | 0.206332076 | 1.153751142 | 0.000546761 | 0.003313727 | 2823.404896 | 3257.609562 | ENSG00000100219 |
| ACTR1B | 0.206300436 | 1.153725839 | 0.009884787 | 0.038656908 | 942.4554664 | 1087.061363 | ENSG00000115073 |
| GOLGA8B | 0.206196344 | 1.1536426 | 0.62220867 | 0.791007118 | 160.7038108 | 185.454256 | ENSG00000215252 |
| UBE2G2 | 0.206170363 | 1.153621824 | 0.000779007 | 0.004508595 | 2314.222541 | 2669.461696 | ENSG00000184787 |
| FAM171B | 0.20616771 | 1.153619703 | 0.113985932 | 0.263157008 | 303.7367119 | 350.2865646 | ENSG00000144369 |
| FOXRED2 | 0.206115725 | 1.153578135 | 0.047832569 | 0.136308747 | 688.7167922 | 795.1086268 | ENSG00000100350 |
| TMEM209 | 0.205838307 | 1.153356333 | 0.103977204 | 0.245205981 | 277.8693422 | 320.4811208 | ENSG00000146842 |
| TJAP1 | 0.205816662 | 1.153339029 | 0.018235617 | 0.063750393 | 741.37279 | 855.506252 | ENSG00000137221 |
| MMACHC | 0.205782481 | 1.153311704 | 0.264947348 | 0.472566175 | 132.2633705 | 152.7005538 | ENSG00000132763 |
| ATE1 | 0.205624064 | 1.15318507 | 0.084774699 | 0.211070559 | 414.0325412 | 477.6515135 | ENSG00000107669 |
| NA | 0.205577289 | 1.153147682 | 0.389632805 | 0.601355258 | 84.78104834 | 97.83033599 | ENSG00000272163 |
| KAZN | 0.205516941 | 1.153099447 | 0.057903437 | 0.157954438 | 399.9648815 | 461.2007509 | ENSG00000189337 |
| CCDC97 | 0.205304421 | 1.152929599 | 0.029869037 | 0.094580277 | 606.4287011 | 699.4320033 | ENSG00000142039 |
| MRNIP | 0.205249128 | 1.152885412 | 0.203353565 | 0.394933368 | 302.794909 | 349.1737425 | ENSG00000161010 |
| MLX | 0.205194764 | 1.15284197 | 0.015037065 | 0.054589782 | 798.1518948 | 919.9095393 | ENSG00000108788 |
| FAM3C | 0.205123168 | 1.15278476 | 0.024497158 | 0.081077607 | 1145.517625 | 1320.697466 | ENSG00000196937 |
| FAM216A | 0.205099784 | 1.152766075 | 0.208664251 | 0.401653693 | 171.7897244 | 197.9342983 | ENSG00000204856 |
| CTSB | 0.204987911 | 1.152676688 | 0.009747148 | 0.03818981 | 1071.998659 | 1235.282598 | ENSG00000164733 |
| CHAMP1 | 0.204934709 | 1.152634182 | 0.089713569 | 0.219917023 | 588.1066428 | 678.2960784 | ENSG00000198824 |
| ZNF865 | 0.204912673 | 1.152616576 | 0.097240006 | 0.233309302 | 602.6430533 | 695.2950651 | ENSG00000261221 |
| ATP1A2 | 0.204834822 | 1.15255438 | 0.030669322 | 0.096628318 | 2002.028874 | 2307.456592 | ENSG00000018625 |
| TMEM263 | 0.204686791 | 1.152436126 | 0.151022143 | 0.320491423 | 233.4488285 | 269.2156486 | ENSG00000151135 |
| HECTD1 | 0.204672791 | 1.152424943 | 0.013098878 | 0.048762529 | 1902.020772 | 2191.15655 | ENSG00000092148 |
| TMEM68 | 0.204642274 | 1.152400566 | 0.119725233 | 0.272279747 | 279.7399321 | 322.209386 | ENSG00000167904 |
| ATG16L2 | 0.204629557 | 1.152390408 | 0.267661078 | 0.475391897 | 241.4682052 | 278.3295472 | ENSG00000168010 |
| CREB3L1 | 0.204456151 | 1.152251904 | 0.130166252 | 0.289164766 | 247.7389032 | 285.5589628 | ENSG00000157613 |
| NA | 0.204415417 | 1.152219371 | 0.471942123 | 0.675199014 | 63.22037765 | 72.91777183 | ENSG00000256008 |
| NA | 0.204397692 | 1.152205214 | 0.614909086 | 0.785838998 | 32.63890667 | 37.68305003 | ENSG00000268858 |
| NA | 0.20439608 | 1.152203927 | 0.794329523 | 0.894861791 | 6.616447084 | 7.620404664 | ENSG00000213280 |
| TEAD3 | 0.204336687 | 1.152156494 | 0.022112479 | 0.074663717 | 767.3460279 | 884.4760403 | ENSG00000007866 |
| CAPN7 | 0.204307281 | 1.15213301 | 0.13200785 | 0.292190372 | 302.9627617 | 349.4722646 | ENSG00000131375 |
| NA | 0.20421865 | 1.152062232 | 0.726743829 | 0.856072681 | 14.18568096 | 16.38854747 | ENSG00000258199 |
| ZDBF2 | 0.204180201 | 1.152031529 | 0.123280062 | 0.27795081 | 418.8462305 | 482.0381904 | ENSG00000204186 |
| KCTD11 | 0.204030712 | 1.151912164 | 0.213792764 | 0.408320829 | 178.4130111 | 205.517153 | ENSG00000213859 |
| ZNF764 | 0.203980466 | 1.151872046 | 0.263891885 | 0.471262245 | 141.2204058 | 162.7505152 | ENSG00000169951 |
| C19orf71 | 0.20395055 | 1.15184816 | 0.58794647 | 0.765960892 | 29.37278611 | 33.86683061 | ENSG00000183397 |
| NEURL1B | 0.203921839 | 1.151825238 | 0.094451636 | 0.22873983 | 387.0166928 | 445.8122413 | ENSG00000214357 |
| ZNF414 | 0.20377151 | 1.151705224 | 0.107346514 | 0.251716022 | 301.5921211 | 347.6723246 | ENSG00000133250 |
| TXNDC11 | 0.20373138 | 1.151673189 | 0.149851659 | 0.318687904 | 227.8504339 | 262.5927797 | ENSG00000153066 |
| SAV1 | 0.203632085 | 1.151593926 | 0.007168505 | 0.029886036 | 1512.73931 | 1742.602458 | ENSG00000151748 |
| XPA | 0.203474328 | 1.151468007 | 0.383960726 | 0.596307427 | 84.29294714 | 97.15420409 | ENSG00000136936 |
| NA | 0.203441976 | 1.151442186 | 0.590230481 | 0.768459333 | 37.96388514 | 43.68878775 | ENSG00000272455 |
| SGF29 | 0.203430644 | 1.151433143 | 0.233505798 | 0.434544443 | 161.7687207 | 186.4973907 | ENSG00000176476 |
| NA | 0.203358191 | 1.151375318 | 0.837140337 | 0.918172233 | 4.65155768 | 5.363351681 | ENSG00000228201 |
| CAPN2 | 0.203318681 | 1.151343787 | 0.005168268 | 0.022744351 | 1206.556091 | 1389.204624 | ENSG00000162909 |
| NRBF2 | 0.20331511 | 1.151340937 | 0.059762487 | 0.161901386 | 522.0556144 | 600.9149643 | ENSG00000148572 |
| RHOD | 0.203287284 | 1.151318731 | 0.133996279 | 0.295310973 | 249.2806255 | 287.2310651 | ENSG00000173156 |
| RNF126 | 0.203261426 | 1.151298096 | 0.005921039 | 0.025448003 | 1669.113006 | 1922.17501 | ENSG00000070423 |
| MED13 | 0.203247152 | 1.151286705 | 0.05613168 | 0.154323304 | 522.3215799 | 601.162126 | ENSG00000108510 |
| INTS8 | 0.203223047 | 1.151267468 | 0.005449231 | 0.023733562 | 1322.742166 | 1522.752341 | ENSG00000164941 |
| CPSF6 | 0.203127222 | 1.151191003 | 0.00032617 | 0.002110936 | 3099.293708 | 3567.403462 | ENSG00000111605 |
| TSKU | 0.203070658 | 1.151145869 | 0.014845514 | 0.054081316 | 884.9225468 | 1019.257781 | ENSG00000182704 |
| KMT5C | 0.202925925 | 1.15103039 | 0.032450334 | 0.101059726 | 695.5155319 | 801.1290686 | ENSG00000133247 |
| TMEM198B | 0.202905809 | 1.151014341 | 0.309796026 | 0.520321767 | 144.9046338 | 166.7013525 | ENSG00000182796 |
| NA | 0.202863318 | 1.150980442 | 0.662797895 | 0.819171351 | 21.08769711 | 24.3298616 | ENSG00000228613 |
| LOC101928069 | 0.202861839 | 1.150979262 | 0.763164764 | 0.877513739 | 9.214154701 | 10.62653672 | ENSG00000255517 |
| DENND4A | 0.202349681 | 1.150570735 | 0.339773522 | 0.551398382 | 117.8840564 | 135.5410046 | ENSG00000174485 |

| | | | | | | | |
|---|---|---|---|---|---|---|---|
| ZNF485 | 0.202172997 | 1.150429836 | 0.298015296 | 0.508219151 | 130.2712267 | 150.0090883 | ENSG00000198298 |
| CD22 | 0.202110034 | 1.15037963 | 0.846909849 | 0.924021751 | 4.042460567 | 4.616448073 | ENSG00000012124 |
| SPDYE3 | 0.202005915 | 1.15029661 | 0.543958097 | 0.73244858 | 41.88537595 | 48.11294198 | ENSG00000214300 |
| ZCCHC7 | 0.201791112 | 1.150125354 | 0.063353526 | 0.169106902 | 418.1316223 | 480.7095529 | ENSG00000147905 |
| FERMT2 | 0.201708229 | 1.150059281 | 0.001590518 | 0.008371495 | 2515.603042 | 2892.684665 | ENSG00000073712 |
| NA | 0.20170468 | 1.150056452 | 0.414773756 | 0.625609309 | 67.82716288 | 78.0144124 | ENSG00000228327 |
| ZSCAN23 | 0.201673527 | 1.150031619 | 0.544801228 | 0.733169756 | 38.15101034 | 43.84975178 | ENSG00000187987 |
| NPAS4 | 0.201487289 | 1.14988317 | 0.815285203 | 0.90611235 | 6.923372981 | 7.970541121 | ENSG00000174576 |
| METTL5 | 0.201485069 | 1.149881401 | 0.02192364 | 0.074158687 | 786.8058871 | 904.4879033 | ENSG00000138382 |
| BLOC1S4 | 0.201355342 | 1.149778008 | 0.069526589 | 0.181706288 | 460.1255357 | 529.1925597 | ENSG00000186222 |
| ACTL6B | 0.201219686 | 1.149669901 | 0.656357738 | 0.814302713 | 25.76681469 | 29.53487622 | ENSG00000077080 |
| ADAM28 | 0.201178366 | 1.149636974 | 0.763303713 | 0.877518148 | 9.472443192 | 10.92938007 | ENSG00000042980 |
| TRAF3IP2 | 0.201090964 | 1.149567328 | 0.047045862 | 0.134513414 | 465.3983173 | 534.939771 | ENSG00000056972 |
| SHC1 | 0.201060388 | 1.149542964 | 0.002753643 | 0.013347678 | 6286.084629 | 7226.43661 | ENSG00000160691 |
| H4C4 | 0.200952734 | 1.149457189 | 0.841843848 | 0.92140286 | 5.144413563 | 5.954093685 | ENSG00000277157 |
| ATP8B2 | 0.200943118 | 1.149449527 | 0.001178604 | 0.006466571 | 1784.083811 | 2050.705447 | ENSG00000143515 |
| C19orf84 | 0.200928151 | 1.149437603 | 0.418573738 | 0.629283182 | 71.67181983 | 82.31250533 | ENSG00000262874 |
| FAIM2 | 0.200861791 | 1.149384732 | 0.68626086 | 0.83186293 | 18.22694632 | 20.95117604 | ENSG00000135472 |
| MICB | 0.200688507 | 1.149246687 | 0.075097817 | 0.19277182 | 386.1151348 | 443.4785299 | ENSG00000204516 |
| ZBTB8B | 0.20057305 | 1.149154718 | 0.363968511 | 0.57608415 | 93.77042434 | 107.7785653 | ENSG00000273274 |
| SLC39A9 | 0.20053055 | 1.149120866 | 0.043993589 | 0.127628022 | 535.1432329 | 614.6755957 | ENSG00000029364 |
| CBX4 | 0.200393228 | 1.149011492 | 0.339664576 | 0.55136372 | 107.2634126 | 123.4486582 | ENSG00000141582 |
| GOLGA5 | 0.200376377 | 1.148998072 | 0.045319677 | 0.130506572 | 557.3885876 | 639.9644222 | ENSG00000066455 |
| KIAA1522 | 0.200352834 | 1.148979322 | 0.001646894 | 0.008608248 | 1670.665434 | 1919.918986 | ENSG00000162522 |
| FAM149A | 0.200254584 | 1.148901077 | 0.11907173 | 0.271283583 | 272.2403608 | 312.7128049 | ENSG00000109794 |
| TLCD3A | 0.200249654 | 1.148897151 | 0.047087012 | 0.13461069 | 500.2043662 | 574.9260532 | ENSG00000167695 |
| MKNK2 | 0.200165218 | 1.148829912 | 0.000274327 | 0.001818416 | 3525.267831 | 4049.852305 | ENSG00000099875 |
| NA | 0.200161481 | 1.148826936 | 0.622267227 | 0.791007229 | 30.76178271 | 35.39516169 | ENSG00000279198 |
| KIF27 | 0.200112428 | 1.148787876 | 0.408420809 | 0.619570003 | 78.1732209 | 89.71791555 | ENSG00000165115 |
| SLC25A6 | 0.199945583 | 1.148655028 | 0.000142068 | 0.001018972 | 16041.47363 | 18426.26402 | ENSG00000169100 |
| FUT1 | 0.199925762 | 1.148639247 | 0.14000068 | 0.304072782 | 247.749861 | 284.6745737 | ENSG00000174951 |
| PTCHD4 | 0.199898563 | 1.148617592 | 0.689622582 | 0.83398915 | 19.4335724 | 22.24436643 | ENSG00000244694 |
| NA | 0.199895767 | 1.148615366 | 0.800284249 | 0.898320559 | 6.647287128 | 7.620404664 | ENSG00000259969 |
| C1orf198 | 0.19988989 | 1.148610687 | 0.025621148 | 0.08398809 | 807.7560902 | 927.7033281 | ENSG00000119280 |
| RPL39 | 0.19984162 | 1.148572257 | 0.014992919 | 0.054481677 | 4148.521701 | 4763.945742 | ENSG00000198918 |
| FAM167B | 0.199796186 | 1.148536086 | 0.675000845 | 0.825822166 | 21.48002419 | 24.72010236 | ENSG00000183615 |
| TSC22D3 | 0.199767565 | 1.148513301 | 0.035965205 | 0.109460578 | 628.4323439 | 721.3120463 | ENSG00000157514 |
| TINF2 | 0.199737689 | 1.148489518 | 0.097199705 | 0.233271881 | 381.3180226 | 437.797812 | ENSG00000092330 |
| DUBR | 0.199718637 | 1.148474351 | 0.558530074 | 0.744063237 | 38.22692832 | 43.8528732 | ENSG00000243701 |
| TSPY26P | 0.199698088 | 1.148457993 | 0.128593104 | 0.286647871 | 260.2799427 | 298.9469586 | ENSG00000235217 |
| SH2D7 | 0.199670786 | 1.148436259 | 0.644354506 | 0.805989478 | 24.64798021 | 28.26627835 | ENSG00000183476 |
| NA | 0.199659194 | 1.148427031 | 0.697782511 | 0.838858153 | 17.61428973 | 20.18715227 | ENSG00000279159 |
| NAA35 | 0.199613398 | 1.148390577 | 0.020782013 | 0.070996439 | 770.9000562 | 884.9622514 | ENSG00000135040 |
| NA | 0.199583309 | 1.148366626 | 0.252738177 | 0.458673888 | 149.1908657 | 171.2170441 | ENSG00000225889 |
| COL11A2 | 0.199567629 | 1.148354145 | 0.3901007 | 0.601690159 | 80.0989561 | 91.99040769 | ENSG00000204248 |
| MTIF2 | 0.199557231 | 1.148345869 | 0.016830687 | 0.05982449 | 827.4512312 | 950.1904029 | ENSG00000085760 |
| CD58 | 0.199213924 | 1.148072638 | 0.485245 | 0.68627663 | 55.28787729 | 63.33322617 | ENSG00000116815 |
| ZNF223 | 0.199158189 | 1.148028287 | 0.771056245 | 0.882285172 | 9.224833118 | 10.5994852 | ENSG00000178386 |
| ISM2 | 0.19899019 | 1.147894608 | 0.637066461 | 0.800785418 | 23.69608314 | 27.26287853 | ENSG00000100593 |
| TPD52 | 0.19886968 | 1.147798728 | 0.210928836 | 0.40461057 | 245.8653643 | 282.5919891 | ENSG00000076554 |
| NA | 0.198799548 | 1.147742933 | 0.74415338 | 0.867008242 | 12.81476105 | 14.63606858 | ENSG00000280152 |
| ENOX1 | 0.198784496 | 1.147730958 | 0.646496896 | 0.807960683 | 25.96314376 | 29.85171831 | ENSG00000120658 |
| NUDT13 | 0.198767691 | 1.147717589 | 0.589337754 | 0.767455763 | 30.09456499 | 34.55991472 | ENSG00000166321 |
| MAN1A2 | 0.198701572 | 1.14766499 | 0.068624012 | 0.179870154 | 429.8203776 | 493.0904604 | ENSG00000198162 |
| ZDHHC13 | 0.19861508 | 1.147596187 | 0.08922641 | 0.219092544 | 465.9991787 | 534.7029266 | ENSG00000177054 |
| PIK3CB | 0.198465645 | 1.147477325 | 0.073338806 | 0.189596625 | 724.9632702 | 831.1045142 | ENSG00000051382 |
| ZBED1 | 0.198371809 | 1.147402693 | 0.002658695 | 0.012967319 | 1491.137454 | 1711.146163 | ENSG00000214717 |
| MEF2A | 0.198321786 | 1.147362909 | 0.114598099 | 0.264195014 | 300.4581329 | 344.8836031 | ENSG00000068305 |
| NA | 0.198298903 | 1.14734471 | 0.511574902 | 0.708135455 | 54.00057854 | 61.89418441 | ENSG00000187534 |
| UBALD2 | 0.198292088 | 1.147339291 | 0.079591742 | 0.201109186 | 367.7734473 | 422.1257843 | ENSG00000185262 |
| SNHG1 | 0.198210334 | 1.147274276 | 0.269397625 | 0.477414112 | 1658.857874 | 1903.243197 | ENSG00000255717 |
| NVL | 0.198090006 | 1.147178591 | 0.009271524 | 0.036722469 | 933.2016505 | 1070.743449 | ENSG00000143748 |
| NUDT12 | 0.198049857 | 1.147146667 | 0.180027783 | 0.362802761 | 208.0704765 | 238.5850571 | ENSG00000112874 |
| PIEZO1 | 0.198040109 | 1.147138916 | 0.005037901 | 0.022234345 | 1673.199123 | 1919.740324 | ENSG00000103335 |
| SLC15A4 | 0.198007489 | 1.147112979 | 0.038851036 | 0.116183978 | 697.4798383 | 799.8379584 | ENSG00000139370 |
| RAPGEFL1 | 0.197972804 | 1.1470854 | 0.158899118 | 0.332242011 | 365.1710635 | 419.153484 | ENSG00000108352 |
| ZNF579 | 0.197965043 | 1.14707923 | 0.099698474 | 0.237607986 | 777.6594903 | 892.7465963 | ENSG00000218891 |
| FABP6 | 0.197841531 | 1.14698103 | 0.553372265 | 0.739906204 | 42.32839922 | 48.59464753 | ENSG00000170231 |
| PCSK5 | 0.197824539 | 1.146967521 | 0.152034186 | 0.321987844 | 232.0141436 | 266.0627192 | ENSG00000099139 |
| SIRT5 | 0.197775466 | 1.146928508 | 0.101909488 | 0.241354488 | 325.7519484 | 373.3362159 | ENSG00000124523 |
| NA | 0.19765595 | 1.146833498 | 0.76641032 | 0.879002726 | 9.51040218 | 10.933731 | ENSG00000231064 |
| ARL5B | 0.197536833 | 1.146738813 | 0.078606659 | 0.199179761 | 998.6630028 | 1145.167655 | ENSG00000165997 |
| SH3GLB2 | 0.197498161 | 1.146708074 | 0.037984773 | 0.114099322 | 1362.050835 | 1561.795566 | ENSG00000148341 |
| KCNIP2 | 0.197491672 | 1.146702917 | 0.519354847 | 0.715026338 | 43.74231213 | 50.14338815 | ENSG00000120049 |

| | | | | | | | |
|---|---|---|---|---|---|---|---|
| MYO5A | 0.197462332 | 1.146679596 | 0.037506505 | 0.112986178 | 713.0843877 | 817.0767795 | ENSG00000197535 |
| LMTK3 | 0.197362944 | 1.146600604 | 0.103069924 | 0.243492058 | 354.1313707 | 406.3614841 | ENSG00000142235 |
| SLC26A2 | 0.197340387 | 1.146582676 | 0.174098129 | 0.354331626 | 241.1405067 | 276.4242258 | ENSG00000155850 |
| RITA1 | 0.197315772 | 1.146563114 | 0.036543786 | 0.110828554 | 563.6872028 | 646.4132016 | ENSG00000139405 |
| NA | 0.197239276 | 1.146502321 | 0.838735475 | 0.91909659 | 4.610039219 | 5.28683159 | ENSG00000204092 |
| TRIM69 | 0.197052038 | 1.146353534 | 0.278565638 | 0.487748658 | 134.1137725 | 153.7217362 | ENSG00000185880 |
| CXorf40A | 0.197042652 | 1.146346076 | 0.318149715 | 0.528152019 | 111.4686407 | 127.8052051 | ENSG00000197620 |
| ZNF239 | 0.197033077 | 1.146338468 | 0.233102241 | 0.434092872 | 166.2014234 | 190.6469099 | ENSG00000196793 |
| NA | 0.197003296 | 1.146314805 | 0.175387589 | 0.355933754 | 12241.64602 | 14033.05426 | ENSG00000211459 |
| ADAT2 | 0.196987052 | 1.146301898 | 0.193988807 | 0.382129824 | 225.6791218 | 258.3771232 | ENSG00000189007 |
| SPTBN4 | 0.196928842 | 1.146255648 | 0.375380966 | 0.587433845 | 98.21708562 | 112.4888216 | ENSG00000160460 |
| RBP7 | 0.196755018 | 1.146117549 | 0.587035383 | 0.765497261 | 32.23025676 | 36.91060795 | ENSG00000162444 |
| MAN2B1 | 0.196709369 | 1.146081284 | 0.017502548 | 0.061730031 | 942.5400295 | 1079.918111 | ENSG00000104774 |
| G6PC3 | 0.196706222 | 1.146078784 | 0.004429726 | 0.019979787 | 2117.238891 | 2427.010684 | ENSG00000141349 |
| KLHL22 | 0.196679796 | 1.146057792 | 0.236447323 | 0.43845703 | 155.7168776 | 178.4942176 | ENSG00000099910 |
| LOC257396 | 0.196629957 | 1.146018201 | 0.646866783 | 0.808315966 | 26.75639141 | 30.58490672 | ENSG00000247796 |
| WDR25 | 0.196497861 | 1.145913274 | 0.20354767 | 0.395188419 | 203.9692844 | 233.8746852 | ENSG00000176473 |
| GMIP | 0.196428776 | 1.145858402 | 0.057528197 | 0.157224313 | 559.0498824 | 640.8498525 | ENSG00000089639 |
| PCBP3 | 0.19626826 | 1.14573092 | 0.276117963 | 0.485068627 | 141.3016632 | 161.8313187 | ENSG00000183570 |
| GET1 | 0.196178172 | 1.145659377 | 0.402560506 | 0.61393481 | 89.1503171 | 102.3321672 | ENSG00000182093 |
| XRN1 | 0.196133587 | 1.145623973 | 0.219116408 | 0.415546633 | 178.5728295 | 204.5579975 | ENSG00000114127 |
| LAMTOR2 | 0.196001705 | 1.145519252 | 0.029329217 | 0.093241472 | 710.6331759 | 814.1590768 | ENSG00000116586 |
| NA | 0.195951751 | 1.145479588 | 0.582796246 | 0.76255721 | 38.09764441 | 43.67601851 | ENSG00000280063 |
| SOD1 | 0.195760474 | 1.145327727 | 0.000251354 | 0.001683869 | 3392.57455 | 3885.490321 | ENSG00000142168 |
| SLFNL1-AS1 | 0.195593523 | 1.145195196 | 0.746769846 | 0.86838024 | 11.60101602 | 13.27912735 | ENSG00000281207 |
| EMB | 0.195483589 | 1.145107935 | 0.094463433 | 0.22873983 | 685.0678284 | 784.5389224 | ENSG00000170571 |
| CHML | 0.195480587 | 1.145105552 | 0.076109581 | 0.194760087 | 1084.440851 | 1241.802129 | ENSG00000203668 |
| FAM217B | 0.195435589 | 1.145069837 | 0.118146492 | 0.27002257 | 378.3110797 | 433.2605407 | ENSG00000196227 |
| NA | 0.195077845 | 1.144785929 | 0.003117122 | 0.014880397 | 2616.05139 | 2995.602234 | ENSG00000215030 |
| IGSF8 | 0.195006322 | 1.144729177 | 0.000558814 | 0.003379188 | 2668.359834 | 3054.81593 | ENSG00000162729 |
| LRRC17 | 0.195002207 | 1.144725912 | 0.713569122 | 0.848956862 | 16.1564941 | 18.48849289 | ENSG00000128606 |
| NA | 0.194978414 | 1.144707033 | 0.836382174 | 0.917869805 | 4.655117152 | 5.324476883 | ENSG00000279253 |
| ZNF141 | 0.194914914 | 1.14465665 | 0.128574884 | 0.286641092 | 300.6641989 | 343.925888 | ENSG00000131127 |
| NOTCH2 | 0.194866368 | 1.144618134 | 0.019847916 | 0.068336901 | 1712.235746 | 1959.173365 | ENSG00000134250 |
| AGAP6 | 0.194581669 | 1.144392279 | 0.692129886 | 0.835483064 | 21.3023822 | 24.37431683 | ENSG00000204149 |
| STXBP5 | 0.194535616 | 1.144355749 | 0.290490865 | 0.500038282 | 148.1283718 | 169.5651834 | ENSG00000164506 |
| MIER2 | 0.194533674 | 1.144354208 | 0.013145259 | 0.048896637 | 890.2993917 | 1018.606577 | ENSG00000105556 |
| MRM3 | 0.194531821 | 1.144352738 | 0.090640042 | 0.22152675 | 345.4456149 | 395.1753969 | ENSG00000171861 |
| MICU1 | 0.194406644 | 1.144253452 | 0.02591621 | 0.084676401 | 813.8170337 | 930.8948191 | ENSG00000107745 |
| SENP2 | 0.194404128 | 1.144251456 | 0.020437567 | 0.070009695 | 834.2103098 | 954.7462215 | ENSG00000163904 |
| DCUN1D1 | 0.19434565 | 1.144205076 | 0.096674725 | 0.232307186 | 473.627981 | 541.9554646 | ENSG00000043093 |
| PKD1L1 | 0.194311202 | 1.144177756 | 0.80294062 | 0.899379218 | 7.825158102 | 9.003167918 | ENSG00000158683 |
| NA | 0.194283895 | 1.144156099 | 0.83040751 | 0.914737804 | 4.937126742 | 5.666195032 | ENSG00000235554 |
| TMEM86A | 0.194253574 | 1.144132053 | 0.611714999 | 0.784141186 | 28.37492139 | 32.48406735 | ENSG00000151117 |
| NA | 0.194188877 | 1.144080746 | 0.386885115 | 0.599124673 | 114.1689365 | 130.522041 | ENSG00000224631 |
| SIKE1 | 0.194168585 | 1.144064654 | 0.026376201 | 0.085867016 | 741.8528825 | 848.9352451 | ENSG00000052723 |
| PAG1 | 0.194122376 | 1.144028011 | 0.809335331 | 0.902986417 | 6.636608711 | 7.566585167 | ENSG00000076641 |
| TANC2 | 0.194062405 | 1.143980456 | 0.180999193 | 0.363944821 | 231.6666412 | 265.0998025 | ENSG00000170921 |
| MEPCE | 0.194029531 | 1.143954389 | 0.005067702 | 0.022344979 | 1513.674937 | 1731.824603 | ENSG00000146834 |
| NA | 0.193866672 | 1.143825261 | 0.711398679 | 0.847591497 | 15.69803773 | 17.89434591 | ENSG00000234678 |
| MIA2 | 0.193824579 | 1.143791888 | 0.141193625 | 0.305679424 | 272.8444239 | 312.0637769 | ENSG00000150527 |
| HOMER1 | 0.193767529 | 1.143746659 | 0.028504997 | 0.091147455 | 805.2594439 | 920.7246268 | ENSG00000152413 |
| NA | 0.193741347 | 1.143725903 | 0.795453831 | 0.895554092 | 11.6576621 | 13.42223617 | ENSG00000283877 |
| CHCHD4 | 0.193721415 | 1.143710101 | 0.108608783 | 0.253751932 | 375.5950198 | 429.6486493 | ENSG00000163528 |
| MAPK12 | 0.193540719 | 1.143566861 | 0.138209969 | 0.301361882 | 568.9505279 | 650.6160592 | ENSG00000188130 |
| CLIP2 | 0.193414097 | 1.143466497 | 0.005510559 | 0.023967505 | 1912.220664 | 2186.215803 | ENSG00000106665 |
| PFKM | 0.193369744 | 1.143431344 | 0.001671043 | 0.008727228 | 2730.542864 | 3121.56403 | ENSG00000152556 |
| PIR | 0.193352376 | 1.143417579 | 0.021222656 | 0.07216933 | 846.3189271 | 967.1354747 | ENSG00000087842 |
| NA | 0.193347982 | 1.143414097 | 0.840661571 | 0.920561463 | 4.647998207 | 5.308302679 | ENSG00000237991 |
| ZNF688 | 0.193261518 | 1.143345572 | 0.509773411 | 0.706366636 | 54.68887377 | 62.56825644 | ENSG00000229809 |
| TMPRSS6 | 0.19314573 | 1.143253813 | 0.804846247 | 0.90064726 | 11.06992182 | 12.71704524 | ENSG00000187045 |
| NKD1 | 0.193127729 | 1.143239548 | 0.19989219 | 0.39021749 | 219.3577526 | 250.9196901 | ENSG00000140807 |
| PDIK1L | 0.193047674 | 1.143176111 | 0.502215912 | 0.700181291 | 57.63294082 | 65.80477914 | ENSG00000175087 |
| RFK | 0.193000683 | 1.143138877 | 0.207355045 | 0.399826428 | 251.7262178 | 287.9560758 | ENSG00000135002 |
| NA | 0.192943064 | 1.143093223 | 0.703641158 | 0.842524014 | 16.32137274 | 18.62942625 | ENSG00000283638 |
| MCF2L | 0.192903475 | 1.143061855 | 0.059669841 | 0.161719998 | 539.6055804 | 616.7066949 | ENSG00000126217 |
| CLSTN2 | 0.192896806 | 1.143056571 | 0.525080648 | 0.718717695 | 54.19215293 | 62.07522207 | ENSG00000158258 |
| ZNF852 | 0.192636419 | 1.142850283 | 0.287062176 | 0.496231181 | 165.8832087 | 189.5333395 | ENSG00000178917 |
| CCDC181 | 0.192625924 | 1.14284197 | 0.430530733 | 0.640353855 | 74.90829555 | 85.54212277 | ENSG00000117477 |
| KDM7A-DT | 0.192508784 | 1.142749181 | 0.515633828 | 0.711976214 | 57.15549188 | 65.29880758 | ENSG00000260231 |
| MPV17 | 0.192380851 | 1.14264785 | 0.038554109 | 0.115515523 | 778.2862307 | 889.0338008 | ENSG00000115204 |
| TRIM16 | 0.192283833 | 1.142571012 | 0.316636508 | 0.526735445 | 118.348131 | 135.2050349 | ENSG00000221926 |
| HACD1 | 0.192242522 | 1.142538296 | 0.22609322 | 0.424768251 | 190.2969264 | 217.4528434 | ENSG00000165996 |
| PMP22 | 0.191940507 | 1.142299141 | 0.625995394 | 0.793253855 | 26.45747418 | 30.23883765 | ENSG00000109099 |

| | | | | | | | |
|---|---|---|---|---|---|---|---|
| MFSD2B | 0.1919262 | 1.142287812 | 0.772074499 | 0.882599617 | 9.868329748 | 11.29786619 | ENSG00000205639 |
| POR | 0.191864774 | 1.142239178 | 0.002245958 | 0.011226819 | 3038.011373 | 3470.208315 | ENSG00000127948 |
| NCOA4 | 0.19185757 | 1.142233474 | 0.00758995 | 0.031337168 | 1698.498253 | 1939.699688 | ENSG00000266412 |
| ANKZF1 | 0.191747543 | 1.142146365 | 0.022696352 | 0.076158603 | 887.6913373 | 1013.829891 | ENSG00000163516 |
| MIOS | 0.191723236 | 1.142127123 | 0.016786397 | 0.05971202 | 846.1099382 | 966.2469753 | ENSG00000164654 |
| DLGAP3 | 0.191684787 | 1.142096684 | 0.090176374 | 0.220736336 | 595.7347824 | 680.9727172 | ENSG00000116544 |
| FAM227A | 0.191683644 | 1.142095779 | 0.597515423 | 0.77367698 | 38.14745087 | 43.5580693 | ENSG00000184949 |
| SEPTIN7P2 | 0.191669999 | 1.142084977 | 0.17839169 | 0.360587647 | 224.8671871 | 256.672264 | ENSG00000214765 |
| TCP11L1 | 0.191591264 | 1.14202265 | 0.255921396 | 0.462814344 | 181.3565194 | 207.2446605 | ENSG00000176148 |
| RBBP5 | 0.191584002 | 1.142016902 | 0.0399583 | 0.118593823 | 671.6156703 | 766.6958549 | ENSG00000117222 |
| GNE | 0.191551207 | 1.141990942 | 0.103548848 | 0.244409447 | 385.0877031 | 439.9616762 | ENSG00000159921 |
| TOMM7 | 0.19144103 | 1.141903733 | 0.013503961 | 0.049945614 | 1749.592645 | 1997.837521 | ENSG00000196683 |
| UQCRB | 0.19124299 | 1.141746993 | 0.00356271 | 0.016645664 | 1689.758799 | 1929.345805 | ENSG00000156467 |
| MTG2 | 0.19104337 | 1.141589025 | 0.047292213 | 0.135115497 | 594.941562 | 679.2819816 | ENSG00000101181 |
| TEF | 0.190946968 | 1.141512746 | 0.220142822 | 0.416823461 | 227.7015215 | 259.5318822 | ENSG00000167074 |
| NA | 0.190883677 | 1.141462669 | 0.482183356 | 0.683636087 | 62.10805172 | 70.86048504 | ENSG00000218426 |
| WDR13 | 0.190827451 | 1.141418183 | 0.073222047 | 0.189450385 | 466.7657039 | 532.9566609 | ENSG00000101940 |
| NA | 0.190792887 | 1.141390838 | 0.667263106 | 0.821678649 | 21.79999277 | 24.89913252 | ENSG00000264015 |
| SYT6 | 0.190702747 | 1.141319525 | 0.038868011 | 0.116216337 | 704.3317163 | 803.518571 | ENSG00000134207 |
| TUT1 | 0.19055136 | 1.141199769 | 0.216696797 | 0.412282795 | 201.8303647 | 230.4849544 | ENSG00000149016 |
| ITM2C | 0.190490533 | 1.141151654 | 1.45E-05 | 0.000132886 | 13140.46929 | 14994.98764 | ENSG00000135916 |
| LOC100129215 | 0.190433401 | 1.141106465 | 0.641975979 | 0.804214244 | 27.59146243 | 31.56371402 | ENSG00000226180 |
| GGA3 | 0.190351972 | 1.14104206 | 0.004336795 | 0.019630603 | 1433.745816 | 1635.673791 | ENSG00000125447 |
| CBX5 | 0.190319048 | 1.141016021 | 0.007237831 | 0.03009389 | 3307.772399 | 3773.526591 | ENSG00000094916 |
| GOLPH3L | 0.190300909 | 1.141001675 | 0.027780969 | 0.089508108 | 812.1074581 | 926.1046174 | ENSG00000143457 |
| DNHD1 | 0.190281549 | 1.140986363 | 0.463458735 | 0.667971431 | 102.9350519 | 117.4808923 | ENSG00000179532 |
| KDM1B | 0.190147897 | 1.140880666 | 0.262359433 | 0.469413348 | 204.7551337 | 233.5525033 | ENSG00000165097 |
| ZFHX2 | 0.190056568 | 1.140808446 | 0.05088332 | 0.143102235 | 665.5147084 | 759.6316684 | ENSG00000136367 |
| COG4 | 0.18994886 | 1.14072328 | 0.01328808 | 0.049359831 | 1134.083932 | 1293.316979 | ENSG00000103051 |
| TMEM131L | 0.189893074 | 1.140679171 | 0.043819742 | 0.127315704 | 652.7408299 | 744.3865605 | ENSG00000121210 |
| RHOBTB1 | 0.189706018 | 1.140531283 | 0.172481954 | 0.352028714 | 283.1050546 | 323.0276284 | ENSG00000072422 |
| KMT2B | 0.189582261 | 1.140433451 | 0.0018379 | 0.009431724 | 2176.597629 | 2481.78823 | ENSG00000272333 |
| NFE2L2 | 0.189522321 | 1.14038607 | 0.085580493 | 0.212460577 | 468.1191575 | 534.2074332 | ENSG00000116044 |
| KLHL21 | 0.18949657 | 1.140365715 | 0.023517932 | 0.078469282 | 941.0953545 | 1073.231017 | ENSG00000162413 |
| VPS35L | 0.189465492 | 1.140341149 | 0.017504068 | 0.061730031 | 979.7982316 | 1117.270268 | ENSG00000103544 |
| EFNA5 | 0.189387993 | 1.140279894 | 0.355515534 | 0.567747007 | 121.3719804 | 138.5908139 | ENSG00000184349 |
| ABHD6 | 0.189379351 | 1.140273063 | 0.286164402 | 0.495345072 | 157.7546841 | 179.6399482 | ENSG00000163686 |
| LOC100128568 | 0.189175307 | 1.140111803 | 0.779957952 | 0.887747464 | 10.51538585 | 12.03512199 | ENSG00000266983 |
| PLEKHG1 | 0.18897411 | 1.139952815 | 0.524737367 | 0.718516443 | 48.33574919 | 55.0909606 | ENSG00000120278 |
| FAM86B2 | 0.188910447 | 1.139902513 | 0.746912761 | 0.86838024 | 13.72958886 | 15.59435075 | ENSG00000145002 |
| FANCL | 0.188899144 | 1.139893582 | 0.048553687 | 0.137947401 | 556.3830708 | 634.424522 | ENSG00000115392 |
| POLRMT | 0.188866444 | 1.139867745 | 0.005446245 | 0.023728989 | 3855.922126 | 4396.169415 | ENSG00000099821 |
| RBMS3 | 0.188761721 | 1.139785007 | 0.648651468 | 0.809250856 | 31.20007745 | 35.66594018 | ENSG00000144642 |
| NA | 0.188550514 | 1.139618158 | 0.723824457 | 0.854499273 | 14.29836272 | 16.27003116 | ENSG00000237977 |
| TMEM42 | 0.188530239 | 1.139602143 | 0.302680741 | 0.512817736 | 138.5429158 | 157.71131 | ENSG00000169964 |
| LINC00649 | 0.188504317 | 1.139581666 | 0.378077486 | 0.590532551 | 110.4936074 | 125.7269713 | ENSG00000237945 |
| MOCS2 | 0.188386407 | 1.139488533 | 0.113699197 | 0.262527123 | 321.3435772 | 366.2182099 | ENSG00000164172 |
| NA | 0.188341399 | 1.139452986 | 0.779303624 | 0.887377084 | 9.901534055 | 11.29351527 | ENSG00000223749 |
| TMEM125 | 0.188274313 | 1.139400001 | 0.09974529 | 0.237659577 | 363.7057917 | 414.3135802 | ENSG00000179178 |
| ICE2 | 0.187995853 | 1.139180103 | 0.035464856 | 0.10830117 | 883.4940394 | 1006.221537 | ENSG00000128915 |
| MAP3K11 | 0.187961156 | 1.139152705 | 0.010938813 | 0.042090358 | 1585.256347 | 1806.476855 | ENSG00000173327 |
| MAP3K2 | 0.18794834 | 1.139142586 | 0.194967089 | 0.383696695 | 209.0822736 | 238.1518742 | ENSG00000169967 |
| NA | 0.18789907 | 1.139103683 | 0.798337803 | 0.897150218 | 7.545512776 | 8.616332126 | ENSG00000213683 |
| CRYGS | 0.18777487 | 1.139005624 | 0.845564612 | 0.923288344 | 4.685957196 | 5.321355461 | ENSG00000213139 |
| CD4 | 0.187729537 | 1.138969834 | 0.429964854 | 0.639713721 | 80.63836446 | 91.70447359 | ENSG00000010610 |
| NA | 0.187655823 | 1.13891164 | 0.801138344 | 0.898333333 | 7.600073919 | 8.672327087 | ENSG00000279822 |
| MLH3 | 0.187606014 | 1.13887232 | 0.169886632 | 0.347981017 | 254.8656725 | 290.0313561 | ENSG00000119684 |
| SEC23A | 0.187572759 | 1.138846068 | 0.020571634 | 0.070405093 | 1112.749235 | 1266.748874 | ENSG00000100934 |
| CDCA7L | 0.187437681 | 1.138739444 | 0.002790779 | 0.013505755 | 2621.289034 | 2984.305403 | ENSG00000164649 |
| UBE2J2 | 0.187435771 | 1.138737937 | 0.008785845 | 0.035272256 | 1766.630709 | 2011.686633 | ENSG00000160087 |
| STX16 | 0.187401449 | 1.138710846 | 0.094785403 | 0.229196202 | 958.3911252 | 1091.331324 | ENSG00000124222 |
| LOC100505549 | 0.187371369 | 1.138687104 | 0.747365252 | 0.868553402 | 14.49616633 | 16.50505627 | ENSG00000267040 |
| SORD | 0.187296275 | 1.138627836 | 0.006858504 | 0.028789253 | 1467.250406 | 1671.123485 | ENSG00000140263 |
| IGF1R | 0.187234612 | 1.13857917 | 0.021420349 | 0.072733977 | 2006.548574 | 2283.934588 | ENSG00000140443 |
| FRA10AC1 | 0.187108819 | 1.138479899 | 0.120350481 | 0.273109689 | 345.6980052 | 393.8990431 | ENSG00000148690 |
| TMEM223 | 0.186927772 | 1.138337037 | 0.141061054 | 0.305427426 | 348.0098127 | 396.3378392 | ENSG00000168569 |
| SLC12A7 | 0.186858439 | 1.138282332 | 0.008939107 | 0.035765899 | 1433.521752 | 1632.079637 | ENSG00000113504 |
| NFRKB | 0.186822294 | 1.138253814 | 0.02523191 | 0.083043676 | 926.1701234 | 1054.407376 | ENSG00000170322 |
| TRAM2 | 0.18676527 | 1.138208825 | 0.035894324 | 0.109332953 | 730.2207738 | 831.1131637 | ENSG00000065308 |
| ZNF280C | 0.186743452 | 1.138191611 | 0.236577248 | 0.43859751 | 224.0158711 | 255.001602 | ENSG00000056277 |
| PELI2 | 0.186716111 | 1.138170041 | 0.230042783 | 0.429903311 | 254.0098812 | 288.8850154 | ENSG00000139946 |
| NA | 0.186517557 | 1.138013409 | 0.671271812 | 0.82390724 | 26.45155045 | 30.07983817 | ENSG00000223611 |
| SIRT3 | 0.186414393 | 1.137932036 | 0.114603418 | 0.264195014 | 378.9302187 | 431.2258818 | ENSG00000142082 |
| DICER1-AS1 | 0.18635143 | 1.137882374 | 0.643831728 | 0.805803452 | 31.59240454 | 36.05154647 | ENSG00000235706 |

| | | | | | | | |
|---|---|---|---|---|---|---|---|
| CPSF2 | 0.186342917 | 1.13787566 | 0.009082983 | 0.036197981 | 1888.490135 | 2149.360401 | ENSG00000165934 |
| RNF123 | 0.186315115 | 1.137853732 | 0.037513795 | 0.112990114 | 704.3002653 | 801.4910486 | ENSG00000164068 |
| NUDT9 | 0.18625893 | 1.13780942 | 0.016624214 | 0.059246705 | 885.6297835 | 1007.646856 | ENSG00000170502 |
| TMEM160 | 0.186252799 | 1.137804584 | 0.048115743 | 0.136929748 | 634.0112384 | 721.8386586 | ENSG00000130748 |
| NA | 0.186227994 | 1.137785021 | 0.801829008 | 0.898704449 | 7.28010534 | 8.276789441 | ENSG00000273382 |
| KRBA1 | 0.186174777 | 1.137743053 | 0.137353991 | 0.300104537 | 319.8739079 | 364.0684171 | ENSG00000133619 |
| SYK | 0.186165354 | 1.137735622 | 0.334310232 | 0.546292653 | 115.5953334 | 131.4969489 | ENSG00000165025 |
| ZBED4 | 0.186065054 | 1.137656526 | 0.004366339 | 0.019736729 | 1526.550489 | 1737.041055 | ENSG00000100426 |
| STX18-AS1 | 0.186015468 | 1.137617425 | 0.331540498 | 0.542983463 | 118.7689338 | 135.1411161 | ENSG00000247708 |
| SSX2IP | 0.185908005 | 1.137532689 | 0.072480269 | 0.187917336 | 485.4626753 | 551.9484554 | ENSG00000117155 |
| CACNG5 | 0.185751124 | 1.137408999 | 0.414028774 | 0.625152061 | 90.02512713 | 102.482821 | ENSG00000075429 |
| MRPL1 | 0.185668542 | 1.137343894 | 0.150942399 | 0.320394202 | 404.0647739 | 459.408188 | ENSG00000169288 |
| ERBIN | 0.185541422 | 1.137243684 | 0.074139618 | 0.191143576 | 459.3126851 | 522.4357556 | ENSG00000112851 |
| TSEN2 | 0.185473034 | 1.137189776 | 0.177199939 | 0.358750091 | 262.701509 | 298.5226455 | ENSG00000154743 |
| SURF2 | 0.185444555 | 1.137167329 | 0.174471236 | 0.354785108 | 302.999831 | 344.8805972 | ENSG00000148291 |
| SLC30A6 | 0.185364183 | 1.137103979 | 0.05652848 | 0.155188303 | 575.8857733 | 655.1103618 | ENSG00000152683 |
| FSCN1 | 0.18506528 | 1.136868414 | 0.000446127 | 0.002785161 | 13070.38013 | 14859.96882 | ENSG00000075618 |
| CARNMT1 | 0.184942471 | 1.136771642 | 0.101373435 | 0.240265729 | 543.9024901 | 618.2172825 | ENSG00000156017 |
| ABCC4 | 0.184879422 | 1.136721964 | 0.122710584 | 0.276967775 | 363.4006191 | 413.0487232 | ENSG00000125257 |
| RSU1 | 0.184845617 | 1.136695329 | 0.009352177 | 0.03697987 | 1288.404667 | 1464.604405 | ENSG00000148484 |
| YBX2 | 0.184639492 | 1.136532935 | 0.063335258 | 0.169097703 | 653.4404139 | 743.1146865 | ENSG00000006047 |
| PAXX | 0.184616652 | 1.136514942 | 0.077414158 | 0.197057065 | 554.7301408 | 630.9188292 | ENSG00000148362 |
| KRT19 | 0.184516664 | 1.136436177 | 0.003029849 | 0.014537837 | 2298.331758 | 2612.290998 | ENSG00000171345 |
| FGF18 | 0.184509959 | 1.136430895 | 0.751573138 | 0.870657509 | 16.87323896 | 19.27058278 | ENSG00000156427 |
| EHD3 | 0.184279344 | 1.136249252 | 0.049210551 | 0.139376117 | 629.5150762 | 715.6708 | ENSG00000013016 |
| NA | 0.184184799 | 1.136174792 | 0.637488756 | 0.801102952 | 28.00395114 | 31.8631524 | ENSG00000272556 |
| MYH7B | 0.183854075 | 1.135914364 | 0.762949355 | 0.877431251 | 13.21509681 | 15.04532142 | ENSG00000078814 |
| ZNF879 | 0.183798302 | 1.135870452 | 0.564169166 | 0.748528238 | 46.7415507 | 53.19927232 | ENSG00000234284 |
| TVP23A | 0.183693626 | 1.135788041 | 0.448040425 | 0.654733041 | 82.14657695 | 93.30949273 | ENSG00000166676 |
| TUT1 | 0.183594413 | 1.135709936 | 0.709955926 | 0.846808416 | 17.58820437 | 19.97454421 | ENSG00000255508 |
| ADO | 0.183524947 | 1.135655253 | 0.02865932 | 0.091486014 | 816.9509468 | 927.8821177 | ENSG00000181915 |
| NR1D2 | 0.183395591 | 1.135553431 | 0.114921058 | 0.264636531 | 340.1301196 | 386.2251025 | ENSG00000174738 |
| SLC35F3 | 0.183371947 | 1.135534821 | 0.521328761 | 0.71612552 | 53.39239731 | 60.69463428 | ENSG00000183780 |
| ATP5MC1 | 0.183361699 | 1.135526755 | 0.038697334 | 0.115852751 | 1544.382128 | 1752.977282 | ENSG00000159199 |
| SELENOI | 0.183292165 | 1.135472027 | 0.041352897 | 0.121780719 | 1242.222074 | 1410.703584 | ENSG00000138018 |
| TAF4 | 0.183259061 | 1.135445973 | 0.028533458 | 0.091176709 | 830.3000828 | 942.6545281 | ENSG00000130699 |
| PLS3 | 0.183104562 | 1.135324384 | 0.00131659 | 0.007113498 | 6706.94187 | 7614.37442 | ENSG00000102024 |
| ATF3 | 0.18307906 | 1.135304315 | 0.091105544 | 0.222319188 | 432.6538478 | 491.018964 | ENSG00000162772 |
| HES6 | 0.182952924 | 1.135205059 | 0.022023039 | 0.074441575 | 1877.67222 | 2132.443925 | ENSG00000144485 |
| MEA1 | 0.182896644 | 1.135160775 | 0.014311032 | 0.052468081 | 1220.878403 | 1385.665773 | ENSG00000124733 |
| BCL9L | 0.182886371 | 1.135152692 | 0.009304137 | 0.036828471 | 1459.385483 | 1657.14709 | ENSG00000186174 |
| MRPS36 | 0.18281402 | 1.135095766 | 0.151602825 | 0.321279071 | 336.3823797 | 382.0447909 | ENSG00000134056 |
| PRKCI | 0.182795953 | 1.135081551 | 0.00699133 | 0.029229857 | 2494.223442 | 2830.618996 | ENSG00000163558 |
| USP45 | 0.182792606 | 1.135078918 | 0.090674763 | 0.221582931 | 402.1482948 | 456.3184424 | ENSG00000123552 |
| FASTK | 0.182662156 | 1.134976287 | 0.040899052 | 0.120778513 | 854.6859352 | 970.4492861 | ENSG00000164896 |
| NFAT5 | 0.182607649 | 1.134933407 | 0.237074863 | 0.439066273 | 227.4886158 | 257.8969307 | ENSG00000102908 |
| TRIM66 | 0.182603738 | 1.13493033 | 0.541905111 | 0.731175343 | 51.55798702 | 58.58163608 | ENSG00000166436 |
| GPN3 | 0.182516862 | 1.134861989 | 0.031883121 | 0.099770131 | 876.8648028 | 994.8921227 | ENSG00000111231 |
| PARP4 | 0.182504996 | 1.134852655 | 0.013994448 | 0.051467411 | 1136.075543 | 1289.332395 | ENSG00000102699 |
| KIF7 | 0.182225588 | 1.134632888 | 0.037622536 | 0.113173238 | 719.0271602 | 816.0717846 | ENSG00000166813 |
| REL | 0.18216789 | 1.134587512 | 0.583014807 | 0.762575961 | 43.37014667 | 49.22738575 | ENSG00000162924 |
| DKK3 | 0.18199058 | 1.134448077 | 0.060097982 | 0.162530487 | 561.9013776 | 637.5015938 | ENSG00000050165 |
| FABP3 | 0.181840668 | 1.134330202 | 0.172885681 | 0.352459025 | 278.5658993 | 316.1555248 | ENSG00000121769 |
| LARGE2 | 0.181764216 | 1.134270092 | 0.005489711 | 0.02389334 | 2114.8469 | 2399.421308 | ENSG00000165905 |
| MIB2 | 0.181745548 | 1.134255415 | 0.040324045 | 0.119529053 | 1288.972245 | 1462.661267 | ENSG00000197530 |
| BLVRA | 0.181658845 | 1.134187251 | 0.059457119 | 0.161212879 | 613.4854506 | 695.2999401 | ENSG00000106605 |
| F11 | 0.181631203 | 1.134165521 | 0.846206938 | 0.923574988 | 5.828233445 | 6.652474682 | ENSG00000088926 |
| FBXL2 | 0.181425488 | 1.134003811 | 0.543392197 | 0.732264995 | 69.30423041 | 78.50491512 | ENSG00000153558 |
| NOXA1 | 0.181281561 | 1.133890685 | 0.66938238 | 0.82295975 | 29.3813796 | 33.2521585 | ENSG00000188747 |
| URI1 | 0.181265679 | 1.133878202 | 0.018131014 | 0.063431678 | 1359.47629 | 1540.767474 | ENSG00000105176 |
| SLC16A10 | 0.181169994 | 1.133803002 | 0.207076489 | 0.399574894 | 215.3585905 | 244.1672597 | ENSG00000112394 |
| LOC107985075 | 0.181144078 | 1.133782635 | 0.814572736 | 0.905800278 | 7.879719244 | 8.952469843 | ENSG00000214999 |
| ZNF581 | 0.181116333 | 1.133760831 | 0.040349027 | 0.119552984 | 765.6776647 | 868.4490831 | ENSG00000171425 |
| LIN7A | 0.181095707 | 1.133744622 | 0.262148535 | 0.469187557 | 219.5321406 | 248.9800463 | ENSG00000111052 |
| PRMT2 | 0.181089659 | 1.133739869 | 0.008864763 | 0.035536201 | 1315.110894 | 1491.074801 | ENSG00000160310 |
| OVOL1 | 0.181015985 | 1.133681974 | 0.522254671 | 0.716804556 | 54.21053511 | 61.34231721 | ENSG00000172818 |
| TNFRSF12A | 0.180989646 | 1.133661276 | 0.037465995 | 0.112900161 | 947.3881427 | 1074.345191 | ENSG00000006327 |
| ERI1 | 0.180813726 | 1.133523048 | 0.090930546 | 0.221993119 | 451.2630794 | 511.5012037 | ENSG00000104626 |
| LEMD1 | 0.180807657 | 1.13351828 | 0.711142964 | 0.847591497 | 18.19730148 | 20.64804915 | ENSG00000186007 |
| CUTC | 0.180754666 | 1.133476646 | 0.147149739 | 0.314606907 | 405.8589138 | 459.6338715 | ENSG00000119929 |
| TENM3 | 0.180408208 | 1.133204478 | 0.011012887 | 0.042323641 | 1896.35841 | 2148.703154 | ENSG00000218336 |
| NAA30 | 0.180300172 | 1.133119622 | 0.131088512 | 0.290768916 | 610.5025331 | 692.1335791 | ENSG00000139977 |
| TMEM108 | 0.180296061 | 1.133116392 | 0.20021938 | 0.390694667 | 226.9948958 | 257.3715909 | ENSG00000144868 |
| DCUN1D3 | 0.180241483 | 1.133073527 | 0.279742348 | 0.488880785 | 179.017048 | 202.8858225 | ENSG00000188215 |

| | | | | | | | |
|---|---|---|---|---|---|---|---|
| RNASEH1-AS1 | 0.1801975 | 1.133038984 | 0.163090118 | 0.338120875 | 364.0651671 | 412.6781999 | ENSG00000234171 |
| IRF8 | 0.180135944 | 1.132990641 | 0.596788848 | 0.773266374 | 35.47857995 | 40.17446572 | ENSG00000140968 |
| NA | 0.180116223 | 1.132975154 | 0.577883336 | 0.759450456 | 51.053563 | 57.70902736 | ENSG00000259768 |
| VCL | 0.180103712 | 1.132965329 | 0.000525146 | 0.003206395 | 10603.31965 | 12012.65937 | ENSG00000035403 |
| ZNF850 | 0.180090981 | 1.132955331 | 0.387366879 | 0.599624878 | 106.4825972 | 120.7624896 | ENSG00000267041 |
| MPDZ | 0.180013998 | 1.132894878 | 0.013636613 | 0.050357341 | 1696.259842 | 1921.100371 | ENSG00000107186 |
| NA | 0.180006873 | 1.132889282 | 0.837515159 | 0.918202392 | 8.22132399 | 9.231950188 | ENSG00000230084 |
| CCN1 | 0.179946049 | 1.132841521 | 0.018059377 | 0.063298316 | 1669.805264 | 1891.802629 | ENSG00000142871 |
| WASF1 | 0.179736813 | 1.132677235 | 0.021048153 | 0.071736938 | 870.4044138 | 985.9185082 | ENSG00000112290 |
| PLPPR2 | 0.179728213 | 1.132670483 | 0.137199714 | 0.299920893 | 341.4346309 | 386.8161281 | ENSG00000105520 |
| FGGY | 0.179547004 | 1.132528223 | 0.356169774 | 0.568384838 | 149.5671754 | 169.073095 | ENSG00000172456 |
| ASTN1 | 0.179488291 | 1.132482134 | 0.18515149 | 0.369493871 | 265.1203794 | 300.0320598 | ENSG00000152092 |
| FRAT2 | 0.179436628 | 1.132441581 | 0.022406576 | 0.075454383 | 1385.914978 | 1569.925037 | ENSG00000181274 |
| IDUA | 0.17936654 | 1.132386567 | 0.552162003 | 0.739047459 | 59.19985873 | 67.02082997 | ENSG00000127415 |
| HERC1 | 0.179350377 | 1.13237388 | 0.078082677 | 0.198251055 | 733.5783946 | 830.0848019 | ENSG00000103657 |
| CHROMR | 0.179329507 | 1.132357499 | 0.378115944 | 0.590532551 | 104.2881745 | 118.0392429 | ENSG00000223960 |
| SEC14L2 | 0.179041278 | 1.132131294 | 0.585531241 | 0.764237853 | 51.120582 | 57.97320683 | ENSG00000100003 |
| C12orf66 | 0.178992861 | 1.1320933 | 0.448387103 | 0.655148197 | 95.23119463 | 107.8472866 | ENSG00000174206 |
| FEZ2 | 0.178758555 | 1.131909453 | 0.08553388 | 0.2124007 | 449.3616239 | 508.4442056 | ENSG00000171055 |
| NA | 0.178676843 | 1.131845346 | 0.685792881 | 0.831762616 | 29.17022782 | 33.04106349 | ENSG00000246596 |
| SNHG11 | 0.178635482 | 1.131812897 | 0.450647142 | 0.656976322 | 101.1807295 | 114.6407376 | ENSG00000174365 |
| NA | 0.178579526 | 1.131768999 | 0.859689858 | 0.931886123 | 5.349615452 | 6.057381751 | ENSG00000284728 |
| TMEM131 | 0.178562695 | 1.131755796 | 0.070251248 | 0.183270454 | 611.9295683 | 692.5041883 | ENSG00000075568 |
| PIP4P2 | 0.178542442 | 1.131739909 | 0.174636693 | 0.354968674 | 371.8598423 | 421.1409747 | ENSG00000155099 |
| GUCA1B | 0.178460882 | 1.131675929 | 0.715390888 | 0.849695403 | 17.93901299 | 20.28826488 | ENSG00000112599 |
| DCX | 0.178410608 | 1.131636494 | 0.397336822 | 0.609116478 | 102.4626887 | 116.0838038 | ENSG00000077279 |
| FAM155B | 0.178399619 | 1.131627874 | 0.065454055 | 0.173341723 | 762.471698 | 862.6989253 | ENSG00000130054 |
| ZNF429 | 0.178353935 | 1.131592041 | 0.47533364 | 0.67789426 | 67.16942836 | 76.01290966 | ENSG00000197013 |
| TBC1D23 | 0.178309504 | 1.131557192 | 0.147098978 | 0.314569607 | 425.1791929 | 480.9359488 | ENSG00000036054 |
| CDR2 | 0.178284892 | 1.131537888 | 0.018085586 | 0.063319669 | 975.4701497 | 1103.77106 | ENSG00000140743 |
| PIMREG | 0.178165993 | 1.131444637 | 0.004401565 | 0.019871757 | 1976.775504 | 2236.360477 | ENSG00000129195 |
| GFOD1 | 0.178083723 | 1.131380118 | 0.23637835 | 0.438448644 | 196.6648737 | 222.503536 | ENSG00000145990 |
| TRIM21 | 0.17795731 | 1.131280987 | 0.209092301 | 0.402231649 | 254.0466188 | 287.2922414 | ENSG00000132109 |
| RNASEH2C | 0.177907892 | 1.131242237 | 0.050645021 | 0.142538186 | 609.400328 | 689.445489 | ENSG00000172922 |
| NOA1 | 0.177832613 | 1.131183211 | 0.044266249 | 0.128157 | 825.5563105 | 933.5532331 | ENSG00000084092 |
| LONP1 | 0.177832212 | 1.131182897 | 0.000388914 | 0.002467125 | 4543.714273 | 5140.100999 | ENSG00000196365 |
| NA | 0.177681111 | 1.131064428 | 0.552304935 | 0.739107051 | 49.42969352 | 55.9053989 | ENSG00000214389 |
| MRPL17 | 0.177532255 | 1.130947732 | 0.012579495 | 0.04725756 | 1125.852234 | 1273.538301 | ENSG00000158042 |
| NA | 0.17743707 | 1.130873118 | 0.555821057 | 0.742266897 | 47.25960223 | 53.47913153 | ENSG00000185275 |
| CARD10 | 0.177436761 | 1.130872875 | 0.019880263 | 0.06839839 | 1143.363045 | 1293.579602 | ENSG00000100065 |
| TEPSIN | 0.17736814 | 1.130819087 | 0.329095295 | 0.540807923 | 150.5439882 | 170.3473162 | ENSG00000167302 |
| L1CAM | 0.177354933 | 1.130808736 | 0.037860783 | 0.11376307 | 1120.903846 | 1266.924543 | ENSG00000198910 |
| AMZ2 | 0.177298712 | 1.130764669 | 0.028089836 | 0.090201759 | 813.7832702 | 920.3752386 | ENSG00000196704 |
| FAHD2B | 0.177279124 | 1.130749317 | 0.076483784 | 0.195350252 | 506.2182509 | 572.3728512 | ENSG00000144199 |
| NA | 0.177147719 | 1.130646329 | 0.54231035 | 0.731381201 | 59.81015105 | 67.60409963 | ENSG00000260853 |
| PRKCQ-AS1 | 0.177135287 | 1.130636587 | 0.198897344 | 0.389030323 | 258.5964786 | 292.2760616 | ENSG00000237943 |
| PFKL | 0.177112098 | 1.130618414 | 0.001502292 | 0.007976041 | 4030.915065 | 4557.757086 | ENSG00000141959 |
| ZNF2 | 0.17708527 | 1.130597388 | 0.537211934 | 0.72743395 | 57.89595784 | 65.487202 | ENSG00000275111 |
| ZNF394 | 0.176888312 | 1.130443049 | 0.084958134 | 0.211387252 | 488.4624725 | 551.8410273 | ENSG00000160908 |
| COQ10B | 0.1768675 | 1.130426742 | 0.214009952 | 0.408652977 | 268.8585838 | 303.7848549 | ENSG00000115520 |
| GLMN | 0.176696765 | 1.13029297 | 0.118441994 | 0.270315491 | 386.5961687 | 437.188015 | ENSG00000174842 |
| SGK3 | 0.176507713 | 1.130144865 | 0.299448958 | 0.509697483 | 177.4848351 | 200.6014345 | ENSG00000104205 |
| CEP72 | 0.176417031 | 1.13007383 | 0.091745276 | 0.223518254 | 469.6442259 | 530.8403101 | ENSG00000112877 |
| RSBN1 | 0.176256953 | 1.129948448 | 0.191016591 | 0.377811932 | 279.5003307 | 315.6895943 | ENSG00000081019 |
| HADHB | 0.176249499 | 1.129942609 | 0.020611328 | 0.070502614 | 986.4084233 | 1114.736309 | ENSG00000138029 |
| NDUFA3 | 0.176182883 | 1.129890436 | 0.075631002 | 0.193923343 | 865.4973866 | 977.6534897 | ENSG00000170906 |
| PPFIA1 | 0.176137987 | 1.129855275 | 0.012924497 | 0.048294248 | 1339.598471 | 1513.35167 | ENSG00000131626 |
| ZNF92 | 0.175968082 | 1.129722221 | 0.146561896 | 0.313833491 | 359.5141905 | 406.3515301 | ENSG00000146757 |
| ATF6B | 0.175833046 | 1.129616484 | 0.086226018 | 0.213557818 | 599.788228 | 677.6686301 | ENSG00000213676 |
| DNPH1 | 0.175799752 | 1.129590415 | 0.033138505 | 0.102694564 | 887.7132516 | 1003.134758 | ENSG00000112667 |
| SNAP23 | 0.175788572 | 1.129581661 | 0.195602059 | 0.384625666 | 343.4880982 | 387.5467849 | ENSG00000092531 |
| LOC389641 | 0.175579312 | 1.12941783 | 0.666135332 | 0.821155848 | 29.768952 | 33.52009912 | ENSG00000246582 |
| IFNGR2 | 0.175538066 | 1.129385541 | 0.144507823 | 0.310575878 | 314.9276035 | 355.6781247 | ENSG00000159128 |
| CRYBG2 | 0.175464428 | 1.129327896 | 0.161626455 | 0.336269208 | 299.464107 | 338.3417208 | ENSG00000176092 |
| ME2 | 0.175459467 | 1.129324013 | 0.005645353 | 0.024463627 | 1672.038946 | 1887.99557 | ENSG00000082212 |
| PRX | 0.17541863 | 1.129292046 | 0.156105787 | 0.328072311 | 330.4975263 | 373.1042693 | ENSG00000105227 |
| PCDHAC2 | 0.175417187 | 1.129290917 | 0.340185761 | 0.551773042 | 129.630653 | 146.2784697 | ENSG00000243232 |
| KCNK15 | 0.175366376 | 1.129251145 | 0.80968617 | 0.90305216 | 7.943763596 | 8.972994974 | ENSG00000124249 |
| SEMA6A-AS1 | 0.175349258 | 1.129237746 | 0.748451525 | 0.869205832 | 15.57231329 | 17.57409886 | ENSG00000248445 |
| ZNF710 | 0.175322326 | 1.129216666 | 0.115173052 | 0.265152151 | 637.1441578 | 719.7858477 | ENSG00000140548 |
| SGSH | 0.175307381 | 1.129204968 | 0.131777543 | 0.291885866 | 342.7631165 | 387.1305112 | ENSG00000181523 |
| ZBTB10 | 0.175305198 | 1.129203259 | 0.24155584 | 0.445095329 | 252.8293399 | 285.7908665 | ENSG00000205189 |
| LSAMP | 0.175044723 | 1.128999403 | 0.417126688 | 0.628084636 | 112.7155901 | 127.115315 | ENSG00000185565 |
| THOC7 | 0.175037159 | 1.128993483 | 0.069376895 | 0.181440614 | 616.0799815 | 695.8959493 | ENSG00000163634 |

| | | | | | | | |
|---|---|---|---|---|---|---|---|
| PODXL2 | 0.174989417 | 1.128956123 | 0.013259378 | 0.049272598 | 1429.532325 | 1614.310999 | ENSG00000114631 |
| NA | 0.174849435 | 1.128846588 | 0.888012876 | 0.94955683 | 3.22104262 | 3.667813717 | ENSG00000260464 |
| SERPINI1 | 0.174775925 | 1.128789071 | 0.449485843 | 0.655840234 | 83.64205223 | 94.54434473 | ENSG00000163536 |
| SUGP2 | 0.174775567 | 1.128788791 | 0.038524389 | 0.115444794 | 3402.167741 | 3839.940545 | ENSG00000064607 |
| NINJ2-AS1 | 0.174527076 | 1.128594385 | 0.619475343 | 0.788725553 | 35.83859245 | 40.54295184 | ENSG00000177406 |
| TENT5B | 0.174421998 | 1.128512186 | 0.010861674 | 0.041827631 | 2612.415085 | 2948.394576 | ENSG00000158246 |
| WDR63 | 0.174413069 | 1.128505202 | 0.854322464 | 0.928675516 | 5.605539679 | 6.340929476 | ENSG00000162643 |
| ACADS | 0.174340832 | 1.128448698 | 0.397653166 | 0.609205812 | 134.1743097 | 151.3741643 | ENSG00000122971 |
| ADRA2B | 0.174268716 | 1.128392292 | 0.223425053 | 0.421267916 | 255.7203203 | 288.7773467 | ENSG00000274286 |
| OGDHL | 0.174214056 | 1.128349541 | 0.148255252 | 0.316118937 | 618.3321427 | 697.5218631 | ENSG00000197444 |
| ZNF554 | 0.174170221 | 1.128315258 | 0.310059207 | 0.520717486 | 160.3541974 | 180.8910471 | ENSG00000172006 |
| FADS1 | 0.174167594 | 1.128313203 | 0.004026714 | 0.01848432 | 3326.415604 | 3752.380584 | ENSG00000149485 |
| PDF | 0.1741274 | 1.128281768 | 0.258136744 | 0.464941574 | 328.0498479 | 370.4658883 | ENSG00000258429 |
| ZNF551 | 0.174026536 | 1.128202888 | 0.081110551 | 0.203937258 | 537.5494178 | 606.4896087 | ENSG00000204519 |
| PRMT8 | 0.173964885 | 1.128154678 | 0.739031234 | 0.863544416 | 18.03865207 | 20.31408689 | ENSG00000111218 |
| ZNRD1ASP | 0.173955107 | 1.128147032 | 0.584399999 | 0.76319043 | 49.02076428 | 55.35230219 | ENSG00000204623 |
| NDUFV3 | 0.173796563 | 1.128023062 | 0.042094575 | 0.123459551 | 718.4883105 | 810.5336381 | ENSG00000160194 |
| LINC02731 | 0.173788405 | 1.128016683 | 0.808895577 | 0.902949338 | 7.954442013 | 8.970819511 | ENSG00000204241 |
| THAP7 | 0.173783403 | 1.128012772 | 0.228301633 | 0.427681689 | 271.2908803 | 306.1376283 | ENSG00000184436 |
| ACTL10 | 0.173739341 | 1.127978321 | 0.762290392 | 0.876987748 | 13.26373421 | 14.95820756 | ENSG00000182584 |
| ABHD12 | 0.173598932 | 1.127868548 | 0.010077141 | 0.039319587 | 1506.590829 | 1699.825879 | ENSG00000100997 |
| ALG11 | 0.173250503 | 1.127596186 | 0.340176863 | 0.551773042 | 182.8741372 | 205.9689262 | ENSG00000253710 |
| ENTPD2 | 0.173231618 | 1.127581426 | 0.448567008 | 0.655281496 | 96.22399977 | 108.3540791 | ENSG00000054179 |
| CCSER2 | 0.173097127 | 1.127476315 | 0.180386412 | 0.363318074 | 306.5535538 | 345.5426148 | ENSG00000107771 |
| ATP12A | 0.17295133 | 1.12736238 | 0.61123289 | 0.783844194 | 40.66123184 | 45.72242798 | ENSG00000075673 |
| HS3ST5 | 0.172938008 | 1.127351969 | 0.376861817 | 0.589012283 | 175.8280924 | 198.5288968 | ENSG00000249853 |
| C2orf50 | 0.172876528 | 1.127303929 | 0.79066561 | 0.892831861 | 10.54266642 | 11.88331131 | ENSG00000150873 |
| MSL3P1 | 0.172784123 | 1.127231727 | 0.831244523 | 0.915293173 | 7.354828109 | 8.255318351 | ENSG00000224287 |
| GPR107 | 0.172761927 | 1.127214384 | 0.006332342 | 0.026927782 | 1781.933219 | 2008.356342 | ENSG00000148358 |
| AGL | 0.172673751 | 1.127145492 | 0.13376522 | 0.294939425 | 530.5740756 | 597.6522275 | ENSG00000162688 |
| ZNF24 | 0.172587093 | 1.12707779 | 0.031374677 | 0.098423717 | 1449.788536 | 1634.256089 | ENSG00000172466 |
| LRRC57 | 0.17255424 | 1.127052125 | 0.474448021 | 0.67731814 | 80.86492318 | 91.00658705 | ENSG00000180979 |
| STARD4-AS1 | 0.172537068 | 1.12703871 | 0.520172148 | 0.715550501 | 69.3383506 | 78.12422686 | ENSG00000246859 |
| ABHD13 | 0.172534575 | 1.127036762 | 0.261207671 | 0.468328763 | 206.9095126 | 233.2827864 | ENSG00000139826 |
| POLG2 | 0.172515242 | 1.127021659 | 0.0984319 | 0.235366592 | 476.9631799 | 537.7203375 | ENSG00000256525 |
| LGR4 | 0.172500009 | 1.12700976 | 0.042813419 | 0.125101565 | 1361.317563 | 1534.433763 | ENSG00000205213 |
| NA | 0.172426414 | 1.12695227 | 0.828613489 | 0.913933287 | 7.954442013 | 8.959942193 | ENSG00000278987 |
| SIK1 | 0.1723232 | 1.126871648 | 0.724707349 | 0.854899475 | 21.82874788 | 24.48458279 | ENSG00000142178 |
| TMEM60 | 0.172235835 | 1.12680341 | 0.254960915 | 0.46159878 | 228.1463503 | 257.0490004 | ENSG00000135211 |
| CSRNP2 | 0.172121155 | 1.126713844 | 0.027091777 | 0.087810658 | 950.9784291 | 1071.101015 | ENSG00000110925 |
| ZRANB1 | 0.172095877 | 1.126694102 | 0.263055523 | 0.470251837 | 214.3645907 | 241.49303 | ENSG00000019995 |
| RXYLT1 | 0.172077538 | 1.12667978 | 0.389090311 | 0.600936593 | 113.9311407 | 128.4438073 | ENSG00000118600 |
| SIAH1 | 0.172067892 | 1.126672247 | 0.461612468 | 0.666428519 | 115.2563978 | 130.173302 | ENSG00000196470 |
| NA | 0.172030007 | 1.126642662 | 0.754383181 | 0.872093926 | 13.82063445 | 15.59964764 | ENSG00000276386 |
| SLC25A28 | 0.171929659 | 1.1265643 | 0.221876219 | 0.419180911 | 230.6307914 | 259.9604307 | ENSG00000155287 |
| TMPRSS11E | 0.171850398 | 1.126502408 | 0.783010051 | 0.889773441 | 14.17411282 | 15.87666897 | ENSG00000087128 |
| ABI1 | 0.171800753 | 1.126463645 | 0.058448361 | 0.159140327 | 744.6978686 | 839.2280431 | ENSG00000136754 |
| CDC40 | 0.171789563 | 1.126454908 | 0.086159548 | 0.213451578 | 597.9994276 | 673.5056659 | ENSG00000168438 |
| LAGE3 | 0.171746103 | 1.126420974 | 0.018809665 | 0.065309896 | 1377.379732 | 1552.146887 | ENSG00000196976 |
| ZBED8 | 0.171676738 | 1.126366817 | 0.441360684 | 0.64904972 | 103.9136709 | 117.1116281 | ENSG00000221886 |
| CRCP | 0.171523863 | 1.126247468 | 0.122304487 | 0.276315377 | 394.7276685 | 444.6648593 | ENSG00000241258 |
| DDI2 | 0.171459791 | 1.126197452 | 0.036450699 | 0.110617299 | 1085.735698 | 1222.505266 | ENSG00000197312 |
| PKP2 | 0.171314118 | 1.126083741 | 0.100270889 | 0.238490647 | 453.034004 | 510.0442114 | ENSG00000057294 |
| NA | 0.171267388 | 1.126047268 | 0.851342314 | 0.92641986 | 5.005925775 | 5.612375535 | ENSG00000259088 |
| YTHDF2 | 0.171186981 | 1.12598451 | 0.001832593 | 0.009409603 | 2690.901262 | 3029.833016 | ENSG00000198492 |
| ZC3HC1 | 0.171185666 | 1.125983484 | 0.068598281 | 0.179852637 | 603.1130278 | 678.7807172 | ENSG00000091732 |
| SLC2A10 | 0.171181847 | 1.125980503 | 0.265559988 | 0.473143151 | 195.9326957 | 220.4141413 | ENSG00000197496 |
| STON2 | 0.171157875 | 1.125961794 | 0.28235217 | 0.491435108 | 220.1681872 | 247.6403932 | ENSG00000140022 |
| NTF4 | 0.171157353 | 1.125961387 | 0.73937689 | 0.863587169 | 17.34148402 | 19.59612672 | ENSG00000225950 |
| NA | 0.171042221 | 1.125871535 | 0.4631457 | 0.667825192 | 111.0143548 | 124.7713591 | ENSG00000253507 |
| PHYKPL | 0.171008418 | 1.125845155 | 0.236874467 | 0.438867292 | 252.5025323 | 284.2985426 | ENSG00000175309 |
| CLDN3 | 0.170908166 | 1.125766924 | 0.209377247 | 0.402574761 | 338.8430479 | 381.4828767 | ENSG00000165215 |
| PIK3R5 | 0.17085919 | 1.125728707 | 0.23448557 | 0.435785775 | 215.0021375 | 241.9490815 | ENSG00000141506 |
| TANC1 | 0.170830004 | 1.125705934 | 0.151741495 | 0.321512187 | 327.0472814 | 368.3230007 | ENSG00000115183 |
| LCN12 | 0.170828874 | 1.125705052 | 0.787319128 | 0.891546762 | 13.39301812 | 15.05932016 | ENSG00000184925 |
| ZNF273 | 0.170826474 | 1.12570318 | 0.226562234 | 0.42548009 | 240.840141 | 271.2991362 | ENSG00000198039 |
| SLX4 | 0.17079285 | 1.125676944 | 0.133172468 | 0.294010061 | 370.2573041 | 416.9535696 | ENSG00000188827 |
| HS6ST1 | 0.170781945 | 1.125668435 | 0.004850701 | 0.021518791 | 2456.807819 | 2765.726807 | ENSG00000136720 |
| MLPH | 0.170763496 | 1.12565404 | 0.329767421 | 0.541217709 | 154.1033816 | 173.2944999 | ENSG00000115648 |
| CYP2J2 | 0.170728052 | 1.125626386 | 0.683151863 | 0.830790283 | 29.9029906 | 33.52511246 | ENSG00000134716 |
| DNAJC27 | 0.170627687 | 1.125548081 | 0.567126142 | 0.750748437 | 50.03139236 | 56.40032519 | ENSG00000115137 |
| PYM1 | 0.170560617 | 1.125495756 | 0.029059825 | 0.092568873 | 1235.160494 | 1389.872957 | ENSG00000170473 |
| AP5B1 | 0.170486309 | 1.125437787 | 0.074938929 | 0.192442437 | 739.6442741 | 832.4536269 | ENSG00000254470 |
| NA | 0.170364392 | 1.125342685 | 0.296415087 | 0.506588137 | 223.8017703 | 252.0484162 | ENSG00000196922 |

| | | | | | | | |
|---|---|---|---|---|---|---|---|
| NA | 0.170362074 | 1.125340877 | 0.859598359 | 0.931886123 | 4.736958866 | 5.28683159 | ENSG00000261613 |
| LRIG3 | 0.170333474 | 1.125318568 | 0.209188399 | 0.402375525 | 259.3153084 | 291.892252 | ENSG00000139263 |
| ARHGAP22 | 0.170230221 | 1.125238033 | 0.147084724 | 0.314569607 | 422.2067023 | 474.9769371 | ENSG00000128805 |
| NA | 0.170181936 | 1.125200373 | 0.790384746 | 0.892671866 | 10.65890765 | 11.97695156 | ENSG00000284294 |
| HBEGF | 0.170116752 | 1.125149535 | 0.524897195 | 0.718550866 | 74.17672799 | 83.61300011 | ENSG00000113070 |
| CHPF2 | 0.170099818 | 1.125136329 | 0.070169738 | 0.183113634 | 614.3593959 | 691.5058245 | ENSG00000033100 |
| TMEM120A | 0.170036697 | 1.125087102 | 0.183233097 | 0.366977969 | 307.4502532 | 345.7516929 | ENSG00000189077 |
| SERPINE2 | 0.169956694 | 1.125024714 | 0.018449481 | 0.064319449 | 1339.378195 | 1506.449553 | ENSG00000135919 |
| LOC171391 | 0.169859827 | 1.124949179 | 0.571671077 | 0.754332608 | 58.8926535 | 66.31686854 | ENSG00000255284 |
| CCDC84 | 0.169796607 | 1.124899884 | 0.313032492 | 0.523337342 | 202.2970835 | 227.4454553 | ENSG00000186166 |
| THUMPD3 | 0.169746177 | 1.124860563 | 0.136055096 | 0.298247299 | 384.6983251 | 432.7771971 | ENSG00000134077 |
| LINC02048 | 0.169712886 | 1.124834607 | 0.848315707 | 0.9249161 | 5.329453825 | 5.985212587 | ENSG00000228271 |
| RAPH1 | 0.169707185 | 1.124830161 | 0.44603513 | 0.653422136 | 103.6434826 | 116.474496 | ENSG00000173166 |
| TIPARP | 0.169706897 | 1.124829937 | 0.383497465 | 0.595734937 | 130.673037 | 147.1258235 | ENSG00000163659 |
| FZR1 | 0.169606305 | 1.124751511 | 0.025762793 | 0.084308183 | 994.6685176 | 1118.863296 | ENSG00000105325 |
| BDP1 | 0.169602051 | 1.124748195 | 0.175277829 | 0.355850549 | 519.8833864 | 584.0577508 | ENSG00000145734 |
| PPP1R21 | 0.169579158 | 1.124730347 | 0.141632673 | 0.306384082 | 359.5456154 | 404.3940108 | ENSG00000162869 |
| NA | 0.169544762 | 1.124703532 | 0.374845355 | 0.586783366 | 149.5434799 | 168.399633 | ENSG00000272168 |
| POLR1C | 0.16951545 | 1.124680681 | 0.159461374 | 0.333012059 | 336.0022579 | 378.0499202 | ENSG00000171453 |
| TMEM150C | 0.169513642 | 1.124679272 | 0.523298119 | 0.71758545 | 62.86748409 | 70.71179579 | ENSG00000249242 |
| ZFAT | 0.169388204 | 1.124581488 | 0.296104316 | 0.506240268 | 194.1688639 | 218.2723677 | ENSG00000066827 |
| ATP6V1C2 | 0.169292702 | 1.124507047 | 0.701858219 | 0.841401159 | 21.88183448 | 24.60376152 | ENSG00000143882 |
| TRIM52 | 0.169278896 | 1.124496286 | 0.508877125 | 0.705487044 | 88.64647789 | 99.59011927 | ENSG00000183718 |
| CNP | 0.169219691 | 1.12445014 | 0.008846517 | 0.03547812 | 1605.753784 | 1805.835907 | ENSG00000173786 |
| ARRDC1 | 0.169081571 | 1.124342493 | 0.259621 | 0.466511832 | 246.9821662 | 278.0029714 | ENSG00000197070 |
| PDK1 | 0.169071305 | 1.124334493 | 0.244343587 | 0.448705182 | 216.3413015 | 243.103514 | ENSG00000152256 |
| INTS2 | 0.168955219 | 1.124244027 | 0.113436082 | 0.262055082 | 416.9368437 | 468.6609266 | ENSG00000108506 |
| FBXO32 | 0.168940766 | 1.124232764 | 0.823336839 | 0.910506307 | 8.001884211 | 8.980467323 | ENSG00000156804 |
| USF1 | 0.168914753 | 1.124212494 | 0.021649668 | 0.073350195 | 1352.067965 | 1519.537869 | ENSG00000158773 |
| SP4 | 0.168898813 | 1.124200072 | 0.194661848 | 0.383218045 | 360.2762926 | 405.122542 | ENSG00000105866 |
| CASP10 | 0.16886176 | 1.1241712 | 0.609226757 | 0.782326304 | 44.98661787 | 50.62291824 | ENSG00000003400 |
| TBRG4 | 0.168681076 | 1.124030416 | 0.001517874 | 0.008037351 | 2995.96755 | 3367.4455 | ENSG00000136270 |
| ZC2HC1A | 0.168560822 | 1.123936729 | 0.459841928 | 0.664802287 | 94.00796639 | 105.8540342 | ENSG00000104427 |
| C6orf136 | 0.168521302 | 1.12390594 | 0.253956325 | 0.460177048 | 213.6341922 | 240.2179058 | ENSG00000204564 |
| NDUFA1 | 0.168478503 | 1.123872599 | 0.123391739 | 0.278073149 | 596.2050612 | 670.3691552 | ENSG00000125356 |
| ZNF646 | 0.168430695 | 1.123835357 | 0.039666532 | 0.117987471 | 924.448849 | 1038.984435 | ENSG00000167395 |
| CAVIN1 | 0.168427857 | 1.123833146 | 0.023118473 | 0.077341427 | 1130.831743 | 1271.087988 | ENSG00000177469 |
| TFDP2 | 0.168413454 | 1.123821927 | 0.028572051 | 0.091255646 | 1044.336364 | 1173.607387 | ENSG00000114126 |
| TJP1 | 0.168359694 | 1.12378005 | 0.013382084 | 0.049573299 | 2603.088635 | 2924.601805 | ENSG00000104067 |
| RETSAT | 0.168263996 | 1.123705509 | 0.158576572 | 0.331714505 | 321.970751 | 361.8136349 | ENSG00000042445 |
| NA | 0.168235134 | 1.123683029 | 0.82600183 | 0.91237981 | 7.730553039 | 8.651801956 | ENSG00000229119 |
| NA | 0.167889286 | 1.123413688 | 0.677241436 | 0.827303061 | 30.82938654 | 34.64948759 | ENSG00000280832 |
| FARSA | 0.1676322 | 1.123213515 | 0.002906061 | 0.014010966 | 3813.079483 | 4283.451044 | ENSG00000179115 |
| RASAL1 | 0.167618726 | 1.123203025 | 0.301835228 | 0.5121436 | 198.357516 | 222.6387733 | ENSG00000111344 |
| CACNB1 | 0.167608078 | 1.123194735 | 0.255942042 | 0.462814344 | 233.1065872 | 261.882859 | ENSG00000067191 |
| EP400P1 | 0.167497611 | 1.123108735 | 0.1783858 | 0.360587647 | 318.7447005 | 358.1596519 | ENSG00000185684 |
| CNTN3 | 0.16748018 | 1.123095166 | 0.845270055 | 0.923032554 | 5.587742316 | 6.282759052 | ENSG00000113805 |
| BCAT2 | 0.167434051 | 1.123059256 | 0.057520916 | 0.157224313 | 768.8515968 | 863.8437232 | ENSG00000105552 |
| CDYL2 | 0.167380115 | 1.123017271 | 0.391836049 | 0.603234759 | 115.1140457 | 129.2316456 | ENSG00000166446 |
| CHTF18 | 0.167333019 | 1.122980611 | 0.509587195 | 0.706264067 | 2096.147337 | 2354.072226 | ENSG00000127586 |
| MUS81 | 0.167087737 | 1.122789702 | 0.110593359 | 0.257528432 | 530.6158989 | 595.8942768 | ENSG00000172732 |
| ACACB | 0.166989494 | 1.122713246 | 0.164728294 | 0.340248696 | 409.239801 | 459.0578633 | ENSG00000076555 |
| MBD6 | 0.166948252 | 1.122681152 | 0.143848674 | 0.309758519 | 498.3984573 | 560.0887483 | ENSG00000166987 |
| NA | 0.166806144 | 1.122570571 | 0.856947182 | 0.930171411 | 5.628065569 | 6.320404345 | ENSG00000235513 |
| PDXK | 0.166779283 | 1.122549671 | 0.008330631 | 0.033775003 | 2106.965655 | 2365.41393 | ENSG00000160209 |
| SLC35D2 | 0.166679098 | 1.12247172 | 0.508428876 | 0.705176213 | 72.16022652 | 80.83950679 | ENSG00000130958 |
| SUPT20H | 0.166658851 | 1.122455968 | 0.026097852 | 0.085225305 | 1114.137316 | 1250.146196 | ENSG00000102710 |
| AATK | 0.166602869 | 1.122412412 | 0.430990901 | 0.640685073 | 112.5554661 | 126.3836826 | ENSG00000181409 |
| C10orf88 | 0.166498717 | 1.122331386 | 0.349352362 | 0.561491009 | 145.3260209 | 163.1294271 | ENSG00000119965 |
| AGGF1 | 0.166452288 | 1.122295268 | 0.357703106 | 0.569801407 | 166.3870485 | 186.4785465 | ENSG00000164252 |
| RNF144A-AS1 | 0.166265371 | 1.122149871 | 0.699603539 | 0.839970842 | 22.75633903 | 25.56762412 | ENSG00000228203 |
| FLVCR2 | 0.166102102 | 1.122022885 | 0.512613358 | 0.708949343 | 70.55387507 | 79.2257858 | ENSG00000119686 |
| TMEM181 | 0.166084846 | 1.122009465 | 0.113453129 | 0.262055082 | 561.6660245 | 630.2914176 | ENSG00000146433 |
| TGFBR3 | 0.165964902 | 1.121916186 | 0.438497352 | 0.6466204 | 109.0820587 | 122.2620452 | ENSG00000069702 |
| PROB1 | 0.16593404 | 1.121892186 | 0.344863989 | 0.556903268 | 174.3764481 | 195.577909 | ENSG00000228672 |
| BRF1 | 0.165857978 | 1.12183304 | 0.105836767 | 0.248907516 | 581.2881699 | 652.6579591 | ENSG00000185024 |
| EPHX4 | 0.16554495 | 1.121589657 | 0.799220905 | 0.897822068 | 11.54645488 | 12.9672986 | ENSG00000172031 |
| WDR1 | 0.165432925 | 1.121502569 | 0.000372961 | 0.002377106 | 10802.45837 | 12114.71762 | ENSG00000071127 |
| PRDX6 | 0.165370899 | 1.121454353 | 0.001398208 | 0.007507862 | 4765.96352 | 5345.245985 | ENSG00000117592 |
| SNX8 | 0.16532912 | 1.121421877 | 0.03039834 | 0.096014826 | 1219.555841 | 1367.356006 | ENSG00000106266 |
| NOTCH1 | 0.165130798 | 1.12126773 | 0.019027739 | 0.065921639 | 3000.161261 | 3363.746478 | ENSG00000148400 |
| MRPL35 | 0.165069505 | 1.121220094 | 0.06109079 | 0.1647201 | 719.0721858 | 806.2758901 | ENSG00000132313 |
| GUSBP2 | 0.164979406 | 1.121150074 | 0.655816553 | 0.814050821 | 35.31873533 | 39.52744521 | ENSG00000241549 |
| NA | 0.164780633 | 1.120995613 | 0.568838921 | 0.752014213 | 52.95527162 | 59.33863901 | ENSG00000249855 |

| | | | | | | | |
|---|---|---|---|---|---|---|---|
| PIGK | 0.164768968 | 1.12098655 | 0.237698414 | 0.439841186 | 300.190335 | 336.5260152 | ENSG00000142892 |
| JAGN1 | 0.164699812 | 1.120932816 | 0.074792523 | 0.192249462 | 583.3425531 | 653.7875882 | ENSG00000171135 |
| NSG2 | 0.164690224 | 1.120925367 | 0.854838076 | 0.929013468 | 5.360293869 | 5.989563514 | ENSG00000170091 |
| NOP53 | 0.164664998 | 1.120905767 | 0.00170571 | 0.008881261 | 7893.90962 | 8848.932557 | ENSG00000105373 |
| TEPP | 0.164653639 | 1.120896942 | 0.807829724 | 0.902233387 | 12.21486782 | 13.72602548 | ENSG00000159648 |
| LOC100129917 | 0.164605811 | 1.120859782 | 0.520913972 | 0.715920265 | 72.48611884 | 81.18607032 | ENSG00000249592 |
| GTF2B | 0.164584129 | 1.120842938 | 0.285661376 | 0.49492799 | 229.9537594 | 257.6606464 | ENSG00000137947 |
| NIPAL3 | 0.164547597 | 1.120814556 | 0.296060586 | 0.506211333 | 176.4469264 | 197.7103185 | ENSG00000001461 |
| ZADH2 | 0.164535303 | 1.120805005 | 0.493161681 | 0.692597482 | 79.79469996 | 89.31785903 | ENSG00000180011 |
| NA | 0.164513562 | 1.120788115 | 0.849464899 | 0.925384716 | 6.836776587 | 7.716220381 | ENSG00000280279 |
| ATP6V1B2 | 0.164463896 | 1.120749531 | 0.018749782 | 0.065149887 | 1815.397304 | 2034.122492 | ENSG00000147416 |
| TAOK1 | 0.164428124 | 1.120721742 | 0.113468307 | 0.262058088 | 1265.025396 | 1417.007425 | ENSG00000160551 |
| POLR3C | 0.164370894 | 1.120677286 | 0.07329048 | 0.189575499 | 631.7714851 | 708.039518 | ENSG00000186141 |
| APBA3 | 0.164309362 | 1.120629489 | 0.175846781 | 0.356507403 | 338.5261063 | 379.6346977 | ENSG00000011132 |
| RASD1 | 0.164280003 | 1.120606684 | 0.60066816 | 0.776307564 | 45.26031331 | 50.73072519 | ENSG00000108551 |
| ALDH6A1 | 0.164276706 | 1.120604123 | 0.13820011 | 0.301361882 | 502.757529 | 562.9787598 | ENSG00000119711 |
| CORO2B | 0.164242458 | 1.120577522 | 0.364607571 | 0.576732891 | 146.7218833 | 164.4305414 | ENSG00000103647 |
| NA | 0.16420702 | 1.120549997 | 0.78404057 | 0.890065126 | 11.25732635 | 12.61186525 | ENSG00000227347 |
| OSBP | 0.164190745 | 1.120537356 | 0.012441789 | 0.046852079 | 1650.601889 | 1849.216449 | ENSG00000110048 |
| TOMM70 | 0.164189398 | 1.12053631 | 0.02277768 | 0.076363657 | 1619.376108 | 1814.784632 | ENSG00000154174 |
| POLR2L | 0.164080594 | 1.120451805 | 0.03056143 | 0.09638487 | 1440.949114 | 1614.503303 | ENSG00000177700 |
| UGCG | 0.164070447 | 1.120443924 | 0.16036319 | 0.334414625 | 522.2345027 | 585.2805686 | ENSG00000148154 |
| CREBBP | 0.16404975 | 1.12042785 | 0.027917365 | 0.089800619 | 1757.31613 | 1968.623191 | ENSG00000005339 |
| AGPS | 0.163672591 | 1.120134979 | 0.054156414 | 0.149905916 | 1614.404581 | 1808.554377 | ENSG00000018510 |
| TEC | 0.163634436 | 1.120105355 | 0.578220045 | 0.759450456 | 48.59403768 | 54.45576337 | ENSG00000135605 |
| MED13L | 0.163631176 | 1.120102824 | 0.095313508 | 0.229947402 | 689.5621339 | 771.8766129 | ENSG00000123066 |
| ZNF620 | 0.163608085 | 1.120084896 | 0.247394572 | 0.452330079 | 243.1190497 | 272.1281404 | ENSG00000177842 |
| TWF1 | 0.163607877 | 1.120084735 | 0.181873903 | 0.365120512 | 392.4867188 | 439.9486961 | ENSG00000151239 |
| MRC2 | 0.163565613 | 1.120051922 | 0.093322369 | 0.226586904 | 729.5525879 | 817.2904851 | ENSG00000011028 |
| NDUFB3 | 0.163550059 | 1.120039847 | 0.138266709 | 0.301450797 | 412.1234082 | 461.921381 | ENSG00000119013 |
| NA | 0.163406316 | 1.119928257 | 0.357468077 | 0.569542279 | 150.6041938 | 168.6987449 | ENSG00000237517 |
| ERMARD | 0.163354327 | 1.1198879 | 0.3910951 | 0.602433194 | 129.0817614 | 144.6110448 | ENSG00000130023 |
| SMAD5 | 0.163338005 | 1.119875231 | 0.045546901 | 0.131041015 | 941.6838804 | 1054.381699 | ENSG00000113658 |
| FN3K | 0.163328016 | 1.119867476 | 0.222932419 | 0.420755419 | 261.3729973 | 292.5652624 | ENSG00000167363 |
| GCN1 | 0.163327744 | 1.119867265 | 0.003377469 | 0.015929945 | 7250.765464 | 8118.961306 | ENSG00000089154 |
| CBR4 | 0.163323836 | 1.119864232 | 0.148740839 | 0.316798697 | 373.2372191 | 418.0274576 | ENSG00000145439 |
| MRPS16 | 0.163189071 | 1.119759628 | 0.017747859 | 0.062426581 | 1673.35609 | 1873.693391 | ENSG00000182180 |
| TAF6 | 0.163132349 | 1.119715604 | 0.022046394 | 0.074507184 | 1540.787653 | 1724.949225 | ENSG00000106290 |
| PRDM14 | 0.163059331 | 1.119658934 | 0.008133542 | 0.033098696 | 2169.480576 | 2429.625787 | ENSG00000147596 |
| NA | 0.16305436 | 1.119655076 | 0.738681882 | 0.86329645 | 18.72603142 | 20.89518108 | ENSG00000276570 |
| PROKR2 | 0.162983554 | 1.119600126 | 0.676851325 | 0.827121605 | 31.25107912 | 35.00225101 | ENSG00000101292 |
| TUBA4A | 0.162429024 | 1.119169867 | 0.032199393 | 0.100443597 | 971.8938492 | 1087.524748 | ENSG00000127824 |
| IGSF1 | 0.162407766 | 1.119153376 | 0.031672854 | 0.099243711 | 1072.599189 | 1200.095997 | ENSG00000147255 |
| EXO1 | 0.162362512 | 1.119118271 | 0.048588184 | 0.138024647 | 816.8310944 | 913.885409 | ENSG00000174371 |
| IGSF3 | 0.162307499 | 1.119075598 | 0.014413929 | 0.052753303 | 1786.972934 | 1999.180587 | ENSG00000143061 |
| UNC5D | 0.162152827 | 1.118955627 | 0.064821367 | 0.172124791 | 1296.363201 | 1450.071495 | ENSG00000156687 |
| ZBTB25 | 0.162078185 | 1.118897737 | 0.444650673 | 0.652110582 | 120.1890237 | 134.3459734 | ENSG00000089775 |
| ZSWIM7 | 0.162038078 | 1.118866631 | 0.136511059 | 0.298865216 | 423.1493694 | 473.397615 | ENSG00000214941 |
| IMPA1 | 0.161941259 | 1.118791547 | 0.168442542 | 0.346065861 | 481.6408224 | 539.1907388 | ENSG00000133731 |
| STK10 | 0.161935605 | 1.118787163 | 0.064878515 | 0.172252321 | 683.6514733 | 765.0691038 | ENSG00000072786 |
| NA | 0.161772137 | 1.118660403 | 0.850943459 | 0.926228658 | 6.234798418 | 7.000435675 | ENSG00000260018 |
| ATP6V1G2 | 0.161685522 | 1.118593244 | 0.778031328 | 0.886837375 | 16.78130364 | 18.7392407 | ENSG00000213760 |
| CRTAC1 | 0.161674972 | 1.118585064 | 0.54737402 | 0.735112955 | 59.89051823 | 66.97521787 | ENSG00000095713 |
| ATP5PD | 0.161659392 | 1.118572984 | 0.067756331 | 0.178090232 | 1381.732269 | 1545.716626 | ENSG00000167863 |
| NA | 0.161657671 | 1.11857165 | 0.658474512 | 0.81591045 | 46.80323079 | 52.47092929 | ENSG00000272734 |
| SAMD3 | 0.161639618 | 1.118557653 | 0.707987071 | 0.845910689 | 23.38679298 | 26.18948502 | ENSG00000164483 |
| RPS27A | 0.161601104 | 1.118527792 | 0.00153168 | 0.008093388 | 10347.43299 | 11574.21156 | ENSG00000143947 |
| NA | 0.161593332 | 1.118521767 | 0.313540174 | 0.523815126 | 188.6250305 | 211.0571914 | ENSG00000286619 |
| NA | 0.161515363 | 1.118461319 | 0.696264569 | 0.837886726 | 27.09532641 | 30.2086647 | ENSG00000237506 |
| CERS6 | 0.161482401 | 1.118435765 | 0.142131691 | 0.307111784 | 469.9294901 | 525.2108441 | ENSG00000172292 |
| ACVR2A | 0.161227248 | 1.118237978 | 0.335511398 | 0.547537071 | 167.8101391 | 187.5334224 | ENSG00000121989 |
| NA | 0.161207232 | 1.118222464 | 0.747500678 | 0.868603883 | 19.02464317 | 21.21325268 | ENSG00000269983 |
| NMD3 | 0.161203178 | 1.118219321 | 0.062967412 | 0.168409513 | 927.395943 | 1036.844068 | ENSG00000169251 |
| LOC100130950 | 0.161124766 | 1.118158546 | 0.852624633 | 0.927577406 | 5.645862931 | 6.305459647 | ENSG00000261879 |
| CDC42BPG | 0.161085683 | 1.118128256 | 0.097227028 | 0.233307805 | 584.688609 | 653.8425646 | ENSG00000171219 |
| NA | 0.161071485 | 1.118117252 | 0.759108112 | 0.874877523 | 17.31567799 | 19.3526833 | ENSG00000232499 |
| ZNF215 | 0.161042277 | 1.118094615 | 0.396224825 | 0.608200671 | 141.1089454 | 157.9168246 | ENSG00000149054 |
| PUS7 | 0.160892872 | 1.117978832 | 0.034371542 | 0.105734904 | 1097.752255 | 1227.830818 | ENSG00000091127 |
| CNGA1 | 0.160887153 | 1.1179744 | 0.784085613 | 0.890065126 | 12.16978988 | 13.64326254 | ENSG00000198515 |
| ZIC2 | 0.160861274 | 1.117954346 | 0.008151827 | 0.033149708 | 3941.166561 | 4407.127928 | ENSG00000043355 |
| LEPROT | 0.160793297 | 1.117901671 | 0.224953989 | 0.423271978 | 278.8574183 | 311.8328191 | ENSG00000213625 |
| RFX7 | 0.160668279 | 1.117804803 | 0.123293298 | 0.27795081 | 697.8827909 | 779.8670894 | ENSG00000181827 |
| CLDN9 | 0.160652863 | 1.117792859 | 0.830334045 | 0.914737804 | 8.814708672 | 9.867809834 | ENSG00000213937 |
| NAPSA | 0.160633727 | 1.117778032 | 0.846595579 | 0.923736594 | 5.934991466 | 6.643772827 | ENSG00000131400 |

| | | | | | | | |
|---|---|---|---|---|---|---|---|
| PDSS1 | 0.160633114 | 1.117777557 | 0.250452166 | 0.455942388 | 272.0639658 | 304.2896696 | ENSG00000148459 |
| NA | 0.160560855 | 1.117721573 | 0.688002194 | 0.83320004 | 27.96034775 | 31.19371485 | ENSG00000264112 |
| CCNYL1 | 0.160467575 | 1.117649307 | 0.086319503 | 0.213733292 | 791.9628364 | 885.0858334 | ENSG00000163249 |
| PRSS46P | 0.160362625 | 1.117568006 | 0.863888856 | 0.934234728 | 5.983628872 | 6.692295438 | ENSG00000261603 |
| ZNF462 | 0.160291298 | 1.117512755 | 0.076411252 | 0.195283477 | 1290.550986 | 1441.593236 | ENSG00000148143 |
| RNF43 | 0.160235162 | 1.117469273 | 0.818535527 | 0.907641561 | 9.244994745 | 10.35698773 | ENSG00000108375 |
| SH3BP4 | 0.160180129 | 1.117426647 | 0.036511408 | 0.110748143 | 1455.002665 | 1625.935602 | ENSG00000130147 |
| WDR89 | 0.160129743 | 1.117387621 | 0.234678866 | 0.436082888 | 285.5622412 | 319.3109656 | ENSG00000140006 |
| NPR2 | 0.160073779 | 1.117344278 | 0.278244924 | 0.48756309 | 275.5373476 | 307.9384416 | ENSG00000159899 |
| ACSL3 | 0.160034032 | 1.117313494 | 0.025250285 | 0.083075198 | 2200.102903 | 2458.252055 | ENSG00000123983 |
| MFSD14A | 0.159890269 | 1.117202161 | 0.65688627 | 0.814690826 | 43.48135389 | 48.58705959 | ENSG00000156875 |
| ZNF146 | 0.15988541 | 1.117198398 | 0.026126694 | 0.085304741 | 2285.114862 | 2553.563726 | ENSG00000167635 |
| NA | 0.159870026 | 1.117186485 | 0.518978085 | 0.7148203 | 73.30222344 | 81.93588444 | ENSG00000261505 |
| PRCC | 0.159845738 | 1.117167677 | 0.008616119 | 0.03472373 | 2097.233624 | 2343.173642 | ENSG00000143294 |
| CSTB | 0.15978642 | 1.117121745 | 0.025803862 | 0.084396594 | 1438.00235 | 1606.880449 | ENSG00000160213 |
| TNFAIP1 | 0.159635872 | 1.117005177 | 0.16312209 | 0.338120875 | 428.9022697 | 478.9722726 | ENSG00000109079 |
| ID2 | 0.159621847 | 1.116994318 | 0.245649689 | 0.450136241 | 281.0428909 | 313.5917291 | ENSG00000115738 |
| PRKCZ-AS1 | 0.159594745 | 1.116973334 | 0.567855296 | 0.7511345 | 55.06487803 | 61.4810751 | ENSG00000182873 |
| LOC107984875 | 0.159561806 | 1.116947833 | 0.725017196 | 0.85505213 | 21.99660117 | 24.65445959 | ENSG00000235560 |
| RABIF | 0.159514826 | 1.116911461 | 0.319151166 | 0.529256214 | 176.7277408 | 197.3437243 | ENSG00000183155 |
| GNG4 | 0.159396081 | 1.116819534 | 0.047883552 | 0.13639229 | 1308.254697 | 1460.926709 | ENSG00000168243 |
| FITM2 | 0.159271072 | 1.116722766 | 0.443150845 | 0.650550076 | 108.9803609 | 121.5846838 | ENSG00000197296 |
| ZCCHC10 | 0.1592012 | 1.116668683 | 0.247152893 | 0.452056405 | 300.4990665 | 335.8251026 | ENSG00000155329 |
| USE1 | 0.158919254 | 1.116450473 | 0.323147319 | 0.53410092 | 170.9635779 | 190.7817684 | ENSG00000053501 |
| SELENOT | 0.158903207 | 1.116438056 | 0.051022285 | 0.143376491 | 901.0155297 | 1005.863878 | ENSG00000198843 |
| PTPRG | 0.158887755 | 1.116426098 | 0.087816005 | 0.216276116 | 1763.07185 | 1967.755035 | ENSG00000144724 |
| SNX17 | 0.158816632 | 1.116371061 | 0.010720118 | 0.04134152 | 2020.206492 | 2255.10514 | ENSG00000115234 |
| NA | 0.158778105 | 1.116341249 | 0.794869024 | 0.895116369 | 13.0620918 | 14.64883782 | ENSG00000267980 |
| GAA | 0.158676955 | 1.116262983 | 0.05578056 | 0.153514409 | 1049.615707 | 1171.35657 | ENSG00000171298 |
| HIPK3 | 0.158629852 | 1.116226538 | 0.310219967 | 0.520802212 | 205.5501353 | 229.1794463 | ENSG00000110422 |
| B3GNT2 | 0.158572102 | 1.116181857 | 0.092443574 | 0.224806141 | 714.8859253 | 797.4192157 | ENSG00000170340 |
| SDCBP | 0.158215533 | 1.115906022 | 0.012232714 | 0.046258471 | 1824.820298 | 2036.101905 | ENSG00000137575 |
| FTL | 0.158205256 | 1.115898073 | 0.000634567 | 0.003767305 | 12596.78241 | 14056.51066 | ENSG00000087086 |
| LOC100287896 | 0.158153633 | 1.115858144 | 0.702801948 | 0.842126015 | 26.9230495 | 30.07955462 | ENSG00000254837 |
| MOG | 0.158099501 | 1.115816277 | 0.851091098 | 0.926282558 | 6.2478411 | 6.990787862 | ENSG00000204655 |
| RPS6KL1 | 0.158027066 | 1.115760255 | 0.379998212 | 0.59241652 | 169.9360683 | 189.6683136 | ENSG00000198208 |
| ZCWPW1 | 0.157952543 | 1.115702621 | 0.712262111 | 0.848059389 | 25.10079217 | 27.92143878 | ENSG00000078487 |
| SPEN | 0.157913805 | 1.115672664 | 0.048954184 | 0.138751278 | 1714.420381 | 1911.851659 | ENSG00000065526 |
| SERTM1 | 0.157875004 | 1.115642658 | 0.690862705 | 0.834552267 | 26.43019361 | 29.48663715 | ENSG00000180440 |
| ROR2 | 0.15783463 | 1.115611437 | 0.452174361 | 0.658283762 | 128.6683568 | 143.3984943 | ENSG00000169071 |
| DKKL1 | 0.157709357 | 1.11551457 | 0.656528848 | 0.814354502 | 37.70531732 | 42.03692701 | ENSG00000104901 |
| JADE2 | 0.157688715 | 1.11549861 | 0.100073538 | 0.238201257 | 872.5510884 | 973.547825 | ENSG00000043143 |
| ABCC8 | 0.157672588 | 1.11548614 | 0.611619104 | 0.784141186 | 44.66425887 | 49.84149076 | ENSG00000006071 |
| NA | 0.157659577 | 1.11547608 | 0.630888223 | 0.796625039 | 41.15408772 | 45.88795385 | ENSG00000233762 |
| RHBDD2 | 0.157628081 | 1.115451727 | 0.083888189 | 0.209304855 | 653.0485932 | 728.6050151 | ENSG00000005486 |
| PAM | 0.157492236 | 1.115346701 | 0.057905845 | 0.157954438 | 989.9773255 | 1104.170201 | ENSG00000145730 |
| DRAM2 | 0.157258344 | 1.115165894 | 0.214399333 | 0.409272354 | 280.1378774 | 312.3440352 | ENSG00000156171 |
| CCDC77 | 0.157248686 | 1.115158429 | 0.131584321 | 0.291663135 | 523.4612131 | 583.2745697 | ENSG00000120647 |
| PPP1R14A | 0.157085639 | 1.115032405 | 0.204726647 | 0.396621126 | 364.1822724 | 405.8412599 | ENSG00000167641 |
| SDSL | 0.157027758 | 1.114987671 | 0.791444249 | 0.893312556 | 11.63897501 | 12.96105575 | ENSG00000139410 |
| NA | 0.156998761 | 1.114965261 | 0.847694354 | 0.924450417 | 6.554766997 | 7.342153824 | ENSG00000239911 |
| ASNSD1 | 0.156993196 | 1.11496096 | 0.403876625 | 0.615097586 | 127.146543 | 141.7450681 | ENSG00000138381 |
| BAZ2B | 0.156979009 | 1.114949996 | 0.174900469 | 0.35535184 | 402.9486353 | 448.8973821 | ENSG00000123636 |
| TRMT61A | 0.156977006 | 1.114948448 | 0.154087625 | 0.325133498 | 645.6490438 | 720.5532671 | ENSG00000166166 |
| APBB1 | 0.156972897 | 1.114945273 | 0.062707751 | 0.167810101 | 942.5602162 | 1051.398996 | ENSG00000166313 |
| ZNF747 | 0.156958064 | 1.11493381 | 0.623994587 | 0.792264204 | 47.82273168 | 53.20691263 | ENSG00000169955 |
| FAM107B | 0.156872428 | 1.114867631 | 0.10780136 | 0.252569692 | 609.9890103 | 680.1536444 | ENSG00000065809 |
| KCNB2 | 0.156816554 | 1.114824454 | 0.696364487 | 0.837900181 | 28.331318 | 31.50989453 | ENSG00000182674 |
| QDPR | 0.156794299 | 1.114807257 | 0.064098135 | 0.170492011 | 884.8596198 | 986.6755339 | ENSG00000151552 |
| PGM2L1 | 0.15676891 | 1.114787639 | 0.153812936 | 0.324844252 | 932.939191 | 1040.080809 | ENSG00000165434 |
| FLYWCH1 | 0.156752119 | 1.114774664 | 0.12713991 | 0.284179664 | 589.5504275 | 657.6416935 | ENSG00000059122 |
| PANX1 | 0.156705059 | 1.114738301 | 0.115353204 | 0.265372793 | 683.631818 | 761.9325892 | ENSG00000110218 |
| ATRX | 0.156657129 | 1.114701268 | 0.11968178 | 0.272255747 | 1238.151237 | 1379.418668 | ENSG00000085224 |
| IGDCC4 | 0.156559264 | 1.114625654 | 0.132328416 | 0.29269409 | 547.309184 | 610.2327989 | ENSG00000103742 |
| POLR3E | 0.156557577 | 1.114624351 | 0.010259776 | 0.039842869 | 2386.277964 | 2659.25022 | ENSG00000058600 |
| ADCY6 | 0.156508574 | 1.114586492 | 0.143145046 | 0.308665698 | 456.6147542 | 508.8426241 | ENSG00000174233 |
| RPLP0 | 0.1564587 | 1.114547961 | 0.000283318 | 0.001872102 | 41928.91297 | 46730.70574 | ENSG00000089157 |
| RNF125 | 0.156348775 | 1.114463042 | 0.272619114 | 0.48124995 | 260.0228494 | 289.7247289 | ENSG00000101695 |
| TRMT10B | 0.156341191 | 1.114457184 | 0.36713999 | 0.579491407 | 144.0767073 | 160.6387465 | ENSG00000165275 |
| NLGN1 | 0.156212108 | 1.114357474 | 0.435274991 | 0.643868786 | 115.6128253 | 128.8264601 | ENSG00000169760 |
| NA | 0.156162941 | 1.114319497 | 0.856722636 | 0.930106754 | 5.703983547 | 6.336578549 | ENSG00000227014 |
| SEPTIN4 | 0.155742959 | 1.113995156 | 0.404479133 | 0.615476848 | 122.693958 | 136.6932617 | ENSG00000108387 |
| DCAF10 | 0.155673451 | 1.113941485 | 0.24464665 | 0.448950732 | 308.445195 | 343.4067051 | ENSG00000122741 |
| RNH1 | 0.155612875 | 1.113894714 | 0.064557006 | 0.171519282 | 1248.131163 | 1391.111604 | ENSG00000023191 |

| | | | | | | | |
|---|---|---|---|---|---|---|---|
| NA | 0.155563681 | 1.113856733 | 0.739897372 | 0.863848501 | 27.17925305 | 30.27714535 | ENSG00000274828 |
| SLC36A1 | 0.155502395 | 1.113809416 | 0.163153792 | 0.338143037 | 382.4430858 | 426.0592905 | ENSG00000123643 |
| NA | 0.155498371 | 1.11380631 | 0.713600277 | 0.848956862 | 25.05927371 | 27.8085029 | ENSG00000280088 |
| THOC2 | 0.155446299 | 1.113766109 | 0.041440279 | 0.121957249 | 1176.057631 | 1309.306003 | ENSG00000125676 |
| SRSF5 | 0.155436059 | 1.113758204 | 0.036797176 | 0.11138238 | 4441.023222 | 4946.508895 | ENSG00000100650 |
| STARD9 | 0.155282217 | 1.113639445 | 0.294930461 | 0.505010587 | 218.6107264 | 243.4211341 | ENSG00000159433 |
| NA | 0.155272702 | 1.1136321 | 0.815672035 | 0.906328805 | 9.833930231 | 10.99719831 | ENSG00000259845 |
| GNPDA1 | 0.155198369 | 1.113574723 | 0.017325104 | 0.061253636 | 2241.960513 | 2495.706651 | ENSG00000113552 |
| RMI1 | 0.155149604 | 1.113537083 | 0.243594379 | 0.447846558 | 380.0090872 | 423.2651338 | ENSG00000178966 |
| NA | 0.155120585 | 1.113514685 | 0.598083698 | 0.774094349 | 62.84762795 | 69.95356336 | ENSG00000272994 |
| DCPS | 0.155102336 | 1.113500601 | 0.139557331 | 0.303388994 | 485.4555302 | 540.2691456 | ENSG00000110063 |
| MFAP2 | 0.155094425 | 1.113494494 | 0.020995272 | 0.071634276 | 1860.270148 | 2071.770399 | ENSG00000117122 |
| NA | 0.155036013 | 1.113449412 | 0.207282921 | 0.399768992 | 494.9648156 | 550.9433543 | ENSG00000272888 |
| TRIM71 | 0.155025095 | 1.113440986 | 0.022996841 | 0.076989069 | 4298.949815 | 4786.170663 | ENSG00000206557 |
| PKD2 | 0.154615901 | 1.113125223 | 0.060270388 | 0.162926749 | 844.3639299 | 940.0280227 | ENSG00000118762 |
| XPO1 | 0.1546106 | 1.113121134 | 0.002542006 | 0.012465795 | 6485.662293 | 7218.821483 | ENSG00000082898 |
| ARHGAP11A | 0.154567855 | 1.113088154 | 0.066921317 | 0.1764774 | 1261.736243 | 1403.852671 | ENSG00000198826 |
| RMI2 | 0.154524195 | 1.113054469 | 0.018982778 | 0.065806736 | 1491.541908 | 1660.253128 | ENSG00000175643 |
| CDC20 | 0.154396204 | 1.112955727 | 0.001382447 | 0.007431685 | 4647.634732 | 5172.785009 | ENSG00000117399 |
| ACOT9 | 0.154299562 | 1.112881176 | 0.083502582 | 0.208542121 | 814.903301 | 906.357626 | ENSG00000123130 |
| TIMELESS | 0.154265019 | 1.11285453 | 0.002477282 | 0.012227777 | 3796.412644 | 4224.488597 | ENSG00000111602 |
| XKR5 | 0.154104858 | 1.112730993 | 0.490986892 | 0.690563743 | 97.73577173 | 108.6651298 | ENSG00000275591 |
| NA | 0.153974418 | 1.112630391 | 0.66979395 | 0.82299482 | 34.59428671 | 38.55762331 | ENSG00000226221 |
| MYL6 | 0.153818066 | 1.112509817 | 0.022334089 | 0.075263967 | 2577.841879 | 2867.802711 | ENSG00000092841 |
| EHD2 | 0.153798672 | 1.112494861 | 0.176744994 | 0.357982541 | 381.5516736 | 424.4871439 | ENSG00000024422 |
| NA | 0.153794742 | 1.11249183 | 0.599113644 | 0.775055567 | 60.17903637 | 66.78095947 | ENSG00000170919 |
| CNTROB | 0.153776357 | 1.112477654 | 0.06727413 | 0.177217457 | 861.438227 | 958.2611333 | ENSG00000170037 |
| DTD2 | 0.1536002 | 1.112341826 | 0.572301937 | 0.754795871 | 57.05704801 | 63.42563692 | ENSG00000129480 |
| MRAS | 0.153598575 | 1.112340573 | 0.213337671 | 0.407657787 | 324.9599215 | 361.7750436 | ENSG00000158186 |
| HRH2 | 0.153539132 | 1.112294742 | 0.229112042 | 0.428893682 | 311.3951858 | 346.1818068 | ENSG00000113749 |
| LTO1 | 0.153479918 | 1.11224909 | 0.199431508 | 0.389610777 | 337.1338296 | 375.1219892 | ENSG00000149716 |
| FAM156A | 0.153463807 | 1.112236669 | 0.831537272 | 0.915296977 | 8.694018246 | 9.644607996 | ENSG00000268350 |
| TCF19 | 0.153411272 | 1.112196168 | 0.208772306 | 0.401738843 | 405.6775103 | 451.5593073 | ENSG00000137310 |
| NA | 0.153409189 | 1.112194563 | 0.65946006 | 0.816488772 | 40.90647765 | 45.41901754 | ENSG00000224905 |
| SRFBP1 | 0.153376049 | 1.112169014 | 0.198901701 | 0.389030323 | 321.2810074 | 357.2838062 | ENSG00000151304 |
| TMEM14B | 0.153366586 | 1.11216172 | 0.201960332 | 0.393198028 | 416.5171843 | 462.9838019 | ENSG00000137210 |
| ACYP1 | 0.153269173 | 1.112086627 | 0.382172416 | 0.594654863 | 166.3330454 | 185.0560055 | ENSG00000119640 |
| BCL10 | 0.153249761 | 1.112071664 | 0.312957017 | 0.523257482 | 211.9436087 | 235.9508161 | ENSG00000142867 |
| KIAA0319L | 0.153241448 | 1.112065256 | 0.043479844 | 0.126637394 | 1541.007677 | 1713.012628 | ENSG00000142687 |
| NA | 0.153149055 | 1.111994039 | 0.877591553 | 0.943087245 | 6.014468916 | 6.682647625 | ENSG00000278616 |
| STARD8 | 0.153040023 | 1.111910004 | 0.406792023 | 0.617595253 | 135.6973188 | 150.7964751 | ENSG00000130052 |
| M6PR | 0.153015802 | 1.111891336 | 0.035900348 | 0.109333668 | 1424.020957 | 1582.654829 | ENSG00000003056 |
| SIK3 | 0.152994988 | 1.111875294 | 0.076993033 | 0.196282997 | 838.814121 | 932.375247 | ENSG00000160584 |
| LOC285074 | 0.152962475 | 1.111850237 | 0.611783756 | 0.784141186 | 63.57594095 | 70.78339786 | ENSG00000285793 |
| CNTNAP3C | 0.15293393 | 1.111828239 | 0.745018275 | 0.867396409 | 20.36739278 | 22.58826004 | ENSG00000283378 |
| ASPHD1 | 0.152769092 | 1.111701212 | 0.143817972 | 0.309727723 | 445.9419135 | 495.9124763 | ENSG00000174939 |
| SPATA41 | 0.152683777 | 1.111635473 | 0.823319389 | 0.910506307 | 10.42878945 | 11.65263712 | ENSG00000189419 |
| IMPACT | 0.152559042 | 1.111539365 | 0.089446007 | 0.219415763 | 799.3980579 | 888.5835071 | ENSG00000154059 |
| ZNF317 | 0.152319907 | 1.111355136 | 0.052282509 | 0.146108362 | 887.8345275 | 986.4467289 | ENSG00000130803 |
| CARD8 | 0.152270161 | 1.111316816 | 0.326807583 | 0.538266283 | 212.7057114 | 236.2787465 | ENSG00000105483 |
| DYRK1A | 0.152203911 | 1.111265784 | 0.107851865 | 0.252572711 | 769.8140165 | 854.9600473 | ENSG00000157540 |
| C15orf41 | 0.152082644 | 1.11117238 | 0.333745254 | 0.545760531 | 207.7104896 | 230.9352145 | ENSG00000186073 |
| ATL2 | 0.151828339 | 1.11097653 | 0.029354373 | 0.093294738 | 1358.608598 | 1508.953055 | ENSG00000119787 |
| MED10 | 0.151704529 | 1.110881191 | 0.168906798 | 0.34662375 | 461.6219013 | 512.582564 | ENSG00000133398 |
| MTPAP | 0.151696907 | 1.110875323 | 0.102816172 | 0.243044613 | 585.471457 | 650.4644032 | ENSG00000107951 |
| DMAC2L | 0.151665169 | 1.110850885 | 0.486131002 | 0.687118027 | 105.4719692 | 117.1998559 | ENSG00000125375 |
| SLF2 | 0.151650524 | 1.110839608 | 0.175531667 | 0.35602143 | 501.953096 | 557.6615217 | ENSG00000119906 |
| MALL | 0.151542782 | 1.110756653 | 0.778301846 | 0.886943686 | 14.68713033 | 16.27003116 | ENSG00000144063 |
| MTBP | 0.15153819 | 1.110753117 | 0.319127884 | 0.529256214 | 200.7722683 | 223.039796 | ENSG00000172167 |
| SOCS6 | 0.15146047 | 1.110693281 | 0.172660757 | 0.352203326 | 407.1011596 | 452.2219646 | ENSG00000170677 |
| ECSIT | 0.151435834 | 1.110674315 | 0.045193023 | 0.130280909 | 1110.283759 | 1233.537358 | ENSG00000130159 |
| NA | 0.15137844 | 1.11063013 | 0.547028596 | 0.734791389 | 81.54462493 | 90.62968262 | ENSG00000224546 |
| PRRG3 | 0.151321585 | 1.110586362 | 0.439261648 | 0.647222924 | 123.8273097 | 137.3851163 | ENSG00000130032 |
| NA | 0.151213486 | 1.11050315 | 0.550801911 | 0.738037043 | 79.8729822 | 88.54541695 | ENSG00000230551 |
| GATA3 | 0.151126168 | 1.11043594 | 0.898276329 | 0.955069726 | 3.313562751 | 3.688338848 | ENSG00000107485 |
| BEST1 | 0.151098551 | 1.110414684 | 0.891282693 | 0.951694019 | 3.561172824 | 3.981534386 | ENSG00000167995 |
| PIAS4 | 0.15109183 | 1.110409511 | 0.026661415 | 0.086646159 | 1504.754055 | 1671.318669 | ENSG00000105229 |
| KLHL26 | 0.151065292 | 1.110389085 | 0.194441088 | 0.382900932 | 377.7918074 | 419.2916615 | ENSG00000167487 |
| RMND1 | 0.15105584 | 1.110381811 | 0.199500582 | 0.389614115 | 359.9225093 | 399.600876 | ENSG00000155906 |
| VASP | 0.150786178 | 1.110174282 | 0.02694551 | 0.087418992 | 2782.552951 | 3089.886523 | ENSG00000125753 |
| NA | 0.150749784 | 1.110146277 | 0.835275961 | 0.917410484 | 8.703501455 | 9.629379751 | ENSG00000262333 |
| GNG11 | 0.15073654 | 1.110136086 | 0.616366609 | 0.786917483 | 48.84787698 | 54.17997154 | ENSG00000127920 |
| PIN1 | 0.150672363 | 1.110086704 | 0.030529076 | 0.09632672 | 1954.77023 | 2170.056042 | ENSG00000127445 |
| ACOT13 | 0.150655157 | 1.110073465 | 0.301472637 | 0.511636058 | 256.4821175 | 284.4673876 | ENSG00000112304 |

| | | | | | | | |
|---|---|---|---|---|---|---|---|
| SIRT1 | 0.150537782 | 1.109983154 | 0.070956039 | 0.184853744 | 1399.269549 | 1553.380147 | ENSG00000096717 |
| HECA | 0.150383153 | 1.109864192 | 0.569042921 | 0.752125949 | 71.55855326 | 79.57961078 | ENSG00000112406 |
| RND3 | 0.150329281 | 1.109822749 | 0.082634904 | 0.206810608 | 674.6609027 | 748.6456536 | ENSG00000115963 |
| LATS1 | 0.150301055 | 1.109801036 | 0.27098683 | 0.479399183 | 332.8298265 | 369.3893937 | ENSG00000131023 |
| EIF3G | 0.150248222 | 1.109760395 | 0.008240881 | 0.033490216 | 4235.451729 | 4700.774296 | ENSG00000130811 |
| ARFIP1 | 0.15010339 | 1.109648992 | 0.138583585 | 0.301862862 | 711.3980513 | 789.6994521 | ENSG00000164144 |
| TRMT10C | 0.150059648 | 1.109615348 | 0.13457161 | 0.296175059 | 732.9274985 | 813.4305525 | ENSG00000174173 |
| RABL2B | 0.150028461 | 1.109591362 | 0.26007706 | 0.467011057 | 279.8706644 | 310.236123 | ENSG00000079974 |
| WDR6 | 0.1499832 | 1.109556552 | 0.002136831 | 0.010769697 | 4900.301176 | 5436.91863 | ENSG00000178252 |
| ELL2 | 0.149978728 | 1.109553112 | 0.303824244 | 0.513856425 | 259.2992649 | 287.4038095 | ENSG00000118985 |
| PLTP | 0.149677303 | 1.109321315 | 0.025369871 | 0.083381485 | 2241.401279 | 2486.978016 | ENSG00000100979 |
| EFCC1 | 0.149636246 | 1.109289746 | 0.799610932 | 0.898046533 | 13.42830736 | 14.94326286 | ENSG00000114654 |
| PCK2 | 0.149467291 | 1.109159844 | 0.032323462 | 0.100746102 | 1978.01216 | 2193.171817 | ENSG00000100889 |
| ZNF304 | 0.14936964 | 1.109084771 | 0.311128385 | 0.521747681 | 234.9350743 | 260.2865417 | ENSG00000131845 |
| KCNK2 | 0.149323343 | 1.109049181 | 0.721716339 | 0.853183917 | 26.33322429 | 29.11692152 | ENSG00000082482 |
| C19orf44 | 0.149194214 | 1.108949919 | 0.526903675 | 0.720012728 | 97.09524975 | 107.8079603 | ENSG00000105072 |
| GPR61 | 0.149173276 | 1.108933825 | 0.862967135 | 0.933718002 | 5.700424074 | 6.289285443 | ENSG00000156097 |
| MAGI3 | 0.149167748 | 1.108929576 | 0.348057089 | 0.560124254 | 210.8754186 | 233.7676133 | ENSG00000081026 |
| NCAPD3 | 0.149128254 | 1.108899219 | 0.036902958 | 0.111613127 | 1632.099649 | 1810.061808 | ENSG00000151503 |
| ITM2B | 0.149124961 | 1.108896688 | 0.020125626 | 0.069154374 | 1780.721077 | 1975.135945 | ENSG00000136156 |
| ZNF415 | 0.149112353 | 1.108886997 | 0.369527271 | 0.581680252 | 165.766662 | 183.8040333 | ENSG00000170954 |
| ZNHIT2 | 0.14910859 | 1.108884105 | 0.489680428 | 0.689811643 | 99.6390072 | 110.461118 | ENSG00000174276 |
| SNAI1 | 0.149002076 | 1.108802239 | 0.389248286 | 0.601059189 | 163.9118107 | 181.9063131 | ENSG00000124216 |
| DNAH11 | 0.148992374 | 1.108794782 | 0.49768748 | 0.696725579 | 112.2974569 | 124.3633358 | ENSG00000105877 |
| ZNF546 | 0.148945281 | 1.108758589 | 0.789216716 | 0.891980289 | 16.73714159 | 18.60360423 | ENSG00000187187 |
| COA7 | 0.148924178 | 1.108742371 | 0.02499121 | 0.082467064 | 1438.095508 | 1594.238892 | ENSG00000162377 |
| EIF4A1 | 0.148871851 | 1.108702157 | 0.428649952 | 0.638461578 | 147.3852361 | 163.6482109 | ENSG00000161960 |
| C3orf14 | 0.148805234 | 1.108650964 | 0.464274083 | 0.668593683 | 142.5521715 | 158.296966 | ENSG00000114405 |
| HHIP | 0.148791482 | 1.108640396 | 0.894846363 | 0.953032171 | 3.282722707 | 3.648518091 | ENSG00000164161 |
| FARSB | 0.148670035 | 1.108547074 | 0.031334212 | 0.098345785 | 1291.251459 | 1431.527819 | ENSG00000116120 |
| CCDC88C | 0.148618095 | 1.108507165 | 0.043085988 | 0.125650946 | 1373.318202 | 1521.854121 | ENSG00000015133 |
| FZD3 | 0.148604132 | 1.108496437 | 0.246926209 | 0.451745658 | 594.2034336 | 658.7309847 | ENSG00000104290 |
| SLC25A18 | 0.148602377 | 1.108495088 | 0.797824693 | 0.896947216 | 13.43898578 | 14.94203336 | ENSG00000182902 |
| LHX6 | 0.148592206 | 1.108487273 | 0.654957794 | 0.813380102 | 43.5842731 | 48.44595827 | ENSG00000106852 |
| NA | 0.148587225 | 1.108483445 | 0.761037128 | 0.876406255 | 21.81156091 | 24.30091816 | ENSG00000229344 |
| RAB3IP | 0.148497197 | 1.108414275 | 0.174423203 | 0.354763832 | 407.1293043 | 451.0414266 | ENSG00000127328 |
| NCCRP1 | 0.148459468 | 1.108385289 | 0.528634613 | 0.721282036 | 79.13905037 | 87.72967651 | ENSG00000188505 |
| EEF1A1 | 0.148264374 | 1.108235413 | 0.003222191 | 0.015296439 | 204647.6232 | 226796.2106 | ENSG00000156508 |
| YTHDF3 | 0.148167599 | 1.108161077 | 0.072412378 | 0.187788938 | 1854.763812 | 2055.119265 | ENSG00000185728 |
| SPTSSB | 0.148089909 | 1.108101403 | 0.385011133 | 0.597138171 | 155.5561432 | 172.2491038 | ENSG00000196542 |
| RGMB | 0.148044325 | 1.108066391 | 0.406037859 | 0.617045544 | 139.9787959 | 155.1477251 | ENSG00000174136 |
| PREPL | 0.148025415 | 1.108051867 | 0.115646383 | 0.265852944 | 618.8027514 | 685.2944339 | ENSG00000138078 |
| ATOX1 | 0.147983152 | 1.108019408 | 0.189936654 | 0.376345997 | 410.9214589 | 455.29312 | ENSG00000177556 |
| BLMH | 0.147952613 | 1.107995954 | 0.017167172 | 0.060803084 | 2005.038204 | 2222.167639 | ENSG00000108578 |
| TMEM268 | 0.147896043 | 1.107952508 | 0.200797741 | 0.391580476 | 375.6323945 | 416.0467205 | ENSG00000157693 |
| MED1 | 0.147847943 | 1.10791557 | 0.060581636 | 0.163534048 | 1414.227706 | 1566.39046 | ENSG00000125686 |
| CEP76 | 0.147828591 | 1.107900708 | 0.31737916 | 0.527477286 | 222.6914514 | 246.484375 | ENSG00000101624 |
| GIT1 | 0.147666048 | 1.107775892 | 0.025997002 | 0.084925338 | 2466.493019 | 2732.858986 | ENSG00000108262 |
| NA | 0.147554732 | 1.107690421 | 0.647427034 | 0.808694989 | 41.98590475 | 46.54433863 | ENSG00000264204 |
| SNHG16 | 0.147545547 | 1.10768337 | 0.071120517 | 0.185180057 | 909.0784841 | 1006.590982 | ENSG00000163597 |
| CETN3 | 0.147542017 | 1.10768066 | 0.261370475 | 0.468515166 | 345.5499564 | 382.5564155 | ENSG00000153140 |
| KIAA1107 | 0.147493561 | 1.107643456 | 0.685933148 | 0.831762616 | 37.25486962 | 41.24736477 | ENSG00000189195 |
| MPZL3 | 0.147383628 | 1.107559057 | 0.599298255 | 0.775241287 | 53.57980183 | 59.32899119 | ENSG00000160588 |
| YIPF5 | 0.147336354 | 1.107522765 | 0.15695871 | 0.329068783 | 476.1814749 | 527.378578 | ENSG00000145817 |
| NOB1 | 0.147187261 | 1.107408316 | 0.041330111 | 0.121765858 | 1375.586126 | 1523.144843 | ENSG00000141101 |
| TMEM178A | 0.147075841 | 1.107322794 | 0.224852208 | 0.423236713 | 364.6375527 | 404.087143 | ENSG00000152154 |
| LMO2 | 0.147038148 | 1.107293863 | 0.675566614 | 0.826353747 | 40.03316831 | 44.2571127 | ENSG00000135363 |
| SNORD3A | 0.147004506 | 1.107268043 | 0.779846646 | 0.887674276 | 17.41976626 | 19.35239976 | ENSG00000263934 |
| PRMT3 | 0.146910558 | 1.10719594 | 0.11701034 | 0.268237469 | 659.2412889 | 729.7202463 | ENSG00000185238 |
| DNAJC18 | 0.146860502 | 1.107157525 | 0.239433855 | 0.44239393 | 360.7640889 | 399.2513792 | ENSG00000170464 |
| SMOX | 0.14668756 | 1.107024814 | 0.224217984 | 0.422421849 | 345.2979227 | 382.4255518 | ENSG00000088826 |
| MRPL36 | 0.146539825 | 1.106911458 | 0.085509299 | 0.212367584 | 740.2628022 | 819.5298578 | ENSG00000171421 |
| ZNF251 | 0.146519339 | 1.10689574 | 0.359474586 | 0.571533723 | 225.5658291 | 249.8328353 | ENSG00000198169 |
| ZBBX | 0.146510537 | 1.106888987 | 0.860387599 | 0.932156491 | 6.000231026 | 6.620126274 | ENSG00000169064 |
| SLC25A48 | 0.146448157 | 1.106841127 | 0.648763104 | 0.809250856 | 43.1462577 | 47.67975904 | ENSG00000145832 |
| DSG2 | 0.146400256 | 1.106804379 | 0.01626891 | 0.058222164 | 4070.509843 | 4505.086177 | ENSG00000046604 |
| TMEM101 | 0.146342796 | 1.106760297 | 0.19233885 | 0.379705144 | 417.3172971 | 461.7091815 | ENSG00000091947 |
| NA | 0.146332301 | 1.106752246 | 0.752246424 | 0.871085164 | 19.49733742 | 21.59450804 | ENSG00000216285 |
| NA | 0.146307708 | 1.10673338 | 0.784111753 | 0.890065126 | 16.86167082 | 18.69383951 | ENSG00000272221 |
| TBL3 | 0.146236644 | 1.106678866 | 0.026340921 | 0.085826137 | 1927.08122 | 2133.162468 | ENSG00000183751 |
| CCNG1 | 0.146047453 | 1.106533749 | 0.014568501 | 0.053226181 | 3697.732095 | 4090.793095 | ENSG00000113328 |
| PBX2 | 0.14602422 | 1.106515929 | 0.010656384 | 0.041120411 | 3040.194833 | 3363.512057 | ENSG00000204304 |
| RASGEF1C | 0.145873061 | 1.1064 | 0.73308077 | 0.859729886 | 22.75633903 | 25.19790849 | ENSG00000146090 |
| ELMOD2 | 0.145824437 | 1.10636271 | 0.248426157 | 0.453402881 | 449.2462724 | 496.891218 | ENSG00000179387 |

| | | | | | | | |
|---|---|---|---|---|---|---|---|
| LLGL2 | 0.145814361 | 1.106354984 | 0.029062264 | 0.092568873 | 1776.396783 | 1965.460043 | ENSG00000073350 |
| SCAF1 | 0.145746998 | 1.106303326 | 0.01815614 | 0.063507817 | 3605.764704 | 3989.662762 | ENSG00000126461 |
| TIPIN | 0.145733376 | 1.106292881 | 0.243715821 | 0.448026213 | 300.8442052 | 332.6274547 | ENSG00000075131 |
| CXCL2 | 0.145718933 | 1.106281805 | 0.853711746 | 0.928338837 | 6.891337729 | 7.648402144 | ENSG00000081041 |
| NA | 0.145713401 | 1.106277563 | 0.797118393 | 0.896454171 | 13.45322367 | 14.9183868 | ENSG00000259394 |
| FAM219A | 0.145568238 | 1.106166256 | 0.170377003 | 0.348653062 | 435.6714942 | 482.1500351 | ENSG00000164970 |
| EI24 | 0.145557794 | 1.106158248 | 0.005839114 | 0.025126983 | 3225.006866 | 3567.110332 | ENSG00000149547 |
| NA | 0.145548455 | 1.106151088 | 0.832117218 | 0.915391953 | 10.13017771 | 11.23345293 | ENSG00000279694 |
| DAZAP2 | 0.145500562 | 1.106114368 | 0.270804447 | 0.479166306 | 338.1115594 | 373.7632514 | ENSG00000183283 |
| ZNF461 | 0.14540494 | 1.106041057 | 0.627334542 | 0.793729923 | 58.40928083 | 64.60600701 | ENSG00000197808 |
| RADIL | 0.145324334 | 1.105979262 | 0.309201358 | 0.519692741 | 241.47446 | 267.0914465 | ENSG00000157927 |
| CDC37L1 | 0.145253425 | 1.105924904 | 0.189974604 | 0.376381704 | 410.7130547 | 454.4396913 | ENSG00000106993 |
| FAM185A | 0.145200624 | 1.105884429 | 0.491711866 | 0.691126718 | 104.5583366 | 115.559156 | ENSG00000222011 |
| POGLUT1 | 0.145170804 | 1.105861571 | 0.258734755 | 0.465619007 | 323.3321621 | 357.2081367 | ENSG00000163389 |
| VPS37A | 0.145014244 | 1.105741571 | 0.192834795 | 0.380332648 | 429.4207046 | 474.7311502 | ENSG00000155975 |
| NA | 0.144886193 | 1.105643431 | 0.652749513 | 0.811801095 | 64.896444 | 71.83231445 | ENSG00000278041 |
| EIF5A2 | 0.144873866 | 1.105633984 | 0.522459733 | 0.71692967 | 83.39116202 | 92.1709509 | ENSG00000163577 |
| UNC5B-AS1 | 0.144729927 | 1.105523679 | 0.817232414 | 0.906888832 | 14.24883559 | 15.6718168 | ENSG00000237512 |
| WFS1 | 0.144556228 | 1.105390584 | 0.10600188 | 0.249202727 | 1174.012861 | 1298.437486 | ENSG00000109501 |
| YTHDC1 | 0.144470729 | 1.105325077 | 0.021151015 | 0.072022525 | 2026.01698 | 2239.441433 | ENSG00000083896 |
| C19orf25 | 0.144352997 | 1.105234879 | 0.088826131 | 0.218223166 | 947.1602397 | 1047.492657 | ENSG00000119559 |
| FAM234A | 0.144241398 | 1.105149388 | 0.012258379 | 0.046336967 | 2374.968979 | 2625.129942 | ENSG00000167930 |
| BPNT1 | 0.144200667 | 1.105118187 | 0.280495777 | 0.489390497 | 303.0661617 | 334.8283448 | ENSG00000162813 |
| ARHGEF7 | 0.144183079 | 1.105104714 | 0.067890612 | 0.178393518 | 1130.529825 | 1249.033863 | ENSG00000102606 |
| PPP2R2A | 0.144061868 | 1.10501187 | 0.029295741 | 0.093171113 | 1633.39597 | 1804.71939 | ENSG00000221914 |
| XYLT1 | 0.144018396 | 1.104978574 | 0.094886521 | 0.229328138 | 991.8582627 | 1095.612207 | ENSG00000103489 |
| OGA | 0.143810147 | 1.104819085 | 0.016936538 | 0.060121526 | 2762.772151 | 3051.92373 | ENSG00000198408 |
| ROBO1 | 0.143705974 | 1.104739312 | 0.036660851 | 0.111112208 | 2521.273945 | 2784.602805 | ENSG00000169855 |
| MIIP | 0.143564132 | 1.104630703 | 0.047874611 | 0.136387393 | 1396.422504 | 1542.966455 | ENSG00000116691 |
| NR4A1 | 0.143555438 | 1.104624046 | 0.223905408 | 0.42204389 | 501.0077069 | 553.915289 | ENSG00000123358 |
| TBC1D17 | 0.143553812 | 1.104622801 | 0.120708333 | 0.273698333 | 630.2780947 | 696.3546481 | ENSG00000104946 |
| UBE4A | 0.143443203 | 1.104538115 | 0.15178492 | 0.321568119 | 601.1860974 | 663.6270741 | ENSG00000110344 |
| AQP11 | 0.143361862 | 1.104475841 | 0.674845503 | 0.825790913 | 37.28570967 | 41.24473781 | ENSG00000178301 |
| NPHP4 | 0.142884914 | 1.104110767 | 0.206771728 | 0.399262489 | 446.9228706 | 493.7961887 | ENSG00000131697 |
| MGME1 | 0.142826933 | 1.104066394 | 0.100052874 | 0.23818209 | 1156.997927 | 1276.744798 | ENSG00000125871 |
| NFATC2 | 0.142809007 | 1.104052676 | 0.701670349 | 0.841282746 | 30.6562199 | 33.90352994 | ENSG00000101096 |
| COX4I1 | 0.142688398 | 1.103960382 | 0.012112721 | 0.045905827 | 7286.292342 | 8043.3826 | ENSG00000131143 |
| ZNF43 | 0.142685995 | 1.103958542 | 0.289610454 | 0.498977595 | 265.9854269 | 293.4928046 | ENSG00000198521 |
| UBXN1 | 0.142678075 | 1.103952483 | 0.027270173 | 0.088214664 | 1876.312818 | 2071.860194 | ENSG00000162191 |
| E2F7 | 0.142643598 | 1.1039261 | 0.221255946 | 0.418343867 | 469.5991741 | 518.407357 | ENSG00000165891 |
| KCNA7 | 0.14257268 | 1.103871837 | 0.831210321 | 0.915293173 | 9.612405521 | 10.59853924 | ENSG00000104848 |
| ATXN1L | 0.142559175 | 1.103861504 | 0.083124971 | 0.207839941 | 913.5281462 | 1007.976613 | ENSG00000224470 |
| NA | 0.142526617 | 1.103836592 | 0.864635573 | 0.934774605 | 6.647287128 | 7.337802897 | ENSG00000268670 |
| UHRF1BP1L | 0.14248353 | 1.103803626 | 0.267260946 | 0.474965442 | 451.1418308 | 497.6894618 | ENSG00000111647 |
| MRPL19 | 0.142379215 | 1.103723817 | 0.151305374 | 0.320984272 | 743.2228865 | 820.2629603 | ENSG00000115364 |
| KIFBP | 0.142343922 | 1.103696817 | 0.0904822 | 0.221284168 | 1034.513801 | 1142.029727 | ENSG00000198954 |
| CNRIP1 | 0.1422624 | 1.103634453 | 0.622737844 | 0.79114079 | 58.09404078 | 64.05497018 | ENSG00000119865 |
| ZNF148 | 0.141855604 | 1.103323305 | 0.333031176 | 0.54504669 | 249.8594412 | 275.5248491 | ENSG00000163848 |
| WWTR1 | 0.141780869 | 1.103266152 | 0.356401903 | 0.568515071 | 264.8920668 | 292.1945579 | ENSG00000018408 |
| DOT1L | 0.141727872 | 1.103225625 | 0.103222994 | 0.243792677 | 2267.501053 | 2502.317698 | ENSG00000104885 |
| DSC2 | 0.14170533 | 1.103208387 | 0.183913677 | 0.367963745 | 683.4773908 | 753.939513 | ENSG00000134755 |
| NA | 0.141691918 | 1.103198131 | 0.89908827 | 0.95546587 | 3.333724377 | 3.676515571 | ENSG00000284726 |
| CARS2 | 0.141663587 | 1.103176467 | 0.04555584 | 0.131046768 | 1279.636562 | 1412.200266 | ENSG00000134905 |
| SMYD2 | 0.141600383 | 1.103128139 | 0.424096296 | 0.634226465 | 140.4953729 | 155.0490715 | ENSG00000143499 |
| NA | 0.141573844 | 1.103107846 | 0.899112348 | 0.95546587 | 3.330164905 | 3.672164644 | ENSG00000227638 |
| ASCC3 | 0.141520756 | 1.103067255 | 0.059273801 | 0.160854394 | 1925.822522 | 2123.807288 | ENSG00000112249 |
| TMEM205 | 0.141482737 | 1.103038187 | 0.537852269 | 0.727796205 | 88.88994368 | 98.15571198 | ENSG00000105518 |
| NPHP1 | 0.141388723 | 1.102966308 | 0.711092561 | 0.847591497 | 36.22972433 | 40.06637522 | ENSG00000144061 |
| NSUN3 | 0.141368981 | 1.102951216 | 0.531479625 | 0.723282606 | 97.96352566 | 107.8572179 | ENSG00000178694 |
| KCNJ2-AS1 | 0.141350628 | 1.102937185 | 0.907363398 | 0.960082797 | 3.317122223 | 3.658165904 | ENSG00000267365 |
| FOPNL | 0.141335697 | 1.10292577 | 0.259416262 | 0.466401 | 483.2187498 | 533.3636523 | ENSG00000133393 |
| CDK9 | 0.141305011 | 1.102902311 | 0.03625256 | 0.110175346 | 2227.18381 | 2455.962293 | ENSG00000136807 |
| SYNGR3 | 0.141271534 | 1.102876719 | 0.138297898 | 0.301483989 | 637.7654863 | 703.8547398 | ENSG00000127561 |
| TIGAR | 0.141226076 | 1.102841969 | 0.504586858 | 0.702012878 | 98.70607655 | 108.9687512 | ENSG00000078237 |
| RNASE1 | 0.141211009 | 1.102830452 | 0.873427287 | 0.940645204 | 6.074953795 | 6.669594844 | ENSG00000129538 |
| GFY | 0.140862989 | 1.102564449 | 0.734180545 | 0.860316207 | 27.68515162 | 30.56873252 | ENSG00000261949 |
| SAT2 | 0.140816167 | 1.102528666 | 0.203228909 | 0.39477247 | 543.0323836 | 598.4574828 | ENSG00000141504 |
| PGM3 | 0.140751006 | 1.10247887 | 0.113130099 | 0.261666857 | 643.6182794 | 709.4375095 | ENSG00000013375 |
| TSG101 | 0.140748729 | 1.10247713 | 0.117080577 | 0.268291789 | 962.7118784 | 1061.112418 | ENSG00000074319 |
| YEATS4 | 0.140472993 | 1.102266439 | 0.129564872 | 0.288269628 | 590.6802204 | 651.0778458 | ENSG00000127337 |
| CCDC51 | 0.140390393 | 1.102203332 | 0.14404169 | 0.309962092 | 571.1662546 | 629.6405609 | ENSG00000164051 |
| SPCS1 | 0.140346068 | 1.102169469 | 0.075086748 | 0.192769608 | 892.4805186 | 983.6603937 | ENSG00000114902 |
| TMEM179 | 0.140256788 | 1.102101263 | 0.347295025 | 0.559353678 | 232.841537 | 256.9611086 | ENSG00000258986 |
| NA | 0.140167334 | 1.10203293 | 0.458589346 | 0.663992931 | 130.529489 | 144.0045801 | ENSG00000247134 |

| | | | | | | | |
|---|---|---|---|---|---|---|---|
| SORD2P | 0.140058682 | 1.101949937 | 0.588387955 | 0.766483171 | 65.04677926 | 71.74347662 | ENSG00000259479 |
| PGM2 | 0.139945194 | 1.101863257 | 0.10376978 | 0.24486971 | 703.7356874 | 775.5232094 | ENSG00000169299 |
| PARD6B | 0.139875357 | 1.10180992 | 0.520621786 | 0.715714945 | 125.1451947 | 138.0537758 | ENSG00000124171 |
| TMUB1 | 0.139866279 | 1.101802987 | 0.191085817 | 0.377830139 | 922.7286973 | 1017.469802 | ENSG00000164897 |
| SHISAL1 | 0.139770609 | 1.101729925 | 0.315612729 | 0.525761748 | 283.7468235 | 312.708411 | ENSG00000138944 |
| ARHGAP12 | 0.13976792 | 1.101727872 | 0.131943293 | 0.292081713 | 912.8242165 | 1005.480213 | ENSG00000165322 |
| UQCRC1 | 0.139736146 | 1.101703607 | 0.00320978 | 0.015260999 | 5829.66137 | 6422.424164 | ENSG00000010256 |
| NA | 0.139716049 | 1.10168826 | 0.894413021 | 0.953032171 | 4.211788399 | 4.677739919 | ENSG00000287310 |
| BAIAP2L2 | 0.139650597 | 1.10163828 | 0.557619986 | 0.743624488 | 120.3201393 | 132.6875566 | ENSG00000128298 |
| PEX7 | 0.139639937 | 1.101630141 | 0.423577437 | 0.633654934 | 160.2002765 | 176.4330314 | ENSG00000112357 |
| TRAP1 | 0.139402843 | 1.101449112 | 0.010169473 | 0.039581718 | 3481.044607 | 3834.134217 | ENSG00000126602 |
| ARCN1 | 0.139342157 | 1.101402782 | 0.013958798 | 0.051366309 | 2904.355169 | 3198.702518 | ENSG00000095139 |
| ZNF283 | 0.139331014 | 1.101394274 | 0.583046561 | 0.762575961 | 70.84831695 | 77.87660046 | ENSG00000167637 |
| HECW2 | 0.139314063 | 1.101381334 | 0.479612122 | 0.681705228 | 119.1565062 | 131.1513313 | ENSG00000138411 |
| VPS9D1 | 0.139214917 | 1.101305647 | 0.558991596 | 0.744415506 | 84.74667498 | 93.3673796 | ENSG00000075399 |
| SLC2A4 | 0.139143556 | 1.101251173 | 0.493803653 | 0.693241302 | 118.9326173 | 130.9896322 | ENSG00000181856 |
| NA | 0.139131455 | 1.101241936 | 0.895181803 | 0.953032171 | 3.888260348 | 4.309253795 | ENSG00000230956 |
| ARFIP2 | 0.139063434 | 1.101190016 | 0.056850234 | 0.155867693 | 1156.874085 | 1274.227275 | ENSG00000132254 |
| BUD13 | 0.138993378 | 1.101136544 | 0.129283747 | 0.287851869 | 666.5273703 | 733.7545589 | ENSG00000137656 |
| EPB41L4B | 0.138992897 | 1.101136176 | 0.120170511 | 0.272832423 | 726.2476448 | 799.3698094 | ENSG00000095203 |
| SCN5A | 0.138894162 | 1.10106082 | 0.209590592 | 0.402779925 | 387.2974811 | 426.3625854 | ENSG00000183873 |
| VWCE | 0.138821747 | 1.101005554 | 0.368312221 | 0.580382661 | 285.1731948 | 314.1498094 | ENSG00000167992 |
| RFX3 | 0.138655459 | 1.100878657 | 0.36952036 | 0.581680252 | 236.1034615 | 259.9167831 | ENSG00000080298 |
| LOC651337 | 0.138587449 | 1.100826762 | 0.862110833 | 0.933113427 | 6.660329809 | 7.322858199 | ENSG00000255585 |
| NA | 0.13836973 | 1.100660647 | 0.900271231 | 0.956071542 | 3.330164905 | 3.660341367 | ENSG00000286974 |
| DDX11 | 0.138364695 | 1.100656806 | 0.110368658 | 0.257100296 | 2352.324765 | 2589.397328 | ENSG00000013573 |
| TMEM17 | 0.138283594 | 1.100594935 | 0.766359304 | 0.878997598 | 24.40006465 | 26.87850175 | ENSG00000186889 |
| ARHGEF37 | 0.138279102 | 1.100591508 | 0.867194158 | 0.936739326 | 7.585836029 | 8.374780621 | ENSG00000183111 |
| ETFA | 0.138247416 | 1.100567336 | 0.052570301 | 0.146612739 | 1867.51303 | 2054.602422 | ENSG00000140374 |
| EIF2B4 | 0.138212189 | 1.100540463 | 0.052438285 | 0.146428658 | 1195.624936 | 1315.781214 | ENSG00000115211 |
| DALRD3 | 0.138152705 | 1.100495088 | 0.217078112 | 0.412841907 | 410.7898107 | 452.1363341 | ENSG00000178149 |
| AP1AR | 0.137980503 | 1.100363738 | 0.154543312 | 0.325985752 | 598.488724 | 658.5430348 | ENSG00000138660 |
| METTL1 | 0.137956104 | 1.100345129 | 0.26475793 | 0.472451432 | 314.6526867 | 346.2652524 | ENSG00000037897 |
| S100PBP | 0.137868325 | 1.100278182 | 0.125945066 | 0.282411845 | 765.050161 | 841.3666954 | ENSG00000116497 |
| ZNF518A | 0.137781685 | 1.100212108 | 0.231025339 | 0.431331045 | 414.81272 | 456.3650301 | ENSG00000177853 |
| DPY30 | 0.137777812 | 1.100209154 | 0.052454292 | 0.146451701 | 1477.178761 | 1624.662973 | ENSG00000162961 |
| NA | 0.1376172 | 1.100086677 | 0.761554428 | 0.876689032 | 21.75847431 | 23.99060246 | ENSG00000226396 |
| EFCAB7 | 0.13758816 | 1.100064534 | 0.620273039 | 0.789290233 | 55.56841234 | 61.16640848 | ENSG00000203965 |
| ZKSCAN7-AS1 | 0.13757666 | 1.100055765 | 0.902418911 | 0.957425906 | 3.310003278 | 3.636694814 | ENSG00000236869 |
| PPP3CC | 0.137361573 | 1.099891773 | 0.65362919 | 0.812465428 | 46.232372 | 50.76212764 | ENSG00000120910 |
| RBBP6 | 0.137340721 | 1.099875876 | 0.044115041 | 0.127876062 | 1369.152485 | 1505.820766 | ENSG00000122257 |
| BRPF1 | 0.137247753 | 1.099805002 | 0.111805603 | 0.259514137 | 826.2934957 | 908.6140463 | ENSG00000156983 |
| PEX19 | 0.137164231 | 1.099741332 | 0.179153201 | 0.361714629 | 641.0111409 | 704.9209672 | ENSG00000162735 |
| STK3 | 0.137022499 | 1.099633298 | 0.355529579 | 0.567747007 | 213.9743224 | 235.3402114 | ENSG00000104375 |
| UBE3D | 0.137002426 | 1.099617998 | 0.32015883 | 0.530613338 | 275.6657156 | 302.8984154 | ENSG00000118420 |
| WDR74 | 0.136933854 | 1.099565734 | 0.048872714 | 0.138582763 | 1611.788616 | 1772.161654 | ENSG00000133316 |
| ZNF25 | 0.136852885 | 1.099504024 | 0.845762108 | 0.92340969 | 10.05929375 | 11.04231595 | ENSG00000175395 |
| SREBF1 | 0.136836889 | 1.099491834 | 0.040399122 | 0.119638925 | 2231.362731 | 2454.108385 | ENSG00000072310 |
| CASQ1 | 0.136743233 | 1.09942046 | 0.774942951 | 0.884331605 | 19.68949663 | 21.59791301 | ENSG00000143318 |
| HEATR3 | 0.136658926 | 1.099356215 | 0.195498374 | 0.384501854 | 468.1161311 | 514.3720984 | ENSG00000155393 |
| SLC25A46 | 0.136608711 | 1.099317951 | 0.214734974 | 0.409718068 | 450.9771787 | 495.8994962 | ENSG00000164209 |
| CDK1 | 0.136546537 | 1.099270576 | 0.064026321 | 0.170324986 | 1189.645783 | 1307.300927 | ENSG00000170312 |
| CLASRP | 0.136418987 | 1.099173393 | 0.05636424 | 0.154827444 | 1445.133569 | 1588.639415 | ENSG00000104859 |
| NDUFA6-DT | 0.13638198 | 1.099145198 | 0.763580242 | 0.877522527 | 20.88041028 | 22.95485425 | ENSG00000237037 |
| RGPD8 | 0.136334065 | 1.099108693 | 0.785381143 | 0.890433415 | 17.17571566 | 18.87750414 | ENSG00000169629 |
| RPL7A | 0.136292895 | 1.099077329 | 0.003482198 | 0.016334155 | 25520.97985 | 28048.83761 | ENSG00000148303 |
| PI4KAP2 | 0.136264887 | 1.099055992 | 0.64780195 | 0.808876289 | 359.427036 | 395.1148964 | ENSG00000183506 |
| BRSK1 | 0.136216163 | 1.099018874 | 0.215077045 | 0.4101504 | 602.6753691 | 662.7669119 | ENSG00000160469 |
| EDN1 | 0.136105657 | 1.098934696 | 0.86719691 | 0.936739326 | 6.694729326 | 7.342153824 | ENSG00000078401 |
| ALKBH4 | 0.136017667 | 1.098867674 | 0.277302859 | 0.486459867 | 369.0859411 | 405.7961423 | ENSG00000160993 |
| NPM3 | 0.135967167 | 1.09882921 | 0.044438234 | 0.128497329 | 2601.490845 | 2857.839024 | ENSG00000107833 |
| NA | 0.135955529 | 1.098820346 | 0.895679262 | 0.953435706 | 4.218907344 | 4.67026757 | ENSG00000261348 |
| PRKD2 | 0.135787562 | 1.098692422 | 0.084439036 | 0.210345758 | 1200.910664 | 1319.583959 | ENSG00000105287 |
| IL18BP | 0.135753026 | 1.098666121 | 0.542591183 | 0.731655444 | 115.6828195 | 127.2308482 | ENSG00000137496 |
| ZNF707 | 0.13573146 | 1.098649699 | 0.267273379 | 0.474965442 | 311.3898463 | 342.2259265 | ENSG00000181135 |
| CAMSAP3 | 0.135631524 | 1.098573597 | 0.07832202 | 0.19870414 | 1297.590728 | 1426.280495 | ENSG00000076826 |
| MCOLN1 | 0.135627393 | 1.098570452 | 0.246401148 | 0.451227611 | 346.8500446 | 380.8791546 | ENSG00000090674 |
| UBE2K | 0.135588523 | 1.098540853 | 0.063406974 | 0.169217444 | 3354.119028 | 3684.810153 | ENSG00000078140 |
| SEC14L1 | 0.135555206 | 1.098515484 | 0.035471621 | 0.10830117 | 1916.165065 | 2104.460475 | ENSG00000129657 |
| ZNF653 | 0.135520872 | 1.098489342 | 0.563824963 | 0.748513341 | 86.29788046 | 94.92711314 | ENSG00000161914 |
| DNAJC30 | 0.135520706 | 1.098489215 | 0.669813629 | 0.82299482 | 58.02198776 | 63.68319468 | ENSG00000176410 |
| NDUFS8 | 0.13547789 | 1.098456615 | 0.078234517 | 0.19858318 | 1470.647843 | 1616.156278 | ENSG00000110717 |
| NA | 0.135458277 | 1.098441682 | 0.869256287 | 0.93821255 | 6.37832022 | 6.975843164 | ENSG00000236439 |
| HAGH | 0.135438398 | 1.098426547 | 0.196131119 | 0.385184711 | 450.1415496 | 494.24132 | ENSG00000063854 |

| | | | | | | | |
|---|---|---|---|---|---|---|---|
| ANAPC11 | 0.135312337 | 1.098330572 | 0.027265416 | 0.088214388 | 2196.800168 | 2412.976416 | ENSG00000141552 |
| EIF2D | 0.135301787 | 1.09832254 | 0.12126418 | 0.274457579 | 814.4350298 | 894.0590299 | ENSG00000143486 |
| SLC1A1 | 0.135237545 | 1.098273633 | 0.280825834 | 0.489649698 | 319.0305751 | 350.3387031 | ENSG00000106688 |
| STX1A | 0.135083648 | 1.098156483 | 0.546209655 | 0.734280278 | 139.8684784 | 153.755357 | ENSG00000106089 |
| CREBRF | 0.135026915 | 1.0981133 | 0.723807665 | 0.854499273 | 32.60242222 | 35.71027982 | ENSG00000164463 |
| C8orf58 | 0.134979149 | 1.098076943 | 0.476218332 | 0.678694679 | 145.7136457 | 159.9736599 | ENSG00000241852 |
| ACTR10 | 0.134889539 | 1.09800874 | 0.20297897 | 0.394401261 | 542.25518 | 594.8872181 | ENSG00000131966 |
| NA | 0.134627931 | 1.097809654 | 0.898921658 | 0.95546587 | 4.497357461 | 4.977461848 | ENSG00000267317 |
| SERTAD3 | 0.134617064 | 1.097801385 | 0.249932357 | 0.455244267 | 373.0403058 | 409.1852537 | ENSG00000167565 |
| EEFSEC | 0.134321552 | 1.097576542 | 0.293718103 | 0.503482471 | 363.2138773 | 398.8561478 | ENSG00000132394 |
| CCND2-AS1 | 0.134277401 | 1.097542952 | 0.897189104 | 0.954467117 | 3.94282149 | 4.346899088 | ENSG00000255920 |
| EPHA1-AS1 | 0.134260168 | 1.097529842 | 0.649946836 | 0.810064757 | 57.98522398 | 63.56004389 | ENSG00000229153 |
| TMCC1 | 0.134088438 | 1.097399207 | 0.259471998 | 0.466412401 | 397.366258 | 435.9599956 | ENSG00000172765 |
| PDE9A | 0.134076322 | 1.09738999 | 0.077938308 | 0.19807902 | 954.8257813 | 1047.769678 | ENSG00000160191 |
| ANKS6 | 0.133964167 | 1.097304683 | 0.044112889 | 0.127876062 | 1881.674386 | 2064.827821 | ENSG00000165138 |
| TBL2 | 0.133939366 | 1.09728582 | 0.162693819 | 0.337677347 | 651.0125672 | 713.949397 | ENSG00000106638 |
| FAM189B | 0.133883606 | 1.09724341 | 0.09958625 | 0.237418485 | 1335.600965 | 1466.219019 | ENSG00000160767 |
| CEL | 0.13384077 | 1.097210832 | 0.686720568 | 0.832171942 | 39.39768035 | 43.25153743 | ENSG00000170835 |
| KCNH2 | 0.133800885 | 1.097180499 | 0.257861267 | 0.46471133 | 453.7540807 | 498.262108 | ENSG00000055118 |
| ACAP3 | 0.133773028 | 1.097159313 | 0.166799465 | 0.343325111 | 855.5827124 | 939.1771819 | ENSG00000131584 |
| SGMS1 | 0.133526334 | 1.09697172 | 0.266611777 | 0.474012748 | 391.0627385 | 429.2184198 | ENSG00000198964 |
| H2BC18 | 0.133502037 | 1.096953246 | 0.907011547 | 0.959924795 | 3.357445476 | 3.672164644 | ENSG00000203814 |
| NHS | 0.133493353 | 1.096946643 | 0.357874523 | 0.569904458 | 319.1411196 | 349.7770295 | ENSG00000188158 |
| OCIAD2 | 0.13343801 | 1.096904564 | 0.101075824 | 0.239801122 | 1615.310888 | 1771.717313 | ENSG00000145247 |
| MPL | 0.133400623 | 1.096876139 | 0.908714736 | 0.960165884 | 3.299324861 | 3.610872798 | ENSG00000117400 |
| PTX3 | 0.133293475 | 1.096794677 | 0.761516017 | 0.876689032 | 24.53290803 | 26.89533837 | ENSG00000163661 |
| HGS | 0.133232753 | 1.096748515 | 0.065044223 | 0.172449832 | 1726.971057 | 1894.823538 | ENSG00000185359 |
| KDM4B | 0.133216546 | 1.096736194 | 0.069826557 | 0.182414514 | 1093.452903 | 1199.247371 | ENSG00000127663 |
| TIGD5 | 0.133206309 | 1.096728412 | 0.384191858 | 0.596568265 | 216.7849874 | 237.9339692 | ENSG00000179886 |
| PON2 | 0.133205246 | 1.096727604 | 0.345394926 | 0.557359923 | 266.9669682 | 293.0842571 | ENSG00000105854 |
| NA | 0.133171411 | 1.096701884 | 0.895759925 | 0.953467827 | 4.242628443 | 4.68426631 | ENSG00000246982 |
| ZNF408 | 0.133168332 | 1.096699543 | 0.274448587 | 0.483268716 | 349.2941362 | 383.2748928 | ENSG00000175213 |
| LINC02361 | 0.133137664 | 1.09667623 | 0.715174301 | 0.849673165 | 35.75583486 | 39.22148043 | ENSG00000256576 |
| NA | 0.133048509 | 1.09660846 | 0.614338093 | 0.78538958 | 64.82080536 | 70.96734603 | ENSG00000186076 |
| TTC3 | 0.132976564 | 1.096553775 | 0.162536197 | 0.337566928 | 759.0469809 | 831.9226183 | ENSG00000182670 |
| PPP6R1 | 0.132917718 | 1.096509049 | 0.064903585 | 0.172270444 | 2701.599224 | 2963.087744 | ENSG00000105063 |
| FADS6 | 0.132862202 | 1.096466855 | 0.766238111 | 0.878994404 | 20.87685081 | 22.8708618 | ENSG00000172782 |
| PTCD1 | 0.132782498 | 1.096406281 | 0.731417403 | 0.858577685 | 29.11093815 | 31.94496938 | ENSG00000106246 |
| ADAMTS6 | 0.132654595 | 1.096309082 | 0.815563419 | 0.906261557 | 13.98551308 | 15.31704587 | ENSG00000049192 |
| PPP1R2 | 0.132617663 | 1.096281019 | 0.239227881 | 0.442099866 | 428.416279 | 469.7543508 | ENSG00000184203 |
| FOXN2 | 0.132611733 | 1.096276512 | 0.43968586 | 0.647645852 | 233.9247767 | 256.5502044 | ENSG00000170802 |
| NA | 0.132583994 | 1.096255434 | 0.554803557 | 0.741242673 | 90.21161578 | 98.7954281 | ENSG00000267886 |
| H4C3 | 0.132557851 | 1.096235569 | 0.761637736 | 0.876689032 | 22.0891213 | 24.25144959 | ENSG00000197061 |
| IMMP2L | 0.132531941 | 1.096215881 | 0.490199227 | 0.69013064 | 129.2089604 | 141.5209203 | ENSG00000184903 |
| MARS2 | 0.132527386 | 1.09621242 | 0.140557483 | 0.3047809 | 781.4192547 | 856.6006651 | ENSG00000247626 |
| FREM2 | 0.132503325 | 1.096194138 | 0.585596602 | 0.764254832 | 913.0442867 | 1000.643252 | ENSG00000150893 |
| GNA13 | 0.132421528 | 1.096131989 | 0.135265077 | 0.297205466 | 1323.615832 | 1451.027906 | ENSG00000120063 |
| NA | 0.13238807 | 1.096106568 | 0.909188903 | 0.960400069 | 3.313562751 | 3.622696074 | ENSG00000285668 |
| CARD19 | 0.132324939 | 1.096058605 | 0.256747559 | 0.463633357 | 391.1361881 | 428.6531436 | ENSG00000165233 |
| WDPCP | 0.132279791 | 1.096024304 | 0.378384961 | 0.590824274 | 212.738357 | 233.1283487 | ENSG00000143951 |
| DNAH7 | 0.132227254 | 1.095984393 | 0.912578928 | 0.961482481 | 3.368123893 | 3.680866498 | ENSG00000118997 |
| BRCA1 | 0.132200736 | 1.095964248 | 0.147888382 | 0.315735401 | 748.3973027 | 820.3178843 | ENSG00000012048 |
| SPRED1 | 0.132181332 | 1.095949507 | 0.174962079 | 0.355400544 | 654.4409222 | 717.2191188 | ENSG00000166068 |
| SCO1 | 0.132042976 | 1.09584441 | 0.043908445 | 0.127472864 | 1628.175995 | 1784.591282 | ENSG00000133028 |
| PAIP2 | 0.13203477 | 1.095838177 | 0.078858555 | 0.19955029 | 1897.14878 | 2078.669314 | ENSG00000120727 |
| VENTX | 0.131982437 | 1.095798426 | 0.40459845 | 0.615551577 | 192.278113 | 210.5051133 | ENSG00000151650 |
| ARHGAP32 | 0.131840111 | 1.095690328 | 0.156681367 | 0.328811581 | 621.292863 | 680.8131708 | ENSG00000134909 |
| PRSS23 | 0.131634324 | 1.095534049 | 0.445351456 | 0.65288482 | 150.4289422 | 164.8046079 | ENSG00000150687 |
| KEAP1 | 0.131536886 | 1.095460061 | 0.03974844 | 0.118175267 | 2165.582039 | 2372.747772 | ENSG00000079999 |
| CSTF3-DT | 0.131460654 | 1.095402178 | 0.907722935 | 0.960149634 | 3.606250758 | 3.953536906 | ENSG00000247151 |
| DOCK10 | 0.131307452 | 1.095285862 | 0.645805593 | 0.80736387 | 54.41990686 | 59.63713143 | ENSG00000135905 |
| GALK1 | 0.131301142 | 1.095281071 | 0.150469005 | 0.319640849 | 607.1703105 | 664.8767028 | ENSG00000108479 |
| TGFBR2 | 0.13128264 | 1.095267024 | 0.359053177 | 0.571152228 | 226.2900245 | 247.8440588 | ENSG00000163513 |
| LIMA1 | 0.131181819 | 1.095190486 | 0.091153062 | 0.22236087 | 1110.462239 | 1216.332219 | ENSG00000050405 |
| MAPK6-DT | 0.13103798 | 1.095081299 | 0.898846975 | 0.955461282 | 3.915540919 | 4.304902868 | ENSG00000259438 |
| COG7 | 0.130976761 | 1.095034832 | 0.432943097 | 0.642171723 | 161.8283158 | 177.3472575 | ENSG00000168434 |
| NA | 0.130955228 | 1.095018488 | 0.861033145 | 0.932648138 | 7.848879201 | 8.625979939 | ENSG00000213700 |
| MED14 | 0.130945665 | 1.09501123 | 0.068327943 | 0.179293206 | 1563.947712 | 1712.338742 | ENSG00000180182 |
| FKBP10 | 0.130855522 | 1.094942813 | 0.028721458 | 0.091668874 | 4574.918444 | 5009.60667 | ENSG00000141756 |
| RPUSD2 | 0.130852471 | 1.094940497 | 0.162234354 | 0.337107182 | 813.5519044 | 890.9941603 | ENSG00000166133 |
| RPL13A | 0.130698153 | 1.094823383 | 0.011843284 | 0.044993043 | 48505.56866 | 53104.30707 | ENSG00000142541 |
| TMEM132D | 0.130657298 | 1.09479238 | 0.211828151 | 0.40563523 | 460.12968 | 503.5236847 | ENSG00000151952 |
| PVR | 0.130654459 | 1.094790226 | 0.229751289 | 0.429589414 | 507.0200908 | 555.3655213 | ENSG00000073008 |
| FOXJ3 | 0.130534497 | 1.094699196 | 0.17866167 | 0.360896956 | 745.2403045 | 815.6959808 | ENSG00000198815 |

| | | | | | | | |
|---|---|---|---|---|---|---|---|
| FBXO21 | 0.130511122 | 1.094681459 | 0.073888478 | 0.190652263 | 1336.655801 | 1462.823949 | ENSG00000135108 |
| FHOD1 | 0.130444284 | 1.094630745 | 0.123529132 | 0.278249958 | 1379.237741 | 1510.253794 | ENSG00000135723 |
| NFIL3 | 0.130356075 | 1.09456382 | 0.442960806 | 0.650475342 | 224.3571965 | 245.6494545 | ENSG00000165030 |
| GSTK1 | 0.130298274 | 1.094519967 | 0.304682872 | 0.51480147 | 416.2272439 | 455.6953223 | ENSG00000197448 |
| PTGES2 | 0.130180447 | 1.09443058 | 0.074774563 | 0.192229462 | 2095.110493 | 2293.798086 | ENSG00000148334 |
| LOC150051 | 0.130176315 | 1.094427446 | 0.71777292 | 0.851146028 | 39.40568901 | 43.23943061 | ENSG00000237594 |
| EIF2B3 | 0.130112466 | 1.09437901 | 0.15553613 | 0.327348691 | 783.5448006 | 857.0783127 | ENSG00000070785 |
| PDCD6IP | 0.130095997 | 1.094366518 | 0.097401899 | 0.23357903 | 1440.049492 | 1575.702945 | ENSG00000170248 |
| ALDH1L2 | 0.130035444 | 1.094320586 | 0.509722782 | 0.706348308 | 145.5211554 | 159.0081889 | ENSG00000136010 |
| BCL9 | 0.129963663 | 1.09426614 | 0.09789575 | 0.234376426 | 1573.74076 | 1721.914334 | ENSG00000116128 |
| ZNF565 | 0.129853297 | 1.094182432 | 0.739387924 | 0.863587169 | 30.85046404 | 33.83353624 | ENSG00000196357 |
| RBM26 | 0.129729959 | 1.094088893 | 0.125800135 | 0.282120361 | 1230.506092 | 1346.115591 | ENSG00000139746 |
| H2BC6 | 0.129717875 | 1.094079729 | 0.776409366 | 0.885362652 | 21.86167286 | 23.91748734 | ENSG00000274290 |
| BOP1 | 0.129710012 | 1.094073766 | 0.021461283 | 0.072855712 | 4334.642091 | 4743.189406 | ENSG00000261236 |
| SMG5 | 0.129705743 | 1.094070528 | 0.008096832 | 0.032975774 | 4457.878714 | 4877.617069 | ENSG00000198952 |
| SALL2 | 0.129642862 | 1.094022843 | 0.008066474 | 0.032880502 | 8593.977996 | 9402.352667 | ENSG00000165821 |
| EFNA3 | 0.129633317 | 1.094015605 | 0.620564366 | 0.789501351 | 68.71854891 | 75.14391682 | ENSG00000143590 |
| MTCH2 | 0.129631354 | 1.094014117 | 0.0531798 | 0.147783608 | 1612.066027 | 1763.934755 | ENSG00000109919 |
| HIF1AN | 0.129590657 | 1.093983256 | 0.044457562 | 0.128533537 | 1666.838776 | 1823.182252 | ENSG00000166135 |
| NA | 0.129570685 | 1.093968111 | 0.862418756 | 0.933339725 | 9.341074348 | 10.16753176 | ENSG00000265511 |
| FLT1 | 0.12955887 | 1.093959152 | 0.044305612 | 0.128211997 | 2057.671954 | 2251.288687 | ENSG00000102755 |
| UBE3A | 0.129493868 | 1.093909864 | 0.104671008 | 0.246626573 | 1431.854505 | 1566.478635 | ENSG00000114062 |
| SPTY2D1 | 0.129488406 | 1.093905722 | 0.264516841 | 0.472110438 | 364.734191 | 398.8560525 | ENSG00000179119 |
| TMEM200B | 0.129477743 | 1.093897638 | 0.428853298 | 0.638613353 | 198.3133278 | 216.9603035 | ENSG00000253304 |
| EID2B | 0.129423097 | 1.093856204 | 0.835086841 | 0.917376243 | 10.94447671 | 11.96947922 | ENSG00000176401 |
| KIAA0513 | 0.129422938 | 1.093856083 | 0.306286139 | 0.516539895 | 321.2415221 | 351.1086861 | ENSG00000135709 |
| ACCS | 0.129315396 | 1.093774548 | 0.629909825 | 0.796188143 | 95.46309285 | 104.4010367 | ENSG00000110455 |
| TMEM132A | 0.129290389 | 1.093755589 | 0.035730863 | 0.10893097 | 2991.993685 | 3273.38855 | ENSG00000006118 |
| GPR1 | 0.129275558 | 1.093744345 | 0.839450108 | 0.919708492 | 11.57137119 | 12.67750803 | ENSG00000183671 |
| NA | 0.129171726 | 1.09366563 | 0.834667064 | 0.917141538 | 12.48619899 | 13.71108078 | ENSG00000237753 |
| TIMMDC1 | 0.129098496 | 1.093610118 | 0.052340757 | 0.146242816 | 1698.266224 | 1857.15071 | ENSG00000113845 |
| USP42 | 0.129090974 | 1.093604416 | 0.193603324 | 0.381569482 | 549.8063622 | 601.6568117 | ENSG00000106346 |
| MCPH1 | 0.129081944 | 1.093597571 | 0.334052898 | 0.546107514 | 302.5890706 | 330.7228292 | ENSG00000147316 |
| GOLGA1 | 0.129067267 | 1.093586446 | 0.540494269 | 0.729922359 | 113.4836421 | 124.0806614 | ENSG00000136935 |
| BCDIN3D-AS1 | 0.129059122 | 1.093580271 | 0.903276764 | 0.957851743 | 4.266349541 | 4.687387732 | ENSG00000258057 |
| SRPX | 0.128725118 | 1.093327122 | 0.498711402 | 0.697486842 | 137.3894025 | 150.1411089 | ENSG00000101955 |
| EEF2K | 0.128682889 | 1.09329512 | 0.175448921 | 0.355933754 | 605.7400487 | 662.0025913 | ENSG00000103319 |
| TRIP10 | 0.128676854 | 1.093290546 | 0.051653265 | 0.14472877 | 2043.412388 | 2234.379595 | ENSG00000125733 |
| GUK1 | 0.12862416 | 1.093250615 | 0.058243247 | 0.158764971 | 1488.67129 | 1627.45387 | ENSG00000143774 |
| TNRC6A | 0.12855949 | 1.09320161 | 0.051028745 | 0.143376491 | 2331.976845 | 2548.947582 | ENSG00000090905 |
| FAM91A1 | 0.128521631 | 1.093172923 | 0.211245213 | 0.404929541 | 484.4564703 | 529.294269 | ENSG00000176853 |
| SLC25A24 | 0.128500766 | 1.093157113 | 0.213241119 | 0.407597018 | 532.0989419 | 581.6655558 | ENSG00000085491 |
| BET1L | 0.12843437 | 1.093106805 | 0.211888499 | 0.405668523 | 459.6887155 | 502.3260265 | ENSG00000177951 |
| ANPEP | 0.128372619 | 1.093060018 | 0.606568579 | 0.780476092 | 71.84442781 | 78.61915314 | ENSG00000166825 |
| ISYNA1 | 0.12836877 | 1.093057102 | 0.037491733 | 0.112959698 | 3951.42887 | 4319.861659 | ENSG00000105655 |
| ASB6 | 0.128316655 | 1.093017618 | 0.265983189 | 0.473433051 | 436.6844086 | 477.238216 | ENSG00000148331 |
| AVL9 | 0.128244501 | 1.092962953 | 0.116616995 | 0.267596083 | 846.0232895 | 924.6156127 | ENSG00000105778 |
| VMP1 | 0.128244032 | 1.092962598 | 0.095140017 | 0.229700745 | 1152.185634 | 1258.836138 | ENSG00000062716 |
| MAP3K9 | 0.128212887 | 1.092939003 | 0.198471258 | 0.388608575 | 817.8137542 | 893.0544303 | ENSG00000006432 |
| DIPK1A | 0.128188462 | 1.0929205 | 0.540962954 | 0.730264327 | 116.5582138 | 127.5988399 | ENSG00000154511 |
| CORO1B | 0.128121564 | 1.092869822 | 0.032764406 | 0.101844109 | 2408.657792 | 2632.196735 | ENSG00000172725 |
| NA | 0.12806056 | 1.092823612 | 0.862879617 | 0.933677968 | 7.927161442 | 8.684150363 | ENSG00000251259 |
| LRCH3 | 0.128051314 | 1.092816608 | 0.147951784 | 0.315749738 | 857.1110354 | 936.3105069 | ENSG00000186001 |
| PHLDB3 | 0.127983862 | 1.092765515 | 0.473014072 | 0.676271538 | 167.6517946 | 183.3704942 | ENSG00000176531 |
| ND4 | 0.127970619 | 1.092755485 | 0.656020797 | 0.814152116 | 25737.57317 | 28125.02899 | ENSG00000198886 |
| PELI1 | 0.12791281 | 1.092711699 | 0.095049547 | 0.229570367 | 1210.031025 | 1322.374734 | ENSG00000197329 |
| FGF14-AS2 | 0.127861881 | 1.092673125 | 0.717253951 | 0.851051826 | 37.93779978 | 41.34753141 | ENSG00000272143 |
| IL17RC | 0.127817987 | 1.092639881 | 0.445200641 | 0.65276507 | 171.834523 | 187.6783076 | ENSG00000163702 |
| ACTN1 | 0.127790983 | 1.092619429 | 0.013907141 | 0.051187815 | 7902.674233 | 8635.148075 | ENSG00000072110 |
| MAP3K10 | 0.127729122 | 1.09257258 | 0.204440528 | 0.396206468 | 582.187539 | 636.3055439 | ENSG00000130758 |
| SYNE2 | 0.127641329 | 1.092506096 | 0.148035036 | 0.31582034 | 1377.582187 | 1504.526408 | ENSG00000054654 |
| ABHD10 | 0.127623392 | 1.092492513 | 0.232937172 | 0.433828254 | 466.0465425 | 509.0075172 | ENSG00000144827 |
| NA | 0.127621635 | 1.092491182 | 0.636768332 | 0.800725077 | 83.28737923 | 90.82744131 | ENSG00000273151 |
| RUFY3 | 0.127512819 | 1.092408783 | 0.205911032 | 0.398098878 | 520.6681696 | 569.0961086 | ENSG00000018189 |
| SRP54 | 0.127510795 | 1.09240725 | 0.090006309 | 0.220447418 | 1140.459276 | 1246.023402 | ENSG00000100883 |
| STUB1 | 0.127257349 | 1.092215359 | 0.150930372 | 0.320394202 | 1041.769879 | 1138.535544 | ENSG00000103266 |
| HSPA1A | 0.12720107 | 1.092172752 | 0.154802445 | 0.326313671 | 801.2151771 | 875.2788545 | ENSG00000204389 |
| TMEM150A | 0.127087243 | 1.092086585 | 0.263712679 | 0.471075856 | 377.0801483 | 411.9834121 | ENSG00000168890 |
| KIF26B | 0.127028018 | 1.092041754 | 0.788087203 | 0.891582187 | 24.40629387 | 26.59277856 | ENSG00000162849 |
| S1PR3 | 0.126910892 | 1.0919531 | 0.347777448 | 0.559768847 | 269.1581115 | 293.6646459 | ENSG00000213694 |
| ADAMTS13 | 0.12689005 | 1.091937324 | 0.573428345 | 0.755779923 | 143.9794587 | 157.4097391 | ENSG00000160323 |
| GSDMB | 0.126860746 | 1.091915145 | 0.566257798 | 0.750233591 | 127.6714341 | 139.5128186 | ENSG00000073605 |
| TMEM165 | 0.126848784 | 1.091906092 | 0.068322828 | 0.179293206 | 1512.189198 | 1651.238262 | ENSG00000134851 |
| SCAI | 0.126841396 | 1.0919005 | 0.392531726 | 0.604143117 | 259.4745165 | 283.0637203 | ENSG00000173611 |

| | | | | | | | |
|---|---|---|---|---|---|---|---|
| TCTEX1D1 | 0.126825341 | 1.091888349 | 0.785102714 | 0.890433415 | 20.962252 | 22.82981154 | ENSG00000152760 |
| NA | 0.126630614 | 1.091740982 | 0.919968989 | 0.964147051 | 3.357445476 | 3.651639513 | ENSG00000228835 |
| HAUS8 | 0.126625385 | 1.091737025 | 0.250788673 | 0.456317828 | 391.730768 | 427.6413981 | ENSG00000131351 |
| MIDN | 0.126589139 | 1.091709596 | 0.063647495 | 0.169603915 | 3487.859576 | 3808.769339 | ENSG00000167470 |
| DMTF1 | 0.126544947 | 1.091676157 | 0.121051468 | 0.274239079 | 1013.034082 | 1106.144118 | ENSG00000135164 |
| CEBPA-DT | 0.126506035 | 1.091646712 | 0.609332986 | 0.782366975 | 78.91307646 | 86.00301964 | ENSG00000267296 |
| PKHD1L1 | 0.126392838 | 1.091561063 | 0.899363916 | 0.95553286 | 4.246187915 | 4.658444293 | ENSG00000205038 |
| NA | 0.126298876 | 1.091489972 | 0.876360375 | 0.94241765 | 7.252824769 | 7.91047878 | ENSG00000197149 |
| CA5B | 0.126248637 | 1.091451964 | 0.560496328 | 0.745630681 | 97.72895827 | 106.5138239 | ENSG00000169239 |
| RAB40A | 0.126175143 | 1.091396364 | 0.749258814 | 0.869654826 | 26.75372166 | 29.22143909 | ENSG00000172476 |
| NA | 0.126174619 | 1.091395968 | 0.751650963 | 0.870694255 | 28.22784012 | 30.89304696 | ENSG00000273329 |
| NA | 0.126169565 | 1.091392145 | 0.900825408 | 0.956302469 | 6.160354981 | 6.682647625 | ENSG00000219797 |
| NA | 0.126083662 | 1.091327161 | 0.904110697 | 0.958242075 | 3.922659864 | 4.290904128 | ENSG00000279794 |
| ZNF70 | 0.125969112 | 1.091240513 | 0.679135586 | 0.828368191 | 53.61181093 | 58.53093801 | ENSG00000187792 |
| ZNF777 | 0.125954975 | 1.091229821 | 0.299737747 | 0.509848409 | 405.3990602 | 442.508876 | ENSG00000196453 |
| N4BP2 | 0.125915926 | 1.091200285 | 0.226770327 | 0.425583188 | 627.9881504 | 685.2131411 | ENSG00000078177 |
| SNHG9 | 0.125872571 | 1.091167493 | 0.821069052 | 0.909223 | 15.18946942 | 16.58034686 | ENSG00000255198 |
| NA | 0.125837448 | 1.091140928 | 0.626175703 | 0.793253855 | 88.60760303 | 96.40926503 | ENSG00000234648 |
| FAM81A | 0.125836284 | 1.091140048 | 0.763411726 | 0.877522527 | 26.14698881 | 28.52182859 | ENSG00000157470 |
| INVS | 0.1258339 | 1.091138245 | 0.506559748 | 0.703824371 | 166.6444211 | 181.720473 | ENSG00000119509 |
| NA | 0.125823642 | 1.091130487 | 0.797810427 | 0.896947216 | 18.33370434 | 19.97671967 | ENSG00000270871 |
| SORBS1 | 0.125795245 | 1.09110901 | 0.188185896 | 0.374093752 | 687.0017487 | 749.0241437 | ENSG00000095637 |
| PNPT1 | 0.125778353 | 1.091096235 | 0.113275415 | 0.261836745 | 998.2665065 | 1089.061906 | ENSG00000138035 |
| GCNT2 | 0.125726308 | 1.091056874 | 0.095604353 | 0.230380019 | 987.154813 | 1077.17289 | ENSG00000111846 |
| MRPL32 | 0.125673623 | 1.091017032 | 0.059538965 | 0.161388453 | 1606.111076 | 1752.169633 | ENSG00000106591 |
| NA | 0.125631166 | 1.090984925 | 0.906824655 | 0.95978163 | 3.629971857 | 3.955712369 | ENSG00000260711 |
| ZNF665 | 0.125553241 | 1.090925998 | 0.636867702 | 0.800725077 | 60.20217265 | 65.67086663 | ENSG00000197497 |
| SDHAF2 | 0.125438413 | 1.090839172 | 0.679166229 | 0.828368191 | 53.20735704 | 58.19385433 | ENSG00000167985 |
| GXYLT2 | 0.125418004 | 1.09082374 | 0.664866646 | 0.820559864 | 56.02597898 | 61.08648342 | ENSG00000172986 |
| TMEM74 | 0.125411925 | 1.090819144 | 0.457832124 | 0.663253386 | 160.170911 | 174.6140794 | ENSG00000164841 |
| IRF7 | 0.125384609 | 1.09079849 | 0.442655104 | 0.650193796 | 182.8747738 | 199.5319404 | ENSG00000185507 |
| AMMECR1 | 0.125382498 | 1.090796894 | 0.169882427 | 0.347981017 | 1064.853891 | 1161.597055 | ENSG00000101935 |
| PLAG1 | 0.125368843 | 1.09078657 | 0.456578391 | 0.662631902 | 196.2799449 | 213.8746752 | ENSG00000181690 |
| NA | 0.125293123 | 1.090729322 | 0.732937732 | 0.859722303 | 35.11353344 | 38.3310165 | ENSG00000247679 |
| NA | 0.125277947 | 1.090717848 | 0.881644056 | 0.945637985 | 7.392787098 | 8.051412139 | ENSG00000189423 |
| AAR2 | 0.125275665 | 1.090716123 | 0.169447688 | 0.347337414 | 824.2521308 | 898.571201 | ENSG00000131043 |
| SBF2-AS1 | 0.125187702 | 1.090649623 | 0.836844188 | 0.918172233 | 11.23360525 | 12.24867601 | ENSG00000246273 |
| GALNT1 | 0.125119417 | 1.090598002 | 0.101825886 | 0.241186737 | 1252.114457 | 1365.898075 | ENSG00000141429 |
| NA | 0.125107148 | 1.090588727 | 0.773218236 | 0.883377277 | 24.77014517 | 26.93638863 | ENSG00000272077 |
| FGFRL1 | 0.125083687 | 1.090570992 | 0.151513906 | 0.32116076 | 726.4946701 | 792.1794663 | ENSG00000127418 |
| INAFM2 | 0.124962471 | 1.090479366 | 0.163057951 | 0.338120875 | 822.4153551 | 897.1699858 | ENSG00000259330 |
| GPSM2 | 0.124907578 | 1.090437875 | 0.383085175 | 0.595456969 | 225.6266456 | 245.9443311 | ENSG00000121957 |
| LNX1 | 0.124805428 | 1.090360669 | 0.600565692 | 0.776296479 | 82.58042249 | 90.0207589 | ENSG00000072201 |
| CDC42EP1 | 0.124761054 | 1.090327133 | 0.316277414 | 0.526264103 | 393.6693183 | 429.6338202 | ENSG00000128283 |
| LOC100133091 | 0.12474275 | 1.090313299 | 0.693749577 | 0.836512627 | 49.682948 | 54.31002758 | ENSG00000205485 |
| CHEK2 | 0.124732213 | 1.090305336 | 0.089415853 | 0.219415071 | 1374.427606 | 1498.144096 | ENSG00000183765 |
| FLVCR1 | 0.124528882 | 1.090151681 | 0.078297479 | 0.198689586 | 1480.671289 | 1614.539999 | ENSG00000162769 |
| FKBP5 | 0.124511712 | 1.090138707 | 0.062491752 | 0.167398118 | 1998.713653 | 2178.305649 | ENSG00000096060 |
| SH3YL1 | 0.124495038 | 1.090126108 | 0.266587374 | 0.474012748 | 441.8338823 | 481.5313909 | ENSG00000035115 |
| SLK | 0.124462095 | 1.090101216 | 0.159589487 | 0.33320591 | 898.1343123 | 978.6434243 | ENSG00000065613 |
| MAPK10 | 0.124459761 | 1.090099452 | 0.729284172 | 0.85751636 | 41.86073898 | 45.49922615 | ENSG00000109339 |
| NA | 0.124395263 | 1.090050719 | 0.853418233 | 0.92812329 | 10.39705969 | 11.29039384 | ENSG00000224790 |
| PKIB | 0.124390027 | 1.090046762 | 0.312792624 | 0.523140408 | 379.2629506 | 413.5868229 | ENSG00000135549 |
| OSBPL10 | 0.124285207 | 1.089967567 | 0.076008752 | 0.194660331 | 1430.679559 | 1558.788135 | ENSG00000144645 |
| CCDC43 | 0.124241589 | 1.089934614 | 0.334342096 | 0.546292653 | 282.4876956 | 307.9037794 | ENSG00000180329 |
| MED11 | 0.124106262 | 1.089832381 | 0.612542068 | 0.784474227 | 75.74453563 | 82.68032904 | ENSG00000161920 |
| NFU1 | 0.12405585 | 1.0897943 | 0.275712393 | 0.484659067 | 412.7232497 | 449.6291957 | ENSG00000169599 |
| ACTA2 | 0.123993406 | 1.089747131 | 0.874378985 | 0.941213357 | 7.045537948 | 7.648402144 | ENSG00000107796 |
| NA | 0.123932862 | 1.0897014 | 0.673637245 | 0.825223104 | 60.15442497 | 65.6503415 | ENSG00000268205 |
| SNX16 | 0.123928287 | 1.089697945 | 0.553260908 | 0.739863889 | 124.9102963 | 136.0688464 | ENSG00000104497 |
| PRMT5 | 0.123845012 | 1.089635047 | 0.124072218 | 0.279240118 | 1395.924685 | 1520.995735 | ENSG00000100462 |
| C6orf89 | 0.123803263 | 1.089603515 | 0.289271358 | 0.498666326 | 388.3484068 | 422.7986268 | ENSG00000198663 |
| GAL3ST4 | 0.123789879 | 1.089593407 | 0.572463855 | 0.754956697 | 94.19923588 | 102.7224079 | ENSG00000197093 |
| MARK1 | 0.123716942 | 1.089538323 | 0.190308563 | 0.376687685 | 553.424461 | 602.8922831 | ENSG00000116141 |
| AOC3 | 0.123645163 | 1.089484116 | 0.75952128 | 0.875194299 | 34.56403148 | 37.53341482 | ENSG00000131471 |
| LRRC56 | 0.123442877 | 1.089331366 | 0.683591091 | 0.830895404 | 53.14804122 | 58.02246448 | ENSG00000161328 |
| CISD1 | 0.123369059 | 1.08927563 | 0.302338802 | 0.512445547 | 392.7094132 | 427.9026444 | ENSG00000122873 |
| TDP1 | 0.123363858 | 1.089271703 | 0.103654864 | 0.244629107 | 1136.223819 | 1237.839402 | ENSG00000042088 |
| ANK1 | 0.123342141 | 1.089255306 | 0.36661666 | 0.579183171 | 277.7854418 | 302.4872706 | ENSG00000029534 |
| KIAA1755 | 0.123314015 | 1.089234071 | 0.904203007 | 0.958242075 | 5.509460075 | 6.047733938 | ENSG00000149633 |
| LINC01097 | 0.123304711 | 1.089227046 | 0.807590076 | 0.902019079 | 18.37166333 | 19.97113924 | ENSG00000281202 |
| CCDC115 | 0.1232359 | 1.089175095 | 0.33293879 | 0.54498995 | 350.6549631 | 381.6735622 | ENSG00000136710 |
| REX1BD | 0.12308685 | 1.089062575 | 0.172898089 | 0.352459025 | 847.7319753 | 923.4615294 | ENSG00000006015 |
| EMD | 0.123061737 | 1.089043617 | 0.073433953 | 0.189764668 | 1237.179133 | 1347.440839 | ENSG00000102119 |

| | | | | | | | |
|---|---|---|---|---|---|---|---|
| RABEPK | 0.123030919 | 1.089020354 | 0.107478383 | 0.25192122 | 973.05743 | 1059.595765 | ENSG00000136933 |
| CNOT4 | 0.122981335 | 1.088982926 | 0.339900198 | 0.55150917 | 335.4444412 | 365.5418671 | ENSG00000080802 |
| ZCCHC9 | 0.122971126 | 1.088975221 | 0.202382381 | 0.393732085 | 568.9445513 | 619.3927775 | ENSG00000131732 |
| TMEM14A | 0.122965535 | 1.088971 | 0.477684503 | 0.680219562 | 150.9959105 | 164.2850891 | ENSG00000096092 |
| SSTR3 | 0.122922153 | 1.088938255 | 0.835229247 | 0.917410484 | 15.04030321 | 16.35714502 | ENSG00000278195 |
| GDI1 | 0.122878415 | 1.088905243 | 0.049870515 | 0.140799025 | 2700.531841 | 2940.533186 | ENSG00000203879 |
| NRG1 | 0.122842257 | 1.088877952 | 0.697709852 | 0.838858153 | 47.12083511 | 51.34757275 | ENSG00000157168 |
| NA | 0.122743984 | 1.088803783 | 0.906644161 | 0.959649147 | 4.30430853 | 4.700440514 | ENSG00000236519 |
| PDK3 | 0.122743186 | 1.088803181 | 0.148258693 | 0.316118937 | 1094.676716 | 1191.496729 | ENSG00000067992 |
| CTAGE7P | 0.122701668 | 1.088771847 | 0.869540471 | 0.938304771 | 7.940204123 | 8.651801956 | ENSG00000233122 |
| TSHZ3 | 0.122671632 | 1.08874918 | 0.435220097 | 0.643868786 | 200.7574718 | 218.8400732 | ENSG00000121297 |
| PHF21A | 0.122626342 | 1.088715001 | 0.200458554 | 0.391042454 | 601.6834542 | 655.3572531 | ENSG00000135365 |
| NA | 0.122580124 | 1.088680124 | 0.822022257 | 0.90966346 | 17.18074967 | 18.62289986 | ENSG00000268030 |
| EXOSC2 | 0.122568524 | 1.08867137 | 0.119847914 | 0.272460316 | 1149.064945 | 1250.626775 | ENSG00000130713 |
| VCAN | 0.122557205 | 1.088662829 | 0.067588707 | 0.177872454 | 5217.661784 | 5679.442672 | ENSG00000038427 |
| ZNF227 | 0.122489852 | 1.088612006 | 0.414787088 | 0.625609309 | 228.5603392 | 248.9460169 | ENSG00000131115 |
| COX10-AS1 | 0.122438486 | 1.088573247 | 0.630360451 | 0.796491175 | 82.00244475 | 89.26453399 | ENSG00000236088 |
| DUSP14 | 0.12242987 | 1.088566746 | 0.264214077 | 0.471721696 | 432.8175313 | 471.0882555 | ENSG00000276023 |
| WHRN | 0.122383121 | 1.088531473 | 0.412689348 | 0.624129682 | 275.1361477 | 299.6876421 | ENSG00000095397 |
| SETX | 0.122320681 | 1.088484362 | 0.163597604 | 0.338773684 | 1676.203774 | 1823.616229 | ENSG00000107290 |
| LINC01006 | 0.122275708 | 1.088450431 | 0.541422161 | 0.730652959 | 111.6714783 | 121.5050423 | ENSG00000182648 |
| ZBTB38 | 0.122271535 | 1.088447283 | 0.436759808 | 0.645089622 | 184.3764521 | 200.611933 | ENSG00000177311 |
| SEC31A | 0.122250221 | 1.088431203 | 0.048995861 | 0.138848565 | 1908.631224 | 2077.489372 | ENSG00000138674 |
| MAPK15 | 0.122186759 | 1.088383326 | 0.475477684 | 0.67798911 | 196.6954344 | 214.2428481 | ENSG00000181085 |
| CES3 | 0.12207601 | 1.088299779 | 0.723684042 | 0.854472967 | 45.87652936 | 50.01411011 | ENSG00000172828 |
| MRGBP | 0.122030244 | 1.088265256 | 0.189862793 | 0.376278606 | 750.2185657 | 816.5403515 | ENSG00000101189 |
| RPL31 | 0.122018664 | 1.08825652 | 0.046627707 | 0.133597979 | 11130.81943 | 12112.95934 | ENSG00000071082 |
| CBL | 0.121948574 | 1.088203651 | 0.117103498 | 0.268291789 | 1745.488784 | 1898.590407 | ENSG00000110395 |
| LCOR | 0.121819724 | 1.088106466 | 0.373120955 | 0.585102486 | 313.6693392 | 341.0550659 | ENSG00000196233 |
| TMEM216 | 0.121787117 | 1.088081874 | 0.314322936 | 0.524844267 | 338.8366178 | 368.8275748 | ENSG00000187049 |
| BIRC2 | 0.121710317 | 1.088023952 | 0.200652786 | 0.391338207 | 585.5633405 | 637.0979214 | ENSG00000110330 |
| PRUNE2 | 0.121640712 | 1.08797146 | 0.289858183 | 0.499192516 | 435.084618 | 473.1011394 | ENSG00000106772 |
| LINC00937 | 0.121630867 | 1.087964035 | 0.766278177 | 0.878994404 | 28.18423672 | 30.68601932 | ENSG00000226091 |
| SLC38A4 | 0.121548838 | 1.087902178 | 0.329715916 | 0.541217709 | 330.1060634 | 358.7923472 | ENSG00000139209 |
| RNF113A | 0.121489203 | 1.08785721 | 0.433128558 | 0.64234591 | 220.9908004 | 240.3206267 | ENSG00000125352 |
| MECP2 | 0.121471423 | 1.087843803 | 0.196030233 | 0.385026619 | 774.6885996 | 842.4436095 | ENSG00000169057 |
| HTR3A | 0.121404313 | 1.0877932 | 0.414752906 | 0.625609309 | 205.384698 | 223.4876401 | ENSG00000166736 |
| ITGB4 | 0.121236598 | 1.08766675 | 0.536152121 | 0.726708735 | 119.6793124 | 130.0977279 | ENSG00000132470 |
| MAP4K3 | 0.12119821 | 1.08763781 | 0.113254523 | 0.2618205 | 1149.661557 | 1250.292361 | ENSG00000011566 |
| RTL8B | 0.121036702 | 1.087516057 | 0.57613272 | 0.758153886 | 105.7904632 | 115.1834085 | ENSG00000212747 |
| CEP295 | 0.121028569 | 1.087509926 | 0.445504909 | 0.653059086 | 181.0602458 | 196.9397683 | ENSG00000166004 |
| SMIM13 | 0.121020313 | 1.087503703 | 0.383492909 | 0.595734937 | 258.5042895 | 281.2593568 | ENSG00000224531 |
| NA | 0.121016192 | 1.087500596 | 0.711295836 | 0.847591497 | 40.257947 | 43.82392977 | ENSG00000224592 |
| BCL7B | 0.121013855 | 1.087498835 | 0.221523077 | 0.418786007 | 606.2249737 | 659.3382728 | ENSG00000106635 |
| CD151 | 0.120677125 | 1.087245038 | 0.103973462 | 0.245205981 | 1134.830853 | 1233.955807 | ENSG00000177697 |
| TANGO2 | 0.120676287 | 1.087244407 | 0.522949385 | 0.717284954 | 135.1086882 | 147.0876111 | ENSG00000183597 |
| UBAP2 | 0.12060813 | 1.087193044 | 0.118518673 | 0.270382355 | 1514.790876 | 1646.781421 | ENSG00000137073 |
| VSIG10L | 0.120551127 | 1.087150088 | 0.319139617 | 0.529256214 | 346.5908402 | 376.6239312 | ENSG00000186806 |
| CALML6 | 0.120541036 | 1.087142483 | 0.908497671 | 0.960165884 | 5.202534178 | 5.690787544 | ENSG00000169885 |
| ATM | 0.120411755 | 1.087045069 | 0.206929073 | 0.399331244 | 639.9873655 | 695.4796757 | ENSG00000149311 |
| TFCP2L1 | 0.120279082 | 1.086945106 | 0.553983228 | 0.740460985 | 124.0289521 | 134.5959432 | ENSG00000115112 |
| LDHB | 0.120151498 | 1.086848987 | 0.632314862 | 0.797472043 | 68.71826958 | 74.74525775 | ENSG00000111716 |
| GOLGA3 | 0.120140006 | 1.08684033 | 0.198102062 | 0.388127134 | 859.7196479 | 934.1184663 | ENSG00000090615 |
| GEMIN6 | 0.120131697 | 1.08683407 | 0.357968958 | 0.569904458 | 340.9799355 | 370.244694 | ENSG00000152147 |
| FAM166B | 0.120113299 | 1.08682021 | 0.87181813 | 0.940010585 | 7.64159238 | 8.292963644 | ENSG00000215187 |
| CYP2C8 | 0.12009188 | 1.086804075 | 0.87979229 | 0.944439862 | 7.39634657 | 8.013766846 | ENSG00000138115 |
| COX1 | 0.120042696 | 1.086767024 | 0.563280237 | 0.748016825 | 93505.02081 | 101618.3602 | ENSG00000198804 |
| SNHG15 | 0.119982171 | 1.086721433 | 0.229308066 | 0.429100557 | 548.5769315 | 596.3187149 | ENSG00000232956 |
| TBC1D3 | 0.119863692 | 1.086632191 | 0.769526786 | 0.88150633 | 36.18851135 | 39.32883588 | ENSG00000274512 |
| NA | 0.119860615 | 1.086629873 | 0.907061582 | 0.959924795 | 4.293630113 | 4.676793961 | ENSG00000272744 |
| IL17C | 0.119793718 | 1.086579488 | 0.796803824 | 0.896226338 | 20.50232109 | 22.30036139 | ENSG00000124391 |
| SNX2 | 0.119787291 | 1.086574648 | 0.09533478 | 0.229965171 | 1423.075872 | 1546.184856 | ENSG00000205302 |
| KDM2A | 0.119472192 | 1.086337354 | 0.083239085 | 0.207995508 | 1637.053631 | 1778.702538 | ENSG00000173120 |
| RIC1 | 0.119469556 | 1.08633537 | 0.220685633 | 0.417641865 | 529.2554043 | 574.8212091 | ENSG00000107036 |
| TNS2 | 0.119401305 | 1.086283979 | 0.557497238 | 0.743526508 | 124.5351561 | 135.0428413 | ENSG00000111077 |
| ANP32B | 0.119298102 | 1.086206274 | 0.030989833 | 0.097426835 | 2581.324158 | 2803.809357 | ENSG00000136938 |
| ARMCX2 | 0.119285441 | 1.086196742 | 0.056810078 | 0.155780212 | 2785.736623 | 3025.359212 | ENSG00000184867 |
| DAB2 | 0.119215887 | 1.086144376 | 0.586068451 | 0.764723322 | 114.0812228 | 124.1118529 | ENSG00000153071 |
| LOC101929595 | 0.119178939 | 1.08611656 | 0.910926634 | 0.961009829 | 3.984339952 | 4.326373957 | ENSG00000245293 |
| AACSP1 | 0.119115809 | 1.086069035 | 0.908217096 | 0.960165884 | 5.536740646 | 6.053030824 | ENSG00000250420 |
| LOC100506282 | 0.1190502 | 1.086019645 | 0.802696422 | 0.899212358 | 22.04996711 | 24.01207355 | ENSG00000247137 |
| MAN1A1 | 0.11902599 | 1.086001421 | 0.50401495 | 0.701682443 | 148.671619 | 161.3512942 | ENSG00000111885 |
| STYK1 | 0.118987553 | 1.085972487 | 0.781953529 | 0.88893053 | 25.00087376 | 27.19288483 | ENSG00000060140 |
| NA | 0.118890773 | 1.085899639 | 0.581466349 | 0.761564047 | 110.3044228 | 119.9821237 | ENSG00000224861 |

| | | | | | | | |
|---|---|---|---|---|---|---|---|
| NAGS | 0.118779138 | 1.085815616 | 0.453494784 | 0.6596002 | 174.8470835 | 189.7933133 | ENSG00000161653 |
| FAM126A | 0.118745281 | 1.085790134 | 0.092457998 | 0.224806141 | 1908.162544 | 2071.130647 | ENSG00000122591 |
| MMAA | 0.118652435 | 1.08572026 | 0.606828738 | 0.780598101 | 92.29274642 | 100.0686604 | ENSG00000151611 |
| ABCD1 | 0.118627815 | 1.085701732 | 0.366837015 | 0.57928899 | 311.0494629 | 337.9045658 | ENSG00000101986 |
| MAGED2 | 0.118557907 | 1.085649124 | 0.036393444 | 0.110500747 | 3175.75021 | 3447.053573 | ENSG00000102316 |
| NA | 0.118530359 | 1.085628394 | 0.783831655 | 0.890064386 | 24.08129128 | 26.14871831 | ENSG00000243547 |
| MRPS33 | 0.11845089 | 1.085568595 | 0.23768143 | 0.439841186 | 544.4044987 | 590.5555473 | ENSG00000090263 |
| LINC00476 | 0.118444407 | 1.085563717 | 0.754882371 | 0.87245738 | 31.43375512 | 34.10090976 | ENSG00000175611 |
| C17orf80 | 0.118368901 | 1.085506903 | 0.252270901 | 0.458134047 | 534.8623923 | 580.478439 | ENSG00000141219 |
| MRPS12 | 0.11834352 | 1.085487806 | 0.11843123 | 0.270315491 | 1156.355728 | 1255.578837 | ENSG00000128626 |
| NR2C2AP | 0.118217911 | 1.085393302 | 0.321000025 | 0.531456561 | 357.1756355 | 387.7458886 | ENSG00000184162 |
| ALDOA | 0.118170053 | 1.085357298 | 0.703884679 | 0.842524014 | 48.43391373 | 52.58505172 | ENSG00000149925 |
| GRIA4 | 0.118115079 | 1.085315941 | 0.432116804 | 0.641566127 | 199.4938685 | 216.4268586 | ENSG00000152578 |
| TUBG2 | 0.118060663 | 1.085275005 | 0.501191124 | 0.699401047 | 315.5586684 | 342.565555 | ENSG00000037042 |
| PRDM5 | 0.118056327 | 1.085271743 | 0.376157404 | 0.588252118 | 310.1316081 | 336.6624727 | ENSG00000138738 |
| ZNF681 | 0.118033121 | 1.085254287 | 0.585068114 | 0.763791741 | 121.0164432 | 131.4718619 | ENSG00000196172 |
| BBIP1 | 0.118004966 | 1.085233108 | 0.536619916 | 0.726856257 | 128.9907158 | 139.9631943 | ENSG00000214413 |
| PIP5K1C | 0.117966115 | 1.085203883 | 0.111530479 | 0.259135468 | 1359.952646 | 1476.300281 | ENSG00000186111 |
| SBF2 | 0.117955796 | 1.085196121 | 0.220377559 | 0.417184263 | 760.1886206 | 824.4056697 | ENSG00000133812 |
| EEF2 | 0.117929176 | 1.085176098 | 0.008381117 | 0.033935976 | 82813.33708 | 89866.46138 | ENSG00000167658 |
| SULT1C4 | 0.117922083 | 1.085170763 | 0.809164886 | 0.902949338 | 21.07582348 | 22.90103475 | ENSG00000198075 |
| NA | 0.117828898 | 1.085100673 | 0.821394648 | 0.909334444 | 20.94181104 | 22.75452096 | ENSG00000280433 |
| NUP214 | 0.117749951 | 1.085041296 | 0.146561282 | 0.313833491 | 1053.954999 | 1143.05176 | ENSG00000126883 |
| GALNT17 | 0.117704091 | 1.085006805 | 0.0475481 | 0.135702866 | 1955.286658 | 2121.622052 | ENSG00000185274 |
| ACAP2 | 0.11768695 | 1.084993914 | 0.248073217 | 0.453053109 | 581.1073001 | 630.7059673 | ENSG00000114331 |
| SBNO1 | 0.117678175 | 1.084987314 | 0.179773083 | 0.362560302 | 1150.642489 | 1248.037513 | ENSG00000139697 |
| SSR1 | 0.117654035 | 1.08496916 | 0.10561156 | 0.248563598 | 1632.24482 | 1770.950959 | ENSG00000124783 |
| RAD23B | 0.117482766 | 1.084840366 | 0.032035466 | 0.100129075 | 6145.772921 | 6667.832176 | ENSG00000119318 |
| NA | 0.117429701 | 1.084800464 | 0.904440033 | 0.958385705 | 4.879006127 | 5.323530924 | ENSG00000277406 |
| SLC13A4 | 0.117417862 | 1.084791562 | 0.68217943 | 0.830141023 | 55.99986746 | 60.71733488 | ENSG00000164707 |
| HSPA12B | 0.117399208 | 1.084777536 | 0.488114576 | 0.688710031 | 243.9499509 | 264.608703 | ENSG00000132622 |
| LINC00599 | 0.117335883 | 1.084729922 | 0.817725273 | 0.907141028 | 553.6711819 | 600.6046026 | ENSG00000253230 |
| P3H4 | 0.117299894 | 1.084702863 | 0.185353582 | 0.369786858 | 932.8340121 | 1012.131481 | ENSG00000141696 |
| EPHX3 | 0.11727092 | 1.084681079 | 0.249470924 | 0.454618998 | 480.6160093 | 521.3652726 | ENSG00000105131 |
| RGMA | 0.117213143 | 1.084637641 | 0.165416105 | 0.341204651 | 909.7493904 | 987.2431964 | ENSG00000182175 |
| COL17A1 | 0.117213087 | 1.084637598 | 0.685114722 | 0.831623057 | 53.27287593 | 57.79585768 | ENSG00000065618 |
| NDRG4 | 0.117051619 | 1.084516212 | 0.236977189 | 0.438971478 | 575.280184 | 624.0208513 | ENSG00000103034 |
| NADK2 | 0.11681065 | 1.084335083 | 0.38119056 | 0.593714122 | 349.3291723 | 378.8975669 | ENSG00000152620 |
| LSM14B | 0.116684378 | 1.084240181 | 0.085379026 | 0.212155644 | 1974.761803 | 2140.91372 | ENSG00000149657 |
| INO80 | 0.116602822 | 1.08417889 | 0.175450745 | 0.355933754 | 866.4713549 | 939.1550648 | ENSG00000128908 |
| HSPBAP1 | 0.11660064 | 1.084177251 | 0.488154318 | 0.688710031 | 165.1940493 | 179.1682896 | ENSG00000169087 |
| PRPSAP2 | 0.116565344 | 1.084150726 | 0.140032758 | 0.304107479 | 858.4715556 | 930.7208546 | ENSG00000141127 |
| NA | 0.11652667 | 1.084121664 | 0.71937662 | 0.852033813 | 43.43302197 | 47.03207611 | ENSG00000261377 |
| GALNT13 | 0.116511066 | 1.084109938 | 0.420749521 | 0.630884946 | 215.4748317 | 233.5528298 | ENSG00000144278 |
| TAPBP | 0.116498379 | 1.084100405 | 0.236920349 | 0.438909239 | 732.9324101 | 794.932761 | ENSG00000231925 |
| LOC103344931 | 0.116407644 | 1.084032225 | 0.724327172 | 0.854693478 | 40.5117863 | 43.891748 | ENSG00000260917 |
| PDE6D | 0.116335089 | 1.083977709 | 0.268406031 | 0.476326564 | 504.0946336 | 546.3029964 | ENSG00000156973 |
| NA | 0.116253353 | 1.083916298 | 0.906524564 | 0.959625358 | 4.86832771 | 5.305181257 | ENSG00000263235 |
| ANXA4 | 0.116150368 | 1.083838926 | 0.43403756 | 0.642785392 | 216.9812386 | 235.0188075 | ENSG00000196975 |
| C8orf37 | 0.116096718 | 1.083798622 | 0.707632067 | 0.845853003 | 46.52478068 | 50.47763395 | ENSG00000156172 |
| NOTCH2NLC | 0.116068169 | 1.083777176 | 0.913092531 | 0.961597187 | 4.259230596 | 4.622974464 | ENSG00000286219 |
| NA | 0.115842573 | 1.083607717 | 0.869750725 | 0.938478027 | 8.89062665 | 9.658606736 | ENSG00000226453 |
| CACNA1C-AS2 | 0.1158263 | 1.083595495 | 0.874119396 | 0.941041204 | 8.263732174 | 8.955874812 | ENSG00000256271 |
| NUP42 | 0.11580949 | 1.083582869 | 0.361993822 | 0.573896351 | 319.868569 | 346.6662978 | ENSG00000136243 |
| SHISAL2B | 0.115719798 | 1.083515505 | 0.536082742 | 0.726666852 | 140.2964002 | 152.0196718 | ENSG00000145642 |
| ASAH1 | 0.115667313 | 1.083476088 | 0.191569099 | 0.378547916 | 961.5773582 | 1041.25784 | ENSG00000104763 |
| THTPA | 0.115580829 | 1.083411139 | 0.500046919 | 0.698373943 | 163.1909216 | 177.0279564 | ENSG00000259431 |
| GPR26 | 0.115569397 | 1.083402554 | 0.828924658 | 0.914028945 | 16.4396989 | 17.81593391 | ENSG00000154478 |
| DCAF15 | 0.115563611 | 1.083398209 | 0.049851413 | 0.140771904 | 1972.954646 | 2137.509978 | ENSG00000132017 |
| MTCL1 | 0.115502298 | 1.083352167 | 0.159185562 | 0.332583178 | 1355.484725 | 1468.119652 | ENSG00000168502 |
| LOC105375421 | 0.115501725 | 1.083351736 | 0.913187077 | 0.961597187 | 3.973661534 | 4.299605982 | ENSG00000284523 |
| SLC2A4RG | 0.115421344 | 1.083291378 | 0.303933412 | 0.513995029 | 453.9349772 | 492.0728213 | ENSG00000125520 |
| C11orf95 | 0.115420415 | 1.08329068 | 0.181331112 | 0.364418224 | 1183.596331 | 1282.151618 | ENSG00000188070 |
| HIVEP2 | 0.115405119 | 1.083279195 | 0.504097214 | 0.701693513 | 192.9304825 | 208.6138765 | ENSG00000010818 |
| ZC3H10 | 0.11539619 | 1.083272491 | 0.588487284 | 0.766506819 | 101.9985879 | 110.4471193 | ENSG00000135482 |
| SLC30A7 | 0.115275355 | 1.083181764 | 0.494394603 | 0.693967748 | 213.8829975 | 231.5699173 | ENSG00000162695 |
| NA | 0.115257456 | 1.083168325 | 0.887833762 | 0.94955683 | 8.063564298 | 8.721795656 | ENSG00000241741 |
| RPS18 | 0.115226487 | 1.083145074 | 0.071841949 | 0.186595408 | 23872.52911 | 25856.40869 | ENSG00000231500 |
| MTMR1 | 0.115189276 | 1.083117137 | 0.273599444 | 0.48225924 | 544.1560768 | 589.221402 | ENSG00000063601 |
| SDC3 | 0.115149029 | 1.083086922 | 0.070860409 | 0.184630081 | 1748.923342 | 1893.837318 | ENSG00000162512 |
| TUBGCP4 | 0.114961874 | 1.082946427 | 0.351076858 | 0.563149676 | 422.4006939 | 457.3187208 | ENSG00000137822 |
| MTERF2 | 0.114933096 | 1.082924824 | 0.685081767 | 0.831623057 | 56.12650778 | 60.85826824 | ENSG00000120832 |
| GATA2 | 0.114917153 | 1.082912857 | 0.879826873 | 0.944439862 | 8.639457104 | 9.363235734 | ENSG00000179348 |
| TACO1 | 0.114896546 | 1.08289739 | 0.189659334 | 0.375943765 | 729.2979106 | 789.9677419 | ENSG00000136463 |

| | | | | | | | |
|---|---|---|---|---|---|---|---|
| ASMTL | 0.114823286 | 1.082842401 | 0.104086886 | 0.24540335 | 2226.930051 | 2410.667708 | ENSG00000169093 |
| ZMYM3 | 0.114620871 | 1.082690486 | 0.067250284 | 0.177204077 | 2632.124318 | 2849.452793 | ENSG00000147130 |
| SLC26A11 | 0.114559525 | 1.082644449 | 0.338665026 | 0.550640467 | 410.1641376 | 444.1883256 | ENSG00000181045 |
| RANBP2 | 0.114531787 | 1.082623633 | 0.15267936 | 0.323027222 | 3623.018369 | 3921.37282 | ENSG00000153201 |
| MAGI2 | 0.114462959 | 1.082571984 | 0.322162486 | 0.532799592 | 462.5450427 | 500.7684457 | ENSG00000187391 |
| BBS10 | 0.114459076 | 1.082569071 | 0.433632069 | 0.642497135 | 227.0536012 | 245.5937002 | ENSG00000179941 |
| LDLR | 0.114420136 | 1.082539852 | 0.043169179 | 0.125816349 | 3552.586814 | 3844.895421 | ENSG00000130164 |
| SEMA4A | 0.114405281 | 1.082528705 | 0.351712401 | 0.56379668 | 329.4818643 | 356.7903203 | ENSG00000196189 |
| EIF3F | 0.114398554 | 1.082523658 | 0.027112699 | 0.087855751 | 4810.635351 | 5208.392774 | ENSG00000175390 |
| ZFYVE19 | 0.114374722 | 1.082505776 | 0.366950175 | 0.579419235 | 302.8698849 | 327.643015 | ENSG00000166140 |
| RTN4R | 0.114348184 | 1.082485863 | 0.337482341 | 0.549415519 | 569.1235181 | 616.620695 | ENSG00000040608 |
| FANCG | 0.114290445 | 1.082442541 | 0.113449265 | 0.262055082 | 1171.984383 | 1268.490424 | ENSG00000221829 |
| ICA1 | 0.114231824 | 1.082398559 | 0.333154788 | 0.545201749 | 373.995483 | 404.4814082 | ENSG00000003147 |
| WDR83 | 0.114189669 | 1.082366932 | 0.765798991 | 0.878910348 | 37.952317 | 40.95683915 | ENSG00000123154 |
| NA | 0.114180997 | 1.082360426 | 0.873198806 | 0.940479038 | 8.632338159 | 9.350182953 | ENSG00000287104 |
| DNAJB14 | 0.114140199 | 1.082329819 | 0.351156859 | 0.563192975 | 366.1510524 | 396.3760812 | ENSG00000164031 |
| ERI2 | 0.114122468 | 1.082316517 | 0.528291323 | 0.721043079 | 143.9583812 | 155.7351348 | ENSG00000196678 |
| GIHCG | 0.114051041 | 1.082262934 | 0.76764111 | 0.879862635 | 29.08958132 | 31.47754612 | ENSG00000257698 |
| FER | 0.114018658 | 1.082238641 | 0.538281512 | 0.728027383 | 188.658819 | 204.2043405 | ENSG00000151422 |
| NDUFA4L2 | 0.113988521 | 1.082216033 | 0.91140055 | 0.961009829 | 5.15509198 | 5.610200072 | ENSG00000185633 |
| ACADM | 0.113962106 | 1.082196219 | 0.172634324 | 0.352187449 | 1051.740336 | 1137.959703 | ENSG00000117054 |
| TIMM29 | 0.113944526 | 1.082183032 | 0.339505483 | 0.551295017 | 493.7972386 | 534.4758253 | ENSG00000142444 |
| UNKL | 0.113876071 | 1.082131684 | 0.22723766 | 0.426197558 | 615.9968405 | 666.5677219 | ENSG00000059145 |
| NSMF | 0.113820359 | 1.082089897 | 0.08881265 | 0.218218436 | 2268.745717 | 2455.482249 | ENSG00000165802 |
| ASAH2B | 0.113710803 | 1.082007727 | 0.694056646 | 0.836669205 | 62.38235755 | 67.61788746 | ENSG00000204147 |
| NA | 0.113632533 | 1.081949028 | 0.605825394 | 0.779795526 | 95.26973844 | 103.0149411 | ENSG00000188185 |
| MIEN1 | 0.113555438 | 1.081891212 | 0.598352863 | 0.774336588 | 107.7998202 | 116.5692931 | ENSG00000141741 |
| NA | 0.113530336 | 1.081872388 | 0.849218531 | 0.925311098 | 12.01706421 | 12.97287903 | ENSG00000229944 |
| SLC15A1 | 0.113508468 | 1.081855989 | 0.735738843 | 0.861168893 | 44.84813008 | 48.47149674 | ENSG00000088386 |
| CLDN15 | 0.113425674 | 1.081793904 | 0.666257161 | 0.821189563 | 62.83191552 | 67.96367299 | ENSG00000106404 |
| COL22A1 | 0.113378155 | 1.081758274 | 0.583798621 | 0.763005183 | 111.3185848 | 120.3194909 | ENSG00000169436 |
| GPR158 | 0.113223961 | 1.081642663 | 0.803164313 | 0.89946973 | 22.71273564 | 24.47588094 | ENSG00000151025 |
| DLEU1 | 0.113202884 | 1.08162686 | 0.705727317 | 0.844219917 | 60.6416626 | 65.64343624 | ENSG00000176124 |
| FBXL4 | 0.113200097 | 1.081624771 | 0.467413482 | 0.671109527 | 221.568752 | 239.5927125 | ENSG00000112234 |
| AIMP1 | 0.113189547 | 1.081616861 | 0.218697237 | 0.414936574 | 660.8868724 | 715.0173687 | ENSG00000164022 |
| ADAT1 | 0.113070393 | 1.081527533 | 0.402478015 | 0.613908191 | 296.3934263 | 320.3204403 | ENSG00000065457 |
| ZFAND2A | 0.112910382 | 1.081407586 | 0.627128935 | 0.793729923 | 92.87695338 | 100.4195749 | ENSG00000178381 |
| NA | 0.112757124 | 1.081292714 | 0.626066237 | 0.793253855 | 117.3327744 | 126.9213831 | ENSG00000239883 |
| ERCC6L2 | 0.112720265 | 1.081265089 | 0.390736112 | 0.602224065 | 280.8136886 | 303.3907874 | ENSG00000182150 |
| RBMS1 | 0.112670065 | 1.081227466 | 0.19913929 | 0.389203606 | 1126.348702 | 1217.737479 | ENSG00000153250 |
| PRDX1 | 0.112633315 | 1.081199923 | 0.052540958 | 0.146585313 | 14052.50971 | 15192.62522 | ENSG00000117450 |
| CUL5 | 0.112616864 | 1.081187595 | 0.420330329 | 0.630595667 | 318.2960067 | 344.0396745 | ENSG00000166266 |
| NA | 0.112600987 | 1.081175696 | 0.577831258 | 0.759450456 | 115.9143861 | 125.2107849 | ENSG00000226085 |
| NA | 0.112506893 | 1.081105184 | 0.905319031 | 0.958876702 | 5.171694134 | 5.626374276 | ENSG00000272155 |
| ZNF226 | 0.112455291 | 1.081066515 | 0.463995984 | 0.668542004 | 207.3466127 | 223.9816205 | ENSG00000167380 |
| NA | 0.112407938 | 1.081031033 | 0.807443991 | 0.90192254 | 21.45630309 | 23.20482406 | ENSG00000243517 |
| NRSN2 | 0.112244636 | 1.080908675 | 0.298962164 | 0.50928202 | 640.3504823 | 692.6377017 | ENSG00000125841 |
| NUP58 | 0.112201563 | 1.080876404 | 0.217272849 | 0.413170654 | 1081.056379 | 1168.60468 | ENSG00000139496 |
| SPOCK1 | 0.112104194 | 1.080803457 | 0.342752239 | 0.554597608 | 372.588689 | 402.3554298 | ENSG00000152377 |
| CPSF4 | 0.112101378 | 1.080801347 | 0.119993718 | 0.27264905 | 1141.419488 | 1233.701413 | ENSG00000160917 |
| AP1M2 | 0.11193929 | 1.080679925 | 0.061581807 | 0.165571245 | 2438.014335 | 2634.272194 | ENSG00000129354 |
| RING1 | 0.11187048 | 1.080628383 | 0.184372445 | 0.368608259 | 946.106489 | 1022.502365 | ENSG00000204227 |
| FBXO30 | 0.111782978 | 1.080562843 | 0.169363625 | 0.347315894 | 981.0068374 | 1060.289205 | ENSG00000118496 |
| C1orf210 | 0.111736007 | 1.080527663 | 0.547903535 | 0.735562541 | 134.9838279 | 145.7757875 | ENSG00000253313 |
| NA | 0.111726148 | 1.080520278 | 0.876832731 | 0.942726224 | 8.274410592 | 8.930998754 | ENSG00000231881 |
| FAM83B | 0.111688184 | 1.080491846 | 0.725843873 | 0.855720453 | 43.8618335 | 47.43603207 | ENSG00000168143 |
| CREB1 | 0.111662324 | 1.080472478 | 0.156602688 | 0.328714211 | 1145.943132 | 1237.62488 | ENSG00000118260 |
| PDGFD | 0.111543731 | 1.080383664 | 0.546370085 | 0.734378368 | 172.0990401 | 186.0839549 | ENSG00000170962 |
| NA | 0.111492206 | 1.08034508 | 0.878674002 | 0.943577762 | 8.019681573 | 8.639978679 | ENSG00000226964 |
| KCNK6 | 0.111477757 | 1.08033426 | 0.329355839 | 0.541072433 | 433.4148065 | 468.4810458 | ENSG00000099337 |
| EPB41L5 | 0.111468011 | 1.080326962 | 0.30592142 | 0.516092607 | 533.4507914 | 576.1398427 | ENSG00000115109 |
| NA | 0.111414889 | 1.080287183 | 0.731507279 | 0.858577685 | 44.61592695 | 48.10546963 | ENSG00000255310 |
| NR3C2 | 0.111386406 | 1.080265856 | 0.822210641 | 0.90974033 | 19.046 | 20.49217109 | ENSG00000151623 |
| SNED1 | 0.111251652 | 1.080164959 | 0.787150229 | 0.891474187 | 36.83884759 | 39.85204319 | ENSG00000162804 |
| WAC | 0.111189022 | 1.080118068 | 0.095130717 | 0.229700745 | 2305.164345 | 2490.447125 | ENSG00000095787 |
| CFAP410 | 0.111108033 | 1.080057435 | 0.753060446 | 0.871354394 | 36.30564231 | 39.27529993 | ENSG00000160226 |
| TXN | 0.111007394 | 1.07998209 | 0.092985049 | 0.225913115 | 2639.626523 | 2851.038448 | ENSG00000136810 |
| CAMK2D | 0.110990379 | 1.079969358 | 0.537641392 | 0.727630621 | 182.3771632 | 196.8033538 | ENSG00000145349 |
| PTGS1 | 0.110937683 | 1.079929911 | 0.730563088 | 0.858105398 | 52.08552175 | 56.1461724 | ENSG00000095303 |
| C6orf120 | 0.110758971 | 1.079796145 | 0.278317461 | 0.48756309 | 517.0855882 | 558.404918 | ENSG00000185127 |
| ID1 | 0.110745877 | 1.079786345 | 0.090112057 | 0.220636095 | 1629.021972 | 1759.332925 | ENSG00000125968 |
| TMEM243 | 0.110713501 | 1.079762113 | 0.33036427 | 0.541788157 | 429.4683738 | 463.7416781 | ENSG00000135185 |
| ZYG11B | 0.110676984 | 1.079734783 | 0.392387501 | 0.603980597 | 453.8664058 | 489.6994141 | ENSG00000162378 |
| RPL22 | 0.110656985 | 1.079719815 | 0.106471257 | 0.25018162 | 1353.899251 | 1461.929955 | ENSG00000116251 |

| | | | | | | | |
|---|---|---|---|---|---|---|---|
| ARHGAP26 | 0.110601659 | 1.07967841 | 0.287439809 | 0.496587992 | 502.0319374 | 541.6727176 | ENSG00000145819 |
| OXGR1 | 0.11051838 | 1.079616087 | 0.924867401 | 0.965883673 | 3.704694626 | 3.979358922 | ENSG00000165621 |
| GIT2 | 0.110503229 | 1.079604749 | 0.562714436 | 0.747649264 | 137.7250573 | 148.6420478 | ENSG00000139436 |
| MED9 | 0.110478005 | 1.079585874 | 0.317009284 | 0.527157898 | 401.9036594 | 433.8718508 | ENSG00000141026 |
| ESCO2 | 0.110203972 | 1.079380832 | 0.414861754 | 0.62567022 | 256.93882 | 277.5105041 | ENSG00000171320 |
| MIR570 | 0.110177034 | 1.079360678 | 0.905709023 | 0.959030832 | 6.152956704 | 6.692295438 | ENSG00000207650 |
| NA | 0.11004823 | 1.079264316 | 0.875725575 | 0.942072201 | 8.601498115 | 9.280189252 | ENSG00000228150 |
| ETV7 | 0.110030339 | 1.079250932 | 0.843373697 | 0.922013311 | 14.13467929 | 15.29557478 | ENSG00000010030 |
| IDI1 | 0.109995614 | 1.079224956 | 0.160762526 | 0.33491454 | 1184.753201 | 1278.853895 | ENSG00000067064 |
| PJA2 | 0.109988537 | 1.079219661 | 0.20045973 | 0.391042454 | 1277.18934 | 1377.6992 | ENSG00000198961 |
| PHLDB1 | 0.109900677 | 1.079153939 | 0.103482225 | 0.244343812 | 1562.988312 | 1686.850564 | ENSG00000019144 |
| CDPF1 | 0.109858463 | 1.079122363 | 0.509514634 | 0.706215329 | 191.9634834 | 207.2591537 | ENSG00000205643 |
| SAR1B | 0.109620524 | 1.078944402 | 0.38797162 | 0.600150584 | 296.272125 | 319.4889772 | ENSG00000152700 |
| H4-16 | 0.109486305 | 1.078844028 | 0.828969047 | 0.914028945 | 17.17927513 | 18.52613818 | ENSG00000197837 |
| RARA | 0.109482873 | 1.078841461 | 0.35630532 | 0.568409019 | 357.6524735 | 386.0892981 | ENSG00000131759 |
| PMF1 | 0.109357392 | 1.078747632 | 0.388152685 | 0.600300013 | 361.8625347 | 390.3333606 | ENSG00000160783 |
| INTU | 0.109339674 | 1.078734383 | 0.425684147 | 0.635611886 | 263.737612 | 284.3714468 | ENSG00000164066 |
| ATP2A2 | 0.109212381 | 1.078639208 | 0.047638375 | 0.135898899 | 8111.827839 | 8749.499833 | ENSG00000174437 |
| SLC1A3 | 0.109088479 | 1.078546576 | 0.283730528 | 0.492939376 | 550.9409097 | 594.0355337 | ENSG00000079215 |
| NA | 0.109076907 | 1.078537925 | 0.579578584 | 0.760406596 | 134.1935293 | 144.6546494 | ENSG00000255135 |
| EXOC4 | 0.109029834 | 1.078502734 | 0.16224343 | 0.337107182 | 1283.005168 | 1383.151768 | ENSG00000131558 |
| KTI12 | 0.109003747 | 1.078483233 | 0.817947545 | 0.907309215 | 21.44086999 | 23.11241331 | ENSG00000198841 |
| PLAGL1 | 0.108966161 | 1.078455136 | 0.829895649 | 0.914706392 | 17.3014401 | 18.64342499 | ENSG00000118495 |
| TCHP | 0.108921354 | 1.078421642 | 0.40487462 | 0.615872509 | 275.9510053 | 297.5395302 | ENSG00000139437 |
| LPGAT1 | 0.108913807 | 1.078416 | 0.42569632 | 0.635611886 | 264.433305 | 285.332356 | ENSG00000123684 |
| FAM20C | 0.108906872 | 1.078410817 | 0.175335071 | 0.355890243 | 1194.599615 | 1288.673014 | ENSG00000177706 |
| SECTM1 | 0.108707345 | 1.078261681 | 0.910465 | 0.960811396 | 4.920524588 | 5.323530924 | ENSG00000141574 |
| DCP1A | 0.108613668 | 1.07819167 | 0.334109718 | 0.546110458 | 420.9757448 | 453.7208578 | ENSG00000272886 |
| NA | 0.108588695 | 1.078173006 | 0.878459578 | 0.943401144 | 9.193993074 | 9.928155721 | ENSG00000219507 |
| NA | 0.108586636 | 1.078171468 | 0.9193165 | 0.964136712 | 4.000942105 | 4.299605982 | ENSG00000255200 |
| ERLEC1 | 0.108580793 | 1.078167101 | 0.309425568 | 0.519937763 | 485.4190196 | 523.1072531 | ENSG00000068912 |
| TNFRSF21 | 0.108536473 | 1.07813398 | 0.109398929 | 0.255279849 | 1417.013734 | 1527.839399 | ENSG00000146072 |
| CASP6 | 0.10847046 | 1.078084649 | 0.282423913 | 0.491483191 | 564.265382 | 607.9557973 | ENSG00000138794 |
| SEMA3F | 0.108412161 | 1.078041085 | 0.118482976 | 0.270353041 | 1309.288663 | 1411.301413 | ENSG00000001617 |
| CSTF3 | 0.108387455 | 1.078022624 | 0.187446964 | 0.372938887 | 921.7674946 | 993.2944509 | ENSG00000176102 |
| ZNF7 | 0.108362486 | 1.078003966 | 0.418294976 | 0.629041297 | 277.2208116 | 298.9472422 | ENSG00000147789 |
| CLCN3 | 0.108287348 | 1.077947823 | 0.280681946 | 0.489579621 | 731.1720867 | 787.8710631 | ENSG00000109572 |
| NA | 0.108251595 | 1.07792111 | 0.918814917 | 0.964051188 | 3.980780479 | 4.276905388 | ENSG00000255471 |
| ARHGEF10L | 0.108120988 | 1.07782353 | 0.118969231 | 0.271148194 | 1344.421137 | 1449.050623 | ENSG00000074964 |
| BID | 0.108094494 | 1.077803737 | 0.220075239 | 0.41673728 | 735.2171902 | 792.498926 | ENSG00000015475 |
| NA | 0.108066405 | 1.077782752 | 0.536027382 | 0.726643967 | 157.2425825 | 169.3829592 | ENSG00000214425 |
| TIAM2 | 0.107966032 | 1.07770777 | 0.528206477 | 0.721043079 | 201.0640928 | 216.5211183 | ENSG00000146426 |
| THAP9 | 0.107960775 | 1.077703843 | 0.517092185 | 0.713270264 | 170.1540074 | 183.285912 | ENSG00000168152 |
| NA | 0.107791667 | 1.077577526 | 0.858501109 | 0.930980945 | 15.37895888 | 16.62480209 | ENSG00000287001 |
| MICU2 | 0.107683346 | 1.077496621 | 0.289452567 | 0.498796581 | 835.5553447 | 900.7077731 | ENSG00000165487 |
| NA | 0.10764647 | 1.077469081 | 0.923229366 | 0.965255735 | 3.746213087 | 4.009531866 | ENSG00000188460 |
| CHUK | 0.107584241 | 1.077422607 | 0.309429734 | 0.519937763 | 502.0838288 | 540.5875734 | ENSG00000213341 |
| TMEM128 | 0.107410766 | 1.077293061 | 0.471245872 | 0.674614126 | 229.8594854 | 247.6844196 | ENSG00000132406 |
| FGF11 | 0.107407478 | 1.077290606 | 0.827492842 | 0.913263802 | 17.53600749 | 18.88062556 | ENSG00000161958 |
| RPL18 | 0.107150689 | 1.077098873 | 0.045452517 | 0.130829262 | 14809.9533 | 15951.28782 | ENSG00000063177 |
| KRI1 | 0.107072623 | 1.077040592 | 0.134576552 | 0.296175059 | 1556.448977 | 1676.881517 | ENSG00000129347 |
| BTBD11 | 0.107068534 | 1.077037539 | 0.525188213 | 0.718766931 | 200.4902849 | 215.8177239 | ENSG00000151136 |
| ZFYVE28 | 0.107003913 | 1.076989297 | 0.595583151 | 0.772286996 | 122.399185 | 131.7239346 | ENSG00000159733 |
| MYO19 | 0.106989017 | 1.076978177 | 0.184522013 | 0.36874397 | 1409.844863 | 1518.402841 | ENSG00000278259 |
| CHST9 | 0.106982424 | 1.076973256 | 0.436080982 | 0.644648872 | 293.5389304 | 315.9785116 | ENSG00000154080 |
| BIRC6 | 0.106916257 | 1.076923863 | 0.261770966 | 0.468843082 | 1377.026631 | 1482.316942 | ENSG00000115760 |
| LOC100506411 | 0.106854047 | 1.076877426 | 0.583749713 | 0.763005183 | 130.738809 | 140.6642023 | ENSG00000259153 |
| SFRP5 | 0.106784101 | 1.076825217 | 0.665521692 | 0.82085806 | 74.89405766 | 80.59823883 | ENSG00000120057 |
| LRRC41 | 0.106736551 | 1.076789727 | 0.175772956 | 0.35643426 | 1203.353561 | 1295.76832 | ENSG00000132128 |
| COX18 | 0.106715817 | 1.076774252 | 0.268774207 | 0.476704382 | 598.2716491 | 643.8736718 | ENSG00000163626 |
| C1orf226 | 0.10661856 | 1.076701665 | 0.354061148 | 0.566120124 | 462.2514655 | 497.46273 | ENSG00000239887 |
| ROBO3 | 0.106591797 | 1.076681692 | 0.580737254 | 0.761084181 | 167.6182854 | 180.4687118 | ENSG00000154134 |
| FAAH2 | 0.106557512 | 1.076656105 | 0.687237925 | 0.832541421 | 70.49842421 | 75.86232849 | ENSG00000165591 |
| CAMLG | 0.106477022 | 1.076596038 | 0.228087458 | 0.427494208 | 981.9192231 | 1057.184653 | ENSG00000164615 |
| NAT16 | 0.106416385 | 1.07655079 | 0.920196628 | 0.964276164 | 4.991687885 | 5.382647307 | ENSG00000167011 |
| BLOC1S2 | 0.10640798 | 1.076544518 | 0.244193421 | 0.44864216 | 745.8829108 | 803.249968 | ENSG00000196072 |
| RANGAP1 | 0.106236909 | 1.076416871 | 0.039538545 | 0.117694551 | 6006.526706 | 6466.004344 | ENSG00000100401 |
| NA | 0.105976791 | 1.076222811 | 0.585454637 | 0.764190685 | 147.9110175 | 159.1357859 | ENSG00000185986 |
| NA | 0.105909395 | 1.076172536 | 0.905865461 | 0.959035162 | 5.5533428 | 6.011034604 | ENSG00000279392 |
| PLA2G7 | 0.105828074 | 1.076111877 | 0.688997544 | 0.833924157 | 79.7433928 | 85.77024263 | ENSG00000146070 |
| ATIC | 0.10578772 | 1.076081777 | 0.037161889 | 0.112144817 | 4373.92295 | 4706.258046 | ENSG00000138363 |
| RBM15B | 0.105565862 | 1.07591631 | 0.100023565 | 0.238142339 | 2747.940362 | 2957.320096 | ENSG00000259956 |
| FGF13-AS1 | 0.105488355 | 1.075858509 | 0.756843765 | 0.873724654 | 40.07794076 | 43.09036249 | ENSG00000226031 |
| C17orf67 | 0.105440282 | 1.07582266 | 0.710415815 | 0.847142936 | 62.42059587 | 67.23873492 | ENSG00000214226 |

| | | | | | | | |
|---|---|---|---|---|---|---|---|
| SH2D6 | 0.105418804 | 1.075806644 | 0.918822305 | 0.964051188 | 4.031782149 | 4.316726144 | ENSG00000152292 |
| NA | 0.105287038 | 1.075708391 | 0.823809862 | 0.910816068 | 24.72122844 | 26.63950458 | ENSG00000224152 |
| ISCA2 | 0.105171141 | 1.075621979 | 0.45891177 | 0.664154248 | 297.7776421 | 319.9062265 | ENSG00000165898 |
| CNTLN | 0.105123425 | 1.075586405 | 0.541935518 | 0.731175343 | 204.0591347 | 219.5733139 | ENSG00000044459 |
| EPHX1 | 0.105077102 | 1.07555187 | 0.136256041 | 0.298583818 | 1368.626958 | 1471.524273 | ENSG00000143819 |
| SRP9 | 0.105043804 | 1.075527046 | 0.074405647 | 0.191620161 | 5066.799025 | 5449.932442 | ENSG00000143742 |
| RALGAPA1 | 0.104966418 | 1.075469356 | 0.451965832 | 0.658191107 | 324.5462638 | 349.1618106 | ENSG00000174373 |
| TNKS1BP1 | 0.104917339 | 1.075432771 | 0.042580996 | 0.124530574 | 4464.823541 | 4801.443382 | ENSG00000149115 |
| ANK2 | 0.104894814 | 1.07541598 | 0.325802025 | 0.536984748 | 561.0132461 | 602.9100954 | ENSG00000145362 |
| LRSAM1 | 0.104871793 | 1.075398819 | 0.343140144 | 0.555102163 | 431.1572296 | 463.6594823 | ENSG00000148356 |
| SNTB1 | 0.104764607 | 1.075318925 | 0.367284016 | 0.57955871 | 382.0925565 | 410.6690866 | ENSG00000172164 |
| ZKSCAN8 | 0.104720127 | 1.075285772 | 0.312244008 | 0.522574273 | 471.8765821 | 507.2314347 | ENSG00000198315 |
| PYCR1 | 0.104535226 | 1.075147969 | 0.057283776 | 0.156715067 | 3878.425842 | 4170.587275 | ENSG00000183010 |
| CNTN1 | 0.104490817 | 1.075114874 | 0.367149308 | 0.579491407 | 402.0607826 | 431.9542249 | ENSG00000018236 |
| SLC52A2 | 0.104411159 | 1.075055513 | 0.180386154 | 0.363318074 | 969.8702021 | 1042.829031 | ENSG00000185803 |
| CHAF1A | 0.104360985 | 1.075018125 | 0.037555773 | 0.113058937 | 4664.791759 | 5015.309178 | ENSG00000167670 |
| NA | 0.104297966 | 1.074971168 | 0.891269727 | 0.951694019 | 8.421491866 | 9.023693049 | ENSG00000286447 |
| NA | 0.104262959 | 1.074945085 | 0.92083365 | 0.964660979 | 4.266349541 | 4.57880278 | ENSG00000266680 |
| NXPE3 | 0.104176184 | 1.07488043 | 0.300067236 | 0.51019805 | 608.0786042 | 653.2002809 | ENSG00000144815 |
| ADPRH | 0.104172409 | 1.074877618 | 0.592251661 | 0.769870087 | 121.2340774 | 130.2092234 | ENSG00000144843 |
| RPLP1 | 0.10408776 | 1.074814552 | 0.03043053 | 0.09608436 | 23279.45965 | 25021.12264 | ENSG00000137818 |
| FOXJ2 | 0.103986152 | 1.074738857 | 0.405779366 | 0.616877017 | 339.7771999 | 365.2067183 | ENSG00000065970 |
| METTL8 | 0.103974655 | 1.074730292 | 0.288143637 | 0.497357882 | 639.8415841 | 687.4907849 | ENSG00000123600 |
| LOC101929798 | 0.103934367 | 1.07470028 | 0.882053789 | 0.945771249 | 9.010427352 | 9.671375971 | ENSG00000275585 |
| DCAF4 | 0.103930494 | 1.074697395 | 0.422960747 | 0.633133617 | 332.3521506 | 357.2721938 | ENSG00000119599 |
| FAM172A | 0.103927706 | 1.074695318 | 0.607598178 | 0.781288725 | 113.5474071 | 121.9672413 | ENSG00000113391 |
| EGFL6 | 0.103880676 | 1.074660285 | 0.774887076 | 0.884331605 | 32.99207955 | 35.47932209 | ENSG00000198759 |
| SAMD1 | 0.103854292 | 1.074640631 | 0.251876288 | 0.457549413 | 1738.253047 | 1868.610011 | ENSG00000141858 |
| SYT16 | 0.103845262 | 1.074633906 | 0.801862323 | 0.898704449 | 34.05162437 | 36.53415502 | ENSG00000139973 |
| PTPN23 | 0.103774146 | 1.074580934 | 0.114197766 | 0.263507545 | 1745.506405 | 1875.617312 | ENSG00000076201 |
| RNF34 | 0.103760201 | 1.074570547 | 0.144655164 | 0.310833457 | 1232.544125 | 1324.632048 | ENSG00000170633 |
| ZWILCH | 0.103604314 | 1.074454443 | 0.329743554 | 0.541217709 | 761.0045499 | 817.5696983 | ENSG00000174442 |
| TMEM39A | 0.103580071 | 1.074436389 | 0.210788582 | 0.404382605 | 791.5630593 | 850.4213917 | ENSG00000176142 |
| TGFA | 0.103563058 | 1.074423718 | 0.590786659 | 0.768600583 | 129.0618791 | 138.4981196 | ENSG00000163235 |
| AFG3L2 | 0.103491695 | 1.074370573 | 0.078669712 | 0.199312787 | 2135.4949 | 2294.040779 | ENSG00000141385 |
| GOSR1 | 0.10345912 | 1.074346314 | 0.306400781 | 0.516651321 | 800.7107787 | 860.2941493 | ENSG00000108587 |
| PPP1R9B | 0.103315326 | 1.07423924 | 0.2804273 | 0.489390497 | 631.8500199 | 678.9685445 | ENSG00000108819 |
| TPM2 | 0.103221185 | 1.074169144 | 0.096583212 | 0.232235034 | 2898.277719 | 3113.945202 | ENSG00000198467 |
| PWWP2B | 0.10312127 | 1.074094754 | 0.382069481 | 0.594543684 | 440.3820622 | 473.1763574 | ENSG00000171813 |
| ZNF761 | 0.103099494 | 1.074078542 | 0.317632974 | 0.527758113 | 486.8339001 | 523.0729174 | ENSG00000160336 |
| KDSR | 0.103031657 | 1.074028039 | 0.251120229 | 0.456703152 | 693.3136597 | 744.5444164 | ENSG00000119537 |
| NT5DC2 | 0.102995789 | 1.074001337 | 0.104196137 | 0.245630264 | 7049.681336 | 7571.400898 | ENSG00000168268 |
| MYL12A | 0.10294899 | 1.073966498 | 0.113333022 | 0.26190579 | 1716.932966 | 1843.673207 | ENSG00000101608 |
| CEP128 | 0.102946229 | 1.073964443 | 0.496365808 | 0.695597231 | 235.5156957 | 252.7202401 | ENSG00000100629 |
| GAR1 | 0.102890277 | 1.073922793 | 0.166323018 | 0.342568459 | 1043.914672 | 1121.100486 | ENSG00000109534 |
| ZFYVE16 | 0.102825593 | 1.073874644 | 0.377998977 | 0.590496376 | 395.4749223 | 424.514479 | ENSG00000039319 |
| CITED4 | 0.102775748 | 1.073837542 | 0.545484056 | 0.733722678 | 246.5174806 | 265.1070343 | ENSG00000179862 |
| NRSN2-AS1 | 0.102736963 | 1.073808673 | 0.708953559 | 0.846414716 | 78.70998566 | 84.50514504 | ENSG00000225377 |
| GPANK1 | 0.10261283 | 1.073716285 | 0.324616122 | 0.535450735 | 497.5529354 | 533.9984934 | ENSG00000204438 |
| ZNF234 | 0.102583895 | 1.07369475 | 0.57989155 | 0.760631986 | 145.1365576 | 155.8143248 | ENSG00000263002 |
| AURKB | 0.102543516 | 1.073664699 | 0.1184218 | 0.270315491 | 2289.828986 | 2457.881631 | ENSG00000178999 |
| GJA1 | 0.102537306 | 1.073660077 | 0.048620382 | 0.138053813 | 12362.94247 | 13273.79362 | ENSG00000152661 |
| ZNF713 | 0.102450486 | 1.073595467 | 0.849785282 | 0.925513999 | 14.8635771 | 15.9821325 | ENSG00000178665 |
| TEDC1 | 0.102293925 | 1.073478968 | 0.389795443 | 0.601510618 | 376.0247734 | 404.0222353 | ENSG00000185347 |
| HEXD | 0.102291609 | 1.073477245 | 0.445581475 | 0.653102663 | 377.8223943 | 405.7617997 | ENSG00000169660 |
| MRPL44 | 0.102287985 | 1.073474547 | 0.203631675 | 0.395270242 | 1008.134784 | 1081.775464 | ENSG00000135900 |
| HSD17B8 | 0.102279351 | 1.073468123 | 0.690816995 | 0.834552267 | 74.67370201 | 80.13008052 | ENSG00000204228 |
| NPIPB4 | 0.102171574 | 1.073387932 | 0.692940102 | 0.835910178 | 70.02394993 | 75.29712496 | ENSG00000185864 |
| TFPI2 | 0.102104969 | 1.073338379 | 0.634186256 | 0.799186403 | 96.85417438 | 103.8805289 | ENSG00000105825 |
| AMER1 | 0.101973997 | 1.073240942 | 0.253439089 | 0.45950434 | 770.2059145 | 826.9655363 | ENSG00000184675 |
| CCDC186 | 0.101923516 | 1.073203389 | 0.506777999 | 0.703972234 | 220.090236 | 236.2982827 | ENSG00000165813 |
| SMAD2 | 0.101857067 | 1.07315396 | 0.202823837 | 0.394188346 | 936.257484 | 1004.31932 | ENSG00000175387 |
| RSKR | 0.10179754 | 1.073109681 | 0.663672865 | 0.819648197 | 83.23815701 | 89.40998623 | ENSG00000167524 |
| KCNB1 | 0.101753927 | 1.073077241 | 0.81632915 | 0.906477444 | 26.41031132 | 28.27715566 | ENSG00000158445 |
| ALG13 | 0.101710301 | 1.073044793 | 0.183284946 | 0.366977969 | 1095.01771 | 1174.791606 | ENSG00000101901 |
| RSRC1 | 0.101531349 | 1.0729117 | 0.443560771 | 0.651068331 | 313.7648084 | 336.2967815 | ENSG00000174891 |
| NA | 0.101526965 | 1.07290844 | 0.857335811 | 0.93039751 | 15.53435431 | 16.69980913 | ENSG00000274565 |
| STAT1 | 0.101503075 | 1.072890674 | 0.126411003 | 0.283086896 | 1637.444938 | 1756.739249 | ENSG00000115415 |
| DOLK | 0.101463921 | 1.072861556 | 0.309710713 | 0.520271021 | 522.96874 | 561.3498309 | ENSG00000175283 |
| SHCBP1 | 0.101410278 | 1.072821666 | 0.349881316 | 0.56200635 | 424.9654498 | 455.9942959 | ENSG00000171241 |
| TBCCD1 | 0.101366794 | 1.072789331 | 0.468139928 | 0.671437878 | 288.5211054 | 309.1353944 | ENSG00000113838 |
| WDR81 | 0.101296309 | 1.072736919 | 0.338842536 | 0.550644635 | 609.1847003 | 653.6229653 | ENSG00000167716 |
| RNF111 | 0.101258236 | 1.07270861 | 0.333756334 | 0.545760531 | 550.2709705 | 589.8135617 | ENSG00000157450 |
| BCL7C | 0.101066864 | 1.072566325 | 0.607692744 | 0.781336974 | 117.8146464 | 126.4740859 | ENSG00000099385 |

| | | | | | | | |
|---|---|---|---|---|---|---|---|
| NA | 0.101050977 | 1.072554515 | 0.915540851 | 0.962507751 | 4.624277109 | 4.955990759 | ENSG00000256968 |
| DDX20 | 0.100927654 | 1.072462835 | 0.212975145 | 0.407212272 | 865.2771873 | 927.6719483 | ENSG00000064703 |
| ATP6V1C1 | 0.100830948 | 1.072390949 | 0.342423237 | 0.554269269 | 582.7572032 | 624.6790194 | ENSG00000155097 |
| ZNF446 | 0.100815808 | 1.072379695 | 0.79166605 | 0.893427354 | 32.17333136 | 34.464099 | ENSG00000083838 |
| TBC1D31 | 0.100803418 | 1.072370485 | 0.414893734 | 0.62567022 | 432.427342 | 463.4687968 | ENSG00000156787 |
| ALG2 | 0.100801795 | 1.072369279 | 0.26878201 | 0.476704382 | 720.6420784 | 772.9749481 | ENSG00000119523 |
| RLN1 | 0.100783779 | 1.072355887 | 0.913531066 | 0.961705456 | 5.611463415 | 6.039032084 | ENSG00000107018 |
| WNK4 | 0.100702227 | 1.072295272 | 0.862363552 | 0.933333468 | 13.54218433 | 14.50043211 | ENSG00000126562 |
| ACSL4 | 0.100687373 | 1.072284231 | 0.254007208 | 0.460180941 | 1791.797047 | 1920.918091 | ENSG00000068366 |
| VANGL1 | 0.10068608 | 1.07228327 | 0.34417518 | 0.556247178 | 480.7666239 | 515.3548575 | ENSG00000173218 |
| UBAP1L | 0.100575905 | 1.072201386 | 0.855739163 | 0.929583524 | 14.22600421 | 15.24828168 | ENSG00000246922 |
| NPAT | 0.100252795 | 1.071961279 | 0.507485596 | 0.704540578 | 262.2478329 | 281.2034345 | ENSG00000149308 |
| FBXL7 | 0.100044555 | 1.071806563 | 0.340895491 | 0.552461663 | 477.6111264 | 511.6754314 | ENSG00000183580 |
| DDX55 | 0.099963537 | 1.071746374 | 0.269690523 | 0.477732439 | 753.3017321 | 806.870373 | ENSG00000111364 |
| TPST1 | 0.099952522 | 1.071738192 | 0.413122483 | 0.624495164 | 329.3241307 | 353.1097374 | ENSG00000169902 |
| TONSL | 0.099902448 | 1.071700994 | 0.189663563 | 0.375943765 | 2819.432052 | 3022.587861 | ENSG00000160949 |
| TWNK | 0.09983266 | 1.071649153 | 0.126335717 | 0.282951853 | 1699.102795 | 1820.811097 | ENSG00000107815 |
| TMEM248 | 0.0998123 | 1.07163403 | 0.22091319 | 0.417988738 | 1146.854608 | 1229.133706 | ENSG00000106609 |
| EHBP1 | 0.099779361 | 1.071609563 | 0.243866555 | 0.448172431 | 754.2302391 | 808.0167137 | ENSG00000115504 |
| MKKS | 0.099772789 | 1.071604682 | 0.454953876 | 0.660704703 | 285.5147734 | 305.9494018 | ENSG00000125863 |
| NA | 0.099521485 | 1.071418034 | 0.819909477 | 0.908578247 | 29.63610862 | 31.75506191 | ENSG00000287038 |
| LINC02649 | 0.099371954 | 1.071306991 | 0.932900303 | 0.969708285 | 4.086343292 | 4.356546901 | ENSG00000215244 |
| SNAPC3 | 0.099324225 | 1.071271549 | 0.270706583 | 0.479166306 | 793.97181 | 850.3059969 | ENSG00000164975 |
| HPS3 | 0.099290196 | 1.071246281 | 0.206071227 | 0.398327034 | 1069.833017 | 1145.815489 | ENSG00000163755 |
| RFLNA | 0.099248743 | 1.071215502 | 0.536818486 | 0.727037945 | 217.3193618 | 232.999166 | ENSG00000178882 |
| GTDC1 | 0.099236734 | 1.071206585 | 0.472995005 | 0.676271538 | 257.0212204 | 275.1708562 | ENSG00000121964 |
| RUNX3 | 0.099205646 | 1.071183502 | 0.793536069 | 0.894317058 | 31.71012031 | 33.88735574 | ENSG00000020633 |
| DHX36 | 0.099175581 | 1.07116118 | 0.176805662 | 0.358067016 | 1665.768246 | 1784.55539 | ENSG00000174953 |
| MAP4K5 | 0.099170619 | 1.071157496 | 0.315606741 | 0.525761748 | 499.359812 | 534.8427258 | ENSG00000012983 |
| SCRN2 | 0.099093922 | 1.071100551 | 0.410146897 | 0.621140669 | 529.1237044 | 566.673711 | ENSG00000141295 |
| LDLRAD3 | 0.099077968 | 1.071088706 | 0.364005949 | 0.57608415 | 412.0525242 | 441.56246 | ENSG00000179241 |
| NA | 0.099055266 | 1.071071853 | 0.919923766 | 0.964147051 | 5.308096991 | 5.694192512 | ENSG00000269680 |
| HTRA2 | 0.099054953 | 1.07107162 | 0.447987463 | 0.654716239 | 323.6227646 | 346.6606018 | ENSG00000115317 |
| ROR1 | 0.099037597 | 1.071058735 | 0.221172373 | 0.418269604 | 1654.972989 | 1772.200555 | ENSG00000185483 |
| RPL31P11 | 0.099022013 | 1.071047165 | 0.912750803 | 0.961580883 | 5.287935364 | 5.675842845 | ENSG00000213075 |
| GLE1 | 0.099018371 | 1.071044461 | 0.423321573 | 0.633372517 | 424.8325553 | 454.5383449 | ENSG00000119392 |
| ERCC4 | 0.098996773 | 1.071028428 | 0.670101542 | 0.823185784 | 111.123782 | 118.9099168 | ENSG00000175595 |
| ENPP1 | 0.098908401 | 1.070962824 | 0.354138388 | 0.566195688 | 660.2570033 | 706.852463 | ENSG00000197594 |
| UBL7-AS1 | 0.09883829 | 1.070910779 | 0.708299085 | 0.846168283 | 62.65190927 | 67.06327769 | ENSG00000247240 |
| NA | 0.098602722 | 1.070735932 | 0.916627781 | 0.962979286 | 4.944245687 | 5.296479403 | ENSG00000241061 |
| EIF2B5 | 0.098425305 | 1.070604265 | 0.174585376 | 0.354940771 | 1246.607973 | 1334.423242 | ENSG00000145191 |
| ENTPD5 | 0.098413807 | 1.070595732 | 0.641002413 | 0.803593373 | 133.0910177 | 142.587336 | ENSG00000187097 |
| GLB1 | 0.098251802 | 1.070475519 | 0.242312261 | 0.445966967 | 923.3347697 | 987.9379186 | ENSG00000170266 |
| NA | 0.098246834 | 1.070471832 | 0.888514033 | 0.949931358 | 10.60790598 | 11.37221082 | ENSG00000214772 |
| RPL37A | 0.098196441 | 1.070434441 | 0.114687485 | 0.264292026 | 13450.12462 | 14397.46813 | ENSG00000197756 |
| NA | 0.098163719 | 1.070410163 | 0.366585514 | 0.579182417 | 441.1981412 | 472.453826 | ENSG00000249353 |
| MAP3K20 | 0.0978179 | 1.070153613 | 0.467673609 | 0.671176845 | 280.7653567 | 300.1757357 | ENSG00000091436 |
| METRNL | 0.097795553 | 1.070137036 | 0.406899579 | 0.617708887 | 362.4807834 | 388.0855993 | ENSG00000176845 |
| SIRT7 | 0.097743355 | 1.070098319 | 0.350873131 | 0.563021036 | 449.4141519 | 481.0561164 | ENSG00000187531 |
| NA | 0.097701822 | 1.070067512 | 0.663115528 | 0.819398128 | 130.8254572 | 140.2231908 | ENSG00000273209 |
| GALNT9 | 0.097596254 | 1.069989214 | 0.847193431 | 0.924172673 | 19.66458032 | 21.02958805 | ENSG00000182870 |
| SMG1P1 | 0.097508448 | 1.069924094 | 0.839507931 | 0.919718486 | 22.73409247 | 24.26734025 | ENSG00000237296 |
| ZNF714 | 0.097488224 | 1.069909096 | 0.371920887 | 0.583850869 | 457.3178977 | 489.4331881 | ENSG00000160352 |
| CA13 | 0.097398825 | 1.069842799 | 0.79354517 | 0.894317058 | 33.28860637 | 35.56331453 | ENSG00000185015 |
| SYMPK | 0.097386591 | 1.069833727 | 0.122042918 | 0.275922484 | 2453.030175 | 2624.125042 | ENSG00000125755 |
| TRDMT1 | 0.097372037 | 1.069822934 | 0.565014859 | 0.749213988 | 212.159489 | 226.9743732 | ENSG00000107614 |
| WEE1 | 0.097328815 | 1.069790883 | 0.18724494 | 0.372615458 | 1410.249673 | 1508.723284 | ENSG00000166483 |
| SRSF6 | 0.097142038 | 1.069652393 | 0.21860514 | 0.414826977 | 1425.435383 | 1524.769514 | ENSG00000124193 |
| TPP1 | 0.097135214 | 1.069647334 | 0.424694432 | 0.634622119 | 354.8012582 | 379.5730794 | ENSG00000166340 |
| CYP27C1 | 0.097048285 | 1.069582885 | 0.636800513 | 0.800725077 | 120.22317 | 128.5978887 | ENSG00000186684 |
| NA | 0.097011498 | 1.069555612 | 0.933931256 | 0.970094702 | 4.36954809 | 4.659390252 | ENSG00000240291 |
| HMGXB3 | 0.097004181 | 1.069550187 | 0.206069322 | 0.398327034 | 1055.877149 | 1129.068872 | ENSG00000113716 |
| ZNF444 | 0.09698855 | 1.069538599 | 0.288502578 | 0.497704503 | 894.9121518 | 957.7199458 | ENSG00000167685 |
| NA | 0.096878011 | 1.069456654 | 0.862080691 | 0.933113427 | 14.62189077 | 15.62424015 | ENSG00000273137 |
| METTL4 | 0.096779724 | 1.069383798 | 0.604204291 | 0.778610299 | 163.0832738 | 174.478064 | ENSG00000101574 |
| MYLK2 | 0.096687093 | 1.069315139 | 0.914400377 | 0.962084301 | 5.295054309 | 5.670545959 | ENSG00000101306 |
| NUTF2 | 0.096487405 | 1.069167142 | 0.09597891 | 0.231076201 | 3299.567931 | 3527.109309 | ENSG00000102898 |
| ZSCAN30 | 0.096484312 | 1.069164849 | 0.513308858 | 0.709495556 | 215.311707 | 230.0377727 | ENSG00000186814 |
| DNM1L | 0.096468765 | 1.069153328 | 0.16582625 | 0.341731638 | 1623.667069 | 1735.987774 | ENSG00000087470 |
| NA | 0.096411816 | 1.069111124 | 0.911343526 | 0.961009829 | 5.887549269 | 6.322579809 | ENSG00000276997 |
| GLT8D1 | 0.096396583 | 1.069099836 | 0.303630075 | 0.51363963 | 639.1745672 | 683.037381 | ENSG00000016864 |
| NA | 0.096390307 | 1.069095185 | 0.912568111 | 0.961482481 | 5.893473005 | 6.326930736 | ENSG00000258655 |
| MAGED1 | 0.09634957 | 1.069064998 | 0.147085473 | 0.314569607 | 2686.145799 | 2871.67109 | ENSG00000179222 |
| DVL2 | 0.096252426 | 1.068993015 | 0.136458688 | 0.298785197 | 1678.682089 | 1794.577274 | ENSG00000004975 |

| | | | | | | | |
|---|---|---|---|---|---|---|---|
| SLC25A30 | 0.096244198 | 1.068986918 | 0.613505505 | 0.784868773 | 137.0640688 | 146.2787533 | ENSG00000174032 |
| APP | 0.096217769 | 1.068967335 | 0.033299069 | 0.103123018 | 20383.92886 | 21789.12134 | ENSG00000142192 |
| SRD5A3-AS1 | 0.096090142 | 1.068872774 | 0.876375613 | 0.94241765 | 11.4610537 | 12.29596912 | ENSG00000249700 |
| CLDND1 | 0.096052061 | 1.068844561 | 0.502969538 | 0.700950463 | 395.3026966 | 423.0670619 | ENSG00000080822 |
| ZC3H12C | 0.096009011 | 1.068812667 | 0.748380819 | 0.869205832 | 50.20219473 | 53.65749927 | ENSG00000149289 |
| CCDC127 | 0.095810172 | 1.068665368 | 0.361983143 | 0.573896351 | 461.112722 | 492.7017327 | ENSG00000164366 |
| LDB3 | 0.09573914 | 1.068612753 | 0.881243934 | 0.945351753 | 11.12804244 | 11.92530753 | ENSG00000122367 |
| HTR1B | 0.095692204 | 1.068577988 | 0.837356314 | 0.918185992 | 19.56257698 | 20.8821283 | ENSG00000135312 |
| WDR5B | 0.095669099 | 1.068560875 | 0.778635901 | 0.886993808 | 53.12518427 | 56.70148753 | ENSG00000196981 |
| DGCR8 | 0.095609129 | 1.068516458 | 0.218336652 | 0.414542705 | 1086.531283 | 1161.093789 | ENSG00000128191 |
| ARF6 | 0.0955681 | 1.06848607 | 0.129638031 | 0.288376048 | 2368.004951 | 2530.676713 | ENSG00000165527 |
| KCNK1 | 0.095566606 | 1.068484964 | 0.511857921 | 0.708423367 | 301.9523351 | 322.4077844 | ENSG00000135750 |
| CAPRIN2 | 0.095543319 | 1.068467717 | 0.724565776 | 0.85484005 | 786.8510703 | 840.7318083 | ENSG00000110888 |
| SLC25A19 | 0.095499909 | 1.068435568 | 0.484456849 | 0.685521324 | 266.8237263 | 284.8834933 | ENSG00000125454 |
| SLC12A8 | 0.095465272 | 1.068409917 | 0.528719975 | 0.721294281 | 232.3952597 | 248.4099248 | ENSG00000221955 |
| EML6 | 0.095392098 | 1.068355728 | 0.822924798 | 0.910263856 | 31.18967837 | 33.33085406 | ENSG00000214595 |
| PIGG | 0.095343867 | 1.068320012 | 0.349093417 | 0.561170342 | 596.0194373 | 636.5393598 | ENSG00000174227 |
| HNRNPU | 0.095312316 | 1.068296649 | 0.08712445 | 0.215134053 | 20826.2106 | 22249.1548 | ENSG00000153187 |
| ZNF195 | 0.095269509 | 1.068264951 | 0.30771005 | 0.51808647 | 891.0480227 | 952.1427636 | ENSG00000005801 |
| SLC6A1 | 0.09521964 | 1.068228025 | 0.875238431 | 0.94170908 | 13.78178573 | 14.6780648 | ENSG00000157103 |
| ARSJ | 0.095193729 | 1.068208841 | 0.65011703 | 0.810064757 | 119.0467736 | 127.0730782 | ENSG00000180801 |
| PHLPP2 | 0.095172594 | 1.068193192 | 0.441987132 | 0.649616856 | 326.3272303 | 348.5898401 | ENSG00000040199 |
| CROCCP2 | 0.095167143 | 1.068189156 | 0.399203518 | 0.610689221 | 571.4287397 | 610.5491895 | ENSG00000215908 |
| MATN2 | 0.095102116 | 1.06814101 | 0.433816497 | 0.642616854 | 323.4718707 | 345.6034551 | ENSG00000132561 |
| ZBTB43 | 0.094923156 | 1.06800852 | 0.517315569 | 0.713310786 | 267.7679458 | 285.8163096 | ENSG00000169155 |
| NDRG3 | 0.094836126 | 1.067944095 | 0.292432903 | 0.502202531 | 641.1024135 | 684.582476 | ENSG00000101079 |
| APLP2 | 0.094786633 | 1.067907458 | 0.033178773 | 0.102785617 | 11974.07738 | 12786.52211 | ENSG00000084234 |
| OSBPL2 | 0.094752704 | 1.067882344 | 0.280251724 | 0.489230192 | 694.9357492 | 741.9756799 | ENSG00000130703 |
| SDHA | 0.094629079 | 1.067790841 | 0.800451231 | 0.898320559 | 31.13987191 | 33.2599144 | ENSG00000073578 |
| NA | 0.094594179 | 1.067765011 | 0.840301442 | 0.920427628 | 20.2914748 | 21.65485393 | ENSG00000220472 |
| NA | 0.094582692 | 1.067756509 | 0.920569735 | 0.964598537 | 4.991687885 | 5.324476883 | ENSG00000245768 |
| BASP1 | 0.094483159 | 1.067682846 | 0.1408754 | 0.305165396 | 4873.060899 | 5203.780126 | ENSG00000176788 |
| CNNM4 | 0.094434804 | 1.06764706 | 0.381844655 | 0.594291767 | 454.0167928 | 484.9350251 | ENSG00000158158 |
| ARHGAP33 | 0.094418209 | 1.067634779 | 0.420771718 | 0.630884946 | 569.4365953 | 608.372724 | ENSG00000004777 |
| HS2ST1 | 0.094369795 | 1.067598952 | 0.495301455 | 0.69503403 | 343.9009953 | 367.4490672 | ENSG00000153936 |
| CXADR | 0.094322778 | 1.06756416 | 0.295665602 | 0.505764936 | 1760.996466 | 1880.025045 | ENSG00000154639 |
| NA | 0.094304256 | 1.067550455 | 0.922106358 | 0.964771134 | 4.706118822 | 5.007634792 | ENSG00000262136 |
| INO80E | 0.094244087 | 1.067505932 | 0.250937986 | 0.456459629 | 1225.598175 | 1308.160256 | ENSG00000169592 |
| TSPAN10 | 0.094232216 | 1.067497148 | 0.860226627 | 0.932142521 | 18.59288255 | 19.92507564 | ENSG00000182612 |
| R3HDM4 | 0.094209799 | 1.067480562 | 0.132754038 | 0.293291998 | 1827.88399 | 1951.503868 | ENSG00000198858 |
| SHISA9 | 0.094193313 | 1.067468363 | 0.381879237 | 0.594296612 | 468.759374 | 500.7579176 | ENSG00000237515 |
| ING1 | 0.094165209 | 1.067447569 | 0.298946331 | 0.50928202 | 745.9613945 | 796.1701017 | ENSG00000153487 |
| ZC3H12A | 0.094149494 | 1.067435941 | 0.513874524 | 0.710121498 | 272.6259262 | 291.234733 | ENSG00000163874 |
| NA | 0.094097654 | 1.067397586 | 0.757074023 | 0.873807894 | 50.5165189 | 53.88431699 | ENSG00000246451 |
| CHD6 | 0.094017143 | 1.067338021 | 0.271236914 | 0.47966187 | 1286.025686 | 1371.808238 | ENSG00000124177 |
| LARP4 | 0.094009474 | 1.067332347 | 0.317020505 | 0.527157898 | 1128.283484 | 1203.947208 | ENSG00000161813 |
| UBE2E2 | 0.093883146 | 1.067238891 | 0.359336681 | 0.571533011 | 594.5351724 | 634.22114 | ENSG00000182247 |
| PFDN4 | 0.093848842 | 1.067213515 | 0.358285981 | 0.570123925 | 572.9499681 | 611.877217 | ENSG00000101132 |
| GTF2H3 | 0.093825656 | 1.067196364 | 0.381319237 | 0.59376395 | 438.8966549 | 468.3892022 | ENSG00000111358 |
| NA | 0.093771598 | 1.067156376 | 0.904437094 | 0.958385705 | 8.09084487 | 8.574335906 | ENSG00000267799 |
| JADE1 | 0.093683037 | 1.06709087 | 0.114901074 | 0.264622779 | 2794.965113 | 2981.747838 | ENSG00000077684 |
| ZMYM5 | 0.093596647 | 1.067026974 | 0.570789923 | 0.75364383 | 166.0771473 | 177.2172014 | ENSG00000132950 |
| NSRP1 | 0.093400788 | 1.066882125 | 0.354313962 | 0.566428441 | 564.8419113 | 602.5822736 | ENSG00000126653 |
| LOH12CR2 | 0.093383172 | 1.066869098 | 0.93354966 | 0.969858363 | 3.773493658 | 3.983709849 | ENSG00000205791 |
| C1orf115 | 0.093334074 | 1.066832791 | 0.432643375 | 0.641979267 | 379.1232682 | 404.0953074 | ENSG00000162817 |
| NA | 0.093251399 | 1.066771656 | 0.566565261 | 0.750496917 | 181.5702887 | 193.6695951 | ENSG00000159247 |
| WDR44 | 0.093202014 | 1.06673514 | 0.266436974 | 0.473880417 | 842.2405925 | 898.0916186 | ENSG00000131725 |
| TDRD12 | 0.093145885 | 1.06669364 | 0.838602749 | 0.91904671 | 23.05970546 | 24.6556891 | ENSG00000173809 |
| H2AC12 | 0.093124261 | 1.066677651 | 0.925245765 | 0.965883673 | 4.3553102 | 4.618623537 | ENSG00000274997 |
| NA | 0.093075354 | 1.066641492 | 0.910938606 | 0.961009829 | 6.227679473 | 6.673945771 | ENSG00000280383 |
| ZNF706 | 0.093053383 | 1.066625248 | 0.268510248 | 0.476401356 | 809.8283219 | 864.0450025 | ENSG00000120963 |
| ZNF385A | 0.092931197 | 1.066534916 | 0.479301377 | 0.681494241 | 324.1693698 | 345.9962502 | ENSG00000161642 |
| VPS4B | 0.092796462 | 1.066435315 | 0.345131181 | 0.557034745 | 629.0993596 | 671.243191 | ENSG00000119541 |
| NDUFA10 | 0.092759233 | 1.066407797 | 0.1863133 | 0.371309273 | 1563.345831 | 1666.4479 | ENSG00000130414 |
| BMP2 | 0.092690726 | 1.066357159 | 0.753386579 | 0.871421911 | 63.60291604 | 67.96461894 | ENSG00000125845 |
| PTK2B | 0.092684815 | 1.066352789 | 0.653975999 | 0.81254148 | 108.5782713 | 115.747666 | ENSG00000120899 |
| ICA1L | 0.092680898 | 1.066349895 | 0.819764101 | 0.908474441 | 32.63623692 | 34.95110144 | ENSG00000163596 |
| HRK | 0.092537023 | 1.066243556 | 0.743908897 | 0.866776871 | 55.43940776 | 59.0420385 | ENSG00000135116 |
| ANKRD16 | 0.092532837 | 1.066240463 | 0.539295073 | 0.729040189 | 240.5780138 | 256.7496144 | ENSG00000134461 |
| NA | 0.092527538 | 1.066236547 | 0.077599043 | 0.197447848 | 642530.663 | 685089.2585 | N_noFeature |
| TUBA1C | 0.092495257 | 1.066212689 | 0.099965642 | 0.238064457 | 3553.835367 | 3788.644884 | ENSG00000167553 |
| CAMKV | 0.092459256 | 1.066186083 | 0.108672257 | 0.253868824 | 2259.104679 | 2408.894735 | ENSG00000164076 |
| LRFN1 | 0.092415731 | 1.066153918 | 0.481367954 | 0.683198091 | 272.4624959 | 290.625714 | ENSG00000128011 |
| MAP7D2 | 0.092410552 | 1.06615009 | 0.674162877 | 0.825438662 | 98.66870237 | 105.0234217 | ENSG00000184368 |

| | | | | | | | |
|---|---|---|---|---|---|---|---|
| COX7A2L | 0.092309085 | 1.066075109 | 0.170557729 | 0.348959313 | 1529.014028 | 1629.595631 | ENSG00000115944 |
| MED18 | 0.092263738 | 1.0660416 | 0.465300422 | 0.669506404 | 321.9681074 | 343.3796239 | ENSG00000130772 |
| FBXL12 | 0.092258245 | 1.066037542 | 0.313658916 | 0.52396715 | 735.9303233 | 784.7903162 | ENSG00000127452 |
| SCARB2 | 0.092257328 | 1.066036864 | 0.262867299 | 0.470010315 | 1015.730461 | 1082.581368 | ENSG00000138760 |
| DPF2 | 0.092241864 | 1.066025437 | 0.127686213 | 0.285164869 | 2078.004897 | 2214.811493 | ENSG00000133884 |
| ZNF675 | 0.09222083 | 1.066009895 | 0.562898574 | 0.747720305 | 203.3934439 | 216.9133666 | ENSG00000197372 |
| TMEM63C | 0.092166045 | 1.065969415 | 0.661606491 | 0.818287941 | 115.5303993 | 123.0145293 | ENSG00000165548 |
| LSM6 | 0.09207681 | 1.065903484 | 0.27445364 | 0.483268716 | 874.1197863 | 932.0209502 | ENSG00000164167 |
| NA | 0.092017059 | 1.065859339 | 0.933775201 | 0.969985956 | 4.00806105 | 4.241435558 | ENSG00000231084 |
| DBNDD1 | 0.091993147 | 1.065841673 | 0.282662092 | 0.491655782 | 1351.977203 | 1441.871724 | ENSG00000003249 |
| MAST4 | 0.091928521 | 1.06579393 | 0.648195035 | 0.809119125 | 113.8555282 | 121.2878724 | ENSG00000069020 |
| PPM1K | 0.091911884 | 1.065781639 | 0.622442778 | 0.791034769 | 126.7257401 | 135.0605286 | ENSG00000163644 |
| DYNLL2 | 0.091898823 | 1.06577199 | 0.117782395 | 0.269451307 | 3099.761568 | 3303.914054 | ENSG00000264364 |
| SELENON | 0.09185112 | 1.065736751 | 0.240302716 | 0.443825603 | 1989.883292 | 2120.748089 | ENSG00000162430 |
| TAF1 | 0.091799365 | 1.06569852 | 0.279982599 | 0.488994296 | 868.1340737 | 924.773192 | ENSG00000147133 |
| C3orf38 | 0.091783916 | 1.065687107 | 0.489753088 | 0.689862539 | 421.9430488 | 449.3493232 | ENSG00000179021 |
| JMJD1C | 0.091776922 | 1.065681941 | 0.178248622 | 0.360448144 | 2606.280313 | 2776.802708 | ENSG00000171988 |
| RAMP2-AS1 | 0.091770587 | 1.065677262 | 0.810402605 | 0.903450601 | 38.7005123 | 41.38567117 | ENSG00000197291 |
| UBA52 | 0.091687005 | 1.065615524 | 0.122476504 | 0.276604734 | 9401.298253 | 10018.88468 | ENSG00000221983 |
| GDF11 | 0.091678076 | 1.065608929 | 0.360485136 | 0.572516706 | 551.7139951 | 587.9339802 | ENSG00000135414 |
| DLGAP1 | 0.091594225 | 1.065546996 | 0.495471025 | 0.695165367 | 284.3401826 | 302.8179006 | ENSG00000170579 |
| C16orf91 | 0.091500011 | 1.065477414 | 0.507642902 | 0.70470716 | 265.0094777 | 282.0750246 | ENSG00000174109 |
| NA | 0.091331008 | 1.065352607 | 0.914130366 | 0.961960992 | 5.934991466 | 6.340929476 | ENSG00000249014 |
| SLBP | 0.091326915 | 1.065349584 | 0.259177747 | 0.466162945 | 1072.553273 | 1142.720204 | ENSG00000163950 |
| CASKIN2 | 0.091309039 | 1.065336384 | 0.284497583 | 0.493726967 | 874.8976787 | 932.6052814 | ENSG00000177303 |
| MAPKAPK5-AS1 | 0.09119324 | 1.065250877 | 0.379827721 | 0.59241652 | 551.2540387 | 587.1322815 | ENSG00000234608 |
| NR1H2 | 0.091188477 | 1.06524736 | 0.226732739 | 0.425583188 | 1216.135954 | 1295.806439 | ENSG00000131408 |
| CASP3 | 0.091149846 | 1.065218837 | 0.620867607 | 0.789780732 | 1535.397841 | 1635.648939 | ENSG00000164305 |
| MBOAT1 | 0.091139456 | 1.065211165 | 0.513162356 | 0.709344978 | 258.3595208 | 275.1136746 | ENSG00000172197 |
| C11orf71 | 0.091118581 | 1.065195752 | 0.858513803 | 0.930980945 | 16.84150919 | 17.96339366 | ENSG00000180425 |
| DCAF16 | 0.091104946 | 1.065185686 | 0.311725274 | 0.522197973 | 841.7723219 | 895.9913466 | ENSG00000163257 |
| SPTSSA | 0.091096287 | 1.065179292 | 0.204457217 | 0.396206468 | 1309.511252 | 1394.809999 | ENSG00000165389 |
| TINCR | 0.09101682 | 1.065120621 | 0.860056494 | 0.932067365 | 17.67569049 | 18.87627464 | ENSG00000223573 |
| MPST | 0.09100277 | 1.065110248 | 0.21187887 | 0.405668523 | 2179.701311 | 2322.113223 | ENSG00000128309 |
| VEGFB | 0.090995145 | 1.065104619 | 0.524310187 | 0.71828784 | 397.8405303 | 424.2150836 | ENSG00000173511 |
| THORLNC | 0.090988887 | 1.065099999 | 0.924001545 | 0.965515917 | 4.947805159 | 5.261009573 | ENSG00000226856 |
| NFKB2 | 0.09094949 | 1.065070914 | 0.338394219 | 0.550342303 | 681.8894215 | 726.1521048 | ENSG00000077150 |
| CPS1 | 0.090818433 | 1.064974165 | 0.417960489 | 0.628751794 | 392.3689257 | 417.631709 | ENSG00000021826 |
| HIP1R | 0.090714483 | 1.064897433 | 0.344251089 | 0.55632225 | 630.2264565 | 671.1738824 | ENSG00000130787 |
| SLC25A25 | 0.090676238 | 1.064869204 | 0.355404997 | 0.567706049 | 519.9380249 | 553.6563111 | ENSG00000148339 |
| NA | 0.090579856 | 1.064798066 | 0.635203091 | 0.799988093 | 130.4423084 | 138.929338 | ENSG00000248124 |
| NA | 0.090549971 | 1.064776009 | 0.916476631 | 0.962927628 | 5.876870851 | 6.275286703 | ENSG00000242457 |
| TMEM14C | 0.090545928 | 1.064773025 | 0.208893987 | 0.401932036 | 1699.612735 | 1809.072176 | ENSG00000111843 |
| DLGAP4 | 0.090528695 | 1.064760307 | 0.24873765 | 0.453768404 | 1003.271211 | 1068.147262 | ENSG00000080845 |
| PRRX2 | 0.09046729 | 1.064714989 | 0.673077594 | 0.824962394 | 119.9420763 | 127.7458052 | ENSG00000167157 |
| GTPBP4 | 0.090394164 | 1.064661023 | 0.119043517 | 0.271252028 | 2445.165129 | 2602.986886 | ENSG00000107937 |
| GORASP2 | 0.090290691 | 1.064584666 | 0.152311522 | 0.322358299 | 2229.220272 | 2372.770419 | ENSG00000115806 |
| TNKS | 0.090268497 | 1.064568289 | 0.398944403 | 0.610475104 | 847.5976062 | 901.7364872 | ENSG00000173273 |
| TMEM18 | 0.090189388 | 1.064509915 | 0.490410891 | 0.690222811 | 300.7605579 | 319.8642732 | ENSG00000151353 |
| ARFGAP3 | 0.090161723 | 1.064489503 | 0.449787314 | 0.656111635 | 340.8711704 | 362.7367308 | ENSG00000242247 |
| SDHAF1 | 0.090121917 | 1.064460132 | 0.463448147 | 0.667971431 | 340.5621852 | 362.7777084 | ENSG00000205138 |
| AGFG1 | 0.090116555 | 1.064456176 | 0.30651497 | 0.516741226 | 1426.366891 | 1518.252045 | ENSG00000173744 |
| KCTD9 | 0.089930419 | 1.064318849 | 0.500985985 | 0.699166445 | 320.0269653 | 340.5427359 | ENSG00000104756 |
| FAM214B | 0.089924698 | 1.064314628 | 0.651530721 | 0.810977963 | 137.0288051 | 145.9929574 | ENSG00000005238 |
| LAMP2 | 0.089871497 | 1.064275381 | 0.312929852 | 0.523257482 | 772.1692777 | 821.800016 | ENSG00000005893 |
| TAB2 | 0.089834924 | 1.064248402 | 0.24914796 | 0.454215016 | 1232.653046 | 1312.020736 | ENSG00000055208 |
| LRCH1 | 0.089745353 | 1.064182329 | 0.519832889 | 0.715410267 | 267.6054053 | 284.5874169 | ENSG00000136141 |
| ADA | 0.089737182 | 1.064176302 | 0.499236459 | 0.697674451 | 325.6095957 | 346.3832969 | ENSG00000196839 |
| ZBTB11-AS1 | 0.089732606 | 1.064172927 | 0.823265678 | 0.910506307 | 28.75657006 | 30.64307715 | ENSG00000256628 |
| ARRDC1-AS1 | 0.089709451 | 1.064155847 | 0.374819713 | 0.586783366 | 637.9857647 | 678.9412953 | ENSG00000203993 |
| OTULIN | 0.089695074 | 1.064145243 | 0.435987392 | 0.644560975 | 459.7824303 | 489.2843403 | ENSG00000154124 |
| IFI16 | 0.089656207 | 1.064116574 | 0.89557814 | 0.953381803 | 10.02489423 | 10.64271092 | ENSG00000163565 |
| ZNF611 | 0.089646136 | 1.064109146 | 0.692705325 | 0.835855442 | 98.03913815 | 104.3258187 | ENSG00000213020 |
| ESRRA | 0.089643113 | 1.064106916 | 0.241229763 | 0.444754865 | 1245.221008 | 1325.492243 | ENSG00000173153 |
| RFT1 | 0.089513481 | 1.064011306 | 0.389628908 | 0.601355258 | 462.8018049 | 492.3126488 | ENSG00000163933 |
| NA | 0.08949701 | 1.063999158 | 0.800750737 | 0.898320559 | 39.0779911 | 41.46264276 | ENSG00000264350 |
| EVI5L | 0.089466583 | 1.063976719 | 0.372585319 | 0.584602338 | 531.5521353 | 565.9000691 | ENSG00000142459 |
| MTERF4 | 0.089454306 | 1.063967665 | 0.388605816 | 0.60048837 | 511.2499565 | 544.0516395 | ENSG00000122085 |
| EVI5 | 0.089414703 | 1.063938458 | 0.692712604 | 0.835855442 | 100.9998335 | 107.2782992 | ENSG00000067208 |
| DOCK1 | 0.089319506 | 1.063868256 | 0.262158703 | 0.469187557 | 1386.582444 | 1474.415643 | ENSG00000150760 |
| ACVR2B | 0.089301384 | 1.063854893 | 0.300619453 | 0.510754566 | 1863.478375 | 1982.207652 | ENSG00000114739 |
| PSD2 | 0.08914682 | 1.063740922 | 0.919571388 | 0.964147051 | 5.318775408 | 5.652196292 | ENSG00000146005 |
| LINC00659 | 0.0891336 | 1.063731174 | 0.924731692 | 0.965883673 | 5.615022888 | 5.976510733 | ENSG00000228705 |
| CDK5RAP2 | 0.089046261 | 1.06366678 | 0.194037576 | 0.382186026 | 1481.575641 | 1575.394093 | ENSG00000136861 |

| | | | | | | | |
|---|---|---|---|---|---|---|---|
| NA | 0.089021296 | 1.063648374 | 0.916570293 | 0.962972461 | 6.26207899 | 6.681418121 | ENSG00000275155 |
| CDK19 | 0.088888056 | 1.063550145 | 0.60830078 | 0.78179939 | 174.4005008 | 185.3415178 | ENSG00000155111 |
| UBTD2 | 0.08885909 | 1.063528791 | 0.293342856 | 0.503113236 | 869.0478097 | 924.444411 | ENSG00000168246 |
| NA | 0.088837048 | 1.063512543 | 0.650066951 | 0.810064757 | 145.7056364 | 155.2563101 | ENSG00000286456 |
| DCTN1 | 0.088812545 | 1.06349448 | 0.087609125 | 0.21599195 | 4532.349552 | 4820.161231 | ENSG00000204843 |
| TMEM69 | 0.088522559 | 1.063280736 | 0.360029145 | 0.572222723 | 550.6298134 | 585.4774673 | ENSG00000159596 |
| GRID1 | 0.088441131 | 1.063220724 | 0.89142912 | 0.951700997 | 11.30240428 | 12.02112325 | ENSG00000182771 |
| MSANTD4 | 0.088400971 | 1.063191128 | 0.39266381 | 0.604159691 | 478.3109374 | 508.5176043 | ENSG00000170903 |
| MTMR7 | 0.088394165 | 1.063186112 | 0.689292221 | 0.83398915 | 95.90492091 | 101.8724998 | ENSG00000003987 |
| NA | 0.088267097 | 1.063092475 | 0.814997137 | 0.905897873 | 31.24307046 | 33.1600313 | ENSG00000267787 |
| CLDN5 | 0.088058705 | 1.062938926 | 0.843506697 | 0.922013311 | 21.54050907 | 22.87833415 | ENSG00000184113 |
| CCDC28B | 0.087993451 | 1.062890849 | 0.538212633 | 0.728027383 | 272.2851071 | 289.4237048 | ENSG00000160050 |
| R3HCC1 | 0.087950071 | 1.06285889 | 0.381553889 | 0.593986085 | 530.0237612 | 563.5478628 | ENSG00000104679 |
| AMOT | 0.087879715 | 1.062807059 | 0.459915581 | 0.664802287 | 354.9503727 | 376.983956 | ENSG00000126016 |
| TTLL6 | 0.08773838 | 1.062702945 | 0.752917304 | 0.871306041 | 79.34425226 | 84.33379815 | ENSG00000170703 |
| CRB3 | 0.087705447 | 1.062678687 | 0.498182397 | 0.697160028 | 336.691696 | 357.8705761 | ENSG00000130545 |
| ZNF513 | 0.087700952 | 1.062675375 | 0.41173972 | 0.622872301 | 609.7226347 | 648.142551 | ENSG00000163795 |
| ZNF260 | 0.087643752 | 1.062633244 | 0.531978433 | 0.723643528 | 383.1194806 | 407.3628537 | ENSG00000254004 |
| CC2D1A | 0.087533299 | 1.062551891 | 0.293116598 | 0.502804708 | 1040.27621 | 1106.0095 | ENSG00000132024 |
| PSAP | 0.087524628 | 1.062545505 | 0.05984162 | 0.161953138 | 13087.63619 | 13905.17284 | ENSG00000197746 |
| TULP2 | 0.087498557 | 1.062526304 | 0.845867922 | 0.923418446 | 22.24212631 | 23.62741323 | ENSG00000104804 |
| KDM6B | 0.087473201 | 1.06250763 | 0.579225377 | 0.760157835 | 186.1266565 | 197.8195658 | ENSG00000132510 |
| UMPS | 0.087291327 | 1.062373693 | 0.220858555 | 0.417927235 | 1319.605686 | 1402.147664 | ENSG00000114491 |
| LOC155060 | 0.087250527 | 1.062343649 | 0.846128813 | 0.923543095 | 22.89986083 | 24.33638799 | ENSG00000244560 |
| SEC23IP | 0.087193671 | 1.062301783 | 0.226838916 | 0.425661062 | 1510.110764 | 1603.858937 | ENSG00000107651 |
| FAM124A | 0.087128035 | 1.062253454 | 0.225371447 | 0.423707294 | 1479.088048 | 1571.531642 | ENSG00000150510 |
| RBM45 | 0.086901649 | 1.06208678 | 0.636163143 | 0.80066251 | 168.9257457 | 179.1938281 | ENSG00000155636 |
| GTF3C2 | 0.086850555 | 1.062049166 | 0.142374228 | 0.307495117 | 2051.030618 | 2178.235532 | ENSG00000115207 |
| FAM43B | 0.086811256 | 1.062020236 | 0.708421778 | 0.846207797 | 82.42533255 | 87.48980601 | ENSG00000183114 |
| IGLON5 | 0.08672361 | 1.061955719 | 0.205005135 | 0.396997741 | 1512.926794 | 1607.100391 | ENSG00000142549 |
| NA | 0.086471876 | 1.061770435 | 0.92654374 | 0.966701516 | 5.009485247 | 5.30083033 | ENSG00000274423 |
| RABL2A | 0.086439634 | 1.061746707 | 0.734092486 | 0.860316207 | 91.69821825 | 97.59268391 | ENSG00000144134 |
| NA | 0.086109492 | 1.061503768 | 0.938598628 | 0.97242896 | 4.059062721 | 4.271608502 | ENSG00000287576 |
| TRAK2 | 0.086065184 | 1.061471167 | 0.532461392 | 0.723985867 | 322.4526753 | 342.2808372 | ENSG00000115993 |
| NA | 0.086019296 | 1.061437406 | 0.701003468 | 0.840766866 | 87.26282021 | 92.68159989 | ENSG00000213976 |
| CSMD2 | 0.085973727 | 1.06140388 | 0.4457378 | 0.653197669 | 469.5043159 | 498.3200675 | ENSG00000121904 |
| ZNF182 | 0.085963168 | 1.061396111 | 0.649720952 | 0.809846226 | 155.8313394 | 165.2148067 | ENSG00000147118 |
| COG5 | 0.085851771 | 1.061314159 | 0.454199127 | 0.660217866 | 438.0933404 | 464.614811 | ENSG00000164597 |
| NA | 0.085800265 | 1.06127627 | 0.927795672 | 0.967350062 | 5.291494837 | 5.60707865 | ENSG00000276116 |
| PROSER2 | 0.085702562 | 1.0612044 | 0.555307531 | 0.741653658 | 278.4805243 | 295.5313628 | ENSG00000148426 |
| RPS25 | 0.085688218 | 1.061193849 | 0.151812238 | 0.321589918 | 10154.3116 | 10775.52751 | ENSG00000118181 |
| TMEM63B | 0.085647197 | 1.061163675 | 0.249281777 | 0.454322173 | 1112.20413 | 1180.315355 | ENSG00000137216 |
| UQCC2 | 0.08558826 | 1.061120325 | 0.473146504 | 0.676314272 | 552.7171215 | 586.3035836 | ENSG00000137288 |
| COMMD1 | 0.085535585 | 1.061081583 | 0.520154217 | 0.715550501 | 344.9270047 | 366.231401 | ENSG00000173163 |
| NPLOC4 | 0.085501232 | 1.061056317 | 0.116403906 | 0.267204055 | 3099.399401 | 3288.385197 | ENSG00000182446 |
| PIK3CA | 0.085449941 | 1.061018595 | 0.608771583 | 0.782191546 | 177.4957928 | 188.2064329 | ENSG00000121879 |
| NA | 0.085334566 | 1.060933747 | 0.897912897 | 0.954898359 | 10.05573427 | 10.62748268 | ENSG00000220804 |
| TAZ | 0.085326637 | 1.060927916 | 0.54949164 | 0.736804148 | 361.305836 | 383.1327729 | ENSG00000102125 |
| PCYOX1 | 0.085223992 | 1.060852436 | 0.241140527 | 0.444668216 | 1261.490236 | 1338.558887 | ENSG00000116005 |
| NA | 0.085193085 | 1.060829709 | 0.924212225 | 0.965629205 | 5.315215936 | 5.626374276 | ENSG00000231510 |
| WDR75 | 0.084962032 | 1.060659827 | 0.219270588 | 0.415703519 | 1759.941652 | 1866.168358 | ENSG00000115368 |
| HSPB8 | 0.084943439 | 1.060646158 | 0.592955118 | 0.770254327 | 189.0982834 | 200.3990413 | ENSG00000152137 |
| ARL17A | 0.084913867 | 1.060624417 | 0.881243446 | 0.945351753 | 14.74525094 | 15.57477158 | ENSG00000228696 |
| F3 | 0.084885204 | 1.060603345 | 0.724483666 | 0.854796578 | 79.21941754 | 84.12968103 | ENSG00000117525 |
| RUNX2 | 0.084868742 | 1.060591243 | 0.885250651 | 0.947622636 | 15.30187185 | 16.31079787 | ENSG00000124813 |
| WDR53 | 0.084816056 | 1.060552512 | 0.573692776 | 0.755930071 | 210.3277484 | 223.1312608 | ENSG00000185798 |
| CISD2 | 0.084792329 | 1.06053507 | 0.61409908 | 0.785269226 | 282.2513476 | 299.6463582 | ENSG00000145354 |
| SOS1 | 0.084644452 | 1.06042637 | 0.411752068 | 0.622872301 | 474.1000649 | 502.7748891 | ENSG00000115904 |
| PBX4 | 0.084511682 | 1.060328785 | 0.83262746 | 0.91561881 | 35.56426047 | 37.60463802 | ENSG00000105717 |
| ITPRID2 | 0.084295045 | 1.060169576 | 0.625586906 | 0.793059591 | 166.4780679 | 176.3231216 | ENSG00000138434 |
| PIP4K2A | 0.084261489 | 1.060144918 | 0.350412247 | 0.56252425 | 820.4910677 | 870.0861444 | ENSG00000150867 |
| NDUFB9 | 0.084256212 | 1.06014104 | 0.317703821 | 0.527777749 | 915.6312179 | 970.7345148 | ENSG00000147684 |
| COX17 | 0.084231603 | 1.060122957 | 0.445819893 | 0.653267279 | 473.7329852 | 502.6174457 | ENSG00000138495 |
| IFT27 | 0.084161751 | 1.060071629 | 0.474499285 | 0.67731814 | 368.060186 | 389.9430472 | ENSG00000100360 |
| NUP37 | 0.084110515 | 1.060033982 | 0.450599839 | 0.656958077 | 508.3305788 | 538.9937581 | ENSG00000075188 |
| TENT4B | 0.084072741 | 1.060006228 | 0.499226094 | 0.697674451 | 335.2253069 | 355.5617838 | ENSG00000121274 |
| KCTD14 | 0.083922825 | 1.059896084 | 0.758172816 | 0.874316737 | 57.94579045 | 61.40578452 | ENSG00000151364 |
| LAMB2 | 0.083880487 | 1.05986498 | 0.165595304 | 0.341442003 | 2119.949081 | 2247.004236 | ENSG00000172037 |
| ZNF497 | 0.083863277 | 1.059852338 | 0.848366253 | 0.9249161 | 23.80284116 | 25.23990471 | ENSG00000174586 |
| LINC02210 | 0.083844469 | 1.059838521 | 0.524217725 | 0.718239903 | 385.8871538 | 409.1474834 | ENSG00000204650 |
| SMDT1 | 0.083740533 | 1.059762169 | 0.542719046 | 0.73167113 | 268.2127485 | 284.4803153 | ENSG00000183172 |
| FGF22 | 0.083723783 | 1.059749865 | 0.83725918 | 0.918172233 | 34.28499655 | 36.42315402 | ENSG00000070388 |
| GOLGA6L9 | 0.083710855 | 1.059740369 | 0.668632713 | 0.822292836 | 118.1610058 | 125.2465383 | ENSG00000197978 |
| FBLIM1 | 0.083692847 | 1.059727141 | 0.168182081 | 0.345645038 | 2311.945846 | 2450.66821 | ENSG00000162458 |

| | | | | | | | |
|---|---|---|---|---|---|---|---|
| TBCD | 0.083675348 | 1.059714287 | 0.104375547 | 0.245991789 | 4158.641593 | 4407.13146 | ENSG00000141556 |
| FBXL16 | 0.08361097 | 1.059667 | 0.548917727 | 0.736359583 | 285.6763458 | 302.5652311 | ENSG00000127585 |
| KLC2 | 0.083516898 | 1.059597906 | 0.257747772 | 0.464655567 | 1100.765017 | 1166.456887 | ENSG00000174996 |
| MTRF1 | 0.083492502 | 1.059579988 | 0.6352884 | 0.799988093 | 171.3416409 | 181.5614735 | ENSG00000120662 |
| DEPDC7 | 0.083455064 | 1.059552493 | 0.673180265 | 0.824984381 | 128.8516432 | 136.5301948 | ENSG00000121690 |
| GNL3L | 0.08330676 | 1.05944358 | 0.258659372 | 0.465572078 | 1119.622801 | 1186.231214 | ENSG00000130119 |
| RWDD2B | 0.083292598 | 1.05943318 | 0.718285462 | 0.851388915 | 83.26187811 | 88.24011459 | ENSG00000156253 |
| TP53RK | 0.083285413 | 1.059427904 | 0.456588267 | 0.662631902 | 467.3599521 | 495.2128229 | ENSG00000172315 |
| SMURF2 | 0.083266948 | 1.059414345 | 0.393540859 | 0.605164003 | 584.2111077 | 618.7448111 | ENSG00000108854 |
| FAR2 | 0.08315031 | 1.059328697 | 0.457774285 | 0.663253386 | 445.0976125 | 471.3430278 | ENSG00000064763 |
| NA | 0.083101635 | 1.059292957 | 0.57803507 | 0.759450456 | 243.0126228 | 257.4701062 | ENSG00000231205 |
| GPD2 | 0.083098761 | 1.059290847 | 0.363993172 | 0.57608415 | 793.5813414 | 840.15696 | ENSG00000115159 |
| NSUN4 | 0.08306356 | 1.059265001 | 0.446152157 | 0.653449937 | 462.3074489 | 489.2790864 | ENSG00000117481 |
| SDC1 | 0.083060443 | 1.059262713 | 0.199490567 | 0.389614115 | 2065.236764 | 2188.324244 | ENSG00000115884 |
| FLT4 | 0.082966771 | 1.059193938 | 0.390170187 | 0.601716195 | 571.8767447 | 605.6950034 | ENSG00000037280 |
| LIMD1 | 0.082927565 | 1.059165155 | 0.524178465 | 0.718239903 | 344.793882 | 365.3661263 | ENSG00000144791 |
| RPA3 | 0.082781335 | 1.059057804 | 0.397481147 | 0.609205812 | 517.7738835 | 548.4152596 | ENSG00000106399 |
| MFSD5 | 0.082711822 | 1.059006777 | 0.453428795 | 0.6596002 | 440.7699662 | 467.0646523 | ENSG00000182544 |
| TUBGCP3 | 0.082654006 | 1.058964338 | 0.341551512 | 0.55319042 | 774.794799 | 820.2184231 | ENSG00000126216 |
| TDRP | 0.082645041 | 1.058957758 | 0.265431027 | 0.473026795 | 1113.228411 | 1178.648223 | ENSG00000180190 |
| LPIN3 | 0.082627381 | 1.058944795 | 0.686281411 | 0.83186293 | 131.7165383 | 139.4885096 | ENSG00000132793 |
| REPS1 | 0.082568644 | 1.058901683 | 0.225448546 | 0.423763008 | 1367.168934 | 1447.953728 | ENSG00000135597 |
| CAD | 0.082511884 | 1.058860024 | 0.102901135 | 0.243184575 | 6188.179433 | 6552.087087 | ENSG00000084774 |
| ZNF696 | 0.082304304 | 1.058707682 | 0.413486776 | 0.624700612 | 615.4489427 | 652.0239687 | ENSG00000185730 |
| IGF2BP3 | 0.082257577 | 1.058673392 | 0.272915066 | 0.481592322 | 3296.969921 | 3489.222283 | ENSG00000136231 |
| RASGRP2 | 0.081990045 | 1.058477091 | 0.479547941 | 0.681690845 | 560.661294 | 593.7929339 | ENSG00000068831 |
| ZSCAN25 | 0.081931494 | 1.058434134 | 0.478825268 | 0.681414206 | 387.8929768 | 410.6779993 | ENSG00000197037 |
| ZNF692 | 0.081888851 | 1.05840285 | 0.486354962 | 0.68722884 | 670.5001165 | 709.618844 | ENSG00000171163 |
| OSER1-DT | 0.081868416 | 1.058387858 | 0.748977635 | 0.869571046 | 65.56602599 | 69.38408153 | ENSG00000223891 |
| PRDM4 | 0.081863838 | 1.058384499 | 0.451722179 | 0.657984832 | 436.3822636 | 461.7189449 | ENSG00000110851 |
| DTWD2 | 0.081862639 | 1.05838362 | 0.694707638 | 0.837133343 | 143.9815175 | 152.4126122 | ENSG00000169570 |
| GOLGB1 | 0.081831422 | 1.058360719 | 0.418909073 | 0.629503762 | 1012.224662 | 1070.508463 | ENSG00000173230 |
| NA | 0.081798577 | 1.058336624 | 0.860343956 | 0.932156491 | 21.93136161 | 23.3049907 | ENSG00000226084 |
| SPATA20 | 0.081678325 | 1.058248413 | 0.438708813 | 0.646711102 | 574.5774238 | 608.1868409 | ENSG00000006282 |
| SEC24A | 0.081658086 | 1.058233567 | 0.507791824 | 0.704758477 | 423.1393537 | 447.6658195 | ENSG00000113615 |
| N4BP1 | 0.081628321 | 1.058211735 | 0.32065898 | 0.531150323 | 1136.704548 | 1202.773451 | ENSG00000102921 |
| VPS37C | 0.081606405 | 1.05819566 | 0.393591236 | 0.605192191 | 646.5107594 | 684.4255364 | ENSG00000167987 |
| DRAXIN | 0.081565175 | 1.058165418 | 0.298093733 | 0.508307011 | 1030.994705 | 1091.138815 | ENSG00000162490 |
| PKN3 | 0.081550368 | 1.058154558 | 0.306032597 | 0.516204501 | 967.5275034 | 1023.553909 | ENSG00000160447 |
| MAPK8IP3 | 0.081545259 | 1.05815081 | 0.374530938 | 0.58653427 | 1282.179099 | 1357.064622 | ENSG00000138834 |
| THOP1 | 0.081544933 | 1.058150572 | 0.142911327 | 0.308324747 | 3610.985036 | 3821.676875 | ENSG00000172009 |
| HLA-V | 0.081510437 | 1.05812527 | 0.92307197 | 0.965185435 | 5.993112081 | 6.338754013 | ENSG00000181126 |
| FAT3 | 0.081470085 | 1.058095675 | 0.510759468 | 0.707234478 | 897.1295569 | 948.5697228 | ENSG00000165323 |
| TMEM245 | 0.081411013 | 1.058052352 | 0.301011299 | 0.511112963 | 1431.849752 | 1514.609137 | ENSG00000106771 |
| CNOT10 | 0.081357294 | 1.058012956 | 0.360486955 | 0.572516706 | 826.0758354 | 874.2024849 | ENSG00000182973 |
| DYNC2LI1 | 0.081317674 | 1.057983901 | 0.748432822 | 0.869205832 | 68.00716913 | 71.99779736 | ENSG00000138036 |
| SEH1L | 0.08131372 | 1.057981001 | 0.340989307 | 0.552564189 | 1227.056248 | 1298.247345 | ENSG00000085415 |
| B4GALNT3 | 0.081306945 | 1.057976033 | 0.310960338 | 0.521674158 | 1075.640303 | 1138.018411 | ENSG00000139044 |
| RPS9 | 0.081301729 | 1.057972208 | 0.145674503 | 0.312664915 | 29793.13536 | 31519.5804 | ENSG00000170889 |
| MAGED4B | 0.081290489 | 1.057963965 | 0.805067726 | 0.90066817 | 45.59451977 | 48.30674887 | ENSG00000187243 |
| PFDN6 | 0.081207804 | 1.057903332 | 0.311111699 | 0.521747681 | 1181.937094 | 1250.229876 | ENSG00000204220 |
| VAMP2 | 0.081190615 | 1.057890727 | 0.666418203 | 0.821280851 | 158.8578323 | 168.0682251 | ENSG00000220205 |
| GYG1 | 0.081121804 | 1.057840271 | 0.394858631 | 0.606647027 | 564.5257815 | 596.958431 | ENSG00000163754 |
| TRIM28 | 0.080994839 | 1.057747179 | 0.144802987 | 0.31106864 | 30906.79975 | 32691.94838 | ENSG00000130726 |
| TMEM104 | 0.080993818 | 1.057746431 | 0.396029953 | 0.607952013 | 679.5319524 | 718.6821563 | ENSG00000109066 |
| NA | 0.080974444 | 1.057732227 | 0.691460669 | 0.834978882 | 122.860285 | 129.8502171 | ENSG00000213621 |
| ISY1 | 0.080969807 | 1.057728827 | 0.733616515 | 0.860144526 | 87.56527076 | 92.5435044 | ENSG00000240682 |
| ASTL | 0.080875864 | 1.057659954 | 0.921355448 | 0.964771134 | 6.905575619 | 7.323804157 | ENSG00000188886 |
| IFI27L1 | 0.080874316 | 1.057658819 | 0.631649253 | 0.797106325 | 186.6043586 | 197.4431559 | ENSG00000165948 |
| LONRF1 | 0.0807567 | 1.057572596 | 0.281586473 | 0.490568316 | 1130.963443 | 1196.204065 | ENSG00000154359 |
| RAB39A | 0.080730692 | 1.057553531 | 0.659383204 | 0.816486541 | 139.9980678 | 147.9139453 | ENSG00000179331 |
| TMEM214 | 0.080707909 | 1.057536831 | 0.291846217 | 0.501459414 | 1104.430331 | 1167.672285 | ENSG00000119777 |
| NA | 0.08054765 | 1.057419363 | 0.904696802 | 0.958481852 | 10.36858391 | 10.91973226 | ENSG00000286158 |
| MDH2 | 0.080507129 | 1.057389663 | 0.082179889 | 0.205967777 | 5673.194115 | 5999.193047 | ENSG00000146701 |
| NA | 0.080462184 | 1.057356723 | 0.635281829 | 0.799988093 | 264.4406515 | 279.324799 | ENSG00000279865 |
| ZDHHC8 | 0.080429763 | 1.057332961 | 0.405464034 | 0.616620091 | 874.8081338 | 925.5304191 | ENSG00000099904 |
| PPP1R3F | 0.080350051 | 1.057274543 | 0.682259146 | 0.830184534 | 123.5097315 | 130.4640386 | ENSG00000049769 |
| IVD | 0.08034862 | 1.057273495 | 0.181178673 | 0.364189381 | 2187.297242 | 2312.872641 | ENSG00000128928 |
| PRDX5 | 0.080282892 | 1.057225327 | 0.170318691 | 0.348638065 | 4471.99935 | 4727.83316 | ENSG00000126432 |
| MALSU1 | 0.08027753 | 1.057221397 | 0.334587698 | 0.546552087 | 925.1777724 | 978.5179434 | ENSG00000156928 |
| LDLRAP1 | 0.080191175 | 1.057158118 | 0.401130228 | 0.612513691 | 580.4534044 | 613.8633759 | ENSG00000157978 |
| FAM83G | 0.080020166 | 1.057032816 | 0.370670388 | 0.582905344 | 1034.529181 | 1094.208197 | ENSG00000188522 |
| MRTFA | 0.079985844 | 1.057007669 | 0.285478646 | 0.494732775 | 1046.174386 | 1105.865047 | ENSG00000196588 |
| MAGEL2 | 0.079958667 | 1.056987757 | 0.683890833 | 0.830938582 | 132.2031911 | 139.8051408 | ENSG00000254585 |

| | | | | | | | |
|---|---|---|---|---|---|---|---|
| NA | 0.079950026 | 1.056981427 | 0.93151789 | 0.969133724 | 5.291494837 | 5.574730242 | ENSG00000251654 |
| SCAND1 | 0.079904873 | 1.056948346 | 0.307848478 | 0.518202915 | 1011.158075 | 1069.104074 | ENSG00000171222 |
| SYNDIG1 | 0.079870165 | 1.056922919 | 0.683193557 | 0.830790283 | 124.6609323 | 131.9354287 | ENSG00000101463 |
| NBR1 | 0.079735031 | 1.056823924 | 0.370864097 | 0.583064309 | 931.9516492 | 984.2143934 | ENSG00000188554 |
| SRPRA | 0.079718712 | 1.056811969 | 0.277311241 | 0.486459867 | 1678.442567 | 1773.32404 | ENSG00000182934 |
| NR1D1 | 0.079658449 | 1.056767826 | 0.711990317 | 0.847851541 | 119.4008362 | 126.2031394 | ENSG00000126368 |
| CINP | 0.079606835 | 1.05673002 | 0.427463307 | 0.637365179 | 512.8065015 | 541.6561645 | ENSG00000100865 |
| EXOSC7 | 0.079602252 | 1.056726663 | 0.259626124 | 0.466511832 | 1210.720132 | 1279.202205 | ENSG00000075914 |
| THOC1 | 0.079481318 | 1.056638087 | 0.338376935 | 0.550342303 | 864.5854366 | 913.6465274 | ENSG00000079134 |
| PIGP | 0.079400887 | 1.05657918 | 0.725411392 | 0.855410187 | 87.82683939 | 92.8304571 | ENSG00000185808 |
| TRIP11 | 0.079290353 | 1.056498232 | 0.508153614 | 0.704897972 | 418.0767039 | 441.3245541 | ENSG00000100815 |
| LCK | 0.079265533 | 1.056480057 | 0.217620304 | 0.413599581 | 1872.850265 | 1978.173364 | ENSG00000182866 |
| USPL1 | 0.079153258 | 1.056397841 | 0.462452828 | 0.667386646 | 452.10677 | 477.7163056 | ENSG00000132952 |
| ZNF816 | 0.079141358 | 1.056389127 | 0.695128007 | 0.837250559 | 116.5602732 | 122.9575884 | ENSG00000180257 |
| LINC01012 | 0.079076023 | 1.056341288 | 0.7900166 | 0.892419158 | 50.86821725 | 53.76267926 | ENSG00000281706 |
| SDHAP2 | 0.079069486 | 1.056336502 | 0.710347154 | 0.847142936 | 116.2417791 | 122.6151352 | ENSG00000242086 |
| FAM53C | 0.079038863 | 1.056314079 | 0.267363648 | 0.475081153 | 1614.646802 | 1705.154199 | ENSG00000120709 |
| OGT | 0.079033124 | 1.056309878 | 0.307345968 | 0.517726215 | 2071.819322 | 2188.211005 | ENSG00000147162 |
| CCNT1 | 0.079008655 | 1.056291962 | 0.451124066 | 0.657452882 | 740.5794394 | 781.6361213 | ENSG00000129315 |
| ITM2A | 0.078999778 | 1.056285463 | 0.430209348 | 0.640027063 | 524.9285437 | 554.3955314 | ENSG00000078596 |
| RIF1 | 0.078964189 | 1.056259406 | 0.362946672 | 0.574867178 | 1358.517168 | 1434.716373 | ENSG00000080345 |
| R3HCC1L | 0.078956044 | 1.056253443 | 0.520517173 | 0.715685882 | 397.2467111 | 419.13611 | ENSG00000166024 |
| ZSWIM3 | 0.078953 | 1.056251214 | 0.686032476 | 0.831774737 | 123.3881508 | 130.4481479 | ENSG00000132801 |
| WDCP | 0.078943356 | 1.056244154 | 0.360421582 | 0.572516706 | 716.6554265 | 757.1048782 | ENSG00000163026 |
| ANKRD34B | 0.078883924 | 1.056200642 | 0.761662324 | 0.876689032 | 71.459499 | 75.40956638 | ENSG00000189127 |
| COPB2 | 0.078733475 | 1.056090504 | 0.136404478 | 0.298778204 | 2959.157204 | 3125.113997 | ENSG00000184432 |
| LINC01963 | 0.078595825 | 1.055989746 | 0.799982558 | 0.898237613 | 51.65851582 | 54.59431035 | ENSG00000260804 |
| PMPCB | 0.078532517 | 1.055943407 | 0.232827025 | 0.433708657 | 1913.126568 | 2019.525121 | ENSG00000105819 |
| KIF18A | 0.078520262 | 1.055934438 | 0.508100755 | 0.704897972 | 343.816101 | 362.8911684 | ENSG00000121621 |
| MVK | 0.078518269 | 1.055932979 | 0.434576878 | 0.643382278 | 518.9164652 | 547.8070709 | ENSG00000110921 |
| GSK3B | 0.078495139 | 1.05591605 | 0.353851564 | 0.56592876 | 774.0724091 | 817.1810565 | ENSG00000082701 |
| CAMTA2 | 0.078439991 | 1.055875688 | 0.689635868 | 0.83398915 | 125.4740361 | 132.6200945 | ENSG00000108509 |
| ZNF480 | 0.078430572 | 1.055868795 | 0.446015369 | 0.653422136 | 545.6645942 | 576.3802802 | ENSG00000198464 |
| PLD2 | 0.078425956 | 1.055865416 | 0.315940815 | 0.525964589 | 1122.554715 | 1185.501022 | ENSG00000129219 |
| DDX46 | 0.07832074 | 1.055788415 | 0.132313328 | 0.29269409 | 3500.262631 | 3695.128524 | ENSG00000145833 |
| RRAGA | 0.078315416 | 1.055784518 | 0.379946578 | 0.59241652 | 755.154628 | 797.5178896 | ENSG00000155876 |
| RNF135 | 0.078254162 | 1.055739693 | 0.636116283 | 0.80066251 | 186.6251567 | 197.1327972 | ENSG00000181481 |
| CRK | 0.078121698 | 1.055642762 | 0.242741621 | 0.446626608 | 1757.413047 | 1855.07565 | ENSG00000167193 |
| HELB | 0.078066917 | 1.055602679 | 0.658624484 | 0.8159892 | 160.5086769 | 169.2359509 | ENSG00000127311 |
| INTS14 | 0.078051695 | 1.055591542 | 0.374418589 | 0.586455591 | 704.0950896 | 743.3072936 | ENSG00000138614 |
| NA | 0.078046641 | 1.055587843 | 0.924729111 | 0.965883673 | 6.296478506 | 6.645948291 | ENSG00000154898 |
| NA | 0.078044404 | 1.055586207 | 0.925722905 | 0.966106933 | 5.962272037 | 6.284934516 | ENSG00000261614 |
| NA | 0.077805823 | 1.055411657 | 0.90061367 | 0.956262744 | 11.92810355 | 12.57299045 | ENSG00000280303 |
| ZNF516 | 0.077764737 | 1.0553816 | 0.431278109 | 0.641011104 | 537.2638226 | 567.0200503 | ENSG00000101493 |
| PPP1R26 | 0.077754685 | 1.055374247 | 0.229430544 | 0.429202162 | 1669.190121 | 1761.764047 | ENSG00000196422 |
| FRMD4A | 0.077663031 | 1.055307201 | 0.46889093 | 0.671961924 | 462.8213823 | 488.1648535 | ENSG00000151474 |
| WDR37 | 0.077609137 | 1.05526778 | 0.512081847 | 0.708546306 | 352.2803321 | 371.965276 | ENSG00000047056 |
| ITGA2 | 0.077605967 | 1.055265461 | 0.680180026 | 0.829014226 | 142.4851786 | 150.3027354 | ENSG00000164171 |
| TRIM26 | 0.077527301 | 1.055207922 | 0.273959992 | 0.482659928 | 1180.818032 | 1245.998286 | ENSG00000234127 |
| MTX2 | 0.07752235 | 1.055204301 | 0.348596475 | 0.560514639 | 814.5545506 | 859.4160687 | ENSG00000128654 |
| ZNF320 | 0.07751543 | 1.05519924 | 0.380262704 | 0.592611089 | 724.8873766 | 764.9214332 | ENSG00000182986 |
| RECQL4 | 0.077368777 | 1.055091982 | 0.353957298 | 0.566010113 | 1959.746822 | 2068.553294 | ENSG00000160957 |
| TBL1XR1 | 0.077340274 | 1.055071136 | 0.429763249 | 0.639615333 | 668.6933429 | 705.7432637 | ENSG00000177565 |
| VARS1 | 0.077324624 | 1.055059692 | 0.130913821 | 0.290483786 | 7013.813027 | 7400.188301 | ENSG00000204394 |
| RACGAP1 | 0.077317114 | 1.055054199 | 0.297572562 | 0.507812023 | 1138.284099 | 1200.632761 | ENSG00000161800 |
| PCOLCE | 0.077283279 | 1.055029456 | 0.520989405 | 0.715971832 | 453.146738 | 478.1683554 | ENSG00000106333 |
| GJD3 | 0.077165483 | 1.054943316 | 0.842527851 | 0.92165416 | 30.25112947 | 31.87062475 | ENSG00000183153 |
| SHROOM3 | 0.077135482 | 1.054921379 | 0.379889726 | 0.59241652 | 1295.496173 | 1366.286865 | ENSG00000138771 |
| DSE | 0.077057394 | 1.054864282 | 0.315882643 | 0.525914107 | 1077.210397 | 1136.17045 | ENSG00000111817 |
| PTCH1 | 0.077006154 | 1.054826816 | 0.531753705 | 0.723460345 | 351.014644 | 370.000357 | ENSG00000185920 |
| ABTB2 | 0.076990856 | 1.054815631 | 0.467759552 | 0.671249175 | 651.6474436 | 687.5183106 | ENSG00000166016 |
| GPX4 | 0.076970912 | 1.054801049 | 0.126272444 | 0.282905877 | 4032.137788 | 4253.405838 | ENSG00000167468 |
| GJB3 | 0.076915179 | 1.054760302 | 0.892877559 | 0.952281345 | 12.9224088 | 13.63143927 | ENSG00000188910 |
| EPC2 | 0.076906892 | 1.054754243 | 0.488439787 | 0.688783257 | 430.9790028 | 454.5022853 | ENSG00000135999 |
| ZNF224 | 0.076878671 | 1.054733612 | 0.782147253 | 0.889060484 | 59.77219207 | 62.94516088 | ENSG00000267680 |
| SAMD14 | 0.076800471 | 1.054676442 | 0.694689128 | 0.837133343 | 121.6424218 | 128.4382269 | ENSG00000167100 |
| CMTR2 | 0.076664489 | 1.054577038 | 0.673091719 | 0.824962394 | 183.5927144 | 193.3578389 | ENSG00000180917 |
| TBC1D8 | 0.076610983 | 1.054537927 | 0.424356452 | 0.634226465 | 701.3882602 | 739.4334935 | ENSG00000204634 |
| NA | 0.076543945 | 1.054488927 | 0.818006621 | 0.907321452 | 56.12500708 | 59.25994345 | ENSG00000229152 |
| PIGBOS1 | 0.076427713 | 1.054403974 | 0.841874839 | 0.92140286 | 30.540258 | 32.15975291 | ENSG00000225973 |
| SYT10 | 0.0763586 | 1.054353464 | 0.875441119 | 0.941819843 | 26.10338542 | 27.42601803 | ENSG00000110975 |
| NDUFB4 | 0.076312377 | 1.054319683 | 0.26788218 | 0.475733964 | 1633.262184 | 1722.25225 | ENSG00000065518 |
| NA | 0.076215831 | 1.05424913 | 0.94724877 | 0.976838866 | 4.100581182 | 4.279080851 | ENSG00000270804 |
| IMPAD1 | 0.076103429 | 1.054166995 | 0.477738224 | 0.680244768 | 655.6090301 | 691.4789909 | ENSG00000104331 |

| | | | | | | | |
|---|---|---|---|---|---|---|---|
| MIER3 | 0.076074322 | 1.054145727 | 0.570360663 | 0.75319722 | 377.1696676 | 397.3582505 | ENSG00000155545 |
| POLR2I | 0.075991525 | 1.054085231 | 0.831639276 | 0.915296977 | 48.15218347 | 50.78749815 | ENSG00000105258 |
| SHLD2 | 0.075962991 | 1.054064383 | 0.474965042 | 0.677782852 | 577.173377 | 608.6763844 | ENSG00000122376 |
| UBE2H | 0.075959614 | 1.054061916 | 0.246718973 | 0.451612658 | 2277.952544 | 2401.18773 | ENSG00000186591 |
| TTC23 | 0.075821644 | 1.053961118 | 0.557696236 | 0.743634359 | 288.5412926 | 304.0288225 | ENSG00000103852 |
| NA | 0.075821598 | 1.053961084 | 0.938746071 | 0.972431174 | 5.91482984 | 6.226764092 | ENSG00000189014 |
| ZNF547 | 0.075800925 | 1.053945981 | 0.893132496 | 0.952281345 | 13.24949632 | 13.96880649 | ENSG00000152433 |
| KHNYN | 0.075796303 | 1.053942605 | 0.460000474 | 0.664863689 | 649.1733244 | 683.6248133 | ENSG00000100441 |
| MTERF3 | 0.075792573 | 1.053939879 | 0.476739092 | 0.679283066 | 492.9210069 | 519.5841566 | ENSG00000156469 |
| JRKL | 0.075698808 | 1.053871383 | 0.699742076 | 0.839984052 | 133.1633762 | 140.380203 | ENSG00000183340 |
| HDAC8 | 0.075680641 | 1.053858112 | 0.78870647 | 0.891842679 | 64.56371207 | 67.90143518 | ENSG00000147099 |
| NCOR2 | 0.075622718 | 1.053815802 | 0.223335686 | 0.421263925 | 6294.160653 | 6633.399927 | ENSG00000196498 |
| GRINA | 0.075526232 | 1.053745326 | 0.32365525 | 0.534472665 | 1049.257525 | 1105.732037 | ENSG00000178719 |
| COX7A2 | 0.075494972 | 1.053722494 | 0.360858938 | 0.572914758 | 2039.154984 | 2148.301243 | ENSG00000112695 |
| ARHGAP23 | 0.07546361 | 1.053699588 | 0.248855114 | 0.453807341 | 2056.737958 | 2167.941109 | ENSG00000275832 |
| SLC18B1 | 0.075306334 | 1.053584725 | 0.443830678 | 0.651211433 | 687.8548958 | 724.3600887 | ENSG00000146409 |
| SSC4D | 0.075290586 | 1.053573224 | 0.555026917 | 0.741479028 | 285.7256197 | 300.9684194 | ENSG00000146700 |
| RRP9 | 0.075282397 | 1.053567244 | 0.256173567 | 0.463083931 | 2101.311678 | 2214.293939 | ENSG00000114767 |
| PPP2R3C | 0.075235798 | 1.053533214 | 0.647125133 | 0.808497522 | 192.4181271 | 202.8881136 | ENSG00000092020 |
| EPB41L2 | 0.07522674 | 1.0535266 | 0.168550745 | 0.346212836 | 3037.788551 | 3199.881325 | ENSG00000079819 |
| B3GLCT | 0.075166387 | 1.053482528 | 0.706165262 | 0.844420374 | 127.9163483 | 134.8300224 | ENSG00000187676 |
| TRIP4 | 0.074988195 | 1.053352417 | 0.619804531 | 0.788853523 | 301.9266075 | 317.9988288 | ENSG00000103671 |
| ZNF385B | 0.074939617 | 1.053316949 | 0.823739316 | 0.910791393 | 44.23783776 | 46.55275694 | ENSG00000144331 |
| DVL1 | 0.074903827 | 1.053290819 | 0.281999266 | 0.49092516 | 2267.137987 | 2388.690435 | ENSG00000107404 |
| ANKRD10 | 0.074799602 | 1.053214728 | 0.346103453 | 0.558121741 | 1338.35475 | 1409.779213 | ENSG00000088448 |
| PTOV1-AS1 | 0.074421019 | 1.052938387 | 0.899976243 | 0.95585464 | 14.15603612 | 14.9249132 | ENSG00000268006 |
| SP3 | 0.074393118 | 1.052918024 | 0.322747044 | 0.533532731 | 1293.554472 | 1362.326193 | ENSG00000172845 |
| NA | 0.074362178 | 1.052895443 | 0.150522819 | 0.319683247 | 11007.86461 | 11589.91473 | ENSG00000224078 |
| NRARP | 0.074313584 | 1.052859979 | 0.507189602 | 0.704350311 | 750.8660848 | 790.5400092 | ENSG00000198435 |
| GPBP1L1 | 0.074298499 | 1.052848971 | 0.286423857 | 0.495567068 | 1806.278496 | 1901.665489 | ENSG00000159592 |
| MRPL14 | 0.074290645 | 1.052843239 | 0.317728633 | 0.527777749 | 1663.903477 | 1751.861137 | ENSG00000180992 |
| IP6K2 | 0.074222979 | 1.052793859 | 0.281119836 | 0.490071827 | 1355.859054 | 1427.374035 | ENSG00000068745 |
| NA | 0.074178255 | 1.052761223 | 0.818344405 | 0.907536208 | 47.78624724 | 50.18850579 | ENSG00000275964 |
| NINJ1 | 0.074136391 | 1.052730674 | 0.46340274 | 0.667971431 | 529.3431953 | 557.5477055 | ENSG00000131669 |
| LRRC8A | 0.074084724 | 1.052692973 | 0.202487844 | 0.393819045 | 3954.063907 | 4162.88218 | ENSG00000136802 |
| RPL32 | 0.074040404 | 1.052660635 | 0.19018667 | 0.376548176 | 8610.626675 | 9063.802717 | ENSG00000144713 |
| DTX3 | 0.073765485 | 1.05246006 | 0.522248469 | 0.716804556 | 806.8516098 | 849.3628312 | ENSG00000178498 |
| PSME1 | 0.073687968 | 1.052403512 | 0.424825328 | 0.634767491 | 788.003641 | 828.9402414 | ENSG00000092010 |
| CBFA2T2 | 0.073678731 | 1.052396774 | 0.446063975 | 0.653422136 | 532.7264212 | 560.6801098 | ENSG00000078699 |
| OPTN | 0.073677759 | 1.052396065 | 0.603230354 | 0.77799629 | 250.1266287 | 263.117711 | ENSG00000123240 |
| HCN2 | 0.073677293 | 1.052395725 | 0.735806015 | 0.861168893 | 100.0971581 | 105.4768462 | ENSG00000099822 |
| PHETA1 | 0.073646124 | 1.052372988 | 0.481278867 | 0.683198091 | 484.9386733 | 510.6042563 | ENSG00000198324 |
| TRMT2A | 0.073603446 | 1.052341857 | 0.424362489 | 0.634226465 | 671.5454223 | 707.0103391 | ENSG00000099899 |
| BET1 | 0.073584422 | 1.052327981 | 0.531383629 | 0.723253105 | 394.346273 | 414.8417579 | ENSG00000105829 |
| MBD5 | 0.073565051 | 1.052313851 | 0.667666562 | 0.821869982 | 159.3548581 | 167.8023217 | ENSG00000204406 |
| LTBP3 | 0.073550107 | 1.052302951 | 0.567550534 | 0.751047019 | 300.0640264 | 315.8322517 | ENSG00000168056 |
| PPM1A | 0.073510606 | 1.052274139 | 0.403828505 | 0.615073902 | 893.3570814 | 940.3238023 | ENSG00000100614 |
| H2AZ2 | 0.073482347 | 1.052253528 | 0.213784918 | 0.408320829 | 2979.363427 | 3135.051184 | ENSG00000105968 |
| PLEKHA8 | 0.073346186 | 1.052154221 | 0.409452639 | 0.620487328 | 961.687117 | 1011.367151 | ENSG00000106086 |
| STXBP6 | 0.073320373 | 1.052135397 | 0.680974062 | 0.829598484 | 153.4246213 | 161.5326154 | ENSG00000168952 |
| ZNRF2 | 0.073297117 | 1.052118436 | 0.676568675 | 0.827022925 | 223.7913712 | 235.4056003 | ENSG00000180233 |
| SCMH1 | 0.073285661 | 1.052110082 | 0.525456251 | 0.718994605 | 427.8291497 | 450.1580788 | ENSG00000010803 |
| ARL13B | 0.073256993 | 1.052089175 | 0.718477071 | 0.851388915 | 118.7253043 | 124.8444742 | ENSG00000169379 |
| C1orf50 | 0.073113223 | 1.051984336 | 0.835018256 | 0.917376243 | 43.12134139 | 45.28924504 | ENSG00000164008 |
| ZNF766 | 0.073085442 | 1.051964079 | 0.48272463 | 0.683859246 | 512.0262704 | 538.582804 | ENSG00000196214 |
| TXNDC9 | 0.073062918 | 1.051947655 | 0.577918283 | 0.759450456 | 284.7677728 | 299.5678736 | ENSG00000115514 |
| TTI2 | 0.07297059 | 1.051880336 | 0.573946366 | 0.756172536 | 355.7205358 | 374.0206708 | ENSG00000129696 |
| NA | 0.07294227 | 1.051859657 | 0.895989367 | 0.953604554 | 13.57658385 | 14.28035169 | ENSG00000271780 |
| C2orf72 | 0.072870207 | 1.051807149 | 0.801951841 | 0.898712919 | 54.43770423 | 57.26714257 | ENSG00000204128 |
| NA | 0.072851472 | 1.051793489 | 0.861120037 | 0.932648138 | 29.16905877 | 30.71184134 | ENSG00000274925 |
| ZNF83 | 0.072797346 | 1.05175403 | 0.648190397 | 0.809119125 | 211.2431611 | 222.1632582 | ENSG00000167766 |
| AGO4 | 0.072772828 | 1.051736156 | 0.594702367 | 0.771674743 | 283.0514099 | 297.5115328 | ENSG00000134698 |
| NA | 0.072670544 | 1.051661593 | 0.788445698 | 0.891711403 | 69.76090675 | 73.52054802 | ENSG00000204860 |
| RNF169 | 0.072586854 | 1.051600588 | 0.650759649 | 0.810498975 | 201.9545629 | 212.0738846 | ENSG00000166439 |
| FAM27E3 | 0.072506777 | 1.05154222 | 0.925804003 | 0.966116406 | 6.936415663 | 7.297036182 | ENSG00000274026 |
| RGS11 | 0.072504516 | 1.051540573 | 0.876396017 | 0.94241765 | 23.76368696 | 25.0444168 | ENSG00000076344 |
| NA | 0.072435552 | 1.051490308 | 0.848609582 | 0.925021181 | 32.80139489 | 34.46220708 | ENSG00000226360 |
| GNB1 | 0.07237278 | 1.051444558 | 0.140253137 | 0.304411044 | 10511.78693 | 11051.53962 | ENSG00000078369 |
| LOC100996842 | 0.072254055 | 1.051358034 | 0.698935458 | 0.839656243 | 130.5452014 | 137.2613031 | ENSG00000233223 |
| TWF2 | 0.072198502 | 1.051317551 | 0.433399894 | 0.642497135 | 741.2947348 | 779.4467413 | ENSG00000247596 |
| SLC37A1 | 0.072146983 | 1.051280009 | 0.4103242 | 0.621309535 | 759.6964269 | 798.8208433 | ENSG00000160190 |
| TPPP3 | 0.072062703 | 1.051218597 | 0.922667456 | 0.965083075 | 7.896321398 | 8.324082547 | ENSG00000159713 |
| STAM-AS1 | 0.072043312 | 1.051204467 | 0.930670815 | 0.96853421 | 6.980298388 | 7.337802897 | ENSG00000260589 |
| MTRF1L | 0.071950305 | 1.051136701 | 0.582684987 | 0.762472699 | 294.5281754 | 309.5717714 | ENSG00000112031 |

| | | | | | | | |
|---|---|---|---|---|---|---|---|
| DUSP5 | 0.071908839 | 1.05110649 | 0.387870885 | 0.600150584 | 904.1731395 | 949.9985309 | ENSG00000138166 |
| CAMK1D | 0.071764082 | 1.051001029 | 0.535812483 | 0.726437264 | 368.5198632 | 387.5055237 | ENSG00000183049 |
| MAK16 | 0.071691337 | 1.050948036 | 0.532705608 | 0.724168256 | 431.8642114 | 454.2628366 | ENSG00000198042 |
| TUT4 | 0.071519413 | 1.050822803 | 0.396997819 | 0.608843902 | 912.5179004 | 959.2586465 | ENSG00000134744 |
| PPP1R12C | 0.071492448 | 1.050803163 | 0.388544106 | 0.60048837 | 1795.044992 | 1887.084535 | ENSG00000125503 |
| BRAP | 0.071461962 | 1.050780958 | 0.506648592 | 0.703844245 | 562.4022689 | 590.6028927 | ENSG00000089234 |
| NRGN | 0.071439988 | 1.050764954 | 0.924277299 | 0.96564377 | 7.899880871 | 8.321907083 | ENSG00000154146 |
| TGS1 | 0.071393416 | 1.050731034 | 0.388624244 | 0.60048837 | 1003.81273 | 1054.630344 | ENSG00000137574 |
| FRYL | 0.071284972 | 1.050652056 | 0.552562215 | 0.739346566 | 534.3235432 | 560.7705427 | ENSG00000075539 |
| GNAO1 | 0.071147842 | 1.050552195 | 0.45554947 | 0.661432233 | 1083.28459 | 1138.281661 | ENSG00000087258 |
| KHDC4 | 0.071142652 | 1.050548416 | 0.71407329 | 0.849104018 | 1260.479304 | 1324.291438 | ENSG00000132680 |
| CKMT2-AS1 | 0.071139383 | 1.050546035 | 0.734539055 | 0.860531641 | 119.3809539 | 125.2926019 | ENSG00000247572 |
| NA | 0.071086673 | 1.050507654 | 0.919077447 | 0.964051188 | 12.65316256 | 13.3727676 | ENSG00000232600 |
| DAD1 | 0.071086203 | 1.050507311 | 0.300608089 | 0.510754566 | 1377.572756 | 1446.809962 | ENSG00000129562 |
| ASB1 | 0.071001636 | 1.050445736 | 0.391217168 | 0.602572076 | 828.9677061 | 870.5303 | ENSG00000065802 |
| ZBTB45 | 0.070944873 | 1.050404406 | 0.497603799 | 0.696660074 | 546.8252259 | 574.8325809 | ENSG00000119574 |
| NA | 0.070836494 | 1.0503255 | 0.924564389 | 0.965883673 | 7.85836241 | 8.274613977 | ENSG00000225345 |
| WDR49 | 0.070827552 | 1.05031899 | 0.930949965 | 0.968756341 | 6.343920703 | 6.645948291 | ENSG00000174776 |
| FAM118A | 0.07068929 | 1.050218337 | 0.590488602 | 0.768530555 | 660.5022747 | 694.0814568 | ENSG00000100376 |
| HIC2 | 0.070638841 | 1.050181613 | 0.273355164 | 0.482053626 | 1599.921686 | 1680.079267 | ENSG00000169635 |
| FNTA | 0.070549103 | 1.050116292 | 0.5134119 | 0.709586046 | 457.481886 | 480.6683347 | ENSG00000168522 |
| MASP2 | 0.070511921 | 1.050089227 | 0.903528467 | 0.957906355 | 13.91434979 | 14.6080711 | ENSG00000009724 |
| TESC | 0.070430766 | 1.050030159 | 0.680991894 | 0.829598484 | 182.2523547 | 191.5848779 | ENSG00000088992 |
| NA | 0.070353087 | 1.049973624 | 0.883438479 | 0.946765652 | 21.68967527 | 22.78809887 | ENSG00000267023 |
| SERHL2 | 0.070263185 | 1.049908197 | 0.820541636 | 0.908947936 | 53.20527211 | 55.80908873 | ENSG00000183569 |
| SCAMP1-AS1 | 0.070209006 | 1.049868769 | 0.800554107 | 0.898320559 | 60.38365344 | 63.32641624 | ENSG00000245556 |
| SPRED3 | 0.070149959 | 1.049825801 | 0.712355582 | 0.848103373 | 129.4040942 | 135.8848983 | ENSG00000188766 |
| SBF1 | 0.070040966 | 1.049746491 | 0.210625439 | 0.40411068 | 3239.574656 | 3401.114332 | ENSG00000100241 |
| GCFC2 | 0.070015725 | 1.049728126 | 0.719995564 | 0.852430332 | 126.0878879 | 132.4673379 | ENSG00000005436 |
| GRM2 | 0.069936575 | 1.049670536 | 0.69867855 | 0.839454411 | 135.6543258 | 142.4058038 | ENSG00000164082 |
| MBOAT7 | 0.069914113 | 1.049654193 | 0.301350766 | 0.511505412 | 1935.334021 | 2031.931313 | ENSG00000125505 |
| FAM104A | 0.06987866 | 1.049628399 | 0.573698783 | 0.755930071 | 374.4359144 | 393.1108784 | ENSG00000133193 |
| EPHA1 | 0.069869599 | 1.049621807 | 0.168592097 | 0.346260114 | 4739.346098 | 4974.378737 | ENSG00000146904 |
| LOC100506302 | 0.069868985 | 1.04962136 | 0.887054278 | 0.949069176 | 19.99642253 | 20.95959435 | ENSG00000216895 |
| CTNNA1 | 0.069828221 | 1.049591703 | 0.134073221 | 0.295418971 | 8759.581434 | 9193.478897 | ENSG00000044115 |
| NUAK1 | 0.069821768 | 1.049587009 | 0.476398054 | 0.678899581 | 542.5274265 | 569.3330257 | ENSG00000074590 |
| DNAJC5 | 0.06967029 | 1.049476811 | 0.328576082 | 0.540048678 | 1801.383471 | 1890.14679 | ENSG00000101152 |
| TMEM92 | 0.069669418 | 1.049476177 | 0.59366065 | 0.770905685 | 271.2683021 | 284.6930914 | ENSG00000167105 |
| DIMT1 | 0.069580903 | 1.049411789 | 0.424327878 | 0.634226465 | 1097.250831 | 1151.957714 | ENSG00000086189 |
| FASTKD2 | 0.069349171 | 1.049243242 | 0.404484149 | 0.615476848 | 994.3913928 | 1043.614272 | ENSG00000118246 |
| MLXIP | 0.069322109 | 1.04922356 | 0.350008649 | 0.562160261 | 1490.781079 | 1564.385094 | ENSG00000175727 |
| NA | 0.06925005 | 1.049171156 | 0.894929864 | 0.953032171 | 16.09244975 | 16.92329451 | ENSG00000287016 |
| MGAT5 | 0.069028485 | 1.04901004 | 0.387149024 | 0.599336775 | 1233.973777 | 1293.898606 | ENSG00000152127 |
| STX3 | 0.06901878 | 1.049002983 | 0.295181808 | 0.505268741 | 1598.819175 | 1676.726927 | ENSG00000166900 |
| PLXNC1 | 0.069015606 | 1.049000675 | 0.804995259 | 0.900665647 | 63.39001096 | 66.47733812 | ENSG00000136040 |
| RPL27 | 0.069013389 | 1.048999063 | 0.277120465 | 0.486332123 | 12694.44505 | 13315.85083 | ENSG00000131469 |
| MKRN2 | 0.06901189 | 1.048997973 | 0.479059539 | 0.68145793 | 532.0838143 | 558.2599899 | ENSG00000075975 |
| NA | 0.068990065 | 1.048982104 | 0.937307248 | 0.971904957 | 5.731264118 | 5.977740238 | ENSG00000287601 |
| PSMA1 | 0.068986422 | 1.048979455 | 0.727980639 | 0.856731016 | 121.6163365 | 127.7022006 | ENSG00000129084 |
| EIF2AK2 | 0.068976106 | 1.048971954 | 0.463996826 | 0.668542004 | 917.0976844 | 962.0539968 | ENSG00000055332 |
| SLC7A10 | 0.068888652 | 1.048908369 | 0.766862904 | 0.879325885 | 83.30723537 | 87.30519542 | ENSG00000130876 |
| C21orf59-TCP10L | 0.068711289 | 1.048779426 | 0.92292572 | 0.965130891 | 9.135872459 | 9.621907401 | ENSG00000265590 |
| SACM1L | 0.068688926 | 1.048763169 | 0.339316308 | 0.551111994 | 1405.691753 | 1474.021322 | ENSG00000211456 |
| RPL18A | 0.068686872 | 1.048761676 | 0.188650229 | 0.374568749 | 18356.55569 | 19251.86953 | ENSG00000105640 |
| EPB41L3 | 0.06858502 | 1.048687637 | 0.510681731 | 0.707210126 | 470.3060786 | 493.0455537 | ENSG00000082397 |
| POLR3G | 0.068578465 | 1.048682872 | 0.389105251 | 0.600936593 | 3651.5133 | 3829.706118 | ENSG00000113356 |
| RAB11FIP1 | 0.068577386 | 1.048682088 | 0.469490018 | 0.67271009 | 605.414488 | 634.6856527 | ENSG00000156675 |
| FAM66C | 0.068489182 | 1.048617976 | 0.856208524 | 0.929762405 | 29.74731584 | 31.20894309 | ENSG00000226711 |
| VPS9D1-AS1 | 0.068467097 | 1.048601923 | 0.773580232 | 0.883577034 | 88.5910526 | 93.05349098 | ENSG00000261373 |
| NA | 0.068466901 | 1.048601781 | 0.927750578 | 0.967350062 | 8.877583968 | 9.337413718 | ENSG00000286064 |
| KATNAL1 | 0.068461979 | 1.048598203 | 0.694032673 | 0.836669205 | 178.4948528 | 187.0086591 | ENSG00000102781 |
| NDEL1 | 0.068446135 | 1.048586687 | 0.48557874 | 0.686491579 | 634.7315172 | 665.6092085 | ENSG00000166579 |
| CHRDL1 | 0.068425139 | 1.048571427 | 0.49581981 | 0.695207823 | 784.9149354 | 822.5313851 | ENSG00000101938 |
| NA | 0.068411655 | 1.048561627 | 0.689520315 | 0.83398915 | 158.980887 | 166.6721981 | ENSG00000174028 |
| ZC3H4 | 0.068381218 | 1.048539505 | 0.401409391 | 0.612708101 | 1160.630696 | 1217.406987 | ENSG00000130749 |
| NA | 0.068373211 | 1.048533686 | 0.753368714 | 0.871421911 | 97.06235093 | 101.8264362 | ENSG00000272645 |
| TXLNA | 0.068323378 | 1.048497468 | 0.344876025 | 0.556903268 | 1760.416023 | 1845.114329 | ENSG00000084652 |
| CORO7 | 0.068300053 | 1.048480517 | 0.825776593 | 0.912242817 | 51.7857148 | 54.30132573 | ENSG00000262246 |
| ZNF75D | 0.068237114 | 1.048434776 | 0.708030306 | 0.845910689 | 141.7162369 | 148.5015659 | ENSG00000186376 |
| AURKAIP1 | 0.068234689 | 1.048433014 | 0.321068992 | 0.531456561 | 2801.835856 | 2938.433722 | ENSG00000175756 |
| NA | 0.068030463 | 1.04828461 | 0.882443694 | 0.945904709 | 21.18970045 | 22.19707332 | ENSG00000271533 |
| COL4A5 | 0.067977144 | 1.048245869 | 0.448837425 | 0.655430939 | 892.6607786 | 935.6792021 | ENSG00000188153 |
| BCAM | 0.067925189 | 1.048208119 | 0.321781792 | 0.532309841 | 1824.484109 | 1913.149247 | ENSG00000187244 |
| MASTL | 0.067791494 | 1.048110986 | 0.370613295 | 0.582864097 | 1041.04121 | 1090.89017 | ENSG00000120539 |

| | | | | | | | |
|---|---|---|---|---|---|---|---|
| LINC00237 | 0.067766387 | 1.048092746 | 0.920747215 | 0.964633548 | 11.76469945 | 12.28102442 | ENSG00000225127 |
| VPS37B | 0.067579585 | 1.047957047 | 0.666155369 | 0.821155848 | 225.8259493 | 236.8527245 | ENSG00000139722 |
| UGGT1 | 0.067555908 | 1.047939848 | 0.367579061 | 0.579733637 | 1907.522177 | 1998.889742 | ENSG00000136731 |
| GKAP1 | 0.067550896 | 1.047936207 | 0.766179376 | 0.878994404 | 94.37031758 | 98.72760986 | ENSG00000165113 |
| METTL15 | 0.067475644 | 1.047881548 | 0.623117313 | 0.7914164 | 295.1072705 | 309.0009874 | ENSG00000169519 |
| NPTN-IT1 | 0.067442795 | 1.047857688 | 0.905008517 | 0.958611579 | 13.96535146 | 14.61554345 | ENSG00000281183 |
| ZC3H7A | 0.067433114 | 1.047850657 | 0.420109544 | 0.630464776 | 916.4304411 | 960.1119166 | ENSG00000122299 |
| MGA | 0.067411242 | 1.047834771 | 0.499083665 | 0.697608609 | 991.8150878 | 1038.648001 | ENSG00000174197 |
| VGLL4 | 0.067389643 | 1.047819084 | 0.544032631 | 0.73244858 | 595.6840095 | 623.8079666 | ENSG00000144560 |
| SPRYD7 | 0.067320109 | 1.047768583 | 0.739227894 | 0.863587169 | 123.1182942 | 129.0705136 | ENSG00000123178 |
| PRR12 | 0.067310326 | 1.047761478 | 0.368948505 | 0.581116973 | 2211.917896 | 2318.026546 | ENSG00000126464 |
| CCDC167 | 0.067283621 | 1.047742084 | 0.705481228 | 0.844050998 | 179.1303407 | 187.6842145 | ENSG00000198937 |
| ZSCAN2 | 0.067283341 | 1.04774188 | 0.600249318 | 0.776046304 | 433.2282411 | 453.358163 | ENSG00000176371 |
| NA | 0.067276955 | 1.047737243 | 0.908811545 | 0.960165884 | 12.99120783 | 13.57326884 | ENSG00000274114 |
| RNPC3 | 0.067224236 | 1.047698957 | 0.72492546 | 0.855050735 | 133.6292308 | 139.9930837 | ENSG00000185946 |
| ECE1 | 0.067181426 | 1.047667868 | 0.277833168 | 0.487039764 | 2092.449911 | 2192.707799 | ENSG00000117298 |
| SEPTIN11 | 0.067110687 | 1.0476165 | 0.294456273 | 0.504427315 | 2705.036168 | 2833.435862 | ENSG00000138758 |
| CGNL1 | 0.067036723 | 1.047562792 | 0.299523889 | 0.509733139 | 1963.236548 | 2056.035682 | ENSG00000128849 |
| NA | 0.067015928 | 1.047547692 | 0.863814933 | 0.934208281 | 42.06182273 | 44.1166738 | ENSG00000260261 |
| SREK1IP1 | 0.067009458 | 1.047542995 | 0.603565884 | 0.778237515 | 540.5305797 | 566.1754688 | ENSG00000153006 |
| FIGN | 0.06695779 | 1.047505479 | 0.636794587 | 0.800725077 | 254.8784358 | 266.8924583 | ENSG00000182263 |
| PGAP1 | 0.066929187 | 1.047484712 | 0.575110061 | 0.757335855 | 422.0163232 | 441.9233355 | ENSG00000197121 |
| RPS6KA5 | 0.066883152 | 1.047451287 | 0.857398275 | 0.93039751 | 33.98785935 | 35.53342513 | ENSG00000100784 |
| PLCB3 | 0.066852768 | 1.047429228 | 0.186969927 | 0.372303571 | 4885.272341 | 5117.743328 | ENSG00000149782 |
| U2SURP | 0.066756635 | 1.047359436 | 0.310115069 | 0.520717933 | 2393.792007 | 2506.979303 | ENSG00000163714 |
| USP11 | 0.066703433 | 1.047320813 | 0.174009232 | 0.35422705 | 5027.889449 | 5265.881411 | ENSG00000102226 |
| ZNF264 | 0.06657788 | 1.047229672 | 0.71399173 | 0.849101626 | 192.4646011 | 201.5260434 | ENSG00000083844 |
| UBE2V2 | 0.06657716 | 1.047229149 | 0.414319089 | 0.625302964 | 1257.694522 | 1317.575383 | ENSG00000169139 |
| GLYATL1 | 0.066473538 | 1.047153934 | 0.755546009 | 0.872903853 | 116.4238703 | 121.7398565 | ENSG00000166840 |
| RAB43 | 0.066451432 | 1.04713789 | 0.722525176 | 0.853712567 | 160.8962749 | 168.6529648 | ENSG00000172780 |
| UTP23 | 0.066357083 | 1.047069411 | 0.734733733 | 0.86059955 | 170.6599315 | 178.9549036 | ENSG00000147679 |
| VPS54 | 0.066342043 | 1.047058495 | 0.604123836 | 0.778603606 | 355.4637474 | 372.234614 | ENSG00000143952 |
| CARD6 | 0.066290291 | 1.047020937 | 0.937921022 | 0.972140333 | 6.942339399 | 7.259390889 | ENSG00000132357 |
| OSBPL8 | 0.065989383 | 1.046802579 | 0.56761684 | 0.751047019 | 1015.779578 | 1063.155597 | ENSG00000091039 |
| RPRD1B | 0.065964461 | 1.046784496 | 0.569589518 | 0.752634622 | 445.7491701 | 466.4013717 | ENSG00000101413 |
| BTBD6 | 0.065944117 | 1.046769735 | 0.661981821 | 0.818441256 | 242.6692124 | 253.9250442 | ENSG00000184887 |
| EIF3D | 0.065924491 | 1.046755495 | 0.201604924 | 0.392587026 | 4154.535012 | 4348.280484 | ENSG00000100353 |
| CBWD2 | 0.065820499 | 1.046680046 | 0.524920718 | 0.718550866 | 528.5003426 | 553.1248534 | ENSG00000136682 |
| NIPAL2 | 0.065699781 | 1.046592468 | 0.847619937 | 0.924422644 | 39.92374054 | 41.82053511 | ENSG00000104361 |
| ZNF483 | 0.065656444 | 1.04656103 | 0.678101278 | 0.827792025 | 357.3654037 | 373.962909 | ENSG00000173258 |
| PDLIM4 | 0.065654167 | 1.046559377 | 0.691825336 | 0.835312414 | 256.0963507 | 268.2301696 | ENSG00000131435 |
| NA | 0.06564388 | 1.046551915 | 0.932585665 | 0.969708285 | 8.592014906 | 9.007518845 | ENSG00000278192 |
| TBC1D22A | 0.065535861 | 1.04647356 | 0.519174813 | 0.714901185 | 481.921969 | 504.3468248 | ENSG00000054611 |
| STAM | 0.06540242 | 1.046376772 | 0.569920087 | 0.752843159 | 389.681952 | 407.6264661 | ENSG00000136738 |
| UXT | 0.06529219 | 1.046296826 | 0.775453757 | 0.884700308 | 115.9728378 | 121.5536805 | ENSG00000126756 |
| PCNP | 0.065267523 | 1.046278936 | 0.228244158 | 0.427618147 | 3086.580671 | 3229.05261 | ENSG00000081154 |
| PELP1 | 0.065264401 | 1.046276672 | 0.182566508 | 0.366160632 | 5812.993793 | 6082.468717 | ENSG00000141456 |
| SLC17A7 | 0.06521071 | 1.046237735 | 0.814025467 | 0.905404967 | 62.3165076 | 65.20828875 | ENSG00000104888 |
| NA | 0.065160246 | 1.046201139 | 0.927805755 | 0.967350062 | 8.587260225 | 9.000992454 | ENSG00000269399 |
| WWC2 | 0.065159684 | 1.046200731 | 0.300153409 | 0.51025269 | 2478.642641 | 2592.494785 | ENSG00000151718 |
| NA | 0.06515941 | 1.046200533 | 0.934646739 | 0.970489863 | 7.620235545 | 7.970541121 | ENSG00000257907 |
| GFRA3 | 0.065136071 | 1.046183608 | 0.909699529 | 0.960563356 | 14.29243898 | 14.93238554 | ENSG00000146013 |
| ATP11A | 0.065107391 | 1.046162811 | 0.37949888 | 0.592055873 | 1546.513071 | 1617.479404 | ENSG00000068650 |
| PPP2R3B | 0.065057659 | 1.046126749 | 0.404826925 | 0.615849564 | 926.8767224 | 969.8065306 | ENSG00000167393 |
| GTF2IRD1 | 0.065031626 | 1.046107871 | 0.420082441 | 0.630464776 | 957.9460414 | 1002.289988 | ENSG00000006704 |
| TTC19 | 0.065022336 | 1.046101136 | 0.300222453 | 0.510284964 | 1917.254864 | 2005.682501 | ENSG00000011295 |
| RERE | 0.064813291 | 1.045949567 | 0.282231366 | 0.491283929 | 2611.080249 | 2731.274384 | ENSG00000142599 |
| RFX3-AS1 | 0.064777867 | 1.045923886 | 0.947333555 | 0.976838866 | 6.365277538 | 6.6394219 | ENSG00000232104 |
| IDH3B | 0.064761238 | 1.04591183 | 0.312831593 | 0.523140408 | 1942.466863 | 2032.020711 | ENSG00000101365 |
| VTI1A | 0.064650028 | 1.045831209 | 0.65246345 | 0.811703295 | 252.5526959 | 264.3324894 | ENSG00000151532 |
| NA | 0.064574728 | 1.045776624 | 0.884141749 | 0.947149327 | 23.0905455 | 24.14816153 | ENSG00000235238 |
| NEU1 | 0.064570083 | 1.045773257 | 0.398577563 | 0.610324922 | 968.7902169 | 1012.908601 | ENSG00000204386 |
| C11orf87 | 0.064537649 | 1.045749747 | 0.903876823 | 0.95821903 | 15.78908332 | 16.52028452 | ENSG00000185742 |
| S1PR5 | 0.064499888 | 1.045722376 | 0.642817279 | 0.804960169 | 275.7783968 | 288.4605773 | ENSG00000180739 |
| ZSCAN29 | 0.064473541 | 1.045703279 | 0.440043052 | 0.647767794 | 802.9352047 | 839.5749426 | ENSG00000140265 |
| ZNF396 | 0.064430911 | 1.04567238 | 0.923937876 | 0.965502811 | 10.42522998 | 10.95085116 | ENSG00000186496 |
| TROAP | 0.064421064 | 1.045665243 | 0.529096101 | 0.721546784 | 687.0216821 | 718.8161547 | ENSG00000135451 |
| DGCR2 | 0.064359021 | 1.045620275 | 0.25035818 | 0.455931941 | 3028.559904 | 3166.929195 | ENSG00000070413 |
| SND1 | 0.064332745 | 1.045601231 | 0.138745563 | 0.302180828 | 9008.296817 | 9418.962884 | ENSG00000197157 |
| RUFY1 | 0.064233095 | 1.045529012 | 0.516089076 | 0.71230543 | 564.2164914 | 589.6046219 | ENSG00000176783 |
| CPLANE1 | 0.064225466 | 1.045523483 | 0.747450052 | 0.868598501 | 122.6992713 | 128.1052106 | ENSG00000197603 |
| EFNA2 | 0.064153696 | 1.045471472 | 0.609156464 | 0.782326193 | 320.3362554 | 335.0355833 | ENSG00000099617 |
| BDNF-AS | 0.063969269 | 1.045337833 | 0.897861422 | 0.95489739 | 18.3808672 | 19.26746136 | ENSG00000245573 |
| ACSL1 | 0.063926891 | 1.045307128 | 0.612985087 | 0.78462019 | 312.3213024 | 326.5861745 | ENSG00000151726 |

| | | | | | | | |
|---|---|---|---|---|---|---|---|
| MIR600HG | 0.063854982 | 1.045255027 | 0.658523399 | 0.815917491 | 374.9105634 | 391.7236486 | ENSG00000236901 |
| CACNA1D | 0.063852238 | 1.045253039 | 0.881818424 | 0.945689642 | 25.67282002 | 26.87386728 | ENSG00000157388 |
| GRTP1 | 0.063798501 | 1.045214106 | 0.448858265 | 0.655430939 | 1129.290021 | 1180.587354 | ENSG00000139835 |
| UBN2 | 0.063762578 | 1.045188081 | 0.599060084 | 0.77503937 | 434.8020498 | 454.3151727 | ENSG00000157741 |
| GPR137B | 0.06372676 | 1.045162132 | 0.753335305 | 0.871421911 | 120.6060139 | 125.9680713 | ENSG00000077585 |
| MVD | 0.063695892 | 1.045139771 | 0.364692665 | 0.576819177 | 2542.539463 | 2657.975003 | ENSG00000167508 |
| MFSD6L | 0.063688834 | 1.045134657 | 0.874498683 | 0.941288551 | 29.59578537 | 30.89645193 | ENSG00000185156 |
| NPY1R | 0.063679085 | 1.045127594 | 0.758822678 | 0.874822186 | 119.4124043 | 124.8202606 | ENSG00000164128 |
| NA | 0.063570177 | 1.045048702 | 0.866862302 | 0.936573648 | 34.77429296 | 36.39995896 | ENSG00000180015 |
| FAM50A | 0.063534013 | 1.045022506 | 0.370949045 | 0.583100776 | 1322.79192 | 1382.379498 | ENSG00000071859 |
| FNDC3B | 0.063528861 | 1.045018774 | 0.538965004 | 0.728690872 | 700.5563642 | 731.8606522 | ENSG00000075420 |
| KATNA1 | 0.063525822 | 1.045016573 | 0.656079464 | 0.814171421 | 274.932979 | 287.1802514 | ENSG00000186625 |
| TIFA | 0.063418425 | 1.044938783 | 0.843189905 | 0.921933845 | 44.11656252 | 46.18767578 | ENSG00000145365 |
| TBC1D10B | 0.06338561 | 1.044915015 | 0.453173662 | 0.659387056 | 1022.278492 | 1068.636462 | ENSG00000169221 |
| TUT7 | 0.063313139 | 1.044862527 | 0.588680739 | 0.766705914 | 395.5523149 | 413.0498801 | ENSG00000083223 |
| C11orf80 | 0.063133412 | 1.044732369 | 0.684143499 | 0.831079275 | 210.5913764 | 219.9368297 | ENSG00000173715 |
| NGLY1 | 0.063098961 | 1.044707421 | 0.561099575 | 0.746222921 | 416.033584 | 434.6809922 | ENSG00000151092 |
| ATPAF1 | 0.063089672 | 1.044700695 | 0.377111303 | 0.589304672 | 1348.896578 | 1409.601188 | ENSG00000123472 |
| SRRM3 | 0.063086905 | 1.044698692 | 0.587382587 | 0.765666704 | 454.2444944 | 474.8500157 | ENSG00000177679 |
| CAP1 | 0.062898808 | 1.044562494 | 0.3356128 | 0.547537071 | 3861.223375 | 4032.314652 | ENSG00000131236 |
| SNTB2 | 0.062829468 | 1.044512291 | 0.676412084 | 0.826926971 | 280.267493 | 292.4727361 | ENSG00000168807 |
| MCUB | 0.062769449 | 1.044468837 | 0.606258537 | 0.780183474 | 404.8901602 | 423.1843057 | ENSG00000005059 |
| NA | 0.062765196 | 1.044465758 | 0.947355247 | 0.976838866 | 6.44355978 | 6.706294178 | ENSG00000266903 |
| STAU2 | 0.062718971 | 1.044432293 | 0.524237163 | 0.718239903 | 735.2515636 | 767.6856645 | ENSG00000040341 |
| PDPK1 | 0.062691386 | 1.044412324 | 0.553228421 | 0.739863889 | 512.1627511 | 535.0383816 | ENSG00000140992 |
| N4BP2L2 | 0.062538747 | 1.044301829 | 0.46984572 | 0.673066556 | 739.5315407 | 772.1804881 | ENSG00000244754 |
| DNTTIP2 | 0.062537009 | 1.044300571 | 0.401210806 | 0.612569009 | 1628.842322 | 1701.391411 | ENSG00000067334 |
| MTOR | 0.062530125 | 1.044295588 | 0.27346357 | 0.482117812 | 3850.108609 | 4019.824982 | ENSG00000198793 |
| CUX1 | 0.062435827 | 1.044227332 | 0.421636818 | 0.631702603 | 1108.248598 | 1156.875368 | ENSG00000257923 |
| NA | 0.062395624 | 1.044198234 | 0.933134368 | 0.969789847 | 7.62735449 | 7.955596422 | ENSG00000287825 |
| EFCAB14 | 0.062343329 | 1.044160385 | 0.496586563 | 0.695700092 | 855.6119477 | 893.1853893 | ENSG00000159658 |
| BTN2A3P | 0.062273527 | 1.044109866 | 0.833181569 | 0.915922575 | 60.55237088 | 63.0344502 | ENSG00000124549 |
| UCHL5 | 0.062162371 | 1.044029423 | 0.524031463 | 0.718166487 | 727.9666774 | 759.7891845 | ENSG00000116750 |
| FAM204A | 0.062147983 | 1.044019011 | 0.714983774 | 0.849668508 | 179.5549818 | 187.4968911 | ENSG00000165669 |
| MEI4 | 0.062087153 | 1.043974992 | 0.935882099 | 0.971266332 | 8.823022826 | 9.225423797 | ENSG00000269964 |
| MORC3 | 0.062034284 | 1.043936735 | 0.641319807 | 0.803884545 | 307.9072873 | 321.3268459 | ENSG00000159256 |
| FBXO24 | 0.061992012 | 1.043906147 | 0.926861498 | 0.966739168 | 11.1304067 | 11.66635232 | ENSG00000106336 |
| AFTPH | 0.061917903 | 1.043852525 | 0.600181447 | 0.776046304 | 351.1309369 | 366.7054959 | ENSG00000119844 |
| CNEP1R1 | 0.061884636 | 1.043828455 | 0.699652471 | 0.839970842 | 205.2951007 | 214.0843727 | ENSG00000205423 |
| NA | 0.061748622 | 1.04373005 | 0.877405939 | 0.943020384 | 30.80330118 | 32.12333712 | ENSG00000128692 |
| CTDSPL | 0.061695996 | 1.043691978 | 0.447012246 | 0.653867333 | 923.074117 | 963.1174519 | ENSG00000144677 |
| LAMTOR4 | 0.061604798 | 1.043626004 | 0.455299445 | 0.66115571 | 1401.359729 | 1462.383538 | ENSG00000188186 |
| SLCO4A1 | 0.061472013 | 1.043529954 | 0.536241733 | 0.726726247 | 578.814355 | 604.2084576 | ENSG00000101187 |
| BRINP2 | 0.061462128 | 1.043522804 | 0.875372058 | 0.941799197 | 29.6728724 | 30.90392428 | ENSG00000198797 |
| ADAM15 | 0.061378432 | 1.043462267 | 0.466747887 | 0.670765835 | 936.6560647 | 977.9322147 | ENSG00000143537 |
| DAPK1 | 0.0612867 | 1.043395922 | 0.306910146 | 0.517222747 | 3253.234199 | 3394.237231 | ENSG00000196730 |
| UXS1 | 0.06124901 | 1.043368664 | 0.425636837 | 0.635611886 | 1040.848998 | 1085.863639 | ENSG00000115652 |
| SRPK2 | 0.06114239 | 1.043291558 | 0.427678551 | 0.637542812 | 1158.253521 | 1207.950962 | ENSG00000135250 |
| PGAP4 | 0.061134784 | 1.043286058 | 0.524850612 | 0.718550866 | 586.4008026 | 611.4538498 | ENSG00000165152 |
| ATP13A2 | 0.061096522 | 1.043258389 | 0.371847124 | 0.583850869 | 2280.633191 | 2379.680095 | ENSG00000159363 |
| ATP4A | 0.061058381 | 1.043230809 | 0.899751355 | 0.955743556 | 19.06824656 | 19.94124984 | ENSG00000105675 |
| SPTLC1 | 0.060932986 | 1.043140138 | 0.441903508 | 0.649544501 | 1094.836028 | 1141.963537 | ENSG00000090054 |
| DCTN2 | 0.060903752 | 1.043119 | 0.390785685 | 0.602231765 | 1767.79198 | 1843.230333 | ENSG00000175203 |
| TMCC3 | 0.060875843 | 1.043098822 | 0.816826458 | 0.906598158 | 74.61944635 | 77.70570507 | ENSG00000057704 |
| ANAPC16 | 0.060825965 | 1.04306276 | 0.537330838 | 0.727443977 | 585.4580571 | 610.6246051 | ENSG00000166295 |
| TTC5 | 0.060780652 | 1.043029999 | 0.660392588 | 0.817373256 | 263.6264042 | 275.0798858 | ENSG00000136319 |
| EIF1B | 0.060722255 | 1.04298778 | 0.461140007 | 0.665899145 | 1113.699196 | 1161.188804 | ENSG00000114784 |
| LYSMD4 | 0.060697222 | 1.042969683 | 0.617215112 | 0.787224317 | 390.7159701 | 407.3845787 | ENSG00000183060 |
| HACE1 | 0.060604593 | 1.04290272 | 0.613071513 | 0.78462019 | 365.758752 | 381.203469 | ENSG00000085382 |
| ST6GALNAC5 | 0.060551903 | 1.042864633 | 0.765050988 | 0.878469069 | 138.0580941 | 144.1640311 | ENSG00000117069 |
| LRRC55 | 0.06037969 | 1.042740154 | 0.802374996 | 0.899050996 | 97.04546944 | 101.2759665 | ENSG00000183908 |
| NDUFS1 | 0.060374124 | 1.042736131 | 0.397094657 | 0.608935474 | 2031.075383 | 2117.229959 | ENSG00000023228 |
| MRPS5 | 0.060324178 | 1.042700032 | 0.404080061 | 0.615208962 | 1159.220876 | 1208.786957 | ENSG00000144029 |
| SNUPN | 0.060321589 | 1.042698161 | 0.658829996 | 0.816144261 | 287.5413168 | 299.6351974 | ENSG00000169371 |
| WDR70 | 0.060316661 | 1.0426946 | 0.570347983 | 0.75319722 | 729.0068998 | 759.5017506 | ENSG00000082068 |
| NCBP3 | 0.060221768 | 1.042626019 | 0.37125432 | 0.58329264 | 1614.085375 | 1682.310192 | ENSG00000074356 |
| ZNF768 | 0.060193618 | 1.042605675 | 0.457083128 | 0.662777555 | 974.3975616 | 1016.154204 | ENSG00000169957 |
| NA | 0.060139993 | 1.042566923 | 0.828787487 | 0.914018435 | 71.79042533 | 74.68659287 | ENSG00000274317 |
| EZR | 0.059940059 | 1.04242245 | 0.260733878 | 0.467656881 | 6378.324434 | 6648.97487 | ENSG00000092820 |
| FBXO45 | 0.059900019 | 1.042393519 | 0.537879356 | 0.727796205 | 558.894793 | 582.5224848 | ENSG00000174013 |
| NCK1 | 0.059813582 | 1.042331067 | 0.676139365 | 0.826733047 | 268.8580507 | 280.4738619 | ENSG00000158092 |
| CHD8 | 0.059778625 | 1.042305811 | 0.325050717 | 0.53607395 | 3916.789299 | 4081.650108 | ENSG00000100888 |
| ZNF791 | 0.059609505 | 1.042183634 | 0.609601399 | 0.782504214 | 362.7660476 | 377.9162482 | ENSG00000173875 |
| CREB3 | 0.059559506 | 1.042147516 | 0.529885268 | 0.72210155 | 595.0708198 | 620.3039049 | ENSG00000107175 |

| | | | | | | | |
|---|---|---|---|---|---|---|---|
| SUGT1P4-STRA6LP | 0.059526605 | 1.04212375 | 0.937691917 | 0.972022827 | 8.295767427 | 8.642154143 | ENSG00000254876 |
| RAD51D | 0.059439873 | 1.042061102 | 0.734218497 | 0.860316207 | 166.5281798 | 173.3454815 | ENSG00000185379 |
| TAF15 | 0.059403917 | 1.042035131 | 0.235070807 | 0.436682293 | 10001.42357 | 10421.45036 | ENSG00000270647 |
| ZNF721 | 0.059395066 | 1.042028738 | 0.563780383 | 0.748513341 | 640.2267393 | 667.2607833 | ENSG00000182903 |
| PATL1 | 0.059381996 | 1.042019298 | 0.467090058 | 0.671002262 | 887.0315179 | 923.9218661 | ENSG00000166889 |
| CC2D2A | 0.059276723 | 1.041943265 | 0.680214637 | 0.829014226 | 242.6709924 | 252.774722 | ENSG00000048342 |
| XIAP | 0.059263558 | 1.041933757 | 0.541964123 | 0.731175343 | 731.0539602 | 761.8012083 | ENSG00000101966 |
| TTC9 | 0.059159172 | 1.04185837 | 0.438689099 | 0.646711102 | 1015.976317 | 1058.723367 | ENSG00000133985 |
| TUFM | 0.059132856 | 1.041839366 | 0.197638938 | 0.387340323 | 5668.673264 | 5905.840758 | ENSG00000178952 |
| ETV1 | 0.059054194 | 1.041782562 | 0.464108127 | 0.668593683 | 1369.363636 | 1426.501821 | ENSG00000006468 |
| LCMT2 | 0.058997664 | 1.041741742 | 0.636147031 | 0.80066251 | 307.1039199 | 319.7665656 | ENSG00000168806 |
| RPAIN | 0.05896185 | 1.041715882 | 0.584546353 | 0.763268885 | 540.057782 | 562.2790017 | ENSG00000129197 |
| SENP3 | 0.058958859 | 1.041713722 | 0.465015237 | 0.669294471 | 1055.378394 | 1099.522488 | ENSG00000161956 |
| SLC6A12 | 0.058942443 | 1.041701869 | 0.935987685 | 0.971322531 | 9.633762356 | 10.04856395 | ENSG00000111181 |
| SLC6A3 | 0.058923039 | 1.041687859 | 0.889147349 | 0.95044703 | 26.86403916 | 27.99890483 | ENSG00000142319 |
| FZD4 | 0.058916969 | 1.041683475 | 0.69702372 | 0.838372903 | 226.4382749 | 235.9196246 | ENSG00000174804 |
| NEU3 | 0.05891261 | 1.041680328 | 0.542680559 | 0.73167113 | 620.2439187 | 645.7021037 | ENSG00000162139 |
| CHSY1 | 0.058887842 | 1.041662445 | 0.453922767 | 0.659968547 | 1278.588892 | 1331.653226 | ENSG00000131873 |
| LRWD1 | 0.058722468 | 1.041543048 | 0.503728506 | 0.701470507 | 915.8720657 | 954.0717504 | ENSG00000161036 |
| NA | 0.058688858 | 1.041518783 | 0.853092424 | 0.927913672 | 43.03357594 | 44.81891127 | ENSG00000214654 |
| RNF185 | 0.05867351 | 1.041507703 | 0.563628673 | 0.748426908 | 563.782672 | 587.1141951 | ENSG00000138942 |
| NBL1 | 0.058660579 | 1.041498368 | 0.727977834 | 0.856731016 | 164.5033898 | 171.3393443 | ENSG00000158747 |
| EMC10 | 0.058592945 | 1.041449544 | 0.405840816 | 0.616895027 | 1577.398667 | 1642.625094 | ENSG00000161671 |
| NA | 0.058571104 | 1.041433777 | 0.915125456 | 0.962364642 | 15.37806915 | 15.97995704 | ENSG00000265095 |
| WDR77 | 0.058495735 | 1.041379372 | 0.461390087 | 0.666209327 | 923.9391122 | 961.9362515 | ENSG00000116455 |
| TAX1BP1 | 0.058478793 | 1.041367143 | 0.459786016 | 0.664802287 | 1065.387655 | 1109.323235 | ENSG00000106052 |
| NA | 0.05846745 | 1.041358955 | 0.773476658 | 0.883539046 | 106.4437485 | 110.8949634 | ENSG00000272711 |
| NA | 0.058451253 | 1.041347264 | 0.948211538 | 0.977265239 | 6.374760747 | 6.605181576 | ENSG00000255142 |
| TBCE | 0.05840245 | 1.041312038 | 0.878154605 | 0.94334185 | 41.32341555 | 43.10362618 | ENSG00000285053 |
| CNNM1 | 0.058364912 | 1.041284945 | 0.845154659 | 0.923013286 | 60.76797186 | 63.15419602 | ENSG00000119946 |
| MOSPD1 | 0.058299302 | 1.041237591 | 0.594348762 | 0.771358959 | 437.3807387 | 455.4772361 | ENSG00000101928 |
| SERF2 | 0.058255724 | 1.04120614 | 0.598195557 | 0.774186067 | 626.3479331 | 651.9862938 | ENSG00000140264 |
| ZNF275 | 0.058246967 | 1.04119982 | 0.612607384 | 0.784504675 | 439.4671827 | 457.5219873 | ENSG00000063587 |
| NA | 0.058181683 | 1.041152705 | 0.877024557 | 0.942825113 | 35.45513819 | 36.79019972 | ENSG00000214199 |
| VPS13A | 0.057771264 | 1.040856559 | 0.580173922 | 0.760873925 | 728.3820112 | 757.705268 | ENSG00000197969 |
| FOS | 0.057723341 | 1.040821985 | 0.646119107 | 0.80759543 | 372.7900259 | 387.9359117 | ENSG00000170345 |
| THEM6 | 0.057690629 | 1.040798386 | 0.713032012 | 0.848591123 | 195.5644208 | 203.7297714 | ENSG00000130193 |
| TTLL1 | 0.057677725 | 1.040789076 | 0.793477673 | 0.894317058 | 126.3612784 | 131.4430568 | ENSG00000100271 |
| TRMT61B | 0.057662643 | 1.040778196 | 0.58413523 | 0.763106094 | 464.6249264 | 483.309102 | ENSG00000171103 |
| RAI1 | 0.057626967 | 1.040752459 | 0.771631215 | 0.882491333 | 120.4248124 | 125.2839 | ENSG00000108557 |
| MBTD1 | 0.057484037 | 1.040649355 | 0.49535878 | 0.695062824 | 1336.797238 | 1390.429697 | ENSG00000011258 |
| GMEB1 | 0.057474544 | 1.040642507 | 0.597951572 | 0.774082495 | 508.148487 | 529.0931054 | ENSG00000162419 |
| G2E3 | 0.057430246 | 1.040610555 | 0.549643543 | 0.736903188 | 931.1051118 | 969.3183283 | ENSG00000092140 |
| OIP5-AS1 | 0.05742858 | 1.040609353 | 0.445270832 | 0.652817302 | 1513.497853 | 1575.10388 | ENSG00000247556 |
| TRUB1 | 0.057317822 | 1.040529467 | 0.618631126 | 0.788422785 | 506.3415849 | 526.5529132 | ENSG00000165832 |
| FBXO34 | 0.057276702 | 1.040499811 | 0.624450557 | 0.7923564 | 363.052227 | 377.7725723 | ENSG00000178974 |
| LEPROTL1 | 0.057109635 | 1.040379326 | 0.54066646 | 0.730042795 | 851.7250122 | 886.4649285 | ENSG00000104660 |
| UTP11 | 0.057090136 | 1.040365264 | 0.475131041 | 0.677841249 | 976.8544172 | 1016.415029 | ENSG00000183520 |
| TMEM164 | 0.05704802 | 1.040334893 | 0.435280529 | 0.643868786 | 1177.515505 | 1224.882151 | ENSG00000157600 |
| ESYT2 | 0.056955684 | 1.040268311 | 0.487657044 | 0.6884316 | 1004.646581 | 1044.761852 | ENSG00000117868 |
| SLMAP | 0.056921152 | 1.040243412 | 0.524574754 | 0.718404067 | 968.3264466 | 1007.234564 | ENSG00000163681 |
| C1GALT1 | 0.056865966 | 1.040203621 | 0.505855655 | 0.703208248 | 1015.137634 | 1056.120179 | ENSG00000106392 |
| CEP131 | 0.056757156 | 1.040125171 | 0.531842397 | 0.723515608 | 892.7601889 | 929.2252937 | ENSG00000141577 |
| BDH2 | 0.056723357 | 1.040100803 | 0.745610427 | 0.867742215 | 217.6971461 | 226.3695801 | ENSG00000164039 |
| CNOT11 | 0.056612316 | 1.040020752 | 0.468679093 | 0.67185333 | 1105.65778 | 1150.409395 | ENSG00000158435 |
| TESK2 | 0.056536333 | 1.039965979 | 0.769197807 | 0.881236386 | 123.654143 | 128.5574056 | ENSG00000070759 |
| MRPL9 | 0.056318573 | 1.039809018 | 0.491018795 | 0.690563743 | 846.4254831 | 880.3499137 | ENSG00000143436 |
| PITPNA | 0.056261863 | 1.039768146 | 0.347017904 | 0.559118942 | 2212.995565 | 2300.50774 | ENSG00000174238 |
| ATP5F1A | 0.056112577 | 1.039660559 | 0.225062141 | 0.423336507 | 14282.73165 | 14848.22102 | ENSG00000152234 |
| ARHGEF40 | 0.056069398 | 1.039629444 | 0.376708387 | 0.588869952 | 2288.736389 | 2380.175223 | ENSG00000165801 |
| DHX15 | 0.055973614 | 1.039560422 | 0.282507024 | 0.491542983 | 6007.545294 | 6244.955228 | ENSG00000109606 |
| TRIM14 | 0.055656171 | 1.039331708 | 0.574545659 | 0.756750926 | 524.4235349 | 544.98907 | ENSG00000106785 |
| BZW2 | 0.055633274 | 1.039315213 | 0.469089277 | 0.672186887 | 1315.401044 | 1366.897925 | ENSG00000136261 |
| LMO4 | 0.055522027 | 1.039235073 | 0.391921954 | 0.603313181 | 1673.258868 | 1738.564949 | ENSG00000143013 |
| NA | 0.055378386 | 1.039131608 | 0.937423817 | 0.971904957 | 10.27606378 | 10.69094999 | ENSG00000242477 |
| DARS1 | 0.055338677 | 1.039103008 | 0.448068299 | 0.654733041 | 1089.028819 | 1131.783869 | ENSG00000115866 |
| NUDT14 | 0.055290052 | 1.039067986 | 0.708738223 | 0.846326173 | 332.3453109 | 345.4859707 | ENSG00000183828 |
| HELQ | 0.055177647 | 1.038987032 | 0.797273219 | 0.896487332 | 110.9867426 | 115.186151 | ENSG00000163312 |
| DYNC1LI2 | 0.055146942 | 1.038964919 | 0.410218659 | 0.621199532 | 1466.729835 | 1523.512478 | ENSG00000135720 |
| EEF1D | 0.055082516 | 1.038918523 | 0.451987187 | 0.658191107 | 1386.586184 | 1440.698323 | ENSG00000104529 |
| SLC25A3 | 0.054910266 | 1.038794489 | 0.248079517 | 0.453053109 | 9585.703738 | 9956.660131 | ENSG00000075415 |
| DDIT4 | 0.054871018 | 1.03876623 | 0.831586754 | 0.915296977 | 718.3756756 | 746.0080077 | ENSG00000168209 |
| CDC42 | 0.054859726 | 1.038758099 | 0.413720473 | 0.624848938 | 3611.429783 | 3750.597329 | ENSG00000070831 |
| APBA1 | 0.054851049 | 1.038751852 | 0.911781584 | 0.96109596 | 19.69870051 | 20.48346923 | ENSG00000107282 |

| | | | | | | | |
|---|---|---|---|---|---|---|---|
| PALD1 | 0.054747008 | 1.038676944 | 0.379589772 | 0.592148757 | 1826.891108 | 1897.271052 | ENSG00000107719 |
| ABCB8 | 0.054650855 | 1.038607721 | 0.622782749 | 0.79114079 | 469.8020379 | 487.9889448 | ENSG00000197150 |
| MAPKBP1 | 0.054608523 | 1.038577246 | 0.659395393 | 0.816486541 | 314.7810809 | 326.7539485 | ENSG00000137802 |
| TRIM35 | 0.054583194 | 1.038559012 | 0.603927112 | 0.778509455 | 489.6385891 | 508.7128086 | ENSG00000104228 |
| NA | 0.054566368 | 1.0385469 | 0.873687189 | 0.94079043 | 39.57176286 | 41.13492335 | ENSG00000259485 |
| CALCOCO1 | 0.054523125 | 1.038515771 | 0.727804892 | 0.856731016 | 293.685044 | 304.6895058 | ENSG00000012822 |
| DCAF13 | 0.054497661 | 1.038497441 | 0.361023014 | 0.573030729 | 2266.197809 | 2352.98305 | ENSG00000164934 |
| MED12L | 0.054336566 | 1.038381486 | 0.804957091 | 0.900665647 | 163.476134 | 169.4122588 | ENSG00000144893 |
| NA | 0.054299718 | 1.038354965 | 0.204116775 | 0.395805041 | 503772.4923 | 523094.1002 | N_multimapping |
| ABHD17B | 0.054242446 | 1.038313745 | 0.565129686 | 0.749313637 | 880.1342291 | 913.8624545 | ENSG00000107362 |
| PRIMPOL | 0.054198558 | 1.038282159 | 0.844479256 | 0.922542416 | 72.04042582 | 74.75641861 | ENSG00000164306 |
| RHOBTB2 | 0.054195557 | 1.03828 | 0.752577295 | 0.871085164 | 197.5417681 | 205.4337277 | ENSG00000008853 |
| TNNT1 | 0.054066527 | 1.038187143 | 0.436298828 | 0.644810651 | 1570.413413 | 1630.803444 | ENSG00000105048 |
| TIMM9 | 0.054046688 | 1.038172867 | 0.753009934 | 0.871354394 | 178.7955756 | 185.7366992 | ENSG00000100575 |
| SNIP1 | 0.054008316 | 1.038145255 | 0.644407473 | 0.805989478 | 375.7409326 | 389.795871 | ENSG00000163877 |
| ACTB | 0.053972931 | 1.038119792 | 0.198969906 | 0.389060538 | 96255.34299 | 99925.08932 | ENSG00000075624 |
| ZNF845 | 0.05393642 | 1.03809352 | 0.652543258 | 0.811703295 | 397.395318 | 412.5484047 | ENSG00000213799 |
| AUNIP | 0.053742689 | 1.03795413 | 0.672839341 | 0.824887754 | 336.2207556 | 348.9094247 | ENSG00000127423 |
| GNB4 | 0.053736117 | 1.037949402 | 0.61504108 | 0.785862706 | 508.4312829 | 527.3651034 | ENSG00000114450 |
| NDUFB11 | 0.053727976 | 1.037943545 | 0.426417323 | 0.636390955 | 1622.99268 | 1684.971643 | ENSG00000147123 |
| C1orf61 | 0.053574729 | 1.037833298 | 0.854921952 | 0.929013468 | 66.99387131 | 69.63549176 | ENSG00000125462 |
| SLC35A1 | 0.053554392 | 1.037818668 | 0.894523768 | 0.953032171 | 33.74736822 | 34.96632968 | ENSG00000164414 |
| PAK2 | 0.053429809 | 1.037729052 | 0.489427739 | 0.689558553 | 1403.170294 | 1456.277346 | ENSG00000180370 |
| RHPN2 | 0.053375698 | 1.03769013 | 0.514953951 | 0.711196824 | 835.2253615 | 866.7615846 | ENSG00000131941 |
| NA | 0.053362541 | 1.037680667 | 0.735810058 | 0.861168893 | 210.1041388 | 218.0080632 | ENSG00000214182 |
| PIN4 | 0.053330501 | 1.037657622 | 0.579908887 | 0.760631986 | 639.9227631 | 663.7839613 | ENSG00000102309 |
| RPS19BP1 | 0.053242665 | 1.037594448 | 0.523412283 | 0.717682492 | 1369.150756 | 1421.390576 | ENSG00000187051 |
| GAREM2 | 0.053216889 | 1.03757591 | 0.529309326 | 0.721629123 | 858.6065618 | 891.23796 | ENSG00000157833 |
| FAM162B | 0.053201465 | 1.037564817 | 0.891321796 | 0.951694019 | 28.51132425 | 29.57592648 | ENSG00000183807 |
| KRT8 | 0.053200911 | 1.037564419 | 0.224783686 | 0.423192096 | 8448.807367 | 8765.887197 | ENSG00000170421 |
| TNRC6B | 0.053171947 | 1.037543588 | 0.616639451 | 0.787212602 | 666.0789296 | 690.7372889 | ENSG00000100354 |
| CGGBP1 | 0.053107236 | 1.037497051 | 0.493404221 | 0.692854676 | 1128.349107 | 1170.632143 | ENSG00000163320 |
| ALDH5A1 | 0.05305009 | 1.037455956 | 0.808912462 | 0.902949338 | 103.7179261 | 107.5811426 | ENSG00000112294 |
| LOC112267983 | 0.053050053 | 1.037455929 | 0.912515948 | 0.961482481 | 21.69559901 | 22.54219644 | ENSG00000259826 |
| DNAJC19 | 0.052969174 | 1.03739777 | 0.544238347 | 0.73258071 | 832.987283 | 864.0796943 | ENSG00000205981 |
| RAD54L2 | 0.052894039 | 1.037343745 | 0.562162481 | 0.747162956 | 806.4814775 | 836.0283448 | ENSG00000164080 |
| NAE1 | 0.052885883 | 1.03733788 | 0.408293804 | 0.619476861 | 2164.924409 | 2245.928266 | ENSG00000159593 |
| KCTD2 | 0.052843168 | 1.037307167 | 0.496822076 | 0.695950868 | 1145.854048 | 1188.494183 | ENSG00000180901 |
| SRD5A1 | 0.052806947 | 1.037281124 | 0.598773655 | 0.774828044 | 575.9684013 | 597.1628317 | ENSG00000145545 |
| SEMA6B | 0.052795359 | 1.037272793 | 0.507983959 | 0.704869733 | 3071.724775 | 3186.741142 | ENSG00000167680 |
| NA | 0.052647838 | 1.037166733 | 0.835656729 | 0.917415262 | 73.2962997 | 76.09879402 | ENSG00000204253 |
| NDUFA8 | 0.052568767 | 1.03710989 | 0.459873734 | 0.664802287 | 1423.228599 | 1475.729379 | ENSG00000119421 |
| FAM114A2 | 0.052551934 | 1.037097789 | 0.781969079 | 0.88893053 | 179.4737511 | 185.9972199 | ENSG00000055147 |
| NPW | 0.052431614 | 1.0370113 | 0.840608525 | 0.920561463 | 82.65367072 | 85.77251341 | ENSG00000183971 |
| EXOC1 | 0.052383186 | 1.03697649 | 0.700315549 | 0.84040536 | 253.1970306 | 262.4067287 | ENSG00000090989 |
| SETMAR | 0.05232513 | 1.036934761 | 0.660434128 | 0.817373256 | 404.7456975 | 419.3993435 | ENSG00000170364 |
| MPV17L | 0.052309017 | 1.03692318 | 0.811634036 | 0.904393876 | 93.45373591 | 96.98962418 | ENSG00000156968 |
| NACAD | 0.052182281 | 1.036832094 | 0.784890689 | 0.890433415 | 146.239121 | 151.74802 | ENSG00000136274 |
| RTL6 | 0.052180722 | 1.036830974 | 0.505594053 | 0.702948074 | 1095.873274 | 1135.783782 | ENSG00000188636 |
| TSGA10IP | 0.05208608 | 1.036762959 | 0.863407018 | 0.933980504 | 47.97545736 | 49.72325799 | ENSG00000175513 |
| PRELID3B | 0.052084878 | 1.036762095 | 0.583264657 | 0.762808383 | 751.1996274 | 778.966737 | ENSG00000101166 |
| FAM168B | 0.052045483 | 1.036733785 | 0.380965295 | 0.593545089 | 2143.957047 | 2222.344974 | ENSG00000152102 |
| RAD52 | 0.052001104 | 1.036701895 | 0.742528401 | 0.865919752 | 182.9320047 | 189.5807052 | ENSG00000002016 |
| TPK1 | 0.051959798 | 1.036672213 | 0.9098997 | 0.960615044 | 20.76060958 | 21.54068854 | ENSG00000196511 |
| PDK2 | 0.051917612 | 1.0366419 | 0.754231596 | 0.872025449 | 189.5537907 | 196.5361912 | ENSG00000005882 |
| HUWE1 | 0.051787622 | 1.0365485 | 0.688618108 | 0.83362519 | 8012.344185 | 8304.809685 | ENSG00000086758 |
| PITHD1 | 0.051742506 | 1.036516086 | 0.457832732 | 0.663253386 | 1614.354392 | 1673.480285 | ENSG00000057757 |
| NA | 0.051689023 | 1.036477661 | 0.896630715 | 0.954233365 | 31.06395393 | 32.21480191 | ENSG00000263272 |
| PFDN5 | 0.05168232 | 1.036472846 | 0.48409999 | 0.685221729 | 1607.637671 | 1666.003778 | ENSG00000123349 |
| RAB5A | 0.051660785 | 1.036457374 | 0.495830244 | 0.695207823 | 1193.204974 | 1237.274714 | ENSG00000144566 |
| ND2 | 0.051639605 | 1.036442159 | 0.793864643 | 0.894518722 | 13198.67142 | 13679.95282 | ENSG00000198763 |
| DENND1C | 0.051448032 | 1.03630454 | 0.619651538 | 0.788813199 | 492.5396114 | 510.3555886 | ENSG00000205744 |
| JMJD8 | 0.051407925 | 1.036275731 | 0.597297963 | 0.773554521 | 724.4843967 | 751.0187209 | ENSG00000161999 |
| H3C12 | 0.051397716 | 1.036268398 | 0.919025397 | 0.964051188 | 17.57396648 | 18.17694768 | ENSG00000197153 |
| GPAT4 | 0.05133599 | 1.036224062 | 0.498479308 | 0.697317165 | 1197.185346 | 1240.523976 | ENSG00000158669 |
| RAMAC | 0.051181308 | 1.036112967 | 0.825429498 | 0.911912722 | 95.94229508 | 99.48321532 | ENSG00000169612 |
| CEP170B | 0.051172034 | 1.036106306 | 0.48550934 | 0.686444852 | 1973.686953 | 2045.045659 | ENSG00000099814 |
| GAK | 0.051066924 | 1.036030822 | 0.408773578 | 0.61989223 | 2221.364248 | 2301.463178 | ENSG00000178950 |
| LIMS2 | 0.051032868 | 1.036006366 | 0.864209095 | 0.93447402 | 49.37513238 | 51.13836966 | ENSG00000072163 |
| ZBED9 | 0.050990689 | 1.035976077 | 0.832452761 | 0.915601071 | 79.25765586 | 82.08618207 | ENSG00000232040 |
| IRX4 | 0.050929558 | 1.035932181 | 0.872512913 | 0.940222903 | 46.48918595 | 48.24640299 | ENSG00000113430 |
| TFIP11 | 0.050903517 | 1.035913482 | 0.578601633 | 0.759655996 | 647.8028134 | 671.0944291 | ENSG00000100109 |
| PIBF1 | 0.050857811 | 1.035880664 | 0.778325864 | 0.886943686 | 152.2941664 | 157.7774043 | ENSG00000083535 |
| PLOD1 | 0.05076742 | 1.035815764 | 0.419600436 | 0.630081242 | 1904.622805 | 1973.111313 | ENSG00000083444 |

| | | | | | | | |
|---|---|---|---|---|---|---|---|
| EXOC5 | 0.050700209 | 1.035767509 | 0.506634999 | 0.703844245 | 1375.564873 | 1424.575158 | ENSG00000070367 |
| BNIPL | 0.050621716 | 1.035711158 | 0.916908268 | 0.963119654 | 22.51968671 | 23.22629515 | ENSG00000163141 |
| SENP5 | 0.05054329 | 1.035654857 | 0.62442461 | 0.7923564 | 531.6015851 | 550.1505753 | ENSG00000119231 |
| SERP1 | 0.050538928 | 1.035651726 | 0.353647741 | 0.565746514 | 4054.225102 | 4199.157897 | ENSG00000120742 |
| EDA | 0.050454493 | 1.035591115 | 0.734862246 | 0.860696695 | 223.4847246 | 231.5621317 | ENSG00000158813 |
| FXN | 0.050442001 | 1.035582149 | 0.774601735 | 0.884208808 | 154.6383663 | 160.1744446 | ENSG00000165060 |
| GRHPR | 0.050371641 | 1.035531644 | 0.417556307 | 0.628430894 | 2386.628876 | 2471.470668 | ENSG00000137106 |
| AP3B1 | 0.050271166 | 1.035459528 | 0.542715385 | 0.73167113 | 1381.520054 | 1429.894222 | ENSG00000132842 |
| ZNF597 | 0.050228318 | 1.035428776 | 0.853840475 | 0.92841878 | 72.24832303 | 74.94074565 | ENSG00000167981 |
| SMARCE1 | 0.050186507 | 1.035398768 | 0.890751253 | 0.951299972 | 39.92257148 | 41.3288982 | ENSG00000073584 |
| RPS13 | 0.050162713 | 1.035381692 | 0.348345438 | 0.56034954 | 8948.147089 | 9264.797133 | ENSG00000110700 |
| APLNR | 0.050139188 | 1.035364809 | 0.950914137 | 0.978958432 | 7.41058446 | 7.634403404 | ENSG00000134817 |
| NANS | 0.050118681 | 1.035350092 | 0.564816685 | 0.749003799 | 843.5006107 | 873.0414761 | ENSG00000095380 |
| ZNF549 | 0.050115772 | 1.035348004 | 0.751027138 | 0.870392545 | 186.5074154 | 192.9862314 | ENSG00000121406 |
| AHCY | 0.049993602 | 1.035260333 | 0.304410527 | 0.514571501 | 7447.927258 | 7709.872359 | ENSG00000101444 |
| NFX1 | 0.04980487 | 1.03512491 | 0.548976777 | 0.7363752 | 971.972437 | 1005.948006 | ENSG00000086102 |
| DTX3L | 0.049726502 | 1.035068683 | 0.790785816 | 0.892914208 | 160.4036989 | 165.7486078 | ENSG00000163840 |
| COA4 | 0.049679732 | 1.035035128 | 0.482393869 | 0.683726181 | 1302.952673 | 1348.676542 | ENSG00000181924 |
| GINS1 | 0.049613217 | 1.034987409 | 0.481597992 | 0.683370506 | 1514.395976 | 1566.856908 | ENSG00000101003 |
| POLR1E | 0.049577459 | 1.034961756 | 0.588437396 | 0.766494705 | 699.6537933 | 724.4719966 | ENSG00000137054 |
| CAPZA2 | 0.049575819 | 1.03496058 | 0.47986805 | 0.681940561 | 1327.237997 | 1373.461918 | ENSG00000198898 |
| PIK3CD | 0.049515302 | 1.034917168 | 0.611869595 | 0.784174879 | 626.957413 | 648.8937197 | ENSG00000171608 |
| RABL3 | 0.049423943 | 1.034851633 | 0.695873673 | 0.837581566 | 406.9397625 | 420.857427 | ENSG00000144840 |
| ZNF507 | 0.049293883 | 1.034758345 | 0.614424042 | 0.785406126 | 701.5272799 | 725.5900953 | ENSG00000168813 |
| SPATS2 | 0.049112128 | 1.034627991 | 0.577565274 | 0.759450456 | 742.4628182 | 767.8223126 | ENSG00000123352 |
| POMGNT1 | 0.049103631 | 1.034621897 | 0.502102822 | 0.700181291 | 1128.814326 | 1167.646338 | ENSG00000085998 |
| IWS1 | 0.049027696 | 1.034567442 | 0.412918035 | 0.624336036 | 2015.645395 | 2085.321271 | ENSG00000163166 |
| ZNF416 | 0.048891867 | 1.034470043 | 0.775660659 | 0.884862404 | 153.7297416 | 158.9659091 | ENSG00000083817 |
| NA | 0.04869363 | 1.034327909 | 0.886775856 | 0.948932572 | 40.15385873 | 41.50151756 | ENSG00000231007 |
| RNF11 | 0.048651683 | 1.034297836 | 0.626307528 | 0.793253855 | 699.7839926 | 724.1510965 | ENSG00000123091 |
| TYW5 | 0.048594368 | 1.034256746 | 0.757190342 | 0.873807894 | 198.4402736 | 205.1958647 | ENSG00000162971 |
| SAP18 | 0.048565532 | 1.034236074 | 0.422383781 | 0.632520633 | 4108.211081 | 4248.124699 | ENSG00000150459 |
| C11orf58 | 0.048495824 | 1.034186103 | 0.456782664 | 0.662717707 | 5209.893209 | 5387.800229 | ENSG00000110696 |
| SLC18A2 | 0.048391068 | 1.034111012 | 0.859944801 | 0.932051054 | 71.76550902 | 74.17475733 | ENSG00000165646 |
| DHX32 | 0.048356735 | 1.034086403 | 0.696241785 | 0.837886726 | 313.7354946 | 324.3540702 | ENSG00000089876 |
| EXOC6B | 0.048288189 | 1.034037272 | 0.563988863 | 0.748528238 | 1176.043673 | 1215.473995 | ENSG00000144036 |
| PCGF3 | 0.048283889 | 1.03403419 | 0.601313657 | 0.776412445 | 656.5424683 | 678.699322 | ENSG00000185619 |
| SYNE1 | 0.048236854 | 1.034000479 | 0.877695225 | 0.943087245 | 54.8187425 | 56.57143149 | ENSG00000131018 |
| NA | 0.048129658 | 1.033923653 | 0.940084939 | 0.97320185 | 10.8863561 | 11.25927494 | ENSG00000254610 |
| SNAP29 | 0.048115104 | 1.033913223 | 0.765878827 | 0.878910348 | 188.6463612 | 195.1142173 | ENSG00000099940 |
| PSMB3 | 0.048092941 | 1.03389734 | 0.466360022 | 0.670412501 | 2945.791156 | 3044.936666 | ENSG00000277791 |
| ABHD5 | 0.04807425 | 1.033883945 | 0.734430621 | 0.860499306 | 289.8371842 | 299.6751861 | ENSG00000011198 |
| NSUN5P1 | 0.048058435 | 1.033872611 | 0.915662516 | 0.962507751 | 611.8384971 | 632.5888982 | ENSG00000223705 |
| SNX29 | 0.048002986 | 1.033832876 | 0.740069097 | 0.863848501 | 227.3115842 | 235.019869 | ENSG00000048471 |
| GRK6 | 0.047942167 | 1.033789294 | 0.453129703 | 0.659373898 | 1840.769531 | 1902.739752 | ENSG00000198055 |
| PARM1 | 0.047927477 | 1.033778767 | 0.774358123 | 0.884054753 | 206.8825637 | 213.900141 | ENSG00000169116 |
| SLC39A8 | 0.047820768 | 1.033702307 | 0.626832929 | 0.793550956 | 530.0658383 | 547.9397969 | ENSG00000138821 |
| WDR59 | 0.047767154 | 1.033663893 | 0.499446288 | 0.69783084 | 1399.23393 | 1445.933068 | ENSG00000103091 |
| KIAA2026 | 0.047583774 | 1.033532513 | 0.770603466 | 0.881997379 | 199.1214499 | 205.8395755 | ENSG00000183354 |
| YTHDF1 | 0.047372763 | 1.033381358 | 0.471859911 | 0.675132542 | 1768.293331 | 1827.624034 | ENSG00000149658 |
| FPGS | 0.047345067 | 1.03336152 | 0.486280098 | 0.687181815 | 1581.884203 | 1634.885459 | ENSG00000136877 |
| MRPS26 | 0.047320655 | 1.033344035 | 0.46213852 | 0.667035 | 2103.486193 | 2174.166897 | ENSG00000125901 |
| BTBD3 | 0.047314239 | 1.033339439 | 0.605362113 | 0.779667429 | 934.5647888 | 965.4885617 | ENSG00000132640 |
| KRR1 | 0.047257575 | 1.033298854 | 0.69751611 | 0.838804873 | 482.0862105 | 498.0793237 | ENSG00000111615 |
| PKMYT1 | 0.047204856 | 1.033261096 | 0.54545325 | 0.733722678 | 1444.732751 | 1493.43347 | ENSG00000127564 |
| MAT2B | 0.047059701 | 1.03315714 | 0.558222738 | 0.743916196 | 1143.692342 | 1181.290052 | ENSG00000038274 |
| UBALD1 | 0.047050748 | 1.033150729 | 0.595472152 | 0.772286996 | 1126.322257 | 1164.392527 | ENSG00000153443 |
| AASDHPPT | 0.047049635 | 1.033149932 | 0.635230369 | 0.799988093 | 530.7413441 | 548.2504687 | ENSG00000149313 |
| LSM7 | 0.04699477 | 1.033110643 | 0.524369354 | 0.718312522 | 3433.97802 | 3547.619505 | ENSG00000130332 |
| NA | 0.046994103 | 1.033110165 | 0.651191913 | 0.810631949 | 639.9631125 | 661.0558395 | ENSG00000227063 |
| C9orf16 | 0.046978041 | 1.033098663 | 0.638675019 | 0.802036409 | 553.8464585 | 572.5499168 | ENSG00000171159 |
| GATD3B | 0.046941909 | 1.03307279 | 0.801089355 | 0.898333333 | 149.9888674 | 155.1474416 | ENSG00000280071 |
| ZNF382 | 0.046921162 | 1.033057933 | 0.846475763 | 0.923736594 | 79.67817939 | 82.34513728 | ENSG00000161298 |
| NUDCD1 | 0.046629799 | 1.032849321 | 0.56234114 | 0.747295225 | 1329.621402 | 1373.444923 | ENSG00000120526 |
| CCN2 | 0.046550249 | 1.032792371 | 0.600669135 | 0.776307564 | 876.5682498 | 905.2880163 | ENSG00000118523 |
| PPM1F | 0.046538544 | 1.032783992 | 0.593801157 | 0.770974282 | 767.641054 | 793.1129583 | ENSG00000100034 |
| UFSP2 | 0.046519197 | 1.032770142 | 0.694488861 | 0.837010823 | 369.6885041 | 381.9547235 | ENSG00000109775 |
| FXR1 | 0.046488973 | 1.032748506 | 0.390836776 | 0.602231765 | 4997.650999 | 5160.447774 | ENSG00000114416 |
| ZNF512B | 0.046482311 | 1.032743737 | 0.532476554 | 0.723985867 | 1193.895329 | 1233.375731 | ENSG00000196700 |
| NA | 0.04645819 | 1.03272647 | 0.939311252 | 0.972799059 | 12.60244022 | 13.03104945 | ENSG00000249684 |
| LOC100129534 | 0.046311492 | 1.032621464 | 0.921860733 | 0.964771134 | 21.49426208 | 22.18647955 | ENSG00000269896 |
| AIMP2 | 0.046220821 | 1.032556568 | 0.627784736 | 0.794139912 | 665.8980331 | 687.5114982 | ENSG00000106305 |
| PCP4 | 0.046159564 | 1.032512726 | 0.942130844 | 0.974494871 | 15.12926387 | 15.55424645 | ENSG00000183036 |
| FSD1 | 0.046102849 | 1.032472137 | 0.484652056 | 0.685694792 | 2022.724748 | 2088.821252 | ENSG00000105255 |

| | | | | | | | |
|---|---|---|---|---|---|---|---|
| PICK1 | 0.046064711 | 1.032444844 | 0.644029539 | 0.805941266 | 626.9167326 | 647.3052215 | ENSG00000100151 |
| RAD54B | 0.046037054 | 1.032425052 | 0.849482029 | 0.925384716 | 90.04228795 | 92.83849654 | ENSG00000197275 |
| SNX6 | 0.046014243 | 1.032408728 | 0.561556927 | 0.746568291 | 982.1037314 | 1013.65151 | ENSG00000129515 |
| LOC100631378 | 0.045832302 | 1.032278537 | 0.945046124 | 0.976056661 | 10.25590215 | 10.57366318 | ENSG00000225868 |
| MUC3A | 0.045786817 | 1.032245992 | 0.739491241 | 0.863654427 | 268.882636 | 277.640939 | ENSG00000169894 |
| PKN1 | 0.045775995 | 1.032238249 | 0.396871016 | 0.608821689 | 5142.764417 | 5308.482291 | ENSG00000123143 |
| GCSH | 0.045685248 | 1.032173322 | 0.92800738 | 0.967366576 | 21.83439228 | 22.5294272 | ENSG00000140905 |
| NA | 0.045666674 | 1.032160034 | 0.945909279 | 0.976250041 | 9.997613659 | 10.29976327 | ENSG00000241556 |
| SOCS3 | 0.045609661 | 1.032119245 | 0.669490499 | 0.82295975 | 541.327867 | 558.8529504 | ENSG00000184557 |
| ZBTB22 | 0.045533757 | 1.032064944 | 0.720518119 | 0.852637136 | 329.1971593 | 339.4881139 | ENSG00000236104 |
| NA | 0.045485411 | 1.032030359 | 0.942558696 | 0.97460693 | 10.91363667 | 11.25709948 | ENSG00000231154 |
| IQSEC1 | 0.045481859 | 1.032027818 | 0.560027407 | 0.745321888 | 1166.559511 | 1204.207502 | ENSG00000144711 |
| BCOR | 0.045191212 | 1.031819926 | 0.507721724 | 0.704758477 | 1666.733901 | 1720.158725 | ENSG00000183337 |
| PCDH17 | 0.045076112 | 1.031737609 | 0.931802769 | 0.969216528 | 17.79932999 | 18.29953137 | ENSG00000118946 |
| UBL4A | 0.04495354 | 1.031649956 | 0.587355551 | 0.765666704 | 990.2559247 | 1021.384378 | ENSG00000102178 |
| C19orf12 | 0.044849379 | 1.031575475 | 0.71193501 | 0.847851541 | 356.691347 | 367.6801202 | ENSG00000131943 |
| RPS5 | 0.044805296 | 1.031543954 | 0.373118829 | 0.585102486 | 15925.69144 | 16427.66082 | ENSG00000083845 |
| NRIP3 | 0.044760049 | 1.031511603 | 0.78535269 | 0.890433415 | 184.4986433 | 190.1697233 | ENSG00000175352 |
| ALDH2 | 0.044724236 | 1.031485997 | 0.683554814 | 0.830895404 | 590.952364 | 609.2709962 | ENSG00000111275 |
| NA | 0.044626653 | 1.031416231 | 0.908512072 | 0.960165884 | 33.31708215 | 34.3398343 | ENSG00000279059 |
| CMPK1 | 0.044622478 | 1.031413245 | 0.655436064 | 0.813800794 | 663.3478711 | 684.6406035 | ENSG00000162368 |
| RAB22A | 0.044527193 | 1.031345127 | 0.771721195 | 0.882491333 | 225.1379596 | 232.3401542 | ENSG00000124209 |
| NA | 0.044512357 | 1.031334521 | 0.946596906 | 0.976430116 | 10.68618822 | 11.00684612 | ENSG00000236782 |
| TBXAS1 | 0.0444499 | 1.031289873 | 0.915687658 | 0.962507751 | 25.14675983 | 25.95445991 | ENSG00000059377 |
| EIF2A | 0.044423378 | 1.031270914 | 0.60310005 | 0.777921211 | 1159.953743 | 1196.115184 | ENSG00000144895 |
| SPATA24 | 0.044401556 | 1.031255316 | 0.812625425 | 0.904860327 | 146.2322553 | 150.8647449 | ENSG00000170469 |
| PMPCA | 0.044378378 | 1.031238748 | 0.417936394 | 0.628751794 | 2767.020502 | 2853.566048 | ENSG00000165688 |
| NA | 0.044288797 | 1.031174718 | 0.806428359 | 0.901201229 | 168.9506097 | 174.0202159 | ENSG00000230202 |
| NA | 0.044210035 | 1.031118423 | 0.93826515 | 0.972296826 | 13.16409514 | 13.59379397 | ENSG00000279798 |
| SRBD1 | 0.044140351 | 1.031068621 | 0.664877259 | 0.820559864 | 566.8254878 | 584.3387872 | ENSG00000068784 |
| NME6 | 0.04412923 | 1.031060672 | 0.743319526 | 0.866407964 | 258.3349356 | 266.4862543 | ENSG00000172113 |
| PCSK9 | 0.044126975 | 1.031059061 | 0.469737678 | 0.673013888 | 2257.940182 | 2328.629492 | ENSG00000169174 |
| CDC5L | 0.043932086 | 1.030919788 | 0.539454996 | 0.729040189 | 1356.416312 | 1398.303563 | ENSG00000096401 |
| FXR2 | 0.043920265 | 1.030911341 | 0.534673301 | 0.725637667 | 1304.858604 | 1345.092585 | ENSG00000129245 |
| PCF11 | 0.043893422 | 1.030892159 | 0.602092195 | 0.7770989 | 846.0325718 | 872.197956 | ENSG00000165494 |
| LRRC8B | 0.04388118 | 1.030883412 | 0.70957629 | 0.846496645 | 406.0730652 | 418.867296 | ENSG00000197147 |
| ARMC8 | 0.043873515 | 1.030877935 | 0.634890085 | 0.799859856 | 776.3522592 | 800.3470716 | ENSG00000114098 |
| DOCK5 | 0.043586144 | 1.030672614 | 0.755878027 | 0.872977739 | 271.2736416 | 279.317422 | ENSG00000147459 |
| PGAP3 | 0.043582837 | 1.030670252 | 0.686403287 | 0.831903869 | 482.3842898 | 497.550411 | ENSG00000161395 |
| NA | 0.043527876 | 1.030630988 | 0.942334937 | 0.974513868 | 12.27891217 | 12.65698289 | ENSG00000237758 |
| RBM10 | 0.043484449 | 1.030599965 | 0.478213535 | 0.680667428 | 2278.859083 | 2348.919322 | ENSG00000182872 |
| SLC27A6 | 0.043471489 | 1.030590707 | 0.833337885 | 0.915987775 | 115.5917478 | 119.0759371 | ENSG00000113396 |
| SEC24B | 0.043425009 | 1.030557505 | 0.674885327 | 0.825790913 | 577.6858103 | 595.5555941 | ENSG00000138802 |
| DBR1 | 0.043349575 | 1.030503621 | 0.645997433 | 0.807496791 | 641.5943535 | 660.8666765 | ENSG00000138231 |
| CEP55 | 0.043313103 | 1.03047757 | 0.728119291 | 0.856731016 | 364.0569314 | 375.2900467 | ENSG00000138180 |
| UBE3C | 0.043269731 | 1.030446591 | 0.529457191 | 0.721752405 | 1901.739554 | 1959.514151 | ENSG00000009335 |
| NA | 0.043133026 | 1.030348954 | 0.9559434 | 0.982155003 | 8.435729756 | 8.651801956 | ENSG00000233013 |
| PPOX | 0.043131396 | 1.03034779 | 0.664659309 | 0.820451733 | 573.6394074 | 590.9570872 | ENSG00000143224 |
| SENP7 | 0.043097011 | 1.030323233 | 0.849155062 | 0.925311098 | 86.02682862 | 88.64842147 | ENSG00000138468 |
| H2AJ | 0.043077641 | 1.0303094 | 0.737356328 | 0.862227508 | 327.0852142 | 337.2219075 | ENSG00000246705 |
| PIAS1 | 0.043076765 | 1.030308774 | 0.667532997 | 0.821850228 | 519.368413 | 535.3353086 | ENSG00000033800 |
| PDE6B | 0.043061113 | 1.030297596 | 0.836747633 | 0.918172233 | 147.7855463 | 152.2661843 | ENSG00000133256 |
| SAC3D1 | 0.043026318 | 1.030272748 | 0.703821726 | 0.842524014 | 486.6096533 | 501.4475538 | ENSG00000168061 |
| ZNF671 | 0.042932003 | 1.030205397 | 0.800415468 | 0.898320559 | 164.4500239 | 169.3510623 | ENSG00000083814 |
| FBL | 0.042929378 | 1.030203522 | 0.403463181 | 0.614765384 | 11736.18872 | 12091.01485 | ENSG00000105202 |
| CYBRD1 | 0.042793949 | 1.03010682 | 0.832648674 | 0.91561881 | 127.9427391 | 131.8486937 | ENSG00000071967 |
| BORA | 0.0426612 | 1.030012039 | 0.826337466 | 0.912467781 | 162.8938105 | 167.7231316 | ENSG00000136122 |
| PLXNA3 | 0.04262458 | 1.029985895 | 0.689669502 | 0.83398915 | 504.1563921 | 519.2667676 | ENSG00000130827 |
| TUSC3 | 0.042565716 | 1.02994387 | 0.561249234 | 0.746231726 | 1179.845337 | 1214.775344 | ENSG00000104723 |
| BTG3 | 0.042560856 | 1.029940401 | 0.785608908 | 0.890476216 | 196.9969691 | 202.754369 | ENSG00000154640 |
| TINAGL1 | 0.042533019 | 1.029920529 | 0.789759284 | 0.892242184 | 191.2257127 | 197.1591137 | ENSG00000142910 |
| PURPL | 0.042505264 | 1.029900715 | 0.837009819 | 0.918172233 | 137.9836513 | 142.015774 | ENSG00000250337 |
| UBAP1 | 0.042472904 | 1.029877614 | 0.586263659 | 0.764770961 | 997.1440346 | 1026.712755 | ENSG00000165006 |
| ZNF430 | 0.042447953 | 1.029859802 | 0.826866341 | 0.91285928 | 138.4032845 | 142.6908646 | ENSG00000118620 |
| NGFR | 0.042419515 | 1.029839503 | 0.761898426 | 0.87673587 | 253.9713374 | 261.6327736 | ENSG00000064300 |
| TGFBR3L | 0.042389394 | 1.029818001 | 0.910300759 | 0.960811396 | 32.84499828 | 33.91970414 | ENSG00000260001 |
| WDR20 | 0.042340155 | 1.029782854 | 0.69499358 | 0.837250559 | 488.6326895 | 503.2502363 | ENSG00000140153 |
| NA | 0.042306158 | 1.029758588 | 0.878308772 | 0.94339673 | 62.22309774 | 63.93673741 | ENSG00000213300 |
| ZNF566 | 0.042211453 | 1.029690992 | 0.820628134 | 0.908947936 | 166.6201407 | 171.8345541 | ENSG00000186017 |
| WDR43 | 0.042190892 | 1.029676317 | 0.483702179 | 0.684709966 | 2534.109861 | 2608.960347 | ENSG00000163811 |
| FAM110A | 0.042147791 | 1.029645556 | 0.762045233 | 0.876818633 | 250.90241 | 258.3597625 | ENSG00000125898 |
| NA | 0.04214542 | 1.029643864 | 0.925743768 | 0.966106933 | 25.83297013 | 26.60772326 | ENSG00000287678 |
| RHOBTB3 | 0.042091633 | 1.029605477 | 0.749121327 | 0.869654826 | 278.855054 | 287.1714543 | ENSG00000164292 |
| TBKBP1 | 0.042055071 | 1.029579384 | 0.681366487 | 0.829757233 | 665.3744406 | 685.5208728 | ENSG00000198933 |

| | | | | | | | |
|---|---|---|---|---|---|---|---|
| TEAD4 | 0.042028912 | 1.029560716 | 0.545872658 | 0.733992237 | 1952.597625 | 2010.655803 | ENSG00000197905 |
| ST6GALNAC3 | 0.041991146 | 1.029533765 | 0.722479189 | 0.853711643 | 395.6540389 | 407.080252 | ENSG00000184005 |
| NA | 0.04196783 | 1.029517127 | 0.966216115 | 0.986931074 | 5.815190764 | 6.03685662 | ENSG00000261549 |
| WDR33 | 0.04188244 | 1.029456193 | 0.468749774 | 0.67185333 | 2225.271884 | 2290.797279 | ENSG00000136709 |
| PPP1R11 | 0.041761293 | 1.029369751 | 0.679665039 | 0.828703108 | 482.884239 | 497.1234709 | ENSG00000204619 |
| ZNF28 | 0.041747196 | 1.029359692 | 0.760308017 | 0.875780231 | 280.8330123 | 289.1290689 | ENSG00000198538 |
| MAMDC2 | 0.041622888 | 1.029271003 | 0.784558079 | 0.890365666 | 203.3489246 | 209.1797425 | ENSG00000165072 |
| RAB40B | 0.041598821 | 1.029253832 | 0.904733783 | 0.958481852 | 41.39012965 | 42.5027419 | ENSG00000141542 |
| CBX6 | 0.041596779 | 1.029252376 | 0.594256803 | 0.771358959 | 1012.865819 | 1042.517255 | ENSG00000183741 |
| ZBTB9 | 0.041580315 | 1.02924063 | 0.610810358 | 0.783429128 | 856.2249621 | 881.3350068 | ENSG00000213588 |
| RPS24 | 0.041504165 | 1.029186305 | 0.459067354 | 0.66424263 | 11252.54374 | 11580.76364 | ENSG00000138326 |
| GCC1 | 0.041482648 | 1.029170956 | 0.649681283 | 0.809846226 | 733.2323668 | 754.2036426 | ENSG00000179562 |
| JMJD7-PLA2G4B | 0.041479405 | 1.029168642 | 0.878454452 | 0.943401144 | 69.54947622 | 71.50816796 | ENSG00000168970 |
| ZC3H3 | 0.041446631 | 1.029145263 | 0.668223771 | 0.822110685 | 788.121459 | 811.5166378 | ENSG00000014164 |
| NA | 0.041386036 | 1.029102038 | 0.959998886 | 0.983806109 | 7.134498607 | 7.280861978 | ENSG00000246334 |
| CDKN1B | 0.041381673 | 1.029098925 | 0.615833749 | 0.786503033 | 837.9276905 | 861.9750917 | ENSG00000111276 |
| LINC01176 | 0.041378341 | 1.029096549 | 0.971986393 | 0.989699845 | 3.853860832 | 4.002059517 | ENSG00000281404 |
| TRERF1 | 0.041186994 | 1.028960067 | 0.613182821 | 0.784656293 | 882.9914733 | 908.4856415 | ENSG00000124496 |
| BCL2L12 | 0.041170757 | 1.028948487 | 0.531538517 | 0.723310628 | 2137.145542 | 2199.74852 | ENSG00000126453 |
| SPON2 | 0.041073824 | 1.028879355 | 0.911513434 | 0.961009829 | 35.17017952 | 36.28751754 | ENSG00000159674 |
| NCS1 | 0.041060305 | 1.028869714 | 0.578670186 | 0.759685603 | 1407.680773 | 1448.243614 | ENSG00000107130 |
| ACTR6 | 0.040997906 | 1.028825215 | 0.710709459 | 0.847279094 | 402.2556371 | 413.894396 | ENSG00000075089 |
| PPRC1 | 0.040959896 | 1.028798109 | 0.500916772 | 0.69914168 | 2765.237939 | 2845.211318 | ENSG00000148840 |
| RPF2 | 0.040941464 | 1.028784965 | 0.692797901 | 0.835855442 | 949.7308932 | 977.4588189 | ENSG00000197498 |
| GTF2F2 | 0.040929225 | 1.028776238 | 0.621757087 | 0.790645922 | 871.7540292 | 896.8327209 | ENSG00000188342 |
| STRN4 | 0.040872188 | 1.028735565 | 0.448613046 | 0.655281496 | 3345.087655 | 3441.427251 | ENSG00000090372 |
| ZNF780A | 0.040822143 | 1.028699881 | 0.85550734 | 0.929428529 | 100.7688256 | 103.5064624 | ENSG00000197782 |
| RAB10 | 0.0408113 | 1.02869215 | 0.504492022 | 0.701984367 | 3088.36741 | 3177.206094 | ENSG00000084733 |
| SEZ6L2 | 0.040778484 | 1.028668751 | 0.543184972 | 0.732037979 | 1705.532274 | 1754.455217 | ENSG00000174938 |
| CACNA2D1 | 0.040775564 | 1.028666669 | 0.777495704 | 0.886333839 | 345.4945055 | 355.0626316 | ENSG00000153956 |
| RPS4X | 0.040723227 | 1.028629352 | 0.351924568 | 0.564085129 | 29038.44007 | 29869.10821 | ENSG00000198034 |
| KLC4 | 0.040698331 | 1.028611602 | 0.705805921 | 0.84425797 | 503.4977417 | 517.6767456 | ENSG00000137171 |
| UVRAG | 0.040560214 | 1.028513132 | 0.729447745 | 0.85755378 | 426.3307253 | 438.5793847 | ENSG00000198382 |
| RAB3GAP1 | 0.040392432 | 1.028393526 | 0.61968931 | 0.788813199 | 910.2416103 | 936.1648891 | ENSG00000115839 |
| IFNGR1 | 0.040302249 | 1.028329242 | 0.685871488 | 0.831762616 | 504.8659919 | 519.109828 | ENSG00000027697 |
| BCDIN3D | 0.040267124 | 1.028304206 | 0.930304561 | 0.968394286 | 23.8381304 | 24.61898976 | ENSG00000186666 |
| TMC4 | 0.040209828 | 1.028263368 | 0.883447409 | 0.946765652 | 67.63083381 | 69.42948272 | ENSG00000167608 |
| COIL | 0.040105861 | 1.02818927 | 0.685658099 | 0.831762616 | 588.9243978 | 605.3953338 | ENSG00000121058 |
| TBCA | 0.040056346 | 1.028153982 | 0.498831162 | 0.697509604 | 2647.109572 | 2721.36435 | ENSG00000171530 |
| HNRNPA1 | 0.040018781 | 1.028127211 | 0.339042182 | 0.550859767 | 62070.44454 | 63815.55577 | ENSG00000135486 |
| DNAJA3 | 0.039972311 | 1.028094095 | 0.532494768 | 0.723985867 | 1870.755228 | 1922.902731 | ENSG00000103423 |
| RBM48 | 0.039956226 | 1.028082633 | 0.849422061 | 0.925384716 | 105.0170723 | 107.9547576 | ENSG00000127993 |
| TLE4 | 0.03994432 | 1.028074148 | 0.685819604 | 0.831762616 | 537.0592055 | 552.2078324 | ENSG00000106829 |
| L3HYPDH | 0.039925368 | 1.028060642 | 0.851434634 | 0.926442905 | 107.6788504 | 110.7149872 | ENSG00000126790 |
| SLU7 | 0.039830161 | 1.027992801 | 0.726063066 | 0.855720453 | 508.4686565 | 522.8335938 | ENSG00000164609 |
| RPS6 | 0.039818801 | 1.027984706 | 0.431953801 | 0.641510501 | 22294.20626 | 22917.34782 | ENSG00000137154 |
| RAPGEF2 | 0.039811651 | 1.027979611 | 0.657602507 | 0.81515086 | 1229.832794 | 1263.744852 | ENSG00000109756 |
| SLC38A9 | 0.03980251 | 1.027973098 | 0.735644526 | 0.861168893 | 432.9287646 | 444.8105521 | ENSG00000177058 |
| CARMIL2 | 0.039730653 | 1.027921899 | 0.636711242 | 0.800725077 | 1025.868421 | 1054.44457 | ENSG00000159753 |
| GLT8D2 | 0.039725094 | 1.027917938 | 0.873753499 | 0.940808189 | 73.66162555 | 75.64943688 | ENSG00000120820 |
| NXN | 0.039679294 | 1.027885306 | 0.479544026 | 0.681690845 | 3464.982899 | 3562.163678 | ENSG00000167693 |
| LHFPL2 | 0.039623492 | 1.027845549 | 0.564033547 | 0.748528238 | 1515.35774 | 1557.27912 | ENSG00000145685 |
| CEP57 | 0.039621146 | 1.027843878 | 0.629012496 | 0.79537335 | 942.6855584 | 968.6004541 | ENSG00000166037 |
| RAD18 | 0.039473773 | 1.027738888 | 0.702743437 | 0.842126015 | 497.6368882 | 511.2256954 | ENSG00000070950 |
| PRMT7 | 0.039368965 | 1.027664228 | 0.587563699 | 0.76572348 | 1156.816575 | 1189.035121 | ENSG00000132600 |
| MBD4 | 0.039286175 | 1.027605256 | 0.692321014 | 0.835643765 | 615.7053483 | 632.3587943 | ENSG00000129071 |
| LZIC | 0.039249028 | 1.027578798 | 0.753774773 | 0.871630402 | 337.4633337 | 346.7842767 | ENSG00000162441 |
| RYR3 | 0.039247397 | 1.027577636 | 0.91054496 | 0.960811396 | 45.75375401 | 46.94176819 | ENSG00000198838 |
| ARHGAP42 | 0.039206441 | 1.027548465 | 0.737906751 | 0.862550693 | 513.4129283 | 527.5775006 | ENSG00000165895 |
| ZFC3H1 | 0.039181695 | 1.02753084 | 0.640783968 | 0.80337285 | 851.0135295 | 874.2296544 | ENSG00000133858 |
| NA | 0.039117957 | 1.027485445 | 0.891397987 | 0.951700997 | 55.30892864 | 56.83880501 | ENSG00000183199 |
| INAFM1 | 0.039062655 | 1.027446059 | 0.850158921 | 0.92567733 | 120.6919993 | 124.1196088 | ENSG00000257704 |
| DYNC1LI1 | 0.039042033 | 1.027431373 | 0.580853228 | 0.761150628 | 1499.550299 | 1541.264348 | ENSG00000144635 |
| RPL19 | 0.038969438 | 1.027379675 | 0.409975324 | 0.62103024 | 15693.51225 | 16123.87484 | ENSG00000108298 |
| FCHSD1 | 0.038948561 | 1.027364808 | 0.751749194 | 0.870754633 | 653.3940635 | 671.3070505 | ENSG00000197948 |
| NA | 0.038885019 | 1.02731956 | 0.943765204 | 0.975320449 | 14.49853059 | 14.90561757 | ENSG00000232888 |
| EXOSC4 | 0.038805019 | 1.027262595 | 0.780820505 | 0.888300918 | 239.1032849 | 245.6921858 | ENSG00000178896 |
| VPS28 | 0.038759775 | 1.02723038 | 0.674821746 | 0.825790913 | 666.1975351 | 684.075232 | ENSG00000160948 |
| ACTL6A | 0.038723381 | 1.027204467 | 0.568785295 | 0.752014213 | 1744.983112 | 1792.05685 | ENSG00000136518 |
| SNRPA | 0.038712968 | 1.027197053 | 0.581045687 | 0.761150628 | 2077.196423 | 2134.410166 | ENSG00000077312 |
| POLD1 | 0.038661691 | 1.027160544 | 0.504782868 | 0.70218212 | 3980.709012 | 4089.753939 | ENSG00000062822 |
| ZIC3 | 0.038591389 | 1.027110492 | 0.59383595 | 0.770974282 | 1659.964754 | 1705.49305 | ENSG00000156925 |
| RUVBL1 | 0.038556246 | 1.027085473 | 0.474490041 | 0.67731814 | 3746.836629 | 3847.678898 | ENSG00000175792 |
| TUBG1 | 0.038490683 | 1.027038798 | 0.559290351 | 0.744708333 | 1755.136251 | 1802.746699 | ENSG00000131462 |

| | | | | | | | |
|---|---|---|---|---|---|---|---|
| NA | 0.038433409 | 1.026998026 | 0.94545186 | 0.976189572 | 17.0238797 | 17.51280701 | ENSG00000286536 |
| MYNN | 0.038349893 | 1.026938577 | 0.800018886 | 0.898237613 | 224.0368963 | 230.1022586 | ENSG00000085274 |
| NOC4L | 0.038249041 | 1.02686679 | 0.639129556 | 0.802206613 | 1229.052946 | 1262.319205 | ENSG00000184967 |
| ATF4 | 0.038212513 | 1.026840791 | 0.393173584 | 0.604796215 | 10442.34595 | 10722.19905 | ENSG00000128272 |
| KLC1 | 0.038206635 | 1.026836607 | 0.744753447 | 0.867262195 | 543.4099136 | 558.2168299 | ENSG00000126214 |
| AGPAT2 | 0.038194948 | 1.026828289 | 0.683253086 | 0.830790283 | 994.2433684 | 1021.602326 | ENSG00000169692 |
| RPL13 | 0.038098133 | 1.026759384 | 0.384980768 | 0.597138171 | 18798.02289 | 19301.479 | ENSG00000167526 |
| MARK2 | 0.038076185 | 1.026743764 | 0.570883446 | 0.753714617 | 1456.877269 | 1495.881849 | ENSG00000072518 |
| CYP2T1P | 0.038071455 | 1.026740398 | 0.946355628 | 0.97639456 | 14.19991885 | 14.58319505 | ENSG00000233622 |
| RPS11 | 0.038060219 | 1.026732401 | 0.57348981 | 0.75578193 | 24082.09823 | 24725.19182 | ENSG00000142534 |
| XYLT2 | 0.03805374 | 1.02672779 | 0.612173731 | 0.784199347 | 1229.039929 | 1262.072502 | ENSG00000015532 |
| CCT4 | 0.038046159 | 1.026722395 | 0.438386681 | 0.64658954 | 5304.338669 | 5445.534314 | ENSG00000115484 |
| DOCK6 | 0.038044501 | 1.026721215 | 0.676642367 | 0.827022925 | 1015.575449 | 1042.882138 | ENSG00000130158 |
| RMDN1 | 0.037984515 | 1.026678526 | 0.749687838 | 0.869914275 | 405.4828834 | 415.9618251 | ENSG00000176623 |
| ZNF697 | 0.03791598 | 1.026629755 | 0.685039155 | 0.831623057 | 674.0405162 | 691.9023378 | ENSG00000143067 |
| FIRRE | 0.03789064 | 1.026611723 | 0.768441092 | 0.880529702 | 446.9860776 | 459.1697283 | ENSG00000213468 |
| BRF2 | 0.037879906 | 1.026604085 | 0.81479523 | 0.905887667 | 215.2746377 | 220.9261152 | ENSG00000104221 |
| LRFN2 | 0.03785024 | 1.026582975 | 0.907677663 | 0.960149634 | 44.86798622 | 46.0884551 | ENSG00000156564 |
| ZC3H6 | 0.037842943 | 1.026577783 | 0.876064567 | 0.942329518 | 80.45421392 | 82.69404424 | ENSG00000188177 |
| SURF6 | 0.037814829 | 1.026557778 | 0.504969471 | 0.70232988 | 3099.512003 | 3182.293175 | ENSG00000148296 |
| TNIK | 0.037602004 | 1.026406352 | 0.621830735 | 0.790655706 | 1570.421345 | 1611.094797 | ENSG00000154310 |
| LOC105372321 | 0.037569405 | 1.02638316 | 0.774587972 | 0.884208808 | 277.3551295 | 284.6149629 | ENSG00000268119 |
| TTK | 0.037558356 | 1.026375299 | 0.661990478 | 0.818441256 | 992.2328697 | 1018.663096 | ENSG00000112742 |
| TRAPPC1 | 0.037509493 | 1.026340537 | 0.662220676 | 0.818618673 | 874.4512457 | 897.1654536 | ENSG00000170043 |
| SKAP2 | 0.037497513 | 1.026332015 | 0.877114328 | 0.942832774 | 79.53582665 | 81.71580403 | ENSG00000005020 |
| STXBP4 | 0.037474607 | 1.02631572 | 0.822748659 | 0.910164243 | 179.776812 | 184.4714634 | ENSG00000166263 |
| DAAM2 | 0.037439657 | 1.026290857 | 0.816353164 | 0.906477444 | 185.2794325 | 190.152814 | ENSG00000146122 |
| KCNA5 | 0.037386383 | 1.02625296 | 0.951275232 | 0.979170042 | 11.63897501 | 11.91565972 | ENSG00000130037 |
| SLX4IP | 0.037382842 | 1.026250441 | 0.846063935 | 0.923543095 | 139.5037374 | 143.0209274 | ENSG00000149346 |
| NA | 0.037370152 | 1.026241414 | 0.949438258 | 0.977918816 | 12.90936612 | 13.23495566 | ENSG00000228204 |
| GPC3 | 0.037309553 | 1.026198309 | 0.531195264 | 0.72312089 | 3042.807674 | 3122.605289 | ENSG00000147257 |
| XPNPEP3 | 0.03730441 | 1.026194651 | 0.816146694 | 0.906477444 | 221.0219715 | 226.9702762 | ENSG00000196236 |
| LOC101928674 | 0.037276078 | 1.026174498 | 0.912337457 | 0.961482481 | 48.43210813 | 49.62358581 | ENSG00000266970 |
| SAMD8 | 0.037254365 | 1.026159054 | 0.728655936 | 0.857148832 | 427.6529822 | 438.689011 | ENSG00000156671 |
| ELK3 | 0.037233488 | 1.026144205 | 0.790634818 | 0.892831861 | 275.973278 | 283.1115079 | ENSG00000111145 |
| DYNC1H1 | 0.037228973 | 1.026140993 | 0.809443461 | 0.903021016 | 6741.277724 | 6917.140249 | ENSG00000197102 |
| NPAS2 | 0.037202518 | 1.026122177 | 0.935790602 | 0.971224748 | 21.9930417 | 22.64330904 | ENSG00000170485 |
| SLC12A6 | 0.037148719 | 1.026083913 | 0.804003707 | 0.900089514 | 247.4571474 | 253.5896118 | ENSG00000140199 |
| KRAS | 0.037041548 | 1.026007693 | 0.729499681 | 0.85755378 | 922.3769995 | 946.5467169 | ENSG00000133703 |
| NPEPL1 | 0.036888558 | 1.025898896 | 0.863531698 | 0.934008941 | 152.599339 | 156.5884909 | ENSG00000215440 |
| TMEM258 | 0.03686895 | 1.025884953 | 0.746162944 | 0.867969455 | 773.0436009 | 792.7267099 | ENSG00000134825 |
| BSCL2 | 0.036866629 | 1.025883303 | 0.951969223 | 0.97945433 | 12.55499802 | 12.86306457 | ENSG00000168000 |
| PLEKHM1 | 0.036829753 | 1.025857081 | 0.781064336 | 0.888429637 | 295.3157525 | 302.7484013 | ENSG00000225190 |
| UGT8 | 0.036805988 | 1.025840182 | 0.744472339 | 0.867135375 | 445.9531768 | 457.2387528 | ENSG00000174607 |
| PTGFRN | 0.036757565 | 1.025805751 | 0.658035241 | 0.815526679 | 971.0195711 | 996.0660617 | ENSG00000134247 |
| MAN1B1 | 0.036722132 | 1.025780558 | 0.617210487 | 0.787224317 | 1101.611683 | 1129.886841 | ENSG00000177239 |
| RNASEH2B | 0.036709374 | 1.025771487 | 0.676046865 | 0.826726999 | 714.9900386 | 733.6656914 | ENSG00000136104 |
| QRICH1 | 0.036648517 | 1.025728217 | 0.53554665 | 0.726305098 | 2254.133075 | 2312.453428 | ENSG00000198218 |
| CHAC1 | 0.036617543 | 1.025706196 | 0.685170186 | 0.831636928 | 764.6377496 | 784.624863 | ENSG00000128965 |
| C12orf45 | 0.036555919 | 1.025662384 | 0.757163824 | 0.873807894 | 435.4324514 | 446.683294 | ENSG00000151131 |
| RAP2B | 0.036492659 | 1.025617411 | 0.609771897 | 0.78257168 | 1261.187708 | 1293.687808 | ENSG00000181467 |
| NSMCE3 | 0.036489006 | 1.025614815 | 0.722760467 | 0.853937151 | 482.7721421 | 495.0059108 | ENSG00000185115 |
| NA | 0.036470377 | 1.025601571 | 0.945631857 | 0.976189572 | 16.44206316 | 16.88129829 | ENSG00000280187 |
| UBE2E1 | 0.036466512 | 1.025598823 | 0.582870709 | 0.76255721 | 1999.683041 | 2050.521726 | ENSG00000170142 |
| MOB1A | 0.036447659 | 1.025585421 | 0.617153048 | 0.787224317 | 1556.175229 | 1596.079197 | ENSG00000114978 |
| TANK | 0.036426175 | 1.025570149 | 0.791602004 | 0.893408465 | 264.9133981 | 271.4291923 | ENSG00000136560 |
| CRYGN | 0.036411716 | 1.02555987 | 0.94875584 | 0.977623748 | 14.5946102 | 14.96596346 | ENSG00000127377 |
| IREB2 | 0.036405339 | 1.025555337 | 0.639226997 | 0.802218099 | 1915.098373 | 1963.942329 | ENSG00000136381 |
| TP53I13 | 0.036278037 | 1.025464847 | 0.745057772 | 0.867396409 | 483.6958683 | 496.1891415 | ENSG00000167543 |
| RYBP | 0.036272107 | 1.025460632 | 0.690622083 | 0.834528492 | 1076.601708 | 1104.349135 | ENSG00000163602 |
| NA | 0.036251156 | 1.02544574 | 0.942136571 | 0.974494871 | 18.08461972 | 18.57995768 | ENSG00000272293 |
| NA | 0.036243829 | 1.025440532 | 0.91065798 | 0.960876967 | 48.08102017 | 49.33256573 | ENSG00000232593 |
| ZCCHC24 | 0.036197689 | 1.025407737 | 0.784276772 | 0.890195147 | 286.9088296 | 294.3825335 | ENSG00000165424 |
| MEAF6 | 0.036182239 | 1.025396756 | 0.705729406 | 0.844219917 | 679.5939107 | 697.2432201 | ENSG00000163875 |
| PTTG1 | 0.036061175 | 1.025310713 | 0.617202994 | 0.787224317 | 1499.441048 | 1536.99832 | ENSG00000164611 |
| CCNB3 | 0.036058829 | 1.025309046 | 0.973990962 | 0.990992905 | 4.500916933 | 4.649742439 | ENSG00000147082 |
| ANXA5 | 0.036051613 | 1.025303918 | 0.525782035 | 0.719260683 | 3831.274268 | 3927.434772 | ENSG00000164111 |
| POMP | 0.036032135 | 1.025290076 | 0.53028711 | 0.722388522 | 2867.184627 | 2939.844714 | ENSG00000132963 |
| BAZ2A | 0.036000327 | 1.025267471 | 0.631729618 | 0.797127712 | 3617.482875 | 3707.861305 | ENSG00000076108 |
| ATG7 | 0.035957501 | 1.025237036 | 0.787865007 | 0.891582187 | 287.2403408 | 294.276859 | ENSG00000197548 |
| NA | 0.03588916 | 1.025188471 | 0.949379733 | 0.977911872 | 14.61240756 | 14.97655723 | ENSG00000260805 |
| NA | 0.035725634 | 1.025072275 | 0.939760949 | 0.972992627 | 22.99683016 | 23.63895296 | ENSG00000245205 |
| ZSCAN9 | 0.035667912 | 1.025031263 | 0.842112113 | 0.921502333 | 220.3356833 | 225.7276886 | ENSG00000137185 |
| STX8 | 0.035649349 | 1.025018074 | 0.795517906 | 0.895554092 | 344.6280102 | 352.7772452 | ENSG00000170310 |

| | | | | | | | |
|---|---|---|---|---|---|---|---|
| RINT1 | 0.035612223 | 1.024991697 | 0.746667145 | 0.868329887 | 420.2065237 | 430.6953904 | ENSG00000135249 |
| ZNF85 | 0.035538254 | 1.024939146 | 0.826163462 | 0.912403332 | 189.1757276 | 193.970052 | ENSG00000105750 |
| RTKN2 | 0.035452 | 1.02487787 | 0.766265587 | 0.878994404 | 518.9271169 | 531.8737016 | ENSG00000182010 |
| CRACD | 0.035424499 | 1.024858333 | 0.711985408 | 0.847851541 | 596.063014 | 610.9162649 | ENSG00000109265 |
| DNM2 | 0.035352486 | 1.024807178 | 0.502181961 | 0.700181291 | 4340.940196 | 4448.310449 | ENSG00000079805 |
| PIK3C2B | 0.035275026 | 1.024752157 | 0.718574949 | 0.851388915 | 807.9649027 | 828.2860188 | ENSG00000133056 |
| NA | 0.035230059 | 1.024720216 | 0.801109689 | 0.898333333 | 286.7492388 | 293.5794669 | ENSG00000244716 |
| PDPR | 0.035214519 | 1.024709179 | 0.667078998 | 0.82154232 | 935.1000796 | 957.8542966 | ENSG00000090857 |
| TXNDC17 | 0.035201717 | 1.024700086 | 0.647251537 | 0.80858274 | 1014.588463 | 1039.802614 | ENSG00000129235 |
| NR5A2 | 0.035128448 | 1.024648047 | 0.844110705 | 0.922299853 | 165.4831778 | 169.5468337 | ENSG00000116833 |
| HERC5 | 0.035081627 | 1.024614793 | 0.772821452 | 0.883060194 | 344.4006129 | 353.0687597 | ENSG00000138646 |
| GLRX | 0.035035669 | 1.024582154 | 0.783284372 | 0.889784424 | 399.0725012 | 408.8765463 | ENSG00000173221 |
| COMTD1 | 0.034945901 | 1.024518404 | 0.752146329 | 0.871054359 | 453.5802775 | 465.0959495 | ENSG00000165644 |
| RBM12 | 0.034912001 | 1.024494331 | 0.583911419 | 0.763005183 | 2447.664062 | 2506.958148 | ENSG00000244462 |
| ZC3H8 | 0.034862309 | 1.024459044 | 0.860152174 | 0.932115328 | 128.9070941 | 132.2317457 | ENSG00000144161 |
| RPL23 | 0.034838228 | 1.024441944 | 0.520684664 | 0.715714945 | 14973.93278 | 15339.80068 | ENSG00000125691 |
| DDX42 | 0.034806083 | 1.024419118 | 0.480447521 | 0.682558597 | 4061.979073 | 4161.139972 | ENSG00000198231 |
| TADA2B | 0.03475626 | 1.024383741 | 0.744257111 | 0.867034858 | 565.451645 | 579.2042178 | ENSG00000173011 |
| THRAP3 | 0.034706241 | 1.024348226 | 0.533386756 | 0.724676898 | 4742.322855 | 4857.266651 | ENSG00000054118 |
| CRLF3 | 0.034660804 | 1.024315965 | 0.792337575 | 0.893932102 | 333.38383 | 341.7552138 | ENSG00000176390 |
| HABP4 | 0.034623369 | 1.024289386 | 0.767123442 | 0.879553531 | 401.0772075 | 411.0218203 | ENSG00000130956 |
| UBLCP1 | 0.034524547 | 1.024219227 | 0.730265658 | 0.857920184 | 610.2312025 | 625.3179707 | ENSG00000164332 |
| TRABD | 0.034513342 | 1.024211272 | 0.586657679 | 0.7651264 | 1867.115849 | 1912.602947 | ENSG00000170638 |
| FBXO9 | 0.034455341 | 1.024170096 | 0.731265976 | 0.858455957 | 606.4461668 | 621.2870404 | ENSG00000112146 |
| AHR | 0.034377431 | 1.02411479 | 0.917096829 | 0.96312259 | 43.70791261 | 44.73302691 | ENSG00000106546 |
| COX7C | 0.034330225 | 1.02408128 | 0.615414742 | 0.786233758 | 5005.631675 | 5126.147908 | ENSG00000127184 |
| NA | 0.034317101 | 1.024071964 | 0.769359662 | 0.881368348 | 450.327454 | 460.8261375 | ENSG00000225178 |
| NOG | 0.034310637 | 1.024067376 | 0.956732689 | 0.982324626 | 13.08016849 | 13.3715381 | ENSG00000183691 |
| NA | 0.03425493 | 1.024027835 | 0.899649496 | 0.955743556 | 61.64100187 | 63.10756533 | ENSG00000230707 |
| TSPAN2 | 0.034235414 | 1.024013982 | 0.819274246 | 0.90814076 | 288.0024173 | 294.5597873 | ENSG00000134198 |
| UBQLN1 | 0.034150764 | 1.0239539 | 0.543507968 | 0.732315108 | 3249.40248 | 3326.966969 | ENSG00000135018 |
| AFDN | 0.034080167 | 1.023903795 | 0.545053666 | 0.733457209 | 3863.90293 | 3955.840512 | ENSG00000130396 |
| RHOT2 | 0.034076702 | 1.023901335 | 0.816409413 | 0.906477444 | 4394.35535 | 4499.66213 | ENSG00000140983 |
| OSTM1 | 0.034027201 | 1.023866204 | 0.75805605 | 0.874306125 | 486.0999683 | 497.7804751 | ENSG00000081087 |
| CA12 | 0.03399921 | 1.02384634 | 0.866894751 | 0.936573648 | 120.4666358 | 123.5488248 | ENSG00000074410 |
| NA | 0.033867413 | 1.023752811 | 0.616893155 | 0.787224317 | 1948.534951 | 1994.650357 | ENSG00000198618 |
| TTYH2 | 0.033782971 | 1.023692891 | 0.866461782 | 0.936266565 | 126.0620557 | 129.0323738 | ENSG00000141540 |
| MRPL47 | 0.03363643 | 1.023588916 | 0.684042505 | 0.831069398 | 924.640528 | 946.684018 | ENSG00000136522 |
| SNAPC2 | 0.033535179 | 1.023517081 | 0.74130222 | 0.86502054 | 483.9897515 | 495.4998185 | ENSG00000104976 |
| NA | 0.033534239 | 1.023516414 | 0.979978384 | 0.994673319 | 3.932143073 | 4.044055737 | ENSG00000272097 |
| FADS3 | 0.033525644 | 1.023510316 | 0.748612712 | 0.869254247 | 736.7363866 | 754.2989771 | ENSG00000221968 |
| DDX6 | 0.033356365 | 1.023390229 | 0.624151749 | 0.7923564 | 2019.988247 | 2067.097301 | ENSG00000110367 |
| U2AF1 | 0.033280734 | 1.023336581 | 0.934425056 | 0.970447491 | 27.64393864 | 28.24168584 | ENSG00000160201 |
| LMBR1L | 0.03327668 | 1.023333706 | 0.749578503 | 0.869840852 | 586.9025313 | 600.2149117 | ENSG00000139636 |
| ZNF335 | 0.033114421 | 1.023218619 | 0.743021722 | 0.866170692 | 587.8907107 | 601.9410218 | ENSG00000198026 |
| PPP3CA | 0.033094257 | 1.023204317 | 0.683411578 | 0.830890855 | 1231.74737 | 1260.495154 | ENSG00000138814 |
| NA | 0.033032189 | 1.023160298 | 0.95019393 | 0.978483689 | 17.45772525 | 17.87164532 | ENSG00000285976 |
| FLCN | 0.032978931 | 1.023122528 | 0.770056575 | 0.881899237 | 402.9801119 | 412.2305441 | ENSG00000154803 |
| GHDC | 0.032957418 | 1.023107271 | 0.816573697 | 0.906477444 | 228.6068916 | 233.8727933 | ENSG00000167925 |
| TNFAIP2 | 0.032884308 | 1.023055426 | 0.803027364 | 0.89940498 | 563.7120935 | 576.7538883 | ENSG00000185215 |
| IRF3 | 0.032742115 | 1.022954598 | 0.614019136 | 0.785269226 | 1783.46373 | 1824.805298 | ENSG00000126456 |
| C19orf48 | 0.032695837 | 1.022921784 | 0.632361337 | 0.797472043 | 1957.479405 | 2002.996765 | ENSG00000167747 |
| AP5Z1 | 0.032626107 | 1.022872345 | 0.74508364 | 0.867396409 | 507.5034641 | 519.1728009 | ENSG00000242802 |
| EIF1 | 0.032472623 | 1.02276353 | 0.520544002 | 0.715685882 | 5301.26044 | 5422.116523 | ENSG00000173812 |
| NA | 0.032339329 | 1.022669039 | 0.948661254 | 0.977598373 | 18.8787571 | 19.32251036 | ENSG00000223813 |
| MAPRE1 | 0.032309134 | 1.022647636 | 0.606796064 | 0.780598101 | 4607.712971 | 4711.821807 | ENSG00000101367 |
| UQCRHL | 0.032295355 | 1.022637868 | 0.711940452 | 0.847851541 | 764.4178747 | 781.4321854 | ENSG00000233954 |
| MRPL52 | 0.032279406 | 1.022626563 | 0.697744987 | 0.838858153 | 986.760628 | 1009.282586 | ENSG00000172590 |
| SC5D | 0.032264314 | 1.022615865 | 0.677578727 | 0.82756234 | 1724.63123 | 1763.29438 | ENSG00000109929 |
| MGAT1 | 0.032212102 | 1.022578857 | 0.554780472 | 0.741242673 | 2937.954512 | 3004.659248 | ENSG00000131446 |
| GTF2A2 | 0.032127992 | 1.022519242 | 0.669523622 | 0.82295975 | 1339.560409 | 1369.252211 | ENSG00000140307 |
| TMEM70 | 0.032073187 | 1.022480399 | 0.747848673 | 0.868847874 | 721.7582949 | 738.3532471 | ENSG00000175606 |
| DUSP11 | 0.032056035 | 1.022468243 | 0.808672103 | 0.902949338 | 319.5928404 | 326.8716868 | ENSG00000144048 |
| SLC25A43 | 0.031982519 | 1.022416142 | 0.860333332 | 0.932156491 | 154.6884783 | 158.1616131 | ENSG00000077713 |
| NKAIN4 | 0.031959245 | 1.022399648 | 0.868518124 | 0.93779101 | 121.4532117 | 124.1880894 | ENSG00000101198 |
| CSNK1D | 0.031920894 | 1.02237247 | 0.58742061 | 0.765666704 | 2909.060013 | 2974.468244 | ENSG00000141551 |
| LOC100507250 | 0.031857528 | 1.022327567 | 0.977831785 | 0.993188472 | 3.561172824 | 3.66998918 | ENSG00000247363 |
| PAK4 | 0.031802761 | 1.022288758 | 0.664341564 | 0.820188829 | 2950.508033 | 3017.06045 | ENSG00000130669 |
| ARRB1 | 0.031754141 | 1.022254307 | 0.808771962 | 0.902949338 | 467.8379609 | 477.8980284 | ENSG00000137486 |
| HADHA | 0.031705166 | 1.022219605 | 0.557988365 | 0.743866319 | 4347.093616 | 4443.194315 | ENSG00000084754 |
| HNRNPA1L2 | 0.03168485 | 1.02220521 | 0.880450297 | 0.944837604 | 108.516565 | 110.9630651 | ENSG00000139675 |
| TMEM145 | 0.031675608 | 1.022198662 | 0.882120916 | 0.945771249 | 139.7543221 | 143.0512387 | ENSG00000167619 |
| STK16 | 0.031645204 | 1.02217712 | 0.798098882 | 0.897070233 | 367.0774751 | 375.2357031 | ENSG00000115661 |
| PLK4 | 0.031641396 | 1.022174422 | 0.716162011 | 0.85029043 | 748.6208606 | 765.0462649 | ENSG00000142731 |

| | | | | | | | |
|---|---|---|---|---|---|---|---|
| NBPF9 | 0.031601318 | 1.022146026 | 0.830456958 | 0.914737804 | 211.348139 | 216.0593184 | ENSG00000269713 |
| FRG1-DT | 0.031499614 | 1.022073972 | 0.950865179 | 0.978958432 | 18.53150795 | 18.94967331 | ENSG00000245685 |
| FTH1 | 0.031451048 | 1.022039566 | 0.67205064 | 0.82439555 | 3308.665444 | 3382.638819 | ENSG00000167996 |
| ADNP2 | 0.031444744 | 1.0220351 | 0.665371431 | 0.82085806 | 1405.510273 | 1435.99191 | ENSG00000101544 |
| STARD4 | 0.031417899 | 1.022016083 | 0.773452468 | 0.883539046 | 621.7537092 | 635.5922252 | ENSG00000164211 |
| SEL1L3 | 0.031374312 | 1.021985206 | 0.616871672 | 0.787224317 | 1753.221474 | 1791.468283 | ENSG00000091490 |
| TMEM161A | 0.031338489 | 1.02195983 | 0.676828306 | 0.827121605 | 1115.490997 | 1139.956562 | ENSG00000064545 |
| ACTR2 | 0.031270828 | 1.021911901 | 0.64907712 | 0.809458211 | 5539.856336 | 5660.841151 | ENSG00000138071 |
| SPRYD3 | 0.031220712 | 1.021876403 | 0.830244982 | 0.914737804 | 250.3529336 | 256.10148 | ENSG00000167778 |
| PDPN | 0.031173289 | 1.021842814 | 0.630301896 | 0.796470479 | 2115.296634 | 2161.285593 | ENSG00000162493 |
| ITGB3BP | 0.031087559 | 1.021782094 | 0.771690165 | 0.882491333 | 435.706173 | 445.2898416 | ENSG00000142856 |
| GMDS | 0.030825444 | 1.021596469 | 0.717404253 | 0.851122916 | 780.9196365 | 798.1027676 | ENSG00000112699 |
| WRNIP1 | 0.030775584 | 1.021561163 | 0.724146892 | 0.854559399 | 1140.224403 | 1164.940527 | ENSG00000124535 |
| HDAC5 | 0.030768172 | 1.021555915 | 0.600233642 | 0.776046304 | 2395.89144 | 2447.699236 | ENSG00000108840 |
| ASL | 0.030754147 | 1.021545984 | 0.785448804 | 0.890433415 | 417.061068 | 425.7236043 | ENSG00000126522 |
| ZNF771 | 0.03069837 | 1.02150649 | 0.828847075 | 0.914028945 | 265.2582824 | 271.172907 | ENSG00000179965 |
| FARP1 | 0.030694104 | 1.021503469 | 0.669502231 | 0.82295975 | 2056.474079 | 2100.453884 | ENSG00000152767 |
| ENTPD4 | 0.03068957 | 1.021500259 | 0.750858262 | 0.870392545 | 810.3748492 | 828.1829026 | ENSG00000197217 |
| PMS2CL | 0.030601905 | 1.02143819 | 0.854423086 | 0.92871587 | 171.1658307 | 174.9106799 | ENSG00000187953 |
| ATP11B | 0.03057896 | 1.021421944 | 0.809123905 | 0.902949338 | 396.0956137 | 404.4057911 | ENSG00000058063 |
| BUB1 | 0.030470725 | 1.021345318 | 0.565971229 | 0.750008186 | 3639.774456 | 3717.02725 | ENSG00000169679 |
| RLF | 0.0303633 | 1.02126927 | 0.786234802 | 0.890918291 | 507.5615068 | 517.9803966 | ENSG00000117000 |
| SOX21 | 0.030349534 | 1.021259525 | 0.911965077 | 0.961235724 | 60.36082207 | 61.69368317 | ENSG00000125285 |
| SRSF10 | 0.030201409 | 1.021154675 | 0.603440584 | 0.778147861 | 3695.185586 | 3773.55574 | ENSG00000188529 |
| DTWD1 | 0.030151971 | 1.021119683 | 0.812070433 | 0.904562352 | 349.9812101 | 357.6082362 | ENSG00000104047 |
| FAM43A | 0.030119947 | 1.021097017 | 0.880399962 | 0.944837604 | 122.5092232 | 125.0546233 | ENSG00000185112 |
| RBM42 | 0.030111378 | 1.021090952 | 0.659086062 | 0.816293303 | 2202.744493 | 2249.903544 | ENSG00000126254 |
| CTBP1 | 0.029864661 | 1.020916349 | 0.690645793 | 0.834528492 | 1871.638736 | 1910.422523 | ENSG00000159692 |
| NA | 0.029758336 | 1.020841112 | 0.980177369 | 0.994714902 | 3.533892253 | 3.632343887 | ENSG00000223393 |
| NDUFB6 | 0.029677183 | 1.02078369 | 0.734654578 | 0.860583311 | 767.7270662 | 783.8025008 | ENSG00000165264 |
| TMEM251 | 0.029627433 | 1.02074849 | 0.896731325 | 0.954280691 | 112.7476515 | 114.9895063 | ENSG00000153485 |
| C14orf119 | 0.029570014 | 1.020707865 | 0.739197419 | 0.863587169 | 726.5841894 | 741.6004268 | ENSG00000179933 |
| NUMA1 | 0.02954971 | 1.0206935 | 0.667830102 | 0.821935407 | 1874.393416 | 1913.631668 | ENSG00000137497 |
| SASS6 | 0.0295431 | 1.020688823 | 0.84653018 | 0.923736594 | 227.1461469 | 232.1042558 | ENSG00000156876 |
| CECR2 | 0.029535015 | 1.020683104 | 0.69562039 | 0.837485087 | 3282.293387 | 3349.521173 | ENSG00000099954 |
| CDCA3 | 0.02952226 | 1.020674079 | 0.750276726 | 0.870058771 | 616.8017086 | 629.5359707 | ENSG00000111665 |
| SSH3 | 0.029461382 | 1.02063101 | 0.794488529 | 0.89490134 | 537.5425259 | 548.8423379 | ENSG00000172830 |
| RARS2 | 0.029457946 | 1.02062858 | 0.742891413 | 0.866125715 | 761.0536681 | 776.4041208 | ENSG00000146282 |
| MRPS9 | 0.029442298 | 1.02061751 | 0.754289969 | 0.87203955 | 733.3556747 | 748.1823407 | ENSG00000135972 |
| SMC5 | 0.029413459 | 1.020597108 | 0.787898621 | 0.891582187 | 620.3462793 | 632.8234617 | ENSG00000198887 |
| CIAO1 | 0.029358408 | 1.020558165 | 0.652793827 | 0.811801095 | 1713.622736 | 1748.518949 | ENSG00000144021 |
| NPIPB5 | 0.029343562 | 1.020547662 | 0.93052326 | 0.968525781 | 45.43083635 | 46.41712047 | ENSG00000243716 |
| RHOQ | 0.029299814 | 1.020516716 | 0.778990529 | 0.887181048 | 570.9794355 | 582.2285113 | ENSG00000119729 |
| BTF3L4 | 0.029263314 | 1.020490897 | 0.733777817 | 0.860173437 | 1285.709658 | 1311.950321 | ENSG00000134717 |
| PI4KAP1 | 0.029137573 | 1.020401959 | 0.898016628 | 0.954954898 | 155.7406248 | 159.0658352 | ENSG00000274602 |
| CLTC | 0.029122076 | 1.020390998 | 0.638981393 | 0.802177997 | 7433.599967 | 7584.290669 | ENSG00000141367 |
| DNAAF2 | 0.029042139 | 1.020334461 | 0.732136097 | 0.859102161 | 998.5538027 | 1018.889419 | ENSG00000165506 |
| THAP6 | 0.028770356 | 1.020142263 | 0.925029043 | 0.965883673 | 63.57980591 | 64.79201506 | ENSG00000174796 |
| HDAC7 | 0.028731169 | 1.020114554 | 0.673807595 | 0.825338127 | 2039.775532 | 2081.355409 | ENSG00000061273 |
| AP1G1 | 0.028686192 | 1.020082751 | 0.700436595 | 0.840454689 | 1452.997376 | 1481.835438 | ENSG00000166747 |
| HDAC1 | 0.028685333 | 1.020082145 | 0.626316306 | 0.793253855 | 3917.453665 | 3995.457679 | ENSG00000116478 |
| LYPLAL1 | 0.028673379 | 1.020073692 | 0.907691543 | 0.960149634 | 78.65776263 | 80.24592691 | ENSG00000143353 |
| DISP2 | 0.02865976 | 1.020064062 | 0.926748901 | 0.966701516 | 48.09169859 | 48.98382674 | ENSG00000140323 |
| SMG8 | 0.028556665 | 1.019991171 | 0.775801872 | 0.884937083 | 632.1598432 | 644.4615759 | ENSG00000167447 |
| SEC62 | 0.028524997 | 1.019968782 | 0.731267196 | 0.858455957 | 1521.161308 | 1551.70402 | ENSG00000008952 |
| FBXO16 | 0.028495779 | 1.019948126 | 0.914782238 | 0.96227163 | 70.8930894 | 72.25864449 | ENSG00000214050 |
| PRMT6 | 0.028485217 | 1.019940658 | 0.71791591 | 0.851146028 | 1264.577825 | 1290.148188 | ENSG00000198890 |
| NA | 0.028443093 | 1.019910879 | 0.954563436 | 0.981271052 | 17.95205567 | 18.29518045 | ENSG00000224699 |
| ARFGAP1 | 0.028390387 | 1.019873619 | 0.686637247 | 0.832134019 | 2921.006474 | 2979.30919 | ENSG00000101199 |
| TRAF2 | 0.028371016 | 1.019859925 | 0.695203919 | 0.837250559 | 1321.624117 | 1347.966644 | ENSG00000127191 |
| TRPC5 | 0.028330928 | 1.019831587 | 0.928957435 | 0.967758568 | 51.52269778 | 52.49625685 | ENSG00000072315 |
| KIF5B | 0.028207492 | 1.019744334 | 0.669328086 | 0.82295975 | 1963.588401 | 2002.396467 | ENSG00000170759 |
| TTYH1 | 0.028177957 | 1.019723458 | 0.791417448 | 0.893312556 | 441.9872446 | 450.751636 | ENSG00000167614 |
| CHD3 | 0.028128247 | 1.019688323 | 0.667722215 | 0.821869982 | 1536.59989 | 1566.885086 | ENSG00000170004 |
| NECTIN2 | 0.028031549 | 1.01961998 | 0.635159611 | 0.799988093 | 4325.656074 | 4411.091651 | ENSG00000130202 |
| HERPUD2 | 0.028022168 | 1.01961335 | 0.811872308 | 0.904395006 | 358.4312039 | 365.59389 | ENSG00000122557 |
| INPP5B | 0.027976703 | 1.019581218 | 0.804596355 | 0.900509221 | 477.2731328 | 486.8192411 | ENSG00000204084 |
| WDFY1 | 0.027873242 | 1.019508103 | 0.758874083 | 0.874822186 | 1128.695771 | 1150.927787 | ENSG00000085449 |
| RBM12B | 0.027845887 | 1.019488772 | 0.758261834 | 0.874329877 | 673.6985804 | 686.7688729 | ENSG00000183808 |
| TMEM41A | 0.027843275 | 1.019486926 | 0.83754061 | 0.918202392 | 271.5841526 | 277.0779836 | ENSG00000163900 |
| PHACTR4 | 0.027820053 | 1.019470517 | 0.717560646 | 0.851122916 | 971.4668433 | 990.4144755 | ENSG00000204138 |
| ZSCAN32 | 0.027806433 | 1.019460893 | 0.845857909 | 0.923418446 | 276.7077685 | 281.8394324 | ENSG00000140987 |
| DDX19B | 0.027791993 | 1.019450688 | 0.838121622 | 0.918786024 | 260.3395378 | 265.3875332 | ENSG00000157349 |
| ZSCAN12 | 0.027786467 | 1.019446783 | 0.861732601 | 0.932860229 | 212.2176102 | 216.1688493 | ENSG00000158691 |

| | | | | | | | |
|---|---|---|---|---|---|---|---|
| NA | 0.027772104 | 1.019436634 | 0.930685217 | 0.96853421 | 52.92682199 | 54.0255339 | ENSG00000257086 |
| GPKOW | 0.027669306 | 1.019363998 | 0.78874915 | 0.891842679 | 492.4966184 | 502.0732188 | ENSG00000068394 |
| LRIG2 | 0.027658019 | 1.019356023 | 0.804370561 | 0.900446832 | 416.7106166 | 424.7270144 | ENSG00000198799 |
| LOC648987 | 0.027634377 | 1.019339319 | 0.959669934 | 0.983629129 | 15.88160346 | 16.1602167 | ENSG00000177738 |
| GPRASP1 | 0.027438275 | 1.019200771 | 0.926670685 | 0.966701516 | 54.25266396 | 55.3863316 | ENSG00000198932 |
| NECAP1 | 0.027399377 | 1.019173292 | 0.857691633 | 0.930409931 | 221.3089889 | 225.6477635 | ENSG00000089818 |
| FBLL1 | 0.027347472 | 1.019136625 | 0.870293895 | 0.938742301 | 185.0576023 | 188.6035789 | ENSG00000188573 |
| AKAP8 | 0.027220938 | 1.019047244 | 0.730583023 | 0.858105398 | 1016.920537 | 1036.226928 | ENSG00000105127 |
| ZNF775 | 0.027183483 | 1.019020788 | 0.790141278 | 0.892506611 | 617.8153369 | 630.1132084 | ENSG00000196456 |
| ZNF567 | 0.02717738 | 1.019016478 | 0.872463358 | 0.940222903 | 167.2600523 | 170.3180463 | ENSG00000189042 |
| CTTNBP2NL | 0.027175475 | 1.019015132 | 0.807338079 | 0.901913344 | 558.6904815 | 569.0668589 | ENSG00000143079 |
| YIPF4 | 0.027153744 | 1.018999783 | 0.828138333 | 0.91367299 | 356.0888357 | 363.1109653 | ENSG00000119820 |
| CD2AP | 0.026966501 | 1.018867538 | 0.755011414 | 0.872553121 | 1286.778154 | 1310.772294 | ENSG00000198087 |
| SPTAN1 | 0.026949718 | 1.018855686 | 0.635669277 | 0.800307619 | 6062.499221 | 6175.846553 | ENSG00000197694 |
| NA | 0.026913156 | 1.018829865 | 0.982061488 | 0.995402713 | 3.27204429 | 3.355322553 | ENSG00000207751 |
| NHLRC2 | 0.026867417 | 1.018797565 | 0.836864496 | 0.918172233 | 332.0019262 | 338.3466912 | ENSG00000196865 |
| NA | 0.02685268 | 1.018787158 | 0.953303687 | 0.980402991 | 27.55881679 | 27.99890483 | ENSG00000214106 |
| OSBPL9 | 0.026850434 | 1.018785572 | 0.718567746 | 0.851388915 | 1364.901342 | 1390.559724 | ENSG00000117859 |
| RAB9A | 0.026849403 | 1.018784844 | 0.810949666 | 0.903953201 | 418.4495054 | 426.5448525 | ENSG00000123595 |
| NELFE | 0.026822346 | 1.018765738 | 0.721676165 | 0.853183917 | 1373.23308 | 1398.937544 | ENSG00000204356 |
| AKAP10 | 0.026816606 | 1.018761684 | 0.809643661 | 0.90305216 | 395.3373243 | 402.7253564 | ENSG00000108599 |
| RAB21 | 0.026706892 | 1.018684213 | 0.793783841 | 0.894481046 | 650.9280029 | 663.1672387 | ENSG00000080371 |
| EIF4B | 0.026671433 | 1.018659175 | 0.63066373 | 0.796625039 | 12554.67877 | 12787.81148 | ENSG00000063046 |
| ATG14 | 0.02659182 | 1.018602964 | 0.849964173 | 0.925589517 | 365.0277693 | 371.4389769 | ENSG00000126775 |
| NA | 0.026529175 | 1.018558735 | 0.954775224 | 0.981381925 | 22.08556183 | 22.50209213 | ENSG00000258592 |
| ARAP1 | 0.026507325 | 1.018543308 | 0.663492644 | 0.819600901 | 2515.411851 | 2561.950355 | ENSG00000186635 |
| USP12 | 0.026316811 | 1.018408814 | 0.816560952 | 0.906477444 | 536.7021416 | 546.4229399 | ENSG00000152484 |
| POC5 | 0.025991317 | 1.018179072 | 0.872231628 | 0.94018803 | 205.2019964 | 208.6222948 | ENSG00000152359 |
| NA | 0.0259618 | 1.01815824 | 0.936407063 | 0.97149082 | 578.3016958 | 588.849535 | ENSG00000111788 |
| CCDC18 | 0.025950215 | 1.018150064 | 0.867974475 | 0.937364783 | 194.1223376 | 197.7725563 | ENSG00000122483 |
| PORCN | 0.025936123 | 1.018140119 | 0.843000757 | 0.9217804 | 305.9429821 | 311.7301411 | ENSG00000102312 |
| MYCL | 0.025840093 | 1.018072351 | 0.813085009 | 0.905169301 | 468.4879911 | 476.876068 | ENSG00000116990 |
| TNPO1 | 0.025807965 | 1.01804968 | 0.737638092 | 0.862343405 | 4227.982256 | 4303.926092 | ENSG00000083312 |
| ANKLE1 | 0.025708729 | 1.017979656 | 0.824682275 | 0.911460453 | 400.8924471 | 407.879294 | ENSG00000160117 |
| NA | 0.025660808 | 1.017945842 | 0.94982962 | 0.978215214 | 27.34858089 | 27.87414567 | ENSG00000180581 |
| HCCS | 0.025607334 | 1.017908112 | 0.780066815 | 0.887787612 | 715.5531942 | 728.5932872 | ENSG00000004961 |
| RIC8B | 0.025572895 | 1.017883814 | 0.838332345 | 0.918919053 | 331.3077072 | 337.4643097 | ENSG00000111785 |
| TUBB4B | 0.025567623 | 1.017880094 | 0.589540698 | 0.767667109 | 7199.626931 | 7328.819496 | ENSG00000188229 |
| KARS1 | 0.025529194 | 1.017852982 | 0.605833199 | 0.779795526 | 5233.504667 | 5326.777898 | ENSG00000065427 |
| TRAPPC4 | 0.025523226 | 1.017848771 | 0.788189813 | 0.891582187 | 675.4479728 | 687.5470204 | ENSG00000196655 |
| ENTPD3-AS1 | 0.025510734 | 1.017839957 | 0.956512234 | 0.982211615 | 20.19895467 | 20.54476108 | ENSG00000223797 |
| PACS2 | 0.025469599 | 1.017810937 | 0.756966129 | 0.873742764 | 1721.137818 | 1752.095797 | ENSG00000179364 |
| RNMT | 0.025461895 | 1.017805501 | 0.73626177 | 0.861374273 | 1268.649906 | 1290.744679 | ENSG00000101654 |
| TOR1B | 0.025357858 | 1.017732107 | 0.813893184 | 0.905404967 | 463.4627934 | 471.6384941 | ENSG00000136816 |
| PRIM1 | 0.025315596 | 1.017702295 | 0.865959946 | 0.935884948 | 260.7126186 | 265.6049437 | ENSG00000198056 |
| CLN8 | 0.025157215 | 1.017590576 | 0.772765362 | 0.883060194 | 786.0554572 | 799.952555 | ENSG00000182372 |
| MED29 | 0.024924959 | 1.01742677 | 0.746299278 | 0.867969455 | 1116.587435 | 1136.384027 | ENSG00000063322 |
| SMYD4 | 0.02483152 | 1.017360876 | 0.815524315 | 0.906261557 | 580.3813002 | 590.0152198 | ENSG00000186532 |
| HIPK1 | 0.024830722 | 1.017360314 | 0.795853733 | 0.89563779 | 1116.147205 | 1134.814821 | ENSG00000163349 |
| ZDHHC12 | 0.024796037 | 1.017335855 | 0.854457942 | 0.92871587 | 390.1701311 | 397.3300781 | ENSG00000160446 |
| XRN2 | 0.024793373 | 1.017333976 | 0.647620724 | 0.808876289 | 3504.130633 | 3564.870261 | ENSG00000088930 |
| RC3H2 | 0.024778684 | 1.017323618 | 0.825305187 | 0.911828729 | 539.4781015 | 548.8682193 | ENSG00000056586 |
| TYK2 | 0.024746773 | 1.017301116 | 0.798124122 | 0.897070233 | 683.8226067 | 696.0764199 | ENSG00000105397 |
| MED22 | 0.024700967 | 1.017268817 | 0.711947826 | 0.847851541 | 1890.825001 | 1924.172624 | ENSG00000148297 |
| CPPED1 | 0.024688965 | 1.017260354 | 0.862496742 | 0.933370637 | 250.3727904 | 254.6648972 | ENSG00000103381 |
| RPL34 | 0.024651213 | 1.017233735 | 0.690195824 | 0.834305072 | 6111.571693 | 6216.839585 | ENSG00000109475 |
| INSC | 0.024645646 | 1.01722981 | 0.96351824 | 0.985647688 | 16.00496363 | 16.23550729 | ENSG00000188487 |
| NA | 0.024624849 | 1.017215146 | 0.981880218 | 0.995402617 | 4.22483108 | 4.321077071 | ENSG00000250258 |
| BST2 | 0.024569419 | 1.017176064 | 0.774139164 | 0.883955535 | 997.1274331 | 1013.975155 | ENSG00000130303 |
| WDR3 | 0.024541372 | 1.01715629 | 0.716512058 | 0.850492157 | 1541.931455 | 1568.304483 | ENSG00000065183 |
| TECR | 0.024434107 | 1.017080667 | 0.694042509 | 0.836669205 | 1920.85458 | 1953.940844 | ENSG00000099797 |
| NA | 0.02438769 | 1.017047944 | 0.958517241 | 0.982874414 | 20.96937094 | 21.30878485 | ENSG00000224261 |
| CCDC142 | 0.0243722 | 1.017037024 | 0.899142747 | 0.95546587 | 137.8004166 | 140.1342577 | ENSG00000135637 |
| CHMP1B2P | 0.024270802 | 1.016965546 | 0.936303324 | 0.97143656 | 53.18894929 | 53.99023202 | ENSG00000278530 |
| UBE2G1 | 0.02422424 | 1.016932725 | 0.697769924 | 0.838858153 | 2847.975937 | 2896.309203 | ENSG00000132388 |
| KCTD21 | 0.024190756 | 1.016909122 | 0.937959704 | 0.972140333 | 50.94441456 | 51.69023686 | ENSG00000188997 |
| ATP5F1C | 0.024115226 | 1.016855885 | 0.681468609 | 0.829757233 | 2828.74627 | 2875.81609 | ENSG00000165629 |
| NA | 0.023997633 | 1.016773005 | 0.983563629 | 0.995796876 | 3.275603762 | 3.350971626 | ENSG00000229018 |
| DNAJC13 | 0.023937229 | 1.016730435 | 0.7932288 | 0.894207674 | 718.9530478 | 731.0780382 | ENSG00000138246 |
| TSNAXIP1 | 0.023686398 | 1.016553678 | 0.967075733 | 0.987033408 | 16.07732214 | 16.30115006 | ENSG00000102904 |
| HAUS3 | 0.023667161 | 1.016540124 | 0.872954649 | 0.940412849 | 251.5150666 | 255.5989431 | ENSG00000214367 |
| AFAP1L1 | 0.023595748 | 1.016489807 | 0.899087863 | 0.95546587 | 160.6762253 | 163.1564786 | ENSG00000157510 |
| PROM1 | 0.023561104 | 1.016465397 | 0.755880125 | 0.872977739 | 1267.309851 | 1287.835193 | ENSG00000007062 |
| CXXC4 | 0.023518823 | 1.016435608 | 0.952018199 | 0.97945433 | 29.41666884 | 29.92852195 | ENSG00000168772 |

| | | | | | | | |
|---|---|---|---|---|---|---|---|
| ZNF14 | 0.023502762 | 1.016424293 | 0.921199732 | 0.964771134 | 81.16058585 | 82.53939568 | ENSG00000105708 |
| TMEM219 | 0.02338575 | 1.016341858 | 0.807352135 | 0.901913344 | 673.9767779 | 684.8015379 | ENSG00000149932 |
| ZNRF1 | 0.023245289 | 1.016242911 | 0.841025901 | 0.920847294 | 573.4094194 | 582.8722651 | ENSG00000186187 |
| TBC1D2 | 0.023239211 | 1.01623863 | 0.893470137 | 0.952533789 | 161.7894671 | 164.477383 | ENSG00000095383 |
| NIF3L1 | 0.023233968 | 1.016234937 | 0.791264455 | 0.893303941 | 842.6896697 | 856.4359469 | ENSG00000196290 |
| FASTKD5 | 0.023213404 | 1.016220451 | 0.816758083 | 0.90657555 | 567.2169769 | 576.7085894 | ENSG00000215251 |
| ANKFY1 | 0.02317609 | 1.016194168 | 0.772853688 | 0.883060194 | 1376.674453 | 1398.14594 | ENSG00000185722 |
| SNX1 | 0.023108441 | 1.01614652 | 0.73996573 | 0.863848501 | 1441.253065 | 1464.451092 | ENSG00000028528 |
| GALC | 0.023094409 | 1.016136636 | 0.880423934 | 0.944837604 | 237.9381773 | 241.7239448 | ENSG00000054983 |
| ABCC6P1 | 0.023090152 | 1.016133638 | 0.9823654 | 0.995631107 | 4.266349541 | 4.358722365 | ENSG00000256340 |
| ZNF473 | 0.02306214 | 1.016113908 | 0.880565602 | 0.944856897 | 207.975566 | 211.3531295 | ENSG00000142528 |
| TMEM59 | 0.022996263 | 1.016067511 | 0.771094348 | 0.882285172 | 1486.81697 | 1510.812036 | ENSG00000116209 |
| PARVA | 0.02291865 | 1.016012851 | 0.844997347 | 0.92294823 | 395.0475592 | 401.082481 | ENSG00000197702 |
| ZFP90 | 0.022888842 | 1.015991859 | 0.863812538 | 0.934208281 | 290.3184981 | 295.1365306 | ENSG00000184939 |
| MIR1915 | 0.022864042 | 1.015974394 | 0.981927025 | 0.995402617 | 4.531756977 | 4.630446813 | ENSG00000222071 |
| TMEM184B | 0.022818697 | 1.015942462 | 0.773500188 | 0.883539046 | 1767.160109 | 1796.074012 | ENSG00000198792 |
| TRAF6 | 0.022773748 | 1.01591081 | 0.88414962 | 0.947149327 | 192.376888 | 195.5099229 | ENSG00000175104 |
| RWDD4 | 0.022764395 | 1.015904223 | 0.881842446 | 0.945689642 | 247.4452993 | 251.2441725 | ENSG00000182552 |
| UQCRFS1 | 0.022633215 | 1.015811854 | 0.715967685 | 0.850113155 | 2034.238129 | 2066.537503 | ENSG00000169021 |
| ALDH7A1 | 0.022544612 | 1.01574947 | 0.666475496 | 0.821297862 | 3749.015086 | 3807.496559 | ENSG00000164904 |
| ZNF774 | 0.022539049 | 1.015745554 | 0.95433733 | 0.981119172 | 28.79689331 | 29.26561078 | ENSG00000196391 |
| RASSF3 | 0.022520144 | 1.015732243 | 0.802473146 | 0.899050996 | 763.5333539 | 775.5898505 | ENSG00000153179 |
| EML1 | 0.022494043 | 1.015713867 | 0.811442836 | 0.904255331 | 616.4520429 | 625.9860632 | ENSG00000066629 |
| ST6GALNAC4 | 0.022439813 | 1.015675688 | 0.822775198 | 0.910164243 | 504.9140445 | 512.8753807 | ENSG00000136840 |
| RARS1 | 0.022438562 | 1.015674807 | 0.732772991 | 0.859594158 | 2237.490005 | 2272.78821 | ENSG00000113643 |
| NRDC | 0.022302066 | 1.015578716 | 0.673507414 | 0.825146432 | 3576.144109 | 3632.025784 | ENSG00000078618 |
| PDE4A | 0.022301842 | 1.015578559 | 0.898198018 | 0.955040233 | 162.070587 | 164.4858739 | ENSG00000065989 |
| COL18A1 | 0.022289221 | 1.015569675 | 0.700737624 | 0.840591375 | 7436.277447 | 7552.879846 | ENSG00000182871 |
| ING3 | 0.022272887 | 1.015558176 | 0.872415534 | 0.940222903 | 258.7349664 | 262.608743 | ENSG00000071243 |
| RALB | 0.022230842 | 1.01552858 | 0.820912394 | 0.909161355 | 734.7250939 | 745.7880725 | ENSG00000144118 |
| NA | 0.022189341 | 1.015499367 | 0.977443824 | 0.992954632 | 7.821598629 | 7.983593902 | ENSG00000267890 |
| DNAJB9 | 0.02218261 | 1.01549463 | 0.887920491 | 0.94955683 | 223.5867018 | 227.0978505 | ENSG00000128590 |
| ADSL | 0.022182379 | 1.015494467 | 0.92904554 | 0.967771899 | 84.60551744 | 86.06175716 | ENSG00000239900 |
| PPP4R3A | 0.022174241 | 1.015488739 | 0.721680612 | 0.853183917 | 2081.914949 | 2114.279592 | ENSG00000100796 |
| INTS13 | 0.022092922 | 1.015431502 | 0.78739081 | 0.891546762 | 1112.355076 | 1129.20355 | ENSG00000064102 |
| TYRO3 | 0.022034608 | 1.015390458 | 0.712388869 | 0.848103373 | 3516.197069 | 3569.550923 | ENSG00000092445 |
| FLYWCH2 | 0.021989383 | 1.015358629 | 0.89936969 | 0.95553286 | 225.4792583 | 229.2404546 | ENSG00000162076 |
| PWP1 | 0.021970441 | 1.015345298 | 0.75479681 | 0.872411882 | 1602.328534 | 1627.052779 | ENSG00000136045 |
| CCT8 | 0.021961514 | 1.015339015 | 0.671716742 | 0.824135456 | 6761.944019 | 6864.81619 | ENSG00000156261 |
| KIAA1109 | 0.021953222 | 1.015333179 | 0.853939113 | 0.928472575 | 637.9604394 | 647.1152805 | ENSG00000138688 |
| PAXBP1-AS1 | 0.021948052 | 1.015329541 | 0.958576064 | 0.982881365 | 25.13371714 | 25.51039965 | ENSG00000238197 |
| TTC38 | 0.021913633 | 1.015305318 | 0.785302968 | 0.890433415 | 920.1481266 | 933.84132 | ENSG00000075234 |
| SESN2 | 0.021904862 | 1.015299145 | 0.806334619 | 0.901201229 | 968.4435003 | 982.9587984 | ENSG00000130766 |
| HOOK3 | 0.021894428 | 1.015291802 | 0.889039741 | 0.950385798 | 332.0217306 | 336.8553001 | ENSG00000168172 |
| NQO1 | 0.021889298 | 1.015288192 | 0.775246439 | 0.884624109 | 991.2523934 | 1006.282799 | ENSG00000181019 |
| GPATCH1 | 0.021732708 | 1.015177998 | 0.877458088 | 0.943022763 | 280.268662 | 284.4395486 | ENSG00000076650 |
| RTKN | 0.021668455 | 1.015132787 | 0.750046477 | 0.869965151 | 1374.620374 | 1395.551039 | ENSG00000114993 |
| ITGA4 | 0.021566938 | 1.015061359 | 0.929081522 | 0.967771899 | 89.71225135 | 91.0982198 | ENSG00000115232 |
| RASL10B | 0.021551898 | 1.015050776 | 0.800604093 | 0.898320559 | 1039.241783 | 1054.957403 | ENSG00000270885 |
| CNBP | 0.02152561 | 1.015032281 | 0.636821968 | 0.800725077 | 8060.650099 | 8181.394807 | ENSG00000169714 |
| MED26 | 0.021442567 | 1.014973856 | 0.913956752 | 0.961911055 | 143.6464213 | 145.8224909 | ENSG00000105085 |
| NPRL2 | 0.021354794 | 1.014912108 | 0.864883595 | 0.934989221 | 319.8012445 | 324.3789463 | ENSG00000114388 |
| HK1 | 0.021348264 | 1.014907514 | 0.658982264 | 0.816271812 | 9455.242411 | 9595.24839 | ENSG00000156515 |
| C6orf132 | 0.021330687 | 1.014895149 | 0.770730408 | 0.881997379 | 1690.030206 | 1715.511398 | ENSG00000188112 |
| PSMC2 | 0.021306665 | 1.01487825 | 0.695922285 | 0.837581566 | 2742.829583 | 2783.596689 | ENSG00000161057 |
| ETV4 | 0.021230813 | 1.014824893 | 0.683811299 | 0.830895404 | 3535.515166 | 3588.128571 | ENSG00000175832 |
| ZNF35 | 0.021165735 | 1.014779116 | 0.884342929 | 0.947241575 | 264.5045206 | 268.4682505 | ENSG00000169981 |
| AKAP5 | 0.021156411 | 1.014772558 | 0.928119 | 0.967366576 | 82.01164863 | 83.21195466 | ENSG00000179841 |
| NA | 0.021108391 | 1.014738782 | 0.947788759 | 0.977056885 | 52.83814067 | 53.49046035 | ENSG00000261512 |
| LOC102723582 | 0.021084199 | 1.014721766 | 0.982845326 | 0.995796876 | 4.896803489 | 4.993636052 | ENSG00000272758 |
| RPL8 | 0.021068117 | 1.014710455 | 0.70787872 | 0.845910689 | 50792.07029 | 51538.77182 | ENSG00000161016 |
| MLLT1 | 0.021065817 | 1.014708838 | 0.75969222 | 0.875284459 | 1781.322244 | 1808.198308 | ENSG00000130382 |
| AP1S2 | 0.021026265 | 1.014681019 | 0.795343435 | 0.895543809 | 4607.828098 | 4675.933754 | ENSG00000182287 |
| PPP1R16A | 0.021021407 | 1.014677603 | 0.880367218 | 0.944837604 | 250.7135048 | 254.5077897 | ENSG00000160972 |
| TOP1MT | 0.020992492 | 1.014657266 | 0.735959656 | 0.861271552 | 1974.56303 | 2003.536769 | ENSG00000184428 |
| ACOX1 | 0.020978922 | 1.014647722 | 0.85695095 | 0.930171411 | 366.1234669 | 371.3368887 | ENSG00000161533 |
| OSBPL6 | 0.020977334 | 1.014646605 | 0.898781424 | 0.95544538 | 189.9398891 | 192.5554348 | ENSG00000079156 |
| NUP54 | 0.020956713 | 1.014632103 | 0.774192457 | 0.883955535 | 1416.367315 | 1436.935488 | ENSG00000138750 |
| TNK2 | 0.020952596 | 1.014629207 | 0.835968135 | 0.917543777 | 856.4311578 | 869.4034267 | ENSG00000061938 |
| NA | 0.020866103 | 1.01456838 | 0.919036445 | 0.964051188 | 115.7077358 | 117.4018702 | ENSG00000124224 |
| ATF7IP | 0.020827186 | 1.014541012 | 0.798450437 | 0.897223346 | 1921.41321 | 1948.667711 | ENSG00000171681 |
| RPL15 | 0.020747004 | 1.014484628 | 0.616708496 | 0.787224317 | 19082.66261 | 19358.62063 | ENSG00000174748 |
| IFFO2 | 0.020716159 | 1.014462938 | 0.87117999 | 0.939537097 | 338.9127634 | 343.4448448 | ENSG00000169991 |
| ANKRD40 | 0.020698234 | 1.014450334 | 0.778314385 | 0.886943686 | 1154.75169 | 1171.048248 | ENSG00000154945 |

| | | | | | | | |
|---|---|---|---|---|---|---|---|
| LEMD3 | 0.020682106 | 1.014438993 | 0.826436569 | 0.9124915 | 959.8693089 | 973.2269882 | ENSG00000174106 |
| HSF2 | 0.020668607 | 1.014429501 | 0.885031781 | 0.947622636 | 236.2555768 | 239.6833267 | ENSG00000025156 |
| WNK1 | 0.020659905 | 1.014423382 | 0.788750671 | 0.891842679 | 3785.64799 | 3839.6608 | ENSG00000060237 |
| SEC23B | 0.020622358 | 1.014396982 | 0.727933751 | 0.856731016 | 2244.006151 | 2276.480606 | ENSG00000101310 |
| NA | 0.020588928 | 1.014373477 | 0.967529507 | 0.987242444 | 17.71748828 | 17.91269558 | ENSG00000229546 |
| MRPS25 | 0.02049212 | 1.014305412 | 0.852795906 | 0.927710293 | 801.9038825 | 813.7221233 | ENSG00000131368 |
| FIS1 | 0.020488587 | 1.014302928 | 0.787974854 | 0.891582187 | 1061.575539 | 1077.073995 | ENSG00000214253 |
| NEK9 | 0.020461252 | 1.01428371 | 0.778453355 | 0.886943686 | 1160.771777 | 1177.202435 | ENSG00000119638 |
| VAMP7 | 0.020410634 | 1.014248124 | 0.864958927 | 0.935004026 | 360.3910593 | 365.8048171 | ENSG00000124333 |
| NA | 0.02022618 | 1.014118456 | 0.981312754 | 0.995225356 | 6.227679473 | 6.345280403 | ENSG00000217835 |
| NA | 0.020118157 | 1.014042527 | 0.954363738 | 0.981119172 | 43.02645699 | 43.67913994 | ENSG00000263266 |
| SECISBP2 | 0.0200979 | 1.014028288 | 0.77863825 | 0.886993808 | 1486.610069 | 1507.269732 | ENSG00000187742 |
| NA | 0.020066977 | 1.014006554 | 0.966945101 | 0.987033408 | 19.30784796 | 19.52518706 | ENSG00000264016 |
| HERPUD1 | 0.02004871 | 1.013993715 | 0.857665979 | 0.930409931 | 490.7598145 | 497.6525851 | ENSG00000051108 |
| CYB5R2 | 0.019925905 | 1.013907405 | 0.849017646 | 0.925283873 | 521.4720178 | 528.4223227 | ENSG00000166394 |
| CBLB | 0.019914957 | 1.013899711 | 0.877767779 | 0.943087245 | 421.0323914 | 426.4680786 | ENSG00000114423 |
| GNAQ | 0.019562986 | 1.013652383 | 0.787417083 | 0.891546762 | 1304.782355 | 1322.174008 | ENSG00000156052 |
| FUBP1 | 0.01949957 | 1.013607827 | 0.711366645 | 0.847591497 | 4203.122035 | 4259.811646 | ENSG00000162613 |
| MAPKAPK2 | 0.01949007 | 1.013601152 | 0.788914574 | 0.891843683 | 1158.626576 | 1174.100444 | ENSG00000162889 |
| PRRC2A | 0.019485981 | 1.01359828 | 0.681381616 | 0.829757233 | 14693.35537 | 14892.9265 | ENSG00000204469 |
| POP1 | 0.019482484 | 1.013595823 | 0.854928517 | 0.929013468 | 446.0981987 | 452.3282061 | ENSG00000104356 |
| FLJ20021 | 0.019455646 | 1.013576967 | 0.951691733 | 0.979385231 | 42.84617141 | 43.48561658 | ENSG00000254531 |
| HMSD | 0.019427245 | 1.013557014 | 0.965630177 | 0.986716954 | 24.53618817 | 24.97035572 | ENSG00000221887 |
| NA | 0.019413677 | 1.013547482 | 0.984759134 | 0.996174529 | 4.266349541 | 4.344723624 | ENSG00000231434 |
| SRSF3 | 0.01937189 | 1.013518125 | 0.703877209 | 0.842524014 | 12852.50258 | 13025.12507 | ENSG00000112081 |
| KIF2C | 0.019360347 | 1.013510017 | 0.816654134 | 0.906513449 | 1463.988529 | 1483.086384 | ENSG00000142945 |
| UBIAD1 | 0.019276957 | 1.013451436 | 0.84443872 | 0.922542416 | 667.4561305 | 676.2694684 | ENSG00000120942 |
| CKM | 0.019265371 | 1.013443297 | 0.986000734 | 0.996661342 | 3.592012868 | 3.658165904 | ENSG00000104879 |
| METTL14 | 0.019264919 | 1.013442979 | 0.83716428 | 0.918172233 | 606.3966403 | 614.3296422 | ENSG00000145388 |
| OTUD4 | 0.019241384 | 1.013426447 | 0.806371103 | 0.901201229 | 1281.773066 | 1298.591196 | ENSG00000164164 |
| CBX3P2 | 0.019239284 | 1.013424972 | 0.984876482 | 0.996174529 | 4.579199175 | 4.660619757 | ENSG00000266405 |
| ZSCAN20 | 0.019199625 | 1.013397114 | 0.944119817 | 0.975473396 | 63.50742125 | 64.35686757 | ENSG00000121903 |
| NFKBIL1 | 0.019183537 | 1.013385813 | 0.91524672 | 0.962438571 | 140.9022172 | 142.7021208 | ENSG00000204498 |
| SEC22C | 0.01915849 | 1.01336822 | 0.846127019 | 0.923543095 | 690.8438399 | 699.879142 | ENSG00000093183 |
| SNORD14A | 0.019093839 | 1.013322809 | 0.975307338 | 0.991904738 | 12.40791675 | 12.62803946 | ENSG00000272034 |
| SS18 | 0.019091781 | 1.013321363 | 0.765292516 | 0.878574417 | 1701.205107 | 1724.133397 | ENSG00000141380 |
| DDX39A | 0.019082274 | 1.013314686 | 0.72130012 | 0.853073731 | 3360.175907 | 3405.0677 | ENSG00000123136 |
| CSNK1G3 | 0.019065762 | 1.013303088 | 0.842859729 | 0.9217804 | 596.0787525 | 603.9590549 | ENSG00000151292 |
| FAM110D | 0.019052469 | 1.013293752 | 0.964486254 | 0.986052631 | 30.97471394 | 31.49343678 | ENSG00000197245 |
| JKAMP | 0.019051954 | 1.01329339 | 0.861589987 | 0.932816999 | 447.3834386 | 453.3234712 | ENSG00000050130 |
| SMCR8 | 0.018962902 | 1.013230845 | 0.809847461 | 0.90313851 | 1016.69896 | 1030.385675 | ENSG00000176994 |
| CDV3 | 0.018924754 | 1.013204054 | 0.723081049 | 0.854154341 | 3191.61862 | 3233.609817 | ENSG00000091527 |
| SLC16A13 | 0.018921506 | 1.013201772 | 0.933469049 | 0.969858363 | 97.15576078 | 98.56853775 | ENSG00000174327 |
| TSGA10 | 0.018844576 | 1.013147746 | 0.956949384 | 0.982418568 | 39.43207986 | 39.97708589 | ENSG00000135951 |
| STK38L | 0.018816906 | 1.013128315 | 0.851140123 | 0.926282558 | 1073.36757 | 1087.715441 | ENSG00000211455 |
| LINC00678 | 0.018613804 | 1.012985697 | 0.761623107 | 0.876689032 | 3361.86194 | 3404.688664 | ENSG00000254934 |
| DLC1 | 0.018579357 | 1.01296151 | 0.942365489 | 0.974513868 | 73.71499148 | 74.56516605 | ENSG00000164741 |
| ULBP1 | 0.018534315 | 1.012929886 | 0.979163881 | 0.994113751 | 9.537682752 | 9.699373451 | ENSG00000111981 |
| CRKL | 0.018488662 | 1.012897833 | 0.749922874 | 0.869935742 | 2721.018911 | 2756.031142 | ENSG00000099942 |
| MRPL51 | 0.018420814 | 1.012850199 | 0.770258858 | 0.881997379 | 2303.197898 | 2332.392595 | ENSG00000111639 |
| SCNN1B | 0.018418673 | 1.012848696 | 0.985092908 | 0.99621074 | 4.889684544 | 4.973110921 | ENSG00000168447 |
| PDLIM7 | 0.018413653 | 1.012845171 | 0.80882845 | 0.902949338 | 1553.039413 | 1573.734859 | ENSG00000196923 |
| LATS2 | 0.018401257 | 1.012836468 | 0.853421283 | 0.92812329 | 587.1599812 | 594.754871 | ENSG00000150457 |
| SERINC5 | 0.018252274 | 1.012731881 | 0.803058871 | 0.89940498 | 1746.555091 | 1767.927206 | ENSG00000164300 |
| FTSJ3 | 0.018212057 | 1.012703651 | 0.749193721 | 0.869654826 | 2697.570412 | 2731.85408 | ENSG00000108592 |
| NA | 0.018134324 | 1.012649087 | 0.977361134 | 0.992924042 | 11.79170069 | 11.98782888 | ENSG00000231856 |
| TSC22D4 | 0.018129331 | 1.012645582 | 0.84966199 | 0.925513999 | 723.6686493 | 733.3676934 | ENSG00000166925 |
| SESN3 | 0.018119001 | 1.012638332 | 0.885922206 | 0.948287755 | 475.6528228 | 481.7443122 | ENSG00000149212 |
| TRIM33 | 0.018030219 | 1.012576017 | 0.848947178 | 0.92528236 | 1154.501715 | 1168.692601 | ENSG00000197323 |
| PLCXD2 | 0.017948971 | 1.012518994 | 0.951939716 | 0.97945433 | 65.24959074 | 66.05117603 | ENSG00000240891 |
| RAB33B | 0.017943246 | 1.012514975 | 0.90882262 | 0.960165884 | 227.9055537 | 230.6005899 | ENSG00000172007 |
| SNRNP70 | 0.017859632 | 1.012456295 | 0.742775879 | 0.866044482 | 9655.132159 | 9775.93349 | ENSG00000104852 |
| ANKRD27 | 0.017834729 | 1.012438819 | 0.789400491 | 0.892025074 | 1660.551118 | 1681.068332 | ENSG00000105186 |
| ATP8A2 | 0.017833792 | 1.012438161 | 0.884287236 | 0.947235646 | 339.9142132 | 344.108702 | ENSG00000132932 |
| EYA2 | 0.017818267 | 1.012427266 | 0.962642407 | 0.985098636 | 31.51915631 | 31.89550081 | ENSG00000064655 |
| SBSPON | 0.017812162 | 1.012422982 | 0.951085746 | 0.979047559 | 55.29797089 | 56.06825485 | ENSG00000164764 |
| ROCK2 | 0.017802057 | 1.012415891 | 0.877329945 | 0.942992374 | 624.2635479 | 631.6014743 | ENSG00000134318 |
| MET | 0.017752845 | 1.012381356 | 0.878779976 | 0.943637906 | 376.7865444 | 381.4154147 | ENSG00000105976 |
| ARHGAP45 | 0.01773466 | 1.012368596 | 0.795526442 | 0.895554092 | 1867.421073 | 1891.095649 | ENSG00000180448 |
| SLC12A5 | 0.01771532 | 1.012355024 | 0.966460396 | 0.98694666 | 26.65555713 | 26.94821191 | ENSG00000124140 |
| NA | 0.017685725 | 1.012334258 | 0.985659415 | 0.996476341 | 4.93356727 | 5.012931678 | ENSG00000108958 |
| SHISAL2A | 0.017664504 | 1.012319367 | 0.974824367 | 0.991520341 | 14.73309798 | 14.97438176 | ENSG00000182183 |
| DYNLT1 | 0.017625977 | 1.012292333 | 0.863093703 | 0.933749117 | 1056.24176 | 1068.946528 | ENSG00000146425 |
| NME4 | 0.017543 | 1.012234113 | 0.752432188 | 0.871085164 | 5184.456387 | 5248.086081 | ENSG00000103202 |

| | | | | | | | |
|---|---|---|---|---|---|---|---|
| HCN3 | 0.017511228 | 1.012211821 | 0.911617794 | 0.961009829 | 233.9078697 | 236.9166136 | ENSG00000143630 |
| SMIM24 | 0.017459059 | 1.012175219 | 0.949306983 | 0.977907982 | 72.84704721 | 73.85015935 | ENSG00000095932 |
| TXNL1 | 0.017404293 | 1.012136797 | 0.789765441 | 0.892242184 | 1570.025893 | 1589.030872 | ENSG00000091164 |
| ZNF66 | 0.017363899 | 1.012108458 | 0.929837743 | 0.968086748 | 120.7542642 | 122.119998 | ENSG00000160229 |
| SLC20A2 | 0.017345803 | 1.012095763 | 0.843847803 | 0.922223826 | 726.1515908 | 735.0497568 | ENSG00000168575 |
| NA | 0.017313963 | 1.012073427 | 0.967120263 | 0.987033408 | 25.98241566 | 26.25295233 | ENSG00000206228 |
| LINC01547 | 0.017287833 | 1.012055096 | 0.972607928 | 0.989959077 | 18.72959089 | 18.88715196 | ENSG00000183250 |
| PHF21B | 0.017259748 | 1.012035395 | 0.89153981 | 0.951765365 | 555.5737803 | 562.32078 | ENSG00000056487 |
| EIF3M | 0.01712878 | 1.011943526 | 0.732045746 | 0.859049519 | 3788.120741 | 3833.156069 | ENSG00000149100 |
| COX11 | 0.017081474 | 1.011910345 | 0.856046195 | 0.929646557 | 763.0565415 | 772.2890099 | ENSG00000166260 |
| MED20 | 0.016971956 | 1.011833532 | 0.847482937 | 0.924379996 | 989.6005818 | 1000.598801 | ENSG00000124641 |
| MPLKIP | 0.016971244 | 1.011833033 | 0.855955605 | 0.929646557 | 868.0364149 | 878.087092 | ENSG00000168303 |
| ITGA2B | 0.016958621 | 1.01182418 | 0.923446194 | 0.965309497 | 221.739275 | 224.515959 | ENSG00000005961 |
| RBMX | 0.016894215 | 1.01177901 | 0.709529201 | 0.846496645 | 11330.20508 | 11463.01977 | ENSG00000147274 |
| RHD | 0.016880813 | 1.011769611 | 0.987139537 | 0.997171979 | 4.280587431 | 4.346899088 | ENSG00000187010 |
| TMTC3 | 0.016860061 | 1.011755058 | 0.904687655 | 0.958481852 | 478.6140501 | 484.2924579 | ENSG00000139324 |
| DIRAS1 | 0.016849957 | 1.011747972 | 0.849482099 | 0.925384716 | 929.7531844 | 941.2785545 | ENSG00000176490 |
| ATP5PO | 0.016838153 | 1.011739694 | 0.944758505 | 0.975866344 | 87.6468593 | 88.71529375 | ENSG00000241837 |
| ARMC1 | 0.01683075 | 1.011734502 | 0.850123155 | 0.92567733 | 762.356958 | 770.9197712 | ENSG00000104442 |
| CYP2S1 | 0.016754589 | 1.011681093 | 0.789002031 | 0.891843683 | 7205.611817 | 7290.747263 | ENSG00000167600 |
| TAOK2 | 0.016732248 | 1.011665427 | 0.801574719 | 0.898520796 | 1596.551555 | 1615.655758 | ENSG00000149930 |
| SLC7A6OS | 0.016679626 | 1.011628528 | 0.913915311 | 0.961911055 | 217.5206987 | 220.2618792 | ENSG00000103061 |
| DPYSL3 | 0.016627434 | 1.01159193 | 0.720343414 | 0.852628353 | 19910.98253 | 20141.31932 | ENSG00000113657 |
| CYFIP2 | 0.016574435 | 1.01155477 | 0.799677933 | 0.898068373 | 2004.854691 | 2028.214094 | ENSG00000055163 |
| CDK17 | 0.016509261 | 1.011509074 | 0.905538327 | 0.958927594 | 316.3545591 | 319.819775 | ENSG00000059758 |
| RAP1B | 0.016507231 | 1.01150765 | 0.839730385 | 0.919855472 | 1478.727321 | 1495.535503 | ENSG00000127314 |
| LPAR3 | 0.016507068 | 1.011507536 | 0.885198165 | 0.947622636 | 415.9105293 | 420.5540166 | ENSG00000171517 |
| PRIM2 | 0.016367851 | 1.011409932 | 0.8612053 | 0.932648138 | 642.9060099 | 650.0639475 | ENSG00000146143 |
| SOCS2-AS1 | 0.01634837 | 1.011396275 | 0.969212513 | 0.988047412 | 25.00471257 | 25.23337832 | ENSG00000246985 |
| ATXN7 | 0.016238825 | 1.011319482 | 0.983269183 | 0.995796876 | 8.457976314 | 8.581808255 | ENSG00000163635 |
| TMSB4X | 0.016202901 | 1.0112943 | 0.794406033 | 0.894861791 | 16514.12735 | 16700.19713 | ENSG00000205542 |
| ZNF718 | 0.016125533 | 1.011240068 | 0.91903168 | 0.964051188 | 190.6779647 | 192.6498101 | ENSG00000250312 |
| RXRB | 0.016094074 | 1.011218017 | 0.872940738 | 0.940412849 | 576.8050248 | 583.4128035 | ENSG00000204231 |
| ITGB1BP1 | 0.015934889 | 1.011106447 | 0.895199242 | 0.953032171 | 367.3259487 | 371.2925061 | ENSG00000119185 |
| DGKZ | 0.015830616 | 1.01103337 | 0.786731033 | 0.89126668 | 3258.719683 | 3295.457815 | ENSG00000149091 |
| PTPRD | 0.015807742 | 1.01101734 | 0.892925373 | 0.952281345 | 782.0160954 | 789.9859829 | ENSG00000153707 |
| DNAJC24 | 0.015769865 | 1.010990797 | 0.912856277 | 0.961597187 | 242.9381532 | 245.7167783 | ENSG00000170946 |
| NA | 0.015705904 | 1.010945977 | 0.975658629 | 0.991946116 | 18.42978394 | 18.55631112 | ENSG00000227376 |
| NA | 0.015675857 | 1.010924922 | 0.987758565 | 0.997423832 | 4.902727226 | 4.970935457 | ENSG00000272416 |
| KLHDC8B | 0.015586499 | 1.010862309 | 0.89715228 | 0.954467117 | 367.6414937 | 371.8445842 | ENSG00000185909 |
| DLEC1 | 0.015584927 | 1.010861208 | 0.984886134 | 0.996174529 | 7.855998146 | 7.964298277 | ENSG00000008226 |
| SWAP70 | 0.015581699 | 1.010858945 | 0.83697175 | 0.918172233 | 1352.269353 | 1366.529353 | ENSG00000133789 |
| SEMA3A | 0.015569984 | 1.010850737 | 0.859743284 | 0.931886123 | 896.8737878 | 906.3926311 | ENSG00000075213 |
| DCUN1D4 | 0.015551582 | 1.010837844 | 0.872493319 | 0.940222903 | 711.325641 | 719.2041114 | ENSG00000109184 |
| DGKB | 0.015530146 | 1.010822824 | 0.964055974 | 0.985984136 | 42.30971214 | 42.63977584 | ENSG00000136267 |
| TRIM44 | 0.015527142 | 1.01082072 | 0.818132409 | 0.907407678 | 1658.414082 | 1676.238863 | ENSG00000166326 |
| NMRK2 | 0.01552582 | 1.010819794 | 0.859508073 | 0.931845109 | 829.1006013 | 838.4060075 | ENSG00000077009 |
| SMN1 | 0.015463952 | 1.010776447 | 0.960775897 | 0.984255629 | 47.20267682 | 47.72922761 | ENSG00000205571 |
| TGOLN2 | 0.015454009 | 1.01076948 | 0.821253975 | 0.909334444 | 1688.714687 | 1706.382189 | ENSG00000152291 |
| HIKESHI | 0.015404983 | 1.010735133 | 0.853130267 | 0.927913672 | 825.3475224 | 834.3500855 | ENSG00000149196 |
| PTH2R | 0.01536096 | 1.010704292 | 0.985792649 | 0.996541255 | 5.876870851 | 5.961566034 | ENSG00000144407 |
| PPP1R7 | 0.015326136 | 1.010679895 | 0.843784418 | 0.922210124 | 1019.721842 | 1030.377563 | ENSG00000115685 |
| EHMT2 | 0.015192722 | 1.010586437 | 0.766114624 | 0.878994404 | 5121.7172 | 5175.922513 | ENSG00000204371 |
| TMSB10 | 0.01510828 | 1.010527288 | 0.792848863 | 0.89406764 | 7247.135941 | 7322.741994 | ENSG00000034510 |
| TMEM98 | 0.015085565 | 1.010511378 | 0.819162468 | 0.908070158 | 1713.768291 | 1731.236909 | ENSG00000006042 |
| RPS6KA1 | 0.015045889 | 1.010483587 | 0.815463756 | 0.90625746 | 1955.883778 | 1975.98434 | ENSG00000117676 |
| CLCN7 | 0.015043902 | 1.010482196 | 0.827899029 | 0.913518736 | 1386.17667 | 1400.927968 | ENSG00000103249 |
| PPP1R9A | 0.015043592 | 1.010481979 | 0.913070731 | 0.961597187 | 290.6858572 | 293.4591313 | ENSG00000158528 |
| NA | 0.015029252 | 1.010471934 | 0.988413836 | 0.997507036 | 4.627836581 | 4.689563196 | ENSG00000260669 |
| WWC2-AS2 | 0.015014177 | 1.010461376 | 0.974443208 | 0.991254655 | 22.38180931 | 22.68190029 | ENSG00000251359 |
| RACK1 | 0.014886925 | 1.010372253 | 0.726495773 | 0.855887253 | 24967.34027 | 25226.34094 | ENSG00000204628 |
| PAFAH1B1 | 0.014783351 | 1.010299719 | 0.831505901 | 0.915296977 | 1866.061975 | 1885.373561 | ENSG00000007168 |
| TMEM11 | 0.014741849 | 1.010270656 | 0.8666443 | 0.936410207 | 708.8474292 | 716.1903617 | ENSG00000178307 |
| RPL36AL | 0.014693527 | 1.010236818 | 0.829725853 | 0.914626006 | 3681.144521 | 3718.909439 | ENSG00000165502 |
| NA | 0.014477771 | 1.010085748 | 0.984448137 | 0.995944879 | 8.577777016 | 8.689447249 | ENSG00000241081 |
| SGPL1 | 0.014451555 | 1.010067393 | 0.834979738 | 0.917376243 | 1494.781027 | 1509.587465 | ENSG00000166224 |
| TP53I3 | 0.014340314 | 1.009989514 | 0.891714654 | 0.951844407 | 517.3889285 | 522.5468722 | ENSG00000115129 |
| RHBDD3 | 0.014296976 | 1.009959174 | 0.856043966 | 0.929646557 | 1639.733511 | 1656.888455 | ENSG00000100263 |
| CCDC130 | 0.014213383 | 1.009900657 | 0.91130829 | 0.961009829 | 347.2379219 | 350.9177601 | ENSG00000104957 |
| TMEM184C | 0.014156315 | 1.009860709 | 0.884526818 | 0.947384809 | 731.6453384 | 738.8897478 | ENSG00000164168 |
| MAF | 0.014136241 | 1.009846658 | 0.946945422 | 0.976736266 | 133.6218064 | 135.0107038 | ENSG00000178573 |
| NFXL1 | 0.014127103 | 1.009840262 | 0.917141315 | 0.96312259 | 290.5788715 | 293.6803686 | ENSG00000170448 |
| TMEM175 | 0.013785301 | 1.00960104 | 0.956403073 | 0.982211615 | 71.04464603 | 71.78859427 | ENSG00000127419 |
| PLEKHB2 | 0.013781609 | 1.009598456 | 0.805422721 | 0.900817151 | 2607.081794 | 2632.113709 | ENSG00000115762 |

| | | | | | | | |
|---|---|---|---|---|---|---|---|
| GAD1 | 0.013721586 | 1.009556452 | 0.986548027 | 0.996898042 | 7.21486578 | 7.303562573 | ENSG00000128683 |
| CPOX | 0.01369896 | 1.00954062 | 0.870285805 | 0.938742301 | 868.452872 | 876.8640406 | ENSG00000080819 |
| TMEM235 | 0.013627343 | 1.009490507 | 0.990052641 | 0.99804732 | 3.602691286 | 3.648518091 | ENSG00000204278 |
| HERC6 | 0.01358118 | 1.009458205 | 0.948075102 | 0.977207636 | 105.4458838 | 106.4531945 | ENSG00000138642 |
| TANGO6 | 0.013534236 | 1.009425359 | 0.907155366 | 0.95997025 | 434.1431462 | 438.2851503 | ENSG00000103047 |
| LOC100288175 | 0.013525093 | 1.009418962 | 0.971586379 | 0.989399232 | 35.9681557 | 36.26727596 | ENSG00000217801 |
| KLLN | 0.013523014 | 1.009417507 | 0.984969966 | 0.996205967 | 9.486681082 | 9.601382271 | ENSG00000227268 |
| ARNT2 | 0.013431424 | 1.009353426 | 0.921177785 | 0.964771134 | 345.3759 | 348.2961071 | ENSG00000172379 |
| LYPLA1 | 0.013414914 | 1.009341875 | 0.86345595 | 0.933980504 | 1915.805356 | 1933.708242 | ENSG00000120992 |
| ANO8 | 0.013275231 | 1.009244154 | 0.960831524 | 0.984255629 | 1285.72161 | 1297.726366 | ENSG00000074855 |
| NA | 0.013168555 | 1.009169531 | 0.990990706 | 0.99806433 | 3.962983117 | 4.007356402 | ENSG00000279140 |
| NA | 0.013112841 | 1.00913056 | 0.984358685 | 0.995944879 | 9.826811286 | 9.945275883 | ENSG00000235558 |
| ENTPD1 | 0.013076793 | 1.009105346 | 0.937116427 | 0.971870005 | 180.7746767 | 182.4667666 | ENSG00000138185 |
| ADAMTS2 | 0.012856436 | 1.008951227 | 0.923772804 | 0.965407741 | 291.4865287 | 293.8890773 | ENSG00000087116 |
| NA | 0.012830537 | 1.008933115 | 0.990422266 | 0.99806433 | 3.911981447 | 3.957887833 | ENSG00000185839 |
| RAB25 | 0.012802495 | 1.008913504 | 0.911435661 | 0.961009829 | 437.4910306 | 441.2576521 | ENSG00000132698 |
| ZNF772 | 0.012781846 | 1.008899064 | 0.958809897 | 0.983014381 | 75.65260031 | 76.23299008 | ENSG00000197128 |
| MRPL49 | 0.012721826 | 1.008857092 | 0.869360407 | 0.938217686 | 1142.392969 | 1152.180267 | ENSG00000149792 |
| NA | 0.012677267 | 1.008825933 | 0.991531957 | 0.99806433 | 3.295765389 | 3.332621958 | ENSG00000275719 |
| SLC35G2 | 0.012626937 | 1.00879074 | 0.96268338 | 0.985098636 | 62.26670113 | 62.82619306 | ENSG00000168917 |
| RPS6KA4 | 0.012619116 | 1.008785271 | 0.891150443 | 0.951672479 | 650.6564447 | 656.64749 | ENSG00000162302 |
| NECTIN3 | 0.012597529 | 1.008770176 | 0.902981029 | 0.957841349 | 523.0324533 | 527.4185441 | ENSG00000177707 |
| SCOC | 0.012590216 | 1.008765063 | 0.918054475 | 0.96367399 | 474.2933926 | 478.6418403 | ENSG00000153130 |
| CYP4X1 | 0.012335194 | 1.008586761 | 0.967020866 | 0.987033408 | 54.81014902 | 55.26830975 | ENSG00000186377 |
| TM2D3 | 0.012280669 | 1.008548644 | 0.913326383 | 0.961597187 | 403.9136267 | 407.3429184 | ENSG00000184277 |
| LRP5L | 0.012257946 | 1.008532759 | 0.965111771 | 0.986395147 | 76.04375834 | 76.61925878 | ENSG00000100068 |
| NAF1 | 0.012209391 | 1.008498817 | 0.902079711 | 0.957227361 | 603.358578 | 608.8037182 | ENSG00000145414 |
| BRD3 | 0.012156319 | 1.008461718 | 0.861345848 | 0.932648138 | 2029.565694 | 2046.949801 | ENSG00000169925 |
| SELENOS | 0.012150822 | 1.008457875 | 0.899770436 | 0.955743556 | 681.3781835 | 687.537007 | ENSG00000131871 |
| ZNRD1 | 0.012127298 | 1.008441432 | 0.953824209 | 0.980831483 | 115.7744238 | 116.7353861 | ENSG00000066379 |
| KRIT1 | 0.012091715 | 1.00841656 | 0.967635809 | 0.987242444 | 50.92753307 | 51.33357401 | ENSG00000001631 |
| GDPD3 | 0.012073589 | 1.00840389 | 0.985058087 | 0.99621074 | 11.86761867 | 11.99312577 | ENSG00000102886 |
| DDB2 | 0.012058962 | 1.008393667 | 0.913127105 | 0.961597187 | 598.649765 | 603.4123393 | ENSG00000134574 |
| ZNF529-AS1 | 0.012008937 | 1.008358701 | 0.974458177 | 0.991254655 | 34.58271857 | 34.81205999 | ENSG00000233527 |
| ZNF318 | 0.011853164 | 1.008249831 | 0.895194145 | 0.953032171 | 740.4371907 | 746.2238098 | ENSG00000171467 |
| TEX9 | 0.011823046 | 1.008228783 | 0.971383563 | 0.989388263 | 44.47093061 | 44.93184715 | ENSG00000151575 |
| NA | 0.011795398 | 1.008209461 | 0.983539606 | 0.995796876 | 13.85147449 | 13.99680397 | ENSG00000269054 |
| NA | 0.011773003 | 1.008193811 | 0.990747619 | 0.99806433 | 4.627836581 | 4.675564456 | ENSG00000258682 |
| ETFDH | 0.01174201 | 1.008172153 | 0.941291653 | 0.97409682 | 204.6587231 | 206.1123186 | ENSG00000171503 |
| PAQR3 | 0.011685121 | 1.008132398 | 0.912586648 | 0.961482481 | 540.3075543 | 544.9464114 | ENSG00000163291 |
| PRKCD | 0.011668325 | 1.008120662 | 0.903273625 | 0.957851743 | 711.1817364 | 716.6841312 | ENSG00000163932 |
| MTA1 | 0.01165943 | 1.008114446 | 0.832277825 | 0.915515303 | 3541.702801 | 3571.082298 | ENSG00000182979 |
| GGA1 | 0.011588065 | 1.008064579 | 0.883596655 | 0.946818136 | 1022.653224 | 1030.957187 | ENSG00000100083 |
| PRKCB | 0.0115636 | 1.008047485 | 0.929680011 | 0.968057669 | 322.4321826 | 324.7977313 | ENSG00000166501 |
| WBP4 | 0.011523017 | 1.008019129 | 0.94086959 | 0.973820136 | 206.9062586 | 208.7120786 | ENSG00000120688 |
| KLF6 | 0.011452687 | 1.00796999 | 0.921658542 | 0.964771134 | 442.6829125 | 445.8761171 | ENSG00000067082 |
| WASH2P | 0.01133215 | 1.007885778 | 0.969031288 | 0.987969328 | 54.92344117 | 55.47221596 | ENSG00000146556 |
| IRF9 | 0.011327535 | 1.007882554 | 0.989095721 | 0.997604649 | 7.583471765 | 7.657103998 | ENSG00000213928 |
| GNRHR2 | 0.011268574 | 1.007841364 | 0.984016261 | 0.995916668 | 14.18568096 | 14.3276448 | ENSG00000211451 |
| UBE2Q2 | 0.011094797 | 1.007719974 | 0.922159625 | 0.964771134 | 422.1989992 | 425.4606973 | ENSG00000140367 |
| ACOXL | 0.011059572 | 1.00769537 | 0.919697832 | 0.964147051 | 429.4215681 | 432.8636485 | ENSG00000153093 |
| REEP6 | 0.010980481 | 1.007640127 | 0.922396666 | 0.964906721 | 466.062586 | 469.781257 | ENSG00000115255 |
| MGAT2 | 0.010946051 | 1.00761608 | 0.991363509 | 0.99806433 | 4.647998207 | 4.691738659 | ENSG00000168282 |
| LOC339192 | 0.010945057 | 1.007615386 | 0.990848462 | 0.99806433 | 5.284375892 | 5.334124696 | ENSG00000267121 |
| SCART1 | 0.010800317 | 1.007514301 | 0.990067169 | 0.99804732 | 5.918389312 | 5.975564774 | ENSG00000214279 |
| CALY | 0.010752697 | 1.007481046 | 0.98367988 | 0.995807802 | 19.80189906 | 19.98078705 | ENSG00000130643 |
| GPBP1 | 0.010722465 | 1.007459934 | 0.882102824 | 0.945771249 | 1737.060199 | 1749.389462 | ENSG00000062194 |
| NDUFS7 | 0.010568515 | 1.007352434 | 0.968589914 | 0.987666064 | 66.41205476 | 66.88469904 | ENSG00000115286 |
| COX3 | 0.010566276 | 1.00735087 | 0.947225601 | 0.976838866 | 49090.55549 | 49451.645 | ENSG00000198938 |
| FBXO42 | 0.010478222 | 1.007289389 | 0.914383545 | 0.962084301 | 562.1585232 | 566.2942819 | ENSG00000037637 |
| GAS8 | 0.010459194 | 1.007276104 | 0.955667592 | 0.981978479 | 133.1965805 | 134.1265554 | ENSG00000141013 |
| CABLES2 | 0.010415086 | 1.007245308 | 0.922002745 | 0.964771134 | 445.966856 | 449.2962521 | ENSG00000149679 |
| AP1G2 | 0.010372463 | 1.007215551 | 0.947806022 | 0.977056885 | 290.9364425 | 292.8496109 | ENSG00000213983 |
| MAP2K1 | 0.010326591 | 1.007183526 | 0.89161488 | 0.951791702 | 1512.727132 | 1523.86122 | ENSG00000169032 |
| SGO1 | 0.010272663 | 1.007145878 | 0.960082438 | 0.983838344 | 132.2040547 | 133.2255703 | ENSG00000129810 |
| GLI2 | 0.010231718 | 1.007117295 | 0.916166334 | 0.962708726 | 1134.864012 | 1143.416178 | ENSG00000074047 |
| HBS1L | 0.010212258 | 1.00710371 | 0.885210478 | 0.947622636 | 1370.839762 | 1380.249535 | ENSG00000112339 |
| CHMP6 | 0.010139179 | 1.007052698 | 0.935609104 | 0.971184542 | 319.3235424 | 321.4400654 | ENSG00000176108 |
| NA | 0.010123084 | 1.007041462 | 0.991522623 | 0.99806433 | 4.971526258 | 5.015107141 | ENSG00000230832 |
| GAPDH | 0.010087063 | 1.007016319 | 0.812299178 | 0.904657072 | 38365.30007 | 38635.22404 | ENSG00000111640 |
| SUB1 | 0.010085522 | 1.007015244 | 0.875140916 | 0.94170908 | 2296.599298 | 2312.997872 | ENSG00000113387 |
| ARFGAP2 | 0.010073336 | 1.007006738 | 0.882418539 | 0.945904709 | 1666.626709 | 1677.739102 | ENSG00000149182 |
| DHX37 | 0.010050214 | 1.006990598 | 0.874751747 | 0.941359938 | 2676.406978 | 2695.500194 | ENSG00000150990 |
| SPATA7 | 0.009986689 | 1.006946259 | 0.967110328 | 0.987033408 | 94.27927199 | 94.72214538 | ENSG00000042317 |

| | | | | | | | |
|---|---|---|---|---|---|---|---|
| SERPINF1 | 0.009930501 | 1.006907043 | 0.930630095 | 0.96853421 | 408.5203858 | 411.322014 | ENSG00000132386 |
| GPATCH3 | 0.009890723 | 1.006879281 | 0.935394156 | 0.97102675 | 370.3175358 | 373.1357147 | ENSG00000198746 |
| NAA20 | 0.009825232 | 1.006833575 | 0.913322068 | 0.961597187 | 651.7853472 | 656.3101228 | ENSG00000173418 |
| CLP1 | 0.009797772 | 1.006814411 | 0.926976036 | 0.966805238 | 622.6029147 | 626.6212378 | ENSG00000172409 |
| STAM2 | 0.009790593 | 1.006809401 | 0.956840751 | 0.982382167 | 155.4434353 | 156.3514153 | ENSG00000115145 |
| E4F1 | 0.009764369 | 1.006791101 | 0.904152795 | 0.958242075 | 888.0652299 | 894.113133 | ENSG00000167967 |
| MUCL3 | 0.009755338 | 1.006784798 | 0.992750571 | 0.998354254 | 4.580394383 | 4.618623537 | ENSG00000168631 |
| NA | 0.009744075 | 1.006776938 | 0.962550752 | 0.985098636 | 110.5810935 | 111.3645621 | ENSG00000254285 |
| ARL5A | 0.009706066 | 1.006750414 | 0.939330303 | 0.972799059 | 563.8156475 | 568.1338317 | ENSG00000162980 |
| SIRT4 | 0.009686205 | 1.006736555 | 0.982574174 | 0.99570258 | 26.93136365 | 27.15117216 | ENSG00000089163 |
| TFCP2 | 0.009685237 | 1.00673588 | 0.892210357 | 0.951943084 | 1413.421241 | 1423.270935 | ENSG00000135457 |
| SLC35E1 | 0.009646205 | 1.006708643 | 0.92817778 | 0.967366576 | 526.6340017 | 530.0639686 | ENSG00000127526 |
| GCDH | 0.00964214 | 1.006705806 | 0.901221151 | 0.956531301 | 1195.963592 | 1204.249627 | ENSG00000105607 |
| ZDHHC11 | 0.009637894 | 1.006702843 | 0.966340614 | 0.986931074 | 113.9350056 | 114.7258439 | ENSG00000188818 |
| TMEM151B | 0.009600981 | 1.006677086 | 0.936510712 | 0.971528999 | 410.6499263 | 413.7599164 | ENSG00000178233 |
| ASPSCR1 | 0.00957017 | 1.006655587 | 0.919812871 | 0.964147051 | 1002.631378 | 1009.793541 | ENSG00000169696 |
| CNNM2 | 0.009530588 | 1.006627968 | 0.924153925 | 0.965621715 | 546.7169422 | 550.1175645 | ENSG00000148842 |
| NA | 0.009459944 | 1.006578679 | 0.985829162 | 0.996541255 | 16.55594013 | 16.68921535 | ENSG00000223803 |
| SLC66A1 | 0.009420995 | 1.006551504 | 0.933452759 | 0.969858363 | 386.7643019 | 389.1901843 | ENSG00000040487 |
| SNHG21 | 0.00932862 | 1.006487057 | 0.988895244 | 0.997539083 | 9.557844378 | 9.635906142 | ENSG00000250988 |
| RNF139-AS1 | 0.009315179 | 1.00647768 | 0.987841387 | 0.99745413 | 13.95111357 | 14.05809582 | ENSG00000245149 |
| FUNDC1 | 0.009308323 | 1.006472897 | 0.951104396 | 0.979047559 | 212.00323 | 213.5411348 | ENSG00000069509 |
| EMG1 | 0.009299507 | 1.006466747 | 0.894035338 | 0.952921173 | 1327.439081 | 1335.925279 | ENSG00000126749 |
| ELOB | 0.009165848 | 1.006373506 | 0.892348238 | 0.951982628 | 2818.217682 | 2836.365208 | ENSG00000103363 |
| CDCA4 | 0.009119127 | 1.006340916 | 0.912574511 | 0.961482481 | 1493.105388 | 1502.682353 | ENSG00000170779 |
| INSR | 0.009046816 | 1.006290478 | 0.919958906 | 0.964147051 | 717.1088233 | 721.4343762 | ENSG00000171105 |
| NPM1 | 0.009034628 | 1.006281976 | 0.830447866 | 0.914737804 | 45888.68907 | 46176.43766 | ENSG00000181163 |
| C2orf69 | 0.009009893 | 1.006264724 | 0.943347546 | 0.974995534 | 444.8550876 | 447.9220751 | ENSG00000178074 |
| MIF | 0.008990227 | 1.006251007 | 0.991777842 | 0.99806433 | 6.296478506 | 6.344050899 | ENSG00000240972 |
| TRIAP1 | 0.008979885 | 1.006243794 | 0.910382575 | 0.960811396 | 1104.3612 | 1111.405811 | ENSG00000170855 |
| NA | 0.008928108 | 1.006207681 | 0.9789126 | 0.994072402 | 42.52500763 | 42.84446005 | ENSG00000225031 |
| KCTD1 | 0.00890978 | 1.006194898 | 0.942807469 | 0.974757421 | 339.4083408 | 341.4312353 | ENSG00000134504 |
| ATG4B | 0.008889845 | 1.006180995 | 0.903508267 | 0.957906355 | 1815.980726 | 1827.002143 | ENSG00000168397 |
| ARHGAP44 | 0.008876095 | 1.006171405 | 0.931297005 | 0.968990425 | 524.8888047 | 528.1400045 | ENSG00000006740 |
| FIBP | 0.008812593 | 1.006127118 | 0.904133805 | 0.958242075 | 1318.877705 | 1327.092417 | ENSG00000172500 |
| NXPH3 | 0.008744125 | 1.006079371 | 0.961040869 | 0.984350589 | 168.999552 | 170.0907044 | ENSG00000182575 |
| GATAD1 | 0.008700307 | 1.006048814 | 0.926709211 | 0.966701516 | 678.5995799 | 682.8335903 | ENSG00000157259 |
| NA | 0.008661753 | 1.006021929 | 0.988733234 | 0.997539083 | 12.24451265 | 12.33361441 | ENSG00000279342 |
| RCHY1 | 0.008640766 | 1.006007294 | 0.94657567 | 0.976430116 | 362.8716098 | 365.08984 | ENSG00000163743 |
| YBX3 | 0.008610416 | 1.005986131 | 0.895923878 | 0.953588595 | 5345.400655 | 5378.53367 | ENSG00000060138 |
| COPB1 | 0.008606731 | 1.005983562 | 0.907644205 | 0.960149634 | 1418.970745 | 1427.532876 | ENSG00000129083 |
| NA | 0.008568565 | 1.005956949 | 0.992043679 | 0.998068556 | 6.942339399 | 6.990787862 | ENSG00000243014 |
| HSD3B7 | 0.008562195 | 1.005952507 | 0.988237783 | 0.997507036 | 13.49354692 | 13.59161851 | ENSG00000099377 |
| LOC93622 | 0.008502422 | 1.00591083 | 0.936805078 | 0.971690224 | 442.2007345 | 444.9271468 | ENSG00000170846 |
| ATF7 | 0.008427298 | 1.005858452 | 0.969359245 | 0.988090319 | 104.2570551 | 104.9843789 | ENSG00000170653 |
| TBC1D2B | 0.008260771 | 1.005742354 | 0.929845143 | 0.968086748 | 578.150418 | 581.4483562 | ENSG00000167202 |
| AMDHD2 | 0.008103308 | 1.005632589 | 0.950799209 | 0.978946847 | 324.2811357 | 326.3926941 | ENSG00000162066 |
| ZNF282 | 0.00810067 | 1.00563075 | 0.914977077 | 0.962281868 | 1608.025014 | 1617.259514 | ENSG00000170265 |
| UBE2A | 0.008063532 | 1.005604863 | 0.922936597 | 0.965130891 | 814.0356611 | 818.6547901 | ENSG00000077721 |
| KCNQ2 | 0.007954688 | 1.005528999 | 0.895116772 | 0.953032171 | 2372.683407 | 2385.94312 | ENSG00000075043 |
| NA | 0.007840501 | 1.005449416 | 0.983845629 | 0.995868827 | 31.70389109 | 31.90855359 | ENSG00000279672 |
| TMEM52 | 0.007830392 | 1.00544237 | 0.963078077 | 0.98530416 | 182.6105877 | 183.6493121 | ENSG00000178821 |
| YLPM1 | 0.007787536 | 1.005412504 | 0.908377591 | 0.960165884 | 2481.584649 | 2494.43305 | ENSG00000119596 |
| ACTR3B | 0.007763246 | 1.005395576 | 0.928447272 | 0.96753817 | 828.7852839 | 832.7741778 | ENSG00000133627 |
| GLS2 | 0.007749714 | 1.005386146 | 0.991867655 | 0.99806433 | 7.610752336 | 7.660225421 | ENSG00000135423 |
| NA | 0.00772851 | 1.00537137 | 0.993753057 | 0.998500968 | 4.640879262 | 4.67026757 | ENSG00000232004 |
| ZSCAN26 | 0.00771153 | 1.005359536 | 0.972521426 | 0.989935773 | 108.957835 | 109.4038987 | ENSG00000197062 |
| DMPK | 0.007626583 | 1.005300342 | 0.952527946 | 0.979818632 | 415.9205968 | 418.4541571 | ENSG00000104936 |
| WASF2 | 0.00757182 | 1.005262183 | 0.894635133 | 0.953032171 | 2991.746181 | 3007.870328 | ENSG00000158195 |
| ASB7 | 0.00756827 | 1.005259709 | 0.961395005 | 0.984595698 | 207.5262873 | 208.5460583 | ENSG00000183475 |
| EIF2AK1 | 0.007562209 | 1.005255485 | 0.894030159 | 0.952921173 | 2676.108595 | 2690.14478 | ENSG00000086232 |
| NA | 0.007519291 | 1.005225581 | 0.970447533 | 0.988890135 | 108.3140329 | 108.8396411 | ENSG00000197744 |
| FSTL1 | 0.0074669 | 1.005189077 | 0.897204144 | 0.954467117 | 4084.85501 | 4106.00311 | ENSG00000163430 |
| KCNQ1OT1 | 0.007398554 | 1.005141459 | 0.98352113 | 0.995796876 | 39.83033068 | 39.94424303 | ENSG00000269821 |
| NA | 0.007380668 | 1.005128997 | 0.983759561 | 0.995835083 | 35.66095047 | 35.87580569 | ENSG00000271888 |
| TRAPPC8 | 0.007309359 | 1.005079318 | 0.934311735 | 0.970383167 | 1050.925226 | 1056.162894 | ENSG00000153339 |
| CHST10 | 0.007307924 | 1.005078318 | 0.933278299 | 0.969789847 | 813.7183355 | 818.253715 | ENSG00000115526 |
| NKIRAS2 | 0.007286232 | 1.005063206 | 0.93377413 | 0.969985956 | 877.0537603 | 881.5312001 | ENSG00000168256 |
| NA | 0.007283331 | 1.005061185 | 0.993454275 | 0.998500968 | 8.697577719 | 8.745442209 | ENSG00000205976 |
| NA | 0.007269395 | 1.005051476 | 0.993836563 | 0.998500968 | 4.971526258 | 5.001108401 | ENSG00000235078 |
| NA | 0.007238262 | 1.005029788 | 0.993201202 | 0.998500968 | 5.96583151 | 6.001386791 | ENSG00000226200 |
| MTO1 | 0.007194324 | 1.00499918 | 0.942729249 | 0.974729914 | 515.9786274 | 518.4575833 | ENSG00000135297 |
| FAM83D | 0.007138203 | 1.004960086 | 0.90804991 | 0.960165884 | 1776.967947 | 1785.751054 | ENSG00000101447 |
| LPCAT2 | 0.007126794 | 1.004952139 | 0.975162899 | 0.991811253 | 90.18197095 | 90.56347275 | ENSG00000087253 |

| | | | | | | | |
|---|---|---|---|---|---|---|---|
| GRK1 | 0.007091131 | 1.004927297 | 0.995497831 | 0.998990416 | 3.32304596 | 3.341323813 | ENSG00000185974 |
| NA | 0.006946977 | 1.004826889 | 0.982435292 | 0.995631107 | 44.62749509 | 44.88020312 | ENSG00000226824 |
| DNAJA2 | 0.006876945 | 1.004778114 | 0.905817381 | 0.959035162 | 2918.568864 | 2932.238701 | ENSG00000069345 |
| IL12A | 0.006855051 | 1.004762866 | 0.994127373 | 0.998568983 | 5.948034148 | 5.979915701 | ENSG00000168811 |
| B4GALT6 | 0.006776544 | 1.004708192 | 0.955527064 | 0.981940937 | 409.0654386 | 410.8113221 | ENSG00000118276 |
| MAP7D1 | 0.006670772 | 1.004634533 | 0.921662933 | 0.964771134 | 3225.094958 | 3240.918983 | ENSG00000116871 |
| PPP2R5B | 0.006438006 | 1.004472457 | 0.972483932 | 0.989935773 | 131.5288283 | 132.2028022 | ENSG00000068971 |
| BRAF | 0.006435598 | 1.004470781 | 0.955585671 | 0.981947731 | 444.9170999 | 446.8121705 | ENSG00000157764 |
| GABPA | 0.006420769 | 1.004460456 | 0.951541676 | 0.979284172 | 464.431903 | 466.6368389 | ENSG00000154727 |
| RNF13 | 0.006339069 | 1.004403575 | 0.961689959 | 0.98479096 | 327.1510125 | 328.6297891 | ENSG00000082996 |
| NEMF | 0.006337268 | 1.004402321 | 0.944835765 | 0.975892771 | 618.4626468 | 621.268051 | ENSG00000165525 |
| SNRPD2 | 0.006296768 | 1.004374125 | 0.934660683 | 0.970489863 | 5769.948449 | 5794.77253 | ENSG00000125743 |
| ADGRL1 | 0.006245396 | 1.004338362 | 0.925143324 | 0.965883673 | 4863.738345 | 4885.575699 | ENSG00000072071 |
| EIF2AK3-DT | 0.006233917 | 1.004330371 | 0.991833852 | 0.99806433 | 12.26586949 | 12.32614206 | ENSG00000234028 |
| FANCB | 0.006206854 | 1.004311531 | 0.976901334 | 0.992510313 | 110.9722254 | 111.3817549 | ENSG00000181544 |
| NA | 0.006177568 | 1.004291145 | 0.995537535 | 0.998990416 | 4.335148574 | 4.355600942 | ENSG00000265574 |
| IFI27 | 0.00613638 | 1.004262473 | 0.994264978 | 0.998600974 | 5.958712565 | 5.987388051 | ENSG00000165949 |
| NA | 0.006134342 | 1.004261055 | 0.991056062 | 0.99806433 | 14.91457877 | 14.98620504 | ENSG00000254365 |
| NA | 0.006112714 | 1.004245999 | 0.99383171 | 0.998500968 | 7.300266966 | 7.334681475 | ENSG00000205740 |
| KMT2E | 0.005921511 | 1.004112913 | 0.957470182 | 0.98244088 | 495.5151294 | 497.1327922 | ENSG00000005483 |
| NTNG2 | 0.005883917 | 1.004086749 | 0.987089023 | 0.997171979 | 39.5103621 | 39.68578227 | ENSG00000196358 |
| TRAPPC6B | 0.00587294 | 1.004079109 | 0.976248741 | 0.992155697 | 137.0329494 | 137.4126193 | ENSG00000182400 |
| NA | 0.00581936 | 1.004041819 | 0.994117606 | 0.998568983 | 8.91910243 | 8.958996234 | ENSG00000284968 |
| NA | 0.005807476 | 1.004033548 | 0.972843162 | 0.990091783 | 238.5579528 | 239.747404 | ENSG00000183458 |
| ZNF782 | 0.005791964 | 1.004022753 | 0.979951406 | 0.994673319 | 88.74225201 | 88.99638246 | ENSG00000196597 |
| SPC24 | 0.00578786 | 1.004019897 | 0.954769095 | 0.981381925 | 613.8089263 | 616.0759212 | ENSG00000161888 |
| NA | 0.005758889 | 1.003999735 | 0.991747394 | 0.99806433 | 15.54147325 | 15.61024141 | ENSG00000287630 |
| BBS2 | 0.005600131 | 1.003889259 | 0.966098319 | 0.986897104 | 345.0809 | 346.3169917 | ENSG00000125124 |
| RPL37 | 0.005594566 | 1.003885386 | 0.93411234 | 0.970229433 | 8952.832867 | 8988.220485 | ENSG00000145592 |
| HECTD3 | 0.005592083 | 1.003883658 | 0.948328614 | 0.977308915 | 1038.912941 | 1043.093744 | ENSG00000126107 |
| PRICKLE1 | 0.005583984 | 1.003878023 | 0.968629239 | 0.987666064 | 264.5875052 | 265.700097 | ENSG00000139174 |
| NBN | 0.005573749 | 1.003870901 | 0.956122593 | 0.982211615 | 694.7907523 | 697.5586741 | ENSG00000104320 |
| ALMS1-IT1 | 0.005487097 | 1.003810608 | 0.990500941 | 0.99806433 | 21.48238845 | 21.57209099 | ENSG00000230002 |
| DNMBP | 0.005486892 | 1.003810465 | 0.956274067 | 0.982211615 | 669.9785055 | 672.3243631 | ENSG00000107554 |
| KLK13 | 0.005485847 | 1.003809738 | 0.984258761 | 0.995944879 | 57.91495041 | 58.157155 | ENSG00000167759 |
| NA | 0.005460423 | 1.003792048 | 0.99532425 | 0.998990416 | 4.975085731 | 4.995811515 | ENSG00000273449 |
| PCDHA11 | 0.005437305 | 1.003775964 | 0.971439331 | 0.989388263 | 215.5189938 | 216.1413033 | ENSG00000249158 |
| DERA | 0.005429719 | 1.003770686 | 0.967104523 | 0.987033408 | 286.1146667 | 287.0216511 | ENSG00000023697 |
| MTIF3 | 0.005407645 | 1.003755328 | 0.969792445 | 0.988425189 | 291.2902252 | 292.6543112 | ENSG00000122033 |
| KDM6A | 0.005390058 | 1.003743091 | 0.946160123 | 0.976299526 | 1030.861453 | 1034.213885 | ENSG00000147050 |
| AFMID | 0.005386151 | 1.003740373 | 0.961867965 | 0.984866434 | 514.3199756 | 515.8011362 | ENSG00000183077 |
| MARVELD3 | 0.005299922 | 1.003680382 | 0.952641709 | 0.979882281 | 705.2574751 | 707.9065515 | ENSG00000140832 |
| MRPL20-AS1 | 0.00522717 | 1.00362977 | 0.960582155 | 0.984255629 | 499.56326 | 501.5950565 | ENSG00000224870 |
| MYO9A | 0.005218754 | 1.003623915 | 0.966825506 | 0.987033408 | 314.5053261 | 315.7287527 | ENSG00000066933 |
| NA | 0.00520535 | 1.00361459 | 0.996448173 | 0.999046366 | 3.326605432 | 3.339148349 | ENSG00000269984 |
| TMA16 | 0.005181697 | 1.003598137 | 0.96082434 | 0.984255629 | 721.1624419 | 723.8484843 | ENSG00000198498 |
| BRAT1 | 0.005161291 | 1.003583941 | 0.946516091 | 0.976430116 | 1700.287435 | 1707.035132 | ENSG00000106009 |
| NA | 0.005148928 | 1.003575342 | 0.995686361 | 0.998990416 | 4.960847841 | 4.979637312 | ENSG00000234026 |
| BMPR1A | 0.005099835 | 1.003541191 | 0.941475112 | 0.974126534 | 1633.123723 | 1638.464979 | ENSG00000107779 |
| RHCG | 0.00506985 | 1.003520334 | 0.994678783 | 0.998751 | 8.332531207 | 8.362957345 | ENSG00000140519 |
| CASP8AP2 | 0.005067204 | 1.003518494 | 0.960820574 | 0.984255629 | 510.1497836 | 511.6960721 | ENSG00000118412 |
| SPATA6L | 0.005018476 | 1.0034846 | 0.994041786 | 0.998568983 | 9.595803367 | 9.631555214 | ENSG00000106686 |
| C16orf95 | 0.004981022 | 1.003458549 | 0.992645184 | 0.998301433 | 18.89683379 | 18.85168213 | ENSG00000260456 |
| PFN1 | 0.004928969 | 1.003422344 | 0.913920841 | 0.961911055 | 14054.47188 | 14102.59371 | ENSG00000108518 |
| MAP4 | 0.004899049 | 1.003401534 | 0.916078503 | 0.962669998 | 6587.53712 | 6609.498295 | ENSG00000047849 |
| MYH9 | 0.004865621 | 1.003378285 | 0.920689703 | 0.964633548 | 15358.08806 | 15409.37019 | ENSG00000100345 |
| GPATCH2L | 0.004835323 | 1.003357213 | 0.951385445 | 0.97918437 | 1252.01174 | 1255.602157 | ENSG00000089916 |
| DIPK1B | 0.004825458 | 1.003350353 | 0.967538473 | 0.987242444 | 430.0253786 | 431.4187428 | ENSG00000165716 |
| ATP8 | 0.004796298 | 1.003330073 | 0.986846836 | 0.997067861 | 59.56195616 | 59.89563514 | ENSG00000228253 |
| PCNX4 | 0.004736269 | 1.003288326 | 0.962563912 | 0.985098636 | 565.559877 | 567.6949004 | ENSG00000126773 |
| ZFAND5 | 0.004727459 | 1.0032822 | 0.94225829 | 0.974509737 | 2371.045682 | 2379.27578 | ENSG00000107372 |
| NA | 0.004684522 | 1.003252341 | 0.996123661 | 0.998990416 | 4.624277109 | 4.640094626 | ENSG00000262766 |
| DUSP8 | 0.004667458 | 1.003240475 | 0.988421984 | 0.997507036 | 44.55989127 | 44.71127228 | ENSG00000184545 |
| STAG3 | 0.004665516 | 1.003239124 | 0.988840767 | 0.997539083 | 51.80824069 | 51.97740047 | ENSG00000066923 |
| PGGT1B | 0.004664759 | 1.003238598 | 0.972532613 | 0.989935773 | 313.8208958 | 314.7597814 | ENSG00000164219 |
| EHD4 | 0.004626951 | 1.003212306 | 0.946523078 | 0.976430116 | 1661.33318 | 1666.762807 | ENSG00000103966 |
| UBE2C | 0.004596461 | 1.003191105 | 0.942192314 | 0.974494871 | 2310.024581 | 2316.943907 | ENSG00000175063 |
| SLC38A10 | 0.004559606 | 1.003165477 | 0.951921194 | 0.97945433 | 1199.584796 | 1203.451445 | ENSG00000157637 |
| DR1 | 0.004552488 | 1.003160528 | 0.959136436 | 0.983242406 | 1068.417097 | 1071.713162 | ENSG00000117505 |
| NA | 0.00450365 | 1.00312657 | 0.995946342 | 0.998990416 | 6.623566029 | 6.645002332 | ENSG00000256742 |
| HSPBP1 | 0.004495059 | 1.003120597 | 0.95708577 | 0.98244088 | 1656.070465 | 1661.86912 | ENSG00000133265 |
| MFSD10 | 0.004486178 | 1.003114421 | 0.94753541 | 0.976917957 | 2264.672057 | 2272.593839 | ENSG00000109736 |
| TMF1 | 0.004405075 | 1.003058032 | 0.976853013 | 0.992510313 | 225.6678586 | 226.2468808 | ENSG00000144747 |
| OGFOD2 | 0.004395472 | 1.003051355 | 0.997114632 | 0.999046366 | 4.229585761 | 4.243611022 | ENSG00000111325 |

| | | | | | | | |
|---|---|---|---|---|---|---|---|
| DEPDC1 | 0.004350175 | 1.003019862 | 0.969278002 | 0.988060838 | 555.437299 | 556.7944208 | ENSG00000024526 |
| NA | 0.004330812 | 1.0030064 | 0.994983365 | 0.998897499 | 10.32114171 | 10.3514073 | ENSG00000164845 |
| TBC1D15 | 0.004321257 | 1.002999757 | 0.964489459 | 0.986052631 | 781.2631972 | 782.9847825 | ENSG00000121749 |
| NA | 0.004262034 | 1.002958585 | 0.996110999 | 0.998990416 | 6.000231026 | 6.017560994 | ENSG00000287721 |
| PTPRO | 0.004070734 | 1.002825602 | 0.99195459 | 0.99806433 | 28.05851228 | 28.13716826 | ENSG00000151490 |
| UBE3B | 0.004053113 | 1.002813354 | 0.966520852 | 0.986955029 | 577.7382347 | 579.2181439 | ENSG00000151148 |
| SEC61G | 0.004045308 | 1.002807929 | 0.95653174 | 0.982211615 | 1512.268572 | 1516.72211 | ENSG00000132432 |
| SMC4 | 0.004001259 | 1.002777311 | 0.955288041 | 0.981793467 | 1512.666037 | 1516.660215 | ENSG00000113810 |
| MSL2 | 0.003973571 | 1.002758066 | 0.962506073 | 0.985098636 | 771.8925553 | 773.9671644 | ENSG00000174579 |
| SBDSP1 | 0.003972294 | 1.002757179 | 0.98101767 | 0.995108659 | 189.874649 | 190.6210879 | ENSG00000225648 |
| RGPD3 | 0.003948602 | 1.002740711 | 0.99657215 | 0.999046366 | 5.9800694 | 5.996089905 | ENSG00000153165 |
| BOK | 0.003930264 | 1.002727966 | 0.957941856 | 0.982680048 | 1358.56624 | 1362.657109 | ENSG00000176720 |
| ATXN3 | 0.003875143 | 1.002689655 | 0.975510008 | 0.991946116 | 303.7352112 | 304.6524073 | ENSG00000066427 |
| NA | 0.003827695 | 1.002656679 | 0.983282105 | 0.995796876 | 152.2321814 | 152.4888917 | ENSG00000241743 |
| APCDD1L | 0.003823831 | 1.002653994 | 0.983392735 | 0.995796876 | 142.6343448 | 143.1312364 | ENSG00000198768 |
| LINC01838 | 0.003789417 | 1.002630076 | 0.997119117 | 0.999046366 | 3.949940435 | 3.960063296 | ENSG00000269037 |
| EAF1 | 0.003676325 | 1.002551484 | 0.97460575 | 0.991351373 | 397.5239916 | 398.3589508 | ENSG00000144597 |
| AHSA2P | 0.003669811 | 1.002546957 | 0.991986628 | 0.99806433 | 424.7640873 | 425.8814305 | ENSG00000173209 |
| NA | 0.003628241 | 1.00251807 | 0.996125097 | 0.998990416 | 8.26136791 | 8.281140368 | ENSG00000275029 |
| UCHL1 | 0.003624771 | 1.002515659 | 0.945241202 | 0.976178103 | 11392.29893 | 11421.39789 | ENSG00000154277 |
| AMACR | 0.003594552 | 1.00249466 | 0.993949646 | 0.998549796 | 20.61500285 | 20.66110193 | ENSG00000242110 |
| ZMYND19 | 0.003545815 | 1.002460795 | 0.96591052 | 0.986812019 | 1128.084032 | 1130.663541 | ENSG00000165724 |
| RETREG3 | 0.003518443 | 1.002441775 | 0.970871553 | 0.989151342 | 591.6389047 | 593.0905676 | ENSG00000141699 |
| ZNF77 | 0.003503222 | 1.002431199 | 0.988894232 | 0.997539083 | 73.17797354 | 73.3397939 | ENSG00000175691 |
| DTNBP1 | 0.003490674 | 1.00242248 | 0.98120772 | 0.995201106 | 226.1953933 | 226.6927604 | ENSG00000047579 |
| PSEN1 | 0.003431471 | 1.002381345 | 0.96494406 | 0.986304859 | 998.6615033 | 1000.869579 | ENSG00000080815 |
| VPS39 | 0.003416878 | 1.002371206 | 0.964592867 | 0.986052631 | 1326.24494 | 1329.215392 | ENSG00000166887 |
| CUL3 | 0.003409475 | 1.002366063 | 0.963638228 | 0.985663648 | 1637.842151 | 1641.551823 | ENSG00000036257 |
| REV1 | 0.003388239 | 1.002351308 | 0.973734134 | 0.990838359 | 589.8130885 | 590.9921781 | ENSG00000135945 |
| NA | 0.003381746 | 1.002346797 | 0.996965262 | 0.999046366 | 5.95159362 | 5.963741498 | ENSG00000287686 |
| TRPC4AP | 0.00332254 | 1.002305663 | 0.95805313 | 0.982680048 | 2638.295501 | 2643.599874 | ENSG00000100991 |
| GPRIN2 | 0.003319242 | 1.002303372 | 0.981576392 | 0.995385822 | 296.5449301 | 297.4591537 | ENSG00000204175 |
| SPG11 | 0.003213038 | 1.00222959 | 0.975700762 | 0.991946116 | 732.3928989 | 733.5030761 | ENSG00000104133 |
| ZNF362 | 0.00315835 | 1.002191599 | 0.975611346 | 0.991946116 | 594.8368889 | 596.2963502 | ENSG00000160094 |
| NUDT16P1 | 0.003049745 | 1.002116158 | 0.989099786 | 0.997604649 | 117.4027425 | 117.5162035 | ENSG00000246082 |
| TSTA3 | 0.003048962 | 1.002115615 | 0.962574378 | 0.985098636 | 2099.902696 | 2104.620811 | ENSG00000104522 |
| NCAPG2 | 0.002923455 | 1.002028439 | 0.959754322 | 0.983649986 | 2396.559802 | 2401.088688 | ENSG00000146918 |
| NA | 0.002896795 | 1.002009923 | 0.997540205 | 0.999046366 | 5.325894353 | 5.334124696 | ENSG00000259005 |
| MRPL57 | 0.002880889 | 1.001998875 | 0.972042958 | 0.989704079 | 975.4769364 | 977.7958003 | ENSG00000173141 |
| ZCCHC18 | 0.002817235 | 1.001954666 | 0.990740855 | 0.99806433 | 103.2485119 | 103.6143123 | ENSG00000166707 |
| NUB1 | 0.002741279 | 1.001901916 | 0.973538256 | 0.990692422 | 936.4694732 | 937.7856606 | ENSG00000013374 |
| TLE3 | 0.002681356 | 1.001860303 | 0.97211777 | 0.98972689 | 1303.234228 | 1305.641333 | ENSG00000140332 |
| ADAM10 | 0.002675999 | 1.001856582 | 0.971370848 | 0.989388263 | 2260.148076 | 2263.996595 | ENSG00000137845 |
| MIB1 | 0.002668583 | 1.001851432 | 0.978355811 | 0.993667285 | 1297.413211 | 1299.222523 | ENSG00000101752 |
| SRGAP2C | 0.002667283 | 1.00185053 | 0.988749308 | 0.997539083 | 138.2131585 | 138.4062033 | ENSG00000171943 |
| TNFRSF25 | 0.002640245 | 1.001831754 | 0.989194491 | 0.997646887 | 145.2794685 | 145.7112063 | ENSG00000215788 |
| PDCD6 | 0.002638331 | 1.001830425 | 0.990965147 | 0.99806433 | 87.4834552 | 87.60274189 | ENSG00000249915 |
| ERICH1 | 0.002592243 | 1.001798421 | 0.975813185 | 0.991991818 | 833.7912353 | 835.1929955 | ENSG00000104714 |
| NIN | 0.002589804 | 1.001796728 | 0.981921857 | 0.995402617 | 433.2154772 | 433.6060856 | ENSG00000100503 |
| LINC00634 | 0.00253421 | 1.001758124 | 0.997686226 | 0.999061689 | 6.320199605 | 6.327876695 | ENSG00000205704 |
| NA | 0.002401773 | 1.001666168 | 0.997946846 | 0.999110755 | 5.257095321 | 5.263185037 | ENSG00000270820 |
| PDZD11 | 0.002327519 | 1.001614615 | 0.984418059 | 0.995944879 | 393.1639528 | 393.8371318 | ENSG00000120509 |
| VDAC3 | 0.002259633 | 1.001567486 | 0.967743295 | 0.987242444 | 2763.526628 | 2767.81492 | ENSG00000078668 |
| MED7 | 0.002234748 | 1.001550209 | 0.991401592 | 0.99806433 | 116.4615238 | 116.7039837 | ENSG00000155868 |
| HERC4 | 0.002206942 | 1.001530906 | 0.980838787 | 0.995011844 | 649.0761274 | 649.8616699 | ENSG00000148634 |
| GPR173 | 0.002182246 | 1.001513762 | 0.988909194 | 0.997539083 | 220.4027808 | 220.6269603 | ENSG00000184194 |
| CHFR | 0.002179618 | 1.001511938 | 0.979041051 | 0.994080282 | 1074.044552 | 1075.465821 | ENSG00000072609 |
| MRPL37 | 0.002162651 | 1.001500159 | 0.971892315 | 0.989657412 | 3036.813113 | 3041.953784 | ENSG00000116221 |
| RPS14 | 0.002129924 | 1.001477441 | 0.975532704 | 0.991946116 | 22351.51174 | 22383.82206 | ENSG00000164587 |
| KLHL2 | 0.00212677 | 1.001475252 | 0.988316576 | 0.997507036 | 274.4033332 | 274.5746064 | ENSG00000109466 |
| PSKH1 | 0.002121078 | 1.0014713 | 0.980593496 | 0.994924101 | 815.0789086 | 816.5083819 | ENSG00000159792 |
| ARHGAP19 | 0.001968076 | 1.001365098 | 0.983478857 | 0.995796876 | 676.9407522 | 678.0411703 | ENSG00000213390 |
| SEMA6A | 0.00193093 | 1.001339315 | 0.972287279 | 0.989846105 | 4807.195405 | 4812.96685 | ENSG00000092421 |
| C2orf76 | 0.001924521 | 1.001334866 | 0.993649303 | 0.998500968 | 85.03280271 | 85.0891927 | ENSG00000186132 |
| DNMT3A | 0.001910625 | 1.001325222 | 0.976289612 | 0.992155697 | 4027.006486 | 4032.226612 | ENSG00000119772 |
| GRSF1 | 0.001904162 | 1.001320736 | 0.975715703 | 0.991946116 | 2721.084813 | 2724.761706 | ENSG00000132463 |
| DDX54 | 0.001874883 | 1.001300415 | 0.974395687 | 0.991254655 | 3455.729461 | 3460.293858 | ENSG00000123064 |
| ANKRD19P | 0.001811276 | 1.00125627 | 0.998708724 | 0.999633714 | 3.622852912 | 3.624871538 | ENSG00000187984 |
| RBBP8 | 0.001790597 | 1.001241918 | 0.983367638 | 0.995796876 | 789.1620386 | 790.1344019 | ENSG00000101773 |
| GFM1 | 0.001769499 | 1.001227276 | 0.982971133 | 0.995796876 | 935.4502772 | 936.4255026 | ENSG00000168827 |
| ST6GAL2 | 0.001750274 | 1.001213933 | 0.988161103 | 0.997507036 | 357.1779998 | 357.4973602 | ENSG00000144057 |
| GOLGA7 | 0.001724145 | 1.001195801 | 0.986930872 | 0.997067861 | 615.6459802 | 616.0096684 | ENSG00000147533 |
| H2BC4 | 0.001708252 | 1.001184771 | 0.997799959 | 0.999069626 | 11.9447057 | 11.94895408 | ENSG00000180596 |
| RNF170 | 0.00166318 | 1.001153493 | 0.992408464 | 0.998169676 | 203.2181138 | 203.4416618 | ENSG00000120925 |

| | | | | | | | |
|---|---|---|---|---|---|---|---|
| CIAPIN1 | 0.001643861 | 1.001140087 | 0.981921082 | 0.995402617 | 1990.802764 | 1992.198921 | ENSG00000005194 |
| TBC1D30 | 0.001620933 | 1.001124176 | 0.993813855 | 0.998500968 | 107.0175564 | 107.0550974 | ENSG00000111490 |
| ABCD3 | 0.001597534 | 1.001107939 | 0.986551629 | 0.996898042 | 675.1611568 | 675.6351445 | ENSG00000117528 |
| TTC31 | 0.00158804 | 1.001101352 | 0.988267072 | 0.997507036 | 498.0939224 | 498.7700347 | ENSG00000115282 |
| HMGB2 | 0.001587519 | 1.00110099 | 0.97761918 | 0.993079351 | 4096.027743 | 4099.88725 | ENSG00000164104 |
| ANKHD1 | 0.001577426 | 1.001093986 | 0.997373128 | 0.999046366 | 23.88852168 | 23.78497229 | ENSG00000131503 |
| KIF5A | 0.001503599 | 1.001042759 | 0.986923822 | 0.997067861 | 788.9149355 | 789.4140257 | ENSG00000155980 |
| PSMB8 | 0.001463091 | 1.001014652 | 0.990893532 | 0.99806433 | 305.0859439 | 305.1650209 | ENSG00000204264 |
| TRIM17 | 0.001421333 | 1.000985678 | 0.997022241 | 0.999046366 | 35.21525745 | 35.34758504 | ENSG00000162931 |
| TM9SF3 | 0.001401525 | 1.000971935 | 0.983287144 | 0.995796876 | 2941.932343 | 2945.130151 | ENSG00000077147 |
| SUOX | 0.001379536 | 1.000956679 | 0.989842427 | 0.99804732 | 445.4887705 | 445.8265962 | ENSG00000139531 |
| TOB2 | 0.00136645 | 1.0009476 | 0.983495483 | 0.995796876 | 2043.663837 | 2045.758544 | ENSG00000183864 |
| BAHD1 | 0.001290939 | 1.000895211 | 0.985462098 | 0.996383537 | 1238.226499 | 1239.363287 | ENSG00000140320 |
| C15orf39 | 0.001135032 | 1.000787054 | 0.988249392 | 0.997507036 | 1500.631915 | 1502.387526 | ENSG00000167173 |
| ZNF212 | 0.001084387 | 1.000751922 | 0.991342037 | 0.99806433 | 558.2352006 | 558.4592617 | ENSG00000170260 |
| C11orf49 | 0.001069905 | 1.000741876 | 0.992501922 | 0.998210513 | 399.7258649 | 399.9390212 | ENSG00000149179 |
| CHMP2A | 0.001050045 | 1.000728101 | 0.989942811 | 0.99804732 | 1099.747218 | 1100.286723 | ENSG00000130724 |
| HELLS | 0.000987222 | 1.000684524 | 0.987530607 | 0.997300295 | 2015.423975 | 2016.891645 | ENSG00000119969 |
| CFTR | 0.000961412 | 1.000666622 | 0.998765623 | 0.999633714 | 11.6282966 | 11.62341014 | ENSG00000001626 |
| SRPK1 | 0.0009489 | 1.000657944 | 0.989962942 | 0.99804732 | 1505.52935 | 1505.971776 | ENSG00000096063 |
| OST4 | 0.000945142 | 1.000655337 | 0.991797528 | 0.99806433 | 767.8375846 | 768.1248497 | ENSG00000228474 |
| VPS36 | 0.00093767 | 1.000650155 | 0.993713613 | 0.998500968 | 345.7982291 | 346.1022809 | ENSG00000136100 |
| FNBP4 | 0.000911571 | 1.000632053 | 0.992210569 | 0.998130112 | 1082.453534 | 1083.149717 | ENSG00000109920 |
| TIMP3 | 0.000872454 | 1.000604922 | 0.996776926 | 0.999046366 | 99.02368086 | 99.12721489 | ENSG00000100234 |
| RUNDC1 | 0.000836242 | 1.000579807 | 0.993520545 | 0.998500968 | 488.564196 | 488.9084248 | ENSG00000198863 |
| MELTF-AS1 | 0.000821921 | 1.000569874 | 0.998845595 | 0.999633714 | 15.31758428 | 15.31174899 | ENSG00000228109 |
| DIP2B | 0.000819214 | 1.000567997 | 0.991891485 | 0.99806433 | 1172.102021 | 1172.21219 | ENSG00000066084 |
| NA | 0.000812027 | 1.000563013 | 0.999338151 | 0.999701317 | 4.658676625 | 4.65626883 | ENSG00000213856 |
| STK32C | 0.000713046 | 1.000494368 | 0.994793494 | 0.998759975 | 549.8668209 | 550.4835619 | ENSG00000165752 |
| MMP15 | 0.000672736 | 1.000466414 | 0.991904747 | 0.99806433 | 2034.329986 | 2035.571109 | ENSG00000102996 |
| WDR60 | 0.000524193 | 1.000363409 | 0.996366683 | 0.999046366 | 445.7316265 | 445.5287662 | ENSG00000126870 |
| MPHOSPH10 | 0.000490818 | 1.000340267 | 0.99565327 | 0.998990416 | 772.5598247 | 773.0077253 | ENSG00000124383 |
| CAMK4 | 0.00047859 | 1.000331789 | 0.997754285 | 0.999069626 | 159.6232146 | 159.6375221 | ENSG00000152495 |
| OTX1 | 0.00047381 | 1.000328474 | 0.999137573 | 0.999666917 | 23.87280925 | 23.85279053 | ENSG00000115507 |
| NA | 0.000444762 | 1.000308333 | 0.999419741 | 0.999701317 | 11.94826517 | 11.93713081 | ENSG00000236603 |
| PRRC1 | 0.000301729 | 1.000209165 | 0.99690587 | 0.999046366 | 1262.72348 | 1262.953012 | ENSG00000164244 |
| FBXO38 | 0.000291995 | 1.000202416 | 0.997523421 | 0.999046366 | 614.6233032 | 614.6200952 | ENSG00000145868 |
| PPP2CB | 0.000254163 | 1.000176188 | 0.997029815 | 0.999046366 | 1671.232603 | 1671.667197 | ENSG00000104695 |
| SLC25A14 | 0.000249611 | 1.000173032 | 0.998811869 | 0.999633714 | 195.8582784 | 195.7069238 | ENSG00000102078 |
| ANKRD18A | 0.000208377 | 1.000144446 | 0.999436635 | 0.999701317 | 58.54420915 | 58.61370094 | ENSG00000180071 |
| BPGM | 0.000169156 | 1.000117257 | 0.998952351 | 0.999633714 | 362.1124823 | 362.5458773 | ENSG00000172331 |
| PRKAG2 | 0.000157233 | 1.000108992 | 0.999251021 | 0.999675693 | 250.9171543 | 250.7633605 | ENSG00000106617 |
| BSN | 2.46E-05 | 1.000017025 | 0.999895785 | 0.999948735 | 158.5100245 | 158.5794522 | ENSG00000164061 |
| NA | -3.99E-06 | 0.999997233 | 0.999997251 | 0.999997251 | 3.33728385 | 3.332621958 | ENSG00000287790 |
| INIP | -1.92E-05 | 0.999986697 | 0.999887656 | 0.999948735 | 279.6604285 | 279.4729211 | ENSG00000148153 |
| THRA | -0.000132882 | 0.999907898 | 0.999051455 | 0.999633714 | 500.2173566 | 500.2989126 | ENSG00000126351 |
| SRF | -0.000177642 | 0.999876876 | 0.997671033 | 0.999061689 | 3115.072111 | 3114.869789 | ENSG00000112658 |
| NA | -0.000230207 | 0.999840446 | 0.999750557 | 0.999909399 | 9.377838129 | 9.364181693 | ENSG00000225218 |
| RPS20 | -0.000241459 | 0.999832647 | 0.99615377 | 0.998990416 | 14093.05211 | 14090.75633 | ENSG00000008988 |
| CCDC134 | -0.000253307 | 0.999824436 | 0.999031479 | 0.999633714 | 107.2165291 | 107.1950848 | ENSG00000100147 |
| ATP5MPL | -0.000333692 | 0.999768729 | 0.997343229 | 0.999046366 | 803.5849305 | 803.6190571 | ENSG00000156411 |
| ZNF598 | -0.000384718 | 0.999733369 | 0.996239688 | 0.998990416 | 1385.285055 | 1385.718247 | ENSG00000167962 |
| TOX4 | -0.000400352 | 0.999722536 | 0.995717893 | 0.998990416 | 1231.308465 | 1230.713618 | ENSG00000092203 |
| RNF216 | -0.000406422 | 0.999718329 | 0.995691367 | 0.998990416 | 1103.385634 | 1103.382049 | ENSG00000011275 |
| PIEZO2 | -0.000422523 | 0.999707172 | 0.9968857 | 0.999046366 | 498.8574473 | 498.3037254 | ENSG00000154864 |
| MEIS2 | -0.00047895 | 0.999668072 | 0.999565243 | 0.999777004 | 6.01683318 | 6.005737718 | ENSG00000134138 |
| CDKL2 | -0.000542804 | 0.999623828 | 0.999252214 | 0.999675693 | 14.34936439 | 14.32329387 | ENSG00000138769 |
| SNX14 | -0.000561242 | 0.999611052 | 0.996600599 | 0.999046366 | 302.7346767 | 302.8484 | ENSG00000135317 |
| DDHD1 | -0.000600436 | 0.999583896 | 0.995824288 | 0.998990416 | 408.8916354 | 408.469258 | ENSG00000100523 |
| NFATC4 | -0.000611099 | 0.999576508 | 0.996181045 | 0.998990416 | 593.0908034 | 592.885824 | ENSG00000100968 |
| WDSUB1 | -0.000692058 | 0.999520417 | 0.997430247 | 0.999046366 | 102.2951148 | 102.0978771 | ENSG00000196151 |
| PRKD3 | -0.000761362 | 0.999472404 | 0.993066787 | 0.998500968 | 1086.847998 | 1085.915764 | ENSG00000115825 |
| PPP1CB | -0.000934296 | 0.999352605 | 0.991162328 | 0.99806433 | 1648.676789 | 1647.575163 | ENSG00000213639 |
| NA | -0.001049803 | 0.999272597 | 0.999018299 | 0.999633714 | 5.962272037 | 5.947567294 | ENSG00000285103 |
| DMWD | -0.001062519 | 0.999263789 | 0.991779563 | 0.99806433 | 705.5115932 | 705.2982708 | ENSG00000185800 |
| RDH11 | -0.001064691 | 0.999262285 | 0.986519978 | 0.996898042 | 1799.856902 | 1798.923531 | ENSG00000072042 |
| NA | -0.001110028 | 0.999230883 | 0.998927039 | 0.999633714 | 6.31308066 | 6.296757792 | ENSG00000259287 |
| P3H1 | -0.001120478 | 0.999223645 | 0.988090167 | 0.997507036 | 1565.075496 | 1564.445621 | ENSG00000117385 |
| NA | -0.00114564 | 0.999206218 | 0.998737495 | 0.999633714 | 8.336090679 | 8.314434734 | ENSG00000249492 |
| NA | -0.001148444 | 0.999204276 | 0.998329339 | 0.999440711 | 18.54574584 | 18.50873447 | ENSG00000240731 |
| GLB1L2 | -0.001173504 | 0.99918692 | 0.993697992 | 0.998500968 | 209.5825278 | 209.3561457 | ENSG00000149328 |
| TIE1 | -0.001198981 | 0.999169275 | 0.996212255 | 0.998990416 | 82.27708222 | 82.21820266 | ENSG00000066056 |
| CWC25 | -0.001227333 | 0.999149639 | 0.992147861 | 0.998120198 | 375.8150705 | 375.5613423 | ENSG00000273559 |
| ID4 | -0.001250081 | 0.999133885 | 0.995717293 | 0.998990416 | 110.864857 | 110.538752 | ENSG00000172201 |

| | | | | | | | |
|---|---|---|---|---|---|---|---|
| NEMP1 | -0.001287855 | 0.999107725 | 0.986462002 | 0.996898042 | 1065.416079 | 1064.333712 | ENSG00000166881 |
| HDLBP | -0.00135399 | 0.999061926 | 0.976539177 | 0.992294905 | 10378.47666 | 10368.6581 | ENSG00000115677 |
| MRPS30 | -0.001433149 | 0.99900711 | 0.985185691 | 0.99621074 | 1186.535746 | 1185.597087 | ENSG00000112996 |
| CBX2 | -0.001490496 | 0.9989674 | 0.980175279 | 0.994714902 | 4089.012619 | 4085.485395 | ENSG00000173894 |
| ATL1 | -0.001502683 | 0.998958962 | 0.994498934 | 0.998676612 | 101.1427705 | 101.0095389 | ENSG00000198513 |
| HSH2D | -0.001524702 | 0.998943716 | 0.99746919 | 0.999046366 | 24.60765695 | 24.55022557 | ENSG00000196684 |
| GNB2 | -0.001611213 | 0.998883816 | 0.979418021 | 0.994264868 | 3054.576847 | 3051.940444 | ENSG00000172354 |
| SFT2D1 | -0.001682049 | 0.998834772 | 0.984328557 | 0.995944879 | 941.0344339 | 940.0385439 | ENSG00000198818 |
| GUCY1A2 | -0.00168666 | 0.998831579 | 0.996613558 | 0.999046366 | 31.45600168 | 31.49901721 | ENSG00000152402 |
| USP37 | -0.001713502 | 0.998812996 | 0.989915329 | 0.99804732 | 304.3786823 | 303.7072506 | ENSG00000135913 |
| ZNF652 | -0.001741333 | 0.998793728 | 0.991064167 | 0.99806433 | 221.3439727 | 221.2192154 | ENSG00000198740 |
| PSMC4 | -0.001765481 | 0.998777011 | 0.976183477 | 0.992155697 | 3312.923651 | 3308.920015 | ENSG00000013275 |
| BCAR3 | -0.001804991 | 0.998749658 | 0.990008373 | 0.99804732 | 246.4050264 | 246.0775085 | ENSG00000137936 |
| ORAI2 | -0.001811488 | 0.99874516 | 0.98478476 | 0.996174529 | 860.5859417 | 859.8770906 | ENSG00000160991 |
| TAPT1-AS1 | -0.00184957 | 0.998718797 | 0.997041154 | 0.999046366 | 18.56115279 | 18.50343759 | ENSG00000263327 |
| SRGAP1 | -0.001898718 | 0.998684774 | 0.991595109 | 0.99806433 | 152.7013162 | 152.3722673 | ENSG00000196935 |
| SPIDR | -0.001911533 | 0.998675903 | 0.981236145 | 0.995201106 | 1073.724838 | 1071.741057 | ENSG00000164808 |
| FGF23 | -0.002020222 | 0.998600669 | 0.997195514 | 0.999046366 | 14.54360853 | 14.49825665 | ENSG00000118972 |
| IRS2 | -0.002052717 | 0.998578177 | 0.986508469 | 0.996898042 | 463.7327279 | 463.3244155 | ENSG00000185950 |
| CARNS1 | -0.002070821 | 0.998565646 | 0.996127472 | 0.998990416 | 25.67904924 | 25.59344613 | ENSG00000172508 |
| EXOC3 | -0.002074654 | 0.998562993 | 0.980320448 | 0.994806645 | 1012.672211 | 1011.419662 | ENSG00000180104 |
| PANX2 | -0.002090138 | 0.998552275 | 0.990182532 | 0.99806433 | 219.2554438 | 219.2260583 | ENSG00000073150 |
| TOPBP1 | -0.002126175 | 0.998527333 | 0.977750847 | 0.993159678 | 1132.524716 | 1131.114388 | ENSG00000163781 |
| DENND4C | -0.00216244 | 0.998502233 | 0.986834345 | 0.997067861 | 300.0669754 | 299.3675403 | ENSG00000137145 |
| LSM8 | -0.002162801 | 0.998501984 | 0.982679621 | 0.995756004 | 649.4717597 | 648.892016 | ENSG00000128534 |
| MAP3K21 | -0.002239207 | 0.998449104 | 0.979409466 | 0.994264868 | 794.7138545 | 793.4025677 | ENSG00000143674 |
| AP4S1 | -0.00227416 | 0.998424914 | 0.989598542 | 0.997947801 | 174.6726955 | 174.4658846 | ENSG00000100478 |
| NDFIP1 | -0.002351907 | 0.99837111 | 0.976257359 | 0.992155697 | 1278.588228 | 1276.809756 | ENSG00000131507 |
| TMEM51-AS1 | -0.002394733 | 0.998341474 | 0.997382854 | 0.999046366 | 8.31236958 | 8.281140368 | ENSG00000175147 |
| PER1 | -0.00246515 | 0.998292747 | 0.980405217 | 0.994839209 | 884.4439283 | 883.2808802 | ENSG00000179094 |
| KHDRBS1 | -0.00247576 | 0.998285406 | 0.964547039 | 0.986052631 | 7091.385244 | 7079.914505 | ENSG00000121774 |
| RPS7 | -0.002504614 | 0.99826544 | 0.962847701 | 0.985175209 | 11634.90865 | 11614.24179 | ENSG00000171863 |
| FAM189A1 | -0.002512987 | 0.998259646 | 0.987209894 | 0.997189712 | 203.3085752 | 202.8409885 | ENSG00000104059 |
| NA | -0.002556059 | 0.998229843 | 0.99151572 | 0.99806433 | 87.76218465 | 87.44875575 | ENSG00000260727 |
| CUTA | -0.002573311 | 0.998217906 | 0.965136856 | 0.986395147 | 3223.012507 | 3217.252392 | ENSG00000112514 |
| SPEF2 | -0.002815504 | 0.998050345 | 0.996989802 | 0.999046366 | 8.294572218 | 8.260615237 | ENSG00000152582 |
| MSRA | -0.002819791 | 0.998047379 | 0.991679573 | 0.99806433 | 70.50910263 | 70.36723976 | ENSG00000175806 |
| HSDL2 | -0.002847014 | 0.998028546 | 0.97986339 | 0.99466352 | 420.6252155 | 419.7621539 | ENSG00000119471 |
| P2RY1 | -0.002854014 | 0.998023704 | 0.979078276 | 0.994080282 | 696.8201941 | 694.8700169 | ENSG00000169860 |
| USP27X | -0.002915189 | 0.997981385 | 0.985561681 | 0.996430877 | 186.3042984 | 185.9575147 | ENSG00000273820 |
| MRPL40 | -0.002959232 | 0.997950919 | 0.982989349 | 0.995796876 | 258.2379662 | 257.8489455 | ENSG00000185608 |
| ASS1 | -0.003048644 | 0.997889072 | 0.961843047 | 0.984866434 | 2258.544344 | 2253.022975 | ENSG00000130707 |
| GPC2 | -0.003056999 | 0.997883293 | 0.971017199 | 0.989230603 | 924.2366066 | 922.5332295 | ENSG00000213420 |
| CEND1 | -0.003100618 | 0.997853123 | 0.987497709 | 0.997300295 | 150.4642053 | 150.3117637 | ENSG00000184524 |
| MED19 | -0.003106562 | 0.997849012 | 0.980594243 | 0.994924101 | 362.7325895 | 362.3025292 | ENSG00000156603 |
| GLUD1P3 | -0.003110814 | 0.997846071 | 0.995230821 | 0.998990416 | 18.35862065 | 18.28241121 | ENSG00000250959 |
| TPST2 | -0.003130936 | 0.997832153 | 0.966705757 | 0.987033408 | 2661.674813 | 2656.307871 | ENSG00000128294 |
| C1D | -0.003142 | 0.997824501 | 0.981773503 | 0.995402617 | 303.9501756 | 303.1082582 | ENSG00000197223 |
| CKMT1A | -0.003178493 | 0.997799262 | 0.986185179 | 0.996794429 | 144.6881949 | 144.4695444 | ENSG00000223572 |
| FIP1L1 | -0.003218971 | 0.997771267 | 0.98306359 | 0.995796876 | 209.2412285 | 208.7386357 | ENSG00000145216 |
| RPL38 | -0.003283196 | 0.997726849 | 0.965777215 | 0.986738908 | 7411.853062 | 7394.956191 | ENSG00000172809 |
| ATP6V1E1 | -0.003291412 | 0.997721168 | 0.964005564 | 0.985984136 | 1272.428511 | 1269.519237 | ENSG00000131100 |
| SLC2A12 | -0.003305657 | 0.997711316 | 0.988081439 | 0.997507036 | 102.8959239 | 102.5808122 | ENSG00000146411 |
| SLC25A36 | -0.003310836 | 0.997707735 | 0.974145098 | 0.991096335 | 807.5445301 | 805.0451781 | ENSG00000114120 |
| SPHK2 | -0.003337086 | 0.997689582 | 0.971054112 | 0.989230603 | 844.7926885 | 843.0517646 | ENSG00000063176 |
| NA | -0.003426114 | 0.997628016 | 0.997565122 | 0.999046366 | 3.691651944 | 3.674340108 | ENSG00000231249 |
| MICB-DT | -0.00345552 | 0.997607682 | 0.997866641 | 0.999083422 | 3.344402795 | 3.330446495 | ENSG00000286940 |
| C2orf68 | -0.003601629 | 0.997506655 | 0.966899614 | 0.987033408 | 826.768197 | 824.6423164 | ENSG00000168887 |
| ZBTB44 | -0.003690342 | 0.997445318 | 0.964456044 | 0.986052631 | 1511.056048 | 1506.883457 | ENSG00000196323 |
| NA | -0.003694256 | 0.997442612 | 0.994426658 | 0.998657135 | 19.24736308 | 19.16199783 | ENSG00000285531 |
| ARHGEF12 | -0.003732756 | 0.997415995 | 0.962184262 | 0.985060835 | 1052.850961 | 1049.805829 | ENSG00000196914 |
| NA | -0.003821464 | 0.997354668 | 0.996221746 | 0.998990416 | 6.667448754 | 6.631949551 | ENSG00000263345 |
| PLK1 | -0.003838204 | 0.997343096 | 0.941437118 | 0.974126534 | 3197.171936 | 3188.730088 | ENSG00000166851 |
| PCID2 | -0.00400473 | 0.997227982 | 0.960713324 | 0.984255629 | 1118.699608 | 1115.106855 | ENSG00000126226 |
| PDIA4 | -0.004113975 | 0.997152472 | 0.929343754 | 0.967945949 | 10035.3289 | 10007.24927 | ENSG00000155660 |
| AFAP1 | -0.004193995 | 0.997097165 | 0.98334659 | 0.995796876 | 122.0042144 | 121.631766 | ENSG00000196526 |
| ARHGEF9 | -0.004270602 | 0.997044221 | 0.958356131 | 0.982815939 | 848.1747198 | 845.6491086 | ENSG00000131089 |
| CHIC2 | -0.004364892 | 0.99697906 | 0.981961914 | 0.995402617 | 133.8572896 | 133.530022 | ENSG00000109220 |
| ATRN | -0.004384548 | 0.996965476 | 0.9627207 | 0.985098636 | 853.252953 | 850.0212196 | ENSG00000088812 |
| MAPK14 | -0.00442404 | 0.996938186 | 0.953092298 | 0.980238975 | 1212.009133 | 1207.868905 | ENSG00000112062 |
| HS3ST3A1 | -0.004427119 | 0.996936058 | 0.982450923 | 0.995631107 | 160.5312284 | 160.2274135 | ENSG00000153976 |
| ZXDC | -0.004428095 | 0.996935384 | 0.970458082 | 0.988890135 | 466.6488529 | 464.9116224 | ENSG00000070476 |
| MEX3B | -0.004473061 | 0.996904312 | 0.964883394 | 0.986296233 | 611.8708361 | 610.3387436 | ENSG00000183496 |
| TBK1 | -0.004514579 | 0.996875623 | 0.971560122 | 0.989399232 | 434.9640049 | 434.0248909 | ENSG00000183735 |

| | | | | | | | |
|---|---|---|---|---|---|---|---|
| RCOR1 | -0.004550705 | 0.996850661 | 0.96259123 | 0.985098636 | 789.0597036 | 786.3859148 | ENSG00000089902 |
| OPHN1 | -0.004554502 | 0.996848038 | 0.984017753 | 0.995916668 | 108.704275 | 108.1429411 | ENSG00000079482 |
| ARFGEF1 | -0.004558783 | 0.99684508 | 0.946263602 | 0.976352955 | 2517.30522 | 2508.893187 | ENSG00000066777 |
| DNAJC2 | -0.004620789 | 0.996802237 | 0.959572995 | 0.983627629 | 842.0583249 | 839.8121634 | ENSG00000105821 |
| CACTIN | -0.004641244 | 0.996788104 | 0.957264204 | 0.98244088 | 911.8304426 | 909.3281929 | ENSG00000105298 |
| RPS21 | -0.004653361 | 0.996779732 | 0.944753328 | 0.975866344 | 6709.760398 | 6688.489047 | ENSG00000171858 |
| SATB1 | -0.004700828 | 0.996746937 | 0.961488605 | 0.98463816 | 605.9889062 | 603.8733612 | ENSG00000182568 |
| PRMT9 | -0.004769197 | 0.996699703 | 0.973826509 | 0.990878967 | 250.5767709 | 249.5443992 | ENSG00000164169 |
| PCMTD2 | -0.004785162 | 0.996688673 | 0.956319791 | 0.982211615 | 914.5547138 | 911.0034261 | ENSG00000203880 |
| TGIF1 | -0.004794453 | 0.996682254 | 0.93186351 | 0.969226325 | 2683.223665 | 2674.330727 | ENSG00000177426 |
| NA | -0.004796551 | 0.996680805 | 0.993848171 | 0.998500968 | 12.25519107 | 12.18303324 | ENSG00000213881 |
| TRRAP | -0.004885911 | 0.996619073 | 0.951996555 | 0.97945433 | 2800.031115 | 2789.734601 | ENSG00000196367 |
| IDH2 | -0.005067594 | 0.996493573 | 0.941748972 | 0.974316559 | 1533.386551 | 1527.624826 | ENSG00000182054 |
| PLEKHA1 | -0.00523634 | 0.996377025 | 0.952952619 | 0.980148696 | 672.6041035 | 670.244689 | ENSG00000107679 |
| MOGS | -0.00532611 | 0.996315028 | 0.940519705 | 0.973564713 | 1942.131337 | 1935.799323 | ENSG00000115275 |
| GALR2 | -0.005579569 | 0.996140007 | 0.994252731 | 0.998600974 | 8.285089009 | 8.231671798 | ENSG00000182687 |
| GPR135 | -0.005587814 | 0.996134313 | 0.992270882 | 0.998137616 | 14.60172914 | 14.505729 | ENSG00000181619 |
| RASSF7 | -0.005654331 | 0.996088387 | 0.957420407 | 0.98244088 | 546.9215593 | 544.9390046 | ENSG00000099849 |
| NA | -0.005746009 | 0.996025091 | 0.993521393 | 0.998500968 | 9.637321828 | 9.572438832 | ENSG00000272836 |
| NA | -0.005753945 | 0.996019612 | 0.987672364 | 0.997390118 | 36.16567998 | 36.05417343 | ENSG00000204685 |
| CD8B | -0.005754984 | 0.996018895 | 0.994751352 | 0.998759975 | 6.36408233 | 6.320404345 | ENSG00000172116 |
| SYBU | -0.005781437 | 0.996000632 | 0.992365591 | 0.998169676 | 12.34415173 | 12.25614836 | ENSG00000147642 |
| LUC7L2 | -0.005798027 | 0.995989179 | 0.956184028 | 0.982211615 | 616.1638035 | 613.8638274 | ENSG00000146963 |
| SUGT1 | -0.005841168 | 0.995959396 | 0.932808518 | 0.969708285 | 1807.747046 | 1801.067104 | ENSG00000165416 |
| DPH1 | -0.005861545 | 0.995945329 | 0.967737951 | 0.987242444 | 275.900614 | 275.0074761 | ENSG00000108963 |
| NA | -0.005927478 | 0.995899814 | 0.99382333 | 0.998500968 | 7.69259405 | 7.636578868 | ENSG00000224765 |
| NA | -0.005969791 | 0.995870606 | 0.995606245 | 0.998990416 | 3.688092472 | 3.660341367 | ENSG00000254701 |
| QRFP | -0.005982747 | 0.995861663 | 0.994626394 | 0.998751 | 5.703983547 | 5.661844105 | ENSG00000188710 |
| GPR75 | -0.006042072 | 0.995820712 | 0.991540439 | 0.99806433 | 17.25519311 | 17.14337493 | ENSG00000119737 |
| CHST3 | -0.006148281 | 0.995747404 | 0.958140538 | 0.982701573 | 367.3885447 | 365.9528439 | ENSG00000122863 |
| VPS13C | -0.006271646 | 0.995662262 | 0.971470826 | 0.989388263 | 216.394057 | 215.1236939 | ENSG00000129003 |
| DGUOK-AS1 | -0.006280199 | 0.995656359 | 0.994388668 | 0.998657135 | 6.303597451 | 6.259112499 | ENSG00000237883 |
| GAS6 | -0.006313721 | 0.995633224 | 0.950702649 | 0.978900798 | 584.6443952 | 581.9009301 | ENSG00000183087 |
| PRDX3 | -0.006360397 | 0.995601013 | 0.904912829 | 0.958563988 | 4265.747171 | 4246.169847 | ENSG00000165672 |
| CNN2 | -0.006457739 | 0.99553384 | 0.894887787 | 0.953032171 | 9300.919371 | 9260.349476 | ENSG00000064666 |
| USP10 | -0.00648876 | 0.995512434 | 0.900460347 | 0.956153706 | 4887.473241 | 4865.15314 | ENSG00000103194 |
| HLA-C | -0.006526086 | 0.995486678 | 0.909785787 | 0.960600726 | 5898.674508 | 5872.100346 | ENSG00000204525 |
| PARP8 | -0.00653093 | 0.995483335 | 0.957633088 | 0.982554648 | 397.7755189 | 395.7841097 | ENSG00000151883 |
| DAG1 | -0.00658319 | 0.995447276 | 0.921037143 | 0.964771134 | 2841.630796 | 2828.516739 | ENSG00000173402 |
| BAAT | -0.006610051 | 0.995428742 | 0.995557548 | 0.998990416 | 3.993823161 | 3.965360182 | ENSG00000136881 |
| GPAT3 | -0.006632562 | 0.99541321 | 0.985322251 | 0.996295482 | 34.60260086 | 34.47592228 | ENSG00000138678 |
| MVB12A | -0.006633909 | 0.995412281 | 0.949324174 | 0.977907982 | 497.6911439 | 495.4807338 | ENSG00000141971 |
| VAV2 | -0.006702881 | 0.995364693 | 0.915720242 | 0.962507751 | 1762.774005 | 1754.411728 | ENSG00000160293 |
| NACC1 | -0.006805399 | 0.995293965 | 0.918703638 | 0.964051188 | 4930.553432 | 4907.881249 | ENSG00000160877 |
| MRPL15 | -0.006947965 | 0.995195616 | 0.9229687 | 0.965130891 | 1259.75516 | 1253.659698 | ENSG00000137547 |
| SESTD1 | -0.006970851 | 0.995179829 | 0.9461031 | 0.97629403 | 497.8367768 | 495.3583907 | ENSG00000187231 |
| MIGA1 | -0.00699015 | 0.995166516 | 0.975884553 | 0.992010969 | 94.22621096 | 93.86961028 | ENSG00000180488 |
| LIMS1 | -0.007077273 | 0.995106421 | 0.945691725 | 0.976189572 | 759.8405846 | 756.2650264 | ENSG00000169756 |
| ITPK1 | -0.007080447 | 0.995104231 | 0.894896098 | 0.953032171 | 4025.913544 | 4006.646132 | ENSG00000100605 |
| CEP83-DT | -0.007134596 | 0.995066883 | 0.991000069 | 0.99806433 | 13.46181716 | 13.35753936 | ENSG00000278916 |
| PRR19 | -0.007187599 | 0.995030326 | 0.981525704 | 0.995385822 | 64.0264148 | 63.75553179 | ENSG00000188368 |
| NA | -0.007254088 | 0.994984469 | 0.992883346 | 0.998381448 | 7.052656893 | 6.992963326 | ENSG00000234645 |
| TAP2 | -0.00729879 | 0.99495364 | 0.968219142 | 0.987407847 | 140.2338043 | 139.470895 | ENSG00000204267 |
| NA | -0.0073396 | 0.994925496 | 0.993347394 | 0.998500968 | 5.700424074 | 5.650020829 | ENSG00000217325 |
| CTIF | -0.007367523 | 0.994906239 | 0.944328925 | 0.975636071 | 492.5761481 | 490.1845379 | ENSG00000134030 |
| DRG1 | -0.007456757 | 0.994844704 | 0.91027438 | 0.960811396 | 1831.964558 | 1822.664899 | ENSG00000185721 |
| DHX8 | -0.00747369 | 0.994833028 | 0.917475931 | 0.963339688 | 1441.801449 | 1434.558501 | ENSG00000067596 |
| RELA | -0.007491242 | 0.994820925 | 0.94965021 | 0.978083782 | 538.4957484 | 535.919336 | ENSG00000173039 |
| FAM117A | -0.007501464 | 0.994813876 | 0.938227332 | 0.972296826 | 643.9071281 | 640.7858384 | ENSG00000121104 |
| PTMA | -0.007501465 | 0.994813875 | 0.870187426 | 0.938734693 | 40717.20189 | 40506.39032 | ENSG00000187514 |
| CEMIP2 | -0.007660145 | 0.994704463 | 0.922837722 | 0.965130891 | 1517.6726 | 1509.312658 | ENSG00000135048 |
| RBBP9 | -0.007673241 | 0.994695434 | 0.969136675 | 0.988023435 | 196.9397637 | 196.1915828 | ENSG00000089050 |
| NIBAN2 | -0.007776393 | 0.994624316 | 0.897198038 | 0.954467117 | 2913.279888 | 2898.065624 | ENSG00000136830 |
| TMOD3 | -0.007840449 | 0.994580155 | 0.938083331 | 0.972215109 | 725.2485587 | 721.0622944 | ENSG00000138594 |
| DDX3X | -0.00788046 | 0.994552572 | 0.886368201 | 0.948550092 | 7504.769574 | 7463.73519 | ENSG00000215301 |
| CEP250 | -0.008116175 | 0.994390091 | 0.925097551 | 0.965883673 | 812.9923874 | 808.3292275 | ENSG00000126001 |
| NA | -0.008142429 | 0.994371995 | 0.991010903 | 0.99806433 | 8.656059258 | 8.574335906 | ENSG00000254332 |
| TMLHE | -0.008159678 | 0.994360106 | 0.966283506 | 0.986931074 | 140.305578 | 139.6765252 | ENSG00000185973 |
| TMEM229B | -0.008217304 | 0.994320389 | 0.9840511 | 0.995916668 | 28.62372667 | 28.49600658 | ENSG00000198133 |
| METTL3 | -0.008229922 | 0.994311693 | 0.914846267 | 0.962281155 | 1996.302259 | 1985.024989 | ENSG00000165819 |
| PALB2 | -0.008298387 | 0.994264508 | 0.942434195 | 0.974531554 | 428.805632 | 425.9946663 | ENSG00000083093 |
| PARD6G | -0.008411974 | 0.99418623 | 0.945267088 | 0.976178103 | 376.3064519 | 374.0171503 | ENSG00000178184 |
| FOXQ1 | -0.00844334 | 0.994164615 | 0.993559305 | 0.998500968 | 4.011620523 | 3.971886573 | ENSG00000164379 |
| SLC41A3 | -0.008464365 | 0.994150127 | 0.938930491 | 0.972506023 | 520.7630546 | 517.824153 | ENSG00000114544 |

| | | | | | | | |
|---|---|---|---|---|---|---|---|
| RAP2C | -0.008483392 | 0.994137016 | 0.940752694 | 0.973752514 | 769.1890822 | 765.3526718 | ENSG00000123728 |
| PCBD1 | -0.008651687 | 0.994021053 | 0.927746467 | 0.967350062 | 677.1482411 | 672.6845559 | ENSG00000166228 |
| GRIK3 | -0.008756068 | 0.993949137 | 0.964281188 | 0.986052631 | 121.8372508 | 121.0571982 | ENSG00000163873 |
| PDCD2L | -0.008800841 | 0.993918291 | 0.968455499 | 0.987595556 | 95.17721831 | 94.51540129 | ENSG00000126249 |
| MZF1 | -0.008804031 | 0.993916093 | 0.962991899 | 0.985269369 | 165.8491402 | 165.0894281 | ENSG00000099326 |
| TYW1 | -0.008839571 | 0.993891609 | 0.933527774 | 0.969858363 | 498.257859 | 495.1444805 | ENSG00000198874 |
| PTPRK | -0.008982261 | 0.993793313 | 0.908037034 | 0.960165884 | 1138.001479 | 1130.730463 | ENSG00000152894 |
| RALA | -0.008985181 | 0.993791302 | 0.90946016 | 0.96047171 | 1838.638897 | 1827.689756 | ENSG00000006451 |
| PPP2R5E | -0.009052545 | 0.993744899 | 0.903531157 | 0.957906355 | 1221.75151 | 1214.284347 | ENSG00000154001 |
| PAGR1 | -0.009069466 | 0.993733244 | 0.953880598 | 0.980836061 | 197.9020343 | 196.7254792 | ENSG00000280789 |
| GUCY1B1 | -0.009072121 | 0.993731415 | 0.972822753 | 0.990091783 | 71.30232413 | 70.730429 | ENSG00000061918 |
| MIPEP | -0.009097005 | 0.993714275 | 0.927473958 | 0.96721772 | 548.0128861 | 544.5397128 | ENSG00000027001 |
| CCDC9 | -0.009105605 | 0.993708351 | 0.911512116 | 0.961009829 | 1048.364386 | 1041.833535 | ENSG00000105321 |
| ENO2 | -0.009105639 | 0.993708328 | 0.884887391 | 0.947622636 | 2435.749916 | 2420.813645 | ENSG00000111674 |
| SLC16A3 | -0.00916827 | 0.99366519 | 0.969567145 | 0.988248895 | 116.3710369 | 115.4346078 | ENSG00000141526 |
| RNF182 | -0.009314576 | 0.993564426 | 0.966737797 | 0.987033408 | 105.5618457 | 104.8856098 | ENSG00000180537 |
| NA | -0.009378631 | 0.993520313 | 0.992845835 | 0.998381448 | 4.386150244 | 4.340372697 | ENSG00000204434 |
| COX6C | -0.009487681 | 0.993445218 | 0.91901885 | 0.964051188 | 655.1374793 | 651.1845817 | ENSG00000164919 |
| EIF3L | -0.009534837 | 0.993412747 | 0.873452704 | 0.940645204 | 3452.137711 | 3429.177604 | ENSG00000100129 |
| CDK11B | -0.009548025 | 0.993403666 | 0.886878402 | 0.948988533 | 1612.552122 | 1602.350736 | ENSG00000248333 |
| XPNPEP1 | -0.009619382 | 0.993354532 | 0.908936112 | 0.960186747 | 937.3004267 | 930.7934229 | ENSG00000108039 |
| TMCO3 | -0.009674559 | 0.993316541 | 0.913770792 | 0.961904203 | 976.1058129 | 969.0451862 | ENSG00000150403 |
| C22orf23 | -0.009679907 | 0.993312859 | 0.989321663 | 0.997721863 | 9.048386341 | 8.951523885 | ENSG00000128346 |
| OSMR | -0.009741678 | 0.99327033 | 0.967888471 | 0.987263577 | 83.66160347 | 83.0394509 | ENSG00000145623 |
| DCAF17 | -0.009846906 | 0.993197885 | 0.92860945 | 0.967653781 | 412.4845642 | 409.6594869 | ENSG00000115827 |
| NPTN | -0.009856514 | 0.99319127 | 0.890440882 | 0.95121103 | 1361.965458 | 1352.764139 | ENSG00000156642 |
| LAMTOR5 | -0.009889243 | 0.993168739 | 0.941813544 | 0.974316559 | 291.9680958 | 289.7831102 | ENSG00000134248 |
| PTPRM | -0.009947519 | 0.993128622 | 0.956980247 | 0.982418568 | 165.0905458 | 163.8561845 | ENSG00000173482 |
| PCYT1A | -0.010000789 | 0.993091952 | 0.921677288 | 0.964771134 | 516.6900589 | 513.312631 | ENSG00000161217 |
| NTN4 | -0.01005245 | 0.993056391 | 0.978968983 | 0.994076204 | 30.08744604 | 29.89399807 | ENSG00000074527 |
| SERINC1 | -0.010139359 | 0.992996571 | 0.922094526 | 0.964771134 | 789.0647626 | 783.886312 | ENSG00000111897 |
| PAAF1 | -0.010204504 | 0.992951734 | 0.928835981 | 0.967758568 | 384.7848692 | 382.1199659 | ENSG00000175575 |
| NA | -0.010224938 | 0.992937669 | 0.988929134 | 0.997539083 | 9.07210744 | 8.972049015 | ENSG00000273893 |
| TRIM45 | -0.010421181 | 0.992802614 | 0.948234485 | 0.977265239 | 364.3154736 | 361.7390926 | ENSG00000134253 |
| NA | -0.010442141 | 0.99278819 | 0.939367945 | 0.972799059 | 252.9918287 | 251.0634613 | ENSG00000233836 |
| SUPT6H | -0.010477477 | 0.992763874 | 0.859134134 | 0.931546657 | 3414.622282 | 3390.004425 | ENSG00000109111 |
| PBDC1 | -0.010494714 | 0.992752013 | 0.911648985 | 0.961009829 | 737.5132842 | 732.5489769 | ENSG00000102390 |
| ASTN2 | -0.010582978 | 0.992691278 | 0.922541293 | 0.96500456 | 452.7496563 | 449.3244175 | ENSG00000148219 |
| ISCU | -0.010652022 | 0.992643772 | 0.899154855 | 0.95546587 | 1401.009278 | 1390.609155 | ENSG00000136003 |
| ACAD8 | -0.01070696 | 0.992605972 | 0.879335634 | 0.944127207 | 1414.485943 | 1404.008319 | ENSG00000151498 |
| LRRC40 | -0.010707993 | 0.992605261 | 0.914893203 | 0.962281155 | 784.984214 | 778.8559766 | ENSG00000066557 |
| SMARCAL1 | -0.010773746 | 0.992560023 | 0.919808472 | 0.964147051 | 536.3982432 | 532.4261492 | ENSG00000138375 |
| SLC29A4 | -0.010790049 | 0.992548807 | 0.93668784 | 0.971621985 | 330.2099502 | 327.9139614 | ENSG00000164638 |
| SPIRE1 | -0.010792264 | 0.992547283 | 0.941349168 | 0.974102961 | 311.0760552 | 308.3412703 | ENSG00000134278 |
| ZNF587 | -0.010801042 | 0.992541244 | 0.933272322 | 0.969789847 | 303.8111553 | 301.6045626 | ENSG00000198466 |
| METTL7B | -0.010911922 | 0.992464963 | 0.978892606 | 0.994072402 | 27.06301182 | 26.87916417 | ENSG00000170439 |
| DGKE | -0.01099814 | 0.992405654 | 0.947825371 | 0.977056885 | 179.3035074 | 177.7267889 | ENSG00000153933 |
| TRAPPC10 | -0.011042469 | 0.992375162 | 0.900853636 | 0.956302469 | 725.6928034 | 720.0336851 | ENSG00000160218 |
| NA | -0.011045497 | 0.992373079 | 0.982067426 | 0.995402713 | 20.46227717 | 20.22044664 | ENSG00000238000 |
| SUV39H1 | -0.011145733 | 0.992304133 | 0.927245638 | 0.967033018 | 329.3338933 | 326.7231131 | ENSG00000101945 |
| PACRGL | -0.011146813 | 0.99230339 | 0.939625654 | 0.972905887 | 221.7431138 | 219.8837452 | ENSG00000163138 |
| RAP1GDS1 | -0.011188218 | 0.992274911 | 0.917988937 | 0.963658759 | 468.8566232 | 465.222317 | ENSG00000138698 |
| SLC22A23 | -0.011201116 | 0.99226604 | 0.892050127 | 0.951914471 | 1182.404833 | 1172.629578 | ENSG00000137266 |
| RASGRP1 | -0.011205175 | 0.992263248 | 0.965452972 | 0.986611438 | 69.57884173 | 69.00689355 | ENSG00000172575 |
| WNK3 | -0.011226018 | 0.992248913 | 0.931648692 | 0.969216414 | 352.9243874 | 350.0504779 | ENSG00000196632 |
| GPR176 | -0.011259819 | 0.992225666 | 0.847463251 | 0.924379996 | 3582.786032 | 3554.47299 | ENSG00000166073 |
| KIF20B | -0.011320094 | 0.992184212 | 0.901934369 | 0.957126914 | 1035.847853 | 1027.650687 | ENSG00000138182 |
| SELENOF | -0.011406875 | 0.992124532 | 0.85658616 | 0.930012052 | 2158.181193 | 2140.824125 | ENSG00000183291 |
| RPS19 | -0.011446444 | 0.992097322 | 0.835628621 | 0.917415262 | 17386.52771 | 17249.25167 | ENSG00000105372 |
| CLPB | -0.011488248 | 0.992068575 | 0.918525834 | 0.964008024 | 470.7597802 | 466.8755322 | ENSG00000162129 |
| MFSD14B | -0.011534609 | 0.992036695 | 0.893078005 | 0.952281345 | 1185.457821 | 1175.642332 | ENSG00000148110 |
| RNF224 | -0.011594146 | 0.991995756 | 0.990209544 | 0.99806433 | 5.376896023 | 5.309532184 | ENSG00000233198 |
| GPN2 | -0.011599926 | 0.991991782 | 0.911178282 | 0.961009829 | 534.0413316 | 530.0125557 | ENSG00000142751 |
| MATR3 | -0.011636596 | 0.991966568 | 0.980810678 | 0.995011844 | 22.39009731 | 22.24029904 | ENSG00000015479 |
| NDUFAF6 | -0.011778304 | 0.991869137 | 0.945957083 | 0.976250041 | 193.1882379 | 191.6758053 | ENSG00000156170 |
| ITGA5 | -0.011931707 | 0.991763677 | 0.896776247 | 0.954280691 | 818.9586284 | 812.4132196 | ENSG00000161638 |
| NEIL2 | -0.011963308 | 0.991741953 | 0.910913543 | 0.961009829 | 489.8476303 | 485.5214162 | ENSG00000154328 |
| CYCS | -0.0119684 | 0.991738453 | 0.837472035 | 0.918202392 | 5880.098505 | 5831.985154 | ENSG00000172115 |
| GID4 | -0.011972231 | 0.991735819 | 0.921681967 | 0.964771134 | 365.7791924 | 362.7128733 | ENSG00000141034 |
| UCP2 | -0.012045469 | 0.991685476 | 0.891878581 | 0.951906189 | 739.6112707 | 733.4441706 | ENSG00000175567 |
| NA | -0.012063923 | 0.99167279 | 0.987282185 | 0.997209395 | 8.036283727 | 7.929774406 | ENSG00000286895 |
| ANKRD36C | -0.012149042 | 0.991614284 | 0.976584226 | 0.992294905 | 26.80472333 | 26.59154906 | ENSG00000174501 |
| TMOD2 | -0.012161312 | 0.99160585 | 0.95398918 | 0.980894304 | 103.7170363 | 102.7673147 | ENSG00000128872 |
| CCDC82 | -0.012176115 | 0.991595675 | 0.929963963 | 0.968157081 | 325.9409309 | 323.4735376 | ENSG00000149231 |

| | | | | | | | |
|---|---|---|---|---|---|---|---|
| ATG101 | -0.01234254 | 0.991481295 | 0.900816203 | 0.956302469 | 858.6882728 | 851.5197137 | ENSG00000123395 |
| NA | -0.012428853 | 0.991421979 | 0.991664855 | 0.99806433 | 3.688092472 | 3.641045742 | ENSG00000235297 |
| USP47 | -0.012455992 | 0.991403329 | 0.899323698 | 0.95553286 | 597.3634333 | 592.4602883 | ENSG00000170242 |
| CARMIL1 | -0.012474518 | 0.991390598 | 0.894994152 | 0.953032171 | 946.9175602 | 938.4385748 | ENSG00000079691 |
| INHBE | -0.012495381 | 0.991376261 | 0.918874102 | 0.964051188 | 317.6296787 | 314.8771636 | ENSG00000139269 |
| SSB | -0.012537925 | 0.991347027 | 0.819767697 | 0.908474441 | 4030.707318 | 3995.305163 | ENSG00000138385 |
| NA | -0.012586364 | 0.991313743 | 0.964126291 | 0.986002654 | 62.35210232 | 61.78486441 | ENSG00000287356 |
| MLF2 | -0.012586857 | 0.991313404 | 0.80105081 | 0.898333333 | 5893.937706 | 5843.081775 | ENSG00000089693 |
| SPINDOC | -0.0126351 | 0.991280255 | 0.830379632 | 0.914737804 | 2651.760037 | 2629.201543 | ENSG00000168005 |
| ZNF510 | -0.012798161 | 0.991168222 | 0.926249574 | 0.966474568 | 270.4572315 | 267.8690175 | ENSG00000081386 |
| NA | -0.012813525 | 0.991157667 | 0.990682249 | 0.99806433 | 3.715373043 | 3.662516831 | ENSG00000258670 |
| TRAPPC11 | -0.012851264 | 0.991131739 | 0.895076008 | 0.953032171 | 713.6134748 | 706.9063027 | ENSG00000168538 |
| CACNA1A | -0.012868212 | 0.991120096 | 0.922002648 | 0.964771134 | 293.0881772 | 290.678136 | ENSG00000141837 |
| NA | -0.012933131 | 0.991075498 | 0.988389609 | 0.997507036 | 7.600073919 | 7.505293321 | ENSG00000270184 |
| MXI1 | -0.012983734 | 0.991040737 | 0.948789412 | 0.977623748 | 157.6093829 | 155.9526836 | ENSG00000119950 |
| ALAS1 | -0.013030296 | 0.991008752 | 0.861355774 | 0.932648138 | 1152.513384 | 1142.191387 | ENSG00000023330 |
| GNPDA2 | -0.013075499 | 0.990977702 | 0.945655614 | 0.976189572 | 139.4111911 | 138.1475317 | ENSG00000163281 |
| STRN | -0.013265016 | 0.990847532 | 0.892723375 | 0.952281345 | 719.3729092 | 712.5200234 | ENSG00000115808 |
| HYAL2 | -0.01327729 | 0.990839103 | 0.876159205 | 0.942347141 | 871.3839487 | 863.6980077 | ENSG00000068001 |
| NA | -0.013287236 | 0.990832272 | 0.975469174 | 0.991946116 | 25.14675983 | 24.92741354 | ENSG00000253683 |
| IMMT | -0.013351535 | 0.990788113 | 0.814117506 | 0.905454011 | 2537.274362 | 2513.622911 | ENSG00000132305 |
| AGO2 | -0.013357969 | 0.990783695 | 0.875203338 | 0.94170908 | 2034.566256 | 2015.23978 | ENSG00000123908 |
| TRAF7 | -0.01343349 | 0.990731831 | 0.795848934 | 0.89563779 | 3604.822267 | 3571.361774 | ENSG00000131653 |
| ZNF346 | -0.013440157 | 0.990727253 | 0.896855161 | 0.954310892 | 526.9346199 | 521.9991248 | ENSG00000113761 |
| CCDC71 | -0.013497867 | 0.990687623 | 0.890067021 | 0.951059472 | 585.8287997 | 580.628054 | ENSG00000177352 |
| ARL6 | -0.013613759 | 0.990608044 | 0.953495215 | 0.980546566 | 88.5756195 | 87.83805055 | ENSG00000113966 |
| DAGLB | -0.013614182 | 0.990607753 | 0.897716619 | 0.954850935 | 524.0446077 | 519.278337 | ENSG00000164535 |
| MAU2 | -0.013620617 | 0.990603335 | 0.868701914 | 0.937813209 | 934.3347746 | 925.8543211 | ENSG00000129933 |
| SHANK2 | -0.013653472 | 0.990580776 | 0.874666209 | 0.941359938 | 918.453016 | 909.2699069 | ENSG00000162105 |
| LSR | -0.013659145 | 0.99057688 | 0.824335644 | 0.911237336 | 3779.082068 | 3744.363877 | ENSG00000105699 |
| NCOA6 | -0.01372281 | 0.990533168 | 0.912606485 | 0.961482481 | 316.9337058 | 314.0778809 | ENSG00000198646 |
| PWP2 | -0.013753854 | 0.990511854 | 0.968026098 | 0.987288708 | 44.86681717 | 44.47908504 | ENSG00000241945 |
| BAZ1A | -0.013775487 | 0.990497002 | 0.888219901 | 0.949670658 | 611.5769535 | 605.7693777 | ENSG00000198604 |
| INO80C | -0.013812122 | 0.99047185 | 0.932125245 | 0.969445162 | 181.6734872 | 179.8287418 | ENSG00000153391 |
| BLCAP | -0.013844391 | 0.990449696 | 0.878355558 | 0.94339673 | 1317.652923 | 1305.620508 | ENSG00000166619 |
| SETD2 | -0.013874645 | 0.990428926 | 0.842484353 | 0.92165416 | 1345.746953 | 1332.609945 | ENSG00000181555 |
| ZNF639 | -0.013898924 | 0.990412258 | 0.899837377 | 0.955760904 | 419.5319077 | 415.5266346 | ENSG00000121864 |
| TMEM242 | -0.013938851 | 0.990384849 | 0.914674188 | 0.96227163 | 302.6978874 | 299.6045729 | ENSG00000215712 |
| NGDN | -0.01394406 | 0.990381273 | 0.870979172 | 0.939374167 | 921.3443269 | 912.4825627 | ENSG00000129460 |
| MXD4 | -0.013994261 | 0.990346812 | 0.879171127 | 0.944004249 | 1052.033843 | 1042.187214 | ENSG00000123933 |
| ENPP5 | -0.014005523 | 0.990339081 | 0.943266277 | 0.974964898 | 137.0741886 | 135.7265612 | ENSG00000112796 |
| SHANK3 | -0.014065786 | 0.990297714 | 0.897839658 | 0.95489739 | 590.6509333 | 584.8797998 | ENSG00000251322 |
| POLDIP2 | -0.014159215 | 0.990233585 | 0.797268836 | 0.896487332 | 2907.317181 | 2879.070257 | ENSG00000004142 |
| PTGIS | -0.014176937 | 0.99022142 | 0.87974183 | 0.944439862 | 866.5157963 | 857.649977 | ENSG00000124212 |
| ATXN7L3 | -0.014257999 | 0.990165784 | 0.808301512 | 0.902706923 | 2990.592539 | 2961.112837 | ENSG00000087152 |
| THAP11 | -0.014272837 | 0.9901556 | 0.870131786 | 0.938728293 | 948.0148096 | 939.2650605 | ENSG00000168286 |
| SKI | -0.014561185 | 0.989957719 | 0.813097273 | 0.905169301 | 2252.333473 | 2230.149255 | ENSG00000157933 |
| NA | -0.014691054 | 0.989868609 | 0.988751129 | 0.997539083 | 4.035341622 | 3.969711109 | ENSG00000277491 |
| NDC80 | -0.014735017 | 0.989838445 | 0.874764392 | 0.941359938 | 621.7963973 | 615.525903 | ENSG00000080986 |
| SLC26A6 | -0.014767093 | 0.989816438 | 0.879717454 | 0.944439862 | 1387.767336 | 1373.828865 | ENSG00000225697 |
| ZIC5 | -0.014776514 | 0.989809974 | 0.884949169 | 0.947622636 | 875.1542134 | 866.5564988 | ENSG00000139800 |
| CCDC66 | -0.014843779 | 0.989763826 | 0.917737219 | 0.963484043 | 283.7657638 | 281.1380456 | ENSG00000180376 |
| IRS1 | -0.014867928 | 0.989747259 | 0.858563678 | 0.930981572 | 1275.679171 | 1262.67434 | ENSG00000169047 |
| DHX30 | -0.014879987 | 0.989738986 | 0.788835419 | 0.891843683 | 3398.33422 | 3363.542328 | ENSG00000132153 |
| YY1 | -0.014892042 | 0.989730716 | 0.810014237 | 0.90313851 | 2883.718755 | 2854.438485 | ENSG00000100811 |
| ADAM17 | -0.014896126 | 0.989727914 | 0.887607895 | 0.949392563 | 535.8456881 | 529.9969283 | ENSG00000151694 |
| AURKA | -0.014922724 | 0.989709667 | 0.81135992 | 0.904250935 | 2134.316751 | 2111.738688 | ENSG00000087586 |
| AFAP1L2 | -0.014965358 | 0.98968042 | 0.945497729 | 0.976189572 | 109.495184 | 108.439469 | ENSG00000169129 |
| NA | -0.015055944 | 0.989618281 | 0.965786707 | 0.986738908 | 40.29857574 | 39.80039916 | ENSG00000260007 |
| SMIM12 | -0.015148438 | 0.989554837 | 0.841777645 | 0.92140286 | 1016.337779 | 1005.611113 | ENSG00000163866 |
| ZNF426 | -0.015157654 | 0.989548515 | 0.909420845 | 0.96047171 | 326.3762249 | 323.0514 | ENSG00000130818 |
| PLXNA1 | -0.01520552 | 0.989515685 | 0.778929051 | 0.887164534 | 3773.280176 | 3733.962436 | ENSG00000114554 |
| DESI1 | -0.015235771 | 0.989494936 | 0.867858615 | 0.937346866 | 751.789732 | 744.0331346 | ENSG00000100418 |
| PLEKHA2 | -0.015295264 | 0.989454133 | 0.940081537 | 0.97320185 | 146.596081 | 144.8535423 | ENSG00000169499 |
| B4GALT2 | -0.015297044 | 0.989452912 | 0.810348215 | 0.903443299 | 1914.971786 | 1894.629194 | ENSG00000117411 |
| TMEM138 | -0.015327838 | 0.989431793 | 0.907731615 | 0.960149634 | 296.3302717 | 293.067872 | ENSG00000149483 |
| PSRC1 | -0.015372644 | 0.989401064 | 0.838602552 | 0.91904671 | 1356.523576 | 1342.253294 | ENSG00000134222 |
| OFD1 | -0.015441028 | 0.989354167 | 0.902477263 | 0.957434027 | 340.9390536 | 337.1951396 | ENSG00000046651 |
| MSMO1 | -0.015472465 | 0.989332609 | 0.850587874 | 0.926001729 | 1810.774734 | 1791.53122 | ENSG00000052802 |
| DNAJB12 | -0.01553872 | 0.989287175 | 0.875984533 | 0.942297101 | 549.3609485 | 543.741587 | ENSG00000148719 |
| NA | -0.015561316 | 0.989271681 | 0.967650692 | 0.987242444 | 45.79883194 | 45.14490672 | ENSG00000226562 |
| FBXL15 | -0.015593452 | 0.989249645 | 0.902956191 | 0.957841349 | 319.3487375 | 316.1306488 | ENSG00000107872 |
| NA | -0.015607764 | 0.989239832 | 0.988479835 | 0.997512113 | 4.00806105 | 3.941713629 | ENSG00000270039 |
| ZBED5 | -0.015619389 | 0.989231861 | 0.884156499 | 0.947149327 | 648.3686899 | 641.556062 | ENSG00000236287 |

| | | | | | | | |
|---|---|---|---|---|---|---|---|
| E2F5 | -0.01567879 | 0.989191131 | 0.894156991 | 0.952997053 | 521.1373038 | 516.0059883 | ENSG00000133740 |
| SYPL1 | -0.015745571 | 0.989145343 | 0.842066492 | 0.921502333 | 905.9858869 | 896.179006 | ENSG00000008282 |
| TRIP12 | -0.015763618 | 0.98913297 | 0.813705619 | 0.905396008 | 2694.349701 | 2664.713544 | ENSG00000153827 |
| TNFAIP8 | -0.015779219 | 0.989122274 | 0.913235654 | 0.961597187 | 274.6542234 | 271.6190044 | ENSG00000145779 |
| RPP25 | -0.015825091 | 0.989090824 | 0.861375493 | 0.932648138 | 767.4476485 | 758.5954484 | ENSG00000178718 |
| KCNJ4 | -0.015984434 | 0.988981587 | 0.961051425 | 0.984350589 | 41.58289925 | 41.16169132 | ENSG00000168135 |
| PCDHB3 | -0.01605711 | 0.988931768 | 0.975989144 | 0.992063889 | 17.74093005 | 17.57627432 | ENSG00000113205 |
| ANKMY1 | -0.016067028 | 0.98892497 | 0.938508953 | 0.972389399 | 112.3609426 | 111.1480546 | ENSG00000144504 |
| CBSL | -0.016142167 | 0.988873465 | 0.861050781 | 0.932648138 | 888.5723504 | 878.6290185 | ENSG00000274276 |
| C16orf96 | -0.016153347 | 0.988865802 | 0.967464725 | 0.987242444 | 29.60763284 | 29.24073472 | ENSG00000205832 |
| EIF3A | -0.016153594 | 0.988865633 | 0.764047988 | 0.877785999 | 11557.37466 | 11427.83585 | ENSG00000107581 |
| MAP2K7 | -0.016159947 | 0.988861278 | 0.820072314 | 0.908627926 | 1430.324657 | 1414.960779 | ENSG00000076984 |
| ORC4 | -0.016169719 | 0.988854581 | 0.855753078 | 0.929583524 | 894.2617377 | 884.2079911 | ENSG00000115947 |
| ACBD4 | -0.016174906 | 0.988851025 | 0.9254685 | 0.965963518 | 196.333337 | 194.1077913 | ENSG00000181513 |
| ABHD17A | -0.016220626 | 0.988819688 | 0.856558405 | 0.930012052 | 1595.338776 | 1578.30577 | ENSG00000129968 |
| ARHGEF10 | -0.016224595 | 0.988816968 | 0.957461033 | 0.98244088 | 57.39272902 | 56.58058485 | ENSG00000104728 |
| PRKDC | -0.016272893 | 0.988783865 | 0.917771483 | 0.963484043 | 8078.143475 | 7987.194439 | ENSG00000253729 |
| STRA6 | -0.016347427 | 0.988732783 | 0.903377014 | 0.957904263 | 264.6047183 | 261.5766107 | ENSG00000137868 |
| MIR4292 | -0.016389702 | 0.988703811 | 0.965234704 | 0.986441765 | 32.91527185 | 32.48529686 | ENSG00000265806 |
| RRP1 | -0.016412152 | 0.988688426 | 0.786866869 | 0.891330078 | 2131.677321 | 2107.94188 | ENSG00000160214 |
| LOC100287042 | -0.0164395 | 0.988669684 | 0.967920843 | 0.987263577 | 28.66196499 | 28.30515314 | ENSG00000263843 |
| FLNA | -0.01644172 | 0.988668163 | 0.763253412 | 0.877513739 | 42559.89868 | 42077.38073 | ENSG00000196924 |
| TAF3 | -0.016472904 | 0.988646793 | 0.890166971 | 0.951106265 | 407.9551709 | 403.5707027 | ENSG00000165632 |
| PUF60 | -0.016494047 | 0.988632304 | 0.760466615 | 0.875856073 | 4039.168652 | 3993.534694 | ENSG00000179950 |
| ETNK1 | -0.016557587 | 0.988588763 | 0.89192375 | 0.951906189 | 885.6530689 | 875.1383891 | ENSG00000139163 |
| LMF2 | -0.016605106 | 0.988556202 | 0.844098024 | 0.922299853 | 899.4753604 | 888.9963758 | ENSG00000100258 |
| CCDC144B | -0.016635623 | 0.988535291 | 0.987444451 | 0.997300295 | 4.365988618 | 4.290904128 | ENSG00000154874 |
| IPPK | -0.016661256 | 0.988517728 | 0.922215966 | 0.964771134 | 160.1113158 | 158.27623 | ENSG00000127080 |
| KDM8 | -0.016669694 | 0.988511946 | 0.920756385 | 0.964633548 | 167.3911418 | 165.5667398 | ENSG00000155666 |
| FNDC7 | -0.016720897 | 0.988476863 | 0.970647432 | 0.989029716 | 22.89037762 | 22.62713484 | ENSG00000143107 |
| SEMA6C | -0.016755107 | 0.988453424 | 0.843426302 | 0.922013311 | 839.3265792 | 829.9630095 | ENSG00000143434 |
| B3GNT4 | -0.016766305 | 0.988445752 | 0.970854398 | 0.989151342 | 22.56209489 | 22.30253685 | ENSG00000176383 |
| AAMDC | -0.016781829 | 0.988435116 | 0.958067516 | 0.982680048 | 50.56101202 | 50.07216495 | ENSG00000087884 |
| PTK7 | -0.016968434 | 0.988307275 | 0.73711382 | 0.861997306 | 4924.855162 | 4866.62966 | ENSG00000112655 |
| TTC39B | -0.017044976 | 0.988254842 | 0.917480944 | 0.963339688 | 182.2615068 | 179.979323 | ENSG00000155158 |
| NOSIP | -0.017048909 | 0.988252148 | 0.816556467 | 0.906477444 | 1769.136176 | 1748.658884 | ENSG00000142546 |
| MYO18A | -0.017063153 | 0.988242391 | 0.794596085 | 0.894952975 | 1788.618542 | 1767.635691 | ENSG00000196535 |
| ZNF233 | -0.017165717 | 0.988172137 | 0.967589038 | 0.987242444 | 25.16692145 | 24.85089345 | ENSG00000159915 |
| SHROOM1 | -0.017197829 | 0.988150142 | 0.92614146 | 0.966415155 | 153.9426728 | 152.251124 | ENSG00000164403 |
| ACTR3 | -0.017199425 | 0.988149049 | 0.798119988 | 0.897070233 | 3406.773429 | 3366.428833 | ENSG00000115091 |
| TAF13 | -0.017258414 | 0.988108647 | 0.894515926 | 0.953032171 | 431.723307 | 426.7178804 | ENSG00000197780 |
| POLL | -0.017341576 | 0.98805169 | 0.876180749 | 0.942347141 | 425.9585343 | 420.7326382 | ENSG00000166169 |
| CSNK1A1 | -0.017366588 | 0.988034561 | 0.778137863 | 0.886905278 | 3212.362314 | 3173.552304 | ENSG00000113712 |
| ZNHIT1 | -0.017438292 | 0.987985455 | 0.826645097 | 0.912668381 | 1204.604246 | 1189.733241 | ENSG00000106400 |
| CSNK2A1 | -0.017474826 | 0.987960436 | 0.768692258 | 0.880764064 | 4215.24761 | 4164.031638 | ENSG00000101266 |
| SON | -0.017511464 | 0.987935347 | 0.734465889 | 0.860499306 | 4847.63107 | 4789.087262 | ENSG00000159140 |
| NA | -0.017547147 | 0.987910912 | 0.985177624 | 0.99621074 | 5.373336551 | 5.275008313 | ENSG00000250993 |
| KBTBD8 | -0.017577568 | 0.987890081 | 0.890610094 | 0.95121103 | 349.3632146 | 345.0386078 | ENSG00000163376 |
| HMOX1 | -0.017721555 | 0.98779149 | 0.83310237 | 0.915888826 | 848.6803646 | 838.537586 | ENSG00000100292 |
| USP38 | -0.01772303 | 0.98779048 | 0.901521827 | 0.956745272 | 267.0458614 | 263.6502396 | ENSG00000170185 |
| CIT | -0.017790178 | 0.987744506 | 0.841748798 | 0.92140286 | 1137.108262 | 1122.437456 | ENSG00000122966 |
| FAM193A | -0.017809455 | 0.987731308 | 0.835544864 | 0.917415262 | 747.0426539 | 737.7324477 | ENSG00000125386 |
| RHOC | -0.017900148 | 0.987669218 | 0.885214783 | 0.947622636 | 430.8751945 | 425.3097599 | ENSG00000155366 |
| MARCHF6 | -0.01800788 | 0.987595467 | 0.814170559 | 0.905459691 | 1621.139454 | 1600.227088 | ENSG00000145495 |
| STXBP1 | -0.018042654 | 0.987571663 | 0.806917298 | 0.901640919 | 1330.732203 | 1313.901003 | ENSG00000136854 |
| RHNO1 | -0.018056786 | 0.987561989 | 0.85796017 | 0.930556858 | 637.2441297 | 629.2665373 | ENSG00000171792 |
| CMTM8 | -0.018066925 | 0.987555049 | 0.929564951 | 0.968057669 | 144.2208657 | 142.5574036 | ENSG00000170293 |
| SMC3 | -0.018089508 | 0.987539591 | 0.792442706 | 0.893961668 | 2628.743709 | 2595.775035 | ENSG00000108055 |
| COG8 | -0.018097858 | 0.987533875 | 0.93487897 | 0.970598645 | 96.62850538 | 95.49269554 | ENSG00000213380 |
| FAM222B | -0.018170052 | 0.987484459 | 0.795665635 | 0.895554092 | 1889.996621 | 1866.629792 | ENSG00000173065 |
| TBCB | -0.01831088 | 0.987388071 | 0.781303282 | 0.888626825 | 2113.619293 | 2087.47529 | ENSG00000105254 |
| POU5F1P3 | -0.0184112 | 0.987319413 | 0.938755242 | 0.972431174 | 88.75145589 | 87.60330898 | ENSG00000235602 |
| VDAC1 | -0.018448835 | 0.987293658 | 0.693227871 | 0.836043703 | 5859.444184 | 5785.205489 | ENSG00000213585 |
| NA | -0.018505636 | 0.987254787 | 0.97134827 | 0.989388263 | 17.18875834 | 16.99640963 | ENSG00000259343 |
| FGD5 | -0.018522792 | 0.987243048 | 0.855797678 | 0.929583524 | 750.5204588 | 740.4733911 | ENSG00000154783 |
| CLCC1 | -0.018533653 | 0.987235615 | 0.893061439 | 0.952281345 | 280.1696072 | 276.5607129 | ENSG00000121940 |
| WWC3 | -0.018546303 | 0.987226959 | 0.80971047 | 0.90305216 | 1207.962938 | 1192.272401 | ENSG00000047644 |
| NA | -0.018586699 | 0.987199316 | 0.949281136 | 0.977907982 | 55.15711883 | 54.50428598 | ENSG00000228782 |
| TRIO | -0.018717599 | 0.987109749 | 0.791326317 | 0.893310867 | 1487.445619 | 1468.108532 | ENSG00000038382 |
| FUS | -0.018759901 | 0.987080806 | 0.710677086 | 0.847279094 | 18240.19405 | 18005.38201 | ENSG00000089280 |
| PSMG3-AS1 | -0.018769974 | 0.987073914 | 0.923721763 | 0.965407741 | 124.7142982 | 123.2406846 | ENSG00000230487 |
| SNCB | -0.018831841 | 0.987031587 | 0.898070834 | 0.954958767 | 225.406621 | 222.6383944 | ENSG00000074317 |
| TACC3 | -0.018863171 | 0.987010152 | 0.696776402 | 0.838128812 | 4989.911449 | 4925.182032 | ENSG00000013810 |
| NA | -0.018902358 | 0.986983343 | 0.983017946 | 0.995796876 | 6.440000307 | 6.317282923 | ENSG00000249485 |

| | | | | | | | |
|---|---|---|---|---|---|---|---|
| RBFA | -0.018984961 | 0.986926834 | 0.912881169 | 0.961597187 | 160.7254469 | 158.5206193 | ENSG00000101546 |
| CBLL1 | -0.018987474 | 0.986925115 | 0.851580102 | 0.926547804 | 580.0403314 | 572.5558534 | ENSG00000105879 |
| NRXN3 | -0.019000104 | 0.986916474 | 0.950638715 | 0.978888339 | 58.60677896 | 57.85816812 | ENSG00000021645 |
| PTPRA | -0.019000546 | 0.986916172 | 0.774943274 | 0.884331605 | 1609.070121 | 1588.03381 | ENSG00000132670 |
| CIZ1 | -0.019015643 | 0.986905845 | 0.765478015 | 0.878733956 | 2383.307738 | 2352.746628 | ENSG00000148337 |
| PCDH10 | -0.019027053 | 0.98689804 | 0.945787832 | 0.976226823 | 72.16642959 | 71.05786486 | ENSG00000138650 |
| RAB38 | -0.019038648 | 0.986890108 | 0.911614176 | 0.961009829 | 169.4559758 | 167.2094042 | ENSG00000123892 |
| NA | -0.019103558 | 0.986845707 | 0.957225633 | 0.98244088 | 37.425672 | 36.95790106 | ENSG00000213058 |
| NEFM | -0.019114463 | 0.986838247 | 0.857961904 | 0.930556858 | 546.4856288 | 539.2380521 | ENSG00000104722 |
| NA | -0.019138524 | 0.986821789 | 0.970349265 | 0.988885967 | 17.12707825 | 16.92546997 | ENSG00000237214 |
| RNF146 | -0.019159324 | 0.986807562 | 0.843894622 | 0.922223826 | 581.1176213 | 573.640714 | ENSG00000118518 |
| COL27A1 | -0.019356452 | 0.986672735 | 0.949113185 | 0.977904004 | 784.2678787 | 773.8836406 | ENSG00000196739 |
| APEX1 | -0.019372527 | 0.986661741 | 0.69291998 | 0.835910178 | 9109.965712 | 8987.892767 | ENSG00000100823 |
| ENY2 | -0.019385949 | 0.986652562 | 0.763455014 | 0.877522527 | 2546.867924 | 2512.807814 | ENSG00000120533 |
| AVPI1 | -0.019430087 | 0.986622376 | 0.907997574 | 0.960165884 | 176.749708 | 174.549099 | ENSG00000119986 |
| SULF2 | -0.019450236 | 0.986608597 | 0.838547251 | 0.91904671 | 732.4089412 | 722.5942937 | ENSG00000196562 |
| SART1 | -0.019480955 | 0.98658759 | 0.77758027 | 0.886376737 | 2819.453653 | 2782.432872 | ENSG00000175467 |
| MCMBP | -0.01951329 | 0.986565477 | 0.741628829 | 0.865282909 | 2210.836736 | 2180.752507 | ENSG00000197771 |
| DUSP1 | -0.019517495 | 0.986562602 | 0.907889128 | 0.960165884 | 195.4430938 | 192.8530969 | ENSG00000120129 |
| NRBP1 | -0.019670317 | 0.986458103 | 0.717509121 | 0.851122916 | 3038.753009 | 2997.532781 | ENSG00000115216 |
| BECN1 | -0.019717739 | 0.986425678 | 0.828990615 | 0.914028945 | 659.5132311 | 650.3512993 | ENSG00000126581 |
| QTRT2 | -0.019761953 | 0.986395448 | 0.791469721 | 0.893312556 | 1240.730517 | 1223.938339 | ENSG00000151576 |
| NUP107 | -0.01977089 | 0.986389337 | 0.756963518 | 0.873742764 | 1929.017478 | 1902.805887 | ENSG00000111581 |
| WDFY2 | -0.01977693 | 0.986385208 | 0.825940484 | 0.912370498 | 841.9562187 | 830.0628269 | ENSG00000139668 |
| MRPL45 | -0.019844767 | 0.986338828 | 0.793127632 | 0.894207674 | 1245.569148 | 1228.51729 | ENSG00000278845 |
| CKAP2 | -0.019923557 | 0.986284962 | 0.801192533 | 0.898333333 | 1412.300192 | 1392.437726 | ENSG00000136108 |
| ZMIZ2 | -0.019948985 | 0.986267579 | 0.809645792 | 0.90305216 | 1683.585076 | 1661.163373 | ENSG00000122515 |
| ZNF131 | -0.020006811 | 0.986228048 | 0.835295189 | 0.917410484 | 602.5330163 | 594.4113765 | ENSG00000172262 |
| FAAP100 | -0.020124836 | 0.98614737 | 0.803870568 | 0.899993815 | 1232.359572 | 1215.741665 | ENSG00000185504 |
| IQGAP2 | -0.020224067 | 0.986079543 | 0.791272879 | 0.893303941 | 2059.332283 | 2030.283938 | ENSG00000145703 |
| PIH1D1 | -0.020297407 | 0.986029416 | 0.784885396 | 0.890433415 | 2028.322071 | 1999.906284 | ENSG00000104872 |
| ZNF333 | -0.020351069 | 0.985992742 | 0.921731593 | 0.964771134 | 125.1579063 | 123.4730827 | ENSG00000160961 |
| NKRF | -0.020378193 | 0.985974204 | 0.864172896 | 0.93447402 | 386.1622976 | 380.6586953 | ENSG00000186416 |
| PROKR1 | -0.02039852 | 0.985960312 | 0.942160558 | 0.974494871 | 61.18788442 | 60.44315142 | ENSG00000169618 |
| RDH13 | -0.020438359 | 0.985933086 | 0.892046365 | 0.951914471 | 218.202713 | 215.1070682 | ENSG00000160439 |
| NA | -0.020593434 | 0.985827114 | 0.96805004 | 0.987288708 | 16.78338858 | 16.56540216 | ENSG00000227398 |
| LENG8-AS1 | -0.020651573 | 0.985787387 | 0.961949542 | 0.984896563 | 26.05713843 | 25.64073924 | ENSG00000226696 |
| MDGA2 | -0.020746908 | 0.985722247 | 0.956570613 | 0.982211615 | 35.27129313 | 34.66488379 | ENSG00000139915 |
| TMX1 | -0.020756178 | 0.985715913 | 0.809197181 | 0.902949338 | 1626.407102 | 1603.410787 | ENSG00000139921 |
| SEMA4B | -0.020765221 | 0.985709735 | 0.779216193 | 0.887331031 | 1435.7089 | 1414.788976 | ENSG00000185033 |
| CFDP1 | -0.020854393 | 0.985648811 | 0.839645467 | 0.919815803 | 715.8399334 | 705.3201934 | ENSG00000153774 |
| TUNAR | -0.020911281 | 0.985609945 | 0.793543714 | 0.894317058 | 1059.182398 | 1043.476061 | ENSG00000250366 |
| ANKRD46 | -0.020977564 | 0.985564664 | 0.868215194 | 0.937517522 | 407.4887326 | 401.3057555 | ENSG00000186106 |
| CROT | -0.020982321 | 0.985561414 | 0.934687537 | 0.970489863 | 78.09165851 | 76.82874542 | ENSG00000005469 |
| PTP4A2 | -0.020985087 | 0.985559525 | 0.72887735 | 0.857302489 | 2759.116406 | 2719.608011 | ENSG00000184007 |
| CDH5 | -0.021017718 | 0.985537233 | 0.980798256 | 0.995011844 | 9.35739717 | 9.26306909 | ENSG00000179776 |
| RASA4B | -0.021052121 | 0.985513732 | 0.983631352 | 0.995807802 | 5.022527928 | 4.918345466 | ENSG00000170667 |
| SNF8 | -0.021101438 | 0.985480044 | 0.867614282 | 0.937136566 | 397.4124794 | 391.2760584 | ENSG00000159210 |
| PCED1A | -0.0211422 | 0.985452201 | 0.887935859 | 0.94955683 | 498.858337 | 491.6785264 | ENSG00000132635 |
| EP300 | -0.021210006 | 0.985405886 | 0.73525614 | 0.860997842 | 2902.920302 | 2859.744002 | ENSG00000100393 |
| RPL10 | -0.021248092 | 0.985379872 | 0.673916519 | 0.825366912 | 10156.45869 | 10008.4181 | ENSG00000147403 |
| ZNF221 | -0.021263028 | 0.985369671 | 0.965660723 | 0.986716954 | 20.96700668 | 20.64710319 | ENSG00000159905 |
| WIPF2 | -0.021265697 | 0.985367848 | 0.805219334 | 0.900702903 | 838.7538625 | 826.3384512 | ENSG00000171475 |
| UPF2 | -0.021287773 | 0.98535277 | 0.786549748 | 0.891168226 | 1211.033696 | 1193.353201 | ENSG00000151461 |
| FAM47E | -0.021316154 | 0.985333386 | 0.984225315 | 0.995944879 | 5.111488587 | 5.008580751 | ENSG00000189157 |
| RPL10A | -0.021332237 | 0.985322402 | 0.637242512 | 0.800900096 | 16864.33906 | 16616.29538 | ENSG00000198755 |
| SMAD4 | -0.021390897 | 0.98528234 | 0.787892491 | 0.891582187 | 1028.315383 | 1012.92757 | ENSG00000141646 |
| KLHL17 | -0.021420631 | 0.985262033 | 0.873529401 | 0.94067416 | 692.1335279 | 682.3911947 | ENSG00000187961 |
| NA | -0.021453358 | 0.985239682 | 0.978883959 | 0.994072402 | 10.10081221 | 9.904225622 | ENSG00000214433 |
| ZNF813 | -0.021489301 | 0.985215137 | 0.864597851 | 0.934774605 | 448.4844235 | 441.6918833 | ENSG00000198346 |
| ING5 | -0.021536147 | 0.985183146 | 0.814000706 | 0.905404967 | 1137.101448 | 1120.423043 | ENSG00000168395 |
| ZFP91 | -0.021541819 | 0.985179274 | 0.850689853 | 0.926059366 | 451.7675295 | 445.1530483 | ENSG00000186660 |
| UTP15 | -0.021560591 | 0.985166454 | 0.827478371 | 0.913263802 | 601.7332084 | 592.8789781 | ENSG00000164338 |
| SNX22 | -0.021583292 | 0.985150953 | 0.932694573 | 0.969708285 | 88.51213382 | 87.2226434 | ENSG00000157734 |
| MED23 | -0.021583293 | 0.985150952 | 0.872069405 | 0.940089565 | 330.535817 | 325.4537372 | ENSG00000112282 |
| MYPOP | -0.021620384 | 0.985125625 | 0.889597182 | 0.950712624 | 218.3625838 | 215.2643866 | ENSG00000176182 |
| DIS3 | -0.021735747 | 0.985046854 | 0.772163668 | 0.882599617 | 1206.403671 | 1188.09588 | ENSG00000083520 |
| TMEM129 | -0.021767079 | 0.985025461 | 0.784035826 | 0.890065126 | 1111.673978 | 1095.021426 | ENSG00000168936 |
| RPS15 | -0.021781226 | 0.985015802 | 0.649109619 | 0.809458211 | 14231.11368 | 14018.20694 | ENSG00000115268 |
| BLM | -0.021786744 | 0.985012034 | 0.81002701 | 0.90313851 | 739.9656911 | 728.7742592 | ENSG00000197299 |
| CCNB1IP1 | -0.021851171 | 0.984968047 | 0.778787367 | 0.88705666 | 1753.252106 | 1726.337358 | ENSG00000100814 |
| NGB | -0.021889225 | 0.984942067 | 0.970303827 | 0.988885967 | 13.74591168 | 13.57326884 | ENSG00000165553 |
| NA | -0.021892432 | 0.984939878 | 0.981850154 | 0.995402617 | 7.763478014 | 7.692290282 | ENSG00000270761 |
| VIPR2 | -0.021916958 | 0.984923134 | 0.944504637 | 0.975763473 | 52.87434578 | 52.21176317 | ENSG00000106018 |

| | | | | | | | |
|---|---|---|---|---|---|---|---|
| NEO1 | -0.021919273 | 0.984921553 | 0.754779561 | 0.872411882 | 1455.766444 | 1433.56437 | ENSG00000067141 |
| MPP5 | -0.021950598 | 0.984900168 | 0.840646809 | 0.920561463 | 532.7622691 | 524.529227 | ENSG00000072415 |
| EXOSC6 | -0.021981438 | 0.984879115 | 0.83718009 | 0.918172233 | 541.0447401 | 533.2949741 | ENSG00000223496 |
| TIMM8A | -0.02198316 | 0.984877939 | 0.849796809 | 0.925513999 | 436.2242251 | 429.4465826 | ENSG00000126953 |
| PAOX | -0.021995452 | 0.984869548 | 0.976407034 | 0.992221634 | 10.35643095 | 10.24499781 | ENSG00000148832 |
| FBH1 | -0.021999685 | 0.984866658 | 0.780185416 | 0.88782839 | 1068.520191 | 1052.145477 | ENSG00000134452 |
| CYP4V2 | -0.022081915 | 0.984810525 | 0.941189777 | 0.97409682 | 61.90818875 | 61.14493739 | ENSG00000145476 |
| FYTTD1 | -0.022180367 | 0.984743322 | 0.829859182 | 0.914706392 | 540.6645143 | 532.2351372 | ENSG00000122068 |
| NA | -0.022182792 | 0.984741666 | 0.961168888 | 0.984417509 | 21.94707404 | 21.58268476 | ENSG00000257489 |
| VAT1 | -0.022193919 | 0.984734071 | 0.613012507 | 0.78462019 | 8256.321206 | 8130.511813 | ENSG00000108828 |
| MAP4K1 | -0.022292309 | 0.984666916 | 0.915958702 | 0.962597668 | 103.4658668 | 101.7788596 | ENSG00000104814 |
| SLC25A51 | -0.022309478 | 0.984655198 | 0.936890493 | 0.971725448 | 72.07866414 | 70.7619994 | ENSG00000122696 |
| GPS1 | -0.022361305 | 0.984619826 | 0.6671042 | 0.82154232 | 3571.781043 | 3517.155919 | ENSG00000169727 |
| SNAP91 | -0.022397861 | 0.984594877 | 0.966027503 | 0.986878145 | 20.85521465 | 20.62251068 | ENSG00000065609 |
| CTDSP1 | -0.022399691 | 0.984593629 | 0.834002945 | 0.916612094 | 452.2392823 | 445.1459978 | ENSG00000144579 |
| AKT1 | -0.022433707 | 0.984570414 | 0.755887318 | 0.872977739 | 2126.394423 | 2093.07187 | ENSG00000142208 |
| GMPPA | -0.022467514 | 0.984547343 | 0.778772612 | 0.88705666 | 957.1811661 | 942.084248 | ENSG00000144591 |
| MPP3 | -0.022475931 | 0.984541598 | 0.911526334 | 0.961009829 | 144.0654185 | 141.8382569 | ENSG00000161647 |
| FAM102A | -0.022476137 | 0.984541458 | 0.781074783 | 0.888429637 | 949.1580291 | 934.6126482 | ENSG00000167106 |
| APBA2 | -0.022548087 | 0.984492358 | 0.78341683 | 0.889807352 | 838.6482997 | 825.6737864 | ENSG00000034053 |
| QKI | -0.022623054 | 0.984441202 | 0.761746906 | 0.876689032 | 2242.606249 | 2207.375148 | ENSG00000112531 |
| PLXND1 | -0.022629557 | 0.984436765 | 0.755755045 | 0.872977739 | 1463.587406 | 1440.249383 | ENSG00000004399 |
| PSENEN | -0.022717471 | 0.984376778 | 0.954050499 | 0.980903945 | 33.80993803 | 33.29472182 | ENSG00000205155 |
| ZNF418 | -0.022784363 | 0.984331137 | 0.888111457 | 0.949608474 | 219.128804 | 215.6624786 | ENSG00000196724 |
| SSTR2 | -0.022805205 | 0.984316917 | 0.885138449 | 0.947622636 | 192.3166824 | 189.3273304 | ENSG00000180616 |
| SNCAIP | -0.022837514 | 0.984294874 | 0.922065351 | 0.964771134 | 87.30937269 | 85.81384722 | ENSG00000064692 |
| EPN1 | -0.022850898 | 0.984285742 | 0.729746361 | 0.857642318 | 4429.676578 | 4360.952328 | ENSG00000063245 |
| AHI1 | -0.022854934 | 0.984282989 | 0.89873122 | 0.95544538 | 153.9168668 | 151.4125447 | ENSG00000135541 |
| SLC37A4 | -0.022930688 | 0.984231307 | 0.802423801 | 0.899050996 | 768.1279612 | 755.7115641 | ENSG00000137700 |
| HCK | -0.022985082 | 0.984194199 | 0.962214309 | 0.985060835 | 18.54099116 | 18.24041499 | ENSG00000101336 |
| LMBR1 | -0.022992517 | 0.984189127 | 0.800449545 | 0.898320559 | 989.6981092 | 973.4378426 | ENSG00000105983 |
| NA | -0.023031781 | 0.984162342 | 0.912965546 | 0.961597187 | 111.1379937 | 109.3379724 | ENSG00000288066 |
| NA | -0.023049614 | 0.984150177 | 0.904600494 | 0.958481852 | 236.9762906 | 233.5077551 | ENSG00000237973 |
| MRPL46 | -0.023059751 | 0.984143262 | 0.944555576 | 0.975763473 | 40.47382731 | 39.78762992 | ENSG00000259494 |
| ZRSR2 | -0.023069228 | 0.984136797 | 0.832049008 | 0.915381416 | 501.626872 | 494.0244766 | ENSG00000169249 |
| ZC3HAV1 | -0.023215793 | 0.984036822 | 0.793195692 | 0.894207674 | 979.6077217 | 963.6100215 | ENSG00000105939 |
| DOHH | -0.023223881 | 0.984031306 | 0.82009865 | 0.908627926 | 702.390904 | 691.7873124 | ENSG00000129932 |
| APEH | -0.023310005 | 0.983972564 | 0.629643975 | 0.795905386 | 5412.690103 | 5325.371692 | ENSG00000164062 |
| KAT5 | -0.023323227 | 0.983963546 | 0.886173754 | 0.948401451 | 211.5901047 | 208.1825129 | ENSG00000172977 |
| NA | -0.023411591 | 0.983903281 | 0.984143647 | 0.995944879 | 3.364564421 | 3.283153389 | ENSG00000203321 |
| MRPL22 | -0.023461848 | 0.983869007 | 0.783177781 | 0.889784424 | 845.6947018 | 831.8886545 | ENSG00000082515 |
| HSD17B11 | -0.023466853 | 0.983865593 | 0.910228549 | 0.960811396 | 130.0105478 | 128.0771404 | ENSG00000198189 |
| NOC3L | -0.023610606 | 0.983767564 | 0.817299842 | 0.906910366 | 544.0089426 | 535.3388816 | ENSG00000173145 |
| HDAC2-AS2 | -0.02368147 | 0.983719243 | 0.766792372 | 0.879325885 | 876.9443326 | 862.5716251 | ENSG00000228624 |
| PGAP6 | -0.023751264 | 0.983671655 | 0.652894575 | 0.81187292 | 3488.952652 | 3432.485343 | ENSG00000129925 |
| RECQL5 | -0.023777561 | 0.983653725 | 0.803687475 | 0.899842175 | 575.4720377 | 566.0056053 | ENSG00000108469 |
| HEBP2 | -0.023798313 | 0.983639576 | 0.784994136 | 0.890433415 | 839.5381851 | 825.8999417 | ENSG00000051620 |
| CHDH | -0.023945479 | 0.983539243 | 0.846560519 | 0.923736594 | 358.4727485 | 352.7994217 | ENSG00000016391 |
| TEX2 | -0.024080029 | 0.983447519 | 0.827022918 | 0.91297877 | 464.4141318 | 456.6921824 | ENSG00000136478 |
| AATF | -0.024093448 | 0.983438372 | 0.69797306 | 0.838980412 | 2063.697127 | 2029.905145 | ENSG00000275700 |
| UBA2 | -0.024154714 | 0.98339661 | 0.671745675 | 0.824135456 | 4040.014229 | 3972.222857 | ENSG00000126261 |
| NXNL2 | -0.024173896 | 0.983383534 | 0.963480898 | 0.985647688 | 21.05357692 | 20.67822209 | ENSG00000130045 |
| ISCA1 | -0.024257065 | 0.983326846 | 0.790429385 | 0.892671866 | 731.8576075 | 719.624736 | ENSG00000135070 |
| NUDT3 | -0.024263563 | 0.983322416 | 0.876735501 | 0.942675355 | 267.1535086 | 262.9797835 | ENSG00000272325 |
| AK3 | -0.024273358 | 0.98331574 | 0.842510132 | 0.92165416 | 526.2279948 | 517.29285 | ENSG00000147853 |
| FBXO8 | -0.024419758 | 0.983215962 | 0.905360364 | 0.958876702 | 113.2301083 | 111.2301551 | ENSG00000164117 |
| NA | -0.024429848 | 0.983209085 | 0.943081045 | 0.974880154 | 40.1381463 | 39.54825388 | ENSG00000227615 |
| FARP2 | -0.02445787 | 0.983189988 | 0.816151963 | 0.906477444 | 481.3771956 | 473.3218729 | ENSG00000006607 |
| RRBP1 | -0.024525112 | 0.983144164 | 0.655730181 | 0.814050821 | 5795.165685 | 5696.965505 | ENSG00000125844 |
| NA | -0.024526699 | 0.983143083 | 0.93564901 | 0.971184542 | 76.03544419 | 74.79181581 | ENSG00000244313 |
| ZNF367 | -0.024588343 | 0.983101075 | 0.892778018 | 0.952281345 | 182.8365355 | 179.7511601 | ENSG00000165244 |
| ATP6AP2 | -0.024640604 | 0.983065464 | 0.765133313 | 0.878472002 | 1249.321388 | 1228.233415 | ENSG00000182220 |
| CASC3 | -0.024656139 | 0.983054878 | 0.63160875 | 0.797106325 | 3723.246622 | 3659.917143 | ENSG00000108349 |
| TSPAN9 | -0.024673105 | 0.983043317 | 0.742666029 | 0.865969866 | 1240.505965 | 1219.264152 | ENSG00000011105 |
| DNAJC10 | -0.02473274 | 0.983002684 | 0.718819428 | 0.85151831 | 2225.299522 | 2187.584019 | ENSG00000077232 |
| SHISA5 | -0.024734274 | 0.983001638 | 0.643483107 | 0.805526878 | 3101.430698 | 3049.04235 | ENSG00000164054 |
| INTS10 | -0.024755943 | 0.982986874 | 0.746950403 | 0.86838024 | 1084.991673 | 1066.13566 | ENSG00000104613 |
| ZNF300 | -0.024769552 | 0.982977601 | 0.889927763 | 0.951012098 | 196.8920428 | 193.7418798 | ENSG00000145908 |
| CTSO | -0.024829098 | 0.982937031 | 0.957406414 | 0.98244088 | 20.58772228 | 20.19992151 | ENSG00000256043 |
| NA | -0.024846747 | 0.982925006 | 0.963613109 | 0.985663648 | 14.88017926 | 14.64136547 | ENSG00000258741 |
| RBL2 | -0.024929474 | 0.982868645 | 0.843485411 | 0.922013311 | 334.7808097 | 329.0161304 | ENSG00000103479 |
| MYOZ3 | -0.025081823 | 0.982764859 | 0.828720018 | 0.913997403 | 364.0562948 | 357.7599313 | ENSG00000164591 |
| NA | -0.025108533 | 0.982746665 | 0.908197808 | 0.960165884 | 123.7054496 | 121.7185963 | ENSG00000279207 |
| CHID1 | -0.025159191 | 0.982712158 | 0.694948108 | 0.837250559 | 1738.707535 | 1708.954542 | ENSG00000177830 |

| | | | | | | | |
|---|---|---|---|---|---|---|---|
| LOC554206 | -0.025180787 | 0.982697447 | 0.966326414 | 0.986931074 | 15.31282959 | 15.05525278 | ENSG00000262587 |
| ANO4 | -0.025500063 | 0.982479995 | 0.925360183 | 0.965920134 | 64.72444642 | 63.59066834 | ENSG00000151572 |
| KLHDC2 | -0.025547475 | 0.982447708 | 0.781389307 | 0.888626825 | 730.3185806 | 717.5781907 | ENSG00000165516 |
| HFM1 | -0.025550837 | 0.982445419 | 0.962661505 | 0.985098636 | 15.93972407 | 15.66216899 | ENSG00000162669 |
| LOC112267871 | -0.025559277 | 0.982439671 | 0.957794515 | 0.982666888 | 19.58629808 | 19.2114664 | ENSG00000228549 |
| CRISPLD1 | -0.025582045 | 0.982424167 | 0.827145277 | 0.913049207 | 431.4286125 | 423.9774409 | ENSG00000121005 |
| MAFG | -0.025678392 | 0.98235856 | 0.781702071 | 0.888776749 | 1026.303585 | 1008.090152 | ENSG00000197063 |
| ROBO2 | -0.025686991 | 0.982352705 | 0.914061581 | 0.96194221 | 92.9151917 | 91.06520898 | ENSG00000185008 |
| CLOCK | -0.02575576 | 0.98230588 | 0.8630155 | 0.933718002 | 298.6634616 | 293.1525565 | ENSG00000134852 |
| ZNF114 | -0.025787661 | 0.98228416 | 0.859695547 | 0.931886123 | 241.5313854 | 237.3675362 | ENSG00000178150 |
| RPL36 | -0.025923694 | 0.982191544 | 0.692038574 | 0.835483064 | 5719.376386 | 5617.75374 | ENSG00000130255 |
| N6AMT1 | -0.026054676 | 0.982102375 | 0.868966987 | 0.93800752 | 225.9448342 | 222.0950611 | ENSG00000156239 |
| NSMCE2 | -0.026075296 | 0.982088338 | 0.851364399 | 0.92641986 | 270.8193551 | 265.8449822 | ENSG00000156831 |
| CHAC2 | -0.026144742 | 0.982041065 | 0.873858806 | 0.940814295 | 200.6144825 | 197.0483533 | ENSG00000143942 |
| TRIM23 | -0.026252424 | 0.981967768 | 0.872984193 | 0.940412849 | 197.5568439 | 194.0496936 | ENSG00000113595 |
| EDNRA | -0.026295365 | 0.981938541 | 0.9075756 | 0.960149634 | 93.4525407 | 91.77418375 | ENSG00000151617 |
| NOC2L | -0.026700358 | 0.98166293 | 0.62605974 | 0.793253855 | 5434.115361 | 5334.621916 | ENSG00000188976 |
| AGPAT1 | -0.026754029 | 0.981626411 | 0.784925014 | 0.890433415 | 555.0794754 | 544.9153878 | ENSG00000204310 |
| DHX9 | -0.026789306 | 0.981602409 | 0.613739112 | 0.785102156 | 7582.785473 | 7442.909415 | ENSG00000135829 |
| PPP3CB | -0.026812073 | 0.981586918 | 0.79470069 | 0.894980174 | 577.2118691 | 566.3712106 | ENSG00000107758 |
| ZNF738 | -0.02681905 | 0.981582171 | 0.764221781 | 0.877932255 | 921.4971049 | 904.7274379 | ENSG00000172687 |
| BFAR | -0.027014946 | 0.981448896 | 0.687915033 | 0.83320004 | 1847.794883 | 1812.933987 | ENSG00000103429 |
| NA | -0.027018634 | 0.981446387 | 0.981039609 | 0.995108659 | 3.746213087 | 3.644167164 | ENSG00000253476 |
| TIAM1 | -0.027049952 | 0.981425082 | 0.81621321 | 0.906477444 | 604.6724696 | 592.8813941 | ENSG00000156299 |
| NA | -0.027209759 | 0.981316376 | 0.958903026 | 0.983056492 | 17.27535474 | 16.93606375 | ENSG00000272078 |
| PTGER4 | -0.027377787 | 0.981202091 | 0.967839467 | 0.987263577 | 10.52130959 | 10.36446008 | ENSG00000171522 |
| TRIQK | -0.027411724 | 0.98117901 | 0.876548966 | 0.942528453 | 189.0671379 | 185.3619273 | ENSG00000205133 |
| RARRES2 | -0.027425899 | 0.981169369 | 0.653422804 | 0.812465428 | 4640.493185 | 4553.426911 | ENSG00000106538 |
| DYNLRB1 | -0.027485169 | 0.981129061 | 0.966399277 | 0.986937613 | 11.51325057 | 11.32803914 | ENSG00000125971 |
| PRKACA | -0.027521546 | 0.981104322 | 0.781574659 | 0.88874409 | 1232.595005 | 1209.172965 | ENSG00000072062 |
| MPV17L2 | -0.027566231 | 0.981073935 | 0.762331526 | 0.876987748 | 854.2849361 | 838.3233922 | ENSG00000254858 |
| TMEM107 | -0.027600199 | 0.981050836 | 0.923241426 | 0.965255735 | 62.59259345 | 61.38932677 | ENSG00000179029 |
| MRGPRF | -0.027634337 | 0.981027622 | 0.892135377 | 0.951916865 | 112.5385846 | 110.481076 | ENSG00000172935 |
| UTRN | -0.027642089 | 0.981022351 | 0.767432698 | 0.879694476 | 897.2425697 | 879.5485805 | ENSG00000152818 |
| MAP3K12 | -0.02764759 | 0.98101861 | 0.834671533 | 0.917141538 | 321.8204157 | 315.7898039 | ENSG00000139625 |
| RRAGC | -0.02771804 | 0.980970706 | 0.894231936 | 0.953020709 | 121.7951475 | 119.5851457 | ENSG00000116954 |
| ERLIN1 | -0.027797129 | 0.98091693 | 0.723140658 | 0.854154341 | 995.8523123 | 976.924421 | ENSG00000107566 |
| GJA3 | -0.027807228 | 0.980910064 | 0.959616386 | 0.983627629 | 14.84577974 | 14.56389942 | ENSG00000121743 |
| SNAI3-AS1 | -0.027832537 | 0.980892856 | 0.921353821 | 0.964771134 | 65.82815329 | 64.525136 | ENSG00000260630 |
| RCC1L | -0.027844876 | 0.980884467 | 0.702078793 | 0.841612162 | 2288.148823 | 2244.862724 | ENSG00000274523 |
| DDX41 | -0.027897697 | 0.980848555 | 0.653789251 | 0.81254148 | 1944.090096 | 1907.037005 | ENSG00000183258 |
| COX6B1 | -0.02802799 | 0.980759976 | 0.677859645 | 0.827602494 | 2712.20834 | 2659.622573 | ENSG00000126267 |
| MRPL27 | -0.028075052 | 0.980727983 | 0.748279577 | 0.86918808 | 861.2414878 | 844.6652879 | ENSG00000108826 |
| TFEB | -0.028106132 | 0.980706855 | 0.916435849 | 0.962927628 | 70.29647631 | 69.10053381 | ENSG00000112561 |
| RAD51AP1 | -0.028144888 | 0.980680511 | 0.761293582 | 0.876553329 | 713.0959814 | 699.2314591 | ENSG00000111247 |
| GOLIM4 | -0.028174713 | 0.980660237 | 0.689646307 | 0.83398915 | 1259.426598 | 1235.193454 | ENSG00000173905 |
| PRKAA2 | -0.02818015 | 0.980656541 | 0.861384635 | 0.932648138 | 228.6756645 | 224.1119171 | ENSG00000162409 |
| LOC101929243 | -0.028192898 | 0.980647876 | 0.960854476 | 0.984255629 | 15.66483343 | 15.35592067 | ENSG00000272288 |
| MSRB1 | -0.028219098 | 0.980630067 | 0.720050631 | 0.852442087 | 959.4562609 | 941.1125771 | ENSG00000198736 |
| CDH1 | -0.028247336 | 0.980610874 | 0.639444539 | 0.802386055 | 4112.859294 | 4032.807096 | ENSG00000039068 |
| FAM199X | -0.028271037 | 0.980594764 | 0.74991732 | 0.869935742 | 1206.37789 | 1182.834353 | ENSG00000123575 |
| YWHAH | -0.028323576 | 0.980559054 | 0.630279681 | 0.796470479 | 3094.456732 | 3035.088785 | ENSG00000128245 |
| RSPH3 | -0.028406097 | 0.980502969 | 0.92824733 | 0.967383192 | 45.36943559 | 44.4740717 | ENSG00000130363 |
| DNAJC14 | -0.028463153 | 0.980464192 | 0.813433808 | 0.90530915 | 338.6325594 | 331.8928688 | ENSG00000135392 |
| NA | -0.028465422 | 0.98046265 | 0.955833197 | 0.982095209 | 16.92691038 | 16.57410401 | ENSG00000275457 |
| SSBP3 | -0.028585387 | 0.980381125 | 0.623000942 | 0.791321818 | 3401.499967 | 3335.244908 | ENSG00000157216 |
| SH3GL1 | -0.028652675 | 0.9803354 | 0.635267076 | 0.799988093 | 3727.292813 | 3654.469917 | ENSG00000141985 |
| IL11 | -0.028728027 | 0.980284198 | 0.944022815 | 0.975469288 | 28.54928324 | 27.99360794 | ENSG00000095752 |
| ZNF324B | -0.028753073 | 0.98026718 | 0.923795779 | 0.965407741 | 69.98688124 | 68.44024961 | ENSG00000249471 |
| ACER3 | -0.028842694 | 0.980206287 | 0.847585321 | 0.924422644 | 259.1154454 | 254.1113787 | ENSG00000078124 |
| NA | -0.028863649 | 0.98019205 | 0.856875373 | 0.930171411 | 266.1968319 | 260.9034322 | ENSG00000282048 |
| ESF1 | -0.028910671 | 0.980160103 | 0.755449138 | 0.872845334 | 828.0158347 | 811.7547093 | ENSG00000089048 |
| MAPKAP1 | -0.028942456 | 0.980138509 | 0.638473266 | 0.80191325 | 2197.45882 | 2153.20793 | ENSG00000119487 |
| ARMCX4 | -0.02909557 | 0.980034492 | 0.804695303 | 0.900543454 | 400.4158879 | 392.1255447 | ENSG00000196440 |
| DCUN1D2 | -0.029130382 | 0.980010844 | 0.787732299 | 0.891582187 | 445.3135451 | 436.1964384 | ENSG00000150401 |
| CPTP | -0.029138288 | 0.980005474 | 0.844324747 | 0.92248035 | 328.8057731 | 322.6394475 | ENSG00000224051 |
| MIEF2 | -0.029184514 | 0.979974073 | 0.842543681 | 0.92165416 | 235.0127979 | 230.1480387 | ENSG00000177427 |
| PHAX | -0.029201688 | 0.979962407 | 0.75106788 | 0.870392545 | 1125.357521 | 1102.297175 | ENSG00000164902 |
| MAP1LC3A | -0.029203128 | 0.97996143 | 0.902386753 | 0.957425906 | 80.82281991 | 79.25936372 | ENSG00000101460 |
| ENC1 | -0.029280421 | 0.979908929 | 0.783798419 | 0.890064386 | 568.0519968 | 556.3292159 | ENSG00000171617 |
| ABCE1 | -0.029283347 | 0.979906942 | 0.667624343 | 0.821869982 | 2012.36692 | 1972.055024 | ENSG00000164163 |
| GALNT11 | -0.029394344 | 0.979831553 | 0.778380971 | 0.886943686 | 560.2365483 | 548.8305443 | ENSG00000178234 |
| OSCP1 | -0.029435447 | 0.979803637 | 0.952202661 | 0.979537382 | 21.96695634 | 21.57833384 | ENSG00000116885 |
| CD320 | -0.029463222 | 0.979784774 | 0.677148908 | 0.827271129 | 1561.356816 | 1530.142305 | ENSG00000167775 |

| | | | | | | | |
|---|---|---|---|---|---|---|---|
| KLF15 | -0.02948935 | 0.97976703 | 0.831824002 | 0.915381416 | 382.1676882 | 374.8409958 | ENSG00000163884 |
| NA | -0.029517652 | 0.97974781 | 0.938687582 | 0.972431174 | 37.12378011 | 36.30492125 | ENSG00000257135 |
| LOC100506405 | -0.029518692 | 0.979747103 | 0.938906166 | 0.972506023 | 31.18019516 | 30.52238537 | ENSG00000228496 |
| FAM174A | -0.029591609 | 0.979697586 | 0.904132694 | 0.958242075 | 82.44282442 | 80.79561865 | ENSG00000174132 |
| NOP56 | -0.02967187 | 0.979643085 | 0.626807675 | 0.793550956 | 5072.991018 | 4969.820758 | ENSG00000101361 |
| PRKACB | -0.029769911 | 0.979576513 | 0.796201544 | 0.895922431 | 517.9645937 | 507.5076351 | ENSG00000142875 |
| RUVBL2 | -0.029799173 | 0.979556645 | 0.632359772 | 0.797472043 | 4095.232915 | 4011.06322 | ENSG00000183207 |
| HNF1B | -0.029804858 | 0.979552785 | 0.980107681 | 0.994714902 | 3.422685036 | 3.320798682 | ENSG00000275410 |
| ZFR | -0.029810217 | 0.979549146 | 0.663198326 | 0.819398128 | 2163.144586 | 2119.131263 | ENSG00000056097 |
| ZC3H12B | -0.029858478 | 0.979516379 | 0.92512762 | 0.965883673 | 44.79473799 | 43.85315675 | ENSG00000102053 |
| PMS2P3 | -0.029889154 | 0.979495552 | 0.898656051 | 0.955419666 | 97.16346455 | 95.18694168 | ENSG00000127957 |
| FAM234B | -0.029956584 | 0.979449773 | 0.730124852 | 0.857920184 | 718.2122508 | 703.3506172 | ENSG00000084444 |
| NEK6 | -0.029975309 | 0.97943706 | 0.792470865 | 0.893961668 | 420.9282764 | 412.2875803 | ENSG00000119408 |
| PAPOLA | -0.029984362 | 0.979430914 | 0.601032414 | 0.776307564 | 4436.848197 | 4345.038974 | ENSG00000090060 |
| LMNTD2 | -0.029990098 | 0.97942702 | 0.90884028 | 0.960165884 | 83.20139323 | 81.60714644 | ENSG00000185522 |
| TNIP2 | -0.030004962 | 0.979416929 | 0.712658051 | 0.848263411 | 1428.618718 | 1399.698013 | ENSG00000168884 |
| C6orf226 | -0.030023247 | 0.979404516 | 0.937184255 | 0.971870005 | 33.14361002 | 32.5465887 | ENSG00000221821 |
| SF3A2 | -0.030078378 | 0.97936709 | 0.670127798 | 0.823185784 | 2040.527235 | 1998.837753 | ENSG00000104897 |
| MAK | -0.030114041 | 0.97934288 | 0.968976581 | 0.987966889 | 8.42713627 | 8.302611457 | ENSG00000111837 |
| SLC7A6 | -0.03013514 | 0.979328558 | 0.809235772 | 0.902949338 | 343.4314777 | 336.046907 | ENSG00000103064 |
| NUMB | -0.030146711 | 0.979320703 | 0.754024831 | 0.871839768 | 674.5684604 | 660.8027577 | ENSG00000133961 |
| CDKL1 | -0.03020973 | 0.979277926 | 0.949911047 | 0.978245726 | 22.30144214 | 21.76729535 | ENSG00000100490 |
| TXLNG | -0.030219817 | 0.979271079 | 0.630869064 | 0.796625039 | 2699.886443 | 2643.292446 | ENSG00000086712 |
| SLC66A3 | -0.030241705 | 0.979256222 | 0.844743654 | 0.922724502 | 200.6231021 | 196.4153315 | ENSG00000162976 |
| SLC25A5 | -0.030251618 | 0.979249493 | 0.53981237 | 0.729314394 | 6474.997028 | 6340.195702 | ENSG00000005022 |
| CDC16 | -0.03033544 | 0.9791926 | 0.666611983 | 0.821412456 | 1441.871164 | 1412.167822 | ENSG00000130177 |
| PEX10 | -0.030365737 | 0.979172037 | 0.792290197 | 0.893932102 | 463.5971108 | 454.2020392 | ENSG00000157911 |
| NUDT16 | -0.030405272 | 0.979145204 | 0.720791457 | 0.852731108 | 1028.650175 | 1006.910306 | ENSG00000198585 |
| MUL1 | -0.03041893 | 0.979135934 | 0.728648592 | 0.857148832 | 742.1618678 | 726.5378267 | ENSG00000090432 |
| TPCN2 | -0.03044494 | 0.979118283 | 0.830805676 | 0.915015174 | 287.2166197 | 281.2566572 | ENSG00000162341 |
| STK35 | -0.030487108 | 0.979089664 | 0.67871116 | 0.828161805 | 1414.987295 | 1385.707021 | ENSG00000125834 |
| COPS7A | -0.030626437 | 0.978995113 | 0.706763284 | 0.845081969 | 840.4085982 | 822.6782481 | ENSG00000111652 |
| PEX13 | -0.030657192 | 0.978974243 | 0.813993365 | 0.905404967 | 468.846425 | 458.7114677 | ENSG00000162928 |
| AAAS | -0.030675569 | 0.978961773 | 0.666742719 | 0.821519949 | 1387.764283 | 1358.295918 | ENSG00000094914 |
| SH3GL3 | -0.030721771 | 0.978930422 | 0.835919034 | 0.917543777 | 238.665575 | 233.4852225 | ENSG00000140600 |
| CCDC65 | -0.030873638 | 0.978827379 | 0.943833688 | 0.97533785 | 25.46792361 | 24.94888463 | ENSG00000139537 |
| ZC2HC1C | -0.030906056 | 0.978805385 | 0.940117625 | 0.97320185 | 31.62324458 | 31.03398032 | ENSG00000119703 |
| TSC22D2 | -0.031078841 | 0.978688165 | 0.739591527 | 0.863664729 | 659.9084805 | 645.8120338 | ENSG00000196428 |
| TMEM106C | -0.03110282 | 0.978671899 | 0.789919902 | 0.892363303 | 416.2951009 | 407.3659455 | ENSG00000134291 |
| NA | -0.031120826 | 0.978659684 | 0.95801759 | 0.982680048 | 13.97127519 | 13.67031407 | ENSG00000285128 |
| CHMP5 | -0.03115264 | 0.978638103 | 0.713550453 | 0.848956862 | 806.691739 | 789.1323996 | ENSG00000086065 |
| TRMT44 | -0.03128265 | 0.978549916 | 0.800744219 | 0.898320559 | 321.4942179 | 314.642422 | ENSG00000155275 |
| GOLPH3 | -0.031384232 | 0.978481018 | 0.647680924 | 0.808876289 | 1974.716671 | 1932.174963 | ENSG00000113384 |
| METAP1 | -0.031396714 | 0.978472552 | 0.643804955 | 0.805803452 | 2316.696426 | 2266.371465 | ENSG00000164024 |
| NFYA | -0.031404578 | 0.978467218 | 0.719146267 | 0.851852051 | 822.1193608 | 804.5384715 | ENSG00000001167 |
| WTAP | -0.031422086 | 0.978455344 | 0.658836074 | 0.816144261 | 1405.387522 | 1375.524347 | ENSG00000146457 |
| MEGF6 | -0.031497035 | 0.978404514 | 0.910404131 | 0.960811396 | 976.2640595 | 955.2839416 | ENSG00000162591 |
| GOLGA2 | -0.031551509 | 0.978367571 | 0.695305779 | 0.837266443 | 1177.702526 | 1152.348159 | ENSG00000167110 |
| PREB | -0.031560805 | 0.978361268 | 0.626278886 | 0.793253855 | 1593.442837 | 1558.774347 | ENSG00000138073 |
| CEBPZ | -0.031591429 | 0.9783405 | 0.653561355 | 0.812465428 | 2557.237246 | 2502.094073 | ENSG00000115816 |
| ELP5 | -0.031628509 | 0.978315356 | 0.703920408 | 0.842524014 | 813.0084308 | 795.4442177 | ENSG00000170291 |
| ZNF557 | -0.031646625 | 0.978303071 | 0.89050545 | 0.95121103 | 96.65250581 | 94.53157549 | ENSG00000130544 |
| HSP90AA1 | -0.031692642 | 0.978271867 | 0.494753515 | 0.694419928 | 42794.40016 | 41864.31143 | ENSG00000080824 |
| ARMT1 | -0.031752343 | 0.978231385 | 0.729105105 | 0.85741017 | 1000.986099 | 979.6064793 | ENSG00000146476 |
| LOC652276 | -0.031787878 | 0.97820729 | 0.908284959 | 0.960165884 | 78.98040095 | 77.27084112 | ENSG00000215154 |
| SLC35F6 | -0.031801154 | 0.978198289 | 0.857483117 | 0.93039751 | 184.0265856 | 180.1494833 | ENSG00000213699 |
| FST | -0.031905219 | 0.978127731 | 0.857627543 | 0.930409931 | 151.1400427 | 147.7910781 | ENSG00000134363 |
| SHROOM4 | -0.031908866 | 0.978125259 | 0.854154902 | 0.928574107 | 164.3883438 | 160.8820219 | ENSG00000158352 |
| R3HDM1 | -0.032047119 | 0.97803153 | 0.696695042 | 0.838084329 | 1195.518666 | 1169.542239 | ENSG00000048991 |
| VPS41 | -0.032080459 | 0.978008928 | 0.72059875 | 0.852637136 | 888.6007733 | 868.9108079 | ENSG00000006715 |
| IFNAR1 | -0.032083991 | 0.978006534 | 0.832028427 | 0.915381416 | 226.619197 | 221.751619 | ENSG00000142166 |
| G6PD | -0.032085231 | 0.978005694 | 0.667004546 | 0.82154232 | 1344.944936 | 1315.954634 | ENSG00000160211 |
| SEPTIN1 | -0.032107651 | 0.977990495 | 0.957866042 | 0.982680048 | 12.5917618 | 12.32179114 | ENSG00000180096 |
| ATG16L1 | -0.032215348 | 0.977917491 | 0.758127684 | 0.874316737 | 837.3965974 | 818.4847157 | ENSG00000085978 |
| BARD1 | -0.032226372 | 0.977910019 | 0.792324744 | 0.893932102 | 341.6593579 | 333.8907557 | ENSG00000138376 |
| RARG | -0.032258777 | 0.977888054 | 0.667863321 | 0.821935407 | 1235.137792 | 1208.371273 | ENSG00000172819 |
| IST1 | -0.032298236 | 0.977861307 | 0.565718174 | 0.749830693 | 3381.706072 | 3306.159765 | ENSG00000182149 |
| CPSF3 | -0.032301412 | 0.977859155 | 0.608987258 | 0.782305701 | 1868.733238 | 1827.408282 | ENSG00000119203 |
| NAA25 | -0.03237563 | 0.977808851 | 0.698796993 | 0.839543305 | 1018.090678 | 995.7185522 | ENSG00000111300 |
| FLOT2 | -0.032429942 | 0.977772041 | 0.559770302 | 0.745137248 | 4384.883994 | 4286.958849 | ENSG00000132589 |
| TASOR2 | -0.032545056 | 0.977694027 | 0.671239409 | 0.82390724 | 1549.82447 | 1515.010486 | ENSG00000108021 |
| KAT6A | -0.032625007 | 0.977639847 | 0.78317243 | 0.889784424 | 628.7441992 | 614.2507358 | ENSG00000083168 |
| CMIP | -0.032663726 | 0.97761361 | 0.625035923 | 0.792680372 | 1605.065463 | 1568.841712 | ENSG00000153815 |
| VPS35 | -0.032670612 | 0.977608943 | 0.64847475 | 0.809147384 | 1591.186176 | 1555.244447 | ENSG00000069329 |

| | | | | | | | |
|---|---|---|---|---|---|---|---|
| SLC25A33 | -0.032690567 | 0.977595421 | 0.715192197 | 0.849673165 | 716.7903548 | 700.9614483 | ENSG00000171612 |
| EXOC2 | -0.032698749 | 0.977589877 | 0.663450482 | 0.819600901 | 1262.941472 | 1234.34965 | ENSG00000112685 |
| ARFGEF2 | -0.032721579 | 0.977574407 | 0.732782968 | 0.859594158 | 829.2769446 | 810.1281558 | ENSG00000124198 |
| IDE | -0.032754235 | 0.977552279 | 0.721442444 | 0.853073731 | 1062.560972 | 1038.441309 | ENSG00000119912 |
| NA | -0.03277309 | 0.977539504 | 0.95731869 | 0.98244088 | 13.21273254 | 12.9147086 | ENSG00000267248 |
| INTS1 | -0.032784441 | 0.977531813 | 0.534796247 | 0.725704328 | 5847.071179 | 5716.215115 | ENSG00000164880 |
| PEX14 | -0.032785819 | 0.977530879 | 0.696572503 | 0.838043685 | 824.3938197 | 805.8468699 | ENSG00000142655 |
| LMBRD2 | -0.032790024 | 0.977528029 | 0.889257251 | 0.950456913 | 83.47775841 | 81.54564368 | ENSG00000164187 |
| UTP4 | -0.032912403 | 0.977445113 | 0.55093333 | 0.738132515 | 2744.287937 | 2682.124886 | ENSG00000141076 |
| CAPN10 | -0.033009253 | 0.977379498 | 0.758441859 | 0.874484066 | 1007.163082 | 984.891729 | ENSG00000142330 |
| ISOC2 | -0.033037394 | 0.977360434 | 0.617028836 | 0.787224317 | 1814.337836 | 1773.537208 | ENSG00000063241 |
| NACC2 | -0.033064707 | 0.97734193 | 0.742946526 | 0.866136499 | 645.3047181 | 630.7819203 | ENSG00000148411 |
| DEF8 | -0.033133018 | 0.977295655 | 0.638741202 | 0.802036409 | 1421.270068 | 1389.328152 | ENSG00000140995 |
| PHF12 | -0.033201878 | 0.97724901 | 0.633784684 | 0.798733651 | 1302.174326 | 1272.558831 | ENSG00000109118 |
| ZNF189 | -0.033234684 | 0.977226788 | 0.813760588 | 0.905396008 | 267.9482314 | 261.6402459 | ENSG00000136870 |
| IER3IP1 | -0.033243425 | 0.977220867 | 0.894457189 | 0.953032171 | 95.88153087 | 93.8505982 | ENSG00000134049 |
| SLC25A17 | -0.033276063 | 0.977198759 | 0.706112711 | 0.844411002 | 698.9844645 | 683.1258197 | ENSG00000100372 |
| TCERG1 | -0.033316251 | 0.977171538 | 0.595160137 | 0.772037516 | 2670.549042 | 2609.451328 | ENSG00000113649 |
| GPR108 | -0.033338413 | 0.977156528 | 0.720899003 | 0.852804916 | 667.6889184 | 652.4458026 | ENSG00000125734 |
| FAT1 | -0.033379793 | 0.977128501 | 0.709604892 | 0.846496645 | 2519.290316 | 2460.953852 | ENSG00000083857 |
| NA | -0.033387137 | 0.977123527 | 0.370721317 | 0.582936892 | 2277604.932 | 2225501.429 | N_ambiguous |
| NA | -0.03351495 | 0.977036964 | 0.719389947 | 0.852033813 | 887.0606046 | 866.4565361 | ENSG00000237550 |
| S100A1 | -0.03356962 | 0.976999941 | 0.955040991 | 0.981601672 | 13.29101478 | 12.98035138 | ENSG00000160678 |
| CPAMD8 | -0.033577352 | 0.976994705 | 0.828007582 | 0.913585136 | 228.6697669 | 223.4211467 | ENSG00000160111 |
| PPP1R16B | -0.03362259 | 0.97696407 | 0.738306595 | 0.862911254 | 663.0984566 | 647.4839775 | ENSG00000101445 |
| ZNF30 | -0.033639439 | 0.97695266 | 0.865045786 | 0.935004026 | 122.7630363 | 119.8505546 | ENSG00000168661 |
| ACTN4 | -0.033642712 | 0.976950444 | 0.531763465 | 0.723460345 | 9264.437317 | 9050.430585 | ENSG00000130402 |
| MEX3C | -0.033662521 | 0.97693703 | 0.59564265 | 0.772289098 | 1828.395228 | 1786.422341 | ENSG00000176624 |
| MMADHC | -0.033682656 | 0.976923395 | 0.649296031 | 0.809530306 | 1599.945284 | 1563.299591 | ENSG00000168288 |
| BICD2 | -0.033698137 | 0.976912912 | 0.692806208 | 0.835855442 | 1019.553254 | 996.1602261 | ENSG00000185963 |
| APOL2 | -0.033807997 | 0.976838524 | 0.765903816 | 0.878910348 | 402.9815864 | 393.4412677 | ENSG00000128335 |
| ARHGEF26 | -0.033834435 | 0.976820624 | 0.772727733 | 0.883060194 | 360.1241518 | 351.7289387 | ENSG00000114790 |
| NA | -0.03388438 | 0.976786807 | 0.939550375 | 0.972905887 | 32.17124642 | 31.37692797 | ENSG00000244625 |
| NOP58 | -0.0338958 | 0.976779075 | 0.562726325 | 0.747649264 | 2286.610436 | 2233.481214 | ENSG00000055044 |
| CWF19L1 | -0.033901347 | 0.976775319 | 0.760375672 | 0.875804743 | 424.8764636 | 414.8284646 | ENSG00000095485 |
| PRDM2 | -0.033968479 | 0.976729869 | 0.742531346 | 0.865919752 | 532.3171871 | 519.7099343 | ENSG00000116731 |
| ABHD2 | -0.034034079 | 0.976685458 | 0.711241562 | 0.847591497 | 654.0821049 | 638.8540458 | ENSG00000140526 |
| SP110 | -0.03412381 | 0.976624713 | 0.890310561 | 0.951205869 | 85.45924997 | 83.48880804 | ENSG00000135899 |
| SECISBP2L | -0.034181457 | 0.97658569 | 0.793139086 | 0.894207674 | 455.9463151 | 444.9614895 | ENSG00000138593 |
| PCNX1 | -0.034325029 | 0.976488508 | 0.653894524 | 0.81254148 | 1034.300207 | 1009.83238 | ENSG00000100731 |
| HEATR6 | -0.034382462 | 0.976449635 | 0.730154524 | 0.857920184 | 599.8356702 | 585.5154485 | ENSG00000068097 |
| KBTBD6 | -0.034447166 | 0.976405843 | 0.792596566 | 0.894035655 | 369.0345828 | 359.9969107 | ENSG00000165572 |
| C5orf22 | -0.034449437 | 0.976404306 | 0.73176493 | 0.858773347 | 590.3359459 | 576.4776614 | ENSG00000082213 |
| AP2A1 | -0.034457304 | 0.976398982 | 0.546097588 | 0.734181885 | 3585.972193 | 3501.737436 | ENSG00000196961 |
| PTPN5 | -0.034462015 | 0.976395794 | 0.804883443 | 0.90064726 | 264.7055526 | 258.3132044 | ENSG00000110786 |
| TDRD3 | -0.034465302 | 0.976393569 | 0.8055797 | 0.900892559 | 251.9726333 | 246.003731 | ENSG00000083544 |
| NDUFAB1 | -0.034488571 | 0.976377821 | 0.604444858 | 0.778857884 | 1513.216351 | 1477.276491 | ENSG00000004779 |
| RPL9 | -0.034605555 | 0.976298652 | 0.548708261 | 0.736171889 | 9997.891537 | 9760.158834 | ENSG00000163682 |
| DLEU2 | -0.034622852 | 0.976286948 | 0.959794455 | 0.983649986 | 10.12305877 | 9.90763059 | ENSG00000231607 |
| SFPQ | -0.034661241 | 0.97626097 | 0.482430165 | 0.683726181 | 10619.37442 | 10367.17391 | ENSG00000116560 |
| TOM1L1 | -0.034829585 | 0.976147059 | 0.708038669 | 0.845910689 | 718.5205995 | 700.9978641 | ENSG00000141198 |
| TTC21B | -0.03487639 | 0.976115391 | 0.771804323 | 0.882509363 | 356.3580825 | 347.7468372 | ENSG00000123607 |
| ATP23 | -0.034894458 | 0.976103166 | 0.857028464 | 0.930171411 | 123.0566141 | 120.1203872 | ENSG00000166896 |
| NOL9 | -0.035097934 | 0.975965507 | 0.726074042 | 0.855720453 | 565.2766204 | 551.6908453 | ENSG00000162408 |
| PUM1 | -0.035192521 | 0.975901523 | 0.537299145 | 0.727443977 | 2836.280292 | 2767.55817 | ENSG00000134644 |
| NUP85 | -0.035192789 | 0.975901341 | 0.646490211 | 0.807960683 | 1432.607964 | 1397.905507 | ENSG00000125450 |
| EVA1A | -0.035231092 | 0.975875432 | 0.918140518 | 0.963710742 | 43.78708458 | 42.7807092 | ENSG00000115363 |
| CTU1 | -0.035307304 | 0.975823881 | 0.786942966 | 0.891346363 | 337.0988452 | 329.1992279 | ENSG00000142544 |
| FAM3A | -0.03542944 | 0.975741273 | 0.749829966 | 0.869935742 | 416.3440694 | 406.2225153 | ENSG00000071889 |
| GSTM1 | -0.035448213 | 0.975728577 | 0.891814696 | 0.95189739 | 79.42461943 | 77.55306403 | ENSG00000134184 |
| DAXX | -0.03550068 | 0.975693093 | 0.597423464 | 0.773610951 | 2330.061131 | 2274.002432 | ENSG00000204209 |
| WARS2 | -0.035505112 | 0.975690095 | 0.857974902 | 0.930556858 | 120.4968654 | 117.5810159 | ENSG00000116874 |
| ZNF160 | -0.035576354 | 0.975641916 | 0.766036615 | 0.878994404 | 397.5267136 | 387.6629148 | ENSG00000170949 |
| GPATCH8 | -0.035601807 | 0.975624703 | 0.615262341 | 0.786092234 | 1580.880189 | 1541.842634 | ENSG00000186566 |
| POGLUT3 | -0.035618128 | 0.975613666 | 0.783607838 | 0.889920261 | 324.4845837 | 316.8229549 | ENSG00000178202 |
| LINC00294 | -0.0358415 | 0.975462624 | 0.867948245 | 0.937364783 | 97.97984848 | 95.55833831 | ENSG00000280798 |
| ITPRIP | -0.035955396 | 0.975385617 | 0.697765665 | 0.838858153 | 646.9757249 | 630.8545612 | ENSG00000148841 |
| NEK8 | -0.035963984 | 0.975379811 | 0.786032005 | 0.890741939 | 284.1192683 | 277.3000715 | ENSG00000160602 |
| SNU13 | -0.035998299 | 0.975356612 | 0.525940362 | 0.719293069 | 2753.973947 | 2686.648442 | ENSG00000100138 |
| NDUFA11 | -0.036014283 | 0.975345806 | 0.88711716 | 0.949082686 | 88.96675138 | 86.94183823 | ENSG00000174886 |
| NA | -0.036044213 | 0.975325571 | 0.956483884 | 0.982211615 | 10.49402901 | 10.25247016 | ENSG00000262583 |
| MANEA | -0.036168792 | 0.975241354 | 0.887522375 | 0.94935486 | 92.90629273 | 90.52873797 | ENSG00000172469 |
| LRP8 | -0.036190398 | 0.975226749 | 0.69154019 | 0.835021515 | 1303.068663 | 1270.506907 | ENSG00000157193 |
| ESS2 | -0.036211661 | 0.975212375 | 0.732725295 | 0.859594158 | 545.5456571 | 532.4429858 | ENSG00000100056 |

| | | | | | | | |
|---|---|---|---|---|---|---|---|
| DSTN | -0.036230653 | 0.975199538 | 0.569295982 | 0.752342899 | 2487.869381 | 2426.734061 | ENSG00000125868 |
| FNBP1L | -0.036334199 | 0.975129548 | 0.587526941 | 0.76572348 | 2906.807419 | 2833.847268 | ENSG00000137942 |
| OSBPL1A | -0.036369242 | 0.975105862 | 0.648448477 | 0.809147384 | 1181.979327 | 1152.385673 | ENSG00000141447 |
| EIF4A2 | -0.036378788 | 0.97509941 | 0.503517577 | 0.701351929 | 6884.507551 | 6712.036812 | ENSG00000156976 |
| SOCS4 | -0.036396713 | 0.975087295 | 0.744784658 | 0.867262195 | 417.1180451 | 406.7209621 | ENSG00000180008 |
| GRPEL1 | -0.036414758 | 0.975075098 | 0.626366892 | 0.793253855 | 1075.687998 | 1049.032884 | ENSG00000109519 |
| PTOV1 | -0.036441631 | 0.975056936 | 0.720223067 | 0.852539339 | 990.2780745 | 966.0340446 | ENSG00000104960 |
| PKIA | -0.036474298 | 0.975034858 | 0.774910994 | 0.884331605 | 350.3767656 | 341.5754057 | ENSG00000171033 |
| MRPS10 | -0.036505456 | 0.9750138 | 0.686222451 | 0.83186293 | 749.9837196 | 731.0297991 | ENSG00000048544 |
| TRNP1 | -0.036554305 | 0.974980787 | 0.677728256 | 0.82756234 | 984.6452542 | 960.2287746 | ENSG00000253368 |
| CDK14 | -0.036558326 | 0.97497807 | 0.704223285 | 0.842791935 | 805.1983231 | 784.4524412 | ENSG00000058091 |
| RXRA | -0.036566857 | 0.974972305 | 0.612811172 | 0.784606033 | 1617.801332 | 1576.820724 | ENSG00000186350 |
| ST13 | -0.036569616 | 0.97497044 | 0.417553941 | 0.628430894 | 7310.919179 | 7127.863257 | ENSG00000100380 |
| ADGRG6 | -0.03658983 | 0.974956779 | 0.873847589 | 0.940814295 | 90.15469038 | 87.81412045 | ENSG00000112414 |
| OGDH | -0.036610217 | 0.974943002 | 0.484250585 | 0.685332156 | 3884.290257 | 3786.812114 | ENSG00000105953 |
| SALL3 | -0.036667226 | 0.974904478 | 0.683776151 | 0.830895404 | 808.5624061 | 788.3230473 | ENSG00000256463 |
| NOP14 | -0.036863601 | 0.974771786 | 0.661904461 | 0.818441256 | 1142.423022 | 1113.861818 | ENSG00000087269 |
| PARG | -0.036875029 | 0.974764064 | 0.635339562 | 0.799999175 | 1227.210781 | 1195.90818 | ENSG00000227345 |
| FOXK1 | -0.036981327 | 0.974692246 | 0.783349528 | 0.889784424 | 385.7791488 | 375.6150759 | ENSG00000164916 |
| CSNK2B | -0.037025185 | 0.974662616 | 0.915926006 | 0.962597668 | 48.51695065 | 47.37495113 | ENSG00000204435 |
| GADD45GIP1 | -0.037048796 | 0.974646665 | 0.586541876 | 0.765028202 | 1981.574777 | 1931.730253 | ENSG00000179271 |
| FAM169A | -0.037062363 | 0.9746375 | 0.733874246 | 0.860179688 | 580.5820262 | 565.4769628 | ENSG00000198780 |
| C20orf96 | -0.037134941 | 0.974588469 | 0.799976881 | 0.898237613 | 556.3207809 | 542.2507466 | ENSG00000196476 |
| GTPBP1 | -0.037233707 | 0.974521752 | 0.533553668 | 0.724818824 | 2488.587044 | 2425.408189 | ENSG00000100226 |
| NDC1 | -0.037313555 | 0.974467817 | 0.624453153 | 0.7923564 | 1808.416628 | 1762.373008 | ENSG00000058804 |
| NDUFS5 | -0.037443301 | 0.974380184 | 0.530426971 | 0.722526929 | 5262.470752 | 5127.34753 | ENSG00000168653 |
| SMC1A | -0.037469246 | 0.974362662 | 0.479359346 | 0.681525352 | 4591.390432 | 4472.970444 | ENSG00000072501 |
| GALK2 | -0.037511415 | 0.974334182 | 0.860478154 | 0.932201121 | 117.3849713 | 114.2420356 | ENSG00000156958 |
| ARVCF | -0.03752341 | 0.974326081 | 0.619264491 | 0.788725553 | 1679.940126 | 1637.429026 | ENSG00000099889 |
| RBM26-AS1 | -0.037540884 | 0.97431428 | 0.97332409 | 0.990527857 | 4.451389804 | 4.295255055 | ENSG00000227354 |
| HACD3 | -0.037625053 | 0.974257439 | 0.488961808 | 0.689210609 | 4325.293062 | 4214.096763 | ENSG00000074696 |
| NA | -0.037685764 | 0.974216442 | 0.921823435 | 0.964771134 | 35.10849943 | 34.31836321 | ENSG00000266066 |
| GTSE1 | -0.037704509 | 0.974203783 | 0.64714051 | 0.808497522 | 1265.527892 | 1232.46405 | ENSG00000075218 |
| PCCB | -0.037725339 | 0.974189718 | 0.602852116 | 0.777760774 | 2033.746437 | 1980.494715 | ENSG00000114054 |
| CLPTM1L | -0.037760116 | 0.974166234 | 0.550136882 | 0.737355218 | 2710.039022 | 2640.511412 | ENSG00000049656 |
| DOK4 | -0.037888327 | 0.974079665 | 0.657648204 | 0.815153999 | 860.1891922 | 837.9948721 | ENSG00000125170 |
| CNOT2 | -0.037931377 | 0.974050599 | 0.609927022 | 0.782664389 | 1192.825866 | 1161.437597 | ENSG00000111596 |
| LARGE1 | -0.038128666 | 0.973917407 | 0.575484638 | 0.75767062 | 1514.589435 | 1474.876191 | ENSG00000133424 |
| VPS33B | -0.038184019 | 0.97388004 | 0.934524699 | 0.970489863 | 25.04267155 | 24.36779044 | ENSG00000184056 |
| SAFB | -0.038187705 | 0.973877552 | 0.480770155 | 0.682708803 | 5043.490664 | 4912.137871 | ENSG00000160633 |
| BMP4 | -0.038198891 | 0.973870001 | 0.871342846 | 0.939605415 | 88.91219024 | 86.7424509 | ENSG00000125378 |
| SLC2A11 | -0.038252852 | 0.973833576 | 0.863416584 | 0.933980504 | 116.7710933 | 113.6891498 | ENSG00000133460 |
| FLII | -0.03840866 | 0.97372841 | 0.429254895 | 0.639110587 | 5971.516198 | 5814.98424 | ENSG00000177731 |
| SRA1 | -0.038491691 | 0.973672371 | 0.648358409 | 0.809147384 | 993.9363135 | 967.8659635 | ENSG00000213523 |
| C16orf72 | -0.038550133 | 0.973632929 | 0.667108895 | 0.82154232 | 1001.699337 | 975.4507898 | ENSG00000182831 |
| SGO2 | -0.03855902 | 0.973626931 | 0.722157195 | 0.853559651 | 512.5755192 | 499.0216426 | ENSG00000163535 |
| LINC02387 | -0.038568693 | 0.973620403 | 0.943707206 | 0.975313883 | 19.75473619 | 19.15793045 | ENSG00000256232 |
| GFER | -0.038602813 | 0.973597378 | 0.717584277 | 0.851122916 | 530.0186749 | 516.2245557 | ENSG00000127554 |
| IQCC | -0.038640915 | 0.973571665 | 0.889321705 | 0.950472009 | 65.70659871 | 64.11559961 | ENSG00000160051 |
| DCBLD2 | -0.038655443 | 0.973561861 | 0.669625345 | 0.82299482 | 1149.603385 | 1119.041671 | ENSG00000057019 |
| NA | -0.038699587 | 0.973532072 | 0.919414246 | 0.964136712 | 36.5120394 | 35.51942639 | ENSG00000251687 |
| ATG12 | -0.038702711 | 0.973529964 | 0.699547144 | 0.839970842 | 600.8210765 | 584.8863035 | ENSG00000145782 |
| SLAIN2 | -0.038782615 | 0.973476046 | 0.685498812 | 0.831762616 | 646.6952416 | 629.5949488 | ENSG00000109171 |
| MTREX | -0.038825892 | 0.973446845 | 0.549807042 | 0.737017745 | 2717.964397 | 2645.115947 | ENSG00000039123 |
| RNF14 | -0.038931053 | 0.973375891 | 0.757652 | 0.874053636 | 335.5844041 | 326.5443665 | ENSG00000013561 |
| TAF1A-AS1 | -0.038938788 | 0.973370673 | 0.952117384 | 0.979503013 | 10.52842853 | 10.25682109 | ENSG00000225265 |
| TLCD4 | -0.038984179 | 0.973340048 | 0.919328157 | 0.964136712 | 30.97855274 | 30.18501815 | ENSG00000152078 |
| MED15 | -0.038987634 | 0.973337717 | 0.59711511 | 0.77347684 | 1436.785125 | 1398.534441 | ENSG00000099917 |
| NA | -0.039016722 | 0.973318092 | 0.929677167 | 0.968057669 | 25.57793562 | 24.98530042 | ENSG00000273270 |
| PLRG1 | -0.03914709 | 0.973230144 | 0.578704638 | 0.759685603 | 1644.300019 | 1599.848031 | ENSG00000171566 |
| BICDL1 | -0.039163525 | 0.973219056 | 0.736082756 | 0.861271552 | 649.1974806 | 632.0533896 | ENSG00000135127 |
| YIPF3 | -0.039204999 | 0.973191079 | 0.611724238 | 0.784141186 | 1178.118626 | 1146.724643 | ENSG00000137207 |
| NA | -0.039276694 | 0.973142718 | 0.932816338 | 0.969708285 | 23.38915724 | 22.74704861 | ENSG00000227486 |
| NR2F6 | -0.039343273 | 0.973097809 | 0.560261024 | 0.745422674 | 2606.021672 | 2536.414292 | ENSG00000160113 |
| ZNF680 | -0.03939171 | 0.973065139 | 0.829661604 | 0.914608569 | 157.376926 | 153.2334089 | ENSG00000173041 |
| MPC1 | -0.039427894 | 0.973040734 | 0.792631135 | 0.894035655 | 218.3862787 | 212.5362945 | ENSG00000060762 |
| ZUP1 | -0.039490664 | 0.972998399 | 0.782829767 | 0.889742909 | 269.6533572 | 262.5493431 | ENSG00000153975 |
| NA | -0.039511353 | 0.972984446 | 0.903068567 | 0.957841349 | 50.51888317 | 49.08862785 | ENSG00000233966 |
| LIPE | -0.039566562 | 0.972947212 | 0.797028868 | 0.896426072 | 240.5273176 | 234.1504771 | ENSG00000079435 |
| GRK2 | -0.039599433 | 0.972925045 | 0.526741088 | 0.719946837 | 2224.962795 | 2164.939423 | ENSG00000173020 |
| ARHGEF5 | -0.039624773 | 0.972907956 | 0.713454143 | 0.848943449 | 557.8666214 | 542.2599593 | ENSG00000050327 |
| EMP2 | -0.039784664 | 0.972800136 | 0.73815717 | 0.862790008 | 463.8783091 | 451.4223257 | ENSG00000213853 |
| APC | -0.039933739 | 0.972699621 | 0.719837522 | 0.852321847 | 490.6438003 | 476.9776094 | ENSG00000134982 |
| POC1A | -0.039982939 | 0.97266645 | 0.621179857 | 0.790018291 | 947.8131166 | 921.7635097 | ENSG00000164087 |

| | | | | | | | |
|---|---|---|---|---|---|---|---|
| ARMC10 | -0.040130806 | 0.972566763 | 0.624929618 | 0.79259878 | 927.2701662 | 901.8312749 | ENSG00000170632 |
| SNRNP25 | -0.040222275 | 0.972505103 | 0.564180616 | 0.748528238 | 1451.379351 | 1411.692454 | ENSG00000161981 |
| HMGN3 | -0.040269675 | 0.972473151 | 0.670347966 | 0.823292434 | 861.7668664 | 838.5405824 | ENSG00000118418 |
| PRADC1 | -0.040328421 | 0.972433554 | 0.831251957 | 0.915293173 | 139.2958658 | 135.4767593 | ENSG00000135617 |
| EYA1 | -0.040343043 | 0.972423698 | 0.936212845 | 0.97143656 | 19.41313144 | 18.90550162 | ENSG00000104313 |
| CROCCP3 | -0.040382043 | 0.972397411 | 0.883829766 | 0.947014192 | 96.66054064 | 94.15414692 | ENSG00000080947 |
| PROSER1 | -0.040399954 | 0.972385339 | 0.53463353 | 0.725637667 | 2159.158851 | 2099.596682 | ENSG00000120685 |
| MEAK7 | -0.040416291 | 0.972374328 | 0.752520427 | 0.871085164 | 471.3519709 | 457.796316 | ENSG00000140950 |
| CXCL6 | -0.040480173 | 0.972331272 | 0.842641616 | 0.921654519 | 109.5429055 | 106.4954743 | ENSG00000124875 |
| FAM110B | -0.040549577 | 0.972284497 | 0.853717796 | 0.928338837 | 109.0115058 | 106.0953022 | ENSG00000169122 |
| SYNJ2 | -0.040567499 | 0.972272419 | 0.721382557 | 0.853073731 | 568.0449551 | 552.4037289 | ENSG00000078269 |
| MAPK8 | -0.040569077 | 0.972271355 | 0.683131392 | 0.830790283 | 812.3659992 | 789.4788046 | ENSG00000107643 |
| QRSL1 | -0.040570815 | 0.972270184 | 0.688353816 | 0.833414711 | 658.6419538 | 640.5506453 | ENSG00000130348 |
| DOP1B | -0.040574776 | 0.972267515 | 0.776481566 | 0.885391521 | 250.7493789 | 243.769263 | ENSG00000142197 |
| SNX30 | -0.040575401 | 0.972267093 | 0.735069586 | 0.860886148 | 401.5940643 | 390.1286764 | ENSG00000148158 |
| MRPS7 | -0.040823294 | 0.972100047 | 0.579616253 | 0.760406596 | 1561.890095 | 1517.588063 | ENSG00000125445 |
| BTBD7 | -0.04083928 | 0.972089276 | 0.717521427 | 0.851122916 | 596.2079842 | 579.3044368 | ENSG00000011114 |
| TP53 | -0.040861778 | 0.972074116 | 0.451910307 | 0.658191107 | 3558.407805 | 3458.519273 | ENSG00000141510 |
| MCU | -0.04094121 | 0.972020597 | 0.626372774 | 0.793253855 | 906.6611394 | 881.2375921 | ENSG00000156026 |
| HMOX2 | -0.040968837 | 0.972001983 | 0.603423852 | 0.778147861 | 1095.840602 | 1065.13696 | ENSG00000103415 |
| NA | -0.040972152 | 0.97199975 | 0.890498063 | 0.95121103 | 56.76524973 | 55.26395882 | ENSG00000235363 |
| RNF41 | -0.041017652 | 0.971969096 | 0.583660488 | 0.762956206 | 1211.001558 | 1176.621538 | ENSG00000181852 |
| ADRM1 | -0.041057395 | 0.97194232 | 0.547872603 | 0.735562541 | 1569.860604 | 1525.981899 | ENSG00000130706 |
| NA | -0.041068183 | 0.971935053 | 0.809364632 | 0.902986417 | 177.4952342 | 172.6847458 | ENSG00000277072 |
| DAAM1 | -0.041124592 | 0.971897051 | 0.718355387 | 0.851388915 | 447.8967093 | 435.1066598 | ENSG00000100592 |
| ZNF586 | -0.041134312 | 0.971890503 | 0.758847563 | 0.874822186 | 279.8045607 | 271.9116329 | ENSG00000083828 |
| RRAS2 | -0.041269892 | 0.971799172 | 0.482515183 | 0.683747222 | 3056.522765 | 2970.264553 | ENSG00000133818 |
| SIX1 | -0.041300816 | 0.971778341 | 0.958647781 | 0.982901533 | 7.494511106 | 7.322858199 | ENSG00000126778 |
| PPARA | -0.041324279 | 0.971762537 | 0.813777699 | 0.905396008 | 149.5259619 | 145.2946491 | ENSG00000186951 |
| CCDC174 | -0.0413428 | 0.971750062 | 0.757439938 | 0.873915764 | 312.3471345 | 303.699422 | ENSG00000154781 |
| BTAF1 | -0.041343539 | 0.971749564 | 0.615777777 | 0.786503033 | 1302.01087 | 1265.268335 | ENSG00000095564 |
| MED27 | -0.041375085 | 0.971728316 | 0.71509956 | 0.849670014 | 567.3749125 | 550.7591725 | ENSG00000160563 |
| PLPP5 | -0.041389159 | 0.971718837 | 0.666907129 | 0.82154232 | 571.1653911 | 555.0397368 | ENSG00000147535 |
| CDS2 | -0.041413211 | 0.971702637 | 0.563047519 | 0.747865551 | 1454.831708 | 1413.207291 | ENSG00000101290 |
| BCLAF3 | -0.041417935 | 0.971699455 | 0.805373209 | 0.900817151 | 172.3816606 | 167.6024828 | ENSG00000173681 |
| WBP1L | -0.041462067 | 0.971669731 | 0.644614801 | 0.806145123 | 684.2914366 | 665.0667259 | ENSG00000166272 |
| CTNND2 | -0.041522153 | 0.971629264 | 0.711420324 | 0.847591497 | 416.6750219 | 404.8140456 | ENSG00000169862 |
| PSMB2 | -0.041614512 | 0.971567063 | 0.46658318 | 0.670631202 | 2806.530741 | 2727.115724 | ENSG00000126067 |
| NA | -0.04167651 | 0.971525313 | 0.942913564 | 0.974760382 | 14.07447374 | 13.6292638 | ENSG00000272631 |
| PNPLA3 | -0.041742137 | 0.971481119 | 0.682765172 | 0.830639714 | 589.7386451 | 572.4459234 | ENSG00000100344 |
| ARL6IP6 | -0.041774974 | 0.971459008 | 0.650533254 | 0.810373005 | 876.5917956 | 851.623147 | ENSG00000177917 |
| RRP8 | -0.041815088 | 0.971431997 | 0.770651472 | 0.881997379 | 312.8683883 | 303.9578531 | ENSG00000132275 |
| G3BP1 | -0.04184369 | 0.971412738 | 0.490734193 | 0.690477988 | 7101.1824 | 6897.629709 | ENSG00000145907 |
| NA | -0.041950926 | 0.971340536 | 0.893056094 | 0.952281345 | 52.00871405 | 50.39988436 | ENSG00000276291 |
| UBE2B | -0.041962822 | 0.971332526 | 0.73579358 | 0.861168893 | 348.1990229 | 338.4238214 | ENSG00000119048 |
| CSGALNACT2 | -0.042108457 | 0.971234479 | 0.801586775 | 0.898520796 | 169.3800317 | 164.4793475 | ENSG00000169826 |
| MDFIC | -0.042318796 | 0.971092887 | 0.764605864 | 0.87816748 | 261.1870929 | 253.8622822 | ENSG00000135272 |
| BTBD2 | -0.042422625 | 0.971023001 | 0.510614632 | 0.707210126 | 2820.182788 | 2738.930442 | ENSG00000133243 |
| RNF181 | -0.042502722 | 0.970969092 | 0.685444545 | 0.831762616 | 672.4670647 | 652.8888645 | ENSG00000168894 |
| MTRR | -0.042512489 | 0.970962519 | 0.636738845 | 0.800725077 | 788.2699637 | 765.0998735 | ENSG00000124275 |
| RAB8B | -0.042557043 | 0.970932533 | 0.674476389 | 0.825608414 | 572.3295822 | 555.7632344 | ENSG00000166128 |
| ABCC5 | -0.042589409 | 0.970910752 | 0.635765846 | 0.800375842 | 1250.134923 | 1212.946134 | ENSG00000114770 |
| FAM200A | -0.042624519 | 0.970887123 | 0.822471438 | 0.909975576 | 161.6693871 | 156.9996653 | ENSG00000221909 |
| OSTF1 | -0.042854715 | 0.970732221 | 0.744492082 | 0.867135375 | 282.4639484 | 274.2700094 | ENSG00000134996 |
| KPNA5 | -0.042890698 | 0.97070801 | 0.812915925 | 0.905077069 | 169.7311713 | 165.0298173 | ENSG00000196911 |
| CCDC85C | -0.043033041 | 0.97061224 | 0.597092834 | 0.77347684 | 925.1626715 | 897.759317 | ENSG00000205476 |
| THSD7B | -0.04304347 | 0.970605224 | 0.842460852 | 0.92165416 | 109.9933532 | 106.6716666 | ENSG00000144229 |
| TCEAL8 | -0.043092451 | 0.970572272 | 0.497530194 | 0.696660074 | 2112.462578 | 2050.47126 | ENSG00000180964 |
| PNP | -0.043133502 | 0.970544655 | 0.529950973 | 0.72213898 | 1455.677431 | 1412.694108 | ENSG00000198805 |
| FXYD5 | -0.043157319 | 0.970528632 | 0.617211537 | 0.787224317 | 870.6274393 | 845.1444618 | ENSG00000089327 |
| CNOT6 | -0.043253887 | 0.970463672 | 0.636537977 | 0.800725077 | 849.3575469 | 823.7740157 | ENSG00000113300 |
| PITPNM2 | -0.043294656 | 0.970436248 | 0.800650139 | 0.898320559 | 161.944557 | 157.17238 | ENSG00000090975 |
| FAM49B | -0.043404835 | 0.970362138 | 0.581176234 | 0.761236869 | 1809.587199 | 1755.535879 | ENSG00000153310 |
| PIK3R4 | -0.043490439 | 0.970304562 | 0.590337909 | 0.768493238 | 1007.115362 | 977.0246603 | ENSG00000196455 |
| RUNDC3B | -0.043503067 | 0.970296069 | 0.957169237 | 0.98244088 | 7.160304638 | 6.98766644 | ENSG00000105784 |
| LSM10 | -0.043515828 | 0.970287486 | 0.621848515 | 0.790655706 | 827.9369676 | 803.827911 | ENSG00000181817 |
| NA | -0.043543996 | 0.970268543 | 0.873117046 | 0.940444614 | 65.26238024 | 63.39640994 | ENSG00000227827 |
| FDPS | -0.043583375 | 0.970242059 | 0.53356798 | 0.724818824 | 1815.398447 | 1760.946603 | ENSG00000160752 |
| SLC7A8 | -0.043724308 | 0.970147283 | 0.395378829 | 0.607298022 | 6277.56785 | 6089.895238 | ENSG00000092068 |
| TWISTNB | -0.043764285 | 0.970120401 | 0.703398692 | 0.842499163 | 641.9233501 | 623.0909524 | ENSG00000105849 |
| MAGOHB | -0.04377166 | 0.970115441 | 0.689604108 | 0.83398915 | 459.242106 | 445.6832038 | ENSG00000111196 |
| TMEM191C | -0.043785454 | 0.970106166 | 0.947263375 | 0.976838866 | 11.3368038 | 10.98319957 | ENSG00000206140 |
| QSOX2 | -0.043799589 | 0.970096661 | 0.553301441 | 0.739863889 | 1462.09679 | 1418.397183 | ENSG00000165661 |
| SLC30A9 | -0.043809312 | 0.970090123 | 0.523565236 | 0.717778069 | 1619.481594 | 1570.934489 | ENSG00000014824 |

| | | | | | | | |
|---|---|---|---|---|---|---|---|
| NUS1 | -0.043917118 | 0.970017636 | 0.572046724 | 0.754564671 | 1977.66318 | 1918.125578 | ENSG00000153989 |
| GRN | -0.044030625 | 0.96994132 | 0.496029572 | 0.695334407 | 2648.658699 | 2569.202557 | ENSG00000030582 |
| TOMM5 | -0.044065533 | 0.969917852 | 0.919711152 | 0.964147051 | 23.62875865 | 22.86651088 | ENSG00000175768 |
| TRMT6 | -0.044076344 | 0.969910584 | 0.686848984 | 0.832230404 | 580.7086915 | 563.4033862 | ENSG00000089195 |
| NA | -0.044118346 | 0.969882346 | 0.903091599 | 0.957841349 | 36.23831781 | 35.06354286 | ENSG00000254536 |
| DISP3 | -0.044186691 | 0.969836401 | 0.771428706 | 0.882491333 | 211.6488619 | 205.3767867 | ENSG00000204624 |
| PRDM11 | -0.044227005 | 0.969809301 | 0.733836182 | 0.860179688 | 300.0141937 | 290.9572172 | ENSG00000019485 |
| SLC22A4 | -0.044344554 | 0.969730285 | 0.951392851 | 0.97918437 | 8.532699083 | 8.295139108 | ENSG00000197208 |
| PRKCZ | -0.044495104 | 0.969629096 | 0.592612253 | 0.76996783 | 901.9970716 | 874.5414605 | ENSG00000067606 |
| EPDR1 | -0.044576787 | 0.969574199 | 0.787619007 | 0.891577351 | 195.9534416 | 190.0196366 | ENSG00000086289 |
| KDM3A | -0.04468943 | 0.969498499 | 0.655888583 | 0.814050821 | 589.7478751 | 571.414737 | ENSG00000115548 |
| IQCB1 | -0.044777752 | 0.969439148 | 0.614014511 | 0.785269226 | 828.1323808 | 803.2008056 | ENSG00000173226 |
| PCDHB9 | -0.044783913 | 0.969435008 | 0.9155096 | 0.962507751 | 28.83129283 | 27.96154308 | ENSG00000177839 |
| RUNDC3A | -0.044817222 | 0.969412626 | 0.848519638 | 0.924976528 | 91.24271097 | 88.42588204 | ENSG00000108309 |
| MRPL50 | -0.044885338 | 0.969366857 | 0.580342631 | 0.760936651 | 940.9173291 | 912.1649653 | ENSG00000136897 |
| MALT1 | -0.044891902 | 0.969362446 | 0.746311253 | 0.867969455 | 311.9073391 | 302.3871766 | ENSG00000172175 |
| FEM1B | -0.044897485 | 0.969358695 | 0.588954586 | 0.767009679 | 1374.982778 | 1332.435378 | ENSG00000169018 |
| NDUFC1 | -0.044914969 | 0.969346948 | 0.641879982 | 0.804214244 | 610.3451056 | 591.9686944 | ENSG00000109390 |
| NA | -0.044926805 | 0.969338995 | 0.904817665 | 0.958516946 | 37.13209427 | 35.90503268 | ENSG00000224086 |
| TMEM19 | -0.044994679 | 0.969293392 | 0.816204377 | 0.906477444 | 129.5494216 | 125.6811187 | ENSG00000139291 |
| PPP1CC | -0.045025563 | 0.969272642 | 0.382603958 | 0.595081184 | 7085.842446 | 6867.622258 | ENSG00000186298 |
| SVIP | -0.045175111 | 0.969172174 | 0.820917099 | 0.909161355 | 125.7560457 | 121.9003691 | ENSG00000198168 |
| ATP1B2 | -0.045178648 | 0.969169798 | 0.734674299 | 0.860583311 | 310.7194007 | 301.1824679 | ENSG00000129244 |
| GPR21 | -0.045300229 | 0.969088126 | 0.59169344 | 0.769356281 | 801.3798293 | 776.4186867 | ENSG00000011454 |
| ZNF638 | -0.045363483 | 0.969045638 | 0.456871402 | 0.662717707 | 2330.036086 | 2257.844318 | ENSG00000075292 |
| ASTE1 | -0.045372438 | 0.969039623 | 0.880695213 | 0.944942282 | 60.13960284 | 58.14918819 | ENSG00000034533 |
| PEX11G | -0.045372879 | 0.969039326 | 0.91130848 | 0.961009829 | 31.29051265 | 30.29889999 | ENSG00000104883 |
| B3GALT1 | -0.045392927 | 0.969025861 | 0.744736041 | 0.867262195 | 315.2039425 | 305.3190796 | ENSG00000172318 |
| MICU3 | -0.045459342 | 0.968981252 | 0.872996195 | 0.940412849 | 62.8924004 | 60.88577126 | ENSG00000155970 |
| CYYR1 | -0.045806138 | 0.968748356 | 0.607959988 | 0.781467763 | 666.0567348 | 645.2791786 | ENSG00000166265 |
| FAM241B | -0.045810759 | 0.968745253 | 0.689261644 | 0.83398915 | 468.8681135 | 453.9921737 | ENSG00000171224 |
| TICRR | -0.045895536 | 0.968688328 | 0.591526265 | 0.769255138 | 865.8388945 | 838.6944529 | ENSG00000140534 |
| RELN | -0.045898417 | 0.968686394 | 0.964577328 | 0.986052631 | 6.511163604 | 6.25164015 | ENSG00000189056 |
| VWC2 | -0.046068286 | 0.968572344 | 0.947299505 | 0.976838866 | 10.34130334 | 10.00779723 | ENSG00000188730 |
| BTF3 | -0.046095262 | 0.968554233 | 0.383388974 | 0.595734937 | 11071.14139 | 10722.61644 | ENSG00000145741 |
| LSM1 | -0.046095448 | 0.968554108 | 0.741450649 | 0.865140285 | 350.7844996 | 339.7219298 | ENSG00000175324 |
| ARF4 | -0.046101572 | 0.968549997 | 0.483428446 | 0.68442107 | 2110.713395 | 2044.513194 | ENSG00000168374 |
| NA | -0.046140785 | 0.968523671 | 0.822146484 | 0.909722644 | 114.2416261 | 110.6625652 | ENSG00000250462 |
| PLP2 | -0.046148353 | 0.968518591 | 0.845104545 | 0.923011933 | 89.8264338 | 86.95912635 | ENSG00000102007 |
| C11orf1 | -0.046346702 | 0.968385443 | 0.813922672 | 0.905404967 | 122.101463 | 118.2800594 | ENSG00000137720 |
| PPP2R3A | -0.046392186 | 0.968354913 | 0.732512362 | 0.859490273 | 297.9746077 | 288.5353008 | ENSG00000073711 |
| MAP2K4 | -0.046435425 | 0.968325891 | 0.611673536 | 0.784141186 | 722.7214557 | 699.6853351 | ENSG00000065559 |
| TTC7B | -0.046473412 | 0.968300395 | 0.706960723 | 0.845211019 | 322.5469493 | 312.2342207 | ENSG00000165914 |
| NPIPB11 | -0.046532506 | 0.968260733 | 0.929714691 | 0.968057669 | 17.20299623 | 16.68486443 | ENSG00000254206 |
| CTCF | -0.046557239 | 0.968244134 | 0.403255961 | 0.614548807 | 2717.802597 | 2631.465531 | ENSG00000102974 |
| LINC01970 | -0.046594313 | 0.968219253 | 0.956376918 | 0.982211615 | 7.238586879 | 7.03713501 | ENSG00000265692 |
| SHOC2 | -0.046598136 | 0.968216687 | 0.735812616 | 0.861168893 | 391.2138856 | 378.7028343 | ENSG00000108061 |
| PXMP4 | -0.046614523 | 0.968205689 | 0.872589826 | 0.940252132 | 59.43562133 | 57.52006585 | ENSG00000101417 |
| LHX8 | -0.046647873 | 0.968183308 | 0.949313337 | 0.977907982 | 9.279394261 | 8.988223219 | ENSG00000162624 |
| SIRT6 | -0.046678574 | 0.968162705 | 0.617273904 | 0.787224317 | 644.7074947 | 624.435358 | ENSG00000077463 |
| CFAP46 | -0.046687183 | 0.968156928 | 0.818486459 | 0.907640445 | 119.2760015 | 115.5391254 | ENSG00000171811 |
| KIF16B | -0.046715155 | 0.968138157 | 0.771289806 | 0.882455349 | 244.606464 | 236.5622512 | ENSG00000089177 |
| RBM44 | -0.04673151 | 0.968127182 | 0.942900476 | 0.974760382 | 10.29742061 | 9.95709916 | ENSG00000177483 |
| CBR1 | -0.046804147 | 0.968078439 | 0.57156952 | 0.75430401 | 839.9207763 | 812.8690602 | ENSG00000159228 |
| RPL11 | -0.046843257 | 0.968052196 | 0.4114181 | 0.622616461 | 15399.59347 | 14908.13507 | ENSG00000142676 |
| SPCS2 | -0.046866722 | 0.968036451 | 0.459276166 | 0.664325244 | 1972.308452 | 1909.494684 | ENSG00000118363 |
| TGFBRAP1 | -0.046873356 | 0.968032 | 0.572721175 | 0.755190573 | 836.2525662 | 809.3064991 | ENSG00000135966 |
| RAB32 | -0.04687503 | 0.968030876 | 0.820555273 | 0.908947936 | 129.3626019 | 125.2705637 | ENSG00000118508 |
| MRPS6 | -0.046970876 | 0.967966567 | 0.674379005 | 0.825608414 | 463.9298694 | 449.1839262 | ENSG00000243927 |
| SMIM15 | -0.046982884 | 0.96795851 | 0.701485409 | 0.84111441 | 409.1600954 | 396.3176273 | ENSG00000188725 |
| POLR3K | -0.047046695 | 0.967915699 | 0.567280566 | 0.750900224 | 1017.823641 | 984.8779576 | ENSG00000161980 |
| PRKRIP1 | -0.047217116 | 0.967801368 | 0.619361924 | 0.788725553 | 669.2490229 | 647.6178806 | ENSG00000128563 |
| PKP4 | -0.04722627 | 0.967795227 | 0.443812817 | 0.651211433 | 2331.440437 | 2255.979931 | ENSG00000144283 |
| MLST8 | -0.047327741 | 0.967727161 | 0.498282086 | 0.697247866 | 1334.040068 | 1291.010793 | ENSG00000167965 |
| HAUS5 | -0.047331213 | 0.967724831 | 0.628965243 | 0.795366855 | 760.1432889 | 735.8672142 | ENSG00000249115 |
| RNF220 | -0.047343559 | 0.96771655 | 0.502136839 | 0.700181291 | 1517.951198 | 1469.000196 | ENSG00000187147 |
| BAIAP2-DT | -0.047365149 | 0.967702069 | 0.689499921 | 0.83398915 | 364.3270412 | 352.7127593 | ENSG00000226137 |
| RALBP1 | -0.047381317 | 0.967691224 | 0.431732274 | 0.6413842 | 2306.794568 | 2232.244355 | ENSG00000017797 |
| PLEKHO2 | -0.047386473 | 0.967687765 | 0.627505954 | 0.793893612 | 596.2157396 | 577.1209637 | ENSG00000241839 |
| KLHL5 | -0.047414271 | 0.96766912 | 0.707781484 | 0.845910689 | 327.506933 | 317.0526831 | ENSG00000109790 |
| SLC16A4 | -0.047443443 | 0.967649553 | 0.956372239 | 0.982211615 | 6.860497685 | 6.670824349 | ENSG00000168679 |
| KIN | -0.047482404 | 0.967623422 | 0.660532435 | 0.817400463 | 455.2604358 | 440.5754024 | ENSG00000151657 |
| PRKD1 | -0.047482731 | 0.967623202 | 0.776880889 | 0.885739893 | 185.8793257 | 179.8883526 | ENSG00000184304 |
| RER1 | -0.047494625 | 0.967615225 | 0.517287473 | 0.713310786 | 1153.779072 | 1116.608491 | ENSG00000157916 |

| | | | | | | | |
|---|---|---|---|---|---|---|---|
| SCAF11 | -0.047547642 | 0.967579667 | 0.590283353 | 0.768475191 | 1470.121653 | 1422.191263 | ENSG00000139218 |
| EXOC3-AS1 | -0.047550268 | 0.967577906 | 0.925038823 | 0.965883673 | 22.29404386 | 21.63773376 | ENSG00000221990 |
| BRPF3 | -0.04755747 | 0.967573076 | 0.528054279 | 0.720973828 | 1263.760578 | 1222.710728 | ENSG00000096070 |
| LSM11 | -0.047570776 | 0.967564152 | 0.7921933 | 0.893932102 | 143.2816802 | 138.5936518 | ENSG00000155858 |
| STRIP2 | -0.047654623 | 0.967507921 | 0.739984246 | 0.863848501 | 230.4709467 | 223.0317566 | ENSG00000128578 |
| GNPTAB | -0.047687794 | 0.967485676 | 0.551090672 | 0.738214452 | 1164.951303 | 1126.727625 | ENSG00000111670 |
| PIK3C2A | -0.0477064 | 0.967473198 | 0.543546836 | 0.732315108 | 1854.950484 | 1794.17698 | ENSG00000011405 |
| NA | -0.047796537 | 0.967412754 | 0.958263818 | 0.982774639 | 6.180237276 | 6.019736458 | ENSG00000234791 |
| OGFR | -0.047817175 | 0.967398915 | 0.545321997 | 0.73371373 | 1320.755919 | 1278.257724 | ENSG00000060491 |
| TMEM134 | -0.047822535 | 0.967395321 | 0.712989884 | 0.848591123 | 352.8852332 | 341.495408 | ENSG00000172663 |
| ZNF787 | -0.047863729 | 0.967367699 | 0.522970428 | 0.717284954 | 1663.006427 | 1609.397077 | ENSG00000142409 |
| POLB | -0.047947649 | 0.96731143 | 0.648330825 | 0.809147384 | 486.2220815 | 470.1864198 | ENSG00000070501 |
| PAPOLG | -0.048017115 | 0.967264855 | 0.681162515 | 0.829598484 | 419.4251241 | 405.3663643 | ENSG00000115421 |
| TRAPPC3 | -0.048061984 | 0.967234772 | 0.583453559 | 0.762833684 | 956.1728499 | 924.5373889 | ENSG00000054116 |
| PLD3 | -0.04807199 | 0.967228064 | 0.402420912 | 0.613870672 | 4816.378085 | 4659.326014 | ENSG00000105223 |
| C4orf48 | -0.048124584 | 0.967192804 | 0.77165301 | 0.882491333 | 231.2899999 | 223.9772695 | ENSG00000243449 |
| RFC4 | -0.048162141 | 0.967167626 | 0.502232236 | 0.700181291 | 1379.384335 | 1334.198949 | ENSG00000163918 |
| ABL1 | -0.048223215 | 0.967126683 | 0.387836836 | 0.600150584 | 3919.795402 | 3790.658475 | ENSG00000097007 |
| AHCYL1 | -0.04823199 | 0.967120801 | 0.388742742 | 0.600524064 | 3613.837318 | 3494.668956 | ENSG00000168710 |
| NA | -0.048310124 | 0.967068425 | 0.917022511 | 0.96312259 | 24.09224903 | 23.22629515 | ENSG00000224831 |
| MYADM | -0.048337699 | 0.967049941 | 0.595847813 | 0.772365019 | 802.1787731 | 775.8495111 | ENSG00000179820 |
| COA8 | -0.048373582 | 0.967025888 | 0.819157948 | 0.908070158 | 116.1344624 | 112.3585976 | ENSG00000256053 |
| CSNK1E | -0.04838434 | 0.967018677 | 0.542779198 | 0.731699989 | 1433.359771 | 1386.081691 | ENSG00000213923 |
| NA | -0.048410339 | 0.967001251 | 0.947414169 | 0.976846287 | 9.268715843 | 8.962401202 | ENSG00000176933 |
| SETD3 | -0.048533456 | 0.966918732 | 0.503751285 | 0.701470507 | 1727.758557 | 1670.314426 | ENSG00000183576 |
| APPL1 | -0.048562685 | 0.966899143 | 0.604760723 | 0.7791033 | 890.9782816 | 861.4935314 | ENSG00000157500 |
| DSCC1 | -0.048610289 | 0.966867239 | 0.549216683 | 0.736487756 | 927.1453582 | 896.3459496 | ENSG00000136982 |
| TRIM37 | -0.048764114 | 0.966764153 | 0.492674551 | 0.692222223 | 1666.526537 | 1610.882924 | ENSG00000108395 |
| PPARD | -0.048822883 | 0.966724773 | 0.57505044 | 0.75731015 | 709.8153698 | 686.2492385 | ENSG00000112033 |
| FAM160A2 | -0.048831832 | 0.966718776 | 0.630772505 | 0.796625039 | 531.6274429 | 513.9209447 | ENSG00000051009 |
| LOC101927751 | -0.048874337 | 0.966690295 | 0.964225727 | 0.986050947 | 5.815190764 | 5.650966787 | ENSG00000259583 |
| STARD5 | -0.048909569 | 0.966666688 | 0.925267294 | 0.965883673 | 16.756108 | 16.22056259 | ENSG00000172345 |
| RNF166 | -0.048977935 | 0.966620881 | 0.687209055 | 0.832541421 | 442.501509 | 428.1278335 | ENSG00000158717 |
| RILPL2 | -0.048984076 | 0.966616766 | 0.816257917 | 0.906477444 | 112.2630836 | 108.3516927 | ENSG00000150977 |
| PHACTR3 | -0.048993522 | 0.966610437 | 0.936164155 | 0.97143656 | 13.67355317 | 13.27695188 | ENSG00000087495 |
| NTMT1 | -0.049072354 | 0.966557621 | 0.611930792 | 0.784174879 | 781.2463145 | 755.0553043 | ENSG00000148335 |
| DDX31 | -0.04921414 | 0.966462634 | 0.627096743 | 0.793729923 | 627.6175124 | 606.2980928 | ENSG00000125485 |
| NIP7 | -0.04922218 | 0.966457248 | 0.53141965 | 0.723253105 | 1245.182517 | 1203.692785 | ENSG00000132603 |
| CHCHD1 | -0.049230314 | 0.966451799 | 0.552182136 | 0.739047459 | 883.419927 | 854.0547245 | ENSG00000172586 |
| SEC24B-AS1 | -0.049284024 | 0.96641582 | 0.923433283 | 0.965309497 | 18.21509884 | 17.60332584 | ENSG00000247950 |
| GRIPAP1 | -0.049306405 | 0.966400828 | 0.481994074 | 0.68352167 | 1380.759932 | 1334.106212 | ENSG00000068400 |
| USP19 | -0.049471895 | 0.966289979 | 0.457710897 | 0.663253386 | 2204.696927 | 2129.905473 | ENSG00000172046 |
| BCR | -0.049489972 | 0.966277871 | 0.39945658 | 0.610828948 | 4329.838681 | 4184.132965 | ENSG00000186716 |
| MAP3K5 | -0.049508024 | 0.966265781 | 0.835450135 | 0.917415262 | 87.11601827 | 84.09371674 | ENSG00000197442 |
| KIF11 | -0.049549387 | 0.966238077 | 0.482775296 | 0.683859246 | 1654.529634 | 1598.452383 | ENSG00000138160 |
| ATP2A1 | -0.049569095 | 0.966224878 | 0.864518593 | 0.93475516 | 58.46236743 | 56.55620325 | ENSG00000196296 |
| LASP1 | -0.049632991 | 0.966182086 | 0.329686095 | 0.541217709 | 5538.994583 | 5351.735299 | ENSG00000002834 |
| BNIP2 | -0.049660815 | 0.966163453 | 0.623882218 | 0.792228058 | 576.5022949 | 557.0987046 | ENSG00000140299 |
| FOXN3 | -0.049716513 | 0.966126152 | 0.578310342 | 0.759450456 | 2009.173842 | 1940.331803 | ENSG00000053254 |
| TASOR | -0.04972527 | 0.966120288 | 0.624199987 | 0.7923564 | 835.0542537 | 806.6429452 | ENSG00000163946 |
| ZNRD2 | -0.049744002 | 0.966107744 | 0.580455749 | 0.760979301 | 728.832894 | 704.029587 | ENSG00000173465 |
| TSPAN15 | -0.049767969 | 0.966091694 | 0.681656797 | 0.829825863 | 351.6400639 | 339.7095394 | ENSG00000099282 |
| MFSD12 | -0.04977592 | 0.96608637 | 0.583940371 | 0.763005183 | 959.2066694 | 927.3300325 | ENSG00000161091 |
| NKD2 | -0.049833303 | 0.966047945 | 0.691011812 | 0.83465034 | 326.5568159 | 315.550952 | ENSG00000145506 |
| LOC100506100 | -0.049856932 | 0.966032123 | 0.937773721 | 0.972054269 | 12.80764211 | 12.32302064 | ENSG00000223478 |
| PDCL | -0.049874215 | 0.96602055 | 0.643969577 | 0.805922164 | 490.3259167 | 473.7108841 | ENSG00000136940 |
| ZDHHC20 | -0.049891322 | 0.966009096 | 0.56388313 | 0.748513341 | 1587.490457 | 1533.570328 | ENSG00000180776 |
| IKBKB | -0.049965616 | 0.96595935 | 0.611418416 | 0.783992177 | 681.4407539 | 658.3363563 | ENSG00000104365 |
| IGFBP2 | -0.049998346 | 0.965937436 | 0.384362776 | 0.59668648 | 5830.845838 | 5631.783523 | ENSG00000115457 |
| GPR37 | -0.050011223 | 0.965928815 | 0.840761023 | 0.920610652 | 116.5385853 | 112.3356135 | ENSG00000170775 |
| CLUHP3 | -0.050012209 | 0.965928154 | 0.798777766 | 0.897431025 | 205.290957 | 198.2390765 | ENSG00000131797 |
| HNRNPK | -0.050098345 | 0.965870486 | 0.310375004 | 0.521016173 | 18317.29848 | 17691.86275 | ENSG00000165119 |
| SMIM26 | -0.050240684 | 0.965775196 | 0.883144601 | 0.94654857 | 53.5009092 | 51.57930849 | ENSG00000232388 |
| DIXDC1 | -0.050291527 | 0.96574116 | 0.70309603 | 0.842350348 | 334.1174563 | 322.7193296 | ENSG00000150764 |
| HMG20A | -0.050312345 | 0.965727225 | 0.535635675 | 0.72637367 | 1150.775943 | 1110.807164 | ENSG00000140382 |
| GCC2 | -0.050317958 | 0.965723468 | 0.571669949 | 0.754332608 | 723.6137577 | 698.6786257 | ENSG00000135968 |
| NA | -0.050331195 | 0.965714607 | 0.716937517 | 0.850836738 | 283.3704626 | 273.3435811 | ENSG00000132967 |
| GSDME | -0.05034574 | 0.965704871 | 0.786999187 | 0.891356582 | 155.3669587 | 150.0973161 | ENSG00000105928 |
| NUDT16L1 | -0.05037634 | 0.965684389 | 0.558491051 | 0.744063237 | 946.1216683 | 913.862214 | ENSG00000168101 |
| CPLANE2 | -0.050466937 | 0.965623748 | 0.849197113 | 0.925311098 | 76.01289214 | 73.33610539 | ENSG00000132881 |
| SOS2 | -0.05049561 | 0.965604557 | 0.750905722 | 0.870392545 | 232.7246859 | 224.5001933 | ENSG00000100485 |
| INTS6 | -0.050509756 | 0.965595089 | 0.614160787 | 0.785269226 | 612.4010673 | 591.3375418 | ENSG00000102786 |
| IL1R1 | -0.050519248 | 0.965588737 | 0.945556106 | 0.976189572 | 8.870465024 | 8.564688093 | ENSG00000115594 |
| TMEM50B | -0.050578992 | 0.965548751 | 0.770711948 | 0.881997379 | 206.6405463 | 199.4045543 | ENSG00000142188 |

| | | | | | | | |
|---|---|---|---|---|---|---|---|
| NRF1 | -0.050581683 | 0.965546949 | 0.652523682 | 0.811703295 | 430.9066443 | 415.9882666 | ENSG00000106459 |
| SPG7 | -0.050589555 | 0.965541681 | 0.474171952 | 0.677209204 | 2248.599738 | 2171.002682 | ENSG00000197912 |
| NA | -0.050594037 | 0.965538682 | 0.935779319 | 0.971224748 | 15.92340125 | 15.30087167 | ENSG00000287492 |
| SYN2 | -0.050611722 | 0.965526846 | 0.7761495 | 0.885173217 | 169.284563 | 163.3265234 | ENSG00000157152 |
| SH2B3 | -0.050656816 | 0.965496667 | 0.521056739 | 0.715972717 | 1061.859965 | 1025.222877 | ENSG00000111252 |
| DENND11 | -0.050663467 | 0.965492216 | 0.540288341 | 0.729834265 | 935.8337059 | 903.2017205 | ENSG00000257093 |
| HEATR1 | -0.050664959 | 0.965491218 | 0.440000607 | 0.647767794 | 2882.510526 | 2782.510405 | ENSG00000119285 |
| TRIM3 | -0.050714431 | 0.96545811 | 0.699496144 | 0.839970842 | 323.0377202 | 311.8267872 | ENSG00000110171 |
| MAEA | -0.050846806 | 0.965369528 | 0.375621306 | 0.587680921 | 2455.116826 | 2370.418246 | ENSG00000090316 |
| RNF19A | -0.050865049 | 0.965357321 | 0.789034909 | 0.891843683 | 148.4432808 | 143.0915836 | ENSG00000034677 |
| LOC645513 | -0.050870521 | 0.965353659 | 0.80691522 | 0.901640919 | 132.8632899 | 128.3606882 | ENSG00000245958 |
| SULF1 | -0.050891966 | 0.96533931 | 0.764031382 | 0.877785999 | 300.215785 | 289.3059236 | ENSG00000137573 |
| NA | -0.050922534 | 0.965318857 | 0.909563608 | 0.960527246 | 29.73218822 | 28.66588337 | ENSG00000269352 |
| CCNE1 | -0.050949262 | 0.965300973 | 0.521103786 | 0.715972717 | 1217.831878 | 1175.341022 | ENSG00000105173 |
| FAM98A | -0.051129675 | 0.965180267 | 0.406736645 | 0.617595253 | 2441.824538 | 2356.670852 | ENSG00000119812 |
| GMPS | -0.051160659 | 0.965159539 | 0.360311011 | 0.572516706 | 3542.07105 | 3418.440768 | ENSG00000163655 |
| ATP6V0B | -0.051165111 | 0.96515656 | 0.446101114 | 0.653425854 | 1813.206746 | 1749.583546 | ENSG00000117410 |
| EXOSC3 | -0.051271645 | 0.965085292 | 0.641781757 | 0.804214244 | 437.4154437 | 422.0307036 | ENSG00000107371 |
| IKZF5 | -0.051435709 | 0.964975548 | 0.784709572 | 0.890433415 | 171.0875223 | 165.133178 | ENSG00000095574 |
| BNIP3 | -0.051477324 | 0.964947713 | 0.580716402 | 0.761084181 | 807.6192523 | 778.8615797 | ENSG00000176171 |
| SDK2 | -0.051486271 | 0.964941729 | 0.693307959 | 0.836086897 | 300.4649469 | 289.7562267 | ENSG00000069188 |
| SERPINE3 | -0.051507481 | 0.964927543 | 0.955331573 | 0.981793467 | 6.207517847 | 6.024087385 | ENSG00000253309 |
| SLC35A5 | -0.051544807 | 0.964902578 | 0.608721618 | 0.782180564 | 506.3042363 | 488.6468426 | ENSG00000138459 |
| GALNT7 | -0.05167142 | 0.964817901 | 0.593496302 | 0.770745267 | 1261.221549 | 1216.706889 | ENSG00000109586 |
| TNKS2 | -0.051682066 | 0.964810782 | 0.493683035 | 0.693175027 | 2214.368797 | 2135.846119 | ENSG00000107854 |
| KCNG3 | -0.05175934 | 0.964759106 | 0.772001247 | 0.882599617 | 163.1485134 | 157.544482 | ENSG00000171126 |
| SUPT7L | -0.051783976 | 0.964742631 | 0.67132511 | 0.82390724 | 454.0485231 | 438.1913851 | ENSG00000119760 |
| FBXO5 | -0.051787127 | 0.964740524 | 0.555125566 | 0.741515512 | 737.5592268 | 711.4523486 | ENSG00000112029 |
| SUMO3 | -0.051832787 | 0.964709991 | 0.378279733 | 0.590739499 | 3167.045433 | 3054.978575 | ENSG00000184900 |
| OTUD7A | -0.051843173 | 0.964703046 | 0.786611142 | 0.891184319 | 149.0588859 | 143.7718984 | ENSG00000169918 |
| NOP16 | -0.05185082 | 0.964697933 | 0.57545351 | 0.75767062 | 707.3538118 | 682.4242219 | ENSG00000048162 |
| YEATS2 | -0.051901989 | 0.964663718 | 0.483425838 | 0.68442107 | 2041.093465 | 1968.204135 | ENSG00000163872 |
| MRE11 | -0.051939215 | 0.964638827 | 0.597376585 | 0.773603292 | 612.556436 | 591.21952 | ENSG00000020922 |
| NDUFS3 | -0.051982694 | 0.964609755 | 0.480748094 | 0.682708803 | 1647.738441 | 1589.686689 | ENSG00000213619 |
| NA | -0.051999724 | 0.964598369 | 0.925274108 | 0.965883673 | 22.74538128 | 22.05133753 | ENSG00000259877 |
| SLC11A2 | -0.052058741 | 0.964558911 | 0.492981066 | 0.692446816 | 1195.27596 | 1152.637861 | ENSG00000110911 |
| SFSWAP | -0.052112271 | 0.964523122 | 0.428364124 | 0.638154973 | 1568.748052 | 1513.288049 | ENSG00000061936 |
| RFWD3 | -0.05219283 | 0.964469265 | 0.459078359 | 0.66424263 | 2688.732619 | 2592.811918 | ENSG00000168411 |
| KCNK5 | -0.052268735 | 0.964418523 | 0.420259005 | 0.630538755 | 1813.311218 | 1748.461126 | ENSG00000164626 |
| NA | -0.052542828 | 0.964235313 | 0.939620236 | 0.972905887 | 10.39942395 | 10.00127084 | ENSG00000225022 |
| ASAP3 | -0.052643743 | 0.964167869 | 0.479032448 | 0.68145793 | 1816.691599 | 1751.694257 | ENSG00000088280 |
| SCAMP1 | -0.052665819 | 0.964153115 | 0.558883045 | 0.744323435 | 819.6251567 | 790.3305093 | ENSG00000085365 |
| NA | -0.052696333 | 0.964132723 | 0.919808364 | 0.964147051 | 17.81921229 | 17.16796744 | ENSG00000206120 |
| STK19 | -0.052835706 | 0.964039587 | 0.727379961 | 0.856608292 | 214.4908994 | 206.8858222 | ENSG00000204344 |
| SEC16A | -0.052860479 | 0.964023032 | 0.466068042 | 0.670237274 | 2086.307224 | 2010.819421 | ENSG00000148396 |
| SMARCC1 | -0.052919764 | 0.963983418 | 0.32643871 | 0.537799445 | 8265.630396 | 7967.315972 | ENSG00000173473 |
| NA | -0.052920424 | 0.963982977 | 0.893253157 | 0.952356228 | 36.2706324 | 34.83069321 | ENSG00000288075 |
| OTUD3 | -0.053019371 | 0.963916865 | 0.620829452 | 0.789780732 | 483.1567398 | 465.5269569 | ENSG00000169914 |
| NA | -0.053046839 | 0.963898513 | 0.910496528 | 0.960811396 | 27.69702525 | 26.76889821 | ENSG00000273253 |
| AOX1 | -0.053065047 | 0.963886348 | 0.722417811 | 0.853692532 | 243.1386004 | 234.3977246 | ENSG00000138356 |
| RAB5IF | -0.053077679 | 0.963877908 | 0.71136489 | 0.847591497 | 255.7986025 | 246.6543244 | ENSG00000101084 |
| COL4A2 | -0.053159507 | 0.963823239 | 0.364454042 | 0.576538329 | 3320.458442 | 3200.119168 | ENSG00000134871 |
| TRIM31 | -0.053230405 | 0.963775876 | 0.958506065 | 0.982874414 | 7.301462175 | 7.047728781 | ENSG00000204616 |
| KIF4A | -0.053445136 | 0.963632438 | 0.500752643 | 0.698998707 | 1182.649773 | 1139.243108 | ENSG00000090889 |
| ANAPC5 | -0.053470582 | 0.963615442 | 0.385681782 | 0.597653274 | 1961.472846 | 1889.83242 | ENSG00000089053 |
| RAB18 | -0.053512364 | 0.963587535 | 0.523127462 | 0.71744823 | 941.6592179 | 906.9565754 | ENSG00000099246 |
| TWSG1 | -0.053519448 | 0.963582803 | 0.654896488 | 0.813380102 | 592.8627172 | 571.1739502 | ENSG00000128791 |
| ZBTB20 | -0.053593044 | 0.963533649 | 0.94558202 | 0.976189572 | 7.603633391 | 7.339978361 | ENSG00000181722 |
| SBK1 | -0.05360165 | 0.963527902 | 0.411200301 | 0.622386607 | 4566.439243 | 4400.719789 | ENSG00000188322 |
| BRCC3 | -0.053645762 | 0.963498442 | 0.630809894 | 0.796625039 | 460.6456722 | 443.7426664 | ENSG00000185515 |
| RIC8A | -0.053684913 | 0.963472295 | 0.373002367 | 0.585013678 | 2001.758352 | 1928.614307 | ENSG00000177963 |
| AUP1 | -0.053886748 | 0.963337514 | 0.548495603 | 0.736043449 | 1008.891346 | 972.151634 | ENSG00000115307 |
| NAGK | -0.053930673 | 0.963308184 | 0.5309391 | 0.72296467 | 834.1301964 | 803.4633041 | ENSG00000124357 |
| HNRNPC | -0.053968345 | 0.96328303 | 0.224983008 | 0.423271978 | 22091.18711 | 21279.12031 | ENSG00000092199 |
| CD3EAP | -0.054007645 | 0.963256789 | 0.752634187 | 0.871085164 | 194.7195349 | 187.6328111 | ENSG00000117877 |
| MIR762HG | -0.054012872 | 0.9632533 | 0.94554916 | 0.976189572 | 7.256384241 | 7.00913753 | ENSG00000260083 |
| NA | -0.054101142 | 0.963194366 | 0.885112997 | 0.947622636 | 34.05042916 | 32.78501879 | ENSG00000213430 |
| NLE1 | -0.054127 | 0.963177102 | 0.443821054 | 0.651211433 | 1501.003314 | 1446.173011 | ENSG00000073536 |
| YES1 | -0.054156843 | 0.963157178 | 0.432240647 | 0.641583323 | 1725.877646 | 1662.342698 | ENSG00000176105 |
| WDYHV1 | -0.054210918 | 0.963121078 | 0.713741453 | 0.849017847 | 252.949726 | 243.3624692 | ENSG00000156795 |
| SEL1L | -0.054268883 | 0.963082382 | 0.632024357 | 0.797206785 | 669.5998054 | 644.463214 | ENSG00000071537 |
| TOPORS | -0.054316223 | 0.963050781 | 0.584405444 | 0.76319043 | 659.5905713 | 634.8287912 | ENSG00000197579 |
| ATPAF2 | -0.054346534 | 0.963030547 | 0.739188311 | 0.863587169 | 211.7152966 | 204.0533508 | ENSG00000171953 |
| SEC24D | -0.054425232 | 0.962978016 | 0.376620895 | 0.58878192 | 2120.244718 | 2041.705272 | ENSG00000150961 |

| | | | | | | | |
|---|---|---|---|---|---|---|---|
| ERC2 | -0.054439815 | 0.962968282 | 0.783239379 | 0.889784424 | 120.6059877 | 116.0657376 | ENSG00000187672 |
| NA | -0.054466629 | 0.962950385 | 0.935074928 | 0.970748723 | 10.64822923 | 10.21482487 | ENSG00000242588 |
| KBTBD4 | -0.054480136 | 0.962941369 | 0.866432621 | 0.936266565 | 49.54534993 | 47.8291107 | ENSG00000123444 |
| SMAD6 | -0.054483489 | 0.962939131 | 0.73073869 | 0.858155597 | 198.3489225 | 191.0095321 | ENSG00000137834 |
| ST3GAL2 | -0.05450413 | 0.962925354 | 0.476854623 | 0.679293925 | 1270.872473 | 1223.202177 | ENSG00000157350 |
| LDB1 | -0.054570555 | 0.96288102 | 0.347499206 | 0.55952193 | 2612.177901 | 2515.67524 | ENSG00000198728 |
| COBL | -0.054628083 | 0.962842625 | 0.528065131 | 0.720973828 | 904.4118506 | 870.6956806 | ENSG00000106078 |
| TSTD2 | -0.054707425 | 0.962789675 | 0.692761528 | 0.835855442 | 355.6658706 | 342.0063702 | ENSG00000136925 |
| WDR54 | -0.054778595 | 0.96274218 | 0.545468164 | 0.733722678 | 733.811182 | 706.2804401 | ENSG00000005448 |
| ZNF222 | -0.054908239 | 0.96265567 | 0.878248836 | 0.943389413 | 37.57247394 | 36.2274552 | ENSG00000159885 |
| NA | -0.054917045 | 0.962649794 | 0.855320289 | 0.929332239 | 55.00172341 | 52.98465675 | ENSG00000261578 |
| QPRT | -0.054931943 | 0.962639853 | 0.313498941 | 0.523792578 | 4132.356421 | 3978.457782 | ENSG00000103485 |
| GLA | -0.054955275 | 0.962624285 | 0.55505899 | 0.741479028 | 666.1670785 | 641.1652971 | ENSG00000102393 |
| UBXN7 | -0.055111254 | 0.962520215 | 0.533081862 | 0.724314769 | 1335.345113 | 1284.700495 | ENSG00000163960 |
| GIPC1 | -0.055143156 | 0.962498931 | 0.454291905 | 0.660251086 | 2435.150196 | 2344.206389 | ENSG00000123159 |
| NUP93 | -0.055183462 | 0.962472042 | 0.295850107 | 0.505988885 | 3735.439139 | 3594.713155 | ENSG00000102900 |
| APAF1 | -0.05526127 | 0.962420134 | 0.676429327 | 0.826926971 | 379.748435 | 365.2669783 | ENSG00000120868 |
| PLAC8 | -0.055267811 | 0.962415771 | 0.946030205 | 0.976272154 | 6.909135092 | 6.671770308 | ENSG00000145287 |
| RABGGTB | -0.055273934 | 0.962411686 | 0.530657422 | 0.72273658 | 959.4851201 | 923.9299915 | ENSG00000137955 |
| PER2 | -0.055390505 | 0.962333926 | 0.785776689 | 0.890559489 | 135.7334716 | 130.657351 | ENSG00000132326 |
| CYP27B1 | -0.055453562 | 0.962291865 | 0.865045821 | 0.935004026 | 45.2116759 | 43.56743356 | ENSG00000111012 |
| RPL35 | -0.055471237 | 0.962280076 | 0.401412861 | 0.612708101 | 18199.35959 | 17512.61156 | ENSG00000136942 |
| NRXN2 | -0.055498329 | 0.962262005 | 0.609430301 | 0.782399133 | 499.6183792 | 480.9282358 | ENSG00000110076 |
| COX5A | -0.055521959 | 0.962246245 | 0.321261613 | 0.531728795 | 3093.870487 | 2976.978406 | ENSG00000178741 |
| RTN2 | -0.05552474 | 0.96224439 | 0.701279184 | 0.84099452 | 287.6009637 | 276.9077506 | ENSG00000125744 |
| GIGYF1 | -0.055540878 | 0.962233626 | 0.405653923 | 0.616809527 | 2905.877418 | 2796.592629 | ENSG00000146830 |
| CRACR2B | -0.055557561 | 0.962222499 | 0.700508917 | 0.840477185 | 298.417631 | 287.3700339 | ENSG00000177685 |
| FBXO2 | -0.055576349 | 0.962209969 | 0.527566467 | 0.720657702 | 956.9919031 | 920.8588361 | ENSG00000116661 |
| CLASP2 | -0.055578631 | 0.962208447 | 0.516096118 | 0.71230543 | 1360.742913 | 1308.727694 | ENSG00000163539 |
| HSPA4 | -0.055592356 | 0.962199293 | 0.259730432 | 0.466654858 | 11797.27942 | 11350.98228 | ENSG00000170606 |
| GTPBP3 | -0.055613778 | 0.962185005 | 0.478251316 | 0.680667428 | 1197.489654 | 1152.725252 | ENSG00000130299 |
| DBN1 | -0.055674695 | 0.962144378 | 0.287112426 | 0.496231181 | 14167.2007 | 13631.80371 | ENSG00000113758 |
| NA | -0.055782591 | 0.962072424 | 0.913469848 | 0.961694619 | 17.54668591 | 16.85765174 | ENSG00000207425 |
| MOAP1 | -0.055855614 | 0.96202373 | 0.601659142 | 0.77664613 | 619.5648274 | 595.9259425 | ENSG00000165943 |
| AKIRIN1 | -0.055906663 | 0.96198969 | 0.336429601 | 0.548375022 | 4860.406778 | 4676.338223 | ENSG00000174574 |
| NELFA | -0.055915872 | 0.961983549 | 0.419883601 | 0.630376166 | 1659.939227 | 1597.447292 | ENSG00000185049 |
| BVES | -0.055936928 | 0.961969509 | 0.813523608 | 0.90530915 | 82.07688819 | 78.93476573 | ENSG00000112276 |
| PPP1R8 | -0.055976858 | 0.961942885 | 0.462811999 | 0.667551225 | 1048.158242 | 1008.306814 | ENSG00000117751 |
| SLC39A14 | -0.056003375 | 0.961925205 | 0.301883579 | 0.512145775 | 7145.073907 | 6872.215036 | ENSG00000104635 |
| NAT14 | -0.056006172 | 0.961923339 | 0.528383117 | 0.721043079 | 1163.338948 | 1119.56995 | ENSG00000090971 |
| RAB11FIP4 | -0.05602299 | 0.961912126 | 0.386274853 | 0.598424988 | 2183.824166 | 2100.562752 | ENSG00000131242 |
| YIPF6 | -0.056090898 | 0.961866849 | 0.555848835 | 0.742266897 | 653.4376669 | 628.1270589 | ENSG00000181704 |
| PTRH2 | -0.056152726 | 0.961825629 | 0.56896876 | 0.752125949 | 641.4274684 | 616.7089001 | ENSG00000141378 |
| FAM102B | -0.056156963 | 0.961822804 | 0.642276565 | 0.804443091 | 388.4453756 | 373.5915553 | ENSG00000162636 |
| ICE1 | -0.056189087 | 0.961801388 | 0.40863691 | 0.619748478 | 2048.293254 | 1969.593003 | ENSG00000164151 |
| YAE1 | -0.056278617 | 0.961741703 | 0.675950736 | 0.826662974 | 277.587612 | 266.9092223 | ENSG00000241127 |
| TMTC1 | -0.056305689 | 0.961723656 | 0.634961599 | 0.79989659 | 375.3554451 | 360.7957419 | ENSG00000133687 |
| RTN4 | -0.056310182 | 0.961720661 | 0.416510265 | 0.627456633 | 6313.057269 | 6071.741949 | ENSG00000115310 |
| RPA2 | -0.05634056 | 0.961700411 | 0.449430369 | 0.65582771 | 1175.707764 | 1130.566377 | ENSG00000117748 |
| TMEM50A | -0.056357485 | 0.961689128 | 0.40371577 | 0.614998476 | 1608.678607 | 1546.584576 | ENSG00000183726 |
| NA | -0.056390427 | 0.96166717 | 0.830776841 | 0.915015174 | 69.01095759 | 66.41966215 | ENSG00000182912 |
| ADAT3 | -0.056410915 | 0.961653513 | 0.757366696 | 0.87388465 | 144.6146412 | 139.2010625 | ENSG00000213638 |
| COA5 | -0.056419571 | 0.961647744 | 0.618949787 | 0.788624184 | 485.5578129 | 466.8063595 | ENSG00000183513 |
| FBXO31 | -0.05641972 | 0.961647644 | 0.597073355 | 0.77347684 | 529.7588863 | 509.3420466 | ENSG00000103264 |
| TSSC4 | -0.056458319 | 0.961621915 | 0.578233109 | 0.759450456 | 719.4781654 | 692.2599896 | ENSG00000184281 |
| TRMT112 | -0.056484678 | 0.961604346 | 0.338139086 | 0.550211669 | 4036.750745 | 3881.111423 | ENSG00000173113 |
| MED4 | -0.056538801 | 0.961568272 | 0.613955705 | 0.785269226 | 406.2853343 | 390.5869463 | ENSG00000136146 |
| ELP3 | -0.056562435 | 0.96155252 | 0.563891941 | 0.748513341 | 665.7744203 | 639.8046447 | ENSG00000134014 |
| ADK | -0.056633005 | 0.961505487 | 0.575347417 | 0.757595591 | 506.7777947 | 487.1134982 | ENSG00000156110 |
| SF1 | -0.056717173 | 0.961449393 | 0.291219301 | 0.50083567 | 6104.100765 | 5869.259115 | ENSG00000168066 |
| LTN1 | -0.056726438 | 0.961443219 | 0.609363825 | 0.782366975 | 570.0135793 | 547.7581265 | ENSG00000198862 |
| RHEBL1 | -0.056785782 | 0.961403672 | 0.787543883 | 0.891546762 | 117.5252129 | 112.7878811 | ENSG00000167550 |
| NA | -0.056792376 | 0.961399278 | 0.800909138 | 0.898333333 | 95.80411278 | 92.108924 | ENSG00000215158 |
| SDAD1 | -0.056821952 | 0.961379568 | 0.471566975 | 0.674832351 | 1183.799882 | 1138.200214 | ENSG00000198301 |
| MFN1 | -0.056853675 | 0.961358429 | 0.526738162 | 0.719946837 | 683.1102855 | 656.7525747 | ENSG00000171109 |
| GRIA1 | -0.056878376 | 0.961341969 | 0.894768205 | 0.953032171 | 26.99568733 | 25.90376183 | ENSG00000155511 |
| LHFPL4 | -0.056878705 | 0.96134175 | 0.533804837 | 0.724879869 | 785.5860431 | 755.1621653 | ENSG00000156959 |
| SLCO4C1 | -0.056893946 | 0.961331595 | 0.766878334 | 0.879325885 | 163.1734297 | 156.8123848 | ENSG00000173930 |
| ZNF525 | -0.056925878 | 0.961310317 | 0.561264254 | 0.746231726 | 581.2851958 | 558.6923949 | ENSG00000203326 |
| CRBN | -0.056942947 | 0.961298943 | 0.670361015 | 0.823292434 | 311.0242155 | 298.8727093 | ENSG00000113851 |
| PIK3CD-AS2 | -0.056990402 | 0.961267324 | 0.834624107 | 0.917141538 | 75.5413931 | 72.56690032 | ENSG00000231789 |
| HIGD1A | -0.057014952 | 0.961250966 | 0.755609539 | 0.872914862 | 208.0912479 | 200.3327362 | ENSG00000181061 |
| FAM83F | -0.057039675 | 0.961234494 | 0.512741574 | 0.709022818 | 876.4376222 | 842.7103433 | ENSG00000133477 |
| WDR82 | -0.057124397 | 0.961178047 | 0.336019727 | 0.548038047 | 3623.395373 | 3482.242143 | ENSG00000164091 |

| | | | | | | | |
|---|---|---|---|---|---|---|---|
| NELFB | -0.057159671 | 0.961154546 | 0.335133459 | 0.547159624 | 3272.684982 | 3146.203439 | ENSG00000188986 |
| DNAJC25 | -0.057203912 | 0.961125073 | 0.7767689 | 0.885665681 | 123.8483872 | 118.9558124 | ENSG00000059769 |
| FANCC | -0.057204254 | 0.961124845 | 0.649498924 | 0.809729811 | 314.700155 | 302.5098033 | ENSG00000158169 |
| GNB5 | -0.057245252 | 0.961097532 | 0.59554506 | 0.772286996 | 512.7673473 | 492.5483136 | ENSG00000069966 |
| ZNF765 | -0.0572774 | 0.961076116 | 0.685940037 | 0.831762616 | 245.0296829 | 235.6265017 | ENSG00000196417 |
| SIGIRR | -0.057362777 | 0.961019242 | 0.502528711 | 0.700542904 | 848.9804771 | 816.1359667 | ENSG00000185187 |
| NA | -0.0575643 | 0.960885011 | 0.881277078 | 0.945351753 | 31.43967886 | 30.21178612 | ENSG00000286358 |
| STAT5B | -0.057598926 | 0.960861949 | 0.397608864 | 0.609205812 | 1533.168333 | 1472.892441 | ENSG00000173757 |
| RSL1D1 | -0.05768358 | 0.96080557 | 0.290999603 | 0.500640202 | 4566.640271 | 4387.669471 | ENSG00000171490 |
| KCNS3 | -0.057745195 | 0.960764537 | 0.570318687 | 0.75319722 | 519.5887425 | 499.4172232 | ENSG00000170745 |
| MYDGF | -0.057756672 | 0.960756894 | 0.498559703 | 0.697377971 | 1969.509402 | 1891.775871 | ENSG00000074842 |
| FER1L4 | -0.057769021 | 0.96074867 | 0.868676875 | 0.937813209 | 46.16357297 | 44.36787312 | ENSG00000088340 |
| DUS1L | -0.057778093 | 0.960742629 | 0.437174551 | 0.645246932 | 1916.730104 | 1841.752776 | ENSG00000169718 |
| TEDC2 | -0.057876722 | 0.96067695 | 0.509917162 | 0.706513984 | 1045.22521 | 1004.790969 | ENSG00000162062 |
| ARSK | -0.05792899 | 0.960642146 | 0.646790096 | 0.808273621 | 376.4804821 | 361.6529114 | ENSG00000164291 |
| FBXO27 | -0.057940736 | 0.960634325 | 0.702383573 | 0.841924073 | 217.6303797 | 209.09161 | ENSG00000161243 |
| PAFAH1B3 | -0.057982625 | 0.960606433 | 0.580999862 | 0.761150628 | 559.2150927 | 537.4336684 | ENSG00000079462 |
| GPR153 | -0.057986729 | 0.960603701 | 0.650153642 | 0.810064757 | 356.826938 | 342.709481 | ENSG00000158292 |
| PTS | -0.058289062 | 0.960402416 | 0.655248585 | 0.813621508 | 339.6710006 | 326.0616424 | ENSG00000150787 |
| NA | -0.058318484 | 0.96038283 | 0.947890542 | 0.977070732 | 6.942339399 | 6.679242657 | ENSG00000259448 |
| TFG | -0.058333397 | 0.960372902 | 0.417376002 | 0.628359837 | 1338.364922 | 1285.67567 | ENSG00000114354 |
| POLR2D | -0.05839051 | 0.960334885 | 0.436858991 | 0.645089622 | 1409.467532 | 1353.6209 | ENSG00000144231 |
| PHF23 | -0.058391234 | 0.960334403 | 0.400536646 | 0.611935487 | 1509.286065 | 1449.660375 | ENSG00000040633 |
| ZNF449 | -0.058419767 | 0.960315409 | 0.762426159 | 0.877043188 | 132.182724 | 127.0638819 | ENSG00000173275 |
| PPCDC | -0.058544113 | 0.960232643 | 0.677262659 | 0.827303061 | 258.6415304 | 248.1988297 | ENSG00000138621 |
| FAM98C | -0.058581507 | 0.960207755 | 0.781709388 | 0.888776749 | 112.8576635 | 108.4274778 | ENSG00000130244 |
| NFIC | -0.058585701 | 0.960204963 | 0.813555816 | 0.90530915 | 83.3137695 | 80.15032211 | ENSG00000141905 |
| EVC | -0.058609136 | 0.960189367 | 0.650757075 | 0.810498975 | 318.8484839 | 306.0329951 | ENSG00000072840 |
| IFT43 | -0.058729124 | 0.960109511 | 0.714990199 | 0.849668508 | 210.4712702 | 202.1713103 | ENSG00000119650 |
| FGFR2 | -0.058759616 | 0.960089219 | 0.375734177 | 0.587736266 | 1627.061279 | 1562.184741 | ENSG00000066468 |
| TMEM167B | -0.058833292 | 0.960040191 | 0.679848364 | 0.828799713 | 254.4027925 | 244.2420558 | ENSG00000215717 |
| CUL4A | -0.058845113 | 0.960032324 | 0.315658824 | 0.52577323 | 2899.379038 | 2782.859651 | ENSG00000139842 |
| OXSR1 | -0.058870349 | 0.960015531 | 0.429370949 | 0.639232981 | 1553.993012 | 1491.460898 | ENSG00000172939 |
| CIDECP1 | -0.058920145 | 0.959982396 | 0.835580143 | 0.917415262 | 57.675349 | 55.43986756 | ENSG00000186162 |
| NA | -0.059118989 | 0.959850092 | 0.872090499 | 0.940089565 | 36.88270417 | 35.42550259 | ENSG00000228502 |
| VAMP3 | -0.059396468 | 0.959665498 | 0.538118158 | 0.727964146 | 603.7867791 | 579.663529 | ENSG00000049245 |
| TPT1 | -0.059412541 | 0.959654807 | 0.346519984 | 0.558650324 | 5331.15449 | 5115.882157 | ENSG00000133112 |
| ZNF93 | -0.059416555 | 0.959652137 | 0.539538411 | 0.729048576 | 634.535266 | 609.2443735 | ENSG00000184635 |
| SUV39H2 | -0.059423054 | 0.959647814 | 0.551133063 | 0.738218873 | 673.0349477 | 645.9636132 | ENSG00000152455 |
| PRR14 | -0.059446375 | 0.959632301 | 0.689471957 | 0.83398915 | 227.43675 | 218.415025 | ENSG00000156858 |
| CTDP1 | -0.059646353 | 0.959499292 | 0.419599608 | 0.630081242 | 1747.225489 | 1676.786096 | ENSG00000060069 |
| METAP2 | -0.059673125 | 0.959481487 | 0.303561864 | 0.51363963 | 3343.158004 | 3207.446856 | ENSG00000111142 |
| CHCHD5 | -0.059779103 | 0.959411007 | 0.667439087 | 0.821822412 | 281.4924483 | 270.0749209 | ENSG00000125611 |
| VPS51 | -0.059811338 | 0.959389571 | 0.36727244 | 0.57955871 | 2118.171238 | 2032.469561 | ENSG00000149823 |
| TBC1D8B | -0.059825527 | 0.959380135 | 0.703531187 | 0.842524014 | 211.6879899 | 203.1064701 | ENSG00000133138 |
| MAT2A | -0.059840453 | 0.95937021 | 0.691200946 | 0.834825394 | 4024.775923 | 3861.273881 | ENSG00000168906 |
| APEX2 | -0.059933068 | 0.959308624 | 0.526030945 | 0.719293069 | 645.1309934 | 618.8885823 | ENSG00000169188 |
| ANLN | -0.059951183 | 0.959296579 | 0.388273141 | 0.600387969 | 1452.390922 | 1393.026306 | ENSG00000011426 |
| GLB1L | -0.059962093 | 0.959289324 | 0.855142347 | 0.929192361 | 48.56319764 | 46.65973352 | ENSG00000163521 |
| TRAPPC9 | -0.059970073 | 0.959284018 | 0.6079154 | 0.781467763 | 480.3476534 | 460.6869079 | ENSG00000167632 |
| NA | -0.060027913 | 0.95924556 | 0.933205341 | 0.969789847 | 8.652499785 | 8.278964904 | ENSG00000261790 |
| ZNRF3 | -0.060100997 | 0.959196968 | 0.533619841 | 0.724837136 | 682.8113427 | 654.5752882 | ENSG00000183579 |
| TAC3 | -0.060103979 | 0.959194985 | 0.780087246 | 0.887787612 | 114.7857891 | 110.0861055 | ENSG00000166863 |
| SMKR1 | -0.060112706 | 0.959189183 | 0.889222813 | 0.950456913 | 24.54478166 | 23.50889691 | ENSG00000240204 |
| CA8 | -0.060113816 | 0.959188445 | 0.762328847 | 0.876987748 | 130.3643055 | 124.8990288 | ENSG00000178538 |
| PSME4 | -0.060142025 | 0.95916969 | 0.366681007 | 0.579187924 | 2889.47636 | 2770.952785 | ENSG00000068878 |
| COPE | -0.060144824 | 0.959167829 | 0.49177365 | 0.691162118 | 803.4621285 | 770.9400987 | ENSG00000105669 |
| POLR1D | -0.060147737 | 0.959165892 | 0.344989079 | 0.556999337 | 2284.215383 | 2190.976805 | ENSG00000186184 |
| HDAC2 | -0.060185892 | 0.959140526 | 0.265349397 | 0.472940557 | 7379.060893 | 7077.235484 | ENSG00000196591 |
| BLOC1S6 | -0.060233511 | 0.959108868 | 0.68565853 | 0.831762616 | 440.3153219 | 422.4003966 | ENSG00000104164 |
| CDKAL1 | -0.060261229 | 0.959090441 | 0.606151967 | 0.780099489 | 385.5226659 | 369.5216049 | ENSG00000145996 |
| ELF2 | -0.060285404 | 0.95907437 | 0.560609784 | 0.745729081 | 617.7669021 | 591.9971434 | ENSG00000109381 |
| PLEKHF1 | -0.060292709 | 0.959069514 | 0.603592528 | 0.778237515 | 426.3790311 | 409.1953959 | ENSG00000166289 |
| TCF20 | -0.06032472 | 0.959048234 | 0.346782608 | 0.558942677 | 2254.548714 | 2161.615678 | ENSG00000100207 |
| NCR3LG1 | -0.060362624 | 0.959023037 | 0.861778107 | 0.932860229 | 41.96602246 | 40.1785331 | ENSG00000188211 |
| ZNF324 | -0.060412612 | 0.958989808 | 0.772010561 | 0.882599617 | 112.0916964 | 107.6429289 | ENSG00000083812 |
| COG1 | -0.0604199 | 0.958984964 | 0.678538367 | 0.828057969 | 278.2857215 | 267.1854133 | ENSG00000166685 |
| NECTIN1 | -0.060499657 | 0.958931949 | 0.287821889 | 0.497120576 | 3574.568292 | 3428.004149 | ENSG00000110400 |
| TFDP1 | -0.060561215 | 0.958891034 | 0.331450827 | 0.542977869 | 2635.576926 | 2526.962581 | ENSG00000198176 |
| ADH5 | -0.060586915 | 0.958873952 | 0.256559916 | 0.463427453 | 6077.750187 | 5827.004619 | ENSG00000197894 |
| NA | -0.060630215 | 0.958845174 | 0.91325593 | 0.961597187 | 15.27962529 | 14.64789186 | ENSG00000273716 |
| CCDC112 | -0.060684228 | 0.958809276 | 0.631435558 | 0.797092269 | 309.9038803 | 297.2379164 | ENSG00000164221 |
| SHMT1 | -0.06076605 | 0.9587549 | 0.475821768 | 0.678283068 | 991.6407515 | 950.4151405 | ENSG00000176974 |
| FOXK2 | -0.060797163 | 0.958734223 | 0.39759511 | 0.609205812 | 1531.998269 | 1468.622071 | ENSG00000141568 |

| | | | | | | | |
|---|---|---|---|---|---|---|---|
| NUP188 | -0.060811556 | 0.958724658 | 0.283181223 | 0.492186485 | 3211.56971 | 3079.347048 | ENSG00000095319 |
| KNTC1 | -0.060909428 | 0.958659621 | 0.502082404 | 0.700181291 | 1100.560401 | 1054.905836 | ENSG00000184445 |
| DDX17 | -0.060916983 | 0.9586546 | 0.294495226 | 0.504448284 | 7367.403788 | 7062.258862 | ENSG00000100201 |
| TOMM22 | -0.0609323 | 0.958644423 | 0.323492227 | 0.53439037 | 2069.477003 | 1983.923245 | ENSG00000100216 |
| NPR1 | -0.060973207 | 0.958617241 | 0.525263917 | 0.718812252 | 594.3499312 | 569.7125047 | ENSG00000169418 |
| FAM161B | -0.060992741 | 0.958604262 | 0.938335074 | 0.972315933 | 7.287224285 | 6.989841904 | ENSG00000156050 |
| SCAF8 | -0.061085233 | 0.958542807 | 0.51193372 | 0.708476352 | 875.4788337 | 838.7985784 | ENSG00000213079 |
| TMEM254 | -0.061221861 | 0.958452034 | 0.725001158 | 0.85505213 | 175.1504761 | 167.8172664 | ENSG00000133678 |
| POFUT2 | -0.061309954 | 0.958393511 | 0.478985396 | 0.68145793 | 880.9247558 | 843.8947901 | ENSG00000186866 |
| TUBGCP6 | -0.061364152 | 0.958357508 | 0.507756804 | 0.704758477 | 1068.80111 | 1024.730676 | ENSG00000128159 |
| RAB39B | -0.061414352 | 0.958324161 | 0.661121784 | 0.817849134 | 283.6015222 | 271.7614306 | ENSG00000155961 |
| PLIN2 | -0.06142676 | 0.95831592 | 0.510671798 | 0.707210126 | 730.4297622 | 699.6768012 | ENSG00000147872 |
| TAMM41 | -0.061465488 | 0.958290195 | 0.699606892 | 0.839970842 | 228.3521888 | 218.8575722 | ENSG00000144559 |
| COQ8B | -0.061480308 | 0.95828035 | 0.489053947 | 0.689237596 | 960.073315 | 919.8426038 | ENSG00000123815 |
| CCDC117 | -0.061505175 | 0.958263834 | 0.446770888 | 0.653682173 | 1084.058411 | 1038.336426 | ENSG00000159873 |
| KLK4 | -0.061536573 | 0.958242979 | 0.91691439 | 0.963119654 | 13.18425676 | 12.66568475 | ENSG00000167749 |
| GMPPB | -0.061550716 | 0.958233585 | 0.569326575 | 0.752342899 | 603.6672833 | 578.8118242 | ENSG00000173540 |
| GID8 | -0.06157153 | 0.958219761 | 0.304818394 | 0.514984378 | 2486.857813 | 2382.745321 | ENSG00000101193 |
| NA | -0.061719286 | 0.958121628 | 0.943250471 | 0.974964898 | 6.620006557 | 6.35710368 | ENSG00000260751 |
| C7orf26 | -0.061749255 | 0.958101725 | 0.528370857 | 0.721043079 | 771.9005895 | 739.9818831 | ENSG00000146576 |
| ETAA1 | -0.061942442 | 0.957973437 | 0.572863194 | 0.75529058 | 463.8364851 | 444.5641256 | ENSG00000143971 |
| STK11IP | -0.061971054 | 0.957954438 | 0.504652213 | 0.702052084 | 620.4085698 | 594.4202893 | ENSG00000144589 |
| RPL5 | -0.061983446 | 0.95794621 | 0.177268121 | 0.358849658 | 19991.56296 | 19150.14136 | ENSG00000122406 |
| MSN | -0.062021145 | 0.957921179 | 0.219998574 | 0.416675667 | 7931.426703 | 7596.796986 | ENSG00000147065 |
| BRMS1 | -0.062071437 | 0.957887786 | 0.461822498 | 0.666680774 | 1030.053103 | 986.2890839 | ENSG00000174744 |
| PAPSS1 | -0.062124648 | 0.957852457 | 0.30329284 | 0.513486816 | 2649.565128 | 2537.444174 | ENSG00000138801 |
| PPP4R3B | -0.062187896 | 0.957810465 | 0.438328247 | 0.646553851 | 1204.613117 | 1153.428494 | ENSG00000275052 |
| CACNG8 | -0.062286898 | 0.95774474 | 0.679776426 | 0.828765513 | 237.6460741 | 227.5653691 | ENSG00000142408 |
| NA | -0.062307864 | 0.957730822 | 0.94180884 | 0.974316559 | 6.258519517 | 6.015385531 | ENSG00000199476 |
| ANKRD11 | -0.062341505 | 0.957708489 | 0.269461263 | 0.477415889 | 3906.185238 | 3740.737699 | ENSG00000167522 |
| COMMD2 | -0.062418339 | 0.957657485 | 0.499093986 | 0.697608609 | 742.0378705 | 710.7655073 | ENSG00000114744 |
| BRIP1 | -0.062422604 | 0.957654655 | 0.617976158 | 0.787824768 | 394.3240265 | 377.5464373 | ENSG00000136492 |
| MED21 | -0.062483413 | 0.95761429 | 0.546725638 | 0.734660145 | 495.7589001 | 474.8856308 | ENSG00000152944 |
| VTA1 | -0.062509123 | 0.957597225 | 0.438544564 | 0.6466204 | 1105.964323 | 1059.195359 | ENSG00000009844 |
| CCDC40 | -0.062518321 | 0.95759112 | 0.701842717 | 0.841401159 | 194.8559639 | 186.7540978 | ENSG00000141519 |
| TCF7L1 | -0.062596912 | 0.957538957 | 0.335654576 | 0.547537071 | 3756.739065 | 3598.00145 | ENSG00000152284 |
| PARP10 | -0.06262494 | 0.957520354 | 0.794397182 | 0.894861791 | 85.44145261 | 81.85333242 | ENSG00000178685 |
| ING2 | -0.062651953 | 0.957502426 | 0.63972442 | 0.802577272 | 291.1692298 | 278.69655 | ENSG00000168556 |
| GPR63 | -0.06266693 | 0.957492485 | 0.652274306 | 0.811636045 | 287.8251843 | 275.6503862 | ENSG00000112218 |
| RASD2 | -0.062676295 | 0.95748627 | 0.72593143 | 0.855720453 | 172.8445406 | 165.310128 | ENSG00000100302 |
| PRKAR1A | -0.06271102 | 0.957463224 | 0.491385475 | 0.690822206 | 1444.692429 | 1383.260571 | ENSG00000108946 |
| PPP2R5C | -0.062726114 | 0.957453207 | 0.433426378 | 0.642497135 | 1203.442722 | 1152.291182 | ENSG00000078304 |
| MAP3K4 | -0.062734121 | 0.957447893 | 0.373262889 | 0.585276457 | 1344.176528 | 1286.85359 | ENSG00000085511 |
| TTC28 | -0.062749393 | 0.957437758 | 0.607484127 | 0.781281513 | 379.4622294 | 362.9859226 | ENSG00000100154 |
| SNX5 | -0.062842755 | 0.957375801 | 0.345517903 | 0.557510732 | 1910.422616 | 1828.71115 | ENSG00000089006 |
| ATXN2 | -0.062884011 | 0.957348424 | 0.467471812 | 0.671142252 | 954.0326121 | 913.3316499 | ENSG00000204842 |
| CKAP2L | -0.062886302 | 0.957346903 | 0.665028385 | 0.820639117 | 235.8988706 | 225.7806145 | ENSG00000169607 |
| EIF5AL1 | -0.062893938 | 0.957341836 | 0.869944905 | 0.938580298 | 37.47547846 | 35.85963149 | ENSG00000253626 |
| ZBED3-AS1 | -0.062918927 | 0.957325254 | 0.890617603 | 0.95121103 | 21.49662634 | 20.58581134 | ENSG00000250802 |
| SERTAD2 | -0.062945799 | 0.957307423 | 0.695715656 | 0.837492999 | 181.2002081 | 173.4716381 | ENSG00000179833 |
| SNX32 | -0.063028298 | 0.957252682 | 0.892082727 | 0.951914471 | 23.57183324 | 22.53444054 | ENSG00000172803 |
| TSEN54 | -0.063047841 | 0.957239714 | 0.422311165 | 0.632462042 | 1168.137614 | 1118.417818 | ENSG00000182173 |
| CYBC1 | -0.063079023 | 0.957219026 | 0.422436797 | 0.632549866 | 978.1669377 | 936.4312213 | ENSG00000178927 |
| SRRM2 | -0.063107102 | 0.957200395 | 0.259283589 | 0.466251245 | 13012.76542 | 12455.79009 | ENSG00000167978 |
| MIR25 | -0.063128854 | 0.957185963 | 0.892277859 | 0.951961322 | 24.24972939 | 23.17152969 | ENSG00000207547 |
| PGM1 | -0.063253816 | 0.957103059 | 0.464132849 | 0.668593683 | 1330.960413 | 1273.138412 | ENSG00000079739 |
| KIAA0586 | -0.063258671 | 0.957099838 | 0.569024392 | 0.752125949 | 433.6707313 | 414.9588768 | ENSG00000100578 |
| SPEG | -0.063274546 | 0.957089306 | 0.534316921 | 0.725262349 | 1181.003987 | 1130.827083 | ENSG00000072195 |
| CYB561D2 | -0.063300989 | 0.957071764 | 0.789250232 | 0.891980289 | 85.94854638 | 82.27325167 | ENSG00000114395 |
| TBP | -0.063316646 | 0.957061377 | 0.554722474 | 0.741242673 | 512.5921213 | 490.792779 | ENSG00000112592 |
| CTDSP2 | -0.063331407 | 0.957051585 | 0.248795492 | 0.453807341 | 3438.88544 | 3290.947407 | ENSG00000175215 |
| SLC38A1 | -0.063347654 | 0.957040807 | 0.34982551 | 0.561964508 | 6824.676227 | 6531.220118 | ENSG00000111371 |
| HSF1 | -0.063351179 | 0.957038469 | 0.324531571 | 0.53535803 | 2206.714316 | 2112.22444 | ENSG00000185122 |
| HIBCH | -0.06348179 | 0.95695183 | 0.485294738 | 0.686295576 | 681.6429549 | 652.0347305 | ENSG00000198130 |
| PGBD1 | -0.063484782 | 0.956949845 | 0.721112106 | 0.853003578 | 199.7584124 | 191.1018475 | ENSG00000137338 |
| DGKQ | -0.063511996 | 0.956931794 | 0.586111795 | 0.764723322 | 620.3596792 | 593.9689379 | ENSG00000145214 |
| HEBP1 | -0.063527755 | 0.956921341 | 0.505187153 | 0.702537509 | 729.8858524 | 698.424747 | ENSG00000013583 |
| GSKIP | -0.063716304 | 0.956796287 | 0.521723445 | 0.716459224 | 708.2611367 | 677.6863041 | ENSG00000100744 |
| JAG1 | -0.063800398 | 0.956740518 | 0.542364378 | 0.731401834 | 637.1734727 | 609.1385834 | ENSG00000101384 |
| PLEKHA7 | -0.063812597 | 0.956732427 | 0.576699536 | 0.758688313 | 453.5968547 | 433.6580132 | ENSG00000166689 |
| ZNF117 | -0.063817932 | 0.956728889 | 0.743284686 | 0.866407964 | 150.1854764 | 143.5964842 | ENSG00000152926 |
| GDI2 | -0.063818351 | 0.956728612 | 0.215346013 | 0.410457151 | 9078.186723 | 8684.625658 | ENSG00000057608 |
| CAMSAP1 | -0.063909668 | 0.956668056 | 0.335395069 | 0.547444761 | 1795.568306 | 1717.713234 | ENSG00000130559 |
| PLA2G15 | -0.063942707 | 0.956646148 | 0.709736484 | 0.846600146 | 186.3763515 | 178.5149536 | ENSG00000103066 |

| | | | | | | | |
|---|---|---|---|---|---|---|---|
| SNORD100 | -0.064029788 | 0.956588406 | 0.909383696 | 0.96047171 | 14.96202097 | 14.30087683 | ENSG00000221500 |
| SLC46A3 | -0.064057993 | 0.956569705 | 0.944064187 | 0.975469288 | 5.860268697 | 5.628549739 | ENSG00000139508 |
| SMIM30 | -0.064079107 | 0.956555706 | 0.610627258 | 0.783380531 | 418.8042045 | 400.8943499 | ENSG00000214194 |
| NFKBIB | -0.064105021 | 0.956538525 | 0.44981082 | 0.656111635 | 1031.472939 | 986.9068478 | ENSG00000104825 |
| KLHL23 | -0.064137207 | 0.956517184 | 0.418693601 | 0.629340071 | 1268.929348 | 1214.180512 | ENSG00000213160 |
| RFC3 | -0.064178034 | 0.956490116 | 0.413816678 | 0.624944259 | 1060.063158 | 1013.550891 | ENSG00000133119 |
| RPL35A | -0.064231085 | 0.956454945 | 0.237285305 | 0.439326763 | 9897.26581 | 9466.07971 | ENSG00000182899 |
| USP44 | -0.064243423 | 0.956446765 | 0.343442493 | 0.555395742 | 4184.992335 | 4002.457169 | ENSG00000136014 |
| XKR8 | -0.064277887 | 0.956423917 | 0.584152567 | 0.763106094 | 414.4186391 | 396.5458424 | ENSG00000158156 |
| GSR | -0.064293031 | 0.956413877 | 0.30245549 | 0.512522914 | 2078.85718 | 1988.331552 | ENSG00000104687 |
| RPL4 | -0.064346547 | 0.956378401 | 0.155671565 | 0.327451271 | 33362.01191 | 31905.74942 | ENSG00000174444 |
| ZSCAN21 | -0.06436006 | 0.956369443 | 0.712121552 | 0.847945493 | 224.4324257 | 214.6038619 | ENSG00000166529 |
| HJURP | -0.064453928 | 0.956307219 | 0.363473033 | 0.575516747 | 1437.84098 | 1374.540252 | ENSG00000123485 |
| UBXN2B | -0.064471724 | 0.956295423 | 0.540699422 | 0.730042795 | 600.8136782 | 574.5014862 | ENSG00000215114 |
| PITPNB | -0.064482399 | 0.956288347 | 0.440926688 | 0.648613531 | 1839.806648 | 1759.568985 | ENSG00000180957 |
| POLR2M | -0.064484151 | 0.956287186 | 0.763725784 | 0.877575992 | 200.3693145 | 191.7072803 | ENSG00000255529 |
| PKHD1 | -0.064598845 | 0.956211164 | 0.932721042 | 0.969708285 | 8.019681573 | 7.651523566 | ENSG00000170927 |
| PNPLA8 | -0.064624655 | 0.956194057 | 0.677902197 | 0.827602494 | 204.0374991 | 195.0029624 | ENSG00000135241 |
| BMP8A | -0.064628165 | 0.956191731 | 0.877725477 | 0.943087245 | 26.77032382 | 25.62891596 | ENSG00000183682 |
| WWP1 | -0.064639399 | 0.956184285 | 0.419974203 | 0.630411957 | 1134.816083 | 1085.15909 | ENSG00000123124 |
| PCMT1 | -0.064641744 | 0.956182731 | 0.378936829 | 0.591374433 | 1444.719911 | 1381.495454 | ENSG00000120265 |
| KCNMB3 | -0.064705596 | 0.956140413 | 0.908617299 | 0.960165884 | 16.60457753 | 15.83590226 | ENSG00000171121 |
| ANO5 | -0.064770903 | 0.956097132 | 0.669778664 | 0.82299482 | 236.2226518 | 225.6959596 | ENSG00000171714 |
| PAK6 | -0.06477486 | 0.95609451 | 0.668285713 | 0.822110685 | 209.0327465 | 199.7871118 | ENSG00000137843 |
| C16orf70 | -0.064841804 | 0.956050146 | 0.600937411 | 0.776307564 | 372.102419 | 356.0000957 | ENSG00000125149 |
| MEMO1 | -0.064946158 | 0.955980995 | 0.757207635 | 0.873807894 | 119.6276998 | 114.4812436 | ENSG00000162959 |
| DAZAP1 | -0.065035906 | 0.955921526 | 0.175814258 | 0.356479736 | 8935.45295 | 8541.978658 | ENSG00000071626 |
| RCE1 | -0.065090663 | 0.955885245 | 0.482635377 | 0.683859246 | 861.1899537 | 823.2554655 | ENSG00000173653 |
| MADD | -0.065130221 | 0.955859036 | 0.358276532 | 0.570123925 | 1417.223691 | 1354.660264 | ENSG00000110514 |
| DDX24 | -0.065313874 | 0.955737364 | 0.241886199 | 0.445443284 | 3365.084306 | 3215.786062 | ENSG00000089737 |
| P3H3 | -0.065358102 | 0.955708065 | 0.411026219 | 0.622172984 | 1160.62719 | 1109.43692 | ENSG00000110811 |
| ZMYM6 | -0.065429711 | 0.955660629 | 0.821471847 | 0.909334444 | 65.57050134 | 62.73501182 | ENSG00000163867 |
| PTCD3 | -0.065555765 | 0.955577132 | 0.281813878 | 0.490709708 | 2326.134601 | 2222.233499 | ENSG00000132300 |
| HDAC4 | -0.065565625 | 0.955570602 | 0.594773985 | 0.771714648 | 422.3096739 | 403.606624 | ENSG00000068024 |
| C3orf52 | -0.065586217 | 0.955556963 | 0.615929154 | 0.786518498 | 292.2818613 | 279.2011241 | ENSG00000114529 |
| YJEFN3 | -0.065642935 | 0.955519397 | 0.798334181 | 0.897150218 | 113.0118637 | 108.0536221 | ENSG00000250067 |
| ERCC5 | -0.065753606 | 0.955446101 | 0.841736204 | 0.92140286 | 44.39501263 | 42.50056644 | ENSG00000134899 |
| ZFP64 | -0.065903341 | 0.955346941 | 0.512491996 | 0.708885325 | 624.6381032 | 596.9670805 | ENSG00000020256 |
| ZNF71 | -0.065911196 | 0.95534174 | 0.63545406 | 0.800090001 | 280.6138257 | 268.0434859 | ENSG00000197951 |
| BAG6 | -0.065982298 | 0.955294658 | 0.24376591 | 0.448050818 | 6903.682544 | 6595.71846 | ENSG00000204463 |
| ERCC6 | -0.066019512 | 0.955270016 | 0.686396959 | 0.831903869 | 198.5304033 | 189.4691668 | ENSG00000225830 |
| MAST1 | -0.066024342 | 0.955266819 | 0.583925194 | 0.763005183 | 375.9423213 | 359.226855 | ENSG00000105613 |
| ARIH2 | -0.066068941 | 0.955237288 | 0.334055282 | 0.546107514 | 2033.309447 | 1942.488527 | ENSG00000177479 |
| PDIA5 | -0.066158316 | 0.955178113 | 0.468726758 | 0.67185333 | 719.6124583 | 687.2024347 | ENSG00000065485 |
| LUC7L3 | -0.066173125 | 0.955168308 | 0.42769552 | 0.637542812 | 3065.216076 | 2928.304787 | ENSG00000108848 |
| IRGQ | -0.06621167 | 0.955142789 | 0.578039251 | 0.759450456 | 422.1586236 | 403.0331701 | ENSG00000167378 |
| AUTS2 | -0.06625248 | 0.955115771 | 0.307450257 | 0.517738517 | 1920.502654 | 1834.836372 | ENSG00000158321 |
| L3MBTL4 | -0.066269755 | 0.955104335 | 0.857469496 | 0.93039751 | 35.88158545 | 34.25054498 | ENSG00000154655 |
| NAA50 | -0.066362999 | 0.955042606 | 0.272568878 | 0.481206251 | 4486.826472 | 4285.478144 | ENSG00000121579 |
| WAPL | -0.066421434 | 0.955003924 | 0.340404626 | 0.551995651 | 1788.81154 | 1707.907356 | ENSG00000062650 |
| CKAP5 | -0.066432032 | 0.954996909 | 0.246822798 | 0.45171514 | 4589.896511 | 4382.534328 | ENSG00000175216 |
| JMJD6 | -0.066568405 | 0.954906641 | 0.544438264 | 0.732785733 | 549.5487108 | 524.9840396 | ENSG00000070495 |
| CKS2 | -0.066624076 | 0.954869793 | 0.665410988 | 0.82085806 | 3454.237509 | 3298.162097 | ENSG00000123975 |
| ANKRD37 | -0.066681239 | 0.954831959 | 0.90849831 | 0.960165884 | 13.95111357 | 13.3189481 | ENSG00000186352 |
| NA | -0.066690132 | 0.954826074 | 0.684269248 | 0.831119741 | 198.584659 | 189.5157975 | ENSG00000248641 |
| CDK6 | -0.066694694 | 0.954823054 | 0.625771581 | 0.793240456 | 355.8177577 | 339.7625379 | ENSG00000105810 |
| UBASH3B | -0.066767281 | 0.954775015 | 0.61615224 | 0.786750173 | 283.1184027 | 270.3024737 | ENSG00000154127 |
| HOOK1 | -0.066789182 | 0.954760521 | 0.515867244 | 0.71214568 | 692.9557845 | 661.2443886 | ENSG00000134709 |
| TMED9 | -0.066856069 | 0.954716257 | 0.231651638 | 0.432316371 | 4080.84715 | 3896.592883 | ENSG00000184840 |
| ZNF720 | -0.066969258 | 0.954641356 | 0.658343249 | 0.815801329 | 290.5416988 | 276.9913212 | ENSG00000197302 |
| SRCAP | -0.067002589 | 0.954619301 | 0.831643147 | 0.915296977 | 50.15978655 | 47.84746037 | ENSG00000080603 |
| ZNF76 | -0.067088175 | 0.954562671 | 0.564807144 | 0.749003799 | 390.3993078 | 372.5584043 | ENSG00000065029 |
| SLC34A3 | -0.067164274 | 0.954512321 | 0.934722802 | 0.970489863 | 6.987417333 | 6.66741938 | ENSG00000198569 |
| DIAPH1 | -0.067313269 | 0.954413749 | 0.19704814 | 0.38658364 | 4175.251336 | 3984.732943 | ENSG00000131504 |
| RNF103 | -0.067313299 | 0.954413729 | 0.789014908 | 0.891843683 | 74.82050395 | 71.41764913 | ENSG00000239305 |
| ACTL8 | -0.067321933 | 0.954408017 | 0.847194986 | 0.924172673 | 40.20458107 | 38.38434154 | ENSG00000117148 |
| SERF1A | -0.067341629 | 0.954394987 | 0.907886561 | 0.960165884 | 14.28532004 | 13.62273741 | ENSG00000205572 |
| EBLN3P | -0.067375009 | 0.954372906 | 0.433285598 | 0.642497135 | 830.9397139 | 793.3435372 | ENSG00000281649 |
| UQCRH | -0.067446818 | 0.954325403 | 0.244866804 | 0.449070278 | 4404.323213 | 4202.81821 | ENSG00000173660 |
| PET100 | -0.067518121 | 0.954278239 | 0.908686366 | 0.960165884 | 12.86784766 | 12.29596912 | ENSG00000229833 |
| SF3B3 | -0.067533034 | 0.954268374 | 0.139178293 | 0.302913686 | 7866.909138 | 7506.971297 | ENSG00000189091 |
| NLN | -0.067584284 | 0.954234476 | 0.230390636 | 0.430358771 | 3821.243376 | 3646.378668 | ENSG00000123213 |
| RAB4B | -0.067662791 | 0.954182551 | 0.938396357 | 0.972326084 | 5.876870851 | 5.626374276 | ENSG00000167578 |
| FAM207A | -0.067768137 | 0.954112879 | 0.476993544 | 0.67944057 | 1311.024575 | 1251.363107 | ENSG00000160256 |

| | | | | | | | |
|---|---|---|---|---|---|---|---|
| MIR589 | -0.067822223 | 0.95407711 | 0.937376153 | 0.971904957 | 5.904151422 | 5.652196292 | ENSG00000207973 |
| MYL12B | -0.067828222 | 0.954073143 | 0.329774539 | 0.541217709 | 2537.947244 | 2420.568539 | ENSG00000118680 |
| CCDC122 | -0.067834368 | 0.954069078 | 0.849893867 | 0.925566327 | 43.29898337 | 41.26543089 | ENSG00000151773 |
| DNA2 | -0.067845984 | 0.954061396 | 0.482228975 | 0.68364944 | 815.4691764 | 778.2686193 | ENSG00000138346 |
| PIGO | -0.067850022 | 0.954058726 | 0.498469176 | 0.697317165 | 587.7954436 | 560.7397206 | ENSG00000165282 |
| ATP5MF | -0.067968613 | 0.953980305 | 0.615913282 | 0.786518498 | 326.2047865 | 311.0908765 | ENSG00000241468 |
| SCAF4 | -0.068048961 | 0.953927177 | 0.366361041 | 0.578876192 | 1569.48959 | 1497.482065 | ENSG00000156304 |
| AARS2 | -0.068091957 | 0.953898747 | 0.413364027 | 0.624700612 | 872.8737796 | 832.8229113 | ENSG00000124608 |
| ZC3H13 | -0.068117838 | 0.953881635 | 0.357958798 | 0.569904458 | 2346.396736 | 2237.536899 | ENSG00000123200 |
| ANKRD6 | -0.068134091 | 0.953870889 | 0.724351025 | 0.854693478 | 168.106692 | 160.2797402 | ENSG00000135299 |
| TMEM218 | -0.068300283 | 0.953761014 | 0.590744201 | 0.768598294 | 343.8232461 | 327.9360425 | ENSG00000150433 |
| ADIPOR2 | -0.068313119 | 0.953752528 | 0.317465241 | 0.527552069 | 1600.260316 | 1526.304509 | ENSG00000006831 |
| TMED8 | -0.068341539 | 0.95373374 | 0.543595413 | 0.732315108 | 444.6567003 | 423.8673429 | ENSG00000100580 |
| NA | -0.068378917 | 0.95370903 | 0.929567421 | 0.968057669 | 7.310945384 | 6.95966896 | ENSG00000260196 |
| SMAGP | -0.068424199 | 0.953679097 | 0.761304088 | 0.876553329 | 98.9009055 | 94.2356374 | ENSG00000170545 |
| SNX25 | -0.068505753 | 0.953625188 | 0.595576036 | 0.772286996 | 333.7079167 | 318.3888586 | ENSG00000109762 |
| CLNS1A | -0.068631718 | 0.953541929 | 0.229265995 | 0.429064346 | 2399.96024 | 2288.428447 | ENSG00000074201 |
| TPRG1L | -0.068644668 | 0.953533369 | 0.438585884 | 0.646630839 | 727.9993492 | 694.2229737 | ENSG00000158109 |
| ULBP3 | -0.068714537 | 0.953487191 | 0.785390428 | 0.890433415 | 77.53417346 | 73.98246349 | ENSG00000131019 |
| RPL30 | -0.068717998 | 0.953484904 | 0.332104281 | 0.543859638 | 2933.307678 | 2796.660286 | ENSG00000156482 |
| SPPL2A | -0.068821294 | 0.953416637 | 0.703603439 | 0.842524014 | 285.8101562 | 272.6727758 | ENSG00000138600 |
| DDX10 | -0.068851671 | 0.953396562 | 0.386231085 | 0.598406271 | 1105.668914 | 1053.968254 | ENSG00000178105 |
| TMEM143 | -0.068986928 | 0.953307183 | 0.648415093 | 0.809147384 | 251.8830622 | 239.8425667 | ENSG00000161558 |
| FAF2 | -0.069067284 | 0.953254087 | 0.380154984 | 0.592537433 | 1265.924414 | 1206.685182 | ENSG00000113194 |
| NA | -0.069108148 | 0.953227086 | 0.886102312 | 0.948401451 | 19.8410271 | 18.91609539 | ENSG00000231993 |
| BRD1 | -0.069114541 | 0.953222862 | 0.408556251 | 0.619725687 | 1256.87547 | 1197.8119 | ENSG00000100425 |
| TMEM140 | -0.069254531 | 0.953130372 | 0.694908088 | 0.837250559 | 173.0082496 | 165.0063387 | ENSG00000146859 |
| ATP5F1B | -0.069257047 | 0.95312871 | 0.107928285 | 0.252662163 | 20288.21096 | 19336.49882 | ENSG00000110955 |
| RTCA | -0.069269866 | 0.953120241 | 0.422919032 | 0.633121358 | 768.6406726 | 732.6658122 | ENSG00000137996 |
| ZNHIT3 | -0.069273681 | 0.953117721 | 0.473151398 | 0.676314272 | 582.5027791 | 555.241919 | ENSG00000273611 |
| NA | -0.069304551 | 0.953097326 | 0.93758846 | 0.972022292 | 6.207517847 | 5.923920741 | ENSG00000230897 |
| PDIA3 | -0.069549659 | 0.952935413 | 0.178586055 | 0.360821402 | 9236.125196 | 8801.23946 | ENSG00000167004 |
| ERCC3 | -0.069577676 | 0.952916907 | 0.285068952 | 0.49430972 | 1671.095974 | 1592.656376 | ENSG00000163161 |
| SLC8A1 | -0.069587405 | 0.952910481 | 0.732770438 | 0.859594158 | 124.0408257 | 117.9950712 | ENSG00000183023 |
| INTS11 | -0.069608192 | 0.952896751 | 0.262317936 | 0.469383573 | 3607.399375 | 3437.967424 | ENSG00000127054 |
| GRIK5 | -0.06963839 | 0.952876806 | 0.314713878 | 0.524986448 | 1457.225792 | 1388.639586 | ENSG00000105737 |
| CEP44 | -0.069642479 | 0.952874105 | 0.666019303 | 0.821155848 | 193.7614092 | 184.6007844 | ENSG00000164118 |
| VPS16 | -0.069661437 | 0.952861584 | 0.525946089 | 0.719293069 | 424.1057418 | 403.9859874 | ENSG00000215305 |
| NA | -0.069707105 | 0.952831422 | 0.931780941 | 0.969216528 | 6.612887612 | 6.30110872 | ENSG00000259972 |
| TRA2B | -0.069722156 | 0.952821481 | 0.206923813 | 0.399331244 | 6958.078149 | 6630.165073 | ENSG00000136527 |
| PSORS1C2 | -0.069917067 | 0.952692762 | 0.932728382 | 0.969708285 | 6.616447084 | 6.303284183 | ENSG00000204538 |
| TTYH3 | -0.070112017 | 0.952564034 | 0.217874153 | 0.414022276 | 6851.234081 | 6526.401904 | ENSG00000136295 |
| MSI2 | -0.070162681 | 0.952530583 | 0.418273521 | 0.629041297 | 1009.580427 | 961.6121549 | ENSG00000153944 |
| FBXO44 | -0.070208585 | 0.952500275 | 0.568800659 | 0.752014213 | 694.4604636 | 661.7068977 | ENSG00000132879 |
| MPP7 | -0.070209525 | 0.952499655 | 0.670563934 | 0.823434547 | 238.0828681 | 226.6568687 | ENSG00000150054 |
| RNFT2 | -0.070257712 | 0.952467841 | 0.626856932 | 0.793550956 | 228.8628159 | 218.0070017 | ENSG00000135119 |
| ARHGAP10 | -0.070322367 | 0.952425157 | 0.68675667 | 0.832171942 | 170.4882139 | 162.2569864 | ENSG00000071205 |
| MIS12 | -0.070366409 | 0.952396082 | 0.537524055 | 0.727576103 | 419.0330758 | 399.0956395 | ENSG00000167842 |
| LETM1 | -0.070394265 | 0.952377694 | 0.270402955 | 0.478859697 | 2254.605874 | 2147.180206 | ENSG00000168924 |
| NDUFS6 | -0.07040135 | 0.952373016 | 0.264945914 | 0.472566175 | 3181.331616 | 3029.979609 | ENSG00000145494 |
| ZSWIM1 | -0.070418503 | 0.952361693 | 0.63280774 | 0.797821895 | 231.7439814 | 220.6605382 | ENSG00000168612 |
| SEC24C | -0.070515009 | 0.952297989 | 0.297110482 | 0.507363365 | 1397.354799 | 1330.665173 | ENSG00000176986 |
| TMEM208 | -0.070543476 | 0.952279198 | 0.471247527 | 0.674614126 | 571.1715941 | 543.8943437 | ENSG00000168701 |
| FAU | -0.070629057 | 0.95222271 | 0.315341399 | 0.525592986 | 1999.723976 | 1904.263713 | ENSG00000149806 |
| CCNL1 | -0.07075078 | 0.952142373 | 0.398786847 | 0.610475104 | 856.9719628 | 815.8444725 | ENSG00000163660 |
| TMEFF2 | -0.070863364 | 0.952068074 | 0.842384109 | 0.92165416 | 37.70887679 | 35.9320842 | ENSG00000144339 |
| ZNF202 | -0.07093586 | 0.952020233 | 0.383097811 | 0.595456969 | 979.964227 | 933.0957318 | ENSG00000166261 |
| PHYHIP | -0.070954979 | 0.952007617 | 0.640138933 | 0.802937284 | 255.2651441 | 243.0837239 | ENSG00000168490 |
| INCENP | -0.070956195 | 0.952006814 | 0.265201835 | 0.472796815 | 2347.968357 | 2234.527418 | ENSG00000149503 |
| CTSK | -0.071023946 | 0.951962107 | 0.736076966 | 0.861271552 | 112.3289335 | 107.0273108 | ENSG00000143387 |
| FAM120A | -0.071045026 | 0.951948198 | 0.344145017 | 0.556247178 | 1470.617031 | 1399.552845 | ENSG00000048828 |
| PPP1R37 | -0.071229617 | 0.951826405 | 0.388253156 | 0.600387969 | 884.5251863 | 842.2720314 | ENSG00000104866 |
| HMCES | -0.071256624 | 0.951808588 | 0.323051602 | 0.533989453 | 1438.650524 | 1369.032467 | ENSG00000183624 |
| CHST14 | -0.07133056 | 0.95175981 | 0.537354289 | 0.727443977 | 425.9366443 | 405.7295965 | ENSG00000169105 |
| RPS29 | -0.071335765 | 0.951756376 | 0.347614988 | 0.559556672 | 3973.012078 | 3781.184646 | ENSG00000213741 |
| SZRD1 | -0.071396698 | 0.951716179 | 0.228129705 | 0.427530959 | 2479.623214 | 2359.65477 | ENSG00000055070 |
| MAST2 | -0.071400628 | 0.951713587 | 0.323485722 | 0.53439037 | 1494.875605 | 1423.179071 | ENSG00000086015 |
| MAFG-DT | -0.071417433 | 0.951702501 | 0.781356782 | 0.888626825 | 85.75847211 | 81.79979647 | ENSG00000265688 |
| WASL | -0.071495748 | 0.95165084 | 0.541161714 | 0.730458078 | 676.4621854 | 644.2221999 | ENSG00000106299 |
| PDXDC1 | -0.07154977 | 0.951615206 | 0.190390284 | 0.376770461 | 2849.959821 | 2711.873978 | ENSG00000179889 |
| BEGAIN | -0.071556977 | 0.951610452 | 0.776083632 | 0.885151552 | 77.86390458 | 74.09076489 | ENSG00000183092 |
| LYRM4 | -0.071560749 | 0.951607964 | 0.442472323 | 0.650077004 | 657.1314037 | 625.162966 | ENSG00000214113 |
| CHPF | -0.071561189 | 0.951607674 | 0.290248275 | 0.499711795 | 2535.574754 | 2413.54595 | ENSG00000123989 |
| DYNLL1 | -0.07161402 | 0.951572827 | 0.17557668 | 0.356074484 | 4611.741395 | 4388.708578 | ENSG00000088986 |

| | | | | | | | |
|---|---|---|---|---|---|---|---|
| SETBP1 | -0.071615374 | 0.951571934 | 0.757275408 | 0.873832707 | 92.66343734 | 88.12935417 | ENSG00000152217 |
| MTF2 | -0.071659374 | 0.951542913 | 0.275752486 | 0.484659067 | 1969.48116 | 1873.896161 | ENSG00000143033 |
| ACSF3 | -0.07171626 | 0.951505394 | 0.557285074 | 0.743506084 | 358.583904 | 341.0133962 | ENSG00000176715 |
| DGLUCY | -0.07173759 | 0.951491326 | 0.543939258 | 0.73244858 | 362.2657673 | 344.6739485 | ENSG00000133943 |
| RSL24D1 | -0.071821527 | 0.951435969 | 0.35003667 | 0.562160261 | 2633.418674 | 2505.450267 | ENSG00000137876 |
| HPS1 | -0.071824879 | 0.951433759 | 0.428165832 | 0.638042432 | 808.7429193 | 769.5139417 | ENSG00000107521 |
| FOXO3B | -0.071855999 | 0.951413236 | 0.621382046 | 0.790222218 | 313.432713 | 297.8794815 | ENSG00000240445 |
| TMEM135 | -0.071909955 | 0.951377654 | 0.541262021 | 0.73050678 | 564.99728 | 537.4511744 | ENSG00000166575 |
| TOR1A | -0.07192422 | 0.951368248 | 0.323349312 | 0.534294493 | 1176.340303 | 1119.231855 | ENSG00000136827 |
| LRRC2 | -0.071958761 | 0.95134547 | 0.731137121 | 0.858455957 | 104.6294999 | 99.49910598 | ENSG00000163827 |
| CANT1 | -0.072027867 | 0.951299901 | 0.362574437 | 0.574479255 | 1079.75479 | 1027.229676 | ENSG00000171302 |
| KSR1 | -0.072050222 | 0.95128516 | 0.441670309 | 0.649302802 | 877.0388865 | 833.7023596 | ENSG00000141068 |
| ANKRD39 | -0.072082654 | 0.951263776 | 0.792840762 | 0.89406764 | 63.128163 | 60.07343579 | ENSG00000213337 |
| GPALPP1 | -0.072117402 | 0.951240864 | 0.520222817 | 0.715550501 | 429.9829176 | 409.2334927 | ENSG00000133114 |
| RNF25 | -0.072123282 | 0.951236987 | 0.512874351 | 0.70906535 | 495.7612905 | 471.7548552 | ENSG00000163481 |
| ZNF621 | -0.072136069 | 0.951228556 | 0.574274408 | 0.75655195 | 355.6344981 | 338.1542724 | ENSG00000172888 |
| UBE2Z | -0.072157775 | 0.951214244 | 0.211493168 | 0.405240309 | 2444.54421 | 2325.062158 | ENSG00000159202 |
| COA1 | -0.072201262 | 0.951185573 | 0.395519062 | 0.607422671 | 1988.638214 | 1891.017313 | ENSG00000106603 |
| NA | -0.072235218 | 0.951163186 | 0.848502489 | 0.924976528 | 31.3744393 | 29.85635278 | ENSG00000260947 |
| PLPPR3 | -0.072275349 | 0.951136728 | 0.410084528 | 0.621140669 | 1615.680389 | 1537.186753 | ENSG00000129951 |
| CCT6A | -0.072289813 | 0.951127192 | 0.149734565 | 0.318510617 | 10111.15134 | 9616.644645 | ENSG00000146731 |
| RBPJ | -0.072294754 | 0.951123934 | 0.343441981 | 0.555395742 | 2357.29166 | 2241.478244 | ENSG00000168214 |
| RAB15 | -0.072361421 | 0.951079984 | 0.256359845 | 0.463198973 | 2581.280229 | 2455.46067 | ENSG00000139998 |
| SCRIB | -0.072363545 | 0.951078583 | 0.213624797 | 0.40808257 | 4323.730975 | 4113.006748 | ENSG00000180900 |
| SLC25A10 | -0.072488979 | 0.950995896 | 0.748587758 | 0.869254247 | 108.948936 | 103.7115255 | ENSG00000183048 |
| SLC39A10 | -0.07249194 | 0.950993944 | 0.347460897 | 0.55952193 | 2719.062003 | 2585.759911 | ENSG00000196950 |
| NSFL1C | -0.072509306 | 0.950982498 | 0.235902086 | 0.437795882 | 2672.710338 | 2541.775155 | ENSG00000088833 |
| PDCD10 | -0.072524266 | 0.950972636 | 0.371308661 | 0.58329264 | 1181.942128 | 1123.351086 | ENSG00000114209 |
| SF3B1 | -0.072527291 | 0.950970642 | 0.119463571 | 0.27197945 | 8531.18696 | 8112.338812 | ENSG00000115524 |
| RPL7 | -0.072533321 | 0.950966668 | 0.105809108 | 0.248873459 | 27608.82298 | 26254.67332 | ENSG00000147604 |
| NA | -0.072574587 | 0.950939467 | 0.826362312 | 0.912467781 | 47.74653437 | 45.46318922 | ENSG00000235944 |
| CCDC12 | -0.072579023 | 0.950936543 | 0.336855311 | 0.548737388 | 1199.480603 | 1140.874457 | ENSG00000160799 |
| TBC1D10A | -0.07265396 | 0.95088715 | 0.747324672 | 0.868553402 | 93.02700931 | 88.50843407 | ENSG00000099992 |
| WDR46 | -0.072694927 | 0.950860149 | 0.21216804 | 0.406080211 | 2302.714428 | 2189.52559 | ENSG00000227057 |
| POM121 | -0.072757848 | 0.95081868 | 0.277830239 | 0.487039764 | 1960.880629 | 1864.476719 | ENSG00000196313 |
| NA | -0.072786624 | 0.950799715 | 0.840667355 | 0.920561463 | 37.08554179 | 35.28506369 | ENSG00000179611 |
| AK2 | -0.072903498 | 0.950722693 | 0.257354501 | 0.464241068 | 2506.099096 | 2382.493272 | ENSG00000004455 |
| CNKSR3 | -0.072931346 | 0.950704342 | 0.779658468 | 0.887571291 | 80.09125234 | 76.29210646 | ENSG00000153721 |
| DYM | -0.072947005 | 0.950694023 | 0.407183837 | 0.61809073 | 909.7891032 | 865.0529711 | ENSG00000141627 |
| SPG21 | -0.073005087 | 0.950655749 | 0.303374307 | 0.513486816 | 1248.206367 | 1186.728032 | ENSG00000090487 |
| PTBP1 | -0.073057547 | 0.950621182 | 0.161831987 | 0.336511459 | 19583.59264 | 18617.07508 | ENSG00000011304 |
| NA | -0.073067477 | 0.950614638 | 0.928180162 | 0.967366576 | 6.99809575 | 6.638475942 | ENSG00000282978 |
| POU5F1 | -0.073074502 | 0.950610009 | 0.083711428 | 0.208974265 | 52398.47322 | 49809.68208 | ENSG00000204531 |
| HCRTR1 | -0.073112341 | 0.950585078 | 0.94128351 | 0.97409682 | 5.629260778 | 5.36429764 | ENSG00000121764 |
| BMERB1 | -0.073165167 | 0.950550271 | 0.551441966 | 0.738527873 | 333.3556074 | 317.0357739 | ENSG00000166780 |
| CD99 | -0.073196625 | 0.950529545 | 0.291587233 | 0.501194584 | 1651.484707 | 1569.649363 | ENSG00000002586 |
| SCRN1 | -0.073216856 | 0.950516216 | 0.198856193 | 0.389030323 | 3019.332298 | 2869.373111 | ENSG00000136193 |
| FNBP1 | -0.0732383 | 0.950502087 | 0.378759642 | 0.591195623 | 818.2079642 | 777.6734014 | ENSG00000187239 |
| RNF19B | -0.073311674 | 0.950453747 | 0.483255111 | 0.68438491 | 603.1782407 | 573.5190037 | ENSG00000116514 |
| WDFY3 | -0.073313762 | 0.950452371 | 0.449302928 | 0.65577601 | 776.2356608 | 737.3953937 | ENSG00000163625 |
| FAM219B | -0.073330915 | 0.950441071 | 0.657251775 | 0.81498357 | 293.5249462 | 278.8236955 | ENSG00000178761 |
| NDNF | -0.073377462 | 0.950410407 | 0.82757127 | 0.913263802 | 40.1574182 | 38.11479255 | ENSG00000173376 |
| CAPZA1 | -0.073389072 | 0.950402758 | 0.304159267 | 0.514238833 | 1956.789149 | 1859.202897 | ENSG00000116489 |
| H3P47 | -0.073449904 | 0.950362685 | 0.888848899 | 0.950235576 | 17.49568424 | 16.65809645 | ENSG00000179101 |
| HPRT1 | -0.073571753 | 0.950282421 | 0.3268874 | 0.538350794 | 1307.798474 | 1242.941245 | ENSG00000165704 |
| PRRG4 | -0.073605097 | 0.950260458 | 0.831358268 | 0.915296977 | 43.57982391 | 41.42036299 | ENSG00000135378 |
| HEG1 | -0.073620484 | 0.950250323 | 0.574906842 | 0.757173842 | 288.0318345 | 273.6056578 | ENSG00000173706 |
| TAPT1 | -0.073620726 | 0.950250164 | 0.71454919 | 0.84957753 | 130.2812685 | 123.7188696 | ENSG00000169762 |
| DEPDC1B | -0.073624786 | 0.95024749 | 0.33226105 | 0.544069187 | 1057.405322 | 1004.989536 | ENSG00000035499 |
| IFNLR1 | -0.073646156 | 0.950233414 | 0.595558131 | 0.772286996 | 293.5202171 | 278.8470155 | ENSG00000185436 |
| RPS3A | -0.073661334 | 0.950223417 | 0.219368038 | 0.415703519 | 6104.820802 | 5799.874219 | ENSG00000145425 |
| NA | -0.073767467 | 0.950153516 | 0.928099114 | 0.967366576 | 6.667448754 | 6.326930736 | ENSG00000287878 |
| PAIP1 | -0.073869453 | 0.950086351 | 0.240963715 | 0.444524739 | 2074.410902 | 1971.359444 | ENSG00000172239 |
| UBE2M | -0.073940375 | 0.950039646 | 0.21931997 | 0.415703519 | 2838.178597 | 2696.59458 | ENSG00000130725 |
| MAPT | -0.073944194 | 0.950037131 | 0.829424713 | 0.9144008 | 40.55775396 | 38.60274095 | ENSG00000186868 |
| NA | -0.073969187 | 0.950020673 | 0.937344575 | 0.971904957 | 5.236933694 | 4.997986979 | ENSG00000213453 |
| TOGARAM1 | -0.074069472 | 0.949954637 | 0.689810506 | 0.83399945 | 143.9631359 | 136.7604175 | ENSG00000198718 |
| TMEM170A | -0.074105821 | 0.949930703 | 0.502812002 | 0.700834354 | 566.9284332 | 538.4034246 | ENSG00000166822 |
| CBWD5 | -0.074110795 | 0.949927428 | 0.72046702 | 0.852637136 | 114.007338 | 108.2722621 | ENSG00000147996 |
| RELT | -0.074139659 | 0.949908424 | 0.704270946 | 0.842795565 | 142.2345933 | 135.2429637 | ENSG00000054967 |
| PSMD12 | -0.074182838 | 0.949879994 | 0.292554619 | 0.502308753 | 2012.12335 | 1911.206288 | ENSG00000197170 |
| SAP130 | -0.074227895 | 0.949850328 | 0.277949491 | 0.487198453 | 1450.450591 | 1377.621906 | ENSG00000136715 |
| NA | -0.074280346 | 0.949815796 | 0.937649565 | 0.972022827 | 5.209653123 | 4.969989499 | ENSG00000202058 |
| ADD1 | -0.074384025 | 0.94974754 | 0.188660757 | 0.374568749 | 3470.796872 | 3295.800298 | ENSG00000087274 |

| | | | | | | | |
|---|---|---|---|---|---|---|---|
| TFB1M | -0.074451635 | 0.949703032 | 0.564083044 | 0.748528238 | 341.030073 | 323.6924839 | ENSG00000029639 |
| PSMD7 | -0.074490999 | 0.94967712 | 0.313226312 | 0.523568676 | 1997.465099 | 1896.427555 | ENSG00000103035 |
| ACIN1 | -0.074496493 | 0.949673503 | 0.11869595 | 0.270688686 | 6045.329198 | 5741.351733 | ENSG00000100813 |
| MANBAL | -0.07451983 | 0.949658142 | 0.373973941 | 0.585953529 | 1141.874229 | 1083.957975 | ENSG00000101363 |
| UBA6 | -0.074726025 | 0.949522423 | 0.43909556 | 0.647129674 | 729.0677663 | 692.296435 | ENSG00000033178 |
| EWSR1 | -0.074737293 | 0.949515007 | 0.1231294 | 0.277647608 | 9972.19297 | 9469.022487 | ENSG00000182944 |
| PGAM5 | -0.074783675 | 0.949484481 | 0.221801361 | 0.419081411 | 2360.958395 | 2241.291504 | ENSG00000247077 |
| HDGFL2 | -0.074846918 | 0.949442859 | 0.172344125 | 0.351823456 | 4387.493457 | 4166.438323 | ENSG00000167674 |
| AP1S1 | -0.074877429 | 0.94942278 | 0.314055242 | 0.524490029 | 1514.259242 | 1437.454317 | ENSG00000106367 |
| MGRN1 | -0.07500211 | 0.949340732 | 0.276965265 | 0.48615011 | 1879.954714 | 1785.439835 | ENSG00000102858 |
| SPATA33 | -0.075017508 | 0.9493306 | 0.643037092 | 0.805122663 | 207.5859342 | 197.3376923 | ENSG00000167523 |
| ZNF544 | -0.075046689 | 0.949311399 | 0.432829004 | 0.642153801 | 610.2407386 | 579.3508698 | ENSG00000198131 |
| ITIH5 | -0.075050116 | 0.949309143 | 0.626006469 | 0.793253855 | 215.4745524 | 204.5676453 | ENSG00000123243 |
| FAM149B1 | -0.075147829 | 0.94924485 | 0.630728672 | 0.796625039 | 278.8028577 | 264.5688596 | ENSG00000138286 |
| MARCHF9 | -0.075151212 | 0.949242623 | 0.745433066 | 0.86764274 | 100.9989432 | 96.05527211 | ENSG00000139266 |
| VDAC2 | -0.075164875 | 0.949233634 | 0.13691976 | 0.299627263 | 4072.144593 | 3865.147438 | ENSG00000165637 |
| SNRPD1 | -0.075293708 | 0.949148871 | 0.242202115 | 0.445851149 | 2973.969754 | 2822.739818 | ENSG00000167088 |
| BSN-DT | -0.075299967 | 0.949144753 | 0.925082022 | 0.965883673 | 7.318064329 | 6.919848203 | ENSG00000226913 |
| TSC1 | -0.075341172 | 0.949117645 | 0.394191749 | 0.605868901 | 824.6156244 | 782.4164052 | ENSG00000165699 |
| GNA12 | -0.075433837 | 0.949056684 | 0.154937671 | 0.326525824 | 3777.740108 | 3585.004635 | ENSG00000146535 |
| KANSL1 | -0.075487809 | 0.94902118 | 0.353248406 | 0.565470134 | 1034.628744 | 981.6379573 | ENSG00000120071 |
| SRP68 | -0.075496259 | 0.949015622 | 0.161206905 | 0.335692183 | 3173.739175 | 3011.801688 | ENSG00000167881 |
| NUBP2 | -0.075557268 | 0.94897549 | 0.309682624 | 0.520270114 | 1920.526446 | 1822.498387 | ENSG00000095906 |
| HSD17B10 | -0.075567936 | 0.948968473 | 0.276258601 | 0.485225417 | 2386.793573 | 2264.532832 | ENSG00000072506 |
| KIAA1328 | -0.07556846 | 0.948968129 | 0.795634139 | 0.895554092 | 61.24364077 | 58.24086389 | ENSG00000150477 |
| DDX1 | -0.075680132 | 0.948894676 | 0.16894493 | 0.346645482 | 4879.863656 | 4630.551174 | ENSG00000079785 |
| PMAIP1 | -0.075705726 | 0.948877843 | 0.265920324 | 0.473407364 | 2385.142501 | 2262.646382 | ENSG00000141682 |
| EIF2AK3 | -0.075807333 | 0.948811017 | 0.553786673 | 0.740250659 | 319.5269381 | 303.1480063 | ENSG00000172071 |
| NA | -0.07582269 | 0.948800917 | 0.882219027 | 0.945771249 | 18.19610627 | 17.26813408 | ENSG00000248334 |
| SPACA6 | -0.075900118 | 0.948749998 | 0.822994488 | 0.910287625 | 72.18483734 | 68.56940264 | ENSG00000182310 |
| CTBP1-AS | -0.075922126 | 0.948735525 | 0.885006519 | 0.947622636 | 17.53008375 | 16.65034056 | ENSG00000280927 |
| SEPTIN9 | -0.075927869 | 0.948731748 | 0.125383428 | 0.281419781 | 10796.34773 | 10242.94383 | ENSG00000184640 |
| C3orf33 | -0.075938578 | 0.948724706 | 0.922966458 | 0.965130891 | 8.476053008 | 7.997592643 | ENSG00000174928 |
| TIMM8B | -0.075981202 | 0.948696676 | 0.291040003 | 0.500664098 | 1389.006171 | 1317.882033 | ENSG00000150779 |
| INSIG1 | -0.075995149 | 0.948687505 | 0.32857373 | 0.540048678 | 1336.587106 | 1267.742037 | ENSG00000186480 |
| GAB2 | -0.076157234 | 0.948580927 | 0.585372162 | 0.764135846 | 290.3427779 | 275.3507596 | ENSG00000033327 |
| IL4R | -0.076302717 | 0.948485276 | 0.366782734 | 0.579251708 | 886.796138 | 841.2346318 | ENSG00000077238 |
| NA | -0.076329883 | 0.948467416 | 0.936291204 | 0.97143656 | 6.574928623 | 6.226764092 | ENSG00000231563 |
| UTP18 | -0.076600147 | 0.948289754 | 0.229693763 | 0.429529871 | 2040.294597 | 1934.567201 | ENSG00000011260 |
| ATN1 | -0.076673789 | 0.94824135 | 0.279382795 | 0.488473282 | 2443.303975 | 2317.749739 | ENSG00000111676 |
| NA | -0.076677068 | 0.948239195 | 0.71500831 | 0.849668508 | 123.1337273 | 116.8442546 | ENSG00000232533 |
| EED | -0.076702367 | 0.948222566 | 0.382960855 | 0.595391104 | 729.4277532 | 691.7316736 | ENSG00000074266 |
| NA | -0.076709347 | 0.948217979 | 0.122351512 | 0.276388554 | 2436977.556 | 2310785.995 | N_unmapped |
| CRIPT | -0.076799183 | 0.948158936 | 0.639268212 | 0.802218099 | 209.2569665 | 198.6681491 | ENSG00000119878 |
| C2CD5 | -0.076811202 | 0.948151037 | 0.348488788 | 0.560425772 | 1153.453355 | 1093.183165 | ENSG00000111731 |
| ZNF330 | -0.076873307 | 0.948110222 | 0.398254809 | 0.610078039 | 1006.934336 | 954.0634047 | ENSG00000109445 |
| NA | -0.077007161 | 0.948022259 | 0.835055005 | 0.917376243 | 38.38646745 | 36.46248032 | ENSG00000284610 |
| DCLRE1B | -0.077013332 | 0.948018205 | 0.600949191 | 0.776307564 | 268.3886111 | 254.3911653 | ENSG00000118655 |
| BOD1L1 | -0.077129848 | 0.947941643 | 0.457868601 | 0.663254471 | 948.8702199 | 898.7393835 | ENSG00000038219 |
| PSMB1 | -0.077168447 | 0.947916282 | 0.152284797 | 0.322337861 | 3722.371429 | 3528.349142 | ENSG00000008018 |
| SGCE | -0.077198119 | 0.947896786 | 0.5210931 | 0.715972717 | 367.4463342 | 348.0782451 | ENSG00000127990 |
| MRPL20 | -0.077208806 | 0.947889764 | 0.302338592 | 0.512445547 | 1299.874515 | 1232.086747 | ENSG00000242485 |
| OARD1 | -0.07723058 | 0.947875458 | 0.43915401 | 0.647141256 | 601.4452489 | 570.3007877 | ENSG00000124596 |
| NA | -0.077255541 | 0.947859058 | 0.845602076 | 0.923288344 | 29.05874127 | 27.56789735 | ENSG00000287126 |
| CENPA | -0.077286056 | 0.94783901 | 0.612152046 | 0.784199347 | 291.1203131 | 275.9071887 | ENSG00000115163 |
| KIAA2013 | -0.077291853 | 0.947835201 | 0.32831528 | 0.539854946 | 1546.066032 | 1465.965878 | ENSG00000116685 |
| KDM5C | -0.077349023 | 0.947797642 | 0.09556006 | 0.230332065 | 6356.275264 | 6024.110539 | ENSG00000126012 |
| HNRNPA0 | -0.077376918 | 0.947779317 | 0.097890678 | 0.234376426 | 13157.18985 | 12470.63436 | ENSG00000177733 |
| GARNL3 | -0.077386428 | 0.947773069 | 0.706102266 | 0.844411002 | 127.6820864 | 121.1047022 | ENSG00000136895 |
| EIF1AD | -0.077445969 | 0.947733954 | 0.420664011 | 0.630845789 | 617.117228 | 584.6891345 | ENSG00000175376 |
| TLK2 | -0.077510646 | 0.947691468 | 0.311234488 | 0.521809598 | 1063.634476 | 1008.071505 | ENSG00000146872 |
| CYP2U1 | -0.077516576 | 0.947687572 | 0.678399162 | 0.828048616 | 180.2263694 | 170.9688506 | ENSG00000155016 |
| SLC15A3 | -0.077548281 | 0.947666746 | 0.577311831 | 0.759282257 | 430.9524373 | 408.5428676 | ENSG00000110446 |
| EGF | -0.07764349 | 0.947604208 | 0.751364525 | 0.870629467 | 78.08426024 | 73.98965229 | ENSG00000138798 |
| RPL22L1 | -0.0776593 | 0.947593823 | 0.29800225 | 0.508219151 | 1203.622754 | 1140.979898 | ENSG00000163584 |
| POLR2J | -0.077663885 | 0.947590812 | 0.473004894 | 0.676271538 | 465.0691449 | 440.6337837 | ENSG00000005075 |
| EARS2 | -0.077721207 | 0.947553162 | 0.233253725 | 0.434160909 | 1584.529152 | 1501.373081 | ENSG00000103356 |
| VAPB | -0.077731505 | 0.947546399 | 0.289394588 | 0.498742179 | 1224.78159 | 1160.412957 | ENSG00000124164 |
| PPP2R1B | -0.077752849 | 0.947532381 | 0.277429806 | 0.486558495 | 2002.408052 | 1896.992861 | ENSG00000137713 |
| KCNA2 | -0.077884539 | 0.947445893 | 0.876934138 | 0.942781578 | 22.56000996 | 21.29601562 | ENSG00000177301 |
| RSPO4 | -0.077996065 | 0.947372654 | 0.667033913 | 0.82154232 | 298.058255 | 282.6537457 | ENSG00000101282 |
| DTX2 | -0.078073781 | 0.947321622 | 0.436949027 | 0.645089622 | 638.604676 | 605.2360414 | ENSG00000091073 |
| BUB1B | -0.078084866 | 0.947314344 | 0.185956456 | 0.370832911 | 3276.86201 | 3103.48506 | ENSG00000156970 |
| TNFRSF11A | -0.078101026 | 0.947303732 | 0.608089016 | 0.781580417 | 221.8089638 | 210.0882725 | ENSG00000141655 |

| | | | | | | | |
|---|---|---|---|---|---|---|---|
| PCDHGB7 | -0.078127037 | 0.947286653 | 0.623980307 | 0.792264204 | 206.1975224 | 195.3718274 | ENSG00000254122 |
| PARP9 | -0.078186288 | 0.947247749 | 0.709192595 | 0.846486042 | 109.0974918 | 103.3035021 | ENSG00000138496 |
| CLMP | -0.078281136 | 0.947185476 | 0.485138371 | 0.686177213 | 481.4638437 | 456.127546 | ENSG00000166250 |
| CHRNA5 | -0.078299923 | 0.947173141 | 0.612936421 | 0.78462019 | 216.715578 | 205.2131528 | ENSG00000169684 |
| KIAA0825 | -0.078307423 | 0.947168218 | 0.844037974 | 0.922299853 | 31.79137721 | 30.18936908 | ENSG00000185261 |
| TMPRSS13 | -0.078337525 | 0.947148455 | 0.93292076 | 0.969708285 | 5.925508257 | 5.612375535 | ENSG00000137747 |
| NA | -0.078381625 | 0.947119503 | 0.92677421 | 0.966701516 | 8.371379919 | 7.983593902 | ENSG00000233672 |
| AP3D1 | -0.078411733 | 0.947099738 | 0.100343507 | 0.238633311 | 7262.639978 | 6878.669003 | ENSG00000065000 |
| PCDHB4 | -0.078413394 | 0.947098647 | 0.914995791 | 0.962281868 | 9.44516262 | 8.994466064 | ENSG00000081818 |
| NA | -0.078473643 | 0.947059096 | 0.936546752 | 0.971528999 | 4.902727226 | 4.665916643 | ENSG00000235545 |
| LCA5 | -0.078572606 | 0.946994134 | 0.770636599 | 0.881997379 | 67.20858256 | 63.60154565 | ENSG00000135338 |
| RAB7A | -0.078687778 | 0.946918537 | 0.120589794 | 0.27354231 | 4323.680275 | 4093.932075 | ENSG00000075785 |
| LOC729870 | -0.078692008 | 0.946915761 | 0.887269953 | 0.949163187 | 17.54432164 | 16.61799215 | ENSG00000287642 |
| HMGXB4 | -0.078715011 | 0.946900663 | 0.308685793 | 0.519149631 | 1013.619406 | 959.9670971 | ENSG00000100281 |
| BUD23 | -0.078735037 | 0.946887519 | 0.186239532 | 0.371201431 | 2292.461117 | 2170.948514 | ENSG00000071462 |
| CELF3 | -0.07889316 | 0.946783744 | 0.869337549 | 0.938217686 | 21.34242612 | 20.25497051 | ENSG00000159409 |
| CCL2 | -0.078989425 | 0.94672057 | 0.830115354 | 0.914737804 | 34.72476583 | 32.89048232 | ENSG00000108691 |
| OSBPL3 | -0.079012947 | 0.946705135 | 0.512221959 | 0.708615611 | 395.3815642 | 374.4035346 | ENSG00000070882 |
| NDUFA12 | -0.07912203 | 0.946633557 | 0.311142303 | 0.521747681 | 1014.328675 | 960.1820689 | ENSG00000184752 |
| GTF3C6 | -0.079169517 | 0.946602399 | 0.295381739 | 0.505462499 | 1287.979693 | 1219.59139 | ENSG00000155115 |
| PCBP2 | -0.079231176 | 0.946561943 | 0.257914129 | 0.464762245 | 2072.60024 | 1962.004783 | ENSG00000197111 |
| PRKRA | -0.079379067 | 0.946464915 | 0.394822792 | 0.606641317 | 832.5269949 | 787.7442668 | ENSG00000180228 |
| PAFAH1B2 | -0.07942386 | 0.94643553 | 0.257044652 | 0.463948031 | 1856.34441 | 1756.422289 | ENSG00000168092 |
| DCTN4 | -0.079493872 | 0.946389602 | 0.361258423 | 0.573333011 | 1046.846054 | 990.4080514 | ENSG00000132912 |
| UBE2W | -0.079636987 | 0.946295725 | 0.729660708 | 0.857636302 | 152.2248087 | 144.1071628 | ENSG00000104343 |
| OXA1L | -0.079753123 | 0.946219552 | 0.22170151 | 0.418934663 | 2267.524697 | 2145.98085 | ENSG00000155463 |
| MB21D2 | -0.079796314 | 0.946191225 | 0.499709245 | 0.698106902 | 398.6031132 | 377.127632 | ENSG00000180611 |
| NA | -0.07985597 | 0.9461521 | 0.699099429 | 0.839692979 | 119.3106803 | 112.8342283 | ENSG00000213866 |
| BCL2 | -0.079861208 | 0.946148665 | 0.759893993 | 0.875410118 | 83.13851793 | 78.62925246 | ENSG00000171791 |
| MTMR12 | -0.079907993 | 0.946117983 | 0.274814408 | 0.483588343 | 1280.712937 | 1211.341303 | ENSG00000150712 |
| ZCCHC17 | -0.080052057 | 0.946023511 | 0.460430418 | 0.665128773 | 662.0326041 | 626.1113668 | ENSG00000121766 |
| LSM4 | -0.080148301 | 0.945960402 | 0.155702176 | 0.327479184 | 5105.519084 | 4829.71078 | ENSG00000130520 |
| EMC8 | -0.080151691 | 0.945958179 | 0.43688958 | 0.645089622 | 683.5412599 | 646.9434821 | ENSG00000131148 |
| DUSP3 | -0.080169599 | 0.945946437 | 0.468404363 | 0.671664116 | 620.50465 | 586.9373607 | ENSG00000108861 |
| AKTIP | -0.080185686 | 0.94593589 | 0.577264085 | 0.759282257 | 249.1189753 | 235.5528195 | ENSG00000166971 |
| NA | -0.080227025 | 0.945908785 | 0.906648668 | 0.959649147 | 10.56994699 | 10.02274193 | ENSG00000214770 |
| LRRC14 | -0.080311225 | 0.945853581 | 0.481418575 | 0.683218591 | 568.0214866 | 537.8040667 | ENSG00000160959 |
| NRSN1 | -0.080335417 | 0.945837721 | 0.933231261 | 0.969789847 | 5.249976376 | 4.977461848 | ENSG00000152954 |
| TMEM25 | -0.080349731 | 0.945828336 | 0.556147704 | 0.742544206 | 278.2767708 | 263.329752 | ENSG00000149582 |
| PCCA | -0.080351469 | 0.945827196 | 0.311115985 | 0.521747681 | 962.7445763 | 910.5122312 | ENSG00000175198 |
| LPIN1 | -0.080363011 | 0.94581963 | 0.527418596 | 0.720507826 | 391.6062132 | 370.2726188 | ENSG00000134324 |
| KPNA2 | -0.080377526 | 0.945810114 | 0.059822908 | 0.161932333 | 11417.3971 | 10798.23665 | ENSG00000182481 |
| NA | -0.080415752 | 0.945785054 | 0.721621553 | 0.853183917 | 102.5104364 | 96.99520461 | ENSG00000267002 |
| GLS | -0.080458865 | 0.94575679 | 0.398424035 | 0.610274365 | 915.1724811 | 865.654096 | ENSG00000115419 |
| SLC16A7 | -0.080511774 | 0.945722107 | 0.821340665 | 0.909334444 | 39.14384105 | 37.04574997 | ENSG00000118596 |
| BRD4 | -0.080525378 | 0.945713189 | 0.203832675 | 0.395475152 | 2546.444125 | 2408.830753 | ENSG00000141867 |
| B3GAT1 | -0.080560604 | 0.945690098 | 0.275326307 | 0.484218412 | 1289.099191 | 1219.513997 | ENSG00000109956 |
| ERN1 | -0.080585046 | 0.945674076 | 0.711997754 | 0.847851541 | 108.1527399 | 102.3196815 | ENSG00000178607 |
| ZFAND2B | -0.080589305 | 0.945671285 | 0.624360356 | 0.7923564 | 200.2835561 | 189.5561127 | ENSG00000158552 |
| ZHX2 | -0.080756407 | 0.945561758 | 0.393786491 | 0.605435767 | 744.8470882 | 704.1534432 | ENSG00000178764 |
| TFAM | -0.080797598 | 0.945534761 | 0.290032491 | 0.499385813 | 2033.268278 | 1922.470785 | ENSG00000108064 |
| KIFC1 | -0.080825502 | 0.945516473 | 0.184916887 | 0.369146229 | 2228.60431 | 2106.880659 | ENSG00000237649 |
| MAD2L1 | -0.080846225 | 0.945502892 | 0.165870141 | 0.34178479 | 2257.095524 | 2134.02155 | ENSG00000164109 |
| RPF1 | -0.080863525 | 0.945491553 | 0.32361423 | 0.53445166 | 964.216025 | 911.7448125 | ENSG00000117133 |
| DENND5A | -0.080889971 | 0.945474222 | 0.24566159 | 0.450136241 | 1704.570433 | 1611.09869 | ENSG00000184014 |
| RRS1 | -0.080967245 | 0.945423581 | 0.300867438 | 0.511018736 | 1308.217497 | 1237.302849 | ENSG00000179041 |
| RAE1 | -0.081058013 | 0.945364101 | 0.343316495 | 0.555334647 | 838.2640075 | 792.7067316 | ENSG00000101146 |
| HDDC3 | -0.081234792 | 0.945248269 | 0.681157709 | 0.829598484 | 123.4029735 | 116.5901018 | ENSG00000184508 |
| ZNF45 | -0.081271287 | 0.945224358 | 0.498379137 | 0.697317165 | 369.7774653 | 349.2575537 | ENSG00000124459 |
| SYNGAP1 | -0.081286234 | 0.945214565 | 0.666186325 | 0.821155848 | 2328.260965 | 2200.840176 | ENSG00000197283 |
| CTTNBP2 | -0.081340595 | 0.945178951 | 0.467246364 | 0.67102263 | 450.8603538 | 425.8884678 | ENSG00000077063 |
| DCAKD | -0.081364456 | 0.945163318 | 0.381159944 | 0.593714122 | 718.7356151 | 679.4951271 | ENSG00000172992 |
| NA | -0.081375972 | 0.945155773 | 0.253322727 | 0.459425689 | 1376.911768 | 1301.004476 | ENSG00000188856 |
| PSMA7 | -0.081467959 | 0.945095511 | 0.122567036 | 0.276709919 | 7171.786428 | 6778.363117 | ENSG00000101182 |
| PRTFDC1 | -0.081589296 | 0.945016028 | 0.613515109 | 0.784868773 | 210.4152345 | 198.8674935 | ENSG00000099256 |
| P2RX3 | -0.0816778 | 0.944958057 | 0.923533327 | 0.965347146 | 6.316640132 | 5.954093685 | ENSG00000109991 |
| LMAN2 | -0.08175192 | 0.94490951 | 0.159138213 | 0.332557834 | 3225.222945 | 3047.90298 | ENSG00000169223 |
| PSME3IP1 | -0.081782601 | 0.944889415 | 0.264298757 | 0.471810381 | 1177.116618 | 1112.067429 | ENSG00000172775 |
| RCL1 | -0.081940597 | 0.944785942 | 0.424268714 | 0.634226465 | 492.3163589 | 465.2115123 | ENSG00000120158 |
| HSPA4L | -0.082001861 | 0.944745822 | 0.645430288 | 0.807001522 | 153.6066608 | 144.9989945 | ENSG00000164070 |
| DPP9-AS1 | -0.082041006 | 0.944720189 | 0.931328843 | 0.968990425 | 5.615022888 | 5.306127216 | ENSG00000205790 |
| CLPP | -0.082074675 | 0.944698142 | 0.295913625 | 0.506051689 | 1141.055993 | 1078.482297 | ENSG00000125656 |
| MEDAG | -0.082125115 | 0.944665113 | 0.842626249 | 0.921654519 | 34.95635857 | 32.91384532 | ENSG00000102802 |
| SIN3B | -0.082127608 | 0.94466348 | 0.202559624 | 0.393877523 | 2206.602883 | 2084.639792 | ENSG00000127511 |

| | | | | | | | |
|---|---|---|---|---|---|---|---|
| NA | -0.082191677 | 0.94462153 | 0.751414387 | 0.870633824 | 98.66692235 | 93.34841048 | ENSG00000214223 |
| TSC2 | -0.082209786 | 0.944609673 | 0.24775694 | 0.452766578 | 2173.183185 | 2053.508002 | ENSG00000103197 |
| ANGEL1 | -0.082226845 | 0.944598504 | 0.339658467 | 0.55136372 | 769.0122794 | 726.4763669 | ENSG00000013523 |
| ZBTB1 | -0.082240912 | 0.944589293 | 0.525192739 | 0.718766931 | 349.7718389 | 330.1412406 | ENSG00000126804 |
| POLE3 | -0.082367509 | 0.944506409 | 0.337509806 | 0.549415519 | 992.4997005 | 937.2108295 | ENSG00000148229 |
| PPIF | -0.08257221 | 0.944372404 | 0.173315689 | 0.353081638 | 2580.744529 | 2437.623716 | ENSG00000108179 |
| NUTM2G | -0.08269886 | 0.944289504 | 0.831695268 | 0.915300998 | 32.72074838 | 30.92634132 | ENSG00000188152 |
| EIF5B | -0.082741375 | 0.944261677 | 0.150740759 | 0.320074121 | 4035.426815 | 3811.234976 | ENSG00000158417 |
| SNORA5C | -0.082747978 | 0.944257356 | 0.929351123 | 0.967945949 | 6.01683318 | 5.674896887 | ENSG00000201772 |
| RBSN | -0.082784565 | 0.944233409 | 0.393873605 | 0.605478472 | 664.7175185 | 627.7804859 | ENSG00000131381 |
| ORAI3 | -0.082799086 | 0.944223906 | 0.832929034 | 0.915782079 | 36.27955694 | 34.32053868 | ENSG00000175938 |
| NUP43 | -0.082854344 | 0.94418774 | 0.265674942 | 0.473175215 | 1249.14756 | 1179.403922 | ENSG00000120253 |
| DDR1 | -0.082859972 | 0.944184057 | 0.100092779 | 0.23821703 | 6496.241117 | 6134.226752 | ENSG00000204580 |
| RPSA | -0.08289506 | 0.944161094 | 0.093301215 | 0.226586904 | 18505.40601 | 17471.52714 | ENSG00000168028 |
| FAM72B | -0.082896754 | 0.944159985 | 0.527896114 | 0.720939647 | 309.1820753 | 291.9368751 | ENSG00000188610 |
| G3BP2 | -0.08290526 | 0.944154418 | 0.165213261 | 0.34091482 | 5549.715373 | 5239.433958 | ENSG00000138757 |
| USP6NL | -0.082917488 | 0.944146416 | 0.355753296 | 0.568056227 | 719.2561093 | 679.1391993 | ENSG00000148429 |
| CSNK1G2 | -0.082980995 | 0.944104856 | 0.189137651 | 0.37523527 | 4393.545536 | 4148.651945 | ENSG00000133275 |
| MRPL42 | -0.082986461 | 0.944101279 | 0.247491056 | 0.452455817 | 1633.75237 | 1542.483652 | ENSG00000198015 |
| HDX | -0.083008281 | 0.944087 | 0.700249332 | 0.84040536 | 105.7338433 | 99.85170145 | ENSG00000165259 |
| NA | -0.083163359 | 0.943985524 | 0.832375228 | 0.915569118 | 33.60354093 | 31.72004358 | ENSG00000253341 |
| IFFO1 | -0.083182479 | 0.943973013 | 0.490784782 | 0.690491739 | 409.241606 | 386.5425547 | ENSG00000010295 |
| NA | -0.08321105 | 0.943954319 | 0.926631997 | 0.966701516 | 5.635184514 | 5.319179997 | ENSG00000230626 |
| TRIM74 | -0.083286407 | 0.943905015 | 0.763044877 | 0.877487669 | 64.03350759 | 60.52061747 | ENSG00000174353 |
| RALGAPA2 | -0.08338119 | 0.943843003 | 0.36877075 | 0.580952094 | 627.4745487 | 592.1450546 | ENSG00000188559 |
| HPS5 | -0.083395867 | 0.943833401 | 0.49633411 | 0.695597231 | 448.7895699 | 423.2384611 | ENSG00000110756 |
| C1QBP | -0.083499413 | 0.943765663 | 0.092967066 | 0.225898488 | 6577.046419 | 6207.856874 | ENSG00000108561 |
| SLC25A34 | -0.083513624 | 0.943756366 | 0.921268382 | 0.964771134 | 6.718450425 | 6.31510746 | ENSG00000162461 |
| RNF114 | -0.083609192 | 0.943693851 | 0.165935248 | 0.341881645 | 2159.025241 | 2037.198945 | ENSG00000124226 |
| TRUB2 | -0.083638808 | 0.943674479 | 0.356653576 | 0.568724376 | 677.7909253 | 639.8118765 | ENSG00000167112 |
| C14orf28 | -0.083661686 | 0.943659514 | 0.789417649 | 0.892025074 | 58.21445187 | 54.99145637 | ENSG00000179476 |
| ZNF321P | -0.083684872 | 0.943644349 | 0.877887439 | 0.943162151 | 18.71179353 | 17.62139196 | ENSG00000213801 |
| NA | -0.083715898 | 0.943624055 | 0.796367075 | 0.896001919 | 48.84401202 | 46.11985755 | ENSG00000249896 |
| RIC3 | -0.08371842 | 0.943622406 | 0.690867967 | 0.834552267 | 190.4090239 | 179.7474622 | ENSG00000166405 |
| CXorf40B | -0.083777866 | 0.943583525 | 0.349667619 | 0.561787696 | 684.7237559 | 645.9263671 | ENSG00000197021 |
| DCK | -0.083787858 | 0.943576989 | 0.581771302 | 0.761901835 | 256.115622 | 241.9696066 | ENSG00000156136 |
| ANKRD9 | -0.08379984 | 0.943569153 | 0.275253532 | 0.484135508 | 1076.922769 | 1016.187278 | ENSG00000156381 |
| SMIM19 | -0.08380748 | 0.943564156 | 0.579560082 | 0.760406596 | 217.8222595 | 205.6586535 | ENSG00000176209 |
| DHX16 | -0.083811657 | 0.943561425 | 0.224191554 | 0.422414197 | 1524.329701 | 1438.375782 | ENSG00000204560 |
| RABEP1 | -0.083866208 | 0.943525747 | 0.281849521 | 0.490709708 | 1098.32451 | 1036.302554 | ENSG00000029725 |
| FKBPL | -0.083867711 | 0.943524764 | 0.626412219 | 0.793253855 | 158.6965132 | 149.677964 | ENSG00000204315 |
| CBFB | -0.083911233 | 0.943496301 | 0.353090505 | 0.565381906 | 667.7354703 | 630.1476893 | ENSG00000067955 |
| NA | -0.084009804 | 0.94343184 | 0.770721376 | 0.881997379 | 56.03546219 | 52.86162155 | ENSG00000254510 |
| NA | -0.084040789 | 0.943411577 | 0.582453931 | 0.762275987 | 245.3253717 | 231.7448504 | ENSG00000277200 |
| GMFB | -0.084049774 | 0.943405702 | 0.407599823 | 0.61857302 | 1321.627578 | 1246.943153 | ENSG00000197045 |
| XRRA1 | -0.084128019 | 0.943354538 | 0.770749417 | 0.881997379 | 58.40691657 | 55.0359116 | ENSG00000166435 |
| CYSTM1 | -0.084156164 | 0.943336135 | 0.535341039 | 0.72613054 | 309.938533 | 292.4141571 | ENSG00000120306 |
| ELOVL1 | -0.084158117 | 0.943334857 | 0.400339336 | 0.611782518 | 552.730242 | 521.3853463 | ENSG00000066322 |
| NA | -0.0841582 | 0.943334803 | 0.894592954 | 0.953032171 | 10.90651772 | 10.29446638 | ENSG00000275088 |
| MYT1 | -0.084174687 | 0.943324022 | 0.733681346 | 0.860167136 | 79.4910542 | 74.94888042 | ENSG00000196132 |
| LINC00491 | -0.084218806 | 0.943295176 | 0.784527883 | 0.890365666 | 51.34475031 | 48.40178659 | ENSG00000250682 |
| COPS5 | -0.084262006 | 0.94326693 | 0.32037099 | 0.530859537 | 988.178328 | 931.7432665 | ENSG00000121022 |
| SEMA3E | -0.084344797 | 0.943212801 | 0.611976138 | 0.784174879 | 224.7491664 | 211.6634023 | ENSG00000170381 |
| AKIRIN2 | -0.084365016 | 0.943199582 | 0.235606889 | 0.437506007 | 1733.642469 | 1635.036514 | ENSG00000135334 |
| FBXW7 | -0.084365407 | 0.943199326 | 0.47705983 | 0.679483739 | 419.8939528 | 396.2712075 | ENSG00000109670 |
| PSMB5 | -0.084375641 | 0.943192636 | 0.106278297 | 0.249790372 | 3954.689242 | 3730.193603 | ENSG00000100804 |
| PPHLN1 | -0.084388321 | 0.943184346 | 0.23118049 | 0.431578049 | 1409.282012 | 1329.203954 | ENSG00000134283 |
| CSTF1 | -0.084390942 | 0.943182633 | 0.279983867 | 0.488994296 | 1000.570052 | 943.9857426 | ENSG00000101138 |
| MSX1 | -0.084436437 | 0.94315289 | 0.937134797 | 0.971870005 | 4.208228926 | 3.999884053 | ENSG00000163132 |
| DLEU7 | -0.084457728 | 0.943138971 | 0.71367592 | 0.848993371 | 111.4028163 | 105.2533608 | ENSG00000186047 |
| PHF8 | -0.084582589 | 0.943057349 | 0.300226435 | 0.510284964 | 957.2345576 | 902.6390713 | ENSG00000172943 |
| MMAB | -0.084618072 | 0.943034155 | 0.454244704 | 0.660233298 | 408.5144621 | 385.1510466 | ENSG00000139428 |
| ERAP1 | -0.08463693 | 0.943021828 | 0.544662428 | 0.733035202 | 288.2919031 | 271.6489891 | ENSG00000164307 |
| ZNF749 | -0.084649244 | 0.943013779 | 0.55201643 | 0.738930414 | 246.8015234 | 232.7810501 | ENSG00000186230 |
| SARAF | -0.084668902 | 0.943000929 | 0.233430361 | 0.434446869 | 1564.702135 | 1476.003426 | ENSG00000133872 |
| PAN3 | -0.08475742 | 0.942943072 | 0.424203872 | 0.634226465 | 551.9752327 | 520.2044388 | ENSG00000152520 |
| FNIP2 | -0.084768182 | 0.942936039 | 0.593954587 | 0.771075299 | 221.5073774 | 208.6423255 | ENSG00000052795 |
| EIF4G1 | -0.084834089 | 0.942892963 | 0.077290157 | 0.196794474 | 20719.99801 | 19536.87078 | ENSG00000114867 |
| PSMG1 | -0.08485551 | 0.942878963 | 0.315754157 | 0.525792898 | 968.1712526 | 912.7567985 | ENSG00000183527 |
| CGRRF1 | -0.084871493 | 0.942868517 | 0.582012351 | 0.761962097 | 206.2239133 | 194.3979381 | ENSG00000100532 |
| RAVER1 | -0.084894248 | 0.942853646 | 0.284746541 | 0.493977441 | 1048.316001 | 988.6635823 | ENSG00000161847 |
| SRSF9 | -0.084922592 | 0.942835122 | 0.251588485 | 0.457246515 | 3830.632649 | 3612.210235 | ENSG00000111786 |
| CALU | -0.084946625 | 0.942819416 | 0.140107948 | 0.304200805 | 4648.76826 | 4382.273349 | ENSG00000128595 |
| IKBKG | -0.084990129 | 0.942790986 | 0.690028612 | 0.83420974 | 108.2796334 | 101.9998859 | ENSG00000269335 |

| | | | | | | | |
|---|---|---|---|---|---|---|---|
| ACO2 | -0.08499749 | 0.942786176 | 0.182148082 | 0.365476681 | 1740.636805 | 1640.837261 | ENSG00000100412 |
| BBX | -0.085000442 | 0.942784247 | 0.399654598 | 0.611082267 | 782.8088361 | 737.5361162 | ENSG00000114439 |
| TLCD3B | -0.085018472 | 0.942772465 | 0.514517182 | 0.710801535 | 423.807766 | 399.7816074 | ENSG00000149926 |
| NA | -0.085040659 | 0.942757966 | 0.925047332 | 0.965883673 | 5.652981876 | 5.326652347 | ENSG00000255458 |
| SLCO5A1 | -0.085102494 | 0.94271756 | 0.702780758 | 0.842126015 | 105.5241922 | 99.46023118 | ENSG00000137571 |
| CADPS2 | -0.085171947 | 0.942672177 | 0.470788392 | 0.674161267 | 454.6700781 | 428.6311554 | ENSG00000081803 |
| NA | -0.0851738 | 0.942670966 | 0.932943067 | 0.969708285 | 4.569715966 | 4.330724884 | ENSG00000213755 |
| COPG1 | -0.085271868 | 0.94260689 | 0.158281146 | 0.331133204 | 2411.944231 | 2273.346004 | ENSG00000181789 |
| COPS8 | -0.085376069 | 0.942538812 | 0.267018003 | 0.474620031 | 1189.567728 | 1121.1575 | ENSG00000198612 |
| ANKRD52 | -0.085414645 | 0.942513609 | 0.183086509 | 0.366944518 | 1628.762209 | 1535.160029 | ENSG00000139645 |
| NDUFA5 | -0.085475313 | 0.942473976 | 0.336230967 | 0.548193198 | 1114.786761 | 1050.737584 | ENSG00000128609 |
| LMAN1 | -0.08552525 | 0.942441354 | 0.253728555 | 0.459940849 | 1935.451555 | 1823.736565 | ENSG00000074695 |
| HP1BP3 | -0.085581206 | 0.942404801 | 0.39688662 | 0.608821689 | 1485.529166 | 1400.31802 | ENSG00000127483 |
| GGA2 | -0.085590784 | 0.942398545 | 0.180819162 | 0.363839553 | 2642.76296 | 2490.204428 | ENSG00000103365 |
| CPEB1 | -0.085626148 | 0.942375445 | 0.858217377 | 0.930766391 | 20.44420048 | 19.30293119 | ENSG00000214575 |
| CREB3L2 | -0.085680852 | 0.942339712 | 0.327093113 | 0.538494422 | 1121.364692 | 1056.399281 | ENSG00000182158 |
| SNORA14B | -0.08577448 | 0.942278558 | 0.945831199 | 0.976226823 | 3.902498238 | 3.704513051 | ENSG00000207181 |
| PACSIN2 | -0.085866442 | 0.942218496 | 0.182647293 | 0.366283755 | 2384.568926 | 2246.941482 | ENSG00000100266 |
| EEPD1 | -0.085951755 | 0.94216278 | 0.485473326 | 0.686444852 | 348.5305602 | 328.2801041 | ENSG00000122547 |
| ABCG1 | -0.086001992 | 0.942129973 | 0.663118293 | 0.819398128 | 127.7129264 | 120.2664065 | ENSG00000160179 |
| CAPN12 | -0.086023735 | 0.942115774 | 0.764449457 | 0.878140389 | 65.30626296 | 61.64470906 | ENSG00000182472 |
| CHERP | -0.086023847 | 0.942115701 | 0.217347866 | 0.413271692 | 1730.3344 | 1630.715131 | ENSG00000085872 |
| PPIA | -0.086031383 | 0.94211078 | 0.043973386 | 0.127603744 | 24449.80887 | 23034.1466 | ENSG00000196262 |
| NA | -0.086043485 | 0.942102877 | 0.930075793 | 0.968220129 | 4.906286698 | 4.63479774 | ENSG00000268189 |
| NA | -0.086116502 | 0.942055197 | 0.847986547 | 0.924715668 | 22.20180306 | 20.9048289 | ENSG00000250132 |
| NA | -0.086211622 | 0.941993087 | 0.931094265 | 0.968853115 | 4.967966786 | 4.691738659 | ENSG00000228818 |
| MIER1 | -0.086213208 | 0.941992052 | 0.445726424 | 0.653197669 | 545.9685187 | 514.4966466 | ENSG00000198160 |
| HNRNPH1 | -0.086221721 | 0.941986493 | 0.070843517 | 0.184611539 | 10525.21501 | 9914.867973 | ENSG00000169045 |
| LRRCC1 | -0.086283162 | 0.941946377 | 0.582258496 | 0.762178672 | 246.6369246 | 231.9992438 | ENSG00000133739 |
| ETV6 | -0.086336322 | 0.941911669 | 0.439998484 | 0.647767794 | 419.7931453 | 395.2598409 | ENSG00000139083 |
| RBPMS | -0.086371492 | 0.941888708 | 0.176045916 | 0.356796214 | 2229.5825 | 2100.357221 | ENSG00000157110 |
| SLC2A6 | -0.086382314 | 0.941881643 | 0.413537848 | 0.624700612 | 636.4521284 | 600.0003594 | ENSG00000160326 |
| ARID4A | -0.086404047 | 0.941867454 | 0.559984595 | 0.745321888 | 247.1540859 | 232.6136323 | ENSG00000032219 |
| LSM2 | -0.086544748 | 0.941775601 | 0.238638231 | 0.441312475 | 1176.632991 | 1108.310725 | ENSG00000204392 |
| KDM7A | -0.086575412 | 0.941755585 | 0.51226644 | 0.708625235 | 317.1676367 | 298.6388184 | ENSG00000006459 |
| PRSS8 | -0.086579957 | 0.941752618 | 0.157398117 | 0.329724174 | 2046.909316 | 1927.777567 | ENSG00000052344 |
| FAM184A | -0.086615908 | 0.94172915 | 0.582388626 | 0.762243343 | 243.4903255 | 229.1240411 | ENSG00000111879 |
| SYNCRIP | -0.086659825 | 0.941700483 | 0.144347026 | 0.310371579 | 3950.264653 | 3719.890827 | ENSG00000135316 |
| TBC1D20 | -0.086760556 | 0.941634735 | 0.380391508 | 0.592762904 | 582.5253573 | 548.6667424 | ENSG00000125875 |
| NA | -0.086809401 | 0.941602855 | 0.870670885 | 0.939095303 | 16.53458329 | 15.55982688 | ENSG00000225200 |
| EEF1AKMT2 | -0.086894039 | 0.941547615 | 0.800387688 | 0.898320559 | 43.33190835 | 40.91578889 | ENSG00000203791 |
| JARID2 | -0.086911433 | 0.941536263 | 0.095973502 | 0.231076201 | 8239.245915 | 7757.194145 | ENSG00000008083 |
| LRIG1 | -0.086913904 | 0.941534651 | 0.118571471 | 0.270470133 | 2938.312747 | 2766.28496 | ENSG00000144749 |
| NA | -0.087085497 | 0.941422672 | 0.877695819 | 0.943087245 | 16.45037732 | 15.47800991 | ENSG00000213178 |
| NA | -0.08709526 | 0.941416301 | 0.878021014 | 0.943251997 | 15.42879149 | 14.54025287 | ENSG00000238207 |
| NCAPH2 | -0.087109727 | 0.941406861 | 0.189100235 | 0.375200455 | 1529.814725 | 1440.274702 | ENSG00000025770 |
| METTL2A | -0.087204293 | 0.941345156 | 0.327162927 | 0.538522781 | 762.8382458 | 717.7427479 | ENSG00000087995 |
| STAT2 | -0.087247794 | 0.941316772 | 0.381019175 | 0.593545089 | 607.8449788 | 572.0974382 | ENSG00000170581 |
| DNASE2 | -0.08725552 | 0.941311731 | 0.453579715 | 0.659622125 | 440.0238559 | 414.3315737 | ENSG00000105612 |
| WDR91 | -0.087379311 | 0.941230965 | 0.545610702 | 0.7337885 | 255.6743788 | 240.8133776 | ENSG00000105875 |
| ORMDL1 | -0.087449503 | 0.941185172 | 0.45959992 | 0.664609277 | 363.0830671 | 341.7756233 | ENSG00000128699 |
| NA | -0.087514316 | 0.941142891 | 0.871947855 | 0.940043115 | 15.9361646 | 15.00455471 | ENSG00000233426 |
| ERP44 | -0.087627201 | 0.941069253 | 0.373714373 | 0.585708628 | 701.5271046 | 660.4443185 | ENSG00000023318 |
| RPS3 | -0.087678745 | 0.941035631 | 0.074234554 | 0.191321714 | 38675.48712 | 36394.2615 | ENSG00000149273 |
| PPTC7 | -0.087850025 | 0.940923916 | 0.427847429 | 0.637657464 | 594.9495446 | 559.7645886 | ENSG00000196850 |
| SPAG7 | -0.088063971 | 0.940784391 | 0.232339043 | 0.433268018 | 1455.16388 | 1369.507376 | ENSG00000091640 |
| NA | -0.088071172 | 0.940779695 | 0.904092453 | 0.958242075 | 10.17525564 | 9.582086645 | ENSG00000205940 |
| GTPBP2 | -0.088094427 | 0.940764531 | 0.262253983 | 0.469313605 | 1304.81439 | 1227.500494 | ENSG00000172432 |
| NDUFAF4 | -0.088125558 | 0.940744231 | 0.386384966 | 0.598546476 | 614.4083643 | 578.3332604 | ENSG00000123545 |
| TRIT1 | -0.088170685 | 0.940714805 | 0.384711846 | 0.596983008 | 560.7890517 | 527.5358012 | ENSG00000043514 |
| ZBED3 | -0.08819645 | 0.940698005 | 0.681048277 | 0.829598484 | 104.0545229 | 97.94696039 | ENSG00000132846 |
| LENG8 | -0.088217456 | 0.940684309 | 0.723711681 | 0.854472967 | 3466.997267 | 3261.417052 | ENSG00000167615 |
| BICRA | -0.088220493 | 0.940682328 | 0.441647837 | 0.649302802 | 596.7828114 | 561.7138278 | ENSG00000063169 |
| PYGB | -0.088227364 | 0.940677848 | 0.118839805 | 0.270904935 | 2684.173835 | 2524.692729 | ENSG00000100994 |
| NA | -0.08831793 | 0.940618798 | 0.928966233 | 0.967758568 | 5.302173254 | 4.988339166 | ENSG00000253919 |
| NFE2L1 | -0.088327601 | 0.940612493 | 0.08189016 | 0.205481957 | 6632.400012 | 6238.007598 | ENSG00000082641 |
| POM121C | -0.088345547 | 0.940600793 | 0.306406926 | 0.516651321 | 1359.046184 | 1278.483213 | ENSG00000272391 |
| TDG | -0.088386534 | 0.940574071 | 0.299633701 | 0.509828132 | 1265.370845 | 1190.329643 | ENSG00000139372 |
| CHCHD7 | -0.088414144 | 0.94055607 | 0.343959304 | 0.556041042 | 721.6029006 | 678.8422629 | ENSG00000170791 |
| KIF22 | -0.088421057 | 0.940551563 | 0.144181334 | 0.310085922 | 3049.591995 | 2867.994933 | ENSG00000079616 |
| SRP19 | -0.088452669 | 0.940530954 | 0.668085701 | 0.822070092 | 122.2067465 | 115.0618863 | ENSG00000153037 |
| ITGAE | -0.08851202 | 0.940492263 | 0.486556253 | 0.687410402 | 397.2238797 | 373.351088 | ENSG00000083457 |
| SLC49A3 | -0.088569488 | 0.9404548 | 0.660575697 | 0.817400463 | 124.9109066 | 117.6507261 | ENSG00000169026 |
| ADGRL2 | -0.08862931 | 0.940415804 | 0.228883926 | 0.428561664 | 2176.503259 | 2046.323642 | ENSG00000117114 |

| | | | | | | | |
|---|---|---|---|---|---|---|---|
| ZNF3 | -0.088743997 | 0.940341049 | 0.405997122 | 0.617033288 | 856.9007483 | 805.2666036 | ENSG00000166526 |
| HGH1 | -0.088810192 | 0.940297905 | 0.303011717 | 0.513237777 | 952.5483728 | 895.9865239 | ENSG00000235173 |
| NEK1 | -0.088835558 | 0.940281372 | 0.517567231 | 0.713553596 | 277.6098847 | 260.9776183 | ENSG00000137601 |
| CARS1 | -0.088837288 | 0.940280244 | 0.19591795 | 0.384886142 | 1651.053026 | 1552.191793 | ENSG00000110619 |
| ACOX3 | -0.088889122 | 0.940246462 | 0.531037187 | 0.722993099 | 250.7126151 | 235.6773677 | ENSG00000087008 |
| PRKAB1 | -0.088904014 | 0.940236756 | 0.471289979 | 0.674623759 | 394.6529202 | 370.8190509 | ENSG00000111725 |
| DHDH | -0.088905838 | 0.940235568 | 0.835725794 | 0.91743775 | 28.04218946 | 26.33476931 | ENSG00000104808 |
| ARAF | -0.088963927 | 0.940197711 | 0.286760249 | 0.496012759 | 935.7471613 | 880.0556269 | ENSG00000078061 |
| RTEL1 | -0.088989529 | 0.940181026 | 0.831094335 | 0.915279712 | 27.82274968 | 26.12166679 | ENSG00000258366 |
| LYPLA2 | -0.088999918 | 0.940174256 | 0.090198855 | 0.22076275 | 3662.608439 | 3443.649485 | ENSG00000011009 |
| CCDC6 | -0.08900028 | 0.94017402 | 0.294025867 | 0.503827103 | 1298.438284 | 1221.096082 | ENSG00000108091 |
| HNRNPUL2 | -0.089090448 | 0.940115261 | 0.324339495 | 0.535228186 | 822.0428325 | 772.5710218 | ENSG00000214753 |
| ABLIM1 | -0.089105638 | 0.940105363 | 0.200828271 | 0.391599577 | 1516.88764 | 1425.91663 | ENSG00000099204 |
| FAM171A2 | -0.089181034 | 0.940056234 | 0.324432329 | 0.535287833 | 961.8592175 | 904.574575 | ENSG00000161682 |
| WASHC3 | -0.089239885 | 0.940017888 | 0.742506557 | 0.865919752 | 63.43745316 | 59.62436219 | ENSG00000120860 |
| NOL11 | -0.089437576 | 0.939889087 | 0.207209016 | 0.399708096 | 2065.537819 | 1941.420154 | ENSG00000130935 |
| MBD1 | -0.089465938 | 0.93987061 | 0.256772853 | 0.463634701 | 1069.620469 | 1005.201692 | ENSG00000141644 |
| PRKAR2B | -0.089466626 | 0.939870161 | 0.311316929 | 0.521901483 | 847.5099704 | 796.6128168 | ENSG00000005249 |
| PIP4K2B | -0.089481448 | 0.939860506 | 0.110706244 | 0.257727738 | 2488.586024 | 2338.906459 | ENSG00000276293 |
| NA | -0.089533072 | 0.939826875 | 0.930345647 | 0.968394286 | 4.569715966 | 4.311429258 | ENSG00000240449 |
| ARL8B | -0.089600113 | 0.939783203 | 0.320824813 | 0.531378407 | 758.8530673 | 713.282913 | ENSG00000134108 |
| TMBIM1 | -0.089677883 | 0.939732544 | 0.361730547 | 0.573767965 | 597.9967584 | 561.7599141 | ENSG00000135926 |
| PROZ | -0.089699829 | 0.939718249 | 0.91151695 | 0.961009829 | 8.134448263 | 7.69569525 | ENSG00000126231 |
| C2orf42 | -0.089700047 | 0.939718107 | 0.655864573 | 0.814050821 | 117.6776331 | 110.5286527 | ENSG00000115998 |
| CHKB | -0.089822781 | 0.939638166 | 0.919985696 | 0.964147051 | 6.394922374 | 5.983037123 | ENSG00000100288 |
| CCK | -0.089824732 | 0.939636895 | 0.913981041 | 0.961911055 | 8.179805529 | 7.627877013 | ENSG00000187094 |
| ZC3H14 | -0.08983726 | 0.939628736 | 0.211267384 | 0.404930939 | 1639.345733 | 1539.896128 | ENSG00000100722 |
| CHP1 | -0.089916693 | 0.939577003 | 0.185738277 | 0.370436938 | 1716.999226 | 1612.913341 | ENSG00000187446 |
| NOL6 | -0.089930913 | 0.939567742 | 0.139885289 | 0.30389206 | 2346.784969 | 2205.101506 | ENSG00000165271 |
| REST | -0.090004735 | 0.939519665 | 0.40447927 | 0.615476848 | 622.8929329 | 584.9665644 | ENSG00000084093 |
| POLR2K | -0.090015385 | 0.93951273 | 0.433821671 | 0.642616854 | 695.2278774 | 653.4418324 | ENSG00000147669 |
| PSORS1C1 | -0.090048908 | 0.9394909 | 0.92795272 | 0.967366576 | 4.566156493 | 4.307078331 | ENSG00000204540 |
| SUN2 | -0.090120188 | 0.939444482 | 0.549583872 | 0.7368755 | 248.9303755 | 233.8975967 | ENSG00000100242 |
| TMEM127 | -0.090141471 | 0.939430624 | 0.328173902 | 0.539669465 | 814.4522673 | 765.1867992 | ENSG00000135956 |
| ZNF331 | -0.090144827 | 0.939428438 | 0.818214278 | 0.907445186 | 1148.211492 | 1078.704962 | ENSG00000130844 |
| WIZ | -0.090178576 | 0.939406463 | 0.315819613 | 0.525855528 | 1290.573465 | 1212.991462 | ENSG00000011451 |
| CEP63 | -0.090223683 | 0.939377092 | 0.468299004 | 0.671564029 | 331.2205004 | 311.0805233 | ENSG00000182923 |
| NA | -0.090278063 | 0.939341684 | 0.869414454 | 0.938222398 | 18.72219262 | 17.63009382 | ENSG00000255306 |
| CCNI | -0.090341029 | 0.939300688 | 0.164554877 | 0.340038572 | 2877.134193 | 2703.263605 | ENSG00000118816 |
| CABP1 | -0.090432868 | 0.939240896 | 0.690825907 | 0.834552267 | 100.0061386 | 94.02577189 | ENSG00000157782 |
| JOSD1 | -0.090460604 | 0.939222839 | 0.338652106 | 0.550640467 | 754.6406429 | 708.8003907 | ENSG00000100221 |
| RAD9A | -0.090487866 | 0.939205091 | 0.458959752 | 0.664172791 | 380.4509419 | 357.422775 | ENSG00000172613 |
| RPL41 | -0.090581883 | 0.939143888 | 0.156789847 | 0.328887732 | 9684.525957 | 9095.389482 | ENSG00000229117 |
| REEP2 | -0.090602522 | 0.939130452 | 0.274875518 | 0.483650811 | 934.6891166 | 878.1576528 | ENSG00000132563 |
| GOLGA4 | -0.090628221 | 0.939113723 | 0.278090164 | 0.487399791 | 910.1389966 | 854.5235749 | ENSG00000144674 |
| RHPN1 | -0.090759487 | 0.939028281 | 0.70932794 | 0.846496645 | 917.8061412 | 861.9799637 | ENSG00000158106 |
| DGUOK | -0.090830146 | 0.938982291 | 0.265167542 | 0.47279405 | 945.0395203 | 887.1698788 | ENSG00000114956 |
| PTEN | -0.090830861 | 0.938981825 | 0.355837571 | 0.568142758 | 724.5283317 | 680.4794289 | ENSG00000171862 |
| KAT6B | -0.090843761 | 0.93897343 | 0.391433679 | 0.602856388 | 546.4423564 | 513.0420931 | ENSG00000156650 |
| CENPI | -0.090911615 | 0.938929268 | 0.464280517 | 0.668593683 | 340.401704 | 319.5322029 | ENSG00000102384 |
| SEPTIN7 | -0.090997362 | 0.938873464 | 0.159824194 | 0.333585312 | 2675.640549 | 2512.53783 | ENSG00000122545 |
| CMSS1 | -0.091014231 | 0.938862486 | 0.27968982 | 0.488880785 | 888.9201315 | 834.4219711 | ENSG00000184220 |
| SEM1 | -0.091022756 | 0.938856938 | 0.259572203 | 0.466503717 | 2230.028387 | 2093.52935 | ENSG00000127922 |
| SLC47A1 | -0.091037269 | 0.938847494 | 0.78818093 | 0.891582187 | 39.93441895 | 37.5315229 | ENSG00000142494 |
| FAM186B | -0.091285277 | 0.938686114 | 0.932922747 | 0.969708285 | 4.222466816 | 3.985885313 | ENSG00000135436 |
| CPEB2 | -0.091320331 | 0.938663307 | 0.842962684 | 0.9217804 | 25.04948501 | 23.52724658 | ENSG00000137449 |
| PDIA6 | -0.091353811 | 0.938641524 | 0.06177247 | 0.165889351 | 10139.20475 | 9516.266656 | ENSG00000143870 |
| SNRNP40 | -0.091390469 | 0.938617674 | 0.151360534 | 0.321029166 | 2118.031505 | 1987.542935 | ENSG00000060688 |
| TCEA1 | -0.091428002 | 0.938593255 | 0.158873188 | 0.332224576 | 3481.519437 | 3267.782042 | ENSG00000187735 |
| AFG3L1P | -0.091470006 | 0.938565928 | 0.592558817 | 0.769952585 | 184.7984502 | 173.4683487 | ENSG00000223959 |
| NUDT4 | -0.091497727 | 0.938547894 | 0.268233623 | 0.476089471 | 1403.247075 | 1317.12528 | ENSG00000173598 |
| GGCT | -0.091511322 | 0.93853905 | 0.42017097 | 0.630506855 | 408.3383202 | 383.4373699 | ENSG00000006625 |
| ZFP69 | -0.091513718 | 0.938537491 | 0.67189158 | 0.824260912 | 107.078931 | 100.521139 | ENSG00000187815 |
| MRPL12 | -0.091516921 | 0.938535407 | 0.685897833 | 0.831762616 | 98.16905861 | 92.24747099 | ENSG00000262814 |
| EDC3 | -0.091588047 | 0.938489138 | 0.253368719 | 0.459455015 | 1020.951351 | 957.8471078 | ENSG00000179151 |
| SMARCC2 | -0.091614743 | 0.938471772 | 0.284180531 | 0.493356131 | 2208.637305 | 2072.982588 | ENSG00000139613 |
| HNRNPDL | -0.09177912 | 0.938364851 | 0.042581941 | 0.124530574 | 7786.550192 | 7306.860698 | ENSG00000152795 |
| PGAP2 | -0.09179853 | 0.938352227 | 0.358280882 | 0.570123925 | 675.6297062 | 634.43211 | ENSG00000148985 |
| TMED2 | -0.091814636 | 0.938341751 | 0.224444346 | 0.422763962 | 3062.333207 | 2873.638847 | ENSG00000086598 |
| CEP192 | -0.091840637 | 0.93832484 | 0.318566345 | 0.528518441 | 700.9798629 | 657.487045 | ENSG00000101639 |
| FAM189A2 | -0.091866222 | 0.9383082 | 0.564696448 | 0.748949534 | 286.9993171 | 269.3797567 | ENSG00000135063 |
| TIAL1 | -0.091869309 | 0.938306192 | 0.156860803 | 0.328933604 | 1810.585525 | 1698.975424 | ENSG00000151923 |
| SYCE2 | -0.091929924 | 0.93826677 | 0.871914423 | 0.940043115 | 18.14985928 | 17.07822661 | ENSG00000161860 |
| CCDC63 | -0.091990363 | 0.938227464 | 0.868673618 | 0.937813209 | 15.85788236 | 14.86456731 | ENSG00000173093 |

| | | | | | | | |
|---|---|---|---|---|---|---|---|
| NXT1 | -0.091995758 | 0.938223955 | 0.438534804 | 0.6466204 | 381.3535651 | 357.6487194 | ENSG00000132661 |
| MAPKAPK5 | -0.092133264 | 0.938134536 | 0.31174265 | 0.522197973 | 856.5509317 | 803.9048397 | ENSG00000089022 |
| NDUFB8 | -0.092146916 | 0.938125658 | 0.832059168 | 0.915381416 | 24.43802364 | 22.95797567 | ENSG00000166136 |
| ZNF571 | -0.092152514 | 0.938122018 | 0.804517467 | 0.900504526 | 37.01526822 | 34.81234354 | ENSG00000180479 |
| ATG9A | -0.092156185 | 0.938119631 | 0.305869705 | 0.516092607 | 668.8806957 | 627.5196912 | ENSG00000198925 |
| NHSL2 | -0.09226405 | 0.938049494 | 0.57824024 | 0.759450456 | 212.1250639 | 198.7288309 | ENSG00000204131 |
| GATAD2A | -0.092309654 | 0.938019842 | 0.106622568 | 0.250386381 | 3995.194094 | 3747.172364 | ENSG00000167491 |
| TSR3 | -0.092332259 | 0.938005145 | 0.283192922 | 0.492186485 | 1151.40027 | 1080.601495 | ENSG00000007520 |
| TTBK2 | -0.092435973 | 0.937937715 | 0.59057921 | 0.768530555 | 186.2025483 | 174.5843579 | ENSG00000128881 |
| ZNF451 | -0.092457108 | 0.937923975 | 0.374235785 | 0.586315149 | 491.5542568 | 461.1086237 | ENSG00000112200 |
| SH3BGRL | -0.092478124 | 0.937910312 | 0.352982112 | 0.56525627 | 650.3181201 | 610.3298738 | ENSG00000131171 |
| NA | -0.092540147 | 0.937869991 | 0.848356648 | 0.9249161 | 22.32724817 | 21.01341385 | ENSG00000269929 |
| NA | -0.092591104 | 0.937836866 | 0.847194213 | 0.924172673 | 20.86380813 | 19.55441405 | ENSG00000249936 |
| CD82 | -0.092613399 | 0.937822373 | 0.655525363 | 0.813858161 | 119.7691628 | 112.3850821 | ENSG00000085117 |
| PCDHA10 | -0.09263522 | 0.937808188 | 0.538099221 | 0.727964146 | 220.5753364 | 206.9458845 | ENSG00000250120 |
| NA | -0.092661449 | 0.937791138 | 0.859448434 | 0.931833946 | 20.9859731 | 19.6707549 | ENSG00000246792 |
| ATG5 | -0.092676864 | 0.937781118 | 0.377865323 | 0.590336418 | 732.4506873 | 686.4688542 | ENSG00000057663 |
| CIC | -0.092704592 | 0.937763095 | 0.216299459 | 0.411751274 | 1749.248935 | 1640.869725 | ENSG00000079432 |
| YWHAZ | -0.092807671 | 0.937696094 | 0.085593659 | 0.212465332 | 11938.48612 | 11194.38872 | ENSG00000164924 |
| MRPL18 | -0.092850085 | 0.937668527 | 0.215098203 | 0.4101504 | 1414.734086 | 1326.197246 | ENSG00000112110 |
| H2AC13 | -0.09309291 | 0.937510718 | 0.869913893 | 0.938580298 | 14.547168 | 13.65726128 | ENSG00000196747 |
| POLR1B | -0.093161379 | 0.937466226 | 0.180398919 | 0.363318074 | 1662.048782 | 1557.953412 | ENSG00000125630 |
| NA | -0.093231134 | 0.937420901 | 0.93237116 | 0.969647522 | 4.531756977 | 4.257609762 | ENSG00000242338 |
| KBTBD2 | -0.093291061 | 0.937381962 | 0.362261241 | 0.574079344 | 564.2805613 | 529.1974777 | ENSG00000170852 |
| TPMT | -0.093350164 | 0.937343562 | 0.355269121 | 0.567571047 | 629.4773448 | 589.8948842 | ENSG00000137364 |
| MICALL2 | -0.093411718 | 0.937303569 | 0.614939301 | 0.785838998 | 184.6555132 | 173.0440786 | ENSG00000164877 |
| RAB1A | -0.093429877 | 0.937291772 | 0.111253598 | 0.258714961 | 3318.439237 | 3109.867492 | ENSG00000138069 |
| ASAP1 | -0.093532938 | 0.937224818 | 0.320220976 | 0.530657523 | 800.0112999 | 749.4302322 | ENSG00000153317 |
| SHQ1 | -0.093671209 | 0.937134996 | 0.314688191 | 0.524986448 | 695.1261022 | 651.4331111 | ENSG00000144736 |
| PHETA2 | -0.093869195 | 0.937006399 | 0.691331738 | 0.834876575 | 93.02105942 | 87.2159061 | ENSG00000177096 |
| RALGAPB | -0.0938698 | 0.937006006 | 0.272033023 | 0.480664637 | 1277.861924 | 1196.677457 | ENSG00000170471 |
| IDNK | -0.093875102 | 0.937002562 | 0.918444747 | 0.963976494 | 5.696864602 | 5.321355461 | ENSG00000148057 |
| COL4A1 | -0.093876156 | 0.937001878 | 0.141044982 | 0.305427426 | 2850.229476 | 2670.487044 | ENSG00000187498 |
| TMEM80 | -0.093893352 | 0.936990709 | 0.593764655 | 0.770974282 | 225.5878486 | 211.6328208 | ENSG00000177042 |
| STK25 | -0.093937463 | 0.936962061 | 0.098936745 | 0.236329423 | 4305.976439 | 4034.901129 | ENSG00000115694 |
| LOC107984813 | -0.094048778 | 0.93688977 | 0.919026259 | 0.964051188 | 5.298613782 | 4.955990759 | ENSG00000259804 |
| SEPT5-GP1BB | -0.094114324 | 0.936847205 | 0.914748659 | 0.96227163 | 6.694729326 | 6.231115019 | ENSG00000284874 |
| PLP1 | -0.09412944 | 0.936837389 | 0.169708009 | 0.347757786 | 1451.52829 | 1360.150317 | ENSG00000123560 |
| GTF2IP12 | -0.094212323 | 0.936783569 | 0.673939062 | 0.825366912 | 100.2798341 | 94.06407959 | ENSG00000283050 |
| ELOVL5 | -0.094213632 | 0.936782719 | 0.140812639 | 0.305099436 | 3779.237181 | 3539.910358 | ENSG00000012660 |
| OLMALINC | -0.094280172 | 0.936739514 | 0.32607273 | 0.537312565 | 669.7178266 | 627.0238189 | ENSG00000235823 |
| SAR1A | -0.094287758 | 0.936734588 | 0.15014255 | 0.319162772 | 2016.320901 | 1888.653656 | ENSG00000079332 |
| PARD3 | -0.094337223 | 0.936702472 | 0.187244928 | 0.372615458 | 1622.37273 | 1519.677603 | ENSG00000148498 |
| POLR2G | -0.094340319 | 0.936700461 | 0.244277331 | 0.448705182 | 1288.292265 | 1206.692919 | ENSG00000168002 |
| NA | -0.094368137 | 0.9366824 | 0.920211016 | 0.964276164 | 5.356734397 | 5.009810256 | ENSG00000218418 |
| PPM1J | -0.094448875 | 0.936629982 | 0.506247945 | 0.703598207 | 253.8450025 | 237.7959893 | ENSG00000155367 |
| PPA1 | -0.094452223 | 0.936627808 | 0.107638421 | 0.252265027 | 2750.419432 | 2576.002489 | ENSG00000180817 |
| CDK13 | -0.094457931 | 0.936624102 | 0.247649375 | 0.452657613 | 1262.649472 | 1182.890088 | ENSG00000065883 |
| RB1 | -0.094528928 | 0.936578011 | 0.442896258 | 0.650446833 | 356.8592264 | 334.063543 | ENSG00000139687 |
| DNAJC16 | -0.094544319 | 0.93656802 | 0.462752226 | 0.667551225 | 348.0196276 | 325.7715253 | ENSG00000116138 |
| NA | -0.094557925 | 0.936559186 | 0.83188609 | 0.915381416 | 23.34527452 | 21.90822871 | ENSG00000253771 |
| MFSD2A | -0.094610829 | 0.936524844 | 0.359633387 | 0.571738068 | 505.0442443 | 473.0775358 | ENSG00000168389 |
| MIS18BP1 | -0.094788759 | 0.936409348 | 0.281789215 | 0.490709708 | 1080.700179 | 1011.966312 | ENSG00000129534 |
| SPINT1-AS1 | -0.094797304 | 0.936403801 | 0.814496763 | 0.90576913 | 34.68116244 | 32.37330694 | ENSG00000261183 |
| DYNLT3 | -0.094830159 | 0.936382476 | 0.741243841 | 0.865005867 | 72.04668061 | 67.6079561 | ENSG00000165169 |
| MAGT1 | -0.094917166 | 0.936326006 | 0.257011816 | 0.463948031 | 1087.490962 | 1018.059279 | ENSG00000102158 |
| SNRK | -0.094938552 | 0.936312126 | 0.626635646 | 0.793430384 | 178.6783662 | 167.2163821 | ENSG00000163788 |
| FKBP8 | -0.094975597 | 0.936288085 | 0.057756214 | 0.157687741 | 8220.211071 | 7696.967085 | ENSG00000105701 |
| C12orf49 | -0.095068783 | 0.93622761 | 0.202873955 | 0.394245177 | 1575.54064 | 1474.689886 | ENSG00000111412 |
| EDF1 | -0.095235314 | 0.936119547 | 0.149755278 | 0.3185188 | 4388.096846 | 4107.780329 | ENSG00000107223 |
| PURG | -0.095248838 | 0.936110772 | 0.609166358 | 0.782326193 | 143.2001179 | 133.9436258 | ENSG00000172733 |
| SRGAP2B | -0.095299193 | 0.936078099 | 0.561235869 | 0.746231726 | 217.918924 | 203.7954868 | ENSG00000196369 |
| C20orf194 | -0.095332178 | 0.936056697 | 0.536601968 | 0.726856257 | 229.9338771 | 214.977223 | ENSG00000088854 |
| RNF149 | -0.095396585 | 0.936014909 | 0.449577635 | 0.655923489 | 367.2710821 | 343.85743 | ENSG00000163162 |
| C5orf15 | -0.095504152 | 0.935945123 | 0.399104588 | 0.610587342 | 623.4033842 | 583.8618114 | ENSG00000113583 |
| BIN1 | -0.095506177 | 0.935943809 | 0.199159462 | 0.389203606 | 1407.305834 | 1317.362748 | ENSG00000136717 |
| AGK | -0.095542972 | 0.935919939 | 0.350283671 | 0.562428154 | 583.6856847 | 546.1125218 | ENSG00000006530 |
| DELE1 | -0.095556383 | 0.935911239 | 0.278959011 | 0.488061971 | 909.1087132 | 850.8094274 | ENSG00000081791 |
| NA | -0.095562834 | 0.935907054 | 0.924877559 | 0.965883673 | 5.009485247 | 4.688617237 | ENSG00000279300 |
| MRM1 | -0.095567496 | 0.93590403 | 0.347201249 | 0.559318911 | 533.1047898 | 498.8589249 | ENSG00000278619 |
| GRAMD2A | -0.095636016 | 0.93585958 | 0.823590411 | 0.910733394 | 24.77223011 | 23.18023155 | ENSG00000175318 |
| DFFA | -0.095678609 | 0.935831951 | 0.178769921 | 0.361077001 | 1553.305432 | 1453.548038 | ENSG00000160049 |
| ZNF354A | -0.095683927 | 0.935828501 | 0.546672471 | 0.734640975 | 216.2171033 | 202.5881081 | ENSG00000169131 |
| HNRNPAB | -0.095688569 | 0.93582549 | 0.064007374 | 0.170298572 | 17285.74097 | 16177.19972 | ENSG00000197451 |

| | | | | | | | |
|---|---|---|---|---|---|---|---|
| SIN3A | -0.095736114 | 0.93579465 | 0.110108252 | 0.256651981 | 2535.605641 | 2372.486061 | ENSG00000169375 |
| SLC35C2 | -0.095749296 | 0.935786099 | 0.252660485 | 0.458591633 | 983.1280132 | 919.9313167 | ENSG00000080189 |
| NA | -0.095798947 | 0.935753895 | 0.789753162 | 0.892242184 | 35.52246268 | 33.21479676 | ENSG00000259917 |
| POLDIP3 | -0.095823653 | 0.93573787 | 0.09788442 | 0.234376426 | 3451.264674 | 3228.834624 | ENSG00000100227 |
| SDF2 | -0.095856748 | 0.935716405 | 0.331251761 | 0.542840116 | 638.7033475 | 597.233066 | ENSG00000132581 |
| SCD5 | -0.095953598 | 0.935653591 | 0.786881391 | 0.891330078 | 43.49350685 | 40.60062781 | ENSG00000145284 |
| RBM27 | -0.095979207 | 0.935636983 | 0.752782026 | 0.871202878 | 342.952631 | 320.9669234 | ENSG00000091009 |
| BCLAF1 | -0.096001027 | 0.935622832 | 0.23893406 | 0.441643313 | 3008.149902 | 2814.258977 | ENSG00000029363 |
| FGD1 | -0.09607586 | 0.935574302 | 0.316244651 | 0.526264103 | 694.3334911 | 649.7634678 | ENSG00000102302 |
| AK9 | -0.096125492 | 0.935542117 | 0.821065582 | 0.909223 | 28.98521371 | 27.12724206 | ENSG00000155085 |
| TSPO | -0.096142795 | 0.935530896 | 0.489657151 | 0.689811643 | 305.1176998 | 285.458272 | ENSG00000100300 |
| NKTR | -0.096159808 | 0.935519864 | 0.291807301 | 0.501459414 | 1112.519396 | 1040.864409 | ENSG00000114857 |
| NYAP1 | -0.096196217 | 0.935496255 | 0.55102587 | 0.738180007 | 204.9641744 | 191.7875546 | ENSG00000166924 |
| NA | -0.096229656 | 0.935474572 | 0.746967425 | 0.86838024 | 59.72472372 | 55.85687629 | ENSG00000259295 |
| PHRF1 | -0.096301963 | 0.935427688 | 0.125009307 | 0.280843278 | 2713.887918 | 2538.937518 | ENSG00000070047 |
| FZD8 | -0.096422661 | 0.935349432 | 0.413479429 | 0.624700612 | 420.9298027 | 393.8229746 | ENSG00000177283 |
| PRKX | -0.09644717 | 0.935333542 | 0.179674679 | 0.362426894 | 2290.335624 | 2141.745944 | ENSG00000183943 |
| PTK2 | -0.0964521 | 0.935330346 | 0.09474284 | 0.229196202 | 3206.971644 | 2999.036513 | ENSG00000169398 |
| WDR47 | -0.096554713 | 0.935263822 | 0.625353219 | 0.792816565 | 148.4121353 | 138.6677855 | ENSG00000085433 |
| NUDT5 | -0.096712092 | 0.935161802 | 0.215679155 | 0.410880115 | 1071.763838 | 1002.378417 | ENSG00000165609 |
| PWWP3A | -0.096742099 | 0.935142352 | 0.274753509 | 0.483571296 | 853.3343133 | 798.0449034 | ENSG00000160953 |
| SSNA1 | -0.096820616 | 0.935091459 | 0.143870887 | 0.30977103 | 2083.998488 | 1948.657942 | ENSG00000176101 |
| LOC100507373 | -0.096837071 | 0.935080794 | 0.737837673 | 0.862523334 | 66.51583813 | 62.03010443 | ENSG00000267169 |
| PSMG4 | -0.096842527 | 0.935077257 | 0.330175385 | 0.541601209 | 769.1154785 | 719.252627 | ENSG00000180822 |
| NA | -0.096878047 | 0.935054236 | 0.927955827 | 0.967366576 | 4.228390553 | 3.969711109 | ENSG00000286585 |
| UNK | -0.097007874 | 0.934970095 | 0.35470076 | 0.566806892 | 508.3424001 | 475.4837725 | ENSG00000132478 |
| NA | -0.097033605 | 0.934953419 | 0.931802115 | 0.969216528 | 4.638514998 | 4.342548161 | ENSG00000275263 |
| SLC19A1 | -0.097037526 | 0.934950878 | 0.197217904 | 0.38677353 | 1113.981972 | 1041.65732 | ENSG00000173638 |
| FOXP2 | -0.097064952 | 0.934933105 | 0.921434706 | 0.964771134 | 4.613598691 | 4.321077071 | ENSG00000128573 |
| TMEM53 | -0.097076895 | 0.934925365 | 0.590452911 | 0.768530555 | 153.4690627 | 143.4272698 | ENSG00000126106 |
| SLC35E2B | -0.097116843 | 0.934899478 | 0.168979496 | 0.346678738 | 1457.714809 | 1362.961202 | ENSG00000189339 |
| GOLT1B | -0.097146189 | 0.934880461 | 0.418408125 | 0.629161353 | 387.2880234 | 362.3284239 | ENSG00000111711 |
| MSH6 | -0.097165557 | 0.93486791 | 0.0570908 | 0.15639103 | 6372.726016 | 5957.269311 | ENSG00000116062 |
| PTPRJ | -0.097173153 | 0.934862988 | 0.346160884 | 0.558166691 | 563.4629304 | 526.505874 | ENSG00000149177 |
| RASGEF1A | -0.097183514 | 0.934856274 | 0.389442811 | 0.601255569 | 466.2603634 | 436.0056803 | ENSG00000198915 |
| NA | -0.097200531 | 0.934845248 | 0.921054352 | 0.964771134 | 4.951364632 | 4.626095886 | ENSG00000244265 |
| SS18L1 | -0.097248539 | 0.93481414 | 0.487171346 | 0.688022052 | 484.935978 | 453.5323087 | ENSG00000184402 |
| DNAJC17 | -0.097266283 | 0.934802642 | 0.500264033 | 0.698623549 | 254.2568547 | 237.4267252 | ENSG00000104129 |
| UFD1 | -0.097560269 | 0.934612172 | 0.225262295 | 0.423586423 | 1195.580112 | 1117.186414 | ENSG00000070010 |
| MAPK1IP1L | -0.097604203 | 0.93458371 | 0.202191717 | 0.393486251 | 1348.250517 | 1260.327137 | ENSG00000168175 |
| MON1B | -0.097631002 | 0.93456635 | 0.212780981 | 0.406923426 | 955.402817 | 892.7894431 | ENSG00000103111 |
| TACC1 | -0.097662939 | 0.934545662 | 0.447799362 | 0.654542644 | 397.5984094 | 371.1432497 | ENSG00000147526 |
| LRP1 | -0.097685244 | 0.934531213 | 0.504890254 | 0.702279771 | 6125.693761 | 5724.330723 | ENSG00000123384 |
| ULK3 | -0.097738179 | 0.934496924 | 0.307374095 | 0.517727416 | 704.5543839 | 658.3873379 | ENSG00000140474 |
| ILRUN | -0.09785397 | 0.934421925 | 0.078559215 | 0.199086256 | 2988.842841 | 2792.716389 | ENSG00000196821 |
| EYA3 | -0.097854941 | 0.934421296 | 0.299511069 | 0.509733139 | 715.1347823 | 668.0836197 | ENSG00000158161 |
| ALKBH1 | -0.097879939 | 0.934405104 | 0.419463098 | 0.629995276 | 351.3672582 | 328.2407348 | ENSG00000100601 |
| ZDHHC7 | -0.097921861 | 0.934377953 | 0.180754092 | 0.363761831 | 1500.083837 | 1401.43746 | ENSG00000153786 |
| DYSF | -0.097922927 | 0.934377262 | 0.243077764 | 0.447070858 | 858.1317571 | 801.682027 | ENSG00000135636 |
| TNFSF15 | -0.098066902 | 0.93428402 | 0.928924643 | 0.967758568 | 3.888260348 | 3.658165904 | ENSG00000181634 |
| B3GNTL1 | -0.098085706 | 0.934271843 | 0.592444622 | 0.769952585 | 141.3390374 | 132.1468073 | ENSG00000175711 |
| MRPS23 | -0.098116265 | 0.934252053 | 0.155659916 | 0.327451271 | 1522.707735 | 1422.337581 | ENSG00000181610 |
| NA | -0.09813948 | 0.93423702 | 0.917172182 | 0.96312259 | 4.971526258 | 4.640094626 | ENSG00000253973 |
| L2HGDH | -0.098179052 | 0.934211395 | 0.280649522 | 0.489568283 | 801.8456578 | 748.7278791 | ENSG00000087299 |
| PAN2 | -0.098179435 | 0.934211147 | 0.44678104 | 0.653682173 | 506.991032 | 473.5311189 | ENSG00000135473 |
| UBXN11 | -0.098201896 | 0.934196603 | 0.558185774 | 0.743916196 | 209.5893413 | 195.961004 | ENSG00000158062 |
| YWHAG | -0.09821095 | 0.93419074 | 0.065426012 | 0.173291758 | 7793.154068 | 7279.865528 | ENSG00000170027 |
| GLUD1 | -0.098262873 | 0.934157119 | 0.18184709 | 0.365120512 | 2136.604634 | 1995.635174 | ENSG00000148672 |
| DCAF1 | -0.098441617 | 0.934041388 | 0.232884021 | 0.433772042 | 1112.621295 | 1038.78366 | ENSG00000145041 |
| PPP5C | -0.098447013 | 0.934037894 | 0.136368877 | 0.298778204 | 3462.353606 | 3233.545545 | ENSG00000011485 |
| APPL2 | -0.098476615 | 0.934018729 | 0.437298156 | 0.645322083 | 353.2040851 | 329.8522804 | ENSG00000136044 |
| ACP1 | -0.098477362 | 0.934018246 | 0.086580487 | 0.214098794 | 2664.943606 | 2489.050574 | ENSG00000143727 |
| PXDN | -0.098496083 | 0.934006126 | 0.09102482 | 0.22217962 | 6443.255767 | 6017.629097 | ENSG00000130508 |
| CIAO2A | -0.098522358 | 0.933989115 | 0.235991434 | 0.437918663 | 1026.523913 | 958.79919 | ENSG00000166797 |
| C11orf24 | -0.098603571 | 0.93393654 | 0.285391559 | 0.494673527 | 775.7437981 | 724.9750576 | ENSG00000171067 |
| CREBL2 | -0.09860573 | 0.933935142 | 0.322057771 | 0.532673061 | 537.391632 | 502.0188322 | ENSG00000111269 |
| C20orf27 | -0.098633058 | 0.933917452 | 0.137584689 | 0.300379984 | 2389.009559 | 2231.756252 | ENSG00000101220 |
| TTC34 | -0.098762736 | 0.93383351 | 0.92868755 | 0.967681769 | 4.65155768 | 4.347845047 | ENSG00000215912 |
| RHOH | -0.098773016 | 0.933826856 | 0.501467724 | 0.699683633 | 247.924807 | 231.6642629 | ENSG00000168421 |
| LINC00926 | -0.098802749 | 0.93380761 | 0.865646321 | 0.935599541 | 14.25328478 | 13.31459718 | ENSG00000247982 |
| LMTK2 | -0.098913595 | 0.933735866 | 0.250436877 | 0.455942388 | 904.7676671 | 844.4044502 | ENSG00000164715 |
| RPS8 | -0.098945041 | 0.933715514 | 0.060308087 | 0.162984918 | 17937.6512 | 16748.16237 | ENSG00000142937 |
| NA | -0.098993391 | 0.933684222 | 0.917077083 | 0.96312259 | 4.995247357 | 4.658444293 | ENSG00000286181 |
| RAF1 | -0.099116396 | 0.933604619 | 0.098399343 | 0.235366592 | 2703.007644 | 2523.765672 | ENSG00000132155 |

| | | | | | | | |
|---|---|---|---|---|---|---|---|
| KCNN1 | -0.099160894 | 0.933575824 | 0.517795289 | 0.713763798 | 235.4184464 | 219.8542347 | ENSG00000105642 |
| SRRD | -0.099231681 | 0.933530018 | 0.502760139 | 0.700813789 | 228.8592302 | 213.6569811 | ENSG00000100104 |
| CD72 | -0.099294784 | 0.933489187 | 0.780217101 | 0.88782839 | 44.25680418 | 41.2895719 | ENSG00000137101 |
| NA | -0.099326731 | 0.933468516 | 0.460969306 | 0.665754462 | 270.5239973 | 252.5688083 | ENSG00000184319 |
| SNX13 | -0.099382945 | 0.933432144 | 0.434690366 | 0.643487883 | 407.0190124 | 379.9665142 | ENSG00000071189 |
| MON2 | -0.099399644 | 0.93342134 | 0.359722756 | 0.571832007 | 497.7599435 | 464.3005232 | ENSG00000061987 |
| PSMD3 | -0.09944414 | 0.933392552 | 0.0481533 | 0.137015983 | 7405.72519 | 6912.673228 | ENSG00000108344 |
| SPATA6 | -0.099475205 | 0.933372454 | 0.557345287 | 0.743522876 | 187.7274152 | 175.2796175 | ENSG00000132122 |
| KCMF1 | -0.099476069 | 0.933371895 | 0.164410807 | 0.339885648 | 2130.314158 | 1988.213569 | ENSG00000176407 |
| LIG4 | -0.099558138 | 0.9333188 | 0.631332856 | 0.797026605 | 115.2810092 | 107.6979779 | ENSG00000174405 |
| RPL26 | -0.099603619 | 0.933289378 | 0.304602056 | 0.514710972 | 646.9866827 | 603.6542267 | ENSG00000161970 |
| KRT18 | -0.099785422 | 0.933171776 | 0.084969556 | 0.211387252 | 3248.169315 | 3031.338921 | ENSG00000111057 |
| RP2 | -0.099793513 | 0.933166542 | 0.530775209 | 0.722844871 | 248.4102646 | 231.9519507 | ENSG00000102218 |
| SOCS5 | -0.099794878 | 0.933165659 | 0.399431891 | 0.610828948 | 370.1013245 | 345.3809157 | ENSG00000171150 |
| KIAA1549 | -0.099823159 | 0.933147367 | 0.319091076 | 0.529256214 | 670.2759476 | 625.196458 | ENSG00000122778 |
| TMEM41B | -0.099879609 | 0.933110855 | 0.36706514 | 0.579491407 | 518.5208574 | 483.8928232 | ENSG00000166471 |
| CHD1L | -0.099893524 | 0.933101855 | 0.922203316 | 0.964771134 | 4.582758647 | 4.279080851 | ENSG00000131778 |
| NA | -0.099907729 | 0.933092668 | 0.919794357 | 0.964147051 | 4.596996537 | 4.293079591 | ENSG00000275882 |
| LINC00106 | -0.099912192 | 0.933089781 | 0.824577155 | 0.911397611 | 25.01895046 | 23.31681398 | ENSG00000236871 |
| ZNF569 | -0.099973037 | 0.933050429 | 0.590969819 | 0.768732954 | 150.2447917 | 140.3361993 | ENSG00000196437 |
| CHD1 | -0.099994069 | 0.933036827 | 0.242604953 | 0.446418643 | 1287.074986 | 1200.735773 | ENSG00000153922 |
| SIAH2 | -0.100046159 | 0.93300314 | 0.326480294 | 0.537821036 | 517.2448742 | 482.6888495 | ENSG00000181788 |
| TLN1 | -0.100110829 | 0.932961318 | 0.0570109 | 0.156217476 | 4224.25255 | 3940.694256 | ENSG00000137076 |
| LYN | -0.100366815 | 0.932795792 | 0.219194509 | 0.415653007 | 1021.340476 | 952.6057727 | ENSG00000254087 |
| PIK3AP1 | -0.100384868 | 0.932784119 | 0.632023214 | 0.797206785 | 113.6147316 | 105.9664756 | ENSG00000155629 |
| PGS1 | -0.10044089 | 0.932747899 | 0.223309011 | 0.421260403 | 1133.200781 | 1057.165895 | ENSG00000087157 |
| GTF2I | -0.10057583 | 0.93266066 | 0.139800056 | 0.303741837 | 1616.055985 | 1507.555789 | ENSG00000263001 |
| MAP2K2 | -0.100682385 | 0.932591778 | 0.102386983 | 0.242363771 | 3223.642714 | 3006.473072 | ENSG00000126934 |
| ABRACL | -0.100724899 | 0.932564296 | 0.144305595 | 0.310317828 | 1482.103811 | 1382.425308 | ENSG00000146386 |
| VRTN | -0.100791049 | 0.932521537 | 0.048722866 | 0.138324011 | 5587.067087 | 5210.604136 | ENSG00000133980 |
| NA | -0.101010961 | 0.932379402 | 0.457051517 | 0.662777555 | 286.279597 | 267.0415265 | ENSG00000284691 |
| LARP1B | -0.101067763 | 0.932342694 | 0.260185742 | 0.467028585 | 731.0421133 | 681.6310141 | ENSG00000138709 |
| ARAP3 | -0.101068276 | 0.932342362 | 0.090505041 | 0.221311368 | 2605.129111 | 2428.458661 | ENSG00000120318 |
| MMS19 | -0.101081458 | 0.932333843 | 0.084062053 | 0.209600193 | 2476.823131 | 2309.158501 | ENSG00000155229 |
| PRB3 | -0.101122637 | 0.932307232 | 0.827682069 | 0.913332702 | 22.0404839 | 20.58798681 | ENSG00000197870 |
| OXNAD1 | -0.101179188 | 0.932270688 | 0.390878068 | 0.602246252 | 432.4154678 | 403.2219113 | ENSG00000154814 |
| TMEM51 | -0.101192522 | 0.932262071 | 0.433513516 | 0.642497135 | 328.5024073 | 306.4658945 | ENSG00000171729 |
| DOCK11 | -0.101214166 | 0.932248085 | 0.36151041 | 0.573611502 | 457.2855569 | 426.3459167 | ENSG00000147251 |
| CDKN2AIP | -0.101232363 | 0.932236327 | 0.343409798 | 0.555395742 | 502.9886142 | 469.1778184 | ENSG00000168564 |
| FAM210A | -0.101286821 | 0.932201138 | 0.405796303 | 0.616877017 | 365.4040785 | 340.4119448 | ENSG00000177150 |
| CEP85 | -0.101385896 | 0.932137122 | 0.226962152 | 0.425807692 | 1223.971435 | 1140.543498 | ENSG00000130695 |
| TCP1 | -0.101466099 | 0.932085304 | 0.027665152 | 0.089232518 | 7368.182396 | 6867.946734 | ENSG00000120438 |
| RPLP2 | -0.101484358 | 0.932073507 | 0.074239106 | 0.191321714 | 10822.20799 | 10087.62188 | ENSG00000177600 |
| PANK2 | -0.101494157 | 0.932067177 | 0.271909086 | 0.4805251 | 713.7840496 | 665.4373875 | ENSG00000125779 |
| SPTBN2 | -0.101534213 | 0.932041299 | 0.120678972 | 0.273690691 | 2161.967579 | 2015.119567 | ENSG00000173898 |
| ATL3 | -0.101550146 | 0.932031006 | 0.435459391 | 0.644005602 | 308.4312364 | 287.3624233 | ENSG00000184743 |
| ATMIN | -0.101641073 | 0.931972265 | 0.213331329 | 0.407657787 | 1311.698181 | 1222.48017 | ENSG00000166454 |
| DIS3L | -0.101642002 | 0.931971665 | 0.21354916 | 0.40799167 | 1093.916647 | 1018.89767 | ENSG00000166938 |
| BEX3 | -0.101679206 | 0.931947632 | 0.02606001 | 0.085116445 | 8174.392769 | 7618.498217 | ENSG00000166681 |
| ZFAND4 | -0.101683925 | 0.931944583 | 0.510774215 | 0.707234478 | 212.0779266 | 197.7124939 | ENSG00000172671 |
| PURB | -0.101743234 | 0.931906272 | 0.244219434 | 0.448646305 | 808.5774826 | 753.3357113 | ENSG00000146676 |
| SKA1 | -0.101758316 | 0.93189653 | 0.456596714 | 0.662631902 | 330.6309801 | 308.2192967 | ENSG00000154839 |
| HSPA14 | -0.101807979 | 0.931864451 | 0.410673903 | 0.621689511 | 425.9119562 | 396.5331888 | ENSG00000284024 |
| GOLGA2P7 | -0.101808028 | 0.93186442 | 0.542061905 | 0.731177781 | 202.888662 | 189.1142279 | ENSG00000225151 |
| LXN | -0.101833318 | 0.931848084 | 0.690120475 | 0.834267391 | 71.61133495 | 66.68514376 | ENSG00000079257 |
| CERS4 | -0.101856973 | 0.931832806 | 0.155984962 | 0.32785487 | 1208.105238 | 1125.655999 | ENSG00000090661 |
| LSM14A | -0.101910071 | 0.93179851 | 0.066927781 | 0.1764774 | 4718.886507 | 4396.946047 | ENSG00000257103 |
| NA | -0.101941475 | 0.931778227 | 0.722311268 | 0.853620043 | 58.71145205 | 54.62637522 | ENSG00000255121 |
| LCLAT1 | -0.101948134 | 0.931773927 | 0.545854008 | 0.733992237 | 180.8876378 | 168.4493422 | ENSG00000172954 |
| LINC02610 | -0.101966284 | 0.931762204 | 0.79714879 | 0.896454171 | 36.36640652 | 33.80241734 | ENSG00000186235 |
| NAPG | -0.102029168 | 0.931721592 | 0.3702854 | 0.582445426 | 411.0451763 | 383.1789818 | ENSG00000134265 |
| EBNA1BP2 | -0.102143419 | 0.931647809 | 0.155636272 | 0.327449977 | 1423.847868 | 1326.396917 | ENSG00000117395 |
| CEPT1 | -0.102147398 | 0.931645239 | 0.489395168 | 0.689558553 | 264.2174247 | 246.2653132 | ENSG00000134255 |
| ATAD2B | -0.102223961 | 0.931595799 | 0.408980368 | 0.620117475 | 343.0125066 | 319.4606008 | ENSG00000119778 |
| ARMCX1 | -0.102239748 | 0.931585605 | 0.312288043 | 0.522595648 | 675.0955339 | 628.5679977 | ENSG00000126947 |
| FKBP9P1 | -0.102242968 | 0.931583526 | 0.92148231 | 0.964771134 | 4.682397724 | 4.363073292 | ENSG00000176826 |
| NA | -0.102248254 | 0.931580112 | 0.765156789 | 0.878472002 | 42.82036538 | 39.86698789 | ENSG00000225177 |
| PCDHA4 | -0.102297101 | 0.931548571 | 0.658119522 | 0.815577609 | 106.4404684 | 99.05693764 | ENSG00000204967 |
| HMGA2 | -0.102332096 | 0.931525975 | 0.181605794 | 0.364737363 | 3438.468856 | 3203.164974 | ENSG00000149948 |
| EPO | -0.102350067 | 0.931514372 | 0.894014608 | 0.952921173 | 8.165288307 | 7.636578868 | ENSG00000130427 |
| TSPAN31 | -0.102354854 | 0.931511281 | 0.490691218 | 0.690477988 | 245.2108588 | 228.2211641 | ENSG00000135452 |
| ILF3-DT | -0.102383468 | 0.931492806 | 0.42006524 | 0.630464776 | 396.5264323 | 369.3432801 | ENSG00000267100 |
| NKAPD1 | -0.102532255 | 0.931396744 | 0.353303383 | 0.56548304 | 499.0297231 | 464.9168037 | ENSG00000150776 |
| PIAS2 | -0.102533963 | 0.931395642 | 0.287280492 | 0.496369165 | 731.1046831 | 680.8799073 | ENSG00000078043 |

| | | | | | | | |
|---|---|---|---|---|---|---|---|
| GRB14 | -0.102633162 | 0.931331602 | 0.566963851 | 0.750638834 | 165.165879 | 153.7436587 | ENSG00000115290 |
| RELCH | -0.102638871 | 0.931327917 | 0.286865942 | 0.496104699 | 601.9227501 | 560.7981316 | ENSG00000134444 |
| AGPAT5 | -0.102660992 | 0.931313636 | 0.256284904 | 0.463107876 | 1793.03594 | 1670.171841 | ENSG00000155189 |
| NA | -0.102692117 | 0.931293545 | 0.921977612 | 0.964771134 | 5.278452155 | 4.899995798 | ENSG00000212440 |
| LINC00960 | -0.102765622 | 0.931246096 | 0.748478964 | 0.869205832 | 47.31268883 | 44.1560001 | ENSG00000242516 |
| NAB1 | -0.102885909 | 0.931168455 | 0.330471628 | 0.541843783 | 466.9711851 | 434.7121111 | ENSG00000138386 |
| SMARCD2 | -0.102906163 | 0.931155383 | 0.066261073 | 0.175208675 | 2884.08505 | 2685.746543 | ENSG00000108604 |
| MFHAS1 | -0.1029721 | 0.931112826 | 0.156371387 | 0.328376446 | 1658.893512 | 1544.102627 | ENSG00000147324 |
| KIAA1549L | -0.103037983 | 0.931070307 | 0.619248175 | 0.788725553 | 113.6850052 | 105.7681498 | ENSG00000110427 |
| TM2D2 | -0.103084843 | 0.931040065 | 0.169890653 | 0.347981017 | 1167.547816 | 1087.045377 | ENSG00000169490 |
| ATP5PF | -0.103094946 | 0.931033545 | 0.355878224 | 0.568159629 | 512.6766072 | 477.2413804 | ENSG00000154723 |
| TMCC2 | -0.103147076 | 0.930999904 | 0.439171953 | 0.647141256 | 298.6545888 | 278.1868469 | ENSG00000133069 |
| NEDD8 | -0.103203563 | 0.930963452 | 0.535739148 | 0.726437264 | 212.1707784 | 197.5240995 | ENSG00000129559 |
| KPNA6 | -0.103257642 | 0.930928556 | 0.156530166 | 0.32863504 | 1741.63322 | 1621.193186 | ENSG00000025800 |
| NA | -0.10325977 | 0.930927183 | 0.925504046 | 0.965963518 | 3.919100392 | 3.667813717 | ENSG00000249465 |
| SS18L2 | -0.103315495 | 0.930891226 | 0.296781062 | 0.506984201 | 874.232162 | 814.3060421 | ENSG00000008324 |
| FBXL20 | -0.103328106 | 0.930883089 | 0.557878427 | 0.743772262 | 152.610881 | 142.0438441 | ENSG00000108306 |
| PPP4R2 | -0.103382828 | 0.930847781 | 0.291651365 | 0.501259194 | 911.9523543 | 849.030807 | ENSG00000163605 |
| MARK3 | -0.103447401 | 0.930806118 | 0.126210865 | 0.282854985 | 1846.988564 | 1719.390225 | ENSG00000075413 |
| NA | -0.10351059 | 0.930765351 | 0.817589287 | 0.907131026 | 26.47051687 | 24.66722883 | ENSG00000268297 |
| STMN3 | -0.10353811 | 0.930747596 | 0.186091593 | 0.370984876 | 2103.135259 | 1958.375592 | ENSG00000197457 |
| STRADB | -0.103596516 | 0.930709917 | 0.333451002 | 0.545539816 | 730.8975527 | 680.3606493 | ENSG00000082146 |
| NUP50 | -0.103688272 | 0.930650725 | 0.095315238 | 0.229947402 | 2533.906237 | 2358.194621 | ENSG00000093000 |
| NA | -0.103715786 | 0.930632976 | 0.704878175 | 0.843415341 | 71.95708341 | 66.99668896 | ENSG00000230074 |
| NA | -0.103806323 | 0.930574576 | 0.923389103 | 0.965309497 | 3.911981447 | 3.660341367 | ENSG00000213087 |
| LSM12 | -0.103810619 | 0.930571805 | 0.283421577 | 0.492464654 | 971.5465205 | 904.4938626 | ENSG00000161654 |
| ZFYVE21 | -0.103931917 | 0.930493568 | 0.244704295 | 0.449012884 | 827.3367171 | 770.2025593 | ENSG00000100711 |
| HHEX | -0.103942283 | 0.930486882 | 0.880911893 | 0.945121072 | 10.33774386 | 9.586437572 | ENSG00000152804 |
| CCDC57 | -0.104047928 | 0.930418748 | 0.681879462 | 0.82993643 | 94.36739465 | 87.93776569 | ENSG00000176155 |
| FN3KRP | -0.104270194 | 0.930275415 | 0.270787324 | 0.479166306 | 656.9569896 | 610.9663958 | ENSG00000141560 |
| HIRIP3 | -0.104293154 | 0.930260611 | 0.324477559 | 0.53531569 | 547.4129144 | 509.3282587 | ENSG00000149929 |
| SAMD4B | -0.104464594 | 0.930150071 | 0.054159813 | 0.149905916 | 3272.959542 | 3044.807724 | ENSG00000179134 |
| SIK2 | -0.104492943 | 0.930131794 | 0.302296884 | 0.512445547 | 549.3600332 | 510.70438 | ENSG00000170145 |
| NA | -0.104541711 | 0.930100353 | 0.753167191 | 0.871381633 | 51.22642415 | 47.62548804 | ENSG00000259943 |
| COX7B | -0.10456537 | 0.9300851 | 0.262784409 | 0.469982842 | 1042.051149 | 969.3458352 | ENSG00000131174 |
| PDZD8 | -0.104568256 | 0.93008324 | 0.243992498 | 0.448360253 | 870.2037135 | 809.3370282 | ENSG00000165650 |
| NA | -0.104604058 | 0.930060159 | 0.3892339 | 0.601059189 | 523.8687719 | 487.4519699 | ENSG00000281398 |
| SLC39A7 | -0.104609745 | 0.930056493 | 0.053005763 | 0.147468156 | 3620.29987 | 3367.154829 | ENSG00000112473 |
| STK33 | -0.104737859 | 0.929973905 | 0.457640376 | 0.663253386 | 278.1068849 | 258.5123082 | ENSG00000130413 |
| CCDC61 | -0.104761062 | 0.929958949 | 0.402854059 | 0.614207252 | 375.9203017 | 349.6209538 | ENSG00000104983 |
| LOC729732 | -0.104824832 | 0.929917844 | 0.65698949 | 0.814765334 | 103.5969563 | 96.46083643 | ENSG00000283674 |
| NA | -0.10484303 | 0.929906114 | 0.748826252 | 0.869448741 | 43.81319609 | 40.78667881 | ENSG00000230555 |
| CDCA5 | -0.104899612 | 0.929869644 | 0.078057992 | 0.198215031 | 2666.871914 | 2479.995943 | ENSG00000146670 |
| DPM3 | -0.104922151 | 0.929855117 | 0.473685604 | 0.676821717 | 352.378445 | 327.7756683 | ENSG00000179085 |
| C1orf109 | -0.104947537 | 0.929838755 | 0.273652216 | 0.482262235 | 750.5095522 | 698.2228054 | ENSG00000116922 |
| TSEN34 | -0.104969483 | 0.929824611 | 0.3275782 | 0.53897145 | 497.7949267 | 462.9877537 | ENSG00000170892 |
| HLA-B | -0.104992822 | 0.929809568 | 0.293861452 | 0.503682475 | 554.8380423 | 515.9712965 | ENSG00000234745 |
| ACTR5 | -0.105085044 | 0.929750134 | 0.291382271 | 0.500979078 | 552.3334396 | 513.3331561 | ENSG00000101442 |
| DRAP1 | -0.10510392 | 0.929737969 | 0.147909893 | 0.315735401 | 3019.545883 | 2807.410482 | ENSG00000175550 |
| HILPDA | -0.105112483 | 0.929732451 | 0.4210198 | 0.631006395 | 301.1561651 | 279.8297949 | ENSG00000135245 |
| ZNF550 | -0.105147511 | 0.929709878 | 0.53060563 | 0.722718162 | 173.5790567 | 161.3402489 | ENSG00000251369 |
| HSPA9 | -0.105223709 | 0.929660775 | 0.023240762 | 0.077693865 | 12298.77983 | 11433.83495 | ENSG00000113013 |
| RNF44 | -0.105225837 | 0.929659404 | 0.052365405 | 0.146290039 | 3968.879931 | 3689.926508 | ENSG00000146083 |
| KLHL4 | -0.105396897 | 0.929549181 | 0.266972113 | 0.474608713 | 805.3407269 | 748.4130016 | ENSG00000102271 |
| AMPD2 | -0.105409592 | 0.929541001 | 0.232456832 | 0.433361033 | 941.0974133 | 875.2359553 | ENSG00000116337 |
| ADGRV1 | -0.10541664 | 0.929536461 | 0.254410994 | 0.460779862 | 878.9123987 | 816.4451255 | ENSG00000164199 |
| GTF3C4 | -0.105442876 | 0.929519556 | 0.137178033 | 0.299917687 | 1799.82779 | 1672.808511 | ENSG00000125484 |
| PPP1R18 | -0.10576135 | 0.929314388 | 0.225490837 | 0.423763008 | 838.6021045 | 779.7609994 | ENSG00000146112 |
| ZNF622 | -0.105839969 | 0.929263747 | 0.179862128 | 0.362662428 | 1280.55988 | 1189.78856 | ENSG00000173545 |
| MCUR1 | -0.105840514 | 0.929263396 | 0.260646522 | 0.467589016 | 738.9604798 | 687.1028352 | ENSG00000050393 |
| GTF2F1 | -0.105888052 | 0.929232776 | 0.098744814 | 0.235960496 | 2724.175498 | 2531.811253 | ENSG00000125651 |
| PSTK | -0.105995689 | 0.929163451 | 0.654802537 | 0.813335037 | 89.58355225 | 83.12238179 | ENSG00000179988 |
| PAXIP1 | -0.106026271 | 0.929143755 | 0.21746276 | 0.413406909 | 897.7533008 | 833.7416133 | ENSG00000157212 |
| FGF12-AS2 | -0.106058985 | 0.929122686 | 0.871326757 | 0.939605415 | 11.73713955 | 10.95085116 | ENSG00000230126 |
| ATP1B3 | -0.10609604 | 0.929098822 | 0.05053471 | 0.142291338 | 3376.076251 | 3136.419764 | ENSG00000069849 |
| TIMM44 | -0.106261057 | 0.928992557 | 0.092079257 | 0.224116094 | 2917.302868 | 2710.575316 | ENSG00000104980 |
| ME1 | -0.10627943 | 0.928980726 | 0.517946352 | 0.713860533 | 208.7222867 | 194.1479683 | ENSG00000065833 |
| LARS1 | -0.106383133 | 0.928913952 | 0.022061212 | 0.07453059 | 6293.602504 | 5845.824325 | ENSG00000133706 |
| EID2 | -0.106449802 | 0.928871026 | 0.256588675 | 0.463435073 | 652.0403555 | 605.7014142 | ENSG00000176396 |
| DHPS | -0.106453111 | 0.928868896 | 0.750257848 | 0.870058771 | 48.79687531 | 45.34450496 | ENSG00000095059 |
| EZH1 | -0.106528795 | 0.928820169 | 0.496482652 | 0.69565773 | 372.573893 | 345.9463731 | ENSG00000108799 |
| VEZT | -0.106777697 | 0.928659937 | 0.240936533 | 0.444518018 | 829.1601964 | 769.9408685 | ENSG00000028203 |
| C1orf220 | -0.106815193 | 0.928635802 | 0.887981899 | 0.94955683 | 8.961789946 | 8.321907083 | ENSG00000213057 |
| ENTPD6 | -0.106834546 | 0.928623344 | 0.145921296 | 0.313008142 | 1164.979778 | 1082.009346 | ENSG00000197586 |

| | | | | | | | |
|---|---|---|---|---|---|---|---|
| NA | -0.1068375 | 0.928621443 | 0.684603918 | 0.831377081 | 70.76291577 | 65.60304839 | ENSG00000227838 |
| RPAP3 | -0.106880419 | 0.928593818 | 0.232659797 | 0.433525433 | 772.9811863 | 717.7419699 | ENSG00000005175 |
| POFUT1 | -0.10708437 | 0.928462553 | 0.152078534 | 0.32204565 | 1070.84723 | 994.2967892 | ENSG00000101346 |
| NA | -0.107145298 | 0.928423343 | 0.900911487 | 0.956310125 | 8.283893801 | 7.705343063 | ENSG00000235997 |
| PXYLP1 | -0.107185679 | 0.928397357 | 0.355480468 | 0.567747007 | 443.5372293 | 411.6243856 | ENSG00000155893 |
| NA | -0.107236871 | 0.928364415 | 0.857378638 | 0.93039751 | 12.86428818 | 11.95765594 | ENSG00000271119 |
| FRG1 | -0.107241984 | 0.928361125 | 0.435795942 | 0.644328377 | 364.827854 | 338.7820199 | ENSG00000109536 |
| MARK4 | -0.107367946 | 0.928280073 | 0.141869974 | 0.306721748 | 1585.040468 | 1471.681215 | ENSG00000007047 |
| TBC1D24 | -0.107453119 | 0.928225271 | 0.420880638 | 0.630920054 | 314.7852769 | 292.1980058 | ENSG00000162065 |
| UPRT | -0.107460783 | 0.92822034 | 0.401142141 | 0.612513691 | 349.0470848 | 323.8163697 | ENSG00000094841 |
| MTX3 | -0.107502793 | 0.928193312 | 0.338841681 | 0.550644635 | 463.1550557 | 430.1554418 | ENSG00000177034 |
| SMAP1 | -0.107600884 | 0.928130204 | 0.464260217 | 0.668593683 | 260.9349813 | 242.2970425 | ENSG00000112305 |
| WDR61 | -0.107639181 | 0.928105567 | 0.284980697 | 0.494247449 | 604.2790513 | 560.6201201 | ENSG00000140395 |
| PABPC4 | -0.107667185 | 0.928087552 | 0.039133848 | 0.116789303 | 4622.829506 | 4290.692768 | ENSG00000090621 |
| BLOC1S5 | -0.107723137 | 0.928051559 | 0.510372582 | 0.70698938 | 205.9662352 | 191.1908103 | ENSG00000188428 |
| DMKN | -0.107768513 | 0.92802237 | 0.104077815 | 0.24540335 | 2648.386524 | 2457.621534 | ENSG00000161249 |
| GLOD4 | -0.107883032 | 0.927948708 | 0.111384342 | 0.258987109 | 1667.179439 | 1546.699572 | ENSG00000167699 |
| PHF13 | -0.107955815 | 0.927901895 | 0.152283691 | 0.322337861 | 1306.155029 | 1211.821938 | ENSG00000116273 |
| RBBP7 | -0.108020538 | 0.927860268 | 0.022391149 | 0.07541588 | 6909.817308 | 6411.481408 | ENSG00000102054 |
| ZBTB2 | -0.108059271 | 0.927835357 | 0.367008719 | 0.579463227 | 472.1604752 | 438.3369826 | ENSG00000181472 |
| RAP1GAP2 | -0.108130957 | 0.927789255 | 0.174500905 | 0.354807235 | 1577.806838 | 1463.831 | ENSG00000132359 |
| TMEM161B | -0.108139268 | 0.92778391 | 0.51288551 | 0.70906535 | 191.0287994 | 177.0868618 | ENSG00000164180 |
| SHC4 | -0.108178597 | 0.927758619 | 0.88049759 | 0.944837604 | 8.910788276 | 8.262790701 | ENSG00000185634 |
| SEMA4D | -0.10821556 | 0.927734849 | 0.165156086 | 0.340834081 | 1241.449705 | 1151.713391 | ENSG00000187764 |
| TCEANC2 | -0.108215694 | 0.927734763 | 0.463354654 | 0.667971431 | 239.0813438 | 221.669802 | ENSG00000116205 |
| GHRLOS | -0.108288939 | 0.927687663 | 0.854760474 | 0.928991226 | 13.23881791 | 12.28102442 | ENSG00000240288 |
| AP3S2 | -0.108352942 | 0.927646508 | 0.490142991 | 0.69013064 | 218.1899758 | 202.3453998 | ENSG00000157823 |
| NA | -0.10843564 | 0.927593336 | 0.88034941 | 0.944837604 | 9.244994745 | 8.560337166 | ENSG00000235636 |
| ZNF532 | -0.10847322 | 0.927569174 | 0.051587392 | 0.144565649 | 4375.910392 | 4058.572993 | ENSG00000074657 |
| ZFP1 | -0.108487208 | 0.92756018 | 0.611036418 | 0.783661987 | 112.884944 | 104.6244791 | ENSG00000184517 |
| TRO | -0.108537385 | 0.92752792 | 0.166615701 | 0.343021641 | 1212.640074 | 1124.853216 | ENSG00000067445 |
| PSMA6 | -0.108590394 | 0.92749384 | 0.801137176 | 0.898333333 | 31.20127266 | 29.03247758 | ENSG00000100902 |
| CHST1 | -0.108655967 | 0.927451685 | 0.73332577 | 0.859899685 | 58.94690916 | 54.72285334 | ENSG00000175264 |
| SAAL1 | -0.108723145 | 0.9274085 | 0.215212535 | 0.410285556 | 943.1165066 | 874.8391655 | ENSG00000166788 |
| IGFBP4 | -0.108732366 | 0.927402573 | 0.032622419 | 0.101545142 | 4913.542358 | 4556.35478 | ENSG00000141753 |
| TMEM260 | -0.108737191 | 0.927399471 | 0.432925475 | 0.642171723 | 274.2340309 | 254.492845 | ENSG00000070269 |
| H3C1 | -0.108909067 | 0.927288991 | 0.915666876 | 0.962507751 | 5.056927445 | 4.673388992 | ENSG00000275714 |
| BABAM2 | -0.108916497 | 0.927284216 | 0.528990871 | 0.72146 | 228.4542438 | 212.0734331 | ENSG00000158019 |
| ATP5MD | -0.108979272 | 0.927243868 | 0.213167211 | 0.407496991 | 1258.430409 | 1167.152915 | ENSG00000173915 |
| GRM4 | -0.108987911 | 0.927238316 | 0.207622554 | 0.400179464 | 762.8441951 | 707.3231662 | ENSG00000124493 |
| SNAPC4 | -0.109025653 | 0.927214059 | 0.18092482 | 0.363879543 | 1060.345855 | 983.6388929 | ENSG00000165684 |
| MBTPS1 | -0.109043289 | 0.927202724 | 0.118084375 | 0.269945942 | 1560.699057 | 1446.77733 | ENSG00000140943 |
| NA | -0.109062436 | 0.927190419 | 0.877131533 | 0.942832774 | 9.319717514 | 8.623804476 | ENSG00000231233 |
| PXDC1 | -0.109115019 | 0.927156626 | 0.759026886 | 0.874877523 | 40.40118947 | 37.54892661 | ENSG00000168994 |
| MZT2A | -0.109171549 | 0.927120297 | 0.15357404 | 0.324521175 | 1786.066749 | 1656.457361 | ENSG00000173272 |
| POLR3F | -0.109239689 | 0.927076509 | 0.44671666 | 0.653682173 | 293.7788367 | 272.4198955 | ENSG00000132664 |
| NA | -0.109255602 | 0.927066284 | 0.894280116 | 0.953020709 | 8.690458774 | 8.061059952 | ENSG00000279619 |
| SLIRP | -0.109283762 | 0.927048188 | 0.176564623 | 0.357693941 | 1472.302501 | 1365.155055 | ENSG00000119705 |
| AP5S1 | -0.109291978 | 0.927042909 | 0.718493799 | 0.851388915 | 55.71638333 | 51.67652167 | ENSG00000125843 |
| YJU2 | -0.109345385 | 0.927008592 | 0.347233994 | 0.559323949 | 527.012038 | 488.8892877 | ENSG00000105248 |
| EPHX2 | -0.109359454 | 0.926999552 | 0.446534248 | 0.653604036 | 255.1373091 | 236.360069 | ENSG00000120915 |
| MRPL13 | -0.109411582 | 0.926966058 | 0.224521557 | 0.422867219 | 764.5703467 | 709.0024776 | ENSG00000172172 |
| SGSM3 | -0.109454613 | 0.926938409 | 0.905778388 | 0.959035162 | 6.826098169 | 6.37110242 | ENSG00000100359 |
| AHDC1 | -0.109480297 | 0.926921908 | 0.201044402 | 0.391899622 | 913.1714133 | 846.3197124 | ENSG00000126705 |
| AASDH | -0.109552092 | 0.926875781 | 0.498968922 | 0.697608609 | 202.1420192 | 187.4301344 | ENSG00000157426 |
| NA | -0.109614303 | 0.926835813 | 0.914212211 | 0.961993514 | 5.033206346 | 4.647566976 | ENSG00000213113 |
| GLO1 | -0.109633438 | 0.926823521 | 0.059816159 | 0.161932333 | 3317.737219 | 3074.217356 | ENSG00000124767 |
| ARPC4 | -0.10967053 | 0.926799692 | 0.186076832 | 0.370984876 | 1354.682326 | 1255.825929 | ENSG00000241553 |
| PPP1CA | -0.109671975 | 0.926798764 | 0.042644181 | 0.124664915 | 5013.01524 | 4645.707745 | ENSG00000172531 |
| SPN | -0.109683544 | 0.926791332 | 0.772420418 | 0.882839653 | 35.61142334 | 32.98941945 | ENSG00000197471 |
| MICA | -0.109694404 | 0.926784355 | 0.388445089 | 0.600457191 | 341.3275419 | 316.3145243 | ENSG00000204520 |
| PRRC2B | -0.10972395 | 0.926765375 | 0.038468683 | 0.115314456 | 6052.075004 | 5607.956754 | ENSG00000130723 |
| TACSTD2 | -0.109784557 | 0.926726443 | 0.801544515 | 0.898520796 | 30.07173361 | 27.97866324 | ENSG00000184292 |
| STXBP2 | -0.10983147 | 0.926696309 | 0.311439124 | 0.522013656 | 455.3286768 | 421.8864082 | ENSG00000076944 |
| OGFOD1 | -0.109896839 | 0.926654321 | 0.178548538 | 0.360784201 | 1157.390662 | 1072.326532 | ENSG00000087263 |
| PNISR | -0.109906304 | 0.926648241 | 0.551946673 | 0.738889411 | 1998.936224 | 1852.431192 | ENSG00000132424 |
| REXO4 | -0.109955397 | 0.926616709 | 0.091434279 | 0.222833443 | 1596.914185 | 1479.855304 | ENSG00000148300 |
| PRPF6 | -0.109987406 | 0.92659615 | 0.043963549 | 0.127603744 | 3381.221046 | 3133.403229 | ENSG00000101161 |
| UNC119 | -0.110021076 | 0.926574526 | 0.161293816 | 0.335799109 | 955.0366276 | 884.9345078 | ENSG00000109103 |
| PLCG1 | -0.110060461 | 0.926549231 | 0.105384534 | 0.248122046 | 4538.220782 | 4204.843363 | ENSG00000124181 |
| NA | -0.110074941 | 0.926539932 | 0.849722613 | 0.925513999 | 15.70248693 | 14.58972144 | ENSG00000269737 |
| NA | -0.11008024 | 0.926536528 | 0.799211692 | 0.897822068 | 27.23500941 | 25.20197587 | ENSG00000143429 |
| RECK | -0.110094099 | 0.926527628 | 0.438163072 | 0.646360694 | 264.2325267 | 244.8415426 | ENSG00000122707 |
| CENPN | -0.110171511 | 0.926477913 | 0.303090412 | 0.513304854 | 483.906078 | 448.2676927 | ENSG00000166451 |

| | | | | | | | |
|---|---|---|---|---|---|---|---|
| BTRC | -0.110186792 | 0.9264681 | 0.212444589 | 0.406485924 | 784.6252214 | 726.8337648 | ENSG00000166167 |
| ZGRF1 | -0.11028606 | 0.926404354 | 0.317990137 | 0.528072787 | 423.2724764 | 392.0392088 | ENSG00000138658 |
| NA | -0.110292995 | 0.926399901 | 0.35163204 | 0.563763569 | 665.3169054 | 616.6200982 | ENSG00000250251 |
| RAI14 | -0.110294931 | 0.926398658 | 0.170340663 | 0.348638065 | 2371.366673 | 2196.45097 | ENSG00000039560 |
| NA | -0.110325386 | 0.926379102 | 0.714704507 | 0.849628264 | 56.95651921 | 52.6292234 | ENSG00000281091 |
| ZNF420 | -0.110355543 | 0.926359738 | 0.661988575 | 0.818441256 | 81.91704357 | 75.77398512 | ENSG00000197050 |
| KIAA1841 | -0.110359271 | 0.926357344 | 0.442553839 | 0.650095617 | 260.7737144 | 241.6457437 | ENSG00000162929 |
| ZNF542P | -0.110360084 | 0.926356822 | 0.720478405 | 0.852637136 | 62.53032854 | 58.07098709 | ENSG00000240225 |
| NA | -0.110434847 | 0.926308819 | 0.883514839 | 0.946784188 | 8.648940313 | 8.013766846 | ENSG00000253170 |
| SUMO2 | -0.110477823 | 0.926281226 | 0.038727985 | 0.115907766 | 4735.069977 | 4386.05673 | ENSG00000188612 |
| FGD5-AS1 | -0.110582622 | 0.926213942 | 0.140609159 | 0.304798436 | 1571.668983 | 1455.081623 | ENSG00000225733 |
| PLEKHG3 | -0.110690292 | 0.92614482 | 0.124686696 | 0.280410105 | 1327.23154 | 1229.579114 | ENSG00000126822 |
| MAP4K4 | -0.110691807 | 0.926143847 | 0.051272213 | 0.143874553 | 4739.177983 | 4388.930669 | ENSG00000071054 |
| CHD2 | -0.110713144 | 0.92613015 | 0.121121197 | 0.274260186 | 1336.564502 | 1237.639416 | ENSG00000173575 |
| NA | -0.110797117 | 0.926076245 | 0.882173281 | 0.945771249 | 11.13516138 | 10.35169085 | ENSG00000237015 |
| CD63 | -0.110806469 | 0.926070242 | 0.03918797 | 0.116876944 | 6874.223241 | 6365.664652 | ENSG00000135404 |
| SERPINB8 | -0.110807146 | 0.926069808 | 0.883127857 | 0.94654857 | 8.209171032 | 7.613878273 | ENSG00000166401 |
| COMMD8 | -0.110917779 | 0.925998795 | 0.739548839 | 0.863664729 | 44.00624502 | 40.8470247 | ENSG00000169019 |
| SLC35B4 | -0.110941182 | 0.925983774 | 0.188260166 | 0.374162622 | 959.8607415 | 888.4198005 | ENSG00000205060 |
| HSD17B6 | -0.110957597 | 0.925973238 | 0.709126086 | 0.846486042 | 54.224773 | 50.13903723 | ENSG00000025423 |
| FURIN | -0.111002262 | 0.925944571 | 0.163924725 | 0.339274231 | 1225.568912 | 1134.816495 | ENSG00000140564 |
| ZNF16 | -0.111013985 | 0.925937047 | 0.576500884 | 0.758517416 | 154.324627 | 142.9201413 | ENSG00000170631 |
| YBX1 | -0.11102987 | 0.925926852 | 0.061505235 | 0.165475583 | 27546.51166 | 25506.31806 | ENSG00000065978 |
| SMARCD1 | -0.111054504 | 0.925911042 | 0.034932979 | 0.107095668 | 4152.529526 | 3844.925508 | ENSG00000066117 |
| SORT1 | -0.111162654 | 0.925841634 | 0.113299952 | 0.261861412 | 1349.449335 | 1249.229205 | ENSG00000134243 |
| RCAN2 | -0.111170658 | 0.925836498 | 0.454935141 | 0.660704703 | 246.8341173 | 228.3725756 | ENSG00000172348 |
| ATRAID | -0.111183412 | 0.925828313 | 0.299372566 | 0.509613387 | 557.0440861 | 515.6977755 | ENSG00000138085 |
| CALR | -0.111201592 | 0.925816647 | 0.009587148 | 0.037750893 | 28840.44875 | 26700.45114 | ENSG00000179218 |
| TPBG | -0.11122889 | 0.925799129 | 0.200079411 | 0.390461886 | 1064.584567 | 985.7859112 | ENSG00000146242 |
| LRRC8C | -0.111257955 | 0.925780477 | 0.665903902 | 0.821129297 | 80.89250866 | 75.01603624 | ENSG00000171488 |
| RRAGD | -0.111261444 | 0.925778238 | 0.339334406 | 0.551111994 | 386.8443636 | 358.1226261 | ENSG00000025039 |
| NA | -0.111276523 | 0.925768562 | 0.923621984 | 0.965386383 | 3.592012868 | 3.346620699 | ENSG00000233916 |
| DNMT1 | -0.111331364 | 0.925733372 | 0.018587177 | 0.064727798 | 7035.948476 | 6513.399332 | ENSG00000130816 |
| PDLIM1 | -0.111380164 | 0.925702059 | 0.0451124 | 0.130088207 | 2969.374232 | 2748.865355 | ENSG00000107438 |
| MGST1 | -0.111383557 | 0.925699882 | 0.063035786 | 0.168496931 | 3598.290232 | 3331.652312 | ENSG00000008394 |
| ZNF19 | -0.11142001 | 0.925676493 | 0.811734779 | 0.904395006 | 24.03265387 | 22.29317258 | ENSG00000157429 |
| IFRD2 | -0.111420753 | 0.925676016 | 0.112678981 | 0.260809237 | 1682.596206 | 1558.183859 | ENSG00000214706 |
| ALG12 | -0.11151606 | 0.925614866 | 0.402105575 | 0.613587894 | 289.7568171 | 268.378437 | ENSG00000182858 |
| VPS25 | -0.111615806 | 0.925550872 | 0.084163053 | 0.209824325 | 1926.664913 | 1782.864269 | ENSG00000131475 |
| ZNF48 | -0.111704613 | 0.925493901 | 0.216218968 | 0.411663999 | 1036.370736 | 959.7049978 | ENSG00000180035 |
| POLA2 | -0.111803734 | 0.925430316 | 0.61793334 | 0.787824768 | 125.4719773 | 116.1881534 | ENSG00000014138 |
| MRPL11 | -0.11181903 | 0.925420505 | 0.075638832 | 0.193923343 | 1879.892906 | 1739.616011 | ENSG00000174547 |
| NSF | -0.111821434 | 0.925418963 | 0.308387855 | 0.51877645 | 523.6021686 | 484.7700664 | ENSG00000073969 |
| YARS2 | -0.111844072 | 0.925404441 | 0.206899248 | 0.399331244 | 750.2123894 | 694.3254541 | ENSG00000139131 |
| RNFT1 | -0.111933484 | 0.92534709 | 0.784964841 | 0.890433415 | 28.62609094 | 26.52590629 | ENSG00000189050 |
| NA | -0.111970852 | 0.925323123 | 0.787544775 | 0.891546762 | 26.55355379 | 24.54152371 | ENSG00000268573 |
| SLC25A29 | -0.111971366 | 0.925322793 | 0.209223973 | 0.402391224 | 1905.035294 | 1763.141771 | ENSG00000197119 |
| RAPGEF1 | -0.111997709 | 0.925305897 | 0.064000595 | 0.170298572 | 3263.481306 | 3020.291185 | ENSG00000107263 |
| ERGIC3 | -0.112006213 | 0.925300443 | 0.051547617 | 0.144475622 | 2845.855244 | 2632.868042 | ENSG00000125991 |
| HNRNPA2B1 | -0.11205945 | 0.9252663 | 0.006645401 | 0.02806996 | 43326.73424 | 40088.50463 | ENSG00000122566 |
| GYG2 | -0.112132568 | 0.925219407 | 0.045307316 | 0.130490874 | 3241.965402 | 2999.000787 | ENSG00000056998 |
| SNRPB | -0.112201494 | 0.925175204 | 0.04893369 | 0.13871401 | 3267.670895 | 3022.652341 | ENSG00000125835 |
| MRPL4 | -0.112308807 | 0.925106389 | 0.207463882 | 0.399954614 | 883.5842208 | 817.7852005 | ENSG00000105364 |
| SPRY1 | -0.112380755 | 0.925060254 | 0.272427931 | 0.481047357 | 499.2236884 | 461.7966219 | ENSG00000164056 |
| TRAM1 | -0.112536971 | 0.924960094 | 0.155826713 | 0.327558713 | 1706.69875 | 1578.582027 | ENSG00000067167 |
| RAB2A | -0.112567765 | 0.924940351 | 0.133526627 | 0.29461974 | 1563.18273 | 1445.737494 | ENSG00000104388 |
| MHENCR | -0.112654 | 0.924885066 | 0.741795271 | 0.865381993 | 47.0567646 | 43.62314498 | ENSG00000232442 |
| TMEM126B | -0.112735786 | 0.924832636 | 0.175111593 | 0.355608499 | 890.8014515 | 823.6638224 | ENSG00000171204 |
| CKMT1B | -0.112738027 | 0.924831199 | 0.587808953 | 0.765834569 | 163.105521 | 150.8296015 | ENSG00000237289 |
| TAB3 | -0.11288866 | 0.924734642 | 0.407284334 | 0.618193591 | 287.1365574 | 265.5467733 | ENSG00000157625 |
| COG3 | -0.11289723 | 0.924729149 | 0.343143376 | 0.555102163 | 384.8033031 | 355.9703742 | ENSG00000136152 |
| ZBTB12 | -0.112941194 | 0.924700969 | 0.192027527 | 0.37925537 | 936.1785908 | 865.3731253 | ENSG00000204366 |
| CLSTN1 | -0.112971128 | 0.924681783 | 0.05572893 | 0.153417033 | 3626.70787 | 3353.367163 | ENSG00000171603 |
| ELAC2 | -0.112996981 | 0.924665213 | 0.038565119 | 0.115530182 | 3371.108771 | 3117.627432 | ENSG00000006744 |
| NA | -0.11300549 | 0.92465976 | 0.910449494 | 0.960811396 | 4.70255935 | 4.335075811 | ENSG00000267605 |
| ZNF407 | -0.113036973 | 0.924639581 | 0.521976931 | 0.716650988 | 206.7268104 | 191.2399954 | ENSG00000215421 |
| PLEKHG6 | -0.113175214 | 0.924550985 | 0.261991796 | 0.468977732 | 631.0021861 | 583.0216464 | ENSG00000008323 |
| NPRL3 | -0.113195407 | 0.924538045 | 0.259565115 | 0.466503717 | 644.1038149 | 595.7248085 | ENSG00000103148 |
| MFN2 | -0.113381338 | 0.9244189 | 0.061512388 | 0.165475583 | 2650.727696 | 2449.859748 | ENSG00000116688 |
| VIRMA | -0.113390737 | 0.924412878 | 0.137060524 | 0.299790131 | 1084.524344 | 1002.305332 | ENSG00000164944 |
| RRP12 | -0.113497225 | 0.924344648 | 0.146927482 | 0.314380863 | 2060.742817 | 1905.2777 | ENSG00000052749 |
| THOC3 | -0.113531131 | 0.924322924 | 0.07725143 | 0.196748921 | 2299.321231 | 2124.925101 | ENSG00000051596 |
| GNA11 | -0.113576386 | 0.92429393 | 0.245285034 | 0.44964161 | 636.3100032 | 588.1297314 | ENSG00000088256 |
| TRNT1 | -0.113638257 | 0.924254292 | 0.297589795 | 0.507812023 | 519.7746208 | 480.1767704 | ENSG00000072756 |

| | | | | | | | |
|---|---|---|---|---|---|---|---|
| SNRPA1 | -0.113776015 | 0.924166042 | 0.074553966 | 0.191845162 | 2367.81228 | 2187.972855 | ENSG00000131876 |
| THAP9-AS1 | -0.113826879 | 0.92413346 | 0.460194547 | 0.664999857 | 279.649523 | 258.4918987 | ENSG00000251022 |
| RDH5 | -0.113872586 | 0.924104182 | 0.915689926 | 0.962507751 | 5.016604192 | 4.616448073 | ENSG00000135437 |
| SRC | -0.113886195 | 0.924095465 | 0.099886696 | 0.237918452 | 1356.311307 | 1253.586174 | ENSG00000197122 |
| MLLT10 | -0.113891066 | 0.924092346 | 0.198845679 | 0.389030323 | 938.2326946 | 866.6686699 | ENSG00000078403 |
| EIF4EBP2 | -0.113921491 | 0.924072858 | 0.036159359 | 0.109945176 | 6901.899514 | 6377.822353 | ENSG00000148730 |
| GTPBP6 | -0.11402928 | 0.924003819 | 0.080887505 | 0.203484819 | 2032.61118 | 1878.319582 | ENSG00000178605 |
| TBC1D25 | -0.114064531 | 0.923981242 | 0.536591501 | 0.726856257 | 177.4880896 | 163.785601 | ENSG00000068354 |
| DOK1 | -0.114094068 | 0.923962326 | 0.397198414 | 0.609003171 | 281.4782366 | 259.949626 | ENSG00000115325 |
| GAB1 | -0.114139328 | 0.92393334 | 0.424582539 | 0.634505124 | 280.4311502 | 258.8127652 | ENSG00000109458 |
| SLC29A1 | -0.11418299 | 0.923905378 | 0.022338187 | 0.075264345 | 4662.364777 | 4307.559589 | ENSG00000112759 |
| TCP11L2 | -0.114247711 | 0.923863931 | 0.828250408 | 0.913692911 | 16.51798114 | 15.26880681 | ENSG00000166046 |
| SLC35E4 | -0.114251402 | 0.923861568 | 0.502207382 | 0.700181291 | 206.6369862 | 190.9655284 | ENSG00000100036 |
| SYT12 | -0.114269197 | 0.923850173 | 0.53008639 | 0.722219282 | 175.7660295 | 162.192784 | ENSG00000173227 |
| RHOA | -0.114270031 | 0.923849638 | 0.010690792 | 0.041236847 | 9203.845876 | 8502.682517 | ENSG00000067560 |
| NA | -0.114289449 | 0.923837203 | 0.679007801 | 0.828363199 | 63.54154143 | 58.80918885 | ENSG00000279041 |
| GOSR2 | -0.114377554 | 0.923780787 | 0.31122292 | 0.521809598 | 440.2969665 | 406.8185542 | ENSG00000108433 |
| PSMD11 | -0.114381963 | 0.923777964 | 0.035445841 | 0.10830117 | 3116.743198 | 2878.913905 | ENSG00000108671 |
| CCAR1 | -0.114436129 | 0.923743281 | 0.102435834 | 0.242418637 | 2483.347615 | 2294.096508 | ENSG00000060339 |
| GRB2 | -0.114471333 | 0.923720741 | 0.040758735 | 0.12049604 | 2727.745293 | 2519.408139 | ENSG00000177885 |
| RBM14 | -0.114529369 | 0.923683583 | 0.095374145 | 0.230030745 | 1933.412736 | 1786.136617 | ENSG00000239306 |
| TAF1C | -0.114578734 | 0.923651977 | 0.183246839 | 0.366977969 | 1553.445193 | 1435.249792 | ENSG00000103168 |
| GNS | -0.114616881 | 0.923627555 | 0.270504922 | 0.478950446 | 884.8995419 | 817.3462035 | ENSG00000135677 |
| PTPN12 | -0.114778225 | 0.923524267 | 0.075010778 | 0.192600753 | 2172.826427 | 2007.335772 | ENSG00000127947 |
| TMEM115 | -0.114858003 | 0.923473199 | 0.3070661 | 0.517393228 | 466.3816387 | 430.60381 | ENSG00000126062 |
| SLC9A3R1 | -0.114916114 | 0.923436003 | 0.042882126 | 0.125205464 | 2857.060782 | 2638.662866 | ENSG00000109062 |
| PSMG2 | -0.115022743 | 0.923367755 | 0.237271848 | 0.439326763 | 628.1927937 | 580.2647694 | ENSG00000128789 |
| NUP98 | -0.115096897 | 0.923320295 | 0.055339821 | 0.152612809 | 3124.188208 | 2884.167712 | ENSG00000110713 |
| STX4 | -0.115155791 | 0.923282604 | 0.219582426 | 0.415929199 | 793.7747202 | 733.2859491 | ENSG00000103496 |
| SLC25A22 | -0.115240526 | 0.923228377 | 0.161669037 | 0.336298878 | 1203.844662 | 1111.848272 | ENSG00000177542 |
| DENND10 | -0.115305358 | 0.92318689 | 0.204829346 | 0.396738687 | 784.442546 | 723.8705749 | ENSG00000119979 |
| SIMC1 | -0.11531469 | 0.923180918 | 0.328405913 | 0.539956961 | 377.5471982 | 348.6900067 | ENSG00000170085 |
| PDCD2 | -0.115415274 | 0.923116557 | 0.093579651 | 0.227094797 | 1377.928624 | 1272.037137 | ENSG00000071994 |
| NAIF1 | -0.115459585 | 0.923088205 | 0.427475105 | 0.637365179 | 247.5057842 | 228.5229889 | ENSG00000171169 |
| STX5 | -0.115575315 | 0.92301416 | 0.294421451 | 0.504413418 | 480.7790556 | 443.8697166 | ENSG00000162236 |
| NIFK | -0.115587939 | 0.923006083 | 0.226707906 | 0.425583188 | 688.5275309 | 635.4252293 | ENSG00000155438 |
| RAB35 | -0.115615084 | 0.922988717 | 0.11579044 | 0.266095758 | 1628.335098 | 1503.197656 | ENSG00000111737 |
| MIR1244-2 | -0.115707429 | 0.922929639 | 0.640089341 | 0.80292841 | 97.11571686 | 89.6878582 | ENSG00000283498 |
| NA | -0.11576008 | 0.922895958 | 0.640632162 | 0.803289183 | 87.97450548 | 81.06471614 | ENSG00000259429 |
| ZZZ3 | -0.115815161 | 0.922860723 | 0.263634849 | 0.470981376 | 735.505734 | 678.6358414 | ENSG00000036549 |
| ARHGEF6 | -0.115876355 | 0.922821579 | 0.311528709 | 0.52207114 | 399.3765047 | 368.393278 | ENSG00000129675 |
| NA | -0.115977987 | 0.922756572 | 0.52930846 | 0.721629123 | 142.7117112 | 131.6212136 | ENSG00000182484 |
| XRCC1 | -0.116023264 | 0.922727613 | 0.271050185 | 0.47946635 | 651.6812072 | 601.3073806 | ENSG00000073050 |
| ATP9B | -0.116073333 | 0.92269559 | 0.318139054 | 0.528152019 | 415.0066587 | 383.0780371 | ENSG00000166377 |
| MCRIP1 | -0.116223231 | 0.922599726 | 0.117119275 | 0.268291789 | 1888.633144 | 1742.839547 | ENSG00000225663 |
| ADM2 | -0.116285977 | 0.922559601 | 0.34699965 | 0.559118942 | 436.3188552 | 402.9686318 | ENSG00000128165 |
| KATNBL1 | -0.116336119 | 0.922527537 | 0.345163918 | 0.557034745 | 357.6160152 | 329.9343809 | ENSG00000134152 |
| RBM47 | -0.116438873 | 0.922461834 | 0.167906451 | 0.345258686 | 1608.986473 | 1483.999994 | ENSG00000163694 |
| EXTL1 | -0.116603245 | 0.92235674 | 0.78814159 | 0.891582187 | 25.24995837 | 23.27481776 | ENSG00000158008 |
| CD44 | -0.116610901 | 0.922351845 | 0.626976552 | 0.793649185 | 99.35165811 | 91.8284977 | ENSG00000026508 |
| NA | -0.116689397 | 0.922301662 | 0.914706308 | 0.96227163 | 3.922659864 | 3.622696074 | ENSG00000188512 |
| WDR86-AS1 | -0.116694479 | 0.922298413 | 0.791225411 | 0.893303941 | 26.69560105 | 24.68340303 | ENSG00000243836 |
| SURF4 | -0.116704469 | 0.922292027 | 0.025895997 | 0.084653956 | 6397.74151 | 5900.420922 | ENSG00000148248 |
| OPA1 | -0.116756283 | 0.922258903 | 0.090800095 | 0.221803996 | 1956.937451 | 1804.385081 | ENSG00000198836 |
| DSCAML1 | -0.116794297 | 0.922234603 | 0.866096963 | 0.935979464 | 12.27298843 | 11.32179629 | ENSG00000177103 |
| DBF4B | -0.116808942 | 0.922225241 | 0.182089421 | 0.365414231 | 901.718395 | 831.4981505 | ENSG00000161692 |
| PISD | -0.116849052 | 0.922199602 | 0.148505488 | 0.316458791 | 983.6490393 | 907.100106 | ENSG00000241878 |
| OXR1 | -0.116907135 | 0.922162475 | 0.421038474 | 0.631006395 | 255.0107205 | 235.2582991 | ENSG00000164830 |
| PSMC3IP | -0.116974811 | 0.922119218 | 0.41996122 | 0.630411957 | 326.1979202 | 300.9276957 | ENSG00000131470 |
| NDUFAF2 | -0.116981896 | 0.922114689 | 0.327117184 | 0.538494422 | 485.9382402 | 447.910483 | ENSG00000164182 |
| COX6A1 | -0.117036776 | 0.922079612 | 0.568093519 | 0.751344359 | 117.7437624 | 108.6077374 | ENSG00000111775 |
| CAPZB | -0.117144771 | 0.922010592 | 0.018430297 | 0.064304723 | 7848.705087 | 7236.226126 | ENSG00000077549 |
| RPS26 | -0.117149495 | 0.922007572 | 0.059825358 | 0.161932333 | 3513.669529 | 3239.826252 | ENSG00000197728 |
| SPECC1L | -0.117181794 | 0.921986931 | 0.198271468 | 0.388277622 | 736.7702013 | 679.3954846 | ENSG00000100014 |
| DHDDS | -0.117187047 | 0.921983574 | 0.155816496 | 0.327558713 | 866.3340623 | 798.6045991 | ENSG00000117682 |
| PLXNB1 | -0.117200597 | 0.921974915 | 0.09748403 | 0.233716633 | 3699.185926 | 3411.270595 | ENSG00000164050 |
| ITCH | -0.117282907 | 0.921922315 | 0.236834094 | 0.438858749 | 705.7299168 | 650.349648 | ENSG00000078747 |
| NA | -0.117332461 | 0.921890649 | 0.743548645 | 0.866517504 | 38.56558398 | 35.62678184 | ENSG00000224043 |
| EPHB4 | -0.117373699 | 0.921864298 | 0.026204615 | 0.085470207 | 4705.655732 | 4338.207788 | ENSG00000196411 |
| BICD1 | -0.117388265 | 0.921854991 | 0.236393121 | 0.438448644 | 688.1544752 | 634.4184042 | ENSG00000151746 |
| NA | -0.117397055 | 0.921849374 | 0.916685963 | 0.962986839 | 3.599131813 | 3.334797422 | ENSG00000241511 |
| MTF1 | -0.117488286 | 0.921791081 | 0.371908605 | 0.583850869 | 325.9672701 | 300.2872312 | ENSG00000188786 |
| GAL3ST3 | -0.117499692 | 0.921783794 | 0.368813061 | 0.580952094 | 291.5369717 | 268.8154764 | ENSG00000175229 |
| LPCAT1 | -0.117696859 | 0.921657826 | 0.027544564 | 0.088921327 | 7215.184763 | 6649.729306 | ENSG00000153395 |

| | | | | | | | |
|---|---|---|---|---|---|---|---|
| NA | -0.117703239 | 0.92165375 | 0.631469284 | 0.797092269 | 82.45734165 | 75.9706299 | ENSG00000250986 |
| FLJ16779 | -0.117722256 | 0.921641601 | 0.852139468 | 0.927102999 | 13.76518358 | 12.64951055 | ENSG00000275620 |
| ZDHHC5 | -0.11783083 | 0.921572243 | 0.061688014 | 0.165762399 | 1922.781005 | 1771.93801 | ENSG00000156599 |
| RPL24 | -0.11791385 | 0.921519213 | 0.041831204 | 0.122839728 | 15810.24152 | 14568.58809 | ENSG00000114391 |
| CRLF1 | -0.117929025 | 0.92150952 | 0.55739574 | 0.743522876 | 130.0689739 | 119.8723093 | ENSG00000006016 |
| SF3B6 | -0.117969147 | 0.921483893 | 0.139308506 | 0.303022476 | 1025.02849 | 944.7562928 | ENSG00000115128 |
| PAQR8 | -0.118027304 | 0.921446747 | 0.399801057 | 0.611207234 | 301.7739336 | 278.1550656 | ENSG00000170915 |
| KHDRBS3 | -0.118030606 | 0.921444638 | 0.285717065 | 0.494933661 | 655.3344978 | 603.9896067 | ENSG00000131773 |
| MBLAC1 | -0.118035113 | 0.92144176 | 0.767211667 | 0.879601283 | 31.00138412 | 28.48976373 | ENSG00000214309 |
| API5 | -0.118053496 | 0.921430019 | 0.10111776 | 0.239840353 | 3225.497836 | 2971.861747 | ENSG00000166181 |
| KIAA1671 | -0.118087465 | 0.921408323 | 0.230120559 | 0.42998187 | 866.5122885 | 798.1920436 | ENSG00000197077 |
| SLC38A2 | -0.118122261 | 0.9213861 | 0.061116882 | 0.164766384 | 6559.84101 | 6043.680838 | ENSG00000134294 |
| FCF1 | -0.118150268 | 0.921368213 | 0.102534649 | 0.242530917 | 1235.263108 | 1138.060553 | ENSG00000119616 |
| ALG1L2 | -0.118237097 | 0.921312763 | 0.909901081 | 0.960615044 | 4.671719306 | 4.285607242 | ENSG00000251287 |
| GAPVD1 | -0.118303928 | 0.921270085 | 0.181386163 | 0.364451291 | 889.9661011 | 819.6316625 | ENSG00000165219 |
| SH3GL2 | -0.118322214 | 0.921258408 | 0.602656358 | 0.777614465 | 113.8466292 | 105.0758437 | ENSG00000107295 |
| PSMC5 | -0.118341592 | 0.921246033 | 0.102846074 | 0.24308487 | 1790.283397 | 1649.973713 | ENSG00000087191 |
| NA | -0.118396056 | 0.921211256 | 0.883989218 | 0.947131305 | 7.201823099 | 6.664297958 | ENSG00000278867 |
| PPP1R1B | -0.118459114 | 0.921170992 | 0.366726369 | 0.57921113 | 311.1899839 | 286.5779697 | ENSG00000131771 |
| MRO | -0.118464249 | 0.921167713 | 0.763924261 | 0.877750649 | 30.64345655 | 28.15551793 | ENSG00000134042 |
| C3orf18 | -0.118464455 | 0.921167581 | 0.514622799 | 0.710843444 | 201.401196 | 185.3015088 | ENSG00000088543 |
| IMP3 | -0.118476226 | 0.921160066 | 0.351452351 | 0.563571168 | 346.4413947 | 318.9896046 | ENSG00000177971 |
| STIL | -0.118497561 | 0.921146444 | 0.2384513 | 0.441046059 | 632.8609018 | 582.7605157 | ENSG00000123473 |
| HDAC6 | -0.118498728 | 0.921145699 | 0.242827014 | 0.446696684 | 835.7336243 | 770.2226759 | ENSG00000094631 |
| NA | -0.118626292 | 0.921064254 | 0.909649339 | 0.960563356 | 5.013044719 | 4.590626057 | ENSG00000279668 |
| KLF8 | -0.118661124 | 0.921042016 | 0.520367172 | 0.715666089 | 182.2657028 | 167.8750579 | ENSG00000102349 |
| WDR41 | -0.118722959 | 0.921002541 | 0.229525046 | 0.42933642 | 810.1748566 | 745.8116988 | ENSG00000164253 |
| CNKSR2 | -0.118738527 | 0.920992602 | 0.36089761 | 0.57292799 | 362.5703545 | 334.0380045 | ENSG00000149970 |
| MAZ | -0.118777346 | 0.920967821 | 0.329762155 | 0.541217709 | 522.878942 | 481.7602161 | ENSG00000103495 |
| ALG8 | -0.118793815 | 0.920957308 | 0.158643535 | 0.331781081 | 946.336965 | 871.375383 | ENSG00000159063 |
| OCLN | -0.118826665 | 0.920936338 | 0.319512216 | 0.529761915 | 438.5192028 | 403.6204118 | ENSG00000197822 |
| SLC12A2 | -0.118829476 | 0.920934544 | 0.403032411 | 0.61430727 | 316.0716853 | 291.137833 | ENSG00000064651 |
| IFT88 | -0.118951837 | 0.920856439 | 0.597240036 | 0.773554521 | 93.37692821 | 85.97530571 | ENSG00000032742 |
| ANKS3 | -0.118957292 | 0.920852957 | 0.506429338 | 0.703694948 | 318.4846064 | 293.4109056 | ENSG00000168096 |
| PKD1 | -0.118968162 | 0.920846019 | 0.12900816 | 0.287335665 | 1176.835829 | 1083.908928 | ENSG00000008710 |
| HINT3 | -0.118969857 | 0.920844937 | 0.433374338 | 0.642497135 | 229.6622921 | 211.6050639 | ENSG00000111911 |
| TTC28-AS1 | -0.118973743 | 0.920842456 | 0.51673506 | 0.712978856 | 148.488943 | 136.7651473 | ENSG00000235954 |
| GART | -0.118979942 | 0.9208385 | 0.024317566 | 0.080567937 | 4319.832111 | 3977.313679 | ENSG00000159131 |
| NA | -0.119066843 | 0.920783035 | 0.911388207 | 0.961009829 | 5.098445906 | 4.67026757 | ENSG00000237840 |
| FAM131B | -0.119077463 | 0.920776256 | 0.140223064 | 0.304380755 | 1148.119147 | 1056.775852 | ENSG00000159784 |
| MICAL3 | -0.119107462 | 0.92075711 | 0.156701309 | 0.328811581 | 873.8152253 | 804.4166026 | ENSG00000243156 |
| CYP20A1 | -0.119142508 | 0.920734743 | 0.421073865 | 0.631009359 | 238.8954394 | 220.1643824 | ENSG00000119004 |
| FOCAD | -0.119194921 | 0.920701294 | 0.133291237 | 0.294203485 | 1272.974733 | 1171.701258 | ENSG00000188352 |
| PTGER1 | -0.119203749 | 0.92069566 | 0.854148322 | 0.928574107 | 11.56661651 | 10.66730343 | ENSG00000160951 |
| TOP3A | -0.119314164 | 0.920625198 | 0.160403281 | 0.334424372 | 1147.723695 | 1056.630848 | ENSG00000177302 |
| FAM177A1 | -0.119381787 | 0.920582047 | 0.318506303 | 0.528511689 | 446.2871033 | 411.0139261 | ENSG00000151327 |
| GUCD1 | -0.119398262 | 0.920571534 | 0.0874333 | 0.215755634 | 1404.262458 | 1293.066231 | ENSG00000138867 |
| LRP10 | -0.119463175 | 0.920530115 | 0.25569273 | 0.462568943 | 615.5313954 | 566.7812641 | ENSG00000197324 |
| NA | -0.119465636 | 0.920528545 | 0.850191861 | 0.92567733 | 12.57515965 | 11.55899687 | ENSG00000231566 |
| CDCA8 | -0.119477622 | 0.920520897 | 0.095300029 | 0.229947402 | 1269.364897 | 1168.263426 | ENSG00000134690 |
| SKA2 | -0.119501211 | 0.920505846 | 0.156698479 | 0.328811581 | 839.9038426 | 772.9663189 | ENSG00000182628 |
| VPS52 | -0.119595184 | 0.920445889 | 0.236822497 | 0.438858749 | 652.9893564 | 600.8484053 | ENSG00000223501 |
| EBAG9 | -0.119599213 | 0.920443318 | 0.416709038 | 0.62765594 | 265.6785004 | 244.7213024 | ENSG00000147654 |
| GLRX3 | -0.11964479 | 0.920414241 | 0.066855399 | 0.176358474 | 2478.609941 | 2281.588882 | ENSG00000108010 |
| PSMA3-AS1 | -0.119773333 | 0.920332236 | 0.467915129 | 0.671349859 | 210.6806681 | 194.0576604 | ENSG00000257621 |
| NA | -0.119773831 | 0.920331918 | 0.693032201 | 0.835967879 | 53.00034955 | 48.75220659 | ENSG00000249565 |
| DCUN1D5 | -0.119809808 | 0.920308968 | 0.0566375 | 0.155439327 | 2112.517698 | 1944.590051 | ENSG00000137692 |
| ACP2 | -0.119816793 | 0.920304512 | 0.367863811 | 0.579853662 | 323.462693 | 297.7288706 | ENSG00000134575 |
| NAMPT | -0.119830009 | 0.920296081 | 0.120085293 | 0.272737313 | 1655.761455 | 1523.624811 | ENSG00000105835 |
| TRIR | -0.119835897 | 0.920292326 | 0.058124861 | 0.158465136 | 3723.915992 | 3427.547323 | ENSG00000123144 |
| VSIG10 | -0.119964419 | 0.920210345 | 0.109042845 | 0.254608571 | 1328.561552 | 1223.102663 | ENSG00000176834 |
| UBB | -0.120026949 | 0.920170462 | 0.046943575 | 0.134281976 | 6685.183175 | 6150.969712 | ENSG00000170315 |
| LIFR-AS1 | -0.120060002 | 0.920149381 | 0.919158091 | 0.964082229 | 3.991458897 | 3.674340108 | ENSG00000244968 |
| PELO | -0.120114273 | 0.920114767 | 0.165014113 | 0.340652768 | 881.5015639 | 811.06295 | ENSG00000152684 |
| ZEB2 | -0.120220858 | 0.920046793 | 0.862640114 | 0.933472298 | 11.93046781 | 10.97884864 | ENSG00000169554 |
| MAFA-AS1 | -0.12033525 | 0.919973845 | 0.872810339 | 0.940412849 | 10.15389881 | 9.390003709 | ENSG00000254338 |
| NA | -0.120410699 | 0.919925734 | 0.749280046 | 0.869654826 | 36.55353171 | 33.61345582 | ENSG00000226415 |
| NA | -0.120520543 | 0.919855695 | 0.832956779 | 0.915782079 | 17.09178901 | 15.69763882 | ENSG00000234769 |
| NA | -0.1205548 | 0.919833853 | 0.91753355 | 0.963341363 | 3.94282149 | 3.627047002 | ENSG00000258611 |
| ADPGK | -0.120638111 | 0.919780737 | 0.191563875 | 0.378547916 | 729.2225257 | 670.6769689 | ENSG00000159322 |
| SUZ12 | -0.120695654 | 0.919744051 | 0.226418888 | 0.425281481 | 849.9525358 | 781.6410394 | ENSG00000178691 |
| RNF145 | -0.120750329 | 0.919709196 | 0.072289268 | 0.187550876 | 1949.535564 | 1792.790635 | ENSG00000145860 |
| EHMT1 | -0.120756057 | 0.919705545 | 0.035418164 | 0.108266758 | 3411.168557 | 3137.306733 | ENSG00000181090 |
| MAP1LC3B | -0.120772922 | 0.919694794 | 0.174222078 | 0.354507482 | 875.6188478 | 805.1419195 | ENSG00000140941 |

| | | | | | | | |
|---|---|---|---|---|---|---|---|
| KIDINS220 | -0.120777777 | 0.919691698 | 0.093830621 | 0.227557632 | 1312.097418 | 1206.622084 | ENSG00000134313 |
| ZNF555 | -0.120824602 | 0.919661849 | 0.576106896 | 0.758153886 | 103.6731536 | 95.34573024 | ENSG00000186300 |
| SNAP25 | -0.120840358 | 0.919651805 | 0.78331614 | 0.889784424 | 54.19004185 | 49.85454354 | ENSG00000132639 |
| MTA3 | -0.120918615 | 0.919601921 | 0.033114988 | 0.102638529 | 3063.274525 | 2816.432393 | ENSG00000057935 |
| RBP4 | -0.120926662 | 0.919596792 | 0.835312326 | 0.917410484 | 15.22506414 | 14.02480145 | ENSG00000138207 |
| KCTD10 | -0.120959973 | 0.919575559 | 0.11310246 | 0.261661148 | 1349.51475 | 1240.481755 | ENSG00000110906 |
| CPSF1 | -0.120974184 | 0.919566501 | 0.036466534 | 0.110647573 | 4725.227688 | 4345.876327 | ENSG00000071894 |
| TIMM21 | -0.121047726 | 0.919519627 | 0.140568792 | 0.3047809 | 848.1747198 | 779.7839406 | ENSG00000075336 |
| ZCCHC3 | -0.121049348 | 0.919518593 | 0.036405657 | 0.110516127 | 2607.19656 | 2397.675827 | ENSG00000247315 |
| SNRNP27 | -0.121098115 | 0.919487511 | 0.543623272 | 0.732315108 | 182.8457132 | 168.2696496 | ENSG00000124380 |
| MRPL16 | -0.121153481 | 0.919452225 | 0.125730049 | 0.282030168 | 1534.798555 | 1410.521499 | ENSG00000166902 |
| B2M | -0.121285351 | 0.919368186 | 0.169397752 | 0.347337414 | 823.3168098 | 756.6450428 | ENSG00000166710 |
| AKR1A1 | -0.121292767 | 0.91936346 | 0.024919114 | 0.082272285 | 3424.496005 | 3148.601153 | ENSG00000117448 |
| NA | -0.121310744 | 0.919352004 | 0.638236116 | 0.801722623 | 101.4496964 | 93.3975955 | ENSG00000196295 |
| USP54 | -0.121313428 | 0.919350294 | 0.223752319 | 0.421839127 | 605.9168532 | 556.7921727 | ENSG00000166348 |
| C3orf70 | -0.121346678 | 0.919329106 | 0.669743253 | 0.82299482 | 62.31266879 | 57.31538163 | ENSG00000187068 |
| CASKIN1 | -0.121467264 | 0.919252268 | 0.462989204 | 0.667710361 | 202.6431631 | 186.4841699 | ENSG00000167971 |
| PDHA1 | -0.121562515 | 0.919191578 | 0.04090999 | 0.120791927 | 2443.538843 | 2245.913621 | ENSG00000131828 |
| CEP57L1 | -0.121581828 | 0.919179273 | 0.56962704 | 0.752634622 | 108.9421226 | 100.0714983 | ENSG00000183137 |
| BCL6 | -0.121600715 | 0.91916724 | 0.708859646 | 0.846409611 | 51.79758843 | 47.60903029 | ENSG00000113916 |
| PCDHB11 | -0.121615682 | 0.919157704 | 0.886179085 | 0.948401451 | 7.658194534 | 7.048958286 | ENSG00000197479 |
| PPP4C | -0.121664821 | 0.919126398 | 0.048813378 | 0.138456089 | 4638.971215 | 4263.630039 | ENSG00000149923 |
| ZNF773 | -0.121692131 | 0.919108999 | 0.627286402 | 0.793729923 | 85.45660638 | 78.58557523 | ENSG00000152439 |
| RLN1 | -0.121764949 | 0.91906261 | 0.790201725 | 0.892521506 | 23.17475148 | 21.31966217 | ENSG00000107014 |
| LSM3 | -0.121837067 | 0.919016668 | 0.082029456 | 0.205699944 | 1617.431836 | 1486.184809 | ENSG00000170860 |
| CHGA | -0.121865826 | 0.918998348 | 0.082278858 | 0.20618846 | 1758.87626 | 1616.615089 | ENSG00000100604 |
| WDHD1 | -0.1219003 | 0.918976389 | 0.100924367 | 0.239519362 | 1588.088415 | 1459.160558 | ENSG00000198554 |
| ARMC9 | -0.121944175 | 0.918948442 | 0.183789564 | 0.367793358 | 760.4756636 | 698.5998348 | ENSG00000135931 |
| NIPBL | -0.121991844 | 0.918918078 | 0.135644358 | 0.297761676 | 1299.021699 | 1193.419368 | ENSG00000164190 |
| PEX1 | -0.122012062 | 0.918905201 | 0.268031717 | 0.47591002 | 440.5430503 | 404.7466062 | ENSG00000127980 |
| PRDM15 | -0.122035448 | 0.918890305 | 0.367142162 | 0.579491407 | 349.1577601 | 320.7204242 | ENSG00000141956 |
| DDOST | -0.122045152 | 0.918884125 | 0.009636972 | 0.037876795 | 6687.653698 | 6145.00521 | ENSG00000244038 |
| RTBDN | -0.122102108 | 0.918847849 | 0.86873732 | 0.937813209 | 8.679780357 | 7.949070032 | ENSG00000132026 |
| SP2 | -0.122149234 | 0.918817835 | 0.365378716 | 0.577614017 | 332.437525 | 305.7907812 | ENSG00000167182 |
| TNS3 | -0.12218016 | 0.918798139 | 0.169088184 | 0.346864039 | 887.9404475 | 815.6392508 | ENSG00000136205 |
| MTMR4 | -0.122206841 | 0.918781147 | 0.224097362 | 0.422320995 | 803.972809 | 738.2157913 | ENSG00000108389 |
| STAT3 | -0.122209614 | 0.918779381 | 0.055140915 | 0.152219878 | 3015.052881 | 2769.377909 | ENSG00000168610 |
| EXTL3 | -0.122211977 | 0.918777877 | 0.049375376 | 0.139714412 | 3908.431092 | 3590.285336 | ENSG00000012232 |
| SLC35E3 | -0.122339581 | 0.918696616 | 0.312182975 | 0.522564748 | 350.2706703 | 321.9124067 | ENSG00000175782 |
| UTP6 | -0.122342062 | 0.918695036 | 0.095666191 | 0.230458288 | 1722.685229 | 1582.181877 | ENSG00000108651 |
| SRM | -0.122368664 | 0.918678096 | 0.076299432 | 0.195060888 | 3348.02563 | 3076.269693 | ENSG00000116649 |
| XPO7 | -0.122489463 | 0.918601177 | 0.035427882 | 0.108278938 | 3361.167673 | 3087.550732 | ENSG00000130227 |
| SLC25A16 | -0.122509034 | 0.918588716 | 0.601101704 | 0.776307564 | 87.05433818 | 79.9480969 | ENSG00000122912 |
| NA | -0.1225118 | 0.918586955 | 0.907335573 | 0.960082797 | 3.973661534 | 3.648518091 | ENSG00000197670 |
| MND1 | -0.122520216 | 0.918581596 | 0.34510281 | 0.557034745 | 311.344489 | 286.2021268 | ENSG00000121211 |
| DHRS3 | -0.122556981 | 0.918558187 | 0.390271227 | 0.601716195 | 249.5792372 | 229.4318025 | ENSG00000162496 |
| DCLRE1A | -0.122625747 | 0.918514405 | 0.406404256 | 0.617304301 | 335.183458 | 307.4649731 | ENSG00000198924 |
| MRFAP1L1 | -0.122686889 | 0.918475479 | 0.100442458 | 0.238808487 | 1479.82124 | 1359.150932 | ENSG00000178988 |
| LRRC3B | -0.122695836 | 0.918469783 | 0.692091553 | 0.835483064 | 51.59327626 | 47.49079752 | ENSG00000179796 |
| OCIAD1 | -0.122707146 | 0.918462583 | 0.065616436 | 0.173649999 | 1875.26568 | 1722.050107 | ENSG00000109180 |
| CYP7B1 | -0.122763776 | 0.918426531 | 0.871739409 | 0.939979373 | 10.1624923 | 9.27706783 | ENSG00000172817 |
| MAGOH | -0.12288772 | 0.918347631 | 0.118338892 | 0.27026605 | 1547.715903 | 1420.927104 | ENSG00000162385 |
| SLC35E2A | -0.122925866 | 0.91832335 | 0.458812012 | 0.664077828 | 204.089696 | 187.4254273 | ENSG00000215790 |
| ATR | -0.12295004 | 0.918307962 | 0.276681257 | 0.485832221 | 567.6296415 | 520.7446936 | ENSG00000175054 |
| THEMIS2 | -0.12302729 | 0.918258793 | 0.32937086 | 0.541072433 | 335.0028925 | 307.7842445 | ENSG00000130775 |
| NA | -0.123047029 | 0.918246229 | 0.604686874 | 0.779063421 | 90.03105087 | 82.79421088 | ENSG00000205746 |
| NR2C2 | -0.123054439 | 0.918241513 | 0.222675946 | 0.420355381 | 722.8296889 | 663.576861 | ENSG00000177463 |
| RBM7 | -0.123089038 | 0.918219491 | 0.381348316 | 0.59376395 | 318.1557906 | 292.3636996 | ENSG00000076053 |
| PPIH | -0.123090055 | 0.918218844 | 0.413066678 | 0.624460791 | 237.5924288 | 218.2045924 | ENSG00000171960 |
| ZAN | -0.123108879 | 0.918206863 | 0.903116649 | 0.957841349 | 5.060486917 | 4.612097146 | ENSG00000146839 |
| RAB28 | -0.123149957 | 0.91818072 | 0.315232956 | 0.525481011 | 388.6811643 | 356.8289413 | ENSG00000157869 |
| NA | -0.12319093 | 0.918154643 | 0.912210322 | 0.961440559 | 4.011620523 | 3.681812457 | ENSG00000287202 |
| SPECC1 | -0.123327086 | 0.918067996 | 0.074904373 | 0.192383452 | 1680.245753 | 1542.449457 | ENSG00000128487 |
| TFAP4 | -0.123341082 | 0.918059089 | 0.202372937 | 0.393732085 | 615.4123804 | 564.9303198 | ENSG00000090447 |
| HSF2BP | -0.123359576 | 0.91804732 | 0.787452031 | 0.891546762 | 25.22356753 | 23.21447187 | ENSG00000160207 |
| C2orf49 | -0.123421978 | 0.918007612 | 0.324895525 | 0.535864802 | 389.1019149 | 357.3305525 | ENSG00000135974 |
| NA | -0.123593345 | 0.917898575 | 0.909351822 | 0.96047171 | 4.000942105 | 3.66998918 | ENSG00000242952 |
| MKLN1 | -0.123601299 | 0.917893515 | 0.248335934 | 0.45329829 | 617.3503209 | 566.3853772 | ENSG00000128585 |
| MRPS11 | -0.123617763 | 0.91788304 | 0.240798732 | 0.444437456 | 550.847142 | 505.5020353 | ENSG00000181991 |
| NA | -0.123631797 | 0.917874111 | 0.796582914 | 0.896137983 | 19.88966451 | 18.24259045 | ENSG00000286651 |
| ANG | -0.123668415 | 0.917850814 | 0.908247676 | 0.960165884 | 4.00806105 | 3.676515571 | ENSG00000214274 |
| RIOK3 | -0.123676235 | 0.917845839 | 0.321367423 | 0.531857312 | 344.1781985 | 315.9812971 | ENSG00000101782 |
| RAD21 | -0.123681104 | 0.917842741 | 0.043861122 | 0.12739423 | 3746.753598 | 3438.675127 | ENSG00000164754 |
| NA | -0.123832101 | 0.917746682 | 0.915886125 | 0.962597668 | 3.970102062 | 3.641045742 | ENSG00000183298 |

| | | | | | | | |
|---|---|---|---|---|---|---|---|
| TM9SF1 | -0.123856313 | 0.91773128 | 0.675851982 | 0.826595731 | 59.80600676 | 54.9497437 | ENSG00000100926 |
| CENPC | -0.123940997 | 0.917677413 | 0.408350167 | 0.619512604 | 263.1606019 | 241.6273214 | ENSG00000145241 |
| STK24 | -0.123973565 | 0.917656697 | 0.122102474 | 0.27599105 | 2122.953843 | 1947.938215 | ENSG00000102572 |
| LZTS3 | -0.124080083 | 0.917588946 | 0.211097671 | 0.404769977 | 1280.431869 | 1175.238046 | ENSG00000088899 |
| TIMM22 | -0.124081848 | 0.917587823 | 0.163073418 | 0.338120875 | 775.7318733 | 711.6666806 | ENSG00000177370 |
| SPCS3 | -0.124139863 | 0.917550926 | 0.231828879 | 0.432574685 | 1163.13502 | 1067.415405 | ENSG00000129128 |
| FASN | -0.124169926 | 0.917531806 | 0.035539964 | 0.108463513 | 34942.34317 | 32060.97212 | ENSG00000169710 |
| UBXN2A | -0.124170446 | 0.917531475 | 0.263189501 | 0.47031924 | 515.09169 | 472.8905389 | ENSG00000173960 |
| RUBCN | -0.124232914 | 0.917491747 | 0.450456568 | 0.656799898 | 228.3219074 | 209.2166098 | ENSG00000145016 |
| NA | -0.124233697 | 0.917491249 | 0.73938352 | 0.863587169 | 38.47364867 | 35.20353026 | ENSG00000284946 |
| MPRIP | -0.124353755 | 0.917414901 | 0.017218657 | 0.060939718 | 3630.730291 | 3330.763012 | ENSG00000133030 |
| TIMM17B | -0.124375292 | 0.917401205 | 0.148226208 | 0.316118937 | 1010.663666 | 927.3613194 | ENSG00000126768 |
| PFN2 | -0.124464472 | 0.917344498 | 0.036788626 | 0.11137435 | 2447.32271 | 2244.727123 | ENSG00000070087 |
| NA | -0.124498835 | 0.917322649 | 0.8818306 | 0.945689642 | 8.408449184 | 7.700046177 | ENSG00000231305 |
| PAWR | -0.124537491 | 0.91729807 | 0.155143038 | 0.326812746 | 2202.575216 | 2020.719348 | ENSG00000177425 |
| GRM3 | -0.124571811 | 0.917276249 | 0.717745282 | 0.851146028 | 41.47614123 | 38.0749718 | ENSG00000198822 |
| CCT5 | -0.124601579 | 0.917257322 | 0.008227087 | 0.033447538 | 11668.7451 | 10702.32075 | ENSG00000150753 |
| LLPH | -0.124640932 | 0.917232302 | 0.451331922 | 0.657517808 | 222.6900024 | 204.3762974 | ENSG00000139233 |
| ANAPC1 | -0.124695204 | 0.917197798 | 0.063587178 | 0.169519847 | 2153.793647 | 1974.882788 | ENSG00000153107 |
| CLTA | -0.124709425 | 0.917188757 | 0.015503051 | 0.055990653 | 4110.774447 | 3770.669337 | ENSG00000122705 |
| APBB2 | -0.124775728 | 0.917146606 | 0.129416725 | 0.28804182 | 875.6365929 | 803.0287534 | ENSG00000163697 |
| TM6SF2 | -0.124811172 | 0.917124074 | 0.820625968 | 0.908947936 | 15.23098788 | 13.9784543 | ENSG00000213996 |
| GPR27 | -0.124872519 | 0.917085076 | 0.222196659 | 0.41961839 | 513.7088709 | 471.1136784 | ENSG00000170837 |
| NFE2L3 | -0.124902067 | 0.917066294 | 0.023763139 | 0.079091803 | 3811.897827 | 3495.43996 | ENSG00000050344 |
| SAMHD1 | -0.12496303 | 0.917027542 | 0.112465768 | 0.260399869 | 1083.790184 | 994.0478379 | ENSG00000101347 |
| N4BP3 | -0.12497739 | 0.917018415 | 0.032110423 | 0.100248857 | 2587.922476 | 2373.280178 | ENSG00000145911 |
| ENTR1 | -0.124985816 | 0.917013059 | 0.06803577 | 0.1786755 | 1976.994236 | 1813.007188 | ENSG00000165689 |
| UMAD1 | -0.124985856 | 0.917013033 | 0.626551561 | 0.793377111 | 75.92245694 | 69.65921095 | ENSG00000219545 |
| EIF4A3 | -0.125014811 | 0.916994629 | 0.01938896 | 0.066953092 | 5188.198838 | 4757.890693 | ENSG00000141543 |
| UFL1 | -0.125020941 | 0.916990733 | 0.305403051 | 0.515649228 | 339.2096213 | 310.9494487 | ENSG00000014123 |
| RPS16 | -0.125115885 | 0.916930387 | 0.028442864 | 0.091010417 | 20390.68376 | 18696.3536 | ENSG00000105193 |
| NA | -0.125192702 | 0.916881567 | 0.908612329 | 0.960165884 | 3.973661534 | 3.638870278 | ENSG00000262904 |
| ARPC3 | -0.12525694 | 0.916840742 | 0.160513594 | 0.334543563 | 850.9976157 | 780.3968161 | ENSG00000111229 |
| RBL1 | -0.12537953 | 0.916762839 | 0.446468878 | 0.653604036 | 179.4022823 | 164.5135878 | ENSG00000080839 |
| DGKA | -0.125392727 | 0.916754452 | 0.426109435 | 0.636082261 | 205.9763032 | 188.7706179 | ENSG00000065357 |
| GBE1 | -0.125397366 | 0.916751505 | 0.468745026 | 0.67185333 | 191.1661176 | 175.4762623 | ENSG00000114480 |
| NA | -0.12563981 | 0.916597458 | 0.903165523 | 0.957841349 | 4.685957196 | 4.267257575 | ENSG00000260518 |
| KIF23 | -0.12580476 | 0.916492665 | 0.100683814 | 0.239192486 | 1358.949617 | 1245.364819 | ENSG00000137807 |
| LURAP1 | -0.12582804 | 0.916477876 | 0.824969803 | 0.911618182 | 14.48192844 | 13.3027739 | ENSG00000171357 |
| STAR | -0.125865864 | 0.916453849 | 0.648748064 | 0.809250856 | 68.36181656 | 62.55208224 | ENSG00000147465 |
| ST6GAL1 | -0.125867047 | 0.916453097 | 0.031155285 | 0.097849253 | 2730.824977 | 2502.329184 | ENSG00000073849 |
| BAG4 | -0.12591369 | 0.916423469 | 0.276542856 | 0.485634354 | 595.1541355 | 545.507999 | ENSG00000156735 |
| TLNRD1 | -0.125977214 | 0.916383118 | 0.129684154 | 0.288376048 | 1308.496661 | 1199.543193 | ENSG00000140406 |
| MYLK3 | -0.126061987 | 0.916329273 | 0.850433971 | 0.925887555 | 11.88066135 | 10.87243915 | ENSG00000140795 |
| NA | -0.126116297 | 0.916294778 | 0.73544437 | 0.861096497 | 33.27764862 | 30.53203318 | ENSG00000229939 |
| TMEM203 | -0.126180278 | 0.916254143 | 0.113451566 | 0.262055082 | 990.9756966 | 908.1811601 | ENSG00000187713 |
| MRPL43 | -0.126310986 | 0.916171134 | 0.130804853 | 0.290310218 | 890.1623523 | 815.2875716 | ENSG00000055950 |
| MRPS17 | -0.126352391 | 0.916144841 | 0.565964249 | 0.750008186 | 110.6653256 | 101.3993282 | ENSG00000239789 |
| RAB11FIP5 | -0.126415424 | 0.916104814 | 0.168182804 | 0.345645038 | 706.3790834 | 647.4184933 | ENSG00000135631 |
| POP7 | -0.126419252 | 0.916102383 | 0.073424815 | 0.189764668 | 1537.469081 | 1408.816893 | ENSG00000172336 |
| ZCCHC8 | -0.126457739 | 0.916077945 | 0.244878248 | 0.449070278 | 622.7583886 | 570.2654631 | ENSG00000033030 |
| CACNG7 | -0.126539347 | 0.916026127 | 0.080842757 | 0.203399343 | 1361.027241 | 1246.929938 | ENSG00000105605 |
| GNAI3 | -0.126564359 | 0.916010246 | 0.073704898 | 0.190282569 | 2144.195225 | 1963.975571 | ENSG00000065135 |
| KXD1 | -0.126668577 | 0.915944078 | 0.147755575 | 0.315580642 | 799.8093519 | 732.6259188 | ENSG00000105700 |
| CEP95 | -0.126734331 | 0.915902332 | 0.327432988 | 0.538826418 | 397.9104211 | 364.5232094 | ENSG00000258890 |
| PAPSS2 | -0.126742625 | 0.915897067 | 0.274336669 | 0.483152849 | 507.4207588 | 464.7312245 | ENSG00000198682 |
| LNPK | -0.126781757 | 0.915872224 | 0.519190646 | 0.714901185 | 139.587664 | 127.7639439 | ENSG00000144320 |
| ADCY1 | -0.126835379 | 0.915838184 | 0.136243586 | 0.298583818 | 1235.311875 | 1131.460015 | ENSG00000164742 |
| DPM2 | -0.126839864 | 0.915835336 | 0.133882377 | 0.295163284 | 1005.304237 | 920.6207622 | ENSG00000136908 |
| CKB | -0.126906147 | 0.91579326 | 0.021878114 | 0.074017947 | 7378.733074 | 6758.264419 | ENSG00000166165 |
| C12orf43 | -0.126959469 | 0.915759413 | 0.222674462 | 0.420355381 | 536.3290602 | 491.150744 | ENSG00000157895 |
| ALKBH7 | -0.126986854 | 0.91574203 | 0.140545338 | 0.3047809 | 1113.360922 | 1019.835041 | ENSG00000125652 |
| ZGPAT | -0.127027744 | 0.915716076 | 0.486966394 | 0.687938387 | 172.6838057 | 158.1807931 | ENSG00000197114 |
| GALM | -0.127081957 | 0.915681667 | 0.292307145 | 0.502066434 | 388.1969019 | 355.3384664 | ENSG00000143891 |
| HCFC1 | -0.127091607 | 0.915675542 | 0.044668866 | 0.12904567 | 4908.736961 | 4495.086989 | ENSG00000172534 |
| CASC4 | -0.127122969 | 0.915655636 | 0.224594724 | 0.422914571 | 532.4216064 | 487.4949215 | ENSG00000166734 |
| PKIG | -0.12714015 | 0.915644732 | 0.263572515 | 0.470914564 | 421.0433486 | 385.558902 | ENSG00000168734 |
| SNRPG | -0.127158348 | 0.915633182 | 0.082723829 | 0.206973967 | 1940.651569 | 1776.923357 | ENSG00000143977 |
| TMED7 | -0.127189975 | 0.91561311 | 0.331411852 | 0.542962899 | 495.7414332 | 454.2853263 | ENSG00000134970 |
| NA | -0.127280497 | 0.915555661 | 0.8161333 | 0.906477444 | 17.00491329 | 15.62982058 | ENSG00000283103 |
| TSPEAR | -0.127313156 | 0.915534936 | 0.773884572 | 0.883817739 | 29.13021005 | 26.60649376 | ENSG00000175894 |
| SORCS1 | -0.12744399 | 0.915451913 | 0.492130533 | 0.691560764 | 155.4132057 | 142.3806442 | ENSG00000108018 |
| HINFP | -0.127483238 | 0.915427008 | 0.182917361 | 0.36670853 | 640.4571113 | 586.0103224 | ENSG00000172273 |
| ZNF33A | -0.127508211 | 0.915411163 | 0.192753539 | 0.380212094 | 590.1953993 | 540.2880624 | ENSG00000189180 |

| | | | | | | | |
|---|---|---|---|---|---|---|---|
| SMN1 | -0.127522274 | 0.915402239 | 0.487275558 | 0.688066316 | 158.8724013 | 145.5414975 | ENSG00000172062 |
| CD83 | -0.127534554 | 0.915394448 | 0.43068226 | 0.640528782 | 237.8501587 | 217.8290854 | ENSG00000112149 |
| PRPF4 | -0.127543848 | 0.91538855 | 0.061203056 | 0.164928612 | 1581.661789 | 1447.568857 | ENSG00000136875 |
| NARS1 | -0.127575648 | 0.915368373 | 0.01266363 | 0.047469759 | 5683.701109 | 5202.463115 | ENSG00000134440 |
| LINC01278 | -0.127581511 | 0.915364653 | 0.312404902 | 0.522611975 | 358.0652677 | 327.6878061 | ENSG00000235437 |
| RMND5A | -0.127643346 | 0.915325421 | 0.13256079 | 0.293002167 | 1712.087066 | 1567.067726 | ENSG00000153561 |
| SMARCAD1 | -0.127726283 | 0.915272803 | 0.159215192 | 0.332608286 | 1393.020286 | 1274.902352 | ENSG00000163104 |
| PPP1R14B | -0.127731921 | 0.915269226 | 0.035829818 | 0.109206925 | 6351.449381 | 5813.116295 | ENSG00000173457 |
| CHKB-CPT1B | -0.127769648 | 0.915245292 | 0.881589076 | 0.945632722 | 6.905575619 | 6.341875435 | ENSG00000254413 |
| GINS4 | -0.127775644 | 0.915241488 | 0.1536043 | 0.324548804 | 772.5185345 | 706.6614845 | ENSG00000147536 |
| KYAT3 | -0.12783242 | 0.91520547 | 0.177749618 | 0.359670153 | 748.93282 | 685.6734615 | ENSG00000137944 |
| MORF4L1 | -0.127846916 | 0.915196274 | 0.017140204 | 0.060741743 | 10429.15469 | 9543.696222 | ENSG00000185787 |
| HSPA13 | -0.127851339 | 0.915193468 | 0.205262545 | 0.397211104 | 765.6796979 | 700.6065915 | ENSG00000155304 |
| WWC1 | -0.127976643 | 0.915113984 | 0.079741173 | 0.201325141 | 1589.192356 | 1454.340563 | ENSG00000113645 |
| E2F1 | -0.127986913 | 0.915107469 | 0.135977311 | 0.298230426 | 1092.079086 | 999.5264822 | ENSG00000101412 |
| NUDT2 | -0.128118873 | 0.91502377 | 0.444037451 | 0.651464207 | 197.7540372 | 180.9089452 | ENSG00000164978 |
| ASXL2 | -0.128124205 | 0.915020389 | 0.187108717 | 0.372462118 | 829.9139327 | 759.0358701 | ENSG00000143970 |
| SPATA13 | -0.128307914 | 0.91490388 | 0.29351631 | 0.503273608 | 390.1949956 | 356.8199559 | ENSG00000182957 |
| GLG1 | -0.128331331 | 0.914889029 | 0.032927766 | 0.102175627 | 5123.118251 | 4686.064093 | ENSG00000090863 |
| ARRB2 | -0.128409893 | 0.914839211 | 0.082700584 | 0.206943227 | 1777.926988 | 1626.7826 | ENSG00000141480 |
| FBXW2 | -0.128530452 | 0.914762765 | 0.112790855 | 0.261004203 | 1084.518114 | 992.0277543 | ENSG00000119402 |
| DPY19L1 | -0.128547496 | 0.914751958 | 0.303346071 | 0.513486816 | 374.2852736 | 342.6239231 | ENSG00000173852 |
| LNX2 | -0.128572317 | 0.914736221 | 0.259839552 | 0.466762099 | 420.4064122 | 384.5679449 | ENSG00000139517 |
| TRIM59 | -0.128590202 | 0.91472488 | 0.317167584 | 0.527263189 | 394.4106484 | 360.8508635 | ENSG00000213186 |
| NA | -0.128716233 | 0.914644975 | 0.433896778 | 0.642632302 | 404.2125701 | 369.8950223 | ENSG00000214826 |
| NA | -0.128719787 | 0.914642722 | 0.873037759 | 0.940412849 | 11.26325008 | 10.30222228 | ENSG00000253570 |
| POLG | -0.128742106 | 0.914628573 | 0.052810635 | 0.147055269 | 1870.350623 | 1711.044651 | ENSG00000140521 |
| SLC3A2 | -0.128755107 | 0.91462033 | 0.00545414 | 0.023749468 | 9322.684837 | 8526.610209 | ENSG00000168003 |
| PLEKHJ1 | -0.128920389 | 0.914515553 | 0.062509424 | 0.167399639 | 2073.247548 | 1896.382368 | ENSG00000104886 |
| RGS12 | -0.128939735 | 0.91450329 | 0.168687411 | 0.346342873 | 612.9496272 | 560.6122915 | ENSG00000159788 |
| SLC2A3 | -0.128948256 | 0.914497889 | 0.030573712 | 0.096407505 | 12874.60376 | 11772.61703 | ENSG00000059804 |
| ERCC6L | -0.128972551 | 0.914482489 | 0.186621706 | 0.371806194 | 772.703321 | 706.5421152 | ENSG00000186871 |
| MRFAP1 | -0.12901725 | 0.914454156 | 0.006679246 | 0.028174571 | 10275.67767 | 9396.676225 | ENSG00000179010 |
| CDKN3 | -0.129023622 | 0.914450117 | 0.215314518 | 0.410438546 | 505.0714993 | 461.6889829 | ENSG00000100526 |
| SLC39A6 | -0.129035275 | 0.91444273 | 0.126230591 | 0.282854985 | 1094.855657 | 1001.230119 | ENSG00000141424 |
| GFRA2 | -0.129116968 | 0.914390951 | 0.488005271 | 0.688684766 | 140.3304943 | 128.3797729 | ENSG00000168546 |
| PINX1 | -0.129190389 | 0.914344418 | 0.329681719 | 0.541217709 | 305.0174498 | 279.0674951 | ENSG00000254093 |
| FHOD3 | -0.12927397 | 0.914291448 | 0.330380324 | 0.541788157 | 331.132735 | 302.6954325 | ENSG00000134775 |
| NA | -0.129286449 | 0.914283539 | 0.881993502 | 0.945771249 | 6.540529107 | 6.011034604 | ENSG00000232150 |
| VMA21 | -0.129428185 | 0.914193721 | 0.312401168 | 0.522611975 | 574.5516684 | 525.620664 | ENSG00000160131 |
| WDR12 | -0.129468991 | 0.914167864 | 0.049566909 | 0.1401304 | 1899.947676 | 1736.449564 | ENSG00000138442 |
| RAB40C | -0.129517917 | 0.914136862 | 0.133013053 | 0.293692447 | 779.5237747 | 712.4145825 | ENSG00000197562 |
| TCF4 | -0.129559469 | 0.914110534 | 0.161345066 | 0.33586878 | 1178.749639 | 1077.088957 | ENSG00000196628 |
| ZRANB3 | -0.129567248 | 0.914105605 | 0.491683046 | 0.691126718 | 164.6626502 | 150.3068028 | ENSG00000121988 |
| TAFA4 | -0.129579232 | 0.914098012 | 0.456357421 | 0.662488269 | 171.0691145 | 156.2809701 | ENSG00000163377 |
| GPX8 | -0.129589452 | 0.914091536 | 0.117113605 | 0.268291789 | 958.3647332 | 876.034753 | ENSG00000164294 |
| TPI1 | -0.129619753 | 0.914072338 | 0.006794399 | 0.028545546 | 13375.7858 | 12225.91597 | ENSG00000111669 |
| TRIM52-AS1 | -0.129643929 | 0.914057021 | 0.508325765 | 0.705084985 | 134.897868 | 123.3982437 | ENSG00000248275 |
| NA | -0.129661621 | 0.914045811 | 0.869027337 | 0.938019048 | 8.319488525 | 7.57623298 | ENSG00000220157 |
| SFMBT1 | -0.129665529 | 0.914043335 | 0.296405906 | 0.506588137 | 405.6341862 | 370.7872696 | ENSG00000163935 |
| ATP6V0A1 | -0.129697594 | 0.91402302 | 0.155452801 | 0.327282738 | 805.5479882 | 736.1390637 | ENSG00000033627 |
| KIAA1958 | -0.129701003 | 0.91402086 | 0.180897871 | 0.363879543 | 642.1370943 | 586.7240996 | ENSG00000165185 |
| IK | -0.129728913 | 0.914003178 | 0.078011366 | 0.198161867 | 1527.169219 | 1395.460089 | ENSG00000113141 |
| COMMD7 | -0.129789162 | 0.913965009 | 0.331123717 | 0.542677375 | 297.2409291 | 271.692782 | ENSG00000149600 |
| ZNF174 | -0.129835276 | 0.913935796 | 0.438458804 | 0.6466204 | 220.673832 | 201.9203516 | ENSG00000103343 |
| SEMA5A | -0.129840668 | 0.91393238 | 0.450073867 | 0.656361128 | 296.1578908 | 270.3305962 | ENSG00000112902 |
| ATP2A1-AS1 | -0.129922495 | 0.913880545 | 0.582861063 | 0.76255721 | 85.04912553 | 77.7501603 | ENSG00000260442 |
| TMEM39B | -0.129989195 | 0.913838294 | 0.220661979 | 0.417638954 | 487.785771 | 445.8655663 | ENSG00000121775 |
| ZNF230 | -0.12999443 | 0.913834978 | 0.681868051 | 0.82993643 | 57.43986573 | 52.4486802 | ENSG00000159882 |
| CAPN15 | -0.130034162 | 0.913809812 | 0.071087705 | 0.185120147 | 2236.875873 | 2044.691205 | ENSG00000103326 |
| CLEC16A | -0.130088672 | 0.913775285 | 0.246972153 | 0.451769576 | 442.7784073 | 404.4970153 | ENSG00000038532 |
| RSPRY1 | -0.130097161 | 0.913769909 | 0.193779865 | 0.381837725 | 574.8425258 | 525.2072711 | ENSG00000159579 |
| ZNF792 | -0.130105841 | 0.913764411 | 0.492727029 | 0.692244453 | 139.5579931 | 127.5860706 | ENSG00000180884 |
| KNOP1 | -0.130226873 | 0.913687756 | 0.080689117 | 0.203105771 | 1272.182934 | 1162.541533 | ENSG00000103550 |
| RAB11B-AS1 | -0.130296716 | 0.913643524 | 0.857549159 | 0.930409931 | 10.19897674 | 9.348007489 | ENSG00000269386 |
| RPL27A | -0.130341401 | 0.913615225 | 0.027679722 | 0.089263724 | 14060.26345 | 12845.67276 | ENSG00000166441 |
| RBM28 | -0.130416546 | 0.91356764 | 0.066587176 | 0.175898561 | 1427.485491 | 1303.695678 | ENSG00000106344 |
| SLC29A2 | -0.130492237 | 0.913519711 | 0.159490917 | 0.333036927 | 806.7194291 | 737.0094016 | ENSG00000174669 |
| LSS | -0.130503192 | 0.913512774 | 0.02165497 | 0.073354995 | 5473.214253 | 5000.526698 | ENSG00000160285 |
| SFXN2 | -0.130567132 | 0.913472288 | 0.328447022 | 0.539977541 | 367.6201636 | 335.8125645 | ENSG00000156398 |
| ARL1 | -0.13060886 | 0.913445868 | 0.138139946 | 0.301308012 | 943.6971273 | 862.2510718 | ENSG00000120805 |
| NOP53-AS1 | -0.130633811 | 0.91343007 | 0.919376018 | 0.964136712 | 3.584893923 | 3.275681039 | ENSG00000269656 |
| UQCRC2 | -0.130667728 | 0.913408596 | 0.029173038 | 0.092843414 | 3494.628181 | 3191.262396 | ENSG00000140740 |
| SOD2 | -0.13067764 | 0.91340232 | 0.177690291 | 0.359588635 | 753.6294556 | 688.3061035 | ENSG00000112096 |

| | | | | | | | |
|---|---|---|---|---|---|---|---|
| KHDRBS2 | -0.130702153 | 0.913386801 | 0.416406565 | 0.627350458 | 189.9745935 | 173.6085041 | ENSG00000112232 |
| OAZ2 | -0.130769689 | 0.913344044 | 0.010016708 | 0.039124206 | 6181.825583 | 5646.259101 | ENSG00000180304 |
| MAP3K1 | -0.130772077 | 0.913342533 | 0.328831481 | 0.540421419 | 301.9580057 | 275.7303113 | ENSG00000095015 |
| UBE2D3 | -0.130827346 | 0.913307543 | 0.01703172 | 0.060413982 | 6097.9467 | 5568.498765 | ENSG00000109332 |
| SYNE3 | -0.130844767 | 0.913296515 | 0.905481775 | 0.958927594 | 3.622852912 | 3.311150869 | ENSG00000176438 |
| MTHFD1L | -0.130929953 | 0.913242589 | 0.024097498 | 0.079951028 | 2880.496546 | 2630.297238 | ENSG00000120254 |
| NEGR1 | -0.130947244 | 0.913231644 | 0.679272438 | 0.828368191 | 47.18012478 | 43.114955 | ENSG00000172260 |
| HS3ST3B1 | -0.130979178 | 0.91321143 | 0.43158926 | 0.641354345 | 181.2464813 | 165.5104613 | ENSG00000125430 |
| SMG7-AS1 | -0.131009735 | 0.913192088 | 0.824950701 | 0.911618182 | 14.11451766 | 12.90941172 | ENSG00000232860 |
| KCNMB4 | -0.131019774 | 0.913185734 | 0.281521731 | 0.490500774 | 378.0094933 | 345.2713122 | ENSG00000135643 |
| DENR | -0.131038832 | 0.913173671 | 0.037543629 | 0.113043915 | 2574.608908 | 2351.41059 | ENSG00000139726 |
| UBC | -0.131147968 | 0.913104594 | 0.007911735 | 0.032361516 | 10312.25273 | 9416.056977 | ENSG00000150991 |
| SRSF7 | -0.131237078 | 0.913048196 | 0.005397275 | 0.023556168 | 7039.739935 | 6427.592553 | ENSG00000115875 |
| UBFD1 | -0.131251388 | 0.91303914 | 0.041941889 | 0.12312647 | 3188.50731 | 2911.04513 | ENSG00000103353 |
| NA | -0.131290076 | 0.913014656 | 0.585998881 | 0.764689667 | 83.97534283 | 76.70041334 | ENSG00000267106 |
| APPBP2 | -0.131437606 | 0.912921296 | 0.30877749 | 0.519257604 | 343.6351789 | 313.486928 | ENSG00000062725 |
| ELP6 | -0.131513147 | 0.912873495 | 0.153997508 | 0.325040918 | 827.8828617 | 755.4618349 | ENSG00000163832 |
| NA | -0.131532946 | 0.912860968 | 0.609862092 | 0.782634248 | 76.61726073 | 69.84136253 | ENSG00000233328 |
| LZTS2 | -0.131692089 | 0.912760276 | 0.023657113 | 0.07883617 | 3249.848808 | 2967.044409 | ENSG00000107816 |
| MAGI1 | -0.131713088 | 0.91274699 | 0.048495635 | 0.137859222 | 2150.651138 | 1962.815297 | ENSG00000151276 |
| NA | -0.131767444 | 0.912712602 | 0.828376709 | 0.913778864 | 13.46626635 | 12.33266845 | ENSG00000229994 |
| NAP1L1 | -0.131804965 | 0.912688865 | 0.001580832 | 0.008334452 | 14415.00208 | 13156.34106 | ENSG00000187109 |
| PIK3IP1 | -0.131828014 | 0.912674284 | 0.735383793 | 0.861093931 | 33.08223542 | 30.19372 | ENSG00000100100 |
| ATP5PB | -0.131839116 | 0.91266726 | 0.058606393 | 0.159485011 | 3092.046191 | 2821.266337 | ENSG00000116459 |
| NA | -0.131844116 | 0.912664097 | 0.434257747 | 0.643061049 | 195.1933988 | 177.9179258 | ENSG00000244198 |
| RAC3 | -0.131861684 | 0.912652984 | 0.067702965 | 0.178011494 | 1766.192674 | 1612.474221 | ENSG00000169750 |
| DDX21 | -0.131879552 | 0.91264168 | 0.015823249 | 0.056929331 | 7050.491192 | 6434.806437 | ENSG00000165732 |
| ACTR8 | -0.131892489 | 0.912633496 | 0.173823365 | 0.353886831 | 671.4774613 | 612.7345545 | ENSG00000113812 |
| RREB1 | -0.131907198 | 0.912624192 | 0.226010693 | 0.42465545 | 755.6165398 | 689.2875175 | ENSG00000124782 |
| DEK | -0.131916681 | 0.912618193 | 0.035008194 | 0.107253389 | 4712.810436 | 4301.076705 | ENSG00000124795 |
| CYGB | -0.132060423 | 0.912527269 | 0.641892771 | 0.804214244 | 59.31462542 | 54.10685642 | ENSG00000161544 |
| WSB2 | -0.132106619 | 0.91249805 | 0.100730339 | 0.239221889 | 974.0061254 | 888.5576554 | ENSG00000176871 |
| KHSRP | -0.132139837 | 0.91247704 | 0.009122815 | 0.036262759 | 13045.50903 | 11903.90936 | ENSG00000088247 |
| ICK | -0.1321487 | 0.912471434 | 0.416931914 | 0.627891483 | 235.1468882 | 214.7313206 | ENSG00000112144 |
| NAA15 | -0.132225005 | 0.912423174 | 0.052475505 | 0.14648927 | 2889.39284 | 2636.502924 | ENSG00000164134 |
| SSH2 | -0.132237563 | 0.912415232 | 0.359395337 | 0.571533011 | 259.6456499 | 236.903307 | ENSG00000141298 |
| PABPC1 | -0.132278669 | 0.912389236 | 0.017154429 | 0.060780749 | 26788.69549 | 24442.40293 | ENSG00000070756 |
| ASCC1 | -0.132309463 | 0.912369761 | 0.179892503 | 0.36268495 | 566.8788799 | 517.2380025 | ENSG00000138303 |
| ZNF143 | -0.132439733 | 0.912287382 | 0.164611083 | 0.340071206 | 828.8603633 | 756.0275625 | ENSG00000166478 |
| RPL29 | -0.132615546 | 0.912176213 | 0.016035078 | 0.057581753 | 17365.44468 | 15840.22467 | ENSG00000162244 |
| NA | -0.132646007 | 0.912156954 | 0.887045623 | 0.949069176 | 6.509689063 | 5.958444612 | ENSG00000227627 |
| LOC105372482 | -0.13266531 | 0.912144749 | 0.743900048 | 0.866776871 | 34.25771598 | 31.31469017 | ENSG00000268516 |
| IPO13 | -0.132671606 | 0.912140768 | 0.166604765 | 0.343021641 | 659.3883958 | 601.5743846 | ENSG00000117408 |
| PRKCG | -0.132679329 | 0.912135885 | 0.468034991 | 0.671389351 | 219.4179326 | 200.2309315 | ENSG00000126583 |
| MRPL39 | -0.13283849 | 0.912035262 | 0.126027025 | 0.282528533 | 1087.5608 | 991.7046397 | ENSG00000154719 |
| TMEM222 | -0.132913128 | 0.911988079 | 0.241687146 | 0.44519938 | 487.009204 | 444.274335 | ENSG00000186501 |
| ZNF543 | -0.133160759 | 0.911831555 | 0.617727299 | 0.787696154 | 80.4423403 | 73.44119005 | ENSG00000178229 |
| ECHS1 | -0.13319352 | 0.911810849 | 0.022628189 | 0.075970372 | 2990.747421 | 2726.607273 | ENSG00000127884 |
| COTL1 | -0.133353371 | 0.911709826 | 0.028572657 | 0.091255646 | 2830.305744 | 2580.381812 | ENSG00000103187 |
| AMBRA1 | -0.133391499 | 0.911685731 | 0.079409647 | 0.200729645 | 1111.823092 | 1013.380731 | ENSG00000110497 |
| RN7SL1 | -0.133400896 | 0.911679793 | 0.679918242 | 0.828801912 | 385.9394487 | 351.6949616 | ENSG00000276168 |
| NA | -0.133403166 | 0.911678358 | 0.908805553 | 0.960165884 | 4.440711387 | 4.030056997 | ENSG00000271815 |
| HCG18 | -0.133422231 | 0.911666311 | 0.243007052 | 0.446984335 | 508.4933207 | 463.2524573 | ENSG00000231074 |
| CHD9 | -0.133474567 | 0.911633239 | 0.244850714 | 0.449070278 | 491.1123509 | 447.3435125 | ENSG00000177200 |
| TRAIP | -0.133495028 | 0.91162031 | 0.284862311 | 0.494087503 | 367.6800119 | 334.9749539 | ENSG00000183763 |
| ATG9B | -0.133524576 | 0.91160164 | 0.737454078 | 0.862288419 | 36.36348359 | 33.24714516 | ENSG00000181652 |
| NDUFA2 | -0.133530021 | 0.911598198 | 0.398934421 | 0.610475104 | 245.5251568 | 223.6910949 | ENSG00000131495 |
| YPEL5 | -0.13359875 | 0.911554772 | 0.210055303 | 0.403426666 | 485.7535571 | 442.693035 | ENSG00000119801 |
| PANK1 | -0.133625421 | 0.91153792 | 0.18837269 | 0.374346864 | 667.8721787 | 608.6942599 | ENSG00000152782 |
| IL12RB2 | -0.133640487 | 0.911528401 | 0.677781664 | 0.82756234 | 46.15853895 | 42.10720426 | ENSG00000081985 |
| ADAMTSL1 | -0.133647395 | 0.911524036 | 0.729292811 | 0.85751636 | 34.21383325 | 31.25434428 | ENSG00000178031 |
| EEF1E1 | -0.133690486 | 0.911496811 | 0.614434213 | 0.785406126 | 92.74205064 | 84.73811029 | ENSG00000124802 |
| STARD3 | -0.133713702 | 0.911482143 | 0.070161709 | 0.183113634 | 1453.923439 | 1325.540703 | ENSG00000131748 |
| ZNF134 | -0.133740047 | 0.911465499 | 0.076935887 | 0.196190904 | 1214.404437 | 1107.274199 | ENSG00000213762 |
| C14orf132 | -0.133832427 | 0.911407137 | 0.059894922 | 0.162074165 | 1292.013893 | 1177.587903 | ENSG00000227051 |
| RNASET2 | -0.13385063 | 0.911395637 | 0.372128672 | 0.58398288 | 260.8077823 | 237.8854466 | ENSG00000026297 |
| EXOSC8 | -0.133883189 | 0.911375069 | 0.074374213 | 0.191591463 | 1370.416035 | 1249.490306 | ENSG00000120699 |
| LSM5 | -0.13388705 | 0.91137263 | 0.164542753 | 0.340038572 | 680.0742564 | 620.0868506 | ENSG00000106355 |
| MCCC1 | -0.133914083 | 0.911355553 | 0.212663073 | 0.406739126 | 527.4123226 | 480.4344664 | ENSG00000078070 |
| RIPK2 | -0.133916284 | 0.911354163 | 0.335588984 | 0.547537071 | 339.1894597 | 308.8785717 | ENSG00000104312 |
| AVPR1A | -0.134014883 | 0.91129188 | 0.915504669 | 0.962507751 | 4.066181665 | 3.697040702 | ENSG00000166148 |
| MAP6D1 | -0.134046783 | 0.91127173 | 0.721377303 | 0.853073731 | 40.37154464 | 36.87390862 | ENSG00000180834 |
| TRAF3 | -0.134104772 | 0.911235102 | 0.180593492 | 0.363593614 | 537.8305115 | 490.1234141 | ENSG00000131323 |
| CXorf56 | -0.134178529 | 0.911188517 | 0.224890627 | 0.423243133 | 552.8100767 | 503.8047008 | ENSG00000018610 |

| | | | | | | | |
|---|---|---|---|---|---|---|---|
| CLIP1 | -0.134232919 | 0.911154165 | 0.20665653 | 0.399172401 | 478.4832405 | 435.8155619 | ENSG00000130779 |
| ZNF655 | -0.134268799 | 0.911131505 | 0.248590009 | 0.453542877 | 511.9652791 | 466.5708172 | ENSG00000197343 |
| KLHL13 | -0.134322938 | 0.911097314 | 0.185280942 | 0.369680993 | 606.5348226 | 552.4148897 | ENSG00000003096 |
| NEIL1 | -0.134352055 | 0.911078927 | 0.56630023 | 0.750233591 | 104.78965 | 95.48136672 | ENSG00000140398 |
| FAM120C | -0.134411599 | 0.911041325 | 0.442428104 | 0.650077004 | 174.4521129 | 159.0073382 | ENSG00000184083 |
| KIF14 | -0.13442561 | 0.911032477 | 0.181092256 | 0.364054423 | 644.4201467 | 586.9857677 | ENSG00000118193 |
| SHISA7 | -0.134530186 | 0.910966442 | 0.721249048 | 0.853073731 | 33.14627977 | 30.17007345 | ENSG00000187902 |
| MEN1 | -0.134557976 | 0.910948894 | 0.053736023 | 0.14895124 | 1728.900073 | 1575.264933 | ENSG00000133895 |
| NA | -0.134623562 | 0.910907483 | 0.772159611 | 0.882599617 | 21.5132285 | 19.60170716 | ENSG00000235587 |
| GYS1 | -0.134630744 | 0.910902948 | 0.059790347 | 0.161930404 | 1470.804434 | 1340.010866 | ENSG00000104812 |
| MFF | -0.134712808 | 0.910851136 | 0.143081104 | 0.308594775 | 710.1092791 | 646.8912912 | ENSG00000168958 |
| GHET1 | -0.134771649 | 0.910813987 | 0.844650365 | 0.922675968 | 10.19422206 | 9.306011269 | ENSG00000281189 |
| MINDY2 | -0.134783482 | 0.910806516 | 0.262402475 | 0.469445883 | 453.2451552 | 412.8868859 | ENSG00000128923 |
| LARS2 | -0.134887905 | 0.910740594 | 0.023815971 | 0.079225754 | 2275.91555 | 2072.519455 | ENSG00000011376 |
| F2RL1 | -0.135088665 | 0.910613867 | 0.257982797 | 0.464808695 | 437.8273214 | 398.6815412 | ENSG00000164251 |
| PSMF1 | -0.135104537 | 0.910603849 | 0.039550116 | 0.117696807 | 1796.131689 | 1635.279325 | ENSG00000125818 |
| B3GALT5 | -0.135187003 | 0.910551799 | 0.132238458 | 0.292563645 | 819.7544156 | 745.9758999 | ENSG00000183778 |
| ZDHHC18 | -0.135243593 | 0.910516084 | 0.087033476 | 0.215021874 | 1070.626063 | 974.7138832 | ENSG00000204160 |
| SGK1 | -0.135265645 | 0.910502166 | 0.315598497 | 0.525761748 | 335.8506757 | 305.5731598 | ENSG00000118515 |
| USP46 | -0.135315761 | 0.910470538 | 0.076759834 | 0.195946131 | 1087.37439 | 989.7412111 | ENSG00000109189 |
| USP39 | -0.135327491 | 0.910463136 | 0.024105909 | 0.079964884 | 2645.054194 | 2408.077112 | ENSG00000168883 |
| TPRA1 | -0.135360426 | 0.910442351 | 0.203715721 | 0.395352111 | 510.3366288 | 464.9044336 | ENSG00000163870 |
| SMPD2 | -0.13536879 | 0.910437073 | 0.480179472 | 0.68228044 | 129.816583 | 118.1661775 | ENSG00000135587 |
| KIF3C | -0.135374992 | 0.910433159 | 0.051003894 | 0.143355937 | 2115.500562 | 1925.884639 | ENSG00000084731 |
| MKI67 | -0.135384743 | 0.910427005 | 0.044172638 | 0.127984084 | 3120.294759 | 2840.001239 | ENSG00000148773 |
| NOL4L | -0.135451752 | 0.910384719 | 0.176011262 | 0.356764267 | 635.6345498 | 578.8163861 | ENSG00000197183 |
| PCDHGA6 | -0.135483579 | 0.910364636 | 0.798497897 | 0.897223346 | 17.48264156 | 15.93048847 | ENSG00000253731 |
| CHEK1 | -0.135607598 | 0.910286381 | 0.080350767 | 0.2023772 | 1786.944126 | 1626.123407 | ENSG00000149554 |
| NA | -0.135697462 | 0.910229682 | 0.901524073 | 0.956745272 | 3.653692956 | 3.322974145 | ENSG00000263884 |
| HPN-AS1 | -0.13570312 | 0.910226113 | 0.770296486 | 0.881997379 | 24.92879459 | 22.70024996 | ENSG00000227392 |
| LOC389831 | -0.135739581 | 0.910203109 | 0.498468668 | 0.697317165 | 118.816376 | 108.1176136 | ENSG00000276256 |
| ARL9 | -0.135803994 | 0.910162471 | 0.812474715 | 0.904745857 | 15.66007874 | 14.20723657 | ENSG00000196503 |
| ARHGAP17 | -0.135837921 | 0.910141068 | 0.057074897 | 0.156370149 | 1363.954653 | 1241.424589 | ENSG00000140750 |
| CEP41 | -0.135875798 | 0.910117173 | 0.488960138 | 0.689210609 | 163.9189297 | 149.0893545 | ENSG00000106477 |
| NPC2 | -0.135944522 | 0.91007382 | 0.112174899 | 0.260056834 | 1006.503771 | 915.8269353 | ENSG00000119655 |
| CELF1 | -0.136006696 | 0.9100346 | 0.032779393 | 0.101844109 | 2634.793539 | 2397.095024 | ENSG00000149187 |
| SNN | -0.136017907 | 0.910027529 | 0.017185802 | 0.060840479 | 2549.559171 | 2320.189511 | ENSG00000184602 |
| TMEM230 | -0.13606869 | 0.909995496 | 0.104741104 | 0.246760947 | 1290.12924 | 1173.800418 | ENSG00000089063 |
| ARHGAP35 | -0.136460695 | 0.909748268 | 0.048799227 | 0.138456089 | 3566.740692 | 3244.501537 | ENSG00000160007 |
| TMEM30A | -0.136483617 | 0.909733814 | 0.125022475 | 0.280843278 | 1129.090489 | 1027.345028 | ENSG00000112697 |
| ATP11C | -0.136507708 | 0.909718623 | 0.068609105 | 0.179856045 | 1891.535492 | 1720.410194 | ENSG00000101974 |
| RBX1 | -0.136530149 | 0.909704472 | 0.054845305 | 0.151558909 | 1989.288269 | 1809.215684 | ENSG00000100387 |
| FAM163A | -0.13658653 | 0.909668921 | 0.735477178 | 0.861096497 | 28.79452905 | 26.17983721 | ENSG00000143340 |
| NA | -0.13670297 | 0.909595505 | 0.887292926 | 0.949163187 | 5.831792917 | 5.34282655 | ENSG00000268584 |
| NFYB | -0.136724727 | 0.909581788 | 0.098302277 | 0.235200621 | 899.2689627 | 818.2124538 | ENSG00000120837 |
| P4HB | -0.13680864 | 0.909528884 | 0.005229416 | 0.022961523 | 7626.152975 | 6935.95169 | ENSG00000185624 |
| FUBP3 | -0.136853816 | 0.909500404 | 0.034916427 | 0.107062304 | 2771.012083 | 2520.114988 | ENSG00000107164 |
| GRK3 | -0.136928058 | 0.909453602 | 0.139285494 | 0.303007321 | 819.5946483 | 745.3528821 | ENSG00000100077 |
| TRMT1 | -0.13695137 | 0.909438906 | 0.027187194 | 0.088041131 | 2830.597105 | 2574.848439 | ENSG00000104907 |
| FUT10 | -0.136990851 | 0.909414019 | 0.376260561 | 0.588339792 | 227.0732042 | 206.491682 | ENSG00000172728 |
| AIF1L | -0.137036749 | 0.909385087 | 0.00567073 | 0.024534187 | 5642.152533 | 5130.270462 | ENSG00000126878 |
| NA | -0.137065924 | 0.909366697 | 0.90074192 | 0.956302469 | 4.015179995 | 3.634519351 | ENSG00000279212 |
| DDX19A | -0.137115297 | 0.909335577 | 0.314371848 | 0.524872614 | 371.2740835 | 337.3851923 | ENSG00000168872 |
| CSRP2 | -0.137130958 | 0.909325705 | 0.020260811 | 0.069517698 | 2476.704369 | 2251.841112 | ENSG00000175183 |
| ZC3H7B | -0.137175103 | 0.909297882 | 0.028494893 | 0.091130576 | 3782.774279 | 3440.24736 | ENSG00000100403 |
| PTPN1 | -0.137318257 | 0.909207659 | 0.116894721 | 0.268004953 | 1150.543486 | 1045.975757 | ENSG00000196396 |
| NA | -0.137349011 | 0.909188278 | 0.747318708 | 0.868553402 | 27.04432474 | 24.5091753 | ENSG00000260290 |
| SNAPC1 | -0.137399854 | 0.909156237 | 0.503314777 | 0.701172881 | 120.3625475 | 109.539914 | ENSG00000023608 |
| DCTD | -0.137443488 | 0.90912874 | 0.043718681 | 0.127097798 | 1516.107358 | 1378.065946 | ENSG00000129187 |
| SPATA2 | -0.137454215 | 0.909121981 | 0.204342256 | 0.396160918 | 504.0810322 | 458.4231606 | ENSG00000158480 |
| RANBP3 | -0.137535504 | 0.909070757 | 0.027783772 | 0.089508108 | 2484.947867 | 2259.492921 | ENSG00000031823 |
| ETF1 | -0.137748173 | 0.90893676 | 0.024621716 | 0.081461302 | 4843.82205 | 4402.712436 | ENSG00000120705 |
| VPS26B | -0.13782358 | 0.908889253 | 0.145628847 | 0.312664915 | 824.3253779 | 749.2053064 | ENSG00000151502 |
| EXTL2 | -0.137848192 | 0.908873748 | 0.147338473 | 0.314867834 | 733.2116715 | 666.6307674 | ENSG00000162694 |
| EMC2 | -0.13786415 | 0.908863695 | 0.144035345 | 0.309962092 | 635.3916159 | 577.672452 | ENSG00000104412 |
| PDE8B | -0.137907012 | 0.908836693 | 0.761239784 | 0.876553329 | 24.06112965 | 21.88770358 | ENSG00000113231 |
| CTNNBL1 | -0.137987783 | 0.908785812 | 0.061284907 | 0.165078515 | 1515.395595 | 1377.011061 | ENSG00000132792 |
| AGFG2 | -0.138050046 | 0.908746592 | 0.534198996 | 0.725216459 | 110.3091775 | 100.4255342 | ENSG00000106351 |
| SFXN1 | -0.138089018 | 0.908722044 | 0.032122315 | 0.100263819 | 2140.948072 | 1945.089882 | ENSG00000164466 |
| FKBP1B | -0.138175785 | 0.908667393 | 0.604653139 | 0.779063421 | 66.50813437 | 60.44873185 | ENSG00000119782 |
| ZNF142 | -0.13818246 | 0.908663189 | 0.157450451 | 0.329797223 | 670.119993 | 609.2960175 | ENSG00000115568 |
| ZNF219 | -0.138301042 | 0.908588505 | 0.067634515 | 0.177943414 | 2500.819532 | 2272.941497 | ENSG00000165804 |
| TBPL1 | -0.138396716 | 0.908528252 | 0.340450193 | 0.552022143 | 304.0490029 | 276.2586999 | ENSG00000028839 |
| CBX3 | -0.138473002 | 0.908480214 | 0.015053906 | 0.054629711 | 3860.773132 | 3507.563624 | ENSG00000122565 |

| | | | | | | | |
|---|---|---|---|---|---|---|---|
| DEAF1 | -0.138475372 | 0.908478721 | 0.051695531 | 0.144761286 | 1271.671416 | 1155.455549 | ENSG00000177030 |
| PCGF1 | -0.138536441 | 0.908440266 | 0.41364005 | 0.624777441 | 177.7434029 | 161.385818 | ENSG00000115289 |
| MFSD9 | -0.138554175 | 0.908429099 | 0.265530121 | 0.473143151 | 383.8279648 | 348.5637345 | ENSG00000135953 |
| BMF | -0.138594731 | 0.908403562 | 0.17434579 | 0.354644576 | 528.4843776 | 479.917291 | ENSG00000104081 |
| NA | -0.13867375 | 0.908353809 | 0.517852194 | 0.713790138 | 110.2872103 | 100.1231424 | ENSG00000251363 |
| TTLL5 | -0.138694654 | 0.908340648 | 0.170571534 | 0.348959313 | 697.5510265 | 633.8417968 | ENSG00000119685 |
| CMTM6 | -0.138737633 | 0.908313588 | 0.15535263 | 0.327108308 | 933.6120531 | 848.2486444 | ENSG00000091317 |
| ST7L | -0.138755411 | 0.908302395 | 0.380006261 | 0.59241652 | 235.4818804 | 213.8769889 | ENSG00000007341 |
| EIF4E3 | -0.13882395 | 0.908259244 | 0.361958662 | 0.573896351 | 229.1442406 | 208.2871031 | ENSG00000163412 |
| EIPR1 | -0.138856267 | 0.908238899 | 0.183112795 | 0.366944518 | 572.3420407 | 519.7788665 | ENSG00000032389 |
| GPD1L | -0.138916987 | 0.908200674 | 0.441314086 | 0.64904972 | 179.0004458 | 162.5386422 | ENSG00000152642 |
| PECR | -0.138921569 | 0.90819779 | 0.3478066 | 0.559768847 | 266.7996736 | 242.18912 | ENSG00000115425 |
| NA | -0.139005492 | 0.908144961 | 0.90215263 | 0.957250951 | 3.657252428 | 3.315501796 | ENSG00000234789 |
| NA | -0.139030676 | 0.908129108 | 0.689568408 | 0.83398915 | 47.05323128 | 42.78411417 | ENSG00000175741 |
| IRF1 | -0.139060054 | 0.908110615 | 0.597835067 | 0.773984729 | 70.52539929 | 64.14983994 | ENSG00000125347 |
| HNRNPLL | -0.139097167 | 0.908087255 | 0.07011871 | 0.183051125 | 1898.175989 | 1724.174973 | ENSG00000143889 |
| PUS3 | -0.139138193 | 0.908061432 | 0.253109434 | 0.459188499 | 367.9309021 | 333.8593532 | ENSG00000110060 |
| UBN1 | -0.139159424 | 0.908048069 | 0.053218149 | 0.147857736 | 1677.581181 | 1523.668218 | ENSG00000118900 |
| APMAP | -0.139227536 | 0.9080052 | 0.026680497 | 0.086693252 | 2006.032815 | 1821.884418 | ENSG00000101474 |
| STT3A | -0.139245682 | 0.907993779 | 0.009508194 | 0.037486898 | 3571.55021 | 3242.856837 | ENSG00000134910 |
| TOMM40 | -0.139304842 | 0.907956546 | 0.0274413 | 0.088631598 | 5674.620079 | 5153.024489 | ENSG00000130204 |
| OAF | -0.139431945 | 0.907876557 | 0.660941058 | 0.817732696 | 48.32863025 | 43.89922035 | ENSG00000184232 |
| ZNF287 | -0.139528338 | 0.9078159 | 0.540116462 | 0.729673037 | 101.555233 | 92.17530182 | ENSG00000141040 |
| ZBTB46 | -0.139532557 | 0.907813245 | 0.202401434 | 0.393732085 | 533.5362455 | 483.923207 | ENSG00000130584 |
| FABP5 | -0.139604187 | 0.907768174 | 0.069485225 | 0.181623319 | 1068.444351 | 970.0121178 | ENSG00000164687 |
| TTLL10 | -0.1396684 | 0.907727771 | 0.830252596 | 0.914737804 | 12.8749666 | 11.72357678 | ENSG00000162571 |
| ARL3 | -0.13972475 | 0.907692316 | 0.10836012 | 0.253359027 | 822.7583565 | 746.6287547 | ENSG00000138175 |
| NA | -0.139869814 | 0.907601052 | 0.306105592 | 0.516281514 | 317.5780928 | 288.1223797 | ENSG00000249152 |
| GSTP1 | -0.139897114 | 0.907583878 | 0.000961168 | 0.005420025 | 15080.93389 | 13687.43153 | ENSG00000084207 |
| HNRNPA3 | -0.139914602 | 0.907572876 | 0.001508291 | 0.008000103 | 15427.71704 | 14002.34879 | ENSG00000170144 |
| KCTD5 | -0.140005989 | 0.907515388 | 0.094832391 | 0.229280464 | 928.9866076 | 843.3846559 | ENSG00000167977 |
| ZNF263 | -0.140022018 | 0.907505305 | 0.109639183 | 0.255779614 | 911.7447109 | 827.5746186 | ENSG00000006194 |
| ASB9 | -0.140099907 | 0.907456312 | 0.906463036 | 0.959614016 | 3.984339952 | 3.599049521 | ENSG00000102048 |
| NA | -0.14017832 | 0.907406991 | 0.861336581 | 0.932648138 | 7.709196204 | 6.969316773 | ENSG00000272894 |
| PNPLA5 | -0.14020832 | 0.907388122 | 0.357754895 | 0.569806965 | 209.1747938 | 189.7706127 | ENSG00000100341 |
| MTHFSD | -0.140227796 | 0.907375873 | 0.439830968 | 0.647758546 | 146.5596483 | 132.9511033 | ENSG00000103248 |
| C12orf65 | -0.140238329 | 0.907369248 | 0.288396292 | 0.497657526 | 331.3862949 | 300.6302742 | ENSG00000130921 |
| DLG3 | -0.140258107 | 0.907356809 | 0.063997126 | 0.170298572 | 1230.064692 | 1116.012001 | ENSG00000082458 |
| RNF6 | -0.140273296 | 0.907347257 | 0.293136343 | 0.502804708 | 289.5290893 | 262.8806784 | ENSG00000127870 |
| NA | -0.140413404 | 0.907259143 | 0.900358511 | 0.95609933 | 3.688092472 | 3.339148349 | ENSG00000259416 |
| HMGA1 | -0.140432666 | 0.90724703 | 0.004282115 | 0.019458071 | 54012.12992 | 49003.31223 | ENSG00000137309 |
| SLCO2A1 | -0.140538884 | 0.907180237 | 0.697089811 | 0.838398998 | 36.60425405 | 33.26426533 | ENSG00000174640 |
| LRRC34 | -0.140712026 | 0.90707137 | 0.663546308 | 0.81961358 | 52.13771863 | 47.29464721 | ENSG00000171757 |
| MXRA7 | -0.140717345 | 0.907068026 | 0.143942799 | 0.309855209 | 892.4380854 | 809.0829411 | ENSG00000182534 |
| CCNB1 | -0.140748387 | 0.907048509 | 0.00989547 | 0.038682664 | 5705.767543 | 5174.7685 | ENSG00000134057 |
| TMEM120B | -0.140753849 | 0.907045074 | 0.115862666 | 0.266188132 | 711.6770868 | 645.6585421 | ENSG00000188735 |
| NA | -0.140754256 | 0.907044819 | 0.218772348 | 0.41497748 | 639.6125053 | 579.677623 | ENSG00000218283 |
| PTPRF | -0.140786457 | 0.907024574 | 0.005566292 | 0.02414873 | 10122.43408 | 9181.05276 | ENSG00000142949 |
| DNAJC21 | -0.140854636 | 0.906981711 | 0.068882931 | 0.180398579 | 1046.679369 | 949.2337291 | ENSG00000168724 |
| TSPAN3 | -0.140859329 | 0.90697876 | 0.013008463 | 0.048531177 | 2906.124157 | 2635.517416 | ENSG00000140391 |
| ZSCAN16-AS1 | -0.141020123 | 0.906877679 | 0.681448203 | 0.829757233 | 41.33884865 | 37.4926481 | ENSG00000269293 |
| NINL | -0.141028518 | 0.906872403 | 0.067881629 | 0.178393518 | 1867.164994 | 1693.551575 | ENSG00000101004 |
| KIAA1586 | -0.141031314 | 0.906870645 | 0.404019762 | 0.615208962 | 248.0887954 | 225.2103976 | ENSG00000168116 |
| NA | -0.141046625 | 0.906861021 | 0.764612543 | 0.87816748 | 21.24426159 | 19.24599027 | ENSG00000260276 |
| SPATC1L | -0.141062978 | 0.906850741 | 0.096288066 | 0.231643329 | 936.4042848 | 849.5779803 | ENSG00000160284 |
| IFT22 | -0.141116063 | 0.906817374 | 0.336970661 | 0.548877949 | 242.2682918 | 219.8991844 | ENSG00000128581 |
| RRP7BP | -0.141143186 | 0.906800326 | 0.388024285 | 0.600150584 | 257.9811523 | 234.0601168 | ENSG00000182841 |
| STRN3 | -0.141163665 | 0.906787454 | 0.164821666 | 0.340404327 | 633.7659153 | 574.3736486 | ENSG00000196792 |
| HAP1 | -0.141189849 | 0.906770996 | 0.416769749 | 0.62769732 | 166.1765071 | 150.7861649 | ENSG00000173805 |
| RAB36 | -0.141249801 | 0.906733316 | 0.688355979 | 0.833414711 | 42.06894168 | 38.1222649 | ENSG00000100228 |
| FAIM | -0.141424092 | 0.90662378 | 0.517982313 | 0.713860533 | 111.1884367 | 100.9938592 | ENSG00000158234 |
| ANO6 | -0.141431264 | 0.906619273 | 0.06183492 | 0.165991822 | 1260.815517 | 1142.621079 | ENSG00000177119 |
| SP1 | -0.141522701 | 0.906561814 | 0.113179658 | 0.261679461 | 1387.764563 | 1257.391364 | ENSG00000185591 |
| LAMTOR1 | -0.141580014 | 0.9065258 | 0.069375164 | 0.181440614 | 1046.764491 | 949.0825512 | ENSG00000149357 |
| ASCL2 | -0.141616291 | 0.906503006 | 0.821316563 | 0.909334444 | 12.45891842 | 11.34203788 | ENSG00000183734 |
| CYTH1 | -0.141634613 | 0.906491494 | 0.193333577 | 0.381236776 | 451.4522378 | 409.1015448 | ENSG00000108669 |
| SWI5 | -0.141659418 | 0.906475908 | 0.261776154 | 0.468843082 | 332.7799939 | 301.6817451 | ENSG00000175854 |
| FBXO6 | -0.141707638 | 0.906445611 | 0.750483746 | 0.870196202 | 25.60165672 | 23.27046683 | ENSG00000116663 |
| ZMYM2 | -0.141725433 | 0.90643443 | 0.018440515 | 0.064304723 | 2548.868512 | 2309.998181 | ENSG00000121741 |
| NA | -0.141780101 | 0.906400083 | 0.689144059 | 0.83398915 | 46.4545071 | 42.25031308 | ENSG00000260257 |
| TLE1 | -0.141785126 | 0.906396926 | 0.009608197 | 0.03780225 | 3587.651278 | 3252.129788 | ENSG00000196781 |
| NA | -0.141789096 | 0.906394432 | 0.841246573 | 0.921035511 | 9.912212473 | 9.000046496 | ENSG00000234782 |
| PSMC6 | -0.141846409 | 0.906358425 | 0.037566578 | 0.113058937 | 1558.439979 | 1412.521887 | ENSG00000100519 |
| ATF2 | -0.141877371 | 0.906338974 | 0.261544645 | 0.468691468 | 447.1705068 | 405.2389352 | ENSG00000115966 |

| | | | | | | | |
|---|---|---|---|---|---|---|---|
| DDX18 | -0.141960846 | 0.906286534 | 0.018080761 | 0.063314513 | 3221.799091 | 2919.764888 | ENSG00000088205 |
| MAFF | -0.141993318 | 0.906266136 | 0.483461852 | 0.68442107 | 131.0424809 | 118.6435321 | ENSG00000185022 |
| INPP5K | -0.141997462 | 0.906263533 | 0.246879255 | 0.451745658 | 382.2699191 | 346.3804161 | ENSG00000132376 |
| NA | -0.142003148 | 0.906259961 | 0.90886552 | 0.960165884 | 3.704694626 | 3.350971626 | ENSG00000227973 |
| NCBP2 | -0.142012041 | 0.906254374 | 0.021421572 | 0.072733977 | 2218.639874 | 2010.567944 | ENSG00000114503 |
| TMUB2 | -0.142015719 | 0.906252064 | 0.246338669 | 0.451212164 | 353.7034234 | 320.5513254 | ENSG00000168591 |
| MST1 | -0.142046152 | 0.906232947 | 0.657358942 | 0.815009428 | 60.74541981 | 54.99781481 | ENSG00000173531 |
| CSE1L | -0.142082843 | 0.9062099 | 0.016566318 | 0.059096131 | 7080.09316 | 6416.465057 | ENSG00000124207 |
| TBC1D5 | -0.142166188 | 0.90615755 | 0.168654714 | 0.346313394 | 620.2238612 | 561.9262476 | ENSG00000131374 |
| DNMT3B | -0.142182218 | 0.906147481 | 0.001716313 | 0.008919253 | 40028.04834 | 36270.62298 | ENSG00000088305 |
| NA | -0.142272222 | 0.906090952 | 0.900281718 | 0.956071542 | 3.677414055 | 3.322974145 | ENSG00000251666 |
| IBTK | -0.142282148 | 0.906084718 | 0.170783478 | 0.34929273 | 678.6296081 | 615.0969757 | ENSG00000005700 |
| RASSF2 | -0.142382328 | 0.906021802 | 0.308000794 | 0.5183669 | 303.9626341 | 275.1250761 | ENSG00000101265 |
| SMIM1 | -0.142404773 | 0.906007706 | 0.802864278 | 0.899347048 | 15.68707998 | 14.25888061 | ENSG00000235169 |
| RPTOR | -0.142432037 | 0.905990585 | 0.0733157 | 0.189584112 | 1006.838282 | 911.9988497 | ENSG00000141564 |
| SLC12A9 | -0.142551701 | 0.905915441 | 0.068148538 | 0.178921887 | 1235.945278 | 1120.134491 | ENSG00000146828 |
| APBB3 | -0.142556563 | 0.905912388 | 0.440912824 | 0.648613531 | 159.4927355 | 144.5305527 | ENSG00000113108 |
| NA | -0.142582311 | 0.90589622 | 0.894862844 | 0.953032171 | 4.066181665 | 3.660341367 | ENSG00000224713 |
| MTFR1 | -0.142692913 | 0.905826774 | 0.120017556 | 0.27264905 | 701.1118427 | 634.8819913 | ENSG00000066855 |
| SREBF2 | -0.142812417 | 0.905751744 | 0.005113786 | 0.022521885 | 7929.821252 | 7182.694487 | ENSG00000198911 |
| IGBP1 | -0.142833877 | 0.90573827 | 0.147918903 | 0.315735401 | 557.1920048 | 504.6550807 | ENSG00000089289 |
| ADORA2B | -0.142844337 | 0.905731704 | 0.571954562 | 0.754495803 | 87.89889241 | 79.73861026 | ENSG00000170425 |
| EPOR | -0.143061585 | 0.905595325 | 0.352169882 | 0.564290534 | 255.9026652 | 231.7547091 | ENSG00000187266 |
| PCDHA8 | -0.14310902 | 0.90556555 | 0.817047685 | 0.906790406 | 12.79904862 | 11.61905921 | ENSG00000204962 |
| DOCK9 | -0.143170743 | 0.905526808 | 0.15396609 | 0.325040918 | 559.42375 | 506.4487778 | ENSG00000088387 |
| PCOLCE2 | -0.143175902 | 0.90552357 | 0.333373646 | 0.54546537 | 272.7142241 | 247.05331 | ENSG00000163710 |
| SLC7A5 | -0.143188056 | 0.905515941 | 0.015169812 | 0.054955286 | 9899.410814 | 8964.332482 | ENSG00000103257 |
| KLHDC10 | -0.143200319 | 0.905508244 | 0.212062306 | 0.405918978 | 449.086785 | 406.7351288 | ENSG00000128607 |
| NA | -0.143243714 | 0.905481008 | 0.897269738 | 0.954483129 | 3.653692956 | 3.297152129 | ENSG00000258645 |
| NA | -0.143258083 | 0.905471989 | 0.788643496 | 0.891842679 | 18.34290822 | 16.67956754 | ENSG00000272476 |
| PLPPR5 | -0.143408375 | 0.905377667 | 0.63175062 | 0.797127712 | 61.32192301 | 55.50406991 | ENSG00000117598 |
| CNIH1 | -0.143496172 | 0.905322571 | 0.077360715 | 0.196947575 | 1031.341064 | 933.7083105 | ENSG00000100528 |
| NA | -0.143607421 | 0.905252762 | 0.759129764 | 0.874877523 | 27.24685688 | 24.66722883 | ENSG00000279821 |
| SOBP | -0.143608487 | 0.905252093 | 0.409382018 | 0.620469288 | 174.108397 | 157.6932439 | ENSG00000112320 |
| PRELID1 | -0.143613839 | 0.905248735 | 0.069360436 | 0.181440614 | 3656.022523 | 3310.062727 | ENSG00000169230 |
| ZNF607 | -0.14362758 | 0.905240113 | 0.294785095 | 0.504899013 | 289.3322015 | 262.097359 | ENSG00000198182 |
| UQCRQ | -0.143735706 | 0.905172271 | 0.313939266 | 0.524342711 | 251.9895147 | 227.951994 | ENSG00000164405 |
| NA | -0.143774048 | 0.905148214 | 0.905560747 | 0.958927594 | 3.660811901 | 3.304624478 | ENSG00000248161 |
| ELOVL6 | -0.143822893 | 0.90511757 | 0.016522337 | 0.058983807 | 2451.172277 | 2218.880141 | ENSG00000170522 |
| NA | -0.143847818 | 0.905101932 | 0.781586542 | 0.88874409 | 19.12072277 | 17.23361021 | ENSG00000228944 |
| NUP155 | -0.143882243 | 0.905080335 | 0.056502684 | 0.155162597 | 1766.926582 | 1599.031131 | ENSG00000113569 |
| CYTH3 | -0.143936222 | 0.905046472 | 0.168915985 | 0.34662375 | 536.1254113 | 485.0900299 | ENSG00000008256 |
| HSD17B4 | -0.143949552 | 0.90503811 | 0.003659102 | 0.017028622 | 7474.236002 | 6763.849358 | ENSG00000133835 |
| PHTF2 | -0.143971364 | 0.905024427 | 0.226682525 | 0.425579032 | 389.6937995 | 352.5687045 | ENSG00000006576 |
| NA | -0.143996007 | 0.905008968 | 0.901154054 | 0.956513844 | 3.715373043 | 3.353147089 | ENSG00000175730 |
| KNDC1 | -0.144050767 | 0.904974617 | 0.2255721 | 0.423873544 | 466.7712966 | 422.370879 | ENSG00000171798 |
| GGN | -0.144183088 | 0.904891619 | 0.714768733 | 0.849651131 | 34.13199154 | 30.98886268 | ENSG00000179168 |
| NA | -0.144190029 | 0.904887265 | 0.837270004 | 0.918172233 | 9.901534055 | 8.968644047 | ENSG00000213763 |
| NA | -0.144208653 | 0.904875584 | 0.689612799 | 0.83398915 | 37.90695974 | 34.28289338 | ENSG00000178715 |
| HNRNPD | -0.144235594 | 0.904858686 | 0.010217768 | 0.039736932 | 19439.45668 | 17590.86576 | ENSG00000138668 |
| SELENOW | -0.144288141 | 0.904825729 | 0.017673471 | 0.062234477 | 3242.368357 | 2933.294012 | ENSG00000178980 |
| CHMP7 | -0.14434518 | 0.904789956 | 0.034967012 | 0.107182605 | 1385.481509 | 1253.498114 | ENSG00000147457 |
| ABCD4 | -0.144395964 | 0.904758108 | 0.082012029 | 0.205683555 | 1153.164637 | 1043.011466 | ENSG00000119688 |
| RNF5 | -0.144448517 | 0.904725151 | 0.026237954 | 0.085549682 | 2229.905768 | 2017.263688 | ENSG00000204308 |
| ISG20 | -0.144565809 | 0.904651599 | 0.763146052 | 0.877513739 | 20.85312971 | 18.83862934 | ENSG00000172183 |
| HSPA12A | -0.14473604 | 0.904544861 | 0.377826576 | 0.590324719 | 208.6157824 | 188.62091 | ENSG00000165868 |
| BNC2 | -0.144794148 | 0.904508429 | 0.224039583 | 0.422254243 | 432.4412221 | 390.8372723 | ENSG00000173068 |
| ANKRD13B | -0.144840074 | 0.904479635 | 0.15343855 | 0.324380054 | 1010.137257 | 913.9203977 | ENSG00000198720 |
| SIGMAR1 | -0.145292166 | 0.904196247 | 0.015816998 | 0.056917686 | 2687.222041 | 2430.418936 | ENSG00000147955 |
| AGAP1 | -0.145296344 | 0.904193628 | 0.057906567 | 0.157954438 | 1386.045224 | 1252.804093 | ENSG00000157985 |
| ZNF213 | -0.145370712 | 0.90414702 | 0.334538579 | 0.546519123 | 243.5656842 | 220.4740357 | ENSG00000085644 |
| LARP4B | -0.145475405 | 0.90408141 | 0.047381725 | 0.13533029 | 1233.745438 | 1115.175765 | ENSG00000107929 |
| SH3BP1 | -0.145516124 | 0.904055894 | 0.527925569 | 0.720939647 | 94.22504191 | 85.28997749 | ENSG00000100092 |
| NEXN | -0.145558041 | 0.904029627 | 0.503269624 | 0.701172881 | 107.3725349 | 97.17641023 | ENSG00000162614 |
| NA | -0.145577699 | 0.904017309 | 0.806295662 | 0.901201229 | 13.55642222 | 12.24867601 | ENSG00000225093 |
| STT3B | -0.145629531 | 0.903984831 | 0.020161558 | 0.06925264 | 2798.415766 | 2529.460124 | ENSG00000163527 |
| TSPYL1 | -0.145637783 | 0.90397966 | 0.088106057 | 0.216788302 | 1053.895989 | 952.3628534 | ENSG00000189241 |
| ZNF431 | -0.145805498 | 0.903874577 | 0.150725994 | 0.320074121 | 555.0417696 | 501.7090836 | ENSG00000196705 |
| SMNDC1 | -0.145816923 | 0.90386742 | 0.149489561 | 0.318101141 | 673.4875588 | 608.8717997 | ENSG00000119953 |
| ARL10 | -0.145905298 | 0.903812053 | 0.281009388 | 0.48992451 | 279.1924889 | 252.1766756 | ENSG00000175414 |
| H2BU1 | -0.145922966 | 0.903800985 | 0.868182309 | 0.937517522 | 6.633049238 | 5.998265368 | ENSG00000196890 |
| FKBP14 | -0.145974381 | 0.903768776 | 0.454828028 | 0.660692258 | 175.0330391 | 158.3905633 | ENSG00000106080 |
| NA | -0.146025299 | 0.903736879 | 0.535836436 | 0.726437264 | 97.95404245 | 88.50703661 | ENSG00000234742 |
| NCOA1 | -0.146171261 | 0.90364545 | 0.188647224 | 0.374568749 | 466.4910926 | 421.3693055 | ENSG00000084676 |

| | | | | | | | |
|---|---|---|---|---|---|---|---|
| RAP1A | -0.146178626 | 0.903640837 | 0.155212045 | 0.326921646 | 599.2834723 | 541.618206 | ENSG00000116473 |
| USP13 | -0.146210359 | 0.90362096 | 0.146528458 | 0.313833491 | 670.0450176 | 605.6574308 | ENSG00000058056 |
| TMEM100 | -0.146278122 | 0.903578518 | 0.805731104 | 0.900901829 | 13.97839414 | 12.60874383 | ENSG00000166292 |
| MDH1B | -0.146393914 | 0.903506 | 0.698511847 | 0.839414332 | 35.61826295 | 32.2200988 | ENSG00000138400 |
| CD164 | -0.146403459 | 0.903500022 | 0.064781676 | 0.172043586 | 1567.305434 | 1416.54462 | ENSG00000135535 |
| NA | -0.146427542 | 0.90348494 | 0.690567284 | 0.834528492 | 41.63034145 | 37.67122675 | ENSG00000230724 |
| SLC35G1 | -0.146567631 | 0.903397214 | 0.37707123 | 0.589290812 | 173.781615 | 156.9771756 | ENSG00000176273 |
| TMBIM6 | -0.146639802 | 0.903352022 | 0.003707982 | 0.017228777 | 11516.10844 | 10402.07138 | ENSG00000139644 |
| NECAP2 | -0.146654965 | 0.903342528 | 0.099445986 | 0.237215794 | 1033.026764 | 932.6170094 | ENSG00000157191 |
| ATP6V0A2 | -0.146744013 | 0.903286772 | 0.101404585 | 0.240279246 | 800.6467611 | 723.0595415 | ENSG00000185344 |
| VWA1 | -0.146760614 | 0.903276378 | 0.128972887 | 0.287290986 | 1217.89366 | 1100.723005 | ENSG00000179403 |
| SLC44A1 | -0.146779313 | 0.903264671 | 0.023736217 | 0.079043988 | 1987.369327 | 1795.077505 | ENSG00000070214 |
| MCM10 | -0.146831056 | 0.903232275 | 0.066524279 | 0.175781587 | 1125.671618 | 1016.333495 | ENSG00000065328 |
| AAK1 | -0.146969134 | 0.903145833 | 0.285032882 | 0.49429256 | 325.3140103 | 293.9774206 | ENSG00000115977 |
| ULK1 | -0.147038606 | 0.903102343 | 0.026920147 | 0.087351714 | 1882.794162 | 1700.559803 | ENSG00000177169 |
| EIF2S1 | -0.147059386 | 0.903089335 | 0.030795913 | 0.096930136 | 2094.120838 | 1891.649879 | ENSG00000134001 |
| SLC16A1 | -0.147071672 | 0.903081645 | 0.00507549 | 0.022374098 | 9704.763356 | 8763.87497 | ENSG00000155380 |
| CDH4 | -0.147161294 | 0.903025546 | 0.226774881 | 0.425583188 | 475.9787413 | 430.0028531 | ENSG00000179242 |
| STIM2 | -0.147219584 | 0.902989061 | 0.176993559 | 0.35840911 | 437.2745916 | 394.7661441 | ENSG00000109689 |
| RETREG1 | -0.147307717 | 0.9029339 | 0.697412138 | 0.83873325 | 33.77553851 | 30.4716873 | ENSG00000154153 |
| CFL1 | -0.147360783 | 0.902900689 | 0.001169445 | 0.006421914 | 9392.286559 | 8480.939984 | ENSG00000172757 |
| CCT7 | -0.147414048 | 0.902867353 | 0.000692616 | 0.004076053 | 14963.53284 | 13509.74734 | ENSG00000135624 |
| SF3B2 | -0.147433297 | 0.902855307 | 0.001378039 | 0.007410099 | 7427.186642 | 6705.570243 | ENSG00000087365 |
| NA | -0.147438642 | 0.902851962 | 0.899763816 | 0.955743556 | 3.677414055 | 3.306799942 | ENSG00000205274 |
| NA | -0.147440892 | 0.902850554 | 0.89304978 | 0.952281345 | 4.409871343 | 3.943889093 | ENSG00000259790 |
| LRRC47 | -0.147501311 | 0.902812744 | 0.013075889 | 0.048705752 | 2330.213012 | 2103.971245 | ENSG00000130764 |
| DNAAF3 | -0.147513018 | 0.902805418 | 0.676112696 | 0.826733047 | 42.98821867 | 38.92204205 | ENSG00000167646 |
| FAM83H-AS1 | -0.147535731 | 0.902791205 | 0.632491196 | 0.797582548 | 48.84640244 | 44.13017809 | ENSG00000203499 |
| EIF4E2 | -0.147667522 | 0.902708738 | 0.027254296 | 0.088203674 | 2243.837583 | 2025.585529 | ENSG00000135930 |
| AP4B1 | -0.147737033 | 0.902665246 | 0.412715461 | 0.624129682 | 177.0987372 | 159.9209019 | ENSG00000134262 |
| IL3RA | -0.147901336 | 0.902562451 | 0.727239389 | 0.856496155 | 26.10074183 | 23.57141827 | ENSG00000185291 |
| ZNF609 | -0.147956443 | 0.902527976 | 0.020901852 | 0.071328419 | 2394.035649 | 2160.033475 | ENSG00000180357 |
| TRAPPC2L | -0.147985365 | 0.902509883 | 0.063587475 | 0.169519847 | 1020.925798 | 921.2394291 | ENSG00000167515 |
| HPCAL1 | -0.148074622 | 0.902454047 | 0.313276233 | 0.523574846 | 230.9723961 | 208.448278 | ENSG00000115756 |
| RAD1 | -0.148191897 | 0.902380691 | 0.120258406 | 0.272933542 | 740.4339617 | 668.2453187 | ENSG00000113456 |
| NGEF | -0.148334718 | 0.902291364 | 0.518169233 | 0.714018677 | 89.46019208 | 80.70559428 | ENSG00000066248 |
| LAP3 | -0.148346357 | 0.902284085 | 0.042143959 | 0.123585195 | 1887.75658 | 1703.165203 | ENSG00000002549 |
| DAW1 | -0.148358202 | 0.902276676 | 0.849046558 | 0.925283873 | 8.783868629 | 7.959947349 | ENSG00000123977 |
| PAF1 | -0.148458127 | 0.902214184 | 0.071969908 | 0.186902051 | 1122.453192 | 1012.563571 | ENSG00000006712 |
| PPIL1 | -0.148499451 | 0.902188342 | 0.049265059 | 0.139485852 | 1358.069267 | 1225.376487 | ENSG00000137168 |
| MAP7 | -0.148662445 | 0.902086419 | 0.01693149 | 0.060114908 | 2354.919951 | 2123.920805 | ENSG00000135525 |
| VAMP1 | -0.148678916 | 0.902076121 | 0.578085864 | 0.759450456 | 133.2526162 | 120.2504862 | ENSG00000139190 |
| PPP2R2D | -0.148697568 | 0.902064458 | 0.091137361 | 0.22236087 | 743.6872934 | 670.8514372 | ENSG00000175470 |
| NA | -0.148723609 | 0.902048176 | 0.87423258 | 0.941109404 | 6.320199605 | 5.71254218 | ENSG00000231409 |
| KDM5A | -0.148734796 | 0.902041181 | 0.097638129 | 0.233907913 | 887.7215146 | 800.2968774 | ENSG00000073614 |
| NA | -0.14880468 | 0.901997487 | 0.261886433 | 0.46892247 | 300.3922829 | 271.0285686 | ENSG00000185596 |
| MTMR9 | -0.148814535 | 0.901991326 | 0.192149145 | 0.379455883 | 474.3126906 | 427.9490439 | ENSG00000104643 |
| ZNF157 | -0.148923962 | 0.901922914 | 0.778438485 | 0.886943686 | 18.10981536 | 16.27655755 | ENSG00000147117 |
| NA | -0.149022316 | 0.901861428 | 0.652449039 | 0.811703295 | 47.87373335 | 43.11032052 | ENSG00000272923 |
| CDC23 | -0.149044911 | 0.901847304 | 0.085553687 | 0.212421955 | 900.047869 | 811.278396 | ENSG00000094880 |
| AKAP12 | -0.149108328 | 0.901807662 | 0.025138906 | 0.082824187 | 3655.373893 | 3295.881122 | ENSG00000131016 |
| HENMT1 | -0.149133643 | 0.901791838 | 0.440641033 | 0.64836974 | 144.4732043 | 130.4901441 | ENSG00000162639 |
| UGT3A2 | -0.149181044 | 0.901762209 | 0.149663243 | 0.318394768 | 582.4577535 | 525.2542807 | ENSG00000168671 |
| CYP26C1 | -0.149190798 | 0.901756112 | 0.893776842 | 0.95280698 | 4.066181665 | 3.638870278 | ENSG00000187553 |
| SLC39A13 | -0.149405258 | 0.901622074 | 0.145678506 | 0.312664915 | 538.4007599 | 485.5073651 | ENSG00000165915 |
| SET | -0.149500614 | 0.901562482 | 0.005428182 | 0.023680116 | 22951.27506 | 20692.35388 | ENSG00000119335 |
| EPCAM | -0.149586394 | 0.901508879 | 0.028530408 | 0.091176709 | 3417.178372 | 3081.324222 | ENSG00000119888 |
| EEA1 | -0.149620503 | 0.901487565 | 0.298910767 | 0.50928202 | 314.3866688 | 283.1349435 | ENSG00000102189 |
| DHX34 | -0.149630443 | 0.901481354 | 0.042622375 | 0.124620459 | 1580.963633 | 1425.835677 | ENSG00000134815 |
| PP2D1 | -0.149634807 | 0.901478627 | 0.861038842 | 0.932648138 | 6.578488096 | 5.926096204 | ENSG00000183977 |
| DNAJC11 | -0.149640775 | 0.901474898 | 0.016395547 | 0.058597635 | 1881.888635 | 1696.506027 | ENSG00000007923 |
| RTCB | -0.149656538 | 0.901465049 | 0.007621084 | 0.031431354 | 3387.756161 | 3053.477853 | ENSG00000100220 |
| GNG5 | -0.149684445 | 0.901447611 | 0.505710734 | 0.703058541 | 121.94017 | 109.8848262 | ENSG00000174021 |
| MSL1 | -0.149747257 | 0.901408365 | 0.069348666 | 0.181440614 | 1859.569194 | 1676.769824 | ENSG00000188895 |
| MDH1 | -0.149747297 | 0.90140834 | 0.002711145 | 0.013175497 | 4217.547343 | 3801.422973 | ENSG00000014641 |
| TELO2 | -0.149766964 | 0.901396051 | 0.083832389 | 0.209220915 | 2143.336588 | 1932.704475 | ENSG00000100726 |
| BPTF | -0.149866938 | 0.90133359 | 0.018397132 | 0.064231804 | 3580.242554 | 3226.495001 | ENSG00000171634 |
| GNL2 | -0.149869616 | 0.901331917 | 0.029496264 | 0.093652355 | 2759.911219 | 2487.102796 | ENSG00000134697 |
| PITPNM1 | -0.150035228 | 0.901228456 | 0.04304545 | 0.12558525 | 2188.656216 | 1973.100109 | ENSG00000110697 |
| RABL6 | -0.150107411 | 0.901183365 | 0.015045093 | 0.054608224 | 3236.050167 | 2917.078538 | ENSG00000196642 |
| FAM168A | -0.150110803 | 0.901181247 | 0.114465938 | 0.263974752 | 786.0589905 | 708.2658936 | ENSG00000054965 |
| TMEM116 | -0.150123713 | 0.901173183 | 0.466935307 | 0.670914738 | 130.513751 | 117.5135766 | ENSG00000198270 |
| AKT2 | -0.150146661 | 0.901158849 | 0.009339969 | 0.03695481 | 3048.168982 | 2747.019394 | ENSG00000105221 |
| SMU1 | -0.15015911 | 0.901151072 | 0.030995113 | 0.097427215 | 1821.515661 | 1640.930457 | ENSG00000122692 |

| | | | | | | | |
|---|---|---|---|---|---|---|---|
| NORAD | -0.150218735 | 0.901113829 | 0.013400532 | 0.049621382 | 3948.301743 | 3557.780786 | ENSG00000260032 |
| RFC5 | -0.150318838 | 0.901051307 | 0.127154787 | 0.284179664 | 580.5971794 | 523.2077759 | ENSG00000111445 |
| CHST4 | -0.150324741 | 0.90104762 | 0.383460796 | 0.595734937 | 241.1981209 | 217.3883348 | ENSG00000140835 |
| RAC1 | -0.150378542 | 0.901014019 | 0.009507607 | 0.037486898 | 5562.446119 | 5011.732437 | ENSG00000136238 |
| YWHAQ | -0.15040036 | 0.901000393 | 0.003619767 | 0.016887055 | 10730.61278 | 9668.599676 | ENSG00000134308 |
| ZNF684 | -0.150520493 | 0.90092537 | 0.499076648 | 0.697608609 | 100.2045009 | 90.17815001 | ENSG00000117010 |
| NA | -0.150619783 | 0.900863368 | 0.897670721 | 0.954850935 | 3.650133483 | 3.271330112 | ENSG00000241478 |
| ZNF101 | -0.150646498 | 0.900846687 | 0.129985802 | 0.288865836 | 627.2734911 | 564.78492 | ENSG00000181896 |
| CYHR1 | -0.150666038 | 0.900834485 | 0.138055196 | 0.301163495 | 1033.723223 | 931.7500171 | ENSG00000187954 |
| DTYMK | -0.150667468 | 0.900833592 | 0.034640635 | 0.106337514 | 1633.072974 | 1470.960134 | ENSG00000168393 |
| METTL17 | -0.150719872 | 0.900800871 | 0.120196953 | 0.272859653 | 772.6893886 | 696.152205 | ENSG00000165792 |
| DPP3 | -0.150789837 | 0.900757187 | 0.012768951 | 0.047800106 | 2373.90112 | 2138.497233 | ENSG00000254986 |
| DLG5 | -0.150875172 | 0.900703909 | 0.011309332 | 0.043285613 | 2670.568138 | 2405.194467 | ENSG00000151208 |
| RGS19 | -0.150875191 | 0.900703897 | 0.315141756 | 0.525468132 | 241.292953 | 217.508008 | ENSG00000171700 |
| MARCHF8 | -0.150892181 | 0.90069329 | 0.278369309 | 0.487572287 | 392.2081658 | 352.8682585 | ENSG00000165406 |
| KLF1 | -0.150982498 | 0.900636906 | 0.81314712 | 0.905169301 | 12.44468053 | 11.22257561 | ENSG00000105610 |
| REEP1 | -0.150997754 | 0.900627382 | 0.709445701 | 0.846496645 | 29.12161657 | 26.23148124 | ENSG00000068615 |
| LIN52 | -0.151012574 | 0.90061813 | 0.323773633 | 0.53453945 | 220.4736386 | 198.5695909 | ENSG00000205659 |
| KCTD7 | -0.151029922 | 0.900607301 | 0.291321552 | 0.500936139 | 267.1383554 | 240.6679253 | ENSG00000243335 |
| TMEM233 | -0.15107032 | 0.900582082 | 0.86005856 | 0.932067365 | 6.663889282 | 5.996089905 | ENSG00000224982 |
| LYAR | -0.151143025 | 0.900536699 | 0.026278546 | 0.085667243 | 1652.727006 | 1488.432224 | ENSG00000145220 |
| ANKRD17 | -0.151166352 | 0.900522138 | 0.045612861 | 0.131150865 | 2814.637292 | 2534.008194 | ENSG00000132466 |
| DPPA4 | -0.151178707 | 0.900514426 | 0.001958739 | 0.00997325 | 12795.14843 | 11521.44408 | ENSG00000121570 |
| OSGEP | -0.151240327 | 0.900475965 | 0.134401301 | 0.295927314 | 541.2371792 | 487.2609579 | ENSG00000092094 |
| ATP13A1 | -0.151251996 | 0.900468681 | 0.040976466 | 0.120950385 | 2145.938212 | 1933.0054 | ENSG00000105726 |
| BMPR2 | -0.151278991 | 0.900451832 | 0.181942606 | 0.365180797 | 427.7692235 | 384.9513757 | ENSG00000204217 |
| KPNA4 | -0.151323627 | 0.900423973 | 0.056751739 | 0.15564284 | 2055.767986 | 1850.92082 | ENSG00000186432 |
| USP6 | -0.151423814 | 0.900361446 | 0.803645507 | 0.899842175 | 16.74098039 | 15.06367109 | ENSG00000129204 |
| STYXL1 | -0.151476141 | 0.90032879 | 0.163125194 | 0.338120875 | 458.3095332 | 412.7550762 | ENSG00000127952 |
| TSPAN4 | -0.15147836 | 0.900327406 | 0.067454979 | 0.177570015 | 1618.491178 | 1457.71237 | ENSG00000214063 |
| POLR3H | -0.151541136 | 0.90028823 | 0.066388662 | 0.175521472 | 946.0188276 | 851.9498908 | ENSG00000100413 |
| PHLPP1 | -0.151555921 | 0.900279004 | 0.032838572 | 0.10194911 | 1556.817637 | 1401.317262 | ENSG00000081913 |
| NA | -0.151591039 | 0.90025709 | 0.688745128 | 0.833725514 | 33.23168096 | 29.88312076 | ENSG00000260565 |
| EPB41L4A | -0.151615555 | 0.900241792 | 0.670897764 | 0.823737356 | 37.88056889 | 34.13108271 | ENSG00000129595 |
| MFSD13A | -0.151682047 | 0.900200302 | 0.299319309 | 0.509612123 | 262.3548702 | 236.3661439 | ENSG00000138111 |
| NA | -0.151830566 | 0.900107635 | 0.670943346 | 0.823739767 | 40.60519615 | 36.59934629 | ENSG00000272655 |
| SNX12 | -0.151910627 | 0.900057686 | 0.034601314 | 0.106268632 | 1949.228534 | 1754.006196 | ENSG00000147164 |
| BYSL | -0.15192578 | 0.900048232 | 0.040930709 | 0.120834209 | 1210.976388 | 1090.155838 | ENSG00000112578 |
| ZNF383 | -0.15195238 | 0.900031637 | 0.666309399 | 0.821200352 | 43.37014667 | 38.97725901 | ENSG00000188283 |
| RALGPS1 | -0.15202998 | 0.899983227 | 0.227089495 | 0.426004284 | 408.1325347 | 367.1000876 | ENSG00000136828 |
| HDAC3 | -0.152107509 | 0.899934865 | 0.019789664 | 0.06814876 | 1619.136176 | 1457.308021 | ENSG00000171720 |
| CHD4 | -0.152217739 | 0.899866107 | 0.284205526 | 0.493356131 | 268.6679509 | 241.9411577 | ENSG00000111642 |
| PSMC1 | -0.152312861 | 0.899806778 | 0.093635252 | 0.227182774 | 1637.670067 | 1473.166684 | ENSG00000100764 |
| DNAJB2 | -0.15236132 | 0.899776554 | 0.072420801 | 0.187788938 | 1081.735593 | 973.5857148 | ENSG00000135924 |
| NA | -0.152371738 | 0.899770057 | 0.695010471 | 0.837250559 | 33.46624835 | 30.05808353 | ENSG00000287271 |
| SHLD3 | -0.152438828 | 0.899728216 | 0.722067128 | 0.853545174 | 25.45368572 | 22.91285802 | ENSG00000253251 |
| MAPK3 | -0.152561771 | 0.899651546 | 0.062080298 | 0.166532164 | 903.4614275 | 812.9641408 | ENSG00000102882 |
| CIAO3 | -0.152590697 | 0.899633509 | 0.20362758 | 0.395270242 | 380.3221905 | 342.1224704 | ENSG00000103245 |
| NA | -0.152610222 | 0.899621333 | 0.893097393 | 0.952281345 | 3.684532999 | 3.294976665 | ENSG00000235272 |
| ARHGEF35 | -0.152646997 | 0.899598401 | 0.36955886 | 0.581680252 | 201.3766107 | 181.0666723 | ENSG00000213214 |
| HYDIN | -0.152657168 | 0.89959206 | 0.759838902 | 0.875400053 | 19.56613645 | 17.59027306 | ENSG00000157423 |
| C1QTNF1 | -0.152884289 | 0.89945045 | 0.515974064 | 0.712241079 | 91.28956835 | 82.1609782 | ENSG00000173918 |
| IFITM3 | -0.152892076 | 0.899445595 | 0.004744216 | 0.021125802 | 3175.696894 | 2856.567378 | ENSG00000142089 |
| SOX8 | -0.152941041 | 0.899415068 | 0.16631772 | 0.342568459 | 432.2846832 | 388.9194058 | ENSG00000005513 |
| HTT | -0.152941817 | 0.899414584 | 0.027257434 | 0.088203674 | 3168.470941 | 2849.224263 | ENSG00000197386 |
| SCHIP1 | -0.153047305 | 0.899348823 | 0.514490027 | 0.710801535 | 107.2506493 | 96.49893323 | ENSG00000283154 |
| TBCEL | -0.15306492 | 0.899337842 | 0.475152897 | 0.677841249 | 120.2204741 | 108.1617853 | ENSG00000154114 |
| DNALI1 | -0.153081149 | 0.899327725 | 0.387729065 | 0.600037973 | 248.4983106 | 223.1891477 | ENSG00000163879 |
| PHF20 | -0.153142547 | 0.899289452 | 0.034112564 | 0.105143751 | 1560.647899 | 1402.914126 | ENSG00000025293 |
| MLYCD | -0.153200973 | 0.899253034 | 0.156249259 | 0.328261238 | 503.3115324 | 452.4931649 | ENSG00000103150 |
| DDAH2 | -0.153277893 | 0.89920509 | 0.038692439 | 0.115852751 | 1370.443265 | 1232.315572 | ENSG00000213722 |
| NA | -0.153384008 | 0.899138953 | 0.836885224 | 0.918172233 | 8.852667661 | 7.98794483 | ENSG00000229582 |
| ANKIB1 | -0.153400853 | 0.899128454 | 0.063185126 | 0.168811317 | 887.568282 | 797.9504625 | ENSG00000001629 |
| RNF20 | -0.153638009 | 0.898980664 | 0.029496649 | 0.093652355 | 1539.874403 | 1384.114161 | ENSG00000155827 |
| SEC13 | -0.153647929 | 0.898974483 | 0.006847302 | 0.028748621 | 2799.819254 | 2516.968127 | ENSG00000157020 |
| ARHGAP40 | -0.153836009 | 0.898857294 | 0.85699886 | 0.930171411 | 6.698288798 | 6.011034604 | ENSG00000124143 |
| LPAR6 | -0.153852976 | 0.898846723 | 0.733928027 | 0.860189337 | 34.57323536 | 31.09621812 | ENSG00000139679 |
| GFM2 | -0.153919641 | 0.898805189 | 0.129694773 | 0.288376048 | 536.6223853 | 482.1761406 | ENSG00000164347 |
| ARHGEF17 | -0.154005551 | 0.898751668 | 0.220033607 | 0.416700228 | 343.1213496 | 308.4138613 | ENSG00000110237 |
| HAUS4 | -0.154022756 | 0.898740951 | 0.465304292 | 0.669506404 | 115.7779833 | 104.0226918 | ENSG00000092036 |
| DMAC2 | -0.154135825 | 0.898670516 | 0.025175372 | 0.082929861 | 1432.641524 | 1287.685571 | ENSG00000105341 |
| CRAMP1 | -0.154224228 | 0.89861545 | 0.199774364 | 0.390027795 | 366.590874 | 329.5001794 | ENSG00000007545 |
| CLIC1 | -0.154310667 | 0.898561612 | 0.002869885 | 0.013857782 | 4604.157621 | 4136.475066 | ENSG00000213719 |
| NEURL4 | -0.154342928 | 0.898541518 | 0.04560789 | 0.131150865 | 1208.446868 | 1086.217362 | ENSG00000215041 |

| | | | | | | | |
|---|---|---|---|---|---|---|---|
| GSPT2 | -0.154351735 | 0.898536033 | 0.072002034 | 0.186959771 | 785.5246162 | 705.9000354 | ENSG00000189369 |
| CAPRIN1 | -0.154372502 | 0.898523099 | 0.001778937 | 0.009196615 | 7419.55695 | 6666.211507 | ENSG00000135387 |
| DNAL4 | -0.154398354 | 0.898506999 | 0.436326995 | 0.644810651 | 185.3861649 | 166.2509111 | ENSG00000100246 |
| ARL2BP | -0.154417913 | 0.898494817 | 0.795684518 | 0.895554092 | 13.33489751 | 11.95765594 | ENSG00000102931 |
| GTF2H1 | -0.154433241 | 0.898485271 | 0.23535013 | 0.437158179 | 440.7151252 | 396.4098701 | ENSG00000110768 |
| CHORDC1 | -0.154478674 | 0.898456977 | 0.046925799 | 0.134271775 | 1888.044033 | 1696.580893 | ENSG00000110172 |
| SLC23A2 | -0.154532329 | 0.898423563 | 0.105207012 | 0.247734966 | 658.7499331 | 591.9646996 | ENSG00000089057 |
| PRSS35 | -0.154644596 | 0.898353652 | 0.544011268 | 0.73244858 | 84.26683562 | 75.74865756 | ENSG00000146250 |
| DHRS4 | -0.154654709 | 0.898347355 | 0.364930589 | 0.577106435 | 188.9455577 | 169.5800555 | ENSG00000157326 |
| RETREG2 | -0.154673042 | 0.89833594 | 0.04162261 | 0.12234132 | 1264.328942 | 1135.648452 | ENSG00000144567 |
| CRNKL1 | -0.154708105 | 0.898314107 | 0.094769182 | 0.229196202 | 795.615283 | 714.2358982 | ENSG00000101343 |
| HMGCR | -0.154752375 | 0.898286542 | 0.01722534 | 0.060951947 | 5075.821706 | 4558.921959 | ENSG00000113161 |
| HMGN1 | -0.154852448 | 0.898224234 | 0.002516212 | 0.01237143 | 7788.720168 | 6996.222276 | ENSG00000205581 |
| LGMN | -0.154883002 | 0.898205212 | 0.022654233 | 0.076044293 | 1582.283982 | 1420.821568 | ENSG00000100600 |
| TRIM24 | -0.154922072 | 0.898180887 | 0.014923764 | 0.054293061 | 3274.483519 | 2941.052969 | ENSG00000122779 |
| RAD17 | -0.154946606 | 0.898165613 | 0.174935973 | 0.355385741 | 464.6816503 | 417.4569142 | ENSG00000152942 |
| GCA | -0.155009396 | 0.898126523 | 0.165786882 | 0.341687795 | 490.7099552 | 440.8819342 | ENSG00000115271 |
| USP22 | -0.155062564 | 0.898093425 | 0.002366354 | 0.011738533 | 6245.60132 | 5608.446297 | ENSG00000124422 |
| SLC37A3 | -0.155071332 | 0.898087967 | 0.058963569 | 0.160265831 | 1067.334975 | 958.4214053 | ENSG00000157800 |
| SDHAP1 | -0.155120749 | 0.898057205 | 0.383243005 | 0.595633623 | 228.4084776 | 205.1362241 | ENSG00000185485 |
| NAB2 | -0.155168811 | 0.898027288 | 0.125799784 | 0.282120361 | 688.3883537 | 618.284336 | ENSG00000166886 |
| TRMT13 | -0.155412481 | 0.897875624 | 0.42666963 | 0.636616573 | 144.9559666 | 130.147648 | ENSG00000122435 |
| MITD1 | -0.155461303 | 0.89784524 | 0.189930728 | 0.376345997 | 372.8392737 | 334.7544946 | ENSG00000158411 |
| NA | -0.155471649 | 0.897838802 | 0.890072814 | 0.951059472 | 3.715373043 | 3.313326333 | ENSG00000233589 |
| SNHG17 | -0.15550973 | 0.897815103 | 0.183372136 | 0.367113622 | 446.7583498 | 401.3440335 | ENSG00000196756 |
| FSIP1 | -0.155530034 | 0.897802467 | 0.822044829 | 0.90966346 | 9.599362839 | 8.611981199 | ENSG00000150667 |
| CNPPD1 | -0.155540758 | 0.897795794 | 0.096643591 | 0.232270166 | 718.2976264 | 644.4779611 | ENSG00000115649 |
| NA | -0.155617953 | 0.897747756 | 0.724114472 | 0.854559399 | 28.78857916 | 25.82847125 | ENSG00000231503 |
| SERINC2 | -0.15562843 | 0.897741237 | 0.069897868 | 0.182550302 | 972.1204068 | 873.1374597 | ENSG00000168528 |
| PCBD2 | -0.155723173 | 0.897682283 | 0.402176309 | 0.613595236 | 168.2973506 | 151.0602757 | ENSG00000132570 |
| CHMP1B | -0.155959239 | 0.897535409 | 0.10987254 | 0.256165792 | 886.9748712 | 796.3297769 | ENSG00000255112 |
| ANK3 | -0.155982449 | 0.897520969 | 0.09873399 | 0.235960496 | 828.5617521 | 743.2453487 | ENSG00000151150 |
| HSP90AB1 | -0.156119617 | 0.89743564 | 0.00012575 | 0.000912331 | 82402.39687 | 73949.98158 | ENSG00000096384 |
| OAZ1 | -0.156139523 | 0.897423257 | 0.001216709 | 0.006654372 | 5175.978231 | 4644.941403 | ENSG00000104904 |
| MIR22HG | -0.156162326 | 0.897409072 | 0.692384259 | 0.835666692 | 34.14086436 | 30.54820739 | ENSG00000186594 |
| HOMER2 | -0.156213031 | 0.897377533 | 0.135573366 | 0.29767504 | 497.4373569 | 446.4708741 | ENSG00000103942 |
| SPPL3 | -0.156346387 | 0.897294587 | 0.162609839 | 0.337645609 | 653.5694708 | 586.7704171 | ENSG00000157837 |
| ADGRB3 | -0.156358692 | 0.897286934 | 0.724023472 | 0.854559399 | 25.95038041 | 23.25646809 | ENSG00000135298 |
| AP2S1 | -0.156361839 | 0.897284977 | 0.026205898 | 0.085470207 | 2841.264321 | 2549.288342 | ENSG00000042753 |
| HIF1A | -0.156414992 | 0.897251919 | 0.021675485 | 0.0734053 | 2714.385153 | 2435.679824 | ENSG00000100644 |
| ANTXR2 | -0.156421349 | 0.897247965 | 0.394511407 | 0.606245474 | 161.8861054 | 145.0963233 | ENSG00000163297 |
| RPUSD1 | -0.156430549 | 0.897242243 | 0.11142425 | 0.25901612 | 913.7037533 | 820.3672803 | ENSG00000007376 |
| ZKSCAN2 | -0.156511535 | 0.897191878 | 0.283776842 | 0.492974488 | 252.5520855 | 226.530923 | ENSG00000155592 |
| GREM1 | -0.156526837 | 0.897182362 | 0.855475206 | 0.929428529 | 6.650846601 | 5.951918221 | ENSG00000166923 |
| NA | -0.156544809 | 0.897171185 | 0.890587211 | 0.95121103 | 3.688092472 | 3.285328852 | ENSG00000181359 |
| ZCCHC2 | -0.156619291 | 0.897124868 | 0.285386288 | 0.494673527 | 367.3226941 | 329.5647379 | ENSG00000141664 |
| ZNF192P1 | -0.156639277 | 0.89711244 | 0.802012646 | 0.898712919 | 12.91648506 | 11.57176611 | ENSG00000226314 |
| PPWD1 | -0.156694195 | 0.897078292 | 0.076090228 | 0.194760087 | 813.1074851 | 729.4378328 | ENSG00000113593 |
| HNRNPUL1 | -0.156700751 | 0.897074215 | 0.001125286 | 0.006217387 | 6941.362634 | 6227.389889 | ENSG00000105323 |
| DCBLD1 | -0.156791349 | 0.897017882 | 0.251978944 | 0.457691868 | 264.0913953 | 236.8521574 | ENSG00000164465 |
| ANP32AP1 | -0.156817599 | 0.897001561 | 0.799902164 | 0.898237613 | 13.75450516 | 12.29719862 | ENSG00000259516 |
| KLF13 | -0.15682218 | 0.896998713 | 0.024226986 | 0.080338083 | 1347.889205 | 1208.856341 | ENSG00000169926 |
| CIP2A | -0.15683792 | 0.896988927 | 0.086881476 | 0.214674432 | 781.5847693 | 701.0689288 | ENSG00000163507 |
| VPS26C | -0.156883882 | 0.896960351 | 0.036063373 | 0.109688646 | 1337.121708 | 1199.303356 | ENSG00000157538 |
| FAM98B | -0.15690497 | 0.89694724 | 0.057318018 | 0.156786033 | 1272.970283 | 1141.886638 | ENSG00000171262 |
| RHOG | -0.156975625 | 0.896903313 | 0.175438803 | 0.355933754 | 448.2081362 | 402.3404125 | ENSG00000177105 |
| APTX | -0.157006811 | 0.896883925 | 0.079614509 | 0.201112894 | 820.633928 | 735.7621795 | ENSG00000137074 |
| NCKAP1 | -0.157053754 | 0.896854743 | 0.018817098 | 0.065323692 | 3621.065512 | 3247.670198 | ENSG00000061676 |
| LRPPRC | -0.157061162 | 0.896850138 | 0.004457412 | 0.020090268 | 6232.137364 | 5588.76981 | ENSG00000138095 |
| BRD8 | -0.157127359 | 0.896808987 | 0.040807488 | 0.120621289 | 1549.965986 | 1389.89737 | ENSG00000112983 |
| SEC61B | -0.157168803 | 0.896783225 | 0.023491039 | 0.078413304 | 1669.626025 | 1497.700267 | ENSG00000106803 |
| UGGT2 | -0.157173377 | 0.896780382 | 0.303749298 | 0.513775683 | 347.981895 | 312.3095543 | ENSG00000102595 |
| GJC1 | -0.157209397 | 0.896757992 | 0.014604418 | 0.053347085 | 1776.071222 | 1592.895729 | ENSG00000182963 |
| PQBP1 | -0.1572296 | 0.896745434 | 0.075830209 | 0.194324331 | 1228.004255 | 1101.180392 | ENSG00000102103 |
| OSER1 | -0.15725569 | 0.896729217 | 0.102722014 | 0.24288284 | 607.9220136 | 544.9868219 | ENSG00000132823 |
| PLCE1 | -0.157268583 | 0.896721204 | 0.698572954 | 0.839434349 | 30.00560433 | 26.95663022 | ENSG00000138193 |
| CNNM3 | -0.157309678 | 0.896695661 | 0.080542836 | 0.202779823 | 977.8660128 | 877.3459571 | ENSG00000168763 |
| METTL6 | -0.157334218 | 0.896680408 | 0.286678463 | 0.495916714 | 262.7098226 | 235.7311872 | ENSG00000206562 |
| ATP6V0A4 | -0.157462735 | 0.896600535 | 0.799393637 | 0.897909287 | 14.46204614 | 12.90070986 | ENSG00000105929 |
| LRP5 | -0.157588455 | 0.896522406 | 0.025363237 | 0.083374191 | 2202.393991 | 1974.236723 | ENSG00000162337 |
| NR3C1 | -0.157642934 | 0.896488552 | 0.273336515 | 0.482053626 | 250.3235681 | 224.4753172 | ENSG00000113580 |
| NA | -0.157682595 | 0.896463908 | 0.854180004 | 0.928574107 | 6.725569369 | 6.013210067 | ENSG00000273306 |
| ANKRD44 | -0.157705975 | 0.896449379 | 0.752538627 | 0.871085164 | 18.86095974 | 16.89964796 | ENSG00000065413 |
| BAP1 | -0.157721427 | 0.896439778 | 0.003449029 | 0.016218854 | 3123.61399 | 2799.985405 | ENSG00000163930 |

| | | | | | | | |
|---|---|---|---|---|---|---|---|
| EIF3B | -0.157736988 | 0.896430109 | 0.001193961 | 0.006543226 | 11531.9267 | 10337.05777 | ENSG00000106263 |
| REXO2 | -0.157787493 | 0.896398728 | 0.126180734 | 0.282839545 | 585.616505 | 525.2971705 | ENSG00000076043 |
| EPS15 | -0.157794832 | 0.896394168 | 0.087354549 | 0.215589474 | 782.2481227 | 700.9908003 | ENSG00000085832 |
| RNF217 | -0.157865992 | 0.896349955 | 0.546879918 | 0.734786085 | 83.5284546 | 74.87945381 | ENSG00000146373 |
| CYP3A5 | -0.157920626 | 0.896316012 | 0.831483086 | 0.915296977 | 8.863346079 | 7.962122813 | ENSG00000106258 |
| FHAD1 | -0.157961659 | 0.896290519 | 0.661742271 | 0.818402279 | 36.94438425 | 33.12333197 | ENSG00000142621 |
| ZNF384 | -0.157962428 | 0.896290041 | 0.112586508 | 0.26062714 | 848.1992266 | 760.1594078 | ENSG00000126746 |
| EEF1AKMT1 | -0.158126637 | 0.89618803 | 0.540974834 | 0.730264327 | 76.01617228 | 68.19512518 | ENSG00000150456 |
| HNRNPF | -0.158159089 | 0.896167872 | 0.00142534 | 0.007629691 | 9268.52603 | 8305.874858 | ENSG00000169813 |
| NA | -0.158161736 | 0.896166228 | 0.77400887 | 0.883906242 | 17.75516794 | 15.94231175 | ENSG00000234405 |
| TMEM231 | -0.158170733 | 0.896160639 | 0.426599414 | 0.6365621 | 130.8880014 | 117.2921713 | ENSG00000205084 |
| RPL14 | -0.158176979 | 0.896156759 | 0.001241906 | 0.006764754 | 18329.43721 | 16425.79171 | ENSG00000188846 |
| CMTM7 | -0.158203177 | 0.896140486 | 0.223083098 | 0.420989977 | 360.1188646 | 322.9191388 | ENSG00000153551 |
| LRBA | -0.158262344 | 0.896103735 | 0.076844943 | 0.196110371 | 2120.758725 | 1899.585986 | ENSG00000198589 |
| CENPT | -0.158362894 | 0.896041282 | 0.605818834 | 0.779795526 | 1063.778277 | 953.2618614 | ENSG00000102901 |
| ZFR2 | -0.158494485 | 0.895959556 | 0.244036766 | 0.448397969 | 323.9255468 | 290.2080858 | ENSG00000105278 |
| ZNF790 | -0.158542967 | 0.895929448 | 0.617995028 | 0.787824768 | 46.74302525 | 41.8731251 | ENSG00000197863 |
| ANAPC13 | -0.158555109 | 0.895921907 | 0.136662339 | 0.299152794 | 490.2414314 | 439.3175139 | ENSG00000129055 |
| TEKT2 | -0.158657058 | 0.895858598 | 0.739019402 | 0.863544416 | 20.76060958 | 18.63907406 | ENSG00000092850 |
| ZNF385C | -0.158720684 | 0.89581909 | 0.431766869 | 0.6413842 | 121.032435 | 108.4074041 | ENSG00000187595 |
| NA | -0.158742264 | 0.895805691 | 0.811858422 | 0.904395006 | 12.24807213 | 10.97761913 | ENSG00000214391 |
| MYD88 | -0.158775964 | 0.895784766 | 0.140862744 | 0.305165396 | 677.1154713 | 606.8097604 | ENSG00000172936 |
| DDX27 | -0.158776768 | 0.895784266 | 0.016712149 | 0.05949273 | 1857.153298 | 1664.05232 | ENSG00000124228 |
| CRAT | -0.158954475 | 0.895673933 | 0.271363117 | 0.479795193 | 320.1553594 | 286.7152146 | ENSG00000095321 |
| MTPN | -0.159029266 | 0.895627502 | 0.031705888 | 0.099330741 | 2076.263843 | 1859.544563 | ENSG00000105887 |
| NCOR1 | -0.159053926 | 0.895612193 | 0.012547879 | 0.04717633 | 2613.747542 | 2340.547555 | ENSG00000141027 |
| NDUFB7 | -0.159059743 | 0.895608582 | 0.065396739 | 0.17323852 | 1452.347466 | 1300.875431 | ENSG00000099795 |
| PTBP3 | -0.15917169 | 0.895539089 | 0.035835689 | 0.109207194 | 1538.364995 | 1377.501369 | ENSG00000119314 |
| RPL21 | -0.159270131 | 0.895477985 | 0.001509677 | 0.008002316 | 11997.41792 | 10742.91233 | ENSG00000122026 |
| POLE | -0.159358289 | 0.895423267 | 0.018775465 | 0.065203136 | 2286.616438 | 2047.816289 | ENSG00000177084 |
| XPO4 | -0.159412439 | 0.895389659 | 0.113161915 | 0.261679461 | 1048.158546 | 938.5594345 | ENSG00000132953 |
| EIF3K | -0.159461307 | 0.89535933 | 0.005432703 | 0.023690735 | 3370.207949 | 3016.99078 | ENSG00000178982 |
| TCEANC | -0.159522351 | 0.895321446 | 0.651045436 | 0.810534186 | 40.47619157 | 36.2274552 | ENSG00000176896 |
| AP2M1 | -0.15956163 | 0.89529707 | 0.000549295 | 0.003325885 | 12696.55063 | 11367.01531 | ENSG00000161203 |
| TTF1 | -0.159619141 | 0.895261381 | 0.086026352 | 0.213203104 | 647.025252 | 579.2659314 | ENSG00000125482 |
| CCDC47 | -0.159622226 | 0.895259467 | 0.032558248 | 0.101362102 | 1989.687389 | 1781.461155 | ENSG00000108588 |
| APLP1 | -0.159731926 | 0.895191395 | 0.110174106 | 0.256710426 | 822.5938095 | 736.4160756 | ENSG00000105290 |
| NKAPP1 | -0.15973522 | 0.895189351 | 0.762941737 | 0.877431251 | 18.88943552 | 16.89406753 | ENSG00000233382 |
| SYN3 | -0.159753491 | 0.895178014 | 0.173050023 | 0.352665222 | 445.7562373 | 398.7983039 | ENSG00000185666 |
| SRI | -0.15988614 | 0.895095711 | 0.052216811 | 0.14596976 | 897.9510521 | 803.5015891 | ENSG00000075142 |
| SMCHD1 | -0.160003982 | 0.8950226 | 0.095204738 | 0.229827621 | 998.6069172 | 893.2188719 | ENSG00000101596 |
| DROSHA | -0.160008033 | 0.895020087 | 0.007687454 | 0.031663591 | 2599.366318 | 2326.222112 | ENSG00000113360 |
| MAPK7 | -0.16001113 | 0.895018166 | 0.091410182 | 0.22283223 | 747.3499103 | 669.0902135 | ENSG00000166484 |
| SETD7 | -0.160065755 | 0.894984278 | 0.355125506 | 0.567389609 | 184.6323502 | 165.2312645 | ENSG00000145391 |
| CFL2 | -0.160076324 | 0.894977722 | 0.35383879 | 0.56592876 | 215.6904321 | 193.2576723 | ENSG00000165410 |
| ZNF561 | -0.160087376 | 0.894970866 | 0.089764168 | 0.220012499 | 762.9654186 | 682.6859853 | ENSG00000171469 |
| TARS3 | -0.16010173 | 0.894961962 | 0.210027587 | 0.403414461 | 319.9548861 | 286.3301754 | ENSG00000185418 |
| NA | -0.160189049 | 0.894907796 | 0.685809242 | 0.831762616 | 38.67145228 | 34.5580228 | ENSG00000248927 |
| GRIP1 | -0.160365447 | 0.894798382 | 0.204071414 | 0.395757718 | 331.3571826 | 296.3927381 | ENSG00000155974 |
| MITF | -0.160446433 | 0.894748154 | 0.751538813 | 0.870657509 | 19.64294416 | 17.64532206 | ENSG00000187098 |
| NA | -0.160468634 | 0.894734385 | 0.566794956 | 0.750531324 | 61.90877357 | 55.36552293 | ENSG00000280206 |
| STAU1 | -0.16049701 | 0.894716787 | 0.005964461 | 0.025605559 | 3281.261208 | 2935.556127 | ENSG00000124214 |
| COPZ1 | -0.160500472 | 0.89471464 | 0.028475197 | 0.091083011 | 1956.209315 | 1749.555097 | ENSG00000111481 |
| NA | -0.160590579 | 0.89465876 | 0.663320641 | 0.819495637 | 35.53522603 | 31.83855989 | ENSG00000124097 |
| TRIM41 | -0.160614566 | 0.894643885 | 0.082574887 | 0.206794351 | 720.4363446 | 644.722509 | ENSG00000146063 |
| NA | -0.160634722 | 0.894631386 | 0.867194751 | 0.936739326 | 5.580623371 | 5.014161183 | ENSG00000280374 |
| TLCD2 | -0.160660824 | 0.8946152 | 0.727538826 | 0.856710154 | 28.33607268 | 25.33326142 | ENSG00000185561 |
| NDUFA4 | -0.160666861 | 0.894611456 | 0.012749463 | 0.047765297 | 3283.277509 | 2937.765021 | ENSG00000189043 |
| TTC30A | -0.160745898 | 0.894562448 | 0.679275076 | 0.828368191 | 34.7719287 | 31.1725273 | ENSG00000197557 |
| DGKD | -0.160893801 | 0.894470743 | 0.062510048 | 0.167399639 | 850.4246458 | 760.7682683 | ENSG00000077044 |
| EMC3 | -0.16095261 | 0.894434282 | 0.094160965 | 0.228252342 | 587.072164 | 524.9942116 | ENSG00000125037 |
| STAT5A | -0.16102432 | 0.894389825 | 0.256492983 | 0.463395196 | 262.297029 | 234.6138329 | ENSG00000126561 |
| SPON1 | -0.161025881 | 0.894388857 | 0.400144032 | 0.61158304 | 136.5623139 | 122.0571931 | ENSG00000262655 |
| RAB8A | -0.161036801 | 0.894382087 | 0.210961377 | 0.404631892 | 406.6346678 | 363.9048784 | ENSG00000167461 |
| UFM1 | -0.161046534 | 0.894376053 | 0.123576822 | 0.278324184 | 499.2762164 | 446.7071288 | ENSG00000120686 |
| HS3ST1 | -0.161063541 | 0.89436551 | 0.683216667 | 0.830790283 | 31.57727692 | 28.20281104 | ENSG00000002587 |
| NA | -0.161099997 | 0.89434291 | 0.74826421 | 0.86918808 | 20.12214697 | 17.93104525 | ENSG00000263301 |
| RUSC2 | -0.161142416 | 0.894316615 | 0.22169176 | 0.418934663 | 307.0481897 | 274.5507192 | ENSG00000198853 |
| SNRPB2 | -0.161176494 | 0.89429549 | 0.023323223 | 0.077929771 | 1520.720066 | 1360.271431 | ENSG00000125870 |
| ARPIN | -0.161179796 | 0.894293443 | 0.371982493 | 0.583899043 | 159.9802525 | 142.9137829 | ENSG00000242498 |
| WDR86 | -0.161189809 | 0.894287236 | 0.247379064 | 0.452330079 | 365.8212695 | 327.1875532 | ENSG00000187260 |
| CCDC78 | -0.16127073 | 0.894237077 | 0.516892859 | 0.71314448 | 135.5458139 | 121.4450229 | ENSG00000162004 |
| BMI1 | -0.161284058 | 0.894228816 | 0.667466887 | 0.821822412 | 35.31133705 | 31.63247822 | ENSG00000168283 |
| LY6G5C | -0.161332824 | 0.89419859 | 0.777379694 | 0.886255087 | 14.45464787 | 12.95888029 | ENSG00000204428 |

| | | | | | | | |
|---|---|---|---|---|---|---|---|
| KAT14 | -0.161482366 | 0.894105907 | 0.11960583 | 0.27216434 | 557.3005428 | 498.0292584 | ENSG00000149474 |
| TSR2 | -0.161504097 | 0.894092439 | 0.045476445 | 0.130878188 | 1342.44699 | 1200.588181 | ENSG00000158526 |
| COPS9 | -0.161551656 | 0.894062965 | 0.067694395 | 0.178011494 | 996.8596608 | 891.3587376 | ENSG00000172428 |
| RPS6KB1 | -0.161555812 | 0.89406039 | 0.146557094 | 0.313833491 | 442.868537 | 395.8108777 | ENSG00000108443 |
| MTNR1A | -0.161801359 | 0.893908234 | 0.824804101 | 0.911541751 | 10.3591007 | 9.215775984 | ENSG00000168412 |
| SAP30BP | -0.161801573 | 0.893908101 | 0.014307017 | 0.052463693 | 1604.92113 | 1434.787013 | ENSG00000161526 |
| MINPP1 | -0.161849941 | 0.893878133 | 0.111191322 | 0.258633836 | 706.5759705 | 631.9123476 | ENSG00000107789 |
| NCDN | -0.161962569 | 0.893808352 | 0.052082428 | 0.145736651 | 1201.315422 | 1074.066169 | ENSG00000020129 |
| LRP2 | -0.162010731 | 0.893778514 | 0.716582312 | 0.850522089 | 27.71955114 | 24.75507774 | ENSG00000081479 |
| AP1S3 | -0.162114612 | 0.89371416 | 0.481294232 | 0.683198091 | 92.4691932 | 82.58545928 | ENSG00000152056 |
| ELP1 | -0.162145064 | 0.893695296 | 0.027018773 | 0.087626572 | 1516.274321 | 1354.844011 | ENSG00000070061 |
| ALG1 | -0.162295041 | 0.893602396 | 0.10194659 | 0.241412083 | 546.2555367 | 488.1299508 | ENSG00000033011 |
| PDZD4 | -0.162645664 | 0.893385247 | 0.004135582 | 0.018905948 | 7066.360945 | 6313.073244 | ENSG00000067840 |
| LOC441242 | -0.162663102 | 0.893374448 | 0.727958194 | 0.856731016 | 21.16241987 | 18.90644758 | ENSG00000272693 |
| PCGF6 | -0.162790402 | 0.893295623 | 0.142250728 | 0.30726765 | 420.1306057 | 375.3159641 | ENSG00000156374 |
| SDF4 | -0.162838984 | 0.893265542 | 0.0088489 | 0.035480144 | 2875.09979 | 2568.739788 | ENSG00000078808 |
| OSBPL7 | -0.162841279 | 0.893264121 | 0.412761187 | 0.624148852 | 184.213659 | 164.5126848 | ENSG00000006025 |
| BCAS3 | -0.162854644 | 0.893255846 | 0.220947183 | 0.418011175 | 316.380034 | 282.453821 | ENSG00000141376 |
| REPIN1 | -0.162914918 | 0.893218528 | 0.008317606 | 0.033729436 | 2615.996044 | 2337.259981 | ENSG00000214022 |
| WDR55 | -0.162925963 | 0.89321169 | 0.115228813 | 0.265163842 | 502.0983716 | 448.5908932 | ENSG00000120314 |
| ATP2C1 | -0.163006321 | 0.893161939 | 0.015291543 | 0.055343195 | 1891.504704 | 1689.451706 | ENSG00000017260 |
| RPA1 | -0.163055429 | 0.893131537 | 0.00605102 | 0.025918236 | 4628.016725 | 4132.578412 | ENSG00000132383 |
| USP33 | -0.16313031 | 0.893085182 | 0.026593047 | 0.086438845 | 1579.690951 | 1410.125845 | ENSG00000077254 |
| MSANTD3 | -0.163135959 | 0.893081685 | 0.163225738 | 0.338180807 | 437.955463 | 390.9528781 | ENSG00000066697 |
| MCM3AP | -0.163141604 | 0.893078191 | 0.010513852 | 0.040670648 | 1866.681446 | 1667.243013 | ENSG00000160294 |
| CALM2 | -0.163148367 | 0.893074004 | 0.002540419 | 0.012461248 | 4428.455965 | 3954.973841 | ENSG00000143933 |
| NOP9 | -0.16317166 | 0.893059585 | 0.053605681 | 0.148720917 | 981.220023 | 876.0650642 | ENSG00000196943 |
| KIF1B | -0.163213084 | 0.893033943 | 0.019115479 | 0.06616885 | 2457.650841 | 2194.072934 | ENSG00000054523 |
| PPP6R2 | -0.163214096 | 0.893033316 | 0.062857481 | 0.168186954 | 1067.51587 | 953.6458327 | ENSG00000100239 |
| NA | -0.163224526 | 0.89302686 | 0.636741459 | 0.800725077 | 44.37868981 | 39.67301304 | ENSG00000278635 |
| CSPP1 | -0.163347247 | 0.892950899 | 0.26181727 | 0.468843082 | 268.1495939 | 239.6472897 | ENSG00000104218 |
| ANKLE2 | -0.163444066 | 0.892890976 | 0.012986938 | 0.048467996 | 1670.834126 | 1491.806265 | ENSG00000176915 |
| RNF144B | -0.16346461 | 0.892878261 | 0.456860347 | 0.662717707 | 105.6644339 | 94.2284486 | ENSG00000137393 |
| NA | -0.163536864 | 0.892833544 | 0.771600943 | 0.882491333 | 17.3782478 | 15.53495082 | ENSG00000249572 |
| HNRNPL | -0.163660992 | 0.892756729 | 0.00027314 | 0.001811821 | 9987.613651 | 8915.96801 | ENSG00000104824 |
| CASZ1 | -0.163952812 | 0.892576166 | 0.441410238 | 0.64907205 | 141.2794684 | 126.174153 | ENSG00000130940 |
| TYW1B | -0.163988829 | 0.892553883 | 0.601811821 | 0.77679012 | 53.78797838 | 48.12353575 | ENSG00000277149 |
| NA | -0.163996949 | 0.892548859 | 0.709368545 | 0.846496645 | 26.11525905 | 23.23187558 | ENSG00000260588 |
| SLC45A4 | -0.16399738 | 0.892548592 | 0.060954882 | 0.164424074 | 891.632963 | 795.7412494 | ENSG00000022567 |
| KLHL35 | -0.16400986 | 0.892540871 | 0.770639485 | 0.881997379 | 15.94684302 | 14.19418379 | ENSG00000149243 |
| KCNK12 | -0.164133795 | 0.892464201 | 0.36832863 | 0.580382661 | 207.4290642 | 185.3329112 | ENSG00000184261 |
| DLX3 | -0.164169575 | 0.892442067 | 0.856481788 | 0.930005667 | 5.962272037 | 5.323530924 | ENSG00000064195 |
| VHL | -0.16423801 | 0.892399735 | 0.067190762 | 0.177112263 | 786.0859656 | 701.6817746 | ENSG00000134086 |
| RPS15A | -0.164263739 | 0.89238382 | 0.11751303 | 0.268932812 | 661.9651043 | 590.90665 | ENSG00000134419 |
| LHX1-DT | -0.164290979 | 0.892366971 | 0.768071372 | 0.880225765 | 17.57752595 | 15.70605713 | ENSG00000277268 |
| PDP2 | -0.164325437 | 0.892345657 | 0.082054198 | 0.205734668 | 646.7168778 | 577.0194293 | ENSG00000172840 |
| CTNS | -0.164329327 | 0.892343251 | 0.24173018 | 0.44519938 | 290.1844595 | 259.1781728 | ENSG00000040531 |
| LRATD2 | -0.164414432 | 0.892290613 | 0.072576352 | 0.188088982 | 835.8166351 | 745.7826827 | ENSG00000168672 |
| PCDHA6 | -0.164429572 | 0.892281249 | 0.420664073 | 0.630845789 | 114.8723593 | 102.45454 | ENSG00000081842 |
| DUS3L | -0.164432519 | 0.892279427 | 0.040351111 | 0.119552984 | 1122.184303 | 1001.7113 | ENSG00000141994 |
| GPR19 | -0.164544833 | 0.892209965 | 0.29913288 | 0.509389038 | 245.513361 | 219.2554009 | ENSG00000183150 |
| CUL2 | -0.164569724 | 0.892194572 | 0.074121334 | 0.191143576 | 805.7215382 | 718.8703601 | ENSG00000108094 |
| PRPF31 | -0.16464166 | 0.892150086 | 0.041396203 | 0.121865518 | 1117.476988 | 997.4567394 | ENSG00000105618 |
| CWC22 | -0.164660295 | 0.892138562 | 0.055879269 | 0.153689898 | 909.3908524 | 811.5499524 | ENSG00000163510 |
| STX1B | -0.164721234 | 0.89210088 | 0.235835966 | 0.437749338 | 295.8709512 | 264.0710751 | ENSG00000099365 |
| ZNF154 | -0.164912488 | 0.891982624 | 0.249179183 | 0.454215016 | 246.5058607 | 219.839122 | ENSG00000179909 |
| TAP1 | -0.164967175 | 0.891948813 | 0.172208063 | 0.351697769 | 400.909939 | 357.6084471 | ENSG00000168394 |
| SPINT2 | -0.165077676 | 0.891880498 | 0.02952372 | 0.093706798 | 2687.141108 | 2396.421247 | ENSG00000167642 |
| NOM1 | -0.165280733 | 0.891754976 | 0.040868414 | 0.120706911 | 991.9234761 | 884.3402226 | ENSG00000146909 |
| AGPAT4 | -0.165399184 | 0.891681763 | 0.080805082 | 0.203358737 | 698.013881 | 622.1265227 | ENSG00000026652 |
| KLHDC4 | -0.165469634 | 0.891638222 | 0.031384928 | 0.098439523 | 1284.637376 | 1145.75691 | ENSG00000104731 |
| RNF8 | -0.165506198 | 0.891615624 | 0.038492209 | 0.115366666 | 982.2087612 | 876.0532612 | ENSG00000112130 |
| NUP35 | -0.165658244 | 0.891521661 | 0.056930883 | 0.156066152 | 815.0319734 | 726.4539498 | ENSG00000163002 |
| ATP5MC2 | -0.165818596 | 0.891422576 | 0.004031333 | 0.018501028 | 3007.970351 | 2681.269054 | ENSG00000135390 |
| PAXBP1 | -0.166023392 | 0.891296044 | 0.051086028 | 0.143458682 | 926.9223851 | 826.2762993 | ENSG00000159086 |
| POLR2B | -0.166198546 | 0.891187841 | 0.004501172 | 0.020253665 | 2993.529207 | 2667.637967 | ENSG00000047315 |
| SPTLC2 | -0.166199504 | 0.891187249 | 0.02578353 | 0.084359313 | 1447.82774 | 1290.026172 | ENSG00000100596 |
| UBE2N | -0.166280331 | 0.891137322 | 0.006624158 | 0.028011022 | 3834.347921 | 3416.80955 | ENSG00000177889 |
| RNF4 | -0.166336889 | 0.891102387 | 0.011363664 | 0.043433097 | 2011.28411 | 1791.775455 | ENSG00000063978 |
| CD34 | -0.16634028 | 0.891100293 | 0.739792521 | 0.863846025 | 19.69038635 | 17.58715164 | ENSG00000174059 |
| PSMC3 | -0.166425843 | 0.891047445 | 0.001629975 | 0.008533982 | 4427.688699 | 3945.143039 | ENSG00000165916 |
| TSPAN13 | -0.166473133 | 0.891018238 | 0.140153554 | 0.304264843 | 675.7485649 | 602.1312425 | ENSG00000106537 |
| TK1 | -0.166475157 | 0.891016988 | 0.009228529 | 0.036598229 | 2084.710076 | 1857.892224 | ENSG00000167900 |
| RBFOX2 | -0.166505844 | 0.890998036 | 0.00181995 | 0.009352315 | 3722.635972 | 3316.707042 | ENSG00000100320 |

| | | | | | | | |
|---|---|---|---|---|---|---|---|
| MYH10 | -0.166572987 | 0.890956569 | 0.002249069 | 0.011239396 | 10247.33064 | 9129.170286 | ENSG00000133026 |
| CYB5R4 | -0.166627035 | 0.890923192 | 0.279108483 | 0.488187802 | 251.6884864 | 224.0399588 | ENSG00000065615 |
| PARP14 | -0.166769538 | 0.890835195 | 0.164264995 | 0.339700442 | 380.6502201 | 339.0287301 | ENSG00000173193 |
| CCDC180 | -0.1667921 | 0.890821264 | 0.807455885 | 0.90192254 | 11.96962201 | 10.66106059 | ENSG00000197816 |
| ZKSCAN4 | -0.166836717 | 0.890793714 | 0.317481031 | 0.527552069 | 181.0427278 | 161.2468922 | ENSG00000187626 |
| ELOA | -0.166839669 | 0.890791892 | 0.041390966 | 0.121865518 | 1246.02715 | 1110.024527 | ENSG00000011007 |
| SNRPF | -0.166873029 | 0.890771294 | 0.014738635 | 0.053754174 | 2350.646937 | 2093.892143 | ENSG00000139343 |
| MANBA | -0.166891052 | 0.890760166 | 0.061329739 | 0.165128617 | 738.610152 | 657.9581997 | ENSG00000109323 |
| FBXW5 | -0.166904669 | 0.890751758 | 0.021247256 | 0.072233022 | 1820.940454 | 1622.661062 | ENSG00000159069 |
| PPM1E | -0.166917219 | 0.89074401 | 0.212343974 | 0.406375754 | 366.2753284 | 326.2380752 | ENSG00000175175 |
| CCNJ | -0.167032394 | 0.890672901 | 0.058897834 | 0.160143573 | 917.5388497 | 817.2276443 | ENSG00000107443 |
| MPZ | -0.167111356 | 0.890624154 | 0.564134371 | 0.748528238 | 69.18590425 | 61.49195242 | ENSG00000158887 |
| GLYR1 | -0.167121419 | 0.890617942 | 0.001689144 | 0.008812011 | 3290.325586 | 2930.644858 | ENSG00000140632 |
| NA | -0.167176629 | 0.89058386 | 0.851113608 | 0.926282558 | 5.985993136 | 5.32882781 | ENSG00000279833 |
| INTS3 | -0.167194038 | 0.890573113 | 0.031629527 | 0.099157306 | 2083.098265 | 1855.246644 | ENSG00000143624 |
| TUSC2 | -0.167221736 | 0.890556015 | 0.064539901 | 0.171497964 | 982.2938563 | 875.1619036 | ENSG00000114383 |
| RPGRIP1L | -0.167338439 | 0.890483979 | 0.272304305 | 0.480918994 | 233.115233 | 207.4670843 | ENSG00000103494 |
| NHSL1 | -0.167421712 | 0.890432582 | 0.186397801 | 0.371399291 | 406.1287698 | 361.5629824 | ENSG00000135540 |
| DLAT | -0.167543625 | 0.89035734 | 0.01229263 | 0.046402492 | 2349.853313 | 2091.942782 | ENSG00000150768 |
| PPIL4 | -0.167563681 | 0.890344963 | 0.099214139 | 0.236872188 | 577.2631239 | 513.8894899 | ENSG00000131013 |
| RPARP-AS1 | -0.167581861 | 0.890333743 | 0.394533062 | 0.606245474 | 154.1950376 | 137.455888 | ENSG00000269609 |
| FMNL3 | -0.167610821 | 0.890315871 | 0.053915357 | 0.149338739 | 824.4564157 | 734.1248846 | ENSG00000161791 |
| ATXN2L | -0.167654156 | 0.890289128 | 0.003269582 | 0.015479084 | 3947.937594 | 3515.578541 | ENSG00000168488 |
| ERICH5 | -0.167682141 | 0.890271859 | 0.650853065 | 0.810506282 | 39.0516264 | 34.73364798 | ENSG00000177459 |
| CHMP2B | -0.167702008 | 0.890259599 | 0.063966338 | 0.170285352 | 771.9500655 | 686.9369828 | ENSG00000083937 |
| UBE2D1 | -0.167715445 | 0.890251308 | 0.110839688 | 0.257943007 | 682.8058012 | 608.2604372 | ENSG00000072401 |
| CTTN | -0.16778839 | 0.890206296 | 0.001372499 | 0.007386618 | 5224.903884 | 4651.577281 | ENSG00000085733 |
| RESF1 | -0.167882137 | 0.890148452 | 0.046079816 | 0.132271975 | 1288.308413 | 1146.655276 | ENSG00000174718 |
| NOL7 | -0.168037172 | 0.8900528 | 0.036961771 | 0.11172081 | 2114.800137 | 1882.903557 | ENSG00000225921 |
| STN1 | -0.168114061 | 0.890005366 | 0.344691301 | 0.556747794 | 208.6709546 | 185.4781955 | ENSG00000107960 |
| LTA4H | -0.168122803 | 0.889999973 | 0.003334285 | 0.015738061 | 2842.561058 | 2529.731052 | ENSG00000111144 |
| RAB3A | -0.168134306 | 0.889992877 | 0.316205049 | 0.526264103 | 200.454664 | 178.3700255 | ENSG00000105649 |
| KLHDC3 | -0.168160969 | 0.889976429 | 0.004047945 | 0.018563729 | 5299.58271 | 4715.852457 | ENSG00000124702 |
| METTL2B | -0.168221874 | 0.889938858 | 0.099832716 | 0.237837875 | 517.8658188 | 461.037895 | ENSG00000165055 |
| TMED10 | -0.168319672 | 0.889878532 | 0.003227521 | 0.015306812 | 3116.44024 | 2773.115875 | ENSG00000170348 |
| APOO | -0.168427333 | 0.889812128 | 0.094950953 | 0.229420259 | 629.0533146 | 559.4677904 | ENSG00000184831 |
| MFAP1 | -0.168461615 | 0.889790984 | 0.020422504 | 0.06997079 | 1620.767976 | 1441.758376 | ENSG00000140259 |
| NA | -0.168507572 | 0.88976264 | 0.78814843 | 0.891582187 | 11.98978363 | 10.65983108 | ENSG00000146677 |
| SMARCB1 | -0.168512283 | 0.889759734 | 0.001409808 | 0.007557034 | 3724.798131 | 3314.507612 | ENSG00000099956 |
| TMEM9B-AS1 | -0.168545964 | 0.889738962 | 0.668442413 | 0.822178087 | 32.56446323 | 28.96806431 | ENSG00000254860 |
| A4GALT | -0.168592183 | 0.889710459 | 0.288500191 | 0.497704503 | 212.7537378 | 189.2086032 | ENSG00000128274 |
| ANO10 | -0.168646873 | 0.889676732 | 0.078274552 | 0.198658099 | 971.394615 | 864.345911 | ENSG00000160746 |
| ABCA1 | -0.168650715 | 0.889674363 | 0.33962098 | 0.55136372 | 217.181712 | 193.0379481 | ENSG00000165029 |
| NA | -0.168696865 | 0.889645904 | 0.664956983 | 0.820604628 | 35.81039601 | 31.83297946 | ENSG00000233757 |
| NBPF15 | -0.168721633 | 0.88963063 | 0.271928607 | 0.4805251 | 303.2853227 | 269.6494213 | ENSG00000266338 |
| EXOSC10 | -0.168761441 | 0.889606084 | 0.054349525 | 0.150342872 | 1237.119032 | 1100.767662 | ENSG00000171824 |
| VAPA | -0.168812449 | 0.889574631 | 0.014285717 | 0.05239576 | 1796.253243 | 1597.946338 | ENSG00000101558 |
| RNF216P1 | -0.168822736 | 0.889568288 | 0.146952457 | 0.31439868 | 416.0400926 | 370.102742 | ENSG00000196204 |
| INPP5E | -0.168932886 | 0.889500372 | 0.120805445 | 0.273780251 | 591.1547202 | 526.2291065 | ENSG00000148384 |
| AP3M1 | -0.168986201 | 0.889467501 | 0.042054285 | 0.123379707 | 982.825335 | 874.2341506 | ENSG00000185009 |
| NA | -0.169093913 | 0.889401096 | 0.874609898 | 0.941354606 | 5.274892683 | 4.71226379 | ENSG00000230415 |
| PDCD11 | -0.169122161 | 0.889383681 | 0.001751657 | 0.009077948 | 4461.3016 | 3967.56638 | ENSG00000148843 |
| SYVN1 | -0.169140101 | 0.889372622 | 0.123036498 | 0.277511292 | 727.8559315 | 647.7441528 | ENSG00000162298 |
| CTPS1 | -0.169173737 | 0.889351886 | 0.00255014 | 0.012495951 | 2897.678158 | 2577.454409 | ENSG00000171793 |
| ARMCX5 | -0.169294003 | 0.889277752 | 0.303480223 | 0.513550538 | 186.1966245 | 165.6031555 | ENSG00000125962 |
| SH3BP5 | -0.169368995 | 0.889231528 | 0.47338136 | 0.676540562 | 99.34929384 | 88.42784659 | ENSG00000131370 |
| DERL1 | -0.169376873 | 0.889226672 | 0.062620206 | 0.167623329 | 832.869594 | 740.61625 | ENSG00000136986 |
| KDM1A | -0.169394595 | 0.889215749 | 0.000709702 | 0.004159754 | 8246.177886 | 7333.00094 | ENSG00000004487 |
| NA | -0.169401579 | 0.889211444 | 0.840464271 | 0.92055259 | 7.855998146 | 7.018785343 | ENSG00000262902 |
| NA | -0.169444746 | 0.889184839 | 0.418627235 | 0.629290403 | 112.7520745 | 100.300775 | ENSG00000284648 |
| IFT52 | -0.169495436 | 0.889153597 | 0.099891735 | 0.237918452 | 553.4315276 | 492.1172765 | ENSG00000101052 |
| USP36 | -0.169595291 | 0.889092057 | 0.018205804 | 0.063657954 | 1345.503564 | 1196.682221 | ENSG00000055483 |
| NRAS | -0.169604536 | 0.889086359 | 0.027188556 | 0.088041131 | 1750.60541 | 1556.06024 | ENSG00000213281 |
| MTRNR2L8 | -0.169648924 | 0.889059005 | 0.872500002 | 0.940222903 | 4.844606611 | 4.342548161 | ENSG00000255823 |
| B4GALT7 | -0.16980095 | 0.888965324 | 0.13447288 | 0.296015893 | 408.8860427 | 363.5301495 | ENSG00000027847 |
| CCNY | -0.169820696 | 0.888953157 | 0.025756654 | 0.084308183 | 1287.317004 | 1144.112028 | ENSG00000108100 |
| FAM13B | -0.17004447 | 0.888815284 | 0.283950745 | 0.493112905 | 219.6966876 | 195.4059724 | ENSG00000031003 |
| SRP72 | -0.170171856 | 0.888736807 | 0.006660784 | 0.028102973 | 2375.606371 | 2110.689748 | ENSG00000174780 |
| MNS1 | -0.170180192 | 0.888731672 | 0.567523301 | 0.751047019 | 57.61661799 | 51.11443956 | ENSG00000138587 |
| TMEM43 | -0.170188902 | 0.888726306 | 0.058358429 | 0.158987151 | 781.9957579 | 694.8618594 | ENSG00000170876 |
| CALCR | -0.170269294 | 0.888676785 | 0.709253235 | 0.846496645 | 26.37563247 | 23.42272901 | ENSG00000004948 |
| HSP90B1 | -0.170321501 | 0.888644627 | 0.000406527 | 0.002563359 | 14817.16167 | 13167.17404 | ENSG00000166598 |
| SOWAHB | -0.170392521 | 0.888600882 | 0.766309974 | 0.878994404 | 14.54953226 | 12.92558592 | ENSG00000186212 |
| KDM4A | -0.170431605 | 0.88857681 | 0.026348243 | 0.085835185 | 1025.180378 | 911.010111 | ENSG00000066135 |

| | | | | | | | |
|---|---|---|---|---|---|---|---|
| EXOSC9 | -0.170502852 | 0.888532929 | 0.04023373 | 0.119298789 | 1003.649099 | 892.2379024 | ENSG00000123737 |
| TRAF4 | -0.170536233 | 0.88851237 | 0.001972843 | 0.010029585 | 3288.034753 | 2921.007941 | ENSG00000076604 |
| KIAA1324L | -0.170560116 | 0.888497661 | 0.052284899 | 0.146108362 | 820.6007493 | 729.007626 | ENSG00000164659 |
| PLEKHM2 | -0.170610087 | 0.888466887 | 0.009710293 | 0.038116582 | 1861.979219 | 1654.127223 | ENSG00000116786 |
| NOMO3 | -0.170678472 | 0.888424774 | 0.415842139 | 0.626739688 | 125.3797104 | 111.2480103 | ENSG00000103226 |
| CEP78 | -0.170737224 | 0.888388594 | 0.041611372 | 0.12232733 | 852.8684064 | 757.7314891 | ENSG00000148019 |
| VPS33A | -0.170794873 | 0.888353096 | 0.261348694 | 0.468515166 | 254.6133333 | 226.260596 | ENSG00000139719 |
| NA | -0.170800854 | 0.888349413 | 0.439889722 | 0.647767794 | 128.8137104 | 114.5797293 | ENSG00000279457 |
| STOML2 | -0.17096363 | 0.888249188 | 0.002127091 | 0.010729195 | 4290.689235 | 3810.637637 | ENSG00000165283 |
| CHMP4A | -0.170972758 | 0.888243568 | 0.731256904 | 0.858455957 | 20.48096426 | 18.17789364 | ENSG00000254505 |
| PPIB | -0.171008917 | 0.888221306 | 0.008420842 | 0.034074911 | 2150.158971 | 1910.099801 | ENSG00000166794 |
| YIF1B | -0.171010194 | 0.88822052 | 0.017603539 | 0.062011527 | 1488.677188 | 1322.157521 | ENSG00000167645 |
| MAP3K14 | -0.17104581 | 0.888198592 | 0.365443367 | 0.577667866 | 154.0779066 | 136.9365371 | ENSG00000006062 |
| STIMATE | -0.171061427 | 0.888188978 | 0.794224872 | 0.894817844 | 11.23004578 | 9.990393526 | ENSG00000213533 |
| NA | -0.171109747 | 0.88815923 | 0.782999154 | 0.889773441 | 12.36787283 | 10.95085116 | ENSG00000267390 |
| SSR3 | -0.171165555 | 0.888124874 | 0.010033894 | 0.039158935 | 2452.941674 | 2178.561934 | ENSG00000114850 |
| KDM4D | -0.171195987 | 0.888106141 | 0.642184951 | 0.804381694 | 38.02737083 | 33.86370918 | ENSG00000186280 |
| RCC1 | -0.171252704 | 0.888071227 | 0.001586463 | 0.008357143 | 3447.453882 | 3061.372378 | ENSG00000180198 |
| MYLPF | -0.171301183 | 0.888041386 | 0.848695179 | 0.92506109 | 5.611463415 | 4.991460588 | ENSG00000180209 |
| TOX2 | -0.171322378 | 0.888028339 | 0.501714443 | 0.699924448 | 76.26853704 | 67.80270895 | ENSG00000124191 |
| RANBP9 | -0.171377275 | 0.887994548 | 0.106468976 | 0.25018162 | 702.7304512 | 623.775598 | ENSG00000010017 |
| ARID2 | -0.171410083 | 0.887974355 | 0.059989742 | 0.16230749 | 760.1358389 | 674.6634307 | ENSG00000189079 |
| CSF1 | -0.171442644 | 0.887954314 | 0.359443875 | 0.571533011 | 151.7203847 | 134.8765804 | ENSG00000184371 |
| UQCR11 | -0.171457156 | 0.887945383 | 0.267639883 | 0.475391897 | 270.7936008 | 240.523544 | ENSG00000127540 |
| NMT1 | -0.171490156 | 0.887925072 | 0.000978197 | 0.005506184 | 3632.856091 | 3225.443999 | ENSG00000136448 |
| NA | -0.171506797 | 0.88791483 | 0.460227946 | 0.664999857 | 90.15944506 | 80.02773841 | ENSG00000228409 |
| SCFD1 | -0.171552282 | 0.887886837 | 0.072798424 | 0.188638618 | 786.2835154 | 698.4899477 | ENSG00000092108 |
| BRD2 | -0.171567354 | 0.887877561 | 0.006129059 | 0.026204953 | 3609.650352 | 3204.672032 | ENSG00000204256 |
| SORBS3 | -0.171606461 | 0.887853493 | 0.053048354 | 0.147564908 | 970.8948666 | 862.5148224 | ENSG00000120896 |
| ARL6IP1 | -0.171774894 | 0.887749843 | 0.031663948 | 0.09923227 | 1537.059087 | 1364.600112 | ENSG00000170540 |
| PGLS | -0.171782199 | 0.887745348 | 0.172785156 | 0.352380959 | 403.8407357 | 358.7544613 | ENSG00000130313 |
| PSMA3 | -0.171806676 | 0.887730287 | 0.004142444 | 0.018932367 | 2460.599869 | 2184.781661 | ENSG00000100567 |
| PARN | -0.171907541 | 0.887668224 | 0.022886628 | 0.076674468 | 1247.985452 | 1108.262581 | ENSG00000140694 |
| NA | -0.172052668 | 0.887578934 | 0.703954801 | 0.842524014 | 23.60859702 | 20.93159687 | ENSG00000233325 |
| THAP3 | -0.172105929 | 0.887546168 | 0.270256271 | 0.478644816 | 222.031963 | 197.1651886 | ENSG00000041988 |
| ARSA | -0.172166635 | 0.887508822 | 0.234879115 | 0.436369117 | 263.8019875 | 234.2641909 | ENSG00000100299 |
| SUCLA2 | -0.17217459 | 0.887503928 | 0.025826044 | 0.084454519 | 1102.965416 | 978.737602 | ENSG00000136143 |
| GDF1 | -0.172194829 | 0.887491478 | 0.312084564 | 0.52250938 | 181.6702326 | 161.4264168 | ENSG00000223802 |
| SCN4B | -0.172239048 | 0.887464277 | 0.381263468 | 0.59376395 | 170.3503109 | 150.8997202 | ENSG00000177098 |
| DMRTA1 | -0.172243907 | 0.887461287 | 0.356295627 | 0.568409019 | 143.9960608 | 127.8988453 | ENSG00000176399 |
| GTF3A | -0.172274789 | 0.887442291 | 0.031971373 | 0.099963474 | 1229.871241 | 1091.471169 | ENSG00000122034 |
| PREP | -0.17235227 | 0.887394631 | 0.022209536 | 0.074924453 | 1868.822888 | 1658.093506 | ENSG00000085377 |
| KLHL15 | -0.172367876 | 0.887385032 | 0.245773535 | 0.450231178 | 324.8489681 | 287.9369481 | ENSG00000174010 |
| NA | -0.172390158 | 0.887371327 | 0.624449693 | 0.7923564 | 41.28812631 | 36.60246772 | ENSG00000181524 |
| FDFT1 | -0.172501577 | 0.887302798 | 0.000311324 | 0.002030163 | 12471.08132 | 11064.74461 | ENSG00000079459 |
| KDM3B | -0.172578045 | 0.887255769 | 0.001527769 | 0.008077243 | 3301.901024 | 2929.188136 | ENSG00000120733 |
| SLC25A1 | -0.172624947 | 0.887226925 | 0.003649563 | 0.016996791 | 5133.544753 | 4555.230922 | ENSG00000100075 |
| ABCF1 | -0.172697177 | 0.887182506 | 0.001441336 | 0.00769786 | 5726.977624 | 5081.642937 | ENSG00000204574 |
| RAB2B | -0.172779941 | 0.887131612 | 0.293114075 | 0.502804708 | 242.351888 | 214.7126874 | ENSG00000129472 |
| IL15RA | -0.172841733 | 0.887093616 | 0.716325473 | 0.85043104 | 21.65291149 | 19.25998901 | ENSG00000134470 |
| GMNN | -0.172846843 | 0.887090474 | 0.004864564 | 0.021570157 | 1985.21098 | 1761.040316 | ENSG00000112312 |
| AMMECR1L | -0.172861478 | 0.887081475 | 0.060400872 | 0.163162705 | 769.5917061 | 682.447374 | ENSG00000144233 |
| RAB3B | -0.172920005 | 0.887045489 | 0.010123065 | 0.039425464 | 1566.577376 | 1389.524753 | ENSG00000169213 |
| FAM95C | -0.172925074 | 0.887042372 | 0.861650365 | 0.932828889 | 5.264214265 | 4.689563196 | ENSG00000283486 |
| NLGN2 | -0.172942426 | 0.887031704 | 0.040005155 | 0.11871423 | 2740.790833 | 2431.709717 | ENSG00000169992 |
| RASL11B | -0.173028849 | 0.886978568 | 0.683459624 | 0.830895404 | 33.55462419 | 29.87347294 | ENSG00000128045 |
| DTD1 | -0.173055296 | 0.886962309 | 0.135896498 | 0.298176527 | 403.666296 | 358.0439735 | ENSG00000125821 |
| RANBP10 | -0.173153857 | 0.886901716 | 0.088416 | 0.217413563 | 580.5563236 | 515.1659389 | ENSG00000141084 |
| NA | -0.173201163 | 0.886872635 | 0.842861251 | 0.9217804 | 6.371201275 | 5.622023348 | ENSG00000286681 |
| STARD7 | -0.173302394 | 0.886810407 | 0.008963021 | 0.03581605 | 3943.840644 | 3497.355267 | ENSG00000084090 |
| NA | -0.173315765 | 0.886802188 | 0.590154298 | 0.768413122 | 50.50820475 | 44.71250178 | ENSG00000179967 |
| STAG2 | -0.173342959 | 0.886785473 | 0.021581149 | 0.073157422 | 3568.816911 | 3164.382329 | ENSG00000101972 |
| PGK1 | -0.173455355 | 0.886716388 | 0.002318391 | 0.011524826 | 9735.605039 | 8631.751358 | ENSG00000102144 |
| PPP3R1 | -0.173470011 | 0.88670738 | 0.218011702 | 0.414158635 | 450.9407715 | 400.2902918 | ENSG00000221823 |
| OLA1 | -0.173506239 | 0.886685114 | 0.008812119 | 0.035362698 | 2135.550401 | 1893.932754 | ENSG00000138430 |
| FAF1 | -0.173514061 | 0.886680307 | 0.053089879 | 0.147658669 | 1074.797503 | 952.5874003 | ENSG00000185104 |
| ANP32A | -0.173522932 | 0.886674855 | 0.000481605 | 0.002982981 | 7647.209136 | 6780.257927 | ENSG00000140350 |
| ZNF470 | -0.173533027 | 0.886668651 | 0.333601392 | 0.545601654 | 181.1652238 | 160.7611193 | ENSG00000197016 |
| NA | -0.173598961 | 0.886628129 | 0.824394406 | 0.911248953 | 7.831081838 | 6.9736677 | ENSG00000272438 |
| ZNF711 | -0.17372419 | 0.886551171 | 0.141690939 | 0.306440122 | 403.4596457 | 357.7337304 | ENSG00000147180 |
| ZNF875 | -0.173769996 | 0.886523023 | 0.115794278 | 0.266095758 | 514.9866608 | 456.3410703 | ENSG00000181666 |
| LPAR4 | -0.173790254 | 0.886510575 | 0.433301586 | 0.642497135 | 102.7613005 | 91.19866999 | ENSG00000147145 |
| WDR36 | -0.173793647 | 0.88650849 | 0.087330074 | 0.215557242 | 786.3617727 | 696.8993561 | ENSG00000134987 |
| PRPS2 | -0.173795543 | 0.886507325 | 0.055265027 | 0.152495621 | 777.2528747 | 689.2016237 | ENSG00000101911 |

| | | | | | | | |
|---|---|---|---|---|---|---|---|
| GLIS2 | -0.173926215 | 0.886427033 | 0.172621382 | 0.352187449 | 363.3297869 | 322.3798299 | ENSG00000126603 |
| VBP1 | -0.173941851 | 0.886417426 | 0.041117532 | 0.121271996 | 1172.051453 | 1038.95444 | ENSG00000155959 |
| COLGALT1 | -0.174016115 | 0.886371798 | 0.003635053 | 0.016936733 | 2598.561047 | 2303.145774 | ENSG00000130309 |
| ROCK1 | -0.174054688 | 0.8863481 | 0.035020529 | 0.107253389 | 1304.51748 | 1155.742109 | ENSG00000067900 |
| ZNF529 | -0.174094904 | 0.886323393 | 0.352695514 | 0.564941033 | 142.681787 | 126.5667802 | ENSG00000186020 |
| IFT20 | -0.174273715 | 0.886213547 | 0.278895184 | 0.48800877 | 201.5628724 | 178.6450823 | ENSG00000109083 |
| GCH1 | -0.174420066 | 0.886123651 | 0.313127872 | 0.523450462 | 199.0380813 | 176.4470301 | ENSG00000131979 |
| TOP2B | -0.174497568 | 0.886076049 | 0.006414397 | 0.02723991 | 4299.480713 | 3809.949211 | ENSG00000077097 |
| NABP2 | -0.174519616 | 0.886062508 | 0.021818426 | 0.073842469 | 1148.81764 | 1018.135624 | ENSG00000139579 |
| DBI | -0.174526367 | 0.886058362 | 0.007898311 | 0.032334619 | 1994.274494 | 1766.972791 | ENSG00000155368 |
| LYPD3 | -0.174554569 | 0.886041041 | 0.497198 | 0.696298423 | 81.39573806 | 72.05407587 | ENSG00000124466 |
| NA | -0.174639785 | 0.885988707 | 0.559097141 | 0.744503562 | 62.72930179 | 55.47316192 | ENSG00000223546 |
| ZNF250 | -0.17465892 | 0.885976956 | 0.274644848 | 0.483425106 | 201.3781114 | 178.4324743 | ENSG00000196150 |
| NDST1 | -0.174701348 | 0.8859509 | 0.003829995 | 0.017706111 | 2710.677153 | 2401.752545 | ENSG00000070614 |
| ABCF3 | -0.174736305 | 0.885929434 | 0.019110173 | 0.06616885 | 1324.681477 | 1173.435212 | ENSG00000161204 |
| RPUSD4 | -0.174796897 | 0.885892226 | 0.021785542 | 0.073757611 | 1117.321847 | 989.8394562 | ENSG00000165526 |
| UBAC2-AS1 | -0.174866944 | 0.885849215 | 0.674016866 | 0.825366912 | 29.18329666 | 25.81447251 | ENSG00000228889 |
| SLC24A4 | -0.17489274 | 0.885833376 | 0.7078616 | 0.845910689 | 21.69679422 | 19.26216447 | ENSG00000140090 |
| SLC31A1 | -0.174970432 | 0.885785673 | 0.006221249 | 0.026551026 | 1959.005845 | 1734.757926 | ENSG00000136868 |
| PPIEL | -0.175100003 | 0.885706123 | 0.592294099 | 0.769872259 | 50.79140955 | 45.00401631 | ENSG00000243970 |
| SLC27A4 | -0.17510403 | 0.88570365 | 0.046113123 | 0.132327355 | 887.705574 | 786.4736614 | ENSG00000167114 |
| SH3KBP1 | -0.17513083 | 0.885687197 | 0.08350519 | 0.208542121 | 592.2148243 | 524.3178687 | ENSG00000147010 |
| RRP1B | -0.175219527 | 0.885632747 | 0.004303456 | 0.019517476 | 3419.215901 | 3028.564636 | ENSG00000160208 |
| CACUL1 | -0.175227864 | 0.885627629 | 0.087707803 | 0.216133088 | 719.6000266 | 636.8280029 | ENSG00000151893 |
| LINC02701 | -0.175262764 | 0.885606205 | 0.796278039 | 0.895955122 | 10.86619447 | 9.642432532 | ENSG00000250508 |
| NA | -0.17537192 | 0.885539202 | 0.729468571 | 0.85755378 | 18.74115904 | 16.64286821 | ENSG00000215302 |
| NA | -0.175378438 | 0.885535201 | 0.831502035 | 0.915296977 | 8.203247295 | 7.272160124 | ENSG00000227470 |
| DNASE1 | -0.175413201 | 0.885513863 | 0.210301638 | 0.403817634 | 266.8738639 | 236.3846615 | ENSG00000213918 |
| PRDX2 | -0.175415262 | 0.885512599 | 0.004961456 | 0.021937978 | 2404.161228 | 2129.107453 | ENSG00000167815 |
| PPP2CA | -0.175535435 | 0.88543884 | 0.006362182 | 0.02703641 | 2285.207148 | 2023.52021 | ENSG00000113575 |
| PARP6 | -0.175606668 | 0.885395123 | 0.21634 | 0.411769896 | 359.9551294 | 318.774297 | ENSG00000137817 |
| GABPB1-IT1 | -0.175622303 | 0.885385528 | 0.365130341 | 0.577318025 | 179.9912178 | 159.5598155 | ENSG00000285410 |
| PUM3 | -0.175654437 | 0.885365807 | 0.008147806 | 0.033140494 | 1728.882277 | 1530.749126 | ENSG00000080608 |
| DIDO1 | -0.175687686 | 0.885345403 | 0.008025671 | 0.032759256 | 2942.201137 | 2605.399886 | ENSG00000101191 |
| CCDC24 | -0.175709122 | 0.885332248 | 0.342740201 | 0.554597608 | 157.6393326 | 139.4570642 | ENSG00000159214 |
| GNAZ | -0.175709214 | 0.885332192 | 0.251859124 | 0.457549413 | 280.4427178 | 248.5005592 | ENSG00000128266 |
| PLCXD1 | -0.17582234 | 0.885262773 | 0.002010982 | 0.010211721 | 3212.632459 | 2844.101218 | ENSG00000182378 |
| H3-3B | -0.175861567 | 0.885238703 | 0.00048527 | 0.003002729 | 14126.12378 | 12505.17747 | ENSG00000132475 |
| ZNF343 | -0.17588978 | 0.885221392 | 0.391616808 | 0.60304007 | 116.101232 | 102.7436681 | ENSG00000088876 |
| NA | -0.176043087 | 0.885127329 | 0.775449886 | 0.884700308 | 16.18260562 | 14.3503454 | ENSG00000256673 |
| EIF3H | -0.176060337 | 0.885116746 | 0.001166728 | 0.006412588 | 3729.496802 | 3300.796806 | ENSG00000147677 |
| DDX56 | -0.176064195 | 0.885114379 | 0.010906038 | 0.041989914 | 1567.794681 | 1387.437883 | ENSG00000136271 |
| PHB | -0.176068102 | 0.885111982 | 0.004960711 | 0.021937978 | 5910.02066 | 5230.57662 | ENSG00000167085 |
| DHCR7 | -0.17608666 | 0.885100597 | 0.00060901 | 0.003645373 | 5199.457491 | 4602.157934 | ENSG00000172893 |
| MRPS2 | -0.176194273 | 0.885034578 | 0.081743281 | 0.205173028 | 801.2387485 | 709.6870278 | ENSG00000122140 |
| PSME2 | -0.176312594 | 0.884961996 | 0.028563228 | 0.091255646 | 1268.139454 | 1121.779288 | ENSG00000100911 |
| PLEKHB1 | -0.176336349 | 0.884947425 | 0.250593786 | 0.456097114 | 209.2848058 | 185.2779778 | ENSG00000021300 |
| MED30 | -0.17633946 | 0.884945516 | 0.285345244 | 0.494673527 | 200.9919874 | 177.8361518 | ENSG00000164758 |
| USH1G | -0.176460323 | 0.884871382 | 0.496030984 | 0.695334407 | 71.28424743 | 63.03227474 | ENSG00000182040 |
| GRWD1 | -0.176530827 | 0.88482814 | 0.003209677 | 0.015260999 | 2633.198326 | 2330.12151 | ENSG00000105447 |
| CCNT2 | -0.176601057 | 0.884785068 | 0.081702548 | 0.205098049 | 836.2527938 | 739.4245151 | ENSG00000082258 |
| MYBBP1A | -0.176625336 | 0.884770178 | 0.001092392 | 0.006055127 | 4879.447466 | 4317.907282 | ENSG00000132382 |
| PRR11 | -0.176808973 | 0.884657565 | 0.085914171 | 0.213036911 | 623.8920713 | 551.9005226 | ENSG00000068489 |
| CLDN10 | -0.176920694 | 0.88458906 | 0.110062807 | 0.256577719 | 536.8417728 | 474.5744215 | ENSG00000134873 |
| NA | -0.177060577 | 0.884503296 | 0.648737561 | 0.809250856 | 32.2990558 | 28.50470843 | ENSG00000259570 |
| ZDHHC2 | -0.177115652 | 0.88446953 | 0.142252638 | 0.30726765 | 387.5050728 | 342.8755739 | ENSG00000104219 |
| FAM92A | -0.177136494 | 0.884456753 | 0.225030839 | 0.423319791 | 237.4488809 | 210.032772 | ENSG00000188343 |
| EIF2S3B | -0.177170121 | 0.884436138 | 0.68295622 | 0.830790283 | 25.1740404 | 22.26829652 | ENSG00000180574 |
| CDC42EP3 | -0.17718815 | 0.884425085 | 0.085718721 | 0.2126919 | 556.4842106 | 492.0774128 | ENSG00000163171 |
| UBXN4 | -0.177255497 | 0.8843838 | 0.007400656 | 0.030635992 | 1807.234432 | 1597.895166 | ENSG00000144224 |
| LIN28A | -0.177261918 | 0.884379864 | 0.001433854 | 0.00766644 | 23215.76897 | 20530.89791 | ENSG00000131914 |
| TBC1D14 | -0.177298634 | 0.884357357 | 0.042497584 | 0.124428972 | 849.8881348 | 751.2007046 | ENSG00000132405 |
| H3-5 | -0.177399436 | 0.884295568 | 0.755647984 | 0.872914862 | 14.2639632 | 12.60004198 | ENSG00000188375 |
| THAP4 | -0.177474531 | 0.88424954 | 0.00870838 | 0.035005906 | 1743.930894 | 1542.348503 | ENSG00000176946 |
| UPF1 | -0.177537391 | 0.884211013 | 0.0011549 | 0.006360538 | 4428.621814 | 3915.238625 | ENSG00000005007 |
| POLR2H | -0.177600726 | 0.884172197 | 0.012770335 | 0.047800106 | 1441.896788 | 1274.752249 | ENSG00000163882 |
| URB1 | -0.177602607 | 0.884171044 | 0.01115803 | 0.042768297 | 1991.438659 | 1760.463823 | ENSG00000142207 |
| ZNF703 | -0.177725611 | 0.884095663 | 0.590735838 | 0.768598294 | 45.80417086 | 40.56659839 | ENSG00000183779 |
| UPF3A | -0.177784383 | 0.884059648 | 0.016733462 | 0.059557375 | 1646.357786 | 1455.637425 | ENSG00000169062 |
| MIR4664 | -0.177862789 | 0.884011603 | 0.48422399 | 0.685332156 | 76.25310394 | 67.49333921 | ENSG00000265660 |
| MBLAC2 | -0.177884637 | 0.883998216 | 0.475557687 | 0.678008978 | 91.64037697 | 81.00576771 | ENSG00000176055 |
| PSMB6 | -0.177888535 | 0.883995828 | 0.003579021 | 0.016705342 | 2999.611986 | 2652.009614 | ENSG00000142507 |
| FKBP4 | -0.177938754 | 0.883965057 | 4.63E-05 | 0.000375736 | 9856.183551 | 8712.415134 | ENSG00000004478 |
| MAP3K6 | -0.178008209 | 0.883922501 | 0.196741164 | 0.386101723 | 269.2346137 | 238.0329063 | ENSG00000142733 |

| | | | | | | | |
|---|---|---|---|---|---|---|---|
| DMAP1 | -0.178080402 | 0.883878271 | 0.02469462 | 0.081659588 | 1009.655507 | 892.5796206 | ENSG00000178028 |
| CWF19L2 | -0.178214383 | 0.88379619 | 0.118355896 | 0.2702722 | 427.87893 | 378.0775911 | ENSG00000152404 |
| C5orf30 | -0.178232827 | 0.883784891 | 0.135155048 | 0.297032827 | 400.3100452 | 353.991005 | ENSG00000181751 |
| MIR5587 | -0.178236066 | 0.883782907 | 0.848219937 | 0.9249161 | 5.673143503 | 5.011985719 | ENSG00000266124 |
| RHEB | -0.178270991 | 0.883761513 | 0.003142679 | 0.014979681 | 2635.429006 | 2329.606879 | ENSG00000106615 |
| CHCHD3 | -0.178400968 | 0.883681895 | 0.003261816 | 0.015453936 | 3000.868451 | 2652.379293 | ENSG00000106554 |
| KIF21A | -0.178408303 | 0.883677403 | 0.046638856 | 0.133597979 | 1213.39709 | 1072.150034 | ENSG00000139116 |
| GIN1 | -0.1784809 | 0.883632937 | 0.548474747 | 0.736043449 | 66.47048086 | 58.64924341 | ENSG00000145723 |
| CZIB | -0.178510415 | 0.883614859 | 0.048810726 | 0.138456089 | 799.6654462 | 706.6295353 | ENSG00000162384 |
| THOC6 | -0.178640121 | 0.883535421 | 0.01632334 | 0.058372709 | 1167.93592 | 1031.638477 | ENSG00000131652 |
| NA | -0.178786095 | 0.883446028 | 0.812785391 | 0.90498509 | 8.298131691 | 7.334681475 | ENSG00000224837 |
| MANF | -0.178808642 | 0.883432222 | 0.004609938 | 0.020634908 | 2376.502206 | 2099.252423 | ENSG00000145050 |
| UNC5CL | -0.178818963 | 0.883425902 | 0.683772966 | 0.830895404 | 33.5750913 | 29.74485732 | ENSG00000124602 |
| NA | -0.178885074 | 0.88338542 | 0.79269622 | 0.894055669 | 11.21700309 | 9.9011042 | ENSG00000236675 |
| FIZ1 | -0.178926349 | 0.883360147 | 0.16306852 | 0.338120875 | 448.8735477 | 396.8681923 | ENSG00000179943 |
| NDUFA6 | -0.179029873 | 0.883296761 | 0.018710484 | 0.065049243 | 1932.623216 | 1706.848301 | ENSG00000184983 |
| PNMA8B | -0.179038759 | 0.883291321 | 0.466282562 | 0.670403211 | 81.04937863 | 71.55496661 | ENSG00000204851 |
| RSF1 | -0.179084453 | 0.883263345 | 0.028289259 | 0.090703336 | 1409.383021 | 1244.535436 | ENSG00000048649 |
| LRP4 | -0.179161627 | 0.883216098 | 0.045530496 | 0.131013778 | 962.3629282 | 849.5259442 | ENSG00000134569 |
| RPGR | -0.179203873 | 0.883190236 | 0.297251626 | 0.507512608 | 164.9191586 | 145.7116578 | ENSG00000156313 |
| NA | -0.179220087 | 0.88318031 | 0.741664519 | 0.865282909 | 16.45630105 | 14.53590194 | ENSG00000260401 |
| PCBP1 | -0.179254524 | 0.883159229 | 0.350733668 | 0.562848855 | 191.8294965 | 169.5984778 | ENSG00000169564 |
| ZFYVE1 | -0.179283188 | 0.883141682 | 0.134447758 | 0.295995093 | 388.2046051 | 342.9042338 | ENSG00000165861 |
| TPM4 | -0.17935187 | 0.883099639 | 3.40E-05 | 0.000285006 | 9009.780193 | 7956.414078 | ENSG00000167460 |
| CMTM4 | -0.179358876 | 0.883095351 | 0.047276684 | 0.135091568 | 1249.469563 | 1102.77303 | ENSG00000183723 |
| MCM8 | -0.179384757 | 0.883079509 | 0.230055751 | 0.429903311 | 238.1641512 | 210.5275303 | ENSG00000125885 |
| FAXC | -0.179396647 | 0.883072231 | 0.24567404 | 0.450136241 | 218.2724018 | 192.8750195 | ENSG00000146267 |
| SPINT1 | -0.179399786 | 0.88307031 | 0.001520808 | 0.008047199 | 3016.345777 | 2664.073214 | ENSG00000166145 |
| AK4 | -0.179413667 | 0.883061813 | 0.010246574 | 0.039807972 | 3386.689421 | 2989.755496 | ENSG00000162433 |
| RAB3D | -0.179474218 | 0.883024751 | 0.244545482 | 0.448895939 | 220.8359892 | 195.0135562 | ENSG00000105514 |
| ZIK1 | -0.179608068 | 0.88294283 | 0.054959762 | 0.151786356 | 710.3213206 | 627.2712344 | ENSG00000171649 |
| FDX1 | -0.179621538 | 0.882934586 | 0.105543987 | 0.248466491 | 534.1613849 | 471.8258472 | ENSG00000137714 |
| AOPEP | -0.179742308 | 0.882860678 | 0.288756729 | 0.497979727 | 208.6033502 | 184.1001394 | ENSG00000148120 |
| IGF2R | -0.179762374 | 0.882848398 | 0.011974961 | 0.045465851 | 4211.288868 | 3717.28737 | ENSG00000197081 |
| ATAT1 | -0.179817062 | 0.882814932 | 0.160311788 | 0.334344353 | 365.7472095 | 323.0553424 | ENSG00000137343 |
| DPY19L2P2 | -0.17984663 | 0.88279684 | 0.725881641 | 0.855720453 | 19.08248445 | 16.85112535 | ENSG00000170629 |
| OPA3 | -0.179884297 | 0.882773791 | 0.043398425 | 0.126419752 | 785.9860739 | 693.7865967 | ENSG00000125741 |
| TXNL4A | -0.179916075 | 0.882754346 | 0.008475782 | 0.034245859 | 2078.281948 | 1835.282651 | ENSG00000141759 |
| SINHCAF | -0.180000972 | 0.882702402 | 0.005504438 | 0.023946397 | 6622.893546 | 5845.332414 | ENSG00000139146 |
| HIVEP3 | -0.180019881 | 0.882690832 | 0.696650855 | 0.838084329 | 24.79119652 | 21.88240669 | ENSG00000127124 |
| POU5F1B | -0.180056104 | 0.88266867 | 0.288720959 | 0.497979727 | 173.4548068 | 153.005289 | ENSG00000212993 |
| HARS1 | -0.1800798 | 0.882654173 | 0.007359079 | 0.030490613 | 1609.276186 | 1420.753015 | ENSG00000170445 |
| MOV10 | -0.180152769 | 0.88260953 | 0.036677715 | 0.111127651 | 1176.746589 | 1038.643214 | ENSG00000155363 |
| IPO7 | -0.180152909 | 0.882609445 | 0.002824801 | 0.01366104 | 5547.89876 | 4896.15879 | ENSG00000205339 |
| ORC6 | -0.180216821 | 0.882570346 | 0.011985846 | 0.045488888 | 1268.971654 | 1119.753833 | ENSG00000091651 |
| NA | -0.180293872 | 0.882523211 | 0.758022125 | 0.874306125 | 18.07986504 | 15.99641479 | ENSG00000188599 |
| DCTN6 | -0.180341651 | 0.882493985 | 0.488861751 | 0.689172452 | 79.25974079 | 69.96545927 | ENSG00000104671 |
| NDUFB2 | -0.180346062 | 0.882491286 | 0.18012322 | 0.36295636 | 377.4860506 | 333.2330164 | ENSG00000090266 |
| PRC1 | -0.180401762 | 0.882457215 | 0.809943463 | 0.90313851 | 8.308810108 | 7.332506011 | ENSG00000198901 |
| EPN2 | -0.180444863 | 0.882430852 | 0.066793702 | 0.176288107 | 602.9805155 | 532.0756862 | ENSG00000072134 |
| MFSD6 | -0.180447855 | 0.882429022 | 0.494883673 | 0.69449938 | 76.71687365 | 67.60142971 | ENSG00000151690 |
| ZNF280B | -0.180449545 | 0.882427988 | 0.25717424 | 0.46400454 | 227.311331 | 200.6187429 | ENSG00000275004 |
| PPP2R1A | -0.180479061 | 0.882409935 | 0.000114467 | 0.000837875 | 8868.238795 | 7825.523919 | ENSG00000105568 |
| LGALSL | -0.180530477 | 0.882378487 | 0.101230513 | 0.239997826 | 437.9094948 | 386.3730138 | ENSG00000119862 |
| ERLIN2 | -0.180613238 | 0.882327871 | 0.032091737 | 0.100223657 | 1252.293497 | 1104.27869 | ENSG00000147475 |
| MROH1 | -0.180616011 | 0.882326174 | 0.149508379 | 0.318101141 | 514.8600461 | 454.4305083 | ENSG00000179832 |
| FBXO17 | -0.18071041 | 0.882268444 | 0.262827048 | 0.469982842 | 234.7524245 | 207.0109602 | ENSG00000269190 |
| PML | -0.180778042 | 0.882227085 | 0.046693201 | 0.133687249 | 852.14968 | 752.2324284 | ENSG00000140464 |
| TBC1D16 | -0.180790848 | 0.882219254 | 0.000616646 | 0.003679417 | 6468.056246 | 5706.054931 | ENSG00000167291 |
| SRRM4 | -0.180886748 | 0.882160613 | 0.480745905 | 0.682708803 | 77.051996 | 67.96272703 | ENSG00000139767 |
| TCF3 | -0.180970084 | 0.882109657 | 0.00058858 | 0.003535409 | 7277.913075 | 6420.37015 | ENSG00000071564 |
| TPRKB | -0.18104333 | 0.882064873 | 0.149495297 | 0.318101141 | 432.5992867 | 381.7212544 | ENSG00000144034 |
| SLC51A | -0.181096856 | 0.882032148 | 0.466862687 | 0.670879763 | 80.13188108 | 70.77554664 | ENSG00000163959 |
| RHOT1 | -0.181127477 | 0.882013427 | 0.049424195 | 0.139789716 | 677.963787 | 597.9824715 | ENSG00000126858 |
| OSBP2 | -0.1811447 | 0.882002897 | 0.321568371 | 0.532049998 | 148.9758751 | 131.4798287 | ENSG00000184792 |
| CAPN1 | -0.181218519 | 0.881957769 | 0.000698301 | 0.004105668 | 4558.425124 | 4020.339869 | ENSG00000014216 |
| MED16 | -0.181289465 | 0.881914399 | 0.004408216 | 0.019897027 | 2285.778059 | 2016.335588 | ENSG00000175221 |
| SNX10 | -0.181320344 | 0.881895522 | 0.375636558 | 0.587680921 | 121.660804 | 107.2548636 | ENSG00000086300 |
| CDH3 | -0.181328993 | 0.881890236 | 0.002126965 | 0.010729195 | 3804.206736 | 3354.211184 | ENSG00000062038 |
| SYCP2L | -0.181461698 | 0.88180912 | 0.681534097 | 0.829783471 | 25.21199939 | 22.20700468 | ENSG00000153157 |
| MBD2 | -0.181484618 | 0.88179511 | 0.184656702 | 0.368858755 | 424.1301256 | 374.2297716 | ENSG00000134046 |
| NA | -0.181570364 | 0.881742703 | 0.400944384 | 0.612310747 | 148.4157209 | 130.948414 | ENSG00000254615 |
| MRPL33 | -0.181573638 | 0.881740702 | 0.148115436 | 0.315956174 | 392.4294368 | 346.0223987 | ENSG00000243147 |
| SUMF1 | -0.181623437 | 0.881710267 | 0.183990937 | 0.368057883 | 271.1108479 | 239.0421701 | ENSG00000144455 |

| | | | | | | | |
|---|---|---|---|---|---|---|---|
| INTS9 | -0.181694646 | 0.881666747 | 0.114755153 | 0.26436509 | 439.4969059 | 387.4428344 | ENSG00000104299 |
| NA | -0.181795559 | 0.881605079 | 0.442180852 | 0.649844397 | 88.20640313 | 77.73928298 | ENSG00000269837 |
| WIPI2 | -0.181813843 | 0.881593906 | 0.004730411 | 0.021089193 | 1904.859431 | 1679.502692 | ENSG00000157954 |
| RNF138 | -0.181815233 | 0.881593057 | 0.138385105 | 0.301639278 | 734.2712675 | 647.7069887 | ENSG00000134758 |
| CREM | -0.181840039 | 0.881577899 | 0.157925372 | 0.330608653 | 327.2868567 | 288.5448533 | ENSG00000095794 |
| BRK1 | -0.18190986 | 0.881535235 | 0.00688356 | 0.028875174 | 3938.15254 | 3471.673436 | ENSG00000254999 |
| YRDC | -0.182012658 | 0.881472424 | 0.140588456 | 0.304788543 | 354.0177731 | 312.1915621 | ENSG00000196449 |
| UROD | -0.182025006 | 0.881464879 | 0.006458086 | 0.027406956 | 1658.975685 | 1462.409791 | ENSG00000126088 |
| PHF6 | -0.182136154 | 0.881396972 | 0.011135568 | 0.042708204 | 1985.908505 | 1750.145803 | ENSG00000156531 |
| NA | -0.182173811 | 0.881373967 | 0.846599574 | 0.923736594 | 5.267773738 | 4.651917903 | ENSG00000257556 |
| COQ3 | -0.182243883 | 0.881331159 | 0.177654085 | 0.359553895 | 280.2846794 | 247.0643553 | ENSG00000132423 |
| ACP6 | -0.182366313 | 0.881256371 | 0.159863084 | 0.333629611 | 338.8439905 | 298.3985917 | ENSG00000162836 |
| PIGW | -0.182478447 | 0.881187877 | 0.167700525 | 0.344916621 | 323.4253182 | 285.227954 | ENSG00000277161 |
| PEPD | -0.18256224 | 0.881136699 | 0.01229537 | 0.046402492 | 2296.688667 | 2024.061347 | ENSG00000124299 |
| SLC7A1 | -0.182636623 | 0.88109127 | 0.011147434 | 0.042736356 | 3687.729647 | 3248.661226 | ENSG00000139514 |
| NEDD1 | -0.182656247 | 0.881079285 | 0.17009208 | 0.34828027 | 406.7390616 | 358.4505135 | ENSG00000139350 |
| DMXL1 | -0.182667781 | 0.881072241 | 0.105901662 | 0.249029122 | 661.3843534 | 582.2203335 | ENSG00000172869 |
| FLT3 | -0.182672518 | 0.881069348 | 0.76371759 | 0.877575992 | 15.37450968 | 13.47650717 | ENSG00000122025 |
| TMED4 | -0.182721503 | 0.881039433 | 0.045960944 | 0.132011015 | 975.8722132 | 859.9878846 | ENSG00000158604 |
| UBL3 | -0.182744109 | 0.881025628 | 0.073690063 | 0.190270281 | 555.8579004 | 489.7711748 | ENSG00000122042 |
| SNAI3 | -0.182898905 | 0.880931102 | 0.355413747 | 0.567706049 | 136.753609 | 120.6568582 | ENSG00000185669 |
| DHODH | -0.183020911 | 0.880856606 | 0.127010609 | 0.283991871 | 406.0288776 | 357.5883306 | ENSG00000102967 |
| COL9A3 | -0.183113257 | 0.880800225 | 0.049211541 | 0.139376117 | 1835.201853 | 1617.09594 | ENSG00000092758 |
| EGFL8 | -0.183291272 | 0.88069155 | 0.765061201 | 0.878469069 | 17.50427772 | 15.38703957 | ENSG00000241404 |
| AKAP8L | -0.18331596 | 0.880676479 | 0.014267087 | 0.052347763 | 1228.917103 | 1081.997239 | ENSG00000011243 |
| LRRC8D | -0.183496715 | 0.880566146 | 0.044935379 | 0.129696567 | 790.044578 | 695.7273747 | ENSG00000171492 |
| PUM2 | -0.18352504 | 0.880548858 | 0.014456093 | 0.052887121 | 1890.951668 | 1664.845778 | ENSG00000055917 |
| CPNE3 | -0.183560082 | 0.88052747 | 0.07687639 | 0.196137614 | 896.855049 | 789.2628415 | ENSG00000085719 |
| BMS1 | -0.183822922 | 0.880367064 | 0.000747344 | 0.004350675 | 3394.470544 | 2988.285013 | ENSG00000165733 |
| PAM16 | -0.183870432 | 0.880338073 | 0.556981375 | 0.743258428 | 63.20169056 | 55.70924859 | ENSG00000217930 |
| VEZF1 | -0.183998907 | 0.88025968 | 0.038416313 | 0.115175752 | 933.8247062 | 821.9403595 | ENSG00000136451 |
| ZNF32 | -0.184042191 | 0.880233271 | 0.310606036 | 0.521172366 | 156.1020596 | 137.4367603 | ENSG00000169740 |
| ZNF519 | -0.184102707 | 0.880196349 | 0.2575492 | 0.464425636 | 185.8209258 | 163.5864246 | ENSG00000175322 |
| ENGASE | -0.184117758 | 0.880187166 | 0.126852544 | 0.283739227 | 883.677658 | 777.9731796 | ENSG00000167280 |
| HAUS1 | -0.184133345 | 0.880177657 | 0.023637856 | 0.078799808 | 968.1749429 | 852.1647173 | ENSG00000152240 |
| ACVR1B | -0.184151102 | 0.880166824 | 0.01707866 | 0.060569107 | 1024.830739 | 901.9825185 | ENSG00000135503 |
| LYPD1 | -0.18415815 | 0.880162524 | 0.720757213 | 0.852731108 | 19.92762349 | 17.49474089 | ENSG00000150551 |
| NSMAF | -0.184246693 | 0.880108507 | 0.058068566 | 0.158334516 | 605.3368161 | 532.6396602 | ENSG00000035681 |
| C1QTNF2 | -0.184265277 | 0.88009717 | 0.661370265 | 0.818102931 | 26.40291304 | 23.26734541 | ENSG00000145861 |
| RBMXL1 | -0.184268905 | 0.880094957 | 0.082439803 | 0.206536971 | 563.9795859 | 496.1080369 | ENSG00000213516 |
| CMTM3 | -0.184326511 | 0.880059816 | 0.124811996 | 0.280537317 | 455.2711921 | 401.0176163 | ENSG00000140931 |
| AP3S1 | -0.184345107 | 0.880048472 | 0.022291765 | 0.075161575 | 1025.41197 | 902.6755503 | ENSG00000177879 |
| TRMU | -0.184346414 | 0.880047675 | 0.021113272 | 0.071932913 | 1281.258601 | 1127.77376 | ENSG00000100416 |
| TRAPPC13 | -0.184352186 | 0.880044154 | 0.299669316 | 0.509842796 | 176.7983199 | 155.4558654 | ENSG00000113597 |
| CXXC1 | -0.184415511 | 0.880005526 | 0.003844444 | 0.017764212 | 2080.55066 | 1830.911706 | ENSG00000154832 |
| USP48 | -0.184450661 | 0.879984086 | 0.003488308 | 0.01635875 | 2080.984077 | 1831.454094 | ENSG00000090686 |
| DMAC1 | -0.184464633 | 0.879975564 | 0.022768164 | 0.076345309 | 1063.322691 | 935.6385806 | ENSG00000137038 |
| NA | -0.184486175 | 0.879962424 | 0.369349627 | 0.581613339 | 127.9486629 | 112.6568065 | ENSG00000240356 |
| ZFAND3 | -0.184499695 | 0.879954178 | 0.010573115 | 0.040871423 | 1594.872437 | 1402.896532 | ENSG00000156639 |
| DHX33 | -0.184801051 | 0.879770389 | 0.001712377 | 0.008908605 | 2284.374774 | 2009.658614 | ENSG00000005100 |
| PACS1 | -0.184889882 | 0.87971622 | 0.029593547 | 0.093881091 | 847.8714313 | 746.0708126 | ENSG00000175115 |
| UBE2I | -0.185046057 | 0.879620994 | 0.001811102 | 0.009316986 | 3172.879337 | 2790.867978 | ENSG00000103275 |
| INPP5F | -0.185050963 | 0.879618003 | 0.00423606 | 0.019281272 | 1869.775543 | 1644.593185 | ENSG00000198825 |
| NA | -0.18506012 | 0.87961242 | 0.625839993 | 0.793253855 | 36.98117419 | 32.55841198 | ENSG00000218175 |
| RBM22 | -0.185111581 | 0.879581045 | 0.036299861 | 0.110283604 | 751.8934636 | 661.1024796 | ENSG00000086589 |
| NA | -0.185150638 | 0.879557233 | 0.779599734 | 0.887571291 | 11.38424599 | 9.989447567 | ENSG00000248873 |
| DNAJB4 | -0.185331803 | 0.87944679 | 0.292746879 | 0.502547478 | 171.5910566 | 151.0224195 | ENSG00000162616 |
| DPY19L2 | -0.185376108 | 0.879419783 | 0.581064718 | 0.761150628 | 44.57677276 | 39.13342061 | ENSG00000177990 |
| FKBP15 | -0.185433857 | 0.879384582 | 0.085058921 | 0.211499372 | 469.3408595 | 412.7460908 | ENSG00000119321 |
| NHP2 | -0.185482002 | 0.879355235 | 0.011479485 | 0.043831394 | 1845.463113 | 1623.099971 | ENSG00000145912 |
| NAT8L | -0.185488999 | 0.879350971 | 0.042336926 | 0.124035502 | 1005.665041 | 884.6762447 | ENSG00000185818 |
| NAXD | -0.185544869 | 0.879316918 | 0.03540035 | 0.108229823 | 786.0762292 | 691.490501 | ENSG00000213995 |
| USP41 | -0.185579189 | 0.879296 | 0.788419464 | 0.891711403 | 10.13966092 | 8.939700608 | ENSG00000161133 |
| KLK14 | -0.185726192 | 0.879206409 | 0.812205979 | 0.904606623 | 9.472443192 | 8.37572658 | ENSG00000129437 |
| PHF2 | -0.185927705 | 0.879083611 | 0.118491508 | 0.270353041 | 438.1408343 | 385.1081044 | ENSG00000197724 |
| NA | -0.185928781 | 0.879082956 | 0.726096461 | 0.855720453 | 22.27388223 | 19.64152791 | ENSG00000262691 |
| PIGZ | -0.186079729 | 0.878990983 | 0.814710377 | 0.905846657 | 7.521791677 | 6.634125014 | ENSG00000119227 |
| SETD4 | -0.186090292 | 0.878984547 | 0.124696005 | 0.280410105 | 587.0993923 | 516.186901 | ENSG00000185917 |
| NA | -0.186293621 | 0.878860675 | 0.80597234 | 0.901064862 | 8.359811778 | 7.334681475 | ENSG00000228420 |
| DDX5 | -0.186390844 | 0.87880145 | 0.000108761 | 0.000801386 | 9613.143091 | 8448.223935 | ENSG00000108654 |
| ATXN10 | -0.186393633 | 0.878799752 | 0.003741563 | 0.017348247 | 2013.81943 | 1769.701271 | ENSG00000130638 |
| RIMKLB | -0.1864124 | 0.87878832 | 0.018272397 | 0.06384352 | 1178.381644 | 1035.450293 | ENSG00000166532 |
| FAM214A | -0.186518089 | 0.878723944 | 0.481346895 | 0.683198091 | 69.75409329 | 61.27232351 | ENSG00000047346 |
| FYCO1 | -0.186528403 | 0.878717662 | 0.151413313 | 0.321032941 | 360.0633614 | 316.2869354 | ENSG00000163820 |

| | | | | | | | |
|---|---|---|---|---|---|---|---|
| GALNT14 | -0.186548483 | 0.878705432 | 0.121116222 | 0.274260186 | 433.0221745 | 380.6924115 | ENSG00000158089 |
| EXOSC5 | -0.186745505 | 0.878585439 | 0.016571377 | 0.059103014 | 1194.13508 | 1048.804816 | ENSG00000077348 |
| LDHA | -0.186909944 | 0.878485303 | 0.000214596 | 0.001462521 | 9466.064554 | 8315.308578 | ENSG00000134333 |
| GRIA3 | -0.186921385 | 0.878478337 | 0.280055986 | 0.489029772 | 205.3392628 | 180.2124562 | ENSG00000125675 |
| ASF1B | -0.18701892 | 0.878418948 | 0.018295896 | 0.063901978 | 968.8098199 | 851.0661642 | ENSG00000105011 |
| BACE2 | -0.187062779 | 0.878392244 | 0.141450833 | 0.306025775 | 330.4616784 | 290.3720556 | ENSG00000182240 |
| CHCHD2 | -0.187126586 | 0.878353396 | 0.003540715 | 0.016559286 | 3419.491446 | 3004.097849 | ENSG00000106153 |
| EHD1 | -0.187199345 | 0.878309099 | 0.032241468 | 0.100525034 | 1495.186421 | 1313.391234 | ENSG00000110047 |
| NA | -0.187213551 | 0.878300451 | 0.62920659 | 0.795425152 | 32.90103396 | 28.85484489 | ENSG00000127589 |
| MYC | -0.187219585 | 0.878296777 | 0.01135971 | 0.043433097 | 1325.617381 | 1164.34349 | ENSG00000136997 |
| SPATA5L1 | -0.187274458 | 0.878263372 | 0.135995214 | 0.298230426 | 356.9051679 | 313.4325414 | ENSG00000171763 |
| CENPK | -0.187324156 | 0.878233118 | 0.079845095 | 0.201506699 | 540.6977442 | 474.9919247 | ENSG00000123219 |
| SYAP1 | -0.187426681 | 0.878170708 | 0.004222754 | 0.019234613 | 1791.979216 | 1574.004984 | ENSG00000169895 |
| DNAAF5 | -0.187429134 | 0.878169215 | 0.01502043 | 0.054560652 | 1476.951824 | 1297.342978 | ENSG00000164818 |
| PES1 | -0.187530029 | 0.878107803 | 0.005086227 | 0.022410966 | 4493.593533 | 3946.220083 | ENSG00000100029 |
| TMEM170B | -0.187546423 | 0.878097824 | 0.352788531 | 0.5650421 | 191.9252451 | 168.4157213 | ENSG00000205269 |
| SH3D19 | -0.187587745 | 0.878072674 | 0.020866061 | 0.071244903 | 1459.911265 | 1281.670056 | ENSG00000109686 |
| NA | -0.187623171 | 0.878051113 | 0.785007005 | 0.890433415 | 10.54147121 | 9.256542699 | ENSG00000213228 |
| WASH8P | -0.187736346 | 0.877982235 | 0.229008578 | 0.428752552 | 300.1135791 | 263.8507112 | ENSG00000226210 |
| SETD5 | -0.187738915 | 0.877980671 | 0.000826293 | 0.004744467 | 3767.63729 | 3307.734961 | ENSG00000168137 |
| P4HA1 | -0.187766582 | 0.877963835 | 0.087902163 | 0.216403642 | 524.444002 | 460.1932408 | ENSG00000122884 |
| C2orf16 | -0.187840346 | 0.877918946 | 0.757940781 | 0.874279969 | 13.20797786 | 11.59664216 | ENSG00000221843 |
| PSMD14 | -0.187843903 | 0.877916782 | 0.003080699 | 0.014728862 | 2410.430242 | 2116.136693 | ENSG00000115233 |
| DENND1A | -0.187893118 | 0.877886834 | 0.024327053 | 0.080585231 | 968.4453843 | 850.0240575 | ENSG00000119522 |
| ZNF618 | -0.187934588 | 0.877861599 | 0.025211628 | 0.082991387 | 1272.491536 | 1116.557446 | ENSG00000157657 |
| PSPH | -0.188208537 | 0.87769492 | 0.020592515 | 0.070451021 | 1091.242742 | 957.272372 | ENSG00000146733 |
| BABAM1 | -0.18822094 | 0.877687375 | 0.311477079 | 0.52203094 | 232.5746294 | 204.2995797 | ENSG00000105393 |
| SLC43A3 | -0.188345789 | 0.877611424 | 0.164989294 | 0.340638769 | 300.3425026 | 263.6493366 | ENSG00000134802 |
| COPS4 | -0.188371267 | 0.877595926 | 0.048391042 | 0.137609521 | 813.3769839 | 713.9713626 | ENSG00000138663 |
| ARNTL2 | -0.188378319 | 0.877591636 | 0.158641874 | 0.331781081 | 309.2621631 | 271.6264041 | ENSG00000029153 |
| UST | -0.188393101 | 0.877582644 | 0.052407039 | 0.146363047 | 771.6538448 | 676.8881056 | ENSG00000111962 |
| WFIKKN1 | -0.188411752 | 0.877571299 | 0.313285452 | 0.523574846 | 202.2339283 | 177.6632489 | ENSG00000127578 |
| DEDD2 | -0.188556828 | 0.877483056 | 0.008441835 | 0.034128968 | 1319.112447 | 1157.591152 | ENSG00000160570 |
| MFSD1 | -0.18856041 | 0.877480877 | 0.32441562 | 0.535287833 | 153.4136374 | 134.8115047 | ENSG00000118855 |
| NA | -0.188626542 | 0.877440655 | 0.824235208 | 0.911179646 | 7.205382571 | 6.354928216 | ENSG00000226179 |
| THAP1 | -0.18863491 | 0.877435565 | 0.448686525 | 0.655281496 | 94.78077309 | 83.26605771 | ENSG00000131931 |
| COPS3 | -0.188649327 | 0.877426797 | 0.004738135 | 0.021108675 | 1672.22752 | 1467.063041 | ENSG00000141030 |
| ILF3 | -0.188652064 | 0.877425133 | 2.15E-05 | 0.000189837 | 22807.09588 | 20011.13902 | ENSG00000129351 |
| WNT4 | -0.188677988 | 0.877409366 | 0.710964865 | 0.847530076 | 19.26040577 | 16.86294862 | ENSG00000162552 |
| C19orf53 | -0.188690391 | 0.877401823 | 0.004621082 | 0.020667237 | 2057.837567 | 1805.478582 | ENSG00000104979 |
| VPS26A | -0.188880822 | 0.877286016 | 0.055485069 | 0.152901727 | 886.5559251 | 778.0814544 | ENSG00000122958 |
| ARPC1A | -0.188933551 | 0.877253953 | 0.368017812 | 0.580038089 | 111.9840486 | 98.30986608 | ENSG00000241685 |
| MED8 | -0.188982254 | 0.877224339 | 0.225471134 | 0.423763008 | 212.3353515 | 186.3364266 | ENSG00000159479 |
| ADGRL3 | -0.189009261 | 0.877207918 | 0.207888377 | 0.40048679 | 249.6693675 | 218.8702461 | ENSG00000150471 |
| P2RX5 | -0.189034201 | 0.877192754 | 0.595666561 | 0.772289098 | 37.06538016 | 32.50676795 | ENSG00000083454 |
| OSTC | -0.189108383 | 0.87714765 | 0.005179316 | 0.022768013 | 1995.49096 | 1750.397487 | ENSG00000198856 |
| NDUFV1 | -0.189115279 | 0.877143457 | 0.003015738 | 0.014476921 | 2645.460661 | 2321.155433 | ENSG00000167792 |
| ARL15 | -0.189348713 | 0.877001543 | 0.320166216 | 0.530613338 | 203.718472 | 178.8248475 | ENSG00000185305 |
| PNPO | -0.189349176 | 0.877001262 | 0.522635959 | 0.716982427 | 56.89720339 | 49.83619387 | ENSG00000108439 |
| AP5M1 | -0.189402628 | 0.87696877 | 0.05291923 | 0.147292507 | 739.5398804 | 648.4485752 | ENSG00000053770 |
| NA | -0.189528315 | 0.876892372 | 0.452419005 | 0.658542694 | 79.26477481 | 69.50448976 | ENSG00000240489 |
| GNL3 | -0.189550741 | 0.876878741 | 0.000449848 | 0.002805608 | 3316.935253 | 2908.899506 | ENSG00000163938 |
| GPX2 | -0.189718667 | 0.87677668 | 0.819033827 | 0.908070158 | 7.531274886 | 6.615775347 | ENSG00000176153 |
| H2AZ1 | -0.189742758 | 0.87676204 | 0.000749023 | 0.004359103 | 4987.497133 | 4373.195481 | ENSG00000164032 |
| NA | -0.189776709 | 0.876741407 | 0.601102632 | 0.776307564 | 38.68657989 | 33.95734944 | ENSG00000288398 |
| UNC13B | -0.189828003 | 0.876710236 | 0.004621538 | 0.020667237 | 2483.546895 | 2176.747576 | ENSG00000198722 |
| FBRSL1 | -0.189903613 | 0.87666429 | 0.033661333 | 0.103990558 | 3105.524372 | 2723.403614 | ENSG00000112787 |
| PHKB | -0.189942049 | 0.876640934 | 0.016682625 | 0.059398824 | 1005.779069 | 881.4764643 | ENSG00000102893 |
| NA | -0.190008304 | 0.876600676 | 0.622572837 | 0.791044071 | 37.1590955 | 32.56276291 | ENSG00000271976 |
| YME1L1 | -0.190084003 | 0.876554681 | 0.001187405 | 0.006512968 | 3497.306898 | 3065.658773 | ENSG00000136758 |
| SMAD7 | -0.190172389 | 0.876500981 | 0.026307191 | 0.085745823 | 1062.849717 | 931.2889484 | ENSG00000101665 |
| EDC4 | -0.190239341 | 0.876460306 | 0.007336004 | 0.030415026 | 1690.174034 | 1481.86777 | ENSG00000038358 |
| GTF3C3 | -0.190248627 | 0.876454664 | 0.01924667 | 0.066533656 | 1077.204143 | 943.7328193 | ENSG00000119041 |
| UNQ6494 | -0.190285043 | 0.876432542 | 0.811679736 | 0.904393876 | 8.383532877 | 7.329384589 | ENSG00000237372 |
| ERH | -0.190331672 | 0.876404215 | 0.002683071 | 0.013062592 | 3285.006446 | 2878.623316 | ENSG00000100632 |
| THUMPD3-AS1 | -0.190504309 | 0.876299348 | 0.300021343 | 0.510168792 | 188.6223607 | 165.4827903 | ENSG00000206573 |
| SCGB3A2 | -0.19051687 | 0.876291719 | 0.038832312 | 0.116146375 | 1077.102138 | 944.1707227 | ENSG00000164265 |
| RNF167 | -0.190537045 | 0.876279464 | 0.001307806 | 0.007072715 | 2225.523612 | 1950.411083 | ENSG00000108523 |
| CROCC | -0.190569044 | 0.876260029 | 0.01427988 | 0.052384525 | 2267.379089 | 1987.489544 | ENSG00000058453 |
| NCAPD2 | -0.190573141 | 0.876257541 | 0.000281332 | 0.001860932 | 5183.315829 | 4541.859853 | ENSG00000010292 |
| NA | -0.190636392 | 0.876219124 | 0.747634851 | 0.868652896 | 14.00211524 | 12.22502946 | ENSG00000283692 |
| WWP2 | -0.190675255 | 0.876195521 | 0.011590113 | 0.044155596 | 1446.023332 | 1267.26639 | ENSG00000198373 |
| LRPAP1 | -0.190685838 | 0.876189093 | 0.001633849 | 0.008551897 | 2195.001788 | 1923.510804 | ENSG00000163956 |
| DPP9 | -0.190787486 | 0.876127362 | 0.014423431 | 0.052777852 | 1357.522207 | 1189.790064 | ENSG00000142002 |

| | | | | | | | |
|---|---|---|---|---|---|---|---|
| LEO1 | -0.190829822 | 0.876101652 | 0.022603027 | 0.07592639 | 1002.80205 | 878.4211996 | ENSG00000166477 |
| NA | -0.1909244 | 0.87604422 | 0.025198059 | 0.082987771 | 842.6178437 | 738.1751105 | ENSG00000232445 |
| USP1 | -0.190988467 | 0.876005318 | 0.014818647 | 0.054004274 | 1309.518248 | 1147.414477 | ENSG00000162607 |
| SEC61A1 | -0.191021127 | 0.875985487 | 0.000877702 | 0.004994093 | 5144.785611 | 4506.73574 | ENSG00000058262 |
| HYOU1 | -0.191038411 | 0.875974992 | 4.59E-05 | 0.000373131 | 9578.307369 | 8390.087985 | ENSG00000149428 |
| LRRC3 | -0.191079244 | 0.8759502 | 0.156996246 | 0.329101354 | 300.151207 | 263.0164828 | ENSG00000160233 |
| NA | -0.191091914 | 0.875942507 | 0.803665721 | 0.899842175 | 8.694018246 | 7.57623298 | ENSG00000261211 |
| NAIP | -0.191137524 | 0.875914815 | 0.850877583 | 0.926210338 | 4.909846171 | 4.313604722 | ENSG00000249437 |
| ODF2 | -0.191247648 | 0.875847957 | 0.085037512 | 0.211473982 | 512.9360909 | 449.161321 | ENSG00000136811 |
| FNDC3A | -0.19126466 | 0.87583763 | 0.092176928 | 0.224296004 | 553.362501 | 484.8515045 | ENSG00000102531 |
| KANSL2 | -0.191280936 | 0.875827748 | 0.065376754 | 0.173209876 | 596.1093138 | 521.8088384 | ENSG00000139620 |
| CCDC136 | -0.191425482 | 0.875740002 | 0.120167094 | 0.272832423 | 652.0489234 | 571.0188135 | ENSG00000128596 |
| EEF1AKMT4 | -0.19144174 | 0.875730133 | 0.601428181 | 0.776474614 | 49.92313422 | 43.64149464 | ENSG00000284753 |
| SLC33A1 | -0.191467383 | 0.875714568 | 0.467964199 | 0.671349859 | 95.47733016 | 83.70253002 | ENSG00000169359 |
| SPTBN1 | -0.191533931 | 0.875674175 | 0.009680073 | 0.038013762 | 5837.89492 | 5111.303836 | ENSG00000115306 |
| SERPING1 | -0.191627809 | 0.875617195 | 0.055358395 | 0.152641742 | 715.3109753 | 626.4980143 | ENSG00000149131 |
| ZNF341 | -0.191658056 | 0.875598837 | 0.183501453 | 0.367333575 | 245.2188674 | 214.5829149 | ENSG00000131061 |
| NCK2 | -0.191804394 | 0.875510027 | 0.010216203 | 0.039736932 | 1587.159277 | 1390.224334 | ENSG00000071051 |
| NA | -0.19183909 | 0.875488972 | 0.805464571 | 0.900817151 | 7.579912292 | 6.648123755 | ENSG00000217261 |
| CALM1 | -0.191904164 | 0.875449482 | 0.000284493 | 0.001878549 | 6252.604723 | 5474.390738 | ENSG00000198668 |
| PYDC1 | -0.192092912 | 0.875334955 | 0.783120476 | 0.889784424 | 9.884931902 | 8.679799436 | ENSG00000169900 |
| SLTM | -0.192093474 | 0.875334614 | 0.00040936 | 0.002580362 | 3758.09574 | 3290.122627 | ENSG00000137776 |
| DPY19L4 | -0.19214488 | 0.875303424 | 0.184554674 | 0.36874397 | 249.6255104 | 218.5921632 | ENSG00000156162 |
| VCPKMT | -0.192169185 | 0.875288678 | 0.341709787 | 0.553351855 | 131.6080264 | 115.34638 | ENSG00000100483 |
| SIM1 | -0.192273788 | 0.875225218 | 0.843590258 | 0.92205128 | 4.916965116 | 4.316726144 | ENSG00000112246 |
| MIS18A | -0.192333392 | 0.875189059 | 0.012366591 | 0.046643314 | 1846.58322 | 1616.575429 | ENSG00000159055 |
| NA | -0.19236162 | 0.875171935 | 0.619461622 | 0.788725553 | 34.83597305 | 30.54915335 | ENSG00000264070 |
| RUBCNL | -0.19247353 | 0.875104051 | 0.42787373 | 0.637657464 | 82.29365822 | 72.05161686 | ENSG00000102445 |
| TOP2A | -0.192544691 | 0.875060887 | 0.000437176 | 0.00273561 | 7800.202356 | 6825.143642 | ENSG00000131747 |
| FMNL2 | -0.192715682 | 0.874957179 | 0.008096474 | 0.032975774 | 1327.511413 | 1161.677982 | ENSG00000157827 |
| CDYL | -0.192799954 | 0.874906072 | 0.006099896 | 0.026103907 | 1767.751201 | 1546.544218 | ENSG00000153046 |
| MYO1B | -0.192833355 | 0.874885817 | 0.004774733 | 0.021231653 | 2053.353078 | 1796.004341 | ENSG00000128641 |
| NA | -0.192849798 | 0.874875845 | 0.685065322 | 0.831623057 | 24.72003323 | 21.67727097 | ENSG00000228506 |
| CDC34 | -0.192885403 | 0.874854254 | 0.000666683 | 0.003935699 | 2770.913342 | 2424.369319 | ENSG00000099804 |
| PPP1R3E | -0.192902034 | 0.874844169 | 0.565213323 | 0.749341839 | 49.54237528 | 43.34300222 | ENSG00000235194 |
| ITGA6 | -0.192911814 | 0.874838239 | 0.006682998 | 0.028184104 | 2436.741423 | 2130.981711 | ENSG00000091409 |
| ATP5MG | -0.19292938 | 0.874827586 | 0.387915208 | 0.600150584 | 119.7204993 | 104.7028911 | ENSG00000167283 |
| NA | -0.192940627 | 0.874820767 | 0.351002156 | 0.563088321 | 139.0206441 | 121.4692889 | ENSG00000204789 |
| CELF2 | -0.192953999 | 0.874812658 | 0.097044608 | 0.232958869 | 550.3591449 | 481.3889944 | ENSG00000048740 |
| GSPT1 | -0.193052042 | 0.87475321 | 0.00062037 | 0.003693469 | 4838.156338 | 4232.505785 | ENSG00000103342 |
| MIR924HG | -0.193086276 | 0.874732453 | 0.507957097 | 0.704869733 | 58.83333768 | 51.49692442 | ENSG00000267374 |
| NA | -0.193095896 | 0.87472662 | 0.83372283 | 0.916357563 | 5.676702975 | 4.939816555 | ENSG00000238186 |
| NA | -0.193201395 | 0.874662657 | 0.832662825 | 0.91561881 | 5.703983547 | 4.963463108 | ENSG00000231381 |
| TMEM250 | -0.193252867 | 0.874631451 | 0.006830881 | 0.028692433 | 1442.536524 | 1261.873419 | ENSG00000238227 |
| ANKRD28 | -0.19328253 | 0.874613468 | 0.025666351 | 0.084082043 | 1029.680379 | 900.3016714 | ENSG00000206560 |
| TBC1D22B | -0.193411485 | 0.874535295 | 0.075203148 | 0.192989734 | 481.4332313 | 420.9135602 | ENSG00000065491 |
| NA | -0.193474833 | 0.874496895 | 0.306831693 | 0.517146011 | 182.2018855 | 159.1886165 | ENSG00000204959 |
| GRM8 | -0.193475999 | 0.874496188 | 0.542082081 | 0.731177781 | 50.98949249 | 44.51389245 | ENSG00000179603 |
| JMY | -0.193503823 | 0.874479323 | 0.167904027 | 0.345258686 | 287.8829732 | 251.892749 | ENSG00000152409 |
| ARF5 | -0.193513024 | 0.874473746 | 0.004210439 | 0.019201677 | 2141.074224 | 1872.254762 | ENSG00000004059 |
| STIP1 | -0.193550225 | 0.874451197 | 5.93E-05 | 0.000468484 | 12144.96546 | 10619.42386 | ENSG00000168439 |
| GSE1 | -0.19359918 | 0.874421525 | 0.002150704 | 0.010807888 | 2668.676166 | 2333.237601 | ENSG00000131149 |
| GLI4 | -0.193613795 | 0.874412666 | 0.548258527 | 0.735934486 | 56.01886003 | 49.02631741 | ENSG00000250571 |
| NA | -0.193614059 | 0.874412507 | 0.680487697 | 0.829255013 | 23.06326493 | 20.16880261 | ENSG00000280649 |
| TRIM27 | -0.19362357 | 0.874406742 | 0.002413148 | 0.011951032 | 2197.066927 | 1921.498984 | ENSG00000204713 |
| ELOVL7 | -0.193647247 | 0.874392392 | 0.487095227 | 0.688003853 | 77.91729667 | 68.00612071 | ENSG00000164181 |
| IZUMO1 | -0.193719211 | 0.874348776 | 0.74448515 | 0.867135375 | 14.8386608 | 13.00399793 | ENSG00000182264 |
| NXT2 | -0.193746474 | 0.874332254 | 0.190270864 | 0.376652545 | 386.9413324 | 338.4565783 | ENSG00000101888 |
| NA | -0.193810853 | 0.874293239 | 0.856052614 | 0.929646557 | 5.65534614 | 4.925817815 | ENSG00000286408 |
| SETD6 | -0.193824061 | 0.874285234 | 0.041585577 | 0.122289572 | 695.0896695 | 608.0114563 | ENSG00000103037 |
| FAM161A | -0.193967253 | 0.874198463 | 0.139202131 | 0.302930657 | 300.8816055 | 263.0491148 | ENSG00000170264 |
| SEZ6 | -0.1940474 | 0.8741499 | 0.458662889 | 0.663997598 | 73.68504116 | 64.46043918 | ENSG00000063015 |
| SLAIN1 | -0.19419183 | 0.874062392 | 0.063189155 | 0.168811317 | 563.4380403 | 492.2223182 | ENSG00000139737 |
| PSMG3 | -0.194342605 | 0.873971049 | 0.088806614 | 0.218218436 | 765.676444 | 669.4942257 | ENSG00000157778 |
| PRPS1 | -0.194379884 | 0.873948466 | 0.000211794 | 0.001447082 | 6933.246173 | 6058.704064 | ENSG00000147224 |
| MED25 | -0.194500114 | 0.873875636 | 0.009666078 | 0.037966697 | 1489.865509 | 1302.180096 | ENSG00000104973 |
| USP14 | -0.19457498 | 0.873830289 | 0.006241783 | 0.0266146 | 2220.321051 | 1940.111336 | ENSG00000101557 |
| CTDSPL2 | -0.194631386 | 0.873796126 | 0.052837859 | 0.147109388 | 723.5253295 | 631.918264 | ENSG00000137770 |
| WNT5A | -0.194654124 | 0.873782354 | 0.490128226 | 0.69013064 | 62.90516375 | 55.01661598 | ENSG00000114251 |
| SEC31B | -0.194761053 | 0.873717594 | 0.612926031 | 0.78462019 | 55.43200948 | 48.53356659 | ENSG00000075826 |
| CNMD | -0.194886482 | 0.873641635 | 0.000550921 | 0.003334661 | 3305.421742 | 2887.590751 | ENSG00000136110 |
| GRB10 | -0.19491769 | 0.873622737 | 0.003718711 | 0.017259961 | 2999.36929 | 2619.6998 | ENSG00000106070 |
| UNC5A | -0.194937078 | 0.873610997 | 0.200388617 | 0.390984506 | 312.8580933 | 273.2982526 | ENSG00000113763 |
| TXNDC15 | -0.194938653 | 0.873610043 | 0.045569635 | 0.131066488 | 616.2875215 | 538.5509273 | ENSG00000113621 |

| | | | | | | | |
|---|---|---|---|---|---|---|---|
| NA | -0.194980991 | 0.873584406 | 0.758976957 | 0.874877523 | 13.36217808 | 11.65481259 | ENSG00000250397 |
| OTUD5 | -0.195000449 | 0.873572624 | 0.003497017 | 0.016387384 | 1590.836588 | 1389.998558 | ENSG00000068308 |
| RALY | -0.195026631 | 0.873556771 | 0.000506634 | 0.003111474 | 5092.583559 | 4449.194255 | ENSG00000125970 |
| RMST | -0.195103858 | 0.873510011 | 0.395824994 | 0.607686774 | 119.9058712 | 104.6858865 | ENSG00000255794 |
| DDA1 | -0.195214968 | 0.873442739 | 0.008670681 | 0.034906374 | 1214.036923 | 1060.426493 | ENSG00000130311 |
| MXD1 | -0.195219612 | 0.873439927 | 0.637141585 | 0.800826545 | 28.14271826 | 24.58446589 | ENSG00000059728 |
| TMEM87A | -0.195455185 | 0.873297318 | 0.127933076 | 0.285475 | 374.7979862 | 327.3684526 | ENSG00000103978 |
| LOC100505501 | -0.195561457 | 0.873232992 | 0.376527149 | 0.588684097 | 101.1916872 | 88.3035819 | ENSG00000167912 |
| C15orf61 | -0.195586807 | 0.873217648 | 0.695587077 | 0.837485087 | 22.75781357 | 19.77979135 | ENSG00000189227 |
| JUP | -0.195630642 | 0.873191117 | 0.002499354 | 0.012312014 | 3186.873605 | 2783.417428 | ENSG00000173801 |
| STX10 | -0.195673164 | 0.87316538 | 0.045104282 | 0.130084661 | 944.1546167 | 824.4967392 | ENSG00000104915 |
| NAV2 | -0.195735594 | 0.873127597 | 0.024274959 | 0.080455003 | 1306.887154 | 1140.608329 | ENSG00000166833 |
| GCAT | -0.195748741 | 0.87311964 | 0.044575498 | 0.128827932 | 691.9300015 | 604.3920496 | ENSG00000100116 |
| UNC50 | -0.195997542 | 0.872969078 | 0.062978742 | 0.168415966 | 484.3197881 | 422.7184182 | ENSG00000115446 |
| LINC01816 | -0.196000594 | 0.872967232 | 0.699030677 | 0.839692979 | 20.47028584 | 17.86322701 | ENSG00000231327 |
| RASGEF1B | -0.19604528 | 0.872940193 | 0.495977468 | 0.695334407 | 61.42868103 | 53.60462574 | ENSG00000138670 |
| HSPA8 | -0.196076943 | 0.872921035 | 0.000133614 | 0.001003463 | 46074.29342 | 40219.18071 | ENSG00000109971 |
| DPP7 | -0.196086404 | 0.87291531 | 0.147452886 | 0.315041041 | 308.7073223 | 269.5010156 | ENSG00000176978 |
| NAA60 | -0.196102885 | 0.872905338 | 0.028336322 | 0.090807983 | 766.0003031 | 668.8718571 | ENSG00000122390 |
| PDCD5 | -0.196226172 | 0.872830747 | 0.022959754 | 0.076878539 | 1164.737605 | 1016.417079 | ENSG00000105185 |
| NA | -0.196276764 | 0.872800139 | 0.356112827 | 0.568384838 | 118.5476885 | 103.4338417 | ENSG00000242474 |
| SNRPC | -0.196371447 | 0.87274286 | 0.00099465 | 0.005590464 | 2422.900399 | 2114.5043 | ENSG00000124562 |
| RWDD3 | -0.196513952 | 0.872656657 | 0.682041056 | 0.830032766 | 23.27914523 | 20.30538504 | ENSG00000122481 |
| SRP14 | -0.19652474 | 0.872650131 | 0.000580102 | 0.003492262 | 5871.86587 | 5124.125026 | ENSG00000140319 |
| ARHGAP39 | -0.196538773 | 0.872641643 | 0.086160518 | 0.213451578 | 569.4650705 | 497.1996644 | ENSG00000147799 |
| NUDT8 | -0.196542877 | 0.872639161 | 0.395556474 | 0.607422671 | 102.8630245 | 89.91701934 | ENSG00000167799 |
| SNX3 | -0.196737421 | 0.872521495 | 0.003424296 | 0.016126643 | 1601.550693 | 1397.708608 | ENSG00000112335 |
| PLAC1 | -0.196753174 | 0.872511968 | 0.797741589 | 0.896947216 | 8.366930723 | 7.264687775 | ENSG00000170965 |
| PRR5 | -0.196761866 | 0.872506711 | 0.198680802 | 0.388978534 | 230.4623794 | 201.2287079 | ENSG00000186654 |
| NDOR1 | -0.19678587 | 0.872492195 | 0.040714602 | 0.120403268 | 789.6401241 | 689.1416996 | ENSG00000188566 |
| SNHG3 | -0.196821786 | 0.872470474 | 0.312095155 | 0.52250938 | 1203.153395 | 1049.717438 | ENSG00000242125 |
| ST3GAL4 | -0.196900413 | 0.872422926 | 0.127291673 | 0.284451928 | 413.9267252 | 361.2545359 | ENSG00000110080 |
| TMPO | -0.197074292 | 0.872317784 | 0.00201514 | 0.010230088 | 3156.065446 | 2753.254959 | ENSG00000120802 |
| BACE1 | -0.197118986 | 0.872290761 | 0.336123747 | 0.54816036 | 131.5703729 | 114.785033 | ENSG00000186318 |
| DDX52 | -0.197189468 | 0.872248147 | 0.04289854 | 0.125234029 | 918.4345881 | 801.5081258 | ENSG00000278053 |
| PAICS | -0.197202543 | 0.872240241 | 4.69E-05 | 0.000380338 | 10413.15262 | 9082.540854 | ENSG00000128050 |
| ADNP | -0.197282781 | 0.872191732 | 0.00331885 | 0.015688729 | 2495.316934 | 2175.748171 | ENSG00000101126 |
| ADCY5 | -0.197284855 | 0.872190478 | 0.052560296 | 0.146612739 | 549.623102 | 479.5043497 | ENSG00000173175 |
| MYO9B | -0.197352304 | 0.872149702 | 0.002781503 | 0.013468895 | 2346.547556 | 2047.120027 | ENSG00000099331 |
| THAP12 | -0.197424239 | 0.872106216 | 0.033753866 | 0.104259365 | 896.9221459 | 782.0537925 | ENSG00000137492 |
| NA | -0.197459563 | 0.872084863 | 0.684127437 | 0.831079275 | 24.17826061 | 21.12055838 | ENSG00000280273 |
| NA | -0.197565027 | 0.872021114 | 0.695669151 | 0.837490401 | 20.22387097 | 17.64220064 | ENSG00000180211 |
| ZNF112 | -0.197638363 | 0.871976789 | 0.574397612 | 0.756661475 | 40.3208223 | 35.16806043 | ENSG00000062370 |
| UCKL1-AS1 | -0.197734152 | 0.871918895 | 0.746189677 | 0.867969455 | 12.57871912 | 10.97355175 | ENSG00000280213 |
| FANCI | -0.197748403 | 0.871910282 | 0.0042692 | 0.019408726 | 1787.429715 | 1558.079654 | ENSG00000140525 |
| SLITRK5 | -0.197773635 | 0.871895033 | 0.293421738 | 0.503173226 | 140.3554106 | 122.4368401 | ENSG00000165300 |
| XRCC4 | -0.197774391 | 0.871894576 | 0.25628264 | 0.463107876 | 166.1981432 | 144.8216454 | ENSG00000152422 |
| SAT1 | -0.197845958 | 0.871851325 | 0.059754372 | 0.161901386 | 693.2854115 | 604.0618712 | ENSG00000130066 |
| LOC102724250 | -0.197965095 | 0.871779331 | 0.322269697 | 0.532891388 | 142.4854841 | 124.1924404 | ENSG00000271254 |
| PSMA4 | -0.19798501 | 0.871767298 | 0.000319311 | 0.002075082 | 5293.656975 | 4614.330064 | ENSG00000041357 |
| SCN2A | -0.197995441 | 0.871760994 | 0.627317208 | 0.793729923 | 28.19847461 | 24.56517026 | ENSG00000136531 |
| AP2B1 | -0.198064857 | 0.87171905 | 0.000442249 | 0.002765521 | 5471.573813 | 4769.123153 | ENSG00000006125 |
| CALCOCO2 | -0.198069221 | 0.871716413 | 0.118667803 | 0.270657182 | 384.1479334 | 334.810206 | ENSG00000136436 |
| THUMPD1 | -0.198088957 | 0.871704488 | 0.029603684 | 0.093885868 | 1097.456313 | 956.4328991 | ENSG00000066654 |
| PPID | -0.19809329 | 0.87170187 | 0.009637172 | 0.037876795 | 1179.045074 | 1027.941321 | ENSG00000171497 |
| CCDC85B | -0.198093654 | 0.87170165 | 0.031503107 | 0.098793784 | 1621.645837 | 1414.031778 | ENSG00000175602 |
| DTX1 | -0.1981845 | 0.871646761 | 0.108354648 | 0.253359027 | 342.8725181 | 298.7755688 | ENSG00000135144 |
| NANOGP8 | -0.198258514 | 0.871602045 | 0.249758673 | 0.455015679 | 170.394804 | 148.4781303 | ENSG00000255192 |
| HSPA5 | -0.198260534 | 0.871600824 | 0.000151024 | 0.001074231 | 21031.85461 | 18331.17045 | ENSG00000044574 |
| NA | -0.198271867 | 0.871593978 | 0.488293472 | 0.688779985 | 67.42624231 | 58.90623407 | ENSG00000261188 |
| SYT17 | -0.198338425 | 0.871553768 | 0.528841926 | 0.721408536 | 67.04373007 | 58.47182163 | ENSG00000103528 |
| UBL7 | -0.198468124 | 0.871475418 | 0.009120704 | 0.036262001 | 1224.227001 | 1066.835336 | ENSG00000138629 |
| ZNF254 | -0.198523801 | 0.871441787 | 0.166436488 | 0.342764785 | 277.2039301 | 241.655108 | ENSG00000213096 |
| TMED5 | -0.198600522 | 0.871395446 | 0.151080529 | 0.320579302 | 357.469265 | 311.7275438 | ENSG00000117500 |
| ITGA1 | -0.198662834 | 0.87135781 | 0.304547205 | 0.514710396 | 147.9178048 | 128.8052726 | ENSG00000213949 |
| SEC11A | -0.198707032 | 0.871331115 | 0.000506088 | 0.003109554 | 3546.789698 | 3090.02216 | ENSG00000140612 |
| MEST | -0.198730779 | 0.871316773 | 0.002492441 | 0.012296174 | 2174.095545 | 1894.303182 | ENSG00000106484 |
| STARD10 | -0.198748273 | 0.871306208 | 0.098419363 | 0.235366592 | 355.863726 | 310.0536879 | ENSG00000214530 |
| PMS1 | -0.198777566 | 0.871288517 | 0.025390815 | 0.083411847 | 901.7982024 | 786.158448 | ENSG00000064933 |
| ASPHD2 | -0.198836082 | 0.871253177 | 0.193861433 | 0.381958599 | 232.237169 | 202.3378548 | ENSG00000128203 |
| PRPF19 | -0.198840408 | 0.871250565 | 0.000167481 | 0.001175789 | 4305.19387 | 3750.748134 | ENSG00000110107 |
| INPP4A | -0.19893655 | 0.871192506 | 0.022616651 | 0.075945137 | 899.2780893 | 783.1999652 | ENSG00000040933 |
| CDK7 | -0.198949975 | 0.8711844 | 0.016236252 | 0.058160398 | 1135.267012 | 988.6600093 | ENSG00000134058 |
| NA | -0.198953923 | 0.871182015 | 0.656311926 | 0.81429937 | 27.17332932 | 23.70364977 | ENSG00000273142 |

| | | | | | | | |
|---|---|---|---|---|---|---|---|
| PAK1 | -0.198984271 | 0.87116369 | 0.000192059 | 0.001331019 | 4247.179894 | 3699.374805 | ENSG00000149269 |
| GABRA5 | -0.199036379 | 0.871132226 | 0.075778401 | 0.194228434 | 474.8918124 | 413.5746434 | ENSG00000186297 |
| FRRS1L | -0.19904776 | 0.871125353 | 0.541275159 | 0.73050678 | 51.39547265 | 44.83101809 | ENSG00000260230 |
| NDUFAF5 | -0.19914972 | 0.87106379 | 0.200073301 | 0.390461886 | 261.7846224 | 227.8583537 | ENSG00000101247 |
| NRM | -0.199158987 | 0.871058195 | 0.075624394 | 0.193923343 | 500.7345962 | 436.5192828 | ENSG00000137404 |
| MICAL1 | -0.199203624 | 0.871031245 | 0.059403745 | 0.161160714 | 1570.784157 | 1368.46849 | ENSG00000135596 |
| TRABD2B | -0.199216223 | 0.871023638 | 0.286285029 | 0.495500079 | 154.9212657 | 135.0804866 | ENSG00000269113 |
| GON7 | -0.199307073 | 0.87096879 | 0.301264564 | 0.511405061 | 139.127707 | 121.253843 | ENSG00000170270 |
| NA | -0.199331657 | 0.870953948 | 0.711548019 | 0.847690156 | 17.59532331 | 15.33009865 | ENSG00000271032 |
| RDH14 | -0.199505851 | 0.870848794 | 0.178482238 | 0.360688825 | 242.4248563 | 211.1457754 | ENSG00000240857 |
| TIMM10 | -0.199561097 | 0.870815446 | 0.018261855 | 0.063818491 | 1082.078978 | 942.4016478 | ENSG00000134809 |
| NA | -0.199606727 | 0.870787904 | 0.177508267 | 0.359297281 | 405.2192031 | 352.7974344 | ENSG00000224114 |
| GRPEL2 | -0.199706752 | 0.870727533 | 0.1109407 | 0.25814627 | 404.8088772 | 352.4969869 | ENSG00000164284 |
| CCDC150 | -0.19984618 | 0.870643386 | 0.236392447 | 0.438448644 | 265.5999394 | 231.418687 | ENSG00000144395 |
| NA | -0.199955388 | 0.870577483 | 0.073640874 | 0.190220131 | 454.6614068 | 395.6712692 | ENSG00000184809 |
| ESRP1 | -0.199988759 | 0.870557346 | 0.000862063 | 0.004918446 | 2822.622886 | 2457.097216 | ENSG00000104413 |
| TLN2 | -0.200125987 | 0.870474544 | 0.013872724 | 0.051099356 | 1074.956684 | 935.6634661 | ENSG00000171914 |
| PTPN2 | -0.20017807 | 0.870443119 | 0.039808525 | 0.11831665 | 744.7331334 | 647.9627202 | ENSG00000175354 |
| UBR2 | -0.200217157 | 0.870419536 | 0.055586968 | 0.153115503 | 700.1080287 | 608.8615621 | ENSG00000024048 |
| ZC4H2 | -0.200242923 | 0.870403991 | 0.048370758 | 0.137572555 | 580.2404476 | 504.8981452 | ENSG00000126970 |
| ILVBL | -0.200256819 | 0.870395608 | 0.011047499 | 0.042433725 | 1399.508417 | 1218.606864 | ENSG00000105135 |
| ST3GAL5 | -0.200321931 | 0.870356325 | 0.496911062 | 0.695999808 | 64.23781976 | 55.85375487 | ENSG00000115525 |
| KLHL25 | -0.200531427 | 0.870229949 | 0.040388011 | 0.119638925 | 800.3976497 | 696.935597 | ENSG00000183655 |
| PLS1 | -0.200544695 | 0.870221945 | 0.086452414 | 0.213879248 | 553.8953491 | 481.9290384 | ENSG00000120756 |
| NA | -0.200714789 | 0.870119352 | 0.834678931 | 0.917141538 | 4.947805159 | 4.307078331 | ENSG00000275318 |
| UBE2L3 | -0.200758843 | 0.870092783 | 0.000155626 | 0.001102398 | 3780.01769 | 3288.968437 | ENSG00000185651 |
| NUP160 | -0.200766352 | 0.870088254 | 0.000419689 | 0.002639305 | 3534.571097 | 3074.777399 | ENSG00000030066 |
| TALDO1 | -0.200773981 | 0.870083653 | 0.000143822 | 0.001029208 | 5197.571489 | 4521.914644 | ENSG00000177156 |
| XRCC6 | -0.200802001 | 0.870066755 | 2.09E-05 | 0.000184925 | 16389.72781 | 14259.33746 | ENSG00000196419 |
| CLSTN3 | -0.200820862 | 0.87005538 | 0.020273446 | 0.069535784 | 954.6078934 | 830.9569294 | ENSG00000139182 |
| CLGN | -0.200836512 | 0.870045942 | 0.26600996 | 0.473433051 | 168.7460182 | 146.7394392 | ENSG00000153132 |
| BAZ1B | -0.20091117 | 0.870000919 | 0.000159971 | 0.001130208 | 3579.906846 | 3114.482025 | ENSG00000009954 |
| AQR | -0.200935132 | 0.869986469 | 0.003134962 | 0.014958 | 2017.294922 | 1755.113871 | ENSG00000021776 |
| RTL8A | -0.200971385 | 0.869964608 | 0.051849975 | 0.145150724 | 576.9088337 | 502.1469237 | ENSG00000203950 |
| PAK1IP1 | -0.2010303 | 0.869929082 | 0.009190452 | 0.0364702 | 1179.95273 | 1026.660237 | ENSG00000111845 |
| IL6ST | -0.201051862 | 0.86991608 | 0.378800419 | 0.591210406 | 102.1533724 | 88.97178995 | ENSG00000134352 |
| AP1M1 | -0.201113721 | 0.869878781 | 0.022611776 | 0.075942272 | 792.2830849 | 688.9407766 | ENSG00000072958 |
| PIP4K2C | -0.201275988 | 0.869780947 | 0.037459142 | 0.112897526 | 641.4188488 | 557.8802703 | ENSG00000166908 |
| CENPO | -0.20132579 | 0.869750923 | 0.056712247 | 0.155557122 | 611.4639139 | 531.9098768 | ENSG00000138092 |
| SYNJ1 | -0.201425541 | 0.869690788 | 0.517172874 | 0.713270264 | 49.92610888 | 43.40909649 | ENSG00000159082 |
| NA | -0.201458257 | 0.869671067 | 0.489229828 | 0.68943402 | 58.51484364 | 50.83741822 | ENSG00000255857 |
| ZBTB21 | -0.201461078 | 0.869669366 | 0.119305952 | 0.271732614 | 368.3439495 | 319.9885379 | ENSG00000173276 |
| TPD52L1 | -0.201494135 | 0.869649439 | 0.412960394 | 0.624350096 | 104.210477 | 90.46642753 | ENSG00000111907 |
| IFT81 | -0.201522074 | 0.869632598 | 0.137077197 | 0.299791879 | 312.744164 | 272.2154651 | ENSG00000122970 |
| NA | -0.201633699 | 0.869565315 | 0.815044255 | 0.905897873 | 7.300266966 | 6.358049639 | ENSG00000287620 |
| PHLDA1 | -0.201664795 | 0.869546573 | 0.078368332 | 0.19876255 | 461.2212856 | 401.0732847 | ENSG00000139289 |
| PHKG2 | -0.201670428 | 0.869543178 | 0.128914915 | 0.287209334 | 406.1299912 | 353.3139701 | ENSG00000156873 |
| ATG3 | -0.201707349 | 0.869520925 | 0.027367905 | 0.088415385 | 737.1141442 | 640.8634724 | ENSG00000144848 |
| SRSF2 | -0.201764814 | 0.869486291 | 0.002282229 | 0.011371649 | 2618.120499 | 2277.026367 | ENSG00000161547 |
| CAST | -0.201851747 | 0.8694339 | 0.084007541 | 0.2095196 | 438.3958427 | 381.0168509 | ENSG00000153113 |
| ACAA2 | -0.201882003 | 0.869415666 | 0.003015349 | 0.014476921 | 1806.91223 | 1570.796554 | ENSG00000167315 |
| DLGAP5 | -0.201903769 | 0.869402549 | 0.0015323 | 0.008094402 | 2832.882725 | 2462.252572 | ENSG00000126787 |
| ADGRB1 | -0.202053819 | 0.86931213 | 0.04494241 | 0.129697038 | 590.5378676 | 513.5819394 | ENSG00000181790 |
| NA | -0.202276584 | 0.869177911 | 0.440126783 | 0.647819643 | 70.6407508 | 61.34014174 | ENSG00000223804 |
| ADAM9 | -0.202436037 | 0.869081851 | 0.057243239 | 0.156626856 | 506.0263448 | 439.79118 | ENSG00000168615 |
| ZNF326 | -0.20253118 | 0.869024538 | 0.03756588 | 0.113058937 | 884.9972957 | 769.2751195 | ENSG00000162664 |
| TTL | -0.202594235 | 0.868986558 | 0.009205321 | 0.036513859 | 1120.982737 | 974.1201877 | ENSG00000114999 |
| HMGB1 | -0.202701163 | 0.868922153 | 0.000218472 | 0.001487326 | 15557.05388 | 13517.53948 | ENSG00000189403 |
| EIF4H | -0.202781002 | 0.868874068 | 4.50E-06 | 4.61E-05 | 10567.71825 | 9181.738252 | ENSG00000106682 |
| C22orf46 | -0.202840991 | 0.86883794 | 0.128869135 | 0.287209334 | 340.9170863 | 295.8686574 | ENSG00000184208 |
| NA | -0.202974472 | 0.868757558 | 0.800298388 | 0.898320559 | 7.225544198 | 6.284934516 | ENSG00000267742 |
| TMEM97 | -0.203008768 | 0.868736905 | 0.001594324 | 0.008382183 | 2316.687072 | 2012.640657 | ENSG00000109084 |
| CD59 | -0.203125746 | 0.868666468 | 0.063604431 | 0.169537006 | 491.7057356 | 427.0580559 | ENSG00000085063 |
| NET1 | -0.203126246 | 0.868666167 | 0.004288481 | 0.019477624 | 1855.064893 | 1611.099547 | ENSG00000173848 |
| PKM | -0.203141242 | 0.868657138 | 9.68E-05 | 0.000720194 | 20250.48864 | 17589.57976 | ENSG00000067225 |
| ARL16 | -0.203196088 | 0.868624116 | 0.071158894 | 0.185228907 | 433.4874187 | 376.3971135 | ENSG00000214087 |
| NA | -0.203532875 | 0.868421365 | 0.630249105 | 0.796470479 | 28.7343235 | 24.88418782 | ENSG00000286067 |
| NARF | -0.203589508 | 0.868387276 | 0.008790759 | 0.035284483 | 1200.209476 | 1042.025397 | ENSG00000141562 |
| MAPK9 | -0.203599332 | 0.868381363 | 0.012283507 | 0.046399988 | 1027.23011 | 892.020574 | ENSG00000050748 |
| NA | -0.203602276 | 0.86837959 | 0.801198217 | 0.898333333 | 7.699712995 | 6.676121235 | ENSG00000254331 |
| ZMIZ1 | -0.203692798 | 0.868325106 | 0.003853418 | 0.017796967 | 1493.449209 | 1296.551902 | ENSG00000108175 |
| LDLRAD2 | -0.203865835 | 0.868220965 | 0.812396932 | 0.904712588 | 6.878295048 | 5.996089905 | ENSG00000187942 |
| CAND1 | -0.203882977 | 0.868210649 | 0.001737014 | 0.009011951 | 4911.602202 | 4263.632243 | ENSG00000111530 |
| DBF4 | -0.203966984 | 0.868160095 | 0.028378574 | 0.090891106 | 844.1273025 | 733.242061 | ENSG00000006634 |

| | | | | | | | |
|---|---|---|---|---|---|---|---|
| CERCAM | -0.203984734 | 0.868149414 | 0.260526261 | 0.46750455 | 152.9023999 | 132.729131 | ENSG00000167123 |
| PDE7A | -0.204158819 | 0.868044664 | 0.004107844 | 0.018801246 | 1382.5019 | 1199.882074 | ENSG00000205268 |
| SPDL1 | -0.204158888 | 0.868044622 | 0.025524056 | 0.08380073 | 938.3110536 | 814.7140427 | ENSG00000040275 |
| ERRFI1 | -0.204239476 | 0.867996135 | 0.005176524 | 0.022761035 | 1721.055768 | 1493.242436 | ENSG00000116285 |
| TUBGCP2 | -0.204257954 | 0.867985018 | 0.001078879 | 0.005989013 | 1866.18175 | 1619.937634 | ENSG00000130640 |
| TAF1D | -0.204368669 | 0.86791841 | 0.01628659 | 0.058263357 | 1103.55471 | 957.6682159 | ENSG00000166012 |
| PNN | -0.204379684 | 0.867911783 | 0.171783406 | 0.351020301 | 4533.519294 | 3934.807736 | ENSG00000100941 |
| ADGRA3 | -0.204479078 | 0.867851991 | 0.003480385 | 0.016329708 | 1787.503853 | 1550.966975 | ENSG00000152990 |
| PHF10 | -0.204627898 | 0.867762473 | 0.045021397 | 0.129885286 | 1044.270645 | 906.1616342 | ENSG00000130024 |
| NA | -0.204661123 | 0.867742489 | 0.842953953 | 0.9217804 | 5.022527928 | 4.356546901 | ENSG00000286136 |
| MCRS1 | -0.204722458 | 0.867705598 | 0.001428379 | 0.007643789 | 1931.077558 | 1675.571163 | ENSG00000187778 |
| NA | -0.204762907 | 0.867681271 | 0.8418695 | 0.92140286 | 4.909846171 | 4.255434298 | ENSG00000214646 |
| YKT6 | -0.20477701 | 0.867672789 | 0.000494819 | 0.003051815 | 3516.271204 | 3051.336343 | ENSG00000106636 |
| PAQR4 | -0.204823933 | 0.867644568 | 0.131212469 | 0.291009687 | 363.7500049 | 315.958313 | ENSG00000162073 |
| KIAA0100 | -0.204857449 | 0.867624412 | 0.001376039 | 0.007401453 | 2645.427406 | 2295.075001 | ENSG00000007202 |
| BIRC5 | -0.204891791 | 0.867603759 | 0.000465216 | 0.002894508 | 3862.33433 | 3350.346277 | ENSG00000089685 |
| AKNA | -0.205001268 | 0.867537925 | 0.007609097 | 0.031388772 | 1154.431519 | 1001.796123 | ENSG00000106948 |
| EGR2 | -0.205045551 | 0.867511296 | 0.636550522 | 0.800725077 | 25.27367947 | 21.88675762 | ENSG00000122877 |
| DENND2C | -0.205066363 | 0.867498782 | 0.443029826 | 0.650491235 | 73.38434448 | 63.54895566 | ENSG00000175984 |
| DERL2 | -0.205092078 | 0.86748332 | 0.048799167 | 0.138456089 | 524.6198378 | 455.1626648 | ENSG00000072849 |
| GYPC | -0.205121731 | 0.86746549 | 0.0533826 | 0.148167313 | 508.0769926 | 440.8954815 | ENSG00000136732 |
| NUTM1 | -0.205187999 | 0.867425645 | 0.649640185 | 0.809846226 | 25.17759987 | 21.80181922 | ENSG00000184507 |
| UTP3 | -0.205371421 | 0.867315368 | 0.0076068 | 0.031386152 | 1436.18195 | 1246.214741 | ENSG00000132467 |
| NA | -0.205379533 | 0.867310492 | 0.493104008 | 0.692567989 | 67.14331684 | 58.29496694 | ENSG00000254612 |
| ZNF770 | -0.205381986 | 0.867309017 | 0.01209461 | 0.04584639 | 1573.647473 | 1364.409216 | ENSG00000198146 |
| TMEM201 | -0.205385758 | 0.86730675 | 0.032328376 | 0.100746102 | 789.9787007 | 685.6515186 | ENSG00000188807 |
| VRK2 | -0.205447623 | 0.867269559 | 0.125570724 | 0.28171456 | 346.4526312 | 300.3930736 | ENSG00000028116 |
| NKAP | -0.205451522 | 0.867267215 | 0.264853418 | 0.472532527 | 161.7075992 | 140.3500301 | ENSG00000101882 |
| NA | -0.205523829 | 0.867223749 | 0.796740206 | 0.89620816 | 7.634473435 | 6.603952071 | ENSG00000270055 |
| CCNF | -0.205594286 | 0.867181398 | 0.006574722 | 0.027826898 | 1192.475745 | 1034.293411 | ENSG00000162063 |
| INPP1 | -0.205606201 | 0.867174236 | 0.490250537 | 0.690151427 | 55.40828838 | 48.08276903 | ENSG00000151689 |
| ERGIC2 | -0.20573786 | 0.867095102 | 0.085326349 | 0.212052652 | 382.898262 | 331.9126159 | ENSG00000087502 |
| TRA2A | -0.205774327 | 0.867073185 | 0.001913107 | 0.009767243 | 1747.708148 | 1515.551038 | ENSG00000164548 |
| NA | -0.205850979 | 0.867027117 | 0.551221925 | 0.738285536 | 44.10616343 | 38.23983526 | ENSG00000213713 |
| RRN3 | -0.205869366 | 0.867016067 | 0.026366083 | 0.085863681 | 799.4374914 | 693.0395548 | ENSG00000085721 |
| NA | -0.205923031 | 0.866983816 | 0.413535287 | 0.624700612 | 76.9416785 | 66.76601478 | ENSG00000262539 |
| NANOG | -0.205970081 | 0.866955543 | 0.004779336 | 0.021247118 | 1633.103971 | 1415.559758 | ENSG00000111704 |
| SRRT | -0.205984962 | 0.8669466 | 1.30E-05 | 0.000120519 | 6707.007031 | 5815.063144 | ENSG00000087087 |
| GABRA4 | -0.206039256 | 0.866913975 | 0.805179383 | 0.900702903 | 7.674796688 | 6.638475942 | ENSG00000109158 |
| ZNF100 | -0.206056808 | 0.866903427 | 0.314704835 | 0.524986448 | 145.6178454 | 126.1836329 | ENSG00000197020 |
| CEBPG | -0.206066299 | 0.866897724 | 0.008923065 | 0.03570928 | 1205.53481 | 1045.071053 | ENSG00000153879 |
| AQP4 | -0.206121302 | 0.866864675 | 0.659953409 | 0.816938918 | 25.38043749 | 21.96734509 | ENSG00000171885 |
| RNPS1 | -0.206155765 | 0.866843967 | 5.11E-05 | 0.000410184 | 7298.430695 | 6326.822481 | ENSG00000205937 |
| CDC123 | -0.206194768 | 0.866820533 | 0.001202298 | 0.006579368 | 3181.540377 | 2757.935003 | ENSG00000151465 |
| ZFX | -0.206195493 | 0.866820097 | 0.147660061 | 0.315447993 | 417.1456306 | 361.6040826 | ENSG00000005889 |
| MAP3K3 | -0.206226555 | 0.866801434 | 0.016373594 | 0.058530251 | 875.8341433 | 759.1989596 | ENSG00000198909 |
| AGTPBP1 | -0.206232347 | 0.866797954 | 0.017746385 | 0.062426581 | 1088.045297 | 943.225605 | ENSG00000135049 |
| FANK1 | -0.206374224 | 0.866712716 | 0.809004875 | 0.902949338 | 6.909135092 | 6.00885914 | ENSG00000203780 |
| CD24 | -0.206375251 | 0.866712099 | 0.000388076 | 0.002462638 | 32103.6162 | 27825.3414 | ENSG00000272398 |
| LINC01128 | -0.206394075 | 0.86670079 | 0.333022091 | 0.54504669 | 110.2267254 | 95.49799242 | ENSG00000228794 |
| NFATC3 | -0.206538779 | 0.866613863 | 0.07538079 | 0.193419323 | 407.6438225 | 353.1285385 | ENSG00000072736 |
| DICER1 | -0.206597867 | 0.866578371 | 0.034436905 | 0.105884233 | 1127.176446 | 976.3611688 | ENSG00000100697 |
| ICMT | -0.206688123 | 0.866524158 | 0.000205031 | 0.001406215 | 4012.544144 | 3476.68199 | ENSG00000116237 |
| ADD2 | -0.206697025 | 0.866518812 | 0.000110178 | 0.000810245 | 7090.25305 | 6143.429501 | ENSG00000075340 |
| PNO1 | -0.206714978 | 0.866508029 | 0.02539236 | 0.083411847 | 672.8490439 | 583.2524792 | ENSG00000115946 |
| ACLY | -0.206778985 | 0.866469586 | 2.66E-05 | 0.000230272 | 14243.51836 | 12340.88188 | ENSG00000131473 |
| ARHGAP1 | -0.20678394 | 0.86646661 | 0.019577181 | 0.067515534 | 931.9995521 | 807.6377291 | ENSG00000175220 |
| SAPCD2 | -0.20687072 | 0.866414492 | 0.011897985 | 0.045191765 | 1128.480476 | 978.1640028 | ENSG00000186193 |
| ZFP41 | -0.206921308 | 0.866384112 | 0.179200533 | 0.361714629 | 271.6576801 | 235.3376141 | ENSG00000181638 |
| LOC100506258 | -0.206962609 | 0.86635931 | 0.750054149 | 0.869965151 | 13.44165553 | 11.5944667 | ENSG00000251364 |
| BORCS5 | -0.20704056 | 0.866312501 | 0.234301599 | 0.43559615 | 184.6688608 | 159.9523567 | ENSG00000165714 |
| PTDSS1 | -0.207209371 | 0.866211138 | 0.000807719 | 0.004656217 | 1941.620861 | 1681.863264 | ENSG00000156471 |
| KIF2A | -0.207217599 | 0.866206198 | 0.022183392 | 0.074863002 | 1035.482859 | 896.6735831 | ENSG00000068796 |
| IFIH1 | -0.207236953 | 0.866194578 | 0.664623665 | 0.820451733 | 24.63254711 | 21.32808048 | ENSG00000115267 |
| KCP | -0.207292778 | 0.866161061 | 0.261522995 | 0.468691468 | 166.1554557 | 143.9852119 | ENSG00000135253 |
| SP6 | -0.207310406 | 0.866150478 | 0.467623089 | 0.671155346 | 62.15549392 | 53.7694892 | ENSG00000189120 |
| NA | -0.207359264 | 0.866121145 | 0.838745599 | 0.91909659 | 6.521562689 | 5.673667382 | ENSG00000225972 |
| ZNF200 | -0.207382057 | 0.866107462 | 0.279822707 | 0.488938917 | 148.1268972 | 128.4842905 | ENSG00000010539 |
| USP30 | -0.207471778 | 0.8660536 | 0.341978403 | 0.553644419 | 115.5769257 | 99.93663985 | ENSG00000135093 |
| CEP104 | -0.207641748 | 0.865951573 | 0.031055549 | 0.0975847 | 647.6919373 | 560.9037631 | ENSG00000116198 |
| GABPB1 | -0.207642314 | 0.865951233 | 0.061022722 | 0.164583562 | 523.1587359 | 453.007743 | ENSG00000104064 |
| GPR157 | -0.207648714 | 0.865947392 | 0.334114918 | 0.546110458 | 117.4344984 | 101.7591124 | ENSG00000180758 |
| NCLN | -0.207699523 | 0.865916895 | 0.002352546 | 0.011676173 | 4395.500861 | 3806.965124 | ENSG00000125912 |
| POC1B | -0.207723114 | 0.865902736 | 0.172761063 | 0.352369875 | 218.1721784 | 188.9115512 | ENSG00000139323 |

| | | | | | | | |
|---|---|---|---|---|---|---|---|
| TRIP13 | -0.207755232 | 0.865883459 | 0.006649987 | 0.02806996 | 1178.813735 | 1020.59801 | ENSG00000071539 |
| SASH1 | -0.207804561 | 0.865853853 | 0.068353592 | 0.179335592 | 568.5370187 | 491.9285556 | ENSG00000111961 |
| FBXO33 | -0.20780488 | 0.865853662 | 0.159173438 | 0.332583178 | 282.2466197 | 244.4958297 | ENSG00000165355 |
| NA | -0.207863486 | 0.865818489 | 0.694019416 | 0.836669205 | 28.96978061 | 25.15640672 | ENSG00000174403 |
| NMNAT3 | -0.207865346 | 0.865817373 | 0.529514406 | 0.721752405 | 48.87515755 | 42.30942946 | ENSG00000163864 |
| NA | -0.208068656 | 0.865695367 | 0.799494471 | 0.897969138 | 9.697806707 | 8.400602638 | ENSG00000267108 |
| PIK3R1 | -0.208115655 | 0.865667166 | 0.172195112 | 0.351697769 | 247.883263 | 214.4453139 | ENSG00000145675 |
| NA | -0.208237115 | 0.865594288 | 0.645943288 | 0.807482557 | 24.17617567 | 20.92752949 | ENSG00000234518 |
| POMT1 | -0.208337943 | 0.865533795 | 0.039222271 | 0.116919808 | 570.7119426 | 493.8512901 | ENSG00000130714 |
| MLLT3 | -0.208372344 | 0.865513157 | 0.253260233 | 0.459356464 | 151.1329237 | 130.8595762 | ENSG00000171843 |
| RFC1 | -0.208436851 | 0.865474458 | 0.001350764 | 0.00727794 | 1956.769091 | 1693.200654 | ENSG00000035928 |
| ITPRIPL1 | -0.20844004 | 0.865472545 | 0.087841604 | 0.216310952 | 399.789857 | 345.8431147 | ENSG00000198885 |
| NA | -0.208483424 | 0.86544652 | 0.498838481 | 0.697509604 | 51.70387309 | 44.68980119 | ENSG00000229132 |
| NA | -0.208509687 | 0.865430765 | 0.308263393 | 0.518716402 | 159.3299162 | 137.8374269 | ENSG00000267383 |
| LRRC37A | -0.208514334 | 0.865427977 | 0.708744952 | 0.846326173 | 17.98053145 | 15.51253378 | ENSG00000176681 |
| MTMR14 | -0.208519736 | 0.865424737 | 0.001889151 | 0.009663221 | 1627.221045 | 1408.35567 | ENSG00000163719 |
| LKAAEAR1 | -0.208541809 | 0.865411496 | 0.804617029 | 0.900509221 | 6.885413993 | 5.975564774 | ENSG00000171695 |
| DNPEP | -0.208608515 | 0.865371483 | 0.005252426 | 0.023043772 | 1486.602235 | 1286.233803 | ENSG00000123992 |
| ATP5MC3 | -0.208665779 | 0.865337135 | 7.19E-05 | 0.000553634 | 5092.758807 | 4406.773062 | ENSG00000154518 |
| GMEB2 | -0.208723472 | 0.865302531 | 0.04523752 | 0.13034949 | 782.5170634 | 676.8426911 | ENSG00000101216 |
| TUB | -0.208750901 | 0.86528608 | 0.005681293 | 0.024557385 | 1360.959053 | 1177.474822 | ENSG00000166402 |
| TCF12 | -0.208771344 | 0.865273819 | 0.00462128 | 0.020667237 | 1753.389762 | 1517.182907 | ENSG00000140262 |
| TCAP | -0.208843998 | 0.865230245 | 0.652619598 | 0.811744787 | 22.65461502 | 19.55318454 | ENSG00000173991 |
| LDB2 | -0.208905032 | 0.865193641 | 0.042023814 | 0.123309467 | 822.7526866 | 711.273727 | ENSG00000169744 |
| TRAK1 | -0.208910792 | 0.865190187 | 0.007365551 | 0.030510732 | 1185.914088 | 1025.628724 | ENSG00000182606 |
| RTF2 | -0.209032107 | 0.865117437 | 0.019715013 | 0.067916456 | 978.6821905 | 846.1417438 | ENSG00000022277 |
| SCN4A | -0.209032433 | 0.865117242 | 0.126844354 | 0.283739227 | 279.4395403 | 241.7251017 | ENSG00000007314 |
| AACS | -0.209068467 | 0.865095634 | 0.00479176 | 0.021289151 | 1517.744218 | 1312.588158 | ENSG00000081760 |
| SCPEP1 | -0.209134678 | 0.865055932 | 0.019184754 | 0.066331761 | 873.9761878 | 755.5903779 | ENSG00000121064 |
| NA | -0.209256273 | 0.864983026 | 0.801415105 | 0.898520796 | 6.894897202 | 5.979915701 | ENSG00000263503 |
| HS6ST2 | -0.209352121 | 0.864925561 | 0.062182414 | 0.166758718 | 472.7315616 | 408.7979234 | ENSG00000171004 |
| MZT1 | -0.209551588 | 0.864805985 | 0.057859677 | 0.157901735 | 599.732776 | 518.9174186 | ENSG00000204899 |
| LOC102724064 | -0.20963943 | 0.864753331 | 0.548402812 | 0.736023531 | 47.28805185 | 40.92666621 | ENSG00000255468 |
| GNAI2 | -0.209663957 | 0.864738629 | 6.19E-05 | 0.000485469 | 7034.798279 | 6083.244641 | ENSG00000114353 |
| OTUD6B-AS1 | -0.209666782 | 0.864736936 | 0.164867532 | 0.34046182 | 282.1976768 | 244.1122833 | ENSG00000253738 |
| ADD3 | -0.209792098 | 0.864661826 | 0.006626912 | 0.028016396 | 1456.118291 | 1258.693059 | ENSG00000148700 |
| ZC3H18 | -0.209797099 | 0.864658828 | 0.000277534 | 0.001837739 | 3673.225626 | 3176.336348 | ENSG00000158545 |
| GPRASP2 | -0.209833403 | 0.86463707 | 0.184102633 | 0.368146783 | 257.3555047 | 222.7400741 | ENSG00000158301 |
| MKX | -0.209891564 | 0.864602214 | 0.575988267 | 0.758153886 | 38.298702 | 33.21697222 | ENSG00000150051 |
| MAPKAPK3 | -0.20996294 | 0.86455944 | 0.000863835 | 0.004927068 | 2130.173175 | 1841.549908 | ENSG00000114738 |
| EIF2S2 | -0.209978456 | 0.864550141 | 3.51E-05 | 0.000292973 | 5343.505138 | 4620.234244 | ENSG00000125977 |
| SPAG9 | -0.209996686 | 0.864539217 | 0.010113814 | 0.039399727 | 1233.48359 | 1066.240869 | ENSG00000008294 |
| ID3 | -0.210044424 | 0.864510611 | 0.000395883 | 0.002507128 | 3368.16167 | 2911.881299 | ENSG00000117318 |
| ATP6V1FNB | -0.210056287 | 0.864503502 | 0.762930377 | 0.877431251 | 11.60813497 | 10.0572658 | ENSG00000272899 |
| SLC25A39 | -0.210068231 | 0.864496345 | 3.20E-05 | 0.00027006 | 5733.649158 | 4957.202441 | ENSG00000013306 |
| IMPDH1 | -0.21023801 | 0.864394615 | 0.005040464 | 0.022240459 | 1449.94449 | 1253.910604 | ENSG00000106348 |
| SCP2 | -0.210361493 | 0.864320633 | 0.023005824 | 0.077005491 | 678.6031132 | 586.5606766 | ENSG00000116171 |
| NA | -0.210377602 | 0.864310982 | 0.83594014 | 0.917543777 | 4.593437065 | 3.979358922 | ENSG00000253838 |
| MPHOSPH8 | -0.210406942 | 0.864293405 | 0.011527108 | 0.043995441 | 899.7190532 | 777.7142634 | ENSG00000196199 |
| FCHO1 | -0.210527954 | 0.864220912 | 0.036425291 | 0.110557958 | 715.1178997 | 618.5120044 | ENSG00000130475 |
| AGBL3 | -0.210567006 | 0.864197519 | 0.578306949 | 0.759450456 | 36.83762623 | 31.89767628 | ENSG00000146856 |
| KDM2B | -0.210609618 | 0.864171994 | 0.000233355 | 0.001578405 | 2863.169831 | 2473.990453 | ENSG00000089094 |
| PIKFYVE | -0.210640812 | 0.864153309 | 0.0285318 | 0.091176709 | 1069.812447 | 923.8507882 | ENSG00000115020 |
| POLD2 | -0.210805915 | 0.864054421 | 0.00013964 | 0.001003463 | 3483.687165 | 3010.308273 | ENSG00000106628 |
| KCNN2 | -0.210833527 | 0.864037883 | 0.13883424 | 0.302339095 | 322.1294521 | 278.5631616 | ENSG00000080709 |
| ABI2 | -0.210848324 | 0.864029021 | 0.001915266 | 0.009775623 | 1687.653642 | 1457.798248 | ENSG00000138443 |
| LOC100288123 | -0.210858156 | 0.864023133 | 0.624219331 | 0.7923564 | 26.4444315 | 22.83539197 | ENSG00000267244 |
| DCAF7 | -0.21092795 | 0.863981334 | 0.000656239 | 0.003881328 | 3039.241112 | 2625.341824 | ENSG00000136485 |
| NA | -0.210996383 | 0.863940353 | 0.637941015 | 0.80145796 | 23.42711623 | 20.26338882 | ENSG00000246889 |
| NA | -0.211168294 | 0.863837413 | 0.707193216 | 0.845435456 | 17.57961088 | 15.24393075 | ENSG00000229358 |
| WBP11 | -0.211189765 | 0.863824556 | 0.000123679 | 0.000898337 | 3458.655199 | 2987.53132 | ENSG00000084463 |
| NA | -0.211256231 | 0.86378476 | 0.612257774 | 0.784216498 | 29.59814963 | 25.52033101 | ENSG00000277459 |
| FAM83H | -0.211264027 | 0.863780093 | 0.001799673 | 0.009268294 | 2272.530752 | 1963.524358 | ENSG00000180921 |
| NA | -0.211406776 | 0.863694629 | 0.609249296 | 0.782326304 | 46.2202452 | 39.96469553 | ENSG00000274471 |
| NA | -0.211434191 | 0.863678217 | 0.595186349 | 0.772037516 | 34.52195436 | 29.86600059 | ENSG00000165121 |
| NA | -0.211568491 | 0.863597822 | 0.459270353 | 0.664325244 | 64.26718526 | 55.50986126 | ENSG00000227946 |
| GNPNAT1 | -0.2116088 | 0.863573693 | 0.022137413 | 0.074725119 | 748.0695787 | 645.8803761 | ENSG00000100522 |
| ARPP19 | -0.211681178 | 0.86353037 | 0.004424173 | 0.01995951 | 2276.721105 | 1966.247097 | ENSG00000128989 |
| NA | -0.211735606 | 0.863497792 | 0.522222465 | 0.716804556 | 45.71195621 | 39.44591178 | ENSG00000232640 |
| RCBTB1 | -0.211775656 | 0.863473821 | 0.013869352 | 0.051096902 | 963.5871496 | 832.0487023 | ENSG00000136144 |
| MARS1 | -0.211811808 | 0.863452184 | 2.76E-06 | 2.93E-05 | 9421.483403 | 8134.542934 | ENSG00000166986 |
| OGFOD3 | -0.211910968 | 0.863392839 | 0.028408301 | 0.090930637 | 689.5623609 | 595.0465011 | ENSG00000181396 |
| CACNG4 | -0.211968634 | 0.863358329 | 0.26557148 | 0.473143151 | 135.9671754 | 117.3558066 | ENSG00000075461 |
| ACBD5 | -0.211985541 | 0.863348211 | 0.094132888 | 0.228232069 | 362.7363766 | 313.0048435 | ENSG00000107897 |

| | | | | | | | |
|---|---|---|---|---|---|---|---|
| FAM200B | -0.212158985 | 0.863244424 | 0.194465852 | 0.382909772 | 194.9900025 | 168.4309199 | ENSG00000237765 |
| CDIP1 | -0.212320411 | 0.863147839 | 0.00406409 | 0.018628724 | 1748.134088 | 1509.596232 | ENSG00000089486 |
| CLTB | -0.212520195 | 0.863028319 | 0.030209583 | 0.09551913 | 741.186756 | 639.724522 | ENSG00000175416 |
| LRRC59 | -0.21257259 | 0.862996976 | 7.66E-05 | 0.000586185 | 3210.287394 | 2770.697324 | ENSG00000108829 |
| CYP2D7 | -0.212724223 | 0.862906277 | 0.64273441 | 0.80490977 | 34.12545741 | 29.43236615 | ENSG00000205702 |
| NA | -0.212741296 | 0.862896065 | 0.800814973 | 0.89833031 | 9.161957822 | 7.912654243 | ENSG00000255129 |
| PDE5A | -0.212742281 | 0.862895476 | 0.099998939 | 0.238113727 | 339.5998634 | 292.8068139 | ENSG00000138735 |
| LGI2 | -0.21285427 | 0.862828496 | 0.174287811 | 0.35456483 | 234.6753107 | 202.5487389 | ENSG00000153012 |
| DYRK1B | -0.213005529 | 0.862738038 | 0.153691469 | 0.324669522 | 265.6972143 | 229.1975124 | ENSG00000105204 |
| C11orf68 | -0.213030285 | 0.862723234 | 0.046686617 | 0.133687249 | 703.0245085 | 606.8029638 | ENSG00000175573 |
| F2R | -0.213042043 | 0.862716203 | 0.025101775 | 0.082730718 | 641.8289988 | 553.8742161 | ENSG00000181104 |
| RNF157 | -0.2130576 | 0.8627069 | 0.135207794 | 0.29711417 | 327.4983401 | 282.6132196 | ENSG00000141576 |
| LOC389906 | -0.213061534 | 0.862704548 | 0.086259513 | 0.213612759 | 442.3057909 | 381.8362502 | ENSG00000285756 |
| STK40 | -0.213075849 | 0.862695987 | 0.009180927 | 0.03644772 | 958.9939925 | 827.2072987 | ENSG00000196182 |
| UACA | -0.213083035 | 0.862691691 | 0.053649122 | 0.148797718 | 469.3551235 | 404.8487374 | ENSG00000137831 |
| NA | -0.213197727 | 0.862623111 | 0.83737979 | 0.918185992 | 4.647998207 | 4.016058257 | ENSG00000287817 |
| USP15 | -0.213205235 | 0.862618622 | 0.056633169 | 0.155439327 | 458.8756117 | 395.7259393 | ENSG00000135655 |
| SLC24A3 | -0.213277059 | 0.862575677 | 0.277322051 | 0.486459867 | 126.8535495 | 109.4530837 | ENSG00000185052 |
| MREG | -0.213290362 | 0.862567724 | 0.062243989 | 0.166829084 | 452.3796019 | 390.2635778 | ENSG00000118242 |
| VSTM5 | -0.213369419 | 0.862520458 | 0.788961813 | 0.891843683 | 7.726993566 | 6.634125014 | ENSG00000214376 |
| NIPBL-DT | -0.213428189 | 0.862485323 | 0.311391886 | 0.521980806 | 120.0618247 | 103.5699297 | ENSG00000285967 |
| WNK2 | -0.213474454 | 0.862457664 | 0.002141968 | 0.010778328 | 2448.474449 | 2112.180341 | ENSG00000165238 |
| FUCA2 | -0.213480649 | 0.862453961 | 0.01410842 | 0.051846177 | 905.4673034 | 780.9984784 | ENSG00000001036 |
| DNAJC9 | -0.213581197 | 0.862393855 | 0.032099054 | 0.100229934 | 553.0834149 | 476.8489212 | ENSG00000213551 |
| EDEM1 | -0.213588869 | 0.862389269 | 0.090586061 | 0.221452138 | 380.7854539 | 328.257855 | ENSG00000134109 |
| POLE4 | -0.213596883 | 0.862384478 | 0.046950696 | 0.134281976 | 488.0396365 | 420.9104388 | ENSG00000115350 |
| GPC6 | -0.213693214 | 0.862326898 | 0.121103345 | 0.274260186 | 354.2351024 | 305.1293401 | ENSG00000183098 |
| TMEM126A | -0.213812996 | 0.862255304 | 0.07889372 | 0.199612526 | 377.9190326 | 325.6335454 | ENSG00000171202 |
| POSTN | -0.21385511 | 0.862230134 | 0.678850938 | 0.828278844 | 20.04030525 | 17.22736737 | ENSG00000133110 |
| ZRANB2 | -0.213900659 | 0.862202912 | 0.003922921 | 0.018070306 | 1812.526868 | 1563.153984 | ENSG00000132485 |
| MLEC | -0.213978752 | 0.862156243 | 6.94E-05 | 0.000537872 | 6644.627546 | 5728.277028 | ENSG00000110917 |
| SENP6 | -0.214074133 | 0.862099245 | 0.003622423 | 0.016887055 | 1484.846645 | 1280.337131 | ENSG00000112701 |
| GNL1 | -0.214129821 | 0.862065969 | 0.001111448 | 0.006149924 | 2100.923366 | 1810.834282 | ENSG00000204590 |
| NA | -0.214162465 | 0.862046463 | 0.673818546 | 0.825338127 | 20.04386472 | 17.22301644 | ENSG00000225234 |
| STK26 | -0.214205437 | 0.862020786 | 0.000591049 | 0.003547983 | 4196.185494 | 3617.15173 | ENSG00000134602 |
| LPAL2 | -0.214249539 | 0.861994436 | 0.683758128 | 0.830895404 | 21.48002419 | 18.4993702 | ENSG00000213071 |
| ARHGDIA | -0.214309892 | 0.861958376 | 0.001034627 | 0.005773689 | 3766.771114 | 3247.263014 | ENSG00000141522 |
| GLRX5 | -0.214317135 | 0.861954048 | 0.000118782 | 0.000866097 | 3016.611971 | 2600.244168 | ENSG00000182512 |
| GRAMD1B | -0.214335192 | 0.86194326 | 0.050923352 | 0.143193492 | 456.0756251 | 392.9379231 | ENSG00000023171 |
| TMED3 | -0.214350871 | 0.861933893 | 0.006406223 | 0.02721132 | 1129.013552 | 972.8514219 | ENSG00000166557 |
| CELSR3 | -0.214353037 | 0.861932598 | 0.010421565 | 0.040382182 | 1500.50868 | 1293.170352 | ENSG00000008300 |
| PEX6 | -0.214387826 | 0.861911814 | 0.015720744 | 0.05660367 | 847.1765752 | 730.3052399 | ENSG00000124587 |
| LAMTOR3 | -0.214393339 | 0.861908521 | 0.089394118 | 0.219390242 | 366.7509456 | 316.2230595 | ENSG00000109270 |
| FAM228B | -0.214444117 | 0.861878185 | 0.730929646 | 0.858326474 | 12.3714323 | 10.63400907 | ENSG00000219626 |
| LOC728554 | -0.21447945 | 0.861857077 | 0.048613356 | 0.138053813 | 554.0332003 | 477.5254522 | ENSG00000170089 |
| NA | -0.214555008 | 0.86181194 | 0.31601084 | 0.526034792 | 119.5105689 | 103.1266474 | ENSG00000276529 |
| NCKIPSD | -0.214564733 | 0.861806131 | 0.011072105 | 0.042516614 | 967.1981557 | 833.6005549 | ENSG00000213672 |
| CPXM2 | -0.214629105 | 0.861767679 | 0.20576002 | 0.397929132 | 183.8506974 | 158.3148212 | ENSG00000121898 |
| NA | -0.214721229 | 0.861712652 | 0.696436036 | 0.837932883 | 16.97554778 | 14.60466614 | ENSG00000269982 |
| THSD1 | -0.214800694 | 0.861665189 | 0.784115091 | 0.890065126 | 11.66061118 | 9.981975218 | ENSG00000136114 |
| CCDC86 | -0.214887053 | 0.861613612 | 0.003074682 | 0.014707539 | 1520.003424 | 1310.1049 | ENSG00000110104 |
| CTH | -0.214986556 | 0.861554188 | 0.11812619 | 0.270008847 | 345.8281277 | 297.5873475 | ENSG00000116761 |
| GTF3C5 | -0.215060258 | 0.861510176 | 0.000195642 | 0.001352877 | 2939.006478 | 2531.725629 | ENSG00000148308 |
| UBP1 | -0.215176953 | 0.861440493 | 0.00199604 | 0.010144031 | 1892.611516 | 1630.566913 | ENSG00000153560 |
| ELOC | -0.215212898 | 0.861419031 | 0.00339557 | 0.016007323 | 1460.919372 | 1258.646682 | ENSG00000154582 |
| CRYZL2P | -0.215227638 | 0.86141023 | 0.690886425 | 0.834552267 | 16.97910726 | 14.59936925 | ENSG00000242193 |
| MON1A | -0.215260007 | 0.861390903 | 0.155302207 | 0.327038601 | 309.3301242 | 266.8243995 | ENSG00000164077 |
| MOSMO | -0.215261667 | 0.861389912 | 0.293995845 | 0.503821373 | 126.3123617 | 108.7042882 | ENSG00000185716 |
| SPAG5 | -0.215420937 | 0.861294822 | 0.001780885 | 0.009201779 | 1743.233448 | 1501.314175 | ENSG00000076382 |
| TRIB1 | -0.21543245 | 0.861287948 | 0.010281335 | 0.039910176 | 1064.826506 | 917.1214575 | ENSG00000173334 |
| STK4 | -0.215480479 | 0.861259276 | 0.008988993 | 0.035897046 | 918.4555099 | 791.0910757 | ENSG00000101109 |
| GUF1 | -0.215575891 | 0.861202319 | 0.032677593 | 0.101649868 | 645.7160901 | 556.1602851 | ENSG00000151806 |
| ZNF668 | -0.21561084 | 0.861181457 | 0.216308414 | 0.411751274 | 166.6918888 | 143.6292841 | ENSG00000167394 |
| LRIF1 | -0.215669099 | 0.861146681 | 0.276169986 | 0.485114891 | 162.0456701 | 139.766266 | ENSG00000121931 |
| IKZF4 | -0.215707108 | 0.861123993 | 0.133448326 | 0.294515793 | 256.9370406 | 221.15764 | ENSG00000123411 |
| GLCE | -0.215741857 | 0.861103253 | 0.099593817 | 0.237418485 | 397.7668208 | 342.4363591 | ENSG00000138604 |
| SDF2L1 | -0.215749995 | 0.861098396 | 0.018834567 | 0.065360307 | 1666.535589 | 1435.659558 | ENSG00000128228 |
| FBLN5 | -0.215768345 | 0.861087443 | 0.517678712 | 0.713655192 | 45.40858979 | 39.04725271 | ENSG00000140092 |
| B9D2 | -0.215787102 | 0.861076247 | 0.505537121 | 0.702948074 | 53.49201023 | 46.06698401 | ENSG00000123810 |
| RRM1 | -0.215900096 | 0.861008809 | 0.000545917 | 0.003309678 | 2107.68166 | 1814.484123 | ENSG00000167325 |
| ZNF121 | -0.215912382 | 0.861001477 | 0.003321169 | 0.015691839 | 1494.683038 | 1286.492735 | ENSG00000197961 |
| KIAA1143 | -0.216053918 | 0.860917013 | 0.009059153 | 0.0361455 | 910.8322735 | 784.0722904 | ENSG00000163807 |
| RND2 | -0.216195145 | 0.860832741 | 0.000396593 | 0.002509097 | 3328.066796 | 2864.665271 | ENSG00000108830 |
| ATG2A | -0.216454342 | 0.860678096 | 0.012158805 | 0.046034289 | 902.7550811 | 777.3541003 | ENSG00000110046 |

| | | | | | | | |
|---|---|---|---|---|---|---|---|
| PSMD2 | -0.216569363 | 0.86060948 | 1.70E-06 | 1.88E-05 | 11029.69185 | 9491.93429 | ENSG00000175166 |
| RBM11 | -0.216737408 | 0.860509242 | 0.534708699 | 0.725637667 | 47.18754921 | 40.59052849 | ENSG00000185272 |
| CNOT9 | -0.216839525 | 0.860448335 | 0.001403701 | 0.007533078 | 1775.909903 | 1527.811451 | ENSG00000144580 |
| CTAGE15 | -0.216865545 | 0.860432817 | 0.787529128 | 0.891546762 | 7.310945384 | 6.270935776 | ENSG00000271079 |
| PNRC2 | -0.216933175 | 0.860392482 | 0.001152936 | 0.006351574 | 2441.483998 | 2100.760449 | ENSG00000189266 |
| TMEM33 | -0.217001988 | 0.860351445 | 0.032988154 | 0.102278984 | 1098.319554 | 945.1220754 | ENSG00000109133 |
| SPRY4 | -0.21712462 | 0.860278316 | 0.000176531 | 0.001231543 | 4092.564208 | 3521.026914 | ENSG00000187678 |
| ZBTB11 | -0.217402557 | 0.860112599 | 0.16126179 | 0.33576945 | 262.2650199 | 225.5819098 | ENSG00000066422 |
| SPTB | -0.217462907 | 0.86007662 | 0.432021 | 0.641532588 | 65.50554112 | 56.3088604 | ENSG00000070182 |
| NA | -0.21752806 | 0.860037779 | 0.838901275 | 0.919213843 | 4.242628443 | 3.66998918 | ENSG00000258477 |
| ECD | -0.217588396 | 0.860001812 | 0.004567948 | 0.020485801 | 1152.631684 | 991.2199057 | ENSG00000122882 |
| FAM76B | -0.217618214 | 0.859984037 | 0.103504112 | 0.244364939 | 308.8523703 | 265.6879902 | ENSG00000077458 |
| ZMYND11 | -0.217688412 | 0.859942193 | 0.034535345 | 0.106135066 | 618.8270568 | 531.9015311 | ENSG00000015171 |
| NA | -0.217735517 | 0.859914116 | 0.838340289 | 0.918919053 | 4.63731979 | 3.988060777 | ENSG00000259687 |
| EXT1 | -0.217866573 | 0.859836004 | 0.014316549 | 0.052468081 | 835.3407653 | 718.3245475 | ENSG00000182197 |
| KLHL7 | -0.217872781 | 0.859832304 | 0.001514059 | 0.008020479 | 1781.215941 | 1531.185008 | ENSG00000122550 |
| ARHGEF25 | -0.217936656 | 0.859794236 | 0.472865784 | 0.676213109 | 64.26359964 | 55.2907268 | ENSG00000240771 |
| NA | -0.21798426 | 0.859765866 | 0.793590959 | 0.894317058 | 6.98385786 | 6.005737718 | ENSG00000228169 |
| ZNF17 | -0.218069442 | 0.859715104 | 0.475627514 | 0.678057341 | 52.48941698 | 45.16715581 | ENSG00000186272 |
| LOC102724159 | -0.218088096 | 0.859703988 | 0.014960923 | 0.05439681 | 932.4705376 | 801.4805072 | ENSG00000275464 |
| GOLGA2P5 | -0.218208025 | 0.859632525 | 0.319754258 | 0.530116685 | 152.7416394 | 131.3355334 | ENSG00000238105 |
| L3MBTL1 | -0.218215523 | 0.859628057 | 0.496122844 | 0.695402896 | 91.34946899 | 78.57937534 | ENSG00000185513 |
| SMG6 | -0.218263798 | 0.859599293 | 0.024231174 | 0.080338083 | 794.5076595 | 682.4539957 | ENSG00000070366 |
| ARID3B | -0.218283172 | 0.85958775 | 8.22E-05 | 0.000624364 | 5340.467467 | 4591.029025 | ENSG00000179361 |
| AVIL | -0.218590081 | 0.859404907 | 0.663704539 | 0.819648197 | 20.06611128 | 17.28090332 | ENSG00000135407 |
| SNX33 | -0.218606923 | 0.859394874 | 0.006774733 | 0.028475593 | 981.2588717 | 843.183304 | ENSG00000173548 |
| CRY1 | -0.218629629 | 0.859381349 | 0.025691722 | 0.084146406 | 604.5174831 | 519.5400575 | ENSG00000008405 |
| UBR1 | -0.218743823 | 0.859313328 | 0.034232431 | 0.105382147 | 512.6179018 | 440.4159514 | ENSG00000159459 |
| EPS15L1 | -0.218874488 | 0.859235504 | 0.016427243 | 0.058688703 | 923.6787911 | 793.3987339 | ENSG00000127527 |
| NA | -0.218898947 | 0.859220937 | 0.745200428 | 0.867425425 | 12.01114047 | 10.30193873 | ENSG00000271833 |
| NA | -0.218966178 | 0.859180897 | 0.629113471 | 0.795425152 | 23.62519918 | 20.27426614 | ENSG00000225159 |
| HAUS6 | -0.219018892 | 0.859149504 | 0.031997332 | 0.100028076 | 1284.57819 | 1103.59358 | ENSG00000147874 |
| RNASEH1 | -0.21911592 | 0.859091725 | 0.025010646 | 0.082502366 | 603.6524618 | 518.6140284 | ENSG00000171865 |
| NOP10 | -0.219266677 | 0.859001957 | 0.001601721 | 0.008409371 | 1638.669388 | 1407.867313 | ENSG00000182117 |
| VAC14 | -0.219534021 | 0.858842791 | 0.004143366 | 0.018932367 | 1413.533923 | 1214.179117 | ENSG00000103043 |
| MAF1 | -0.219611497 | 0.85879667 | 0.000347112 | 0.002229665 | 2842.891528 | 2441.778156 | ENSG00000179632 |
| ALOX12-AS1 | -0.219646678 | 0.858775729 | 0.702566611 | 0.842036584 | 16.28341375 | 13.97940026 | ENSG00000215067 |
| NA | -0.219658185 | 0.858768879 | 0.773754058 | 0.883722128 | 10.48126567 | 8.953699348 | ENSG00000234911 |
| IQCH | -0.219734317 | 0.858723562 | 0.785077675 | 0.890433415 | 7.365506526 | 6.30110872 | ENSG00000103599 |
| CCAR2 | -0.21975212 | 0.858712965 | 3.06E-05 | 0.000260499 | 3675.558129 | 3156.135145 | ENSG00000158941 |
| BIN3 | -0.219819971 | 0.858672581 | 0.09190209 | 0.223857985 | 332.810529 | 285.6347478 | ENSG00000147439 |
| ARMH1 | -0.219979623 | 0.858577563 | 0.82718342 | 0.913049207 | 4.624277109 | 3.967535646 | ENSG00000198520 |
| AGAP4 | -0.220026806 | 0.858549484 | 0.372877812 | 0.584915481 | 104.8433214 | 89.88249547 | ENSG00000188234 |
| TIMM50 | -0.220030044 | 0.858547557 | 0.000306027 | 0.002001152 | 3374.84033 | 2897.287207 | ENSG00000105197 |
| CERT1 | -0.220146581 | 0.858478209 | 0.006077986 | 0.02602784 | 1327.716107 | 1140.161233 | ENSG00000113163 |
| LOC442497 | -0.220329141 | 0.858369583 | 0.703199662 | 0.842421061 | 15.39319677 | 13.24025255 | ENSG00000249574 |
| NA | -0.220388083 | 0.858334514 | 0.708028491 | 0.845910689 | 13.54218433 | 11.63740888 | ENSG00000258674 |
| NUBP1 | -0.220394563 | 0.858330659 | 0.118293574 | 0.27026605 | 297.7129868 | 255.463234 | ENSG00000103274 |
| GET3 | -0.220479946 | 0.858279862 | 0.000911951 | 0.005176495 | 2366.270838 | 2030.37753 | ENSG00000198356 |
| RMDN3 | -0.220490077 | 0.858273835 | 0.010651276 | 0.041118017 | 862.2696339 | 740.0415963 | ENSG00000137824 |
| STRCP1 | -0.220502379 | 0.858266517 | 0.750874388 | 0.870392545 | 11.63661075 | 9.982921177 | ENSG00000166763 |
| LOC283922 | -0.220620314 | 0.858196359 | 0.520387966 | 0.715666089 | 49.35288582 | 42.32030678 | ENSG00000214331 |
| TMEM237 | -0.220626251 | 0.858192827 | 0.03264024 | 0.101567134 | 648.9288186 | 557.1319466 | ENSG00000155755 |
| SLC39A3 | -0.220656936 | 0.858174575 | 0.086429683 | 0.213879248 | 478.8468898 | 411.2911584 | ENSG00000141873 |
| RAMP1 | -0.220718127 | 0.858138177 | 0.720127466 | 0.852479609 | 14.6183313 | 12.49987533 | ENSG00000132329 |
| RNF165 | -0.220736774 | 0.858127085 | 0.135107622 | 0.297005724 | 289.5032838 | 248.2446098 | ENSG00000141622 |
| NA | -0.220760492 | 0.858112978 | 0.806410785 | 0.901201229 | 6.169558858 | 5.32882781 | ENSG00000280332 |
| MOB2 | -0.220830263 | 0.858071479 | 0.11290039 | 0.261225664 | 436.8792631 | 374.9373583 | ENSG00000182208 |
| NA | -0.220906411 | 0.85802619 | 0.84522586 | 0.923032554 | 4.228390553 | 3.6419917 | ENSG00000267676 |
| NA | -0.220937801 | 0.858007521 | 0.785746119 | 0.890559489 | 7.399906043 | 6.324755273 | ENSG00000276571 |
| RASA1 | -0.221177652 | 0.857864887 | 0.00449561 | 0.020243109 | 1033.577844 | 886.6483525 | ENSG00000145715 |
| NA | -0.221385587 | 0.857741253 | 0.268168994 | 0.47603598 | 128.3671015 | 110.0795791 | ENSG00000226259 |
| NDUFAF8 | -0.221409217 | 0.857727204 | 0.041133043 | 0.1212988 | 609.4715169 | 523.1676419 | ENSG00000224877 |
| LRRC20 | -0.221524119 | 0.857658894 | 0.0152902 | 0.055343195 | 709.7845041 | 608.5012537 | ENSG00000172731 |
| MARF1 | -0.221526725 | 0.857657345 | 0.04718469 | 0.134849102 | 465.5727053 | 399.1029009 | ENSG00000166783 |
| ADGRB2 | -0.221531767 | 0.857654347 | 0.002936996 | 0.014127653 | 1225.818023 | 1051.579087 | ENSG00000121753 |
| POP4 | -0.221545843 | 0.857645979 | 0.049247713 | 0.13945765 | 438.9927345 | 376.5909204 | ENSG00000105171 |
| ARHGEF18 | -0.221562658 | 0.857635983 | 0.024850142 | 0.082101982 | 611.4337371 | 524.3884093 | ENSG00000104880 |
| CSNK2A2 | -0.221578531 | 0.857626547 | 0.011980405 | 0.045477378 | 923.1627983 | 791.5786453 | ENSG00000070770 |
| LTV1 | -0.221596223 | 0.85761603 | 0.11961684 | 0.27216434 | 284.4152103 | 244.1543225 | ENSG00000135521 |
| CD81 | -0.221609012 | 0.857608428 | 0.000582646 | 0.003504226 | 3390.922981 | 2908.146623 | ENSG00000110651 |
| ZZEF1 | -0.221666317 | 0.857574363 | 0.032040244 | 0.100129075 | 609.979728 | 522.9593715 | ENSG00000074755 |
| UBAC2 | -0.221686784 | 0.857562197 | 0.005790434 | 0.024977694 | 1164.584041 | 998.2796257 | ENSG00000134882 |
| GUSB | -0.221771037 | 0.857512117 | 0.013658053 | 0.050416795 | 941.6720324 | 807.8691516 | ENSG00000169919 |

| | | | | | | | |
|---|---|---|---|---|---|---|---|
| RPL23A | -0.221780593 | 0.857506437 | 0.002140443 | 0.010778187 | 1614.633199 | 1385.124216 | ENSG00000198242 |
| PALM2AKAP2 | -0.221810187 | 0.857488847 | 0.001604694 | 0.008417959 | 2101.561679 | 1802.020861 | ENSG00000157654 |
| CDC42BPB | -0.221822923 | 0.857481278 | 6.87E-05 | 0.000532998 | 3691.236896 | 3164.543154 | ENSG00000198752 |
| RBP1 | -0.221868085 | 0.857454435 | 0.085753806 | 0.212723056 | 361.7364531 | 309.9256917 | ENSG00000114115 |
| NA | -0.221989429 | 0.857382319 | 0.489961837 | 0.690105108 | 48.67707461 | 41.6858446 | ENSG00000259985 |
| DUSP7 | -0.222058917 | 0.857341024 | 0.018441852 | 0.064304723 | 672.3052125 | 576.4576307 | ENSG00000164086 |
| PPP1R26-AS1 | -0.222085133 | 0.857325445 | 0.552369209 | 0.739140686 | 39.45788589 | 33.8972871 | ENSG00000225361 |
| PSMB7 | -0.222089418 | 0.857322898 | 8.51E-05 | 0.000643319 | 3918.368342 | 3359.210296 | ENSG00000136930 |
| CCDC58 | -0.22209787 | 0.857317876 | 0.020645057 | 0.070605197 | 678.6496395 | 581.8335837 | ENSG00000160124 |
| NA | -0.222209949 | 0.857251276 | 0.683785184 | 0.830895404 | 20.50440602 | 17.66584719 | ENSG00000262519 |
| NKILA | -0.222292554 | 0.857202193 | 0.662609304 | 0.819045471 | 23.71149009 | 20.38285109 | ENSG00000278709 |
| NCL | -0.222477514 | 0.857092303 | 9.97E-08 | 1.39E-06 | 36626.26674 | 31392.33953 | ENSG00000115053 |
| SF3B5 | -0.222581029 | 0.857030808 | 0.001126543 | 0.006222511 | 1704.708362 | 1460.924781 | ENSG00000169976 |
| NA | -0.222622781 | 0.857006005 | 0.717312276 | 0.851067559 | 13.9961915 | 11.98224845 | ENSG00000260526 |
| ATF1 | -0.222786308 | 0.856908871 | 0.059192212 | 0.160725365 | 449.9597371 | 385.455571 | ENSG00000123268 |
| ENPP4 | -0.222794327 | 0.856904108 | 0.351300187 | 0.563375002 | 98.82084324 | 84.70715934 | ENSG00000001561 |
| NA | -0.2228566 | 0.856867121 | 0.116332307 | 0.267104634 | 64555.29303 | 55315.58204 | ENSG00000210082 |
| SMARCA4 | -0.222900925 | 0.856840795 | 2.02E-06 | 2.20E-05 | 10578.35899 | 9064.293865 | ENSG00000127616 |
| MADCAM1 | -0.222904765 | 0.856838514 | 0.623323547 | 0.791625097 | 24.0290944 | 20.6405768 | ENSG00000099866 |
| HSBP1 | -0.222943353 | 0.856815597 | 0.001901739 | 0.009717377 | 3502.892925 | 3001.209423 | ENSG00000230989 |
| COL7A1 | -0.222958574 | 0.856806557 | 0.511661938 | 0.708204024 | 282.2427553 | 241.8773771 | ENSG00000114270 |
| HLA-A | -0.222988151 | 0.856788992 | 5.16E-05 | 0.000413402 | 6183.442102 | 5297.346373 | ENSG00000206503 |
| NA | -0.223180426 | 0.856674811 | 0.104285941 | 0.245811283 | 306.8848112 | 263.1681255 | ENSG00000210196 |
| TCEAL9 | -0.223328328 | 0.856586991 | 0.00183931 | 0.009436393 | 1372.274778 | 1175.580157 | ENSG00000185222 |
| HCN4 | -0.223527785 | 0.856468573 | 0.025667619 | 0.084082043 | 639.6742883 | 547.9679623 | ENSG00000138622 |
| TXNDC12 | -0.223528466 | 0.856468169 | 0.227155979 | 0.426086677 | 161.3318743 | 138.1337009 | ENSG00000117862 |
| NA | -0.223642604 | 0.856400413 | 0.702864191 | 0.842126015 | 13.56234596 | 11.61593779 | ENSG00000239377 |
| NA | -0.223809773 | 0.856301185 | 0.798235041 | 0.897141508 | 7.041978476 | 6.02285788 | ENSG00000262692 |
| PYURF | -0.223836876 | 0.856285098 | 0.020543714 | 0.070335034 | 763.1883911 | 653.315982 | ENSG00000145337 |
| AGAP9 | -0.223850711 | 0.856276887 | 0.500502907 | 0.698853782 | 55.65500873 | 47.70730502 | ENSG00000204172 |
| GHRHR | -0.223926202 | 0.856232082 | 0.609653456 | 0.782504214 | 31.47435771 | 26.87046231 | ENSG00000106128 |
| AGER | -0.224035683 | 0.856167108 | 0.382906136 | 0.595355045 | 111.8995633 | 95.99419117 | ENSG00000204305 |
| MIR4442 | -0.224155259 | 0.856096149 | 0.609982932 | 0.782665336 | 30.18944938 | 25.80671661 | ENSG00000283971 |
| RFTN1 | -0.224176304 | 0.856083661 | 0.300115783 | 0.510234657 | 110.5396012 | 94.64545733 | ENSG00000131378 |
| PTGES3 | -0.224225893 | 0.856054235 | 1.53E-05 | 0.00013964 | 9024.533424 | 7725.418516 | ENSG00000110958 |
| METTL9 | -0.224295153 | 0.85601314 | 0.000692609 | 0.004076053 | 2462.639428 | 2108.383744 | ENSG00000197006 |
| CTBP2 | -0.224332517 | 0.85599097 | 0.000122141 | 0.000888878 | 3728.670348 | 3191.468316 | ENSG00000175029 |
| SERPINA10 | -0.224503691 | 0.855889414 | 0.830194168 | 0.914737804 | 4.655117152 | 3.977183459 | ENSG00000140093 |
| IRF2 | -0.224580903 | 0.855843609 | 0.177856805 | 0.359809937 | 228.3927913 | 195.2533837 | ENSG00000168310 |
| C8orf33 | -0.224612001 | 0.855825161 | 0.002795934 | 0.013524903 | 1207.370747 | 1033.597364 | ENSG00000182307 |
| PNKD | -0.224631879 | 0.855813369 | 0.022460046 | 0.075553611 | 696.4499115 | 596.2468722 | ENSG00000127838 |
| TCTN1 | -0.224673183 | 0.855788867 | 0.335597485 | 0.547537071 | 92.00097429 | 78.76538339 | ENSG00000204852 |
| PGRMC2 | -0.224691674 | 0.855777899 | 0.001757266 | 0.009104519 | 1476.823248 | 1264.102161 | ENSG00000164040 |
| CELF4 | -0.224700223 | 0.855772828 | 0.518727396 | 0.714579245 | 45.46431999 | 38.89622003 | ENSG00000101489 |
| ALDH18A1 | -0.224757761 | 0.855738698 | 3.87E-05 | 0.000320082 | 4493.488203 | 3844.531724 | ENSG00000059573 |
| NUP62 | -0.224764583 | 0.855734652 | 0.000385391 | 0.002447247 | 2368.764183 | 2027.352982 | ENSG00000213024 |
| KLF3 | -0.224842536 | 0.855688416 | 0.105926979 | 0.249057643 | 306.7765786 | 262.7450419 | ENSG00000109787 |
| CARM1 | -0.224950216 | 0.855624551 | 9.38E-05 | 0.000701222 | 5000.898674 | 4278.902344 | ENSG00000142453 |
| FAM229A | -0.22499366 | 0.855598786 | 0.663955152 | 0.819904077 | 20.29859374 | 17.37577308 | ENSG00000225828 |
| ZC3H15 | -0.225004304 | 0.855592473 | 0.000257064 | 0.001719682 | 2249.472592 | 1924.530503 | ENSG00000065548 |
| VAMP8 | -0.225010465 | 0.85558882 | 0.096226666 | 0.23152511 | 364.5740153 | 311.8160255 | ENSG00000118640 |
| NA | -0.22505143 | 0.855564525 | 0.656272633 | 0.81429937 | 19.40837676 | 16.65157006 | ENSG00000240376 |
| SART3 | -0.225098181 | 0.855536801 | 0.001168052 | 0.00641613 | 2707.686755 | 2315.958778 | ENSG00000075856 |
| RPP30 | -0.225191259 | 0.855481607 | 0.009369234 | 0.037016314 | 818.132098 | 700.1995274 | ENSG00000148688 |
| EIF5A | -0.225225688 | 0.855461191 | 0.00024199 | 0.00163039 | 19666.97176 | 16824.76322 | ENSG00000132507 |
| ZNF276 | -0.225245845 | 0.855449239 | 0.280325591 | 0.48927438 | 150.9722149 | 129.3867886 | ENSG00000158805 |
| ZNF732 | -0.225255674 | 0.855443411 | 0.186768721 | 0.371981365 | 178.5182684 | 152.78833 | ENSG00000186777 |
| TMEM240 | -0.225272658 | 0.85543334 | 0.384332449 | 0.59668648 | 79.05481824 | 67.60265921 | ENSG00000205090 |
| CBX8 | -0.225336773 | 0.855395325 | 0.521918774 | 0.71662324 | 40.43706353 | 34.59973547 | ENSG00000141570 |
| CD9 | -0.225349923 | 0.855387528 | 0.000218204 | 0.001486036 | 2700.149356 | 2309.087752 | ENSG00000010278 |
| ZNF682 | -0.225351656 | 0.8553865 | 0.205882957 | 0.398085353 | 176.4638079 | 150.9193424 | ENSG00000197124 |
| ARSG | -0.22536767 | 0.855377006 | 0.446380099 | 0.653581034 | 63.57622028 | 54.34963743 | ENSG00000141337 |
| TMEM18-DT | -0.225408302 | 0.855352915 | 0.806321489 | 0.901201229 | 6.622370821 | 5.671491918 | ENSG00000233296 |
| GTF2E2 | -0.225460174 | 0.855322161 | 0.012373536 | 0.046650873 | 836.5063799 | 715.1037045 | ENSG00000197265 |
| ERCC8 | -0.225583376 | 0.855249123 | 0.264879189 | 0.472533864 | 151.0389552 | 129.2266322 | ENSG00000049167 |
| AZI2 | -0.225589784 | 0.855245324 | 0.22481965 | 0.423217612 | 164.2454068 | 140.3681689 | ENSG00000163512 |
| NA | -0.225852328 | 0.855089699 | 0.672310538 | 0.82439555 | 17.81445761 | 15.2364584 | ENSG00000228205 |
| PIGA | -0.225853429 | 0.855089047 | 0.250811705 | 0.456317828 | 148.738638 | 127.3578555 | ENSG00000165195 |
| NCKAP5-AS2 | -0.225883266 | 0.855071362 | 0.759642028 | 0.87528003 | 10.10673594 | 8.59051011 | ENSG00000226953 |
| CASP2 | -0.225961431 | 0.855025036 | 0.050762577 | 0.142805194 | 575.1028736 | 491.5667841 | ENSG00000106144 |
| NBR2 | -0.225986472 | 0.855010195 | 0.474449931 | 0.67731814 | 49.40241295 | 42.24596215 | ENSG00000198496 |
| ARHGEF19 | -0.226050987 | 0.854971961 | 0.003315339 | 0.01567606 | 1466.907022 | 1254.499078 | ENSG00000142632 |
| LINGO2 | -0.226074341 | 0.854958121 | 0.44030935 | 0.648008267 | 59.72235945 | 50.98220805 | ENSG00000174482 |
| CHMP4B | -0.226181542 | 0.854894595 | 0.001223754 | 0.006681296 | 2293.989413 | 1961.041647 | ENSG00000101421 |

| | | | | | | | |
|---|---|---|---|---|---|---|---|
| MMP14 | -0.22631005 | 0.854818449 | 0.000612336 | 0.003659512 | 2134.107508 | 1824.141292 | ENSG00000157227 |
| VLDLR | -0.226324298 | 0.854810007 | 0.048852144 | 0.138545239 | 485.5050318 | 414.7368318 | ENSG00000147852 |
| ZNF417 | -0.226354752 | 0.854791963 | 0.252847744 | 0.458828638 | 141.9063367 | 121.4533983 | ENSG00000173480 |
| TIMM23 | -0.226418582 | 0.854754144 | 0.000163159 | 0.001149294 | 2427.787388 | 2074.89634 | ENSG00000265354 |
| SBDS | -0.226440537 | 0.854741137 | 0.089795724 | 0.220061283 | 318.2150809 | 271.9688573 | ENSG00000126524 |
| GPR62 | -0.226899009 | 0.854469553 | 0.756230619 | 0.8732639 | 9.316158041 | 7.939422219 | ENSG00000180929 |
| NA | -0.227003262 | 0.854407809 | 0.763586322 | 0.877522527 | 8.508977984 | 7.290509791 | ENSG00000223612 |
| PEX5 | -0.2270651 | 0.854371187 | 0.000279022 | 0.001846945 | 1922.703535 | 1642.590518 | ENSG00000139197 |
| COMMD5 | -0.227298749 | 0.854232831 | 0.017632008 | 0.062100049 | 770.6182731 | 658.2267598 | ENSG00000170619 |
| ACP4 | -0.227397335 | 0.854174459 | 0.833299435 | 0.915987775 | 5.033206346 | 4.273783965 | ENSG00000142513 |
| STK38 | -0.227436792 | 0.854151098 | 0.027616344 | 0.089105527 | 552.1268149 | 471.8020327 | ENSG00000112079 |
| IFT140 | -0.227446902 | 0.854145112 | 0.029952192 | 0.094811791 | 629.3086023 | 537.1618486 | ENSG00000187535 |
| ZFP36L1 | -0.227486763 | 0.854121513 | 0.000523403 | 0.003199889 | 2359.940933 | 2016.234727 | ENSG00000185650 |
| BANF1 | -0.227565317 | 0.854075008 | 0.000185644 | 0.001290354 | 4699.213277 | 4012.786969 | ENSG00000175334 |
| NEPRO | -0.227573994 | 0.854069871 | 0.026835993 | 0.087123558 | 701.2130592 | 599.301273 | ENSG00000163608 |
| UBE4B | -0.227629028 | 0.854037292 | 0.000420104 | 0.002641031 | 3358.242529 | 2867.968033 | ENSG00000130939 |
| FARS2 | -0.227630011 | 0.85403671 | 0.076224285 | 0.194947945 | 359.0414962 | 306.6897361 | ENSG00000145982 |
| WDR5 | -0.227819363 | 0.853924626 | 0.000144062 | 0.001030537 | 2543.113656 | 2171.884147 | ENSG00000196363 |
| ARPC2 | -0.227830446 | 0.853918066 | 0.000127171 | 0.000920513 | 4178.06858 | 3567.720619 | ENSG00000163466 |
| CSDE1 | -0.227858002 | 0.853901756 | 3.54E-06 | 3.68E-05 | 13425.28927 | 11463.67007 | ENSG00000009307 |
| CHKB-DT | -0.227870292 | 0.853894482 | 0.468567074 | 0.671795414 | 54.0774124 | 46.25766948 | ENSG00000205559 |
| QARS1 | -0.227884624 | 0.853885999 | 5.18E-06 | 5.25E-05 | 6153.532079 | 5254.102691 | ENSG00000172053 |
| TUBA1B | -0.227916549 | 0.853867104 | 1.26E-05 | 0.000117299 | 4392.256087 | 3750.189734 | ENSG00000123416 |
| XPO6 | -0.227921322 | 0.853864279 | 4.89E-05 | 0.000394298 | 4228.668421 | 3610.70774 | ENSG00000169180 |
| VKORC1L1 | -0.228108283 | 0.853753633 | 0.003567236 | 0.01666269 | 1540.669202 | 1315.555279 | ENSG00000196715 |
| UBOX5 | -0.228140049 | 0.853734834 | 0.284197919 | 0.493356131 | 139.212244 | 119.0329949 | ENSG00000185019 |
| NA | -0.228386718 | 0.853588877 | 0.814997502 | 0.905897873 | 7.471069339 | 6.354928216 | ENSG00000108785 |
| NA | -0.228424015 | 0.85356681 | 0.468896703 | 0.671961924 | 62.84762795 | 53.52375472 | ENSG00000228222 |
| CEP83 | -0.228432938 | 0.853561531 | 0.073398273 | 0.189724389 | 369.0103292 | 314.7553149 | ENSG00000173588 |
| TMEM62 | -0.22857218 | 0.853479153 | 0.548596632 | 0.736074416 | 40.62179831 | 34.6272385 | ENSG00000137842 |
| NOLC1 | -0.228657862 | 0.853428466 | 1.32E-05 | 0.00012216 | 6709.412352 | 5726.175864 | ENSG00000166197 |
| KIAA0930 | -0.228720738 | 0.853391273 | 0.00048515 | 0.003002729 | 2382.460561 | 2032.919768 | ENSG00000100364 |
| SOX12 | -0.228778251 | 0.853357253 | 0.000856986 | 0.004892435 | 4121.790265 | 3517.991513 | ENSG00000177732 |
| FAM126B | -0.228790978 | 0.853349725 | 0.402981601 | 0.61430727 | 69.46376955 | 59.22882455 | ENSG00000155744 |
| ZNF584 | -0.228799262 | 0.853344825 | 0.310418038 | 0.521021071 | 98.8424794 | 84.40263498 | ENSG00000171574 |
| PHF20L1 | -0.228851629 | 0.853313851 | 0.041553288 | 0.122232686 | 472.8655996 | 403.7285453 | ENSG00000129292 |
| HAUS2 | -0.228945461 | 0.853258354 | 0.038309897 | 0.114911435 | 459.5585675 | 392.2805924 | ENSG00000137814 |
| NA | -0.229041754 | 0.853201405 | 0.532216704 | 0.723764128 | 41.21785274 | 35.20636813 | ENSG00000240253 |
| CNOT8 | -0.229085293 | 0.853175656 | 0.005800299 | 0.025014536 | 989.7856994 | 844.3410461 | ENSG00000155508 |
| IGHMBP2 | -0.229119138 | 0.853155641 | 0.029512283 | 0.093686244 | 534.8606129 | 456.11832 | ENSG00000132740 |
| GRM7 | -0.22912938 | 0.853149585 | 0.825123845 | 0.911703624 | 4.266349541 | 3.648518091 | ENSG00000196277 |
| PSIP1 | -0.229184809 | 0.853116807 | 2.41E-05 | 0.000211028 | 8138.471599 | 6943.170507 | ENSG00000164985 |
| NA | -0.229226245 | 0.853092305 | 0.317286763 | 0.527414886 | 92.91460688 | 79.26371464 | ENSG00000213707 |
| ZNHIT6 | -0.229571158 | 0.852888376 | 0.020125532 | 0.069154374 | 602.0057871 | 513.5877604 | ENSG00000117174 |
| CTNNB1 | -0.229630549 | 0.852853266 | 1.98E-06 | 2.17E-05 | 10782.44935 | 9195.294105 | ENSG00000168036 |
| GSK3A | -0.229640771 | 0.852847223 | 0.003739874 | 0.017348247 | 1141.829384 | 973.954115 | ENSG00000105723 |
| CCNJL | -0.229690556 | 0.852817793 | 0.002285808 | 0.01138383 | 1123.396853 | 958.0862432 | ENSG00000135083 |
| DDX51 | -0.229702318 | 0.85281084 | 0.017503981 | 0.061730031 | 953.6381472 | 813.6740319 | ENSG00000185163 |
| H2AC16 | -0.229748091 | 0.852783783 | 0.803361033 | 0.899636688 | 6.945898872 | 5.905571074 | ENSG00000276903 |
| CLCN6 | -0.229860449 | 0.85271737 | 0.014028984 | 0.051574336 | 740.9249864 | 631.8878208 | ENSG00000011021 |
| PIGU | -0.229888413 | 0.852700842 | 0.018091022 | 0.063326961 | 703.9259893 | 599.9787727 | ENSG00000101464 |
| RAB11A | -0.229942075 | 0.852669126 | 0.000583747 | 0.00350861 | 2646.718822 | 2256.614686 | ENSG00000103769 |
| PTPN18 | -0.229948125 | 0.85266555 | 0.067613794 | 0.177913683 | 352.5712145 | 300.6522397 | ENSG00000072135 |
| HSDL1 | -0.230278631 | 0.852470236 | 0.002494025 | 0.012300215 | 1223.600107 | 1042.674267 | ENSG00000103160 |
| NA | -0.23033242 | 0.852438454 | 0.785014913 | 0.890433415 | 7.458026658 | 6.320404345 | ENSG00000263551 |
| ARL2 | -0.230668043 | 0.852240169 | 0.051033983 | 0.143376491 | 479.6673668 | 408.8466999 | ENSG00000213465 |
| SVBP | -0.230771925 | 0.852178805 | 0.373497798 | 0.585549432 | 83.05489677 | 70.71179579 | ENSG00000177868 |
| RTRAF | -0.230806163 | 0.852158581 | 6.92E-05 | 0.000536921 | 3248.308944 | 2768.7028 | ENSG00000087302 |
| SLC2A9 | -0.230816526 | 0.85215246 | 0.78543391 | 0.890433415 | 6.609328139 | 5.630725203 | ENSG00000109667 |
| NSUN5P2 | -0.230900138 | 0.852103075 | 0.558214593 | 0.743916196 | 36.37591588 | 30.9543388 | ENSG00000106133 |
| CNN3 | -0.231050342 | 0.852014364 | 4.97E-06 | 5.05E-05 | 4698.6871 | 4003.165418 | ENSG00000117519 |
| NA | -0.231050857 | 0.85201406 | 0.825143677 | 0.911703624 | 4.269909014 | 3.644167164 | ENSG00000235254 |
| CDADC1 | -0.231080281 | 0.851996683 | 0.415376006 | 0.626197483 | 64.35584044 | 54.91911925 | ENSG00000102543 |
| C9orf135 | -0.231105887 | 0.851981562 | 0.229172659 | 0.428932177 | 177.6443742 | 151.5266374 | ENSG00000204711 |
| UPK1A-AS1 | -0.231124162 | 0.851970769 | 0.793257144 | 0.894207674 | 6.203958375 | 5.305181257 | ENSG00000226510 |
| PPAT | -0.231140989 | 0.851960832 | 0.005173979 | 0.022755147 | 1764.954896 | 1503.488736 | ENSG00000128059 |
| ERCC2 | -0.231182319 | 0.851936426 | 0.00104625 | 0.005827848 | 2625.828704 | 2237.047069 | ENSG00000104884 |
| PCDHB2 | -0.231194577 | 0.851929187 | 0.218757219 | 0.41497748 | 170.1377363 | 145.1737894 | ENSG00000112852 |
| HINT1 | -0.231219694 | 0.851914355 | 0.000287634 | 0.001895314 | 3236.279924 | 2756.663673 | ENSG00000169567 |
| TYMS | -0.23127778 | 0.851880056 | 0.088936339 | 0.218437088 | 304.2227282 | 259.2379516 | ENSG00000176890 |
| SEC22B | -0.231421375 | 0.851795271 | 0.015214566 | 0.055106841 | 722.1989812 | 614.9752223 | ENSG00000265808 |
| NA | -0.231578788 | 0.851702336 | 0.348177694 | 0.560232732 | 86.90725691 | 74.06153791 | ENSG00000174977 |
| PRR5-ARHGAP8 | -0.231600593 | 0.851689463 | 0.84332034 | 0.922013311 | 4.328029629 | 3.69146027 | ENSG00000248405 |
| SF3A1 | -0.231813822 | 0.851563594 | 5.92E-06 | 5.94E-05 | 4402.570929 | 3748.927032 | ENSG00000099995 |

| | | | | | | | |
|---|---|---|---|---|---|---|---|
| OLFM1 | -0.231856702 | 0.851538284 | 0.001100841 | 0.006096594 | 1450.10403 | 1235.142064 | ENSG00000130558 |
| LINC01670 | -0.231964954 | 0.851474391 | 0.653991554 | 0.81254148 | 18.35742544 | 15.58782436 | ENSG00000279094 |
| NFYC | -0.232013876 | 0.851445518 | 0.015386437 | 0.05560139 | 788.4174801 | 671.6528227 | ENSG00000066136 |
| SLC35F5 | -0.232169379 | 0.851353749 | 0.081161297 | 0.204037686 | 406.8989329 | 346.2339749 | ENSG00000115084 |
| LOC101927060 | -0.232184629 | 0.85134475 | 0.14796975 | 0.315752399 | 202.9213083 | 172.695056 | ENSG00000262879 |
| CEP89 | -0.232221243 | 0.851323144 | 0.032790057 | 0.101844109 | 471.2704858 | 401.1680819 | ENSG00000121289 |
| GLCCI1 | -0.232366032 | 0.851237709 | 0.087792473 | 0.21624636 | 334.3354216 | 284.6633699 | ENSG00000106415 |
| EIF3I | -0.232411075 | 0.851211133 | 3.37E-06 | 3.52E-05 | 5616.535651 | 4780.677417 | ENSG00000084623 |
| L3MBTL2 | -0.232428194 | 0.851201033 | 0.019724515 | 0.067936799 | 553.7539128 | 471.3413468 | ENSG00000100395 |
| MAPK1 | -0.232827906 | 0.850965232 | 0.000560783 | 0.00338784 | 1859.247447 | 1581.80407 | ENSG00000100030 |
| BANCR | -0.232881022 | 0.850933903 | 0.756284662 | 0.8732639 | 9.017546297 | 7.662400884 | ENSG00000278910 |
| NUDT10 | -0.232930682 | 0.850904613 | 0.029811661 | 0.094430262 | 579.6461203 | 493.1775016 | ENSG00000122824 |
| NA | -0.23303584 | 0.850842593 | 0.823667401 | 0.910765202 | 4.269909014 | 3.636694814 | ENSG00000281501 |
| NA | -0.233125523 | 0.850789703 | 0.659078518 | 0.816293303 | 17.14843509 | 14.60589564 | ENSG00000228532 |
| CARHSP1 | -0.233150414 | 0.850775024 | 0.00081102 | 0.004673821 | 1576.641905 | 1341.158911 | ENSG00000153048 |
| MACO1 | -0.233161704 | 0.850768366 | 0.073860181 | 0.19063134 | 319.1687574 | 271.7166394 | ENSG00000204178 |
| NEIL3 | -0.233343632 | 0.850661088 | 0.141212184 | 0.305684559 | 220.5047579 | 187.4582272 | ENSG00000109674 |
| RNF144A | -0.233358495 | 0.850652325 | 0.033894564 | 0.10464261 | 480.1978501 | 408.4387289 | ENSG00000151692 |
| DSC3 | -0.233407602 | 0.85062337 | 0.398946212 | 0.610475104 | 80.49453717 | 68.45614026 | ENSG00000134762 |
| QSER1 | -0.233503877 | 0.850566608 | 0.003772273 | 0.017481566 | 1851.440695 | 1574.594586 | ENSG00000060749 |
| STRBP | -0.23358468 | 0.85051897 | 0.000378978 | 0.002410081 | 2441.013851 | 2075.892854 | ENSG00000165209 |
| RP9 | -0.233621342 | 0.850497357 | 0.179351954 | 0.361904225 | 181.6577748 | 154.5724949 | ENSG00000164610 |
| MTAP | -0.233661636 | 0.850473603 | 0.015876656 | 0.0571106 | 708.3030386 | 602.6312251 | ENSG00000099810 |
| SCFD2 | -0.233678269 | 0.850463798 | 0.064296826 | 0.170900148 | 328.0110259 | 278.9777113 | ENSG00000184178 |
| FAM111A-DT | -0.233748225 | 0.85042256 | 0.500754753 | 0.698998707 | 39.79593117 | 33.80553876 | ENSG00000245571 |
| NA | -0.233788956 | 0.850398551 | 0.520000417 | 0.715549976 | 42.72339605 | 36.28222066 | ENSG00000277531 |
| CORO1C | -0.233812977 | 0.850384392 | 0.000152253 | 0.001081752 | 2916.871441 | 2479.846513 | ENSG00000110880 |
| SIGLEC10 | -0.233830428 | 0.850374106 | 0.56467873 | 0.748949534 | 31.66267811 | 26.86734089 | ENSG00000142512 |
| NA | -0.233957963 | 0.850298935 | 0.247701146 | 0.452708424 | 176.105374 | 149.8198433 | ENSG00000260404 |
| POLR2E | -0.234012625 | 0.850266719 | 3.46E-05 | 0.000289647 | 5744.412736 | 4884.932938 | ENSG00000099817 |
| CD276 | -0.23406182 | 0.850237726 | 2.69E-05 | 0.000232393 | 4367.099875 | 3713.44445 | ENSG00000103855 |
| PRCP | -0.234080919 | 0.850226471 | 0.002224446 | 0.011136973 | 1517.536373 | 1289.710275 | ENSG00000137509 |
| FAM131A | -0.23413563 | 0.850194228 | 0.05587044 | 0.153689898 | 383.4638074 | 326.240389 | ENSG00000175182 |
| FICD | -0.234233174 | 0.850136746 | 0.229853487 | 0.429695417 | 136.7280823 | 116.2483837 | ENSG00000198855 |
| EPC1 | -0.234235009 | 0.850135665 | 0.150991625 | 0.32046267 | 251.8908177 | 214.0476734 | ENSG00000120616 |
| ELOF1 | -0.23427578 | 0.85011164 | 0.002376346 | 0.011785002 | 1463.09173 | 1244.419036 | ENSG00000130165 |
| GABARAPL2 | -0.234398043 | 0.850039599 | 0.008987956 | 0.035897046 | 828.7743773 | 704.6270663 | ENSG00000034713 |
| P2RX2 | -0.234437661 | 0.850016257 | 0.520821961 | 0.715845904 | 49.02193334 | 41.71556604 | ENSG00000187848 |
| SVIL | -0.234543616 | 0.849953832 | 0.019624215 | 0.067652608 | 635.4609742 | 539.7903935 | ENSG00000197321 |
| STC2 | -0.234562084 | 0.849942952 | 0.000784319 | 0.004532394 | 1939.885909 | 1648.238931 | ENSG00000113739 |
| FAM104B | -0.234632763 | 0.849901313 | 0.215798379 | 0.410987533 | 145.6872287 | 123.9122976 | ENSG00000182518 |
| ANKRD12 | -0.234642933 | 0.849895322 | 0.08262915 | 0.206810608 | 323.1768184 | 274.6235078 | ENSG00000101745 |
| TM2D1 | -0.234705278 | 0.849858595 | 0.208769161 | 0.401738843 | 155.2696839 | 132.0889204 | ENSG00000162604 |
| AMN1 | -0.234705619 | 0.849858395 | 0.433638787 | 0.642497135 | 68.93234429 | 58.43729776 | ENSG00000151743 |
| MPZL2 | -0.234801607 | 0.849801852 | 0.554287572 | 0.740815343 | 30.02932543 | 25.53310025 | ENSG00000149573 |
| MOCS1 | -0.234806755 | 0.84979882 | 0.008259528 | 0.033558776 | 741.7629804 | 630.2316817 | ENSG00000124615 |
| SLF1 | -0.23487031 | 0.849761384 | 0.052704971 | 0.146909055 | 407.1598906 | 346.1360267 | ENSG00000133302 |
| CAMKMT | -0.234878355 | 0.849756646 | 0.081993099 | 0.205663391 | 318.6921726 | 270.7318728 | ENSG00000143919 |
| JAML | -0.234941642 | 0.84971937 | 0.810997039 | 0.903953201 | 6.316640132 | 5.379525885 | ENSG00000160593 |
| NA | -0.234964126 | 0.849706128 | 0.820405857 | 0.908914972 | 4.736958866 | 4.007356402 | ENSG00000236154 |
| NA | -0.235082704 | 0.849636292 | 0.835423244 | 0.917415262 | 3.919100392 | 3.353147089 | ENSG00000178660 |
| UNC119B | -0.235115128 | 0.849617196 | 0.00039984 | 0.002526255 | 2119.804235 | 1800.478007 | ENSG00000175970 |
| DOLPP1 | -0.235140811 | 0.849602071 | 0.015116091 | 0.0548072 | 797.8565894 | 677.7540724 | ENSG00000167130 |
| ERAL1 | -0.235443757 | 0.849423686 | 0.002165912 | 0.010872249 | 1318.085932 | 1119.577472 | ENSG00000132591 |
| BCAP31 | -0.235446219 | 0.849422236 | 1.42E-05 | 0.000130502 | 3962.80429 | 3365.750384 | ENSG00000185825 |
| IARS1 | -0.235527688 | 0.849374271 | 6.40E-06 | 6.37E-05 | 8031.969431 | 6821.482621 | ENSG00000196305 |
| MYO1E | -0.235618737 | 0.849320668 | 0.00197299 | 0.010029585 | 1324.094678 | 1124.201686 | ENSG00000157483 |
| SLC43A2 | -0.235656797 | 0.849298262 | 0.022683072 | 0.076127565 | 542.1437185 | 460.613981 | ENSG00000167703 |
| GANAB | -0.235661698 | 0.849295377 | 8.83E-07 | 1.03E-05 | 12232.17288 | 10388.13811 | ENSG00000089597 |
| ZNF580 | -0.235729305 | 0.849255579 | 0.125344119 | 0.281364993 | 234.7296187 | 199.5679773 | ENSG00000213015 |
| DKC1 | -0.235758279 | 0.849238523 | 2.81E-05 | 0.000240853 | 4531.809249 | 3847.914992 | ENSG00000130826 |
| PAIP2B | -0.235872285 | 0.849171416 | 0.009114273 | 0.036251693 | 918.2442546 | 779.2914304 | ENSG00000124374 |
| MAST3 | -0.235995211 | 0.849099065 | 0.089706493 | 0.219917023 | 317.1881561 | 269.2459168 | ENSG00000099308 |
| ARGLU1 | -0.236058771 | 0.849061657 | 0.006230899 | 0.026580195 | 3017.28699 | 2562.017278 | ENSG00000134884 |
| USP28 | -0.236074279 | 0.849052531 | 0.000122481 | 0.000891005 | 2136.609136 | 1814.150583 | ENSG00000048028 |
| CENPX | -0.236097268 | 0.849039002 | 0.000418849 | 0.0026349 | 1775.365382 | 1507.468246 | ENSG00000169689 |
| PLEKHH2 | -0.236198877 | 0.848979206 | 0.35037652 | 0.562514713 | 99.68316925 | 84.63314121 | ENSG00000152527 |
| TMPRSS3 | -0.236208733 | 0.848973406 | 0.832855819 | 0.915777709 | 3.888260348 | 3.322974145 | ENSG00000160183 |
| NSA2 | -0.236216171 | 0.848969029 | 0.000534778 | 0.003255698 | 1494.913792 | 1269.161229 | ENSG00000164346 |
| NRIP2 | -0.236233914 | 0.848958588 | 0.50883322 | 0.705487044 | 43.01224526 | 36.5215967 | ENSG00000053702 |
| PPIL3 | -0.236373917 | 0.848876207 | 0.032766128 | 0.101844109 | 482.7475045 | 409.9576957 | ENSG00000240344 |
| ADPRHL2 | -0.23645056 | 0.848831111 | 0.012287336 | 0.046399988 | 675.4117416 | 573.4434068 | ENSG00000116863 |
| FAM174C | -0.236458329 | 0.84882654 | 0.052394691 | 0.146350205 | 604.1085266 | 513.152402 | ENSG00000228300 |
| WDR45B | -0.236508096 | 0.84879726 | 6.33E-05 | 0.000495048 | 2605.68616 | 2211.303552 | ENSG00000141580 |

| | | | | | | | |
|---|---|---|---|---|---|---|---|
| RCN2 | -0.236612426 | 0.848735881 | 0.000159156 | 0.001124871 | 2740.08398 | 2325.908902 | ENSG00000117906 |
| THOC5 | -0.236710428 | 0.848678228 | 0.002137903 | 0.010772223 | 1456.158569 | 1235.814798 | ENSG00000100296 |
| COPRS | -0.236716528 | 0.84867464 | 0.011006404 | 0.042307337 | 817.4869972 | 693.6602721 | ENSG00000172301 |
| DCHS2 | -0.236797139 | 0.848627221 | 0.508861012 | 0.705487044 | 45.49162671 | 38.60084903 | ENSG00000197410 |
| ZNF836 | -0.236840202 | 0.848601891 | 0.384703544 | 0.596983008 | 70.13810622 | 59.57226666 | ENSG00000196267 |
| RAB6B | -0.236849144 | 0.848596631 | 0.03502986 | 0.107253389 | 438.1037389 | 371.905141 | ENSG00000154917 |
| NA | -0.23694765 | 0.848538691 | 0.69451732 | 0.837010823 | 12.92952774 | 10.97137629 | ENSG00000225850 |
| PARK7 | -0.237137767 | 0.848426879 | 6.28E-06 | 6.25E-05 | 6279.405272 | 5327.44009 | ENSG00000116288 |
| KCND3 | -0.237164307 | 0.848411272 | 0.737633681 | 0.862343405 | 10.13017771 | 8.623804476 | ENSG00000171385 |
| FUCA1 | -0.237170519 | 0.848407619 | 0.1427744 | 0.308112734 | 223.1653409 | 189.2611206 | ENSG00000179163 |
| LIMK2 | -0.23719009 | 0.848396109 | 0.003433805 | 0.016159335 | 1182.815313 | 1003.697133 | ENSG00000182541 |
| LIN54 | -0.237220358 | 0.84837831 | 0.145212555 | 0.311806607 | 191.5462661 | 162.4669675 | ENSG00000189308 |
| KMT5B | -0.237236572 | 0.848368775 | 0.004867286 | 0.021576643 | 913.1856512 | 774.8855961 | ENSG00000110066 |
| SRSF11 | -0.237337854 | 0.848309219 | 4.51E-05 | 0.000367346 | 3680.181439 | 3121.781409 | ENSG00000116754 |
| NA | -0.23736426 | 0.848293692 | 0.836410986 | 0.917869805 | 4.277027959 | 3.627047002 | ENSG00000263766 |
| FTO | -0.237399404 | 0.848273029 | 0.00722768 | 0.030064643 | 805.3397855 | 682.868661 | ENSG00000140718 |
| LINC00662 | -0.237407857 | 0.848268058 | 0.122120429 | 0.275998599 | 268.1741536 | 227.1537299 | ENSG00000261824 |
| PHC1 | -0.237414142 | 0.848264363 | 0.001424195 | 0.007625724 | 14741.99615 | 12505.16701 | ENSG00000111752 |
| PRKCA | -0.237423946 | 0.848258598 | 0.041751999 | 0.12264528 | 569.0448007 | 482.5630061 | ENSG00000154229 |
| TTC37 | -0.237469863 | 0.848231601 | 0.001209479 | 0.006616748 | 1432.665629 | 1215.278026 | ENSG00000198677 |
| RBPMS2 | -0.237708935 | 0.84809105 | 0.000149633 | 0.001065543 | 3621.340587 | 3071.669004 | ENSG00000166831 |
| YAF2 | -0.237746912 | 0.848068726 | 0.627316197 | 0.793729923 | 20.78789015 | 17.66896861 | ENSG00000015153 |
| RNF122 | -0.237859097 | 0.848002782 | 0.074905773 | 0.192383452 | 308.4259231 | 261.6512389 | ENSG00000133874 |
| LOC105374042 | -0.237866118 | 0.847998655 | 0.775689547 | 0.884862404 | 6.667448754 | 5.640373016 | ENSG00000239482 |
| ENDOD1 | -0.237929806 | 0.847961221 | 0.252382683 | 0.458248915 | 117.0365269 | 99.16703565 | ENSG00000149218 |
| ENOPH1 | -0.238028019 | 0.847903497 | 0.000397838 | 0.002515287 | 1935.435738 | 1640.909408 | ENSG00000145293 |
| RPGRIP1 | -0.238043082 | 0.847894644 | 0.699438502 | 0.839970842 | 12.87140713 | 10.90790898 | ENSG00000092200 |
| COASY | -0.238170005 | 0.847820053 | 0.000329871 | 0.002131428 | 2111.501023 | 1790.455351 | ENSG00000068120 |
| BTN2A1 | -0.238248584 | 0.847773876 | 0.067196683 | 0.177112263 | 317.5576257 | 269.1706262 | ENSG00000112763 |
| MMP17 | -0.238296688 | 0.847745609 | 0.011113036 | 0.042656351 | 1198.947275 | 1016.736657 | ENSG00000198598 |
| MYO3A | -0.238330605 | 0.84772568 | 0.312113911 | 0.52250938 | 93.27965339 | 79.12516766 | ENSG00000095777 |
| ACOT8 | -0.23837491 | 0.847699646 | 0.043887387 | 0.127431319 | 391.2429723 | 331.559569 | ENSG00000101473 |
| GPATCH11 | -0.238435974 | 0.847663767 | 0.341912059 | 0.553584467 | 87.09082263 | 73.83131522 | ENSG00000152133 |
| NDN | -0.238482411 | 0.847636483 | 0.013063233 | 0.048687419 | 729.7454093 | 618.9032864 | ENSG00000182636 |
| CLPTM1 | -0.238555651 | 0.847593453 | 0.00021 | 0.001435861 | 2566.312206 | 2175.855276 | ENSG00000104853 |
| HNRNPM | -0.238569313 | 0.847585426 | 7.03E-06 | 6.96E-05 | 15054.5476 | 12760.6491 | ENSG00000099783 |
| NA | -0.238572879 | 0.847583331 | 0.52889499 | 0.721428811 | 35.71670682 | 30.31195277 | ENSG00000257354 |
| RSBN1L | -0.238782104 | 0.847460421 | 0.039543126 | 0.117694551 | 490.0137292 | 415.4979748 | ENSG00000187257 |
| ELAVL1 | -0.238814241 | 0.847441543 | 2.11E-05 | 0.000186781 | 3196.933888 | 2709.419328 | ENSG00000066044 |
| CCDC25 | -0.238815824 | 0.847440613 | 0.005492176 | 0.02389856 | 834.4151806 | 706.9911685 | ENSG00000147419 |
| STK39 | -0.238854142 | 0.847418106 | 0.015918544 | 0.057228576 | 753.0837674 | 637.715504 | ENSG00000198648 |
| IFT122 | -0.238856775 | 0.847416559 | 0.014689979 | 0.053628505 | 630.2128034 | 533.9608911 | ENSG00000163913 |
| SLCO3A1 | -0.238971589 | 0.847349121 | 0.034231175 | 0.105382147 | 429.7142561 | 364.278777 | ENSG00000176463 |
| NA | -0.238987794 | 0.847339604 | 0.821506749 | 0.909334444 | 4.70255935 | 3.96223876 | ENSG00000286788 |
| CNOT1 | -0.239422019 | 0.847084608 | 4.72E-05 | 0.000381953 | 6431.545346 | 5447.157332 | ENSG00000125107 |
| CPSF7 | -0.239523153 | 0.847025229 | 9.85E-05 | 0.000731925 | 2154.32248 | 1825.003297 | ENSG00000149532 |
| PRSS16 | -0.239526166 | 0.847023461 | 0.135341881 | 0.297270462 | 244.7384433 | 207.3990255 | ENSG00000112812 |
| SUCLG1 | -0.23954665 | 0.847011434 | 0.00926417 | 0.036716441 | 711.608236 | 602.6462127 | ENSG00000163541 |
| ZNF22 | -0.239548232 | 0.847010505 | 0.056680171 | 0.155491724 | 329.0172048 | 278.7156347 | ENSG00000165512 |
| UTS2R | -0.239592771 | 0.846984357 | 0.75518812 | 0.872703931 | 8.185449933 | 6.947845683 | ENSG00000181408 |
| VPS50 | -0.239663479 | 0.846942846 | 0.219375233 | 0.415703519 | 128.5817866 | 108.8514644 | ENSG00000004766 |
| SLC16A1-AS1 | -0.239682623 | 0.846931607 | 0.27724151 | 0.48645414 | 124.9269246 | 105.8511237 | ENSG00000226419 |
| DCP2 | -0.239730761 | 0.846903349 | 0.007198652 | 0.029977188 | 1448.724081 | 1226.636864 | ENSG00000172795 |
| DAB2IP | -0.239831524 | 0.8468442 | 0.000750451 | 0.004363382 | 1975.351704 | 1672.958337 | ENSG00000136848 |
| NA | -0.239986476 | 0.84675325 | 0.716471508 | 0.850492157 | 10.97531676 | 9.272716903 | ENSG00000268087 |
| LOC150776 | -0.240043166 | 0.846719978 | 0.05737839 | 0.156901149 | 357.9828417 | 303.0706559 | ENSG00000152117 |
| DEF6 | -0.240302972 | 0.846567511 | 0.132042152 | 0.292232044 | 220.1320338 | 186.499777 | ENSG00000023892 |
| IDH1 | -0.240308822 | 0.846564078 | 7.40E-07 | 8.75E-06 | 10284.14353 | 8705.652778 | ENSG00000138413 |
| AFG1L | -0.240326999 | 0.846553412 | 0.29184752 | 0.501459414 | 94.54795957 | 79.97826984 | ENSG00000135537 |
| NA | -0.240370696 | 0.846527772 | 0.580536936 | 0.760980082 | 28.50984971 | 24.18486086 | ENSG00000224858 |
| ZSWIM8 | -0.240399434 | 0.846510909 | 0.013475178 | 0.049862443 | 634.1692519 | 536.849641 | ENSG00000214655 |
| VILL | -0.24041658 | 0.846500849 | 0.301898127 | 0.512145775 | 100.0263003 | 84.76554067 | ENSG00000136059 |
| C1orf174 | -0.240461052 | 0.846474755 | 0.013876708 | 0.051104062 | 776.4491501 | 657.2186529 | ENSG00000198912 |
| FBXO25 | -0.240557461 | 0.846418191 | 0.16169434 | 0.336299297 | 190.2936724 | 160.9719307 | ENSG00000147364 |
| MAP1A | -0.240587693 | 0.846400454 | 0.261117369 | 0.468211309 | 152.7840732 | 129.4271335 | ENSG00000166963 |
| LCMT1 | -0.241246716 | 0.846013907 | 0.00197204 | 0.010029585 | 1080.15955 | 914.0194801 | ENSG00000205629 |
| SMAP2 | -0.241338073 | 0.845960336 | 0.176625795 | 0.357779484 | 172.1337451 | 145.7732332 | ENSG00000084070 |
| MCC | -0.241369913 | 0.845941666 | 0.027956089 | 0.089894559 | 594.2698945 | 502.872007 | ENSG00000171444 |
| EIF4G3 | -0.24141048 | 0.845917879 | 8.67E-05 | 0.000653989 | 2791.977417 | 2361.098381 | ENSG00000075151 |
| ANAPC7 | -0.24141215 | 0.8459169 | 0.002772963 | 0.013434428 | 1022.68409 | 864.9458493 | ENSG00000196510 |
| GBF1 | -0.241436077 | 0.845902871 | 0.002526937 | 0.012411234 | 1273.24164 | 1077.116924 | ENSG00000107862 |
| KIAA0753 | -0.241510725 | 0.845859103 | 0.032221162 | 0.100494904 | 541.726755 | 457.9336335 | ENSG00000198920 |
| PRPF4B | -0.241631142 | 0.845788505 | 0.000729035 | 0.004259848 | 1888.93316 | 1597.859446 | ENSG00000112739 |
| EDRF1 | -0.241668884 | 0.845766378 | 0.028129632 | 0.090283498 | 466.7953226 | 394.7760754 | ENSG00000107938 |

| | | | | | | | |
|---|---|---|---|---|---|---|---|
| WRAP53 | -0.241801582 | 0.845688589 | 0.007387869 | 0.030589764 | 756.2221298 | 639.6122485 | ENSG00000141499 |
| CWC27 | -0.241927103 | 0.845615014 | 0.002196095 | 0.01100378 | 1161.068329 | 981.9572881 | ENSG00000153015 |
| BRI3BP | -0.242013244 | 0.845564525 | 0.064987135 | 0.172379782 | 370.469956 | 313.5047105 | ENSG00000184992 |
| SAE1 | -0.242101134 | 0.845513014 | 1.50E-07 | 2.01E-06 | 8123.976398 | 6868.573393 | ENSG00000142230 |
| MTFR1L | -0.242183937 | 0.845464488 | 0.142061647 | 0.307030695 | 186.8119509 | 157.8632886 | ENSG00000117640 |
| FBXO11 | -0.242192465 | 0.84545949 | 0.013680815 | 0.050477939 | 599.2775225 | 506.518277 | ENSG00000138081 |
| PEMT | -0.242267258 | 0.84541566 | 0.233982712 | 0.435174661 | 122.8502431 | 103.9362404 | ENSG00000133027 |
| JADE3 | -0.242294862 | 0.845399485 | 0.022057699 | 0.07453059 | 613.2117807 | 518.2013277 | ENSG00000102221 |
| RABGGTA | -0.242428952 | 0.845320913 | 0.110141355 | 0.256665781 | 253.0075667 | 214.0291558 | ENSG00000100949 |
| BSDC1 | -0.242485689 | 0.84528767 | 0.002552197 | 0.012502784 | 1051.845392 | 889.3381369 | ENSG00000160058 |
| ZFP82 | -0.242539352 | 0.845256229 | 0.047570787 | 0.135747102 | 400.7207544 | 338.6339275 | ENSG00000181007 |
| XPOT | -0.242611245 | 0.845214109 | 2.41E-06 | 2.60E-05 | 5421.26719 | 4581.566737 | ENSG00000184575 |
| SPAG16 | -0.242706651 | 0.845158216 | 0.154690487 | 0.326192361 | 207.2217257 | 175.0763737 | ENSG00000144451 |
| ELMOD1 | -0.242792338 | 0.845108021 | 0.55296707 | 0.739709489 | 31.74660476 | 26.81380494 | ENSG00000110675 |
| MAD2L2 | -0.242801317 | 0.845102761 | 2.12E-05 | 0.000187448 | 4431.084359 | 3744.266636 | ENSG00000116670 |
| AQP5 | -0.242816499 | 0.845093868 | 0.45705585 | 0.662777555 | 50.38751432 | 42.48845962 | ENSG00000161798 |
| SLC25A37 | -0.243003443 | 0.844984368 | 0.009384387 | 0.037052928 | 900.1662736 | 760.5923072 | ENSG00000147454 |
| EIF2S3 | -0.243071484 | 0.844944517 | 3.58E-07 | 4.51E-06 | 11632.46808 | 9828.133654 | ENSG00000130741 |
| CCDC137 | -0.243128492 | 0.84491113 | 0.000760877 | 0.004417204 | 1654.714089 | 1398.568774 | ENSG00000185298 |
| HADH | -0.243140468 | 0.844904116 | 0.003062431 | 0.014656898 | 1253.87768 | 1058.857141 | ENSG00000138796 |
| CDK2AP1 | -0.243218831 | 0.844858225 | 0.00322507 | 0.015302874 | 3069.492328 | 2593.343059 | ENSG00000111328 |
| ZDHHC23 | -0.243254184 | 0.844837522 | 0.072039354 | 0.18700525 | 363.0003095 | 306.785123 | ENSG00000184307 |
| BCL2L2 | -0.243269945 | 0.844828292 | 0.044066996 | 0.127815271 | 415.6619772 | 351.1866466 | ENSG00000129473 |
| HSPB11 | -0.24338969 | 0.844758174 | 0.073187526 | 0.189420012 | 398.412734 | 336.5904941 | ENSG00000081870 |
| MED12 | -0.243472557 | 0.844709653 | 0.001272964 | 0.00691001 | 1244.647584 | 1051.573936 | ENSG00000184634 |
| NTHL1 | -0.24353119 | 0.844675324 | 0.000998306 | 0.005607676 | 1259.49982 | 1064.005637 | ENSG00000065057 |
| LOC152048 | -0.243709289 | 0.844571056 | 0.81571992 | 0.906328805 | 4.331589101 | 3.650693555 | ENSG00000270194 |
| LYPD6B | -0.243754753 | 0.844544441 | 0.397652404 | 0.609205812 | 59.29326859 | 50.0348032 | ENSG00000150556 |
| DNAJB5 | -0.243811001 | 0.844511515 | 0.001699716 | 0.008857698 | 1387.122156 | 1171.080145 | ENSG00000137094 |
| C17orf58 | -0.244040241 | 0.844377335 | 0.045915014 | 0.131919221 | 394.273609 | 332.9445803 | ENSG00000186665 |
| NA | -0.244110349 | 0.844336303 | 0.768082625 | 0.880225765 | 7.825158102 | 6.631949551 | ENSG00000251400 |
| FKBP1A | -0.24420463 | 0.844281127 | 0.000869057 | 0.004947889 | 2940.643088 | 2483.342149 | ENSG00000088832 |
| MACROD2 | -0.244239668 | 0.844260623 | 0.636864494 | 0.800725077 | 23.62311424 | 19.83578631 | ENSG00000172264 |
| NA | -0.244322959 | 0.844211882 | 0.759153719 | 0.874877523 | 8.203247295 | 6.929496016 | ENSG00000279500 |
| ADAMTSL3 | -0.244383041 | 0.844176726 | 0.556896215 | 0.743249825 | 29.16193982 | 24.62958353 | ENSG00000156218 |
| SMIM4 | -0.244396609 | 0.844168786 | 0.099026633 | 0.236514224 | 258.6115801 | 218.360754 | ENSG00000168273 |
| SHPRH | -0.244399105 | 0.844167326 | 0.037292585 | 0.112521243 | 550.9989524 | 464.6791813 | ENSG00000146414 |
| ZNF334 | -0.244472733 | 0.844124245 | 0.135486894 | 0.297519767 | 251.0669576 | 211.8019196 | ENSG00000198185 |
| PLA2G12A | -0.244493681 | 0.844111988 | 0.081260011 | 0.204199725 | 295.2185295 | 249.4492889 | ENSG00000123739 |
| PRKAR2A | -0.244532216 | 0.844089442 | 0.0097252 | 0.038159235 | 798.5282301 | 673.7450419 | ENSG00000114302 |
| C1QL2 | -0.244553706 | 0.844076869 | 0.637410855 | 0.801058361 | 19.19753047 | 16.1763909 | ENSG00000144119 |
| CEBPB | -0.244659937 | 0.844014719 | 0.043087928 | 0.125650946 | 418.9376066 | 353.7793429 | ENSG00000172216 |
| CDCA2 | -0.244768475 | 0.843951223 | 0.004041258 | 0.018537565 | 988.6763475 | 834.2789873 | ENSG00000184661 |
| TOE1 | -0.244773908 | 0.843948045 | 0.15223243 | 0.322299265 | 239.1190752 | 201.9648068 | ENSG00000132773 |
| DHX57 | -0.244791871 | 0.843937537 | 0.010766844 | 0.041504766 | 868.8526497 | 733.1034281 | ENSG00000163214 |
| SDHAF3 | -0.244809979 | 0.843926945 | 0.034108717 | 0.105143751 | 449.540435 | 379.2244559 | ENSG00000196636 |
| C9orf40 | -0.24481604 | 0.843923399 | 0.057122386 | 0.156454861 | 328.5972917 | 277.4566846 | ENSG00000135045 |
| ZNF700 | -0.244841217 | 0.843908672 | 0.291447005 | 0.500999152 | 100.7436299 | 84.9952689 | ENSG00000196757 |
| VCP | -0.244848367 | 0.843904489 | 5.69E-09 | 9.78E-08 | 15006.72061 | 12663.82125 | ENSG00000165280 |
| SLC25A21-AS1 | -0.244849419 | 0.843903874 | 0.766750329 | 0.879325885 | 7.903440343 | 6.695416861 | ENSG00000258708 |
| PIP5KL1 | -0.244898509 | 0.843875159 | 0.263832152 | 0.471244699 | 165.2803402 | 139.6755066 | ENSG00000167103 |
| LPAR2 | -0.244987825 | 0.843822917 | 0.01546676 | 0.055870267 | 843.3519051 | 711.9658581 | ENSG00000064547 |
| MTG1 | -0.244994877 | 0.843818793 | 0.385640846 | 0.597653274 | 77.76309645 | 65.69935857 | ENSG00000148824 |
| SERTM2 | -0.245060827 | 0.84378022 | 0.623777668 | 0.792148562 | 20.4074367 | 17.2401366 | ENSG00000260802 |
| PKN2 | -0.245077188 | 0.843770651 | 0.023369445 | 0.078050447 | 613.1956849 | 517.1458323 | ENSG00000065243 |
| MRPL21 | -0.24522788 | 0.843682522 | 0.0031366 | 0.014962032 | 1089.171068 | 918.7235661 | ENSG00000197345 |
| HHAT | -0.245250045 | 0.84366956 | 0.770487976 | 0.881997379 | 9.048386341 | 7.605176419 | ENSG00000054392 |
| NR2C1 | -0.245484374 | 0.843532539 | 0.024658369 | 0.081553992 | 491.5136287 | 414.5131355 | ENSG00000120798 |
| C12orf73 | -0.245494675 | 0.843526516 | 0.251840926 | 0.457549413 | 133.7481674 | 112.9724191 | ENSG00000204954 |
| CCDC59 | -0.245508818 | 0.843518247 | 0.030602168 | 0.096465021 | 588.2393816 | 496.5959126 | ENSG00000133773 |
| NT5C3B | -0.2456075 | 0.843460551 | 0.000367935 | 0.002348245 | 1504.619607 | 1269.05092 | ENSG00000141698 |
| MCM7 | -0.24571582 | 0.843397225 | 8.41E-09 | 1.40E-07 | 12209.8267 | 10297.4303 | ENSG00000166508 |
| CRADD | -0.245717199 | 0.843396419 | 0.351706742 | 0.56379668 | 70.0942235 | 59.15837935 | ENSG00000169372 |
| CCDC32 | -0.245756925 | 0.843373196 | 0.432166786 | 0.641566127 | 55.88245718 | 47.14349894 | ENSG00000128891 |
| GIGYF2 | -0.2458393 | 0.843325042 | 0.001769 | 0.009150252 | 1653.898773 | 1394.095873 | ENSG00000204120 |
| SAMD12 | -0.245944705 | 0.84326343 | 0.297627075 | 0.507812023 | 119.3272825 | 100.4901881 | ENSG00000177570 |
| CWC15 | -0.2459496 | 0.843260569 | 0.01047568 | 0.04056453 | 826.4843813 | 697.043971 | ENSG00000150316 |
| ZNF500 | -0.246029129 | 0.843214085 | 0.13280662 | 0.293373846 | 223.9274947 | 188.8998959 | ENSG00000103199 |
| CNOT7 | -0.246082112 | 0.843183119 | 0.000800179 | 0.004614163 | 1853.285653 | 1562.667663 | ENSG00000198791 |
| ECHDC3 | -0.246254228 | 0.843082531 | 0.053263431 | 0.147923514 | 391.6441978 | 330.2126544 | ENSG00000134463 |
| ALAD | -0.246326088 | 0.843040539 | 0.040860422 | 0.120702186 | 394.7445238 | 332.8833314 | ENSG00000148218 |
| MAPK6 | -0.246480332 | 0.842950411 | 9.26E-05 | 0.000694209 | 2804.677156 | 2363.893595 | ENSG00000069956 |
| NA | -0.246520626 | 0.842926868 | 0.175902972 | 0.356583044 | 178.349246 | 150.3637866 | ENSG00000078319 |
| TKT | -0.246644639 | 0.842854414 | 3.02E-08 | 4.58E-07 | 23535.24479 | 19836.73833 | ENSG00000163931 |

| | | | | | | | |
|---|---|---|---|---|---|---|---|
| VTI1B | -0.246674323 | 0.842837072 | 0.019452931 | 0.067148348 | 484.2257934 | 408.1535728 | ENSG00000100568 |
| ZNF660 | -0.246675061 | 0.842836641 | 0.164100094 | 0.339508138 | 162.2087693 | 136.7974957 | ENSG00000144792 |
| NUDT11 | -0.246702833 | 0.842820416 | 0.007194418 | 0.029966164 | 813.9178936 | 685.8274797 | ENSG00000196368 |
| NA | -0.246819183 | 0.842752447 | 0.829421833 | 0.9144008 | 3.95705938 | 3.346620699 | ENSG00000244055 |
| KAT2A | -0.246951488 | 0.842675165 | 0.321964805 | 0.53256594 | 1165.322351 | 982.0913131 | ENSG00000108773 |
| IP6K1 | -0.246983087 | 0.842656708 | 0.000856945 | 0.004892435 | 1244.184627 | 1048.525682 | ENSG00000176095 |
| CLDN12 | -0.247037927 | 0.842624678 | 0.012649404 | 0.047444685 | 686.0635303 | 577.9277484 | ENSG00000157224 |
| RCOR2 | -0.247041078 | 0.842622837 | 0.000358302 | 0.002292186 | 2759.721744 | 2325.340999 | ENSG00000167771 |
| C2CD2 | -0.24704957 | 0.842617877 | 0.039150543 | 0.116820668 | 376.4125995 | 317.1035921 | ENSG00000157617 |
| NELL2 | -0.247218395 | 0.842519279 | 0.002112634 | 0.010667673 | 1283.884785 | 1081.267864 | ENSG00000184613 |
| VGF | -0.247298546 | 0.842472473 | 0.194662997 | 0.383218045 | 238.7833681 | 201.5017048 | ENSG00000128564 |
| ADGRA1 | -0.247361993 | 0.842435424 | 0.741179718 | 0.864984487 | 9.786488034 | 8.267141628 | ENSG00000197177 |
| GAS1 | -0.247470768 | 0.842371909 | 0.202631119 | 0.39393542 | 137.0516104 | 115.4068939 | ENSG00000180447 |
| NA | -0.247612926 | 0.842288909 | 0.475986019 | 0.678429646 | 42.17775848 | 35.59348747 | ENSG00000236565 |
| CPE | -0.247688925 | 0.842244539 | 0.018833708 | 0.065360307 | 530.8027448 | 447.0325571 | ENSG00000109472 |
| SPIN1 | -0.247721079 | 0.842225768 | 6.94E-05 | 0.000537872 | 2759.05246 | 2323.218907 | ENSG00000106723 |
| DHRS4-AS1 | -0.247851115 | 0.842149858 | 0.132882799 | 0.293473471 | 190.415558 | 160.4476095 | ENSG00000215256 |
| LINC00539 | -0.24786959 | 0.842139074 | 0.750908346 | 0.870392545 | 8.291012746 | 6.981140049 | ENSG00000224429 |
| LOC400684 | -0.247874613 | 0.842136142 | 0.639132715 | 0.802206613 | 19.69514103 | 16.63199089 | ENSG00000267213 |
| CDK2 | -0.24788097 | 0.842132431 | 0.047446574 | 0.13547453 | 393.3075008 | 331.0780517 | ENSG00000123374 |
| TOMM34 | -0.247920338 | 0.842109451 | 0.001803424 | 0.009282548 | 1165.16843 | 981.1880627 | ENSG00000025772 |
| GPHN | -0.248170961 | 0.841963174 | 0.064891883 | 0.172263594 | 342.5917555 | 288.6717879 | ENSG00000171723 |
| MPHOSPH6 | -0.248272631 | 0.841903841 | 0.029455232 | 0.093552314 | 637.0759436 | 536.6450724 | ENSG00000135698 |
| NA | -0.248401852 | 0.841828436 | 0.693547867 | 0.836322807 | 12.67479873 | 10.65765562 | ENSG00000228274 |
| OIP5 | -0.248409158 | 0.841824173 | 0.298009947 | 0.508219151 | 88.18952164 | 74.25078297 | ENSG00000104147 |
| MZT2B | -0.248410228 | 0.841823548 | 0.360370881 | 0.572516706 | 82.72452911 | 69.55953876 | ENSG00000152082 |
| DRP2 | -0.248440178 | 0.841806072 | 0.596549772 | 0.77306268 | 23.33251117 | 19.61012546 | ENSG00000102385 |
| NOL10 | -0.248461489 | 0.841793638 | 0.007742105 | 0.031847015 | 779.8106169 | 656.6961282 | ENSG00000115761 |
| ERI3 | -0.248629392 | 0.841695674 | 0.000304037 | 0.001988828 | 1603.532185 | 1350.091581 | ENSG00000117419 |
| TDP2 | -0.248645526 | 0.841686261 | 0.002267072 | 0.011314391 | 1193.132538 | 1004.252784 | ENSG00000111802 |
| RNGTT | -0.248923955 | 0.841523838 | 0.021223732 | 0.07216933 | 689.5322614 | 580.1203881 | ENSG00000111880 |
| NA | -0.248961916 | 0.841501696 | 0.562568134 | 0.747544273 | 26.77388329 | 22.52261726 | ENSG00000243176 |
| GPT2 | -0.249023473 | 0.841465791 | 2.82E-05 | 0.000241969 | 2774.450637 | 2334.750203 | ENSG00000166123 |
| ASPRV1 | -0.249064974 | 0.841441586 | 0.685287613 | 0.831721353 | 14.51157327 | 12.19703198 | ENSG00000244617 |
| CRMP1 | -0.249138503 | 0.841398702 | 3.17E-07 | 4.04E-06 | 6144.766108 | 5170.392476 | ENSG00000072832 |
| ZNF767P | -0.249217014 | 0.841352914 | 0.24753111 | 0.452485239 | 196.5035545 | 165.4898111 | ENSG00000133624 |
| NA | -0.249306847 | 0.841300527 | 0.665561879 | 0.82085806 | 16.2525737 | 13.68838019 | ENSG00000228393 |
| CIB2 | -0.249357933 | 0.841270737 | 0.001927567 | 0.009825129 | 1095.552188 | 921.6772262 | ENSG00000136425 |
| DMC1 | -0.249384349 | 0.841255333 | 0.293056683 | 0.502804708 | 90.1947343 | 75.86951729 | ENSG00000100206 |
| LRRFIP2 | -0.249470931 | 0.841204848 | 0.016628994 | 0.059252558 | 605.471997 | 509.5478876 | ENSG00000093167 |
| TMEM9B | -0.249516047 | 0.841178542 | 0.136933073 | 0.299627263 | 243.4387651 | 204.7260777 | ENSG00000175348 |
| MARCHF11 | -0.249574414 | 0.841144511 | 0.796087707 | 0.895847715 | 5.958712565 | 5.024754954 | ENSG00000183654 |
| LZTS1 | -0.249849583 | 0.840984093 | 0.037323176 | 0.112595556 | 483.8150841 | 406.9954291 | ENSG00000061337 |
| TTLL12 | -0.249931062 | 0.840936598 | 9.30E-06 | 8.95E-05 | 6259.011954 | 5263.88325 | ENSG00000100304 |
| ARID5A | -0.249935189 | 0.840934192 | 0.092155494 | 0.224272746 | 250.2126403 | 210.3968119 | ENSG00000196843 |
| GOT1 | -0.249968276 | 0.840914907 | 0.000961032 | 0.005420025 | 1477.75379 | 1242.408096 | ENSG00000120053 |
| NA | -0.250072098 | 0.840854393 | 0.752561306 | 0.871085164 | 8.356252306 | 7.02313627 | ENSG00000279531 |
| LAPTM4A | -0.250097548 | 0.84083956 | 4.33E-06 | 4.44E-05 | 4938.452894 | 4151.756418 | ENSG00000068697 |
| C5orf51 | -0.25013777 | 0.840816118 | 0.053177605 | 0.147783608 | 464.6999802 | 390.5376028 | ENSG00000205765 |
| HSD17B12 | -0.250204711 | 0.840777105 | 0.029130293 | 0.092723004 | 521.9097533 | 438.9867254 | ENSG00000149084 |
| CDC26 | -0.250282125 | 0.84073199 | 0.044361807 | 0.128295975 | 350.4702278 | 294.6131123 | ENSG00000176386 |
| ISOC1 | -0.250339615 | 0.840698489 | 0.003526935 | 0.016498928 | 1049.154222 | 882.1224232 | ENSG00000066583 |
| KAT7 | -0.250345971 | 0.840694785 | 6.15E-06 | 6.15E-05 | 4629.748126 | 3892.018425 | ENSG00000136504 |
| ZNF74 | -0.250565645 | 0.840566785 | 8.53E-05 | 0.000644692 | 1900.9653 | 1597.936922 | ENSG00000185252 |
| PIGT | -0.250565798 | 0.840566696 | 0.00298998 | 0.014367879 | 1031.904525 | 867.5434339 | ENSG00000124155 |
| SLC27A1 | -0.250590706 | 0.840552184 | 0.094900683 | 0.229328138 | 240.6040735 | 202.1739373 | ENSG00000130304 |
| XXYLT1 | -0.250741578 | 0.840464287 | 0.007700956 | 0.031712289 | 732.9002964 | 616.0994092 | ENSG00000173950 |
| SCAMP2 | -0.25094678 | 0.840344751 | 0.001590372 | 0.008371495 | 974.2338526 | 818.7427773 | ENSG00000140497 |
| NA | -0.250982587 | 0.840323895 | 0.743362894 | 0.866407964 | 8.694018246 | 7.276511051 | ENSG00000226015 |
| LOC101929295 | -0.251031797 | 0.840295232 | 0.745196413 | 0.867425425 | 8.298131691 | 6.964965846 | ENSG00000245552 |
| FJX1 | -0.251090038 | 0.84026131 | 0.025541443 | 0.083814101 | 526.7075025 | 442.8826293 | ENSG00000179431 |
| CDK20 | -0.251165671 | 0.840217261 | 0.289810053 | 0.499184864 | 91.71657428 | 77.07135848 | ENSG00000156345 |
| CLK1 | -0.251270568 | 0.840156171 | 0.070268447 | 0.183290004 | 309.9030168 | 260.3823574 | ENSG00000013441 |
| SOGA1 | -0.251331773 | 0.84012053 | 0.006708273 | 0.02826545 | 935.1857084 | 785.1654108 | ENSG00000149639 |
| FOXP4 | -0.251343937 | 0.840113446 | 0.015714692 | 0.056595802 | 850.4401823 | 714.8518428 | ENSG00000137166 |
| LAMA1 | -0.251362036 | 0.840102907 | 0.000204578 | 0.001404584 | 2742.435704 | 2303.211236 | ENSG00000101680 |
| DPM1 | -0.251368113 | 0.840099368 | 0.001720776 | 0.008937194 | 1081.778638 | 909.0913712 | ENSG00000000419 |
| PBRM1 | -0.251391555 | 0.840085717 | 0.005616644 | 0.024355986 | 1215.175997 | 1020.722819 | ENSG00000163939 |
| TLK1 | -0.25165418 | 0.839932804 | 0.003215132 | 0.015274912 | 1154.52574 | 969.8874313 | ENSG00000198586 |
| PKD1P5-LOC105376752 | -0.251778775 | 0.839860268 | 0.079775415 | 0.201384669 | 296.1170867 | 248.927311 | ENSG00000254681 |
| HDHD2 | -0.251992721 | 0.83973573 | 0.725503411 | 0.855465279 | 10.32826066 | 8.668922118 | ENSG00000167220 |
| ANO1 | -0.252053434 | 0.839700392 | 0.170384915 | 0.348653062 | 164.9434901 | 138.6679964 | ENSG00000131620 |
| CLK3 | -0.25210949 | 0.839667766 | 0.003686318 | 0.017134164 | 830.5915495 | 697.4021469 | ENSG00000179335 |
| CALML4 | -0.25217949 | 0.839627026 | 0.751059779 | 0.870392545 | 7.950882541 | 6.690119975 | ENSG00000129007 |

| | | | | | | | |
|---|---|---|---|---|---|---|---|
| SH3D21 | -0.25227492 | 0.839571489 | 0.12888333 | 0.287209334 | 282.291672 | 237.0525336 | ENSG00000214193 |
| LOC728485 | -0.252342718 | 0.839532035 | 0.747125461 | 0.868435054 | 8.356252306 | 7.006962066 | ENSG00000267260 |
| ADCY3 | -0.252379068 | 0.839510883 | 0.005169116 | 0.022744351 | 807.7457428 | 678.2420707 | ENSG00000138031 |
| FAAP20 | -0.252411074 | 0.839492258 | 0.001586087 | 0.008357143 | 1013.284284 | 850.8297416 | ENSG00000162585 |
| CTAGE8 | -0.252423982 | 0.839484747 | 0.779709172 | 0.887571291 | 6.300037978 | 5.275008313 | ENSG00000244693 |
| MRPS18A | -0.252491052 | 0.839445721 | 0.035259995 | 0.107853095 | 527.7583766 | 442.9303942 | ENSG00000096080 |
| CCND2 | -0.252511492 | 0.839433828 | 0.000895617 | 0.005086834 | 1343.55075 | 1127.630691 | ENSG00000118971 |
| C1QL3 | -0.252521775 | 0.839427845 | 0.4741112 | 0.677209204 | 49.3549446 | 41.39577048 | ENSG00000165985 |
| CMAS | -0.252527032 | 0.839424786 | 0.000819432 | 0.004709364 | 1575.013865 | 1322.090646 | ENSG00000111726 |
| PPP2R5D | -0.252596195 | 0.839384545 | 0.000614848 | 0.00367101 | 1431.773885 | 1202.177731 | ENSG00000112640 |
| GTF2IP4 | -0.252603225 | 0.839380455 | 0.046103332 | 0.132319364 | 581.5788265 | 487.9797484 | ENSG00000233369 |
| LINC00174 | -0.252644323 | 0.839356543 | 0.47982602 | 0.681932148 | 52.12495528 | 43.71801473 | ENSG00000179406 |
| NA | -0.252644964 | 0.839356171 | 0.783610485 | 0.889920261 | 5.876870851 | 4.939816555 | ENSG00000178429 |
| MANSC1 | -0.252708077 | 0.839319453 | 0.348477866 | 0.560425772 | 90.53692386 | 75.88306453 | ENSG00000111261 |
| SLC43A1 | -0.252716757 | 0.839314403 | 0.020271858 | 0.069535784 | 584.242611 | 490.3220663 | ENSG00000149150 |
| GARS1 | -0.252843693 | 0.839240559 | 2.04E-05 | 0.000181119 | 4816.724801 | 4041.71134 | ENSG00000106105 |
| ZNF32-AS2 | -0.252845696 | 0.839239393 | 0.651886458 | 0.811279351 | 16.25613318 | 13.64449205 | ENSG00000230565 |
| FOXP1 | -0.252946153 | 0.839180958 | 0.311701227 | 0.522197973 | 86.006667 | 72.11023879 | ENSG00000114861 |
| CENPH | -0.253040829 | 0.839125889 | 0.009146844 | 0.036335327 | 678.2606705 | 569.4642159 | ENSG00000153044 |
| TUBB | -0.253044508 | 0.839123749 | 5.18E-11 | 1.21E-09 | 32781.29884 | 27507.33481 | ENSG00000196230 |
| OS9 | -0.253252834 | 0.839002588 | 0.000229178 | 0.001552934 | 2014.728685 | 1689.948465 | ENSG00000135506 |
| MSH3 | -0.253544781 | 0.838832823 | 0.032091554 | 0.100223657 | 419.6122749 | 351.9868323 | ENSG00000113318 |
| CCNA2 | -0.253604206 | 0.838798272 | 0.000111778 | 0.000821327 | 2643.618112 | 2217.421813 | ENSG00000145386 |
| SLC35C1 | -0.253694497 | 0.838745777 | 0.028215137 | 0.09052716 | 482.3226358 | 404.9032623 | ENSG00000181830 |
| RNF40 | -0.253699031 | 0.838743141 | 0.000190616 | 0.00132199 | 1844.676277 | 1547.662268 | ENSG00000103549 |
| WDR19 | -0.253713543 | 0.838734705 | 0.120740313 | 0.273698333 | 323.1444783 | 271.146182 | ENSG00000157796 |
| NA | -0.25382623 | 0.838669194 | 0.81712362 | 0.906821388 | 3.953499908 | 3.320798682 | ENSG00000228252 |
| PTPA | -0.253849787 | 0.838655501 | 1.16E-06 | 1.33E-05 | 5815.443654 | 4876.878639 | ENSG00000119383 |
| TMEM167A | -0.253935392 | 0.838605739 | 0.000164741 | 0.001158712 | 3306.495888 | 2773.153171 | ENSG00000174695 |
| SGTA | -0.254020918 | 0.838556026 | 6.80E-06 | 6.74E-05 | 3267.368013 | 2740.322356 | ENSG00000104969 |
| TIGD3 | -0.25404325 | 0.838543046 | 0.566689162 | 0.750531324 | 24.16313299 | 20.26773975 | ENSG00000173825 |
| NA | -0.254103116 | 0.83850825 | 0.698041634 | 0.839009436 | 12.64039921 | 10.56931225 | ENSG00000226243 |
| ZPR1 | -0.254167923 | 0.838470584 | 0.000327519 | 0.002117984 | 1585.289449 | 1329.114066 | ENSG00000109917 |
| MAL2 | -0.254232126 | 0.838433272 | 0.002728396 | 0.013243306 | 1829.139557 | 1533.874446 | ENSG00000147676 |
| NA | -0.254252492 | 0.838421436 | 0.713982464 | 0.849101626 | 9.905093528 | 8.316610197 | ENSG00000260296 |
| PGLYRP1 | -0.254266613 | 0.83841323 | 0.776291923 | 0.885282184 | 5.942110411 | 4.989285125 | ENSG00000008438 |
| TBC1D9 | -0.254324927 | 0.838379341 | 0.351100041 | 0.563149676 | 76.18905959 | 63.87951295 | ENSG00000109436 |
| NA | -0.25445441 | 0.838304099 | 0.672320456 | 0.82439555 | 13.81471071 | 11.5944667 | ENSG00000260167 |
| C19orf73 | -0.254490478 | 0.838283142 | 0.579281982 | 0.760179295 | 29.01513788 | 24.29533773 | ENSG00000221916 |
| CCM2L | -0.254499172 | 0.83827809 | 0.241716389 | 0.44519938 | 126.6860273 | 106.3923971 | ENSG00000101331 |
| ANKS1B | -0.254685724 | 0.838169701 | 0.441839917 | 0.649501582 | 44.77813584 | 37.50693039 | ENSG00000185046 |
| LMNB1 | -0.254779052 | 0.838115482 | 3.66E-05 | 0.000304835 | 4773.817632 | 4001.15402 | ENSG00000113368 |
| MAIP1 | -0.254781981 | 0.83811378 | 0.084734192 | 0.210997523 | 234.7183299 | 196.7585627 | ENSG00000162972 |
| SLC24A1 | -0.254877794 | 0.838058121 | 0.194880109 | 0.383565488 | 152.2950567 | 127.4933763 | ENSG00000074621 |
| MRPL54 | -0.255067731 | 0.837947794 | 0.029366432 | 0.09331736 | 470.0163396 | 393.7553149 | ENSG00000183617 |
| DIABLO | -0.255112776 | 0.837921631 | 0.618831241 | 0.78857139 | 24.12489467 | 20.29668318 | ENSG00000184047 |
| C17orf97 | -0.255151545 | 0.837899115 | 0.77437298 | 0.884054753 | 5.96583151 | 5.005459328 | ENSG00000187624 |
| CDC14A | -0.255403432 | 0.837752835 | 0.156439134 | 0.328480437 | 163.1562171 | 136.5479774 | ENSG00000079335 |
| ST8SIA1 | -0.255470421 | 0.837713936 | 0.357073685 | 0.569121372 | 73.54124003 | 61.73009896 | ENSG00000111728 |
| PRMT1 | -0.255569146 | 0.837656612 | 1.71E-06 | 1.89E-05 | 15728.86477 | 13175.51343 | ENSG00000126457 |
| MOB3B | -0.255632023 | 0.837620105 | 0.02118659 | 0.07210466 | 503.2854209 | 421.3015826 | ENSG00000120162 |
| C12orf76 | -0.255698161 | 0.837581707 | 0.184441189 | 0.36866764 | 161.5020663 | 135.1526558 | ENSG00000174456 |
| USP31 | -0.255719433 | 0.837569357 | 0.034139277 | 0.105208918 | 397.6200973 | 332.8132948 | ENSG00000103404 |
| MIR1915HG | -0.255743107 | 0.837555613 | 0.096743618 | 0.232413589 | 223.7083343 | 187.5157352 | ENSG00000204682 |
| POLA1 | -0.255757914 | 0.837547017 | 5.27E-05 | 0.000421104 | 2311.900333 | 1936.26297 | ENSG00000101868 |
| SOX21-AS1 | -0.255763517 | 0.837543765 | 0.465639272 | 0.669835287 | 44.38669848 | 37.26632484 | ENSG00000227640 |
| ATAD3C | -0.25576414 | 0.837543403 | 0.604106785 | 0.778603606 | 21.41358942 | 17.97181196 | ENSG00000215915 |
| NA | -0.255786411 | 0.837530474 | 0.813478513 | 0.90530915 | 3.977221007 | 3.334797422 | ENSG00000231977 |
| CRB2 | -0.255798155 | 0.837523656 | 0.080202505 | 0.202111596 | 323.7601351 | 271.3027091 | ENSG00000148204 |
| RNF168 | -0.255811061 | 0.837516164 | 0.000608974 | 0.003645373 | 1199.266224 | 1004.272143 | ENSG00000163961 |
| TSR1 | -0.25587697 | 0.837477903 | 8.45E-05 | 0.000639931 | 3556.545228 | 2978.844835 | ENSG00000167721 |
| RAB14 | -0.25592658 | 0.837449105 | 0.013266654 | 0.04928994 | 934.9851103 | 783.1954557 | ENSG00000119396 |
| ALG3 | -0.256304975 | 0.837229485 | 0.00102671 | 0.005738209 | 1040.521378 | 871.2703413 | ENSG00000214160 |
| C10orf25 | -0.25631842 | 0.837221683 | 0.552996367 | 0.739709489 | 26.17546459 | 21.90728275 | ENSG00000165511 |
| SPATA5 | -0.256360235 | 0.837197417 | 0.100134863 | 0.238287158 | 249.527599 | 208.8298896 | ENSG00000145375 |
| AGAP3 | -0.256518105 | 0.837105809 | 0.000262642 | 0.001752027 | 1830.88028 | 1532.967644 | ENSG00000133612 |
| ACOT7 | -0.256593782 | 0.8370619 | 1.25E-05 | 0.000116206 | 2744.163355 | 2296.898666 | ENSG00000097021 |
| EPHB2 | -0.256633278 | 0.837038985 | 0.002272886 | 0.011337015 | 891.1051751 | 746.0876492 | ENSG00000133216 |
| RFESD | -0.256706682 | 0.836996397 | 0.717872307 | 0.851146028 | 9.939493044 | 8.325312052 | ENSG00000175449 |
| AIP | -0.256725212 | 0.836985647 | 0.007848021 | 0.032170584 | 678.1037232 | 567.5827652 | ENSG00000110711 |
| LINC01126 | -0.256750665 | 0.83697088 | 0.726953372 | 0.856266103 | 9.541242224 | 8.013766846 | ENSG00000279873 |
| LOC145694 | -0.256779148 | 0.836954357 | 0.767410762 | 0.879694476 | 7.937839859 | 6.647177796 | ENSG00000259426 |
| COX14 | -0.257100103 | 0.83676818 | 0.398665991 | 0.610410835 | 70.24074611 | 58.92647565 | ENSG00000178449 |
| COL9A1 | -0.257388818 | 0.836600741 | 0.042845533 | 0.125137318 | 495.8312331 | 414.9024006 | ENSG00000112280 |

| | | | | | | | |
|---|---|---|---|---|---|---|---|
| NA | -0.257392442 | 0.83659864 | 0.742614436 | 0.865963175 | 11.18616305 | 9.393125132 | ENSG00000286548 |
| ANKRD13D | -0.257414463 | 0.836585871 | 0.027869854 | 0.089708914 | 469.2501449 | 392.7910531 | ENSG00000172932 |
| RAB6A | -0.257466686 | 0.836555588 | 0.004584158 | 0.020543858 | 1135.529014 | 950.0852753 | ENSG00000175582 |
| BTBD10 | -0.25749128 | 0.836541328 | 0.016034723 | 0.057581753 | 522.046741 | 436.7893559 | ENSG00000148925 |
| NA | -0.257539779 | 0.836513206 | 0.721411796 | 0.853073731 | 9.930009835 | 8.310083807 | ENSG00000249042 |
| CCT2 | -0.25757361 | 0.83649359 | 7.98E-08 | 1.13E-06 | 8772.928402 | 7338.607701 | ENSG00000166226 |
| ZNF140 | -0.257678618 | 0.836432707 | 0.131757442 | 0.291875577 | 174.9194421 | 146.215286 | ENSG00000196387 |
| MRPL34 | -0.25776736 | 0.836381259 | 0.000720276 | 0.004213883 | 1221.530239 | 1021.760495 | ENSG00000130312 |
| MRPS22 | -0.257851905 | 0.836332246 | 0.010128851 | 0.039439865 | 602.55905 | 503.7841529 | ENSG00000175110 |
| CXXC5 | -0.258022061 | 0.836233613 | 8.98E-05 | 0.000675738 | 2937.611354 | 2456.691253 | ENSG00000171604 |
| TECPR1 | -0.258051335 | 0.836216645 | 0.047919555 | 0.136453678 | 383.2896994 | 320.5950456 | ENSG00000205356 |
| PRRT3 | -0.258088867 | 0.83619489 | 0.145697233 | 0.312669572 | 217.849845 | 182.3279134 | ENSG00000163704 |
| PIF1 | -0.258139032 | 0.836165815 | 0.008738382 | 0.035104095 | 1842.466475 | 1540.844412 | ENSG00000140451 |
| EML4 | -0.258147922 | 0.836160663 | 4.72E-05 | 0.000381871 | 2245.833521 | 1877.949202 | ENSG00000143924 |
| LOC101928718 | -0.258173901 | 0.836145605 | 0.798684258 | 0.897379355 | 5.942110411 | 4.968759994 | ENSG00000284830 |
| FAM118B | -0.258178798 | 0.836142767 | 0.002437987 | 0.012052719 | 900.8604397 | 753.1439416 | ENSG00000197798 |
| PABPN1 | -0.258203022 | 0.836128728 | 0.000297334 | 0.001948356 | 2089.366388 | 1747.594773 | ENSG00000100836 |
| MKRN1 | -0.258245129 | 0.836104325 | 4.03E-05 | 0.000331445 | 2629.130996 | 2197.744094 | ENSG00000133606 |
| KRTDAP | -0.258290377 | 0.836078102 | 0.781985856 | 0.88893053 | 6.749290468 | 5.605849145 | ENSG00000188508 |
| UBA6-AS1 | -0.258291433 | 0.83607749 | 0.30834364 | 0.518758988 | 79.00737604 | 66.08787536 | ENSG00000248049 |
| CPNE2 | -0.258331506 | 0.836054267 | 0.014313962 | 0.052468081 | 696.322051 | 581.8095809 | ENSG00000140848 |
| WASHC2A | -0.258340965 | 0.836048785 | 0.001589419 | 0.00837038 | 932.0688058 | 779.1311717 | ENSG00000099290 |
| CLSPN | -0.258386901 | 0.836022166 | 0.003250378 | 0.015407477 | 1011.374794 | 845.0060695 | ENSG00000092853 |
| TMEM187 | -0.25839352 | 0.83601833 | 0.207870917 | 0.40048679 | 127.3893984 | 106.4874348 | ENSG00000177854 |
| NA | -0.258529091 | 0.835939773 | 0.337874221 | 0.549828063 | 80.47231677 | 67.21487746 | ENSG00000204745 |
| PPIL2 | -0.258558376 | 0.835922805 | 0.00461543 | 0.020654596 | 848.3788305 | 709.2271495 | ENSG00000100023 |
| LRFN4 | -0.25868681 | 0.835848391 | 0.018679149 | 0.064964223 | 880.4404472 | 736.4335317 | ENSG00000173621 |
| PEX11A | -0.25874623 | 0.835813965 | 0.375174024 | 0.587200518 | 57.54398016 | 48.09553827 | ENSG00000166821 |
| CCDC160 | -0.258808011 | 0.835778174 | 0.305318055 | 0.51555392 | 95.01295006 | 79.48937549 | ENSG00000203952 |
| TSNARE1 | -0.258997303 | 0.835668521 | 0.062230013 | 0.166829084 | 335.8459204 | 280.7288881 | ENSG00000171045 |
| GNG2 | -0.259000153 | 0.83566687 | 0.043691912 | 0.127059095 | 327.3790975 | 273.6585313 | ENSG00000186469 |
| CYP51A1 | -0.259011954 | 0.835660035 | 0.08840026 | 0.217403165 | 263.3740656 | 220.0858548 | ENSG00000001630 |
| HNRNPR | -0.2591449 | 0.835583031 | 3.63E-08 | 5.45E-07 | 6347.167028 | 5303.555042 | ENSG00000125944 |
| PI4KA | -0.259182235 | 0.835561408 | 0.00020372 | 0.00140095 | 1618.204213 | 1352.603901 | ENSG00000241973 |
| LAPTM4B | -0.259209957 | 0.835545352 | 1.84E-07 | 2.42E-06 | 11099.14149 | 9273.987251 | ENSG00000104341 |
| RIMKLA | -0.259281212 | 0.835504085 | 0.146062758 | 0.313151159 | 245.5373097 | 204.8137157 | ENSG00000177181 |
| EEF1B2 | -0.259521525 | 0.835364925 | 2.29E-07 | 2.98E-06 | 7131.323869 | 5956.940454 | ENSG00000114942 |
| RAP2A | -0.259672364 | 0.83527759 | 0.003121903 | 0.014899455 | 878.4412557 | 733.7983744 | ENSG00000125249 |
| CDC7 | -0.259674781 | 0.83527619 | 0.000932439 | 0.005277295 | 1075.910744 | 898.8703035 | ENSG00000097046 |
| NA | -0.259693272 | 0.835265484 | 0.769153707 | 0.881236386 | 6.756409413 | 5.598376795 | ENSG00000276007 |
| CDK2AP2 | -0.259804941 | 0.835200835 | 0.001881634 | 0.009632599 | 923.256182 | 770.8665414 | ENSG00000167797 |
| NA | -0.259977754 | 0.835100796 | 0.609149077 | 0.782326193 | 19.42261465 | 16.23550729 | ENSG00000269883 |
| NUP153 | -0.260089895 | 0.835035886 | 0.001705631 | 0.008881261 | 2339.949292 | 1953.759431 | ENSG00000124789 |
| NA | -0.26032927 | 0.834897347 | 0.535152187 | 0.726030823 | 29.43327099 | 24.58541185 | ENSG00000224934 |
| TSPOAP1 | -0.260363446 | 0.834877569 | 0.165425988 | 0.341204651 | 274.5824235 | 229.3352947 | ENSG00000005379 |
| PRPF18 | -0.260556985 | 0.834765577 | 0.272751973 | 0.481439481 | 114.9438281 | 95.85219626 | ENSG00000165630 |
| ZNF830 | -0.260580835 | 0.834751777 | 0.037798193 | 0.113629237 | 425.8363176 | 355.118218 | ENSG00000198783 |
| STX7 | -0.260583433 | 0.834750274 | 0.010969022 | 0.042180814 | 618.5744388 | 516.2006029 | ENSG00000079950 |
| RANBP6 | -0.260653085 | 0.834709974 | 0.039561437 | 0.117711948 | 502.1259833 | 419.5054171 | ENSG00000137040 |
| TENT4A | -0.260824433 | 0.834610842 | 0.00565253 | 0.024481515 | 733.9767489 | 612.9051366 | ENSG00000112941 |
| ZNF592 | -0.260906102 | 0.834563597 | 0.000336113 | 0.002167119 | 1485.026626 | 1239.471684 | ENSG00000166716 |
| USP27X-AS1 | -0.260922187 | 0.834554293 | 0.655023483 | 0.813395481 | 13.94399462 | 11.62993653 | ENSG00000234390 |
| NA | -0.26100064 | 0.834508911 | 0.674602526 | 0.825709294 | 14.7674975 | 12.37684014 | ENSG00000258559 |
| NA | -0.261092629 | 0.834455703 | 0.811788812 | 0.904395006 | 4.000942105 | 3.336972886 | ENSG00000272518 |
| C19orf47 | -0.261099608 | 0.834451666 | 0.044189106 | 0.12801216 | 377.8437244 | 315.5263595 | ENSG00000160392 |
| ASB3 | -0.261115974 | 0.8344422 | 0.028905581 | 0.092163077 | 411.0902281 | 343.1355907 | ENSG00000115239 |
| NA | -0.261164666 | 0.834414037 | 0.18879092 | 0.374704836 | 130.4660033 | 108.8331147 | ENSG00000261409 |
| NATD1 | -0.261253936 | 0.834362408 | 0.035446874 | 0.10830117 | 381.5427752 | 318.1504582 | ENSG00000274180 |
| EMC1 | -0.26129764 | 0.834337133 | 0.000203855 | 0.00140095 | 1617.789004 | 1349.595392 | ENSG00000127463 |
| DPY19L2P3 | -0.261322444 | 0.834322788 | 0.459613299 | 0.664609277 | 40.13962084 | 33.4392281 | ENSG00000227855 |
| KRBOX4 | -0.261345606 | 0.834309393 | 0.19567805 | 0.384660843 | 134.8895539 | 112.6913304 | ENSG00000147121 |
| RAB5B | -0.261403291 | 0.834276035 | 0.000742667 | 0.004328788 | 1342.389028 | 1119.691479 | ENSG00000111540 |
| ATG2B | -0.261711439 | 0.834097859 | 0.049650287 | 0.140303108 | 353.6929981 | 294.8440701 | ENSG00000066739 |
| ZNF708 | -0.26179418 | 0.834050024 | 0.125061697 | 0.280897972 | 236.9673399 | 197.6357629 | ENSG00000182141 |
| CPD | -0.261914064 | 0.83398072 | 0.004433424 | 0.019989078 | 1503.044856 | 1253.064212 | ENSG00000108582 |
| TMPO-AS1 | -0.261968124 | 0.83394947 | 0.337347581 | 0.549275142 | 66.73677802 | 55.64500327 | ENSG00000257167 |
| ZNF419 | -0.26210265 | 0.83387171 | 0.151336223 | 0.321013656 | 172.2736818 | 143.5115458 | ENSG00000105136 |
| SLC25A26 | -0.262139444 | 0.833850444 | 0.013331604 | 0.049463133 | 561.8838857 | 468.294405 | ENSG00000144741 |
| HNRNPH3 | -0.262201497 | 0.833814579 | 1.90E-05 | 0.000170533 | 4781.548125 | 3987.246246 | ENSG00000096746 |
| SYNRG | -0.262269879 | 0.833775058 | 0.003520072 | 0.016473313 | 827.32754 | 689.5482396 | ENSG00000275066 |
| PAXIP1-AS1 | -0.262390603 | 0.833705292 | 0.236165141 | 0.438154896 | 184.5357381 | 154.0849683 | ENSG00000273344 |
| TSPAN17 | -0.262391522 | 0.83370476 | 0.000103705 | 0.000767122 | 1989.324324 | 1658.727846 | ENSG00000048140 |
| RNF214 | -0.262590521 | 0.833589771 | 0.028230596 | 0.090533227 | 474.3790743 | 395.0342526 | ENSG00000167257 |
| UBE2O | -0.262754055 | 0.833495286 | 2.80E-05 | 0.000240104 | 2687.028888 | 2239.518688 | ENSG00000175931 |

| | | | | | | | |
|---|---|---|---|---|---|---|---|
| SDR39U1 | -0.262938729 | 0.8333886 | 0.299111933 | 0.509389038 | 91.30500145 | 76.22818765 | ENSG00000100445 |
| NPM2 | -0.262974254 | 0.833368079 | 0.532916634 | 0.724314769 | 27.50753579 | 22.92033037 | ENSG00000158806 |
| GLB1L3 | -0.26301409 | 0.833345068 | 0.000612291 | 0.003659512 | 1520.805876 | 1267.96723 | ENSG00000166105 |
| SUN1 | -0.263018625 | 0.833342449 | 0.000237059 | 0.001601162 | 2166.060101 | 1805.157876 | ENSG00000164828 |
| SNHG26 | -0.263113917 | 0.833287407 | 0.179995072 | 0.362775553 | 140.1125552 | 116.7868192 | ENSG00000228649 |
| ZNF385D | -0.263189883 | 0.833243531 | 0.297428793 | 0.507723289 | 89.25351565 | 74.50436866 | ENSG00000151789 |
| NA | -0.263192011 | 0.833242302 | 0.574439279 | 0.756663582 | 21.93639562 | 18.25441373 | ENSG00000253954 |
| SYF2 | -0.263263401 | 0.833201071 | 0.008288405 | 0.033639917 | 682.2686792 | 568.7416212 | ENSG00000117614 |
| BNIP1 | -0.263392931 | 0.833126267 | 0.233936448 | 0.435131471 | 108.9649278 | 90.7186884 | ENSG00000113734 |
| LIPT2 | -0.263422789 | 0.833109024 | 0.444497104 | 0.65193599 | 42.174199 | 35.1786542 | ENSG00000175536 |
| DHX29 | -0.263434957 | 0.833101998 | 0.005375337 | 0.023476696 | 1099.967451 | 916.2282807 | ENSG00000067248 |
| NFKB1 | -0.263476395 | 0.833078069 | 0.044170757 | 0.127984084 | 328.6598621 | 273.6772598 | ENSG00000109320 |
| ERGIC1 | -0.263556144 | 0.83303202 | 1.06E-05 | 9.98E-05 | 3016.253634 | 2512.278959 | ENSG00000113719 |
| SLC35F2 | -0.263598768 | 0.833007408 | 0.005814114 | 0.025056795 | 630.9909491 | 525.5050285 | ENSG00000110660 |
| MCF2L2 | -0.263611384 | 0.833000124 | 0.314529942 | 0.524936499 | 70.89192035 | 59.01309506 | ENSG00000053524 |
| LSMEM1 | -0.263616851 | 0.832996968 | 0.767672578 | 0.879862635 | 5.972950455 | 4.967814035 | ENSG00000181016 |
| RNF10 | -0.263688812 | 0.832955419 | 9.92E-07 | 1.15E-05 | 3965.536934 | 3302.85841 | ENSG00000022840 |
| MRPL58 | -0.263706406 | 0.832945261 | 0.008765823 | 0.035206843 | 591.5387331 | 492.8554353 | ENSG00000167862 |
| HMGN5 | -0.263876059 | 0.832847317 | 0.008958418 | 0.03581605 | 731.2390522 | 609.404886 | ENSG00000198157 |
| MYCN | -0.264096074 | 0.832720316 | 0.000736049 | 0.004296841 | 1808.844933 | 1506.594531 | ENSG00000134323 |
| FBXO3 | -0.264104689 | 0.832715343 | 0.024259976 | 0.080419455 | 418.0429665 | 348.3526395 | ENSG00000110429 |
| NA | -0.264139122 | 0.832695468 | 0.531700556 | 0.723460345 | 29.19989881 | 24.32674018 | ENSG00000269825 |
| LOC220729 | -0.264168646 | 0.832678428 | 0.077649254 | 0.197548992 | 309.6633898 | 257.6613815 | ENSG00000214135 |
| THYN1 | -0.264189076 | 0.832666636 | 0.000567486 | 0.00342614 | 1171.858886 | 975.4465045 | ENSG00000151500 |
| ZNF570 | -0.264283208 | 0.832612309 | 0.303325687 | 0.513486816 | 80.73533379 | 67.157653 | ENSG00000171827 |
| PLA2G4B | -0.26431205 | 0.832595664 | 0.813190488 | 0.905169301 | 3.997382633 | 3.322974145 | ENSG00000243708 |
| ASRGL1 | -0.264318719 | 0.832591815 | 0.084255257 | 0.20999875 | 249.8282963 | 207.8182823 | ENSG00000162174 |
| TBRG1 | -0.264434572 | 0.832524958 | 0.027933449 | 0.089837054 | 392.99849 | 327.0317478 | ENSG00000154144 |
| GPR68 | -0.264458013 | 0.832511431 | 0.67134152 | 0.82390724 | 14.75236989 | 12.22597542 | ENSG00000119714 |
| SARS1 | -0.264592466 | 0.832433848 | 2.66E-05 | 0.000230272 | 3841.447667 | 3196.891483 | ENSG00000031698 |
| CLASP1 | -0.264594063 | 0.832432927 | 0.000527513 | 0.003216689 | 1419.178487 | 1180.95158 | ENSG00000074054 |
| KNSTRN | -0.264613578 | 0.832421667 | 0.000539371 | 0.003277614 | 1171.489974 | 975.2561455 | ENSG00000128944 |
| TARS1 | -0.264637992 | 0.83240758 | 6.68E-08 | 9.59E-07 | 8604.354084 | 7162.277472 | ENSG00000113407 |
| ATP9A | -0.264646369 | 0.832402747 | 0.068006089 | 0.178622392 | 256.6828958 | 213.7702006 | ENSG00000054793 |
| NA | -0.26484423 | 0.832288593 | 0.811459629 | 0.904255331 | 3.980780479 | 3.306799942 | ENSG00000258982 |
| IGF2BP1 | -0.264922672 | 0.832243341 | 4.88E-07 | 6.01E-06 | 9038.399749 | 7521.677226 | ENSG00000159217 |
| FAM171A1 | -0.265047481 | 0.832171347 | 0.00044271 | 0.002767488 | 1537.88526 | 1279.638582 | ENSG00000148468 |
| IL17RA | -0.265078858 | 0.832153248 | 0.036561938 | 0.1108658 | 331.3957781 | 275.8180876 | ENSG00000177663 |
| IRF5 | -0.265149087 | 0.832112741 | 0.724751951 | 0.854899475 | 9.92289089 | 8.24226557 | ENSG00000128604 |
| AMPH | -0.265150652 | 0.832111838 | 0.0394408 | 0.117474864 | 325.9014457 | 271.3198723 | ENSG00000078053 |
| FRMPD3 | -0.265173101 | 0.83209889 | 0.240999519 | 0.444547365 | 100.4865105 | 83.53002626 | ENSG00000147234 |
| RGS7BP | -0.265189855 | 0.832089227 | 0.413408256 | 0.624700612 | 52.88382899 | 44.06794028 | ENSG00000186479 |
| TIMP2 | -0.265192166 | 0.832087894 | 0.018839361 | 0.065364932 | 542.6286174 | 451.9213397 | ENSG00000035862 |
| ZNF460 | -0.265295722 | 0.832028169 | 0.496153488 | 0.695402896 | 34.28054735 | 28.44776751 | ENSG00000197714 |
| NRCAM | -0.265418184 | 0.831957546 | 0.229129381 | 0.428893682 | 123.136397 | 102.3036752 | ENSG00000091129 |
| SUPT16H | -0.265470591 | 0.831927325 | 1.92E-07 | 2.52E-06 | 9765.709862 | 8123.869814 | ENSG00000092201 |
| ZBTB14 | -0.265497983 | 0.83191153 | 0.080838651 | 0.203399343 | 227.7478202 | 189.4217058 | ENSG00000198081 |
| KANK2 | -0.265517958 | 0.831900012 | 0.000488409 | 0.003020169 | 1185.022727 | 985.9229748 | ENSG00000197256 |
| R3HDM2 | -0.265686736 | 0.831802695 | 0.108953281 | 0.254462368 | 215.6536689 | 179.3467527 | ENSG00000179912 |
| NUTM2A-AS1 | -0.265720156 | 0.831783426 | 0.029767283 | 0.094337159 | 387.7918371 | 322.7114058 | ENSG00000223482 |
| NSG1 | -0.265726113 | 0.831779992 | 0.001189907 | 0.006524796 | 1079.098609 | 897.7334427 | ENSG00000168824 |
| NA | -0.26584607 | 0.831710834 | 0.195214127 | 0.38402279 | 148.2170275 | 123.1613266 | ENSG00000280511 |
| NUDC | -0.265972994 | 0.831637666 | 1.12E-07 | 1.54E-06 | 6483.413053 | 5392.160712 | ENSG00000090273 |
| GPR143 | -0.266162632 | 0.831528357 | 0.024094066 | 0.079951028 | 403.7588417 | 335.964922 | ENSG00000101850 |
| NSDHL | -0.266197167 | 0.831508452 | 0.000868535 | 0.004946406 | 1332.579857 | 1107.520793 | ENSG00000147383 |
| GSS | -0.266223312 | 0.831493383 | 0.002313254 | 0.011505348 | 900.4630275 | 748.3608764 | ENSG00000100983 |
| AQP3 | -0.266291777 | 0.831453925 | 0.304586241 | 0.514710972 | 75.09778501 | 62.46858426 | ENSG00000165272 |
| TNFSF12 | -0.266466145 | 0.831353439 | 0.526899014 | 0.720012728 | 29.84548037 | 24.86489219 | ENSG00000239697 |
| IPO11 | -0.266501935 | 0.831332815 | 0.005180812 | 0.022769287 | 895.6283124 | 744.5396733 | ENSG00000086200 |
| HECTD2 | -0.26669519 | 0.831221462 | 0.090435781 | 0.221256603 | 214.6000217 | 178.3557432 | ENSG00000165338 |
| MED28 | -0.26670118 | 0.831218011 | 3.19E-05 | 0.000269646 | 2226.174378 | 1850.044098 | ENSG00000118579 |
| PCDHA2 | -0.266737977 | 0.83119681 | 0.539301515 | 0.729040189 | 27.0428502 | 22.46785181 | ENSG00000204969 |
| GDE1 | -0.266887805 | 0.831110493 | 0.000866401 | 0.004938719 | 1107.206544 | 919.9859735 | ENSG00000006007 |
| U2AF1L5 | -0.266935829 | 0.831082827 | 0.528697922 | 0.721294281 | 27.62614128 | 22.92997819 | ENSG00000275895 |
| ZBTB49 | -0.266994258 | 0.831049169 | 0.25130449 | 0.456862274 | 107.1871897 | 89.04244606 | ENSG00000168826 |
| BCAS2 | -0.267121074 | 0.830976121 | 0.000535748 | 0.003259537 | 1194.725795 | 992.4291036 | ENSG00000116752 |
| SPRED2 | -0.267307688 | 0.830868641 | 0.008917365 | 0.035701437 | 536.2256346 | 445.6091427 | ENSG00000198369 |
| PRR29-AS1 | -0.267346722 | 0.830846161 | 0.808884243 | 0.902949338 | 3.977221007 | 3.294976665 | ENSG00000264954 |
| NA | -0.267357691 | 0.830839844 | 0.785323367 | 0.890433415 | 5.600784998 | 4.663741179 | ENSG00000258702 |
| BRSK2 | -0.267382174 | 0.830825744 | 0.004297666 | 0.019500583 | 907.6602511 | 754.5860024 | ENSG00000174672 |
| KIF13A | -0.267446818 | 0.830788518 | 0.001263552 | 0.006868791 | 997.8483478 | 828.6198131 | ENSG00000137177 |
| TMEM255B | -0.267525284 | 0.830743333 | 0.295297911 | 0.505410645 | 75.55771592 | 62.74995652 | ENSG00000184497 |
| ERICH6-AS1 | -0.267552956 | 0.830727399 | 0.716809982 | 0.850738845 | 9.523444862 | 7.911424738 | ENSG00000240137 |
| NRP1 | -0.267559106 | 0.830723858 | 0.46760857 | 0.671155346 | 43.03952583 | 35.71463075 | ENSG00000099250 |

| | | | | | | | |
|---|---|---|---|---|---|---|---|
| NACA | -0.267587151 | 0.83070771 | 9.27E-06 | 8.93E-05 | 4889.328815 | 4061.95227 | ENSG00000196531 |
| SLC35A4 | -0.267821861 | 0.830572574 | 7.75E-06 | 7.62E-05 | 2276.245411 | 1890.83766 | ENSG00000176087 |
| NA | -0.267835271 | 0.830564854 | 0.020226309 | 0.069442678 | 545.4622641 | 452.7895975 | ENSG00000254277 |
| MAML1 | -0.267911505 | 0.830520967 | 0.000166328 | 0.001169 | 1754.868024 | 1457.670179 | ENSG00000161021 |
| TOM1L2 | -0.268053589 | 0.830439177 | 0.289699316 | 0.499085165 | 85.22201283 | 70.85206673 | ENSG00000175662 |
| MTURN | -0.268055809 | 0.830437899 | 0.142560614 | 0.307767153 | 163.6395898 | 135.9026808 | ENSG00000180354 |
| CDIPT | -0.268067946 | 0.830430913 | 0.009236134 | 0.0366207 | 946.0731611 | 786.0679494 | ENSG00000103502 |
| NA | -0.268196783 | 0.830356756 | 0.753442024 | 0.871432669 | 7.187585209 | 5.996089905 | ENSG00000269038 |
| CCDC69 | -0.268294042 | 0.83030078 | 0.046001977 | 0.132108778 | 313.5222579 | 260.186543 | ENSG00000198624 |
| DCAF12 | -0.268312374 | 0.830290229 | 0.002909717 | 0.014021435 | 852.8716865 | 708.016112 | ENSG00000198876 |
| JPH4 | -0.268316366 | 0.830287932 | 8.66E-05 | 0.000653116 | 3295.109932 | 2736.438838 | ENSG00000092051 |
| DNTTIP1 | -0.268479162 | 0.830194246 | 0.004550623 | 0.020424975 | 813.1616367 | 675.0592949 | ENSG00000101457 |
| ECT2 | -0.268611547 | 0.830118069 | 0.000367037 | 0.002343304 | 1597.348667 | 1325.924467 | ENSG00000114346 |
| JPH3 | -0.268719997 | 0.83005567 | 0.000452359 | 0.002820336 | 1310.47955 | 1088.141807 | ENSG00000154118 |
| LARP1 | -0.268721635 | 0.830054727 | 3.01E-08 | 4.57E-07 | 9558.124564 | 7933.286585 | ENSG00000155506 |
| ATAD2 | -0.268732686 | 0.830048369 | 0.005680388 | 0.024557385 | 642.5122345 | 533.3391028 | ENSG00000156802 |
| MERTK | -0.268751739 | 0.830037407 | 0.095294117 | 0.229947402 | 229.6293416 | 190.3693941 | ENSG00000153208 |
| ZNF286A | -0.268791568 | 0.830014493 | 0.027142369 | 0.087936807 | 477.8700247 | 396.9041339 | ENSG00000187607 |
| XPO5 | -0.268807889 | 0.830005103 | 5.15E-06 | 5.23E-05 | 3197.692249 | 2653.542078 | ENSG00000124571 |
| NA | -0.268809087 | 0.830004413 | 0.448647096 | 0.655281496 | 43.13201981 | 35.74480369 | ENSG00000279348 |
| PDHX | -0.268880212 | 0.829963495 | 0.004591317 | 0.020566183 | 850.6894173 | 705.6739527 | ENSG00000110435 |
| DDX23 | -0.268966472 | 0.829913873 | 2.05E-06 | 2.24E-05 | 4849.020818 | 4023.988502 | ENSG00000174243 |
| CADM1 | -0.269244702 | 0.829753835 | 0.028842305 | 0.092023471 | 398.5942671 | 330.5419595 | ENSG00000182985 |
| TADA3 | -0.269272196 | 0.829738023 | 0.000925974 | 0.005246631 | 1272.289465 | 1055.821228 | ENSG00000171148 |
| LINC01801 | -0.269277707 | 0.829734853 | 0.581070087 | 0.761150628 | 21.54050907 | 17.83958046 | ENSG00000267767 |
| ZNF436 | -0.269430008 | 0.829647265 | 0.449173154 | 0.655637271 | 40.85784024 | 33.86465514 | ENSG00000125945 |
| NA | -0.269618727 | 0.829538746 | 0.719858801 | 0.852321847 | 9.192797866 | 7.64622668 | ENSG00000184441 |
| SERPINB4 | -0.269634625 | 0.829529605 | 0.821793129 | 0.9095419 | 3.953499908 | 3.269154649 | ENSG00000206073 |
| EXOC6 | -0.269673257 | 0.829507392 | 0.149997062 | 0.318925299 | 142.0907666 | 117.8406336 | ENSG00000138190 |
| SLC10A4 | -0.269678267 | 0.829504511 | 0.293545328 | 0.503277669 | 83.81138007 | 69.62440353 | ENSG00000145248 |
| GATC | -0.269724579 | 0.829477884 | 0.069482933 | 0.181623319 | 241.7914539 | 200.5318399 | ENSG00000257218 |
| LRRFIP1 | -0.269802242 | 0.829433233 | 5.15E-05 | 0.000412884 | 1758.529493 | 1458.253323 | ENSG00000124831 |
| PCDHA7 | -0.269889316 | 0.829383174 | 0.580514739 | 0.760980082 | 20.73688848 | 17.20996366 | ENSG00000204963 |
| NA | -0.270061476 | 0.829284207 | 0.384835738 | 0.597077137 | 53.36183659 | 44.27328691 | ENSG00000233230 |
| SSU72 | -0.270122311 | 0.829249239 | 6.48E-06 | 6.44E-05 | 3707.720816 | 3074.938253 | ENSG00000160075 |
| MAML3 | -0.270188881 | 0.829210976 | 0.114185439 | 0.263507545 | 178.4817839 | 147.9472397 | ENSG00000196782 |
| CD40 | -0.270330275 | 0.829129712 | 0.030735995 | 0.096773804 | 382.7096367 | 317.1284682 | ENSG00000101017 |
| TMEM147 | -0.270351636 | 0.829117436 | 0.000361014 | 0.002307189 | 1279.952971 | 1061.510625 | ENSG00000105677 |
| NA | -0.270490949 | 0.829037376 | 0.817603432 | 0.907131026 | 3.946380963 | 3.259506836 | ENSG00000283907 |
| HERC2 | -0.270526358 | 0.829017029 | 0.000418829 | 0.0026349 | 4336.833025 | 3594.420512 | ENSG00000128731 |
| SNORA55 | -0.270794124 | 0.828863177 | 0.835637915 | 0.917415262 | 3.633531329 | 3.022306258 | ENSG00000201457 |
| LDAH | -0.270920343 | 0.828790664 | 0.013602408 | 0.050280188 | 508.4508858 | 421.1988616 | ENSG00000118961 |
| OTUB1 | -0.271046154 | 0.828718392 | 5.41E-05 | 0.000431515 | 2069.21513 | 1715.056296 | ENSG00000167770 |
| MID1IP1 | -0.271067361 | 0.82870621 | 3.14E-07 | 4.01E-06 | 4490.257292 | 3721.084817 | ENSG00000165175 |
| NA | -0.271197288 | 0.828631582 | 0.29748829 | 0.507759941 | 79.92042439 | 66.14169486 | ENSG00000237094 |
| NA | -0.271269425 | 0.82859015 | 0.641699527 | 0.804214244 | 14.83985601 | 12.32179114 | ENSG00000240914 |
| OBI1 | -0.271349563 | 0.828544125 | 0.153695818 | 0.324669522 | 176.0353013 | 145.8754597 | ENSG00000152193 |
| SLC6A9 | -0.271362278 | 0.828536823 | 0.009136434 | 0.036301609 | 575.6360783 | 477.0343824 | ENSG00000196517 |
| AXIN1 | -0.271403447 | 0.82851318 | 1.11E-05 | 0.000104682 | 2064.306225 | 1710.624017 | ENSG00000103126 |
| BNIP3L | -0.2714102 | 0.828509302 | 0.007258857 | 0.03013487 | 592.0985307 | 490.3626948 | ENSG00000104765 |
| PHB2 | -0.27142153 | 0.828502795 | 2.46E-09 | 4.46E-08 | 7954.854305 | 6590.430834 | ENSG00000215021 |
| NUDT21 | -0.271433724 | 0.828495792 | 2.99E-05 | 0.00025544 | 4643.577448 | 3846.948453 | ENSG00000167005 |
| RPRM | -0.271556673 | 0.82842519 | 0.043884285 | 0.127431319 | 318.0439474 | 263.2576554 | ENSG00000177519 |
| TERF2IP | -0.271674276 | 0.828357663 | 0.038277909 | 0.114833727 | 324.3641464 | 268.8092336 | ENSG00000166848 |
| FANCA | -0.271770855 | 0.828302211 | 0.008941398 | 0.035767485 | 1250.189919 | 1035.695853 | ENSG00000187741 |
| NA | -0.271771788 | 0.828301676 | 0.542824265 | 0.731708511 | 26.55355379 | 22.00716585 | ENSG00000203279 |
| CHCHD6 | -0.271902773 | 0.828226476 | 0.002522744 | 0.012397089 | 841.3464844 | 696.6896516 | ENSG00000159685 |
| PHACTR1 | -0.271921986 | 0.828215446 | 0.146116485 | 0.313177825 | 143.0305107 | 118.4217278 | ENSG00000112137 |
| RIOK1 | -0.2719433 | 0.82820321 | 0.00343985 | 0.016183748 | 746.3050847 | 617.7764296 | ENSG00000124784 |
| RAB1B | -0.271961169 | 0.828192952 | 0.001252385 | 0.006812007 | 895.1772282 | 741.4778001 | ENSG00000174903 |
| LCORL | -0.272023081 | 0.828157412 | 0.107424752 | 0.251826767 | 192.3389028 | 159.3552039 | ENSG00000178177 |
| PRPF39 | -0.272102461 | 0.828111846 | 0.008903704 | 0.035669589 | 635.9873137 | 526.4891756 | ENSG00000185246 |
| AAGAB | -0.272129746 | 0.828096185 | 0.001888756 | 0.009663221 | 955.0879609 | 790.8261612 | ENSG00000103591 |
| KIAA1191 | -0.272165848 | 0.828075463 | 3.70E-05 | 0.000307481 | 1821.093435 | 1507.72702 | ENSG00000122203 |
| ARID1A | -0.27219618 | 0.828058053 | 3.72E-05 | 0.000309031 | 4117.293872 | 3408.998234 | ENSG00000117713 |
| FEN1 | -0.27225959 | 0.828021659 | 1.52E-05 | 0.000138389 | 2363.184117 | 1957.183996 | ENSG00000168496 |
| SEMA6A-AS2 | -0.272266427 | 0.828017734 | 0.729085667 | 0.85741017 | 7.607192864 | 6.296757792 | ENSG00000249167 |
| DNAJC1 | -0.272482857 | 0.827893526 | 0.024377568 | 0.080738404 | 395.3109596 | 327.1972307 | ENSG00000136770 |
| SCT | -0.272485413 | 0.82789206 | 0.608999214 | 0.782305701 | 18.4757516 | 15.32697723 | ENSG00000070031 |
| C9orf85 | -0.272584512 | 0.827835194 | 0.077289156 | 0.196794474 | 251.3842559 | 208.2260221 | ENSG00000155621 |
| DSTNP2 | -0.27278811 | 0.827718375 | 0.524743383 | 0.718516443 | 26.52627322 | 21.9620482 | ENSG00000248593 |
| SKP1 | -0.272801244 | 0.827710839 | 0.000832605 | 0.004774901 | 1573.76476 | 1302.784782 | ENSG00000113558 |
| NSUN2 | -0.2728197 | 0.827700251 | 3.18E-05 | 0.000269542 | 3714.732711 | 3074.761926 | ENSG00000037474 |
| BRD7 | -0.273087351 | 0.827546709 | 4.74E-06 | 4.83E-05 | 2340.281258 | 1936.570019 | ENSG00000166164 |

| | | | | | | | |
|---|---|---|---|---|---|---|---|
| EIF1AX | -0.273166481 | 0.82750132 | 0.000276131 | 0.00182909 | 2988.200562 | 2472.75812 | ENSG00000173674 |
| AADAT | -0.273273895 | 0.827439712 | 0.067562382 | 0.177827955 | 248.7023684 | 205.9362419 | ENSG00000109576 |
| PHACTR2 | -0.273336058 | 0.82740406 | 0.022914912 | 0.076755609 | 545.4451283 | 451.2595721 | ENSG00000112419 |
| NCBP1 | -0.273336627 | 0.827403733 | 8.78E-05 | 0.000661411 | 1900.75481 | 1572.79426 | ENSG00000136937 |
| CHSY3 | -0.273413171 | 0.827359836 | 0.34160125 | 0.553223531 | 62.42179108 | 51.62676955 | ENSG00000198108 |
| WDFY3-AS2 | -0.273473003 | 0.827325524 | 0.382767991 | 0.595287285 | 55.47973101 | 45.87272561 | ENSG00000180769 |
| NA | -0.273530336 | 0.827292646 | 0.63158894 | 0.797106325 | 14.82561812 | 12.28197038 | ENSG00000257576 |
| NREP | -0.273554632 | 0.827278715 | 0.062703769 | 0.167810101 | 297.5718298 | 245.9072529 | ENSG00000134986 |
| GPN1 | -0.273852949 | 0.827107669 | 0.015365638 | 0.055547488 | 675.3246394 | 558.3695505 | ENSG00000198522 |
| ZNF821 | -0.273857413 | 0.82710511 | 0.320918583 | 0.531440498 | 75.90760866 | 62.67220692 | ENSG00000102984 |
| GNAS | -0.273918788 | 0.827069924 | 5.16E-11 | 1.21E-09 | 17549.6487 | 14515.17547 | ENSG00000087460 |
| ABHD15 | -0.273937499 | 0.827059198 | 0.53529988 | 0.72613054 | 24.45226153 | 20.22574353 | ENSG00000168792 |
| NA | -0.273951945 | 0.827050917 | 0.764992763 | 0.878469069 | 5.642303459 | 4.67026757 | ENSG00000275198 |
| NA | -0.274030134 | 0.827006094 | 0.641678491 | 0.804214244 | 16.5594996 | 13.72507952 | ENSG00000230091 |
| NA | -0.274057577 | 0.826990363 | 0.795026676 | 0.89524052 | 5.223891013 | 4.342548161 | ENSG00000279072 |
| NPIPB3 | -0.274099473 | 0.826966348 | 0.556366754 | 0.742700653 | 24.05757018 | 19.91949521 | ENSG00000169246 |
| TRMT2B | -0.274182876 | 0.826918542 | 0.012399092 | 0.046728567 | 456.4172816 | 377.5199755 | ENSG00000188917 |
| MYBL1 | -0.274221704 | 0.826896287 | 0.29505611 | 0.505134134 | 77.21804369 | 63.86941363 | ENSG00000185697 |
| SH2D5 | -0.274398358 | 0.826795042 | 0.002710618 | 0.013175497 | 727.3768522 | 601.415639 | ENSG00000189410 |
| NA | -0.274471415 | 0.826753175 | 0.557649457 | 0.743624488 | 25.600767 | 21.0990873 | ENSG00000267565 |
| SNRPN | -0.274505908 | 0.826733408 | 0.29658313 | 0.506829465 | 85.34092381 | 70.41905175 | ENSG00000128739 |
| CNDP2 | -0.274613192 | 0.826671932 | 5.41E-09 | 9.33E-08 | 6478.18195 | 5355.40176 | ENSG00000133313 |
| NA | -0.274702729 | 0.826620628 | 0.787687549 | 0.891582187 | 5.222695804 | 4.340372697 | ENSG00000235084 |
| NA | -0.274717821 | 0.82661198 | 0.488615944 | 0.688980221 | 35.02604732 | 28.89930012 | ENSG00000237863 |
| IFT172 | -0.274757886 | 0.826589025 | 0.030758108 | 0.096827283 | 365.3961215 | 302.203555 | ENSG00000138002 |
| SERBP1 | -0.274763828 | 0.826585621 | 4.24E-08 | 6.28E-07 | 22197.87566 | 18348.22186 | ENSG00000142864 |
| LRP11 | -0.274815887 | 0.826555794 | 0.586962394 | 0.765470949 | 20.82584914 | 17.20154535 | ENSG00000120256 |
| ABHD17C | -0.274895191 | 0.82651036 | 0.008364246 | 0.033882193 | 549.1931214 | 453.9972597 | ENSG00000136379 |
| TUBE1 | -0.27491029 | 0.82650171 | 0.064235664 | 0.170761614 | 270.3021416 | 223.2856258 | ENSG00000074935 |
| CAB39 | -0.275054535 | 0.826419078 | 0.001817324 | 0.009341362 | 1245.080339 | 1028.977877 | ENSG00000135932 |
| PPP6C | -0.275088066 | 0.826399871 | 0.001286383 | 0.006970827 | 952.6245701 | 787.4333613 | ENSG00000119414 |
| SELENOK | -0.275164768 | 0.826355936 | 0.006094926 | 0.026088547 | 623.6334256 | 515.1258346 | ENSG00000113811 |
| EIF5 | -0.275183106 | 0.826345432 | 7.69E-07 | 9.07E-06 | 4395.92507 | 3632.818118 | ENSG00000100664 |
| TEP1 | -0.275187524 | 0.826342902 | 0.054051669 | 0.1496724 | 355.1979833 | 293.174733 | ENSG00000129566 |
| ZNF274 | -0.275335962 | 0.826257884 | 0.012626082 | 0.047394862 | 557.3984542 | 460.5189433 | ENSG00000171606 |
| LOC101928517 | -0.275481683 | 0.826174431 | 0.736250585 | 0.861374273 | 7.681915632 | 6.340929476 | ENSG00000269072 |
| SETD1A | -0.275555502 | 0.826132159 | 0.000482026 | 0.002984613 | 1773.952566 | 1466.025947 | ENSG00000099381 |
| HSD17B14 | -0.275621869 | 0.826094156 | 0.145188766 | 0.311790976 | 185.7102772 | 153.3730401 | ENSG00000087076 |
| LINC01980 | -0.275649279 | 0.826078461 | 0.298172456 | 0.508395345 | 73.6153524 | 60.79480093 | ENSG00000225548 |
| CLDN7 | -0.275766906 | 0.826011112 | 0.000171639 | 0.001201857 | 1463.219079 | 1208.689955 | ENSG00000181885 |
| BTN3A3 | -0.275888725 | 0.825941368 | 0.566578477 | 0.750496917 | 29.42823698 | 24.41073262 | ENSG00000111801 |
| MORN2 | -0.276369816 | 0.825665989 | 0.277211223 | 0.486446194 | 79.20281539 | 65.47443277 | ENSG00000188010 |
| VPS53 | -0.276442786 | 0.825624229 | 0.002445842 | 0.012082064 | 798.8820919 | 659.4636716 | ENSG00000141252 |
| ZHX3 | -0.276518745 | 0.82558076 | 0.162653901 | 0.337677347 | 136.6299439 | 112.686485 | ENSG00000174306 |
| NA | -0.276554681 | 0.825560196 | 0.821720502 | 0.90951774 | 3.633531329 | 3.007078013 | ENSG00000258515 |
| NA | -0.276570334 | 0.825551239 | 0.820075864 | 0.908627926 | 3.657252428 | 3.027603144 | ENSG00000274080 |
| TAOK3 | -0.27661504 | 0.825525658 | 0.062237902 | 0.166829084 | 283.0110605 | 233.4337464 | ENSG00000135090 |
| KIF24 | -0.276776133 | 0.825433483 | 0.135788559 | 0.298008942 | 166.6293452 | 137.5106105 | ENSG00000186638 |
| NA | -0.276808346 | 0.825415053 | 0.760561396 | 0.875911816 | 5.642303459 | 4.65626883 | ENSG00000286129 |
| TRAM1L1 | -0.276955537 | 0.825330844 | 0.523303512 | 0.71758545 | 25.76297588 | 21.29166469 | ENSG00000174599 |
| SLC4A1AP | -0.276973174 | 0.825320755 | 0.008137364 | 0.033105151 | 615.1493895 | 507.8801066 | ENSG00000163798 |
| TMED1 | -0.277201884 | 0.825189927 | 0.079816164 | 0.201460607 | 229.6290361 | 189.2932581 | ENSG00000099203 |
| RAB5C | -0.277274842 | 0.825148198 | 0.027709081 | 0.08932844 | 385.9048727 | 318.3427521 | ENSG00000108774 |
| KLF9 | -0.277281193 | 0.825144566 | 0.540459602 | 0.729922359 | 24.61950443 | 20.30103411 | ENSG00000119138 |
| PCMTD1 | -0.277441431 | 0.825052923 | 0.034101793 | 0.105143751 | 333.8321666 | 275.3962335 | ENSG00000168300 |
| PRXL2B | -0.277500283 | 0.825019267 | 3.97E-05 | 0.000327169 | 2035.148402 | 1679.378499 | ENSG00000157870 |
| NPDC1 | -0.277513803 | 0.825011536 | 0.340487605 | 0.552035407 | 528.7089549 | 436.2962622 | ENSG00000107281 |
| MBIP | -0.277598187 | 0.824963282 | 0.018432734 | 0.064304723 | 427.8166912 | 353.1800146 | ENSG00000151332 |
| NA | -0.277706199 | 0.82490152 | 0.733362156 | 0.859899685 | 7.620235545 | 6.273111239 | ENSG00000250220 |
| HEATR5B | -0.277786129 | 0.824855819 | 0.013098783 | 0.048762529 | 474.133854 | 390.9394262 | ENSG00000008869 |
| LINC01134 | -0.277857756 | 0.824814868 | 0.704471099 | 0.842981669 | 9.221273646 | 7.608581387 | ENSG00000236423 |
| ADAL | -0.27797412 | 0.824748343 | 0.141268655 | 0.305771753 | 193.4124585 | 159.5942964 | ENSG00000168803 |
| BRIX1 | -0.278008514 | 0.824728681 | 0.012881816 | 0.048153819 | 585.6630569 | 483.443341 | ENSG00000113460 |
| ANAPC15 | -0.278086425 | 0.824684144 | 0.15797345 | 0.330672644 | 148.7166707 | 122.6723596 | ENSG00000110200 |
| CDCP1 | -0.278087077 | 0.824683772 | 0.016589521 | 0.059156466 | 453.3809738 | 373.9234577 | ENSG00000163814 |
| BEND4 | -0.278104199 | 0.824673984 | 0.002075689 | 0.010506401 | 1258.581459 | 1037.725524 | ENSG00000188848 |
| NA | -0.278311512 | 0.824555488 | 0.631644558 | 0.797106325 | 14.89085768 | 12.27761945 | ENSG00000227885 |
| RTTN | -0.278410424 | 0.824498958 | 0.022545316 | 0.075759484 | 383.9699859 | 316.5700746 | ENSG00000176225 |
| COG6 | -0.278523819 | 0.824434155 | 0.036491897 | 0.110706742 | 320.8006094 | 264.4820816 | ENSG00000133103 |
| GTF2IP1 | -0.278572271 | 0.824406468 | 0.090529795 | 0.221343239 | 203.3619934 | 167.618657 | ENSG00000277053 |
| RPN1 | -0.278652786 | 0.82436046 | 2.94E-07 | 3.76E-06 | 5129.42889 | 4228.430209 | ENSG00000163902 |
| TJP2 | -0.278837877 | 0.824254705 | 0.256098522 | 0.46299259 | 84.47206367 | 69.56311169 | ENSG00000119139 |
| PFDN1 | -0.279015757 | 0.824153083 | 0.003657142 | 0.017025845 | 611.2124656 | 503.6950745 | ENSG00000113068 |
| NA | -0.279025768 | 0.824147365 | 0.644659976 | 0.806145123 | 16.61286554 | 13.70360843 | ENSG00000274840 |

| | | | | | | | |
|---|---|---|---|---|---|---|---|
| SSBP1 | -0.279199701 | 0.82404801 | 1.03E-05 | 9.80E-05 | 3459.92599 | 2851.340261 | ENSG00000106028 |
| TRPM7 | -0.279280101 | 0.824002088 | 0.004524676 | 0.020335198 | 823.535638 | 678.4743306 | ENSG00000092439 |
| RPL7L1 | -0.279424487 | 0.823919626 | 1.14E-06 | 1.31E-05 | 2900.890965 | 2390.189764 | ENSG00000146223 |
| UBE2V1 | -0.27946523 | 0.823896358 | 0.452138588 | 0.658283762 | 38.30493123 | 31.59890031 | ENSG00000244687 |
| MTM1 | -0.279486463 | 0.823884232 | 0.061379247 | 0.165214807 | 345.7288197 | 284.6347827 | ENSG00000171100 |
| GTPBP8 | -0.279639648 | 0.823796757 | 0.111596491 | 0.259225059 | 171.9623323 | 141.674412 | ENSG00000163607 |
| GMPR2 | -0.279640621 | 0.823796201 | 7.62E-05 | 0.000583803 | 1477.956626 | 1217.488208 | ENSG00000100938 |
| SEMA4F | -0.279743401 | 0.823737515 | 0.064743601 | 0.171966654 | 239.9395256 | 197.5070747 | ENSG00000135622 |
| MRPS18C | -0.279771555 | 0.82372144 | 0.213846089 | 0.408381373 | 118.5432648 | 97.83139754 | ENSG00000163319 |
| IMPA2 | -0.279804436 | 0.823702666 | 3.15E-05 | 0.00026687 | 2468.932708 | 2033.558239 | ENSG00000141401 |
| TCF15 | -0.279949617 | 0.82361978 | 0.039227106 | 0.116919808 | 336.4396623 | 277.3378848 | ENSG00000125878 |
| PRELID3A | -0.280092881 | 0.823537996 | 0.14054192 | 0.3047809 | 173.9684347 | 143.4548158 | ENSG00000141391 |
| CNOT6L | -0.280171932 | 0.823492872 | 0.057853267 | 0.157901735 | 258.4962286 | 212.8677978 | ENSG00000138767 |
| EME2 | -0.280296778 | 0.823421613 | 0.330892215 | 0.542345033 | 565.5112919 | 465.7622991 | ENSG00000197774 |
| ELK1 | -0.280501299 | 0.823304891 | 0.001274043 | 0.006913878 | 1055.558376 | 868.6786957 | ENSG00000126767 |
| MORN4 | -0.280536846 | 0.823284605 | 0.038976336 | 0.116448047 | 331.120251 | 272.6281229 | ENSG00000171160 |
| GATB | -0.280632537 | 0.82323 | 0.014939542 | 0.054329531 | 433.0171149 | 356.2377478 | ENSG00000059691 |
| HMGCS1 | -0.280635554 | 0.823228279 | 3.52E-08 | 5.31E-07 | 5849.08027 | 4814.781054 | ENSG00000112972 |
| RRP7A | -0.280691859 | 0.823196151 | 1.18E-05 | 0.000110074 | 2012.998719 | 1657.246166 | ENSG00000189306 |
| RPP21 | -0.280702882 | 0.823189861 | 0.597271454 | 0.773554521 | 20.27487264 | 16.70198459 | ENSG00000241370 |
| NPHP3 | -0.280873358 | 0.823092594 | 0.499753983 | 0.698117758 | 27.86782762 | 22.92997819 | ENSG00000113971 |
| FAAH | -0.280893041 | 0.823081365 | 0.041035543 | 0.121086911 | 301.1108078 | 247.8470123 | ENSG00000117480 |
| SMCO4 | -0.280961615 | 0.823042243 | 0.003154508 | 0.015024689 | 741.3335579 | 610.1668296 | ENSG00000166002 |
| ERBB2 | -0.281132736 | 0.822944626 | 5.98E-05 | 0.000471428 | 13463.22099 | 11079.96273 | ENSG00000141736 |
| CTSD | -0.281184671 | 0.822915002 | 0.008134024 | 0.033098696 | 690.9992871 | 568.7819364 | ENSG00000117984 |
| MRPS15 | -0.281189928 | 0.822912003 | 1.01E-05 | 9.61E-05 | 1959.2007 | 1612.116204 | ENSG00000116898 |
| TNFRSF13C | -0.281206639 | 0.822902472 | 0.55378228 | 0.740250659 | 23.01462752 | 18.99791237 | ENSG00000159958 |
| UBE2D2 | -0.281231144 | 0.822888494 | 2.19E-05 | 0.000192911 | 3387.300142 | 2787.977314 | ENSG00000131508 |
| USB1 | -0.281242422 | 0.822882061 | 4.52E-05 | 0.000368024 | 1701.484369 | 1400.462311 | ENSG00000103005 |
| EPS8L1 | -0.281394049 | 0.822795581 | 0.002525926 | 0.012409498 | 766.7006989 | 631.0173446 | ENSG00000131037 |
| SLC25A42 | -0.281399001 | 0.822792757 | 0.164337542 | 0.339776055 | 136.5086425 | 112.285934 | ENSG00000181035 |
| RANBP17 | -0.281446777 | 0.82276551 | 0.168868944 | 0.346602543 | 168.9800014 | 139.001886 | ENSG00000204764 |
| DOCK7 | -0.281503181 | 0.822733344 | 0.004231664 | 0.019265906 | 754.2859955 | 620.4637323 | ENSG00000116641 |
| NA | -0.281553365 | 0.822704726 | 0.669232683 | 0.82295975 | 12.80972704 | 10.53384242 | ENSG00000274422 |
| PSAT1 | -0.281599752 | 0.822678273 | 3.00E-06 | 3.17E-05 | 8760.006998 | 7206.317296 | ENSG00000135069 |
| NA | -0.281633644 | 0.822658947 | 0.165105578 | 0.340767086 | 140.8167899 | 115.7974182 | ENSG00000230330 |
| NA | -0.281791997 | 0.822568655 | 0.812134671 | 0.904580553 | 3.650133483 | 3.006132054 | ENSG00000270171 |
| ZNF69 | -0.281921253 | 0.822494962 | 0.191059421 | 0.377817504 | 109.2608959 | 89.81060985 | ENSG00000198429 |
| RASA3 | -0.281921887 | 0.822494601 | 0.002256996 | 0.011272094 | 750.3781833 | 617.4971602 | ENSG00000185989 |
| PPP2R2B | -0.282112918 | 0.822385699 | 0.000158659 | 0.001121778 | 1480.132692 | 1217.319306 | ENSG00000156475 |
| ELOVL3 | -0.282168234 | 0.822354168 | 0.807219727 | 0.901913344 | 3.657252428 | 3.01142894 | ENSG00000119915 |
| DNAH9 | -0.28225265 | 0.822306051 | 0.622206708 | 0.791007118 | 14.97981833 | 12.30467097 | ENSG00000007174 |
| DIAPH3 | -0.282266693 | 0.822298047 | 0.003266849 | 0.01547002 | 666.3861605 | 547.8711053 | ENSG00000139734 |
| NTM | -0.282364616 | 0.822242235 | 0.041190854 | 0.121393459 | 302.6694377 | 248.8224146 | ENSG00000182667 |
| EIF4E | -0.282436639 | 0.822201188 | 9.06E-05 | 0.000681283 | 2127.935454 | 1749.445625 | ENSG00000151247 |
| ARHGEF28 | -0.282439721 | 0.822199431 | 0.547435635 | 0.735112974 | 22.17096302 | 18.21676844 | ENSG00000214944 |
| THY1 | -0.282468636 | 0.822182953 | 6.57E-08 | 9.47E-07 | 6280.976012 | 5163.909581 | ENSG00000154096 |
| FBXW11 | -0.282490316 | 0.822170597 | 0.000922412 | 0.005228017 | 1523.043825 | 1251.99434 | ENSG00000072803 |
| NA | -0.282709034 | 0.822045963 | 0.254874033 | 0.461485726 | 81.61131288 | 67.03745568 | ENSG00000283567 |
| FAM111A | -0.282800288 | 0.821993968 | 0.018745669 | 0.065149887 | 447.8842515 | 367.8771212 | ENSG00000166801 |
| PNMA8A | -0.282831513 | 0.821976177 | 0.000347387 | 0.002230578 | 1066.212275 | 876.2338044 | ENSG00000182013 |
| ZDHHC16 | -0.282847412 | 0.821967119 | 0.000292444 | 0.001922316 | 1218.252349 | 1001.333103 | ENSG00000171307 |
| HYKK | -0.282849889 | 0.821965708 | 0.60324065 | 0.77799629 | 21.12565609 | 17.41559383 | ENSG00000188266 |
| ZNF214 | -0.282877744 | 0.821949838 | 0.462728266 | 0.667551225 | 33.17356034 | 27.26287853 | ENSG00000149050 |
| VWA8 | -0.28307258 | 0.821838841 | 0.008598423 | 0.034678696 | 527.0725496 | 432.999242 | ENSG00000102763 |
| DPAGT1 | -0.283089521 | 0.82182919 | 0.000325263 | 0.002107238 | 1046.151555 | 859.9108997 | ENSG00000172269 |
| SRCIN1 | -0.283180408 | 0.821777418 | 0.177882079 | 0.359822522 | 119.4239986 | 98.21556341 | ENSG00000277363 |
| ARL6IP4 | -0.283344663 | 0.821683862 | 0.690496046 | 0.834528492 | 9.299555887 | 7.63222794 | ENSG00000182196 |
| CDR2L | -0.283402257 | 0.82165106 | 0.000954515 | 0.005390555 | 1418.982182 | 1166.0746 | ENSG00000109089 |
| NRIP1 | -0.283425589 | 0.821637772 | 0.119461495 | 0.27197945 | 198.2557659 | 162.7029385 | ENSG00000180530 |
| FNDC5 | -0.283464691 | 0.821615503 | 0.478848182 | 0.681414206 | 31.16003353 | 25.60403991 | ENSG00000160097 |
| NA | -0.283931421 | 0.821349743 | 0.677757848 | 0.82756234 | 13.85058477 | 11.33768695 | ENSG00000285804 |
| NA | -0.284011137 | 0.82130436 | 0.578585808 | 0.759655996 | 18.59911177 | 15.25480807 | ENSG00000269044 |
| NA | -0.28406318 | 0.821274734 | 0.605629864 | 0.779795526 | 16.59270391 | 13.61961599 | ENSG00000278864 |
| CACNA2D2 | -0.284064733 | 0.82127385 | 0.000787388 | 0.004548737 | 885.37508 | 727.2954396 | ENSG00000007402 |
| PFAS | -0.284127772 | 0.821237965 | 5.95E-09 | 1.02E-07 | 5744.898023 | 4718.024081 | ENSG00000178921 |
| CNTNAP3B | -0.284132461 | 0.821235296 | 0.398966573 | 0.610475104 | 45.68823511 | 37.52187509 | ENSG00000154529 |
| ATG13 | -0.284317262 | 0.821130107 | 0.00024468 | 0.00164499 | 1099.255357 | 902.3684084 | ENSG00000175224 |
| EIF3J | -0.284371332 | 0.821099333 | 0.000534945 | 0.003255698 | 1677.066942 | 1377.454465 | ENSG00000104131 |
| ACTR1A | -0.284411893 | 0.821076248 | 1.42E-06 | 1.60E-05 | 2490.951581 | 2045.324009 | ENSG00000138107 |
| NA | -0.284483533 | 0.821035477 | 0.205715884 | 0.397884522 | 114.9835666 | 94.51220723 | ENSG00000229320 |
| LOC100128770 | -0.284516356 | 0.821016798 | 0.448669689 | 0.655281496 | 37.70175785 | 30.99973999 | ENSG00000205890 |
| NA | -0.284660281 | 0.820934896 | 0.30510805 | 0.515335437 | 70.07642614 | 57.59970736 | ENSG00000237004 |
| TSPYL2 | -0.28496084 | 0.820763887 | 0.003510045 | 0.016436199 | 637.6364044 | 523.197393 | ENSG00000184205 |

| | | | | | | | |
|---|---|---|---|---|---|---|---|
| GINS3 | -0.285160328 | 0.820650404 | 0.052530963 | 0.146585313 | 243.3281421 | 199.7118212 | ENSG00000181938 |
| FBXW8 | -0.285163348 | 0.820648686 | 0.009382803 | 0.037052928 | 475.7414262 | 390.4563232 | ENSG00000174989 |
| SUMO1 | -0.285267329 | 0.820589541 | 1.03E-05 | 9.80E-05 | 4331.307398 | 3554.582162 | ENSG00000116030 |
| ATAD3A | -0.285273136 | 0.820586238 | 5.73E-05 | 0.000454717 | 1893.647792 | 1554.326358 | ENSG00000197785 |
| DHRS11 | -0.285476647 | 0.820470492 | 0.011132543 | 0.042708204 | 719.4514447 | 590.3094269 | ENSG00000278535 |
| NA | -0.285615449 | 0.820391558 | 0.458149661 | 0.663610704 | 34.83152385 | 28.56193289 | ENSG00000257122 |
| RAB31 | -0.285701911 | 0.820342392 | 0.001929978 | 0.009834766 | 741.267176 | 607.7751587 | ENSG00000168461 |
| ARAP2 | -0.285914953 | 0.820221262 | 0.514866853 | 0.71112854 | 26.30830798 | 21.52356838 | ENSG00000047365 |
| TOP1 | -0.285920952 | 0.820217851 | 3.58E-05 | 0.000298905 | 3884.472931 | 3185.94881 | ENSG00000198900 |
| TARDBP | -0.285925045 | 0.820215525 | 8.36E-06 | 8.15E-05 | 2868.314296 | 2352.817449 | ENSG00000120948 |
| FBLN1 | -0.286060057 | 0.82013877 | 2.26E-08 | 3.50E-07 | 5613.583679 | 4603.852882 | ENSG00000077942 |
| GAS2L3 | -0.286127339 | 0.820100522 | 0.144668074 | 0.310833457 | 159.4897608 | 130.6386225 | ENSG00000139354 |
| STMP1 | -0.2862502 | 0.820030684 | 0.001088986 | 0.006038021 | 903.2981013 | 740.9036838 | ENSG00000243317 |
| MRPS31P5 | -0.286331478 | 0.819984488 | 0.724747689 | 0.854899475 | 7.689034577 | 6.277462167 | ENSG00000243406 |
| DACH1 | -0.286348478 | 0.819974825 | 0.734140449 | 0.860316207 | 7.33110701 | 6.011034604 | ENSG00000276644 |
| ST14 | -0.286390894 | 0.819950718 | 1.62E-05 | 0.000147174 | 1944.313478 | 1594.459562 | ENSG00000149418 |
| NBPF8 | -0.286776033 | 0.819731854 | 0.374789112 | 0.586783366 | 55.44032363 | 45.48153889 | ENSG00000270231 |
| ERO1A | -0.286826593 | 0.819703128 | 0.010602649 | 0.040955415 | 521.3059701 | 427.0453163 | ENSG00000197930 |
| EMSY | -0.286888071 | 0.819668198 | 0.00843665 | 0.034116944 | 571.5381675 | 468.2600921 | ENSG00000158636 |
| CHMP4C | -0.286935467 | 0.81964127 | 0.56040897 | 0.745566988 | 21.95419299 | 17.99016163 | ENSG00000164695 |
| PSMA5 | -0.286952668 | 0.819631498 | 4.43E-07 | 5.50E-06 | 3488.342309 | 2859.095091 | ENSG00000143106 |
| HTRA4 | -0.286967889 | 0.81962285 | 0.565780773 | 0.749861036 | 20.28555106 | 16.60739838 | ENSG00000169495 |
| PTPN13 | -0.286999968 | 0.819604626 | 1.03E-05 | 9.80E-05 | 2631.945401 | 2156.885493 | ENSG00000163629 |
| CAMTA1 | -0.287046426 | 0.819578233 | 0.000386724 | 0.002454886 | 1364.692652 | 1118.296223 | ENSG00000171735 |
| C3orf67 | -0.287133084 | 0.819529005 | 0.197272633 | 0.386823141 | 108.9673182 | 89.17787162 | ENSG00000163689 |
| TRIM25 | -0.287258981 | 0.819457492 | 0.000301447 | 0.001973248 | 1054.70322 | 864.2963471 | ENSG00000121060 |
| NUP88 | -0.287416061 | 0.819368274 | 3.28E-05 | 0.000275993 | 1742.849025 | 1427.572726 | ENSG00000108559 |
| SNHG20 | -0.287452853 | 0.819347379 | 0.332633238 | 0.544584196 | 69.48543187 | 56.91314964 | ENSG00000234912 |
| DNAI1 | -0.287636675 | 0.819242988 | 0.703792549 | 0.842524014 | 9.395635491 | 7.683871974 | ENSG00000122735 |
| FERMT1 | -0.287714616 | 0.81919873 | 0.004103582 | 0.018791499 | 602.41319 | 493.3238771 | ENSG00000101311 |
| NA | -0.287859438 | 0.8191165 | 0.275756509 | 0.484659067 | 95.85722554 | 78.60659482 | ENSG00000227232 |
| UBL5 | -0.287874918 | 0.819107712 | 4.92E-05 | 0.000396109 | 2471.799505 | 2024.453788 | ENSG00000198258 |
| NTN5 | -0.287894459 | 0.819096617 | 0.742105621 | 0.865637101 | 6.885413993 | 5.659668642 | ENSG00000142233 |
| NHLRC4 | -0.287953391 | 0.819063159 | 0.730246563 | 0.857920184 | 7.747155192 | 6.319458387 | ENSG00000257108 |
| TCOF1 | -0.287984214 | 0.81904566 | 3.84E-10 | 7.85E-09 | 8491.984767 | 6955.656888 | ENSG00000070814 |
| SEMA3B | -0.288014309 | 0.819028574 | 0.266570937 | 0.474012748 | 89.0479827 | 72.92668459 | ENSG00000012171 |
| NA | -0.288061016 | 0.819002059 | 0.523965936 | 0.718166487 | 27.14129407 | 22.22317888 | ENSG00000238072 |
| FAHD1 | -0.288134001 | 0.818960628 | 0.02149708 | 0.07296412 | 373.2523467 | 305.7771613 | ENSG00000180185 |
| BTN1A1 | -0.288174804 | 0.818937466 | 0.67544648 | 0.826260317 | 10.53554748 | 8.64650507 | ENSG00000124557 |
| FUT11 | -0.288267513 | 0.818884842 | 0.127004837 | 0.283991871 | 146.0267485 | 119.482352 | ENSG00000196968 |
| PATZ1 | -0.288329623 | 0.818849588 | 2.21E-05 | 0.000194154 | 2313.003026 | 1894.481784 | ENSG00000100105 |
| MCAT | -0.288376298 | 0.818823097 | 0.011048272 | 0.042433725 | 442.0534 | 362.1186537 | ENSG00000100294 |
| NA | -0.288393972 | 0.818813066 | 0.830013084 | 0.914737804 | 3.695211417 | 3.02542768 | ENSG00000287972 |
| ANKS1A | -0.288509361 | 0.818747578 | 7.61E-06 | 7.49E-05 | 1863.612441 | 1525.469315 | ENSG00000064999 |
| GABBR1 | -0.288554707 | 0.818721844 | 0.165651437 | 0.341483177 | 1551.009208 | 1269.888056 | ENSG00000204681 |
| LINS1 | -0.288554801 | 0.818721791 | 0.245544396 | 0.45002969 | 85.23032698 | 69.86455758 | ENSG00000140471 |
| ATG4C | -0.288567275 | 0.818714712 | 0.011215391 | 0.042953288 | 428.3196413 | 350.6801377 | ENSG00000125703 |
| NA | -0.288634091 | 0.818676795 | 0.665947708 | 0.821129699 | 14.48312365 | 11.84566602 | ENSG00000228302 |
| NUTM2B-AS1 | -0.288883231 | 0.81853543 | 0.130478277 | 0.289755675 | 163.3549366 | 133.7303553 | ENSG00000225484 |
| TATDN2 | -0.289078054 | 0.818424901 | 0.00949051 | 0.037432806 | 512.6395374 | 419.607485 | ENSG00000157014 |
| GPR83 | -0.289291924 | 0.818303584 | 0.785476847 | 0.890433415 | 4.858844501 | 4.005180939 | ENSG00000123901 |
| PSMD13 | -0.289340415 | 0.81827608 | 2.36E-05 | 0.000207005 | 3130.004004 | 2560.498153 | ENSG00000185627 |
| CASTOR3 | -0.289464537 | 0.818205683 | 0.163601589 | 0.338773684 | 135.3192296 | 110.7100692 | ENSG00000239521 |
| TMEM91 | -0.289592609 | 0.818133052 | 0.659348072 | 0.816486541 | 12.5680407 | 10.28169715 | ENSG00000142046 |
| TMEM189 | -0.289634544 | 0.818109272 | 0.035581608 | 0.108555521 | 314.3401425 | 257.0621056 | ENSG00000240849 |
| CHMP1A | -0.289701241 | 0.818071451 | 1.48E-05 | 0.000135653 | 2071.218815 | 1694.457 | ENSG00000131165 |
| RPN2 | -0.289767672 | 0.818033782 | 2.88E-07 | 3.69E-06 | 5103.065445 | 4174.569173 | ENSG00000118705 |
| CDC37 | -0.289779752 | 0.818026932 | 2.39E-06 | 2.58E-05 | 5191.720784 | 4247.821988 | ENSG00000105401 |
| THG1L | -0.289813305 | 0.818007908 | 0.027317102 | 0.088321087 | 308.6406082 | 252.4595609 | ENSG00000113272 |
| IGSF22 | -0.289864955 | 0.817978623 | 0.747929677 | 0.86888853 | 6.925737245 | 5.681139731 | ENSG00000179057 |
| STARD3NL | -0.28990861 | 0.817953872 | 0.002667051 | 0.013001359 | 657.8315726 | 537.9390931 | ENSG00000010270 |
| AMFR | -0.29009253 | 0.817849603 | 0.009431495 | 0.037223361 | 544.3597524 | 444.8694279 | ENSG00000159461 |
| CCDC17 | -0.290231921 | 0.817770587 | 0.651583377 | 0.810990053 | 12.1804683 | 9.98509664 | ENSG00000159588 |
| NA | -0.290285512 | 0.81774021 | 0.049938979 | 0.140950174 | 274.5815338 | 224.3538904 | ENSG00000255769 |
| SUPV3L1 | -0.290337014 | 0.817711018 | 0.0019985 | 0.010153799 | 810.4081058 | 662.7921739 | ENSG00000156502 |
| NA | -0.290430268 | 0.817658165 | 0.647364463 | 0.80867032 | 12.19351098 | 9.994744453 | ENSG00000251689 |
| AAMP | -0.290507961 | 0.817614133 | 2.82E-07 | 3.62E-06 | 3915.628437 | 3201.880527 | ENSG00000127837 |
| HPF1 | -0.290698157 | 0.817506351 | 0.012470734 | 0.046933004 | 471.2398989 | 385.1238069 | ENSG00000056050 |
| TNFRSF1A | -0.291080717 | 0.817289601 | 0.000205068 | 0.001406215 | 1121.963109 | 917.2589133 | ENSG00000067182 |
| CTNNBIP1 | -0.291117081 | 0.817269001 | 0.000510591 | 0.003132721 | 996.3749432 | 814.5969465 | ENSG00000178585 |
| TCEAL4 | -0.291167246 | 0.817240583 | 0.000127068 | 0.000920124 | 1279.043327 | 1045.043972 | ENSG00000133142 |
| PCGF5 | -0.291179314 | 0.817233747 | 0.201448202 | 0.392476389 | 129.9521478 | 106.2690354 | ENSG00000180628 |
| FBXO36 | -0.291615946 | 0.816986449 | 0.381695887 | 0.594109191 | 55.86374394 | 45.63022815 | ENSG00000153832 |
| BRD3OS | -0.291642235 | 0.816971562 | 0.026209053 | 0.085470207 | 342.3728743 | 279.897046 | ENSG00000235106 |

| | | | | | | | |
|---|---|---|---|---|---|---|---|
| CNPY3 | -0.29177572 | 0.816895975 | 6.16E-06 | 6.16E-05 | 2215.034513 | 1809.356903 | ENSG00000137161 |
| PLBD2 | -0.291907936 | 0.816821114 | 0.00016231 | 0.001144167 | 1110.204919 | 906.7713024 | ENSG00000151176 |
| NA | -0.292071808 | 0.816728339 | 0.594377314 | 0.771358959 | 20.24995634 | 16.50070534 | ENSG00000235381 |
| SNHG18 | -0.292203549 | 0.816653762 | 0.190864455 | 0.377550564 | 104.8320582 | 85.62706117 | ENSG00000250786 |
| CEBPD | -0.292205238 | 0.816652806 | 0.386988164 | 0.599235116 | 47.12200416 | 38.53738172 | ENSG00000221869 |
| LRRIQ1 | -0.292245117 | 0.816630232 | 0.779162797 | 0.88732373 | 4.848166083 | 3.985885313 | ENSG00000133640 |
| BMT2 | -0.292345627 | 0.816573341 | 0.284556504 | 0.493783844 | 66.61697732 | 54.46228976 | ENSG00000164603 |
| LINC01465 | -0.292475529 | 0.816499819 | 0.68111313 | 0.829598484 | 10.13017771 | 8.302611457 | ENSG00000221949 |
| PPP4R4 | -0.292500558 | 0.816485654 | 0.139765756 | 0.303702254 | 135.4826337 | 110.6201175 | ENSG00000119698 |
| LINC01664 | -0.292507003 | 0.816482006 | 0.770500044 | 0.881997379 | 5.329453825 | 4.356546901 | ENSG00000235478 |
| PPAN | -0.292566254 | 0.816448474 | 0.117303264 | 0.268615513 | 142.3039771 | 116.1904968 | ENSG00000130810 |
| PPIP5K2 | -0.292613858 | 0.816421535 | 0.009581188 | 0.037735294 | 571.5704298 | 466.3236423 | ENSG00000145725 |
| ADRA2A | -0.292732605 | 0.816354339 | 0.518348595 | 0.714213717 | 28.95109353 | 23.61681945 | ENSG00000150594 |
| KPNB1 | -0.292836284 | 0.816295674 | 3.60E-07 | 4.53E-06 | 11014.06303 | 8990.517815 | ENSG00000108424 |
| E2F3 | -0.292930537 | 0.816242346 | 0.000120645 | 0.000878664 | 1312.429871 | 1070.945987 | ENSG00000112242 |
| NA | -0.29296351 | 0.816223691 | 0.246590731 | 0.451421671 | 77.60889623 | 63.37928977 | ENSG00000264577 |
| NA | -0.293265105 | 0.816053077 | 0.130053463 | 0.288982192 | 162.8380542 | 133.0116601 | ENSG00000204791 |
| NCBP2AS2 | -0.29333052 | 0.816016076 | 0.006690313 | 0.028208654 | 868.7763739 | 709.2284674 | ENSG00000270170 |
| APIP | -0.293451458 | 0.815947674 | 0.033575334 | 0.103826787 | 319.3541025 | 260.8138164 | ENSG00000149089 |
| BCCIP | -0.293536145 | 0.815899779 | 1.67E-06 | 1.84E-05 | 2211.281792 | 1803.991469 | ENSG00000107949 |
| PPME1 | -0.293540043 | 0.815897574 | 0.000628742 | 0.003739777 | 866.3821155 | 706.6038812 | ENSG00000214517 |
| GALNS | -0.293626125 | 0.815848893 | 0.007547644 | 0.031169309 | 461.7333873 | 376.8732386 | ENSG00000141012 |
| ABT1 | -0.293707243 | 0.815803023 | 0.000178214 | 0.001241909 | 1362.390632 | 1111.338192 | ENSG00000146109 |
| ELMO2 | -0.293757036 | 0.815774866 | 0.006530284 | 0.027663618 | 505.7568454 | 412.5323259 | ENSG00000062598 |
| SAP30L | -0.293804058 | 0.815748278 | 0.009105699 | 0.036235188 | 546.4937159 | 445.6580013 | ENSG00000164576 |
| PNKP | -0.293843861 | 0.815725772 | 0.000445624 | 0.002782938 | 1311.145599 | 1069.782605 | ENSG00000039650 |
| RPS6KA3 | -0.293870183 | 0.81571089 | 0.01898412 | 0.065806736 | 370.9900407 | 302.5483516 | ENSG00000177189 |
| RRAGB | -0.294178451 | 0.815536611 | 0.118744257 | 0.270766154 | 145.5807761 | 118.6925063 | ENSG00000083750 |
| CALN1 | -0.294322276 | 0.815455313 | 0.004868312 | 0.021576643 | 554.2141748 | 451.9722783 | ENSG00000183166 |
| PRKAR1B | -0.294395636 | 0.815413849 | 0.000224708 | 0.001524131 | 2025.416676 | 1651.682887 | ENSG00000188191 |
| USP2 | -0.294402222 | 0.815410126 | 0.118331808 | 0.27026605 | 160.0769424 | 130.5927698 | ENSG00000036672 |
| CBS | -0.294403951 | 0.815409149 | 0.000524738 | 0.003205976 | 998.5624485 | 814.5348673 | ENSG00000160200 |
| CTSA | -0.29444237 | 0.815387435 | 0.003549282 | 0.016595243 | 645.5244896 | 526.5533054 | ENSG00000064601 |
| KLF10 | -0.294555606 | 0.815323438 | 0.005545821 | 0.02408205 | 536.2701022 | 437.1087523 | ENSG00000155090 |
| TAF9 | -0.294570675 | 0.815314922 | 0.014991252 | 0.054481677 | 517.52869 | 422.0793848 | ENSG00000273841 |
| TACC2 | -0.294674942 | 0.815256 | 0.000206583 | 0.001416086 | 1121.490441 | 914.4375598 | ENSG00000138162 |
| INPP5A | -0.294795439 | 0.81518791 | 0.008045944 | 0.032825158 | 555.1800048 | 452.3064086 | ENSG00000068383 |
| ECHDC2 | -0.294939729 | 0.815106384 | 0.07063339 | 0.184165618 | 218.5944558 | 178.0127956 | ENSG00000121310 |
| KDELR2 | -0.29495161 | 0.815099671 | 0.00015759 | 0.001114636 | 1236.579012 | 1008.217146 | ENSG00000136240 |
| COMMD10 | -0.295030146 | 0.815055301 | 0.032870388 | 0.102014343 | 298.9668019 | 243.5234332 | ENSG00000145781 |
| ACE | -0.295060168 | 0.81503834 | 0.336192457 | 0.548193198 | 55.30064064 | 45.03028983 | ENSG00000159640 |
| NA | -0.295091535 | 0.81502062 | 0.074385579 | 0.191594607 | 217.272121 | 176.9150205 | ENSG00000231698 |
| NA | -0.295260679 | 0.814925071 | 0.580713891 | 0.761084181 | 18.68067416 | 15.26228042 | ENSG00000213144 |
| NA | -0.295333532 | 0.81488392 | 0.763219366 | 0.877513739 | 5.66958403 | 4.594976984 | ENSG00000242262 |
| BAG1 | -0.295346893 | 0.814876374 | 0.00100939 | 0.005653123 | 1223.207726 | 996.9702838 | ENSG00000107262 |
| MEGF8 | -0.295412813 | 0.814839141 | 1.83E-06 | 2.01E-05 | 3393.387662 | 2764.903007 | ENSG00000105429 |
| FBXO7 | -0.295450867 | 0.814817648 | 2.47E-05 | 0.000215467 | 1395.594832 | 1137.231213 | ENSG00000100225 |
| INPPL1 | -0.295470508 | 0.814806555 | 1.17E-06 | 1.33E-05 | 2402.279348 | 1957.664043 | ENSG00000165458 |
| IPO8 | -0.295539242 | 0.814767737 | 7.67E-05 | 0.000586339 | 1351.131299 | 1100.598602 | ENSG00000133704 |
| ABCB9 | -0.295645113 | 0.814707948 | 0.188184897 | 0.374093752 | 112.2562695 | 91.56743966 | ENSG00000150967 |
| GPLD1 | -0.295648417 | 0.814706082 | 0.140542157 | 0.3047809 | 136.2331676 | 110.8778432 | ENSG00000112293 |
| NA | -0.295760925 | 0.81464255 | 0.654582797 | 0.813115577 | 11.84982131 | 9.687550175 | ENSG00000226608 |
| MBP | -0.295827719 | 0.814604834 | 0.003044688 | 0.014597344 | 623.8258374 | 508.0858617 | ENSG00000197971 |
| NF1 | -0.295995327 | 0.814510201 | 0.008620694 | 0.034734758 | 656.7832381 | 534.7109028 | ENSG00000196712 |
| NELFCD | -0.296090145 | 0.814456671 | 2.72E-05 | 0.000234206 | 1412.498579 | 1150.42167 | ENSG00000101158 |
| SIX2 | -0.296132292 | 0.814432878 | 0.644229742 | 0.805989478 | 13.37374622 | 10.92502914 | ENSG00000170577 |
| COPS2 | -0.296222951 | 0.814381701 | 0.00094991 | 0.005369901 | 1134.952258 | 924.4116408 | ENSG00000166200 |
| CEP70 | -0.296272568 | 0.814353693 | 0.019590021 | 0.067547478 | 414.9286819 | 338.1803053 | ENSG00000114107 |
| RPL6 | -0.296289154 | 0.814344331 | 8.40E-11 | 1.90E-09 | 19148.34796 | 15592.91202 | ENSG00000089009 |
| NUDCD3 | -0.296312253 | 0.814331292 | 3.01E-05 | 0.000257156 | 1652.772591 | 1345.658662 | ENSG00000015676 |
| C7orf50 | -0.296365469 | 0.814301255 | 9.23E-06 | 8.91E-05 | 2064.314486 | 1681.480159 | ENSG00000146540 |
| SPAST | -0.296369069 | 0.814299223 | 0.004581187 | 0.020535417 | 573.6408814 | 467.0891065 | ENSG00000021574 |
| ANKDD1A | -0.296493569 | 0.814228955 | 0.31461728 | 0.524986448 | 57.41230583 | 46.78059325 | ENSG00000166839 |
| MELK | -0.296501733 | 0.814224347 | 0.000141882 | 0.001018025 | 1129.157787 | 919.7263262 | ENSG00000165304 |
| FDXACB1 | -0.296689851 | 0.814118184 | 0.566820806 | 0.750531324 | 18.68659789 | 15.24828168 | ENSG00000255561 |
| MESD | -0.296704275 | 0.814110045 | 3.24E-05 | 0.00027321 | 1359.684465 | 1107.008338 | ENSG00000117899 |
| BEX2 | -0.296839423 | 0.814033785 | 0.000293867 | 0.001930993 | 1038.045504 | 844.9517985 | ENSG00000133134 |
| UBTD1 | -0.296950737 | 0.813970979 | 0.008663646 | 0.034893321 | 620.6793156 | 505.6854594 | ENSG00000165886 |
| DERL3 | -0.296957578 | 0.813967119 | 0.154912457 | 0.326509124 | 119.1055046 | 96.97222047 | ENSG00000099958 |
| TIMM13 | -0.29700694 | 0.81393927 | 3.98E-06 | 4.10E-05 | 4307.12954 | 3506.241265 | ENSG00000099800 |
| MARVELD1 | -0.29707616 | 0.813900218 | 0.001353887 | 0.007292684 | 1338.003437 | 1088.949617 | ENSG00000155254 |
| CALM3 | -0.297080679 | 0.813897669 | 2.48E-09 | 4.50E-08 | 9650.828995 | 7854.776445 | ENSG00000160014 |
| PBX3 | -0.297150844 | 0.813858086 | 0.004294554 | 0.019491144 | 563.5252988 | 458.7583094 | ENSG00000167081 |
| ZNF345 | -0.297509346 | 0.813655872 | 0.342772745 | 0.554597608 | 59.22210529 | 48.21433813 | ENSG00000251247 |

| | | | | | | | |
|---|---|---|---|---|---|---|---|
| HDDC2 | -0.297658955 | 0.813571499 | 8.21E-05 | 0.000623922 | 1284.849776 | 1045.257075 | ENSG00000111906 |
| STAMBP | -0.297735042 | 0.813528593 | 0.00026005 | 0.001735967 | 983.0740363 | 799.5499012 | ENSG00000124356 |
| NA | -0.297892922 | 0.81343957 | 0.519226087 | 0.714901185 | 22.8547829 | 18.58525456 | ENSG00000224094 |
| WASHC1 | -0.297914474 | 0.813427419 | 0.165372372 | 0.341182746 | 112.2209803 | 91.38595049 | ENSG00000181404 |
| CRY2 | -0.297927023 | 0.813420343 | 0.007804029 | 0.03203197 | 509.8289508 | 414.5632235 | ENSG00000121671 |
| USP5 | -0.297966357 | 0.813398166 | 5.49E-08 | 7.97E-07 | 4307.639308 | 3503.45082 | ENSG00000111667 |
| CARMIL3 | -0.298065609 | 0.813342209 | 0.069848218 | 0.182445863 | 340.0995066 | 276.7554361 | ENSG00000186648 |
| HECTD4 | -0.298108979 | 0.813317759 | 0.000970047 | 0.005465198 | 762.7471735 | 620.3732362 | ENSG00000173064 |
| ACSS1 | -0.298393891 | 0.813157156 | 0.360142349 | 0.572354478 | 49.03617123 | 39.87474379 | ENSG00000154930 |
| KIT | -0.298493127 | 0.813101225 | 0.003622328 | 0.016887055 | 554.0341168 | 450.3182851 | ENSG00000157404 |
| NA | -0.298549987 | 0.813069179 | 0.188244165 | 0.374162622 | 108.8641196 | 88.51327945 | ENSG00000235552 |
| CNIH2 | -0.298574388 | 0.813055427 | 0.086731171 | 0.214415258 | 218.6033804 | 177.8270711 | ENSG00000174871 |
| SCAP | -0.298600614 | 0.813040647 | 4.76E-07 | 5.87E-06 | 3670.06436 | 2984.475692 | ENSG00000114650 |
| PRXL2A | -0.298604456 | 0.813038482 | 8.01E-05 | 0.000609985 | 1405.553063 | 1143.103825 | ENSG00000122378 |
| PPIP5K1 | -0.298606365 | 0.813037406 | 0.111953049 | 0.259717416 | 177.6138141 | 144.3066884 | ENSG00000168781 |
| STAG3L5P | -0.298717174 | 0.812974962 | 0.244786252 | 0.449070278 | 116.8752077 | 95.12113095 | ENSG00000242294 |
| LOC374443 | -0.298757731 | 0.812952108 | 0.678145356 | 0.827792325 | 11.82017647 | 9.628433792 | ENSG00000256594 |
| GPR3 | -0.298760693 | 0.812950439 | 0.041415876 | 0.121904429 | 241.7629781 | 196.5049044 | ENSG00000181773 |
| RFNG | -0.29876996 | 0.812945217 | 0.012955252 | 0.048380448 | 390.0716355 | 317.1904224 | ENSG00000169733 |
| NA | -0.298878173 | 0.812884243 | 0.751011211 | 0.870392545 | 5.318775408 | 4.318901608 | ENSG00000278291 |
| ZNF350-AS1 | -0.299118951 | 0.812748588 | 0.664138385 | 0.820076723 | 10.17761991 | 8.288612717 | ENSG00000269235 |
| NDE1 | -0.299295645 | 0.812649052 | 0.012155179 | 0.046029787 | 434.3566622 | 353.0807939 | ENSG00000072864 |
| ANKRD54 | -0.299329863 | 0.812629778 | 0.063208933 | 0.168817627 | 220.1981631 | 179.1100466 | ENSG00000100124 |
| GK5 | -0.299377887 | 0.812602728 | 0.107216917 | 0.251555734 | 160.4944652 | 130.312795 | ENSG00000175066 |
| MSL3 | -0.299409252 | 0.812585062 | 0.03279219 | 0.101844109 | 294.1281452 | 238.9423824 | ENSG00000005302 |
| NA | -0.299505671 | 0.812530756 | 0.774158828 | 0.883955535 | 6.158880441 | 5.042158663 | ENSG00000270504 |
| RFXAP | -0.299538131 | 0.812512475 | 0.103961054 | 0.245205981 | 161.9353532 | 131.6005205 | ENSG00000133111 |
| C2CD2L | -0.299561657 | 0.812499226 | 0.336689702 | 0.548688178 | 51.00581531 | 41.40447233 | ENSG00000172375 |
| TBCK | -0.29963329 | 0.812458884 | 0.059720328 | 0.161833605 | 272.1496986 | 221.1479922 | ENSG00000145348 |
| ZBTB8A | -0.299819569 | 0.812353987 | 0.156963283 | 0.329068783 | 126.529716 | 102.9029511 | ENSG00000160062 |
| NA | -0.299954608 | 0.812277953 | 0.385654553 | 0.597653274 | 44.01692344 | 35.81886477 | ENSG00000286431 |
| TSPAN6 | -0.300071606 | 0.812212083 | 3.03E-05 | 0.000258237 | 1725.141832 | 1400.627352 | ENSG00000000003 |
| GPR50-AS1 | -0.300160903 | 0.812161811 | 0.807290922 | 0.901913344 | 3.684532999 | 2.981255996 | ENSG00000234696 |
| COA3 | -0.300186768 | 0.812147251 | 0.000272592 | 0.001808821 | 1058.495602 | 859.8271182 | ENSG00000183978 |
| EIF6 | -0.300273899 | 0.812098203 | 5.56E-05 | 0.000442605 | 1511.674602 | 1227.824901 | ENSG00000242372 |
| NA | -0.300497581 | 0.811972302 | 0.753297455 | 0.871421911 | 5.754985217 | 4.64227009 | ENSG00000255320 |
| NSD3 | -0.300525047 | 0.811956843 | 0.000879879 | 0.005003466 | 1243.884359 | 1009.539708 | ENSG00000147548 |
| AP4M1 | -0.300541159 | 0.811947776 | 0.053811619 | 0.149095132 | 234.3835392 | 190.238036 | ENSG00000221838 |
| PCNA | -0.300564851 | 0.811934442 | 2.09E-08 | 3.26E-07 | 5710.947881 | 4636.702368 | ENSG00000132646 |
| SAMD10 | -0.300566069 | 0.811933756 | 0.206891545 | 0.399331244 | 101.3901012 | 82.47954425 | ENSG00000130590 |
| MRPS28 | -0.300613349 | 0.811907148 | 0.188563591 | 0.374552717 | 103.2188671 | 83.84440934 | ENSG00000147586 |
| CDK16 | -0.300628347 | 0.811898707 | 2.46E-08 | 3.78E-07 | 4492.706581 | 3648.08055 | ENSG00000102225 |
| ZNF527 | -0.300643498 | 0.811890181 | 0.329335797 | 0.541072433 | 53.06530978 | 43.12677827 | ENSG00000189164 |
| TRIM16L | -0.300913758 | 0.811738104 | 0.156265212 | 0.328261238 | 117.0400864 | 94.97071773 | ENSG00000108448 |
| NA | -0.300969337 | 0.811706833 | 0.497966034 | 0.696960538 | 26.20630464 | 21.26489671 | ENSG00000283544 |
| NFKBIE | -0.301022677 | 0.811676823 | 0.02642429 | 0.086008742 | 385.1505522 | 312.5594382 | ENSG00000146232 |
| PLEKHA3 | -0.301075518 | 0.811647094 | 0.064362118 | 0.17104962 | 234.9834057 | 190.8408848 | ENSG00000116095 |
| SLC25A23 | -0.301205422 | 0.811574014 | 0.000557018 | 0.003369404 | 1514.507494 | 1229.571882 | ENSG00000125648 |
| GPM6B | -0.301469756 | 0.81142533 | 1.74E-05 | 0.000156821 | 2004.722353 | 1626.572125 | ENSG00000046653 |
| KPNA3 | -0.301488892 | 0.811414567 | 0.000185216 | 0.001287854 | 1540.465753 | 1250.228495 | ENSG00000102753 |
| NA | -0.301688396 | 0.811302368 | 0.709159469 | 0.846486042 | 8.134448263 | 6.626652665 | ENSG00000228960 |
| VAV3 | -0.301786912 | 0.811246969 | 0.030643588 | 0.096565539 | 273.717149 | 221.9381215 | ENSG00000134215 |
| GNG7 | -0.301826745 | 0.811224571 | 0.014719239 | 0.053704177 | 459.4126297 | 372.786946 | ENSG00000176533 |
| NA | -0.301860417 | 0.811205637 | 0.718490279 | 0.851388915 | 7.777995236 | 6.259112499 | ENSG00000260448 |
| CHST6 | -0.301898411 | 0.811184274 | 0.021585987 | 0.07316069 | 521.8229043 | 423.1356246 | ENSG00000183196 |
| NECAB2 | -0.301938902 | 0.811161507 | 0.070780537 | 0.184498336 | 223.4167897 | 181.1684902 | ENSG00000103154 |
| ZKSCAN7 | -0.302079981 | 0.811082189 | 0.121749403 | 0.275423751 | 165.2008633 | 133.7730866 | ENSG00000196345 |
| THAP10 | -0.302095056 | 0.811073714 | 0.208182394 | 0.400930504 | 99.00705255 | 80.37134848 | ENSG00000129028 |
| HLA-DPB2 | -0.302149113 | 0.811043324 | 0.015559113 | 0.056139443 | 406.8517183 | 329.6300245 | ENSG00000224557 |
| GUSBP1 | -0.302285875 | 0.810966444 | 0.215503015 | 0.41069697 | 83.57620228 | 67.74671399 | ENSG00000183666 |
| ATXN7L1 | -0.302375064 | 0.810916311 | 0.00031531 | 0.002053323 | 905.2240647 | 733.8585524 | ENSG00000146776 |
| FAM122A | -0.302390297 | 0.810907749 | 0.469787044 | 0.673033556 | 27.85122546 | 22.57331534 | ENSG00000187866 |
| RABEP2 | -0.302477171 | 0.81085892 | 0.00608007 | 0.026030861 | 527.7216384 | 427.8456832 | ENSG00000177548 |
| ZNF358 | -0.302488979 | 0.810852283 | 0.077774608 | 0.197841254 | 195.3366669 | 158.542374 | ENSG00000198816 |
| KIF1A | -0.302540404 | 0.810823381 | 2.29E-08 | 3.54E-07 | 14126.203 | 11453.56496 | ENSG00000130294 |
| IFITM2 | -0.302613359 | 0.810782379 | 0.000166458 | 0.001169476 | 1068.644136 | 866.2493702 | ENSG00000185201 |
| WRN | -0.302614491 | 0.810781744 | 0.00083201 | 0.004772936 | 750.7553572 | 608.5178794 | ENSG00000165392 |
| C1orf52 | -0.302687792 | 0.81074055 | 0.00265327 | 0.012950892 | 612.8704291 | 497.006227 | ENSG00000162642 |
| LOC105375304 | -0.302696538 | 0.810735635 | 0.753097507 | 0.871354394 | 5.738383063 | 4.618623537 | ENSG00000225792 |
| PPP2R2C | -0.302800587 | 0.810677166 | 0.018770402 | 0.065197542 | 342.98103 | 277.9120011 | ENSG00000074211 |
| NA | -0.302942849 | 0.81059723 | 0.799316881 | 0.897876474 | 3.691651944 | 2.978134574 | ENSG00000254027 |
| NONO | -0.303050855 | 0.810536548 | 1.33E-11 | 3.33E-10 | 15765.78675 | 12778.05882 | ENSG00000147140 |
| MRPS34 | -0.30328019 | 0.810407713 | 3.35E-05 | 0.000281322 | 2024.181906 | 1640.886342 | ENSG00000074071 |
| DUSP9 | -0.303350982 | 0.810367948 | 0.261558876 | 0.468691468 | 73.90923562 | 59.92457858 | ENSG00000130829 |

| | | | | | | | |
|---|---|---|---|---|---|---|---|
| ZNF488 | -0.303394061 | 0.81034375 | 0.038945806 | 0.116375246 | 246.7392329 | 199.9410979 | ENSG00000265763 |
| NA | -0.303423941 | 0.810326968 | 0.648006109 | 0.80899024 | 11.91862034 | 9.671375971 | ENSG00000225259 |
| PHF5A | -0.303545817 | 0.810258516 | 7.60E-06 | 7.48E-05 | 1589.774322 | 1288.101327 | ENSG00000100410 |
| FBXW9 | -0.303853294 | 0.810085846 | 0.091582767 | 0.223166523 | 180.5902213 | 146.3289998 | ENSG00000132004 |
| NA | -0.303911884 | 0.810052948 | 0.75685791 | 0.873724654 | 5.724145173 | 4.602449333 | ENSG00000279641 |
| UBE2L6 | -0.303942532 | 0.81003574 | 3.34E-05 | 0.000280997 | 1274.869933 | 1032.716957 | ENSG00000156587 |
| NPIPB15 | -0.303946866 | 0.810033306 | 0.577309769 | 0.759282257 | 16.82490704 | 13.65726128 | ENSG00000196436 |
| ZNF582 | -0.303984155 | 0.81001237 | 0.587348869 | 0.765666704 | 17.22435307 | 13.95263229 | ENSG00000018869 |
| SMARCA5-AS1 | -0.304035371 | 0.809983615 | 0.264803609 | 0.472488299 | 70.54230693 | 57.14410737 | ENSG00000245112 |
| NA | -0.304055845 | 0.80997212 | 0.44653415 | 0.653604036 | 32.75275748 | 26.52032585 | ENSG00000237854 |
| GDF6 | -0.304121889 | 0.809935042 | 0.639723158 | 0.802577272 | 12.2753527 | 9.924750753 | ENSG00000156466 |
| CFAP221 | -0.304127642 | 0.809931812 | 0.764984062 | 0.878469069 | 4.944245687 | 4.021355143 | ENSG00000163075 |
| METTL25 | -0.304138379 | 0.809925784 | 0.284139933 | 0.493356131 | 64.66424087 | 52.29838257 | ENSG00000127720 |
| NAP1L4 | -0.304181397 | 0.809901634 | 6.97E-10 | 1.37E-08 | 5104.355048 | 4134.42931 | ENSG00000205531 |
| SETDB2 | -0.304303542 | 0.809833067 | 0.097059927 | 0.232966029 | 166.6029026 | 134.7499293 | ENSG00000136169 |
| RAPGEF3 | -0.30433576 | 0.809814983 | 0.050111747 | 0.141303229 | 261.9023376 | 212.2602621 | ENSG00000079337 |
| GNAL | -0.304351601 | 0.809806091 | 0.00566967 | 0.024534187 | 554.6144326 | 448.9698348 | ENSG00000141404 |
| NRTN | -0.304473344 | 0.809737757 | 0.258725933 | 0.465619007 | 91.51704295 | 74.248156 | ENSG00000171119 |
| PSMD8 | -0.304481599 | 0.809733124 | 9.17E-08 | 1.29E-06 | 3123.23023 | 2528.958415 | ENSG00000099341 |
| ASIC3 | -0.304489124 | 0.809728901 | 0.60794266 | 0.781467763 | 16.52657463 | 13.33606827 | ENSG00000213199 |
| AJM1 | -0.30451372 | 0.809715096 | 0.328099177 | 0.539669465 | 52.79870714 | 42.77229089 | ENSG00000232434 |
| KCTD6 | -0.304532019 | 0.809704826 | 0.072520455 | 0.187995715 | 203.879434 | 165.2143552 | ENSG00000168301 |
| SNX7 | -0.304602739 | 0.809665135 | 0.009664843 | 0.037966697 | 440.7222191 | 356.7872715 | ENSG00000162627 |
| SIPA1L3 | -0.304620452 | 0.809655195 | 0.000298703 | 0.001956646 | 1019.440494 | 825.3536846 | ENSG00000105738 |
| PRKCSH | -0.304701317 | 0.809609814 | 1.48E-10 | 3.21E-09 | 7458.415978 | 6038.838828 | ENSG00000130175 |
| DDB1 | -0.304969261 | 0.809459463 | 5.22E-11 | 1.22E-09 | 9985.318218 | 8082.147645 | ENSG00000167986 |
| WSB1 | -0.305009797 | 0.80943672 | 0.095001997 | 0.229514227 | 1605.379282 | 1299.546176 | ENSG00000109046 |
| CCNQ | -0.305014204 | 0.809434247 | 0.021270629 | 0.072299464 | 398.9761939 | 323.2292436 | ENSG00000262919 |
| RAB23 | -0.305026346 | 0.809427435 | 0.024137588 | 0.080055908 | 289.5109609 | 234.2533863 | ENSG00000112210 |
| ACO1 | -0.305027964 | 0.809426527 | 2.41E-06 | 2.59E-05 | 2163.648655 | 1750.891942 | ENSG00000122729 |
| CSTF2T | -0.305230902 | 0.809312677 | 0.000100476 | 0.000744698 | 1331.930157 | 1078.130028 | ENSG00000177613 |
| SPIN4 | -0.305306519 | 0.809270258 | 0.055650007 | 0.153222098 | 251.7985502 | 204.0196775 | ENSG00000186767 |
| OTUB2 | -0.305436867 | 0.809197144 | 0.203544755 | 0.395188419 | 107.9398086 | 87.26509112 | ENSG00000089723 |
| QRFPR | -0.305539332 | 0.809139674 | 0.481517868 | 0.683308156 | 29.03382497 | 23.48742582 | ENSG00000186867 |
| TMSB15B | -0.305589801 | 0.809111369 | 0.672162162 | 0.82439555 | 9.877812957 | 8.021239196 | ENSG00000269226 |
| ASIC2 | -0.305724631 | 0.809035755 | 0.482717979 | 0.683859246 | 28.36780245 | 23.00091785 | ENSG00000108684 |
| NA | -0.305783834 | 0.809002556 | 0.026203104 | 0.085470207 | 390.2762008 | 315.6977721 | ENSG00000264538 |
| C17orf100 | -0.305839224 | 0.808971497 | 0.4190537 | 0.629503762 | 39.46408896 | 31.92567376 | ENSG00000256806 |
| SLC66A2 | -0.305914008 | 0.808929563 | 0.016039492 | 0.057586657 | 462.9097052 | 374.5948825 | ENSG00000122490 |
| DLL3 | -0.305948056 | 0.808910473 | 0.039444433 | 0.117474864 | 301.3865365 | 243.5622354 | ENSG00000090932 |
| NA | -0.306072787 | 0.80884054 | 0.68430399 | 0.831119741 | 9.064988495 | 7.302333068 | ENSG00000271869 |
| TTC7A | -0.306124794 | 0.808811383 | 0.005066488 | 0.022344844 | 565.1600737 | 457.0061867 | ENSG00000068724 |
| PCGF2 | -0.306170945 | 0.80878551 | 0.001365836 | 0.007352856 | 700.7071351 | 566.8419661 | ENSG00000277258 |
| DCLK3 | -0.306190405 | 0.8087746 | 0.273980113 | 0.482659928 | 70.5933086 | 57.06569537 | ENSG00000163673 |
| ZNF33B | -0.30627679 | 0.808726174 | 0.010962673 | 0.042168603 | 393.0406188 | 317.8903165 | ENSG00000196693 |
| NA | -0.30628544 | 0.808721325 | 0.450863768 | 0.657139944 | 34.13466129 | 27.615474 | ENSG00000285728 |
| NIPA2 | -0.306314317 | 0.808705138 | 5.32E-06 | 5.38E-05 | 1936.099195 | 1566.067243 | ENSG00000140157 |
| CERK | -0.306343707 | 0.808688664 | 6.34E-07 | 7.60E-06 | 2454.984133 | 1984.865448 | ENSG00000100422 |
| LIMK1 | -0.306411092 | 0.808650893 | 1.19E-07 | 1.63E-06 | 3205.226478 | 2592.18441 | ENSG00000106683 |
| SYNPO2 | -0.306469936 | 0.80861791 | 0.117760582 | 0.269434044 | 140.7186515 | 113.6363489 | ENSG00000172403 |
| DUT | -0.306484022 | 0.808610016 | 2.62E-06 | 2.80E-05 | 2041.532901 | 1650.88575 | ENSG00000128951 |
| NA | -0.306493634 | 0.808604628 | 0.750966537 | 0.870392545 | 5.376896023 | 4.335075811 | ENSG00000228434 |
| PLAUR | -0.306533334 | 0.808582377 | 0.016993203 | 0.060300007 | 339.6752228 | 274.6848426 | ENSG00000011422 |
| COPS7B | -0.306581734 | 0.808555251 | 1.97E-05 | 0.000175547 | 1597.395727 | 1291.375182 | ENSG00000144524 |
| JAM3 | -0.306658053 | 0.80851248 | 1.62E-05 | 0.000147174 | 1389.63378 | 1123.792235 | ENSG00000166086 |
| PGAM1 | -0.306717497 | 0.808479166 | 1.81E-06 | 1.99E-05 | 13417.6708 | 10848.12416 | ENSG00000171314 |
| PNMA2 | -0.306743139 | 0.808464797 | 0.156591345 | 0.328714211 | 118.341012 | 95.7496862 | ENSG00000240694 |
| CYB561D1 | -0.306885758 | 0.808384879 | 0.101052732 | 0.23977646 | 150.9466882 | 122.0618275 | ENSG00000174151 |
| JPT2 | -0.306908636 | 0.80837206 | 5.00E-10 | 1.01E-08 | 4720.226746 | 3815.549622 | ENSG00000206053 |
| FBXW4P1 | -0.306932779 | 0.808358533 | 0.601054329 | 0.776307564 | 15.25234472 | 12.34231627 | ENSG00000230701 |
| AJAP1 | -0.306946834 | 0.808350657 | 0.027862813 | 0.08970154 | 318.7287088 | 257.7993614 | ENSG00000196581 |
| EPS8 | -0.306955512 | 0.808345795 | 0.191884358 | 0.379091547 | 106.4170528 | 85.90067753 | ENSG00000151491 |
| NA | -0.306981659 | 0.808331145 | 0.014410776 | 0.052751987 | 395.4319288 | 319.8695701 | ENSG00000259781 |
| ADCY2 | -0.307091338 | 0.808269695 | 2.16E-05 | 0.000190818 | 1981.870339 | 1601.208703 | ENSG00000078295 |
| GS1-124K5.11 | -0.307210495 | 0.80820294 | 0.017685299 | 0.062254428 | 330.2748064 | 266.958312 | ENSG00000229180 |
| FLRT2 | -0.307226858 | 0.808193774 | 0.15549695 | 0.327339193 | 109.5236336 | 88.58372465 | ENSG00000185070 |
| SDCBP2 | -0.307306912 | 0.808148929 | 0.197226851 | 0.38677353 | 87.10061133 | 70.380009 | ENSG00000125775 |
| ERP29 | -0.307333606 | 0.808133976 | 5.48E-08 | 7.97E-07 | 4029.040611 | 3256.443396 | ENSG00000089248 |
| KAT8 | -0.307352061 | 0.808123638 | 0.000743441 | 0.004330626 | 986.304231 | 797.0598962 | ENSG00000103510 |
| LINC01224 | -0.307372745 | 0.808112052 | 0.003193221 | 0.015195367 | 541.6621258 | 437.7207021 | ENSG00000269416 |
| KISS1R | -0.307587886 | 0.807991552 | 0.44014688 | 0.647819643 | 32.49446899 | 26.24983091 | ENSG00000116014 |
| RNF26 | -0.307810466 | 0.807866904 | 2.13E-05 | 0.000188296 | 1825.145839 | 1474.969475 | ENSG00000173456 |
| MRPL3 | -0.307972585 | 0.807776127 | 2.84E-06 | 3.01E-05 | 2549.274062 | 2059.688077 | ENSG00000114686 |
| SCARB1 | -0.307985991 | 0.807768621 | 1.18E-05 | 0.000110359 | 1377.256652 | 1112.506855 | ENSG00000073060 |

| | | | | | | | |
|---|---|---|---|---|---|---|---|
| NA | -0.308091389 | 0.807709611 | 0.716441796 | 0.850492157 | 7.392787098 | 5.933568554 | ENSG00000275325 |
| NA | -0.308115873 | 0.807695903 | 0.340093992 | 0.551728807 | 58.09554147 | 46.83441275 | ENSG00000280287 |
| NA | -0.308118035 | 0.807694693 | 0.626264688 | 0.793253855 | 12.36787283 | 9.961450087 | ENSG00000276488 |
| P2RY11 | -0.308142306 | 0.807681105 | 0.323473548 | 0.53439037 | 52.08997095 | 42.10067787 | ENSG00000244165 |
| FGF13 | -0.308250782 | 0.807620377 | 0.000237735 | 0.001605155 | 1195.574442 | 965.106469 | ENSG00000129682 |
| OTUD6B | -0.308314888 | 0.807584492 | 0.020113707 | 0.069138578 | 481.282921 | 388.7565796 | ENSG00000155100 |
| NA | -0.308514304 | 0.807472872 | 0.761714314 | 0.876689032 | 4.967966786 | 4.023530606 | ENSG00000229759 |
| NA | -0.308681352 | 0.807379381 | 0.792349985 | 0.893932102 | 3.708254098 | 2.973783647 | ENSG00000236875 |
| MYO10 | -0.30868797 | 0.807375677 | 5.56E-08 | 8.06E-07 | 4159.265515 | 3357.465161 | ENSG00000145555 |
| ZNF395 | -0.308866546 | 0.807275747 | 0.019631257 | 0.067652608 | 355.3904475 | 286.6996074 | ENSG00000186918 |
| GSTM2 | -0.308871853 | 0.807272777 | 0.388006685 | 0.600150584 | 41.45389467 | 33.41000111 | ENSG00000213366 |
| UQCR10 | -0.309057046 | 0.807169157 | 0.108297334 | 0.25330631 | 144.6407527 | 116.8114117 | ENSG00000184076 |
| KLF12 | -0.309092338 | 0.807149412 | 0.051681948 | 0.144744712 | 248.0606257 | 200.0835014 | ENSG00000118922 |
| MPDU1 | -0.309108767 | 0.807140221 | 0.001521572 | 0.008048986 | 690.2071823 | 557.1879939 | ENSG00000129255 |
| NA | -0.309122517 | 0.807132528 | 0.725966559 | 0.855720453 | 7.383303889 | 5.923920741 | ENSG00000237950 |
| LINC01285 | -0.309137666 | 0.807124053 | 0.756320061 | 0.8732639 | 4.920524588 | 3.981534386 | ENSG00000203650 |
| SCARF2 | -0.309285683 | 0.807041248 | 0.024807521 | 0.081989855 | 393.4706511 | 317.693935 | ENSG00000244486 |
| MYADML2 | -0.309307185 | 0.80702922 | 0.788078485 | 0.891582187 | 3.711813571 | 2.973783647 | ENSG00000185105 |
| HLA-DPB1 | -0.309474572 | 0.806935591 | 0.025749985 | 0.084307987 | 299.4753958 | 241.8221469 | ENSG00000223865 |
| CDC6 | -0.309486924 | 0.806928682 | 3.20E-06 | 3.36E-05 | 3067.569453 | 2475.23496 | ENSG00000094804 |
| PPIG | -0.309649887 | 0.806837539 | 1.71E-05 | 0.000154867 | 2026.611203 | 1634.717817 | ENSG00000138398 |
| NXPH2 | -0.309698644 | 0.806810271 | 0.27078176 | 0.479166306 | 62.08522034 | 50.12409253 | ENSG00000144227 |
| PRORP | -0.309828324 | 0.806737752 | 0.215731195 | 0.410901021 | 89.57167863 | 72.15563997 | ENSG00000100890 |
| RRM2 | -0.309829507 | 0.806737091 | 3.05E-06 | 3.21E-05 | 3851.89779 | 3107.37053 | ENSG00000171848 |
| NADSYN1 | -0.309909028 | 0.806692625 | 0.001406444 | 0.007543507 | 696.6800042 | 561.8166871 | ENSG00000172890 |
| PTPRG-AS1 | -0.309952347 | 0.806668403 | 0.457631311 | 0.663253386 | 29.29122373 | 23.6003617 | ENSG00000241472 |
| ATP6AP1 | -0.309978906 | 0.806653554 | 5.03E-06 | 5.12E-05 | 1646.677221 | 1328.536663 | ENSG00000071553 |
| ETV5 | -0.310023497 | 0.806628622 | 0.000259271 | 0.001732319 | 996.0907196 | 803.0671767 | ENSG00000244405 |
| ORC1 | -0.310027368 | 0.806626457 | 7.35E-05 | 0.000564964 | 1453.884259 | 1172.498481 | ENSG00000085840 |
| BAG5 | -0.310171116 | 0.80654609 | 0.00024242 | 0.001632121 | 919.9749856 | 741.9257169 | ENSG00000166170 |
| E2F4 | -0.310635402 | 0.806286571 | 2.98E-05 | 0.000254031 | 2303.863901 | 1858.14579 | ENSG00000205250 |
| FGD4 | -0.310703574 | 0.806248472 | 0.028379684 | 0.090891106 | 476.6906791 | 384.386601 | ENSG00000139132 |
| CABLES1 | -0.310795849 | 0.806196905 | 0.0068871 | 0.028883608 | 439.7979075 | 354.7487859 | ENSG00000134508 |
| FYN | -0.310894906 | 0.806141553 | 1.52E-05 | 0.000138389 | 1722.317137 | 1388.347906 | ENSG00000010810 |
| PDZD7 | -0.311037824 | 0.806061698 | 0.397473785 | 0.609205812 | 41.60189182 | 33.56710868 | ENSG00000186862 |
| SEC11C | -0.311107855 | 0.806022572 | 0.00101984 | 0.005704882 | 719.0638717 | 579.5814284 | ENSG00000166562 |
| CYB5A | -0.311159775 | 0.805993565 | 3.28E-05 | 0.000276232 | 1704.393654 | 1373.465838 | ENSG00000166347 |
| SMC2 | -0.311237172 | 0.805950326 | 0.00026659 | 0.001775232 | 1349.375859 | 1087.564104 | ENSG00000136824 |
| VEGFA | -0.311304312 | 0.80591282 | 0.000536481 | 0.003262945 | 919.1456107 | 741.1326473 | ENSG00000112715 |
| CST1 | -0.311619321 | 0.80573687 | 0.444290974 | 0.651684275 | 30.50941796 | 24.57046715 | ENSG00000170373 |
| TNPO2 | -0.311627735 | 0.805732171 | 1.32E-06 | 1.50E-05 | 2802.386862 | 2258.386854 | ENSG00000105576 |
| AIFM3 | -0.311673926 | 0.805706374 | 0.727039303 | 0.856313911 | 9.57088706 | 7.73769147 | ENSG00000183773 |
| KANSL3 | -0.311696212 | 0.805693928 | 5.04E-05 | 0.000405193 | 1210.732032 | 975.3723708 | ENSG00000114982 |
| NA | -0.311724985 | 0.80567786 | 0.489362469 | 0.689558553 | 29.6645844 | 23.84531818 | ENSG00000223703 |
| NCOA7 | -0.311807494 | 0.805631784 | 0.063579917 | 0.169519847 | 186.5047456 | 150.2534777 | ENSG00000111912 |
| FOXN4 | -0.311947593 | 0.805553553 | 0.654467054 | 0.813025281 | 12.38063618 | 10.03144379 | ENSG00000139445 |
| PPM1G | -0.31197396 | 0.80553883 | 7.20E-11 | 1.64E-09 | 8596.301624 | 6925.025841 | ENSG00000115241 |
| NA | -0.312045397 | 0.805498944 | 0.749856906 | 0.869935742 | 4.937126742 | 3.983709849 | ENSG00000280486 |
| UBA1 | -0.312152944 | 0.8054389 | 9.62E-14 | 3.10E-12 | 19180.61705 | 15448.65706 | ENSG00000130985 |
| BSG | -0.312161842 | 0.805433932 | 3.29E-11 | 7.90E-10 | 20943.29115 | 16868.36837 | ENSG00000172270 |
| NSD2 | -0.312495683 | 0.805247576 | 1.21E-09 | 2.31E-08 | 5132.816055 | 4132.567287 | ENSG00000109685 |
| PDCL3 | -0.312651462 | 0.805160632 | 0.000380665 | 0.002419679 | 771.7582635 | 621.4572961 | ENSG00000115539 |
| UIMC1 | -0.312699466 | 0.805133841 | 0.000590572 | 0.003546247 | 837.9826606 | 674.8392871 | ENSG00000087206 |
| DPP10-AS1 | -0.312862259 | 0.805042995 | 0.306745002 | 0.517082867 | 55.67606008 | 44.91037606 | ENSG00000235026 |
| KLHL9 | -0.31293357 | 0.805003204 | 0.001598023 | 0.00839631 | 698.0253712 | 561.6241097 | ENSG00000198642 |
| GM2A | -0.313025961 | 0.804951652 | 0.000568373 | 0.003430399 | 838.3583599 | 674.7347695 | ENSG00000196743 |
| USP35 | -0.313171093 | 0.80487068 | 0.030732045 | 0.096773804 | 263.5662249 | 212.0717091 | ENSG00000118369 |
| TSPAN7 | -0.313388481 | 0.80474941 | 0.364044916 | 0.57608415 | 48.14242093 | 38.75594909 | ENSG00000156298 |
| ACAD9 | -0.313420611 | 0.804731488 | 0.001491282 | 0.007924271 | 606.1570389 | 487.8201295 | ENSG00000177646 |
| EPHB6 | -0.313442352 | 0.804719361 | 0.000526804 | 0.003213402 | 768.3613584 | 618.5696077 | ENSG00000106123 |
| PRPF40A | -0.313484205 | 0.804696016 | 5.84E-08 | 8.46E-07 | 3704.15399 | 2980.506133 | ENSG00000196504 |
| NA | -0.313495124 | 0.804689925 | 0.391516802 | 0.602935237 | 42.97784574 | 34.62338203 | ENSG00000276791 |
| RFX2 | -0.313611439 | 0.804625051 | 0.141707259 | 0.306440122 | 119.6185221 | 96.30060745 | ENSG00000087903 |
| PRPF8 | -0.313633052 | 0.804612998 | 1.94E-09 | 3.58E-08 | 11439.32636 | 9203.883309 | ENSG00000174231 |
| APCDD1 | -0.313728204 | 0.804559932 | 0.137743498 | 0.300657184 | 119.4278374 | 96.02432116 | ENSG00000154856 |
| LINGO1 | -0.313859214 | 0.804486873 | 3.95E-05 | 0.000325585 | 2158.571466 | 1737.154399 | ENSG00000169783 |
| SNX11 | -0.313866236 | 0.804482958 | 0.012410719 | 0.046763055 | 523.2461964 | 420.7545381 | ENSG00000002919 |
| CDH23 | -0.313926721 | 0.80444923 | 0.037086034 | 0.112023312 | 244.5365221 | 196.6221912 | ENSG00000107736 |
| CLN6 | -0.313974274 | 0.804422715 | 0.000890798 | 0.005060985 | 946.7120784 | 761.9737457 | ENSG00000128973 |
| NA | -0.314193972 | 0.804300225 | 0.462725119 | 0.667551225 | 30.24401052 | 24.32674018 | ENSG00000215859 |
| C4orf33 | -0.314490064 | 0.804135171 | 0.40979405 | 0.620855246 | 33.39388985 | 26.84804526 | ENSG00000151470 |
| LINC00877 | -0.314531088 | 0.804112305 | 0.543831279 | 0.732438575 | 19.86238393 | 15.98742939 | ENSG00000241163 |
| RAD51 | -0.314550953 | 0.804101233 | 0.002534241 | 0.012436619 | 846.0615016 | 680.4144594 | ENSG00000051180 |
| NA | -0.314711126 | 0.804011964 | 0.631822676 | 0.797156893 | 12.86428818 | 10.37845882 | ENSG00000214535 |

| | | | | | | | |
|---|---|---|---|---|---|---|---|
| EIF2AK4 | -0.31480178 | 0.803961444 | 7.09E-09 | 1.20E-07 | 3710.265431 | 2982.429827 | ENSG00000128829 |
| CSK | -0.31481341 | 0.803954963 | 4.15E-05 | 0.000340379 | 1680.978339 | 1351.927304 | ENSG00000103653 |
| ITGA9 | -0.314895513 | 0.803909212 | 0.051449941 | 0.144266092 | 194.4117455 | 156.3304386 | ENSG00000144668 |
| SLC9A1 | -0.315057846 | 0.80381876 | 0.00215891 | 0.010840473 | 862.8088141 | 693.7874997 | ENSG00000090020 |
| SLC35F1 | -0.315113325 | 0.80378785 | 0.030081934 | 0.095174623 | 261.8857355 | 210.4539208 | ENSG00000196376 |
| TRMT9B | -0.315195724 | 0.803741943 | 0.379981851 | 0.59241652 | 41.76913472 | 33.5714596 | ENSG00000250305 |
| ANAPC4 | -0.315469887 | 0.803589219 | 0.001681807 | 0.008778589 | 614.2164071 | 493.4527466 | ENSG00000053900 |
| ZMYND8 | -0.315514477 | 0.803564382 | 2.81E-07 | 3.61E-06 | 3262.243633 | 2620.980786 | ENSG00000101040 |
| ALKBH6 | -0.315617026 | 0.803507266 | 0.401323455 | 0.612691466 | 35.97646985 | 28.84207565 | ENSG00000239382 |
| CCDC152 | -0.315768717 | 0.803422786 | 0.073584144 | 0.190100762 | 217.8480656 | 174.9728747 | ENSG00000198865 |
| CHAF1B | -0.315774182 | 0.803419742 | 0.000648283 | 0.003841486 | 813.7013511 | 653.3892127 | ENSG00000159259 |
| CEP97 | -0.31589677 | 0.803351478 | 0.079479775 | 0.200880027 | 183.1772243 | 147.0261084 | ENSG00000182504 |
| NA | -0.316202294 | 0.803181367 | 0.779679201 | 0.887571291 | 4.544799659 | 3.674340108 | ENSG00000287742 |
| CARD9 | -0.316203837 | 0.803180508 | 0.479630255 | 0.681705228 | 25.19064255 | 20.21174479 | ENSG00000187796 |
| IRF4 | -0.31630645 | 0.803123383 | 0.340208912 | 0.551773042 | 47.08760465 | 37.88506433 | ENSG00000137265 |
| WLS | -0.316472381 | 0.803031018 | 0.402593456 | 0.61393481 | 43.99139674 | 35.30000839 | ENSG00000116729 |
| NA | -0.31660037 | 0.80295978 | 0.769903143 | 0.881776995 | 4.937126742 | 3.964414224 | ENSG00000279838 |
| SLC25A38 | -0.316812744 | 0.802841587 | 0.000113524 | 0.00083258 | 1088.843118 | 873.7546635 | ENSG00000144659 |
| TMEM74B | -0.316899412 | 0.802793359 | 0.44664326 | 0.653662273 | 32.3797023 | 25.95540587 | ENSG00000125895 |
| SAMM50 | -0.316961692 | 0.802758704 | 7.33E-05 | 0.000563727 | 1464.812849 | 1176.232204 | ENSG00000100347 |
| DPH3 | -0.317008188 | 0.802732833 | 0.001760029 | 0.00911383 | 652.3122971 | 523.919915 | ENSG00000154813 |
| LINC00545 | -0.317024363 | 0.802723833 | 0.545878791 | 0.733992237 | 18.23406526 | 14.64136547 | ENSG00000236094 |
| STRIP1 | -0.317085138 | 0.802690018 | 3.23E-05 | 0.000272448 | 1206.060287 | 967.965613 | ENSG00000143093 |
| ACACA | -0.317168523 | 0.802643625 | 3.45E-08 | 5.20E-07 | 5979.297625 | 4798.557777 | ENSG00000278540 |
| LAS1L | -0.317385746 | 0.802522782 | 8.19E-08 | 1.16E-06 | 2564.666798 | 2058.434278 | ENSG00000001497 |
| SUFU | -0.317543508 | 0.80243503 | 0.003778304 | 0.017492835 | 514.1735303 | 412.5018694 | ENSG00000107882 |
| NA | -0.317620476 | 0.802392221 | 0.06725984 | 0.17720453 | 182.2033862 | 146.1078579 | ENSG00000254682 |
| HEY1 | -0.317737652 | 0.802327053 | 0.278311851 | 0.48756309 | 74.58387778 | 59.76406605 | ENSG00000164683 |
| YWHAE | -0.317759081 | 0.802315136 | 7.78E-09 | 1.31E-07 | 22039.86098 | 17681.93241 | ENSG00000108953 |
| CABIN1 | -0.317841166 | 0.802269488 | 4.70E-08 | 6.91E-07 | 3325.014374 | 2667.155504 | ENSG00000099991 |
| FRMPD1 | -0.317872572 | 0.802252023 | 0.633542274 | 0.798559516 | 12.37707671 | 9.977624291 | ENSG00000070601 |
| TIMM10B | -0.317943079 | 0.802212817 | 0.00565337 | 0.024481515 | 411.2118088 | 329.8030227 | ENSG00000132286 |
| HUS1 | -0.317952673 | 0.802207482 | 0.035942125 | 0.109407969 | 279.2657894 | 223.9576904 | ENSG00000136273 |
| WDR7 | -0.318234963 | 0.80205053 | 0.037530595 | 0.11302269 | 316.259954 | 253.4418256 | ENSG00000091157 |
| CENPBD1P1 | -0.318273903 | 0.802028883 | 0.006546042 | 0.027717938 | 424.2979266 | 340.4690966 | ENSG00000213753 |
| GOT2 | -0.318308475 | 0.802009664 | 4.46E-09 | 7.79E-08 | 4766.991992 | 3822.659984 | ENSG00000125166 |
| TP73-AS1 | -0.318316799 | 0.802005036 | 1.00E-04 | 0.000741686 | 1142.196185 | 915.9521983 | ENSG00000227372 |
| TBC1D12 | -0.318352443 | 0.801985222 | 0.278902762 | 0.48800877 | 60.0138784 | 48.09459231 | ENSG00000108239 |
| FAM92B | -0.318392909 | 0.801962727 | 0.804515881 | 0.900504526 | 3.704694626 | 2.94796163 | ENSG00000153789 |
| LRTOMT | -0.318414787 | 0.801950566 | 0.508840991 | 0.705487044 | 21.52390691 | 17.25725676 | ENSG00000184154 |
| MNAT1 | -0.318660738 | 0.80181386 | 0.006415863 | 0.027240014 | 400.2412206 | 320.8580252 | ENSG00000020426 |
| GRK5 | -0.318668178 | 0.801809726 | 0.183102579 | 0.366944518 | 90.25076998 | 72.28910098 | ENSG00000198873 |
| GMCL1 | -0.318710022 | 0.80178647 | 0.002681648 | 0.013062284 | 708.3293771 | 568.0398126 | ENSG00000087338 |
| ZNF207 | -0.318820901 | 0.801724851 | 4.17E-05 | 0.000341143 | 2683.295628 | 2151.560914 | ENSG00000010244 |
| LINC01534 | -0.31882214 | 0.801724163 | 0.592807987 | 0.770116175 | 14.98337781 | 12.01242139 | ENSG00000225975 |
| NA | -0.318901028 | 0.801680325 | 0.794629149 | 0.894952975 | 4.548359131 | 3.66346279 | ENSG00000272338 |
| PODXL | -0.3189852 | 0.801633553 | 3.48E-11 | 8.32E-10 | 24770.19913 | 19856.24724 | ENSG00000128567 |
| DNAJC4 | -0.319051851 | 0.801596519 | 0.07321771 | 0.189450385 | 168.2531885 | 134.8799854 | ENSG00000110011 |
| ABCC6 | -0.319071736 | 0.801585471 | 0.561880294 | 0.746840467 | 18.14866408 | 14.53902336 | ENSG00000091262 |
| ECEL1 | -0.319208238 | 0.801509631 | 0.035721267 | 0.10893097 | 259.7393914 | 208.1881659 | ENSG00000171551 |
| NA | -0.319266214 | 0.801477423 | 0.604800338 | 0.7791033 | 17.04551587 | 13.63891162 | ENSG00000284617 |
| TTF2 | -0.319335981 | 0.801438665 | 1.15E-06 | 1.32E-05 | 1919.812555 | 1538.257656 | ENSG00000116830 |
| CYB561A3 | -0.319367208 | 0.801421318 | 0.034021613 | 0.10497954 | 256.2377614 | 205.4202234 | ENSG00000162144 |
| CDK12 | -0.319437631 | 0.801382199 | 7.97E-05 | 0.000607579 | 1931.872176 | 1547.672028 | ENSG00000167258 |
| PYGL | -0.31943814 | 0.801381916 | 3.19E-05 | 0.000270024 | 1262.10785 | 1011.010953 | ENSG00000100504 |
| GPC1 | -0.319625897 | 0.801277629 | 1.68E-05 | 0.000151968 | 1772.187793 | 1420.461856 | ENSG00000063660 |
| MTCH1 | -0.319676686 | 0.801249421 | 9.35E-12 | 2.38E-10 | 5742.864737 | 4601.414897 | ENSG00000137409 |
| ESCO1 | -0.319802518 | 0.801179539 | 0.006513128 | 0.027609521 | 722.1622953 | 578.5042808 | ENSG00000141446 |
| KCNJ8 | -0.319827411 | 0.801165715 | 0.620261039 | 0.789290233 | 14.82442291 | 11.86713711 | ENSG00000121361 |
| TXNRD1 | -0.319990324 | 0.80107525 | 2.00E-08 | 3.14E-07 | 3582.665795 | 2869.704622 | ENSG00000198431 |
| FTSJ1 | -0.320017465 | 0.80106018 | 0.000529162 | 0.003223621 | 1008.492537 | 808.1688938 | ENSG00000068438 |
| HEXIM2 | -0.320060482 | 0.801036295 | 0.443173245 | 0.650550076 | 30.20368727 | 24.14910749 | ENSG00000168517 |
| NA | -0.32012167 | 0.801002322 | 0.753516345 | 0.871465257 | 5.009485247 | 4.01170733 | ENSG00000273226 |
| LMNB2 | -0.320146393 | 0.800988596 | 4.42E-11 | 1.04E-09 | 12977.89852 | 10395.2012 | ENSG00000176619 |
| CYLD | -0.320411243 | 0.800841564 | 0.042712874 | 0.124846405 | 314.0747868 | 251.5111672 | ENSG00000083799 |
| POLR3A | -0.320466651 | 0.800810807 | 3.69E-05 | 0.000306558 | 1478.263045 | 1183.821055 | ENSG00000148606 |
| NA | -0.320567835 | 0.800754644 | 0.67276984 | 0.824856094 | 8.308810108 | 6.648123755 | ENSG00000279759 |
| SFXN4 | -0.320674565 | 0.800695407 | 0.012182788 | 0.046115846 | 332.232603 | 265.9488373 | ENSG00000183605 |
| RTN3 | -0.320697933 | 0.800682437 | 5.15E-08 | 7.52E-07 | 3697.407246 | 2960.477865 | ENSG00000133318 |
| SOWAHD | -0.32070347 | 0.800679364 | 0.196219287 | 0.385257474 | 85.10991589 | 68.06696105 | ENSG00000187808 |
| COX8A | -0.320742522 | 0.800657691 | 0.038655458 | 0.115764087 | 279.0827818 | 223.6355943 | ENSG00000176340 |
| CNR1 | -0.320761611 | 0.800647097 | 0.495759752 | 0.695207823 | 29.1192523 | 23.3438655 | ENSG00000118432 |
| NTF3 | -0.320767239 | 0.800643974 | 0.558320106 | 0.743993452 | 17.0573895 | 13.61961599 | ENSG00000185652 |
| GSTZ1 | -0.320897124 | 0.800571896 | 0.008601642 | 0.034680191 | 457.5568108 | 365.9561333 | ENSG00000100577 |

| | | | | | | | |
|---|---|---|---|---|---|---|---|
| BCAT1 | -0.320993703 | 0.800518305 | 5.53E-05 | 0.000440403 | 4961.126052 | 3971.128908 | ENSG00000060982 |
| TRNAU1AP | -0.321038836 | 0.800493262 | 0.004527075 | 0.020341139 | 490.8682741 | 392.646125 | ENSG00000180098 |
| UPP1 | -0.321051387 | 0.800486298 | 0.000490496 | 0.0030301 | 744.9525982 | 596.4773355 | ENSG00000183696 |
| NA | -0.321121251 | 0.800447534 | 0.680573124 | 0.829255013 | 9.499723763 | 7.619175159 | ENSG00000271862 |
| ZNF155 | -0.321246758 | 0.800377903 | 0.353489377 | 0.565636916 | 43.44814958 | 34.81669447 | ENSG00000204920 |
| CRYM | -0.321438776 | 0.800271382 | 0.029443087 | 0.093544589 | 270.2366227 | 216.0622289 | ENSG00000103316 |
| SLC22A15 | -0.32149228 | 0.800241703 | 0.132229418 | 0.292563645 | 128.7997518 | 103.1238096 | ENSG00000163393 |
| TAF8 | -0.321497861 | 0.800238608 | 0.004665411 | 0.020843691 | 464.5872729 | 371.5424033 | ENSG00000137413 |
| TLL2 | -0.321529385 | 0.800221122 | 0.063623422 | 0.169563693 | 174.2320627 | 139.3822681 | ENSG00000095587 |
| RNF130 | -0.321623872 | 0.800168715 | 9.49E-05 | 0.000708482 | 1093.906734 | 875.5698317 | ENSG00000113269 |
| TRIM36 | -0.321635674 | 0.800162169 | 0.029055483 | 0.092568873 | 303.0039235 | 242.4163368 | ENSG00000152503 |
| SCYL1 | -0.321815374 | 0.800062508 | 2.15E-06 | 2.33E-05 | 1754.361769 | 1403.781808 | ENSG00000142186 |
| ZDHHC3 | -0.322036371 | 0.799939961 | 1.61E-06 | 1.79E-05 | 1616.053342 | 1292.787653 | ENSG00000163812 |
| MARCHF5 | -0.322171697 | 0.79986493 | 0.004539234 | 0.020386074 | 564.3531991 | 451.2565663 | ENSG00000198060 |
| AP3M2 | -0.322180478 | 0.799860061 | 0.012527096 | 0.047116951 | 455.1409674 | 364.0087765 | ENSG00000070718 |
| CHM | -0.322401389 | 0.799737593 | 0.001063207 | 0.005915928 | 647.4584866 | 517.9301406 | ENSG00000188419 |
| C1S | -0.322454515 | 0.799708144 | 0.05924763 | 0.160806481 | 191.9222704 | 153.4496328 | ENSG00000182326 |
| HCFC2 | -0.322488465 | 0.799689325 | 0.480346593 | 0.682466552 | 31.29615706 | 24.99350782 | ENSG00000111727 |
| ZNF211 | -0.322503311 | 0.799681096 | 0.028859898 | 0.092064049 | 334.3041982 | 267.4660505 | ENSG00000121417 |
| SLC1A5 | -0.322589985 | 0.799633054 | 1.90E-10 | 4.07E-09 | 7384.979431 | 5905.013782 | ENSG00000105281 |
| EGLN1 | -0.322604599 | 0.799624955 | 0.185569301 | 0.370178119 | 90.38096977 | 72.22222871 | ENSG00000135766 |
| PAFAH2 | -0.322638746 | 0.799606028 | 0.038905303 | 0.116274599 | 230.1530375 | 183.8566233 | ENSG00000158006 |
| MTMR2 | -0.322649559 | 0.799600036 | 9.50E-06 | 9.12E-05 | 1503.290563 | 1202.024981 | ENSG00000087053 |
| PTTG1IP | -0.323154447 | 0.799320255 | 2.19E-08 | 3.41E-07 | 3724.752035 | 2976.793342 | ENSG00000183255 |
| NA | -0.323208032 | 0.799290567 | 0.650937187 | 0.810506282 | 10.72859641 | 8.609805735 | ENSG00000227543 |
| ECHDC1 | -0.323233484 | 0.799276466 | 0.000135617 | 0.000976783 | 1014.574836 | 811.1020155 | ENSG00000093144 |
| CCZ1 | -0.3233755 | 0.79919779 | 0.012560687 | 0.047205687 | 343.9060037 | 274.8436312 | ENSG00000122674 |
| ZFYVE27 | -0.323455547 | 0.799153449 | 0.01342392 | 0.04969824 | 587.361825 | 469.4748837 | ENSG00000155256 |
| TAF4B | -0.323744546 | 0.79899338 | 0.006968405 | 0.029153373 | 430.3308038 | 343.8790893 | ENSG00000141384 |
| LOC100287015 | -0.324008488 | 0.798847216 | 0.053719536 | 0.148949264 | 195.9181785 | 156.5843079 | ENSG00000246089 |
| NA | -0.324037627 | 0.798831082 | 0.765910913 | 0.878910348 | 4.566156493 | 3.666867758 | ENSG00000243759 |
| YIPF1 | -0.324290637 | 0.798691 | 0.002659544 | 0.012968107 | 497.5713693 | 397.4930067 | ENSG00000058799 |
| ZNF740 | -0.324337153 | 0.798665249 | 0.000570049 | 0.003437223 | 712.6132453 | 569.3650381 | ENSG00000139651 |
| GLTP | -0.324351727 | 0.798657181 | 0.002027141 | 0.010282718 | 500.6503647 | 399.8198425 | ENSG00000139433 |
| UBA5 | -0.324497254 | 0.798576624 | 0.00033108 | 0.002138322 | 804.9857479 | 642.9808645 | ENSG00000081307 |
| LINC00511 | -0.324650807 | 0.798491632 | 0.573114001 | 0.75549755 | 16.96575909 | 13.59161851 | ENSG00000227036 |
| ZMYM4 | -0.324766568 | 0.798427564 | 1.59E-06 | 1.78E-05 | 2024.448842 | 1615.982802 | ENSG00000146463 |
| CRELD2 | -0.324950984 | 0.798325509 | 0.000114063 | 0.000835884 | 1046.919301 | 836.1918537 | ENSG00000184164 |
| LOC102724450 | -0.324951677 | 0.798325126 | 0.746224274 | 0.867969455 | 5.033206346 | 4.01170733 | ENSG00000226944 |
| ORC2 | -0.325026418 | 0.798283768 | 0.000115519 | 0.000844594 | 1163.082746 | 928.2463779 | ENSG00000115942 |
| SNRPD3 | -0.325167508 | 0.798205703 | 0.002549477 | 0.012495946 | 756.8701273 | 604.1049157 | ENSG00000100028 |
| VKORC1 | -0.325190752 | 0.798192843 | 0.605637266 | 0.779795526 | 12.95680832 | 10.35046134 | ENSG00000167397 |
| HSPH1 | -0.325363124 | 0.798097481 | 1.18E-07 | 1.62E-06 | 4782.686267 | 3816.947732 | ENSG00000120694 |
| SLC6A8 | -0.325441722 | 0.798054002 | 4.92E-07 | 6.05E-06 | 3096.035986 | 2471.276343 | ENSG00000130821 |
| NA | -0.325555211 | 0.797991226 | 0.606313547 | 0.7802011 | 13.70439322 | 10.98755049 | ENSG00000286511 |
| TEX13C | -0.325628065 | 0.797950929 | 0.67342048 | 0.825137853 | 10.72623214 | 8.581808255 | ENSG00000282815 |
| DBNL | -0.325764529 | 0.797875455 | 4.56E-05 | 0.00037076 | 1352.408756 | 1079.295915 | ENSG00000136279 |
| NA | -0.325901854 | 0.797799512 | 0.484877854 | 0.685911482 | 26.6745497 | 21.24096661 | ENSG00000232472 |
| DVL3 | -0.325951218 | 0.797772214 | 3.47E-07 | 4.38E-06 | 2352.604434 | 1877.421709 | ENSG00000161202 |
| RFC2 | -0.325964833 | 0.797764686 | 5.47E-07 | 6.63E-06 | 1758.431737 | 1402.701067 | ENSG00000049541 |
| MBD3 | -0.32606993 | 0.797706572 | 0.000355522 | 0.002276718 | 1023.693879 | 816.9825198 | ENSG00000071655 |
| TTC1 | -0.326427935 | 0.797508646 | 1.51E-05 | 0.000137753 | 1494.996725 | 1192.086749 | ENSG00000113312 |
| DHRS7 | -0.326429232 | 0.797507929 | 0.000856535 | 0.004892435 | 795.6811074 | 634.1703693 | ENSG00000100612 |
| RGPD5 | -0.326778014 | 0.797315149 | 0.651901729 | 0.811279351 | 9.585124949 | 7.650577608 | ENSG00000015568 |
| GNAI1 | -0.327333537 | 0.797008195 | 0.013125903 | 0.048834255 | 611.0671376 | 486.9022219 | ENSG00000127955 |
| ARMC6 | -0.327681855 | 0.796815792 | 3.77E-06 | 3.91E-05 | 1727.432157 | 1376.973089 | ENSG00000105676 |
| ZNF12 | -0.327712864 | 0.796798665 | 0.007828938 | 0.032106298 | 367.1139334 | 292.5281116 | ENSG00000164631 |
| SLC15A2 | -0.327722494 | 0.796793347 | 0.292672931 | 0.50246621 | 59.06406627 | 47.17707685 | ENSG00000163406 |
| NA | -0.327725142 | 0.796791884 | 0.594981156 | 0.771930416 | 15.0213368 | 11.94025223 | ENSG00000256338 |
| TMX2 | -0.327908912 | 0.796690396 | 9.57E-05 | 0.000713957 | 1125.531175 | 896.8504511 | ENSG00000213593 |
| SCAMP5 | -0.328028073 | 0.796624595 | 1.48E-05 | 0.000135609 | 2002.126601 | 1594.680385 | ENSG00000198794 |
| GATD1 | -0.328123705 | 0.796571791 | 0.007315351 | 0.030349388 | 1034.16055 | 824.1499054 | ENSG00000177225 |
| SMIM7 | -0.328268283 | 0.796491967 | 0.039796595 | 0.118299811 | 228.6395117 | 182.0262269 | ENSG00000214046 |
| PMEL | -0.328341381 | 0.796451612 | 3.56E-06 | 3.70E-05 | 2042.452407 | 1626.242095 | ENSG00000185664 |
| ZNF827 | -0.328449464 | 0.796391946 | 0.004925683 | 0.021800214 | 597.2912768 | 475.2330544 | ENSG00000151612 |
| GABRQ | -0.328631099 | 0.796291687 | 0.007877892 | 0.032265014 | 375.4301935 | 299.1513593 | ENSG00000268089 |
| RBM17 | -0.328631709 | 0.79629135 | 2.32E-08 | 3.57E-07 | 2668.521381 | 2124.846112 | ENSG00000134453 |
| HNRNPH2 | -0.328641972 | 0.796285686 | 9.63E-06 | 9.24E-05 | 1420.044423 | 1130.598231 | ENSG00000126945 |
| IGF2BP2 | -0.328846697 | 0.796172697 | 1.68E-09 | 3.13E-08 | 2988.290415 | 2379.170287 | ENSG00000073792 |
| PNCK | -0.328907631 | 0.79613907 | 0.207951278 | 0.400567104 | 83.1838752 | 66.2645621 | ENSG00000130822 |
| ECRG4 | -0.32917533 | 0.795991357 | 0.441088659 | 0.648801256 | 30.93970403 | 24.60839599 | ENSG00000119147 |
| NA | -0.329221348 | 0.795965967 | 0.619652105 | 0.788813199 | 13.69874881 | 10.84973856 | ENSG00000256238 |
| XKR6 | -0.329272073 | 0.795937981 | 0.039004268 | 0.116510679 | 238.0372315 | 189.5503214 | ENSG00000171044 |
| KIF26A | -0.329298217 | 0.795923558 | 2.13E-05 | 0.000187923 | 4063.676855 | 3234.835374 | ENSG00000066735 |

| | | | | | | | |
|---|---|---|---|---|---|---|---|
| PCDHB13 | -0.329417834 | 0.795857569 | 0.338740954 | 0.550644635 | 43.92023345 | 34.92527942 | ENSG00000187372 |
| PCSK7 | -0.329425295 | 0.795853453 | 8.90E-05 | 0.000669853 | 918.356455 | 731.0795942 | ENSG00000160613 |
| SPINK2 | -0.329552268 | 0.795783412 | 0.614139685 | 0.785269226 | 12.56092176 | 10.01744505 | ENSG00000128040 |
| GSN | -0.329569446 | 0.795773937 | 1.36E-05 | 0.000125017 | 1246.849633 | 991.96433 | ENSG00000148180 |
| DPH2 | -0.329726372 | 0.795687383 | 2.02E-05 | 0.000179245 | 1211.09891 | 963.7943391 | ENSG00000132768 |
| PITPNC1 | -0.329805266 | 0.795643872 | 4.01E-05 | 0.00033059 | 1151.598608 | 916.2366491 | ENSG00000154217 |
| DRD4 | -0.330011195 | 0.79553031 | 0.289387743 | 0.498742179 | 54.06909824 | 43.01696382 | ENSG00000069696 |
| ZNF606 | -0.330424917 | 0.795302209 | 0.061444312 | 0.165342809 | 178.4604532 | 141.8503208 | ENSG00000166704 |
| RPS23 | -0.331034917 | 0.794966011 | 3.22E-06 | 3.37E-05 | 2287.337344 | 1818.313468 | ENSG00000186468 |
| ZNF133 | -0.331130132 | 0.794913546 | 0.016998581 | 0.060307758 | 279.3475788 | 221.9453103 | ENSG00000125846 |
| MOCOS | -0.331249943 | 0.794847534 | 0.000686611 | 0.004043232 | 628.6805644 | 499.5351724 | ENSG00000075643 |
| NUDCD2 | -0.331314689 | 0.794811863 | 9.01E-05 | 0.000677279 | 982.107011 | 780.7003219 | ENSG00000170584 |
| H1-10 | -0.331432804 | 0.794746793 | 2.46E-06 | 2.65E-05 | 2328.2333 | 1850.641201 | ENSG00000184897 |
| PSMD10 | -0.33144021 | 0.794742714 | 9.13E-06 | 8.82E-05 | 2285.920076 | 1815.995996 | ENSG00000101843 |
| PRSS12 | -0.331554099 | 0.794679978 | 0.04879295 | 0.138456089 | 205.4086984 | 163.2775493 | ENSG00000164099 |
| TBC1D1 | -0.331823441 | 0.79453163 | 2.81E-07 | 3.61E-06 | 2228.353498 | 1770.293424 | ENSG00000065882 |
| NME1 | -0.331841922 | 0.794521452 | 0.002333714 | 0.011591842 | 552.4968699 | 439.0320016 | ENSG00000239672 |
| AP2A2 | -0.331903359 | 0.794487618 | 3.25E-06 | 3.39E-05 | 1634.931158 | 1298.813372 | ENSG00000183020 |
| OGG1 | -0.331921195 | 0.794477796 | 0.001197478 | 0.006556789 | 608.7796361 | 483.8428305 | ENSG00000114026 |
| PCYT2 | -0.332038589 | 0.794413151 | 1.70E-05 | 0.000153838 | 1241.242924 | 986.2004273 | ENSG00000185813 |
| ZNF445 | -0.33207645 | 0.794392303 | 0.001941792 | 0.009892297 | 630.6877841 | 500.6650124 | ENSG00000185219 |
| NA | -0.332110553 | 0.794373525 | 0.565230315 | 0.749341839 | 15.40743466 | 12.26797164 | ENSG00000269069 |
| BUB3 | -0.332132574 | 0.7943614 | 1.29E-09 | 2.43E-08 | 5237.676151 | 4160.579827 | ENSG00000154473 |
| LONP2 | -0.332198864 | 0.794324901 | 0.000852964 | 0.004878322 | 647.2038361 | 514.067742 | ENSG00000102910 |
| RNF24 | -0.332332066 | 0.794251566 | 0.007512336 | 0.031030286 | 358.6322621 | 284.8050813 | ENSG00000101236 |
| ZW10 | -0.332349407 | 0.794242019 | 0.001249844 | 0.006800143 | 688.1720199 | 546.4908011 | ENSG00000086827 |
| RBM23 | -0.332494495 | 0.794162148 | 0.000209229 | 0.00143163 | 998.217642 | 792.6823367 | ENSG00000100461 |
| MOB3C | -0.332558041 | 0.794127169 | 0.157149867 | 0.329350265 | 92.89355553 | 73.73315608 | ENSG00000142961 |
| GLT1D1 | -0.332622357 | 0.794091767 | 0.018567919 | 0.064672657 | 338.1343902 | 268.4164182 | ENSG00000151948 |
| NA | -0.332725312 | 0.7940351 | 0.417967089 | 0.628751794 | 30.97262901 | 24.65228413 | ENSG00000274021 |
| TNFRSF8 | -0.332767959 | 0.794011629 | 4.93E-05 | 0.000396196 | 1112.243387 | 883.5467312 | ENSG00000120949 |
| LOC399900 | -0.332784936 | 0.794002285 | 0.67160482 | 0.824069717 | 7.961560958 | 6.322579809 | ENSG00000279491 |
| LRRN1 | -0.332827425 | 0.793978901 | 1.33E-06 | 1.51E-05 | 2706.619156 | 2148.707051 | ENSG00000175928 |
| CELSR1 | -0.332863234 | 0.793959194 | 6.50E-05 | 0.000507384 | 972.5366832 | 771.8798726 | ENSG00000075275 |
| CRYBB2P1 | -0.332902312 | 0.793937688 | 0.01751629 | 0.061761602 | 284.4501946 | 225.9391624 | ENSG00000100058 |
| EMC9 | -0.33294812 | 0.79391248 | 0.006494422 | 0.027554968 | 448.1644805 | 355.9732121 | ENSG00000100908 |
| RPIA | -0.33296812 | 0.793901474 | 0.001200835 | 0.006573267 | 592.4401611 | 470.2902547 | ENSG00000153574 |
| MBNL3 | -0.333035572 | 0.793864357 | 0.176408967 | 0.357416945 | 90.4862271 | 71.80739543 | ENSG00000076770 |
| HTR7 | -0.333078311 | 0.793840839 | 0.256269339 | 0.463107876 | 62.27115033 | 49.50942042 | ENSG00000148680 |
| ARPC5L | -0.333150412 | 0.793801167 | 8.48E-07 | 9.95E-06 | 1603.002792 | 1272.620858 | ENSG00000136950 |
| MAP3K8 | -0.333195221 | 0.793776512 | 0.184143918 | 0.368190353 | 80.31928561 | 63.8028249 | ENSG00000107968 |
| CCDC126 | -0.333195795 | 0.793776196 | 0.322457915 | 0.533101437 | 47.62197899 | 37.8968876 | ENSG00000169193 |
| AHSA1 | -0.333200928 | 0.793773372 | 2.63E-10 | 5.51E-09 | 3798.571217 | 3015.452541 | ENSG00000100591 |
| CNPY2 | -0.333214142 | 0.793766102 | 0.408797344 | 0.61989223 | 30.51061317 | 24.19015775 | ENSG00000257727 |
| NA | -0.333227629 | 0.793758681 | 0.524865086 | 0.718550866 | 22.65817449 | 17.968407 | ENSG00000262801 |
| RSRC2 | -0.333331581 | 0.79370149 | 3.50E-07 | 4.43E-06 | 2085.052442 | 1654.646544 | ENSG00000111011 |
| FAM117B | -0.33336493 | 0.793683143 | 0.002231465 | 0.011163236 | 541.1470234 | 429.2894321 | ENSG00000138439 |
| SLC2A13 | -0.33375973 | 0.793465978 | 0.084036586 | 0.209564364 | 162.2832127 | 128.7750996 | ENSG00000151229 |
| ELMO1 | -0.334222529 | 0.793211484 | 0.046320938 | 0.132842963 | 189.6027074 | 150.4192871 | ENSG00000155849 |
| HSPD1 | -0.334281582 | 0.793179017 | 2.81E-12 | 7.63E-11 | 28435.07768 | 22553.51887 | ENSG00000144381 |
| CTSC | -0.334406289 | 0.793110457 | 2.44E-12 | 6.66E-11 | 7882.085539 | 6250.977804 | ENSG00000109861 |
| CELF5 | -0.33445014 | 0.793086351 | 0.031052868 | 0.0975847 | 290.7111045 | 230.6658038 | ENSG00000161082 |
| PLPPR1 | -0.334469065 | 0.793075948 | 0.318423935 | 0.528431801 | 70.45657411 | 55.97444664 | ENSG00000148123 |
| ZNF695 | -0.33457705 | 0.793016589 | 0.228082548 | 0.427494208 | 68.55013696 | 54.31388405 | ENSG00000197472 |
| CCDC141 | -0.334579468 | 0.793015259 | 0.145887209 | 0.312970571 | 104.019513 | 82.43442661 | ENSG00000163492 |
| GRAMD2B | -0.334729452 | 0.792932821 | 0.073313816 | 0.189584112 | 156.7639384 | 124.2064391 | ENSG00000155324 |
| HTATSF1 | -0.334736241 | 0.79292909 | 1.06E-07 | 1.47E-06 | 2071.247902 | 1642.152467 | ENSG00000102241 |
| AEBP1 | -0.334826484 | 0.792879492 | 2.80E-07 | 3.60E-06 | 1923.327226 | 1525.317023 | ENSG00000106624 |
| LOC101927974 | -0.334868589 | 0.792856352 | 0.305938932 | 0.516092607 | 47.334325 | 37.53681979 | ENSG00000241764 |
| SQLE | -0.33488047 | 0.792849823 | 8.02E-09 | 1.35E-07 | 4301.24823 | 3409.955754 | ENSG00000104549 |
| ESPL1 | -0.335105171 | 0.792726345 | 7.91E-07 | 9.30E-06 | 1934.500067 | 1533.791295 | ENSG00000135476 |
| EPM2AIP1 | -0.33510553 | 0.792726148 | 0.036844798 | 0.111472928 | 233.6267242 | 185.0687747 | ENSG00000178567 |
| TEAD2 | -0.33522516 | 0.792660417 | 5.20E-08 | 7.58E-07 | 2760.73437 | 2188.622094 | ENSG00000074219 |
| ACAN | -0.335250102 | 0.792646714 | 0.454036763 | 0.660032659 | 30.12985423 | 23.95948356 | ENSG00000157766 |
| CNOT2-DT | -0.335278694 | 0.792631005 | 0.595839866 | 0.772365019 | 13.43097711 | 10.59636377 | ENSG00000257815 |
| ST3GAL1 | -0.335364503 | 0.792583862 | 0.045289774 | 0.130460248 | 223.1778249 | 176.8100085 | ENSG00000008513 |
| SWSAP1 | -0.335383162 | 0.792573611 | 0.447339314 | 0.654123697 | 31.42427191 | 24.91559027 | ENSG00000173928 |
| KATNB1 | -0.335527822 | 0.792494143 | 1.33E-06 | 1.51E-05 | 1597.011259 | 1265.9087 | ENSG00000140854 |
| ZNF350 | -0.335539625 | 0.79248766 | 0.16270097 | 0.337677347 | 90.00049016 | 71.31152319 | ENSG00000256683 |
| CRNDE | -0.335615574 | 0.792445941 | 0.205467472 | 0.397566926 | 95.68166848 | 75.82723753 | ENSG00000245694 |
| SYNJ2BP | -0.33584929 | 0.792317575 | 0.003804514 | 0.017596927 | 417.5376266 | 330.8122865 | ENSG00000213463 |
| NA | -0.335865997 | 0.7923084 | 0.444827639 | 0.652319456 | 27.14960822 | 21.47722123 | ENSG00000280058 |
| DPY19L3 | -0.335868831 | 0.792306844 | 0.000185734 | 0.001290503 | 922.2931257 | 730.8442426 | ENSG00000178904 |
| LRRC42 | -0.335950063 | 0.792262234 | 1.62E-05 | 0.000147059 | 1154.435864 | 914.6529331 | ENSG00000116212 |

| | | | | | | | |
|---|---|---|---|---|---|---|---|
| MEX3D | -0.335968358 | 0.792252187 | 2.44E-06 | 2.62E-05 | 2242.166582 | 1776.795683 | ENSG00000181588 |
| ZNF180 | -0.336009883 | 0.792229384 | 0.057193277 | 0.156535512 | 180.0324308 | 142.7789244 | ENSG00000167384 |
| NA | -0.336058517 | 0.792202678 | 0.650122766 | 0.810064757 | 9.17263624 | 7.280861978 | ENSG00000213104 |
| NA | -0.336136172 | 0.792160038 | 0.703851539 | 0.842524014 | 6.31308066 | 5.003283865 | ENSG00000280422 |
| PPT1 | -0.336238301 | 0.792103962 | 1.39E-07 | 1.88E-06 | 3628.234661 | 2873.126902 | ENSG00000131238 |
| RWDD1 | -0.336565599 | 0.791924281 | 0.000283138 | 0.001872102 | 709.0553264 | 561.6980754 | ENSG00000111832 |
| PRSS36 | -0.336674592 | 0.791864455 | 0.564496241 | 0.748848511 | 16.0322442 | 12.68715584 | ENSG00000178226 |
| ZNF271P | -0.336701282 | 0.791849806 | 0.00333195 | 0.015734902 | 476.016648 | 377.150995 | ENSG00000257267 |
| LINC01018 | -0.336747396 | 0.791824496 | 0.519601102 | 0.715241508 | 22.55113714 | 17.90427727 | ENSG00000250056 |
| TRPC1 | -0.336999939 | 0.7916859 | 0.081183003 | 0.204065088 | 164.6021391 | 130.1472691 | ENSG00000144935 |
| CCDC189 | -0.337007441 | 0.791681783 | 0.682372168 | 0.830268565 | 8.859786606 | 7.05113375 | ENSG00000196118 |
| ZNF614 | -0.337462678 | 0.79143201 | 0.003657604 | 0.017025845 | 500.2665788 | 395.7497538 | ENSG00000142556 |
| GBA2 | -0.337576036 | 0.791369826 | 4.04E-08 | 6.01E-07 | 3061.149233 | 2422.578306 | ENSG00000070610 |
| NA | -0.337673749 | 0.791316229 | 0.454754653 | 0.660692258 | 26.46578834 | 20.96082386 | ENSG00000248367 |
| NA | -0.337819492 | 0.791236293 | 0.286097551 | 0.495345072 | 52.86750617 | 41.86101828 | ENSG00000268222 |
| ZNF426-DT | -0.337832654 | 0.791229075 | 0.532051384 | 0.723643528 | 18.4757516 | 14.64259497 | ENSG00000278611 |
| SRPRB | -0.338117607 | 0.791072811 | 0.000519816 | 0.003180022 | 628.1696318 | 496.7593357 | ENSG00000144867 |
| NA | -0.338320696 | 0.790961459 | 0.708654082 | 0.846326173 | 7.583471765 | 6.014439572 | ENSG00000268230 |
| AKR1B1 | -0.338367356 | 0.790935878 | 2.77E-10 | 5.79E-09 | 4264.367762 | 3372.508082 | ENSG00000085662 |
| STRAP | -0.338638133 | 0.790787442 | 1.24E-07 | 1.69E-06 | 4279.09351 | 3383.847062 | ENSG00000023734 |
| AMD1 | -0.338657988 | 0.79077656 | 3.14E-05 | 0.000266283 | 4087.675738 | 3232.761564 | ENSG00000123505 |
| CMC2 | -0.338734362 | 0.790734698 | 0.143798225 | 0.309720516 | 110.6119335 | 87.51054205 | ENSG00000103121 |
| HSPE1 | -0.338898666 | 0.790644649 | 0.011133808 | 0.042708204 | 542.4354639 | 429.1604806 | ENSG00000115541 |
| KIF1C | -0.338993241 | 0.790592821 | 3.59E-10 | 7.38E-09 | 4718.96525 | 3731.103603 | ENSG00000129250 |
| ARTN | -0.339105896 | 0.790531088 | 0.026535411 | 0.086296062 | 223.757251 | 176.9700695 | ENSG00000117407 |
| DBP | -0.339121057 | 0.790522781 | 0.111737702 | 0.259440091 | 162.5053217 | 128.7297414 | ENSG00000105516 |
| LOXL1-AS1 | -0.339184523 | 0.790488005 | 0.208624085 | 0.401653693 | 83.0990844 | 65.84225648 | ENSG00000261801 |
| TBC1D13 | -0.339229039 | 0.790463615 | 0.000358475 | 0.002292515 | 949.365978 | 750.7302755 | ENSG00000107021 |
| NA | -0.339304217 | 0.790422425 | 0.016253568 | 0.058189314 | 513.0758506 | 406.0275648 | ENSG00000284976 |
| TMEM38B | -0.339306263 | 0.790421304 | 0.04566399 | 0.131257908 | 322.0273971 | 254.4264015 | ENSG00000095209 |
| ILKAP | -0.339326033 | 0.790410472 | 0.002760409 | 0.013377039 | 604.1050206 | 477.7395437 | ENSG00000132323 |
| SERGEF | -0.33941238 | 0.790363167 | 0.036959367 | 0.11172081 | 216.8107417 | 171.2913888 | ENSG00000129158 |
| TSPAN14 | -0.339473545 | 0.790329659 | 4.30E-07 | 5.34E-06 | 1851.002121 | 1463.247359 | ENSG00000108219 |
| C9orf78 | -0.339607477 | 0.790256293 | 6.26E-06 | 6.24E-05 | 1212.643529 | 958.521361 | ENSG00000136819 |
| NA | -0.339785757 | 0.790158643 | 0.679937941 | 0.828801912 | 8.050521617 | 6.352752753 | ENSG00000242083 |
| PTPN9 | -0.339934039 | 0.790077434 | 3.21E-06 | 3.37E-05 | 1291.930272 | 1020.661142 | ENSG00000169410 |
| WASHC2C | -0.340153881 | 0.789957049 | 6.69E-05 | 0.000520332 | 920.820099 | 727.2122252 | ENSG00000172661 |
| METRN | -0.340160122 | 0.789953632 | 0.006144131 | 0.026257504 | 392.4241234 | 310.0120705 | ENSG00000103260 |
| NA | -0.340243609 | 0.789907919 | 0.495681762 | 0.695207823 | 20.15507194 | 15.92613754 | ENSG00000273373 |
| GPRC5B | -0.340258448 | 0.789899794 | 2.27E-08 | 3.50E-07 | 3097.902423 | 2447.157804 | ENSG00000167191 |
| EXOC7 | -0.340302227 | 0.789875825 | 2.52E-06 | 2.70E-05 | 1792.603059 | 1415.699171 | ENSG00000182473 |
| ARL17A | -0.340321006 | 0.789865544 | 0.295643199 | 0.505764936 | 52.2180858 | 41.30480015 | ENSG00000185829 |
| RNF208 | -0.340464033 | 0.789787241 | 0.154765477 | 0.32630859 | 114.1639542 | 90.36202555 | ENSG00000212864 |
| SOX7 | -0.34059037 | 0.789718083 | 0.65080667 | 0.810504087 | 9.697806707 | 7.638754331 | ENSG00000171056 |
| TIAF1 | -0.340801735 | 0.789602392 | 0.528036647 | 0.720973828 | 18.87044295 | 14.88197102 | ENSG00000221995 |
| TM9SF4 | -0.340879097 | 0.789560052 | 5.89E-06 | 5.92E-05 | 1851.607304 | 1461.421817 | ENSG00000101337 |
| TM9SF2 | -0.340931152 | 0.789531564 | 8.07E-08 | 1.14E-06 | 2833.759574 | 2237.261878 | ENSG00000125304 |
| NA | -0.340983281 | 0.789503036 | 0.665597006 | 0.82085806 | 10.03409811 | 7.962122813 | ENSG00000277476 |
| HYAL3 | -0.341027848 | 0.789478647 | 0.346623113 | 0.558768886 | 37.46007151 | 29.56845413 | ENSG00000186792 |
| FRRS1 | -0.341075753 | 0.789452433 | 0.237051639 | 0.439066273 | 66.8518502 | 52.77762911 | ENSG00000156869 |
| CADPS | -0.34125941 | 0.789351941 | 0.025536271 | 0.083811693 | 237.0844192 | 186.9802101 | ENSG00000163618 |
| COQ7 | -0.341449439 | 0.789247976 | 0.02528532 | 0.083175975 | 224.4825893 | 177.2128505 | ENSG00000167186 |
| ND1 | -0.341483896 | 0.789229126 | 0.125255735 | 0.281225158 | 18126.08508 | 14305.78771 | ENSG00000198888 |
| ANKRD26 | -0.341485208 | 0.789228408 | 0.05860078 | 0.159485011 | 166.9151674 | 131.6806135 | ENSG00000107890 |
| DPP4 | -0.341557471 | 0.789188878 | 0.201030584 | 0.391899622 | 85.33202484 | 67.42797998 | ENSG00000197635 |
| CAMK2B | -0.34163853 | 0.789144538 | 0.584295109 | 0.763175369 | 15.61856028 | 12.3251961 | ENSG00000058404 |
| PTPRU | -0.342098937 | 0.788892739 | 1.77E-07 | 2.34E-06 | 1836.100001 | 1448.606141 | ENSG00000060656 |
| NUFIP1 | -0.342299718 | 0.788782955 | 0.002134224 | 0.010759431 | 473.0600975 | 372.9974083 | ENSG00000083635 |
| MSH2 | -0.342372355 | 0.788743243 | 1.40E-11 | 3.49E-10 | 6737.081483 | 5313.107017 | ENSG00000095002 |
| NA | -0.342582851 | 0.788628169 | 0.164624356 | 0.340071206 | 84.17078217 | 66.4510646 | ENSG00000269958 |
| LRRTM4 | -0.342648155 | 0.788592473 | 0.180692752 | 0.363677141 | 101.944891 | 80.18529748 | ENSG00000176204 |
| SSRP1 | -0.342717718 | 0.78855445 | 1.92E-14 | 6.61E-13 | 8231.77139 | 6491.372716 | ENSG00000149136 |
| CCNL2 | -0.342736798 | 0.788544021 | 0.093574088 | 0.227094797 | 2136.998236 | 1685.202021 | ENSG00000221978 |
| CCNB2 | -0.342764517 | 0.788528871 | 3.79E-07 | 4.75E-06 | 1660.901395 | 1309.346361 | ENSG00000157456 |
| PODNL1 | -0.342810911 | 0.788503514 | 0.4685626 | 0.671795414 | 23.96385484 | 18.94532238 | ENSG00000132000 |
| IRS4 | -0.342826926 | 0.788494761 | 0.05039227 | 0.14193259 | 216.4566268 | 170.5392836 | ENSG00000133124 |
| LUC7L | -0.342838851 | 0.788488243 | 2.05E-05 | 0.000181881 | 2048.817176 | 1615.821624 | ENSG00000007392 |
| DCLRE1C | -0.34284545 | 0.788484637 | 0.059423225 | 0.161190404 | 155.2441833 | 122.3634414 | ENSG00000152457 |
| FBXL19 | -0.343085345 | 0.788353537 | 1.17E-07 | 1.60E-06 | 3373.60333 | 2660.323277 | ENSG00000099364 |
| AKR7A2 | -0.343114689 | 0.788337502 | 0.000309197 | 0.002019084 | 717.2307083 | 565.7365507 | ENSG00000053371 |
| NA | -0.343126876 | 0.788330842 | 0.060612458 | 0.163570488 | 160.2029462 | 126.2561809 | ENSG00000226067 |
| RTL8C | -0.343151116 | 0.788317597 | 9.27E-05 | 0.000694355 | 974.9325202 | 768.9661943 | ENSG00000134590 |
| MACC1 | -0.343271144 | 0.788252014 | 0.031840571 | 0.099663378 | 247.6223309 | 194.9754164 | ENSG00000183742 |
| PCDHGB1 | -0.343341137 | 0.788213772 | 0.458214938 | 0.663654353 | 26.41595572 | 20.80031133 | ENSG00000254221 |

| | | | | | | | |
|---|---|---|---|---|---|---|---|
| TUBD1 | -0.34334534 | 0.788211476 | 0.020807485 | 0.071057749 | 282.111412 | 222.5698411 | ENSG00000108423 |
| HIP1 | -0.343360149 | 0.788203385 | 1.02E-07 | 1.43E-06 | 3508.80421 | 2765.335439 | ENSG00000127946 |
| RBMX2 | -0.343443978 | 0.788157587 | 0.00305805 | 0.014646534 | 473.24656 | 372.9194181 | ENSG00000134597 |
| NCOA2 | -0.343468192 | 0.788144359 | 0.003199451 | 0.015219554 | 738.2955491 | 581.5347944 | ENSG00000140396 |
| FERMT3 | -0.343718087 | 0.788007853 | 0.405459592 | 0.616620091 | 29.45462783 | 23.19858121 | ENSG00000149781 |
| COMT | -0.34372355 | 0.788004869 | 0.063221835 | 0.168826974 | 185.3520442 | 145.9743969 | ENSG00000093010 |
| FANCD2 | -0.343811197 | 0.787956998 | 5.74E-07 | 6.94E-06 | 1845.935608 | 1454.063983 | ENSG00000144554 |
| TMEM159 | -0.343862594 | 0.787928927 | 0.00175856 | 0.009108723 | 469.0404421 | 369.5096137 | ENSG00000011638 |
| DNAJB1 | -0.343949799 | 0.787881301 | 2.19E-09 | 4.02E-08 | 3430.337353 | 2702.219911 | ENSG00000132002 |
| SH2B1 | -0.343967516 | 0.787871626 | 0.000426293 | 0.002672825 | 1227.965309 | 967.9227271 | ENSG00000178188 |
| DPYSL2 | -0.344308402 | 0.787685486 | 2.49E-07 | 3.23E-06 | 2743.679807 | 2161.021865 | ENSG00000092964 |
| CDC42EP4 | -0.344337006 | 0.787669869 | 0.005133966 | 0.02260022 | 455.696087 | 358.9871727 | ENSG00000179604 |
| MFSD4B | -0.344485015 | 0.787589065 | 0.130355524 | 0.289517119 | 112.747346 | 88.89916928 | ENSG00000173214 |
| CDK5RAP3 | -0.34464235 | 0.787503178 | 0.199105222 | 0.389203606 | 88.37812138 | 69.67682557 | ENSG00000108465 |
| CAPNS1 | -0.344756505 | 0.787440868 | 4.95E-09 | 8.56E-08 | 2950.257704 | 2323.595132 | ENSG00000126247 |
| UGDH | -0.344764256 | 0.787436637 | 6.83E-05 | 0.000530585 | 1838.498935 | 1447.837658 | ENSG00000109814 |
| KCNQ4 | -0.344781522 | 0.787427213 | 0.084702184 | 0.210945634 | 132.4783611 | 104.3296025 | ENSG00000117013 |
| NA | -0.345006601 | 0.787304374 | 0.473217653 | 0.676357782 | 26.13661588 | 20.49217109 | ENSG00000232611 |
| TNC | -0.345142128 | 0.787230418 | 0.112103623 | 0.259955418 | 136.5623139 | 107.5594309 | ENSG00000041982 |
| NOMO2 | -0.345490067 | 0.787040583 | 4.43E-06 | 4.54E-05 | 1495.961288 | 1177.550827 | ENSG00000185164 |
| MZF1-AS1 | -0.345680131 | 0.786936903 | 0.428330022 | 0.638154973 | 27.47906001 | 21.64397661 | ENSG00000267858 |
| NT5M | -0.345732278 | 0.786908459 | 0.058901629 | 0.160143573 | 192.2674602 | 151.4373481 | ENSG00000205309 |
| SEC1P | -0.345737299 | 0.78690572 | 0.603076483 | 0.777921211 | 12.70088409 | 9.973273364 | ENSG00000232871 |
| NA | -0.345754956 | 0.786896089 | 0.500383426 | 0.698738613 | 19.75562591 | 15.55235453 | ENSG00000257851 |
| NA | -0.345828624 | 0.786855909 | 0.733257347 | 0.859883569 | 4.65155768 | 3.658165904 | ENSG00000233885 |
| RBM5 | -0.345880526 | 0.786827602 | 0.000739757 | 0.004314487 | 619.4050345 | 487.4329149 | ENSG00000003756 |
| EBP | -0.346034526 | 0.786743617 | 0.012379932 | 0.046665673 | 349.5607389 | 275.210341 | ENSG00000147155 |
| CCDC151 | -0.346175451 | 0.78666677 | 0.312448006 | 0.522637784 | 54.59070924 | 42.8750845 | ENSG00000198003 |
| FZD1 | -0.34653639 | 0.786469984 | 0.266193193 | 0.473656617 | 58.6986887 | 46.04674242 | ENSG00000157240 |
| SOCS7 | -0.346668444 | 0.786397999 | 2.09E-06 | 2.27E-05 | 1549.643243 | 1218.477325 | ENSG00000274211 |
| FAM86DP | -0.34671017 | 0.786375255 | 0.03918606 | 0.116876944 | 191.6580582 | 150.6125996 | ENSG00000244026 |
| LINC02289 | -0.34686181 | 0.786292604 | 0.555884603 | 0.742266897 | 15.63635765 | 12.27544399 | ENSG00000258819 |
| RABGEF1 | -0.347199134 | 0.786108779 | 0.378631875 | 0.591045046 | 34.68680684 | 27.30922568 | ENSG00000154710 |
| CISH | -0.347347739 | 0.78602781 | 0.232361053 | 0.433268018 | 61.5564904 | 48.45655204 | ENSG00000114737 |
| LOC101927420 | -0.347348493 | 0.786027399 | 0.475258277 | 0.677887112 | 23.69252367 | 18.67794886 | ENSG00000224046 |
| C9orf129 | -0.347394135 | 0.786002532 | 0.052734064 | 0.146909055 | 166.2974768 | 130.7319792 | ENSG00000204352 |
| PDCD7 | -0.347403807 | 0.785997262 | 0.000208432 | 0.001427204 | 737.394095 | 579.563267 | ENSG00000090470 |
| ZWINT | -0.347630955 | 0.785873519 | 1.68E-08 | 2.68E-07 | 2157.026257 | 1695.028634 | ENSG00000122952 |
| MGAT4C | -0.347639691 | 0.785868761 | 0.253984959 | 0.460180941 | 62.51105722 | 49.04540213 | ENSG00000182050 |
| NA | -0.347834273 | 0.785762775 | 0.74373036 | 0.86667579 | 5.029646873 | 3.92024254 | ENSG00000255992 |
| VOPP1 | -0.347856084 | 0.785750895 | 1.59E-06 | 1.77E-05 | 1629.124864 | 1280.346232 | ENSG00000154978 |
| NA | -0.347950872 | 0.785699272 | 0.595846259 | 0.772365019 | 12.61311864 | 9.877457647 | ENSG00000253816 |
| LOC101927151 | -0.347955774 | 0.785696602 | 0.202478946 | 0.393819045 | 90.25669372 | 70.93208711 | ENSG00000267575 |
| RNASEK | -0.348001667 | 0.785671609 | 0.511972513 | 0.708478121 | 19.08039952 | 14.94855975 | ENSG00000219200 |
| NA | -0.348089768 | 0.785623632 | 0.699686593 | 0.839970842 | 5.907710895 | 4.65626883 | ENSG00000250461 |
| SELENOV | -0.348114182 | 0.785610337 | 0.190018944 | 0.37643006 | 79.48746857 | 62.44682962 | ENSG00000186838 |
| LRP3 | -0.348195017 | 0.78556632 | 9.06E-06 | 8.77E-05 | 1815.494507 | 1426.604265 | ENSG00000130881 |
| NF2 | -0.348425652 | 0.785440746 | 0.000112166 | 0.000823259 | 781.7249593 | 613.8086104 | ENSG00000186575 |
| RAB11B | -0.348450317 | 0.785427318 | 6.09E-06 | 6.10E-05 | 1732.388381 | 1361.295902 | ENSG00000185236 |
| MGAT4A | -0.34859176 | 0.785350318 | 0.189184152 | 0.3752881 | 90.01264312 | 70.60907481 | ENSG00000071073 |
| CCDC96 | -0.348713573 | 0.785284011 | 0.720541619 | 0.852637136 | 5.95159362 | 4.686441774 | ENSG00000173013 |
| MAPRE2 | -0.348750891 | 0.785263698 | 1.86E-06 | 2.05E-05 | 1255.123433 | 985.495352 | ENSG00000166974 |
| NA | -0.348783293 | 0.785246061 | 0.284472443 | 0.493726967 | 57.69014555 | 45.24868924 | ENSG00000264164 |
| UBXN6 | -0.348958725 | 0.785150581 | 0.000128139 | 0.000926458 | 956.328375 | 751.1375864 | ENSG00000167671 |
| PARL | -0.3490315 | 0.785110976 | 0.303641664 | 0.51363963 | 64.8881037 | 50.98610748 | ENSG00000175193 |
| TTC21A | -0.349159551 | 0.785041294 | 0.384964608 | 0.597138171 | 32.58342965 | 25.5405726 | ENSG00000168026 |
| DCAF11 | -0.349177461 | 0.785031548 | 9.72E-06 | 9.31E-05 | 1567.064737 | 1229.651124 | ENSG00000100897 |
| ZNF432 | -0.349208593 | 0.785014609 | 0.095439511 | 0.230129617 | 122.5186802 | 96.12136638 | ENSG00000256087 |
| RIPOR1 | -0.349419438 | 0.78489989 | 0.043624179 | 0.126920754 | 195.1821879 | 153.1609562 | ENSG00000039523 |
| HIRA | -0.349500873 | 0.784855586 | 0.010490915 | 0.040606873 | 325.4308364 | 255.6261329 | ENSG00000100084 |
| TMOD4 | -0.349585082 | 0.784809776 | 0.674008237 | 0.825366912 | 8.019681573 | 6.26563889 | ENSG00000163157 |
| GAS6-DT | -0.349743886 | 0.784723394 | 0.567717913 | 0.751047019 | 13.9333162 | 10.91320587 | ENSG00000272695 |
| FCN3 | -0.350039452 | 0.784562643 | 0.737560984 | 0.862343405 | 4.627836581 | 3.622696074 | ENSG00000142748 |
| NA | -0.35012471 | 0.784516279 | 0.055823835 | 0.153611121 | 171.2076285 | 134.3763572 | ENSG00000220785 |
| CHPT1 | -0.350211181 | 0.784469259 | 0.011764999 | 0.044758664 | 315.4895633 | 247.7627363 | ENSG00000111666 |
| KCNT1 | -0.3504543 | 0.784337074 | 0.670143407 | 0.823185784 | 9.703451111 | 7.654928535 | ENSG00000107147 |
| NA | -0.3505336 | 0.784293963 | 0.707595427 | 0.845853003 | 5.931431994 | 4.662795221 | ENSG00000230896 |
| BZW1 | -0.35055949 | 0.784279888 | 1.20E-10 | 2.65E-09 | 3535.283138 | 2772.428151 | ENSG00000082153 |
| TCTA | -0.350742664 | 0.784180317 | 0.108578613 | 0.253744229 | 110.8939431 | 86.95099159 | ENSG00000145022 |
| CATSPER2P1 | -0.350795976 | 0.78415134 | 0.565499124 | 0.749592964 | 13.593186 | 10.6694789 | ENSG00000205771 |
| SEMA7A | -0.351015952 | 0.784031785 | 5.65E-05 | 0.000449187 | 785.922867 | 616.2227709 | ENSG00000138623 |
| KCNK10 | -0.351088705 | 0.783992248 | 0.430752008 | 0.640582067 | 33.29900545 | 26.09612832 | ENSG00000100433 |
| CENPV | -0.351097126 | 0.783987672 | 1.09E-06 | 1.26E-05 | 2271.80475 | 1781.692709 | ENSG00000166582 |
| CDCA7 | -0.351198331 | 0.783932678 | 6.03E-08 | 8.72E-07 | 2172.108638 | 1702.53167 | ENSG00000144354 |

| | | | | | | | |
|---|---|---|---|---|---|---|---|
| UBTF | -0.351296701 | 0.783879227 | 4.73E-09 | 8.21E-08 | 3469.025211 | 2719.829427 | ENSG00000108312 |
| PSME3 | -0.351387919 | 0.783829665 | 6.13E-11 | 1.42E-09 | 6250.606782 | 4898.970512 | ENSG00000131467 |
| ACYP2 | -0.351401222 | 0.783822438 | 0.236813262 | 0.438858749 | 58.08811704 | 45.50111806 | ENSG00000170634 |
| ORMDL3 | -0.351403934 | 0.783820965 | 1.21E-05 | 0.000112595 | 1590.54329 | 1247.05535 | ENSG00000172057 |
| CDC25C | -0.351503797 | 0.783766711 | 0.003771409 | 0.017481566 | 379.7863678 | 297.578573 | ENSG00000158402 |
| ENTPD7 | -0.35154732 | 0.783743066 | 0.011611752 | 0.044215979 | 318.5575236 | 249.3909075 | ENSG00000198018 |
| NA | -0.351670284 | 0.783676269 | 0.537180021 | 0.72743395 | 15.67431663 | 12.24432509 | ENSG00000264107 |
| ZNF511 | -0.351811091 | 0.783599786 | 0.129469195 | 0.288107825 | 106.1071778 | 83.11207157 | ENSG00000198546 |
| AGO3 | -0.351948927 | 0.783524924 | 0.011087116 | 0.0425656 | 323.4211745 | 253.4039397 | ENSG00000126070 |
| NA | -0.351983017 | 0.78350641 | 0.597077519 | 0.77347684 | 11.88658509 | 9.330887327 | ENSG00000261143 |
| C18orf32 | -0.351995341 | 0.783499717 | 0.433497552 | 0.642497135 | 30.53906279 | 23.91408237 | ENSG00000177576 |
| SMG9 | -0.352015415 | 0.783488815 | 0.001482931 | 0.007884334 | 523.8319819 | 410.5704033 | ENSG00000105771 |
| TMEM198 | -0.352120781 | 0.783431596 | 0.014874357 | 0.054165489 | 259.3677846 | 203.1533117 | ENSG00000188760 |
| TRIB2 | -0.352148212 | 0.7834167 | 3.83E-05 | 0.000317368 | 1922.808234 | 1506.610187 | ENSG00000071575 |
| ALKBH8 | -0.352166543 | 0.783406746 | 0.008361289 | 0.033877482 | 312.6809838 | 244.8780311 | ENSG00000137760 |
| CACHD1 | -0.352513261 | 0.783218495 | 4.16E-06 | 4.28E-05 | 1720.388426 | 1347.567498 | ENSG00000158966 |
| USP16 | -0.352528738 | 0.783210093 | 0.006700537 | 0.028239154 | 374.1945074 | 293.0561213 | ENSG00000156256 |
| APTR | -0.352595838 | 0.783173667 | 0.037608895 | 0.113150227 | 182.3071951 | 142.8172321 | ENSG00000214293 |
| SPIN2B | -0.352796492 | 0.783064749 | 0.083093712 | 0.207789294 | 130.5606084 | 102.289225 | ENSG00000186787 |
| NA | -0.352835188 | 0.783043745 | 0.494086807 | 0.693587256 | 22.85361384 | 17.88034717 | ENSG00000235316 |
| FAM122C | -0.352853436 | 0.783033841 | 0.278663422 | 0.487770759 | 52.60209873 | 41.20832202 | ENSG00000156500 |
| CAPN5 | -0.353027751 | 0.782939236 | 3.01E-06 | 3.18E-05 | 1159.61384 | 907.9893178 | ENSG00000149260 |
| SMPD4 | -0.3530662 | 0.78291837 | 9.54E-08 | 1.34E-06 | 2846.732398 | 2229.396711 | ENSG00000136699 |
| IER2 | -0.353074611 | 0.782913806 | 1.79E-05 | 0.000160808 | 1537.597624 | 1203.855035 | ENSG00000160888 |
| NA | -0.353155977 | 0.782869652 | 0.388603462 | 0.60048837 | 30.95246738 | 24.283798 | ENSG00000261342 |
| VPS18 | -0.353165704 | 0.782864374 | 5.27E-06 | 5.34E-05 | 1333.53328 | 1044.088789 | ENSG00000104142 |
| P4HTM | -0.353435484 | 0.782717993 | 0.031763717 | 0.099478906 | 252.5354055 | 197.4218527 | ENSG00000178467 |
| PHKA1 | -0.353822144 | 0.782508244 | 0.006900797 | 0.028928202 | 375.5066184 | 293.5783101 | ENSG00000067177 |
| ODF2L | -0.353911805 | 0.782459614 | 0.018931499 | 0.065660488 | 300.2357185 | 235.0660482 | ENSG00000122417 |
| SGSM2 | -0.353980592 | 0.782422307 | 0.003306816 | 0.015639673 | 794.4119894 | 621.7076783 | ENSG00000141258 |
| NEFL | -0.354168502 | 0.782320404 | 0.001789415 | 0.009233089 | 457.5221837 | 357.8020002 | ENSG00000277586 |
| TMEM200C | -0.354204184 | 0.782301055 | 0.022454158 | 0.075547259 | 228.9808365 | 179.2927652 | ENSG00000206432 |
| DXO | -0.354278276 | 0.78226088 | 0.052171574 | 0.145899627 | 160.4647942 | 125.4629302 | ENSG00000204348 |
| HAT1 | -0.354317484 | 0.782239621 | 1.09E-05 | 0.00010315 | 1831.347302 | 1432.738181 | ENSG00000128708 |
| URGCP | -0.354322607 | 0.782236843 | 0.001428999 | 0.007644654 | 463.0556442 | 362.1475971 | ENSG00000106608 |
| FAM13A | -0.354570516 | 0.782102437 | 0.00830098 | 0.033676479 | 339.8291175 | 265.7612006 | ENSG00000138640 |
| WDR48 | -0.354683378 | 0.782041256 | 2.08E-05 | 0.000183782 | 1135.59204 | 887.6885672 | ENSG00000114742 |
| NOS1 | -0.354772464 | 0.781992966 | 0.076638346 | 0.195715369 | 179.0354302 | 139.8132326 | ENSG00000089250 |
| PJA1 | -0.354947743 | 0.781897965 | 3.53E-07 | 4.45E-06 | 1806.757977 | 1412.587603 | ENSG00000181191 |
| SHMT2 | -0.354991611 | 0.78187419 | 1.05E-12 | 3.01E-11 | 7892.421856 | 6170.134654 | ENSG00000182199 |
| CLCNKB | -0.355021364 | 0.781858065 | 0.346819714 | 0.558942677 | 37.39958663 | 29.25596296 | ENSG00000184908 |
| CDK5RAP1 | -0.355027076 | 0.781854969 | 0.000190013 | 0.001318778 | 876.4357125 | 685.2149737 | ENSG00000101391 |
| TXNL4B | -0.355035695 | 0.781850299 | 0.061406106 | 0.165263547 | 153.8771278 | 120.4297569 | ENSG00000140830 |
| POLK | -0.355064552 | 0.78183466 | 0.087944752 | 0.216452058 | 131.3754922 | 102.5911224 | ENSG00000122008 |
| NA | -0.35508693 | 0.781822533 | 0.52141926 | 0.716145653 | 21.84890951 | 17.05770148 | ENSG00000247373 |
| GK | -0.35520225 | 0.781760041 | 0.03278378 | 0.101844109 | 210.8825637 | 164.7977027 | ENSG00000198814 |
| ADSS1 | -0.35537778 | 0.781664932 | 0.028415727 | 0.090938996 | 233.0953245 | 182.2076434 | ENSG00000185100 |
| SPRY2 | -0.355427786 | 0.781637839 | 0.001624374 | 0.008509463 | 922.0766368 | 720.3968516 | ENSG00000136158 |
| MDM1 | -0.355496164 | 0.781600793 | 0.041690948 | 0.122523117 | 201.2565569 | 157.3246421 | ENSG00000111554 |
| CDO1 | -0.355575177 | 0.781557988 | 0.004554152 | 0.020433633 | 358.0773945 | 279.8802094 | ENSG00000129596 |
| DNAJA1 | -0.355664478 | 0.781509612 | 4.00E-11 | 9.49E-10 | 6957.054617 | 5437.345362 | ENSG00000086061 |
| BAG2 | -0.35568437 | 0.781498836 | 0.002141303 | 0.010778187 | 455.5314621 | 355.8144962 | ENSG00000112208 |
| NA | -0.355770597 | 0.781452129 | 0.709137656 | 0.846486042 | 5.996671554 | 4.691738659 | ENSG00000275720 |
| DNAAF1 | -0.355817274 | 0.781426846 | 0.672232639 | 0.82439555 | 7.211306308 | 5.654371756 | ENSG00000154099 |
| SPOP | -0.355830898 | 0.781419467 | 0.000629914 | 0.003745567 | 605.2953232 | 473.0698526 | ENSG00000121067 |
| NA | -0.356068231 | 0.781290929 | 0.674931572 | 0.825790913 | 7.232663142 | 5.670545959 | ENSG00000243485 |
| B3GNT5 | -0.356105065 | 0.781270981 | 0.11112867 | 0.258544124 | 105.7293679 | 82.53192333 | ENSG00000176597 |
| SLC4A7 | -0.356383892 | 0.781120001 | 0.001304557 | 0.007057968 | 1234.471205 | 963.822396 | ENSG00000033867 |
| GRID2 | -0.356445629 | 0.781086576 | 0.003139832 | 0.014973671 | 492.4286835 | 384.3715837 | ENSG00000152208 |
| GPI | -0.356717681 | 0.780939298 | 1.22E-12 | 3.45E-11 | 6486.024518 | 5065.73294 | ENSG00000105220 |
| WDR4 | -0.356814043 | 0.780887139 | 3.85E-06 | 3.99E-05 | 1441.415527 | 1125.212937 | ENSG00000160193 |
| NA | -0.35691141 | 0.780834439 | 0.752437021 | 0.871085164 | 4.262790069 | 3.344445235 | ENSG00000223751 |
| TRMT12 | -0.357052764 | 0.780757937 | 0.002319316 | 0.011526392 | 436.5960851 | 340.8097009 | ENSG00000183665 |
| ADAMTS17 | -0.357060361 | 0.780753826 | 0.265912336 | 0.473407364 | 53.44103471 | 41.7331377 | ENSG00000140470 |
| ARHGAP21 | -0.357106111 | 0.780729068 | 0.000440081 | 0.002752872 | 776.2340115 | 605.8066239 | ENSG00000107863 |
| EIF4G2 | -0.357263995 | 0.780643631 | 1.03E-11 | 2.61E-10 | 17750.09989 | 13856.22581 | ENSG00000110321 |
| CCZ1B | -0.357399663 | 0.780570225 | 0.004337755 | 0.019630603 | 376.2515853 | 293.7206409 | ENSG00000146574 |
| IDH3A | -0.357429331 | 0.780554173 | 0.001787585 | 0.009226167 | 435.2432936 | 339.6524829 | ENSG00000166411 |
| CLIC4 | -0.357446182 | 0.780545056 | 1.60E-07 | 2.13E-06 | 6582.14997 | 5137.324213 | ENSG00000169504 |
| NA | -0.357579996 | 0.780472662 | 0.741866303 | 0.865411398 | 4.71679724 | 3.66998918 | ENSG00000274767 |
| SUSD5 | -0.357605402 | 0.780458918 | 0.128121519 | 0.285765311 | 96.10272453 | 74.95852823 | ENSG00000173705 |
| NA | -0.357993297 | 0.780249105 | 0.680243815 | 0.829014226 | 7.174542527 | 5.610200072 | ENSG00000272619 |
| NA | -0.358296574 | 0.780085102 | 0.31806606 | 0.528112378 | 48.572707 | 37.93046551 | ENSG00000236397 |
| RNF121 | -0.358684766 | 0.77987523 | 0.000519774 | 0.003180022 | 672.3350587 | 524.3390992 | ENSG00000137522 |

| | | | | | | | |
|---|---|---|---|---|---|---|---|
| NA | -0.358750698 | 0.77983959 | 0.477413694 | 0.679936472 | 24.14622535 | 18.88280103 | ENSG00000234825 |
| PPM1B | -0.35875224 | 0.779838756 | 0.013781485 | 0.050822758 | 481.8763057 | 375.898433 | ENSG00000138032 |
| NA | -0.359322547 | 0.779530541 | 0.682464042 | 0.830326854 | 8.920271485 | 6.950967106 | ENSG00000265800 |
| SLC35B1 | -0.359340639 | 0.779520766 | 2.69E-05 | 0.000231965 | 1053.073095 | 821.2172111 | ENSG00000121073 |
| B4GALT1 | -0.359567811 | 0.779398029 | 0.000527913 | 0.003218086 | 552.1223401 | 430.1368812 | ENSG00000086062 |
| IPO5 | -0.359680426 | 0.779337193 | 4.70E-10 | 9.51E-09 | 11648.18555 | 9077.043616 | ENSG00000065150 |
| ZNF319 | -0.359734112 | 0.779308192 | 0.024864916 | 0.082136423 | 216.4875186 | 168.7861423 | ENSG00000166188 |
| LPCAT3 | -0.359749951 | 0.779299637 | 0.0005617 | 0.003392297 | 596.3400157 | 464.7869359 | ENSG00000111684 |
| CLTRN | -0.359780839 | 0.779282952 | 0.664359466 | 0.820188829 | 8.474578468 | 6.62542316 | ENSG00000147003 |
| SLC44A2 | -0.359830292 | 0.77925624 | 2.48E-10 | 5.21E-09 | 3126.13276 | 2436.14518 | ENSG00000129353 |
| RUFY2 | -0.360123732 | 0.779097758 | 0.069161265 | 0.181027096 | 133.9219188 | 104.3233597 | ENSG00000204130 |
| TMEM81 | -0.360214629 | 0.779048672 | 0.223905625 | 0.42204389 | 60.17992609 | 46.83346679 | ENSG00000174529 |
| CHST8 | -0.36024548 | 0.779032013 | 0.003496646 | 0.016387384 | 493.824241 | 384.4403706 | ENSG00000124302 |
| NA | -0.360383185 | 0.778957658 | 0.714616696 | 0.84957753 | 8.173881792 | 6.341875435 | ENSG00000226912 |
| SUPT3H | -0.360406517 | 0.77894506 | 0.013476193 | 0.049862443 | 254.3509011 | 198.1575001 | ENSG00000196284 |
| CCT6P3 | -0.360503816 | 0.778892528 | 0.428376521 | 0.638154973 | 25.57585069 | 19.89367319 | ENSG00000234585 |
| ST8SIA6 | -0.360507803 | 0.778890376 | 0.305290755 | 0.51555392 | 42.89717308 | 33.39940734 | ENSG00000148488 |
| ZNF589 | -0.360592693 | 0.778844546 | 1.15E-07 | 1.58E-06 | 2632.682205 | 2050.473534 | ENSG00000164048 |
| ASNS | -0.360657752 | 0.778809425 | 0.387078435 | 0.599276623 | 31.10310813 | 24.22222261 | ENSG00000070669 |
| NA | -0.360864991 | 0.778697559 | 0.550008015 | 0.737234819 | 2492.590127 | 1941.011109 | ENSG00000248527 |
| CASP7 | -0.360930575 | 0.778662161 | 0.077181888 | 0.196651371 | 124.7451383 | 97.19135492 | ENSG00000165806 |
| ANO2 | -0.360961222 | 0.77864562 | 0.161797296 | 0.336476371 | 77.24176479 | 60.13283572 | ENSG00000047617 |
| NA | -0.361236203 | 0.778497222 | 0.092447989 | 0.224806141 | 139.6455053 | 108.8606607 | ENSG00000260708 |
| KIF18B | -0.361272972 | 0.778477381 | 9.20E-08 | 1.29E-06 | 1603.502131 | 1248.192554 | ENSG00000186185 |
| GABRG3 | -0.361343163 | 0.778439508 | 0.708923467 | 0.846414716 | 5.529621701 | 4.330724884 | ENSG00000182256 |
| SMARCA5 | -0.361473549 | 0.778369158 | 7.28E-11 | 1.66E-09 | 4601.866662 | 3581.604676 | ENSG00000153147 |
| LSG1 | -0.361473583 | 0.77836914 | 3.61E-08 | 5.42E-07 | 1746.412256 | 1359.45839 | ENSG00000041802 |
| CCNH | -0.361603178 | 0.778299223 | 0.013523526 | 0.050008183 | 275.2455493 | 214.1727161 | ENSG00000134480 |
| DACH2 | -0.361607174 | 0.778297067 | 0.274985388 | 0.483753986 | 57.85830433 | 44.9235968 | ENSG00000126733 |
| GPC5 | -0.361687889 | 0.778253525 | 0.603866826 | 0.77848488 | 10.61738919 | 8.236968684 | ENSG00000179399 |
| FMR1 | -0.361759266 | 0.778215022 | 0.000203046 | 0.001397929 | 672.0593301 | 522.8720897 | ENSG00000102081 |
| SENP1 | -0.3620998 | 0.778031353 | 0.00250556 | 0.012335113 | 407.0898969 | 316.6182707 | ENSG00000079387 |
| ZBED5-AS1 | -0.362112793 | 0.778024346 | 0.591388677 | 0.769171843 | 13.29101478 | 10.34705637 | ENSG00000247271 |
| LINC01572 | -0.362222789 | 0.77796503 | 0.34967499 | 0.561787696 | 37.64274751 | 29.24726111 | ENSG00000261008 |
| SLC1A4 | -0.362371701 | 0.777884734 | 0.202621502 | 0.39393542 | 76.03485937 | 59.14918304 | ENSG00000115902 |
| MT1X | -0.362499396 | 0.777815885 | 0.26849421 | 0.476401356 | 57.83219282 | 44.98566665 | ENSG00000187193 |
| ALKBH5 | -0.36252721 | 0.77780089 | 1.31E-06 | 1.49E-05 | 1352.177522 | 1051.762235 | ENSG00000091542 |
| NNT | -0.362591573 | 0.77776619 | 0.002930275 | 0.014102507 | 396.3473158 | 308.0840391 | ENSG00000112992 |
| CCDC107 | -0.362770768 | 0.777669591 | 0.34563277 | 0.557553162 | 41.5407704 | 32.21763979 | ENSG00000159884 |
| KIF15 | -0.362906456 | 0.777596453 | 0.000283236 | 0.001872102 | 691.0178446 | 536.9597913 | ENSG00000163808 |
| MCFD2 | -0.36322538 | 0.777424576 | 1.52E-07 | 2.03E-06 | 1751.833672 | 1361.599867 | ENSG00000180398 |
| MED6 | -0.363332525 | 0.777366841 | 0.006722539 | 0.028312922 | 314.8866954 | 244.9477412 | ENSG00000133997 |
| LINGO3 | -0.36335444 | 0.777355033 | 0.516233816 | 0.712443409 | 16.21580992 | 12.60221744 | ENSG00000220008 |
| JDP2 | -0.363409291 | 0.777325478 | 0.000321644 | 0.002088088 | 693.769167 | 539.40956 | ENSG00000140044 |
| SHKBP1 | -0.363449826 | 0.777303638 | 3.11E-07 | 3.97E-06 | 1574.099498 | 1223.854465 | ENSG00000160410 |
| WAS | -0.363487814 | 0.777283171 | 0.42025439 | 0.630538755 | 25.11591978 | 19.50371598 | ENSG00000015285 |
| MARCHF4 | -0.363571146 | 0.777238276 | 0.152938733 | 0.323504477 | 85.42929965 | 66.48965585 | ENSG00000144583 |
| SH3BGRL2 | -0.36361758 | 0.77721326 | 0.001715092 | 0.008915361 | 495.049021 | 384.7149102 | ENSG00000198478 |
| BMP2K | -0.363825914 | 0.777101034 | 0.027810345 | 0.089578435 | 203.9022653 | 158.3507426 | ENSG00000138756 |
| SCNN1D | -0.364124251 | 0.776940352 | 0.371158578 | 0.583284492 | 43.87279125 | 34.11975389 | ENSG00000162572 |
| TC2N | -0.364253621 | 0.776870685 | 0.0047265 | 0.021082574 | 405.6489828 | 314.9072935 | ENSG00000165929 |
| TMEM132B | -0.364441078 | 0.776769749 | 1.31E-07 | 1.77E-06 | 2080.811366 | 1615.848279 | ENSG00000139364 |
| NA | -0.364672342 | 0.776645243 | 0.616840671 | 0.787224317 | 9.854091858 | 7.686047437 | ENSG00000287225 |
| FAM155A | -0.364698155 | 0.776631347 | 0.011929485 | 0.045302298 | 289.05848 | 224.6265608 | ENSG00000204442 |
| RGL2 | -0.365042406 | 0.776446052 | 0.000514142 | 0.003151434 | 1020.63085 | 792.7789867 | ENSG00000237441 |
| NA | -0.365102079 | 0.776413937 | 0.532180904 | 0.723764128 | 17.53008375 | 13.61403556 | ENSG00000260806 |
| NA | -0.365158355 | 0.776383652 | 0.71000368 | 0.846811892 | 6.01683318 | 4.664970684 | ENSG00000240207 |
| NA | -0.365158704 | 0.776383464 | 0.782888955 | 0.889742909 | 3.860979777 | 3.02542768 | ENSG00000223349 |
| DNAJC7 | -0.365161105 | 0.776382172 | 3.92E-06 | 4.05E-05 | 1383.68613 | 1074.663619 | ENSG00000168259 |
| TMEM65 | -0.365502699 | 0.776198366 | 0.007383966 | 0.030580307 | 390.7572865 | 303.6014309 | ENSG00000164983 |
| TENM4 | -0.365554116 | 0.776170703 | 0.016314622 | 0.058352583 | 284.932041 | 220.9227828 | ENSG00000149256 |
| LAMP1 | -0.3656181 | 0.77613628 | 7.57E-13 | 2.22E-11 | 4607.170239 | 3576.079609 | ENSG00000185896 |
| RAPGEF4 | -0.365711016 | 0.776086295 | 0.264399753 | 0.471946062 | 54.52666489 | 42.27006021 | ENSG00000091428 |
| FAM136A | -0.36590735 | 0.775980686 | 8.36E-08 | 1.18E-06 | 2307.441291 | 1791.123266 | ENSG00000035141 |
| POLR2F | -0.36596214 | 0.775951217 | 1.04E-08 | 1.71E-07 | 2059.666463 | 1598.503398 | ENSG00000100142 |
| NFATC2IP | -0.366019933 | 0.775920134 | 0.00021577 | 0.001469991 | 1138.49177 | 883.568878 | ENSG00000176953 |
| MAN1C1 | -0.366033686 | 0.775912737 | 0.004969735 | 0.021969441 | 344.0014724 | 266.9398468 | ENSG00000117643 |
| ITGA7 | -0.366297122 | 0.775771068 | 0.092259619 | 0.224468296 | 1648.267434 | 1278.747661 | ENSG00000135424 |
| ATP6V1G1 | -0.366328089 | 0.775754417 | 2.72E-07 | 3.51E-06 | 2009.483818 | 1559.178019 | ENSG00000136888 |
| BORCS7 | -0.366370457 | 0.775731635 | 0.376275775 | 0.588339792 | 32.48114698 | 25.26667268 | ENSG00000166275 |
| TGFBR1 | -0.366412887 | 0.775708821 | 0.000582026 | 0.003501615 | 852.5146487 | 661.1094872 | ENSG00000106799 |
| PTRHD1 | -0.366570307 | 0.775624184 | 0.066613647 | 0.175943877 | 132.6002467 | 102.8606714 | ENSG00000184924 |
| POMT2 | -0.366871223 | 0.775462422 | 0.000162188 | 0.001143734 | 717.1512576 | 556.2561438 | ENSG00000009830 |
| ZNF605 | -0.367339738 | 0.775210632 | 0.02391141 | 0.079501227 | 223.8266349 | 173.4982381 | ENSG00000196458 |

| | | | | | | | |
|---|---|---|---|---|---|---|---|
| CDC25A | -0.367442997 | 0.775155149 | 2.50E-07 | 3.24E-06 | 2139.672087 | 1658.612551 | ENSG00000164045 |
| METTL22 | -0.367470433 | 0.775140408 | 0.001640122 | 0.008579976 | 663.7517164 | 514.2806039 | ENSG00000067365 |
| RAD54L | -0.367573052 | 0.775085274 | 1.95E-05 | 0.000174366 | 983.2258466 | 761.6498921 | ENSG00000085999 |
| MCM6 | -0.367914257 | 0.774901984 | 8.45E-13 | 2.45E-11 | 5338.757327 | 4136.738548 | ENSG00000076003 |
| ANTXR1 | -0.367929271 | 0.774893919 | 3.34E-06 | 3.49E-05 | 1173.857057 | 909.7278706 | ENSG00000169604 |
| CRTAP | -0.367952534 | 0.774881425 | 4.33E-10 | 8.80E-09 | 2617.145057 | 2028.267086 | ENSG00000170275 |
| MAPK4 | -0.368006131 | 0.774852638 | 0.02243765 | 0.07550517 | 207.4975584 | 160.7910087 | ENSG00000141639 |
| CPT1B | -0.368112723 | 0.774795391 | 0.493740991 | 0.693204863 | 17.61548494 | 13.64326254 | ENSG00000205560 |
| TESK1 | -0.368115118 | 0.774794105 | 0.007064732 | 0.02949756 | 323.9196748 | 251.1802966 | ENSG00000107140 |
| LDLRAD4 | -0.368147783 | 0.774776563 | 0.275931942 | 0.484922271 | 44.18208141 | 34.19672548 | ENSG00000168675 |
| LAMB1 | -0.368425701 | 0.774627325 | 2.05E-08 | 3.20E-07 | 2458.574216 | 1904.165181 | ENSG00000091136 |
| NA | -0.368689115 | 0.774485903 | 0.729955617 | 0.857822764 | 4.266349541 | 3.308975405 | ENSG00000217624 |
| NBDY | -0.368791174 | 0.774431117 | 0.000148 | 0.001055902 | 661.8084358 | 512.4775949 | ENSG00000204272 |
| ANKH | -0.368807205 | 0.774422511 | 0.005434805 | 0.02369259 | 337.6801037 | 261.5017716 | ENSG00000154122 |
| MDC1 | -0.368975856 | 0.774331986 | 3.94E-07 | 4.94E-06 | 1540.836115 | 1193.039167 | ENSG00000137337 |
| DDX39B | -0.368990353 | 0.774324205 | 0.08960771 | 0.219743098 | 1447.020893 | 1120.50667 | ENSG00000198563 |
| PCNX3 | -0.369068823 | 0.77428209 | 1.39E-07 | 1.88E-06 | 1866.099122 | 1445.232278 | ENSG00000197136 |
| NA | -0.369176961 | 0.774224056 | 0.667723121 | 0.821869982 | 7.3453449 | 5.687666122 | ENSG00000272721 |
| WDR11 | -0.369228884 | 0.774196192 | 1.33E-06 | 1.51E-05 | 1205.976666 | 933.4438178 | ENSG00000120008 |
| ECI2 | -0.36944441 | 0.774080542 | 0.011176847 | 0.042823038 | 276.9171403 | 214.174608 | ENSG00000198721 |
| LIG1 | -0.36945228 | 0.77407632 | 2.49E-07 | 3.23E-06 | 2210.102927 | 1710.75996 | ENSG00000105486 |
| GABBR2 | -0.369542176 | 0.774028087 | 0.333570255 | 0.545597979 | 44.58836705 | 34.53163369 | ENSG00000136928 |
| SIVA1 | -0.369618921 | 0.773986914 | 2.08E-07 | 2.72E-06 | 1638.457371 | 1268.455165 | ENSG00000184990 |
| RFFL | -0.369913785 | 0.773828739 | 0.013676218 | 0.050473983 | 246.7573096 | 190.8324665 | ENSG00000092871 |
| EMILIN3 | -0.369988037 | 0.773788913 | 0.009520101 | 0.037523674 | 266.8990595 | 206.5080671 | ENSG00000183798 |
| HLA-DMA | -0.370242586 | 0.773652398 | 0.308685427 | 0.519149631 | 40.42874938 | 31.29728646 | ENSG00000204257 |
| POLQ | -0.370359305 | 0.773589809 | 4.88E-05 | 0.000393889 | 737.1962914 | 570.3477019 | ENSG00000051341 |
| NA | -0.370461499 | 0.773535013 | 0.709482727 | 0.846496645 | 5.607903943 | 4.353425479 | ENSG00000272502 |
| NA | -0.37062403 | 0.773447874 | 0.459204948 | 0.664323996 | 22.32605296 | 17.32289954 | ENSG00000229212 |
| USP3 | -0.370669742 | 0.773423367 | 0.014179079 | 0.052065313 | 250.1278239 | 193.2641987 | ENSG00000140455 |
| DNAJC3-DT | -0.370761467 | 0.773374196 | 0.244841828 | 0.449070278 | 54.38075267 | 42.07920678 | ENSG00000247400 |
| SIX5 | -0.370777229 | 0.773365746 | 0.03740172 | 0.112760453 | 171.4151946 | 132.5302381 | ENSG00000177045 |
| TAS1R3 | -0.370999329 | 0.773246697 | 0.624611592 | 0.7923564 | 9.037707924 | 6.964019887 | ENSG00000169962 |
| GABRB3 | -0.371167959 | 0.773156321 | 2.66E-09 | 4.78E-08 | 3992.53961 | 3086.266536 | ENSG00000166206 |
| BEND6 | -0.371267524 | 0.773102965 | 0.300904564 | 0.511018736 | 44.5601706 | 34.35986497 | ENSG00000151917 |
| XDH | -0.371280646 | 0.773095933 | 0.299746134 | 0.509848409 | 42.95173422 | 33.23286288 | ENSG00000158125 |
| PPP4R1 | -0.371531255 | 0.772961651 | 1.30E-08 | 2.11E-07 | 1835.03326 | 1418.183164 | ENSG00000154845 |
| ZNF26 | -0.371646748 | 0.772899776 | 0.036017998 | 0.109585935 | 182.6704616 | 141.2841713 | ENSG00000198393 |
| SDHD | -0.371737309 | 0.77285126 | 0.013624669 | 0.050333212 | 403.0522422 | 311.7090786 | ENSG00000204370 |
| PDXDC2P-NPIPB14P | -0.371927999 | 0.772749114 | 0.392557066 | 0.604143117 | 28.8834897 | 22.32958837 | ENSG00000196696 |
| CTR9 | -0.372021039 | 0.772699281 | 1.68E-07 | 2.22E-06 | 1606.199913 | 1241.187536 | ENSG00000198730 |
| KRT10 | -0.372224138 | 0.77259051 | 0.000469872 | 0.002919888 | 573.6370164 | 443.3414233 | ENSG00000186395 |
| ZNF423 | -0.372257084 | 0.772572867 | 5.78E-07 | 6.96E-06 | 1506.959688 | 1164.360376 | ENSG00000102935 |
| ABCA17P | -0.37228074 | 0.772560199 | 0.677662606 | 0.82756234 | 7.658194534 | 5.881924521 | ENSG00000238098 |
| NA | -0.372310247 | 0.772544399 | 0.060363038 | 0.16308383 | 131.9131467 | 101.8497992 | ENSG00000228709 |
| CYB5B | -0.372568754 | 0.772405984 | 3.54E-08 | 5.32E-07 | 2753.820785 | 2127.049784 | ENSG00000103018 |
| SRSF1 | -0.3726594 | 0.772357454 | 0.000921837 | 0.005226328 | 1455.915764 | 1124.834653 | ENSG00000136450 |
| NPY5R | -0.372706277 | 0.772332359 | 0.409407879 | 0.620469288 | 26.5879533 | 20.48034781 | ENSG00000164129 |
| ALYREF | -0.37271403 | 0.772328208 | 4.20E-10 | 8.54E-09 | 2939.222035 | 2269.917988 | ENSG00000183684 |
| ALDH4A1 | -0.372862162 | 0.772248912 | 2.65E-07 | 3.42E-06 | 1495.767804 | 1155.103448 | ENSG00000159423 |
| CEP162 | -0.373567734 | 0.771871324 | 0.066864002 | 0.176358474 | 170.0707173 | 131.4043702 | ENSG00000135315 |
| SLC25A20 | -0.373607316 | 0.771850147 | 0.110773748 | 0.257821326 | 97.58543646 | 75.22088841 | ENSG00000178537 |
| GAS7 | -0.373654812 | 0.771824737 | 0.00546428 | 0.023788135 | 326.6309022 | 251.9514663 | ENSG00000007237 |
| MKS1 | -0.373722818 | 0.771788355 | 0.004895214 | 0.021680611 | 346.8177562 | 267.6020955 | ENSG00000011143 |
| SARM1 | -0.373740753 | 0.771778761 | 0.12519781 | 0.281136819 | 94.86141959 | 73.27973156 | ENSG00000004139 |
| ATAD3B | -0.373953318 | 0.771665056 | 5.70E-07 | 6.89E-06 | 2662.378418 | 2055.17555 | ENSG00000160072 |
| C9orf163 | -0.374067503 | 0.771603983 | 0.647070775 | 0.808497522 | 7.338225955 | 5.652196292 | ENSG00000196366 |
| ENOX2 | -0.374104541 | 0.771584175 | 0.002547349 | 0.012488756 | 536.6060364 | 413.7586213 | ENSG00000165675 |
| DONSON | -0.374188072 | 0.771539502 | 0.001101672 | 0.006099404 | 507.9710982 | 392.0098865 | ENSG00000159147 |
| HSD17B1 | -0.374230953 | 0.77151657 | 0.422799621 | 0.633042959 | 24.95963463 | 19.21704683 | ENSG00000108786 |
| BEST2 | -0.374502766 | 0.771371225 | 0.215539791 | 0.410702144 | 58.14743287 | 44.80150756 | ENSG00000039987 |
| LRRC61 | -0.374583542 | 0.771328037 | 0.668452335 | 0.822178087 | 7.356023318 | 5.666195032 | ENSG00000127399 |
| UBE2E3 | -0.374622057 | 0.771307446 | 3.96E-06 | 4.09E-05 | 1259.917168 | 972.2155099 | ENSG00000170035 |
| SCRN3 | -0.374724773 | 0.771252533 | 0.082575388 | 0.206794351 | 119.3023923 | 91.99570458 | ENSG00000144306 |
| CETN2 | -0.374934619 | 0.771140359 | 0.000396037 | 0.00250726 | 528.3052349 | 407.3312107 | ENSG00000147400 |
| OTULINL | -0.375061578 | 0.771072501 | 0.027828425 | 0.089621385 | 215.4277206 | 166.2931182 | ENSG00000145569 |
| IQCD | -0.375183406 | 0.77100739 | 0.425411444 | 0.635391895 | 24.646785 | 18.99356144 | ENSG00000166578 |
| FOXO6 | -0.375216396 | 0.77098976 | 0.0772371 | 0.196738958 | 131.0084125 | 101.1497372 | ENSG00000204060 |
| PDE3B | -0.375482301 | 0.770847671 | 0.006655619 | 0.028087455 | 376.7912985 | 290.546524 | ENSG00000152270 |
| NA | -0.37568219 | 0.770740876 | 0.749554777 | 0.869840852 | 4.685957196 | 3.585050781 | ENSG00000287262 |
| COPG2 | -0.375796082 | 0.770680033 | 1.00E-05 | 9.54E-05 | 1040.006328 | 801.5235447 | ENSG00000158623 |
| H4C1 | -0.375818444 | 0.770668087 | 0.690597315 | 0.834528492 | 5.594861261 | 4.321077071 | ENSG00000278637 |
| IRF2BPL | -0.375824729 | 0.770664729 | 8.84E-07 | 1.03E-05 | 2045.855695 | 1576.519212 | ENSG00000119669 |
| LRGUK | -0.375947648 | 0.770599071 | 0.325536896 | 0.536688283 | 34.41903514 | 26.50755662 | ENSG00000155530 |

| | | | | | | | |
|---|---|---|---|---|---|---|---|
| METTL16 | -0.376216627 | 0.770455412 | 8.92E-08 | 1.26E-06 | 1699.378856 | 1309.102539 | ENSG00000127804 |
| SF3A3 | -0.376248182 | 0.770438561 | 3.18E-12 | 8.57E-11 | 5066.993875 | 3903.506153 | ENSG00000183431 |
| SMIM20 | -0.37625448 | 0.770435198 | 3.35E-05 | 0.000281292 | 806.0734374 | 620.9364322 | ENSG00000250317 |
| NA | -0.376287396 | 0.77041762 | 0.327375925 | 0.538779463 | 58.05699767 | 44.82309424 | ENSG00000242622 |
| DSP | -0.376356218 | 0.770380869 | 6.70E-09 | 1.14E-07 | 2418.486079 | 1862.527829 | ENSG00000096696 |
| ACBD7 | -0.376709238 | 0.770192384 | 0.00335995 | 0.015855238 | 338.6568648 | 260.8717033 | ENSG00000176244 |
| PWWP2A | -0.377151362 | 0.769956389 | 1.09E-07 | 1.50E-06 | 1363.679204 | 1050.149882 | ENSG00000170234 |
| NAPA | -0.377156699 | 0.769953541 | 0.000825892 | 0.004743606 | 463.2427432 | 356.6873158 | ENSG00000105402 |
| DCHS1 | -0.377174301 | 0.769944147 | 2.60E-05 | 0.000225576 | 840.5213061 | 646.957837 | ENSG00000166341 |
| HSD17B7 | -0.377285712 | 0.769884691 | 6.38E-05 | 0.000498821 | 764.9967161 | 588.850078 | ENSG00000132196 |
| VASH1 | -0.377338973 | 0.769856269 | 0.001152616 | 0.006351574 | 424.9221257 | 327.0713577 | ENSG00000071246 |
| CKAP4 | -0.377515876 | 0.769761875 | 4.13E-09 | 7.22E-08 | 2682.27139 | 2065.149799 | ENSG00000136026 |
| LZTFL1 | -0.377557403 | 0.769739719 | 0.06745438 | 0.177570015 | 131.938063 | 101.4912445 | ENSG00000163818 |
| POMGNT2 | -0.377574372 | 0.769730665 | 0.000271224 | 0.001801638 | 606.6395474 | 467.1574192 | ENSG00000144647 |
| NA | -0.377626223 | 0.769703001 | 0.661821711 | 0.818441256 | 6.88185452 | 5.312653607 | ENSG00000265399 |
| NA | -0.377630357 | 0.769700795 | 0.727823136 | 0.856731016 | 4.335148574 | 3.334797422 | ENSG00000273026 |
| NA | -0.377651469 | 0.769689532 | 0.753797603 | 0.871630402 | 4.342267519 | 3.341323813 | ENSG00000269473 |
| KPTN | -0.377666275 | 0.769681632 | 0.00830304 | 0.033677601 | 290.1562375 | 223.1997415 | ENSG00000118162 |
| ZMAT2 | -0.377714494 | 0.769655908 | 6.99E-07 | 8.30E-06 | 1661.437698 | 1278.507839 | ENSG00000146007 |
| ZFYVE9 | -0.377760981 | 0.769631109 | 0.00010011 | 0.000742566 | 747.5091184 | 575.4176269 | ENSG00000157077 |
| NOX4 | -0.377880079 | 0.769567576 | 0.384822383 | 0.597077137 | 26.75372166 | 20.61163336 | ENSG00000086991 |
| NA | -0.378181524 | 0.769406795 | 0.640781423 | 0.80337285 | 10.21913837 | 7.836134152 | ENSG00000272267 |
| NA | -0.378430169 | 0.769274201 | 0.228859139 | 0.428557743 | 59.50411488 | 45.78003131 | ENSG00000287222 |
| TMEM30B | -0.378489276 | 0.769242685 | 0.001626362 | 0.008517428 | 515.9367767 | 397.0736116 | ENSG00000182107 |
| XRCC5 | -0.378546501 | 0.769212173 | 9.54E-15 | 3.47E-13 | 14161.44926 | 10892.65019 | ENSG00000079246 |
| TXN2 | -0.378612391 | 0.769177043 | 1.39E-06 | 1.56E-05 | 1108.326853 | 852.6515088 | ENSG00000100348 |
| LINC00863 | -0.378667276 | 0.769147781 | 0.160501169 | 0.334543563 | 92.99558445 | 71.68965713 | ENSG00000224914 |
| EXD2 | -0.378765634 | 0.769095345 | 0.000123362 | 0.000896726 | 677.5832818 | 520.7949402 | ENSG00000081177 |
| BHLHE40-AS1 | -0.378887705 | 0.769030272 | 0.710385576 | 0.847142936 | 6.552682065 | 5.001108401 | ENSG00000235831 |
| KEL | -0.378967111 | 0.768987946 | 0.352127604 | 0.564270666 | 32.84171814 | 25.24425564 | ENSG00000197993 |
| WRAP73 | -0.378985051 | 0.768978383 | 0.0003193 | 0.002075082 | 640.420042 | 492.5043232 | ENSG00000116213 |
| ATP6V1E2 | -0.379100384 | 0.768916912 | 0.143771114 | 0.309708159 | 78.58896359 | 60.49290354 | ENSG00000250565 |
| TASP1 | -0.379237384 | 0.768843898 | 0.045358755 | 0.130582748 | 174.9407989 | 134.3557892 | ENSG00000089123 |
| GRHL2 | -0.379248007 | 0.768838237 | 0.000247262 | 0.001661762 | 687.1367805 | 528.2372177 | ENSG00000083307 |
| NA | -0.379478511 | 0.768715407 | 0.196774248 | 0.386126524 | 66.47876886 | 51.01644838 | ENSG00000206791 |
| PSMD6 | -0.379581371 | 0.768660602 | 8.06E-11 | 1.83E-09 | 2551.643835 | 1961.342596 | ENSG00000163636 |
| U2AF2 | -0.379687283 | 0.768604175 | 5.78E-12 | 1.51E-10 | 8991.922097 | 6912.033357 | ENSG00000063244 |
| EPB41 | -0.379858599 | 0.76851291 | 2.59E-10 | 5.44E-09 | 2181.684915 | 1676.701643 | ENSG00000159023 |
| NA | -0.379878616 | 0.768502247 | 0.730222778 | 0.857920184 | 4.358869673 | 3.346620699 | ENSG00000260633 |
| STK36 | -0.380033881 | 0.768419544 | 0.00033906 | 0.002180908 | 701.8740483 | 539.7302062 | ENSG00000163482 |
| MYB | -0.38006948 | 0.768400583 | 0.123909408 | 0.278973435 | 84.15062055 | 64.70946304 | ENSG00000118513 |
| HTRA3 | -0.380311103 | 0.768271903 | 0.012132655 | 0.045962927 | 233.8251382 | 179.6217142 | ENSG00000170801 |
| EIF4EBP1 | -0.380526571 | 0.768157169 | 2.33E-09 | 4.25E-08 | 3040.3381 | 2335.618286 | ENSG00000187840 |
| ABHD4 | -0.380573582 | 0.768132138 | 2.01E-05 | 0.000178938 | 872.8013949 | 670.383971 | ENSG00000100439 |
| GJC3 | -0.380867144 | 0.767975854 | 0.488799837 | 0.6891648 | 16.83202598 | 12.93305827 | ENSG00000176402 |
| MRPL30 | -0.381112433 | 0.767845292 | 0.000222533 | 0.001511162 | 676.4330486 | 519.459345 | ENSG00000185414 |
| STX2 | -0.381128482 | 0.767836751 | 0.00042408 | 0.002660714 | 486.1458842 | 373.2282934 | ENSG00000111450 |
| ZNF175 | -0.381234202 | 0.767780486 | 0.059225771 | 0.160770258 | 143.9560169 | 110.5279903 | ENSG00000105497 |
| WASF3 | -0.381376657 | 0.767704677 | 0.000736457 | 0.004297893 | 525.4640615 | 403.0477359 | ENSG00000132970 |
| UCK1 | -0.381463474 | 0.767658481 | 7.34E-06 | 7.25E-05 | 1250.953364 | 960.2988113 | ENSG00000130717 |
| WDR62 | -0.381565802 | 0.767604034 | 1.04E-08 | 1.71E-07 | 1893.643702 | 1453.932952 | ENSG00000075702 |
| LOC100130548 | -0.381571119 | 0.767601205 | 0.723937922 | 0.854526387 | 4.320910684 | 3.308975405 | ENSG00000235138 |
| NA | -0.381602204 | 0.767584666 | 0.656161085 | 0.814219206 | 8.661982994 | 6.638475942 | ENSG00000285416 |
| FNDC4 | -0.381638076 | 0.767565581 | 0.019116614 | 0.06616885 | 209.3121125 | 160.719334 | ENSG00000115226 |
| DCAF5 | -0.382160692 | 0.76728758 | 1.79E-05 | 0.000160939 | 850.0456151 | 652.0890444 | ENSG00000139990 |
| ZNF90 | -0.382188743 | 0.767272662 | 5.28E-05 | 0.000422194 | 690.3225077 | 529.8122224 | ENSG00000213988 |
| NA | -0.382510842 | 0.767101378 | 0.591823961 | 0.769473013 | 12.08230377 | 9.215775984 | ENSG00000250539 |
| HIGD2A | -0.382553671 | 0.767078605 | 0.04399568 | 0.127628022 | 147.9222802 | 113.5519049 | ENSG00000146066 |
| UTP20 | -0.382640527 | 0.767032426 | 1.17E-06 | 1.34E-05 | 1757.944422 | 1347.837382 | ENSG00000120800 |
| CCDC22 | -0.382797398 | 0.766949027 | 0.005512809 | 0.023971772 | 289.8351255 | 222.365724 | ENSG00000101997 |
| MVB12B | -0.383213624 | 0.76672779 | 0.004072332 | 0.018657445 | 308.3301228 | 236.4745179 | ENSG00000196814 |
| LOC101929709 | -0.383277023 | 0.766694097 | 0.14467705 | 0.310833457 | 81.95027403 | 62.81531575 | ENSG00000251136 |
| ALPL | -0.383299896 | 0.766681941 | 3.73E-14 | 1.26E-12 | 5681.543552 | 4356.16716 | ENSG00000162551 |
| CDC25B | -0.383316771 | 0.766672974 | 2.88E-05 | 0.000246542 | 896.7908288 | 687.7244852 | ENSG00000101224 |
| PI4K2A | -0.383380309 | 0.766639209 | 0.000160574 | 0.001133623 | 568.6320071 | 435.8807532 | ENSG00000155252 |
| STOX1 | -0.383695651 | 0.766471657 | 0.165379235 | 0.341182746 | 75.51472292 | 57.86753238 | ENSG00000165730 |
| AXIN2 | -0.383729915 | 0.766453454 | 0.007816344 | 0.032068576 | 271.8726707 | 208.3962551 | ENSG00000168646 |
| DZIP1 | -0.383868772 | 0.766379687 | 4.06E-06 | 4.18E-05 | 1066.211717 | 816.9601028 | ENSG00000134874 |
| C16orf87 | -0.383969677 | 0.766326087 | 0.001112528 | 0.006154098 | 425.7895113 | 326.2006409 | ENSG00000155330 |
| ESR2 | -0.38423482 | 0.766185262 | 0.614137434 | 0.785269226 | 9.448722093 | 7.255039962 | ENSG00000140009 |
| LY6G5B | -0.384493076 | 0.76604812 | 0.32817126 | 0.539669465 | 32.07961602 | 24.58881682 | ENSG00000240053 |
| KHK | -0.384518519 | 0.76603461 | 0.009808971 | 0.038392212 | 274.7055299 | 210.5497365 | ENSG00000138030 |
| NA | -0.38484508 | 0.765861234 | 0.752409112 | 0.871085164 | 4.342267519 | 3.318623218 | ENSG00000261360 |
| LLGL1 | -0.384857655 | 0.765854558 | 5.95E-10 | 1.18E-08 | 3306.457397 | 2532.798383 | ENSG00000131899 |

| | | | | | | | |
|---|---|---|---|---|---|---|---|
| POLD3 | -0.384862714 | 0.765851873 | 3.95E-05 | 0.000325585 | 779.6240242 | 597.0389029 | ENSG00000077514 |
| STOML1 | -0.384984471 | 0.765787241 | 0.278648612 | 0.487770759 | 50.43020184 | 38.55109692 | ENSG00000067221 |
| NA | -0.385191304 | 0.765677461 | 0.683077575 | 0.830790283 | 8.283893801 | 6.351806794 | ENSG00000232439 |
| HACD2 | -0.385302854 | 0.765618261 | 6.52E-05 | 0.000509144 | 878.067136 | 672.1359554 | ENSG00000206527 |
| NA | -0.385428426 | 0.765551625 | 0.700816972 | 0.840633163 | 5.628065569 | 4.304902868 | ENSG00000264304 |
| AGO1 | -0.38542913 | 0.765551251 | 2.62E-09 | 4.71E-08 | 2511.217398 | 1922.090856 | ENSG00000092847 |
| SREK1 | -0.385455525 | 0.765537245 | 4.92E-05 | 0.000396109 | 707.7232814 | 541.7779702 | ENSG00000153914 |
| WDR24 | -0.385490018 | 0.765518942 | 0.000509559 | 0.003127926 | 482.3386276 | 369.3882595 | ENSG00000127580 |
| NA | -0.385549282 | 0.765487497 | 0.691312266 | 0.834876575 | 6.499010646 | 5.003283865 | ENSG00000273783 |
| AMIGO1 | -0.385668529 | 0.765424227 | 0.058842698 | 0.160029428 | 124.4574843 | 95.25644092 | ENSG00000181754 |
| ZNF578 | -0.385881023 | 0.765311496 | 0.296010831 | 0.506172088 | 38.61213646 | 29.48446169 | ENSG00000258405 |
| ESRG | -0.38600631 | 0.765245038 | 5.05E-09 | 8.71E-08 | 8044.694481 | 6156.250419 | ENSG00000265992 |
| SMYD5 | -0.386324615 | 0.765076219 | 4.54E-07 | 5.62E-06 | 1569.25469 | 1201.139218 | ENSG00000135632 |
| MSANTD2 | -0.386434409 | 0.765017996 | 0.011634526 | 0.044289061 | 276.4738109 | 211.7358979 | ENSG00000120458 |
| MDK | -0.386498916 | 0.764983791 | 4.76E-13 | 1.42E-11 | 6831.229441 | 5226.30649 | ENSG00000110492 |
| CMTR1 | -0.386622215 | 0.764918415 | 7.70E-07 | 9.07E-06 | 1303.130926 | 996.7162895 | ENSG00000137200 |
| WNT11 | -0.386623959 | 0.76491749 | 0.763480725 | 0.877522527 | 3.87758193 | 2.980310038 | ENSG00000085741 |
| CAMK2A | -0.386803523 | 0.764822291 | 0.518608626 | 0.714519874 | 24.23546534 | 18.50996398 | ENSG00000070808 |
| BEX1 | -0.386894813 | 0.764773897 | 1.60E-06 | 1.78E-05 | 1312.244525 | 1003.557324 | ENSG00000133169 |
| AK6 | -0.386928818 | 0.764755871 | 0.013907582 | 0.051187815 | 237.2682898 | 181.591152 | ENSG00000085231 |
| RNPEPL1 | -0.387213334 | 0.764605067 | 0.005333633 | 0.023328432 | 357.8280828 | 273.7573958 | ENSG00000142327 |
| EFTUD2 | -0.387244554 | 0.764588521 | 2.14E-10 | 4.55E-09 | 4506.954835 | 3445.775259 | ENSG00000108883 |
| NA | -0.387280444 | 0.7645695 | 0.579503254 | 0.760406596 | 11.36644863 | 8.684150363 | ENSG00000225689 |
| FAM221A | -0.387293668 | 0.764562492 | 0.0166401 | 0.059280947 | 231.2837456 | 176.6862382 | ENSG00000188732 |
| NA | -0.387359908 | 0.764527389 | 0.728028339 | 0.856731016 | 5.609099151 | 4.283431778 | ENSG00000284624 |
| MEGF11 | -0.387400963 | 0.764505633 | 0.138438136 | 0.301685229 | 77.68600942 | 59.40863271 | ENSG00000157890 |
| DHCR24 | -0.387481068 | 0.764463185 | 7.28E-14 | 2.37E-12 | 14087.31883 | 10769.09381 | ENSG00000116133 |
| MIGA2 | -0.387562957 | 0.764419795 | 0.000155054 | 0.001099588 | 726.9493144 | 555.7167623 | ENSG00000148343 |
| MTMR8 | -0.387587263 | 0.764406917 | 0.087084071 | 0.21509431 | 110.1081199 | 84.13325395 | ENSG00000102043 |
| DDRGK1 | -0.387686189 | 0.764354502 | 4.26E-07 | 5.30E-06 | 1293.660034 | 989.2017999 | ENSG00000198171 |
| NA | -0.387977625 | 0.764200112 | 0.552999555 | 0.739709489 | 13.47813998 | 10.32993621 | ENSG00000272853 |
| ABITRAM | -0.388021584 | 0.764176828 | 0.04742218 | 0.135425353 | 162.5340768 | 124.3381762 | ENSG00000119328 |
| IRX3 | -0.388022441 | 0.764176374 | 0.277556806 | 0.486690834 | 54.11684593 | 41.38472521 | ENSG00000177508 |
| AKIP1 | -0.388105605 | 0.764132324 | 0.002115252 | 0.010678035 | 397.6503008 | 303.6284824 | ENSG00000166452 |
| HCG11 | -0.388147935 | 0.764109904 | 0.009598447 | 0.037787509 | 303.2707788 | 231.6796591 | ENSG00000228223 |
| AP1B1 | -0.388263842 | 0.764048518 | 6.65E-11 | 1.53E-09 | 2914.604256 | 2227.446335 | ENSG00000100280 |
| SLC10A7 | -0.388268875 | 0.764045852 | 0.300410997 | 0.510506764 | 36.93637559 | 28.15986886 | ENSG00000120519 |
| RBM6 | -0.388611764 | 0.763864281 | 1.90E-09 | 3.52E-08 | 2604.953528 | 1989.877794 | ENSG00000004534 |
| PANK4 | -0.388649076 | 0.763844525 | 2.19E-06 | 2.36E-05 | 963.8898539 | 736.0475762 | ENSG00000157881 |
| NA | -0.388798942 | 0.763765182 | 0.640263192 | 0.803013745 | 8.315929053 | 6.359279144 | ENSG00000224738 |
| DIP2C | -0.388865979 | 0.763729694 | 0.000594426 | 0.00356712 | 457.005048 | 349.0651872 | ENSG00000151240 |
| NA | -0.388947641 | 0.763686465 | 0.257833917 | 0.464706387 | 43.0750944 | 32.89672516 | ENSG00000261663 |
| LINC02139 | -0.389092334 | 0.763609876 | 0.695248597 | 0.837250972 | 5.202534178 | 3.997708589 | ENSG00000278214 |
| ZNF789 | -0.389301109 | 0.76349938 | 0.142164744 | 0.30714806 | 78.40303361 | 59.92240312 | ENSG00000198556 |
| ADAM23 | -0.389410535 | 0.763441472 | 1.31E-06 | 1.49E-05 | 1020.703741 | 779.1223542 | ENSG00000114948 |
| BOD1 | -0.389441445 | 0.763425116 | 1.11E-11 | 2.80E-10 | 3387.861314 | 2585.931089 | ENSG00000145919 |
| RSPO1 | -0.3894685 | 0.763410799 | 0.736390787 | 0.861418484 | 3.908421974 | 3.001781127 | ENSG00000169218 |
| CTPS2 | -0.389507313 | 0.763390261 | 1.72E-08 | 2.73E-07 | 2152.573646 | 1643.011182 | ENSG00000047230 |
| C2CD3 | -0.389541591 | 0.763372124 | 0.000505078 | 0.003104948 | 544.6418393 | 415.8471652 | ENSG00000168014 |
| FAHD2A | -0.389558531 | 0.76336316 | 0.0001203 | 0.00087683 | 706.0231896 | 538.9145384 | ENSG00000115042 |
| RND1 | -0.3899165 | 0.763173774 | 0.017500094 | 0.061730031 | 224.5852292 | 171.5208334 | ENSG00000172602 |
| EFHC1 | -0.390115537 | 0.763068492 | 0.016083604 | 0.057712113 | 227.1653932 | 173.1663357 | ENSG00000096093 |
| CCDC81 | -0.390278979 | 0.76298205 | 0.584179232 | 0.763106094 | 12.17126443 | 9.218897406 | ENSG00000149201 |
| ZNF763 | -0.390547534 | 0.762840035 | 0.657201693 | 0.81497498 | 7.832277047 | 6.003562254 | ENSG00000197054 |
| POLR1A | -0.391101888 | 0.762546971 | 1.66E-07 | 2.20E-06 | 2839.209536 | 2164.605883 | ENSG00000068654 |
| ADAMTS15 | -0.391107487 | 0.762544012 | 0.013104401 | 0.048764483 | 253.2421341 | 193.1817193 | ENSG00000166106 |
| TARBP2 | -0.391213407 | 0.762488029 | 2.03E-07 | 2.66E-06 | 1391.519445 | 1061.381821 | ENSG00000139546 |
| SPHK1 | -0.391274892 | 0.762455534 | 0.255322413 | 0.462164648 | 44.35972339 | 33.80241734 | ENSG00000176170 |
| LRRC23 | -0.391280415 | 0.762452615 | 0.475996442 | 0.678429646 | 20.52159299 | 15.58129797 | ENSG00000010626 |
| RCAN1 | -0.391287347 | 0.762448951 | 0.024915643 | 0.082272285 | 179.5508381 | 136.9138365 | ENSG00000159200 |
| GOLGA6L4 | -0.39142547 | 0.762375959 | 0.571144866 | 0.753848952 | 11.78458175 | 9.031165398 | ENSG00000184206 |
| RAN | -0.391558556 | 0.762305634 | 1.94E-20 | 1.19E-18 | 20006.71862 | 15250.66532 | ENSG00000132341 |
| SIRPA | -0.391617125 | 0.762274688 | 0.007255394 | 0.030127115 | 267.1822381 | 203.5772983 | ENSG00000198053 |
| ANKHD1-EIF4EBP3 | -0.39167101 | 0.762246217 | 0.680533276 | 0.829255013 | 6.46936581 | 4.951639832 | ENSG00000254996 |
| YTHDF3-AS1 | -0.39172185 | 0.762219356 | 0.455688433 | 0.661567962 | 17.84293339 | 13.60561725 | ENSG00000270673 |
| TBC1D4 | -0.391855907 | 0.762148533 | 5.23E-07 | 6.39E-06 | 1238.455194 | 944.0922771 | ENSG00000136111 |
| ARMH4 | -0.391929983 | 0.762109401 | 0.006159154 | 0.02631575 | 304.67971 | 232.2568015 | ENSG00000139971 |
| CENPU | -0.392027227 | 0.762058033 | 4.35E-07 | 5.40E-06 | 1224.377719 | 933.4051742 | ENSG00000151725 |
| CDKN2AIPNL | -0.392073651 | 0.762033511 | 7.29E-07 | 8.63E-06 | 1290.900598 | 983.7821268 | ENSG00000237190 |
| NA | -0.392116814 | 0.762010713 | 0.657600363 | 0.81515086 | 6.9186183 | 5.263185037 | ENSG00000259863 |
| NA | -0.392388239 | 0.761867364 | 0.633561939 | 0.798559516 | 8.237646812 | 6.273111239 | ENSG00000273004 |
| NA | -0.392463473 | 0.761827635 | 0.578540959 | 0.759655996 | 10.83535443 | 8.260615237 | ENSG00000233221 |
| SETD9 | -0.392466638 | 0.761825964 | 0.078335357 | 0.198705603 | 121.856802 | 92.73759485 | ENSG00000155542 |
| ZNF518B | -0.392650911 | 0.761728663 | 0.002621682 | 0.012809907 | 413.5324891 | 314.7273604 | ENSG00000178163 |

| | | | | | | | |
|---|---|---|---|---|---|---|---|
| TAF6L | -0.392711673 | 0.761696582 | 0.001464923 | 0.007802347 | 356.8212674 | 271.7472639 | ENSG00000162227 |
| RBM39 | -0.393128466 | 0.76147656 | 2.77E-15 | 1.09E-13 | 4892.818005 | 3725.986988 | ENSG00000131051 |
| PXK | -0.393312475 | 0.761379444 | 0.034556241 | 0.106182005 | 185.3331039 | 140.9223795 | ENSG00000168297 |
| MTRNR2L2 | -0.393555069 | 0.761251426 | 0.351593101 | 0.563749 | 30.87774461 | 23.54654221 | ENSG00000269028 |
| GPRC5C | -0.393789673 | 0.761127645 | 0.009086881 | 0.036203745 | 308.0128234 | 234.4792151 | ENSG00000170412 |
| COQ4 | -0.394063213 | 0.760983346 | 0.000203242 | 0.001398769 | 560.095748 | 426.4464099 | ENSG00000167113 |
| NA | -0.394101848 | 0.760962968 | 0.703366206 | 0.842499163 | 5.649422404 | 4.286553201 | ENSG00000248559 |
| TPD52L2 | -0.394176395 | 0.760923648 | 5.43E-12 | 1.42E-10 | 2910.329288 | 2214.866535 | ENSG00000101150 |
| THBS4 | -0.394183777 | 0.760919755 | 0.001434235 | 0.00766644 | 419.6739294 | 319.309212 | ENSG00000113296 |
| HK2 | -0.39451185 | 0.760746739 | 3.12E-06 | 3.28E-05 | 2443.814696 | 1858.408826 | ENSG00000159399 |
| NOL8 | -0.39457994 | 0.760710836 | 0.00028688 | 0.001891004 | 578.9808128 | 440.543832 | ENSG00000198000 |
| DOP1A | -0.39465062 | 0.760673568 | 0.021848388 | 0.073930624 | 222.058607 | 168.7512396 | ENSG00000083097 |
| RAB34 | -0.394666692 | 0.760665094 | 1.01E-12 | 2.89E-11 | 3408.845487 | 2593.316562 | ENSG00000109113 |
| IL17RD | -0.394753798 | 0.760619168 | 2.12E-08 | 3.30E-07 | 2201.594996 | 1673.973633 | ENSG00000144730 |
| ABLIM3 | -0.394803297 | 0.760593072 | 0.449442532 | 0.65582771 | 19.69542037 | 14.94855975 | ENSG00000173210 |
| ARF3 | -0.394963344 | 0.760508699 | 2.35E-11 | 5.70E-10 | 4376.657845 | 3328.134998 | ENSG00000134287 |
| WDR73 | -0.394982439 | 0.760498634 | 0.076133067 | 0.19479379 | 119.8862938 | 91.19498147 | ENSG00000177082 |
| CPEB3 | -0.395007835 | 0.760485247 | 0.404077895 | 0.615208962 | 24.48902531 | 18.64002002 | ENSG00000107864 |
| MPPED2 | -0.395115691 | 0.760428395 | 0.139174125 | 0.302913686 | 85.05093113 | 64.82580388 | ENSG00000066382 |
| BANP | -0.395442184 | 0.760256323 | 0.000197442 | 0.001364323 | 684.1068766 | 520.2390485 | ENSG00000172530 |
| LINC02570 | -0.395462814 | 0.760245452 | 0.017889649 | 0.062855074 | 197.392297 | 150.1394279 | ENSG00000237923 |
| SOX2-OT | -0.39549253 | 0.760229793 | 0.299026469 | 0.509345619 | 35.34810083 | 26.88692006 | ENSG00000242808 |
| COL14A1 | -0.395513668 | 0.760218655 | 1.10E-05 | 0.000103649 | 987.0141886 | 749.9279964 | ENSG00000187955 |
| MYL6B | -0.395516088 | 0.760217379 | 3.07E-06 | 3.23E-05 | 1208.949942 | 918.7733409 | ENSG00000196465 |
| TRMT11 | -0.395575374 | 0.76018614 | 0.001508518 | 0.008000103 | 407.4756893 | 309.9278148 | ENSG00000066651 |
| ALDH3A2 | -0.395861653 | 0.760035308 | 1.09E-08 | 1.79E-07 | 1990.372736 | 1512.606092 | ENSG00000072210 |
| NA | -0.395877742 | 0.760026832 | 0.751920951 | 0.870846753 | 3.90605771 | 2.978134574 | ENSG00000213609 |
| PCAT1 | -0.395934368 | 0.759997002 | 0.421962001 | 0.632089504 | 21.43494626 | 16.32820158 | ENSG00000253438 |
| RIMS2 | -0.396002515 | 0.759961104 | 0.001003255 | 0.005630392 | 429.2964024 | 326.2231735 | ENSG00000176406 |
| BEND7 | -0.396144696 | 0.759886211 | 0.051211214 | 0.14374861 | 130.0591852 | 98.89436524 | ENSG00000165626 |
| ESYT1 | -0.396331318 | 0.759787921 | 5.23E-10 | 1.05E-08 | 2465.461541 | 1873.326366 | ENSG00000139641 |
| NA | -0.396348489 | 0.759778878 | 0.72297323 | 0.854135093 | 3.915540919 | 2.987782387 | ENSG00000250859 |
| SUSD4 | -0.396361383 | 0.759772088 | 0.135322811 | 0.297270462 | 79.62417691 | 60.38308908 | ENSG00000143502 |
| AOC2 | -0.396452314 | 0.759724202 | 0.217066066 | 0.412841907 | 63.08931429 | 47.93116927 | ENSG00000131480 |
| SLC38A7 | -0.396482722 | 0.759708189 | 4.37E-05 | 0.000356944 | 722.9421425 | 549.3340772 | ENSG00000103042 |
| NA | -0.396649339 | 0.759620456 | 0.531387211 | 0.723253105 | 12.22435103 | 9.288891107 | ENSG00000272716 |
| C4orf46 | -0.396708957 | 0.759589065 | 0.000196463 | 0.001358057 | 503.5680409 | 382.4561762 | ENSG00000205208 |
| ZNF710-AS1 | -0.396716036 | 0.759585338 | 0.653717736 | 0.812522011 | 6.970815179 | 5.281534704 | ENSG00000259291 |
| GRK4 | -0.396956029 | 0.759458991 | 0.344466042 | 0.556431546 | 30.88722782 | 23.47342708 | ENSG00000125388 |
| ATP1A1 | -0.39698245 | 0.759445083 | 1.31E-12 | 3.70E-11 | 6972.541934 | 5294.783552 | ENSG00000163399 |
| MID1 | -0.397103359 | 0.759381439 | 5.16E-07 | 6.30E-06 | 1253.300565 | 951.337363 | ENSG00000101871 |
| MSANTD1 | -0.397277787 | 0.759289631 | 0.636219255 | 0.80067977 | 7.018257377 | 5.31482907 | ENSG00000188981 |
| MGAT4B | -0.39743503 | 0.759206879 | 1.81E-09 | 3.35E-08 | 1841.215049 | 1398.077852 | ENSG00000161013 |
| CDC45 | -0.397501597 | 0.75917185 | 0.000371682 | 0.002369756 | 592.810216 | 449.9638407 | ENSG00000093009 |
| NA | -0.39750791 | 0.759168528 | 0.676682988 | 0.827022925 | 5.731264118 | 4.335075811 | ENSG00000224550 |
| TSC22D1 | -0.397529622 | 0.759157103 | 7.14E-10 | 1.40E-08 | 2027.645299 | 1539.189572 | ENSG00000102804 |
| DCTN5 | -0.397540098 | 0.75915159 | 2.66E-09 | 4.78E-08 | 1837.604908 | 1394.780183 | ENSG00000166847 |
| FRMD4B | -0.397898839 | 0.758962843 | 0.035208542 | 0.107748065 | 152.3739493 | 115.6697485 | ENSG00000114541 |
| ASCC2 | -0.397917252 | 0.758953156 | 3.26E-06 | 3.41E-05 | 936.6797341 | 711.0623187 | ENSG00000100325 |
| TMEM158 | -0.398210754 | 0.758798771 | 0.035398988 | 0.108229823 | 331.5527777 | 251.8382898 | ENSG00000249992 |
| ZNF404 | -0.39833872 | 0.758731469 | 0.423238967 | 0.633337985 | 22.36548649 | 16.93171282 | ENSG00000176222 |
| DENND2A | -0.398366544 | 0.758716835 | 0.0017986 | 0.00926529 | 379.136973 | 287.7621963 | ENSG00000146966 |
| MT1G | -0.398382726 | 0.758708325 | 0.000732125 | 0.004276582 | 450.6551251 | 342.0328976 | ENSG00000125144 |
| MTDH | -0.398604731 | 0.758591583 | 5.97E-05 | 0.000471219 | 1632.02607 | 1237.788079 | ENSG00000147649 |
| PUS10 | -0.398698074 | 0.758542503 | 0.282632561 | 0.491655782 | 38.90126499 | 29.43716858 | ENSG00000162927 |
| SRSF8 | -0.398822433 | 0.75847712 | 0.000873504 | 0.004971708 | 519.9383815 | 394.7144274 | ENSG00000263465 |
| PCDHB8 | -0.398953041 | 0.758408458 | 0.583435913 | 0.762833684 | 9.612405521 | 7.278686515 | ENSG00000120322 |
| UBBP4 | -0.399028148 | 0.758368976 | 0.039697566 | 0.118042598 | 171.8134455 | 130.2771572 | ENSG00000263563 |
| NA | -0.399038609 | 0.758363477 | 0.671175327 | 0.82390724 | 5.703983547 | 4.304902868 | ENSG00000251050 |
| PPP1R32 | -0.399066356 | 0.758348892 | 0.461446145 | 0.666239331 | 17.99357413 | 13.62491288 | ENSG00000162148 |
| RBM4 | -0.399107422 | 0.758327306 | 0.10907096 | 0.254642735 | 111.0861285 | 84.28366717 | ENSG00000173933 |
| TSN | -0.399236725 | 0.758259343 | 2.76E-10 | 5.77E-09 | 2513.530095 | 1905.744843 | ENSG00000211460 |
| KLHL18 | -0.399269178 | 0.758242287 | 1.31E-05 | 0.000121367 | 855.5182847 | 648.5783679 | ENSG00000114648 |
| RBBP8NL | -0.399281947 | 0.758235575 | 0.515657523 | 0.711976214 | 14.42973156 | 10.97449771 | ENSG00000130701 |
| SCLT1 | -0.39929546 | 0.758228474 | 0.01901037 | 0.065885638 | 191.6657614 | 145.384222 | ENSG00000151466 |
| GET4 | -0.399409807 | 0.758168379 | 0.012776534 | 0.047807579 | 210.1429875 | 159.3266393 | ENSG00000239857 |
| MAN2C1 | -0.39966433 | 0.758034634 | 0.000420447 | 0.002641433 | 762.3003643 | 577.9136903 | ENSG00000140400 |
| MAFA | -0.399691289 | 0.758020469 | 0.102461913 | 0.24242186 | 86.19735167 | 65.37332016 | ENSG00000182759 |
| KHDC1 | -0.399927675 | 0.757896277 | 0.077936609 | 0.19807902 | 156.0905176 | 118.3950027 | ENSG00000135314 |
| CASTOR2 | -0.399972172 | 0.757872902 | 0.00066573 | 0.003931893 | 404.8758967 | 306.821684 | ENSG00000274070 |
| CERS5 | -0.400051473 | 0.757831245 | 8.12E-06 | 7.94E-05 | 816.7735586 | 618.850034 | ENSG00000139624 |
| SH3RF3 | -0.400075103 | 0.757818832 | 0.254223719 | 0.46052901 | 41.59594193 | 31.55066124 | ENSG00000172985 |
| USP7 | -0.400480951 | 0.757605678 | 2.28E-13 | 7.06E-12 | 5387.285515 | 4081.108027 | ENSG00000187555 |
| NA | -0.400615244 | 0.75753516 | 0.717994672 | 0.851182007 | 3.946380963 | 2.999605664 | ENSG00000272505 |

| | | | | | | | |
|---|---|---|---|---|---|---|---|
| NA | -0.400875767 | 0.757398376 | 0.717807619 | 0.851146028 | 3.949940435 | 3.001781127 | ENSG00000258427 |
| PSKH2 | -0.400961909 | 0.757353154 | 0.67139986 | 0.823925294 | 5.731264118 | 4.318901608 | ENSG00000147613 |
| CA11 | -0.400991671 | 0.75733753 | 3.15E-06 | 3.30E-05 | 1049.207387 | 794.8859397 | ENSG00000063180 |
| CCDC28A | -0.401300678 | 0.757175335 | 0.029729006 | 0.094231667 | 170.2459171 | 128.7388947 | ENSG00000024862 |
| C15orf62 | -0.4014285 | 0.757108253 | 0.458520021 | 0.663948476 | 17.93545352 | 13.5443254 | ENSG00000188277 |
| PRKCE | -0.401438625 | 0.757102939 | 0.311679157 | 0.522197973 | 33.23732536 | 25.19355756 | ENSG00000171132 |
| CCDC124 | -0.40151003 | 0.757065468 | 5.12E-08 | 7.49E-07 | 2733.162302 | 2069.208255 | ENSG00000007080 |
| PA2G4 | -0.401596583 | 0.757020051 | 4.75E-13 | 1.42E-11 | 3065.165127 | 2320.364998 | ENSG00000170515 |
| HEY2 | -0.40184306 | 0.756890728 | 0.021167245 | 0.072055023 | 182.7549731 | 138.3614646 | ENSG00000135547 |
| TPGS2 | -0.401862722 | 0.756880413 | 4.49E-11 | 1.06E-09 | 2422.347824 | 1833.495183 | ENSG00000134779 |
| NA | -0.402348581 | 0.75662556 | 0.736536845 | 0.861482616 | 4.824444984 | 3.679636993 | ENSG00000287574 |
| CBX1 | -0.402414354 | 0.756591066 | 3.16E-18 | 1.61E-16 | 7465.888952 | 5648.569649 | ENSG00000108468 |
| ATP10A | -0.40243192 | 0.756581854 | 0.001844624 | 0.009458665 | 354.3341566 | 268.0307596 | ENSG00000206190 |
| ATP6V1F | -0.402542193 | 0.756524027 | 0.002066695 | 0.010469296 | 360.7964552 | 273.0991918 | ENSG00000128524 |
| NA | -0.402645697 | 0.756469753 | 0.71869436 | 0.851423561 | 3.94282149 | 2.989957851 | ENSG00000269978 |
| TSPAN33 | -0.402717429 | 0.756432142 | 7.68E-06 | 7.56E-05 | 965.7780653 | 730.797276 | ENSG00000158457 |
| LMAN2L | -0.402936326 | 0.756317378 | 3.35E-05 | 0.000281292 | 692.0186316 | 523.2562259 | ENSG00000114988 |
| DEPTOR | -0.40298542 | 0.756291642 | 0.312312125 | 0.522595648 | 36.10694897 | 27.24575837 | ENSG00000155792 |
| TP53TG5 | -0.403267981 | 0.756143532 | 0.621121622 | 0.78999743 | 8.346769097 | 6.292406865 | ENSG00000124251 |
| ZKSCAN5 | -0.403308645 | 0.756122219 | 4.66E-05 | 0.000377864 | 669.7317585 | 506.2625158 | ENSG00000196652 |
| LRFN3 | -0.403564616 | 0.755988075 | 0.001410166 | 0.007557034 | 423.2633504 | 320.0949474 | ENSG00000126243 |
| PEX3 | -0.403694522 | 0.755920007 | 0.020227915 | 0.069442678 | 198.6051517 | 150.2827047 | ENSG00000034693 |
| CTDNEP1 | -0.403747158 | 0.755892428 | 0.000242178 | 0.001631071 | 600.9162135 | 454.452009 | ENSG00000175826 |
| NA | -0.4037485 | 0.755891725 | 0.405929186 | 0.616979693 | 23.38707231 | 17.65279441 | ENSG00000259556 |
| CDKN1C | -0.403887437 | 0.755818933 | 0.005876418 | 0.025279308 | 306.9909582 | 232.2450736 | ENSG00000129757 |
| ANXA11 | -0.403922347 | 0.755800644 | 2.34E-10 | 4.93E-09 | 1877.431399 | 1419.000441 | ENSG00000122359 |
| HTRA1 | -0.404115775 | 0.755699318 | 0.00015515 | 0.001099852 | 642.9089846 | 485.7010861 | ENSG00000166033 |
| NA | -0.404283112 | 0.755611669 | 0.286368169 | 0.495561521 | 37.00161515 | 27.91396643 | ENSG00000280018 |
| RB1CC1 | -0.404348649 | 0.755577346 | 2.17E-06 | 2.35E-05 | 900.3774243 | 680.3953841 | ENSG00000023287 |
| KCTD17 | -0.404438063 | 0.755530519 | 4.14E-05 | 0.000339383 | 896.8476234 | 677.8938918 | ENSG00000100379 |
| NA | -0.404542956 | 0.755475589 | 0.677110483 | 0.827271129 | 5.295054309 | 4.009531866 | ENSG00000269846 |
| MACROH2A1 | -0.404688132 | 0.75539957 | 4.72E-10 | 9.54E-09 | 2739.985555 | 2069.897838 | ENSG00000113648 |
| FSIP2 | -0.404927726 | 0.755274129 | 0.517158325 | 0.713270264 | 13.64418767 | 10.2891695 | ENSG00000188738 |
| ZNF205 | -0.40507559 | 0.755196723 | 0.003404815 | 0.016042895 | 355.3976175 | 268.6054953 | ENSG00000122386 |
| PIGF | -0.405270173 | 0.755094873 | 0.094178484 | 0.228252342 | 88.42436837 | 66.77566259 | ENSG00000151665 |
| NA | -0.405323353 | 0.75506704 | 0.394921565 | 0.606694359 | 24.10737664 | 18.19312189 | ENSG00000260051 |
| SRRM1 | -0.405366649 | 0.75504438 | 6.95E-11 | 1.59E-09 | 3201.740997 | 2417.871178 | ENSG00000133226 |
| EXOC3L2 | -0.405781499 | 0.754827297 | 0.200990989 | 0.391876403 | 51.0200532 | 38.49850692 | ENSG00000283632 |
| CASP9 | -0.406014035 | 0.754705642 | 0.000316453 | 0.002060051 | 471.9098126 | 355.9893863 | ENSG00000132906 |
| ZNF136 | -0.406046081 | 0.754688879 | 0.06152968 | 0.165475583 | 116.8455367 | 88.1247197 | ENSG00000196646 |
| NA | -0.406315112 | 0.754548159 | 0.556949963 | 0.743258428 | 17.24659962 | 13.08174753 | ENSG00000251188 |
| IFT46 | -0.406435359 | 0.754485271 | 0.008663743 | 0.034893321 | 232.7030236 | 175.4557371 | ENSG00000118096 |
| SMPDL3A | -0.406688365 | 0.754352968 | 0.316537421 | 0.526632824 | 33.45437473 | 25.23243236 | ENSG00000172594 |
| NA | -0.406726793 | 0.754332875 | 0.214741299 | 0.409718068 | 47.87581829 | 36.11764074 | ENSG00000238123 |
| NA | -0.406783587 | 0.75430318 | 0.639117841 | 0.802206613 | 6.605768667 | 4.989285125 | ENSG00000214784 |
| OLFM2 | -0.406903818 | 0.754240321 | 7.53E-11 | 1.71E-09 | 2051.909519 | 1547.770382 | ENSG00000105088 |
| HMGN2 | -0.407023324 | 0.754177845 | 5.41E-11 | 1.26E-09 | 6555.976789 | 4944.625489 | ENSG00000198830 |
| HRAS | -0.407086351 | 0.754144898 | 1.36E-05 | 0.000125344 | 791.5865267 | 597.2787937 | ENSG00000174775 |
| NA | -0.40721954 | 0.754075279 | 0.566715868 | 0.750531324 | 11.04648005 | 8.307908343 | ENSG00000286389 |
| CTC-338M12.4 | -0.407333596 | 0.754015666 | 0.279374062 | 0.488473282 | 41.56777164 | 31.34987645 | ENSG00000233937 |
| EIF3C | -0.407344962 | 0.754009726 | 1.19E-07 | 1.62E-06 | 1205.667629 | 908.7958846 | ENSG00000184110 |
| ELMOD3 | -0.40740049 | 0.753980705 | 0.028450103 | 0.091018159 | 175.5938049 | 132.3304719 | ENSG00000115459 |
| PHTF1 | -0.407583663 | 0.753884982 | 5.40E-05 | 0.000431089 | 614.6203024 | 463.1752021 | ENSG00000116793 |
| PPP1R13L | -0.408001467 | 0.753666688 | 0.002655637 | 0.012959094 | 318.2355218 | 239.8756205 | ENSG00000104881 |
| NAP1L5 | -0.408140139 | 0.75359425 | 0.232033719 | 0.432828604 | 46.57578235 | 35.08624345 | ENSG00000177432 |
| MAP9 | -0.408214377 | 0.753555472 | 0.064224029 | 0.170754722 | 137.0454073 | 103.2357565 | ENSG00000164114 |
| LRRC1 | -0.408258375 | 0.753532491 | 0.005259038 | 0.023054068 | 289.6444152 | 218.0682935 | ENSG00000137269 |
| RIBC1 | -0.408310803 | 0.753505108 | 0.487024236 | 0.687968636 | 14.97981833 | 11.2602209 | ENSG00000158423 |
| AKAP17A | -0.408373663 | 0.753472277 | 5.11E-10 | 1.03E-08 | 2290.786628 | 1726.55169 | ENSG00000197976 |
| GNB3 | -0.408670374 | 0.753317331 | 0.065351666 | 0.173187274 | 134.0606859 | 100.9406068 | ENSG00000111664 |
| SPRYD4 | -0.408757024 | 0.753272087 | 0.01688668 | 0.059986909 | 189.7014568 | 142.8592284 | ENSG00000176422 |
| PEX2 | -0.408868904 | 0.753213674 | 0.000506157 | 0.003109554 | 458.8924671 | 345.4338619 | ENSG00000164751 |
| MLLT6 | -0.408871434 | 0.753212353 | 0.205601359 | 0.397732391 | 386.074279 | 290.8638738 | ENSG00000275023 |
| ZBTB8OS | -0.40892896 | 0.75318232 | 0.003047478 | 0.014607009 | 450.4680005 | 339.3071394 | ENSG00000176261 |
| SNRNP200 | -0.409099602 | 0.753093239 | 1.04E-18 | 5.63E-17 | 12385.08878 | 9326.725329 | ENSG00000144028 |
| TBC1D32 | -0.409208084 | 0.753036613 | 0.119971054 | 0.272641801 | 77.99794317 | 58.65955363 | ENSG00000146350 |
| NAT2 | -0.409216209 | 0.753032372 | 0.393813765 | 0.605435767 | 25.94417734 | 19.57030471 | ENSG00000156006 |
| CIB1 | -0.409562445 | 0.752851672 | 0.012907527 | 0.048240382 | 227.8284667 | 171.5872542 | ENSG00000185043 |
| GRIN3B | -0.409676688 | 0.752792058 | 0.268178278 | 0.47603598 | 46.70242266 | 35.21724545 | ENSG00000116032 |
| SIPA1L1 | -0.409859311 | 0.752696772 | 3.60E-09 | 6.32E-08 | 1641.617066 | 1235.888945 | ENSG00000197555 |
| RANBP1 | -0.409984421 | 0.752631501 | 1.68E-14 | 5.85E-13 | 7987.148163 | 6011.404727 | ENSG00000099901 |
| ADAMTS9 | -0.410274853 | 0.752480002 | 0.398933651 | 0.610475104 | 25.99634807 | 19.58960033 | ENSG00000163638 |
| TMEM199 | -0.410346629 | 0.752442566 | 0.112213239 | 0.260113786 | 88.66633403 | 66.63595873 | ENSG00000244045 |
| B3GAT2 | -0.410351259 | 0.752440152 | 0.386147351 | 0.598325626 | 26.78308717 | 20.20521839 | ENSG00000112309 |

| | | | | | | | |
|---|---|---|---|---|---|---|---|
| ZNF277 | -0.410447166 | 0.752390133 | 0.010932786 | 0.042075741 | 305.0239839 | 229.5424246 | ENSG00000198839 |
| NA | -0.410735898 | 0.752239569 | 0.372064224 | 0.583930264 | 26.90525213 | 20.30008815 | ENSG00000260285 |
| SERTAD4 | -0.410876157 | 0.75216644 | 0.182960279 | 0.366755639 | 57.08076911 | 42.94216769 | ENSG00000082497 |
| ITGB8 | -0.411427911 | 0.751878831 | 0.266050121 | 0.473459908 | 43.30401739 | 32.49834964 | ENSG00000105855 |
| CCND3 | -0.411523577 | 0.751828976 | 1.91E-05 | 0.000171076 | 877.1252018 | 659.6887357 | ENSG00000112576 |
| MLLT11 | -0.411832911 | 0.75166779 | 0.001226404 | 0.006691893 | 422.8235811 | 318.0126892 | ENSG00000213190 |
| ANAPC10 | -0.411850427 | 0.751658664 | 0.442984557 | 0.650475342 | 21.48803286 | 16.19474057 | ENSG00000164162 |
| PHC2 | -0.412056326 | 0.751551396 | 0.00142935 | 0.007644654 | 336.2142732 | 252.6762364 | ENSG00000134686 |
| FADS2 | -0.412193605 | 0.751479886 | 2.80E-13 | 8.60E-12 | 11069.90823 | 8319.310555 | ENSG00000134824 |
| STX18 | -0.412376241 | 0.75138476 | 4.77E-05 | 0.000385707 | 593.4525953 | 445.8208048 | ENSG00000168818 |
| NA | -0.412881954 | 0.75112142 | 0.240356737 | 0.443881966 | 46.46666006 | 34.97909892 | ENSG00000239569 |
| MFSD11 | -0.412911156 | 0.751106217 | 0.016134821 | 0.057873903 | 181.5702887 | 136.2946026 | ENSG00000092931 |
| SCLY | -0.413377113 | 0.750863666 | 0.073131475 | 0.189319795 | 112.8704007 | 84.87658464 | ENSG00000132330 |
| LINC00665 | -0.413390885 | 0.750856498 | 3.44E-08 | 5.19E-07 | 1917.50797 | 1440.237353 | ENSG00000232677 |
| TTC33 | -0.413454714 | 0.750823279 | 0.020132663 | 0.06916597 | 195.8063865 | 146.9853417 | ENSG00000113638 |
| VIPR1 | -0.413554317 | 0.750771444 | 0.129674218 | 0.288376048 | 80.60724508 | 60.60025898 | ENSG00000114812 |
| NA | -0.413615408 | 0.750739653 | 0.577727191 | 0.759450456 | 9.330395931 | 7.00166518 | ENSG00000260293 |
| TTC9C | -0.413684557 | 0.750703671 | 0.000436113 | 0.002729863 | 442.9023261 | 332.3098776 | ENSG00000162222 |
| H4C5 | -0.413715986 | 0.750687317 | 0.416167376 | 0.627090153 | 19.43921681 | 14.61554345 | ENSG00000276966 |
| ZNF846 | -0.413938135 | 0.750571734 | 0.115615843 | 0.265826622 | 77.32569143 | 58.05074551 | ENSG00000196605 |
| TAF11 | -0.413983042 | 0.750548371 | 0.000171613 | 0.001201857 | 478.9571038 | 359.4638876 | ENSG00000064995 |
| CTAGE4 | -0.414207471 | 0.750431623 | 0.584469841 | 0.763221749 | 9.746444113 | 7.276511051 | ENSG00000225932 |
| P2RY2 | -0.414246737 | 0.750411199 | 0.164194259 | 0.339628541 | 62.75154835 | 47.01155098 | ENSG00000175591 |
| GHITM | -0.414375434 | 0.75034426 | 9.82E-11 | 2.20E-09 | 2772.772766 | 2080.227845 | ENSG00000165678 |
| TSSC2 | -0.414398371 | 0.750332331 | 9.07E-09 | 1.50E-07 | 1888.916708 | 1417.203971 | ENSG00000223756 |
| ZNF571-AS1 | -0.414476406 | 0.750291747 | 0.670444686 | 0.82334165 | 5.329453825 | 3.995533126 | ENSG00000267470 |
| TMEM92-AS1 | -0.41469766 | 0.75017669 | 0.712456177 | 0.848130037 | 3.984339952 | 2.985606923 | ENSG00000251179 |
| ZNF573 | -0.414894282 | 0.750074457 | 0.254323201 | 0.460665035 | 39.47952206 | 29.6179227 | ENSG00000189144 |
| STBD1 | -0.414949363 | 0.75004582 | 0.459535486 | 0.664598534 | 17.67449528 | 13.30371986 | ENSG00000118804 |
| NA | -0.414988772 | 0.750025332 | 0.482310552 | 0.683662446 | 18.26965999 | 13.72290406 | ENSG00000261094 |
| ATP2B1 | -0.415019787 | 0.750009208 | 7.03E-05 | 0.000542839 | 875.7175704 | 656.6569566 | ENSG00000070961 |
| ENOSF1 | -0.415159375 | 0.749936644 | 0.001246446 | 0.006784375 | 446.0445278 | 334.6434936 | ENSG00000132199 |
| NA | -0.415210187 | 0.749910232 | 2.47E-06 | 2.65E-05 | 931.9350465 | 698.8371213 | ENSG00000179899 |
| DAP | -0.415458956 | 0.749780933 | 1.36E-06 | 1.53E-05 | 1029.802849 | 772.4575524 | ENSG00000112977 |
| FBXL17 | -0.415662864 | 0.749674968 | 0.099298692 | 0.236959123 | 84.61202599 | 63.38174878 | ENSG00000145743 |
| UGT3A1 | -0.415668245 | 0.749672172 | 0.16141233 | 0.335971768 | 67.47574387 | 50.47640445 | ENSG00000145626 |
| YAP1 | -0.415786282 | 0.749610838 | 1.09E-08 | 1.79E-07 | 2229.767052 | 1671.28141 | ENSG00000137693 |
| DOCK8 | -0.415818285 | 0.74959421 | 0.271887194 | 0.4805251 | 43.51753343 | 32.53571138 | ENSG00000107099 |
| NA | -0.415894062 | 0.749554839 | 0.431191063 | 0.640932171 | 18.65011344 | 13.96133414 | ENSG00000277182 |
| NA | -0.416026314 | 0.74948613 | 0.503194831 | 0.701160952 | 15.55571114 | 11.67628368 | ENSG00000236969 |
| ST7 | -0.416069926 | 0.749463474 | 0.012585446 | 0.047270516 | 210.5626213 | 157.709586 | ENSG00000004866 |
| NA | -0.416094704 | 0.749450603 | 0.602976865 | 0.777868577 | 9.285317997 | 6.948791642 | ENSG00000258317 |
| BBOX1-AS1 | -0.416382232 | 0.749301252 | 0.197505379 | 0.387118741 | 55.01507157 | 41.23045552 | ENSG00000254560 |
| NA | -0.416510262 | 0.74923476 | 0.633448072 | 0.798522585 | 7.579912292 | 5.703840325 | ENSG00000244586 |
| CPNE1 | -0.416568309 | 0.749204615 | 1.70E-08 | 2.71E-07 | 1361.345937 | 1019.662966 | ENSG00000214078 |
| TOX | -0.416980099 | 0.748990799 | 0.007624477 | 0.031438484 | 246.5464894 | 184.5412891 | ENSG00000198846 |
| ZNF530 | -0.41720113 | 0.748876057 | 0.030874571 | 0.097129148 | 157.9166137 | 118.2871528 | ENSG00000183647 |
| SERPINH1 | -0.417271971 | 0.748839286 | 2.16E-19 | 1.23E-17 | 16275.72304 | 12187.95232 | ENSG00000149257 |
| WDR76 | -0.417664014 | 0.748635821 | 0.000753029 | 0.004377022 | 397.7745769 | 297.5667497 | ENSG00000092470 |
| MAGEH1 | -0.417743996 | 0.748594319 | 0.009745252 | 0.03818981 | 266.0705226 | 199.3039361 | ENSG00000187601 |
| PRPSAP1 | -0.417776098 | 0.748577661 | 3.41E-06 | 3.56E-05 | 830.6147113 | 621.8819451 | ENSG00000161542 |
| NA | -0.417797605 | 0.748566502 | 0.321457843 | 0.531913727 | 34.1524325 | 25.55116637 | ENSG00000215835 |
| BAK1 | -0.417817485 | 0.748556187 | 4.99E-07 | 6.14E-06 | 1168.793771 | 874.8402794 | ENSG00000030110 |
| PASK | -0.417842441 | 0.748543239 | 3.14E-08 | 4.77E-07 | 1712.290028 | 1281.618112 | ENSG00000115687 |
| TNPO3 | -0.417844607 | 0.748542115 | 5.63E-14 | 1.86E-12 | 3713.719286 | 2779.540586 | ENSG00000064419 |
| TP53INP2 | -0.417851894 | 0.748538334 | 0.03471215 | 0.106522417 | 144.9370002 | 108.5691462 | ENSG00000078804 |
| GCLM | -0.417935224 | 0.748495099 | 0.000496101 | 0.003056912 | 461.0252615 | 345.2053632 | ENSG00000023909 |
| NA | -0.418117748 | 0.748400409 | 0.35346822 | 0.565636916 | 29.75590932 | 22.20454567 | ENSG00000206341 |
| BTNL9 | -0.418136401 | 0.748390733 | 0.397555151 | 0.609205812 | 25.73361038 | 19.20058908 | ENSG00000165810 |
| CCNDBP1 | -0.418185806 | 0.748365105 | 0.00028605 | 0.001886192 | 473.3587086 | 354.4434836 | ENSG00000166946 |
| NA | -0.418209228 | 0.748352955 | 0.262677619 | 0.469893609 | 37.28570967 | 27.94943626 | ENSG00000248103 |
| CFAP77 | -0.418245296 | 0.748334246 | 0.559610531 | 0.745082126 | 11.51205537 | 8.626925898 | ENSG00000188523 |
| ATP1A3 | -0.418380701 | 0.748264014 | 0.276945996 | 0.48615011 | 36.81865981 | 27.62389231 | ENSG00000105409 |
| CDK4 | -0.418447109 | 0.748229572 | 1.83E-12 | 5.10E-11 | 2413.68716 | 1805.790747 | ENSG00000135446 |
| BBS5 | -0.41849913 | 0.748202592 | 0.480754193 | 0.682708803 | 15.45131738 | 11.56429376 | ENSG00000163093 |
| MMS22L | -0.418622563 | 0.748138581 | 0.000259253 | 0.001732319 | 478.8421095 | 358.4803505 | ENSG00000146263 |
| CIR1 | -0.418734442 | 0.748080566 | 0.003456896 | 0.016243714 | 292.7729633 | 219.1142566 | ENSG00000138433 |
| DNAJC8 | -0.418859779 | 0.748015578 | 1.67E-11 | 4.13E-10 | 3830.686703 | 2864.756137 | ENSG00000126698 |
| TPPP | -0.418933057 | 0.747977586 | 0.173271344 | 0.353029381 | 64.56223753 | 48.35063701 | ENSG00000171368 |
| PPP1R12A | -0.419394063 | 0.747738611 | 3.68E-08 | 5.52E-07 | 1304.104206 | 974.9213826 | ENSG00000058272 |
| CLINT1 | -0.419394258 | 0.74773851 | 6.10E-11 | 1.41E-09 | 2225.641886 | 1663.993613 | ENSG00000113282 |
| CASD1 | -0.419402595 | 0.747734189 | 0.001088106 | 0.006034912 | 379.7187901 | 283.9553841 | ENSG00000127995 |
| NA | -0.419426775 | 0.747721657 | 0.467163084 | 0.67102263 | 15.54503272 | 11.62993653 | ENSG00000235459 |
| AZGP1 | -0.419480361 | 0.747693885 | 0.578324052 | 0.759450456 | 8.887067177 | 6.664297958 | ENSG00000160862 |

| | | | | | | | |
|---|---|---|---|---|---|---|---|
| ARG2 | -0.419494034 | 0.747686799 | 0.000350523 | 0.002247751 | 531.3566704 | 397.3530396 | ENSG00000081181 |
| MAPK8IP1 | -0.419506661 | 0.747680255 | 9.51E-08 | 1.33E-06 | 1256.102077 | 939.5263592 | ENSG00000121653 |
| LIN28B | -0.419562965 | 0.747651075 | 7.91E-06 | 7.76E-05 | 2331.441299 | 1743.015581 | ENSG00000187772 |
| NA | -0.419746613 | 0.74755591 | 0.30039061 | 0.510506764 | 34.70935889 | 25.98463285 | ENSG00000264575 |
| TUBA1A | -0.419800504 | 0.747527985 | 2.04E-15 | 8.16E-14 | 5935.284418 | 4436.371138 | ENSG00000167552 |
| C7orf31 | -0.419840548 | 0.747507237 | 0.286469188 | 0.495600093 | 36.09271108 | 26.92238989 | ENSG00000153790 |
| SEC63 | -0.420095823 | 0.747374983 | 2.22E-08 | 3.43E-07 | 2198.922694 | 1643.393285 | ENSG00000025796 |
| SLC4A8 | -0.420172332 | 0.747335348 | 0.002682316 | 0.013062284 | 336.3159455 | 251.0803706 | ENSG00000050438 |
| ZFP30 | -0.420400331 | 0.747217251 | 0.1686435 | 0.346313394 | 57.2916154 | 42.79347843 | ENSG00000120784 |
| UQCC3 | -0.420450209 | 0.747191418 | 0.09233135 | 0.224590115 | 95.26175534 | 71.28759309 | ENSG00000204922 |
| COQ10A | -0.420697735 | 0.747063233 | 0.011823267 | 0.044944124 | 210.7856206 | 157.4511253 | ENSG00000135469 |
| NA | -0.420707251 | 0.747058305 | 0.689163707 | 0.83398915 | 4.855285028 | 3.648518091 | ENSG00000213261 |
| MARCKSL1 | -0.420778264 | 0.747021534 | 2.77E-10 | 5.78E-09 | 12123.35514 | 9057.287813 | ENSG00000175130 |
| TMEM178B | -0.421030418 | 0.746890981 | 0.000496131 | 0.003056912 | 430.7453513 | 321.7428134 | ENSG00000261115 |
| GABRA3 | -0.421045461 | 0.746883193 | 0.008691726 | 0.034972874 | 209.7957384 | 156.7132071 | ENSG00000011677 |
| RBM41 | -0.421228886 | 0.74678824 | 0.000249047 | 0.001671375 | 507.5321669 | 378.9182076 | ENSG00000089682 |
| DHX40 | -0.421373458 | 0.746713408 | 5.66E-06 | 5.71E-05 | 931.1521724 | 694.9270078 | ENSG00000108406 |
| SLC35D1 | -0.421426825 | 0.746685787 | 0.074735684 | 0.192155668 | 103.7674532 | 77.52958543 | ENSG00000116704 |
| UHRF2 | -0.4214687 | 0.746664114 | 5.54E-07 | 6.71E-06 | 1069.878784 | 798.7072248 | ENSG00000147854 |
| NA | -0.421938615 | 0.746420951 | 0.586531429 | 0.765028202 | 8.872829288 | 6.634125014 | ENSG00000258526 |
| PSMA2 | -0.42215279 | 0.746310149 | 0.143713515 | 0.309644007 | 67.02443202 | 50.11539067 | ENSG00000106588 |
| ATCAY | -0.422384518 | 0.746190285 | 5.12E-08 | 7.49E-07 | 1355.163413 | 1010.884354 | ENSG00000167654 |
| ZBTB17 | -0.422415702 | 0.746174156 | 8.61E-05 | 0.000649855 | 572.0867007 | 426.9393286 | ENSG00000116809 |
| DACT3 | -0.422534782 | 0.746112569 | 0.165969467 | 0.341914845 | 60.4936916 | 45.15287352 | ENSG00000197380 |
| RBM24 | -0.422536111 | 0.746111882 | 0.600080004 | 0.776007344 | 8.005443683 | 5.951918221 | ENSG00000112183 |
| G0S2 | -0.422580925 | 0.746088706 | 0.417867121 | 0.628751794 | 21.65527576 | 16.1602167 | ENSG00000123689 |
| NA | -0.422632748 | 0.746061907 | 0.007833018 | 0.032116054 | 217.5191724 | 162.244428 | ENSG00000234664 |
| DHX38 | -0.422664415 | 0.746045531 | 4.25E-12 | 1.12E-10 | 2358.337602 | 1759.271475 | ENSG00000140829 |
| VPS4A | -0.422710601 | 0.746021647 | 1.00E-05 | 9.54E-05 | 671.5593553 | 501.033993 | ENSG00000132612 |
| TWIST2 | -0.422792905 | 0.745979089 | 0.123312546 | 0.27796102 | 74.71939033 | 55.84287755 | ENSG00000233608 |
| PROM2 | -0.423026272 | 0.745858431 | 0.096839439 | 0.232555036 | 98.99879071 | 73.83878757 | ENSG00000155066 |
| C10orf82 | -0.423027383 | 0.745857857 | 0.041722018 | 0.122576277 | 149.6787137 | 111.7395746 | ENSG00000165863 |
| ADRA1D | -0.423290315 | 0.745721936 | 0.504274909 | 0.701785679 | 12.85716924 | 9.584262108 | ENSG00000171873 |
| RHPN1-AS1 | -0.423377316 | 0.745676967 | 0.367836974 | 0.579853662 | 24.0528155 | 17.97398743 | ENSG00000254389 |
| KIF3A | -0.423378984 | 0.745676105 | 0.003142301 | 0.014979681 | 303.8428851 | 226.671194 | ENSG00000131437 |
| NA | -0.423472903 | 0.745627563 | 0.179966816 | 0.362757318 | 61.03574356 | 45.40558589 | ENSG00000275234 |
| TEX10 | -0.423531282 | 0.745597392 | 8.11E-08 | 1.15E-06 | 2068.748872 | 1542.54738 | ENSG00000136891 |
| PDE8A | -0.423981968 | 0.745364509 | 0.049337744 | 0.139649775 | 120.5627154 | 89.98389161 | ENSG00000073417 |
| COMMD4 | -0.424150256 | 0.745277569 | 8.79E-09 | 1.46E-07 | 1678.71031 | 1251.645627 | ENSG00000140365 |
| TMSB15A | -0.424227547 | 0.745237642 | 0.000202092 | 0.001392886 | 642.9300104 | 479.0910521 | ENSG00000158164 |
| NA | -0.424259915 | 0.745220923 | 0.347615508 | 0.559556672 | 27.63770942 | 20.64927865 | ENSG00000232931 |
| NA | -0.424341086 | 0.745178995 | 0.465715263 | 0.669842555 | 246.7190261 | 183.8861339 | ENSG00000198744 |
| PEBP1 | -0.424370174 | 0.745163971 | 4.20E-20 | 2.54E-18 | 8295.359736 | 6181.775302 | ENSG00000089220 |
| ATP8A1 | -0.424954913 | 0.74486201 | 0.001486703 | 0.007902166 | 324.6102826 | 241.6358123 | ENSG00000124406 |
| EPN3 | -0.424995851 | 0.744840874 | 0.087662772 | 0.216087737 | 88.07711922 | 65.5342842 | ENSG00000049283 |
| PIGH | -0.425022085 | 0.74482733 | 0.068754315 | 0.180136687 | 95.73383921 | 71.3522899 | ENSG00000100564 |
| PCBP4 | -0.425512924 | 0.744573965 | 0.004742185 | 0.021121738 | 335.8717526 | 250.2956241 | ENSG00000090097 |
| NA | -0.426151097 | 0.744244677 | 0.238739005 | 0.441455607 | 42.9078515 | 31.99878888 | ENSG00000271576 |
| PIK3C3 | -0.426189139 | 0.744225053 | 2.17E-05 | 0.00019103 | 632.0338656 | 470.205221 | ENSG00000078142 |
| LHB | -0.42638218 | 0.744125478 | 0.668103351 | 0.822070092 | 6.165999386 | 4.594976984 | ENSG00000104826 |
| ARRDC4 | -0.426410774 | 0.744110729 | 0.036394739 | 0.110500747 | 133.9658015 | 99.66028093 | ENSG00000140450 |
| DHRS13 | -0.426744589 | 0.743938575 | 0.000296347 | 0.001943236 | 550.5085898 | 409.7695646 | ENSG00000167536 |
| TVP23C | -0.426778475 | 0.743921101 | 0.291415433 | 0.500990482 | 31.72168845 | 23.63053465 | ENSG00000175106 |
| NA | -0.426893015 | 0.743862041 | 0.344160016 | 0.556247178 | 27.28008734 | 20.27303663 | ENSG00000215102 |
| TMEM67 | -0.427040726 | 0.743785885 | 0.018114987 | 0.063397791 | 169.9654338 | 126.439562 | ENSG00000164953 |
| LINC01574 | -0.42710571 | 0.743752383 | 0.624531658 | 0.7923564 | 7.645151852 | 5.696367976 | ENSG00000248859 |
| CHRNA7 | -0.427131681 | 0.743738994 | 0.073124129 | 0.189319795 | 94.44684595 | 70.24531848 | ENSG00000175344 |
| RCC2 | -0.427611126 | 0.743491871 | 1.99E-23 | 1.57E-21 | 14946.05573 | 11111.83826 | ENSG00000179051 |
| MAGEF1 | -0.427618735 | 0.74348795 | 4.55E-07 | 5.63E-06 | 1777.289746 | 1321.925304 | ENSG00000177383 |
| ITPKA | -0.427771416 | 0.74340927 | 0.526779691 | 0.719947493 | 13.04576897 | 9.692847061 | ENSG00000137825 |
| NDRG2 | -0.427888116 | 0.743349138 | 0.000540547 | 0.00328239 | 400.9588557 | 297.9727132 | ENSG00000165795 |
| LRRC10B | -0.42800843 | 0.743287149 | 0.282865808 | 0.491799004 | 33.04308122 | 24.57576404 | ENSG00000204950 |
| FPGT | -0.428040811 | 0.743270466 | 0.280494145 | 0.489390497 | 37.78924397 | 28.1051034 | ENSG00000254685 |
| FRMD5 | -0.42804739 | 0.743267077 | 0.004551141 | 0.020424975 | 335.8364634 | 249.4319148 | ENSG00000171877 |
| NA | -0.428237528 | 0.743169125 | 0.674083949 | 0.825395538 | 4.906286698 | 3.662516831 | ENSG00000261634 |
| C11orf74 | -0.428285577 | 0.743144375 | 0.035733511 | 0.10893097 | 150.3562526 | 111.6328816 | ENSG00000166352 |
| UBR5 | -0.428333288 | 0.743119799 | 2.31E-08 | 3.56E-07 | 4214.924123 | 3131.431866 | ENSG00000104517 |
| RHOF | -0.428714356 | 0.74292354 | 0.105597945 | 0.248562531 | 83.43685034 | 62.11816425 | ENSG00000139725 |
| CDC14B | -0.428831246 | 0.742863349 | 1.21E-10 | 2.68E-09 | 1867.721109 | 1387.225473 | ENSG00000081377 |
| NA | -0.428852228 | 0.742852545 | 0.346990572 | 0.559118942 | 25.48808524 | 18.93974195 | ENSG00000223509 |
| GEMIN5 | -0.428866374 | 0.742845261 | 2.80E-09 | 4.99E-08 | 1589.74178 | 1180.837719 | ENSG00000082516 |
| CH17-340M24.3 | -0.428870676 | 0.742843046 | 0.527126462 | 0.720160837 | 11.17548463 | 8.328433474 | ENSG00000197180 |
| PCM1 | -0.428881347 | 0.742837552 | 1.62E-08 | 2.59E-07 | 2065.426788 | 1533.856924 | ENSG00000078674 |
| UBE2J1 | -0.429008227 | 0.742772225 | 0.000140282 | 0.001007312 | 575.5296514 | 427.6536729 | ENSG00000198833 |

| | | | | | | | |
|---|---|---|---|---|---|---|---|
| ZNF630 | -0.429105537 | 0.742722126 | 0.357406049 | 0.569491499 | 30.02251197 | 22.26706702 | ENSG00000221994 |
| MED31 | -0.429141434 | 0.742703646 | 0.20853381 | 0.401525387 | 45.19654829 | 33.53069289 | ENSG00000108590 |
| CDH7 | -0.429273679 | 0.742635569 | 0.317380164 | 0.527477286 | 28.09171659 | 20.84883394 | ENSG00000081138 |
| EFCAB11 | -0.42927965 | 0.742632495 | 0.118317301 | 0.27026605 | 74.7896639 | 55.59290774 | ENSG00000140025 |
| BCL7A | -0.429355642 | 0.74259338 | 0.00021333 | 0.001455994 | 531.9236904 | 395.0140337 | ENSG00000110987 |
| ZNF23 | -0.429369528 | 0.742586232 | 0.259818753 | 0.466762099 | 44.89793654 | 33.34636585 | ENSG00000167377 |
| SNTA1 | -0.429404102 | 0.742568436 | 0.003943743 | 0.018138721 | 246.1879771 | 182.7860677 | ENSG00000101400 |
| DNAH5 | -0.429514284 | 0.742511727 | 0.487122029 | 0.688003853 | 14.2876843 | 10.58548646 | ENSG00000039139 |
| FREM1 | -0.429532476 | 0.742502364 | 0.144097286 | 0.3100432 | 63.04779583 | 46.76999948 | ENSG00000164946 |
| ZNF862 | -0.42972337 | 0.742404124 | 0.136046385 | 0.298247299 | 65.77359215 | 48.7755696 | ENSG00000106479 |
| FOXRED1 | -0.429732239 | 0.74239956 | 2.64E-07 | 3.42E-06 | 1324.238401 | 983.1855114 | ENSG00000110074 |
| ACAA1 | -0.429790964 | 0.742369342 | 6.00E-05 | 0.000472813 | 587.4084548 | 436.3434467 | ENSG00000060971 |
| IL4I1 | -0.430043639 | 0.742239333 | 0.610813369 | 0.783429128 | 7.645151852 | 5.680193772 | ENSG00000104951 |
| AUH | -0.43010597 | 0.742207266 | 0.00074079 | 0.004319177 | 422.4999752 | 313.6253297 | ENSG00000148090 |
| KIF5C | -0.430123138 | 0.742198434 | 2.35E-09 | 4.28E-08 | 2198.353537 | 1631.48576 | ENSG00000168280 |
| MBNL2 | -0.430156245 | 0.742181402 | 0.413510241 | 0.624700612 | 22.40077573 | 16.66244738 | ENSG00000139793 |
| CENPQ | -0.43023494 | 0.742140919 | 0.019628148 | 0.067652608 | 194.4568234 | 144.1980605 | ENSG00000031691 |
| XRCC2 | -0.430367681 | 0.742072639 | 0.000285851 | 0.001885536 | 567.8002935 | 421.7563952 | ENSG00000196584 |
| SMG1P7 | -0.430442512 | 0.742034149 | 0.091189888 | 0.222410053 | 84.48126754 | 62.7378497 | ENSG00000261556 |
| BAG3 | -0.430461851 | 0.742024203 | 9.16E-05 | 0.000687493 | 638.3592288 | 473.9584676 | ENSG00000151929 |
| CENPW | -0.430491701 | 0.74200885 | 5.29E-06 | 5.36E-05 | 691.4882001 | 513.0353261 | ENSG00000203760 |
| FOXD2-AS1 | -0.430559158 | 0.741974156 | 0.674456563 | 0.825608414 | 6.32731855 | 4.703561936 | ENSG00000237424 |
| NA | -0.430798099 | 0.74185128 | 0.126673355 | 0.283501308 | 64.81577134 | 48.07624264 | ENSG00000268362 |
| TTC14 | -0.4309527 | 0.741771786 | 0.00249478 | 0.012300215 | 329.6873711 | 244.7313493 | ENSG00000163728 |
| NA | -0.431193067 | 0.74164821 | 0.453490869 | 0.6596002 | 17.79549119 | 13.18425759 | ENSG00000198225 |
| GTF3C1 | -0.431309897 | 0.741588154 | 3.62E-14 | 1.23E-12 | 5340.723055 | 3960.142534 | ENSG00000077235 |
| NA | -0.431394301 | 0.741544769 | 0.074455558 | 0.191722554 | 102.289191 | 75.83387951 | ENSG00000235501 |
| PRDM10 | -0.431438161 | 0.741522225 | 0.000324546 | 0.002104035 | 478.5351319 | 354.7404402 | ENSG00000170325 |
| NA | -0.432157581 | 0.741152547 | 0.000659342 | 0.003898249 | 355.3886674 | 263.4031506 | ENSG00000260834 |
| EIF3E | -0.432164759 | 0.741148859 | 7.57E-17 | 3.37E-15 | 6168.604902 | 4571.717043 | ENSG00000104408 |
| TCTN2 | -0.432203807 | 0.741128799 | 0.022112413 | 0.074663717 | 153.0954227 | 113.4766143 | ENSG00000168778 |
| PUDP | -0.432297842 | 0.741080494 | 1.19E-05 | 0.000111426 | 804.0283372 | 595.9807579 | ENSG00000130021 |
| TOLLIP | -0.432323162 | 0.741067488 | 2.87E-06 | 3.04E-05 | 863.0459995 | 639.7895547 | ENSG00000078902 |
| GFAP | -0.432346225 | 0.741055641 | 0.137582622 | 0.300379984 | 63.24320902 | 46.8891782 | ENSG00000131095 |
| TBC1D7 | -0.432347992 | 0.741054734 | 0.000122681 | 0.000892115 | 540.3893187 | 400.3599294 | ENSG00000145979 |
| NA | -0.432705417 | 0.740871162 | 0.184544309 | 0.36874397 | 51.8542345 | 38.40798809 | ENSG00000271369 |
| AMN | -0.432770968 | 0.7408375 | 0.228163446 | 0.427551764 | 42.60923976 | 31.60760216 | ENSG00000166126 |
| LOC100507437 | -0.432880773 | 0.740781116 | 0.224900442 | 0.423243133 | 53.54957218 | 39.76710479 | ENSG00000251602 |
| KDELR1 | -0.433160029 | 0.74063774 | 2.29E-14 | 7.88E-13 | 3552.794585 | 2631.696173 | ENSG00000105438 |
| THADA | -0.433454756 | 0.740486451 | 4.62E-06 | 4.72E-05 | 722.8750194 | 535.1914648 | ENSG00000115970 |
| CCDC91 | -0.433541638 | 0.740441859 | 0.01626206 | 0.058208681 | 192.2558921 | 142.2933195 | ENSG00000123106 |
| NA | -0.433653711 | 0.740384341 | 0.353431495 | 0.565636916 | 23.72336371 | 17.58403021 | ENSG00000271524 |
| ADAMTS8 | -0.43373942 | 0.740340357 | 1.05E-05 | 9.96E-05 | 610.4053117 | 451.8884968 | ENSG00000134917 |
| BTBD17 | -0.433741247 | 0.74033942 | 0.000308443 | 0.002014856 | 491.4288111 | 363.9565951 | ENSG00000204347 |
| GCHFR | -0.434084268 | 0.740163415 | 0.62922194 | 0.795425152 | 7.679551368 | 5.6845447 | ENSG00000137880 |
| RPL21P28 | -0.434296494 | 0.740054542 | 0.423147862 | 0.633337985 | 17.47196314 | 12.94488155 | ENSG00000220749 |
| NA | -0.434362906 | 0.740020476 | 0.676903344 | 0.82713165 | 4.971526258 | 3.690514311 | ENSG00000267096 |
| NA | -0.43481936 | 0.739786378 | 0.497504126 | 0.696660074 | 14.90153609 | 11.0510178 | ENSG00000255182 |
| BUD31 | -0.434869154 | 0.739760845 | 3.02E-06 | 3.18E-05 | 806.3581168 | 596.6086211 | ENSG00000106245 |
| SNHG19 | -0.434915101 | 0.739737285 | 0.225403517 | 0.423725406 | 421.2944664 | 311.6777323 | ENSG00000260260 |
| ISL2 | -0.435250594 | 0.739565282 | 0.66992765 | 0.823081366 | 5.832988126 | 4.344723624 | ENSG00000159556 |
| ERG28 | -0.435388624 | 0.739494528 | 1.38E-07 | 1.87E-06 | 1232.5979 | 911.6713185 | ENSG00000133935 |
| TTC4 | -0.435696653 | 0.739336655 | 0.560119023 | 0.74535213 | 8.979587309 | 6.624477201 | ENSG00000243725 |
| AVEN | -0.435819664 | 0.739273619 | 0.000332268 | 0.002145262 | 416.1385882 | 307.8337131 | ENSG00000169857 |
| RTN1 | -0.435836196 | 0.739265147 | 0.090456079 | 0.221277598 | 88.94508964 | 65.61487167 | ENSG00000139970 |
| QPCTL | -0.435953202 | 0.739205193 | 9.44E-05 | 0.000705344 | 556.4049608 | 411.1350397 | ENSG00000011478 |
| CDH24 | -0.436101205 | 0.739129364 | 3.61E-05 | 0.000301327 | 662.8935596 | 490.1525887 | ENSG00000139880 |
| TMA7 | -0.436220814 | 0.739068088 | 0.027740875 | 0.089415673 | 138.3326799 | 102.2412695 | ENSG00000232112 |
| WIPF3 | -0.436344757 | 0.739004597 | 7.76E-05 | 0.000592882 | 472.652058 | 349.2693769 | ENSG00000122574 |
| ITSN1 | -0.436546303 | 0.738901364 | 1.13E-13 | 3.60E-12 | 3197.630847 | 2362.548636 | ENSG00000205726 |
| VMAC | -0.436562147 | 0.738893249 | 0.467212421 | 0.67102263 | 14.84222027 | 10.98849645 | ENSG00000187650 |
| C2orf78 | -0.43658351 | 0.738882308 | 0.638841917 | 0.802056216 | 6.814530028 | 5.007634792 | ENSG00000187833 |
| FIBCD1 | -0.43678485 | 0.738779198 | 0.046640798 | 0.133597979 | 131.3429244 | 97.08979082 | ENSG00000130720 |
| ZNF10 | -0.436831622 | 0.738755247 | 0.09873896 | 0.235960496 | 76.43933941 | 56.49519495 | ENSG00000256223 |
| SLC25A12 | -0.436962136 | 0.738688419 | 0.003609383 | 0.016842897 | 271.2384041 | 200.4316733 | ENSG00000115840 |
| KIAA0895L | -0.437127035 | 0.738603992 | 0.003576499 | 0.016701825 | 614.5254441 | 454.061498 | ENSG00000196123 |
| TPBGL | -0.437134379 | 0.738600232 | 0.414704355 | 0.625609309 | 19.32297558 | 14.32982026 | ENSG00000261594 |
| VEGFC | -0.437138588 | 0.738598077 | 0.307462932 | 0.517738517 | 28.763689 | 21.25496535 | ENSG00000150630 |
| KBTBD11 | -0.437190887 | 0.738571303 | 0.000885871 | 0.005034507 | 356.39899 | 263.1858651 | ENSG00000176595 |
| C6orf52 | -0.437348944 | 0.738490392 | 0.434932377 | 0.643656292 | 18.47219213 | 13.67220598 | ENSG00000137434 |
| ADAMTS3 | -0.437643127 | 0.73833982 | 0.088918394 | 0.21842142 | 104.2810555 | 76.98452816 | ENSG00000156140 |
| XYLB | -0.437654769 | 0.738333862 | 0.087712152 | 0.216133088 | 79.52634344 | 58.77371902 | ENSG00000093217 |
| WHAMM | -0.437951009 | 0.73818227 | 0.02691869 | 0.087351714 | 137.9270308 | 101.8970923 | ENSG00000156232 |
| SLC4A3 | -0.437987573 | 0.738163561 | 0.002520289 | 0.012388251 | 364.8827468 | 269.4005856 | ENSG00000114923 |

| | | | | | | | |
|---|---|---|---|---|---|---|---|
| ZMPSTE24 | -0.438200826 | 0.738054457 | 9.09E-06 | 8.79E-05 | 1084.096193 | 800.2751954 | ENSG00000084073 |
| YIF1A | -0.438224643 | 0.738042273 | 1.97E-09 | 3.63E-08 | 1351.243649 | 997.2304685 | ENSG00000174851 |
| CHRDL2 | -0.43854503 | 0.73787839 | 0.451182915 | 0.657452882 | 14.82917759 | 10.95737755 | ENSG00000054938 |
| PARP12 | -0.438730269 | 0.737783654 | 0.020345084 | 0.069730838 | 179.5615427 | 132.4521096 | ENSG00000059378 |
| NA | -0.438741003 | 0.737778165 | 0.378538171 | 0.59094762 | 21.17309829 | 15.63417151 | ENSG00000262049 |
| PTPN11 | -0.438875716 | 0.737709277 | 1.08E-14 | 3.91E-13 | 3569.352786 | 2632.906515 | ENSG00000179295 |
| TTC27 | -0.439924784 | 0.737173041 | 5.83E-07 | 7.02E-06 | 829.199019 | 611.2749876 | ENSG00000018699 |
| INSYN1 | -0.439998234 | 0.737135511 | 0.00727603 | 0.030199524 | 220.2399609 | 162.4077784 | ENSG00000205363 |
| PREX2 | -0.440023879 | 0.737122408 | 7.28E-08 | 1.04E-06 | 1187.873891 | 875.147694 | ENSG00000046889 |
| TRAF3IP1 | -0.440096595 | 0.737085256 | 4.89E-05 | 0.000394298 | 533.6738168 | 393.2523062 | ENSG00000204104 |
| EPG5 | -0.440768935 | 0.736741832 | 0.000700225 | 0.004115703 | 436.1927485 | 321.0613643 | ENSG00000152223 |
| CEP152 | -0.440785772 | 0.736733233 | 0.004168598 | 0.019031999 | 269.1631972 | 198.3617852 | ENSG00000103995 |
| CREBZF | -0.440875656 | 0.736687334 | 1.82E-06 | 2.00E-05 | 1303.943751 | 960.711723 | ENSG00000137504 |
| MROH6 | -0.441018554 | 0.736614369 | 0.119440896 | 0.27197945 | 71.34000378 | 52.54790089 | ENSG00000204839 |
| PRR36 | -0.441130037 | 0.73655745 | 8.59E-05 | 0.00064833 | 739.4343949 | 544.9708156 | ENSG00000183248 |
| NA | -0.441307798 | 0.736466701 | 0.210439713 | 0.404000608 | 43.72690518 | 32.19522274 | ENSG00000246465 |
| S1PR1 | -0.441326371 | 0.73645722 | 0.305926132 | 0.516092607 | 32.13921117 | 23.61776541 | ENSG00000170989 |
| RAB11FIP2 | -0.441438979 | 0.736399739 | 0.018482863 | 0.064412044 | 205.6225716 | 151.4451766 | ENSG00000107560 |
| TM7SF2 | -0.441771601 | 0.736229977 | 5.65E-07 | 6.84E-06 | 1083.262649 | 797.8912008 | ENSG00000149809 |
| AJUBA | -0.441980322 | 0.736123472 | 4.91E-05 | 0.000395813 | 848.9023951 | 625.0211391 | ENSG00000129474 |
| CCDC34 | -0.44200514 | 0.736110808 | 1.81E-05 | 0.000162345 | 586.9369302 | 432.0995521 | ENSG00000109881 |
| MRTO4 | -0.442093321 | 0.736065817 | 4.83E-11 | 1.13E-09 | 1815.105103 | 1336.282087 | ENSG00000053372 |
| WDR45 | -0.442136694 | 0.736043688 | 0.037376837 | 0.112703427 | 122.5720723 | 90.15299041 | ENSG00000196998 |
| PPP1R1C | -0.442405453 | 0.735906584 | 0.255378582 | 0.462222016 | 35.26773366 | 25.92863789 | ENSG00000150722 |
| LRRC8E | -0.44242236 | 0.73589796 | 0.454476937 | 0.660469176 | 16.62206941 | 12.27884896 | ENSG00000171017 |
| ZNF256 | -0.442464834 | 0.735876295 | 0.013011068 | 0.04853131 | 225.9525124 | 166.064336 | ENSG00000152454 |
| NAPB | -0.442480001 | 0.735868558 | 0.003708498 | 0.017228777 | 246.9865898 | 181.750603 | ENSG00000125814 |
| PLAGL2 | -0.442601436 | 0.735806622 | 1.71E-08 | 2.73E-07 | 1130.604294 | 832.0896276 | ENSG00000126003 |
| LANCL2 | -0.442674648 | 0.735769283 | 5.78E-05 | 0.000457627 | 519.3313699 | 382.30486 | ENSG00000132434 |
| DGCR6L | -0.442863733 | 0.735672856 | 3.96E-06 | 4.09E-05 | 1328.92789 | 977.7723444 | ENSG00000128185 |
| FAM3B | -0.442876261 | 0.735666468 | 0.344320448 | 0.556339122 | 29.26274795 | 21.47627527 | ENSG00000183844 |
| NA | -0.442957678 | 0.735624953 | 0.175331634 | 0.355890243 | 50.04827384 | 36.78055191 | ENSG00000227473 |
| ECPAS | -0.443196126 | 0.735503379 | 9.23E-15 | 3.37E-13 | 4619.360341 | 3396.997839 | ENSG00000136813 |
| STAMBPL1 | -0.443244248 | 0.735478846 | 0.028904584 | 0.092163077 | 133.9489201 | 98.51159683 | ENSG00000138134 |
| NA | -0.443270164 | 0.735465634 | 0.631150301 | 0.796902683 | 6.371201275 | 4.679915383 | ENSG00000175886 |
| SAFB2 | -0.443385489 | 0.735406846 | 1.43E-09 | 2.68E-08 | 2485.815329 | 1828.749217 | ENSG00000130254 |
| ZNF585A | -0.443661798 | 0.735266012 | 0.125002782 | 0.280843278 | 66.2854406 | 48.68438835 | ENSG00000196967 |
| ALG14 | -0.443731097 | 0.735230695 | 0.157661654 | 0.330202987 | 55.54616578 | 40.76426176 | ENSG00000172339 |
| RNLS | -0.443904686 | 0.735142235 | 0.033309664 | 0.103123018 | 140.4369729 | 103.3846567 | ENSG00000184719 |
| NA | -0.444085366 | 0.735050173 | 0.472370592 | 0.675760823 | 13.54930327 | 9.966746973 | ENSG00000259891 |
| PRPF38B | -0.44416362 | 0.735010304 | 4.88E-12 | 1.28E-10 | 2445.218519 | 1797.554892 | ENSG00000134186 |
| CLK4 | -0.44465393 | 0.734760548 | 0.006209774 | 0.026508042 | 218.2810214 | 160.4553654 | ENSG00000113240 |
| FANCM | -0.444742042 | 0.734715674 | 0.004208304 | 0.019196574 | 228.6866484 | 167.9563078 | ENSG00000187790 |
| SLC27A5 | -0.445074765 | 0.734546249 | 0.009876164 | 0.038631183 | 233.327196 | 171.4274041 | ENSG00000083807 |
| GOLM1 | -0.44535527 | 0.734403444 | 2.11E-15 | 8.40E-14 | 4060.990054 | 2982.325811 | ENSG00000135052 |
| EXOSC1 | -0.44539951 | 0.734380924 | 1.99E-05 | 0.00017746 | 703.5158387 | 516.5189284 | ENSG00000171311 |
| TMEM87B | -0.445713132 | 0.734221297 | 0.011602272 | 0.044193002 | 199.6433402 | 146.4316779 | ENSG00000153214 |
| ZFP14 | -0.44578708 | 0.734183664 | 0.023946948 | 0.079577355 | 165.885904 | 121.9163753 | ENSG00000142065 |
| PJVK | -0.446042163 | 0.734053865 | 0.449995122 | 0.656329772 | 15.40534973 | 11.26457183 | ENSG00000204311 |
| RMC1 | -0.446074087 | 0.734037622 | 0.007759655 | 0.031905308 | 218.797319 | 160.7432641 | ENSG00000141452 |
| HEPH | -0.446097989 | 0.734025461 | 4.25E-11 | 1.01E-09 | 1669.795302 | 1225.533282 | ENSG00000089472 |
| HAPLN3 | -0.446156798 | 0.73399554 | 1.48E-12 | 4.16E-11 | 1982.109816 | 1455.061068 | ENSG00000140511 |
| DNAJB11 | -0.446308095 | 0.733918569 | 1.55E-09 | 2.91E-08 | 1324.489674 | 972.2363709 | ENSG00000090520 |
| TMEM94 | -0.446314093 | 0.733915518 | 3.32E-08 | 5.03E-07 | 1450.592158 | 1065.031371 | ENSG00000177728 |
| KLRG1 | -0.446351005 | 0.73389674 | 0.172250706 | 0.351708788 | 53.00894304 | 38.82622633 | ENSG00000139187 |
| CD99P1 | -0.446492081 | 0.733824979 | 0.090815751 | 0.221803996 | 90.66801337 | 66.51859929 | ENSG00000223773 |
| ZNF664 | -0.446787638 | 0.733674659 | 3.53E-10 | 7.27E-09 | 1714.23562 | 1257.220555 | ENSG00000179195 |
| NA | -0.446893373 | 0.733620891 | 0.52780153 | 0.720874514 | 11.70274003 | 8.621629012 | ENSG00000226332 |
| URAHP | -0.446967267 | 0.733583316 | 0.546345977 | 0.734378368 | 10.02489423 | 7.350855679 | ENSG00000222019 |
| NFIX | -0.447110062 | 0.733510711 | 0.502000072 | 0.700181291 | 12.28247164 | 9.021517585 | ENSG00000008441 |
| PTPRT | -0.447142302 | 0.73349432 | 0.077801159 | 0.197855494 | 115.3554265 | 84.58798062 | ENSG00000196090 |
| STAP2 | -0.44719313 | 0.733468478 | 0.001734937 | 0.009004687 | 305.5367227 | 224.0545247 | ENSG00000178078 |
| C1orf54 | -0.447363999 | 0.733381613 | 0.035916374 | 0.109364837 | 125.9600785 | 92.46028997 | ENSG00000118292 |
| MAP2K3 | -0.447566137 | 0.733278865 | 2.06E-06 | 2.24E-05 | 927.0683235 | 679.7040037 | ENSG00000034152 |
| CSAD | -0.447890242 | 0.73311415 | 0.201598501 | 0.392587026 | 62.36989968 | 45.77428293 | ENSG00000139631 |
| NRDE2 | -0.44813558 | 0.732989491 | 0.006988612 | 0.029224964 | 197.4151022 | 144.6457367 | ENSG00000119720 |
| ZNF493 | -0.448206357 | 0.732953532 | 0.027331944 | 0.088353947 | 155.3313639 | 113.8486734 | ENSG00000196268 |
| SEPTIN3 | -0.448339724 | 0.732885779 | 1.28E-09 | 2.42E-08 | 1697.375859 | 1243.646014 | ENSG00000100167 |
| CAAP1 | -0.448347929 | 0.732881611 | 5.49E-05 | 0.000436885 | 665.8642701 | 487.8517296 | ENSG00000120159 |
| LOXL1 | -0.44841107 | 0.732849537 | 0.004836538 | 0.021461568 | 323.8609171 | 237.4010711 | ENSG00000129038 |
| COQ2 | -0.448557222 | 0.732775299 | 0.000840803 | 0.004816064 | 391.2468623 | 286.945131 | ENSG00000173085 |
| UNC45A | -0.448952182 | 0.732574718 | 1.57E-13 | 4.98E-12 | 3244.49718 | 2376.53447 | ENSG00000140553 |
| ARL6IP5 | -0.449063964 | 0.732517959 | 9.47E-06 | 9.10E-05 | 616.1665 | 451.267203 | ENSG00000144746 |
| NA | -0.449128836 | 0.732485022 | 0.142013084 | 0.306988991 | 58.54184488 | 42.92287206 | ENSG00000272849 |

| | | | | | | | |
|---|---|---|---|---|---|---|---|
| THAP2 | -0.449198671 | 0.732449566 | 0.143001231 | 0.308461251 | 61.26052226 | 44.88795901 | ENSG00000173451 |
| DPH7 | -0.449275468 | 0.732410578 | 1.92E-06 | 2.10E-05 | 871.3083624 | 638.3551904 | ENSG00000148399 |
| C22orf39 | -0.449325683 | 0.732385086 | 0.000839803 | 0.004813256 | 343.5768568 | 251.7406548 | ENSG00000242259 |
| UBAC1 | -0.449557166 | 0.732267582 | 1.42E-11 | 3.53E-10 | 1749.946428 | 1281.414635 | ENSG00000130560 |
| NEK4 | -0.449639202 | 0.732225945 | 8.67E-06 | 8.44E-05 | 755.3227351 | 552.8766373 | ENSG00000114904 |
| NA | -0.449925897 | 0.73208045 | 0.638734583 | 0.802036409 | 5.893473005 | 4.332900348 | ENSG00000236989 |
| PER3 | -0.450012604 | 0.732036452 | 0.027545129 | 0.088921327 | 158.6544361 | 116.124076 | ENSG00000049246 |
| SYT13 | -0.450279081 | 0.731901252 | 7.88E-05 | 0.000601475 | 481.4602581 | 352.4564311 | ENSG00000019505 |
| RIDA | -0.450723002 | 0.731676079 | 6.14E-05 | 0.000482241 | 568.6978048 | 415.9888337 | ENSG00000132541 |
| SPC25 | -0.450866791 | 0.731603159 | 0.000152346 | 0.001082009 | 590.0665694 | 431.7496992 | ENSG00000152253 |
| DPCD | -0.450941861 | 0.731565091 | 0.000466554 | 0.002901175 | 359.392 | 262.8563694 | ENSG00000166171 |
| C12orf75 | -0.451089152 | 0.731490406 | 0.00019928 | 0.001375511 | 455.3959745 | 333.2340874 | ENSG00000235162 |
| PGD | -0.451244648 | 0.731411569 | 7.02E-25 | 6.16E-23 | 10636.90083 | 7779.757433 | ENSG00000142657 |
| ATP6V1H | -0.451279968 | 0.731393663 | 8.02E-06 | 7.86E-05 | 621.6208141 | 454.7502179 | ENSG00000047249 |
| PPP1R13B | -0.451300717 | 0.731383144 | 0.001921032 | 0.00979711 | 274.065314 | 200.423255 | ENSG00000088808 |
| STIM1 | -0.451517984 | 0.731273008 | 1.93E-05 | 0.000172981 | 547.6080489 | 400.3222411 | ENSG00000167323 |
| GREB1 | -0.451620357 | 0.731221119 | 7.40E-07 | 8.75E-06 | 948.7946068 | 693.3406445 | ENSG00000196208 |
| C9orf116 | -0.451651407 | 0.731205381 | 0.056176022 | 0.154400258 | 100.6270832 | 73.61709878 | ENSG00000160345 |
| NT5C3A | -0.451797811 | 0.731131183 | 0.000202319 | 0.00139343 | 408.45929 | 298.7480658 | ENSG00000122643 |
| NA | -0.45192899 | 0.731064707 | 0.527074624 | 0.720142112 | 13.64390834 | 10.03361925 | ENSG00000279089 |
| FGF12 | -0.451934695 | 0.731061816 | 0.005813765 | 0.025056795 | 233.3162906 | 170.5004814 | ENSG00000114279 |
| USP49 | -0.452492933 | 0.730778993 | 4.08E-06 | 4.20E-05 | 697.7428291 | 509.8058243 | ENSG00000164663 |
| 2-Mar | -0.452778131 | 0.730634543 | 0.007490334 | 0.030966498 | 213.1772366 | 155.6977004 | ENSG00000117791 |
| HTR7P1 | -0.452965644 | 0.730539586 | 0.089674187 | 0.219877568 | 83.31763446 | 60.94320664 | ENSG00000183935 |
| MARCKS | -0.453000393 | 0.73052199 | 1.58E-10 | 3.41E-09 | 2994.484539 | 2187.643303 | ENSG00000277443 |
| NUP62CL | -0.453131437 | 0.730455638 | 0.233173804 | 0.434097722 | 39.99164985 | 29.13867616 | ENSG00000198088 |
| NA | -0.453232198 | 0.730404623 | 0.282684608 | 0.491655782 | 29.93444103 | 21.8985809 | ENSG00000255836 |
| TRIL | -0.453444862 | 0.730296964 | 0.005394624 | 0.023550041 | 200.8790263 | 146.6998294 | ENSG00000255690 |
| NA | -0.453969602 | 0.730031387 | 0.641759817 | 0.804214244 | 6.86761663 | 5.045280085 | ENSG00000267199 |
| WDR35 | -0.45411816 | 0.729956218 | 1.43E-06 | 1.60E-05 | 762.4722834 | 556.3941237 | ENSG00000118965 |
| SAP30 | -0.454191011 | 0.729919359 | 0.001228605 | 0.006701964 | 330.4922392 | 241.3856746 | ENSG00000164105 |
| ANGPTL4 | -0.454294507 | 0.729866997 | 0.278737387 | 0.48785501 | 40.51798937 | 29.48758311 | ENSG00000167772 |
| FAM53B | -0.454365366 | 0.72983115 | 1.05E-05 | 9.96E-05 | 641.2697082 | 467.9492819 | ENSG00000189319 |
| NA | -0.454454687 | 0.729785966 | 0.602778205 | 0.777718549 | 7.272986395 | 5.308302679 | ENSG00000236643 |
| FILIP1 | -0.45460248 | 0.729711209 | 0.590857949 | 0.768640378 | 7.276545868 | 5.310478143 | ENSG00000118407 |
| SLC22A31 | -0.454666954 | 0.729678599 | 0.044775379 | 0.129294042 | 173.0290222 | 126.3838935 | ENSG00000259803 |
| HMGN4 | -0.454691546 | 0.729666161 | 2.31E-08 | 3.56E-07 | 989.795182 | 722.3720082 | ENSG00000182952 |
| UHRF1BP1 | -0.454798058 | 0.729612293 | 1.92E-07 | 2.51E-06 | 981.3345104 | 715.7205091 | ENSG00000065060 |
| MORC4 | -0.454948488 | 0.72953622 | 0.006305388 | 0.026855492 | 198.0799556 | 144.5149456 | ENSG00000133131 |
| NXF1 | -0.455073378 | 0.729473069 | 1.42E-07 | 1.91E-06 | 1066.55073 | 778.11432 | ENSG00000162231 |
| TERF1 | -0.455216664 | 0.729400622 | 5.83E-14 | 1.92E-12 | 10898.91297 | 7949.542714 | ENSG00000147601 |
| NA | -0.455324993 | 0.729345855 | 0.69311112 | 0.836009676 | 4.538875922 | 3.330446495 | ENSG00000236762 |
| ZFP62 | -0.455603262 | 0.729205191 | 8.62E-06 | 8.40E-05 | 623.6467732 | 454.8763222 | ENSG00000196670 |
| EIF3CL | -0.455609863 | 0.729201855 | 0.001310691 | 0.00708426 | 357.5013008 | 260.7894348 | ENSG00000205609 |
| NA | -0.455677716 | 0.72916756 | 0.543056054 | 0.731968709 | 9.496164291 | 6.93167148 | ENSG00000215006 |
| CHRNA3 | -0.455839577 | 0.729085757 | 0.03599536 | 0.109534704 | 111.887969 | 81.63058208 | ENSG00000080644 |
| BLOC1S1 | -0.456077508 | 0.728965525 | 0.425637823 | 0.635611886 | 17.30619478 | 12.59039416 | ENSG00000135441 |
| LINC01918 | -0.456079032 | 0.728964755 | 0.159063447 | 0.332461334 | 56.581684 | 41.28645048 | ENSG00000226508 |
| NA | -0.456128671 | 0.728939674 | 0.702543158 | 0.842036584 | 4.528197505 | 3.31767726 | ENSG00000253520 |
| MINCR | -0.456171956 | 0.728917804 | 0.167671454 | 0.344894391 | 58.96974054 | 43.03457844 | ENSG00000253716 |
| NA | -0.456239267 | 0.728883796 | 0.370165095 | 0.582353201 | 22.66501411 | 16.53210779 | ENSG00000269867 |
| USP25 | -0.456239648 | 0.728883603 | 3.66E-06 | 3.81E-05 | 999.5082732 | 728.4957248 | ENSG00000155313 |
| TAX1BP3 | -0.456253726 | 0.728876491 | 0.65034059 | 0.81019078 | 5.934991466 | 4.33602177 | ENSG00000213977 |
| N4BP2L1 | -0.45635915 | 0.728823231 | 0.302259596 | 0.512445547 | 28.18187246 | 20.50616983 | ENSG00000139597 |
| PPFIA2 | -0.45651334 | 0.728745341 | 0.36666517 | 0.579187924 | 21.92215773 | 15.9737142 | ENSG00000139220 |
| PABPC1L | -0.45665343 | 0.728674581 | 0.16243918 | 0.337476778 | 880.4740555 | 641.6218899 | ENSG00000101104 |
| NA | -0.456659915 | 0.728671305 | 0.414567712 | 0.625528224 | 18.10834082 | 13.18331163 | ENSG00000251495 |
| NA | -0.456697854 | 0.728652144 | 0.683284865 | 0.830790283 | 4.559037549 | 3.343499276 | ENSG00000235649 |
| ZNF670-ZNF695 | -0.45685614 | 0.728572204 | 0.557428884 | 0.743522876 | 9.554284906 | 6.968370814 | ENSG00000135747 |
| NA | -0.45705102 | 0.728473795 | 0.382391018 | 0.594896975 | 19.63137601 | 14.28347312 | ENSG00000260000 |
| SPSB4 | -0.457139855 | 0.728428939 | 4.69E-06 | 4.79E-05 | 710.403948 | 517.2638768 | ENSG00000175093 |
| OSBPL11 | -0.457162018 | 0.728417749 | 2.44E-06 | 2.62E-05 | 850.5680891 | 619.3884993 | ENSG00000144909 |
| DCTN3 | -0.457210988 | 0.728393025 | 0.000397242 | 0.002512361 | 351.7242961 | 256.104107 | ENSG00000137100 |
| TAF1B | -0.45737096 | 0.728312262 | 0.000681283 | 0.004014362 | 363.9824879 | 265.1407575 | ENSG00000115750 |
| NA | -0.457459925 | 0.728267351 | 0.526074343 | 0.719295827 | 9.561403851 | 6.971492236 | ENSG00000215210 |
| PRR7 | -0.458046997 | 0.72797106 | 0.008378137 | 0.033931184 | 200.9603094 | 146.4063503 | ENSG00000131188 |
| NA | -0.458142641 | 0.7279228 | 0.116759076 | 0.267758973 | 64.22180184 | 46.81388762 | ENSG00000274372 |
| FGFR1OP | -0.458422323 | 0.727781698 | 0.006778114 | 0.028483462 | 206.9605404 | 150.7345209 | ENSG00000213066 |
| NIPSNAP3A | -0.458631305 | 0.727676283 | 0.016175488 | 0.05799774 | 179.790745 | 130.6825106 | ENSG00000136783 |
| ACPP | -0.459045986 | 0.727467153 | 0.397122017 | 0.608935474 | 18.74263358 | 13.58632163 | ENSG00000014257 |
| TMX4 | -0.459100258 | 0.727439788 | 0.003176305 | 0.015124692 | 286.1114383 | 208.1199688 | ENSG00000125827 |
| CD200 | -0.459270903 | 0.72735375 | 1.55E-08 | 2.50E-07 | 1095.353112 | 796.3890479 | ENSG00000091972 |
| DDX49 | -0.459400123 | 0.727288605 | 2.11E-06 | 2.29E-05 | 1036.390366 | 753.9369814 | ENSG00000105671 |
| TMEM234 | -0.459409163 | 0.727284047 | 0.150257316 | 0.319262984 | 61.2830743 | 44.59419638 | ENSG00000160055 |

| | | | | | | | |
|---|---|---|---|---|---|---|---|
| LINC02449 | -0.459513849 | 0.727231275 | 0.538414633 | 0.728155282 | 9.548361169 | 6.94567022 | ENSG00000215241 |
| NA | -0.459573505 | 0.727201205 | 0.460731705 | 0.665462199 | 15.87212025 | 11.51482519 | ENSG00000269688 |
| NA | -0.459961656 | 0.727005581 | 0.382546529 | 0.59504087 | 18.2791432 | 13.27042549 | ENSG00000255389 |
| ENO1 | -0.460431019 | 0.726769097 | 9.36E-33 | 1.28E-30 | 55998.47062 | 40697.73654 | ENSG00000074800 |
| TNRC6C | -0.46048235 | 0.726743239 | 0.000171417 | 0.001201189 | 684.7381435 | 497.1595601 | ENSG00000078687 |
| ARID1B | -0.460737974 | 0.726614482 | 5.54E-12 | 1.45E-10 | 1985.50878 | 1442.432564 | ENSG00000049618 |
| MBOAT2 | -0.460745913 | 0.726610484 | 0.000491191 | 0.0030334 | 395.9684142 | 287.9578951 | ENSG00000143797 |
| H2BC20P | -0.460938856 | 0.726513315 | 0.214388272 | 0.409272354 | 44.32857787 | 32.29538938 | ENSG00000261716 |
| ITFG2 | -0.461367936 | 0.726297271 | 0.001407504 | 0.007547052 | 311.3789147 | 226.3605221 | ENSG00000111203 |
| NME5 | -0.461490946 | 0.726235347 | 0.686206046 | 0.83186293 | 6.434966294 | 4.710088327 | ENSG00000112981 |
| CLDN6 | -0.461755509 | 0.726102181 | 2.20E-23 | 1.72E-21 | 14454.78987 | 10496.10078 | ENSG00000184697 |
| ANKMY2 | -0.461818819 | 0.726070318 | 0.000697938 | 0.004104817 | 317.5730588 | 230.4851223 | ENSG00000106524 |
| NA | -0.461943058 | 0.726007794 | 0.638236585 | 0.801722623 | 5.925508257 | 4.304902868 | ENSG00000277351 |
| NA | -0.462084518 | 0.725936611 | 0.67440156 | 0.825608414 | 5.005925775 | 3.616169684 | ENSG00000261033 |
| SCNN1A | -0.462154862 | 0.725901216 | 3.24E-09 | 5.72E-08 | 1674.054687 | 1215.44644 | ENSG00000111319 |
| STAG1 | -0.462213446 | 0.72587174 | 6.15E-05 | 0.000482495 | 568.7283916 | 412.6803098 | ENSG00000118007 |
| C1R | -0.462282812 | 0.72583684 | 0.04055724 | 0.120031888 | 122.3315551 | 88.67469498 | ENSG00000159403 |
| NA | -0.462673395 | 0.72564036 | 0.685529846 | 0.831762616 | 4.613598691 | 3.364970366 | ENSG00000179978 |
| MCM3 | -0.463178535 | 0.725386331 | 1.33E-25 | 1.26E-23 | 11307.89164 | 8202.210285 | ENSG00000112118 |
| LYPD6 | -0.463326091 | 0.725312144 | 0.008423619 | 0.034078845 | 184.652538 | 133.8586874 | ENSG00000187123 |
| ADRA2C | -0.46341087 | 0.725269523 | 1.53E-06 | 1.71E-05 | 709.0828863 | 514.3045864 | ENSG00000184160 |
| YARS1 | -0.463742142 | 0.725103005 | 3.04E-22 | 2.18E-20 | 6840.887889 | 4960.120532 | ENSG00000134684 |
| CCDC88B | -0.463748271 | 0.725099925 | 0.000718692 | 0.004207224 | 493.6415905 | 358.1544933 | ENSG00000168071 |
| TMEM200A | -0.464068062 | 0.724939215 | 0.000204453 | 0.001404546 | 435.4823096 | 315.9546245 | ENSG00000164484 |
| GUSBP1 | -0.464110754 | 0.724917763 | 0.336806185 | 0.548704693 | 26.20154996 | 19.01096515 | ENSG00000215190 |
| TRIM39 | -0.464392331 | 0.724776292 | 0.002414527 | 0.011952383 | 280.649979 | 203.4376374 | ENSG00000204599 |
| LINC02609 | -0.46443062 | 0.724757057 | 0.615806333 | 0.786503033 | 6.436440835 | 4.630446813 | ENSG00000233593 |
| AGBL5 | -0.464457322 | 0.724743643 | 2.20E-08 | 3.41E-07 | 1131.211665 | 819.7205433 | ENSG00000084693 |
| RARB | -0.464493708 | 0.724725364 | 0.667077193 | 0.82154232 | 4.596996537 | 3.348796162 | ENSG00000077092 |
| C4orf3 | -0.464664708 | 0.724639469 | 1.62E-07 | 2.15E-06 | 903.3401261 | 654.6131444 | ENSG00000164096 |
| DGKG | -0.46482297 | 0.724559981 | 0.517134318 | 0.713270264 | 11.98978363 | 8.692852217 | ENSG00000058866 |
| NA | -0.464959316 | 0.724491508 | 0.456741026 | 0.662717707 | 15.93469006 | 11.58264342 | ENSG00000214222 |
| PLOD3 | -0.465021806 | 0.724460127 | 2.02E-10 | 4.30E-09 | 1675.637188 | 1213.75807 | ENSG00000106397 |
| ECM1 | -0.465043507 | 0.72444923 | 0.404412458 | 0.615476848 | 17.02298998 | 12.31431879 | ENSG00000143369 |
| ZNF354B | -0.465398859 | 0.724270812 | 0.021214072 | 0.07216933 | 191.674381 | 138.9416557 | ENSG00000178338 |
| L3MBTL3 | -0.465628602 | 0.724155484 | 0.000118308 | 0.00086298 | 423.598648 | 306.5610049 | ENSG00000198945 |
| LRRC37A2 | -0.465801145 | 0.724068882 | 0.041072422 | 0.1211768 | 105.1072281 | 76.09472664 | ENSG00000238083 |
| GDAP1 | -0.46584219 | 0.724048282 | 4.60E-07 | 5.68E-06 | 828.1412019 | 599.4860516 | ENSG00000104381 |
| FXYD7 | -0.465954954 | 0.723991692 | 0.28814205 | 0.497357882 | 38.62578953 | 28.07201995 | ENSG00000221946 |
| NA | -0.466074366 | 0.723931769 | 0.619495162 | 0.788725553 | 5.996671554 | 4.342548161 | ENSG00000278356 |
| DHFR | -0.466289292 | 0.723823929 | 1.25E-08 | 2.03E-07 | 1418.486657 | 1026.800086 | ENSG00000228716 |
| NA | -0.466383578 | 0.723776626 | 0.478173816 | 0.680667428 | 11.94234144 | 8.642154143 | ENSG00000283078 |
| RSRP1 | -0.466703559 | 0.723616114 | 0.149281437 | 0.317725677 | 700.7114814 | 507.0779993 | ENSG00000117616 |
| HESX1 | -0.46675142 | 0.723592109 | 0.020728334 | 0.070838688 | 137.4021396 | 99.42722036 | ENSG00000163666 |
| RPS10 | -0.466845382 | 0.723544983 | 0.426992034 | 0.63691977 | 16.41241833 | 11.87801443 | ENSG00000124614 |
| NA | -0.466905913 | 0.723514626 | 0.071242804 | 0.185383326 | 92.7106002 | 67.15880987 | ENSG00000272086 |
| CEP126 | -0.467035237 | 0.723449773 | 0.221534139 | 0.418786007 | 36.26203891 | 26.18201267 | ENSG00000110318 |
| GINM1 | -0.46709116 | 0.723421731 | 0.00220675 | 0.011054236 | 258.9152776 | 187.4289049 | ENSG00000055211 |
| ERBB3 | -0.467196579 | 0.723368871 | 2.69E-09 | 4.82E-08 | 1253.313938 | 906.5178847 | ENSG00000065361 |
| CHRNB1 | -0.467262934 | 0.723335602 | 0.001405629 | 0.007541279 | 327.0745881 | 236.793084 | ENSG00000170175 |
| TRPC3 | -0.467265351 | 0.72333439 | 0.456808868 | 0.662717707 | 13.34201645 | 9.631555214 | ENSG00000138741 |
| GRIK1 | -0.467334005 | 0.723299969 | 0.526012756 | 0.719293069 | 9.615964993 | 6.941319293 | ENSG00000171189 |
| NOTO | -0.467335068 | 0.723299436 | 0.549168692 | 0.736475695 | 9.199916811 | 6.680472162 | ENSG00000214513 |
| CS | -0.467397242 | 0.723268266 | 5.20E-17 | 2.36E-15 | 3023.725165 | 2186.991022 | ENSG00000062485 |
| EGR1 | -0.467742762 | 0.723095067 | 0.001496398 | 0.007946983 | 645.256842 | 466.4377257 | ENSG00000120738 |
| EFL1 | -0.467768159 | 0.723082337 | 8.29E-09 | 1.39E-07 | 1093.029684 | 790.1423258 | ENSG00000140598 |
| ZNF730 | -0.467889295 | 0.723021626 | 0.106493563 | 0.250202903 | 78.41045804 | 56.60016402 | ENSG00000183850 |
| CACNB2 | -0.468004166 | 0.72296406 | 0.129392777 | 0.288022465 | 54.16161837 | 39.18818607 | ENSG00000165995 |
| NYNRIN | -0.46812013 | 0.72290595 | 8.24E-10 | 1.60E-08 | 1350.625401 | 976.073524 | ENSG00000205978 |
| C12orf10 | -0.4682653 | 0.722833212 | 4.89E-06 | 4.98E-05 | 734.6374842 | 531.3171905 | ENSG00000139637 |
| LOC102723313 | -0.468433608 | 0.722748889 | 0.560017349 | 0.745321888 | 8.219849449 | 5.940094945 | ENSG00000248896 |
| L1TD1 | -0.468656148 | 0.722637412 | 7.81E-19 | 4.31E-17 | 14976.60119 | 10822.95591 | ENSG00000240563 |
| NA | -0.468822669 | 0.722554007 | 0.120779939 | 0.273755299 | 61.00935271 | 44.01941767 | ENSG00000286132 |
| MCM2 | -0.468922076 | 0.722504223 | 1.79E-13 | 5.61E-12 | 4884.525996 | 3528.960442 | ENSG00000073111 |
| CCDC148 | -0.468928346 | 0.722501082 | 0.543124008 | 0.732008057 | 9.564963323 | 6.891850723 | ENSG00000153237 |
| NA | -0.468949131 | 0.722490673 | 0.47135875 | 0.674671062 | 12.85836445 | 9.302889847 | ENSG00000255224 |
| PAK5 | -0.469112338 | 0.722408945 | 0.396745801 | 0.608754018 | 17.86309501 | 12.88671112 | ENSG00000101349 |
| ZNF236-DT | -0.469297467 | 0.72231625 | 0.428203666 | 0.638048464 | 20.21527749 | 14.66160705 | ENSG00000264278 |
| NA | -0.469780327 | 0.722074537 | 0.659415024 | 0.816486541 | 5.491662713 | 3.99023624 | ENSG00000237877 |
| ARPC1B | -0.469825338 | 0.722052009 | 0.261957919 | 0.468977732 | 30.75941845 | 22.20142425 | ENSG00000130429 |
| CANX | -0.470033295 | 0.721947936 | 7.30E-18 | 3.61E-16 | 11691.88344 | 8440.652178 | ENSG00000127022 |
| NUP210 | -0.47007895 | 0.72192509 | 1.41E-17 | 6.68E-16 | 4421.139279 | 3191.454079 | ENSG00000132182 |
| SRSF4 | -0.470176922 | 0.721876067 | 5.37E-15 | 2.02E-13 | 3954.069818 | 2854.265965 | ENSG00000116350 |
| CNTN4 | -0.470611126 | 0.721658838 | 0.350634154 | 0.562784804 | 22.00252491 | 15.90031553 | ENSG00000144619 |

| | | | | | | | |
|---|---|---|---|---|---|---|---|
| ANKRD13A | -0.471149502 | 0.721389584 | 0.000472506 | 0.002933816 | 316.5994994 | 228.5273398 | ENSG00000076513 |
| OVOS2 | -0.471341043 | 0.721293814 | 0.083595047 | 0.208711325 | 79.31132728 | 57.11582635 | ENSG00000177359 |
| NA | -0.471434374 | 0.721247154 | 0.495505481 | 0.695165367 | 17.09298422 | 12.30684644 | ENSG00000271043 |
| SPANXA2-OT1 | -0.471575602 | 0.721176553 | 0.652470421 | 0.811703295 | 4.596996537 | 3.325149609 | ENSG00000277215 |
| VAT1L | -0.471700357 | 0.721114193 | 9.01E-14 | 2.91E-12 | 2679.532754 | 1932.083992 | ENSG00000171724 |
| ZSCAN5A | -0.471967103 | 0.720980876 | 0.096330229 | 0.231715244 | 73.68890612 | 53.08198552 | ENSG00000131848 |
| LARP6 | -0.47225227 | 0.720838379 | 2.70E-05 | 0.000233069 | 482.0020568 | 347.627207 | ENSG00000166173 |
| KCTD15 | -0.472301824 | 0.72081362 | 1.49E-08 | 2.41E-07 | 1220.262289 | 879.9907395 | ENSG00000153885 |
| NA | -0.472683926 | 0.720622735 | 0.572877007 | 0.75529058 | 8.349133361 | 6.015385531 | ENSG00000230454 |
| LINC02805 | -0.472885504 | 0.720522055 | 0.650895229 | 0.810506282 | 4.620717636 | 3.339148349 | ENSG00000232265 |
| NAP1L2 | -0.472966993 | 0.720481358 | 9.67E-06 | 9.27E-05 | 527.7282248 | 380.3503668 | ENSG00000186462 |
| TOM1 | -0.473045218 | 0.720442293 | 2.10E-06 | 2.28E-05 | 653.8507915 | 471.1326905 | ENSG00000100284 |
| WNT2B | -0.473162481 | 0.720383738 | 0.371456834 | 0.583413366 | 21.54022974 | 15.55765142 | ENSG00000134245 |
| SH2B2 | -0.473529029 | 0.720200732 | 0.000145352 | 0.001038186 | 446.3188854 | 321.7415113 | ENSG00000160999 |
| PLXDC1 | -0.473713879 | 0.72010846 | 0.463459342 | 0.667971431 | 14.38732338 | 10.3514073 | ENSG00000161381 |
| NA | -0.473836543 | 0.720047236 | 0.58481955 | 0.763520028 | 6.922177773 | 4.995811515 | ENSG00000260095 |
| MTA2 | -0.473898954 | 0.720016087 | 7.84E-20 | 4.61E-18 | 4374.660388 | 3149.872031 | ENSG00000149480 |
| NA | -0.474017305 | 0.719957023 | 0.249492787 | 0.454618998 | 33.1234484 | 23.90755598 | ENSG00000241506 |
| NA | -0.474040026 | 0.719945685 | 0.492327054 | 0.691785447 | 10.58654915 | 7.616053737 | ENSG00000250616 |
| PIGN | -0.47437095 | 0.719780564 | 0.000166956 | 0.001172545 | 432.6683651 | 311.2427168 | ENSG00000197563 |
| RASA2 | -0.474679744 | 0.719626518 | 0.000755429 | 0.004388275 | 321.1531719 | 230.9840207 | ENSG00000155903 |
| NA | -0.474879768 | 0.719526752 | 0.01213881 | 0.045977023 | 197.7356811 | 142.4511324 | ENSG00000273015 |
| SCN1A | -0.475397208 | 0.719268731 | 0.273648729 | 0.482262235 | 29.91811821 | 21.45575014 | ENSG00000144285 |
| SORL1 | -0.475693652 | 0.719120951 | 8.07E-09 | 1.35E-07 | 2370.045781 | 1703.564955 | ENSG00000137642 |
| SKP2 | -0.475861374 | 0.719037354 | 2.34E-10 | 4.93E-09 | 1720.8134 | 1237.154219 | ENSG00000145604 |
| FBXO15 | -0.47588748 | 0.719024343 | 0.426416356 | 0.636390955 | 16.14108715 | 11.62123468 | ENSG00000141665 |
| SLIT3 | -0.476193907 | 0.718871639 | 0.114761187 | 0.26436509 | 67.62404592 | 48.67228153 | ENSG00000184347 |
| SMO | -0.476330315 | 0.718803673 | 3.55E-15 | 1.37E-13 | 2734.475414 | 1965.955471 | ENSG00000128602 |
| CHST11 | -0.47644982 | 0.718744133 | 1.04E-05 | 9.89E-05 | 503.2400636 | 361.6095177 | ENSG00000171310 |
| NA | -0.476700362 | 0.718619325 | 0.357369835 | 0.569481845 | 23.1809807 | 16.60087199 | ENSG00000269918 |
| DIP2A | -0.476762913 | 0.718588168 | 5.76E-07 | 6.96E-06 | 803.5421651 | 577.343714 | ENSG00000160305 |
| H1-2 | -0.476886675 | 0.718526527 | 0.017385503 | 0.06142661 | 153.8202024 | 110.6994755 | ENSG00000187837 |
| FCHSD2 | -0.47697434 | 0.718482867 | 0.023594182 | 0.078681994 | 154.4900643 | 110.8544802 | ENSG00000137478 |
| SUDS3 | -0.477140055 | 0.718400343 | 3.81E-12 | 1.02E-10 | 2232.561063 | 1603.647938 | ENSG00000111707 |
| DUSP22 | -0.477329571 | 0.718305979 | 1.38E-06 | 1.56E-05 | 750.990126 | 539.1345759 | ENSG00000112679 |
| VPS29 | -0.477796503 | 0.718073535 | 6.58E-05 | 0.000512787 | 403.8255558 | 290.0803731 | ENSG00000111237 |
| NAT10 | -0.477802318 | 0.71807064 | 1.03E-17 | 5.01E-16 | 3429.314396 | 2462.167583 | ENSG00000135372 |
| ADNP-AS1 | -0.477806023 | 0.718068797 | 0.520497732 | 0.715685882 | 9.268715843 | 6.664297958 | ENSG00000259456 |
| EFEMP1 | -0.477884426 | 0.718029774 | 0.002312528 | 0.011504763 | 230.601452 | 165.5763876 | ENSG00000115380 |
| ATP6V1A | -0.477948161 | 0.717998054 | 8.15E-07 | 9.56E-06 | 904.5740589 | 649.325891 | ENSG00000114573 |
| PPT2 | -0.478335155 | 0.717805481 | 0.028584341 | 0.091277525 | 117.6186228 | 84.35534188 | ENSG00000221988 |
| NA | -0.478380896 | 0.717782724 | 0.604211487 | 0.778610299 | 6.990976805 | 5.024754954 | ENSG00000251593 |
| FAM218A | -0.478594102 | 0.717676655 | 0.467971912 | 0.671349859 | 13.45825768 | 9.615381011 | ENSG00000250486 |
| NA | -0.47861965 | 0.717663946 | 0.288934492 | 0.498222049 | 27.27505333 | 19.62535371 | ENSG00000255753 |
| GFOD2 | -0.478790457 | 0.717578984 | 1.00E-07 | 1.40E-06 | 923.389278 | 662.5785996 | ENSG00000141098 |
| SALL1 | -0.479138303 | 0.71740599 | 1.48E-08 | 2.39E-07 | 1101.339463 | 789.9785668 | ENSG00000103449 |
| ZBTB40 | -0.479184398 | 0.717383069 | 9.51E-06 | 9.13E-05 | 727.5960127 | 522.0825204 | ENSG00000184677 |
| NA | -0.479726575 | 0.717113521 | 0.510156224 | 0.706793361 | 10.15389881 | 7.302333068 | ENSG00000234630 |
| EMC7 | -0.4798989 | 0.71702787 | 8.52E-07 | 9.99E-06 | 768.4998206 | 550.8220728 | ENSG00000134153 |
| UBE2F | -0.479942894 | 0.717006005 | 0.017743416 | 0.062426581 | 135.5976797 | 97.14283231 | ENSG00000184182 |
| PTGR1 | -0.480022996 | 0.716966196 | 1.18E-05 | 0.000110359 | 604.870552 | 433.4477485 | ENSG00000106853 |
| ZNF322 | -0.480430951 | 0.716763486 | 0.001546778 | 0.008166314 | 263.2963421 | 188.7549381 | ENSG00000181315 |
| ABCF2 | -0.48107137 | 0.716445382 | 8.35E-05 | 0.000633496 | 400.9351602 | 287.2924296 | ENSG00000033050 |
| NA | -0.481200872 | 0.716381074 | 0.553029297 | 0.739709489 | 8.356252306 | 5.968092425 | ENSG00000286966 |
| CCDC92 | -0.481555817 | 0.716204845 | 0.280027127 | 0.489024624 | 26.8948792 | 19.24163934 | ENSG00000119242 |
| CTXN1 | -0.481559726 | 0.716202904 | 2.72E-05 | 0.000234206 | 806.0224863 | 577.6383797 | ENSG00000178531 |
| PKNOX1 | -0.48172983 | 0.716118464 | 3.82E-06 | 3.96E-05 | 610.6917711 | 437.2382615 | ENSG00000160199 |
| KDM4C | -0.481788737 | 0.716089224 | 5.19E-07 | 6.34E-06 | 716.9961154 | 513.454415 | ENSG00000107077 |
| RAVER2 | -0.481823904 | 0.716071769 | 1.15E-07 | 1.57E-06 | 822.644558 | 588.8537665 | ENSG00000162437 |
| SARS2 | -0.482003487 | 0.71598264 | 0.457722186 | 0.663253386 | 12.46247789 | 8.937525145 | ENSG00000104835 |
| IL18R1 | -0.48234278 | 0.715814275 | 0.636666946 | 0.800725077 | 6.514723076 | 4.620799 | ENSG00000115604 |
| NA | -0.482407071 | 0.715782377 | 0.648863815 | 0.809258562 | 4.613598691 | 3.299327592 | ENSG00000188451 |
| NA | -0.482499164 | 0.715736687 | 0.424184733 | 0.634226465 | 16.65290946 | 11.96200687 | ENSG00000288060 |
| GULP1 | -0.482529237 | 0.715721768 | 1.60E-07 | 2.13E-06 | 1929.662372 | 1380.673297 | ENSG00000144366 |
| EFHD1 | -0.482643678 | 0.715664996 | 0.014237633 | 0.05226 | 148.1096847 | 106.0929587 | ENSG00000115468 |
| TRIM8 | -0.482833787 | 0.715570696 | 8.26E-06 | 8.06E-05 | 577.4245471 | 413.3298549 | ENSG00000171206 |
| EFS | -0.482999335 | 0.71548859 | 8.42E-05 | 0.000637938 | 1094.300535 | 783.4317799 | ENSG00000100842 |
| MAX | -0.483022662 | 0.715477021 | 6.16E-06 | 6.16E-05 | 573.3858992 | 410.2210745 | ENSG00000125952 |
| MAP2K5 | -0.483474297 | 0.715253076 | 0.001272161 | 0.006907637 | 276.8145521 | 198.1097555 | ENSG00000137764 |
| NA | -0.483490968 | 0.715244811 | 0.529490105 | 0.721752405 | 9.292436942 | 6.642826869 | ENSG00000235790 |
| NA | -0.483909152 | 0.715037518 | 0.040478599 | 0.119836705 | 103.4747913 | 74.09416986 | ENSG00000180385 |
| SHLD1 | -0.484119027 | 0.714933507 | 0.074895319 | 0.192383452 | 84.09898233 | 60.05536967 | ENSG00000171984 |
| ZNF785 | -0.484141369 | 0.714922435 | 0.029309618 | 0.09319955 | 110.4497246 | 78.87470339 | ENSG00000197162 |
| TAGLN3 | -0.484236749 | 0.714875171 | 0.040313558 | 0.119516727 | 107.8428132 | 76.97771822 | ENSG00000144834 |

| | | | | | | | |
|---|---|---|---|---|---|---|---|
| SEPTIN2 | -0.484516806 | 0.714736413 | 3.78E-28 | 4.08E-26 | 9240.60876 | 6604.62783 | ENSG00000168385 |
| OXCT1 | -0.484617289 | 0.714686633 | 7.50E-08 | 1.07E-06 | 876.4515284 | 626.356135 | ENSG00000083720 |
| ZNF689 | -0.484618449 | 0.714686059 | 3.47E-06 | 3.62E-05 | 594.7153088 | 425.126903 | ENSG00000156853 |
| CCDC93 | -0.484747656 | 0.714622054 | 6.42E-05 | 0.000501724 | 603.1242126 | 430.9235532 | ENSG00000125633 |
| NA | -0.484863586 | 0.714564632 | 0.388381903 | 0.600408663 | 16.2525737 | 11.59976359 | ENSG00000260005 |
| DZANK1 | -0.484965231 | 0.714514289 | 0.004767837 | 0.021210976 | 232.3806646 | 165.9886665 | ENSG00000089091 |
| EPHA4 | -0.485259284 | 0.71436867 | 0.010183389 | 0.039627714 | 176.4866393 | 125.876323 | ENSG00000116106 |
| IL22RA1 | -0.485280932 | 0.714357951 | 0.64347937 | 0.805526878 | 4.644438735 | 3.313326333 | ENSG00000142677 |
| SEMA4G | -0.48539097 | 0.714303468 | 0.020555659 | 0.070363172 | 122.2212637 | 87.27719794 | ENSG00000095539 |
| GNA14 | -0.485393243 | 0.714302342 | 0.000682522 | 0.004020406 | 302.7195497 | 216.082965 | ENSG00000156049 |
| SPAG4 | -0.485475448 | 0.714261642 | 0.249753752 | 0.455015679 | 40.20961508 | 28.74892986 | ENSG00000061656 |
| NA | -0.485569734 | 0.714214964 | 0.117800291 | 0.269459605 | 61.13123834 | 43.61640767 | ENSG00000253636 |
| LRIG2-DT | -0.485671621 | 0.714164526 | 0.672350556 | 0.82439555 | 4.698999877 | 3.355322553 | ENSG00000238198 |
| CCDC27 | -0.485713701 | 0.714143696 | 0.559714622 | 0.745115652 | 8.425051338 | 5.996089905 | ENSG00000162592 |
| TSPAN5 | -0.486143717 | 0.713930867 | 4.41E-06 | 4.52E-05 | 692.4175446 | 494.2244404 | ENSG00000168785 |
| WASH5P | -0.48637482 | 0.713816512 | 0.232285862 | 0.433213362 | 31.14343138 | 22.21542299 | ENSG00000282458 |
| DNAL1 | -0.486418026 | 0.713795135 | 0.006968265 | 0.029153373 | 178.7706331 | 127.5930485 | ENSG00000119661 |
| NA | -0.486613515 | 0.713698421 | 0.285099154 | 0.494316702 | 27.98526406 | 19.95089765 | ENSG00000228623 |
| FLJ13224 | -0.487073572 | 0.713470868 | 0.658023949 | 0.815526679 | 4.70255935 | 3.353147089 | ENSG00000177340 |
| PNPLA6 | -0.487201989 | 0.713407363 | 1.43E-14 | 5.04E-13 | 2785.256834 | 1987.064215 | ENSG00000032444 |
| HDGFL3 | -0.487416086 | 0.713301501 | 9.52E-15 | 3.47E-13 | 2341.858775 | 1670.309383 | ENSG00000166503 |
| FHL1 | -0.487566909 | 0.713226934 | 3.27E-10 | 6.77E-09 | 1211.836707 | 864.351607 | ENSG00000022267 |
| NA | -0.487692283 | 0.713164956 | 0.641831395 | 0.804214244 | 4.655117152 | 3.313326333 | ENSG00000256995 |
| SGPP1 | -0.487896935 | 0.713063797 | 0.003114144 | 0.01486994 | 223.900188 | 159.8568675 | ENSG00000126821 |
| POU4F1 | -0.48830005 | 0.712864583 | 0.628473555 | 0.794869303 | 5.625701305 | 4.023530606 | ENSG00000152192 |
| EFNB3 | -0.488503539 | 0.712764042 | 2.88E-06 | 3.05E-05 | 587.0226363 | 418.5214379 | ENSG00000108947 |
| NA | -0.489066634 | 0.712485898 | 0.436573805 | 0.644973505 | 13.5457438 | 9.671375971 | ENSG00000234618 |
| STAC2 | -0.489182865 | 0.712428499 | 0.001656461 | 0.008655857 | 238.703865 | 170.0248508 | ENSG00000141750 |
| PARD6A | -0.489617646 | 0.712213829 | 0.004244891 | 0.019312158 | 213.1593869 | 151.7820924 | ENSG00000102981 |
| NA | -0.489698328 | 0.712174 | 0.30274556 | 0.5128588 | 26.1318612 | 18.53796146 | ENSG00000234743 |
| NUP205 | -0.489707496 | 0.712169474 | 5.08E-15 | 1.92E-13 | 5615.174768 | 3998.100312 | ENSG00000155561 |
| RET | -0.489733698 | 0.71215654 | 0.000157471 | 0.001114218 | 416.5502127 | 296.8587209 | ENSG00000165731 |
| PRKY | -0.489747517 | 0.712149719 | 0.502901249 | 0.700907017 | 10.19422206 | 7.257215425 | ENSG00000099725 |
| DLST | -0.48996243 | 0.71204364 | 8.63E-15 | 3.18E-13 | 2880.195746 | 2050.385062 | ENSG00000119689 |
| HMG20B | -0.490031433 | 0.712009584 | 1.72E-13 | 5.39E-12 | 1786.530878 | 1272.178334 | ENSG00000064961 |
| PREX1 | -0.490056274 | 0.711997325 | 2.18E-06 | 2.36E-05 | 837.0538742 | 595.8154523 | ENSG00000124126 |
| NKX1-2 | -0.49045347 | 0.711801328 | 2.62E-05 | 0.000226924 | 506.2591584 | 360.283768 | ENSG00000229544 |
| PGRMC1 | -0.490508572 | 0.711774142 | 1.51E-15 | 6.10E-14 | 6511.558211 | 4635.088316 | ENSG00000101856 |
| KIF9 | -0.490797049 | 0.711631832 | 0.05296008 | 0.147384484 | 89.4053516 | 63.65129777 | ENSG00000088727 |
| JRK | -0.491050438 | 0.711506855 | 4.61E-09 | 8.01E-08 | 973.5102943 | 692.7241235 | ENSG00000234616 |
| PCDHB16 | -0.491109985 | 0.711477488 | 0.050035797 | 0.141139064 | 96.37110662 | 68.65240617 | ENSG00000272674 |
| USF2 | -0.49127346 | 0.711396874 | 1.63E-14 | 5.69E-13 | 2717.990306 | 1933.697067 | ENSG00000105698 |
| PHF14 | -0.491635852 | 0.7112182 | 2.81E-09 | 5.01E-08 | 1342.901278 | 954.9267124 | ENSG00000106443 |
| NAGPA | -0.491673878 | 0.711199454 | 0.002333064 | 0.011591663 | 232.1467338 | 165.2093419 | ENSG00000103174 |
| NA | -0.492626908 | 0.710729798 | 0.356068425 | 0.568367188 | 18.24830315 | 12.9685281 | ENSG00000276248 |
| TUG1 | -0.492679381 | 0.710703948 | 8.60E-12 | 2.20E-10 | 1882.072838 | 1337.425663 | ENSG00000253352 |
| PPM1M | -0.492726703 | 0.710680637 | 0.041770723 | 0.122681198 | 119.0981063 | 84.59857439 | ENSG00000164088 |
| WARS2-AS1 | -0.492816858 | 0.710636227 | 0.192393759 | 0.379705144 | 40.77007479 | 29.00041272 | ENSG00000231365 |
| PDAP1 | -0.492872765 | 0.710608689 | 1.37E-17 | 6.52E-16 | 6468.64288 | 4596.915529 | ENSG00000106244 |
| NA | -0.492918878 | 0.710585977 | 0.212519239 | 0.406587563 | 40.1574182 | 28.50499198 | ENSG00000256663 |
| EFEMP2 | -0.492964315 | 0.710563597 | 0.094112976 | 0.22821309 | 88.97590353 | 63.16327674 | ENSG00000172638 |
| SNW1 | -0.493229088 | 0.710433202 | 1.27E-09 | 2.41E-08 | 1109.42878 | 788.0014027 | ENSG00000100603 |
| DDX58 | -0.493298782 | 0.710398883 | 0.023236678 | 0.077693865 | 116.7293217 | 82.92500197 | ENSG00000107201 |
| NA | -0.493461987 | 0.710318523 | 0.50968962 | 0.706348308 | 10.35554123 | 7.355206606 | ENSG00000273188 |
| NA | -0.493642258 | 0.710229772 | 0.642581612 | 0.804771784 | 5.577063899 | 3.963184719 | ENSG00000237082 |
| WDR90 | -0.493782377 | 0.710160795 | 5.46E-05 | 0.00043569 | 794.2397636 | 564.4552695 | ENSG00000161996 |
| NA | -0.493892886 | 0.7101064 | 0.601518492 | 0.776517651 | 7.952077749 | 5.646899406 | ENSG00000181227 |
| LINC00638 | -0.494009471 | 0.710049018 | 0.423223546 | 0.633337985 | 14.46769055 | 10.29134496 | ENSG00000258701 |
| NA | -0.494119076 | 0.709995076 | 0.051420487 | 0.144204916 | 105.0099795 | 74.61418312 | ENSG00000186301 |
| SLITRK3 | -0.49421933 | 0.70994574 | 0.108047562 | 0.252816969 | 61.24956451 | 43.5664876 | ENSG00000121871 |
| FUT8 | -0.494400533 | 0.709856575 | 9.14E-06 | 8.83E-05 | 480.1530776 | 340.7522655 | ENSG00000033170 |
| NA | -0.494415426 | 0.709849248 | 0.49801606 | 0.696978902 | 12.16414548 | 8.579632792 | ENSG00000182165 |
| PAQR5 | -0.494431208 | 0.709841482 | 0.001749007 | 0.009066703 | 218.9307211 | 155.3905061 | ENSG00000137819 |
| ZNF626 | -0.494635874 | 0.709740789 | 0.03116767 | 0.097871873 | 132.8653487 | 94.29605592 | ENSG00000188171 |
| BCL2L1 | -0.494817409 | 0.709651488 | 1.14E-06 | 1.31E-05 | 689.1292559 | 489.085395 | ENSG00000171552 |
| VPS11 | -0.494840672 | 0.709640045 | 4.62E-05 | 0.000374593 | 475.7468181 | 337.5945337 | ENSG00000160695 |
| SLC16A14 | -0.494874833 | 0.709623242 | 0.043973798 | 0.127603744 | 96.9013111 | 68.80639231 | ENSG00000163053 |
| YTHDC2 | -0.495315486 | 0.709406529 | 8.73E-08 | 1.23E-06 | 777.0695889 | 551.1635503 | ENSG00000047188 |
| RPS6KA2 | -0.495510669 | 0.70931056 | 2.18E-07 | 2.84E-06 | 771.6606321 | 547.1613228 | ENSG00000071242 |
| NAA40 | -0.495544035 | 0.709294155 | 7.15E-07 | 8.48E-06 | 947.8810265 | 672.1739929 | ENSG00000110583 |
| ADHFE1 | -0.495622915 | 0.709255376 | 0.68828398 | 0.833414711 | 4.170269938 | 2.975959111 | ENSG00000147576 |
| NA | -0.49582585 | 0.709155616 | 0.683800248 | 0.830895404 | 4.231950025 | 3.023252217 | ENSG00000228748 |
| NA | -0.495873214 | 0.709132335 | 0.013069168 | 0.048699929 | 159.7883987 | 113.225415 | ENSG00000280128 |
| CCDC183-AS1 | -0.495975857 | 0.709081884 | 0.607547067 | 0.781288725 | 5.584182844 | 3.960063296 | ENSG00000228544 |

| | | | | | | | |
|---|---|---|---|---|---|---|---|
| ZDHHC14 | -0.496239427 | 0.708952352 | 0.002112446 | 0.010667673 | 210.0481031 | 148.8849968 | ENSG00000175048 |
| PRRG2 | -0.496271628 | 0.708936528 | 0.192395718 | 0.379705144 | 49.44154099 | 34.97832091 | ENSG00000126460 |
| CDS1 | -0.496579874 | 0.708785073 | 4.62E-05 | 0.000374593 | 412.261896 | 292.3800848 | ENSG00000163624 |
| HERC2P9 | -0.496587 | 0.708781572 | 7.70E-05 | 0.000588863 | 440.3173557 | 312.0728052 | ENSG00000206149 |
| COA6-AS1 | -0.496782501 | 0.708685531 | 0.616850876 | 0.787224317 | 5.673143503 | 4.02570607 | ENSG00000231663 |
| NA | -0.496803648 | 0.708675143 | 0.210563597 | 0.404074132 | 36.61996648 | 25.92334101 | ENSG00000250994 |
| BHLHE40 | -0.497121999 | 0.708518781 | 0.116617197 | 0.267596083 | 70.41625144 | 49.74472908 | ENSG00000134107 |
| NUDT15 | -0.497308511 | 0.70842719 | 1.82E-06 | 2.00E-05 | 795.8614697 | 563.8075304 | ENSG00000136159 |
| PPP6R3 | -0.497334496 | 0.70841443 | 4.36E-11 | 1.03E-09 | 2036.845931 | 1442.856076 | ENSG00000110075 |
| ANAPC2 | -0.497770125 | 0.708200553 | 5.52E-10 | 1.10E-08 | 1367.366685 | 968.784425 | ENSG00000176248 |
| NA | -0.497824463 | 0.70817388 | 0.672142677 | 0.82439555 | 4.627836581 | 3.261682299 | ENSG00000274213 |
| NA | -0.498087268 | 0.708044888 | 0.483459159 | 0.68442107 | 13.72366512 | 9.718669077 | ENSG00000282381 |
| TP53BP1 | -0.49816007 | 0.70800916 | 2.81E-15 | 1.10E-13 | 2682.557494 | 1899.174446 | ENSG00000067369 |
| URM1 | -0.498653782 | 0.70776691 | 1.21E-09 | 2.31E-08 | 1294.543917 | 916.5006808 | ENSG00000167118 |
| GANC | -0.498805373 | 0.707692545 | 0.013061362 | 0.048687419 | 147.4347376 | 104.2654728 | ENSG00000214013 |
| NA | -0.498884857 | 0.707653557 | 0.664222242 | 0.820126646 | 5.696864602 | 4.034407924 | ENSG00000254554 |
| ZDHHC22 | -0.499011818 | 0.707591284 | 1.31E-12 | 3.70E-11 | 1517.961397 | 1074.242164 | ENSG00000177108 |
| ABCC1 | -0.499103198 | 0.707546467 | 4.50E-13 | 1.35E-11 | 2607.165853 | 1844.31584 | ENSG00000103222 |
| ABCA3 | -0.49919961 | 0.707499185 | 7.07E-06 | 7.00E-05 | 476.1708226 | 336.904571 | ENSG00000167972 |
| SPDYE5 | -0.499467557 | 0.707367795 | 0.680691146 | 0.829345309 | 5.122167005 | 3.585050781 | ENSG00000170092 |
| NA | -0.499653842 | 0.707276464 | 0.330356755 | 0.541788157 | 21.49662634 | 15.17734202 | ENSG00000272578 |
| NA | -0.499667968 | 0.707269538 | 0.375470376 | 0.587518275 | 18.41910552 | 13.00834886 | ENSG00000272449 |
| NA | -0.500126324 | 0.707044869 | 0.087986578 | 0.216526783 | 62.0487359 | 43.85315675 | ENSG00000237638 |
| ACVR2B-AS1 | -0.500152515 | 0.707032033 | 0.172572943 | 0.352176359 | 41.40408821 | 29.29672968 | ENSG00000229589 |
| ADCY4 | -0.50065964 | 0.706783546 | 0.564174761 | 0.748528238 | 7.04909742 | 4.953815295 | ENSG00000129467 |
| RIMS4 | -0.500715207 | 0.706756324 | 4.33E-13 | 1.30E-11 | 2059.209559 | 1455.267086 | ENSG00000101098 |
| RPL26L1 | -0.500909434 | 0.706661182 | 3.89E-06 | 4.02E-05 | 589.5171459 | 416.6351621 | ENSG00000037241 |
| FRY | -0.501075841 | 0.706579677 | 0.0069304 | 0.029032966 | 179.0621259 | 126.4801178 | ENSG00000073910 |
| HSBP1L1 | -0.50119445 | 0.706521589 | 0.388860419 | 0.600656717 | 15.53079483 | 10.99067192 | ENSG00000226742 |
| NA | -0.501614706 | 0.70631581 | 0.35461434 | 0.566764709 | 17.87377343 | 12.63021492 | ENSG00000275807 |
| LOC101930370 | -0.501709014 | 0.70626964 | 0.15722945 | 0.329443933 | 46.64930991 | 32.99036541 | ENSG00000245213 |
| NA | -0.501729103 | 0.706259805 | 0.617024805 | 0.787224317 | 5.696864602 | 4.023530606 | ENSG00000213695 |
| EFHD2 | -0.501818301 | 0.70621614 | 7.92E-11 | 1.80E-09 | 1339.01251 | 945.9226212 | ENSG00000142634 |
| PLAC9 | -0.502027317 | 0.706113832 | 0.076929824 | 0.196190904 | 64.83712818 | 45.7793689 | ENSG00000189129 |
| LOC100507283 | -0.502315862 | 0.705972621 | 0.315080297 | 0.525450848 | 21.62710546 | 15.23428294 | ENSG00000245498 |
| FAM122B | -0.502580357 | 0.705843204 | 1.42E-08 | 2.30E-07 | 1065.699257 | 752.4810437 | ENSG00000156504 |
| NA | -0.50279471 | 0.705738338 | 0.000490348 | 0.0030301 | 378.2915291 | 267.0645333 | ENSG00000166492 |
| RIMS3 | -0.503044682 | 0.705616068 | 1.64E-08 | 2.62E-07 | 1050.158038 | 740.9326929 | ENSG00000117016 |
| SOCS2 | -0.503605339 | 0.705341906 | 0.000193036 | 0.001336324 | 340.3136331 | 240.1174556 | ENSG00000120833 |
| KCNJ10 | -0.50364043 | 0.70532475 | 0.276098015 | 0.485068627 | 26.35903032 | 18.62602128 | ENSG00000177807 |
| FIGNL2 | -0.503813526 | 0.70524013 | 0.012568675 | 0.047226308 | 159.0666193 | 112.2190617 | ENSG00000261308 |
| NPEPPS | -0.503991202 | 0.70515328 | 4.94E-09 | 8.55E-08 | 1696.345374 | 1195.894798 | ENSG00000141279 |
| NA | -0.504196241 | 0.70505307 | 0.019865363 | 0.068372038 | 134.3705865 | 94.87350455 | ENSG00000211772 |
| LOC100506142 | -0.504492426 | 0.704908337 | 0.653479325 | 0.812465428 | 4.231950025 | 3.001781127 | ENSG00000250116 |
| MORF4L2 | -0.504558327 | 0.704876138 | 8.82E-23 | 6.58E-21 | 6898.447173 | 4861.85996 | ENSG00000123562 |
| EIF2B1 | -0.504580693 | 0.704865211 | 1.34E-11 | 3.36E-10 | 1350.480657 | 952.0988981 | ENSG00000111361 |
| RARA-AS1 | -0.504847774 | 0.704734734 | 0.654968028 | 0.813380102 | 4.215347871 | 2.987782387 | ENSG00000265666 |
| FAM151B | -0.505288711 | 0.704519375 | 0.556544164 | 0.742884968 | 7.565674402 | 5.352474363 | ENSG00000152380 |
| NCAM2 | -0.505444663 | 0.704443223 | 0.22879344 | 0.428477202 | 30.62329493 | 21.51921745 | ENSG00000154654 |
| PIGS | -0.505506565 | 0.704412998 | 9.79E-12 | 2.48E-10 | 1271.416993 | 895.6010332 | ENSG00000087111 |
| TMEM117 | -0.50565506 | 0.704340497 | 0.28540973 | 0.494673527 | 28.61777678 | 20.15915479 | ENSG00000139173 |
| JPT1 | -0.505704355 | 0.704316431 | 8.07E-21 | 5.10E-19 | 4493.852534 | 3164.800303 | ENSG00000189159 |
| AKT1S1 | -0.505945097 | 0.704198912 | 1.79E-09 | 3.32E-08 | 1482.554641 | 1044.538168 | ENSG00000204673 |
| NA | -0.506035297 | 0.704154886 | 0.471403933 | 0.674684598 | 10.79027649 | 7.6182292 | ENSG00000230979 |
| NA | -0.506210744 | 0.704069258 | 0.36407989 | 0.576091222 | 17.8286955 | 12.5343992 | ENSG00000214975 |
| BICC1 | -0.506346471 | 0.704003023 | 0.18455748 | 0.36874397 | 37.24566575 | 26.16801393 | ENSG00000122870 |
| NA | -0.506811095 | 0.703776333 | 0.455559918 | 0.661432233 | 15.71108041 | 11.06284108 | ENSG00000161132 |
| MGC12916 | -0.507427261 | 0.703475819 | 0.021542992 | 0.073080548 | 127.9967416 | 90.11780413 | ENSG00000266709 |
| EIF2B2 | -0.507591173 | 0.703395898 | 8.12E-08 | 1.15E-06 | 719.8998847 | 506.6015344 | ENSG00000119718 |
| ABTB1 | -0.507654906 | 0.703364825 | 0.033814232 | 0.104428743 | 104.856669 | 73.87938633 | ENSG00000114626 |
| STAG3L4 | -0.507694796 | 0.703345378 | 0.025202551 | 0.082987771 | 132.8727214 | 93.28416517 | ENSG00000106610 |
| PCSK4 | -0.50807384 | 0.70316061 | 0.026874485 | 0.087233526 | 135.9354456 | 95.65152706 | ENSG00000115257 |
| CATSPERG | -0.5081774 | 0.703110137 | 0.432726111 | 0.642051587 | 12.28603111 | 8.630330866 | ENSG00000099338 |
| FBN3 | -0.508208493 | 0.703094984 | 2.00E-08 | 3.14E-07 | 1923.496276 | 1352.67516 | ENSG00000142449 |
| MISP3 | -0.508250276 | 0.703074621 | 0.128160789 | 0.285819143 | 48.7245168 | 34.27107011 | ENSG00000141854 |
| CRYBG1 | -0.508293764 | 0.703053428 | 1.62E-06 | 1.80E-05 | 612.143032 | 430.1537905 | ENSG00000112297 |
| NA | -0.508398986 | 0.703002154 | 0.070219278 | 0.183212362 | 69.830901 | 49.16893178 | ENSG00000273117 |
| NA | -0.508619107 | 0.7028949 | 0.287460944 | 0.496587992 | 24.09077449 | 16.95564292 | ENSG00000229036 |
| PGAM4 | -0.508977466 | 0.702720326 | 0.58030257 | 0.760936651 | 8.02560531 | 5.62107739 | ENSG00000226784 |
| TTLL2 | -0.509415179 | 0.702507153 | 0.663193461 | 0.819398128 | 4.774917855 | 3.336972886 | ENSG00000120440 |
| ZNF514 | -0.509451144 | 0.702489641 | 0.004880307 | 0.021624731 | 218.8945939 | 153.9077872 | ENSG00000144026 |
| C13orf42 | -0.509455774 | 0.702487386 | 3.11E-09 | 5.52E-08 | 1329.436746 | 933.9948843 | ENSG00000226792 |
| GNB1L | -0.510389911 | 0.702032677 | 0.000429025 | 0.002689058 | 390.9143579 | 274.4040172 | ENSG00000185838 |
| C17orf75 | -0.510607916 | 0.701926601 | 6.38E-07 | 7.63E-06 | 738.2718797 | 518.4780787 | ENSG00000108666 |

| | | | | | | | |
|---|---|---|---|---|---|---|---|
| NA | -0.510708379 | 0.701877724 | 0.130671211 | 0.290104314 | 53.00034955 | 37.19822306 | ENSG00000279673 |
| NA | -0.510846469 | 0.701810546 | 0.429599702 | 0.639472718 | 13.28981958 | 9.328711863 | ENSG00000250838 |
| SSBP2 | -0.510852742 | 0.701807494 | 5.97E-05 | 0.000471201 | 379.1526599 | 266.0588627 | ENSG00000145687 |
| LZTR1 | -0.510964831 | 0.70175297 | 0.005294949 | 0.023198386 | 168.3451238 | 118.2448731 | ENSG00000099949 |
| B4GAT1 | -0.510996869 | 0.701737386 | 2.16E-08 | 3.36E-07 | 762.5941946 | 535.2341234 | ENSG00000174684 |
| NA | -0.511088297 | 0.701692916 | 0.188575443 | 0.374552717 | 37.21691063 | 26.11201897 | ENSG00000225285 |
| ARMC2 | -0.511189265 | 0.70164381 | 0.096735023 | 0.232413589 | 57.77646262 | 40.50653605 | ENSG00000118690 |
| NA | -0.511422259 | 0.701530504 | 0.430858685 | 0.640589378 | 14.695139 | 10.36446008 | ENSG00000273796 |
| GRHL1 | -0.51147173 | 0.701506448 | 0.055151647 | 0.152227251 | 85.74809918 | 60.25977034 | ENSG00000134317 |
| EZH2 | -0.511524277 | 0.701480898 | 8.08E-11 | 1.83E-09 | 1598.865804 | 1121.757037 | ENSG00000106462 |
| MIR1244-3 | -0.511713374 | 0.701388959 | 0.267664392 | 0.475391897 | 30.40769395 | 21.29601562 | ENSG00000283429 |
| TPX2 | -0.511820811 | 0.701336729 | 6.90E-21 | 4.39E-19 | 6162.240181 | 4321.078076 | ENSG00000088325 |
| HSPA1L | -0.512165165 | 0.701169348 | 0.093246467 | 0.226489972 | 60.48390291 | 42.4853382 | ENSG00000204390 |
| SLC12A1 | -0.512184702 | 0.701159853 | 0.576101134 | 0.758153886 | 7.494511106 | 5.25883411 | ENSG00000074803 |
| LOC100130111 | -0.512189656 | 0.701157446 | 0.246935157 | 0.451745658 | 29.74495157 | 20.84136159 | ENSG00000256802 |
| CCDC200 | -0.512487255 | 0.701012826 | 0.662698393 | 0.81910198 | 4.774917855 | 3.327325073 | ENSG00000236383 |
| PCTP | -0.512513113 | 0.701000261 | 5.39E-06 | 5.46E-05 | 490.1284703 | 343.6174642 | ENSG00000141179 |
| ZNF793 | -0.512640427 | 0.700938403 | 0.015310323 | 0.05538473 | 164.3705464 | 115.24801 | ENSG00000188227 |
| NA | -0.512646252 | 0.700935572 | 0.473499666 | 0.676607233 | 11.88658509 | 8.353309532 | ENSG00000225808 |
| PIANP | -0.51311491 | 0.700707911 | 0.004704513 | 0.020998519 | 235.4513714 | 165.1353761 | ENSG00000139200 |
| EFR3B | -0.513190113 | 0.700671386 | 1.43E-05 | 0.000131278 | 449.7643762 | 315.2000805 | ENSG00000084710 |
| NA | -0.513255015 | 0.700639866 | 0.638441061 | 0.80191325 | 4.266349541 | 3.001781127 | ENSG00000232909 |
| NA | -0.513457152 | 0.700541706 | 0.436835569 | 0.645089622 | 12.32991384 | 8.616332126 | ENSG00000261420 |
| NA | -0.513619243 | 0.700463002 | 0.541740631 | 0.731030498 | 9.457315578 | 6.578130054 | ENSG00000228010 |
| NA | -0.513695917 | 0.700425776 | 0.677329457 | 0.827331142 | 4.252111651 | 2.985606923 | ENSG00000274425 |
| MAOA | -0.513750752 | 0.700399155 | 0.003677335 | 0.017096621 | 196.7823101 | 137.8136647 | ENSG00000189221 |
| ATPSCKMT | -0.513880281 | 0.700336274 | 0.003930354 | 0.018094765 | 180.2827106 | 126.2728066 | ENSG00000150756 |
| MYZAP | -0.51395369 | 0.700300639 | 0.545406378 | 0.733722678 | 7.559750666 | 5.30083033 | ENSG00000263155 |
| EPB41L1 | -0.514209629 | 0.700176415 | 1.20E-08 | 1.95E-07 | 918.3686335 | 643.2573187 | ENSG00000088367 |
| TCF7 | -0.514221153 | 0.700170822 | 0.000487188 | 0.003013607 | 275.981566 | 193.2687605 | ENSG00000081059 |
| CCDC154 | -0.514386272 | 0.700090691 | 0.528557149 | 0.721228451 | 9.447526884 | 6.638475942 | ENSG00000197599 |
| NA | -0.514423329 | 0.700072708 | 0.370989423 | 0.583115709 | 17.05383002 | 11.88548677 | ENSG00000230383 |
| MYBL2 | -0.514499378 | 0.700035806 | 8.30E-24 | 6.81E-22 | 6508.754195 | 4556.732104 | ENSG00000101057 |
| ZER1 | -0.514744221 | 0.699917012 | 1.19E-07 | 1.63E-06 | 687.4665116 | 481.2546531 | ENSG00000160445 |
| NA | -0.514860561 | 0.699860572 | 0.619412879 | 0.788725553 | 5.213212595 | 3.66998918 | ENSG00000242571 |
| ARMC5 | -0.514969041 | 0.69980795 | 0.000296838 | 0.001945775 | 297.1238243 | 208.0149271 | ENSG00000140691 |
| NA | -0.515053906 | 0.699766786 | 0.648336882 | 0.809147384 | 4.262790069 | 2.992133314 | ENSG00000261625 |
| TET1 | -0.515060045 | 0.699763808 | 3.35E-07 | 4.25E-06 | 1361.038478 | 951.9589271 | ENSG00000138336 |
| FOXI3 | -0.515190561 | 0.699700505 | 0.007244628 | 0.030108891 | 171.2168585 | 119.8433658 | ENSG00000214336 |
| TCN2 | -0.515213777 | 0.699689246 | 2.33E-12 | 6.41E-11 | 1339.550445 | 937.0926921 | ENSG00000185339 |
| CFAP20 | -0.515585685 | 0.699508898 | 1.91E-06 | 2.10E-05 | 559.2058626 | 391.1404945 | ENSG00000070761 |
| PKD1P6-NPIPP1 | -0.515776824 | 0.699416229 | 0.20941227 | 0.402601111 | 33.74945315 | 23.62618372 | ENSG00000270580 |
| EID1 | -0.515838297 | 0.699386427 | 3.39E-15 | 1.31E-13 | 2886.237595 | 2019.024311 | ENSG00000255302 |
| CACFD1 | -0.515887278 | 0.699362682 | 8.64E-07 | 1.01E-05 | 665.1379898 | 465.3003501 | ENSG00000160325 |
| ATP7B | -0.516023149 | 0.699296821 | 0.000283971 | 0.001875761 | 314.7677588 | 219.9924981 | ENSG00000123191 |
| ZNF512 | -0.516070176 | 0.699274026 | 2.02E-05 | 0.000179245 | 456.9240699 | 319.6353956 | ENSG00000243943 |
| ATP6V0E1 | -0.516368485 | 0.699129451 | 1.16E-12 | 3.29E-11 | 1434.384811 | 1002.843243 | ENSG00000113732 |
| NUSAP1 | -0.516530103 | 0.699051135 | 0.644070193 | 0.805941266 | 4.293630113 | 3.01142894 | ENSG00000137804 |
| C1orf216 | -0.516541361 | 0.69904568 | 7.65E-05 | 0.000585467 | 342.3126688 | 239.4268305 | ENSG00000142686 |
| SPICE1 | -0.516850866 | 0.698895729 | 0.569788126 | 0.752794793 | 6.663889282 | 4.649742439 | ENSG00000163611 |
| CUL7 | -0.516861436 | 0.698890608 | 1.22E-05 | 0.00011333 | 590.9624581 | 412.7433685 | ENSG00000044090 |
| HSPA14 | -0.51698374 | 0.698831362 | 5.23E-05 | 0.000418269 | 408.0335316 | 285.2498766 | ENSG00000187522 |
| MED24 | -0.517006388 | 0.698820391 | 1.54E-19 | 8.95E-18 | 2790.784904 | 1950.353931 | ENSG00000008838 |
| TKFC | -0.517168512 | 0.698741866 | 5.06E-07 | 6.21E-06 | 645.1937908 | 450.8061609 | ENSG00000149476 |
| BRINP1 | -0.517177325 | 0.698737597 | 4.44E-08 | 6.55E-07 | 919.8878561 | 642.9140648 | ENSG00000078725 |
| GSTM4 | -0.517226128 | 0.698713961 | 0.000554278 | 0.003353907 | 299.4083506 | 209.3271296 | ENSG00000168765 |
| NA | -0.517305422 | 0.698675559 | 0.51092945 | 0.707345698 | 8.516096929 | 5.963741498 | ENSG00000236540 |
| FBXL14 | -0.517370674 | 0.698643959 | 5.12E-05 | 0.000410534 | 369.6591648 | 258.2086868 | ENSG00000171823 |
| ANKRD42 | -0.517613971 | 0.698526149 | 0.032319872 | 0.100746102 | 108.9928187 | 76.2368895 | ENSG00000137494 |
| NAA10 | -0.517641937 | 0.698512609 | 9.97E-05 | 0.000739926 | 384.88456 | 269.1234487 | ENSG00000102030 |
| FAM8A1 | -0.517804432 | 0.698433938 | 0.000866666 | 0.004938743 | 265.2624523 | 185.4162413 | ENSG00000137414 |
| ARHGAP18 | -0.517816565 | 0.698428064 | 0.019079423 | 0.066088574 | 128.6837899 | 89.78072045 | ENSG00000146376 |
| MIF4GD | -0.518159513 | 0.698262058 | 0.001006184 | 0.005638513 | 251.590348 | 175.5416215 | ENSG00000125457 |
| TSPEAR-AS1 | -0.518390903 | 0.698150074 | 0.083144665 | 0.207861662 | 70.66566711 | 49.24495741 | ENSG00000235890 |
| LOC101927989 | -0.518529447 | 0.698083033 | 0.171990112 | 0.351366644 | 42.38891025 | 29.60704538 | ENSG00000265778 |
| NA | -0.518791361 | 0.697956311 | 0.01021295 | 0.039734562 | 173.0174279 | 120.8059263 | ENSG00000247092 |
| NA | -0.518820944 | 0.697942 | 0.516307613 | 0.712493188 | 9.51040218 | 6.659001072 | ENSG00000184616 |
| C5 | -0.518996277 | 0.697857183 | 0.533745919 | 0.724879869 | 8.097963815 | 5.612375535 | ENSG00000106804 |
| CCDC173 | -0.519077621 | 0.697817836 | 0.385415943 | 0.597584572 | 17.00015861 | 11.87583896 | ENSG00000154479 |
| FEZF1-AS1 | -0.519168564 | 0.69777385 | 0.002496559 | 0.012303631 | 288.0825569 | 201.199145 | ENSG00000230316 |
| PNMA1 | -0.519477162 | 0.697624609 | 2.26E-15 | 8.95E-14 | 2144.669595 | 1496.260994 | ENSG00000176903 |
| NA | -0.519656776 | 0.697537761 | 0.557770578 | 0.743680977 | 6.691169853 | 4.65626883 | ENSG00000243230 |
| RHBDL3 | -0.520018263 | 0.697363005 | 1.59E-06 | 1.77E-05 | 546.0767774 | 381.019142 | ENSG00000141314 |
| NA | -0.520136293 | 0.697305955 | 0.470347405 | 0.673734128 | 10.94091724 | 7.621350622 | ENSG00000281849 |

| | | | | | | | |
|---|---|---|---|---|---|---|---|
| TES | -0.520437247 | 0.697160508 | 1.63E-06 | 1.81E-05 | 534.2844146 | 372.3265733 | ENSG00000135269 |
| NA | -0.520896421 | 0.696938654 | 0.183262762 | 0.366977969 | 46.82222336 | 32.70369627 | ENSG00000231793 |
| NCAPG | -0.520946592 | 0.696914418 | 1.15E-08 | 1.88E-07 | 1207.098954 | 841.0741919 | ENSG00000109805 |
| ABHD16A | -0.520999771 | 0.69688873 | 0.456896737 | 0.662717707 | 10.97175728 | 7.637524826 | ENSG00000204427 |
| MPP2 | -0.521025426 | 0.696876338 | 8.77E-18 | 4.29E-16 | 2428.928494 | 1692.481056 | ENSG00000108852 |
| NA | -0.521043593 | 0.696867562 | 0.583496559 | 0.762833684 | 6.162439913 | 4.318901608 | ENSG00000227440 |
| PCYT1B | -0.52123344 | 0.696775866 | 3.59E-11 | 8.54E-10 | 1152.4799 | 802.8656865 | ENSG00000102230 |
| TTLL4 | -0.521506151 | 0.696644168 | 1.04E-14 | 3.79E-13 | 2045.676962 | 1425.439529 | ENSG00000135912 |
| MYO3B | -0.521574907 | 0.696610968 | 0.519577241 | 0.715241508 | 8.999748935 | 6.235465946 | ENSG00000071909 |
| NA | -0.521602361 | 0.696597712 | 0.335058524 | 0.547084577 | 18.07038183 | 12.61404072 | ENSG00000269119 |
| ZBTB32 | -0.521849789 | 0.696478253 | 0.400662502 | 0.612078253 | 15.77751518 | 10.9499052 | ENSG00000011590 |
| NA | -0.521921901 | 0.696443441 | 0.64428877 | 0.805989478 | 4.324470156 | 3.017955331 | ENSG00000260306 |
| TREX2 | -0.522305064 | 0.696258498 | 0.481289253 | 0.683198091 | 10.48454581 | 7.327209126 | ENSG00000183479 |
| POU6F1 | -0.522514251 | 0.69615755 | 0.02380973 | 0.079218947 | 101.9615187 | 71.04292016 | ENSG00000184271 |
| DDN | -0.522632072 | 0.696100699 | 0.47510599 | 0.677841249 | 10.06285322 | 6.983315513 | ENSG00000181418 |
| H3C2 | -0.522834003 | 0.696003274 | 0.625267148 | 0.792803919 | 5.291494837 | 3.697986661 | ENSG00000286522 |
| MXD3 | -0.523228145 | 0.695813153 | 0.004894622 | 0.021680611 | 160.2797539 | 111.4962991 | ENSG00000213347 |
| CENPM | -0.52330562 | 0.695775787 | 1.55E-08 | 2.49E-07 | 721.9652774 | 502.2799629 | ENSG00000100162 |
| COX16 | -0.524119137 | 0.695383559 | 0.258985894 | 0.465982147 | 30.42640719 | 21.12926024 | ENSG00000133983 |
| LOC112694756 | -0.524238387 | 0.695326083 | 0.494859378 | 0.69449938 | 9.599362839 | 6.692295438 | ENSG00000285043 |
| RFLNB | -0.524869447 | 0.695022001 | 1.00E-10 | 2.25E-09 | 1164.844876 | 809.5280925 | ENSG00000183688 |
| ULK4 | -0.525181344 | 0.69487176 | 0.004541029 | 0.020389284 | 173.7638438 | 120.7067782 | ENSG00000168038 |
| RELB | -0.525908712 | 0.694521513 | 0.000672683 | 0.0039674 | 289.037097 | 200.7692085 | ENSG00000104856 |
| GATD3A | -0.526032337 | 0.694462002 | 0.257813975 | 0.464706387 | 29.23874752 | 20.37972967 | ENSG00000160221 |
| NA | -0.526127569 | 0.694416162 | 0.482276568 | 0.683662446 | 9.57564174 | 6.662122495 | ENSG00000283959 |
| ELMSAN1 | -0.526197024 | 0.694382732 | 0.000258703 | 0.001730028 | 305.6798872 | 212.1247936 | ENSG00000156030 |
| GIPC2 | -0.526471814 | 0.694250485 | 0.304038856 | 0.51412731 | 21.96959993 | 15.21063638 | ENSG00000137960 |
| PIGQ | -0.526526651 | 0.694224097 | 4.47E-05 | 0.000364682 | 464.8459186 | 322.8688399 | ENSG00000007541 |
| ZNF674-AS1 | -0.52653497 | 0.694220094 | 0.063209396 | 0.168817627 | 73.75770515 | 51.17289353 | ENSG00000230844 |
| PLPP6 | -0.526794647 | 0.69409515 | 0.002186047 | 0.010959251 | 201.2138432 | 139.7949259 | ENSG00000205808 |
| PRPF38A | -0.527372249 | 0.693817315 | 3.38E-14 | 1.15E-12 | 1631.453912 | 1131.822816 | ENSG00000134748 |
| ADCK2 | -0.527527929 | 0.69374245 | 0.006278259 | 0.026752013 | 154.0838042 | 106.9395345 | ENSG00000133597 |
| UTF1 | -0.527671973 | 0.693673188 | 0.000135802 | 0.000977741 | 482.1041381 | 334.5861106 | ENSG00000171794 |
| MSI1 | -0.527698854 | 0.693660263 | 2.03E-12 | 5.61E-11 | 1497.242637 | 1038.632123 | ENSG00000135097 |
| NA | -0.527883732 | 0.693571378 | 0.132719191 | 0.293249319 | 46.31245984 | 32.17904854 | ENSG00000260719 |
| SYS1 | -0.528000586 | 0.693515203 | 0.000721584 | 0.004218919 | 252.4904054 | 175.0483762 | ENSG00000204070 |
| NA | -0.528190237 | 0.693424042 | 0.601150215 | 0.776307564 | 5.27133321 | 3.664692295 | ENSG00000231006 |
| LOC101928725 | -0.528238229 | 0.693400975 | 0.645050166 | 0.806579646 | 4.320910684 | 2.995254736 | ENSG00000245975 |
| NA | -0.528397494 | 0.693324432 | 0.631295117 | 0.797026605 | 4.324470156 | 2.9974302 | ENSG00000261220 |
| NA | -0.528398494 | 0.693323952 | 0.573722304 | 0.755930071 | 6.238357891 | 4.344723624 | ENSG00000273674 |
| TOR2A | -0.528780156 | 0.693140559 | 0.000154243 | 0.001094654 | 365.3367795 | 253.2359548 | ENSG00000160404 |
| ZNF484 | -0.528790941 | 0.693135377 | 0.016790207 | 0.059714322 | 120.5751738 | 83.51574397 | ENSG00000127081 |
| GALNT18 | -0.529260757 | 0.692909693 | 0.000812505 | 0.004678099 | 251.4172331 | 174.3688393 | ENSG00000110328 |
| PCDH1 | -0.53014874 | 0.692483336 | 4.44E-17 | 2.02E-15 | 5527.714632 | 3827.855528 | ENSG00000156453 |
| NA | -0.530182889 | 0.692466945 | 0.605557035 | 0.779795526 | 5.236933694 | 3.630168424 | ENSG00000197320 |
| LHFPL5 | -0.530202543 | 0.692457512 | 0.267543454 | 0.475311206 | 23.99380516 | 16.57722543 | ENSG00000197753 |
| SLC27A2 | -0.530286039 | 0.692417437 | 0.066785572 | 0.176288107 | 80.61881323 | 55.94332774 | ENSG00000140284 |
| ZDHHC6 | -0.530377737 | 0.692373428 | 6.80E-07 | 8.11E-06 | 688.2244433 | 476.4508816 | ENSG00000023041 |
| ZNF84 | -0.530541473 | 0.692294853 | 4.70E-06 | 4.80E-05 | 518.4630684 | 359.0147187 | ENSG00000198040 |
| ADAMTS7 | -0.5305629 | 0.692284571 | 2.54E-16 | 1.09E-14 | 2395.640376 | 1658.692439 | ENSG00000136378 |
| NA | -0.530590066 | 0.692271535 | 0.523548234 | 0.717778069 | 9.106506956 | 6.263463426 | ENSG00000242615 |
| CNPY4 | -0.530827484 | 0.69215762 | 0.002435327 | 0.012042722 | 183.2398203 | 126.8795549 | ENSG00000166997 |
| TNIP1 | -0.531405886 | 0.691880178 | 1.60E-07 | 2.14E-06 | 817.3645267 | 565.5022833 | ENSG00000145901 |
| PARPBP | -0.531451013 | 0.691858537 | 1.76E-05 | 0.000158953 | 412.6814775 | 285.6367124 | ENSG00000185480 |
| ZNF266 | -0.531746391 | 0.6917169 | 0.000106968 | 0.000789714 | 328.0154757 | 226.8115602 | ENSG00000174652 |
| TNFAIP3 | -0.53196148 | 0.69161378 | 0.11050822 | 0.257361911 | 51.40882082 | 35.54524841 | ENSG00000118503 |
| RBBP4 | -0.532295739 | 0.691453558 | 6.89E-18 | 3.43E-16 | 4023.835132 | 2781.628962 | ENSG00000162521 |
| SCD | -0.532313766 | 0.691444919 | 6.39E-23 | 4.86E-21 | 29963.57974 | 20717.88981 | ENSG00000099194 |
| LRRC37B | -0.532758009 | 0.691232037 | 0.002048932 | 0.010384885 | 193.0963026 | 133.5435693 | ENSG00000185158 |
| ZNF433 | -0.532903888 | 0.691162147 | 0.362395551 | 0.574243999 | 16.29765164 | 11.23127746 | ENSG00000197647 |
| LOX | -0.533182534 | 0.691028667 | 0.280803042 | 0.489649698 | 32.26910548 | 22.34604612 | ENSG00000113083 |
| CREG2 | -0.53321275 | 0.691014194 | 0.41747568 | 0.628430894 | 12.93901095 | 8.921350941 | ENSG00000175874 |
| NUCB1 | -0.533544113 | 0.690855498 | 8.64E-19 | 4.74E-17 | 2427.916444 | 1677.427454 | ENSG00000104805 |
| SFXN5 | -0.53376575 | 0.690749372 | 3.96E-10 | 8.09E-09 | 946.6346603 | 653.9425929 | ENSG00000144040 |
| AGBL2 | -0.533766373 | 0.690749074 | 0.363919866 | 0.57608415 | 20.78552589 | 14.40075992 | ENSG00000165923 |
| ENKD1 | -0.533996829 | 0.690638742 | 0.001445888 | 0.007719985 | 223.1178987 | 154.0395242 | ENSG00000124074 |
| BAD | -0.534012651 | 0.690631168 | 0.186733993 | 0.371951424 | 32.65194935 | 22.57331534 | ENSG00000002330 |
| LRRC45 | -0.534114201 | 0.690582557 | 6.60E-08 | 9.50E-07 | 909.851367 | 628.7542596 | ENSG00000169683 |
| SOAT1 | -0.534320957 | 0.690483595 | 0.013322365 | 0.049438567 | 124.2605966 | 85.76655411 | ENSG00000057252 |
| FGFR1 | -0.534624965 | 0.69033811 | 1.58E-32 | 2.07E-30 | 9861.498191 | 6807.876783 | ENSG00000077782 |
| MICOS10 | -0.534790968 | 0.690258681 | 0.05500484 | 0.15188864 | 72.56412174 | 50.04851839 | ENSG00000173436 |
| SSBP3-AS1 | -0.534968265 | 0.690173858 | 0.542279242 | 0.731381201 | 7.174542527 | 4.967814035 | ENSG00000198711 |
| NA | -0.535135937 | 0.69009365 | 0.426016567 | 0.635993903 | 13.54218433 | 9.375059011 | ENSG00000223969 |
| NA | -0.535729533 | 0.68980977 | 0.521238202 | 0.716053207 | 9.150110349 | 6.340929476 | ENSG00000273729 |

| | | | | | | | |
|---|---|---|---|---|---|---|---|
| MGAT5B | -0.53593561 | 0.689711243 | 0.000148458 | 0.001058371 | 303.7918834 | 209.5871034 | ENSG00000167889 |
| SLC1A2 | -0.536156737 | 0.689605537 | 0.21954376 | 0.415897673 | 39.71289424 | 27.33143182 | ENSG00000110436 |
| CASC2 | -0.536312397 | 0.689531136 | 0.59966116 | 0.775551364 | 5.260654793 | 3.624871538 | ENSG00000177640 |
| NLK | -0.536633152 | 0.689377849 | 1.08E-07 | 1.50E-06 | 660.6496097 | 455.2420954 | ENSG00000087095 |
| NA | -0.536737004 | 0.689328226 | 0.454840352 | 0.660692258 | 10.57231126 | 7.285212905 | ENSG00000213467 |
| OCRL | -0.536758368 | 0.689318018 | 6.46E-10 | 1.28E-08 | 1258.519196 | 867.2437619 | ENSG00000122126 |
| RBM12B-AS1 | -0.536990596 | 0.689207069 | 0.474968402 | 0.677782852 | 9.670526136 | 6.657771567 | ENSG00000279331 |
| NA | -0.537006902 | 0.689199279 | 0.500931219 | 0.69914168 | 8.717739345 | 5.993914441 | ENSG00000255639 |
| FKBP7 | -0.537044948 | 0.689181105 | 0.102427298 | 0.242418637 | 50.17163402 | 34.5502669 | ENSG00000079150 |
| PCDHB10 | -0.537155923 | 0.689128093 | 0.080116672 | 0.202023809 | 64.56577085 | 44.49336732 | ENSG00000120324 |
| MINDY3 | -0.53722747 | 0.689093918 | 0.00076378 | 0.00442997 | 246.5224367 | 169.7792319 | ENSG00000148481 |
| REC8 | -0.537430012 | 0.688997182 | 7.76E-07 | 9.13E-06 | 590.9509934 | 407.2125264 | ENSG00000100918 |
| NUTM2A | -0.537622515 | 0.688905254 | 0.462026641 | 0.666924491 | 10.54147121 | 7.255039962 | ENSG00000184923 |
| NA | -0.53764249 | 0.688895715 | 0.388524287 | 0.60048837 | 13.48406371 | 9.31253766 | ENSG00000251141 |
| HEMK1 | -0.537674956 | 0.688880213 | 0.006173321 | 0.026358394 | 170.8983122 | 117.7926054 | ENSG00000114735 |
| MCM5 | -0.538011247 | 0.688719654 | 7.04E-25 | 6.16E-23 | 8812.009594 | 6069.05396 | ENSG00000100297 |
| LRG1 | -0.538197625 | 0.688630686 | 0.340884918 | 0.552461663 | 19.83866284 | 13.68866373 | ENSG00000171236 |
| VIM-AS1 | -0.538381696 | 0.68854283 | 0.561784869 | 0.746766189 | 6.269197935 | 4.323252535 | ENSG00000229124 |
| HYPK | -0.538723376 | 0.688379779 | 0.583022564 | 0.762575961 | 5.322334881 | 3.662516831 | ENSG00000242028 |
| PCDHB14 | -0.538750594 | 0.688366792 | 0.097566943 | 0.233856038 | 50.62060718 | 34.84658386 | ENSG00000120327 |
| ZNF780B | -0.539263122 | 0.688122288 | 0.011612987 | 0.044215979 | 129.5592103 | 89.13492945 | ENSG00000128000 |
| UCHL3 | -0.539775907 | 0.687877749 | 0.582195736 | 0.762149347 | 5.291494837 | 3.634519351 | ENSG00000118939 |
| ZNF280A | -0.539953452 | 0.6877931 | 0.523596012 | 0.717778069 | 7.225544198 | 4.981812775 | ENSG00000169548 |
| NA | -0.540155546 | 0.68769676 | 0.56020145 | 0.745395927 | 8.592014906 | 5.888450911 | ENSG00000263535 |
| NA | -0.540622699 | 0.687474116 | 0.647993845 | 0.80899024 | 4.30430853 | 2.945786167 | ENSG00000279573 |
| NA | -0.540986904 | 0.687300587 | 0.417195797 | 0.628138613 | 13.05881166 | 8.941876072 | ENSG00000225616 |
| PIP5K1B | -0.541179175 | 0.687208995 | 0.022308161 | 0.075197985 | 102.9300179 | 70.75095413 | ENSG00000107242 |
| ACADSB | -0.541435137 | 0.687087082 | 1.60E-06 | 1.78E-05 | 494.8140964 | 339.7919758 | ENSG00000196177 |
| ERICH2 | -0.541510101 | 0.68705138 | 0.372222889 | 0.584082198 | 14.44752892 | 9.94310042 | ENSG00000204334 |
| NA | -0.541607246 | 0.687005119 | 0.052117648 | 0.145813596 | 82.14304363 | 56.49642445 | ENSG00000206567 |
| SOX9 | -0.5419632 | 0.686835636 | 0.014841986 | 0.054078894 | 120.1683295 | 82.55963727 | ENSG00000125398 |
| UNC79 | -0.54199558 | 0.686820221 | 0.063900628 | 0.170134408 | 72.34941049 | 49.67785681 | ENSG00000133958 |
| CDNF | -0.542054039 | 0.686792391 | 0.551613398 | 0.738582061 | 6.254960045 | 4.295255055 | ENSG00000185267 |
| NA | -0.542183619 | 0.686730708 | 0.530939741 | 0.72296467 | 7.236222615 | 4.977461848 | ENSG00000235027 |
| NMNAT1 | -0.542263163 | 0.686692845 | 0.080135576 | 0.202023809 | 59.96613072 | 41.25873655 | ENSG00000173614 |
| HRC | -0.542479964 | 0.68658966 | 0.06186114 | 0.16602049 | 76.56180986 | 52.51990341 | ENSG00000130528 |
| CCPG1 | -0.542673493 | 0.686497565 | 0.287944971 | 0.497213837 | 24.23785576 | 16.63946324 | ENSG00000260916 |
| USP43 | -0.54276264 | 0.686455146 | 0.000541215 | 0.003285389 | 232.8210443 | 159.7854334 | ENSG00000154914 |
| COL26A1 | -0.542797 | 0.686438797 | 6.00E-20 | 3.54E-18 | 2861.000175 | 1964.310179 | ENSG00000160963 |
| RADX | -0.542850928 | 0.686413138 | 0.000547647 | 0.003317953 | 229.7283959 | 157.5993201 | ENSG00000147231 |
| MCM4 | -0.54290265 | 0.68638853 | 2.54E-25 | 2.36E-23 | 8764.033061 | 6015.790676 | ENSG00000104738 |
| TMEM109 | -0.543749305 | 0.685985837 | 2.03E-10 | 4.32E-09 | 1036.419881 | 711.2847855 | ENSG00000110108 |
| NISCH | -0.54377175 | 0.685975165 | 2.36E-23 | 1.83E-21 | 3407.537975 | 2337.598209 | ENSG00000010322 |
| UBXN8 | -0.543792665 | 0.68596522 | 0.013763829 | 0.050767559 | 129.3338207 | 88.85159263 | ENSG00000104691 |
| NA | -0.543839191 | 0.685943098 | 0.122152384 | 0.276028315 | 43.93655627 | 30.12590177 | ENSG00000244479 |
| NA | -0.54407936 | 0.685828917 | 0.614934067 | 0.785838998 | 5.387574441 | 3.692689775 | ENSG00000232545 |
| PTBP2 | -0.544219784 | 0.685762166 | 8.19E-09 | 1.37E-07 | 816.3833426 | 559.5804724 | ENSG00000117569 |
| PTDSS2 | -0.544284271 | 0.685731513 | 1.59E-12 | 4.45E-11 | 1483.994361 | 1018.045767 | ENSG00000174915 |
| PSMD1 | -0.544378847 | 0.685686561 | 8.05E-28 | 8.50E-26 | 5612.641109 | 3848.234504 | ENSG00000173692 |
| ZNF302 | -0.544604237 | 0.685579446 | 4.21E-06 | 4.33E-05 | 456.9300198 | 313.2685285 | ENSG00000089335 |
| NA | -0.54462597 | 0.685569118 | 0.367489938 | 0.579660284 | 17.46247993 | 11.99965216 | ENSG00000275734 |
| ZNF528 | -0.54497149 | 0.685404947 | 0.557455144 | 0.743522876 | 6.340361231 | 4.346899088 | ENSG00000167555 |
| MIR4453HG | -0.54564806 | 0.685083593 | 0.163081552 | 0.338120875 | 38.36778037 | 26.32077057 | ENSG00000268471 |
| NA | -0.545897754 | 0.684965033 | 0.636933656 | 0.800725077 | 5.304537518 | 3.624871538 | ENSG00000257195 |
| NA | -0.546016867 | 0.684908483 | 0.396392842 | 0.608311297 | 15.62687444 | 10.72112293 | ENSG00000253931 |
| NA | -0.546104666 | 0.684866802 | 0.348388315 | 0.560370781 | 16.05240583 | 10.96702536 | ENSG00000214439 |
| NA | -0.546315172 | 0.684766879 | 0.467212861 | 0.67102263 | 9.210595228 | 6.324755273 | ENSG00000265194 |
| ABHD8 | -0.54654199 | 0.68465923 | 0.001455583 | 0.007756403 | 235.3247572 | 161.1716969 | ENSG00000127220 |
| NA | -0.546713249 | 0.68457796 | 0.430250488 | 0.640037846 | 11.70421457 | 8.002889528 | ENSG00000231043 |
| NA | -0.546804669 | 0.684534582 | 0.358827532 | 0.570841373 | 15.48691211 | 10.61036251 | ENSG00000257740 |
| GALR3 | -0.546838333 | 0.684518609 | 0.415972912 | 0.626847147 | 13.1620102 | 8.967698088 | ENSG00000128310 |
| NCAPH | -0.547033921 | 0.684425814 | 9.60E-10 | 1.85E-08 | 1356.025844 | 927.8840096 | ENSG00000121152 |
| ACSS2 | -0.547178342 | 0.684357303 | 5.87E-10 | 1.17E-08 | 954.1087048 | 652.7483921 | ENSG00000131069 |
| DPYSL4 | -0.547193728 | 0.684350004 | 9.72E-10 | 1.87E-08 | 881.3154325 | 603.4094585 | ENSG00000151640 |
| EPAS1 | -0.547205714 | 0.684344319 | 0.063872682 | 0.170083982 | 70.33173997 | 48.01154583 | ENSG00000116016 |
| NA | -0.547599733 | 0.684157441 | 0.059446995 | 0.161210784 | 66.5475679 | 45.45912184 | ENSG00000273247 |
| SLC16A9 | -0.548335285 | 0.683808715 | 0.000322675 | 0.00209406 | 292.9909024 | 200.2858945 | ENSG00000165449 |
| SLC22A5 | -0.54868234 | 0.683644238 | 0.01357807 | 0.05020005 | 113.9394548 | 77.9055439 | ENSG00000197375 |
| NA | -0.548761347 | 0.6836068 | 0.49900587 | 0.697608609 | 8.212730504 | 5.616726463 | ENSG00000217130 |
| VSTM2B | -0.548786197 | 0.683595025 | 0.236846618 | 0.438858749 | 26.61375933 | 18.21336347 | ENSG00000187135 |
| NA | -0.548825992 | 0.683576169 | 0.362971471 | 0.574867178 | 15.57943224 | 10.65765562 | ENSG00000236801 |
| KCNC1 | -0.548957484 | 0.683513869 | 0.322347862 | 0.53296615 | 18.43067367 | 12.61716214 | ENSG00000129159 |
| NMT2 | -0.549052175 | 0.683469008 | 3.35E-05 | 0.000281668 | 351.3829968 | 240.1843278 | ENSG00000152465 |
| SLC4A4 | -0.549310438 | 0.683346668 | 0.131246711 | 0.291029168 | 41.66830044 | 28.44559205 | ENSG00000080493 |

| | | | | | | | |
|---|---|---|---|---|---|---|---|
| CCDC144NL | -0.549318678 | 0.683342765 | 0.373762907 | 0.585708628 | 13.60386442 | 9.286715643 | ENSG00000205212 |
| CHD7 | -0.550004924 | 0.683017797 | 1.89E-13 | 5.88E-12 | 2992.257894 | 2043.049147 | ENSG00000171316 |
| ARSB | -0.550272879 | 0.682890951 | 2.03E-05 | 0.000180305 | 347.78 | 237.4320221 | ENSG00000113273 |
| TRIM7 | -0.550577353 | 0.682746845 | 0.636340383 | 0.800725077 | 5.418414484 | 3.693635733 | ENSG00000146054 |
| SLC25A13 | -0.550603701 | 0.682734376 | 5.38E-17 | 2.43E-15 | 2216.402946 | 1513.094324 | ENSG00000004864 |
| GORASP1 | -0.550714417 | 0.682681983 | 6.97E-10 | 1.37E-08 | 1050.754319 | 717.1971236 | ENSG00000114745 |
| NA | -0.550768965 | 0.682656171 | 0.431945098 | 0.641510501 | 11.22292683 | 7.683871974 | ENSG00000270878 |
| KCNAB3 | -0.550818062 | 0.68263294 | 0.091997588 | 0.223975049 | 56.50935165 | 38.62979247 | ENSG00000170049 |
| ARHGEF15 | -0.550874399 | 0.682606284 | 0.167119254 | 0.3438709 | 34.04686969 | 23.25552213 | ENSG00000198844 |
| RSPH1 | -0.551019973 | 0.68253741 | 0.585625188 | 0.764254832 | 5.329453825 | 3.620520611 | ENSG00000160188 |
| NA | -0.551027447 | 0.682533874 | 0.368806913 | 0.580952094 | 15.11622119 | 10.27081983 | ENSG00000228501 |
| NDUFA9 | -0.551600597 | 0.682262772 | 0.058504135 | 0.159269221 | 69.67016665 | 47.52286238 | ENSG00000139180 |
| IFRD1 | -0.551883104 | 0.682129185 | 1.48E-11 | 3.68E-10 | 1033.583488 | 705.1241883 | ENSG00000006652 |
| ESPN | -0.552024633 | 0.682062272 | 1.10E-06 | 1.26E-05 | 513.4091156 | 350.3782403 | ENSG00000187017 |
| NA | -0.552108624 | 0.682022565 | 0.673118641 | 0.824962394 | 4.880201335 | 3.344445235 | ENSG00000234241 |
| NR6A1 | -0.55336684 | 0.681428012 | 1.61E-20 | 9.96E-19 | 5702.650349 | 3885.523242 | ENSG00000148200 |
| LINC01668 | -0.553716449 | 0.681262901 | 0.626303453 | 0.793253855 | 4.858844501 | 3.330446495 | ENSG00000283051 |
| NOMO1 | -0.553717962 | 0.681262187 | 7.06E-17 | 3.16E-15 | 2162.20487 | 1472.844043 | ENSG00000103512 |
| KCNQ1 | -0.554101843 | 0.681080937 | 0.041354501 | 0.121780719 | 74.92815169 | 51.0434999 | ENSG00000053918 |
| KIF3B | -0.554229687 | 0.681020586 | 7.75E-11 | 1.76E-09 | 1226.628458 | 835.1853622 | ENSG00000101350 |
| ZNF501 | -0.554731272 | 0.680783855 | 0.010444848 | 0.040461733 | 142.0379849 | 96.73424189 | ENSG00000186446 |
| NA | -0.554820311 | 0.68074184 | 0.289867362 | 0.499192516 | 20.50468536 | 13.97098195 | ENSG00000239881 |
| NA | -0.5548653 | 0.680720612 | 0.388695203 | 0.600524064 | 13.12969562 | 8.955874812 | ENSG00000265750 |
| DUSP16 | -0.555396993 | 0.680469785 | 1.02E-13 | 3.28E-12 | 2569.612955 | 1748.171404 | ENSG00000111266 |
| LINC02728 | -0.555702916 | 0.680325506 | 0.581804944 | 0.761901835 | 5.880430324 | 4.027881533 | ENSG00000251323 |
| UNG | -0.555737576 | 0.680309162 | 7.52E-23 | 5.66E-21 | 4649.267012 | 3162.545297 | ENSG00000076248 |
| NA | -0.556511626 | 0.679944253 | 0.00450322 | 0.020258052 | 146.6133197 | 99.65621354 | ENSG00000280079 |
| SCAMP4 | -0.556532529 | 0.679934402 | 3.53E-09 | 6.20E-08 | 878.7281507 | 597.6501177 | ENSG00000227500 |
| SYT9 | -0.55681456 | 0.679801495 | 0.039212324 | 0.116912653 | 87.60175521 | 59.49090118 | ENSG00000170743 |
| NA | -0.557329559 | 0.679558869 | 0.00474861 | 0.021135401 | 149.3050476 | 101.5696565 | ENSG00000197462 |
| TMTC2 | -0.557390719 | 0.679530062 | 0.014150424 | 0.051970199 | 117.7852809 | 80.06784272 | ENSG00000179104 |
| TAFA5 | -0.557506732 | 0.67947542 | 3.88E-12 | 1.03E-10 | 1113.419427 | 756.7501502 | ENSG00000219438 |
| WTIP | -0.557743171 | 0.679364072 | 2.98E-06 | 3.15E-05 | 440.5288647 | 299.2252095 | ENSG00000142279 |
| PHLDB2 | -0.557760877 | 0.679355735 | 5.96E-08 | 8.63E-07 | 774.1012683 | 525.7297004 | ENSG00000144824 |
| NA | -0.557835924 | 0.679320396 | 0.172860293 | 0.352458068 | 34.2402241 | 23.23282154 | ENSG00000236255 |
| FRMD6-AS1 | -0.557872965 | 0.679302955 | 0.496839269 | 0.695950868 | 8.24713002 | 5.588728982 | ENSG00000273888 |
| NA | -0.55806671 | 0.679211735 | 0.097988743 | 0.234539596 | 53.94632288 | 36.62053384 | ENSG00000243302 |
| AURKC | -0.558256191 | 0.679122535 | 0.242258293 | 0.445911097 | 25.91926104 | 17.66461769 | ENSG00000105146 |
| NA | -0.558281171 | 0.679110775 | 0.536242004 | 0.726726247 | 7.383303889 | 5.005459328 | ENSG00000256897 |
| KCNIP3 | -0.558573883 | 0.678973003 | 0.03501823 | 0.107253389 | 83.80512527 | 56.86680249 | ENSG00000115041 |
| RBCK1 | -0.558983938 | 0.678780047 | 2.20E-09 | 4.02E-08 | 1296.980819 | 880.8818885 | ENSG00000125826 |
| CCP110 | -0.56013296 | 0.678239654 | 4.81E-05 | 0.000388305 | 365.6363077 | 247.9563323 | ENSG00000103540 |
| NPTX1 | -0.56027763 | 0.678171645 | 0.02823141 | 0.090533227 | 111.8641962 | 75.74307713 | ENSG00000171246 |
| SFXN3 | -0.56032745 | 0.678148226 | 0.123101145 | 0.277617043 | 40.14673979 | 27.19941123 | ENSG00000107819 |
| HLA-L | -0.560358753 | 0.678133512 | 0.09843396 | 0.235366592 | 56.53215688 | 38.34624475 | ENSG00000243753 |
| MAP1LC3B2 | -0.560920251 | 0.677869634 | 0.460280047 | 0.664999857 | 9.310234305 | 6.30110872 | ENSG00000258102 |
| PCDHGA8 | -0.561235948 | 0.677721315 | 0.337265351 | 0.549215845 | 16.16717252 | 10.97232225 | ENSG00000253767 |
| NA | -0.561455154 | 0.677618349 | 0.334506223 | 0.546513542 | 16.68641925 | 11.28074603 | ENSG00000239218 |
| NA | -0.561832334 | 0.677441214 | 0.547084462 | 0.734791389 | 6.813055488 | 4.63479774 | ENSG00000073905 |
| MAP2K6 | -0.561833123 | 0.677440844 | 2.04E-09 | 3.74E-08 | 963.8476728 | 652.6012159 | ENSG00000108984 |
| GATA5 | -0.561960276 | 0.67738114 | 0.480766653 | 0.682708803 | 9.854091858 | 6.708469642 | ENSG00000130700 |
| PM20D2 | -0.561969139 | 0.677376978 | 0.001034892 | 0.005773689 | 277.8589687 | 188.1947776 | ENSG00000146281 |
| TEX13B | -0.562015985 | 0.677354984 | 0.266501823 | 0.473951119 | 28.63259949 | 19.35239976 | ENSG00000170925 |
| ZBTB26 | -0.562228418 | 0.677255252 | 0.005195852 | 0.022830074 | 150.6353393 | 102.0616722 | ENSG00000171448 |
| NA | -0.562424804 | 0.677163068 | 0.270442023 | 0.478883976 | 21.65438603 | 14.65536421 | ENSG00000260563 |
| NA | -0.56248151 | 0.677136452 | 0.54709567 | 0.734791389 | 6.854573949 | 4.662795221 | ENSG00000273554 |
| WASHC4 | -0.56250393 | 0.677125929 | 8.52E-09 | 1.42E-07 | 877.9419703 | 594.2951083 | ENSG00000136051 |
| PRAF2 | -0.562603073 | 0.677079398 | 0.206645719 | 0.399172401 | 30.86587099 | 20.9508925 | ENSG00000243279 |
| ARRDC3 | -0.563271278 | 0.676765871 | 0.0296195 | 0.093916093 | 80.94973955 | 54.83200539 | ENSG00000113369 |
| GNG8 | -0.563323538 | 0.676741357 | 0.503936795 | 0.701625359 | 7.85836241 | 5.345002014 | ENSG00000167414 |
| SDAD1P1 | -0.56332778 | 0.676739367 | 0.46615975 | 0.670277667 | 10.20490048 | 6.889675259 | ENSG00000228451 |
| EML3 | -0.563436653 | 0.676688299 | 3.62E-08 | 5.43E-07 | 645.4936239 | 436.7635768 | ENSG00000149499 |
| NA | -0.563437934 | 0.676687698 | 0.291177518 | 0.50081865 | 20.16930983 | 13.67343549 | ENSG00000284428 |
| SIRT2 | -0.563486437 | 0.676664948 | 0.002727026 | 0.013242451 | 204.2883376 | 138.0863351 | ENSG00000068903 |
| NA | -0.563574017 | 0.676623872 | 0.481872623 | 0.683503506 | 10.36502444 | 7.008191571 | ENSG00000229932 |
| YY2 | -0.563589528 | 0.676616597 | 0.530258643 | 0.722388522 | 11.38157625 | 7.746393325 | ENSG00000230797 |
| NA | -0.563734833 | 0.676548454 | 0.486392755 | 0.687230822 | 8.843184452 | 6.011034604 | ENSG00000272604 |
| GTSE1-DT | -0.564213772 | 0.676323894 | 0.367821076 | 0.579853662 | 15.74400539 | 10.69312545 | ENSG00000277232 |
| NA | -0.564386606 | 0.676242875 | 0.034802821 | 0.106783309 | 81.99743689 | 55.48091782 | ENSG00000237238 |
| NA | -0.564657132 | 0.676116082 | 0.244790831 | 0.449070278 | 22.58345173 | 15.24610621 | ENSG00000271147 |
| TTC26 | -0.565247132 | 0.675839636 | 0.003711399 | 0.017238016 | 154.6602818 | 104.4003313 | ENSG00000105948 |
| DHX35 | -0.565261582 | 0.675832867 | 5.43E-07 | 6.59E-06 | 548.7215706 | 370.5552503 | ENSG00000101452 |
| ZFP37 | -0.565304692 | 0.675812672 | 0.054741439 | 0.151360481 | 85.32223615 | 57.76067139 | ENSG00000136866 |
| RORB | -0.565724857 | 0.67561588 | 0.004109105 | 0.018801246 | 154.6499083 | 104.5897873 | ENSG00000198963 |

| | | | | | | | |
|---|---|---|---|---|---|---|---|
| NA | -0.566124598 | 0.675428707 | 0.515669084 | 0.711976214 | 7.392787098 | 4.969989499 | ENSG00000277010 |
| NA | -0.566195972 | 0.675395292 | 0.364026309 | 0.57608415 | 15.18235047 | 10.24717328 | ENSG00000229644 |
| ADGRG2 | -0.566360873 | 0.675318098 | 0.000315128 | 0.002052844 | 258.2319908 | 174.201138 | ENSG00000173698 |
| FAM32A | -0.567163685 | 0.674942411 | 7.98E-16 | 3.33E-14 | 1798.53045 | 1213.864942 | ENSG00000105058 |
| NA | -0.567638648 | 0.674720243 | 0.545959209 | 0.734048103 | 6.402041319 | 4.290904128 | ENSG00000274818 |
| CCDC163 | -0.567642043 | 0.674718656 | 0.116651652 | 0.267610113 | 42.88204547 | 28.91083985 | ENSG00000280670 |
| AHCYL2 | -0.567699223 | 0.674691914 | 1.06E-07 | 1.47E-06 | 598.5782701 | 403.6372914 | ENSG00000158467 |
| NA | -0.567819042 | 0.674635882 | 0.241932718 | 0.445485509 | 25.60997087 | 17.33595232 | ENSG00000272079 |
| SOX18 | -0.568122575 | 0.674493958 | 0.570371906 | 0.75319722 | 5.920753576 | 4.007356402 | ENSG00000203883 |
| EAPP | -0.568360976 | 0.674382509 | 1.00E-05 | 9.57E-05 | 382.1901106 | 257.5351819 | ENSG00000129518 |
| SLC35A3 | -0.568400337 | 0.67436411 | 0.008913751 | 0.035694697 | 133.5619319 | 90.15980035 | ENSG00000117620 |
| DHRSX | -0.568554615 | 0.674291999 | 4.09E-05 | 0.000335606 | 333.0371395 | 224.5555259 | ENSG00000169084 |
| ASAP2 | -0.568766392 | 0.674193026 | 5.27E-07 | 6.42E-06 | 495.6272002 | 334.1332961 | ENSG00000151693 |
| NA | -0.568812583 | 0.67417144 | 0.140658631 | 0.304870682 | 43.83930761 | 29.60080254 | ENSG00000230715 |
| WDR27 | -0.568946236 | 0.674108987 | 0.001290579 | 0.006987553 | 384.969072 | 259.6593542 | ENSG00000184465 |
| CD53 | -0.569099825 | 0.674037225 | 0.472738093 | 0.676081702 | 8.87638876 | 6.003562254 | ENSG00000143119 |
| NA | -0.56918433 | 0.673997745 | 0.594513963 | 0.771483281 | 5.853149752 | 3.951361442 | ENSG00000272008 |
| PDE1C | -0.569299893 | 0.673943758 | 0.599589536 | 0.77551184 | 5.828233445 | 3.932065816 | ENSG00000154678 |
| NA | -0.569383449 | 0.673904727 | 0.252296417 | 0.458136331 | 21.20393834 | 14.29217497 | ENSG00000224825 |
| HDAC9 | -0.569758499 | 0.673729559 | 0.000799382 | 0.004612384 | 212.9524311 | 143.4796489 | ENSG00000048052 |
| NHLH1 | -0.569801704 | 0.673709382 | 0.598053717 | 0.774094349 | 4.889684544 | 3.301503056 | ENSG00000171786 |
| CHN1 | -0.569864771 | 0.673679932 | 4.66E-10 | 9.42E-09 | 929.5102779 | 626.004937 | ENSG00000128656 |
| DTX4 | -0.569880921 | 0.67367239 | 1.40E-10 | 3.05E-09 | 890.714628 | 600.0091863 | ENSG00000110042 |
| SYTL4 | -0.570111424 | 0.673564765 | 0.567749391 | 0.751047019 | 5.846030807 | 3.943889093 | ENSG00000102362 |
| UBXN10 | -0.570209752 | 0.673518859 | 0.155275462 | 0.327018746 | 35.04028521 | 23.53906986 | ENSG00000162543 |
| UHRF1 | -0.570312746 | 0.673470778 | 4.25E-15 | 1.63E-13 | 2512.288588 | 1691.975259 | ENSG00000276043 |
| PPM1L | -0.570657623 | 0.673309804 | 0.000827673 | 0.004750942 | 278.0950107 | 187.1910942 | ENSG00000163590 |
| GPX3 | -0.570762536 | 0.673260843 | 5.02E-09 | 8.68E-08 | 714.980276 | 481.5853986 | ENSG00000211445 |
| AARSD1 | -0.570878666 | 0.673206651 | 0.419002518 | 0.629503762 | 11.77509854 | 7.943773146 | ENSG00000266967 |
| ZNF778 | -0.571077989 | 0.673113646 | 2.07E-06 | 2.26E-05 | 520.6038465 | 350.3410465 | ENSG00000170100 |
| DAZL | -0.571232037 | 0.673041777 | 0.490855657 | 0.690540009 | 7.889202453 | 5.324476883 | ENSG00000092345 |
| HIPK2 | -0.571311265 | 0.673004816 | 2.48E-09 | 4.50E-08 | 896.7904454 | 603.3911385 | ENSG00000064393 |
| MRPL48 | -0.571331508 | 0.672995373 | 9.54E-05 | 0.000712089 | 372.6776497 | 250.8203444 | ENSG00000175581 |
| SVEP1 | -0.572287961 | 0.67254935 | 0.248119517 | 0.453053109 | 22.24332152 | 14.95291068 | ENSG00000165124 |
| LINC00843 | -0.573014935 | 0.672210538 | 0.625135484 | 0.792700169 | 3.970102062 | 2.676237182 | ENSG00000178440 |
| TTN-AS1 | -0.573128097 | 0.672157813 | 0.152523456 | 0.322734506 | 35.51890321 | 23.8334949 | ENSG00000237298 |
| NA | -0.573497484 | 0.671985737 | 0.075976152 | 0.194603231 | 70.30240004 | 47.37741014 | ENSG00000213888 |
| COL2A1 | -0.573619127 | 0.671929079 | 3.41E-06 | 3.56E-05 | 418.6322075 | 281.3686042 | ENSG00000139219 |
| LBH | -0.574090708 | 0.671709478 | 1.42E-12 | 3.99E-11 | 1170.887359 | 786.3897846 | ENSG00000213626 |
| ATP6V0D1 | -0.574209691 | 0.671654083 | 7.21E-13 | 2.12E-11 | 1332.863315 | 895.116889 | ENSG00000159720 |
| FMNL1 | -0.574364961 | 0.6715818 | 0.002733782 | 0.013265024 | 206.5388478 | 138.8651357 | ENSG00000184922 |
| ST3GAL6 | -0.574571136 | 0.671485831 | 0.076829903 | 0.196098489 | 56.85451587 | 38.11573851 | ENSG00000064225 |
| CLIP3 | -0.574600949 | 0.671471956 | 4.14E-05 | 0.000339837 | 343.9145966 | 231.0878985 | ENSG00000105270 |
| RPL13AP20 | -0.574710542 | 0.67142095 | 0.371190758 | 0.583286525 | 15.44686819 | 10.35698773 | ENSG00000234498 |
| FNTB | -0.574816745 | 0.671371525 | 0.05775543 | 0.157687741 | 65.87086696 | 44.17775474 | ENSG00000257365 |
| TIGD1 | -0.575095046 | 0.671242028 | 0.019986285 | 0.068750637 | 116.8936155 | 78.62711995 | ENSG00000221944 |
| NA | -0.575470389 | 0.671067415 | 0.594331368 | 0.771358959 | 4.982204676 | 3.350971626 | ENSG00000214263 |
| NA | -0.575720747 | 0.670950972 | 0.406156723 | 0.617103833 | 13.40250133 | 8.968644047 | ENSG00000225356 |
| WFDC2 | -0.576075077 | 0.670786204 | 6.76E-07 | 8.07E-06 | 444.9176585 | 298.3581085 | ENSG00000101443 |
| C1QL1 | -0.576710456 | 0.670490848 | 0.000212484 | 0.001451273 | 286.0871068 | 191.9079628 | ENSG00000131094 |
| NA | -0.576943912 | 0.670382358 | 0.31709487 | 0.527235132 | 17.8286955 | 11.96418233 | ENSG00000263968 |
| NA | -0.577604966 | 0.670075254 | 0.4489633 | 0.655482947 | 11.51708938 | 7.69569525 | ENSG00000274525 |
| HDAC11 | -0.577616574 | 0.670069863 | 2.94E-05 | 0.000251355 | 362.3282854 | 242.592697 | ENSG00000163517 |
| NA | -0.577916212 | 0.669930708 | 0.60837687 | 0.781843974 | 4.947805159 | 3.316447755 | ENSG00000270558 |
| UBR7 | -0.578288894 | 0.669757671 | 8.80E-09 | 1.46E-07 | 715.6807248 | 479.2609789 | ENSG00000012963 |
| EPOP | -0.578317336 | 0.669744468 | 0.004861451 | 0.021561416 | 150.2447917 | 100.654768 | ENSG00000273604 |
| ADI1 | -0.578393279 | 0.669709214 | 2.32E-09 | 4.23E-08 | 858.6616555 | 574.9757194 | ENSG00000182551 |
| BOLA2B | -0.578582037 | 0.669621596 | 0.298374224 | 0.50864752 | 19.25417654 | 12.93523373 | ENSG00000169627 |
| ARMH3 | -0.578834755 | 0.669504309 | 6.60E-08 | 9.50E-07 | 548.3636686 | 367.2261716 | ENSG00000120029 |
| PMS2P4 | -0.578956232 | 0.669447938 | 0.362963183 | 0.574867178 | 12.89512823 | 8.637803216 | ENSG00000067601 |
| DGKI | -0.579027629 | 0.669414808 | 0.244381396 | 0.448725588 | 23.32183275 | 15.57694704 | ENSG00000157680 |
| SFRP1 | -0.579254395 | 0.669309597 | 1.33E-19 | 7.72E-18 | 14465.82615 | 9681.878026 | ENSG00000104332 |
| NA | -0.579354375 | 0.669263215 | 0.612167881 | 0.784199347 | 3.980780479 | 2.666589369 | ENSG00000232702 |
| FAM222A | -0.579573775 | 0.669161443 | 0.063020303 | 0.168479391 | 64.11654452 | 42.96486828 | ENSG00000139438 |
| KALRN | -0.57963907 | 0.669131158 | 0.000480749 | 0.002979635 | 231.8059409 | 155.0466125 | ENSG00000160145 |
| TGFB1 | -0.580101479 | 0.668916724 | 9.39E-05 | 0.000701785 | 297.0520762 | 198.8282625 | ENSG00000105329 |
| NA | -0.580323346 | 0.668813862 | 0.429857018 | 0.639654068 | 9.950171462 | 6.657771567 | ENSG00000287811 |
| KCNC4 | -0.5806473 | 0.668663698 | 7.06E-06 | 6.99E-05 | 401.7618909 | 268.7657243 | ENSG00000116396 |
| PIK3R3 | -0.580815948 | 0.668585537 | 0.000170828 | 0.001197506 | 275.6491129 | 184.3378344 | ENSG00000117461 |
| AGAP2 | -0.580834881 | 0.668576763 | 0.075610211 | 0.193923343 | 52.67089777 | 35.21724545 | ENSG00000135439 |
| NA | -0.580847392 | 0.668570965 | 0.496546291 | 0.695695282 | 9.404839368 | 6.319458387 | ENSG00000278840 |
| NA | -0.581482549 | 0.668276687 | 0.481286338 | 0.683198091 | 7.937839859 | 5.303005794 | ENSG00000240409 |
| ABAT | -0.58158733 | 0.668228152 | 0.000108853 | 0.000801749 | 290.0648864 | 193.6485755 | ENSG00000183044 |
| UBE2R2 | -0.582260576 | 0.667916391 | 5.66E-20 | 3.35E-18 | 2257.768412 | 1508.344165 | ENSG00000107341 |

| | | | | | | | |
|---|---|---|---|---|---|---|---|
| NA | -0.58266253 | 0.667730326 | 0.343579381 | 0.55552197 | 15.42047734 | 10.33428714 | ENSG00000229097 |
| NA | -0.582718929 | 0.667704224 | 0.487559235 | 0.688363946 | 8.447297896 | 5.678018309 | ENSG00000259319 |
| CACNA2D3 | -0.583211978 | 0.667476071 | 0.027369677 | 0.088415385 | 85.13689098 | 56.83880501 | ENSG00000157445 |
| FBXW4 | -0.583281752 | 0.66744379 | 1.51E-08 | 2.44E-07 | 717.3391941 | 478.9806909 | ENSG00000107829 |
| WDTC1 | -0.583507313 | 0.667339446 | 5.76E-10 | 1.15E-08 | 850.0130474 | 567.0523987 | ENSG00000142784 |
| NA | -0.583561195 | 0.667314522 | 0.601132199 | 0.776307564 | 4.937126742 | 3.28967978 | ENSG00000269925 |
| MPP1 | -0.583615137 | 0.667289572 | 0.000248451 | 0.001667967 | 278.9522771 | 186.0609184 | ENSG00000130830 |
| NA | -0.584003591 | 0.667109924 | 0.487959731 | 0.688671961 | 7.903440343 | 5.263185037 | ENSG00000178162 |
| NA | -0.584525562 | 0.666868606 | 0.501552954 | 0.699750853 | 6.956577289 | 4.632622277 | ENSG00000280020 |
| NA | -0.584931432 | 0.666681024 | 1.25E-07 | 1.70E-06 | 946.8582461 | 631.0480549 | ENSG00000254339 |
| PINLYP | -0.585234686 | 0.666540902 | 0.06177918 | 0.165889351 | 68.39652156 | 45.52258915 | ENSG00000234465 |
| GSTO1 | -0.585372677 | 0.666477152 | 5.90E-09 | 1.01E-07 | 977.4832681 | 651.1017985 | ENSG00000148834 |
| MMD | -0.585496485 | 0.666419959 | 1.41E-07 | 1.90E-06 | 614.6265827 | 409.6523504 | ENSG00000108960 |
| INSYN2A | -0.586374831 | 0.666014351 | 0.402625503 | 0.61393481 | 13.44490952 | 8.977345901 | ENSG00000188916 |
| CPXM1 | -0.586523001 | 0.665945952 | 1.72E-14 | 5.97E-13 | 1859.870223 | 1238.88436 | ENSG00000088882 |
| PLA2R1 | -0.58661071 | 0.665905467 | 0.043705863 | 0.127080096 | 70.23062636 | 46.7330166 | ENSG00000153246 |
| NAALAD2 | -0.586717694 | 0.665856088 | 0.069586078 | 0.181836597 | 51.30206279 | 34.14290598 | ENSG00000077616 |
| RCN1 | -0.587683551 | 0.66541046 | 4.52E-09 | 7.88E-08 | 901.65646 | 600.2311586 | ENSG00000049449 |
| ZNF615 | -0.58812877 | 0.665205144 | 0.002904157 | 0.014008941 | 158.7617527 | 105.5816473 | ENSG00000197619 |
| CDC42EP5 | -0.588204249 | 0.665170343 | 0.015311818 | 0.05538473 | 116.4884995 | 77.34497484 | ENSG00000167617 |
| IGDCC3 | -0.588705118 | 0.664939451 | 2.38E-17 | 1.11E-15 | 2254.985795 | 1499.293597 | ENSG00000174498 |
| NA | -0.58877681 | 0.66490641 | 0.030594689 | 0.096457546 | 89.12598561 | 59.31425741 | ENSG00000213612 |
| BRD9 | -0.589083455 | 0.664765098 | 8.32E-12 | 2.13E-10 | 1108.076906 | 736.4795726 | ENSG00000028310 |
| CHST7 | -0.58916507 | 0.664727493 | 1.36E-07 | 1.84E-06 | 682.9359748 | 454.3041704 | ENSG00000147119 |
| NTRK3 | -0.589306889 | 0.664662153 | 0.001644706 | 0.008599189 | 177.9183751 | 118.4051021 | ENSG00000140538 |
| ZFYVE26 | -0.589327189 | 0.6646528 | 8.84E-12 | 2.25E-10 | 978.1831565 | 649.9617936 | ENSG00000072121 |
| F13A1 | -0.589413446 | 0.664613063 | 0.214209288 | 0.408992257 | 25.03199314 | 16.60087199 | ENSG00000124491 |
| FASTKD1 | -0.589828497 | 0.664421887 | 3.22E-09 | 5.70E-08 | 707.7980565 | 470.4098326 | ENSG00000138399 |
| TNNI3 | -0.589906571 | 0.664385931 | 0.134644292 | 0.296255092 | 42.05470379 | 27.96154308 | ENSG00000129991 |
| CBY1 | -0.590576941 | 0.664077286 | 0.005110567 | 0.02251296 | 157.7286249 | 104.7164383 | ENSG00000100211 |
| GCGR | -0.591501193 | 0.663651986 | 0.484503916 | 0.685536559 | 8.029164782 | 5.31482907 | ENSG00000215644 |
| TSC22D1-AS1 | -0.591540291 | 0.663634001 | 0.24460765 | 0.448950732 | 23.95912631 | 15.93266393 | ENSG00000278156 |
| NA | -0.59156856 | 0.663620998 | 0.3828986 | 0.595355045 | 12.05502319 | 7.979242975 | ENSG00000230373 |
| GLIDR | -0.592100442 | 0.663376384 | 0.137363178 | 0.300104537 | 36.48920803 | 24.25797599 | ENSG00000278175 |
| PTPMT1 | -0.592146273 | 0.66335531 | 0.503307928 | 0.701172881 | 7.025376322 | 4.646621017 | ENSG00000110536 |
| KCND2 | -0.592393545 | 0.663241624 | 3.51E-05 | 0.000293383 | 283.7248563 | 188.2283985 | ENSG00000184408 |
| CHRAC1 | -0.592418823 | 0.663230003 | 1.61E-07 | 2.14E-06 | 678.2208797 | 450.0158409 | ENSG00000104472 |
| NA | -0.59298347 | 0.662970476 | 0.404169097 | 0.615294913 | 11.56425224 | 7.688222901 | ENSG00000278126 |
| RNF7 | -0.593077712 | 0.66292717 | 1.60E-09 | 3.00E-08 | 664.1971289 | 440.3322425 | ENSG00000114125 |
| CD2BP2 | -0.593226458 | 0.662858824 | 2.75E-13 | 8.48E-12 | 2365.226679 | 1568.30387 | ENSG00000169217 |
| SERAC1 | -0.593686697 | 0.662647397 | 6.67E-05 | 0.000519231 | 273.5101671 | 181.3720176 | ENSG00000122335 |
| TYMSOS | -0.593860969 | 0.662567356 | 0.174720561 | 0.355062713 | 30.9002705 | 20.47382142 | ENSG00000176912 |
| GLYCTK | -0.593874671 | 0.662561064 | 0.03792123 | 0.113926572 | 73.25358603 | 48.62453692 | ENSG00000168237 |
| KRT7 | -0.594100015 | 0.662457582 | 0.000153027 | 0.001086431 | 278.8929868 | 184.7904083 | ENSG00000135480 |
| LOC101927556 | -0.594968965 | 0.662058697 | 0.543693167 | 0.732348903 | 6.027511597 | 3.9762375 | ENSG00000260927 |
| MLH1 | -0.595088572 | 0.662003811 | 3.28E-13 | 1.00E-11 | 1205.852032 | 798.0422537 | ENSG00000076242 |
| TMEM35A | -0.595235572 | 0.661936361 | 0.008715189 | 0.035025823 | 116.6477593 | 77.11692762 | ENSG00000126950 |
| NA | -0.595839797 | 0.661659189 | 0.061534551 | 0.165475583 | 54.7520284 | 36.21345646 | ENSG00000286279 |
| HID1 | -0.596197717 | 0.661495058 | 4.70E-05 | 0.000380469 | 265.9343991 | 175.8652735 | ENSG00000167861 |
| EOGT | -0.596460588 | 0.661374539 | 2.32E-05 | 0.000203683 | 311.8151245 | 206.2641292 | ENSG00000163378 |
| NA | -0.596469267 | 0.66137056 | 0.463902521 | 0.668542004 | 8.556420182 | 5.680193772 | ENSG00000236814 |
| TRIM61 | -0.596823319 | 0.661208273 | 0.002508737 | 0.0123434 | 169.4944929 | 112.1510755 | ENSG00000183439 |
| ATXN7L3B | -0.59689671 | 0.661174638 | 2.98E-19 | 1.69E-17 | 1758.399728 | 1162.625934 | ENSG00000253719 |
| FAM78A | -0.597049838 | 0.661104464 | 0.000634513 | 0.003767305 | 202.0165741 | 133.6574511 | ENSG00000126882 |
| ULK2 | -0.597257152 | 0.661009471 | 0.000457218 | 0.002845931 | 204.4057996 | 135.1557772 | ENSG00000083290 |
| LINC01637 | -0.597634476 | 0.660836612 | 0.613258924 | 0.784700506 | 4.066181665 | 2.678412645 | ENSG00000237476 |
| NUBPL | -0.597678142 | 0.660816611 | 0.00064765 | 0.003838941 | 198.1807899 | 130.8270169 | ENSG00000151413 |
| PDE4D | -0.598104763 | 0.660621229 | 0.235435077 | 0.437272957 | 25.75229747 | 17.06205241 | ENSG00000113448 |
| NA | -0.598686652 | 0.660354832 | 0.221929661 | 0.419239936 | 23.67620085 | 15.60494452 | ENSG00000204121 |
| LINC00032 | -0.598848779 | 0.660280626 | 0.534838551 | 0.725709588 | 7.130939134 | 4.696089587 | ENSG00000283945 |
| TCTN3 | -0.598858029 | 0.660276393 | 1.59E-08 | 2.55E-07 | 832.2982806 | 549.5254977 | ENSG00000119977 |
| NA | -0.598892309 | 0.660260704 | 0.572146406 | 0.754643448 | 5.033206346 | 3.306799942 | ENSG00000285859 |
| MAPRE3 | -0.598961727 | 0.660228935 | 0.002929887 | 0.014102507 | 154.5306924 | 102.0884402 | ENSG00000084764 |
| DRG2 | -0.598979507 | 0.660220799 | 3.43E-09 | 6.05E-08 | 657.9199223 | 434.3306675 | ENSG00000108591 |
| POLR2C | -0.599056359 | 0.66018563 | 3.48E-15 | 1.35E-13 | 1525.964171 | 1007.495844 | ENSG00000102978 |
| FUNDC2 | -0.599483731 | 0.659990091 | 1.43E-10 | 3.12E-09 | 793.2364565 | 523.3163609 | ENSG00000165775 |
| AKAP6 | -0.600063521 | 0.659724907 | 0.49185771 | 0.691228817 | 7.514672732 | 4.972164962 | ENSG00000151320 |
| EXT2 | -0.600162911 | 0.659679459 | 3.61E-17 | 1.65E-15 | 2106.724372 | 1389.25616 | ENSG00000151348 |
| NA | -0.600629025 | 0.659466361 | 0.491111113 | 0.690591354 | 8.026800518 | 5.265360501 | ENSG00000230170 |
| NUCB2 | -0.600721218 | 0.65942422 | 7.66E-05 | 0.000586051 | 313.3707268 | 206.7825341 | ENSG00000070081 |
| AASS | -0.601014642 | 0.659290116 | 5.35E-26 | 5.10E-24 | 4244.507132 | 2798.150131 | ENSG00000008311 |
| TRAPPC2 | -0.601265646 | 0.659175421 | 0.00116784 | 0.00641613 | 168.1028271 | 110.702386 | ENSG00000196459 |
| LRP4-AS1 | -0.601562033 | 0.659040014 | 0.552965415 | 0.739709489 | 5.556902273 | 3.686163384 | ENSG00000247675 |
| LOC728752 | -0.602182573 | 0.658756605 | 0.257695539 | 0.464634358 | 19.70253931 | 12.94923247 | ENSG00000267309 |

| | | | | | | | |
|---|---|---|---|---|---|---|---|
| GOLGA8H | -0.602445913 | 0.658636371 | 0.329918874 | 0.541407537 | 15.69447826 | 10.32558528 | ENSG00000261794 |
| SEC61A2 | -0.602801713 | 0.658473957 | 5.09E-07 | 6.24E-06 | 454.895721 | 299.4926783 | ENSG00000065665 |
| NHLRC3 | -0.602980431 | 0.658392391 | 0.00038402 | 0.00243936 | 225.1800367 | 148.1076367 | ENSG00000188811 |
| TRPM3 | -0.603353422 | 0.658222194 | 0.001152804 | 0.006351574 | 192.3922693 | 126.4796663 | ENSG00000083067 |
| WNT3A | -0.603364523 | 0.65821713 | 0.563088745 | 0.747867709 | 5.495222185 | 3.632343887 | ENSG00000154342 |
| AFF2 | -0.604001795 | 0.657926444 | 0.000116711 | 0.000852644 | 244.9848843 | 161.1787904 | ENSG00000155966 |
| FOXM1 | -0.60407747 | 0.657891934 | 5.81E-21 | 3.73E-19 | 3071.948692 | 2021.531542 | ENSG00000111206 |
| NA | -0.604323349 | 0.657779819 | 0.540351744 | 0.729834265 | 6.105793839 | 3.995533126 | ENSG00000271855 |
| TCEAL3 | -0.604431871 | 0.657730342 | 2.24E-06 | 2.42E-05 | 363.3935263 | 239.1118803 | ENSG00000196507 |
| ARL4C | -0.604803078 | 0.657561129 | 8.66E-09 | 1.44E-07 | 614.8249717 | 404.1037257 | ENSG00000188042 |
| NA | -0.60519532 | 0.657382374 | 0.293638388 | 0.503391517 | 16.15057036 | 10.63400907 | ENSG00000277159 |
| NA | -0.605346644 | 0.657313425 | 0.467382101 | 0.671109527 | 7.57635282 | 4.991460588 | ENSG00000226049 |
| RECQL | -0.605602326 | 0.657196943 | 3.73E-08 | 5.59E-07 | 638.6884992 | 419.485584 | ENSG00000004700 |
| ERG | -0.605622736 | 0.657187645 | 0.187069788 | 0.372423883 | 27.84291131 | 18.30823323 | ENSG00000157554 |
| TIA1 | -0.605980271 | 0.657024798 | 3.74E-14 | 1.26E-12 | 1641.3046 | 1078.671681 | ENSG00000116001 |
| ATG4A | -0.606216856 | 0.656917063 | 0.013382998 | 0.049573299 | 94.53344234 | 62.1495667 | ENSG00000101844 |
| ANGPT1 | -0.606677614 | 0.656707295 | 0.063341125 | 0.169097703 | 55.60133731 | 36.59688728 | ENSG00000154188 |
| RNF150 | -0.606687341 | 0.656702867 | 0.000707546 | 0.00414969 | 201.9246387 | 132.4525611 | ENSG00000170153 |
| TEX261 | -0.606810608 | 0.65664676 | 7.14E-07 | 8.48E-06 | 416.3695961 | 273.5868566 | ENSG00000144043 |
| NA | -0.607493646 | 0.656335946 | 0.131282895 | 0.291029168 | 39.97446287 | 26.33041838 | ENSG00000272667 |
| PDSS2 | -0.60789231 | 0.656154604 | 0.00010789 | 0.000795903 | 286.851651 | 188.3060325 | ENSG00000164494 |
| NA | -0.608080433 | 0.656069049 | 0.39973416 | 0.611154437 | 11.62473712 | 7.612932314 | ENSG00000255921 |
| NA | -0.608186989 | 0.656020594 | 0.5691007 | 0.752149676 | 5.060486917 | 3.294976665 | ENSG00000256393 |
| TEX26-AS1 | -0.608828683 | 0.655728869 | 0.512096824 | 0.708546306 | 6.582047568 | 4.326373957 | ENSG00000224743 |
| PARTICL | -0.608923095 | 0.655685958 | 0.076873348 | 0.196137614 | 48.49795808 | 31.80498198 | ENSG00000286532 |
| INSIG2 | -0.609113577 | 0.655599393 | 0.004260427 | 0.01937817 | 136.5726869 | 89.52365715 | ENSG00000125629 |
| PARP3 | -0.609272738 | 0.65552707 | 0.116025783 | 0.266498044 | 40.12301869 | 26.32171653 | ENSG00000041880 |
| GPNMB | -0.609430753 | 0.655455275 | 0.391751358 | 0.603198075 | 10.62094866 | 6.947845683 | ENSG00000136235 |
| SEMA5B | -0.609452725 | 0.655445293 | 1.16E-10 | 2.57E-09 | 798.0087564 | 523.2087648 | ENSG00000082684 |
| ATOH7 | -0.609599792 | 0.655378481 | 0.525511386 | 0.718994605 | 6.520367481 | 4.276905388 | ENSG00000179774 |
| BTG3-AS1 | -0.609649004 | 0.655356125 | 0.44245373 | 0.650077004 | 9.704925652 | 6.360225102 | ENSG00000280594 |
| LINC01003 | -0.609790604 | 0.655291806 | 0.223427029 | 0.421267916 | 22.78242439 | 14.93891194 | ENSG00000261455 |
| NA | -0.610059216 | 0.65516981 | 0.001790635 | 0.009236679 | 148.3540146 | 97.14406182 | ENSG00000188002 |
| NA | -0.610957562 | 0.654761971 | 0.495700826 | 0.695207823 | 6.605768667 | 4.335075811 | ENSG00000270175 |
| NA | -0.611234658 | 0.654636224 | 0.012933126 | 0.048316931 | 100.5431566 | 65.88273965 | ENSG00000287086 |
| NA | -0.611255249 | 0.654626882 | 0.591928514 | 0.769555968 | 4.569715966 | 3.009253477 | ENSG00000272562 |
| BEND3 | -0.611803047 | 0.654378364 | 1.74E-11 | 4.28E-10 | 1153.460403 | 754.8688745 | ENSG00000178409 |
| LAT2 | -0.612180769 | 0.654207059 | 0.319872957 | 0.530266924 | 15.7689217 | 10.35793369 | ENSG00000086730 |
| NA | -0.61264117 | 0.653998318 | 0.591534186 | 0.769255138 | 4.545994867 | 2.987782387 | ENSG00000284642 |
| NDUFAF7 | -0.61267785 | 0.65398169 | 2.64E-11 | 6.38E-10 | 988.0379116 | 646.2225591 | ENSG00000003509 |
| ZNF184 | -0.612696651 | 0.653973168 | 0.001236893 | 0.006739388 | 168.2614765 | 109.9287144 | ENSG00000096654 |
| NA | -0.613132023 | 0.653775844 | 0.429178146 | 0.639046699 | 10.1527036 | 6.662122495 | ENSG00000285077 |
| NA | -0.61381427 | 0.653466748 | 0.269300328 | 0.477354674 | 23.53387425 | 15.41409109 | ENSG00000260362 |
| NA | -0.613964691 | 0.653398618 | 0.450805312 | 0.657105459 | 8.570658071 | 5.586553519 | ENSG00000273391 |
| NA | -0.614244053 | 0.653272107 | 0.538119114 | 0.727964146 | 5.549783328 | 3.632343887 | ENSG00000236015 |
| CDC27 | -0.614482951 | 0.65316394 | 5.14E-08 | 7.50E-07 | 787.5265219 | 514.284658 | ENSG00000004897 |
| CCDC120 | -0.614605433 | 0.65310849 | 4.41E-06 | 4.52E-05 | 398.4650088 | 260.4374064 | ENSG00000147144 |
| GPR6 | -0.614996081 | 0.652931667 | 0.530999936 | 0.722993099 | 6.536969634 | 4.265082111 | ENSG00000146360 |
| ADAM11 | -0.61568021 | 0.652622119 | 0.000149936 | 0.001067298 | 255.4534389 | 166.6943316 | ENSG00000073670 |
| HES1 | -0.615785557 | 0.652574465 | 0.000291439 | 0.001916375 | 220.5098437 | 143.957287 | ENSG00000114315 |
| NA | -0.615847675 | 0.652546368 | 0.414263766 | 0.62526944 | 9.657483454 | 6.282759052 | ENSG00000257303 |
| NA | -0.616301104 | 0.65234131 | 0.449665564 | 0.656001095 | 8.652499785 | 5.632900666 | ENSG00000234297 |
| LOC100506098 | -0.616391339 | 0.65230051 | 0.093029422 | 0.225991849 | 40.66807145 | 26.52467678 | ENSG00000233834 |
| NA | -0.616648352 | 0.652184314 | 0.472478935 | 0.675813429 | 7.634473435 | 4.973110921 | ENSG00000229020 |
| ZNF165 | -0.616913828 | 0.652064314 | 0.023665411 | 0.078849908 | 79.32022567 | 51.71946385 | ENSG00000197279 |
| IQUB | -0.616944456 | 0.652050471 | 0.532538532 | 0.723993246 | 6.53460537 | 4.255434298 | ENSG00000164675 |
| NA | -0.61721875 | 0.651926511 | 0.571489194 | 0.754250711 | 4.566156493 | 2.989957851 | ENSG00000272370 |
| ATP6V1D | -0.617273759 | 0.651901654 | 3.27E-12 | 8.76E-11 | 870.5422651 | 567.4508899 | ENSG00000100554 |
| KCTD8 | -0.617281897 | 0.651897977 | 0.007168851 | 0.029886036 | 143.2932484 | 93.40351184 | ENSG00000183783 |
| PRR5L | -0.61733954 | 0.651871931 | 0.005347884 | 0.023378425 | 133.088043 | 86.66091747 | ENSG00000135362 |
| SLC49A4 | -0.617869569 | 0.651632485 | 1.73E-05 | 0.000155841 | 289.6394068 | 188.8266128 | ENSG00000138463 |
| LOC101928000 | -0.618055861 | 0.651548346 | 0.172830589 | 0.352435554 | 31.77416466 | 20.68474848 | ENSG00000234327 |
| PRRT4 | -0.618126294 | 0.651516538 | 0.108121552 | 0.252957818 | 41.74391293 | 27.25323072 | ENSG00000224940 |
| LONRF2 | -0.618349195 | 0.651415885 | 0.33467402 | 0.546630623 | 13.74591168 | 8.991344641 | ENSG00000170500 |
| TOLLIP-AS1 | -0.618416839 | 0.651385342 | 0.323780607 | 0.53453945 | 14.25803946 | 9.27706783 | ENSG00000255153 |
| TNFSF13 | -0.61847393 | 0.651359566 | 0.067355457 | 0.177382208 | 54.19393296 | 35.29688697 | ENSG00000161955 |
| LINC00862 | -0.618614395 | 0.651296151 | 0.280816944 | 0.489649698 | 17.87257822 | 11.65140762 | ENSG00000203721 |
| CCNC | -0.618822293 | 0.651202303 | 2.02E-08 | 3.17E-07 | 700.2735945 | 456.2536502 | ENSG00000112237 |
| PCSK1N | -0.619317529 | 0.650978802 | 0.011352711 | 0.043417566 | 149.8190071 | 97.5689944 | ENSG00000102109 |
| NBPF1 | -0.619705249 | 0.650803877 | 6.89E-07 | 8.20E-06 | 435.3245243 | 283.4974407 | ENSG00000219481 |
| SLC35A2 | -0.619976741 | 0.650681418 | 6.51E-10 | 1.28E-08 | 679.6941608 | 442.4060097 | ENSG00000102100 |
| CIPC | -0.619983735 | 0.650678264 | 2.97E-07 | 3.80E-06 | 435.4985813 | 283.3412791 | ENSG00000198894 |
| MYCBP | -0.620379979 | 0.650499576 | 0.000288821 | 0.00190181 | 243.5339806 | 158.461503 | ENSG00000214114 |
| NA | -0.620479054 | 0.650454905 | 0.311865445 | 0.522311008 | 15.26182792 | 9.9011042 | ENSG00000235508 |

| | | | | | | | |
|---|---|---|---|---|---|---|---|
| LOC100128398 | -0.620928086 | 0.650252486 | 0.005722116 | 0.024715195 | 134.0686946 | 87.26180174 | ENSG00000176593 |
| WARS1 | -0.621088214 | 0.650180316 | 2.68E-20 | 1.63E-18 | 2874.965521 | 1868.610495 | ENSG00000140105 |
| ZNF107 | -0.621239802 | 0.650112004 | 8.52E-06 | 8.31E-05 | 428.3525918 | 278.5521687 | ENSG00000196247 |
| PTPRZ1 | -0.621253296 | 0.650105923 | 1.54E-22 | 1.13E-20 | 5816.463418 | 3780.778731 | ENSG00000106278 |
| NA | -0.621383098 | 0.650047435 | 0.421006751 | 0.631006395 | 11.80949805 | 7.714044917 | ENSG00000279452 |
| ARHGAP8 | -0.621399325 | 0.650040123 | 0.131616324 | 0.291699834 | 35.14912817 | 22.80521903 | ENSG00000241484 |
| NA | -0.621460305 | 0.650012648 | 0.417625347 | 0.628484714 | 9.179755185 | 5.987388051 | ENSG00000228519 |
| NA | -0.6220322 | 0.649755029 | 0.466993605 | 0.670914738 | 7.69259405 | 4.987109661 | ENSG00000235370 |
| NRG3 | -0.6224056 | 0.64958688 | 0.065616041 | 0.173649999 | 50.02071394 | 32.54876417 | ENSG00000185737 |
| EBLN2 | -0.622604097 | 0.649497511 | 0.26528761 | 0.47290509 | 19.30281395 | 12.55369483 | ENSG00000255423 |
| PDIA2 | -0.622820821 | 0.64939995 | 0.326996649 | 0.538425113 | 15.91836724 | 10.3514073 | ENSG00000185615 |
| RSPO2 | -0.622929194 | 0.64935117 | 0.367736455 | 0.5798366 | 14.32327902 | 9.285769684 | ENSG00000147655 |
| UPF3B | -0.623144015 | 0.649254487 | 7.42E-15 | 2.74E-13 | 1093.819703 | 710.108839 | ENSG00000125351 |
| NGRN | -0.624804906 | 0.648507468 | 1.40E-08 | 2.27E-07 | 560.993415 | 364.00845 | ENSG00000182768 |
| FBN2 | -0.624809667 | 0.648505328 | 3.31E-09 | 5.84E-08 | 930.6058727 | 603.0236937 | ENSG00000138829 |
| NA | -0.625778469 | 0.648069988 | 0.115028608 | 0.264851897 | 40.38250239 | 26.1462593 | ENSG00000213222 |
| CCNI2 | -0.625897741 | 0.648016413 | 0.260266873 | 0.467129814 | 21.15441121 | 13.68743423 | ENSG00000205089 |
| LTBP1 | -0.626304499 | 0.647833735 | 1.09E-14 | 3.92E-13 | 1303.616942 | 844.2121564 | ENSG00000049323 |
| ZNF253 | -0.626396117 | 0.647792596 | 3.52E-06 | 3.66E-05 | 355.2211452 | 230.0515605 | ENSG00000256771 |
| DFFB | -0.626411325 | 0.647785767 | 5.73E-05 | 0.000454576 | 233.6056985 | 151.3240334 | ENSG00000169598 |
| NA | -0.626446779 | 0.647769848 | 0.467347942 | 0.671109527 | 7.665313479 | 4.944167482 | ENSG00000272140 |
| NA | -0.626636022 | 0.647684884 | 0.135420395 | 0.297408322 | 38.58958441 | 24.99494823 | ENSG00000215105 |
| PERP | -0.626796141 | 0.647613003 | 4.06E-11 | 9.62E-10 | 971.9276889 | 629.6029679 | ENSG00000112378 |
| FLJ45513 | -0.626804852 | 0.647609093 | 0.511405694 | 0.707953125 | 7.212501516 | 4.696089587 | ENSG00000204584 |
| NBAS | -0.626913689 | 0.647560239 | 3.48E-10 | 7.18E-09 | 856.8263043 | 554.4182844 | ENSG00000151779 |
| U2AF1L4 | -0.627288058 | 0.647392224 | 0.202683473 | 0.393996642 | 28.21034824 | 18.25100876 | ENSG00000161265 |
| ZNF491 | -0.627336203 | 0.64737062 | 0.521902833 | 0.71662324 | 5.676702975 | 3.674340108 | ENSG00000177599 |
| NA | -0.627412198 | 0.64733652 | 0.338734316 | 0.550644635 | 15.34842432 | 9.900158241 | ENSG00000226239 |
| ARIH1 | -0.628194827 | 0.64698545 | 2.99E-13 | 9.15E-12 | 1289.37731 | 833.9212629 | ENSG00000166233 |
| ZNF844 | -0.628288488 | 0.646943449 | 0.027494597 | 0.088788553 | 69.23276163 | 44.76197035 | ENSG00000223547 |
| MTHFD2L | -0.628771007 | 0.64672711 | 0.01511695 | 0.0548072 | 97.82625865 | 63.37928977 | ENSG00000163738 |
| KLHL3 | -0.629449726 | 0.646422928 | 3.06E-06 | 3.22E-05 | 367.0222774 | 237.0174723 | ENSG00000146021 |
| ST3GAL3 | -0.630117435 | 0.646123819 | 0.24046602 | 0.443996948 | 21.65319083 | 13.98185927 | ENSG00000126091 |
| BMS1P1 | -0.630276125 | 0.646052752 | 0.178393984 | 0.360587647 | 30.68113621 | 19.8022084 | ENSG00000204177 |
| PDRG1 | -0.630450771 | 0.645974548 | 1.15E-10 | 2.55E-09 | 986.3418578 | 637.0939063 | ENSG00000088356 |
| DYNC1I2 | -0.630532048 | 0.645938157 | 4.49E-20 | 2.70E-18 | 1840.257151 | 1188.405744 | ENSG00000077380 |
| DHRS1 | -0.630546673 | 0.64593161 | 9.78E-06 | 9.35E-05 | 293.6139325 | 189.6248769 | ENSG00000157379 |
| LRP2BP | -0.630659474 | 0.645881107 | 0.191569 | 0.378547916 | 24.06349392 | 15.53618033 | ENSG00000109771 |
| C8orf48 | -0.630776424 | 0.645828752 | 0.514568487 | 0.710820414 | 5.676702975 | 3.660341367 | ENSG00000164743 |
| TRAM2-AS1 | -0.63082961 | 0.645804944 | 0.040852117 | 0.120696533 | 65.8017886 | 42.51361922 | ENSG00000225791 |
| OAT | -0.631043647 | 0.64570914 | 1.19E-23 | 9.66E-22 | 2523.838857 | 1629.533245 | ENSG00000065154 |
| MGC32805 | -0.631435905 | 0.6455336 | 0.272423498 | 0.481047357 | 16.95775042 | 10.92190772 | ENSG00000250328 |
| SEPTIN8 | -0.631537043 | 0.645488348 | 5.61E-18 | 2.82E-16 | 1601.971314 | 1033.870938 | ENSG00000164402 |
| LOC105370689 | -0.631946448 | 0.645305198 | 0.440015463 | 0.647767794 | 9.805454452 | 6.362400566 | ENSG00000287346 |
| KCNH4 | -0.631988881 | 0.645286218 | 0.27315761 | 0.481840226 | 18.01938016 | 11.67722964 | ENSG00000089558 |
| RRP36 | -0.632037673 | 0.645264395 | 1.12E-19 | 6.57E-18 | 1631.883639 | 1053.113849 | ENSG00000124541 |
| RCBTB2 | -0.63253255 | 0.645043093 | 8.76E-06 | 8.52E-05 | 312.9069839 | 201.8943616 | ENSG00000136161 |
| CCDC88A | -0.633566133 | 0.644581133 | 5.43E-15 | 2.04E-13 | 1155.605169 | 744.7991659 | ENSG00000115355 |
| NA | -0.63376164 | 0.644493789 | 0.288225429 | 0.497453594 | 16.02868473 | 10.31499151 | ENSG00000279656 |
| ANKRD36B | -0.633984364 | 0.644394299 | 0.035026663 | 0.107253389 | 64.48039582 | 41.59704973 | ENSG00000196912 |
| BRICD5 | -0.634352844 | 0.644229735 | 0.037068035 | 0.111986857 | 88.94658975 | 57.4343924 | ENSG00000182685 |
| SLC32A1 | -0.634400809 | 0.644208317 | 0.506197977 | 0.70358054 | 6.722009897 | 4.313604722 | ENSG00000101438 |
| NA | -0.634557045 | 0.644138556 | 0.052153938 | 0.145888445 | 52.98227286 | 34.17525439 | ENSG00000277534 |
| CBLN1 | -0.636130668 | 0.643436343 | 0.00042121 | 0.002644464 | 245.743758 | 158.346151 | ENSG00000102924 |
| PLPBP | -0.636550191 | 0.643249265 | 9.19E-13 | 2.64E-11 | 840.7077942 | 540.7987114 | ENSG00000147471 |
| AREL1 | -0.636822996 | 0.643127642 | 1.92E-11 | 4.73E-10 | 1102.49267 | 708.5946316 | ENSG00000119682 |
| PEX12 | -0.637207709 | 0.642956167 | 0.034304565 | 0.105580461 | 69.42789549 | 44.64998043 | ENSG00000108733 |
| CYP27A1 | -0.637490525 | 0.642830138 | 0.002744168 | 0.013305165 | 129.4586554 | 83.15813517 | ENSG00000135929 |
| NA | -0.637710122 | 0.642732299 | 0.464898111 | 0.669294471 | 7.245705824 | 4.672443033 | ENSG00000270964 |
| NA | -0.638281345 | 0.642477865 | 0.491111564 | 0.690591354 | 7.201823099 | 4.635743699 | ENSG00000223343 |
| LOC101928424 | -0.638782181 | 0.642254865 | 0.549168526 | 0.736475695 | 5.649422404 | 3.613048261 | ENSG00000255337 |
| ARMC7 | -0.638848749 | 0.642225231 | 3.16E-06 | 3.32E-05 | 326.2397703 | 209.4989709 | ENSG00000125449 |
| NA | -0.638959284 | 0.642176028 | 0.465176294 | 0.669437616 | 8.343209624 | 5.36429764 | ENSG00000260144 |
| RPP40 | -0.640142617 | 0.641649516 | 0.007782977 | 0.031966779 | 105.963656 | 68.02997817 | ENSG00000124787 |
| PHF3 | -0.640271654 | 0.641592128 | 3.04E-18 | 1.56E-16 | 1677.035896 | 1075.566031 | ENSG00000118482 |
| SCG3 | -0.640549968 | 0.641468369 | 2.75E-09 | 4.91E-08 | 1087.844265 | 697.8504495 | ENSG00000104112 |
| SLC20A1 | -0.640675965 | 0.641412349 | 1.90E-26 | 1.90E-24 | 2546.290049 | 1633.234943 | ENSG00000144136 |
| IFT57 | -0.640733364 | 0.64138683 | 2.52E-09 | 4.56E-08 | 584.3297649 | 374.8876991 | ENSG00000114446 |
| ZNF268 | -0.64078328 | 0.641364639 | 1.79E-05 | 0.000160939 | 291.4135854 | 186.8719087 | ENSG00000090612 |
| ALDH16A1 | -0.642070222 | 0.640792771 | 7.72E-14 | 2.51E-12 | 1112.142832 | 712.8302664 | ENSG00000161618 |
| HES7 | -0.642250517 | 0.640712696 | 0.53420631 | 0.725216459 | 6.269197935 | 4.035353883 | ENSG00000179111 |
| LRRN3 | -0.642547003 | 0.640581038 | 0.527677955 | 0.720757861 | 5.66958403 | 3.610872798 | ENSG00000173114 |
| POLR3B | -0.642614182 | 0.64055121 | 1.97E-08 | 3.09E-07 | 536.8474172 | 343.6767485 | ENSG00000013503 |
| ARFRP1 | -0.642917842 | 0.6404164 | 5.56E-11 | 1.29E-09 | 729.7055929 | 467.3855618 | ENSG00000101246 |

| | | | | | | | |
|---|---|---|---|---|---|---|---|
| NA | -0.642928717 | 0.640411572 | 0.510913238 | 0.707345698 | 6.258519517 | 4.027881533 | ENSG00000232186 |
| GDAP2 | -0.6435186 | 0.640149777 | 5.87E-05 | 0.000463958 | 231.4972873 | 148.1455655 | ENSG00000196505 |
| SLC44A3 | -0.643714318 | 0.64006294 | 0.02374302 | 0.079052704 | 67.88172402 | 43.41997381 | ENSG00000143036 |
| FLRT1 | -0.643935646 | 0.639964753 | 0.066471129 | 0.175673862 | 47.64717463 | 30.4716873 | ENSG00000126500 |
| PMM1 | -0.644226521 | 0.639835737 | 0.000618286 | 0.003685713 | 180.8001517 | 115.6682354 | ENSG00000100417 |
| NA | -0.645151991 | 0.639425422 | 0.401464127 | 0.612708101 | 9.80664966 | 6.282759052 | ENSG00000232630 |
| NA | -0.645483478 | 0.639278519 | 0.335632347 | 0.547537071 | 11.96606254 | 7.644051217 | ENSG00000286835 |
| C5orf38 | -0.645502165 | 0.639270239 | 0.124899687 | 0.280667608 | 32.59263353 | 20.83918612 | ENSG00000186493 |
| FBXO43 | -0.645637509 | 0.639210269 | 0.155094497 | 0.326765869 | 28.58368275 | 18.24911685 | ENSG00000156509 |
| SEPTIN5 | -0.645741114 | 0.639164367 | 0.507273952 | 0.704350311 | 13.04668485 | 8.364849262 | ENSG00000184702 |
| PHKA2 | -0.645744219 | 0.639162991 | 1.28E-11 | 3.21E-10 | 757.5773372 | 484.3210357 | ENSG00000044446 |
| GPR160 | -0.64581753 | 0.639130513 | 8.66E-10 | 1.68E-08 | 571.889508 | 365.5439699 | ENSG00000173890 |
| DOCK4 | -0.645994529 | 0.639052105 | 6.66E-11 | 1.53E-09 | 692.8638486 | 442.6654164 | ENSG00000128512 |
| ETHE1 | -0.646306868 | 0.638913767 | 0.023681792 | 0.078890569 | 75.96544995 | 48.53061312 | ENSG00000105755 |
| KCNJ14 | -0.646340411 | 0.638898912 | 0.449114606 | 0.655602485 | 7.28010534 | 4.654093366 | ENSG00000182324 |
| MTHFD1 | -0.646724887 | 0.638728669 | 3.18E-35 | 4.92E-33 | 5371.672593 | 3430.544173 | ENSG00000100714 |
| DUSP18 | -0.647087291 | 0.638568242 | 0.335247927 | 0.547299196 | 11.96250306 | 7.627877013 | ENSG00000167065 |
| MRPL2 | -0.64752431 | 0.638374837 | 0.271102112 | 0.479513289 | 20.15003793 | 12.90070986 | ENSG00000112651 |
| KMT5A | -0.647545704 | 0.63836537 | 1.25E-06 | 1.42E-05 | 335.3424384 | 214.042871 | ENSG00000183955 |
| GLDC | -0.647604889 | 0.638339182 | 3.56E-33 | 4.97E-31 | 4349.416556 | 2776.012206 | ENSG00000178445 |
| NA | -0.647732012 | 0.638282938 | 0.174823314 | 0.3552333 | 28.2127125 | 18.02903643 | ENSG00000275437 |
| TMEM123 | -0.648111651 | 0.638114998 | 1.18E-15 | 4.84E-14 | 1921.463981 | 1226.204881 | ENSG00000152558 |
| GPR182 | -0.648156751 | 0.638095051 | 0.386718687 | 0.598965177 | 11.46697743 | 7.348680215 | ENSG00000166856 |
| NA | -0.648165962 | 0.638090977 | 0.558768225 | 0.744275493 | 4.685957196 | 2.978134574 | ENSG00000233747 |
| VARS2 | -0.648449782 | 0.637965458 | 0.357061294 | 0.569121372 | 10.96463834 | 6.997314253 | ENSG00000137411 |
| NA | -0.648842421 | 0.637791855 | 0.037155428 | 0.112144817 | 64.59783226 | 41.20709251 | ENSG00000279495 |
| E2F8 | -0.649063635 | 0.637694068 | 3.39E-05 | 0.00028411 | 253.2877707 | 161.5111443 | ENSG00000129173 |
| FXYD6 | -0.649781025 | 0.637377049 | 0.002155233 | 0.010827765 | 127.0575562 | 80.98819605 | ENSG00000137726 |
| ZNF576 | -0.650273616 | 0.637159461 | 0.002911047 | 0.014024264 | 125.9965368 | 80.3595252 | ENSG00000124444 |
| ZNF800 | -0.650400396 | 0.637103472 | 0.055283638 | 0.1525053 | 56.14074567 | 35.75785647 | ENSG00000048405 |
| NA | -0.65135371 | 0.636682621 | 0.488367771 | 0.688779985 | 6.231238946 | 3.974062036 | ENSG00000286360 |
| CDK5 | -0.652196721 | 0.636310697 | 1.13E-07 | 1.56E-06 | 416.4825833 | 265.0397402 | ENSG00000164885 |
| NA | -0.652374833 | 0.636232144 | 0.470928093 | 0.674310185 | 7.816843948 | 4.997986979 | ENSG00000229413 |
| NA | -0.652426149 | 0.636209514 | 0.457508433 | 0.663190571 | 7.276545868 | 4.620799 | ENSG00000250308 |
| NA | -0.65268287 | 0.636096313 | 0.284697869 | 0.49393838 | 15.13846775 | 9.654255809 | ENSG00000272927 |
| EXOC3L1 | -0.653310007 | 0.635819863 | 0.036962225 | 0.11172081 | 68.43509094 | 43.541328 | ENSG00000179044 |
| FEM1A | -0.653311714 | 0.635819111 | 0.049740684 | 0.140516502 | 53.79448693 | 34.16749849 | ENSG00000141965 |
| IGSF5 | -0.653669735 | 0.635661345 | 0.08568934 | 0.212646936 | 38.60354297 | 24.55769791 | ENSG00000183067 |
| GLIPR2 | -0.654098165 | 0.635472604 | 0.000318989 | 0.00207442 | 217.6086918 | 138.2113778 | ENSG00000122694 |
| FZD2 | -0.654141999 | 0.635453296 | 1.98E-10 | 4.23E-09 | 1453.752661 | 924.083377 | ENSG00000180340 |
| NA | -0.654525012 | 0.635284616 | 0.501366443 | 0.699594006 | 6.275121671 | 3.991182199 | ENSG00000268947 |
| VN1R1 | -0.65479683 | 0.635164933 | 0.502215191 | 0.700181291 | 6.838251127 | 4.309253795 | ENSG00000178201 |
| IGFLR1 | -0.655920618 | 0.634670364 | 0.465675017 | 0.669835685 | 7.296707494 | 4.618623537 | ENSG00000126246 |
| H2BC8 | -0.656912286 | 0.634234259 | 0.115984673 | 0.266436024 | 31.42544097 | 19.94437126 | ENSG00000273802 |
| SLC25A53 | -0.657512911 | 0.633970269 | 5.47E-05 | 0.00043569 | 250.8792732 | 159.0628387 | ENSG00000269743 |
| CYP2D6 | -0.657714606 | 0.633881643 | 0.218386842 | 0.414542705 | 23.15578506 | 14.71353463 | ENSG00000100197 |
| SLC2A1-AS1 | -0.65811407 | 0.633706154 | 0.446238978 | 0.653521693 | 7.310945384 | 4.616448073 | ENSG00000227533 |
| C1QTNF4 | -0.65878634 | 0.633410927 | 0.001874401 | 0.00960338 | 144.3628868 | 91.56185923 | ENSG00000172247 |
| TMEM266 | -0.659177337 | 0.633239284 | 0.001083025 | 0.006010263 | 141.6405982 | 89.64964582 | ENSG00000169758 |
| C12orf57 | -0.65997478 | 0.632889361 | 0.006742262 | 0.028383329 | 100.7910721 | 63.77823239 | ENSG00000111678 |
| ZNF699 | -0.660001446 | 0.632877662 | 0.130681356 | 0.290104314 | 36.8465252 | 23.27075038 | ENSG00000196110 |
| SLC26A8 | -0.660040473 | 0.632860543 | 0.438995658 | 0.647083438 | 7.854802937 | 4.989285125 | ENSG00000112053 |
| NA | -0.660476058 | 0.632669495 | 0.002094439 | 0.010595629 | 219.6441858 | 139.0791841 | ENSG00000263146 |
| SHC3 | -0.660484651 | 0.632665727 | 3.06E-05 | 0.000260499 | 250.3893925 | 158.4134748 | ENSG00000148082 |
| NPEPPSP1 | -0.660833355 | 0.632512828 | 0.010356011 | 0.040158782 | 90.92391145 | 57.4976488 | ENSG00000274487 |
| TCF24 | -0.661319791 | 0.632299599 | 0.069342851 | 0.181440614 | 41.98590475 | 26.55702519 | ENSG00000261787 |
| GFI1 | -0.661755231 | 0.632108784 | 0.532021359 | 0.723643528 | 6.334437495 | 4.00423498 | ENSG00000162676 |
| KLHL14 | -0.661855355 | 0.632064917 | 0.086390581 | 0.213853207 | 41.9775906 | 26.53460814 | ENSG00000197705 |
| NA | -0.662331912 | 0.631856165 | 0.336261499 | 0.548195628 | 12.64276347 | 7.983593902 | ENSG00000286753 |
| PCDHB7 | -0.662360924 | 0.631843459 | 0.302134143 | 0.512420591 | 13.66910398 | 8.614156663 | ENSG00000113212 |
| CLCNKA | -0.662422932 | 0.631816303 | 0.251531373 | 0.457186716 | 21.7291088 | 13.7369028 | ENSG00000186510 |
| ZNF521 | -0.662977254 | 0.631573588 | 1.92E-05 | 0.000172344 | 272.1057897 | 171.94749 | ENSG00000198795 |
| NA | -0.663285161 | 0.631438809 | 0.211045233 | 0.404751622 | 20.49639735 | 12.94705701 | ENSG00000267523 |
| CUL1 | -0.663392666 | 0.631391758 | 2.20E-26 | 2.18E-24 | 2687.448652 | 1696.953238 | ENSG00000055130 |
| LIME1 | -0.663427295 | 0.631376603 | 0.397511983 | 0.609205812 | 11.06160767 | 7.027487197 | ENSG00000203896 |
| ARHGAP29 | -0.66365954 | 0.631274972 | 0.010550599 | 0.040804434 | 83.60556779 | 52.81309894 | ENSG00000137962 |
| RBM25 | -0.663757218 | 0.631232233 | 3.44E-26 | 3.31E-24 | 3412.778488 | 2153.994266 | ENSG00000119707 |
| NA | -0.663858459 | 0.631187938 | 0.216903605 | 0.412634692 | 19.41193624 | 12.26702568 | ENSG00000228218 |
| MSC | -0.664015857 | 0.631119079 | 0.257514757 | 0.464425636 | 17.87140917 | 11.2763951 | ENSG00000178860 |
| AMER3 | -0.664017643 | 0.631118298 | 0.434449672 | 0.643294814 | 7.876159772 | 4.983988239 | ENSG00000178171 |
| PRR29 | -0.664295467 | 0.630996773 | 0.418181866 | 0.628971371 | 8.879948232 | 5.596201332 | ENSG00000224383 |
| NA | -0.664479796 | 0.630916158 | 0.197717703 | 0.38745448 | 24.74139006 | 15.63105009 | ENSG00000235023 |
| POU3F1 | -0.665011942 | 0.630683484 | 5.83E-11 | 1.35E-09 | 1116.619496 | 704.4661976 | ENSG00000185668 |
| CRIP2 | -0.665241866 | 0.630582979 | 3.73E-05 | 0.000309398 | 227.2893377 | 143.2284495 | ENSG00000182809 |

| | | | | | | | |
|---|---|---|---|---|---|---|---|
| SNORD62A | -0.665714295 | 0.630376521 | 0.601444038 | 0.776474614 | 4.246187915 | 2.696762312 | ENSG00000235284 |
| LINC01090 | -0.666143081 | 0.630189193 | 0.004807066 | 0.021345272 | 138.9755923 | 87.58110284 | ENSG00000231689 |
| KIF6 | -0.666355251 | 0.630096521 | 0.122162809 | 0.276028315 | 33.07837046 | 20.88865469 | ENSG00000164627 |
| KIF20A | -0.666545197 | 0.630013568 | 5.96E-18 | 2.99E-16 | 1716.896587 | 1081.502144 | ENSG00000112984 |
| DUSP26 | -0.667355631 | 0.629659757 | 0.039906452 | 0.118495809 | 54.19838215 | 34.11273304 | ENSG00000133878 |
| NA | -0.667833841 | 0.629451078 | 0.25266857 | 0.458591633 | 17.38536674 | 10.97572722 | ENSG00000227542 |
| HSPA1B | -0.668275113 | 0.62925858 | 4.14E-14 | 1.39E-12 | 1292.906675 | 813.7708662 | ENSG00000204388 |
| RPE | -0.668683605 | 0.629080433 | 4.23E-13 | 1.28E-11 | 1088.656017 | 684.7158714 | ENSG00000197713 |
| COMMD9 | -0.670067962 | 0.62847708 | 1.07E-06 | 1.23E-05 | 364.3199745 | 229.0863005 | ENSG00000110442 |
| PDYN | -0.670233518 | 0.628404964 | 0.120014863 | 0.27264905 | 38.50835309 | 24.28597347 | ENSG00000101327 |
| NA | -0.670563852 | 0.628261095 | 0.006646544 | 0.02806996 | 93.02372917 | 58.45035054 | ENSG00000263823 |
| NA | -0.670601695 | 0.628244615 | 0.222856159 | 0.420653529 | 20.57556932 | 12.97940542 | ENSG00000268262 |
| NA | -0.670797023 | 0.628159562 | 0.295452792 | 0.505492478 | 19.02792331 | 11.98101895 | ENSG00000284636 |
| TMEM191B | -0.671752384 | 0.627743729 | 0.477855285 | 0.680360153 | 8.472214204 | 5.343772509 | ENSG00000278558 |
| CCNE2 | -0.671976329 | 0.627646294 | 0.048345511 | 0.13752146 | 60.54677821 | 37.99873525 | ENSG00000175305 |
| LINC00632 | -0.672384122 | 0.627468908 | 0.008033925 | 0.032783207 | 105.5656845 | 66.32244898 | ENSG00000203930 |
| GAD2 | -0.672445058 | 0.627442406 | 0.462641894 | 0.667551225 | 7.999519947 | 5.025984459 | ENSG00000136750 |
| NA | -0.672922233 | 0.627234912 | 0.591109529 | 0.76886173 | 4.215347871 | 2.656941556 | ENSG00000271275 |
| NA | -0.674190883 | 0.626683588 | 0.584662545 | 0.763367822 | 4.791520009 | 2.980310038 | ENSG00000278009 |
| FNDC11 | -0.674570115 | 0.626518878 | 0.032156115 | 0.100325167 | 67.39598708 | 42.2133302 | ENSG00000125531 |
| TFPT | -0.675149492 | 0.626267322 | 1.33E-05 | 0.000122962 | 333.2841386 | 208.9771087 | ENSG00000105619 |
| NA | -0.675284853 | 0.626208565 | 0.498777579 | 0.697509604 | 5.291494837 | 3.311150869 | ENSG00000272918 |
| ITFG1 | -0.675429435 | 0.626145812 | 5.38E-07 | 6.54E-06 | 352.1833111 | 220.4221082 | ENSG00000129636 |
| PPM1H | -0.675803348 | 0.625983551 | 6.39E-21 | 4.09E-19 | 2009.843501 | 1257.731613 | ENSG00000111110 |
| NA | -0.675836688 | 0.625969084 | 0.335646669 | 0.547537071 | 15.33271189 | 9.62531237 | ENSG00000250046 |
| NA | -0.676030823 | 0.625884857 | 0.512601575 | 0.708949343 | 5.311656463 | 3.322974145 | ENSG00000284526 |
| NA | -0.676175678 | 0.625822018 | 0.040553813 | 0.120031888 | 54.60347259 | 34.24524809 | ENSG00000272686 |
| RGPD1 | -0.678094272 | 0.62499031 | 0.497037862 | 0.69612578 | 5.322334881 | 3.322974145 | ENSG00000187627 |
| DUSP28 | -0.678277196 | 0.624911071 | 0.159978752 | 0.33379723 | 30.27040137 | 18.82368465 | ENSG00000188542 |
| FOXN3-AS1 | -0.679527799 | 0.6243696 | 0.179687742 | 0.362426894 | 23.46863469 | 14.68553715 | ENSG00000258920 |
| MIR4713HG | -0.679580288 | 0.624346884 | 0.465563781 | 0.669803268 | 6.884218784 | 4.304902868 | ENSG00000259240 |
| ZNF613 | -0.679777412 | 0.624261582 | 0.00052494 | 0.003206176 | 160.7218875 | 100.3312315 | ENSG00000176024 |
| GPT | -0.679777614 | 0.624261495 | 0.462215941 | 0.667095762 | 6.963696234 | 4.360897828 | ENSG00000167701 |
| RNF38 | -0.679813432 | 0.624245996 | 4.22E-09 | 7.38E-08 | 659.0474281 | 411.3751281 | ENSG00000137075 |
| SMIM10L1 | -0.679946191 | 0.624188555 | 5.31E-07 | 6.46E-06 | 361.7213249 | 225.8872349 | ENSG00000256537 |
| CCDC15 | -0.680813892 | 0.623813253 | 0.025664393 | 0.084082043 | 66.563001 | 41.45488686 | ENSG00000149548 |
| TTBK1 | -0.68091553 | 0.623769307 | 0.014873226 | 0.054165489 | 78.74168927 | 49.19503734 | ENSG00000146216 |
| LINC01089 | -0.681870708 | 0.623356459 | 0.005127646 | 0.022577663 | 189.8266487 | 118.4600784 | ENSG00000212694 |
| ZNF891 | -0.681877437 | 0.623353551 | 0.013624749 | 0.050333212 | 77.87725275 | 48.48454952 | ENSG00000214029 |
| NA | -0.682103912 | 0.623255705 | 0.348293082 | 0.560313047 | 10.66483139 | 6.631949551 | ENSG00000267500 |
| HM13 | -0.682243617 | 0.623195354 | 5.63E-24 | 4.68E-22 | 2423.343428 | 1510.5623 | ENSG00000101294 |
| KREMEN1 | -0.682390831 | 0.623131766 | 7.05E-05 | 0.000544276 | 218.0055204 | 135.7682739 | ENSG00000183762 |
| FAM111B | -0.682488119 | 0.623089746 | 4.14E-08 | 6.15E-07 | 641.9607243 | 400.231627 | ENSG00000189057 |
| NCAM1 | -0.682620725 | 0.623032477 | 1.27E-05 | 0.000118237 | 249.2891928 | 155.3615627 | ENSG00000149294 |
| TMEM102 | -0.682841054 | 0.622937335 | 0.026747438 | 0.086895815 | 85.80916829 | 53.59547238 | ENSG00000181284 |
| FBRS | -0.683469441 | 0.622666064 | 1.76E-08 | 2.79E-07 | 511.1397169 | 318.5146364 | ENSG00000156860 |
| PSMD9 | -0.683668243 | 0.622580267 | 0.309598371 | 0.520174843 | 14.33009248 | 8.951523885 | ENSG00000110801 |
| NA | -0.684149382 | 0.622372672 | 0.323547899 | 0.534435484 | 13.45113874 | 8.360781881 | ENSG00000270938 |
| GNRH1 | -0.684438614 | 0.622247911 | 0.522587859 | 0.716982427 | 5.338937034 | 3.313326333 | ENSG00000147437 |
| LOC100288748 | -0.684455978 | 0.622240421 | 0.161028924 | 0.335395525 | 25.74072933 | 16.00048217 | ENSG00000261437 |
| MCOLN3 | -0.685065124 | 0.62197775 | 0.131678509 | 0.291769171 | 27.75751012 | 17.27778189 | ENSG00000055732 |
| SLC9A2 | -0.685190356 | 0.621923762 | 0.519625144 | 0.715241508 | 5.889913533 | 3.681812457 | ENSG00000115616 |
| NA | -0.685219341 | 0.621911267 | 0.237400919 | 0.439497731 | 18.63084154 | 11.62341014 | ENSG00000236166 |
| SAP30-DT | -0.685299373 | 0.621876768 | 0.367421925 | 0.579660284 | 10.65178871 | 6.603006112 | ENSG00000272870 |
| NA | -0.685326367 | 0.621865132 | 0.551625584 | 0.738582061 | 4.293630113 | 2.680588109 | ENSG00000285399 |
| LOC106699570 | -0.685952971 | 0.621595097 | 0.524442311 | 0.718312522 | 5.924313049 | 3.702337588 | ENSG00000246777 |
| NA | -0.686473463 | 0.62137088 | 0.510262354 | 0.706888538 | 5.360293869 | 3.320798682 | ENSG00000215630 |
| PCDHA12 | -0.687495207 | 0.620930969 | 0.000143445 | 0.001027677 | 175.2201387 | 108.8085222 | ENSG00000251664 |
| ASGR1 | -0.687700525 | 0.620842607 | 9.59E-05 | 0.000714716 | 201.8312027 | 125.2537271 | ENSG00000141505 |
| NA | -0.688069253 | 0.620683951 | 0.093707062 | 0.22731636 | 33.60829561 | 20.87030503 | ENSG00000227896 |
| CLCF1 | -0.68817322 | 0.620639223 | 0.252235348 | 0.458113536 | 15.47623369 | 9.614435052 | ENSG00000175505 |
| MAGEE1 | -0.688358185 | 0.620559657 | 3.30E-07 | 4.20E-06 | 333.8822524 | 207.2453229 | ENSG00000198934 |
| NA | -0.688366716 | 0.620555988 | 0.487243832 | 0.688066316 | 5.883989796 | 3.66998918 | ENSG00000228110 |
| ZNF616 | -0.688515455 | 0.620492013 | 6.42E-10 | 1.27E-08 | 567.9636459 | 352.4237038 | ENSG00000204611 |
| CYFIP1 | -0.688801821 | 0.620368861 | 5.87E-28 | 6.26E-26 | 2350.576236 | 1458.074105 | ENSG00000273749 |
| ARSI | -0.689343 | 0.620136194 | 0.000311705 | 0.002031947 | 179.2504202 | 111.238857 | ENSG00000183876 |
| NA | -0.689406256 | 0.620109004 | 0.387648718 | 0.599962796 | 8.570658071 | 5.326652347 | ENSG00000266947 |
| ADRB1 | -0.68956978 | 0.620038722 | 0.571099068 | 0.753841189 | 4.246187915 | 2.634240961 | ENSG00000043591 |
| NA | -0.68965211 | 0.620003339 | 0.522634275 | 0.716982427 | 5.400617122 | 3.339148349 | ENSG00000230747 |
| MMP25 | -0.690512086 | 0.619633871 | 7.76E-10 | 1.51E-08 | 499.9552816 | 309.8533546 | ENSG00000008516 |
| FSCN2 | -0.690839603 | 0.61949322 | 0.558740895 | 0.744275493 | 4.30430853 | 2.671886255 | ENSG00000186765 |
| NA | -0.691083343 | 0.619388566 | 0.54034077 | 0.729834265 | 4.277027959 | 2.652590629 | ENSG00000278952 |
| NA | -0.691457414 | 0.619227988 | 0.434870879 | 0.643615717 | 8.128803858 | 5.019458068 | ENSG00000244459 |
| SLC9A7 | -0.691501825 | 0.619208927 | 1.57E-08 | 2.52E-07 | 480.1118123 | 297.1431193 | ENSG00000065923 |

| | | | | | | | |
|---|---|---|---|---|---|---|---|
| CYB5D2 | -0.691838256 | 0.619064547 | 0.004293328 | 0.019491144 | 112.4745141 | 69.6726426 | ENSG00000167740 |
| SV2C | -0.691984102 | 0.619001966 | 0.488320741 | 0.688779985 | 5.927872521 | 3.686163384 | ENSG00000122012 |
| NA | -0.692054578 | 0.618971729 | 0.587683141 | 0.76572348 | 4.228390553 | 2.614945335 | ENSG00000254717 |
| NA | -0.692372196 | 0.618835474 | 0.473815525 | 0.676956135 | 5.907710895 | 3.672164644 | ENSG00000226002 |
| ACAT2 | -0.692625435 | 0.618726858 | 1.87E-26 | 1.88E-24 | 2563.04893 | 1586.004571 | ENSG00000120437 |
| NA | -0.693168994 | 0.618493786 | 0.335924038 | 0.547929302 | 12.94968937 | 8.020009691 | ENSG00000184844 |
| CAVIN4 | -0.693812251 | 0.618218078 | 0.000449803 | 0.002805608 | 158.3697311 | 97.78862332 | ENSG00000170681 |
| NA | -0.693941525 | 0.618162685 | 0.371911418 | 0.583850869 | 9.633762356 | 5.935744017 | ENSG00000225416 |
| NA | -0.694190011 | 0.618056224 | 0.564637054 | 0.748949534 | 4.25330686 | 2.626768612 | ENSG00000276434 |
| RBM15-AS1 | -0.694361564 | 0.617982734 | 0.4877257 | 0.688444565 | 5.91482984 | 3.667813717 | ENSG00000227963 |
| LINC01108 | -0.694405892 | 0.617963746 | 8.99E-06 | 8.71E-05 | 299.6263164 | 185.1338504 | ENSG00000226673 |
| ZNF737 | -0.694544546 | 0.617904358 | 0.000465328 | 0.002894508 | 165.5994191 | 102.2566657 | ENSG00000237440 |
| SLC2A8 | -0.694635552 | 0.617865381 | 6.05E-13 | 1.79E-11 | 834.3354494 | 515.7232913 | ENSG00000136856 |
| HAS3 | -0.695483948 | 0.617502144 | 5.14E-29 | 5.82E-27 | 2595.979226 | 1603.003818 | ENSG00000103044 |
| TCF25 | -0.69565203 | 0.617430206 | 4.87E-19 | 2.72E-17 | 1337.762282 | 825.9762735 | ENSG00000141002 |
| SLC25A35 | -0.695742534 | 0.617391474 | 0.066161461 | 0.174994285 | 40.93019875 | 25.2386752 | ENSG00000125434 |
| ENPEP | -0.696454067 | 0.617087054 | 0.289102553 | 0.49846633 | 13.923833 | 8.566863557 | ENSG00000138792 |
| NA | -0.696623671 | 0.617014513 | 0.530022421 | 0.722184229 | 4.297189585 | 2.650415165 | ENSG00000270607 |
| BAALC | -0.696864402 | 0.616911565 | 0.109851203 | 0.256147669 | 31.76676638 | 19.60577454 | ENSG00000164929 |
| NA | -0.697050706 | 0.616831905 | 0.390279146 | 0.601716195 | 8.65961873 | 5.345002014 | ENSG00000280300 |
| SCYL2 | -0.69722311 | 0.616758197 | 3.62E-10 | 7.42E-09 | 700.2006517 | 431.8981346 | ENSG00000136021 |
| NAA16 | -0.697232352 | 0.616754246 | 5.84E-07 | 7.02E-06 | 483.000734 | 297.7588529 | ENSG00000172766 |
| BCL6B | -0.697792536 | 0.616514813 | 1.82E-07 | 2.40E-06 | 379.3468779 | 233.736778 | ENSG00000161940 |
| NA | -0.699330232 | 0.615858051 | 0.528278566 | 0.721043079 | 4.328029629 | 2.664413905 | ENSG00000232946 |
| PCLAF | -0.700057637 | 0.615547615 | 0.002236491 | 0.011185414 | 116.8523764 | 71.94861234 | ENSG00000166803 |
| TMEM121 | -0.700494272 | 0.615361346 | 0.017462194 | 0.061639914 | 68.50533836 | 42.07703131 | ENSG00000184986 |
| KCTD21-AS1 | -0.700642961 | 0.615297928 | 0.014335371 | 0.05252687 | 78.43743312 | 48.32699046 | ENSG00000246174 |
| NA | -0.701154678 | 0.615079723 | 0.539448179 | 0.729040189 | 5.449254528 | 3.336972886 | ENSG00000260288 |
| CFL1P1 | -0.701540314 | 0.614915333 | 0.434990838 | 0.643690371 | 7.498070578 | 4.618623537 | ENSG00000223820 |
| EEF1AKMT3 | -0.701598385 | 0.614890582 | 0.083568072 | 0.208671564 | 37.61635666 | 23.1027655 | ENSG00000123427 |
| KCNJ11 | -0.701635033 | 0.614874962 | 4.15E-05 | 0.000340379 | 240.0525384 | 147.742882 | ENSG00000187486 |
| ADGRG5 | -0.702012471 | 0.61471412 | 1.63E-07 | 2.16E-06 | 373.5492313 | 229.5541752 | ENSG00000159618 |
| NA | -0.702174663 | 0.614645016 | 0.084817476 | 0.211127828 | 34.03026753 | 20.93500184 | ENSG00000254252 |
| RAB11FIP3 | -0.702239789 | 0.61461727 | 8.36E-21 | 5.26E-19 | 1413.455614 | 868.9240943 | ENSG00000090565 |
| NA | -0.702275201 | 0.614602184 | 0.491163615 | 0.690613124 | 6.472925283 | 4.005180939 | ENSG00000286004 |
| VAMP5 | -0.703475477 | 0.614091067 | 0.107221037 | 0.251555734 | 31.95625584 | 19.67917321 | ENSG00000168899 |
| NA | -0.703645721 | 0.614018607 | 0.297504037 | 0.507759941 | 11.8735424 | 7.294860718 | ENSG00000286277 |
| ING4 | -0.703671146 | 0.614007786 | 5.88E-06 | 5.91E-05 | 340.2614368 | 208.9345454 | ENSG00000111653 |
| ZNF181 | -0.703693362 | 0.613998331 | 0.002709315 | 0.013175254 | 130.1991736 | 79.95869067 | ENSG00000197841 |
| PTN | -0.703734057 | 0.613981011 | 2.40E-13 | 7.43E-12 | 815.9821421 | 501.042273 | ENSG00000105894 |
| NAA38 | -0.703808808 | 0.6139492 | 1.26E-07 | 1.71E-06 | 382.9053548 | 235.1241031 | ENSG00000183011 |
| SEPTIN6 | -0.704481484 | 0.613663004 | 1.39E-14 | 4.92E-13 | 920.4464585 | 565.0378136 | ENSG00000125354 |
| TRAF3IP2-AS1 | -0.705213507 | 0.613351711 | 0.026353132 | 0.085836306 | 62.8366702 | 38.57067609 | ENSG00000231889 |
| ZNF138 | -0.70570947 | 0.613140892 | 5.98E-07 | 7.19E-06 | 307.8438272 | 188.8350311 | ENSG00000197008 |
| SLC2A1 | -0.705755978 | 0.613121126 | 9.07E-29 | 1.01E-26 | 4904.647224 | 3006.449432 | ENSG00000117394 |
| PFKFB4 | -0.70620338 | 0.612931018 | 1.31E-06 | 1.49E-05 | 365.3186255 | 223.7328802 | ENSG00000114268 |
| MXRA5 | -0.706350449 | 0.612868538 | 0.001246586 | 0.006784375 | 150.6969932 | 92.23515325 | ENSG00000101825 |
| NA | -0.706747233 | 0.612700005 | 0.543592475 | 0.732315108 | 4.351750728 | 2.660062978 | ENSG00000261120 |
| NA | -0.70712797 | 0.61253833 | 0.526167469 | 0.719371075 | 4.871887182 | 3.003956591 | ENSG00000233971 |
| CCDC74B | -0.708294649 | 0.612043183 | 8.75E-05 | 0.000659821 | 199.0060722 | 121.9111737 | ENSG00000152076 |
| BARX1 | -0.708502015 | 0.611955217 | 0.049292932 | 0.139543849 | 45.02218644 | 27.54983123 | ENSG00000131668 |
| PINK1-AS | -0.708564599 | 0.611928671 | 0.126686129 | 0.283501308 | 27.73645877 | 16.93294232 | ENSG00000117242 |
| TMEM163 | -0.708610498 | 0.611909203 | 0.084994269 | 0.21139428 | 36.43136674 | 22.31464367 | ENSG00000152128 |
| CHRNB4 | -0.708902419 | 0.611785399 | 0.005520314 | 0.02399888 | 104.6419577 | 64.11626202 | ENSG00000117971 |
| SLC30A3 | -0.709971065 | 0.6113324 | 1.73E-06 | 1.91E-05 | 309.8495985 | 189.507801 | ENSG00000115194 |
| ATP6V0E2 | -0.710212914 | 0.611229926 | 3.64E-12 | 9.73E-11 | 644.1707822 | 393.7963222 | ENSG00000171130 |
| TMEM191A | -0.710547215 | 0.611088309 | 0.012594462 | 0.047294971 | 75.80977519 | 46.32376376 | ENSG00000226287 |
| NASP | -0.711013116 | 0.610890997 | 4.73E-67 | 2.88E-64 | 18000.47091 | 10996.27933 | ENSG00000132780 |
| NA | -0.711030907 | 0.610883464 | 0.143577177 | 0.309430369 | 24.92732005 | 15.26010495 | ENSG00000213740 |
| NA | -0.711524568 | 0.610674468 | 0.415442201 | 0.626247284 | 8.172407252 | 5.019458068 | ENSG00000286311 |
| FAM210B | -0.711569077 | 0.610655628 | 1.20E-09 | 2.29E-08 | 468.5879356 | 286.0310205 | ENSG00000124098 |
| NA | -0.711653949 | 0.610619704 | 0.416612745 | 0.627560955 | 7.589395501 | 4.632622277 | ENSG00000232811 |
| MAGED4 | -0.711836562 | 0.610542419 | 0.085904049 | 0.213036911 | 34.26483492 | 20.96299932 | ENSG00000154545 |
| NA | -0.711889666 | 0.610519946 | 0.02803891 | 0.090099508 | 62.5970688 | 38.29772213 | ENSG00000271109 |
| MYCNOS | -0.712269581 | 0.610359194 | 0.036675048 | 0.111127651 | 65.09068814 | 39.70827196 | ENSG00000233718 |
| NA | -0.712324708 | 0.610335872 | 0.507241552 | 0.704350311 | 4.886125072 | 2.999605664 | ENSG00000232187 |
| SEPHS1 | -0.712407183 | 0.610300982 | 1.82E-51 | 5.83E-49 | 15736.26242 | 9603.894965 | ENSG00000086475 |
| NA | -0.712413711 | 0.610298221 | 0.218576457 | 0.414814229 | 19.15156281 | 11.66105543 | ENSG00000279069 |
| SLITRK4 | -0.71250814 | 0.610258276 | 3.69E-05 | 0.000306558 | 198.4420531 | 121.1863083 | ENSG00000179542 |
| NA | -0.712699081 | 0.610177513 | 0.528310868 | 0.721043079 | 6.014468916 | 3.667813717 | ENSG00000233542 |
| NA | -0.713074774 | 0.610018638 | 0.200631986 | 0.391338055 | 18.49235375 | 11.28821838 | ENSG00000198237 |
| NA | -0.714040654 | 0.609610368 | 0.186386921 | 0.371399291 | 27.9256689 | 17.08910393 | ENSG00000256667 |
| PRKCH | -0.714392602 | 0.609461671 | 1.98E-05 | 0.000176651 | 289.7739779 | 176.5855045 | ENSG00000027075 |
| DLD | -0.714421934 | 0.60944928 | 2.64E-24 | 2.24E-22 | 1841.351094 | 1122.130967 | ENSG00000091140 |

| | | | | | | | |
|---|---|---|---|---|---|---|---|
| NA | -0.714881447 | 0.609255195 | 0.172965299 | 0.352557978 | 24.67170131 | 15.03907858 | ENSG00000256092 |
| OPRL1 | -0.715338372 | 0.609062264 | 0.476840222 | 0.679293925 | 5.9800694 | 3.632343887 | ENSG00000125510 |
| ANKRD63 | -0.716549242 | 0.608551286 | 0.358725391 | 0.570775045 | 9.244994745 | 5.622023348 | ENSG00000230778 |
| PEX26 | -0.717798999 | 0.608024348 | 1.89E-19 | 1.08E-17 | 1536.662791 | 934.2348275 | ENSG00000215193 |
| IQCK | -0.71807985 | 0.607905994 | 0.083169608 | 0.207896498 | 36.58289721 | 22.27236391 | ENSG00000174628 |
| ASF1A | -0.718154319 | 0.607874616 | 1.72E-09 | 3.20E-08 | 463.2858141 | 281.6841214 | ENSG00000111875 |
| TMPRSS5 | -0.719013995 | 0.607512503 | 0.420802872 | 0.630884946 | 8.833980575 | 5.348123436 | ENSG00000166682 |
| LINC01426 | -0.719638745 | 0.60724948 | 0.276750563 | 0.485908738 | 12.54431961 | 7.616053737 | ENSG00000234380 |
| CFAP36 | -0.719860527 | 0.607156136 | 7.45E-06 | 7.34E-05 | 258.3287063 | 156.9388679 | ENSG00000163001 |
| NA | -0.720060921 | 0.607071807 | 0.000157199 | 0.001112706 | 197.6751707 | 119.8027671 | ENSG00000282936 |
| TIGD7 | -0.72011778 | 0.607047882 | 0.054777173 | 0.151417332 | 43.76099921 | 26.62919435 | ENSG00000140993 |
| POGLUT2 | -0.720360029 | 0.606945958 | 6.39E-11 | 1.47E-09 | 629.3582323 | 382.1869108 | ENSG00000134901 |
| NA | -0.721932151 | 0.606284922 | 0.064212068 | 0.170746959 | 37.84172018 | 22.93745054 | ENSG00000123965 |
| TRAPPC5 | -0.722499919 | 0.606046368 | 0.03782393 | 0.113671482 | 50.95837312 | 30.8933305 | ENSG00000181029 |
| ZNF252P-AS1 | -0.722623652 | 0.605994392 | 0.540691835 | 0.730042795 | 4.931203006 | 2.995254736 | ENSG00000255559 |
| UTP14A | -0.723182779 | 0.60575958 | 2.42E-16 | 1.04E-14 | 1077.934975 | 653.108859 | ENSG00000156697 |
| NOVA1 | -0.723223106 | 0.605742648 | 5.50E-05 | 0.000437848 | 203.0233372 | 123.0782801 | ENSG00000139910 |
| NA | -0.723305477 | 0.605708064 | 0.219443147 | 0.415790492 | 18.58459455 | 11.30221712 | ENSG00000264964 |
| H2AW | -0.723521202 | 0.6056175 | 3.81E-08 | 5.69E-07 | 367.722725 | 222.7544517 | ENSG00000181218 |
| NA | -0.723702022 | 0.605541599 | 0.400423667 | 0.611812375 | 7.696153522 | 4.646621017 | ENSG00000228305 |
| EFNB1 | -0.723932783 | 0.60544475 | 3.38E-08 | 5.12E-07 | 444.6985754 | 269.4783876 | ENSG00000090776 |
| MAGIX | -0.725090308 | 0.604959175 | 0.215698462 | 0.410880115 | 16.98503099 | 10.24064689 | ENSG00000269313 |
| POMC | -0.725203547 | 0.604911693 | 0.012129845 | 0.045961502 | 79.95276455 | 48.41332632 | ENSG00000115138 |
| RAB33A | -0.725524795 | 0.604777011 | 0.005258828 | 0.023054068 | 102.3710583 | 62.01052526 | ENSG00000134594 |
| NA | -0.726032539 | 0.604564202 | 0.097979479 | 0.234539596 | 34.96141815 | 21.17655335 | ENSG00000261754 |
| USP4 | -0.727088867 | 0.604121708 | 3.04E-16 | 1.29E-14 | 954.0840684 | 576.1866843 | ENSG00000114316 |
| PCDHGA9 | -0.727300688 | 0.604033016 | 0.390822371 | 0.602231765 | 8.787428101 | 5.338475623 | ENSG00000261934 |
| CAB39L | -0.727518133 | 0.603941982 | 0.010457703 | 0.040503223 | 75.95477153 | 45.86988773 | ENSG00000102547 |
| TTLL7 | -0.72753407 | 0.60393531 | 0.00391361 | 0.018039671 | 96.57989413 | 58.37070902 | ENSG00000137941 |
| LOC283856 | -0.728201308 | 0.603656058 | 0.564501318 | 0.748848511 | 4.848166083 | 2.922139614 | ENSG00000246379 |
| SIAH3 | -0.728696264 | 0.603448993 | 0.393378914 | 0.605013502 | 8.192568878 | 4.944167482 | ENSG00000215475 |
| ZNF823 | -0.72900305 | 0.603320684 | 0.026557893 | 0.086354305 | 57.1872478 | 34.50921664 | ENSG00000197933 |
| EXOG | -0.729363605 | 0.603169923 | 7.43E-05 | 0.000570849 | 219.8152937 | 132.3903233 | ENSG00000157036 |
| RBM3 | -0.730003809 | 0.602902322 | 4.53E-21 | 2.93E-19 | 1691.034937 | 1019.779169 | ENSG00000102317 |
| GNG13 | -0.730303526 | 0.602777084 | 0.34276187 | 0.554597608 | 9.930009835 | 5.991738978 | ENSG00000127588 |
| SRPK3 | -0.730870809 | 0.602540112 | 0.079195744 | 0.200323014 | 36.79493872 | 22.1951814 | ENSG00000184343 |
| LOC728392 | -0.731025892 | 0.602475345 | 1.00E-05 | 9.56E-05 | 224.3716615 | 135.1526558 | ENSG00000286190 |
| STX16-NPEPL1 | -0.731192143 | 0.602405922 | 0.360460078 | 0.572516706 | 11.14611913 | 6.706294178 | ENSG00000254995 |
| MOK | -0.731488712 | 0.6022821 | 0.146279249 | 0.31341997 | 23.73048266 | 14.32669884 | ENSG00000080823 |
| BOLA3-AS1 | -0.731495792 | 0.602279144 | 2.52E-05 | 0.000219435 | 213.1419467 | 128.4700082 | ENSG00000225439 |
| LCA5L | -0.732464546 | 0.601874857 | 0.519216066 | 0.714901185 | 4.926448325 | 2.96196037 | ENSG00000157578 |
| ZAR1 | -0.732579483 | 0.601826908 | 0.140690589 | 0.304904955 | 23.17950616 | 13.95263229 | ENSG00000182223 |
| NA | -0.732678049 | 0.601785792 | 0.000320578 | 0.00208188 | 157.5058789 | 94.74439447 | ENSG00000276710 |
| ZNF440 | -0.733356475 | 0.60150287 | 0.004493046 | 0.020236389 | 115.0713064 | 69.0843596 | ENSG00000171295 |
| NA | -0.733572357 | 0.601412869 | 0.503567188 | 0.701361077 | 4.995247357 | 3.006132054 | ENSG00000225330 |
| KTN1 | -0.733849198 | 0.601297473 | 1.84E-29 | 2.13E-27 | 2679.71154 | 1611.136006 | ENSG00000126777 |
| CDC37L1-DT | -0.734110094 | 0.601188745 | 0.478932042 | 0.68145793 | 4.971526258 | 2.987782387 | ENSG00000273061 |
| NA | -0.734188612 | 0.601156027 | 0.131638096 | 0.291713852 | 23.74352534 | 14.2965259 | ENSG00000231113 |
| LRRC73 | -0.734712019 | 0.600937968 | 0.446427846 | 0.65360027 | 6.571369151 | 3.941713629 | ENSG00000204052 |
| NA | -0.734971065 | 0.600830075 | 0.221239982 | 0.418343867 | 17.16147777 | 10.33334118 | ENSG00000267493 |
| CLCN2 | -0.735271037 | 0.60070516 | 3.73E-06 | 3.87E-05 | 260.675576 | 156.6463348 | ENSG00000114859 |
| DUSP4 | -0.73542755 | 0.600639996 | 2.04E-08 | 3.19E-07 | 397.3194001 | 238.5879676 | ENSG00000120875 |
| LNP1 | -0.735926437 | 0.600432329 | 9.37E-05 | 0.000700784 | 188.201253 | 112.9636219 | ENSG00000206535 |
| REM2 | -0.736408383 | 0.600231782 | 0.009367352 | 0.037016314 | 74.12036124 | 44.4202522 | ENSG00000139890 |
| CCDC153 | -0.736918152 | 0.600019731 | 0.08861004 | 0.217805622 | 33.71533297 | 20.15792529 | ENSG00000248712 |
| ARHGAP6 | -0.736930999 | 0.600014388 | 0.111973482 | 0.259717416 | 27.67566841 | 16.64069274 | ENSG00000047648 |
| MYO5B | -0.737474589 | 0.599788352 | 6.19E-09 | 1.06E-07 | 402.8220728 | 241.6543729 | ENSG00000167306 |
| RTN4RL1 | -0.739109224 | 0.599109151 | 0.374316722 | 0.586393306 | 8.270851119 | 4.941992019 | ENSG00000185924 |
| C19orf54 | -0.739548327 | 0.598926832 | 3.13E-10 | 6.50E-09 | 527.0704647 | 315.7350385 | ENSG00000188493 |
| NTS | -0.739549079 | 0.59892652 | 5.06E-05 | 0.000406829 | 184.7145236 | 110.6008219 | ENSG00000133636 |
| NA | -0.739620216 | 0.598896989 | 7.31E-05 | 0.000562455 | 193.6884665 | 115.9281366 | ENSG00000248605 |
| DNAJB6 | -0.739931707 | 0.598767696 | 3.87E-51 | 1.20E-48 | 8629.556256 | 5166.799878 | ENSG00000105993 |
| MMP2 | -0.740097418 | 0.598698924 | 7.55E-25 | 6.57E-23 | 2038.707699 | 1220.818064 | ENSG00000087245 |
| GRIA2 | -0.740306164 | 0.598612303 | 0.169429256 | 0.347337414 | 25.47501641 | 15.24298479 | ENSG00000120251 |
| CFAP298 | -0.740325523 | 0.598604271 | 0.007853873 | 0.032181743 | 90.44826811 | 54.20390164 | ENSG00000159079 |
| IKZF2 | -0.740432342 | 0.598559951 | 0.108608333 | 0.253751932 | 28.39655756 | 16.97276308 | ENSG00000030419 |
| NA | -0.740901605 | 0.59836529 | 0.265032404 | 0.472628608 | 13.85147449 | 8.293909603 | ENSG00000276564 |
| MYL5 | -0.741110629 | 0.598278602 | 0.00565889 | 0.024494187 | 85.21252962 | 50.94891369 | ENSG00000215375 |
| CCSER1 | -0.741358511 | 0.598175816 | 0.323900733 | 0.534691026 | 10.00829208 | 5.973389311 | ENSG00000184305 |
| NA | -0.741412719 | 0.59815334 | 0.229696699 | 0.429529871 | 15.03437948 | 8.975170438 | ENSG00000276728 |
| ATF7IP2 | -0.741588162 | 0.598080605 | 2.78E-06 | 2.95E-05 | 262.4156606 | 156.9283695 | ENSG00000166669 |
| NA | -0.741622272 | 0.598066464 | 0.005360139 | 0.023426573 | 84.27042125 | 50.39988436 | ENSG00000234771 |
| NA | -0.741718836 | 0.598026435 | 0.491379771 | 0.690822206 | 5.587742316 | 3.360619439 | ENSG00000279812 |
| NA | -0.742013372 | 0.597904356 | 0.482181715 | 0.683636087 | 5.005925775 | 2.985606923 | ENSG00000224888 |

| | | | | | | | |
|---|---|---|---|---|---|---|---|
| TLR2 | -0.742691302 | 0.597623464 | 0.170789996 | 0.34929273 | 22.79457735 | 13.57109338 | ENSG00000137462 |
| PITX3 | -0.74280152 | 0.597577809 | 0.259186961 | 0.466162945 | 13.33489751 | 7.945948609 | ENSG00000107859 |
| NA | -0.743058641 | 0.597471317 | 0.329488404 | 0.541124317 | 11.15768727 | 6.699767788 | ENSG00000280734 |
| LOC100996756 | -0.74385143 | 0.597143085 | 0.34877569 | 0.560755057 | 8.863346079 | 5.30083033 | ENSG00000287978 |
| HTR1A | -0.744014874 | 0.597075438 | 0.077204999 | 0.196683717 | 35.46078259 | 21.12585527 | ENSG00000178394 |
| LINC00458 | -0.744042342 | 0.59706407 | 3.67E-05 | 0.000305275 | 207.39197 | 123.6643877 | ENSG00000234787 |
| IGFBP6 | -0.74416248 | 0.597014352 | 1.12E-05 | 0.000105207 | 227.0058797 | 135.5836633 | ENSG00000167779 |
| GMFG | -0.744190074 | 0.597002934 | 0.521541707 | 0.716261736 | 4.937126742 | 2.93396289 | ENSG00000130755 |
| PRSS53 | -0.744986851 | 0.59667331 | 0.19415948 | 0.382386253 | 17.73025163 | 10.58983738 | ENSG00000151006 |
| EPB41L4A-DT | -0.746253488 | 0.596149681 | 0.232516802 | 0.433430046 | 15.53671857 | 9.24907035 | ENSG00000278921 |
| MAP3K13 | -0.747559012 | 0.595610457 | 0.00086438 | 0.004928691 | 123.7324514 | 73.63951583 | ENSG00000073803 |
| NA | -0.747739685 | 0.595535872 | 0.34183281 | 0.553503611 | 11.12832177 | 6.582480981 | ENSG00000226471 |
| NA | -0.747912788 | 0.59546442 | 0.345570566 | 0.557548068 | 8.897745595 | 5.30083033 | ENSG00000260793 |
| PRC1-AS1 | -0.748341565 | 0.595287471 | 0.490738446 | 0.690477988 | 5.61858236 | 3.358443975 | ENSG00000258725 |
| ZNF577 | -0.748473197 | 0.595233159 | 0.000176983 | 0.001233785 | 151.0644302 | 89.86755077 | ENSG00000161551 |
| ALKBH3 | -0.748503586 | 0.595220621 | 5.87E-05 | 0.000464012 | 192.4426862 | 114.465353 | ENSG00000166199 |
| TLL1 | -0.748764593 | 0.595112946 | 0.125455425 | 0.281514461 | 24.60882601 | 14.63919001 | ENSG00000038295 |
| PGAM1P5 | -0.74905394 | 0.594993602 | 0.052005683 | 0.145543468 | 47.58696908 | 28.31073358 | ENSG00000257150 |
| SFRP2 | -0.74923803 | 0.594917685 | 5.63E-46 | 1.42E-43 | 6993.180566 | 4160.630961 | ENSG00000145423 |
| WWOX | -0.749298683 | 0.594892674 | 3.53E-05 | 0.000294548 | 212.7125248 | 126.3843021 | ENSG00000186153 |
| DNAJC3 | -0.749431672 | 0.594837839 | 2.81E-13 | 8.64E-12 | 756.642705 | 450.2016607 | ENSG00000102580 |
| GDPD2 | -0.75003262 | 0.594590113 | 1.84E-08 | 2.91E-07 | 368.3968085 | 219.1628746 | ENSG00000130055 |
| ZFP69B | -0.750394571 | 0.594440958 | 0.000271608 | 0.001802921 | 139.0307377 | 82.63738686 | ENSG00000187801 |
| SLIT2 | -0.750446813 | 0.594419433 | 3.21E-05 | 0.000271088 | 197.4136538 | 117.3066216 | ENSG00000145147 |
| UBQLN2 | -0.75090125 | 0.594232225 | 2.53E-16 | 1.09E-14 | 1076.62515 | 640.1589345 | ENSG00000188021 |
| CLEC4G | -0.751077612 | 0.594159588 | 0.158227436 | 0.331057514 | 24.57295195 | 14.64477044 | ENSG00000182566 |
| NA | -0.75156856 | 0.59395743 | 0.037635424 | 0.113193976 | 47.38385213 | 28.16857071 | ENSG00000259642 |
| NA | -0.752327185 | 0.593645186 | 0.46633691 | 0.670412501 | 5.5533428 | 3.301503056 | ENSG00000249675 |
| LDOC1 | -0.752615433 | 0.593526589 | 6.92E-15 | 2.57E-13 | 878.9513764 | 522.0223628 | ENSG00000182195 |
| RASGRF1 | -0.753137642 | 0.59331179 | 0.155528281 | 0.327348691 | 20.17879304 | 11.98877484 | ENSG00000058335 |
| EVL | -0.75326261 | 0.593260399 | 3.16E-17 | 1.46E-15 | 1220.760895 | 724.3827072 | ENSG00000196405 |
| STAB2 | -0.753404099 | 0.59320222 | 0.243104418 | 0.447076339 | 15.08154234 | 8.969873552 | ENSG00000136011 |
| CPO | -0.754328047 | 0.592822435 | 0.198783543 | 0.389030323 | 17.31923746 | 10.22882361 | ENSG00000144410 |
| NA | -0.754461351 | 0.592767662 | 0.427007742 | 0.63691977 | 7.328742746 | 4.349074552 | ENSG00000285763 |
| NA | -0.754537093 | 0.592736542 | 0.240854327 | 0.444457861 | 14.59816967 | 8.657098842 | ENSG00000278376 |
| RAD54B | -0.754724173 | 0.592659685 | 0.001061679 | 0.005909504 | 138.6277322 | 82.1011994 | ENSG00000265817 |
| DANCR | -0.755032649 | 0.592532976 | 8.98E-22 | 6.24E-20 | 1488.611215 | 882.2925539 | ENSG00000226950 |
| MAP7D3 | -0.755351046 | 0.592402221 | 1.79E-19 | 1.04E-17 | 1037.91978 | 614.7276389 | ENSG00000129680 |
| SLC6A2 | -0.755684669 | 0.592265244 | 0.474857153 | 0.677776537 | 5.629260778 | 3.341323813 | ENSG00000103546 |
| KIAA1324 | -0.756252368 | 0.592032234 | 0.051068132 | 0.143429756 | 41.4503352 | 24.51040481 | ENSG00000116299 |
| RTN4RL2 | -0.75644622 | 0.591952689 | 5.39E-07 | 6.55E-06 | 348.6444889 | 206.4863125 | ENSG00000186907 |
| SLC16A11 | -0.756850003 | 0.591787036 | 0.283137326 | 0.492186485 | 14.15987493 | 8.368254231 | ENSG00000174326 |
| ONECUT1 | -0.757537079 | 0.591505267 | 0.124307882 | 0.279703844 | 32.03036822 | 18.8622759 | ENSG00000169856 |
| FOXL2NB | -0.757658315 | 0.591455563 | 0.08321801 | 0.207989957 | 30.95986566 | 18.29953137 | ENSG00000206262 |
| NA | -0.758164421 | 0.591248113 | 0.450689921 | 0.656987971 | 5.580623371 | 3.299327592 | ENSG00000273387 |
| GINS2 | -0.758616474 | 0.591062881 | 3.24E-23 | 2.49E-21 | 1586.919775 | 937.5325694 | ENSG00000131153 |
| ERVK3-1 | -0.758646622 | 0.591050529 | 0.190041034 | 0.376434332 | 18.50066791 | 10.93250149 | ENSG00000142396 |
| NA | -0.759202932 | 0.590822661 | 0.389895352 | 0.601542768 | 7.818039157 | 4.632622277 | ENSG00000287430 |
| GFRA1 | -0.759267242 | 0.590796325 | 0.074509393 | 0.191800015 | 35.39793345 | 20.91759813 | ENSG00000151892 |
| ZNF20 | -0.759310701 | 0.590778529 | 0.384965359 | 0.597138171 | 7.318064329 | 4.318901608 | ENSG00000132010 |
| NDUFC2 | -0.759663686 | 0.590634 | 0.018019692 | 0.063206146 | 66.35957855 | 39.30112195 | ENSG00000151366 |
| NA | -0.759696738 | 0.590620469 | 0.481872686 | 0.683503506 | 5.091326961 | 2.987782387 | ENSG00000280114 |
| NA | -0.760756884 | 0.590186619 | 9.56E-16 | 3.95E-14 | 809.6998759 | 478.0061392 | ENSG00000281566 |
| ALDH3A1 | -0.761149485 | 0.590026033 | 0.241528601 | 0.445088566 | 13.51846323 | 7.995417179 | ENSG00000108602 |
| LEAP2 | -0.761874888 | 0.589729436 | 0.330770831 | 0.542193138 | 11.2893616 | 6.659947031 | ENSG00000164406 |
| C20orf204 | -0.762013044 | 0.589672964 | 0.298257045 | 0.508493662 | 11.23952898 | 6.624477201 | ENSG00000196421 |
| NAP1L3 | -0.762230141 | 0.589584237 | 2.03E-25 | 1.90E-23 | 1507.533724 | 888.8936252 | ENSG00000186310 |
| SMIM27 | -0.762292377 | 0.589558804 | 0.237615442 | 0.439808648 | 14.65273081 | 8.622858517 | ENSG00000235453 |
| FLJ37453 | -0.762870578 | 0.589322569 | 0.003741323 | 0.017348247 | 111.8298484 | 66.03455032 | ENSG00000179743 |
| VPS8 | -0.763639504 | 0.589008556 | 3.31E-14 | 1.13E-12 | 755.8081409 | 444.9329679 | ENSG00000156931 |
| TAF5 | -0.763792193 | 0.588946221 | 3.12E-12 | 8.41E-11 | 608.9809736 | 358.5907118 | ENSG00000148835 |
| TGIF2 | -0.76469739 | 0.588576812 | 1.71E-17 | 8.09E-16 | 977.5175173 | 575.3940006 | ENSG00000118707 |
| FOXI2 | -0.764716415 | 0.58856905 | 0.259061503 | 0.466073781 | 14.53293011 | 8.522691873 | ENSG00000186766 |
| TMX3 | -0.765041104 | 0.588436603 | 3.51E-08 | 5.29E-07 | 383.1928079 | 225.5178754 | ENSG00000166479 |
| NA | -0.765056256 | 0.588430423 | 0.190756514 | 0.377376573 | 18.74263358 | 11.02302032 | ENSG00000266777 |
| CUX2 | -0.765806855 | 0.588124357 | 3.66E-07 | 4.61E-06 | 294.6800114 | 173.2121884 | ENSG00000111249 |
| NA | -0.766665032 | 0.587774619 | 0.425294723 | 0.635267804 | 6.200398902 | 3.662516831 | ENSG00000214832 |
| NA | -0.76672248 | 0.587751215 | 0.218233132 | 0.414454214 | 15.86024662 | 9.342710603 | ENSG00000227252 |
| LOC105371899 | -0.767063747 | 0.587612199 | 0.496412467 | 0.695610999 | 5.11504806 | 2.98343146 | ENSG00000267568 |
| LINC01950 | -0.767755696 | 0.587330435 | 0.004050988 | 0.018573174 | 89.97707455 | 52.8230303 | ENSG00000251027 |
| LINC00115 | -0.768302739 | 0.587107772 | 0.188409885 | 0.374381384 | 17.01587104 | 9.975448827 | ENSG00000225880 |
| ELF4 | -0.768308862 | 0.587105281 | 0.000151658 | 0.001078337 | 161.4267849 | 94.71618608 | ENSG00000102034 |
| NA | -0.769491279 | 0.586624293 | 0.399046963 | 0.610548642 | 7.427186614 | 4.346899088 | ENSG00000274292 |
| NA | -0.76954323 | 0.586603169 | 0.1205599 | 0.273519187 | 24.37781809 | 14.26417749 | ENSG00000244932 |

| | | | | | | | |
|---|---|---|---|---|---|---|---|
| TPR | -0.77094159 | 0.586034868 | 1.56E-37 | 2.69E-35 | 3509.767242 | 2056.548833 | ENSG00000047410 |
| CTB-178M22.2 | -0.77194864 | 0.585625939 | 0.135771937 | 0.298007094 | 22.61993617 | 13.28007331 | ENSG00000253978 |
| ZSCAN18 | -0.772643253 | 0.585344046 | 2.04E-08 | 3.19E-07 | 338.5976012 | 198.2819758 | ENSG00000121413 |
| LOC105372401 | -0.774487263 | 0.584596354 | 0.503372489 | 0.701201552 | 4.528197505 | 2.666589369 | ENSG00000286177 |
| DDX11-AS1 | -0.776491667 | 0.583784711 | 0.069394658 | 0.181461938 | 34.13110182 | 19.92167067 | ENSG00000245614 |
| NA | -0.776590446 | 0.583744742 | 0.360548912 | 0.572566952 | 9.715604069 | 5.66307361 | ENSG00000260236 |
| NA | -0.77708997 | 0.583542659 | 0.360716399 | 0.572736607 | 7.978163112 | 4.651917903 | ENSG00000272087 |
| BMP7 | -0.777297497 | 0.583458724 | 2.20E-08 | 3.41E-07 | 390.1632141 | 227.4925375 | ENSG00000101144 |
| NRG2 | -0.778037532 | 0.583159514 | 0.261815801 | 0.468843082 | 11.98266469 | 6.981140049 | ENSG00000158458 |
| XKR7 | -0.778512105 | 0.582967716 | 3.95E-07 | 4.95E-06 | 281.8118327 | 164.1158747 | ENSG00000260903 |
| ZNF610 | -0.778857446 | 0.582828186 | 0.014028056 | 0.051574336 | 69.01212664 | 40.27898329 | ENSG00000167554 |
| NA | -0.779709698 | 0.58248399 | 0.080370145 | 0.202399012 | 30.7748254 | 17.92764028 | ENSG00000240429 |
| DUSP6 | -0.779843985 | 0.582429775 | 4.56E-31 | 5.63E-29 | 2287.081299 | 1331.772784 | ENSG00000139318 |
| NA | -0.780648513 | 0.58210507 | 0.294191726 | 0.504065574 | 10.23218105 | 5.951918221 | ENSG00000271781 |
| ZDHHC15 | -0.781429243 | 0.581790142 | 0.000927531 | 0.005253877 | 114.2549481 | 66.50299218 | ENSG00000102383 |
| C9orf24 | -0.781762349 | 0.581655827 | 0.068513591 | 0.179655536 | 31.94557742 | 18.61542751 | ENSG00000164972 |
| CAMK2N2 | -0.782805419 | 0.581235441 | 0.000568867 | 0.003432288 | 145.1878904 | 84.41178834 | ENSG00000163888 |
| NLRP12 | -0.782929848 | 0.581185314 | 4.45E-05 | 0.000363015 | 185.2162778 | 107.7810244 | ENSG00000142405 |
| LRRC71 | -0.783401114 | 0.580995496 | 0.406472084 | 0.617357674 | 6.292919033 | 3.658165904 | ENSG00000160838 |
| XPNPEP2 | -0.78346724 | 0.580968867 | 0.425146148 | 0.635106861 | 6.323759077 | 3.678691035 | ENSG00000122121 |
| MAGI2-AS3 | -0.783578003 | 0.580924265 | 0.00022809 | 0.001546116 | 127.9439343 | 74.34536918 | ENSG00000234456 |
| NA | -0.783672896 | 0.580886056 | 0.533048885 | 0.724314769 | 4.57208023 | 2.666589369 | ENSG00000226009 |
| NA | -0.783724124 | 0.58086543 | 0.083849031 | 0.209234797 | 33.83393846 | 19.60917951 | ENSG00000286797 |
| ZNF841 | -0.783919903 | 0.58078661 | 0.00023991 | 0.001618106 | 168.7614775 | 98.05932917 | ENSG00000197608 |
| NIPSNAP3B | -0.784000698 | 0.580754085 | 0.278563613 | 0.487748658 | 10.84839711 | 6.31510746 | ENSG00000165028 |
| KCNMA1 | -0.784004384 | 0.580752601 | 0.000749714 | 0.004360439 | 131.1563579 | 76.10560396 | ENSG00000156113 |
| NA | -0.784050354 | 0.580734097 | 0.296274099 | 0.506438845 | 10.31402277 | 5.985212587 | ENSG00000287009 |
| NA | -0.784373023 | 0.580604226 | 0.288099371 | 0.497357882 | 10.90295825 | 6.348401826 | ENSG00000261526 |
| TAF7 | -0.784591002 | 0.580516508 | 2.34E-36 | 3.78E-34 | 3114.542162 | 1808.133981 | ENSG00000178913 |
| LOC101927551 | -0.785240496 | 0.580255221 | 0.28693857 | 0.496139433 | 12.59532128 | 7.307629954 | ENSG00000230922 |
| KCNK13 | -0.785366967 | 0.580204357 | 0.187999429 | 0.373880499 | 16.62354395 | 9.638081605 | ENSG00000152315 |
| MCTP1 | -0.786661484 | 0.579683978 | 0.004024713 | 0.018479629 | 82.85175366 | 48.02147719 | ENSG00000175471 |
| TOB1-AS1 | -0.786896212 | 0.57958967 | 0.11272366 | 0.260880677 | 24.08365554 | 13.98715616 | ENSG00000229980 |
| NA | -0.787755463 | 0.579244577 | 0.507214497 | 0.704350311 | 5.173168675 | 2.964135834 | ENSG00000286299 |
| NA | -0.788315621 | 0.579019716 | 0.446298388 | 0.653521693 | 5.738383063 | 3.299327592 | ENSG00000273891 |
| TCEAL1 | -0.788991273 | 0.578748609 | 0.0070344 | 0.029396912 | 67.10062933 | 38.85516977 | ENSG00000172465 |
| ZNF596 | -0.790138813 | 0.578288448 | 0.258013744 | 0.464808695 | 14.92022318 | 8.679799436 | ENSG00000172748 |
| C16orf86 | -0.790170236 | 0.578275852 | 0.128260072 | 0.286006784 | 21.86047765 | 12.65386147 | ENSG00000159761 |
| DM1-AS | -0.791067205 | 0.577916432 | 0.214491434 | 0.409406785 | 14.93118093 | 8.619453548 | ENSG00000267395 |
| NA | -0.791092525 | 0.577906289 | 0.209817584 | 0.403130403 | 17.76137101 | 10.2126494 | ENSG00000273340 |
| TRMT10A | -0.791110246 | 0.57789919 | 0.032673841 | 0.101649868 | 45.41303898 | 26.31329822 | ENSG00000145331 |
| NUP210L | -0.791214307 | 0.577857508 | 0.41638717 | 0.627350458 | 7.507553787 | 4.367424219 | ENSG00000143552 |
| SLC1A6 | -0.791616356 | 0.577696494 | 1.36E-07 | 1.84E-06 | 293.708843 | 169.6126874 | ENSG00000105143 |
| ENHO | -0.791796064 | 0.577624538 | 4.56E-05 | 0.000370982 | 185.8591897 | 107.478349 | ENSG00000168913 |
| NA | -0.792304496 | 0.577421009 | 0.192735681 | 0.380212094 | 18.88796098 | 10.94120335 | ENSG00000235043 |
| C8G | -0.792716704 | 0.577256051 | 0.414155344 | 0.625189379 | 6.32731855 | 3.644167164 | ENSG00000176919 |
| PARP11 | -0.793188704 | 0.577067224 | 0.00092143 | 0.005225585 | 126.1554917 | 72.74978695 | ENSG00000111224 |
| LINC01315 | -0.794262562 | 0.576637848 | 1.24E-10 | 2.73E-09 | 446.3896659 | 257.3734828 | ENSG00000229891 |
| NOL4 | -0.794928908 | 0.576371575 | 0.135616116 | 0.297734288 | 22.42449683 | 12.93088281 | ENSG00000101746 |
| NA | -0.795061708 | 0.576318522 | 0.479290988 | 0.681494241 | 4.596996537 | 2.654766092 | ENSG00000217289 |
| NA | -0.795705788 | 0.576061287 | 0.012271694 | 0.046368735 | 57.66022139 | 33.21262129 | ENSG00000261971 |
| PDGFRL | -0.795769478 | 0.576035856 | 0.003428629 | 0.016142332 | 80.10015131 | 46.14132864 | ENSG00000104213 |
| NA | -0.796316033 | 0.57581767 | 0.11074147 | 0.257777968 | 28.85264966 | 16.61146576 | ENSG00000267480 |
| SLC47A2 | -0.796583225 | 0.575711037 | 0.491659674 | 0.691126718 | 4.604115482 | 2.654766092 | ENSG00000180638 |
| LUARIS | -0.796884764 | 0.575590719 | 0.307019027 | 0.517360072 | 9.2378758 | 5.32882781 | ENSG00000231638 |
| CD300A | -0.797808963 | 0.575222111 | 0.268644749 | 0.476550449 | 11.50849589 | 6.626652665 | ENSG00000167851 |
| PEF1 | -0.799252599 | 0.574646801 | 2.34E-12 | 6.43E-11 | 541.303614 | 310.9899318 | ENSG00000162517 |
| TPI1P2 | -0.799266779 | 0.574641153 | 0.499251921 | 0.697674451 | 4.63731979 | 2.668764832 | ENSG00000230359 |
| NA | -0.801021818 | 0.573942527 | 0.393157366 | 0.604796215 | 7.47434948 | 4.304902868 | ENSG00000264456 |
| NA | -0.801040219 | 0.573935207 | 0.475471474 | 0.67798911 | 4.640879262 | 2.666589369 | ENSG00000235748 |
| NA | -0.801618486 | 0.573705206 | 0.0326378 | 0.101567134 | 47.82570634 | 27.47860802 | ENSG00000260920 |
| H2BC7 | -0.801846437 | 0.573614565 | 0.133933825 | 0.295207783 | 22.02772055 | 12.60751432 | ENSG00000277224 |
| NA | -0.802109887 | 0.573509827 | 0.271825265 | 0.480477361 | 11.05003953 | 6.324755273 | ENSG00000232450 |
| NA | -0.802795866 | 0.573237197 | 0.012193646 | 0.046147696 | 56.13955046 | 32.20269509 | ENSG00000243199 |
| GABRA2 | -0.803556926 | 0.572934879 | 0.137685343 | 0.300564987 | 23.15606439 | 13.20043179 | ENSG00000151834 |
| ESRRB | -0.804181525 | 0.572686887 | 0.144820704 | 0.311071315 | 19.68802209 | 11.30874351 | ENSG00000119715 |
| NA | -0.804810965 | 0.572437081 | 0.000726767 | 0.00424791 | 102.661051 | 58.76189574 | ENSG00000277013 |
| TEK | -0.804891534 | 0.572405114 | 0.022000585 | 0.074378992 | 48.16494682 | 27.61859542 | ENSG00000120156 |
| B3GNT7 | -0.805226728 | 0.572272137 | 2.95E-35 | 4.61E-33 | 2315.135563 | 1324.644606 | ENSG00000156966 |
| FBXL13 | -0.805800505 | 0.572044583 | 0.258271504 | 0.465051236 | 12.77296326 | 7.336856939 | ENSG00000161040 |
| SMPDL3B | -0.806028471 | 0.571954199 | 1.66E-13 | 5.22E-12 | 601.8399926 | 344.2352577 | ENSG00000130768 |
| ATOH8 | -0.80631257 | 0.57184158 | 0.126470834 | 0.283153729 | 24.93471832 | 14.23088313 | ENSG00000168874 |
| RIPPLY3 | -0.806582889 | 0.571734443 | 0.055173272 | 0.152264685 | 40.59810336 | 23.22099826 | ENSG00000183145 |
| NA | -0.807945149 | 0.57119484 | 0.460714793 | 0.665462199 | 5.191855761 | 2.975959111 | ENSG00000261305 |

| | | | | | | | |
|---|---|---|---|---|---|---|---|
| POPDC2 | -0.808139774 | 0.571117789 | 0.457822569 | 0.663253386 | 4.65155768 | 2.652590629 | ENSG00000121577 |
| NPPC | -0.808291291 | 0.571057811 | 0.101770516 | 0.241085825 | 24.37990302 | 13.93768759 | ENSG00000163273 |
| NA | -0.808563732 | 0.570949982 | 0.303589273 | 0.51363963 | 9.870694012 | 5.644723943 | ENSG00000267419 |
| FAM229B | -0.80898916 | 0.570781642 | 0.000631182 | 0.003751927 | 108.8424835 | 62.16763282 | ENSG00000203778 |
| PYGO1 | -0.809001142 | 0.570776902 | 0.127789294 | 0.28535232 | 27.21456845 | 15.5467741 | ENSG00000171016 |
| ZNF433-AS1 | -0.809008155 | 0.570774127 | 0.007673724 | 0.031620833 | 75.22973867 | 42.91729163 | ENSG00000219665 |
| NA | -0.809688056 | 0.570505201 | 0.007211049 | 0.030015576 | 69.16487848 | 39.51060859 | ENSG00000272106 |
| SHF | -0.810354136 | 0.570241865 | 1.42E-14 | 5.01E-13 | 713.1568758 | 406.7327854 | ENSG00000138606 |
| MFGE8 | -0.811466462 | 0.569802374 | 3.77E-89 | 5.94E-86 | 19799.31405 | 11281.70147 | ENSG00000140545 |
| NA | -0.811496205 | 0.569790627 | 0.17230017 | 0.351771752 | 16.31425379 | 9.272716903 | ENSG00000226074 |
| NA | -0.81223384 | 0.569499373 | 0.409080973 | 0.620172929 | 6.409160263 | 3.622696074 | ENSG00000263603 |
| NA | -0.812269387 | 0.569485342 | 0.44088078 | 0.648613531 | 5.818750236 | 3.339148349 | ENSG00000237080 |
| NA | -0.812526466 | 0.569383872 | 0.112387031 | 0.260261077 | 23.3345961 | 13.32859592 | ENSG00000231982 |
| BMP6 | -0.812716142 | 0.569309018 | 0.001164646 | 0.006406738 | 97.35623414 | 55.4862147 | ENSG00000153162 |
| SHISA6 | -0.812978106 | 0.569205653 | 3.08E-05 | 0.000262028 | 165.0579519 | 93.98226262 | ENSG00000188803 |
| BEND5 | -0.813060995 | 0.56917295 | 0.000797094 | 0.004601993 | 113.9551411 | 64.90733731 | ENSG00000162373 |
| AIFM1 | -0.813062375 | 0.569172406 | 5.15E-46 | 1.33E-43 | 3569.522673 | 2031.45358 | ENSG00000156709 |
| SMARCA2 | -0.813164283 | 0.569132202 | 1.90E-08 | 2.99E-07 | 338.3392349 | 192.385558 | ENSG00000080503 |
| ASXL3 | -0.813768248 | 0.568893993 | 0.28237026 | 0.491435108 | 12.68992634 | 7.219570132 | ENSG00000141431 |
| DEXI | -0.814525033 | 0.56859565 | 0.013220339 | 0.049137198 | 73.23372989 | 41.68728501 | ENSG00000182108 |
| GUSBP3 | -0.814957484 | 0.568425238 | 0.103056123 | 0.243489915 | 22.75277956 | 12.94705701 | ENSG00000253203 |
| NA | -0.815341896 | 0.568273799 | 0.238872257 | 0.441572803 | 13.49118266 | 7.700046177 | ENSG00000270605 |
| CXCL12 | -0.815429486 | 0.568239298 | 2.19E-15 | 8.70E-14 | 815.1633684 | 463.1554323 | ENSG00000107562 |
| NA | -0.815585595 | 0.568177815 | 0.504139332 | 0.701700419 | 4.695440405 | 2.662238442 | ENSG00000273819 |
| FGF21 | -0.81562167 | 0.568163607 | 0.258410957 | 0.465213625 | 11.71133352 | 6.634125014 | ENSG00000105550 |
| LOC101927811 | -0.816061379 | 0.567990467 | 0.30501202 | 0.51521932 | 10.57706594 | 6.027208807 | ENSG00000271133 |
| WDR34 | -0.81626968 | 0.567908465 | 3.35E-22 | 2.38E-20 | 1358.327223 | 771.6591298 | ENSG00000119333 |
| PLCG1-AS1 | -0.816784309 | 0.56770592 | 0.281521586 | 0.490500774 | 11.04856499 | 6.296757792 | ENSG00000226648 |
| XKRX | -0.816936998 | 0.567645839 | 0.023108315 | 0.077321143 | 48.52198466 | 27.49695769 | ENSG00000182489 |
| LOC101929748 | -0.816969456 | 0.567633069 | 0.139440537 | 0.303274737 | 19.32089065 | 10.94555427 | ENSG00000288062 |
| NA | -0.817445366 | 0.567445851 | 0.248260452 | 0.453204315 | 13.51726802 | 7.697870714 | ENSG00000235082 |
| NA | -0.818490442 | 0.567034947 | 0.102462869 | 0.24242186 | 28.85648847 | 16.33879535 | ENSG00000279233 |
| NCR3 | -0.818678256 | 0.566961134 | 0.437199156 | 0.645246932 | 5.298613782 | 3.013604404 | ENSG00000204475 |
| SKIV2L | -0.819030802 | 0.566822604 | 5.83E-25 | 5.15E-23 | 1331.175279 | 754.5710148 | ENSG00000204351 |
| MT1L | -0.819036455 | 0.566820383 | 0.457188492 | 0.662879437 | 5.20609365 | 2.950137094 | ENSG00000260549 |
| HEXA | -0.819515615 | 0.566632158 | 0.004914158 | 0.021759409 | 74.94625454 | 42.42622181 | ENSG00000213614 |
| HPCA | -0.820345013 | 0.566306497 | 0.026375681 | 0.085867016 | 47.42417538 | 26.88909552 | ENSG00000121905 |
| LRAT | -0.820436939 | 0.566270414 | 1.67E-11 | 4.13E-10 | 503.4624775 | 285.2035294 | ENSG00000121207 |
| NA | -0.82089413 | 0.566090991 | 0.283960966 | 0.493112905 | 11.7480973 | 6.62542316 | ENSG00000274104 |
| NA | -0.821340958 | 0.56591569 | 0.356147004 | 0.568384838 | 8.263732174 | 4.693914123 | ENSG00000272121 |
| NA | -0.822549903 | 0.565441665 | 0.231416545 | 0.431933331 | 12.31687116 | 6.943494756 | ENSG00000287710 |
| LCMT1-AS2 | -0.822745776 | 0.5653649 | 0.414452681 | 0.625404624 | 5.84959028 | 3.325149609 | ENSG00000260034 |
| NA | -0.823330504 | 0.565135803 | 0.154693078 | 0.326192361 | 21.18970045 | 11.98536987 | ENSG00000217648 |
| MYO5C | -0.824657382 | 0.564616274 | 4.79E-12 | 1.26E-10 | 518.5861237 | 292.6291813 | ENSG00000128833 |
| NA | -0.824709311 | 0.564595951 | 0.333223525 | 0.545266985 | 9.407203632 | 5.347177478 | ENSG00000225695 |
| NA | -0.824750837 | 0.564579701 | 0.338287423 | 0.550263392 | 8.240011076 | 4.662795221 | ENSG00000278341 |
| NA | -0.825046856 | 0.564463869 | 0.215515328 | 0.41069697 | 13.46390209 | 7.60423046 | ENSG00000260747 |
| SCN2B | -0.825276998 | 0.564373832 | 0.158087327 | 0.330837674 | 18.29694056 | 10.32681479 | ENSG00000149575 |
| ZNF594 | -0.826161396 | 0.564027967 | 1.19E-05 | 0.000111148 | 208.3299596 | 117.7001947 | ENSG00000180626 |
| NA | -0.826163597 | 0.564027106 | 0.196229369 | 0.385257474 | 14.7144109 | 8.264966164 | ENSG00000270157 |
| SPNS1 | -0.826291796 | 0.563976988 | 0.222071126 | 0.419465215 | 12.89156876 | 7.255039962 | ENSG00000169682 |
| NA | -0.826788631 | 0.5637828 | 0.432938711 | 0.642171723 | 5.315215936 | 2.9974302 | ENSG00000239345 |
| BNC1 | -0.827661158 | 0.563441933 | 0.48881983 | 0.6891648 | 4.685957196 | 2.622417685 | ENSG00000169594 |
| AHSP | -0.828328974 | 0.563181179 | 0.337697959 | 0.549635953 | 7.665313479 | 4.304902868 | ENSG00000169877 |
| NA | -0.828739395 | 0.563020987 | 0.285612789 | 0.494889212 | 10.01185155 | 5.614550999 | ENSG00000275888 |
| MACROH2A2 | -0.829066972 | 0.562893162 | 2.80E-32 | 3.63E-30 | 1844.263328 | 1038.013858 | ENSG00000099284 |
| C2orf81 | -0.829165301 | 0.562854798 | 0.478216709 | 0.680667428 | 4.764239437 | 2.668764832 | ENSG00000284308 |
| ACP7 | -0.829417095 | 0.562756572 | 0.2049644 | 0.396959562 | 15.39111184 | 8.644329606 | ENSG00000183760 |
| LOC105375624 | -0.830202107 | 0.562450443 | 0.4398189 | 0.647758546 | 5.260654793 | 2.950137094 | ENSG00000254231 |
| ZSWIM5 | -0.830521101 | 0.562326094 | 1.99E-05 | 0.000177561 | 204.8862238 | 115.1747066 | ENSG00000162415 |
| SAP30L-AS1 | -0.830565192 | 0.562308908 | 0.152696003 | 0.323027222 | 18.9499204 | 10.65670966 | ENSG00000245275 |
| NA | -0.83078286 | 0.562224076 | 0.340579909 | 0.552137658 | 8.24713002 | 4.636973204 | ENSG00000232320 |
| IL1RAPL2 | -0.832722669 | 0.561468633 | 0.258362493 | 0.465170719 | 11.16836569 | 6.268760312 | ENSG00000189108 |
| LOC101928415 | -0.832782099 | 0.561445504 | 0.017254625 | 0.061044134 | 53.18627954 | 29.8744189 | ENSG00000286724 |
| RIN3 | -0.833300868 | 0.561243654 | 0.007502344 | 0.031005381 | 74.07233423 | 41.6788667 | ENSG00000100599 |
| GLDN | -0.834038541 | 0.560956754 | 0.198036828 | 0.388039583 | 15.34366964 | 8.569984979 | ENSG00000186417 |
| NA | -0.834101599 | 0.560932236 | 0.04789139 | 0.136394044 | 34.23874956 | 19.22016825 | ENSG00000255397 |
| TEX14 | -0.834828183 | 0.560649805 | 0.094335549 | 0.228518065 | 26.65319286 | 14.91186041 | ENSG00000121101 |
| NA | -0.834987144 | 0.560588034 | 0.489023345 | 0.689237596 | 5.282011628 | 2.952312557 | ENSG00000259151 |
| KSR2 | -0.835083607 | 0.560550553 | 0.004301324 | 0.01951249 | 84.99367466 | 47.52794836 | ENSG00000171435 |
| CRYBB3 | -0.835180036 | 0.560513088 | 0.258069753 | 0.464865251 | 11.91150139 | 6.697592324 | ENSG00000100053 |
| NA | -0.835445733 | 0.560409869 | 0.463108216 | 0.667822133 | 5.349615452 | 2.993079273 | ENSG00000222881 |
| NA | -0.835547787 | 0.560370228 | 0.137146914 | 0.299909618 | 19.01871943 | 10.64365688 | ENSG00000279738 |
| TMEM88 | -0.835653577 | 0.560329138 | 0.071628634 | 0.186169385 | 27.75870533 | 15.53618033 | ENSG00000167874 |

| | | | | | | | |
|---|---|---|---|---|---|---|---|
| NA | -0.835774467 | 0.560282188 | 0.312807305 | 0.523140408 | 9.458205302 | 5.321355461 | ENSG00000277383 |
| NA | -0.836040001 | 0.560179075 | 0.435543529 | 0.644005602 | 5.842471335 | 3.271330112 | ENSG00000228057 |
| OGFR-AS1 | -0.836363676 | 0.56005341 | 0.267028744 | 0.474620031 | 12.36876255 | 6.943494756 | ENSG00000229873 |
| CRTC3 | -0.836818431 | 0.559876903 | 6.10E-07 | 7.33E-06 | 260.0724027 | 145.6292738 | ENSG00000140577 |
| LINC01273 | -0.838999975 | 0.559030935 | 0.280469842 | 0.489390497 | 15.01658212 | 8.412425914 | ENSG00000231742 |
| C3orf80 | -0.840274833 | 0.558537158 | 0.03106796 | 0.097607458 | 42.6483678 | 23.78809371 | ENSG00000180044 |
| PDE1B | -0.840380713 | 0.558496168 | 0.007705088 | 0.031722388 | 65.41774951 | 36.57134881 | ENSG00000123360 |
| LMF1 | -0.84102492 | 0.558246838 | 7.25E-05 | 0.000558149 | 159.1016036 | 88.92593726 | ENSG00000260807 |
| BBS9 | -0.841532831 | 0.558050338 | 5.39E-10 | 1.08E-08 | 451.1815437 | 251.5776833 | ENSG00000122507 |
| NA | -0.841548287 | 0.55804436 | 0.100826361 | 0.239359626 | 24.36477541 | 13.54744683 | ENSG00000234062 |
| SLC5A6 | -0.841593049 | 0.558027046 | 6.31E-48 | 1.75E-45 | 3976.59353 | 2219.488069 | ENSG00000138074 |
| NA | -0.842001457 | 0.557869098 | 0.419069171 | 0.629503762 | 5.380455496 | 2.989957851 | ENSG00000229539 |
| KIAA0319 | -0.84208063 | 0.557838484 | 0.107242532 | 0.251555734 | 23.26490734 | 12.97600045 | ENSG00000137261 |
| NA | -0.84254929 | 0.557657299 | 0.390915767 | 0.602255202 | 5.962272037 | 3.327325073 | ENSG00000252061 |
| DPF1 | -0.842966778 | 0.557495947 | 1.14E-09 | 2.18E-08 | 365.5757961 | 203.8552656 | ENSG00000011332 |
| STS | -0.843775909 | 0.557183365 | 0.001340196 | 0.00722925 | 118.0086112 | 65.75095964 | ENSG00000101846 |
| NA | -0.844646279 | 0.556847321 | 0.314544952 | 0.524936499 | 9.608846048 | 5.368648567 | ENSG00000123009 |
| THSD7A | -0.844659238 | 0.556842319 | 0.025620266 | 0.08398809 | 42.83193352 | 23.86555976 | ENSG00000005108 |
| LOC102724532 | -0.844841519 | 0.556771968 | 0.080434761 | 0.202534729 | 28.06680029 | 15.65346714 | ENSG00000264920 |
| NA | -0.845044157 | 0.55669377 | 0.3101255 | 0.520717933 | 8.901305067 | 4.949464368 | ENSG00000282787 |
| SLC39A1 | -0.845447509 | 0.55653815 | 0.010605479 | 0.040957968 | 59.08214296 | 32.9256686 | ENSG00000143570 |
| PCDHA5 | -0.845529355 | 0.556506578 | 0.351952082 | 0.564085129 | 9.490240554 | 5.285885631 | ENSG00000204965 |
| CLDND2 | -0.845842975 | 0.556385615 | 0.291182885 | 0.50081865 | 13.23881791 | 7.38944693 | ENSG00000160318 |
| NA | -0.845960788 | 0.556340181 | 0.218706839 | 0.414936574 | 12.53127692 | 6.983315513 | ENSG00000262454 |
| NA | -0.846350735 | 0.556189828 | 0.074580534 | 0.191861243 | 31.018876 | 17.26472911 | ENSG00000273619 |
| TNFAIP8L3 | -0.846537049 | 0.556118005 | 0.255602987 | 0.462495201 | 15.51271814 | 8.675448509 | ENSG00000183578 |
| TSPAN18 | -0.846814945 | 0.556010894 | 1.35E-14 | 4.78E-13 | 585.2511019 | 325.3293999 | ENSG00000157570 |
| NA | -0.847182413 | 0.555869291 | 0.396296888 | 0.608213469 | 6.616447084 | 3.692689775 | ENSG00000229816 |
| RIPOR3 | -0.848516323 | 0.555355574 | 0.00791096 | 0.032361516 | 69.31760415 | 38.58439128 | ENSG00000042062 |
| GPR137 | -0.848795241 | 0.555248216 | 7.38E-14 | 2.40E-12 | 665.7964648 | 369.9801583 | ENSG00000173264 |
| GGT7 | -0.849061255 | 0.555145845 | 8.03E-06 | 7.86E-05 | 1569.372531 | 871.3197934 | ENSG00000131067 |
| COL9A2 | -0.849596925 | 0.554939759 | 4.55E-05 | 0.000369969 | 155.1377308 | 86.10999622 | ENSG00000049089 |
| NA | -0.850076507 | 0.554755316 | 0.367713027 | 0.5798366 | 7.160304638 | 3.991182199 | ENSG00000272599 |
| NA | -0.852648411 | 0.553767231 | 0.408609636 | 0.619748478 | 6.047673224 | 3.346620699 | ENSG00000250012 |
| NA | -0.853365813 | 0.55349193 | 0.361569919 | 0.573611572 | 7.208942044 | 4.009531866 | ENSG00000235489 |
| CAMK2N1 | -0.85403127 | 0.553236685 | 0.000338619 | 0.002179107 | 107.2423351 | 59.35263775 | ENSG00000162545 |
| NA | -0.854044889 | 0.553231463 | 0.015633378 | 0.056385856 | 48.05610387 | 26.59031955 | ENSG00000272841 |
| NA | -0.854215536 | 0.553166028 | 0.24069635 | 0.444335345 | 11.94590091 | 6.584656445 | ENSG00000269906 |
| TGFB3 | -0.854446619 | 0.553077432 | 0.238019559 | 0.440340848 | 11.97910522 | 6.603952071 | ENSG00000119699 |
| MYOM2 | -0.854522284 | 0.553048426 | 0.061664644 | 0.165733956 | 31.22735803 | 17.31883216 | ENSG00000036448 |
| PDS5A | -0.854708484 | 0.552977052 | 1.01E-35 | 1.62E-33 | 2590.880249 | 1432.326848 | ENSG00000121892 |
| NA | -0.854854817 | 0.552920966 | 0.246784722 | 0.45168923 | 16.98147152 | 9.409299335 | ENSG00000273723 |
| ZNF329 | -0.855067278 | 0.552839545 | 3.93E-08 | 5.86E-07 | 284.6491417 | 157.1963101 | ENSG00000181894 |
| PLXNA4 | -0.85533593 | 0.552736608 | 0.014746754 | 0.053773403 | 50.33740238 | 27.78268088 | ENSG00000221866 |
| SELENOP | -0.855552873 | 0.552653497 | 0.041563424 | 0.122243462 | 37.72519962 | 20.8821283 | ENSG00000250722 |
| NA | -0.856191536 | 0.552408898 | 0.210479215 | 0.404013758 | 13.25897953 | 7.332506011 | ENSG00000272205 |
| TRIM9 | -0.856873658 | 0.552147775 | 0.00039026 | 0.002474836 | 115.4079027 | 63.73689858 | ENSG00000100505 |
| NA | -0.856915386 | 0.552131805 | 0.079679309 | 0.201222752 | 25.78433272 | 14.22218127 | ENSG00000249129 |
| ADCYAP1R1 | -0.856989133 | 0.552103582 | 1.10E-17 | 5.30E-16 | 850.2270948 | 469.3744203 | ENSG00000078549 |
| PPP1R14B-AS1 | -0.859239011 | 0.55124325 | 0.374367726 | 0.586424561 | 6.574928623 | 3.618345147 | ENSG00000256940 |
| CYP26A1 | -0.859271387 | 0.551230879 | 0.002495214 | 0.012300215 | 75.31602958 | 41.49064024 | ENSG00000095596 |
| PCDH19 | -0.859304703 | 0.55121815 | 0.02556128 | 0.083864625 | 62.39153585 | 34.28741226 | ENSG00000165194 |
| NA | -0.860207421 | 0.550873351 | 0.356200205 | 0.568385371 | 7.272986395 | 4.021355143 | ENSG00000267394 |
| DCTPP1 | -0.861069028 | 0.550544457 | 7.92E-23 | 5.93E-21 | 1054.497889 | 580.357559 | ENSG00000179958 |
| SCUBE3 | -0.861160796 | 0.550509438 | 0.001690727 | 0.008817837 | 84.59542384 | 46.58633485 | ENSG00000146197 |
| TRABD2A | -0.86168083 | 0.550311038 | 4.17E-05 | 0.000341595 | 170.2305619 | 93.65650776 | ENSG00000186854 |
| PCDHA3 | -0.861875511 | 0.550236782 | 0.085849839 | 0.212933309 | 28.79096957 | 15.80043243 | ENSG00000255408 |
| NA | -0.861953614 | 0.550206995 | 0.345611701 | 0.557553162 | 7.872600299 | 4.354371437 | ENSG00000286379 |
| PRRT1 | -0.863406316 | 0.54965325 | 0.076427862 | 0.195283477 | 32.34502345 | 17.729766 | ENSG00000204314 |
| NA | -0.864296672 | 0.549314138 | 0.090317785 | 0.221025187 | 25.46436414 | 14.0173291 | ENSG00000282022 |
| SHE | -0.865263244 | 0.548946233 | 0.055845117 | 0.153647295 | 35.51386919 | 19.4931222 | ENSG00000169291 |
| FBXL19-AS1 | -0.865310636 | 0.548928201 | 0.107906975 | 0.252643593 | 28.0398252 | 15.4075647 | ENSG00000260852 |
| PAX3 | -0.865864127 | 0.548717644 | 0.384287525 | 0.596667755 | 6.623566029 | 3.624871538 | ENSG00000135903 |
| NA | -0.866207299 | 0.548587137 | 0.365196547 | 0.577374364 | 7.263503186 | 3.989006735 | ENSG00000280145 |
| NA | -0.86641 | 0.548510065 | 0.276044591 | 0.485029968 | 10.2155789 | 5.590904446 | ENSG00000271992 |
| LOC105369760 | -0.867397784 | 0.54813464 | 0.483033216 | 0.684173275 | 4.879006127 | 2.696762312 | ENSG00000258017 |
| LOC730101 | -0.868032993 | 0.547893353 | 1.50E-07 | 2.01E-06 | 273.1768765 | 149.8023443 | ENSG00000216775 |
| C8orf76 | -0.868226677 | 0.547819803 | 0.179182807 | 0.361714629 | 15.69419893 | 8.61850759 | ENSG00000189376 |
| NA | -0.868621607 | 0.54766986 | 1.58E-07 | 2.11E-06 | 266.887212 | 146.2012146 | ENSG00000267321 |
| NA | -0.869324808 | 0.547402979 | 0.226665135 | 0.425579032 | 12.08794817 | 6.638475942 | ENSG00000271933 |
| RIPK4 | -0.869760462 | 0.547237704 | 0.00017367 | 0.001214275 | 126.2666989 | 69.22094204 | ENSG00000183421 |
| NA | -0.869883157 | 0.547191166 | 0.339056101 | 0.550859767 | 7.272986395 | 3.979358922 | ENSG00000183889 |
| TIMM23B | -0.871384583 | 0.546621995 | 0.018768869 | 0.065197542 | 50.19863526 | 27.40454694 | ENSG00000204152 |
| ANKRD36 | -0.8716145 | 0.546534889 | 0.006525155 | 0.027648092 | 71.96806731 | 39.38511439 | ENSG00000135976 |

| | | | | | | | |
|---|---|---|---|---|---|---|---|
| ASXL1 | -0.87214895 | 0.546332461 | 2.03E-17 | 9.55E-16 | 1001.631786 | 547.1219106 | ENSG00000171456 |
| ZNF558 | -0.872174324 | 0.546322852 | 6.91E-07 | 8.22E-06 | 203.7032927 | 111.3088507 | ENSG00000167785 |
| CAPN10-DT | -0.872299855 | 0.546275318 | 0.003796162 | 0.017562597 | 68.13408878 | 37.25667703 | ENSG00000260942 |
| ZNF704 | -0.873161936 | 0.545948989 | 2.66E-05 | 0.000230272 | 150.1051348 | 82.02924115 | ENSG00000164684 |
| NA | -0.873219245 | 0.545927303 | 0.115534854 | 0.265717212 | 23.5647143 | 12.81577146 | ENSG00000212769 |
| ANAPC1P1 | -0.873788383 | 0.545711979 | 0.214731569 | 0.409718068 | 12.11166927 | 6.622301738 | ENSG00000233673 |
| ST8SIA2 | -0.874791445 | 0.545332694 | 0.001035344 | 0.005774506 | 89.38755424 | 48.78304195 | ENSG00000140557 |
| NA | -0.875683076 | 0.544995765 | 0.513456893 | 0.709596299 | 4.810207095 | 2.628944076 | ENSG00000213018 |
| PRSS27 | -0.876258988 | 0.544778251 | 0.064000828 | 0.170298572 | 30.46169643 | 16.65374553 | ENSG00000172382 |
| ALOXE3 | -0.876616533 | 0.544643254 | 0.393297029 | 0.60493683 | 6.085632213 | 3.285328852 | ENSG00000179148 |
| SEC14L1P1 | -0.877425325 | 0.544338006 | 0.236467294 | 0.43845703 | 10.96819781 | 5.957215107 | ENSG00000213693 |
| NA | -0.878810788 | 0.543815513 | 0.371295214 | 0.58329264 | 6.674567699 | 3.606521871 | ENSG00000251468 |
| ZHX1 | -0.878904323 | 0.543780257 | 0.000490202 | 0.0030301 | 95.82279987 | 52.1383645 | ENSG00000165156 |
| LOC101929524 | -0.880044257 | 0.543350763 | 0.460255805 | 0.664999857 | 4.300749058 | 2.34539635 | ENSG00000246214 |
| NA | -0.880115969 | 0.543323755 | 0.315355792 | 0.525592986 | 8.58014128 | 4.674618497 | ENSG00000270587 |
| GALNT16 | -0.881401371 | 0.542839884 | 2.78E-07 | 3.58E-06 | 251.665428 | 136.7379278 | ENSG00000100626 |
| IQGAP1 | -0.881870187 | 0.542663512 | 2.40E-43 | 5.46E-41 | 3367.311908 | 1826.798095 | ENSG00000140575 |
| TFE3 | -0.882636676 | 0.542375277 | 2.92E-48 | 8.36E-46 | 3043.600591 | 1651.030016 | ENSG00000068323 |
| PCDHGA2 | -0.882967191 | 0.542251036 | 0.14686235 | 0.314277107 | 15.89820561 | 8.611981199 | ENSG00000081853 |
| ACSM3 | -0.883333611 | 0.542113331 | 0.138861989 | 0.302364655 | 19.08751846 | 10.3364626 | ENSG00000005187 |
| NA | -0.883558975 | 0.542028654 | 0.006973892 | 0.029169868 | 56.32815019 | 30.51368352 | ENSG00000235859 |
| SNORC | -0.883729392 | 0.541964631 | 0.111744014 | 0.259440091 | 21.52894093 | 11.62776107 | ENSG00000182600 |
| ADGRF3 | -0.883771895 | 0.541948664 | 0.001224242 | 0.006682025 | 85.12832364 | 46.18862174 | ENSG00000173567 |
| NA | -0.885704243 | 0.541223263 | 0.032130254 | 0.100263819 | 40.59095826 | 22.02116459 | ENSG00000260924 |
| NA | -0.886354201 | 0.540979488 | 0.308961025 | 0.519427486 | 8.643016576 | 4.68426631 | ENSG00000279307 |
| CCDC30 | -0.886469546 | 0.540936238 | 0.266226797 | 0.473656617 | 9.768690671 | 5.303005794 | ENSG00000186409 |
| TTLL11 | -0.887270989 | 0.540635822 | 0.090096198 | 0.220625868 | 23.26699228 | 12.60751432 | ENSG00000175764 |
| NA | -0.8882654 | 0.540263304 | 0.420626861 | 0.630845789 | 4.889684544 | 2.650415165 | ENSG00000287780 |
| ZNF438 | -0.889004799 | 0.539986484 | 0.039887292 | 0.118457542 | 34.35499079 | 18.54448785 | ENSG00000183621 |
| STK32B | -0.889136545 | 0.539937175 | 0.000124311 | 0.000902578 | 127.6992989 | 69.03460749 | ENSG00000152953 |
| EPHB1 | -0.889372914 | 0.539848719 | 5.87E-09 | 1.01E-07 | 282.4529906 | 152.4741149 | ENSG00000154928 |
| DLL1 | -0.889850005 | 0.539670224 | 0.000738694 | 0.00430962 | 95.96187247 | 51.73251663 | ENSG00000198719 |
| ULK4P2 | -0.890264576 | 0.539515167 | 0.315213806 | 0.525481011 | 8.67385662 | 4.68426631 | ENSG00000260128 |
| LOC107985164 | -0.891074796 | 0.53921226 | 0.010244413 | 0.039807764 | 51.71663644 | 27.94196391 | ENSG00000267327 |
| NA | -0.892130684 | 0.538817762 | 0.336502831 | 0.548447049 | 7.389227625 | 3.957887833 | ENSG00000244471 |
| NA | -0.892674726 | 0.538614612 | 0.190636656 | 0.377178968 | 20.44595435 | 10.9648499 | ENSG00000230615 |
| PIGL | -0.892936048 | 0.538517059 | 9.10E-09 | 1.51E-07 | 264.5359193 | 142.4748516 | ENSG00000108474 |
| NA | -0.894564878 | 0.537909406 | 0.149272728 | 0.317725677 | 17.38684128 | 9.350182953 | ENSG00000255201 |
| NA | -0.894664015 | 0.537872444 | 0.420411615 | 0.630667515 | 4.947805159 | 2.668764832 | ENSG00000280010 |
| SMAD9 | -0.895886017 | 0.537417044 | 0.047704446 | 0.136025739 | 30.25944362 | 16.2302104 | ENSG00000120693 |
| PCDHGA3 | -0.896357199 | 0.537241554 | 0.244983449 | 0.449219578 | 9.854091858 | 5.30083033 | ENSG00000254245 |
| NA | -0.89698413 | 0.537008143 | 0.29293032 | 0.50277098 | 9.921695682 | 5.337529665 | ENSG00000275202 |
| SNURF | -0.89750595 | 0.536813943 | 0.473070889 | 0.676301569 | 4.351750728 | 2.333573074 | ENSG00000273173 |
| MCF2 | -0.897828315 | 0.536694008 | 0.061142894 | 0.164790003 | 30.38249831 | 16.33879535 | ENSG00000101977 |
| NA | -0.897832161 | 0.536692577 | 0.412147789 | 0.623371056 | 5.615022888 | 3.029778607 | ENSG00000255050 |
| MANEAL | -0.897963399 | 0.536643758 | 0.001015183 | 0.005682195 | 89.28138104 | 47.92710189 | ENSG00000185090 |
| MYLK-AS1 | -0.89820168 | 0.536555131 | 0.417969289 | 0.628751794 | 6.847455004 | 3.697040702 | ENSG00000239523 |
| NA | -0.898597148 | 0.536408072 | 0.392641837 | 0.604159691 | 6.231238946 | 3.364970366 | ENSG00000278434 |
| TOMM20L | -0.898755512 | 0.536349194 | 0.485008538 | 0.686044959 | 4.338708046 | 2.322695756 | ENSG00000196860 |
| LINC01783 | -0.899170216 | 0.536195042 | 0.28162055 | 0.490582426 | 9.227197382 | 4.937641091 | ENSG00000233421 |
| NA | -0.899680798 | 0.536005311 | 0.000274952 | 0.00182192 | 107.2129958 | 57.49641929 | ENSG00000261061 |
| CDHR1 | -0.900220326 | 0.535804898 | 3.50E-11 | 8.36E-10 | 523.3172106 | 280.5859901 | ENSG00000148600 |
| RBM19 | -0.902467065 | 0.534971127 | 8.29E-37 | 1.40E-34 | 1796.900935 | 961.5794207 | ENSG00000122965 |
| PSPN | -0.902662623 | 0.534898616 | 0.025566557 | 0.083867367 | 39.04539718 | 20.94029873 | ENSG00000125650 |
| NA | -0.905458033 | 0.533863184 | 0.409659525 | 0.620751034 | 4.947805159 | 2.636416425 | ENSG00000273271 |
| NA | -0.905719334 | 0.533766499 | 0.42394532 | 0.634104813 | 4.954924104 | 2.640767352 | ENSG00000279481 |
| NA | -0.906222923 | 0.533580215 | 0.395548557 | 0.607422671 | 6.278681143 | 3.367145829 | ENSG00000232546 |
| PRSS22 | -0.906251771 | 0.533569545 | 0.118319262 | 0.27026605 | 18.67383454 | 9.937803534 | ENSG00000005001 |
| ALOX12B | -0.906484257 | 0.533483569 | 0.422097874 | 0.632192747 | 5.656541349 | 3.027603144 | ENSG00000179477 |
| SNORA11F | -0.906661411 | 0.533418065 | 0.117072647 | 0.268291789 | 21.25997402 | 11.29692023 | ENSG00000221164 |
| PRRT2 | -0.907019427 | 0.533285709 | 0.000710981 | 0.004165956 | 170.83104 | 91.19313251 | ENSG00000167371 |
| NA | -0.909729367 | 0.532284932 | 0.147434961 | 0.315038383 | 16.13633247 | 8.581808255 | ENSG00000236924 |
| NTN3 | -0.909786907 | 0.532263703 | 0.413556106 | 0.624700612 | 5.666024558 | 3.023252217 | ENSG00000162068 |
| NA | -0.909845553 | 0.532242067 | 0.052895523 | 0.147248224 | 35.07943941 | 18.65836969 | ENSG00000273162 |
| ZMYND15 | -0.91008261 | 0.532154619 | 0.400714978 | 0.612108903 | 5.569944954 | 2.96196037 | ENSG00000141497 |
| NA | -0.910275075 | 0.53208363 | 0.108894208 | 0.254355859 | 25.13608141 | 13.41788524 | ENSG00000277283 |
| MAN2A2 | -0.910851985 | 0.531870902 | 1.03E-09 | 1.97E-08 | 414.7471756 | 220.7358718 | ENSG00000196547 |
| WNT8B | -0.911642658 | 0.531579488 | 0.261054391 | 0.468142832 | 9.41935659 | 4.991460588 | ENSG00000075290 |
| NA | -0.911848464 | 0.531503662 | 0.257133071 | 0.463974588 | 9.956095198 | 5.275008313 | ENSG00000244171 |
| SLC38A5 | -0.91195052 | 0.531466065 | 9.20E-27 | 9.34E-25 | 1155.092709 | 614.0496378 | ENSG00000017483 |
| YWHAB | -0.912298941 | 0.531337727 | 2.38E-59 | 1.00E-56 | 6499.358099 | 3453.135235 | ENSG00000166913 |
| NA | -0.912607353 | 0.531224153 | 0.008026321 | 0.032759256 | 57.4351372 | 30.46449849 | ENSG00000234009 |
| TBC1D19 | -0.913886414 | 0.53075339 | 0.019367935 | 0.066903868 | 40.00321799 | 21.23472377 | ENSG00000109680 |
| NA | -0.91450903 | 0.530524385 | 0.246926844 | 0.451745658 | 9.966773615 | 5.265360501 | ENSG00000215007 |

| | | | | | | | |
|---|---|---|---|---|---|---|---|
| UBE2QL1 | -0.914523236 | 0.530519161 | 0.029640941 | 0.093968302 | 36.91145928 | 19.54665815 | ENSG00000215218 |
| ARRDC3-AS1 | -0.915450838 | 0.530178166 | 0.380030943 | 0.59241652 | 5.652981876 | 2.9974302 | ENSG00000281357 |
| MMGT1 | -0.915513619 | 0.530155095 | 4.07E-15 | 1.56E-13 | 650.5755188 | 344.8942695 | ENSG00000169446 |
| DUOX2 | -0.915733921 | 0.530074145 | 0.454733417 | 0.660692258 | 4.42054976 | 2.33139761 | ENSG00000140279 |
| NCKAP5 | -0.916539225 | 0.529778344 | 0.084320919 | 0.210106946 | 21.8925129 | 11.57611703 | ENSG00000176771 |
| NA | -0.917026259 | 0.529599528 | 0.445978088 | 0.653422136 | 4.416990288 | 2.32487122 | ENSG00000232303 |
| NA | -0.917054439 | 0.529589183 | 0.367650328 | 0.579797615 | 6.203958375 | 3.283153389 | ENSG00000233461 |
| NA | -0.917208267 | 0.529532719 | 0.372709617 | 0.584700209 | 5.666024558 | 2.999605664 | ENSG00000256591 |
| NPAS3 | -0.91735614 | 0.529478445 | 0.418903285 | 0.629503762 | 5.560461745 | 2.93396289 | ENSG00000151322 |
| NA | -0.917746017 | 0.529335378 | 0.173055261 | 0.352665222 | 15.09458502 | 8.026536081 | ENSG00000271966 |
| LHFPL3 | -0.918312473 | 0.529127582 | 0.345158813 | 0.557034745 | 6.884218784 | 3.650693555 | ENSG00000187416 |
| TAS2R14 | -0.919170167 | 0.528813104 | 0.380043746 | 0.59241652 | 6.231238946 | 3.292801202 | ENSG00000212127 |
| PIM2 | -0.91950486 | 0.528690438 | 6.03E-63 | 3.08E-60 | 5341.16595 | 2823.53961 | ENSG00000102096 |
| NA | -0.920029783 | 0.52849811 | 0.033216413 | 0.102885346 | 33.27172488 | 17.61297365 | ENSG00000261762 |
| DNAJC12 | -0.920045612 | 0.528492311 | 0.060590916 | 0.163535722 | 25.07559653 | 13.25425129 | ENSG00000108176 |
| ATP2B1-AS1 | -0.921329265 | 0.528022289 | 0.300928746 | 0.511018736 | 8.832506035 | 4.690792701 | ENSG00000271614 |
| LOXL3 | -0.922249653 | 0.527685537 | 6.33E-13 | 1.87E-11 | 436.6886046 | 230.3973461 | ENSG00000115318 |
| NA | -0.922401556 | 0.527629979 | 0.370934717 | 0.583100776 | 5.652981876 | 2.973783647 | ENSG00000287787 |
| NA | -0.92326023 | 0.527316034 | 6.10E-05 | 0.000479402 | 130.4230103 | 68.86427919 | ENSG00000274712 |
| NA | -0.923450488 | 0.527246498 | 0.08482006 | 0.211127828 | 24.62158936 | 13.03540038 | ENSG00000287737 |
| ACSM1 | -0.92369161 | 0.527158385 | 0.346268451 | 0.558292469 | 6.851014476 | 3.608697334 | ENSG00000166743 |
| NA | -0.92481174 | 0.52674925 | 0.054115905 | 0.149828304 | 27.06301182 | 14.26105607 | ENSG00000258568 |
| NA | -0.924928868 | 0.526706486 | 0.381313902 | 0.59376395 | 5.731264118 | 3.015779867 | ENSG00000277968 |
| CNTFR | -0.925615228 | 0.526455966 | 0.000203394 | 0.00139931 | 112.8135014 | 59.44722396 | ENSG00000122756 |
| GDF10 | -0.925794174 | 0.526390671 | 0.282510327 | 0.491542983 | 8.158169362 | 4.307078331 | ENSG00000266524 |
| ADPRHL1 | -0.92609374 | 0.526281381 | 0.032046856 | 0.100133171 | 33.31560761 | 17.52150886 | ENSG00000153531 |
| GPR4 | -0.926827583 | 0.52601375 | 0.324321149 | 0.535228186 | 6.970815179 | 3.672164644 | ENSG00000177464 |
| NA | -0.92819096 | 0.525516891 | 0.268076109 | 0.475944093 | 10.20637502 | 5.34282655 | ENSG00000287906 |
| EXOSC10-AS1 | -0.928191984 | 0.525516518 | 0.268834377 | 0.476752485 | 8.863346079 | 4.679915383 | ENSG00000230337 |
| SCN9A | -0.928690574 | 0.525334933 | 0.002144676 | 0.010786204 | 77.21684848 | 40.58305614 | ENSG00000169432 |
| NA | -0.928955359 | 0.525238524 | 0.277522467 | 0.486675809 | 8.185449933 | 4.309253795 | ENSG00000266490 |
| ECEL1P2 | -0.929074502 | 0.52519515 | 0.073933434 | 0.190742199 | 25.40652286 | 13.33294684 | ENSG00000244280 |
| PCDHGB6 | -0.929878255 | 0.524902635 | 1.37E-05 | 0.000126086 | 140.2225672 | 73.64386676 | ENSG00000253305 |
| C16orf54 | -0.931176527 | 0.524430491 | 0.09507816 | 0.229610109 | 21.62354599 | 11.34951023 | ENSG00000185905 |
| NA | -0.931781983 | 0.524210449 | 0.27083641 | 0.479177965 | 8.24713002 | 4.335075811 | ENSG00000213938 |
| NA | -0.931962017 | 0.524145037 | 0.184850327 | 0.369108957 | 14.02583634 | 7.355206606 | ENSG00000287707 |
| ANKRD18B | -0.932143175 | 0.524079225 | 0.000102818 | 0.000761157 | 114.0704926 | 59.69530185 | ENSG00000230453 |
| GPAM | -0.932186537 | 0.524063473 | 2.03E-21 | 1.36E-19 | 752.2546713 | 394.1358649 | ENSG00000119927 |
| NIBAN3 | -0.932365359 | 0.523998519 | 0.389567723 | 0.601355258 | 5.642303459 | 2.936138354 | ENSG00000167483 |
| TCEA2 | -0.932951341 | 0.523785729 | 8.07E-17 | 3.59E-15 | 859.3001181 | 450.4358258 | ENSG00000171703 |
| ATAD1 | -0.933553212 | 0.523567259 | 3.06E-18 | 1.57E-16 | 801.252603 | 419.6894902 | ENSG00000138138 |
| JAM2 | -0.933709949 | 0.523510381 | 1.37E-05 | 0.000126054 | 145.2931221 | 76.02974628 | ENSG00000154721 |
| HERC2P3 | -0.934831514 | 0.523103557 | 0.000612371 | 0.003659512 | 108.3241009 | 56.65977486 | ENSG00000180229 |
| TTLL3 | -0.937007745 | 0.522315077 | 0.037418587 | 0.112793298 | 37.71391081 | 19.70376572 | ENSG00000214021 |
| NA | -0.937388922 | 0.522177094 | 1.82E-07 | 2.40E-06 | 252.2184377 | 131.7191321 | ENSG00000159712 |
| NA | -0.937915224 | 0.521986636 | 0.303255897 | 0.513486816 | 7.685475105 | 4.013882793 | ENSG00000256167 |
| LINC01252 | -0.937956837 | 0.52197158 | 0.363354438 | 0.575377207 | 7.06333531 | 3.686163384 | ENSG00000247157 |
| ADAM33 | -0.939042526 | 0.521578922 | 5.41E-07 | 6.57E-06 | 336.4343234 | 175.6944211 | ENSG00000149451 |
| SAMD13 | -0.93922056 | 0.521514561 | 0.272117082 | 0.480723209 | 8.257808438 | 4.304902868 | ENSG00000203943 |
| LOC105371067 | -0.939501835 | 0.521412894 | 0.120716718 | 0.273698333 | 16.50493846 | 8.600157923 | ENSG00000285567 |
| ATF5 | -0.940986759 | 0.520876495 | 8.02E-08 | 1.14E-06 | 217.94295 | 113.6172641 | ENSG00000169136 |
| GPR137C | -0.940999631 | 0.520871848 | 0.000855534 | 0.004890061 | 77.02115595 | 40.11099841 | ENSG00000180998 |
| NA | -0.943180227 | 0.520085158 | 0.220274876 | 0.417031683 | 11.53697167 | 6.02285788 | ENSG00000272525 |
| RTL5 | -0.944229094 | 0.519707184 | 3.10E-05 | 0.000262713 | 125.8666687 | 65.4100195 | ENSG00000242732 |
| CCDC74A | -0.944561475 | 0.519587463 | 2.21E-09 | 4.04E-08 | 302.2703228 | 157.1725909 | ENSG00000163040 |
| ELAVL4 | -0.94512326 | 0.519385175 | 0.298587772 | 0.508919683 | 7.651075589 | 3.960063296 | ENSG00000162374 |
| H2BC15 | -0.945322659 | 0.519313394 | 0.095673443 | 0.230458288 | 18.55166958 | 9.635906142 | ENSG00000233822 |
| YPEL2 | -0.945609098 | 0.519210297 | 0.001603477 | 0.008413908 | 78.7600453 | 40.87161721 | ENSG00000175155 |
| VWA5B2 | -0.946041007 | 0.519054881 | 0.196273103 | 0.38530328 | 12.20062993 | 6.35710368 | ENSG00000145198 |
| NA | -0.947399194 | 0.518566461 | 0.115234294 | 0.265163842 | 16.67218136 | 8.628155403 | ENSG00000232000 |
| SLC9A6 | -0.950066334 | 0.517608662 | 3.47E-14 | 1.18E-12 | 672.7991857 | 348.0370566 | ENSG00000198689 |
| ADAMTS19-AS1 | -0.95139038 | 0.51713384 | 0.006769018 | 0.028463243 | 55.95123006 | 28.96153792 | ENSG00000249421 |
| SHISA2 | -0.953058305 | 0.516536318 | 2.69E-11 | 6.50E-10 | 436.0341503 | 225.1479489 | ENSG00000180730 |
| TMEM132E | -0.953213731 | 0.516480673 | 0.014706127 | 0.053677081 | 39.86117073 | 20.56093528 | ENSG00000181291 |
| NA | -0.954062283 | 0.516176983 | 0.219336662 | 0.415703519 | 14.92169772 | 7.724638689 | ENSG00000275180 |
| GLYATL2 | -0.955949913 | 0.515502056 | 0.207324942 | 0.399809204 | 11.5096911 | 5.923920741 | ENSG00000156689 |
| EPHA10 | -0.956005487 | 0.515482199 | 0.024065703 | 0.079887643 | 38.14745087 | 19.69317195 | ENSG00000183317 |
| NOL12 | -0.956298877 | 0.51537738 | 0.002182816 | 0.010945956 | 66.45476843 | 34.30001354 | ENSG00000273899 |
| NA | -0.956462785 | 0.51531883 | 0.275158923 | 0.484014183 | 9.75236785 | 5.023808996 | ENSG00000270933 |
| TTC25 | -0.958291901 | 0.514665898 | 0.055885016 | 0.153689898 | 26.29735023 | 13.52597574 | ENSG00000204815 |
| NA | -0.960217808 | 0.51397931 | 0.288390763 | 0.497657526 | 8.379973404 | 4.283431778 | ENSG00000276408 |
| NA | -0.96029704 | 0.513951084 | 0.448913952 | 0.655461573 | 4.53531645 | 2.350693236 | ENSG00000280107 |
| NA | -0.960532781 | 0.513867109 | 0.248523168 | 0.453472125 | 9.030588979 | 4.616448073 | ENSG00000227008 |
| HUNK | -0.960636001 | 0.513830345 | 4.36E-10 | 8.84E-09 | 399.3893198 | 205.1367053 | ENSG00000142149 |

| | | | | | | | |
|---|---|---|---|---|---|---|---|
| NA | -0.961056683 | 0.513680537 | 0.490074177 | 0.69013064 | 4.538875922 | 2.346342309 | ENSG00000268938 |
| NA | -0.961862783 | 0.5133936 | 0.090817464 | 0.221803996 | 21.35813855 | 10.94025739 | ENSG00000288586 |
| TXNDC16 | -0.962749501 | 0.513078152 | 1.02E-09 | 1.96E-08 | 369.3179172 | 189.5507002 | ENSG00000087301 |
| NA | -0.963282766 | 0.512888538 | 0.199436981 | 0.389610777 | 11.5808544 | 5.919569814 | ENSG00000275582 |
| NA | -0.963581312 | 0.512782413 | 0.385323942 | 0.597540044 | 6.510884272 | 3.362794902 | ENSG00000253210 |
| DCLK2 | -0.963598268 | 0.512776386 | 0.179383576 | 0.361929363 | 12.300269 | 6.296757792 | ENSG00000170390 |
| DPYSL5 | -0.963910812 | 0.512665311 | 3.05E-30 | 3.62E-28 | 1101.832317 | 564.8836891 | ENSG00000157851 |
| C5orf49 | -0.965102139 | 0.512242145 | 0.170310428 | 0.348638065 | 13.5457438 | 6.925145089 | ENSG00000215217 |
| SMARCA1 | -0.965991692 | 0.511926398 | 8.52E-58 | 3.28E-55 | 3297.183538 | 1687.541859 | ENSG00000102038 |
| CHTF8 | -0.966194495 | 0.51185444 | 0.22530768 | 0.423629584 | 10.43382347 | 5.324476883 | ENSG00000168802 |
| BBOF1 | -0.966288625 | 0.511821045 | 0.390017402 | 0.60165648 | 6.475289547 | 3.331676 | ENSG00000119636 |
| SHROOM2 | -0.96651012 | 0.511742472 | 2.07E-23 | 1.62E-21 | 891.5333245 | 456.0967333 | ENSG00000146950 |
| HSP90B2P | -0.968512887 | 0.511032558 | 0.307182495 | 0.517497169 | 8.396296226 | 4.311429258 | ENSG00000259706 |
| NKAIN1 | -0.968612044 | 0.510997436 | 1.11E-33 | 1.59E-31 | 2092.464026 | 1069.412666 | ENSG00000084628 |
| C1GALT1C1 | -0.970532886 | 0.510317533 | 3.29E-13 | 1.00E-11 | 369.9008511 | 188.7986153 | ENSG00000171155 |
| FAAP24 | -0.970899889 | 0.510187731 | 0.000219408 | 0.001492083 | 96.15967608 | 48.98817766 | ENSG00000131944 |
| SMIM3 | -0.971653173 | 0.509921413 | 2.21E-20 | 1.35E-18 | 681.2022698 | 347.2777329 | ENSG00000256235 |
| NA | -0.971719878 | 0.509897836 | 0.112324126 | 0.260228509 | 16.92098664 | 8.61850759 | ENSG00000263731 |
| NA | -0.971728234 | 0.509894883 | 0.030483472 | 0.096222038 | 30.59601435 | 15.5694747 | ENSG00000243679 |
| DRC3 | -0.972058226 | 0.509778267 | 0.122824564 | 0.277182172 | 16.36881493 | 8.342432214 | ENSG00000171962 |
| H2AX | -0.972691733 | 0.509554465 | 2.89E-32 | 3.72E-30 | 3545.992291 | 1807.163418 | ENSG00000188486 |
| ATP8B1 | -0.972960096 | 0.509459689 | 0.003551528 | 0.016601634 | 62.65666395 | 31.93409206 | ENSG00000081923 |
| GOPC | -0.973692559 | 0.5092011 | 1.90E-12 | 5.29E-11 | 475.0726823 | 241.7037032 | ENSG00000047932 |
| EDNRB | -0.973756201 | 0.509178638 | 1.09E-08 | 1.78E-07 | 252.263795 | 128.2724174 | ENSG00000136160 |
| IRAK1BP1 | -0.973915708 | 0.509122345 | 7.00E-06 | 6.94E-05 | 138.1200797 | 70.40148008 | ENSG00000146243 |
| DUSP15 | -0.975358637 | 0.508613395 | 0.001888675 | 0.009663221 | 66.44142027 | 33.8357117 | ENSG00000149599 |
| NA | -0.975610382 | 0.508524652 | 0.001221828 | 0.006672707 | 76.39754162 | 38.92733894 | ENSG00000229321 |
| SLC16A2 | -0.976075636 | 0.508360684 | 1.34E-45 | 3.29E-43 | 2406.418421 | 1223.243606 | ENSG00000147100 |
| RBM20 | -0.977483923 | 0.50786469 | 0.190342784 | 0.376715938 | 13.14421284 | 6.655596104 | ENSG00000203867 |
| EPHB3 | -0.980866366 | 0.50667538 | 0.000811655 | 0.004674626 | 79.32022567 | 40.22488024 | ENSG00000182580 |
| INCA1 | -0.98105077 | 0.506610622 | 0.053944506 | 0.149397565 | 29.45790797 | 14.96785537 | ENSG00000196388 |
| AMOTL1 | -0.983209287 | 0.505853213 | 5.55E-40 | 1.08E-37 | 1724.657289 | 872.2746441 | ENSG00000166025 |
| FAM133B | -0.983449113 | 0.505769129 | 1.81E-20 | 1.11E-18 | 789.9899116 | 399.635021 | ENSG00000234545 |
| DNLZ | -0.984288919 | 0.505474802 | 0.295188246 | 0.505268741 | 7.238586879 | 3.672164644 | ENSG00000213221 |
| KIAA0895 | -0.984561046 | 0.505379467 | 0.334881332 | 0.546869227 | 5.873311379 | 2.973783647 | ENSG00000164542 |
| GDF7 | -0.984658161 | 0.505345448 | 0.046614223 | 0.133582639 | 26.36050486 | 13.30589532 | ENSG00000143869 |
| NA | -0.985242292 | 0.505140881 | 0.385498443 | 0.597632758 | 6.576403164 | 3.285328852 | ENSG00000263011 |
| EFCAB6 | -0.985492405 | 0.505053315 | 0.137950325 | 0.300969487 | 14.43448624 | 7.308859458 | ENSG00000186976 |
| NA | -0.986020024 | 0.504868641 | 0.163079732 | 0.338120875 | 14.33484716 | 7.241041222 | ENSG00000261499 |
| H2AC11 | -0.986174246 | 0.504814675 | 0.004570873 | 0.020494051 | 61.168918 | 30.94715 | ENSG00000196787 |
| RNF31 | -0.986386244 | 0.504740499 | 0.261386062 | 0.468515166 | 7.899880871 | 3.999884053 | ENSG00000092098 |
| COL21A1 | -0.986950679 | 0.504543065 | 0.000675752 | 0.003984256 | 76.4144231 | 38.59432264 | ENSG00000124749 |
| HRH3 | -0.987551048 | 0.504333146 | 1.02E-06 | 1.18E-05 | 175.6616875 | 88.69978195 | ENSG00000101180 |
| LOC100128108 | -0.987768581 | 0.504257108 | 0.095042118 | 0.229570367 | 17.80853387 | 9.005343381 | ENSG00000259658 |
| NA | -0.988151649 | 0.504123234 | 0.073982071 | 0.190841607 | 23.83960494 | 12.05224215 | ENSG00000203497 |
| ZNF677 | -0.988279448 | 0.504078579 | 6.58E-11 | 1.51E-09 | 400.3532658 | 201.5824899 | ENSG00000197928 |
| NA | -0.98874959 | 0.503914337 | 0.357000858 | 0.569121372 | 5.233374222 | 2.638591889 | ENSG00000197332 |
| ACSL6 | -0.989181247 | 0.503763588 | 0.272904912 | 0.481592322 | 7.879719244 | 3.974062036 | ENSG00000164398 |
| CYP1B1 | -0.989210605 | 0.503753337 | 0.053332187 | 0.148049146 | 26.20838957 | 13.20913365 | ENSG00000138061 |
| PCBP1-AS1 | -0.989379289 | 0.50369444 | 1.78E-07 | 2.35E-06 | 230.3793425 | 116.2149214 | ENSG00000179818 |
| NA | -0.989750007 | 0.503565026 | 0.357235444 | 0.569315726 | 5.264214265 | 2.654766092 | ENSG00000232842 |
| NA | -0.990286088 | 0.503377945 | 0.19877319 | 0.389030323 | 11.15056833 | 5.614550999 | ENSG00000257511 |
| NA | -0.990311381 | 0.50336912 | 0.336720363 | 0.548688178 | 5.972950455 | 3.015779867 | ENSG00000268157 |
| OR7E14P | -0.990479184 | 0.503310575 | 0.082881508 | 0.207341009 | 19.10976502 | 9.631555214 | ENSG00000184669 |
| ITGB1BP2 | -0.991022944 | 0.50312091 | 0.034564332 | 0.10618959 | 27.72430582 | 13.97098195 | ENSG00000147166 |
| MCTS1 | -0.991260556 | 0.503038053 | 1.08E-19 | 6.35E-18 | 635.956551 | 319.8961272 | ENSG00000232119 |
| CLEC14A | -0.99150831 | 0.502951674 | 0.2930923 | 0.502804708 | 7.218425253 | 3.630168424 | ENSG00000176435 |
| LOC101928445 | -0.991745914 | 0.502868847 | 0.276439421 | 0.485497858 | 7.865481354 | 3.953536906 | ENSG00000244945 |
| NA | -0.9933153 | 0.502322116 | 0.266235901 | 0.473656617 | 8.648940313 | 4.353425479 | ENSG00000280219 |
| YPEL1 | -0.993846819 | 0.502137084 | 2.88E-05 | 0.000246816 | 144.284579 | 72.44876288 | ENSG00000100027 |
| CSTF2 | -0.994136201 | 0.502036374 | 5.69E-39 | 1.03E-36 | 1535.619796 | 770.8113971 | ENSG00000101811 |
| NA | -0.994387541 | 0.501948919 | 0.301158314 | 0.51131662 | 6.574928623 | 3.297152129 | ENSG00000287966 |
| NA | -0.994616625 | 0.501869221 | 0.312088357 | 0.52250938 | 7.841760256 | 3.927714889 | ENSG00000267868 |
| NA | -0.994718745 | 0.501833698 | 0.167921685 | 0.345258686 | 12.49331794 | 6.263463426 | ENSG00000242375 |
| NA | -0.995182936 | 0.501672258 | 0.280610247 | 0.489544989 | 9.35411703 | 4.701386472 | ENSG00000287234 |
| GNG3 | -0.995315765 | 0.50162607 | 0.430828735 | 0.640589378 | 4.671719306 | 2.347571814 | ENSG00000162188 |
| RNF39 | -0.995328266 | 0.501621724 | 0.007093213 | 0.029596846 | 59.83328734 | 29.9837389 | ENSG00000204618 |
| LOC105378083 | -0.9957071 | 0.501490021 | 0.364935401 | 0.577106435 | 5.305732727 | 2.662238442 | ENSG00000271913 |
| NA | -0.996244343 | 0.501303307 | 0.383467235 | 0.595734937 | 4.647998207 | 2.33139761 | ENSG00000285090 |
| IRX5 | -0.9963498 | 0.501266665 | 0.294916502 | 0.505010587 | 6.612887612 | 3.313326333 | ENSG00000176842 |
| NA | -0.996449427 | 0.50123205 | 0.311800162 | 0.522247988 | 6.674567699 | 3.350971626 | ENSG00000228559 |
| NA | -0.996876369 | 0.501083741 | 0.385576577 | 0.597653274 | 5.325894353 | 2.670940296 | ENSG00000240616 |
| KPNA1 | -0.997839536 | 0.500749321 | 8.44E-19 | 4.65E-17 | 638.1672449 | 319.7613117 | ENSG00000114030 |
| NPY2R | -0.998032201 | 0.500682453 | 0.419106854 | 0.629510295 | 4.582758647 | 2.287225926 | ENSG00000185149 |

| | | | | | | | |
|---|---|---|---|---|---|---|---|
| LOC105371414 | -0.998124284 | 0.500650497 | 0.056053345 | 0.154130377 | 22.56209489 | 11.29039384 | ENSG00000260279 |
| NA | -0.998219429 | 0.50061748 | 0.23610434 | 0.438085131 | 9.337514876 | 4.677739919 | ENSG00000254244 |
| GSC2 | -0.998402166 | 0.500554074 | 0.295444928 | 0.505492478 | 6.663889282 | 3.336972886 | ENSG00000063515 |
| NA | -0.998450682 | 0.500537241 | 0.122998038 | 0.277483926 | 15.20014784 | 7.596758111 | ENSG00000282886 |
| NA | -0.999170536 | 0.500287553 | 0.339633392 | 0.55136372 | 5.911270367 | 2.94796163 | ENSG00000233588 |
| TAF2 | -0.999385242 | 0.500213104 | 1.93E-23 | 1.54E-21 | 755.9777475 | 378.0763616 | ENSG00000064313 |
| NA | -1.000262598 | 0.499908999 | 0.3559752 | 0.568266411 | 6.685246117 | 3.343499276 | ENSG00000261570 |
| ST8SIA5 | -1.001727366 | 0.499401699 | 0.000844445 | 0.004835458 | 82.99619134 | 41.49130265 | ENSG00000101638 |
| NA | -1.002086063 | 0.499277548 | 0.386873066 | 0.599124673 | 4.627836581 | 2.30340013 | ENSG00000241889 |
| SMPD1 | -1.003060202 | 0.498940539 | 0.003940037 | 0.018130506 | 52.65429561 | 26.29712402 | ENSG00000166311 |
| BISPR | -1.003230974 | 0.498881483 | 0.114278174 | 0.263638323 | 16.0085231 | 7.974892048 | ENSG00000282851 |
| ME3 | -1.003615549 | 0.498748515 | 0.009106286 | 0.036235188 | 46.59505425 | 23.19640575 | ENSG00000151376 |
| NA | -1.004036397 | 0.498603046 | 0.320520531 | 0.531060733 | 6.003790499 | 2.987782387 | ENSG00000279637 |
| NA | -1.004940034 | 0.498290843 | 0.102492439 | 0.242461444 | 17.35956071 | 8.633452289 | ENSG00000217094 |
| CPNE9 | -1.005569277 | 0.498073557 | 5.48E-05 | 0.000436701 | 108.0750163 | 53.76920565 | ENSG00000144550 |
| NA | -1.006298727 | 0.497821786 | 0.325788607 | 0.536984748 | 6.000231026 | 2.978134574 | ENSG00000256356 |
| NA | -1.006640646 | 0.497703816 | 0.040643977 | 0.120250902 | 26.73711951 | 13.34041919 | ENSG00000288559 |
| NA | -1.006945608 | 0.497598621 | 0.230295019 | 0.430265278 | 8.65961873 | 4.293079591 | ENSG00000275120 |
| NA | -1.008065022 | 0.497212675 | 0.361991241 | 0.573896351 | 6.08207274 | 3.023252217 | ENSG00000245322 |
| NA | -1.010528849 | 0.496364262 | 0.302465866 | 0.512522914 | 8.563539126 | 4.227436818 | ENSG00000234692 |
| LINC00652 | -1.011605172 | 0.495994088 | 0.343513032 | 0.555462249 | 6.089191685 | 3.015779867 | ENSG00000179935 |
| NA | -1.011893612 | 0.495894933 | 0.223123573 | 0.420989977 | 9.422916062 | 4.658444293 | ENSG00000260774 |
| NA | -1.012274075 | 0.495764174 | 0.347505042 | 0.55952193 | 5.353174924 | 2.640767352 | ENSG00000260060 |
| LRRC6 | -1.014786265 | 0.494901642 | 0.003478581 | 0.016326486 | 56.29522522 | 27.78268088 | ENSG00000129295 |
| LOC100505715 | -1.016714509 | 0.49424062 | 0.197151401 | 0.38670588 | 10.75379205 | 5.28683159 | ENSG00000267058 |
| NA | -1.016760319 | 0.494224926 | 0.255282453 | 0.462136611 | 8.101523287 | 3.985885313 | ENSG00000279161 |
| SPACA9 | -1.017320156 | 0.49403318 | 0.147792978 | 0.315624832 | 14.12400087 | 7.000435675 | ENSG00000165698 |
| ZFP42 | -1.018510768 | 0.493625638 | 9.90E-48 | 2.71E-45 | 4504.218823 | 2223.221721 | ENSG00000179059 |
| APLN | -1.019733056 | 0.493207603 | 6.77E-05 | 0.000526043 | 115.3859617 | 56.90321828 | ENSG00000171388 |
| ZNF599 | -1.022390181 | 0.49230006 | 0.003520587 | 0.016473313 | 62.07782207 | 30.64525261 | ENSG00000153896 |
| NA | -1.023579786 | 0.49189429 | 0.071317036 | 0.185487155 | 19.55309377 | 9.607908661 | ENSG00000272070 |
| TMEM99 | -1.023721919 | 0.491845832 | 3.67E-05 | 0.000305275 | 124.4073468 | 61.10417067 | ENSG00000167920 |
| NA | -1.024109719 | 0.49171364 | 0.309751445 | 0.520293163 | 6.749290468 | 3.294976665 | ENSG00000284720 |
| CRELD1 | -1.025573421 | 0.49121502 | 0.044666534 | 0.12904567 | 25.04712075 | 12.33796534 | ENSG00000163703 |
| CCBE1 | -1.025780766 | 0.491144427 | 4.76E-06 | 4.85E-05 | 128.4326204 | 63.0195055 | ENSG00000183287 |
| H2BC9 | -1.026975233 | 0.490737956 | 0.000604757 | 0.003624513 | 88.70579314 | 43.64707508 | ENSG00000275713 |
| SLC25A27 | -1.027431547 | 0.490582764 | 0.004760104 | 0.021181566 | 74.24135716 | 36.46420428 | ENSG00000153291 |
| UCP1 | -1.028076404 | 0.490363532 | 0.237153406 | 0.439168668 | 8.767266475 | 4.321077071 | ENSG00000109424 |
| CCDC157 | -1.028108142 | 0.490352745 | 0.000711295 | 0.004166503 | 75.59300516 | 37.04016954 | ENSG00000187860 |
| DZIP3 | -1.028366431 | 0.490264963 | 1.34E-16 | 5.83E-15 | 593.2435546 | 291.0719926 | ENSG00000198919 |
| NA | -1.02865218 | 0.490167868 | 0.117196477 | 0.268436079 | 15.51775215 | 7.596758111 | ENSG00000267265 |
| E2F2 | -1.033223584 | 0.488617153 | 7.15E-10 | 1.40E-08 | 230.3846558 | 112.5957982 | ENSG00000007968 |
| NA | -1.034877646 | 0.48805727 | 0.203931777 | 0.395527534 | 10.95515513 | 5.361176218 | ENSG00000274825 |
| NA | -1.03624465 | 0.487595038 | 0.163201051 | 0.338180807 | 14.407485 | 7.026257692 | ENSG00000279259 |
| CRHR2 | -1.036456375 | 0.487523486 | 0.030535801 | 0.09632672 | 29.21383122 | 14.26852842 | ENSG00000106113 |
| DYNC2H1 | -1.038630479 | 0.486789354 | 1.14E-12 | 3.26E-11 | 392.7090036 | 191.0914217 | ENSG00000187240 |
| FGFBP3 | -1.039759517 | 0.486408546 | 1.74E-45 | 4.22E-43 | 2150.608327 | 1046.307346 | ENSG00000174721 |
| ACD | -1.0405475 | 0.486142948 | 1.62E-27 | 1.70E-25 | 800.8146144 | 389.4216365 | ENSG00000102977 |
| AMBP | -1.041544618 | 0.485807067 | 0.013350047 | 0.049512104 | 38.8870271 | 18.84610169 | ENSG00000106927 |
| FIG4 | -1.041735128 | 0.48574292 | 6.00E-22 | 4.21E-20 | 697.7078192 | 338.6904169 | ENSG00000112367 |
| MAN2A1 | -1.04218714 | 0.485590755 | 9.08E-33 | 1.25E-30 | 1620.739221 | 786.7594346 | ENSG00000112893 |
| TMEM221 | -1.042400731 | 0.485518869 | 0.006518373 | 0.027625554 | 46.52030533 | 22.56243802 | ENSG00000188051 |
| NA | -1.042521281 | 0.485478301 | 0.177980445 | 0.359982939 | 10.92431509 | 5.298654867 | ENSG00000286868 |
| NA | -1.043380663 | 0.485189198 | 0.239614996 | 0.442598708 | 8.19612835 | 3.988060777 | ENSG00000213757 |
| PLCL2 | -1.045897195 | 0.484343607 | 4.98E-08 | 7.32E-07 | 209.8381465 | 101.5486369 | ENSG00000154822 |
| NA | -1.046003854 | 0.484307801 | 0.206429273 | 0.398815021 | 8.89062665 | 4.309253795 | ENSG00000280828 |
| NA | -1.046404804 | 0.484173222 | 0.20640457 | 0.398808093 | 9.530563807 | 4.60680026 | ENSG00000259577 |
| HLA-F-AS1 | -1.048861963 | 0.483349293 | 0.022207852 | 0.074924453 | 30.29264793 | 14.63171766 | ENSG00000214922 |
| LRCH2 | -1.049310556 | 0.483199024 | 4.64E-08 | 6.84E-07 | 213.5630034 | 103.0238109 | ENSG00000130224 |
| RAB27A | -1.049697545 | 0.483069427 | 7.43E-07 | 8.78E-06 | 141.5857578 | 68.4236239 | ENSG00000069974 |
| C10orf143 | -1.050044983 | 0.482953106 | 0.004068449 | 0.018644177 | 48.08694391 | 23.19735171 | ENSG00000237489 |
| NA | -1.0514064 | 0.482497576 | 0.206837331 | 0.399317419 | 8.928585639 | 4.304902868 | ENSG00000279039 |
| CABP7 | -1.0522415 | 0.482218365 | 0.235741549 | 0.437669991 | 8.260172702 | 3.985885313 | ENSG00000100314 |
| RASL10A | -1.052665405 | 0.482076696 | 0.02473146 | 0.081767089 | 30.17493216 | 14.56172396 | ENSG00000100276 |
| SYNDIG1L | -1.052873598 | 0.482007133 | 0.050079508 | 0.141241265 | 22.7542541 | 10.92938007 | ENSG00000183379 |
| PVALB | -1.053589432 | 0.481768031 | 0.04434403 | 0.128275587 | 27.72814462 | 13.36189028 | ENSG00000100362 |
| HS6ST3 | -1.054389417 | 0.481500961 | 0.004215527 | 0.019210962 | 48.18630366 | 23.20699952 | ENSG00000185352 |
| C6orf118 | -1.05445202 | 0.481480068 | 0.100532473 | 0.238992415 | 15.91600297 | 7.676399624 | ENSG00000112539 |
| DIO3 | -1.055507991 | 0.481127781 | 0.2656897 | 0.473175215 | 6.86761663 | 3.311150869 | ENSG00000197406 |
| TM4SF18 | -1.057770285 | 0.480373914 | 0.000702377 | 0.004123215 | 79.58474339 | 38.2388893 | ENSG00000163762 |
| WNT7B | -1.058682496 | 0.480070271 | 0.199466028 | 0.389614115 | 10.46822298 | 5.017282605 | ENSG00000188064 |
| CFAP43 | -1.060642958 | 0.479418352 | 0.253118998 | 0.459188499 | 7.572793347 | 3.624871538 | ENSG00000197748 |
| NA | -1.061017742 | 0.479293825 | 0.123470126 | 0.278183408 | 15.93113058 | 7.679521046 | ENSG00000269388 |
| NA | -1.063710595 | 0.478400037 | 0.171454834 | 0.350500599 | 11.79050548 | 5.659668642 | ENSG00000279227 |

| | | | | | | | |
|---|---|---|---|---|---|---|---|
| RSPH4A | -1.06467439 | 0.478080547 | 0.1263033 | 0.2829128 | 13.2056136 | 6.324755273 | ENSG00000111834 |
| RGPD6 | -1.064889354 | 0.478009318 | 0.32943454 | 0.541082909 | 5.564021218 | 2.674061718 | ENSG00000183054 |
| LMNTD2-AS1 | -1.06566568 | 0.477752166 | 0.157863715 | 0.330589517 | 16.77654896 | 8.009415919 | ENSG00000254815 |
| NA | -1.066468675 | 0.477486327 | 0.285989731 | 0.49531512 | 6.26207899 | 2.9974302 | ENSG00000259408 |
| NA | -1.067596722 | 0.477113125 | 0.260149386 | 0.467028585 | 7.620235545 | 3.624871538 | ENSG00000225137 |
| NA | -1.069605353 | 0.476449313 | 0.221057155 | 0.418177339 | 9.674085608 | 4.576627316 | ENSG00000226900 |
| NA | -1.069904983 | 0.476350371 | 0.353243424 | 0.565470134 | 4.882565599 | 2.343220887 | ENSG00000254094 |
| DBNDD2 | -1.070359238 | 0.476200408 | 0.032783194 | 0.101844109 | 29.39058348 | 14.03662473 | ENSG00000244274 |
| NA | -1.07057576 | 0.476128945 | 0.283873426 | 0.493051563 | 6.976738915 | 3.318623218 | ENSG00000280434 |
| CDK5R2 | -1.071205532 | 0.475921148 | 0.184922983 | 0.369146229 | 14.80217635 | 7.048012328 | ENSG00000171450 |
| AKR1C3 | -1.071732685 | 0.47574728 | 0.005446924 | 0.023728989 | 43.9579131 | 20.92629999 | ENSG00000196139 |
| PARP2 | -1.072089267 | 0.475629708 | 1.26E-23 | 1.02E-21 | 756.2188235 | 359.7415916 | ENSG00000129484 |
| NA | -1.074405497 | 0.474866702 | 0.383957458 | 0.596307427 | 4.903922434 | 2.341045423 | ENSG00000285803 |
| NA | -1.075670332 | 0.474450561 | 0.361865347 | 0.573885378 | 4.858844501 | 2.313047943 | ENSG00000266648 |
| NA | -1.076408251 | 0.474207948 | 0.39025284 | 0.601716195 | 4.902727226 | 2.334519033 | ENSG00000232470 |
| NA | -1.076744476 | 0.474097445 | 0.350292822 | 0.562428154 | 4.882565599 | 2.32487122 | ENSG00000241255 |
| LOC644656 | -1.076899094 | 0.474046637 | 0.178869158 | 0.361200176 | 10.55214963 | 5.023808996 | ENSG00000268403 |
| RAI2 | -1.077685727 | 0.473788233 | 0.011142113 | 0.042724631 | 39.0797967 | 18.52490868 | ENSG00000131831 |
| NA | -1.078288774 | 0.473590231 | 0.198213609 | 0.388265119 | 9.85765133 | 4.693914123 | ENSG00000241627 |
| NA | -1.078291152 | 0.47358945 | 0.302193933 | 0.512445547 | 5.611463415 | 2.662238442 | ENSG00000263745 |
| CYP39A1 | -1.078648649 | 0.47347211 | 0.002741966 | 0.013297904 | 50.52719732 | 23.89790817 | ENSG00000146233 |
| ZNF428 | -1.081233549 | 0.472624542 | 5.50E-08 | 7.98E-07 | 224.8440758 | 106.3274894 | ENSG00000131116 |
| LINC02043 | -1.08145493 | 0.472552024 | 0.05700856 | 0.156217476 | 20.40268202 | 9.673551435 | ENSG00000232233 |
| NA | -1.081735525 | 0.472460124 | 0.235650431 | 0.437543839 | 10.65771244 | 5.051806476 | ENSG00000248971 |
| NA | -1.082109424 | 0.472337694 | 0.065032658 | 0.172443379 | 20.46080263 | 9.697197988 | ENSG00000285693 |
| NA | -1.083693885 | 0.471819227 | 0.073546757 | 0.190030168 | 20.38519014 | 9.584262108 | ENSG00000267904 |
| RBFADN | -1.084913121 | 0.471420656 | 0.354356206 | 0.566448024 | 4.967966786 | 2.3528687 | ENSG00000261126 |
| NA | -1.085134914 | 0.471348188 | 0.283372278 | 0.492452881 | 6.347480176 | 2.98343146 | ENSG00000268279 |
| SYPL2 | -1.085967797 | 0.471076152 | 0.000241785 | 0.00162959 | 80.82757459 | 38.06437803 | ENSG00000143028 |
| DQX1 | -1.087084931 | 0.470711521 | 0.107784945 | 0.252569692 | 14.10027977 | 6.662122495 | ENSG00000144045 |
| ENTPD3 | -1.088677228 | 0.470192285 | 0.331412938 | 0.542962899 | 6.258519517 | 2.924315077 | ENSG00000168032 |
| ODC1 | -1.089577722 | 0.469898894 | 1.64E-73 | 1.55E-70 | 4426.181025 | 2079.718362 | ENSG00000115758 |
| NA | -1.089730728 | 0.469849061 | 0.11785049 | 0.269541784 | 14.14535771 | 6.666473422 | ENSG00000253333 |
| NA | -1.08973835 | 0.469846579 | 0.10060848 | 0.239082818 | 16.11973032 | 7.554761891 | ENSG00000229692 |
| CHN2 | -1.090022828 | 0.469753941 | 0.326086275 | 0.537312565 | 5.615022888 | 2.628944076 | ENSG00000106069 |
| ICAM5 | -1.091213373 | 0.46936645 | 4.03E-05 | 0.000331401 | 107.0842705 | 50.26162092 | ENSG00000105376 |
| ZNF585B | -1.091677753 | 0.469215393 | 0.001477072 | 0.007859823 | 66.08171325 | 31.0470331 | ENSG00000245680 |
| H2AC8 | -1.092849429 | 0.468834477 | 0.034103157 | 0.105143751 | 26.84832673 | 12.62056711 | ENSG00000277075 |
| ZNF559 | -1.092907369 | 0.468815649 | 1.31E-08 | 2.13E-07 | 188.2418811 | 88.29677196 | ENSG00000188321 |
| LEF1 | -1.095120215 | 0.468097117 | 0.295048548 | 0.505134134 | 5.703983547 | 2.664413905 | ENSG00000138795 |
| DPH5 | -1.0966812 | 0.467590914 | 1.48E-26 | 1.50E-24 | 727.3335543 | 339.9070872 | ENSG00000117543 |
| TBL1X | -1.096893777 | 0.46752202 | 2.73E-55 | 9.72E-53 | 1929.931495 | 902.1796148 | ENSG00000101849 |
| UBE2Q2P1 | -1.096919503 | 0.467513684 | 0.184703275 | 0.368879161 | 9.870694012 | 4.604624797 | ENSG00000189136 |
| MROH8 | -1.096964171 | 0.467499209 | 0.007716716 | 0.031754543 | 38.3654161 | 17.94939492 | ENSG00000101353 |
| NA | -1.097157074 | 0.467436704 | 0.329580014 | 0.541217709 | 4.975085731 | 2.32487122 | ENSG00000260209 |
| H4C8 | -1.098607274 | 0.466967071 | 0.008551075 | 0.034520532 | 39.89765517 | 18.68324574 | ENSG00000158406 |
| DLK2 | -1.098624869 | 0.466961376 | 0.163927972 | 0.339274231 | 10.72651147 | 5.003283865 | ENSG00000171462 |
| LSMEM2 | -1.102537855 | 0.465696565 | 0.346804861 | 0.558942677 | 4.975085731 | 2.310872479 | ENSG00000179564 |
| ZNF443 | -1.102914471 | 0.46557501 | 0.000184891 | 0.001286067 | 81.19259495 | 37.81601658 | ENSG00000180855 |
| SPATA31C2 | -1.10341996 | 0.465411911 | 0.326980265 | 0.538425113 | 4.967966786 | 2.30340013 | ENSG00000177910 |
| ISM1 | -1.104087784 | 0.465196522 | 0.015144497 | 0.054884635 | 30.70010263 | 14.30399825 | ENSG00000101230 |
| NA | -1.104881613 | 0.464940623 | 0.012633512 | 0.047403909 | 34.27935215 | 15.90684192 | ENSG00000284727 |
| NA | -1.105420936 | 0.464766846 | 0.189297605 | 0.375434285 | 10.01065634 | 4.64227009 | ENSG00000273014 |
| NA | -1.10573608 | 0.464665333 | 0.301557461 | 0.511718271 | 5.758544689 | 2.664413905 | ENSG00000268295 |
| NA | -1.106518171 | 0.464413504 | 0.224180596 | 0.422414197 | 7.854802937 | 3.662516831 | ENSG00000228395 |
| LRRC49 | -1.106716444 | 0.464349683 | 3.94E-08 | 5.87E-07 | 202.5663553 | 94.16698879 | ENSG00000137821 |
| OR7E12P | -1.108428707 | 0.463798897 | 8.10E-06 | 7.92E-05 | 110.4082062 | 51.2307804 | ENSG00000189398 |
| NA | -1.108616652 | 0.46373848 | 0.218002908 | 0.414158635 | 7.889202453 | 3.674340108 | ENSG00000260643 |
| PPP2R5A | -1.108985664 | 0.46361988 | 3.64E-29 | 4.17E-27 | 881.216632 | 408.3598225 | ENSG00000066027 |
| NA | -1.111276216 | 0.46288438 | 0.084907498 | 0.21131762 | 15.75824328 | 7.313210386 | ENSG00000286018 |
| NA | -1.111751782 | 0.462731821 | 0.247833411 | 0.452799082 | 8.697577719 | 4.033178419 | ENSG00000284602 |
| SCUBE2 | -1.11201238 | 0.462648244 | 0.097535604 | 0.233810597 | 14.36360228 | 6.624477201 | ENSG00000175356 |
| INHBB | -1.115269693 | 0.461604856 | 0.307724434 | 0.51808647 | 5.78582526 | 2.652590629 | ENSG00000163083 |
| SLC7A11 | -1.116406371 | 0.461241308 | 1.19E-12 | 3.39E-11 | 466.2950678 | 214.9979888 | ENSG00000151012 |
| ABCB7 | -1.118017496 | 0.460726506 | 1.58E-31 | 1.99E-29 | 852.8402099 | 392.9028321 | ENSG00000131269 |
| NA | -1.11834845 | 0.460620827 | 0.019151482 | 0.066253112 | 30.96934887 | 14.22870766 | ENSG00000277103 |
| NA | -1.118679883 | 0.46051502 | 0.330003307 | 0.541451994 | 5.053367972 | 2.313047943 | ENSG00000270108 |
| CLIC2 | -1.120043445 | 0.46007997 | 0.232636119 | 0.433524088 | 7.225544198 | 3.339148349 | ENSG00000155962 |
| NA | -1.121099255 | 0.459743392 | 0.051121687 | 0.143537475 | 19.49733742 | 8.970819511 | ENSG00000271851 |
| LRRC4 | -1.121649491 | 0.459568082 | 1.90E-12 | 5.28E-11 | 296.37418 | 136.2115561 | ENSG00000128594 |
| CEP135 | -1.123004325 | 0.459136705 | 1.15E-10 | 2.56E-09 | 299.3677742 | 137.4175373 | ENSG00000174799 |
| NA | -1.123213508 | 0.459070137 | 0.069153282 | 0.181027096 | 17.33228014 | 7.927598942 | ENSG00000237398 |
| ZNF781 | -1.124462294 | 0.458672941 | 0.339741995 | 0.551394601 | 5.811631291 | 2.690235922 | ENSG00000196381 |
| TMSB15B | -1.126352275 | 0.458072457 | 0.005680809 | 0.024557385 | 50.73923882 | 23.2337675 | ENSG00000158427 |

| | | | | | | | |
|---|---|---|---|---|---|---|---|
| RIT2 | -1.127231652 | 0.45779333 | 0.114077964 | 0.263337288 | 13.78031119 | 6.330052158 | ENSG00000152214 |
| ZNF649 | -1.127405943 | 0.457738028 | 9.38E-22 | 6.49E-20 | 518.8936338 | 237.5022993 | ENSG00000198093 |
| EMC6 | -1.129483449 | 0.457079351 | 0.03992367 | 0.118528298 | 23.27675482 | 10.6318336 | ENSG00000127774 |
| NA | -1.130567405 | 0.456736058 | 0.195799537 | 0.384807823 | 8.717739345 | 3.960063296 | ENSG00000255139 |
| NA | -1.13083865 | 0.456650194 | 0.147905543 | 0.315735401 | 14.59105072 | 6.670540803 | ENSG00000248015 |
| LRRC37A3 | -1.131087863 | 0.456571318 | 0.020255796 | 0.069513123 | 28.36543818 | 12.97287903 | ENSG00000176809 |
| TSPYL4 | -1.131891999 | 0.456316903 | 7.76E-29 | 8.68E-27 | 875.577277 | 399.7051529 | ENSG00000187189 |
| NA | -1.1333193 | 0.455865678 | 2.91E-08 | 4.44E-07 | 160.1857592 | 73.08518962 | ENSG00000278922 |
| P2RX5-TAX1BP3 | -1.13414194 | 0.455605813 | 0.006162854 | 0.026325604 | 37.90695974 | 17.24448753 | ENSG00000257950 |
| SLC25A41 | -1.135162538 | 0.455283621 | 0.16555451 | 0.341432447 | 10.862635 | 4.935465628 | ENSG00000181240 |
| BCORL1 | -1.13535891 | 0.455221654 | 1.30E-16 | 5.67E-15 | 392.670259 | 178.74693 | ENSG00000085185 |
| NPY | -1.136487172 | 0.454865786 | 0.2348198 | 0.436301843 | 8.087285397 | 3.672164644 | ENSG00000122585 |
| SRSF12 | -1.137887381 | 0.45442453 | 4.50E-07 | 5.57E-06 | 154.2578606 | 70.17515683 | ENSG00000154548 |
| LINC00852 | -1.138760991 | 0.454149441 | 0.096611483 | 0.232270166 | 13.82419392 | 6.273111239 | ENSG00000231177 |
| LOC728673 | -1.139245293 | 0.453997012 | 0.146180596 | 0.313244133 | 11.68405295 | 5.284656126 | ENSG00000170161 |
| DLG1 | -1.139355164 | 0.453962438 | 3.23E-40 | 6.35E-38 | 1212.512439 | 550.3990524 | ENSG00000075711 |
| C18orf65 | -1.141361972 | 0.453331409 | 0.173628614 | 0.353642836 | 10.27606378 | 4.662795221 | ENSG00000275763 |
| NA | -1.142025322 | 0.453123016 | 0.023697855 | 0.078930157 | 26.42307467 | 11.99842265 | ENSG00000249637 |
| NA | -1.142414238 | 0.453000881 | 0.034187397 | 0.105340022 | 24.24733897 | 10.98849645 | ENSG00000230562 |
| NA | -1.1434211 | 0.45268484 | 0.270667753 | 0.479148905 | 6.527486426 | 2.950137094 | ENSG00000273486 |
| NA | -1.146131301 | 0.45183524 | 0.280274368 | 0.489230192 | 5.846030807 | 2.652590629 | ENSG00000235772 |
| NA | -1.147684114 | 0.451349178 | 0.253654825 | 0.459851336 | 6.595090249 | 2.973783647 | ENSG00000279722 |
| MAMDC4 | -1.148483828 | 0.451099056 | 0.001005415 | 0.005635878 | 341.5983399 | 154.1798381 | ENSG00000177943 |
| ZSCAN1 | -1.150196019 | 0.450564009 | 0.130245726 | 0.289307285 | 13.89179774 | 6.235465946 | ENSG00000152467 |
| NA | -1.15110589 | 0.450279939 | 0.034382581 | 0.105751635 | 25.17195547 | 11.30968947 | ENSG00000177337 |
| NA | -1.15200033 | 0.450000862 | 0.153180447 | 0.323906925 | 9.595803367 | 4.321077071 | ENSG00000213790 |
| NA | -1.152694529 | 0.449784382 | 0.157910221 | 0.330608653 | 9.557844378 | 4.295255055 | ENSG00000226468 |
| NA | -1.152753748 | 0.44976592 | 0.142921622 | 0.308324747 | 13.40250133 | 6.033735198 | ENSG00000274943 |
| EXTL3-AS1 | -1.152772561 | 0.449760054 | 0.267914937 | 0.475747399 | 8.146601221 | 3.646342627 | ENSG00000246339 |
| NA | -1.153266825 | 0.449605994 | 0.241397464 | 0.444977171 | 6.629489766 | 2.973783647 | ENSG00000230295 |
| GPR155 | -1.153650856 | 0.44948633 | 0.004079141 | 0.018684109 | 42.80849176 | 19.21581732 | ENSG00000163328 |
| SHD | -1.154793928 | 0.449130335 | 0.000444244 | 0.002776154 | 67.95349771 | 30.59673 | ENSG00000105251 |
| RAB7B | -1.156433081 | 0.448620334 | 0.198913254 | 0.389030323 | 9.572082268 | 4.286553201 | ENSG00000276600 |
| LTB4R2 | -1.158897312 | 0.447854711 | 0.060077469 | 0.162498282 | 19.28056739 | 8.62503398 | ENSG00000213906 |
| NA | -1.159417431 | 0.44769328 | 0.140525073 | 0.3047809 | 11.09956666 | 4.989285125 | ENSG00000235912 |
| NA | -1.160282791 | 0.447424825 | 0.033593315 | 0.103848379 | 23.02411073 | 10.33428714 | ENSG00000214189 |
| LRRC46 | -1.16111167 | 0.447167837 | 0.240736992 | 0.444366934 | 6.722009897 | 2.999605664 | ENSG00000141294 |
| RTF1 | -1.161287797 | 0.447113249 | 9.75E-63 | 4.85E-60 | 2309.374757 | 1032.778314 | ENSG00000137815 |
| NA | -1.161715755 | 0.446980638 | 0.173467989 | 0.353353789 | 8.952306737 | 4.01170733 | ENSG00000233859 |
| ROM1 | -1.162615825 | 0.446701862 | 0.112147015 | 0.260024111 | 13.32065962 | 5.921745277 | ENSG00000149489 |
| OLIG1 | -1.164137996 | 0.446230801 | 0.005673222 | 0.024539349 | 36.45864732 | 16.26255881 | ENSG00000184221 |
| ADAMTS19 | -1.16462347 | 0.446080667 | 2.07E-21 | 1.38E-19 | 569.8801511 | 254.1214054 | ENSG00000145808 |
| NA | -1.166455133 | 0.445514678 | 0.260574683 | 0.46750455 | 5.972950455 | 2.662238442 | ENSG00000227038 |
| NA | -1.166969227 | 0.44535595 | 0.042205339 | 0.123692124 | 21.61406278 | 9.610084125 | ENSG00000254461 |
| CASR | -1.167761585 | 0.445111418 | 0.10314377 | 0.243636034 | 13.40458626 | 5.989563514 | ENSG00000036828 |
| NA | -1.168330865 | 0.444935814 | 0.058015954 | 0.158213905 | 22.38328385 | 9.934682112 | ENSG00000219891 |
| SPACA4 | -1.168725003 | 0.444814276 | 0.290843254 | 0.500508005 | 5.897032477 | 2.612769872 | ENSG00000177202 |
| NA | -1.170969095 | 0.444122912 | 0.187060848 | 0.372423883 | 8.240011076 | 3.672164644 | ENSG00000272092 |
| GNA15 | -1.171975585 | 0.443813179 | 0.184019197 | 0.368057883 | 8.24713002 | 3.672164644 | ENSG00000060558 |
| NA | -1.172231322 | 0.443734514 | 0.021524175 | 0.073029832 | 25.43944783 | 11.2990957 | ENSG00000196741 |
| ZNF487 | -1.172454877 | 0.44366576 | 0.065359052 | 0.173187274 | 18.77199908 | 8.359552376 | ENSG00000243660 |
| SAMD15 | -1.172843378 | 0.443546302 | 0.101107323 | 0.239840353 | 14.84577974 | 6.561009892 | ENSG00000100583 |
| NA | -1.173059712 | 0.443479797 | 3.28E-28 | 3.56E-26 | 1312.207205 | 581.6921785 | ENSG00000236673 |
| NA | -1.173460696 | 0.443356553 | 0.320957496 | 0.531456561 | 5.948034148 | 2.628944076 | ENSG00000259051 |
| FOSB | -1.177959267 | 0.441976244 | 0.007709234 | 0.031732538 | 34.52312341 | 15.29245336 | ENSG00000125740 |
| ZFP28 | -1.17798185 | 0.441969326 | 0.076420524 | 0.195283477 | 15.77128596 | 6.989841904 | ENSG00000196867 |
| CXXC5-AS1 | -1.179404676 | 0.441533658 | 0.282859299 | 0.491799004 | 5.257095321 | 2.327046683 | ENSG00000250635 |
| LOC339260 | -1.179602078 | 0.441473248 | 2.80E-21 | 1.84E-19 | 472.3421313 | 208.5185553 | ENSG00000233098 |
| PODN | -1.180196247 | 0.441291466 | 0.27909086 | 0.488187802 | 5.260654793 | 2.327046683 | ENSG00000174348 |
| COBLL1 | -1.180306868 | 0.441257631 | 0.000236952 | 0.001601015 | 84.34575441 | 37.32354931 | ENSG00000082438 |
| NRN1 | -1.182605447 | 0.440555155 | 0.115821301 | 0.266125473 | 11.31544696 | 4.977461848 | ENSG00000124785 |
| NLGN4X | -1.183125964 | 0.440396234 | 1.81E-72 | 1.42E-69 | 4894.629026 | 2155.186244 | ENSG00000146938 |
| KBTBD11-OT1 | -1.185089324 | 0.439797307 | 0.282731363 | 0.491655782 | 5.260654793 | 2.313047943 | ENSG00000253696 |
| NA | -1.185303902 | 0.439731899 | 0.145950285 | 0.313034767 | 10.61382972 | 4.676793961 | ENSG00000248590 |
| DOK3 | -1.1875093 | 0.43906021 | 5.37E-07 | 6.53E-06 | 122.3653959 | 53.76920565 | ENSG00000146094 |
| NA | -1.187856765 | 0.438954477 | 0.281414197 | 0.490403904 | 5.302173254 | 2.329222147 | ENSG00000248476 |
| NA | -1.188569168 | 0.438737775 | 0.284626328 | 0.493859629 | 5.260654793 | 2.30340013 | ENSG00000280007 |
| NSMCE4A | -1.189346517 | 0.438501439 | 1.59E-37 | 2.71E-35 | 982.3048413 | 430.8704914 | ENSG00000107672 |
| C1orf229 | -1.190702135 | 0.438089598 | 0.22169412 | 0.418934663 | 8.263732174 | 3.604346407 | ENSG00000221953 |
| NA | -1.191476839 | 0.437854414 | 0.182364214 | 0.365832609 | 9.906288736 | 4.353425479 | ENSG00000279605 |
| NA | -1.19161783 | 0.437811626 | 0.286794764 | 0.496027028 | 5.27133321 | 2.301224667 | ENSG00000237233 |
| NA | -1.192620202 | 0.437507544 | 0.251665999 | 0.457343379 | 6.795258125 | 2.987782387 | ENSG00000274225 |
| TTI1 | -1.194102554 | 0.437058241 | 1.04E-40 | 2.10E-38 | 1363.702011 | 595.726038 | ENSG00000101407 |
| LOC100129484 | -1.195622367 | 0.436598062 | 0.290600528 | 0.50018146 | 5.311656463 | 2.313047943 | ENSG00000234432 |

| | | | | | | | |
|---|---|---|---|---|---|---|---|
| MMEL1 | -1.196819539 | 0.436235916 | 0.20311524 | 0.394592254 | 8.408449184 | 3.658165904 | ENSG00000142606 |
| HGSNAT | -1.197576563 | 0.436007071 | 1.33E-05 | 0.000123134 | 123.1527198 | 53.65910765 | ENSG00000165102 |
| NA | -1.19948785 | 0.43542983 | 0.05843382 | 0.159123675 | 22.19824359 | 9.681969743 | ENSG00000231160 |
| LOC642361 | -1.199674121 | 0.435373614 | 0.000762872 | 0.004427425 | 54.79207232 | 23.85156102 | ENSG00000272447 |
| CNTF | -1.200203333 | 0.435213938 | 0.18855069 | 0.374552717 | 7.638032907 | 3.322974145 | ENSG00000242689 |
| NA | -1.201054642 | 0.434957202 | 0.201936907 | 0.393192956 | 8.328971734 | 3.596874058 | ENSG00000279601 |
| LINC02523 | -1.201838097 | 0.434721063 | 0.032499762 | 0.101196703 | 20.64940237 | 8.963347161 | ENSG00000224506 |
| MEGF9 | -1.202593895 | 0.434493381 | 0.00085792 | 0.004896288 | 57.94014605 | 25.09244496 | ENSG00000106780 |
| LINC01431 | -1.20442572 | 0.433942044 | 0.274609804 | 0.483408478 | 5.384014968 | 2.329222147 | ENSG00000232645 |
| PCDHGB4 | -1.205034731 | 0.433758901 | 2.61E-06 | 2.79E-05 | 122.8226832 | 53.27066348 | ENSG00000253953 |
| NA | -1.20512058 | 0.433733091 | 0.341022698 | 0.552570884 | 4.631396053 | 2.019852405 | ENSG00000285676 |
| NA | -1.205121235 | 0.433732894 | 0.113170313 | 0.261679461 | 12.28603111 | 5.334124696 | ENSG00000258748 |
| LOC101927278 | -1.206520744 | 0.433312349 | 0.099580003 | 0.237418485 | 17.62377294 | 7.629106518 | ENSG00000287261 |
| NA | -1.209241005 | 0.43249609 | 1.54E-10 | 3.34E-09 | 254.4881681 | 110.0545351 | ENSG00000262001 |
| DLGAP4-AS1 | -1.209500039 | 0.432418443 | 0.096021839 | 0.231120626 | 12.25875054 | 5.291182517 | ENSG00000232907 |
| TMEM255A | -1.210195621 | 0.432210007 | 0.001634558 | 0.008553237 | 46.75578859 | 20.22356806 | ENSG00000125355 |
| LOC101927124 | -1.210656124 | 0.432072069 | 0.275571941 | 0.484515046 | 6.140193355 | 2.624593148 | ENSG00000250365 |
| NA | -1.211226975 | 0.431901139 | 0.067712454 | 0.178011494 | 16.15293463 | 6.990787862 | ENSG00000234373 |
| HIC1 | -1.216169513 | 0.430424018 | 0.23033859 | 0.430304113 | 6.887778257 | 2.959784907 | ENSG00000177374 |
| CEP290 | -1.216370959 | 0.430363921 | 4.69E-11 | 1.10E-09 | 208.2721189 | 89.57530119 | ENSG00000198707 |
| PLS3-AS1 | -1.216506208 | 0.430323577 | 0.208048981 | 0.400714431 | 7.770876291 | 3.334797422 | ENSG00000271826 |
| SHISA8 | -1.216826689 | 0.430227995 | 0.094711131 | 0.229196202 | 13.14985725 | 5.682369236 | ENSG00000234965 |
| NA | -1.216871064 | 0.430214763 | 0.269409701 | 0.477414112 | 6.152956704 | 2.662238442 | ENSG00000267898 |
| ANKRD45 | -1.219457081 | 0.429444297 | 0.338761546 | 0.550644635 | 5.322334881 | 2.263579373 | ENSG00000183831 |
| CDK8 | -1.224990837 | 0.42780023 | 2.62E-25 | 2.42E-23 | 707.4400243 | 302.5576205 | ENSG00000132964 |
| NA | -1.225184529 | 0.427742799 | 0.217567619 | 0.413564618 | 7.045537948 | 3.015779867 | ENSG00000272501 |
| NA | -1.225235131 | 0.427727796 | 0.26068617 | 0.467615722 | 7.059775838 | 3.023252217 | ENSG00000260498 |
| GEMIN7-AS1 | -1.229202254 | 0.426553246 | 0.202031111 | 0.393281894 | 7.018257377 | 2.985606923 | ENSG00000267348 |
| NA | -1.229906277 | 0.426345142 | 0.059192008 | 0.160725365 | 18.62135833 | 7.956542381 | ENSG00000237087 |
| C19orf57 | -1.233722992 | 0.425218717 | 6.13E-05 | 0.000481938 | 90.05713623 | 38.39353785 | ENSG00000132016 |
| SLFN11 | -1.233745711 | 0.425212021 | 0.033355642 | 0.103231942 | 19.51869426 | 8.302611457 | ENSG00000172716 |
| NA | -1.235299071 | 0.424754439 | 0.246490858 | 0.451326338 | 6.275121671 | 2.674061718 | ENSG00000285454 |
| NA | -1.235908067 | 0.424575178 | 0.040733851 | 0.12044133 | 20.26508395 | 8.625979939 | ENSG00000286606 |
| TMEM45A | -1.236283544 | 0.424464692 | 3.93E-15 | 1.51E-13 | 282.3216218 | 119.9003067 | ENSG00000181458 |
| ELP2 | -1.237009217 | 0.42425124 | 1.56E-68 | 1.02E-65 | 1998.193114 | 847.6341106 | ENSG00000134759 |
| BIVM | -1.237740167 | 0.424036345 | 1.68E-05 | 0.000152477 | 92.19193829 | 39.0226602 | ENSG00000134897 |
| DOCK2 | -1.239299566 | 0.423578255 | 3.07E-07 | 3.93E-06 | 118.4014969 | 50.1897353 | ENSG00000134516 |
| BOC | -1.2406452 | 0.423183358 | 1.22E-10 | 2.70E-09 | 189.6050978 | 80.18652698 | ENSG00000144857 |
| ZNF583 | -1.242370244 | 0.422677656 | 0.007474831 | 0.030922713 | 32.91171238 | 13.86769389 | ENSG00000198440 |
| OVOL2 | -1.243255252 | 0.422418448 | 4.02E-12 | 1.07E-10 | 221.482766 | 93.62682927 | ENSG00000125850 |
| NA | -1.243792722 | 0.422261107 | 0.148288163 | 0.316146078 | 10.30098008 | 4.365248755 | ENSG00000214776 |
| CUL4B | -1.244274522 | 0.422120113 | 5.90E-32 | 7.48E-30 | 1566.319497 | 660.8771843 | ENSG00000158290 |
| NA | -1.24509862 | 0.421879058 | 0.012936014 | 0.04831816 | 28.26727365 | 11.88766224 | ENSG00000274964 |
| NA | -1.250248475 | 0.4203758 | 0.148377119 | 0.316264323 | 11.83202394 | 4.98058327 | ENSG00000273008 |
| LAMA4 | -1.255464389 | 0.418858719 | 0.012534822 | 0.047136622 | 27.65314252 | 11.58358938 | ENSG00000112769 |
| CACNA1G | -1.255668869 | 0.418799356 | 2.07E-16 | 8.98E-15 | 340.3771711 | 142.5643089 | ENSG00000006283 |
| FAM133A | -1.256566909 | 0.418538746 | 0.032699092 | 0.101699991 | 21.49306687 | 9.02028808 | ENSG00000179083 |
| HERC2P2 | -1.257573081 | 0.418246948 | 0.000203789 | 0.00140095 | 1843.871717 | 771.2489481 | ENSG00000276550 |
| NA | -1.258189709 | 0.418068222 | 0.272278732 | 0.480918805 | 5.540300119 | 2.32487122 | ENSG00000229237 |
| NA | -1.260881671 | 0.417288865 | 0.278169673 | 0.487493901 | 6.329682814 | 2.628944076 | ENSG00000275956 |
| H2BC21 | -1.26490314 | 0.416127304 | 0.000102261 | 0.000757332 | 80.39372904 | 33.46098273 | ENSG00000184678 |
| ZNF460-AS1 | -1.265173583 | 0.416049306 | 0.24052098 | 0.444055011 | 5.580623371 | 2.329222147 | ENSG00000267871 |
| MIR4500HG | -1.265747376 | 0.415883867 | 0.252939765 | 0.458951524 | 5.577063899 | 2.32487122 | ENSG00000228824 |
| NA | -1.266666344 | 0.415619041 | 0.231025037 | 0.431331045 | 7.924797178 | 3.283153389 | ENSG00000280233 |
| LTB4R | -1.266723492 | 0.415602578 | 2.82E-05 | 0.000241969 | 111.581628 | 46.43936956 | ENSG00000213903 |
| JAKMIP2 | -1.268836717 | 0.414994259 | 1.29E-14 | 4.58E-13 | 283.651608 | 117.6412462 | ENSG00000176049 |
| MIR9-3HG | -1.269556291 | 0.414787324 | 0.17323083 | 0.352984919 | 9.694247235 | 4.030056997 | ENSG00000255571 |
| SLC30A10 | -1.269655185 | 0.414758892 | 0.169838083 | 0.347981017 | 8.036283727 | 3.327325073 | ENSG00000196660 |
| GRIN2B | -1.272783111 | 0.413860622 | 6.57E-05 | 0.000512169 | 74.42700781 | 30.7670583 | ENSG00000273079 |
| NA | -1.272828682 | 0.413847549 | 0.242805491 | 0.446696684 | 7.250460505 | 3.01142894 | ENSG00000279179 |
| LOC105376748 | -1.272987724 | 0.413801929 | 0.138416402 | 0.301672677 | 9.605286576 | 3.967535646 | ENSG00000233654 |
| UBE2E1-AS1 | -1.274036814 | 0.413501132 | 0.190108492 | 0.376488975 | 7.211306308 | 2.989957851 | ENSG00000223791 |
| PRR18 | -1.280159064 | 0.411750109 | 0.156771429 | 0.328887732 | 8.872829288 | 3.664692295 | ENSG00000176381 |
| TSFM | -1.280806907 | 0.411565254 | 0.238872531 | 0.441572803 | 5.615022888 | 2.30340013 | ENSG00000123297 |
| NA | -1.280971671 | 0.411518253 | 0.005404845 | 0.023583757 | 33.02172439 | 13.58726758 | ENSG00000235494 |
| PCDHGA1 | -1.281217245 | 0.411448211 | 0.039320728 | 0.11718036 | 20.13282538 | 8.236968684 | ENSG00000204956 |
| TTC23L | -1.282881901 | 0.410973735 | 0.117369388 | 0.26870177 | 11.30240428 | 4.636973204 | ENSG00000205838 |
| LINC01524 | -1.286989126 | 0.409805393 | 0.311088225 | 0.521747681 | 4.837487666 | 1.994030388 | ENSG00000234948 |
| ADM | -1.289925587 | 0.408972123 | 1.32E-51 | 4.29E-49 | 1142.51727 | 467.3380577 | ENSG00000148926 |
| NA | -1.290000455 | 0.4089509 | 0.226929972 | 0.42578962 | 6.478849019 | 2.666589369 | ENSG00000227848 |
| IGSF11 | -1.291284908 | 0.408586967 | 0.016822689 | 0.059807318 | 23.55640014 | 9.614435052 | ENSG00000144847 |
| TMEM202-AS1 | -1.292871107 | 0.408137985 | 0.192589356 | 0.379967613 | 7.3453449 | 2.995254736 | ENSG00000261423 |
| NA | -1.294715464 | 0.40761655 | 0.248527217 | 0.453472125 | 7.390422834 | 3.013604404 | ENSG00000224195 |
| ZNF528-AS1 | -1.297874693 | 0.406724924 | 0.233827294 | 0.435014132 | 6.567809678 | 2.688060458 | ENSG00000269834 |

| | | | | | | | |
|---|---|---|---|---|---|---|---|
| SPAG1 | -1.2990538 | 0.406392646 | 0.001791578 | 0.009236679 | 41.60898461 | 16.91147123 | ENSG00000104450 |
| NA | -1.300293512 | 0.406043581 | 0.289930577 | 0.499255854 | 4.93356727 | 2.015501478 | ENSG00000215414 |
| CAPS2 | -1.300555217 | 0.405969932 | 0.026768553 | 0.086949454 | 20.48452373 | 8.334959865 | ENSG00000180881 |
| NA | -1.300820698 | 0.405895233 | 0.263146413 | 0.470286743 | 4.913405643 | 2.005853665 | ENSG00000286281 |
| NA | -1.302247228 | 0.405494084 | 0.20108589 | 0.391940038 | 6.520367481 | 2.652590629 | ENSG00000279696 |
| NA | -1.304663444 | 0.404815534 | 0.15021949 | 0.319254452 | 8.18189046 | 3.322974145 | ENSG00000212789 |
| HSPB6 | -1.305589378 | 0.404555803 | 0.168215702 | 0.34567503 | 8.151050417 | 3.301503056 | ENSG00000004776 |
| NA | -1.307971611 | 0.403888336 | 0.264669264 | 0.472337842 | 5.738383063 | 2.299049203 | ENSG00000230185 |
| LOC105374618 | -1.307998271 | 0.403880873 | 0.052536927 | 0.146585313 | 16.43850369 | 6.655596104 | ENSG00000248994 |
| NA | -1.30809574 | 0.403853587 | 0.091955561 | 0.223901594 | 11.47765585 | 4.644445553 | ENSG00000253549 |
| UNC45B | -1.31058815 | 0.403156489 | 0.187829126 | 0.373590945 | 7.372625471 | 2.94796163 | ENSG00000141161 |
| NA | -1.313894562 | 0.402233581 | 0.156155698 | 0.328104179 | 8.308810108 | 3.353147089 | ENSG00000241168 |
| NA | -1.314452287 | 0.402078114 | 0.042528055 | 0.124441017 | 17.32724613 | 6.997314253 | ENSG00000235245 |
| AVP | -1.314856126 | 0.40196558 | 0.206787736 | 0.399262489 | 6.544088579 | 2.624593148 | ENSG00000101200 |
| NA | -1.315791295 | 0.401705106 | 0.02657603 | 0.086398404 | 19.74970218 | 7.927598942 | ENSG00000272931 |
| NA | -1.319367037 | 0.400710707 | 0.014899394 | 0.054242511 | 28.31115637 | 11.35698258 | ENSG00000285679 |
| NA | -1.319400861 | 0.400701312 | 0.142026096 | 0.306988991 | 9.126668583 | 3.632343887 | ENSG00000228060 |
| LOC728975 | -1.319893069 | 0.400564627 | 0.022357466 | 0.075315867 | 23.92264186 | 9.544441352 | ENSG00000254614 |
| FRZB | -1.321041892 | 0.400245783 | 1.10E-10 | 2.46E-09 | 170.1139635 | 68.06790701 | ENSG00000162998 |
| LOC102724474 | -1.321206537 | 0.400200108 | 0.050502023 | 0.142220505 | 15.7689217 | 6.332227622 | ENSG00000283828 |
| KCND1 | -1.324232453 | 0.399361606 | 6.94E-15 | 2.57E-13 | 302.6964646 | 121.0514498 | ENSG00000102057 |
| S100A6 | -1.324260354 | 0.399353883 | 0.003503188 | 0.01641223 | 34.80276874 | 13.91281153 | ENSG00000197956 |
| DRD2 | -1.325589966 | 0.398986001 | 0.070350176 | 0.183477844 | 13.27677689 | 5.281534704 | ENSG00000149295 |
| NA | -1.32694057 | 0.398612658 | 0.195857786 | 0.38481354 | 6.687610381 | 2.662238442 | ENSG00000233514 |
| NA | -1.328898524 | 0.398072048 | 0.286144419 | 0.495345072 | 4.937126742 | 1.952034168 | ENSG00000232732 |
| NA | -1.332609711 | 0.397049364 | 0.259918072 | 0.46681434 | 5.873311379 | 2.347571814 | ENSG00000285669 |
| NA | -1.334497617 | 0.396530126 | 0.178617672 | 0.360846677 | 7.503994315 | 2.98343146 | ENSG00000236264 |
| NA | -1.334938854 | 0.396408869 | 0.14442252 | 0.310446807 | 10.06641269 | 3.978412964 | ENSG00000269318 |
| PAPLN | -1.335146431 | 0.396351837 | 4.10E-10 | 8.36E-09 | 172.8116157 | 68.47621389 | ENSG00000100767 |
| GSAP | -1.335206312 | 0.396335386 | 0.071731318 | 0.186359326 | 14.21296153 | 5.629495698 | ENSG00000186088 |
| LILRB3 | -1.336377905 | 0.396013659 | 0.272942097 | 0.48159502 | 5.894668214 | 2.347571814 | ENSG00000204577 |
| ZNF582-AS1 | -1.337700567 | 0.39565076 | 0.010926625 | 0.042060603 | 26.08085953 | 10.29881731 | ENSG00000267454 |
| MID2 | -1.337937875 | 0.395585685 | 0.000165752 | 0.001165383 | 72.40427712 | 28.66645047 | ENSG00000080561 |
| CP | -1.338761171 | 0.395360002 | 0.038156117 | 0.114541134 | 18.5187446 | 7.290509791 | ENSG00000047457 |
| ZAR1L | -1.339657386 | 0.395114477 | 0.299978698 | 0.510168792 | 5.118607532 | 2.022027868 | ENSG00000189167 |
| DKFZP434A062 | -1.340685674 | 0.394832958 | 0.268624036 | 0.476550449 | 5.040325291 | 1.977856185 | ENSG00000262075 |
| LINC01347 | -1.341263796 | 0.394674771 | 0.188499414 | 0.374519876 | 6.756409413 | 2.652590629 | ENSG00000214837 |
| CHRM4 | -1.341664583 | 0.394565143 | 0.034831054 | 0.106852575 | 18.59911177 | 7.353977101 | ENSG00000180720 |
| C17orf49 | -1.34208325 | 0.394450658 | 0.080350321 | 0.2023772 | 12.6546371 | 4.989285125 | ENSG00000258315 |
| ALDH1L1 | -1.343127377 | 0.394165284 | 0.001972539 | 0.010029585 | 37.83104176 | 14.87762009 | ENSG00000144908 |
| NA | -1.344113259 | 0.393896019 | 0.233150285 | 0.434097722 | 5.934991466 | 2.350693236 | ENSG00000229271 |
| NA | -1.344662874 | 0.393745987 | 5.87E-06 | 5.91E-05 | 92.7791199 | 36.61429099 | ENSG00000182057 |
| NA | -1.347379342 | 0.393005296 | 0.006007731 | 0.02577374 | 28.6913305 | 11.28604292 | ENSG00000285825 |
| MTTP | -1.347697865 | 0.392918536 | 0.154656354 | 0.326187764 | 7.630913962 | 3.003956591 | ENSG00000138823 |
| NA | -1.347894507 | 0.392864984 | 0.162534313 | 0.337566928 | 7.583471765 | 2.978134574 | ENSG00000250299 |
| LINC02328 | -1.350043284 | 0.39228028 | 0.210349711 | 0.403868879 | 5.918389312 | 2.327046683 | ENSG00000258733 |
| LETM2 | -1.351361785 | 0.391921932 | 0.010809838 | 0.041661999 | 24.5519006 | 9.610084125 | ENSG00000165046 |
| MT2A | -1.352850394 | 0.391517746 | 5.76E-34 | 8.36E-32 | 670.1685531 | 262.4112906 | ENSG00000125148 |
| SRRM2-AS1 | -1.3548472 | 0.390976229 | 0.038886003 | 0.116251729 | 16.16361305 | 6.338754013 | ENSG00000205913 |
| NA | -1.35634688 | 0.390570021 | 0.065933564 | 0.174415934 | 13.67146824 | 5.345002014 | ENSG00000280046 |
| NA | -1.35675247 | 0.390460234 | 0.224613881 | 0.422914571 | 5.95159362 | 2.32487122 | ENSG00000270532 |
| NA | -1.361206991 | 0.389256493 | 0.015379659 | 0.055587534 | 24.65982768 | 9.558440092 | ENSG00000220848 |
| NEK10 | -1.365304926 | 0.38815239 | 0.226273904 | 0.425065421 | 6.051232696 | 2.3528687 | ENSG00000163491 |
| NA | -1.3662984 | 0.38788519 | 1.09E-05 | 0.000103142 | 84.09516968 | 32.67352332 | ENSG00000278133 |
| B4GALNT1 | -1.367095268 | 0.387671003 | 0.001578428 | 0.008324101 | 42.97665053 | 16.70851098 | ENSG00000135454 |
| NA | -1.369152585 | 0.387118569 | 0.211145382 | 0.404820359 | 5.976509927 | 2.30340013 | ENSG00000286529 |
| HACL1 | -1.369513293 | 0.387021792 | 1.14E-36 | 1.88E-34 | 715.7034777 | 277.082829 | ENSG00000131373 |
| NA | -1.372534377 | 0.386212194 | 0.090011651 | 0.220447418 | 12.02626808 | 4.672443033 | ENSG00000240668 |
| SPOCK2 | -1.373457399 | 0.385965179 | 6.97E-05 | 0.000539218 | 67.94960718 | 26.17454032 | ENSG00000107742 |
| NA | -1.375256033 | 0.385484289 | 0.192379216 | 0.379705144 | 6.939975135 | 2.690235922 | ENSG00000260269 |
| NA | -1.37566975 | 0.385373761 | 0.207454239 | 0.399954614 | 6.034630542 | 2.315223407 | ENSG00000224126 |
| LIG3 | -1.375888477 | 0.385315339 | 1.78E-91 | 3.36E-88 | 2292.77463 | 883.3355206 | ENSG00000005156 |
| PARP16 | -1.377382864 | 0.384916424 | 1.18E-08 | 1.92E-07 | 134.6621054 | 51.92074309 | ENSG00000138617 |
| PRTG | -1.378045233 | 0.384739742 | 1.79E-07 | 2.36E-06 | 104.2502155 | 40.07117765 | ENSG00000166450 |
| NA | -1.38272392 | 0.383494045 | 0.273742811 | 0.482376877 | 5.196610441 | 2.005853665 | ENSG00000232748 |
| NA | -1.383483983 | 0.38329206 | 0.187134735 | 0.37247465 | 6.919813509 | 2.654766092 | ENSG00000267342 |
| PTGER2 | -1.385879587 | 0.382656129 | 0.000127693 | 0.000923582 | 52.90190569 | 20.24939008 | ENSG00000125384 |
| NA | -1.386094225 | 0.382599204 | 0.160709357 | 0.334840711 | 7.831081838 | 3.017955331 | ENSG00000261572 |
| NA | -1.387103164 | 0.382331729 | 0.16040261 | 0.334424372 | 6.956577289 | 2.662238442 | ENSG00000276436 |
| PDE10A | -1.387467671 | 0.382235142 | 0.000612501 | 0.003659512 | 47.02887364 | 17.96216415 | ENSG00000112541 |
| NA | -1.38855077 | 0.381948288 | 0.050941564 | 0.143223378 | 13.88231453 | 5.298654867 | ENSG00000261324 |
| STEAP3-AS1 | -1.388565823 | 0.381944303 | 0.106276372 | 0.249790372 | 10.42434026 | 3.957887833 | ENSG00000229867 |
| ABCA5 | -1.389215252 | 0.381772409 | 0.000213559 | 0.001457031 | 50.46432203 | 19.25251666 | ENSG00000154265 |
| CTNNAL1 | -1.392649616 | 0.380864673 | 3.67E-65 | 2.10E-62 | 1412.503944 | 537.9463545 | ENSG00000119326 |

| | | | | | | | |
|---|---|---|---|---|---|---|---|
| GPR50 | -1.393006573 | 0.38077045 | 2.75E-12 | 7.47E-11 | 195.1385845 | 74.24643204 | ENSG00000102195 |
| ADAM20P1 | -1.393393798 | 0.380668263 | 0.231970247 | 0.43275295 | 6.174592871 | 2.341045423 | ENSG00000259158 |
| NA | -1.393493185 | 0.38064204 | 0.271274572 | 0.479683548 | 5.278452155 | 2.022027868 | ENSG00000284959 |
| TRH | -1.397021636 | 0.379712228 | 0.257321801 | 0.464226425 | 6.111438243 | 2.336694496 | ENSG00000170893 |
| LHX2 | -1.397347492 | 0.379626473 | 0.122834927 | 0.277182172 | 10.56282805 | 4.028827492 | ENSG00000106689 |
| NA | -1.3980384 | 0.379444713 | 0.104592942 | 0.246473384 | 9.670526136 | 3.672164644 | ENSG00000287837 |
| PDGFA | -1.399284054 | 0.379117234 | 8.96E-41 | 1.84E-38 | 770.0551965 | 291.7934828 | ENSG00000197461 |
| NA | -1.399407732 | 0.379084735 | 0.06850713 | 0.179655536 | 12.2990738 | 4.665916643 | ENSG00000280087 |
| NA | -1.399903458 | 0.378954499 | 0.147311477 | 0.314867834 | 7.912923552 | 3.01142894 | ENSG00000281468 |
| KCNJ12 | -1.400235719 | 0.378867234 | 5.37E-08 | 7.83E-07 | 111.3678326 | 42.24908358 | ENSG00000184185 |
| CALCB | -1.402086035 | 0.378381633 | 7.01E-09 | 1.19E-07 | 126.272902 | 47.79051945 | ENSG00000175868 |
| NA | -1.402825532 | 0.378187732 | 0.249805927 | 0.455057869 | 5.282011628 | 2.003678201 | ENSG00000236972 |
| H2BC10 | -1.402871698 | 0.37817563 | 0.265357593 | 0.472940557 | 5.308096991 | 2.015501478 | ENSG00000278588 |
| PNPLA4 | -1.402988367 | 0.378145048 | 2.74E-26 | 2.67E-24 | 506.8192614 | 191.5404953 | ENSG00000006757 |
| NA | -1.403754723 | 0.377944232 | 0.040331609 | 0.119532711 | 15.82704231 | 5.996089905 | ENSG00000226361 |
| NKX2-4 | -1.404481325 | 0.377753931 | 0.214875675 | 0.409891629 | 5.291494837 | 2.005853665 | ENSG00000125816 |
| LOC729603 | -1.406953119 | 0.377107273 | 0.049079894 | 0.139065837 | 15.90296029 | 6.010088645 | ENSG00000213073 |
| NA | -1.411627855 | 0.375887318 | 0.208654114 | 0.401653693 | 6.187356221 | 2.341045423 | ENSG00000230105 |
| NA | -1.41204136 | 0.375779596 | 0.201357816 | 0.392348571 | 6.19683943 | 2.34539635 | ENSG00000272054 |
| MTSS1 | -1.412679022 | 0.375613541 | 5.49E-08 | 7.97E-07 | 108.7570823 | 40.88996688 | ENSG00000170873 |
| NA | -1.413092102 | 0.375506009 | 0.246566634 | 0.451421317 | 5.349615452 | 2.013326014 | ENSG00000259869 |
| LOC101928521 | -1.414304445 | 0.375190591 | 0.048906403 | 0.138657471 | 14.18212149 | 5.33630016 | ENSG00000242686 |
| LINC02696 | -1.416578901 | 0.374599557 | 0.127663534 | 0.285147941 | 8.012562628 | 3.001781127 | ENSG00000254427 |
| NA | -1.417773374 | 0.374289538 | 0.134762978 | 0.296378112 | 7.985282057 | 2.98343146 | ENSG00000269935 |
| SCML1 | -1.418803547 | 0.374022367 | 0.000174768 | 0.001220146 | 53.19370397 | 19.84855555 | ENSG00000047634 |
| LAMP5 | -1.419637106 | 0.373806327 | 0.001454702 | 0.007753896 | 35.58770224 | 13.29624751 | ENSG00000125869 |
| LOC105376114 | -1.420134984 | 0.373677348 | 0.055604945 | 0.15312564 | 14.28176056 | 5.348123436 | ENSG00000254473 |
| CFAP69 | -1.423192589 | 0.372886226 | 0.001078637 | 0.005989013 | 44.49821118 | 16.61297881 | ENSG00000105792 |
| NA | -1.423398151 | 0.372833099 | 0.001217339 | 0.006655893 | 41.79374555 | 15.60087714 | ENSG00000286707 |
| NA | -1.426778427 | 0.371960563 | 0.174137599 | 0.354373768 | 8.094404342 | 3.008307518 | ENSG00000279744 |
| NA | -1.42762085 | 0.37174343 | 7.87E-12 | 2.02E-10 | 237.3566662 | 88.36442224 | ENSG00000259203 |
| NA | -1.42774344 | 0.371711843 | 0.083723255 | 0.208976165 | 10.75379205 | 3.983709849 | ENSG00000285602 |
| NA | -1.429996527 | 0.371131786 | 0.10129858 | 0.240118448 | 9.810209133 | 3.650693555 | ENSG00000248489 |
| SNORD62B | -1.430805186 | 0.370923818 | 0.1071377 | 0.251434754 | 8.986706254 | 3.336972886 | ENSG00000231587 |
| NA | -1.431754801 | 0.370679747 | 0.071302075 | 0.185487155 | 11.62237286 | 4.297430518 | ENSG00000225506 |
| TFAP2E | -1.432490428 | 0.370490787 | 0.097436841 | 0.233633156 | 10.84987165 | 4.01170733 | ENSG00000116819 |
| TPP2 | -1.432811033 | 0.370408463 | 1.94E-81 | 2.15E-78 | 1863.092078 | 689.9000047 | ENSG00000134900 |
| MAP3K7 | -1.4370619 | 0.36931867 | 9.35E-65 | 5.19E-62 | 1297.395848 | 479.253055 | ENSG00000135341 |
| PRR22 | -1.439570516 | 0.368677042 | 0.001913043 | 0.009767243 | 35.09337182 | 12.93211231 | ENSG00000212123 |
| NA | -1.443365545 | 0.367708506 | 0.148585315 | 0.316529461 | 7.252824769 | 2.680588109 | ENSG00000248126 |
| NA | -1.44342989 | 0.367692106 | 0.17172653 | 0.350980034 | 6.289359561 | 2.30340013 | ENSG00000260992 |
| ADAP2 | -1.445432647 | 0.367182028 | 0.026457396 | 0.086086824 | 17.20536049 | 6.326930736 | ENSG00000184060 |
| FOXB2 | -1.44898717 | 0.366278476 | 0.022767649 | 0.076345309 | 18.17950412 | 6.678296698 | ENSG00000204612 |
| INTS6L | -1.450549709 | 0.365881986 | 1.29E-17 | 6.16E-16 | 296.8510186 | 108.7038367 | ENSG00000165359 |
| CTU2 | -1.450996166 | 0.365768777 | 2.81E-28 | 3.06E-26 | 527.1360609 | 192.9733895 | ENSG00000174177 |
| TMEM121B | -1.452191036 | 0.365465965 | 1.04E-15 | 4.30E-14 | 242.1499918 | 88.47173473 | ENSG00000183307 |
| NA | -1.452713264 | 0.365333698 | 0.161120823 | 0.335549928 | 7.221984725 | 2.634240961 | ENSG00000271918 |
| ZDHHC9 | -1.45291389 | 0.365282897 | 2.30E-80 | 2.41E-77 | 1720.445658 | 628.3858461 | ENSG00000188706 |
| NA | -1.452947637 | 0.365274352 | 0.042406947 | 0.124202109 | 13.65130661 | 4.979637312 | ENSG00000231713 |
| NA | -1.457724763 | 0.364066837 | 0.021135904 | 0.071984047 | 18.31354271 | 6.673945771 | ENSG00000269993 |
| LTK | -1.461820474 | 0.363034743 | 0.240449196 | 0.443996948 | 5.511824339 | 2.019852405 | ENSG00000062524 |
| D21S2088E | -1.466417971 | 0.361879686 | 3.35E-44 | 7.91E-42 | 750.2168386 | 271.5318179 | ENSG00000228592 |
| NA | -1.466472409 | 0.361866031 | 0.122617288 | 0.276790279 | 9.201112019 | 3.344445235 | ENSG00000272764 |
| SHISA3 | -1.467916323 | 0.36150404 | 4.02E-07 | 5.03E-06 | 101.791606 | 36.86236889 | ENSG00000178343 |
| SLITRK6 | -1.471987771 | 0.360485273 | 0.094229239 | 0.228319114 | 11.01772494 | 3.991182199 | ENSG00000184564 |
| SLC5A2 | -1.472463443 | 0.360366437 | 0.066438667 | 0.175629091 | 12.01350473 | 4.326373957 | ENSG00000140675 |
| ZBTB33 | -1.474508592 | 0.359855948 | 5.36E-46 | 1.37E-43 | 830.272243 | 298.7715014 | ENSG00000177485 |
| LOC101928837 | -1.478255523 | 0.358922552 | 0.090635617 | 0.22152675 | 9.272275316 | 3.334797422 | ENSG00000255100 |
| ORM1 | -1.480702902 | 0.358314194 | 0.053470348 | 0.148367252 | 13.79691335 | 4.932344206 | ENSG00000229314 |
| LOC101928963 | -1.481619464 | 0.358086624 | 0.211709659 | 0.405531687 | 5.5533428 | 1.996205852 | ENSG00000272677 |
| NA | -1.481959493 | 0.358002237 | 0.030032107 | 0.095048825 | 15.74756486 | 5.654371756 | ENSG00000262877 |
| GRIN1 | -1.483087695 | 0.357722385 | 5.87E-12 | 1.53E-10 | 156.8713068 | 56.13730259 | ENSG00000176884 |
| THBD | -1.484920431 | 0.357268239 | 0.021569032 | 0.073140022 | 19.46057364 | 6.948791642 | ENSG00000178726 |
| NA | -1.489388362 | 0.356163514 | 0.11996288 | 0.272641801 | 11.0355223 | 3.922418003 | ENSG00000230042 |
| ZNF568 | -1.490993038 | 0.355767582 | 0.000336557 | 0.002169244 | 42.94964929 | 15.2526326 | ENSG00000198453 |
| NA | -1.500886248 | 0.353336269 | 0.188742239 | 0.374651129 | 5.587742316 | 1.966032908 | ENSG00000217275 |
| NA | -1.501798977 | 0.3531128 | 0.195816954 | 0.384807823 | 5.611463415 | 1.975680721 | ENSG00000185065 |
| NA | -1.501890289 | 0.353090451 | 0.178793274 | 0.361085549 | 5.673143503 | 2.005853665 | ENSG00000262410 |
| ARHGAP36 | -1.504821524 | 0.352373778 | 0.004366431 | 0.019736729 | 25.44656678 | 8.977345901 | ENSG00000147256 |
| NA | -1.514564575 | 0.350002086 | 0.048542081 | 0.137935178 | 13.21509681 | 4.612097146 | ENSG00000279329 |
| NMU | -1.518010134 | 0.34916718 | 3.22E-15 | 1.25E-13 | 215.0534185 | 75.07297716 | ENSG00000109255 |
| NA | -1.521257858 | 0.348382037 | 0.047678208 | 0.135971453 | 13.2530558 | 4.59280152 | ENSG00000259529 |
| PPP1R3G | -1.521902706 | 0.348226353 | 0.150178925 | 0.319204163 | 7.558555457 | 2.622417685 | ENSG00000219607 |
| LINC00648 | -1.522328888 | 0.3481235 | 0.003864284 | 0.017842787 | 31.40260959 | 10.95832351 | ENSG00000259129 |

| | | | | | | | |
|---|---|---|---|---|---|---|---|
| NA | -1.525932122 | 0.347255122 | 0.195277197 | 0.38410685 | 5.731264118 | 1.975680721 | ENSG00000275329 |
| FOXL2 | -1.529495492 | 0.346398481 | 0.00310929 | 0.014850518 | 29.80068177 | 10.34611042 | ENSG00000183770 |
| CCT6B | -1.529946794 | 0.346290138 | 0.034533504 | 0.106135066 | 15.24522577 | 5.253537224 | ENSG00000132141 |
| LOC730183 | -1.536507614 | 0.344718919 | 0.02272575 | 0.076243702 | 17.27624446 | 5.975564774 | ENSG00000261840 |
| ZNF878 | -1.541884191 | 0.343436626 | 2.51E-05 | 0.000218398 | 57.98939384 | 19.95184361 | ENSG00000257446 |
| ZNF880 | -1.544512279 | 0.342811573 | 8.29E-13 | 2.41E-11 | 173.0725733 | 59.31622196 | ENSG00000221923 |
| CMPK2 | -1.546812848 | 0.34226535 | 0.013785579 | 0.050827929 | 19.3194161 | 6.620126274 | ENSG00000134326 |
| CHRD | -1.553648511 | 0.340647492 | 0.011536911 | 0.044023956 | 20.56044171 | 7.025311733 | ENSG00000090539 |
| MMUT | -1.556148267 | 0.340057763 | 6.29E-31 | 7.71E-29 | 497.0627749 | 168.9147579 | ENSG00000146085 |
| COL15A1 | -1.55635974 | 0.340007921 | 0.105018321 | 0.247383184 | 8.747104848 | 2.987782387 | ENSG00000204291 |
| SHBG | -1.557943454 | 0.339634883 | 0.036714232 | 0.111184776 | 14.7618531 | 5.015107141 | ENSG00000129214 |
| ANKS4B | -1.557980745 | 0.339626104 | 0.161676331 | 0.336298878 | 6.887778257 | 2.357219627 | ENSG00000175311 |
| TMEM114 | -1.559083982 | 0.339366489 | 0.01817553 | 0.063563868 | 21.57018006 | 7.325979621 | ENSG00000232258 |
| NA | -1.563974349 | 0.338218071 | 0.086347258 | 0.213773985 | 10.82467601 | 3.67746153 | ENSG00000235100 |
| DBX1 | -1.564506149 | 0.338093422 | 0.130136451 | 0.289132573 | 7.834641311 | 2.656941556 | ENSG00000109851 |
| RANGRF | -1.571757924 | 0.336398243 | 0.066543644 | 0.175808159 | 10.79027649 | 3.630168424 | ENSG00000108961 |
| NA | -1.580070856 | 0.334465462 | 0.120599062 | 0.27354231 | 6.953017817 | 2.327046683 | ENSG00000272277 |
| CRHR1 | -1.580498859 | 0.334366251 | 0.126571233 | 0.28334492 | 6.902016147 | 2.301224667 | ENSG00000120088 |
| NA | -1.580628467 | 0.334336213 | 0.233758372 | 0.434928754 | 5.018968456 | 1.684660646 | ENSG00000275017 |
| NA | -1.5807284 | 0.334313055 | 0.063589017 | 0.169519847 | 9.935933572 | 3.322974145 | ENSG00000240093 |
| SLC17A6 | -1.581437376 | 0.334148806 | 0.047841884 | 0.136314723 | 11.92217981 | 3.983709849 | ENSG00000091664 |
| CBLN3 | -1.583293537 | 0.333719169 | 0.052646864 | 0.14679404 | 11.86642346 | 3.94824002 | ENSG00000139899 |
| NA | -1.585434411 | 0.333224317 | 0.164002252 | 0.339342888 | 6.007349971 | 2.003678201 | ENSG00000248996 |
| WASHC5 | -1.587880736 | 0.332659759 | 2.15E-59 | 9.23E-57 | 935.3907344 | 311.1234655 | ENSG00000164961 |
| C1QTNF3 | -1.588868255 | 0.332432133 | 0.113132639 | 0.261666857 | 7.954442013 | 2.634240961 | ENSG00000082196 |
| HINT2 | -1.58955383 | 0.332274197 | 0.171779799 | 0.351020301 | 6.054792169 | 2.015501478 | ENSG00000137133 |
| GPD1 | -1.589729333 | 0.332233779 | 0.044283387 | 0.128186967 | 11.96250306 | 3.960063296 | ENSG00000167588 |
| C2orf15 | -1.589875919 | 0.332200024 | 0.002580969 | 0.012624085 | 27.98023004 | 9.316888587 | ENSG00000273045 |
| DGKK | -1.593455832 | 0.331376722 | 3.12E-39 | 5.83E-37 | 680.19411 | 225.2316578 | ENSG00000274588 |
| CA5BP1 | -1.595333664 | 0.330945678 | 7.92E-13 | 2.31E-11 | 172.5693445 | 57.11242138 | ENSG00000186312 |
| NA | -1.598764392 | 0.330159625 | 0.08424791 | 0.20999875 | 10.06849762 | 3.348796162 | ENSG00000257557 |
| NKX3-2 | -1.603834257 | 0.329001426 | 0.000195272 | 0.001350812 | 43.49706632 | 14.32329387 | ENSG00000109705 |
| HLF | -1.603944489 | 0.328976289 | 0.145672885 | 0.312664915 | 7.138058079 | 2.34539635 | ENSG00000108924 |
| TNKS2-AS1 | -1.605340934 | 0.328658013 | 0.155797385 | 0.327558713 | 6.089191685 | 1.991854925 | ENSG00000228701 |
| ZNF829 | -1.606469129 | 0.328401101 | 0.000218663 | 0.001488093 | 41.47733644 | 13.66161221 | ENSG00000185869 |
| SCN7A | -1.609279786 | 0.327761934 | 0.18941705 | 0.375552859 | 5.056927445 | 1.642664426 | ENSG00000136546 |
| ANKRD61 | -1.618760448 | 0.32561511 | 0.087670863 | 0.216087737 | 9.094353998 | 2.96196037 | ENSG00000157999 |
| MT1E | -1.619796814 | 0.325381287 | 6.84E-19 | 3.79E-17 | 286.1677533 | 93.20691003 | ENSG00000169715 |
| NA | -1.625009224 | 0.324207816 | 0.005899214 | 0.025360038 | 21.54762801 | 6.995138789 | ENSG00000280604 |
| MGC16275 | -1.63143377 | 0.322767278 | 0.068057408 | 0.178707475 | 9.25211369 | 2.987782387 | ENSG00000246731 |
| FLJ38576 | -1.632037325 | 0.322632275 | 0.016681621 | 0.059398824 | 17.39960463 | 5.608024608 | ENSG00000279943 |
| ANOS1 | -1.632241945 | 0.322586519 | 1.40E-94 | 2.94E-91 | 4495.998759 | 1450.089654 | ENSG00000011201 |
| ZGLP1 | -1.635399628 | 0.321881234 | 0.029332676 | 0.093241472 | 15.38490877 | 4.951639832 | ENSG00000220201 |
| NA | -1.638520824 | 0.321185613 | 0.069118781 | 0.180966059 | 10.25234268 | 3.273505576 | ENSG00000203706 |
| NA | -1.642614916 | 0.320275442 | 0.010847003 | 0.041779655 | 18.71179353 | 5.985212587 | ENSG00000286342 |
| METTL7A | -1.644336181 | 0.319893552 | 1.83E-55 | 6.64E-53 | 935.4959406 | 299.2365812 | ENSG00000185432 |
| WNT5B | -1.64767198 | 0.319154749 | 4.38E-06 | 4.49E-05 | 68.62038438 | 21.85005828 | ENSG00000111186 |
| UBA7 | -1.653547163 | 0.317857677 | 0.123671379 | 0.278503935 | 6.258519517 | 1.991854925 | ENSG00000182179 |
| ASGR2 | -1.65390342 | 0.317779195 | 0.181057453 | 0.364023208 | 5.216772068 | 1.666310979 | ENSG00000161944 |
| NA | -1.655194453 | 0.317494949 | 0.08857895 | 0.217757546 | 8.387092349 | 2.654766092 | ENSG00000259172 |
| NA | -1.655417153 | 0.317445943 | 0.044226328 | 0.128080693 | 11.48833427 | 3.65599044 | ENSG00000259404 |
| NA | -1.656895029 | 0.317120923 | 0.146533623 | 0.313833491 | 6.340361231 | 2.019852405 | ENSG00000286650 |
| NA | -1.659498544 | 0.316549156 | 0.030091351 | 0.095188468 | 13.69638455 | 4.337251275 | ENSG00000287431 |
| NA | -1.660329435 | 0.316366899 | 0.14358159 | 0.309430369 | 6.271562199 | 1.980031648 | ENSG00000269752 |
| NA | -1.667401359 | 0.3148199 | 0.049521748 | 0.140023687 | 10.531988 | 3.320798682 | ENSG00000253893 |
| NA | -1.669237107 | 0.314419564 | 0.171632155 | 0.350825115 | 5.318775408 | 1.680309719 | ENSG00000262921 |
| H2BC5 | -1.669884208 | 0.314278567 | 2.79E-09 | 4.98E-08 | 109.7154873 | 34.50732473 | ENSG00000158373 |
| NA | -1.672036418 | 0.313810076 | 0.180593357 | 0.363593614 | 5.244052639 | 1.640488963 | ENSG00000260855 |
| RTP1 | -1.680985005 | 0.311869634 | 3.33E-07 | 4.23E-06 | 77.69902595 | 24.24274774 | ENSG00000175077 |
| NA | -1.686209479 | 0.310742294 | 0.074192485 | 0.191253764 | 8.516096929 | 2.650415165 | ENSG00000248774 |
| NA | -1.686473003 | 0.310685539 | 0.046904929 | 0.134260509 | 12.85864378 | 3.9762375 | ENSG00000237436 |
| PCDHA9 | -1.691125802 | 0.309685169 | 0.002097002 | 0.010604332 | 27.8295893 | 8.644329606 | ENSG00000204961 |
| H3C6 | -1.69771219 | 0.308274574 | 0.037700674 | 0.11337217 | 12.8026081 | 3.929890353 | ENSG00000274750 |
| EMX1 | -1.700789857 | 0.30761764 | 6.66E-14 | 2.18E-12 | 161.7752031 | 49.75532286 | ENSG00000135638 |
| NA | -1.707135538 | 0.306267558 | 0.07605276 | 0.194720224 | 8.703501455 | 2.662238442 | ENSG00000231831 |
| MIR3176 | -1.707263847 | 0.306240321 | 0.008296272 | 0.033664609 | 20.63991916 | 6.319458387 | ENSG00000266235 |
| NA | -1.713747467 | 0.304867134 | 0.03235472 | 0.100811564 | 11.96250306 | 3.636694814 | ENSG00000286808 |
| NA | -1.714314349 | 0.304747365 | 0.012783942 | 0.047825821 | 16.31781326 | 4.953815295 | ENSG00000279692 |
| PCDHB15 | -1.717156633 | 0.304147567 | 4.62E-09 | 8.03E-08 | 108.0293797 | 32.87553762 | ENSG00000113248 |
| KAZALD1 | -1.727194206 | 0.3020388 | 0.004000403 | 0.018381412 | 23.13679249 | 7.006962066 | ENSG00000107821 |
| ST8SIA4 | -1.729050461 | 0.301650429 | 0.108350268 | 0.253359027 | 6.602209194 | 1.994030388 | ENSG00000113532 |
| NA | -1.740493736 | 0.29926724 | 0.030244318 | 0.095608295 | 13.4202987 | 4.023530606 | ENSG00000213642 |
| GLTPD2 | -1.742829809 | 0.298783046 | 0.044341525 | 0.128275587 | 11.05212446 | 3.311150869 | ENSG00000182327 |
| H3C8 | -1.745856715 | 0.298156829 | 7.82E-06 | 7.68E-05 | 52.22400954 | 15.55330049 | ENSG00000273983 |

| | | | | | | | |
|---|---|---|---|---|---|---|---|
| LRRC37A11P | -1.745921087 | 0.298143526 | 0.053181215 | 0.147783608 | 10.03557265 | 2.98343146 | ENSG00000214553 |
| NA | -1.746125095 | 0.298101369 | 0.046914752 | 0.134260509 | 11.10312613 | 3.318623218 | ENSG00000277511 |
| NA | -1.746277918 | 0.298069793 | 0.126448616 | 0.283137552 | 6.687610381 | 1.987503998 | ENSG00000188801 |
| NA | -1.748542774 | 0.297602227 | 0.13356138 | 0.294661994 | 5.580623371 | 1.668486443 | ENSG00000272189 |
| NA | -1.749603802 | 0.297383436 | 0.005443907 | 0.023726791 | 21.07701869 | 6.247289223 | ENSG00000286482 |
| NA | -1.762300609 | 0.29477772 | 0.0786822 | 0.199317685 | 7.845319728 | 2.315223407 | ENSG00000285646 |
| SHLD2P1 | -1.76887929 | 0.293436595 | 0.137236269 | 0.299959367 | 5.68382192 | 1.668486443 | ENSG00000165874 |
| H3C10 | -1.772893665 | 0.292621228 | 2.42E-05 | 0.000211226 | 49.94271103 | 14.62613722 | ENSG00000278828 |
| NOX1 | -1.77418468 | 0.292359489 | 0.11663496 | 0.267604327 | 6.843895532 | 2.019852405 | ENSG00000007952 |
| NA | -1.776336591 | 0.291923733 | 0.02807826 | 0.090179923 | 12.5680407 | 3.686163384 | ENSG00000273284 |
| NA | -1.779497058 | 0.291284925 | 2.40E-18 | 1.24E-16 | 212.8077141 | 61.9934051 | ENSG00000278630 |
| SEC14L4 | -1.781237526 | 0.29093373 | 0.000569502 | 0.003435018 | 34.12990661 | 9.904509168 | ENSG00000133488 |
| ZNF876P | -1.7941464 | 0.28834214 | 0.016657803 | 0.05933282 | 18.43779261 | 5.327881852 | ENSG00000198155 |
| NA | -1.795106761 | 0.288150262 | 0.067384072 | 0.177432821 | 9.17263624 | 2.642942816 | ENSG00000277342 |
| NA | -1.795887753 | 0.287994317 | 0.132854977 | 0.293446345 | 6.956577289 | 2.011150551 | ENSG00000267191 |
| ZNF799 | -1.796181982 | 0.287935588 | 0.000155235 | 0.001100045 | 40.24281939 | 11.54499813 | ENSG00000196466 |
| RDX | -1.797115635 | 0.287749308 | 7.40E-99 | 1.75E-95 | 1695.472876 | 487.7164196 | ENSG00000137710 |
| FAM135B | -1.801708948 | 0.286834617 | 0.029447847 | 0.093544589 | 11.52273378 | 3.299327592 | ENSG00000147724 |
| NA | -1.802074238 | 0.286761999 | 0.010163515 | 0.039566683 | 16.24901423 | 4.668092106 | ENSG00000286340 |
| LRRTM3 | -1.811270891 | 0.284939811 | 0.003420795 | 0.016114172 | 22.08080715 | 6.289285443 | ENSG00000198739 |
| LOC105371050 | -1.81515086 | 0.284174526 | 0.139923178 | 0.303939407 | 7.069259047 | 2.011150551 | ENSG00000260874 |
| PCDHB12 | -1.815535799 | 0.284098713 | 1.14E-08 | 1.87E-07 | 89.64612206 | 25.4018147 | ENSG00000120328 |
| ETV2 | -1.817089301 | 0.283792958 | 0.011581305 | 0.044139 | 16.46489454 | 4.668092106 | ENSG00000105672 |
| MAP3K2-DT | -1.821070235 | 0.283010947 | 0.031812021 | 0.099597151 | 11.77537787 | 3.322974145 | ENSG00000236682 |
| NA | -1.825771693 | 0.282090172 | 0.000799806 | 0.004613418 | 35.51445401 | 10.02586336 | ENSG00000224945 |
| NA | -1.831282719 | 0.281014656 | 0.125574433 | 0.28171456 | 5.945669884 | 1.682485183 | ENSG00000260577 |
| PHGDH | -1.834298137 | 0.280427913 | 0 | 0 | 15325.66663 | 4297.344873 | ENSG00000092621 |
| NA | -1.841826796 | 0.27896832 | 0.000579845 | 0.003492007 | 30.82318347 | 8.602333386 | ENSG00000279413 |
| NA | -1.850203537 | 0.277353236 | 0.116693511 | 0.267673625 | 7.165059318 | 1.994030388 | ENSG00000225883 |
| PHKG1 | -1.855116895 | 0.276410266 | 0.082628156 | 0.206810608 | 7.221984725 | 2.008029128 | ENSG00000164776 |
| NA | -1.856810639 | 0.276085947 | 0.105685256 | 0.248690145 | 6.020392653 | 1.656663166 | ENSG00000236060 |
| LOC105373383 | -1.861063126 | 0.275273354 | 0.034422694 | 0.105857771 | 10.88991557 | 3.01142894 | ENSG00000260081 |
| NA | -1.861587326 | 0.275173353 | 0.030644283 | 0.096565539 | 10.83771869 | 2.987782387 | ENSG00000218690 |
| RNF103-CHMP3 | -1.862111726 | 0.275073349 | 0.117729678 | 0.269395975 | 6.013273708 | 1.642664426 | ENSG00000249884 |
| HMX3 | -1.867096409 | 0.274124578 | 0.078461204 | 0.198917955 | 7.283664813 | 2.001502738 | ENSG00000188620 |
| NA | -1.868799262 | 0.273801212 | 0.032848441 | 0.101962987 | 10.85195658 | 2.964135834 | ENSG00000253200 |
| LOC100128164 | -1.871061814 | 0.273372151 | 0.126016284 | 0.282528533 | 6.061911114 | 1.64483989 | ENSG00000239219 |
| POU3F4 | -1.872617443 | 0.273077538 | 0.002895751 | 0.013971962 | 24.40125986 | 6.691065933 | ENSG00000196767 |
| NA | -1.879451135 | 0.271787096 | 0.051712323 | 0.144786839 | 9.77817388 | 2.674061718 | ENSG00000223461 |
| CXCL16 | -1.884885053 | 0.270765334 | 9.23E-05 | 0.000692282 | 39.18772378 | 10.62095629 | ENSG00000161921 |
| ABALON | -1.886620094 | 0.270439897 | 0.002239082 | 0.011195412 | 24.49494904 | 6.612653925 | ENSG00000281376 |
| LOC101928058 | -1.888194164 | 0.270144991 | 0.076276835 | 0.195055929 | 7.392787098 | 1.989679461 | ENSG00000253764 |
| GRPR | -1.897422963 | 0.268422411 | 3.76E-20 | 2.28E-18 | 230.4386066 | 61.88654411 | ENSG00000126010 |
| SEMA3F-AS1 | -1.902253243 | 0.267525211 | 0.111473998 | 0.259099871 | 6.227679473 | 1.678134256 | ENSG00000235016 |
| NA | -1.904181593 | 0.267167867 | 0.10780854 | 0.252569692 | 6.2478411 | 1.682485183 | ENSG00000236184 |
| NA | -1.908264181 | 0.266412895 | 0.050678417 | 0.142610924 | 10.02252997 | 2.674061718 | ENSG00000270344 |
| NA | -1.916809382 | 0.264839574 | 0.037015715 | 0.111864582 | 10.03557265 | 2.648239701 | ENSG00000262227 |
| TCHH | -1.920246673 | 0.264209332 | 0.045037212 | 0.129911065 | 10.05781921 | 2.678412645 | ENSG00000159450 |
| ZNF442 | -1.92892091 | 0.262625533 | 0.000204607 | 0.001404584 | 33.00067304 | 8.677623972 | ENSG00000198342 |
| RSPH9 | -1.934394665 | 0.261630989 | 0.009939048 | 0.038835315 | 16.54317678 | 4.309253795 | ENSG00000172426 |
| NA | -1.935545422 | 0.261422384 | 9.74E-06 | 9.32E-05 | 58.57710855 | 15.27845462 | ENSG00000238178 |
| DRD5 | -1.938041658 | 0.260970446 | 0.083963802 | 0.209438172 | 7.534834359 | 1.952034168 | ENSG00000169676 |
| LINC00309 | -1.946129335 | 0.259511552 | 0.032793965 | 0.101844109 | 10.23574052 | 2.660062978 | ENSG00000230923 |
| NA | -1.950100426 | 0.258798215 | 0.002827884 | 0.013672451 | 20.48215946 | 5.295533444 | ENSG00000270354 |
| NA | -1.960003523 | 0.257027829 | 0.012598423 | 0.047300439 | 14.23075889 | 3.664692295 | ENSG00000261485 |
| AMH | -1.968164211 | 0.255578041 | 2.31E-19 | 1.32E-17 | 302.3578089 | 77.44440644 | ENSG00000104899 |
| NA | -1.97504486 | 0.254362014 | 0.000782069 | 0.004522159 | 27.30113869 | 6.936968366 | ENSG00000268798 |
| H2BC17 | -1.983064619 | 0.25295197 | 0.042078377 | 0.123431213 | 10.58654915 | 2.696762312 | ENSG00000274641 |
| NKAIN3 | -1.984290779 | 0.252737075 | 0.015103342 | 0.054788057 | 14.547168 | 3.700162124 | ENSG00000185942 |
| ZNF536 | -1.985192958 | 0.252579077 | 0.163451766 | 0.338611956 | 5.308096991 | 1.349468888 | ENSG00000198597 |
| PTENP1-AS | -1.992918736 | 0.251230106 | 0.076088591 | 0.194760087 | 6.589166513 | 1.654487703 | ENSG00000281128 |
| ZNF441 | -1.993032856 | 0.251210234 | 0.005992327 | 0.025719339 | 17.24807416 | 4.346899088 | ENSG00000197044 |
| MAATS1 | -2.000176442 | 0.249969427 | 0.119918344 | 0.272587618 | 5.318775408 | 1.328943757 | ENSG00000183833 |
| NA | -2.01016697 | 0.248244392 | 0.018436924 | 0.064304723 | 13.48909773 | 3.353147089 | ENSG00000274849 |
| GPR85 | -2.011956834 | 0.247936601 | 0.107065348 | 0.251296183 | 5.380455496 | 1.331119221 | ENSG00000164604 |
| NA | -2.012141301 | 0.247904901 | 0.002777615 | 0.013453518 | 156.7292851 | 38.88924214 | ENSG00000225138 |
| NA | -2.024302087 | 0.24582404 | 0.082130384 | 0.205871025 | 6.77657104 | 1.656663166 | ENSG00000228477 |
| ADM5 | -2.029954084 | 0.244862867 | 2.53E-09 | 4.58E-08 | 75.96873009 | 18.63472313 | ENSG00000224420 |
| GLIPR1L1 | -2.032825861 | 0.244375937 | 0.006694814 | 0.028221331 | 14.97625886 | 3.660341367 | ENSG00000173401 |
| FABP7 | -2.034917786 | 0.244021846 | 1.73E-19 | 1.00E-17 | 179.6166881 | 43.89297751 | ENSG00000164434 |
| LINC02724 | -2.054856317 | 0.240672579 | 0.114207533 | 0.263507545 | 5.529621701 | 1.342942497 | ENSG00000181908 |
| NA | -2.061958973 | 0.239490615 | 0.082468024 | 0.206580266 | 6.974374652 | 1.678134256 | ENSG00000238287 |
| LIPE-AS1 | -2.067931307 | 0.238501244 | 0.012037777 | 0.04565845 | 12.54076013 | 2.999605664 | ENSG00000213904 |
| NA | -2.067982375 | 0.238492801 | 1.40E-13 | 4.47E-12 | 118.4216585 | 28.28027709 | ENSG00000206417 |

| | | | | | | | |
|---|---|---|---|---|---|---|---|
| MYO16 | -2.07056198 | 0.238066746 | 0.110612743 | 0.257541814 | 5.536740646 | 1.317120481 | ENSG00000041515 |
| LONRF3 | -2.076470443 | 0.237093753 | 0.121329697 | 0.274572958 | 5.53910491 | 1.307472668 | ENSG00000175556 |
| SLC7A4 | -2.079145591 | 0.236654524 | 0.121187083 | 0.274348845 | 5.666024558 | 1.349468888 | ENSG00000099960 |
| NA | -2.07991568 | 0.236528235 | 0.00451693 | 0.020310054 | 16.80474541 | 3.974062036 | ENSG00000283236 |
| NA | -2.082563265 | 0.236094565 | 0.099306583 | 0.236959123 | 5.673143503 | 1.347293424 | ENSG00000287806 |
| NA | -2.083471864 | 0.235945921 | 0.067334901 | 0.177352804 | 7.083496937 | 1.670661906 | ENSG00000272787 |
| NA | -2.084073195 | 0.235847597 | 0.051679059 | 0.144744712 | 8.41289838 | 1.994030388 | ENSG00000235586 |
| SLC9A3 | -2.085583987 | 0.235600746 | 4.38E-05 | 0.000357915 | 35.29117543 | 8.344607677 | ENSG00000066230 |
| NUTM2B | -2.099680114 | 0.233309974 | 0.086469053 | 0.213879248 | 7.092980146 | 1.640488963 | ENSG00000188199 |
| NA | -2.102697455 | 0.232822525 | 0.137558937 | 0.300379984 | 5.772782579 | 1.347293424 | ENSG00000240882 |
| CNTD2 | -2.109863385 | 0.231668952 | 0.001132301 | 0.006252488 | 24.51394161 | 5.710366716 | ENSG00000105219 |
| NA | -2.110781912 | 0.231521501 | 0.027912906 | 0.089800619 | 10.08301485 | 2.33139761 | ENSG00000261338 |
| CDH9 | -2.113131325 | 0.231144778 | 0.068789429 | 0.180203686 | 7.185220945 | 1.670661906 | ENSG00000113100 |
| SPEF1 | -2.114426539 | 0.230937355 | 0.00848012 | 0.034256055 | 15.76180275 | 3.639816237 | ENSG00000101222 |
| NA | -2.123106279 | 0.229552129 | 0.000665733 | 0.003931893 | 23.11663086 | 5.307356721 | ENSG00000261189 |
| NA | -2.144623124 | 0.226153915 | 0.038812267 | 0.116104809 | 8.818268145 | 2.008029128 | ENSG00000286873 |
| NA | -2.151399661 | 0.22509413 | 0.000540299 | 0.00328194 | 24.95223635 | 5.608024608 | ENSG00000278493 |
| ZBTB16 | -2.176859819 | 0.221156597 | 0.042585038 | 0.124530574 | 9.07210744 | 2.010204592 | ENSG00000109906 |
| NA | -2.17847219 | 0.220909568 | 0.03610015 | 0.109782824 | 8.990265726 | 1.975680721 | ENSG00000269086 |
| ZNF44 | -2.193906553 | 0.218558811 | 8.89E-10 | 1.72E-08 | 79.02845354 | 17.31883216 | ENSG00000197857 |
| NA | -2.212053513 | 0.215826884 | 0.032986721 | 0.102278984 | 9.152474613 | 1.968208372 | ENSG00000272944 |
| NA | -2.217542262 | 0.215007328 | 0.022489823 | 0.075613371 | 10.86975394 | 2.355044163 | ENSG00000284634 |
| MAP6 | -2.243392181 | 0.211189179 | 1.91E-12 | 5.29E-11 | 103.9296627 | 21.94275258 | ENSG00000171533 |
| NA | -2.243664109 | 0.211149376 | 0.061038966 | 0.164603867 | 6.296478506 | 1.331119221 | ENSG00000238082 |
| NA | -2.243727566 | 0.211140089 | 0.002035784 | 0.010323786 | 17.3014401 | 3.646342627 | ENSG00000266865 |
| MIOX | -2.24804879 | 0.210508619 | 0.076657004 | 0.195736548 | 6.333242286 | 1.335470148 | ENSG00000100253 |
| ANKRD53 | -2.251944733 | 0.209940916 | 0.035381994 | 0.108208738 | 7.92360197 | 1.670661906 | ENSG00000144031 |
| DNAJC9-AS1 | -2.261062489 | 0.208618284 | 0.014390135 | 0.052686643 | 12.7981589 | 2.668764832 | ENSG00000236756 |
| NA | -2.273431569 | 0.206837321 | 0.062238545 | 0.166829084 | 6.42932189 | 1.319295944 | ENSG00000260160 |
| NA | -2.288885421 | 0.204633547 | 0.0440782 | 0.127828133 | 8.147490944 | 1.684660646 | ENSG00000269425 |
| TCEAL5 | -2.297785315 | 0.203375061 | 3.65E-11 | 8.67E-10 | 84.82493107 | 17.26160769 | ENSG00000204065 |
| NA | -2.314320048 | 0.201057485 | 0.001177613 | 0.006463009 | 18.13918087 | 3.648518091 | ENSG00000215014 |
| NA | -2.35068415 | 0.196053031 | 0.01508077 | 0.054716685 | 10.17050096 | 2.005853665 | ENSG00000286478 |
| H2BC11 | -2.351179681 | 0.195985703 | 1.35E-12 | 3.81E-11 | 88.08632309 | 17.29613156 | ENSG00000124635 |
| H4C12 | -2.357450825 | 0.195135636 | 0.014552963 | 0.053184865 | 10.22506211 | 2.003678201 | ENSG00000273542 |
| HMX2 | -2.411564076 | 0.187951968 | 3.37E-21 | 2.21E-19 | 158.9939559 | 29.88841764 | ENSG00000188816 |
| ORM2 | -2.414162552 | 0.187613747 | 0.000643349 | 0.003815686 | 21.28786498 | 4.023530606 | ENSG00000228278 |
| H4C11 | -2.419782869 | 0.186884281 | 0.003374537 | 0.015920093 | 14.22719942 | 2.664413905 | ENSG00000197238 |
| NA | -2.431179997 | 0.185413732 | 0.023616756 | 0.078743368 | 8.932145111 | 1.652312239 | ENSG00000283355 |
| RASAL3 | -2.432791697 | 0.185206714 | 0.022521381 | 0.075698038 | 9.034148451 | 1.682485183 | ENSG00000105122 |
| HRAT92 | -2.470717217 | 0.180401443 | 0.001276868 | 0.006927218 | 18.50303217 | 3.362794902 | ENSG00000223855 |
| NA | -2.491094582 | 0.177871271 | 0.008475532 | 0.034245859 | 11.18852731 | 1.994030388 | ENSG00000278012 |
| NA | -2.503743337 | 0.17631861 | 0.008991056 | 0.035897695 | 11.27036903 | 1.980031648 | ENSG00000233967 |
| AQP4-AS1 | -2.511315924 | 0.175395553 | 0.065736743 | 0.173944009 | 5.689745657 | 0.995927462 | ENSG00000260372 |
| NA | -2.521842994 | 0.174120383 | 0.02977906 | 0.094358649 | 7.596514446 | 1.317120481 | ENSG00000237410 |
| FBP1 | -2.555446752 | 0.170111579 | 1.38E-06 | 1.56E-05 | 36.97641951 | 6.275286703 | ENSG00000165140 |
| NA | -2.564358912 | 0.169063964 | 0.013976716 | 0.05141221 | 9.779369089 | 1.656663166 | ENSG00000286964 |
| NA | -2.565161833 | 0.168969899 | 0.024980628 | 0.08244655 | 7.845319728 | 1.331119221 | ENSG00000224080 |
| NA | -2.570466222 | 0.168349784 | 0.095397253 | 0.230057095 | 5.944474675 | 1.005575275 | ENSG00000276744 |
| TBXT | -2.594511074 | 0.165567215 | 0.048457776 | 0.137778546 | 5.993112081 | 0.984104186 | ENSG00000164458 |
| H2BC12 | -2.597873266 | 0.165181811 | 2.65E-39 | 5.01E-37 | 355.9239832 | 58.90037009 | ENSG00000197903 |
| MT1F | -2.63517211 | 0.160966 | 5.45E-48 | 1.54E-45 | 387.6805787 | 62.319895 | ENSG00000198417 |
| DLX4 | -2.658469259 | 0.158387539 | 0.015134298 | 0.054858201 | 10.51419064 | 1.675958792 | ENSG00000108813 |
| ZNF563 | -2.701954146 | 0.153684743 | 1.75E-07 | 2.31E-06 | 43.24353251 | 6.642826869 | ENSG00000188868 |
| SHPK | -2.722041926 | 0.151559698 | 1.16E-18 | 6.21E-17 | 122.5101129 | 18.5603785 | ENSG00000197417 |
| PCDHB5 | -2.728983885 | 0.150832175 | 1.03E-36 | 1.72E-34 | 259.7852812 | 39.17295782 | ENSG00000113209 |
| RHOXF1-AS1 | -2.739474488 | 0.149739372 | 1.59E-08 | 2.56E-07 | 46.53426388 | 6.975843164 | ENSG00000258545 |
| NA | -2.740570934 | 0.149625613 | 0.035575484 | 0.108554372 | 6.656770337 | 0.993751999 | ENSG00000265451 |
| WDR97 | -2.80695799 | 0.142896453 | 0.001597014 | 0.008393991 | 16.31870299 | 2.355044163 | ENSG00000179698 |
| H2AC6 | -2.81722805 | 0.141882833 | 8.98E-69 | 6.06E-66 | 533.0123469 | 75.68613621 | ENSG00000180573 |
| NA | -2.827582709 | 0.140868143 | 0.006141918 | 0.02625399 | 11.71133352 | 1.628665686 | ENSG00000286747 |
| H4C9 | -2.833906864 | 0.140251989 | 7.23E-18 | 3.59E-16 | 111.2180816 | 15.6394684 | ENSG00000276180 |
| MT1M | -2.877259897 | 0.136100107 | 7.08E-05 | 0.000546394 | 22.02179681 | 3.001781127 | ENSG00000205364 |
| NA | -2.892492274 | 0.134670683 | 0.008912134 | 0.035694697 | 9.902729264 | 1.345117961 | ENSG00000272468 |
| LINC00540 | -3.049472367 | 0.120786208 | 2.71E-06 | 2.88E-05 | 30.23333211 | 3.646342627 | ENSG00000276476 |
| CFAP65 | -3.061286146 | 0.119801166 | 0.01250007 | 0.047034036 | 8.370490196 | 1.009926202 | ENSG00000181378 |
| NA | -3.074298993 | 0.11872544 | 0.004214906 | 0.019210962 | 11.13516138 | 1.319295944 | ENSG00000224664 |
| PCDHGC3 | -3.206846377 | 0.108303639 | 1.73E-21 | 1.17E-19 | 129.0345985 | 14.02168003 | ENSG00000240184 |
| ZNF439 | -3.266182829 | 0.103939589 | 0.005345426 | 0.023373088 | 9.619524466 | 1.007750739 | ENSG00000171291 |
| NA | -3.313710847 | 0.100571201 | 0.005333978 | 0.023328432 | 9.902729264 | 0.998102926 | ENSG00000286416 |
| TMED6 | -3.361194093 | 0.097314993 | 0.004378428 | 0.019781488 | 10.21794316 | 0.993751999 | ENSG00000157315 |
| HAS1 | -3.384080347 | 0.095783413 | 4.92E-08 | 7.23E-07 | 38.18393531 | 3.681812457 | ENSG00000105509 |
| DNAJA4 | -3.431462611 | 0.092688707 | 9.36E-19 | 5.11E-17 | 125.8603877 | 11.70617307 | ENSG00000140403 |
| NA | -3.519891234 | 0.087178051 | 0.01237272 | 0.046650873 | 7.64159238 | 0.670383517 | ENSG00000281641 |

| | | | | | | | |
|---|---|---|---|---|---|---|---|
| NA | -3.532160344 | 0.086439807 | 0.012285126 | 0.046399988 | 7.623795017 | 0.646736964 | ENSG00000279412 |
| NA | -3.589474803 | 0.0830731 | 3.11E-09 | 5.51E-08 | 44.01216876 | 3.683987921 | ENSG00000228065 |
| ZNF229 | -3.659489283 | 0.079137797 | 8.77E-15 | 3.21E-13 | 75.34864907 | 5.956269148 | ENSG00000278318 |
| ZNF736 | -3.685067783 | 0.077747075 | 3.88E-24 | 3.26E-22 | 128.3433804 | 10.03361925 | ENSG00000234444 |
| NA | -3.695222571 | 0.077201754 | 0.006615552 | 0.027980896 | 8.583700753 | 0.660735704 | ENSG00000280254 |
| DMRT2 | -3.819823798 | 0.070813891 | 6.07E-05 | 0.000477611 | 18.75687147 | 1.319295944 | ENSG00000173253 |
| CUTALP | -4.072800451 | 0.059424413 | 2.20E-10 | 4.66E-09 | 44.72001523 | 2.664413905 | ENSG00000226752 |
| PKLR | -4.182780684 | 0.055062706 | 1.42E-14 | 5.01E-13 | 66.30412769 | 3.662516831 | ENSG00000143627 |
| NA | -4.227668691 | 0.053375863 | 0.006922909 | 0.029008016 | 6.698288798 | 0.323368482 | ENSG00000285600 |
| NA | -4.299112555 | 0.050797012 | 0.007924373 | 0.032406192 | 7.036054739 | 0.337367222 | ENSG00000260949 |
| NCR1 | -5.20614079 | 0.027089155 | 0.001401005 | 0.007520744 | 6.787249457 | 0 | ENSG00000189430 |
| BMP3 | -5.304442673 | 0.025304843 | 0.001095832 | 0.006072416 | 7.231467934 | 0 | ENSG00000152785 |
| ZNF280D | -5.974772384 | 0.015900629 | 6.31E-08 | 9.11E-07 | 41.72405679 | 0.65856024 | ENSG00000137871 |
| NA | -6.221967891 | 0.013396799 | 8.96E-06 | 8.69E-05 | 13.69282508 | 0 | ENSG00000270325 |
| PSMD5 | -6.595887355 | 0.010338084 | 4.59E-58 | 1.84E-55 | 449.1866511 | 4.618623537 | ENSG00000095261 |
| ZNF572 | -6.732502711 | 0.009404046 | 5.04E-07 | 6.20E-06 | 19.46769259 | 0 | ENSG00000180938 |
| NLRP7 | -7.227593881 | 0.006672329 | 3.03E-08 | 4.60E-07 | 27.50781512 | 0 | ENSG00000167634 |
| NLRP2 | -8.104963942 | 0.003632141 | 1.04E-30 | 1.27E-28 | 3882.973088 | 14.06724917 | ENSG00000022556 |

| **Table S3. Total gene expression using RNA-seq data in 3 biological replicates of SF3B1$^{K700E}$ versus SF3B1$^{WT}$ ES cells (K700E ES line 3).** Average baseMean, log2fold change, fold change (FC) and p-value are shown. | | | | | | | |
|---|---|---|---|---|---|---|---|
| **Gene Symbol** | **log2 FC** | **FC** | **p-value** | **padj** | **WT baseMean** | **MT baseMean** | **Ensembl ID** |
| LOC101928254 | 7.196991406 | 146.7270857 | 1.28E-11 | 3.19E-10 | 0.621806095 | 93.24428294 | ENSG00000219445 |
| NA | 7.109480909 | 138.0915191 | 8.58E-08 | 1.16E-06 | 0 | 24.29430264 | ENSG00000233309 |
| NA | 6.88196224 | 117.9443288 | 4.72E-08 | 6.73E-07 | 0.310903048 | 40.38883132 | ENSG00000272405 |
| FAR2P1 | 6.256714652 | 76.46431187 | 1.87E-06 | 1.98E-05 | 0.337436127 | 26.23736792 | ENSG00000286058 |
| RHOXF1P1 | 6.239128756 | 75.53789921 | 1.30E-05 | 0.000113615 | 0 | 13.30818593 | ENSG00000234493 |
| FOXC2 | 6.16985815 | 71.99666373 | 1.57E-05 | 0.000134348 | 0 | 12.63633215 | ENSG00000176692 |
| PRAC1 | 6.119885244 | 69.54549912 | 2.51E-05 | 0.000204645 | 0 | 12.19352916 | ENSG00000159182 |
| NAP1L6P | 6.025679002 | 65.14935551 | 3.19E-05 | 0.000254084 | 0 | 11.46635008 | ENSG00000204118 |
| NPAP1 | 6.007325678 | 64.32580395 | 5.27E-06 | 5.06E-05 | 0.337436127 | 22.01714987 | ENSG00000185823 |
| NA | 6.000961013 | 64.04264612 | 4.24E-05 | 0.000324999 | 0 | 11.23718768 | ENSG00000272068 |
| C7orf57 | 6.000294458 | 64.01306391 | 5.10E-05 | 0.000382869 | 0 | 11.29025147 | ENSG00000164746 |
| MOV10L1 | 5.97977569 | 63.10908014 | 5.03E-05 | 0.000378724 | 0 | 11.10254153 | ENSG00000073146 |
| MED15P9 | 5.950459683 | 61.83962573 | 8.60E-06 | 7.80E-05 | 0.310903048 | 21.19611583 | ENSG00000223760 |
| NA | 5.947688115 | 61.72093939 | 6.48E-05 | 0.000473854 | 0 | 10.81706615 | ENSG00000257275 |
| KRT80 | 5.857995762 | 58.00059368 | 8.58E-05 | 0.000608087 | 0 | 10.16822024 | ENSG00000167767 |
| KHDC3L | 5.836282162 | 57.13417936 | 8.12E-05 | 0.000577884 | 0 | 10.05223347 | ENSG00000203908 |
| DDX43 | 5.829745098 | 56.87588159 | 0.000167211 | 0.00109586 | 0 | 9.94654392 | ENSG00000080007 |
| NA | 5.745350678 | 53.64421442 | 0.000178982 | 0.001165582 | 0 | 9.448527246 | ENSG00000286168 |
| LOC283299 | 5.729541551 | 53.05958738 | 1.00E-12 | 2.97E-11 | 1.266988357 | 67.51808038 | ENSG00000254951 |
| TBX3 | 5.707734363 | 52.2635915 | 4.33E-07 | 5.17E-06 | 0.618649183 | 33.2060603 | ENSG00000135111 |
| CYTOR | 5.700794743 | 52.01279798 | 4.51E-07 | 5.37E-06 | 0.645182262 | 33.16103224 | ENSG00000222041 |
| NA | 5.549894211 | 46.84730694 | 0.000409685 | 0.002403643 | 0 | 8.229168613 | ENSG00000248215 |
| GDF15 | 5.541707786 | 46.5822295 | 2.69E-279 | 4.99E-275 | 41.52248755 | 1939.484302 | ENSG00000130513 |
| REG1A | 5.503257001 | 45.35711588 | 0.000583842 | 0.003285041 | 0 | 7.978097646 | ENSG00000115386 |
| NA | 5.438842727 | 43.37652959 | 0.000521541 | 0.002978811 | 0 | 7.632948467 | ENSG00000267696 |
| HORMAD1 | 5.437236471 | 43.32826226 | 8.67E-05 | 0.000613474 | 0.307746135 | 14.8381777 | ENSG00000143452 |
| SIX3 | 5.3836572 | 41.74863722 | 4.64E-08 | 6.64E-07 | 0.952928397 | 39.73833646 | ENSG00000138083 |
| KLF18 | 5.361259529 | 41.10549977 | 0.000856612 | 0.004609884 | 0 | 7.224000232 | ENSG00000283039 |
| NA | 5.328579909 | 40.18485326 | 0.001054984 | 0.005520074 | 0 | 7.093041305 | ENSG00000232949 |
| ANXA1 | 5.218125142 | 37.22306998 | 1.43E-18 | 7.95E-17 | 2.50112981 | 94.38756991 | ENSG00000135046 |
| CH25H | 5.190377514 | 36.51399237 | 0.000245802 | 0.001545872 | 0.307746135 | 12.52034537 | ENSG00000138135 |
| NA | 5.132571352 | 35.0798665 | 0.000319225 | 0.00193603 | 0.337436127 | 11.99877117 | ENSG00000287862 |
| MIR205HG | 5.073051214 | 33.66205189 | 3.53E-20 | 2.28E-18 | 9.143817732 | 309.0143782 | ENSG00000230937 |
| NA | 5.070795242 | 33.60945497 | 2.16E-17 | 1.07E-15 | 2.563666707 | 85.70237829 | ENSG00000287839 |
| LOC105373085 | 4.999917985 | 31.99818091 | 3.11E-09 | 5.35E-08 | 1.29667835 | 40.8803496 | ENSG00000286381 |
| NA | 4.994638744 | 31.88130423 | 0.000545964 | 0.003100153 | 0.307746135 | 10.89215012 | ENSG00000271662 |
| LGALS12 | 4.89793122 | 29.81427232 | 1.23E-06 | 1.35E-05 | 0.959242223 | 28.41901676 | ENSG00000133317 |
| ANKRD1 | 4.884622445 | 29.54050215 | 0.00109325 | 0.005696457 | 0.310903048 | 10.07425365 | ENSG00000148677 |
| GDA | 4.880899918 | 29.46437825 | 9.47E-09 | 1.53E-07 | 1.263831445 | 37.45534657 | ENSG00000119125 |
| NA | 4.841430962 | 28.66922425 | 2.84E-10 | 5.75E-09 | 1.634114477 | 45.74798783 | ENSG00000267279 |
| 1-Dec | 4.833458176 | 28.51122608 | 0.001205303 | 0.006165909 | 0.310903048 | 9.717931174 | ENSG00000173077 |
| PITX2 | 4.806506655 | 27.98354158 | 2.97E-22 | 2.28E-20 | 4.118181946 | 115.6707075 | ENSG00000164093 |
| NA | 4.723906324 | 26.42636932 | 1.85E-11 | 4.50E-10 | 1.941860612 | 50.5907947 | ENSG00000226539 |
| ZNF385D-AS1 | 4.630332151 | 24.76674127 | 1.58E-07 | 2.04E-06 | 1.234141452 | 31.29238967 | ENSG00000225542 |
| FAM13A-AS1 | 4.595944733 | 24.18339242 | 0.006194652 | 0.024743897 | 0.310903048 | 8.323573238 | ENSG00000248019 |
| CCR7 | 4.553232871 | 23.47792286 | 0.002939696 | 0.013087757 | 0.310903048 | 8.060667697 | ENSG00000126353 |
| MYL7 | 4.531944526 | 23.13402716 | 1.26E-12 | 3.66E-11 | 2.507443636 | 58.81346493 | ENSG00000106631 |
| MAB21L4 | 4.51351268 | 22.84034725 | 0.003393804 | 0.014779314 | 0.310903048 | 7.797762156 | ENSG00000172478 |
| MAGEA4 | 4.508638073 | 22.76330407 | 1.57E-21 | 1.13E-19 | 5.385170303 | 122.8021265 | ENSG00000147381 |
| NA | 4.437943528 | 21.6747512 | 0.004493682 | 0.018787707 | 0.307746135 | 7.388813921 | ENSG00000286735 |
| PAX7 | 4.387670267 | 20.93246441 | 1.21E-11 | 3.04E-10 | 2.646422858 | 53.69591794 | ENSG00000009709 |
| LOC101928371 | 4.370597002 | 20.68620368 | 0.011469865 | 0.041232369 | 0.337436127 | 7.108013452 | ENSG00000225420 |
| DMRTB1 | 4.317361038 | 19.93678725 | 0.006151112 | 0.024601795 | 0.337436127 | 6.811364769 | ENSG00000143006 |
| NA | 4.256464857 | 19.11276841 | 4.24E-06 | 4.19E-05 | 1.29667835 | 24.49329751 | ENSG00000231240 |
| SCG2 | 4.175350343 | 18.06781751 | 1.56E-13 | 5.04E-12 | 3.886878136 | 69.45878097 | ENSG00000171951 |
| TDRD1 | 4.141973122 | 17.65461092 | 0.000239429 | 0.001508352 | 0.932709143 | 16.67902587 | ENSG00000095627 |
| CER1 | 4.132084733 | 17.53401805 | 1.59E-23 | 1.34E-21 | 6.553617945 | 116.6196745 | ENSG00000147869 |
| TLR8-AS1 | 4.130527382 | 17.51510075 | 0.00173805 | 0.008426108 | 0.621806095 | 11.10622876 | ENSG00000233338 |
| PRAME | 4.096819019 | 17.11060681 | 0.125013819 | 0.267794977 | 0.674872254 | 11.36770858 | ENSG00000185686 |
| MIR4435-2HG | 4.023900439 | 16.26727212 | 1.14E-24 | 1.05E-22 | 8.108035486 | 130.1643973 | ENSG00000172965 |
| AADACL3 | 3.980605373 | 15.78634604 | 2.67E-06 | 2.74E-05 | 1.627800651 | 25.28438726 | ENSG00000188984 |
| LINC01414 | 3.931376967 | 15.25676269 | 0.000565284 | 0.0031942 | 0.932709143 | 14.47805638 | ENSG00000253554 |
| TBX1 | 3.91322193 | 15.06597286 | 8.80E-07 | 9.87E-06 | 1.862261374 | 28.5811309 | ENSG00000184058 |
| NA | 3.871675854 | 14.63829738 | 0.003955658 | 0.016884882 | 0.645182262 | 9.358359501 | ENSG00000271743 |
| COL8A1 | 3.835740133 | 14.27817944 | 3.05E-09 | 5.25E-08 | 2.871412842 | 40.84550716 | ENSG00000144810 |
| NA | 3.829793958 | 14.21945197 | 0.005761136 | 0.023222466 | 0.61549227 | 8.971981107 | ENSG00000286717 |
| VGLL2 | 3.772595074 | 13.66671943 | 0.000779846 | 0.004244008 | 0.92955223 | 12.9325428 | ENSG00000170162 |
| TCL1B | 3.756926121 | 13.51908983 | 3.61E-06 | 3.62E-05 | 1.935546786 | 25.93312154 | ENSG00000213231 |
| OTUD6A | 3.751165934 | 13.46522035 | 0.005762976 | 0.023224832 | 0.648339175 | 8.61185978 | ENSG00000189401 |
| LINC01508 | 3.745612362 | 13.4134864 | 0.000157482 | 0.001038707 | 1.320054516 | 17.26494878 | ENSG00000231107 |
| PANCR | 3.731322375 | 13.28128078 | 3.03E-05 | 0.00024307 | 1.657490643 | 21.33781002 | ENSG00000250103 |

| | | | | | | | |
|---|---|---|---|---|---|---|---|
| NA | 3.728322191 | 13.25369014 | 5.59E-24 | 4.91E-22 | 9.193726977 | 122.3146706 | ENSG00000286699 |
| NKX2-5 | 3.703223862 | 13.0251119 | 6.38E-20 | 4.06E-18 | 7.704905549 | 99.64290063 | ENSG00000183072 |
| NA | 3.685275616 | 12.8640732 | 0.001462812 | 0.007251072 | 0.959242223 | 12.34804561 | ENSG00000254186 |
| NA | 3.671100727 | 12.73829892 | 0.008700601 | 0.03280247 | 0.618649183 | 8.071952618 | ENSG00000286186 |
| GALR1 | 3.661327536 | 12.652298 | 1.19E-07 | 1.56E-06 | 2.587042873 | 32.35062184 | ENSG00000166573 |
| LINC01266 | 3.639666272 | 12.46374979 | 0.0117585 | 0.0420637 | 0.61549227 | 7.839102992 | ENSG00000224957 |
| LINC01833 | 3.631927037 | 12.39706791 | 0.001344997 | 0.006753773 | 0.95608531 | 11.87149947 | ENSG00000259439 |
| PTPRB | 3.620449993 | 12.298837 | 3.50E-17 | 1.70E-15 | 7.430006319 | 90.44274481 | ENSG00000127329 |
| NECTIN4 | 3.618729926 | 12.28418233 | 6.66E-06 | 6.26E-05 | 1.915327532 | 23.49137832 | ENSG00000143217 |
| NA | 3.614535207 | 12.24851724 | 0.000106486 | 0.000736694 | 1.607581397 | 19.60221338 | ENSG00000248213 |
| LINC01139 | 3.61187673 | 12.22596749 | 6.33E-54 | 3.26E-51 | 27.98737008 | 339.9933462 | ENSG00000215808 |
| NA | 3.55510973 | 11.75424305 | 4.54E-06 | 4.44E-05 | 2.243292921 | 26.18766494 | ENSG00000258636 |
| SGCG | 3.554851356 | 11.75213815 | 1.27E-06 | 1.39E-05 | 2.824660509 | 33.50595818 | ENSG00000102683 |
| NA | 3.524534022 | 11.50775114 | 0.004220699 | 0.017828611 | 0.932709143 | 10.94141507 | ENSG00000282057 |
| TEX11 | 3.480067374 | 11.15847042 | 0.015203927 | 0.051628724 | 0.674872254 | 7.235285152 | ENSG00000120498 |
| NA | 3.431506936 | 10.78913229 | 0.01570923 | 0.053024016 | 0.648339175 | 6.909018588 | ENSG00000259205 |
| RGS4 | 3.429717613 | 10.77575921 | 4.05E-05 | 0.000312806 | 1.882480627 | 20.52470009 | ENSG00000117152 |
| CNTN5 | 3.413679458 | 10.65663062 | 0.02375025 | 0.074318273 | 0.61549227 | 6.736718825 | ENSG00000149972 |
| EBF2 | 3.411512531 | 10.64063638 | 0.023722647 | 0.074256975 | 0.61549227 | 6.661085195 | ENSG00000221818 |
| NA | 3.410484609 | 10.63305761 | 1.57E-20 | 1.05E-18 | 10.96682068 | 115.4632388 | ENSG00000280623 |
| NA | 3.39023735 | 10.48487204 | 6.19E-07 | 7.17E-06 | 2.868255929 | 30.06653265 | ENSG00000250131 |
| HCLS1 | 3.383283939 | 10.4344593 | 2.41E-27 | 2.78E-25 | 15.0085603 | 156.3188687 | ENSG00000180353 |
| SERPINE1 | 3.366546271 | 10.3141017 | 3.98E-69 | 3.69E-66 | 42.4879459 | 436.8225074 | ENSG00000106366 |
| LINC00654 | 3.363973239 | 10.295723 | 0.004454447 | 0.018653607 | 0.985775302 | 9.917037658 | ENSG00000205181 |
| TRPC6 | 3.35563858 | 10.23641459 | 3.01E-09 | 5.18E-08 | 4.90234343 | 49.3794718 | ENSG00000137672 |
| DPPA5 | 3.350992619 | 10.20350293 | 2.55E-18 | 1.39E-16 | 9.773215429 | 100.3034695 | ENSG00000203909 |
| LINC-ROR | 3.345755772 | 10.16653232 | 1.57E-26 | 1.66E-24 | 14.02406277 | 142.3345921 | ENSG00000258609 |
| NA | 3.344694402 | 10.15905569 | 0.020169025 | 0.065078813 | 0.645182262 | 6.481299358 | ENSG00000234699 |
| LINC02709 | 3.342162967 | 10.14124567 | 0.005413748 | 0.022037867 | 0.926395317 | 9.579486173 | ENSG00000245522 |
| TRDN | 3.320077854 | 9.987183325 | 1.51E-06 | 1.62E-05 | 2.84172285 | 28.52481792 | ENSG00000186439 |
| FZD10-AS1 | 3.266385038 | 9.622321734 | 0.002512196 | 0.011510762 | 1.237298365 | 12.07710434 | ENSG00000250208 |
| PLA2G2A | 3.238396603 | 9.437446775 | 7.84E-47 | 2.91E-44 | 32.50690053 | 306.6625937 | ENSG00000188257 |
| POT1-AS1 | 3.229308543 | 9.378183726 | 0.004195707 | 0.017750112 | 1.230984539 | 11.80010272 | ENSG00000224897 |
| SP5 | 3.195835317 | 9.163097163 | 0.010064135 | 0.036962107 | 0.959242223 | 8.724047713 | ENSG00000204335 |
| LOC157273 | 3.189878722 | 9.125342585 | 1.46E-08 | 2.28E-07 | 4.992788881 | 46.11223441 | ENSG00000248538 |
| NA | 3.177766349 | 9.049050047 | 0.002821951 | 0.012639449 | 1.320054516 | 11.65742084 | ENSG00000256417 |
| PRDM1 | 3.146191508 | 8.853153945 | 1.46E-05 | 0.000126515 | 2.954168993 | 25.61390302 | ENSG00000057657 |
| MYCT1 | 3.144792264 | 8.844571589 | 0.01140408 | 0.041017506 | 0.985775302 | 8.529178109 | ENSG00000120279 |
| LCP1 | 3.141183664 | 8.822476391 | 2.18E-39 | 5.77E-37 | 28.82163621 | 255.0657545 | ENSG00000136167 |
| PADI3 | 3.133743585 | 8.77709538 | 3.40E-39 | 8.89E-37 | 28.1890816 | 248.3195113 | ENSG00000142619 |
| AGMO | 3.123282119 | 8.713679918 | 0.000489757 | 0.002811138 | 1.918484445 | 16.69878455 | ENSG00000187546 |
| TEX37 | 3.113486281 | 8.654714813 | 0.000129325 | 0.000873829 | 2.196540588 | 19.16645842 | ENSG00000172073 |
| LAPTM5 | 3.111222535 | 8.641145268 | 0.000127885 | 0.000867256 | 2.193383675 | 19.11763151 | ENSG00000162511 |
| CCDC60 | 3.061517111 | 8.348500601 | 0.01552241 | 0.052508185 | 0.92955223 | 7.891617128 | ENSG00000183273 |
| LINC01152 | 3.055829578 | 8.315653114 | 0.000944202 | 0.005012846 | 1.971550604 | 16.06458438 | ENSG00000256124 |
| NLRP10 | 3.054597776 | 8.308556076 | 0.000573834 | 0.00323364 | 1.888794453 | 15.79831802 | ENSG00000182261 |
| CDKN1A | 3.04537355 | 8.255602741 | 1.12E-201 | 1.04E-197 | 227.1284045 | 1877.927819 | ENSG00000124762 |
| IZUMO2 | 3.042124282 | 8.23703022 | 0.001800125 | 0.008685853 | 1.891951366 | 15.69730339 | ENSG00000161652 |
| NA | 3.035022819 | 8.196584228 | 0.000143329 | 0.00095453 | 2.868255929 | 23.55583865 | ENSG00000288271 |
| NA | 3.033791798 | 8.189593243 | 0.000584989 | 0.003290496 | 1.88563754 | 15.58847627 | ENSG00000259345 |
| C1orf87 | 3.025641904 | 8.143460035 | 0.017744764 | 0.058441072 | 0.985775302 | 7.861561213 | ENSG00000162598 |
| NA | 3.011288359 | 8.062841483 | 0.006798421 | 0.026774742 | 1.29667835 | 10.33727081 | ENSG00000229282 |
| NA | 3.006435595 | 8.035766231 | 0.017000365 | 0.056476562 | 0.926395317 | 7.595406479 | ENSG00000237346 |
| UNC93A | 3.001564942 | 8.008682589 | 1.29E-06 | 1.41E-05 | 4.211686612 | 33.35579022 | ENSG00000112494 |
| NPIPA5 | 3.000147677 | 8.000818936 | 0.017922242 | 0.05883295 | 0.926395317 | 7.554065643 | ENSG00000183793 |
| NTSR2 | 2.996283236 | 7.979416409 | 0.021545979 | 0.068767551 | 0.932709143 | 7.546579569 | ENSG00000169006 |
| FOXL1 | 2.991469284 | 7.952835246 | 0.027565714 | 0.083943092 | 0.926395317 | 7.482780512 | ENSG00000176678 |
| LINC01265 | 2.990980223 | 7.950139763 | 0.017823457 | 0.058585823 | 0.982618389 | 7.693060297 | ENSG00000249740 |
| LINC01612 | 2.983982899 | 7.911673551 | 0.002573641 | 0.011740059 | 1.571577579 | 12.48605258 | ENSG00000250266 |
| FLI1 | 2.980427004 | 7.892197198 | 2.67E-10 | 5.42E-09 | 7.036347121 | 55.42231666 | ENSG00000151702 |
| PCDHGB5 | 2.976868111 | 7.872752428 | 1.88E-08 | 2.88E-07 | 5.783264191 | 45.22685166 | ENSG00000276547 |
| LAMP3 | 2.975022855 | 7.862689348 | 2.70E-06 | 2.76E-05 | 3.857188144 | 30.10363661 | ENSG00000078081 |
| POTEF | 2.969451909 | 7.832386231 | 3.30E-05 | 0.000261923 | 3.454058207 | 27.33639127 | ENSG00000196604 |
| PTPRH | 2.965579132 | 7.811389132 | 0.000672491 | 0.00372169 | 2.279296739 | 17.63679306 | ENSG00000080031 |
| TCEA3 | 2.964839269 | 7.807384213 | 0.018493958 | 0.060378195 | 0.985775302 | 7.535294649 | ENSG00000204219 |
| SLC25A3P1 | 2.959895243 | 7.780674589 | 0.012734043 | 0.044927765 | 1.243612191 | 9.838592867 | ENSG00000236253 |
| SNTG1 | 2.957759732 | 7.769165974 | 1.85E-05 | 0.000155374 | 3.208848969 | 24.80404227 | ENSG00000147481 |
| CYP2B6 | 2.948660555 | 7.720319505 | 7.38E-65 | 5.95E-62 | 58.17778998 | 449.148198 | ENSG00000197408 |
| NA | 2.94445783 | 7.697862115 | 0.001029454 | 0.005403543 | 1.941860612 | 14.7893508 | ENSG00000225087 |
| LINC02432 | 2.937502389 | 7.660838931 | 0.026049299 | 0.07999962 | 0.923238405 | 7.216514158 | ENSG00000248810 |
| NA | 2.922663516 | 7.582447039 | 0.018981354 | 0.0617302 | 0.95608531 | 7.246570073 | ENSG00000270265 |
| NA | 2.917455672 | 7.555125243 | 0.00113209 | 0.005854394 | 1.941860612 | 14.53068213 | ENSG00000276975 |
| SRRM5 | 2.912327775 | 7.528319074 | 0.000637002 | 0.003544339 | 2.255920572 | 16.85231331 | ENSG00000226763 |
| NA | 2.894623979 | 7.436501056 | 0.000179621 | 0.001169331 | 2.957325906 | 21.57065965 | ENSG00000282024 |

| | | | | | | | |
|---|---|---|---|---|---|---|---|
| NA | 2.886015923 | 7.392262168 | 0.014771555 | 0.050430501 | 1.230984539 | 9.27567783 | ENSG00000286458 |
| GRAMD1C | 2.867947676 | 7.300259137 | 3.15E-07 | 3.85E-06 | 6.44738793 | 46.79190912 | ENSG00000178075 |
| ODF3L2 | 2.846766558 | 7.193862367 | 0.011187174 | 0.040410671 | 1.29667835 | 9.208079926 | ENSG00000181781 |
| NA | 2.846096777 | 7.190523341 | 0.017369266 | 0.057450939 | 1.349744508 | 9.418471331 | ENSG00000157306 |
| F7 | 2.843432575 | 7.17725697 | 0.027663752 | 0.084162833 | 0.959242223 | 6.860191679 | ENSG00000057593 |
| NA | 2.838659812 | 7.153552207 | 0.06243238 | 0.158842186 | 0.932709143 | 6.778059661 | ENSG00000230183 |
| LRFN5 | 2.829689567 | 7.109211554 | 2.33E-10 | 4.81E-09 | 8.217422414 | 58.62096845 | ENSG00000165379 |
| NA | 2.827755301 | 7.099686399 | 0.029889899 | 0.089351513 | 0.959242223 | 6.770023934 | ENSG00000279267 |
| NTRK2 | 2.815238862 | 7.038357826 | 0.014512938 | 0.049719554 | 1.320054516 | 9.125947908 | ENSG00000148053 |
| PTGDR2 | 2.811500914 | 7.020145411 | 0.005024376 | 0.020670763 | 1.574734492 | 11.15505567 | ENSG00000183134 |
| NA | 2.809789387 | 7.011822071 | 0.006896673 | 0.027086871 | 1.607581397 | 11.24522341 | ENSG00000231464 |
| NA | 2.80229283 | 6.975481615 | 0.003916523 | 0.016744829 | 1.99492677 | 13.61188265 | ENSG00000248587 |
| CBY2 | 2.800287916 | 6.965794517 | 0.045585594 | 0.124577322 | 0.923238405 | 6.556383336 | ENSG00000174015 |
| CREB5 | 2.798450493 | 6.956928498 | 6.56E-12 | 1.72E-10 | 10.39502152 | 72.70147314 | ENSG00000146592 |
| NA | 2.788989618 | 6.91145576 | 0.034089027 | 0.099200192 | 0.926395317 | 6.518841347 | ENSG00000173727 |
| NA | 2.779915833 | 6.868122792 | 0.013451902 | 0.046811407 | 1.263831445 | 8.747055588 | ENSG00000213514 |
| NODAL | 2.762858729 | 6.78739855 | 1.28E-40 | 3.95E-38 | 44.3565211 | 300.5240301 | ENSG00000156574 |
| LINC02617 | 2.761368006 | 6.780388814 | 9.71E-20 | 6.08E-18 | 19.5482123 | 132.0657953 | ENSG00000256288 |
| PRPH2 | 2.760445342 | 6.776053857 | 0.002714137 | 0.012230443 | 1.938703699 | 13.00014071 | ENSG00000112619 |
| KRT81 | 2.760117966 | 6.774516413 | 0.006523611 | 0.025840693 | 1.912170619 | 12.86070802 | ENSG00000205426 |
| LOC102724770 | 2.759365753 | 6.77098514 | 0.034263312 | 0.099582285 | 0.985775302 | 6.548897262 | ENSG00000278817 |
| LNCTAM34A | 2.759041144 | 6.769461827 | 4.67E-05 | 0.000354379 | 3.735271262 | 25.64819581 | ENSG00000234546 |
| SATB2-AS1 | 2.758597412 | 6.767380054 | 0.000313881 | 0.001907365 | 2.861942103 | 19.39930805 | ENSG00000225953 |
| LINC01162 | 2.747079543 | 6.71356724 | 0.006680603 | 0.026355533 | 1.571577579 | 10.62219654 | ENSG00000232790 |
| SERPINB9 | 2.736618 | 6.665060635 | 4.40E-134 | 1.63E-130 | 220.0317079 | 1465.935419 | ENSG00000170542 |
| KRT17 | 2.734469756 | 6.655143416 | 3.55E-05 | 0.00027909 | 5.074267255 | 33.69191599 | ENSG00000128422 |
| NA | 2.728379669 | 6.627109094 | 0.062588224 | 0.159188986 | 0.982618389 | 6.398617687 | ENSG00000227719 |
| FMR1-AS1 | 2.722236534 | 6.598950219 | 0.050050102 | 0.133876257 | 0.982618389 | 6.417938335 | ENSG00000268066 |
| DPEP3 | 2.721935093 | 6.597571555 | 2.85E-16 | 1.24E-14 | 15.27083188 | 100.8295038 | ENSG00000141096 |
| SPAAR | 2.710058917 | 6.543483689 | 0.047521488 | 0.128541048 | 0.985775302 | 6.293477795 | ENSG00000235387 |
| ARHGDIB | 2.695098465 | 6.475979709 | 0.004268561 | 0.017997538 | 1.915327532 | 12.37766349 | ENSG00000111348 |
| SPDYE6 | 2.694582016 | 6.473661883 | 0.037238401 | 0.106164671 | 0.95608531 | 6.181289862 | ENSG00000260097 |
| NA | 2.680841713 | 6.412299063 | 0.046223126 | 0.125720292 | 0.959242223 | 6.121178032 | ENSG00000272320 |
| MCCC2 | 2.679649415 | 6.407001884 | 0.000466187 | 0.002694201 | 2.871412842 | 18.40162574 | ENSG00000131844 |
| CTCFL | 2.675499634 | 6.388599206 | 4.38E-08 | 6.27E-07 | 9.1670962 | 57.68653555 | ENSG00000124092 |
| ARHGAP24 | 2.67496857 | 6.38624796 | 7.16E-06 | 6.69E-05 | 4.747579643 | 30.44922382 | ENSG00000138639 |
| NA | 2.671754778 | 6.37203759 | 0.011007738 | 0.039917319 | 1.627800651 | 10.21704715 | ENSG00000253966 |
| SGCD | 2.664247416 | 6.3389655 | 0.011143142 | 0.040297949 | 1.548201413 | 9.973350641 | ENSG00000170624 |
| NA | 2.662622956 | 6.331831906 | 0.080464829 | 0.192579435 | 0.985775302 | 6.193012816 | ENSG00000257869 |
| MIR2052HG | 2.654346792 | 6.295612707 | 0.00569282 | 0.023007135 | 1.968393691 | 12.20905096 | ENSG00000254349 |
| MRPS9-AS2 | 2.649037445 | 6.272486434 | 0.026394447 | 0.080952288 | 1.266988357 | 7.978097646 | ENSG00000224509 |
| NEAT1 | 2.643037762 | 6.246455437 | 8.09E-16 | 3.40E-14 | 28.18267008 | 175.8984494 | ENSG00000245532 |
| GOLGA6C | 2.635923424 | 6.215728221 | 0.009099223 | 0.034001009 | 1.888794453 | 11.76580993 | ENSG00000167195 |
| P4HA3 | 2.631238134 | 6.195574794 | 0.059524857 | 0.152929539 | 1.012308381 | 6.053580129 | ENSG00000149380 |
| SEMA3C | 2.613419367 | 6.119523695 | 1.33E-05 | 0.000116039 | 4.777269635 | 29.21120581 | ENSG00000075223 |
| NA | 2.611570608 | 6.111686779 | 0.003230848 | 0.014169553 | 2.223073667 | 13.64573741 | ENSG00000229646 |
| MEIOB | 2.607077686 | 6.092683026 | 1.64E-29 | 2.22E-27 | 35.2594557 | 213.9454041 | ENSG00000162039 |
| TMEM232 | 2.603635939 | 6.078165422 | 4.25E-07 | 5.09E-06 | 6.973810223 | 42.58742768 | ENSG00000186952 |
| HAO2 | 2.603346462 | 6.076945956 | 0.004316968 | 0.01817274 | 2.255920572 | 13.5813887 | ENSG00000116882 |
| GCNT4 | 2.602828011 | 6.074762521 | 9.61E-28 | 1.14E-25 | 33.30063045 | 202.4513712 | ENSG00000176928 |
| KCNMB1 | 2.598115599 | 6.054952319 | 2.51E-09 | 4.39E-08 | 10.09674613 | 61.38886662 | ENSG00000145936 |
| LINC02301 | 2.592835277 | 6.032831455 | 2.32E-05 | 0.000190683 | 4.757050382 | 28.79900838 | ENSG00000258743 |
| NA | 2.575005816 | 5.958733875 | 0.02878771 | 0.086677915 | 1.29667835 | 7.663004382 | ENSG00000233695 |
| GPER1 | 2.57417421 | 5.955300098 | 0.019482926 | 0.063139807 | 1.581048318 | 9.482270388 | ENSG00000164850 |
| NA | 2.573795498 | 5.953737019 | 0.003102956 | 0.01371576 | 2.799405206 | 16.85611216 | ENSG00000189229 |
| ERVH48-1 | 2.562599536 | 5.907712161 | 6.25E-38 | 1.43E-35 | 508.809632 | 3004.655753 | ENSG00000233056 |
| CALHM3 | 2.555653183 | 5.87933581 | 0.007856474 | 0.03017484 | 1.909013706 | 11.19215962 | ENSG00000183128 |
| NA | 2.541332058 | 5.82126243 | 0.008771148 | 0.033028132 | 1.968393691 | 11.24098653 | ENSG00000229931 |
| PLAAT4 | 2.540414991 | 5.81756325 | 0.008890344 | 0.033354989 | 1.888794453 | 11.06868677 | ENSG00000133321 |
| NA | 2.531993032 | 5.783701243 | 0.014908585 | 0.050802669 | 1.604424484 | 9.219364847 | ENSG00000233058 |
| NA | 2.516964794 | 5.7237664 | 0.032088437 | 0.094491118 | 1.293521437 | 7.362994886 | ENSG00000228157 |
| ITPRID1 | 2.516180959 | 5.720657451 | 0.000103261 | 0.000717064 | 5.078701945 | 29.18582481 | ENSG00000180347 |
| FGF17 | 2.512064085 | 5.704356252 | 0.011021004 | 0.039949792 | 1.965236778 | 11.03863085 | ENSG00000158815 |
| NA | 2.511322369 | 5.701424293 | 0.037905658 | 0.107641395 | 1.237298365 | 7.156402328 | ENSG00000228577 |
| NA | 2.507101164 | 5.684766784 | 1.11E-09 | 2.05E-08 | 15.3143296 | 86.14788083 | ENSG00000287492 |
| CHL1 | 2.499828773 | 5.656182902 | 0.047203588 | 0.127923863 | 1.323211429 | 7.291160102 | ENSG00000134121 |
| PADI1 | 2.493447885 | 5.631221445 | 7.26E-07 | 8.32E-06 | 7.642368652 | 43.0935917 | ENSG00000142623 |
| ACTA1 | 2.487007762 | 5.60613996 | 9.79E-74 | 1.07E-70 | 191.7435819 | 1075.235985 | ENSG00000143632 |
| MECOM | 2.485036949 | 5.598486844 | 2.03E-06 | 2.13E-05 | 7.020562556 | 39.16903465 | ENSG00000085276 |
| TMEM82 | 2.481021205 | 5.582925114 | 0.001014177 | 0.005333062 | 3.176002064 | 17.71479981 | ENSG00000162460 |
| ABHD12B | 2.478971962 | 5.575000608 | 2.77E-30 | 4.01E-28 | 49.84594459 | 276.9625373 | ENSG00000131969 |
| BAHCC1 | 2.474290075 | 5.556937735 | 2.63E-21 | 1.85E-19 | 27.9609347 | 155.5079087 | ENSG00000266074 |
| CFAP100 | 2.474062269 | 5.556060347 | 0.018853596 | 0.061433288 | 1.604424484 | 8.851757447 | ENSG00000163885 |
| TAGLN | 2.473045198 | 5.552144817 | 8.28E-187 | 5.11E-183 | 515.4779282 | 2861.259272 | ENSG00000149591 |

| | | | | | | | |
|---|---|---|---|---|---|---|---|
| NA | 2.461315797 | 5.507187763 | 5.01E-05 | 0.000377423 | 5.117862675 | 28.16133578 | ENSG00000249626 |
| FAT4 | 2.460795075 | 5.505200373 | 3.44E-27 | 3.89E-25 | 46.4684264 | 256.0427307 | ENSG00000196159 |
| CMKLR1 | 2.460206288 | 5.502954072 | 3.18E-13 | 9.95E-12 | 20.44995367 | 112.2909557 | ENSG00000174600 |
| NA | 2.459825748 | 5.501502747 | 0.03872713 | 0.109662859 | 1.263831445 | 7.006672407 | ENSG00000255239 |
| PCDH11Y | 2.459540151 | 5.500413776 | 0.000162919 | 0.00107114 | 4.77095581 | 26.19883824 | ENSG00000099715 |
| NA | 2.457753252 | 5.493605269 | 0.001805282 | 0.008695193 | 3.11662208 | 17.26451074 | ENSG00000276627 |
| MUC4 | 2.446431355 | 5.450661601 | 6.79E-12 | 1.77E-10 | 17.73595787 | 97.11384111 | ENSG00000145113 |
| PLK2 | 2.444013775 | 5.441535367 | 3.91E-105 | 1.21E-101 | 196.1459384 | 1065.924064 | ENSG00000145632 |
| SLC52A1 | 2.441223187 | 5.431020041 | 0.000241343 | 0.001519894 | 4.191467358 | 22.56834196 | ENSG00000132517 |
| KCNJ2 | 2.436187919 | 5.412097833 | 6.12E-08 | 8.50E-07 | 9.256263875 | 50.04328985 | ENSG00000123700 |
| NA | 2.435488417 | 5.409474371 | 0.045681011 | 0.124782882 | 1.27014527 | 6.878962673 | ENSG00000267723 |
| ZNF436-AS1 | 2.428985547 | 5.385146329 | 0.022169105 | 0.070320417 | 1.601267572 | 8.589289939 | ENSG00000249087 |
| CCDC146 | 2.424730327 | 5.369286274 | 0.002159679 | 0.010167003 | 3.126092818 | 16.9650509 | ENSG00000135205 |
| NA | 2.424531758 | 5.368547311 | 0.026738859 | 0.081819056 | 1.627800651 | 8.638116849 | ENSG00000245146 |
| VAV1 | 2.407988461 | 5.307338113 | 3.13E-16 | 1.36E-14 | 26.74375789 | 140.9233983 | ENSG00000141968 |
| PIWIL2 | 2.40676904 | 5.302854044 | 1.72E-06 | 1.83E-05 | 8.225014016 | 43.68871254 | ENSG00000197181 |
| NA | 2.406276166 | 5.301042718 | 0.001212938 | 0.006198125 | 3.785180508 | 20.12648712 | ENSG00000257951 |
| LEFTY1 | 2.399592028 | 5.276539308 | 1.17E-73 | 1.20E-70 | 129.613626 | 682.6245303 | ENSG00000243709 |
| PAGE4 | 2.39683358 | 5.26646015 | 0.061329354 | 0.156761585 | 1.266988357 | 6.649800274 | ENSG00000101951 |
| LINC00989 | 2.395057447 | 5.259980485 | 0.021340146 | 0.068216326 | 1.876166801 | 9.954579647 | ENSG00000250334 |
| RERG | 2.392401517 | 5.250306028 | 0.007136678 | 0.027840705 | 2.590199786 | 13.49165899 | ENSG00000134533 |
| ZNF648 | 2.385140583 | 5.22394817 | 0.027931448 | 0.0848102 | 1.601267572 | 8.35319112 | ENSG00000179930 |
| ADGRL4 | 2.375969844 | 5.190846563 | 4.67E-07 | 5.54E-06 | 9.484410772 | 49.29222683 | ENSG00000162618 |
| NA | 2.371137662 | 5.173489359 | 5.95E-40 | 1.75E-37 | 91.06364752 | 469.7891568 | ENSG00000251095 |
| LOC100505774 | 2.368655614 | 5.164596414 | 0.03499621 | 0.101200075 | 1.607581397 | 8.259224528 | ENSG00000226833 |
| ANXA3 | 2.366644329 | 5.157401378 | 8.79E-91 | 1.63E-87 | 188.680026 | 972.9615607 | ENSG00000138772 |
| LOC100130691 | 2.363631868 | 5.146643553 | 8.45E-06 | 7.68E-05 | 7.297340922 | 37.63948091 | ENSG00000213963 |
| NA | 2.363553929 | 5.146365524 | 0.012651727 | 0.044698268 | 2.279296739 | 11.56356587 | ENSG00000198857 |
| ZIC1 | 2.361949551 | 5.140645584 | 0.000367602 | 0.002188611 | 4.886558866 | 24.83409819 | ENSG00000152977 |
| LINC02588 | 2.359777765 | 5.132912847 | 0.056283214 | 0.146489691 | 1.263831445 | 6.507556426 | ENSG00000257842 |
| LOC728755 | 2.357605788 | 5.125191061 | 0.002113205 | 0.009971014 | 4.119459722 | 21.17734484 | ENSG00000258932 |
| LINC02582 | 2.352474567 | 5.106994717 | 5.48E-13 | 1.68E-11 | 18.87334004 | 96.22747067 | ENSG00000261780 |
| FAM20A | 2.351150491 | 5.102309774 | 2.92E-06 | 2.97E-05 | 8.26733166 | 42.17738013 | ENSG00000108950 |
| NA | 2.34612312 | 5.084560655 | 0.049662688 | 0.133070443 | 1.293521437 | 6.511355273 | ENSG00000236992 |
| PTP4A1 | 2.34308018 | 5.073847577 | 0.019086215 | 0.061984213 | 1.941860612 | 9.793564804 | ENSG00000112245 |
| NA | 2.342443494 | 5.071608894 | 0.023001081 | 0.072426442 | 1.945017524 | 9.801050878 | ENSG00000267547 |
| MYOCD | 2.338064208 | 5.056237421 | 1.35E-10 | 2.91E-09 | 16.04246341 | 81.67053146 | ENSG00000141052 |
| NA | 2.335907987 | 5.048686119 | 0.01609636 | 0.054074667 | 2.16053677 | 11.09874268 | ENSG00000283696 |
| MAFB | 2.329757553 | 5.027208596 | 8.22E-19 | 4.75E-17 | 30.82603372 | 155.2732356 | ENSG00000204103 |
| GATA4 | 2.328384567 | 5.022426573 | 0.043879849 | 0.120878289 | 1.568420667 | 7.940444037 | ENSG00000136574 |
| NA | 2.324949808 | 5.010483435 | 0.014215824 | 0.048891462 | 2.305829818 | 11.34612642 | ENSG00000256139 |
| ADGRG3 | 2.323344677 | 5.00491191 | 0.008450895 | 0.032050109 | 2.983858985 | 14.616613 | ENSG00000182885 |
| NA | 2.315704288 | 4.978476387 | 0.032288732 | 0.094869904 | 1.601267572 | 7.951728958 | ENSG00000236212 |
| NA | 2.304170476 | 4.938833992 | 0.023022902 | 0.072482837 | 1.915327532 | 9.482270388 | ENSG00000237713 |
| TNFRSF10C | 2.304115879 | 4.938647089 | 5.06E-12 | 1.36E-10 | 17.79533785 | 87.92211877 | ENSG00000173535 |
| LINC00504 | 2.300804064 | 4.927323056 | 0.012120804 | 0.043159841 | 3.010392064 | 14.51939721 | ENSG00000248360 |
| LMX1B | 2.299241207 | 4.921988227 | 0.003180334 | 0.013991052 | 3.176002064 | 15.60301039 | ENSG00000136944 |
| BCAS1 | 2.294721693 | 4.906593297 | 0.00122302 | 0.006238473 | 3.797808159 | 18.72040623 | ENSG00000064787 |
| LINC02525 | 2.293966466 | 4.904025449 | 5.80E-06 | 5.52E-05 | 8.71721393 | 42.33058249 | ENSG00000230269 |
| SMIM38 | 2.292711119 | 4.899760117 | 0.01551002 | 0.052485431 | 2.530819802 | 12.51329733 | ENSG00000284713 |
| PTGIR | 2.292320938 | 4.898435141 | 0.090769374 | 0.210922714 | 1.290364524 | 6.233365965 | ENSG00000160013 |
| PARVG | 2.291335537 | 4.895090514 | 0.000758327 | 0.004139865 | 4.135244286 | 20.23173863 | ENSG00000138964 |
| NA | 2.288276645 | 4.884722636 | 0.060203339 | 0.154415817 | 1.263831445 | 6.200060856 | ENSG00000280202 |
| ITGAX | 2.27851717 | 4.851790208 | 0.044109383 | 0.121438399 | 1.5450445 | 7.625462394 | ENSG00000140678 |
| SERHL | 2.275790812 | 4.842630114 | 0.001434784 | 0.007127386 | 4.218000437 | 20.28480242 | ENSG00000172250 |
| LAIR1 | 2.273449981 | 4.834779122 | 0.070485655 | 0.173860287 | 1.320054516 | 6.278943681 | ENSG00000167613 |
| NA | 2.270030683 | 4.82333389 | 0.078132197 | 0.188383263 | 1.240455278 | 6.034809134 | ENSG00000279532 |
| DCC | 2.268760165 | 4.819088065 | 6.30E-06 | 5.95E-05 | 8.336182383 | 40.04368214 | ENSG00000187323 |
| NA | 2.267312774 | 4.814255715 | 0.017579683 | 0.057954793 | 2.279296739 | 10.88510208 | ENSG00000273355 |
| SCARA5 | 2.263471444 | 4.801454294 | 0.017972024 | 0.058975476 | 2.56682362 | 12.32178854 | ENSG00000168079 |
| NA | 2.256482806 | 4.778251549 | 0.066971083 | 0.167373523 | 1.538730674 | 7.448925751 | ENSG00000224786 |
| PTCHD4 | 2.250503127 | 4.758487652 | 1.50E-10 | 3.20E-09 | 18.68878857 | 88.32683012 | ENSG00000244694 |
| NA | 2.246649029 | 4.745792518 | 1.94E-12 | 5.48E-11 | 34.40200881 | 163.5659171 | ENSG00000223745 |
| INMT | 2.243095542 | 4.734117591 | 0.009946074 | 0.036604553 | 2.901102834 | 13.69825155 | ENSG00000241644 |
| BTG2 | 2.242048334 | 4.73068249 | 1.44E-31 | 2.23E-29 | 60.25684837 | 285.2989328 | ENSG00000159388 |
| NA | 2.232763653 | 4.700335226 | 0.066745561 | 0.166944969 | 1.657490643 | 7.659317155 | ENSG00000259635 |
| PRKN | 2.231954213 | 4.697698791 | 0.000133859 | 0.000900245 | 6.811357136 | 31.78869449 | ENSG00000185345 |
| CDKN2A | 2.224114169 | 4.672239307 | 0.019197765 | 0.062302816 | 2.252763659 | 10.49872369 | ENSG00000147889 |
| SCNN1B | 2.21250335 | 4.634787999 | 0.000817038 | 0.004420018 | 4.70082731 | 21.94955197 | ENSG00000168447 |
| LINC01630 | 2.210555251 | 4.628533779 | 0.037199394 | 0.106092846 | 1.879323714 | 8.807167418 | ENSG00000227115 |
| NA | 2.209272494 | 4.624420198 | 0.084738366 | 0.200048566 | 1.604424484 | 7.419419489 | ENSG00000266217 |
| NA | 2.20638125 | 4.615161872 | 0.024769544 | 0.07688473 | 2.22623058 | 10.26956129 | ENSG00000257883 |
| NA | 2.206014252 | 4.613988 | 0.00127779 | 0.006467113 | 4.478994239 | 20.60303326 | ENSG00000227712 |
| VSIR | 2.205881308 | 4.613562841 | 0.004495112 | 0.018789445 | 3.486905112 | 16.1889333 | ENSG00000107738 |

| | | | | | | | |
|---|---|---|---|---|---|---|---|
| NA | 2.203246863 | 4.605145897 | 0.030182123 | 0.090079786 | 1.941860612 | 8.840472526 | ENSG00000287527 |
| DCDC2 | 2.199750311 | 4.593998262 | 0.074593154 | 0.181765647 | 1.266988357 | 5.832453456 | ENSG00000146038 |
| DNAH6 | 2.19899898 | 4.59160641 | 0.054867687 | 0.143632445 | 1.571577579 | 7.272827141 | ENSG00000115423 |
| NA | 2.194738521 | 4.578066823 | 0.060774738 | 0.15551553 | 1.660647556 | 7.418869836 | ENSG00000266573 |
| LAMA2 | 2.192408707 | 4.57067965 | 1.88E-25 | 1.88E-23 | 68.09166837 | 310.1420459 | ENSG00000196569 |
| TWIST1 | 2.191467574 | 4.567698968 | 0.00973047 | 0.035939522 | 2.924479001 | 13.2480741 | ENSG00000122691 |
| PKD1L2 | 2.189137151 | 4.560326601 | 7.35E-13 | 2.23E-11 | 29.81688225 | 136.4043103 | ENSG00000166473 |
| GABRP | 2.186132905 | 4.550840127 | 4.21E-24 | 3.71E-22 | 47.16549475 | 215.0885794 | ENSG00000094755 |
| NA | 2.181097877 | 4.534985307 | 0.002600837 | 0.011837897 | 4.049331223 | 18.56264058 | ENSG00000237118 |
| LBX2-AS1 | 2.179686091 | 4.530549654 | 0.037702237 | 0.107206698 | 1.888794453 | 8.608060934 | ENSG00000257702 |
| NA | 2.177997022 | 4.52524851 | 4.95E-05 | 0.000373341 | 8.492223946 | 37.99949061 | ENSG00000286811 |
| ARSL | 2.176701295 | 4.521186075 | 2.81E-40 | 8.39E-38 | 85.41522096 | 386.4848622 | ENSG00000157399 |
| CATSPER2 | 2.169967147 | 4.50013146 | 5.13E-05 | 0.000384767 | 8.128352438 | 36.97230205 | ENSG00000166762 |
| LRP1B | 2.16870368 | 4.496192114 | 0.061608675 | 0.157237013 | 1.598110659 | 7.156402328 | ENSG00000168702 |
| DLGAP2 | 2.160786571 | 4.471585838 | 1.45E-07 | 1.88E-06 | 12.0916729 | 54.08653321 | ENSG00000198010 |
| PF4 | 2.159440212 | 4.46741479 | 0.008976986 | 0.033618803 | 3.810435811 | 17.00964093 | ENSG00000163737 |
| SPAG17 | 2.153073958 | 4.447744633 | 0.010050177 | 0.036936345 | 3.490062025 | 15.62276907 | ENSG00000155761 |
| NA | 2.147170433 | 4.429581595 | 0.001731447 | 0.008400692 | 4.38676735 | 19.58289273 | ENSG00000286214 |
| MMP9 | 2.146441827 | 4.427345084 | 1.03E-25 | 1.04E-23 | 63.90422974 | 282.8378605 | ENSG00000100985 |
| COLEC11 | 2.145243913 | 4.423670449 | 0.000314867 | 0.001912103 | 6.570680286 | 29.23702485 | ENSG00000118004 |
| SYTL2 | 2.141757568 | 4.412993334 | 1.86E-22 | 1.44E-20 | 47.5540202 | 209.6327768 | ENSG00000137501 |
| CSMD1 | 2.141417871 | 4.411954372 | 0.000590307 | 0.003314375 | 5.392761905 | 23.85518687 | ENSG00000183117 |
| TMPRSS4 | 2.132601658 | 4.385075424 | 0.045254545 | 0.12381868 | 1.968393691 | 8.4803512 | ENSG00000137648 |
| LINC01091 | 2.120556606 | 4.348616865 | 0.028934508 | 0.087014793 | 2.193383675 | 9.609542088 | ENSG00000249464 |
| LINC00304 | 2.119048209 | 4.344072583 | 0.026070813 | 0.080051921 | 2.249606746 | 9.744737895 | ENSG00000180422 |
| SGCA | 2.118008744 | 4.340943798 | 0.056151868 | 0.146188891 | 1.862261374 | 8.165369556 | ENSG00000108823 |
| SOX1 | 2.115386545 | 4.333060994 | 0.029939229 | 0.089457732 | 2.279296739 | 9.793564804 | ENSG00000182968 |
| SYT4 | 2.114613806 | 4.330740733 | 3.32E-13 | 1.03E-11 | 26.63140945 | 115.4452407 | ENSG00000132872 |
| LINC00472 | 2.11083702 | 4.319418251 | 0.000293017 | 0.001800057 | 6.933371716 | 30.11915841 | ENSG00000233237 |
| NA | 2.105996916 | 4.304951296 | 0.000854484 | 0.004599771 | 6.506767914 | 27.78255508 | ENSG00000242242 |
| PURPL | 2.104499194 | 4.300484464 | 2.94E-36 | 6.06E-34 | 132.6789633 | 570.2258157 | ENSG00000250337 |
| CEACAM1 | 2.101441383 | 4.291379184 | 9.21E-07 | 1.03E-05 | 11.89321599 | 50.8311304 | ENSG00000079385 |
| NA | 2.099689766 | 4.286172063 | 0.005641661 | 0.022840225 | 4.194624271 | 17.86116892 | ENSG00000261068 |
| LINC00607 | 2.093306217 | 4.267248766 | 0.022790003 | 0.071940412 | 2.504286723 | 10.80621926 | ENSG00000235770 |
| LOC105370802 | 2.092167751 | 4.263882704 | 0.050048366 | 0.133876257 | 2.202854414 | 9.470985467 | ENSG00000259200 |
| GBX2 | 2.086103646 | 4.245997865 | 1.43E-19 | 8.88E-18 | 44.57825648 | 188.5220221 | ENSG00000168505 |
| COL6A3 | 2.082282739 | 4.234767427 | 8.80E-36 | 1.79E-33 | 101.3935766 | 428.2823624 | ENSG00000163359 |
| TMCC1-AS1 | 2.073848428 | 4.210082277 | 4.79E-06 | 4.66E-05 | 10.40133535 | 43.99675776 | ENSG00000271270 |
| ZFP57 | 2.07130509 | 4.202666829 | 0.065844198 | 0.165219035 | 9.050215368 | 37.96151059 | ENSG00000204644 |
| VLDLR-AS1 | 2.06597666 | 4.187173386 | 1.70E-12 | 4.85E-11 | 36.39377866 | 152.5259722 | ENSG00000236404 |
| STARD13 | 2.059590125 | 4.168678539 | 7.05E-25 | 6.56E-23 | 54.72185264 | 228.1498196 | ENSG00000133121 |
| DDIT3 | 2.054637692 | 4.154392978 | 2.05E-11 | 4.95E-10 | 24.98782423 | 104.1560801 | ENSG00000175197 |
| SMIM10L2A | 2.05410746 | 4.1528664 | 7.79E-49 | 3.01E-46 | 162.2849564 | 674.4967833 | ENSG00000178947 |
| MT1G | 2.053717528 | 4.151744113 | 1.61E-95 | 3.31E-92 | 433.3383172 | 1799.29288 | ENSG00000125144 |
| NA | 2.051948394 | 4.146656072 | 0.083604316 | 0.198097979 | 1.5450445 | 6.492584279 | ENSG00000286667 |
| TNFSF9 | 2.051215125 | 4.144549016 | 7.01E-17 | 3.26E-15 | 38.73067395 | 160.8616661 | ENSG00000125657 |
| NA | 2.048340723 | 4.136299703 | 0.000466588 | 0.002695678 | 7.119103271 | 29.21858027 | ENSG00000203362 |
| PADI4 | 2.047624349 | 4.134246324 | 0.030491602 | 0.090725855 | 2.507443636 | 10.47246662 | ENSG00000159339 |
| NA | 2.046364812 | 4.130638519 | 0.084481587 | 0.199615213 | 1.627800651 | 6.676606995 | ENSG00000288065 |
| HR | 2.045757303 | 4.128899499 | 0.074879734 | 0.182295218 | 1.862261374 | 7.748935247 | ENSG00000168453 |
| BANK1 | 2.043024641 | 4.121086201 | 3.66E-05 | 0.000287094 | 8.876510104 | 36.60750581 | ENSG00000153064 |
| NA | 2.036518832 | 4.102544072 | 0.089951597 | 0.209442575 | 1.581048318 | 6.500070353 | ENSG00000262089 |
| LEFTY2 | 2.035376606 | 4.099297248 | 6.00E-34 | 1.07E-31 | 129.2469058 | 528.2066216 | ENSG00000143768 |
| CDIPTOSP | 2.031819435 | 4.089202295 | 0.070111898 | 0.173169813 | 1.915327532 | 7.877083014 | ENSG00000214725 |
| TAC1 | 2.031603046 | 4.088589001 | 2.55E-16 | 1.12E-14 | 36.1499449 | 148.0854632 | ENSG00000006128 |
| PDGFB | 2.030012054 | 4.084082625 | 1.38E-40 | 4.20E-38 | 121.7507997 | 497.3027291 | ENSG00000100311 |
| RTL9 | 2.026618021 | 4.074485851 | 0.006647975 | 0.026245288 | 4.132087374 | 16.9241481 | ENSG00000243978 |
| NA | 2.023656081 | 4.066129268 | 0.091187104 | 0.211681085 | 1.657490643 | 6.627780086 | ENSG00000251250 |
| LINC00958 | 2.023036615 | 4.064383723 | 0.03706264 | 0.105777481 | 2.563666707 | 10.32543624 | ENSG00000251381 |
| VCX3A | 2.022890305 | 4.063971557 | 0.055235203 | 0.14424817 | 2.187069849 | 8.964495033 | ENSG00000169059 |
| NA | 2.018035259 | 4.050318218 | 0.045878654 | 0.125119921 | 2.279296739 | 9.174774817 | ENSG00000244349 |
| SP140L | 2.017176842 | 4.047908957 | 9.86E-05 | 0.000687912 | 9.164036985 | 37.14752459 | ENSG00000185404 |
| NA | 2.013554766 | 4.037758893 | 0.065123545 | 0.163860952 | 1.909013706 | 7.658767502 | ENSG00000260455 |
| NA | 2.003883374 | 4.010781504 | 0.0088509 | 0.033247382 | 4.324230453 | 17.58004204 | ENSG00000267317 |
| KCNG2 | 2.002065898 | 4.005731989 | 0.063442641 | 0.160876442 | 2.173164421 | 8.772874623 | ENSG00000178342 |
| ALDH1A1 | 1.997190156 | 3.99221704 | 2.71E-09 | 4.71E-08 | 19.82311153 | 78.99213977 | ENSG00000165092 |
| EBI3 | 1.991304059 | 3.975962251 | 0.000266261 | 0.001655998 | 7.251866366 | 28.98628029 | ENSG00000105246 |
| NA | 1.988153008 | 3.967287661 | 1.14E-32 | 1.84E-30 | 171.7664208 | 680.6264351 | ENSG00000255666 |
| NA | 1.985760765 | 3.960714648 | 0.023735855 | 0.074285771 | 3.232225135 | 12.78325091 | ENSG00000283491 |
| NA | 1.98519984 | 3.959175007 | 0.063119987 | 0.160343454 | 1.941860612 | 7.636747314 | ENSG00000228043 |
| PPP1R3B | 1.984841825 | 3.958192633 | 4.09E-96 | 9.47E-93 | 385.7688207 | 1526.37829 | ENSG00000173281 |
| TAFA2 | 1.984515412 | 3.957297181 | 0.000651114 | 0.00361743 | 6.325471048 | 25.10785062 | ENSG00000198673 |
| HAPLN1 | 1.983031286 | 3.953228333 | 4.68E-11 | 1.07E-09 | 25.98750496 | 102.86511 | ENSG00000145681 |
| NA | 1.981377385 | 3.948698963 | 0.003057482 | 0.013543809 | 5.124176501 | 20.22469059 | ENSG00000285130 |

| | | | | | | | |
|---|---|---|---|---|---|---|---|
| NA | 1.980708725 | 3.946869243 | 0.107825964 | 0.239679349 | 1.627800651 | 6.395368493 | ENSG00000256654 |
| KCNJ5 | 1.978421258 | 3.94061624 | 0.132240519 | 0.278871802 | 1.846476809 | 7.336188165 | ENSG00000120457 |
| DPPA2 | 1.974725296 | 3.930533908 | 5.74E-28 | 7.05E-26 | 83.600892 | 328.1563056 | ENSG00000163530 |
| LINC00622 | 1.973652501 | 3.92761223 | 0.034653152 | 0.100431824 | 2.50112981 | 9.894467817 | ENSG00000260941 |
| LOC400499 | 1.973568949 | 3.927384774 | 5.08E-06 | 4.90E-05 | 15.63164417 | 61.52246219 | ENSG00000188897 |
| PI15 | 1.972420506 | 3.924259665 | 0.044598766 | 0.122415045 | 2.61041904 | 10.1611722 | ENSG00000137558 |
| BMPER | 1.971358833 | 3.92137288 | 0.040894598 | 0.114458937 | 2.805719032 | 11.1212009 | ENSG00000164619 |
| PTGS2 | 1.958252164 | 3.885909128 | 0.072084198 | 0.17709551 | 1.88563754 | 7.411821795 | ENSG00000073756 |
| NA | 1.956173403 | 3.880314002 | 0.07650954 | 0.185203324 | 1.971550604 | 7.527808575 | ENSG00000268364 |
| PCDH10 | 1.955209342 | 3.877721904 | 7.78E-20 | 4.92E-18 | 69.39849387 | 268.224219 | ENSG00000138650 |
| CAPN13 | 1.952283591 | 3.869865945 | 3.54E-14 | 1.24E-12 | 45.35935875 | 175.596456 | ENSG00000162949 |
| AACSP1 | 1.950525041 | 3.865151708 | 0.003411805 | 0.014833842 | 5.322633406 | 20.75331283 | ENSG00000250420 |
| PLD6 | 1.949525534 | 3.862474836 | 0.002254166 | 0.01051566 | 5.683445699 | 22.03603292 | ENSG00000179598 |
| GADD45B | 1.948753769 | 3.86040917 | 6.76E-35 | 1.29E-32 | 106.0718996 | 409.9139788 | ENSG00000099860 |
| DMBX1 | 1.941826983 | 3.841918682 | 0.00025317 | 0.001585755 | 8.528325462 | 32.92850903 | ENSG00000197587 |
| CGB2 | 1.941195999 | 3.840238729 | 0.038141761 | 0.108207469 | 2.527662889 | 9.770994963 | ENSG00000104818 |
| NA | 1.940862859 | 3.839352063 | 0.010726097 | 0.038985937 | 4.77095581 | 18.27336635 | ENSG00000260412 |
| NA | 1.938624668 | 3.833400327 | 0.00046105 | 0.002667924 | 9.63739312 | 37.36616651 | ENSG00000273654 |
| ACTG2 | 1.935680195 | 3.825584514 | 1.18E-11 | 2.97E-10 | 30.51197376 | 116.8336415 | ENSG00000163017 |
| IER5L | 1.935336201 | 3.824672456 | 1.22E-07 | 1.61E-06 | 17.07687018 | 65.54919607 | ENSG00000188483 |
| IP6K3 | 1.934181563 | 3.821612664 | 0.025974879 | 0.079828256 | 3.126092818 | 12.05128531 | ENSG00000161896 |
| DUSP19 | 1.933917716 | 3.820913815 | 4.31E-05 | 0.00033005 | 10.12327921 | 38.85872792 | ENSG00000162999 |
| NA | 1.933874623 | 3.820799686 | 0.051216973 | 0.136269792 | 2.521349063 | 9.684626065 | ENSG00000267055 |
| LHX5 | 1.929855122 | 3.810169349 | 0.091546097 | 0.212197037 | 1.862261374 | 7.167687249 | ENSG00000089116 |
| NA | 1.926865981 | 3.802283175 | 0.140557685 | 0.291822128 | 1.581048318 | 6.065303082 | ENSG00000267521 |
| C2orf78 | 1.916954171 | 3.776249719 | 0.00275844 | 0.012399936 | 6.553520247 | 24.44392095 | ENSG00000187833 |
| CEMIP | 1.916696799 | 3.775576107 | 1.53E-10 | 3.26E-09 | 28.95302383 | 109.312881 | ENSG00000103888 |
| NA | 1.914320849 | 3.769363293 | 0.150493622 | 0.306611936 | 1.601267572 | 6.054018162 | ENSG00000213212 |
| PRSS21 | 1.914319111 | 3.76935875 | 0.121941846 | 0.262768302 | 1.657490643 | 6.121178032 | ENSG00000007038 |
| CTSLP2 | 1.913556532 | 3.767366871 | 0.05313838 | 0.140099822 | 2.619889779 | 9.770994963 | ENSG00000280913 |
| FIBIN | 1.909312884 | 3.756301546 | 0.007112525 | 0.027775714 | 5.074267255 | 19.05003361 | ENSG00000176971 |
| TENM1 | 1.908046845 | 3.753006647 | 1.75E-08 | 2.69E-07 | 37.14687468 | 139.0334286 | ENSG00000009694 |
| ZFHX3 | 1.905883813 | 3.747383981 | 1.24E-13 | 4.02E-12 | 43.61091899 | 163.3425515 | ENSG00000140836 |
| LGR5 | 1.902002093 | 3.73731481 | 0.070683951 | 0.174232791 | 2.166850596 | 8.191626624 | ENSG00000139292 |
| NA | 1.899539013 | 3.730939622 | 0.046359618 | 0.126023637 | 2.50112981 | 9.444728399 | ENSG00000248588 |
| CHRNA9 | 1.883931276 | 3.690794151 | 2.57E-08 | 3.82E-07 | 20.29331075 | 74.94410987 | ENSG00000174343 |
| ZYG11A | 1.882187388 | 3.686335521 | 5.99E-11 | 1.35E-09 | 28.26552393 | 104.222802 | ENSG00000203995 |
| NLRC3 | 1.881782715 | 3.685301657 | 0.019368869 | 0.062825101 | 3.774431993 | 14.03960188 | ENSG00000167984 |
| NA | 1.880635775 | 3.682373013 | 0.00120728 | 0.00617432 | 6.914430239 | 25.65199466 | ENSG00000277883 |
| EMP1 | 1.877682367 | 3.674842369 | 0.006430843 | 0.02551686 | 5.055325778 | 18.56968862 | ENSG00000134531 |
| OVGP1 | 1.876310707 | 3.671350126 | 0.00030661 | 0.001871161 | 8.571920882 | 31.55584486 | ENSG00000085465 |
| IFIT3 | 1.876140215 | 3.670916288 | 0.020317115 | 0.065488245 | 3.454058207 | 12.77152796 | ENSG00000119917 |
| TLR4 | 1.874464081 | 3.666653866 | 0.050750674 | 0.135359475 | 2.563666707 | 9.388415416 | ENSG00000136869 |
| NA | 1.873795115 | 3.66495406 | 0.158711238 | 0.318418931 | 1.554515239 | 5.734799638 | ENSG00000244119 |
| ISLR2 | 1.869331918 | 3.653633488 | 7.90E-11 | 1.76E-09 | 27.89524089 | 102.0927912 | ENSG00000167178 |
| HES4 | 1.868771039 | 3.652213333 | 9.31E-07 | 1.04E-05 | 19.0996078 | 69.85969348 | ENSG00000188290 |
| NA | 1.86782784 | 3.649826386 | 0.063592487 | 0.16112415 | 2.249606746 | 8.146598562 | ENSG00000249916 |
| NA | 1.865378561 | 3.643635292 | 0.11196389 | 0.246584885 | 1.998083683 | 7.126346413 | ENSG00000235660 |
| NA | 1.858972419 | 3.627491965 | 0.160236729 | 0.320611848 | 1.541887587 | 5.701494529 | ENSG00000257758 |
| LOC349160 | 1.858142045 | 3.625404686 | 0.001654311 | 0.00807506 | 6.934649492 | 25.28438726 | ENSG00000234352 |
| GRIK2 | 1.857588952 | 3.624015064 | 5.55E-10 | 1.07E-08 | 27.92620866 | 101.2936657 | ENSG00000164418 |
| PAX5 | 1.852689511 | 3.611728661 | 8.04E-05 | 0.000572805 | 11.80730293 | 42.50320867 | ENSG00000196092 |
| IRGM | 1.849468931 | 3.603675061 | 0.145491035 | 0.299149195 | 1.571577579 | 5.739036518 | ENSG00000237693 |
| ALDH1A3 | 1.849451483 | 3.603631479 | 2.55E-12 | 7.02E-11 | 37.38902471 | 134.5639631 | ENSG00000184254 |
| ALPK2 | 1.849227209 | 3.603071321 | 0.043516185 | 0.120126448 | 2.921322088 | 10.45738285 | ENSG00000198796 |
| SULT2B1 | 1.847950114 | 3.599883238 | 0.001294215 | 0.006537744 | 7.261337105 | 26.26318695 | ENSG00000088002 |
| LINC02204 | 1.843786456 | 3.589508854 | 0.064957519 | 0.163598675 | 2.61041904 | 9.260705682 | ENSG00000280639 |
| GP1BA | 1.843722604 | 3.589349989 | 0.00746565 | 0.028880894 | 5.386448079 | 19.3955092 | ENSG00000185245 |
| NUPR1 | 1.843438292 | 3.588642707 | 0.00910458 | 0.034014169 | 4.724203477 | 17.08516294 | ENSG00000176046 |
| TRPM2 | 1.839368883 | 3.578534487 | 0.009217527 | 0.034384273 | 4.697670398 | 16.96125205 | ENSG00000142185 |
| NA | 1.839166072 | 3.57803146 | 0.122817935 | 0.264226071 | 1.852790635 | 6.75548982 | ENSG00000283031 |
| LINC02188 | 1.837680332 | 3.574348569 | 6.11E-15 | 2.34E-13 | 44.52340888 | 159.2478933 | ENSG00000261175 |
| NA | 1.836456829 | 3.571318563 | 0.000392631 | 0.002315314 | 10.248353 | 36.58581203 | ENSG00000284294 |
| NA | 1.834370363 | 3.566157357 | 2.65E-07 | 3.30E-06 | 19.52483613 | 69.43456217 | ENSG00000256969 |
| KCNK7 | 1.832623722 | 3.561842496 | 0.012743989 | 0.044947115 | 4.135244286 | 14.71426682 | ENSG00000173338 |
| SLC12A5-AS1 | 1.831619636 | 3.559364388 | 0.021979008 | 0.069896865 | 3.830655064 | 13.66450841 | ENSG00000204044 |
| ZNF843 | 1.830312562 | 3.556141085 | 0.016858631 | 0.056096564 | 4.099240469 | 14.68844778 | ENSG00000176723 |
| DNAH1 | 1.83001316 | 3.555403156 | 8.60E-06 | 7.80E-05 | 16.04740176 | 56.94157317 | ENSG00000114841 |
| GUCY1B2 | 1.829858762 | 3.555022675 | 0.071933465 | 0.176795498 | 2.616732866 | 9.249858795 | ENSG00000123201 |
| STK32A | 1.829017594 | 3.552950513 | 0.00020867 | 0.001333614 | 9.474940033 | 33.80607929 | ENSG00000169302 |
| PARD6G-AS1 | 1.825956112 | 3.54541894 | 1.95E-05 | 0.000162651 | 14.57258345 | 51.88176488 | ENSG00000267270 |
| MIR3936HG | 1.825759448 | 3.544935673 | 0.035217222 | 0.101748401 | 3.599351255 | 12.58001917 | ENSG00000233006 |
| ZBBX | 1.82487794 | 3.542770327 | 0.005704974 | 0.023051228 | 5.769358763 | 20.30638457 | ENSG00000169064 |
| SLC9A9 | 1.818380413 | 3.526850476 | 0.046147085 | 0.125638294 | 3.208848969 | 11.33539115 | ENSG00000181804 |

| | | | | | | | |
|---|---|---|---|---|---|---|---|
| LOC101928222 | 1.817250505 | 3.524089356 | 0.022484098 | 0.071088411 | 4.463209675 | 15.76132569 | ENSG00000227210 |
| STMN2 | 1.816737144 | 3.522835586 | 2.43E-09 | 4.25E-08 | 28.61992469 | 100.3412347 | ENSG00000104435 |
| SLC7A14 | 1.815606293 | 3.520075306 | 0.001241216 | 0.0063165 | 7.668901731 | 26.88153891 | ENSG00000013293 |
| MYOM1 | 1.815556339 | 3.519953424 | 0.000725972 | 0.003979639 | 8.247112406 | 29.1214761 | ENSG00000101605 |
| LOC283710 | 1.813911928 | 3.515943599 | 0.150198927 | 0.30631458 | 1.968393691 | 6.898171701 | ENSG00000259772 |
| NA | 1.813614874 | 3.515219734 | 0.130938452 | 0.277075556 | 1.601267572 | 5.641382699 | ENSG00000273243 |
| NA | 1.813552875 | 3.515068673 | 0.00237997 | 0.01099735 | 7.618992485 | 26.80678134 | ENSG00000287238 |
| INPP4B | 1.811242805 | 3.509444783 | 0.013567559 | 0.047108777 | 5.461612628 | 18.9968582 | ENSG00000109452 |
| LINC02078 | 1.807834309 | 3.501163193 | 0.054137345 | 0.142136022 | 2.560509794 | 8.938126345 | ENSG00000267719 |
| ADGRD1 | 1.807629781 | 3.500666875 | 0.012978781 | 0.045523225 | 5.01932196 | 17.70775177 | ENSG00000111452 |
| NA | 1.807260663 | 3.499771333 | 0.152644621 | 0.309836317 | 1.627800651 | 5.671438614 | ENSG00000256424 |
| WNT3 | 1.806913594 | 3.498929494 | 5.67E-25 | 5.31E-23 | 83.83535272 | 293.1096204 | ENSG00000108379 |
| NA | 1.801531008 | 3.485899576 | 0.036713959 | 0.105028451 | 4.367825873 | 15.31048696 | ENSG00000234694 |
| NA | 1.796921798 | 3.474780384 | 0.145045282 | 0.298543277 | 1.607581397 | 5.573784796 | ENSG00000286373 |
| ASB2 | 1.794817211 | 3.469715109 | 0.045320281 | 0.123961938 | 2.84172285 | 9.875696822 | ENSG00000100628 |
| CTNNA3 | 1.79211989 | 3.463234049 | 0.010803851 | 0.039231696 | 5.805362581 | 19.93497833 | ENSG00000183230 |
| S100A2 | 1.790712732 | 3.459857769 | 0.09687862 | 0.221372007 | 2.669799024 | 9.031543283 | ENSG00000196754 |
| BHLHE41 | 1.790681664 | 3.459783263 | 1.20E-05 | 0.000105883 | 16.16428259 | 55.50769787 | ENSG00000123095 |
| ULBP1 | 1.789602508 | 3.457196265 | 0.000411957 | 0.002414677 | 9.170350811 | 31.84077059 | ENSG00000111981 |
| LINC01356 | 1.788596404 | 3.454786133 | 2.24E-33 | 3.74E-31 | 157.7306999 | 545.1718562 | ENSG00000215866 |
| NA | 1.786858879 | 3.450627829 | 0.146339023 | 0.300459449 | 1.607581397 | 5.524957886 | ENSG00000227999 |
| ITGAM | 1.786801513 | 3.450490626 | 0.008116645 | 0.031010645 | 5.763044937 | 19.88702749 | ENSG00000169896 |
| FMN1 | 1.786451528 | 3.449653668 | 1.89E-10 | 3.93E-09 | 35.28598878 | 121.1381499 | ENSG00000248905 |
| NA | 1.786302557 | 3.44929748 | 0.081949216 | 0.19522452 | 2.223073667 | 7.704345218 | ENSG00000275560 |
| FAM163B | 1.785816552 | 3.448135699 | 1.15E-14 | 4.26E-13 | 49.30896917 | 170.4960395 | ENSG00000196990 |
| NA | 1.78028477 | 3.434939693 | 0.149485738 | 0.305296822 | 1.634114477 | 5.555013801 | ENSG00000261659 |
| RBM20 | 1.780186442 | 3.43470559 | 0.00021361 | 0.001361431 | 12.64137366 | 43.11929911 | ENSG00000203867 |
| ZMAT3 | 1.775482463 | 3.423524798 | 1.30E-35 | 2.61E-33 | 192.0310111 | 656.7387528 | ENSG00000172667 |
| LOC730100 | 1.77251063 | 3.41647988 | 0.065558814 | 0.164598627 | 2.504286723 | 8.630630775 | ENSG00000231918 |
| IL1A | 1.772094851 | 3.415495404 | 0.050357192 | 0.134600548 | 3.139998246 | 10.81750418 | ENSG00000115008 |
| PLA2G4C | 1.771083272 | 3.41310139 | 2.24E-06 | 2.33E-05 | 18.4923085 | 63.00913776 | ENSG00000105499 |
| CYP26B1 | 1.767186778 | 3.403895574 | 0.003977542 | 0.016950968 | 7.870417851 | 26.46934148 | ENSG00000003137 |
| LINC01234 | 1.767009593 | 3.403477549 | 3.25E-11 | 7.64E-10 | 35.645621 | 121.3612519 | ENSG00000249550 |
| IER3 | 1.766556267 | 3.402408272 | 2.41E-45 | 8.60E-43 | 237.2288112 | 807.196778 | ENSG00000137331 |
| HSD52 | 1.766511941 | 3.402303735 | 0.084883147 | 0.200308665 | 2.223073667 | 7.572836638 | ENSG00000224609 |
| NA | 1.763746932 | 3.395789266 | 0.09648976 | 0.220673891 | 2.196540588 | 7.490266586 | ENSG00000260808 |
| AATBC | 1.762894806 | 3.393784141 | 0.003861609 | 0.016548224 | 6.411384112 | 21.68708446 | ENSG00000215458 |
| TSPYL5 | 1.762450381 | 3.392738839 | 2.23E-15 | 8.95E-14 | 56.32627712 | 191.3527804 | ENSG00000180543 |
| PRTN3 | 1.762136647 | 3.392001122 | 0.03790738 | 0.107641395 | 3.516595104 | 11.95418114 | ENSG00000196415 |
| CYP2B7P | 1.761063392 | 3.38947867 | 0.104610719 | 0.234671394 | 2.196540588 | 7.471495592 | ENSG00000256612 |
| LOC107984525 | 1.761033358 | 3.389408108 | 0.00315659 | 0.013906405 | 7.767442446 | 26.14296329 | ENSG00000286069 |
| LINC02572 | 1.760141974 | 3.387314575 | 0.084153405 | 0.199042777 | 2.252763659 | 7.595406479 | ENSG00000229536 |
| NA | 1.758395523 | 3.38321655 | 0.05896785 | 0.15187745 | 3.754212739 | 12.72974909 | ENSG00000251396 |
| GPR179 | 1.754939512 | 3.375121672 | 0.046159054 | 0.125638294 | 3.19937823 | 10.79873319 | ENSG00000277399 |
| SEZ6L | 1.75473722 | 3.374648453 | 0.00387791 | 0.016614236 | 6.328627961 | 21.43970072 | ENSG00000100095 |
| NA | 1.75408643 | 3.373126515 | 0.115691333 | 0.252820146 | 1.968393691 | 6.548897262 | ENSG00000278000 |
| PAPPA2 | 1.751085729 | 3.366117948 | 0.00035447 | 0.002117223 | 12.16821292 | 41.28494934 | ENSG00000116183 |
| NA | 1.747159999 | 3.356970825 | 1.78E-07 | 2.27E-06 | 21.32140369 | 71.58020636 | ENSG00000268970 |
| INSL3 | 1.738849583 | 3.337689116 | 0.037475604 | 0.106726004 | 3.493218937 | 11.66490691 | ENSG00000248099 |
| INPP5D | 1.737384241 | 3.334300753 | 4.86E-51 | 2.05E-48 | 243.9148992 | 813.3663054 | ENSG00000168918 |
| NA | 1.734131532 | 3.326791688 | 0.006622153 | 0.026175089 | 7.439477058 | 24.55990773 | ENSG00000253706 |
| ABCC3 | 1.731545341 | 3.320833385 | 0.036443312 | 0.104524967 | 3.483748199 | 11.56725309 | ENSG00000108846 |
| PCDH11X | 1.729736142 | 3.316671534 | 1.72E-23 | 1.45E-21 | 107.4877438 | 355.5163257 | ENSG00000102290 |
| MYLK4 | 1.729526653 | 3.316189967 | 0.095935848 | 0.219623888 | 2.510600548 | 8.402018029 | ENSG00000145949 |
| IL32 | 1.726412013 | 3.309038353 | 0.006934824 | 0.027190615 | 7.565828629 | 24.7101873 | ENSG00000008517 |
| KCNJ6 | 1.726363871 | 3.308927933 | 4.76E-06 | 4.63E-05 | 18.99535462 | 63.08520942 | ENSG00000157542 |
| GPR35 | 1.724500469 | 3.304656842 | 0.134914343 | 0.282862266 | 1.938703699 | 6.406653414 | ENSG00000178623 |
| SPATA32 | 1.72370666 | 3.302839032 | 0.051448094 | 0.136708097 | 3.179158977 | 10.55503667 | ENSG00000184361 |
| TENM2 | 1.72274697 | 3.300642694 | 5.56E-12 | 1.48E-10 | 40.58840294 | 133.6781423 | ENSG00000145934 |
| NA | 1.721843127 | 3.2985755 | 0.000274055 | 0.001699912 | 11.01998453 | 36.54414478 | ENSG00000251271 |
| RALYL | 1.721228877 | 3.297171379 | 0.080927915 | 0.193288399 | 3.314981286 | 10.78376104 | ENSG00000184672 |
| GOLGA6A | 1.720702963 | 3.295969662 | 0.008792219 | 0.033087306 | 5.673974961 | 18.74622526 | ENSG00000159289 |
| NA | 1.720083057 | 3.294553733 | 0.022126149 | 0.070232239 | 5.167771921 | 16.8631602 | ENSG00000250993 |
| HAPLN2 | 1.71488183 | 3.282697541 | 0.007199242 | 0.028016631 | 6.31600031 | 20.84348058 | ENSG00000132702 |
| NA | 1.714745784 | 3.282387999 | 0.05790653 | 0.149831024 | 3.533657445 | 11.52222503 | ENSG00000249863 |
| NA | 1.712814178 | 3.27799619 | 0.006312817 | 0.025129242 | 6.292624143 | 20.76079891 | ENSG00000204054 |
| NA | 1.710559197 | 3.272876574 | 0.039021522 | 0.110382308 | 3.903940477 | 12.63633215 | ENSG00000260725 |
| CD8A | 1.710281766 | 3.27224726 | 0.049543252 | 0.13282723 | 3.176002064 | 10.42363971 | ENSG00000153563 |
| TLX2 | 1.705076879 | 3.260463071 | 0.182507015 | 0.352869269 | 1.862261374 | 6.117490805 | ENSG00000115297 |
| COL4A4 | 1.703257248 | 3.25635333 | 0.000278595 | 0.001724609 | 11.10905451 | 36.14973066 | ENSG00000081052 |
| NA | 1.701676785 | 3.252787971 | 0.059915799 | 0.15372085 | 3.910254302 | 12.58739362 | ENSG00000258704 |
| ALOX5 | 1.701135077 | 3.251566832 | 1.24E-08 | 1.96E-07 | 30.42280608 | 98.72128999 | ENSG00000012779 |
| GSTT2B | 1.700863041 | 3.250953771 | 3.26E-05 | 0.00025857 | 20.2028653 | 65.70458859 | ENSG00000133433 |
| LOC283731 | 1.699064569 | 3.246903638 | 0.011216037 | 0.040474756 | 6.04110108 | 19.54111386 | ENSG00000248540 |

| | | | | | | | |
|---|---|---|---|---|---|---|---|
| CCDC80 | 1.696881986 | 3.24199527 | 8.94E-14 | 2.96E-12 | 49.73675305 | 161.5390306 | ENSG00000091986 |
| MVP | 1.696279316 | 3.240641245 | 1.52E-07 | 1.96E-06 | 27.26386636 | 87.90995778 | ENSG00000013364 |
| NA | 1.695737934 | 3.239425399 | 0.067637769 | 0.168585048 | 2.874569755 | 9.298247671 | ENSG00000258623 |
| PRKAR2A-AS1 | 1.695570877 | 3.239050311 | 0.047590154 | 0.128689219 | 3.569661262 | 11.48891992 | ENSG00000224424 |
| TM6SF1 | 1.695402186 | 3.238671599 | 0.064550869 | 0.162884384 | 3.483748199 | 11.36544707 | ENSG00000136404 |
| ERFL | 1.694927273 | 3.237605652 | 0.159950677 | 0.320246932 | 1.876166801 | 6.102407038 | ENSG00000268041 |
| SLC16A3 | 1.694854391 | 3.2374421 | 6.85E-19 | 3.99E-17 | 111.9079368 | 361.2593843 | ENSG00000141526 |
| C14orf39 | 1.69463354 | 3.236946543 | 0.003803783 | 0.016326845 | 7.033190208 | 22.71862153 | ENSG00000179008 |
| TNRC18 | 1.693389343 | 3.234156166 | 1.08E-33 | 1.87E-31 | 151.2126798 | 489.1132205 | ENSG00000182095 |
| EIF1B-AS1 | 1.692387595 | 3.231911281 | 0.004110493 | 0.017425407 | 7.658153216 | 24.793307 | ENSG00000280739 |
| NA | 1.690719158 | 3.228175824 | 0.100112668 | 0.226860031 | 2.983858985 | 9.523611223 | ENSG00000228863 |
| NA | 1.68980036 | 3.226120576 | 0.167626699 | 0.331446501 | 2.183912936 | 7.13438214 | ENSG00000233750 |
| HTR2C | 1.687099243 | 3.220086052 | 2.53E-09 | 4.42E-08 | 32.18336983 | 103.6651115 | ENSG00000147246 |
| SOX14 | 1.684629977 | 3.214579382 | 0.119965615 | 0.259505452 | 2.243292921 | 7.178972169 | ENSG00000168875 |
| DEUP1 | 1.684302615 | 3.213850043 | 0.079803648 | 0.191392444 | 2.557352881 | 8.184140551 | ENSG00000165325 |
| CHADL | 1.68370552 | 3.212520188 | 0.10784514 | 0.239693228 | 2.497972897 | 8.048944743 | ENSG00000100399 |
| TRADD | 1.682210214 | 3.209192241 | 0.037811396 | 0.107467629 | 3.877407397 | 12.3512948 | ENSG00000102871 |
| PDE11A | 1.681460951 | 3.20752598 | 0.150352862 | 0.306459907 | 1.971550604 | 6.248887766 | ENSG00000128655 |
| NLRP1 | 1.680139095 | 3.20458846 | 0.004037534 | 0.017147491 | 7.69543481 | 24.52985181 | ENSG00000091592 |
| NA | 1.67974404 | 3.203711064 | 0.083038313 | 0.197115979 | 2.871412842 | 9.249858795 | ENSG00000267505 |
| SLC2A1-AS1 | 1.675782736 | 3.194926487 | 0.004360866 | 0.018315045 | 7.030033295 | 22.43358418 | ENSG00000227533 |
| NA | 1.672114663 | 3.186813657 | 0.010739982 | 0.039022707 | 5.967815668 | 19.09506167 | ENSG00000259033 |
| SLFN12 | 1.671693562 | 3.185883611 | 4.99E-21 | 3.47E-19 | 94.76233625 | 301.6356609 | ENSG00000172123 |
| BEST1 | 1.669920233 | 3.181969998 | 0.054150343 | 0.142136022 | 3.424368214 | 11.00108886 | ENSG00000167995 |
| TTC22 | 1.668918159 | 3.179760618 | 0.000696777 | 0.003843784 | 12.44173668 | 39.43909985 | ENSG00000006555 |
| PMFBP1 | 1.665741618 | 3.172767089 | 0.008970795 | 0.033609199 | 8.592140136 | 27.39325391 | ENSG00000118557 |
| NA | 1.663928709 | 3.168782654 | 0.102911047 | 0.23141875 | 2.563666707 | 8.147036595 | ENSG00000230552 |
| GUCA1B | 1.663671408 | 3.168217559 | 1.51E-05 | 0.000130183 | 17.2486963 | 54.60821904 | ENSG00000112599 |
| TUBB8B | 1.662724118 | 3.166137956 | 0.048289123 | 0.130218334 | 3.543128183 | 11.1963965 | ENSG00000173213 |
| RGN | 1.660812026 | 3.161944459 | 0.02564641 | 0.079037339 | 5.197461913 | 16.3273783 | ENSG00000130988 |
| LOC105372273 | 1.659910763 | 3.159969784 | 0.031740133 | 0.093703668 | 4.7930542 | 15.11572898 | ENSG00000267082 |
| MIR124-2HG | 1.659014304 | 3.158006858 | 0.064634376 | 0.162984152 | 4.119459722 | 13.08326041 | ENSG00000254377 |
| KIF25 | 1.658662401 | 3.157236649 | 0.144034246 | 0.296944462 | 1.941860612 | 6.053580129 | ENSG00000125337 |
| AZU1 | 1.657057199 | 3.15372573 | 0.005655136 | 0.022884779 | 8.022122422 | 25.31488121 | ENSG00000172232 |
| TEX41 | 1.656120195 | 3.151678109 | 0.017097493 | 0.056748735 | 5.114705763 | 16.06078553 | ENSG00000226674 |
| NPFFR2 | 1.654246399 | 3.147587314 | 1.91E-17 | 9.49E-16 | 85.77080181 | 269.2870773 | ENSG00000056291 |
| ARID5B | 1.651416305 | 3.141418833 | 0.007078219 | 0.027668854 | 7.109632533 | 22.14062273 | ENSG00000150347 |
| RTL4 | 1.649157446 | 3.136504094 | 0.073400523 | 0.179679077 | 3.139998246 | 9.886981743 | ENSG00000187823 |
| CFAP44 | 1.648632685 | 3.13536344 | 0.041781209 | 0.116534678 | 4.685042746 | 14.86498443 | ENSG00000206530 |
| ARHGAP25 | 1.648411179 | 3.134882086 | 3.93E-05 | 0.000305021 | 15.83325799 | 49.80339218 | ENSG00000163219 |
| KIAA1211L | 1.643709029 | 3.124681253 | 0.00689489 | 0.027086871 | 7.056566374 | 21.95714966 | ENSG00000196872 |
| PPARGC1A | 1.643144758 | 3.123459359 | 4.10E-06 | 4.06E-05 | 21.77759979 | 67.776311 | ENSG00000109819 |
| NA | 1.642925871 | 3.122985501 | 0.035907626 | 0.103292245 | 4.988354192 | 15.74992915 | ENSG00000261326 |
| LOC101927314 | 1.642693677 | 3.122482913 | 0.095893952 | 0.219577481 | 2.587042873 | 8.049382777 | ENSG00000271860 |
| NA | 1.637510648 | 3.111285202 | 0.105571551 | 0.236226382 | 2.616732866 | 8.079438691 | ENSG00000225125 |
| NA | 1.635541765 | 3.107042046 | 0.074794657 | 0.182184803 | 2.871412842 | 8.900584356 | ENSG00000266904 |
| PPP1R3C | 1.632190426 | 3.099832854 | 7.15E-06 | 6.68E-05 | 21.19001607 | 65.96618003 | ENSG00000119938 |
| FRK | 1.631560032 | 3.098478659 | 0.09463112 | 0.217254204 | 2.84172285 | 8.893536316 | ENSG00000111816 |
| PTPRR | 1.63103106 | 3.097342794 | 0.02323887 | 0.072988944 | 5.108391937 | 15.79083195 | ENSG00000153233 |
| DDR2 | 1.630971569 | 3.097215075 | 0.000146269 | 0.000972504 | 14.37728346 | 44.37901089 | ENSG00000162733 |
| GDF3 | 1.630271425 | 3.095712351 | 8.05E-20 | 5.06E-18 | 109.8911297 | 340.2211373 | ENSG00000184344 |
| RGMB-AS1 | 1.629855144 | 3.094819231 | 0.04815521 | 0.129932839 | 3.847717405 | 11.91284031 | ENSG00000246763 |
| MYCBPAP | 1.629794617 | 3.094689395 | 0.00097709 | 0.00516084 | 12.4853321 | 38.69815111 | ENSG00000136449 |
| DNASE1L2 | 1.627904233 | 3.090637033 | 0.145764884 | 0.29958243 | 1.912170619 | 5.922621201 | ENSG00000167968 |
| LINC02615 | 1.627622087 | 3.09003266 | 0.002409938 | 0.011110895 | 9.977888462 | 30.78352611 | ENSG00000251432 |
| XKR9 | 1.62761824 | 3.090024421 | 0.178744524 | 0.347714418 | 1.891951366 | 5.873794292 | ENSG00000221947 |
| NA | 1.626748289 | 3.088161686 | 0.000187075 | 0.001210629 | 15.08815954 | 46.50664854 | ENSG00000267102 |
| ATP8B3 | 1.622697432 | 3.079502783 | 3.69E-11 | 8.60E-10 | 51.93182047 | 159.6616281 | ENSG00000130270 |
| PADI2 | 1.618925568 | 3.071462072 | 1.96E-25 | 1.95E-23 | 139.2729221 | 426.7610676 | ENSG00000117115 |
| NAALADL2 | 1.618650086 | 3.070875634 | 0.0510373 | 0.135922274 | 4.052488136 | 12.59174212 | ENSG00000177694 |
| TMEM154 | 1.617892519 | 3.069263523 | 0.000531351 | 0.003027378 | 14.8252866 | 45.26439365 | ENSG00000170006 |
| NA | 1.615750961 | 3.064710842 | 0.176888012 | 0.344953957 | 1.971550604 | 5.990219105 | ENSG00000234353 |
| XIRP2 | 1.61560971 | 3.064410797 | 0.065850492 | 0.165219035 | 3.52290893 | 10.79124711 | ENSG00000163092 |
| OTOF | 1.614897965 | 3.062899363 | 0.041004844 | 0.114704275 | 4.11186812 | 12.60247739 | ENSG00000115155 |
| NA | 1.614630541 | 3.062331663 | 3.74E-05 | 0.000291987 | 17.53306627 | 53.64395346 | ENSG00000242759 |
| PTH1R | 1.614487798 | 3.062028684 | 0.001269046 | 0.006428121 | 9.845223065 | 30.14497744 | ENSG00000160801 |
| NA | 1.613726438 | 3.060413173 | 0.110604101 | 0.244418315 | 2.844879763 | 8.762027735 | ENSG00000214243 |
| RNF213-AS1 | 1.611960542 | 3.056669439 | 0.017463855 | 0.057685812 | 5.816111096 | 17.68898078 | ENSG00000263069 |
| NA | 1.611564086 | 3.055829576 | 0.140030359 | 0.291020708 | 2.249606746 | 6.833934611 | ENSG00000253854 |
| NA | 1.610908504 | 3.054441277 | 0.000588055 | 0.003304734 | 11.41995756 | 34.88589362 | ENSG00000246263 |
| NA | 1.609307678 | 3.051053924 | 0.128137718 | 0.27235143 | 2.246449834 | 6.811364769 | ENSG00000234807 |
| TNRC6C-AS1 | 1.608804752 | 3.049990507 | 0.001972669 | 0.009384353 | 9.451563867 | 28.97499537 | ENSG00000204282 |
| LPO | 1.608711149 | 3.049792628 | 0.020809488 | 0.066784944 | 5.088172683 | 15.53585052 | ENSG00000167419 |
| NA | 1.607857756 | 3.047989127 | 0.099144902 | 0.225294654 | 3.255601302 | 9.879933702 | ENSG00000267413 |

| | | | | | | | |
|---|---|---|---|---|---|---|---|
| NA | 1.606580628 | 3.045292125 | 0.077776885 | 0.187736319 | 3.563347437 | 10.79873319 | ENSG00000246203 |
| ANGPTL6 | 1.604809335 | 3.041555512 | 0.182250921 | 0.352592753 | 1.971550604 | 5.933906122 | ENSG00000130812 |
| LCN15 | 1.603719489 | 3.039258716 | 0.114171497 | 0.250302169 | 2.829095198 | 8.649401769 | ENSG00000177984 |
| TMC8 | 1.603057628 | 3.037864724 | 4.72E-06 | 4.59E-05 | 24.47530737 | 74.55579638 | ENSG00000167895 |
| GADL1 | 1.600734078 | 3.032975992 | 0.174037104 | 0.340828792 | 2.16053677 | 6.654037154 | ENSG00000144644 |
| NA | 1.600392727 | 3.032258456 | 0.012903302 | 0.045310686 | 6.384851033 | 19.35416836 | ENSG00000215156 |
| MAGEB17 | 1.599226943 | 3.029809199 | 0.013325422 | 0.046475983 | 6.024038739 | 18.29648585 | ENSG00000182798 |
| KCTD19 | 1.598888837 | 3.029099224 | 0.016017695 | 0.053888537 | 6.10679489 | 18.46553641 | ENSG00000168676 |
| NA | 1.598089869 | 3.027422165 | 0.162804159 | 0.324313444 | 2.216759841 | 6.76677474 | ENSG00000271216 |
| NA | 1.595173161 | 3.021307785 | 0.042052586 | 0.11708016 | 4.128930461 | 12.5128593 | ENSG00000283312 |
| HLA-DQA1 | 1.595063585 | 3.021078318 | 0.007014713 | 0.02746898 | 7.943800961 | 23.99743072 | ENSG00000196735 |
| NA | 1.594955217 | 3.020851398 | 0.049011861 | 0.131707385 | 4.049331223 | 12.33632266 | ENSG00000287310 |
| ITPR2 | 1.594785965 | 3.020497024 | 1.12E-54 | 6.12E-52 | 809.8102554 | 2444.801801 | ENSG00000123104 |
| RYR2 | 1.591760015 | 3.014168386 | 4.50E-16 | 1.93E-14 | 104.1568803 | 313.3606712 | ENSG00000198626 |
| NDUFV2-AS1 | 1.591462729 | 3.013547341 | 0.032968231 | 0.096560238 | 4.442990421 | 13.40160287 | ENSG00000266053 |
| NA | 1.591189843 | 3.012977382 | 0.157103804 | 0.316083576 | 2.16053677 | 6.609009092 | ENSG00000267149 |
| LINC01376 | 1.591082788 | 3.012753814 | 0.009427267 | 0.035057175 | 7.112789446 | 21.26696293 | ENSG00000236204 |
| EID3 | 1.589123062 | 3.008664131 | 0.006154184 | 0.024608771 | 7.612678659 | 22.90589344 | ENSG00000255150 |
| LINC02348 | 1.587731066 | 3.005762597 | 0.071356746 | 0.175610933 | 3.202535143 | 9.586972247 | ENSG00000286733 |
| PTH2R | 1.585610767 | 3.001348335 | 0.019141948 | 0.062143432 | 5.650598794 | 17.02461308 | ENSG00000144407 |
| TEX19 | 1.585185432 | 3.000463609 | 0.034190059 | 0.099416151 | 4.972569627 | 15.0518183 | ENSG00000182459 |
| ZNF354C | 1.583879975 | 2.997749794 | 4.22E-06 | 4.17E-05 | 28.80259704 | 85.9436614 | ENSG00000177932 |
| RGS6 | 1.580081997 | 2.989868425 | 1.02E-05 | 9.07E-05 | 22.65724277 | 67.7174101 | ENSG00000182732 |
| DEPP1 | 1.577939742 | 2.98543207 | 0.104671743 | 0.234751463 | 3.436995866 | 10.26532441 | ENSG00000165507 |
| NA | 1.576733409 | 2.982936795 | 0.116738675 | 0.254545248 | 2.583885961 | 7.681775377 | ENSG00000273576 |
| NA | 1.5759152 | 2.981245535 | 0.055059325 | 0.143971436 | 3.850874318 | 11.46255124 | ENSG00000223486 |
| NA | 1.575202361 | 2.979772857 | 0.116167073 | 0.253447973 | 2.904259747 | 8.608060934 | ENSG00000270460 |
| SPATA12 | 1.574695911 | 2.978727008 | 0.077664573 | 0.18754856 | 3.546285096 | 10.55134945 | ENSG00000186451 |
| TFAP2C | 1.571493111 | 2.972121535 | 1.81E-18 | 9.99E-17 | 97.68117784 | 289.7471023 | ENSG00000087510 |
| NA | 1.571477166 | 2.972088686 | 0.024149289 | 0.075299952 | 5.745982597 | 17.02081423 | ENSG00000253348 |
| LINC02482 | 1.571397767 | 2.971925122 | 0.157634739 | 0.316842236 | 2.246449834 | 6.680405842 | ENSG00000251580 |
| EBF3 | 1.570158635 | 2.969373629 | 0.063355798 | 0.160766204 | 4.088491954 | 12.15598717 | ENSG00000108001 |
| FLNC | 1.568934808 | 2.966855799 | 5.33E-62 | 4.11E-59 | 726.7068022 | 2156.208944 | ENSG00000128591 |
| NA | 1.567041406 | 2.962964632 | 0.067701119 | 0.168697407 | 3.486905112 | 10.35604181 | ENSG00000250049 |
| GNGT1 | 1.56464192 | 2.958040735 | 5.39E-05 | 0.000402215 | 19.13245471 | 56.66033467 | ENSG00000127928 |
| PHF24 | 1.56413608 | 2.957003764 | 0.010427593 | 0.038063716 | 8.929576263 | 26.50027346 | ENSG00000122733 |
| LOC107986163 | 1.563163934 | 2.955011887 | 0.017892584 | 0.058766818 | 6.020881826 | 17.80496756 | ENSG00000272970 |
| C3P1 | 1.562270016 | 2.953181479 | 0.160605322 | 0.321106692 | 2.279296739 | 6.699176837 | ENSG00000167798 |
| NME9 | 1.561990723 | 2.952609825 | 0.163608986 | 0.32556535 | 2.305829818 | 6.672370115 | ENSG00000181322 |
| SIGLEC15 | 1.557981294 | 2.944415547 | 0.057722618 | 0.14948037 | 3.794651247 | 11.16210371 | ENSG00000197046 |
| MYO15B | 1.55788832 | 2.944225802 | 8.47E-12 | 2.17E-10 | 120.1722319 | 354.1887073 | ENSG00000266714 |
| NA | 1.554706294 | 2.937739136 | 0.12305576 | 0.264522786 | 2.613575953 | 7.595406479 | ENSG00000261469 |
| LINC01976 | 1.553610201 | 2.935508025 | 0.173047191 | 0.339464081 | 2.276139826 | 6.642863853 | ENSG00000261514 |
| TPCN2 | 1.553069319 | 2.934407678 | 0.015947725 | 0.053711636 | 7.165953302 | 21.25611604 | ENSG00000287725 |
| ABCC11 | 1.552888306 | 2.934039525 | 0.206550997 | 0.385152757 | 1.859104461 | 5.517471813 | ENSG00000121270 |
| PRSS30P | 1.552464916 | 2.933178594 | 0.053475999 | 0.140664798 | 4.830335794 | 14.17859654 | ENSG00000172460 |
| RAB17 | 1.551361344 | 2.930935752 | 8.37E-54 | 4.19E-51 | 392.8720644 | 1151.39302 | ENSG00000124839 |
| NA | 1.551081652 | 2.930367594 | 0.009942455 | 0.036600187 | 7.622149398 | 22.44161991 | ENSG00000279602 |
| SATB2 | 1.550713468 | 2.929619843 | 7.72E-10 | 1.46E-08 | 46.00275957 | 134.7838873 | ENSG00000119042 |
| MIR1281 | 1.54926385 | 2.926677643 | 0.194133142 | 0.368050701 | 1.888794453 | 5.566298722 | ENSG00000284015 |
| GJC2 | 1.54705725 | 2.922204715 | 1.34E-06 | 1.45E-05 | 27.09519715 | 79.02500685 | ENSG00000198835 |
| NA | 1.546447267 | 2.920969446 | 0.086590766 | 0.203328678 | 4.555436564 | 13.22941472 | ENSG00000236154 |
| NA | 1.546149492 | 2.920366615 | 0.022924867 | 0.072260116 | 6.424011764 | 18.716719 | ENSG00000258433 |
| LINC01137 | 1.546085447 | 2.920236974 | 0.006882972 | 0.027055593 | 9.074869311 | 26.34586862 | ENSG00000233621 |
| SSTR1 | 1.545370629 | 2.91879043 | 0.000970733 | 0.005137146 | 13.27906202 | 38.86186549 | ENSG00000139874 |
| PSD2 | 1.543918559 | 2.915854152 | 0.028686339 | 0.086493021 | 5.114705763 | 14.90957446 | ENSG00000146005 |
| PRELP | 1.541781612 | 2.91153833 | 0.119963134 | 0.259505452 | 2.927635913 | 8.495873001 | ENSG00000188783 |
| MUC19 | 1.540982393 | 2.909925854 | 0.012836192 | 0.045169162 | 9.395340795 | 27.60320728 | ENSG00000205592 |
| TEX13B | 1.539599857 | 2.907138605 | 1.59E-05 | 0.0001354 | 27.5342332 | 79.63762486 | ENSG00000170925 |
| PIFO | 1.539489108 | 2.906915445 | 1.04E-10 | 2.27E-09 | 49.86488606 | 144.5622567 | ENSG00000173947 |
| ITGA11 | 1.537859347 | 2.903633462 | 0.115537329 | 0.252639082 | 2.874569755 | 8.35319112 | ENSG00000137809 |
| HSPA2 | 1.536499265 | 2.900897389 | 6.40E-57 | 3.82E-54 | 627.4487701 | 1819.932689 | ENSG00000126803 |
| NA | 1.535788431 | 2.899468433 | 0.135264694 | 0.283404458 | 2.527662889 | 7.385564727 | ENSG00000225603 |
| BBC3 | 1.534528403 | 2.896937186 | 3.73E-43 | 1.21E-40 | 268.820546 | 778.1448667 | ENSG00000105327 |
| EPHA3 | 1.533764649 | 2.895403971 | 0.09916811 | 0.225298083 | 2.868255929 | 8.311850284 | ENSG00000044524 |
| UAP1L1 | 1.531057805 | 2.889976587 | 6.21E-34 | 1.10E-31 | 195.1898531 | 563.6285521 | ENSG00000197355 |
| RHCE | 1.529450057 | 2.886757774 | 0.144379804 | 0.297557577 | 2.616732866 | 7.445126904 | ENSG00000188672 |
| NA | 1.529447332 | 2.886752322 | 0.005941981 | 0.023847356 | 8.634457779 | 24.80360424 | ENSG00000254275 |
| MRC1 | 1.528330279 | 2.884518025 | 0.149736778 | 0.305708465 | 2.583885961 | 7.453162631 | ENSG00000260314 |
| NA | 1.527714662 | 2.883287426 | 0.05205488 | 0.137657972 | 4.492899667 | 12.95555068 | ENSG00000285907 |
| NA | 1.527582087 | 2.883022483 | 0.002993899 | 0.013300319 | 10.49356224 | 30.34832081 | ENSG00000231154 |
| ATP1A1-AS1 | 1.527035226 | 2.881929865 | 0.054515019 | 0.142911084 | 4.357077358 | 12.66258922 | ENSG00000203865 |
| DDIT4 | 1.526571526 | 2.881003725 | 1.39E-08 | 2.18E-07 | 690.8145741 | 1989.75534 | ENSG00000168209 |
| KLF5 | 1.522377578 | 2.872640743 | 2.26E-05 | 0.000186508 | 21.90404906 | 62.90042226 | ENSG00000102554 |

| | | | | | | | |
|---|---|---|---|---|---|---|---|
| CATSPER3 | 1.521243061 | 2.870382622 | 0.107522495 | 0.2391195 | 3.469842771 | 9.999607709 | ENSG00000152705 |
| CFD | 1.51884336 | 2.865612152 | 7.81E-06 | 7.20E-05 | 24.87212348 | 71.1221048 | ENSG00000197766 |
| NA | 1.517640067 | 2.863223058 | 0.093250807 | 0.214963987 | 3.788337421 | 10.93013015 | ENSG00000274922 |
| GZMM | 1.515122219 | 2.858230407 | 0.208868531 | 0.387970471 | 1.918484445 | 5.464846056 | ENSG00000197540 |
| SPDYC | 1.514837917 | 2.857667211 | 0.1236203 | 0.265397735 | 2.861942103 | 8.16157071 | ENSG00000204710 |
| NA | 1.514425973 | 2.856851355 | 0.214387244 | 0.394456848 | 1.905856794 | 5.434790142 | ENSG00000229503 |
| IRX1 | 1.514343679 | 2.856688398 | 0.001813655 | 0.008724182 | 11.35426375 | 32.52672045 | ENSG00000170549 |
| ATXN1 | 1.51339595 | 2.854812412 | 1.19E-05 | 0.000105 | 22.38046441 | 63.69563727 | ENSG00000124788 |
| LYST | 1.513185291 | 2.854395589 | 0.106701224 | 0.237949764 | 3.205692056 | 9.167288744 | ENSG00000143669 |
| FBLN7 | 1.511045637 | 2.850165386 | 0.078374731 | 0.188747045 | 3.821184326 | 10.85829536 | ENSG00000144152 |
| CRLF1 | 1.509807249 | 2.847719896 | 6.63E-17 | 3.10E-15 | 125.0620454 | 355.898682 | ENSG00000006016 |
| NA | 1.509703786 | 2.84751568 | 2.16E-07 | 2.72E-06 | 56.32786316 | 159.7775033 | ENSG00000238178 |
| COL5A2 | 1.508172658 | 2.844495223 | 1.51E-57 | 9.32E-55 | 447.4179493 | 1273.017862 | ENSG00000204262 |
| FAM124B | 1.506737139 | 2.841666284 | 0.000300897 | 0.001842967 | 15.84272873 | 45.0202591 | ENSG00000124019 |
| NA | 1.506333432 | 2.840871217 | 1.02E-05 | 9.07E-05 | 28.06696932 | 79.37483094 | ENSG00000260776 |
| NA | 1.500460274 | 2.829329642 | 0.007046601 | 0.02757055 | 9.276483128 | 26.28999367 | ENSG00000286909 |
| NA | 1.500093858 | 2.82861114 | 0.14468198 | 0.298047773 | 2.930792826 | 8.206710392 | ENSG00000236975 |
| NA | 1.499276819 | 2.827009672 | 0.072626005 | 0.178235284 | 4.069550476 | 11.55228095 | ENSG00000249119 |
| NA | 1.499254058 | 2.826965071 | 0.045844492 | 0.125081969 | 4.813273453 | 13.5969105 | ENSG00000259088 |
| NA | 1.498916962 | 2.826304608 | 0.173835122 | 0.340561507 | 2.249606746 | 6.369111425 | ENSG00000284747 |
| ANGPT4 | 1.496463541 | 2.82150233 | 0.033033719 | 0.0967062 | 5.42560881 | 15.33349484 | ENSG00000101280 |
| NA | 1.495769054 | 2.820144437 | 0.217989674 | 0.399034136 | 1.941860612 | 5.419817994 | ENSG00000285812 |
| DPPA3 | 1.494150142 | 2.816981602 | 5.82E-07 | 6.78E-06 | 33.57680746 | 94.62421839 | ENSG00000187569 |
| GAPLINC | 1.49338847 | 2.815494768 | 0.215483248 | 0.395853556 | 1.945017524 | 5.416019147 | ENSG00000266835 |
| SPSB2 | 1.493004239 | 2.814745021 | 3.32E-16 | 1.44E-14 | 109.45277 | 308.9554915 | ENSG00000111671 |
| LOC101927837 | 1.489334323 | 2.807593995 | 0.096140282 | 0.220037538 | 4.082178128 | 11.47719697 | ENSG00000270049 |
| EOMES | 1.488532452 | 2.806033927 | 0.062828219 | 0.159689825 | 4.396238088 | 12.41140663 | ENSG00000163508 |
| LEPR | 1.488021697 | 2.805040687 | 1.19E-06 | 1.31E-05 | 30.64789377 | 86.0987275 | ENSG00000116678 |
| CASC8 | 1.487566761 | 2.80415629 | 0.170053509 | 0.334974145 | 2.22623058 | 6.260172686 | ENSG00000246228 |
| DACT1 | 1.48751749 | 2.804060525 | 2.78E-52 | 1.32E-49 | 382.060977 | 1071.416498 | ENSG00000165617 |
| TUBA3E | 1.487304285 | 2.803646164 | 0.143846645 | 0.296722729 | 2.533976715 | 7.088804425 | ENSG00000152086 |
| NA | 1.485883189 | 2.800885853 | 0.134121407 | 0.281550136 | 2.871412842 | 8.064466544 | ENSG00000259723 |
| GRIN2C | 1.483965192 | 2.797164677 | 0.101163236 | 0.22851425 | 3.18231589 | 8.889299436 | ENSG00000161509 |
| ONECUT1 | 1.483813308 | 2.796870213 | 4.51E-05 | 0.000343573 | 30.80373993 | 85.6150217 | ENSG00000169856 |
| KCNJ9 | 1.483799006 | 2.796842487 | 0.190118302 | 0.362415879 | 2.187069849 | 6.155032794 | ENSG00000162728 |
| MAP4K3-DT | 1.482140927 | 2.793629941 | 1.18E-10 | 2.56E-09 | 59.9571975 | 167.1713899 | ENSG00000231312 |
| LPAR5 | 1.480535641 | 2.7905232 | 0.126803211 | 0.270445422 | 2.835409024 | 7.955527805 | ENSG00000184574 |
| ACRBP | 1.479186296 | 2.78791446 | 0.003021004 | 0.013401459 | 11.38079683 | 31.76524858 | ENSG00000111644 |
| GBP1P1 | 1.479051642 | 2.78765426 | 0.108797172 | 0.241259487 | 3.549442009 | 9.891218623 | ENSG00000225492 |
| EXPH5 | 1.478425478 | 2.786444615 | 1.17E-06 | 1.29E-05 | 36.37671632 | 101.5022337 | ENSG00000110723 |
| DANT2 | 1.476848114 | 2.783399734 | 0.032137612 | 0.094592076 | 6.061320334 | 16.85231331 | ENSG00000235244 |
| DGKH | 1.47559027 | 2.780974026 | 8.36E-13 | 2.52E-11 | 83.61983347 | 232.2747485 | ENSG00000102780 |
| WDR66 | 1.475570984 | 2.78093685 | 0.00181416 | 0.008724348 | 14.90182662 | 41.54036881 | ENSG00000158023 |
| NA | 1.474814261 | 2.779478574 | 0.212035913 | 0.391655731 | 2.308986731 | 6.365312579 | ENSG00000255176 |
| SH3TC2 | 1.473549941 | 2.777043817 | 0.117112091 | 0.255143107 | 3.238538961 | 8.923154197 | ENSG00000169247 |
| LOXHD1 | 1.470582026 | 2.77133675 | 0.004507284 | 0.018827585 | 11.56209369 | 31.99842462 | ENSG00000167210 |
| SYNPR | 1.469882848 | 2.769993994 | 0.003114116 | 0.013751974 | 11.95575289 | 32.97364871 | ENSG00000163630 |
| FAM71E1 | 1.469159539 | 2.768605579 | 8.09E-17 | 3.75E-15 | 93.92262811 | 259.8620044 | ENSG00000142530 |
| TNFAIP6 | 1.468996201 | 2.768292142 | 0.035059075 | 0.101354686 | 5.428765723 | 14.96544941 | ENSG00000123610 |
| GDNF | 1.468229559 | 2.766821475 | 0.106967061 | 0.23834185 | 3.56650435 | 9.747987089 | ENSG00000168621 |
| SPATC1 | 1.468015377 | 2.766410743 | 0.110193563 | 0.243690961 | 3.447744381 | 9.586972247 | ENSG00000186583 |
| PLAC1 | 1.467987807 | 2.766357879 | 0.013546276 | 0.047090575 | 8.045498589 | 22.09928189 | ENSG00000170965 |
| SLC40A1 | 1.467731624 | 2.765866692 | 0.245772546 | 0.433245993 | 1.862261374 | 5.213663469 | ENSG00000138449 |
| NA | 1.466425196 | 2.763363204 | 0.256713081 | 0.445373046 | 1.945017524 | 5.378915192 | ENSG00000287229 |
| DNAH10 | 1.465817586 | 2.762199623 | 0.02366165 | 0.074103587 | 6.626903358 | 18.40542458 | ENSG00000197653 |
| GPAT2 | 1.46542678 | 2.761451483 | 0.063144855 | 0.160362666 | 4.132087374 | 11.43629417 | ENSG00000186281 |
| HAS2 | 1.46344597 | 2.757662631 | 1.14E-15 | 4.72E-14 | 90.81547677 | 250.1090076 | ENSG00000170961 |
| VPS13B-DT | 1.463112481 | 2.757025252 | 0.003205087 | 0.014076559 | 11.12295994 | 30.70040641 | ENSG00000253948 |
| RBMS2 | 1.462963717 | 2.756740974 | 1.86E-22 | 1.44E-20 | 157.5292815 | 434.782473 | ENSG00000076067 |
| NA | 1.461772516 | 2.754465734 | 0.000329589 | 0.001990422 | 17.23291174 | 47.48924708 | ENSG00000258053 |
| NA | 1.461689959 | 2.754308116 | 0.014048662 | 0.048495275 | 8.803126994 | 24.00871564 | ENSG00000227799 |
| CES1P1 | 1.460776505 | 2.752564757 | 0.17797876 | 0.346656777 | 2.785499778 | 7.783228042 | ENSG00000228695 |
| LINC01694 | 1.458640816 | 2.748493021 | 4.45E-06 | 4.36E-05 | 28.69015089 | 78.89112514 | ENSG00000233922 |
| NA | 1.458615074 | 2.748443981 | 0.054148758 | 0.142136022 | 5.911494898 | 16.04570176 | ENSG00000224846 |
| ANKRD33 | 1.456800638 | 2.744989513 | 0.028336212 | 0.085716945 | 6.464450271 | 17.65143879 | ENSG00000167612 |
| ZP3 | 1.456294559 | 2.744026775 | 0.041238609 | 0.115264119 | 6.506767914 | 17.67345898 | ENSG00000188372 |
| VIM | 1.456079054 | 2.743616913 | 1.35E-27 | 1.58E-25 | 202.7693016 | 556.7636856 | ENSG00000026025 |
| UPK1A | 1.45597696 | 2.743422763 | 0.036069724 | 0.103678081 | 7.254925581 | 19.66926163 | ENSG00000105668 |
| NA | 1.455573171 | 2.742655029 | 0.216292246 | 0.396828398 | 1.88563754 | 5.213663469 | ENSG00000275880 |
| GNRH2 | 1.454556051 | 2.740722099 | 0.224938097 | 0.407766303 | 1.888794453 | 5.213663469 | ENSG00000125787 |
| GATA3 | 1.451253628 | 2.734455583 | 0.142019743 | 0.294034174 | 3.185472803 | 8.652650963 | ENSG00000107485 |
| UNC5C | 1.449417376 | 2.730977402 | 0.001240327 | 0.006313709 | 13.78832427 | 37.54551432 | ENSG00000182168 |
| KIF25-AS1 | 1.44864157 | 2.72950922 | 0.033924552 | 0.098845724 | 6.92705789 | 19.07347951 | ENSG00000229921 |
| GOLGA6B | 1.448253397 | 2.728774913 | 0.182415594 | 0.352766634 | 2.891632095 | 7.925909923 | ENSG00000215186 |

| | | | | | | | |
|---|---|---|---|---|---|---|---|
| NA | 1.447879475 | 2.728067753 | 0.027917905 | 0.084794593 | 11.0433607 | 30.31925258 | ENSG00000234537 |
| NA | 1.446719377 | 2.725874945 | 0.201238637 | 0.378118095 | 2.272982913 | 6.132462953 | ENSG00000273374 |
| ITGA9-AS1 | 1.443778037 | 2.720323143 | 0.000100169 | 0.000697685 | 20.29646766 | 55.3585176 | ENSG00000235257 |
| NDRG1 | 1.442825249 | 2.718527173 | 6.49E-08 | 8.96E-07 | 44.14100186 | 119.9367265 | ENSG00000104419 |
| SCARF1 | 1.440411288 | 2.713982255 | 1.08E-13 | 3.54E-12 | 83.81257791 | 227.4906765 | ENSG00000074660 |
| NA | 1.440357346 | 2.713880781 | 9.56E-08 | 1.29E-06 | 38.7104547 | 105.0628302 | ENSG00000224592 |
| DUSP10 | 1.438375077 | 2.710154459 | 0.002268075 | 0.010567244 | 14.54910959 | 39.1164089 | ENSG00000143507 |
| ALX1 | 1.43576394 | 2.705253781 | 9.47E-05 | 0.000663731 | 23.92748574 | 64.98248218 | ENSG00000180318 |
| HMCN1 | 1.433854585 | 2.701675843 | 2.44E-06 | 2.52E-05 | 231.8525028 | 626.1803762 | ENSG00000143341 |
| AHNAK2 | 1.431910745 | 2.698038145 | 1.01E-09 | 1.87E-08 | 57.97745394 | 156.2533091 | ENSG00000185567 |
| CAV1 | 1.430141767 | 2.69473194 | 4.58E-13 | 1.41E-11 | 75.32606643 | 203.0153855 | ENSG00000105974 |
| LINC00707 | 1.428679679 | 2.692002368 | 0.156324746 | 0.314892168 | 2.557352881 | 6.867677753 | ENSG00000238266 |
| C22orf42 | 1.428091 | 2.690904143 | 0.106606663 | 0.237792085 | 3.457215119 | 9.373331648 | ENSG00000205856 |
| CDH5 | 1.427410796 | 2.689635732 | 0.044371403 | 0.121996674 | 8.994089994 | 24.35386482 | ENSG00000179776 |
| SLC39A4 | 1.426758794 | 2.688420469 | 1.95E-05 | 0.000163014 | 25.28472416 | 67.72759571 | ENSG00000147804 |
| NA | 1.426442411 | 2.687830965 | 0.030967307 | 0.091846387 | 6.014568001 | 16.21475233 | ENSG00000281852 |
| TAC4 | 1.426045238 | 2.687091109 | 0.275851954 | 0.466822508 | 2.243292921 | 6.042844861 | ENSG00000176358 |
| LINC01144 | 1.42604068 | 2.687082619 | 0.226929158 | 0.410452329 | 2.779185952 | 7.595844512 | ENSG00000281912 |
| OAZ3 | 1.425949719 | 2.686913205 | 0.130068211 | 0.275664291 | 3.404148961 | 9.238135841 | ENSG00000143450 |
| KCNK3 | 1.423821832 | 2.682953094 | 0.01395091 | 0.048230959 | 8.590862359 | 23.05237417 | ENSG00000171303 |
| ISLR | 1.423311082 | 2.68200343 | 0.013048623 | 0.045708407 | 10.94472229 | 29.23746288 | ENSG00000129009 |
| FOSL1 | 1.422731838 | 2.680926818 | 1.85E-22 | 1.44E-20 | 154.2670733 | 412.9922953 | ENSG00000175592 |
| CFHR3 | 1.4211858 | 2.67805539 | 0.183639069 | 0.354169252 | 2.593356699 | 6.897733667 | ENSG00000116785 |
| B3GNT3 | 1.416807862 | 2.669941 | 0.000422638 | 0.002470631 | 17.06108561 | 45.67334188 | ENSG00000179913 |
| NA | 1.415560114 | 2.667632838 | 0.233390105 | 0.418301151 | 2.510600548 | 6.770461967 | ENSG00000279852 |
| ARIH2OS | 1.41450533 | 2.665683191 | 0.100209847 | 0.226969242 | 3.77127508 | 10.14240121 | ENSG00000221883 |
| KITLG | 1.414304203 | 2.665311592 | 1.18E-12 | 3.46E-11 | 83.7216288 | 222.7722899 | ENSG00000049130 |
| MASP1 | 1.412981311 | 2.662868731 | 1.03E-06 | 1.14E-05 | 40.87542615 | 109.0053854 | ENSG00000127241 |
| NA | 1.41276716 | 2.662473488 | 0.199321375 | 0.375658161 | 2.580729048 | 6.82264969 | ENSG00000231443 |
| C22orf23 | 1.411386077 | 2.659925939 | 0.01593783 | 0.053688067 | 8.700151589 | 23.02231826 | ENSG00000128346 |
| NA | 1.410861064 | 2.658958137 | 0.008815107 | 0.033146518 | 13.31812505 | 35.26197478 | ENSG00000285804 |
| EMP3 | 1.407175758 | 2.652174604 | 3.49E-07 | 4.23E-06 | 44.08152417 | 116.4850283 | ENSG00000142227 |
| RPSAP52 | 1.407044622 | 2.651933542 | 0.116024635 | 0.253275415 | 3.883721223 | 10.29636801 | ENSG00000241749 |
| NA | 1.406628349 | 2.651168469 | 0.186338251 | 0.357514818 | 2.643265945 | 6.909018588 | ENSG00000260136 |
| PLIN4 | 1.405783401 | 2.649616204 | 2.53E-05 | 0.000206045 | 26.24849876 | 69.69702967 | ENSG00000167676 |
| RNASEL | 1.40460596 | 2.647454629 | 0.004442493 | 0.018619895 | 11.50271371 | 30.43425167 | ENSG00000135828 |
| NA | 1.403707436 | 2.645806283 | 1.95E-05 | 0.00016268 | 28.51379238 | 75.39202582 | ENSG00000123870 |
| PEX5L | 1.402644819 | 2.643858232 | 3.27E-05 | 0.000259466 | 33.52058438 | 88.55412877 | ENSG00000114757 |
| NA | 1.401923576 | 2.642536824 | 0.222672026 | 0.404766902 | 2.252763659 | 5.945191042 | ENSG00000253252 |
| NXPH1 | 1.40188453 | 2.642465306 | 0.176607335 | 0.344619331 | 2.590199786 | 6.770023934 | ENSG00000122584 |
| IL18 | 1.401483855 | 2.641731525 | 0.13094339 | 0.277075556 | 3.433838953 | 9.102940034 | ENSG00000150782 |
| STMN4 | 1.401340751 | 2.641469498 | 0.202221271 | 0.379464314 | 2.223073667 | 5.885079212 | ENSG00000015592 |
| NA | 1.400731945 | 2.640355054 | 0.024608092 | 0.076460416 | 8.362715462 | 22.06652644 | ENSG00000205976 |
| GABRG2 | 1.400628902 | 2.640166476 | 0.025186317 | 0.077930418 | 9.476217809 | 25.07030863 | ENSG00000113327 |
| C2CD4B | 1.400368375 | 2.639689749 | 0.126119535 | 0.269266151 | 4.635035802 | 12.08893892 | ENSG00000205502 |
| NA | 1.399745292 | 2.638549944 | 0.15133371 | 0.307984955 | 2.84172285 | 7.509037581 | ENSG00000245025 |
| NA | 1.398431115 | 2.636147536 | 0.174361107 | 0.341251223 | 3.22591131 | 8.507157921 | ENSG00000285257 |
| GS1-24F4.2 | 1.396971009 | 2.633480924 | 0.186604606 | 0.357951745 | 2.50112981 | 6.665322075 | ENSG00000245857 |
| TGM1 | 1.396163356 | 2.632007055 | 0.000888081 | 0.004759886 | 18.92522612 | 49.99446294 | ENSG00000092295 |
| CD79A | 1.394592622 | 2.629143017 | 0.079080878 | 0.190052514 | 4.446147334 | 11.7024489 | ENSG00000105369 |
| LINC01460 | 1.393869796 | 2.62782608 | 0.108250051 | 0.240362578 | 4.181996619 | 10.92600489 | ENSG00000205334 |
| PRODH | 1.393104705 | 2.626432861 | 1.93E-05 | 0.00016139 | 28.00768704 | 73.44373598 | ENSG00000100033 |
| NA | 1.392242742 | 2.624864121 | 0.004419187 | 0.018538978 | 12.63072284 | 33.31131181 | ENSG00000257474 |
| TAS2R5 | 1.391535489 | 2.623577648 | 0.130396675 | 0.276297241 | 3.463528945 | 9.174774817 | ENSG00000127366 |
| CFP | 1.390284494 | 2.621303668 | 0.124366938 | 0.266753482 | 3.52290893 | 9.189308932 | ENSG00000126759 |
| ARHGAP31-AS1 | 1.389245011 | 2.619415659 | 0.014128692 | 0.048670975 | 8.522011636 | 22.41481319 | ENSG00000241155 |
| THBS1 | 1.387668295 | 2.616554474 | 4.60E-33 | 7.55E-31 | 411.2225299 | 1076.045813 | ENSG00000137801 |
| RDM1 | 1.387409693 | 2.616085501 | 1.43E-09 | 2.59E-08 | 70.01536162 | 182.8511332 | ENSG00000278023 |
| PCDHGB3 | 1.38712447 | 2.615568347 | 0.111425244 | 0.245851843 | 4.132087374 | 10.85124732 | ENSG00000262209 |
| NA | 1.386201751 | 2.613896017 | 1.00E-18 | 5.68E-17 | 126.5812332 | 330.6797512 | ENSG00000243004 |
| NA | 1.384630009 | 2.611049864 | 0.169149852 | 0.333619598 | 3.503967452 | 9.186059738 | ENSG00000235888 |
| LINC00698 | 1.383543345 | 2.609083913 | 4.87E-11 | 1.11E-09 | 74.83131097 | 195.6802882 | ENSG00000244342 |
| LOC101928063 | 1.382097855 | 2.606471084 | 0.252795308 | 0.441468788 | 2.166850596 | 5.674687808 | ENSG00000266921 |
| SNORD101 | 1.380803394 | 2.604133471 | 0.192261599 | 0.365549495 | 2.537133628 | 6.639065007 | ENSG00000206754 |
| TDRD10 | 1.380727391 | 2.603996286 | 0.164188342 | 0.326404241 | 2.891632095 | 7.478981666 | ENSG00000163239 |
| RDH16 | 1.380628882 | 2.603818489 | 0.053935157 | 0.141711681 | 5.455298802 | 14.17056081 | ENSG00000139547 |
| LINC00346 | 1.379607005 | 2.601974828 | 0.20559736 | 0.384011032 | 2.219916754 | 5.764855553 | ENSG00000255874 |
| MAML2 | 1.379301284 | 2.601423502 | 7.28E-15 | 2.73E-13 | 94.07739189 | 244.503017 | ENSG00000184384 |
| SOX3 | 1.379082231 | 2.601028542 | 1.48E-44 | 4.98E-42 | 367.4637396 | 956.1607194 | ENSG00000134595 |
| JUN | 1.377275911 | 2.597773974 | 8.33E-29 | 1.10E-26 | 350.0000801 | 909.2672947 | ENSG00000177606 |
| MUC16 | 1.376981872 | 2.59724457 | 0.182508083 | 0.352869269 | 2.904259747 | 7.48646774 | ENSG00000181143 |
| NA | 1.376814795 | 2.596943803 | 0.103125168 | 0.231843991 | 4.164934279 | 10.7268984 | ENSG00000286592 |
| LMCD1-AS1 | 1.376491192 | 2.596361362 | 0.017979107 | 0.058988275 | 9.124876254 | 23.80647158 | ENSG00000227110 |
| FGF18 | 1.374503083 | 2.592785906 | 0.004346837 | 0.018264401 | 16.22513575 | 42.34274348 | ENSG00000156427 |

| | | | | | | | |
|---|---|---|---|---|---|---|---|
| LINC00672 | 1.374502388 | 2.592784655 | 0.001842809 | 0.008841474 | 16.76724491 | 43.45597453 | ENSG00000263874 |
| NA | 1.374452717 | 2.592695389 | 0.016915529 | 0.056255591 | 10.0904323 | 26.34641828 | ENSG00000262583 |
| DENND2D | 1.373991427 | 2.591866529 | 0.011094556 | 0.040141334 | 9.524849279 | 24.69510353 | ENSG00000162777 |
| NA | 1.372444914 | 2.589089637 | 0.036717671 | 0.105028451 | 6.331784874 | 16.40582309 | ENSG00000277287 |
| GRPR | 1.371217634 | 2.586888073 | 2.45E-23 | 2.04E-21 | 221.5835472 | 573.0350059 | ENSG00000126010 |
| DISP1 | 1.370435728 | 2.585486421 | 6.79E-12 | 1.77E-10 | 85.14535778 | 220.2236865 | ENSG00000154309 |
| CAPS2 | 1.369988147 | 2.584684426 | 0.000632215 | 0.003521931 | 19.69803773 | 51.03034851 | ENSG00000180881 |
| NA | 1.365610503 | 2.576853471 | 0.212600138 | 0.392385079 | 2.613575953 | 6.699176837 | ENSG00000235351 |
| LACC1 | 1.364648741 | 2.575136203 | 1.22E-07 | 1.60E-06 | 43.35052655 | 111.5826593 | ENSG00000179630 |
| DHRS2 | 1.364525965 | 2.574917064 | 2.69E-05 | 0.000217597 | 28.54348237 | 73.26438817 | ENSG00000100867 |
| LGALS1 | 1.364172191 | 2.574285728 | 6.20E-18 | 3.28E-16 | 142.3586741 | 365.8202914 | ENSG00000100097 |
| OTOGL | 1.363586879 | 2.573241532 | 0.089483704 | 0.208720233 | 4.525746572 | 11.53719718 | ENSG00000165899 |
| NA | 1.362546776 | 2.571387036 | 0.02541514 | 0.07842633 | 8.048655502 | 20.59174834 | ENSG00000239407 |
| TRIM22 | 1.361846401 | 2.570139027 | 2.35E-25 | 2.29E-23 | 435.4405192 | 1118.986286 | ENSG00000132274 |
| COL23A1 | 1.361285435 | 2.569139868 | 2.56E-34 | 4.70E-32 | 317.5118733 | 815.3300852 | ENSG00000050767 |
| POU6F2 | 1.360832072 | 2.568332649 | 0.006415827 | 0.025468183 | 14.92254954 | 38.06296326 | ENSG00000106536 |
| NAV3 | 1.36003138 | 2.566907627 | 9.61E-05 | 0.000672478 | 30.12274925 | 77.42372149 | ENSG00000067798 |
| LINC01405 | 1.359690606 | 2.566301379 | 2.17E-18 | 1.19E-16 | 161.2168083 | 413.0226861 | ENSG00000185847 |
| TMEM196 | 1.359676229 | 2.566275805 | 0.139419376 | 0.290239101 | 3.49637585 | 8.986953254 | ENSG00000173452 |
| CREBRF | 1.358784402 | 2.564689907 | 1.54E-05 | 0.000132368 | 31.34792363 | 80.17274549 | ENSG00000164463 |
| CREB3L3 | 1.358468936 | 2.564129163 | 0.150825048 | 0.307060718 | 3.152625897 | 8.075201812 | ENSG00000060566 |
| ARSD | 1.357989262 | 2.563276769 | 8.90E-12 | 2.27E-10 | 84.18729563 | 215.704773 | ENSG00000006756 |
| POMK | 1.355434758 | 2.558742127 | 2.75E-06 | 2.81E-05 | 36.48294634 | 93.61711491 | ENSG00000185900 |
| GUCY2C | 1.354470918 | 2.557033246 | 5.92E-06 | 5.63E-05 | 33.19134122 | 85.13577602 | ENSG00000070019 |
| MIR210HG | 1.352612583 | 2.553741651 | 0.001189186 | 0.006110447 | 21.89448062 | 55.82432847 | ENSG00000247095 |
| MIXL1 | 1.35211046 | 2.55285299 | 0.004217804 | 0.017823251 | 20.96256824 | 53.67291007 | ENSG00000185155 |
| SPTLC3 | 1.350244032 | 2.549552475 | 0.081268617 | 0.193927198 | 4.469523501 | 11.38746726 | ENSG00000172296 |
| MARCHF11 | 1.349780637 | 2.548733688 | 0.058800423 | 0.151551666 | 5.730198032 | 14.64341972 | ENSG00000183654 |
| IL15 | 1.349486873 | 2.548214763 | 0.017115945 | 0.056799809 | 10.43408456 | 26.32286075 | ENSG00000164136 |
| CPN1 | 1.349484229 | 2.548210093 | 0.183437896 | 0.35396543 | 3.099559739 | 7.993069793 | ENSG00000120054 |
| LINC00847 | 1.348398871 | 2.546293763 | 0.063667323 | 0.161241205 | 5.617751889 | 14.40677125 | ENSG00000245060 |
| RASSF9 | 1.347093922 | 2.543991626 | 0.079968967 | 0.191725424 | 6.817670962 | 17.30683926 | ENSG00000198774 |
| LINC01138 | 1.346446787 | 2.54285075 | 2.52E-06 | 2.60E-05 | 34.65353187 | 88.18133708 | ENSG00000274020 |
| DSEL | 1.345132375 | 2.540535062 | 1.39E-17 | 7.08E-16 | 174.2034277 | 442.0415615 | ENSG00000171451 |
| MRVI1 | 1.345028407 | 2.540351984 | 7.62E-11 | 1.70E-09 | 68.95384305 | 175.1274673 | ENSG00000072952 |
| MT1X | 1.343384217 | 2.537458482 | 3.10E-08 | 4.57E-07 | 55.61342421 | 140.8459412 | ENSG00000187193 |
| NA | 1.342559453 | 2.536008275 | 0.047548623 | 0.128595678 | 6.044257993 | 15.2842299 | ENSG00000225465 |
| SP8 | 1.342048362 | 2.535110025 | 7.48E-14 | 2.52E-12 | 107.6160722 | 272.2684013 | ENSG00000164651 |
| APOL6 | 1.341459551 | 2.534075574 | 0.128335698 | 0.272615912 | 3.486905112 | 8.821701532 | ENSG00000221963 |
| PLXDC2 | 1.341252168 | 2.533711335 | 7.92E-42 | 2.49E-39 | 375.1541384 | 950.9591052 | ENSG00000120594 |
| NA | 1.341101626 | 2.533446961 | 3.96E-05 | 0.000306999 | 29.55263384 | 74.59191265 | ENSG00000250903 |
| PYROXD2 | 1.340237755 | 2.531930413 | 6.74E-05 | 0.000489811 | 31.00101676 | 78.58830448 | ENSG00000119943 |
| EGFLAM | 1.340071521 | 2.53163869 | 0.142547737 | 0.294962572 | 3.52290893 | 8.881813362 | ENSG00000164318 |
| TMEM220-AS1 | 1.340016685 | 2.531542465 | 0.01604089 | 0.053927418 | 9.649923073 | 24.39184484 | ENSG00000263400 |
| NUTM2F | 1.337884094 | 2.527803105 | 0.204031285 | 0.382048478 | 2.821503596 | 7.160201175 | ENSG00000130950 |
| LHFPL1 | 1.337417968 | 2.526986519 | 7.45E-07 | 8.51E-06 | 44.559315 | 112.162808 | ENSG00000182508 |
| LOC105373958 | 1.336098532 | 2.524676486 | 0.170546169 | 0.335801838 | 2.865099016 | 7.235285152 | ENSG00000234949 |
| KANK3 | 1.335947703 | 2.524412554 | 2.16E-07 | 2.72E-06 | 44.77808886 | 113.0183581 | ENSG00000186994 |
| PDP1 | 1.335565711 | 2.523744237 | 5.15E-33 | 8.37E-31 | 269.1347036 | 679.2653017 | ENSG00000164951 |
| LINC00540 | 1.335462394 | 2.523563507 | 8.90E-05 | 0.000628563 | 29.07433935 | 73.18484408 | ENSG00000276476 |
| RORA-AS1 | 1.331606034 | 2.516826967 | 0.238431855 | 0.424056808 | 2.537133628 | 6.444195403 | ENSG00000245534 |
| CSRNP3 | 1.331162997 | 2.516054193 | 2.28E-13 | 7.24E-12 | 94.36945116 | 237.5450598 | ENSG00000178662 |
| CHRNE | 1.32937374 | 2.51293567 | 0.150086729 | 0.30615314 | 3.176002064 | 8.011840788 | ENSG00000108556 |
| KCNAB2 | 1.329076164 | 2.512417397 | 2.29E-21 | 1.63E-19 | 180.8263774 | 453.5299805 | ENSG00000069424 |
| RGS20 | 1.327957747 | 2.510470455 | 0.125460333 | 0.268373048 | 3.768118167 | 9.478033508 | ENSG00000147509 |
| CYP2F1 | 1.32750945 | 2.509690483 | 0.124876332 | 0.267629271 | 4.197781184 | 10.51749468 | ENSG00000197446 |
| PTAFR | 1.325969157 | 2.507012444 | 8.00E-06 | 7.35E-05 | 41.36063583 | 103.5886018 | ENSG00000169403 |
| NA | 1.325241559 | 2.505748394 | 0.033323561 | 0.097385153 | 8.300178565 | 20.80225136 | ENSG00000287104 |
| LINC01780 | 1.325212273 | 2.50569753 | 0.007437812 | 0.028797244 | 11.4698668 | 28.77643854 | ENSG00000232650 |
| NA | 1.324993614 | 2.505317788 | 0.008365517 | 0.031773607 | 11.73401752 | 29.39479049 | ENSG00000241634 |
| LINC00954 | 1.324174061 | 2.503894994 | 0.094433811 | 0.216959508 | 5.061639604 | 12.78325091 | ENSG00000228784 |
| MYO1F | 1.324108602 | 2.503781388 | 1.58E-10 | 3.35E-09 | 84.33642201 | 211.0247416 | ENSG00000142347 |
| CD81-AS1 | 1.322982011 | 2.501826964 | 0.083040982 | 0.197115979 | 4.833492707 | 12.02502824 | ENSG00000238184 |
| LOC102723566 | 1.322974174 | 2.501813375 | 0.119801911 | 0.259302653 | 5.385170303 | 13.5747787 | ENSG00000225032 |
| HPCAL4 | 1.322074007 | 2.500252859 | 4.09E-06 | 4.05E-05 | 37.11550095 | 92.73724286 | ENSG00000116983 |
| LOC285097 | 1.320521365 | 2.49756351 | 0.122043137 | 0.262894876 | 4.152306627 | 10.36352788 | ENSG00000280119 |
| SLC8A3 | 1.319211035 | 2.495296125 | 0.157814024 | 0.317053267 | 3.596194342 | 8.863042367 | ENSG00000100678 |
| DUSP27 | 1.318113891 | 2.493399218 | 0.092681739 | 0.214069479 | 4.442990421 | 11.11751368 | ENSG00000198842 |
| TMEM204 | 1.317556598 | 2.492436238 | 0.135877886 | 0.284399858 | 3.797808159 | 9.542382217 | ENSG00000131634 |
| G2E3-AS1 | 1.317294395 | 2.491983291 | 0.102748412 | 0.231145322 | 4.856868873 | 12.01754217 | ENSG00000257636 |
| NA | 1.315726906 | 2.489277221 | 0.231123026 | 0.415523734 | 2.494815984 | 6.278943681 | ENSG00000227107 |
| NA | 1.315642027 | 2.489130771 | 0.032359387 | 0.095032304 | 11.96838054 | 29.77879576 | ENSG00000262580 |
| NOS3 | 1.315334386 | 2.488600045 | 1.30E-07 | 1.70E-06 | 50.82993845 | 126.5668797 | ENSG00000164867 |
| VWC2 | 1.315311936 | 2.488561319 | 0.015841189 | 0.053420778 | 9.945041557 | 24.63879055 | ENSG00000188730 |

| | | | | | | | |
|---|---|---|---|---|---|---|---|
| NA | 1.314634762 | 2.48739351 | 0.031728555 | 0.093684409 | 8.283116224 | 20.58426226 | ENSG00000248740 |
| LOC101927531 | 1.314490379 | 2.487144587 | 0.214672057 | 0.394832976 | 2.527662889 | 6.33905551 | ENSG00000255595 |
| BGN | 1.313777175 | 2.485915357 | 6.82E-10 | 1.30E-08 | 72.98473646 | 181.173673 | ENSG00000182492 |
| POLN | 1.312384536 | 2.483516852 | 0.164960479 | 0.327622966 | 3.569661262 | 8.85219548 | ENSG00000130997 |
| NA | 1.311700849 | 2.482340203 | 0.009527506 | 0.035344836 | 11.11221142 | 27.63607435 | ENSG00000262468 |
| MROH5 | 1.310222869 | 2.479798453 | 0.145067484 | 0.298543277 | 3.483748199 | 8.67197161 | ENSG00000226807 |
| SP140 | 1.309181664 | 2.478009406 | 0.176579366 | 0.344619331 | 3.258758215 | 8.023125708 | ENSG00000079263 |
| CPQ | 1.30889386 | 2.477515117 | 0.002290511 | 0.010653033 | 15.8692618 | 39.40232231 | ENSG00000104324 |
| LINC01140 | 1.308506437 | 2.476849892 | 0.038303761 | 0.108633803 | 7.317560176 | 18.09369213 | ENSG00000267272 |
| MUC5B | 1.303776795 | 2.468743235 | 0.161746509 | 0.32272686 | 3.436995866 | 8.532976956 | ENSG00000117983 |
| APOBEC3D | 1.303612643 | 2.468462354 | 0.083284214 | 0.197566826 | 5.349166485 | 13.315672 | ENSG00000243811 |
| OVCH2 | 1.303188417 | 2.467736606 | 0.131698988 | 0.278048445 | 4.069550476 | 10.14240121 | ENSG00000183378 |
| HPGD | 1.303113313 | 2.467608144 | 0.001305626 | 0.006586409 | 18.08286473 | 44.5662828 | ENSG00000164120 |
| ADGRG1 | 1.302955904 | 2.467338925 | 9.03E-05 | 0.000636206 | 30.50062388 | 75.28958546 | ENSG00000205336 |
| BCHE | 1.302413507 | 2.466411476 | 6.73E-07 | 7.74E-06 | 48.93029778 | 120.3168926 | ENSG00000114200 |
| NA | 1.302210234 | 2.466063987 | 1.88E-06 | 1.99E-05 | 40.7445422 | 100.3708526 | ENSG00000259820 |
| NA | 1.300149827 | 2.462544554 | 0.00261631 | 0.011890801 | 17.50021937 | 43.20253044 | ENSG00000254297 |
| GOLGA6D | 1.300072538 | 2.462412632 | 0.077973784 | 0.188074715 | 5.878647993 | 14.30531858 | ENSG00000140478 |
| SOHLH2 | 1.299575998 | 2.461565277 | 0.13420569 | 0.281694921 | 4.472680413 | 10.99349117 | ENSG00000120669 |
| NA | 1.297361594 | 2.457789899 | 0.012176677 | 0.043283952 | 11.13558759 | 27.33171636 | ENSG00000213513 |
| NA | 1.296741919 | 2.45673444 | 0.057683735 | 0.149400551 | 6.959904795 | 17.11857967 | ENSG00000260193 |
| CCDC169 | 1.295807658 | 2.455144023 | 2.72E-08 | 4.03E-07 | 61.09279824 | 149.7450998 | ENSG00000242715 |
| NA | 1.293844418 | 2.4518053 | 0.13110641 | 0.277262233 | 3.791494334 | 9.33578966 | ENSG00000270177 |
| BCL2L14 | 1.292828581 | 2.450079532 | 0.053963726 | 0.141766649 | 6.580151025 | 16.23732217 | ENSG00000121380 |
| LOC100506691 | 1.29279469 | 2.450021976 | 1.24E-09 | 2.26E-08 | 73.19008591 | 179.0633093 | ENSG00000256546 |
| NA | 1.292712166 | 2.449881835 | 0.124313893 | 0.266670567 | 4.535217311 | 11.0424297 | ENSG00000270977 |
| DDX60L | 1.292523775 | 2.449561945 | 0.027773356 | 0.084413148 | 8.247112406 | 20.20916879 | ENSG00000181381 |
| NA | 1.292105 | 2.448851006 | 0.073916881 | 0.180521211 | 5.481831882 | 13.33020612 | ENSG00000238035 |
| NA | 1.292034653 | 2.448731602 | 0.209771509 | 0.388985007 | 2.812032858 | 6.954046651 | ENSG00000268201 |
| THRB | 1.291989435 | 2.448654853 | 2.71E-08 | 4.03E-07 | 54.39704416 | 133.1452832 | ENSG00000151090 |
| FN1 | 1.291873111 | 2.448457426 | 5.72E-37 | 1.21E-34 | 2564.409727 | 6277.590035 | ENSG00000115414 |
| TCAM1P | 1.290677653 | 2.446429405 | 0.036601239 | 0.104783344 | 8.015808597 | 19.63215767 | ENSG00000240280 |
| NA | 1.289523032 | 2.444472256 | 0.252976127 | 0.441659716 | 2.85878519 | 6.942323697 | ENSG00000249894 |
| WNT8B | 1.289505165 | 2.444441984 | 0.025511078 | 0.078685691 | 9.05780697 | 21.99089281 | ENSG00000075290 |
| NA | 1.28573338 | 2.438059577 | 0.011213248 | 0.040474756 | 11.10274068 | 27.08433262 | ENSG00000233577 |
| PPT2-EGFL8 | 1.285299029 | 2.437325663 | 0.158548609 | 0.318161534 | 4.099240469 | 10.0371497 | ENSG00000258388 |
| NA | 1.28486619 | 2.436594522 | 0.0152774 | 0.051849721 | 10.72930074 | 26.25201365 | ENSG00000280734 |
| GAB3 | 1.284743184 | 2.436386785 | 0.059449835 | 0.152757969 | 6.070791072 | 14.75560765 | ENSG00000160219 |
| PLIN1 | 1.284558755 | 2.436075345 | 0.263641901 | 0.453362311 | 2.279296739 | 5.502388045 | ENSG00000166819 |
| ZNF808 | 1.282919037 | 2.433308158 | 5.03E-07 | 5.94E-06 | 55.14953882 | 134.0059462 | ENSG00000198482 |
| RNASE1 | 1.282124972 | 2.431969224 | 0.091788804 | 0.212492343 | 5.842644175 | 14.11044898 | ENSG00000129538 |
| CR1L | 1.281529858 | 2.430966241 | 0.012420435 | 0.043990129 | 11.70117061 | 28.5966527 | ENSG00000197721 |
| FOXB2 | 1.280095668 | 2.428550806 | 0.001623672 | 0.007950628 | 17.48127789 | 42.52307897 | ENSG00000204612 |
| ARRB1 | 1.279814591 | 2.428077702 | 7.68E-23 | 6.16E-21 | 449.8543852 | 1091.294142 | ENSG00000137486 |
| SULF1 | 1.27916241 | 2.42698032 | 1.41E-16 | 6.33E-15 | 288.6858926 | 699.4362588 | ENSG00000137573 |
| NA | 1.278781318 | 2.426339311 | 0.264457771 | 0.454094232 | 2.521349063 | 6.124976879 | ENSG00000231822 |
| SAMSN1 | 1.277721365 | 2.424557326 | 0.073927732 | 0.180523923 | 6.467607184 | 15.52792641 | ENSG00000155307 |
| ALOX12P2 | 1.276927916 | 2.423224242 | 2.41E-05 | 0.00019718 | 54.16446257 | 131.5327617 | ENSG00000262943 |
| LMCD1 | 1.273374234 | 2.417262642 | 1.66E-18 | 9.24E-17 | 163.7859018 | 396.3506998 | ENSG00000071282 |
| FRMPD4 | 1.272975352 | 2.416594399 | 0.001189527 | 0.006110505 | 20.14664223 | 48.52728253 | ENSG00000169933 |
| PICART1 | 1.272064625 | 2.415069362 | 0.236115699 | 0.42110935 | 2.530819802 | 6.106205885 | ENSG00000246640 |
| SAMD4A | 1.271564795 | 2.414232793 | 2.27E-09 | 4.00E-08 | 82.86230277 | 200.425218 | ENSG00000020577 |
| HEATR4 | 1.271560622 | 2.41422581 | 0.028453663 | 0.085959281 | 8.293864739 | 20.01809803 | ENSG00000187105 |
| FGF8 | 1.271237045 | 2.413684393 | 0.000808251 | 0.004380154 | 22.69580215 | 54.68655221 | ENSG00000107831 |
| CPB2-AS1 | 1.269803276 | 2.411286833 | 0.175129711 | 0.34242579 | 4.016484318 | 9.819821873 | ENSG00000235903 |
| MLKL | 1.268534358 | 2.409166926 | 0.228908666 | 0.412798988 | 3.414897476 | 8.300565364 | ENSG00000168404 |
| TGM2 | 1.268238943 | 2.408673661 | 3.01E-07 | 3.69E-06 | 48.77109931 | 117.4151127 | ENSG00000198959 |
| FOXS1 | 1.267856068 | 2.40803451 | 0.29357477 | 0.486117644 | 2.223073667 | 5.408971107 | ENSG00000179772 |
| MUC12 | 1.265767732 | 2.404551344 | 0.008674238 | 0.03273215 | 15.39274876 | 37.02101733 | ENSG00000205277 |
| ITGB2 | 1.265622965 | 2.404310072 | 0.030217353 | 0.090135796 | 8.587705446 | 20.65565901 | ENSG00000160255 |
| NLGN1 | 1.264573323 | 2.402561436 | 3.80E-13 | 1.18E-11 | 111.1682652 | 267.0171958 | ENSG00000169760 |
| NA | 1.264507071 | 2.402451107 | 0.12333044 | 0.26492888 | 4.132087374 | 9.917037658 | ENSG00000213070 |
| IQSEC3 | 1.263774919 | 2.401232199 | 6.37E-05 | 0.000467007 | 30.26676452 | 72.75866219 | ENSG00000120645 |
| NA | 1.261948934 | 2.398194938 | 0.057526112 | 0.149082712 | 7.222176374 | 17.46036803 | ENSG00000272316 |
| NA | 1.261571275 | 2.397567238 | 0.175740354 | 0.343402395 | 3.768118167 | 9.110975761 | ENSG00000267778 |
| NR2F1-AS1 | 1.260438993 | 2.395686274 | 0.030369672 | 0.090479262 | 8.99211316 | 21.40540793 | ENSG00000237187 |
| POLD4 | 1.259958035 | 2.394887745 | 0.000134249 | 0.000901837 | 27.04213099 | 64.69331958 | ENSG00000175482 |
| MSX2 | 1.258559818 | 2.392567817 | 0.095087989 | 0.218033006 | 5.692916438 | 13.62641677 | ENSG00000120149 |
| SPTY2D1OS | 1.258497347 | 2.392464216 | 0.097849342 | 0.222911248 | 6.487826437 | 15.48746164 | ENSG00000247595 |
| AKAP6 | 1.257942975 | 2.391545062 | 0.053120587 | 0.140087785 | 7.225333287 | 17.3171365 | ENSG00000151320 |
| LINC02175 | 1.257897822 | 2.391470212 | 0.010561127 | 0.038486025 | 12.56493134 | 29.89379485 | ENSG00000262155 |
| ATRNL1 | 1.257855103 | 2.391399401 | 7.86E-16 | 3.31E-14 | 149.2550347 | 357.2032473 | ENSG00000107518 |
| ALG10 | 1.256911575 | 2.389835928 | 0.000656915 | 0.003647471 | 23.07437584 | 55.26400136 | ENSG00000139133 |
| NA | 1.256842503 | 2.389721513 | 0.227235384 | 0.41072563 | 2.874569755 | 6.852705605 | ENSG00000269694 |

| | | | | | | | |
|---|---|---|---|---|---|---|---|
| NA | 1.256225882 | 2.388700343 | 0.038487807 | 0.109089007 | 7.675215557 | 18.31525684 | ENSG00000235499 |
| FOXD1 | 1.255664714 | 2.387771386 | 0.106442167 | 0.237543542 | 5.416138072 | 12.88751474 | ENSG00000251493 |
| ZNF790-AS1 | 1.255515605 | 2.387524611 | 0.060451277 | 0.154858877 | 6.683126429 | 15.99362566 | ENSG00000267254 |
| ZNF280A | 1.255145456 | 2.386912128 | 0.051602676 | 0.136942154 | 6.947277144 | 16.67252748 | ENSG00000169548 |
| GADD45G | 1.25512201 | 2.386873338 | 1.83E-16 | 8.12E-15 | 155.1027149 | 370.4967875 | ENSG00000130222 |
| SAMD11 | 1.253748667 | 2.384602285 | 0.165036827 | 0.327704365 | 4.277380422 | 10.10485922 | ENSG00000187634 |
| NA | 1.253008853 | 2.383379775 | 0.073545089 | 0.179851124 | 6.328627961 | 15.10824291 | ENSG00000265975 |
| LOC101929748 | 1.252844658 | 2.383108534 | 0.007100274 | 0.027739562 | 18.57949934 | 44.3471315 | ENSG00000288062 |
| ELFN1 | 1.252743146 | 2.382940858 | 1.77E-10 | 3.72E-09 | 107.0436717 | 254.5550985 | ENSG00000225968 |
| SNORD116-5 | 1.250948657 | 2.379978692 | 0.250760691 | 0.43924002 | 2.808875945 | 6.694939957 | ENSG00000207191 |
| FLRT3 | 1.250745426 | 2.37964345 | 0.072764945 | 0.178460432 | 6.295781056 | 15.01807516 | ENSG00000125848 |
| MSH2-OT1 | 1.250343509 | 2.378980604 | 0.0835762 | 0.198097979 | 6.020881826 | 14.41469536 | ENSG00000235760 |
| PTPN14 | 1.250242914 | 2.378814729 | 2.20E-38 | 5.29E-36 | 1091.755724 | 2596.455928 | ENSG00000152104 |
| ARL11 | 1.248632063 | 2.37616013 | 0.010065164 | 0.036962107 | 13.03700969 | 30.99661705 | ENSG00000152213 |
| RASSF10-DT | 1.247054301 | 2.373562932 | 0.171214673 | 0.336816152 | 4.208529699 | 9.917037658 | ENSG00000254670 |
| POLR3GL | 1.245712094 | 2.371355721 | 0.012846222 | 0.045187299 | 13.98805895 | 33.14159997 | ENSG00000121851 |
| ZNF91 | 1.244876108 | 2.36998201 | 3.88E-27 | 4.31E-25 | 269.4602112 | 638.6458822 | ENSG00000167232 |
| NPNT | 1.244707262 | 2.369704656 | 0.0578875 | 0.149802698 | 6.639531009 | 15.76077603 | ENSG00000168743 |
| PDE6G | 1.243838217 | 2.368277631 | 2.15E-07 | 2.71E-06 | 51.25200986 | 121.3381324 | ENSG00000185527 |
| ACHE | 1.24297525 | 2.366861439 | 1.41E-08 | 2.21E-07 | 77.41707606 | 183.2852392 | ENSG00000087085 |
| IDH1-AS1 | 1.242511686 | 2.366101045 | 0.022798407 | 0.071940412 | 10.19972153 | 24.12525208 | ENSG00000231908 |
| NDUFA4L2 | 1.242452669 | 2.366004256 | 0.154046312 | 0.311657499 | 4.955507287 | 11.85652732 | ENSG00000185633 |
| NA | 1.241660779 | 2.364705921 | 0.092783936 | 0.214216343 | 6.130171057 | 14.5048631 | ENSG00000255142 |
| MIR34AHG | 1.24156854 | 2.364554738 | 0.016311738 | 0.05464963 | 75.28740936 | 178.1240672 | ENSG00000228526 |
| MEF2C | 1.240534194 | 2.36286007 | 0.006088016 | 0.024381004 | 13.94446353 | 32.99946774 | ENSG00000081189 |
| NA | 1.239583695 | 2.361303846 | 0.058131893 | 0.150267296 | 7.734595541 | 18.25558304 | ENSG00000279583 |
| TPRG1 | 1.238889583 | 2.360168045 | 0.230945024 | 0.41528428 | 2.84172285 | 6.683655036 | ENSG00000188001 |
| TRPV4 | 1.238681959 | 2.359828407 | 1.44E-05 | 0.000124875 | 41.41931675 | 97.36261029 | ENSG00000111199 |
| KIAA1107 | 1.238328885 | 2.359250951 | 3.73E-05 | 0.000290981 | 35.82197951 | 84.62939721 | ENSG00000189195 |
| NA | 1.236953902 | 2.357003501 | 0.199026015 | 0.375292316 | 3.212005882 | 7.535294649 | ENSG00000287778 |
| NA | 1.236943777 | 2.35698696 | 4.80E-05 | 0.000363993 | 34.95684332 | 82.40107135 | ENSG00000188013 |
| TRPV2 | 1.23656729 | 2.356371959 | 0.00025803 | 0.001613469 | 27.12676628 | 63.91015393 | ENSG00000187688 |
| PARP15 | 1.236425047 | 2.356139643 | 0.281449926 | 0.47271345 | 2.223073667 | 5.266289226 | ENSG00000173200 |
| NA | 1.234421669 | 2.352870093 | 0.200019079 | 0.376475436 | 3.602508168 | 8.420789023 | ENSG00000254013 |
| POLH | 1.233708956 | 2.351708029 | 1.75E-11 | 4.26E-10 | 127.6472091 | 300.5165843 | ENSG00000170734 |
| PDZD2 | 1.233682685 | 2.351665205 | 7.20E-18 | 3.78E-16 | 264.98714 | 622.5240919 | ENSG00000133401 |
| NA | 1.233316065 | 2.351067672 | 0.266244141 | 0.456270818 | 2.904259747 | 6.777510008 | ENSG00000232936 |
| FALEC | 1.232397359 | 2.349570992 | 0.220282016 | 0.401801681 | 3.146312072 | 7.479419699 | ENSG00000228126 |
| RNVU1-1 | 1.232179436 | 2.349216109 | 0.261920117 | 0.451186754 | 2.560509794 | 6.046532088 | ENSG00000207340 |
| NA | 1.232003807 | 2.348930141 | 0.077674864 | 0.187548974 | 6.352004128 | 14.97337351 | ENSG00000223374 |
| LPP | 1.230614062 | 2.346668511 | 9.52E-17 | 4.36E-15 | 387.5593828 | 908.7313581 | ENSG00000145012 |
| NA | 1.229821094 | 2.345379034 | 0.327100499 | 0.521301591 | 2.507443636 | 5.918933974 | ENSG00000241170 |
| TOGARAM2 | 1.229712006 | 2.345201697 | 0.043100399 | 0.119344957 | 7.986118604 | 18.73549 | ENSG00000189350 |
| ARID3C | 1.22884749 | 2.343796787 | 0.281392534 | 0.47271345 | 2.246449834 | 5.273775299 | ENSG00000205143 |
| KCNE3 | 1.228844407 | 2.343791778 | 0.000873152 | 0.004689367 | 25.47676954 | 59.40880901 | ENSG00000175538 |
| NOTUM | 1.227921064 | 2.342292202 | 0.048367823 | 0.130278918 | 7.340936343 | 17.16360773 | ENSG00000185269 |
| CCDC121 | 1.227683038 | 2.341905786 | 0.00269764 | 0.012182762 | 17.57034787 | 41.12773334 | ENSG00000176714 |
| MS4A15 | 1.227228296 | 2.341167727 | 0.126764907 | 0.270425965 | 4.892872691 | 11.34612642 | ENSG00000166961 |
| HRK | 1.222755947 | 2.333921354 | 3.17E-07 | 3.86E-06 | 53.30947353 | 124.1883859 | ENSG00000135116 |
| NA | 1.222413965 | 2.333368177 | 0.281932738 | 0.473069106 | 2.246449834 | 5.243719384 | ENSG00000225970 |
| GNLY | 1.221183204 | 2.331378433 | 0.296765852 | 0.48960818 | 2.252763659 | 5.247518231 | ENSG00000115523 |
| TSHZ1 | 1.220743106 | 2.330667347 | 0.072602033 | 0.178225971 | 6.334941787 | 14.79314964 | ENSG00000179981 |
| CABYR | 1.220428922 | 2.330159841 | 7.19E-10 | 1.37E-08 | 78.32570998 | 182.517707 | ENSG00000154040 |
| CXCR2 | 1.220289621 | 2.329934861 | 0.014437638 | 0.04948903 | 12.35582361 | 28.8247158 | ENSG00000180871 |
| VLDLR | 1.21979052 | 2.329128958 | 1.62E-32 | 2.59E-30 | 466.8461164 | 1086.33844 | ENSG00000147852 |
| LINC01772 | 1.219160021 | 2.328111284 | 0.151989767 | 0.309042404 | 5.310005754 | 12.52078341 | ENSG00000226029 |
| NA | 1.219155007 | 2.328103192 | 0.143798702 | 0.296699577 | 4.833492707 | 11.24902226 | ENSG00000284630 |
| KIAA1257 | 1.219095273 | 2.328006801 | 6.02E-05 | 0.000443366 | 33.56290203 | 78.24877762 | ENSG00000114656 |
| CLC | 1.219024241 | 2.327892183 | 0.039474228 | 0.111289221 | 8.273645486 | 19.26031339 | ENSG00000105205 |
| NA | 1.218576126 | 2.327169229 | 0.000427601 | 0.002494539 | 26.88599173 | 62.48095355 | ENSG00000264112 |
| NRK | 1.217660624 | 2.325692928 | 2.26E-11 | 5.41E-10 | 97.76591082 | 227.1092591 | ENSG00000123572 |
| MORN3 | 1.216265459 | 2.323444944 | 0.025880423 | 0.079612906 | 10.46702916 | 24.4256996 | ENSG00000139714 |
| PCDHGA7 | 1.215591325 | 2.322359511 | 0.217497295 | 0.398487037 | 3.258758215 | 7.478981666 | ENSG00000253537 |
| NA | 1.215396049 | 2.32204519 | 0.234717306 | 0.419584351 | 3.777588906 | 8.841022179 | ENSG00000287830 |
| RBM15 | 1.215098548 | 2.321566405 | 2.45E-20 | 1.62E-18 | 362.0901071 | 840.6628218 | ENSG00000162775 |
| HNMT | 1.213578555 | 2.319121741 | 0.001558484 | 0.007680124 | 20.13401457 | 46.66397615 | ENSG00000150540 |
| NA | 1.21306148 | 2.318290695 | 0.019034293 | 0.06188065 | 14.35528277 | 33.5172431 | ENSG00000267284 |
| SPATA18 | 1.212678137 | 2.317674777 | 0.001925415 | 0.009211574 | 19.85280152 | 45.83815557 | ENSG00000163071 |
| GRIN2D | 1.21259954 | 2.317548515 | 2.77E-38 | 6.45E-36 | 541.5927921 | 1253.973236 | ENSG00000105464 |
| LTBR | 1.212009946 | 2.316601584 | 2.27E-05 | 0.000186753 | 48.59967914 | 112.1976504 | ENSG00000111321 |
| GADD45A | 1.211746405 | 2.316178442 | 1.74E-28 | 2.27E-26 | 388.7837452 | 899.9897213 | ENSG00000116717 |
| NA | 1.211569893 | 2.315895077 | 0.002341458 | 0.010848774 | 22.53650597 | 52.07294726 | ENSG00000213062 |
| VENTX | 1.21011003 | 2.313552809 | 8.40E-18 | 4.35E-16 | 184.8870585 | 427.4044028 | ENSG00000151650 |
| LOC100129503 | 1.209041227 | 2.311839477 | 0.236550658 | 0.421663883 | 2.808875945 | 6.548897262 | ENSG00000280351 |

| | | | | | | | |
|---|---|---|---|---|---|---|---|
| WDR72 | 1.204727751 | 2.304937692 | 0.000663053 | 0.003675357 | 28.91829779 | 66.51897235 | ENSG00000166415 |
| RENBP | 1.204532521 | 2.304625801 | 9.12E-21 | 6.22E-19 | 222.5785827 | 513.5151796 | ENSG00000102032 |
| SDR42E2 | 1.204294345 | 2.304245359 | 0.201276778 | 0.378151425 | 3.493218937 | 8.098209686 | ENSG00000183921 |
| PLAC8L1 | 1.204206633 | 2.304105272 | 0.173294267 | 0.339794156 | 4.403829691 | 10.24005503 | ENSG00000173261 |
| NA | 1.203858018 | 2.303548571 | 0.316052819 | 0.509633108 | 2.216759841 | 5.10472473 | ENSG00000230521 |
| KIRREL3 | 1.203808452 | 2.30346943 | 0.01363711 | 0.047322636 | 13.76179119 | 31.53608618 | ENSG00000149571 |
| NA | 1.201470042 | 2.299738848 | 0.311000491 | 0.504735823 | 2.272982913 | 5.194892475 | ENSG00000273002 |
| NA | 1.201219312 | 2.299339205 | 0.263046651 | 0.452632683 | 2.583885961 | 5.922621201 | ENSG00000286496 |
| NA | 1.199592675 | 2.296748164 | 0.091774228 | 0.212492343 | 6.676812603 | 15.30300089 | ENSG00000259448 |
| GDPGP1 | 1.199014501 | 2.295827905 | 0.165435636 | 0.328280318 | 5.10711416 | 11.71753267 | ENSG00000183208 |
| FSTL5 | 1.196570826 | 2.291942461 | 0.189576546 | 0.361743201 | 4.122616635 | 9.417921678 | ENSG00000168843 |
| RNF213 | 1.19628427 | 2.291487268 | 1.85E-07 | 2.36E-06 | 1002.897651 | 2298.042022 | ENSG00000173821 |
| SLC6A16 | 1.195651319 | 2.290482148 | 0.069156522 | 0.171486103 | 7.115946358 | 16.20346741 | ENSG00000063127 |
| WNK4 | 1.195068438 | 2.289556929 | 0.012537783 | 0.044363358 | 13.02122513 | 29.81111318 | ENSG00000126562 |
| DNAJC6 | 1.194769774 | 2.289083 | 1.08E-10 | 2.36E-09 | 91.02340441 | 208.2171028 | ENSG00000116675 |
| SLC25A45 | 1.194557881 | 2.288746819 | 0.173245063 | 0.339780318 | 4.085335041 | 9.425957404 | ENSG00000162241 |
| NA | 1.194258559 | 2.288272013 | 0.235180001 | 0.420127821 | 2.871412842 | 6.571467103 | ENSG00000244558 |
| NA | 1.193670442 | 2.287339384 | 0.219224341 | 0.400604922 | 3.549442009 | 8.120779527 | ENSG00000231249 |
| NEB | 1.193325471 | 2.286792512 | 0.001818515 | 0.008738492 | 26.54539869 | 60.77150235 | ENSG00000183091 |
| SNHG8 | 1.192762659 | 2.285900581 | 1.18E-33 | 2.02E-31 | 541.0086432 | 1238.059346 | ENSG00000269893 |
| FAXDC2 | 1.192594191 | 2.285633665 | 7.25E-05 | 0.000521666 | 36.82038247 | 84.23139904 | ENSG00000170271 |
| PCOLCE-AS1 | 1.192149555 | 2.284929346 | 0.019732383 | 0.063825477 | 12.65725592 | 29.09565706 | ENSG00000224729 |
| NLRP14 | 1.190914414 | 2.282973976 | 0.312290523 | 0.505854968 | 2.276139826 | 5.149864413 | ENSG00000158077 |
| SMIM29 | 1.190328022 | 2.282046236 | 6.00E-09 | 9.97E-08 | 78.50512771 | 178.7957288 | ENSG00000186577 |
| PHLDA2 | 1.190079311 | 2.28165286 | 2.05E-10 | 4.27E-09 | 94.74467255 | 215.7783281 | ENSG00000181649 |
| XCL1 | 1.189916953 | 2.281396101 | 0.304430452 | 0.498217572 | 2.640109032 | 5.97144811 | ENSG00000143184 |
| EFCAB10 | 1.189706773 | 2.28106376 | 0.033461395 | 0.097695887 | 9.94819847 | 22.59459903 | ENSG00000185055 |
| KCNS1 | 1.189652611 | 2.280978125 | 1.37E-06 | 1.48E-05 | 57.11940568 | 130.3396198 | ENSG00000124134 |
| MISP | 1.188375244 | 2.278959433 | 0.078117342 | 0.188382244 | 7.2284902 | 16.46918411 | ENSG00000099812 |
| AMIGO2 | 1.187724025 | 2.277930964 | 9.17E-13 | 2.74E-11 | 126.5608186 | 287.7529486 | ENSG00000139211 |
| SLC66A1L | 1.187684674 | 2.277868833 | 0.040079938 | 0.112637085 | 9.828160724 | 22.34297841 | ENSG00000174899 |
| NA | 1.187560221 | 2.277672343 | 0.150218982 | 0.30632024 | 4.413300429 | 10.12363022 | ENSG00000272720 |
| COL28A1 | 1.186016758 | 2.275236884 | 0.070823383 | 0.174530061 | 7.036347121 | 16.02368158 | ENSG00000215018 |
| NA | 1.185964033 | 2.275153734 | 0.278361341 | 0.469483037 | 2.904259747 | 6.586439251 | ENSG00000272328 |
| PSMD6-AS2 | 1.184865135 | 2.273421415 | 0.089897179 | 0.209342168 | 6.437917191 | 14.50767426 | ENSG00000239653 |
| MINAR1 | 1.184795871 | 2.273312269 | 0.011503044 | 0.041301319 | 15.53172798 | 35.09997225 | ENSG00000169330 |
| NA | 1.183297693 | 2.270952755 | 0.03006915 | 0.089785988 | 10.93081686 | 24.71398615 | ENSG00000225689 |
| NA | 1.183137858 | 2.270701172 | 1.90E-05 | 0.000159498 | 44.52854263 | 101.4556682 | ENSG00000260719 |
| CBR3 | 1.183058574 | 2.270576388 | 0.304311992 | 0.498111673 | 2.223073667 | 5.018355832 | ENSG00000159231 |
| RIPOR2 | 1.182662432 | 2.269953008 | 1.45E-44 | 4.98E-42 | 593.2124165 | 1346.730939 | ENSG00000111913 |
| NR5A1 | 1.181549777 | 2.26820302 | 0.125371268 | 0.268317677 | 5.031949612 | 11.50389207 | ENSG00000136931 |
| PRCD | 1.180097677 | 2.265921178 | 0.086410578 | 0.203085676 | 6.364631779 | 14.41425732 | ENSG00000214140 |
| ELOA-AS1 | 1.179924843 | 2.265649739 | 0.037843827 | 0.1075106 | 9.47178312 | 21.58618145 | ENSG00000236810 |
| NA | 1.179734164 | 2.265350311 | 0.023264153 | 0.073031473 | 11.74160912 | 26.56655726 | ENSG00000287858 |
| TRANK1 | 1.17973326 | 2.265348891 | 3.28E-15 | 1.29E-13 | 247.1664491 | 559.310337 | ENSG00000168016 |
| C2orf74 | 1.17887502 | 2.264001667 | 0.235470171 | 0.420524585 | 3.205692056 | 7.242771226 | ENSG00000237651 |
| RBPJL | 1.178041964 | 2.262694741 | 0.149053232 | 0.304682101 | 4.413300429 | 10.04094854 | ENSG00000124232 |
| PAH | 1.177928854 | 2.262517348 | 0.001481436 | 0.007329668 | 23.61017117 | 53.2869697 | ENSG00000171759 |
| NA | 1.177824725 | 2.262354053 | 0.112494256 | 0.247562163 | 5.422451897 | 12.216099 | ENSG00000267506 |
| NA | 1.177713197 | 2.262179168 | 0.213102541 | 0.392921059 | 4.200938097 | 9.493555308 | ENSG00000240291 |
| TAPBPL | 1.176910869 | 2.260921448 | 0.035656642 | 0.102761654 | 9.806062334 | 22.28385426 | ENSG00000139192 |
| SLIT2 | 1.176368204 | 2.26007117 | 9.12E-17 | 4.20E-15 | 189.819385 | 429.3929341 | ENSG00000145147 |
| RAD51-AS1 | 1.175992583 | 2.259482813 | 0.000285944 | 0.001763035 | 41.8806466 | 94.73969579 | ENSG00000245849 |
| NA | 1.175901043 | 2.259339452 | 0.184611057 | 0.355341294 | 4.164934279 | 9.320267859 | ENSG00000261474 |
| MICB-DT | 1.175655006 | 2.258954177 | 0.257826089 | 0.446392079 | 3.215162795 | 7.239083999 | ENSG00000286940 |
| TSKS | 1.175092253 | 2.258073198 | 0.326554074 | 0.520811738 | 2.580729048 | 5.791112621 | ENSG00000126467 |
| SUCLG2-AS1 | 1.174769555 | 2.257568175 | 0.00333001 | 0.014545937 | 23.02702214 | 52.10570271 | ENSG00000241316 |
| IPP | 1.174555369 | 2.257233035 | 1.25E-08 | 1.97E-07 | 80.57787228 | 181.7026216 | ENSG00000197429 |
| KIAA1217 | 1.174300597 | 2.256834455 | 2.65E-27 | 3.03E-25 | 350.7086187 | 791.6441149 | ENSG00000120549 |
| IL6R | 1.173945549 | 2.256279116 | 1.89E-10 | 3.93E-09 | 99.45012994 | 224.7210177 | ENSG00000160712 |
| IGSF21 | 1.173354034 | 2.255354216 | 7.61E-23 | 6.13E-21 | 388.1431406 | 875.4010145 | ENSG00000117154 |
| NA | 1.173061975 | 2.254897688 | 0.310875697 | 0.504706837 | 2.276139826 | 5.067182742 | ENSG00000228528 |
| IL20RB | 1.172546498 | 2.254092154 | 0.000971218 | 0.005137146 | 26.05507791 | 58.75225379 | ENSG00000174564 |
| HCP5 | 1.171728282 | 2.252814121 | 2.22E-05 | 0.000183256 | 41.42266906 | 93.54104325 | ENSG00000206337 |
| OPRK1 | 1.170955598 | 2.251607873 | 0.047374679 | 0.128274988 | 8.604767787 | 19.3955092 | ENSG00000082556 |
| FBXW12 | 1.170896326 | 2.25151537 | 0.097681715 | 0.222685475 | 6.461293358 | 14.4705703 | ENSG00000164049 |
| NA | 1.16972228 | 2.249683862 | 0.000123212 | 0.000838943 | 36.72499866 | 82.83280422 | ENSG00000248092 |
| CKM | 1.168738737 | 2.248150685 | 0.213581164 | 0.393529505 | 3.454058207 | 7.775192315 | ENSG00000104879 |
| NA | 1.168591944 | 2.247921949 | 0.172525847 | 0.338728182 | 4.079021215 | 9.200593852 | ENSG00000183022 |
| WDR31 | 1.168028263 | 2.247043826 | 0.08899502 | 0.207815761 | 6.731757899 | 15.0705893 | ENSG00000148225 |
| TMEM38A | 1.167703793 | 2.24653851 | 1.31E-16 | 5.92E-15 | 180.1301058 | 404.1311881 | ENSG00000072954 |
| INSM1 | 1.16656685 | 2.244768781 | 0.005367502 | 0.021892887 | 19.05285547 | 42.93246523 | ENSG00000173404 |
| NA | 1.16630207 | 2.244356832 | 0.074458949 | 0.181486345 | 7.74406628 | 17.31757453 | ENSG00000280099 |
| NPC1L1 | 1.165629683 | 2.243311064 | 0.345600618 | 0.539171873 | 2.16053677 | 4.924938894 | ENSG00000015520 |

| | | | | | | | |
|---|---|---|---|---|---|---|---|
| NLRP7 | 1.163799096 | 2.240466407 | 0.002485608 | 0.011422829 | 26.4530741 | 58.98086654 | ENSG00000167634 |
| ADGRE5 | 1.16276057 | 2.238854186 | 0.000322101 | 0.001951561 | 32.29138128 | 72.05967528 | ENSG00000123146 |
| IFITM10 | 1.162518055 | 2.23847787 | 0.276428259 | 0.467541533 | 3.314981286 | 7.344223892 | ENSG00000244242 |
| RAP2C-AS1 | 1.162498307 | 2.238447229 | 0.07012233 | 0.173170854 | 8.278080175 | 18.51381367 | ENSG00000232160 |
| LINC01291 | 1.16156978 | 2.237007015 | 0.209244865 | 0.388474839 | 3.90709739 | 8.683256531 | ENSG00000204792 |
| NA | 1.161050374 | 2.236201781 | 1.36E-12 | 3.91E-11 | 180.0993108 | 403.113398 | ENSG00000258649 |
| KDM6B | 1.161043758 | 2.236191526 | 2.11E-16 | 9.29E-15 | 178.9707229 | 400.0049366 | ENSG00000132510 |
| MYOF | 1.160983441 | 2.236098036 | 0.000142161 | 0.000947431 | 32.45699128 | 72.57420144 | ENSG00000138119 |
| SCN11A | 1.159634697 | 2.234008534 | 0.107609104 | 0.239254694 | 5.677131874 | 12.73018712 | ENSG00000168356 |
| STAC3 | 1.158888638 | 2.232853563 | 0.000703069 | 0.003867818 | 27.12173023 | 60.57966715 | ENSG00000185482 |
| F10 | 1.157005566 | 2.229941041 | 6.48E-05 | 0.000473854 | 35.62352261 | 79.44936526 | ENSG00000126218 |
| P2RY8 | 1.155550601 | 2.227693267 | 0.156877482 | 0.31576534 | 4.733674215 | 10.55459864 | ENSG00000182162 |
| IFI35 | 1.154862859 | 2.226631565 | 0.003814427 | 0.016353596 | 21.98552743 | 49.04235835 | ENSG00000068079 |
| TLE6 | 1.154231113 | 2.225656753 | 0.00811825 | 0.031010645 | 16.57698096 | 36.88549511 | ENSG00000104953 |
| NA | 1.153775562 | 2.224954082 | 0.000387968 | 0.002292921 | 40.6566523 | 90.06958643 | ENSG00000237978 |
| CCN1 | 1.152387785 | 2.222814852 | 2.06E-50 | 8.47E-48 | 1605.64571 | 3569.336539 | ENSG00000142871 |
| ISL1 | 1.152274955 | 2.222641017 | 0.341621955 | 0.535896852 | 2.587042873 | 5.769092433 | ENSG00000016082 |
| LOC101927825 | 1.15111921 | 2.220861169 | 0.157063607 | 0.31603701 | 5.025635786 | 11.27148048 | ENSG00000282556 |
| NA | 1.149968398 | 2.219090335 | 0.289497812 | 0.481388161 | 3.510281278 | 7.824568877 | ENSG00000270171 |
| NA | 1.149755654 | 2.218763126 | 0.001616017 | 0.007927809 | 24.27251347 | 54.06027615 | ENSG00000255495 |
| DPH6 | 1.14960458 | 2.218530797 | 5.23E-12 | 1.40E-10 | 123.1991826 | 273.0452918 | ENSG00000134146 |
| CCN2 | 1.14936526 | 2.218162808 | 1.53E-37 | 3.41E-35 | 842.8899094 | 1869.250869 | ENSG00000118523 |
| LOC101927021 | 1.14907468 | 2.217716083 | 0.020938342 | 0.067128737 | 12.96500206 | 28.8966622 | ENSG00000229043 |
| NA | 1.148487815 | 2.216814134 | 0.001704642 | 0.008287995 | 29.95911608 | 66.78820176 | ENSG00000268903 |
| LOC100506274 | 1.148321093 | 2.216557969 | 0.325189611 | 0.519305897 | 2.510600548 | 5.615125631 | ENSG00000229727 |
| LOC100507291 | 1.146706826 | 2.214079194 | 0.000761899 | 0.004156915 | 26.817141 | 59.34153752 | ENSG00000248932 |
| KLHL11 | 1.14647124 | 2.213717674 | 3.93E-06 | 3.91E-05 | 65.43073872 | 144.7075752 | ENSG00000178502 |
| NEBL | 1.146467796 | 2.213712389 | 1.95E-07 | 2.48E-06 | 75.10305328 | 166.2447779 | ENSG00000078114 |
| PRRT1B | 1.146395837 | 2.213601977 | 0.368162773 | 0.561249392 | 2.769715214 | 6.188775936 | ENSG00000283526 |
| SLC22A7 | 1.145475024 | 2.212189575 | 0.136864264 | 0.285850519 | 5.372542652 | 11.98043821 | ENSG00000137204 |
| METTL27 | 1.145330991 | 2.211968731 | 1.89E-07 | 2.40E-06 | 74.03688199 | 163.647794 | ENSG00000165171 |
| ENTPD1-AS1 | 1.143655513 | 2.209401345 | 0.069060713 | 0.171325291 | 8.309649304 | 18.35659768 | ENSG00000226688 |
| MYOM3 | 1.143521382 | 2.209195941 | 0.35125877 | 0.544467173 | 2.216759841 | 4.92873774 | ENSG00000142661 |
| TGFB1I1 | 1.141282438 | 2.20577011 | 4.74E-10 | 9.24E-09 | 97.06954154 | 214.1443989 | ENSG00000140682 |
| NA | 1.140768259 | 2.204984109 | 0.198824752 | 0.375065441 | 4.069550476 | 9.054551158 | ENSG00000228748 |
| OPCML | 1.140724005 | 2.204916474 | 0.000230928 | 0.001459762 | 36.11138553 | 79.74177707 | ENSG00000183715 |
| NA | 1.140252415 | 2.204195845 | 0.150672666 | 0.306909242 | 6.811357136 | 15.0264373 | ENSG00000272787 |
| NA | 1.139719877 | 2.203382367 | 0.293895903 | 0.486518994 | 2.533976715 | 5.622611705 | ENSG00000230131 |
| FAM71F1 | 1.137651737 | 2.200226025 | 0.001000645 | 0.005268392 | 26.94537171 | 59.08622967 | ENSG00000135248 |
| CCND2-AS1 | 1.136547562 | 2.198542714 | 0.204883482 | 0.383091035 | 3.791494334 | 8.390733108 | ENSG00000255920 |
| GCNA | 1.136515816 | 2.198494337 | 0.057528842 | 0.149082712 | 9.382713144 | 20.79085482 | ENSG00000147174 |
| FAHD2CP | 1.135183393 | 2.19646482 | 9.11E-05 | 0.000641014 | 41.10841371 | 89.96631029 | ENSG00000231584 |
| MFAP3L | 1.134715848 | 2.195753112 | 5.86E-08 | 8.16E-07 | 85.59326321 | 187.4552046 | ENSG00000198948 |
| NA | 1.133596565 | 2.194050246 | 0.011575289 | 0.041514766 | 20.18076691 | 44.36579088 | ENSG00000246792 |
| FOXP3 | 1.133402246 | 2.193754744 | 0.206092987 | 0.384640554 | 4.135244286 | 9.110864141 | ENSG00000049768 |
| LINC00634 | 1.133170323 | 2.193402113 | 0.119333144 | 0.258470352 | 6.077104898 | 13.37198499 | ENSG00000205704 |
| GJB2 | 1.131360438 | 2.190652179 | 0.368032445 | 0.561131404 | 2.229387493 | 4.898681826 | ENSG00000165474 |
| NA | 1.13131605 | 2.190584779 | 0.322582214 | 0.516655613 | 2.219916754 | 4.887396905 | ENSG00000227355 |
| NA | 1.131063316 | 2.190201062 | 0.29076386 | 0.483103307 | 3.132406644 | 6.901532514 | ENSG00000243024 |
| SNHG31 | 1.13064397 | 2.189564531 | 0.282158844 | 0.473132364 | 2.805719032 | 6.192574783 | ENSG00000229267 |
| NA | 1.130550084 | 2.189422046 | 0.033328694 | 0.097385153 | 11.61200294 | 25.34818631 | ENSG00000258072 |
| MNX1 | 1.130528401 | 2.18938914 | 0.362488144 | 0.555419581 | 2.272982913 | 4.973765803 | ENSG00000130675 |
| NA | 1.129948981 | 2.188510008 | 0.110317541 | 0.243872294 | 6.384851033 | 13.91514134 | ENSG00000227214 |
| GMPR | 1.129434747 | 2.187730074 | 0.018717096 | 0.061042171 | 15.73787418 | 34.68072678 | ENSG00000137198 |
| FXYD1 | 1.127952316 | 2.185483242 | 0.329646944 | 0.524075867 | 2.580729048 | 5.641382699 | ENSG00000266964 |
| NA | 1.126723444 | 2.183622464 | 0.016727063 | 0.055788986 | 14.34127965 | 31.19473585 | ENSG00000254365 |
| NA | 1.126652045 | 2.183514399 | 0.096635672 | 0.22092581 | 6.719130247 | 14.59404316 | ENSG00000270728 |
| MDM2 | 1.126304604 | 2.182988612 | 7.30E-18 | 3.82E-16 | 384.7844209 | 839.1398999 | ENSG00000135679 |
| CALCA | 1.125809474 | 2.182239542 | 0.162421917 | 0.323795613 | 5.812954183 | 12.68179825 | ENSG00000110680 |
| OR2A42 | 1.124984449 | 2.180991956 | 0.255166841 | 0.443978805 | 3.969634287 | 8.54046303 | ENSG00000212807 |
| KLHDC8A | 1.123884851 | 2.179330274 | 1.65E-05 | 0.000140093 | 49.40563075 | 108.0014279 | ENSG00000162873 |
| NA | 1.12327088 | 2.178403009 | 0.067494382 | 0.168318202 | 8.915670835 | 19.36870248 | ENSG00000254198 |
| NA | 1.12210681 | 2.176646026 | 8.05E-10 | 1.52E-08 | 98.57038926 | 214.6091105 | ENSG00000177822 |
| DNAH3 | 1.122054548 | 2.176567177 | 0.030448047 | 0.090654474 | 11.82308749 | 25.66272993 | ENSG00000158486 |
| EXD3 | 1.121723893 | 2.176068382 | 4.11E-09 | 6.97E-08 | 120.3473126 | 262.1194388 | ENSG00000187609 |
| ATP6AP1L | 1.12125389 | 2.175359575 | 0.032549248 | 0.095514211 | 12.29003211 | 26.60085006 | ENSG00000205464 |
| NBEAL1 | 1.120510098 | 2.174238341 | 2.30E-05 | 0.00018935 | 51.21778749 | 111.2303504 | ENSG00000144426 |
| EFCC1 | 1.120086739 | 2.173600405 | 0.031509423 | 0.093213621 | 12.91321368 | 28.15721052 | ENSG00000114654 |
| DYNC1I1 | 1.119298732 | 2.172413498 | 0.008196373 | 0.031244668 | 17.18615941 | 37.39198555 | ENSG00000158560 |
| C4orf51 | 1.118951753 | 2.171891078 | 0.246270404 | 0.433793543 | 3.755490516 | 8.191626624 | ENSG00000237136 |
| NA | 1.117652081 | 2.169935381 | 0.260911235 | 0.449960811 | 3.596194342 | 7.771943121 | ENSG00000286216 |
| KBTBD12 | 1.117580363 | 2.169827514 | 0.259585205 | 0.448466833 | 3.900783564 | 8.439560018 | ENSG00000187715 |
| COLQ | 1.115493904 | 2.166691726 | 0.15904386 | 0.318844663 | 6.266091064 | 13.6908771 | ENSG00000206561 |
| NA | 1.115155005 | 2.166182817 | 0.06775302 | 0.168713486 | 9.521692366 | 20.73129264 | ENSG00000223749 |

| | | | | | | | |
|---|---|---|---|---|---|---|---|
| LINC01748 | 1.114840452 | 2.165710571 | 0.186865039 | 0.358265918 | 4.135244286 | 8.960696186 | ENSG00000226476 |
| TMC1 | 1.113657436 | 2.163935408 | 0.3113994 | 0.505117843 | 3.391521309 | 7.449363784 | ENSG00000165091 |
| ZFHX4 | 1.112395911 | 2.162044041 | 0.000543872 | 0.003089217 | 31.13821454 | 67.44354605 | ENSG00000091656 |
| SYCP2 | 1.112310007 | 2.161915308 | 0.26259901 | 0.452072269 | 3.143155159 | 6.811364769 | ENSG00000196074 |
| PCDHGA5 | 1.11123907 | 2.160311077 | 0.129681552 | 0.274907685 | 6.908116413 | 14.95785171 | ENSG00000253485 |
| NA | 1.110753858 | 2.159584636 | 0.290763596 | 0.483103307 | 2.894789008 | 6.237602845 | ENSG00000279684 |
| NA | 1.110564741 | 2.159301563 | 0.191085439 | 0.363947952 | 4.188310445 | 9.017009169 | ENSG00000270441 |
| ISL2 | 1.10966549 | 2.157956062 | 0.183740807 | 0.354291732 | 5.607003374 | 12.21653703 | ENSG00000159556 |
| PPIL6 | 1.108909384 | 2.156825388 | 0.091474716 | 0.212162732 | 7.036347121 | 15.14947212 | ENSG00000185250 |
| CALD1 | 1.108551043 | 2.156289736 | 3.32E-78 | 4.10E-75 | 3601.274319 | 7765.555509 | ENSG00000122786 |
| TMEM9B-AS1 | 1.108528701 | 2.156256343 | 0.000809973 | 0.004388203 | 31.31191981 | 67.51483118 | ENSG00000254860 |
| FOXG1 | 1.108391171 | 2.156050801 | 0.151322057 | 0.307984955 | 5.11154885 | 11.00108886 | ENSG00000176165 |
| ANKRD37 | 1.108227282 | 2.155805888 | 0.025893705 | 0.079615298 | 13.41488433 | 28.9754334 | ENSG00000186352 |
| NA | 1.107032312 | 2.154020995 | 0.279060395 | 0.470063197 | 2.868255929 | 6.170004941 | ENSG00000236372 |
| FAM21EP | 1.106824817 | 2.153711217 | 0.310204277 | 0.504061553 | 2.563666707 | 5.506186892 | ENSG00000235618 |
| FOXL2NB | 1.105644209 | 2.15194948 | 0.002976404 | 0.013232111 | 29.77131 | 64.15907501 | ENSG00000206262 |
| NA | 1.105635715 | 2.151936811 | 0.263581166 | 0.453299928 | 3.19937823 | 6.867677753 | ENSG00000224728 |
| NA | 1.104884965 | 2.150817277 | 0.139864122 | 0.29087092 | 6.133327969 | 13.13544813 | ENSG00000261349 |
| ACER2 | 1.10409007 | 2.149632548 | 1.27E-06 | 1.39E-05 | 79.48340911 | 170.5989893 | ENSG00000177076 |
| CERS3 | 1.103939237 | 2.149407815 | 0.182855008 | 0.353293381 | 5.230308818 | 11.17013943 | ENSG00000154227 |
| PPFIBP2 | 1.103013912 | 2.148029657 | 1.24E-12 | 3.60E-11 | 162.0789079 | 348.0809867 | ENSG00000166387 |
| NA | 1.102591453 | 2.147400749 | 0.004381856 | 0.018386764 | 22.0866237 | 47.38041996 | ENSG00000228956 |
| KMT2C | 1.102544872 | 2.147331417 | 2.45E-14 | 8.69E-13 | 467.7672803 | 1004.19489 | ENSG00000055609 |
| NA | 1.101978059 | 2.146487928 | 2.78E-09 | 4.83E-08 | 96.81613934 | 207.6435641 | ENSG00000232098 |
| LRIG2-DT | 1.101195415 | 2.145323802 | 0.200356729 | 0.376957834 | 4.519432746 | 9.647084077 | ENSG00000238198 |
| LINC02035 | 1.101140263 | 2.14524179 | 0.015457562 | 0.052345146 | 16.60794873 | 35.62523368 | ENSG00000273033 |
| SFR1 | 1.100999942 | 2.145033148 | 5.00E-05 | 0.000376659 | 45.48137333 | 97.44496555 | ENSG00000156384 |
| GJA5 | 1.100488372 | 2.144272668 | 0.001348809 | 0.006765582 | 31.29485747 | 67.1816282 | ENSG00000265107 |
| GOLGA6L7 | 1.100238168 | 2.143900823 | 0.344799529 | 0.538284574 | 2.276139826 | 4.84605607 | ENSG00000261649 |
| NA | 1.099845163 | 2.143316882 | 0.194916995 | 0.369159164 | 4.164934279 | 8.889299436 | ENSG00000277173 |
| DNM3 | 1.099778234 | 2.143217451 | 0.000106053 | 0.00073425 | 41.4068845 | 89.02549062 | ENSG00000197959 |
| NA | 1.099700859 | 2.143102509 | 0.341799736 | 0.536084988 | 2.779185952 | 5.974697304 | ENSG00000281100 |
| NA | 1.098645914 | 2.141535976 | 0.003000238 | 0.013322092 | 28.6857162 | 61.58962206 | ENSG00000237886 |
| CDH8 | 1.098571271 | 2.14142518 | 9.47E-06 | 8.52E-05 | 64.53610793 | 138.1452029 | ENSG00000150394 |
| OXCT2 | 1.098411656 | 2.141188272 | 0.017768186 | 0.058492996 | 16.90938104 | 36.27094201 | ENSG00000198754 |
| ACSM3 | 1.098382958 | 2.14114568 | 0.013964077 | 0.048267475 | 18.35323158 | 39.20756433 | ENSG00000005187 |
| LOC653653 | 1.097656758 | 2.140068178 | 0.038658696 | 0.109523124 | 10.82468454 | 23.1988549 | ENSG00000280852 |
| BDNF | 1.097457203 | 2.139772182 | 1.47E-06 | 1.58E-05 | 64.243545 | 137.6626996 | ENSG00000176697 |
| NA | 1.097338106 | 2.139595546 | 0.010594253 | 0.038553679 | 18.39692469 | 39.46612137 | ENSG00000287315 |
| SERPINB1 | 1.096347104 | 2.138126341 | 1.21E-14 | 4.47E-13 | 184.7115718 | 394.8234076 | ENSG00000021355 |
| NHLRC1 | 1.095610641 | 2.137035155 | 0.047945072 | 0.129478939 | 10.14665537 | 21.72473807 | ENSG00000187566 |
| ONECUT2 | 1.095338164 | 2.136631578 | 0.002494058 | 0.011444633 | 27.05801325 | 58.10775639 | ENSG00000119547 |
| MDN1 | 1.09502249 | 2.136164116 | 4.66E-06 | 4.55E-05 | 4958.188881 | 10591.49696 | ENSG00000112159 |
| SLC44A3-AS1 | 1.095011731 | 2.136148185 | 0.002190846 | 0.010282385 | 27.99684082 | 59.59272011 | ENSG00000224081 |
| CBX7 | 1.095001495 | 2.13613303 | 1.30E-07 | 1.69E-06 | 83.09468896 | 177.2589038 | ENSG00000100307 |
| MYL9 | 1.09472311 | 2.135720877 | 6.26E-50 | 2.52E-47 | 1375.349256 | 2938.323605 | ENSG00000101335 |
| CYSRT1 | 1.094509455 | 2.135404612 | 0.054709733 | 0.143299914 | 10.20287845 | 21.79613482 | ENSG00000197191 |
| CALB1 | 1.093218788 | 2.133495086 | 8.99E-19 | 5.16E-17 | 339.2885634 | 723.2700146 | ENSG00000104327 |
| SLC5A11 | 1.093209853 | 2.133481874 | 0.309253879 | 0.503332882 | 2.587042873 | 5.476130977 | ENSG00000158865 |
| LOC100507336 | 1.093141587 | 2.133380923 | 0.269742884 | 0.460054717 | 3.208848969 | 6.807565923 | ENSG00000228793 |
| POU3F2 | 1.092171288 | 2.131946578 | 0.317226196 | 0.510858486 | 2.930792826 | 6.200060856 | ENSG00000184486 |
| PPP5D1 | 1.090896142 | 2.130063061 | 0.04605518 | 0.125472102 | 10.85121762 | 22.99931038 | ENSG00000230510 |
| MIR100HG | 1.090819127 | 2.129949355 | 0.169637115 | 0.334402711 | 5.839487262 | 12.3512948 | ENSG00000255248 |
| TULP4 | 1.089458162 | 2.127941017 | 1.72E-13 | 5.54E-12 | 228.9238822 | 486.3275935 | ENSG00000130338 |
| NA | 1.089339217 | 2.127765583 | 0.224699946 | 0.407454143 | 3.824341239 | 8.093972806 | ENSG00000259349 |
| NA | 1.088584385 | 2.126652607 | 0.286999453 | 0.478306614 | 3.404148961 | 7.332938971 | ENSG00000268047 |
| CSTF3-DT | 1.088373488 | 2.126341749 | 0.279182178 | 0.47018287 | 3.466685858 | 7.351271932 | ENSG00000247151 |
| NA | 1.08785045 | 2.125571001 | 0.253686105 | 0.442440789 | 3.549442009 | 7.48646774 | ENSG00000226823 |
| LOC112267895 | 1.087541144 | 2.125115338 | 0.083756937 | 0.198434241 | 9.206354629 | 19.65852636 | ENSG00000261241 |
| TFPI2 | 1.086790066 | 2.124009275 | 1.25E-08 | 1.97E-07 | 93.12949955 | 197.7047211 | ENSG00000105825 |
| BTBD19 | 1.085908498 | 2.122711783 | 0.17330712 | 0.339794156 | 5.11154885 | 10.88510208 | ENSG00000222009 |
| PURA | 1.085421086 | 2.121994748 | 3.71E-07 | 4.48E-06 | 87.97522709 | 186.7464701 | ENSG00000185129 |
| RNF180 | 1.085220263 | 2.121699388 | 0.004796624 | 0.019843888 | 23.07555592 | 48.92593353 | ENSG00000164197 |
| PCDHAC1 | 1.085077718 | 2.121489765 | 5.44E-06 | 5.21E-05 | 63.00634433 | 133.9296513 | ENSG00000248383 |
| DLG2 | 1.084507631 | 2.120651615 | 3.29E-09 | 5.63E-08 | 114.0331688 | 241.7782118 | ENSG00000150672 |
| NA | 1.083629326 | 2.119360966 | 0.097511758 | 0.22232537 | 8.637614692 | 18.37200786 | ENSG00000228192 |
| NA | 1.082962779 | 2.118382015 | 0.217829427 | 0.398784596 | 4.403829691 | 9.388415416 | ENSG00000253553 |
| TXNRD2 | 1.081694291 | 2.116520249 | 9.63E-11 | 2.11E-09 | 125.5410163 | 265.7825789 | ENSG00000184470 |
| PDCD1 | 1.081586882 | 2.116362679 | 0.005116541 | 0.020989371 | 20.63074688 | 43.71183203 | ENSG00000188389 |
| SYTL1 | 1.081287384 | 2.115923376 | 1.26E-08 | 1.98E-07 | 112.8832567 | 239.2396908 | ENSG00000142765 |
| NPAS4 | 1.081283903 | 2.11591827 | 0.146954056 | 0.301321659 | 6.655315573 | 14.10296291 | ENSG00000174576 |
| TSPAN32 | 1.080818515 | 2.115235823 | 0.180771094 | 0.350351506 | 5.180399573 | 10.85460814 | ENSG00000064201 |
| LOC101929054 | 1.079739844 | 2.113654899 | 0.040445912 | 0.11345914 | 12.04176365 | 25.50540231 | ENSG00000243224 |
| GPR20 | 1.079453516 | 2.113235449 | 0.011156336 | 0.040337793 | 18.99969161 | 39.99865408 | ENSG00000204882 |

| | | | | | | | |
|---|---|---|---|---|---|---|---|
| FAAH2 | 1.079221859 | 2.112896149 | 4.75E-06 | 4.63E-05 | 67.78411763 | 143.0012213 | ENSG00000165591 |
| IRX4 | 1.078370575 | 2.11164977 | 0.000467838 | 0.002701232 | 44.70164653 | 94.32768132 | ENSG00000113430 |
| C4orf47 | 1.078366689 | 2.111644083 | 0.036426674 | 0.104524967 | 12.80698366 | 27.02477044 | ENSG00000205129 |
| PNMA2 | 1.078217237 | 2.111425344 | 7.90E-10 | 1.49E-08 | 113.7913119 | 240.5244487 | ENSG00000240694 |
| RNF227 | 1.077626681 | 2.110561225 | 0.012149689 | 0.043225528 | 17.89703548 | 37.77836394 | ENSG00000179859 |
| EBF4 | 1.076098682 | 2.108327054 | 0.00120515 | 0.006165909 | 36.10299717 | 75.83395275 | ENSG00000088881 |
| NA | 1.076025888 | 2.108220677 | 0.000199064 | 0.001277956 | 42.32065216 | 89.36965211 | ENSG00000228065 |
| SULT1C2 | 1.075816587 | 2.107914845 | 0.03214426 | 0.094595381 | 13.39466507 | 28.20983628 | ENSG00000198203 |
| MED12L | 1.075301662 | 2.107162625 | 2.81E-07 | 3.48E-06 | 157.2020077 | 330.5940267 | ENSG00000144893 |
| MYH3 | 1.07353077 | 2.104577694 | 0.002600215 | 0.011837897 | 27.19699247 | 57.43276504 | ENSG00000109063 |
| NA | 1.072945716 | 2.103724401 | 0.113394222 | 0.249069508 | 7.287870184 | 15.31808466 | ENSG00000175772 |
| FLNB-AS1 | 1.071907618 | 2.1022112 | 0.306441075 | 0.500491619 | 3.824341239 | 8.064354924 | ENSG00000244161 |
| TEKT3 | 1.071485064 | 2.101595569 | 0.152156419 | 0.309217881 | 5.79273493 | 12.14893913 | ENSG00000125409 |
| TNS4 | 1.071254338 | 2.101259494 | 0.051876982 | 0.137375051 | 11.98228597 | 25.0933165 | ENSG00000131746 |
| NA | 1.070748287 | 2.100522569 | 0.00148016 | 0.007325311 | 34.68322186 | 73.07831874 | ENSG00000281490 |
| CPNE4 | 1.069133529 | 2.098172843 | 0.001719827 | 0.00835087 | 32.74569824 | 68.35343374 | ENSG00000196353 |
| TMEM249 | 1.068422556 | 2.0971391 | 0.320906256 | 0.514727525 | 2.901102834 | 6.01603814 | ENSG00000261587 |
| NA | 1.067114965 | 2.095239213 | 0.260942542 | 0.449972932 | 3.463528945 | 7.306681903 | ENSG00000260495 |
| HIVEP2 | 1.066413516 | 2.094220739 | 1.06E-10 | 2.32E-09 | 185.5194177 | 387.9983848 | ENSG00000010818 |
| NA | 1.06610836 | 2.09377782 | 0.243113169 | 0.42965066 | 4.681885833 | 9.917475691 | ENSG00000271978 |
| FAT3 | 1.064827903 | 2.091920322 | 2.96E-07 | 3.64E-06 | 862.6612855 | 1804.443138 | ENSG00000165323 |
| SPOCD1 | 1.062555297 | 2.088627619 | 0.018259962 | 0.059767784 | 16.31283025 | 34.09155468 | ENSG00000134668 |
| SOX6 | 1.062495647 | 2.088541263 | 0.195605013 | 0.370197417 | 4.863182699 | 10.11983137 | ENSG00000110693 |
| SLC44A5 | 1.060515131 | 2.085676104 | 0.002576884 | 0.011749072 | 31.45858833 | 65.78270697 | ENSG00000137968 |
| CPEB4 | 1.060477424 | 2.085621592 | 6.06E-12 | 1.60E-10 | 145.2783124 | 303.0299072 | ENSG00000113742 |
| KLF7 | 1.060368999 | 2.085464854 | 9.53E-17 | 4.36E-15 | 262.2476032 | 547.1457729 | ENSG00000118263 |
| HES2 | 1.060132731 | 2.085123348 | 0.038656611 | 0.109523124 | 13.23230969 | 27.7374154 | ENSG00000069812 |
| NA | 1.059307914 | 2.083931583 | 0.040597657 | 0.113781488 | 11.78392676 | 24.4810249 | ENSG00000287562 |
| SOX15 | 1.059105429 | 2.083639121 | 0.001313828 | 0.006618778 | 29.89016766 | 62.4139053 | ENSG00000129194 |
| NA | 1.059088208 | 2.083614249 | 0.031513788 | 0.093213621 | 12.76023133 | 26.58576629 | ENSG00000284620 |
| UTS2R | 1.058327878 | 2.082516431 | 0.097070166 | 0.221723497 | 7.870515549 | 16.43967785 | ENSG00000181408 |
| LINC02018 | 1.058057669 | 2.082126423 | 0.03647806 | 0.104579156 | 12.41648137 | 25.86183641 | ENSG00000272690 |
| CD79B | 1.05783167 | 2.081800282 | 0.071345199 | 0.175605833 | 9.990516113 | 20.71956969 | ENSG00000007312 |
| UBE2D4 | 1.057739034 | 2.081666613 | 0.002699012 | 0.012185988 | 26.06898334 | 54.26274344 | ENSG00000078967 |
| SCTR | 1.05567441 | 2.078689696 | 0.087161143 | 0.20446637 | 9.236044621 | 19.17763172 | ENSG00000080293 |
| NMI | 1.054747115 | 2.077354042 | 0.000800271 | 0.004343256 | 35.30640343 | 73.57936982 | ENSG00000123609 |
| LOC101926913 | 1.054573597 | 2.077104207 | 0.257718396 | 0.446378734 | 3.483748199 | 7.257854994 | ENSG00000234350 |
| LINC01275 | 1.0541548 | 2.076501336 | 0.303407922 | 0.497026907 | 3.235382048 | 6.717947831 | ENSG00000237595 |
| DHRS12 | 1.053976434 | 2.076244626 | 0.001856364 | 0.008901903 | 27.64687474 | 57.42472931 | ENSG00000102796 |
| WNT9A | 1.053353924 | 2.075348938 | 6.47E-05 | 0.000473842 | 59.77717841 | 124.3599615 | ENSG00000143816 |
| NA | 1.052967916 | 2.074793732 | 0.00109497 | 0.005703817 | 31.61778681 | 65.55668214 | ENSG00000231806 |
| KLHL31 | 1.05208998 | 2.073531523 | 0.169454527 | 0.334113851 | 5.845801088 | 12.02882709 | ENSG00000124743 |
| SP100 | 1.050842129 | 2.071738809 | 0.355812654 | 0.549369521 | 2.888475183 | 5.997705179 | ENSG00000067066 |
| RUNX2 | 1.050519414 | 2.071275436 | 0.034333373 | 0.099707731 | 14.71166038 | 30.6515795 | ENSG00000124813 |
| NA | 1.050082343 | 2.070648028 | 0.278853105 | 0.469927573 | 3.516595104 | 7.280424835 | ENSG00000279368 |
| IFI27 | 1.049998247 | 2.070527332 | 0.163178786 | 0.324850227 | 5.730198032 | 11.91284031 | ENSG00000165949 |
| CARD14 | 1.049526632 | 2.06985059 | 0.252907402 | 0.441581327 | 4.466366588 | 9.309970625 | ENSG00000141527 |
| CCDC7 | 1.049151634 | 2.069312646 | 0.145066422 | 0.298543277 | 6.973810223 | 14.54240509 | ENSG00000216937 |
| HSPG2 | 1.049144803 | 2.069302848 | 3.63E-24 | 3.22E-22 | 2695.702641 | 5577.676893 | ENSG00000142798 |
| HMGCLL1 | 1.049042013 | 2.069155418 | 0.386499736 | 0.579384926 | 2.540290541 | 5.28506022 | ENSG00000146151 |
| NEFH | 1.048870872 | 2.068909978 | 8.75E-09 | 1.42E-07 | 113.8834411 | 235.5505713 | ENSG00000100285 |
| AKR1C2 | 1.048725517 | 2.068701541 | 0.193283215 | 0.366964874 | 5.289786501 | 11.03863085 | ENSG00000151632 |
| RAG1 | 1.048658499 | 2.068605445 | 0.029424396 | 0.088159029 | 14.75072341 | 30.47548089 | ENSG00000166349 |
| CEBPZOS | 1.04846863 | 2.068333219 | 0.051738722 | 0.13716571 | 10.8076222 | 22.36598628 | ENSG00000218739 |
| QPCT | 1.047678086 | 2.067200159 | 2.71E-05 | 0.00021915 | 67.00872783 | 138.1432191 | ENSG00000115828 |
| PLK3 | 1.046802426 | 2.065945829 | 1.19E-09 | 2.17E-08 | 116.0098532 | 239.3748463 | ENSG00000173846 |
| MYH2 | 1.046630442 | 2.065699562 | 0.001074894 | 0.005611857 | 38.4120816 | 79.14489565 | ENSG00000125414 |
| NA | 1.046532093 | 2.065558747 | 0.021408575 | 0.068423267 | 17.51916085 | 36.19150953 | ENSG00000261799 |
| NA | 1.046325984 | 2.065263674 | 0.266693884 | 0.456661534 | 3.546285096 | 7.306681903 | ENSG00000254701 |
| MSRB3 | 1.045862787 | 2.064600701 | 0.015851978 | 0.053447436 | 18.53915853 | 38.42677181 | ENSG00000174099 |
| LINC01484 | 1.045644795 | 2.064288762 | 1.16E-06 | 1.28E-05 | 79.48912157 | 164.316287 | ENSG00000253686 |
| FCGBP | 1.045637791 | 2.06427874 | 0.052946167 | 0.139727218 | 11.58546986 | 23.84390195 | ENSG00000275395 |
| FBXO48 | 1.045077981 | 2.063477892 | 0.022964613 | 0.072348484 | 17.51855949 | 36.13899539 | ENSG00000204923 |
| EXOC3L2 | 1.044753044 | 2.063013189 | 9.44E-05 | 0.000661886 | 49.05734841 | 101.0907604 | ENSG00000283632 |
| NA | 1.044650181 | 2.062866102 | 0.000517488 | 0.002956571 | 35.57045645 | 73.47846681 | ENSG00000267278 |
| CORO2A | 1.044528777 | 2.062692517 | 1.70E-42 | 5.43E-40 | 1330.248073 | 2743.089042 | ENSG00000106789 |
| DICER1-AS1 | 1.043475937 | 2.061187769 | 0.001987535 | 0.009442955 | 30.37615145 | 62.87168044 | ENSG00000235706 |
| NA | 1.042785385 | 2.06020141 | 0.110541721 | 0.24433873 | 7.390845588 | 15.17952804 | ENSG00000260123 |
| AOC3 | 1.042687368 | 2.060061444 | 0.002705281 | 0.012211318 | 33.23789816 | 68.26524137 | ENSG00000131471 |
| SUCLG2 | 1.042288803 | 2.059492401 | 8.53E-27 | 9.25E-25 | 643.4085378 | 1323.761054 | ENSG00000172340 |
| MGMT | 1.042040659 | 2.059138199 | 1.13E-16 | 5.15E-15 | 261.343885 | 537.815614 | ENSG00000170430 |
| NA | 1.039592458 | 2.055646879 | 0.001280662 | 0.00647957 | 33.86739355 | 69.62662061 | ENSG00000249898 |
| NA | 1.039559932 | 2.055600534 | 0.252184978 | 0.440938768 | 3.847717405 | 7.876644981 | ENSG00000287077 |
| TIE1 | 1.039550642 | 2.055587297 | 2.43E-06 | 2.51E-05 | 79.10741361 | 162.6151577 | ENSG00000066056 |

| | | | | | | | |
|---|---|---|---|---|---|---|---|
| TGFB2 | 1.039449975 | 2.05544387 | 0.193937376 | 0.367867704 | 5.594375723 | 11.56725309 | ENSG00000092969 |
| TMEM155 | 1.039448871 | 2.055442297 | 0.35157967 | 0.544557714 | 2.530819802 | 5.232434464 | ENSG00000164112 |
| LBX2 | 1.038854361 | 2.054595459 | 0.332658805 | 0.52722041 | 2.924479001 | 5.997705179 | ENSG00000179528 |
| NA | 1.038790898 | 2.05450508 | 0.256909676 | 0.445408246 | 4.079021215 | 8.469615933 | ENSG00000246982 |
| SLC2A14 | 1.038652139 | 2.054307487 | 1.69E-08 | 2.62E-07 | 103.6868765 | 212.7860456 | ENSG00000173262 |
| CPEB2 | 1.038582061 | 2.054207703 | 0.010053634 | 0.036941722 | 24.08923977 | 49.46496463 | ENSG00000137449 |
| RND3 | 1.037986403 | 2.053359739 | 4.46E-31 | 6.83E-29 | 648.7214899 | 1331.41028 | ENSG00000115963 |
| SPEF2 | 1.037731207 | 2.052996556 | 0.111876469 | 0.246466002 | 7.976647866 | 16.41385882 | ENSG00000152582 |
| LOC285095 | 1.037199936 | 2.052240681 | 0.227176339 | 0.410699011 | 4.419614255 | 9.144718903 | ENSG00000216921 |
| BTK | 1.037086014 | 2.052078633 | 0.184390232 | 0.355026861 | 5.333381921 | 11.00108886 | ENSG00000010671 |
| CFAP300 | 1.037044909 | 2.052020166 | 0.166122087 | 0.329308055 | 6.431603366 | 13.1728785 | ENSG00000137691 |
| MIR635 | 1.035484005 | 2.049801212 | 0.18465436 | 0.355387738 | 5.064796517 | 10.43112579 | ENSG00000207561 |
| NA | 1.035478675 | 2.049793639 | 0.279238893 | 0.470192936 | 3.490062025 | 7.141430181 | ENSG00000287917 |
| APOA1 | 1.035459929 | 2.049767004 | 0.327304264 | 0.521559388 | 2.924479001 | 5.978934184 | ENSG00000118137 |
| NA | 1.03489286 | 2.048961477 | 0.106370851 | 0.23747017 | 9.012332413 | 18.34856195 | ENSG00000214992 |
| SLC5A3 | 1.034531225 | 2.048447935 | 2.78E-19 | 1.67E-17 | 508.1398557 | 1040.959462 | ENSG00000198743 |
| HIVEP1 | 1.034022407 | 2.047725605 | 9.47E-18 | 4.88E-16 | 316.9196443 | 649.6724956 | ENSG00000095951 |
| COL25A1 | 1.033982759 | 2.04766933 | 5.30E-09 | 8.89E-08 | 111.9726459 | 229.459784 | ENSG00000188517 |
| PTCH2 | 1.033927592 | 2.047591031 | 0.141571823 | 0.293303395 | 6.961182572 | 14.3321253 | ENSG00000117425 |
| SPI1 | 1.033661129 | 2.047212879 | 0.211056876 | 0.390555345 | 4.757050382 | 9.789765958 | ENSG00000066336 |
| DUOX1 | 1.033432975 | 2.04688915 | 0.012998251 | 0.045575027 | 20.20424077 | 41.52908389 | ENSG00000137857 |
| LGSN | 1.033022971 | 2.046307521 | 0.379892179 | 0.572945989 | 2.477753643 | 5.108523577 | ENSG00000146166 |
| MKLN1-AS | 1.032525232 | 2.045601654 | 0.007320168 | 0.028408489 | 23.03205819 | 47.20333366 | ENSG00000236753 |
| NA | 1.031550935 | 2.044220661 | 0.216898238 | 0.397456783 | 5.289786501 | 10.86589306 | ENSG00000248923 |
| NA | 1.031463176 | 2.044096316 | 0.336246817 | 0.530473144 | 2.84172285 | 5.817481309 | ENSG00000280077 |
| PLA2G6 | 1.031021436 | 2.043470528 | 6.61E-09 | 1.09E-07 | 121.9428451 | 249.1593164 | ENSG00000184381 |
| SUGCT | 1.030811885 | 2.043173735 | 0.000130941 | 0.000883137 | 50.6845477 | 103.484338 | ENSG00000175600 |
| ROBO2 | 1.03039337 | 2.042581112 | 5.33E-07 | 6.28E-06 | 89.34549912 | 182.2370181 | ENSG00000185008 |
| FZD5 | 1.030063762 | 2.042114503 | 6.64E-47 | 2.51E-44 | 1572.01474 | 3209.252065 | ENSG00000163251 |
| NEK5 | 1.029447319 | 2.041242124 | 0.133200744 | 0.280214117 | 6.662907176 | 13.57010378 | ENSG00000197168 |
| NA | 1.0291566 | 2.040830833 | 0.110907723 | 0.244826553 | 8.492321644 | 17.46025641 | ENSG00000230207 |
| NTS | 1.028163559 | 2.039426564 | 9.96E-13 | 2.95E-11 | 177.6119137 | 361.9065184 | ENSG00000133636 |
| IFITM1 | 1.027946216 | 2.039119346 | 2.03E-18 | 1.12E-16 | 792.1476353 | 1614.767591 | ENSG00000185885 |
| HLA-DQB1 | 1.027044385 | 2.037845087 | 7.94E-14 | 2.67E-12 | 212.6090978 | 432.8606635 | ENSG00000179344 |
| THORLNC | 1.026506283 | 2.037085146 | 0.206654621 | 0.385248114 | 4.757050382 | 9.725966901 | ENSG00000226856 |
| PPP1R15A | 1.026004548 | 2.036376819 | 1.02E-21 | 7.54E-20 | 354.8841061 | 722.8510645 | ENSG00000087074 |
| IRX2 | 1.025983147 | 2.036346611 | 1.13E-13 | 3.70E-12 | 271.8485866 | 553.9514835 | ENSG00000170561 |
| TMEM64 | 1.024483399 | 2.034230835 | 3.84E-21 | 2.69E-19 | 490.3653459 | 997.1309413 | ENSG00000180694 |
| HHLA3 | 1.024456509 | 2.03419292 | 9.62E-07 | 1.07E-05 | 73.87324882 | 150.2079879 | ENSG00000197568 |
| FZD10 | 1.024386882 | 2.034094749 | 0.084093861 | 0.198927328 | 10.39808074 | 21.00449542 | ENSG00000111432 |
| NA | 1.023971281 | 2.033508866 | 0.358368446 | 0.551615379 | 2.50112981 | 5.108523577 | ENSG00000261693 |
| REPS2 | 1.02345686 | 2.032783909 | 3.84E-07 | 4.62E-06 | 99.15471837 | 201.0069754 | ENSG00000169891 |
| OBSCN-AS1 | 1.023193583 | 2.03241298 | 0.32872853 | 0.523226012 | 2.835409024 | 5.764855553 | ENSG00000162913 |
| ADGRB3 | 1.022910308 | 2.032013953 | 0.006325394 | 0.025168497 | 24.95487963 | 50.53013322 | ENSG00000135298 |
| ADAMTSL5 | 1.022534508 | 2.031484712 | 0.303283857 | 0.496875102 | 3.232225135 | 6.522640194 | ENSG00000185761 |
| C1orf167 | 1.022473522 | 2.031398838 | 0.050486689 | 0.134849444 | 12.12136289 | 24.65799958 | ENSG00000215910 |
| DIAPH2 | 1.022409645 | 2.031308898 | 3.42E-35 | 6.67E-33 | 918.7910674 | 1865.27447 | ENSG00000147202 |
| NA | 1.022065583 | 2.030824518 | 0.250937444 | 0.439408185 | 4.460052762 | 9.084607073 | ENSG00000259687 |
| SORCS2 | 1.019856287 | 2.027716959 | 5.77E-05 | 0.000427957 | 61.97747727 | 125.8401386 | ENSG00000184985 |
| NA | 1.019712282 | 2.027514571 | 0.05482345 | 0.143536916 | 14.87529354 | 30.26575075 | ENSG00000283341 |
| CLMN | 1.019641723 | 2.027415412 | 1.18E-07 | 1.56E-06 | 109.3936006 | 221.5753181 | ENSG00000165959 |
| NA | 1.019586611 | 2.027337964 | 0.366150883 | 0.559228716 | 2.613575953 | 5.273775299 | ENSG00000267632 |
| ZKSCAN7-AS1 | 1.019545753 | 2.027280549 | 0.319083482 | 0.513002538 | 3.18231589 | 6.424986375 | ENSG00000236869 |
| MYH7 | 1.018938245 | 2.026427057 | 0.354981816 | 0.548282644 | 2.927635913 | 5.896364133 | ENSG00000092054 |
| ZFP2 | 1.018324807 | 2.025565598 | 0.065110525 | 0.163850438 | 10.75899073 | 21.78430025 | ENSG00000198939 |
| EDA2R | 1.018217111 | 2.025414396 | 1.58E-07 | 2.04E-06 | 115.9492931 | 235.3204699 | ENSG00000131080 |
| LINC01876 | 1.018142352 | 2.025309444 | 0.007227911 | 0.028101972 | 24.06005344 | 48.61420108 | ENSG00000226383 |
| SLC9A7P1 | 1.016834954 | 2.023474901 | 0.076220481 | 0.184759403 | 10.60600838 | 21.39847151 | ENSG00000227825 |
| NA | 1.016717412 | 2.023310047 | 0.342729757 | 0.536797013 | 2.95101208 | 5.922621201 | ENSG00000235092 |
| PHOSPHO1 | 1.016632786 | 2.023191367 | 0.052706305 | 0.139232991 | 12.28697289 | 24.90494528 | ENSG00000173868 |
| GAN | 1.016477259 | 2.022973272 | 3.64E-08 | 5.29E-07 | 118.0408815 | 238.9942337 | ENSG00000261609 |
| SLFN13 | 1.016413664 | 2.022884101 | 4.62E-11 | 1.06E-09 | 180.0504089 | 364.2350146 | ENSG00000154760 |
| SORBS2 | 1.016213875 | 2.022603984 | 9.41E-19 | 5.38E-17 | 384.3571255 | 778.2032582 | ENSG00000154556 |
| GRID2IP | 1.015343035 | 2.021383467 | 0.00278709 | 0.01251052 | 29.63736682 | 59.88524353 | ENSG00000215045 |
| NA | 1.015195046 | 2.021176129 | 0.356417948 | 0.550074964 | 2.918165175 | 5.862509371 | ENSG00000224376 |
| NA | 1.015023854 | 2.020936308 | 0.078830072 | 0.189548073 | 9.501473112 | 19.22657025 | ENSG00000227012 |
| FAM86EP | 1.014417611 | 2.020087257 | 0.000877747 | 0.004712676 | 43.09071282 | 86.78219261 | ENSG00000251669 |
| NA | 1.014139088 | 2.019697301 | 0.31236578 | 0.505932654 | 3.232225135 | 6.473813284 | ENSG00000259376 |
| NA | 1.013232277 | 2.018428213 | 0.134074867 | 0.28151621 | 7.231647112 | 14.62409907 | ENSG00000253540 |
| TENT5C | 1.012810951 | 2.017838836 | 0.007901837 | 0.030312908 | 22.77728052 | 45.84239245 | ENSG00000183508 |
| GLRA4 | 1.012456318 | 2.017342886 | 0.095574581 | 0.21898619 | 10.02661763 | 20.3443646 | ENSG00000188828 |
| NA | 1.012363684 | 2.017213359 | 0.173646912 | 0.340352355 | 5.994348747 | 12.15653682 | ENSG00000262619 |
| LINC01237 | 1.011724974 | 2.016320495 | 0.19777834 | 0.373471984 | 5.633536454 | 11.44757909 | ENSG00000233806 |
| CASC9 | 1.010964791 | 2.015258338 | 0.001815834 | 0.008730137 | 34.04869042 | 68.47128427 | ENSG00000249395 |

| | | | | | | | |
|---|---|---|---|---|---|---|---|
| CNIH3-AS2 | 1.010733306 | 2.014935009 | 0.064450034 | 0.162702466 | 10.70908149 | 21.66082739 | ENSG00000233384 |
| CD27-AS1 | 1.010211478 | 2.014206332 | 0.014961737 | 0.050965049 | 23.26769899 | 47.01748747 | ENSG00000215039 |
| SHC2 | 1.008989184 | 2.012500559 | 3.93E-08 | 5.67E-07 | 107.6174477 | 216.7638897 | ENSG00000129946 |
| NA | 1.008649375 | 2.012026595 | 0.133066249 | 0.280160156 | 7.29418401 | 14.66543991 | ENSG00000257027 |
| HSD17B14 | 1.008434191 | 2.011726515 | 1.21E-10 | 2.61E-09 | 178.5780484 | 358.7966001 | ENSG00000087076 |
| NA | 1.008316771 | 2.011562789 | 0.303651872 | 0.497294672 | 3.513438191 | 7.111812299 | ENSG00000279989 |
| LINC00659 | 1.007930517 | 2.011024304 | 0.19957812 | 0.375954718 | 5.399075731 | 10.92644292 | ENSG00000228705 |
| ZDHHC21 | 1.007789359 | 2.010827548 | 6.07E-06 | 5.76E-05 | 98.17309213 | 197.7035186 | ENSG00000175893 |
| NPIPB6 | 1.007769302 | 2.010799592 | 0.032676802 | 0.095812664 | 15.33780346 | 30.74499643 | ENSG00000198156 |
| CCDC200 | 1.007669509 | 2.010660508 | 0.319916754 | 0.513584853 | 4.591440382 | 9.197344659 | ENSG00000236383 |
| FAM161B | 1.007500158 | 2.0104245 | 0.152105029 | 0.309147341 | 7.006657128 | 14.07301861 | ENSG00000156050 |
| CST1 | 1.007166016 | 2.00995892 | 0.003697333 | 0.015910463 | 29.33849007 | 59.04956375 | ENSG00000170373 |
| BHLHB9 | 1.006829118 | 2.00948961 | 0.398955383 | 0.590735725 | 2.646422858 | 5.273775299 | ENSG00000198908 |
| C4orf50 | 1.006746397 | 2.009374394 | 0.088235541 | 0.206354261 | 10.48281372 | 21.10214924 | ENSG00000181215 |
| EVPL | 1.006214486 | 2.008633689 | 0.044594682 | 0.122415045 | 14.813358 | 29.8299958 | ENSG00000167880 |
| ZNF490 | 1.006123738 | 2.008507346 | 0.00704235 | 0.027559737 | 23.45540738 | 47.09070769 | ENSG00000188033 |
| NA | 1.006004005 | 2.008340662 | 0.061781256 | 0.157568984 | 10.79183764 | 21.70216823 | ENSG00000218537 |
| RCN3 | 1.005745049 | 2.007980208 | 9.63E-16 | 4.02E-14 | 278.8217355 | 559.4330538 | ENSG00000142552 |
| M1AP | 1.003556919 | 2.004937021 | 1.04E-05 | 9.23E-05 | 86.89239942 | 173.7356973 | ENSG00000159374 |
| LOC105371899 | 1.002947981 | 2.004090948 | 0.259081877 | 0.447931297 | 4.919405771 | 9.7104451 | ENSG00000267568 |
| LOC100506258 | 1.002339768 | 2.003246239 | 0.073550253 | 0.179851124 | 12.92446585 | 25.84350345 | ENSG00000251364 |
| CCDC110 | 1.00181585 | 2.002518888 | 0.062767793 | 0.159558122 | 12.7210706 | 25.39602554 | ENSG00000168491 |
| GCC2-AS1 | 1.001580029 | 2.002191586 | 0.080707537 | 0.192960977 | 11.20571609 | 22.30543642 | ENSG00000214184 |
| FAM157A | 1.000776631 | 2.00107693 | 2.32E-05 | 0.000190166 | 74.77163789 | 149.3878299 | ENSG00000236438 |
| NA | 1.000669079 | 2.000927755 | 6.70E-60 | 4.60E-57 | 2257.106347 | 4516.486455 | ENSG00000249790 |
| CARF | 1.00042858 | 2.000594227 | 0.005076704 | 0.020849023 | 24.74577191 | 49.5743414 | ENSG00000138380 |
| SH2D3A | 1.000344428 | 2.000477535 | 7.37E-27 | 8.08E-25 | 513.8648297 | 1028.313525 | ENSG00000125731 |
| OGFRL1 | 1.000084395 | 2.000117 | 1.01E-08 | 1.62E-07 | 133.311112 | 266.2624456 | ENSG00000119900 |
| LPL | 1.00004294 | 2.000059528 | 2.08E-08 | 3.14E-07 | 108.4407699 | 216.7134138 | ENSG00000175445 |
| PLAAT3 | 0.999749293 | 1.999652476 | 3.84E-27 | 4.29E-25 | 1037.142326 | 2073.386772 | ENSG00000176485 |
| MEF2C-AS1 | 0.999390544 | 1.999155293 | 0.126663701 | 0.270272281 | 9.50768924 | 18.81382317 | ENSG00000248309 |
| NA | 0.999169905 | 1.998849575 | 0.019723182 | 0.063806853 | 22.76780978 | 45.55465556 | ENSG00000204121 |
| CPT1A | 0.998619986 | 1.99808781 | 9.38E-08 | 1.27E-06 | 131.7418195 | 262.7719232 | ENSG00000110090 |
| CCDC113 | 0.996966437 | 1.995799007 | 5.57E-10 | 1.07E-08 | 141.1916019 | 281.7151051 | ENSG00000103021 |
| CAV2 | 0.995913821 | 1.994343368 | 0.009240451 | 0.034438471 | 25.30682255 | 50.39223787 | ENSG00000105971 |
| EFHB | 0.995546904 | 1.993836216 | 0.208614174 | 0.38769228 | 5.121019588 | 10.1456504 | ENSG00000163576 |
| VDR | 0.993366669 | 1.990825359 | 0.107332884 | 0.238869797 | 8.293864739 | 16.49219199 | ENSG00000111424 |
| RFX3-AS1 | 0.992896908 | 1.990177225 | 0.244055901 | 0.43079353 | 6.119422542 | 12.15653682 | ENSG00000232104 |
| ELK4 | 0.992719437 | 1.989932421 | 5.01E-17 | 2.39E-15 | 303.3854138 | 603.3880814 | ENSG00000158711 |
| APCDD1L-DT | 0.989716656 | 1.985794944 | 0.40506028 | 0.596724205 | 2.897945921 | 5.787863427 | ENSG00000231290 |
| TUBB2A | 0.989678461 | 1.985742371 | 3.69E-59 | 2.36E-56 | 2688.5815 | 5338.280746 | ENSG00000137267 |
| NA | 0.989515539 | 1.985518137 | 0.010236618 | 0.037481929 | 25.89893865 | 51.45415727 | ENSG00000259692 |
| CDK10 | 0.989436813 | 1.985409793 | 1.08E-08 | 1.72E-07 | 253.2002517 | 503.265028 | ENSG00000185324 |
| EPCAM-DT | 0.989009776 | 1.9848222 | 0.02892665 | 0.08700528 | 18.72932477 | 37.30236746 | ENSG00000234690 |
| NA | 0.988268383 | 1.983802472 | 0.192284165 | 0.365554899 | 5.819268009 | 11.51517699 | ENSG00000262703 |
| PLD1 | 0.988229267 | 1.983748686 | 0.010009651 | 0.036809301 | 21.65568291 | 42.93147755 | ENSG00000075651 |
| EPHA7 | 0.985626617 | 1.980173191 | 8.08E-10 | 1.52E-08 | 155.8437846 | 308.1027123 | ENSG00000135333 |
| LOC100506100 | 0.985287741 | 1.97970812 | 0.07541714 | 0.183243481 | 12.31656518 | 24.18850148 | ENSG00000223478 |
| SEMA6A-AS1 | 0.984738028 | 1.978953932 | 0.043964123 | 0.121092444 | 14.97255648 | 29.72573197 | ENSG00000248445 |
| NA | 0.98449033 | 1.978614191 | 0.43236471 | 0.6222587 | 2.468282905 | 4.962480882 | ENSG00000283897 |
| LINC01869 | 0.984337563 | 1.978404688 | 0.387585937 | 0.580627041 | 2.785499778 | 5.581270869 | ENSG00000180279 |
| LOC101929140 | 0.984000732 | 1.977942836 | 0.193772295 | 0.36762982 | 5.42560881 | 10.73862136 | ENSG00000278390 |
| NABP1 | 0.983341781 | 1.977039617 | 0.002991025 | 0.013290737 | 29.93248531 | 59.16500088 | ENSG00000173559 |
| DOC2A | 0.982523742 | 1.975918911 | 2.70E-05 | 0.000218866 | 124.3910064 | 246.2382646 | ENSG00000149927 |
| SDK2 | 0.981965759 | 1.975154844 | 9.61E-14 | 3.17E-12 | 288.9115665 | 570.3664247 | ENSG00000069188 |
| PTPRM | 0.981528481 | 1.97455627 | 2.79E-10 | 5.65E-09 | 158.7460675 | 312.9089648 | ENSG00000173482 |
| ADARB2 | 0.981462343 | 1.974465751 | 0.000370714 | 0.002205717 | 60.50935612 | 119.0460075 | ENSG00000185736 |
| EDIL3 | 0.980843935 | 1.973619582 | 2.09E-16 | 9.22E-15 | 735.612604 | 1450.232006 | ENSG00000164176 |
| SCAPER | 0.978456145 | 1.970355766 | 6.20E-07 | 7.17E-06 | 102.4994873 | 201.4455415 | ENSG00000140386 |
| TIGD6 | 0.978354123 | 1.970216434 | 0.003517028 | 0.015240728 | 29.03705776 | 57.16594903 | ENSG00000164296 |
| LOC100507551 | 0.978166522 | 1.969960254 | 0.233106153 | 0.417994335 | 5.544368779 | 10.87761601 | ENSG00000269439 |
| RAD51B | 0.977721642 | 1.969352876 | 0.000154406 | 0.001019875 | 49.41816071 | 97.22416529 | ENSG00000182185 |
| NA | 0.977362787 | 1.968863082 | 0.27989427 | 0.471039716 | 3.850874318 | 7.554065643 | ENSG00000280407 |
| NA | 0.97692185 | 1.968261422 | 0.000260096 | 0.00162493 | 57.86669153 | 113.6309761 | ENSG00000170919 |
| REXO5 | 0.976336803 | 1.967463407 | 1.04E-05 | 9.20E-05 | 79.68196371 | 157.0717552 | ENSG00000005189 |
| AMT | 0.976133034 | 1.967185538 | 0.235140951 | 0.420127821 | 5.31631958 | 10.57760651 | ENSG00000145020 |
| NKAIN2 | 0.97517665 | 1.965881893 | 0.302478086 | 0.495942042 | 3.510281278 | 6.897733667 | ENSG00000188580 |
| NA | 0.975064852 | 1.965729559 | 0.413879628 | 0.605206495 | 2.930792826 | 5.693458802 | ENSG00000277782 |
| CXCL14 | 0.974948379 | 1.965570866 | 0.161150508 | 0.321849517 | 7.661310129 | 15.11236817 | ENSG00000145824 |
| CAHM | 0.974913519 | 1.965523373 | 0.17242272 | 0.338575518 | 7.879986287 | 15.63025514 | ENSG00000270419 |
| TTC32 | 0.97380402 | 1.964012375 | 8.59E-05 | 0.000608515 | 62.62323825 | 123.0123434 | ENSG00000183891 |
| PRKG1 | 0.972975316 | 1.962884544 | 2.21E-09 | 3.90E-08 | 148.0639099 | 290.7073542 | ENSG00000185532 |
| CCDC172 | 0.97293373 | 1.962827963 | 0.008109749 | 0.030990947 | 24.38495961 | 47.99172387 | ENSG00000182645 |
| LINC01270 | 0.972868949 | 1.962739829 | 0.385417188 | 0.57834662 | 3.593037429 | 6.953608617 | ENSG00000203999 |

| | | | | | | | |
|---|---|---|---|---|---|---|---|
| PLAU | 0.971940438 | 1.961477026 | 9.69E-12 | 2.46E-10 | 199.0869011 | 390.4546706 | ENSG00000122861 |
| NA | 0.971685947 | 1.961131053 | 0.236732158 | 0.421747913 | 5.389604992 | 10.55839749 | ENSG00000250999 |
| OR2H2 | 0.971309731 | 1.960619709 | 0.315538566 | 0.509025309 | 4.016484318 | 7.963013879 | ENSG00000204657 |
| ATP10B | 0.970628956 | 1.959694755 | 0.231967386 | 0.416637613 | 5.544368779 | 10.80621926 | ENSG00000118322 |
| SNAI2 | 0.969794338 | 1.958561375 | 0.110747639 | 0.244618847 | 8.551701628 | 16.7546595 | ENSG00000019549 |
| PLBD1 | 0.969048542 | 1.957549165 | 2.24E-10 | 4.63E-09 | 171.2650433 | 335.0158444 | ENSG00000121316 |
| LINC00548 | 0.968947686 | 1.957412322 | 0.28884041 | 0.480553695 | 4.128930461 | 8.139550521 | ENSG00000215483 |
| LINC00894 | 0.968200972 | 1.956399461 | 0.003439159 | 0.014938219 | 60.81148749 | 119.1177794 | ENSG00000235703 |
| NA | 0.968193432 | 1.956389236 | 0.299918495 | 0.493415916 | 4.400672778 | 8.653200616 | ENSG00000287460 |
| LINC00858 | 0.967871478 | 1.955952693 | 0.179442041 | 0.34865555 | 7.003500215 | 13.77344715 | ENSG00000229404 |
| DUSP8 | 0.967605325 | 1.955591886 | 0.00058553 | 0.003291639 | 42.84442121 | 83.85261834 | ENSG00000184545 |
| PDGFRB | 0.966861149 | 1.954583406 | 5.63E-15 | 2.16E-13 | 286.0857033 | 559.1325434 | ENSG00000113721 |
| NA | 0.966282434 | 1.953799513 | 0.343259618 | 0.537082521 | 3.830655064 | 7.539531529 | ENSG00000279529 |
| PGBD4 | 0.965901191 | 1.953283275 | 0.03651096 | 0.104590848 | 19.842053 | 38.80296459 | ENSG00000182405 |
| SESN1 | 0.965142494 | 1.952256335 | 4.09E-17 | 1.98E-15 | 307.3131213 | 599.9306091 | ENSG00000080546 |
| WIPI1 | 0.964773268 | 1.951756761 | 1.70E-14 | 6.14E-13 | 332.0842613 | 647.4794099 | ENSG00000070540 |
| VPS13C | 0.964231053 | 1.951023361 | 6.31E-09 | 1.05E-07 | 208.0852077 | 405.5348874 | ENSG00000129003 |
| CHMP4C | 0.963988492 | 1.950695363 | 0.035962076 | 0.103416775 | 21.10904136 | 40.94217329 | ENSG00000164695 |
| LDHD | 0.963699737 | 1.950304971 | 2.53E-07 | 3.15E-06 | 101.3374512 | 197.3695427 | ENSG00000166816 |
| CLHC1 | 0.96346726 | 1.949990723 | 0.002355115 | 0.010904247 | 38.59928633 | 75.51181144 | ENSG00000162994 |
| SLC5A4-AS1 | 0.963411333 | 1.949915131 | 0.205223514 | 0.383544683 | 8.428409272 | 16.32694027 | ENSG00000242082 |
| NA | 0.963095927 | 1.949488882 | 0.243130254 | 0.42965066 | 8.251547096 | 16.19317018 | ENSG00000227598 |
| STRADA | 0.962646223 | 1.948881299 | 0.008473583 | 0.032127163 | 26.90177629 | 52.32358019 | ENSG00000266173 |
| GPR146 | 0.96195332 | 1.947945509 | 0.423030592 | 0.614022163 | 2.824660509 | 5.53244396 | ENSG00000164849 |
| NRROS | 0.961224746 | 1.946962027 | 6.30E-06 | 5.96E-05 | 77.28992774 | 150.2718986 | ENSG00000174004 |
| ASB16 | 0.960804496 | 1.946394968 | 0.067430783 | 0.168204863 | 12.79623514 | 24.88303672 | ENSG00000161664 |
| LOC441204 | 0.960356509 | 1.945790666 | 0.35407304 | 0.547413224 | 3.126092818 | 6.102407038 | ENSG00000214870 |
| NA | 0.960136705 | 1.945494235 | 0.080267547 | 0.192165505 | 11.93563134 | 23.3073556 | ENSG00000287625 |
| RASGRF2 | 0.959506137 | 1.944644091 | 6.61E-18 | 3.48E-16 | 343.6980079 | 668.4371654 | ENSG00000113319 |
| NA | 0.958848889 | 1.943758371 | 0.010304634 | 0.037677273 | 23.23673123 | 45.19354655 | ENSG00000278730 |
| RASGEF1C | 0.958458993 | 1.943233131 | 0.014696039 | 0.050216884 | 21.88067289 | 42.54521077 | ENSG00000146090 |
| NRP2 | 0.958191735 | 1.942873183 | 5.31E-25 | 5.00E-23 | 662.9841624 | 1288.23642 | ENSG00000118257 |
| AKR1C1 | 0.957899439 | 1.942479589 | 0.050410834 | 0.134705075 | 15.34411729 | 29.81155122 | ENSG00000187134 |
| AFDN-DT | 0.957379493 | 1.941779646 | 0.01660124 | 0.055479152 | 26.61297163 | 51.55093502 | ENSG00000198221 |
| RP1L1 | 0.957242873 | 1.941595773 | 0.145798722 | 0.29958243 | 7.284713271 | 14.19313065 | ENSG00000183638 |
| KAT2B | 0.956520256 | 1.94062351 | 0.06058953 | 0.155170147 | 12.44932828 | 24.1477103 | ENSG00000114166 |
| NTN1 | 0.955080577 | 1.938687909 | 7.61E-06 | 7.04E-05 | 84.4287466 | 163.4999908 | ENSG00000065320 |
| STK17A | 0.954157099 | 1.937447341 | 8.18E-14 | 2.74E-12 | 269.2150996 | 521.5869888 | ENSG00000164543 |
| PRNP | 0.953977831 | 1.93720661 | 2.64E-19 | 1.59E-17 | 450.1711583 | 871.7110677 | ENSG00000171867 |
| KDELR3 | 0.953662296 | 1.936782966 | 1.24E-07 | 1.63E-06 | 116.3782571 | 225.0612688 | ENSG00000100196 |
| NA | 0.953173619 | 1.936127041 | 0.218562946 | 0.399846575 | 6.553520247 | 12.61799919 | ENSG00000256967 |
| PCED1B | 0.952168894 | 1.934779148 | 1.50E-05 | 0.000129585 | 74.41871029 | 144.1539698 | ENSG00000179715 |
| NA | 0.952047446 | 1.934616282 | 0.028137538 | 0.085310181 | 19.00600544 | 36.65677075 | ENSG00000234338 |
| FER1L5 | 0.952039994 | 1.934606289 | 0.157449885 | 0.31664238 | 8.231327842 | 15.98245236 | ENSG00000249715 |
| HS1BP3 | 0.952030767 | 1.934593917 | 4.28E-19 | 2.52E-17 | 371.5319146 | 718.5186182 | ENSG00000118960 |
| DDX60 | 0.951622332 | 1.934046299 | 0.000351849 | 0.002103604 | 46.82796092 | 90.4424184 | ENSG00000137628 |
| PHLDA3 | 0.951542119 | 1.93393877 | 2.35E-21 | 1.67E-19 | 870.1839617 | 1683.822258 | ENSG00000174307 |
| SENP8 | 0.951300656 | 1.933615116 | 0.001663807 | 0.008112866 | 44.29082729 | 85.84247225 | ENSG00000166192 |
| RNF152 | 0.950674692 | 1.932776331 | 0.027108812 | 0.082732541 | 22.12084608 | 42.81125389 | ENSG00000176641 |
| PRKG2 | 0.949905221 | 1.931745747 | 0.024071534 | 0.075120695 | 28.70012529 | 55.23405706 | ENSG00000138669 |
| NA | 0.949795852 | 1.931599308 | 9.59E-08 | 1.29E-06 | 125.5096425 | 242.4606489 | ENSG00000247134 |
| LIF | 0.949466819 | 1.931158823 | 1.34E-05 | 0.000116676 | 80.97223054 | 156.7399779 | ENSG00000128342 |
| NA | 0.948929867 | 1.930440204 | 0.408430174 | 0.600182883 | 3.321295112 | 6.365312579 | ENSG00000287923 |
| LINC00886 | 0.948487578 | 1.929848477 | 0.336207067 | 0.530455587 | 3.966477374 | 7.55786449 | ENSG00000240875 |
| NA | 0.948375552 | 1.929698629 | 0.020615897 | 0.066255426 | 23.81927889 | 45.89176902 | ENSG00000272077 |
| NA | 0.948185052 | 1.92944384 | 0.313481605 | 0.507030986 | 3.77127508 | 7.336737818 | ENSG00000279794 |
| NA | 0.947843857 | 1.928987584 | 0.308800294 | 0.502976258 | 3.768118167 | 7.332938971 | ENSG00000228903 |
| TRPS1 | 0.947451243 | 1.928462701 | 7.73E-06 | 7.13E-05 | 74.41545568 | 143.5731598 | ENSG00000104447 |
| NA | 0.947262239 | 1.928210075 | 0.016898086 | 0.056207667 | 23.47109425 | 45.11086488 | ENSG00000240086 |
| ACAD10 | 0.947039702 | 1.927912669 | 4.64E-11 | 1.06E-09 | 289.1427952 | 557.6866506 | ENSG00000111271 |
| IGSF9B | 0.945945716 | 1.926451301 | 3.10E-22 | 2.36E-20 | 1433.686365 | 2761.532342 | ENSG00000080854 |
| RPH3AL | 0.945795127 | 1.926250227 | 0.009300858 | 0.034642704 | 26.53780708 | 51.07102807 | ENSG00000181031 |
| ANKRD50 | 0.945478526 | 1.925827555 | 7.53E-18 | 3.93E-16 | 608.579692 | 1171.746964 | ENSG00000151458 |
| ILDR1 | 0.945074353 | 1.925288108 | 0.005064776 | 0.020813871 | 30.12462839 | 58.05837983 | ENSG00000145103 |
| JAK1 | 0.945031922 | 1.925231484 | 1.31E-22 | 1.03E-20 | 632.4481286 | 1216.972948 | ENSG00000162434 |
| KC6 | 0.94491842 | 1.925080026 | 0.000685512 | 0.003786966 | 52.73767438 | 101.7568887 | ENSG00000267313 |
| ADAMTSL2 | 0.943991869 | 1.923844066 | 0.003503428 | 0.015188897 | 36.33627782 | 69.96808256 | ENSG00000197859 |
| RHOB | 0.943740841 | 1.923509348 | 8.98E-15 | 3.34E-13 | 348.2771137 | 669.7182043 | ENSG00000143878 |
| EXOC8 | 0.94198769 | 1.921173335 | 1.23E-06 | 1.35E-05 | 99.22819917 | 190.5020882 | ENSG00000116903 |
| NA | 0.941406326 | 1.920399315 | 0.185090523 | 0.355873787 | 7.967177127 | 15.32980761 | ENSG00000235997 |
| NDUFA13 | 0.940678092 | 1.919430192 | 0.008504008 | 0.032218577 | 31.69610827 | 60.85743321 | ENSG00000186010 |
| STPG1 | 0.940644087 | 1.919384951 | 9.94E-10 | 1.85E-08 | 150.658349 | 289.1079411 | ENSG00000001460 |
| HLA-E | 0.940026187 | 1.918563062 | 2.04E-32 | 3.24E-30 | 956.3658463 | 1834.197949 | ENSG00000204592 |
| NEDD9 | 0.939170522 | 1.917425498 | 3.63E-12 | 9.84E-11 | 218.7716121 | 418.8981923 | ENSG00000111859 |

| | | | | | | | |
|---|---|---|---|---|---|---|---|
| OXCT1-AS1 | 0.938806918 | 1.916942308 | 0.401551211 | 0.593157738 | 2.888475183 | 5.524957886 | ENSG00000248668 |
| STRC | 0.938662985 | 1.91675107 | 0.417493611 | 0.608571958 | 2.554195968 | 4.913653973 | ENSG00000242866 |
| MACROD1 | 0.935993943 | 1.913208284 | 3.96E-16 | 1.70E-14 | 355.0128178 | 678.9291759 | ENSG00000133315 |
| LINC00571 | 0.935718356 | 1.912842853 | 0.405247912 | 0.596903085 | 2.533976715 | 4.868625911 | ENSG00000223685 |
| TNN | 0.935695343 | 1.912812341 | 0.026314056 | 0.080759183 | 19.4780838 | 37.23053267 | ENSG00000120332 |
| CPS1 | 0.935333763 | 1.912332996 | 1.46E-17 | 7.40E-16 | 377.2865712 | 720.9151102 | ENSG00000021826 |
| RAC2 | 0.935160591 | 1.912103466 | 0.007000534 | 0.027425047 | 42.09951815 | 80.80403132 | ENSG00000128340 |
| METAP1D | 0.934702486 | 1.911496403 | 0.000119398 | 0.000816877 | 63.82956885 | 122.1086277 | ENSG00000172878 |
| RAI1 | 0.93424466 | 1.910889905 | 5.89E-07 | 6.85E-06 | 115.7943339 | 221.3288189 | ENSG00000108557 |
| NA | 0.934121139 | 1.910726304 | 0.033481349 | 0.097738751 | 19.12486311 | 36.63106333 | ENSG00000261490 |
| CATSPER2P1 | 0.933842899 | 1.910357835 | 0.100227028 | 0.226980417 | 13.07113437 | 25.07878239 | ENSG00000205771 |
| BNIP3 | 0.933743892 | 1.910226739 | 5.41E-18 | 2.87E-16 | 776.593952 | 1482.124583 | ENSG00000176171 |
| NA | 0.933479221 | 1.909876328 | 0.230042536 | 0.413986795 | 5.481831882 | 10.44989678 | ENSG00000263004 |
| GOLGA6L10 | 0.933240256 | 1.909560006 | 0.001208603 | 0.006179378 | 43.67345589 | 83.47102647 | ENSG00000278662 |
| FABP6 | 0.932719495 | 1.908870848 | 0.002014074 | 0.00955435 | 40.70222455 | 77.89767971 | ENSG00000170231 |
| CTBS | 0.932681149 | 1.908820112 | 0.181984541 | 0.352224465 | 7.115946358 | 13.5293126 | ENSG00000117151 |
| NA | 0.932218269 | 1.908207776 | 0.320002194 | 0.513677522 | 3.788337421 | 7.257854994 | ENSG00000267745 |
| TBCE | 0.931801661 | 1.907656822 | 0.266331456 | 0.456336064 | 5.167771921 | 9.75927201 | ENSG00000284770 |
| NA | 0.931411074 | 1.907140424 | 0.440579711 | 0.628759865 | 2.593356699 | 4.943709888 | ENSG00000285725 |
| NA | 0.930899489 | 1.906464264 | 0.278568247 | 0.469697987 | 5.781986415 | 10.98980394 | ENSG00000278616 |
| NA | 0.93052211 | 1.905965639 | 0.441276614 | 0.629512066 | 2.580729048 | 4.91745282 | ENSG00000280138 |
| RGS9 | 0.930235566 | 1.905587119 | 3.27E-10 | 6.57E-09 | 163.1469357 | 311.0541322 | ENSG00000108370 |
| NA | 0.930116088 | 1.905429312 | 0.383599672 | 0.576294675 | 2.924479001 | 5.536242807 | ENSG00000268564 |
| DST | 0.929629154 | 1.904786306 | 0.000339713 | 0.002042907 | 1264.122716 | 2407.863839 | ENSG00000151914 |
| CRIM1 | 0.929133406 | 1.904131884 | 5.08E-15 | 1.96E-13 | 354.1215393 | 674.5429248 | ENSG00000150938 |
| NA | 0.928861242 | 1.903772704 | 0.176744566 | 0.344746767 | 8.356401637 | 15.95652171 | ENSG00000279619 |
| CCDC183 | 0.927852421 | 1.902441934 | 0.126117634 | 0.269266151 | 9.30617312 | 17.68149471 | ENSG00000213213 |
| NA | 0.927507564 | 1.901987236 | 0.002629139 | 0.011931552 | 35.46116722 | 67.61964466 | ENSG00000250519 |
| RIMBP2 | 0.927219231 | 1.901607148 | 0.021637997 | 0.069013704 | 23.31760824 | 44.34994267 | ENSG00000060709 |
| LOC102724057 | 0.926924755 | 1.901219041 | 0.297774402 | 0.49083518 | 4.549122739 | 8.581803865 | ENSG00000230303 |
| CSF2RA | 0.926511467 | 1.900674477 | 0.01790926 | 0.058811167 | 23.40224353 | 44.46256863 | ENSG00000198223 |
| FAM27C | 0.926355664 | 1.900469226 | 0.34138709 | 0.535800519 | 3.483748199 | 6.665322075 | ENSG00000231527 |
| SH2D2A | 0.926333913 | 1.900440573 | 2.16E-06 | 2.25E-05 | 90.80916294 | 172.4424656 | ENSG00000027869 |
| ABI3 | 0.926179029 | 1.900236559 | 0.003334957 | 0.014560683 | 35.6861572 | 68.06233604 | ENSG00000108798 |
| AHNAK | 0.925880261 | 1.899843078 | 4.83E-05 | 0.000365655 | 83.73415875 | 158.5661401 | ENSG00000124942 |
| RGS2 | 0.925875991 | 1.899837455 | 1.22E-09 | 2.23E-08 | 181.7119282 | 344.8641849 | ENSG00000116741 |
| CCDC192 | 0.925700742 | 1.899606689 | 0.436198715 | 0.625590574 | 2.56682362 | 4.876111984 | ENSG00000230561 |
| FOXC1 | 0.925389006 | 1.899196269 | 0.375031263 | 0.568069764 | 3.208848969 | 6.042295208 | ENSG00000054598 |
| NA | 0.925137986 | 1.89886585 | 0.092701071 | 0.214069479 | 10.89037835 | 20.6256031 | ENSG00000188459 |
| NA | 0.925045751 | 1.898744454 | 3.13E-20 | 2.03E-18 | 910.4558325 | 1727.310485 | ENSG00000254339 |
| LOC107985177 | 0.925025725 | 1.898718098 | 0.395365557 | 0.587768127 | 2.95101208 | 5.555013801 | ENSG00000287281 |
| KCNQ5 | 0.924432348 | 1.897937321 | 0.010405141 | 0.037992294 | 25.50340031 | 48.44190132 | ENSG00000185760 |
| LMOD1 | 0.923870241 | 1.897197985 | 0.06322125 | 0.160490706 | 13.7321012 | 25.97782319 | ENSG00000163431 |
| LOC105370941 | 0.923723998 | 1.89700568 | 0.323138386 | 0.517322868 | 4.419614255 | 8.443358864 | ENSG00000261136 |
| USP53 | 0.923610517 | 1.896856469 | 7.33E-14 | 2.48E-12 | 314.9002515 | 596.4993938 | ENSG00000145390 |
| NA | 0.922722189 | 1.895688854 | 0.05157854 | 0.136897703 | 15.61901652 | 29.59009813 | ENSG00000213189 |
| NA | 0.92225896 | 1.895080273 | 0.225368795 | 0.408307453 | 6.067634159 | 11.42077237 | ENSG00000261098 |
| UPP1 | 0.921889812 | 1.894595433 | 3.23E-28 | 4.07E-26 | 716.3181696 | 1357.248219 | ENSG00000183696 |
| NA | 0.92174182 | 1.894401095 | 0.236680933 | 0.421747913 | 5.799048755 | 10.91840719 | ENSG00000280310 |
| NA | 0.921251113 | 1.893756858 | 0.019586094 | 0.063407627 | 26.07086247 | 49.43947201 | ENSG00000203635 |
| C12orf56 | 0.920744551 | 1.893092035 | 0.196042388 | 0.370949425 | 7.136165612 | 13.42037386 | ENSG00000185306 |
| SNORA71B | 0.9205966 | 1.892897905 | 0.42171093 | 0.612922709 | 2.507443636 | 4.759687172 | ENSG00000235408 |
| NA | 0.920497289 | 1.892767608 | 0.098029317 | 0.223216685 | 11.49639988 | 21.74730791 | ENSG00000251611 |
| NA | 0.919101551 | 1.890937332 | 0.078652511 | 0.189194762 | 13.09451054 | 24.81956407 | ENSG00000260267 |
| NA | 0.918700793 | 1.890412131 | 0.439555918 | 0.627927345 | 2.593356699 | 4.887396905 | ENSG00000176857 |
| LOC105372493 | 0.918485539 | 1.890130098 | 0.504170138 | 0.683955792 | 2.769715214 | 5.300032367 | ENSG00000286787 |
| BMPR1B | 0.9183862 | 1.889999955 | 0.085062141 | 0.200552311 | 11.86668291 | 22.36229905 | ENSG00000138696 |
| SMG1 | 0.917556375 | 1.888913157 | 4.72E-05 | 0.000358 | 1692.385263 | 3196.722745 | ENSG00000157106 |
| C8orf88 | 0.917282725 | 1.888554903 | 5.28E-08 | 7.44E-07 | 125.9537146 | 237.8752369 | ENSG00000253250 |
| NA | 0.916787683 | 1.887906981 | 0.07355517 | 0.179851124 | 14.48794817 | 27.42199572 | ENSG00000231770 |
| NA | 0.915608749 | 1.886364861 | 0.048208953 | 0.130021062 | 15.27398879 | 28.78717381 | ENSG00000235726 |
| NA | 0.91472122 | 1.885204748 | 0.086138663 | 0.202600758 | 13.38203742 | 25.34113827 | ENSG00000230699 |
| RASIP1 | 0.91458751 | 1.885030034 | 3.00E-08 | 4.43E-07 | 164.1009465 | 308.6954684 | ENSG00000105538 |
| RRM2B | 0.914526021 | 1.884949694 | 2.77E-15 | 1.09E-13 | 1159.693853 | 2184.960263 | ENSG00000048392 |
| TP73 | 0.914206564 | 1.884532354 | 5.79E-06 | 5.52E-05 | 89.03794838 | 167.6255894 | ENSG00000078900 |
| TDRD5 | 0.914183344 | 1.884502023 | 0.127125992 | 0.270791074 | 9.974731549 | 18.66734244 | ENSG00000162782 |
| USHBP1 | 0.912137757 | 1.881831895 | 5.68E-07 | 6.65E-06 | 110.8401044 | 208.3720573 | ENSG00000130307 |
| FAM167B | 0.911780978 | 1.881366575 | 0.048299794 | 0.130228161 | 20.65284527 | 39.0245293 | ENSG00000183615 |
| BHLHE23 | 0.910564962 | 1.879781481 | 0.17126817 | 0.336824289 | 7.287870184 | 13.70909844 | ENSG00000125533 |
| NPIPB2 | 0.910545756 | 1.879756455 | 0.239283215 | 0.42520375 | 5.76620185 | 10.75315547 | ENSG00000234719 |
| ZADH2 | 0.910520441 | 1.879723472 | 3.29E-05 | 0.000260962 | 76.72750162 | 143.9664746 | ENSG00000180011 |
| NA | 0.910350718 | 1.879502348 | 0.302000687 | 0.49542751 | 4.819587279 | 9.054551158 | ENSG00000279668 |
| UBE2D3-AS1 | 0.910021452 | 1.879073439 | 0.211080883 | 0.390555345 | 6.672377914 | 12.59542935 | ENSG00000246560 |
| NA | 0.909882833 | 1.8788929 | 0.334031663 | 0.528643399 | 3.827498152 | 7.171486096 | ENSG00000255471 |

| | | | | | | | |
|---|---|---|---|---|---|---|---|
| BRCA2 | 0.909040172 | 1.877795781 | 9.73E-10 | 1.82E-08 | 209.3817732 | 393.5070564 | ENSG00000139618 |
| EFHC2 | 0.908954761 | 1.877684615 | 0.002889651 | 0.012892824 | 38.14802858 | 71.78734857 | ENSG00000183690 |
| NA | 0.908060489 | 1.87652107 | 0.185328435 | 0.356056516 | 7.003500215 | 13.1617052 | ENSG00000261235 |
| NA | 0.907487175 | 1.875775505 | 0.098470861 | 0.223988578 | 12.83539588 | 24.0880365 | ENSG00000181097 |
| DEGS2 | 0.907029897 | 1.875181052 | 0.091528879 | 0.212197037 | 12.38363447 | 23.19461802 | ENSG00000168350 |
| LOC339874 | 0.907029851 | 1.875180992 | 0.438485209 | 0.626929327 | 2.954168993 | 5.479929824 | ENSG00000250608 |
| MEIS3 | 0.906378091 | 1.874334041 | 3.16E-22 | 2.40E-20 | 609.4503905 | 1141.668742 | ENSG00000105419 |
| NA | 0.906300877 | 1.874233729 | 0.233520804 | 0.418414015 | 6.367788692 | 11.90111735 | ENSG00000256742 |
| TM7SF3 | 0.905910312 | 1.873726406 | 2.02E-19 | 1.23E-17 | 1007.739668 | 1887.807131 | ENSG00000064115 |
| ABCC2 | 0.905885617 | 1.873694333 | 0.186926184 | 0.35834608 | 8.270488573 | 15.59640038 | ENSG00000023839 |
| SPARCL1 | 0.905451865 | 1.873131084 | 0.151537579 | 0.308332143 | 9.659393812 | 17.96966963 | ENSG00000152583 |
| NPIPB13 | 0.904562962 | 1.871977327 | 0.339550552 | 0.533776924 | 3.513438191 | 6.578953177 | ENSG00000198064 |
| TENT5A | 0.904301859 | 1.871638563 | 0.000309877 | 0.00188675 | 63.27473434 | 118.0773934 | ENSG00000112773 |
| CBLC | 0.90382496 | 1.871019973 | 2.22E-10 | 4.60E-09 | 192.7936756 | 360.2997937 | ENSG00000142273 |
| MRGPRG-AS1 | 0.902991431 | 1.869939289 | 0.374620403 | 0.567534294 | 3.83696889 | 7.148916254 | ENSG00000236301 |
| DMTN | 0.902984448 | 1.869930238 | 0.032189677 | 0.09465142 | 22.6458929 | 42.23325508 | ENSG00000158856 |
| NA | 0.902844663 | 1.869749067 | 0.44537181 | 0.633452623 | 2.977545159 | 5.502388045 | ENSG00000275481 |
| NA | 0.902619223 | 1.869456916 | 0.329422801 | 0.523943798 | 4.442990421 | 8.236216654 | ENSG00000277475 |
| TMSB15B-AS1 | 0.902464308 | 1.869256187 | 0.342804006 | 0.536797013 | 3.490062025 | 6.530126268 | ENSG00000231728 |
| THSD4 | 0.902460233 | 1.869250907 | 1.50E-05 | 0.00012937 | 92.34533806 | 172.6530718 | ENSG00000187720 |
| CPM | 0.902215224 | 1.868933484 | 0.062434732 | 0.158842186 | 14.72103342 | 27.43696787 | ENSG00000135678 |
| NA | 0.902126326 | 1.868818325 | 0.169852239 | 0.334684392 | 8.640771605 | 16.20021822 | ENSG00000262903 |
| ATOH1 | 0.901887169 | 1.868508555 | 0.22673549 | 0.410182097 | 6.858109469 | 12.70013121 | ENSG00000172238 |
| NA | 0.901511746 | 1.868022388 | 0.053338338 | 0.140462171 | 17.04530105 | 31.91161769 | ENSG00000276855 |
| SRMS | 0.901231101 | 1.86765904 | 0.184033158 | 0.354670952 | 7.982961692 | 14.91717215 | ENSG00000125508 |
| ALPG | 0.901037465 | 1.867408383 | 0.040038026 | 0.112553432 | 20.2358099 | 37.78128672 | ENSG00000163286 |
| GBGT1 | 0.900792674 | 1.867091556 | 5.54E-05 | 0.000413038 | 104.8830373 | 195.8269837 | ENSG00000148288 |
| LRRC2-AS1 | 0.900648157 | 1.866904535 | 0.455168689 | 0.642603328 | 2.835409024 | 5.34517205 | ENSG00000268324 |
| NA | 0.900485273 | 1.866693769 | 0.075369375 | 0.183151435 | 13.48057814 | 25.0777947 | ENSG00000230882 |
| NA | 0.900166801 | 1.866281745 | 0.372752063 | 0.566091997 | 3.436995866 | 6.432472449 | ENSG00000259045 |
| ACOXL-AS1 | 0.899271398 | 1.865123805 | 0.284670406 | 0.47570835 | 4.866339612 | 9.035780163 | ENSG00000204581 |
| NA | 0.899003966 | 1.864778099 | 0.260270225 | 0.449147891 | 5.617751889 | 10.5132578 | ENSG00000279406 |
| TP53INP1 | 0.898932508 | 1.864685738 | 1.11E-05 | 9.86E-05 | 105.5598863 | 196.7639098 | ENSG00000164938 |
| NA | 0.898795433 | 1.864508576 | 0.306816603 | 0.500840137 | 4.475837326 | 8.356878347 | ENSG00000287935 |
| P4HA2 | 0.898695346 | 1.86437923 | 4.32E-08 | 6.20E-07 | 139.1499228 | 259.284332 | ENSG00000072682 |
| CEP85L | 0.898685074 | 1.864365956 | 0.000189434 | 0.001223758 | 80.63459901 | 149.8386284 | ENSG00000111860 |
| LINC00471 | 0.898600232 | 1.864256319 | 0.089162352 | 0.208127839 | 14.10681892 | 26.3501055 | ENSG00000181798 |
| SPN | 0.898307906 | 1.863878613 | 0.006389528 | 0.025385533 | 34.24399041 | 63.81684861 | ENSG00000197471 |
| NA | 0.898252254 | 1.863806715 | 0.189289276 | 0.361455537 | 7.281556358 | 13.54384671 | ENSG00000287406 |
| KIAA0825 | 0.897886779 | 1.86333462 | 0.010319347 | 0.037717208 | 30.5695723 | 57.15846296 | ENSG00000185261 |
| PILRB | 0.897337753 | 1.862625653 | 0.012861666 | 0.045224457 | 35.13438191 | 65.5994487 | ENSG00000121716 |
| CRTAC1 | 0.896712191 | 1.861818181 | 0.000299717 | 0.001836347 | 57.58567388 | 107.2994066 | ENSG00000095713 |
| PLIN5 | 0.895817756 | 1.860664257 | 0.092694265 | 0.214069479 | 13.27590511 | 24.85330721 | ENSG00000214456 |
| FUT1 | 0.895555419 | 1.860325948 | 1.10E-12 | 3.24E-11 | 238.224636 | 443.2124684 | ENSG00000174951 |
| SIX4 | 0.894954867 | 1.85955171 | 7.82E-20 | 4.93E-18 | 462.474442 | 860.1367037 | ENSG00000100625 |
| OR52A1 | 0.894473173 | 1.858930938 | 0.153408021 | 0.3108072 | 8.578234708 | 15.99742451 | ENSG00000182070 |
| ADCY8 | 0.894470454 | 1.858927435 | 0.007049699 | 0.027576852 | 32.91003047 | 61.15700468 | ENSG00000155897 |
| NA | 0.893592422 | 1.857796426 | 0.001350768 | 0.006771743 | 55.28999121 | 102.7452127 | ENSG00000223459 |
| NA | 0.893511873 | 1.857692703 | 0.418840857 | 0.610090886 | 3.132406644 | 5.851224451 | ENSG00000279711 |
| LYRM9 | 0.892955444 | 1.856976353 | 0.158414563 | 0.317995832 | 8.518854723 | 15.88099969 | ENSG00000232859 |
| CHRM3 | 0.892678732 | 1.856620216 | 1.96E-08 | 2.98E-07 | 148.0244561 | 274.71658 | ENSG00000133019 |
| PDLIM3 | 0.892430739 | 1.856301097 | 6.67E-05 | 0.000486366 | 68.63031235 | 127.4180813 | ENSG00000154553 |
| NA | 0.891954831 | 1.855688853 | 0.282397431 | 0.47331832 | 5.511521874 | 10.12308056 | ENSG00000227288 |
| MGLL | 0.891634098 | 1.855276351 | 0.093872246 | 0.215914405 | 13.92414658 | 25.7533357 | ENSG00000074416 |
| KMT2A | 0.891557661 | 1.855178057 | 9.85E-14 | 3.24E-12 | 721.9007069 | 1339.636894 | ENSG00000118058 |
| NA | 0.890607922 | 1.853957179 | 0.380318014 | 0.573400641 | 3.205692056 | 5.952677116 | ENSG00000284726 |
| NA | 0.890194974 | 1.853426589 | 0.121208923 | 0.261553869 | 12.47911597 | 23.32656463 | ENSG00000236200 |
| C1QTNF6 | 0.88973059 | 1.852830092 | 0.000305951 | 0.001868365 | 58.98976232 | 109.3778909 | ENSG00000133466 |
| C5orf58 | 0.888597864 | 1.851375921 | 0.323024858 | 0.517230472 | 4.555436564 | 8.402018029 | ENSG00000234511 |
| LGALS9 | 0.888584072 | 1.851358223 | 0.267791181 | 0.457948122 | 6.163017962 | 11.33495312 | ENSG00000168961 |
| OBSCN | 0.888416573 | 1.851143289 | 3.73E-16 | 1.61E-14 | 1192.744682 | 2208.663232 | ENSG00000154358 |
| LINC00960 | 0.888332747 | 1.851035735 | 0.00170945 | 0.008302658 | 45.49221954 | 84.45698583 | ENSG00000242516 |
| PLAT | 0.888073661 | 1.850703347 | 5.72E-07 | 6.68E-06 | 124.3282742 | 229.7626047 | ENSG00000104368 |
| HPN | 0.887941865 | 1.850534286 | 1.16E-10 | 2.51E-09 | 213.7415416 | 395.0439931 | ENSG00000105707 |
| NA | 0.887253782 | 1.849651897 | 0.253433287 | 0.442142686 | 6.401913374 | 11.84904125 | ENSG00000255145 |
| NDUFA6-DT | 0.887037329 | 1.849374407 | 0.034228208 | 0.099511469 | 20.0777915 | 37.20840086 | ENSG00000237037 |
| MAP1B | 0.886987789 | 1.849310903 | 1.08E-17 | 5.55E-16 | 1316.847342 | 2435.116697 | ENSG00000131711 |
| NA | 0.88660036 | 1.848814346 | 0.016160858 | 0.054203054 | 25.98612949 | 47.93529927 | ENSG00000213839 |
| TIRAP | 0.886260448 | 1.848378801 | 0.062109797 | 0.158167489 | 16.70648945 | 30.82442891 | ENSG00000150455 |
| ZNF713 | 0.886079531 | 1.848147024 | 0.065376593 | 0.164252375 | 14.2913704 | 26.41302849 | ENSG00000178665 |
| NA | 0.885108431 | 1.846903427 | 0.286867528 | 0.478258772 | 5.402232644 | 9.980948335 | ENSG00000278200 |
| TRPM6 | 0.885040807 | 1.846816858 | 0.082304618 | 0.195918384 | 14.46951035 | 26.67213519 | ENSG00000119121 |
| LIFR-AS1 | 0.884702311 | 1.846383595 | 0.426558804 | 0.617161661 | 3.83696889 | 7.115611146 | ENSG00000244968 |
| SIK1 | 0.884403956 | 1.846001795 | 0.039443439 | 0.111230066 | 20.99018369 | 38.57325254 | ENSG00000142178 |

| | | | | | | | |
|---|---|---|---|---|---|---|---|
| NA | 0.88409388 | 1.845605079 | 0.074385883 | 0.181427551 | 18.79246303 | 34.58544609 | ENSG00000233290 |
| VWA5A | 0.883856617 | 1.84530158 | 1.45E-19 | 8.98E-18 | 514.8447949 | 950.2214057 | ENSG00000110002 |
| NA | 0.883739239 | 1.845151452 | 0.299166116 | 0.492647249 | 4.439833508 | 8.184140551 | ENSG00000229431 |
| HLA-DRB1 | 0.88362838 | 1.845009673 | 2.99E-11 | 7.05E-10 | 267.7561561 | 493.5074905 | ENSG00000196126 |
| NA | 0.883559747 | 1.844921903 | 0.264325237 | 0.454031876 | 5.375699565 | 9.928322578 | ENSG00000273314 |
| GNMT | 0.883344569 | 1.844646754 | 0.25367571 | 0.442440789 | 6.097324152 | 11.27527932 | ENSG00000124713 |
| SEMA3G | 0.883218463 | 1.844485519 | 4.00E-05 | 0.000309374 | 83.56182896 | 154.1458682 | ENSG00000010319 |
| FBXL2 | 0.882982563 | 1.844183946 | 0.001524894 | 0.007530612 | 66.64022623 | 122.9911993 | ENSG00000153558 |
| MSX1 | 0.882698769 | 1.84382121 | 0.344588011 | 0.538181023 | 4.04617431 | 7.505238734 | ENSG00000163132 |
| LINC01841 | 0.882697148 | 1.843819139 | 0.250951529 | 0.439408185 | 5.730198032 | 10.5320288 | ENSG00000266913 |
| SAR1B | 0.882511502 | 1.843581891 | 1.07E-11 | 2.71E-10 | 284.8837323 | 524.461962 | ENSG00000152700 |
| NR5A2 | 0.882310969 | 1.843325653 | 2.93E-07 | 3.60E-06 | 159.1223561 | 293.2008387 | ENSG00000116833 |
| RIN1 | 0.882012645 | 1.842944526 | 0.006709938 | 0.02645437 | 37.34482793 | 68.83151722 | ENSG00000174791 |
| P2RX4 | 0.880796435 | 1.841391555 | 7.54E-06 | 6.99E-05 | 98.94876756 | 182.1245038 | ENSG00000135124 |
| PHOX2A | 0.880402013 | 1.840888201 | 0.280157055 | 0.471396356 | 5.445828064 | 10.01089263 | ENSG00000165462 |
| RBFOX1 | 0.880043388 | 1.84043065 | 0.014090425 | 0.048613538 | 33.6892536 | 61.89448098 | ENSG00000078328 |
| ARMC12 | 0.879898161 | 1.840245395 | 0.169951085 | 0.334807972 | 9.693518493 | 17.80909282 | ENSG00000157343 |
| MYO15A | 0.879892864 | 1.840238638 | 0.120927339 | 0.261159089 | 13.39782199 | 24.7711752 | ENSG00000091536 |
| ZNF578 | 0.879690332 | 1.839980315 | 0.007320543 | 0.028408489 | 37.1280309 | 68.30635897 | ENSG00000258405 |
| NA | 0.879545252 | 1.839795293 | 0.395946109 | 0.588262452 | 3.185472803 | 5.862509371 | ENSG00000238230 |
| CD86 | 0.878862464 | 1.838924774 | 0.44504513 | 0.633328198 | 2.616732866 | 4.748402251 | ENSG00000114013 |
| CAMK2D | 0.878852899 | 1.838912583 | 7.61E-09 | 1.25E-07 | 175.3693949 | 322.5197094 | ENSG00000145349 |
| LINC01184 | 0.878657182 | 1.838663131 | 0.000542412 | 0.00308376 | 56.32312021 | 103.4577545 | ENSG00000245937 |
| IQCH | 0.878532651 | 1.838504428 | 0.205579738 | 0.384011032 | 7.083099453 | 12.9325428 | ENSG00000103599 |
| ENPP2 | 0.878439407 | 1.838385606 | 0.025298863 | 0.078135166 | 32.9196966 | 60.65343702 | ENSG00000136960 |
| NA | 0.878312125 | 1.838223421 | 0.386297766 | 0.579263023 | 3.146312072 | 5.794911468 | ENSG00000278743 |
| IL10RA | 0.877959052 | 1.837773605 | 0.414128921 | 0.605511629 | 3.400992048 | 6.33905551 | ENSG00000110324 |
| KCTD18 | 0.876966017 | 1.836509064 | 0.000102445 | 0.000711932 | 77.53503927 | 142.0007279 | ENSG00000155729 |
| RAB37 | 0.876812973 | 1.836314255 | 0.000375633 | 0.002232117 | 66.28028575 | 121.9466252 | ENSG00000172794 |
| INKA2-AS1 | 0.876193348 | 1.835525743 | 0.380650249 | 0.573714866 | 3.755490516 | 6.946560577 | ENSG00000227811 |
| NA | 0.876170961 | 1.835497261 | 0.020432491 | 0.065791484 | 28.12516923 | 51.63394311 | ENSG00000267432 |
| NXPH4 | 0.876141181 | 1.835459372 | 0.00051534 | 0.002946115 | 61.93781288 | 113.8735733 | ENSG00000182379 |
| MAOB | 0.876121049 | 1.83543376 | 0.003149808 | 0.013879826 | 45.00298114 | 82.42666715 | ENSG00000069535 |
| USP46-AS1 | 0.875688365 | 1.834883371 | 0.157825264 | 0.317053267 | 8.918827748 | 16.42134489 | ENSG00000248866 |
| PHYH | 0.875520003 | 1.834669253 | 2.22E-11 | 5.31E-10 | 255.003539 | 467.3598523 | ENSG00000107537 |
| NLRP6 | 0.875089126 | 1.83412139 | 0.254686389 | 0.443475905 | 6.083418724 | 11.17382666 | ENSG00000174885 |
| KIFAP3 | 0.874510542 | 1.833385974 | 0.001169042 | 0.006020288 | 54.49380344 | 99.84872874 | ENSG00000075945 |
| NA | 0.874273742 | 1.833085072 | 0.470992444 | 0.656343906 | 2.619889779 | 4.740916177 | ENSG00000260884 |
| NA | 0.874194312 | 1.832984152 | 0.17662345 | 0.344619331 | 9.762466914 | 18.05702621 | ENSG00000234630 |
| ACTA2 | 0.874174338 | 1.832958774 | 0.223343904 | 0.405609148 | 6.775353319 | 12.34760758 | ENSG00000107796 |
| NA | 0.874061522 | 1.832815446 | 0.002655556 | 0.012027879 | 44.86390422 | 82.07904167 | ENSG00000239213 |
| PCDH12 | 0.873435793 | 1.832020685 | 0.102937184 | 0.231449443 | 17.26260173 | 31.71762415 | ENSG00000113555 |
| NA | 0.873235939 | 1.831766916 | 0.440645181 | 0.628804881 | 2.802562119 | 5.176121481 | ENSG00000233538 |
| FAM189A2 | 0.872874835 | 1.831308486 | 3.39E-11 | 7.94E-10 | 275.9523972 | 505.3849387 | ENSG00000135063 |
| UCP3 | 0.872029787 | 1.830236125 | 0.220831952 | 0.40236919 | 6.980124049 | 12.80538272 | ENSG00000175564 |
| RASA4 | 0.871979452 | 1.830172269 | 0.395722694 | 0.588110368 | 3.172845151 | 5.802397541 | ENSG00000105808 |
| TRAF1 | 0.871862768 | 1.830024252 | 0.039299674 | 0.11089832 | 28.52957694 | 52.20467908 | ENSG00000056558 |
| NA | 0.871322228 | 1.829338719 | 0.413122673 | 0.604535668 | 2.868255929 | 5.273775299 | ENSG00000223668 |
| RAB20 | 0.870661842 | 1.82850154 | 5.34E-14 | 1.82E-12 | 311.1614174 | 569.0826629 | ENSG00000139832 |
| HIPK3 | 0.870572855 | 1.82838876 | 2.47E-09 | 4.31E-08 | 197.6517245 | 360.8061725 | ENSG00000110422 |
| SLC25A34 | 0.870505788 | 1.828303765 | 0.240194758 | 0.426292219 | 6.461293358 | 11.76635958 | ENSG00000162461 |
| GNG12 | 0.869120028 | 1.826548457 | 2.77E-12 | 7.58E-11 | 499.6830417 | 912.0882071 | ENSG00000172380 |
| NA | 0.867452118 | 1.824437992 | 0.398148801 | 0.590012783 | 3.152625897 | 5.746084558 | ENSG00000272654 |
| NA | 0.866974725 | 1.823834378 | 0.365297345 | 0.558431292 | 4.389924263 | 8.075751464 | ENSG00000230910 |
| LIPA | 0.866299624 | 1.822981125 | 3.71E-14 | 1.29E-12 | 432.3187104 | 788.214119 | ENSG00000107798 |
| SIM1 | 0.865642546 | 1.822151034 | 0.300535916 | 0.493806087 | 4.72736039 | 8.66068669 | ENSG00000112246 |
| B4GALT6 | 0.865485667 | 1.821952904 | 5.45E-12 | 1.45E-10 | 393.338603 | 715.8690397 | ENSG00000118276 |
| FHIT | 0.865376921 | 1.821815576 | 0.052193777 | 0.13799707 | 17.39536483 | 31.79942976 | ENSG00000189283 |
| ADCY2 | 0.86491333 | 1.821230254 | 3.73E-27 | 4.19E-25 | 1905.702328 | 3469.846274 | ENSG00000078295 |
| SPP1 | 0.864371339 | 1.820546184 | 9.87E-09 | 1.59E-07 | 277.0911549 | 503.9952867 | ENSG00000118785 |
| INHBC | 0.864218489 | 1.82035331 | 0.250516451 | 0.438991135 | 6.127014144 | 11.1287986 | ENSG00000175189 |
| SSC4D | 0.86403113 | 1.820116922 | 8.28E-12 | 2.13E-10 | 274.7403315 | 499.6518509 | ENSG00000146700 |
| RASA4B | 0.862944776 | 1.818746884 | 0.327434184 | 0.521675658 | 4.830335794 | 8.693991798 | ENSG00000170667 |
| ICA1L | 0.862215398 | 1.817827619 | 0.018300227 | 0.059872268 | 31.37908679 | 57.24439383 | ENSG00000163596 |
| TYMP | 0.86157768 | 1.817024258 | 0.456396483 | 0.643650313 | 3.388364396 | 6.218831851 | ENSG00000025708 |
| STAT6 | 0.860883509 | 1.816150183 | 2.58E-19 | 1.56E-17 | 962.9145033 | 1749.340005 | ENSG00000166888 |
| PIEZO2 | 0.860444917 | 1.815598142 | 5.67E-12 | 1.50E-10 | 479.6803322 | 870.757163 | ENSG00000154864 |
| LOC100505797 | 0.86014548 | 1.815221347 | 0.212963974 | 0.392722261 | 7.390845588 | 13.35646318 | ENSG00000263958 |
| RASGRF2-AS1 | 0.860060928 | 1.815114965 | 0.349665012 | 0.543044355 | 3.824341239 | 6.957845497 | ENSG00000251450 |
| QRICH2 | 0.859917766 | 1.814934856 | 0.008474679 | 0.032127163 | 36.88923319 | 67.12520359 | ENSG00000129646 |
| LOC101928335 | 0.859912498 | 1.814928229 | 0.21341923 | 0.393348449 | 8.436000875 | 15.23595264 | ENSG00000236064 |
| HMGCL | 0.859616354 | 1.814555714 | 1.14E-06 | 1.26E-05 | 205.6873616 | 372.3740868 | ENSG00000117305 |
| ZSCAN20 | 0.859121559 | 1.81393349 | 0.00077531 | 0.004222625 | 61.06882071 | 110.8818489 | ENSG00000121903 |
| SSBP3-AS1 | 0.859049297 | 1.813842636 | 0.259443173 | 0.448305031 | 6.897367898 | 12.53108064 | ENSG00000198711 |

| | | | | | | | |
|---|---|---|---|---|---|---|---|
| NA | 0.858239262 | 1.812824498 | 0.407221003 | 0.598955168 | 3.463528945 | 6.335256664 | ENSG00000254595 |
| FBXO36 | 0.857700895 | 1.812148136 | 0.002322688 | 0.01078374 | 53.72147285 | 97.26214531 | ENSG00000153832 |
| GNRH1 | 0.857521249 | 1.811922499 | 0.319338041 | 0.513144724 | 5.134925016 | 9.27567783 | ENSG00000147437 |
| DMXL2 | 0.857298982 | 1.811643369 | 4.02E-07 | 4.84E-06 | 173.003486 | 313.5800944 | ENSG00000104093 |
| LOXL4 | 0.857171929 | 1.811483831 | 0.375035159 | 0.568069764 | 4.333701191 | 7.921673043 | ENSG00000138131 |
| HLA-F | 0.856417347 | 1.810536607 | 0.00364357 | 0.015711942 | 41.7555728 | 75.62399937 | ENSG00000204642 |
| WEE1 | 0.856174016 | 1.810231261 | 8.30E-26 | 8.40E-24 | 1356.017668 | 2454.155845 | ENSG00000166483 |
| SCART1 | 0.856082779 | 1.810116784 | 0.329502435 | 0.523943798 | 5.689759525 | 10.4015079 | ENSG00000214279 |
| IGF1R | 0.855961865 | 1.809965082 | 7.79E-21 | 5.33E-19 | 1929.434266 | 3491.501897 | ENSG00000140443 |
| NLGN4Y | 0.855621615 | 1.809538265 | 0.424830998 | 0.615526713 | 2.894789008 | 5.236233311 | ENSG00000165246 |
| GIPC3 | 0.854843577 | 1.808562652 | 0.000392985 | 0.002316661 | 70.12604897 | 126.9493476 | ENSG00000179855 |
| LINC01310 | 0.853703456 | 1.807133962 | 0.463184439 | 0.649713191 | 2.927635913 | 5.277574146 | ENSG00000205632 |
| NA | 0.853673715 | 1.807096708 | 0.465570136 | 0.651875569 | 2.533976715 | 4.624929397 | ENSG00000228373 |
| ADORA2A | 0.853122101 | 1.806405897 | 0.322475423 | 0.51652921 | 4.714732738 | 8.54046303 | ENSG00000128271 |
| TTC39B | 0.853004463 | 1.806258608 | 1.61E-08 | 2.49E-07 | 175.2586325 | 316.0935206 | ENSG00000155158 |
| ATP7A | 0.85264995 | 1.805814812 | 0.002490469 | 0.011436658 | 55.31377334 | 99.54568485 | ENSG00000165240 |
| PRORSD1P | 0.852000024 | 1.805001485 | 0.188886715 | 0.360909935 | 8.531482375 | 15.42691178 | ENSG00000162997 |
| NFIA | 0.851853887 | 1.804818658 | 0.094455934 | 0.216959508 | 13.49448357 | 24.22235624 | ENSG00000162599 |
| B3GALT4 | 0.851737692 | 1.804673302 | 0.028531075 | 0.086165054 | 23.24432283 | 41.92642079 | ENSG00000235863 |
| NCALD | 0.851678261 | 1.804598962 | 0.013550261 | 0.047091794 | 29.19823307 | 52.78135534 | ENSG00000104490 |
| NFAM1 | 0.851676459 | 1.804596708 | 0.03802512 | 0.107893076 | 23.08374888 | 41.53624355 | ENSG00000235568 |
| ZNF486 | 0.851375006 | 1.804219674 | 7.34E-12 | 1.90E-10 | 348.4550809 | 628.2076681 | ENSG00000256229 |
| BLVRB | 0.850034494 | 1.802544022 | 6.02E-15 | 2.31E-13 | 368.9593785 | 665.0528504 | ENSG00000090013 |
| FGF16 | 0.849781259 | 1.802227651 | 0.013267835 | 0.04630999 | 29.47628921 | 53.27298524 | ENSG00000196468 |
| NLRC5 | 0.849167095 | 1.801460596 | 0.026705164 | 0.081740234 | 28.52591637 | 51.4987473 | ENSG00000140853 |
| FOXO3 | 0.848912253 | 1.801142408 | 2.33E-14 | 8.29E-13 | 350.3271964 | 630.5451878 | ENSG00000118689 |
| FGD2 | 0.848686604 | 1.800860717 | 0.436361353 | 0.625775403 | 3.622727421 | 6.439958523 | ENSG00000146192 |
| BCDIN3D-AS1 | 0.848506487 | 1.800635898 | 0.357973773 | 0.551282586 | 4.102397381 | 7.358758006 | ENSG00000258057 |
| HMBOX1 | 0.848141852 | 1.800180853 | 2.20E-06 | 2.29E-05 | 119.3608383 | 214.6767084 | ENSG00000147421 |
| PXN | 0.846371213 | 1.797972822 | 2.76E-23 | 2.28E-21 | 975.6986371 | 1753.802857 | ENSG00000089159 |
| NA | 0.846011523 | 1.79752461 | 0.3571845 | 0.550616112 | 3.824341239 | 6.867677753 | ENSG00000231066 |
| NA | 0.845832582 | 1.797301672 | 0.412885969 | 0.604466327 | 3.232225135 | 5.753570632 | ENSG00000258626 |
| NA | 0.8453568 | 1.796709044 | 0.241414583 | 0.427638142 | 6.983280962 | 12.61051312 | ENSG00000280145 |
| REV3L | 0.845238645 | 1.796561901 | 4.51E-05 | 0.000343573 | 89.48970979 | 160.5667293 | ENSG00000009413 |
| PLOD2 | 0.844781115 | 1.795992238 | 7.31E-12 | 1.90E-10 | 402.1106646 | 721.2327595 | ENSG00000152952 |
| UHMK1 | 0.844709537 | 1.795903134 | 4.98E-12 | 1.34E-10 | 402.395786 | 722.205874 | ENSG00000152332 |
| CTD-2297D10.2 | 0.844493582 | 1.795634327 | 0.000837815 | 0.004520542 | 131.8740789 | 237.2486033 | ENSG00000250579 |
| LPXN | 0.844137109 | 1.795190702 | 0.100752264 | 0.227835829 | 12.65725592 | 22.79370551 | ENSG00000110031 |
| DDHD2 | 0.844126349 | 1.795177314 | 5.21E-11 | 1.18E-09 | 273.8996539 | 491.5060655 | ENSG00000085788 |
| SARDH | 0.843925612 | 1.79492755 | 1.37E-12 | 3.93E-11 | 296.5595499 | 531.9262266 | ENSG00000123453 |
| BRWD1 | 0.843782479 | 1.794749479 | 1.35E-15 | 5.55E-14 | 614.4721703 | 1102.105442 | ENSG00000185658 |
| G0S2 | 0.843693962 | 1.794639365 | 0.043148899 | 0.119414916 | 20.82288995 | 37.47422919 | ENSG00000123689 |
| STC2 | 0.843148086 | 1.793960451 | 7.10E-31 | 1.06E-28 | 1865.337908 | 3344.880298 | ENSG00000113739 |
| NA | 0.841541477 | 1.79196378 | 0.171735586 | 0.337462611 | 9.934293042 | 17.72597312 | ENSG00000269947 |
| HSPB7 | 0.841206495 | 1.791547749 | 0.364509846 | 0.557636838 | 4.489742754 | 8.053181623 | ENSG00000173641 |
| UGDH-AS1 | 0.840782703 | 1.791021559 | 0.237507241 | 0.422736887 | 7.47685635 | 13.50674276 | ENSG00000249348 |
| TPSG1 | 0.840725038 | 1.790949972 | 0.006580941 | 0.026028842 | 38.52274631 | 69.03376128 | ENSG00000116176 |
| IQCN | 0.840296913 | 1.79041858 | 0.387199134 | 0.58014137 | 3.490062025 | 6.278943681 | ENSG00000130518 |
| MPI | 0.840201355 | 1.790299995 | 1.55E-17 | 7.83E-16 | 518.8627297 | 929.1746934 | ENSG00000178802 |
| MMP23A | 0.839342347 | 1.789234333 | 0.461361001 | 0.647890953 | 2.527662889 | 4.557331494 | ENSG00000215914 |
| CEP19 | 0.838941977 | 1.788737862 | 0.035297366 | 0.101884657 | 27.5792041 | 49.22919222 | ENSG00000174007 |
| NA | 0.838768673 | 1.788523003 | 0.312608262 | 0.506161696 | 5.435079549 | 9.729654128 | ENSG00000287608 |
| NCR3LG1 | 0.838505556 | 1.788196843 | 0.004886296 | 0.020169846 | 40.35394221 | 72.03049543 | ENSG00000188211 |
| NA | 0.838337541 | 1.787988604 | 0.15415462 | 0.311774523 | 10.43102534 | 18.72800392 | ENSG00000257681 |
| NA | 0.837953342 | 1.787512514 | 2.08E-07 | 2.63E-06 | 143.4551141 | 256.3750661 | ENSG00000225889 |
| ATF3 | 0.837806444 | 1.787330516 | 1.51E-14 | 5.50E-13 | 416.0262122 | 743.5725608 | ENSG00000162772 |
| APOBEC3H | 0.837590179 | 1.787062608 | 0.472364401 | 0.657542077 | 2.554195968 | 4.587387409 | ENSG00000100298 |
| GIPR | 0.837518577 | 1.786973917 | 0.241443725 | 0.427648891 | 9.945041557 | 17.83969839 | ENSG00000010310 |
| PGPEP1 | 0.837199197 | 1.786578365 | 3.12E-20 | 2.03E-18 | 586.0614511 | 1047.01395 | ENSG00000130517 |
| JPH2 | 0.836964912 | 1.786288259 | 0.007154925 | 0.027888409 | 36.66867789 | 65.37959585 | ENSG00000149596 |
| GPRIN3 | 0.836760293 | 1.786034927 | 0.001754483 | 0.008492451 | 47.76510475 | 85.28181872 | ENSG00000185477 |
| RPL32P3 | 0.83657138 | 1.78580107 | 0.000213555 | 0.001361431 | 98.99995458 | 177.0069971 | ENSG00000251474 |
| NA | 0.836183111 | 1.785320527 | 0.229111229 | 0.412948314 | 6.99718639 | 12.53163029 | ENSG00000279145 |
| MX2 | 0.836125045 | 1.785248672 | 0.002624321 | 0.011924236 | 54.7299479 | 97.67533042 | ENSG00000183486 |
| MATN1 | 0.83588058 | 1.784946187 | 0.430958125 | 0.620909351 | 2.897945921 | 5.146065566 | ENSG00000162510 |
| LOC105375431 | 0.835635088 | 1.784642482 | 0.325685491 | 0.519873817 | 5.670818048 | 10.19502697 | ENSG00000274993 |
| INHBE | 0.835508606 | 1.784486028 | 4.26E-13 | 1.32E-11 | 305.4191931 | 544.827949 | ENSG00000139269 |
| BHLHE40 | 0.835394778 | 1.78434524 | 0.002586373 | 0.011780741 | 67.7118169 | 120.3736436 | ENSG00000134107 |
| SNAPC5 | 0.835042119 | 1.783909119 | 5.31E-09 | 8.89E-08 | 195.5085432 | 348.8195623 | ENSG00000174446 |
| NA | 0.834812727 | 1.783625496 | 0.017570196 | 0.057944129 | 31.21781378 | 55.54848905 | ENSG00000257261 |
| NA | 0.834721176 | 1.783512313 | 0.100031528 | 0.226727797 | 13.10082437 | 23.3073556 | ENSG00000279393 |
| ALG10B | 0.83444061 | 1.783165501 | 0.000306265 | 0.001869667 | 86.02370034 | 153.0869832 | ENSG00000175548 |
| NA | 0.834156736 | 1.782814668 | 0.390570053 | 0.583196503 | 4.277380422 | 7.539093496 | ENSG00000270069 |
| LIN37 | 0.834058387 | 1.782693137 | 0.404111183 | 0.5957522 | 3.450901294 | 6.181289862 | ENSG00000267796 |

| | | | | | | | |
|---|---|---|---|---|---|---|---|
| PRDM13 | 0.833847982 | 1.782433165 | 0.44092169 | 0.629102587 | 2.891632095 | 5.127294571 | ENSG00000112238 |
| NA | 0.833145851 | 1.781565902 | 0.251583558 | 0.440140615 | 6.805043311 | 12.04759808 | ENSG00000224418 |
| UCHL3 | 0.833062523 | 1.781463004 | 0.309267015 | 0.503332882 | 5.088172683 | 9.107176914 | ENSG00000118939 |
| DEPDC4 | 0.832798563 | 1.781137092 | 0.024376706 | 0.075868661 | 26.81586323 | 47.78188212 | ENSG00000166153 |
| ARSG | 0.832152721 | 1.78033992 | 0.000614361 | 0.003435897 | 61.13451452 | 108.8557671 | ENSG00000141337 |
| C6orf163 | 0.831889166 | 1.780014713 | 0.152632676 | 0.309836317 | 11.60884603 | 20.65240982 | ENSG00000203872 |
| TMEM161B-AS1 | 0.831312164 | 1.779302943 | 6.55E-05 | 0.000478365 | 103.7253381 | 184.374953 | ENSG00000247828 |
| NA | 0.830791688 | 1.778661146 | 0.114192017 | 0.250317545 | 16.58023557 | 29.64629949 | ENSG00000251188 |
| LACTB2 | 0.830626459 | 1.778457452 | 0.003426283 | 0.01488927 | 45.00495798 | 80.0488346 | ENSG00000147592 |
| GRIN2A | 0.83057827 | 1.778398048 | 0.000964075 | 0.005109568 | 72.25600129 | 127.986945 | ENSG00000183454 |
| GRHL3 | 0.830465937 | 1.778259581 | 0.162395882 | 0.323778537 | 10.44680991 | 18.61906518 | ENSG00000158055 |
| ZC3H6 | 0.829834716 | 1.777481711 | 0.000140148 | 0.0009357 | 77.36152941 | 137.6012737 | ENSG00000188177 |
| GREM1 | 0.829641689 | 1.777243908 | 0.308642113 | 0.502897789 | 6.394321771 | 11.26300672 | ENSG00000166923 |
| PTAR1 | 0.828749156 | 1.776144743 | 1.02E-19 | 6.39E-18 | 675.5367291 | 1199.510876 | ENSG00000188647 |
| BDP1 | 0.828128289 | 1.77538054 | 4.79E-11 | 1.09E-09 | 499.9158264 | 887.1611711 | ENSG00000145734 |
| NA | 0.82802265 | 1.775250545 | 0.002080961 | 0.009833894 | 66.35732944 | 118.1336435 | ENSG00000182912 |
| NA | 0.82782965 | 1.775013073 | 0.410347692 | 0.60197753 | 4.224314263 | 7.509475614 | ENSG00000212195 |
| LINC00551 | 0.827747036 | 1.774911432 | 0.209369945 | 0.388490944 | 11.39016987 | 20.06257644 | ENSG00000272274 |
| TNXB | 0.827644531 | 1.774785327 | 3.69E-07 | 4.47E-06 | 154.5439494 | 274.182948 | ENSG00000168477 |
| UTF1 | 0.827185142 | 1.774220283 | 0.008062523 | 0.030829544 | 463.5428832 | 822.3157469 | ENSG00000171794 |
| TTPAL | 0.827141192 | 1.774166233 | 1.84E-11 | 4.46E-10 | 290.0973696 | 514.7757272 | ENSG00000124120 |
| TRIM5 | 0.825912692 | 1.772656119 | 7.42E-10 | 1.41E-08 | 240.7362212 | 426.183292 | ENSG00000132256 |
| RASAL3 | 0.825722014 | 1.772421846 | 0.194913884 | 0.369159164 | 8.687523938 | 15.29171597 | ENSG00000105122 |
| KCNIP4 | 0.825662694 | 1.77234897 | 0.512790268 | 0.691606377 | 2.643265945 | 4.673756307 | ENSG00000185774 |
| RLN1 | 0.825661637 | 1.772347671 | 0.037452347 | 0.10669256 | 22.28508061 | 39.46579495 | ENSG00000107014 |
| ADAMTS5 | 0.825597785 | 1.772269231 | 0.010942811 | 0.039705181 | 33.05669899 | 58.65514963 | ENSG00000154736 |
| ZBTB44-DT | 0.825486556 | 1.772132597 | 0.456149045 | 0.64339927 | 3.26507204 | 5.70854257 | ENSG00000175773 |
| TRIML2 | 0.824983403 | 1.771514659 | 4.17E-12 | 1.13E-10 | 495.4556083 | 878.0985297 | ENSG00000179046 |
| PKP1 | 0.824720873 | 1.771192322 | 0.182879934 | 0.353293381 | 9.560853097 | 16.84059036 | ENSG00000081277 |
| NA | 0.824667951 | 1.771127351 | 0.006559761 | 0.025950611 | 329.7423452 | 583.8291674 | ENSG00000274090 |
| NA | 0.824479779 | 1.770896356 | 0.464900118 | 0.651183403 | 2.557352881 | 4.538560499 | ENSG00000231767 |
| FNIP1 | 0.824411148 | 1.770812114 | 3.94E-06 | 3.92E-05 | 158.4405687 | 280.4632771 | ENSG00000217128 |
| EMBP1 | 0.824381093 | 1.770775224 | 0.009598528 | 0.035572708 | 38.71864766 | 68.57912371 | ENSG00000231752 |
| KCNH3 | 0.824229186 | 1.770588782 | 0.248420121 | 0.436542952 | 9.21898228 | 16.28571105 | ENSG00000135519 |
| APOBEC3F | 0.824138869 | 1.770477941 | 0.0003933 | 0.002317782 | 74.45787102 | 131.8015757 | ENSG00000128394 |
| NA | 0.823518794 | 1.769717148 | 0.000290699 | 0.001788783 | 75.36650494 | 133.5783386 | ENSG00000234261 |
| IFI27L2 | 0.823472637 | 1.769660529 | 8.21E-06 | 7.49E-05 | 123.2706866 | 218.5572251 | ENSG00000119632 |
| NA | 0.823246129 | 1.769382708 | 0.028616645 | 0.086325017 | 25.60953263 | 45.16294098 | ENSG00000233396 |
| FBXO32 | 0.823084983 | 1.769185082 | 0.241084565 | 0.427380328 | 7.69543481 | 13.52876295 | ENSG00000156804 |
| NOCT | 0.823078436 | 1.769177054 | 3.04E-13 | 9.53E-12 | 384.1734837 | 679.4607923 | ENSG00000151014 |
| OXCT2P1 | 0.822796857 | 1.768831788 | 0.157412677 | 0.316601906 | 11.60568911 | 20.3852674 | ENSG00000237624 |
| APLNR | 0.822682621 | 1.768691733 | 0.280536454 | 0.471906212 | 7.125417097 | 12.57297113 | ENSG00000134817 |
| ZNF850 | 0.820944927 | 1.766562666 | 2.44E-05 | 0.000199227 | 102.3885144 | 180.9525463 | ENSG00000267041 |
| NT5C2 | 0.820837016 | 1.766430535 | 2.59E-18 | 1.41E-16 | 616.894475 | 1089.765745 | ENSG00000076685 |
| EPPK1 | 0.820500607 | 1.766018684 | 3.36E-08 | 4.89E-07 | 269.6652902 | 475.9098312 | ENSG00000261150 |
| LOC105371083 | 0.820427936 | 1.765929729 | 0.247174389 | 0.434972478 | 6.659750263 | 11.80390157 | ENSG00000263080 |
| NA | 0.820020318 | 1.765430855 | 0.336176613 | 0.530452695 | 5.339695747 | 9.516125149 | ENSG00000230457 |
| NHSL2 | 0.819632867 | 1.764956794 | 9.42E-09 | 1.52E-07 | 203.9740387 | 359.7202978 | ENSG00000204131 |
| ADORA2A-AS1 | 0.819598218 | 1.764914406 | 0.391423972 | 0.584063433 | 4.43036277 | 7.812734304 | ENSG00000178803 |
| SPATA17 | 0.818449144 | 1.763509252 | 0.358211592 | 0.551476183 | 4.45246116 | 7.835304145 | ENSG00000162814 |
| GGT1 | 0.818092723 | 1.763073627 | 0.056878556 | 0.147728022 | 20.14160618 | 35.46009358 | ENSG00000100031 |
| KCNB1 | 0.818007186 | 1.762969097 | 0.044248316 | 0.12169436 | 25.39844807 | 44.70615352 | ENSG00000158445 |
| MAN2B2 | 0.817832228 | 1.762755311 | 5.09E-09 | 8.55E-08 | 197.9350121 | 349.0423379 | ENSG00000013288 |
| SMG1P5 | 0.817000295 | 1.761739108 | 0.027018373 | 0.082497286 | 26.34260479 | 46.45738359 | ENSG00000183604 |
| PLCB1 | 0.816834147 | 1.761536229 | 1.23E-05 | 0.000108238 | 132.134276 | 232.3047013 | ENSG00000182621 |
| NA | 0.816804672 | 1.761500241 | 0.033489314 | 0.097746605 | 25.08005112 | 44.27463545 | ENSG00000224411 |
| NA | 0.816580199 | 1.761226186 | 0.318129562 | 0.511874164 | 5.147552668 | 9.058350005 | ENSG00000278918 |
| NA | 0.816339841 | 1.760932783 | 0.236330696 | 0.421371152 | 7.612678659 | 13.42406109 | ENSG00000287200 |
| NA | 0.816274909 | 1.76085353 | 0.42285655 | 0.613865692 | 3.457215119 | 6.091122117 | ENSG00000234975 |
| TRPV6 | 0.816184393 | 1.760743056 | 0.046751941 | 0.126941178 | 27.89386542 | 49.13241447 | ENSG00000165125 |
| GPR37 | 0.81615633 | 1.760708806 | 0.000290177 | 0.001786166 | 112.0676389 | 196.846145 | ENSG00000170775 |
| NA | 0.815994626 | 1.760511469 | 0.312936746 | 0.506547568 | 5.064796517 | 8.930640271 | ENSG00000257556 |
| SIPA1L2 | 0.814534303 | 1.758730349 | 2.68E-13 | 8.47E-12 | 394.078395 | 692.6157674 | ENSG00000116991 |
| FDXR | 0.813584276 | 1.757572591 | 9.22E-23 | 7.34E-21 | 813.0745608 | 1428.908957 | ENSG00000161513 |
| NA | 0.813548085 | 1.757528501 | 0.502033176 | 0.682511764 | 3.444587468 | 6.125414912 | ENSG00000279283 |
| IDS | 0.813418583 | 1.757370745 | 1.48E-06 | 1.59E-05 | 145.7969477 | 256.0555127 | ENSG00000010404 |
| RPL39L | 0.812511366 | 1.756265996 | 2.33E-16 | 1.02E-14 | 486.5911019 | 854.1881518 | ENSG00000163923 |
| LGI4 | 0.811917259 | 1.755542909 | 0.004904348 | 0.020230843 | 61.28168671 | 107.8153585 | ENSG00000153902 |
| NA | 0.811911012 | 1.755535307 | 0.367732581 | 0.560766469 | 4.108711207 | 7.235285152 | ENSG00000255130 |
| SUGP1 | 0.811635562 | 1.755200159 | 3.03E-24 | 2.70E-22 | 842.8411278 | 1479.078147 | ENSG00000105705 |
| FAM183A | 0.811112852 | 1.754564339 | 0.298832396 | 0.492141428 | 5.99750566 | 10.51705665 | ENSG00000186973 |
| LOC101929494 | 0.810574142 | 1.753909298 | 0.001174538 | 0.006043553 | 73.72805347 | 129.7945284 | ENSG00000276170 |
| NA | 0.810571621 | 1.753906233 | 0.026403936 | 0.080967992 | 26.92525016 | 47.35785012 | ENSG00000272556 |
| XG | 0.808634063 | 1.751552295 | 0.115582117 | 0.252707202 | 13.46479357 | 23.5664623 | ENSG00000124343 |

| | | | | | | | |
|---|---|---|---|---|---|---|---|
| LONRF2 | 0.808080957 | 1.750880908 | 0.123397906 | 0.265043086 | 13.21652512 | 23.17964587 | ENSG00000170500 |
| ACP5 | 0.808036418 | 1.750826855 | 0.000122331 | 0.000834173 | 91.84535115 | 160.5832388 | ENSG00000102575 |
| CCDC162P | 0.807355752 | 1.750001007 | 0.080271142 | 0.192165505 | 19.16845853 | 33.63053034 | ENSG00000203799 |
| HEPHL1 | 0.806882613 | 1.749427179 | 0.513804269 | 0.692278836 | 2.812032858 | 4.97756465 | ENSG00000181333 |
| ADGRD2 | 0.8067136 | 1.749222245 | 0.054130763 | 0.142136022 | 24.02730423 | 42.23490404 | ENSG00000180264 |
| NA | 0.806582658 | 1.749063488 | 0.503703775 | 0.683776217 | 2.590199786 | 4.512303431 | ENSG00000265174 |
| NA | 0.806140608 | 1.748527648 | 0.254176962 | 0.442963405 | 8.111192399 | 14.09547683 | ENSG00000233013 |
| NA | 0.80496797 | 1.747107003 | 0.482987028 | 0.666361675 | 2.838565937 | 5.022592712 | ENSG00000278876 |
| PIM1 | 0.804737485 | 1.746827907 | 2.66E-27 | 3.03E-25 | 1173.975911 | 2050.172103 | ENSG00000137193 |
| ITGA10 | 0.804640197 | 1.746710114 | 0.42811962 | 0.618358482 | 3.179158977 | 5.573784796 | ENSG00000143127 |
| NA | 0.804150581 | 1.746117423 | 0.3022578 | 0.49571242 | 6.676812603 | 11.65362199 | ENSG00000243014 |
| PKHD1 | 0.804108587 | 1.746066599 | 0.246017428 | 0.433575448 | 7.711219375 | 13.42406109 | ENSG00000170927 |
| ARL4D | 0.80410365 | 1.746060623 | 2.86E-07 | 3.53E-06 | 219.8393468 | 383.4187151 | ENSG00000175906 |
| RPS27L | 0.80337176 | 1.745175059 | 1.00E-17 | 5.13E-16 | 657.6742317 | 1147.550635 | ENSG00000185088 |
| ENPP6 | 0.803168446 | 1.744929134 | 0.471247143 | 0.656501394 | 2.924479001 | 5.037126827 | ENSG00000164303 |
| TMPRSS11E | 0.802970201 | 1.744689375 | 0.163150079 | 0.324827969 | 13.6290281 | 23.62613609 | ENSG00000087128 |
| NA | 0.802724623 | 1.744392416 | 0.287339103 | 0.478786563 | 6.989594788 | 12.24290572 | ENSG00000233483 |
| VCL | 0.802670873 | 1.744327426 | 9.96E-40 | 2.84E-37 | 10195.56786 | 17784.01797 | ENSG00000035403 |
| SBF2-AS1 | 0.802402395 | 1.744002847 | 0.149753407 | 0.305708739 | 10.80130837 | 18.82510809 | ENSG00000246273 |
| ZSCAN31 | 0.801887548 | 1.743380585 | 0.000711919 | 0.003913021 | 68.96025457 | 120.3812413 | ENSG00000235109 |
| WNT9B | 0.801433279 | 1.742831724 | 0.283626613 | 0.474563117 | 7.222176374 | 12.71185416 | ENSG00000158955 |
| RLIM | 0.800789034 | 1.742053624 | 2.38E-21 | 1.68E-19 | 736.3836417 | 1282.229526 | ENSG00000131263 |
| PLS3-AS1 | 0.800688722 | 1.741932501 | 0.287706373 | 0.4792262 | 7.473601739 | 12.98940544 | ENSG00000271826 |
| ZBTB34 | 0.80065719 | 1.74189443 | 3.21E-10 | 6.46E-09 | 352.8008986 | 614.2975055 | ENSG00000177125 |
| NA | 0.800649253 | 1.741884847 | 0.063624165 | 0.161160349 | 27.6297147 | 48.18801919 | ENSG00000286067 |
| NA | 0.800475015 | 1.741674488 | 0.037409608 | 0.106619971 | 25.60015959 | 44.78134912 | ENSG00000205918 |
| KIFC3 | 0.800363497 | 1.741539864 | 2.03E-07 | 2.57E-06 | 169.2392238 | 294.9413822 | ENSG00000140859 |
| GAD1 | 0.800214034 | 1.741359451 | 0.261980999 | 0.451186754 | 6.937806405 | 12.1452519 | ENSG00000128683 |
| VSNL1 | 0.800185353 | 1.741324833 | 1.78E-17 | 8.91E-16 | 1251.203459 | 2178.036616 | ENSG00000163032 |
| NA | 0.800151028 | 1.741283403 | 0.004490194 | 0.018777358 | 50.33706212 | 87.72681113 | ENSG00000260077 |
| LGR6 | 0.799761144 | 1.74081289 | 0.058800477 | 0.151551666 | 23.26198653 | 40.45208072 | ENSG00000133067 |
| EPB41L4A | 0.799446602 | 1.740433393 | 0.012417176 | 0.043990129 | 36.42484413 | 63.45562798 | ENSG00000129595 |
| OLFML3 | 0.79897493 | 1.739864471 | 3.85E-19 | 2.27E-17 | 772.453792 | 1343.844775 | ENSG00000116774 |
| NA | 0.798813257 | 1.739669507 | 0.131183129 | 0.277329544 | 12.78676441 | 22.27256934 | ENSG00000224616 |
| IFIT1 | 0.798780196 | 1.739629641 | 0.040841008 | 0.114359759 | 27.58296237 | 47.79955381 | ENSG00000185745 |
| TRIM56 | 0.797985073 | 1.738671131 | 1.15E-18 | 6.51E-17 | 709.9898725 | 1234.735787 | ENSG00000169871 |
| PLXND1 | 0.79763614 | 1.738250664 | 8.88E-27 | 9.57E-25 | 1407.322271 | 2445.328481 | ENSG00000004399 |
| TMEM8B | 0.797269063 | 1.737808442 | 8.24E-09 | 1.34E-07 | 222.4545163 | 386.4810634 | ENSG00000137103 |
| BATF2 | 0.797156967 | 1.737673421 | 0.031633093 | 0.093444256 | 29.18056937 | 50.80867218 | ENSG00000168062 |
| LYPD5 | 0.796707187 | 1.737131761 | 0.298688509 | 0.492019081 | 6.067634159 | 10.48320189 | ENSG00000159871 |
| NA | 0.796247583 | 1.736578446 | 0.475776129 | 0.660526949 | 3.241695874 | 5.622611705 | ENSG00000103832 |
| ESPNL | 0.796156553 | 1.736468876 | 0.035259381 | 0.101822588 | 24.40833578 | 42.37236136 | ENSG00000144488 |
| PAQR9-AS1 | 0.795939366 | 1.736207484 | 0.357658865 | 0.55102655 | 4.77095581 | 8.29307929 | ENSG00000241570 |
| LYPD1 | 0.795908287 | 1.736170083 | 0.063759835 | 0.161371677 | 19.16086692 | 33.17925358 | ENSG00000150551 |
| JPX | 0.79587448 | 1.736129399 | 0.274801323 | 0.465810108 | 6.388007946 | 11.09874268 | ENSG00000225470 |
| DLL3 | 0.795264381 | 1.735395365 | 3.16E-09 | 5.43E-08 | 289.8130898 | 502.1959313 | ENSG00000090932 |
| CCDC167 | 0.794923532 | 1.734985412 | 3.06E-06 | 3.10E-05 | 172.2425279 | 298.6905564 | ENSG00000198937 |
| ADA2 | 0.794373493 | 1.73432406 | 9.91E-19 | 5.65E-17 | 903.3617739 | 1565.877622 | ENSG00000093072 |
| ATAD3C | 0.794351144 | 1.734297194 | 0.069771376 | 0.172534048 | 20.59158614 | 35.8545077 | ENSG00000215915 |
| EML2 | 0.79433298 | 1.734275359 | 8.61E-12 | 2.20E-10 | 357.3482626 | 619.5584585 | ENSG00000125746 |
| C1orf74 | 0.794171905 | 1.734081741 | 1.61E-05 | 0.000137296 | 107.3753954 | 186.125745 | ENSG00000162757 |
| MOG | 0.793634596 | 1.733436029 | 0.310513759 | 0.504406959 | 6.008254175 | 10.47246662 | ENSG00000204655 |
| NA | 0.793119666 | 1.732817438 | 0.397322579 | 0.589448225 | 4.115025033 | 7.11886034 | ENSG00000242085 |
| NA | 0.792751662 | 1.732375485 | 0.224877542 | 0.407696407 | 7.979804779 | 13.82552325 | ENSG00000286207 |
| JSRP1 | 0.792531137 | 1.7321107 | 0.351374331 | 0.544467173 | 5.481831882 | 9.406636757 | ENSG00000167476 |
| COLCA2 | 0.792513619 | 1.732089669 | 0.452889152 | 0.640214136 | 3.205692056 | 5.547527727 | ENSG00000214290 |
| NA | 0.792434308 | 1.731994451 | 0.234769742 | 0.419637612 | 8.03602785 | 13.90440607 | ENSG00000286966 |
| GABRE | 0.792260417 | 1.731785703 | 0.004317178 | 0.01817274 | 70.8095952 | 122.7438156 | ENSG00000102287 |
| TNNC1 | 0.792079305 | 1.731568313 | 0.04856857 | 0.130781617 | 27.6298124 | 48.04902454 | ENSG00000114854 |
| PLCB2 | 0.791960272 | 1.731425452 | 9.12E-08 | 1.23E-06 | 245.3264815 | 424.8903238 | ENSG00000137841 |
| KATNAL2 | 0.791822424 | 1.731260023 | 1.50E-05 | 0.00012937 | 129.3778875 | 223.9463259 | ENSG00000167216 |
| MAP3K15 | 0.791258481 | 1.730583414 | 0.004268759 | 0.017997538 | 46.55759408 | 80.6643754 | ENSG00000180815 |
| RPL13P5 | 0.791131598 | 1.730431218 | 0.1874105 | 0.359163086 | 9.809219247 | 16.94573025 | ENSG00000240370 |
| LHPP | 0.791093242 | 1.730385213 | 1.72E-16 | 7.73E-15 | 794.7853917 | 1374.959051 | ENSG00000107902 |
| CAPN6 | 0.79108805 | 1.730378986 | 1.07E-06 | 1.19E-05 | 165.3692353 | 285.9259733 | ENSG00000077274 |
| PABPC1L2B | 0.790490649 | 1.729662607 | 0.28533683 | 0.476607138 | 7.767442446 | 13.44338174 | ENSG00000184388 |
| NA | 0.790172783 | 1.729281556 | 0.564770367 | 0.732937653 | 2.640109032 | 4.534761653 | ENSG00000280089 |
| RASAL2-AS1 | 0.789676149 | 1.728686369 | 0.046020323 | 0.125432451 | 23.75042817 | 40.93991178 | ENSG00000224687 |
| BASP1-AS1 | 0.789246989 | 1.728172212 | 0.343945096 | 0.537587178 | 5.359915 | 9.294448824 | ENSG00000215196 |
| AKAP13 | 0.788859285 | 1.727707853 | 1.41E-10 | 3.02E-09 | 694.2815075 | 1199.582359 | ENSG00000170776 |
| CSRP1 | 0.788456559 | 1.727225633 | 0.282627462 | 0.47357539 | 7.036347121 | 12.05833335 | ENSG00000159176 |
| SERINC5 | 0.788345266 | 1.727092396 | 5.80E-20 | 3.72E-18 | 1679.431859 | 2899.634045 | ENSG00000164300 |
| FAM214A | 0.787935228 | 1.726601597 | 0.000746459 | 0.004082287 | 67.07136242 | 115.8318339 | ENSG00000047346 |
| ACOT11 | 0.786743232 | 1.725175619 | 9.63E-05 | 0.000673587 | 93.9117819 | 161.8471874 | ENSG00000162390 |

| | | | | | | | |
|---|---|---|---|---|---|---|---|
| NA | 0.786090434 | 1.724395179 | 0.195069755 | 0.369339769 | 9.918508477 | 17.02841192 | ENSG00000287009 |
| NA | 0.785903408 | 1.724171649 | 0.001803909 | 0.008695193 | 64.99821189 | 112.423014 | ENSG00000179277 |
| TMPRSS13 | 0.785901717 | 1.724169628 | 0.363528886 | 0.556600113 | 5.696073351 | 9.804300072 | ENSG00000137747 |
| SYNM | 0.784972426 | 1.723059387 | 0.000827605 | 0.004471964 | 83.77262043 | 143.9739607 | ENSG00000182253 |
| AMOTL2 | 0.784691973 | 1.722724465 | 1.13E-21 | 8.26E-20 | 1054.71444 | 1816.731369 | ENSG00000114019 |
| NA | 0.784662267 | 1.722688993 | 0.238123761 | 0.42363081 | 13.11670663 | 22.80966534 | ENSG00000226450 |
| LOC100268168 | 0.783563264 | 1.721377198 | 0.477180354 | 0.661534945 | 2.84172285 | 4.936223814 | ENSG00000204758 |
| DNAH2 | 0.783542487 | 1.721352408 | 0.300113575 | 0.493470097 | 7.35484177 | 12.70816694 | ENSG00000183914 |
| MPP1 | 0.783327155 | 1.721095503 | 6.79E-09 | 1.12E-07 | 268.2332705 | 461.5262282 | ENSG00000130830 |
| NA | 0.782951549 | 1.720647475 | 0.09929436 | 0.225557258 | 15.71755723 | 27.01348552 | ENSG00000259366 |
| SGSM3 | 0.782912559 | 1.720600973 | 0.330103481 | 0.524400272 | 6.56436646 | 11.35372412 | ENSG00000100359 |
| NA | 0.782091563 | 1.719622108 | 0.131205126 | 0.277344412 | 12.93531207 | 22.29838838 | ENSG00000276805 |
| ZNF292 | 0.78193453 | 1.719434943 | 1.14E-09 | 2.09E-08 | 354.3236342 | 608.64756 | ENSG00000188994 |
| NA | 0.781499569 | 1.718916624 | 0.173759914 | 0.340537045 | 10.85121762 | 18.6491211 | ENSG00000246877 |
| OR2A20P | 0.78118233 | 1.718538687 | 0.252715694 | 0.44137134 | 7.879986287 | 13.64193857 | ENSG00000170356 |
| AASS | 0.78030313 | 1.717491703 | 2.34E-26 | 2.46E-24 | 4081.332894 | 7010.317738 | ENSG00000008311 |
| NA | 0.779883629 | 1.716992371 | 0.488731114 | 0.671520682 | 2.871412842 | 4.962480882 | ENSG00000257605 |
| C12orf60 | 0.779764707 | 1.716850844 | 0.048854787 | 0.131323367 | 24.1833458 | 41.48372941 | ENSG00000182993 |
| C18orf25 | 0.779512539 | 1.716550783 | 2.47E-11 | 5.88E-10 | 361.0823538 | 619.8742131 | ENSG00000152242 |
| TUBB6 | 0.779492076 | 1.716526436 | 1.09E-27 | 1.29E-25 | 2936.01691 | 5040.449442 | ENSG00000176014 |
| PTPDC1 | 0.779469212 | 1.716499233 | 2.83E-05 | 0.000227174 | 127.8904415 | 219.1193269 | ENSG00000158079 |
| CLYBL | 0.77917738 | 1.716152049 | 2.61E-05 | 0.000212197 | 109.6591268 | 188.1665757 | ENSG00000125246 |
| NA | 0.779169754 | 1.716142978 | 0.034377375 | 0.099788609 | 27.72972859 | 47.72501948 | ENSG00000247627 |
| MACF1 | 0.778343105 | 1.715159928 | 3.78E-05 | 0.000294289 | 4522.165416 | 7756.245084 | ENSG00000127603 |
| SV2B | 0.778312113 | 1.715123084 | 0.133702917 | 0.281046253 | 15.38012111 | 26.37581292 | ENSG00000185518 |
| CGREF1 | 0.77830303 | 1.715112285 | 0.002671943 | 0.012083812 | 56.47788399 | 96.80839225 | ENSG00000138028 |
| NA | 0.778299426 | 1.715108001 | 0.096954991 | 0.221519207 | 24.59604417 | 42.14633653 | ENSG00000260400 |
| PPP1R15B | 0.777786511 | 1.714498344 | 1.43E-14 | 5.24E-13 | 971.5246154 | 1665.979025 | ENSG00000158615 |
| ATP1B1 | 0.777771753 | 1.714480806 | 4.56E-16 | 1.95E-14 | 762.857213 | 1307.766655 | ENSG00000143153 |
| LLPH-DT | 0.777695823 | 1.714390574 | 0.118419277 | 0.257090649 | 14.20861425 | 24.4289488 | ENSG00000239335 |
| LRRC15 | 0.777602598 | 1.714279796 | 0.253443611 | 0.442142686 | 7.601930144 | 13.04896761 | ENSG00000172061 |
| PYY2 | 0.777477231 | 1.714130836 | 0.298648183 | 0.492019081 | 6.755134065 | 11.59405982 | ENSG00000237575 |
| SLC48A1 | 0.776704905 | 1.713213446 | 2.39E-08 | 3.59E-07 | 241.5597388 | 413.6633762 | ENSG00000211584 |
| RAB24 | 0.776522234 | 1.712996535 | 0.002276581 | 0.010598883 | 60.84137288 | 104.5054662 | ENSG00000169228 |
| NA | 0.776269326 | 1.712696269 | 0.167731162 | 0.331542141 | 15.95083788 | 27.42568295 | ENSG00000253878 |
| AGPAT3 | 0.775653972 | 1.711965907 | 5.49E-06 | 5.25E-05 | 152.4776163 | 261.0062706 | ENSG00000160216 |
| CDH2 | 0.775519408 | 1.711806235 | 1.08E-15 | 4.47E-14 | 552.4811784 | 945.2782765 | ENSG00000170558 |
| MBTPS2 | 0.774430914 | 1.710515188 | 1.23E-05 | 0.000108229 | 148.5921887 | 254.3528947 | ENSG00000012174 |
| NA | 0.773848445 | 1.709824729 | 0.392273551 | 0.584907321 | 5.455298802 | 9.305845365 | ENSG00000287806 |
| OMA1 | 0.773601233 | 1.709531769 | 9.02E-05 | 0.00063583 | 108.5279608 | 185.0736848 | ENSG00000162600 |
| RILPL1 | 0.773191138 | 1.709045893 | 9.65E-07 | 1.08E-05 | 165.4602594 | 282.8603674 | ENSG00000188026 |
| LINC01852 | 0.773135532 | 1.708980023 | 0.463565264 | 0.650001408 | 3.810435811 | 6.511355273 | ENSG00000236914 |
| NA | 0.77309195 | 1.708928396 | 1.09E-05 | 9.67E-05 | 133.6799218 | 228.1047916 | ENSG00000204789 |
| CEP350 | 0.772479153 | 1.708202668 | 3.04E-10 | 6.11E-09 | 531.1366842 | 907.2707022 | ENSG00000135837 |
| SPATS2L | 0.772387158 | 1.708093747 | 3.48E-16 | 1.50E-14 | 898.7524069 | 1534.881117 | ENSG00000196141 |
| KIF17 | 0.772373003 | 1.708076988 | 5.85E-10 | 1.13E-08 | 283.4078316 | 484.3099226 | ENSG00000117245 |
| C1QTNF12 | 0.77213528 | 1.707795559 | 0.063431046 | 0.160876442 | 22.80203216 | 39.03851376 | ENSG00000184163 |
| LINC01503 | 0.771641021 | 1.707210579 | 0.010577817 | 0.038515255 | 40.01344687 | 68.44491559 | ENSG00000233901 |
| EPHA1-AS1 | 0.770734129 | 1.706137747 | 0.004822094 | 0.019935911 | 55.75616171 | 95.01939695 | ENSG00000229153 |
| EHBP1L1 | 0.770676491 | 1.706069585 | 7.44E-06 | 6.91E-05 | 143.802021 | 245.4351884 | ENSG00000173442 |
| NA | 0.769770602 | 1.704998656 | 0.342902833 | 0.536880932 | 5.955188016 | 10.17190747 | ENSG00000228981 |
| LHX6 | 0.76975169 | 1.704976306 | 0.014017355 | 0.048406495 | 41.90549593 | 71.70411724 | ENSG00000106852 |
| TMPRSS9 | 0.769738508 | 1.704960728 | 0.145213078 | 0.298809725 | 12.96500206 | 22.20117259 | ENSG00000178297 |
| TTC39C | 0.769456493 | 1.704627478 | 6.41E-11 | 1.44E-09 | 323.0628263 | 550.9831337 | ENSG00000168234 |
| TMEM51-AS1 | 0.769209081 | 1.704335171 | 0.257663718 | 0.446353805 | 7.99243243 | 13.66070956 | ENSG00000175147 |
| TRIM68 | 0.769090818 | 1.704195466 | 1.52E-05 | 0.000130469 | 160.7086511 | 273.6102854 | ENSG00000167333 |
| LINGO2 | 0.769058879 | 1.704157738 | 0.002559799 | 0.01168267 | 57.42893325 | 97.70396062 | ENSG00000174482 |
| ZNF503 | 0.769012614 | 1.70410309 | 0.533940785 | 0.708785167 | 2.947855167 | 4.939473008 | ENSG00000165655 |
| SNHG32 | 0.768858105 | 1.703920594 | 1.27E-12 | 3.68E-11 | 585.2847835 | 997.7681053 | ENSG00000204387 |
| APBA1 | 0.768571709 | 1.703582375 | 0.092396248 | 0.213555289 | 18.93913155 | 32.25621722 | ENSG00000107282 |
| GAS5 | 0.768068678 | 1.702988483 | 2.48E-20 | 1.63E-18 | 3694.378444 | 6291.818238 | ENSG00000234741 |
| NA | 0.768005786 | 1.702914246 | 0.490121362 | 0.6727825 | 3.122935905 | 5.307080408 | ENSG00000286415 |
| ZBTB42 | 0.767835342 | 1.70271307 | 0.000203458 | 0.001304353 | 85.74752334 | 145.939819 | ENSG00000179627 |
| NA | 0.767693115 | 1.702545218 | 0.509975366 | 0.689214193 | 2.785499778 | 4.778458166 | ENSG00000277369 |
| LOC100506405 | 0.767207493 | 1.701972224 | 0.030952249 | 0.091831118 | 29.98367233 | 51.01840231 | ENSG00000228496 |
| STK17B | 0.766813008 | 1.701506907 | 3.97E-05 | 0.000307325 | 147.9158659 | 251.2659526 | ENSG00000081320 |
| NA | 0.766563734 | 1.701212941 | 0.002157061 | 0.010157256 | 65.10287103 | 110.4295845 | ENSG00000250320 |
| LOC100129434 | 0.76462641 | 1.698929999 | 0.496028126 | 0.677678408 | 2.901102834 | 4.936223814 | ENSG00000233251 |
| DPEP1 | 0.764595678 | 1.698893808 | 0.003116271 | 0.013758209 | 59.58493764 | 101.0732003 | ENSG00000015413 |
| NA | 0.76420958 | 1.698439207 | 0.35831635 | 0.551580999 | 5.435079549 | 9.193107779 | ENSG00000255458 |
| AKAP7 | 0.764172807 | 1.698395915 | 9.94E-09 | 1.60E-07 | 248.7855758 | 421.9550382 | ENSG00000118507 |
| TNFRSF14 | 0.764065125 | 1.698269153 | 0.443723686 | 0.632030034 | 3.454058207 | 5.911336281 | ENSG00000157873 |
| PAQR7 | 0.763962055 | 1.698147829 | 3.24E-05 | 0.000257881 | 121.7153973 | 206.7709952 | ENSG00000182749 |
| CACNA1F | 0.763662731 | 1.697795541 | 0.1175638 | 0.255652641 | 17.12668173 | 28.85433368 | ENSG00000102001 |

| | | | | | | | |
|---|---|---|---|---|---|---|---|
| NA | 0.763373461 | 1.697455155 | 0.259011624 | 0.447893398 | 7.995589343 | 13.59311166 | ENSG00000224738 |
| NA | 0.762901953 | 1.696900476 | 0.080590144 | 0.192804663 | 20.27624841 | 34.57927412 | ENSG00000228613 |
| LINC01836 | 0.762898337 | 1.696896223 | 0.13321283 | 0.280214117 | 15.18807572 | 25.92618512 | ENSG00000267530 |
| APLN | 0.762041001 | 1.695888125 | 6.35E-05 | 0.000466009 | 110.9512728 | 187.794405 | ENSG00000171388 |
| DHTKD1 | 0.761968014 | 1.695802331 | 2.02E-25 | 2.00E-23 | 1431.237009 | 2426.357706 | ENSG00000181192 |
| ASNS | 0.761922895 | 1.695749298 | 0.027393318 | 0.083518302 | 29.90595223 | 50.76733135 | ENSG00000070669 |
| C11orf65 | 0.761903547 | 1.695726556 | 0.416611502 | 0.607751088 | 4.075864302 | 6.969130418 | ENSG00000166323 |
| DCAF4L1 | 0.760920994 | 1.694572068 | 0.486863791 | 0.669985548 | 2.844879763 | 4.838569996 | ENSG00000182308 |
| LINC02328 | 0.760742287 | 1.694362173 | 0.352308989 | 0.545516575 | 5.689759525 | 9.669653918 | ENSG00000258733 |
| PLEKHA2 | 0.760221238 | 1.693750342 | 0.000123887 | 0.000842922 | 140.9702499 | 238.2678904 | ENSG00000169499 |
| CD101 | 0.760189656 | 1.693713265 | 0.161036211 | 0.321655904 | 11.75739368 | 19.91295814 | ENSG00000134256 |
| XIAP | 0.7601615 | 1.69368021 | 6.83E-12 | 1.78E-10 | 702.9585987 | 1190.085451 | ENSG00000101966 |
| CNN1 | 0.760121932 | 1.693633759 | 2.97E-10 | 5.99E-09 | 321.0194635 | 543.9885735 | ENSG00000130176 |
| WLS | 0.760120191 | 1.693631715 | 0.013208511 | 0.046145807 | 42.30339443 | 71.56198501 | ENSG00000116729 |
| UTRN | 0.759908079 | 1.693382727 | 9.28E-11 | 2.05E-09 | 862.7632763 | 1460.462735 | ENSG00000152818 |
| RNF32 | 0.759891598 | 1.693363383 | 0.131621093 | 0.27797007 | 14.58521111 | 24.71767337 | ENSG00000105982 |
| SERPINA5 | 0.759754579 | 1.693202565 | 0.010480132 | 0.038228453 | 44.14415877 | 74.71268597 | ENSG00000188488 |
| NA | 0.759593451 | 1.693013469 | 0.017681672 | 0.058249576 | 52.52205744 | 89.05752191 | ENSG00000279407 |
| CRYGB | 0.758799183 | 1.692081646 | 0.351110323 | 0.544376671 | 5.852114914 | 9.789327924 | ENSG00000182187 |
| C17orf113 | 0.758727851 | 1.691997986 | 0.008610496 | 0.032522301 | 45.18743492 | 76.24603856 | ENSG00000267221 |
| FOSL2 | 0.758492299 | 1.691721752 | 0.000347964 | 0.002082816 | 117.6332191 | 199.0856358 | ENSG00000075426 |
| SYDE2 | 0.758479368 | 1.691706589 | 0.000196085 | 0.001261451 | 109.9963675 | 185.5773641 | ENSG00000097096 |
| CALHM1 | 0.758065684 | 1.691221573 | 0.281328574 | 0.47271345 | 7.506546342 | 12.78281288 | ENSG00000185933 |
| ADAMTS16 | 0.757612666 | 1.690690598 | 1.06E-06 | 1.18E-05 | 205.3387784 | 347.3631171 | ENSG00000145536 |
| TRIB3 | 0.757221165 | 1.690231861 | 8.61E-11 | 1.91E-09 | 336.4565067 | 568.7986846 | ENSG00000101255 |
| SIL1 | 0.757135974 | 1.690132057 | 0.001969659 | 0.009377256 | 70.29737143 | 118.7940605 | ENSG00000120725 |
| LOC102606465 | 0.757029944 | 1.690007845 | 0.073750014 | 0.180161166 | 20.04178769 | 33.92630295 | ENSG00000233184 |
| NFATC1 | 0.756851453 | 1.68979877 | 3.17E-05 | 0.000252504 | 150.5746082 | 254.5302672 | ENSG00000131196 |
| ESRRB | 0.756674155 | 1.689591117 | 0.089399883 | 0.208581675 | 18.92966081 | 31.99754856 | ENSG00000119715 |
| NEK7 | 0.756327538 | 1.689185231 | 0.001122949 | 0.005814155 | 123.1757088 | 207.508328 | ENSG00000151414 |
| PIK3R1 | 0.756065804 | 1.688878806 | 2.68E-07 | 3.33E-06 | 238.3628184 | 401.9773965 | ENSG00000145675 |
| PCDHGC5 | 0.756049032 | 1.688859172 | 0.441521885 | 0.629813486 | 3.483748199 | 5.911336281 | ENSG00000240764 |
| ZNF285 | 0.755974604 | 1.688772047 | 0.022266768 | 0.070545689 | 37.23732013 | 62.84162453 | ENSG00000267508 |
| FLNB | 0.754916537 | 1.687533962 | 1.38E-21 | 9.98E-20 | 4006.035308 | 6759.520988 | ENSG00000136068 |
| ZNF8 | 0.754682283 | 1.687259975 | 6.55E-06 | 6.17E-05 | 163.7196067 | 276.4482662 | ENSG00000278129 |
| PCLO | 0.754586671 | 1.687148159 | 0.000493916 | 0.002833253 | 89.23818672 | 150.542067 | ENSG00000186472 |
| GRM7 | 0.754543553 | 1.687097735 | 0.410001662 | 0.601708037 | 4.102397381 | 6.965331571 | ENSG00000196277 |
| NA | 0.754107898 | 1.686588355 | 0.464379281 | 0.650601375 | 3.463528945 | 5.866308218 | ENSG00000288546 |
| UNC5D | 0.753791882 | 1.686218956 | 5.17E-21 | 3.58E-19 | 1246.535975 | 2101.213997 | ENSG00000156687 |
| RGS7 | 0.75329018 | 1.685632668 | 0.040551414 | 0.113686265 | 27.48442166 | 46.31145252 | ENSG00000182901 |
| SESN2 | 0.753216978 | 1.685547142 | 6.96E-19 | 4.03E-17 | 931.2259866 | 1568.60206 | ENSG00000130766 |
| NA | 0.752531212 | 1.68474613 | 0.282019022 | 0.473069106 | 7.545707073 | 12.77901403 | ENSG00000213700 |
| TTLL1 | 0.752291351 | 1.684466049 | 6.41E-05 | 0.000469268 | 121.504215 | 204.3995494 | ENSG00000100271 |
| IKBIP | 0.751384179 | 1.683407183 | 5.15E-06 | 4.96E-05 | 219.619979 | 369.4645658 | ENSG00000166130 |
| LINC00910 | 0.751195026 | 1.683186485 | 0.034002696 | 0.098987443 | 35.21733346 | 59.55952662 | ENSG00000188825 |
| HHEX | 0.751043012 | 1.68300914 | 0.228129745 | 0.411808562 | 9.941884644 | 16.63823468 | ENSG00000152804 |
| GIHCG | 0.750876067 | 1.682814398 | 0.039704434 | 0.111819174 | 27.97168322 | 46.96679681 | ENSG00000257698 |
| SNCG | 0.750804495 | 1.682730916 | 0.000362255 | 0.002159547 | 88.84067155 | 149.3910791 | ENSG00000173267 |
| NA | 0.750509029 | 1.682386326 | 0.385575263 | 0.578503353 | 5.280315762 | 8.956897339 | ENSG00000278997 |
| SQSTM1 | 0.750330366 | 1.682177992 | 2.78E-25 | 2.70E-23 | 1373.784789 | 2311.511616 | ENSG00000161011 |
| TLR3 | 0.749923563 | 1.681703728 | 0.16737176 | 0.331099179 | 12.10430055 | 20.36693444 | ENSG00000164342 |
| FAM166B | 0.749531985 | 1.68124734 | 0.3098981 | 0.503769234 | 7.347250168 | 12.32135051 | ENSG00000215187 |
| NA | 0.749304739 | 1.680982539 | 0.130655028 | 0.276654894 | 14.55364198 | 24.54537361 | ENSG00000279756 |
| SLC9B2 | 0.748933853 | 1.68055045 | 0.003479564 | 0.015096031 | 56.32943403 | 94.67783183 | ENSG00000164038 |
| NA | 0.748372258 | 1.679896393 | 0.39485121 | 0.587286119 | 4.472680413 | 7.478981666 | ENSG00000259562 |
| LINC02021 | 0.747880644 | 1.679324048 | 0.109421133 | 0.242266309 | 17.59560317 | 29.46987447 | ENSG00000249846 |
| NA | 0.747606679 | 1.679005177 | 0.336864531 | 0.531250771 | 5.769358763 | 9.673341145 | ENSG00000287721 |
| NA | 0.747343693 | 1.678699143 | 0.337123107 | 0.53144847 | 6.367788692 | 10.69728052 | ENSG00000268521 |
| NA | 0.747293316 | 1.678640525 | 0.111137608 | 0.245304802 | 17.70814701 | 29.84920483 | ENSG00000233871 |
| LDLRAD2 | 0.746608169 | 1.677843516 | 0.314841952 | 0.508473455 | 6.61299793 | 11.17762551 | ENSG00000187942 |
| COLCA1 | 0.746245514 | 1.677421804 | 0.427106665 | 0.617568648 | 4.816430366 | 8.094522459 | ENSG00000196167 |
| RASAL2 | 0.74594245 | 1.677069468 | 1.95E-13 | 6.25E-12 | 574.0972121 | 962.5457822 | ENSG00000075391 |
| ZNF805 | 0.74593445 | 1.677060169 | 0.000310655 | 0.001890861 | 92.29039277 | 154.6744904 | ENSG00000204524 |
| TSGA10 | 0.745437236 | 1.676482282 | 0.016674252 | 0.055682988 | 37.915447 | 63.50087928 | ENSG00000135951 |
| NA | 0.745066247 | 1.67605123 | 0.444817965 | 0.633150769 | 3.513438191 | 5.892565286 | ENSG00000230982 |
| NA | 0.7450071 | 1.675982517 | 0.264117291 | 0.453800859 | 8.532760151 | 14.29783251 | ENSG00000258526 |
| KLHL29 | 0.744782987 | 1.675722185 | 1.89E-08 | 2.88E-07 | 252.4409395 | 423.5944471 | ENSG00000119771 |
| GALNT9 | 0.744757448 | 1.675692521 | 0.122345687 | 0.263332361 | 18.91062164 | 31.69863837 | ENSG00000182870 |
| NPTN-IT1 | 0.744448081 | 1.675333229 | 0.155126309 | 0.313158807 | 13.42751198 | 22.49369601 | ENSG00000281183 |
| PTGES3L | 0.743681031 | 1.674442728 | 0.248287424 | 0.43639252 | 8.264174747 | 13.87435016 | ENSG00000267060 |
| NA | 0.743452009 | 1.674176938 | 0.523489241 | 0.699924224 | 2.927635913 | 4.883598058 | ENSG00000237227 |
| NA | 0.743121024 | 1.67379289 | 0.445828794 | 0.633889072 | 3.486905112 | 5.843738377 | ENSG00000286366 |
| SOCS1 | 0.743014146 | 1.673668895 | 2.18E-06 | 2.27E-05 | 376.7236715 | 630.6420453 | ENSG00000185338 |
| ZBTB4 | 0.742809897 | 1.673431963 | 3.38E-12 | 9.20E-11 | 450.9032457 | 754.8983874 | ENSG00000174282 |

| | | | | | | | |
|---|---|---|---|---|---|---|---|
| ADRB2 | 0.74279981 | 1.673420263 | 0.091533567 | 0.212197037 | 18.15615015 | 30.40788298 | ENSG00000169252 |
| PTPN6 | 0.741667393 | 1.672107259 | 1.61E-10 | 3.41E-09 | 328.652572 | 549.0775617 | ENSG00000111679 |
| TRPV1 | 0.741344887 | 1.67173351 | 0.028815468 | 0.086727166 | 46.63454007 | 78.12359292 | ENSG00000196689 |
| LINC02609 | 0.740473118 | 1.670723647 | 0.346763356 | 0.540170237 | 6.189551041 | 10.24330422 | ENSG00000233593 |
| SLA | 0.74036771 | 1.670601583 | 0.094523844 | 0.217034802 | 18.21237322 | 30.4191679 | ENSG00000155926 |
| PKHD1L1 | 0.740334051 | 1.670562607 | 0.429959888 | 0.62009778 | 4.082178128 | 6.841420684 | ENSG00000205038 |
| FKBP9P1 | 0.740285867 | 1.670506813 | 0.42591858 | 0.616651967 | 4.502370406 | 7.471495592 | ENSG00000176826 |
| USH1C | 0.739929301 | 1.670093994 | 0.032301689 | 0.094892931 | 31.03396136 | 51.9072575 | ENSG00000006611 |
| TMEM54 | 0.739839786 | 1.669990373 | 0.014376423 | 0.049352216 | 49.24563552 | 82.18786878 | ENSG00000121900 |
| GABRG1 | 0.739408893 | 1.669491668 | 0.430489862 | 0.620475861 | 4.082178128 | 6.837621838 | ENSG00000163285 |
| LOC105372233 | 0.73915873 | 1.669202204 | 0.36378352 | 0.556819635 | 5.405389557 | 9.054551158 | ENSG00000267751 |
| CACNA1C | 0.73915526 | 1.669198188 | 0.003880256 | 0.016620447 | 60.36919682 | 100.7128558 | ENSG00000151067 |
| MTHFD2 | 0.738930487 | 1.668938147 | 1.97E-30 | 2.88E-28 | 2355.312616 | 3929.984501 | ENSG00000065911 |
| NA | 0.737941902 | 1.667794923 | 0.514329583 | 0.692724692 | 3.113465167 | 5.247518231 | ENSG00000215452 |
| AZIN2 | 0.737840861 | 1.667678122 | 0.098995626 | 0.225016548 | 17.85156092 | 29.69100114 | ENSG00000142920 |
| LINC01238 | 0.737658698 | 1.667467564 | 0.193078278 | 0.366688472 | 11.22781448 | 18.70163523 | ENSG00000237940 |
| NA | 0.737183187 | 1.666918058 | 0.433300036 | 0.623072609 | 4.449304247 | 7.479419699 | ENSG00000261821 |
| PLB1 | 0.73704908 | 1.666763116 | 0.466233371 | 0.652459173 | 3.463528945 | 5.776140473 | ENSG00000163803 |
| PRR16 | 0.737036908 | 1.666749054 | 0.273962399 | 0.46515382 | 7.989275517 | 13.25512214 | ENSG00000184838 |
| CEP112 | 0.736911203 | 1.666603833 | 0.016710659 | 0.055745331 | 43.66280507 | 73.08167956 | ENSG00000154240 |
| ZNF814 | 0.736660589 | 1.666314348 | 9.11E-09 | 1.48E-07 | 259.9429535 | 433.5823318 | ENSG00000204514 |
| KBTBD3 | 0.73604458 | 1.665603009 | 0.072984953 | 0.17885801 | 24.69072892 | 41.05264937 | ENSG00000182359 |
| MGARP | 0.735953703 | 1.665498094 | 0.388592741 | 0.581422849 | 4.747579643 | 7.92547189 | ENSG00000137463 |
| NA | 0.735797848 | 1.66531818 | 0.455970256 | 0.643266615 | 3.77127508 | 6.346541584 | ENSG00000255062 |
| ZMAT4 | 0.735780063 | 1.66529765 | 6.33E-07 | 7.31E-06 | 188.870989 | 314.5892362 | ENSG00000165061 |
| PCDHGC3 | 0.73557442 | 1.665060293 | 5.63E-05 | 0.00041853 | 124.0768488 | 206.1757224 | ENSG00000240184 |
| NA | 0.735464406 | 1.664933328 | 0.530095059 | 0.705701898 | 2.805719032 | 4.669519427 | ENSG00000214192 |
| SH3RF3 | 0.734547692 | 1.663875736 | 0.022374047 | 0.070825037 | 39.99766231 | 66.47438232 | ENSG00000172985 |
| RIN2 | 0.734506656 | 1.663828409 | 0.117603265 | 0.255708461 | 19.37766395 | 32.32491443 | ENSG00000132669 |
| TNFRSF10B | 0.734225463 | 1.663504147 | 1.50E-17 | 7.55E-16 | 1831.110484 | 3045.949321 | ENSG00000120889 |
| MRTFB | 0.733837892 | 1.663057317 | 1.73E-08 | 2.66E-07 | 276.5640109 | 459.9990475 | ENSG00000186260 |
| TRIM72 | 0.732997238 | 1.662088541 | 0.026393446 | 0.080952288 | 35.5128579 | 58.9514719 | ENSG00000177238 |
| GCSAM | 0.732873769 | 1.661946302 | 0.14610115 | 0.300104036 | 15.40359497 | 25.67032762 | ENSG00000174500 |
| NA | 0.732449886 | 1.661458072 | 0.383605665 | 0.576294675 | 5.200618826 | 8.570518945 | ENSG00000228242 |
| FGF13 | 0.732388683 | 1.66138759 | 9.95E-19 | 5.66E-17 | 1149.636298 | 1909.136222 | ENSG00000129682 |
| ZDHHC11B | 0.732295861 | 1.661280701 | 0.311954072 | 0.505553847 | 6.666064088 | 11.1100276 | ENSG00000206077 |
| NA | 0.732238607 | 1.661214773 | 0.49917626 | 0.68011443 | 3.782023595 | 6.32777059 | ENSG00000272848 |
| CR2 | 0.732018118 | 1.660960907 | 2.12E-05 | 0.0001758 | 128.9590707 | 214.0990444 | ENSG00000117322 |
| TOR4A | 0.731641704 | 1.660527602 | 3.51E-06 | 3.53E-05 | 211.7781613 | 351.0726163 | ENSG00000198113 |
| PABPC5 | 0.731478095 | 1.660339301 | 0.011484254 | 0.041252789 | 43.37390271 | 71.98535574 | ENSG00000174740 |
| EPB41L4A-DT | 0.731308993 | 1.660144699 | 0.142900567 | 0.29539934 | 14.94098735 | 24.82661211 | ENSG00000278921 |
| SLFN5 | 0.730646615 | 1.659382659 | 0.001025926 | 0.005388073 | 89.0485992 | 147.4874825 | ENSG00000166750 |
| SOX4 | 0.730640389 | 1.659375498 | 2.73E-34 | 4.97E-32 | 5295.344605 | 8785.773259 | ENSG00000124766 |
| EDN1 | 0.73047949 | 1.659190444 | 0.347413485 | 0.54072072 | 6.437917191 | 10.63716869 | ENSG00000078401 |
| C19orf71 | 0.730116022 | 1.658772485 | 0.051720855 | 0.137137959 | 28.24214776 | 47.02084828 | ENSG00000183397 |
| ITPR1 | 0.730020847 | 1.658663059 | 0.00014003 | 0.000935252 | 169.442326 | 280.8988491 | ENSG00000150995 |
| GTSF1 | 0.729687092 | 1.658279386 | 0.392022422 | 0.584626937 | 5.088172683 | 8.386496228 | ENSG00000170627 |
| NA | 0.729139024 | 1.657649539 | 0.172429822 | 0.338575518 | 149.7602682 | 248.3297944 | ENSG00000235688 |
| RHBDF1 | 0.72880527 | 1.657266102 | 1.31E-11 | 3.25E-10 | 548.906406 | 910.3922974 | ENSG00000007384 |
| ZBTB32 | 0.728760499 | 1.657214673 | 0.183584706 | 0.354138107 | 15.17091569 | 25.11197588 | ENSG00000011590 |
| CASP10 | 0.728671054 | 1.657111931 | 0.015468925 | 0.05236549 | 43.25386497 | 71.63228246 | ENSG00000003400 |
| CRISPLD2 | 0.728425305 | 1.656829683 | 0.000166136 | 0.001090739 | 101.6636352 | 168.4212424 | ENSG00000103196 |
| SLC25A14 | 0.728324017 | 1.656713365 | 4.08E-06 | 4.04E-05 | 188.3259335 | 311.4376592 | ENSG00000102078 |
| NA | 0.728164705 | 1.656530429 | 0.283249837 | 0.474305963 | 8.558015454 | 14.21949934 | ENSG00000231551 |
| NA | 0.727592101 | 1.655873085 | 0.159282488 | 0.319184953 | 13.7321012 | 22.71113546 | ENSG00000279249 |
| MAPK6-DT | 0.727352684 | 1.655598314 | 0.452249903 | 0.639700809 | 3.764961254 | 6.290228601 | ENSG00000259438 |
| NR4A2 | 0.727135786 | 1.655349426 | 0.395507925 | 0.587932618 | 6.028473429 | 9.988434408 | ENSG00000153234 |
| VEPH1 | 0.727036158 | 1.655235116 | 0.014520332 | 0.049735693 | 55.63494389 | 92.01766148 | ENSG00000197415 |
| MYO5A | 0.726775424 | 1.654935998 | 3.04E-14 | 1.07E-12 | 685.6798594 | 1134.16034 | ENSG00000197535 |
| SUSD2 | 0.726471186 | 1.654587039 | 2.45E-07 | 3.06E-06 | 206.8867845 | 342.3191655 | ENSG00000099994 |
| GALNT10 | 0.726303213 | 1.654394406 | 3.30E-12 | 9.01E-11 | 593.3359945 | 980.6834341 | ENSG00000164574 |
| HACD4 | 0.726259129 | 1.654343854 | 0.059852537 | 0.15360107 | 24.25663121 | 40.03195918 | ENSG00000188921 |
| TMC7 | 0.725978601 | 1.654022203 | 0.003330949 | 0.014546609 | 66.29213929 | 109.5531135 | ENSG00000170537 |
| TXNIP | 0.725674746 | 1.653673876 | 0.001280951 | 0.00647957 | 107.8738114 | 177.7923839 | ENSG00000265972 |
| LOC651337 | 0.725414079 | 1.653375116 | 0.366597811 | 0.559588522 | 6.405070286 | 10.53962649 | ENSG00000255585 |
| ESYT2 | 0.725303171 | 1.653248017 | 4.25E-18 | 2.27E-16 | 966.025706 | 1596.217492 | ENSG00000117868 |
| ZNF502 | 0.72525234 | 1.653189768 | 0.130059868 | 0.275664291 | 16.87841327 | 27.80402562 | ENSG00000196653 |
| APC2 | 0.724782043 | 1.652650941 | 0.009089126 | 0.033970127 | 50.14431768 | 82.84712354 | ENSG00000115266 |
| TRAPPC6A | 0.724238595 | 1.652028522 | 6.80E-14 | 2.30E-12 | 654.6915528 | 1082.124975 | ENSG00000007255 |
| ZCCHC4 | 0.724173048 | 1.651953466 | 4.31E-05 | 0.000329986 | 124.9609265 | 206.3917764 | ENSG00000168228 |
| FGF11 | 0.724075077 | 1.651841288 | 0.111988575 | 0.246595728 | 16.86262871 | 27.85340218 | ENSG00000161958 |
| NA | 0.723619623 | 1.65131989 | 7.96E-12 | 2.06E-10 | 392.5853116 | 648.024112 | ENSG00000224032 |
| ROCK1P1 | 0.723425077 | 1.651097227 | 0.465666172 | 0.651911536 | 3.804121985 | 6.320284516 | ENSG00000263006 |
| NA | 0.723149304 | 1.650781647 | 0.24150171 | 0.427662256 | 9.613919255 | 15.83217279 | ENSG00000241556 |

| | | | | | | | |
|---|---|---|---|---|---|---|---|
| MAN1A1 | 0.722828138 | 1.650414199 | 8.39E-06 | 7.64E-05 | 142.9566004 | 235.8723862 | ENSG00000111885 |
| NA | 0.722800439 | 1.650382512 | 0.23727296 | 0.422603982 | 10.17950228 | 16.78840264 | ENSG00000223551 |
| NA | 0.722701178 | 1.650268965 | 0.472581386 | 0.657668074 | 3.572818175 | 5.855023298 | ENSG00000258670 |
| NA | 0.722533198 | 1.650076828 | 0.047675831 | 0.128902093 | 36.44241013 | 60.26556157 | ENSG00000266869 |
| NA | 0.722490034 | 1.65002746 | 0.005225843 | 0.021399863 | 80.08352278 | 132.1762714 | ENSG00000273151 |
| VPS13D | 0.722307698 | 1.649818933 | 1.77E-08 | 2.71E-07 | 722.9989057 | 1192.456952 | ENSG00000048707 |
| IL1R1 | 0.722156776 | 1.649646353 | 0.26369476 | 0.453369079 | 8.528325462 | 14.10676175 | ENSG00000115594 |
| SH3BP5-AS1 | 0.722034847 | 1.64950694 | 0.25210882 | 0.44090874 | 9.203197716 | 15.2133828 | ENSG00000224660 |
| ADAMTS20 | 0.721959798 | 1.649421134 | 5.05E-05 | 0.000379408 | 134.9671294 | 222.294908 | ENSG00000173157 |
| DMD | 0.721587837 | 1.648995929 | 0.00019192 | 0.001238522 | 131.165082 | 215.9455065 | ENSG00000198947 |
| LOC283177 | 0.721460381 | 1.648850254 | 0.141875325 | 0.293767989 | 16.96748325 | 28.04166177 | ENSG00000255545 |
| NFASC | 0.721359909 | 1.648735428 | 7.29E-09 | 1.20E-07 | 313.4558223 | 517.158275 | ENSG00000163531 |
| FGF13-AS1 | 0.721274926 | 1.648638311 | 0.018952332 | 0.061668278 | 38.53853087 | 63.47819782 | ENSG00000226031 |
| RNF43 | 0.721190135 | 1.64854142 | 0.253048992 | 0.441715576 | 8.889137755 | 14.62409907 | ENSG00000108375 |
| SMTN | 0.721042218 | 1.648372406 | 2.03E-08 | 3.07E-07 | 260.9381018 | 430.0843947 | ENSG00000183963 |
| NA | 0.720921073 | 1.648233995 | 0.450599115 | 0.638193795 | 4.188310445 | 6.875163826 | ENSG00000271727 |
| ENG | 0.719973308 | 1.647151559 | 0.001752952 | 0.008487257 | 78.97602599 | 130.1712305 | ENSG00000106991 |
| NA | 0.719672072 | 1.646807668 | 0.25924685 | 0.448174714 | 8.962423168 | 14.70678074 | ENSG00000231233 |
| AK5 | 0.719327994 | 1.646414958 | 0.002219736 | 0.010378902 | 75.59139722 | 124.1347724 | ENSG00000154027 |
| GRIP2 | 0.719308117 | 1.646392274 | 0.000577015 | 0.003250579 | 191.0850956 | 315.1564771 | ENSG00000144596 |
| NA | 0.719204539 | 1.646274075 | 0.082985122 | 0.197033852 | 23.57228822 | 38.7754966 | ENSG00000259363 |
| MAP3K20 | 0.719033211 | 1.646078583 | 1.65E-07 | 2.12E-06 | 269.9710595 | 443.6367865 | ENSG00000091436 |
| ASH1L | 0.718542749 | 1.645519073 | 5.70E-09 | 9.51E-08 | 426.9882301 | 702.2610196 | ENSG00000116539 |
| NA | 0.717888715 | 1.64477326 | 0.220994058 | 0.402624976 | 13.1323935 | 21.53311766 | ENSG00000232022 |
| TPO | 0.717837388 | 1.644714744 | 0.172783105 | 0.339053687 | 17.88944388 | 29.4326589 | ENSG00000115705 |
| NA | 0.717777743 | 1.644646749 | 0.411322782 | 0.602835398 | 4.747579643 | 7.79396331 | ENSG00000273381 |
| TSSK6 | 0.717057404 | 1.64382578 | 0.020685485 | 0.066456019 | 40.00081922 | 65.94729742 | ENSG00000178093 |
| NA | 0.717034419 | 1.643799591 | 0.095800921 | 0.219396301 | 20.6294691 | 34.01267185 | ENSG00000243517 |
| LINC00624 | 0.71676651 | 1.643494366 | 0.329714223 | 0.524137869 | 6.976967136 | 11.51897584 | ENSG00000278811 |
| FTCDNL1 | 0.716709838 | 1.643429807 | 0.199149431 | 0.375448638 | 11.13558759 | 18.32654176 | ENSG00000226124 |
| NA | 0.716568103 | 1.643268358 | 0.337869862 | 0.531959964 | 6.358317954 | 10.49872369 | ENSG00000259972 |
| FAM160A1 | 0.716565383 | 1.643265261 | 1.45E-12 | 4.16E-11 | 600.950575 | 987.4039388 | ENSG00000164142 |
| NA | 0.71617845 | 1.642824593 | 0.442774706 | 0.631223609 | 4.185153532 | 6.837621838 | ENSG00000213862 |
| ALS2CL | 0.715619866 | 1.642188646 | 0.06605756 | 0.165553823 | 157.1391852 | 258.1966826 | ENSG00000178038 |
| C16orf96 | 0.715222233 | 1.641736091 | 0.048153977 | 0.129932839 | 28.47147473 | 46.61839844 | ENSG00000205832 |
| PWRN1 | 0.714926392 | 1.641399469 | 0.320940505 | 0.514737913 | 7.44706866 | 12.19027996 | ENSG00000259905 |
| HELB | 0.714895841 | 1.641364711 | 1.30E-05 | 0.000113402 | 154.3404564 | 253.26917 | ENSG00000127311 |
| RASEF | 0.714782569 | 1.641235846 | 0.001154638 | 0.005956042 | 82.54676963 | 135.1594188 | ENSG00000165105 |
| RORA | 0.71466572 | 1.641102921 | 0.016064894 | 0.053998321 | 45.29809962 | 74.37513448 | ENSG00000069667 |
| LCN2 | 0.714082512 | 1.640439641 | 0.295757429 | 0.488644305 | 7.890734802 | 12.9513138 | ENSG00000148346 |
| APOBEC3B-AS1 | 0.71408121 | 1.640438161 | 0.541611527 | 0.714474773 | 2.930792826 | 4.759687172 | ENSG00000249310 |
| IGIP | 0.713642389 | 1.639939269 | 0.388076677 | 0.581174299 | 6.309686484 | 10.38609772 | ENSG00000182700 |
| DOCK3 | 0.713447691 | 1.639717966 | 1.39E-05 | 0.000120708 | 250.4949299 | 411.0208549 | ENSG00000088538 |
| RTL10 | 0.713285664 | 1.639533823 | 5.08E-14 | 1.74E-12 | 511.6821946 | 838.9179064 | ENSG00000215012 |
| DLGAP1 | 0.713063332 | 1.639281175 | 1.10E-08 | 1.75E-07 | 273.4063716 | 447.8693888 | ENSG00000170579 |
| PMEPA1 | 0.712712302 | 1.638882362 | 5.36E-10 | 1.04E-08 | 499.5876805 | 818.7157538 | ENSG00000124225 |
| PARD6B | 0.712406596 | 1.63853512 | 0.002585794 | 0.011780741 | 120.3280629 | 197.0360762 | ENSG00000124171 |
| LTBP4 | 0.712323151 | 1.638440351 | 2.33E-22 | 1.79E-20 | 1712.942078 | 2807.572237 | ENSG00000090006 |
| CD37 | 0.711703216 | 1.637736454 | 0.164541114 | 0.326930211 | 18.04044939 | 29.5002568 | ENSG00000104894 |
| GAL3ST4 | 0.711555187 | 1.637568422 | 0.000409916 | 0.002404237 | 90.57667905 | 148.4379869 | ENSG00000197093 |
| NA | 0.71107666 | 1.637025347 | 0.324628432 | 0.518767319 | 8.342496209 | 13.58887478 | ENSG00000227001 |
| GAS2 | 0.711017377 | 1.63695808 | 0.360210275 | 0.553439213 | 6.719130247 | 10.97472018 | ENSG00000148935 |
| PRMT8 | 0.710372211 | 1.636226204 | 0.146021533 | 0.299973741 | 17.34408011 | 28.32246224 | ENSG00000111218 |
| SP2-AS1 | 0.709932067 | 1.635727093 | 0.504005721 | 0.683882985 | 3.232225135 | 5.292546294 | ENSG00000234494 |
| NA | 0.709866431 | 1.635652677 | 0.3437652 | 0.537484749 | 6.361474866 | 10.43861186 | ENSG00000213280 |
| IL13RA1 | 0.709602196 | 1.635353128 | 2.50E-12 | 6.90E-11 | 529.3123284 | 864.8202705 | ENSG00000131724 |
| NA | 0.709598966 | 1.635349467 | 0.526021056 | 0.702433961 | 2.927635913 | 4.729631257 | ENSG00000250938 |
| TMEM45B | 0.709491719 | 1.635227903 | 0.51176424 | 0.690876133 | 3.552598922 | 5.764855553 | ENSG00000151715 |
| PRR14L | 0.709414565 | 1.635140455 | 1.16E-12 | 3.39E-11 | 869.6918073 | 1422.036284 | ENSG00000183530 |
| SMIM1 | 0.709240608 | 1.634943305 | 0.162606541 | 0.3240591 | 15.08194341 | 24.75901421 | ENSG00000235169 |
| NA | 0.709143272 | 1.634833002 | 0.003006007 | 0.01334451 | 83.75339332 | 137.0869623 | ENSG00000176654 |
| STARD8 | 0.708881904 | 1.634536852 | 4.94E-05 | 0.000372941 | 130.4793635 | 213.0487364 | ENSG00000130052 |
| CAPN11 | 0.708872498 | 1.634526195 | 0.06678079 | 0.167010546 | 27.51095474 | 44.84058489 | ENSG00000137225 |
| SLC12A4 | 0.708869843 | 1.634523187 | 6.57E-20 | 4.17E-18 | 1143.739821 | 1869.615142 | ENSG00000124067 |
| NA | 0.707994985 | 1.633532304 | 0.539879138 | 0.713042111 | 2.874569755 | 4.650748432 | ENSG00000285407 |
| RASGRP4 | 0.707885186 | 1.633407986 | 0.009056959 | 0.033877232 | 50.0476561 | 81.88362241 | ENSG00000171777 |
| CFAP45 | 0.707070261 | 1.632485594 | 0.563025304 | 0.731313406 | 2.81518977 | 4.662471386 | ENSG00000213085 |
| WIPF1 | 0.706945673 | 1.632344622 | 5.81E-05 | 0.000430885 | 147.5988444 | 241.1114307 | ENSG00000115935 |
| NA | 0.706934589 | 1.632332081 | 0.225716779 | 0.408778071 | 12.87277517 | 21.12103185 | ENSG00000203644 |
| SLFNL1-AS1 | 0.70692258 | 1.632318494 | 0.208131369 | 0.387222123 | 11.15580684 | 18.18754711 | ENSG00000281207 |
| ERN1 | 0.706684888 | 1.632049583 | 0.000388225 | 0.002293708 | 103.9890078 | 169.7044001 | ENSG00000178607 |
| NA | 0.706057188 | 1.631339652 | 0.022648545 | 0.071571719 | 40.46776383 | 66.07825634 | ENSG00000253320 |
| FGD6 | 0.705983805 | 1.631256675 | 3.36E-12 | 9.15E-11 | 535.0080935 | 872.1856385 | ENSG00000180263 |
| AZIN1 | 0.705827316 | 1.631079742 | 9.44E-18 | 4.87E-16 | 4567.199771 | 7448.074494 | ENSG00000155096 |

| | | | | | | | |
|---|---|---|---|---|---|---|---|
| VPS13B | 0.70546575 | 1.630671015 | 1.57E-08 | 2.44E-07 | 323.0259507 | 526.6728628 | ENSG00000132549 |
| VCPIP1 | 0.705060217 | 1.630212707 | 1.10E-08 | 1.75E-07 | 306.3908675 | 499.3145714 | ENSG00000175073 |
| NA | 0.705049139 | 1.63020019 | 0.355019964 | 0.548282644 | 6.573837199 | 10.81001811 | ENSG00000280279 |
| MUC3A | 0.704887242 | 1.630017262 | 7.88E-08 | 1.07E-06 | 258.543202 | 421.6462604 | ENSG00000169894 |
| CYP1B1-AS1 | 0.704852635 | 1.629978161 | 0.49687216 | 0.678123082 | 3.176002064 | 5.187406401 | ENSG00000232973 |
| FAM50B | 0.704502326 | 1.629582425 | 0.002881677 | 0.012863439 | 66.21066092 | 107.9558502 | ENSG00000145945 |
| NA | 0.704137137 | 1.629169981 | 0.397128325 | 0.589301551 | 5.967815668 | 9.797363651 | ENSG00000230897 |
| NA | 0.704013179 | 1.629030007 | 0.212707774 | 0.392423238 | 13.9678397 | 22.87313799 | ENSG00000154608 |
| TUBA3D | 0.703954139 | 1.628963343 | 0.452039054 | 0.639455292 | 5.203775739 | 8.379010155 | ENSG00000075886 |
| SDCBP2-AS1 | 0.703800886 | 1.628790313 | 0.241520494 | 0.427662256 | 10.4215546 | 17.07019079 | ENSG00000234684 |
| DTNA | 0.703381536 | 1.628316939 | 2.12E-05 | 0.000176273 | 172.5384205 | 281.2254505 | ENSG00000134769 |
| NA | 0.703159055 | 1.628065852 | 0.475972668 | 0.660700826 | 3.804121985 | 6.203859703 | ENSG00000244055 |
| PSD3 | 0.703127362 | 1.628030087 | 8.35E-09 | 1.36E-07 | 343.5661393 | 559.8069767 | ENSG00000156011 |
| CLTRN | 0.702623164 | 1.627461217 | 0.320856595 | 0.514692413 | 8.148571691 | 13.29646297 | ENSG00000147003 |
| CD2BP2-DT | 0.702586386 | 1.627419729 | 0.120201495 | 0.259894366 | 19.47304775 | 31.76612465 | ENSG00000260219 |
| KLLN | 0.702257896 | 1.627049221 | 0.303548532 | 0.497169362 | 9.120441565 | 14.91717215 | ENSG00000227268 |
| TMEM106A | 0.702188842 | 1.626971345 | 0.16002135 | 0.320284634 | 14.00384352 | 22.8124765 | ENSG00000184988 |
| SYT12 | 0.701888117 | 1.626632244 | 2.62E-05 | 0.000212565 | 169.0184731 | 274.6523429 | ENSG00000173227 |
| NA | 0.70187375 | 1.626616045 | 0.102267346 | 0.23047855 | 22.08218901 | 35.98210581 | ENSG00000263826 |
| NA | 0.701803918 | 1.626537313 | 0.400999727 | 0.592731097 | 5.395918818 | 8.844709406 | ENSG00000280322 |
| NA | 0.701453981 | 1.626142831 | 0.439047192 | 0.627394037 | 5.608281151 | 9.231087801 | ENSG00000268584 |
| RRAS | 0.701334772 | 1.62600847 | 1.10E-07 | 1.46E-06 | 264.0402171 | 429.7348341 | ENSG00000126458 |
| NFAT5 | 0.701150622 | 1.625800934 | 5.41E-07 | 6.36E-06 | 218.745079 | 355.633014 | ENSG00000102908 |
| XKR4 | 0.700579088 | 1.625156989 | 0.111474641 | 0.24587302 | 19.19499161 | 31.08254792 | ENSG00000206579 |
| NA | 0.700419788 | 1.624977552 | 0.142955286 | 0.295443049 | 22.62241903 | 36.65665913 | ENSG00000259048 |
| LMO7 | 0.699818299 | 1.624300206 | 4.32E-07 | 5.17E-06 | 224.7110155 | 364.6963251 | ENSG00000136153 |
| TMIGD2 | 0.699742114 | 1.624214434 | 0.205665862 | 0.384061571 | 12.84308518 | 20.96315458 | ENSG00000167664 |
| CRYBG3 | 0.699685224 | 1.624150387 | 9.99E-05 | 0.000695833 | 146.4698431 | 237.7906201 | ENSG00000080200 |
| APOLD1 | 0.699586265 | 1.624038985 | 0.476807559 | 0.661190767 | 4.125773548 | 6.653599121 | ENSG00000178878 |
| LINC00639 | 0.699398612 | 1.623827759 | 0.493736943 | 0.675874798 | 3.854031231 | 6.256373839 | ENSG00000259070 |
| CRTC1 | 0.699136623 | 1.623532903 | 2.34E-09 | 4.13E-08 | 350.9214621 | 569.4663015 | ENSG00000105662 |
| TMEM59L | 0.699022602 | 1.623404595 | 2.35E-09 | 4.13E-08 | 357.871122 | 581.0035474 | ENSG00000105696 |
| ZNF460 | 0.698066699 | 1.622329314 | 0.036979505 | 0.105588992 | 32.96309662 | 53.39265925 | ENSG00000197714 |
| LOC103611081 | 0.697974942 | 1.622226134 | 0.189113199 | 0.361230962 | 13.19620817 | 21.32327591 | ENSG00000255455 |
| KIAA2026 | 0.697845066 | 1.622080103 | 4.48E-06 | 4.39E-05 | 191.4585355 | 310.7959016 | ENSG00000183354 |
| XKR5 | 0.697830479 | 1.622063702 | 0.002208137 | 0.010337361 | 93.98328587 | 152.3339766 | ENSG00000275591 |
| NA | 0.697755445 | 1.621979342 | 0.004420632 | 0.01854069 | 68.98235296 | 112.0662535 | ENSG00000246695 |
| IL18BP | 0.696906595 | 1.621025285 | 0.004689162 | 0.019451417 | 111.2326812 | 180.8003719 | ENSG00000137496 |
| PHYHD1 | 0.696401082 | 1.620457386 | 0.000819705 | 0.004433153 | 90.71240366 | 146.8021939 | ENSG00000175287 |
| SEMA6A | 0.69638387 | 1.620438053 | 1.49E-26 | 1.58E-24 | 4622.351362 | 7489.517945 | ENSG00000092421 |
| KLHL24 | 0.696364532 | 1.620416333 | 2.87E-07 | 3.54E-06 | 319.4169333 | 516.7580463 | ENSG00000114796 |
| NA | 0.696162675 | 1.620189625 | 0.323812383 | 0.517909796 | 7.056566374 | 11.41752317 | ENSG00000269378 |
| NA | 0.696127747 | 1.620150401 | 0.470792559 | 0.656213378 | 3.768118167 | 6.113691958 | ENSG00000284731 |
| SLC44A4 | 0.696102093 | 1.620121592 | 0.043101424 | 0.119344957 | 31.67598671 | 51.33674477 | ENSG00000204385 |
| MYH6 | 0.695974734 | 1.619978576 | 0.361762316 | 0.554809321 | 6.649001748 | 10.8287891 | ENSG00000197616 |
| NA | 0.695885652 | 1.619878551 | 0.335072751 | 0.529337273 | 8.273645486 | 13.49220864 | ENSG00000280211 |
| NPAS3 | 0.695702465 | 1.619672878 | 0.412229447 | 0.603734538 | 5.346009572 | 8.724485747 | ENSG00000151322 |
| NA | 0.695121449 | 1.61902072 | 0.505804694 | 0.685426936 | 3.780745819 | 6.124976879 | ENSG00000255122 |
| NA | 0.694800914 | 1.61866105 | 0.470344723 | 0.655885118 | 3.877407397 | 6.260172686 | ENSG00000277715 |
| ITGB1 | 0.694636936 | 1.618477081 | 7.89E-24 | 6.80E-22 | 5692.774782 | 9212.569901 | ENSG00000150093 |
| TINCR | 0.694373972 | 1.618182104 | 0.153582854 | 0.31102112 | 16.99411403 | 27.51909989 | ENSG00000223573 |
| NA | 0.694246187 | 1.618038782 | 0.118586299 | 0.257332686 | 19.70119465 | 31.97216755 | ENSG00000241860 |
| TNC | 0.694106633 | 1.617882274 | 7.94E-05 | 0.000566857 | 131.3160875 | 212.2832424 | ENSG00000041982 |
| ZBTB37 | 0.694082001 | 1.617854651 | 0.000314111 | 0.001908138 | 157.8220324 | 254.9456425 | ENSG00000185278 |
| P2RY1 | 0.69401375 | 1.617778115 | 5.59E-09 | 9.33E-08 | 670.0521614 | 1083.921546 | ENSG00000169860 |
| GJB7 | 0.693913351 | 1.617665536 | 0.241168739 | 0.427440453 | 11.32447606 | 18.21760302 | ENSG00000164411 |
| ANKAR | 0.693161978 | 1.616823256 | 0.250531861 | 0.438991135 | 12.37100682 | 20.03686903 | ENSG00000151687 |
| GEM | 0.693134229 | 1.616792158 | 0.002669778 | 0.012080645 | 81.3816743 | 131.4038007 | ENSG00000164949 |
| NA | 0.692919657 | 1.61655171 | 0.222610011 | 0.404733563 | 12.11820598 | 19.67349851 | ENSG00000249684 |
| BCL2L13 | 0.692826084 | 1.616446865 | 2.82E-14 | 9.97E-13 | 748.047914 | 1208.357083 | ENSG00000099968 |
| ADGRL3 | 0.69263758 | 1.616235672 | 5.38E-07 | 6.33E-06 | 240.0714211 | 387.4933914 | ENSG00000150471 |
| ZNF185 | 0.69251383 | 1.616097042 | 5.75E-08 | 8.03E-07 | 282.6516915 | 456.3960822 | ENSG00000147394 |
| NA | 0.69244213 | 1.616016725 | 0.027543076 | 0.08389209 | 55.86101625 | 90.28695454 | ENSG00000280287 |
| LOC100507071 | 0.69227857 | 1.615833527 | 0.420090586 | 0.611384442 | 5.627222628 | 9.201031886 | ENSG00000254064 |
| TPCN1 | 0.692272921 | 1.6158272 | 7.73E-10 | 1.46E-08 | 499.2468095 | 806.279436 | ENSG00000186815 |
| OTUD1 | 0.692130568 | 1.615667772 | 0.000324667 | 0.001965824 | 111.2843719 | 179.6178622 | ENSG00000165312 |
| KDM6A | 0.691933203 | 1.615446758 | 2.84E-14 | 1.00E-12 | 991.2385354 | 1600.857568 | ENSG00000147050 |
| NA | 0.691920848 | 1.615432924 | 0.403380719 | 0.595101349 | 5.399075731 | 8.769625429 | ENSG00000255050 |
| CUTALP | 0.691685433 | 1.615169344 | 0.016570232 | 0.055385516 | 43.00046277 | 69.41210395 | ENSG00000226752 |
| SLC30A1 | 0.691389215 | 1.614837747 | 1.52E-06 | 1.63E-05 | 483.5426988 | 780.824612 | ENSG00000170385 |
| FRMD6 | 0.69120841 | 1.614635381 | 3.46E-07 | 4.19E-06 | 249.3313077 | 402.7072123 | ENSG00000139926 |
| SMAD7 | 0.69113156 | 1.614549374 | 5.99E-17 | 2.82E-15 | 1022.003776 | 1650.453486 | ENSG00000101665 |
| NA | 0.690268993 | 1.613584345 | 0.330994346 | 0.525254238 | 8.257860921 | 13.40583975 | ENSG00000269399 |
| NA | 0.690059405 | 1.613349949 | 0.296588554 | 0.489542614 | 9.388929271 | 15.04801946 | ENSG00000235493 |

| | | | | | | | |
|---|---|---|---|---|---|---|---|
| KMT2D | 0.68981057 | 1.613071704 | 0.001386006 | 0.006914735 | 1917.450356 | 3093.072537 | ENSG00000167548 |
| TES | 0.689458988 | 1.612678649 | 5.22E-12 | 1.40E-10 | 513.7502114 | 827.793854 | ENSG00000135269 |
| ZSCAN23 | 0.689359899 | 1.612567889 | 0.042552864 | 0.118118134 | 36.68574024 | 59.29001107 | ENSG00000187987 |
| UBR4 | 0.688910945 | 1.61206615 | 0.002821594 | 0.012639449 | 4228.930687 | 6817.24014 | ENSG00000127481 |
| TGFBR2 | 0.688805747 | 1.611948606 | 1.25E-06 | 1.37E-05 | 217.5850197 | 350.4724857 | ENSG00000163513 |
| DCAKD | 0.688032675 | 1.61108507 | 1.38E-14 | 5.07E-13 | 691.1098877 | 1113.270741 | ENSG00000172992 |
| MBOAT1 | 0.687986471 | 1.611033474 | 2.15E-07 | 2.71E-06 | 248.4304759 | 400.0380672 | ENSG00000172197 |
| NHS | 0.687789086 | 1.610813073 | 3.44E-07 | 4.18E-06 | 306.8806847 | 493.7432142 | ENSG00000188158 |
| NA | 0.687400972 | 1.61037979 | 0.532912763 | 0.707927547 | 3.447744381 | 5.600041864 | ENSG00000237846 |
| ZNF516 | 0.687209427 | 1.610165996 | 5.33E-11 | 1.20E-09 | 516.6036749 | 832.401908 | ENSG00000101493 |
| NA | 0.68678502 | 1.609692393 | 0.134621372 | 0.282445706 | 17.26260173 | 27.7031226 | ENSG00000224699 |
| SERINC3 | 0.686329463 | 1.609184183 | 5.73E-14 | 1.95E-12 | 745.1834164 | 1198.126999 | ENSG00000132824 |
| CXCL1 | 0.686136314 | 1.608968759 | 0.50161879 | 0.682148554 | 3.777588906 | 6.072351123 | ENSG00000163739 |
| SLC2A3 | 0.685817657 | 1.608613416 | 6.87E-23 | 5.59E-21 | 12379.75373 | 19912.25716 | ENSG00000059804 |
| SPART | 0.685441149 | 1.608193662 | 1.74E-16 | 7.79E-15 | 886.6379738 | 1425.472683 | ENSG00000133104 |
| NA | 0.685419504 | 1.608169534 | 0.567265195 | 0.734612098 | 2.788656691 | 4.538560499 | ENSG00000267422 |
| CTH | 0.684417809 | 1.607053334 | 4.74E-08 | 6.75E-07 | 332.5379769 | 533.6768038 | ENSG00000116761 |
| MED13 | 0.684252416 | 1.606869109 | 2.18E-09 | 3.86E-08 | 502.2571412 | 806.5403175 | ENSG00000108510 |
| MMP19 | 0.684010487 | 1.606599672 | 0.163139306 | 0.324827969 | 17.02182718 | 27.29472402 | ENSG00000123342 |
| ARHGEF39 | 0.683745083 | 1.606304142 | 4.10E-06 | 4.06E-05 | 205.4029014 | 329.6031861 | ENSG00000137135 |
| MIR647 | 0.683650771 | 1.606199139 | 0.476116927 | 0.660802091 | 4.125773548 | 6.665322075 | ENSG00000207554 |
| FZD6 | 0.683608287 | 1.606151841 | 3.92E-05 | 0.000304838 | 141.1101235 | 226.430907 | ENSG00000164930 |
| ZBTB10 | 0.683446662 | 1.605971913 | 6.29E-06 | 5.95E-05 | 243.1078274 | 390.6013258 | ENSG00000205189 |
| OR2A7 | 0.682900461 | 1.605364012 | 0.370185979 | 0.563441482 | 6.411384112 | 10.2507903 | ENSG00000243896 |
| ZNF793-AS1 | 0.68260154 | 1.60503142 | 0.2643733 | 0.454072351 | 10.20603536 | 16.43631704 | ENSG00000266916 |
| MYEF2 | 0.682209723 | 1.604595574 | 3.98E-15 | 1.56E-13 | 954.3818786 | 1530.57183 | ENSG00000104177 |
| NA | 0.681916301 | 1.604269258 | 0.13622364 | 0.284930476 | 17.70311096 | 28.43465018 | ENSG00000278974 |
| NMNAT1 | 0.68190742 | 1.604259381 | 0.020874375 | 0.066943395 | 57.66033476 | 92.78825993 | ENSG00000173614 |
| NA | 0.681635587 | 1.603957135 | 0.385963462 | 0.578945192 | 5.733354945 | 9.159253017 | ENSG00000164616 |
| NA | 0.681026563 | 1.603280178 | 0.001085505 | 0.00566247 | 88.16274008 | 141.4430374 | ENSG00000279088 |
| FRYL | 0.68078314 | 1.603009683 | 1.91E-08 | 2.91E-07 | 513.7895827 | 822.7895774 | ENSG00000075539 |
| ALG1L | 0.680707292 | 1.602925408 | 0.067810776 | 0.168789272 | 26.51255178 | 42.4892242 | ENSG00000189366 |
| TTN | 0.680493245 | 1.602687607 | 0.008900139 | 0.033378224 | 86.33195014 | 138.469964 | ENSG00000155657 |
| ATG16L2 | 0.680457027 | 1.602647372 | 0.113137031 | 0.248651932 | 232.1852186 | 372.2479321 | ENSG00000168010 |
| C1RL-AS1 | 0.680161775 | 1.602319419 | 0.115017077 | 0.251768809 | 24.20356505 | 38.93045108 | ENSG00000205885 |
| ID2 | 0.680092813 | 1.602242829 | 2.74E-07 | 3.39E-06 | 270.2452596 | 432.5870226 | ENSG00000115738 |
| NA | 0.679591236 | 1.601685879 | 0.451212243 | 0.638719431 | 4.763364208 | 7.655518308 | ENSG00000279481 |
| NWD2 | 0.679236492 | 1.60129209 | 0.524882171 | 0.701381843 | 3.811713587 | 6.083636043 | ENSG00000174145 |
| TTC23L | 0.678415967 | 1.600381623 | 0.256060608 | 0.444941858 | 10.86700218 | 17.4190272 | ENSG00000205838 |
| FGF2 | 0.678337313 | 1.600294373 | 1.89E-15 | 7.67E-14 | 876.6686464 | 1403.500221 | ENSG00000138685 |
| CDHR3 | 0.678170675 | 1.600109543 | 0.000223276 | 0.001417189 | 127.4330653 | 204.2777255 | ENSG00000128536 |
| ACVR1C | 0.678087169 | 1.600016928 | 0.167636749 | 0.331446501 | 15.05215572 | 24.01251449 | ENSG00000123612 |
| TEPP | 0.677690961 | 1.599577576 | 0.2556036 | 0.444446675 | 11.74348826 | 18.86319973 | ENSG00000159648 |
| MED13L | 0.677656817 | 1.599539719 | 2.14E-09 | 3.79E-08 | 663.0491724 | 1060.292014 | ENSG00000123066 |
| NA | 0.677412214 | 1.599268546 | 0.443580794 | 0.631972743 | 4.747579643 | 7.61797632 | ENSG00000232732 |
| NA | 0.676972572 | 1.598781265 | 0.437324755 | 0.626236344 | 5.412981159 | 8.668172763 | ENSG00000235513 |
| NA | 0.676839559 | 1.598633868 | 0.048892161 | 0.131404772 | 39.96098207 | 63.93597297 | ENSG00000205534 |
| PCDHA13 | 0.675308923 | 1.596938687 | 0.501359934 | 0.681946647 | 4.638192715 | 7.351709965 | ENSG00000239389 |
| ASPDH | 0.675161993 | 1.596776056 | 0.546897561 | 0.718573613 | 3.235382048 | 5.131093418 | ENSG00000204653 |
| SCN1B | 0.674953057 | 1.596544822 | 0.022201818 | 0.070388043 | 44.48287267 | 70.78444169 | ENSG00000105711 |
| SLC6A17 | 0.674771712 | 1.596344152 | 0.296860495 | 0.489677146 | 8.554858541 | 13.64898661 | ENSG00000197106 |
| TMEM169 | 0.674752979 | 1.596323424 | 0.019107433 | 0.062042252 | 48.58714919 | 77.35970764 | ENSG00000163449 |
| PRELID2 | 0.674712193 | 1.596278295 | 0.021159623 | 0.067741612 | 47.86186403 | 76.33141977 | ENSG00000186314 |
| COQ6 | 0.674630342 | 1.596187734 | 0.277017685 | 0.468111395 | 12.33550666 | 19.57909388 | ENSG00000119723 |
| GNA14 | 0.674449273 | 1.595987413 | 6.41E-07 | 7.39E-06 | 291.0801008 | 463.8279572 | ENSG00000156049 |
| LOC101928517 | 0.673966861 | 1.595453831 | 0.342917551 | 0.536880932 | 7.387688675 | 11.76635958 | ENSG00000269072 |
| FES | 0.673887877 | 1.595366487 | 0.000526526 | 0.003006356 | 100.9217913 | 161.2095148 | ENSG00000182511 |
| NA | 0.673716669 | 1.595177171 | 0.309661717 | 0.503645563 | 9.004740811 | 14.38095221 | ENSG00000276900 |
| CDH6 | 0.673340138 | 1.594760899 | 0.04469653 | 0.122545191 | 34.19417886 | 54.7096717 | ENSG00000113361 |
| CNTN4 | 0.672330309 | 1.593645021 | 0.11733997 | 0.255405638 | 21.15716917 | 33.82864913 | ENSG00000144619 |
| KNL1 | 0.672327333 | 1.593641733 | 5.83E-08 | 8.13E-07 | 351.8509166 | 560.4304013 | ENSG00000137812 |
| LOC101927572 | 0.672114741 | 1.593406915 | 0.456705513 | 0.643890155 | 5.570901858 | 8.855994327 | ENSG00000267698 |
| GLIPR1 | 0.671850248 | 1.593114818 | 0.470143525 | 0.655738783 | 4.079021215 | 6.530126268 | ENSG00000139278 |
| NA | 0.671813587 | 1.593074336 | 0.562044176 | 0.730738379 | 4.174405017 | 6.654148774 | ENSG00000261360 |
| NUAK2 | 0.671540752 | 1.59277309 | 2.64E-06 | 2.71E-05 | 211.449309 | 337.0846928 | ENSG00000163545 |
| FYTTD1 | 0.671170281 | 1.592364133 | 8.90E-11 | 1.96E-09 | 519.8862903 | 827.1694015 | ENSG00000122068 |
| LCAT | 0.670998496 | 1.592174538 | 0.00062056 | 0.003463249 | 104.602621 | 166.6329084 | ENSG00000213398 |
| NA | 0.670892314 | 1.592057358 | 0.208851497 | 0.387970471 | 13.71631664 | 21.86373273 | ENSG00000286989 |
| ZNF124 | 0.670867058 | 1.592029488 | 0.000134832 | 0.000905098 | 153.7076912 | 245.1636974 | ENSG00000196418 |
| NA | 0.670849597 | 1.592010219 | 0.031710543 | 0.093646144 | 39.33532001 | 62.60497606 | ENSG00000224905 |
| NA | 0.670769025 | 1.591921311 | 0.258866616 | 0.447684413 | 10.46702916 | 16.74012538 | ENSG00000251615 |
| NA | 0.670627928 | 1.591765627 | 0.328769747 | 0.523226012 | 8.359558549 | 13.32315808 | ENSG00000261211 |
| EIF2AK3-DT | 0.67030364 | 1.591407871 | 0.244843611 | 0.431937129 | 11.7933975 | 18.81437282 | ENSG00000234028 |
| LINC01844 | 0.670087308 | 1.591169258 | 0.014532011 | 0.049744639 | 54.23586885 | 86.0875542 | ENSG00000236714 |

| | | | | | | | |
|---|---|---|---|---|---|---|---|
| COQ5 | 0.669821768 | 1.590876417 | 1.52E-10 | 3.24E-09 | 489.1837595 | 777.6292815 | ENSG00000110871 |
| AKAP9 | 0.6697631 | 1.590811725 | 1.08E-09 | 2.01E-08 | 486.4932376 | 773.6804343 | ENSG00000127914 |
| CUX1 | 0.669751773 | 1.590799235 | 6.14E-15 | 2.34E-13 | 1065.633794 | 1694.742558 | ENSG00000257923 |
| ZNF697 | 0.669561381 | 1.590589311 | 1.56E-10 | 3.31E-09 | 648.1315234 | 1030.176926 | ENSG00000143067 |
| KCTD13 | 0.669410796 | 1.590423298 | 7.51E-05 | 0.000539196 | 164.4392771 | 261.9312908 | ENSG00000174943 |
| ZNF652 | 0.669363163 | 1.590370789 | 8.82E-06 | 7.97E-05 | 212.8377105 | 338.5299159 | ENSG00000198740 |
| DSG2-AS1 | 0.669264414 | 1.590261935 | 0.53741996 | 0.711203176 | 3.70558127 | 5.97144811 | ENSG00000264859 |
| CENPF | 0.668816534 | 1.58976832 | 1.26E-18 | 7.07E-17 | 2814.319726 | 4473.568281 | ENSG00000117724 |
| ZNF81 | 0.668364782 | 1.589270592 | 0.000893663 | 0.004786299 | 109.2275847 | 173.3339087 | ENSG00000197779 |
| CDC42BPA | 0.6682704 | 1.589166625 | 2.74E-12 | 7.52E-11 | 784.9235349 | 1247.243477 | ENSG00000143776 |
| NA | 0.668152778 | 1.589037066 | 0.064720434 | 0.163094745 | 38.72002313 | 61.49649126 | ENSG00000260912 |
| AGBL2 | 0.66805044 | 1.588924352 | 0.127140949 | 0.270791812 | 19.98428684 | 31.83328452 | ENSG00000165923 |
| PEAK1 | 0.66772682 | 1.588567971 | 1.37E-10 | 2.94E-09 | 835.288205 | 1326.167692 | ENSG00000173517 |
| SIRT5 | 0.667593528 | 1.588421208 | 4.24E-08 | 6.10E-07 | 313.2319147 | 497.0050525 | ENSG00000124523 |
| LINC00205 | 0.667372385 | 1.588177746 | 2.63E-06 | 2.70E-05 | 246.7036536 | 392.2422804 | ENSG00000223768 |
| LINC00339 | 0.667077491 | 1.587853147 | 0.16582184 | 0.328875547 | 19.6511877 | 31.21873141 | ENSG00000218510 |
| ICAM1 | 0.666979823 | 1.587745657 | 0.007076015 | 0.027668854 | 63.46244273 | 100.5468396 | ENSG00000090339 |
| TIGD4 | 0.666903863 | 1.587662061 | 0.129523453 | 0.274698213 | 24.05323595 | 38.26195812 | ENSG00000169989 |
| NPAS1 | 0.666685555 | 1.587421835 | 0.000658605 | 0.003654668 | 115.6567302 | 183.4804436 | ENSG00000130751 |
| NA | 0.666186573 | 1.586872891 | 0.330495688 | 0.524839952 | 8.098564747 | 12.79029895 | ENSG00000286447 |
| DUSP5 | 0.665839651 | 1.586491346 | 6.25E-15 | 2.37E-13 | 869.4062799 | 1378.568085 | ENSG00000138166 |
| NUFIP2 | 0.665764823 | 1.586409062 | 4.75E-18 | 2.53E-16 | 2047.416166 | 3247.653259 | ENSG00000108256 |
| NA | 0.66560071 | 1.586228611 | 0.357547538 | 0.55094663 | 8.08465932 | 12.77152796 | ENSG00000231305 |
| SNX30 | 0.665422742 | 1.586032949 | 3.93E-08 | 5.67E-07 | 386.1528365 | 611.679025 | ENSG00000148158 |
| MYRIP | 0.665006507 | 1.585575425 | 0.053297051 | 0.140413298 | 35.68733728 | 56.70536273 | ENSG00000170011 |
| ZNF641 | 0.664513982 | 1.585034214 | 0.000106841 | 0.000738875 | 128.8434676 | 204.2119511 | ENSG00000167528 |
| SYT15 | 0.6643574 | 1.584862193 | 0.510475217 | 0.689712463 | 3.516595104 | 5.585069716 | ENSG00000204176 |
| ITGAL | 0.664156687 | 1.584641715 | 0.534923042 | 0.709580851 | 3.202535143 | 5.029640753 | ENSG00000005844 |
| DOK6 | 0.66398204 | 1.584449897 | 0.180009464 | 0.349313726 | 15.12238191 | 24.03128548 | ENSG00000206052 |
| LOC339192 | 0.663893833 | 1.584353026 | 0.495778085 | 0.677486619 | 5.081858857 | 8.087036385 | ENSG00000267121 |
| MYO9A | 0.663526665 | 1.583949857 | 6.81E-06 | 6.39E-05 | 302.4120556 | 478.9736136 | ENSG00000066933 |
| TMEM200B | 0.663524155 | 1.583947101 | 1.56E-05 | 0.000133417 | 190.6932929 | 302.2748708 | ENSG00000253304 |
| IGSF22 | 0.663517569 | 1.58393987 | 0.365958408 | 0.55902688 | 6.659750263 | 10.52454272 | ENSG00000179057 |
| NMRK1 | 0.663307365 | 1.583709104 | 0.019528348 | 0.063264885 | 50.6694622 | 80.53809138 | ENSG00000106733 |
| CD7 | 0.663247755 | 1.583643669 | 0.233382316 | 0.418301151 | 13.16030205 | 20.92941144 | ENSG00000173762 |
| CA5B | 0.663001309 | 1.583373169 | 0.001382561 | 0.006903648 | 93.97431879 | 148.6859203 | ENSG00000169239 |
| NA | 0.662740178 | 1.583086601 | 0.524783076 | 0.701350557 | 3.549442009 | 5.622611705 | ENSG00000214252 |
| CHAC1 | 0.662625029 | 1.58296025 | 5.35E-12 | 1.43E-10 | 735.2207412 | 1164.176909 | ENSG00000128965 |
| BAIAP3 | 0.662497746 | 1.582820599 | 9.00E-06 | 8.13E-05 | 211.3638996 | 334.9861636 | ENSG00000007516 |
| LOC102724301 | 0.662131735 | 1.582419089 | 0.202286061 | 0.379547465 | 13.07429129 | 20.68571493 | ENSG00000254854 |
| ADPRM | 0.662090022 | 1.582373337 | 0.00226189 | 0.010545845 | 81.29062749 | 128.6414535 | ENSG00000170222 |
| ADCY7 | 0.662001706 | 1.582276474 | 0.002257115 | 0.010526766 | 87.77915298 | 138.9067779 | ENSG00000121281 |
| NPTX2 | 0.661580542 | 1.581814629 | 1.31E-11 | 3.26E-10 | 741.2225461 | 1172.905368 | ENSG00000106236 |
| NA | 0.661173536 | 1.581368438 | 0.318332487 | 0.512037619 | 8.81713012 | 13.95648218 | ENSG00000225173 |
| RAB19 | 0.660718794 | 1.580870064 | 0.08650354 | 0.203226849 | 28.62821536 | 45.31756906 | ENSG00000146955 |
| NXNL2 | 0.660384034 | 1.580503286 | 0.16919641 | 0.333675916 | 20.24773849 | 31.97976525 | ENSG00000130045 |
| HBQ1 | 0.6602978 | 1.580408817 | 0.087943756 | 0.205827703 | 29.20266776 | 46.30071725 | ENSG00000086506 |
| MGA | 0.66024904 | 1.580355403 | 1.88E-11 | 4.57E-10 | 953.7160259 | 1506.49264 | ENSG00000174197 |
| EVI5 | 0.65984408 | 1.579911865 | 0.002226222 | 0.010398386 | 97.1149184 | 153.0422815 | ENSG00000067208 |
| ZNF699 | 0.659640521 | 1.579688961 | 0.081841892 | 0.194998924 | 35.43000406 | 55.89495233 | ENSG00000196110 |
| SNORD19 | 0.65962716 | 1.579674331 | 0.507416681 | 0.686841029 | 3.490062025 | 5.524957886 | ENSG00000212493 |
| PRKAB2 | 0.659444057 | 1.579473856 | 9.45E-11 | 2.08E-09 | 694.1484141 | 1095.220548 | ENSG00000131791 |
| AFAP1L1 | 0.659390952 | 1.579415717 | 0.000237845 | 0.001500415 | 154.5001585 | 243.5755205 | ENSG00000157510 |
| PTPRG | 0.658938515 | 1.578920481 | 1.98E-13 | 6.32E-12 | 1695.327984 | 2675.823232 | ENSG00000144724 |
| NBPF20 | 0.658555446 | 1.578501297 | 0.281768809 | 0.473034702 | 9.859128493 | 15.59596235 | ENSG00000162825 |
| NA | 0.658514402 | 1.578456389 | 0.404499587 | 0.596229947 | 6.434760279 | 10.05928151 | ENSG00000268049 |
| RHBDD1 | 0.65841146 | 1.578343765 | 5.31E-08 | 7.47E-07 | 332.525447 | 524.1716287 | ENSG00000144468 |
| HRH2 | 0.658277172 | 1.578196858 | 9.79E-06 | 8.78E-05 | 299.4221911 | 472.67269 | ENSG00000113749 |
| LOC155060 | 0.658080027 | 1.577981211 | 0.152452629 | 0.309550819 | 22.01965211 | 34.88731934 | ENSG00000244560 |
| ZNF425 | 0.657668808 | 1.577531494 | 6.83E-05 | 0.000495341 | 177.534464 | 279.8517584 | ENSG00000204947 |
| TTC3 | 0.657550914 | 1.577402588 | 4.23E-12 | 1.14E-10 | 729.8583609 | 1150.944944 | ENSG00000182670 |
| PCDHAC2 | 0.657325573 | 1.577156225 | 0.000631035 | 0.003517472 | 124.6487456 | 196.1532185 | ENSG00000243232 |
| CBX4 | 0.657081301 | 1.576889209 | 0.001251189 | 0.0063533 | 103.1342142 | 162.8855493 | ENSG00000141582 |
| NKILA | 0.656880796 | 1.576670069 | 0.121787675 | 0.262527654 | 22.80203216 | 35.96994482 | ENSG00000278709 |
| ARL6IP4 | 0.656828772 | 1.576613215 | 0.348734505 | 0.542098912 | 8.942203914 | 14.17111046 | ENSG00000182196 |
| STEAP1B | 0.656375842 | 1.576118319 | 0.007270385 | 0.028243412 | 72.92613059 | 114.9160688 | ENSG00000105889 |
| AKR7A3 | 0.656209725 | 1.57593685 | 0.403136415 | 0.594898719 | 5.706821866 | 9.02829409 | ENSG00000162482 |
| GFPT1 | 0.656009526 | 1.575718176 | 1.13E-13 | 3.69E-12 | 1811.21223 | 2852.751011 | ENSG00000198380 |
| ACSF2 | 0.655765826 | 1.575452029 | 1.96E-06 | 2.07E-05 | 244.1125593 | 384.5710171 | ENSG00000167107 |
| NA | 0.655756084 | 1.57544139 | 0.172927902 | 0.339265983 | 20.9605914 | 33.21073521 | ENSG00000264015 |
| NA | 0.65556628 | 1.575234136 | 0.464202149 | 0.650549915 | 5.018044184 | 7.951728958 | ENSG00000286540 |
| ITGA3 | 0.654971763 | 1.574585135 | 2.74E-15 | 1.09E-13 | 782.4428197 | 1232.09567 | ENSG00000005884 |
| NA | 0.654871905 | 1.574476151 | 0.342790292 | 0.536797013 | 7.711219375 | 12.12268206 | ENSG00000226964 |
| NA | 0.654727581 | 1.574318653 | 0.418810291 | 0.610090886 | 5.958344929 | 9.470985467 | ENSG00000272054 |

| | | | | | | | |
|---|---|---|---|---|---|---|---|
| MT1F | 0.654444075 | 1.574009311 | 3.31E-07 | 4.03E-06 | 372.7812545 | 585.9249736 | ENSG00000198417 |
| CDK6 | 0.654133135 | 1.573670106 | 3.38E-06 | 3.41E-05 | 342.1419066 | 538.275419 | ENSG00000105810 |
| LOC101927556 | 0.654094324 | 1.573627771 | 0.433345541 | 0.623089702 | 5.795891842 | 9.02405721 | ENSG00000260927 |
| MAVS | 0.65407474 | 1.573606411 | 1.76E-17 | 8.79E-16 | 1036.429037 | 1630.842098 | ENSG00000088888 |
| HAAO | 0.654038207 | 1.573566563 | 0.068881615 | 0.171018332 | 28.66865386 | 45.10717765 | ENSG00000162882 |
| VSIG1 | 0.653664117 | 1.573158591 | 0.089860639 | 0.209342168 | 24.40202196 | 38.41548689 | ENSG00000101842 |
| IQCG | 0.653596087 | 1.57308441 | 0.016831724 | 0.056017088 | 67.94509755 | 106.7820693 | ENSG00000114473 |
| USP32 | 0.653441295 | 1.572915638 | 1.83E-15 | 7.42E-14 | 1582.214364 | 2488.033439 | ENSG00000170832 |
| GCNT1 | 0.653105304 | 1.572549363 | 1.71E-05 | 0.000144647 | 180.8051735 | 284.354529 | ENSG00000187210 |
| RGPD8 | 0.652986298 | 1.57241965 | 0.167135974 | 0.330773961 | 16.51444407 | 25.93268351 | ENSG00000169629 |
| NKPD1 | 0.652893304 | 1.572318298 | 0.268804058 | 0.45904491 | 11.28088064 | 17.7002657 | ENSG00000179846 |
| FAT1 | 0.652892201 | 1.572317096 | 1.32E-10 | 2.85E-09 | 2422.489502 | 3808.550866 | ENSG00000083857 |
| VMO1 | 0.652475743 | 1.571863286 | 0.573921651 | 0.739436906 | 3.400992048 | 5.359706164 | ENSG00000182853 |
| NA | 0.652335934 | 1.571710966 | 0.564948058 | 0.732937653 | 2.874569755 | 4.527275579 | ENSG00000243746 |
| NA | 0.652006651 | 1.571352278 | 0.214912755 | 0.395041446 | 15.28977335 | 24.0049168 | ENSG00000273145 |
| NA | 0.651920221 | 1.571258143 | 0.310872443 | 0.504706837 | 9.756153089 | 15.46120457 | ENSG00000273295 |
| ICAM3 | 0.651910318 | 1.571247357 | 0.010727759 | 0.038985937 | 62.68902976 | 98.33348587 | ENSG00000076662 |
| NA | 0.651512439 | 1.570814085 | 0.496754869 | 0.678121437 | 4.505527319 | 7.104326225 | ENSG00000287262 |
| SHC3 | 0.649874656 | 1.56903187 | 1.55E-06 | 1.66E-05 | 240.7591164 | 377.7730871 | ENSG00000148082 |
| ZNF674 | 0.64939829 | 1.568513873 | 0.054265765 | 0.142365465 | 35.57035875 | 55.64680414 | ENSG00000251192 |
| PDLIM5 | 0.649143033 | 1.56823638 | 2.29E-12 | 6.35E-11 | 783.7524566 | 1228.421148 | ENSG00000163110 |
| PDGFC | 0.648962045 | 1.568039655 | 0.128199983 | 0.272452527 | 25.49383188 | 39.70894181 | ENSG00000145431 |
| ATF7 | 0.648859146 | 1.56792782 | 0.001143415 | 0.005904722 | 100.2469191 | 157.1648457 | ENSG00000170653 |
| MPPE1 | 0.64879533 | 1.567858465 | 0.007503406 | 0.029008788 | 97.18307006 | 152.317619 | ENSG00000154889 |
| MEIOC | 0.648489356 | 1.567525982 | 0.002687633 | 0.01214053 | 99.37910699 | 155.4544068 | ENSG00000180336 |
| KLK14 | 0.648481654 | 1.567517614 | 0.315348976 | 0.508896641 | 9.107813914 | 14.30531858 | ENSG00000129437 |
| FCSK | 0.648449527 | 1.567482707 | 0.000103844 | 0.000720574 | 143.9220587 | 225.4192402 | ENSG00000157353 |
| FMO5 | 0.648342024 | 1.56736591 | 0.282263787 | 0.473179904 | 11.40101608 | 17.8951353 | ENSG00000131781 |
| FAM49A | 0.648154293 | 1.56716197 | 0.000165192 | 0.001084928 | 143.3956364 | 224.6361862 | ENSG00000197872 |
| FAM107A | 0.648114248 | 1.56711847 | 0.437536487 | 0.626268718 | 6.480234835 | 10.13860236 | ENSG00000168309 |
| NA | 0.64792358 | 1.566911372 | 0.427838611 | 0.618144747 | 5.45214189 | 8.491636121 | ENSG00000273759 |
| MTMR10 | 0.647536743 | 1.566491285 | 1.06E-05 | 9.42E-05 | 195.6583686 | 306.4272274 | ENSG00000166912 |
| NA | 0.647344925 | 1.566283021 | 0.554429832 | 0.724617623 | 3.625884334 | 5.65266762 | ENSG00000239268 |
| ZNF582 | 0.64707264 | 1.565987439 | 0.194035658 | 0.367978807 | 16.55991862 | 25.9669763 | ENSG00000018869 |
| SNORD83A | 0.646911047 | 1.565812045 | 0.336155388 | 0.530452695 | 9.487567685 | 14.91326168 | ENSG00000209482 |
| LOC286059 | 0.646738556 | 1.565624846 | 0.25000813 | 0.43853283 | 11.49639988 | 18.00776127 | ENSG00000246130 |
| SLC35F6 | 0.646452874 | 1.565314851 | 0.000123793 | 0.000842593 | 176.9409951 | 276.8217191 | ENSG00000213699 |
| RAPH1 | 0.646400322 | 1.565257834 | 0.002072382 | 0.009798348 | 99.66357466 | 156.0181979 | ENSG00000173166 |
| NA | 0.646104928 | 1.564937378 | 0.359676761 | 0.552940368 | 7.754814795 | 12.08514007 | ENSG00000241741 |
| KLHL15 | 0.645675237 | 1.564471348 | 3.27E-07 | 3.99E-06 | 312.3661771 | 487.9986754 | ENSG00000174010 |
| ZNF57 | 0.64558182 | 1.564370049 | 0.48084616 | 0.664578876 | 4.161777366 | 6.481299358 | ENSG00000171970 |
| IL1RAP | 0.645446579 | 1.564223409 | 0.000805272 | 0.004366563 | 109.1124853 | 170.4017465 | ENSG00000196083 |
| NGF | 0.645409617 | 1.564183334 | 0.228505721 | 0.412177619 | 13.26643437 | 20.74158988 | ENSG00000134259 |
| LOC105370295 | 0.64522202 | 1.563979952 | 0.271598945 | 0.46245428 | 11.37004831 | 17.80496756 | ENSG00000287376 |
| FUOM | 0.645176586 | 1.5639307 | 1.01E-05 | 9.05E-05 | 257.5957907 | 402.6408337 | ENSG00000148803 |
| PPIAP46 | 0.644987363 | 1.563725589 | 0.468931815 | 0.654734441 | 4.783583461 | 7.445126904 | ENSG00000260266 |
| NA | 0.644428765 | 1.563120247 | 0.164696581 | 0.327133901 | 18.47652393 | 28.9754334 | ENSG00000244041 |
| NA | 0.644427786 | 1.563119186 | 0.22541696 | 0.408314889 | 13.27590511 | 20.73779103 | ENSG00000280294 |
| NA | 0.6442506 | 1.562927222 | 0.001100302 | 0.005726763 | 205.513972 | 321.5657184 | ENSG00000244879 |
| NA | 0.643007823 | 1.561581453 | 0.439177879 | 0.627518544 | 5.70050804 | 8.904383203 | ENSG00000232057 |
| NA | 0.64288287 | 1.561446208 | 0.1972077 | 0.372887933 | 19.83889609 | 31.09145971 | ENSG00000260528 |
| CCDC33 | 0.642879472 | 1.561442531 | 0.206242295 | 0.384827816 | 16.96560411 | 26.33739486 | ENSG00000140481 |
| DFFA | 0.642642281 | 1.561185838 | 4.79E-16 | 2.04E-14 | 1493.557492 | 2331.702324 | ENSG00000160049 |
| NHLH1 | 0.64262046 | 1.561162226 | 0.495935681 | 0.677602057 | 4.70082731 | 7.419307869 | ENSG00000171786 |
| ZDHHC1 | 0.642170886 | 1.56067581 | 0.01894568 | 0.06165746 | 51.03333371 | 79.62221468 | ENSG00000159714 |
| NA | 0.641786784 | 1.560260352 | 0.641546863 | 0.789751316 | 2.977545159 | 4.632415471 | ENSG00000260329 |
| OXSM | 0.641504855 | 1.559955479 | 0.183805976 | 0.354380521 | 18.63444463 | 28.91813274 | ENSG00000151093 |
| SRR | 0.641394302 | 1.559835944 | 0.069412087 | 0.171897865 | 32.5708129 | 50.90946358 | ENSG00000167720 |
| ABRAXAS1 | 0.641315972 | 1.559751257 | 1.45E-06 | 1.56E-05 | 285.0469595 | 444.5356444 | ENSG00000163322 |
| RICTOR | 0.641308495 | 1.559743173 | 6.73E-05 | 0.000489811 | 181.7560273 | 282.9648057 | ENSG00000164327 |
| UEVLD | 0.641163141 | 1.559586034 | 0.008122646 | 0.031016626 | 76.26448804 | 118.7976361 | ENSG00000151116 |
| SNORA33 | 0.64102137 | 1.559432784 | 0.304157536 | 0.497923481 | 11.043263 | 17.09221098 | ENSG00000200534 |
| NA | 0.640868195 | 1.559267223 | 0.458238025 | 0.645314456 | 5.071110343 | 7.921673043 | ENSG00000232093 |
| PPL | 0.640700979 | 1.559086507 | 9.09E-16 | 3.81E-14 | 966.9892101 | 1507.592158 | ENSG00000118898 |
| NA | 0.640690143 | 1.559074796 | 0.193239263 | 0.366919015 | 16.04867953 | 25.01063483 | ENSG00000233830 |
| ATF6 | 0.640597248 | 1.558974411 | 2.66E-10 | 5.42E-09 | 457.066572 | 712.3716212 | ENSG00000118217 |
| GPC6 | 0.640465758 | 1.558832329 | 1.58E-07 | 2.03E-06 | 340.619577 | 530.2552648 | ENSG00000183098 |
| DBH-AS1 | 0.640464614 | 1.558831094 | 0.357714647 | 0.551066683 | 8.554858541 | 13.43958289 | ENSG00000225756 |
| NA | 0.640445845 | 1.558810814 | 0.116018413 | 0.253275415 | 27.30312479 | 42.39014467 | ENSG00000260025 |
| PCYOX1L | 0.640366778 | 1.558725385 | 5.72E-10 | 1.10E-08 | 459.6476093 | 715.89384 | ENSG00000145882 |
| TOB1 | 0.640023211 | 1.558354231 | 1.10E-07 | 1.46E-06 | 330.3224797 | 514.8854304 | ENSG00000141232 |
| SLC16A10 | 0.640004975 | 1.558334533 | 4.93E-06 | 4.76E-05 | 207.0756729 | 322.7636292 | ENSG00000112394 |
| PNLIPRP2 | 0.639956427 | 1.558282095 | 0.407785643 | 0.599547764 | 6.497297176 | 10.08228938 | ENSG00000266200 |
| KIF13B | 0.639720425 | 1.558027206 | 8.00E-06 | 7.35E-05 | 269.921323 | 420.070977 | ENSG00000197892 |

| | | | | | | | |
|---|---|---|---|---|---|---|---|
| PPARG | 0.639603874 | 1.557901342 | 0.556352178 | 0.72605487 | 3.454058207 | 5.374678312 | ENSG00000132170 |
| THSD7A | 0.63941935 | 1.557702096 | 0.046880632 | 0.127169874 | 41.18505143 | 64.41719402 | ENSG00000005108 |
| CYP2W1 | 0.639216473 | 1.557483062 | 0.573194686 | 0.738929561 | 3.26507204 | 5.010869759 | ENSG00000073067 |
| ZBTB5 | 0.638862772 | 1.557101264 | 3.03E-05 | 0.00024261 | 323.6490346 | 504.425875 | ENSG00000168795 |
| NUDT18 | 0.638696534 | 1.556921855 | 0.069810235 | 0.172574767 | 31.14137146 | 48.39588557 | ENSG00000275074 |
| RAB12 | 0.638551307 | 1.556765137 | 4.44E-06 | 4.36E-05 | 313.5292054 | 487.6632822 | ENSG00000206418 |
| TMEM192 | 0.638361226 | 1.55656004 | 8.02E-06 | 7.35E-05 | 239.8332021 | 372.6543376 | ENSG00000170088 |
| TCAF1 | 0.63832832 | 1.556524538 | 2.06E-07 | 2.60E-06 | 348.1955528 | 541.3857123 | ENSG00000198420 |
| GRID2 | 0.638243792 | 1.556433342 | 1.83E-09 | 3.27E-08 | 473.5081139 | 736.590943 | ENSG00000152208 |
| RAB40A | 0.638210767 | 1.556397715 | 0.094960824 | 0.217799628 | 25.72523338 | 40.11887774 | ENSG00000172476 |
| CHTF8 | 0.63801638 | 1.556188022 | 0.362946099 | 0.555845562 | 10.03411153 | 15.64522729 | ENSG00000168802 |
| NA | 0.637720977 | 1.555869413 | 0.341530055 | 0.53583425 | 8.247112406 | 12.88795277 | ENSG00000241081 |
| NA | 0.637707744 | 1.555855142 | 0.538566637 | 0.712018978 | 3.510281278 | 5.438588988 | ENSG00000231203 |
| B3GALT5 | 0.637595361 | 1.555733949 | 3.20E-12 | 8.74E-11 | 788.2439582 | 1225.913049 | ENSG00000183778 |
| PLEKHN1 | 0.637572375 | 1.555709161 | 0.072458481 | 0.177920705 | 32.86337583 | 51.33773246 | ENSG00000187583 |
| FAM160B2 | 0.637294417 | 1.555409458 | 6.29E-08 | 8.72E-07 | 330.3607686 | 514.0818935 | ENSG00000158863 |
| NA | 0.637180378 | 1.555286515 | 0.231473884 | 0.415993113 | 15.09447336 | 23.41629434 | ENSG00000188383 |
| NA | 0.636926924 | 1.555013305 | 0.481798947 | 0.665443003 | 4.717889651 | 7.411821795 | ENSG00000268189 |
| LOC648987 | 0.636728615 | 1.554799571 | 0.187267963 | 0.358964157 | 15.27083188 | 23.75004698 | ENSG00000177738 |
| H6PD | 0.636512484 | 1.554566663 | 8.45E-07 | 9.53E-06 | 279.7337218 | 435.1466164 | ENSG00000049239 |
| NA | 0.636294482 | 1.554331774 | 0.49190824 | 0.674535907 | 5.021201097 | 7.816533151 | ENSG00000279072 |
| AKR1E2 | 0.63590381 | 1.553910929 | 0.032728287 | 0.095948446 | 44.17078955 | 68.73755063 | ENSG00000165568 |
| AOC2 | 0.635651027 | 1.553638684 | 0.01516825 | 0.051526456 | 60.66175975 | 94.22786917 | ENSG00000131480 |
| LOC100506207 | 0.635636107 | 1.553622615 | 0.469920327 | 0.655539918 | 5.280315762 | 8.270509449 | ENSG00000285219 |
| CTHRC1 | 0.635613765 | 1.553598557 | 0.156146767 | 0.314602042 | 19.16530161 | 29.67591738 | ENSG00000164932 |
| MFSD14C | 0.635509458 | 1.553486235 | 4.68E-08 | 6.68E-07 | 508.946311 | 790.490708 | ENSG00000196312 |
| A2ML1 | 0.635445969 | 1.553417872 | 0.000174493 | 0.001137948 | 152.1773641 | 236.1004493 | ENSG00000166535 |
| SMG1P3 | 0.635405698 | 1.55337451 | 0.045217505 | 0.123753874 | 58.62056167 | 90.94307172 | ENSG00000180747 |
| NA | 0.635333205 | 1.553296459 | 0.557125191 | 0.726552089 | 3.149468985 | 4.943709888 | ENSG00000231173 |
| OVCH1-AS1 | 0.635301809 | 1.553262656 | 0.572350681 | 0.738457885 | 2.891632095 | 4.47844867 | ENSG00000257599 |
| NA | 0.634584944 | 1.552491042 | 0.258402007 | 0.447074219 | 16.7191171 | 25.98260972 | ENSG00000263412 |
| LOC389831 | 0.634468778 | 1.55236604 | 0.001647895 | 0.008047983 | 114.2462302 | 177.6735146 | ENSG00000276256 |
| RAB25 | 0.633776585 | 1.551621406 | 9.45E-09 | 1.52E-07 | 420.6787163 | 652.1754096 | ENSG00000132698 |
| NTF4 | 0.633719367 | 1.55155987 | 0.193335629 | 0.367003516 | 16.67374024 | 26.01960206 | ENSG00000225950 |
| CACNA1B | 0.633536021 | 1.551362701 | 1.10E-07 | 1.47E-06 | 362.4265586 | 562.6194104 | ENSG00000148408 |
| ZNF79 | 0.63312887 | 1.550924945 | 0.002756807 | 0.012395599 | 102.2134336 | 158.9268513 | ENSG00000196152 |
| NA | 0.633113835 | 1.550908781 | 0.002399101 | 0.011071947 | 98.56339902 | 152.8595729 | ENSG00000267002 |
| MARVELD2 | 0.632797839 | 1.550569121 | 5.71E-07 | 6.67E-06 | 290.6215219 | 450.1532547 | ENSG00000152939 |
| LINC02593 | 0.632720815 | 1.55048634 | 0.316679667 | 0.510333097 | 8.915670835 | 13.84429424 | ENSG00000223764 |
| RPS6KA5 | 0.631747095 | 1.549440221 | 0.073601219 | 0.179916247 | 32.68188357 | 50.41370841 | ENSG00000100784 |
| CHD9 | 0.631719792 | 1.549410898 | 6.04E-09 | 1.00E-07 | 472.2371946 | 731.2273264 | ENSG00000177200 |
| CYB5RL | 0.631646459 | 1.549332143 | 1.54E-05 | 0.00013172 | 203.1983632 | 315.2180373 | ENSG00000215883 |
| APP | 0.630647683 | 1.548259912 | 7.72E-29 | 1.03E-26 | 19600.40333 | 30345.7315 | ENSG00000142192 |
| EPHA2 | 0.630415259 | 1.548010502 | 6.77E-16 | 2.86E-14 | 1180.218197 | 1827.203979 | ENSG00000142627 |
| SLC16A5 | 0.630098303 | 1.547670446 | 4.64E-06 | 4.53E-05 | 265.5753099 | 411.64819 | ENSG00000170190 |
| KLHL28 | 0.629874264 | 1.547430123 | 0.002048405 | 0.009693084 | 105.2958333 | 162.6436763 | ENSG00000179454 |
| SNX32 | 0.629864723 | 1.54741989 | 0.152435408 | 0.309550819 | 22.6635566 | 35.1192929 | ENSG00000172803 |
| NUP50-DT | 0.629796385 | 1.547346593 | 0.048688423 | 0.131052203 | 51.87005769 | 80.20302464 | ENSG00000226328 |
| SLC2A1 | 0.629738507 | 1.547284518 | 1.48E-20 | 9.96E-19 | 4716.173946 | 7295.666243 | ENSG00000117394 |
| TRAF3IP2 | 0.629716437 | 1.547260848 | 1.11E-09 | 2.05E-08 | 447.5100559 | 692.1098266 | ENSG00000056972 |
| PKP3 | 0.629676097 | 1.547217585 | 7.23E-07 | 8.28E-06 | 336.2740298 | 520.6396708 | ENSG00000184363 |
| XPC | 0.629440629 | 1.546965078 | 4.56E-10 | 8.91E-09 | 662.5632636 | 1025.059317 | ENSG00000154767 |
| NA | 0.62942698 | 1.546950443 | 0.253934169 | 0.442748437 | 13.00859748 | 20.0964312 | ENSG00000234129 |
| SMAD3 | 0.629255428 | 1.546766505 | 6.81E-08 | 9.35E-07 | 434.9329255 | 672.6444496 | ENSG00000166949 |
| NA | 0.629212878 | 1.546720885 | 0.390786186 | 0.583440565 | 6.725444073 | 10.40106987 | ENSG00000277270 |
| TMEM91 | 0.62911922 | 1.546620478 | 0.255373291 | 0.444254606 | 12.08535907 | 18.73917722 | ENSG00000142046 |
| SLCO1A2 | 0.628462126 | 1.54591621 | 0.050984759 | 0.13584697 | 41.97415126 | 64.68940911 | ENSG00000084453 |
| INKA2 | 0.628414311 | 1.545864975 | 3.34E-05 | 0.000264771 | 293.4273538 | 453.2443222 | ENSG00000197852 |
| AMZ2P1 | 0.628370352 | 1.545817872 | 0.00113591 | 0.005870873 | 118.3933054 | 182.7388336 | ENSG00000214174 |
| CLEC2D | 0.628317202 | 1.545760924 | 0.151579349 | 0.308383276 | 19.49198923 | 30.02475378 | ENSG00000069493 |
| TFRC | 0.628264008 | 1.545703932 | 5.86E-17 | 2.77E-15 | 3046.886472 | 4708.384741 | ENSG00000072274 |
| NA | 0.628207185 | 1.545643053 | 0.284265317 | 0.475245659 | 11.15896376 | 17.2462894 | ENSG00000258815 |
| SNHG29 | 0.62800417 | 1.545425566 | 2.02E-18 | 1.12E-16 | 3273.871328 | 5059.022858 | ENSG00000175061 |
| CEL | 0.627937831 | 1.545354505 | 0.047771487 | 0.129123048 | 37.88260009 | 58.55012136 | ENSG00000170835 |
| MELTF | 0.627220031 | 1.544585819 | 0.008152918 | 0.031123827 | 82.41400654 | 126.8659015 | ENSG00000163975 |
| ABHD1 | 0.627192954 | 1.544556829 | 0.481891076 | 0.665443003 | 4.439833508 | 6.860191679 | ENSG00000143994 |
| PCDHGB2 | 0.626154433 | 1.543445384 | 0.139955732 | 0.29093086 | 26.92387468 | 41.43858973 | ENSG00000253910 |
| NA | 0.626127166 | 1.543416213 | 0.56562654 | 0.733304582 | 3.149468985 | 4.898681826 | ENSG00000226070 |
| ZNF300 | 0.625689489 | 1.54294805 | 7.17E-05 | 0.000517568 | 189.3164366 | 292.3833258 | ENSG00000145908 |
| FBLN2 | 0.625478249 | 1.542722148 | 6.52E-15 | 2.47E-13 | 1122.42413 | 1730.57988 | ENSG00000163520 |
| TMEM171 | 0.62545726 | 1.542699704 | 0.140116767 | 0.29114756 | 27.09509945 | 41.74316252 | ENSG00000157111 |
| NA | 0.625446893 | 1.542688619 | 0.474301676 | 0.659368981 | 5.667661135 | 8.818452338 | ENSG00000258655 |
| EPS8L2 | 0.625205382 | 1.54243039 | 4.92E-10 | 9.58E-09 | 563.1401553 | 868.8527644 | ENSG00000177106 |
| MORN1 | 0.62514165 | 1.542362254 | 0.004150596 | 0.017579331 | 91.01571511 | 140.3897034 | ENSG00000116151 |

| | | | | | | | |
|---|---|---|---|---|---|---|---|
| PROK2 | 0.624982668 | 1.542192298 | 0.151695185 | 0.308585068 | 24.4442419 | 37.51513199 | ENSG00000163421 |
| LOXL2 | 0.624590663 | 1.541773314 | 3.76E-14 | 1.31E-12 | 906.6400372 | 1397.258762 | ENSG00000134013 |
| LOC101927322 | 0.624382938 | 1.54155134 | 0.555954541 | 0.725587033 | 3.827498152 | 5.907649054 | ENSG00000267226 |
| NA | 0.623538639 | 1.540649452 | 0.050534542 | 0.134932524 | 37.4263063 | 57.59746711 | ENSG00000286388 |
| JUND | 0.623513842 | 1.540622971 | 2.02E-11 | 4.88E-10 | 1011.726477 | 1558.159778 | ENSG00000130522 |
| NA | 0.622605447 | 1.539653221 | 0.481282457 | 0.665033106 | 5.280315762 | 8.17285563 | ENSG00000237877 |
| CTSB | 0.622111837 | 1.539126528 | 5.20E-14 | 1.78E-12 | 1030.791351 | 1585.560931 | ENSG00000164733 |
| GOLGA4 | 0.621583517 | 1.538562998 | 2.30E-12 | 6.37E-11 | 875.1381465 | 1346.869192 | ENSG00000144674 |
| SVEP1 | 0.621493972 | 1.538467505 | 0.149796367 | 0.305729089 | 21.3870975 | 32.81924387 | ENSG00000165124 |
| ALMS1-IT1 | 0.621418513 | 1.538387039 | 0.176240986 | 0.344126678 | 20.65727995 | 31.6369892 | ENSG00000230002 |
| PTK6 | 0.620672175 | 1.537591404 | 0.453022343 | 0.640291705 | 5.727041119 | 8.761589702 | ENSG00000101213 |
| MSH5 | 0.619651795 | 1.53650429 | 0.394675869 | 0.587072437 | 6.994029477 | 10.77996219 | ENSG00000204410 |
| RIPOR1 | 0.61932432 | 1.536155561 | 0.000146519 | 0.00097333 | 187.6701981 | 288.3535088 | ENSG00000039523 |
| NA | 0.619317067 | 1.536147838 | 0.591191463 | 0.752964718 | 2.894789008 | 4.41833684 | ENSG00000228285 |
| AFAP1 | 0.619043396 | 1.535856467 | 0.001364948 | 0.006829899 | 117.3085084 | 180.4143913 | ENSG00000196526 |
| IL2RB | 0.618942899 | 1.535749484 | 0.061836286 | 0.157665941 | 36.7969086 | 56.30258647 | ENSG00000100385 |
| GOLGA3 | 0.618728124 | 1.535520874 | 5.46E-09 | 9.13E-08 | 826.6825351 | 1270.049647 | ENSG00000090615 |
| RLN1 | 0.61870544 | 1.53549673 | 0.45428108 | 0.641557992 | 5.395918818 | 8.30436421 | ENSG00000107018 |
| DLGAP4-AS1 | 0.618647052 | 1.535434588 | 0.267859739 | 0.45798281 | 11.78708368 | 18.04106638 | ENSG00000232907 |
| LRRK1 | 0.618570331 | 1.535352937 | 0.008240107 | 0.031392013 | 102.3989924 | 157.2642517 | ENSG00000154237 |
| MAFF | 0.618413677 | 1.535186231 | 0.001120134 | 0.005809655 | 126.0078632 | 193.3096153 | ENSG00000185022 |
| NA | 0.618136229 | 1.534891024 | 0.20565898 | 0.384061571 | 17.3112332 | 26.51448116 | ENSG00000284606 |
| LOC105373159 | 0.617992733 | 1.534738366 | 0.056610729 | 0.147156135 | 60.05070217 | 92.30946073 | ENSG00000177788 |
| OGFOD2 | 0.617899834 | 1.534639543 | 0.561272914 | 0.730111749 | 4.065115787 | 6.290228601 | ENSG00000111325 |
| RBM33 | 0.616798039 | 1.533467976 | 1.55E-15 | 6.36E-14 | 1321.13048 | 2025.71696 | ENSG00000184863 |
| ENTPD3-AS1 | 0.616704385 | 1.533368433 | 0.16428634 | 0.32652901 | 19.4231385 | 29.84540598 | ENSG00000223797 |
| CAP2 | 0.61661806 | 1.533276685 | 5.73E-07 | 6.68E-06 | 294.7213638 | 451.836824 | ENSG00000112186 |
| DHRS7B | 0.615958011 | 1.532575355 | 9.87E-06 | 8.83E-05 | 232.6068841 | 356.8151571 | ENSG00000109016 |
| TEP1 | 0.615707901 | 1.532309686 | 7.91E-06 | 7.28E-05 | 341.5419436 | 523.118773 | ENSG00000129566 |
| SYTL3 | 0.615552149 | 1.532144268 | 0.562130374 | 0.730738379 | 3.179158977 | 4.894882979 | ENSG00000164674 |
| NA | 0.615450974 | 1.532036824 | 0.582435376 | 0.746119523 | 2.868255929 | 4.362023857 | ENSG00000237804 |
| DENND3 | 0.615284062 | 1.531859585 | 0.011617472 | 0.041639572 | 113.1600577 | 173.7316123 | ENSG00000105339 |
| ZNF106 | 0.615227013 | 1.531799012 | 1.42E-10 | 3.04E-09 | 1538.832666 | 2356.70336 | ENSG00000103994 |
| NA | 0.614908352 | 1.531460707 | 0.458850352 | 0.645816342 | 5.484988795 | 8.398219182 | ENSG00000251050 |
| SAT1 | 0.614323415 | 1.530839907 | 7.69E-09 | 1.26E-07 | 666.6552958 | 1019.919048 | ENSG00000130066 |
| SPINK5 | 0.614214932 | 1.5307248 | 0.026510029 | 0.081226134 | 60.7774605 | 93.25754324 | ENSG00000133710 |
| MSANTD1 | 0.61417817 | 1.530685795 | 0.402447009 | 0.594196834 | 6.748820239 | 10.28464506 | ENSG00000188981 |
| MIR17HG | 0.61416155 | 1.530668161 | 0.003962912 | 0.016900279 | 109.0185746 | 167.2610567 | ENSG00000215417 |
| NA | 0.614115813 | 1.530619636 | 0.368290564 | 0.561340239 | 9.702989232 | 14.6874601 | ENSG00000215086 |
| NID2 | 0.613707271 | 1.530186257 | 5.69E-05 | 0.000422481 | 206.3350092 | 315.5833344 | ENSG00000087303 |
| CCDC191 | 0.613646671 | 1.530121983 | 0.347658401 | 0.540879779 | 9.182978463 | 14.06542092 | ENSG00000163617 |
| LIFR | 0.613547312 | 1.530016607 | 4.21E-05 | 0.000323119 | 197.3391377 | 301.5993927 | ENSG00000113594 |
| NCEH1 | 0.612940817 | 1.529373537 | 0.016703167 | 0.055739377 | 71.30948442 | 108.6930001 | ENSG00000144959 |
| ATF7IP | 0.612548328 | 1.528957524 | 2.86E-18 | 1.54E-16 | 1847.555108 | 2823.706342 | ENSG00000171681 |
| GALNT12 | 0.61151421 | 1.527861967 | 1.52E-05 | 0.000130624 | 316.9990708 | 484.1836067 | ENSG00000119514 |
| BCL6 | 0.611050715 | 1.527371189 | 0.042842493 | 0.118779769 | 49.8042283 | 76.1835536 | ENSG00000113916 |
| NA | 0.611023544 | 1.527342423 | 0.525236117 | 0.701716439 | 3.854031231 | 5.873794292 | ENSG00000286001 |
| MALAT1 | 0.610324736 | 1.526602793 | 3.90E-11 | 9.04E-10 | 1022.211673 | 1560.911748 | ENSG00000251562 |
| KLHL8 | 0.610311859 | 1.526589167 | 3.98E-05 | 0.000308119 | 209.4072391 | 319.1688111 | ENSG00000145332 |
| NR2F2 | 0.610275089 | 1.526550259 | 0.500305406 | 0.680962061 | 4.439833508 | 6.75125294 | ENSG00000185551 |
| NSUN7 | 0.610267275 | 1.526541992 | 0.000204537 | 0.001310366 | 157.9266013 | 241.0184518 | ENSG00000179299 |
| NA | 0.610084994 | 1.526349129 | 0.19896316 | 0.375211967 | 18.47652393 | 28.29653159 | ENSG00000280239 |
| NA | 0.609937396 | 1.526192981 | 0.533017829 | 0.708006407 | 3.88056431 | 5.903850207 | ENSG00000277491 |
| CD70 | 0.609650106 | 1.525889093 | 0.138396438 | 0.288336285 | 25.0364557 | 38.37144651 | ENSG00000125726 |
| ACTG1 | 0.609527056 | 1.525758953 | 8.15E-30 | 1.15E-27 | 53058.21798 | 80953.41052 | ENSG00000184009 |
| PLGRKT | 0.609499058 | 1.525729343 | 0.000255518 | 0.001599921 | 172.6279942 | 263.2731346 | ENSG00000107020 |
| ZNF528 | 0.609219369 | 1.525433586 | 0.431103115 | 0.621021696 | 6.097324152 | 9.268191756 | ENSG00000167555 |
| LOC339803 | 0.609048432 | 1.525252855 | 0.030488149 | 0.090725855 | 58.07558869 | 88.70203522 | ENSG00000212978 |
| TBC1D8-AS1 | 0.608842965 | 1.525035647 | 0.569110227 | 0.735971767 | 3.202535143 | 4.894882979 | ENSG00000272902 |
| SIAE | 0.608363859 | 1.52452928 | 0.005506431 | 0.022356231 | 93.30250575 | 141.8961376 | ENSG00000110013 |
| NIN | 0.608354819 | 1.524519727 | 9.14E-07 | 1.03E-05 | 416.5699074 | 634.6259647 | ENSG00000100503 |
| NIPA1 | 0.608348044 | 1.524512568 | 1.04E-08 | 1.67E-07 | 426.5099204 | 650.5224947 | ENSG00000170113 |
| GDPD5 | 0.608313772 | 1.524476353 | 0.002501789 | 0.011471586 | 109.5440274 | 166.7186245 | ENSG00000158555 |
| TMEM241 | 0.607772913 | 1.523904942 | 0.002144608 | 0.010101186 | 107.5101353 | 163.8827936 | ENSG00000134490 |
| ZNF221 | 0.60683041 | 1.522909709 | 0.179845418 | 0.349178188 | 20.15926988 | 30.65906557 | ENSG00000159905 |
| NA | 0.606752777 | 1.522827762 | 0.475168886 | 0.660029989 | 5.944439501 | 9.144718903 | ENSG00000213225 |
| KLF11 | 0.606390473 | 1.522445383 | 6.91E-05 | 0.000500957 | 205.0179162 | 311.5240281 | ENSG00000172059 |
| NA | 0.606127844 | 1.522168261 | 0.178725825 | 0.347714418 | 21.23549063 | 32.26761376 | ENSG00000258592 |
| NEDD4 | 0.606119634 | 1.522159599 | 0.00013723 | 0.000918867 | 184.1245894 | 279.9458366 | ENSG00000069869 |
| VPS33B-DT | 0.605505028 | 1.521511278 | 0.625493176 | 0.777994129 | 3.136841333 | 4.78594424 | ENSG00000214432 |
| IRAK1 | 0.605458953 | 1.521462687 | 6.98E-15 | 2.63E-13 | 1030.999655 | 1569.191098 | ENSG00000184216 |
| NA | 0.605445839 | 1.521448857 | 0.393566452 | 0.586305751 | 7.886300113 | 12.03251431 | ENSG00000279500 |
| H1-3 | 0.605276326 | 1.521270101 | 0.55647488 | 0.726110513 | 4.485308065 | 6.792593775 | ENSG00000124575 |
| DENND5B | 0.605152888 | 1.521139946 | 8.36E-05 | 0.000593534 | 222.7249581 | 338.5150469 | ENSG00000170456 |

| | | | | | | | |
|---|---|---|---|---|---|---|---|
| AFF4 | 0.605046321 | 1.521027589 | 5.19E-10 | 1.01E-08 | 960.2713949 | 1460.121879 | ENSG00000072364 |
| BLOC1S3 | 0.604908784 | 1.520882591 | 0.001067718 | 0.005580677 | 116.7689547 | 177.7276375 | ENSG00000189114 |
| GALNT17 | 0.604452417 | 1.520401567 | 1.95E-16 | 8.66E-15 | 1880.118005 | 2857.867484 | ENSG00000185274 |
| ARMCX3 | 0.604174877 | 1.520109106 | 1.11E-07 | 1.47E-06 | 361.8807138 | 550.0685308 | ENSG00000102401 |
| IRAK2 | 0.604033939 | 1.519960613 | 0.166895114 | 0.330403123 | 21.77128596 | 33.1352132 | ENSG00000134070 |
| OCEL1 | 0.603783039 | 1.519696298 | 0.000128727 | 0.000871378 | 220.2465281 | 334.1743989 | ENSG00000099330 |
| NA | 0.603776701 | 1.519689622 | 0.586559559 | 0.749312221 | 3.768118167 | 5.71982749 | ENSG00000249465 |
| RGP1 | 0.603357114 | 1.519247707 | 4.79E-09 | 8.09E-08 | 514.2791142 | 781.288225 | ENSG00000107185 |
| DDTL | 0.603216027 | 1.519099141 | 0.102269105 | 0.23047855 | 29.09969235 | 44.37619973 | ENSG00000099974 |
| SDK1 | 0.602791685 | 1.518652391 | 4.25E-06 | 4.19E-05 | 397.8822239 | 603.6049141 | ENSG00000146555 |
| NA | 0.602751284 | 1.518609864 | 0.178856274 | 0.347840733 | 20.13085766 | 30.61078831 | ENSG00000205959 |
| NA | 0.602613629 | 1.518464972 | 0.392549893 | 0.585225202 | 8.039184763 | 12.20536373 | ENSG00000286403 |
| CEACAM19 | 0.602308291 | 1.518143632 | 0.252598963 | 0.44125063 | 14.15998278 | 21.54440258 | ENSG00000186567 |
| PAG1 | 0.602272696 | 1.518106176 | 0.421976424 | 0.613212407 | 6.38169412 | 9.695910986 | ENSG00000076641 |
| NA | 0.601926716 | 1.517742155 | 0.378618817 | 0.571992985 | 8.618673215 | 13.06773861 | ENSG00000287808 |
| CDA | 0.601741925 | 1.517547764 | 0.005413391 | 0.022037867 | 86.74720407 | 131.6006457 | ENSG00000158825 |
| FBXO24 | 0.601604785 | 1.517403516 | 0.319111254 | 0.513002688 | 10.70276766 | 16.34246207 | ENSG00000106336 |
| NA | 0.601363455 | 1.51714971 | 0.143889417 | 0.296770363 | 26.13595492 | 39.74802114 | ENSG00000274828 |
| NPBWR1 | 0.601065957 | 1.51683689 | 0.48159932 | 0.665363104 | 6.534676468 | 9.969663414 | ENSG00000183729 |
| LINC01600 | 0.600886409 | 1.516648127 | 0.406488802 | 0.598163038 | 6.980124049 | 10.57336963 | ENSG00000164385 |
| CLGN | 0.600644044 | 1.516393359 | 0.000372787 | 0.002217343 | 162.2622567 | 245.7348227 | ENSG00000153132 |
| SCRN1 | 0.600181887 | 1.515907671 | 1.49E-19 | 9.17E-18 | 2903.275369 | 4400.093692 | ENSG00000136193 |
| AEBP2 | 0.599870762 | 1.515580794 | 3.48E-10 | 6.94E-09 | 785.0897084 | 1189.672363 | ENSG00000139154 |
| KIAA0556 | 0.599823671 | 1.515531324 | 3.25E-06 | 3.29E-05 | 304.8207631 | 462.2420107 | ENSG00000047578 |
| SLC22A18 | 0.599812559 | 1.515519651 | 0.010462961 | 0.038173329 | 73.84228105 | 112.0483585 | ENSG00000110628 |
| KIF12 | 0.5998116 | 1.515518644 | 0.082145927 | 0.195642843 | 34.31609574 | 52.16691385 | ENSG00000136883 |
| NA | 0.599725279 | 1.515427968 | 0.552311559 | 0.723161339 | 4.108711207 | 6.256373839 | ENSG00000287005 |
| ZNF575 | 0.599620334 | 1.515317736 | 0.539669281 | 0.712815739 | 3.850874318 | 5.806196388 | ENSG00000176472 |
| ZSWIM9 | 0.599556937 | 1.51525115 | 1.16E-06 | 1.28E-05 | 368.341714 | 557.632224 | ENSG00000185453 |
| ZNF18 | 0.598830511 | 1.514488382 | 0.000161715 | 0.001064906 | 158.7367922 | 240.5428933 | ENSG00000154957 |
| SLC45A1 | 0.598004544 | 1.513621561 | 0.012796332 | 0.045063124 | 71.49601273 | 108.2440251 | ENSG00000162426 |
| ITGB5 | 0.597748595 | 1.513353052 | 6.36E-24 | 5.51E-22 | 6817.6943 | 10316.76943 | ENSG00000082781 |
| CCND2 | 0.597744243 | 1.513348487 | 6.96E-10 | 1.32E-08 | 1291.909029 | 1954.914692 | ENSG00000118971 |
| RAPGEF6 | 0.597379207 | 1.512965623 | 0.054664362 | 0.143240312 | 50.63030147 | 76.46562789 | ENSG00000158987 |
| VWDE | 0.597105446 | 1.512678554 | 0.119663229 | 0.259059816 | 199.179609 | 301.4070255 | ENSG00000146530 |
| CLDN11 | 0.596886945 | 1.512449472 | 0.006107427 | 0.024453458 | 82.60634501 | 124.9385728 | ENSG00000013297 |
| ZNF658 | 0.596804668 | 1.512363219 | 0.108003604 | 0.239901583 | 29.80741151 | 45.03914172 | ENSG00000274349 |
| C18orf65 | 0.596777771 | 1.512335023 | 0.356617559 | 0.550199771 | 9.882504659 | 14.98097121 | ENSG00000275763 |
| NA | 0.596325207 | 1.511860688 | 0.577048186 | 0.741833328 | 3.513438191 | 5.352658124 | ENSG00000263884 |
| TMEM119 | 0.596314657 | 1.511849632 | 0.624976965 | 0.777404233 | 3.093245913 | 4.688290421 | ENSG00000183160 |
| NBPF26 | 0.596137438 | 1.51166393 | 0.018597769 | 0.060685044 | 67.34172927 | 101.785622 | ENSG00000273136 |
| GEMIN8 | 0.596088589 | 1.511612746 | 0.000607837 | 0.003403521 | 163.6794764 | 247.4939056 | ENSG00000046647 |
| TNFRSF12A | 0.59598102 | 1.511500042 | 1.58E-11 | 3.89E-10 | 910.9801871 | 1377.153476 | ENSG00000006327 |
| CAVIN3 | 0.595957743 | 1.511475656 | 0.001709196 | 0.008302658 | 163.1946575 | 246.7327283 | ENSG00000170955 |
| ZNF391 | 0.595196931 | 1.510678783 | 0.000971721 | 0.005138338 | 127.5877314 | 192.4901815 | ENSG00000124613 |
| KCNMB4 | 0.594850114 | 1.510315667 | 7.26E-08 | 9.91E-07 | 363.4750662 | 549.0034654 | ENSG00000135643 |
| NMUR1 | 0.594500879 | 1.509950106 | 0.306229151 | 0.50027775 | 12.41332446 | 18.79549021 | ENSG00000171596 |
| NA | 0.594469099 | 1.509916845 | 0.06798661 | 0.169136085 | 49.08881984 | 73.89748904 | ENSG00000259768 |
| EFR3A | 0.594414091 | 1.509859275 | 5.19E-11 | 1.18E-09 | 655.9237175 | 990.3850564 | ENSG00000132294 |
| ASAH2B | 0.593983165 | 1.509408355 | 0.032670307 | 0.095808777 | 59.98332462 | 90.6261147 | ENSG00000204147 |
| ST8SIA4 | 0.59383 | 1.509248116 | 0.434990443 | 0.624534268 | 6.348847215 | 9.575687326 | ENSG00000113532 |
| ANXA2 | 0.593201327 | 1.508590584 | 1.18E-11 | 2.97E-10 | 818.0717843 | 1233.357157 | ENSG00000182718 |
| GRK1 | 0.593175675 | 1.508563761 | 0.612624165 | 0.768999741 | 3.196221317 | 4.816000155 | ENSG00000185974 |
| NA | 0.593129951 | 1.508515951 | 0.416564191 | 0.607751088 | 7.897048628 | 11.91229065 | ENSG00000278238 |
| EMILIN1 | 0.592930684 | 1.508307607 | 0.000619518 | 0.003459518 | 143.3188858 | 216.5190939 | ENSG00000138080 |
| TBX10 | 0.592877498 | 1.508252003 | 0.315960252 | 0.509528175 | 10.93397377 | 16.40582309 | ENSG00000167800 |
| OR4D1 | 0.59271557 | 1.508082726 | 0.609995991 | 0.767318155 | 2.897945921 | 4.313196947 | ENSG00000141194 |
| NA | 0.592120437 | 1.507460748 | 0.579489965 | 0.743734299 | 3.18231589 | 4.789743086 | ENSG00000186676 |
| NA | 0.592014928 | 1.507350506 | 0.455628725 | 0.643007903 | 6.097324152 | 9.121711028 | ENSG00000176700 |
| FOXK1 | 0.591991947 | 1.507326496 | 2.70E-06 | 2.76E-05 | 370.9545909 | 558.6608383 | ENSG00000164916 |
| CCDC8 | 0.591984508 | 1.507318723 | 9.48E-11 | 2.08E-09 | 681.2615078 | 1026.559069 | ENSG00000169515 |
| NA | 0.591802189 | 1.50712825 | 0.594936906 | 0.755905214 | 3.708738183 | 5.671438614 | ENSG00000285589 |
| P4HA1 | 0.591791021 | 1.507116583 | 3.73E-08 | 5.40E-07 | 504.279225 | 759.2642814 | ENSG00000122884 |
| LOC105376805 | 0.591770547 | 1.507095195 | 0.459601064 | 0.646298468 | 5.763044937 | 8.690742605 | ENSG00000238142 |
| CD99L2 | 0.591717806 | 1.507040101 | 2.12E-07 | 2.68E-06 | 483.3056152 | 728.3296089 | ENSG00000102181 |
| C1orf220 | 0.591558756 | 1.506873966 | 0.408555664 | 0.600213036 | 8.618673215 | 13.04571842 | ENSG00000213057 |
| CDKL5 | 0.591518474 | 1.506831893 | 4.40E-05 | 0.000335863 | 279.0247022 | 420.0295246 | ENSG00000008086 |
| CNNM2 | 0.591506225 | 1.5068191 | 5.45E-08 | 7.66E-07 | 525.7042805 | 791.5213662 | ENSG00000148842 |
| HECTD1 | 0.591258388 | 1.506560269 | 3.61E-11 | 8.42E-10 | 1828.926759 | 2754.917152 | ENSG00000092148 |
| TAPT1-AS1 | 0.59125837 | 1.50656025 | 0.219638772 | 0.401023051 | 17.84840401 | 26.8858874 | ENSG00000263327 |
| GABARAPL1 | 0.591233319 | 1.50653409 | 5.67E-12 | 1.50E-10 | 1149.412202 | 1731.107244 | ENSG00000139112 |
| NRG1 | 0.591220447 | 1.506520648 | 0.06362203 | 0.161160349 | 45.30511251 | 68.49797938 | ENSG00000157168 |
| LRIG2 | 0.59111247 | 1.506407899 | 3.89E-07 | 4.69E-06 | 400.6820199 | 603.1920638 | ENSG00000198799 |
| AKAP3 | 0.591092197 | 1.506386731 | 0.358746698 | 0.551942958 | 9.474940033 | 14.26397775 | ENSG00000111254 |

| | | | | | | | |
|---|---|---|---|---|---|---|---|
| TSC22D3 | 0.590813989 | 1.506096268 | 1.78E-09 | 3.20E-08 | 604.2772894 | 909.5941738 | ENSG00000157514 |
| OGDHL | 0.590784024 | 1.506064987 | 2.05E-07 | 2.60E-06 | 594.559523 | 895.4446229 | ENSG00000197444 |
| NDUFAF3 | 0.590783149 | 1.506064074 | 0.000727717 | 0.003986849 | 134.4144672 | 202.5126855 | ENSG00000178057 |
| NA | 0.590654396 | 1.505929671 | 0.515650794 | 0.693461169 | 4.694513485 | 7.07003343 | ENSG00000241255 |
| KLC3 | 0.590639615 | 1.505914243 | 0.012884562 | 0.045270611 | 82.37453754 | 124.0190089 | ENSG00000104892 |
| ZFYVE28 | 0.59059319 | 1.505865784 | 0.001759649 | 0.008513007 | 117.6938769 | 177.3050312 | ENSG00000159733 |
| NA | 0.590442385 | 1.505708384 | 0.499215554 | 0.68011443 | 4.83664962 | 7.257854994 | ENSG00000255992 |
| RUNX1T1 | 0.590173852 | 1.505428148 | 0.000324895 | 0.001966557 | 179.4304819 | 269.7674223 | ENSG00000079102 |
| ANK1 | 0.590091104 | 1.505341805 | 1.96E-05 | 0.000163557 | 267.0972865 | 402.1873902 | ENSG00000029534 |
| NA | 0.58954234 | 1.50476932 | 0.021911287 | 0.069729359 | 76.19180399 | 115.009088 | ENSG00000282508 |
| NA | 0.589321214 | 1.504538697 | 0.408564688 | 0.600213036 | 8.323554731 | 12.43342682 | ENSG00000254332 |
| CTNNA2 | 0.589145959 | 1.504355941 | 6.37E-05 | 0.00046727 | 192.5996533 | 289.534999 | ENSG00000066032 |
| NA | 0.58871829 | 1.503910059 | 0.30675055 | 0.500827092 | 12.00575983 | 18.16552692 | ENSG00000237753 |
| NA | 0.588357374 | 1.503533876 | 0.026909824 | 0.082220027 | 53.88137038 | 81.00247653 | ENSG00000254510 |
| NA | 0.587948541 | 1.503107863 | 0.541112643 | 0.714167763 | 4.357077358 | 6.616495166 | ENSG00000242338 |
| NIPSNAP2 | 0.587935819 | 1.503094609 | 6.84E-10 | 1.30E-08 | 549.9464276 | 826.5932035 | ENSG00000146729 |
| CLIP2 | 0.587915142 | 1.503073066 | 2.05E-17 | 1.01E-15 | 1838.730618 | 2763.595378 | ENSG00000106665 |
| TFPI | 0.58789797 | 1.503055175 | 0.012769225 | 0.045001065 | 74.26700572 | 111.9136008 | ENSG00000003436 |
| USP34 | 0.587864053 | 1.50301984 | 3.30E-11 | 7.72E-10 | 1287.88024 | 1935.400483 | ENSG00000115464 |
| NA | 0.587310264 | 1.502443006 | 0.523273747 | 0.699838045 | 4.83664962 | 7.242771226 | ENSG00000286757 |
| LPAR6 | 0.587276605 | 1.502407953 | 0.121115564 | 0.26144373 | 33.24430968 | 50.06316015 | ENSG00000139679 |
| ZNF77 | 0.587252519 | 1.50238287 | 0.028318746 | 0.085691371 | 70.36168977 | 105.9730846 | ENSG00000175691 |
| ZNF264 | 0.587245806 | 1.50237588 | 0.001199051 | 0.0061492 | 185.0775929 | 277.9360411 | ENSG00000083844 |
| PJA2 | 0.587199499 | 1.502327658 | 1.04E-12 | 3.06E-11 | 1228.110247 | 1843.89053 | ENSG00000198961 |
| LOC257396 | 0.58714501 | 1.502270918 | 0.15529942 | 0.313415096 | 25.72701482 | 38.5319117 | ENSG00000247796 |
| NA | 0.586970795 | 1.502089519 | 0.303288445 | 0.496875102 | 11.90268673 | 17.78988379 | ENSG00000266389 |
| SFMBT2 | 0.586940645 | 1.502058129 | 0.001943033 | 0.009269173 | 137.6944293 | 206.4245319 | ENSG00000198879 |
| RAET1K | 0.586703338 | 1.501811077 | 0.47245167 | 0.657542077 | 6.097324152 | 9.193545812 | ENSG00000218358 |
| SERTM2 | 0.586606685 | 1.501710467 | 0.201829313 | 0.378882254 | 19.62031763 | 29.49244431 | ENSG00000260802 |
| NA | 0.586457451 | 1.501555136 | 0.362852219 | 0.555747709 | 9.859128493 | 14.80823341 | ENSG00000203706 |
| TMEM217 | 0.586138072 | 1.501222764 | 0.525630017 | 0.702129759 | 4.132087374 | 6.211345777 | ENSG00000172738 |
| ZNF311 | 0.585952191 | 1.501029355 | 0.014131828 | 0.048670975 | 67.94459388 | 101.8186007 | ENSG00000197935 |
| C16orf74 | 0.585915678 | 1.500991366 | 0.160476287 | 0.320917944 | 25.51099192 | 38.19023496 | ENSG00000154102 |
| KAZN | 0.585842634 | 1.500915372 | 1.36E-07 | 1.77E-06 | 384.5828297 | 577.3910492 | ENSG00000189337 |
| FAM83B | 0.585564806 | 1.50062636 | 0.056872096 | 0.147728022 | 42.17270586 | 63.26791804 | ENSG00000168143 |
| OSER1-DT | 0.585224253 | 1.500272174 | 0.016827971 | 0.056014658 | 63.04540737 | 94.55380932 | ENSG00000223891 |
| HELZ | 0.585127955 | 1.500172036 | 5.55E-07 | 6.51E-06 | 400.5530901 | 600.6617934 | ENSG00000198265 |
| ARR3 | 0.585056379 | 1.500097611 | 0.534238287 | 0.709027742 | 4.449304247 | 6.725433905 | ENSG00000120500 |
| C5AR2 | 0.584781779 | 1.499812111 | 0.495214661 | 0.677066136 | 5.028792699 | 7.576635484 | ENSG00000134830 |
| FGF21 | 0.5847419 | 1.499770654 | 0.373789302 | 0.567063185 | 11.26066138 | 16.92046087 | ENSG00000105550 |
| SIDT2 | 0.584654421 | 1.499679717 | 2.12E-06 | 2.21E-05 | 386.1258224 | 579.1307627 | ENSG00000149577 |
| SARS2 | 0.584560526 | 1.499582117 | 0.330395974 | 0.524726562 | 11.98238367 | 18.05290095 | ENSG00000104835 |
| INPP5B | 0.584536424 | 1.499557064 | 5.01E-07 | 5.91E-06 | 458.906953 | 688.2047103 | ENSG00000204084 |
| LINC01607 | 0.58453425 | 1.499554805 | 0.645612134 | 0.79292218 | 2.977545159 | 4.403364692 | ENSG00000272138 |
| NA | 0.584122237 | 1.499126615 | 0.256427684 | 0.445149075 | 19.64053688 | 29.60985681 | ENSG00000259605 |
| INO80D | 0.583833291 | 1.498826397 | 0.000129991 | 0.000877685 | 218.1853061 | 326.5788723 | ENSG00000114933 |
| SUMF2 | 0.583745422 | 1.498735112 | 7.04E-11 | 1.57E-09 | 1059.272485 | 1586.417031 | ENSG00000129103 |
| PLSCR1 | 0.583727622 | 1.49871662 | 2.50E-08 | 3.72E-07 | 547.545695 | 820.2742875 | ENSG00000188313 |
| NA | 0.583672834 | 1.498659705 | 0.054705057 | 0.143299914 | 46.96959339 | 70.42487002 | ENSG00000249896 |
| SLC4A7 | 0.583649334 | 1.498635294 | 4.99E-09 | 8.39E-08 | 1187.054178 | 1777.769132 | ENSG00000033867 |
| CPEB1 | 0.583107964 | 1.498073038 | 0.20709106 | 0.385751327 | 19.65759923 | 29.34541393 | ENSG00000214575 |
| NA | 0.582644439 | 1.497591798 | 0.360014788 | 0.553230585 | 9.454720779 | 14.24195756 | ENSG00000259845 |
| MYH9 | 0.582608395 | 1.497554383 | 9.16E-21 | 6.22E-19 | 14767.71811 | 22115.75372 | ENSG00000100345 |
| SGSM1 | 0.582561927 | 1.497506149 | 0.002669813 | 0.012080645 | 115.2161232 | 172.6017686 | ENSG00000167037 |
| USF3 | 0.582283913 | 1.4972176 | 0.000152956 | 0.001011373 | 211.048171 | 315.4596468 | ENSG00000176542 |
| USE1 | 0.582209823 | 1.497140712 | 0.000263543 | 0.001642957 | 164.3891724 | 245.8410619 | ENSG00000053501 |
| WHRN | 0.582150888 | 1.497079555 | 4.19E-05 | 0.000322477 | 264.5477206 | 396.6270889 | ENSG00000095397 |
| RLF | 0.58207622 | 1.497002074 | 2.35E-07 | 2.94E-06 | 488.0616582 | 729.7910461 | ENSG00000117000 |
| C6orf99 | 0.581984848 | 1.496907265 | 0.193826875 | 0.367695732 | 20.50567308 | 30.72242659 | ENSG00000203711 |
| PLEKHA8P1 | 0.581872963 | 1.496791181 | 0.00432207 | 0.018185074 | 107.5675385 | 161.3011083 | ENSG00000134297 |
| TBC1D2 | 0.581458705 | 1.49636145 | 0.000363403 | 0.002165694 | 155.5658262 | 232.8948611 | ENSG00000095383 |
| ZNF397 | 0.581405178 | 1.496305934 | 6.34E-08 | 8.79E-07 | 696.4838589 | 1041.885708 | ENSG00000186812 |
| MARCHF1 | 0.581129753 | 1.496020301 | 0.131013975 | 0.277115066 | 27.83576321 | 41.55132731 | ENSG00000145416 |
| AHR | 0.581040189 | 1.49592743 | 0.086298881 | 0.202874615 | 42.02859289 | 62.65953691 | ENSG00000106546 |
| MAP3K2 | 0.580906213 | 1.495788517 | 0.000103428 | 0.000717952 | 201.0421634 | 300.3915422 | ENSG00000169967 |
| NA | 0.580818514 | 1.495697593 | 0.572807643 | 0.738883513 | 4.168091191 | 6.226429544 | ENSG00000265574 |
| ZNF14 | 0.58063212 | 1.495504364 | 0.011481289 | 0.041252789 | 78.0382808 | 116.7734265 | ENSG00000105708 |
| MAGI3 | 0.579941334 | 1.494788463 | 0.00026619 | 0.001655998 | 202.7780506 | 302.8486956 | ENSG00000081026 |
| NA | 0.579054077 | 1.493869451 | 0.516301252 | 0.694018523 | 4.806959628 | 7.141430181 | ENSG00000229970 |
| NA | 0.578989074 | 1.493802144 | 0.556708827 | 0.726225955 | 4.244533517 | 6.320284516 | ENSG00000206549 |
| KTN1-AS1 | 0.578958248 | 1.493770227 | 0.034477953 | 0.099970961 | 63.48444342 | 94.82887585 | ENSG00000186615 |
| ZNF600 | 0.578781501 | 1.493587233 | 0.017230497 | 0.057057358 | 82.7611088 | 123.894477 | ENSG00000189190 |
| KLF4 | 0.578592712 | 1.493391797 | 0.053429078 | 0.140581281 | 52.77555733 | 78.58121616 | ENSG00000136826 |
| DPP6 | 0.57828247 | 1.493070688 | 0.000756727 | 0.00413356 | 140.924392 | 210.3062108 | ENSG00000130226 |

| | | | | | | | |
|---|---|---|---|---|---|---|---|
| CNTNAP2 | 0.578007954 | 1.492786614 | 9.31E-13 | 2.77E-11 | 1435.002924 | 2141.429071 | ENSG00000174469 |
| LINC00638 | 0.577836263 | 1.492608972 | 0.27131476 | 0.462055294 | 13.91161663 | 20.74907595 | ENSG00000258701 |
| NA | 0.577831883 | 1.49260444 | 0.541311764 | 0.71427255 | 4.15546354 | 6.192574783 | ENSG00000274099 |
| EAF2 | 0.57773744 | 1.492506734 | 0.014399044 | 0.049402422 | 78.96793073 | 118.1578136 | ENSG00000145088 |
| PIK3CB | 0.577720319 | 1.492489021 | 6.26E-07 | 7.24E-06 | 697.097878 | 1039.252653 | ENSG00000051382 |
| DYRK1A | 0.57744307 | 1.492202231 | 5.03E-10 | 9.76E-09 | 740.2394317 | 1104.170661 | ENSG00000157540 |
| GAL3ST1 | 0.57727032 | 1.492023563 | 6.78E-05 | 0.00049287 | 209.6512682 | 312.3944387 | ENSG00000128242 |
| TGFBR3L | 0.576842096 | 1.491580763 | 0.110850085 | 0.244757623 | 31.58060291 | 47.13584738 | ENSG00000260001 |
| NA | 0.576546859 | 1.491275553 | 0.080787079 | 0.19307643 | 62.39894733 | 93.36410885 | ENSG00000278041 |
| ALG13 | 0.576521576 | 1.491249419 | 1.12E-12 | 3.28E-11 | 1052.931913 | 1570.207377 | ENSG00000101901 |
| CLCN5 | 0.576489785 | 1.491216558 | 1.59E-06 | 1.70E-05 | 323.5581832 | 482.064984 | ENSG00000171365 |
| CDX4 | 0.576456282 | 1.491181929 | 0.333478959 | 0.528129457 | 10.59969456 | 15.73072012 | ENSG00000131264 |
| DRAM1 | 0.576424763 | 1.491149351 | 0.003591264 | 0.015508036 | 122.841625 | 182.9387759 | ENSG00000136048 |
| PLLP | 0.575846848 | 1.490552146 | 0.009686354 | 0.035805123 | 84.70938094 | 126.4246358 | ENSG00000102934 |
| NA | 0.575569713 | 1.490265845 | 0.238934077 | 0.42474623 | 17.62972785 | 26.24061711 | ENSG00000270871 |
| NA | 0.575558085 | 1.490253834 | 0.250526076 | 0.438991135 | 15.83957181 | 23.72422795 | ENSG00000237356 |
| NA | 0.575540367 | 1.490235532 | 0.615255721 | 0.770795561 | 3.22591131 | 4.778458166 | ENSG00000235215 |
| C2orf92 | 0.575263371 | 1.489949435 | 0.283534786 | 0.474495145 | 14.74884427 | 22.00261576 | ENSG00000228486 |
| LNCPRESS1 | 0.575182577 | 1.489865998 | 0.000101611 | 0.000706778 | 239.2997895 | 356.4439741 | ENSG00000232301 |
| LSMEM2 | 0.574670653 | 1.48933743 | 0.519032933 | 0.696125362 | 4.783583461 | 7.081318351 | ENSG00000179564 |
| NMB | 0.574332755 | 1.488988649 | 0.000307563 | 0.00187574 | 171.3265729 | 255.4600654 | ENSG00000197696 |
| MICOS13 | 0.573900452 | 1.48854254 | 2.20E-05 | 0.000182395 | 280.4521668 | 417.407821 | ENSG00000174917 |
| VSTM5 | 0.573672411 | 1.48830727 | 0.446771695 | 0.635005107 | 7.430006319 | 11.00488771 | ENSG00000214376 |
| MARCHF10 | 0.573608234 | 1.488241066 | 0.571676215 | 0.737943374 | 3.483748199 | 5.206177396 | ENSG00000173838 |
| NA | 0.573387573 | 1.488013457 | 0.080240796 | 0.192142496 | 45.29564177 | 67.6732581 | ENSG00000273599 |
| TRHDE | 0.572908528 | 1.487519446 | 0.133185385 | 0.280214117 | 29.94017461 | 44.54751181 | ENSG00000072657 |
| MC5R | 0.57274314 | 1.487348928 | 0.543864466 | 0.716162275 | 4.396238088 | 6.578953177 | ENSG00000176136 |
| TMEM256 | 0.572433165 | 1.487029393 | 0.157023661 | 0.315990937 | 22.84307203 | 33.98586513 | ENSG00000205544 |
| TSPAN31 | 0.572289819 | 1.48688165 | 6.52E-05 | 0.000476215 | 235.7845699 | 350.6415766 | ENSG00000135452 |
| HPSE2 | 0.572174254 | 1.48676255 | 0.382389275 | 0.575026598 | 10.40755148 | 15.34802895 | ENSG00000172987 |
| ARHGAP5 | 0.572001481 | 1.48658451 | 4.58E-07 | 5.45E-06 | 388.6685481 | 577.2342225 | ENSG00000100852 |
| TCIRG1 | 0.571999211 | 1.486582172 | 0.000107994 | 0.000746018 | 273.853796 | 407.3803645 | ENSG00000110719 |
| SLC17A9 | 0.571957825 | 1.486539527 | 3.66E-05 | 0.000287094 | 713.410971 | 1061.545248 | ENSG00000101194 |
| PLSCR4 | 0.57177463 | 1.486350777 | 0.252517953 | 0.44125063 | 18.36407779 | 27.18859644 | ENSG00000114698 |
| OLFM3 | 0.571534842 | 1.486103753 | 0.544344502 | 0.716539843 | 4.383610437 | 6.552696109 | ENSG00000118733 |
| LRRC29 | 0.571382317 | 1.485946647 | 0.443683558 | 0.632021449 | 6.92705789 | 10.33347197 | ENSG00000125122 |
| TMEM68 | 0.570736694 | 1.485281817 | 6.70E-05 | 0.000488188 | 268.9809472 | 399.207795 | ENSG00000167904 |
| NA | 0.570631913 | 1.485173947 | 0.039702317 | 0.111819174 | 54.76722949 | 81.18215075 | ENSG00000281091 |
| NEDD4L | 0.570452453 | 1.484989215 | 1.39E-12 | 3.99E-11 | 1266.798919 | 1880.339712 | ENSG00000049759 |
| ETV7 | 0.570421042 | 1.484956883 | 0.287807275 | 0.479308118 | 13.58996507 | 20.20536994 | ENSG00000010030 |
| CYTL1 | 0.570279032 | 1.48481072 | 0.003657274 | 0.015767366 | 110.2070461 | 163.5419929 | ENSG00000170891 |
| PIBF1 | 0.570047222 | 1.484572163 | 0.001468203 | 0.007271959 | 146.4430018 | 217.2596046 | ENSG00000083535 |
| BEND6 | 0.569591731 | 1.484103524 | 0.061434318 | 0.156921671 | 42.8487582 | 63.56131753 | ENSG00000151917 |
| NA | 0.569405308 | 1.483911762 | 0.580281538 | 0.74434377 | 3.454058207 | 5.146065566 | ENSG00000233916 |
| MARCHF2 | 0.569299009 | 1.48380243 | 3.70E-05 | 0.000289271 | 230.0199389 | 341.3293041 | ENSG00000099785 |
| TCP11L2 | 0.56921057 | 1.483711474 | 0.241345227 | 0.427637901 | 15.88316723 | 23.52468343 | ENSG00000166046 |
| DNAJC12 | 0.569178154 | 1.483678137 | 0.150444258 | 0.30654506 | 24.11006039 | 35.82390213 | ENSG00000108176 |
| KCNH5 | 0.569076295 | 1.483573389 | 0.097277855 | 0.22201056 | 32.3350744 | 47.98748699 | ENSG00000140015 |
| HELZ2 | 0.568882781 | 1.483374405 | 1.20E-09 | 2.19E-08 | 645.4790047 | 957.7790151 | ENSG00000130589 |
| BCAS3 | 0.568843603 | 1.483334123 | 3.87E-06 | 3.87E-05 | 304.22086 | 450.9162639 | ENSG00000141376 |
| TSHZ3 | 0.568819756 | 1.483309604 | 0.00029456 | 0.001807742 | 193.0328641 | 286.7639548 | ENSG00000121297 |
| MXD1 | 0.568746021 | 1.483233795 | 0.133830835 | 0.281194955 | 27.06235024 | 40.09575824 | ENSG00000059728 |
| CERS4 | 0.568676101 | 1.483161912 | 1.80E-12 | 5.10E-11 | 1161.659421 | 1722.638137 | ENSG00000090661 |
| NA | 0.568674167 | 1.483159924 | 0.064571234 | 0.162888012 | 52.64722893 | 78.26963561 | ENSG00000286279 |
| TTLL2 | 0.568602862 | 1.48308662 | 0.603460644 | 0.762512924 | 4.591440382 | 6.811802803 | ENSG00000120440 |
| APOBEC3G | 0.568593436 | 1.483076931 | 0.222217699 | 0.404264274 | 19.50273774 | 28.98671833 | ENSG00000239713 |
| CDH11 | 0.568528373 | 1.483010048 | 0.000292636 | 0.001798911 | 188.415973 | 279.2407809 | ENSG00000140937 |
| CCDC186 | 0.568442855 | 1.482922143 | 0.000892236 | 0.004780774 | 211.6205564 | 313.497985 | ENSG00000165813 |
| ZBTB41 | 0.567969018 | 1.482435174 | 0.041770828 | 0.116523258 | 62.12532588 | 91.77242763 | ENSG00000177888 |
| C16orf58 | 0.567805198 | 1.482266851 | 3.99E-07 | 4.80E-06 | 354.3811577 | 525.3592196 | ENSG00000140688 |
| NA | 0.567687848 | 1.482146287 | 0.417476129 | 0.608571958 | 8.840506286 | 13.15410751 | ENSG00000285852 |
| IRF2BP2 | 0.567641192 | 1.482098356 | 4.25E-07 | 5.09E-06 | 440.1796801 | 652.7007911 | ENSG00000168264 |
| FAM86C2P | 0.56756876 | 1.482023948 | 0.02890623 | 0.086957972 | 67.65410549 | 100.2687387 | ENSG00000160172 |
| CSPG4 | 0.567515613 | 1.481969353 | 0.001199798 | 0.006151331 | 153.9854543 | 227.8887376 | ENSG00000173546 |
| COL13A1 | 0.567001219 | 1.48144105 | 0.003906144 | 0.016715878 | 115.0192372 | 170.4049957 | ENSG00000197467 |
| HECW1 | 0.56688259 | 1.48131924 | 0.004562651 | 0.019033123 | 118.6136501 | 176.051603 | ENSG00000002746 |
| NA | 0.566537334 | 1.480964783 | 0.314147862 | 0.507759937 | 11.81045984 | 17.44484623 | ENSG00000280435 |
| SNORA73B | 0.566366604 | 1.480789535 | 0.196758187 | 0.372113902 | 20.06388608 | 29.74406493 | ENSG00000200087 |
| CCDC171 | 0.565909149 | 1.480320075 | 0.054949157 | 0.143766005 | 48.48988625 | 71.88868961 | ENSG00000164989 |
| NELL1 | 0.565682599 | 1.480087634 | 0.482704881 | 0.666104703 | 6.348847215 | 9.33155278 | ENSG00000165973 |
| ZNF160 | 0.565620289 | 1.480023711 | 1.32E-06 | 1.44E-05 | 382.2368697 | 565.4623523 | ENSG00000170949 |
| CRYGS | 0.565384156 | 1.479781488 | 0.575040361 | 0.740467702 | 4.505527319 | 6.582202371 | ENSG00000213139 |
| NA | 0.565223143 | 1.479616345 | 0.269184939 | 0.459483668 | 17.03761175 | 25.23979723 | ENSG00000229546 |
| RGL3 | 0.564623559 | 1.479001545 | 0.041326961 | 0.115482879 | 584.3677687 | 864.5833393 | ENSG00000205517 |

| | | | | | | | |
|---|---|---|---|---|---|---|---|
| ZNF468 | 0.564328153 | 1.478698735 | 0.000242538 | 0.001526904 | 196.7754338 | 290.5265807 | ENSG00000204604 |
| SUCLA2 | 0.564131501 | 1.478497189 | 3.35E-10 | 6.70E-09 | 1060.553003 | 1567.16163 | ENSG00000136143 |
| ALMS1 | 0.563904994 | 1.47826508 | 1.51E-11 | 3.73E-10 | 993.0530597 | 1467.840003 | ENSG00000116127 |
| SPON1 | 0.563867042 | 1.478226193 | 0.00163991 | 0.008017433 | 131.3160875 | 193.9118958 | ENSG00000262655 |
| NA | 0.563593853 | 1.477946302 | 0.042110096 | 0.117152283 | 53.87317742 | 79.61830421 | ENSG00000238278 |
| MYEOV | 0.563558854 | 1.477910448 | 0.325116479 | 0.51923385 | 12.24012286 | 18.06407425 | ENSG00000172927 |
| PYDC1 | 0.563329867 | 1.47767589 | 0.367623364 | 0.560738329 | 9.504630025 | 14.054136 | ENSG00000169900 |
| VSTM2A | 0.563263345 | 1.477607756 | 0.389065513 | 0.581715087 | 8.906200096 | 13.19544834 | ENSG00000170419 |
| NCOA3 | 0.56296946 | 1.477306791 | 8.84E-08 | 1.20E-06 | 492.3694277 | 726.9776021 | ENSG00000124151 |
| TNFRSF10D | 0.562792299 | 1.477125391 | 5.52E-06 | 5.27E-05 | 321.183698 | 474.9804568 | ENSG00000173530 |
| MSS51 | 0.562467045 | 1.476792412 | 0.370585822 | 0.563911136 | 9.845223065 | 14.58699512 | ENSG00000166343 |
| CFAP77 | 0.562362747 | 1.476685652 | 0.368686407 | 0.5616478 | 11.06989378 | 16.43642866 | ENSG00000188523 |
| H1-5 | 0.562340282 | 1.476662659 | 0.612781169 | 0.768999774 | 3.503967452 | 5.176121481 | ENSG00000184357 |
| NA | 0.562202259 | 1.476521393 | 0.595131114 | 0.755943716 | 3.549442009 | 5.236233311 | ENSG00000260645 |
| NA | 0.562066943 | 1.47638291 | 0.638962732 | 0.78834611 | 2.874569755 | 4.276092992 | ENSG00000241429 |
| CTDSPL | 0.561896094 | 1.476208082 | 4.43E-11 | 1.02E-09 | 887.5853026 | 1309.416473 | ENSG00000144677 |
| USP45 | 0.561880618 | 1.476192247 | 3.16E-06 | 3.21E-05 | 386.6882258 | 570.6124484 | ENSG00000123552 |
| LINC02361 | 0.561315513 | 1.475614134 | 0.110735684 | 0.244618847 | 34.38434511 | 50.77861627 | ENSG00000256576 |
| ELOVL3 | 0.561205424 | 1.475501538 | 0.580506873 | 0.744472349 | 3.516595104 | 5.187406401 | ENSG00000119915 |
| CCDC50 | 0.561159207 | 1.47545427 | 2.71E-06 | 2.78E-05 | 387.701331 | 571.6133227 | ENSG00000152492 |
| IL17D | 0.561119732 | 1.475413899 | 0.312463726 | 0.506047072 | 17.29111165 | 25.46318541 | ENSG00000172458 |
| AFF3 | 0.559984135 | 1.474253005 | 0.242607315 | 0.429094759 | 22.13200056 | 32.46704665 | ENSG00000144218 |
| BACH1 | 0.559983272 | 1.474252123 | 7.64E-09 | 1.25E-07 | 584.4728111 | 861.1384625 | ENSG00000156273 |
| NA | 0.559907433 | 1.474174627 | 0.040223411 | 0.112912566 | 61.29233753 | 90.13927133 | ENSG00000137970 |
| PABPC3 | 0.559261521 | 1.47351477 | 0.361872147 | 0.554842685 | 9.871756144 | 14.57950904 | ENSG00000151846 |
| INE1 | 0.558536082 | 1.47277402 | 0.227093096 | 0.410668693 | 19.18108618 | 28.21341188 | ENSG00000224975 |
| PHLPP2 | 0.558527825 | 1.472765591 | 7.74E-06 | 7.14E-05 | 313.7888988 | 461.9125663 | ENSG00000040199 |
| BMPR2 | 0.558500808 | 1.472738011 | 2.54E-06 | 2.62E-05 | 411.3246108 | 605.1679962 | ENSG00000204217 |
| ZEB1 | 0.558468536 | 1.472705067 | 0.44758579 | 0.635762991 | 7.625306311 | 11.19595847 | ENSG00000148516 |
| FANK1 | 0.558318844 | 1.472552268 | 0.478187082 | 0.662286607 | 6.642687922 | 9.853676634 | ENSG00000203780 |
| LIMCH1 | 0.558247083 | 1.472479025 | 0.160196343 | 0.320565647 | 26.84051717 | 39.55628911 | ENSG00000064042 |
| NA | 0.558163431 | 1.472393648 | 0.546316004 | 0.718354974 | 7.186074858 | 10.56252275 | ENSG00000108785 |
| NA | 0.558089394 | 1.472318089 | 0.428445868 | 0.618684513 | 7.728281715 | 11.32735543 | ENSG00000286895 |
| MDM4 | 0.558041322 | 1.47226903 | 1.80E-05 | 0.000151874 | 354.0907821 | 521.4422921 | ENSG00000198625 |
| ZNF284 | 0.557895994 | 1.472120731 | 0.31406718 | 0.507759937 | 14.53963885 | 21.30081769 | ENSG00000186026 |
| NA | 0.557469491 | 1.471685593 | 0.550785236 | 0.721876151 | 4.383610437 | 6.470014438 | ENSG00000235649 |
| PIDD1 | 0.557290221 | 1.471502732 | 2.10E-07 | 2.65E-06 | 535.8226667 | 788.4793263 | ENSG00000177595 |
| ABCG2 | 0.557182127 | 1.471392484 | 0.111612172 | 0.246059236 | 38.48556241 | 56.58327532 | ENSG00000118777 |
| NA | 0.557123186 | 1.471332371 | 0.25680516 | 0.445394336 | 17.67332327 | 25.96328907 | ENSG00000205041 |
| VIL1 | 0.556959605 | 1.471165554 | 0.001326164 | 0.006668237 | 137.6529084 | 202.3774897 | ENSG00000127831 |
| LNX1 | 0.556894642 | 1.47109931 | 0.013403438 | 0.046677811 | 79.40508765 | 116.8670582 | ENSG00000072201 |
| BTBD11 | 0.556885325 | 1.471089809 | 0.000351477 | 0.002102058 | 192.7754331 | 283.3935127 | ENSG00000151136 |
| PPFIA4 | 0.556749586 | 1.470951405 | 0.286180082 | 0.477519715 | 134.1519025 | 197.411299 | ENSG00000143847 |
| SPATA7 | 0.556501318 | 1.470698296 | 0.01942636 | 0.062978513 | 90.66051758 | 132.8551329 | ENSG00000042317 |
| CFAP58 | 0.556500379 | 1.470697339 | 0.373467113 | 0.566807307 | 11.91343525 | 17.44473461 | ENSG00000120051 |
| NSMCE1 | 0.556301431 | 1.470494544 | 0.000287534 | 0.001771519 | 194.052471 | 285.1615871 | ENSG00000169189 |
| AARS1 | 0.555937258 | 1.4701234 | 6.22E-18 | 3.29E-16 | 10657.42275 | 15667.06009 | ENSG00000090861 |
| NA | 0.555710985 | 1.469892844 | 0.597651414 | 0.757703763 | 3.536814357 | 5.187406401 | ENSG00000244009 |
| ND6 | 0.55570911 | 1.469890933 | 0.028960731 | 0.08706173 | 3740.834468 | 5498.643351 | ENSG00000198695 |
| MAPT | 0.555660834 | 1.469841748 | 0.07514789 | 0.182756987 | 38.99798158 | 57.37210356 | ENSG00000186868 |
| TTC9B | 0.555469108 | 1.469646427 | 0.661157716 | 0.802744652 | 2.877726668 | 4.249835924 | ENSG00000174521 |
| GIT2 | 0.555374217 | 1.469549766 | 0.002544889 | 0.011623207 | 132.4332504 | 194.5810501 | ENSG00000139436 |
| URB1-AS1 | 0.555273696 | 1.469447378 | 0.509125579 | 0.688316689 | 5.419294985 | 7.974298799 | ENSG00000256073 |
| AMN1 | 0.555211873 | 1.46938441 | 0.047338518 | 0.128214538 | 66.28818562 | 97.24023674 | ENSG00000151743 |
| UST | 0.554961243 | 1.469129166 | 3.15E-09 | 5.42E-08 | 741.9857445 | 1089.690119 | ENSG00000111962 |
| BCL11B | 0.554565098 | 1.468725817 | 0.023639588 | 0.074059519 | 71.15669746 | 104.4663869 | ENSG00000127152 |
| AEN | 0.553943897 | 1.468093544 | 6.92E-19 | 4.02E-17 | 2938.199366 | 4312.658213 | ENSG00000181026 |
| TECPR2 | 0.553891403 | 1.468040127 | 3.22E-10 | 6.46E-09 | 706.3373348 | 1036.535382 | ENSG00000196663 |
| NA | 0.553643221 | 1.467787606 | 0.152741798 | 0.309931737 | 26.28450258 | 38.56521681 | ENSG00000248121 |
| SUSD3 | 0.553590091 | 1.467733554 | 0.013662567 | 0.047393227 | 95.49548346 | 140.1789856 | ENSG00000157303 |
| PAPPA | 0.553223393 | 1.467360539 | 0.550569217 | 0.721811549 | 4.685042746 | 6.927789582 | ENSG00000182752 |
| SYNPO | 0.553105478 | 1.467240613 | 0.01262946 | 0.044637943 | 89.56921133 | 131.2632058 | ENSG00000171992 |
| MAGEL2 | 0.552796227 | 1.466926134 | 0.005498927 | 0.022330655 | 127.1132929 | 186.8141083 | ENSG00000254585 |
| ARFGAP3 | 0.552640763 | 1.466768067 | 8.16E-06 | 7.45E-05 | 327.7628795 | 480.2376823 | ENSG00000242247 |
| SSH1 | 0.552466917 | 1.466591331 | 1.63E-09 | 2.94E-08 | 653.5386193 | 958.2511011 | ENSG00000084112 |
| MPHOSPH9 | 0.552302321 | 1.466424019 | 1.83E-06 | 1.94E-05 | 437.6369242 | 641.0282378 | ENSG00000051825 |
| SEC31B | 0.552056907 | 1.466174589 | 0.115685663 | 0.252820146 | 53.29882271 | 78.45420798 | ENSG00000075826 |
| ANO7 | 0.551944698 | 1.466060558 | 0.024909596 | 0.077206571 | 63.10794426 | 92.44950597 | ENSG00000146205 |
| NA | 0.551627487 | 1.465738245 | 0.549798208 | 0.721157819 | 6.267970201 | 9.208079926 | ENSG00000225972 |
| SATL1 | 0.551052289 | 1.465153977 | 0.368543224 | 0.561632964 | 10.20919227 | 14.9997422 | ENSG00000184788 |
| NA | 0.550892197 | 1.464991401 | 0.379869539 | 0.572945989 | 10.41711991 | 15.24288906 | ENSG00000233221 |
| PTCD2 | 0.550870002 | 1.464968864 | 0.000841795 | 0.004540694 | 150.3474459 | 220.1944034 | ENSG00000049883 |
| DBT | 0.550670781 | 1.464766581 | 3.02E-07 | 3.70E-06 | 559.6573766 | 819.3445062 | ENSG00000137992 |
| CCNB1IP1 | 0.55065939 | 1.464755016 | 1.37E-11 | 3.40E-10 | 1685.895182 | 2468.314142 | ENSG00000100814 |

| | | | | | | | |
|---|---|---|---|---|---|---|---|
| NFKBIZ | 0.550254442 | 1.464343933 | 0.004639886 | 0.019277215 | 119.4424143 | 175.0923387 | ENSG00000144802 |
| NA | 0.549914971 | 1.463999409 | 0.390247103 | 0.58296444 | 10.27488608 | 15.0673401 | ENSG00000236782 |
| HERC1 | 0.549606724 | 1.463686643 | 3.11E-05 | 0.000248067 | 705.3886457 | 1032.376557 | ENSG00000103657 |
| CADM2 | 0.549281304 | 1.463356526 | 0.004963715 | 0.020457527 | 102.4275774 | 149.8817214 | ENSG00000175161 |
| KDM7A | 0.549087657 | 1.463160119 | 0.000139386 | 0.000931281 | 304.9825172 | 445.7097114 | ENSG00000006459 |
| ZDBF2 | 0.549012654 | 1.463084054 | 7.45E-06 | 6.92E-05 | 402.7549524 | 588.5120898 | ENSG00000204186 |
| NA | 0.548598072 | 1.462663673 | 0.538960823 | 0.712472208 | 4.717889651 | 6.946560577 | ENSG00000214018 |
| ART5 | 0.548454239 | 1.462517857 | 0.0031123 | 0.013747226 | 116.5957531 | 170.4992887 | ENSG00000167311 |
| NA | 0.548348038 | 1.4624102 | 0.355493389 | 0.548968044 | 11.30119759 | 16.63118664 | ENSG00000230658 |
| BTN3A3 | 0.548058798 | 1.462117037 | 0.23294706 | 0.417789899 | 28.29658939 | 41.66158015 | ENSG00000111801 |
| NHLRC2 | 0.548048086 | 1.462106181 | 5.85E-05 | 0.000432843 | 319.228631 | 466.5957522 | ENSG00000196865 |
| EXOC3L1 | 0.547811103 | 1.461866029 | 0.043998115 | 0.121168063 | 65.7986616 | 96.37763862 | ENSG00000179044 |
| INTS4P2 | 0.547708247 | 1.46176181 | 0.599029246 | 0.758920946 | 3.797808159 | 5.577583642 | ENSG00000273024 |
| KLF6 | 0.547457939 | 1.461508216 | 7.88E-07 | 8.95E-06 | 425.664515 | 622.0333953 | ENSG00000067082 |
| GPC5 | 0.54710321 | 1.461148905 | 0.368200678 | 0.561249392 | 10.20919227 | 14.94711644 | ENSG00000179399 |
| LOC101929705 | 0.547002818 | 1.461047232 | 0.365920369 | 0.55902688 | 10.43102534 | 15.32600876 | ENSG00000237686 |
| NA | 0.54685662 | 1.460899182 | 0.268797028 | 0.45904491 | 15.8692618 | 23.13830504 | ENSG00000268869 |
| GXYLT2 | 0.546654237 | 1.46069426 | 0.049348061 | 0.132399677 | 53.87062187 | 78.49583495 | ENSG00000172986 |
| SPTBN4 | 0.546571837 | 1.460610834 | 0.015027158 | 0.051169088 | 94.43880555 | 138.0946557 | ENSG00000160460 |
| NA | 0.546378837 | 1.460415451 | 0.397510445 | 0.58963254 | 10.7046468 | 15.74244307 | ENSG00000237015 |
| OSBPL5 | 0.546299384 | 1.460335024 | 5.99E-05 | 0.000441693 | 302.2974372 | 441.7158304 | ENSG00000021762 |
| DECR2 | 0.546231622 | 1.460266435 | 0.010662219 | 0.038778173 | 95.04039239 | 138.7121316 | ENSG00000242612 |
| ADCY10P1 | 0.546188614 | 1.460222904 | 0.091546752 | 0.212197037 | 62.94864809 | 92.14866068 | ENSG00000161912 |
| MIR4479 | 0.545905989 | 1.459936873 | 0.337720853 | 0.531944773 | 15.30683569 | 22.41167562 | ENSG00000266507 |
| AR | 0.545819851 | 1.459849708 | 7.59E-05 | 0.000544175 | 254.0298877 | 370.2946148 | ENSG00000169083 |
| TEAD1 | 0.54573403 | 1.459762869 | 8.66E-11 | 1.92E-09 | 1318.617617 | 1924.039375 | ENSG00000187079 |
| NA | 0.545653395 | 1.459681283 | 0.496833933 | 0.678123082 | 6.944120231 | 10.11190726 | ENSG00000267249 |
| CLDN19 | 0.545538539 | 1.459565079 | 0.002492308 | 0.011439432 | 146.5912564 | 213.5956916 | ENSG00000164007 |
| TUNAR | 0.545457988 | 1.459483588 | 3.98E-11 | 9.20E-10 | 1018.472403 | 1485.530338 | ENSG00000250366 |
| RRNAD1 | 0.545382966 | 1.459407695 | 0.00013729 | 0.000918935 | 233.6683276 | 340.9330665 | ENSG00000143303 |
| NA | 0.544825079 | 1.458843455 | 0.586799598 | 0.749424538 | 3.797808159 | 5.566298722 | ENSG00000234286 |
| SLC5A5 | 0.544775546 | 1.458793368 | 0.117216764 | 0.255257374 | 37.47749332 | 54.5325854 | ENSG00000105641 |
| LOC105378083 | 0.54474485 | 1.45876233 | 0.567782565 | 0.734939214 | 5.100800335 | 7.498190693 | ENSG00000271913 |
| MYCBP2 | 0.544640481 | 1.458656802 | 2.71E-05 | 0.000219159 | 670.8442827 | 978.4432745 | ENSG00000005810 |
| NA | 0.544604403 | 1.458620326 | 0.119195503 | 0.258306372 | 30.90563296 | 45.05835074 | ENSG00000265519 |
| MYADM | 0.544268723 | 1.45828098 | 4.78E-09 | 8.07E-08 | 771.3193037 | 1125.054701 | ENSG00000179820 |
| GUSBP2 | 0.544212686 | 1.458224338 | 0.175107905 | 0.342419278 | 33.96277735 | 49.65833717 | ENSG00000241549 |
| FITM2 | 0.544130495 | 1.458141265 | 0.006508414 | 0.025786009 | 104.7914117 | 152.6878228 | ENSG00000197296 |
| NA | 0.54398718 | 1.457996423 | 0.149368624 | 0.305158519 | 30.04177454 | 43.86634849 | ENSG00000267787 |
| LOC343052 | 0.543964174 | 1.457973173 | 0.656034084 | 0.79911618 | 3.26507204 | 4.703374188 | ENSG00000231827 |
| RUFY3 | 0.54395964 | 1.45796859 | 1.58E-07 | 2.03E-06 | 500.6411038 | 730.1355743 | ENSG00000018189 |
| RPL12 | 0.543916901 | 1.4579254 | 3.64E-22 | 2.75E-20 | 13610.20527 | 19841.31727 | ENSG00000197958 |
| TEK | 0.543907995 | 1.457916401 | 0.066188625 | 0.165753274 | 46.31238484 | 67.52535166 | ENSG00000120156 |
| ELL2 | 0.543899119 | 1.457907431 | 4.94E-05 | 0.000372941 | 249.3334951 | 363.1134928 | ENSG00000118985 |
| NA | 0.543734035 | 1.457740615 | 0.581889767 | 0.74582662 | 4.244533517 | 6.121178032 | ENSG00000220884 |
| SRGAP1 | 0.543599796 | 1.457604982 | 0.002235097 | 0.010434909 | 146.8339101 | 213.7230665 | ENSG00000196935 |
| MBD5 | 0.543383193 | 1.457386158 | 0.001064716 | 0.005566557 | 153.225368 | 223.6116972 | ENSG00000204406 |
| NA | 0.543262098 | 1.457263835 | 0.343761057 | 0.537484749 | 11.55577987 | 16.76594442 | ENSG00000229944 |
| CENPE | 0.543206924 | 1.457208104 | 4.10E-10 | 8.08E-09 | 740.8960842 | 1079.309979 | ENSG00000138778 |
| IMMP2L | 0.54301342 | 1.457012667 | 0.004518409 | 0.018861304 | 124.245518 | 180.8977707 | ENSG00000184903 |
| LIN7C | 0.542852192 | 1.456849848 | 0.00015374 | 0.001015833 | 292.0276849 | 424.8547311 | ENSG00000148943 |
| CNTNAP5 | 0.542591638 | 1.456586761 | 0.606184908 | 0.764778101 | 3.758647429 | 5.502388045 | ENSG00000155052 |
| ARHGEF3 | 0.542386162 | 1.456379321 | 0.006815589 | 0.026836653 | 101.329356 | 147.8046628 | ENSG00000163947 |
| NA | 0.542137463 | 1.456128285 | 0.469887608 | 0.655539918 | 7.139322525 | 10.33347197 | ENSG00000252051 |
| IAH1 | 0.542132291 | 1.456123065 | 0.000721177 | 0.003958037 | 181.4875396 | 264.534152 | ENSG00000134330 |
| ZMAT5 | 0.542074709 | 1.456064949 | 0.014140028 | 0.048675945 | 110.0507115 | 159.8283769 | ENSG00000100319 |
| SYCE2 | 0.541988831 | 1.455978277 | 0.274467509 | 0.465457116 | 17.45031012 | 25.5285218 | ENSG00000161860 |
| NA | 0.54176142 | 1.455748791 | 0.333791875 | 0.528382698 | 14.83485503 | 21.73634941 | ENSG00000258011 |
| ADA | 0.541693205 | 1.45567996 | 1.39E-05 | 0.000120708 | 313.0986479 | 455.7455244 | ENSG00000196839 |
| LMF1 | 0.541431085 | 1.455415504 | 0.008523983 | 0.032267872 | 110.4049243 | 160.7886204 | ENSG00000103227 |
| PSD4 | 0.541379752 | 1.45536372 | 0.132171033 | 0.278813786 | 38.41731305 | 56.06323845 | ENSG00000125637 |
| CEBPB-AS1 | 0.541317782 | 1.455301207 | 0.455694617 | 0.643051929 | 7.820606303 | 11.47383616 | ENSG00000277449 |
| C17orf107 | 0.541142703 | 1.455124609 | 0.293012885 | 0.485570987 | 19.76816623 | 28.88201647 | ENSG00000205710 |
| CBX5 | 0.540845771 | 1.45482515 | 4.75E-13 | 1.46E-11 | 3180.644129 | 4626.076331 | ENSG00000094916 |
| NA | 0.5408094 | 1.454788473 | 0.12062458 | 0.260653439 | 40.44183211 | 59.00128649 | ENSG00000260261 |
| RGPD3 | 0.540798519 | 1.454777501 | 0.530536939 | 0.706137998 | 5.74913951 | 8.386934262 | ENSG00000153165 |
| ZC3H12B | 0.540717395 | 1.4546957 | 0.072717156 | 0.17836683 | 43.07374818 | 62.6008508 | ENSG00000102053 |
| PPP1R35 | 0.540456139 | 1.454432295 | 0.001297804 | 0.006554085 | 190.4035036 | 277.5351773 | ENSG00000160813 |
| ND5 | 0.540375836 | 1.45435134 | 0.010297099 | 0.03765715 | 16596.39258 | 24137.12164 | ENSG00000198786 |
| NA | 0.539904139 | 1.45387591 | 0.276679165 | 0.467693512 | 16.93907103 | 24.6505135 | ENSG00000279159 |
| ZNF678 | 0.539779158 | 1.453749966 | 0.000397891 | 0.002341857 | 180.9455282 | 263.2798478 | ENSG00000181450 |
| RUNX1 | 0.53960447 | 1.45357395 | 0.472103898 | 0.657299702 | 7.056566374 | 10.20196339 | ENSG00000159216 |
| RGS5 | 0.539595492 | 1.453564904 | 0.000385924 | 0.002282294 | 211.2614052 | 306.6552905 | ENSG00000143248 |
| PTGR2 | 0.539561231 | 1.453530385 | 0.397762631 | 0.589864987 | 9.229730795 | 13.41657501 | ENSG00000140043 |

| | | | | | | | |
|---|---|---|---|---|---|---|---|
| ERMP1 | 0.539168745 | 1.453135006 | 2.09E-09 | 3.70E-08 | 811.8004467 | 1179.164934 | ENSG00000099219 |
| JAK2 | 0.539058607 | 1.453024075 | 0.105305214 | 0.235772751 | 40.05764365 | 58.14810954 | ENSG00000096968 |
| NA | 0.538746264 | 1.45270953 | 0.29384781 | 0.486482833 | 14.25852349 | 20.78336875 | ENSG00000258744 |
| NA | 0.538615359 | 1.452577722 | 0.14741064 | 0.302157576 | 27.72647398 | 40.26100996 | ENSG00000260086 |
| ADAMTS6 | 0.538249018 | 1.452208919 | 0.358218505 | 0.551476183 | 13.44773123 | 19.36826444 | ENSG00000049192 |
| OCA2 | 0.538061455 | 1.452020131 | 0.436451401 | 0.625856114 | 9.253106962 | 13.43154716 | ENSG00000104044 |
| RAB3IL1 | 0.538047979 | 1.452006568 | 0.000106984 | 0.000739589 | 236.3327823 | 343.0461298 | ENSG00000167994 |
| NA | 0.537983768 | 1.451941944 | 0.026786137 | 0.081933275 | 66.41670942 | 96.41755374 | ENSG00000234160 |
| PPP1R16B | 0.537746277 | 1.451702951 | 1.15E-06 | 1.27E-05 | 637.5963804 | 924.8758299 | ENSG00000101445 |
| STOX2 | 0.537574493 | 1.451530104 | 4.69E-08 | 6.70E-07 | 786.7749349 | 1141.920623 | ENSG00000173320 |
| HASPIN | 0.537111224 | 1.451064072 | 4.27E-05 | 0.000327184 | 300.9589449 | 436.9182664 | ENSG00000177602 |
| TMEM141 | 0.537081699 | 1.451034377 | 0.048313972 | 0.130228791 | 58.19938471 | 84.4898529 | ENSG00000244187 |
| NA | 0.536688891 | 1.450639351 | 0.628315553 | 0.780510951 | 3.579132001 | 5.157350486 | ENSG00000233799 |
| BAX | 0.5364253 | 1.450374333 | 3.59E-18 | 1.92E-16 | 2748.446298 | 3986.002449 | ENSG00000087088 |
| HMCN2 | 0.536240169 | 1.45018823 | 1.29E-05 | 0.000112997 | 384.0170513 | 557.2525446 | ENSG00000148357 |
| NA | 0.53602572 | 1.449972682 | 0.592744976 | 0.75444118 | 4.055645048 | 5.941392196 | ENSG00000261348 |
| NA | 0.535979504 | 1.449926234 | 0.612004326 | 0.768689989 | 4.066393563 | 5.945191042 | ENSG00000264281 |
| AFF1 | 0.535911806 | 1.449858198 | 1.03E-08 | 1.65E-07 | 898.4132191 | 1302.448903 | ENSG00000172493 |
| CCNG1 | 0.535861505 | 1.449807648 | 9.04E-13 | 2.71E-11 | 3555.565401 | 5153.320123 | ENSG00000113328 |
| DGAT2 | 0.53581124 | 1.449757136 | 0.016029628 | 0.053911317 | 84.97461403 | 123.1277806 | ENSG00000062282 |
| SLC7A2 | 0.535497227 | 1.449441621 | 6.64E-06 | 6.25E-05 | 498.3422794 | 721.5565329 | ENSG00000003989 |
| DTNB | 0.535467418 | 1.449411672 | 6.10E-08 | 8.47E-07 | 623.8651283 | 904.5801036 | ENSG00000138101 |
| PATJ | 0.535377256 | 1.449321094 | 2.32E-08 | 3.48E-07 | 1020.151 | 1477.991706 | ENSG00000132849 |
| CHKB | 0.535354246 | 1.449297978 | 0.526215316 | 0.702433961 | 6.15039031 | 8.844271373 | ENSG00000100288 |
| BBS9 | 0.535353298 | 1.449297026 | 7.48E-06 | 6.94E-05 | 433.8268193 | 628.1583319 | ENSG00000122507 |
| LOC107985206 | 0.535065716 | 1.449008157 | 0.507589091 | 0.68692604 | 6.341255613 | 9.170537938 | ENSG00000286391 |
| FAM172A | 0.534669702 | 1.448610464 | 0.005344468 | 0.02182295 | 109.1827115 | 158.0052004 | ENSG00000113391 |
| GSTO2 | 0.534175728 | 1.448114549 | 0.008372949 | 0.031787022 | 135.2161947 | 196.022403 | ENSG00000065621 |
| ACTN4 | 0.534150807 | 1.448089536 | 1.30E-14 | 4.76E-13 | 8908.39895 | 12900.83167 | ENSG00000130402 |
| CASP8 | 0.534108003 | 1.448046571 | 0.072622239 | 0.178235284 | 47.25890171 | 68.22236319 | ENSG00000064012 |
| ELAVL2 | 0.534007462 | 1.447945662 | 0.000366896 | 0.002185809 | 261.5171093 | 378.4290524 | ENSG00000107105 |
| LINC00032 | 0.533956366 | 1.44789438 | 0.528048843 | 0.704021174 | 6.858109469 | 9.921274538 | ENSG00000283945 |
| NUTM2D | 0.53373851 | 1.447675756 | 0.333137305 | 0.527678558 | 13.03513056 | 18.85896285 | ENSG00000214562 |
| NAT1 | 0.533609165 | 1.44754597 | 0.049088046 | 0.131835653 | 54.29209192 | 78.40610524 | ENSG00000171428 |
| TMEM132D | 0.533575248 | 1.44751194 | 6.39E-06 | 6.03E-05 | 442.4368936 | 640.4223019 | ENSG00000151952 |
| ZNF654 | 0.533545482 | 1.447482074 | 0.022797639 | 0.071940412 | 89.06448146 | 128.7299808 | ENSG00000175105 |
| GRAMD2A | 0.533533973 | 1.447470528 | 0.213943034 | 0.39403957 | 23.81937659 | 34.6355871 | ENSG00000175318 |
| SMG1P6 | 0.533507497 | 1.447443964 | 0.630240762 | 0.781643622 | 3.549442009 | 5.10472473 | ENSG00000254634 |
| NA | 0.533154506 | 1.447089854 | 0.629988056 | 0.781581472 | 3.146312072 | 4.606158403 | ENSG00000207751 |
| MPL | 0.532972676 | 1.446907482 | 0.63799553 | 0.787529817 | 3.172845151 | 4.636214318 | ENSG00000117400 |
| KLHL36 | 0.532902291 | 1.446836893 | 3.19E-08 | 4.69E-07 | 884.1102283 | 1278.586517 | ENSG00000135686 |
| CNR1 | 0.532059911 | 1.445992341 | 0.19820333 | 0.374045465 | 27.99566074 | 40.55028419 | ENSG00000118432 |
| PTER | 0.531989312 | 1.445921583 | 0.006208158 | 0.024779852 | 113.6698987 | 164.1904008 | ENSG00000165983 |
| NA | 0.531861653 | 1.445793644 | 0.165435861 | 0.328280318 | 29.60264078 | 42.94743738 | ENSG00000280142 |
| NA | 0.531433782 | 1.445364917 | 0.295443835 | 0.488252879 | 15.71765493 | 22.84253242 | ENSG00000249602 |
| IFT80 | 0.531396758 | 1.445327826 | 0.024169031 | 0.075344352 | 86.66464331 | 125.3435074 | ENSG00000068885 |
| NA | 0.531325658 | 1.445256597 | 0.637670813 | 0.787276235 | 3.149468985 | 4.598672329 | ENSG00000282390 |
| KCNV2 | 0.531236102 | 1.445166885 | 0.576134907 | 0.741070468 | 4.45246116 | 6.402416534 | ENSG00000168263 |
| BIRC6 | 0.531123083 | 1.445053676 | 6.14E-07 | 7.12E-06 | 1324.13331 | 1913.35056 | ENSG00000115760 |
| NIPSNAP1 | 0.53105682 | 1.444987307 | 1.70E-17 | 8.53E-16 | 3532.027094 | 5102.606295 | ENSG00000184117 |
| ADAMTSL4 | 0.530815361 | 1.444745485 | 0.010143158 | 0.037189574 | 130.601085 | 188.4222184 | ENSG00000143382 |
| NA | 0.530601735 | 1.444531571 | 0.162635562 | 0.324082088 | 30.74317987 | 44.42111618 | ENSG00000136149 |
| GMFG | 0.530510877 | 1.4444406 | 0.54775302 | 0.719340642 | 4.747579643 | 6.867677753 | ENSG00000130755 |
| GAB2 | 0.530135346 | 1.444064664 | 9.27E-05 | 0.000651654 | 279.180165 | 402.7876953 | ENSG00000033327 |
| NOL3 | 0.530115218 | 1.444044517 | 0.052016976 | 0.137627786 | 54.53098733 | 78.7821864 | ENSG00000140939 |
| CNGA4 | 0.530012055 | 1.443941261 | 0.668949465 | 0.807760075 | 2.930792826 | 4.227266083 | ENSG00000132259 |
| TTC7B | 0.529802564 | 1.443731604 | 1.50E-05 | 0.00012937 | 310.1439002 | 447.7438692 | ENSG00000165914 |
| VASH2 | 0.529648341 | 1.443577279 | 6.98E-11 | 1.56E-09 | 1393.208665 | 2010.729172 | ENSG00000143494 |
| NA | 0.529492171 | 1.443421021 | 0.568627395 | 0.735753989 | 5.383291167 | 7.7451364 | ENSG00000243824 |
| NOSTRIN | 0.529405206 | 1.443334015 | 0.548889913 | 0.720425299 | 5.349166485 | 7.696309491 | ENSG00000163072 |
| ITSN2 | 0.529295014 | 1.443223778 | 0.000127308 | 0.000863978 | 224.3066078 | 323.5736015 | ENSG00000198399 |
| NA | 0.529263463 | 1.443192217 | 0.421774673 | 0.612967285 | 9.696675406 | 13.87011328 | ENSG00000234232 |
| MYSM1 | 0.529088267 | 1.443016971 | 2.60E-09 | 4.53E-08 | 992.5230147 | 1431.334039 | ENSG00000162601 |
| BMP7 | 0.528808625 | 1.442737294 | 4.04E-05 | 0.000312216 | 375.1724937 | 540.4111008 | ENSG00000101144 |
| IFI27L1 | 0.528630307 | 1.442558981 | 0.003395718 | 0.014780702 | 179.4370662 | 259.1973421 | ENSG00000165948 |
| NA | 0.528338563 | 1.442267294 | 0.080671383 | 0.19289942 | 85.30680354 | 122.9743231 | ENSG00000248131 |
| NA | 0.528034963 | 1.441963816 | 0.607947148 | 0.766065623 | 4.058801961 | 5.90005136 | ENSG00000279058 |
| CLCN3 | 0.5280096 | 1.441938467 | 4.02E-06 | 3.99E-05 | 703.0411448 | 1013.055741 | ENSG00000109572 |
| ERCC6 | 0.527881528 | 1.441810467 | 0.000488854 | 0.002806824 | 190.8998451 | 275.3132382 | ENSG00000225830 |
| SPOCK1 | 0.527805694 | 1.441734682 | 1.02E-05 | 9.07E-05 | 358.2651872 | 516.0676935 | ENSG00000152377 |
| NAA80 | 0.527760039 | 1.441689058 | 0.032040944 | 0.094396256 | 75.40616933 | 108.6498358 | ENSG00000243477 |
| INTS6-AS1 | 0.527623457 | 1.441552577 | 0.558744772 | 0.727844785 | 5.289786501 | 7.711831292 | ENSG00000236778 |
| CHODL | 0.527361591 | 1.441290943 | 0.231294356 | 0.415751101 | 22.59145127 | 32.45207451 | ENSG00000154645 |
| TAB1 | 0.527313173 | 1.441242573 | 8.01E-11 | 1.78E-09 | 919.1027672 | 1324.110584 | ENSG00000100324 |

| | | | | | | | |
|---|---|---|---|---|---|---|---|
| ATM | 0.52707372 | 1.441003381 | 0.000106406 | 0.000736416 | 615.3987384 | 887.5109726 | ENSG00000149311 |
| MOB1B | 0.526446673 | 1.440377205 | 0.000907774 | 0.004854206 | 288.5702592 | 415.5656269 | ENSG00000173542 |
| NA | 0.526358666 | 1.440289342 | 0.285584376 | 0.476805768 | 19.35428778 | 28.00043256 | ENSG00000253981 |
| NLRP12 | 0.526324862 | 1.440255595 | 0.001193709 | 0.006128149 | 178.0943346 | 256.9728236 | ENSG00000142405 |
| APOBEC3C | 0.526275847 | 1.440206663 | 5.42E-06 | 5.20E-05 | 357.6847891 | 515.3760097 | ENSG00000244509 |
| DPP10 | 0.526155611 | 1.44008664 | 0.001131178 | 0.005851313 | 188.7959222 | 272.059444 | ENSG00000175497 |
| CENPP | 0.526049816 | 1.43998104 | 0.021030409 | 0.067377285 | 93.41642507 | 134.1676852 | ENSG00000188312 |
| ABO | 0.525550423 | 1.439482673 | 0.321090561 | 0.514905321 | 14.95677191 | 21.48797798 | ENSG00000175164 |
| SIGLEC10 | 0.525351099 | 1.439283807 | 0.203643288 | 0.381441162 | 30.44490447 | 43.9230995 | ENSG00000142512 |
| ATXN2-AS | 0.525337057 | 1.439269798 | 0.656923939 | 0.79954022 | 2.954168993 | 4.238551003 | ENSG00000258099 |
| XRN1 | 0.525073324 | 1.439006714 | 0.000999973 | 0.005266697 | 171.7106636 | 247.1019762 | ENSG00000114127 |
| CCSAP | 0.525034501 | 1.438967991 | 8.67E-05 | 0.000613474 | 396.7466604 | 570.6352874 | ENSG00000154429 |
| LRRC75A | 0.524855488 | 1.438789452 | 2.34E-05 | 0.000191749 | 317.9984358 | 457.2895006 | ENSG00000181350 |
| TNFRSF21 | 0.524696406 | 1.43863081 | 1.08E-11 | 2.72E-10 | 1362.536944 | 1960.328404 | ENSG00000146072 |
| POT1 | 0.524334617 | 1.438270085 | 0.000303471 | 0.001855666 | 295.1470731 | 423.6936693 | ENSG00000128513 |
| C5orf24 | 0.524289096 | 1.438224704 | 0.013981242 | 0.048299785 | 163.9112687 | 235.6927833 | ENSG00000181904 |
| GVQW3 | 0.524160531 | 1.438096543 | 0.030469663 | 0.090704261 | 127.4910698 | 183.7109604 | ENSG00000179240 |
| LCOR | 0.524150339 | 1.438086384 | 0.000100235 | 0.000697877 | 301.6162512 | 433.2585013 | ENSG00000196233 |
| TNKS | 0.524074417 | 1.438010706 | 7.80E-08 | 1.06E-06 | 815.0231185 | 1171.115981 | ENSG00000173273 |
| NA | 0.523502227 | 1.437440487 | 0.096855549 | 0.221346579 | 40.89121072 | 58.86499138 | ENSG00000225031 |
| SPDYA | 0.523286916 | 1.437225977 | 0.628440021 | 0.780570128 | 3.877407397 | 5.558812648 | ENSG00000163806 |
| NIT2 | 0.523242478 | 1.437181708 | 0.000391392 | 0.002309473 | 284.345186 | 408.3138557 | ENSG00000114021 |
| SUN2 | 0.523190642 | 1.437130071 | 0.000121051 | 0.000826661 | 239.3645737 | 344.0760582 | ENSG00000100242 |
| GRM4 | 0.52307281 | 1.437012698 | 2.73E-08 | 4.04E-07 | 733.5131233 | 1054.59569 | ENSG00000124493 |
| CEP44 | 0.52303601 | 1.436976044 | 0.00061612 | 0.003444694 | 186.3159963 | 267.6720392 | ENSG00000164118 |
| C5orf47 | 0.522930397 | 1.436870853 | 0.585618008 | 0.748592604 | 4.135244286 | 5.978934184 | ENSG00000185056 |
| NA | 0.522916775 | 1.436857286 | 0.566625907 | 0.734240688 | 4.860025786 | 6.965331571 | ENSG00000242861 |
| CAPG | 0.522910453 | 1.436850989 | 5.24E-08 | 7.39E-07 | 659.8948401 | 947.6924804 | ENSG00000042493 |
| LRP6 | 0.52288799 | 1.436828617 | 2.29E-08 | 3.44E-07 | 1116.73107 | 1604.001842 | ENSG00000070018 |
| PCDH18 | 0.522521499 | 1.436463664 | 3.29E-08 | 4.82E-07 | 817.5570953 | 1173.366112 | ENSG00000189184 |
| PYCR2 | 0.522460168 | 1.436402599 | 4.11E-06 | 4.06E-05 | 423.0533066 | 608.2238403 | ENSG00000143811 |
| LINC01097 | 0.522183109 | 1.436126775 | 0.273602114 | 0.464871193 | 17.66573167 | 25.23880954 | ENSG00000281202 |
| DNHD1 | 0.521845378 | 1.435790621 | 0.029659959 | 0.088750029 | 98.97973532 | 142.3679375 | ENSG00000179532 |
| SLC17A5 | 0.521792502 | 1.435737998 | 0.000137725 | 0.000921516 | 253.7298309 | 364.0657005 | ENSG00000119899 |
| MMAA | 0.521731444 | 1.435677236 | 0.022370058 | 0.070825037 | 88.74844466 | 127.1059108 | ENSG00000151611 |
| LINC00491 | 0.521602519 | 1.435548944 | 0.076299381 | 0.184853951 | 49.37396393 | 70.96565325 | ENSG00000250682 |
| SPEN | 0.521557997 | 1.435504643 | 2.31E-08 | 3.48E-07 | 1648.516876 | 2366.439977 | ENSG00000065526 |
| PGM1 | 0.521423078 | 1.435370402 | 1.10E-07 | 1.46E-06 | 1279.807953 | 1835.818561 | ENSG00000079739 |
| RHBDD2 | 0.521189342 | 1.435137872 | 5.84E-07 | 6.80E-06 | 627.9501551 | 900.9667435 | ENSG00000005486 |
| MICU3 | 0.521180495 | 1.435129072 | 0.041745588 | 0.116470379 | 60.47542683 | 86.6954973 | ENSG00000155970 |
| RHOD | 0.521147579 | 1.435096329 | 0.000157033 | 0.001036117 | 239.6933359 | 344.0932918 | ENSG00000173156 |
| NA | 0.521142301 | 1.435091079 | 0.583389417 | 0.746670773 | 4.899186517 | 7.006672407 | ENSG00000287516 |
| MOSPD1 | 0.52105051 | 1.434999775 | 1.29E-06 | 1.41E-05 | 420.5692165 | 603.7774858 | ENSG00000101928 |
| NETO1 | 0.52104815 | 1.434997427 | 0.000401476 | 0.002358466 | 257.9280931 | 369.4844277 | ENSG00000166342 |
| FAM169A | 0.520730248 | 1.434681256 | 1.59E-07 | 2.04E-06 | 558.2690879 | 800.7662011 | ENSG00000198780 |
| KDR | 0.52072773 | 1.434678752 | 1.76E-10 | 3.71E-09 | 3170.844239 | 4547.780447 | ENSG00000128052 |
| NEK3 | 0.520473397 | 1.434425855 | 1.40E-05 | 0.000121498 | 517.0213494 | 741.4436578 | ENSG00000136098 |
| TMEM42 | 0.520178834 | 1.434133009 | 0.004298819 | 0.018111913 | 133.2205688 | 190.9123505 | ENSG00000169964 |
| HIBADH | 0.520172985 | 1.434127195 | 1.64E-07 | 2.10E-06 | 512.9449436 | 736.0800412 | ENSG00000106049 |
| COLEC12 | 0.520141619 | 1.434096016 | 0.093706274 | 0.215665229 | 48.1266161 | 69.11578168 | ENSG00000158270 |
| DAW1 | 0.520100046 | 1.434054692 | 0.486308242 | 0.669482059 | 8.446847088 | 12.1109591 | ENSG00000123977 |
| NA | 0.520045078 | 1.434000054 | 0.680510951 | 0.815245229 | 3.503967452 | 4.992098764 | ENSG00000213985 |
| LIMS2 | 0.519338024 | 1.433297433 | 0.071471069 | 0.175822244 | 47.4763001 | 68.11116295 | ENSG00000072163 |
| ZNF34 | 0.519337125 | 1.43329654 | 0.01641574 | 0.054928564 | 84.16845185 | 120.7043703 | ENSG00000196378 |
| WDR88 | 0.518815047 | 1.432777957 | 0.445252284 | 0.633428449 | 8.956109342 | 12.83912586 | ENSG00000166359 |
| STYK1 | 0.518711996 | 1.432675618 | 0.229508751 | 0.4134467 | 24.03677497 | 34.42443125 | ENSG00000060140 |
| FOXN2 | 0.51870619 | 1.432669852 | 0.000969022 | 0.005131616 | 224.932638 | 322.2770719 | ENSG00000170802 |
| DBH | 0.51854032 | 1.432505144 | 0.452595617 | 0.639896801 | 9.620233081 | 13.84853112 | ENSG00000123454 |
| SH2D3C | 0.51848851 | 1.432453701 | 0.020518412 | 0.065976447 | 87.2206579 | 125.2288347 | ENSG00000095370 |
| PTTG1 | 0.518432432 | 1.432398023 | 1.69E-11 | 4.13E-10 | 1441.81323 | 2064.498589 | ENSG00000164611 |
| LOC105376199 | 0.518341219 | 1.432307463 | 0.652221252 | 0.796943154 | 3.113465167 | 4.497219664 | ENSG00000228317 |
| NA | 0.518090855 | 1.432058924 | 0.571846049 | 0.738012302 | 4.789897287 | 6.841420684 | ENSG00000214263 |
| VWA3A | 0.517935942 | 1.431905161 | 0.668144786 | 0.807209144 | 3.215162795 | 4.624929397 | ENSG00000175267 |
| ZNF681 | 0.517744758 | 1.43171542 | 0.015164156 | 0.051522031 | 116.3570531 | 166.6919294 | ENSG00000196172 |
| PHF11 | 0.517578515 | 1.431550451 | 0.042451338 | 0.117889286 | 73.64184731 | 105.4249269 | ENSG00000136147 |
| NA | 0.517498592 | 1.431471147 | 0.646360632 | 0.793354169 | 3.19937823 | 4.606158403 | ENSG00000232385 |
| NA | 0.517496982 | 1.43146955 | 0.537324552 | 0.711203176 | 6.408227199 | 9.216115653 | ENSG00000272669 |
| FAM174A | 0.517416901 | 1.431390094 | 0.024214087 | 0.075425864 | 79.27370003 | 113.6193647 | ENSG00000174132 |
| LOC101930085 | 0.517394427 | 1.431367797 | 0.143934441 | 0.296784442 | 38.74705987 | 55.51953244 | ENSG00000271122 |
| NA | 0.517328212 | 1.431302103 | 0.430112677 | 0.620204481 | 8.912513922 | 12.76772911 | ENSG00000176933 |
| C17orf50 | 0.517294341 | 1.4312685 | 0.648267286 | 0.794589371 | 3.436995866 | 4.985050724 | ENSG00000270806 |
| PTGER4 | 0.517290885 | 1.431265072 | 0.40073703 | 0.592616303 | 10.11696538 | 14.53448098 | ENSG00000171522 |
| CISD3 | 0.517076019 | 1.431051923 | 0.000195949 | 0.001261014 | 258.6078885 | 370.425176 | ENSG00000277972 |
| NT5DC3 | 0.517043059 | 1.431019229 | 0.000421184 | 0.002463307 | 305.8664447 | 437.975176 | ENSG00000111696 |

| | | | | | | | |
|---|---|---|---|---|---|---|---|
| PYGL | 0.517028162 | 1.431004453 | 7.01E-11 | 1.57E-09 | 1213.596969 | 1736.107387 | ENSG00000100504 |
| ZFP36 | 0.51690385 | 1.430881155 | 0.022013126 | 0.069981351 | 91.49114578 | 131.214817 | ENSG00000128016 |
| RYR1 | 0.516628872 | 1.430608453 | 4.34E-09 | 7.35E-08 | 1005.220363 | 1438.937155 | ENSG00000196218 |
| PCED1A | 0.516440882 | 1.430422051 | 0.000203927 | 0.001306908 | 479.6817077 | 686.4978731 | ENSG00000132635 |
| SERPINE2 | 0.516375996 | 1.430357719 | 2.02E-11 | 4.88E-10 | 1287.871108 | 1841.497654 | ENSG00000135919 |
| PAAF1 | 0.516350502 | 1.430332442 | 9.66E-06 | 8.67E-05 | 369.9938604 | 529.5768412 | ENSG00000175575 |
| ZNF236 | 0.516326774 | 1.430308918 | 0.000409315 | 0.002402231 | 219.7671512 | 314.5566955 | ENSG00000130856 |
| FUT4 | 0.516227175 | 1.430210178 | 0.002492287 | 0.011439432 | 143.9638727 | 205.7757976 | ENSG00000196371 |
| OXER1 | 0.516009845 | 1.429994745 | 0.388411616 | 0.581300136 | 11.94953676 | 17.13723904 | ENSG00000162881 |
| KIAA1109 | 0.515908607 | 1.429894401 | 7.62E-05 | 0.000545776 | 613.4625077 | 877.1139389 | ENSG00000138688 |
| UNC45B | 0.515821987 | 1.429808553 | 0.494061407 | 0.676137792 | 7.089413279 | 10.13860236 | ENSG00000141161 |
| YPEL1 | 0.515664735 | 1.429652713 | 0.007990192 | 0.030590832 | 138.7372244 | 198.5995653 | ENSG00000100027 |
| CD19 | 0.515095938 | 1.42908917 | 0.524133582 | 0.7006341 | 5.733354945 | 8.165369556 | ENSG00000177455 |
| NA | 0.515039946 | 1.429033707 | 0.492077141 | 0.674617778 | 7.317560176 | 10.50252254 | ENSG00000253930 |
| NA | 0.514961277 | 1.428955785 | 0.315486144 | 0.509025309 | 15.51604111 | 22.26888211 | ENSG00000236924 |
| ADM2 | 0.514894222 | 1.42888937 | 2.27E-05 | 0.000187056 | 419.5415144 | 599.8979653 | ENSG00000128165 |
| PPP1R1C | 0.514773517 | 1.428769826 | 0.130517045 | 0.27652068 | 33.91286811 | 48.32123963 | ENSG00000150722 |
| SERTAD1 | 0.514690479 | 1.428687591 | 0.001681281 | 0.008189453 | 178.2497748 | 254.9238455 | ENSG00000197019 |
| PPP1R32 | 0.514639732 | 1.428637337 | 0.293506325 | 0.486047733 | 17.30176246 | 24.7444801 | ENSG00000162148 |
| NA | 0.514636584 | 1.42863422 | 0.649537807 | 0.795357672 | 3.232225135 | 4.636214318 | ENSG00000261770 |
| NA | 0.514629272 | 1.428626979 | 0.439202037 | 0.627518544 | 8.998426986 | 12.84292471 | ENSG00000286138 |
| NA | 0.514622182 | 1.428619959 | 0.061514829 | 0.157040749 | 55.52179868 | 79.40901211 | ENSG00000274652 |
| NA | 0.514561676 | 1.428560045 | 0.665739561 | 0.805826608 | 3.090089 | 4.448392755 | ENSG00000188985 |
| GPCPD1 | 0.51436142 | 1.428361764 | 0.000758089 | 0.004139781 | 181.3984696 | 259.0071156 | ENSG00000125772 |
| ROBO3 | 0.514320545 | 1.428321296 | 0.023794089 | 0.074405205 | 161.1726341 | 230.566995 | ENSG00000154134 |
| CRYZL1 | 0.514295819 | 1.428296816 | 0.000496295 | 0.002846019 | 211.5468802 | 301.8976904 | ENSG00000205758 |
| LOC101927752 | 0.514155448 | 1.428157853 | 0.025049426 | 0.077571607 | 89.9793315 | 128.1791151 | ENSG00000253982 |
| NA | 0.51414663 | 1.428149124 | 0.012771425 | 0.045001065 | 106.0515826 | 151.4349442 | ENSG00000251363 |
| NA | 0.513832206 | 1.427837904 | 0.319603873 | 0.51336128 | 17.99883081 | 25.83601737 | ENSG00000255306 |
| ERO1B | 0.513539392 | 1.427548135 | 0.058315929 | 0.150575012 | 62.67186972 | 89.12128069 | ENSG00000086619 |
| PPDPF | 0.513509763 | 1.427518817 | 7.69E-06 | 7.10E-05 | 792.7640596 | 1131.569911 | ENSG00000125534 |
| NDUFAF1 | 0.513451077 | 1.42746075 | 6.28E-05 | 0.000460838 | 323.3161308 | 461.0447436 | ENSG00000137806 |
| VCX3B | 0.513395533 | 1.427405794 | 0.604956361 | 0.763569953 | 4.688199659 | 6.781746888 | ENSG00000205642 |
| CIT | 0.513189613 | 1.42720207 | 5.56E-09 | 9.28E-08 | 1093.407983 | 1560.066121 | ENSG00000122966 |
| TMPPE | 0.512516522 | 1.426536362 | 0.227714934 | 0.411351714 | 21.26833754 | 30.40039691 | ENSG00000188167 |
| LY75 | 0.512133323 | 1.426157506 | 0.201781754 | 0.378831343 | 27.86733234 | 39.60044111 | ENSG00000054219 |
| HECA | 0.512028069 | 1.426053462 | 0.05125948 | 0.136343744 | 68.80676856 | 98.23649333 | ENSG00000112406 |
| ZFAS1 | 0.511922899 | 1.425949509 | 0.005609973 | 0.022736766 | 125.4836131 | 179.0819687 | ENSG00000177410 |
| NA | 0.511815145 | 1.42584301 | 0.560539906 | 0.729465427 | 5.038263437 | 7.205229238 | ENSG00000213598 |
| NA | 0.511327353 | 1.425360997 | 0.266763839 | 0.456696932 | 19.27331307 | 27.34680012 | ENSG00000225234 |
| LOC101928673 | 0.510792659 | 1.424832825 | 0.088117932 | 0.206157251 | 47.16983174 | 67.20332197 | ENSG00000203739 |
| TBC1D8 | 0.510716251 | 1.424757366 | 5.53E-07 | 6.50E-06 | 674.4122304 | 961.0405588 | ENSG00000204634 |
| TTLL6 | 0.510613245 | 1.424655644 | 0.079868275 | 0.191522655 | 76.29468169 | 108.9260016 | ENSG00000170703 |
| NA | 0.510465241 | 1.424509498 | 0.417804412 | 0.608964059 | 9.617076168 | 13.63021561 | ENSG00000213080 |
| NA | 0.509952952 | 1.424003756 | 0.448174994 | 0.635975449 | 11.33847918 | 16.27910104 | ENSG00000279227 |
| NA | 0.509693677 | 1.423747864 | 0.124692703 | 0.267297543 | 41.85794685 | 59.53260828 | ENSG00000269001 |
| ATAD5 | 0.509489464 | 1.423546347 | 0.000162046 | 0.001066536 | 318.1645268 | 452.6322623 | ENSG00000176208 |
| DEF6 | 0.50939814 | 1.423456238 | 0.00056061 | 0.003173591 | 211.6640541 | 301.1273067 | ENSG00000023892 |
| FHL3 | 0.509350038 | 1.423408778 | 1.56E-05 | 0.000133425 | 376.6340678 | 536.0514903 | ENSG00000183386 |
| SLC12A9-AS1 | 0.509222861 | 1.423283306 | 0.319847833 | 0.513518689 | 15.3396826 | 21.84864896 | ENSG00000236305 |
| LANCL1 | 0.509193997 | 1.423254831 | 9.36E-10 | 1.75E-08 | 902.4170984 | 1283.743892 | ENSG00000115365 |
| INTS12 | 0.508841375 | 1.422907003 | 0.017298978 | 0.057263666 | 101.7027959 | 144.8597815 | ENSG00000138785 |
| B3GNTL1 | 0.508757737 | 1.422824515 | 0.01666395 | 0.055658615 | 135.9032886 | 193.9009776 | ENSG00000175711 |
| LOC100507053 | 0.508139039 | 1.42221447 | 0.192944297 | 0.366509133 | 28.15298008 | 40.10803085 | ENSG00000246090 |
| PLPP1 | 0.507943468 | 1.422021688 | 1.46E-05 | 0.000126254 | 450.478927 | 640.0261038 | ENSG00000067113 |
| NA | 0.507844416 | 1.421924058 | 0.014117777 | 0.048662614 | 93.68876875 | 133.1720899 | ENSG00000197258 |
| SLC7A5 | 0.507815775 | 1.42189583 | 8.62E-15 | 3.22E-13 | 9518.763611 | 13534.45045 | ENSG00000103257 |
| ARHGAP4 | 0.507373239 | 1.421459741 | 5.79E-08 | 8.09E-07 | 695.8469521 | 989.3772288 | ENSG00000089820 |
| FCHO2 | 0.506975009 | 1.421067427 | 0.012164339 | 0.043256689 | 123.3125006 | 175.2247947 | ENSG00000157107 |
| TBC1D3 | 0.506850961 | 1.420945243 | 0.166881987 | 0.330403123 | 34.79439023 | 49.46204185 | ENSG00000274512 |
| NA | 0.506817842 | 1.420912624 | 0.563759584 | 0.732061945 | 5.055325778 | 7.19025709 | ENSG00000270820 |
| NA | 0.506764274 | 1.420859866 | 0.16828809 | 0.332308913 | 31.99763827 | 45.56852841 | ENSG00000229939 |
| PRRC2B | 0.506689414 | 1.420786141 | 1.45E-10 | 3.09E-09 | 5819.469093 | 8268.056014 | ENSG00000130723 |
| TMEM147-AS1 | 0.506539291 | 1.420638305 | 0.00857067 | 0.032398286 | 260.6042286 | 370.7066067 | ENSG00000236144 |
| FAM126A | 0.506358081 | 1.420459877 | 1.85E-10 | 3.87E-09 | 1834.809383 | 2604.975231 | ENSG00000122591 |
| CUL9 | 0.506224741 | 1.420328599 | 0.006382693 | 0.025363815 | 177.497611 | 252.3364976 | ENSG00000112659 |
| RPUSD3 | 0.5060829 | 1.420188963 | 1.06E-10 | 2.31E-09 | 1248.326732 | 1771.984822 | ENSG00000156990 |
| DOCK1 | 0.506060095 | 1.420166514 | 1.68E-08 | 2.60E-07 | 1333.307457 | 1893.082293 | ENSG00000150760 |
| TRIM58 | 0.505924421 | 1.420032965 | 0.400683924 | 0.59258505 | 11.81993058 | 16.73957573 | ENSG00000162722 |
| RGS16 | 0.505821481 | 1.419931645 | 0.000224216 | 0.001422667 | 312.0293272 | 442.9915162 | ENSG00000143333 |
| BCAR1 | 0.505700918 | 1.41981299 | 1.33E-11 | 3.30E-10 | 2734.450595 | 3882.829421 | ENSG00000050820 |
| NA | 0.505663582 | 1.419776246 | 0.484553123 | 0.667859565 | 7.291027097 | 10.38229888 | ENSG00000250299 |
| TULP2 | 0.505466938 | 1.419582739 | 0.235158818 | 0.420127821 | 21.38837528 | 30.35157 | ENSG00000104804 |
| NA | 0.505412654 | 1.419529326 | 0.298735287 | 0.492025221 | 16.13784721 | 22.91717836 | ENSG00000227398 |

| | | | | | | | |
|---|---|---|---|---|---|---|---|
| SLC17A7 | 0.505169833 | 1.419290424 | 0.054996626 | 0.143827778 | 59.92374924 | 84.90617559 | ENSG00000104888 |
| PIWIL4 | 0.504781384 | 1.418908328 | 0.53599793 | 0.710391197 | 6.708381732 | 9.470547434 | ENSG00000134627 |
| SPRED1 | 0.504740586 | 1.418868204 | 4.07E-06 | 4.04E-05 | 629.2721114 | 892.2155308 | ENSG00000166068 |
| NA | 0.504711518 | 1.418839616 | 0.640501684 | 0.789444284 | 3.427525127 | 4.924938894 | ENSG00000237438 |
| MKNK1 | 0.50448657 | 1.418618405 | 0.001546861 | 0.007630963 | 194.3822178 | 275.6388435 | ENSG00000079277 |
| SHROOM4 | 0.50427274 | 1.418408159 | 0.001863339 | 0.008931792 | 158.0695115 | 224.3507108 | ENSG00000158352 |
| COL6A2 | 0.504250794 | 1.418386583 | 8.59E-14 | 2.87E-12 | 3166.050522 | 4490.943384 | ENSG00000142173 |
| NA | 0.504125002 | 1.418262915 | 0.582333784 | 0.746040945 | 4.816430366 | 6.788794928 | ENSG00000279300 |
| GREM2 | 0.503703216 | 1.417848333 | 0.633103474 | 0.783736436 | 3.883721223 | 5.47233213 | ENSG00000180875 |
| ZNF333 | 0.503616293 | 1.41776291 | 0.006903795 | 0.027097616 | 120.3492668 | 170.706869 | ENSG00000160961 |
| NA | 0.503506555 | 1.417655072 | 0.535130401 | 0.709650959 | 6.259777238 | 8.911869277 | ENSG00000227627 |
| ENDOG | 0.503438259 | 1.417587963 | 0.004542918 | 0.018955076 | 169.7291613 | 240.6722117 | ENSG00000167136 |
| CEP250 | 0.503436295 | 1.417586033 | 3.08E-08 | 4.54E-07 | 781.7387611 | 1108.4409 | ENSG00000126001 |
| SLC26A2 | 0.503247848 | 1.417400878 | 0.000915833 | 0.004887433 | 231.8764652 | 328.6778798 | ENSG00000155850 |
| EPB41L2 | 0.503224771 | 1.417378206 | 2.29E-15 | 9.18E-14 | 2921.003856 | 4140.382736 | ENSG00000079819 |
| SH3RF1 | 0.503126373 | 1.417281538 | 0.000168313 | 0.001102301 | 248.6133437 | 352.3187732 | ENSG00000154447 |
| CITED1 | 0.503037968 | 1.417194693 | 0.670432234 | 0.808475039 | 3.821184326 | 5.47656901 | ENSG00000125931 |
| CLPSL2 | 0.502795212 | 1.416956247 | 0.324087601 | 0.518260535 | 15.09575114 | 21.32327591 | ENSG00000196748 |
| RAB11FIP1 | 0.502759904 | 1.41692157 | 8.08E-07 | 9.16E-06 | 582.137795 | 825.0659431 | ENSG00000156675 |
| MAMLD1 | 0.502650242 | 1.416813871 | 0.469398838 | 0.655190112 | 8.875232328 | 12.54616441 | ENSG00000013619 |
| ABHD14B | 0.50264589 | 1.416809597 | 0.044241545 | 0.121693796 | 71.73037575 | 101.609197 | ENSG00000114779 |
| STX12 | 0.502514601 | 1.41668067 | 0.003018391 | 0.01339307 | 168.1143716 | 237.7830938 | ENSG00000117758 |
| NA | 0.502503346 | 1.416669618 | 0.649839827 | 0.795523587 | 3.119778992 | 4.440906681 | ENSG00000272129 |
| CSNK1G1 | 0.50241332 | 1.416581218 | 0.002133131 | 0.010049681 | 207.8969054 | 294.6944994 | ENSG00000169118 |
| ISG15 | 0.502398545 | 1.416566711 | 3.38E-05 | 0.000267236 | 316.6836127 | 448.3953115 | ENSG00000187608 |
| IGFBP3 | 0.501952269 | 1.416128585 | 0.077142335 | 0.18648181 | 53.05677039 | 75.01126974 | ENSG00000146674 |
| NA | 0.50180887 | 1.415987834 | 0.281711548 | 0.472981414 | 17.73595787 | 25.17164967 | ENSG00000267325 |
| GRIA4 | 0.501786176 | 1.41596556 | 0.000730815 | 0.00400264 | 191.8262404 | 271.4333828 | ENSG00000152578 |
| NA | 0.501723736 | 1.415904279 | 0.487906127 | 0.670934275 | 7.364312509 | 10.43861186 | ENSG00000282221 |
| ZNF619 | 0.501416471 | 1.415602752 | 0.025321022 | 0.078190575 | 89.52323311 | 126.7244221 | ENSG00000177873 |
| DES | 0.501404305 | 1.415590814 | 0.09544996 | 0.218781791 | 45.05930191 | 63.91764001 | ENSG00000175084 |
| SZT2 | 0.501318531 | 1.415506653 | 0.000376329 | 0.002234997 | 255.669211 | 362.0315201 | ENSG00000198198 |
| FREM2 | 0.501301723 | 1.415490163 | 5.37E-05 | 0.000400652 | 877.9848678 | 1241.985724 | ENSG00000150893 |
| NA | 0.500507664 | 1.414711292 | 0.397045228 | 0.589225419 | 13.11029511 | 18.60453107 | ENSG00000272145 |
| PDCD4 | 0.50034403 | 1.414550841 | 0.000214693 | 0.001367394 | 243.6266732 | 344.4395403 | ENSG00000150593 |
| NA | 0.500298303 | 1.414506006 | 0.297211663 | 0.490169155 | 19.96209075 | 28.22155923 | ENSG00000272419 |
| NA | 0.500195584 | 1.414405298 | 0.010395917 | 0.037973575 | 151.1947457 | 214.1397643 | ENSG00000214425 |
| IDH3G | 0.500095662 | 1.414307339 | 2.61E-06 | 2.68E-05 | 504.095643 | 713.6187029 | ENSG00000067829 |
| LINC00664 | 0.499816645 | 1.414033839 | 0.073981969 | 0.180625034 | 50.94614287 | 72.07552349 | ENSG00000268658 |
| NA | 0.499739503 | 1.413958232 | 0.132936443 | 0.279950464 | 40.03050921 | 56.80301655 | ENSG00000230724 |
| SLC6A6 | 0.499626685 | 1.413847665 | 1.21E-13 | 3.94E-12 | 3254.019947 | 4600.257935 | ENSG00000131389 |
| NA | 0.499507871 | 1.413731231 | 0.540236425 | 0.713361492 | 7.414221755 | 10.47626547 | ENSG00000272894 |
| CACTIN-AS1 | 0.499296569 | 1.413524187 | 0.507883425 | 0.687132274 | 7.622149398 | 10.70812741 | ENSG00000226800 |
| COL6A4P1 | 0.498601683 | 1.412843515 | 0.644432289 | 0.792088868 | 3.146312072 | 4.459677675 | ENSG00000230524 |
| HSD11B1L | 0.498555828 | 1.412798609 | 0.064208374 | 0.162196811 | 75.63294075 | 106.6839775 | ENSG00000167733 |
| ZNF548 | 0.498506994 | 1.412750788 | 0.024013883 | 0.074991291 | 84.55579723 | 119.4213242 | ENSG00000188785 |
| ALDH1L2 | 0.498444201 | 1.4126893 | 0.010209267 | 0.037402365 | 139.9262973 | 197.5179988 | ENSG00000136010 |
| UNC80 | 0.498310459 | 1.412558345 | 0.451012263 | 0.638580864 | 8.575077795 | 12.15273797 | ENSG00000144406 |
| SHARPIN | 0.498265139 | 1.412513973 | 3.28E-08 | 4.81E-07 | 689.7110307 | 974.3270821 | ENSG00000179526 |
| PPP1R17 | 0.498128085 | 1.412379793 | 0.24514411 | 0.432220409 | 20.93849301 | 29.61255635 | ENSG00000106341 |
| C12orf66 | 0.498074781 | 1.412327609 | 0.025543114 | 0.078771393 | 91.57142143 | 129.3339018 | ENSG00000174206 |
| FAM114A2 | 0.497848098 | 1.412105715 | 0.002002751 | 0.009505502 | 172.5693132 | 243.7912481 | ENSG00000055147 |
| ADAM8 | 0.49778799 | 1.412046883 | 0.021656599 | 0.069061151 | 85.53841562 | 120.8358789 | ENSG00000151651 |
| ZNF37A | 0.49769767 | 1.411958484 | 0.061585899 | 0.157200532 | 58.98986002 | 83.30489869 | ENSG00000075407 |
| DNER | 0.497588397 | 1.411851544 | 0.069108509 | 0.171420918 | 57.72781001 | 81.32015772 | ENSG00000187957 |
| PPP1R12B | 0.497406859 | 1.411673898 | 0.001289358 | 0.006514985 | 249.106145 | 352.0786926 | ENSG00000077157 |
| FAM162A | 0.497039652 | 1.411314632 | 1.81E-07 | 2.31E-06 | 723.5832801 | 1020.62713 | ENSG00000114023 |
| BDNF-AS | 0.49697385 | 1.411250263 | 0.314178858 | 0.507759937 | 17.67214319 | 24.97254319 | ENSG00000245573 |
| CTNND1 | 0.496830832 | 1.411110369 | 0.014656939 | 0.05012024 | 103.4271604 | 145.9065139 | ENSG00000198561 |
| GALNT11 | 0.496510441 | 1.410797027 | 2.20E-06 | 2.29E-05 | 538.7148322 | 760.016922 | ENSG00000178234 |
| RPL23P8 | 0.496103065 | 1.410398715 | 0.647031322 | 0.793861944 | 3.202535143 | 4.519789505 | ENSG00000219451 |
| NA | 0.495949653 | 1.410248745 | 0.552379145 | 0.723161339 | 5.653755707 | 8.041896703 | ENSG00000232024 |
| IFT74 | 0.495593811 | 1.409900948 | 0.001971658 | 0.009381952 | 178.4037644 | 251.1550787 | ENSG00000096872 |
| FEM1C | 0.495534194 | 1.409842689 | 0.000413159 | 0.00242096 | 326.8812371 | 460.2124331 | ENSG00000145780 |
| SNORD46 | 0.495511709 | 1.409820715 | 0.437511288 | 0.626268718 | 11.48249445 | 16.27910104 | ENSG00000200913 |
| NA | 0.495471381 | 1.409781307 | 0.440425945 | 0.628637233 | 10.37480227 | 14.75604569 | ENSG00000230979 |
| CKMT2 | 0.495389241 | 1.409701043 | 0.218824816 | 0.400167651 | 26.20796256 | 36.93041156 | ENSG00000131730 |
| CPNE7 | 0.495270563 | 1.409585084 | 0.314067814 | 0.507759937 | 349.1054946 | 492.2485063 | ENSG00000178773 |
| DNAH5 | 0.495254625 | 1.409569511 | 0.354373069 | 0.547662123 | 13.7396928 | 19.36925213 | ENSG00000039139 |
| EDEM1 | 0.495218603 | 1.409534317 | 5.82E-05 | 0.000430985 | 366.1352217 | 515.8905988 | ENSG00000134109 |
| EFCAB13 | 0.495209498 | 1.409525422 | 0.481766276 | 0.665443003 | 7.618992485 | 10.77996219 | ENSG00000178852 |
| C3orf62 | 0.494950614 | 1.409272513 | 0.232450504 | 0.417101137 | 23.69114588 | 33.49858372 | ENSG00000188315 |
| PLEKHM3 | 0.49459161 | 1.408921869 | 0.013273781 | 0.046322021 | 124.2564619 | 175.2450946 | ENSG00000178385 |
| AMDHD1 | 0.494491979 | 1.408824573 | 0.551331488 | 0.722322422 | 5.42560881 | 7.663004382 | ENSG00000139344 |

| | | | | | | | |
|---|---|---|---|---|---|---|---|
| DGAT1 | 0.494245118 | 1.408583529 | 8.11E-06 | 7.41E-05 | 489.693901 | 690.2716236 | ENSG00000185000 |
| MARS2 | 0.493860859 | 1.408208405 | 7.29E-07 | 8.34E-06 | 751.3814535 | 1058.666123 | ENSG00000247626 |
| LIN28B-AS1 | 0.493841856 | 1.408189857 | 0.465246476 | 0.651570061 | 8.664147771 | 12.2128498 | ENSG00000203809 |
| CFAP99 | 0.493801962 | 1.408150917 | 0.677322304 | 0.813471461 | 3.718208921 | 5.300032367 | ENSG00000206113 |
| UBASH3B | 0.493778822 | 1.408128331 | 0.000118789 | 0.000813912 | 272.2386004 | 383.3090035 | ENSG00000154127 |
| YOD1 | 0.49360453 | 1.407958226 | 0.00716207 | 0.027908049 | 146.7469146 | 206.3438969 | ENSG00000180667 |
| IQCA1 | 0.493551705 | 1.407906674 | 0.005677552 | 0.022955442 | 134.1097802 | 188.9511671 | ENSG00000132321 |
| IL3RA | 0.493423012 | 1.407781089 | 0.2196228 | 0.401023051 | 25.09583569 | 35.44176062 | ENSG00000185291 |
| MCEE | 0.49330293 | 1.407663919 | 0.029510581 | 0.088345842 | 92.61510355 | 130.4992492 | ENSG00000124370 |
| MC1R | 0.49310593 | 1.407471715 | 0.369623788 | 0.562770664 | 13.49448357 | 18.93784567 | ENSG00000258839 |
| FAM114A1 | 0.49304712 | 1.407414342 | 0.002773476 | 0.012458458 | 174.2121017 | 244.8817977 | ENSG00000197712 |
| ABLIM2 | 0.492985778 | 1.407354502 | 0.060404721 | 0.154780577 | 61.51870298 | 86.59871955 | ENSG00000163995 |
| NA | 0.492093888 | 1.406484729 | 0.415377149 | 0.606714483 | 13.47742122 | 19.00219438 | ENSG00000253477 |
| FSTL1 | 0.492060115 | 1.406451804 | 2.32E-15 | 9.25E-14 | 3927.789572 | 5523.750722 | ENSG00000163430 |
| SPOCK3 | 0.491630568 | 1.40603311 | 0.604623432 | 0.763311161 | 4.191467358 | 5.873794292 | ENSG00000196104 |
| JADE1 | 0.49157155 | 1.405975593 | 4.74E-14 | 1.63E-12 | 2687.521974 | 3777.864131 | ENSG00000077684 |
| LRCH1 | 0.491527009 | 1.405932186 | 0.000260126 | 0.00162493 | 257.3209892 | 361.5605963 | ENSG00000136141 |
| FUT9 | 0.49148494 | 1.405891189 | 0.043803358 | 0.120703456 | 72.39339447 | 101.7435168 | ENSG00000172461 |
| NA | 0.491366663 | 1.405775935 | 0.366144867 | 0.559228716 | 13.55711816 | 19.06500575 | ENSG00000287670 |
| PVT1 | 0.491306989 | 1.405717789 | 0.001962177 | 0.009346438 | 186.2043469 | 261.8423969 | ENSG00000249859 |
| TMEM245 | 0.491272141 | 1.405683835 | 2.98E-09 | 5.14E-08 | 1376.795099 | 1934.347116 | ENSG00000106771 |
| NA | 0.49123694 | 1.405649537 | 0.645358744 | 0.792859536 | 3.804121985 | 5.307080408 | ENSG00000280106 |
| STX16 | 0.491123901 | 1.405539405 | 0.000121491 | 0.000829358 | 921.5313485 | 1295.740406 | ENSG00000124222 |
| ZNF267 | 0.491119234 | 1.405534858 | 0.000911939 | 0.004872263 | 233.0865541 | 327.204822 | ENSG00000185947 |
| TNS1 | 0.490782859 | 1.405207185 | 0.0031741 | 0.013966943 | 158.7070045 | 222.7171794 | ENSG00000079308 |
| GABRG3 | 0.490711265 | 1.405137454 | 0.582783884 | 0.746399526 | 5.31631958 | 7.490266586 | ENSG00000182256 |
| WDR5B | 0.490574009 | 1.405003777 | 0.123070865 | 0.264524576 | 51.08630217 | 71.7877866 | ENSG00000196981 |
| AKAP11 | 0.49055538 | 1.404985635 | 5.43E-06 | 5.21E-05 | 464.3121245 | 651.9519812 | ENSG00000023516 |
| CAPS | 0.490530323 | 1.404961233 | 0.496609397 | 0.678082046 | 7.906519366 | 11.17013943 | ENSG00000105519 |
| HYLS1 | 0.490450887 | 1.404883877 | 0.003107322 | 0.013731786 | 162.5914248 | 228.5442337 | ENSG00000198331 |
| NA | 0.490318367 | 1.404754836 | 0.46211943 | 0.648661127 | 8.85629085 | 12.49777553 | ENSG00000265194 |
| SLC52A3 | 0.490275562 | 1.404713157 | 0.001800531 | 0.008685853 | 178.2382521 | 250.1243146 | ENSG00000101276 |
| FAM160B1 | 0.490135287 | 1.404576582 | 0.003404829 | 0.014813402 | 201.6972072 | 283.5738481 | ENSG00000151553 |
| NA | 0.489773941 | 1.404224828 | 0.048742148 | 0.131134702 | 67.73982315 | 95.02644499 | ENSG00000242071 |
| ZYX | 0.489707588 | 1.404160246 | 4.30E-09 | 7.28E-08 | 2923.903252 | 4106.066151 | ENSG00000159840 |
| NA | 0.489279081 | 1.403743246 | 0.600197437 | 0.759725903 | 4.919405771 | 6.830135764 | ENSG00000271335 |
| MYO6 | 0.489204943 | 1.403671111 | 6.48E-08 | 8.96E-07 | 774.6283319 | 1086.687674 | ENSG00000196586 |
| AMOT | 0.489177329 | 1.403644244 | 4.13E-05 | 0.000317875 | 341.3053932 | 478.7277756 | ENSG00000126016 |
| NEXMIF | 0.489076821 | 1.403546461 | 0.378754271 | 0.572023464 | 13.12735745 | 18.32230488 | ENSG00000050030 |
| STAM2 | 0.48898047 | 1.403452727 | 0.008440051 | 0.032015535 | 149.4722376 | 209.3179496 | ENSG00000115145 |
| TMEM181 | 0.48877802 | 1.403255798 | 4.43E-06 | 4.35E-05 | 540.0527911 | 757.9856157 | ENSG00000146433 |
| RBP7 | 0.48868532 | 1.403165634 | 0.176587549 | 0.344619331 | 30.99154602 | 43.34367498 | ENSG00000162444 |
| RNF170 | 0.48860029 | 1.403082937 | 0.003192154 | 0.014036388 | 195.4151136 | 274.3109926 | ENSG00000120925 |
| NAT9 | 0.488498889 | 1.402984324 | 2.43E-05 | 0.000198946 | 720.0798762 | 1011.135567 | ENSG00000109065 |
| KCNH8 | 0.488471908 | 1.402958086 | 0.138446491 | 0.288408144 | 39.12295767 | 54.66734318 | ENSG00000183960 |
| DAPK3 | 0.488439679 | 1.402926744 | 2.35E-10 | 4.85E-09 | 1978.497476 | 2776.883496 | ENSG00000167657 |
| ADAMTS14 | 0.488413385 | 1.402901176 | 0.077685532 | 0.187550301 | 58.23707226 | 81.68930246 | ENSG00000138316 |
| MKI67 | 0.487785273 | 1.402290522 | 7.61E-07 | 8.66E-06 | 3000.351277 | 4207.483064 | ENSG00000148773 |
| RNF141 | 0.487777838 | 1.402283295 | 0.000136406 | 0.000914999 | 433.5560087 | 607.7429683 | ENSG00000110315 |
| TSEN15 | 0.487772403 | 1.402278012 | 4.03E-07 | 4.84E-06 | 634.1314155 | 889.1250222 | ENSG00000198860 |
| NA | 0.487762159 | 1.402268055 | 0.562568647 | 0.730892646 | 5.736511858 | 8.086924765 | ENSG00000226200 |
| ZNF839 | 0.487047337 | 1.401573436 | 0.014213652 | 0.048891462 | 116.0491116 | 162.8871982 | ENSG00000022976 |
| FEZ2 | 0.486917882 | 1.401447676 | 6.13E-06 | 5.82E-05 | 432.0893985 | 605.1894122 | ENSG00000171055 |
| LOC100507250 | 0.486803177 | 1.401336255 | 0.652463325 | 0.797031264 | 3.424368214 | 4.838569996 | ENSG00000247363 |
| STX17 | 0.48678386 | 1.401317492 | 0.000255927 | 0.001601942 | 283.3095015 | 396.6042556 | ENSG00000136874 |
| CRNDE | 0.486643015 | 1.401180694 | 0.035400129 | 0.102127704 | 91.99725112 | 128.9182404 | ENSG00000245694 |
| NA | 0.486630325 | 1.401168369 | 0.017812575 | 0.058585823 | 145.1330194 | 203.8179121 | ENSG00000263072 |
| SLC7A3 | 0.486545291 | 1.401085785 | 1.64E-11 | 4.01E-10 | 2750.818078 | 3854.151073 | ENSG00000165349 |
| TMEFF1 | 0.486543513 | 1.401084058 | 0.006837338 | 0.026905143 | 242.0974359 | 338.9659025 | ENSG00000241697 |
| TMEM101 | 0.486079128 | 1.40063314 | 8.01E-06 | 7.35E-05 | 401.277368 | 562.1361024 | ENSG00000091947 |
| SHTN1 | 0.485963184 | 1.400520581 | 0.000425947 | 0.002487241 | 307.9785379 | 430.6649899 | ENSG00000187164 |
| ZNF398 | 0.485721765 | 1.400286239 | 8.20E-11 | 1.82E-09 | 1470.450269 | 2058.982989 | ENSG00000197024 |
| PRKAG1 | 0.485609655 | 1.400177428 | 4.66E-05 | 0.000353974 | 402.3324976 | 563.0186252 | ENSG00000181929 |
| NA | 0.485572275 | 1.40014115 | 0.182749864 | 0.353226195 | 32.50502139 | 45.51612589 | ENSG00000261167 |
| CEP295 | 0.485368272 | 1.399943179 | 0.003455337 | 0.01499443 | 174.0967917 | 244.1037534 | ENSG00000166004 |
| SCO2 | 0.485347025 | 1.399922562 | 0.659354398 | 0.801500257 | 3.229068223 | 4.508504585 | ENSG00000284194 |
| MAP4K2 | 0.485100827 | 1.399683684 | 1.27E-06 | 1.39E-05 | 519.3302983 | 727.2358814 | ENSG00000168067 |
| SGK1 | 0.484954645 | 1.399541866 | 0.00026618 | 0.001655998 | 322.9442769 | 451.1839157 | ENSG00000118515 |
| ZFAND1 | 0.484918555 | 1.399506857 | 0.000105071 | 0.000727998 | 373.7589944 | 523.0811907 | ENSG00000104231 |
| DYRK4 | 0.484797203 | 1.399389142 | 0.194586287 | 0.368721219 | 31.06355366 | 43.29484807 | ENSG00000010219 |
| TEX13C | 0.484685288 | 1.399280591 | 0.50354208 | 0.683776217 | 10.3109876 | 14.50073784 | ENSG00000282815 |
| NA | 0.484548733 | 1.399148151 | 0.664923714 | 0.805385316 | 3.169688238 | 4.42962176 | ENSG00000206147 |
| FAM178B | 0.484326939 | 1.398933069 | 0.653853169 | 0.797703095 | 3.208848969 | 4.47844867 | ENSG00000168754 |
| NR1I2 | 0.484325064 | 1.398931251 | 0.297503693 | 0.490493821 | 19.09517311 | 26.66800993 | ENSG00000144852 |

| | | | | | | | |
|---|---|---|---|---|---|---|---|
| PARS2 | 0.484302518 | 1.398909389 | 0.000292737 | 0.001798934 | 267.5563086 | 374.4836148 | ENSG00000162396 |
| GOLGB1 | 0.484252314 | 1.398860709 | 1.65E-05 | 0.000140093 | 973.3201157 | 1361.599339 | ENSG00000173230 |
| STX8 | 0.483911889 | 1.398530667 | 0.000400003 | 0.002351301 | 331.3864939 | 463.2105216 | ENSG00000170310 |
| LGALS3BP | 0.48318847 | 1.397829569 | 6.82E-10 | 1.30E-08 | 1198.728981 | 1674.951896 | ENSG00000108679 |
| ACTN2 | 0.482723482 | 1.397379113 | 0.113484136 | 0.24917219 | 45.79543329 | 64.13171864 | ENSG00000077522 |
| LPIN2 | 0.482619651 | 1.397278548 | 8.74E-05 | 0.000617807 | 351.214431 | 490.9176578 | ENSG00000101577 |
| TMEM63A | 0.482601606 | 1.397261071 | 0.073253263 | 0.179405512 | 1592.958532 | 2225.981314 | ENSG00000196187 |
| KRBA2 | 0.482592359 | 1.397252115 | 0.448238688 | 0.635989014 | 10.43102534 | 14.65470464 | ENSG00000184619 |
| NA | 0.482546837 | 1.397208028 | 0.584012363 | 0.747158497 | 7.714376287 | 10.84799813 | ENSG00000264769 |
| GNAS-AS1 | 0.482414538 | 1.397079906 | 0.445361685 | 0.633452623 | 9.825003812 | 13.73915435 | ENSG00000235590 |
| TTC17 | 0.482318995 | 1.396987387 | 1.42E-07 | 1.84E-06 | 938.6436962 | 1311.468495 | ENSG00000052841 |
| TRIM32 | 0.481943591 | 1.396623924 | 5.00E-05 | 0.000376765 | 416.6548358 | 582.3264288 | ENSG00000119401 |
| NA | 0.481672911 | 1.396361912 | 0.460188968 | 0.646882239 | 10.20603536 | 14.17392162 | ENSG00000248590 |
| DYNC1H1 | 0.48160739 | 1.396298497 | 0.029949497 | 0.089471951 | 6482.312811 | 9051.141854 | ENSG00000197102 |
| SLC25A16 | 0.48155717 | 1.396249892 | 0.030432717 | 0.090623389 | 83.71027892 | 116.9615745 | ENSG00000122912 |
| ZNF225 | 0.481459875 | 1.396155733 | 0.036551982 | 0.104674664 | 96.94386639 | 135.086708 | ENSG00000256294 |
| DHX58 | 0.481174799 | 1.395879881 | 0.063559177 | 0.16106177 | 62.60617591 | 87.26578679 | ENSG00000108771 |
| ZBTB18 | 0.481144187 | 1.395850262 | 0.001398159 | 0.006965988 | 258.0109469 | 360.3075348 | ENSG00000179456 |
| SEC22A | 0.481143019 | 1.395849133 | 0.108708556 | 0.241149537 | 54.26763337 | 75.9045766 | ENSG00000121542 |
| B9D1 | 0.480945472 | 1.395658013 | 0.000171736 | 0.001121946 | 302.111608 | 421.7865776 | ENSG00000108641 |
| FAM151A | 0.48089193 | 1.395606218 | 0.528992357 | 0.704790766 | 7.033190208 | 9.740501015 | ENSG00000162391 |
| GTF2IP12 | 0.480846172 | 1.395561954 | 0.020734369 | 0.066578448 | 96.42257784 | 134.6607409 | ENSG00000283050 |
| SAMD8 | 0.480750877 | 1.395469776 | 1.87E-05 | 0.000156931 | 411.2121647 | 573.5250582 | ENSG00000156671 |
| PDE12 | 0.480705744 | 1.39542612 | 0.000150158 | 0.000994293 | 363.3881458 | 507.1050135 | ENSG00000174840 |
| ELF1 | 0.480677434 | 1.395398738 | 0.005356155 | 0.021856227 | 189.2746977 | 263.6585882 | ENSG00000120690 |
| SPDYE5 | 0.480645558 | 1.395367907 | 0.62163447 | 0.775152125 | 4.925719596 | 6.792593775 | ENSG00000170092 |
| USP9X | 0.480574449 | 1.395299133 | 1.58E-07 | 2.04E-06 | 4666.689654 | 6510.847954 | ENSG00000124486 |
| GPR89A | 0.4805551 | 1.395280419 | 0.014868903 | 0.050695408 | 124.079908 | 172.7572325 | ENSG00000117262 |
| NA | 0.480382022 | 1.39511304 | 0.219238343 | 0.400604922 | 32.7539889 | 45.79009311 | ENSG00000229852 |
| MATN3 | 0.480312027 | 1.395045355 | 0.016345354 | 0.054742462 | 124.3615119 | 173.7484164 | ENSG00000132031 |
| TSPEAR | 0.480014774 | 1.394757949 | 0.249514789 | 0.438009765 | 28.00946847 | 38.96711701 | ENSG00000175894 |
| C3 | 0.479923599 | 1.394669807 | 0.000593329 | 0.003329326 | 257.6677006 | 359.1277649 | ENSG00000125730 |
| NA | 0.479815144 | 1.394564966 | 0.40859436 | 0.600213036 | 14.00256574 | 19.50769713 | ENSG00000254978 |
| ORMDL2 | 0.47967747 | 1.394431891 | 0.004719801 | 0.019559666 | 177.8217955 | 247.8445654 | ENSG00000123353 |
| OSGEPL1 | 0.479517847 | 1.394277617 | 0.042329978 | 0.117640399 | 78.85213227 | 109.822629 | ENSG00000128694 |
| RINL | 0.479388283 | 1.394152406 | 0.001865571 | 0.008939116 | 182.5200673 | 254.2206217 | ENSG00000187994 |
| CORO7 | 0.479309009 | 1.394075802 | 0.090064768 | 0.209653406 | 49.79603534 | 69.30316521 | ENSG00000262246 |
| NIFK-AS1 | 0.478872817 | 1.393654374 | 0.06005087 | 0.154046065 | 67.77088862 | 94.62239491 | ENSG00000236859 |
| LOC101927483 | 0.478818229 | 1.393601643 | 0.648079447 | 0.794567224 | 5.494459533 | 7.572398604 | ENSG00000286595 |
| FBN1 | 0.47870109 | 1.393488494 | 0.002610525 | 0.011870333 | 184.0290102 | 256.3936138 | ENSG00000166147 |
| PXMP4 | 0.478696945 | 1.39348449 | 0.086534239 | 0.203273206 | 57.14832156 | 79.59573437 | ENSG00000101417 |
| ARHGEF33 | 0.478661883 | 1.393450625 | 0.669465917 | 0.807856359 | 3.179158977 | 4.41833684 | ENSG00000214694 |
| IRF6 | 0.478277771 | 1.393079673 | 2.98E-05 | 0.000239292 | 387.6437324 | 539.8206548 | ENSG00000117595 |
| LAMC1 | 0.477907298 | 1.392721987 | 2.23E-12 | 6.22E-11 | 3991.408922 | 5558.212041 | ENSG00000135862 |
| TNRC6B | 0.477801999 | 1.392620339 | 8.03E-06 | 7.35E-05 | 640.4811723 | 891.6105741 | ENSG00000100354 |
| NA | 0.4777664 | 1.392585976 | 0.623601374 | 0.776735743 | 4.916248858 | 6.76253786 | ENSG00000283538 |
| NA | 0.4772904 | 1.392126585 | 1.99E-08 | 3.01E-07 | 11770.46138 | 16385.83991 | ENSG00000211459 |
| NBPF14 | 0.477196072 | 1.392035566 | 0.31902044 | 0.512945679 | 17.40483557 | 24.34301793 | ENSG00000270629 |
| ESD | 0.477192617 | 1.392032232 | 1.80E-08 | 2.76E-07 | 775.4557054 | 1079.176821 | ENSG00000139684 |
| ADAT2 | 0.477187134 | 1.392026942 | 0.001194139 | 0.006128149 | 217.0034567 | 301.9813597 | ENSG00000189007 |
| KIAA1671 | 0.476896063 | 1.391746121 | 5.26E-08 | 7.43E-07 | 833.2014496 | 1159.888956 | ENSG00000197077 |
| LINC01012 | 0.476833208 | 1.391685488 | 0.106954342 | 0.23834185 | 48.91333314 | 68.22021331 | ENSG00000281706 |
| SHROOM3 | 0.476810502 | 1.391663585 | 2.94E-10 | 5.92E-09 | 1245.687518 | 1732.973888 | ENSG00000138771 |
| TAOK1 | 0.476472582 | 1.391337656 | 3.71E-05 | 0.000290243 | 1216.435294 | 1691.507277 | ENSG00000160551 |
| CMBL | 0.47641365 | 1.391280823 | 0.000128647 | 0.000871151 | 503.9215318 | 700.7577624 | ENSG00000164237 |
| CASK | 0.476303573 | 1.391174673 | 1.36E-08 | 2.14E-07 | 1022.021289 | 1421.985387 | ENSG00000147044 |
| PRICKLE3 | 0.476296984 | 1.391168319 | 0.003552889 | 0.015370971 | 185.9857684 | 258.9433166 | ENSG00000012211 |
| CBFA2T2 | 0.476249256 | 1.391122297 | 5.73E-06 | 5.47E-05 | 512.2433429 | 713.0819333 | ENSG00000078699 |
| TRIM61 | 0.476208585 | 1.39108308 | 0.012693312 | 0.044819556 | 162.9821975 | 226.7158412 | ENSG00000183439 |
| DNAH7 | 0.476185069 | 1.391060406 | 0.676524918 | 0.812987929 | 3.238538961 | 4.482247516 | ENSG00000118997 |
| NA | 0.476079293 | 1.39095842 | 0.410778081 | 0.602279634 | 12.75579664 | 17.79412067 | ENSG00000286615 |
| NA | 0.47589612 | 1.390781826 | 0.556512347 | 0.726110513 | 5.988034921 | 8.390733108 | ENSG00000280383 |
| NA | 0.47584724 | 1.390734706 | 0.052654619 | 0.139116281 | 69.98321377 | 97.45745295 | ENSG00000204055 |
| NA | 0.475451328 | 1.390353105 | 0.630526081 | 0.78189284 | 4.244533517 | 5.855023298 | ENSG00000237821 |
| SMG1P1 | 0.47525641 | 1.390165272 | 0.28266541 | 0.473596159 | 21.86035594 | 30.30643032 | ENSG00000237296 |
| HLA-DMA | 0.475195443 | 1.390106526 | 0.141411053 | 0.293134152 | 38.875967 | 54.02642138 | ENSG00000204257 |
| MYO1D | 0.474993742 | 1.38991219 | 4.03E-06 | 4.00E-05 | 613.1517551 | 852.3381686 | ENSG00000176658 |
| NA | 0.474905644 | 1.389827318 | 0.597641909 | 0.757703763 | 4.777269635 | 6.676606995 | ENSG00000229759 |
| C21orf91 | 0.474691225 | 1.389620772 | 0.000812495 | 0.004399293 | 322.6137182 | 447.8628501 | ENSG00000154642 |
| S100A14 | 0.474663602 | 1.389594166 | 0.298555664 | 0.491947962 | 22.85509832 | 31.72379613 | ENSG00000189334 |
| NA | 0.474641503 | 1.389572881 | 0.589902598 | 0.751995212 | 5.17724266 | 7.13014526 | ENSG00000240087 |
| SEMA6A-AS2 | 0.474220093 | 1.389167047 | 0.526174736 | 0.702433961 | 7.314403263 | 10.22877011 | ENSG00000249167 |
| ENTPD5 | 0.474061793 | 1.389014629 | 0.010430863 | 0.038063716 | 127.9720176 | 177.9275084 | ENSG00000187097 |
| CDK1 | 0.473770919 | 1.388734606 | 1.82E-08 | 2.80E-07 | 1143.911143 | 1587.564523 | ENSG00000170312 |

| | | | | | | | |
|---|---|---|---|---|---|---|---|
| ZBTB20 | 0.473634858 | 1.38860364 | 0.519751604 | 0.696786707 | 7.31124635 | 10.09682349 | ENSG00000181722 |
| LMLN | 0.473629075 | 1.388598074 | 0.06640925 | 0.166215941 | 61.25021528 | 85.18460293 | ENSG00000185621 |
| CILP2 | 0.473556695 | 1.388528409 | 0.000402949 | 0.002366369 | 344.7116167 | 478.1926232 | ENSG00000160161 |
| ZNF404 | 0.473547466 | 1.388519527 | 0.285409284 | 0.476642249 | 21.50713525 | 29.73234198 | ENSG00000176222 |
| GFRA1 | 0.473295277 | 1.388276829 | 0.217747769 | 0.398730689 | 34.03360491 | 47.38454522 | ENSG00000151892 |
| PRKCE | 0.472978511 | 1.387972045 | 0.195191476 | 0.36952797 | 31.95719977 | 44.45822013 | ENSG00000171132 |
| SNHG5 | 0.472755051 | 1.387757078 | 4.46E-11 | 1.03E-09 | 3605.044715 | 5002.872129 | ENSG00000203875 |
| SETD7 | 0.472734933 | 1.387737726 | 0.005431816 | 0.022096168 | 177.5419579 | 246.5973495 | ENSG00000145391 |
| CCM2 | 0.472527889 | 1.387538583 | 5.56E-07 | 6.52E-06 | 604.0756605 | 838.3225707 | ENSG00000136280 |
| CDON | 0.472413441 | 1.387428515 | 0.000277385 | 0.001717696 | 336.7500543 | 466.9981852 | ENSG00000064309 |
| NA | 0.472330468 | 1.387348723 | 0.5753026 | 0.740616683 | 6.230087246 | 8.728284593 | ENSG00000227848 |
| NA | 0.472155537 | 1.387180513 | 0.344040126 | 0.537651269 | 16.78618638 | 23.29238346 | ENSG00000285976 |
| TMC4 | 0.472120413 | 1.387146741 | 0.07162472 | 0.176153472 | 65.03392261 | 90.1170279 | ENSG00000167608 |
| DCP1B | 0.472097918 | 1.387125113 | 0.000291428 | 0.001792675 | 308.0047628 | 426.6317173 | ENSG00000151065 |
| LINC01090 | 0.472026667 | 1.387056608 | 0.017865865 | 0.058710273 | 133.6306139 | 184.9051839 | ENSG00000231689 |
| NA | 0.471791987 | 1.386830996 | 0.101211698 | 0.228595859 | 49.85739216 | 69.20606104 | ENSG00000271369 |
| CARD11 | 0.471774001 | 1.386813707 | 3.31E-10 | 6.63E-09 | 1158.189195 | 1606.533484 | ENSG00000198286 |
| ZNF888 | 0.471676021 | 1.386719525 | 0.013925241 | 0.048160183 | 114.1431571 | 158.1784879 | ENSG00000213793 |
| ZNF43 | 0.471404669 | 1.386458726 | 0.001120481 | 0.005809655 | 255.755049 | 354.0682512 | ENSG00000198521 |
| NA | 0.471325628 | 1.386382768 | 0.389741191 | 0.582349503 | 14.07712893 | 19.55707369 | ENSG00000232648 |
| SPRY4 | 0.47088451 | 1.385958933 | 8.23E-15 | 3.07E-13 | 3935.199104 | 5453.97094 | ENSG00000187678 |
| BACH2 | 0.470763267 | 1.385842463 | 0.089528097 | 0.208771232 | 53.19081126 | 73.7928988 | ENSG00000112182 |
| NA | 0.470620833 | 1.385705649 | 0.093619167 | 0.215545047 | 76.85069628 | 106.6524245 | ENSG00000237094 |
| ARHGAP31 | 0.470582088 | 1.385668435 | 0.003953065 | 0.016877701 | 186.2577964 | 257.9344525 | ENSG00000031081 |
| RBM34 | 0.470572077 | 1.38565882 | 0.646960636 | 0.793827767 | 3.791494334 | 5.303831214 | ENSG00000188739 |
| SCN8A | 0.470150221 | 1.3852537 | 5.77E-05 | 0.000427957 | 483.5221039 | 669.8137316 | ENSG00000196876 |
| RNF157-AS1 | 0.470118482 | 1.385223225 | 0.580394941 | 0.744431822 | 6.266091064 | 8.652650963 | ENSG00000267128 |
| TMEM168 | 0.469945301 | 1.385056953 | 0.001317231 | 0.006630517 | 218.2452875 | 302.141052 | ENSG00000146802 |
| LOC441242 | 0.469822892 | 1.38493944 | 0.287217153 | 0.478626391 | 20.34953382 | 28.18715482 | ENSG00000272693 |
| FRAS1 | 0.469807138 | 1.384924316 | 0.016071506 | 0.05401075 | 2677.737814 | 3708.379144 | ENSG00000138759 |
| NA | 0.469775986 | 1.384894412 | 0.67387738 | 0.810752571 | 3.754212739 | 5.243719384 | ENSG00000213609 |
| HLTF | 0.469557318 | 1.384684521 | 5.95E-06 | 5.65E-05 | 847.7866607 | 1173.068444 | ENSG00000071794 |
| CNTNAP3 | 0.469498873 | 1.384628428 | 0.064791252 | 0.163224285 | 64.70921184 | 89.70862932 | ENSG00000106714 |
| TADA2A | 0.469445755 | 1.384577448 | 0.002283882 | 0.010624866 | 277.9571554 | 384.7466692 | ENSG00000276234 |
| CYS1 | 0.469152547 | 1.384296081 | 0.55532978 | 0.725293281 | 6.348847215 | 8.742818708 | ENSG00000205795 |
| EME1 | 0.468986638 | 1.384136896 | 0.012816732 | 0.045126391 | 149.8336513 | 207.6918817 | ENSG00000154920 |
| CYP1B1 | 0.46872168 | 1.383882716 | 0.276598103 | 0.467615547 | 25.19890879 | 34.8403159 | ENSG00000138061 |
| JUNB | 0.468246967 | 1.38342743 | 0.046902721 | 0.127169874 | 90.66565133 | 125.1632835 | ENSG00000171223 |
| ADCK5 | 0.468139306 | 1.383324195 | 0.011171789 | 0.04037004 | 145.8653925 | 202.2040277 | ENSG00000173137 |
| MICB | 0.467849784 | 1.383046616 | 7.73E-05 | 0.000553001 | 371.2777157 | 513.0732124 | ENSG00000204516 |
| GALNT2 | 0.467808688 | 1.38300722 | 1.46E-11 | 3.61E-10 | 3148.573506 | 4353.685989 | ENSG00000143641 |
| SYT1 | 0.467600471 | 1.382807632 | 1.42E-05 | 0.000122775 | 582.21668 | 804.6971675 | ENSG00000067715 |
| PLCG2 | 0.467210059 | 1.382433477 | 0.000410115 | 0.00240464 | 267.3084461 | 369.7735189 | ENSG00000197943 |
| MYL12A | 0.46685466 | 1.382092965 | 5.36E-10 | 1.04E-08 | 1650.941119 | 2281.897921 | ENSG00000101608 |
| LYNX1 | 0.466593483 | 1.381842781 | 0.030255767 | 0.090212416 | 86.21328787 | 119.0685774 | ENSG00000180155 |
| LAMA5 | 0.466180857 | 1.381447616 | 3.74E-10 | 7.38E-09 | 3443.611696 | 4758.465088 | ENSG00000130702 |
| NA | 0.466176227 | 1.381443183 | 0.137211081 | 0.286446291 | 56.48369416 | 78.31846252 | ENSG00000244627 |
| MAT1A | 0.46610927 | 1.38137907 | 0.051180284 | 0.136213596 | 90.14101048 | 124.1959836 | ENSG00000151224 |
| SEC24A | 0.465946954 | 1.381223661 | 0.000207677 | 0.001328189 | 406.8635734 | 561.4470375 | ENSG00000113615 |
| DUOX2 | 0.465814144 | 1.381096516 | 0.641252423 | 0.789699164 | 4.250847342 | 5.813682462 | ENSG00000140279 |
| RALY-AS1 | 0.465454119 | 1.380751906 | 0.415480258 | 0.606817265 | 12.88214821 | 17.78283575 | ENSG00000285230 |
| IER5 | 0.465208925 | 1.380517259 | 6.77E-06 | 6.35E-05 | 498.7474839 | 688.7947987 | ENSG00000162783 |
| PPARA | 0.464597239 | 1.37993206 | 0.007193416 | 0.028003103 | 143.7768634 | 198.4206152 | ENSG00000186951 |
| TRMT2B | 0.464516055 | 1.37985441 | 1.68E-05 | 0.000142371 | 438.8655486 | 606.0109246 | ENSG00000188917 |
| CFDP1 | 0.463984037 | 1.37934566 | 6.38E-06 | 6.02E-05 | 688.3084685 | 948.734538 | ENSG00000153774 |
| LIMA1 | 0.463810712 | 1.379179956 | 8.20E-09 | 1.34E-07 | 1067.778877 | 1472.593057 | ENSG00000050405 |
| GNGT2 | 0.463780376 | 1.379150955 | 0.622162343 | 0.775568796 | 4.747579643 | 6.563981029 | ENSG00000167083 |
| ZNF852 | 0.463504909 | 1.378887646 | 0.012721939 | 0.044903526 | 159.5069505 | 220.2234003 | ENSG00000178917 |
| BRI3 | 0.46347475 | 1.378858821 | 0.000252945 | 0.001585342 | 632.2264234 | 871.6236164 | ENSG00000164713 |
| FAHD2B | 0.463367281 | 1.378756112 | 7.01E-06 | 6.55E-05 | 486.749917 | 670.8621843 | ENSG00000144199 |
| SYNJ2 | 0.463002363 | 1.37840741 | 0.000147642 | 0.000980439 | 546.1831575 | 753.4085573 | ENSG00000078269 |
| NMNAT2 | 0.46274584 | 1.378162341 | 0.000115348 | 0.000792955 | 352.5382437 | 485.7358738 | ENSG00000157064 |
| PLCD1 | 0.4625604 | 1.377985207 | 5.63E-05 | 0.000418415 | 368.6052634 | 507.6438701 | ENSG00000187091 |
| GPNMB | 0.462532503 | 1.377958561 | 0.484408961 | 0.667762226 | 10.21234918 | 14.15602669 | ENSG00000136235 |
| SHISA9 | 0.462219068 | 1.377659224 | 2.30E-05 | 0.000189315 | 450.7306455 | 621.2130854 | ENSG00000237515 |
| PARVB | 0.46187483 | 1.377330543 | 0.00198435 | 0.009430241 | 235.474441 | 323.9314126 | ENSG00000188677 |
| TMEM182 | 0.461644141 | 1.377110323 | 0.099969726 | 0.226727797 | 57.4890123 | 79.34890028 | ENSG00000170417 |
| RNPC3 | 0.461371974 | 1.376850553 | 0.013048324 | 0.045708407 | 128.4920284 | 177.0350776 | ENSG00000185946 |
| HERC5 | 0.461240165 | 1.376724766 | 0.000341315 | 0.002050543 | 331.1585273 | 455.6880718 | ENSG00000138646 |
| NAGLU | 0.460418327 | 1.375940731 | 0.001635638 | 0.007998659 | 287.2549703 | 395.0797829 | ENSG00000108784 |
| SLC13A4 | 0.46036094 | 1.375886001 | 0.103487658 | 0.232517912 | 53.84980125 | 74.1008324 | ENSG00000164707 |
| KIAA0355 | 0.460340662 | 1.375866662 | 3.37E-06 | 3.40E-05 | 620.0207859 | 853.1320611 | ENSG00000166398 |
| LRRC7 | 0.460236079 | 1.375766927 | 0.513574434 | 0.692196991 | 8.598453961 | 11.86781224 | ENSG00000033122 |
| PLEKHH1 | 0.46000478 | 1.375546375 | 5.91E-05 | 0.000436538 | 648.7801482 | 892.9105536 | ENSG00000054690 |

| | | | | | | | |
|---|---|---|---|---|---|---|---|
| C12orf29 | 0.459885354 | 1.375432513 | 0.002474586 | 0.01138064 | 205.7467264 | 282.6771805 | ENSG00000133641 |
| TLCD5 | 0.459735973 | 1.375290104 | 0.000766264 | 0.004177042 | 239.8904173 | 330.0173589 | ENSG00000181264 |
| NA | 0.459642342 | 1.375200851 | 0.655314481 | 0.798737248 | 3.761804341 | 5.213663469 | ENSG00000213087 |
| BCS1L | 0.459619874 | 1.375179434 | 3.15E-05 | 0.000251029 | 515.1899203 | 708.555047 | ENSG00000074582 |
| LETMD1 | 0.459525481 | 1.375089461 | 9.04E-05 | 0.000636695 | 353.4217201 | 485.8330895 | ENSG00000050426 |
| ZNF611 | 0.459216748 | 1.374795226 | 0.039222339 | 0.110747536 | 94.27396966 | 129.3410614 | ENSG00000213020 |
| PROCR | 0.458906463 | 1.374499577 | 0.021117135 | 0.067631759 | 127.4993605 | 175.1576349 | ENSG00000101000 |
| PGF | 0.458612858 | 1.374219879 | 0.031911505 | 0.094134624 | 98.60964769 | 135.6854531 | ENSG00000119630 |
| C1RL | 0.458231096 | 1.373856285 | 0.006652508 | 0.026255872 | 180.995648 | 248.5290267 | ENSG00000139178 |
| KRCC1 | 0.458137618 | 1.37376727 | 0.020997297 | 0.067282832 | 131.4336674 | 180.1896487 | ENSG00000172086 |
| ELMO1 | 0.458114598 | 1.37374535 | 0.003139027 | 0.013845484 | 182.316672 | 250.5951268 | ENSG00000155849 |
| ITPRIPL2 | 0.457777841 | 1.373424725 | 0.029280853 | 0.087814131 | 102.2493398 | 140.30906 | ENSG00000205730 |
| PROSER3 | 0.45776826 | 1.373415604 | 0.012465618 | 0.044133289 | 138.1047675 | 189.923142 | ENSG00000167595 |
| MXI1 | 0.457567423 | 1.373224424 | 0.01744499 | 0.057633768 | 151.5522057 | 207.5484982 | ENSG00000119950 |
| LINC00106 | 0.457287231 | 1.372957751 | 0.357071853 | 0.550616112 | 24.05689653 | 33.10570694 | ENSG00000236871 |
| SNAPC3 | 0.456811517 | 1.372505107 | 1.83E-06 | 1.94E-05 | 763.4694432 | 1047.967815 | ENSG00000164975 |
| MFAP3 | 0.45662375 | 1.372326486 | 0.001812866 | 0.008722654 | 230.8475981 | 316.4417041 | ENSG00000037749 |
| SLC39A8 | 0.456620916 | 1.372323791 | 2.47E-05 | 0.000201733 | 509.6872678 | 699.0669161 | ENSG00000138821 |
| DCAF16 | 0.456510763 | 1.372219015 | 1.93E-06 | 2.04E-05 | 809.4241127 | 1110.403964 | ENSG00000163257 |
| NETO2 | 0.456452346 | 1.372163452 | 0.000310966 | 0.001892135 | 381.4184706 | 523.2283956 | ENSG00000171208 |
| PPP1R14C | 0.456445011 | 1.372156476 | 0.496843761 | 0.678123082 | 9.028116978 | 12.31755166 | ENSG00000198729 |
| NA | 0.456428249 | 1.372140534 | 0.225228689 | 0.408093511 | 30.93847986 | 42.57234391 | ENSG00000255026 |
| PLEKHG4 | 0.456251332 | 1.371972279 | 0.000281099 | 0.001737209 | 388.1978452 | 532.9021522 | ENSG00000196155 |
| LOC102724788 | 0.456105966 | 1.371834047 | 0.003969305 | 0.016919755 | 196.7710968 | 270.026967 | ENSG00000277196 |
| TUSC1 | 0.456066985 | 1.371796981 | 0.080958009 | 0.193335361 | 61.42401824 | 84.43810321 | ENSG00000198680 |
| MAP1LC3A | 0.456025113 | 1.371757167 | 0.046177901 | 0.125638294 | 77.7147501 | 106.7440893 | ENSG00000101460 |
| ARHGAP9 | 0.456022765 | 1.371754934 | 0.220233004 | 0.401751821 | 30.75086917 | 42.32441052 | ENSG00000123329 |
| RGPD5 | 0.456020965 | 1.371753223 | 0.487089504 | 0.67015937 | 9.217103144 | 12.62124838 | ENSG00000015568 |
| SNX18 | 0.455287678 | 1.371056171 | 7.77E-05 | 0.000555818 | 467.8515921 | 641.0095784 | ENSG00000178996 |
| CUEDC2 | 0.455240225 | 1.371011075 | 3.14E-06 | 3.18E-05 | 880.0859943 | 1206.819694 | ENSG00000107874 |
| C15orf40 | 0.454995718 | 1.370778736 | 0.007915074 | 0.030347122 | 155.7883357 | 213.7023688 | ENSG00000169609 |
| C3orf35 | 0.454989457 | 1.370772787 | 0.624478026 | 0.777188921 | 5.385170303 | 7.44187771 | ENSG00000198590 |
| NA | 0.45498274 | 1.370766406 | 0.556598038 | 0.726171198 | 7.576674842 | 10.43492463 | ENSG00000219102 |
| SLC25A11 | 0.454956041 | 1.370741038 | 6.46E-08 | 8.94E-07 | 1367.599576 | 1873.681424 | ENSG00000108528 |
| CYFIP2 | 0.454580178 | 1.370383967 | 6.50E-09 | 1.07E-07 | 1927.740847 | 2642.075173 | ENSG00000055163 |
| FZD9 | 0.45449963 | 1.370307459 | 0.023053448 | 0.072524499 | 125.2031742 | 171.4886089 | ENSG00000188763 |
| GAREM1 | 0.454394772 | 1.370207865 | 0.092357371 | 0.213515219 | 78.89898231 | 107.8776916 | ENSG00000141441 |
| SUGP2 | 0.454302347 | 1.370120088 | 5.71E-08 | 8.00E-07 | 3271.370725 | 4482.064183 | ENSG00000064607 |
| SDE2 | 0.454285778 | 1.370104352 | 0.000104231 | 0.000722887 | 385.1335152 | 527.2227492 | ENSG00000143751 |
| BAMBI | 0.454050112 | 1.369880562 | 2.01E-06 | 2.11E-05 | 591.1346359 | 809.4375828 | ENSG00000095739 |
| ADAM28 | 0.453931985 | 1.369768402 | 0.504740623 | 0.684288598 | 9.107813914 | 12.57665836 | ENSG00000042980 |
| PMS2P4 | 0.453655179 | 1.369505613 | 0.423451727 | 0.614248596 | 12.39941903 | 17.0326488 | ENSG00000067601 |
| AHRR | 0.45361311 | 1.369465679 | 0.496919561 | 0.678123082 | 12.45701758 | 17.13723904 | ENSG00000286169 |
| ATL3 | 0.453541697 | 1.369397892 | 0.000305385 | 0.001865524 | 296.5754322 | 405.7407239 | ENSG00000184743 |
| FBXO25 | 0.453508957 | 1.369366816 | 0.003759873 | 0.01615708 | 182.9750833 | 250.7227249 | ENSG00000147364 |
| ZNF701 | 0.453402642 | 1.369265909 | 0.064722232 | 0.163094745 | 72.25364113 | 98.99884126 | ENSG00000167562 |
| TMEM267 | 0.453356855 | 1.369222453 | 0.05889803 | 0.151718707 | 72.83756427 | 99.7375285 | ENSG00000151881 |
| ACVRL1 | 0.453335571 | 1.369202252 | 0.531921674 | 0.707117795 | 8.643928518 | 11.84904125 | ENSG00000139567 |
| ARHGEF5 | 0.453233185 | 1.369105086 | 3.21E-05 | 0.000255081 | 536.423547 | 734.0239035 | ENSG00000050327 |
| TPD52L1 | 0.453123925 | 1.369001403 | 0.035266614 | 0.101827608 | 100.2061875 | 137.0660414 | ENSG00000111907 |
| WASF2 | 0.452679896 | 1.36858012 | 2.27E-11 | 5.42E-10 | 2876.680993 | 3937.047189 | ENSG00000158195 |
| ZBTB3 | 0.452640544 | 1.368542791 | 2.84E-05 | 0.000228453 | 454.0238367 | 621.7572781 | ENSG00000185670 |
| NA | 0.452595833 | 1.368500378 | 0.620676296 | 0.774862891 | 5.419294985 | 7.36624408 | ENSG00000230626 |
| MORF4L2 | 0.452536152 | 1.368443768 | 4.41E-15 | 1.72E-13 | 6633.27505 | 9076.576092 | ENSG00000123562 |
| TMEM86B | 0.452424918 | 1.368338262 | 0.659243739 | 0.801470873 | 5.186713398 | 7.111812299 | ENSG00000180089 |
| GPR158 | 0.452413273 | 1.368327218 | 0.307531859 | 0.501731365 | 21.84141446 | 29.76228627 | ENSG00000151025 |
| NA | 0.452383459 | 1.368298941 | 0.671425289 | 0.809097911 | 3.52290893 | 4.808514081 | ENSG00000231720 |
| BBS4 | 0.452306687 | 1.368226129 | 8.01E-06 | 7.35E-05 | 607.6681117 | 830.6754775 | ENSG00000140463 |
| ANO9 | 0.452162244 | 1.368089149 | 0.26010305 | 0.449026633 | 265.0150712 | 362.704021 | ENSG00000185101 |
| INPP1 | 0.452142166 | 1.368070109 | 0.113735039 | 0.249538862 | 53.27544655 | 72.96817752 | ENSG00000151689 |
| FKBP9 | 0.452019656 | 1.367953941 | 0.001061983 | 0.005553838 | 261.403671 | 357.0546881 | ENSG00000122642 |
| NA | 0.45190288 | 1.36784322 | 0.448001414 | 0.635957225 | 13.05407203 | 17.83882232 | ENSG00000271780 |
| RELL1 | 0.451888941 | 1.367830004 | 0.077961331 | 0.188074715 | 71.45497286 | 97.52274906 | ENSG00000181826 |
| RGS3 | 0.451438432 | 1.36740294 | 0.000312371 | 0.001898811 | 411.8306935 | 562.7920047 | ENSG00000138835 |
| FGGY | 0.451199843 | 1.367176821 | 0.025418526 | 0.07842633 | 143.8163324 | 196.1726423 | ENSG00000172456 |
| RO60 | 0.451079075 | 1.367062379 | 0.002298694 | 0.010685726 | 303.9321379 | 415.4718751 | ENSG00000116747 |
| SGF29 | 0.451041344 | 1.367026627 | 0.008604236 | 0.032505281 | 155.5439232 | 212.9063809 | ENSG00000176476 |
| RAD23A | 0.450816958 | 1.366814027 | 1.48E-09 | 2.67E-08 | 1234.331495 | 1687.057001 | ENSG00000179262 |
| ATRX | 0.450734668 | 1.366736067 | 3.43E-06 | 3.45E-05 | 1190.604409 | 1626.550594 | ENSG00000085224 |
| NA | 0.450519057 | 1.366531823 | 0.506961456 | 0.686492557 | 11.25128834 | 15.52511525 | ENSG00000226332 |
| PTGER1 | 0.450397814 | 1.366416986 | 0.468502067 | 0.654300152 | 11.12295994 | 15.22466772 | ENSG00000160951 |
| FIGNL1 | 0.450296985 | 1.366321491 | 0.01168656 | 0.041830573 | 163.3304048 | 223.2608854 | ENSG00000132436 |
| IPMK | 0.450190961 | 1.366221083 | 0.014523776 | 0.049738296 | 178.5190592 | 243.6862911 | ENSG00000151151 |
| SNX19 | 0.450173655 | 1.366204695 | 5.67E-07 | 6.64E-06 | 662.7190196 | 905.0990008 | ENSG00000120451 |

| | | | | | | | |
|---|---|---|---|---|---|---|---|
| SORCS1 | 0.450087113 | 1.366122744 | 0.01167218 | 0.04178717 | 149.4395861 | 204.3281527 | ENSG00000108018 |
| SPAG4 | 0.450002001 | 1.366042151 | 0.189619125 | 0.3617872 | 38.66232689 | 52.79632748 | ENSG00000061656 |
| NA | 0.449728357 | 1.365783071 | 0.448541252 | 0.63620292 | 12.63831445 | 17.32517223 | ENSG00000237350 |
| RAB3IP | 0.449588216 | 1.365650408 | 6.88E-05 | 0.000498966 | 391.4794009 | 534.2955853 | ENSG00000127328 |
| MMP24OS | 0.449568767 | 1.365631998 | 0.016126811 | 0.054143282 | 230.4120121 | 314.4962255 | ENSG00000126005 |
| VIPAS39 | 0.449198306 | 1.36528137 | 6.71E-05 | 0.000488553 | 385.6052853 | 526.1669933 | ENSG00000151445 |
| SLC16A7 | 0.449032295 | 1.365124276 | 0.17947936 | 0.34868663 | 37.63876633 | 51.44188466 | ENSG00000118596 |
| CHMP3 | 0.449028628 | 1.365120807 | 0.060901134 | 0.155760418 | 90.86302586 | 123.9469509 | ENSG00000115561 |
| NA | 0.448924499 | 1.36502228 | 0.661877415 | 0.803355337 | 4.128930461 | 5.682723535 | ENSG00000285399 |
| NOXA1 | 0.448910894 | 1.365009407 | 0.258441415 | 0.447074219 | 28.2515208 | 38.44185558 | ENSG00000188747 |
| ELAVL3 | 0.448810575 | 1.364914493 | 0.325984012 | 0.520215918 | 19.81236301 | 26.94913681 | ENSG00000196361 |
| EIF3J-DT | 0.448722544 | 1.364831211 | 0.014725906 | 0.050309666 | 171.8225084 | 234.7154496 | ENSG00000179523 |
| NA | 0.448688407 | 1.364798917 | 0.678041235 | 0.813965688 | 3.463528945 | 4.718346336 | ENSG00000198658 |
| REST | 0.448659221 | 1.364771307 | 4.44E-06 | 4.35E-05 | 598.9552196 | 817.1422544 | ENSG00000084093 |
| CHRNA10 | 0.448592892 | 1.364708562 | 0.612390711 | 0.76895735 | 5.458455715 | 7.381327847 | ENSG00000129749 |
| ZNF595 | 0.448544287 | 1.364662585 | 0.096562238 | 0.220812401 | 65.51735085 | 89.36684095 | ENSG00000272602 |
| RGPD2 | 0.448270236 | 1.364403382 | 0.266023377 | 0.456018986 | 32.27234211 | 43.84267934 | ENSG00000185304 |
| GPR156 | 0.448148128 | 1.364287906 | 0.138865794 | 0.289119144 | 49.36637232 | 67.52286691 | ENSG00000175697 |
| FNDC3B | 0.447920877 | 1.364073021 | 4.83E-06 | 4.69E-05 | 673.6399371 | 919.073633 | ENSG00000075420 |
| DAGLA | 0.447908706 | 1.364061514 | 0.00122405 | 0.006239422 | 247.7944562 | 338.2663978 | ENSG00000134780 |
| TMEM47 | 0.44770919 | 1.363872885 | 0.000589291 | 0.003310678 | 554.9125479 | 756.3082982 | ENSG00000147027 |
| FAM234A | 0.447691011 | 1.3638557 | 6.63E-12 | 1.74E-10 | 2283.645491 | 3114.643337 | ENSG00000167930 |
| MCTP2 | 0.447644566 | 1.363811794 | 0.143453004 | 0.296240444 | 47.85921078 | 65.18472624 | ENSG00000140563 |
| PRIMPOL | 0.447345933 | 1.363529519 | 0.070009453 | 0.172984295 | 69.27096223 | 94.28699331 | ENSG00000164306 |
| NA | 0.447308025 | 1.363493691 | 0.711599064 | 0.836373711 | 3.119778992 | 4.324919901 | ENSG00000277969 |
| NA | 0.44723806 | 1.363427569 | 0.349737744 | 0.543111801 | 17.19247323 | 23.45383633 | ENSG00000282718 |
| AIF1 | 0.447150052 | 1.363344399 | 0.234341366 | 0.419316749 | 34.09879506 | 46.53940399 | ENSG00000204472 |
| NA | 0.446974765 | 1.363178763 | 0.374211061 | 0.567389377 | 16.57194491 | 22.53828604 | ENSG00000172974 |
| SLC12A5 | 0.446740566 | 1.36295749 | 0.269729289 | 0.460054717 | 25.6329088 | 34.90141542 | ENSG00000124140 |
| ERV3-1 | 0.446703016 | 1.362922016 | 0.227423601 | 0.410959013 | 44.41096274 | 60.79104624 | ENSG00000213462 |
| CCDC9B | 0.446576002 | 1.362802031 | 0.39671155 | 0.588966027 | 14.56311272 | 19.85653354 | ENSG00000188549 |
| PCIF1 | 0.446136273 | 1.362386716 | 7.43E-06 | 6.90E-05 | 608.7905061 | 828.7932969 | ENSG00000100982 |
| PLD5 | 0.44612007 | 1.362371415 | 0.105278344 | 0.235759356 | 52.45834046 | 71.46758039 | ENSG00000180287 |
| NA | 0.445926921 | 1.362189032 | 0.31223467 | 0.505808704 | 22.08040757 | 30.1224076 | ENSG00000230409 |
| LOC100288637 | 0.445800781 | 1.362069936 | 0.007705024 | 0.029689171 | 167.4197837 | 228.1662175 | ENSG00000187951 |
| SLC6A15 | 0.445561765 | 1.361844296 | 2.21E-05 | 0.000183027 | 449.4552688 | 612.0799863 | ENSG00000072041 |
| EP400 | 0.445409681 | 1.361700742 | 2.61E-06 | 2.69E-05 | 2280.291718 | 3105.121061 | ENSG00000183495 |
| ALDH3B1 | 0.445383316 | 1.361675858 | 0.326068506 | 0.520261164 | 23.02386523 | 31.36378642 | ENSG00000006534 |
| PCDHB3 | 0.445360859 | 1.361654663 | 0.373060868 | 0.566421735 | 17.05665092 | 23.2397577 | ENSG00000113205 |
| ZDHHC19 | 0.445235691 | 1.361536531 | 0.097107516 | 0.221758342 | 58.57588387 | 79.602456 | ENSG00000163958 |
| STEAP2 | 0.445134921 | 1.361441433 | 0.012717565 | 0.044896638 | 188.8566776 | 257.2546831 | ENSG00000157214 |
| ZNF573 | 0.445127497 | 1.361434427 | 0.181684465 | 0.351790627 | 37.96219933 | 51.77673661 | ENSG00000189144 |
| GZF1 | 0.445096665 | 1.361405332 | 4.47E-05 | 0.000341307 | 407.1668622 | 554.3449502 | ENSG00000125812 |
| PDGFA | 0.444998287 | 1.3613125 | 5.69E-07 | 6.65E-06 | 740.4480811 | 1007.498626 | ENSG00000197461 |
| MDGA1 | 0.444900218 | 1.361219966 | 0.280197467 | 0.471421556 | 29.5279799 | 40.16379418 | ENSG00000112139 |
| USP24 | 0.444773347 | 1.361100265 | 3.93E-06 | 3.91E-05 | 1496.008404 | 2036.012825 | ENSG00000162402 |
| ZBTB43 | 0.444773009 | 1.361099946 | 0.00167742 | 0.008173703 | 257.4674849 | 350.2825058 | ENSG00000169155 |
| NA | 0.444696598 | 1.361027859 | 0.522765316 | 0.699561903 | 9.143817732 | 12.39968368 | ENSG00000184616 |
| AGO2 | 0.444638931 | 1.360973457 | 5.07E-06 | 4.89E-05 | 1956.349489 | 2662.303324 | ENSG00000123908 |
| NA | 0.444550394 | 1.360889939 | 0.04139538 | 0.11563255 | 91.60910899 | 124.6845791 | ENSG00000188185 |
| APOBEC3B | 0.444526262 | 1.360867175 | 0.048362234 | 0.130278918 | 82.52674578 | 112.3736374 | ENSG00000179750 |
| NA | 0.443847389 | 1.360226957 | 0.653826806 | 0.797703095 | 4.102397381 | 5.630097779 | ENSG00000266680 |
| SLC16A12 | 0.443833792 | 1.360214138 | 0.073212349 | 0.179367836 | 72.47940523 | 98.75101949 | ENSG00000152779 |
| L1CAM | 0.443583545 | 1.359978219 | 1.26E-06 | 1.38E-05 | 1077.795325 | 1465.622573 | ENSG00000198910 |
| CFAP97 | 0.443495202 | 1.359894943 | 5.77E-05 | 0.000427957 | 579.6364546 | 787.5734834 | ENSG00000164323 |
| NA | 0.443469102 | 1.359870341 | 0.642370988 | 0.790289007 | 4.522589659 | 6.094920964 | ENSG00000286788 |
| ASB12 | 0.443151388 | 1.3595709 | 0.504281061 | 0.684010728 | 8.601610874 | 11.68747675 | ENSG00000198881 |
| ATXN3 | 0.442641628 | 1.359090596 | 0.001739432 | 0.008430604 | 292.0574726 | 396.6024003 | ENSG00000066427 |
| PTH2 | 0.442512614 | 1.358969064 | 0.66401367 | 0.804615646 | 3.774431993 | 5.146065566 | ENSG00000142538 |
| MANEA | 0.442380682 | 1.358844794 | 0.06603452 | 0.165546299 | 89.33877933 | 121.1600585 | ENSG00000172469 |
| NA | 0.442052744 | 1.358535951 | 0.553792811 | 0.723938212 | 7.298618699 | 9.905752737 | ENSG00000187952 |
| CCDC102B | 0.441949411 | 1.358438649 | 0.523475843 | 0.699924224 | 9.282796954 | 12.65185395 | ENSG00000150636 |
| ARRDC3-AS1 | 0.441935711 | 1.358425749 | 0.618144857 | 0.772952151 | 5.435079549 | 7.423106716 | ENSG00000281357 |
| TMEM132C | 0.441918691 | 1.358409724 | 0.313427905 | 0.506988374 | 21.69800055 | 29.40227657 | ENSG00000181234 |
| RCSD1 | 0.44180833 | 1.358305814 | 0.573799808 | 0.739401074 | 6.781667144 | 9.129197102 | ENSG00000198771 |
| RAB11B-AS1 | 0.441691293 | 1.358195628 | 0.508352214 | 0.687521886 | 9.804784558 | 13.3189212 | ENSG00000269386 |
| HEATR5A | 0.441599404 | 1.358109123 | 0.0051144 | 0.020985232 | 200.9491398 | 272.5538767 | ENSG00000129493 |
| BTD | 0.441306906 | 1.357833802 | 0.002040617 | 0.009660483 | 269.5060917 | 365.6670891 | ENSG00000169814 |
| CHSY1 | 0.441148798 | 1.357685003 | 2.60E-08 | 3.87E-07 | 1229.412901 | 1668.893132 | ENSG00000131873 |
| KIF5B | 0.44096938 | 1.357516168 | 1.08E-07 | 1.44E-06 | 1888.089332 | 2562.283396 | ENSG00000170759 |
| CCDC90B | 0.440891306 | 1.357442705 | 7.12E-05 | 0.000514312 | 474.5066372 | 643.5309774 | ENSG00000137500 |
| NDFIP2 | 0.440890877 | 1.357442301 | 0.000648174 | 0.003604338 | 393.9241721 | 534.8778295 | ENSG00000102471 |
| NA | 0.440589527 | 1.357158789 | 0.654903106 | 0.798340843 | 4.135244286 | 5.65266762 | ENSG00000279800 |
| TCEA1 | 0.440383417 | 1.356964912 | 2.85E-09 | 4.94E-08 | 3347.690217 | 4542.089427 | ENSG00000187735 |

| | | | | | | | |
|---|---|---|---|---|---|---|---|
| NA | 0.440147295 | 1.35674284 | 0.606374842 | 0.764943597 | 5.802205668 | 7.816533151 | ENSG00000224126 |
| FLYWCH1 | 0.440135691 | 1.356731927 | 4.18E-05 | 0.000321973 | 566.8761104 | 769.3152411 | ENSG00000059122 |
| ZKSCAN1 | 0.440083095 | 1.356682466 | 8.44E-06 | 7.68E-05 | 742.4430078 | 1007.576264 | ENSG00000106261 |
| TGDS | 0.439859646 | 1.356472355 | 0.003063196 | 0.01355618 | 220.0495444 | 298.1557303 | ENSG00000088451 |
| CLCN1 | 0.439484622 | 1.35611979 | 0.415067762 | 0.606501568 | 14.76472653 | 20.15329384 | ENSG00000188037 |
| AGAP1 | 0.439418144 | 1.356057303 | 1.30E-08 | 2.04E-07 | 1332.765551 | 1806.810747 | ENSG00000157985 |
| NA | 0.439406723 | 1.356046568 | 0.584852085 | 0.747954646 | 6.619311756 | 9.020808016 | ENSG00000261172 |
| C2orf88 | 0.438972532 | 1.355638516 | 0.518065921 | 0.695432301 | 9.888818485 | 13.38663072 | ENSG00000187699 |
| CCHCR1 | 0.438958566 | 1.355625393 | 5.21E-05 | 0.000389877 | 827.726736 | 1122.145206 | ENSG00000204536 |
| NA | 0.438729677 | 1.355410335 | 0.62766004 | 0.780008796 | 4.753893469 | 6.421187528 | ENSG00000241061 |
| SLC66A3 | 0.438682401 | 1.35536592 | 0.009333844 | 0.034737642 | 192.9119319 | 261.4309724 | ENSG00000162976 |
| NOTCH1 | 0.438500455 | 1.355194999 | 4.09E-09 | 6.94E-08 | 2884.821206 | 3909.577767 | ENSG00000148400 |
| LGALS7 | 0.438499872 | 1.35519445 | 0.659037043 | 0.801429864 | 3.824341239 | 5.187406401 | ENSG00000205076 |
| TECTA | 0.438448586 | 1.355146276 | 0.256080135 | 0.444941858 | 28.73246854 | 39.02397964 | ENSG00000109927 |
| TRG-AS1 | 0.438391016 | 1.3550922 | 0.524877756 | 0.701381843 | 8.710900104 | 11.77384565 | ENSG00000281103 |
| NR2E3 | 0.43830045 | 1.355007137 | 0.575106969 | 0.740467702 | 6.411384112 | 8.690742605 | ENSG00000278570 |
| WBP1 | 0.438244165 | 1.354954274 | 0.079137843 | 0.190115461 | 78.50768326 | 106.3882048 | ENSG00000239779 |
| NA | 0.438214562 | 1.354926472 | 0.612419307 | 0.76895735 | 5.660069533 | 7.741887206 | ENSG00000276997 |
| FLJ45513 | 0.438182505 | 1.354896365 | 0.585933448 | 0.748944175 | 6.933371716 | 9.463499393 | ENSG00000204584 |
| REL | 0.43816218 | 1.354877277 | 0.166989054 | 0.330553789 | 41.7050622 | 56.40772636 | ENSG00000162924 |
| EXTL1 | 0.438121229 | 1.354838819 | 0.328386773 | 0.522789937 | 24.2787296 | 33.01162873 | ENSG00000158008 |
| NA | 0.437644374 | 1.354391078 | 0.449416674 | 0.637054581 | 12.49164592 | 16.89734138 | ENSG00000274114 |
| NA | 0.437571328 | 1.354322504 | 0.436186528 | 0.625590574 | 12.94921749 | 17.59132696 | ENSG00000271643 |
| LOC100506688 | 0.437472191 | 1.354229443 | 0.705087349 | 0.832415373 | 3.490062025 | 4.703374188 | ENSG00000215246 |
| NA | 0.437454978 | 1.354213286 | 0.58869331 | 0.75096991 | 6.053728732 | 8.17285563 | ENSG00000154898 |
| UVSSA | 0.437315167 | 1.354082056 | 0.18075884 | 0.350351506 | 323.3601549 | 438.035042 | ENSG00000163945 |
| RNFT2 | 0.437242537 | 1.354013889 | 0.001890231 | 0.009050257 | 220.0608942 | 298.0760117 | ENSG00000135119 |
| S100A10 | 0.437196429 | 1.353970615 | 1.59E-05 | 0.000135516 | 591.293233 | 799.949873 | ENSG00000197747 |
| BIK | 0.437190797 | 1.35396533 | 0.433248317 | 0.623046579 | 12.62756593 | 17.1222669 | ENSG00000100290 |
| AP3B2 | 0.437020435 | 1.353805455 | 0.002684832 | 0.012130833 | 251.4288417 | 340.3149378 | ENSG00000103723 |
| ZNF736 | 0.436945222 | 1.353734878 | 0.020664354 | 0.066399642 | 123.4071103 | 167.2865409 | ENSG00000234444 |
| DDAH1 | 0.436919375 | 1.353710625 | 2.17E-05 | 0.000179867 | 747.9050113 | 1011.715403 | ENSG00000153904 |
| ARRDC2 | 0.436833556 | 1.353630101 | 0.058590081 | 0.151156538 | 92.17775123 | 124.7903802 | ENSG00000105643 |
| NA | 0.436806188 | 1.353604424 | 0.350031287 | 0.543382519 | 21.34477986 | 28.97934387 | ENSG00000231160 |
| SOS1 | 0.436770068 | 1.353570534 | 5.92E-05 | 0.000437647 | 455.8729667 | 616.7286477 | ENSG00000115904 |
| ITGA9 | 0.436536759 | 1.353351656 | 0.00662415 | 0.026177399 | 186.9366223 | 253.0394579 | ENSG00000144668 |
| SLC25A5-AS1 | 0.436529862 | 1.353345186 | 0.381534148 | 0.574486465 | 16.70333253 | 22.60263475 | ENSG00000224281 |
| LYRM1 | 0.43646323 | 1.353282682 | 0.003281264 | 0.014366869 | 225.3876692 | 304.4652796 | ENSG00000102897 |
| BRWD3 | 0.436273532 | 1.353104753 | 1.55E-06 | 1.66E-05 | 815.6246902 | 1103.447223 | ENSG00000165288 |
| PANK3 | 0.436109363 | 1.352950787 | 1.46E-06 | 1.57E-05 | 1467.642302 | 1984.965897 | ENSG00000120137 |
| NA | 0.435940504 | 1.352792441 | 0.618881763 | 0.773508301 | 5.11154885 | 6.916504662 | ENSG00000144158 |
| RIPK1 | 0.435651199 | 1.352521192 | 0.000159307 | 0.001049626 | 428.5961796 | 579.0510751 | ENSG00000137275 |
| MST1L | 0.435350426 | 1.352239247 | 0.172845409 | 0.339140039 | 47.89777015 | 64.91631 | ENSG00000186715 |
| CCS | 0.435290934 | 1.352183487 | 7.99E-06 | 7.34E-05 | 628.7300773 | 850.734845 | ENSG00000173992 |
| NA | 0.435203745 | 1.352101771 | 0.002095308 | 0.009891602 | 305.3652551 | 413.2404923 | ENSG00000249152 |
| RNF212 | 0.435099805 | 1.352004361 | 0.101263373 | 0.228684704 | 59.87078078 | 81.01701065 | ENSG00000178222 |
| MADCAM1 | 0.43506444 | 1.351971219 | 0.293479501 | 0.486046741 | 23.10534361 | 31.28914048 | ENSG00000099866 |
| LINC00173 | 0.435001038 | 1.351911806 | 0.490049961 | 0.672761479 | 15.78956487 | 21.42560464 | ENSG00000196668 |
| GRID1 | 0.434945788 | 1.351860033 | 0.483420183 | 0.666793903 | 10.86700218 | 14.62789792 | ENSG00000182771 |
| GUSBP11 | 0.434940325 | 1.351854914 | 0.339722064 | 0.533877978 | 23.82884733 | 32.38915152 | ENSG00000228315 |
| C14orf28 | 0.434883379 | 1.351801555 | 0.128275134 | 0.272549735 | 55.97758882 | 75.73574927 | ENSG00000179476 |
| NIPAL3 | 0.434640698 | 1.351574183 | 0.009848407 | 0.036281131 | 169.6644521 | 229.1687492 | ENSG00000001461 |
| DSTNP2 | 0.434329631 | 1.351282794 | 0.265247055 | 0.454978548 | 25.50655723 | 34.46241127 | ENSG00000248593 |
| MIB1 | 0.434171547 | 1.351134735 | 7.80E-06 | 7.19E-05 | 1247.555484 | 1684.305108 | ENSG00000101752 |
| NANOS1 | 0.433928972 | 1.350907574 | 0.122029938 | 0.262894876 | 71.88839415 | 97.14605536 | ENSG00000188613 |
| NA | 0.433862892 | 1.350845699 | 0.620583381 | 0.774799082 | 5.088172683 | 6.871476599 | ENSG00000223784 |
| ACVR2B | 0.433781078 | 1.350769096 | 2.24E-05 | 0.000184802 | 1791.858852 | 2419.681803 | ENSG00000114739 |
| TPM1 | 0.433674701 | 1.350669501 | 2.06E-09 | 3.67E-08 | 3964.671417 | 5356.017665 | ENSG00000140416 |
| KAZALD1 | 0.433637552 | 1.350634722 | 0.322848168 | 0.516992219 | 22.24907679 | 30.10418626 | ENSG00000107821 |
| DNAJC14 | 0.433612513 | 1.350611281 | 0.000454565 | 0.002635255 | 325.6143691 | 439.9401583 | ENSG00000135392 |
| ERFE | 0.433601101 | 1.350600597 | 0.277571831 | 0.468699553 | 31.75874286 | 42.79595533 | ENSG00000178752 |
| NA | 0.433072081 | 1.350105438 | 0.476603315 | 0.661031643 | 12.88973981 | 17.38517244 | ENSG00000225356 |
| HLA-DRA | 0.433018227 | 1.350055042 | 0.20026897 | 0.376830974 | 42.15870274 | 56.7867303 | ENSG00000204287 |
| C11orf54 | 0.433005127 | 1.350042783 | 0.167757324 | 0.331543426 | 52.76736437 | 71.00089345 | ENSG00000182919 |
| NA | 0.432939552 | 1.349981421 | 0.65351885 | 0.797566245 | 4.102397381 | 5.573784796 | ENSG00000217624 |
| PROX1 | 0.432828725 | 1.34987772 | 0.462897314 | 0.64955624 | 11.22150065 | 15.10064521 | ENSG00000117707 |
| LRCH3 | 0.432756791 | 1.349810416 | 1.31E-06 | 1.43E-05 | 824.154429 | 1112.485259 | ENSG00000186001 |
| TOMM20 | 0.432673149 | 1.349732161 | 1.15E-09 | 2.11E-08 | 4271.290867 | 5763.672698 | ENSG00000173726 |
| C2orf16 | 0.432079095 | 1.349176501 | 0.463078762 | 0.649713191 | 12.70085134 | 17.24292859 | ENSG00000221843 |
| IZUMO4 | 0.431980271 | 1.349084085 | 0.594615318 | 0.75561195 | 8.954831566 | 12.14948878 | ENSG00000099840 |
| OSGIN1 | 0.431971618 | 1.349075994 | 0.37761105 | 0.570898376 | 17.20322175 | 23.13450619 | ENSG00000140961 |
| TNRC6C | 0.431915094 | 1.349023139 | 0.000487876 | 0.002802074 | 658.4451719 | 887.681056 | ENSG00000078687 |
| FERMT2 | 0.431694922 | 1.348817278 | 1.22E-08 | 1.93E-07 | 2418.891368 | 3261.506486 | ENSG00000073712 |
| FBXL18 | 0.431423845 | 1.348563864 | 8.28E-05 | 0.000588802 | 420.9026239 | 567.2645591 | ENSG00000155034 |

| | | | | | | | |
|---|---|---|---|---|---|---|---|
| ACYP2 | 0.431415038 | 1.348555631 | 0.116374447 | 0.25381078 | 55.85735567 | 75.22589803 | ENSG00000170634 |
| STARD9 | 0.431343563 | 1.348488822 | 0.023498878 | 0.073705909 | 210.213093 | 283.9361999 | ENSG00000159433 |
| TEN1-CDK3 | 0.431311082 | 1.348458463 | 0.537488048 | 0.711203176 | 7.982961692 | 10.78744827 | ENSG00000261408 |
| NA | 0.431268497 | 1.34841866 | 0.703708572 | 0.831491391 | 3.943101207 | 5.303831214 | ENSG00000270804 |
| HOOK3 | 0.431151938 | 1.348309722 | 0.00372727 | 0.016031844 | 319.263139 | 429.7974533 | ENSG00000168172 |
| NA | 0.431121615 | 1.348281384 | 0.649327817 | 0.795321709 | 4.74442273 | 6.364874545 | ENSG00000226790 |
| MIA2 | 0.431103622 | 1.348264568 | 0.001204443 | 0.006165909 | 262.3561934 | 353.5335686 | ENSG00000150527 |
| NA | 0.430852007 | 1.348029443 | 0.317895936 | 0.511803621 | 21.25255297 | 28.72761163 | ENSG00000185641 |
| SKIL | 0.430713484 | 1.347900015 | 2.89E-05 | 0.000232292 | 1531.371879 | 2063.82256 | ENSG00000136603 |
| GFOD1 | 0.430696814 | 1.347884441 | 0.004579697 | 0.019076609 | 189.1020974 | 254.9646367 | ENSG00000145990 |
| PPIC | 0.430323977 | 1.347536151 | 0.082762081 | 0.196630228 | 68.67135222 | 92.58090293 | ENSG00000168938 |
| PIGK | 0.430167392 | 1.347389902 | 0.003504759 | 0.015191115 | 288.65282 | 388.5478961 | ENSG00000142892 |
| SNAI3-AS1 | 0.430114681 | 1.347340674 | 0.101655735 | 0.229458945 | 63.30126742 | 85.23722869 | ENSG00000260630 |
| CAPN7 | 0.430082979 | 1.347311067 | 0.004377318 | 0.018375814 | 291.3051733 | 392.6052843 | ENSG00000131375 |
| CDKL2 | 0.430065012 | 1.347294289 | 0.433697975 | 0.623499711 | 13.79907279 | 18.53258467 | ENSG00000138769 |
| RIC1 | 0.43003086 | 1.347262395 | 3.09E-05 | 0.000246856 | 508.90361 | 685.3809203 | ENSG00000107036 |
| DISC1 | 0.429913336 | 1.34715265 | 0.208661196 | 0.387740787 | 39.63241532 | 53.38831075 | ENSG00000162946 |
| LOC100134368 | 0.429866397 | 1.34710882 | 0.332387303 | 0.526940918 | 23.87569736 | 32.36614364 | ENSG00000236829 |
| RC3H2 | 0.429612187 | 1.346871474 | 5.80E-05 | 0.000429679 | 518.7341157 | 698.6121585 | ENSG00000056586 |
| TBX22 | 0.429504772 | 1.346771197 | 0.521382541 | 0.698417462 | 8.921984661 | 11.99877117 | ENSG00000122145 |
| APOOL | 0.42935959 | 1.346635675 | 0.015667819 | 0.052893872 | 147.7401838 | 199.0445266 | ENSG00000155008 |
| NA | 0.429291364 | 1.346571993 | 0.467631313 | 0.653527277 | 12.58387281 | 16.88236923 | ENSG00000285184 |
| FAM86B3P | 0.428931362 | 1.346236019 | 0.100129084 | 0.226869492 | 72.84575724 | 98.28707237 | ENSG00000173295 |
| FAM229B | 0.428680677 | 1.346002115 | 0.044230491 | 0.121693796 | 104.6556871 | 141.1477028 | ENSG00000203778 |
| TRPV3 | 0.428570903 | 1.345899702 | 0.458101406 | 0.645234077 | 13.338442 | 17.87950188 | ENSG00000167723 |
| TTC28 | 0.428503418 | 1.345836747 | 0.000292188 | 0.00179675 | 364.876306 | 490.5763789 | ENSG00000100154 |
| TRIM21 | 0.4284346 | 1.34577255 | 0.001600702 | 0.007863085 | 244.2827017 | 328.7274796 | ENSG00000132109 |
| OTX1 | 0.428387635 | 1.345728741 | 0.309786626 | 0.50373621 | 22.95679595 | 30.84265025 | ENSG00000115507 |
| PIGV | 0.428141107 | 1.345498803 | 0.010276152 | 0.037587963 | 160.5636511 | 216.1209925 | ENSG00000060642 |
| CARD8 | 0.42811622 | 1.345475592 | 0.006833988 | 0.026897672 | 204.5275954 | 275.3180247 | ENSG00000105483 |
| TUBGCP4 | 0.428010398 | 1.345376905 | 0.00065087 | 0.00361716 | 406.1447375 | 546.2505394 | ENSG00000137822 |
| ZNF780A | 0.427809931 | 1.345189973 | 0.05687806 | 0.147728022 | 96.89308533 | 130.2055233 | ENSG00000197782 |
| TTC39A | 0.427563696 | 1.3449604 | 0.041806721 | 0.116588288 | 105.9787032 | 142.2599462 | ENSG00000085831 |
| NA | 0.427447448 | 1.344852032 | 0.582762485 | 0.746399526 | 7.17216943 | 9.556916332 | ENSG00000263551 |
| CGN | 0.427444279 | 1.344849078 | 1.60E-08 | 2.48E-07 | 1815.480418 | 2442.63649 | ENSG00000143375 |
| IBA57 | 0.427340925 | 1.344752736 | 0.003993398 | 0.01700681 | 233.4500196 | 313.8994959 | ENSG00000181873 |
| SMIM14 | 0.427261758 | 1.344678946 | 0.008803673 | 0.033116961 | 194.7014115 | 261.4287737 | ENSG00000163683 |
| NA | 0.427026331 | 1.344459532 | 0.233986447 | 0.418924344 | 38.11518168 | 51.46967907 | ENSG00000236017 |
| ZNF853 | 0.426863883 | 1.344308154 | 0.001945279 | 0.00927548 | 242.2601595 | 325.7106603 | ENSG00000236609 |
| CCNA1 | 0.426791437 | 1.34424065 | 0.266384541 | 0.45634373 | 29.14822612 | 39.18499449 | ENSG00000133101 |
| NA | 0.426683484 | 1.344140068 | 0.009630026 | 0.035660913 | 179.6884916 | 241.5405756 | ENSG00000250182 |
| LOH12CR2 | 0.426380353 | 1.343857675 | 0.711901344 | 0.83656984 | 3.629041247 | 4.834771149 | ENSG00000205791 |
| NEU4 | 0.426126727 | 1.343621445 | 0.598694771 | 0.758663264 | 6.100481064 | 8.142799715 | ENSG00000204099 |
| IRAK4 | 0.426049016 | 1.343549072 | 0.220522329 | 0.402037429 | 33.86305656 | 45.51557624 | ENSG00000198001 |
| SH3TC1 | 0.426023171 | 1.343525004 | 0.681701724 | 0.816192491 | 4.333701191 | 5.873794292 | ENSG00000125089 |
| ZNF83 | 0.425980088 | 1.343484883 | 0.00977037 | 0.036043797 | 203.1229056 | 273.1697926 | ENSG00000167766 |
| SNORA66 | 0.425915481 | 1.34342472 | 0.716620744 | 0.839556455 | 3.490062025 | 4.658234506 | ENSG00000207523 |
| FAM91A1 | 0.425792411 | 1.343310124 | 6.89E-05 | 0.000499584 | 465.8366567 | 625.6328479 | ENSG00000176853 |
| COL6A1 | 0.425774376 | 1.343293331 | 2.16E-14 | 7.68E-13 | 6443.343685 | 8655.680638 | ENSG00000142156 |
| GAD2 | 0.425515835 | 1.343052625 | 0.567438866 | 0.734702435 | 7.691000121 | 10.30341605 | ENSG00000136750 |
| NAPSA | 0.425013768 | 1.342585315 | 0.607930559 | 0.766065623 | 5.706821866 | 7.711831292 | ENSG00000131400 |
| ACOT13 | 0.424968061 | 1.342542781 | 0.004267807 | 0.017997538 | 246.6245353 | 331.4460499 | ENSG00000112304 |
| NA | 0.424959145 | 1.342534484 | 0.575805563 | 0.740853659 | 7.433163232 | 9.917037658 | ENSG00000229119 |
| CYP2D7 | 0.424878634 | 1.342459564 | 0.299927951 | 0.493415916 | 32.81327119 | 44.08042712 | ENSG00000205702 |
| ZNF275 | 0.42459908 | 1.342199459 | 0.000178967 | 0.001165582 | 422.5810103 | 566.9841967 | ENSG00000063587 |
| SSPN | 0.424391772 | 1.342006605 | 0.51798914 | 0.695379597 | 11.27782142 | 15.11897817 | ENSG00000123096 |
| SNTB2 | 0.424140588 | 1.341772971 | 0.00314191 | 0.013854903 | 269.484497 | 361.5402251 | ENSG00000168807 |
| BMPR1A | 0.424099971 | 1.341735197 | 1.22E-08 | 1.93E-07 | 1570.340976 | 2106.234663 | ENSG00000107779 |
| CHEK2 | 0.423957734 | 1.34160292 | 6.75E-08 | 9.28E-07 | 1321.590337 | 1772.531308 | ENSG00000183765 |
| GOLGA6L4 | 0.423838291 | 1.341491851 | 0.538150479 | 0.711875025 | 11.33088758 | 15.35650271 | ENSG00000184206 |
| NA | 0.423408521 | 1.341092288 | 0.660571384 | 0.802356197 | 4.158620453 | 5.59255579 | ENSG00000261220 |
| FAM72B | 0.423373092 | 1.341059355 | 0.001641376 | 0.008022485 | 297.2998077 | 398.3618808 | ENSG00000188610 |
| KCTD20 | 0.423132033 | 1.340835297 | 7.59E-06 | 7.02E-05 | 866.0122555 | 1160.15949 | ENSG00000112078 |
| TMEM263 | 0.423129676 | 1.340833106 | 0.00549133 | 0.022308372 | 224.473691 | 300.7474267 | ENSG00000151135 |
| HMGB3 | 0.422990476 | 1.340703741 | 6.76E-12 | 1.77E-10 | 6260.50286 | 8393.133144 | ENSG00000029993 |
| SBNO1 | 0.422974102 | 1.340688524 | 8.31E-07 | 9.39E-06 | 1106.412228 | 1482.459459 | ENSG00000139697 |
| NSD1 | 0.422798193 | 1.340525063 | 5.88E-08 | 8.19E-07 | 3662.544574 | 4909.384326 | ENSG00000165671 |
| GALNT6 | 0.422761075 | 1.340490574 | 0.007719235 | 0.029737746 | 206.7974214 | 277.3169649 | ENSG00000139629 |
| UBE2S | 0.422651711 | 1.340388962 | 3.05E-11 | 7.16E-10 | 6781.137796 | 9088.831949 | ENSG00000108106 |
| KIF2C | 0.422355031 | 1.340113348 | 1.72E-06 | 1.83E-05 | 1407.728859 | 1885.822359 | ENSG00000142945 |
| ALPK3 | 0.422186568 | 1.339956873 | 2.36E-07 | 2.95E-06 | 1662.552326 | 2228.071085 | ENSG00000136383 |
| GPA33 | 0.422138024 | 1.339911787 | 0.677273132 | 0.813471461 | 4.185153532 | 5.615125631 | ENSG00000143167 |
| NA | 0.421824018 | 1.339620184 | 0.413185764 | 0.604535668 | 15.01930881 | 20.14580777 | ENSG00000268218 |
| POLR2A | 0.421659083 | 1.339467042 | 3.45E-09 | 5.89E-08 | 5592.256963 | 7491.119103 | ENSG00000181222 |

| | | | | | | | |
|---|---|---|---|---|---|---|---|
| SLC47A2 | 0.421653802 | 1.339462138 | 0.669443424 | 0.807856359 | 4.425928081 | 5.97144811 | ENSG00000180638 |
| LRP4 | 0.421596641 | 1.339409069 | 1.55E-06 | 1.66E-05 | 925.3730976 | 1239.486778 | ENSG00000134569 |
| NA | 0.421513498 | 1.33933188 | 0.58177265 | 0.745786021 | 6.748820239 | 9.035780163 | ENSG00000230530 |
| EDEM3 | 0.421496116 | 1.339315744 | 0.00248938 | 0.011434491 | 287.0196902 | 384.0005757 | ENSG00000116406 |
| R3HDM2 | 0.421386689 | 1.339214161 | 0.005994524 | 0.024022164 | 207.3543304 | 277.8823647 | ENSG00000179912 |
| GAMT | 0.421255965 | 1.33909282 | 0.000224918 | 0.001426148 | 413.3470402 | 553.6481619 | ENSG00000130005 |
| SLC35D2 | 0.421161885 | 1.339005499 | 0.106900464 | 0.238250746 | 69.38656528 | 92.96089455 | ENSG00000130958 |
| NA | 0.421044683 | 1.338896725 | 0.093698675 | 0.215665229 | 84.39008953 | 112.8618033 | ENSG00000260727 |
| F12 | 0.420964623 | 1.338822426 | 0.187693274 | 0.359481983 | 45.97612879 | 61.36608198 | ENSG00000131187 |
| MRS2 | 0.420847768 | 1.338713989 | 2.43E-08 | 3.63E-07 | 1323.04668 | 1770.358937 | ENSG00000124532 |
| NA | 0.420847004 | 1.338713281 | 0.594465374 | 0.755547815 | 7.886300113 | 10.65648934 | ENSG00000227470 |
| NA | 0.420769078 | 1.338640973 | 0.661395601 | 0.802980877 | 4.105554294 | 5.513672966 | ENSG00000242611 |
| LRRC23 | 0.420752029 | 1.338625154 | 0.403605517 | 0.595338213 | 19.73266608 | 26.34586862 | ENSG00000010626 |
| RAD51C | 0.42055621 | 1.338443472 | 0.008329148 | 0.031672635 | 182.4501342 | 244.4473653 | ENSG00000108384 |
| NR1D2 | 0.420183326 | 1.338097578 | 0.001045255 | 0.005474247 | 327.0489443 | 437.1593976 | ENSG00000174738 |
| BTG3 | 0.420167596 | 1.338082989 | 0.015212059 | 0.051646876 | 189.4293637 | 252.9920164 | ENSG00000154640 |
| TCEAL7 | 0.420130443 | 1.338048531 | 0.089401793 | 0.208581675 | 80.36245073 | 107.3577663 | ENSG00000182916 |
| NA | 0.420115144 | 1.338034342 | 0.71649766 | 0.839556455 | 4.346328843 | 5.847537224 | ENSG00000237609 |
| MRI1 | 0.420083343 | 1.338004848 | 0.009893469 | 0.036432657 | 188.2105032 | 251.5759732 | ENSG00000037757 |
| MAP1LC3C | 0.419853678 | 1.337791865 | 0.521449174 | 0.698456239 | 11.00913832 | 14.616613 | ENSG00000197769 |
| ERCC1 | 0.419848983 | 1.337787512 | 2.04E-05 | 0.000169568 | 537.2462527 | 718.9089474 | ENSG00000012061 |
| NA | 0.419435488 | 1.337404139 | 0.301492753 | 0.494895641 | 28.39315327 | 37.97703239 | ENSG00000236333 |
| TPK1 | 0.419323633 | 1.337300452 | 0.346726914 | 0.540170237 | 19.96218845 | 26.69795422 | ENSG00000196511 |
| NA | 0.419071216 | 1.337066496 | 0.234650414 | 0.419534371 | 38.74823995 | 51.85930666 | ENSG00000260007 |
| PEAR1 | 0.419069868 | 1.337065246 | 0.361798481 | 0.554809321 | 20.14348531 | 26.91951893 | ENSG00000187800 |
| EGF | 0.419005387 | 1.337005488 | 0.085916579 | 0.202217662 | 75.07957942 | 100.3676034 | ENSG00000138798 |
| ASAH1 | 0.418996901 | 1.336997623 | 8.55E-07 | 9.63E-06 | 924.6255186 | 1235.679936 | ENSG00000104763 |
| LAMC3 | 0.418907356 | 1.336914642 | 2.05E-07 | 2.60E-06 | 990.6985833 | 1325.060467 | ENSG00000050555 |
| THPO | 0.418886004 | 1.336894855 | 0.628314798 | 0.780510951 | 5.40854647 | 7.235285152 | ENSG00000090534 |
| FAH | 0.418885281 | 1.336894186 | 0.000699167 | 0.003853981 | 319.6327457 | 427.4868612 | ENSG00000103876 |
| SH3YL1 | 0.418855707 | 1.336866781 | 0.000336004 | 0.002024547 | 424.8571501 | 567.6102664 | ENSG00000035115 |
| NA | 0.418728443 | 1.336748857 | 0.267303797 | 0.45741008 | 290.6097887 | 388.6792478 | ENSG00000234072 |
| TAF10 | 0.418677587 | 1.336701736 | 0.049311826 | 0.132321613 | 94.87871342 | 126.6447748 | ENSG00000166337 |
| MCHR1 | 0.418549653 | 1.336583207 | 0.16913682 | 0.333619598 | 45.22985026 | 60.3952064 | ENSG00000128285 |
| ALDH2 | 0.418350684 | 1.336398885 | 9.91E-05 | 0.000690819 | 568.2480587 | 758.8790342 | ENSG00000111275 |
| ZNF70 | 0.418216128 | 1.336274249 | 0.158188343 | 0.317610524 | 51.55471995 | 68.91733646 | ENSG00000187792 |
| SCYL3 | 0.41804386 | 1.336114698 | 0.046886176 | 0.127169874 | 104.0616919 | 138.9625413 | ENSG00000000457 |
| LINC02731 | 0.417802065 | 1.335890784 | 0.586485233 | 0.749312221 | 7.648682477 | 10.28888194 | ENSG00000204241 |
| ENPP1 | 0.417798013 | 1.335887032 | 0.000158635 | 0.001045567 | 634.8623607 | 847.3152721 | ENSG00000197594 |
| PHLDA1 | 0.417765541 | 1.335856965 | 0.000798453 | 0.004334659 | 443.4792828 | 591.9857678 | ENSG00000139289 |
| TCP11L1 | 0.417764439 | 1.335855944 | 0.017314138 | 0.057303614 | 174.3741715 | 233.2746295 | ENSG00000176148 |
| TMF1 | 0.417704587 | 1.335800526 | 0.004867982 | 0.020103204 | 216.9923023 | 289.5332469 | ENSG00000144747 |
| NA | 0.417567437 | 1.335673544 | 0.191942432 | 0.36509247 | 46.034231 | 61.26071885 | ENSG00000272923 |
| NA | 0.417412965 | 1.335530539 | 0.271173193 | 0.461984003 | 27.94199323 | 37.29009485 | ENSG00000287126 |
| CYSTM1 | 0.4173618 | 1.335483175 | 0.001558936 | 0.007680309 | 298.0320831 | 397.7742376 | ENSG00000120306 |
| STOML1 | 0.417220822 | 1.33535268 | 0.215967886 | 0.396390255 | 48.49738016 | 64.52415739 | ENSG00000067221 |
| ZNF587 | 0.416943881 | 1.335096369 | 0.006827454 | 0.026877662 | 292.12249 | 390.6196277 | ENSG00000198466 |
| POU2F2 | 0.416888712 | 1.335045315 | 0.056077067 | 0.146035178 | 95.21930646 | 127.250568 | ENSG00000028277 |
| OSCAR | 0.416515098 | 1.334699624 | 0.182359021 | 0.352765066 | 49.00437995 | 65.67365661 | ENSG00000170909 |
| LINC00649 | 0.416387227 | 1.334581331 | 0.038960901 | 0.110244479 | 106.248664 | 141.5543493 | ENSG00000237945 |
| SLCO5A1 | 0.416211361 | 1.334418654 | 0.054417114 | 0.142694792 | 101.4613449 | 135.4865614 | ENSG00000137571 |
| SH3PXD2A | 0.416120306 | 1.334334435 | 0.000138563 | 0.000926455 | 545.748451 | 728.2365812 | ENSG00000107957 |
| RPRD1A | 0.415952145 | 1.334178914 | 2.48E-09 | 4.34E-08 | 2318.549629 | 3092.897225 | ENSG00000141425 |
| TFCP2L1 | 0.415767724 | 1.334008375 | 0.038507683 | 0.109128655 | 119.2679123 | 158.8874456 | ENSG00000115112 |
| WWC2 | 0.415740734 | 1.333983418 | 1.30E-09 | 2.36E-08 | 2383.348663 | 3178.242992 | ENSG00000151718 |
| FARP2 | 0.415740722 | 1.333983407 | 5.56E-05 | 0.000413894 | 462.8718368 | 617.2904005 | ENSG00000006607 |
| PCNX2 | 0.415713653 | 1.333958378 | 8.14E-07 | 9.23E-06 | 1299.683447 | 1733.723812 | ENSG00000135749 |
| TMSB4X | 0.415585907 | 1.333840266 | 1.72E-08 | 2.65E-07 | 15879.42338 | 21180.59458 | ENSG00000205542 |
| BOD1L1 | 0.415533902 | 1.333792186 | 0.00014629 | 0.000972504 | 912.408013 | 1216.819388 | ENSG00000038219 |
| DUS2 | 0.414964194 | 1.333265586 | 0.002383671 | 0.011011704 | 275.3482322 | 366.8370711 | ENSG00000167264 |
| PPP1R3D | 0.41491682 | 1.333221806 | 0.090922328 | 0.211225174 | 75.08982427 | 100.2852481 | ENSG00000132825 |
| PIPOX | 0.41490038 | 1.333206615 | 1.40E-05 | 0.000121127 | 786.8764296 | 1048.52298 | ENSG00000179761 |
| ELN | 0.414687978 | 1.333010346 | 0.013328836 | 0.046477982 | 160.2319274 | 213.8830307 | ENSG00000049540 |
| PELI1 | 0.4146212 | 1.332948647 | 5.06E-07 | 5.96E-06 | 1163.499509 | 1550.906352 | ENSG00000197329 |
| EMID1 | 0.414607686 | 1.332936161 | 0.036941564 | 0.105513169 | 122.005382 | 162.5867507 | ENSG00000186998 |
| NA | 0.414461913 | 1.332801484 | 0.706927476 | 0.833366865 | 3.602508168 | 4.740916177 | ENSG00000188460 |
| LRRC75B | 0.414326199 | 1.332676114 | 0.00689954 | 0.027092388 | 196.9412392 | 262.7925664 | ENSG00000178026 |
| IDNK | 0.414279334 | 1.332632824 | 0.63296841 | 0.783663094 | 5.478674969 | 7.287910909 | ENSG00000148057 |
| MYLIP | 0.41410131 | 1.332468392 | 2.51E-05 | 0.000204481 | 544.6876089 | 725.790498 | ENSG00000007944 |
| YPEL4 | 0.414092736 | 1.332460472 | 0.415746908 | 0.606967561 | 19.65632145 | 26.33133451 | ENSG00000166793 |
| LINC00662 | 0.413909205 | 1.332290976 | 0.00522217 | 0.021389548 | 257.8709982 | 343.0130395 | ENSG00000261824 |
| NA | 0.413467539 | 1.331883171 | 0.374463519 | 0.567389377 | 22.48975364 | 29.76553547 | ENSG00000259556 |
| CADM1 | 0.413308784 | 1.331736618 | 0.000969065 | 0.005131616 | 383.2631812 | 510.0269901 | ENSG00000182985 |
| CLIP1 | 0.413207388 | 1.331643023 | 0.00017442 | 0.001137876 | 460.089494 | 612.9345805 | ENSG00000130779 |

| | | | | | | | |
|---|---|---|---|---|---|---|---|
| TDRP | 0.413194634 | 1.331631251 | 6.75E-08 | 9.28E-07 | 1070.428592 | 1425.400499 | ENSG00000180190 |
| ZDHHC11 | 0.41272249 | 1.331195526 | 0.115631705 | 0.252785801 | 109.5504389 | 146.2136117 | ENSG00000188818 |
| PTGFRN | 0.412578269 | 1.331062458 | 2.23E-05 | 0.000184322 | 933.703026 | 1242.698266 | ENSG00000134247 |
| ZNF724 | 0.412345683 | 1.330847886 | 0.079153022 | 0.190127281 | 75.66350256 | 100.7124177 | ENSG00000196081 |
| NAGA | 0.412205489 | 1.330718567 | 2.61E-05 | 0.000211922 | 528.6452431 | 703.2730296 | ENSG00000198951 |
| LAMP2 | 0.411984656 | 1.33051489 | 7.88E-05 | 0.000563047 | 742.4744641 | 987.1966213 | ENSG00000005893 |
| RIMBP3 | 0.411879422 | 1.330417842 | 0.515480182 | 0.693363165 | 9.815533073 | 13.08282238 | ENSG00000275793 |
| NA | 0.411829731 | 1.330372019 | 0.578717081 | 0.74299961 | 7.982961692 | 10.5884534 | ENSG00000230438 |
| PAQR3 | 0.411795624 | 1.330340568 | 0.001323001 | 0.006655944 | 519.5252901 | 690.6515891 | ENSG00000163291 |
| APLF | 0.411631345 | 1.330189092 | 0.107037883 | 0.238442323 | 61.99266048 | 82.46487041 | ENSG00000169621 |
| PCDHGA9 | 0.411598067 | 1.330158409 | 0.56694068 | 0.734545862 | 8.450004 | 11.27852852 | ENSG00000261934 |
| GLS2 | 0.411546638 | 1.330110992 | 0.589902794 | 0.751995212 | 7.317560176 | 9.767307736 | ENSG00000135423 |
| PTPRJ | 0.411339503 | 1.329920036 | 0.000183982 | 0.001194784 | 541.8147979 | 720.3055584 | ENSG00000149177 |
| MACROD2 | 0.41111575 | 1.32971379 | 0.382940095 | 0.575621389 | 22.71780284 | 30.08910249 | ENSG00000172264 |
| TXNDC5 | 0.411075181 | 1.329676398 | 0.039155586 | 0.110626467 | 256.7032723 | 341.0439488 | ENSG00000239264 |
| BATF3 | 0.410961896 | 1.329571992 | 0.170671845 | 0.336013595 | 46.29975719 | 61.52373601 | ENSG00000123685 |
| TMEM88 | 0.410622012 | 1.329258795 | 0.287694112 | 0.4792262 | 26.69078943 | 35.52020541 | ENSG00000167874 |
| ALDH6A1 | 0.410575462 | 1.329215906 | 0.000207555 | 0.001327863 | 483.4409338 | 642.1316499 | ENSG00000119711 |
| ADAM9 | 0.410464069 | 1.329113279 | 5.69E-05 | 0.000422907 | 486.5767905 | 646.6752202 | ENSG00000168615 |
| PPP3CC | 0.410456413 | 1.329106225 | 0.185279439 | 0.356036197 | 44.45633959 | 59.06029902 | ENSG00000120910 |
| NPIPA1 | 0.41037527 | 1.329031473 | 0.051476506 | 0.136744383 | 109.6035051 | 145.8511886 | ENSG00000183426 |
| GALK2 | 0.410233307 | 1.328900701 | 0.041989335 | 0.116952747 | 112.8685771 | 149.7798307 | ENSG00000156958 |
| CLDN1 | 0.41022103 | 1.328889393 | 0.126656021 | 0.270272281 | 63.31527055 | 83.99073692 | ENSG00000163347 |
| EXOC4 | 0.410221027 | 1.32888939 | 3.09E-06 | 3.14E-05 | 1233.691514 | 1639.003388 | ENSG00000131558 |
| ADGRE2 | 0.410118532 | 1.328794984 | 0.301263288 | 0.494738064 | 26.98778705 | 35.92904202 | ENSG00000127507 |
| AGK | 0.410002106 | 1.328687754 | 0.000103258 | 0.000717064 | 561.2403418 | 745.4032403 | ENSG00000006530 |
| ARFGEF3 | 0.409970964 | 1.328659073 | 0.000663127 | 0.003675357 | 825.1462774 | 1095.907675 | ENSG00000112379 |
| NA | 0.409924847 | 1.328616602 | 0.665509816 | 0.805653751 | 4.475837326 | 5.97144811 | ENSG00000256995 |
| DBIL5P | 0.409705521 | 1.328414634 | 0.624954375 | 0.777404233 | 6.597213365 | 8.855994327 | ENSG00000231784 |
| WDR83 | 0.40966469 | 1.328377037 | 0.270183663 | 0.460679301 | 36.49665637 | 48.33296258 | ENSG00000123154 |
| NECAB1 | 0.409653293 | 1.328366544 | 0.065361039 | 0.164235561 | 109.4393683 | 145.1383918 | ENSG00000123119 |
| FADD | 0.409609033 | 1.328325792 | 0.000380812 | 0.002257826 | 399.7191171 | 530.7623133 | ENSG00000168040 |
| CHD6 | 0.409596877 | 1.3283146 | 6.97E-06 | 6.52E-05 | 1236.607718 | 1642.204828 | ENSG00000124177 |
| LOC101929331 | 0.409564442 | 1.328284737 | 0.721227301 | 0.842401487 | 3.172845151 | 4.249835924 | ENSG00000224307 |
| HEY2 | 0.409398593 | 1.328132049 | 0.013132778 | 0.045968445 | 175.7290271 | 233.4716489 | ENSG00000135547 |
| PCDHA10 | 0.409330002 | 1.328068907 | 0.004596363 | 0.019134995 | 212.0975504 | 281.7486334 | ENSG00000250120 |
| SERPINB6 | 0.409303443 | 1.328044457 | 0.000503276 | 0.002883373 | 428.3993087 | 568.5339472 | ENSG00000124570 |
| TTC21A | 0.409088199 | 1.327846334 | 0.311095737 | 0.504837468 | 31.33341684 | 41.72494118 | ENSG00000168026 |
| BTN3A2 | 0.408891113 | 1.327664949 | 0.102057058 | 0.230208097 | 71.20650901 | 94.61859607 | ENSG00000186470 |
| NA | 0.408796076 | 1.327577493 | 0.330807148 | 0.525154576 | 208.8150479 | 277.3340728 | ENSG00000227671 |
| VAV2 | 0.408652613 | 1.327445484 | 1.76E-08 | 2.70E-07 | 1694.996622 | 2249.858728 | ENSG00000160293 |
| CCDC144NL | 0.408484809 | 1.327291094 | 0.465356086 | 0.65162509 | 13.08060511 | 17.27579566 | ENSG00000205212 |
| SNHG16 | 0.408447709 | 1.327256962 | 3.92E-06 | 3.91E-05 | 874.1367746 | 1160.055419 | ENSG00000163597 |
| HPS4 | 0.408378514 | 1.327193306 | 8.48E-05 | 0.000601286 | 733.5484885 | 973.1386554 | ENSG00000100099 |
| SCG5 | 0.408349905 | 1.327166986 | 0.000971195 | 0.005137146 | 327.6016065 | 434.5334487 | ENSG00000166922 |
| ASPH | 0.408200173 | 1.327029252 | 4.55E-07 | 5.42E-06 | 1401.031143 | 1858.457283 | ENSG00000198363 |
| NA | 0.408183934 | 1.327014315 | 0.676935705 | 0.813270653 | 4.194624271 | 5.555013801 | ENSG00000272800 |
| LTC4S | 0.408029121 | 1.326871923 | 0.509700772 | 0.68889332 | 11.4698668 | 15.23595264 | ENSG00000213316 |
| CKAP5 | 0.407942246 | 1.326792025 | 2.12E-10 | 4.39E-09 | 4413.411602 | 5855.015284 | ENSG00000175216 |
| NCMAP | 0.407923257 | 1.326774562 | 0.620923674 | 0.774962921 | 6.526987168 | 8.593088786 | ENSG00000184454 |
| SLC46A3 | 0.407858683 | 1.326715178 | 0.650734532 | 0.795936006 | 5.633536454 | 7.493953813 | ENSG00000139508 |
| ZNF623 | 0.407852695 | 1.326709671 | 3.06E-05 | 0.000245109 | 622.6296114 | 826.0060068 | ENSG00000183309 |
| ELK3 | 0.407784774 | 1.326647212 | 0.003618379 | 0.015610576 | 265.3538828 | 351.9173111 | ENSG00000111145 |
| DSP | 0.407743123 | 1.326608912 | 4.62E-05 | 0.000350776 | 2325.515916 | 3084.486592 | ENSG00000096696 |
| ACBD4 | 0.407630339 | 1.326505207 | 0.014239566 | 0.048954945 | 188.7835051 | 250.5212537 | ENSG00000181513 |
| C18orf54 | 0.407586185 | 1.32646461 | 0.003659167 | 0.015771859 | 362.4635092 | 480.556686 | ENSG00000166845 |
| ZNF562 | 0.407504077 | 1.326389119 | 0.003058209 | 0.013543809 | 296.8425444 | 393.0239716 | ENSG00000171466 |
| LYSMD1 | 0.407281652 | 1.326184641 | 0.058559591 | 0.151119946 | 101.0183552 | 134.1636003 | ENSG00000163155 |
| NA | 0.40724854 | 1.326154204 | 0.682980512 | 0.816931965 | 4.717889651 | 6.305200749 | ENSG00000261634 |
| BBX | 0.407117694 | 1.326033933 | 3.24E-05 | 0.000257408 | 752.7202541 | 997.2446497 | ENSG00000114439 |
| RHOXF1-AS1 | 0.406895928 | 1.325830115 | 0.175208563 | 0.34254383 | 44.74396417 | 59.32981457 | ENSG00000258545 |
| ADAMTS1 | 0.40659557 | 1.325554116 | 0.000188686 | 0.001219779 | 477.6470188 | 632.6607845 | ENSG00000154734 |
| NFS1 | 0.40657848 | 1.325538414 | 0.005252775 | 0.021505396 | 231.0984448 | 306.5541014 | ENSG00000244005 |
| SEPTIN10 | 0.406414193 | 1.325387477 | 1.10E-06 | 1.22E-05 | 1401.833915 | 1856.886291 | ENSG00000186522 |
| TIGAR | 0.406226205 | 1.325214785 | 0.059741846 | 0.15335947 | 94.91165802 | 125.7448579 | ENSG00000078237 |
| KDSR | 0.406112179 | 1.325110049 | 1.09E-05 | 9.67E-05 | 666.6645786 | 882.9464319 | ENSG00000119537 |
| ATP2A3 | 0.406070699 | 1.325071951 | 5.11E-06 | 4.92E-05 | 989.3012372 | 1311.457161 | ENSG00000074370 |
| NA | 0.405526482 | 1.324572197 | 0.556718375 | 0.726225955 | 8.234484755 | 10.89215012 | ENSG00000232346 |
| KCNA3 | 0.405440024 | 1.324492821 | 0.496272617 | 0.677862531 | 12.07904525 | 16.06502241 | ENSG00000177272 |
| GPSM2 | 0.405226029 | 1.324296373 | 0.005388622 | 0.021969361 | 216.9504883 | 287.0021004 | ENSG00000121957 |
| NA | 0.405097651 | 1.324178536 | 0.307634457 | 0.501782934 | 30.64020447 | 40.51990186 | ENSG00000254469 |
| CHST14 | 0.405057419 | 1.32414161 | 0.000427221 | 0.002493108 | 409.5520056 | 542.6192702 | ENSG00000169105 |
| IL34 | 0.40504028 | 1.324125879 | 0.050710869 | 0.135272775 | 96.84582933 | 128.2739578 | ENSG00000157368 |
| IGFBP2 | 0.405009898 | 1.324097994 | 0.010321662 | 0.037717208 | 5606.583045 | 7423.172921 | ENSG00000115457 |

| | | | | | | | |
|---|---|---|---|---|---|---|---|
| METTL23 | 0.40497566 | 1.324066572 | 0.023587027 | 0.073919846 | 145.4737101 | 192.68885 | ENSG00000181038 |
| NEK6 | 0.404929712 | 1.324024403 | 0.000428807 | 0.002500789 | 404.7529232 | 535.8340509 | ENSG00000119408 |
| ZNF552 | 0.404802849 | 1.32390798 | 0.048456733 | 0.130499433 | 109.9820562 | 145.5458429 | ENSG00000178935 |
| ZSWIM6 | 0.404763383 | 1.323871764 | 0.002963383 | 0.013184913 | 267.443667 | 354.033887 | ENSG00000130449 |
| NDUFB5 | 0.404740627 | 1.323850882 | 0.000250094 | 0.001570206 | 702.1203637 | 929.5666538 | ENSG00000136521 |
| TSSK4 | 0.404708107 | 1.323821041 | 0.393688254 | 0.586309208 | 23.85843962 | 31.62008195 | ENSG00000139908 |
| ZNF519 | 0.404597494 | 1.323719547 | 0.009484438 | 0.035220312 | 178.6756045 | 236.6720613 | ENSG00000175322 |
| ADCK1 | 0.404146595 | 1.323305897 | 0.005597581 | 0.022706404 | 238.2606398 | 315.3101317 | ENSG00000063761 |
| PPP1R2 | 0.404019918 | 1.323189708 | 0.00131554 | 0.006623802 | 411.9487619 | 544.6931509 | ENSG00000184203 |
| HEXB | 0.40365225 | 1.322852538 | 1.26E-05 | 0.000110278 | 942.9791863 | 1246.806831 | ENSG00000049860 |
| TAFA3 | 0.403309359 | 1.322538168 | 0.635200509 | 0.785377333 | 6.951711833 | 9.189308932 | ENSG00000184599 |
| CACNA1D | 0.40329192 | 1.322522182 | 0.337701291 | 0.531944773 | 24.6876697 | 32.71074317 | ENSG00000157388 |
| ZC3H10 | 0.40307809 | 1.322326178 | 0.049940522 | 0.133660309 | 98.0755361 | 129.6574688 | ENSG00000135482 |
| NA | 0.402715979 | 1.321994321 | 0.614537724 | 0.770209997 | 8.579512484 | 11.3424392 | ENSG00000265800 |
| COBL | 0.402474877 | 1.321773408 | 8.71E-06 | 7.89E-05 | 869.6384555 | 1149.43135 | ENSG00000106078 |
| FBXL20 | 0.40211127 | 1.32144032 | 0.0237848 | 0.074394733 | 146.7449378 | 193.6595023 | ENSG00000108306 |
| RAB4B | 0.402011403 | 1.32134885 | 0.638010622 | 0.787529817 | 5.650598794 | 7.475182819 | ENSG00000167578 |
| RSU1 | 0.40191799 | 1.321263296 | 1.15E-07 | 1.52E-06 | 1238.870816 | 1637.47609 | ENSG00000148484 |
| GABBR2 | 0.401909933 | 1.321255918 | 0.242994235 | 0.429533157 | 42.86967651 | 56.56114351 | ENSG00000136928 |
| NA | 0.401728198 | 1.321089491 | 0.564376186 | 0.732605984 | 9.882504659 | 13.04516877 | ENSG00000242477 |
| ERCC5 | 0.40165302 | 1.321020652 | 0.202423488 | 0.379737156 | 42.68650051 | 56.53587413 | ENSG00000134899 |
| ZDHHC24 | 0.401263895 | 1.320664392 | 0.021269333 | 0.068019185 | 159.0389236 | 209.9759306 | ENSG00000174165 |
| ALDH5A1 | 0.401260685 | 1.320661454 | 0.060221681 | 0.154441489 | 99.73252308 | 131.580449 | ENSG00000112294 |
| ITM2B | 0.401026817 | 1.320447385 | 2.58E-06 | 2.66E-05 | 1712.243717 | 2260.395456 | ENSG00000136156 |
| RHBDF2 | 0.400880915 | 1.320313853 | 0.000463874 | 0.00268251 | 485.07014 | 640.8648986 | ENSG00000129667 |
| NA | 0.400759146 | 1.320202419 | 0.400877939 | 0.592682812 | 19.70623069 | 26.10217211 | ENSG00000267523 |
| GPR180 | 0.400452251 | 1.319921611 | 0.012521508 | 0.044314231 | 212.1810056 | 279.7198118 | ENSG00000152749 |
| TBILA | 0.400397034 | 1.319871093 | 0.643845462 | 0.791682598 | 6.676812603 | 8.874765321 | ENSG00000261488 |
| TK2 | 0.400364282 | 1.31984113 | 0.232088079 | 0.416773613 | 38.35901544 | 50.67391441 | ENSG00000166548 |
| DNASE1L1 | 0.400209994 | 1.319699988 | 0.256202044 | 0.444990476 | 33.77073197 | 44.48794963 | ENSG00000013563 |
| UQCC1 | 0.40019192 | 1.319683455 | 0.002531095 | 0.011577322 | 875.3300942 | 1156.002606 | ENSG00000101019 |
| NA | 0.400154352 | 1.319649091 | 0.438818268 | 0.627185333 | 16.34893176 | 21.64954247 | ENSG00000283103 |
| ITPRID2 | 0.400086337 | 1.319586878 | 0.029075802 | 0.087354664 | 160.0801252 | 210.7694966 | ENSG00000138434 |
| PFKFB2 | 0.400072731 | 1.319574433 | 0.000633556 | 0.003527285 | 455.5468956 | 600.7685822 | ENSG00000123836 |
| RC3H1 | 0.400069217 | 1.319571219 | 0.02280015 | 0.071940412 | 202.1364312 | 266.0927337 | ENSG00000135870 |
| EIF4B | 0.400064872 | 1.319567245 | 1.55E-10 | 3.30E-09 | 12072.16092 | 15929.01674 | ENSG00000063046 |
| SLC38A4 | 0.399987265 | 1.319496263 | 0.001617811 | 0.00793451 | 317.4215029 | 418.3588348 | ENSG00000139209 |
| FAM189A1 | 0.399835028 | 1.319357034 | 0.008539391 | 0.032306405 | 195.4970956 | 257.9689685 | ENSG00000104059 |
| E2F7 | 0.399781127 | 1.319307742 | 0.000317107 | 0.001924449 | 451.5385891 | 595.3414611 | ENSG00000165891 |
| MECR | 0.399703216 | 1.319236496 | 4.19E-05 | 0.000322085 | 546.206218 | 720.8291164 | ENSG00000116353 |
| RBM24 | 0.399598137 | 1.319140413 | 0.604004708 | 0.762992316 | 7.698591723 | 10.07425365 | ENSG00000112183 |
| GSK3B | 0.399584704 | 1.31912813 | 3.30E-05 | 0.000261609 | 744.3225043 | 981.0818703 | ENSG00000082701 |
| CHD2 | 0.399389986 | 1.318950102 | 1.68E-05 | 0.000142458 | 1285.193266 | 1695.176585 | ENSG00000173575 |
| NA | 0.399278223 | 1.318847929 | 0.529834953 | 0.705507653 | 10.13718464 | 13.37903303 | ENSG00000213228 |
| JMJD1C | 0.399272082 | 1.318842316 | 1.87E-08 | 2.86E-07 | 2506.059735 | 3304.07476 | ENSG00000171988 |
| NPL | 0.399218157 | 1.318793021 | 0.149129919 | 0.30477163 | 57.51238846 | 75.95395317 | ENSG00000135838 |
| FA2H | 0.399188069 | 1.318765517 | 0.367439251 | 0.560549764 | 24.84303485 | 32.77834107 | ENSG00000103089 |
| CCDC122 | 0.399152543 | 1.318733043 | 0.252681932 | 0.441353965 | 41.6349337 | 54.70021025 | ENSG00000151773 |
| NGFR | 0.399108176 | 1.318692489 | 0.006245789 | 0.024899862 | 244.2007423 | 322.4573361 | ENSG00000064300 |
| ZMYM1 | 0.399082645 | 1.318669153 | 0.00769049 | 0.029645495 | 209.6369568 | 276.3873899 | ENSG00000197056 |
| NA | 0.398965782 | 1.31856234 | 0.583667243 | 0.746923199 | 9.004740811 | 11.87909716 | ENSG00000259424 |
| TOMM7 | 0.398719709 | 1.318337459 | 3.33E-07 | 4.05E-06 | 1682.3369 | 2218.281542 | ENSG00000196683 |
| LENG1 | 0.398484994 | 1.318122994 | 0.03313424 | 0.096954532 | 140.2019706 | 184.6898315 | ENSG00000105617 |
| CRYBA4 | 0.398455223 | 1.318095793 | 0.371717157 | 0.564936924 | 24.26482417 | 32.04367592 | ENSG00000196431 |
| GPC4 | 0.398371049 | 1.318018892 | 2.62E-10 | 5.36E-09 | 15612.40821 | 20577.09769 | ENSG00000076716 |
| SMAD6 | 0.398358818 | 1.318007718 | 0.008349001 | 0.031735106 | 190.724862 | 251.299298 | ENSG00000137834 |
| PLEKHA8 | 0.398283226 | 1.31793866 | 7.26E-06 | 6.77E-05 | 924.7216992 | 1217.812833 | ENSG00000106086 |
| PFKL | 0.398138396 | 1.317806361 | 7.27E-11 | 1.62E-09 | 3875.961359 | 5107.437777 | ENSG00000141959 |
| CA10 | 0.397987642 | 1.317668664 | 0.600352689 | 0.759741754 | 7.67837247 | 10.14608844 | ENSG00000154975 |
| CEPT1 | 0.397985954 | 1.317667123 | 0.010587638 | 0.038537173 | 254.0619227 | 334.5597811 | ENSG00000134255 |
| ANKRA2 | 0.397933521 | 1.317619235 | 0.091606671 | 0.21230937 | 80.02561597 | 105.3702142 | ENSG00000164331 |
| NA | 0.397886387 | 1.317576188 | 0.606093573 | 0.764778101 | 7.664467042 | 10.12363022 | ENSG00000229976 |
| ERC1 | 0.397861157 | 1.317553146 | 0.000303822 | 0.001857174 | 1035.325193 | 1363.771214 | ENSG00000082805 |
| NA | 0.397850205 | 1.317543144 | 0.107939458 | 0.239816582 | 78.40285136 | 103.3265723 | ENSG00000224546 |
| GDPD3 | 0.397691045 | 1.317397799 | 0.523071023 | 0.699675423 | 11.40920904 | 15.02176239 | ENSG00000102886 |
| BMP8B | 0.397664951 | 1.317373972 | 0.295971081 | 0.488855289 | 32.30528671 | 42.56050934 | ENSG00000116985 |
| CRYGN | 0.397348898 | 1.317085405 | 0.474390589 | 0.659443124 | 14.03353351 | 18.55559254 | ENSG00000127377 |
| DOCK6 | 0.397155257 | 1.316908635 | 0.000326678 | 0.001976704 | 976.5053104 | 1286.750956 | ENSG00000130158 |
| IMPDH2 | 0.397143693 | 1.316898079 | 6.37E-10 | 1.22E-08 | 3983.471465 | 5245.514847 | ENSG00000178035 |
| B4GALT5 | 0.397129606 | 1.31688522 | 3.14E-08 | 4.62E-07 | 2341.330759 | 3082.127861 | ENSG00000158470 |
| FBXO22 | 0.396843159 | 1.316623779 | 3.70E-07 | 4.47E-06 | 1467.467387 | 1931.760398 | ENSG00000167196 |
| MLF1 | 0.396607004 | 1.316408278 | 0.000484529 | 0.002787884 | 467.0225726 | 614.1185067 | ENSG00000178053 |
| CIAO2B | 0.396435491 | 1.316251788 | 4.95E-06 | 4.78E-05 | 1073.567267 | 1413.311377 | ENSG00000166595 |
| NA | 0.396219576 | 1.316054811 | 0.116291419 | 0.253659545 | 83.90745805 | 110.283765 | ENSG00000213976 |

| | | | | | | | |
|---|---|---|---|---|---|---|---|
| LINC01535 | 0.395765446 | 1.315640611 | 0.602113348 | 0.76138151 | 8.10172166 | 10.58834178 | ENSG00000226686 |
| PIR | 0.395762575 | 1.315637992 | 1.79E-05 | 0.000150948 | 813.7908562 | 1069.741179 | ENSG00000087842 |
| COQ10B | 0.395353265 | 1.315264782 | 0.009397844 | 0.034968809 | 258.5271244 | 339.4030345 | ENSG00000115520 |
| BOLA3 | 0.395288496 | 1.315205736 | 0.00685278 | 0.026948745 | 210.5044758 | 277.0308367 | ENSG00000163170 |
| ZNF75D | 0.395254169 | 1.315174442 | 0.030187398 | 0.090081024 | 136.2658823 | 179.4018485 | ENSG00000186376 |
| SMYD3 | 0.395000402 | 1.314943127 | 0.002797453 | 0.012541845 | 296.9498416 | 390.6562534 | ENSG00000185420 |
| RMND5B | 0.394982595 | 1.314926896 | 1.45E-06 | 1.56E-05 | 1567.856616 | 2061.819768 | ENSG00000145916 |
| LMNA | 0.394701695 | 1.314670898 | 4.86E-06 | 4.71E-05 | 1189.350793 | 1564.046427 | ENSG00000160789 |
| CALCR | 0.394678304 | 1.314649584 | 0.337021384 | 0.531367421 | 25.36126418 | 33.38953336 | ENSG00000004948 |
| MROH8 | 0.394630183 | 1.314605734 | 0.227234166 | 0.41072563 | 36.89051097 | 48.50906119 | ENSG00000101353 |
| SRD5A3 | 0.394163315 | 1.314180386 | 0.001747249 | 0.008464066 | 344.3927161 | 452.7393372 | ENSG00000128039 |
| PBK | 0.394089295 | 1.314112961 | 0.051472794 | 0.136744383 | 154.0616261 | 201.9062395 | ENSG00000168078 |
| ASMTL | 0.394055156 | 1.314081865 | 2.45E-08 | 3.66E-07 | 2141.321296 | 2813.645143 | ENSG00000169093 |
| SLC35B2 | 0.393904479 | 1.313944628 | 0.00011775 | 0.00080739 | 638.1430743 | 838.2729625 | ENSG00000157593 |
| KCNIP1 | 0.393899614 | 1.313940197 | 0.364536955 | 0.557636838 | 21.1963299 | 27.92100008 | ENSG00000182132 |
| LDB3 | 0.393879182 | 1.313921589 | 0.532303134 | 0.707394694 | 10.69833297 | 14.10676175 | ENSG00000122367 |
| MTCL1 | 0.393813512 | 1.313861782 | 7.38E-07 | 8.43E-06 | 1303.398122 | 1712.301465 | ENSG00000168502 |
| ERCC6L2 | 0.393647476 | 1.313710582 | 0.002729012 | 0.012291498 | 270.0191873 | 354.6269696 | ENSG00000182150 |
| TRMT10B | 0.393361543 | 1.313450238 | 0.030132069 | 0.08995937 | 138.5370838 | 182.1842807 | ENSG00000165275 |
| B4GALT4 | 0.393236594 | 1.313336488 | 0.000302551 | 0.001850653 | 473.8827566 | 621.8946867 | ENSG00000121578 |
| ATP12A | 0.392818943 | 1.31295634 | 0.299531294 | 0.492985792 | 39.10085928 | 51.25818836 | ENSG00000075673 |
| SDCBP | 0.392632805 | 1.312786952 | 6.86E-08 | 9.42E-07 | 1754.677792 | 2302.788029 | ENSG00000137575 |
| TAF12 | 0.392507618 | 1.312673042 | 0.002930872 | 0.013051607 | 327.7196901 | 429.9842646 | ENSG00000120656 |
| NA | 0.392148527 | 1.312346355 | 0.276033 | 0.467043603 | 38.11951867 | 49.95323372 | ENSG00000260265 |
| TMEM191C | 0.392009492 | 1.312219888 | 0.547792811 | 0.719341926 | 10.89984909 | 14.32832646 | ENSG00000206140 |
| NA | 0.391845118 | 1.312070389 | 0.691476607 | 0.823321819 | 4.185153532 | 5.457359983 | ENSG00000228878 |
| BCKDHB | 0.391826783 | 1.312053713 | 0.000611809 | 0.003423695 | 416.6648853 | 546.5821421 | ENSG00000083123 |
| ZNFX1 | 0.391802633 | 1.31203175 | 0.002514988 | 0.011517862 | 335.7367613 | 440.6759857 | ENSG00000124201 |
| TMEM120A | 0.39172167 | 1.311958122 | 0.002336942 | 0.010833638 | 295.6375893 | 387.6296865 | ENSG00000189077 |
| ESRG | 0.391716361 | 1.311953295 | 4.09E-09 | 6.94E-08 | 7735.258465 | 10148.00093 | ENSG00000265992 |
| NA | 0.391341124 | 1.311612107 | 0.522199718 | 0.699117944 | 11.48249445 | 15.01807516 | ENSG00000283078 |
| NA | 0.391313119 | 1.311586647 | 0.310525704 | 0.504406959 | 31.49586992 | 41.22060063 | ENSG00000237854 |
| NA | 0.391309724 | 1.31158356 | 0.187642007 | 0.359420934 | 51.92815989 | 68.18635854 | ENSG00000187534 |
| TJP1 | 0.391091884 | 1.311385532 | 6.00E-08 | 8.35E-07 | 2502.989963 | 3281.430888 | ENSG00000104067 |
| SKIDA1 | 0.391084586 | 1.311378899 | 0.143650616 | 0.296483354 | 56.54811019 | 74.1605062 | ENSG00000180592 |
| TFR2 | 0.391037224 | 1.311335849 | 0.048139956 | 0.129929509 | 116.6645062 | 153.0896114 | ENSG00000106327 |
| SNX8 | 0.390846995 | 1.311162952 | 9.24E-07 | 1.03E-05 | 1172.685907 | 1537.17962 | ENSG00000106266 |
| LOC102546294 | 0.390742648 | 1.311068122 | 0.71531939 | 0.838672165 | 4.591440382 | 6.008990099 | ENSG00000247199 |
| CBWD3 | 0.390399991 | 1.310756764 | 0.518297745 | 0.695519378 | 12.92130894 | 16.86739708 | ENSG00000196873 |
| BICRAL | 0.390295794 | 1.310662099 | 0.005887846 | 0.023681716 | 252.9743747 | 331.7909356 | ENSG00000112624 |
| SPATA25 | 0.390040781 | 1.310430445 | 0.52605362 | 0.702433961 | 11.34351523 | 14.95091529 | ENSG00000149634 |
| C8orf31 | 0.389569614 | 1.310002545 | 0.583150319 | 0.74651941 | 7.986118604 | 10.49872369 | ENSG00000177335 |
| GSDMB | 0.389298754 | 1.30975662 | 0.127659437 | 0.271599539 | 122.7562156 | 161.2058679 | ENSG00000073605 |
| B3GNT4 | 0.389233397 | 1.309697286 | 0.3718351 | 0.565023509 | 21.69612142 | 28.33319751 | ENSG00000176383 |
| DLEU1 | 0.389210071 | 1.309676112 | 0.180007101 | 0.349313726 | 58.30739616 | 76.4678265 | ENSG00000176124 |
| ALKAL2 | 0.389139802 | 1.309612323 | 0.298705012 | 0.492019081 | 34.1530413 | 44.67945842 | ENSG00000189292 |
| GAP43 | 0.389020721 | 1.309504231 | 0.00091575 | 0.004887433 | 383.8178954 | 502.2087535 | ENSG00000172020 |
| NFE2L2 | 0.388880954 | 1.309377373 | 0.000742763 | 0.004066876 | 450.1108162 | 589.5365788 | ENSG00000116044 |
| CDK18 | 0.388791326 | 1.30929603 | 4.61E-07 | 5.47E-06 | 1204.67203 | 1577.782244 | ENSG00000117266 |
| AMZ2 | 0.388693702 | 1.309207436 | 1.55E-05 | 0.000132842 | 782.4992382 | 1024.271396 | ENSG00000196704 |
| NA | 0.388658685 | 1.309175659 | 0.000422703 | 0.002470631 | 521.3210534 | 682.0629293 | ENSG00000230623 |
| C6orf52 | 0.388406664 | 1.308946983 | 0.424544167 | 0.61539978 | 17.76249095 | 23.35662055 | ENSG00000137434 |
| RBM43 | 0.388343166 | 1.308889372 | 0.437688402 | 0.62636992 | 17.36429936 | 22.63973871 | ENSG00000184898 |
| FER1L4 | 0.388296052 | 1.308846629 | 0.274199471 | 0.465250018 | 44.39064578 | 58.28528072 | ENSG00000088340 |
| SUSD1 | 0.388241891 | 1.308797494 | 0.056406532 | 0.146707642 | 104.7024395 | 136.8392521 | ENSG00000106868 |
| METRNL | 0.388130328 | 1.308696289 | 0.002535469 | 0.01158999 | 348.5417681 | 455.785176 | ENSG00000176845 |
| GGCX | 0.387843376 | 1.308436016 | 8.70E-05 | 0.000615434 | 575.7249225 | 753.1986837 | ENSG00000115486 |
| ZRANB3 | 0.387730356 | 1.308333517 | 0.036499684 | 0.104590848 | 158.3309113 | 206.7192052 | ENSG00000121988 |
| NA | 0.387518243 | 1.308141173 | 0.570404825 | 0.73702475 | 11.2089707 | 14.72935059 | ENSG00000283877 |
| PICALM | 0.387468304 | 1.308095892 | 9.37E-05 | 0.000657263 | 1112.622825 | 1454.996631 | ENSG00000073921 |
| TIPARP | 0.387276176 | 1.307921701 | 0.051647815 | 0.137022703 | 125.6416315 | 164.5779816 | ENSG00000163659 |
| ITGA2B | 0.387265856 | 1.307912345 | 0.057623095 | 0.149264353 | 213.2134356 | 279.4331431 | ENSG00000005961 |
| ZNF462 | 0.387213974 | 1.307865311 | 3.31E-05 | 0.000262489 | 1240.943884 | 1622.886544 | ENSG00000148143 |
| COMMD6 | 0.386660781 | 1.307363914 | 0.00070092 | 0.003859431 | 420.2925359 | 549.8756366 | ENSG00000188243 |
| EBPL | 0.386580756 | 1.307291397 | 4.86E-06 | 4.71E-05 | 1461.729237 | 1910.216259 | ENSG00000123179 |
| SMARCD3 | 0.38651232 | 1.307229386 | 0.037023324 | 0.105681549 | 127.5485707 | 166.689333 | ENSG00000082014 |
| TMEM106B | 0.38635301 | 1.307085042 | 0.011698292 | 0.04186448 | 347.1436852 | 453.5104048 | ENSG00000106460 |
| NA | 0.386327594 | 1.307062016 | 0.553576118 | 0.723938212 | 9.220260057 | 12.06257023 | ENSG00000286504 |
| CES2 | 0.385537615 | 1.306346502 | 0.002195144 | 0.010293291 | 362.6983009 | 474.2468909 | ENSG00000172831 |
| NA | 0.385487125 | 1.306300784 | 0.394028102 | 0.586565079 | 21.08704067 | 27.68110241 | ENSG00000226084 |
| CCDC159 | 0.385198854 | 1.306039793 | 0.323729559 | 0.517881475 | 31.07745908 | 40.59899948 | ENSG00000183401 |
| NRSN2-AS1 | 0.385182254 | 1.306024765 | 0.153453015 | 0.310864376 | 75.67545379 | 98.80309559 | ENSG00000225377 |
| SPRY4-AS1 | 0.384688437 | 1.305577806 | 0.465162537 | 0.651501735 | 14.79116192 | 19.2415424 | ENSG00000231185 |
| ACADVL | 0.384651557 | 1.305544431 | 0.00034026 | 0.002045531 | 2471.584493 | 3227.73756 | ENSG00000072778 |

| | | | | | | | |
|---|---|---|---|---|---|---|---|
| NA | 0.384471213 | 1.305381242 | 4.04E-05 | 0.000312216 | 810.2190302 | 1058.17587 | ENSG00000232445 |
| MTFR2 | 0.38445408 | 1.30536574 | 0.039148714 | 0.110623915 | 120.380836 | 157.0718668 | ENSG00000146410 |
| ACSS3 | 0.384408053 | 1.305324095 | 0.000279508 | 0.001729683 | 534.0674996 | 696.3873051 | ENSG00000111058 |
| NA | 0.384244881 | 1.305176468 | 0.682519674 | 0.816696987 | 4.816430366 | 6.297714675 | ENSG00000213236 |
| IQCH-AS1 | 0.384084002 | 1.305030932 | 0.588295684 | 0.750721028 | 7.870515549 | 10.30341605 | ENSG00000259673 |
| PRDX4 | 0.383955222 | 1.304914446 | 6.99E-08 | 9.58E-07 | 2112.188508 | 2755.172326 | ENSG00000123131 |
| RBAK | 0.383891224 | 1.304856561 | 0.017386941 | 0.057493232 | 248.5641335 | 323.9741389 | ENSG00000146587 |
| DYRK2 | 0.383835056 | 1.304805761 | 0.000270505 | 0.00167958 | 712.9178313 | 930.215978 | ENSG00000127334 |
| ERCC4 | 0.383758628 | 1.304736639 | 0.067463042 | 0.168262685 | 106.8490708 | 139.1603337 | ENSG00000175595 |
| NA | 0.38348914 | 1.304492943 | 0.582153495 | 0.745978891 | 8.26733166 | 10.83627517 | ENSG00000133519 |
| LIAS | 0.383392051 | 1.304405158 | 0.016521322 | 0.055251928 | 207.5902643 | 270.964546 | ENSG00000121897 |
| ZNF721 | 0.382991215 | 1.304042795 | 0.000356794 | 0.002129552 | 615.613972 | 802.3577699 | ENSG00000182903 |
| KIF27 | 0.382951871 | 1.304007232 | 0.113360805 | 0.249055139 | 75.16549248 | 98.09881277 | ENSG00000165115 |
| ABCA7 | 0.382861329 | 1.303925397 | 0.001965167 | 0.009358274 | 500.8754442 | 653.8828 | ENSG00000064687 |
| EEF2K | 0.382772418 | 1.30384504 | 8.85E-05 | 0.000624785 | 582.4627762 | 759.2619486 | ENSG00000103319 |
| SLC22A3 | 0.382743149 | 1.303818589 | 0.308436263 | 0.5026871 | 38.28532407 | 50.05413674 | ENSG00000146477 |
| COX20 | 0.382591281 | 1.303681347 | 0.518217215 | 0.695519378 | 11.89952982 | 15.45696769 | ENSG00000203667 |
| TUT7 | 0.38258 | 1.303671153 | 0.001255088 | 0.006366121 | 380.344317 | 495.3283652 | ENSG00000083223 |
| LINC01970 | 0.382476669 | 1.303577783 | 0.645531215 | 0.792913747 | 6.961182572 | 9.129746755 | ENSG00000265692 |
| NA | 0.382350206 | 1.30346352 | 0.5530085 | 0.723664454 | 9.524849279 | 12.44146255 | ENSG00000213326 |
| FCGRT | 0.382260817 | 1.303382759 | 0.001091743 | 0.005691802 | 421.0742621 | 549.3126099 | ENSG00000104870 |
| NA | 0.382214282 | 1.303340719 | 0.736434948 | 0.852322051 | 3.185472803 | 4.159668179 | ENSG00000213399 |
| CECR2 | 0.382065419 | 1.303206241 | 1.19E-05 | 0.000105105 | 3156.17627 | 4112.728171 | ENSG00000099954 |
| ZCRB1 | 0.382057388 | 1.303198987 | 0.002887579 | 0.01288668 | 415.1402402 | 541.1172558 | ENSG00000139168 |
| NA | 0.382021907 | 1.303166937 | 0.701303834 | 0.830275647 | 6.000662573 | 7.812845924 | ENSG00000235008 |
| NTPCR | 0.381906645 | 1.303062827 | 1.78E-05 | 0.000149948 | 968.8219694 | 1261.507646 | ENSG00000135778 |
| FAM222B | 0.381785305 | 1.302953236 | 7.78E-07 | 8.85E-06 | 1817.285825 | 2368.019026 | ENSG00000173065 |
| PRKAA1 | 0.381554087 | 1.302744431 | 0.002624953 | 0.011924236 | 344.1915833 | 447.9767902 | ENSG00000132356 |
| PSTPIP1 | 0.381394429 | 1.302600268 | 0.634243001 | 0.784402761 | 6.944120231 | 8.994001295 | ENSG00000140368 |
| TMTC2 | 0.381384629 | 1.30259142 | 0.10017842 | 0.22695353 | 113.2598535 | 147.9191525 | ENSG00000179104 |
| TRIM52 | 0.381355489 | 1.30256511 | 0.198760677 | 0.374982735 | 85.23866705 | 111.2104172 | ENSG00000183718 |
| NA | 0.381349218 | 1.302559448 | 0.5954403 | 0.755953244 | 9.093810788 | 11.80390157 | ENSG00000228010 |
| PIK3IP1 | 0.381334546 | 1.302546202 | 0.336139377 | 0.530452695 | 31.80737433 | 41.2877605 | ENSG00000100100 |
| PRRC2C | 0.38133359 | 1.302545338 | 3.35E-05 | 0.000264895 | 4356.664982 | 5674.165969 | ENSG00000117523 |
| SYN3 | 0.381318306 | 1.302531539 | 0.001413051 | 0.007031302 | 428.6287334 | 558.1911656 | ENSG00000185666 |
| SLC39A11 | 0.381250639 | 1.302470448 | 0.002682721 | 0.012127206 | 387.3198109 | 503.7061644 | ENSG00000133195 |
| ASTN2 | 0.381196585 | 1.302421649 | 0.000462631 | 0.002676158 | 435.3368295 | 566.739959 | ENSG00000148219 |
| HSDL2 | 0.381185979 | 1.302412074 | 0.002331823 | 0.010820734 | 404.4595861 | 526.1086018 | ENSG00000119471 |
| ECM1 | 0.381098197 | 1.30233283 | 0.466334485 | 0.65250214 | 16.36905332 | 21.3575687 | ENSG00000143369 |
| ATG4A | 0.380964532 | 1.302212175 | 0.083875889 | 0.198614506 | 90.89705284 | 118.318502 | ENSG00000101844 |
| DPP8 | 0.38095859 | 1.302206811 | 0.000837795 | 0.004520542 | 406.9023281 | 530.0692038 | ENSG00000074603 |
| NA | 0.380866054 | 1.302123289 | 0.48197755 | 0.665447848 | 16.19594942 | 21.136004 | ENSG00000279716 |
| SNRNP35 | 0.380864445 | 1.302121837 | 0.021480581 | 0.068594262 | 250.2828833 | 326.5338216 | ENSG00000184209 |
| ATP2C2 | 0.380446974 | 1.301745098 | 0.13586679 | 0.284399858 | 64.64045881 | 84.25039327 | ENSG00000064270 |
| KRTDAP | 0.380072485 | 1.30140724 | 0.649414327 | 0.795321709 | 6.49098335 | 8.447046091 | ENSG00000188508 |
| ZCCHC7 | 0.380034233 | 1.301372735 | 0.00064912 | 0.003608517 | 402.0545392 | 522.8778474 | ENSG00000147905 |
| PLPPR4 | 0.379746743 | 1.301113432 | 0.252859201 | 0.441538763 | 40.27118606 | 52.22559996 | ENSG00000117600 |
| ATP2B3 | 0.37968774 | 1.30106022 | 0.496327941 | 0.67788814 | 14.2212419 | 18.52889744 | ENSG00000067842 |
| VGLL3 | 0.379442135 | 1.300838747 | 0.116047511 | 0.253276563 | 72.40474434 | 94.12185321 | ENSG00000206538 |
| CIART | 0.37943862 | 1.300835577 | 0.059384467 | 0.152653498 | 116.8694723 | 152.2938467 | ENSG00000159208 |
| SELENOV | 0.379367392 | 1.300771354 | 0.158637345 | 0.318305138 | 76.43631417 | 99.57001527 | ENSG00000186838 |
| FAM193B | 0.379364622 | 1.300768857 | 0.101395515 | 0.228955223 | 910.6961183 | 1184.921185 | ENSG00000146067 |
| SRPK2 | 0.379361358 | 1.300765914 | 4.29E-06 | 4.23E-05 | 1113.734125 | 1447.932413 | ENSG00000135250 |
| FBN2 | 0.379189812 | 1.300611254 | 0.002082557 | 0.009838928 | 894.8549979 | 1163.380066 | ENSG00000138829 |
| HNF1B | 0.378849698 | 1.300304672 | 0.752095481 | 0.862102644 | 3.29160512 | 4.276092992 | ENSG00000275410 |
| TWF1 | 0.378820727 | 1.300278561 | 0.027486581 | 0.083733776 | 377.3857958 | 490.2911578 | ENSG00000151239 |
| GTF2H5 | 0.3787687 | 1.30023167 | 0.001103568 | 0.005740535 | 396.1015008 | 514.9525903 | ENSG00000272047 |
| SAMD12 | 0.378601228 | 1.300080745 | 0.069292185 | 0.171669745 | 114.7466981 | 149.0711993 | ENSG00000177570 |
| SLC46A1 | 0.37858973 | 1.300070383 | 0.265747932 | 0.455631098 | 39.77770837 | 51.64792757 | ENSG00000076351 |
| GRAMD4 | 0.378587147 | 1.300068055 | 0.000226106 | 0.001432213 | 610.0093514 | 793.5131242 | ENSG00000075240 |
| NA | 0.378478122 | 1.299969813 | 0.22787617 | 0.411482554 | 51.2461997 | 66.63748415 | ENSG00000235152 |
| ZBTB6 | 0.378389822 | 1.29989025 | 0.02928859 | 0.087823123 | 182.0777766 | 236.2259688 | ENSG00000186130 |
| ASCC3 | 0.378384004 | 1.299885008 | 4.40E-07 | 5.25E-06 | 1851.790155 | 2406.292953 | ENSG00000112249 |
| RNF215 | 0.378354424 | 1.299858356 | 0.008876151 | 0.033321973 | 233.2259392 | 302.9179425 | ENSG00000099999 |
| BBS2 | 0.37827871 | 1.299790141 | 0.004003422 | 0.017041668 | 331.8051228 | 431.2371431 | ENSG00000125124 |
| MRM2 | 0.378242206 | 1.299757253 | 0.586393265 | 0.749312221 | 10.42974757 | 13.62696642 | ENSG00000122687 |
| EHHADH | 0.377890894 | 1.299440786 | 0.293154338 | 0.485725238 | 31.382146 | 40.79343105 | ENSG00000113790 |
| DCAF10 | 0.377696072 | 1.299265322 | 0.005614868 | 0.022751632 | 296.5837228 | 384.8505663 | ENSG00000122741 |
| SEMA3E | 0.377593227 | 1.299172705 | 0.01674432 | 0.055816408 | 216.1085329 | 280.4698242 | ENSG00000170381 |
| USP51 | 0.377575812 | 1.299157022 | 0.116194358 | 0.253477664 | 87.15674553 | 113.3720842 | ENSG00000247746 |
| SLX4 | 0.377389884 | 1.298989603 | 0.002221201 | 0.010381035 | 356.0186245 | 462.6297745 | ENSG00000188827 |
| TTBK2 | 0.377115729 | 1.29874278 | 0.022067371 | 0.0701228 | 179.0497208 | 232.6999915 | ENSG00000128881 |
| WARS2-AS1 | 0.376833251 | 1.298488513 | 0.244053879 | 0.43079353 | 39.20265461 | 50.86454713 | ENSG00000231365 |
| NA | 0.376703582 | 1.29837181 | 0.008878548 | 0.033324222 | 243.2659586 | 315.6023371 | ENSG00000233836 |

| | | | | | | | |
|---|---|---|---|---|---|---|---|
| NA | 0.376582591 | 1.298262928 | 0.505653869 | 0.685272653 | 12.96500206 | 16.86739708 | ENSG00000251141 |
| S1PR2 | 0.376449271 | 1.29814296 | 0.005902275 | 0.02371402 | 244.1892947 | 317.039788 | ENSG00000267534 |
| METRN | 0.376344919 | 1.298049067 | 0.011816078 | 0.042212644 | 377.3296856 | 489.3245678 | ENSG00000103260 |
| USP28 | 0.376283754 | 1.297994036 | 1.19E-07 | 1.56E-06 | 2054.442696 | 2667.209664 | ENSG00000048028 |
| PDXK | 0.376226649 | 1.297942659 | 3.92E-08 | 5.66E-07 | 2025.93067 | 2630.198076 | ENSG00000160209 |
| SEMA5A | 0.376103778 | 1.297832121 | 0.019460163 | 0.063077064 | 284.7900096 | 369.367047 | ENSG00000112902 |
| AQP11 | 0.375929422 | 1.297675282 | 0.272207269 | 0.463277275 | 35.85166951 | 46.61146201 | ENSG00000178301 |
| CDC42EP5 | 0.375613462 | 1.297391113 | 0.10751439 | 0.2391195 | 112.0121149 | 144.8012783 | ENSG00000167617 |
| FAM95C | 0.375444714 | 1.29723937 | 0.668522807 | 0.807507932 | 5.061639604 | 6.597724171 | ENSG00000283486 |
| MTR | 0.375230941 | 1.297047165 | 0.000193575 | 0.001248339 | 651.0832497 | 844.588724 | ENSG00000116984 |
| DDX50 | 0.375127021 | 1.296953739 | 3.34E-05 | 0.000264267 | 865.8921275 | 1122.336433 | ENSG00000107625 |
| EXTL3 | 0.374815985 | 1.296674154 | 9.77E-10 | 1.82E-08 | 3758.170039 | 4872.348803 | ENSG00000012232 |
| MED7 | 0.374796207 | 1.296656378 | 0.056835064 | 0.147697851 | 111.9852736 | 145.1141101 | ENSG00000155868 |
| IMMP1L | 0.374728982 | 1.296595959 | 0.107182672 | 0.238650126 | 80.69357303 | 104.7149815 | ENSG00000148950 |
| TENT2 | 0.374663735 | 1.296537321 | 0.001048809 | 0.005489589 | 435.8592982 | 564.4912132 | ENSG00000164329 |
| NT5DC1 | 0.374420033 | 1.296318327 | 0.000198732 | 0.001276263 | 640.8102276 | 830.2790964 | ENSG00000178425 |
| CMC1 | 0.374335297 | 1.29624219 | 0.004646888 | 0.019301978 | 288.0918822 | 373.1778872 | ENSG00000187118 |
| NA | 0.374131496 | 1.29605909 | 0.434509854 | 0.624209861 | 20.82151448 | 26.96477023 | ENSG00000260563 |
| SMUG1 | 0.373980533 | 1.295923478 | 0.002539478 | 0.011604215 | 332.6242808 | 431.0524591 | ENSG00000123415 |
| IRS2 | 0.373955352 | 1.29590086 | 0.005928326 | 0.02379805 | 445.8884788 | 577.582031 | ENSG00000185950 |
| MYT1L | 0.373876141 | 1.29582971 | 0.742706453 | 0.856105567 | 4.152306627 | 5.34093517 | ENSG00000186487 |
| GPSM1 | 0.373707166 | 1.295677946 | 0.000277029 | 0.00171606 | 570.4283185 | 739.8070291 | ENSG00000160360 |
| WDR54 | 0.37370235 | 1.295673621 | 7.10E-05 | 0.000512962 | 705.6132297 | 914.2882384 | ENSG00000005448 |
| UGCG | 0.373688773 | 1.295661428 | 0.001119911 | 0.005809655 | 502.1483631 | 650.5189904 | ENSG00000148154 |
| CFAP70 | 0.373447025 | 1.295444336 | 0.739274361 | 0.85432791 | 3.19937823 | 4.140897185 | ENSG00000156042 |
| CNTRL | 0.37344366 | 1.295441314 | 0.008061246 | 0.030829544 | 242.5303309 | 313.8577086 | ENSG00000119397 |
| ETS1 | 0.373416406 | 1.295416842 | 3.91E-06 | 3.90E-05 | 1481.860591 | 1918.547189 | ENSG00000134954 |
| NA | 0.373362994 | 1.295368884 | 0.152880267 | 0.310131724 | 65.21857179 | 84.67898857 | ENSG00000228327 |
| FYCO1 | 0.373351041 | 1.295358151 | 0.003299908 | 0.01443486 | 346.220056 | 449.0256415 | ENSG00000163820 |
| CHMP2A | 0.373327841 | 1.295337321 | 9.83E-06 | 8.80E-05 | 1057.490679 | 1369.743063 | ENSG00000130724 |
| NA | 0.373253262 | 1.295270361 | 0.74733787 | 0.858933404 | 3.850874318 | 4.928188088 | ENSG00000270112 |
| ECI1 | 0.3728466 | 1.294905305 | 1.95E-06 | 2.06E-05 | 1680.62675 | 2175.735632 | ENSG00000167969 |
| F8 | 0.37281578 | 1.294877643 | 0.344937263 | 0.538376668 | 27.00800631 | 34.95717875 | ENSG00000185010 |
| PPM1D | 0.372663721 | 1.294741172 | 0.001354043 | 0.006782656 | 395.3707137 | 511.3807801 | ENSG00000170836 |
| NA | 0.372638534 | 1.294718567 | 0.488375007 | 0.671262814 | 16.09425179 | 20.96370424 | ENSG00000260628 |
| PTP4A3 | 0.372550501 | 1.294639566 | 0.000158544 | 0.001045339 | 809.1111578 | 1046.869851 | ENSG00000184489 |
| DUSP13 | 0.372090435 | 1.29422678 | 0.403869664 | 0.595585642 | 25.80030024 | 33.15624571 | ENSG00000079393 |
| LINC02614 | 0.372038741 | 1.294180407 | 0.624188874 | 0.77710194 | 6.967496398 | 9.065836078 | ENSG00000241288 |
| HIF3A | 0.371966088 | 1.294115234 | 0.00726032 | 0.0282146 | 257.6778704 | 333.462009 | ENSG00000124440 |
| NA | 0.371876441 | 1.294034822 | 0.700608403 | 0.829664311 | 5.075545032 | 6.586439251 | ENSG00000276141 |
| BCL2L11 | 0.371812021 | 1.293977042 | 0.019672279 | 0.063664401 | 231.9955409 | 300.637867 | ENSG00000153094 |
| MKRN2 | 0.37175187 | 1.293923092 | 0.001181735 | 0.006077211 | 511.6260692 | 662.6281034 | ENSG00000075975 |
| MME | 0.371532539 | 1.293726395 | 0.018445006 | 0.060260839 | 227.1678509 | 293.3586128 | ENSG00000196549 |
| TAF1 | 0.371497957 | 1.293695383 | 5.27E-05 | 0.000393878 | 834.7584605 | 1079.848157 | ENSG00000147133 |
| LOC105369779 | 0.371340712 | 1.293554387 | 0.138552884 | 0.28858946 | 71.16734828 | 91.79598515 | ENSG00000258056 |
| NA | 0.371268779 | 1.293489891 | 0.345790097 | 0.539240527 | 29.78531312 | 38.70140031 | ENSG00000254602 |
| LRAT | 0.371196156 | 1.29342478 | 0.000990591 | 0.005223219 | 484.1053355 | 626.2450993 | ENSG00000121207 |
| PMAIP1 | 0.370998497 | 1.293247585 | 1.14E-05 | 0.000100933 | 2293.484363 | 2964.761806 | ENSG00000141682 |
| JMJD4 | 0.370909089 | 1.293167441 | 0.291442315 | 0.483761659 | 32.96003741 | 42.68789266 | ENSG00000081692 |
| NAXE | 0.370861287 | 1.293124593 | 8.25E-07 | 9.34E-06 | 1358.864495 | 1756.892691 | ENSG00000163382 |
| PRIM2 | 0.370807555 | 1.293076434 | 0.000114719 | 0.000788946 | 618.1848622 | 798.8638867 | ENSG00000146143 |
| NA | 0.370743715 | 1.293019216 | 0.57131861 | 0.737588404 | 10.16243994 | 13.17299012 | ENSG00000214770 |
| GUCY2D | 0.370691129 | 1.292972086 | 0.729053135 | 0.848129016 | 3.493218937 | 4.538560499 | ENSG00000132518 |
| ZFP1 | 0.370546236 | 1.292842236 | 0.095903729 | 0.219577481 | 108.5425653 | 140.0526528 | ENSG00000184517 |
| NA | 0.370497183 | 1.29279828 | 0.637088067 | 0.786713987 | 6.653436437 | 8.619345854 | ENSG00000259863 |
| ZNF784 | 0.370342953 | 1.292660081 | 0.01352846 | 0.047042491 | 274.6700827 | 355.0558998 | ENSG00000179922 |
| ZBTB21 | 0.370103036 | 1.292445132 | 0.003257725 | 0.014277292 | 354.1930585 | 457.2736524 | ENSG00000173276 |
| SNX21 | 0.369944973 | 1.292303539 | 0.012321218 | 0.043730649 | 215.165541 | 278.1249619 | ENSG00000124104 |
| LOC101927989 | 0.369914816 | 1.292276526 | 0.279043951 | 0.470063197 | 40.75589207 | 52.77810614 | ENSG00000265778 |
| ATP11B | 0.369811142 | 1.292183664 | 0.005339703 | 0.021808296 | 380.8589084 | 491.5325684 | ENSG00000058063 |
| PPCS | 0.369274133 | 1.291702769 | 0.003446794 | 0.014964365 | 340.4004949 | 439.3147265 | ENSG00000127125 |
| HPS6 | 0.368717649 | 1.291204623 | 0.000567147 | 0.003203749 | 507.2209391 | 654.5258943 | ENSG00000166189 |
| DLL1 | 0.368672448 | 1.291164169 | 0.091246637 | 0.211775304 | 92.27323273 | 119.1409618 | ENSG00000198719 |
| TACC1 | 0.368565514 | 1.29106847 | 0.004018801 | 0.01708246 | 382.3226624 | 492.9986898 | ENSG00000147526 |
| RPS6KB2 | 0.368518533 | 1.291026427 | 0.000263378 | 0.001642829 | 542.4303128 | 700.017292 | ENSG00000175634 |
| NA | 0.368517539 | 1.291025538 | 0.408501736 | 0.600213036 | 21.91795449 | 28.34503208 | ENSG00000224063 |
| VAMP4 | 0.368516616 | 1.291024712 | 0.051567305 | 0.136887485 | 128.5311891 | 165.724741 | ENSG00000117533 |
| KAT6A | 0.368328893 | 1.290856735 | 0.002447126 | 0.011259937 | 604.6053072 | 780.1923392 | ENSG00000083168 |
| AP4E1 | 0.367944794 | 1.290513107 | 0.021034085 | 0.067377416 | 211.5365975 | 272.8868649 | ENSG00000081014 |
| COG7 | 0.367939598 | 1.290508459 | 0.02856324 | 0.086234091 | 155.6032055 | 200.9564367 | ENSG00000168434 |
| CYP2R1 | 0.367896906 | 1.290470271 | 0.029695679 | 0.088842569 | 177.5108174 | 228.8688514 | ENSG00000186104 |
| NA | 0.367865399 | 1.290442088 | 0.750632752 | 0.861278241 | 3.400992048 | 4.410850766 | ENSG00000286970 |
| SNX2 | 0.367855925 | 1.290433614 | 2.70E-06 | 2.76E-05 | 1368.384 | 1765.736739 | ENSG00000205302 |
| NA | 0.367835937 | 1.290415736 | 0.305224368 | 0.499164249 | 35.57035875 | 45.9404843 | ENSG00000273230 |

| | | | | | | | |
|---|---|---|---|---|---|---|---|
| NA | 0.367519265 | 1.290132521 | 0.488281552 | 0.671201534 | 22.91498197 | 29.59857189 | ENSG00000285796 |
| NA | 0.367506828 | 1.290121399 | 0.211980634 | 0.391615515 | 58.78361611 | 75.65339402 | ENSG00000270872 |
| NA | 0.367312613 | 1.289947735 | 0.376140243 | 0.569228825 | 26.90809012 | 34.72531681 | ENSG00000228623 |
| PLCD4 | 0.367266939 | 1.289906897 | 0.207950954 | 0.386964532 | 50.62704686 | 65.27489399 | ENSG00000115556 |
| ARID3A | 0.367169861 | 1.289820103 | 4.85E-07 | 5.74E-06 | 2046.127154 | 2638.51499 | ENSG00000116017 |
| LNPEP | 0.367099286 | 1.289757008 | 0.265604154 | 0.455426717 | 56.14162794 | 72.06595042 | ENSG00000113441 |
| NA | 0.367019004 | 1.289685239 | 0.350739657 | 0.544026727 | 26.12836332 | 33.67088349 | ENSG00000273142 |
| NA | 0.366894979 | 1.289574373 | 0.482998961 | 0.666361675 | 14.84432577 | 19.16265957 | ENSG00000267424 |
| FAM177A1 | 0.366396015 | 1.289128443 | 0.001686206 | 0.008209124 | 429.1288029 | 553.4208145 | ENSG00000151327 |
| PLXNB3 | 0.36637455 | 1.289109263 | 0.252401823 | 0.441168072 | 45.83083574 | 59.15437723 | ENSG00000198753 |
| ARCN1 | 0.366363815 | 1.289099671 | 6.49E-08 | 8.96E-07 | 2792.669177 | 3599.221269 | ENSG00000095139 |
| ACTR1B | 0.366258806 | 1.289005846 | 0.000127688 | 0.000866242 | 906.2324499 | 1168.506081 | ENSG00000115073 |
| FGF4 | 0.366144808 | 1.288903996 | 0.378643678 | 0.571992985 | 24.31295198 | 31.4543922 | ENSG00000075388 |
| ZNF155 | 0.366141468 | 1.288901012 | 0.256004899 | 0.444941858 | 41.77716753 | 54.03076988 | ENSG00000204920 |
| GAA | 0.366131884 | 1.28889245 | 1.65E-05 | 0.000139957 | 1009.258788 | 1300.132259 | ENSG00000171298 |
| AAMDC | 0.366090804 | 1.28885575 | 0.22946814 | 0.413430786 | 48.61515544 | 62.87124241 | ENSG00000087884 |
| NA | 0.365918072 | 1.288701445 | 0.755953444 | 0.864333115 | 3.738428175 | 4.853542143 | ENSG00000230956 |
| ZNF529-AS1 | 0.36574376 | 1.288545749 | 0.30570393 | 0.499683979 | 33.2550582 | 42.70991284 | ENSG00000233527 |
| LINC02585 | 0.365587218 | 1.28840594 | 0.672402443 | 0.809585672 | 5.375699565 | 6.965331571 | ENSG00000228350 |
| ORAI3 | 0.36553326 | 1.288357754 | 0.324492271 | 0.518683894 | 34.88040099 | 45.06627485 | ENSG00000175938 |
| WDR63 | 0.365496781 | 1.288325178 | 0.703059418 | 0.831263 | 5.388327216 | 6.969130418 | ENSG00000162643 |
| RIPK4 | 0.365373623 | 1.288215202 | 0.055533434 | 0.14486371 | 121.4104021 | 156.6108425 | ENSG00000183421 |
| ALDOC | 0.365189321 | 1.288050646 | 6.97E-05 | 0.000504923 | 833.3624069 | 1073.065932 | ENSG00000109107 |
| MYH7B | 0.364965835 | 1.287851131 | 0.53255432 | 0.707603539 | 12.70716517 | 16.44391473 | ENSG00000078814 |
| SYNPR-AS1 | 0.364927918 | 1.287817284 | 0.609033704 | 0.7667212 | 8.588983223 | 11.07997169 | ENSG00000241359 |
| ABL2 | 0.364755319 | 1.287663224 | 0.000940456 | 0.004999474 | 516.9466281 | 665.1884355 | ENSG00000143322 |
| WDR37 | 0.364623977 | 1.287546001 | 0.003124965 | 0.013790024 | 338.7339243 | 436.6655465 | ENSG00000047056 |
| RBM26 | 0.364616331 | 1.287539177 | 9.50E-06 | 8.54E-05 | 1183.179808 | 1522.801112 | ENSG00000139746 |
| N4BP2 | 0.36448029 | 1.287417772 | 0.007445693 | 0.028815719 | 603.8471076 | 776.6310403 | ENSG00000078177 |
| PNLDC1 | 0.364412008 | 1.287356841 | 0.773278167 | 0.874965662 | 3.126092818 | 4.020673525 | ENSG00000146453 |
| AVL9 | 0.364384361 | 1.287332171 | 5.91E-05 | 0.000436538 | 813.5058697 | 1046.722557 | ENSG00000105778 |
| SACS | 0.364307635 | 1.287263709 | 0.000201757 | 0.001293898 | 1067.748984 | 1374.04448 | ENSG00000151835 |
| COX10-AS1 | 0.364083953 | 1.287064142 | 0.147736814 | 0.302625343 | 78.85331235 | 101.5765532 | ENSG00000236088 |
| NA | 0.364083574 | 1.287063804 | 0.401059691 | 0.592731097 | 23.54132045 | 30.19336632 | ENSG00000249456 |
| ZNF627 | 0.364004916 | 1.286993632 | 0.007418599 | 0.028734862 | 260.685301 | 335.1994291 | ENSG00000198551 |
| DNMBP | 0.363990918 | 1.286981146 | 0.000258725 | 0.001617269 | 644.2167745 | 828.5228969 | ENSG00000107554 |
| RGS10 | 0.363943319 | 1.286938685 | 0.090251749 | 0.209930461 | 94.68341343 | 121.795796 | ENSG00000148908 |
| LOC101928059 | 0.363925651 | 1.286922924 | 0.713713723 | 0.837849642 | 5.575434246 | 7.220313005 | ENSG00000284738 |
| IGFBP5 | 0.363908381 | 1.286907519 | 0.443150928 | 0.631553963 | 22.65210903 | 28.96327242 | ENSG00000115461 |
| NA | 0.363840627 | 1.286847083 | 0.69980388 | 0.828998907 | 5.164615008 | 6.64655108 | ENSG00000219470 |
| PUS7L | 0.363801072 | 1.286811802 | 0.099643366 | 0.226266863 | 101.8002542 | 131.278178 | ENSG00000129317 |
| ZBED8 | 0.363706206 | 1.286727188 | 0.101600084 | 0.229361261 | 99.91314353 | 128.7434558 | ENSG00000221886 |
| NA | 0.363659176 | 1.286685243 | 0.660517502 | 0.802356197 | 6.289467231 | 8.075201812 | ENSG00000235510 |
| ATP2A1 | 0.363637513 | 1.286665923 | 0.204898653 | 0.383091035 | 56.21383098 | 72.51887614 | ENSG00000196296 |
| SYNJ1 | 0.363403396 | 1.286457143 | 0.211123365 | 0.390584441 | 48.0058793 | 61.72284249 | ENSG00000159082 |
| NA | 0.363267817 | 1.286336252 | 0.03391031 | 0.098819762 | 316.6866719 | 407.9027434 | ENSG00000229191 |
| S100A11 | 0.363191915 | 1.286268578 | 1.60E-05 | 0.000136765 | 1137.43045 | 1462.717369 | ENSG00000163191 |
| MYH11 | 0.363191493 | 1.286268202 | 0.436617548 | 0.62586147 | 19.7839508 | 25.41903341 | ENSG00000133392 |
| HES7 | 0.363186273 | 1.286263548 | 0.66884544 | 0.807687085 | 6.027195652 | 7.707594412 | ENSG00000179111 |
| XPR1 | 0.363136422 | 1.286219103 | 1.96E-06 | 2.07E-05 | 1373.84211 | 1766.279537 | ENSG00000143324 |
| MIR570 | 0.363115761 | 1.286200683 | 0.686037454 | 0.819372224 | 5.914749509 | 7.681775377 | ENSG00000207650 |
| TSPO | 0.362957538 | 1.286059631 | 0.015177248 | 0.051547573 | 293.3829691 | 376.8398333 | ENSG00000100300 |
| PTPRS | 0.362935702 | 1.286040166 | 1.15E-08 | 1.83E-07 | 3663.444503 | 4712.465216 | ENSG00000105426 |
| LOC729867 | 0.362855584 | 1.28596875 | 0.68886244 | 0.821581497 | 5.153866493 | 6.627780086 | ENSG00000227741 |
| TMTC3 | 0.362798775 | 1.285918113 | 0.005603411 | 0.02272508 | 460.2096143 | 591.6181604 | ENSG00000139324 |
| NA | 0.362756507 | 1.285880439 | 0.740608868 | 0.855070377 | 4.115025033 | 5.239482504 | ENSG00000277879 |
| ANKRD13C | 0.362739728 | 1.285865483 | 0.00395705 | 0.016886938 | 375.043376 | 481.7038835 | ENSG00000118454 |
| ST6GALNAC5 | 0.362687294 | 1.28581875 | 0.072634677 | 0.178235284 | 132.7502567 | 170.9874462 | ENSG00000117069 |
| HSD17B8 | 0.362668173 | 1.285801709 | 0.162335878 | 0.323728552 | 71.80749449 | 92.28458066 | ENSG00000204228 |
| PPARGC1B | 0.362611413 | 1.285751122 | 0.007537784 | 0.029123471 | 296.2986538 | 380.8930709 | ENSG00000155846 |
| NSUN6 | 0.362547409 | 1.285694082 | 0.029250547 | 0.087765849 | 164.8422116 | 211.6792098 | ENSG00000241058 |
| TP53I11 | 0.362535897 | 1.285683823 | 1.91E-05 | 0.000160178 | 1127.592127 | 1450.223536 | ENSG00000175274 |
| CENPA | 0.362042946 | 1.285244596 | 0.011028938 | 0.039962922 | 279.9270449 | 359.1873984 | ENSG00000115163 |
| SBNO2 | 0.362027659 | 1.285230977 | 2.88E-06 | 2.94E-05 | 1334.510127 | 1716.060054 | ENSG00000064932 |
| SH2B3 | 0.362019408 | 1.285223627 | 6.55E-06 | 6.17E-05 | 1021.026607 | 1312.232594 | ENSG00000111252 |
| GPR107 | 0.361762549 | 1.284994825 | 6.99E-07 | 8.02E-06 | 1713.435646 | 2201.424862 | ENSG00000148358 |
| USP20 | 0.361652962 | 1.284897221 | 0.003783151 | 0.016245812 | 427.0723617 | 548.8980309 | ENSG00000136878 |
| PGGHG | 0.361618932 | 1.284866912 | 0.331560428 | 0.52576497 | 1419.295132 | 1823.77967 | ENSG00000142102 |
| TSHZ2 | 0.361443213 | 1.284710426 | 0.732377374 | 0.850760404 | 3.460372032 | 4.459677675 | ENSG00000182463 |
| ZNRD1ASP | 0.361351358 | 1.284628633 | 0.229622643 | 0.413467995 | 47.13570706 | 60.65854997 | ENSG00000204623 |
| POLM | 0.361220569 | 1.284512179 | 0.009699787 | 0.03584048 | 228.2295875 | 293.0491821 | ENSG00000122678 |
| C20orf194 | 0.361215856 | 1.284507983 | 0.021833223 | 0.069541189 | 221.1019982 | 283.9024165 | ENSG00000088854 |
| SMOX | 0.361207745 | 1.284500761 | 0.003664777 | 0.015788699 | 332.0280005 | 426.7019434 | ENSG00000088826 |
| NA | 0.36108925 | 1.284395263 | 0.703261076 | 0.831299338 | 4.757050382 | 6.091122117 | ENSG00000217716 |

| | | | | | | | |
|---|---|---|---|---|---|---|---|
| NEMP2 | 0.360960824 | 1.284280934 | 0.133138668 | 0.280214117 | 72.91272882 | 93.64998199 | ENSG00000189362 |
| PLPP2 | 0.360958953 | 1.284279269 | 2.14E-06 | 2.23E-05 | 1537.80198 | 1975.041008 | ENSG00000141934 |
| SAT2 | 0.360942967 | 1.284265038 | 0.001203605 | 0.00616573 | 522.1657976 | 669.9421739 | ENSG00000141504 |
| KCNT2 | 0.360572096 | 1.283934937 | 0.063219746 | 0.160490706 | 140.5141515 | 180.4333855 | ENSG00000162687 |
| VPS37A | 0.360424627 | 1.283803703 | 0.001810377 | 0.008715203 | 412.903299 | 529.7181687 | ENSG00000155975 |
| SRFBP1 | 0.360221576 | 1.283623028 | 0.003882563 | 0.016626482 | 308.9281966 | 396.7680187 | ENSG00000151304 |
| GABRB2 | 0.360180246 | 1.283586255 | 0.599156993 | 0.758989523 | 8.522011636 | 10.95226195 | ENSG00000145864 |
| LOC100288175 | 0.360060875 | 1.283480054 | 0.335258373 | 0.529499674 | 34.58901814 | 44.40614403 | ENSG00000217801 |
| HLCS | 0.36004245 | 1.283463662 | 0.003406725 | 0.014818174 | 316.453083 | 406.327961 | ENSG00000159267 |
| PLCH2 | 0.359858034 | 1.28329961 | 0.593561156 | 0.7550489 | 8.879667017 | 11.42880809 | ENSG00000149527 |
| SULT1C4 | 0.359712935 | 1.283170549 | 0.458607559 | 0.645589587 | 20.26805545 | 26.01156633 | ENSG00000198075 |
| NTRK1 | 0.359679527 | 1.283140835 | 0.7386458 | 0.853862439 | 3.457215119 | 4.448392755 | ENSG00000198400 |
| HCN1 | 0.359546739 | 1.283022739 | 0.283518331 | 0.474495145 | 40.35413761 | 51.9072575 | ENSG00000164588 |
| NR3C2 | 0.359307916 | 1.282810365 | 0.458532015 | 0.645532273 | 18.31407084 | 23.43875256 | ENSG00000151623 |
| RGMB | 0.359279858 | 1.282785417 | 0.058614251 | 0.151176806 | 134.595764 | 172.5437035 | ENSG00000174136 |
| SEC1P | 0.359185273 | 1.282701318 | 0.572107177 | 0.738297898 | 12.21486756 | 15.67104632 | ENSG00000232871 |
| NA | 0.358932683 | 1.28247676 | 0.04870659 | 0.131077094 | 146.3839301 | 187.517515 | ENSG00000241743 |
| NA | 0.358845598 | 1.282399349 | 0.303829227 | 0.497497207 | 38.04693232 | 48.95230222 | ENSG00000259485 |
| ZNF197 | 0.358402118 | 1.282005204 | 0.005890045 | 0.023685418 | 305.7973608 | 392.2163095 | ENSG00000186448 |
| XRCC3 | 0.358283127 | 1.281899471 | 0.001364436 | 0.006829179 | 422.7078881 | 541.633096 | ENSG00000126215 |
| NA | 0.358225249 | 1.281848044 | 0.746929224 | 0.858676729 | 4.058801961 | 5.251205458 | ENSG00000230847 |
| LDB2 | 0.358101171 | 1.281737804 | 0.000303918 | 0.001857174 | 791.1421897 | 1013.078806 | ENSG00000169744 |
| SPRY1 | 0.35808056 | 1.281719494 | 0.000468518 | 0.002703457 | 480.0314632 | 615.0733596 | ENSG00000164056 |
| ICK | 0.358002551 | 1.28165019 | 0.045877721 | 0.125119921 | 226.0907658 | 289.5199866 | ENSG00000112144 |
| KLHL12 | 0.357818534 | 1.281486725 | 4.93E-05 | 0.000372832 | 888.8071997 | 1138.387615 | ENSG00000117153 |
| PEX11G | 0.357802724 | 1.281472682 | 0.344215622 | 0.537826027 | 30.08852687 | 38.47191149 | ENSG00000104883 |
| ZNF28 | 0.357696775 | 1.281378576 | 0.012345709 | 0.04380788 | 270.0263955 | 346.2814793 | ENSG00000198538 |
| B3GNT8 | 0.357499873 | 1.281203703 | 0.538420932 | 0.711982108 | 13.33971978 | 17.06964114 | ENSG00000177191 |
| NEK1 | 0.357475837 | 1.281182357 | 0.0107719 | 0.039131006 | 266.929895 | 341.8574799 | ENSG00000137601 |
| NA | 0.357445735 | 1.281155626 | 0.241396579 | 0.427638142 | 46.2326879 | 59.33080225 | ENSG00000232593 |
| EPRS1 | 0.357347407 | 1.281068311 | 9.62E-08 | 1.29E-06 | 4318.540156 | 5531.327037 | ENSG00000136628 |
| ZBTB38 | 0.357214415 | 1.280950223 | 0.024411711 | 0.075952097 | 177.2910362 | 226.993822 | ENSG00000177311 |
| LPCAT2 | 0.357167634 | 1.280908688 | 0.127025853 | 0.270671093 | 86.7162363 | 110.7819965 | ENSG00000087253 |
| ZNF461 | 0.357092449 | 1.280841936 | 0.196591034 | 0.37187367 | 56.16382403 | 71.74826924 | ENSG00000197808 |
| SBF2 | 0.357062693 | 1.280815519 | 9.51E-05 | 0.000665904 | 730.9679096 | 935.6340439 | ENSG00000133812 |
| NPTXR | 0.356945951 | 1.28071188 | 0.003378572 | 0.014716439 | 426.9293386 | 546.3815612 | ENSG00000221890 |
| NA | 0.356909386 | 1.280679421 | 0.293411105 | 0.486036914 | 36.45000174 | 46.67526107 | ENSG00000178715 |
| PSTPIP2 | 0.356738921 | 1.280528108 | 0.218448274 | 0.399715699 | 58.40386234 | 74.50521734 | ENSG00000152229 |
| DACH2 | 0.356728409 | 1.280518777 | 0.21487066 | 0.395041446 | 55.63424483 | 71.01674167 | ENSG00000126733 |
| TCHP | 0.356516355 | 1.280330575 | 0.010626991 | 0.038665223 | 265.340556 | 339.9593798 | ENSG00000139437 |
| GALE | 0.356457657 | 1.280278484 | 0.002827642 | 0.012661881 | 496.94694 | 636.3295159 | ENSG00000117308 |
| NOTCH4 | 0.356413001 | 1.280238856 | 0.320206529 | 0.513916506 | 35.00171651 | 44.95788576 | ENSG00000204301 |
| SENP5 | 0.356404128 | 1.280230982 | 0.000698177 | 0.003850046 | 511.1738193 | 654.1577775 | ENSG00000119231 |
| PKIG | 0.356239812 | 1.280085178 | 0.001252746 | 0.006359112 | 404.8596569 | 518.2927617 | ENSG00000168734 |
| SNX16 | 0.356199857 | 1.280049727 | 0.067145221 | 0.167718266 | 120.1103714 | 153.6664077 | ENSG00000104497 |
| PDGFD | 0.356159909 | 1.280014283 | 0.049881159 | 0.133520711 | 165.4821472 | 211.9035857 | ENSG00000170962 |
| TATDN3 | 0.356032568 | 1.279901307 | 0.016947024 | 0.056340114 | 206.7004515 | 264.5690658 | ENSG00000203705 |
| HHLA1 | 0.355871752 | 1.279758645 | 0.115349907 | 0.252348346 | 84.25426722 | 107.628596 | ENSG00000132297 |
| EIF3F | 0.355842206 | 1.279732436 | 9.68E-08 | 1.30E-06 | 4625.657157 | 5918.9129 | ENSG00000175390 |
| MAP2K1 | 0.355823699 | 1.279716019 | 9.48E-07 | 1.06E-05 | 1454.56526 | 1861.668222 | ENSG00000169032 |
| MAST4 | 0.355721037 | 1.279624958 | 0.083212026 | 0.197463546 | 109.4752744 | 139.8959865 | ENSG00000069020 |
| SKA3 | 0.355454465 | 1.279388539 | 3.03E-05 | 0.00024307 | 924.4177938 | 1182.611791 | ENSG00000165480 |
| UBE4A | 0.355315344 | 1.279265172 | 0.000683087 | 0.003776948 | 578.0714014 | 739.1790553 | ENSG00000110344 |
| RBM12B | 0.35512184 | 1.2790936 | 0.001031057 | 0.005410423 | 647.8077973 | 829.1232281 | ENSG00000183808 |
| GRIN3B | 0.355015342 | 1.278999182 | 0.312106056 | 0.505732964 | 44.90070479 | 57.52998083 | ENSG00000116032 |
| RPL13AP5 | 0.354950367 | 1.278941581 | 0.020322193 | 0.065493222 | 198.9359706 | 254.1449881 | ENSG00000236552 |
| TMEM184A | 0.35492222 | 1.278916629 | 0.434363088 | 0.624117023 | 20.05125842 | 25.71579371 | ENSG00000164855 |
| NANP | 0.354854347 | 1.278856463 | 0.040545401 | 0.113686265 | 171.1090768 | 218.866879 | ENSG00000170191 |
| OCIAD2 | 0.354730583 | 1.278746759 | 0.000131297 | 0.000885213 | 1553.254454 | 1986.734617 | ENSG00000145247 |
| GRAMD1A | 0.354727973 | 1.278744446 | 5.66E-08 | 7.94E-07 | 3576.957827 | 4574.003152 | ENSG00000089351 |
| AGAP2 | 0.354725587 | 1.278742331 | 0.216115247 | 0.396620313 | 50.64598833 | 64.77632766 | ENSG00000135439 |
| HSPA4L | 0.35470227 | 1.278721663 | 0.047478279 | 0.128461667 | 147.7040823 | 188.4910902 | ENSG00000164070 |
| NA | 0.354336237 | 1.278397274 | 0.682310495 | 0.816552123 | 5.680288787 | 7.299195829 | ENSG00000226002 |
| NA | 0.354333584 | 1.278394923 | 0.633673829 | 0.783961787 | 7.334622517 | 9.403387563 | ENSG00000287825 |
| SLC25A4 | 0.354307722 | 1.278372006 | 9.64E-07 | 1.08E-05 | 1399.15472 | 1788.566944 | ENSG00000151729 |
| PHC3 | 0.354264273 | 1.278333507 | 0.011605618 | 0.041613179 | 229.4182544 | 293.2443781 | ENSG00000173889 |
| ORAI1 | 0.35424103 | 1.278312912 | 0.001098885 | 0.005722601 | 520.7054936 | 665.3912377 | ENSG00000276045 |
| NA | 0.354098968 | 1.278187043 | 0.648203657 | 0.794567224 | 7.817351692 | 9.909551584 | ENSG00000244459 |
| GCNT2 | 0.354040895 | 1.278135593 | 0.000239276 | 0.001507903 | 949.1923839 | 1212.404594 | ENSG00000111846 |
| CPNE8 | 0.353952359 | 1.278057158 | 0.078438018 | 0.188850379 | 108.7655784 | 138.8882302 | ENSG00000139117 |
| BTNL8 | 0.35380417 | 1.277925887 | 0.665850133 | 0.805855189 | 6.947277144 | 8.885500589 | ENSG00000113303 |
| NA | 0.353785175 | 1.277909061 | 0.640425813 | 0.789444284 | 8.756374661 | 11.07193596 | ENSG00000242615 |
| TRPT1 | 0.353626364 | 1.277768398 | 0.014734885 | 0.050324498 | 212.3742311 | 271.2912505 | ENSG00000149743 |
| CXCL5 | 0.353496451 | 1.277653341 | 0.316367826 | 0.51000794 | 69.66549324 | 88.76678168 | ENSG00000163735 |

| | | | | | | | |
|---|---|---|---|---|---|---|---|
| PHPT1 | 0.353392428 | 1.277561222 | 2.52E-05 | 0.000205066 | 1281.825301 | 1638.633004 | ENSG00000054148 |
| RABL2A | 0.353299602 | 1.277479023 | 0.146509955 | 0.300677168 | 88.16797153 | 113.0240206 | ENSG00000144134 |
| EEF2 | 0.353272582 | 1.277455098 | 1.67E-11 | 4.08E-10 | 79630.31972 | 101723.3488 | ENSG00000167658 |
| NA | 0.353216912 | 1.277405805 | 0.07597577 | 0.184329137 | 106.3315179 | 135.7899318 | ENSG00000254285 |
| RACGAP1 | 0.35315543 | 1.277351368 | 5.63E-06 | 5.37E-05 | 1094.529761 | 1397.788483 | ENSG00000161800 |
| WDFY3 | 0.353045566 | 1.277254099 | 0.002417469 | 0.011137305 | 746.4076811 | 953.4231782 | ENSG00000163625 |
| HLA-G | 0.352962518 | 1.277180577 | 0.254233432 | 0.44298754 | 46.0418226 | 58.8054292 | ENSG00000204632 |
| FAM120AOS | 0.352700931 | 1.276949021 | 9.27E-05 | 0.000651654 | 737.9023711 | 942.0249607 | ENSG00000188938 |
| ELF2 | 0.352699911 | 1.276948119 | 0.001302171 | 0.006574352 | 594.0243064 | 757.8347864 | ENSG00000109381 |
| AGRN | 0.352525789 | 1.276794011 | 6.02E-10 | 1.16E-08 | 8020.550316 | 10240.92792 | ENSG00000188157 |
| NUAK1 | 0.352519494 | 1.276788439 | 0.001000864 | 0.005268392 | 521.668276 | 666.0772193 | ENSG00000074590 |
| ZPLD2P | 0.352492463 | 1.276764517 | 0.312616431 | 0.506161696 | 36.66108629 | 46.90062462 | ENSG00000236155 |
| SYT17 | 0.352299782 | 1.276594008 | 0.285482447 | 0.476721476 | 64.45711005 | 82.2054373 | ENSG00000103528 |
| LINC00909 | 0.352282892 | 1.276579063 | 0.144755105 | 0.298132138 | 77.76328387 | 99.393367 | ENSG00000264247 |
| ZNF2 | 0.352175892 | 1.276484387 | 0.219706518 | 0.401067724 | 55.6729019 | 70.91540062 | ENSG00000275111 |
| ZC3H12A | 0.352100318 | 1.276417522 | 0.013485138 | 0.046909448 | 262.1326015 | 334.773024 | ENSG00000163874 |
| LATS2 | 0.352011244 | 1.276338716 | 0.001212366 | 0.00619691 | 564.5812397 | 720.1663171 | ENSG00000150457 |
| NTNG1 | 0.351983679 | 1.27631433 | 0.461814901 | 0.648316025 | 17.57666169 | 22.41861204 | ENSG00000162631 |
| NA | 0.351974579 | 1.276306279 | 0.569620694 | 0.736315119 | 15.9766719 | 20.46513791 | ENSG00000274840 |
| MACC1 | 0.351964942 | 1.276297754 | 0.018307263 | 0.059879913 | 238.1056806 | 303.4143102 | ENSG00000183742 |
| NA | 0.351949324 | 1.276283937 | 0.445013021 | 0.633328198 | 22.46322056 | 28.72804966 | ENSG00000247735 |
| SSBP4 | 0.35183355 | 1.276181522 | 0.004095275 | 0.017372813 | 767.3895292 | 978.8369487 | ENSG00000130511 |
| SYNE1 | 0.35176325 | 1.276119337 | 0.256651201 | 0.445307373 | 52.71282504 | 67.19984954 | ENSG00000131018 |
| ARL4A | 0.351658106 | 1.276026336 | 0.00717964 | 0.027967097 | 310.6477431 | 396.5680361 | ENSG00000122644 |
| NA | 0.351600356 | 1.275975259 | 0.390451727 | 0.583176107 | 24.61616573 | 31.3867943 | ENSG00000267565 |
| STRN3 | 0.351539609 | 1.275921533 | 0.000456547 | 0.002645918 | 609.4156418 | 777.0393449 | ENSG00000196792 |
| ZBED4 | 0.351466455 | 1.275856837 | 8.22E-06 | 7.49E-05 | 1467.842872 | 1873.629858 | ENSG00000100426 |
| ATXN1L | 0.351385316 | 1.275785084 | 5.46E-05 | 0.000407366 | 878.4029255 | 1120.411223 | ENSG00000224470 |
| CBR4 | 0.351063259 | 1.275500318 | 0.003148919 | 0.013879205 | 358.8923975 | 457.4789309 | ENSG00000145439 |
| SYCE3 | 0.350938489 | 1.275390012 | 0.706679112 | 0.833271761 | 4.892872691 | 6.192574783 | ENSG00000217442 |
| UBR3 | 0.350421789 | 1.274933315 | 0.003194024 | 0.014041282 | 409.4090879 | 521.260499 | ENSG00000144357 |
| HUWE1 | 0.35035152 | 1.274871218 | 0.045769407 | 0.124966972 | 7704.515742 | 9822.179013 | ENSG00000086758 |
| SLC24A1 | 0.35033365 | 1.274855427 | 0.055298079 | 0.144392024 | 146.4420323 | 186.5651872 | ENSG00000074621 |
| SAMD3 | 0.350312883 | 1.274837076 | 0.423121438 | 0.614105931 | 22.48541665 | 28.76515362 | ENSG00000164483 |
| MLXIP | 0.350238393 | 1.274771255 | 4.01E-05 | 0.000310051 | 1433.428957 | 1827.432216 | ENSG00000175727 |
| ERC2 | 0.350225002 | 1.274759422 | 0.073988629 | 0.180625034 | 115.9719702 | 147.7472505 | ENSG00000187672 |
| PHF7 | 0.349953682 | 1.274519708 | 0.086064038 | 0.202502333 | 112.0622196 | 142.7821817 | ENSG00000010318 |
| LOC652276 | 0.349948937 | 1.274515516 | 0.149641072 | 0.305580391 | 75.94334014 | 96.85374673 | ENSG00000215154 |
| DPY19L1 | 0.349947638 | 1.274514368 | 0.010429996 | 0.038063716 | 359.8815477 | 458.384032 | ENSG00000173852 |
| ANKMY2 | 0.349798175 | 1.274382336 | 0.00540242 | 0.022002567 | 305.359039 | 389.0964289 | ENSG00000106524 |
| NRBP2 | 0.349762632 | 1.27435094 | 0.067910232 | 0.169014133 | 196.9238838 | 251.3490497 | ENSG00000185189 |
| NA | 0.34974135 | 1.274332142 | 0.678387271 | 0.814170153 | 5.76620185 | 7.374279807 | ENSG00000271714 |
| DAP3 | 0.349596689 | 1.274204369 | 1.21E-05 | 0.000106015 | 1416.478829 | 1804.347032 | ENSG00000132676 |
| SUSD6 | 0.349581386 | 1.274190853 | 0.095966882 | 0.219667802 | 101.8043958 | 129.465522 | ENSG00000100647 |
| DNAI1 | 0.349550759 | 1.274163804 | 0.615112753 | 0.770668544 | 9.034430804 | 11.50389207 | ENSG00000122735 |
| ZBTB39 | 0.349450628 | 1.274075372 | 4.35E-05 | 0.000332404 | 1001.611173 | 1276.390932 | ENSG00000166860 |
| TSPAN2 | 0.349403824 | 1.274034039 | 0.024818625 | 0.076986674 | 276.9408183 | 352.09947 | ENSG00000134198 |
| RASSF10 | 0.349320548 | 1.273960502 | 0.186832645 | 0.35824087 | 60.49752522 | 77.05644895 | ENSG00000189431 |
| NA | 0.349215834 | 1.273868038 | 0.339465767 | 0.533734182 | 42.39641807 | 54.19211114 | ENSG00000269981 |
| ANXA6 | 0.349108536 | 1.2737733 | 1.44E-08 | 2.25E-07 | 2601.962144 | 3314.047476 | ENSG00000197043 |
| HLA-DOA | 0.349106998 | 1.273771941 | 0.001600663 | 0.007863085 | 438.7877082 | 558.8950104 | ENSG00000204252 |
| UGP2 | 0.349046058 | 1.273718138 | 6.21E-06 | 5.88E-05 | 7640.763495 | 9730.682159 | ENSG00000169764 |
| NA | 0.349023942 | 1.273698613 | 0.775466119 | 0.876264422 | 3.268228953 | 4.133411111 | ENSG00000288086 |
| HK1 | 0.349005492 | 1.273682324 | 1.68E-09 | 3.03E-08 | 9091.811236 | 11579.11233 | ENSG00000156515 |
| HRAT92 | 0.348977273 | 1.273657411 | 0.485645381 | 0.66875218 | 17.79218094 | 22.7791714 | ENSG00000223855 |
| FAM13C | 0.348968033 | 1.273649254 | 0.616991986 | 0.772278752 | 8.293864739 | 10.5132578 | ENSG00000148541 |
| ND4L | 0.348948122 | 1.273631676 | 0.081445688 | 0.194274696 | 3396.20956 | 4325.692154 | ENSG00000212907 |
| CTTNBP2NL | 0.348860433 | 1.273554265 | 0.002031373 | 0.009619671 | 537.1998312 | 683.6995496 | ENSG00000143079 |
| SPTY2D1 | 0.348842772 | 1.273538675 | 0.009641003 | 0.03567673 | 350.7115802 | 446.0022433 | ENSG00000179119 |
| NIT1 | 0.348736053 | 1.273444472 | 0.00172801 | 0.008386212 | 477.29806 | 607.5505523 | ENSG00000158793 |
| THY1 | 0.348693368 | 1.273406795 | 1.12E-08 | 1.78E-07 | 6039.512459 | 7691.319976 | ENSG00000154096 |
| PAN2 | 0.348626718 | 1.273347967 | 0.007290555 | 0.028309897 | 487.4925728 | 620.8930798 | ENSG00000135473 |
| PLEKHG1 | 0.348512605 | 1.273247253 | 0.249558685 | 0.438045332 | 46.47789714 | 59.16891134 | ENSG00000120278 |
| PHF19 | 0.348480305 | 1.273218748 | 0.005389943 | 0.021969917 | 329.9848482 | 419.9110615 | ENSG00000119403 |
| ADORA2B | 0.34826464 | 1.273028431 | 0.149264744 | 0.304979909 | 84.52244666 | 107.673624 | ENSG00000170425 |
| ZNF555 | 0.348112672 | 1.272894343 | 0.088737277 | 0.207266124 | 99.68755218 | 126.8330344 | ENSG00000186300 |
| NA | 0.347891414 | 1.27269914 | 0.697022396 | 0.827075873 | 5.141238842 | 6.518841347 | ENSG00000233597 |
| GPR78 | 0.347730484 | 1.272557181 | 0.599458604 | 0.759267803 | 11.51534136 | 14.69213501 | ENSG00000155269 |
| CEP120 | 0.347715562 | 1.272544019 | 0.007201426 | 0.028016631 | 335.010078 | 426.2686732 | ENSG00000168944 |
| ZKSCAN2 | 0.34766792 | 1.272501997 | 0.016133511 | 0.054143282 | 242.8356192 | 309.2320551 | ENSG00000155592 |
| STXBP5 | 0.347517396 | 1.272369237 | 0.052932986 | 0.139712327 | 142.4303735 | 181.2036173 | ENSG00000164506 |
| MTMR3 | 0.347394022 | 1.272260433 | 0.167168938 | 0.330803876 | 65.49031411 | 83.25596016 | ENSG00000100330 |
| DALRD3 | 0.347283463 | 1.272162939 | 0.0031967 | 0.014049713 | 395.0018062 | 502.1811824 | ENSG00000178149 |
| RFK | 0.347269707 | 1.272150809 | 0.040919413 | 0.1145101 | 242.0429565 | 307.7039581 | ENSG00000135002 |

| | | | | | | | |
|---|---|---|---|---|---|---|---|
| NA | 0.34726963 | 1.272150741 | 0.547497584 | 0.719107096 | 12.88540282 | 16.435879 | ENSG00000233406 |
| NA | 0.347198228 | 1.272087781 | 0.154438816 | 0.312110894 | 84.59476256 | 107.5399656 | ENSG00000259429 |
| TMEM44 | 0.347126876 | 1.272024868 | 0.028126373 | 0.08529028 | 198.6660097 | 252.7939464 | ENSG00000145014 |
| NA | 0.347119887 | 1.272018706 | 0.678787208 | 0.814439186 | 5.673974961 | 7.246570073 | ENSG00000235280 |
| SHB | 0.34706524 | 1.271970525 | 0.172717279 | 0.338996299 | 80.31381926 | 102.039401 | ENSG00000107338 |
| NA | 0.347006227 | 1.271918497 | 0.480457248 | 0.6642839 | 17.68910784 | 22.44866795 | ENSG00000263571 |
| SUCO | 0.346874141 | 1.271802051 | 6.21E-05 | 0.000456006 | 826.3870484 | 1050.509536 | ENSG00000094975 |
| NA | 0.346751213 | 1.271693689 | 0.641402956 | 0.789727124 | 8.187732422 | 10.48375154 | ENSG00000236540 |
| KSR1 | 0.346703623 | 1.271651741 | 0.000721415 | 0.003958174 | 843.3312684 | 1072.500948 | ENSG00000141068 |
| TMEM108 | 0.346363188 | 1.271351703 | 0.017201563 | 0.056992092 | 218.2657021 | 277.5281921 | ENSG00000144868 |
| NA | 0.346301259 | 1.27129713 | 0.395708137 | 0.588110368 | 27.41251172 | 35.02152746 | ENSG00000224858 |
| TST | 0.345954906 | 1.270991962 | 0.002909956 | 0.012964693 | 421.5884401 | 535.6004452 | ENSG00000128311 |
| NA | 0.345892416 | 1.27093691 | 0.251278266 | 0.439855628 | 45.11868189 | 57.34204764 | ENSG00000245248 |
| RNF169 | 0.34578264 | 1.270840207 | 0.033323941 | 0.097385153 | 194.1945094 | 246.4450945 | ENSG00000166439 |
| PXDC1 | 0.345755807 | 1.270816571 | 0.335121302 | 0.529337273 | 38.84509693 | 49.43925721 | ENSG00000168994 |
| LAMC2 | 0.345716061 | 1.27078156 | 0.012269645 | 0.043555948 | 263.8700747 | 334.842557 | ENSG00000058085 |
| AGO4 | 0.345602531 | 1.270681562 | 0.013340023 | 0.046500655 | 272.1663223 | 345.7756502 | ENSG00000134698 |
| LDLRAD4 | 0.345584935 | 1.270666065 | 0.30568814 | 0.499683979 | 42.48478899 | 54.11757681 | ENSG00000168675 |
| NA | 0.34540802 | 1.270510255 | 0.395951985 | 0.588262452 | 44.4368194 | 56.68389219 | ENSG00000274471 |
| CMTM1 | 0.345375353 | 1.270481487 | 0.59496845 | 0.755905214 | 9.510943851 | 12.12268206 | ENSG00000089505 |
| TTC30B | 0.345361388 | 1.270469189 | 0.404851956 | 0.596509035 | 30.31992838 | 38.70531077 | ENSG00000196659 |
| NA | 0.345277859 | 1.270395633 | 0.68547932 | 0.818973783 | 5.458455715 | 6.927789582 | ENSG00000231120 |
| ZNF526 | 0.345253059 | 1.270373796 | 0.001381942 | 0.006903648 | 454.375975 | 577.3329757 | ENSG00000167625 |
| KCNF1 | 0.345244302 | 1.270366085 | 0.273039139 | 0.464192108 | 45.00170336 | 57.00405813 | ENSG00000162975 |
| TBC1D10C | 0.3452123 | 1.270337906 | 0.613750427 | 0.769585982 | 8.578234708 | 10.86209421 | ENSG00000175463 |
| GPR89B | 0.345068078 | 1.27021092 | 0.082941559 | 0.19695565 | 116.2310849 | 147.5144008 | ENSG00000188092 |
| NA | 0.345039361 | 1.270185637 | 0.540041497 | 0.713205722 | 13.39150816 | 17.05510703 | ENSG00000271936 |
| RYK | 0.344941396 | 1.270099389 | 0.000124968 | 0.000849029 | 880.4425149 | 1117.55115 | ENSG00000163785 |
| PABPC4L | 0.344900385 | 1.270063285 | 0.209094215 | 0.388311842 | 54.97968952 | 69.86619186 | ENSG00000254535 |
| NA | 0.344892402 | 1.270056256 | 0.342281654 | 0.536395153 | 32.92265812 | 41.80663516 | ENSG00000236255 |
| ZNF554 | 0.344763626 | 1.269942895 | 0.067094144 | 0.167635867 | 154.1876694 | 195.8170843 | ENSG00000172006 |
| SCARB2 | 0.344754874 | 1.269935191 | 7.23E-05 | 0.000521219 | 976.6761065 | 1239.627747 | ENSG00000138760 |
| NA | 0.34467297 | 1.269863098 | 0.213974814 | 0.394042844 | 62.75038658 | 79.97941322 | ENSG00000227827 |
| ABCA11P | 0.344454509 | 1.269670822 | 0.734703779 | 0.851647998 | 4.105554294 | 5.247518231 | ENSG00000251595 |
| HERC4 | 0.344235345 | 1.269477957 | 0.000388905 | 0.002296993 | 624.125348 | 792.1577106 | ENSG00000148634 |
| FRA10AC1 | 0.344151464 | 1.269404149 | 0.008479732 | 0.032133173 | 332.4009594 | 421.9815185 | ENSG00000148690 |
| PAQR6 | 0.343652228 | 1.268964956 | 0.271252114 | 0.462013743 | 85.70896397 | 109.0654343 | ENSG00000160781 |
| YIPF2 | 0.343602366 | 1.268921099 | 0.000337847 | 0.002033665 | 828.3866652 | 1051.743796 | ENSG00000130733 |
| TMEM37 | 0.343549188 | 1.268874328 | 0.01813652 | 0.059420587 | 222.7617587 | 282.9988124 | ENSG00000171227 |
| KLRG2 | 0.343413495 | 1.268754988 | 0.002327142 | 0.010801716 | 457.8149477 | 581.2088174 | ENSG00000188883 |
| SLC25A6 | 0.342944629 | 1.26834272 | 2.04E-09 | 3.62E-08 | 15424.97074 | 19564.16293 | ENSG00000169100 |
| YBX2 | 0.342540318 | 1.267987319 | 0.003543632 | 0.01533828 | 628.3040975 | 797.5505876 | ENSG00000006047 |
| SLC6A3 | 0.342417248 | 1.267879158 | 0.399527396 | 0.591440955 | 25.83008793 | 32.74459793 | ENSG00000142319 |
| MPND | 0.342356174 | 1.267825485 | 0.017605067 | 0.058028155 | 283.3716098 | 359.2974767 | ENSG00000008382 |
| NFIL3 | 0.342321736 | 1.267795222 | 0.039392069 | 0.111125208 | 215.7176472 | 273.4287474 | ENSG00000165030 |
| C9orf72 | 0.342226578 | 1.267711603 | 0.032165037 | 0.094628066 | 243.8086313 | 309.2216059 | ENSG00000147894 |
| RSPO1 | 0.342194387 | 1.267683317 | 0.746387062 | 0.858586011 | 3.758647429 | 4.778458166 | ENSG00000169218 |
| ERMAP | 0.342115766 | 1.267614235 | 0.309878523 | 0.503769234 | 35.62983643 | 45.21231755 | ENSG00000164010 |
| FMO4 | 0.342082317 | 1.267584846 | 0.565270822 | 0.733099812 | 13.5665889 | 17.25366386 | ENSG00000076258 |
| POLR3D | 0.342077024 | 1.267580195 | 2.35E-06 | 2.43E-05 | 1326.439892 | 1681.477594 | ENSG00000168495 |
| CCDC81 | 0.341959564 | 1.267476996 | 0.621016408 | 0.77502647 | 11.70422983 | 14.83872736 | ENSG00000149201 |
| NEURL1 | 0.341462361 | 1.267040255 | 0.535041821 | 0.709590766 | 18.39692469 | 23.40533584 | ENSG00000107954 |
| GXYLT1 | 0.341458567 | 1.267036923 | 0.013355911 | 0.046547283 | 343.1049298 | 434.6727295 | ENSG00000151233 |
| NUDT6 | 0.341441427 | 1.26702187 | 0.28072735 | 0.472098785 | 44.22385571 | 56.14255931 | ENSG00000170917 |
| TRAM2 | 0.341436456 | 1.267017504 | 0.00046844 | 0.002703457 | 702.1296245 | 889.2011652 | ENSG00000065308 |
| OGA | 0.341242573 | 1.266847242 | 1.62E-07 | 2.08E-06 | 2656.588064 | 3365.112395 | ENSG00000198408 |
| GFPT2 | 0.341150757 | 1.26676662 | 7.52E-07 | 8.57E-06 | 2719.603503 | 3445.345874 | ENSG00000131459 |
| PLEKHA4 | 0.341101411 | 1.266723292 | 0.014364214 | 0.049319442 | 341.7947445 | 432.2352232 | ENSG00000105559 |
| ZFC3H1 | 0.340974515 | 1.266611879 | 0.001379177 | 0.006893655 | 818.2941211 | 1037.055557 | ENSG00000133858 |
| SOWAHC | 0.340881909 | 1.266530579 | 0.003701753 | 0.015925788 | 450.1388377 | 569.6486837 | ENSG00000198142 |
| MTHFR | 0.340873185 | 1.266522919 | 0.013317713 | 0.046457838 | 250.6378627 | 317.1170219 | ENSG00000177000 |
| COPS6 | 0.340620541 | 1.266301146 | 3.33E-06 | 3.36E-05 | 2129.199241 | 2696.994019 | ENSG00000168090 |
| ATP5ME | 0.340528224 | 1.266220119 | 0.499677681 | 0.680407475 | 25.03202101 | 31.81137595 | ENSG00000169020 |
| DDB2 | 0.34040596 | 1.266112815 | 0.001803551 | 0.008695193 | 575.6331168 | 728.4615152 | ENSG00000134574 |
| NA | 0.340106795 | 1.265850295 | 0.385439595 | 0.57834662 | 44.7148529 | 56.52046395 | ENSG00000249264 |
| CLCF1 | 0.339841121 | 1.265617208 | 0.514720384 | 0.692846128 | 14.88032959 | 18.84387906 | ENSG00000175505 |
| TMEM120B | 0.339832651 | 1.265609778 | 0.001928214 | 0.009222583 | 684.3066485 | 866.9517246 | ENSG00000188735 |
| NA | 0.339823941 | 1.265602137 | 0.704232138 | 0.831811477 | 5.5178357 | 6.965331571 | ENSG00000273891 |
| RNF19B | 0.339688884 | 1.265483664 | 0.000860137 | 0.004625735 | 579.9960642 | 734.0486232 | ENSG00000116514 |
| PTPN3 | 0.339628882 | 1.265431034 | 0.053353793 | 0.140462954 | 148.1700422 | 187.7203087 | ENSG00000070159 |
| NA | 0.339461211 | 1.265283973 | 0.740063189 | 0.854707161 | 3.768118167 | 4.778458166 | ENSG00000234206 |
| H2BC18 | 0.339250482 | 1.265099171 | 0.76685366 | 0.871099885 | 3.229068223 | 4.043243366 | ENSG00000203814 |
| PYCR1 | 0.33899924 | 1.264878876 | 1.18E-07 | 1.56E-06 | 3729.318494 | 4718.137292 | ENSG00000183010 |
| LPAR1 | 0.338850539 | 1.26474851 | 0.199375423 | 0.375721819 | 65.78217798 | 82.95551263 | ENSG00000198121 |

| | | | | | | | |
|---|---|---|---|---|---|---|---|
| PINK1 | 0.338711057 | 1.264626238 | 0.041902367 | 0.11674962 | 186.0868647 | 235.2517324 | ENSG00000158828 |
| SDHB | 0.338653077 | 1.264575415 | 5.51E-05 | 0.000410285 | 881.0319777 | 1114.310465 | ENSG00000117118 |
| C1QL4 | 0.338609496 | 1.264537215 | 0.432202127 | 0.622133674 | 24.48793502 | 31.09470891 | ENSG00000186897 |
| FOXN3 | 0.3382248 | 1.26420007 | 0.000182298 | 0.001185515 | 1931.958419 | 2440.984 | ENSG00000053254 |
| KPNA2 | 0.338176078 | 1.264157377 | 1.43E-09 | 2.59E-08 | 10978.4303 | 13877.72907 | ENSG00000182481 |
| RELA-DT | 0.338098438 | 1.264089347 | 0.516865 | 0.694373435 | 14.99908956 | 18.95281782 | ENSG00000285533 |
| LINC-PINT | 0.338048185 | 1.264045316 | 0.737844673 | 0.853154481 | 4.085335041 | 5.194892475 | ENSG00000231721 |
| AIPL1 | 0.337710342 | 1.263749343 | 0.562237597 | 0.730738379 | 12.94793972 | 16.36079503 | ENSG00000129221 |
| LINC02076 | 0.337630434 | 1.263679348 | 0.604953439 | 0.763569953 | 10.04367997 | 12.75275697 | ENSG00000220161 |
| ACAP2 | 0.337610512 | 1.263661898 | 0.001833155 | 0.008804274 | 558.7519222 | 705.8953945 | ENSG00000114331 |
| RNF19A | 0.337557927 | 1.26361584 | 0.061373867 | 0.156853731 | 142.7365487 | 180.2825563 | ENSG00000034677 |
| DUBR | 0.337516325 | 1.263579402 | 0.314088307 | 0.507759937 | 36.75774787 | 46.35972977 | ENSG00000243701 |
| NYX | 0.337224363 | 1.263323715 | 0.735476892 | 0.852171088 | 4.108711207 | 5.217462316 | ENSG00000188937 |
| AGO3 | 0.337157799 | 1.263265428 | 0.007950566 | 0.030470595 | 310.9800455 | 392.6653876 | ENSG00000126070 |
| MRPS31 | 0.336978488 | 1.263108428 | 0.003939857 | 0.016829066 | 459.8592952 | 580.1541206 | ENSG00000102738 |
| SPTBN5 | 0.336883498 | 1.263025265 | 0.273931628 | 0.46515382 | 70.33515669 | 89.13998034 | ENSG00000137877 |
| CCSER2 | 0.336731859 | 1.262892517 | 0.013612167 | 0.047253776 | 294.7651546 | 371.8279674 | ENSG00000107771 |
| PID1 | 0.336681341 | 1.262848296 | 0.187861151 | 0.3596177 | 67.46196241 | 85.19869902 | ENSG00000153823 |
| BMP2K | 0.336646733 | 1.262818002 | 0.032003703 | 0.094316526 | 196.0645578 | 247.7785049 | ENSG00000138756 |
| SMURF1 | 0.336384156 | 1.262588185 | 0.00112673 | 0.005831562 | 667.999839 | 842.6662758 | ENSG00000198742 |
| LUZP1 | 0.336351442 | 1.262559555 | 3.32E-06 | 3.36E-05 | 1667.398784 | 2105.459948 | ENSG00000169641 |
| NAPRT | 0.336341129 | 1.26255053 | 0.004968725 | 0.020469071 | 481.0878328 | 608.0830222 | ENSG00000147813 |
| MOB3A | 0.336307078 | 1.262520732 | 0.080480764 | 0.192592703 | 127.8374731 | 161.686937 | ENSG00000172081 |
| FARP1 | 0.336198383 | 1.262425614 | 1.03E-05 | 9.18E-05 | 1977.366567 | 2496.5273 | ENSG00000152767 |
| MN1 | 0.336066361 | 1.262310094 | 0.050035692 | 0.133876257 | 194.1403609 | 244.5822975 | ENSG00000169184 |
| GPC2 | 0.336060343 | 1.262304829 | 0.000120658 | 0.000824888 | 888.6934832 | 1122.071467 | ENSG00000213420 |
| HSPA12A | 0.335982984 | 1.262237144 | 0.029418233 | 0.088154814 | 200.6009552 | 253.1302869 | ENSG00000165868 |
| ZNF682 | 0.335889377 | 1.262155249 | 0.051533972 | 0.136838199 | 169.6858515 | 213.7636428 | ENSG00000197124 |
| CBWD6 | 0.335776997 | 1.262056936 | 0.388918876 | 0.581598708 | 32.41211809 | 40.92114079 | ENSG00000215126 |
| DCP2 | 0.335623969 | 1.261923075 | 0.001119975 | 0.005809655 | 1393.046918 | 1756.957809 | ENSG00000172795 |
| ATP1A4 | 0.335569295 | 1.261875253 | 0.744127534 | 0.857157223 | 3.768118167 | 4.759687172 | ENSG00000132681 |
| KHNYN | 0.335191885 | 1.261545189 | 0.003049374 | 0.01351437 | 624.2349229 | 787.3050276 | ENSG00000100441 |
| STK10 | 0.335158747 | 1.261516212 | 0.000376479 | 0.002234997 | 657.3651101 | 829.1180177 | ENSG00000072786 |
| AP5B1 | 0.335003521 | 1.261380487 | 0.000225945 | 0.00143217 | 711.1946171 | 897.246463 | ENSG00000254470 |
| FAM98C | 0.334978412 | 1.261358534 | 0.102178703 | 0.230358848 | 108.5160322 | 136.9797841 | ENSG00000130244 |
| TLE2 | 0.334813202 | 1.261214098 | 0.006717434 | 0.026478291 | 361.5497869 | 456.4884485 | ENSG00000065717 |
| SLC7A6OS | 0.334695826 | 1.261111491 | 0.030220343 | 0.090135796 | 209.150492 | 264.0221106 | ENSG00000103061 |
| FTL | 0.334689863 | 1.261106279 | 1.81E-10 | 3.80E-09 | 12112.42601 | 15274.44342 | ENSG00000087086 |
| CXorf38 | 0.334682176 | 1.261099559 | 0.030024588 | 0.089681824 | 186.8241762 | 235.6245644 | ENSG00000185753 |
| MGAT5 | 0.334669405 | 1.261088396 | 4.07E-05 | 0.000313765 | 1186.532356 | 1495.41677 | ENSG00000152127 |
| ZNF749 | 0.334664089 | 1.26108375 | 0.047712297 | 0.128981872 | 237.3122438 | 299.8876087 | ENSG00000186230 |
| PTPRVP | 0.334658975 | 1.261079279 | 0.742983723 | 0.85637191 | 4.741265818 | 6.03155994 | ENSG00000243323 |
| CHP2 | 0.334651754 | 1.261072967 | 0.332811061 | 0.527387158 | 36.42662557 | 45.80061358 | ENSG00000166869 |
| DIPK1A | 0.334521202 | 1.260958855 | 0.106219881 | 0.237276036 | 112.074539 | 141.3515556 | ENSG00000154511 |
| MDFI | 0.334340448 | 1.260800881 | 1.45E-05 | 0.000125712 | 2026.762779 | 2555.397366 | ENSG00000112559 |
| CENPL | 0.333958176 | 1.26046685 | 0.039770016 | 0.111969844 | 184.6268388 | 232.6056985 | ENSG00000120334 |
| CBWD1 | 0.333897725 | 1.260414036 | 0.014179467 | 0.048793584 | 253.000517 | 318.7350315 | ENSG00000172785 |
| MPV17 | 0.333822359 | 1.260348193 | 0.000702562 | 0.00386618 | 748.3835687 | 943.4119131 | ENSG00000115204 |
| LOC101929551 | 0.333692561 | 1.260234806 | 0.72941837 | 0.848386575 | 5.141238842 | 6.436271296 | ENSG00000226506 |
| PSPC1 | 0.33368461 | 1.260227861 | 3.03E-07 | 3.71E-06 | 2452.37428 | 3090.551136 | ENSG00000121390 |
| KLHDC7A | 0.333676753 | 1.260220998 | 0.001443353 | 0.007164197 | 510.879287 | 643.6639117 | ENSG00000179023 |
| PCDHA8 | 0.333553497 | 1.260113336 | 0.565267407 | 0.733099812 | 12.30719214 | 15.57719135 | ENSG00000204962 |
| NA | 0.333525311 | 1.260088717 | 0.422565924 | 0.613587969 | 26.10498716 | 32.84181371 | ENSG00000280058 |
| TSTD2 | 0.333344147 | 1.259930494 | 0.013641949 | 0.047330566 | 342.0071819 | 430.3806138 | ENSG00000136925 |
| ANKEF1 | 0.333322083 | 1.259911225 | 0.03581376 | 0.103102242 | 179.1539966 | 225.7864096 | ENSG00000132623 |
| CXADR | 0.333194298 | 1.259799635 | 0.001159708 | 0.005978867 | 1693.311417 | 2132.544111 | ENSG00000154639 |
| BCL9L | 0.333097618 | 1.259715214 | 2.71E-05 | 0.000218949 | 1403.269444 | 1767.67196 | ENSG00000186174 |
| LINC00605 | 0.332946861 | 1.259583585 | 0.392926295 | 0.585550852 | 30.46324459 | 38.20889433 | ENSG00000251533 |
| NA | 0.332780506 | 1.259438352 | 0.730231025 | 0.849118629 | 4.419614255 | 5.611326784 | ENSG00000275882 |
| ZSCAN22 | 0.332774095 | 1.259432756 | 0.265345228 | 0.455066942 | 56.72507007 | 71.51998291 | ENSG00000182318 |
| HARS2 | 0.332766911 | 1.259426484 | 0.004439645 | 0.018612166 | 411.0705095 | 517.2107573 | ENSG00000112855 |
| LENG9 | 0.332441349 | 1.259142311 | 0.193601508 | 0.367381012 | 71.25967287 | 89.49637416 | ENSG00000275183 |
| INTU | 0.332391091 | 1.259098449 | 0.01990703 | 0.064300609 | 253.600127 | 319.3818307 | ENSG00000164066 |
| CCDC125 | 0.332344266 | 1.259057583 | 0.184188747 | 0.35487945 | 67.2635055 | 84.69633384 | ENSG00000183323 |
| USP40 | 0.332336836 | 1.259051099 | 0.075249328 | 0.18297419 | 134.1722194 | 168.8050329 | ENSG00000085982 |
| CTSK | 0.332283753 | 1.259004773 | 0.097886994 | 0.222961478 | 108.0067699 | 135.9960863 | ENSG00000143387 |
| LOC103344931 | 0.332141605 | 1.25888073 | 0.314806602 | 0.508464101 | 38.95428846 | 49.03823309 | ENSG00000260917 |
| EXOC6B | 0.332019956 | 1.258774585 | 3.01E-05 | 0.000241339 | 1130.837934 | 1423.224902 | ENSG00000144036 |
| NA | 0.331816476 | 1.258597058 | 0.584126833 | 0.747231695 | 11.43070607 | 14.37671533 | ENSG00000225808 |
| ALG11 | 0.331810888 | 1.258592184 | 0.072074412 | 0.177094943 | 175.8545819 | 221.0941055 | ENSG00000253710 |
| TRMT5 | 0.331686582 | 1.258483745 | 0.001950846 | 0.009297529 | 516.5551563 | 649.3798287 | ENSG00000126814 |
| ADAMTS9 | 0.330967745 | 1.257856849 | 0.464619209 | 0.650888315 | 24.99857275 | 31.4500437 | ENSG00000163638 |
| FGFRL1 | 0.330917239 | 1.257812814 | 0.000251841 | 0.001580103 | 698.561316 | 878.634111 | ENSG00000127418 |
| UBE2C | 0.330831044 | 1.257737668 | 1.99E-06 | 2.10E-05 | 2221.246642 | 2793.026611 | ENSG00000175063 |

| | | | | | | | |
|---|---|---|---|---|---|---|---|
| TBL1XR1 | 0.330801057 | 1.257711526 | 0.004212396 | 0.017812582 | 642.9729366 | 808.2319535 | ENSG00000177565 |
| SERPINI1 | 0.330800368 | 1.257710925 | 0.16244391 | 0.323804628 | 80.42380755 | 101.2448388 | ENSG00000163536 |
| ENTPD4 | 0.330765334 | 1.257680383 | 0.00783491 | 0.030108276 | 779.2132705 | 981.0116711 | ENSG00000197217 |
| CENPC | 0.330667122 | 1.257594769 | 0.026074968 | 0.080051921 | 253.0311014 | 318.5262891 | ENSG00000145241 |
| NA | 0.330611966 | 1.257546691 | 0.339049308 | 0.533305599 | 37.19697932 | 46.79862231 | ENSG00000288398 |
| SFI1 | 0.330573677 | 1.257513316 | 0.029743278 | 0.088941902 | 284.9298082 | 358.5645721 | ENSG00000198089 |
| DYNC2LI1 | 0.330513827 | 1.257461149 | 0.191644072 | 0.36467997 | 65.39049561 | 82.32262656 | ENSG00000138036 |
| MRPL10 | 0.330428419 | 1.25738671 | 7.61E-06 | 7.04E-05 | 1571.629266 | 1976.080701 | ENSG00000159111 |
| RILP | 0.330344892 | 1.257313913 | 0.06504279 | 0.163746682 | 147.6231076 | 185.3980163 | ENSG00000167705 |
| FRAT1 | 0.330185815 | 1.257175285 | 0.147562419 | 0.302368366 | 87.94604076 | 110.8295496 | ENSG00000165879 |
| CCDC71L | 0.330099081 | 1.257099706 | 0.004081024 | 0.017316322 | 407.9955135 | 512.8858374 | ENSG00000253276 |
| COL4A2 | 0.329958231 | 1.256976982 | 4.53E-08 | 6.49E-07 | 3192.772791 | 4012.930369 | ENSG00000134871 |
| RPL28 | 0.329958178 | 1.256976936 | 4.10E-07 | 4.92E-06 | 13070.60058 | 16430.25786 | ENSG00000108107 |
| TMTC4 | 0.329891103 | 1.256918496 | 0.000564127 | 0.003190582 | 792.3456792 | 996.1749355 | ENSG00000125247 |
| AKT3 | 0.329874737 | 1.256904238 | 0.01811643 | 0.059376726 | 270.7705995 | 339.8870583 | ENSG00000117020 |
| CPLANE1 | 0.329537138 | 1.256610149 | 0.098229828 | 0.223550052 | 117.9857408 | 148.1154075 | ENSG00000197603 |
| ARHGEF37 | 0.329508189 | 1.256584935 | 0.693331618 | 0.82436945 | 7.295461786 | 9.242372721 | ENSG00000183111 |
| ATP13A3 | 0.329476832 | 1.256557623 | 0.000383173 | 0.002268921 | 1248.005103 | 1567.177987 | ENSG00000133657 |
| NA | 0.329186099 | 1.256304426 | 0.366437265 | 0.559481692 | 35.57982949 | 44.68000808 | ENSG00000280018 |
| NA | 0.32913017 | 1.256255724 | 0.759769587 | 0.866398174 | 3.741585088 | 4.699575342 | ENSG00000277959 |
| FAR2 | 0.329095493 | 1.256225529 | 0.004458295 | 0.01865659 | 427.9984261 | 537.3791349 | ENSG00000064763 |
| NA | 0.329091035 | 1.256221647 | 0.223261112 | 0.405519546 | 80.74979611 | 101.715651 | ENSG00000267106 |
| NA | 0.328850083 | 1.256011856 | 0.514275856 | 0.692702668 | 17.75302021 | 22.31336052 | ENSG00000217165 |
| CRACR2A | 0.328601353 | 1.25579533 | 0.197941093 | 0.373626664 | 63.77650269 | 79.9967585 | ENSG00000130038 |
| ZNF483 | 0.328543715 | 1.255745161 | 0.014735678 | 0.050324498 | 343.6320058 | 432.0446307 | ENSG00000173258 |
| FOXJ1 | 0.328438154 | 1.255653282 | 0.103599046 | 0.232711758 | 115.8881317 | 145.6770251 | ENSG00000129654 |
| NA | 0.328401003 | 1.255620948 | 0.764925543 | 0.870029457 | 4.036703571 | 5.085953736 | ENSG00000278383 |
| LOC283922 | 0.328386492 | 1.255608319 | 0.309686381 | 0.503645563 | 47.45598314 | 59.55204054 | ENSG00000214331 |
| REEP3 | 0.328185703 | 1.255433579 | 0.0434967 | 0.12011937 | 352.4949036 | 442.3477514 | ENSG00000165476 |
| NA | 0.32811095 | 1.255368531 | 0.406855012 | 0.598559358 | 30.59788682 | 38.46843906 | ENSG00000272977 |
| SVIL | 0.328092373 | 1.255352366 | 0.002197913 | 0.010299906 | 611.0382337 | 767.1758409 | ENSG00000197321 |
| IQSEC2 | 0.328047151 | 1.255313018 | 3.49E-05 | 0.000274614 | 1449.895964 | 1820.883322 | ENSG00000124313 |
| BRAF | 0.327747667 | 1.255052459 | 0.003411928 | 0.014833842 | 427.8038478 | 536.7072811 | ENSG00000157764 |
| ACAT1 | 0.327698291 | 1.255009506 | 0.003127084 | 0.013796089 | 435.7954835 | 546.7604938 | ENSG00000075239 |
| PGBD2 | 0.327601374 | 1.2549252 | 0.085420886 | 0.201218951 | 129.8849623 | 162.9903628 | ENSG00000185220 |
| TAFA4 | 0.327588642 | 1.254914124 | 0.069457039 | 0.171963232 | 164.4991381 | 206.2220731 | ENSG00000163377 |
| NA | 0.327549073 | 1.254879707 | 0.607603331 | 0.7658678 | 11.47618063 | 14.37302811 | ENSG00000231711 |
| GNG7 | 0.32751248 | 1.254847877 | 0.00428975 | 0.018077815 | 441.7506715 | 554.6743946 | ENSG00000176533 |
| NA | 0.32739095 | 1.254742176 | 0.707187509 | 0.833461357 | 5.650598794 | 7.081318351 | ENSG00000178429 |
| STK3 | 0.327388615 | 1.254740145 | 0.02771021 | 0.084276506 | 205.7474254 | 258.1819479 | ENSG00000104375 |
| RBM26-AS1 | 0.327320939 | 1.254681288 | 0.779232638 | 0.878699058 | 4.280537334 | 5.371429118 | ENSG00000227354 |
| THBS3 | 0.327262069 | 1.25463009 | 0.256821478 | 0.445394336 | 57.99649311 | 73.03665149 | ENSG00000169231 |
| ANP32B | 0.327049991 | 1.254445672 | 1.23E-06 | 1.35E-05 | 2482.086601 | 3112.791124 | ENSG00000136938 |
| NA | 0.327030631 | 1.254428838 | 0.632923041 | 0.783663094 | 9.265734613 | 11.57853801 | ENSG00000272836 |
| TSTD3 | 0.327019538 | 1.254419193 | 0.575747361 | 0.740853659 | 11.78076985 | 14.7705798 | ENSG00000279170 |
| WDR17 | 0.327003422 | 1.25440518 | 0.292031937 | 0.484515256 | 44.68448649 | 56.10121848 | ENSG00000150627 |
| FBXL4 | 0.326914034 | 1.25432746 | 0.031481962 | 0.093178967 | 213.0522603 | 266.8408824 | ENSG00000112234 |
| LOC150051 | 0.326752442 | 1.254186975 | 0.349203357 | 0.5425547 | 37.89028939 | 47.56356661 | ENSG00000237594 |
| CCL2 | 0.326742185 | 1.254178058 | 0.356925394 | 0.550583042 | 33.3909782 | 41.85502404 | ENSG00000108691 |
| NA | 0.326704247 | 1.254145078 | 0.389008446 | 0.581676687 | 29.6462362 | 37.16293477 | ENSG00000280832 |
| CETN3 | 0.326517587 | 1.253982824 | 0.016206182 | 0.054325445 | 332.2760735 | 416.0947904 | ENSG00000153140 |
| PUDP | 0.32649684 | 1.25396479 | 0.001119058 | 0.005809655 | 773.0900373 | 969.1482208 | ENSG00000130021 |
| CATSPERG | 0.326432313 | 1.253908706 | 0.575516569 | 0.740696842 | 11.81361676 | 14.80063572 | ENSG00000099338 |
| NA | 0.326283397 | 1.253779283 | 0.662522518 | 0.803690999 | 7.740909367 | 9.677139991 | ENSG00000242083 |
| GTPBP10 | 0.326266877 | 1.253764927 | 0.039680395 | 0.111785445 | 199.5769136 | 250.11248 | ENSG00000105793 |
| NUMBL | 0.326262052 | 1.253760733 | 0.005148888 | 0.02111739 | 489.2743253 | 613.1496234 | ENSG00000105245 |
| CNKSR1 | 0.325974025 | 1.253510451 | 0.023234802 | 0.072988783 | 255.7158883 | 320.8453239 | ENSG00000142675 |
| STN1 | 0.325922304 | 1.253465513 | 0.055147098 | 0.144119615 | 200.6487148 | 251.2095683 | ENSG00000107960 |
| STX18-AS1 | 0.325846731 | 1.253399854 | 0.098047212 | 0.223216685 | 114.1994779 | 143.058522 | ENSG00000247708 |
| CHST3 | 0.325803874 | 1.253362621 | 0.007780617 | 0.029941691 | 353.262717 | 442.6295714 | ENSG00000122863 |
| DIS3L2 | 0.325698806 | 1.253271345 | 0.008806643 | 0.033121412 | 312.1401047 | 390.8879005 | ENSG00000144535 |
| ZNF574 | 0.325434482 | 1.253041748 | 7.41E-05 | 0.000532211 | 942.6168404 | 1181.33561 | ENSG00000105732 |
| NA | 0.325341697 | 1.252961163 | 0.649887916 | 0.795523587 | 8.039184763 | 10.06351839 | ENSG00000228420 |
| NA | 0.325309675 | 1.252933352 | 0.299355989 | 0.492872338 | 42.73640975 | 53.62507084 | ENSG00000232573 |
| DNAJB14 | 0.325291078 | 1.252917201 | 0.014867202 | 0.050695408 | 352.0792589 | 440.7142603 | ENSG00000164031 |
| MAGI2 | 0.325233221 | 1.252866956 | 0.00360589 | 0.015563941 | 444.7538628 | 557.3371126 | ENSG00000187391 |
| EDNRB | 0.325225895 | 1.252860594 | 0.035476119 | 0.102289109 | 242.570574 | 303.5057921 | ENSG00000136160 |
| AHSA2P | 0.32516215 | 1.252805238 | 0.484410442 | 0.667762226 | 408.4290477 | 511.8048156 | ENSG00000173209 |
| DIPK2A | 0.325099314 | 1.252750674 | 0.016805987 | 0.055971638 | 399.0964014 | 499.6343622 | ENSG00000181744 |
| LRRC28 | 0.325000292 | 1.252664692 | 0.0687918 | 0.170841111 | 146.2993925 | 183.0190844 | ENSG00000168904 |
| CEP72 | 0.324787674 | 1.252480093 | 0.002839142 | 0.012704165 | 451.587897 | 566.0311531 | ENSG00000112877 |
| OTUD3 | 0.324698651 | 1.25240281 | 0.003094453 | 0.013687969 | 464.5827996 | 581.8138377 | ENSG00000169914 |
| FAM120B | 0.32458682 | 1.252305734 | 0.001736631 | 0.008421435 | 488.5975739 | 611.6758246 | ENSG00000112584 |
| CD109 | 0.324577392 | 1.25229755 | 0.134021234 | 0.28143548 | 114.2144657 | 142.5884743 | ENSG00000156535 |

| | | | | | | | |
|---|---|---|---|---|---|---|---|
| LOC105369306 | 0.324264238 | 1.252025753 | 0.396206027 | 0.588546073 | 29.6714915 | 37.11399624 | ENSG00000233393 |
| DDX19B | 0.324210539 | 1.251979152 | 0.017017817 | 0.056504518 | 250.3332735 | 313.3338729 | ENSG00000157349 |
| SLC16A8 | 0.323895895 | 1.251706132 | 0.538423504 | 0.711982108 | 16.07846722 | 20.26223258 | ENSG00000100156 |
| BOLA1 | 0.323779087 | 1.251604791 | 0.102665773 | 0.231035385 | 120.7074259 | 150.7925967 | ENSG00000178096 |
| NA | 0.323626582 | 1.251472493 | 0.006895304 | 0.027086871 | 656.9317413 | 822.1747635 | ENSG00000197582 |
| DTNBP1 | 0.323547577 | 1.251403963 | 0.028210238 | 0.08546414 | 217.5028423 | 272.0407762 | ENSG00000047579 |
| XAB2 | 0.323469831 | 1.251336526 | 6.68E-07 | 7.69E-06 | 1990.689953 | 2490.890569 | ENSG00000076924 |
| MMRN2 | 0.323214219 | 1.251114839 | 0.343195746 | 0.537047446 | 37.13887712 | 46.52866872 | ENSG00000173269 |
| TPM4 | 0.322995814 | 1.250925451 | 2.42E-09 | 4.24E-08 | 8663.360327 | 10836.85445 | ENSG00000167460 |
| CNIH3 | 0.322964085 | 1.250897939 | 0.160807921 | 0.321338438 | 96.11523767 | 119.945598 | ENSG00000143786 |
| LINC01278 | 0.322956094 | 1.250891011 | 0.012888832 | 0.04527703 | 344.2971369 | 431.1600124 | ENSG00000235437 |
| SERTAD2 | 0.322948864 | 1.250884742 | 0.044219825 | 0.121688231 | 174.232614 | 217.8530136 | ENSG00000179833 |
| POC1B | 0.322931635 | 1.250869803 | 0.035471336 | 0.102289109 | 209.7814758 | 262.0548266 | ENSG00000139323 |
| CLCC1 | 0.322897488 | 1.250840198 | 0.015734207 | 0.053079323 | 269.3988543 | 337.1181011 | ENSG00000121940 |
| ZNF561 | 0.322838034 | 1.250788651 | 0.000690375 | 0.003811559 | 733.6434211 | 916.8222674 | ENSG00000171469 |
| PPM1N | 0.322797556 | 1.250753558 | 0.077244826 | 0.186680871 | 172.3512158 | 215.7932289 | ENSG00000213889 |
| CKAP2L | 0.322668672 | 1.250641826 | 0.034290052 | 0.099644376 | 226.825499 | 283.29867 | ENSG00000169607 |
| NA | 0.32266226 | 1.250636267 | 0.743721335 | 0.856902351 | 4.164934279 | 5.213663469 | ENSG00000224272 |
| MED23 | 0.322591028 | 1.25057452 | 0.013802586 | 0.047789486 | 317.8184393 | 397.2307549 | ENSG00000112282 |
| PPFIA1 | 0.322389799 | 1.2504001 | 3.67E-05 | 0.000287811 | 1288.099795 | 1611.219785 | ENSG00000131626 |
| NID1 | 0.322373173 | 1.250385691 | 0.000934046 | 0.004970315 | 1245.559656 | 1556.851747 | ENSG00000116962 |
| DDX28 | 0.32188372 | 1.249961553 | 0.018492944 | 0.060378195 | 328.6548419 | 411.2887384 | ENSG00000182810 |
| NA | 0.321750299 | 1.249845961 | 0.61274653 | 0.768999774 | 12.12767672 | 15.15315935 | ENSG00000253816 |
| METTL5 | 0.321723595 | 1.249822827 | 0.000549557 | 0.003118641 | 756.5744581 | 945.3923685 | ENSG00000138382 |
| TBC1D5 | 0.321565602 | 1.249685963 | 0.007763195 | 0.029882254 | 596.3618105 | 744.6564199 | ENSG00000131374 |
| ARHGEF10 | 0.321324889 | 1.249477471 | 0.29095787 | 0.483338994 | 55.18742177 | 68.91766288 | ENSG00000104728 |
| SRP14-AS1 | 0.321266507 | 1.249426909 | 0.365525105 | 0.558595177 | 35.60636257 | 44.38324777 | ENSG00000248508 |
| HERC6 | 0.321239852 | 1.249403825 | 0.125054793 | 0.267794977 | 101.3918929 | 126.8941339 | ENSG00000138642 |
| NA | 0.321229025 | 1.249394449 | 0.751901989 | 0.86205208 | 3.850874318 | 4.789743086 | ENSG00000231414 |
| PARM1 | 0.321072446 | 1.249258857 | 0.055500214 | 0.144797432 | 198.9170518 | 248.3801325 | ENSG00000169116 |
| TRPM4 | 0.321047762 | 1.249237483 | 0.032611112 | 0.095657172 | 242.283032 | 302.9969285 | ENSG00000130529 |
| TOX3 | 0.320838585 | 1.249056368 | 0.0279856 | 0.084918975 | 268.7440286 | 335.3455834 | ENSG00000103460 |
| ZNF66 | 0.320679573 | 1.248918707 | 0.150442777 | 0.30654506 | 116.1128286 | 145.2142889 | ENSG00000160229 |
| LINC00667 | 0.320672094 | 1.248912232 | 0.150923536 | 0.307217662 | 124.5578192 | 155.8812903 | ENSG00000263753 |
| KATNA1 | 0.320575702 | 1.24882879 | 0.01842766 | 0.060214782 | 264.3610946 | 329.9189322 | ENSG00000186625 |
| DNAJC9-AS1 | 0.32055717 | 1.248812748 | 0.630235512 | 0.781643622 | 11.59178368 | 14.52743294 | ENSG00000227540 |
| MBP | 0.320475311 | 1.248741892 | 0.001306098 | 0.006587 | 599.8562619 | 749.0152835 | ENSG00000197971 |
| NA | 0.320073129 | 1.248393827 | 0.650427142 | 0.795936006 | 8.015808597 | 9.984635562 | ENSG00000249492 |
| NA | 0.319998482 | 1.248329236 | 0.736652633 | 0.852520756 | 4.806959628 | 6.039046014 | ENSG00000237017 |
| SLC25A18 | 0.319932451 | 1.248272102 | 0.576223963 | 0.74109588 | 12.92268441 | 16.18469642 | ENSG00000182902 |
| NARS2 | 0.319873302 | 1.248220925 | 0.000328892 | 0.001987505 | 1007.891853 | 1257.740098 | ENSG00000137513 |
| LINC00877 | 0.319745299 | 1.248110182 | 0.507553537 | 0.68692604 | 19.09833003 | 23.86658341 | ENSG00000241163 |
| NA | 0.31954055 | 1.247933061 | 0.619352608 | 0.773807075 | 10.93081686 | 13.54384671 | ENSG00000230793 |
| HACE1 | 0.319511349 | 1.247907802 | 0.011908559 | 0.042510252 | 351.6933268 | 438.8747749 | ENSG00000085382 |
| NARF | 0.319507591 | 1.247904552 | 0.000169281 | 0.001107469 | 1154.07329 | 1440.366968 | ENSG00000141562 |
| CLPB | 0.319451497 | 1.247856032 | 0.003567437 | 0.015426709 | 452.6696348 | 564.8321657 | ENSG00000162129 |
| YES1 | 0.31934065 | 1.247760159 | 0.000367908 | 0.002189727 | 1659.524766 | 2069.797919 | ENSG00000176105 |
| ADAM17 | 0.319253534 | 1.247684816 | 0.003318773 | 0.014503687 | 515.2558997 | 642.4391455 | ENSG00000151694 |
| ZNF766 | 0.319027733 | 1.247489552 | 0.004614538 | 0.019197722 | 492.3488025 | 613.8230774 | ENSG00000196214 |
| LOC283683 | 0.319014555 | 1.247478157 | 0.64510642 | 0.792759739 | 9.488845461 | 11.91284031 | ENSG00000274253 |
| CCND1 | 0.318982924 | 1.247450807 | 0.00072529 | 0.003977076 | 1943.536281 | 2424.350552 | ENSG00000110092 |
| PET100 | 0.318877537 | 1.247359685 | 0.576258565 | 0.74109588 | 12.37288595 | 15.4381967 | ENSG00000229833 |
| THNSL1 | 0.31878883 | 1.247282991 | 0.091659733 | 0.212361099 | 144.3861395 | 179.9126471 | ENSG00000185875 |
| LOC171391 | 0.318521861 | 1.247052205 | 0.259828828 | 0.448762231 | 56.62643165 | 70.77838134 | ENSG00000255284 |
| URI1 | 0.318396774 | 1.246944085 | 6.61E-05 | 0.000482968 | 1307.204785 | 1629.597701 | ENSG00000105176 |
| DCDC2C | 0.318231862 | 1.246801557 | 0.762872353 | 0.868653694 | 3.768118167 | 4.680804347 | ENSG00000214866 |
| MGAT2 | 0.318148283 | 1.246729329 | 0.735895332 | 0.852173027 | 4.469523501 | 5.59255579 | ENSG00000168282 |
| HMGN3-AS1 | 0.318102698 | 1.246689937 | 0.650957427 | 0.796096664 | 10.10621687 | 12.59130409 | ENSG00000270362 |
| CDKN1B | 0.318050591 | 1.246644909 | 0.000297832 | 0.001826008 | 805.7202977 | 1003.841547 | ENSG00000111276 |
| DIP2B | 0.317978855 | 1.246582924 | 0.00011897 | 0.000814851 | 1127.052153 | 1404.207034 | ENSG00000066084 |
| RASSF8 | 0.317943397 | 1.246552285 | 0.009543144 | 0.035391363 | 370.6018587 | 461.6144919 | ENSG00000123094 |
| FSIP2 | 0.317769836 | 1.24640233 | 0.59075449 | 0.752666784 | 13.12104362 | 16.26314121 | ENSG00000188738 |
| NA | 0.317719933 | 1.246359218 | 0.584617441 | 0.747777753 | 14.03984734 | 17.57679285 | ENSG00000178464 |
| SALL1 | 0.317634604 | 1.246285503 | 0.000419458 | 0.002454763 | 1058.991107 | 1320.10595 | ENSG00000103449 |
| NA | 0.317618386 | 1.246271494 | 0.412562136 | 0.604174038 | 29.1495039 | 36.30424712 | ENSG00000185495 |
| NA | 0.31748622 | 1.246157327 | 0.681293204 | 0.815861488 | 7.529922509 | 9.470985467 | ENSG00000272438 |
| XBP1 | 0.317399058 | 1.246082042 | 2.60E-06 | 2.67E-05 | 2714.819191 | 3382.432382 | ENSG00000100219 |
| PRKY | 0.317356912 | 1.24604564 | 0.640745426 | 0.789444284 | 9.802905421 | 12.30670477 | ENSG00000099725 |
| CSRNP1 | 0.31731524 | 1.246009648 | 0.12008965 | 0.259723505 | 135.3623595 | 168.4050678 | ENSG00000144655 |
| CYP11A1 | 0.317247912 | 1.245951501 | 0.41841938 | 0.609668615 | 32.8790627 | 40.83422224 | ENSG00000140459 |
| GTF2A1 | 0.317053771 | 1.245783847 | 0.196734422 | 0.372106926 | 653.7991693 | 814.2789547 | ENSG00000165417 |
| MTUS2 | 0.316982286 | 1.24572212 | 0.594828585 | 0.755831139 | 14.5194196 | 17.96587078 | ENSG00000132938 |
| NA | 0.316921065 | 1.245669258 | 0.529621441 | 0.705324695 | 17.49834023 | 21.84485011 | ENSG00000248334 |
| NOD2 | 0.316903101 | 1.245653748 | 0.610657034 | 0.767784917 | 11.12295994 | 13.89692 | ENSG00000167207 |

| | | | | | | | |
|---|---|---|---|---|---|---|---|
| TIAM1 | 0.316803668 | 1.245567898 | 0.004709127 | 0.019525492 | 581.44111 | 724.1200145 | ENSG00000156299 |
| PSAP | 0.316649197 | 1.245434542 | 4.79E-08 | 6.81E-07 | 12584.47766 | 15671.9168 | ENSG00000197746 |
| TERF2 | 0.316552413 | 1.245350993 | 0.00072759 | 0.003986849 | 647.3175968 | 806.3309625 | ENSG00000132604 |
| CREB3L2 | 0.316491408 | 1.245298334 | 0.000499328 | 0.002861641 | 1078.235624 | 1342.728908 | ENSG00000182158 |
| MMP23B | 0.31625786 | 1.245096758 | 0.4381082 | 0.626620376 | 26.91883863 | 33.47232666 | ENSG00000189409 |
| PLOD1 | 0.316167925 | 1.245019143 | 4.43E-06 | 4.35E-05 | 1831.390021 | 2280.578504 | ENSG00000083444 |
| ARHGAP44 | 0.316129066 | 1.244985609 | 0.002562528 | 0.011692245 | 504.7111353 | 628.2184437 | ENSG00000006740 |
| PCDHGA10 | 0.316096342 | 1.24495737 | 0.028385609 | 0.085823636 | 243.2136892 | 302.9443112 | ENSG00000253846 |
| ARHGEF12 | 0.316095332 | 1.244956498 | 0.001866605 | 0.008941756 | 1012.377154 | 1259.733351 | ENSG00000196914 |
| SSR1 | 0.315917461 | 1.244803016 | 4.27E-05 | 0.000326826 | 1569.489016 | 1953.228466 | ENSG00000124783 |
| RBMS1 | 0.315851826 | 1.244746386 | 0.000539689 | 0.003070162 | 1083.023627 | 1347.536817 | ENSG00000153250 |
| ZSWIM4 | 0.315655059 | 1.244576628 | 0.006912123 | 0.027124558 | 401.7194859 | 499.7666755 | ENSG00000132003 |
| EEF1AKMT2 | 0.315427596 | 1.244380416 | 0.347275682 | 0.540692833 | 41.66472139 | 51.86536701 | ENSG00000203791 |
| ADRB1 | 0.315335093 | 1.244300632 | 0.754315358 | 0.863474302 | 4.082178128 | 5.078467662 | ENSG00000043591 |
| NA | 0.315245684 | 1.24422352 | 0.707000564 | 0.833400016 | 6.550461033 | 8.176654477 | ENSG00000073905 |
| NA | 0.315230204 | 1.24421017 | 0.693476481 | 0.82436945 | 6.441074104 | 8.011840788 | ENSG00000279278 |
| APAF1 | 0.315112273 | 1.244108468 | 0.009001632 | 0.033697483 | 365.1412534 | 454.005977 | ENSG00000120868 |
| LCORL | 0.31510122 | 1.244098937 | 0.074216921 | 0.181086943 | 184.947408 | 229.7446695 | ENSG00000178177 |
| NA | 0.315082162 | 1.244082502 | 0.653528038 | 0.797566245 | 9.448309256 | 11.66870576 | ENSG00000286099 |
| CLN8 | 0.314887558 | 1.2439147 | 0.000376848 | 0.002235756 | 755.8241583 | 940.1730424 | ENSG00000182372 |
| ASPM | 0.314820748 | 1.243857097 | 0.000534416 | 0.003042971 | 1022.60743 | 1271.124897 | ENSG00000066279 |
| FBXO46 | 0.314741645 | 1.243788898 | 0.003295316 | 0.014419354 | 511.1541862 | 636.3439468 | ENSG00000177051 |
| ATP11A | 0.314559587 | 1.24363195 | 0.000437582 | 0.002545555 | 1487.042914 | 1848.738934 | ENSG00000068650 |
| TBC1D23 | 0.314508298 | 1.243587739 | 0.025917381 | 0.079673702 | 408.8439707 | 507.825774 | ENSG00000036054 |
| NSUN3 | 0.314324018 | 1.243428902 | 0.153597513 | 0.31102112 | 94.19930878 | 116.9229332 | ENSG00000178694 |
| KCTD12 | 0.314321632 | 1.243426845 | 0.033615311 | 0.098052592 | 252.9151826 | 313.9277913 | ENSG00000178695 |
| RBMS3 | 0.314269552 | 1.24338196 | 0.43194617 | 0.6220157 | 29.99955459 | 37.23703106 | ENSG00000144642 |
| PDHB | 0.314075683 | 1.243214886 | 3.61E-05 | 0.000283712 | 2120.728018 | 2636.158191 | ENSG00000168291 |
| OPN3 | 0.314071572 | 1.243211343 | 0.565742435 | 0.733403531 | 16.85503711 | 20.95555689 | ENSG00000054277 |
| RGS14 | 0.313955089 | 1.24311097 | 0.008834469 | 0.033192387 | 365.6273123 | 454.7406909 | ENSG00000169220 |
| GPBP1L1 | 0.313947979 | 1.243104844 | 1.66E-05 | 0.000140693 | 1736.824916 | 2158.809428 | ENSG00000159592 |
| ARHGEF1 | 0.31392089 | 1.243081503 | 8.22E-05 | 0.00058442 | 1291.409388 | 1604.933418 | ENSG00000076928 |
| SMIM10L2B | 0.313898205 | 1.243061957 | 0.092180042 | 0.213185038 | 122.5445296 | 152.2972075 | ENSG00000196972 |
| LOC101927468 | 0.313726794 | 1.242914274 | 0.275407304 | 0.466410725 | 58.11327625 | 72.35774968 | ENSG00000227733 |
| WSCD1 | 0.31341954 | 1.242649595 | 8.49E-06 | 7.71E-05 | 1795.37432 | 2230.585426 | ENSG00000179314 |
| RAB31 | 0.313411432 | 1.242642612 | 0.001804673 | 0.008695193 | 712.7713209 | 884.9390483 | ENSG00000168461 |
| ANK2 | 0.313145671 | 1.242413724 | 0.002370182 | 0.010963053 | 539.4463561 | 670.1061518 | ENSG00000145362 |
| FRS2 | 0.313129525 | 1.242399819 | 0.034590039 | 0.10026459 | 320.832567 | 397.8496162 | ENSG00000166225 |
| LATS1 | 0.313029668 | 1.242313829 | 0.016107338 | 0.054101741 | 320.0353719 | 397.1460978 | ENSG00000131023 |
| SH3KBP1 | 0.312894818 | 1.242197714 | 0.002185928 | 0.010264503 | 569.4419797 | 707.2240585 | ENSG00000147010 |
| DCP1A | 0.312862451 | 1.242169845 | 0.010237052 | 0.037481929 | 404.7926853 | 502.2346355 | ENSG00000272886 |
| JAG1 | 0.312813268 | 1.2421275 | 0.004266482 | 0.017997538 | 612.6932438 | 760.9135813 | ENSG00000101384 |
| RAP1GAP2 | 0.312742414 | 1.242066497 | 0.000801893 | 0.004349514 | 1517.134907 | 1885.077629 | ENSG00000132359 |
| SOX5 | 0.312664941 | 1.2419998 | 0.537527468 | 0.711203176 | 16.33936333 | 20.23542586 | ENSG00000134532 |
| EMC4 | 0.312552599 | 1.241903089 | 0.000421012 | 0.002463078 | 973.9518206 | 1209.401872 | ENSG00000128463 |
| RXYLT1 | 0.312287932 | 1.24167528 | 0.128321223 | 0.272615912 | 109.5499353 | 136.0675946 | ENSG00000118600 |
| NA | 0.312263713 | 1.241654435 | 0.106293703 | 0.237412328 | 123.033069 | 152.8301782 | ENSG00000240356 |
| SH3PXD2B | 0.312192123 | 1.241592823 | 9.54E-06 | 8.57E-05 | 2429.003208 | 3016.269625 | ENSG00000174705 |
| NPIPB11 | 0.312187929 | 1.241589214 | 0.542697969 | 0.715337754 | 16.54097715 | 20.62983998 | ENSG00000254206 |
| ELAC1 | 0.312143853 | 1.241551282 | 0.469345765 | 0.655182226 | 22.27117518 | 27.70735948 | ENSG00000141642 |
| NA | 0.311997712 | 1.241425522 | 0.397415167 | 0.589538394 | 30.2307607 | 37.54182709 | ENSG00000286358 |
| RBBP6 | 0.31192578 | 1.241363628 | 3.70E-05 | 0.0002893 | 1316.505599 | 1634.046715 | ENSG00000122257 |
| FBXO34 | 0.311823736 | 1.241275827 | 0.01402037 | 0.048407887 | 349.0889133 | 433.8613088 | ENSG00000178974 |
| TNFRSF10A | 0.31180674 | 1.241261204 | 0.061965451 | 0.157908383 | 192.334999 | 238.8928927 | ENSG00000104689 |
| LZIC | 0.311430803 | 1.240937798 | 0.022252572 | 0.070512767 | 324.4809406 | 402.465116 | ENSG00000162441 |
| FLCN | 0.311349568 | 1.240867927 | 0.009333215 | 0.034737642 | 387.4838349 | 480.7426757 | ENSG00000154803 |
| CNN2 | 0.311260101 | 1.240790978 | 2.07E-08 | 3.13E-07 | 8943.276705 | 11097.49622 | ENSG00000064666 |
| CRABP2 | 0.311214242 | 1.240751537 | 0.000133866 | 0.000900245 | 1426.405027 | 1768.86213 | ENSG00000143320 |
| CBL | 0.311160541 | 1.240705354 | 0.000141426 | 0.000942871 | 1678.392634 | 2081.561571 | ENSG00000110395 |
| OSBPL10 | 0.311089123 | 1.240643937 | 0.000129019 | 0.000873015 | 1375.676576 | 1705.620987 | ENSG00000144645 |
| NA | 0.310895466 | 1.240477413 | 0.73483631 | 0.851745268 | 4.780426548 | 5.96016319 | ENSG00000267096 |
| NA | 0.310822187 | 1.240414407 | 0.146020213 | 0.299973741 | 153.2079149 | 190.25887 | ENSG00000267383 |
| BARD1 | 0.310743002 | 1.240346326 | 0.022890194 | 0.072175375 | 328.5262204 | 406.8262722 | ENSG00000138376 |
| POU5F1P4 | 0.310487368 | 1.240126566 | 0.446604279 | 0.634882752 | 28.90113775 | 35.74870653 | ENSG00000237872 |
| RCAN3 | 0.310367253 | 1.24002332 | 0.028964208 | 0.08706173 | 244.4615407 | 302.8225502 | ENSG00000117602 |
| CCDC88C | 0.31026892 | 1.239938804 | 5.08E-05 | 0.000381401 | 1320.536853 | 1637.195464 | ENSG00000015133 |
| PIM3 | 0.310066809 | 1.23976511 | 0.000937033 | 0.004984781 | 1097.280234 | 1360.158726 | ENSG00000198355 |
| ADGRG6 | 0.310057553 | 1.239757156 | 0.16139894 | 0.322172101 | 86.68970322 | 107.4642203 | ENSG00000112414 |
| NA | 0.309951952 | 1.239666413 | 0.460547292 | 0.647189591 | 26.05625799 | 32.1553142 | ENSG00000237506 |
| NA | 0.309929731 | 1.239647319 | 0.740553821 | 0.855060088 | 5.438236462 | 6.683655036 | ENSG00000259834 |
| LARGE1 | 0.309912521 | 1.239632531 | 1.80E-05 | 0.000151637 | 1456.347862 | 1805.515088 | ENSG00000133424 |
| ZNF287 | 0.309804481 | 1.239539702 | 0.163756488 | 0.325720408 | 97.65602023 | 121.1662707 | ENSG00000141040 |
| CDKN3 | 0.309780863 | 1.23951941 | 0.005718405 | 0.023085366 | 485.6527779 | 601.3653117 | ENSG00000100526 |
| GAL3ST2 | 0.309690093 | 1.239441425 | 0.74534072 | 0.857749036 | 4.410143516 | 5.476130977 | ENSG00000154252 |

| | | | | | | | |
|---|---|---|---|---|---|---|---|
| SNORD104 | 0.309544394 | 1.23931626 | 0.422204172 | 0.613399085 | 39.04671074 | 48.68278667 | ENSG00000199753 |
| TRMO | 0.309473095 | 1.239255014 | 0.26652897 | 0.456479547 | 58.66820845 | 72.61075574 | ENSG00000136932 |
| PPP1R13L | 0.309457157 | 1.239241323 | 0.015492435 | 0.052435498 | 305.9991852 | 379.0589528 | ENSG00000104881 |
| SIX2 | 0.309394778 | 1.239187742 | 0.613362198 | 0.769307369 | 12.86014752 | 15.94804795 | ENSG00000170577 |
| VAMP7 | 0.309361525 | 1.23915918 | 0.04970351 | 0.133160574 | 346.5311545 | 428.9000212 | ENSG00000124333 |
| BPHL | 0.309126808 | 1.238957593 | 0.016409546 | 0.054917756 | 349.196609 | 432.4184101 | ENSG00000137274 |
| PCDH17 | 0.309075172 | 1.23891325 | 0.555517082 | 0.725307064 | 17.11721099 | 21.16226107 | ENSG00000118946 |
| TRIP11 | 0.30903537 | 1.238879071 | 0.009544711 | 0.035391363 | 402.0157618 | 497.5755652 | ENSG00000100815 |
| ADARB1 | 0.309033004 | 1.238877039 | 0.023109148 | 0.072640009 | 323.5931046 | 401.4771586 | ENSG00000197381 |
| ARHGAP40 | 0.308928878 | 1.238787627 | 0.697728273 | 0.827575567 | 6.441074104 | 7.963013879 | ENSG00000124143 |
| SNHG12 | 0.308927945 | 1.238786826 | 0.371895299 | 0.565064432 | 307.779119 | 381.4111669 | ENSG00000197989 |
| ITPKC | 0.308892007 | 1.238755967 | 0.009811431 | 0.036173676 | 344.9035644 | 427.0387305 | ENSG00000086544 |
| EPS15 | 0.308745522 | 1.238630196 | 0.000396643 | 0.002335255 | 752.1709441 | 931.7469659 | ENSG00000085832 |
| WSCD2 | 0.30852373 | 1.23843979 | 0.31498325 | 0.508572137 | 48.19033308 | 59.55105286 | ENSG00000075035 |
| KIAA0232 | 0.308484868 | 1.238406431 | 0.002000606 | 0.009497754 | 638.4740465 | 790.1856315 | ENSG00000170871 |
| CCDC57 | 0.308410236 | 1.238342368 | 0.212206061 | 0.391891902 | 90.73341967 | 112.7093654 | ENSG00000176155 |
| TFF3 | 0.308296855 | 1.238245051 | 0.609862912 | 0.767202824 | 11.15580684 | 13.82552325 | ENSG00000160180 |
| SBF1 | 0.30829662 | 1.23824485 | 1.19E-06 | 1.31E-05 | 3114.972428 | 3857.70378 | ENSG00000100241 |
| GCC1 | 0.3081988 | 1.238160895 | 0.001072699 | 0.005603552 | 705.0460984 | 872.5661084 | ENSG00000179562 |
| FAN1 | 0.308145576 | 1.238115217 | 0.009820327 | 0.036199273 | 652.6195377 | 808.0323553 | ENSG00000198690 |
| NRAV | 0.30804297 | 1.238027164 | 0.118224709 | 0.256818654 | 118.5374184 | 146.6036371 | ENSG00000248008 |
| COX2 | 0.308021608 | 1.238008833 | 0.114682547 | 0.251198653 | 27471.38662 | 34009.88914 | ENSG00000198712 |
| RAB30 | 0.307985608 | 1.237977941 | 0.1922335 | 0.365533567 | 80.86925513 | 100.1420166 | ENSG00000137502 |
| SLC25A40 | 0.307906138 | 1.23790975 | 0.098531594 | 0.224099221 | 135.7908424 | 168.0306272 | ENSG00000075303 |
| PPIP5K1 | 0.307740121 | 1.237767306 | 0.089889416 | 0.209342168 | 170.7864557 | 211.4002731 | ENSG00000168781 |
| SARS1 | 0.30767312 | 1.237709824 | 3.78E-05 | 0.000294289 | 3693.810794 | 4571.117694 | ENSG00000031698 |
| FADS3 | 0.307619143 | 1.237663517 | 0.011351833 | 0.040877187 | 708.3969933 | 877.6105242 | ENSG00000221968 |
| NUDT9 | 0.307615372 | 1.237660282 | 0.000457638 | 0.002650585 | 851.5760055 | 1053.376482 | ENSG00000170502 |
| ZNF273 | 0.307532027 | 1.237588784 | 0.045841729 | 0.125081969 | 231.5802643 | 286.4345104 | ENSG00000198039 |
| NA | 0.307517369 | 1.23757621 | 0.685169519 | 0.818704911 | 9.488845461 | 11.77395727 | ENSG00000285761 |
| SPOPL | 0.307503106 | 1.237563975 | 0.117851373 | 0.256127738 | 124.8928725 | 154.5959424 | ENSG00000144228 |
| UPK1A-AS1 | 0.307387414 | 1.237464736 | 0.734125589 | 0.851336652 | 5.964658755 | 7.468134778 | ENSG00000226510 |
| GALNT3 | 0.307273369 | 1.237366919 | 0.042027345 | 0.117027465 | 347.0839895 | 429.0110469 | ENSG00000115339 |
| ZNF429 | 0.307039762 | 1.237166576 | 0.277425249 | 0.468543792 | 64.58729495 | 79.6390103 | ENSG00000197013 |
| TRIM16 | 0.306931438 | 1.237073687 | 0.113538585 | 0.249238912 | 113.7976257 | 140.8764351 | ENSG00000221926 |
| NA | 0.306889601 | 1.237037814 | 0.441858383 | 0.63014799 | 25.75808029 | 31.82524879 | ENSG00000287064 |
| TNFAIP1 | 0.306869751 | 1.237020794 | 0.008157359 | 0.031134133 | 412.4247864 | 510.0485094 | ENSG00000109079 |
| STAG3 | 0.30678483 | 1.236947981 | 0.32886762 | 0.523240833 | 49.82068928 | 61.6675172 | ENSG00000066923 |
| GTF2H3 | 0.306704325 | 1.236878959 | 0.010515627 | 0.038335293 | 422.0282503 | 521.5521464 | ENSG00000111358 |
| TPP1 | 0.306614129 | 1.236801632 | 0.015715135 | 0.053034288 | 341.1515239 | 421.8580457 | ENSG00000166340 |
| EPHA6 | 0.306601065 | 1.236790433 | 0.475073614 | 0.659978814 | 22.6711482 | 28.06748081 | ENSG00000080224 |
| DGCR8 | 0.306578255 | 1.236770878 | 0.008884616 | 0.033340247 | 1044.770503 | 1293.114829 | ENSG00000128191 |
| LMO4 | 0.306577982 | 1.236770645 | 1.33E-05 | 0.000115747 | 1608.94369 | 1989.911408 | ENSG00000143013 |
| NA | 0.306537597 | 1.236736025 | 0.244149681 | 0.43088177 | 85.208278 | 105.2708485 | ENSG00000234648 |
| EIF5A2 | 0.30645093 | 1.236661733 | 0.21937742 | 0.40078276 | 80.18609222 | 98.88423992 | ENSG00000163577 |
| BTBD1 | 0.30627225 | 1.236508579 | 0.016366097 | 0.054792132 | 692.7509999 | 856.3442444 | ENSG00000064726 |
| NA | 0.306257296 | 1.236495763 | 0.360396516 | 0.553627965 | 48.57452154 | 60.22801958 | ENSG00000246451 |
| ABCC10 | 0.306120013 | 1.236378107 | 0.009212885 | 0.034383246 | 443.826791 | 549.1597424 | ENSG00000124574 |
| TTC5 | 0.306084615 | 1.236347771 | 0.030766562 | 0.091382606 | 253.496242 | 313.4080808 | ENSG00000136319 |
| NA | 0.306070508 | 1.236335682 | 0.316915362 | 0.510624126 | 47.34806939 | 58.46700165 | ENSG00000265415 |
| ZNF192P1 | 0.30604512 | 1.236313926 | 0.61711469 | 0.772310229 | 12.41836051 | 15.42366258 | ENSG00000226314 |
| FXYD5 | 0.305946984 | 1.236229831 | 0.000425323 | 0.002484379 | 837.147535 | 1034.620501 | ENSG00000089327 |
| NA | 0.305691284 | 1.236010744 | 0.749835394 | 0.86068305 | 4.714732738 | 5.851224451 | ENSG00000235545 |
| HS3ST4 | 0.30565166 | 1.235976797 | 0.174676692 | 0.34175634 | 86.1823201 | 106.6319014 | ENSG00000182601 |
| GAS2L3 | 0.305636387 | 1.235963713 | 0.08254136 | 0.196231601 | 153.3596194 | 189.2723207 | ENSG00000139354 |
| APH1B | 0.305622778 | 1.235952053 | 0.014550975 | 0.04979464 | 329.9182826 | 407.2783134 | ENSG00000138613 |
| SCAMP1-AS1 | 0.305617223 | 1.235947295 | 0.282221345 | 0.473151553 | 58.063367 | 71.87751631 | ENSG00000245556 |
| SLC25A25 | 0.305437381 | 1.235793235 | 0.002725663 | 0.012279397 | 499.9526118 | 617.7517513 | ENSG00000148339 |
| ZRANB1 | 0.305366613 | 1.235732618 | 0.05328235 | 0.140394523 | 206.1169344 | 254.5696729 | ENSG00000019995 |
| ZNF250 | 0.305193262 | 1.235584143 | 0.045959798 | 0.125285909 | 193.6341578 | 239.2447635 | ENSG00000196150 |
| FIGN | 0.305125543 | 1.235526147 | 0.029939911 | 0.089457732 | 245.0783108 | 302.6054372 | ENSG00000182263 |
| PTPRN | 0.305014119 | 1.235430727 | 0.139709029 | 0.290580983 | 112.1210959 | 138.2152771 | ENSG00000054356 |
| ADRA1A | 0.304985675 | 1.235406371 | 0.651201629 | 0.796243605 | 8.889137755 | 10.98231787 | ENSG00000120907 |
| THAP8 | 0.304918809 | 1.235349113 | 0.156441655 | 0.315059181 | 125.1943048 | 154.8788299 | ENSG00000161277 |
| TUBB2B | 0.304877888 | 1.235314074 | 7.72E-07 | 8.78E-06 | 10163.10068 | 12554.46872 | ENSG00000137285 |
| RFX7 | 0.304817822 | 1.235262643 | 0.001656653 | 0.00808436 | 671.0559761 | 828.811613 | ENSG00000181827 |
| GRK6 | 0.304402007 | 1.234906666 | 8.70E-06 | 7.88E-05 | 1770.008606 | 2185.775559 | ENSG00000198055 |
| TMEM175 | 0.304326852 | 1.234842336 | 0.22372766 | 0.406128153 | 68.3118177 | 84.43821483 | ENSG00000127419 |
| NUDCD1 | 0.304169039 | 1.234707267 | 0.000699546 | 0.003854157 | 1278.487161 | 1578.255548 | ENSG00000120526 |
| NFIB | 0.304163177 | 1.234702251 | 0.347861528 | 0.541104957 | 39.43080151 | 48.64002012 | ENSG00000147862 |
| NA | 0.304163062 | 1.234702153 | 0.42283024 | 0.613865692 | 29.41681153 | 36.32301811 | ENSG00000259802 |
| CACNG6 | 0.304161304 | 1.234700648 | 0.51171496 | 0.690859926 | 20.09357607 | 24.80035505 | ENSG00000130433 |
| CHRFAM7A | 0.304159739 | 1.234699309 | 0.576274621 | 0.74109588 | 13.91793045 | 17.22371956 | ENSG00000166664 |
| CENPJ | 0.304153402 | 1.234693885 | 0.003230524 | 0.014169553 | 576.9263073 | 712.2216051 | ENSG00000151849 |

| | | | | | | | |
|---|---|---|---|---|---|---|---|
| MPG | 0.304141929 | 1.234684066 | 0.013558585 | 0.047108777 | 365.1603903 | 450.8296803 | ENSG00000103152 |
| FRS3 | 0.303931899 | 1.234504332 | 0.260630919 | 0.44960289 | 70.9854727 | 87.73115963 | ENSG00000137218 |
| LRRC8C-DT | 0.303911847 | 1.234487174 | 0.822683354 | 0.904849165 | 3.374361271 | 4.11084127 | ENSG00000231999 |
| NA | 0.303893879 | 1.234471799 | 0.428807953 | 0.61915913 | 35.69986724 | 44.0236761 | ENSG00000257135 |
| ZNF862 | 0.303771439 | 1.234367035 | 0.274249569 | 0.465250018 | 63.24820126 | 78.17891821 | ENSG00000106479 |
| FAM135A | 0.303764288 | 1.234360917 | 0.007781867 | 0.029941691 | 508.7220275 | 627.9671579 | ENSG00000082269 |
| RPL7A | 0.303752818 | 1.234351103 | 2.88E-08 | 4.26E-07 | 24539.95157 | 30290.30026 | ENSG00000148303 |
| SLC22A15 | 0.303608548 | 1.234227673 | 0.114036437 | 0.250065229 | 123.8475218 | 152.6338829 | ENSG00000163393 |
| NA | 0.303268359 | 1.233936676 | 0.758603404 | 0.865494336 | 4.70082731 | 5.817481309 | ENSG00000287780 |
| NA | 0.303047758 | 1.23374801 | 0.732301071 | 0.850725088 | 5.339695747 | 6.64655108 | ENSG00000249675 |
| ZNF251 | 0.303036612 | 1.233738479 | 0.045797357 | 0.125007598 | 216.899474 | 267.7586229 | ENSG00000198169 |
| PBXIP1 | 0.302708479 | 1.233457903 | 0.00137138 | 0.006860231 | 635.7590434 | 784.0551524 | ENSG00000163346 |
| ANXA7 | 0.302584259 | 1.233351704 | 0.000296677 | 0.001820129 | 954.8149538 | 1177.460765 | ENSG00000138279 |
| COTL1 | 0.302564781 | 1.233335052 | 2.02E-05 | 0.000168205 | 2721.447507 | 3355.935994 | ENSG00000103187 |
| ISYNA1 | 0.3023976 | 1.233192141 | 6.21E-06 | 5.88E-05 | 3799.505141 | 4686.223773 | ENSG00000105655 |
| NA | 0.302365126 | 1.233164383 | 0.093349691 | 0.215004743 | 143.7883109 | 177.5444908 | ENSG00000272168 |
| LRRC58 | 0.302329229 | 1.2331337 | 0.041217295 | 0.115221921 | 437.2514202 | 538.749457 | ENSG00000163428 |
| ZNF786 | 0.302169077 | 1.232996818 | 0.012386521 | 0.043895177 | 377.3330153 | 465.727572 | ENSG00000197362 |
| XPNPEP2 | 0.301883499 | 1.232752774 | 0.716724468 | 0.839556455 | 6.080261811 | 7.448925751 | ENSG00000122121 |
| MFF-DT | 0.3017371 | 1.232627685 | 0.648045509 | 0.794567224 | 9.474940033 | 11.71373382 | ENSG00000236432 |
| ZNF217 | 0.3016844 | 1.232582659 | 0.000353677 | 0.002113166 | 1198.707943 | 1476.641142 | ENSG00000171940 |
| SLC22A20P | 0.301662093 | 1.232563601 | 0.176817602 | 0.344852933 | 109.5875251 | 135.0373717 | ENSG00000197847 |
| LOC101926933 | 0.301616387 | 1.232524554 | 0.595324293 | 0.755953244 | 16.0568725 | 19.78568644 | ENSG00000257285 |
| DTWD2 | 0.301408963 | 1.232347359 | 0.154207882 | 0.311780178 | 138.4462324 | 170.4148549 | ENSG00000169570 |
| MARF1 | 0.301386424 | 1.232328107 | 0.005397661 | 0.0219917 | 447.6717348 | 551.376623 | ENSG00000166783 |
| HLA-DPA1 | 0.301244123 | 1.232206561 | 0.433627358 | 0.623446548 | 31.68348062 | 38.89539384 | ENSG00000231389 |
| RHOC | 0.301064204 | 1.232052902 | 0.013864022 | 0.047984272 | 414.311935 | 510.1048223 | ENSG00000155366 |
| KTI12 | 0.301046663 | 1.232037923 | 0.522546115 | 0.699420069 | 20.61811922 | 25.41903341 | ENSG00000198841 |
| NFKBIL1 | 0.301013775 | 1.232009837 | 0.106824987 | 0.238111163 | 135.4860578 | 167.069213 | ENSG00000204498 |
| LGALS8 | 0.300999122 | 1.231997324 | 0.100602584 | 0.227608434 | 144.6973508 | 178.053904 | ENSG00000116977 |
| CMYA5 | 0.300944539 | 1.231950713 | 0.534606793 | 0.709313645 | 18.23131469 | 22.45235518 | ENSG00000164309 |
| RECK | 0.300890563 | 1.231904623 | 0.035947861 | 0.103391938 | 254.0805711 | 313.0529608 | ENSG00000122707 |
| EIF4ENIF1 | 0.300422714 | 1.231505195 | 0.026702373 | 0.081740234 | 301.0051185 | 370.1894749 | ENSG00000184708 |
| NA | 0.300145209 | 1.231268336 | 0.788645496 | 0.884433126 | 4.466366588 | 5.491214744 | ENSG00000269696 |
| NA | 0.300076398 | 1.23120961 | 0.76653605 | 0.870835943 | 4.56175039 | 5.603840711 | ENSG00000273001 |
| RBPJ | 0.299814452 | 1.230986084 | 0.000213811 | 0.001362244 | 2266.678535 | 2788.790598 | ENSG00000168214 |
| AHCYL1 | 0.29977472 | 1.230952182 | 1.31E-06 | 1.43E-05 | 3474.924487 | 4276.716069 | ENSG00000168710 |
| NT5E | 0.299611786 | 1.23081317 | 0.613344733 | 0.769307369 | 12.00575983 | 14.81571948 | ENSG00000135318 |
| NA | 0.299494906 | 1.230713459 | 0.381083565 | 0.574027769 | 42.24599128 | 52.14445563 | ENSG00000244479 |
| DSC3 | 0.299408856 | 1.230640055 | 0.224166146 | 0.40676477 | 77.40196791 | 95.23975918 | ENSG00000134762 |
| NA | 0.299335121 | 1.23057716 | 0.380812168 | 0.573865576 | 53.97131217 | 66.56922498 | ENSG00000229152 |
| NQO2 | 0.299270179 | 1.230521768 | 0.000137036 | 0.000918209 | 1157.67086 | 1424.275433 | ENSG00000124588 |
| SMIM10 | 0.29907028 | 1.230351279 | 0.474043756 | 0.659208206 | 31.26319064 | 38.48407248 | ENSG00000184785 |
| PRR13 | 0.298972378 | 1.23026779 | 0.014082878 | 0.048596547 | 421.3215611 | 518.1897717 | ENSG00000205352 |
| PCSK1 | 0.298865611 | 1.230176747 | 0.473227651 | 0.658405892 | 25.59256799 | 31.514066 | ENSG00000175426 |
| AP3S2 | 0.298800971 | 1.23012163 | 0.048767639 | 0.131184241 | 209.7972603 | 257.9531203 | ENSG00000157823 |
| NA | 0.298776587 | 1.230100839 | 0.740991811 | 0.855139616 | 6.034787254 | 7.490704619 | ENSG00000268947 |
| PRDM5 | 0.29875249 | 1.230080293 | 0.029012431 | 0.087192539 | 298.2109221 | 366.4733826 | ENSG00000138738 |
| CALU | 0.298624142 | 1.229970865 | 2.47E-05 | 0.000201733 | 4470.062762 | 5496.827911 | ENSG00000128595 |
| SYTL5 | 0.298556207 | 1.229912949 | 0.70184273 | 0.830701395 | 8.365872375 | 10.239617 | ENSG00000147041 |
| DENND6A | 0.298540247 | 1.229899342 | 0.044621517 | 0.122436047 | 219.9862333 | 270.3639773 | ENSG00000174839 |
| ASIC4 | 0.298436498 | 1.229810899 | 0.72094192 | 0.842237882 | 6.127014144 | 7.478981666 | ENSG00000072182 |
| GJB5 | 0.298332146 | 1.229721949 | 0.682664673 | 0.816765029 | 8.539073977 | 10.46822974 | ENSG00000189280 |
| MALL | 0.298217344 | 1.229624098 | 0.5966931 | 0.75696458 | 14.12260349 | 17.36271421 | ENSG00000144063 |
| CRACD | 0.298191092 | 1.229601723 | 0.003295586 | 0.014419354 | 573.1536416 | 704.5962056 | ENSG00000109265 |
| AIP | 0.298177248 | 1.229589924 | 0.001765755 | 0.008538092 | 652.0403045 | 801.8007641 | ENSG00000110711 |
| CCNK | 0.298145804 | 1.229563125 | 0.012056657 | 0.042972707 | 511.6576309 | 628.8177301 | ENSG00000090061 |
| MARCHF3 | 0.298101353 | 1.229525241 | 0.085256779 | 0.200908983 | 156.9195017 | 193.1668218 | ENSG00000173926 |
| PPIE | 0.298031247 | 1.229465495 | 0.001677607 | 0.008173703 | 655.2464025 | 805.5259999 | ENSG00000084072 |
| NT5M | 0.298009336 | 1.229446823 | 0.06694837 | 0.167339325 | 184.8729425 | 227.659289 | ENSG00000205309 |
| MGAT3 | 0.297780562 | 1.22925188 | 0.00664509 | 0.026245288 | 425.051969 | 522.4734624 | ENSG00000128268 |
| NA | 0.297660355 | 1.229149462 | 0.104663273 | 0.234751463 | 186.2431168 | 229.1204091 | ENSG00000248605 |
| NA | 0.297649094 | 1.229139868 | 0.431596532 | 0.621587546 | 29.86363458 | 36.82582132 | ENSG00000235174 |
| HVCN1 | 0.297367774 | 1.228900213 | 0.18903912 | 0.361126681 | 90.18292216 | 110.5571826 | ENSG00000122986 |
| TVP23B | 0.297333377 | 1.228870914 | 0.064711107 | 0.163094745 | 231.5607215 | 284.654985 | ENSG00000171928 |
| C11orf86 | 0.29710001 | 1.228672151 | 0.643989688 | 0.791807408 | 11.06230218 | 13.69076548 | ENSG00000173237 |
| NA | 0.296987831 | 1.228576618 | 0.690152916 | 0.82232685 | 7.897048628 | 9.759710043 | ENSG00000217130 |
| PBLD | 0.296987669 | 1.22857648 | 0.444621133 | 0.632967822 | 34.36728277 | 42.40004415 | ENSG00000108187 |
| LOC100507283 | 0.296944023 | 1.228539312 | 0.512862609 | 0.691653613 | 20.7949814 | 25.52428492 | ENSG00000245498 |
| PITPNM2 | 0.296771777 | 1.228392643 | 0.081170214 | 0.193767226 | 155.7156516 | 191.2714842 | ENSG00000090975 |
| ZFPM1 | 0.296656358 | 1.228294373 | 0.121499504 | 0.262109784 | 129.9980098 | 159.5392143 | ENSG00000179588 |
| LINC01123 | 0.296569516 | 1.228220439 | 0.692121981 | 0.823657812 | 7.258180192 | 8.938126345 | ENSG00000204588 |
| SLC8A2 | 0.29653503 | 1.22819108 | 0.084464331 | 0.199614524 | 198.6599664 | 244.2757183 | ENSG00000118160 |
| TENM3 | 0.296252953 | 1.227950966 | 0.000837024 | 0.004518906 | 1823.460005 | 2238.501067 | ENSG00000218336 |

| | | | | | | | |
|---|---|---|---|---|---|---|---|
| ZNF41 | 0.296174926 | 1.227884556 | 0.187974185 | 0.359759763 | 95.24446406 | 116.9882696 | ENSG00000147124 |
| NA | 0.29603545 | 1.227765852 | 0.326221743 | 0.520441414 | 78.84010598 | 97.03909199 | ENSG00000237238 |
| DDI2 | 0.295971514 | 1.227711443 | 0.000852174 | 0.004590002 | 1043.996512 | 1281.061207 | ENSG00000197312 |
| MTRES1 | 0.29592655 | 1.227673179 | 0.073283279 | 0.179423024 | 172.7790974 | 212.0551793 | ENSG00000130349 |
| MED1 | 0.295868973 | 1.227624184 | 0.000227831 | 0.001441657 | 1359.859666 | 1668.551461 | ENSG00000125686 |
| ADCY9 | 0.295677649 | 1.227461394 | 0.008777255 | 0.033044414 | 419.1427593 | 514.3531209 | ENSG00000162104 |
| HDAC4 | 0.295558719 | 1.227360211 | 0.020487895 | 0.065914187 | 406.0540815 | 498.4269816 | ENSG00000068024 |
| ARFGEF1 | 0.295507295 | 1.227316463 | 2.15E-05 | 0.000178051 | 2420.520274 | 2969.734648 | ENSG00000066777 |
| HIPK1 | 0.295384343 | 1.22721187 | 0.001385673 | 0.006914735 | 1073.249314 | 1316.716903 | ENSG00000163349 |
| CNGA1 | 0.295349353 | 1.227182107 | 0.617348386 | 0.772320958 | 11.70117061 | 14.36543041 | ENSG00000198515 |
| CFLAR | 0.295347873 | 1.227180848 | 0.226590399 | 0.410079724 | 92.42286277 | 113.293313 | ENSG00000003402 |
| BORCS6 | 0.295331531 | 1.227166948 | 0.377828299 | 0.571087073 | 40.33194152 | 49.52540287 | ENSG00000196544 |
| IFFO2 | 0.295028996 | 1.226909636 | 0.021809766 | 0.069489807 | 325.8903507 | 399.4062317 | ENSG00000169991 |
| GAS2L1 | 0.294831335 | 1.226741551 | 0.036361909 | 0.104402044 | 280.7533135 | 344.3129928 | ENSG00000185340 |
| TUSC8 | 0.294696891 | 1.226627237 | 0.795575845 | 0.889390356 | 3.625884334 | 4.380794851 | ENSG00000237361 |
| ZNF304 | 0.294667999 | 1.226602673 | 0.055605246 | 0.144989819 | 225.9064022 | 276.6047177 | ENSG00000131845 |
| NA | 0.294656547 | 1.226592936 | 0.710673119 | 0.835868485 | 8.386189327 | 10.35972904 | ENSG00000198580 |
| LOC100130357 | 0.294462365 | 1.226427851 | 0.741153837 | 0.855273346 | 5.455298802 | 6.725433905 | ENSG00000215022 |
| POMZP3 | 0.29439424 | 1.22636994 | 0.608063411 | 0.766083016 | 13.21652512 | 16.30112123 | ENSG00000146707 |
| NIPAL1 | 0.294342315 | 1.226325802 | 0.203645209 | 0.381441162 | 103.1531557 | 126.4226605 | ENSG00000163293 |
| C22orf15 | 0.294184406 | 1.226191582 | 0.467347758 | 0.653229513 | 29.77012992 | 36.69106354 | ENSG00000169314 |
| TIMP3 | 0.294142685 | 1.226156123 | 0.176074042 | 0.343909401 | 95.21496947 | 116.6825975 | ENSG00000100234 |
| NCOA4 | 0.294130481 | 1.226145751 | 0.000129069 | 0.000873015 | 1633.184988 | 2001.583672 | ENSG00000266412 |
| SPATA5 | 0.294112307 | 1.226130305 | 0.050848235 | 0.135580663 | 239.942183 | 293.9517985 | ENSG00000145375 |
| SGPP1 | 0.294045177 | 1.226073253 | 0.060363087 | 0.154738034 | 215.2895325 | 263.915545 | ENSG00000126821 |
| PTPRE | 0.293853578 | 1.225910433 | 0.507441147 | 0.686841029 | 24.95941202 | 30.67035049 | ENSG00000132334 |
| AMN | 0.293679124 | 1.225762203 | 0.427259263 | 0.617692916 | 40.9682544 | 50.43434316 | ENSG00000166126 |
| MGST2 | 0.293672161 | 1.225756287 | 0.244382837 | 0.43120635 | 66.61506863 | 81.65121082 | ENSG00000085871 |
| BFSP1 | 0.29367145 | 1.225755683 | 0.305191436 | 0.499154436 | 58.87406156 | 71.88682586 | ENSG00000125864 |
| RAB3C | 0.293666185 | 1.225751209 | 0.065532084 | 0.164553807 | 227.6842237 | 278.541794 | ENSG00000152932 |
| ZFAND4 | 0.293499955 | 1.225609984 | 0.062259151 | 0.158482504 | 203.9269781 | 249.9375923 | ENSG00000172671 |
| GPATCH2L | 0.293466973 | 1.225581965 | 0.000762662 | 0.004159855 | 1203.900368 | 1475.56707 | ENSG00000089916 |
| NA | 0.293406263 | 1.225530392 | 0.779705495 | 0.878986213 | 4.144715025 | 5.048411747 | ENSG00000246575 |
| SEMG1 | 0.293320165 | 1.225457257 | 0.497970738 | 0.678931121 | 23.19125667 | 28.51776988 | ENSG00000124233 |
| CDK11A | 0.293134928 | 1.225299922 | 0.664899954 | 0.805385316 | 9.910916875 | 12.15653682 | ENSG00000008128 |
| RPS6KL1 | 0.293067497 | 1.225242654 | 0.127851279 | 0.271898521 | 163.3989624 | 200.5729097 | ENSG00000198208 |
| CAP1 | 0.293046823 | 1.225225096 | 3.17E-05 | 0.000252534 | 3712.853306 | 4548.866741 | ENSG00000131236 |
| RPL3L | 0.292978231 | 1.225166845 | 0.617214801 | 0.772310229 | 13.24937203 | 16.30871892 | ENSG00000140986 |
| LRATD1 | 0.292756253 | 1.224978351 | 0.143496226 | 0.296263679 | 110.5046451 | 135.2646703 | ENSG00000162981 |
| TBX6 | 0.292378071 | 1.224657282 | 0.459790577 | 0.646469319 | 37.67289101 | 46.20642425 | ENSG00000149922 |
| SPARC | 0.29234359 | 1.224628013 | 1.57E-05 | 0.000133988 | 3904.500993 | 4780.792037 | ENSG00000113140 |
| KCNH6 | 0.292300607 | 1.224591528 | 0.370154246 | 0.563439454 | 52.05451147 | 63.80165323 | ENSG00000173826 |
| UNC93B1 | 0.292256597 | 1.224554171 | 0.022894551 | 0.072176833 | 416.8083817 | 510.2846795 | ENSG00000110057 |
| RBKS | 0.2922399 | 1.224539999 | 0.588524177 | 0.750857513 | 15.4503473 | 18.98667258 | ENSG00000171174 |
| HJURP | 0.292124125 | 1.224441735 | 0.00038582 | 0.002282294 | 1382.581009 | 1693.156343 | ENSG00000123485 |
| EXOSC7 | 0.292015563 | 1.2243496 | 0.000287243 | 0.001770459 | 1164.17614 | 1425.81339 | ENSG00000075914 |
| SDC2 | 0.291982769 | 1.224321769 | 0.000992888 | 0.005233842 | 1258.083096 | 1539.080181 | ENSG00000169439 |
| MAP3K1 | 0.291932138 | 1.224278803 | 0.031807651 | 0.093873089 | 290.3555898 | 355.187248 | ENSG00000095015 |
| RIOK3 | 0.291748579 | 1.224123044 | 0.026504511 | 0.081222654 | 330.9460672 | 404.6847533 | ENSG00000101782 |
| PKP2 | 0.29162941 | 1.224021933 | 0.007325985 | 0.028423656 | 435.6199968 | 533.039346 | ENSG00000057294 |
| KDM1B | 0.291568243 | 1.223970039 | 0.076101401 | 0.184543159 | 196.8817616 | 241.2826798 | ENSG00000165097 |
| ITGB8 | 0.291530348 | 1.223937889 | 0.39808567 | 0.590012783 | 41.64114982 | 50.75517036 | ENSG00000105855 |
| PDPR | 0.291359723 | 1.223793145 | 0.000896095 | 0.00479729 | 899.1501472 | 1100.261394 | ENSG00000090857 |
| NR4A1 | 0.290916737 | 1.223417432 | 0.016160898 | 0.054203054 | 481.7446581 | 589.8952031 | ENSG00000123358 |
| NA | 0.290909817 | 1.223411563 | 0.406785159 | 0.598504099 | 39.57421542 | 48.45307462 | ENSG00000233762 |
| ARMCX2 | 0.290760203 | 1.223284697 | 1.90E-05 | 0.000159003 | 2678.598975 | 3276.096892 | ENSG00000184867 |
| FUT10 | 0.29072696 | 1.223256509 | 0.053217497 | 0.140263523 | 218.3351542 | 267.2522441 | ENSG00000172728 |
| CKMT1B | 0.290571436 | 1.223124648 | 0.129592508 | 0.274781781 | 156.8300636 | 191.771253 | ENSG00000237289 |
| DEPTOR | 0.290560354 | 1.223115253 | 0.478075041 | 0.662180914 | 34.71912798 | 42.47457847 | ENSG00000155792 |
| ATP5PD | 0.2903251 | 1.222915821 | 0.000148653 | 0.000985735 | 1328.631757 | 1624.365879 | ENSG00000167863 |
| YTHDC1 | 0.290284256 | 1.2228812 | 1.12E-05 | 9.88E-05 | 1948.125332 | 2382.292483 | ENSG00000083896 |
| NDRG3 | 0.289918473 | 1.222571188 | 0.002168474 | 0.010198047 | 616.4578218 | 753.6179779 | ENSG00000101079 |
| F3 | 0.289752407 | 1.222430468 | 0.219490311 | 0.40094948 | 76.17286251 | 93.18472077 | ENSG00000117525 |
| NA | 0.289490817 | 1.222208837 | 0.369802807 | 0.56299698 | 55.44219945 | 67.9999627 | ENSG00000261971 |
| SYT6 | 0.289473034 | 1.222193772 | 0.002668234 | 0.012080645 | 677.2484136 | 827.3145999 | ENSG00000134207 |
| ROBO1 | 0.289347771 | 1.222087658 | 0.000116208 | 0.000797408 | 2424.383821 | 2962.626221 | ENSG00000169855 |
| NA | 0.289337941 | 1.222079331 | 0.758378658 | 0.86529526 | 5.277158849 | 6.481299358 | ENSG00000224669 |
| NA | 0.28923471 | 1.22199189 | 0.752744948 | 0.862657174 | 4.803802715 | 5.862509371 | ENSG00000276603 |
| ZNF691 | 0.289192069 | 1.221955772 | 0.021833397 | 0.069541189 | 354.6956236 | 433.5257956 | ENSG00000164011 |
| SLFNL1 | 0.289141102 | 1.221912604 | 0.804712911 | 0.895189973 | 3.440152779 | 4.257321998 | ENSG00000171790 |
| AAAS | 0.289081046 | 1.22186174 | 0.000222523 | 0.001413378 | 1334.415856 | 1630.553413 | ENSG00000094914 |
| SH3GLB2 | 0.288990034 | 1.221784662 | 0.00151109 | 0.007464431 | 1309.688056 | 1600.380086 | ENSG00000148341 |
| NECAP1 | 0.288696431 | 1.221536042 | 0.05187398 | 0.137375051 | 212.8008349 | 260.1863359 | ENSG00000089818 |
| SYT7 | 0.28859819 | 1.221452863 | 0.005357398 | 0.021856484 | 525.7692002 | 641.6765828 | ENSG00000011347 |

| | | | | | | | |
|---|---|---|---|---|---|---|---|
| NA | 0.28858969 | 1.221445667 | 0.110765926 | 0.244630085 | 143.5628173 | 175.6013139 | ENSG00000197462 |
| ZBED1 | 0.288472633 | 1.221346566 | 0.000266457 | 0.001656109 | 1433.788273 | 1751.410929 | ENSG00000214717 |
| NME2 | 0.28835973 | 1.221250988 | 0.47781571 | 0.661871178 | 31.9007813 | 38.9226386 | ENSG00000243678 |
| NA | 0.288289651 | 1.221191667 | 0.307552068 | 0.501731365 | 53.18439974 | 64.90359936 | ENSG00000183199 |
| ABR | 0.288095763 | 1.221027559 | 7.39E-05 | 0.000531242 | 1677.563953 | 2047.86119 | ENSG00000159842 |
| TRIM6 | 0.287930371 | 1.220887587 | 0.051836977 | 0.137308351 | 212.3527341 | 259.0702534 | ENSG00000121236 |
| PIGX | 0.287752559 | 1.220737122 | 0.063673486 | 0.161241205 | 204.7977517 | 250.065302 | ENSG00000163964 |
| NOD1 | 0.287746541 | 1.220732031 | 0.224673491 | 0.407446035 | 82.79711262 | 100.9106481 | ENSG00000106100 |
| STX3 | 0.287441422 | 1.220473882 | 6.00E-05 | 0.000441693 | 1537.350166 | 1875.822496 | ENSG00000166900 |
| YBEY | 0.287365977 | 1.220410059 | 0.023246668 | 0.072988944 | 318.6388751 | 389.1309365 | ENSG00000182362 |
| MAP3K9 | 0.287290774 | 1.220346445 | 0.009545776 | 0.035391363 | 786.3988494 | 959.3958457 | ENSG00000006432 |
| NMRK2 | 0.28719368 | 1.220264318 | 0.00142685 | 0.007091775 | 797.2152582 | 973.2727922 | ENSG00000077009 |
| NA | 0.287070964 | 1.220160527 | 0.000662232 | 0.003672595 | 1151.194534 | 1403.994278 | ENSG00000196205 |
| PDE4A | 0.286713291 | 1.219858062 | 0.100541417 | 0.227525595 | 155.836193 | 190.068197 | ENSG00000065989 |
| PPM1F | 0.286637481 | 1.219793963 | 0.003302077 | 0.014440938 | 738.1275339 | 900.8772313 | ENSG00000100034 |
| IFT27 | 0.286366312 | 1.219564713 | 0.022319704 | 0.070701314 | 353.91392 | 431.6389317 | ENSG00000100360 |
| FAM227A | 0.286343591 | 1.219545505 | 0.457632622 | 0.644706827 | 36.68258332 | 44.9018992 | ENSG00000184949 |
| TUBA1C | 0.286326911 | 1.219531405 | 5.49E-06 | 5.25E-05 | 3417.211921 | 4167.206 | ENSG00000167553 |
| BNIP3L | 0.286256311 | 1.219471728 | 0.007000424 | 0.027425047 | 569.343514 | 693.592815 | ENSG00000104765 |
| SLC13A5 | 0.286224 | 1.219444417 | 0.471224675 | 0.656501394 | 27.49832708 | 33.40038025 | ENSG00000141485 |
| ARL5B | 0.286206413 | 1.219429551 | 0.024491575 | 0.076162225 | 960.264119 | 1170.225699 | ENSG00000165997 |
| WBP2 | 0.28610019 | 1.21933977 | 0.000129111 | 0.000873015 | 1505.074141 | 1834.428903 | ENSG00000132471 |
| SSH2 | 0.28598157 | 1.219239519 | 0.054116902 | 0.142136022 | 249.6617536 | 304.5832333 | ENSG00000141298 |
| LYRM2 | 0.285954714 | 1.219216822 | 0.046737757 | 0.126921259 | 243.5464953 | 296.818673 | ENSG00000083099 |
| SPAG9 | 0.285828231 | 1.219109937 | 0.00051024 | 0.002918763 | 1186.065471 | 1445.384327 | ENSG00000008294 |
| ASIC3 | 0.28577473 | 1.219064728 | 0.629463395 | 0.781306895 | 15.89254027 | 19.38477393 | ENSG00000213199 |
| ARL17A | 0.285615618 | 1.218930288 | 0.609206384 | 0.766741215 | 14.17882656 | 17.22371956 | ENSG00000228696 |
| NA | 0.285278602 | 1.218645577 | 0.270573785 | 0.46113237 | 76.80394395 | 93.22832311 | ENSG00000230551 |
| C19orf25 | 0.28524901 | 1.21862058 | 0.001805207 | 0.008695193 | 910.727221 | 1110.222377 | ENSG00000119559 |
| PI4K2B | 0.285228431 | 1.218603197 | 0.074443345 | 0.181486345 | 401.7860137 | 489.3185703 | ENSG00000038210 |
| TRIM69 | 0.285202868 | 1.218581605 | 0.129321565 | 0.274332825 | 128.9603484 | 156.9700877 | ENSG00000185880 |
| PKN3 | 0.285161288 | 1.218546485 | 0.001705782 | 0.00829136 | 930.3393534 | 1134.044013 | ENSG00000160447 |
| LYSMD3 | 0.285107944 | 1.218501429 | 0.1804765 | 0.350036777 | 186.455464 | 227.0659514 | ENSG00000176018 |
| NA | 0.285062116 | 1.218462724 | 0.699408863 | 0.828667383 | 7.917267881 | 9.673341145 | ENSG00000254910 |
| HCN2 | 0.284972285 | 1.218386858 | 0.207830523 | 0.386856574 | 96.24887258 | 117.4788002 | ENSG00000099822 |
| UBA7 | 0.284872407 | 1.218302512 | 0.736982241 | 0.852689235 | 6.017724914 | 7.385564727 | ENSG00000182179 |
| KRT8 | 0.284827798 | 1.218264841 | 2.94E-07 | 3.62E-06 | 8124.005333 | 9896.712404 | ENSG00000170421 |
| ANKRD53 | 0.284414165 | 1.217915604 | 0.694207772 | 0.824772234 | 7.618992485 | 9.28696275 | ENSG00000144031 |
| RSBN1 | 0.284400351 | 1.217903943 | 0.055118747 | 0.144065839 | 268.7497411 | 326.7958737 | ENSG00000081019 |
| GRB14 | 0.284376094 | 1.217883466 | 0.114804053 | 0.251421282 | 158.8163914 | 193.7530712 | ENSG00000115290 |
| STAU2 | 0.284319126 | 1.217835376 | 0.005309385 | 0.021703604 | 706.9789768 | 860.2346033 | ENSG00000040341 |
| ZZEF1 | 0.284291993 | 1.217812472 | 0.007826267 | 0.03008754 | 586.535679 | 714.652055 | ENSG00000074755 |
| PCAT6 | 0.284256575 | 1.217782575 | 0.321803746 | 0.51576536 | 62.75619675 | 76.67050859 | ENSG00000228288 |
| GNAQ | 0.284250308 | 1.217777285 | 0.000167078 | 0.001095377 | 1254.623115 | 1527.279498 | ENSG00000156052 |
| MICU1 | 0.284248344 | 1.217775627 | 0.002180185 | 0.010242723 | 782.5396919 | 952.3316175 | ENSG00000107745 |
| EPB41L5 | 0.284226981 | 1.217757595 | 0.012606207 | 0.044579934 | 512.9611866 | 624.5065798 | ENSG00000115109 |
| ALG1L2 | 0.284116472 | 1.21766432 | 0.775083825 | 0.875992658 | 4.492899667 | 5.438588988 | ENSG00000251287 |
| GPR157 | 0.283991573 | 1.217558907 | 0.160591908 | 0.321106692 | 112.9154272 | 137.3533403 | ENSG00000180758 |
| LYPLA1 | 0.283968098 | 1.217539095 | 0.00790288 | 0.030312908 | 1842.157745 | 2241.995636 | ENSG00000120992 |
| NA | 0.283632062 | 1.217255536 | 0.805628685 | 0.895843063 | 3.176002064 | 3.900999518 | ENSG00000225536 |
| NA | 0.283476275 | 1.2171241 | 0.578500548 | 0.742824523 | 18.00632471 | 21.7992724 | ENSG00000276570 |
| CERS6 | 0.283462415 | 1.217112407 | 0.010321441 | 0.037717208 | 451.8631045 | 549.8071626 | ENSG00000172292 |
| IPCEF1 | 0.283154077 | 1.216852309 | 0.742523786 | 0.856105567 | 7.189231771 | 8.716999673 | ENSG00000074706 |
| RAB33B | 0.282899026 | 1.216637203 | 0.063870524 | 0.161541525 | 219.1464275 | 266.3316924 | ENSG00000172007 |
| CD55 | 0.282814817 | 1.216566191 | 0.033524432 | 0.097818298 | 300.6655927 | 366.0283097 | ENSG00000196352 |
| NA | 0.282711145 | 1.216478772 | 0.733988125 | 0.851336652 | 5.991191834 | 7.344223892 | ENSG00000278434 |
| FGFR1OP2 | 0.282694845 | 1.216465028 | 0.011782547 | 0.042133459 | 423.0116654 | 514.6191725 | ENSG00000111790 |
| TMEM134 | 0.28265036 | 1.21642752 | 0.025690795 | 0.079121501 | 339.3265215 | 413.0389724 | ENSG00000172663 |
| MRPL2 | 0.28261719 | 1.216399552 | 0.588082667 | 0.750562088 | 19.37332695 | 23.65987924 | ENSG00000112651 |
| SMAD1 | 0.282459566 | 1.21626666 | 0.056986726 | 0.147905333 | 235.4529439 | 285.9095754 | ENSG00000170365 |
| RPP14 | 0.282434688 | 1.216245686 | 0.079517483 | 0.190804899 | 196.5809306 | 238.9541751 | ENSG00000163684 |
| KRT18 | 0.282362528 | 1.216184854 | 6.38E-05 | 0.00046738 | 3123.238595 | 3798.944435 | ENSG00000111057 |
| ORC5 | 0.282151696 | 1.216007137 | 0.022036624 | 0.070044039 | 359.2803442 | 436.4209336 | ENSG00000164815 |
| DAD1 | 0.282019732 | 1.215895913 | 0.000133988 | 0.000900414 | 1324.610087 | 1610.131734 | ENSG00000129562 |
| PIGP | 0.281996063 | 1.215875965 | 0.222858482 | 0.404946971 | 84.45094269 | 102.5947183 | ENSG00000185808 |
| GNA13 | 0.281839936 | 1.215744392 | 0.00290648 | 0.012952318 | 1272.71424 | 1546.871723 | ENSG00000120063 |
| BICD1 | 0.281558066 | 1.215506886 | 0.003985343 | 0.016976406 | 661.6925952 | 804.9304326 | ENSG00000151746 |
| NA | 0.281324507 | 1.215310122 | 0.584980584 | 0.747954646 | 16.07531031 | 19.5903788 | ENSG00000234176 |
| ADGRV1 | 0.281244144 | 1.215242427 | 0.023767499 | 0.074359693 | 845.1436956 | 1027.530647 | ENSG00000164199 |
| CYC1 | 0.281160178 | 1.215171701 | 2.35E-06 | 2.43E-05 | 4570.167739 | 5553.147474 | ENSG00000179091 |
| NA | 0.281050853 | 1.215079621 | 0.677044495 | 0.81334863 | 10.0499938 | 12.216099 | ENSG00000219201 |
| ZNF330 | 0.281032262 | 1.215063963 | 0.001609851 | 0.007903838 | 968.2400005 | 1176.160245 | ENSG00000109445 |
| NA | 0.280847565 | 1.214908418 | 0.814798208 | 0.900075694 | 3.235382048 | 3.957312501 | ENSG00000286156 |
| LOC440173 | 0.280827124 | 1.214891205 | 0.420940182 | 0.612090485 | 36.92208009 | 44.78036144 | ENSG00000269994 |

| | | | | | | | |
|---|---|---|---|---|---|---|---|
| WDR47 | 0.280778659 | 1.214850393 | 0.115466384 | 0.252513742 | 142.709512 | 173.1810412 | ENSG00000085433 |
| NA | 0.280681932 | 1.214768944 | 0.773206144 | 0.874965662 | 4.691356572 | 5.685972729 | ENSG00000277406 |
| TMCC1 | 0.280643134 | 1.214736277 | 0.014248497 | 0.048967486 | 382.0904716 | 464.1953816 | ENSG00000172765 |
| JMJD7-PLA2G4B | 0.280614319 | 1.214712015 | 0.381665903 | 0.574586291 | 66.87468696 | 81.42457344 | ENSG00000168970 |
| SECISBP2L | 0.280545875 | 1.214654389 | 0.022870378 | 0.072137437 | 438.4267982 | 532.1111882 | ENSG00000138593 |
| ACRV1 | 0.280536382 | 1.214646396 | 0.698030109 | 0.827774653 | 8.187732422 | 9.977149488 | ENSG00000134940 |
| NA | 0.280397848 | 1.214529765 | 0.684548185 | 0.818173473 | 9.385772358 | 11.33864035 | ENSG00000226121 |
| KCTD7 | 0.280382327 | 1.214516699 | 0.051527525 | 0.136838199 | 256.8711069 | 312.3387156 | ENSG00000243335 |
| TFB1M | 0.280319215 | 1.21446357 | 0.03927235 | 0.110838089 | 327.9271891 | 397.6746084 | ENSG00000029639 |
| BOK | 0.280226553 | 1.21438557 | 0.000540352 | 0.003072993 | 1306.328133 | 1587.348295 | ENSG00000176720 |
| IGF2BP2 | 0.280090535 | 1.214271083 | 9.00E-06 | 8.13E-05 | 2873.396852 | 3489.529912 | ENSG00000073792 |
| RAD54L2 | 0.280088773 | 1.214269599 | 0.007737521 | 0.029795804 | 775.4849069 | 941.3623283 | ENSG00000164080 |
| PLXNA1 | 0.280040178 | 1.214228699 | 7.81E-06 | 7.20E-05 | 3628.178824 | 4405.87529 | ENSG00000114554 |
| TAZ | 0.279901072 | 1.214111628 | 0.084191925 | 0.199057678 | 347.4213657 | 422.128183 | ENSG00000102125 |
| SWT1 | 0.279819401 | 1.214042899 | 0.208027266 | 0.387067296 | 102.223408 | 124.148256 | ENSG00000116668 |
| KDM4B | 0.279808127 | 1.214033412 | 0.000444487 | 0.002582484 | 1051.418423 | 1276.276737 | ENSG00000127663 |
| COL5A1 | 0.279717204 | 1.213956903 | 0.003962597 | 0.016900279 | 687.275828 | 834.0132355 | ENSG00000130635 |
| TMC6 | 0.279277673 | 1.213587115 | 0.000555682 | 0.003149546 | 1351.204883 | 1640.774045 | ENSG00000141524 |
| NBR1 | 0.27911158 | 1.213447406 | 0.008357436 | 0.031760657 | 896.1315773 | 1086.193925 | ENSG00000188554 |
| NA | 0.278912648 | 1.213280096 | 0.739967843 | 0.854707161 | 6.93149258 | 8.488386927 | ENSG00000261812 |
| SLC25A21-AS1 | 0.278873922 | 1.213247529 | 0.709896563 | 0.835062275 | 7.598773232 | 9.211878773 | ENSG00000258708 |
| CCDC61 | 0.278837235 | 1.213216677 | 0.029129867 | 0.087474554 | 361.4801167 | 438.6338896 | ENSG00000104983 |
| ACTC1 | 0.278225028 | 1.212701959 | 0.086968699 | 0.204061003 | 243.3316373 | 295.6083378 | ENSG00000159251 |
| SLC30A7 | 0.278169054 | 1.212654908 | 0.097349671 | 0.222065083 | 205.6570777 | 249.0459889 | ENSG00000162695 |
| NA | 0.277969382 | 1.212487086 | 0.807111374 | 0.896619594 | 3.454058207 | 4.231064929 | ENSG00000285877 |
| NA | 0.277832004 | 1.212371635 | 0.590260413 | 0.752140701 | 15.93307648 | 19.3279113 | ENSG00000230629 |
| PEG10 | 0.277805897 | 1.212349696 | 0.000293276 | 0.001801056 | 2635.248417 | 3194.466576 | ENSG00000242265 |
| MRPS27 | 0.277797627 | 1.212342747 | 0.000268944 | 0.001670447 | 1717.058118 | 2080.760527 | ENSG00000113048 |
| OIP5-AS1 | 0.277748878 | 1.212301782 | 0.013002086 | 0.045579772 | 1455.305518 | 1763.358991 | ENSG00000247556 |
| GATA2 | 0.277680417 | 1.212244255 | 0.702586215 | 0.831159372 | 8.306492391 | 10.08977545 | ENSG00000179348 |
| NA | 0.27766801 | 1.212233831 | 0.662084487 | 0.803380688 | 11.72454678 | 14.2757007 | ENSG00000251689 |
| S100A4 | 0.277559764 | 1.212142879 | 0.207153153 | 0.385828213 | 89.9017091 | 108.8703012 | ENSG00000196154 |
| MAP3K5 | 0.27746981 | 1.212067303 | 0.27745265 | 0.468547373 | 83.76965891 | 101.3413933 | ENSG00000197442 |
| RPP25L | 0.277320285 | 1.211941687 | 0.020504435 | 0.065942947 | 378.9102076 | 459.2587198 | ENSG00000164967 |
| SVOPL | 0.277297681 | 1.211922699 | 0.467098907 | 0.653162956 | 31.55712904 | 38.11883821 | ENSG00000157703 |
| SLC47A1 | 0.277208447 | 1.211847741 | 0.409549858 | 0.601140181 | 38.39955165 | 46.49817478 | ENSG00000142494 |
| INA | 0.277205296 | 1.211845094 | 0.026706066 | 0.081740234 | 389.3592875 | 471.8776664 | ENSG00000148798 |
| LHX8 | 0.276993035 | 1.211666811 | 0.692647492 | 0.823967294 | 8.921984661 | 10.85884502 | ENSG00000162624 |
| NA | 0.276907981 | 1.211595379 | 0.584918695 | 0.747954646 | 17.38589409 | 21.15097615 | ENSG00000188599 |
| HAGH | 0.276250126 | 1.21104303 | 0.015163368 | 0.051522031 | 432.8337229 | 524.4158035 | ENSG00000063854 |
| TRRAP | 0.275910722 | 1.210758157 | 0.007109704 | 0.027770548 | 2692.421829 | 3259.651825 | ENSG00000196367 |
| DCAF4 | 0.275780658 | 1.210649007 | 0.046235927 | 0.125720292 | 319.5620384 | 387.2793531 | ENSG00000119599 |
| SDHAF4 | 0.27573862 | 1.210613731 | 0.112521076 | 0.247591787 | 169.4299914 | 205.3084897 | ENSG00000154079 |
| ETV3 | 0.275710032 | 1.210589743 | 0.011247901 | 0.040550204 | 693.8626157 | 839.5310967 | ENSG00000117036 |
| ZBTB24 | 0.275696473 | 1.210578365 | 0.015107964 | 0.051415901 | 443.0537387 | 536.0475883 | ENSG00000112365 |
| RPL3 | 0.275686235 | 1.210569774 | 5.89E-07 | 6.85E-06 | 52770.86578 | 63881.86827 | ENSG00000100316 |
| RAPGEF1 | 0.275443291 | 1.210365937 | 3.87E-05 | 0.000301237 | 3137.953397 | 3798.431375 | ENSG00000107263 |
| VPS35L | 0.27534477 | 1.210283284 | 0.001347719 | 0.006761947 | 942.109014 | 1140.185195 | ENSG00000103544 |
| KIAA0319L | 0.275318987 | 1.210261654 | 0.000453459 | 0.002629663 | 1481.770281 | 1792.784955 | ENSG00000142687 |
| ITCH | 0.27524582 | 1.210200277 | 0.002994934 | 0.013301725 | 678.591942 | 821.0300424 | ENSG00000078747 |
| SBK3 | 0.275073357 | 1.210055616 | 0.68904587 | 0.821592842 | 10.82468454 | 12.99970267 | ENSG00000231274 |
| SLC41A3 | 0.275000263 | 1.20999431 | 0.01019844 | 0.037384867 | 500.7322635 | 606.0474789 | ENSG00000114544 |
| NA | 0.274993551 | 1.20998868 | 0.100808311 | 0.22793004 | 187.6953331 | 226.9152656 | ENSG00000244198 |
| CHMP1B2P | 0.274940851 | 1.209944481 | 0.374273572 | 0.567389377 | 51.14577985 | 61.9942218 | ENSG00000278530 |
| CMTM4 | 0.274918084 | 1.209925388 | 0.004661509 | 0.019345597 | 1201.431176 | 1453.095792 | ENSG00000183723 |
| RIC3 | 0.274757276 | 1.209790533 | 0.133628894 | 0.280929892 | 183.0861539 | 221.6707189 | ENSG00000166405 |
| PIKFYVE | 0.274620352 | 1.209675719 | 0.009214502 | 0.034383246 | 1028.714435 | 1243.767671 | ENSG00000115020 |
| IL23A | 0.274588663 | 1.209649148 | 0.567989052 | 0.735133316 | 19.31690849 | 23.24724377 | ENSG00000110944 |
| AJM1 | 0.274373125 | 1.209468441 | 0.37356558 | 0.566909396 | 50.76928069 | 61.63789932 | ENSG00000232434 |
| HINT3 | 0.274278681 | 1.209389267 | 0.11520731 | 0.252095901 | 220.8307443 | 266.8402296 | ENSG00000111911 |
| NA | 0.27401908 | 1.209171667 | 0.643369537 | 0.791307386 | 12.94162589 | 15.76121407 | ENSG00000179066 |
| CEP63 | 0.27395353 | 1.209116729 | 0.026416727 | 0.080978721 | 318.4858927 | 384.9544324 | ENSG00000182923 |
| CLUAP1 | 0.273902332 | 1.209073821 | 0.225634065 | 0.408668207 | 90.51710367 | 109.2491935 | ENSG00000103351 |
| ZCCHC14 | 0.273872626 | 1.209048926 | 0.00343803 | 0.014936815 | 814.7000013 | 984.5191375 | ENSG00000140948 |
| PHACTR4 | 0.273411621 | 1.208662643 | 0.001247311 | 0.006337499 | 934.1090651 | 1128.753965 | ENSG00000204138 |
| CTC-338M12.4 | 0.273407783 | 1.208659427 | 0.479603412 | 0.663554019 | 39.96975375 | 48.4504867 | ENSG00000233937 |
| RBBP8NL | 0.273407565 | 1.208659245 | 0.625583898 | 0.778047089 | 13.87561281 | 16.8034864 | ENSG00000130701 |
| KCNA6 | 0.273371132 | 1.208628722 | 0.371161613 | 0.564401422 | 48.71497393 | 59.04152802 | ENSG00000130035 |
| REEP6 | 0.273351111 | 1.20861195 | 0.013420815 | 0.046720772 | 448.1459628 | 541.9154254 | ENSG00000115255 |
| PCK2 | 0.27327571 | 1.208548784 | 0.000655161 | 0.003638821 | 1901.959608 | 2297.546964 | ENSG00000100889 |
| NA | 0.273150726 | 1.208444089 | 0.581103352 | 0.745134186 | 17.5469717 | 21.26740096 | ENSG00000276248 |
| ZBTB7A | 0.272916225 | 1.20824768 | 0.020189714 | 0.065122896 | 443.0694331 | 535.3633503 | ENSG00000178951 |
| TSPAN11 | 0.272892229 | 1.208227584 | 0.117435137 | 0.255552756 | 161.8298427 | 195.2618785 | ENSG00000110900 |
| KNDC1 | 0.272849554 | 1.208191845 | 0.013098309 | 0.045865117 | 448.8317941 | 542.2236939 | ENSG00000171798 |

| | | | | | | | |
|---|---|---|---|---|---|---|---|
| GPSM3 | 0.272811134 | 1.208159671 | 0.247263617 | 0.435088187 | 80.38948747 | 96.9462876 | ENSG00000213654 |
| ERMARD | 0.272707141 | 1.208072587 | 0.176408642 | 0.34438148 | 124.1192641 | 150.2511925 | ENSG00000130023 |
| NA | 0.27268383 | 1.208053067 | 0.740810999 | 0.855075943 | 6.616154843 | 8.053181623 | ENSG00000265399 |
| GYG1 | 0.272471109 | 1.207874956 | 0.010574568 | 0.038512291 | 542.8252498 | 655.0125144 | ENSG00000163754 |
| PRKAA2 | 0.272352269 | 1.207775463 | 0.092096489 | 0.213071568 | 219.8896468 | 265.1407408 | ENSG00000162409 |
| COL26A1 | 0.272311019 | 1.20774093 | 2.75E-05 | 0.000221248 | 2750.975817 | 3322.718262 | ENSG00000160963 |
| PLAA | 0.272286416 | 1.207720334 | 0.00328305 | 0.014371293 | 1310.50759 | 1581.86701 | ENSG00000137055 |
| NR1H3 | 0.272192331 | 1.207641576 | 0.038422736 | 0.108921227 | 294.0068271 | 355.2956858 | ENSG00000025434 |
| NA | 0.272059848 | 1.207530683 | 0.702588426 | 0.831159372 | 8.548544716 | 10.34844411 | ENSG00000226453 |
| NA | 0.272030122 | 1.207505803 | 0.177985642 | 0.346656777 | 123.8558125 | 149.8494039 | ENSG00000279457 |
| CACNA2D1 | 0.271913074 | 1.20740784 | 0.07850927 | 0.188914428 | 332.2216319 | 400.3913779 | ENSG00000153956 |
| RANBP2 | 0.271883272 | 1.207382898 | 0.001361747 | 0.006817562 | 3483.789009 | 4205.321712 | ENSG00000153201 |
| TEAD3 | 0.271803806 | 1.207316396 | 0.006173994 | 0.024677337 | 737.8348959 | 891.1483876 | ENSG00000007866 |
| SLC9A8 | 0.271728474 | 1.207253356 | 0.093320432 | 0.215004287 | 196.4403805 | 237.0390074 | ENSG00000197818 |
| SGTB | 0.271461111 | 1.207029646 | 0.055587589 | 0.144964172 | 236.4432516 | 285.4698471 | ENSG00000197860 |
| VPS4B | 0.271325488 | 1.206916183 | 0.008283919 | 0.031526521 | 604.898697 | 729.961634 | ENSG00000119541 |
| RAPGEF3 | 0.271276074 | 1.206874845 | 0.128626264 | 0.273107936 | 251.8303856 | 304.5439165 | ENSG00000079337 |
| RHBDL1 | 0.271218708 | 1.206826857 | 0.326271039 | 0.520449904 | 60.60681445 | 73.14471417 | ENSG00000103269 |
| KIAA0040 | 0.27115638 | 1.206774721 | 0.026679959 | 0.081706272 | 373.9720936 | 450.7685807 | ENSG00000235750 |
| TDGF1P3 | 0.271048614 | 1.20668458 | 0.007821871 | 0.030076877 | 503.3669305 | 607.4483352 | ENSG00000225366 |
| NA | 0.270946999 | 1.206599592 | 0.6027549 | 0.761880879 | 17.86744319 | 21.6420564 | ENSG00000264964 |
| ZNF543 | 0.2709386 | 1.206592567 | 0.284582854 | 0.475647815 | 77.35333644 | 93.49645322 | ENSG00000178229 |
| TMEM178A | 0.270728541 | 1.206416897 | 0.035318768 | 0.10193056 | 350.6106793 | 423.5538791 | ENSG00000152154 |
| SALL4 | 0.270618002 | 1.206324466 | 3.41E-05 | 0.000268916 | 3042.845717 | 3670.010172 | ENSG00000101115 |
| ACTN1 | 0.27046613 | 1.206197484 | 1.02E-05 | 9.09E-05 | 7598.747026 | 9165.724886 | ENSG00000072110 |
| PHLPP1 | 0.270378238 | 1.206124001 | 0.000509212 | 0.002913779 | 1496.943684 | 1804.808964 | ENSG00000081913 |
| NANOG | 0.27035515 | 1.206104699 | 0.000492091 | 0.002823661 | 1570.292488 | 1893.524689 | ENSG00000111704 |
| SH3RF2 | 0.270150909 | 1.205933964 | 0.65858426 | 0.800984361 | 13.94436584 | 16.80392444 | ENSG00000156463 |
| LINC01023 | 0.270148335 | 1.205931813 | 0.713021099 | 0.837541981 | 8.923262437 | 10.78744827 | ENSG00000272523 |
| NUDT16P1 | 0.270127942 | 1.205914767 | 0.209421227 | 0.388490944 | 112.891352 | 136.0075944 | ENSG00000246082 |
| TTC31 | 0.270004033 | 1.205811199 | 0.01925814 | 0.06247687 | 478.9301075 | 577.9857723 | ENSG00000115282 |
| GLG1 | 0.269973995 | 1.205786093 | 0.00012206 | 0.000832634 | 4926.178969 | 5939.06832 | ENSG00000090863 |
| FBXL3 | 0.269957285 | 1.205772127 | 0.335668966 | 0.52992237 | 94.45290637 | 113.661653 | ENSG00000005812 |
| GGNBP2 | 0.269841413 | 1.205675288 | 0.003146083 | 0.013870004 | 748.1408626 | 901.902436 | ENSG00000278311 |
| NDUFB1 | 0.269806139 | 1.205645809 | 0.038860637 | 0.110011157 | 425.9662177 | 514.119261 | ENSG00000183648 |
| SLC16A4 | 0.269792119 | 1.205634093 | 0.750872198 | 0.861339687 | 6.597213365 | 8.008041941 | ENSG00000168679 |
| NA | 0.269378762 | 1.205288707 | 0.831495186 | 0.909306807 | 3.575975088 | 4.241800197 | ENSG00000266717 |
| ARFIP1 | 0.269314382 | 1.205234923 | 0.014138019 | 0.048675945 | 684.0387244 | 824.2652082 | ENSG00000164144 |
| ATG4D | 0.269289905 | 1.205214475 | 0.007971047 | 0.030530149 | 576.4853019 | 694.7162804 | ENSG00000130734 |
| LOC101928053 | 0.269210735 | 1.205148339 | 0.82187723 | 0.904391889 | 3.397835135 | 4.129612264 | ENSG00000246308 |
| FLNA | 0.268915786 | 1.20490198 | 2.92E-06 | 2.97E-05 | 40923.53276 | 49308.8023 | ENSG00000196924 |
| RAB32 | 0.26890948 | 1.204896713 | 0.171716939 | 0.337462611 | 124.385294 | 149.7386014 | ENSG00000118508 |
| C6orf89 | 0.268752643 | 1.204765734 | 0.03118416 | 0.092356538 | 373.4203183 | 449.324623 | ENSG00000198663 |
| MIB2 | 0.268537008 | 1.204585676 | 0.002462417 | 0.011327487 | 1239.419803 | 1493.86187 | ENSG00000197530 |
| ZEB1-AS1 | 0.268219665 | 1.204320738 | 0.623021105 | 0.776326016 | 17.04718019 | 20.65620867 | ENSG00000237036 |
| CLU | 0.268120363 | 1.204237846 | 2.40E-05 | 0.000196295 | 3728.564782 | 4490.294542 | ENSG00000120885 |
| PKD1P6-NPIPP1 | 0.268055229 | 1.204183478 | 0.474126513 | 0.659273823 | 32.44939968 | 39.15493857 | ENSG00000270580 |
| NATD1 | 0.267744828 | 1.203924422 | 0.02297653 | 0.072367607 | 366.8758628 | 441.3809298 | ENSG00000274180 |
| CHCHD6 | 0.267736192 | 1.203917215 | 0.010112301 | 0.037091112 | 809.0183894 | 974.5231231 | ENSG00000159685 |
| TMOD3 | 0.267663073 | 1.203856199 | 0.005741328 | 0.023162773 | 697.3731302 | 838.8386569 | ENSG00000138594 |
| SIK1B | 0.267469724 | 1.203694871 | 0.268107141 | 0.458313339 | 75.79616796 | 91.10634807 | ENSG00000275993 |
| C8orf34 | 0.267362678 | 1.203605561 | 0.619130218 | 0.773695794 | 14.66924503 | 17.68518193 | ENSG00000165084 |
| XRCC4 | 0.267213645 | 1.203481233 | 0.128223383 | 0.272471013 | 159.8084805 | 191.9900779 | ENSG00000152422 |
| WNK1 | 0.267183771 | 1.203456313 | 0.001266636 | 0.006418214 | 3640.160215 | 4380.625279 | ENSG00000060237 |
| LOC100505549 | 0.26704209 | 1.203338132 | 0.635078082 | 0.785278349 | 13.93687193 | 16.77343049 | ENSG00000267040 |
| NA | 0.267030266 | 1.20332827 | 0.799194211 | 0.892074001 | 4.463209675 | 5.325851402 | ENSG00000232004 |
| SNAP23 | 0.266990282 | 1.203294921 | 0.06508549 | 0.163809677 | 330.2913317 | 397.2296958 | ENSG00000092531 |
| NA | 0.266863107 | 1.203188854 | 0.468100488 | 0.654035013 | 35.85166951 | 43.34141347 | ENSG00000248103 |
| ATF5 | 0.266679924 | 1.203036091 | 0.100581238 | 0.227587921 | 209.5623936 | 252.3878008 | ENSG00000169136 |
| GLTP | 0.266662342 | 1.20302143 | 0.015420762 | 0.052259718 | 481.4002469 | 578.7031148 | ENSG00000139433 |
| EEF1A1 | 0.266453158 | 1.20284701 | 1.87E-05 | 0.000156931 | 196781.4016 | 236696.4541 | ENSG00000156508 |
| SCRN2 | 0.266361626 | 1.202770697 | 0.013832113 | 0.047882776 | 508.7844592 | 611.6238601 | ENSG00000141295 |
| RASL11B | 0.266093008 | 1.202546772 | 0.517620131 | 0.695085604 | 32.26316446 | 38.97091585 | ENSG00000128045 |
| MATK | 0.266075131 | 1.202531871 | 0.00962417 | 0.035646355 | 644.4901476 | 775.5528757 | ENSG00000007264 |
| NDUFS4 | 0.266060921 | 1.202520027 | 0.015164168 | 0.051522031 | 531.1866911 | 638.7880547 | ENSG00000164258 |
| FOXL2 | 0.266044495 | 1.202506336 | 0.499304831 | 0.68011443 | 28.65730399 | 34.47369619 | ENSG00000183770 |
| MAPK12 | 0.266001256 | 1.202470296 | 0.069455996 | 0.171963232 | 547.0669573 | 658.338281 | ENSG00000188130 |
| SLC30A5 | 0.265943873 | 1.202422469 | 0.015919285 | 0.053635345 | 521.7747014 | 627.0620165 | ENSG00000145740 |
| SCRN3 | 0.265778987 | 1.202285051 | 0.192075446 | 0.365307985 | 114.7176095 | 137.619057 | ENSG00000144306 |
| NA | 0.265744969 | 1.202256702 | 0.382379888 | 0.575026598 | 50.01017911 | 59.92983356 | ENSG00000276291 |
| STON2 | 0.265698933 | 1.202218339 | 0.079513593 | 0.190804899 | 211.7042972 | 254.5599253 | ENSG00000140022 |
| COX15 | 0.265685369 | 1.202207037 | 0.051391694 | 0.136597397 | 282.6544425 | 339.3693945 | ENSG00000014919 |
| SOX18 | 0.265582481 | 1.202121302 | 0.765271165 | 0.870214234 | 5.694194214 | 6.848906758 | ENSG00000203883 |
| OSCP1 | 0.265576227 | 1.202116091 | 0.60125922 | 0.760560905 | 21.1186098 | 25.53980672 | ENSG00000116885 |

| | | | | | | | |
|---|---|---|---|---|---|---|---|
| DEPDC1 | 0.265494274 | 1.202047806 | 0.050905659 | 0.135694734 | 534.1029772 | 641.1869593 | ENSG00000024526 |
| MVB12A | 0.2654305 | 1.201994671 | 0.012833375 | 0.045167826 | 478.5623573 | 575.3656287 | ENSG00000141971 |
| NA | 0.265169597 | 1.201777317 | 0.565092472 | 0.733022388 | 20.16370457 | 24.23407919 | ENSG00000224261 |
| ANKRD29 | 0.265103037 | 1.201721873 | 0.386347861 | 0.579287376 | 56.98280926 | 68.67831486 | ENSG00000154065 |
| OSBPL3 | 0.265097037 | 1.201716876 | 0.03238918 | 0.095089666 | 380.1671692 | 456.7848337 | ENSG00000070882 |
| TRIM71 | 0.265078925 | 1.201701789 | 0.00084467 | 0.004553552 | 4133.705208 | 4966.577988 | ENSG00000206557 |
| ZSCAN16 | 0.265063936 | 1.201689303 | 0.422459531 | 0.613572983 | 40.38185077 | 48.55408925 | ENSG00000196812 |
| RNH1 | 0.264953224 | 1.20159709 | 0.002978605 | 0.013238721 | 1200.107423 | 1442.554319 | ENSG00000023191 |
| MCM9 | 0.264863404 | 1.201522283 | 0.140385033 | 0.291528986 | 143.8376188 | 172.9114141 | ENSG00000111877 |
| BSPRY | 0.264847407 | 1.20150896 | 0.132064092 | 0.278619907 | 160.1081314 | 192.528488 | ENSG00000119411 |
| TIGD2 | 0.264815573 | 1.201482449 | 0.246132592 | 0.433592002 | 116.3334816 | 140.0039863 | ENSG00000180346 |
| SOGA1 | 0.264710871 | 1.201395255 | 0.005538119 | 0.022479961 | 899.2488758 | 1079.803304 | ENSG00000149639 |
| HMX3 | 0.264647118 | 1.201342166 | 0.750575523 | 0.861278241 | 7.003500215 | 8.462129859 | ENSG00000188620 |
| CEP128 | 0.264592991 | 1.201297095 | 0.077792971 | 0.187736319 | 226.4669492 | 271.8052271 | ENSG00000100629 |
| ENTPD8 | 0.264576971 | 1.201283756 | 0.705721017 | 0.832939297 | 9.209511542 | 11.09125661 | ENSG00000188833 |
| FGF14-AS2 | 0.2645722 | 1.201279783 | 0.463973929 | 0.650328428 | 36.47969173 | 43.89260555 | ENSG00000272143 |
| FLOT1 | 0.264295584 | 1.201049477 | 5.17E-05 | 0.000387446 | 2909.122861 | 3494.294489 | ENSG00000137312 |
| KCTD11 | 0.264112559 | 1.200897118 | 0.10701279 | 0.23841508 | 171.5519688 | 205.8803879 | ENSG00000213859 |
| PLXNA2 | 0.264063696 | 1.200856445 | 0.366896201 | 0.559951761 | 62.09002112 | 74.74654073 | ENSG00000076356 |
| NA | 0.264054556 | 1.200848837 | 0.598923068 | 0.758900673 | 18.15171546 | 21.83367681 | ENSG00000223813 |
| SAPCD2 | 0.264033067 | 1.20083095 | 0.001682285 | 0.00819219 | 1085.093756 | 1303.608279 | ENSG00000186193 |
| ZFYVE16 | 0.263842299 | 1.200672175 | 0.029241587 | 0.087753168 | 380.2692501 | 456.0337707 | ENSG00000039319 |
| M6PR | 0.263803553 | 1.200639929 | 0.00120482 | 0.006165909 | 1369.285892 | 1644.106833 | ENSG00000003056 |
| CDC42SE1 | 0.26369903 | 1.200552946 | 0.000699306 | 0.003853981 | 1365.768989 | 1639.51461 | ENSG00000197622 |
| CCPG1 | 0.26369679 | 1.200551082 | 0.547571051 | 0.719152627 | 23.30685973 | 27.86413745 | ENSG00000260916 |
| TMEM86A | 0.263619355 | 1.200486646 | 0.544887036 | 0.716897589 | 27.28290554 | 32.81577144 | ENSG00000151117 |
| WWTR1 | 0.263588502 | 1.200460973 | 0.088233636 | 0.206354261 | 254.7076086 | 305.4349442 | ENSG00000018408 |
| ASXL2 | 0.263515443 | 1.200400182 | 0.014807864 | 0.050543084 | 798.01667 | 957.3923563 | ENSG00000143970 |
| NA | 0.263456815 | 1.200351402 | 0.817009582 | 0.901565698 | 3.477434373 | 4.201009014 | ENSG00000250982 |
| SYN2 | 0.26336156 | 1.20027215 | 0.146808373 | 0.301122884 | 162.7726239 | 195.1800814 | ENSG00000157152 |
| CDK5R1 | 0.263301771 | 1.200222409 | 0.051633819 | 0.137005182 | 321.0072645 | 385.1713222 | ENSG00000176749 |
| MYO1C | 0.263284202 | 1.200207792 | 0.00020782 | 0.001328642 | 1974.867038 | 2370.260166 | ENSG00000197879 |
| NA | 0.263200123 | 1.200137847 | 0.697985458 | 0.827774653 | 9.670142327 | 11.51094011 | ENSG00000220804 |
| CHROMR | 0.262954658 | 1.199933669 | 0.214765619 | 0.394926677 | 100.2809461 | 120.1634755 | ENSG00000223960 |
| ZNF415 | 0.262924583 | 1.199908655 | 0.1337489 | 0.281076629 | 159.3971576 | 191.0531687 | ENSG00000170954 |
| RSKR | 0.26287389 | 1.199866494 | 0.273721145 | 0.464894568 | 80.03636448 | 96.25372773 | ENSG00000167524 |
| TMEM30A | 0.262836911 | 1.19983574 | 0.005104485 | 0.020953827 | 1085.673943 | 1302.376679 | ENSG00000112697 |
| ZXDB | 0.262814118 | 1.199816784 | 0.104950446 | 0.235205751 | 188.8304528 | 226.3855525 | ENSG00000198455 |
| CCT4 | 0.262806483 | 1.199810434 | 1.19E-05 | 0.000105073 | 5100.431892 | 6118.401075 | ENSG00000115484 |
| CPPED1 | 0.262721492 | 1.199739754 | 0.071685869 | 0.176257084 | 240.742054 | 288.5246061 | ENSG00000103381 |
| EGFL7 | 0.262717664 | 1.19973657 | 0.005659266 | 0.022896492 | 895.8180884 | 1074.648901 | ENSG00000172889 |
| PIGBOS1 | 0.262608548 | 1.199645834 | 0.49626215 | 0.677862531 | 29.36818006 | 35.2389669 | ENSG00000225973 |
| UBAP1L | 0.262588718 | 1.199629345 | 0.641117265 | 0.789671608 | 13.68031282 | 16.37576718 | ENSG00000246922 |
| MCAM | 0.262492788 | 1.19954958 | 0.012203514 | 0.043354404 | 504.0517545 | 604.4208839 | ENSG00000076706 |
| HBP1 | 0.262423581 | 1.199492038 | 0.036493513 | 0.104590848 | 359.3804709 | 430.5261068 | ENSG00000105856 |
| BST2 | 0.262234883 | 1.19933516 | 0.002160699 | 0.01016922 | 958.8004856 | 1149.544431 | ENSG00000130303 |
| MEF2A | 0.262196282 | 1.199303071 | 0.089579167 | 0.208864044 | 288.9049444 | 346.0511858 | ENSG00000068305 |
| GTF2H2C | 0.262041938 | 1.199174772 | 0.105432017 | 0.235971139 | 246.7500976 | 295.7925435 | ENSG00000183474 |
| TOM1L2 | 0.261887955 | 1.199046788 | 0.291974698 | 0.484463657 | 81.94665596 | 98.07806641 | ENSG00000175662 |
| RNF185 | 0.261872023 | 1.199033547 | 0.01325834 | 0.046285568 | 542.1042266 | 649.5790383 | ENSG00000138942 |
| TJP3 | 0.261758679 | 1.198939349 | 0.04066927 | 0.113953779 | 337.6751492 | 404.5257138 | ENSG00000105289 |
| CARNMT1 | 0.261685705 | 1.198878706 | 0.064203221 | 0.162196811 | 522.9924569 | 626.4199636 | ENSG00000156017 |
| PTGR1 | 0.261638306 | 1.198839319 | 0.026023271 | 0.079946181 | 581.6355003 | 697.6375646 | ENSG00000106853 |
| ROCK2 | 0.261612006 | 1.198817464 | 0.022096008 | 0.070167013 | 600.2725232 | 719.0982088 | ENSG00000134318 |
| ESRRG | 0.261588417 | 1.198797863 | 0.494154515 | 0.676205048 | 30.36026919 | 36.31118354 | ENSG00000196482 |
| NA | 0.261473222 | 1.198702146 | 0.671696473 | 0.809284433 | 12.48217518 | 14.98477005 | ENSG00000267750 |
| ACACB | 0.26143935 | 1.198674003 | 0.041622217 | 0.116213644 | 393.5049273 | 471.4305552 | ENSG00000076555 |
| LINC00367 | 0.261359955 | 1.198608039 | 0.823046955 | 0.904942493 | 3.232225135 | 3.882228524 | ENSG00000233780 |
| NA | 0.261246785 | 1.198514019 | 0.137517401 | 0.286975096 | 175.2063404 | 210.0640516 | ENSG00000204959 |
| GBAP1 | 0.261231514 | 1.198501333 | 0.443534736 | 0.631972743 | 47.59387999 | 57.16661031 | ENSG00000160766 |
| SPTSSB | 0.261056562 | 1.198356003 | 0.136838545 | 0.28582955 | 149.5745366 | 179.1041005 | ENSG00000196542 |
| LRP8 | 0.261039173 | 1.198341559 | 0.006790073 | 0.026750263 | 1252.964692 | 1501.577801 | ENSG00000157193 |
| C6orf62 | 0.260910064 | 1.198234322 | 0.003009481 | 0.013356732 | 924.4499643 | 1107.580874 | ENSG00000112308 |
| AMFR | 0.260777479 | 1.198124208 | 0.027414018 | 0.083526452 | 523.429554 | 626.4922598 | ENSG00000159461 |
| KCNG1 | 0.260418701 | 1.197826289 | 0.017542353 | 0.057893512 | 619.0651664 | 741.8930793 | ENSG00000026559 |
| CEP170 | 0.260383765 | 1.197797283 | 0.002174435 | 0.010220891 | 962.622181 | 1152.910699 | ENSG00000143702 |
| TMEM179B | 0.260378058 | 1.197792544 | 0.458111321 | 0.645234077 | 42.91652654 | 51.39261972 | ENSG00000185475 |
| TRAP1 | 0.260365732 | 1.197782312 | 3.60E-05 | 0.000282911 | 3347.243928 | 4009.580952 | ENSG00000126602 |
| MYL12B | 0.26026622 | 1.197699695 | 0.000884449 | 0.004744535 | 2440.428096 | 2921.76023 | ENSG00000118680 |
| KDM7A-DT | 0.26013374 | 1.197589718 | 0.357656061 | 0.55102655 | 54.96065035 | 65.75155175 | ENSG00000260231 |
| NA | 0.260127275 | 1.197584351 | 0.707367746 | 0.833517498 | 9.974731549 | 11.98043821 | ENSG00000226525 |
| SGIP1 | 0.260100653 | 1.197562253 | 0.690523016 | 0.822503443 | 10.20603536 | 12.28044771 | ENSG00000118473 |
| NA | 0.26005938 | 1.197527993 | 0.75669083 | 0.864566204 | 5.709978779 | 6.830135764 | ENSG00000259553 |
| ZNF702P | 0.260036025 | 1.197508606 | 0.282782293 | 0.473749165 | 82.56580881 | 98.63729422 | ENSG00000242779 |

| | | | | | | | |
|---|---|---|---|---|---|---|---|
| PYCARD | 0.259982163 | 1.197463899 | 0.2561918 | 0.444990476 | 102.3257821 | 122.2970618 | ENSG00000103490 |
| AK7 | 0.259604883 | 1.197150791 | 0.360853907 | 0.554034205 | 64.4790881 | 76.92439072 | ENSG00000140057 |
| AP1M2 | 0.259402885 | 1.196983185 | 0.000185443 | 0.001201748 | 2344.275246 | 2805.683681 | ENSG00000129354 |
| ZNF430 | 0.259360357 | 1.1969479 | 0.185141796 | 0.355882395 | 133.077448 | 159.3290377 | ENSG00000118620 |
| TATDN1 | 0.259358905 | 1.196946696 | 0.031637137 | 0.093444256 | 340.9934078 | 408.0558567 | ENSG00000147687 |
| PCP4 | 0.259349637 | 1.196939006 | 0.667336018 | 0.807126452 | 14.55038737 | 17.28708058 | ENSG00000183036 |
| ZNF646 | 0.259332196 | 1.196924537 | 0.002262913 | 0.010545845 | 888.919127 | 1063.965414 | ENSG00000167395 |
| ALS2 | 0.259249736 | 1.196856125 | 0.004245441 | 0.017923199 | 1136.946218 | 1360.824186 | ENSG00000003393 |
| NA | 0.258996199 | 1.196645811 | 0.338174055 | 0.532203699 | 59.2696976 | 70.90890223 | ENSG00000230707 |
| SMC5 | 0.258942439 | 1.19660122 | 0.021448547 | 0.068521758 | 596.4931755 | 713.0772979 | ENSG00000198887 |
| PLEKHH3 | 0.258876298 | 1.196546363 | 0.011575931 | 0.041514766 | 532.2668655 | 636.696207 | ENSG00000068137 |
| NA | 0.258423771 | 1.196171103 | 0.713334907 | 0.837723303 | 9.018646239 | 10.77996219 | ENSG00000225218 |
| GVQW2 | 0.258102557 | 1.195904807 | 0.752002359 | 0.86205208 | 6.405070286 | 7.674289303 | ENSG00000279968 |
| NDUFV2 | 0.258012632 | 1.195830267 | 0.774354966 | 0.875489229 | 6.180080302 | 7.385564727 | ENSG00000178127 |
| RBM4B | 0.257840628 | 1.195687704 | 0.02521308 | 0.077947148 | 407.1812713 | 486.7256404 | ENSG00000173914 |
| SEC23A | 0.257780973 | 1.195638264 | 0.004333821 | 0.018217971 | 1069.956845 | 1278.361465 | ENSG00000100934 |
| BLZF1 | 0.257748357 | 1.195611233 | 0.134010184 | 0.28143548 | 175.3947479 | 209.4723545 | ENSG00000117475 |
| PRR3 | 0.257345394 | 1.19527733 | 0.13080189 | 0.276934227 | 161.1709504 | 192.4897435 | ENSG00000204576 |
| LOC285074 | 0.257336871 | 1.195270269 | 0.357968639 | 0.551282586 | 61.13017753 | 73.12170629 | ENSG00000285793 |
| ACTB | 0.257299901 | 1.19523964 | 7.33E-06 | 6.83E-05 | 92554.35189 | 110623.9409 | ENSG00000075624 |
| NA | 0.257216772 | 1.195170772 | 0.727656467 | 0.846909144 | 10.08086387 | 11.96871526 | ENSG00000234911 |
| NA | 0.257091866 | 1.1950673 | 0.75340174 | 0.863119536 | 6.020881826 | 7.197743164 | ENSG00000275155 |
| ZNF837 | 0.257046061 | 1.195029358 | 0.267501416 | 0.457663736 | 93.69971266 | 111.9149862 | ENSG00000152475 |
| CNTNAP3C | 0.257007655 | 1.194997546 | 0.596419067 | 0.756844835 | 19.58421612 | 23.32981383 | ENSG00000283378 |
| METTL4 | 0.256935368 | 1.194937671 | 0.13113927 | 0.277300084 | 156.8120916 | 187.5361744 | ENSG00000101574 |
| NA | 0.256862465 | 1.194877289 | 0.696161661 | 0.826246173 | 10.55294222 | 12.6668261 | ENSG00000268087 |
| NA | 0.256844737 | 1.194862606 | 0.747531857 | 0.858996555 | 7.397159414 | 8.769075776 | ENSG00000224765 |
| ETS2 | 0.256737648 | 1.194773917 | 0.004019456 | 0.01708246 | 741.9866012 | 886.5328878 | ENSG00000157557 |
| PAK2 | 0.256700699 | 1.194743318 | 0.007698298 | 0.029669423 | 1349.220487 | 1611.274128 | ENSG00000180370 |
| TMEM187 | 0.256411627 | 1.194503952 | 0.197631302 | 0.373422219 | 122.4914635 | 146.2470998 | ENSG00000177854 |
| NA | 0.256383952 | 1.194481038 | 0.777809282 | 0.877841751 | 5.723884207 | 6.82264969 | ENSG00000287686 |
| PHLDB3 | 0.256184564 | 1.194315966 | 0.132328925 | 0.278956357 | 161.2064506 | 192.5878354 | ENSG00000176531 |
| RANBP17 | 0.255711302 | 1.193924247 | 0.140995868 | 0.29240432 | 162.4832179 | 193.6601636 | ENSG00000204764 |
| NA | 0.255602664 | 1.193834345 | 0.648094755 | 0.794567224 | 18.29512937 | 21.73558496 | ENSG00000269983 |
| NA | 0.255469946 | 1.193724525 | 0.694696085 | 0.825246586 | 10.83731219 | 13.00437759 | ENSG00000236364 |
| ZNF581 | 0.255407317 | 1.193672706 | 0.009255541 | 0.034487776 | 736.2371534 | 878.7252346 | ENSG00000171425 |
| VEGFA | 0.255366314 | 1.193638781 | 0.029725961 | 0.088918814 | 883.7901551 | 1055.8721259 | ENSG00000112715 |
| PRKX | 0.255318537 | 1.193599252 | 0.000359166 | 0.002142513 | 2202.253046 | 2627.975945 | ENSG00000183943 |
| TCERG1L | 0.255175672 | 1.19348106 | 0.125637947 | 0.268547308 | 217.3872392 | 258.9440407 | ENSG00000176769 |
| ACBD3 | 0.255085147 | 1.193406175 | 0.009181212 | 0.034286639 | 712.1597445 | 849.5053318 | ENSG00000182827 |
| COG4 | 0.254977626 | 1.193317236 | 0.002760845 | 0.01240473 | 1090.47495 | 1301.307609 | ENSG00000103051 |
| YBX3 | 0.254884512 | 1.19324022 | 0.004928492 | 0.020321394 | 5139.756793 | 6132.544486 | ENSG00000060138 |
| CHST4 | 0.254759264 | 1.193136633 | 0.108483704 | 0.240766013 | 231.9100338 | 276.8950913 | ENSG00000140835 |
| NA | 0.25471062 | 1.193096404 | 0.718997311 | 0.840949657 | 11.54818826 | 13.84104505 | ENSG00000271833 |
| PTBP3 | 0.254529943 | 1.192946995 | 0.02048838 | 0.065914187 | 1479.209064 | 1763.565472 | ENSG00000119314 |
| TRMT9B | 0.254471448 | 1.192898628 | 0.518198848 | 0.695519378 | 40.16061906 | 47.90177936 | ENSG00000250305 |
| NA | 0.254311684 | 1.192766533 | 0.530602993 | 0.70614339 | 33.6815643 | 39.95643717 | ENSG00000237863 |
| SRPX | 0.25430778 | 1.192763306 | 0.20410458 | 0.3821194 | 132.1028272 | 157.2578246 | ENSG00000101955 |
| CDKAL1 | 0.254287748 | 1.192746744 | 0.053377102 | 0.140484408 | 370.7019855 | 441.5435448 | ENSG00000145996 |
| NA | 0.254176583 | 1.192654842 | 0.815630888 | 0.900673432 | 3.480591286 | 4.1709531 | ENSG00000231050 |
| NA | 0.254043993 | 1.192545237 | 0.823056321 | 0.904942493 | 3.450901294 | 4.133411111 | ENSG00000224358 |
| GPR39 | 0.25402644 | 1.192530727 | 0.758225999 | 0.86529526 | 6.050571819 | 7.209028084 | ENSG00000183840 |
| ILK | 0.254010364 | 1.192517439 | 0.775981282 | 0.876419078 | 5.848958001 | 6.954046651 | ENSG00000166333 |
| POGLUT3 | 0.253843486 | 1.192379508 | 0.083903832 | 0.19862927 | 312.0015163 | 371.7938979 | ENSG00000178202 |
| NEMP1 | 0.253705348 | 1.192265343 | 0.001334147 | 0.006702922 | 1024.449404 | 1221.480633 | ENSG00000166881 |
| ZNF101 | 0.253527945 | 1.192118743 | 0.008855332 | 0.03325729 | 603.1559849 | 718.6393915 | ENSG00000181896 |
| PPP1R3E | 0.253445206 | 1.192050376 | 0.47955674 | 0.663554019 | 47.63865548 | 56.976304 | ENSG00000235194 |
| RETSAT | 0.253427476 | 1.192035727 | 0.043400874 | 0.12002265 | 309.5948759 | 368.9462726 | ENSG00000042445 |
| ZNF776 | 0.253398946 | 1.192012154 | 0.086863636 | 0.203840292 | 259.2253576 | 309.4078272 | ENSG00000152443 |
| FBF1 | 0.253356392 | 1.191976995 | 0.460260164 | 0.646933254 | 49.47250464 | 59.13279507 | ENSG00000188878 |
| PRDM2 | 0.253177786 | 1.191829437 | 0.018738713 | 0.061091169 | 511.849992 | 610.0078174 | ENSG00000116731 |
| PRKCZ-AS1 | 0.252692833 | 1.191428878 | 0.378263864 | 0.57165219 | 52.94866124 | 62.9453387 | ENSG00000182873 |
| CCDC27 | 0.252662893 | 1.191404153 | 0.740063695 | 0.854707161 | 8.10172166 | 9.590771093 | ENSG00000162592 |
| HSD17B11 | 0.252477146 | 1.191250769 | 0.191779955 | 0.364858314 | 125.0084756 | 148.9660594 | ENSG00000198189 |
| NA | 0.252456887 | 1.191234042 | 0.000355652 | 0.002123599 | 8464.055823 | 10082.4292 | ENSG00000240342 |
| NHSL1 | 0.252398916 | 1.191186176 | 0.057220381 | 0.148420559 | 390.5183772 | 464.7760313 | ENSG00000135540 |
| MSN | 0.252293063 | 1.19109878 | 8.00E-06 | 7.35E-05 | 7626.505521 | 9083.001453 | ENSG00000147065 |
| HMMR | 0.252182463 | 1.191007471 | 0.010705909 | 0.038921795 | 807.478118 | 961.3322457 | ENSG00000072571 |
| LY6E | 0.252050934 | 1.190898893 | 0.000386743 | 0.002286412 | 1575.028153 | 1875.548203 | ENSG00000160932 |
| LRP1 | 0.251916688 | 1.190788083 | 0.201351735 | 0.378253909 | 5890.292406 | 7013.982097 | ENSG00000123384 |
| NA | 0.251816847 | 1.190705678 | 0.837089155 | 0.912409457 | 3.572818175 | 4.264808071 | ENSG00000233589 |
| CRLF3 | 0.2517826 | 1.190677412 | 0.074023306 | 0.180668552 | 320.5524438 | 381.5815139 | ENSG00000176390 |
| PGM3 | 0.251741138 | 1.190643193 | 0.007858851 | 0.03017484 | 618.8761954 | 736.60625 | ENSG00000013375 |
| NPIPB4 | 0.251686401 | 1.190598021 | 0.346515445 | 0.540053587 | 67.33057479 | 80.37229001 | ENSG00000185864 |

| | | | | | | | |
|---|---|---|---|---|---|---|---|
| CDK9 | 0.251588624 | 1.190517332 | 0.000343041 | 0.002058245 | 2141.59079 | 2549.722836 | ENSG00000136807 |
| MORC3 | 0.251554697 | 1.190489336 | 0.080620386 | 0.192832468 | 296.0610588 | 351.9826072 | ENSG00000159256 |
| ZNF518B | 0.251509821 | 1.190452305 | 0.075457673 | 0.183274384 | 397.6464627 | 473.5164627 | ENSG00000178163 |
| DECR1 | 0.251242078 | 1.190231395 | 0.051527368 | 0.136838199 | 388.0539503 | 461.2867996 | ENSG00000104325 |
| TMEM209 | 0.251238942 | 1.190228808 | 0.060660242 | 0.155286869 | 267.1839736 | 317.9015016 | ENSG00000146842 |
| RHEBL1 | 0.251132635 | 1.190141108 | 0.242512363 | 0.429008698 | 113.0087364 | 134.4332274 | ENSG00000167550 |
| E2F6 | 0.251095728 | 1.190110662 | 0.0373466 | 0.106456764 | 431.9382727 | 514.0935133 | ENSG00000169016 |
| PEX16 | 0.250798374 | 1.189865393 | 0.027130502 | 0.082771476 | 444.0432949 | 528.5105733 | ENSG00000121680 |
| FBLIM1 | 0.250574685 | 1.18968092 | 0.000204637 | 0.001310554 | 2223.033663 | 2645.293935 | ENSG00000162458 |
| ZNF620 | 0.250504591 | 1.18962312 | 0.080884652 | 0.193228076 | 233.7738586 | 277.93769 | ENSG00000177842 |
| RSPH3 | 0.250470418 | 1.189594942 | 0.426868627 | 0.617429303 | 43.62670355 | 51.86960389 | ENSG00000130363 |
| ZNF235 | 0.250431443 | 1.189562804 | 0.434231435 | 0.62397623 | 45.15774493 | 53.72696154 | ENSG00000159917 |
| NA | 0.250328291 | 1.189477754 | 0.468954338 | 0.654734441 | 39.95209005 | 47.46255197 | ENSG00000279483 |
| ULBP3 | 0.250269369 | 1.189429176 | 0.313247941 | 0.50683992 | 74.55137569 | 88.63075009 | ENSG00000131019 |
| CDR2 | 0.250242598 | 1.189407104 | 0.002124092 | 0.01001728 | 937.9612171 | 1115.704497 | ENSG00000140743 |
| NA | 0.250237551 | 1.189402943 | 0.768310689 | 0.871924202 | 6.391164859 | 7.599205325 | ENSG00000268670 |
| HOOK1 | 0.2502342 | 1.189400181 | 0.033103178 | 0.096878937 | 666.3068482 | 791.5664741 | ENSG00000134709 |
| ZNF669 | 0.250193276 | 1.189366442 | 0.111151667 | 0.24530662 | 214.2834402 | 254.7451187 | ENSG00000188295 |
| GID4 | 0.249940391 | 1.189157981 | 0.042073766 | 0.11709187 | 351.720228 | 418.0283313 | ENSG00000141034 |
| RASSF1 | 0.249716014 | 1.18897305 | 0.094362052 | 0.216878259 | 275.8538339 | 328.3627321 | ENSG00000068028 |
| DDX25 | 0.24969058 | 1.188952089 | 0.103562706 | 0.232658326 | 194.2339632 | 231.0793939 | ENSG00000109832 |
| LINC02453 | 0.249688329 | 1.188950233 | 0.822360289 | 0.904601199 | 3.427525127 | 4.092070275 | ENSG00000245017 |
| NA | 0.249683424 | 1.188946191 | 0.702963555 | 0.831263 | 12.32287901 | 14.62453711 | ENSG00000280152 |
| NA | 0.249487672 | 1.18878488 | 0.616913575 | 0.772278752 | 20.64021761 | 24.69609122 | ENSG00000278600 |
| BRMS1L | 0.249369944 | 1.188687876 | 0.202698694 | 0.380052362 | 142.8583527 | 169.9551447 | ENSG00000100916 |
| NA | 0.249362239 | 1.188681527 | 0.826725484 | 0.906810387 | 4.357077358 | 5.221149543 | ENSG00000250794 |
| RPL17 | 0.249319955 | 1.188646689 | 0.406094397 | 0.597630106 | 55.19393099 | 65.75590025 | ENSG00000265681 |
| EXOC1 | 0.249094739 | 1.188461147 | 0.073737196 | 0.180153598 | 243.4643178 | 289.2163301 | ENSG00000090989 |
| RYR3 | 0.249027952 | 1.18840613 | 0.5153396 | 0.693278953 | 43.99561111 | 52.40388874 | ENSG00000198838 |
| NA | 0.248862587 | 1.18826992 | 0.807954974 | 0.896899705 | 4.363391183 | 5.236233311 | ENSG00000242797 |
| NA | 0.248862587 | 1.18826992 | 0.807954974 | 0.896899705 | 4.363391183 | 5.236233311 | ENSG00000268938 |
| RPL13A | 0.248847554 | 1.188257538 | 0.000115749 | 0.000794849 | 46641.77049 | 55422.36607 | ENSG00000142541 |
| ZSCAN29 | 0.248822987 | 1.188237304 | 0.004496591 | 0.018791391 | 772.0585694 | 917.3956316 | ENSG00000140265 |
| OGT | 0.248775129 | 1.188197888 | 0.175970558 | 0.343743498 | 1992.171018 | 2367.403364 | ENSG00000147162 |
| BLOC1S2 | 0.248767478 | 1.188191587 | 0.010227541 | 0.037461905 | 717.1988047 | 852.0944318 | ENSG00000196072 |
| ANKRD6 | 0.248730066 | 1.188160775 | 0.163685622 | 0.325649347 | 161.6414579 | 191.8455725 | ENSG00000135299 |
| USP12 | 0.248653495 | 1.188097715 | 0.043500659 | 0.12011937 | 516.0716152 | 612.464349 | ENSG00000152484 |
| PIP4P2 | 0.248541165 | 1.188005212 | 0.058180754 | 0.150348408 | 357.5544913 | 425.246486 | ENSG00000155099 |
| PITRM1 | 0.24853309 | 1.187998563 | 0.007261446 | 0.0282146 | 659.7502461 | 783.5177215 | ENSG00000107959 |
| AOC1 | 0.24850972 | 1.187979318 | 0.824638131 | 0.905655553 | 3.886878136 | 4.636214318 | ENSG00000002726 |
| MAPK11 | 0.248487588 | 1.187961094 | 0.436941697 | 0.625978002 | 48.65431617 | 58.06316636 | ENSG00000185386 |
| FSTL3 | 0.248285245 | 1.18779449 | 0.145396502 | 0.299021169 | 206.5746787 | 245.8299289 | ENSG00000070404 |
| HAX1 | 0.24810795 | 1.187648529 | 0.001756934 | 0.008502094 | 1230.501434 | 1461.027548 | ENSG00000143575 |
| MAN1A2 | 0.247858535 | 1.187443224 | 0.057829432 | 0.149673327 | 413.3045271 | 490.2130712 | ENSG00000198162 |
| PLEKHB2 | 0.247814975 | 1.187407372 | 8.93E-05 | 0.000629771 | 2506.855343 | 2976.333226 | ENSG00000115762 |
| SYNE2 | 0.24775823 | 1.187360669 | 0.034859932 | 0.100904886 | 1324.634816 | 1573.056695 | ENSG00000054654 |
| PYROXD1 | 0.247749104 | 1.187353158 | 0.400007547 | 0.592009891 | 55.25449106 | 65.71868467 | ENSG00000121350 |
| PTK2 | 0.247713418 | 1.187323789 | 0.000164569 | 0.001081217 | 3083.68998 | 3660.188686 | ENSG00000169398 |
| COL20A1 | 0.247257542 | 1.186948666 | 0.785379891 | 0.882568409 | 7.984840828 | 9.512326302 | ENSG00000101203 |
| ITGA5 | 0.247250894 | 1.186943196 | 0.010501373 | 0.03829086 | 787.4470038 | 934.693001 | ENSG00000161638 |
| ZNF132 | 0.24723164 | 1.186927356 | 0.577381696 | 0.742025373 | 24.58143969 | 29.23421369 | ENSG00000131849 |
| AXL | 0.247031661 | 1.186762842 | 0.013155078 | 0.046019539 | 650.3351745 | 771.5609037 | ENSG00000167601 |
| RNF123 | 0.246775063 | 1.186551782 | 0.007962021 | 0.030501882 | 677.2240301 | 803.5740144 | ENSG00000164068 |
| NA | 0.246659599 | 1.186456822 | 0.530534393 | 0.706137998 | 29.41996844 | 34.9013038 | ENSG00000243679 |
| ATP10D | 0.246569236 | 1.186382511 | 0.315191727 | 0.508775777 | 75.65412952 | 89.93680403 | ENSG00000145246 |
| OSGIN2 | 0.24644467 | 1.18628008 | 0.123245785 | 0.264808413 | 233.0422822 | 276.2332083 | ENSG00000164823 |
| SENP7 | 0.24640729 | 1.186249345 | 0.290080615 | 0.482184226 | 82.72006893 | 98.04899818 | ENSG00000138468 |
| RGPD4 | 0.246397676 | 1.18624144 | 0.820346337 | 0.903565639 | 3.513438191 | 4.178439173 | ENSG00000196862 |
| TESMIN | 0.246267226 | 1.186134183 | 0.016791178 | 0.05594141 | 523.3309004 | 620.8474618 | ENSG00000132749 |
| ZNF446 | 0.246180595 | 1.18606296 | 0.543123592 | 0.715644295 | 30.93404517 | 36.7393408 | ENSG00000083838 |
| FBXO38 | 0.246143849 | 1.186032751 | 0.01395008 | 0.048230959 | 590.9936799 | 700.4727653 | ENSG00000145868 |
| PHF21B | 0.246141399 | 1.186030737 | 0.037931963 | 0.107678205 | 534.2146718 | 633.9182582 | ENSG00000056487 |
| PRSS23 | 0.24610439 | 1.186000313 | 0.172305916 | 0.338403926 | 144.6426009 | 171.7081269 | ENSG00000150687 |
| MPPED1 | 0.246015853 | 1.185927531 | 0.744951841 | 0.857520491 | 7.820606303 | 9.33578966 | ENSG00000186732 |
| GCC2 | 0.245889116 | 1.185823355 | 0.008820902 | 0.033161583 | 695.7928786 | 824.6884044 | ENSG00000135968 |
| BICDL2 | 0.245690819 | 1.185660376 | 0.084680743 | 0.199958324 | 277.0517761 | 328.2884839 | ENSG00000162069 |
| AURKA | 0.245478481 | 1.185485881 | 0.001153758 | 0.005953159 | 2052.263421 | 2432.177615 | ENSG00000087586 |
| CHID1 | 0.245410459 | 1.185429988 | 0.000571917 | 0.003224796 | 1671.861648 | 1982.087117 | ENSG00000177830 |
| AMPD3 | 0.244899794 | 1.18501046 | 0.253230182 | 0.441978359 | 109.8506685 | 130.197814 | ENSG00000133805 |
| ERVMER34-1 | 0.244763437 | 1.184898463 | 0.120636948 | 0.260653439 | 209.2348342 | 247.9282432 | ENSG00000226887 |
| FRRS1 | 0.244711226 | 1.184855583 | 0.372158631 | 0.565329733 | 64.27699327 | 76.09575898 | ENSG00000156869 |
| TKT | 0.244683214 | 1.184832578 | 6.21E-06 | 5.88E-05 | 22630.22617 | 26812.35793 | ENSG00000163931 |
| HPS3 | 0.244639861 | 1.184796974 | 0.00260313 | 0.011842517 | 1028.711075 | 1218.737335 | ENSG00000163755 |
| UBE2W | 0.244525239 | 1.184702846 | 0.57976237 | 0.743877865 | 146.368634 | 173.3240177 | ENSG00000104343 |

| | | | | | | | |
|---|---|---|---|---|---|---|---|
| METTL15 | 0.244521471 | 1.184699751 | 0.121037368 | 0.261335809 | 283.7734619 | 335.6474079 | ENSG00000169519 |
| ARL8A | 0.244427938 | 1.184622948 | 0.010652138 | 0.038749113 | 737.980745 | 874.4998641 | ENSG00000143862 |
| TEX45 | 0.244318671 | 1.184533229 | 0.741891573 | 0.855704472 | 7.906519366 | 9.422158558 | ENSG00000198723 |
| ANKRD19P | 0.244308668 | 1.184525017 | 0.824441654 | 0.905596573 | 3.483748199 | 4.133411111 | ENSG00000187984 |
| SLC12A6 | 0.244300516 | 1.184518323 | 0.117490756 | 0.255633748 | 237.9480682 | 281.3006058 | ENSG00000140199 |
| ABCC4 | 0.244289131 | 1.184508976 | 0.064889913 | 0.163450619 | 349.4358952 | 413.5330383 | ENSG00000125257 |
| CALB2 | 0.243988908 | 1.184262507 | 0.492459078 | 0.674850354 | 33.43645276 | 39.60467799 | ENSG00000172137 |
| PGAP6 | 0.243933387 | 1.184216932 | 5.61E-05 | 0.000417408 | 3354.809982 | 3972.967004 | ENSG00000129925 |
| MECP2 | 0.243804411 | 1.184111069 | 0.011223919 | 0.040487371 | 744.9192657 | 881.4533208 | ENSG00000169057 |
| COX7A2L | 0.243759353 | 1.184074087 | 0.001502779 | 0.007425356 | 1470.242875 | 1739.99879 | ENSG00000115944 |
| ZNF433 | 0.243758277 | 1.184073205 | 0.679984299 | 0.815136731 | 15.6708049 | 18.42689512 | ENSG00000197647 |
| RPS28 | 0.243712754 | 1.184035842 | 0.005281013 | 0.02160669 | 6447.796542 | 7635.256975 | ENSG00000233927 |
| LOC100287896 | 0.243393409 | 1.183773781 | 0.565406708 | 0.733122144 | 25.88768647 | 30.62857162 | ENSG00000254837 |
| NA | 0.24332845 | 1.183720482 | 0.651206462 | 0.796243605 | 17.85165862 | 21.2899708 | ENSG00000239415 |
| ZKSCAN3 | 0.24330426 | 1.183700634 | 0.118539987 | 0.25726231 | 192.1031164 | 227.5691213 | ENSG00000189298 |
| FAM168B | 0.243220026 | 1.183631524 | 0.000265639 | 0.001654352 | 2061.532974 | 2439.436697 | ENSG00000152102 |
| NA | 0.242971425 | 1.183427582 | 0.845861863 | 0.917815337 | 3.294762033 | 3.889714598 | ENSG00000254016 |
| IGFBP7 | 0.242965399 | 1.183422638 | 0.77208944 | 0.874541661 | 6.053728732 | 7.137631334 | ENSG00000163453 |
| CSGALNACT2 | 0.242827827 | 1.183309796 | 0.178758312 | 0.347714418 | 162.8701573 | 192.372331 | ENSG00000169826 |
| SLC43A1 | 0.24275771 | 1.183252286 | 0.032205579 | 0.09467061 | 561.7705224 | 665.0425934 | ENSG00000149150 |
| PRKDC | 0.242489905 | 1.183032662 | 0.191538506 | 0.364586081 | 7767.877848 | 9189.529158 | ENSG00000253729 |
| NA | 0.242449784 | 1.182999762 | 0.67084705 | 0.808839398 | 14.01143512 | 16.59364465 | ENSG00000267939 |
| C12orf45 | 0.242387621 | 1.18294879 | 0.042082063 | 0.11709187 | 418.6974844 | 495.0655712 | ENSG00000151131 |
| ZNF318 | 0.242288525 | 1.182867538 | 0.009524016 | 0.035338963 | 711.9644823 | 841.8009153 | ENSG00000171467 |
| NEURL1B | 0.242173717 | 1.182773411 | 0.056002955 | 0.145883171 | 372.1264135 | 440.3226347 | ENSG00000214357 |
| SELENOH | 0.242050626 | 1.182672501 | 0.231243693 | 0.415700349 | 172.2350114 | 203.7297911 | ENSG00000211450 |
| PLCD3 | 0.241928468 | 1.182572365 | 0.024895258 | 0.077184462 | 648.9949234 | 767.6524753 | ENSG00000161714 |
| MYBPC2 | 0.24185946 | 1.1825158 | 0.257726278 | 0.446378734 | 97.23633162 | 114.8037693 | ENSG00000086967 |
| FN3K | 0.24183136 | 1.182492768 | 0.087419133 | 0.204910359 | 251.3240849 | 296.8776856 | ENSG00000167363 |
| IGSF10 | 0.241828651 | 1.182490547 | 0.611135193 | 0.768125571 | 22.75400205 | 26.89271221 | ENSG00000152580 |
| ACAP1 | 0.241698077 | 1.182383529 | 0.028549322 | 0.086206116 | 530.7996767 | 627.6648156 | ENSG00000072818 |
| DCTN2 | 0.241684511 | 1.182372411 | 0.002384417 | 0.011012407 | 1699.843351 | 2008.80369 | ENSG00000175203 |
| BBS10 | 0.241680418 | 1.182369056 | 0.141755041 | 0.293589338 | 218.323609 | 258.3195655 | ENSG00000179941 |
| SLC38A6 | 0.241610793 | 1.182311996 | 0.43395138 | 0.623718873 | 47.60147159 | 56.2918512 | ENSG00000139974 |
| MRPS31P5 | 0.241521675 | 1.182238964 | 0.753241733 | 0.86304292 | 7.394002501 | 8.758340508 | ENSG00000243406 |
| HMG20A | 0.241407341 | 1.182145275 | 0.003375315 | 0.014709166 | 1106.53643 | 1307.29015 | ENSG00000140382 |
| PLEC | 0.241354507 | 1.182101984 | 0.001854802 | 0.008896716 | 4601.526216 | 5439.517321 | ENSG00000178209 |
| PRR29-AS1 | 0.241333488 | 1.182084762 | 0.814146485 | 0.89991352 | 3.824341239 | 4.538560499 | ENSG00000264954 |
| GGACT | 0.241327384 | 1.18207976 | 0.73712805 | 0.852804699 | 9.981143073 | 11.80715076 | ENSG00000134864 |
| UBN2 | 0.241322065 | 1.182075402 | 0.049743986 | 0.133249753 | 418.0787602 | 494.0788305 | ENSG00000157741 |
| BBS7 | 0.241255344 | 1.182020736 | 0.088718587 | 0.207266124 | 296.3217217 | 350.2864247 | ENSG00000138686 |
| CKS2 | 0.241187668 | 1.181965289 | 0.009801239 | 0.036143288 | 3321.601015 | 3925.276627 | ENSG00000123975 |
| CHKB-DT | 0.24113257 | 1.181920149 | 0.421442754 | 0.61262902 | 51.99455276 | 61.62237752 | ENSG00000205559 |
| GOLGA1 | 0.241085626 | 1.181881691 | 0.249137338 | 0.437637326 | 109.1232338 | 129.0697224 | ENSG00000136935 |
| TMBIM1 | 0.24106637 | 1.181865917 | 0.021155017 | 0.067741376 | 575.0130543 | 679.8721857 | ENSG00000135926 |
| CKMT2-AS1 | 0.240957062 | 1.181776374 | 0.286316731 | 0.477588268 | 114.7983888 | 135.7806222 | ENSG00000247572 |
| SFT2D1 | 0.240938495 | 1.181761165 | 0.013489805 | 0.046916878 | 904.8692658 | 1069.962444 | ENSG00000198818 |
| ZNF16 | 0.240877298 | 1.181711037 | 0.195422548 | 0.369889859 | 148.3846897 | 175.3988466 | ENSG00000170631 |
| NANS | 0.240704407 | 1.181569431 | 0.018905671 | 0.061548868 | 811.076184 | 958.1062525 | ENSG00000095380 |
| ANTKMT | 0.240674476 | 1.181544918 | 0.086553673 | 0.203293091 | 368.5409299 | 435.6280289 | ENSG00000103254 |
| UHRF1BP1 | 0.240429146 | 1.181344014 | 0.010097356 | 0.037065638 | 943.6135747 | 1113.877106 | ENSG00000065060 |
| IQGAP2 | 0.240308007 | 1.181244824 | 0.000343799 | 0.002062129 | 1980.159282 | 2338.910596 | ENSG00000145703 |
| PHIP | 0.240303182 | 1.181240873 | 0.042289947 | 0.117564475 | 1023.522362 | 1208.4314 | ENSG00000146247 |
| JAK3 | 0.240289536 | 1.1812297 | 0.339681401 | 0.533877978 | 73.68120344 | 87.22185804 | ENSG00000105639 |
| RBL2 | 0.240234401 | 1.181184558 | 0.070485928 | 0.173860287 | 321.9018952 | 380.4827457 | ENSG00000103479 |
| NA | 0.240101084 | 1.181075412 | 0.720645467 | 0.842146475 | 10.6749568 | 12.63253331 | ENSG00000272153 |
| NA | 0.240002096 | 1.180994377 | 0.815437117 | 0.90056677 | 5.103957248 | 6.076588003 | ENSG00000237938 |
| ZNF714 | 0.239761297 | 1.180797275 | 0.040237948 | 0.112927035 | 439.7268515 | 518.889268 | ENSG00000160352 |
| TRIM35 | 0.239682825 | 1.18073305 | 0.0279166 | 0.084794593 | 470.8097148 | 556.0281443 | ENSG00000104228 |
| SNAPC1 | 0.239446888 | 1.18053997 | 0.22832437 | 0.411950354 | 115.7332702 | 136.723815 | ENSG00000023608 |
| ZNF587B | 0.239399067 | 1.180500839 | 0.081843996 | 0.194998924 | 308.4641983 | 364.5321727 | ENSG00000269343 |
| BCAS4 | 0.23929985 | 1.180419656 | 0.04618397 | 0.125638294 | 348.6427667 | 411.4350588 | ENSG00000124243 |
| CKS1B | 0.239262939 | 1.180389456 | 0.000903266 | 0.00483289 | 3228.343021 | 3810.049009 | ENSG00000173207 |
| ANXA5 | 0.23920182 | 1.180339451 | 0.000345164 | 0.002069475 | 3684.0182 | 4347.054203 | ENSG00000164111 |
| CRKL | 0.23914966 | 1.180296777 | 0.000182051 | 0.001184322 | 2616.39889 | 3088.541898 | ENSG00000099942 |
| NALCN | 0.23914465 | 1.180292678 | 0.687487355 | 0.820416628 | 12.18074288 | 14.30531858 | ENSG00000102452 |
| RPL27 | 0.239126953 | 1.1802782 | 0.001686838 | 0.008210046 | 12206.74933 | 14406.97755 | ENSG00000131469 |
| ASB1 | 0.238985585 | 1.180162551 | 0.009577238 | 0.035500906 | 797.0973852 | 940.6425321 | ENSG00000065802 |
| DCST2 | 0.238498358 | 1.179764054 | 0.822949476 | 0.904942493 | 3.847717405 | 4.54984542 | ENSG00000163354 |
| ZNF585A | 0.238468338 | 1.179739505 | 0.37909009 | 0.572276273 | 63.73606419 | 74.98490105 | ENSG00000196967 |
| OSBPL6 | 0.238453198 | 1.179727126 | 0.16864533 | 0.332872476 | 182.636174 | 215.18601 | ENSG00000079156 |
| NOTCH2 | 0.238373594 | 1.179662033 | 0.011930686 | 0.042564646 | 1646.422093 | 1941.509036 | ENSG00000134250 |
| AGTRAP | 0.238353016 | 1.179645207 | 0.002389952 | 0.01103522 | 1082.110243 | 1276.341635 | ENSG00000177674 |
| GPX1 | 0.238352032 | 1.179644402 | 0.001783089 | 0.008608432 | 4076.733493 | 4808.650053 | ENSG00000233276 |

| | | | | | | | |
|---|---|---|---|---|---|---|---|
| FXR1 | 0.238263587 | 1.179572086 | 0.00031644 | 0.001921026 | 4805.554787 | 5667.069267 | ENSG00000114416 |
| PRPF40B | 0.238245907 | 1.17955763 | 0.171197796 | 0.336816152 | 183.6229188 | 217.0572892 | ENSG00000110844 |
| NA | 0.238194845 | 1.179515882 | 0.56849344 | 0.735683383 | 26.18774331 | 30.94822818 | ENSG00000143429 |
| MTRNR2L8 | 0.238126196 | 1.179459758 | 0.811430984 | 0.898880668 | 4.658509667 | 5.543728881 | ENSG00000255823 |
| MRC2 | 0.238035967 | 1.179385994 | 0.009321157 | 0.034704363 | 701.5071944 | 827.3391451 | ENSG00000011028 |
| NA | 0.237825427 | 1.179213893 | 0.42339748 | 0.614217979 | 50.92216534 | 59.98994539 | ENSG00000249855 |
| ALG1L9P | 0.237551911 | 1.17899035 | 0.535802863 | 0.710188819 | 28.62317931 | 33.7685373 | ENSG00000248671 |
| NBEAL2 | 0.237538839 | 1.178979668 | 0.006274923 | 0.024999875 | 1836.414469 | 2166.100509 | ENSG00000160796 |
| TET3 | 0.23752072 | 1.178964862 | 0.044648321 | 0.122475261 | 549.4463203 | 647.2754053 | ENSG00000187605 |
| TRIOBP | 0.237487101 | 1.178937388 | 0.00417448 | 0.017668376 | 1297.027929 | 1529.871395 | ENSG00000100106 |
| SLC15A4 | 0.237444795 | 1.178902817 | 0.022710923 | 0.071719925 | 670.653583 | 789.9778365 | ENSG00000139370 |
| ZNF449 | 0.237329966 | 1.178808988 | 0.269259849 | 0.45951754 | 127.0957269 | 149.6017567 | ENSG00000173275 |
| SLC5A12 | 0.237274553 | 1.178763712 | 0.431275263 | 0.621221398 | 47.23552555 | 55.59037954 | ENSG00000148942 |
| NPHP1 | 0.23717696 | 1.178683976 | 0.503796497 | 0.683776217 | 34.83620421 | 41.19533125 | ENSG00000144061 |
| DGLUCY | 0.237149505 | 1.178661545 | 0.046760839 | 0.126946739 | 348.3399589 | 410.5660739 | ENSG00000133943 |
| SLC19A3 | 0.236809878 | 1.178384107 | 0.269328749 | 0.45951754 | 122.9259671 | 144.506167 | ENSG00000135917 |
| SURF1 | 0.236807142 | 1.178381873 | 0.073061849 | 0.17902278 | 434.7515833 | 511.6266349 | ENSG00000148290 |
| AKNA | 0.236777147 | 1.178357373 | 0.003978531 | 0.016951286 | 1110.029348 | 1308.118066 | ENSG00000106948 |
| SLC24A2 | 0.236772803 | 1.178353825 | 0.292196893 | 0.484702158 | 96.87296377 | 114.0839647 | ENSG00000155886 |
| MIR1915 | 0.236668407 | 1.17826856 | 0.8177558 | 0.901961189 | 4.357077358 | 5.168635407 | ENSG00000222071 |
| SNHG6 | 0.23662566 | 1.178233649 | 0.01674178 | 0.055816408 | 986.871273 | 1163.554934 | ENSG00000245910 |
| PITPNA | 0.236612107 | 1.17822258 | 0.001090414 | 0.005686478 | 2127.918828 | 2507.562452 | ENSG00000174238 |
| PDE1C | 0.236609253 | 1.17822025 | 0.809026122 | 0.897335896 | 5.605124238 | 6.669120922 | ENSG00000154678 |
| C18orf21 | 0.236525248 | 1.178151647 | 0.296681077 | 0.489580869 | 102.2456041 | 120.4603474 | ENSG00000141428 |
| FSCN1 | 0.236239223 | 1.177918092 | 0.00056367 | 0.00318897 | 12567.86411 | 14803.35158 | ENSG00000075618 |
| OSBPL8 | 0.236194974 | 1.177881964 | 0.053046572 | 0.139932419 | 976.7535708 | 1149.663059 | ENSG00000091039 |
| SEL1L | 0.236187995 | 1.177876267 | 0.028461801 | 0.085969854 | 643.8581118 | 757.8138168 | ENSG00000071537 |
| CRIPT | 0.236175273 | 1.17786588 | 0.159970763 | 0.32025255 | 201.2035342 | 236.9245177 | ENSG00000119878 |
| GPR176 | 0.236004001 | 1.177726056 | 0.000342478 | 0.002056867 | 3445.007216 | 4056.433313 | ENSG00000166073 |
| ABRAXAS2 | 0.235993288 | 1.177717311 | 0.05544023 | 0.144722426 | 342.3898069 | 402.841094 | ENSG00000165660 |
| FAM229A | 0.235938823 | 1.17767285 | 0.619344236 | 0.773807075 | 19.5185223 | 22.91337952 | ENSG00000225828 |
| WWC2-AS2 | 0.235879613 | 1.177624518 | 0.630110362 | 0.781586498 | 21.5198606 | 25.45723667 | ENSG00000251359 |
| PIK3CA | 0.235822325 | 1.177577756 | 0.166146521 | 0.329308055 | 170.6736036 | 200.8694065 | ENSG00000121879 |
| GLUD1 | 0.235649992 | 1.177437101 | 0.001244787 | 0.006327726 | 2054.452144 | 2418.099245 | ENSG00000148672 |
| NA | 0.235584777 | 1.177383877 | 0.522227861 | 0.699117944 | 36.99526781 | 43.46433667 | ENSG00000284946 |
| SLC32A1 | 0.235579806 | 1.177379821 | 0.77349594 | 0.875038864 | 6.464450271 | 7.576635484 | ENSG00000101438 |
| NA | 0.23543201 | 1.177259211 | 0.082299828 | 0.195918384 | 282.9588363 | 333.0980888 | ENSG00000281406 |
| ROMO1 | 0.235401609 | 1.177234403 | 0.018553006 | 0.060550545 | 964.1211945 | 1134.869785 | ENSG00000125995 |
| CASP3 | 0.235214123 | 1.177081425 | 0.33132257 | 0.52552268 | 1476.354239 | 1737.624068 | ENSG00000164305 |
| MTMR6 | 0.235059292 | 1.176955106 | 0.14784744 | 0.302785022 | 252.9675572 | 297.6149471 | ENSG00000139505 |
| ARHGAP11A | 0.23498953 | 1.176898196 | 0.006467469 | 0.025645716 | 1213.25161 | 1426.896444 | ENSG00000198826 |
| NA | 0.234844085 | 1.176779553 | 0.364791833 | 0.557934629 | 64.80912803 | 76.28726777 | ENSG00000229335 |
| NA | 0.234843067 | 1.176778723 | 0.521704406 | 0.698697121 | 37.2058487 | 43.93394639 | ENSG00000278768 |
| GEN1 | 0.234839606 | 1.1767759 | 0.113384774 | 0.249069508 | 211.8903218 | 249.5031515 | ENSG00000178295 |
| ZNF732 | 0.234758075 | 1.176709399 | 0.153359468 | 0.310756099 | 171.6575975 | 202.1043985 | ENSG00000186777 |
| ENPP5 | 0.234742791 | 1.176696933 | 0.241733484 | 0.427957626 | 131.7969602 | 154.9484661 | ENSG00000112796 |
| TDRKH | 0.234696603 | 1.176659261 | 0.026642089 | 0.08160378 | 475.7078341 | 559.6713513 | ENSG00000182134 |
| ZNF347 | 0.234411813 | 1.17642701 | 0.173831128 | 0.340561507 | 188.7434574 | 222.1398418 | ENSG00000197937 |
| ASS1 | 0.234383145 | 1.176403634 | 0.00130954 | 0.006600764 | 2171.732053 | 2553.979193 | ENSG00000130707 |
| COG5 | 0.234374213 | 1.17639635 | 0.050161802 | 0.134116971 | 421.2485613 | 495.0549391 | ENSG00000164597 |
| NA | 0.234231387 | 1.176279893 | 0.063139169 | 0.160362666 | 524.4973864 | 617.0710747 | ENSG00000254277 |
| INVS | 0.23420201 | 1.176255941 | 0.197805821 | 0.373471984 | 160.2411051 | 188.2640148 | ENSG00000119509 |
| GHR | 0.234002561 | 1.176093338 | 0.819389771 | 0.90291228 | 4.069550476 | 4.808514081 | ENSG00000112964 |
| COPA | 0.233936198 | 1.17603924 | 0.000208915 | 0.001334721 | 4196.902915 | 4934.942597 | ENSG00000122218 |
| WDR18 | 0.233910639 | 1.176018405 | 0.001555299 | 0.007666467 | 1699.578148 | 1998.549445 | ENSG00000065268 |
| SH3BP4 | 0.233755735 | 1.175892141 | 0.004570251 | 0.019048581 | 1399.033938 | 1645.07867 | ENSG00000130147 |
| CCDC124 | 0.233596211 | 1.175762126 | 0.002513559 | 0.011514161 | 2628.156132 | 3090.331452 | ENSG00000007080 |
| PHC1 | 0.233566751 | 1.175738116 | 0.008174945 | 0.031188642 | 14175.38612 | 16667.46781 | ENSG00000111752 |
| ATP6V1G2 | 0.233502766 | 1.175685972 | 0.66047032 | 0.802356197 | 16.13774951 | 18.88521992 | ENSG00000213760 |
| NA | 0.233292712 | 1.175514807 | 0.00484223 | 0.020005773 | 2515.463433 | 2957.001232 | ENSG00000215030 |
| AFMID | 0.233190316 | 1.175431377 | 0.046241245 | 0.125720292 | 494.5636984 | 580.9236685 | ENSG00000183077 |
| GMDS | 0.233166183 | 1.175411715 | 0.013189616 | 0.046097746 | 750.8898397 | 882.6090005 | ENSG00000112699 |
| RCOR1 | 0.233163323 | 1.175409385 | 0.01453368 | 0.049744639 | 758.7183459 | 891.6700331 | ENSG00000089902 |
| CTSH | 0.233078139 | 1.175339985 | 0.069187053 | 0.171523922 | 343.628789 | 403.8646356 | ENSG00000103811 |
| RAB4A | 0.233074523 | 1.175337039 | 0.024484703 | 0.07615363 | 622.878248 | 732.0159398 | ENSG00000168118 |
| NKD1 | 0.232988804 | 1.175267207 | 0.143801364 | 0.296699577 | 210.9229844 | 247.9117337 | ENSG00000140807 |
| DIP2C | 0.232868505 | 1.175169212 | 0.061960762 | 0.157908383 | 439.4254192 | 516.1738756 | ENSG00000151240 |
| PLEKHO2 | 0.232815647 | 1.175126157 | 0.031492856 | 0.093191299 | 573.2990323 | 674.2686715 | ENSG00000241839 |
| SYT14 | 0.232716757 | 1.17504561 | 0.229102043 | 0.412948314 | 197.4809656 | 231.7537953 | ENSG00000143469 |
| FAM66B | 0.232586994 | 1.174939926 | 0.699893785 | 0.829029936 | 12.53849595 | 14.80063572 | ENSG00000215374 |
| NADK | 0.232572234 | 1.174927905 | 0.001468096 | 0.007271959 | 1753.061161 | 2059.602481 | ENSG00000008130 |
| CACNA1H | 0.2324588 | 1.174835528 | 0.006535825 | 0.025873688 | 1848.369801 | 2172.086923 | ENSG00000196557 |
| TMEM87B | 0.232445087 | 1.174824362 | 0.225029661 | 0.407850249 | 191.965821 | 224.9615364 | ENSG00000153214 |
| P2RX7 | 0.232208646 | 1.174631837 | 0.817414778 | 0.901837342 | 4.075864302 | 4.808514081 | ENSG00000089041 |

| | | | | | | | |
|---|---|---|---|---|---|---|---|
| DCXR | 0.232001516 | 1.174463206 | 0.009457027 | 0.035139643 | 916.9113226 | 1076.609303 | ENSG00000169738 |
| LIG4 | 0.231913755 | 1.174391763 | 0.260629093 | 0.44960289 | 110.8453359 | 130.3219481 | ENSG00000174405 |
| NDEL1 | 0.23191223 | 1.174390522 | 0.019601861 | 0.063447587 | 610.3304993 | 717.1532572 | ENSG00000166579 |
| MOSPD2 | 0.231791183 | 1.174291991 | 0.195311076 | 0.369716626 | 168.1605452 | 197.1469903 | ENSG00000130150 |
| ZNF665 | 0.231709464 | 1.174225477 | 0.422029965 | 0.613242131 | 57.88838396 | 67.9490488 | ENSG00000197497 |
| PFKP | 0.231669793 | 1.174193189 | 0.000401172 | 0.002357425 | 3543.9947 | 4161.528264 | ENSG00000067057 |
| MEDAG | 0.231645897 | 1.17417374 | 0.564211177 | 0.732480414 | 33.61449501 | 39.24791748 | ENSG00000102802 |
| GRB7 | 0.231515831 | 1.174067887 | 0.02800354 | 0.084959504 | 530.9722844 | 623.63292 | ENSG00000141738 |
| FAM227B | 0.231435653 | 1.174002641 | 0.473253658 | 0.658405892 | 46.47967858 | 54.3786186 | ENSG00000166262 |
| HNF4G | 0.231268044 | 1.173866256 | 0.573168748 | 0.738929561 | 26.65350784 | 31.31919639 | ENSG00000164749 |
| CNTNAP1 | 0.231131176 | 1.173754896 | 0.104562278 | 0.234591123 | 339.8143245 | 398.6924246 | ENSG00000108797 |
| ZFAND6 | 0.230984438 | 1.173635519 | 0.001355131 | 0.006786274 | 2120.160015 | 2487.459577 | ENSG00000086666 |
| NA | 0.230967377 | 1.173621639 | 0.852085495 | 0.92181605 | 3.427525127 | 4.020673525 | ENSG00000287916 |
| NKIRAS1 | 0.230958094 | 1.173614088 | 0.357948966 | 0.551282586 | 77.24464857 | 90.78409514 | ENSG00000197885 |
| NA | 0.230918056 | 1.173581518 | 0.771601285 | 0.874349365 | 7.109632533 | 8.323135205 | ENSG00000189423 |
| BTG1 | 0.230885785 | 1.173555267 | 0.167503148 | 0.331253024 | 222.9156355 | 262.0123235 | ENSG00000133639 |
| EFCAB14 | 0.230776765 | 1.173466588 | 0.010887106 | 0.039518201 | 822.73312 | 965.5568441 | ENSG00000159658 |
| SNCA | 0.230625146 | 1.17334327 | 0.166145419 | 0.329308055 | 192.6806281 | 226.3476128 | ENSG00000145335 |
| FRMPD2 | 0.230451651 | 1.173202175 | 0.844429314 | 0.91704846 | 3.552598922 | 4.159668179 | ENSG00000170324 |
| CAMKK1 | 0.23020711 | 1.173003331 | 0.150781213 | 0.307060718 | 181.6860942 | 213.217787 | ENSG00000004660 |
| COX10 | 0.229815293 | 1.172684802 | 0.045919804 | 0.125195299 | 524.5794736 | 615.3264059 | ENSG00000006695 |
| EPB41L4B | 0.229792965 | 1.172666652 | 0.013885178 | 0.048039555 | 698.3241044 | 818.3872173 | ENSG00000095203 |
| CORO1B | 0.229775984 | 1.17265285 | 0.000475932 | 0.002742824 | 2316.071485 | 2715.556779 | ENSG00000172725 |
| DUSP18 | 0.229754954 | 1.172635757 | 0.70307968 | 0.831263 | 11.50271371 | 13.49545784 | ENSG00000167065 |
| NA | 0.229699011 | 1.172590287 | 0.611925551 | 0.768649825 | 25.44589947 | 29.91691435 | ENSG00000248367 |
| VAMP5 | 0.229698192 | 1.172589621 | 0.537176515 | 0.711203176 | 30.727493 | 36.06759864 | ENSG00000168899 |
| POLL | 0.229690287 | 1.172583196 | 0.042724766 | 0.118524294 | 409.5842663 | 480.0453377 | ENSG00000166169 |
| FHDC1 | 0.22955808 | 1.172475747 | 0.008028073 | 0.030716827 | 841.1839454 | 986.2314889 | ENSG00000137460 |
| COX1 | 0.229453343 | 1.17239063 | 0.315277878 | 0.50882621 | 89901.96331 | 105400.1802 | ENSG00000198804 |
| CLPX | 0.229432254 | 1.172373493 | 0.004480994 | 0.018743117 | 1113.683524 | 1305.377355 | ENSG00000166855 |
| FIRRE | 0.229417604 | 1.172361587 | 0.448652633 | 0.636263517 | 429.7925584 | 504.1141913 | ENSG00000213468 |
| KIRREL1 | 0.22921691 | 1.172198511 | 0.002446546 | 0.011259937 | 1487.479237 | 1743.517998 | ENSG00000183853 |
| CARS2 | 0.229174522 | 1.172164071 | 0.003545439 | 0.015342324 | 1230.428021 | 1442.459794 | ENSG00000134905 |
| DNASE1 | 0.229115046 | 1.172115749 | 0.134993025 | 0.282995218 | 256.6108122 | 301.3160446 | ENSG00000213918 |
| INTS4 | 0.229006185 | 1.172027308 | 0.03988883 | 0.112253201 | 543.2935925 | 637.2830462 | ENSG00000149262 |
| NDUFV3 | 0.229001317 | 1.172023354 | 0.016479217 | 0.055121064 | 690.8683391 | 809.8988622 | ENSG00000160194 |
| INTS2 | 0.228936495 | 1.171970695 | 0.073230793 | 0.179389311 | 400.9089641 | 469.5742481 | ENSG00000108506 |
| ITGAV | 0.228910688 | 1.17194973 | 0.030818895 | 0.091508714 | 722.8177292 | 846.2552165 | ENSG00000138448 |
| GSTO1 | 0.228858055 | 1.171906976 | 0.016384134 | 0.054842611 | 939.9190654 | 1100.733273 | ENSG00000148834 |
| SDHA | 0.228811845 | 1.171869439 | 0.541805445 | 0.714567859 | 29.94323382 | 35.07740241 | ENSG00000073578 |
| DENND5A | 0.228756295 | 1.171824319 | 0.00237921 | 0.010996578 | 1639.049121 | 1920.136649 | ENSG00000184014 |
| GNS | 0.228719958 | 1.171794804 | 0.035717611 | 0.102889368 | 850.8365897 | 996.7890732 | ENSG00000135677 |
| CREB3L1 | 0.228681722 | 1.171763749 | 0.105401588 | 0.235931526 | 238.2108283 | 279.2692366 | ENSG00000157613 |
| VRK1 | 0.228515658 | 1.171628878 | 0.005258406 | 0.021523697 | 1042.12145 | 1220.553749 | ENSG00000100749 |
| MAPKAP1 | 0.228480426 | 1.171600266 | 0.000944151 | 0.005012846 | 2112.984831 | 2474.70584 | ENSG00000119487 |
| NA | 0.228315941 | 1.171466697 | 0.832059498 | 0.909480985 | 3.546285096 | 4.140897185 | ENSG00000235297 |
| NBPF11 | 0.228171161 | 1.171349141 | 0.134703199 | 0.282515457 | 268.400662 | 314.4728517 | ENSG00000263956 |
| SECISBP2 | 0.228162403 | 1.171342031 | 0.002523118 | 0.011552241 | 1429.452009 | 1674.09918 | ENSG00000187742 |
| NA | 0.228136706 | 1.171321167 | 0.832975926 | 0.910227923 | 3.519752017 | 4.11084127 | ENSG00000240036 |
| JHY | 0.227990976 | 1.171202855 | 0.677480534 | 0.81350335 | 15.78009413 | 18.48430741 | ENSG00000109944 |
| CACNA1A | 0.227929199 | 1.171152705 | 0.101127264 | 0.228460838 | 281.8173126 | 330.4804618 | ENSG00000141837 |
| TRIQK | 0.227923976 | 1.171148465 | 0.206352359 | 0.38491693 | 181.8054329 | 212.5362485 | ENSG00000205133 |
| IDUA | 0.22790637 | 1.171134173 | 0.4387182 | 0.627165685 | 56.92460935 | 66.75390897 | ENSG00000127415 |
| CLEC18A | 0.227826258 | 1.171069143 | 0.808285598 | 0.897105531 | 5.121019588 | 6.039046014 | ENSG00000157322 |
| TMEM41A | 0.227794101 | 1.17104304 | 0.179237114 | 0.348418427 | 261.1367541 | 305.6829808 | ENSG00000163900 |
| MRPS18B | 0.227767905 | 1.171021776 | 0.010850054 | 0.039391755 | 918.217667 | 1075.165346 | ENSG00000204568 |
| HPN-AS1 | 0.227761278 | 1.171016398 | 0.609600921 | 0.767081504 | 23.97226124 | 28.10122395 | ENSG00000227392 |
| ZNRF3 | 0.227632855 | 1.170912163 | 0.01811767 | 0.059376726 | 656.56212 | 768.2849149 | ENSG00000183579 |
| IL6ST | 0.227264791 | 1.170613475 | 0.342951971 | 0.536880932 | 98.22801478 | 114.87429 | ENSG00000134352 |
| KIAA1191 | 0.227043107 | 1.170433613 | 0.006304681 | 0.025107639 | 1751.073157 | 2048.391898 | ENSG00000122203 |
| DCAF12L1 | 0.227020335 | 1.170415139 | 0.232578532 | 0.417290466 | 130.2779451 | 152.3601305 | ENSG00000198889 |
| PARD3B | 0.22692882 | 1.170340898 | 0.426504327 | 0.617131017 | 65.66214023 | 76.75566656 | ENSG00000116117 |
| NA | 0.226903824 | 1.170320621 | 0.457136054 | 0.644203135 | 50.35668001 | 58.95851994 | ENSG00000225077 |
| NES | 0.226841664 | 1.170270197 | 5.12E-05 | 0.000384341 | 6093.649321 | 7131.671829 | ENSG00000132688 |
| NEK11 | 0.226734687 | 1.170183424 | 0.72473042 | 0.845161619 | 11.85405526 | 13.76117454 | ENSG00000114670 |
| NA | 0.226733428 | 1.170182403 | 0.582164599 | 0.745978891 | 26.27691098 | 30.86565813 | ENSG00000228175 |
| SEPTIN11 | 0.226542166 | 1.170027279 | 0.000860306 | 0.004625735 | 2601.03426 | 3042.365824 | ENSG00000138758 |
| HACD3 | 0.226525052 | 1.170013399 | 0.001751352 | 0.008481722 | 4158.978154 | 4865.105798 | ENSG00000074696 |
| B3GNT2 | 0.226481221 | 1.169977853 | 0.022889763 | 0.072175375 | 687.3998119 | 803.6438822 | ENSG00000170340 |
| CHAMP1 | 0.226479527 | 1.169976479 | 0.063373687 | 0.160789585 | 565.4676099 | 661.7722411 | ENSG00000198824 |
| SORT1 | 0.226375328 | 1.16989198 | 0.002240774 | 0.010458452 | 1297.561935 | 1517.934613 | ENSG00000134243 |
| NA | 0.226341959 | 1.169864921 | 0.58818971 | 0.750637478 | 27.1407694 | 31.77262304 | ENSG00000273329 |
| FHL2 | 0.226207676 | 1.169756038 | 0.033259232 | 0.097243511 | 633.3177746 | 740.4198845 | ENSG00000115641 |
| PTOV1-AS1 | 0.226163448 | 1.169720178 | 0.709355116 | 0.834795081 | 13.60890654 | 15.98234074 | ENSG00000268006 |

| | | | | | | | |
|---|---|---|---|---|---|---|---|
| XK | 0.226144567 | 1.16970487 | 0.590475311 | 0.752362808 | 27.54517711 | 32.12850748 | ENSG00000047597 |
| TINF2 | 0.22607868 | 1.169651451 | 0.07036799 | 0.173661827 | 366.6519778 | 428.7805394 | ENSG00000092330 |
| WDR1 | 0.225860463 | 1.169474547 | 2.72E-05 | 0.000219928 | 10387.19606 | 12147.5782 | ENSG00000071127 |
| KIF1B | 0.225784833 | 1.169413242 | 0.004016199 | 0.017080364 | 2363.211228 | 2762.816017 | ENSG00000054523 |
| ATE1 | 0.225688522 | 1.169335177 | 0.05824832 | 0.150455131 | 398.1098519 | 465.4263113 | ENSG00000107669 |
| MRPL28 | 0.225668428 | 1.169318891 | 0.001708129 | 0.008300594 | 1931.203781 | 2258.616505 | ENSG00000086504 |
| RBM44 | 0.225501297 | 1.169183437 | 0.747518779 | 0.858996555 | 9.901446137 | 11.63540065 | ENSG00000177483 |
| STYXL1 | 0.225396092 | 1.16909818 | 0.042856266 | 0.118800185 | 440.6811554 | 515.3329168 | ENSG00000127952 |
| NT5C | 0.225364407 | 1.169072504 | 0.179832897 | 0.349178188 | 210.6490925 | 246.6210186 | ENSG00000125458 |
| TANC2 | 0.225341762 | 1.169054155 | 0.172087189 | 0.338062404 | 222.7516116 | 260.45073 | ENSG00000170921 |
| ZNF875 | 0.225318631 | 1.169035411 | 0.053925277 | 0.14170581 | 495.1900595 | 579.0551778 | ENSG00000181666 |
| ARHGAP32 | 0.225300235 | 1.169020504 | 0.026353234 | 0.080866033 | 597.3981714 | 698.2350183 | ENSG00000134909 |
| ARNT | 0.225212012 | 1.16894902 | 0.012733755 | 0.044927765 | 768.5152458 | 898.1680652 | ENSG00000143437 |
| KDM3A | 0.225152727 | 1.168900985 | 0.042701545 | 0.11849535 | 567.0734176 | 662.4602695 | ENSG00000115548 |
| RPS18 | 0.225142353 | 1.168892579 | 0.001607704 | 0.007895385 | 22955.42637 | 26831.69618 | ENSG00000231500 |
| GATM | 0.225140839 | 1.168891352 | 0.12156184 | 0.262162806 | 318.1308081 | 371.5701119 | ENSG00000171766 |
| RPL31 | 0.22510594 | 1.168863077 | 0.000437514 | 0.002545555 | 10703.11305 | 12510.51285 | ENSG00000071082 |
| NA | 0.224884047 | 1.168683315 | 0.255160786 | 0.443978805 | 118.3985368 | 138.3896639 | ENSG00000232533 |
| UBE2Q2 | 0.224750565 | 1.16857519 | 0.054537112 | 0.142948781 | 405.9635157 | 474.1553604 | ENSG00000140367 |
| PPP1R21 | 0.22467382 | 1.168513029 | 0.064657635 | 0.163020624 | 345.7176113 | 404.2167212 | ENSG00000162869 |
| NA | 0.224594526 | 1.168448806 | 0.763449344 | 0.868883644 | 7.645525565 | 8.926841424 | ENSG00000284707 |
| CYTH2 | 0.224519134 | 1.168387747 | 0.045531287 | 0.124447261 | 1460.597462 | 1707.093141 | ENSG00000105443 |
| NA | 0.224394607 | 1.168286902 | 0.799346079 | 0.892098527 | 9.040842327 | 10.52030584 | ENSG00000278840 |
| HOMER1 | 0.224157482 | 1.168094895 | 0.058817465 | 0.1515533 | 774.3081535 | 903.521487 | ENSG00000152413 |
| GSTA4 | 0.22401143 | 1.167976648 | 0.164221415 | 0.326434976 | 228.9925224 | 267.6426445 | ENSG00000170899 |
| NSRP1 | 0.22398155 | 1.167952458 | 0.033432852 | 0.09762793 | 543.125502 | 634.0937838 | ENSG00000126653 |
| FAM72C | 0.223843818 | 1.167840961 | 0.430854298 | 0.620904561 | 58.71989914 | 68.42895575 | ENSG00000263513 |
| NA | 0.223819265 | 1.167821086 | 0.62112057 | 0.775052679 | 23.18178593 | 27.17505001 | ENSG00000260051 |
| COX18 | 0.223573795 | 1.167622402 | 0.02845048 | 0.085959281 | 575.2784828 | 671.1497064 | ENSG00000163626 |
| MTUS1 | 0.223524764 | 1.167582719 | 0.366356204 | 0.559450102 | 84.88101167 | 99.01012618 | ENSG00000129422 |
| PRKAB1 | 0.223417851 | 1.167496197 | 0.079135743 | 0.190115461 | 379.4747461 | 442.5390773 | ENSG00000111725 |
| ANKRD11 | 0.223406089 | 1.167486679 | 0.00116473 | 0.00600142 | 3755.975971 | 4386.049635 | ENSG00000167522 |
| CSNK1E | 0.223244198 | 1.167355678 | 0.008673118 | 0.03273215 | 1378.237206 | 1609.354209 | ENSG00000213923 |
| VAMP3 | 0.223227371 | 1.167342063 | 0.036830903 | 0.105278215 | 580.5731925 | 677.5293383 | ENSG00000049245 |
| KRTCAP3 | 0.222979316 | 1.167141368 | 0.065141492 | 0.163883862 | 451.1758 | 526.8921829 | ENSG00000157992 |
| BAIAP2L1 | 0.222866206 | 1.167049866 | 0.026783342 | 0.081933275 | 709.8223083 | 828.5793617 | ENSG00000006453 |
| FGD5 | 0.222764035 | 1.166967219 | 0.021494864 | 0.068628048 | 721.672485 | 841.7216976 | ENSG00000154783 |
| NA | 0.222672708 | 1.166893348 | 0.74233427 | 0.855995979 | 9.567166922 | 11.21516749 | ENSG00000287811 |
| ITPR3 | 0.222536298 | 1.166783021 | 0.013435681 | 0.046763739 | 2100.853637 | 2452.183842 | ENSG00000096433 |
| RNASET2 | 0.222501259 | 1.166754684 | 0.150302059 | 0.306423756 | 250.784223 | 292.9569761 | ENSG00000026297 |
| ALG9 | 0.22244451 | 1.16670879 | 0.551295892 | 0.722322422 | 32.19284057 | 37.4323387 | ENSG00000086848 |
| SNX9 | 0.222306553 | 1.16659723 | 0.013235365 | 0.046214066 | 779.0341384 | 909.0565768 | ENSG00000130340 |
| ZNF215 | 0.222169012 | 1.166486016 | 0.278590157 | 0.469697987 | 135.6758408 | 158.3548097 | ENSG00000149054 |
| NA | 0.222124885 | 1.166450338 | 0.832238851 | 0.909583375 | 3.824341239 | 4.452191601 | ENSG00000232599 |
| ODF2 | 0.222075553 | 1.166410452 | 0.058675775 | 0.151293379 | 493.2141719 | 575.6011065 | ENSG00000136811 |
| NA | 0.221937798 | 1.166299083 | 0.169851437 | 0.334684392 | 295.0794099 | 343.8049735 | ENSG00000210196 |
| CLIC6 | 0.221910399 | 1.166276934 | 0.406037529 | 0.597593867 | 66.14614718 | 77.06002456 | ENSG00000159212 |
| TNFSF4 | 0.221844146 | 1.166223376 | 0.786133486 | 0.882911566 | 6.917587152 | 8.037659823 | ENSG00000117586 |
| NA | 0.221796789 | 1.166185095 | 0.000311965 | 0.001896969 | 10584.65823 | 12344.8117 | ENSG00000224078 |
| TMEM79 | 0.221754485 | 1.1661509 | 0.558867042 | 0.727952896 | 30.91322456 | 36.00368797 | ENSG00000163472 |
| TUBGCP5 | 0.221735231 | 1.166135337 | 0.068901962 | 0.171045936 | 373.5697604 | 435.5038863 | ENSG00000275835 |
| P3H4 | 0.221713192 | 1.166117522 | 0.016155229 | 0.054203054 | 896.9474957 | 1046.242037 | ENSG00000141696 |
| CHCHD10 | 0.221309414 | 1.165791198 | 0.60101297 | 0.7603532 | 530.1548775 | 618.0042641 | ENSG00000250479 |
| RAD51D | 0.221283576 | 1.165770319 | 0.210847208 | 0.390276855 | 160.1286589 | 186.3502325 | ENSG00000185379 |
| ZNF597 | 0.221248729 | 1.165742162 | 0.410174704 | 0.601819028 | 69.46645761 | 81.13723431 | ENSG00000167981 |
| RPL36A | 0.221211911 | 1.165712411 | 0.594339281 | 0.755547815 | 26.96125397 | 31.51819126 | ENSG00000241343 |
| ARL13B | 0.221164593 | 1.165674179 | 0.295578357 | 0.488431639 | 114.1672097 | 132.8599194 | ENSG00000169379 |
| BET1L | 0.221074206 | 1.16560115 | 0.041550333 | 0.116030417 | 442.0148222 | 515.0703376 | ENSG00000177951 |
| LOC100130548 | 0.221066349 | 1.165594802 | 0.827524068 | 0.907212296 | 4.15546354 | 4.84605607 | ENSG00000235138 |
| LACTB | 0.220939389 | 1.165492232 | 0.145788725 | 0.29958243 | 279.247332 | 325.1235349 | ENSG00000103642 |
| UBA6 | 0.220874628 | 1.165439916 | 0.023786726 | 0.074394733 | 701.0368823 | 816.7880903 | ENSG00000033178 |
| TEF | 0.220869092 | 1.165435443 | 0.188039885 | 0.359787707 | 218.9514358 | 254.8475506 | ENSG00000167074 |
| NA | 0.220843932 | 1.165415119 | 0.789768779 | 0.885301473 | 6.417697938 | 7.418869836 | ENSG00000251468 |
| LONP2 | 0.220728165 | 1.165321605 | 0.024819002 | 0.076986674 | 622.3069978 | 725.1893881 | ENSG00000102910 |
| CEP164 | 0.220718333 | 1.165313664 | 0.109158509 | 0.241829287 | 330.3821528 | 385.5721718 | ENSG00000110274 |
| PSEN2 | 0.220624417 | 1.165237807 | 0.059378081 | 0.152653498 | 411.5342222 | 479.1898993 | ENSG00000143801 |
| SNORD15B | 0.220505517 | 1.165141778 | 0.825833611 | 0.906340611 | 4.108711207 | 4.789743086 | ENSG00000207445 |
| APC | 0.220465964 | 1.165109835 | 0.049654801 | 0.133070443 | 471.7912433 | 549.1702142 | ENSG00000134982 |
| LYG1 | 0.220421959 | 1.165074297 | 0.799303388 | 0.892098527 | 6.361474866 | 7.479419699 | ENSG00000144214 |
| NA | 0.220357252 | 1.165022043 | 0.030554827 | 0.090844221 | 852.9472848 | 993.2645844 | ENSG00000237550 |
| STAT5A | 0.219944901 | 1.164689104 | 0.134109623 | 0.281550136 | 252.2114172 | 293.6145734 | ENSG00000126561 |
| NA | 0.219899762 | 1.164652664 | 0.827376157 | 0.907201323 | 4.138401199 | 4.819799001 | ENSG00000236519 |
| DCAF17 | 0.21970889 | 1.164498588 | 0.08086462 | 0.193228076 | 396.6270513 | 461.2347235 | ENSG00000115827 |
| ETV5 | 0.219637294 | 1.164440799 | 0.014226601 | 0.048919446 | 957.803864 | 1114.693219 | ENSG00000244405 |

| | | | | | | | |
|---|---|---|---|---|---|---|---|
| CC2D1B | 0.219461802 | 1.164299163 | 0.099697194 | 0.226333634 | 379.3761902 | 442.454724 | ENSG00000154222 |
| CTC1 | 0.219297669 | 1.164166711 | 0.037235597 | 0.106164671 | 479.2997665 | 557.8815915 | ENSG00000178971 |
| RBSN | 0.21918106 | 1.164072618 | 0.022382808 | 0.070840671 | 639.1702803 | 744.0906795 | ENSG00000131381 |
| ND3 | 0.218865341 | 1.1638179 | 0.007258152 | 0.028213628 | 4844.722084 | 5637.51565 | ENSG00000198840 |
| SLIT3 | 0.218849611 | 1.163805211 | 0.434528738 | 0.624209861 | 65.02031028 | 75.78066572 | ENSG00000184347 |
| ZNF791 | 0.21869923 | 1.163683907 | 0.07340725 | 0.179679077 | 348.8169756 | 405.5721114 | ENSG00000173875 |
| NA | 0.218555545 | 1.163568016 | 0.665116641 | 0.805385316 | 17.38777323 | 20.26179455 | ENSG00000272293 |
| ZNF816 | 0.218480061 | 1.163507138 | 0.309543325 | 0.503605687 | 112.0792819 | 130.1520215 | ENSG00000180257 |
| NA | 0.218440708 | 1.1634754 | 0.505081964 | 0.684647782 | 42.91139279 | 50.09266641 | ENSG00000226824 |
| MRPS33 | 0.218320226 | 1.16337824 | 0.041007366 | 0.114704275 | 523.4815151 | 608.5396577 | ENSG00000090263 |
| NA | 0.218259853 | 1.163329557 | 0.775188278 | 0.876003875 | 8.810816294 | 10.26956129 | ENSG00000255129 |
| TRAPPC12 | 0.21824959 | 1.163321282 | 0.010889147 | 0.039518201 | 951.9700815 | 1107.690792 | ENSG00000171853 |
| B3GAT3 | 0.218052213 | 1.163162137 | 0.051429536 | 0.136678381 | 540.2134401 | 629.0121072 | ENSG00000149541 |
| NA | 0.2176878 | 1.162868369 | 0.608614324 | 0.766514251 | 26.17511565 | 30.45682151 | ENSG00000259495 |
| SEL1L3 | 0.217618899 | 1.162812833 | 0.001570471 | 0.007730972 | 1685.816605 | 1959.927294 | ENSG00000091490 |
| TMED8 | 0.217523274 | 1.162735762 | 0.071211807 | 0.175324075 | 427.5623742 | 496.5260929 | ENSG00000100580 |
| ALDOA | 0.217501462 | 1.162718183 | 0.507689268 | 0.68692604 | 46.57022173 | 54.39501647 | ENSG00000149925 |
| NA | 0.217469504 | 1.162692427 | 0.750973316 | 0.861402366 | 9.546947669 | 11.13628467 | ENSG00000249042 |
| TAPT1 | 0.217428262 | 1.16265919 | 0.262594632 | 0.452072269 | 125.2737086 | 145.3684302 | ENSG00000169762 |
| RRAGC | 0.217370456 | 1.162612606 | 0.323256995 | 0.517468056 | 117.1096455 | 136.0698562 | ENSG00000116954 |
| NA | 0.217235291 | 1.162503687 | 0.792632275 | 0.887223492 | 6.712816421 | 7.748935247 | ENSG00000287750 |
| MAP3K10 | 0.217216441 | 1.162488498 | 0.034676766 | 0.10048455 | 559.8064279 | 650.9437239 | ENSG00000130758 |
| C11orf1 | 0.217093894 | 1.162389756 | 0.272361349 | 0.463454393 | 117.4064477 | 136.457437 | ENSG00000137720 |
| INSC | 0.217092256 | 1.162388436 | 0.682639476 | 0.816765029 | 15.38959184 | 17.83122463 | ENSG00000188487 |
| FAM72A | 0.217059932 | 1.162362393 | 0.340056411 | 0.53430023 | 97.55294713 | 113.1717752 | ENSG00000196550 |
| CREBBP | 0.217021431 | 1.162331374 | 0.015862604 | 0.053468548 | 1689.754459 | 1964.009422 | ENSG00000005339 |
| ISM2 | 0.216929415 | 1.162257242 | 0.620572139 | 0.774799082 | 22.78369204 | 26.54442545 | ENSG00000100593 |
| TRMT10A | 0.216889808 | 1.162225334 | 0.529500571 | 0.705315768 | 43.66596198 | 50.68757246 | ENSG00000145331 |
| PAWR | 0.216732335 | 1.162098482 | 0.238668172 | 0.42435494 | 2117.879778 | 2460.988919 | ENSG00000177425 |
| UXT | 0.216717823 | 1.162086792 | 0.292556819 | 0.485125532 | 111.5121128 | 129.6570308 | ENSG00000126756 |
| MUS81 | 0.216657345 | 1.162038079 | 0.102077526 | 0.230214756 | 510.2224618 | 593.5558527 | ENSG00000172732 |
| CHKA | 0.216531231 | 1.161936503 | 0.040547283 | 0.113686265 | 645.5915864 | 750.0043889 | ENSG00000110721 |
| LINC02693 | 0.216503151 | 1.161913888 | 0.095352977 | 0.218586528 | 306.9541881 | 356.933955 | ENSG00000212719 |
| NMRAL1 | 0.216357805 | 1.161796835 | 0.013717595 | 0.047548514 | 827.047301 | 960.7719504 | ENSG00000153406 |
| BBOF1 | 0.216307563 | 1.161756376 | 0.807424082 | 0.89663259 | 6.226930333 | 7.295396982 | ENSG00000119636 |
| LOC112268124 | 0.216286845 | 1.161739693 | 0.773269565 | 0.874965662 | 7.698591723 | 8.911869277 | ENSG00000258738 |
| SENP2 | 0.21621256 | 1.161679876 | 0.018729683 | 0.061072475 | 802.1311011 | 931.6072697 | ENSG00000163904 |
| PDLIM1 | 0.216078525 | 1.161571954 | 0.001470203 | 0.00727992 | 2855.224797 | 3316.152549 | ENSG00000107438 |
| CARS1 | 0.21607203 | 1.161566725 | 0.005628812 | 0.022793182 | 1587.560606 | 1844.258585 | ENSG00000110619 |
| NA | 0.216063448 | 1.161559815 | 0.754622366 | 0.863503792 | 13.63987431 | 15.93786233 | ENSG00000258199 |
| TAF6 | 0.215773028 | 1.161326012 | 0.003394786 | 0.01478012 | 1481.55761 | 1720.170521 | ENSG00000106290 |
| PDGFRA | 0.215725472 | 1.161287732 | 0.711960051 | 0.836585785 | 13.51480052 | 15.72692127 | ENSG00000134853 |
| NBPF3 | 0.215678215 | 1.161249693 | 0.042066078 | 0.11709187 | 513.1190322 | 595.8367069 | ENSG00000142794 |
| SND1 | 0.215621836 | 1.161204313 | 7.90E-05 | 0.000564145 | 8661.95705 | 10058.24739 | ENSG00000197157 |
| DACH1 | 0.215615034 | 1.161198839 | 0.785042735 | 0.882418854 | 7.050252548 | 8.135313641 | ENSG00000276644 |
| NA | 0.215411944 | 1.161035386 | 0.744092088 | 0.857157223 | 9.531163105 | 11.09874268 | ENSG00000234782 |
| RERE | 0.215382211 | 1.161011459 | 0.006595721 | 0.026081737 | 2510.654693 | 2914.484598 | ENSG00000142599 |
| ZXDC | 0.215309283 | 1.160952771 | 0.084995933 | 0.200498234 | 448.7050744 | 520.3156573 | ENSG00000070476 |
| NA | 0.215247642 | 1.160903169 | 0.462510241 | 0.649062225 | 52.88622204 | 61.58103668 | ENSG00000261578 |
| ZNF165 | 0.215205332 | 1.160869124 | 0.396696716 | 0.588966027 | 76.27395878 | 88.40527492 | ENSG00000197279 |
| DRAM2 | 0.215167185 | 1.160838429 | 0.111791889 | 0.246338232 | 269.3677889 | 312.4136477 | ENSG00000156171 |
| MTBP | 0.215114997 | 1.160796438 | 0.18269478 | 0.353156556 | 193.0541657 | 224.1856823 | ENSG00000172167 |
| MRPL41 | 0.215029189 | 1.160727399 | 0.020873491 | 0.066943395 | 1514.199904 | 1757.781603 | ENSG00000182154 |
| LOC283856 | 0.214981778 | 1.160689254 | 0.829939062 | 0.908528105 | 4.66166658 | 5.427304068 | ENSG00000246379 |
| COL11A1 | 0.214975061 | 1.16068385 | 0.125246926 | 0.268144453 | 303.7499618 | 352.4486816 | ENSG00000060718 |
| NFU1 | 0.214939698 | 1.1606554 | 0.069162546 | 0.171486103 | 396.8638721 | 460.4022216 | ENSG00000169599 |
| NA | 0.214934797 | 1.160651457 | 0.782188971 | 0.8805327 | 7.030033295 | 8.105257726 | ENSG00000260196 |
| RPL18 | 0.214850616 | 1.160583736 | 0.000244204 | 0.001536868 | 14240.79308 | 16527.07277 | ENSG00000063177 |
| PHF1 | 0.21465616 | 1.160427315 | 0.045744417 | 0.124919261 | 762.9723502 | 886.2387415 | ENSG00000112511 |
| SIRT1 | 0.214616231 | 1.160395199 | 0.020477101 | 0.065912225 | 1345.479172 | 1560.923812 | ENSG00000096717 |
| UNC13A | 0.214540652 | 1.16033441 | 0.025787181 | 0.079352418 | 1754.005107 | 2036.20957 | ENSG00000130477 |
| STX6 | 0.214536896 | 1.160331389 | 0.022348567 | 0.070780647 | 930.0616728 | 1078.94761 | ENSG00000135823 |
| LSM11 | 0.214468639 | 1.160276492 | 0.247856409 | 0.435800274 | 137.7717661 | 160.0701383 | ENSG00000155858 |
| NMD3 | 0.214269915 | 1.160116681 | 0.011664823 | 0.041785045 | 891.7374963 | 1034.260053 | ENSG00000169251 |
| MAP4 | 0.214242973 | 1.160095017 | 0.000224447 | 0.001423647 | 6334.248668 | 7348.015165 | ENSG00000047849 |
| CXCL3 | 0.214097609 | 1.159978133 | 0.512380023 | 0.691254284 | 41.64440443 | 48.25732895 | ENSG00000163734 |
| CA3 | 0.214048621 | 1.159938745 | 0.772656568 | 0.874752298 | 8.567486193 | 9.935808652 | ENSG00000164879 |
| ORC3 | 0.213876811 | 1.159800617 | 0.016544855 | 0.055310669 | 831.6252646 | 964.1309558 | ENSG00000135336 |
| AP1S2 | 0.213784503 | 1.159726412 | 0.301872709 | 0.495324452 | 4430.615272 | 5138.093056 | ENSG00000182287 |
| NA | 0.213650661 | 1.159618826 | 0.809097867 | 0.897335896 | 6.44738793 | 7.415070989 | ENSG00000243926 |
| LOC152048 | 0.213621176 | 1.159595127 | 0.831684017 | 0.909313528 | 4.164934279 | 4.819799001 | ENSG00000270194 |
| PKD1 | 0.213472577 | 1.159475694 | 0.015351319 | 0.052062459 | 1131.578508 | 1312.928739 | ENSG00000008710 |
| MLLT6 | 0.213409063 | 1.15942465 | 0.517876433 | 0.695278654 | 371.2216129 | 430.6127166 | ENSG00000275023 |
| PAK1 | 0.213318154 | 1.159351592 | 0.002169343 | 0.010199542 | 4083.894509 | 4734.187217 | ENSG00000149269 |

| | | | | | | | |
|---|---|---|---|---|---|---|---|
| SPICE1 | 0.213268537 | 1.159311721 | 0.821324702 | 0.904213366 | 6.408227199 | 7.373841773 | ENSG00000163611 |
| PTPN4 | 0.213204334 | 1.159260131 | 0.067710155 | 0.168697407 | 463.0696924 | 536.2664132 | ENSG00000088179 |
| GNB4 | 0.213183314 | 1.159243241 | 0.081577914 | 0.194565057 | 488.8932711 | 565.9143129 | ENSG00000114450 |
| GDI1 | 0.213111116 | 1.159185229 | 0.00808077 | 0.030886572 | 2596.740659 | 3011.027566 | ENSG00000203879 |
| C19orf81 | 0.213022867 | 1.159114324 | 0.386890018 | 0.57981884 | 89.23009146 | 103.6620855 | ENSG00000235034 |
| ADAMTS4 | 0.212967789 | 1.159070073 | 0.184817283 | 0.355553614 | 178.8897482 | 207.4270007 | ENSG00000158859 |
| GSTM3 | 0.212963917 | 1.159066963 | 0.530067874 | 0.705701898 | 43.87567106 | 50.84957498 | ENSG00000134202 |
| NA | 0.212860103 | 1.158983561 | 0.723338912 | 0.844046387 | 13.56333429 | 15.71618601 | ENSG00000285979 |
| SPRYD7 | 0.212817319 | 1.158949191 | 0.283481841 | 0.474492229 | 118.387187 | 137.3060507 | ENSG00000123178 |
| CDAN1 | 0.212732152 | 1.158880776 | 0.105981088 | 0.236885369 | 391.6967016 | 454.3420801 | ENSG00000140326 |
| TMOD1 | 0.212719312 | 1.158870462 | 0.698642539 | 0.82815737 | 14.60858727 | 16.97253697 | ENSG00000136842 |
| MTG2 | 0.212487892 | 1.158684585 | 0.039982733 | 0.112466228 | 572.0573219 | 662.8313493 | ENSG00000101181 |
| MRPS36 | 0.212421795 | 1.158631501 | 0.119202244 | 0.258306372 | 323.4573574 | 374.6311461 | ENSG00000134056 |
| NDUFA10 | 0.212402121 | 1.158615701 | 0.0061935 | 0.024743897 | 1503.264863 | 1740.845572 | ENSG00000130414 |
| TSPAN18 | 0.212371387 | 1.158591019 | 0.050522559 | 0.134925809 | 562.7532159 | 651.8427479 | ENSG00000157570 |
| TMCC3 | 0.212305927 | 1.158538451 | 0.411203891 | 0.602708812 | 71.75177508 | 83.00152838 | ENSG00000057704 |
| SELENOO | 0.212291223 | 1.158526643 | 0.061417691 | 0.156900824 | 483.6805509 | 560.4231872 | ENSG00000073169 |
| IGF2BP3 | 0.212256963 | 1.158499132 | 0.007784872 | 0.029947036 | 3170.254362 | 3672.064629 | ENSG00000136231 |
| ARRDC1 | 0.21200798 | 1.158299213 | 0.184280052 | 0.354925334 | 237.4742159 | 275.5192811 | ENSG00000197070 |
| CDC42BPG | 0.211909978 | 1.158220533 | 0.041686625 | 0.116375947 | 562.2011549 | 651.444154 | ENSG00000171219 |
| LCTL | 0.211862466 | 1.15818239 | 0.283830407 | 0.474732675 | 119.8841037 | 138.832132 | ENSG00000188501 |
| CGGBP1 | 0.211792597 | 1.158126301 | 0.025125235 | 0.077764154 | 1084.964961 | 1256.16316 | ENSG00000163320 |
| GNL3L | 0.211777769 | 1.158114398 | 0.015158158 | 0.051522031 | 1076.563318 | 1246.925469 | ENSG00000130119 |
| TRAF6 | 0.21175157 | 1.158093367 | 0.209399212 | 0.388490944 | 184.9764216 | 214.0681125 | ENSG00000175104 |
| CUTC | 0.21164816 | 1.15801036 | 0.111970346 | 0.246584885 | 390.2568199 | 451.8155683 | ENSG00000119929 |
| FAM86B2 | 0.211642396 | 1.158005733 | 0.733027558 | 0.851101774 | 13.20379977 | 15.27349463 | ENSG00000145002 |
| SSR4 | 0.211617732 | 1.157985936 | 0.00402638 | 0.017104039 | 1464.549959 | 1695.969572 | ENSG00000180879 |
| NA | 0.211502515 | 1.157893461 | 0.755751909 | 0.864316049 | 10.02789541 | 11.64993477 | ENSG00000260948 |
| IGSF5 | 0.211419117 | 1.157826528 | 0.535282206 | 0.709650959 | 37.11865786 | 43.02913137 | ENSG00000183067 |
| PODXL2 | 0.21133647 | 1.157760202 | 0.010110239 | 0.03709089 | 1374.556827 | 1591.145803 | ENSG00000114631 |
| LINC02210 | 0.211294493 | 1.157726516 | 0.117007858 | 0.254982203 | 371.0433753 | 430.0929172 | ENSG00000204650 |
| CD46 | 0.211265806 | 1.157703495 | 0.042752803 | 0.118566579 | 1483.814433 | 1718.036571 | ENSG00000117335 |
| MPDZ | 0.211177529 | 1.157632659 | 0.011791617 | 0.042157761 | 1631.061785 | 1888.336582 | ENSG00000107186 |
| RPA3 | 0.211065202 | 1.157542531 | 0.09344514 | 0.215171106 | 497.8660595 | 575.776695 | ENSG00000106399 |
| NA | 0.211004055 | 1.15749347 | 0.693808106 | 0.824614559 | 15.90022957 | 18.47302249 | ENSG00000225200 |
| NA | 0.210967432 | 1.157464088 | 0.816798623 | 0.901479525 | 5.08501577 | 5.903850207 | ENSG00000263050 |
| DENND4A | 0.210934792 | 1.157437901 | 0.348942792 | 0.542244881 | 113.3468716 | 131.0146112 | ENSG00000174485 |
| EVI5L | 0.210930532 | 1.157434484 | 0.05426848 | 0.142365465 | 511.1106885 | 592.1175887 | ENSG00000142459 |
| HELLS | 0.210929758 | 1.157433862 | 0.003449298 | 0.014971731 | 1937.922574 | 2242.684424 | ENSG00000119969 |
| COL1A2 | 0.210853375 | 1.157372584 | 0.003225702 | 0.014153686 | 2237.761665 | 2589.317551 | ENSG00000164692 |
| TRIM62 | 0.21081447 | 1.157341374 | 0.130835157 | 0.276973021 | 306.4766904 | 354.9485612 | ENSG00000116525 |
| NUDT3 | 0.210812865 | 1.157340086 | 0.188047023 | 0.359787707 | 256.8804648 | 297.6748844 | ENSG00000272325 |
| TBC1D22A | 0.210765901 | 1.157302412 | 0.049245251 | 0.132181242 | 463.3894875 | 536.1938458 | ENSG00000054611 |
| CLEC11A | 0.210685226 | 1.157237698 | 0.245085736 | 0.432199718 | 142.2521358 | 164.8293875 | ENSG00000105472 |
| C16orf95 | 0.210556148 | 1.157134164 | 0.708542177 | 0.834156177 | 18.17183701 | 20.91443929 | ENSG00000260456 |
| EHD2 | 0.210475516 | 1.157069494 | 0.073656458 | 0.180027535 | 366.8849276 | 424.6691484 | ENSG00000024422 |
| VRK3 | 0.210458834 | 1.157056115 | 0.050419307 | 0.134708294 | 504.7316855 | 584.168774 | ENSG00000105053 |
| PHYHIPL | 0.210454057 | 1.157052284 | 0.822352861 | 0.904601199 | 5.138081929 | 5.96016319 | ENSG00000165443 |
| ERRFI1 | 0.210336971 | 1.156958384 | 0.005086635 | 0.020885178 | 1654.909183 | 1913.748409 | ENSG00000116285 |
| RALGAPA1 | 0.210251742 | 1.156890037 | 0.109553136 | 0.242500637 | 312.0608963 | 361.2800191 | ENSG00000174373 |
| CKMT1A | 0.21022295 | 1.156866949 | 0.292967596 | 0.485546045 | 139.1203305 | 161.3199909 | ENSG00000223572 |
| BRSK1 | 0.210177138 | 1.156830214 | 0.131340433 | 0.277535467 | 579.4962953 | 671.2298178 | ENSG00000160469 |
| TLL2 | 0.210052655 | 1.156730401 | 0.204110395 | 0.3821194 | 167.5323276 | 193.7737773 | ENSG00000095587 |
| FBXO10 | 0.209961231 | 1.156657101 | 0.255509324 | 0.444407828 | 142.0384957 | 164.4615568 | ENSG00000147912 |
| NA | 0.209926724 | 1.156629436 | 0.662010455 | 0.803380688 | 20.68696995 | 23.87720706 | ENSG00000235587 |
| ELOVL7 | 0.20990241 | 1.156609944 | 0.393646005 | 0.586309208 | 74.92156102 | 86.5758233 | ENSG00000164181 |
| ND4 | 0.209731464 | 1.156472904 | 0.456191882 | 0.643410726 | 24744.93365 | 28616.8931 | ENSG00000198886 |
| RPS27 | 0.209657024 | 1.156413234 | 0.000908552 | 0.004856969 | 9751.838959 | 11278.2886 | ENSG00000177954 |
| ZBTB7C | 0.209655983 | 1.156412399 | 0.776095935 | 0.876495159 | 9.841968454 | 11.27852852 | ENSG00000184828 |
| ZNF383 | 0.209638672 | 1.156398523 | 0.528421425 | 0.704224631 | 41.7050622 | 48.11844592 | ENSG00000188283 |
| SPINK1 | 0.209601213 | 1.156368499 | 0.77040977 | 0.873601287 | 8.88282393 | 10.26956129 | ENSG00000164266 |
| ZNF675 | 0.20948167 | 1.156272685 | 0.232304531 | 0.417039061 | 195.5641875 | 225.8703651 | ENSG00000197372 |
| DDX6 | 0.209459926 | 1.156255258 | 0.022863859 | 0.072129151 | 1942.337708 | 2244.95779 | ENSG00000110367 |
| CEBPA-DT | 0.209398455 | 1.156205993 | 0.412701538 | 0.604282691 | 75.88070554 | 87.54344969 | ENSG00000267296 |
| PRAG1 | 0.209358237 | 1.156173762 | 0.012418481 | 0.043990129 | 1061.142721 | 1227.448712 | ENSG00000275342 |
| PLEKHA7 | 0.209284501 | 1.156114671 | 0.143807217 | 0.296699577 | 436.155424 | 504.6794223 | ENSG00000166689 |
| PGAP1 | 0.209124508 | 1.155986467 | 0.123562978 | 0.265305401 | 405.7898105 | 468.3894769 | ENSG00000197121 |
| ZNF860 | 0.209110873 | 1.155975541 | 0.450143455 | 0.637871361 | 66.47490933 | 76.91376707 | ENSG00000197385 |
| CEP126 | 0.209106581 | 1.155972103 | 0.562918907 | 0.731226453 | 34.86895342 | 40.24972504 | ENSG00000110318 |
| EFNB2 | 0.208752733 | 1.155688613 | 0.21541682 | 0.395770751 | 161.875415 | 186.9656617 | ENSG00000125266 |
| MMGT1 | 0.208625865 | 1.155586989 | 0.124972776 | 0.267743086 | 625.5510387 | 722.2917243 | ENSG00000169446 |
| NA | 0.208593716 | 1.155561238 | 0.724521254 | 0.845024034 | 13.28221893 | 15.33685565 | ENSG00000287535 |
| SRGAP2B | 0.208586625 | 1.155555558 | 0.198602727 | 0.374722889 | 209.5393257 | 242.1708653 | ENSG00000196369 |
| TMEM128 | 0.208470263 | 1.155462359 | 0.152369961 | 0.309550819 | 221.0214142 | 255.3765077 | ENSG00000132406 |

| | | | | | | | |
|---|---|---|---|---|---|---|---|
| NA | 0.208467084 | 1.155459813 | 0.58188118 | 0.74582662 | 33.75820202 | 39.18119564 | ENSG00000266066 |
| CHTOP | 0.208440177 | 1.155438264 | 0.004661564 | 0.019345597 | 1677.5514 | 1938.761314 | ENSG00000160679 |
| MRRF | 0.208329884 | 1.155349934 | 0.087678912 | 0.205363446 | 381.6057882 | 441.0494789 | ENSG00000148187 |
| SRXN1 | 0.208304804 | 1.15532985 | 0.466180277 | 0.652459173 | 73.21799446 | 84.56559815 | ENSG00000271303 |
| ASH2L | 0.208301941 | 1.155327557 | 0.007399102 | 0.028677323 | 1302.230682 | 1504.310172 | ENSG00000129691 |
| DMTF1 | 0.208280516 | 1.1553104 | 0.043060891 | 0.119278217 | 974.0631546 | 1126.178152 | ENSG00000135164 |
| TTC19 | 0.208151122 | 1.155206786 | 0.003810136 | 0.016346542 | 1843.546185 | 2129.172453 | ENSG00000011295 |
| SRPRA | 0.208025723 | 1.15510638 | 0.008544871 | 0.032320537 | 1613.887156 | 1863.707524 | ENSG00000182934 |
| KXD1 | 0.207855521 | 1.154970114 | 0.025655003 | 0.079050677 | 769.0530179 | 888.2936825 | ENSG00000105700 |
| NFE2L1 | 0.207811501 | 1.154934874 | 0.000443108 | 0.002575278 | 6377.481636 | 7365.472986 | ENSG00000082641 |
| CPSF6 | 0.20768468 | 1.154833353 | 0.000705748 | 0.003881406 | 2980.151907 | 3441.71463 | ENSG00000111605 |
| OSTF1 | 0.207655128 | 1.154809698 | 0.124841496 | 0.267585555 | 271.6061434 | 313.7126536 | ENSG00000134996 |
| REXO1 | 0.20764507 | 1.154801647 | 0.023244021 | 0.072988944 | 967.9341258 | 1117.756804 | ENSG00000079313 |
| ZNF765 | 0.207614314 | 1.154777029 | 0.163625835 | 0.32556535 | 235.6092786 | 271.9499557 | ENSG00000196417 |
| PCBP3 | 0.207561955 | 1.154735119 | 0.266800117 | 0.456716853 | 135.8689685 | 156.9000051 | ENSG00000183570 |
| F8A1 | 0.207306612 | 1.154530761 | 0.448510204 | 0.63620292 | 76.90721245 | 88.66361716 | ENSG00000277203 |
| SPATA24 | 0.206970738 | 1.154262005 | 0.291324501 | 0.483761659 | 140.6142631 | 162.5337901 | ENSG00000170469 |
| RBP5 | 0.20695002 | 1.15424543 | 0.677370412 | 0.813476526 | 20.48092144 | 23.67908826 | ENSG00000139194 |
| PDE4B | 0.206932998 | 1.154231811 | 0.256055919 | 0.444941858 | 146.5477586 | 169.0444925 | ENSG00000184588 |
| SOCS4 | 0.206900346 | 1.154205688 | 0.074863001 | 0.182295218 | 401.0796101 | 463.0779222 | ENSG00000180008 |
| ARFGEF2 | 0.206803746 | 1.154128407 | 0.035335364 | 0.101962577 | 797.4072961 | 919.6046913 | ENSG00000124198 |
| MED9 | 0.206647696 | 1.154003577 | 0.087325167 | 0.204767795 | 386.4508036 | 445.5958201 | ENSG00000141026 |
| TMEM219 | 0.206471514 | 1.153862659 | 0.039808933 | 0.112062389 | 648.0627105 | 747.3566028 | ENSG00000149932 |
| LOC101927974 | 0.206346705 | 1.153762841 | 0.507666625 | 0.68692604 | 45.51549801 | 52.59788227 | ENSG00000241764 |
| CHP1 | 0.206322639 | 1.153743595 | 0.007284735 | 0.028293226 | 1650.970081 | 1904.076986 | ENSG00000187446 |
| CALCOCO1 | 0.206215341 | 1.15365779 | 0.21689703 | 0.397456783 | 282.4068656 | 325.8502449 | ENSG00000012822 |
| UGGT2 | 0.206199631 | 1.153645228 | 0.153984963 | 0.311567392 | 334.6170952 | 386.1064164 | ENSG00000102595 |
| CHST2 | 0.205992187 | 1.153479358 | 0.046153717 | 0.125638294 | 807.1576313 | 930.7907587 | ENSG00000175040 |
| TMEM131 | 0.205908566 | 1.153412503 | 0.061470262 | 0.156991848 | 588.3894996 | 678.0940684 | ENSG00000075568 |
| AP1G2 | 0.205803923 | 1.153328845 | 0.247841453 | 0.435800274 | 279.7522725 | 322.9410821 | ENSG00000213983 |
| CHIC1 | 0.205778213 | 1.153308292 | 0.323396495 | 0.517601955 | 115.5455618 | 133.0879825 | ENSG00000204116 |
| PRRC2A | 0.205697317 | 1.153243624 | 0.000127977 | 0.000867566 | 14128.40176 | 16293.35181 | ENSG00000204469 |
| ISCU | 0.205639024 | 1.153197028 | 0.016793892 | 0.05594141 | 1347.165316 | 1553.856027 | ENSG00000136003 |
| NA | 0.205581652 | 1.153151169 | 0.523264029 | 0.699838045 | 42.79263282 | 49.3527767 | ENSG00000185834 |
| SERTAD4 | 0.205554661 | 1.153129596 | 0.465887243 | 0.652171765 | 54.88736494 | 63.21952916 | ENSG00000082497 |
| SNX10 | 0.205412021 | 1.153015591 | 0.317335807 | 0.510990603 | 116.981723 | 134.588468 | ENSG00000086300 |
| RCCD1 | 0.205193794 | 1.152841195 | 0.173869867 | 0.340573253 | 473.0007308 | 545.8428911 | ENSG00000166965 |
| SNHG15 | 0.205159286 | 1.15281362 | 0.115430675 | 0.252474543 | 527.4731576 | 608.8777278 | ENSG00000232956 |
| AIMP1 | 0.205096469 | 1.152763426 | 0.035561617 | 0.102519679 | 635.4654207 | 732.562895 | ENSG00000164022 |
| NA | 0.204990901 | 1.152679077 | 0.395928663 | 0.588262452 | 92.12162586 | 106.4042763 | ENSG00000215158 |
| PLEKHG2 | 0.204853894 | 1.152569617 | 0.010202373 | 0.037391889 | 1136.822144 | 1310.358767 | ENSG00000090924 |
| N4BP1 | 0.204838212 | 1.152557088 | 0.017569278 | 0.057944129 | 1092.978643 | 1259.298323 | ENSG00000102921 |
| NA | 0.204779276 | 1.152510006 | 0.70664883 | 0.833271761 | 17.11415177 | 19.78579806 | ENSG00000275765 |
| LMBR1L | 0.204510717 | 1.152295485 | 0.062098669 | 0.158160881 | 564.3435395 | 650.1109501 | ENSG00000139636 |
| LONP1 | 0.204492678 | 1.152281077 | 0.000622959 | 0.003475596 | 4369.015565 | 5033.989151 | ENSG00000196365 |
| CPVL | 0.204415977 | 1.152219818 | 0.070953718 | 0.174828 | 530.6367199 | 610.6138564 | ENSG00000106066 |
| WDR60 | 0.20435938 | 1.152174617 | 0.127002486 | 0.270652417 | 428.6039818 | 494.0327922 | ENSG00000126870 |
| RSAD1 | 0.204334807 | 1.152154992 | 0.02206897 | 0.0701228 | 856.6712661 | 987.4631117 | ENSG00000136444 |
| CDKN2D | 0.204297874 | 1.152125498 | 0.5739872 | 0.739436906 | 33.57237277 | 38.81738709 | ENSG00000129355 |
| RSRC1 | 0.204276396 | 1.152108345 | 0.136656924 | 0.285578919 | 301.7114396 | 347.5011728 | ENSG00000174891 |
| SPRYD3 | 0.204134538 | 1.151995066 | 0.166885853 | 0.330403123 | 240.7215265 | 277.3287592 | ENSG00000167778 |
| SH3BP2 | 0.204008125 | 1.151894129 | 0.050539634 | 0.134932524 | 802.5552774 | 924.6594127 | ENSG00000087266 |
| MUTYH | 0.203978951 | 1.151870837 | 0.672725756 | 0.809789244 | 18.83733623 | 21.67591116 | ENSG00000132781 |
| PIP4P1 | 0.203811732 | 1.151737334 | 0.033222329 | 0.097166269 | 619.9941551 | 714.0498543 | ENSG00000165782 |
| RAB5A | 0.203461481 | 1.151457754 | 0.03523519 | 0.101779962 | 1147.314488 | 1320.622697 | ENSG00000144566 |
| MTERF1 | 0.203437425 | 1.151438554 | 0.242653907 | 0.429116199 | 178.8366821 | 205.5901259 | ENSG00000127989 |
| BCL2L2 | 0.203159414 | 1.151216691 | 0.117122993 | 0.255143107 | 399.6911859 | 460.5676252 | ENSG00000129473 |
| IGF2R | 0.202916983 | 1.151023256 | 0.017080427 | 0.056702244 | 4049.452805 | 4660.480198 | ENSG00000197081 |
| GPR173 | 0.202807796 | 1.150936147 | 0.208332455 | 0.387432465 | 211.9222969 | 243.9184082 | ENSG00000184194 |
| LYRM4 | 0.202761956 | 1.150899578 | 0.035698967 | 0.102853088 | 631.8681365 | 727.0582455 | ENSG00000214113 |
| NA | 0.202693864 | 1.15084526 | 0.82310999 | 0.904942493 | 5.637971143 | 6.511355273 | ENSG00000235001 |
| ITPKB | 0.202593551 | 1.150765242 | 0.153731067 | 0.31122355 | 283.1790083 | 325.6570469 | ENSG00000143772 |
| GORAB | 0.202557523 | 1.150736505 | 0.21560655 | 0.395955923 | 197.7008445 | 227.7783018 | ENSG00000120370 |
| DAB1 | 0.202418057 | 1.150625268 | 0.085034564 | 0.200513407 | 423.3746651 | 486.6701548 | ENSG00000173406 |
| ADAM10 | 0.202388562 | 1.150601744 | 0.005617602 | 0.022757736 | 2173.273677 | 2499.785083 | ENSG00000137845 |
| ZNF571 | 0.202261102 | 1.150500094 | 0.595022291 | 0.7559218 | 35.5907734 | 41.10136466 | ENSG00000180479 |
| MANBA | 0.202256172 | 1.150496163 | 0.038731686 | 0.109662859 | 710.2207477 | 816.6416011 | ENSG00000109323 |
| GLUL | 0.202166082 | 1.150424322 | 0.001221913 | 0.00623539 | 5506.651958 | 6334.30821 | ENSG00000135821 |
| SLC35G2 | 0.202153498 | 1.150414287 | 0.461838593 | 0.648316025 | 59.87256222 | 68.88359332 | ENSG00000168917 |
| LRTOMT | 0.202147762 | 1.150409713 | 0.655985033 | 0.79911618 | 20.69644069 | 23.83272865 | ENSG00000184154 |
| ZNF331 | 0.202008584 | 1.150298737 | 0.665157712 | 0.805385316 | 1104.058601 | 1270.132415 | ENSG00000130844 |
| SLC9A3R2 | 0.201964861 | 1.150263876 | 0.150234757 | 0.30632024 | 339.2881801 | 390.2073497 | ENSG00000065054 |
| NA | 0.201963166 | 1.150262525 | 0.514784139 | 0.692846128 | 57.50913385 | 66.3003707 | ENSG00000260853 |
| LOC100133091 | 0.201962896 | 1.15026231 | 0.540586774 | 0.713722415 | 47.77151628 | 55.1337037 | ENSG00000205485 |

| | | | | | | | |
|---|---|---|---|---|---|---|---|
| HAGHL | 0.201943072 | 1.150246504 | 0.125324699 | 0.268270364 | 546.2904699 | 628.9501803 | ENSG00000103253 |
| CYB5R3 | 0.20194045 | 1.150244414 | 0.003735097 | 0.016058058 | 2034.851219 | 2341.270702 | ENSG00000100243 |
| SPAG16 | 0.201827499 | 1.150154363 | 0.227156121 | 0.410699011 | 199.2646124 | 228.9962347 | ENSG00000144451 |
| GINM1 | 0.201741604 | 1.150085887 | 0.194881417 | 0.369159164 | 248.957289 | 286.1002081 | ENSG00000055211 |
| OSBPL2 | 0.201523611 | 1.149912121 | 0.061729636 | 0.157480672 | 668.2236263 | 768.932622 | ENSG00000130703 |
| BUB3 | 0.201503093 | 1.149895767 | 0.001489791 | 0.007363141 | 5036.381857 | 5791.872331 | ENSG00000154473 |
| TOR1AIP2 | 0.201325897 | 1.149754542 | 0.092984059 | 0.214535858 | 501.166609 | 575.8977778 | ENSG00000169905 |
| AGL | 0.201237885 | 1.149684403 | 0.108614881 | 0.241028282 | 510.1836093 | 585.6500731 | ENSG00000162688 |
| TNFAIP8L1 | 0.201213868 | 1.149665264 | 0.214982778 | 0.395129989 | 205.4467672 | 236.2318462 | ENSG00000185361 |
| MSRB2 | 0.201033134 | 1.149521249 | 0.176648569 | 0.344632061 | 222.1404563 | 255.2713678 | ENSG00000148450 |
| ZNF93 | 0.201016637 | 1.149508104 | 0.067778128 | 0.168730667 | 610.1272244 | 701.1588352 | ENSG00000184635 |
| ZNF148 | 0.200977702 | 1.149477081 | 0.200533117 | 0.377174832 | 240.2581221 | 275.7103116 | ENSG00000163848 |
| ADRA2A | 0.200959312 | 1.149462429 | 0.610356438 | 0.767487918 | 27.83892012 | 31.92670146 | ENSG00000150594 |
| INCENP | 0.200906778 | 1.149420574 | 0.005327358 | 0.021762674 | 2257.719945 | 2594.242903 | ENSG00000149503 |
| RABAC1 | 0.200903875 | 1.149418261 | 0.66100151 | 0.802702466 | 27.23555184 | 31.44365693 | ENSG00000105404 |
| CINP | 0.20079334 | 1.149330199 | 0.079746469 | 0.191304821 | 493.0904736 | 566.2102459 | ENSG00000100865 |
| INAFM1 | 0.200739834 | 1.149287574 | 0.343890678 | 0.537587178 | 116.0517648 | 133.396689 | ENSG00000257704 |
| ILDR2 | 0.200634683 | 1.149203812 | 0.636873976 | 0.786554426 | 23.91919508 | 27.44825279 | ENSG00000143195 |
| ZNF224 | 0.200546043 | 1.149133206 | 0.475381914 | 0.660127999 | 57.47313003 | 65.95390742 | ENSG00000267680 |
| RRP7BP | 0.200525096 | 1.149116521 | 0.314103418 | 0.507759937 | 248.0621697 | 285.5995725 | ENSG00000182841 |
| FGF23 | 0.20049596 | 1.149093315 | 0.716660445 | 0.839556455 | 13.98362426 | 16.03821569 | ENSG00000118972 |
| HMGXB3 | 0.200457726 | 1.149062862 | 0.01396701 | 0.048268612 | 1015.291967 | 1166.381111 | ENSG00000113716 |
| NA | 0.200432794 | 1.149043005 | 0.803461493 | 0.89448314 | 7.552020899 | 8.739569514 | ENSG00000262902 |
| SPRY2 | 0.200358512 | 1.148983844 | 0.075757112 | 0.183900761 | 886.6000914 | 1018.405686 | ENSG00000136158 |
| ZBTB7B | 0.200336894 | 1.148966627 | 0.045086388 | 0.123449713 | 714.4439796 | 821.5604655 | ENSG00000160685 |
| NA | 0.200212726 | 1.148867744 | 0.83783848 | 0.91290408 | 4.505527319 | 5.157350486 | ENSG00000231680 |
| PNPLA7 | 0.199981362 | 1.148683515 | 0.695017007 | 0.825416199 | 18.18140545 | 20.95610654 | ENSG00000130653 |
| NA | 0.199961151 | 1.148667423 | 0.523010269 | 0.699675423 | 47.23878016 | 54.18699819 | ENSG00000212802 |
| ABHD11 | 0.199939172 | 1.148649924 | 0.142925327 | 0.295414074 | 275.0735284 | 315.905699 | ENSG00000106077 |
| IMPACT | 0.199918354 | 1.148633349 | 0.049830629 | 0.133404719 | 768.657269 | 882.3338852 | ENSG00000154059 |
| ANAPC16 | 0.199805626 | 1.148543602 | 0.051831535 | 0.137308351 | 562.9613162 | 646.2682071 | ENSG00000166295 |
| ZNF770 | 0.199723068 | 1.148477878 | 0.018325141 | 0.059922046 | 1513.154869 | 1736.868365 | ENSG00000198146 |
| SNTG2 | 0.199680573 | 1.14844405 | 0.198154982 | 0.373992305 | 192.1264926 | 220.6214614 | ENSG00000172554 |
| RBM18 | 0.199676778 | 1.148441029 | 0.264034446 | 0.453742644 | 182.5556878 | 209.6794967 | ENSG00000119446 |
| RANGAP1 | 0.199667911 | 1.148433971 | 0.000748864 | 0.004093484 | 5775.630079 | 6633.50777 | ENSG00000100401 |
| HDAC10 | 0.199500171 | 1.148300451 | 0.811630947 | 0.898880668 | 6.709659508 | 7.741887206 | ENSG00000100429 |
| FUT11 | 0.199461204 | 1.148269437 | 0.275807912 | 0.466822508 | 140.4162121 | 161.0287413 | ENSG00000196968 |
| CDV3 | 0.199438452 | 1.148251328 | 0.010206702 | 0.037400359 | 3068.896158 | 3522.612409 | ENSG00000091527 |
| CNEP1R1 | 0.199387479 | 1.148210759 | 0.212760747 | 0.392447024 | 197.4091685 | 226.3783928 | ENSG00000205423 |
| NA | 0.199364093 | 1.148192147 | 0.599712675 | 0.759411523 | 35.46560191 | 40.83889715 | ENSG00000228502 |
| PATL1 | 0.199259379 | 1.148108811 | 0.031788415 | 0.093831262 | 852.927336 | 979.0091921 | ENSG00000166889 |
| MIEN1 | 0.199239227 | 1.148092774 | 0.344743889 | 0.538243049 | 103.6584642 | 118.8963892 | ENSG00000141741 |
| CAMLG | 0.199181431 | 1.148046782 | 0.040671428 | 0.113953779 | 944.1952878 | 1083.630278 | ENSG00000164615 |
| NA | 0.199150674 | 1.148022306 | 0.870619029 | 0.932827673 | 3.563347437 | 4.054528287 | ENSG00000227973 |
| TXNDC12 | 0.198970897 | 1.147879258 | 0.261949182 | 0.451186754 | 155.1336827 | 178.2233926 | ENSG00000117862 |
| IFITM2 | 0.198884372 | 1.147810416 | 0.016736753 | 0.055811259 | 1027.577737 | 1179.276661 | ENSG00000185201 |
| NPC1 | 0.198763391 | 1.147714168 | 0.018968367 | 0.061709621 | 929.4703687 | 1066.535479 | ENSG00000141458 |
| DIO3 | 0.198615193 | 1.147596277 | 0.808953295 | 0.897335896 | 6.603527191 | 7.621663547 | ENSG00000197406 |
| DET1 | 0.198540132 | 1.147536571 | 0.809361358 | 0.897454647 | 6.388007946 | 7.362994886 | ENSG00000140543 |
| NA | 0.198456211 | 1.147469821 | 0.822923568 | 0.904942493 | 5.819268009 | 6.687891916 | ENSG00000229666 |
| SETD1B | 0.19840979 | 1.1474329 | 0.087286787 | 0.204703697 | 447.8026414 | 513.9038195 | ENSG00000139718 |
| RPAP1 | 0.198238078 | 1.147296339 | 0.005574773 | 0.022618835 | 1721.443893 | 1975.122678 | ENSG00000103932 |
| PPP1R26-AS1 | 0.198169216 | 1.147241578 | 0.571430279 | 0.737681186 | 37.93892086 | 43.70478399 | ENSG00000225361 |
| MYLK-AS1 | 0.198164691 | 1.14723798 | 0.807650101 | 0.896776118 | 6.583307938 | 7.595406479 | ENSG00000239523 |
| NA | 0.198072756 | 1.147164875 | 0.757101328 | 0.864710144 | 12.21358978 | 13.93391234 | ENSG00000280173 |
| C9orf16 | 0.198066305 | 1.147159745 | 0.059911733 | 0.15372085 | 532.5476953 | 611.0111304 | ENSG00000171159 |
| EYA3 | 0.198062363 | 1.147156611 | 0.046904784 | 0.127169874 | 687.625794 | 788.7052311 | ENSG00000158161 |
| AQP4 | 0.198057791 | 1.147152975 | 0.667441027 | 0.80717496 | 24.40380339 | 27.84167922 | ENSG00000171885 |
| SATB1 | 0.19805764 | 1.147152855 | 0.058652913 | 0.151255473 | 582.6864134 | 668.7809291 | ENSG00000182568 |
| WNK3 | 0.1978758 | 1.147008275 | 0.120785901 | 0.260944852 | 339.3612475 | 389.3928543 | ENSG00000196632 |
| THOC2 | 0.197850595 | 1.146988236 | 0.015314241 | 0.051946219 | 1130.846225 | 1296.455077 | ENSG00000125676 |
| ZNF644 | 0.197825823 | 1.146968542 | 0.047899744 | 0.129401553 | 910.2810744 | 1043.805318 | ENSG00000122482 |
| KCNQ1OT1 | 0.197800917 | 1.146948741 | 0.561746072 | 0.730570307 | 38.29963546 | 43.97093872 | ENSG00000269821 |
| SDC3 | 0.197684697 | 1.14685635 | 0.004666963 | 0.019363667 | 1681.697122 | 1928.297464 | ENSG00000162512 |
| COA5 | 0.197371585 | 1.146607472 | 0.078488902 | 0.188899274 | 466.8987766 | 535.322711 | ENSG00000183513 |
| UXS1 | 0.19734443 | 1.14658589 | 0.01660794 | 0.055491538 | 1000.842367 | 1147.786814 | ENSG00000115652 |
| MRPL38 | 0.197216909 | 1.146484546 | 0.631882063 | 0.782788727 | 29.86541602 | 34.19245769 | ENSG00000204316 |
| NA | 0.197206879 | 1.146476576 | 0.626853476 | 0.779190626 | 30.04620923 | 34.41347274 | ENSG00000259612 |
| ABCA17P | 0.197095797 | 1.146388305 | 0.802894422 | 0.894151534 | 7.364312509 | 8.363926387 | ENSG00000238098 |
| UGGT1 | 0.197033661 | 1.146338932 | 0.018066654 | 0.059254533 | 1834.157301 | 2102.235188 | ENSG00000136731 |
| NA | 0.197003217 | 1.146314742 | 0.714655556 | 0.838366554 | 17.13121411 | 19.67294885 | ENSG00000228205 |
| NA | 0.196990856 | 1.146304921 | 0.840255251 | 0.914223597 | 4.525746572 | 5.16483656 | ENSG00000262136 |
| ZNF518A | 0.196918091 | 1.146247106 | 0.127924514 | 0.272023055 | 398.8655035 | 457.2561008 | ENSG00000177853 |
| NA | 0.196896968 | 1.146230324 | 0.779867811 | 0.878986213 | 8.951674653 | 10.21704715 | ENSG00000250421 |

| | | | | | | | |
|---|---|---|---|---|---|---|---|
| NA | 0.196869651 | 1.14620862 | 0.835789833 | 0.911983321 | 4.866339612 | 5.562499875 | ENSG00000256393 |
| STXBP1 | 0.196701006 | 1.146074641 | 0.012357051 | 0.043829515 | 1279.56233 | 1466.830647 | ENSG00000136854 |
| EPM2A | 0.196652367 | 1.146036002 | 0.414810877 | 0.606314808 | 76.28846557 | 87.36300258 | ENSG00000112425 |
| RUBCN | 0.196610648 | 1.146002863 | 0.200620927 | 0.377225146 | 219.5437473 | 251.41941 | ENSG00000145016 |
| LINC00926 | 0.196513943 | 1.145926048 | 0.732842765 | 0.851034324 | 13.7068459 | 15.6518373 | ENSG00000247982 |
| SS18 | 0.196445539 | 1.145871716 | 0.011612049 | 0.041628184 | 1635.786793 | 1873.805478 | ENSG00000141380 |
| OCRL | 0.196252321 | 1.145718262 | 0.022679568 | 0.071633114 | 1210.12076 | 1385.934982 | ENSG00000122126 |
| SHISA4 | 0.196166261 | 1.145649919 | 0.413200488 | 0.604535668 | 103.1939775 | 118.2373577 | ENSG00000198892 |
| TGOLN2 | 0.196122752 | 1.145615369 | 0.007571587 | 0.029229703 | 1623.790128 | 1860.299835 | ENSG00000152291 |
| SEMA3A | 0.196108012 | 1.145603664 | 0.056117252 | 0.146119296 | 862.3824476 | 987.1063419 | ENSG00000075213 |
| NA | 0.19605477 | 1.145561387 | 0.762327894 | 0.868300466 | 10.12643612 | 11.58982294 | ENSG00000253893 |
| MRPL23 | 0.195942662 | 1.145472372 | 0.197428482 | 0.373191177 | 210.060306 | 240.6810119 | ENSG00000214026 |
| EIF3M | 0.195941097 | 1.145471129 | 0.002275921 | 0.010598472 | 3642.49009 | 4171.769638 | ENSG00000149100 |
| RNF113A | 0.195939204 | 1.145469626 | 0.235768632 | 0.420773791 | 212.4934947 | 243.5525613 | ENSG00000125352 |
| ZNF706 | 0.195935503 | 1.145466688 | 0.036271783 | 0.10416195 | 778.6882237 | 891.9910291 | ENSG00000120963 |
| RIMKLA | 0.195931647 | 1.145463626 | 0.264806216 | 0.454521061 | 236.1085816 | 269.7917124 | ENSG00000177181 |
| BCKDK | 0.195891811 | 1.145431998 | 0.015426278 | 0.052268854 | 987.6039619 | 1131.212115 | ENSG00000103507 |
| NUDT1 | 0.195825169 | 1.145379089 | 0.031586314 | 0.093338766 | 860.9593724 | 986.4395072 | ENSG00000106268 |
| ZNF169 | 0.195692826 | 1.145274024 | 0.501987708 | 0.682500013 | 63.05309667 | 72.51920255 | ENSG00000175787 |
| NA | 0.195662341 | 1.145249824 | 0.824390678 | 0.905596573 | 5.455298802 | 6.248887766 | ENSG00000271013 |
| MIR4500HG | 0.195525368 | 1.145141096 | 0.827059877 | 0.907078985 | 5.363071913 | 6.143747873 | ENSG00000228824 |
| FAR1 | 0.195405604 | 1.145046038 | 0.029222912 | 0.087725535 | 908.1477396 | 1039.692367 | ENSG00000197601 |
| CAMKMT | 0.195385551 | 1.145030122 | 0.137526319 | 0.286975096 | 306.4439337 | 350.4466667 | ENSG00000143919 |
| NUDT12 | 0.195313183 | 1.144972686 | 0.24858994 | 0.436799957 | 200.0676403 | 228.6992596 | ENSG00000112874 |
| NA | 0.195147168 | 1.144840939 | 0.591153258 | 0.752964718 | 41.7631644 | 47.58872437 | ENSG00000261377 |
| NBPF12 | 0.195123926 | 1.144822495 | 0.308140718 | 0.502293761 | 163.4328917 | 187.0227391 | ENSG00000268043 |
| NCK2 | 0.194884054 | 1.144632165 | 0.014128837 | 0.048670975 | 1526.11119 | 1747.318557 | ENSG00000071051 |
| NA | 0.194726561 | 1.144507217 | 0.855077232 | 0.923059911 | 3.847717405 | 4.362023857 | ENSG00000255200 |
| PSTK | 0.194664423 | 1.144457924 | 0.395314297 | 0.587768127 | 86.13990476 | 98.43168935 | ENSG00000179988 |
| C12orf4 | 0.194500422 | 1.144327833 | 0.34676516 | 0.540170237 | 129.4621168 | 147.9485068 | ENSG00000047621 |
| ZYG11B | 0.19433556 | 1.144197074 | 0.084541694 | 0.199731772 | 436.4172896 | 499.0721402 | ENSG00000162378 |
| LURAP1L | 0.194194841 | 1.144085476 | 0.861654409 | 0.927350492 | 3.910254302 | 4.433420607 | ENSG00000153714 |
| MINDY2 | 0.194106434 | 1.144015369 | 0.127750667 | 0.271715728 | 435.8213023 | 498.4924925 | ENSG00000128923 |
| C8orf48 | 0.194096594 | 1.144007567 | 0.824873917 | 0.905753491 | 5.458455715 | 6.241401692 | ENSG00000164743 |
| SLC35E2A | 0.194074384 | 1.143989955 | 0.266073733 | 0.456063126 | 196.2401422 | 224.8516905 | ENSG00000215790 |
| UQCC2 | 0.194031862 | 1.143956237 | 0.155207522 | 0.313288632 | 531.4852748 | 608.3362748 | ENSG00000137288 |
| STK19 | 0.193906827 | 1.143857097 | 0.206582689 | 0.385152757 | 206.2441577 | 236.1432158 | ENSG00000204344 |
| COL12A1 | 0.193857975 | 1.143818366 | 0.523591443 | 0.700010373 | 48.19170855 | 55.10634733 | ENSG00000111799 |
| TNK2 | 0.193795572 | 1.143768891 | 0.060621593 | 0.15523081 | 823.4795642 | 942.7276122 | ENSG00000061938 |
| STAT1 | 0.193755828 | 1.143737383 | 0.006538698 | 0.025873688 | 1574.498476 | 1800.562203 | ENSG00000115415 |
| PRSS53 | 0.193754094 | 1.143736008 | 0.721369407 | 0.842461284 | 17.04718019 | 19.62847044 | ENSG00000151006 |
| OPRD1 | 0.193703095 | 1.143695578 | 0.496939015 | 0.678123082 | 63.9276059 | 73.29532016 | ENSG00000116329 |
| NA | 0.193561056 | 1.143582982 | 0.800011767 | 0.892525635 | 7.879986287 | 9.073322152 | ENSG00000213757 |
| LOC101927060 | 0.193514458 | 1.143546045 | 0.222221022 | 0.404264274 | 195.1197246 | 223.1769299 | ENSG00000262879 |
| CISD1 | 0.193482514 | 1.143520726 | 0.112212176 | 0.24705874 | 377.6002478 | 431.881649 | ENSG00000122873 |
| SACM1L | 0.193457719 | 1.143501073 | 0.017519309 | 0.057827759 | 1351.651406 | 1544.925552 | ENSG00000211456 |
| PKM | 0.193166037 | 1.143269904 | 0.002189967 | 0.010280862 | 19472.17285 | 22260.62645 | ENSG00000067225 |
| MAPRE1 | 0.193051955 | 1.143179504 | 0.001935068 | 0.009245823 | 4430.580299 | 5064.340019 | ENSG00000101367 |
| ZKSCAN8 | 0.192917424 | 1.143072907 | 0.090902815 | 0.211206314 | 453.7361144 | 518.9065907 | ENSG00000198315 |
| HEXD | 0.192753641 | 1.142943146 | 0.214272762 | 0.394411696 | 363.2898383 | 415.8759429 | ENSG00000169660 |
| LOC554206 | 0.192433433 | 1.142689497 | 0.724378827 | 0.844911087 | 14.72546811 | 16.77343049 | ENSG00000262587 |
| POFUT2 | 0.192393804 | 1.142658109 | 0.032973855 | 0.096561451 | 847.0745971 | 967.6456256 | ENSG00000186866 |
| NANOGP8 | 0.192352992 | 1.142625785 | 0.264459542 | 0.454094232 | 163.8438086 | 187.2405134 | ENSG00000255192 |
| UPF1 | 0.192293535 | 1.142578696 | 0.001379822 | 0.006895017 | 4258.353782 | 4864.750044 | ENSG00000005007 |
| STAT5B | 0.192242362 | 1.142538169 | 0.022392246 | 0.070858442 | 1474.223687 | 1684.412702 | ENSG00000173757 |
| IL11RA | 0.192234092 | 1.142531619 | 0.621491294 | 0.775152125 | 33.65375344 | 38.43860638 | ENSG00000137070 |
| MYLK | 0.192127949 | 1.142447563 | 0.1003142 | 0.22712233 | 401.2582537 | 458.4943251 | ENSG00000065534 |
| MTFMT | 0.191952377 | 1.142308539 | 0.188318437 | 0.360158282 | 332.7505421 | 380.1934311 | ENSG00000103707 |
| WDR78 | 0.191734744 | 1.142136233 | 0.845320966 | 0.917497001 | 4.872653438 | 5.543728881 | ENSG00000152763 |
| LOC100506411 | 0.191690664 | 1.142101336 | 0.346296421 | 0.53984842 | 125.7198326 | 143.6074526 | ENSG00000259153 |
| WHAMM | 0.191664238 | 1.142080417 | 0.324728313 | 0.518837462 | 132.6211542 | 151.7844335 | ENSG00000156232 |
| EPAS1 | 0.191663749 | 1.142080029 | 0.477364281 | 0.661721941 | 67.63043623 | 76.98450255 | ENSG00000116016 |
| GPX8 | 0.191651794 | 1.142070566 | 0.029499836 | 0.088327941 | 921.5108808 | 1052.025615 | ENSG00000164294 |
| RAB26 | 0.191627849 | 1.142051611 | 0.467670794 | 0.653533174 | 71.3316805 | 81.65968458 | ENSG00000167964 |
| FAM221A | 0.191557683 | 1.141996068 | 0.218684902 | 0.39995125 | 222.3973987 | 253.7866158 | ENSG00000188732 |
| CTSV | 0.19155526 | 1.14199415 | 0.012792729 | 0.045058997 | 1488.224741 | 1700.114858 | ENSG00000136943 |
| GCK | 0.191498104 | 1.141948908 | 0.264502962 | 0.454126716 | 160.8189098 | 183.4582002 | ENSG00000106633 |
| SOWAHB | 0.191481142 | 1.141935481 | 0.736915842 | 0.85266564 | 13.99121587 | 15.91050596 | ENSG00000186212 |
| COLGALT1 | 0.191448121 | 1.141909345 | 0.004517305 | 0.018860945 | 2498.637771 | 2853.280218 | ENSG00000130309 |
| FANCB | 0.191440949 | 1.141903668 | 0.378557697 | 0.571960962 | 106.7093925 | 121.60721 | ENSG00000181544 |
| PBX2 | 0.191369638 | 1.141847226 | 0.003937014 | 0.016824677 | 2923.358347 | 3338.031838 | ENSG00000204304 |
| DNAJC21 | 0.191306204 | 1.141797022 | 0.021472501 | 0.068580276 | 1006.433609 | 1149.194049 | ENSG00000168724 |
| SLC36A4 | 0.191225984 | 1.141733534 | 0.167470063 | 0.331233232 | 336.7654329 | 384.0663098 | ENSG00000180773 |
| SEMA4C | 0.191128755 | 1.141656591 | 0.004215408 | 0.017821253 | 2314.415453 | 2641.732725 | ENSG00000168758 |

| | | | | | | | |
|---|---|---|---|---|---|---|---|
| NA | 0.191117955 | 1.141648045 | 0.814126487 | 0.89991352 | 7.417378668 | 8.382697382 | ENSG00000228929 |
| AIM2 | 0.191044972 | 1.141590292 | 0.706144513 | 0.833185791 | 17.2486963 | 19.6955187 | ENSG00000163568 |
| G2E3 | 0.191004707 | 1.141558432 | 0.14261845 | 0.295010085 | 895.3030759 | 1021.530192 | ENSG00000092140 |
| PFKFB1 | 0.190975858 | 1.141535605 | 0.852291598 | 0.921931393 | 4.247690429 | 4.808514081 | ENSG00000158571 |
| TDRD7 | 0.190904017 | 1.141478762 | 0.499783569 | 0.680501665 | 55.66531029 | 63.65148527 | ENSG00000196116 |
| C2orf76 | 0.190752781 | 1.141359108 | 0.4599269 | 0.646562892 | 81.76832061 | 93.34314769 | ENSG00000186132 |
| BEX4 | 0.190714315 | 1.141328677 | 0.030796379 | 0.091456512 | 945.4763171 | 1079.060883 | ENSG00000102409 |
| KPNA5 | 0.190601723 | 1.141239608 | 0.311695779 | 0.505327089 | 163.200295 | 186.3461072 | ENSG00000196911 |
| NCOR2 | 0.190565289 | 1.141210787 | 0.005991227 | 0.024014144 | 6052.117911 | 6906.962809 | ENSG00000196498 |
| MIEF1 | 0.190539424 | 1.141190328 | 0.016644647 | 0.05560416 | 1169.553237 | 1334.432547 | ENSG00000100335 |
| ARSA | 0.190479815 | 1.141143177 | 0.182388458 | 0.352766634 | 253.6566432 | 289.8482688 | ENSG00000100299 |
| NA | 0.190465177 | 1.141131599 | 0.313019836 | 0.506593577 | 135.63323 | 154.7282155 | ENSG00000258634 |
| FIBCD1 | 0.190351963 | 1.141042053 | 0.36409735 | 0.55719431 | 126.2854383 | 144.0038018 | ENSG00000130720 |
| PCSK6 | 0.190338419 | 1.141031341 | 0.439721088 | 0.628007655 | 75.22111419 | 85.71278714 | ENSG00000140479 |
| SNX13 | 0.190186247 | 1.140910994 | 0.114413609 | 0.250684695 | 391.3699011 | 446.4928142 | ENSG00000071189 |
| KDF1 | 0.1898497 | 1.140644878 | 0.126976896 | 0.270629001 | 335.5418446 | 382.6594963 | ENSG00000175707 |
| ICE2 | 0.189806744 | 1.140610915 | 0.037551524 | 0.106876528 | 849.5356792 | 968.4506511 | ENSG00000128915 |
| NA | 0.189805075 | 1.140609595 | 0.732962387 | 0.851101774 | 14.26799423 | 16.34246207 | ENSG00000240914 |
| HSCB | 0.189709321 | 1.140533894 | 0.539127611 | 0.712557309 | 55.02003033 | 62.47215338 | ENSG00000100209 |
| LRG1 | 0.189676454 | 1.140507911 | 0.70794844 | 0.833950537 | 19.07495386 | 21.67167428 | ENSG00000171236 |
| TRIP4 | 0.189605422 | 1.140451759 | 0.18627021 | 0.357421273 | 290.3148808 | 331.0080963 | ENSG00000103671 |
| COX7A2 | 0.189579025 | 1.140430892 | 0.024138011 | 0.075277453 | 1960.824281 | 2235.407049 | ENSG00000112695 |
| CNTROB | 0.189570145 | 1.140423873 | 0.038995198 | 0.110324683 | 828.3031954 | 944.9516525 | ENSG00000170037 |
| DHRS7 | 0.189501761 | 1.140369818 | 0.051908294 | 0.1374187 | 765.1023177 | 872.087086 | ENSG00000100612 |
| ZNF426 | 0.189352962 | 1.140252207 | 0.149839423 | 0.305769441 | 313.8200846 | 357.8885444 | ENSG00000130818 |
| PODXL | 0.189229411 | 1.14015456 | 0.000528099 | 0.003011624 | 23817.57152 | 27155.47772 | ENSG00000128567 |
| UBE2G2 | 0.18922053 | 1.140147542 | 0.015953562 | 0.053721533 | 2225.255076 | 2537.824986 | ENSG00000184787 |
| HADHB | 0.189158749 | 1.140098718 | 0.043446067 | 0.120096115 | 948.4725408 | 1081.045845 | ENSG00000138029 |
| FLT1 | 0.189091397 | 1.140045494 | 0.014394456 | 0.049395825 | 1978.564584 | 2255.150558 | ENSG00000102755 |
| ACVR2A | 0.189001919 | 1.139974789 | 0.283477912 | 0.474492229 | 161.3597412 | 183.7738431 | ENSG00000121989 |
| PRRG1 | 0.188982665 | 1.139959576 | 0.398612753 | 0.590322714 | 92.36821057 | 105.291881 | ENSG00000130962 |
| NA | 0.188938306 | 1.139924525 | 0.518605984 | 0.69575413 | 52.3724274 | 59.8406535 | ENSG00000214176 |
| GPR21 | 0.188902616 | 1.139896326 | 0.049482611 | 0.132683843 | 770.5651178 | 878.4596529 | ENSG00000011454 |
| DUSP14 | 0.188847106 | 1.139852467 | 0.1137209 | 0.249538862 | 416.1854107 | 474.6917322 | ENSG00000276023 |
| PCNX4 | 0.188833977 | 1.139842094 | 0.07111242 | 0.175149181 | 543.8084544 | 619.9260257 | ENSG00000126773 |
| RRAS2 | 0.188818806 | 1.139830108 | 0.028789751 | 0.086677915 | 2939.054858 | 3348.970369 | ENSG00000133818 |
| GM2A | 0.188793764 | 1.139810323 | 0.050861175 | 0.135595657 | 806.1116194 | 918.9112956 | ENSG00000196743 |
| TRIO | 0.188409471 | 1.13950675 | 0.046387311 | 0.126080425 | 1430.285418 | 1630.46002 | ENSG00000038382 |
| IFT140 | 0.188387285 | 1.139489227 | 0.080208516 | 0.1921075 | 605.1251679 | 688.8051589 | ENSG00000187535 |
| VMAC | 0.1883596 | 1.13946736 | 0.734564217 | 0.851605263 | 14.27242892 | 16.32369107 | ENSG00000187650 |
| LENG8-AS1 | 0.188303196 | 1.139422812 | 0.683670649 | 0.817493665 | 25.05657726 | 28.56290956 | ENSG00000226696 |
| GTF2E1 | 0.188208982 | 1.139348406 | 0.020310121 | 0.065486739 | 1150.082933 | 1309.597247 | ENSG00000153767 |
| CDC42SE2 | 0.188202753 | 1.139343487 | 0.078221514 | 0.18847602 | 558.2466812 | 635.5126382 | ENSG00000158985 |
| YIPF5 | 0.188102554 | 1.139264359 | 0.105072039 | 0.235411458 | 457.8785443 | 521.1738037 | ENSG00000145817 |
| KLHL21 | 0.188034303 | 1.139210464 | 0.029804854 | 0.089111657 | 904.8946641 | 1031.014335 | ENSG00000162413 |
| SMDT1 | 0.187973087 | 1.139162127 | 0.177682573 | 0.346284838 | 257.8993952 | 293.9991368 | ENSG00000183172 |
| ETFA | 0.187867131 | 1.139078465 | 0.018905528 | 0.061548868 | 1795.753968 | 2044.796572 | ENSG00000140374 |
| EDEM2 | 0.187793552 | 1.139020373 | 0.110728332 | 0.244618847 | 413.4233097 | 471.0896429 | ENSG00000088298 |
| EPSTI1 | 0.18775728 | 1.138991736 | 0.745209529 | 0.857657316 | 14.76916122 | 16.88616808 | ENSG00000133106 |
| TEC | 0.187607987 | 1.138873877 | 0.555338133 | 0.725293281 | 46.72626329 | 53.20852491 | ENSG00000135605 |
| HDAC2-AS2 | 0.187601354 | 1.138868641 | 0.02860876 | 0.08631528 | 843.2328102 | 960.4259655 | ENSG00000228624 |
| LRCH4 | 0.187466504 | 1.138762195 | 0.182120675 | 0.352377555 | 258.5710129 | 294.7315405 | ENSG00000077454 |
| TSC22D2 | 0.187418054 | 1.138723953 | 0.091111664 | 0.211587352 | 634.55033 | 721.8949773 | ENSG00000196428 |
| THAP6 | 0.187310902 | 1.13863938 | 0.51410107 | 0.692567904 | 61.13068119 | 69.63378027 | ENSG00000174796 |
| FNIP2 | 0.18730895 | 1.13863784 | 0.241204455 | 0.427440453 | 212.990227 | 242.0704717 | ENSG00000052795 |
| SLC22A23 | 0.187287658 | 1.138621036 | 0.028189799 | 0.08542825 | 1136.943181 | 1294.367547 | ENSG00000137266 |
| ELMO3 | 0.187224739 | 1.138571379 | 0.2067649 | 0.385414931 | 247.309382 | 281.6034668 | ENSG00000102890 |
| PACS2 | 0.187088986 | 1.138464248 | 0.023032195 | 0.072498536 | 1654.93755 | 1884.863416 | ENSG00000179364 |
| NKAPP1 | 0.187079861 | 1.138457047 | 0.713645016 | 0.837849642 | 18.16118619 | 20.68191608 | ENSG00000233382 |
| MTFP1 | 0.186808445 | 1.138242888 | 0.706260805 | 0.833246982 | 22.67568059 | 25.89200394 | ENSG00000242114 |
| NA | 0.186804179 | 1.138239522 | 0.819332943 | 0.90291228 | 7.328308691 | 8.36447604 | ENSG00000250220 |
| PAIP2 | 0.186674711 | 1.138137381 | 0.029034227 | 0.087243897 | 1824.272022 | 2076.380578 | ENSG00000120727 |
| MEF2D | 0.186639869 | 1.138109894 | 0.095498244 | 0.218838336 | 468.9213036 | 533.4098762 | ENSG00000116604 |
| NEU1 | 0.186609583 | 1.138086002 | 0.026537161 | 0.081295827 | 931.5446016 | 1059.679198 | ENSG00000204386 |
| CSNK1A1 | 0.186607154 | 1.138084086 | 0.007427515 | 0.028763387 | 3088.922822 | 3515.589177 | ENSG00000113712 |
| DNAL4 | 0.186565517 | 1.138051241 | 0.321616119 | 0.51555381 | 178.2631167 | 202.5317914 | ENSG00000100246 |
| GATD3B | 0.186480664 | 1.137984308 | 0.337756912 | 0.531949916 | 144.221905 | 164.5527122 | ENSG00000280071 |
| NA | 0.186312263 | 1.137851482 | 0.746657663 | 0.858611443 | 12.93531207 | 14.74432273 | ENSG00000259394 |
| NA | 0.186016644 | 1.137618352 | 0.801129222 | 0.893214577 | 8.320397819 | 9.512326302 | ENSG00000261790 |
| NA | 0.186008893 | 1.137612241 | 0.403950501 | 0.595657458 | 90.9829659 | 103.510595 | ENSG00000229638 |
| TMLHE | 0.185964042 | 1.137576875 | 0.326976231 | 0.521267668 | 134.9124772 | 153.3953548 | ENSG00000185973 |
| ZNF394 | 0.185951559 | 1.137567032 | 0.090738342 | 0.210877045 | 469.6835999 | 534.1024763 | ENSG00000160908 |
| TMEM242 | 0.185948027 | 1.137564247 | 0.201743437 | 0.378831343 | 291.0638126 | 331.1849594 | ENSG00000215712 |
| RAPGEF2 | 0.185632504 | 1.137315484 | 0.023523451 | 0.073758017 | 1182.569787 | 1344.291958 | ENSG00000109756 |

| | | | | | | | |
|---|---|---|---|---|---|---|---|
| ADSS2 | 0.185458534 | 1.137178347 | 0.012231509 | 0.043437209 | 1452.942473 | 1652.458017 | ENSG00000035687 |
| UFL1 | 0.185448298 | 1.137170279 | 0.137837759 | 0.287398473 | 326.1717591 | 370.6036477 | ENSG00000014123 |
| MYO3A | 0.185426401 | 1.137153019 | 0.43600336 | 0.625455593 | 89.69387916 | 101.735043 | ENSG00000095777 |
| ADAT1 | 0.185334449 | 1.137080543 | 0.185989552 | 0.357030592 | 284.9954043 | 323.885334 | ENSG00000065457 |
| ALDH1B1 | 0.185193411 | 1.136969387 | 0.00619722 | 0.024748819 | 2966.619706 | 3372.398983 | ENSG00000137124 |
| UBE2E2 | 0.184950976 | 1.136778344 | 0.087943148 | 0.205827703 | 571.6980504 | 649.7807815 | ENSG00000182247 |
| GPR83 | 0.18487507 | 1.136718535 | 0.859997686 | 0.926192962 | 4.671137318 | 5.288309414 | ENSG00000123901 |
| ITM2C | 0.184865468 | 1.136710969 | 0.000279928 | 0.001730739 | 12635.31027 | 14362.58439 | ENSG00000135916 |
| SRF | 0.184812871 | 1.136669529 | 0.005607437 | 0.022731462 | 2995.331233 | 3404.350493 | ENSG00000112658 |
| NA | 0.184596038 | 1.136498703 | 0.807849707 | 0.896899705 | 7.278399445 | 8.300565364 | ENSG00000254893 |
| MFSD3 | 0.184537906 | 1.13645291 | 0.04527814 | 0.12386495 | 1214.080231 | 1379.839648 | ENSG00000167700 |
| TIMMDC1 | 0.184446931 | 1.136381249 | 0.011669856 | 0.04178717 | 1632.999075 | 1855.672174 | ENSG00000113845 |
| RNF139 | 0.184334805 | 1.136292933 | 0.117527794 | 0.255634331 | 424.894146 | 482.6044532 | ENSG00000170881 |
| CST6 | 0.184172756 | 1.136165307 | 0.835227163 | 0.911720374 | 5.763044937 | 6.533925114 | ENSG00000175315 |
| GPX4 | 0.184156253 | 1.136152311 | 0.001396304 | 0.006958618 | 3877.11418 | 4405.070304 | ENSG00000167468 |
| SCN2A | 0.183825958 | 1.135892226 | 0.647298691 | 0.794137415 | 27.11413862 | 30.7036556 | ENSG00000136531 |
| TSG101 | 0.18367859 | 1.135776204 | 0.066340616 | 0.166089014 | 925.7310985 | 1050.745818 | ENSG00000074319 |
| STK38L | 0.183592477 | 1.135708412 | 0.085861261 | 0.202127856 | 1032.096658 | 1171.825138 | ENSG00000211455 |
| NEU3 | 0.183370466 | 1.135533655 | 0.068766068 | 0.170800095 | 596.4076457 | 677.0979637 | ENSG00000162139 |
| FUBP1 | 0.183362449 | 1.135527346 | 0.001489175 | 0.007362061 | 4041.541536 | 4589.059428 | ENSG00000162613 |
| ACYP1 | 0.183244133 | 1.135434224 | 0.325359004 | 0.519510828 | 159.945077 | 181.8437033 | ENSG00000119640 |
| PHKB | 0.183200306 | 1.135399732 | 0.042076992 | 0.11709187 | 967.1148626 | 1097.421835 | ENSG00000102893 |
| RPL32 | 0.183181397 | 1.135384851 | 0.002025826 | 0.009607641 | 8279.692249 | 9400.550667 | ENSG00000144713 |
| PITPNM3 | 0.183107487 | 1.135326686 | 0.403780072 | 0.59554439 | 101.1745922 | 114.9959394 | ENSG00000091622 |
| SLC8B1 | 0.182995867 | 1.13523885 | 0.152981915 | 0.310249131 | 344.5509451 | 391.2915528 | ENSG00000089060 |
| SOCS2-AS1 | 0.182966551 | 1.135215782 | 0.671030583 | 0.808859269 | 24.04426888 | 27.25718203 | ENSG00000246985 |
| TRIM66 | 0.182894398 | 1.135159008 | 0.554720987 | 0.724742618 | 49.57302219 | 56.3749709 | ENSG00000166436 |
| RIF1 | 0.182845378 | 1.135120438 | 0.068680866 | 0.170611335 | 1306.315573 | 1481.932628 | ENSG00000080345 |
| PPCDC | 0.182828751 | 1.135107356 | 0.234064387 | 0.419001939 | 248.7022183 | 282.3191377 | ENSG00000138621 |
| BVES | 0.182806255 | 1.135089657 | 0.455232556 | 0.642644542 | 78.92226077 | 89.7649423 | ENSG00000112276 |
| LRRFIP1 | 0.182635419 | 1.134955253 | 0.011053398 | 0.040028073 | 1690.940466 | 1918.506599 | ENSG00000124831 |
| RMDN1 | 0.182575804 | 1.134908356 | 0.13761629 | 0.287098189 | 389.8999386 | 442.1891903 | ENSG00000176623 |
| RAB21 | 0.18247462 | 1.134828761 | 0.152417207 | 0.309550819 | 625.9168417 | 709.7479733 | ENSG00000080371 |
| LYSMD4 | 0.182433536 | 1.134796445 | 0.160919969 | 0.321493015 | 375.7048238 | 426.5585374 | ENSG00000183060 |
| BTN2A2 | 0.182404636 | 1.134773713 | 0.281435423 | 0.47271345 | 203.6225994 | 231.1183616 | ENSG00000124508 |
| RGL1 | 0.182384151 | 1.1347576 | 0.45036068 | 0.637977385 | 87.51056758 | 99.15898004 | ENSG00000143344 |
| NA | 0.182204869 | 1.134616594 | 0.737411605 | 0.852943558 | 15.90970031 | 18.10921394 | ENSG00000228648 |
| NA | 0.182198922 | 1.134611917 | 0.873509191 | 0.934704533 | 3.546285096 | 3.964360542 | ENSG00000181359 |
| NA | 0.182171303 | 1.134590196 | 0.744844676 | 0.857450386 | 14.03669042 | 15.9485976 | ENSG00000278376 |
| NA | 0.182078809 | 1.134517457 | 0.397848578 | 0.589910045 | 97.54605459 | 110.704651 | ENSG00000196295 |
| TAMM41 | 0.182014161 | 1.13446662 | 0.238596308 | 0.424267866 | 219.5647633 | 249.0035974 | ENSG00000144559 |
| LOC100507557 | 0.181862774 | 1.134347584 | 0.809506882 | 0.897492586 | 7.556455588 | 8.60057486 | ENSG00000235652 |
| LRP12 | 0.181817485 | 1.134311975 | 0.280717385 | 0.472098785 | 195.0154715 | 220.9170192 | ENSG00000147650 |
| NA | 0.181786475 | 1.134287593 | 0.791480556 | 0.886486471 | 10.70266996 | 12.09262614 | ENSG00000226471 |
| YEATS2 | 0.18171809 | 1.134233828 | 0.027872131 | 0.084699471 | 1962.613679 | 2225.60548 | ENSG00000163872 |
| FAM102B | 0.181716952 | 1.134232934 | 0.197724867 | 0.373471984 | 373.5162656 | 423.3295174 | ENSG00000162636 |
| MICALL1 | 0.181534619 | 1.134089594 | 0.031870727 | 0.094029298 | 1069.018197 | 1212.513725 | ENSG00000100139 |
| PTPMT1 | 0.181420861 | 1.134000173 | 0.822012106 | 0.904456123 | 6.755134065 | 7.663004382 | ENSG00000110536 |
| NA | 0.181205387 | 1.133830817 | 0.744594441 | 0.857268814 | 14.89611415 | 16.93119614 | ENSG00000273066 |
| PHYKPL | 0.181051069 | 1.133709543 | 0.291250115 | 0.483694412 | 242.7957594 | 275.7064901 | ENSG00000175309 |
| MSANTD4 | 0.181020458 | 1.133685489 | 0.10089369 | 0.228040811 | 459.9279127 | 521.5715073 | ENSG00000170903 |
| EIF1 | 0.18097923 | 1.133653092 | 0.001024417 | 0.005381671 | 5097.428851 | 5778.624653 | ENSG00000173812 |
| EIF3D | 0.180898522 | 1.133589675 | 0.00326711 | 0.014311656 | 3994.847202 | 4528.038112 | ENSG00000100353 |
| PLP1 | 0.180887251 | 1.133580818 | 0.015381386 | 0.052145345 | 1395.716356 | 1582.052204 | ENSG00000123560 |
| LINC01775 | 0.180804527 | 1.133515821 | 0.862479095 | 0.927914759 | 4.433519683 | 4.988299917 | ENSG00000267201 |
| ZSCAN16-AS1 | 0.180777991 | 1.133494971 | 0.604359785 | 0.76323532 | 39.74801838 | 45.18281128 | ENSG00000269293 |
| RUNDC1 | 0.180623379 | 1.133373503 | 0.093291449 | 0.215004191 | 469.7814264 | 532.7413119 | ENSG00000198863 |
| FBXO8 | 0.180621099 | 1.133371712 | 0.388173502 | 0.581225374 | 108.8767468 | 123.3376223 | ENSG00000164117 |
| GBE1 | 0.180566627 | 1.13332892 | 0.303239337 | 0.496875102 | 183.8108226 | 208.4571121 | ENSG00000114480 |
| COA1 | 0.180519866 | 1.133292186 | 0.036067448 | 0.103678081 | 1912.233743 | 2166.225541 | ENSG00000106603 |
| SERF1A | 0.180417179 | 1.133211525 | 0.829758664 | 0.908426196 | 6.020881826 | 6.811364769 | ENSG00000172058 |
| ATP4A | 0.180394814 | 1.133193957 | 0.716083315 | 0.839249283 | 18.3343878 | 20.88850864 | ENSG00000105675 |
| PURB | 0.180159656 | 1.133009263 | 0.083887184 | 0.198615874 | 777.4888234 | 881.3987458 | ENSG00000146676 |
| GLB1L | 0.180113697 | 1.13297317 | 0.553400852 | 0.723938212 | 46.6965733 | 52.91991196 | ENSG00000163521 |
| PRKCQ-AS1 | 0.180107578 | 1.132968364 | 0.219703782 | 0.401067724 | 248.6529104 | 281.87397 | ENSG00000237943 |
| LARP4 | 0.180082036 | 1.132948306 | 0.059286585 | 0.152528813 | 1084.923703 | 1228.334524 | ENSG00000161813 |
| RBM48 | 0.180020569 | 1.132900038 | 0.39626322 | 0.588563426 | 100.9823514 | 114.4333507 | ENSG00000127993 |
| VMP1 | 0.180006458 | 1.132888957 | 0.040894954 | 0.114458937 | 1107.90943 | 1254.508302 | ENSG00000062716 |
| RTN4IP1 | 0.180004257 | 1.132887228 | 0.166818929 | 0.330358161 | 288.845369 | 327.3097874 | ENSG00000130347 |
| WDR91 | 0.179952818 | 1.132846836 | 0.21660341 | 0.397254684 | 245.8380137 | 278.8580268 | ENSG00000105875 |
| TCAIM | 0.179848109 | 1.132764619 | 0.348279263 | 0.541618381 | 184.0729965 | 208.2787117 | ENSG00000179152 |
| KLHL26 | 0.17970046 | 1.132648694 | 0.137918202 | 0.287533853 | 363.2714755 | 411.1003185 | ENSG00000167487 |
| PIK3R5 | 0.179579481 | 1.132553719 | 0.254656639 | 0.443475905 | 206.7349822 | 234.1473016 | ENSG00000141506 |
| ALG6 | 0.179344847 | 1.13236954 | 0.155517858 | 0.313739889 | 384.5870464 | 435.1552731 | ENSG00000088035 |

| | | | | | | | |
|---|---|---|---|---|---|---|---|
| SLC25A37 | 0.179338368 | 1.132364455 | 0.10775253 | 0.239544846 | 865.5509709 | 980.7002332 | ENSG00000147454 |
| HDHD5 | 0.179263977 | 1.132306067 | 0.006432728 | 0.025518873 | 2023.327444 | 2290.865642 | ENSG00000069998 |
| SLC39A9 | 0.179187727 | 1.132246224 | 0.084957538 | 0.200433174 | 514.5680165 | 582.2834072 | ENSG00000029364 |
| ACVR1 | 0.179176437 | 1.132237363 | 0.131847389 | 0.278321127 | 380.6528372 | 430.6350456 | ENSG00000115170 |
| AGBL3 | 0.179054006 | 1.132141282 | 0.658180762 | 0.80059869 | 35.42328427 | 40.32959555 | ENSG00000146856 |
| RBM12 | 0.178994166 | 1.132094324 | 0.015610278 | 0.052738041 | 2353.567183 | 2663.233825 | ENSG00000244462 |
| RETREG3 | 0.178951533 | 1.13206087 | 0.081246145 | 0.193898538 | 568.8996373 | 643.92747 | ENSG00000141699 |
| NA | 0.178847587 | 1.131979308 | 0.089874654 | 0.209342168 | 1261.747134 | 1428.223915 | ENSG00000236673 |
| ERLEC1 | 0.17884509 | 1.131977349 | 0.130883142 | 0.277023924 | 466.7686667 | 527.7222545 | ENSG00000068912 |
| PKLR | 0.178730723 | 1.131887617 | 0.515218013 | 0.693216018 | 63.75322422 | 72.35340118 | ENSG00000143627 |
| TMEM229B | 0.178657711 | 1.131830336 | 0.668568936 | 0.807511024 | 27.52052317 | 31.28957851 | ENSG00000198133 |
| THEM4 | 0.178516464 | 1.13171953 | 0.284140875 | 0.475123329 | 179.4406291 | 202.8804045 | ENSG00000159445 |
| NA | 0.17843917 | 1.131658898 | 0.807325297 | 0.896619594 | 7.989275517 | 9.065836078 | ENSG00000279759 |
| HCFC1R1 | 0.178260678 | 1.131518897 | 0.130994393 | 0.277115066 | 373.0916387 | 422.2794813 | ENSG00000103145 |
| HMGB2 | 0.178120036 | 1.131408595 | 0.006496828 | 0.025745607 | 3938.618662 | 4455.651119 | ENSG00000164104 |
| SDC4 | 0.1780092 | 1.131321677 | 0.059656063 | 0.153181694 | 936.9838826 | 1059.533656 | ENSG00000124145 |
| NRBF2 | 0.177981373 | 1.131299856 | 0.097292376 | 0.222012766 | 501.9914422 | 567.5324744 | ENSG00000148572 |
| GTSE1 | 0.177960186 | 1.131283242 | 0.038894714 | 0.110090812 | 1216.877457 | 1375.864312 | ENSG00000075218 |
| TMEM161A | 0.177825243 | 1.131177433 | 0.027736448 | 0.084342467 | 1072.611806 | 1212.756434 | ENSG00000064545 |
| CLOCK | 0.177772145 | 1.1311358 | 0.237927812 | 0.423363486 | 287.1776335 | 324.2523514 | ENSG00000134852 |
| SEMA3F | 0.177651138 | 1.13104093 | 0.022157514 | 0.070307716 | 1258.94243 | 1423.404218 | ENSG00000001617 |
| NA | 0.177645655 | 1.131036631 | 0.736817048 | 0.852627858 | 15.81619565 | 17.93636452 | ENSG00000213178 |
| SEC16A | 0.177628895 | 1.131023492 | 0.024171658 | 0.075344352 | 2006.095908 | 2268.952173 | ENSG00000148396 |
| DCK | 0.177483299 | 1.130909356 | 0.354565404 | 0.547900216 | 246.2667671 | 278.2406936 | ENSG00000156136 |
| RIMS1 | 0.177317648 | 1.130779511 | 0.593554014 | 0.7550489 | 42.94877208 | 48.6099642 | ENSG00000079841 |
| NXN | 0.17721456 | 1.130698714 | 0.00433232 | 0.018215793 | 3331.754938 | 3767.459382 | ENSG00000167693 |
| SOS2 | 0.177195424 | 1.130683716 | 0.285091896 | 0.476304295 | 223.7718198 | 252.6676221 | ENSG00000100485 |
| MPZL3 | 0.177094173 | 1.130604366 | 0.549315568 | 0.720779805 | 51.51931749 | 58.3137993 | ENSG00000160588 |
| NA | 0.177061411 | 1.130578692 | 0.786518204 | 0.883102122 | 11.45911829 | 13.0344335 | ENSG00000225259 |
| NA | 0.176881759 | 1.130437915 | 0.67992918 | 0.815136731 | 41.95463107 | 47.56499233 | ENSG00000261786 |
| LARP4B | 0.176790724 | 1.130366585 | 0.022146072 | 0.070283442 | 1186.314649 | 1340.558194 | ENSG00000107929 |
| DMXL1 | 0.176647543 | 1.130254407 | 0.096237392 | 0.22020541 | 635.9709172 | 718.0564946 | ENSG00000172869 |
| ZNF283 | 0.176548631 | 1.130176919 | 0.501094188 | 0.681735278 | 68.12716853 | 76.95163547 | ENSG00000167637 |
| MKNK2 | 0.176541665 | 1.130171462 | 0.003679889 | 0.015839077 | 3389.712617 | 3830.600109 | ENSG00000099875 |
| NA | 0.176487695 | 1.130129184 | 0.875688986 | 0.935624952 | 4.436676596 | 5.078905695 | ENSG00000179978 |
| DNAJC27 | 0.176412726 | 1.130070459 | 0.569714305 | 0.736317339 | 48.10451771 | 54.48799538 | ENSG00000115137 |
| ZNF195 | 0.176394761 | 1.130056387 | 0.089724042 | 0.209122911 | 856.756142 | 967.864625 | ENSG00000005801 |
| MROH1 | 0.1762904 | 1.129974644 | 0.180090552 | 0.349434494 | 495.0678343 | 560.0945878 | ENSG00000179832 |
| RAB3GAP1 | 0.176244213 | 1.129938469 | 0.051091005 | 0.136032277 | 875.2350035 | 988.4534506 | ENSG00000115839 |
| PPM1K | 0.176164675 | 1.129876176 | 0.400912485 | 0.592686607 | 121.852595 | 137.4958073 | ENSG00000163644 |
| PDZD8 | 0.176156258 | 1.129869584 | 0.07455816 | 0.181704265 | 836.7325741 | 944.8146155 | ENSG00000165650 |
| GPATCH2 | 0.176152096 | 1.129866324 | 0.249215204 | 0.437732619 | 265.5569471 | 299.6891945 | ENSG00000092978 |
| TMEM251 | 0.176136844 | 1.12985438 | 0.435045794 | 0.624541154 | 108.4085244 | 122.214715 | ENSG00000153485 |
| TTC8 | 0.176115353 | 1.129837549 | 0.368343216 | 0.561374324 | 145.5411627 | 164.408493 | ENSG00000165533 |
| ZBTB1 | 0.176065535 | 1.129798535 | 0.20328785 | 0.381002833 | 336.3308468 | 380.0971401 | ENSG00000126804 |
| LOC100129917 | 0.175785811 | 1.1295795 | 0.581278604 | 0.745255786 | 69.70190302 | 78.93400331 | ENSG00000249592 |
| C1orf43 | 0.175672268 | 1.129490603 | 0.018484934 | 0.060370002 | 2050.769932 | 2316.567251 | ENSG00000143612 |
| ITGB4 | 0.175645804 | 1.129469885 | 0.391847582 | 0.584554337 | 115.0758663 | 130.0029444 | ENSG00000132470 |
| SETX | 0.175611374 | 1.12944293 | 0.07650696 | 0.185203324 | 1611.782887 | 1819.850889 | ENSG00000107290 |
| MCOLN2 | 0.17554476 | 1.129390781 | 0.636277485 | 0.786236877 | 33.26462663 | 37.66205075 | ENSG00000153898 |
| TEX264 | 0.175416135 | 1.129290094 | 0.255565737 | 0.444422535 | 205.0762138 | 231.5704742 | ENSG00000164081 |
| PTEN | 0.175241206 | 1.129153174 | 0.096633606 | 0.22092581 | 696.6648171 | 786.3982138 | ENSG00000171862 |
| FAM210A | 0.175238949 | 1.129151407 | 0.209138859 | 0.388355837 | 351.355387 | 396.0457604 | ENSG00000177150 |
| ABCB8 | 0.1752065 | 1.129126011 | 0.141430281 | 0.293141225 | 451.7255234 | 509.8511147 | ENSG00000197150 |
| ANO6 | 0.175114163 | 1.129053745 | 0.039585971 | 0.111557628 | 1212.356108 | 1367.980132 | ENSG00000177119 |
| CD164L2 | 0.175105342 | 1.129046842 | 0.628226056 | 0.780510951 | 45.80745958 | 51.8407589 | ENSG00000174950 |
| TEX2 | 0.175096138 | 1.129039639 | 0.107224428 | 0.238685749 | 446.5547673 | 504.3622102 | ENSG00000136478 |
| BTBD9 | 0.17504732 | 1.129001435 | 0.358171358 | 0.551476183 | 135.5922878 | 152.8473003 | ENSG00000183826 |
| PDZD4 | 0.174955176 | 1.128929329 | 0.040694591 | 0.113984214 | 6794.702762 | 7671.886491 | ENSG00000067840 |
| C2orf68 | 0.174885125 | 1.128874514 | 0.056583436 | 0.147105821 | 794.9975813 | 897.0340963 | ENSG00000168887 |
| MTX2 | 0.174695673 | 1.128726283 | 0.048003 | 0.129578737 | 783.245825 | 883.866992 | ENSG00000128654 |
| GRHPR | 0.174528395 | 1.128595416 | 0.011105903 | 0.040171119 | 2294.886813 | 2590.375776 | ENSG00000137106 |
| SGCB | 0.174501964 | 1.12857474 | 0.162233928 | 0.323560058 | 628.4919408 | 709.3683484 | ENSG00000163069 |
| NPAT | 0.174373747 | 1.128474444 | 0.318071188 | 0.511863454 | 252.1572686 | 284.1630916 | ENSG00000149308 |
| NA | 0.174277964 | 1.128399525 | 0.874094104 | 0.934994283 | 4.994668017 | 5.701494529 | ENSG00000232748 |
| MARK4 | 0.174267927 | 1.128391675 | 0.026986346 | 0.082426655 | 1524.075937 | 1719.508494 | ENSG00000007047 |
| BMP4 | 0.174262107 | 1.128387123 | 0.456443407 | 0.643667511 | 85.49303876 | 96.59040315 | ENSG00000125378 |
| MYLK3 | 0.174116007 | 1.128272858 | 0.781793125 | 0.880354443 | 11.42311447 | 12.9668356 | ENSG00000140795 |
| TTC25 | 0.174024655 | 1.128201418 | 0.677106413 | 0.813370294 | 25.28482185 | 28.5999019 | ENSG00000204815 |
| SP4 | 0.174015713 | 1.128194425 | 0.191938181 | 0.36509247 | 346.4217675 | 390.5523957 | ENSG00000105866 |
| NA | 0.173986491 | 1.128171573 | 0.819225002 | 0.90291228 | 8.370307064 | 9.456013319 | ENSG00000262333 |
| TMEM143 | 0.173899401 | 1.128103472 | 0.23940683 | 0.42534185 | 242.1978331 | 273.1221279 | ENSG00000161558 |
| CEP68 | 0.173780543 | 1.128010536 | 0.107914559 | 0.239790006 | 557.4842348 | 629.3900118 | ENSG00000011523 |
| REV1 | 0.173760933 | 1.127995204 | 0.095692092 | 0.21917415 | 567.1429447 | 639.4923376 | ENSG00000135945 |

| | | | | | | | |
|---|---|---|---|---|---|---|---|
| NA | 0.173712683 | 1.127957479 | 0.624510338 | 0.777188921 | 49.26142009 | 55.7119173 | ENSG00000259943 |
| BCL10 | 0.173693456 | 1.127942447 | 0.318345175 | 0.512037619 | 203.7944104 | 229.7148284 | ENSG00000142867 |
| DGKQ | 0.173667291 | 1.12792199 | 0.19164686 | 0.36467997 | 596.4927921 | 673.6744754 | ENSG00000145214 |
| PSAT1 | 0.173660849 | 1.127916954 | 0.006846189 | 0.026934253 | 8423.360476 | 9500.101371 | ENSG00000135069 |
| BCL7C | 0.173624644 | 1.127888649 | 0.398567829 | 0.590322714 | 113.2864843 | 127.9664622 | ENSG00000099385 |
| DTWD1 | 0.173516309 | 1.127803957 | 0.188181307 | 0.359970311 | 336.5317463 | 379.5531706 | ENSG00000104047 |
| FAM89A | 0.173341591 | 1.127667381 | 0.674978583 | 0.811873412 | 29.13126148 | 32.84929979 | ENSG00000182118 |
| CD72 | 0.173206838 | 1.127562058 | 0.616419502 | 0.771888321 | 42.5580744 | 48.06724588 | ENSG00000137101 |
| DOK7 | 0.173036108 | 1.127428629 | 0.814766663 | 0.900075694 | 9.260698564 | 10.46866777 | ENSG00000175920 |
| NORAD | 0.17299594 | 1.127397239 | 0.012925833 | 0.045372613 | 3796.521838 | 4279.558168 | ENSG00000260032 |
| PGK1 | 0.17290413 | 1.127325497 | 0.021792187 | 0.069457676 | 9361.46069 | 10551.66952 | ENSG00000102144 |
| G6PC3 | 0.1727738 | 1.127223661 | 0.023892444 | 0.074674966 | 2035.85568 | 2295.058687 | ENSG00000141349 |
| CLCN7 | 0.172709904 | 1.127173738 | 0.024401744 | 0.075933835 | 1332.87572 | 1503.104847 | ENSG00000103249 |
| AK3 | 0.172549717 | 1.127048592 | 0.181888545 | 0.352075436 | 505.9970651 | 569.6262884 | ENSG00000147853 |
| SEC14L1 | 0.172203505 | 1.126778159 | 0.017774483 | 0.058503333 | 1842.494866 | 2076.116246 | ENSG00000129657 |
| FECH | 0.172147797 | 1.126734651 | 0.072700456 | 0.178349472 | 779.2375336 | 878.093823 | ENSG00000066926 |
| ATP6V1E1 | 0.172043454 | 1.126653163 | 0.028211374 | 0.08546414 | 1223.494491 | 1378.357416 | ENSG00000131100 |
| TXNRD3 | 0.172021999 | 1.126636408 | 0.351570529 | 0.544557714 | 144.8872088 | 163.283101 | ENSG00000197763 |
| NA | 0.171981668 | 1.126604913 | 0.736380833 | 0.852322051 | 17.72195474 | 19.8716173 | ENSG00000227376 |
| CYP26A1 | 0.171925609 | 1.126561137 | 0.518695869 | 0.695809231 | 72.42043121 | 81.76113724 | ENSG00000095596 |
| CDT1 | 0.171840129 | 1.12649439 | 0.037670078 | 0.107148132 | 2228.70945 | 2510.456227 | ENSG00000167513 |
| LMAN1 | 0.171783989 | 1.126450555 | 0.038180245 | 0.10830007 | 1861.046208 | 2095.40729 | ENSG00000074695 |
| NA | 0.171697555 | 1.12638307 | 0.753695356 | 0.863295824 | 17.00801945 | 19.19640271 | ENSG00000231369 |
| CEP57L1 | 0.171612285 | 1.126316497 | 0.436170398 | 0.625590574 | 104.7510709 | 117.739842 | ENSG00000183137 |
| NA | 0.171555396 | 1.126272085 | 0.857146707 | 0.924108956 | 4.839806533 | 5.397248153 | ENSG00000271943 |
| PHF20 | 0.171542487 | 1.126262007 | 0.026860155 | 0.08212242 | 1500.669823 | 1689.38392 | ENSG00000025293 |
| DCUN1D3 | 0.171165021 | 1.125967371 | 0.354381776 | 0.547662123 | 172.1385452 | 193.7082176 | ENSG00000188215 |
| RNF112 | 0.171106861 | 1.125921981 | 0.600799137 | 0.760238357 | 43.39549744 | 48.95923864 | ENSG00000128482 |
| LNPK | 0.170974809 | 1.125818928 | 0.366589058 | 0.559588522 | 134.2178894 | 150.9309301 | ENSG00000144320 |
| RPL22 | 0.170959297 | 1.125806823 | 0.028239477 | 0.085503133 | 1301.839413 | 1465.222473 | ENSG00000116251 |
| ZNF414 | 0.170941357 | 1.125792823 | 0.203928012 | 0.381893683 | 289.992515 | 326.7018443 | ENSG00000133250 |
| RNF41 | 0.170920297 | 1.125776389 | 0.043206043 | 0.11955521 | 1164.443853 | 1310.536561 | ENSG00000181852 |
| FBXO17 | 0.170864681 | 1.125732992 | 0.27425662 | 0.465250018 | 225.7257067 | 253.8122231 | ENSG00000269190 |
| CADM4 | 0.17079405 | 1.12567788 | 0.132206555 | 0.278856982 | 561.7015135 | 632.7546713 | ENSG00000105767 |
| TMEM167B | 0.170720628 | 1.125620593 | 0.233089268 | 0.417994335 | 244.6190554 | 275.4510303 | ENSG00000215717 |
| DPY19L4 | 0.170707293 | 1.125610189 | 0.243564772 | 0.430235347 | 240.0231804 | 270.2105602 | ENSG00000156162 |
| GOLGA6L9 | 0.170546493 | 1.125484737 | 0.398000531 | 0.589934568 | 113.6193881 | 127.7137423 | ENSG00000197978 |
| SCAF8 | 0.170511772 | 1.125457651 | 0.055496494 | 0.144797432 | 841.819657 | 946.9494531 | ENSG00000213079 |
| SELENOS | 0.170503736 | 1.125451383 | 0.077890984 | 0.187948379 | 655.1794158 | 737.9299128 | ENSG00000131871 |
| H2AZ2 | 0.170436714 | 1.125399099 | 0.013171274 | 0.046051006 | 2864.861641 | 3224.160469 | ENSG00000105968 |
| XPO1 | 0.170421055 | 1.125386884 | 0.007005209 | 0.027437563 | 6236.347197 | 7017.090602 | ENSG00000082898 |
| PLK5 | 0.170339453 | 1.125323232 | 0.777910418 | 0.877902435 | 13.52742817 | 15.197861 | ENSG00000185988 |
| ZNF316 | 0.170237467 | 1.125243684 | 0.063544825 | 0.16106177 | 767.4691058 | 863.9336626 | ENSG00000205903 |
| CCDC141 | 0.170193793 | 1.125209621 | 0.435334983 | 0.624883718 | 100.0204559 | 112.3887212 | ENSG00000163492 |
| ARID4B | 0.170014318 | 1.125069651 | 0.114377609 | 0.250635446 | 508.70486 | 572.1358217 | ENSG00000054267 |
| TMEM186 | 0.169892196 | 1.124974419 | 0.308646938 | 0.502897789 | 222.1382841 | 250.0040196 | ENSG00000184857 |
| NA | 0.169783087 | 1.124889342 | 0.682200871 | 0.816506824 | 28.30280552 | 31.90848012 | ENSG00000224934 |
| SEPSECS | 0.169747432 | 1.124861542 | 0.539212264 | 0.71257188 | 64.95639791 | 72.99713413 | ENSG00000109618 |
| ZNF212 | 0.169672476 | 1.1248031 | 0.118938499 | 0.257885604 | 536.7861029 | 603.9706435 | ENSG00000170260 |
| DYNC1LI2 | 0.169555472 | 1.124711882 | 0.022233981 | 0.070477955 | 1410.351875 | 1585.861316 | ENSG00000135720 |
| SMIM13 | 0.169272423 | 1.124491241 | 0.311627043 | 0.505327089 | 248.5565419 | 279.2403513 | ENSG00000224531 |
| RPL37A | 0.169220728 | 1.124450949 | 0.00653316 | 0.025872988 | 12933.28514 | 14543.09097 | ENSG00000197756 |
| ASL | 0.169103151 | 1.124359311 | 0.189774372 | 0.362008872 | 401.0337448 | 450.9985563 | ENSG00000126522 |
| NA | 0.16909444 | 1.124352522 | 0.664005356 | 0.804615646 | 33.76441815 | 38.10886739 | ENSG00000247679 |
| STYX | 0.169091027 | 1.124349863 | 0.21556888 | 0.395955923 | 519.2263907 | 583.2257868 | ENSG00000198252 |
| SPIRE1 | 0.168938988 | 1.124231379 | 0.247578542 | 0.435547443 | 299.1274559 | 335.9808745 | ENSG00000134278 |
| RPS3 | 0.168868342 | 1.124176328 | 0.007211045 | 0.028048164 | 37189.20072 | 41806.99541 | ENSG00000149273 |
| TLN2 | 0.16880618 | 1.124127891 | 0.061064498 | 0.156127661 | 1033.635209 | 1161.858127 | ENSG00000171914 |
| MOCS2 | 0.168798331 | 1.124121776 | 0.187897168 | 0.3596495 | 308.988952 | 347.2175209 | ENSG00000164172 |
| TLCD2 | 0.16870055 | 1.12404559 | 0.691981784 | 0.823650512 | 27.24552624 | 30.5464396 | ENSG00000185561 |
| NA | 0.168636591 | 1.123995758 | 0.323499742 | 0.517691704 | 185.0112453 | 207.8865994 | ENSG00000189343 |
| SULT4A1 | 0.16862059 | 1.123983292 | 0.313848568 | 0.507580224 | 205.7913291 | 231.3545721 | ENSG00000130540 |
| KIZ | 0.168498877 | 1.123888471 | 0.156020227 | 0.314415449 | 447.9688302 | 503.8888504 | ENSG00000088970 |
| ASXL3 | 0.168494903 | 1.123885376 | 0.791396347 | 0.886482527 | 12.20105983 | 13.78049519 | ENSG00000141431 |
| NA | 0.168423741 | 1.12382994 | 0.803229458 | 0.894332258 | 9.190570065 | 10.35224296 | ENSG00000226468 |
| CHPF2 | 0.168345345 | 1.123768873 | 0.109624301 | 0.242600215 | 590.7374138 | 664.5165591 | ENSG00000033100 |
| UBXN1 | 0.16832885 | 1.123756024 | 0.014576496 | 0.049872766 | 1804.172305 | 2027.948552 | ENSG00000162191 |
| NA | 0.168249695 | 1.12369437 | 0.705821197 | 0.832963392 | 27.68543411 | 31.15806993 | ENSG00000231503 |
| CHST12 | 0.168216819 | 1.123668764 | 0.147671592 | 0.302525178 | 409.723734 | 460.3146502 | ENSG00000136213 |
| NDUFB4 | 0.168156648 | 1.123621899 | 0.024547759 | 0.076319914 | 1570.485419 | 1764.43763 | ENSG00000065518 |
| PIP5K1C | 0.167768517 | 1.12331965 | 0.034442943 | 0.099900704 | 1307.640153 | 1469.455219 | ENSG00000186111 |
| ZSCAN12 | 0.167759728 | 1.123312807 | 0.314307125 | 0.50786627 | 204.0561184 | 228.8160108 | ENSG00000158691 |
| YIPF6 | 0.167725822 | 1.123286407 | 0.114080285 | 0.250131787 | 628.3185968 | 705.1307766 | ENSG00000181704 |
| ZNF845 | 0.167421458 | 1.123049453 | 0.169872459 | 0.334688652 | 382.1197556 | 429.1500734 | ENSG00000213799 |

| | | | | | | | |
|---|---|---|---|---|---|---|---|
| PCNT | 0.167391296 | 1.123025974 | 0.0810712 | 0.193555794 | 1071.569049 | 1203.534218 | ENSG00000160299 |
| GCN1 | 0.167387634 | 1.123023124 | 0.019686253 | 0.0636985 | 6972.044978 | 7829.97504 | ENSG00000089154 |
| EID2B | 0.167367931 | 1.123007787 | 0.805352674 | 0.895650273 | 10.52325223 | 11.84904125 | ENSG00000176401 |
| LPIN1 | 0.167357324 | 1.12299953 | 0.189174408 | 0.361310642 | 376.5540027 | 422.858571 | ENSG00000134324 |
| ADGRA2 | 0.167350253 | 1.122994026 | 0.52913622 | 0.704931764 | 95.83509182 | 107.6202338 | ENSG00000020181 |
| MXRA8 | 0.167330274 | 1.122978474 | 0.325076818 | 0.519215249 | 162.2988618 | 182.1116815 | ENSG00000162576 |
| NOA1 | 0.167262681 | 1.122925862 | 0.073256767 | 0.179405512 | 793.8099443 | 890.7579637 | ENSG00000084092 |
| RACK1 | 0.167251423 | 1.122917099 | 0.001173664 | 0.006040737 | 24007.51504 | 26957.68016 | ENSG00000204628 |
| RARS2 | 0.167222407 | 1.122894515 | 0.086222827 | 0.202747257 | 731.7906531 | 821.6615345 | ENSG00000146282 |
| PFDN2 | 0.167196425 | 1.122874293 | 0.056448461 | 0.146796095 | 1188.093297 | 1334.256386 | ENSG00000143256 |
| ATRN | 0.167146172 | 1.122835181 | 0.13170321 | 0.278048445 | 820.4602878 | 920.3757991 | ENSG00000088812 |
| NA | 0.1671401 | 1.122830454 | 0.536270003 | 0.71065553 | 75.93565084 | 85.27520871 | ENSG00000187653 |
| LEMD1 | 0.166998769 | 1.122720464 | 0.736172316 | 0.852230962 | 17.49706246 | 19.66177555 | ENSG00000186007 |
| EGFR | 0.166987742 | 1.122711882 | 0.334929693 | 0.529296293 | 153.153165 | 171.8913852 | ENSG00000146648 |
| GALNT1 | 0.1669834 | 1.122708504 | 0.094742904 | 0.217456964 | 1203.976299 | 1351.073195 | ENSG00000141429 |
| FKTN | 0.166827452 | 1.122587151 | 0.298003028 | 0.491168352 | 214.1638084 | 240.1999025 | ENSG00000106692 |
| TMA16 | 0.166783623 | 1.122553047 | 0.154761622 | 0.312592841 | 693.4633191 | 778.0794271 | ENSG00000198498 |
| BZW2 | 0.166744581 | 1.12252267 | 0.048314081 | 0.130228791 | 1264.803068 | 1419.380725 | ENSG00000136261 |
| MIGA1 | 0.166681964 | 1.12247395 | 0.495328918 | 0.677172395 | 90.60586538 | 101.6077713 | ENSG00000180488 |
| QDPR | 0.166551144 | 1.122372172 | 0.072892862 | 0.178726856 | 850.8491116 | 955.6160659 | ENSG00000151552 |
| NEK10 | 0.166489108 | 1.12232391 | 0.850955649 | 0.921239019 | 5.819268009 | 6.488785432 | ENSG00000163491 |
| SGO2 | 0.166476344 | 1.122313981 | 0.197762507 | 0.373471984 | 492.8639952 | 552.7318069 | ENSG00000163535 |
| TMPRSS5 | 0.166437348 | 1.122283645 | 0.844325814 | 0.917016342 | 8.495380859 | 9.52741007 | ENSG00000166682 |
| RPAIN | 0.166362849 | 1.122225693 | 0.148716585 | 0.304128123 | 519.3025777 | 582.8466889 | ENSG00000129197 |
| RECQL5 | 0.166324069 | 1.122195528 | 0.100468001 | 0.227442766 | 553.351253 | 621.2083476 | ENSG00000108469 |
| MYO5B | 0.166299857 | 1.122176695 | 0.164640002 | 0.32705657 | 387.327891 | 434.924502 | ENSG00000167306 |
| NA | 0.166284357 | 1.122164639 | 0.796611086 | 0.890279177 | 12.1870567 | 13.72462024 | ENSG00000228274 |
| AASDHPPT | 0.166265291 | 1.122149809 | 0.135992616 | 0.284576598 | 510.3413195 | 572.5094979 | ENSG00000149313 |
| TRAF3IP2-AS1 | 0.166263394 | 1.122148333 | 0.578986109 | 0.743190563 | 60.41664821 | 67.92400266 | ENSG00000231889 |
| AK4 | 0.166252855 | 1.122140136 | 0.048725077 | 0.131107807 | 3256.497332 | 3652.683684 | ENSG00000162433 |
| CCDC14 | 0.166236217 | 1.122127195 | 0.206091943 | 0.384640554 | 703.2839713 | 789.7205165 | ENSG00000175455 |
| VKORC1 | 0.166171486 | 1.122076848 | 0.786160007 | 0.882911566 | 12.45879902 | 13.91514134 | ENSG00000167397 |
| CERS2 | 0.166152643 | 1.122062193 | 0.022095631 | 0.070167013 | 2100.797406 | 2357.485876 | ENSG00000143418 |
| MKRN1 | 0.166128273 | 1.122043239 | 0.016167803 | 0.054206596 | 2528.097133 | 2835.92738 | ENSG00000133606 |
| SLC22A17 | 0.16608642 | 1.122010689 | 0.046448586 | 0.12622846 | 1104.286883 | 1239.287473 | ENSG00000092096 |
| ELMOD2 | 0.165876711 | 1.121847606 | 0.165006531 | 0.327679315 | 431.9783279 | 484.1302081 | ENSG00000179387 |
| ACTR3C | 0.165804348 | 1.121791338 | 0.8206065 | 0.903761083 | 8.648363207 | 9.70339706 | ENSG00000106526 |
| NA | 0.16579732 | 1.121785873 | 0.26846624 | 0.458721609 | 233.6648624 | 262.0139238 | ENSG00000231205 |
| ETV4 | 0.165629154 | 1.121655121 | 0.00538499 | 0.021959384 | 3399.558539 | 3813.410986 | ENSG00000175832 |
| NA | 0.165557377 | 1.121599318 | 0.54972784 | 0.721116554 | 62.33127668 | 69.94551272 | ENSG00000186076 |
| SCARF2 | 0.165430454 | 1.121500648 | 0.241488667 | 0.427662256 | 378.3297571 | 424.6327142 | ENSG00000244486 |
| LRRC57 | 0.165425316 | 1.121496654 | 0.549570955 | 0.720961784 | 77.75559457 | 86.83664184 | ENSG00000180979 |
| FNDC1 | 0.165411053 | 1.121485566 | 0.777340226 | 0.877472669 | 15.82556869 | 17.72652277 | ENSG00000164694 |
| ZNF252P-AS1 | 0.165378996 | 1.121460647 | 0.886523614 | 0.942188695 | 4.739988041 | 5.363943044 | ENSG00000255559 |
| RNF6 | 0.165367474 | 1.121451691 | 0.223932166 | 0.406387203 | 278.393042 | 312.5508189 | ENSG00000127870 |
| SPG11 | 0.165213316 | 1.121331865 | 0.132020914 | 0.278560522 | 704.2261663 | 789.2002218 | ENSG00000104133 |
| ANKRD46 | 0.165204752 | 1.121325209 | 0.227925015 | 0.411530661 | 391.8192198 | 438.6393516 | ENSG00000186106 |
| LINC01426 | 0.165157192 | 1.121288244 | 0.78273732 | 0.880989463 | 12.06198291 | 13.58562558 | ENSG00000234380 |
| TET1 | 0.16514108 | 1.121275721 | 0.067065929 | 0.167587964 | 1308.707204 | 1467.052003 | ENSG00000138336 |
| KIF16B | 0.165119714 | 1.121259116 | 0.30273031 | 0.49618001 | 235.2146272 | 263.5040004 | ENSG00000089177 |
| RFTN1 | 0.165097434 | 1.1212418 | 0.444327308 | 0.632840393 | 106.2853669 | 119.2232054 | ENSG00000131378 |
| HIKESHI | 0.164601202 | 1.120856202 | 0.068299765 | 0.169778412 | 793.6177634 | 890.0703423 | ENSG00000149196 |
| COL4A1 | 0.164441291 | 1.120731971 | 0.013004523 | 0.045579772 | 2740.614861 | 3071.025543 | ENSG00000187498 |
| CAPRIN2 | 0.164318644 | 1.120636699 | 0.513281827 | 0.691998545 | 756.5864697 | 848.0390225 | ENSG00000110888 |
| PRKRIP1 | 0.164307358 | 1.120627932 | 0.109579177 | 0.242529313 | 643.5136854 | 721.3788199 | ENSG00000128563 |
| EPHX4 | 0.164243404 | 1.120578256 | 0.810882726 | 0.898642696 | 11.10274068 | 12.36214169 | ENSG00000172031 |
| RNF125 | 0.164127981 | 1.120488608 | 0.260212668 | 0.449102778 | 250.0243473 | 279.8108556 | ENSG00000101695 |
| SLC25A20 | 0.164119469 | 1.120481997 | 0.458182221 | 0.6452849 | 93.83533957 | 105.1013599 | ENSG00000178537 |
| C11orf74 | 0.164070068 | 1.12044363 | 0.39287819 | 0.585526245 | 144.5810487 | 161.6057524 | ENSG00000166352 |
| WWP1 | 0.164029948 | 1.120412472 | 0.043514545 | 0.120126448 | 1091.186856 | 1222.483062 | ENSG00000123124 |
| C3orf67 | 0.163904667 | 1.120315182 | 0.443022793 | 0.631468464 | 104.7775063 | 117.2460622 | ENSG00000163689 |
| SRGAP2C | 0.163888661 | 1.120302753 | 0.427386182 | 0.617780027 | 132.9047274 | 148.6199226 | ENSG00000171943 |
| USP42 | 0.163788236 | 1.120224771 | 0.144909136 | 0.298383061 | 528.6652518 | 592.229393 | ENSG00000106346 |
| LEMD2 | 0.163676618 | 1.120138106 | 0.034847896 | 0.100899589 | 1339.483757 | 1500.681946 | ENSG00000161904 |
| MRPS35 | 0.16360504 | 1.120082533 | 0.093668429 | 0.21563168 | 794.8625405 | 890.3985037 | ENSG00000061794 |
| PTMS | 0.163554494 | 1.12004329 | 0.014107278 | 0.048651104 | 7160.174325 | 8019.27562 | ENSG00000159335 |
| C1orf53 | 0.163514598 | 1.120012317 | 0.854961434 | 0.923041604 | 5.481831882 | 6.083636043 | ENSG00000203724 |
| FZD3 | 0.163463909 | 1.119972967 | 0.18828947 | 0.360140046 | 571.3588403 | 639.5412987 | ENSG00000104290 |
| BRIP1 | 0.163321229 | 1.119862208 | 0.177810511 | 0.346461314 | 379.1672704 | 424.1940994 | ENSG00000136492 |
| WDR7 | 0.163228675 | 1.119790367 | 0.227349533 | 0.410891882 | 304.10092 | 340.0594696 | ENSG00000091157 |
| USP8 | 0.163126301 | 1.11971091 | 0.066102064 | 0.165603629 | 806.4023556 | 902.676355 | ENSG00000138592 |
| ZNF280C | 0.162914642 | 1.119546648 | 0.339955433 | 0.534186858 | 215.3909596 | 241.0670639 | ENSG00000056277 |
| ATG4B | 0.162847237 | 1.119494343 | 0.083497201 | 0.197970721 | 1746.160486 | 1955.516876 | ENSG00000168397 |
| ARHGAP42 | 0.162837801 | 1.11948702 | 0.173367601 | 0.339876785 | 493.6768394 | 552.3453084 | ENSG00000165895 |

| | | | | | | | |
|---|---|---|---|---|---|---|---|
| DPP7 | 0.162713175 | 1.119390319 | 0.222733225 | 0.404838442 | 296.8395829 | 332.5135916 | ENSG00000176978 |
| FAM20B | 0.162619268 | 1.119317458 | 0.116097656 | 0.253326343 | 900.5591966 | 1007.049356 | ENSG00000116199 |
| POGZ | 0.162320627 | 1.119085781 | 0.065992519 | 0.165508166 | 1277.182711 | 1429.606269 | ENSG00000143442 |
| BCO2 | 0.162197368 | 1.118990174 | 0.87728137 | 0.936413113 | 4.383610437 | 4.966279729 | ENSG00000197580 |
| RPS2 | 0.162130404 | 1.118938236 | 0.006242076 | 0.024895769 | 55144.68137 | 61702.80557 | ENSG00000140988 |
| CCL25 | 0.162070619 | 1.118891869 | 0.841266724 | 0.914864605 | 7.225333287 | 8.063916891 | ENSG00000131142 |
| FBXO16 | 0.162000479 | 1.118837473 | 0.573248151 | 0.738947088 | 68.17213942 | 76.31429773 | ENSG00000214050 |
| ARPC5 | 0.161999136 | 1.118836431 | 0.034019631 | 0.099013794 | 2245.343834 | 2511.417186 | ENSG00000162704 |
| FAM174B | 0.161983796 | 1.118824535 | 0.197752189 | 0.373471984 | 440.4595927 | 492.6303264 | ENSG00000185442 |
| NA | 0.16190102 | 1.118760343 | 0.85214627 | 0.921827991 | 7.225235589 | 8.053181623 | ENSG00000267466 |
| ARAP1 | 0.161836918 | 1.118710635 | 0.020858828 | 0.066922269 | 2418.687319 | 2706.330066 | ENSG00000186635 |
| ZNF146 | 0.161817823 | 1.118695828 | 0.07568882 | 0.18375905 | 2197.203102 | 2457.531877 | ENSG00000167635 |
| PC | 0.1617976 | 1.118680147 | 0.083790347 | 0.198488022 | 716.1338287 | 800.715184 | ENSG00000173599 |
| LYRM7 | 0.161705615 | 1.118608823 | 0.396922265 | 0.589098354 | 166.6538647 | 185.8866831 | ENSG00000186687 |
| EP300 | 0.161616773 | 1.11853994 | 0.071415952 | 0.175733306 | 2791.334601 | 3121.77707 | ENSG00000100393 |
| RALBP1 | 0.161576078 | 1.11850839 | 0.016982958 | 0.056439331 | 2218.090295 | 2480.83514 | ENSG00000017797 |
| APRT | 0.161546591 | 1.118485529 | 0.036651176 | 0.104877705 | 1951.670144 | 2182.527747 | ENSG00000198931 |
| CSRNP2 | 0.161518035 | 1.118463391 | 0.0644963 | 0.162768849 | 914.4151689 | 1022.110564 | ENSG00000110925 |
| NUSAP1 | 0.161494656 | 1.118445266 | 0.881882291 | 0.939373634 | 4.128930461 | 4.673756307 | ENSG00000137804 |
| ARHGEF17 | 0.16143486 | 1.11839891 | 0.243043315 | 0.429578943 | 329.9222362 | 369.4297955 | ENSG00000110237 |
| NA | 0.161319441 | 1.118309439 | 0.584969206 | 0.747954646 | 57.43336794 | 64.25150426 | ENSG00000259295 |
| CAVIN1 | 0.160974593 | 1.118042161 | 0.065211572 | 0.16397114 | 1087.356201 | 1215.255772 | ENSG00000177469 |
| RBM46 | 0.160959242 | 1.118030265 | 0.865322352 | 0.929978419 | 5.531741128 | 6.132462953 | ENSG00000151962 |
| MIEF2 | 0.160628499 | 1.117773981 | 0.28318752 | 0.474256573 | 225.9741706 | 252.4922879 | ENSG00000177427 |
| PRKCI | 0.160399651 | 1.117596689 | 0.046693726 | 0.126820269 | 2398.360898 | 2679.224179 | ENSG00000163558 |
| LTB4R | 0.160340202 | 1.117550637 | 0.542189512 | 0.714870913 | 107.2925415 | 120.133245 | ENSG00000213903 |
| NUF2 | 0.160332505 | 1.117544674 | 0.180255292 | 0.34964433 | 400.9197126 | 447.7156452 | ENSG00000143228 |
| NA | 0.16022487 | 1.117461301 | 0.859064129 | 0.925476317 | 5.481831882 | 6.064865049 | ENSG00000217325 |
| MPHOSPH8 | 0.1601054 | 1.117368768 | 0.063857364 | 0.161530284 | 865.1332964 | 966.9259242 | ENSG00000196199 |
| ABHD6 | 0.160090882 | 1.117357523 | 0.394624406 | 0.587043001 | 151.6939358 | 169.1746786 | ENSG00000163686 |
| MAGED2 | 0.160081917 | 1.11735058 | 0.012967662 | 0.045502204 | 3053.671484 | 3411.585349 | ENSG00000102316 |
| SPTAN1 | 0.160078308 | 1.117347785 | 0.024065621 | 0.075120695 | 5829.448523 | 6513.692085 | ENSG00000197694 |
| NA | 0.160064331 | 1.11733696 | 0.762315248 | 0.868300466 | 21.86686516 | 24.56045738 | ENSG00000259877 |
| LINC00240 | 0.159936891 | 1.117238265 | 0.824387684 | 0.905596573 | 9.842066152 | 10.99740164 | ENSG00000224843 |
| RPS6KA2 | 0.159877054 | 1.117191928 | 0.088488744 | 0.20684202 | 742.0017018 | 828.4446753 | ENSG00000071242 |
| YLPM1 | 0.15980207 | 1.117133863 | 0.047239939 | 0.127984948 | 2386.174797 | 2665.994852 | ENSG00000119596 |
| NA | 0.159672113 | 1.117033237 | 0.286642959 | 0.478013371 | 275.7346831 | 307.9792797 | ENSG00000244716 |
| ARHGEF18 | 0.159646242 | 1.117013206 | 0.110676378 | 0.244548879 | 587.9170153 | 656.6164279 | ENSG00000104880 |
| ZNF565 | 0.159626281 | 1.116997751 | 0.695649166 | 0.825955263 | 29.66084069 | 33.2736582 | ENSG00000196357 |
| RNF11 | 0.159604141 | 1.116980609 | 0.241186043 | 0.427440453 | 672.8690121 | 751.1935079 | ENSG00000123091 |
| SLC35A5 | 0.159584494 | 1.116965398 | 0.181828346 | 0.351995672 | 486.8367019 | 543.1775817 | ENSG00000138459 |
| EPHX1 | 0.159453318 | 1.116863843 | 0.040518214 | 0.11364476 | 1316.011437 | 1469.379751 | ENSG00000143819 |
| MTIF3 | 0.159254309 | 1.116709791 | 0.269235471 | 0.45951754 | 280.0935267 | 312.732205 | ENSG00000122033 |
| AIG1 | 0.159199115 | 1.116667069 | 0.172624567 | 0.338886107 | 427.1513444 | 476.9460601 | ENSG00000146416 |
| NA | 0.159175567 | 1.116648843 | 0.647483176 | 0.794206035 | 36.6313963 | 40.87611272 | ENSG00000280063 |
| DGKD | 0.159140789 | 1.116621925 | 0.070341546 | 0.173619684 | 817.7169026 | 913.2630966 | ENSG00000077044 |
| LRRC27 | 0.159116122 | 1.116602833 | 0.629856073 | 0.781581472 | 45.71573635 | 51.12789071 | ENSG00000148814 |
| MAPK10 | 0.159056708 | 1.11655685 | 0.665114286 | 0.805385316 | 40.2553038 | 44.84102292 | ENSG00000109339 |
| OLIG3 | 0.158921242 | 1.116452012 | 0.883559835 | 0.939962378 | 4.043017397 | 4.568616414 | ENSG00000177468 |
| PTGIS | 0.158830573 | 1.116381849 | 0.080117061 | 0.191995035 | 833.216242 | 930.0243576 | ENSG00000124212 |
| AMACR | 0.158769335 | 1.116334462 | 0.736006468 | 0.852173027 | 19.82311153 | 22.12988746 | ENSG00000242110 |
| CRCP | 0.158615998 | 1.11621582 | 0.217600048 | 0.3985571 | 379.5433862 | 424.0288795 | ENSG00000241258 |
| RIBC2 | 0.158604892 | 1.116207227 | 0.756536335 | 0.864466365 | 20.39500838 | 22.88062406 | ENSG00000128408 |
| RBBP9 | 0.158538721 | 1.116156032 | 0.410562286 | 0.602171734 | 189.3698709 | 211.3477187 | ENSG00000089050 |
| CD2AP | 0.158429155 | 1.116071268 | 0.067245044 | 0.167831894 | 1237.300246 | 1380.130474 | ENSG00000198087 |
| NEMF | 0.158325269 | 1.115990905 | 0.114954898 | 0.251685044 | 594.6822965 | 663.6278953 | ENSG00000165525 |
| GLB1 | 0.158316116 | 1.115983824 | 0.086798749 | 0.203739616 | 887.8381034 | 989.934289 | ENSG00000170266 |
| MRPL17 | 0.158248569 | 1.115931575 | 0.046820856 | 0.127072447 | 1082.561094 | 1208.417681 | ENSG00000158042 |
| SLC37A4 | 0.158172621 | 1.11587283 | 0.101085927 | 0.22839529 | 738.5955987 | 823.6055557 | ENSG00000137700 |
| NA | 0.158108831 | 1.115823493 | 0.396434055 | 0.58874269 | 162.4635774 | 181.2019054 | ENSG00000230202 |
| BCAT1 | 0.158019121 | 1.11575411 | 0.04682072 | 0.127072447 | 4770.335823 | 5321.399431 | ENSG00000060982 |
| PCNP | 0.157778013 | 1.115567657 | 0.011653017 | 0.041750826 | 2967.924509 | 3310.204143 | ENSG00000081154 |
| ZNF135 | 0.157727501 | 1.115528599 | 0.650531436 | 0.795936006 | 38.7982469 | 43.39728842 | ENSG00000176293 |
| CEP104 | 0.157673783 | 1.115487064 | 0.109187911 | 0.241865513 | 622.7784748 | 694.789206 | ENSG00000116198 |
| TAB2 | 0.157661754 | 1.115477763 | 0.067750926 | 0.168713486 | 1185.249959 | 1321.755177 | ENSG00000055208 |
| CCDC97 | 0.157585794 | 1.115419033 | 0.112641035 | 0.2477675 | 583.1051173 | 650.4883135 | ENSG00000142039 |
| DPF2 | 0.157576177 | 1.115411598 | 0.017821538 | 0.058585823 | 1998.122898 | 2228.629699 | ENSG00000133884 |
| PGM2L1 | 0.157565063 | 1.115403005 | 0.187317833 | 0.359022617 | 897.074095 | 1000.021284 | ENSG00000165434 |
| ARHGAP26 | 0.157524212 | 1.115371422 | 0.15596628 | 0.314340912 | 482.7313582 | 537.8770399 | ENSG00000145819 |
| INO80C | 0.157474372 | 1.11533289 | 0.353956315 | 0.547324073 | 174.6874347 | 194.7750352 | ENSG00000153391 |
| TRIP10 | 0.157467528 | 1.1153276 | 0.029308802 | 0.087855303 | 1964.864435 | 2192.084049 | ENSG00000125733 |
| PTPRD | 0.157456411 | 1.115319005 | 0.165344264 | 0.3282003 | 751.9779968 | 838.0561356 | ENSG00000153707 |
| ETFRF1 | 0.157430891 | 1.115299276 | 0.642058145 | 0.790074341 | 47.17298865 | 52.36729415 | ENSG00000205707 |
| SEC24B-AS1 | 0.157393673 | 1.115270504 | 0.773531712 | 0.875038864 | 17.51284702 | 19.50032268 | ENSG00000247950 |

| | | | | | | | |
|---|---|---|---|---|---|---|---|
| NA | 0.157352681 | 1.115238817 | 0.884794804 | 0.940944853 | 4.066393563 | 4.587387409 | ENSG00000286585 |
| APTR | 0.157311326 | 1.115206849 | 0.414385039 | 0.605838314 | 175.2979886 | 195.9523288 | ENSG00000214293 |
| PTPN5 | 0.157286428 | 1.115187602 | 0.371051675 | 0.564342011 | 254.5240645 | 284.0431802 | ENSG00000110786 |
| CAMTA2 | 0.15726861 | 1.115173829 | 0.424274598 | 0.61515336 | 120.6495191 | 134.563087 | ENSG00000108509 |
| EZR | 0.157180388 | 1.115105638 | 0.010018299 | 0.036833793 | 6133.15576 | 6839.195221 | ENSG00000092820 |
| ZSCAN21 | 0.157149248 | 1.115081569 | 0.332495539 | 0.527067424 | 215.8135871 | 240.6748399 | ENSG00000166529 |
| WDFY2 | 0.157131304 | 1.1150677 | 0.089051722 | 0.207921971 | 809.591895 | 902.6146345 | ENSG00000139668 |
| ZFAND2A | 0.157122867 | 1.115061179 | 0.491600406 | 0.674213551 | 89.30515831 | 99.42568443 | ENSG00000178381 |
| USP4 | 0.157101205 | 1.115044436 | 0.073440513 | 0.179713027 | 917.4121366 | 1022.858642 | ENSG00000114316 |
| GGH | 0.157083238 | 1.11503055 | 0.112935757 | 0.24823901 | 592.8898705 | 660.7618969 | ENSG00000137563 |
| PIEZO1 | 0.15703727 | 1.114995023 | 0.032939569 | 0.096506794 | 1608.849809 | 1794.479371 | ENSG00000103335 |
| REEP5 | 0.156897963 | 1.114887364 | 0.067401845 | 0.168177948 | 1077.148038 | 1200.496907 | ENSG00000129625 |
| CRLS1 | 0.156886642 | 1.114878615 | 0.401391252 | 0.593015979 | 301.7641823 | 336.2504698 | ENSG00000088766 |
| ZNF629 | 0.156845848 | 1.114847091 | 0.084062639 | 0.198887961 | 835.4567164 | 931.6338935 | ENSG00000102870 |
| NA | 0.156734639 | 1.114761157 | 0.832920788 | 0.910221337 | 8.359558549 | 9.290761597 | ENSG00000226015 |
| MB21D2 | 0.156726759 | 1.114755069 | 0.197397872 | 0.373171375 | 383.2862265 | 427.0526033 | ENSG00000180611 |
| ABI1 | 0.156672047 | 1.114712794 | 0.125662978 | 0.268569839 | 716.0753053 | 798.0716977 | ENSG00000136754 |
| HEYL | 0.156671775 | 1.114712584 | 0.415614526 | 0.606869899 | 130.6527605 | 145.7559079 | ENSG00000163909 |
| MAFK | 0.1566362 | 1.114685096 | 0.108149877 | 0.240197701 | 628.8430271 | 701.2273495 | ENSG00000198517 |
| BACE1 | 0.156515657 | 1.114591964 | 0.464717577 | 0.650976915 | 126.5041145 | 140.8450651 | ENSG00000186318 |
| PNMA8B | 0.156498442 | 1.114578664 | 0.535045374 | 0.709590766 | 77.93205078 | 86.69505927 | ENSG00000204851 |
| ARHGEF35 | 0.156448006 | 1.1145397 | 0.323111196 | 0.517322868 | 193.6380888 | 215.8130589 | ENSG00000213214 |
| DYNLT1 | 0.156387197 | 1.114492723 | 0.118875685 | 0.257839901 | 1015.674945 | 1131.631727 | ENSG00000146425 |
| STK32C | 0.156288169 | 1.114416226 | 0.149103343 | 0.304750923 | 528.7328998 | 589.4787772 | ENSG00000165752 |
| GSTZ1 | 0.15627204 | 1.114403767 | 0.205567238 | 0.384011032 | 439.9796449 | 489.8499727 | ENSG00000100577 |
| NTAN1 | 0.156213622 | 1.114358644 | 0.255693512 | 0.444561309 | 277.9435657 | 309.9667202 | ENSG00000157045 |
| TCF20 | 0.156168681 | 1.114323931 | 0.034387863 | 0.09979583 | 2167.899578 | 2415.23386 | ENSG00000100207 |
| APLP2 | 0.156136725 | 1.114299249 | 0.003378456 | 0.014716439 | 11513.76194 | 12829.43201 | ENSG00000084234 |
| COQ9 | 0.156103671 | 1.114273719 | 0.118671165 | 0.257456557 | 621.8347991 | 692.9336005 | ENSG00000088682 |
| ABHD17B | 0.155988677 | 1.114184906 | 0.167228823 | 0.330887051 | 846.3045213 | 942.2646662 | ENSG00000107362 |
| RFX3 | 0.155907401 | 1.114122139 | 0.356683203 | 0.550255243 | 227.0291647 | 252.6164849 | ENSG00000080298 |
| CNBP | 0.155854458 | 1.114081254 | 0.005069945 | 0.020830496 | 7750.800699 | 8634.297841 | ENSG00000169714 |
| FAM3C | 0.155788572 | 1.114030377 | 0.111559844 | 0.245991187 | 1101.496407 | 1226.894422 | ENSG00000196937 |
| NA | 0.155787618 | 1.11402964 | 0.868694256 | 0.931788838 | 5.799048755 | 6.406215381 | ENSG00000226564 |
| NA | 0.155656248 | 1.113928203 | 0.609664711 | 0.767109691 | 51.87244049 | 57.77020491 | ENSG00000243302 |
| USP38 | 0.155561875 | 1.113855338 | 0.30933797 | 0.503404156 | 256.7750467 | 285.6163907 | ENSG00000170185 |
| NA | 0.155264652 | 1.113625886 | 0.828755742 | 0.907724296 | 9.996829939 | 11.12131252 | ENSG00000224790 |
| UBE2V1 | 0.155200109 | 1.113576066 | 0.663533007 | 0.804258763 | 36.82985321 | 40.9544459 | ENSG00000244687 |
| TGFA | 0.155165845 | 1.113549619 | 0.429338712 | 0.619564961 | 124.1033819 | 138.0363673 | ENSG00000163235 |
| ZNF473 | 0.155041144 | 1.113453372 | 0.350549289 | 0.543916327 | 199.9811259 | 222.6733538 | ENSG00000142528 |
| OSBP | 0.155041036 | 1.113453288 | 0.044215514 | 0.121688231 | 1587.154064 | 1766.319778 | ENSG00000110048 |
| NA | 0.155003606 | 1.113424401 | 0.823072865 | 0.904942493 | 9.821846899 | 10.92600489 | ENSG00000225507 |
| FOPNL | 0.154784961 | 1.113255671 | 0.347768032 | 0.541004927 | 464.6395711 | 516.8954718 | ENSG00000133393 |
| GREB1 | 0.154671197 | 1.113167889 | 0.122265913 | 0.26327923 | 912.3356971 | 1015.318441 | ENSG00000196208 |
| UBALD2 | 0.154657468 | 1.113157296 | 0.290178582 | 0.482260567 | 353.6356533 | 393.1156364 | ENSG00000185262 |
| MGME1 | 0.154647986 | 1.113149979 | 0.084636999 | 0.199905969 | 1112.532846 | 1237.483958 | ENSG00000125871 |
| BTBD7 | 0.154629862 | 1.113135995 | 0.171670704 | 0.337442447 | 573.3026703 | 637.9407149 | ENSG00000011114 |
| SLC9A5 | 0.154580414 | 1.113097843 | 0.340600792 | 0.534974159 | 249.1646531 | 277.440661 | ENSG00000135740 |
| SNX24 | 0.154570868 | 1.113090478 | 0.362329131 | 0.555267746 | 196.4923643 | 218.8559205 | ENSG00000064652 |
| NA | 0.154565381 | 1.113086245 | 0.896096329 | 0.946766059 | 3.794651247 | 4.264808071 | ENSG00000272505 |
| NUDT19 | 0.154507586 | 1.113041655 | 0.640309493 | 0.789444284 | 43.41631806 | 48.3363234 | ENSG00000213965 |
| C1orf122 | 0.154377548 | 1.112941335 | 0.187504439 | 0.359259308 | 526.7440164 | 585.9392929 | ENSG00000197982 |
| RPS4X | 0.154308632 | 1.112888173 | 0.007363899 | 0.028546854 | 27922.30647 | 31074.13125 | ENSG00000198034 |
| C17orf67 | 0.154148569 | 1.112764708 | 0.612192126 | 0.768828355 | 60.02366543 | 66.94889019 | ENSG00000214226 |
| GPAA1 | 0.154120182 | 1.112742812 | 0.057570313 | 0.149148474 | 1145.188482 | 1274.409529 | ENSG00000197858 |
| AKR1A1 | 0.15411293 | 1.112737219 | 0.011568689 | 0.041512473 | 3292.822703 | 3664.58158 | ENSG00000117448 |
| RPS12 | 0.154069612 | 1.112703809 | 0.006312213 | 0.025129242 | 13549.84172 | 15077.32277 | ENSG00000112306 |
| NA | 0.154050409 | 1.112688998 | 0.901499668 | 0.949934326 | 3.536814357 | 3.968597422 | ENSG00000251666 |
| NSUN5P2 | 0.15404647 | 1.11268596 | 0.754552872 | 0.863477583 | 34.97696487 | 39.04753717 | ENSG00000106133 |
| RPL39 | 0.15402155 | 1.112666741 | 0.085752366 | 0.201897148 | 3989.209938 | 4437.612689 | ENSG00000198918 |
| EIF3G | 0.153968688 | 1.112625972 | 0.012629836 | 0.044637943 | 4072.607848 | 4532.193109 | ENSG00000130811 |
| DSCAML1 | 0.153905021 | 1.112576872 | 0.805379705 | 0.895650273 | 11.79971133 | 13.13913536 | ENSG00000177103 |
| NA | 0.153894832 | 1.112569014 | 0.59687934 | 0.757087441 | 59.95531837 | 66.75358255 | ENSG00000287356 |
| GRIPAP1 | 0.153803714 | 1.112498749 | 0.075459524 | 0.183274384 | 1327.689307 | 1477.578237 | ENSG00000068400 |
| INSR | 0.153803286 | 1.112498419 | 0.150061622 | 0.30613562 | 689.5349805 | 766.8279032 | ENSG00000171105 |
| SMAD5 | 0.153801606 | 1.112497124 | 0.073840231 | 0.180357781 | 905.4885164 | 1007.694859 | ENSG00000113658 |
| ARHGAP19 | 0.153795695 | 1.112492566 | 0.146940292 | 0.301321659 | 650.9071691 | 723.7858549 | ENSG00000213390 |
| DOT1L | 0.153732997 | 1.112444219 | 0.059849095 | 0.15360107 | 2180.291331 | 2426.775752 | ENSG00000104885 |
| NA | 0.153666614 | 1.112393033 | 0.813328115 | 0.899577686 | 10.62938455 | 11.81518649 | ENSG00000234589 |
| OSBP2 | 0.153547629 | 1.112301293 | 0.437089797 | 0.626141776 | 143.2486596 | 159.5670002 | ENSG00000184792 |
| MED4 | 0.153533688 | 1.112290544 | 0.233174001 | 0.418075548 | 390.6712619 | 435.0098204 | ENSG00000136146 |
| MAMSTR | 0.153272491 | 1.112089185 | 0.692589547 | 0.823958834 | 30.30730073 | 33.81356537 | ENSG00000176909 |
| NA | 0.153118322 | 1.111970352 | 0.814342519 | 0.899952335 | 10.46702916 | 11.634851 | ENSG00000254610 |
| PLPP5 | 0.153110195 | 1.111964088 | 0.21122801 | 0.390708577 | 549.2014268 | 611.4311093 | ENSG00000147535 |

| | | | | | | | |
|---|---|---|---|---|---|---|---|
| C21orf58 | 0.153090386 | 1.111948819 | 0.652274473 | 0.796943154 | 313.517292 | 348.7494655 | ENSG00000160298 |
| IER2 | 0.153027892 | 1.111900654 | 0.066445409 | 0.166283987 | 1478.52554 | 1644.0594 | ENSG00000160888 |
| OTUD4 | 0.152931359 | 1.111826258 | 0.078276844 | 0.188560311 | 1232.499984 | 1369.39487 | ENSG00000164164 |
| MAP4K3 | 0.152857015 | 1.111768965 | 0.070300407 | 0.173541256 | 1105.46275 | 1228.372226 | ENSG00000011566 |
| SLC25A26 | 0.152825092 | 1.111744365 | 0.143157414 | 0.295695923 | 540.2863347 | 600.4259178 | ENSG00000144741 |
| CDC40 | 0.152739559 | 1.111678455 | 0.184271516 | 0.354925334 | 575.0171807 | 638.7717367 | ENSG00000168438 |
| NA | 0.152734747 | 1.111674747 | 0.503811806 | 0.683776217 | 109.7853807 | 122.1086277 | ENSG00000224631 |
| KMT2B | 0.152721315 | 1.111664397 | 0.064768204 | 0.163188402 | 2092.942209 | 2326.346909 | ENSG00000272333 |
| RNF114 | 0.152702612 | 1.111649986 | 0.034298907 | 0.099654482 | 2076.02102 | 2307.685572 | ENSG00000124226 |
| YIPF4 | 0.152702323 | 1.111649763 | 0.259931737 | 0.448814496 | 342.392851 | 380.5305446 | ENSG00000119820 |
| KRR1 | 0.152546891 | 1.111530003 | 0.219118492 | 0.400546616 | 463.5597049 | 514.8521966 | ENSG00000111615 |
| EVC | 0.152506143 | 1.111498609 | 0.269377632 | 0.459558637 | 306.5854911 | 340.9830959 | ENSG00000072840 |
| PYCR3 | 0.152461763 | 1.111464418 | 0.251522683 | 0.440117202 | 294.2639876 | 327.0770493 | ENSG00000104524 |
| ETV3L | 0.152366027 | 1.111390664 | 0.840040772 | 0.914174887 | 7.737752454 | 8.551747951 | ENSG00000253831 |
| H1-2 | 0.152292779 | 1.111334238 | 0.430190344 | 0.620237094 | 147.9028323 | 164.5966494 | ENSG00000187837 |
| HPS5 | 0.152251509 | 1.111302448 | 0.221187256 | 0.402739425 | 431.5452149 | 478.9168795 | ENSG00000110756 |
| FGF19 | 0.15218664 | 1.111252481 | 0.240427633 | 0.426582975 | 340.3423927 | 377.7064366 | ENSG00000162344 |
| PML | 0.152137967 | 1.111214991 | 0.118892688 | 0.257846605 | 819.3655087 | 910.6339211 | ENSG00000140464 |
| TOGARAM1 | 0.152082306 | 1.111172119 | 0.416246169 | 0.607457204 | 138.4264191 | 153.8316594 | ENSG00000198718 |
| NA | 0.151859611 | 1.111000612 | 0.845852053 | 0.917815337 | 7.631620137 | 8.435323138 | ENSG00000240409 |
| PLXNB2 | 0.151852315 | 1.110994993 | 0.025128387 | 0.077764154 | 3618.988231 | 4021.325992 | ENSG00000196576 |
| NUMB | 0.151777082 | 1.110937059 | 0.131176592 | 0.277329544 | 648.6137942 | 720.8203162 | ENSG00000133961 |
| U2AF1 | 0.151776039 | 1.110936256 | 0.715435402 | 0.838729455 | 26.5814025 | 29.43233248 | ENSG00000160201 |
| NA | 0.151716139 | 1.110890131 | 0.359192447 | 0.552379092 | 186.095441 | 206.5217477 | ENSG00000213553 |
| U2AF1L5 | 0.151691227 | 1.110870949 | 0.717406364 | 0.839899927 | 26.56561794 | 29.41356149 | ENSG00000275895 |
| HOMEZ | 0.15167475 | 1.110858262 | 0.392784597 | 0.585480917 | 176.1505198 | 195.727363 | ENSG00000215271 |
| ST6GAL1 | 0.1515722 | 1.110779302 | 0.028950435 | 0.087048565 | 2625.820704 | 2915.76172 | ENSG00000073849 |
| CSNK1G3 | 0.151230902 | 1.110516557 | 0.1788912 | 0.347844922 | 573.1623382 | 635.8602721 | ENSG00000151292 |
| NA | 0.151208338 | 1.110499188 | 0.847089317 | 0.918589821 | 9.815435375 | 10.88130324 | ENSG00000287906 |
| FAM217B | 0.151194078 | 1.110488211 | 0.226363175 | 0.409748518 | 363.765344 | 403.9175475 | ENSG00000196227 |
| CNOT6 | 0.151069099 | 1.110392016 | 0.139614125 | 0.290514 | 816.724881 | 906.0857875 | ENSG00000113300 |
| TMEM18-DT | 0.15105166 | 1.110378593 | 0.87035411 | 0.932705584 | 6.369066468 | 7.130583293 | ENSG00000233296 |
| RNF216 | 0.151028598 | 1.110360844 | 0.062861213 | 0.15975178 | 1060.955516 | 1178.311623 | ENSG00000011275 |
| MRPL24 | 0.151011508 | 1.11034769 | 0.184681217 | 0.355402521 | 415.7532898 | 461.7116363 | ENSG00000143314 |
| UBE3C | 0.150882918 | 1.110248727 | 0.027752926 | 0.084378728 | 1828.610158 | 2030.129826 | ENSG00000009335 |
| NA | 0.150845728 | 1.110220108 | 0.827757599 | 0.907253657 | 9.342176938 | 10.34095804 | ENSG00000260236 |
| ANKS3 | 0.150824438 | 1.110203725 | 0.312172186 | 0.5057959 | 306.2411398 | 340.3773514 | ENSG00000168096 |
| SH3GLB1 | 0.150797841 | 1.110183257 | 0.125730839 | 0.268621944 | 788.8799703 | 875.6451575 | ENSG00000097033 |
| TTYH3 | 0.150690487 | 1.110100649 | 0.01206732 | 0.042994176 | 6587.75553 | 7312.809888 | ENSG00000136295 |
| NA | 0.150585865 | 1.11002015 | 0.814043259 | 0.89991352 | 10.46071533 | 11.59730901 | ENSG00000242858 |
| TMEM18 | 0.150491636 | 1.109947652 | 0.274302281 | 0.465262054 | 289.2058882 | 320.8170915 | ENSG00000151353 |
| NA | 0.150363265 | 1.109848892 | 0.032067067 | 0.094443194 | 617824.1515 | 685692.1862 | N_noFeature |
| R3HCC1L | 0.150358548 | 1.109845264 | 0.249681623 | 0.438219621 | 381.9838508 | 423.3319393 | ENSG00000166024 |
| RPLP0 | 0.150310599 | 1.109808378 | 0.009603435 | 0.035583779 | 40317.1868 | 44743.56896 | ENSG00000089157 |
| SYCE1L | 0.150293672 | 1.109795357 | 0.594138617 | 0.755368716 | 65.32470411 | 72.54100795 | ENSG00000205078 |
| RTTN | 0.150214905 | 1.109734767 | 0.209573973 | 0.388735389 | 369.2056702 | 409.49232 | ENSG00000176225 |
| TOPBP1 | 0.150199178 | 1.109722669 | 0.071152327 | 0.175200906 | 1088.98104 | 1208.719738 | ENSG00000163781 |
| CPSF2 | 0.150194396 | 1.109718991 | 0.083452223 | 0.197914715 | 1815.853865 | 2014.568259 | ENSG00000165934 |
| MDGA2 | 0.150079557 | 1.10963066 | 0.678939346 | 0.814568995 | 33.91602502 | 37.60562615 | ENSG00000139915 |
| NA | 0.150053089 | 1.109610304 | 0.891373451 | 0.944688675 | 3.735271262 | 4.189724094 | ENSG00000215283 |
| TOPORS | 0.149926289 | 1.109512783 | 0.138827063 | 0.28907098 | 634.2473268 | 703.4015123 | ENSG00000197579 |
| CLTC | 0.149711421 | 1.109347549 | 0.023091547 | 0.072600293 | 7147.832844 | 7928.127534 | ENSG00000141367 |
| NA | 0.149626435 | 1.109282202 | 0.809937593 | 0.897702628 | 12.90236746 | 14.31704154 | ENSG00000261884 |
| ZBTB45 | 0.149516302 | 1.109197525 | 0.201766013 | 0.378831343 | 525.7997469 | 583.1896076 | ENSG00000119574 |
| ZNF525 | 0.149474239 | 1.109165186 | 0.152499971 | 0.309610523 | 558.9283936 | 619.7689128 | ENSG00000203326 |
| NA | 0.149283569 | 1.109018606 | 0.895992604 | 0.946766059 | 3.516595104 | 3.938541507 | ENSG00000274080 |
| CRISPLD1 | 0.149259136 | 1.108999823 | 0.209938771 | 0.389139432 | 414.8296681 | 460.3614304 | ENSG00000121005 |
| ASB16-AS1 | 0.149250578 | 1.108993245 | 0.564801763 | 0.732937653 | 74.96269859 | 83.27191999 | ENSG00000267080 |
| GOLGA5 | 0.149171593 | 1.108932532 | 0.165357419 | 0.3282003 | 535.9668472 | 593.9268436 | ENSG00000066455 |
| EIF4A2 | 0.149126627 | 1.108897969 | 0.024115687 | 0.075220489 | 6619.882603 | 7339.281758 | ENSG00000156976 |
| SEC14L5 | 0.149094358 | 1.108873166 | 0.301427391 | 0.494876009 | 231.8808248 | 257.113092 | ENSG00000103184 |
| ARMC1 | 0.14907826 | 1.108860793 | 0.111831728 | 0.246396725 | 733.047667 | 812.281123 | ENSG00000104442 |
| HERPUD1 | 0.149053748 | 1.108841953 | 0.178761004 | 0.347714418 | 471.885372 | 522.9761625 | ENSG00000051108 |
| HARBI1 | 0.14902218 | 1.10881769 | 0.564209056 | 0.732480414 | 76.06988711 | 84.3849278 | ENSG00000180423 |
| NA | 0.148873096 | 1.108703115 | 0.888791262 | 0.943202869 | 4.108711207 | 4.598672329 | ENSG00000205871 |
| PPP2CB | 0.14875317 | 1.108610956 | 0.051325903 | 0.136461668 | 1606.997552 | 1781.517075 | ENSG00000104695 |
| ZDHHC17 | 0.148533133 | 1.108441885 | 0.222376097 | 0.404427298 | 393.6800679 | 436.3291657 | ENSG00000186908 |
| OLFM2 | 0.148512163 | 1.108425774 | 0.025685624 | 0.079118723 | 1973.028325 | 2187.031806 | ENSG00000105088 |
| PSEN1 | 0.148468276 | 1.108392056 | 0.076362208 | 0.184981984 | 960.2633601 | 1063.939769 | ENSG00000080815 |
| PLK1 | 0.148397641 | 1.10833779 | 0.014347854 | 0.049272398 | 3074.245189 | 3407.351647 | ENSG00000166851 |
| NPIPB12 | 0.148313588 | 1.108273219 | 0.834064515 | 0.911041473 | 9.173507724 | 10.13436548 | ENSG00000169203 |
| KBTBD6 | 0.148223333 | 1.108203888 | 0.254878564 | 0.44368548 | 354.8611133 | 392.9706217 | ENSG00000165572 |
| NA | 0.148207326 | 1.108191592 | 0.730187446 | 0.849118629 | 28.04684777 | 31.21873141 | ENSG00000274925 |
| IKZF3 | 0.148104525 | 1.108112629 | 0.847170379 | 0.918589821 | 10.27488608 | 11.45930204 | ENSG00000161405 |

| | | | | | | | |
|---|---|---|---|---|---|---|---|
| GOLGA7 | 0.147970264 | 1.10800951 | 0.214159563 | 0.394320817 | 592.0006365 | 656.1530707 | ENSG00000147533 |
| THAP9-AS1 | 0.147852018 | 1.107918699 | 0.349186722 | 0.5425547 | 268.8849846 | 297.9036088 | ENSG00000251022 |
| ELP4 | 0.147784617 | 1.10786694 | 0.45602029 | 0.643266615 | 182.5372273 | 201.7324024 | ENSG00000109911 |
| NA | 0.147751911 | 1.107841825 | 0.651488108 | 0.796430353 | 42.59417592 | 47.230252 | ENSG00000244398 |
| ATXN2 | 0.147710524 | 1.107810044 | 0.108698983 | 0.241149537 | 917.3279374 | 1016.027916 | ENSG00000204842 |
| TUSC3 | 0.14767317 | 1.107781361 | 0.101939043 | 0.229986393 | 1134.494884 | 1255.735934 | ENSG00000104723 |
| EPOP | 0.14766121 | 1.107772177 | 0.429445995 | 0.619597701 | 144.4658364 | 160.2002212 | ENSG00000273604 |
| PRSS12 | 0.147603816 | 1.107728108 | 0.345849258 | 0.539287409 | 197.512843 | 218.7052028 | ENSG00000164099 |
| ASB3 | 0.147574713 | 1.107705763 | 0.206332012 | 0.38491693 | 395.2863715 | 437.8630595 | ENSG00000115239 |
| MANSC1 | 0.147554735 | 1.107690424 | 0.591985238 | 0.753820295 | 87.05851308 | 96.24134351 | ENSG00000111261 |
| OPLAH | 0.147552674 | 1.107688841 | 0.137755219 | 0.287323343 | 812.4947414 | 900.5867864 | ENSG00000178814 |
| GOT1 | 0.147497056 | 1.107646139 | 0.082233624 | 0.195801387 | 1420.947947 | 1573.088966 | ENSG00000120053 |
| CCNY | 0.147287969 | 1.107485622 | 0.07292699 | 0.178763237 | 1237.80796 | 1370.220078 | ENSG00000108100 |
| NA | 0.147252915 | 1.107458713 | 0.853425591 | 0.922280546 | 7.459696311 | 8.202911545 | ENSG00000198416 |
| ADAD2 | 0.147238245 | 1.107447452 | 0.883057489 | 0.939798306 | 4.413300429 | 4.936223814 | ENSG00000140955 |
| NA | 0.147143574 | 1.107374782 | 0.378787783 | 0.572023464 | 171.4924911 | 189.7940065 | ENSG00000078319 |
| MAN2B1 | 0.14710466 | 1.107344914 | 0.081331623 | 0.194052561 | 906.3021951 | 1003.725823 | ENSG00000104774 |
| PKP4 | 0.147085165 | 1.10732995 | 0.029449214 | 0.088219127 | 2241.794084 | 2481.687521 | ENSG00000144283 |
| UACA | 0.147054321 | 1.107306276 | 0.181080004 | 0.350840142 | 451.3028506 | 499.9179427 | ENSG00000137831 |
| PDCD2L | 0.147038526 | 1.107294153 | 0.512297982 | 0.691244234 | 91.52003901 | 101.2072968 | ENSG00000126249 |
| NHLH2 | 0.146851477 | 1.107150599 | 0.516652747 | 0.694239253 | 91.92148521 | 101.9130054 | ENSG00000177551 |
| ZNF783 | 0.146811418 | 1.107119858 | 0.342747997 | 0.536797013 | 238.7345973 | 264.449933 | ENSG00000204946 |
| MEGF6 | 0.146766862 | 1.107085666 | 0.5660368 | 0.733733812 | 938.7091869 | 1039.509782 | ENSG00000162591 |
| FBXO21 | 0.146735108 | 1.107061299 | 0.100661591 | 0.227686347 | 1285.290604 | 1421.954549 | ENSG00000135108 |
| FZR1 | 0.146725916 | 1.107054246 | 0.08260588 | 0.196309447 | 956.4373125 | 1058.746741 | ENSG00000105325 |
| RBM47 | 0.146655015 | 1.106999841 | 0.06958689 | 0.172238704 | 1547.114595 | 1711.961531 | ENSG00000163694 |
| PMP22 | 0.146531715 | 1.106905235 | 0.737294401 | 0.852943558 | 25.44086342 | 28.02895113 | ENSG00000109099 |
| FRMD8 | 0.146521255 | 1.10689721 | 0.188787325 | 0.360757221 | 542.5724338 | 601.1502312 | ENSG00000126391 |
| L2HGDH | 0.1463586 | 1.106772421 | 0.106780311 | 0.238068842 | 771.0232681 | 853.0004493 | ENSG00000087299 |
| ADGRL2 | 0.146354957 | 1.106769626 | 0.080632979 | 0.192832468 | 2092.790129 | 2315.16011 | ENSG00000117114 |
| LRRC9 | 0.146280118 | 1.106712214 | 0.796226634 | 0.889956847 | 15.9766719 | 17.7002657 | ENSG00000131951 |
| NA | 0.14621606 | 1.106663076 | 0.582300231 | 0.746040945 | 68.88301549 | 76.24278936 | ENSG00000216866 |
| RPS6KA6 | 0.146210272 | 1.106658636 | 0.338740414 | 0.532864949 | 243.3851094 | 269.3884268 | ENSG00000072133 |
| TMEM185A | 0.146167926 | 1.106626154 | 0.278824903 | 0.469922774 | 323.891861 | 358.7285013 | ENSG00000269556 |
| BBIP1 | 0.146150872 | 1.106613073 | 0.454357783 | 0.641605116 | 124.0332534 | 137.0270737 | ENSG00000214413 |
| RAP1GAP | 0.146024437 | 1.106516095 | 0.254017681 | 0.442852384 | 402.2463514 | 445.3405525 | ENSG00000076864 |
| CXCR4 | 0.14602092 | 1.106513398 | 0.748003515 | 0.859325429 | 21.71821981 | 23.97865973 | ENSG00000121966 |
| TCF4 | 0.146010863 | 1.106505685 | 0.075034777 | 0.182553763 | 1133.436019 | 1253.987252 | ENSG00000196628 |
| IVNS1ABP | 0.145931281 | 1.106444649 | 0.013361936 | 0.046559525 | 5492.637811 | 6076.440316 | ENSG00000116679 |
| MLXIPL | 0.14580824 | 1.106350289 | 0.616927855 | 0.772278752 | 107.3155344 | 119.0642776 | ENSG00000009950 |
| PSORS1C1 | 0.145768919 | 1.106320136 | 0.894883347 | 0.946203969 | 4.389924263 | 4.891195752 | ENSG00000204540 |
| LARP1B | 0.1457523 | 1.106307392 | 0.11955201 | 0.258882691 | 702.9410705 | 777.2975756 | ENSG00000138709 |
| NA | 0.145676411 | 1.1062492 | 0.866671512 | 0.930273692 | 5.961501842 | 6.657836001 | ENSG00000214832 |
| PGM2 | 0.145553727 | 1.10615513 | 0.137988854 | 0.287648794 | 676.6718715 | 748.2544242 | ENSG00000169299 |
| NA | 0.145447874 | 1.106073973 | 0.373917541 | 0.56711847 | 254.292039 | 281.0546478 | ENSG00000279865 |
| C2CD2 | 0.145428369 | 1.106059018 | 0.264421949 | 0.454094232 | 361.9424539 | 399.737571 | ENSG00000157617 |
| NA | 0.14537839 | 1.106020702 | 0.478368899 | 0.662348428 | 169.319447 | 187.4655421 | ENSG00000260404 |
| RPL23 | 0.145353098 | 1.106001313 | 0.015069065 | 0.05130236 | 14398.54795 | 15924.46358 | ENSG00000125691 |
| TTC12 | 0.145309258 | 1.105967705 | 0.533048527 | 0.708006407 | 95.21231622 | 105.4129808 | ENSG00000149292 |
| NA | 0.145280127 | 1.105945373 | 0.3708467 | 0.564139796 | 190.0341075 | 209.9048602 | ENSG00000233476 |
| NA | 0.145207728 | 1.105889875 | 0.878001724 | 0.937000286 | 5.325790319 | 5.941392196 | ENSG00000229237 |
| PACSIN3 | 0.145095967 | 1.105804208 | 0.065011229 | 0.163689459 | 1110.036331 | 1227.346392 | ENSG00000165912 |
| VPS72 | 0.1449154 | 1.105665815 | 0.075278629 | 0.1830029 | 1671.09149 | 1847.104896 | ENSG00000163159 |
| CDC20 | 0.14464529 | 1.105458825 | 0.024720226 | 0.076757357 | 4468.939976 | 4939.351687 | ENSG00000117399 |
| RBFOX3 | 0.144572286 | 1.105402887 | 0.235602422 | 0.420544675 | 438.9872323 | 485.7221125 | ENSG00000167281 |
| VAMP8 | 0.144526666 | 1.105367933 | 0.26855081 | 0.458781521 | 350.5671742 | 387.3868661 | ENSG00000118640 |
| SEC61G | 0.144463449 | 1.105319499 | 0.082169005 | 0.195672664 | 1454.150367 | 1607.293075 | ENSG00000132432 |
| CREBL2 | 0.144462285 | 1.105318607 | 0.209239185 | 0.388474839 | 516.7269672 | 570.6601272 | ENSG00000111269 |
| DYSF | 0.144433259 | 1.105296369 | 0.126844084 | 0.270483072 | 825.1310266 | 911.9369406 | ENSG00000135636 |
| POGK | 0.144363895 | 1.105243228 | 0.057942255 | 0.149902533 | 1294.864709 | 1431.052171 | ENSG00000143157 |
| SHLD2 | 0.144154365 | 1.10508272 | 0.235592491 | 0.420544675 | 554.9694926 | 613.0589689 | ENSG00000122376 |
| BUB1B | 0.144030857 | 1.104988119 | 0.034148939 | 0.099325074 | 3150.926614 | 3481.471468 | ENSG00000156970 |
| PTPN21 | 0.143964186 | 1.104937055 | 0.349921609 | 0.543306283 | 224.5221271 | 248.273822 | ENSG00000070778 |
| TRUB1 | 0.14395258 | 1.104928166 | 0.237498537 | 0.422736887 | 486.8756672 | 537.2137312 | ENSG00000165832 |
| NPM3 | 0.143936091 | 1.104915538 | 0.059242875 | 0.152437518 | 2501.532515 | 2763.644087 | ENSG00000107833 |
| CX3CL1 | 0.143816254 | 1.104823762 | 0.645345846 | 0.792859536 | 46.13602633 | 50.99974294 | ENSG00000006210 |
| ZFP3 | 0.143797841 | 1.104809661 | 0.756093277 | 0.864333115 | 26.67500487 | 29.6585721 | ENSG00000180787 |
| SFT2D2 | 0.143751047 | 1.104773827 | 0.401049214 | 0.592731097 | 178.0278666 | 196.736545 | ENSG00000213064 |
| PPP1R9A | 0.143704905 | 1.104738494 | 0.431988045 | 0.6220157 | 279.5142489 | 308.382843 | ENSG00000158528 |
| WDR70 | 0.143659354 | 1.104703614 | 0.215040856 | 0.395197543 | 700.9904686 | 773.5646305 | ENSG00000082068 |
| FAM216A | 0.143451916 | 1.104544785 | 0.400184121 | 0.59215821 | 165.1885171 | 182.3712262 | ENSG00000204856 |
| SLC34A2 | 0.14341679 | 1.104517893 | 0.654784592 | 0.798301383 | 45.06551804 | 49.65267457 | ENSG00000157765 |
| SELENOM | 0.143247334 | 1.104388166 | 0.430562753 | 0.620532664 | 164.2719833 | 181.5680871 | ENSG00000198832 |
| ZNF92 | 0.143139349 | 1.104305506 | 0.349990408 | 0.543367585 | 345.6862375 | 381.2705057 | ENSG00000146757 |

| | | | | | | | |
|---|---|---|---|---|---|---|---|
| NA | 0.143137453 | 1.104304055 | 0.631720487 | 0.782745522 | 50.8349745 | 56.19843426 | ENSG00000268222 |
| TXN | 0.14307687 | 1.104257683 | 0.050217336 | 0.134246085 | 2538.193014 | 2803.075086 | ENSG00000136810 |
| SYMPK | 0.143034315 | 1.104225111 | 0.039235585 | 0.110768064 | 2358.756602 | 2604.745572 | ENSG00000125755 |
| FAM186B | 0.142925667 | 1.104141957 | 0.899499391 | 0.948527635 | 4.058801961 | 4.515990658 | ENSG00000135436 |
| NDUFS2 | 0.142921791 | 1.104138989 | 0.037099544 | 0.105855696 | 2077.067062 | 2293.533227 | ENSG00000158864 |
| JARID2 | 0.142782458 | 1.104032359 | 0.013702725 | 0.047505855 | 7922.498457 | 8746.341818 | ENSG00000008083 |
| LGMN | 0.142760332 | 1.104015427 | 0.053167149 | 0.140150753 | 1521.452898 | 1679.131426 | ENSG00000100600 |
| PPP1R3F | 0.14265322 | 1.103933463 | 0.475209803 | 0.660037357 | 118.7624083 | 130.9589595 | ENSG00000049769 |
| FBXW9 | 0.142631559 | 1.103916889 | 0.430947444 | 0.620909351 | 173.6454363 | 191.246939 | ENSG00000132004 |
| SYNE3 | 0.142526167 | 1.103836248 | 0.901158288 | 0.949787537 | 3.483748199 | 3.878429677 | ENSG00000176438 |
| NA | 0.142502335 | 1.103818014 | 0.89407691 | 0.94574241 | 4.072707389 | 4.531074426 | ENSG00000230593 |
| SETD2 | 0.142431607 | 1.1037639 | 0.087511207 | 0.205048383 | 1294.014658 | 1428.133212 | ENSG00000181555 |
| SNX25 | 0.142243466 | 1.103619969 | 0.272681718 | 0.463871772 | 320.8720209 | 354.3576773 | ENSG00000109762 |
| PACS1 | 0.14222288 | 1.103604221 | 0.150177696 | 0.306304987 | 815.2595636 | 899.2131084 | ENSG00000175115 |
| IL11 | 0.142049561 | 1.103471647 | 0.720889394 | 0.842237882 | 27.45157475 | 30.26140225 | ENSG00000095752 |
| FBXW7 | 0.141910071 | 1.103364961 | 0.246029621 | 0.433575448 | 403.7560911 | 445.6352258 | ENSG00000109670 |
| CIDECP1 | 0.141896007 | 1.103354205 | 0.617971472 | 0.772868503 | 55.45620257 | 61.28058915 | ENSG00000186162 |
| NOB1 | 0.141722743 | 1.103221702 | 0.081818318 | 0.194987875 | 1322.725975 | 1458.906297 | ENSG00000141101 |
| KMT5B | 0.141600009 | 1.103127853 | 0.126798883 | 0.270445422 | 878.0728704 | 968.4300482 | ENSG00000110066 |
| NA | 0.14153619 | 1.103079056 | 0.645744791 | 0.792972261 | 47.52936625 | 52.29688509 | ENSG00000214389 |
| MED14 | 0.141466202 | 1.103025545 | 0.085540287 | 0.201449006 | 1503.795773 | 1658.500247 | ENSG00000180182 |
| MOV10 | 0.141311816 | 1.102907513 | 0.128094875 | 0.272291594 | 1131.485913 | 1248.58662 | ENSG00000155363 |
| PCNX1 | 0.141301666 | 1.102899754 | 0.122340295 | 0.263332361 | 994.5338836 | 1096.753048 | ENSG00000100731 |
| APPL2 | 0.141092747 | 1.102740053 | 0.278990208 | 0.470063197 | 339.6145746 | 374.3658758 | ENSG00000136044 |
| MDH2 | 0.141059097 | 1.102714332 | 0.01980078 | 0.064013198 | 5455.059942 | 6014.632965 | ENSG00000146701 |
| LINC02367 | 0.141047608 | 1.102705551 | 0.866754893 | 0.930279181 | 6.302094882 | 7.006672407 | ENSG00000260423 |
| HROB | 0.141037279 | 1.102697656 | 0.274517261 | 0.465474106 | 325.1988069 | 358.3563935 | ENSG00000125319 |
| PCSK5 | 0.140684552 | 1.102428089 | 0.348224914 | 0.541579304 | 223.097218 | 245.7095533 | ENSG00000099139 |
| PFKFB3 | 0.14067022 | 1.102417137 | 0.100830819 | 0.22793004 | 1032.37966 | 1138.006899 | ENSG00000170525 |
| ZC3HAV1L | 0.140662958 | 1.102411588 | 0.611837374 | 0.768591155 | 66.86974861 | 73.84662386 | ENSG00000146858 |
| PRDX5 | 0.140411056 | 1.102219118 | 0.028754464 | 0.08661756 | 4300.15307 | 4739.504503 | ENSG00000126432 |
| CC2D2A | 0.140315153 | 1.10214585 | 0.342017171 | 0.536244502 | 233.3384831 | 256.9815835 | ENSG00000048342 |
| WDR59 | 0.140282679 | 1.102121042 | 0.094386382 | 0.216907279 | 1345.445214 | 1483.209927 | ENSG00000103091 |
| TAGLN2 | 0.14025347 | 1.102098729 | 0.093025055 | 0.21460373 | 1577.195289 | 1737.289521 | ENSG00000158710 |
| NGLY1 | 0.140244161 | 1.102091618 | 0.229588509 | 0.4134467 | 400.0388895 | 440.7545421 | ENSG00000151092 |
| IL27RA | 0.140229196 | 1.102080185 | 0.216136029 | 0.396620313 | 437.2707903 | 482.0216049 | ENSG00000104998 |
| MPP5 | 0.139958503 | 1.101873422 | 0.215661028 | 0.395955923 | 512.2862393 | 563.9132144 | ENSG00000072415 |
| RWDD2A | 0.139884358 | 1.101816794 | 0.545667698 | 0.717771832 | 108.4995712 | 119.605785 | ENSG00000013392 |
| FBXL7 | 0.139769292 | 1.101728919 | 0.20877779 | 0.387918553 | 459.2464562 | 505.5413902 | ENSG00000183580 |
| PTPN23 | 0.139646554 | 1.101635193 | 0.053663613 | 0.141118246 | 1678.383156 | 1848.532519 | ENSG00000076201 |
| CLEC2L | 0.139633126 | 1.101624939 | 0.796957536 | 0.890505274 | 15.38643493 | 16.93499499 | ENSG00000236279 |
| NELFE | 0.139459486 | 1.101492358 | 0.074871095 | 0.182295218 | 1320.458434 | 1454.534463 | ENSG00000204356 |
| LAMA4 | 0.139404591 | 1.101450447 | 0.740391655 | 0.854997645 | 26.58781403 | 29.36885984 | ENSG00000112769 |
| FER | 0.139376733 | 1.101429178 | 0.433075808 | 0.622850174 | 181.4088122 | 199.7093952 | ENSG00000151422 |
| PNPO | 0.139372718 | 1.101426113 | 0.666860782 | 0.806709596 | 54.7122842 | 60.37786112 | ENSG00000108439 |
| TMEM177 | 0.139293053 | 1.101365295 | 0.141283305 | 0.292934868 | 918.2663962 | 1011.907273 | ENSG00000144120 |
| ZNF710 | 0.1392344 | 1.10132052 | 0.236176666 | 0.421177555 | 612.6596675 | 675.0228487 | ENSG00000140548 |
| FCHSD2 | 0.139153399 | 1.101258686 | 0.473466068 | 0.658574174 | 148.5512843 | 163.3471233 | ENSG00000137478 |
| XRRA1 | 0.139087623 | 1.101208479 | 0.635856997 | 0.785926881 | 56.15938934 | 61.87735895 | ENSG00000166435 |
| NINJ1 | 0.13904293 | 1.101174365 | 0.199421413 | 0.375770279 | 508.9861556 | 560.8762874 | ENSG00000131669 |
| ZBTB22 | 0.139002225 | 1.101143297 | 0.350066781 | 0.543382519 | 316.5481737 | 348.7409514 | ENSG00000236104 |
| ANLN | 0.138929953 | 1.101088136 | 0.115434837 | 0.252474543 | 1396.543956 | 1536.663038 | ENSG00000011426 |
| CFL2 | 0.138799065 | 1.100988244 | 0.479577079 | 0.663554019 | 207.393957 | 228.2899849 | ENSG00000165410 |
| ENTPD2 | 0.138762761 | 1.10096054 | 0.553786276 | 0.723938212 | 92.52909278 | 101.672558 | ENSG00000054179 |
| CDCA4 | 0.1387415 | 1.100944315 | 0.14419912 | 0.297251305 | 1435.7388 | 1581.495296 | ENSG00000170779 |
| TMEM234 | 0.138720606 | 1.10092837 | 0.633342882 | 0.783917255 | 58.92594764 | 65.0435817 | ENSG00000160055 |
| CD74 | 0.138565285 | 1.10080985 | 0.165996588 | 0.329151681 | 583.7519607 | 642.2882131 | ENSG00000019582 |
| NA | 0.138562754 | 1.100807919 | 0.657134318 | 0.799693201 | 46.80458475 | 51.61897096 | ENSG00000259985 |
| SLC4A11 | 0.138258376 | 1.100575697 | 0.33154494 | 0.52576497 | 563.503373 | 620.8859604 | ENSG00000088836 |
| KIAA1324L | 0.138107243 | 1.100460409 | 0.146171512 | 0.300215293 | 789.0588769 | 867.723038 | ENSG00000164659 |
| KIFC1 | 0.13810267 | 1.100456921 | 0.039854926 | 0.112174822 | 2142.93831 | 2357.805844 | ENSG00000237649 |
| SYDE1 | 0.138074344 | 1.100435315 | 0.256893038 | 0.445408246 | 423.4971455 | 466.4953275 | ENSG00000105137 |
| MIER2 | 0.137933611 | 1.100327974 | 0.119867833 | 0.259384753 | 856.0757977 | 941.7633608 | ENSG00000105556 |
| SLC25A15 | 0.137887431 | 1.100292753 | 0.188442741 | 0.360358829 | 634.1496579 | 697.7343021 | ENSG00000102743 |
| NA | 0.137811932 | 1.100235175 | 0.891071457 | 0.944638699 | 4.813273453 | 5.330088282 | ENSG00000261033 |
| NA | 0.137783974 | 1.100213853 | 0.561599078 | 0.730433496 | 86.69158236 | 95.48845702 | ENSG00000228409 |
| MINK1 | 0.137722034 | 1.100166619 | 0.051195434 | 0.136232041 | 2392.783811 | 2633.428642 | ENSG00000141503 |
| TRAPPC6B | 0.137656692 | 1.100116791 | 0.495507173 | 0.677275366 | 131.7621364 | 144.9950889 | ENSG00000182400 |
| NA | 0.137649715 | 1.100111471 | 0.886857642 | 0.942266356 | 5.505208048 | 5.974697304 | ENSG00000279641 |
| WDPCP | 0.137637637 | 1.100102261 | 0.401238424 | 0.592837448 | 204.5530461 | 225.0889516 | ENSG00000143951 |
| NTSR1 | 0.137626864 | 1.100094046 | 0.78069225 | 0.879518677 | 18.16246397 | 19.98424327 | ENSG00000101188 |
| H1-0 | 0.137569657 | 1.100050425 | 0.077009542 | 0.186257974 | 3243.255048 | 3567.19108 | ENSG00000189060 |
| NKX3-1 | 0.137501882 | 1.099998748 | 0.706801843 | 0.833271761 | 33.78020271 | 37.1366777 | ENSG00000167034 |
| PRDX1 | 0.137451108 | 1.099960035 | 0.041205356 | 0.115205923 | 13512.62968 | 14862.71064 | ENSG00000117450 |

| | | | | | | | |
|---|---|---|---|---|---|---|---|
| ZSCAN32 | 0.13733398 | 1.099870737 | 0.350604433 | 0.543956351 | 266.0701782 | 292.2878219 | ENSG00000140987 |
| ZNF551 | 0.137036648 | 1.099644083 | 0.212199118 | 0.391891902 | 516.8715839 | 568.3301658 | ENSG00000204519 |
| BSN | 0.137008176 | 1.099622381 | 0.450499616 | 0.638101635 | 152.4088633 | 167.7584121 | ENSG00000164061 |
| NA | 0.1369496 | 1.099577736 | 0.675102711 | 0.811910561 | 45.00180106 | 49.67243325 | ENSG00000272734 |
| NFYA | 0.136907901 | 1.099545954 | 0.150829868 | 0.307060718 | 790.5150468 | 869.8241727 | ENSG00000001167 |
| SPRTN | 0.13686697 | 1.099514759 | 0.379772411 | 0.572904271 | 199.0453575 | 218.9493374 | ENSG00000010072 |
| FUS | 0.136724147 | 1.099405916 | 0.020485945 | 0.065914187 | 17539.0068 | 19283.81 | ENSG00000089280 |
| USP54 | 0.136703071 | 1.099389855 | 0.200567711 | 0.377199309 | 582.6149094 | 640.5578326 | ENSG00000166348 |
| CSNK2A3 | 0.136639436 | 1.099341363 | 0.89399551 | 0.94574241 | 4.377296611 | 4.84605607 | ENSG00000254598 |
| WDR41 | 0.136633487 | 1.09933683 | 0.222201213 | 0.404264274 | 779.0510804 | 855.5295802 | ENSG00000164253 |
| DOP1B | 0.136602073 | 1.099312893 | 0.34355408 | 0.537290635 | 241.1099543 | 265.0571117 | ENSG00000142197 |
| NUDT14 | 0.13658461 | 1.099299586 | 0.412019906 | 0.603523033 | 319.5600616 | 351.8829957 | ENSG00000183828 |
| FCN3 | 0.136418466 | 1.099172996 | 0.89123599 | 0.944651026 | 4.449304247 | 4.924938894 | ENSG00000142748 |
| OTULIN | 0.136356651 | 1.0991259 | 0.227819338 | 0.411460107 | 442.1049594 | 485.8914492 | ENSG00000154124 |
| KIAA0586 | 0.136242855 | 1.099039208 | 0.245759921 | 0.433245993 | 416.992971 | 458.0801608 | ENSG00000100578 |
| GPR63 | 0.136223205 | 1.099024238 | 0.342457075 | 0.536616455 | 276.7474049 | 304.4267902 | ENSG00000112218 |
| TBC1D2B | 0.136221374 | 1.099022844 | 0.216280355 | 0.396828398 | 555.9231126 | 611.5714894 | ENSG00000167202 |
| SYT3 | 0.13618957 | 1.098998616 | 0.307470132 | 0.501730184 | 322.5146965 | 354.8562436 | ENSG00000213023 |
| AASDH | 0.13616356 | 1.098978803 | 0.41183131 | 0.603342147 | 194.3654485 | 213.4149181 | ENSG00000157426 |
| NET1 | 0.13614314 | 1.098963248 | 0.066147428 | 0.165672491 | 1783.763286 | 1959.43051 | ENSG00000173848 |
| ECHDC2 | 0.136122449 | 1.098947487 | 0.393846139 | 0.586497216 | 210.1863871 | 230.8654269 | ENSG00000121310 |
| KHDRBS2 | 0.136077442 | 1.098913204 | 0.408326589 | 0.600105067 | 182.6687126 | 200.7004675 | ENSG00000112232 |
| NA | 0.136071697 | 1.098908828 | 0.847897427 | 0.918948068 | 8.885980843 | 9.831106793 | ENSG00000250138 |
| GARS1 | 0.136033348 | 1.098879618 | 0.05307655 | 0.139991573 | 4631.504996 | 5088.436241 | ENSG00000106105 |
| OXLD1 | 0.135938861 | 1.098807651 | 0.230377954 | 0.414487336 | 463.782643 | 510.0015547 | ENSG00000204237 |
| PLCB4 | 0.135809459 | 1.098709098 | 0.405816105 | 0.597457755 | 219.7306589 | 241.2479406 | ENSG00000101333 |
| POLI | 0.135790898 | 1.098694963 | 0.309431701 | 0.50351248 | 308.3848096 | 338.5070196 | ENSG00000101751 |
| NA | 0.135788371 | 1.098693039 | 0.841984189 | 0.915376288 | 9.924822303 | 10.83583714 | ENSG00000164845 |
| ANKRD12 | 0.135777411 | 1.098684691 | 0.341147396 | 0.535546635 | 310.7469602 | 340.988678 | ENSG00000101745 |
| PPP3CB-AS1 | 0.13571183 | 1.09863475 | 0.886943419 | 0.942266356 | 5.052168865 | 5.585069716 | ENSG00000221817 |
| NA | 0.13565967 | 1.09859503 | 0.838810801 | 0.913533869 | 19.66499543 | 21.5979044 | ENSG00000230615 |
| SYT5 | 0.135640676 | 1.098580566 | 0.726413633 | 0.84637897 | 34.86085815 | 38.39335508 | ENSG00000129990 |
| CCDC144B | 0.135632783 | 1.098574556 | 0.910241397 | 0.95534368 | 4.197781184 | 4.632415471 | ENSG00000154874 |
| UBE3A | 0.135617002 | 1.098562539 | 0.115128786 | 0.251953823 | 1376.801668 | 1512.07896 | ENSG00000114062 |
| ZNF121 | 0.135514461 | 1.098484461 | 0.153983322 | 0.311567392 | 1437.236198 | 1579.337866 | ENSG00000197961 |
| DOCK5 | 0.13535955 | 1.098366516 | 0.373058397 | 0.566421735 | 260.8444018 | 285.9615399 | ENSG00000147459 |
| COL7A1 | 0.135323101 | 1.098338766 | 0.731036635 | 0.849522415 | 271.3854154 | 298.2301791 | ENSG00000114270 |
| NA | 0.135247825 | 1.098281459 | 0.882086792 | 0.939373634 | 5.363071913 | 5.930107275 | ENSG00000237082 |
| WDR13 | 0.135163664 | 1.098217392 | 0.239927382 | 0.425930474 | 448.816904 | 493.3373893 | ENSG00000101940 |
| LIMS1 | 0.135137504 | 1.098197478 | 0.221134142 | 0.402736883 | 730.6113289 | 802.0257207 | ENSG00000169756 |
| NA | 0.134870736 | 1.09799443 | 0.720275611 | 0.841873505 | 35.21270337 | 38.51259106 | ENSG00000250994 |
| CRIP3 | 0.13483067 | 1.097963938 | 0.573637082 | 0.73924278 | 165.1731385 | 181.6217408 | ENSG00000146215 |
| CMTR2 | 0.134681981 | 1.097850783 | 0.473929328 | 0.659098534 | 176.5298827 | 193.5822685 | ENSG00000180917 |
| CTNNA1 | 0.134555231 | 1.097754335 | 0.017665572 | 0.05821723 | 8422.791896 | 9245.619173 | ENSG00000044115 |
| KLHL42 | 0.134329497 | 1.097582586 | 0.300216942 | 0.49358829 | 340.1986932 | 373.7749834 | ENSG00000087448 |
| PARVA | 0.134259376 | 1.09752924 | 0.307933748 | 0.502044694 | 379.8697581 | 416.2711038 | ENSG00000197702 |
| NUDT7 | 0.134172015 | 1.097462782 | 0.846479005 | 0.918323688 | 11.18233992 | 12.33296184 | ENSG00000140876 |
| CEP55 | 0.134128077 | 1.097429359 | 0.309959754 | 0.503769234 | 350.0501324 | 383.8761152 | ENSG00000138180 |
| C1QL3 | 0.134071594 | 1.097386394 | 0.70989755 | 0.835062275 | 47.46307108 | 51.92872804 | ENSG00000165985 |
| NA | 0.134047981 | 1.097368433 | 0.899878577 | 0.948765547 | 4.555436564 | 5.022592712 | ENSG00000261613 |
| FKBP2 | 0.134014561 | 1.097343013 | 0.895212803 | 0.946203969 | 4.433519683 | 4.894882979 | ENSG00000173486 |
| POGLUT1 | 0.133962209 | 1.097303194 | 0.342249702 | 0.536395153 | 310.9057679 | 340.7771159 | ENSG00000163389 |
| RPS6 | 0.133945888 | 1.09729078 | 0.022248134 | 0.070510758 | 21437.21914 | 23522.33033 | ENSG00000137154 |
| FGR | 0.13391633 | 1.097268299 | 0.890146578 | 0.943928176 | 5.53489804 | 5.985982225 | ENSG00000000938 |
| NA | 0.133910071 | 1.097263538 | 0.808074501 | 0.896924949 | 16.43474713 | 18.0002752 | ENSG00000234769 |
| MTRNR2L2 | 0.133584265 | 1.09701577 | 0.754828466 | 0.863526374 | 29.68737377 | 32.72615335 | ENSG00000269028 |
| GOSR1 | 0.133518358 | 1.096965656 | 0.222303448 | 0.404374532 | 769.939012 | 844.1733713 | ENSG00000108587 |
| WDR44 | 0.133517768 | 1.096965207 | 0.168311045 | 0.332318835 | 809.868072 | 887.5195798 | ENSG00000131725 |
| TEX30 | 0.133361092 | 1.096846083 | 0.575806461 | 0.740853659 | 80.78452215 | 88.70682175 | ENSG00000151287 |
| EIF2B3 | 0.133291594 | 1.096793247 | 0.214272205 | 0.394411696 | 753.4419537 | 826.6536736 | ENSG00000070785 |
| LOC107985075 | 0.133213895 | 1.096734179 | 0.866378772 | 0.930273692 | 7.575397065 | 8.35319112 | ENSG00000214999 |
| PSMD4 | 0.133023251 | 1.096589262 | 0.037613169 | 0.107035541 | 4163.39168 | 4565.457271 | ENSG00000159352 |
| TRAPPC3 | 0.132996951 | 1.096569271 | 0.156573563 | 0.315222074 | 919.4377002 | 1008.404581 | ENSG00000054116 |
| DNAJC18 | 0.132977835 | 1.096554741 | 0.321463566 | 0.515353839 | 346.8935528 | 380.1183387 | ENSG00000170464 |
| ZZZ3 | 0.132960613 | 1.096541651 | 0.209419953 | 0.388490944 | 707.215512 | 774.9727665 | ENSG00000036549 |
| CCDC151 | 0.1329413 | 1.096526972 | 0.654855057 | 0.798334778 | 52.49246514 | 57.61612649 | ENSG00000198003 |
| ARMC9 | 0.132874337 | 1.096476078 | 0.148156494 | 0.303283902 | 731.248213 | 801.8080354 | ENSG00000135931 |
| SLC35A1 | 0.132797565 | 1.096417732 | 0.739419813 | 0.854442722 | 32.44930198 | 35.51260771 | ENSG00000164414 |
| EDA | 0.132682489 | 1.09633028 | 0.376366427 | 0.569481211 | 214.8912432 | 235.673941 | ENSG00000158813 |
| TMCO6 | 0.132554891 | 1.09623332 | 0.445849679 | 0.633889072 | 300.7604129 | 330.1165903 | ENSG00000113119 |
| CRYL1 | 0.132506642 | 1.096196658 | 0.765984015 | 0.870752437 | 29.46040695 | 32.12113303 | ENSG00000165475 |
| TMEM185B | 0.132455743 | 1.096157985 | 0.29746745 | 0.490493821 | 409.4953917 | 449.401173 | ENSG00000226479 |
| MHENCR | 0.132432414 | 1.09614026 | 0.720129366 | 0.841791517 | 45.24828808 | 49.87577662 | ENSG00000232442 |
| P3H2 | 0.132223059 | 1.095981206 | 0.307784734 | 0.501890045 | 395.6390885 | 433.3186216 | ENSG00000090530 |

| | | | | | | | |
|---|---|---|---|---|---|---|---|
| MTIF2 | 0.132190977 | 1.095956834 | 0.142199756 | 0.294373985 | 795.6296775 | 871.7233403 | ENSG00000085760 |
| NA | 0.132096851 | 1.095885333 | 0.56675354 | 0.734354734 | 93.54781269 | 102.2833123 | ENSG00000213442 |
| CPAMD8 | 0.132093922 | 1.095883108 | 0.385105508 | 0.577938906 | 219.875065 | 241.1397747 | ENSG00000160111 |
| POLR1E | 0.132027224 | 1.095832445 | 0.161264366 | 0.321972827 | 672.7459303 | 737.3569866 | ENSG00000137054 |
| TCFL5 | 0.131839741 | 1.095690047 | 0.245960859 | 0.433536715 | 570.7625227 | 625.1237377 | ENSG00000101190 |
| SULT1A1 | 0.131807642 | 1.095665669 | 0.602731607 | 0.761880879 | 74.53223881 | 81.54986977 | ENSG00000196502 |
| C1orf159 | 0.131751901 | 1.095623337 | 0.389456575 | 0.582018075 | 246.4868565 | 270.3211394 | ENSG00000131591 |
| ITGB3BP | 0.131646823 | 1.095543541 | 0.264244506 | 0.453935274 | 418.9572004 | 458.6895925 | ENSG00000142856 |
| MAPK8IP3 | 0.131623144 | 1.09552556 | 0.246069239 | 0.433592002 | 1232.873544 | 1351.66171 | ENSG00000138834 |
| FAS | 0.131611973 | 1.095517077 | 0.771477666 | 0.874346286 | 22.45384752 | 24.59744971 | ENSG00000026103 |
| NA | 0.131457109 | 1.095399486 | 0.820621305 | 0.903761083 | 13.94130662 | 15.35226583 | ENSG00000232888 |
| CDK17 | 0.131413022 | 1.095366013 | 0.351902996 | 0.545012963 | 304.1900877 | 332.7112321 | ENSG00000059758 |
| UQCRB | 0.131372697 | 1.095335396 | 0.106117124 | 0.23708321 | 1624.799745 | 1779.368546 | ENSG00000156467 |
| CD200 | 0.131326532 | 1.095300347 | 0.127622288 | 0.271567334 | 1053.251009 | 1153.838013 | ENSG00000091972 |
| TAC3 | 0.131324888 | 1.095299099 | 0.542764794 | 0.715374961 | 110.367545 | 121.0766526 | ENSG00000166863 |
| KLC2 | 0.131316907 | 1.09529304 | 0.120944701 | 0.261166152 | 1058.454168 | 1159.613182 | ENSG00000174996 |
| SOCS6 | 0.131284733 | 1.095268613 | 0.259064382 | 0.447931297 | 391.4468471 | 429.012951 | ENSG00000170677 |
| STK11 | 0.131261468 | 1.095250951 | 0.079334559 | 0.1904646 | 2166.083273 | 2372.652552 | ENSG00000118046 |
| ZNF254 | 0.131250254 | 1.095242438 | 0.425757646 | 0.616580107 | 266.537709 | 291.6362361 | ENSG00000213096 |
| CCNT1 | 0.131217104 | 1.095217272 | 0.20448176 | 0.382541509 | 712.1233651 | 779.1007701 | ENSG00000129315 |
| NA | 0.131197673 | 1.095202521 | 0.811347772 | 0.898880668 | 14.90370576 | 16.27062728 | ENSG00000277011 |
| ANTXR2 | 0.131153163 | 1.095168733 | 0.471415338 | 0.656550632 | 155.666727 | 170.1767094 | ENSG00000163297 |
| ADAM22 | 0.131051788 | 1.09509178 | 0.275076318 | 0.466037706 | 409.821959 | 448.6815513 | ENSG00000008277 |
| ZCCHC18 | 0.130855754 | 1.094942989 | 0.628490775 | 0.780570128 | 99.27820612 | 109.0795304 | ENSG00000166707 |
| NA | 0.130836108 | 1.094928079 | 0.435640978 | 0.62514666 | 215.1858806 | 235.6049174 | ENSG00000196922 |
| EI24 | 0.13083371 | 1.094926258 | 0.03117037 | 0.092330451 | 3101.002987 | 3395.005983 | ENSG00000149547 |
| LIMD1 | 0.130679689 | 1.094809371 | 0.346752969 | 0.540170237 | 331.5225191 | 363.0009784 | ENSG00000144791 |
| CREB3 | 0.130610925 | 1.09475719 | 0.188538004 | 0.360466615 | 572.1906637 | 626.4344582 | ENSG00000107175 |
| XYLT2 | 0.130551768 | 1.094712301 | 0.097830704 | 0.222911248 | 1181.788286 | 1293.922339 | ENSG00000015532 |
| LOC107986257 | 0.130487374 | 1.09466344 | 0.898187371 | 0.947822814 | 4.505527319 | 4.954994809 | ENSG00000286176 |
| TEX9 | 0.130446364 | 1.094632323 | 0.707958895 | 0.833950537 | 42.75850814 | 47.00207729 | ENSG00000151575 |
| DSE | 0.130363942 | 1.094569788 | 0.136164517 | 0.284838957 | 1035.802368 | 1133.32462 | ENSG00000111817 |
| LRRC3-DT | 0.130220577 | 1.094461023 | 0.90739975 | 0.953927791 | 3.844560492 | 4.227266083 | ENSG00000229356 |
| NA | 0.130197677 | 1.094443651 | 0.784115966 | 0.881898496 | 20.92143067 | 23.00409691 | ENSG00000226396 |
| PARP4 | 0.130174173 | 1.09442582 | 0.119957117 | 0.259505452 | 1092.383949 | 1196.106445 | ENSG00000102699 |
| MNT | 0.130150392 | 1.09440778 | 0.384560733 | 0.577495433 | 252.7714831 | 277.053733 | ENSG00000070444 |
| NR1D1 | 0.130082327 | 1.094356148 | 0.525026099 | 0.70152359 | 114.8142711 | 125.4522229 | ENSG00000126368 |
| RASSF3 | 0.130044226 | 1.094327247 | 0.163052997 | 0.324716931 | 734.1781189 | 803.2488068 | ENSG00000153179 |
| TNK1 | 0.130032704 | 1.094318508 | 0.43091679 | 0.620909351 | 190.4988874 | 208.7585473 | ENSG00000174292 |
| DOCK2 | 0.129809657 | 1.094149334 | 0.52759546 | 0.703541422 | 113.8519697 | 124.4995687 | ENSG00000134516 |
| CHST8 | 0.129686613 | 1.094056021 | 0.303023826 | 0.496573261 | 474.8405257 | 518.798439 | ENSG00000124302 |
| MTHFD1L | 0.129625651 | 1.094009792 | 0.043532983 | 0.120148101 | 2769.734299 | 3029.611436 | ENSG00000120254 |
| DUS4L | 0.129583758 | 1.093978025 | 0.740407479 | 0.854997645 | 34.1081681 | 37.33958303 | ENSG00000105865 |
| UBE2H | 0.129483813 | 1.09390224 | 0.055464544 | 0.144765506 | 2190.373833 | 2396.079038 | ENSG00000186591 |
| SEMA4F | 0.129191538 | 1.093680649 | 0.41313607 | 0.604535668 | 230.7166165 | 252.0644655 | ENSG00000135622 |
| CHCHD5 | 0.129141257 | 1.093642533 | 0.368674616 | 0.5616478 | 270.6790491 | 296.1226492 | ENSG00000125611 |
| HMBS | 0.129136706 | 1.093639083 | 0.146294026 | 0.30040034 | 1018.801857 | 1113.904655 | ENSG00000256269 |
| SPG21 | 0.128968406 | 1.09351151 | 0.118094909 | 0.256566762 | 1200.205663 | 1311.87936 | ENSG00000090487 |
| ANG | 0.128888871 | 1.093451227 | 0.906596305 | 0.953407621 | 3.854031231 | 4.231064929 | ENSG00000214274 |
| STAG3L4 | 0.128874609 | 1.093440418 | 0.558067557 | 0.727371596 | 127.7713884 | 139.5680081 | ENSG00000106610 |
| GET1 | 0.128870516 | 1.093437315 | 0.628676704 | 0.78073535 | 85.72118565 | 93.56543656 | ENSG00000182093 |
| NA | 0.128846602 | 1.093419191 | 0.756000968 | 0.864333115 | 34.09544275 | 37.30225584 | ENSG00000214199 |
| BCL3 | 0.128824488 | 1.093402431 | 0.68101688 | 0.815688694 | 48.46019626 | 52.82550733 | ENSG00000069399 |
| SMIM19 | 0.128693921 | 1.09330348 | 0.432872943 | 0.622845053 | 209.4454151 | 228.6421737 | ENSG00000176209 |
| NA | 0.128679894 | 1.09329285 | 0.908145339 | 0.954386806 | 3.490062025 | 3.833401615 | ENSG00000267150 |
| SLC35E3 | 0.128675857 | 1.093289791 | 0.299424048 | 0.492940602 | 336.800159 | 368.3692615 | ENSG00000175782 |
| C1orf116 | 0.128648688 | 1.093269202 | 0.588090139 | 0.750562088 | 99.71458892 | 109.0667081 | ENSG00000182795 |
| MOCOS | 0.128531216 | 1.093180186 | 0.210741361 | 0.390119896 | 604.5132233 | 660.3702855 | ENSG00000075643 |
| ERLIN1 | 0.128479633 | 1.0931411 | 0.140635962 | 0.291919244 | 957.5760701 | 1046.415602 | ENSG00000107566 |
| HPSE | 0.128317748 | 1.093018446 | 0.562235277 | 0.730738379 | 93.45884041 | 102.1905566 | ENSG00000173083 |
| CEBPB | 0.128158185 | 1.092897564 | 0.310971525 | 0.50473301 | 402.8360096 | 440.4373074 | ENSG00000172216 |
| RNF126 | 0.128152209 | 1.092893037 | 0.100909637 | 0.228040811 | 1604.939998 | 1754.266718 | ENSG00000070423 |
| NA | 0.128120496 | 1.092869013 | 0.874922661 | 0.935237434 | 6.672377914 | 7.325452897 | ENSG00000260269 |
| PEX11A | 0.127938239 | 1.092730959 | 0.652820422 | 0.797217744 | 55.3297533 | 60.44772053 | ENSG00000166821 |
| SIKE1 | 0.127824529 | 1.092644836 | 0.191535815 | 0.364586081 | 713.3287935 | 779.2856774 | ENSG00000052723 |
| VPS26C | 0.127753352 | 1.09259093 | 0.094269411 | 0.216719086 | 1285.727783 | 1404.386431 | ENSG00000157538 |
| ZNF484 | 0.127633795 | 1.09250039 | 0.53898634 | 0.712472208 | 115.93529 | 126.5251008 | ENSG00000127081 |
| GRM1 | 0.127624656 | 1.09249347 | 0.834441783 | 0.911185059 | 12.78360749 | 14.02083089 | ENSG00000152822 |
| FKRP | 0.12756759 | 1.092450257 | 0.20960456 | 0.388753219 | 622.2552845 | 680.3939832 | ENSG00000181027 |
| PAXIP1-AS1 | 0.127552716 | 1.092438994 | 0.822094073 | 0.904469402 | 177.4340668 | 193.9450752 | ENSG00000273344 |
| GALC | 0.127511958 | 1.092408131 | 0.402815626 | 0.594646341 | 228.7902322 | 249.6809619 | ENSG00000054983 |
| BEST4 | 0.127486499 | 1.092388854 | 0.728202152 | 0.847450627 | 32.85380739 | 35.90278495 | ENSG00000142959 |
| FLVCR1-DT | 0.127484059 | 1.092387007 | 0.452451394 | 0.639762821 | 177.3428246 | 193.9476144 | ENSG00000198468 |
| MRPS25 | 0.127470053 | 1.092376402 | 0.233687501 | 0.418631756 | 771.0538752 | 842.9107713 | ENSG00000131368 |

| | | | | | | | |
|---|---|---|---|---|---|---|---|
| MCF2L2 | 0.127397841 | 1.092321725 | 0.62831679 | 0.780510951 | 68.16642696 | 74.48743403 | ENSG00000053524 |
| BBS12 | 0.127337018 | 1.092275675 | 0.681863661 | 0.816316087 | 51.57997525 | 56.17867558 | ENSG00000181004 |
| OSTM1 | 0.127320004 | 1.092262794 | 0.294104533 | 0.486777407 | 467.4030103 | 510.1754144 | ENSG00000081087 |
| MTF1 | 0.127305995 | 1.092252187 | 0.33535463 | 0.52957373 | 313.4310706 | 342.17681 | ENSG00000188786 |
| TADA2B | 0.127272925 | 1.092227151 | 0.250456035 | 0.438991135 | 543.7013525 | 594.1985408 | ENSG00000173011 |
| ACTR2 | 0.127236636 | 1.092199678 | 0.096224463 | 0.220203013 | 5327.012819 | 5817.246489 | ENSG00000138071 |
| SSUH2 | 0.127216217 | 1.09218422 | 0.885340483 | 0.941108896 | 5.769358763 | 6.32777059 | ENSG00000125046 |
| ZNF3 | 0.127091576 | 1.092089865 | 0.232946192 | 0.417789899 | 823.9591441 | 899.6492704 | ENSG00000166526 |
| TRIM16L | 0.127082896 | 1.092083294 | 0.553453534 | 0.723938212 | 112.5387326 | 122.6347249 | ENSG00000108448 |
| MSL2 | 0.127074742 | 1.092077122 | 0.19249091 | 0.365835371 | 742.2202651 | 811.1966027 | ENSG00000174579 |
| C14orf93 | 0.127073265 | 1.092076004 | 0.606080665 | 0.764778101 | 79.84559688 | 87.40164387 | ENSG00000100802 |
| BICD2 | 0.127068809 | 1.09207263 | 0.167119711 | 0.330773961 | 980.3360253 | 1071.334939 | ENSG00000185963 |
| RN7SL1 | 0.127046713 | 1.092055905 | 0.759166328 | 0.866029958 | 371.1382405 | 405.3344845 | ENSG00000276168 |
| CEP95 | 0.127022987 | 1.092037945 | 0.335126986 | 0.529337273 | 382.6110837 | 418.2738202 | ENSG00000258890 |
| VDAC1 | 0.126889564 | 1.091936956 | 0.045042217 | 0.12336522 | 5634.1331 | 6151.687719 | ENSG00000213585 |
| ABLIM3 | 0.126881514 | 1.091930863 | 0.797951668 | 0.891359341 | 18.94031163 | 20.6256031 | ENSG00000173210 |
| NPRL3 | 0.126811594 | 1.091877945 | 0.234642578 | 0.419534371 | 619.320275 | 676.3188831 | ENSG00000103148 |
| NA | 0.126755668 | 1.091835619 | 0.874923605 | 0.935237434 | 6.669221001 | 7.314167977 | ENSG00000214653 |
| NA | 0.126373167 | 1.09154618 | 0.895177204 | 0.946203969 | 5.756731112 | 6.305200749 | ENSG00000261251 |
| DUSP3 | 0.126080428 | 1.091324715 | 0.229832562 | 0.413725398 | 596.6394758 | 650.9583612 | ENSG00000108861 |
| SAV1 | 0.12606438 | 1.091312575 | 0.134683451 | 0.282506007 | 1454.57549 | 1587.425242 | ENSG00000151748 |
| GTPBP1 | 0.126047422 | 1.091299748 | 0.05942607 | 0.152718078 | 2392.873332 | 2611.299779 | ENSG00000100226 |
| ZNF670 | 0.126010121 | 1.091271533 | 0.514796647 | 0.692846128 | 206.1261945 | 224.687784 | ENSG00000277462 |
| SETDB1 | 0.126001501 | 1.091265013 | 0.226723526 | 0.410182097 | 617.0651437 | 673.455298 | ENSG00000143379 |
| NA | 0.125967659 | 1.091239414 | 0.849495549 | 0.920078138 | 11.17474832 | 12.1109591 | ENSG00000225506 |
| RPS27A | 0.125959332 | 1.091233116 | 0.02001668 | 0.064609743 | 9949.685382 | 10856.88772 | ENSG00000143947 |
| RPL9 | 0.125893787 | 1.09118354 | 0.049766303 | 0.133271009 | 9613.704269 | 10489.25648 | ENSG00000163682 |
| NRG3 | 0.12588771 | 1.091178943 | 0.692019304 | 0.823650512 | 48.09504698 | 52.6691674 | ENSG00000185737 |
| TAF8 | 0.12588652 | 1.091178043 | 0.26344801 | 0.453128031 | 446.7293596 | 487.2848598 | ENSG00000137413 |
| LOC400684 | 0.125880992 | 1.091173862 | 0.812528407 | 0.899366839 | 18.93597464 | 20.72336854 | ENSG00000267213 |
| SGO1 | 0.125749835 | 1.091074667 | 0.543967722 | 0.716179733 | 127.1216586 | 138.5717112 | ENSG00000129810 |
| NA | 0.125724418 | 1.091055445 | 0.787178417 | 0.883573449 | 26.20292651 | 28.66425061 | ENSG00000279821 |
| NA | 0.125660275 | 1.091006937 | 0.522426457 | 0.699310412 | 142.5165948 | 155.1770788 | ENSG00000280511 |
| RNF2 | 0.125595635 | 1.090958056 | 0.299872977 | 0.493415916 | 610.0005646 | 665.3256865 | ENSG00000121481 |
| TMEM268 | 0.125591352 | 1.090954817 | 0.309703706 | 0.503645563 | 361.1937926 | 393.6887774 | ENSG00000157693 |
| LSM12 | 0.125566567 | 1.090936074 | 0.209904529 | 0.389121223 | 934.2142273 | 1019.700566 | ENSG00000161654 |
| HABP4 | 0.125244369 | 1.090692463 | 0.290174486 | 0.482260567 | 385.6541121 | 420.6358759 | ENSG00000130956 |
| LAMTOR4 | 0.125212411 | 1.090668302 | 0.155817622 | 0.314142171 | 1347.519386 | 1470.033584 | ENSG00000188186 |
| KLF12 | 0.125173909 | 1.090639195 | 0.401230418 | 0.592837448 | 238.5259706 | 260.128812 | ENSG00000118922 |
| CSNK2A2 | 0.125137184 | 1.090611433 | 0.185118117 | 0.355873787 | 887.6668786 | 967.3734247 | ENSG00000070770 |
| KBTBD7 | 0.125074344 | 1.090563929 | 0.299484802 | 0.492966815 | 378.2530217 | 412.8023239 | ENSG00000120696 |
| VPS54 | 0.125060506 | 1.090553469 | 0.390591248 | 0.583196503 | 341.7846799 | 372.3484423 | ENSG00000143952 |
| NA | 0.12490359 | 1.09043486 | 0.751390037 | 0.861720373 | 29.57926462 | 32.36614364 | ENSG00000279198 |
| HDLBP | 0.12474823 | 1.090317441 | 0.026420542 | 0.080978721 | 9979.418261 | 10881.28346 | ENSG00000115677 |
| EXOC5 | 0.124628972 | 1.090227315 | 0.220042095 | 0.401561657 | 1322.681417 | 1441.143889 | ENSG00000070367 |
| SWAP70 | 0.124453742 | 1.090094904 | 0.136812192 | 0.285806713 | 1300.297984 | 1416.577833 | ENSG00000133789 |
| ZNF738 | 0.124345163 | 1.090012865 | 0.249844054 | 0.438338668 | 886.0709176 | 966.5781729 | ENSG00000172687 |
| LFNG | 0.124293908 | 1.089974141 | 0.265935237 | 0.455910063 | 514.4043834 | 560.2880627 | ENSG00000106003 |
| ZNF239 | 0.124238051 | 1.089931941 | 0.513321486 | 0.691998545 | 159.8073005 | 174.6054954 | ENSG00000196793 |
| SLC35B3 | 0.124238017 | 1.089931916 | 0.494694945 | 0.676705004 | 175.0162719 | 190.5160727 | ENSG00000124786 |
| ZNF346 | 0.124181519 | 1.089889233 | 0.266659399 | 0.456644673 | 506.6851438 | 552.6034048 | ENSG00000113761 |
| POU2F1 | 0.124179206 | 1.089887486 | 0.234988291 | 0.419939869 | 979.0317101 | 1066.673541 | ENSG00000143190 |
| MAN2C1 | 0.12415636 | 1.089870227 | 0.316735104 | 0.510378058 | 732.9805227 | 799.4802096 | ENSG00000140400 |
| PLEKHF1 | 0.124131107 | 1.08985115 | 0.322620599 | 0.516672443 | 409.9911167 | 447.1742326 | ENSG00000166289 |
| UCP2 | 0.124117966 | 1.089841223 | 0.189553054 | 0.361735616 | 711.1858001 | 774.9468762 | ENSG00000175567 |
| MPRIP | 0.123868261 | 1.089652607 | 0.055930853 | 0.145715831 | 3491.126217 | 3803.652109 | ENSG00000133030 |
| SRRD | 0.123796258 | 1.089598225 | 0.42454064 | 0.61539978 | 220.0647276 | 240.0748209 | ENSG00000100104 |
| APBB2 | 0.123777372 | 1.089583961 | 0.190821546 | 0.363557415 | 841.9736088 | 916.7935885 | ENSG00000163697 |
| SPPL2B | 0.123626123 | 1.089469737 | 0.361764625 | 0.554809321 | 722.9984775 | 788.5816812 | ENSG00000005206 |
| ADCY4 | 0.12360495 | 1.089453748 | 0.883006774 | 0.939798298 | 6.778510231 | 7.404335721 | ENSG00000129467 |
| ESRP2 | 0.123603923 | 1.089452973 | 0.148700513 | 0.304128123 | 1158.748449 | 1263.24052 | ENSG00000103067 |
| GNPTG | 0.123559953 | 1.089419769 | 0.33291606 | 0.527418263 | 382.1916656 | 416.8667517 | ENSG00000090581 |
| RDH12 | 0.123337197 | 1.089251573 | 0.865813333 | 0.930066343 | 11.50281141 | 12.56113655 | ENSG00000139988 |
| COPG1 | 0.123306643 | 1.089228505 | 0.081613577 | 0.194625068 | 2319.188483 | 2526.086967 | ENSG00000181789 |
| C1GALT1 | 0.123246087 | 1.089182787 | 0.243901741 | 0.430644458 | 976.1193551 | 1062.556839 | ENSG00000106392 |
| CYP20A1 | 0.123156273 | 1.089114982 | 0.419842247 | 0.611197394 | 229.7046989 | 250.5767938 | ENSG00000119004 |
| AFG3L2 | 0.123087135 | 1.08906279 | 0.078236269 | 0.188487067 | 2053.403614 | 2235.514745 | ENSG00000141385 |
| RBIS | 0.123068914 | 1.089049035 | 0.662394374 | 0.803667015 | 58.93226147 | 64.22904604 | ENSG00000176731 |
| TBC1D8B | 0.123014824 | 1.089008205 | 0.450273527 | 0.6379748 | 203.5478256 | 221.4336953 | ENSG00000133138 |
| CYBA | 0.122945703 | 1.088956031 | 0.08387045 | 0.198614506 | 2916.952457 | 3176.17989 | ENSG00000051523 |
| NA | 0.122886667 | 1.088911471 | 0.902327072 | 0.950373924 | 4.492899667 | 4.906167899 | ENSG00000237493 |
| GSKIP | 0.12287048 | 1.088899253 | 0.234620189 | 0.419534371 | 681.0449812 | 741.5762254 | ENSG00000100744 |
| NCOA5 | 0.122829866 | 1.0888686 | 0.154916489 | 0.312861652 | 898.7684014 | 979.1395469 | ENSG00000124160 |
| EHD3 | 0.12276617 | 1.088820526 | 0.225406081 | 0.408314889 | 605.3083136 | 659.4574834 | ENSG00000013016 |

| | | | | | | | |
|---|---|---|---|---|---|---|---|
| CEBPG | 0.122624182 | 1.088713372 | 0.125607727 | 0.268544655 | 1159.190326 | 1262.500965 | ENSG00000153879 |
| MAP3K14 | 0.12258899 | 1.088686815 | 0.53825875 | 0.711948165 | 148.1471697 | 161.6162015 | ENSG00000006062 |
| KDM5A | 0.122581487 | 1.088681153 | 0.263261881 | 0.452876898 | 853.6053425 | 928.7896433 | ENSG00000073614 |
| NA | 0.1225612 | 1.088665844 | 0.913473713 | 0.95711017 | 4.108711207 | 4.485934743 | ENSG00000287733 |
| PAPSS2 | 0.122243997 | 1.088426507 | 0.277996628 | 0.469124055 | 487.8988445 | 531.1237403 | ENSG00000198682 |
| FBXO4 | 0.122129842 | 1.088340387 | 0.838760737 | 0.913533026 | 13.15389053 | 14.3508963 | ENSG00000151876 |
| AKTIP | 0.122042391 | 1.088274418 | 0.415352948 | 0.606714483 | 239.5458706 | 260.9257876 | ENSG00000166971 |
| RCL1 | 0.122021708 | 1.088258817 | 0.276828787 | 0.467834845 | 473.384536 | 514.8012198 | ENSG00000120158 |
| NA | 0.121942326 | 1.088198938 | 0.912029166 | 0.956299421 | 4.241376604 | 4.624929397 | ENSG00000259790 |
| KIF7 | 0.121936862 | 1.088194817 | 0.191302219 | 0.364285968 | 691.3783981 | 752.7150267 | ENSG00000166813 |
| GATAD1 | 0.121807378 | 1.088097155 | 0.216657513 | 0.397262598 | 652.5081436 | 709.6245323 | ENSG00000157259 |
| TOMM70 | 0.121692733 | 1.088010691 | 0.15771381 | 0.316966791 | 1557.110109 | 1693.570323 | ENSG00000154174 |
| DLG4 | 0.121654721 | 1.087982024 | 0.177619979 | 0.346199252 | 947.3248685 | 1031.199712 | ENSG00000132535 |
| ADAMTS10 | 0.121620898 | 1.087956518 | 0.539055357 | 0.712512622 | 171.281602 | 186.8002354 | ENSG00000142303 |
| NA | 0.121579772 | 1.087925504 | 0.379554416 | 0.572761891 | 260.1210934 | 283.1630764 | ENSG00000184319 |
| NA | 0.12152882 | 1.087887083 | 0.31652798 | 0.510177371 | 639.7270166 | 696.7781612 | ENSG00000250251 |
| CDC5L | 0.121420651 | 1.08780552 | 0.136161052 | 0.284838957 | 1304.256509 | 1418.074631 | ENSG00000096401 |
| KLF3 | 0.121369938 | 1.087767282 | 0.397195783 | 0.589354467 | 294.9746531 | 321.1652666 | ENSG00000109787 |
| EIF3A | 0.121238766 | 1.087668385 | 0.059600806 | 0.153061014 | 11113.18742 | 12086.77874 | ENSG00000107581 |
| EARS2 | 0.12121101 | 1.08764746 | 0.115778762 | 0.252958101 | 1523.601394 | 1656.463733 | ENSG00000103356 |
| ZBTB48 | 0.121156704 | 1.087606519 | 0.351519512 | 0.544557714 | 351.6455671 | 382.5183602 | ENSG00000204859 |
| PREPL | 0.121071291 | 1.087542131 | 0.243577182 | 0.430235347 | 595.0149071 | 646.4352823 | ENSG00000138078 |
| NA | 0.1210225 | 1.087505351 | 0.880656682 | 0.9386446 | 7.328308691 | 7.98558372 | ENSG00000257907 |
| WIPF2 | 0.120857183 | 1.087380742 | 0.182530801 | 0.352876382 | 806.5149972 | 877.0255628 | ENSG00000171475 |
| LOC101928445 | 0.120791021 | 1.087330876 | 0.87379127 | 0.934875449 | 7.562769414 | 8.24425238 | ENSG00000244945 |
| XDH | 0.120746157 | 1.087297063 | 0.719399841 | 0.841168169 | 41.30065448 | 45.00627464 | ENSG00000158125 |
| EIF1B | 0.120743384 | 1.087294974 | 0.174363282 | 0.341251223 | 1070.911584 | 1163.817057 | ENSG00000114784 |
| NA | 0.120676036 | 1.087244217 | 0.397283333 | 0.589437183 | 389.6619901 | 423.8046152 | ENSG00000224114 |
| MANBAL | 0.120619451 | 1.087201574 | 0.178772563 | 0.347714418 | 1098.006684 | 1193.273273 | ENSG00000101363 |
| GPS2 | 0.12061376 | 1.087197286 | 0.782085762 | 0.8805327 | 23.83200424 | 25.97077515 | ENSG00000132522 |
| KRBA1 | 0.120541494 | 1.087142828 | 0.351067515 | 0.544355837 | 307.5779484 | 334.458154 | ENSG00000133619 |
| PRDM4 | 0.120466759 | 1.087086514 | 0.313019274 | 0.506593577 | 419.6129358 | 456.2208596 | ENSG00000110851 |
| ASB13 | 0.120385474 | 1.087025266 | 0.244152388 | 0.43088177 | 613.0200366 | 666.1264525 | ENSG00000196372 |
| AKAP10 | 0.120237732 | 1.086913953 | 0.309041979 | 0.503187564 | 380.1378625 | 412.9200225 | ENSG00000108599 |
| VPS39 | 0.120213746 | 1.086895882 | 0.12864323 | 0.273112671 | 1275.243542 | 1385.666136 | ENSG00000166887 |
| CACNG7 | 0.120195439 | 1.08688209 | 0.15935463 | 0.31926048 | 1308.691404 | 1423.247569 | ENSG00000105605 |
| ING3 | 0.120130651 | 1.086833282 | 0.408797099 | 0.600404094 | 248.7856735 | 270.0748147 | ENSG00000071243 |
| SUPT20H | 0.120095682 | 1.086806939 | 0.152838061 | 0.310093116 | 1071.299576 | 1163.670361 | ENSG00000102710 |
| HUS1 | 0.120017074 | 1.086747724 | 0.425143731 | 0.615858592 | 268.5177608 | 291.6800936 | ENSG00000136273 |
| NA | 0.119992124 | 1.08672893 | 0.808529588 | 0.897215147 | 18.32807397 | 19.96178505 | ENSG00000198134 |
| NPIPB5 | 0.119891871 | 1.086653415 | 0.726601951 | 0.846456835 | 43.68174655 | 47.67706864 | ENSG00000243716 |
| IPO7 | 0.119864824 | 1.086633044 | 0.097080295 | 0.221723497 | 5334.613886 | 5795.472327 | ENSG00000205339 |
| COX5B | 0.119853525 | 1.086624533 | 0.105611774 | 0.236249791 | 1904.7372 | 2069.354784 | ENSG00000135940 |
| APELA | 0.119813431 | 1.086594335 | 0.526902907 | 0.702967066 | 219.7660688 | 238.5735307 | ENSG00000248329 |
| SMCHD1 | 0.119802217 | 1.086585889 | 0.273631321 | 0.464871193 | 960.2125942 | 1042.249347 | ENSG00000101596 |
| CDC42 | 0.119768081 | 1.08656018 | 0.16883974 | 0.333157972 | 3472.657104 | 3771.863714 | ENSG00000070831 |
| IRX5 | 0.119654634 | 1.086474741 | 0.889857902 | 0.943784055 | 6.358317954 | 6.923990736 | ENSG00000176842 |
| ITGA2 | 0.119388729 | 1.08627451 | 0.531627638 | 0.70682831 | 137.0033892 | 148.729411 | ENSG00000164171 |
| ATG7 | 0.119285632 | 1.086196886 | 0.384476659 | 0.577415963 | 276.200643 | 299.9100351 | ENSG00000197548 |
| USP37 | 0.119238984 | 1.086161765 | 0.394049952 | 0.586565079 | 292.680767 | 317.5412687 | ENSG00000135913 |
| NA | 0.119143937 | 1.08609021 | 0.745381783 | 0.857749036 | 33.67722731 | 36.64928467 | ENSG00000246067 |
| RNF103 | 0.119078624 | 1.086041042 | 0.649858472 | 0.795523587 | 71.94529364 | 78.13681293 | ENSG00000239305 |
| TMEM30B | 0.119042403 | 1.086013776 | 0.341945653 | 0.536223092 | 496.1098926 | 538.6896002 | ENSG00000182107 |
| SYAP1 | 0.118808686 | 1.085837855 | 0.100206002 | 0.226969242 | 1723.088403 | 1870.892445 | ENSG00000169895 |
| SPC24 | 0.118684169 | 1.085744142 | 0.263120695 | 0.452718062 | 590.2281441 | 640.6780647 | ENSG00000161888 |
| ENDOD1 | 0.11868177 | 1.085742337 | 0.561149268 | 0.730002144 | 112.5355757 | 122.125973 | ENSG00000149218 |
| STMN1 | 0.118607028 | 1.085686089 | 0.066868876 | 0.167207562 | 5687.492671 | 6173.55922 | ENSG00000117632 |
| SIRT3 | 0.118591759 | 1.085674598 | 0.344196665 | 0.537826027 | 364.3705688 | 395.6942647 | ENSG00000142082 |
| LOC400622 | 0.118558432 | 1.085649519 | 0.915338814 | 0.958169563 | 3.457215119 | 3.762004864 | ENSG00000267665 |
| NAPG | 0.118496479 | 1.085602899 | 0.377307501 | 0.570485987 | 395.2370636 | 428.568682 | ENSG00000134265 |
| VGLL4 | 0.118480402 | 1.085590801 | 0.256511821 | 0.44519057 | 572.7929422 | 621.601336 | ENSG00000144560 |
| MAFG | 0.118469459 | 1.085582568 | 0.192833838 | 0.36635403 | 986.8606374 | 1071.460507 | ENSG00000197063 |
| PPA2 | 0.11845242 | 1.085569746 | 0.280753685 | 0.472100236 | 512.4503988 | 555.8355362 | ENSG00000138777 |
| GNB1 | 0.118444693 | 1.085563932 | 0.04451404 | 0.122280008 | 10107.70329 | 10971.40565 | ENSG00000078369 |
| TMEM106C | 0.118314697 | 1.085466121 | 0.328004432 | 0.522275161 | 400.2977111 | 434.2511109 | ENSG00000134291 |
| NA | 0.118303402 | 1.085457622 | 0.853535293 | 0.922307785 | 12.45248519 | 13.41982421 | ENSG00000184844 |
| MTPAP | 0.118284796 | 1.085443623 | 0.250051353 | 0.438536316 | 562.9609329 | 610.8222096 | ENSG00000107951 |
| NUDT15 | 0.118268979 | 1.085431723 | 0.219951869 | 0.401476063 | 765.2646427 | 830.4264532 | ENSG00000136159 |
| KLF9 | 0.118257084 | 1.085422774 | 0.788660099 | 0.884433126 | 23.67398584 | 25.6026181 | ENSG00000119138 |
| BTBD2 | 0.118256628 | 1.085422431 | 0.162780815 | 0.324301798 | 2711.691459 | 2942.617935 | ENSG00000133243 |
| PRRG3 | 0.118165163 | 1.085353619 | 0.574355155 | 0.739808078 | 119.07271 | 129.2945363 | ENSG00000130032 |
| PRKAR1A | 0.118150121 | 1.085342303 | 0.146480066 | 0.300677168 | 1389.139249 | 1507.604136 | ENSG00000108946 |
| SCHIP1 | 0.118089951 | 1.085297037 | 0.875946059 | 0.935667514 | 8.939047001 | 9.710994753 | ENSG00000151967 |
| ALCAM | 0.118084584 | 1.085293 | 0.362160278 | 0.555144293 | 500.5646841 | 543.0471724 | ENSG00000170017 |

| | | | | | | | |
|---|---|---|---|---|---|---|---|
| METTL2B | 0.118057037 | 1.085272278 | 0.279237311 | 0.470192936 | 497.9534457 | 540.3497236 | ENSG00000165055 |
| BABAM2 | 0.118017903 | 1.085242839 | 0.503713194 | 0.683776217 | 219.6529463 | 238.7010575 | ENSG00000158019 |
| RXRB | 0.117996655 | 1.085226856 | 0.275844755 | 0.466822508 | 554.6252542 | 602.5089025 | ENSG00000204231 |
| CAPN12 | 0.117964505 | 1.085202672 | 0.755830068 | 0.864333115 | 62.793982 | 68.35662004 | ENSG00000182472 |
| ELFN2 | 0.117881094 | 1.085139932 | 0.642132058 | 0.790099951 | 75.47441869 | 81.88417206 | ENSG00000166897 |
| IRF5 | 0.117856915 | 1.085121745 | 0.861456455 | 0.927245133 | 9.540633843 | 10.37101396 | ENSG00000128604 |
| NA | 0.117699966 | 1.085003703 | 0.893099164 | 0.94538237 | 5.789578017 | 6.290228601 | ENSG00000261461 |
| SGPL1 | 0.117676677 | 1.084986188 | 0.135993036 | 0.284576598 | 1437.29857 | 1559.241686 | ENSG00000166224 |
| NA | 0.117675961 | 1.084985649 | 0.892823955 | 0.945253009 | 5.686602612 | 6.181289862 | ENSG00000189014 |
| PER2 | 0.117656922 | 1.084971331 | 0.570988032 | 0.737373586 | 130.5219516 | 141.5693214 | ENSG00000132326 |
| TTC21B | 0.117591026 | 1.084921776 | 0.349283255 | 0.542633349 | 342.6526799 | 371.8624121 | ENSG00000123607 |
| NA | 0.117580954 | 1.084914201 | 0.912238427 | 0.956464729 | 3.764961254 | 4.092070275 | ENSG00000260367 |
| LOC101929709 | 0.117472375 | 1.084832552 | 0.66772451 | 0.80717496 | 78.79769064 | 85.66224838 | ENSG00000251136 |
| EMILIN2 | 0.117367026 | 1.084753338 | 0.509624492 | 0.688840456 | 199.8433268 | 216.8726137 | ENSG00000132205 |
| TCF7L2 | 0.117365542 | 1.084752222 | 0.30576771 | 0.499744157 | 419.9006581 | 455.7864272 | ENSG00000148737 |
| CDCA8 | 0.117339412 | 1.084732575 | 0.139655358 | 0.290567176 | 1220.559339 | 1323.837922 | ENSG00000134690 |
| SON | 0.11726894 | 1.084679591 | 0.106345575 | 0.237454745 | 4661.217715 | 5055.488712 | ENSG00000159140 |
| PRR11 | 0.117198339 | 1.084626511 | 0.265115281 | 0.454798835 | 599.8922133 | 650.2914685 | ENSG00000068489 |
| LINC01456 | 0.117185938 | 1.084617188 | 0.867652515 | 0.931048617 | 10.64644689 | 11.44334221 | ENSG00000225882 |
| SCAI | 0.117130258 | 1.084575328 | 0.452988066 | 0.640291705 | 249.5035398 | 270.302981 | ENSG00000173611 |
| SMG8 | 0.117113088 | 1.084562421 | 0.261960447 | 0.451186754 | 607.8719728 | 659.3918609 | ENSG00000167447 |
| NIPSNAP3A | 0.117070637 | 1.084530508 | 0.533375302 | 0.70829298 | 172.8755862 | 187.2696933 | ENSG00000136783 |
| SNAP91 | 0.116952253 | 1.084441518 | 0.822033203 | 0.904456123 | 20.05135612 | 21.8823921 | ENSG00000065609 |
| RPL13 | 0.116898952 | 1.084401453 | 0.043724267 | 0.120539277 | 18075.41023 | 19600.0725 | ENSG00000167526 |
| MED18 | 0.116863215 | 1.084374592 | 0.371923007 | 0.565064432 | 309.5861042 | 335.5032375 | ENSG00000130772 |
| DELE1 | 0.11679513 | 1.084323418 | 0.256280566 | 0.445063056 | 874.1717711 | 948.4147959 | ENSG00000081791 |
| DCTN4 | 0.116663345 | 1.084224374 | 0.184916706 | 0.355634142 | 1006.584945 | 1091.260696 | ENSG00000132912 |
| PRMT2 | 0.116655156 | 1.08421822 | 0.125048712 | 0.267794977 | 1264.560933 | 1370.933425 | ENSG00000160310 |
| PRR12 | 0.116595374 | 1.084173293 | 0.170980631 | 0.336550023 | 2126.826905 | 2305.611221 | ENSG00000126464 |
| CMIP | 0.11631375 | 1.083961675 | 0.174134666 | 0.340947804 | 1543.376332 | 1672.981807 | ENSG00000153815 |
| SIRT4 | 0.116305068 | 1.083955152 | 0.788917955 | 0.88450822 | 25.89272252 | 28.10078592 | ENSG00000089163 |
| TRAFD1 | 0.116234951 | 1.083902472 | 0.228987676 | 0.412805912 | 713.9806205 | 774.1729487 | ENSG00000135148 |
| CELF1 | 0.116189012 | 1.083867958 | 0.117770405 | 0.256011809 | 2533.52199 | 2746.18891 | ENSG00000149187 |
| ANKRD10 | 0.116171864 | 1.083855076 | 0.246524222 | 0.434116858 | 1286.909054 | 1395.657998 | ENSG00000088448 |
| DEAF1 | 0.116091357 | 1.083794595 | 0.149153043 | 0.30478528 | 1222.774513 | 1324.912752 | ENSG00000177030 |
| DAZAP2 | 0.115963703 | 1.083698701 | 0.410656686 | 0.60224015 | 325.1061141 | 351.8488371 | ENSG00000183283 |
| TREML2 | 0.115947223 | 1.083686322 | 0.877135401 | 0.93639635 | 9.051590842 | 9.819383839 | ENSG00000112195 |
| MTFR1 | 0.115917638 | 1.0836641 | 0.257373018 | 0.445975272 | 674.1577232 | 729.9006377 | ENSG00000066855 |
| ST13 | 0.115798633 | 1.083574714 | 0.058166884 | 0.150336777 | 7029.853026 | 7616.346744 | ENSG00000100380 |
| AKR1C3 | 0.115739018 | 1.08352994 | 0.73324025 | 0.851101774 | 42.26493275 | 45.71044584 | ENSG00000196139 |
| RNF122 | 0.115738199 | 1.083529325 | 0.423303256 | 0.614180691 | 296.564879 | 321.1068666 | ENSG00000133874 |
| NA | 0.115660622 | 1.083471062 | 0.335100673 | 0.529337273 | 424.234863 | 459.2824603 | ENSG00000249353 |
| CD22 | 0.115635642 | 1.083452302 | 0.920187845 | 0.960669268 | 3.886878136 | 4.212293935 | ENSG00000012124 |
| RPS19 | 0.115607888 | 1.08343146 | 0.058891228 | 0.151718707 | 16718.34277 | 18113.8369 | ENSG00000105372 |
| HPAT5 | 0.115584895 | 1.083414192 | 0.234614829 | 0.419534371 | 928.1165954 | 1006.236639 | ENSG00000280707 |
| CDK19 | 0.115412036 | 1.083284389 | 0.501181788 | 0.681804409 | 167.6934052 | 181.5571286 | ENSG00000155111 |
| IQCE | 0.115237455 | 1.083153309 | 0.484988032 | 0.668210541 | 187.8306744 | 203.6015232 | ENSG00000106012 |
| POM121C | 0.1151896 | 1.08311738 | 0.183108263 | 0.353550219 | 1306.752633 | 1415.747744 | ENSG00000272391 |
| C15orf56 | 0.115138027 | 1.083078662 | 0.918398213 | 0.959665811 | 3.463528945 | 3.75451879 | ENSG00000176753 |
| NA | 0.115073768 | 1.083030422 | 0.914979463 | 0.958037973 | 3.774431993 | 4.092070275 | ENSG00000279970 |
| C17orf97 | 0.115060032 | 1.08302011 | 0.898217326 | 0.947822814 | 5.736511858 | 6.215144624 | ENSG00000187624 |
| FBXO45 | 0.115049445 | 1.083012162 | 0.355015343 | 0.548282644 | 537.39916 | 581.2652646 | ENSG00000174013 |
| HIP1R | 0.114986801 | 1.082965137 | 0.249299596 | 0.437756392 | 605.9959263 | 656.6204903 | ENSG00000130787 |
| SELENON | 0.114935478 | 1.082926612 | 0.18470672 | 0.355414695 | 1913.31176 | 2071.645881 | ENSG00000162430 |
| RAP2B | 0.114933503 | 1.08292513 | 0.166592231 | 0.330050285 | 1212.710185 | 1312.840935 | ENSG00000181467 |
| TET2 | 0.114696697 | 1.082747392 | 0.768100307 | 0.871908897 | 33.19124352 | 35.84212347 | ENSG00000168769 |
| COPB2 | 0.114636312 | 1.082702074 | 0.06815271 | 0.169458322 | 2845.3929 | 3080.134369 | ENSG00000184432 |
| ABCA2 | 0.114598814 | 1.082673933 | 0.255848435 | 0.444788941 | 734.3638957 | 795.4506469 | ENSG00000107331 |
| CLDN23 | 0.114546871 | 1.082634953 | 0.564239931 | 0.732480414 | 129.2524077 | 140.0061362 | ENSG00000253958 |
| ETFBKMT | 0.114473271 | 1.082579722 | 0.851766169 | 0.921524381 | 15.00974038 | 16.14715443 | ENSG00000139160 |
| XYLT1 | 0.114348869 | 1.082486377 | 0.202871986 | 0.380318511 | 953.7140041 | 1032.224877 | ENSG00000103489 |
| MAP11 | 0.114293228 | 1.082444629 | 0.451245939 | 0.638719431 | 248.2956458 | 269.0903927 | ENSG00000146826 |
| C12orf49 | 0.11426837 | 1.082425979 | 0.134220775 | 0.281694921 | 1514.964861 | 1639.052964 | ENSG00000111412 |
| CIRBP | 0.114221432 | 1.082390762 | 0.218911708 | 0.400247568 | 2642.308119 | 2860.792766 | ENSG00000099622 |
| RPS25 | 0.114198933 | 1.082373883 | 0.060904055 | 0.155760418 | 9763.940457 | 10567.99169 | ENSG00000118181 |
| FAM110C | 0.114110828 | 1.082307784 | 0.523419878 | 0.699924224 | 154.9541672 | 167.7810935 | ENSG00000184731 |
| ATP6V1C2 | 0.11384152 | 1.082105768 | 0.819651327 | 0.902946927 | 21.04019064 | 22.66544612 | ENSG00000143882 |
| MOB3B | 0.113812061 | 1.082083673 | 0.330027 | 0.524365231 | 483.9371851 | 522.9542052 | ENSG00000120162 |
| AOX1 | 0.113790977 | 1.082067859 | 0.468038205 | 0.653997293 | 233.7993843 | 252.7609274 | ENSG00000138356 |
| RHOQ | 0.113606775 | 1.081929711 | 0.294721673 | 0.48753762 | 549.0265567 | 593.7965375 | ENSG00000119729 |
| ZNF528-AS1 | 0.113602968 | 1.081926855 | 0.89415857 | 0.94574241 | 6.31600031 | 6.837621838 | ENSG00000269834 |
| DCBLD1 | 0.113561083 | 1.081895445 | 0.422420484 | 0.613569064 | 253.9390363 | 274.4667828 | ENSG00000164465 |
| WDR26 | 0.113525701 | 1.081868912 | 0.141556146 | 0.293303395 | 1486.336375 | 1607.308582 | ENSG00000162923 |
| HIF1AN | 0.113524857 | 1.081868279 | 0.14263907 | 0.295019812 | 1602.759874 | 1734.667428 | ENSG00000166135 |

| | | | | | | | |
|---|---|---|---|---|---|---|---|
| NA | 0.113523615 | 1.081867348 | 0.757753078 | 0.865109988 | 33.58500042 | 36.35252437 | ENSG00000214485 |
| RAC1 | 0.113431158 | 1.081798017 | 0.171283529 | 0.336824289 | 5348.470506 | 5784.828914 | ENSG00000136238 |
| CA2 | 0.113374405 | 1.081755462 | 0.576945789 | 0.741753139 | 116.8553715 | 126.3189463 | ENSG00000104267 |
| NAIP | 0.113302121 | 1.081701263 | 0.905973185 | 0.952968616 | 4.721046564 | 5.108523577 | ENSG00000249437 |
| MOSPD3 | 0.113281134 | 1.081685528 | 0.454289796 | 0.641557992 | 227.6452584 | 246.2994983 | ENSG00000106330 |
| CCDC149 | 0.113219086 | 1.081639008 | 0.544588632 | 0.716707097 | 172.1235574 | 186.3274394 | ENSG00000181982 |
| MMACHC | 0.113138614 | 1.081578677 | 0.562450512 | 0.730874136 | 127.1766039 | 137.7218237 | ENSG00000132763 |
| MFSD12 | 0.113005377 | 1.081478794 | 0.251550492 | 0.440124311 | 922.3096316 | 997.6190359 | ENSG00000161091 |
| COA8 | 0.112944655 | 1.081433276 | 0.594538273 | 0.755565849 | 111.6642234 | 120.6587926 | ENSG00000256053 |
| TLN1 | 0.112936493 | 1.081427158 | 0.09839798 | 0.223877752 | 4061.830051 | 4392.399485 | ENSG00000137076 |
| CLCN4 | 0.112934416 | 1.081425601 | 0.490947235 | 0.673766461 | 267.9248253 | 289.6220207 | ENSG00000073464 |
| SMCR8 | 0.11292968 | 1.081422051 | 0.184839276 | 0.355559018 | 977.611063 | 1057.813121 | ENSG00000176994 |
| ZNF688 | 0.112876812 | 1.081382423 | 0.703403172 | 0.831365258 | 52.58478973 | 56.88047365 | ENSG00000229809 |
| NA | 0.112816295 | 1.081337063 | 0.898692338 | 0.948216088 | 5.76620185 | 6.233915618 | ENSG00000272462 |
| FAM214B | 0.112709423 | 1.081256962 | 0.573465596 | 0.73914336 | 131.7572958 | 142.5869856 | ENSG00000005238 |
| SLC25A33 | 0.112520275 | 1.081115211 | 0.241831606 | 0.428063718 | 689.2256483 | 744.9995795 | ENSG00000171612 |
| ACSF3 | 0.112395228 | 1.081021509 | 0.382367844 | 0.575026598 | 344.7986121 | 372.9548255 | ENSG00000176715 |
| IKBKG | 0.112306632 | 1.080955125 | 0.62152526 | 0.775152125 | 104.117915 | 112.7764137 | ENSG00000269335 |
| NA | 0.112304708 | 1.080953683 | 0.915758337 | 0.95827165 | 3.830655064 | 4.140897185 | ENSG00000251661 |
| C1orf61 | 0.11228474 | 1.080938722 | 0.732560917 | 0.850813638 | 64.41990352 | 69.95811174 | ENSG00000125462 |
| GOLGA2P7 | 0.112118601 | 1.08081425 | 0.511927543 | 0.690995927 | 195.0966189 | 210.9673293 | ENSG00000225151 |
| APMAP | 0.112059639 | 1.080770078 | 0.113288431 | 0.248925641 | 1928.885685 | 2084.846864 | ENSG00000101474 |
| NA | 0.111978421 | 1.080709237 | 0.803959902 | 0.894704967 | 22.44625592 | 24.25989823 | ENSG00000253771 |
| CALCRL | 0.111894609 | 1.080646456 | 0.864098252 | 0.92922524 | 11.14505833 | 12.04011201 | ENSG00000064989 |
| FRMD4A | 0.111570065 | 1.080403385 | 0.326710632 | 0.520989637 | 445.0269957 | 480.6310459 | ENSG00000151474 |
| FAM156A | 0.111535766 | 1.0803777 | 0.879811461 | 0.938030898 | 8.359558549 | 9.02829409 | ENSG00000268350 |
| VDAC2 | 0.111529224 | 1.0803728 | 0.093322941 | 0.215004287 | 3915.570902 | 4229.12931 | ENSG00000165637 |
| NUTF2 | 0.111429715 | 1.080298285 | 0.091696239 | 0.212410709 | 3172.761407 | 3426.930853 | ENSG00000102898 |
| NA | 0.111229056 | 1.08014804 | 0.916953001 | 0.959042743 | 3.818027413 | 4.12212619 | ENSG00000267219 |
| SDHAF2 | 0.111082694 | 1.080038465 | 0.728461773 | 0.847646315 | 51.1586029 | 55.2411051 | ENSG00000167985 |
| ZNF469 | 0.111056115 | 1.080018568 | 0.718039065 | 0.840266197 | 53.58574824 | 57.86307219 | ENSG00000225614 |
| MCPH1 | 0.111024503 | 1.079994903 | 0.491068959 | 0.673883613 | 290.955027 | 313.528041 | ENSG00000147316 |
| PRDM11 | 0.111011366 | 1.079985068 | 0.449521843 | 0.637107184 | 288.4813998 | 311.5062363 | ENSG00000019485 |
| PSMA6 | 0.110895839 | 1.07989859 | 0.785856077 | 0.882841791 | 29.99827682 | 32.3877258 | ENSG00000100902 |
| LOC100289495 | 0.11071037 | 1.079759769 | 0.873530658 | 0.934704533 | 9.303016208 | 10.04094854 | ENSG00000286760 |
| ERICH2 | 0.110671469 | 1.079730655 | 0.695265136 | 0.825605076 | 57.22298245 | 61.89612994 | ENSG00000239467 |
| ROR2 | 0.110658526 | 1.079720969 | 0.586538757 | 0.749312221 | 123.7280628 | 133.4190356 | ENSG00000169071 |
| TLE4 | 0.11063253 | 1.079701513 | 0.289201502 | 0.480981752 | 516.4069994 | 557.389412 | ENSG00000106829 |
| CEP83 | 0.110563131 | 1.079649577 | 0.364990977 | 0.558171313 | 354.8220729 | 382.8276708 | ENSG00000173588 |
| RELN | 0.11054179 | 1.079633606 | 0.914506335 | 0.957916954 | 6.262836453 | 6.672370115 | ENSG00000189056 |
| NA | 0.110493182 | 1.079597231 | 0.778625808 | 0.878442338 | 31.12746603 | 33.7164612 | ENSG00000287763 |
| SNRNP48 | 0.110457431 | 1.079570478 | 0.444789724 | 0.633150769 | 491.7947269 | 530.6869658 | ENSG00000168566 |
| ZC3H13 | 0.11044827 | 1.079563624 | 0.108426825 | 0.240668595 | 2256.197487 | 2435.063707 | ENSG00000123200 |
| FNBP1L | 0.110370365 | 1.079505329 | 0.102490141 | 0.230796777 | 2795.062671 | 3016.262035 | ENSG00000137942 |
| ANO5 | 0.110324856 | 1.079471277 | 0.512074613 | 0.69104346 | 227.1477293 | 244.6761525 | ENSG00000171714 |
| NDUFA8 | 0.11030874 | 1.079459218 | 0.193144138 | 0.366775968 | 1368.524701 | 1476.986761 | ENSG00000119421 |
| SPCS1 | 0.110266237 | 1.079427417 | 0.205182287 | 0.383506303 | 858.1581333 | 926.0992594 | ENSG00000114902 |
| CBFA2T3 | 0.11010244 | 1.079304871 | 0.780327181 | 0.879291341 | 32.27747586 | 34.82242098 | ENSG00000129993 |
| DOK5 | 0.110044898 | 1.079261824 | 0.474805026 | 0.659821275 | 241.3727295 | 260.0742193 | ENSG00000101134 |
| RMND1 | 0.109949308 | 1.079190316 | 0.379766097 | 0.572904271 | 346.0828582 | 373.5119747 | ENSG00000155906 |
| TOX4 | 0.109852242 | 1.079117709 | 0.164068143 | 0.326200275 | 1183.969757 | 1278.022273 | ENSG00000092203 |
| PPTC7 | 0.109803931 | 1.079081574 | 0.322313553 | 0.516359183 | 572.0720015 | 616.9373901 | ENSG00000196850 |
| RPL4 | 0.1097612 | 1.079049613 | 0.057805514 | 0.149632319 | 32079.53875 | 34614.57264 | ENSG00000174444 |
| ZNF320 | 0.10975206 | 1.079042778 | 0.252549062 | 0.44125063 | 697.025915 | 752.4978105 | ENSG00000182986 |
| OST4 | 0.109736644 | 1.079031248 | 0.296366 | 0.48934053 | 738.3351458 | 797.206927 | ENSG00000228474 |
| DGKE | 0.109687528 | 1.078994513 | 0.537992906 | 0.711717382 | 172.4124749 | 186.1385673 | ENSG00000153933 |
| VTA1 | 0.109641699 | 1.078960238 | 0.239944971 | 0.425930474 | 1063.445973 | 1146.857923 | ENSG00000009844 |
| ZNF582-AS1 | 0.109575871 | 1.078911008 | 0.804059255 | 0.894704967 | 25.07995342 | 27.03974259 | ENSG00000267454 |
| PLEKHG5 | 0.109503272 | 1.078856716 | 0.27249018 | 0.463610599 | 769.8495965 | 831.3840777 | ENSG00000171680 |
| JRKL | 0.109436091 | 1.078806479 | 0.621402558 | 0.775152125 | 128.0408683 | 138.4178963 | ENSG00000183340 |
| COL4A6 | 0.109425032 | 1.078798209 | 0.204240052 | 0.382276682 | 1112.44349 | 1200.37549 | ENSG00000197565 |
| NA | 0.109348766 | 1.078741182 | 0.794557743 | 0.888681022 | 35.10529328 | 37.94730289 | ENSG00000251687 |
| FAM83D | 0.109338423 | 1.078733448 | 0.11531529 | 0.252302395 | 1708.655513 | 1842.87935 | ENSG00000101447 |
| TMEM60 | 0.109330017 | 1.078727163 | 0.47558914 | 0.660316812 | 219.3763558 | 236.7138402 | ENSG00000135211 |
| WASL | 0.109272238 | 1.078683961 | 0.419794258 | 0.611191421 | 650.4371501 | 701.4965612 | ENSG00000106299 |
| CDNF | 0.10911664 | 1.078567629 | 0.89688864 | 0.947229702 | 6.014568001 | 6.481299358 | ENSG00000185267 |
| MTFR1L | 0.109115557 | 1.078566819 | 0.521623546 | 0.698639312 | 179.6328699 | 193.9476144 | ENSG00000117640 |
| MCU | 0.108930124 | 1.078428197 | 0.27659124 | 0.467615547 | 871.7987817 | 939.4423104 | ENSG00000156026 |
| NOP53 | 0.108928684 | 1.078427121 | 0.074132561 | 0.180904921 | 7590.479137 | 8185.761514 | ENSG00000105373 |
| PRKCQ | 0.108813453 | 1.078340989 | 0.348504883 | 0.54178741 | 487.535454 | 526.2058897 | ENSG00000065675 |
| NA | 0.108789454 | 1.07832305 | 0.602013275 | 0.761306908 | 227.8412801 | 245.617625 | ENSG00000237973 |
| ATP5F1A | 0.108676627 | 1.078238722 | 0.066045108 | 0.165550451 | 13733.75109 | 14807.63307 | ENSG00000152234 |
| CUL5 | 0.108659864 | 1.078226194 | 0.438085997 | 0.626620376 | 306.0598429 | 329.5691165 | ENSG00000166266 |
| ZSCAN26 | 0.108618123 | 1.078194999 | 0.646056634 | 0.793142577 | 104.7667578 | 112.6838325 | ENSG00000197062 |

| | | | | | | | |
|---|---|---|---|---|---|---|---|
| TRAK1 | 0.108483695 | 1.078094539 | 0.175941889 | 0.343723723 | 1140.3256 | 1228.836643 | ENSG00000182606 |
| POLDIP3 | 0.108472654 | 1.078086288 | 0.102515989 | 0.230796777 | 3318.599147 | 3577.264719 | ENSG00000100227 |
| EGR2 | 0.10846984 | 1.078084186 | 0.811559932 | 0.898880668 | 24.30210577 | 26.1694436 | ENSG00000122877 |
| STXBP3 | 0.108445013 | 1.078065633 | 0.379756002 | 0.572904271 | 606.3228093 | 653.1853412 | ENSG00000116266 |
| DNAJC30 | 0.108393424 | 1.078027084 | 0.752002937 | 0.86205208 | 55.7934433 | 60.27315926 | ENSG00000176410 |
| SPIN1 | 0.108390896 | 1.078025194 | 0.133867874 | 0.281240894 | 2652.975762 | 2859.026508 | ENSG00000106723 |
| NA | 0.108200619 | 1.077883023 | 0.892711618 | 0.945188068 | 6.748820239 | 7.265341067 | ENSG00000225518 |
| SLC2A4 | 0.108200526 | 1.077882954 | 0.606755159 | 0.765267106 | 114.3586764 | 123.3796244 | ENSG00000181856 |
| NA | 0.108191393 | 1.07787613 | 0.847804568 | 0.918948068 | 14.06006659 | 15.13449997 | ENSG00000273137 |
| GGA3 | 0.108176154 | 1.077864745 | 0.146553393 | 0.300733014 | 1378.622845 | 1485.814365 | ENSG00000125447 |
| ICOSLG | 0.108108354 | 1.077814091 | 0.754301246 | 0.863474302 | 42.40834667 | 45.53770804 | ENSG00000160223 |
| NF1 | 0.107982881 | 1.077720357 | 0.342676302 | 0.536797013 | 631.5418171 | 680.7959378 | ENSG00000196712 |
| MBOAT7 | 0.107899102 | 1.077657774 | 0.200847106 | 0.377497238 | 1860.886461 | 2005.010842 | ENSG00000125505 |
| NA | 0.107860287 | 1.077628781 | 0.906785657 | 0.953498546 | 6.258499462 | 6.739968019 | ENSG00000253210 |
| ZFAND3 | 0.107776566 | 1.077566247 | 0.14683509 | 0.301144358 | 1533.569804 | 1652.028752 | ENSG00000156639 |
| AP3M2 | 0.107772599 | 1.077563284 | 0.406016228 | 0.597593867 | 437.635376 | 471.5735635 | ENSG00000070718 |
| TSPAN9 | 0.107764808 | 1.077557464 | 0.199294781 | 0.375658161 | 1192.826122 | 1285.217294 | ENSG00000011105 |
| GOLPH3L | 0.107666562 | 1.077484087 | 0.308258118 | 0.502440943 | 780.8977904 | 840.7631575 | ENSG00000143457 |
| STK4 | 0.107479347 | 1.077344273 | 0.258376332 | 0.447074219 | 883.1377646 | 952.0719301 | ENSG00000101109 |
| AURKB | 0.1071336 | 1.077086114 | 0.183016131 | 0.353519643 | 2201.799008 | 2371.462873 | ENSG00000178999 |
| CCNB1 | 0.107112093 | 1.077070058 | 0.103706625 | 0.232896957 | 5486.503655 | 5908.740539 | ENSG00000134057 |
| OVOL1 | 0.106993472 | 1.076981503 | 0.726617473 | 0.846456835 | 52.12839824 | 55.93866629 | ENSG00000172818 |
| SLC25A28 | 0.106982933 | 1.076973636 | 0.478374666 | 0.662348428 | 221.7633558 | 239.0087763 | ENSG00000155287 |
| LRBA | 0.106701668 | 1.076763692 | 0.251070327 | 0.439533148 | 2039.254425 | 2194.968226 | ENSG00000198589 |
| TGIF1 | 0.106678056 | 1.076746068 | 0.121650234 | 0.262292396 | 2580.077773 | 2778.962857 | ENSG00000177426 |
| LOC101927420 | 0.106652809 | 1.076727226 | 0.812027515 | 0.899212401 | 22.78053513 | 24.63174251 | ENSG00000224046 |
| NA | 0.106580909 | 1.076673566 | 0.64677687 | 0.793707376 | 85.57569721 | 92.1344127 | ENSG00000139239 |
| OLFML2B | 0.10637281 | 1.076518274 | 0.569738154 | 0.736317339 | 177.1939687 | 191.2022776 | ENSG00000162745 |
| ATP5IF1 | 0.106370544 | 1.076516583 | 0.207551447 | 0.386453539 | 2037.747 | 2194.259248 | ENSG00000130770 |
| NA | 0.106347304 | 1.076499242 | 0.866216685 | 0.930189756 | 11.41680064 | 12.27241198 | ENSG00000286277 |
| TSPAN12 | 0.106343191 | 1.076496173 | 0.445139282 | 0.633364921 | 316.4602687 | 341.1367363 | ENSG00000106025 |
| NDUFA1 | 0.106325737 | 1.07648315 | 0.306944021 | 0.501004007 | 573.2895616 | 617.5670044 | ENSG00000125356 |
| ARHGAP12 | 0.106243078 | 1.076421474 | 0.242955919 | 0.429506391 | 877.711983 | 944.4550926 | ENSG00000165322 |
| NDUFS1 | 0.106065659 | 1.076289107 | 0.153060375 | 0.310374288 | 1952.986884 | 2101.423114 | ENSG00000023228 |
| WBP4 | 0.105979273 | 1.076224663 | 0.499272099 | 0.68011443 | 198.9474182 | 214.2833936 | ENSG00000120688 |
| NA | 0.105978109 | 1.076223794 | 0.528130032 | 0.704021174 | 208.9288469 | 224.865237 | ENSG00000231698 |
| LOC100507373 | 0.105880321 | 1.076150848 | 0.733413922 | 0.851164285 | 63.96085877 | 68.84301693 | ENSG00000267169 |
| ATXN2L | 0.105820672 | 1.076106355 | 0.094649603 | 0.217269724 | 3796.09661 | 4085.5634 | ENSG00000168488 |
| CBWD5 | 0.10566594 | 1.075990947 | 0.622408655 | 0.775781707 | 109.6262799 | 118.0007092 | ENSG00000147996 |
| RGPD1 | 0.105611339 | 1.075950226 | 0.907346503 | 0.953925922 | 5.117862675 | 5.494901971 | ENSG00000187627 |
| ESF1 | 0.105610475 | 1.075949581 | 0.267408279 | 0.457546626 | 796.1771909 | 856.4074055 | ENSG00000089048 |
| CCT8 | 0.105533968 | 1.075892524 | 0.091855262 | 0.212619637 | 6502.051159 | 6994.3277 | ENSG00000156261 |
| GPR19 | 0.105497192 | 1.075865099 | 0.479348908 | 0.663418777 | 236.069233 | 253.8149227 | ENSG00000183150 |
| NA | 0.105461908 | 1.075838787 | 0.759741517 | 0.866398174 | 38.61053851 | 41.59679341 | ENSG00000231007 |
| ADAM12 | 0.105452664 | 1.075831894 | 0.908740689 | 0.954598925 | 6.847458652 | 7.358758006 | ENSG00000148848 |
| RAB2B | 0.105264043 | 1.075691246 | 0.551542225 | 0.722473927 | 233.0367577 | 250.4140184 | ENSG00000129472 |
| HIC2 | 0.105205541 | 1.075647627 | 0.183045401 | 0.353536331 | 1538.420344 | 1654.198956 | ENSG00000169635 |
| TNFRSF1B | 0.105205244 | 1.075647406 | 0.639743451 | 0.789046724 | 92.68414966 | 99.64301225 | ENSG00000028137 |
| CDCA7L | 0.10517296 | 1.075623336 | 0.110882743 | 0.244800567 | 2520.534777 | 2710.613379 | ENSG00000164649 |
| RALGPS1 | 0.105070885 | 1.075547235 | 0.427949704 | 0.618257059 | 392.4352157 | 422.0820154 | ENSG00000136828 |
| GPKOW | 0.104997359 | 1.075492422 | 0.334157501 | 0.528670131 | 473.567787 | 509.3768788 | ENSG00000068394 |
| SOCS5 | 0.104975209 | 1.075475909 | 0.398853523 | 0.590632086 | 355.8674235 | 382.8707723 | ENSG00000171150 |
| ILRUN | 0.104932042 | 1.07544373 | 0.106386623 | 0.237476776 | 2873.934136 | 3090.217079 | ENSG00000196821 |
| RAI14 | 0.104906008 | 1.075424324 | 0.143479038 | 0.296261195 | 2280.153648 | 2452.187774 | ENSG00000039560 |
| DARS2 | 0.104843364 | 1.075377629 | 0.306757279 | 0.500827092 | 727.8876896 | 783.3556561 | ENSG00000117593 |
| CCDC84 | 0.104763655 | 1.075318215 | 0.541934149 | 0.714666944 | 194.5199192 | 209.2547405 | ENSG00000186166 |
| ADAMTS15 | 0.10476106 | 1.075316281 | 0.492155526 | 0.674641507 | 243.5019902 | 262.2644938 | ENSG00000166106 |
| NA | 0.104668548 | 1.075247329 | 0.623452926 | 0.776603033 | 170.6649296 | 184.0100939 | ENSG00000277072 |
| POLQ | 0.104614756 | 1.075207239 | 0.270349182 | 0.460858117 | 708.847552 | 762.115666 | ENSG00000051341 |
| TBC1D9B | 0.104528612 | 1.075143039 | 0.150710152 | 0.306951864 | 1607.935358 | 1728.543422 | ENSG00000197226 |
| NA | 0.104480784 | 1.075107397 | 0.874583448 | 0.935130738 | 9.874913057 | 10.59214063 | ENSG00000255618 |
| TIPRL | 0.104292702 | 1.074967246 | 0.370557929 | 0.563911136 | 518.1238474 | 556.3291012 | ENSG00000143155 |
| NA | 0.104219057 | 1.074912374 | 0.909880177 | 0.95507273 | 6.861266382 | 7.359307659 | ENSG00000246334 |
| FURIN | 0.104175445 | 1.07487988 | 0.193485721 | 0.367198889 | 1178.448553 | 1267.249359 | ENSG00000140564 |
| SRRM2 | 0.103986566 | 1.074739165 | 0.199061286 | 0.375320638 | 12512.22191 | 13448.10989 | ENSG00000167978 |
| LRRC8B | 0.103956824 | 1.074717009 | 0.447954861 | 0.635957225 | 390.4549533 | 419.3310958 | ENSG00000197147 |
| TAF3 | 0.103892543 | 1.074669125 | 0.421372536 | 0.612623045 | 392.2767536 | 421.4748049 | ENSG00000165632 |
| IGFBPL1 | 0.103887328 | 1.07466524 | 0.293035043 | 0.485570987 | 1278.662491 | 1373.171695 | ENSG00000137142 |
| GPR108 | 0.103875887 | 1.074656718 | 0.278444551 | 0.46955051 | 642.0137095 | 689.7819203 | ENSG00000125734 |
| THOC7 | 0.103874439 | 1.074655639 | 0.347381664 | 0.54072072 | 592.3791426 | 636.3189233 | ENSG00000163634 |
| SEPTIN7P2 | 0.10371365 | 1.074535875 | 0.526762925 | 0.702967066 | 216.2237299 | 232.5327015 | ENSG00000214765 |
| ENDOV | 0.103363439 | 1.074275065 | 0.61699884 | 0.772278752 | 134.3472025 | 144.2527229 | ENSG00000173818 |
| GUCY1B1 | 0.103286292 | 1.074217621 | 0.70810962 | 0.83401045 | 68.56314537 | 73.37971368 | ENSG00000061918 |
| FAF1 | 0.103268342 | 1.074204256 | 0.238269221 | 0.42383574 | 1033.461872 | 1109.659462 | ENSG00000185104 |

| | | | | | | | |
|---|---|---|---|---|---|---|---|
| UBAP2 | 0.103239954 | 1.074183119 | 0.226655885 | 0.410158189 | 1456.517757 | 1563.980921 | ENSG00000137073 |
| UROS | 0.10312879 | 1.074100352 | 0.277720701 | 0.468710105 | 674.8063555 | 724.7012222 | ENSG00000188690 |
| BRD4 | 0.103122982 | 1.074096028 | 0.143069897 | 0.295598758 | 2448.52288 | 2630.426211 | ENSG00000141867 |
| MRE11 | 0.103116855 | 1.074091467 | 0.325381319 | 0.519510828 | 589.0004217 | 633.3017382 | ENSG00000020922 |
| FAM122C | 0.103031706 | 1.074028075 | 0.741896793 | 0.855704472 | 50.58029452 | 54.44351697 | ENSG00000156500 |
| CYTH3 | 0.102945613 | 1.073963984 | 0.354042156 | 0.547411139 | 515.5045816 | 553.2544635 | ENSG00000008256 |
| AGFG1 | 0.102941618 | 1.073961011 | 0.301569365 | 0.494940359 | 1371.470579 | 1472.476601 | ENSG00000173744 |
| TMEM184C | 0.102867814 | 1.073906072 | 0.301629308 | 0.494988276 | 703.5076457 | 755.2560953 | ENSG00000164168 |
| BCL11A | 0.102792662 | 1.073850131 | 0.419949667 | 0.61127363 | 356.2929223 | 383.0893026 | ENSG00000119866 |
| SNHG1 | 0.10264706 | 1.07374176 | 0.676657783 | 0.813094875 | 1595.02343 | 1712.901075 | ENSG00000255717 |
| NA | 0.102630646 | 1.073729544 | 0.250547546 | 0.438991135 | 62070.32285 | 66646.60622 | ENSG00000210082 |
| BRI3BP | 0.102535688 | 1.073658873 | 0.450306806 | 0.6379748 | 356.215999 | 382.4212074 | ENSG00000184992 |
| CD99P1 | 0.102533111 | 1.073656955 | 0.686822376 | 0.819887022 | 87.18062536 | 93.7186792 | ENSG00000223773 |
| ADAMTS13 | 0.102470007 | 1.073609995 | 0.689721604 | 0.822077183 | 138.4391444 | 149.0982696 | ENSG00000160323 |
| TRAF2 | 0.102329116 | 1.073505153 | 0.230304375 | 0.414413718 | 1270.830857 | 1364.906191 | ENSG00000127191 |
| PARP9 | 0.102320147 | 1.073498479 | 0.628486023 | 0.780570128 | 104.9052333 | 112.4445961 | ENSG00000138496 |
| REX1BD | 0.102222701 | 1.073425972 | 0.319325427 | 0.513144724 | 815.1463957 | 875.3682534 | ENSG00000006015 |
| TRAPPC8 | 0.102190361 | 1.07340191 | 0.27440713 | 0.465397305 | 1010.52074 | 1084.855071 | ENSG00000153339 |
| TMCO4 | 0.102152988 | 1.073374105 | 0.650513118 | 0.795936006 | 103.969871 | 111.5643263 | ENSG00000162542 |
| GGA1 | 0.101979397 | 1.073244959 | 0.24496006 | 0.432101433 | 983.3343836 | 1055.717961 | ENSG00000100083 |
| JAGN1 | 0.101943141 | 1.073217988 | 0.316665437 | 0.510333097 | 560.9202385 | 601.6333387 | ENSG00000171135 |
| EVC2 | 0.101831047 | 1.073134605 | 0.53069339 | 0.70614339 | 190.1069119 | 204.1071376 | ENSG00000173040 |
| COX19 | 0.101685596 | 1.073026418 | 0.513691606 | 0.692267885 | 277.0256338 | 297.9065634 | ENSG00000240230 |
| NR2C2 | 0.101667252 | 1.073012775 | 0.402999099 | 0.594775068 | 695.0390236 | 746.2540675 | ENSG00000177463 |
| OXR1 | 0.101601241 | 1.072963679 | 0.485265523 | 0.668376071 | 245.1991453 | 263.132929 | ENSG00000164830 |
| SPTLC1 | 0.101585229 | 1.072951771 | 0.273670529 | 0.464871193 | 1052.733967 | 1128.770832 | ENSG00000090054 |
| AURKC | 0.101501549 | 1.072889539 | 0.831463947 | 0.909306807 | 24.92085265 | 26.78801035 | ENSG00000105146 |
| LINGO3 | 0.101362577 | 1.072786195 | 0.851389474 | 0.921493373 | 15.59120566 | 16.67577667 | ENSG00000220008 |
| ATP6 | 0.101334383 | 1.07276523 | 0.277669207 | 0.468699553 | 23591.58175 | 25308.26495 | ENSG00000198899 |
| PALLD | 0.101221877 | 1.072681576 | 0.460983362 | 0.647605976 | 349.4143232 | 374.3557531 | ENSG00000129116 |
| MT1L | 0.101199545 | 1.072664971 | 0.918454131 | 0.959670135 | 5.005416532 | 5.355907317 | ENSG00000260549 |
| GLMN | 0.101145104 | 1.072624494 | 0.463908423 | 0.65028579 | 371.7312433 | 398.5650094 | ENSG00000174842 |
| DNM1 | 0.101126379 | 1.072610573 | 0.711012308 | 0.836108249 | 755.3802741 | 810.4999268 | ENSG00000106976 |
| ANKRD16 | 0.101114631 | 1.072601838 | 0.518383439 | 0.695556263 | 231.3244043 | 248.5853397 | ENSG00000134461 |
| ZNF761 | 0.101113917 | 1.072601307 | 0.358685581 | 0.551935738 | 468.1106317 | 501.9673185 | ENSG00000160336 |
| UNC5B | 0.101104286 | 1.072594147 | 0.215320459 | 0.395632932 | 1533.540617 | 1645.204565 | ENSG00000107731 |
| FAM241B | 0.101022273 | 1.072533175 | 0.398581533 | 0.590322714 | 450.8557118 | 483.5544482 | ENSG00000171224 |
| JAZF1 | 0.101012117 | 1.072525625 | 0.647437739 | 0.794202863 | 125.4640703 | 134.2726103 | ENSG00000153814 |
| CDC37L1-DT | 0.10100418 | 1.072519724 | 0.914921384 | 0.958037973 | 4.780426548 | 5.108523577 | ENSG00000273061 |
| ZNF746 | 0.101000914 | 1.072517296 | 0.346396113 | 0.539958415 | 607.2757151 | 650.8421194 | ENSG00000181220 |
| SMC6 | 0.100981398 | 1.072502788 | 0.256562277 | 0.445236447 | 1329.105234 | 1425.074432 | ENSG00000163029 |
| ERBIN | 0.10094142 | 1.072473069 | 0.383786786 | 0.576520043 | 441.6579409 | 473.3789248 | ENSG00000112851 |
| YTHDF3 | 0.100932822 | 1.072466677 | 0.257482901 | 0.446123969 | 1783.500353 | 1911.940673 | ENSG00000185728 |
| MYO3B | 0.100908456 | 1.072448564 | 0.887969586 | 0.942849507 | 8.654677033 | 9.249420762 | ENSG00000071909 |
| ZNF485 | 0.100881426 | 1.072428471 | 0.636439761 | 0.786332546 | 125.2542861 | 134.2208606 | ENSG00000198298 |
| ARMC10 | 0.100821278 | 1.072383761 | 0.311135396 | 0.504837468 | 891.6282821 | 955.959766 | ENSG00000170632 |
| MYCL | 0.10074967 | 1.072330534 | 0.365933793 | 0.55902688 | 450.4800094 | 482.7182901 | ENSG00000116990 |
| COX4I1 | 0.100650096 | 1.072256525 | 0.149410349 | 0.305176482 | 7006.293463 | 7513.084048 | ENSG00000131143 |
| CLDN4 | 0.100621041 | 1.072234931 | 0.450664186 | 0.638237184 | 292.2234886 | 313.526481 | ENSG00000189143 |
| LOC106660606 | 0.100600421 | 1.072219606 | 0.906321903 | 0.953173135 | 6.322314136 | 6.758739013 | ENSG00000260778 |
| RAP1A | 0.100571761 | 1.072198306 | 0.339093814 | 0.533330341 | 576.2460157 | 617.2574932 | ENSG00000116473 |
| WDR89 | 0.100533804 | 1.072170097 | 0.539455547 | 0.712722396 | 274.5808248 | 294.0999995 | ENSG00000140006 |
| FAM83F | 0.10051679 | 1.072157453 | 0.262734354 | 0.45226326 | 842.7255474 | 903.4226399 | ENSG00000133477 |
| KANSL1 | 0.100431422 | 1.072094013 | 0.242665743 | 0.429116199 | 994.855648 | 1066.469133 | ENSG00000120071 |
| NA | 0.100287167 | 1.071986819 | 0.930177372 | 0.965658848 | 3.56650435 | 3.810831773 | ENSG00000267892 |
| FAM86B1 | 0.100281685 | 1.071982746 | 0.774991865 | 0.875992658 | 39.15590228 | 41.96776162 | ENSG00000186523 |
| ACOX1 | 0.100218961 | 1.07193614 | 0.44808439 | 0.635975449 | 352.0530341 | 377.1901902 | ENSG00000161533 |
| KPNA6 | 0.100210352 | 1.071929744 | 0.208603672 | 0.38769228 | 1674.672711 | 1795.043941 | ENSG00000025800 |
| RASA3 | 0.100140606 | 1.071877924 | 0.292465223 | 0.485060443 | 721.5229375 | 773.4080758 | ENSG00000185989 |
| MOXD1 | 0.100057733 | 1.071816353 | 0.692980864 | 0.824106983 | 79.90369908 | 85.51324263 | ENSG00000079931 |
| VAT1 | 0.099841388 | 1.071655637 | 0.065163538 | 0.163894832 | 7938.892871 | 8507.376751 | ENSG00000108828 |
| HSPA4 | 0.099827624 | 1.071645413 | 0.104856635 | 0.235080789 | 11343.67223 | 12155.28186 | ENSG00000170606 |
| CBX8 | 0.099817569 | 1.071637943 | 0.775177698 | 0.876003875 | 38.88100305 | 41.65646721 | ENSG00000141570 |
| CMTM3 | 0.099791711 | 1.071618737 | 0.419231972 | 0.610612603 | 437.7538125 | 469.2947786 | ENSG00000140931 |
| TXLNA | 0.099760331 | 1.071595428 | 0.35239942 | 0.545516575 | 1692.769098 | 1814.606626 | ENSG00000084652 |
| FOCAD | 0.099752941 | 1.071589939 | 0.220740633 | 0.402242344 | 1224.022191 | 1311.059091 | ENSG00000188352 |
| NA | 0.099658568 | 1.071519844 | 0.929371237 | 0.965200409 | 3.460372032 | 3.69440696 | ENSG00000241511 |
| KMT2E | 0.099602782 | 1.071478411 | 0.412892182 | 0.604466327 | 476.4652671 | 510.3781283 | ENSG00000005483 |
| NA | 0.099557707 | 1.071444935 | 0.907938646 | 0.954223696 | 6.077104898 | 6.488785432 | ENSG00000231409 |
| SOX11 | 0.099540694 | 1.0714323 | 0.262794297 | 0.452324433 | 1692.78734 | 1813.914549 | ENSG00000176887 |
| HS6ST2 | 0.099507197 | 1.071407423 | 0.41558097 | 0.606868713 | 454.5464256 | 486.5423419 | ENSG00000171004 |
| NAIF1 | 0.0994675 | 1.071377943 | 0.513810031 | 0.692278836 | 237.9958877 | 255.2394884 | ENSG00000171169 |
| NUP133 | 0.099429419 | 1.071349664 | 0.207024486 | 0.38570385 | 1767.086442 | 1892.570251 | ENSG00000069248 |
| IL17RD | 0.099373395 | 1.071308061 | 0.198546227 | 0.374654426 | 2116.959502 | 2267.061438 | ENSG00000144730 |

| | | | | | | | |
|---|---|---|---|---|---|---|---|
| ACTN3 | 0.099010859 | 1.071038884 | 0.440310561 | 0.628529445 | 341.9126925 | 366.3193445 | ENSG00000248746 |
| NA | 0.098986803 | 1.071021026 | 0.896497021 | 0.947031963 | 8.362715462 | 8.926841424 | ENSG00000284602 |
| ARMCX6 | 0.09893689 | 1.070983972 | 0.503078671 | 0.683503536 | 266.0184875 | 285.0433303 | ENSG00000198960 |
| UBA6-AS1 | 0.098915932 | 1.070968414 | 0.707555205 | 0.833682656 | 75.97252647 | 81.44907838 | ENSG00000248049 |
| COPG2 | 0.098906056 | 1.070961083 | 0.264162117 | 0.453835806 | 1000.00499 | 1070.430928 | ENSG00000158623 |
| LCLAT1 | 0.098873477 | 1.070936899 | 0.550349282 | 0.721625298 | 173.9355187 | 186.2685385 | ENSG00000172954 |
| NA | 0.098799169 | 1.07088174 | 0.911191727 | 0.955762377 | 5.742825684 | 6.124976879 | ENSG00000227038 |
| LPCAT4 | 0.098641772 | 1.070764914 | 0.282363198 | 0.473303746 | 828.3039695 | 887.4530501 | ENSG00000176454 |
| ZNF24 | 0.098595002 | 1.070730202 | 0.252117774 | 0.44090874 | 1394.030168 | 1492.233823 | ENSG00000172466 |
| NA | 0.098589841 | 1.070726372 | 0.875189792 | 0.935364718 | 16.39370726 | 17.59925107 | ENSG00000284617 |
| NA | 0.098508986 | 1.070666365 | 0.924229692 | 0.962449326 | 4.685042746 | 4.999584838 | ENSG00000271737 |
| NA | 0.098465116 | 1.070633809 | 0.862892058 | 0.928197411 | 15.2518904 | 16.39508782 | ENSG00000227252 |
| POU6F1 | 0.098461887 | 1.070631412 | 0.660368989 | 0.802312608 | 98.04090775 | 105.2295077 | ENSG00000184271 |
| DSG2 | 0.098454346 | 1.070625816 | 0.10635113 | 0.237454745 | 3913.98136 | 4189.946595 | ENSG00000046604 |
| TAF4 | 0.098386456 | 1.070575436 | 0.271596374 | 0.46245428 | 798.374348 | 854.728497 | ENSG00000130699 |
| MUCL3 | 0.098373297 | 1.070565672 | 0.92895999 | 0.964935518 | 4.402551914 | 4.699575342 | ENSG00000168631 |
| TUBE1 | 0.098337448 | 1.070539069 | 0.501338065 | 0.681946647 | 259.9173074 | 278.057802 | ENSG00000074935 |
| DLGAP4 | 0.098319924 | 1.070526066 | 0.246991821 | 0.434733752 | 964.7065233 | 1032.996145 | ENSG00000080845 |
| PSMD5 | 0.098305233 | 1.070515165 | 0.40258748 | 0.594356887 | 431.9260358 | 462.3822876 | ENSG00000095261 |
| SMURF2 | 0.098241961 | 1.070468217 | 0.381853892 | 0.574734451 | 561.7601194 | 600.7116877 | ENSG00000108854 |
| MLH3 | 0.098103703 | 1.070365635 | 0.491371298 | 0.674098864 | 245.0687423 | 262.1239618 | ENSG00000119684 |
| TEPSIN | 0.097943059 | 1.070246457 | 0.693587947 | 0.824405759 | 144.7563248 | 155.3056017 | ENSG00000167302 |
| FZD4 | 0.097878114 | 1.070198279 | 0.527435141 | 0.70347459 | 217.7328683 | 233.2748527 | ENSG00000174804 |
| SLU7 | 0.097809303 | 1.070147236 | 0.409055165 | 0.600556752 | 488.9299362 | 523.2079128 | ENSG00000164609 |
| STOM | 0.097800101 | 1.07014041 | 0.350648441 | 0.543979092 | 617.8937498 | 660.9703044 | ENSG00000148175 |
| ZNF747 | 0.09773958 | 1.070095518 | 0.75877324 | 0.865634817 | 45.98432175 | 49.1503094 | ENSG00000169955 |
| TMED5 | 0.09765284 | 1.070031183 | 0.543931217 | 0.716179733 | 343.7136045 | 367.494359 | ENSG00000117500 |
| EAF1 | 0.097509896 | 1.069925168 | 0.420773855 | 0.61194465 | 382.2490687 | 408.6365551 | ENSG00000144597 |
| EHD4 | 0.097485561 | 1.069907121 | 0.20681871 | 0.385476465 | 1597.468967 | 1708.762052 | ENSG00000103966 |
| LENG8 | 0.097417294 | 1.069856495 | 0.779797324 | 0.878986213 | 3333.690493 | 3566.760015 | ENSG00000167615 |
| DNAJC16 | 0.097400556 | 1.069844083 | 0.447512144 | 0.635716262 | 334.6324366 | 357.8175857 | ENSG00000116138 |
| UQCRHL | 0.097370062 | 1.06982147 | 0.308898716 | 0.50308689 | 735.0426689 | 786.4728287 | ENSG00000233954 |
| IMPA1 | 0.097366152 | 1.06981857 | 0.502453808 | 0.682933329 | 463.1284558 | 495.2963423 | ENSG00000133731 |
| CYTB | 0.097252058 | 1.069733968 | 0.621197207 | 0.775052679 | 21025.41105 | 22491.64084 | ENSG00000198727 |
| SPRED2 | 0.097238147 | 1.069723654 | 0.361218596 | 0.554482811 | 515.6039942 | 551.4967351 | ENSG00000198369 |
| ADAM19 | 0.097115918 | 1.069633028 | 0.272957407 | 0.464170345 | 786.8340444 | 841.5648164 | ENSG00000135074 |
| RHOU | 0.097114641 | 1.069632081 | 0.587719366 | 0.750329471 | 189.6849155 | 202.7780758 | ENSG00000116574 |
| PPM1A | 0.097078014 | 1.069604925 | 0.364521672 | 0.557636838 | 859.0080485 | 918.3600399 | ENSG00000100614 |
| SYNGR1 | 0.097067973 | 1.069597481 | 0.399489649 | 0.591432315 | 579.6161225 | 620.8204721 | ENSG00000100321 |
| SLC44A3 | 0.096995436 | 1.069543704 | 0.715458762 | 0.838729455 | 65.27163795 | 69.79468349 | ENSG00000143036 |
| GPATCH8 | 0.096879399 | 1.069457684 | 0.226762759 | 0.41019139 | 1520.092135 | 1625.501815 | ENSG00000186566 |
| C6orf120 | 0.096847672 | 1.069434165 | 0.413831428 | 0.605206495 | 497.2094296 | 531.0514272 | ENSG00000185127 |
| IGSF3 | 0.096821788 | 1.069414978 | 0.194832792 | 0.369112846 | 1718.284728 | 1837.068462 | ENSG00000143061 |
| MFSD14B | 0.096792736 | 1.069393443 | 0.30492195 | 0.498845743 | 1139.857227 | 1218.321315 | ENSG00000148110 |
| BSN-DT | 0.096667041 | 1.069300276 | 0.906035339 | 0.952979909 | 7.036347121 | 7.493953813 | ENSG00000226913 |
| GLT8D1 | 0.096656147 | 1.069292201 | 0.330105643 | 0.524400272 | 614.6106459 | 656.8382159 | ENSG00000016864 |
| NDE1 | 0.096635933 | 1.06927722 | 0.451890192 | 0.639387192 | 417.6451661 | 446.4290067 | ENSG00000072864 |
| FAM218A | 0.09662241 | 1.069267197 | 0.876498385 | 0.936003742 | 12.94152819 | 13.78049519 | ENSG00000250486 |
| HLA-B | 0.096602315 | 1.069252303 | 0.3672495 | 0.560352534 | 533.5136346 | 570.1640859 | ENSG00000234745 |
| NA | 0.096578235 | 1.069234456 | 0.922552839 | 0.961675721 | 4.896029604 | 5.213663469 | ENSG00000280114 |
| COIL | 0.096483697 | 1.069164393 | 0.342251695 | 0.536395153 | 566.2781993 | 605.114105 | ENSG00000121058 |
| ISG20 | 0.09646439 | 1.069150085 | 0.84034279 | 0.91422735 | 20.05125842 | 21.48472879 | ENSG00000172183 |
| MIF4GD | 0.096464328 | 1.069150039 | 0.516707582 | 0.694262601 | 241.9236103 | 258.2825245 | ENSG00000125457 |
| SH3GL3 | 0.096369332 | 1.069079642 | 0.527645926 | 0.703553208 | 229.4932235 | 245.0125418 | ENSG00000140600 |
| EGLN3 | 0.096359406 | 1.069072287 | 0.411551248 | 0.60307486 | 409.7699827 | 438.280155 | ENSG00000129521 |
| LINC02610 | 0.096324878 | 1.069046701 | 0.812224243 | 0.899322786 | 34.9732066 | 37.36204125 | ENSG00000186235 |
| KLK6 | 0.096308603 | 1.069034641 | 0.882063822 | 0.939373634 | 10.59969456 | 11.27852852 | ENSG00000167755 |
| JMY | 0.096231465 | 1.068977483 | 0.501510326 | 0.682098008 | 276.8132715 | 295.7518555 | ENSG00000152409 |
| EMB | 0.096228934 | 1.068975608 | 0.37856076 | 0.571960962 | 658.7101343 | 704.0207 | ENSG00000170571 |
| PCDH8 | 0.096211365 | 1.06896259 | 0.861922005 | 0.927530769 | 16.32683337 | 17.50495806 | ENSG00000136099 |
| VPS13A | 0.09618127 | 1.068940292 | 0.41012412 | 0.601792451 | 700.4122951 | 748.5773702 | ENSG00000197969 |
| ECH1 | 0.096170543 | 1.068932344 | 0.371570472 | 0.564806621 | 573.8478687 | 613.678014 | ENSG00000104823 |
| GEMIN2 | 0.096103461 | 1.068882642 | 0.456874291 | 0.643981147 | 395.0381182 | 422.1940917 | ENSG00000092208 |
| SLC25A51 | 0.095996084 | 1.06880309 | 0.724998521 | 0.845367888 | 69.31130303 | 73.90563638 | ENSG00000122696 |
| DDX11 | 0.095917647 | 1.068744983 | 0.353196887 | 0.546358303 | 2261.858399 | 2418.402972 | ENSG00000013573 |
| CZIB | 0.09589077 | 1.068725073 | 0.322126588 | 0.516148888 | 768.9376627 | 822.1507998 | ENSG00000162384 |
| SLC2A10 | 0.095836586 | 1.068684935 | 0.550813236 | 0.721876151 | 188.4018722 | 201.2564462 | ENSG00000197496 |
| TNNI1 | 0.095818761 | 1.06867173 | 0.824606777 | 0.905655553 | 26.77995711 | 28.62572093 | ENSG00000159173 |
| NFATC3 | 0.095815411 | 1.068669249 | 0.433048321 | 0.622850174 | 391.9690452 | 418.5007437 | ENSG00000072736 |
| SEC24B | 0.095749973 | 1.068620777 | 0.4365406 | 0.62586147 | 555.4590393 | 593.1484871 | ENSG00000138802 |
| OAS3 | 0.095679699 | 1.068568726 | 0.84922233 | 0.920007404 | 21.48187995 | 22.86835145 | ENSG00000111331 |
| TSC1 | 0.095533843 | 1.068460699 | 0.290057369 | 0.482184226 | 792.9195302 | 846.876795 | ENSG00000165699 |
| ARHGEF4 | 0.095440422 | 1.068391514 | 0.380953623 | 0.573941228 | 596.483607 | 637.6882041 | ENSG00000136002 |
| ROR1 | 0.09540307 | 1.068363853 | 0.179774222 | 0.3491131 | 1591.339435 | 1700.220347 | ENSG00000185483 |

| | | | | | | | |
|---|---|---|---|---|---|---|---|
| PNPLA5 | 0.095380957 | 1.068347478 | 0.548583298 | 0.720124856 | 201.1312334 | 215.0866443 | ENSG00000100341 |
| SELENOT | 0.095334817 | 1.068313311 | 0.274527758 | 0.465474106 | 866.3813432 | 925.3414747 | ENSG00000198843 |
| SKAP2 | 0.095233278 | 1.068238124 | 0.697039232 | 0.827075873 | 76.47745173 | 81.72216953 | ENSG00000005020 |
| NUP155 | 0.095231077 | 1.068236494 | 0.239463771 | 0.425361465 | 1698.963047 | 1814.137888 | ENSG00000113569 |
| EPG5 | 0.095195001 | 1.068209782 | 0.495638052 | 0.677345204 | 419.4349087 | 447.9622879 | ENSG00000152223 |
| CHKB-CPT1B | 0.095191953 | 1.068207526 | 0.915576343 | 0.95827165 | 6.639531009 | 7.179410203 | ENSG00000254413 |
| MARCHF7 | 0.095187902 | 1.068204526 | 0.25532883 | 0.444218955 | 1452.973035 | 1551.67511 | ENSG00000136536 |
| CAB39L | 0.095129803 | 1.068161509 | 0.714382502 | 0.838210008 | 73.03720126 | 77.79816214 | ENSG00000102547 |
| ADSS1 | 0.094908939 | 1.067997995 | 0.529407628 | 0.705242649 | 224.1391187 | 239.2141579 | ENSG00000185100 |
| CUL7 | 0.094736172 | 1.067870107 | 0.463128293 | 0.649713191 | 568.2535007 | 606.9659838 | ENSG00000044090 |
| BTF3L4 | 0.09467973 | 1.06782833 | 0.272088754 | 0.463118099 | 1236.288885 | 1319.574906 | ENSG00000134717 |
| COQ8A | 0.094677821 | 1.067826917 | 0.292872866 | 0.485432453 | 925.4613631 | 988.9746673 | ENSG00000163050 |
| SMAD2 | 0.094669513 | 1.067820768 | 0.30561949 | 0.499634086 | 900.2614023 | 960.6007012 | ENSG00000175387 |
| EFHD1 | 0.094634816 | 1.067795087 | 0.610379406 | 0.767487918 | 142.4132134 | 152.1510532 | ENSG00000115468 |
| PMS2 | 0.094615493 | 1.067780785 | 0.485288485 | 0.668376071 | 274.2449747 | 293.0243508 | ENSG00000122512 |
| SOD1 | 0.094569562 | 1.067746791 | 0.142872939 | 0.29539934 | 3262.178673 | 3483.617087 | ENSG00000142168 |
| NDUFA3 | 0.094514676 | 1.06770617 | 0.403263119 | 0.595022582 | 832.2580595 | 888.9487894 | ENSG00000170906 |
| INF2 | 0.09441718 | 1.067634018 | 0.179324243 | 0.34853141 | 1742.537323 | 1860.759227 | ENSG00000203485 |
| SRBD1 | 0.09440577 | 1.067625575 | 0.357881152 | 0.551277366 | 545.0314565 | 581.5850731 | ENSG00000068784 |
| IFNAR1 | 0.094192407 | 1.067467693 | 0.549567278 | 0.720961784 | 217.8991774 | 232.5557722 | ENSG00000142166 |
| CTSO | 0.094118394 | 1.067412931 | 0.843327119 | 0.916298794 | 19.79657845 | 21.16226107 | ENSG00000256043 |
| RSL24D1 | 0.094094205 | 1.067395034 | 0.37229266 | 0.565486982 | 2532.234842 | 2701.868096 | ENSG00000137876 |
| ZNF524 | 0.094047021 | 1.067360125 | 0.790766466 | 0.886205447 | 43.45547879 | 46.30297876 | ENSG00000171443 |
| ENAH | 0.094040059 | 1.067354975 | 0.129287218 | 0.274291357 | 6239.429199 | 6658.754594 | ENSG00000154380 |
| ABHD18 | 0.094012432 | 1.067334535 | 0.555591168 | 0.725317088 | 222.1430118 | 236.8012681 | ENSG00000164074 |
| PPP1R11 | 0.093934543 | 1.067276913 | 0.383562076 | 0.576294675 | 464.3171606 | 495.5206384 | ENSG00000204619 |
| NEK2 | 0.09385388 | 1.067217242 | 0.392598565 | 0.585250687 | 561.6945385 | 598.8350497 | ENSG00000117650 |
| JDP2 | 0.093840174 | 1.067207103 | 0.315718205 | 0.509182144 | 667.0989694 | 711.8816402 | ENSG00000140044 |
| TNRC6A | 0.093818365 | 1.06719097 | 0.219185638 | 0.400590338 | 2242.318334 | 2393.353738 | ENSG00000090905 |
| NA | 0.093799324 | 1.067176886 | 0.925525294 | 0.963290836 | 4.49605658 | 4.770972092 | ENSG00000270060 |
| RD3 | 0.09376034 | 1.067148049 | 0.933002229 | 0.967187875 | 3.510281278 | 3.724462875 | ENSG00000198570 |
| GTF2H2 | 0.093733486 | 1.067128185 | 0.506906532 | 0.686468338 | 278.6247292 | 297.3218027 | ENSG00000145736 |
| ATF4 | 0.093627355 | 1.067049686 | 0.085416806 | 0.201218951 | 10040.87049 | 10714.23122 | ENSG00000128272 |
| MRPL19 | 0.093340412 | 1.066837477 | 0.475096436 | 0.659978814 | 714.6514036 | 761.8082184 | ENSG00000115364 |
| RPL15 | 0.093288884 | 1.066799374 | 0.069918037 | 0.172781452 | 18349.06931 | 19573.86426 | ENSG00000174748 |
| AHCTF1 | 0.093254125 | 1.066773672 | 0.220563104 | 0.402037429 | 1465.528939 | 1562.892012 | ENSG00000153207 |
| VPS41 | 0.093191018 | 1.06672701 | 0.295202878 | 0.48807227 | 854.4500489 | 911.3566972 | ENSG00000006715 |
| MAD2L1 | 0.093095095 | 1.066656087 | 0.32497585 | 0.519098716 | 2170.324371 | 2313.791442 | ENSG00000164109 |
| FAM83H-AS1 | 0.093032912 | 1.066610113 | 0.770722443 | 0.873834328 | 46.96703784 | 50.23122303 | ENSG00000203499 |
| PRKD2 | 0.093026414 | 1.066605309 | 0.285858703 | 0.477177808 | 1154.709227 | 1231.460872 | ENSG00000105287 |
| ARFGAP1 | 0.092923869 | 1.066529498 | 0.310771088 | 0.50467286 | 2808.709152 | 2996.686882 | ENSG00000101199 |
| NA | 0.0927818 | 1.066424477 | 0.918334501 | 0.959665811 | 5.42560881 | 5.753570632 | ENSG00000217241 |
| C1orf115 | 0.092690391 | 1.066356911 | 0.469626351 | 0.655376462 | 364.554053 | 388.5218941 | ENSG00000162817 |
| RPS24 | 0.092562215 | 1.066262175 | 0.102244 | 0.230478022 | 10819.98746 | 11536.77292 | ENSG00000138326 |
| KDM5B | 0.092426914 | 1.066162182 | 0.189498876 | 0.361715592 | 4816.25185 | 5133.855406 | ENSG00000117139 |
| POLR2D | 0.092422384 | 1.066158835 | 0.257524487 | 0.446154312 | 1355.305853 | 1445.273645 | ENSG00000144231 |
| NA | 0.092413459 | 1.066152239 | 0.875168792 | 0.935364718 | 14.05059585 | 15.03348534 | ENSG00000277142 |
| DNAJB2 | 0.092309177 | 1.066075177 | 0.285104125 | 0.476304295 | 1040.16415 | 1109.06141 | ENSG00000135924 |
| NA | 0.092306889 | 1.066073486 | 0.80063767 | 0.8929372 | 44.03792875 | 46.96062483 | ENSG00000226562 |
| COL18A1 | 0.092270577 | 1.066046654 | 0.240762172 | 0.426972165 | 7150.206495 | 7622.581809 | ENSG00000182871 |
| NA | 0.092201137 | 1.065995344 | 0.852886261 | 0.922280546 | 28.05256023 | 30.04874934 | ENSG00000246596 |
| ACP4 | 0.092151288 | 1.065958512 | 0.92290134 | 0.961876704 | 4.839806533 | 5.127294571 | ENSG00000142513 |
| FAM53A | 0.092020739 | 1.065862058 | 0.780924755 | 0.879590362 | 48.29784087 | 51.69829182 | ENSG00000174137 |
| FBL | 0.092016273 | 1.065858759 | 0.104160617 | 0.233774869 | 11285.11151 | 12028.82682 | ENSG00000105202 |
| ELL | 0.091836574 | 1.065726006 | 0.441954793 | 0.630236988 | 470.9386219 | 502.3528527 | ENSG00000105656 |
| WSB2 | 0.09176912 | 1.065676178 | 0.288943228 | 0.480638465 | 936.5511231 | 997.634326 | ENSG00000176871 |
| NA | 0.091726939 | 1.065645021 | 0.91202309 | 0.956299421 | 6.437917191 | 6.818850843 | ENSG00000141028 |
| SOCS7 | 0.091724145 | 1.065642957 | 0.216631221 | 0.397254684 | 1490.085605 | 1587.700452 | ENSG00000274211 |
| RPTOR | 0.091722439 | 1.065641697 | 0.278709533 | 0.469845709 | 968.1431283 | 1031.464289 | ENSG00000141564 |
| OPHN1 | 0.091680367 | 1.065610621 | 0.672648242 | 0.809789244 | 104.527261 | 111.305108 | ENSG00000079482 |
| TMEM14C | 0.091602048 | 1.065552774 | 0.281867284 | 0.473069106 | 1634.311937 | 1740.401964 | ENSG00000111843 |
| SGK3 | 0.091448847 | 1.065439629 | 0.599243194 | 0.75904684 | 170.6597959 | 182.1134336 | ENSG00000104205 |
| THTPA | 0.091334123 | 1.065354907 | 0.620883354 | 0.774962921 | 156.9151647 | 167.4966058 | ENSG00000259431 |
| CHTF18 | 0.091329508 | 1.0653515 | 0.766536726 | 0.870835943 | 2015.497698 | 2147.454598 | ENSG00000127586 |
| USP13 | 0.091092419 | 1.065176436 | 0.407642756 | 0.599385231 | 644.2617529 | 686.1751476 | ENSG00000058056 |
| D2HGDH | 0.091038766 | 1.065136824 | 0.595536491 | 0.755953244 | 195.1110507 | 207.9221031 | ENSG00000180902 |
| SMC1A | 0.090966927 | 1.065083786 | 0.159738787 | 0.319926375 | 4414.86868 | 4701.364222 | ENSG00000072501 |
| NA | 0.090928147 | 1.065055157 | 0.858527621 | 0.925221009 | 20.97143761 | 22.45235518 | ENSG00000229344 |
| MFAP4 | 0.090894757 | 1.065030508 | 0.721758357 | 0.842703123 | 74.96081945 | 80.03528817 | ENSG00000166482 |
| TUT1 | 0.090819095 | 1.064974654 | 0.577938388 | 0.742308391 | 194.0627385 | 206.7964878 | ENSG00000149016 |
| NA | 0.090811011 | 1.064968687 | 0.931943322 | 0.96667654 | 3.850874318 | 4.073299281 | ENSG00000260593 |
| RNASEH2C | 0.090799997 | 1.064960556 | 0.367504875 | 0.560603733 | 585.966864 | 623.7214473 | ENSG00000172922 |
| RABL2B | 0.090722067 | 1.064903032 | 0.535168654 | 0.709650959 | 269.1173482 | 286.6336572 | ENSG00000079974 |
| FAM107B | 0.090652288 | 1.064851527 | 0.392064395 | 0.584642489 | 586.5211198 | 624.3662711 | ENSG00000065809 |

| | | | | | | | |
|---|---|---|---|---|---|---|---|
| NA | 0.090624226 | 1.064830814 | 0.756946294 | 0.864633317 | 59.83330379 | 63.50021801 | ENSG00000213300 |
| NSMCE2 | 0.090616357 | 1.064825006 | 0.51775398 | 0.695164609 | 260.4040879 | 277.2737603 | ENSG00000156831 |
| TYMSOS | 0.090587978 | 1.064804061 | 0.836386128 | 0.9121504 | 29.71202771 | 31.59251079 | ENSG00000176912 |
| HEMK1 | 0.090502195 | 1.064740749 | 0.622416855 | 0.775781707 | 164.3336258 | 175.2200485 | ENSG00000114735 |
| TRIM37 | 0.090384777 | 1.064654096 | 0.219166423 | 0.400590338 | 1602.45548 | 1705.540933 | ENSG00000108395 |
| KCMF1 | 0.090354591 | 1.06463182 | 0.233515774 | 0.418414015 | 2048.387381 | 2180.339031 | ENSG00000176407 |
| MYRF | 0.09033343 | 1.064616204 | 0.551724712 | 0.722610827 | 255.2803174 | 271.77236 | ENSG00000124920 |
| ADNP2 | 0.09032439 | 1.064609533 | 0.247753521 | 0.435784736 | 1351.471733 | 1438.270843 | ENSG00000101544 |
| ZNF131 | 0.090225618 | 1.064536648 | 0.409606151 | 0.601175198 | 579.3630285 | 616.4585909 | ENSG00000172262 |
| P3H1 | 0.090202684 | 1.064519727 | 0.252596285 | 0.44125063 | 1504.876105 | 1602.804387 | ENSG00000117385 |
| LINC01670 | 0.090192265 | 1.064512038 | 0.86800072 | 0.931260512 | 17.65310402 | 18.80688675 | ENSG00000279094 |
| ZFP82 | 0.090140435 | 1.064473796 | 0.483650565 | 0.667012395 | 385.3134214 | 409.6730532 | ENSG00000181007 |
| DDHD1 | 0.090080178 | 1.064429337 | 0.46035345 | 0.647015306 | 393.1786078 | 418.2387143 | ENSG00000100523 |
| VWCE | 0.090022347 | 1.064386669 | 0.587574643 | 0.750214139 | 274.2007029 | 292.1220692 | ENSG00000167992 |
| RPS9 | 0.089989195 | 1.064362211 | 0.192360077 | 0.36566171 | 28648.49688 | 30492.16731 | ENSG00000170889 |
| ADK | 0.089955341 | 1.064337235 | 0.400794173 | 0.592653519 | 487.2958595 | 518.7097457 | ENSG00000156110 |
| NECTIN2 | 0.089885106 | 1.064285421 | 0.170238357 | 0.335302626 | 4159.321175 | 4426.6817 | ENSG00000130202 |
| IRGQ | 0.089799662 | 1.06422239 | 0.474577844 | 0.65965395 | 405.9370577 | 432.4020525 | ENSG00000167378 |
| ZBTB25 | 0.089737444 | 1.064176496 | 0.691885328 | 0.823649732 | 115.577612 | 123.0306764 | ENSG00000089775 |
| BORCS5 | 0.089679244 | 1.064133566 | 0.587276516 | 0.749936816 | 177.5679123 | 188.9986799 | ENSG00000165714 |
| HELQ | 0.089407102 | 1.063932853 | 0.685561659 | 0.81901507 | 106.7263572 | 113.3976884 | ENSG00000163312 |
| BECN1 | 0.08939506 | 1.063923972 | 0.362145432 | 0.555144293 | 634.1582795 | 674.6823659 | ENSG00000126581 |
| FBXO30 | 0.08918495 | 1.063769037 | 0.341009408 | 0.53542778 | 943.300176 | 1003.967155 | ENSG00000118496 |
| DNM2 | 0.089100346 | 1.063706656 | 0.127214985 | 0.270856121 | 4174.07023 | 4440.297414 | ENSG00000079805 |
| RNASEH1-AS1 | 0.089063951 | 1.063679822 | 0.49487698 | 0.676904055 | 350.0761391 | 372.3177738 | ENSG00000234171 |
| KCNK10 | 0.08893214 | 1.063582644 | 0.814346639 | 0.899952335 | 32.01657975 | 34.10608879 | ENSG00000100433 |
| MTRF1L | 0.088923154 | 1.063576019 | 0.518991046 | 0.696119558 | 283.2043387 | 300.9581445 | ENSG00000112031 |
| ANKFY1 | 0.088738764 | 1.063440093 | 0.322058583 | 0.516084538 | 1323.762509 | 1407.510133 | ENSG00000185722 |
| DAG1 | 0.088556779 | 1.063305957 | 0.228986135 | 0.412805912 | 2732.387787 | 2904.580582 | ENSG00000173402 |
| MID1 | 0.088459113 | 1.063233976 | 0.279368255 | 0.470368027 | 1205.126437 | 1280.607056 | ENSG00000101871 |
| GALNT13 | 0.088444471 | 1.063223186 | 0.599773827 | 0.759411523 | 207.1881191 | 219.9329865 | ENSG00000144278 |
| UBL7-AS1 | 0.088382311 | 1.063177377 | 0.767474394 | 0.871585629 | 60.24284525 | 64.1432268 | ENSG00000247240 |
| RYBP | 0.088358013 | 1.063159471 | 0.463525216 | 0.649994428 | 1035.192986 | 1099.973169 | ENSG00000163602 |
| PLEKHA5 | 0.088239703 | 1.063072289 | 0.301123533 | 0.494596204 | 1196.346611 | 1271.211258 | ENSG00000052126 |
| SEC22B | 0.088205926 | 1.0630474 | 0.371207003 | 0.564401422 | 694.4252976 | 738.1443807 | ENSG00000265808 |
| FOXK2 | 0.088183689 | 1.063031015 | 0.242308013 | 0.428688116 | 1473.077247 | 1566.043638 | ENSG00000141568 |
| TWSG1 | 0.08818264 | 1.063030241 | 0.456770745 | 0.643894208 | 570.0814867 | 605.560993 | ENSG00000128791 |
| RPS6KC1 | 0.088151671 | 1.063007423 | 0.461325727 | 0.647890508 | 452.5201403 | 480.5462772 | ENSG00000136643 |
| DR1 | 0.088049557 | 1.062932186 | 0.395308212 | 0.587768127 | 1027.333136 | 1091.166042 | ENSG00000117505 |
| PPFIBP1 | 0.088032439 | 1.062919574 | 0.411151393 | 0.602690656 | 522.1003969 | 554.5418581 | ENSG00000110841 |
| MICAL3 | 0.088005083 | 1.062899419 | 0.359936792 | 0.553202463 | 840.2237936 | 893.5965663 | ENSG00000243156 |
| MAPK13 | 0.08797659 | 1.062878427 | 0.541746055 | 0.714559918 | 291.8731317 | 310.2940861 | ENSG00000156711 |
| FSD1L | 0.087973546 | 1.062876185 | 0.806110152 | 0.896221316 | 39.72404085 | 42.33200821 | ENSG00000106701 |
| ANKRD49 | 0.087789596 | 1.062740672 | 0.737965341 | 0.853187551 | 69.07713541 | 73.59109277 | ENSG00000168876 |
| NOC3L | 0.087783969 | 1.062736527 | 0.432913617 | 0.622850174 | 523.0968078 | 555.6921931 | ENSG00000173145 |
| SLC25A24 | 0.087745219 | 1.062707983 | 0.494409739 | 0.676364785 | 511.6400723 | 543.085765 | ENSG00000085491 |
| NA | 0.087726298 | 1.062694045 | 0.832032862 | 0.909480985 | 33.19006344 | 35.30221631 | ENSG00000165121 |
| GUF1 | 0.08763666 | 1.06262802 | 0.468192419 | 0.654064845 | 620.8758726 | 659.1874408 | ENSG00000151806 |
| BTAF1 | 0.087581359 | 1.062587288 | 0.447123333 | 0.635261315 | 1251.958156 | 1331.122046 | ENSG00000095564 |
| PPP2R3B | 0.087550134 | 1.06256429 | 0.326873811 | 0.521187132 | 891.2319923 | 947.3367648 | ENSG00000167393 |
| NA | 0.087528602 | 1.062548432 | 0.892059514 | 0.944983483 | 15.55708098 | 16.63542352 | ENSG00000256673 |
| C2orf27A | 0.087354799 | 1.062420433 | 0.794750591 | 0.888735818 | 46.47671706 | 49.32217113 | ENSG00000287151 |
| SKI | 0.087235698 | 1.06233273 | 0.209330699 | 0.388490944 | 2165.719228 | 2300.729138 | ENSG00000157933 |
| TARS2 | 0.08714975 | 1.062269443 | 0.277323683 | 0.468457633 | 1164.108898 | 1236.227218 | ENSG00000143374 |
| YIF1B | 0.087100541 | 1.062233211 | 0.247595115 | 0.435547443 | 1431.461285 | 1520.562475 | ENSG00000167645 |
| PLIN2 | 0.086823802 | 1.062029472 | 0.359131723 | 0.552331537 | 702.3494581 | 745.5132211 | ENSG00000147872 |
| CAMKK2 | 0.086768952 | 1.061989095 | 0.300471281 | 0.493745352 | 1078.352444 | 1144.904369 | ENSG00000110931 |
| ARHGEF11 | 0.086689258 | 1.061930432 | 0.360838647 | 0.554034205 | 945.8893464 | 1004.854608 | ENSG00000132694 |
| EBLN3P | 0.086584663 | 1.061853446 | 0.357484336 | 0.550895043 | 798.9948385 | 848.7806149 | ENSG00000281649 |
| S1PR3 | 0.086573502 | 1.061845231 | 0.539483247 | 0.712722396 | 258.8103143 | 274.5423049 | ENSG00000213694 |
| RBPMS | 0.086552187 | 1.061829543 | 0.206551635 | 0.385152757 | 2143.832783 | 2276.418907 | ENSG00000157110 |
| OSBPL9 | 0.086551924 | 1.061829349 | 0.304710623 | 0.498588036 | 1312.396234 | 1393.413302 | ENSG00000117859 |
| TECPR1 | 0.086443166 | 1.061749306 | 0.478539739 | 0.662527492 | 368.5567296 | 391.4795976 | ENSG00000205356 |
| LOC107987373 | 0.086435453 | 1.06174363 | 0.64410777 | 0.79190006 | 148.7100769 | 157.9808071 | ENSG00000276345 |
| RNF168 | 0.086402776 | 1.061719582 | 0.286192349 | 0.477519715 | 1153.155479 | 1223.73418 | ENSG00000163961 |
| CLASP2 | 0.086375256 | 1.061699329 | 0.34089346 | 0.535297754 | 1308.475441 | 1388.572702 | ENSG00000163539 |
| DCPS | 0.086360436 | 1.061688423 | 0.452466335 | 0.639762821 | 466.7946211 | 495.0270331 | ENSG00000110063 |
| MOB1A | 0.086112117 | 1.061505699 | 0.31961493 | 0.51336128 | 1496.356408 | 1587.724 | ENSG00000114978 |
| SLC38A2 | 0.08605655 | 1.061464815 | 0.261763644 | 0.451053122 | 6307.616334 | 6693.930493 | ENSG00000134294 |
| ZNF496 | 0.086041977 | 1.061454093 | 0.183837193 | 0.354403841 | 2374.746842 | 2520.643099 | ENSG00000162714 |
| NA | 0.085941748 | 1.061380353 | 0.631704216 | 0.782745522 | 152.8732297 | 162.2973649 | ENSG00000174028 |
| MZT1 | 0.08591156 | 1.061358144 | 0.544532366 | 0.716707097 | 576.6708229 | 611.5733036 | ENSG00000204899 |
| TMEM248 | 0.085771553 | 1.061255148 | 0.330858166 | 0.525190583 | 1102.733616 | 1170.091446 | ENSG00000106609 |
| TOP2A | 0.085737569 | 1.06123015 | 0.225050421 | 0.407850249 | 7500.350989 | 7958.138995 | ENSG00000131747 |

| | | | | | | | |
|---|---|---|---|---|---|---|---|
| CPT2 | 0.085701212 | 1.061203406 | 0.550152021 | 0.721465527 | 255.5309308 | 270.9785304 | ENSG00000157184 |
| NA | 0.085675734 | 1.061184666 | 0.861776721 | 0.927428275 | 28.52198534 | 30.38250198 | ENSG00000223703 |
| DDIAS | 0.085628271 | 1.061149754 | 0.508657993 | 0.687784872 | 367.1592481 | 389.318503 | ENSG00000165490 |
| CHFR | 0.085624329 | 1.061146855 | 0.290986714 | 0.483343589 | 1032.749079 | 1096.209184 | ENSG00000072609 |
| GOLT1B | 0.085529856 | 1.06107737 | 0.791140584 | 0.886400725 | 372.3910301 | 394.9458207 | ENSG00000111711 |
| PTK7 | 0.085516053 | 1.061067218 | 0.134816534 | 0.282689177 | 4735.513329 | 5024.163246 | ENSG00000112655 |
| NA | 0.085503731 | 1.061058155 | 0.094782177 | 0.217520168 | 484409.616 | 513986.6228 | N_multimapping |
| CYYR1 | 0.085416704 | 1.060994152 | 0.433835931 | 0.623649666 | 640.4492199 | 678.7903155 | ENSG00000166265 |
| TNNT1 | 0.085381457 | 1.06096823 | 0.251070141 | 0.439533148 | 1510.042261 | 1602.593566 | ENSG00000105048 |
| ATR | 0.085362646 | 1.060954397 | 0.446679773 | 0.634923147 | 545.8194665 | 578.6273067 | ENSG00000175054 |
| METAP1 | 0.085292679 | 1.060902945 | 0.260845085 | 0.449888591 | 2227.678884 | 2362.671226 | ENSG00000164024 |
| AVPR1B | 0.085266535 | 1.06088372 | 0.896194224 | 0.946766059 | 13.80104962 | 14.76353176 | ENSG00000198049 |
| BPGM | 0.085240998 | 1.060864941 | 0.537042736 | 0.711203176 | 348.1882392 | 369.4513374 | ENSG00000172331 |
| NA | 0.085182891 | 1.060822214 | 0.740809484 | 0.855075943 | 84.64339403 | 89.69679475 | ENSG00000215492 |
| LMBRD2 | 0.085134832 | 1.060786876 | 0.730536961 | 0.849314518 | 80.26757059 | 84.93199463 | ENSG00000164187 |
| ATP6AP2 | 0.08509462 | 1.060757309 | 0.357148202 | 0.550616112 | 1201.274082 | 1273.689197 | ENSG00000182220 |
| C15orf41 | 0.085048993 | 1.060723763 | 0.600291746 | 0.759741754 | 199.7191474 | 211.9947411 | ENSG00000186073 |
| SRGAP2 | 0.08501974 | 1.060702254 | 0.374440776 | 0.567389377 | 844.8443604 | 895.3103971 | ENSG00000266028 |
| LINC00678 | 0.084887248 | 1.060604848 | 0.219258462 | 0.400604922 | 3232.648822 | 3428.023212 | ENSG00000254934 |
| PEX1 | 0.084806874 | 1.060545762 | 0.47124553 | 0.656501394 | 423.6123578 | 448.8515493 | ENSG00000127980 |
| NUDT17 | 0.084798303 | 1.060539462 | 0.839027524 | 0.913662522 | 27.77963784 | 29.51110369 | ENSG00000186364 |
| HBS1L | 0.084795416 | 1.060537339 | 0.287784682 | 0.479308118 | 1318.138819 | 1397.168665 | ENSG00000112339 |
| NCAPD2 | 0.084487953 | 1.060311344 | 0.21847071 | 0.39971729 | 4983.963408 | 5284.556187 | ENSG00000010292 |
| XPNPEP3 | 0.084437307 | 1.060274123 | 0.604786179 | 0.763459136 | 212.5158862 | 225.5588961 | ENSG00000196236 |
| H3C6 | 0.084412408 | 1.060255824 | 0.888146861 | 0.942849507 | 12.31034906 | 13.09030845 | ENSG00000274750 |
| NA | 0.084352608 | 1.060211877 | 0.91780008 | 0.959419441 | 7.109632533 | 7.478981666 | ENSG00000277010 |
| MTOR | 0.084137794 | 1.060054256 | 0.279012615 | 0.470063197 | 3702.110656 | 3924.498086 | ENSG00000198793 |
| FLJ20021 | 0.084134313 | 1.060051468 | 0.800632885 | 0.8929372 | 41.19767908 | 43.74557517 | ENSG00000254531 |
| MZF1 | 0.084118891 | 1.060040137 | 0.712999293 | 0.837541981 | 159.4668051 | 169.5132665 | ENSG00000099326 |
| KDM5C | 0.084110946 | 1.060034299 | 0.132001442 | 0.278551146 | 6111.885875 | 6479.087577 | ENSG00000126012 |
| DTX2 | 0.084092645 | 1.060020852 | 0.387940345 | 0.581110996 | 614.0479342 | 650.8749151 | ENSG00000091073 |
| ZNF768 | 0.084001977 | 1.059954236 | 0.338024872 | 0.532145841 | 936.9310345 | 993.8064125 | ENSG00000169957 |
| CD164 | 0.083982031 | 1.059939581 | 0.381398881 | 0.574329449 | 1507.038558 | 1597.24993 | ENSG00000135535 |
| SLC2A13 | 0.083979472 | 1.059937701 | 0.649356449 | 0.795321709 | 156.0405578 | 165.3303269 | ENSG00000151229 |
| NA | 0.083936598 | 1.059906203 | 0.937172317 | 0.969033102 | 4.188310445 | 4.403364692 | ENSG00000265458 |
| ZNF154 | 0.083854076 | 1.059845578 | 0.56182252 | 0.730570307 | 237.0295576 | 251.1496797 | ENSG00000179909 |
| TFCP2 | 0.083799836 | 1.059805732 | 0.304666129 | 0.498559247 | 1359.053241 | 1440.097658 | ENSG00000135457 |
| ASB6 | 0.083773056 | 1.05978606 | 0.511495728 | 0.690614251 | 419.9118876 | 444.9639132 | ENSG00000148331 |
| NIPBL | 0.083687513 | 1.059723223 | 0.33739457 | 0.531831175 | 1249.094297 | 1323.048066 | ENSG00000164190 |
| TRIAP1 | 0.083680851 | 1.05971833 | 0.297506087 | 0.490493821 | 1061.906077 | 1125.22266 | ENSG00000170855 |
| NA | 0.08366283 | 1.059705093 | 0.901432141 | 0.949917177 | 12.9447828 | 13.63726365 | ENSG00000260747 |
| ALDH3A2 | 0.08360352 | 1.059661528 | 0.242768079 | 0.429237444 | 1913.823593 | 2027.806761 | ENSG00000072210 |
| KCNJ4 | 0.083352532 | 1.059477193 | 0.806289607 | 0.896253917 | 39.98375688 | 42.29347853 | ENSG00000168135 |
| NFX1 | 0.083344765 | 1.05947149 | 0.330612058 | 0.524947056 | 934.5966349 | 990.1732796 | ENSG00000086102 |
| XPNPEP1 | 0.083319838 | 1.059453184 | 0.343425359 | 0.537179981 | 901.2629621 | 954.5096278 | ENSG00000108039 |
| COG3 | 0.083245869 | 1.059398866 | 0.495837392 | 0.67751771 | 369.9996932 | 392.073954 | ENSG00000136152 |
| RPS6KB1 | 0.083239011 | 1.059393829 | 0.548728598 | 0.720264578 | 425.8466837 | 450.3632795 | ENSG00000108443 |
| HTRA2 | 0.08320552 | 1.059369237 | 0.538285712 | 0.711948165 | 311.1892282 | 329.6231596 | ENSG00000115317 |
| NA | 0.0831806 | 1.059350938 | 0.883482892 | 0.939962378 | 17.06927858 | 18.18429791 | ENSG00000234405 |
| CNST | 0.082977885 | 1.059202098 | 0.47885195 | 0.662811206 | 409.2997835 | 433.6662873 | ENSG00000162852 |
| SLC35F5 | 0.082877823 | 1.059128637 | 0.530858836 | 0.706211432 | 391.2476161 | 413.8968553 | ENSG00000115084 |
| INPP5J | 0.082841905 | 1.059102269 | 0.681587717 | 0.816108711 | 123.5557556 | 131.0039875 | ENSG00000185133 |
| EIF3H | 0.082814357 | 1.059082045 | 0.17967885 | 0.348999285 | 3586.105478 | 3797.196806 | ENSG00000147677 |
| MAPKAPK5-AS1 | 0.082798253 | 1.059070224 | 0.475270389 | 0.660072042 | 530.0649208 | 561.7158063 | ENSG00000234608 |
| SPATS2 | 0.082757797 | 1.059040526 | 0.380383295 | 0.573445561 | 713.9252618 | 755.9550193 | ENSG00000123352 |
| BLVRA | 0.082679681 | 1.058983185 | 0.428326212 | 0.618608121 | 589.9032606 | 624.2093815 | ENSG00000106605 |
| SLIT1 | 0.082638226 | 1.058952755 | 0.839353145 | 0.913691856 | 32.12280977 | 34.12117256 | ENSG00000187122 |
| TMEM254 | 0.082505502 | 1.058855339 | 0.64802343 | 0.794567224 | 168.409415 | 178.1280003 | ENSG00000133678 |
| RBBP8 | 0.082488724 | 1.058843025 | 0.384966151 | 0.577916949 | 758.8085209 | 803.3145897 | ENSG00000101773 |
| RPL7 | 0.082365872 | 1.058752864 | 0.129538749 | 0.274699219 | 26547.66866 | 28107.046 | ENSG00000147604 |
| STX5 | 0.082176077 | 1.058613588 | 0.472907725 | 0.658072811 | 462.3021873 | 489.5082875 | ENSG00000162236 |
| PRICKLE2 | 0.081936391 | 1.058437727 | 0.793669847 | 0.888170326 | 55.95095804 | 59.03641508 | ENSG00000163637 |
| TARBP1 | 0.081917773 | 1.058424068 | 0.50219883 | 0.682636825 | 516.1344603 | 546.5812032 | ENSG00000059588 |
| WWC3 | 0.081876661 | 1.058393907 | 0.337676852 | 0.531944773 | 1161.509828 | 1228.714771 | ENSG00000047644 |
| OXSR1 | 0.081872351 | 1.058390745 | 0.296618602 | 0.489542614 | 1494.245044 | 1580.641073 | ENSG00000172939 |
| IVD | 0.081857267 | 1.058379679 | 0.242276935 | 0.428674053 | 2103.214412 | 2226.496208 | ENSG00000128928 |
| IMMT | 0.081852955 | 1.058376515 | 0.219990084 | 0.401506274 | 2439.720302 | 2581.556653 | ENSG00000132305 |
| WASHC3 | 0.081843648 | 1.058369688 | 0.764892924 | 0.870029457 | 60.99741444 | 64.56224903 | ENSG00000120860 |
| NLRX1 | 0.08183072 | 1.058360204 | 0.444557277 | 0.632925532 | 538.5462083 | 569.8532296 | ENSG00000160703 |
| SORD2P | 0.081590998 | 1.058184358 | 0.767034851 | 0.871187027 | 62.54689363 | 66.29949464 | ENSG00000259479 |
| NA | 0.081477285 | 1.058100956 | 0.932715087 | 0.967047518 | 5.097643422 | 5.348421244 | ENSG00000253919 |
| TMEM129 | 0.081462412 | 1.058090048 | 0.384723219 | 0.57759904 | 1068.922536 | 1131.46752 | ENSG00000168936 |
| ZNF317 | 0.081391777 | 1.058038245 | 0.348784931 | 0.542131827 | 853.7073332 | 903.2826979 | ENSG00000130803 |
| DBF4B | 0.081334966 | 1.057996582 | 0.413112496 | 0.604535668 | 867.0386123 | 917.7950873 | ENSG00000161692 |

| | | | | | | | |
|---|---|---|---|---|---|---|---|
| SERINC1 | 0.081288242 | 1.057962317 | 0.436861423 | 0.625959769 | 758.7222618 | 802.7871297 | ENSG00000111897 |
| GLI3 | 0.081263425 | 1.057944119 | 0.472455371 | 0.657542077 | 439.929044 | 465.3980076 | ENSG00000106571 |
| DYDC2 | 0.081247077 | 1.05793213 | 0.789704229 | 0.885282657 | 64.62457655 | 68.13566788 | ENSG00000133665 |
| LOC220729 | 0.081178549 | 1.05788188 | 0.614538979 | 0.770209997 | 297.7529824 | 315.1374426 | ENSG00000214135 |
| NA | 0.081118874 | 1.057838123 | 0.903162983 | 0.950822081 | 11.26066138 | 11.81843568 | ENSG00000281344 |
| RPS11 | 0.081078588 | 1.057808584 | 0.303887851 | 0.497549243 | 23157.04588 | 24495.49508 | ENSG00000142534 |
| CARD19 | 0.081063003 | 1.057797157 | 0.514915296 | 0.692859017 | 376.1049922 | 398.1265867 | ENSG00000165233 |
| RAB10 | 0.081034407 | 1.057776191 | 0.254304437 | 0.44298754 | 2969.630195 | 3140.609262 | ENSG00000084733 |
| GABPB2 | 0.080979818 | 1.057736167 | 0.805001572 | 0.895390934 | 44.91391116 | 47.3719462 | ENSG00000143458 |
| CWF19L2 | 0.080939977 | 1.057706957 | 0.487161143 | 0.670208181 | 411.4347718 | 434.9972842 | ENSG00000152404 |
| NA | 0.08092402 | 1.057695258 | 0.751095543 | 0.86148925 | 106.7432467 | 112.8924004 | ENSG00000253507 |
| DHRS4L2 | 0.08083413 | 1.057629358 | 0.772178116 | 0.874575239 | 70.48447846 | 74.78788156 | ENSG00000187630 |
| RAB6A | 0.080792622 | 1.057598929 | 0.459233593 | 0.646127338 | 1091.863592 | 1154.170728 | ENSG00000175582 |
| MRPL35 | 0.080726382 | 1.057550372 | 0.415237437 | 0.606606026 | 691.4346962 | 730.8091484 | ENSG00000132313 |
| NA | 0.080637621 | 1.057485309 | 0.799012428 | 0.89199472 | 47.61655709 | 50.32452835 | ENSG00000228232 |
| CST3 | 0.080563861 | 1.057431245 | 0.535778535 | 0.710188819 | 341.4372693 | 361.1291898 | ENSG00000101439 |
| PDLIM7 | 0.080350015 | 1.057274517 | 0.370790518 | 0.564129983 | 1493.312809 | 1580.010667 | ENSG00000196923 |
| TUBB4B | 0.08011537 | 1.057102572 | 0.165282705 | 0.328122286 | 6922.856903 | 7317.978195 | ENSG00000188229 |
| PTPN18 | 0.080027356 | 1.057038084 | 0.525969647 | 0.702433961 | 339.0193767 | 358.2905562 | ENSG00000072135 |
| GPX7 | 0.079983339 | 1.057005834 | 0.636731124 | 0.786535234 | 174.2604249 | 184.0617948 | ENSG00000116157 |
| COCH | 0.079887284 | 1.056935461 | 0.609188773 | 0.766741215 | 221.1596944 | 233.8143218 | ENSG00000100473 |
| MIA3 | 0.079749506 | 1.056834527 | 0.352873245 | 0.54602018 | 962.5833884 | 1017.122591 | ENSG00000154305 |
| POMGNT1 | 0.079721933 | 1.056814329 | 0.323897393 | 0.51800106 | 1085.423728 | 1147.099765 | ENSG00000085998 |
| ARID2 | 0.07939016 | 1.056571324 | 0.416132899 | 0.607339723 | 730.9175944 | 771.8245737 | ENSG00000189079 |
| NA | 0.079370921 | 1.056557234 | 0.931201886 | 0.966296058 | 5.478674969 | 5.734799638 | ENSG00000213695 |
| NA | 0.079366015 | 1.056553641 | 0.783105339 | 0.88124313 | 60.79086227 | 64.31855252 | ENSG00000256008 |
| ACTRT3 | 0.079344184 | 1.056537654 | 0.788309191 | 0.884326033 | 52.73136055 | 55.75183242 | ENSG00000184378 |
| ZNF572 | 0.079296978 | 1.056503083 | 0.875750722 | 0.935624952 | 18.71729848 | 19.86456926 | ENSG00000180938 |
| GPAT4 | 0.079210424 | 1.056439701 | 0.323746513 | 0.517881475 | 1151.165565 | 1215.867761 | ENSG00000158669 |
| GSTK1 | 0.079144099 | 1.056391134 | 0.565091881 | 0.733022388 | 400.2194123 | 423.3493651 | ENSG00000197448 |
| FAM204A | 0.079143596 | 1.056390766 | 0.667855573 | 0.807175429 | 172.654219 | 182.1589711 | ENSG00000165669 |
| CLDN15 | 0.079088241 | 1.056350234 | 0.776956666 | 0.877253417 | 60.41476907 | 63.91764001 | ENSG00000106404 |
| MAP1S | 0.079069782 | 1.056336718 | 0.295770448 | 0.488644305 | 1857.387071 | 1961.679703 | ENSG00000130479 |
| GCH1 | 0.079028518 | 1.056306505 | 0.638122085 | 0.787571068 | 191.3898576 | 202.4274159 | ENSG00000131979 |
| ARL8B | 0.078987529 | 1.056276495 | 0.471424183 | 0.656550632 | 729.6688559 | 770.1589163 | ENSG00000134108 |
| MKLN1 | 0.078947033 | 1.056246846 | 0.476158163 | 0.660809838 | 593.6110141 | 626.5175291 | ENSG00000128585 |
| LLGL2 | 0.078851238 | 1.056176713 | 0.257854408 | 0.446392079 | 1708.112704 | 1804.09151 | ENSG00000073350 |
| KLC4 | 0.078820873 | 1.056154484 | 0.468589004 | 0.654372258 | 484.1418581 | 511.249097 | ENSG00000137171 |
| DDX10 | 0.078670651 | 1.056044516 | 0.376525701 | 0.569675701 | 1063.172269 | 1122.343178 | ENSG00000178105 |
| ABTB2 | 0.078616873 | 1.056005152 | 0.436657746 | 0.62586147 | 626.6005083 | 661.6649513 | ENSG00000166016 |
| C1GALT1C1 | 0.078579534 | 1.055977821 | 0.526227539 | 0.702433961 | 355.6779337 | 375.6044435 | ENSG00000171155 |
| NA | 0.078577763 | 1.055976525 | 0.843031521 | 0.91613874 | 31.38086823 | 33.22863014 | ENSG00000268858 |
| ZBTB9 | 0.078397306 | 1.055844449 | 0.369050916 | 0.5619446 | 823.3054455 | 869.1903788 | ENSG00000213588 |
| GART | 0.078290535 | 1.05576631 | 0.187775703 | 0.359602695 | 4153.750755 | 4384.596111 | ENSG00000159131 |
| RPS3A | 0.078282761 | 1.055760621 | 0.274595867 | 0.465547 | 5870.20894 | 6196.118688 | ENSG00000145425 |
| NA | 0.0781726 | 1.055680009 | 0.938766357 | 0.96979109 | 4.916248858 | 5.146065566 | ENSG00000223508 |
| FOXO1 | 0.078091372 | 1.055620572 | 0.531087511 | 0.706419294 | 629.7102378 | 664.1189982 | ENSG00000150907 |
| C10orf105 | 0.078031458 | 1.055576734 | 0.920362084 | 0.96068883 | 7.734595541 | 8.086486732 | ENSG00000214688 |
| EP400P1 | 0.077964255 | 1.055527565 | 0.646826422 | 0.793715632 | 306.4852667 | 324.134407 | ENSG00000185684 |
| ECE1 | 0.077934629 | 1.05550589 | 0.278314181 | 0.469483037 | 2011.977608 | 2123.854334 | ENSG00000117298 |
| PITPNB | 0.077900513 | 1.05548093 | 0.379172785 | 0.572325792 | 1769.093274 | 1866.997139 | ENSG00000180957 |
| NA | 0.077796471 | 1.055404815 | 0.823830175 | 0.905349068 | 556.0561089 | 587.0480504 | ENSG00000111788 |
| PDCL | 0.077688644 | 1.055325938 | 0.48574198 | 0.66875218 | 471.4835949 | 497.6750828 | ENSG00000136940 |
| ENO3 | 0.077677282 | 1.055317626 | 0.585069384 | 0.747994479 | 259.2867296 | 273.5987856 | ENSG00000108515 |
| SLC20A2 | 0.077608947 | 1.055267641 | 0.444360898 | 0.632840393 | 698.2272322 | 736.6001409 | ENSG00000168575 |
| TBX19 | 0.077521523 | 1.055203696 | 0.898503643 | 0.948070966 | 12.46511284 | 13.16539243 | ENSG00000143178 |
| PTCH1 | 0.077415246 | 1.055125967 | 0.553074098 | 0.723664454 | 337.5272029 | 355.8027888 | ENSG00000185920 |
| UFSP2 | 0.077346136 | 1.055075423 | 0.548413209 | 0.719971981 | 355.4802509 | 374.8863507 | ENSG00000109775 |
| GOLIM4 | 0.077291874 | 1.055035742 | 0.318026871 | 0.511836577 | 1211.00271 | 1277.578666 | ENSG00000173905 |
| SESTD1 | 0.077273358 | 1.055022201 | 0.511375855 | 0.690531168 | 478.694444 | 504.5228499 | ENSG00000187231 |
| CCDC130 | 0.077151355 | 1.054932985 | 0.577898898 | 0.742308391 | 333.8877063 | 352.8444098 | ENSG00000104957 |
| SEPTIN1 | 0.077049215 | 1.054858301 | 0.90256605 | 0.950449757 | 12.10873524 | 12.78650011 | ENSG00000180096 |
| GLP1R | 0.07698345 | 1.054810217 | 0.754323167 | 0.863474302 | 82.72756284 | 87.22510723 | ENSG00000112164 |
| SLMAP | 0.07693888 | 1.054777631 | 0.3882479 | 0.581249696 | 931.0958842 | 981.8119093 | ENSG00000163681 |
| UHRF1BP1L | 0.076827044 | 1.054695869 | 0.532232583 | 0.707378895 | 433.7834193 | 457.3412503 | ENSG00000111647 |
| NA | 0.076779015 | 1.054660757 | 0.895775877 | 0.946647856 | 13.50079739 | 14.23447149 | ENSG00000268575 |
| CAMSAP2 | 0.076761313 | 1.054647816 | 0.486645867 | 0.669847345 | 652.5212522 | 687.515016 | ENSG00000118200 |
| LINC00294 | 0.076734113 | 1.054627932 | 0.734124329 | 0.851336652 | 94.21203413 | 99.47158856 | ENSG00000280798 |
| ACO2 | 0.076683382 | 1.054590848 | 0.274198541 | 0.465250018 | 1673.721174 | 1765.041926 | ENSG00000100412 |
| SLC3A2 | 0.076613204 | 1.05453955 | 0.154589497 | 0.312347296 | 8964.225331 | 9452.700886 | ENSG00000168003 |
| TALDO1 | 0.076592116 | 1.054524136 | 0.204540804 | 0.382541509 | 4997.820777 | 5269.895298 | ENSG00000177156 |
| NA | 0.07650971 | 1.054463904 | 0.925122668 | 0.963054568 | 8.573800019 | 9.077120999 | ENSG00000284968 |
| NFRKB | 0.076302324 | 1.054312336 | 0.422162727 | 0.613386954 | 890.5369235 | 939.5404199 | ENSG00000170322 |
| MYOZ1 | 0.076300195 | 1.054310781 | 0.889001106 | 0.943307179 | 15.92676265 | 16.76214557 | ENSG00000177791 |

| | | | | | | | |
|---|---|---|---|---|---|---|---|
| C6orf47 | 0.076264358 | 1.054284591 | 0.629311883 | 0.781223472 | 218.3776446 | 230.5738912 | ENSG00000204439 |
| NA | 0.076214841 | 1.054248406 | 0.919516316 | 0.96040098 | 8.290707826 | 8.788396423 | ENSG00000179131 |
| SORBS1 | 0.076182151 | 1.054224519 | 0.466198366 | 0.652459173 | 660.597155 | 695.8721579 | ENSG00000095637 |
| PKIB | 0.076072313 | 1.054144259 | 0.625619713 | 0.778047089 | 364.6878833 | 383.9716904 | ENSG00000135549 |
| C3orf52 | 0.07589165 | 1.054012261 | 0.577453402 | 0.742035609 | 281.0454856 | 295.9897149 | ENSG00000114529 |
| SIN3B | 0.07588701 | 1.054008872 | 0.324689389 | 0.518819996 | 2121.731284 | 2237.167535 | ENSG00000127511 |
| PAFAH2 | 0.075774605 | 1.053926753 | 0.624163567 | 0.77710194 | 221.308648 | 232.9728678 | ENSG00000158006 |
| TLE5 | 0.07577315 | 1.05392569 | 0.289396596 | 0.481263033 | 3058.779448 | 3224.171376 | ENSG00000104964 |
| KCNJ14 | 0.075712157 | 1.053881135 | 0.928089212 | 0.964548443 | 7.000343303 | 7.434391636 | ENSG00000182324 |
| TDGF1 | 0.07571164 | 1.053880757 | 0.207951157 | 0.386964532 | 5441.233059 | 5733.636766 | ENSG00000241186 |
| TMEM170B | 0.075608585 | 1.053805479 | 0.675842709 | 0.812484191 | 184.5448794 | 194.1272483 | ENSG00000205269 |
| ZCCHC9 | 0.075571202 | 1.053778173 | 0.459285555 | 0.646150409 | 547.0756912 | 576.5953877 | ENSG00000131732 |
| ZFR | 0.075435208 | 1.053678844 | 0.331108992 | 0.525273821 | 2079.967345 | 2191.195471 | ENSG00000056097 |
| NA | 0.075341127 | 1.053610134 | 0.890432027 | 0.944068828 | 16.93779326 | 17.92931648 | ENSG00000234287 |
| PDE6B | 0.075107365 | 1.05343943 | 0.69087172 | 0.822760165 | 142.1052719 | 149.6573455 | ENSG00000133256 |
| CDC73 | 0.075099284 | 1.053433529 | 0.34153226 | 0.53583425 | 1105.040581 | 1163.907886 | ENSG00000134371 |
| LOC105372321 | 0.075070556 | 1.053412553 | 0.611480206 | 0.768345443 | 266.691676 | 281.2028891 | ENSG00000268119 |
| GARNL3 | 0.075007131 | 1.053366242 | 0.713687477 | 0.837849642 | 122.7726765 | 129.5876094 | ENSG00000136895 |
| RABEPK | 0.074876877 | 1.053271143 | 0.377225997 | 0.570455828 | 935.6553647 | 985.5857806 | ENSG00000136933 |
| DCAF8 | 0.074845522 | 1.053248252 | 0.631211394 | 0.782480902 | 288.949254 | 304.4955905 | ENSG00000132716 |
| MRPL42 | 0.074828043 | 1.053235492 | 0.450377474 | 0.637977385 | 1570.956994 | 1653.663285 | ENSG00000198015 |
| PLIN3 | 0.074729569 | 1.053163604 | 0.292184434 | 0.484702158 | 1627.180431 | 1713.469926 | ENSG00000105355 |
| DNAJA2 | 0.074664775 | 1.053116305 | 0.285383493 | 0.476642125 | 2806.376556 | 2954.585709 | ENSG00000069345 |
| ADO | 0.07461503 | 1.053079994 | 0.479389739 | 0.663418777 | 785.5412971 | 826.5213913 | ENSG00000181915 |
| PIGA | 0.074603898 | 1.053071868 | 0.733512962 | 0.851224605 | 143.0218882 | 150.3176508 | ENSG00000165195 |
| NCOR1 | 0.074492422 | 1.052990501 | 0.276739896 | 0.467727269 | 2513.243006 | 2646.420369 | ENSG00000141027 |
| PROB1 | 0.074485351 | 1.05298534 | 0.676127309 | 0.812720899 | 167.6796725 | 176.6101695 | ENSG00000228672 |
| ZNF407 | 0.074091671 | 1.052698042 | 0.675377285 | 0.812188074 | 198.7865133 | 209.4181285 | ENSG00000215421 |
| ZNF493 | 0.07408974 | 1.052696634 | 0.743510734 | 0.856819495 | 149.3599869 | 156.9151602 | ENSG00000196268 |
| KIAA1958 | 0.074067971 | 1.052680749 | 0.492998269 | 0.675364364 | 617.4403651 | 649.7863953 | ENSG00000165185 |
| PFKM | 0.074044562 | 1.052663669 | 0.280614234 | 0.471994212 | 2625.569887 | 2763.460342 | ENSG00000152556 |
| RHNO1 | 0.074000069 | 1.052631205 | 0.469872444 | 0.655539918 | 612.7407179 | 644.6138579 | ENSG00000171792 |
| HSPA13 | 0.073899309 | 1.05255769 | 0.467193516 | 0.653162956 | 736.2488866 | 774.5420045 | ENSG00000155304 |
| MRPL22 | 0.073889614 | 1.052550617 | 0.410696901 | 0.602251473 | 813.1933958 | 855.8458533 | ENSG00000082515 |
| NA | 0.073881339 | 1.05254458 | 0.830195876 | 0.90859717 | 42.3225313 | 44.73620944 | ENSG00000286431 |
| NA | 0.073712165 | 1.052421163 | 0.871259772 | 0.933190511 | 22.20104668 | 23.38623843 | ENSG00000235238 |
| PCDHGA11 | 0.073672785 | 1.052392436 | 0.841907147 | 0.915346223 | 34.88167877 | 36.72761785 | ENSG00000253873 |
| ZBTB47 | 0.073643702 | 1.052371222 | 0.746892285 | 0.858676729 | 96.21327472 | 101.0040651 | ENSG00000114853 |
| TC2N | 0.073624257 | 1.052357038 | 0.536920712 | 0.711203176 | 390.0519363 | 410.1065466 | ENSG00000165929 |
| B4GALNT3 | 0.073555564 | 1.052306931 | 0.392423786 | 0.585084262 | 1034.266644 | 1088.59902 | ENSG00000139044 |
| NA | 0.073504816 | 1.052269916 | 0.914357802 | 0.9578199 | 10.7046468 | 11.27852852 | ENSG00000224664 |
| ANKRD33B | 0.073499503 | 1.052266042 | 0.564131391 | 0.732480414 | 369.4004665 | 388.3509798 | ENSG00000164236 |
| KDM2A | 0.073400212 | 1.052193623 | 0.329854991 | 0.524185173 | 1574.088304 | 1656.778517 | ENSG00000173120 |
| ARL14EP | 0.073363663 | 1.052166967 | 0.558296862 | 0.727516983 | 444.5749034 | 467.3461539 | ENSG00000152219 |
| ATP8A2 | 0.073259158 | 1.052090754 | 0.576763684 | 0.741570456 | 326.8481045 | 343.9551733 | ENSG00000132932 |
| FBXO33 | 0.073153097 | 1.052013411 | 0.603131234 | 0.762304575 | 271.388264 | 285.5247343 | ENSG00000165355 |
| NA | 0.073084686 | 1.051963527 | 0.873419206 | 0.934693202 | 21.01050065 | 22.13737353 | ENSG00000225630 |
| TANK | 0.073074514 | 1.051956111 | 0.614821489 | 0.770459869 | 254.7311954 | 267.8108106 | ENSG00000136560 |
| UBE3D | 0.073065378 | 1.051949449 | 0.612551838 | 0.768991559 | 265.0699938 | 278.4877828 | ENSG00000118420 |
| ZNF417 | 0.072937577 | 1.051856266 | 0.72675541 | 0.846511109 | 136.4455931 | 143.8592964 | ENSG00000173480 |
| PMM2 | 0.072886737 | 1.0518192 | 0.586395725 | 0.749312221 | 284.7587562 | 299.6364974 | ENSG00000140650 |
| ARMT1 | 0.072860548 | 1.051800106 | 0.48819147 | 0.671127481 | 962.5033454 | 1012.179399 | ENSG00000146476 |
| NUDT2 | 0.072849096 | 1.051791758 | 0.713608899 | 0.837849642 | 190.1586776 | 200.3854145 | ENSG00000164978 |
| MTNR1A | 0.072768642 | 1.051733104 | 0.915668389 | 0.95827165 | 9.960826121 | 10.49872369 | ENSG00000168412 |
| ATP2C1 | 0.072725614 | 1.051701737 | 0.309038049 | 0.503187564 | 1818.769467 | 1912.566472 | ENSG00000017260 |
| NA | 0.072615928 | 1.051621781 | 0.815395225 | 0.90056677 | 57.84428488 | 60.83366089 | ENSG00000268205 |
| NA | 0.072518375 | 1.051550674 | 0.735075248 | 0.85191875 | 102.3511351 | 107.5271433 | ENSG00000272711 |
| SORCS3 | 0.072517727 | 1.051550202 | 0.831548762 | 0.909311745 | 38.90437921 | 40.8648278 | ENSG00000156395 |
| INO80 | 0.072510795 | 1.051545149 | 0.43024289 | 0.620264601 | 833.1639726 | 876.3672958 | ENSG00000128908 |
| VANGL1 | 0.072408319 | 1.051470459 | 0.529559079 | 0.705324695 | 462.2829754 | 485.7516817 | ENSG00000173218 |
| PLAAT5 | 0.072386369 | 1.051454462 | 0.779196131 | 0.878699058 | 69.56608071 | 73.01265593 | ENSG00000168004 |
| RAB3B | 0.072368156 | 1.051441188 | 0.394126251 | 0.586584433 | 1506.343129 | 1584.161635 | ENSG00000169213 |
| NA | 0.072329846 | 1.051413268 | 0.855183229 | 0.923101397 | 34.87832646 | 36.56291578 | ENSG00000288075 |
| CREB1 | 0.072206318 | 1.051323247 | 0.390834061 | 0.583465041 | 1101.907084 | 1157.920804 | ENSG00000118260 |
| SYN1 | 0.072181127 | 1.05130489 | 0.779831573 | 0.878986213 | 85.39627948 | 89.52967927 | ENSG00000008056 |
| NA | 0.072158693 | 1.051288542 | 0.633600117 | 0.783961787 | 217.0440906 | 228.0855825 | ENSG00000224578 |
| UFM1 | 0.072156866 | 1.051287211 | 0.543773624 | 0.716144415 | 480.0727961 | 505.0730402 | ENSG00000120686 |
| AGER | 0.072146328 | 1.051279532 | 0.763561205 | 0.868904241 | 107.5885545 | 113.459226 | ENSG00000204305 |
| IP6K2 | 0.072117979 | 1.051258874 | 0.389155671 | 0.581802953 | 1303.733627 | 1371.064807 | ENSG00000068745 |
| RMI2 | 0.072033628 | 1.051197411 | 0.393591031 | 0.586305751 | 1434.18439 | 1506.980464 | ENSG00000175643 |
| ATAT1 | 0.071950051 | 1.051136516 | 0.634700599 | 0.784863951 | 351.6778353 | 370.3211757 | ENSG00000137343 |
| CYREN | 0.071918382 | 1.051113442 | 0.680116339 | 0.815242286 | 173.8407363 | 182.961386 | ENSG00000122783 |
| GNAI1 | 0.071821367 | 1.051042762 | 0.498449867 | 0.679334498 | 587.584863 | 617.5784638 | ENSG00000127955 |
| CCNI | 0.071816144 | 1.051038957 | 0.400611982 | 0.592545385 | 2766.480478 | 2907.224992 | ENSG00000118816 |

| | | | | | | | |
|---|---|---|---|---|---|---|---|
| DGUOK | 0.071655767 | 1.050922125 | 0.459469854 | 0.646251652 | 908.7218239 | 955.5733397 | ENSG00000114956 |
| PCYOX1 | 0.071552541 | 1.050846933 | 0.448344578 | 0.636021309 | 1212.987858 | 1273.984531 | ENSG00000116005 |
| DVL2 | 0.071429163 | 1.05075707 | 0.315685804 | 0.509182144 | 1614.144023 | 1696.450029 | ENSG00000004975 |
| DSTYK | 0.071425053 | 1.050754076 | 0.526274335 | 0.702433961 | 463.7186329 | 487.2090029 | ENSG00000133059 |
| GSR | 0.071409256 | 1.050742571 | 0.326980571 | 0.521267668 | 1998.940763 | 2100.188895 | ENSG00000104687 |
| ZFAT | 0.071405683 | 1.050739969 | 0.676425606 | 0.812987929 | 186.7065963 | 196.3736125 | ENSG00000066827 |
| FAM83G | 0.071364626 | 1.050710066 | 0.449275946 | 0.636903807 | 994.7486734 | 1045.535994 | ENSG00000188522 |
| MSTO1 | 0.071280157 | 1.05064855 | 0.612945355 | 0.769113472 | 385.4754837 | 405.3897695 | ENSG00000125459 |
| GINS4 | 0.07125096 | 1.050627287 | 0.472424754 | 0.657542077 | 742.8251218 | 780.6491754 | ENSG00000147536 |
| ZFP90 | 0.071231236 | 1.050612923 | 0.609336572 | 0.766800925 | 279.1481149 | 293.4324229 | ENSG00000184939 |
| H4C4 | 0.071052523 | 1.050482787 | 0.951763211 | 0.976472701 | 4.946036548 | 5.240032157 | ENSG00000277157 |
| SLC12A2 | 0.071001364 | 1.050445537 | 0.614533023 | 0.770209997 | 303.9123246 | 319.1605521 | ENSG00000064651 |
| CCDC28A | 0.070991187 | 1.050438127 | 0.703435307 | 0.831365258 | 163.7059118 | 171.9222055 | ENSG00000024862 |
| FOXJ2 | 0.070987249 | 1.05043526 | 0.583887883 | 0.747118958 | 326.7184007 | 342.866344 | ENSG00000065970 |
| ANKZF1 | 0.070941156 | 1.0504017 | 0.516165139 | 0.693936217 | 853.5587105 | 897.2922245 | ENSG00000163516 |
| HEATR6 | 0.070907287 | 1.050377041 | 0.482168631 | 0.665612499 | 576.7673269 | 605.7080637 | ENSG00000068097 |
| B3GALNT2 | 0.070865073 | 1.050346306 | 0.525765709 | 0.702258694 | 520.4155013 | 545.981748 | ENSG00000162885 |
| WAC | 0.070839696 | 1.050327831 | 0.382572756 | 0.575209171 | 2216.484315 | 2327.716835 | ENSG00000095787 |
| TMEM271 | 0.070828116 | 1.050319401 | 0.957259291 | 0.979400701 | 4.0682727 | 4.324919901 | ENSG00000273238 |
| SLC51B | 0.070814569 | 1.050309538 | 0.951189881 | 0.976154665 | 3.546285096 | 3.68312204 | ENSG00000186198 |
| WDFY1 | 0.070663594 | 1.050199631 | 0.519600006 | 0.696633863 | 1085.299902 | 1139.225046 | ENSG00000085449 |
| ACSL4 | 0.070559574 | 1.050123914 | 0.44829217 | 0.635995643 | 1722.935007 | 1808.245122 | ENSG00000068366 |
| LOC389906 | 0.070545457 | 1.050113637 | 0.574292761 | 0.739779123 | 425.2832955 | 446.4209794 | ENSG00000285756 |
| PIP5K1B | 0.070504115 | 1.050083546 | 0.748105217 | 0.859335736 | 98.9735192 | 103.8102781 | ENSG00000107242 |
| NCAPD3 | 0.070492496 | 1.050075089 | 0.394110999 | 0.586584433 | 1569.312335 | 1647.795154 | ENSG00000151503 |
| LIPT1 | 0.070484412 | 1.050069205 | 0.836598623 | 0.912220332 | 43.22535505 | 45.40393796 | ENSG00000144182 |
| DCTN1 | 0.070388652 | 1.049999508 | 0.220209163 | 0.401751821 | 4358.099135 | 4576.342183 | ENSG00000204843 |
| POM121 | 0.070378102 | 1.04999183 | 0.329984934 | 0.524343345 | 1885.497831 | 1979.756239 | ENSG00000196313 |
| ATG16L1 | 0.070334102 | 1.049959807 | 0.557418947 | 0.726781763 | 805.2019108 | 845.6014272 | ENSG00000085978 |
| LOC112267983 | 0.070283005 | 1.04992262 | 0.887652513 | 0.942577728 | 20.86332846 | 21.91580883 | ENSG00000259826 |
| RPL35 | 0.070225485 | 1.049880761 | 0.382262731 | 0.575026598 | 17500.22952 | 18373.70131 | ENSG00000136942 |
| ZNRD1 | 0.070216563 | 1.049874268 | 0.737556215 | 0.852980587 | 111.327366 | 116.8788928 | ENSG00000066379 |
| PTPRK | 0.070153966 | 1.049828717 | 0.397935009 | 0.589931808 | 1094.263325 | 1148.358775 | ENSG00000152894 |
| GPN3 | 0.070151364 | 1.049826823 | 0.461309908 | 0.647890508 | 843.1716261 | 884.8209116 | ENSG00000111231 |
| AP1S1 | 0.070138933 | 1.049817777 | 0.360422738 | 0.553627965 | 1456.067281 | 1528.581355 | ENSG00000106367 |
| UTP11 | 0.070123811 | 1.049806773 | 0.419864316 | 0.611197394 | 939.2918096 | 985.9161405 | ENSG00000183520 |
| ACADS | 0.070114176 | 1.049799762 | 0.75477643 | 0.863520379 | 129.0070257 | 135.4568404 | ENSG00000122971 |
| PREP | 0.069993167 | 1.049711712 | 0.376911258 | 0.570049493 | 1796.932082 | 1886.195812 | ENSG00000085377 |
| PPP2R2A | 0.06994527 | 1.049676862 | 0.425948672 | 0.616651967 | 1570.599978 | 1647.583153 | ENSG00000221914 |
| FLOT2 | 0.06989493 | 1.049640237 | 0.303436735 | 0.497030177 | 4216.366469 | 4425.829071 | ENSG00000132589 |
| POLA2 | 0.069888469 | 1.049635536 | 0.930716917 | 0.966002539 | 7.367469422 | 7.779429195 | ENSG00000285816 |
| DLC1 | 0.069823062 | 1.04958795 | 0.794729634 | 0.888735818 | 70.88297831 | 74.47267668 | ENSG00000164741 |
| SETD5 | 0.069721663 | 1.049514183 | 0.377275769 | 0.570484548 | 3622.742151 | 3802.823201 | ENSG00000168137 |
| NA | 0.069710839 | 1.049506309 | 0.933108233 | 0.967210907 | 9.801529947 | 10.20956108 | ENSG00000225913 |
| SLC38A3 | 0.069639692 | 1.049454553 | 0.912579017 | 0.956487804 | 11.4995568 | 12.058883 | ENSG00000188338 |
| STX7 | 0.06961569 | 1.049437094 | 0.480417658 | 0.6642839 | 594.7902103 | 623.9509361 | ENSG00000079950 |
| BRD3 | 0.069597136 | 1.049423597 | 0.35256272 | 0.545624369 | 1951.542168 | 2047.928135 | ENSG00000169925 |
| LINC01128 | 0.06955223 | 1.049390933 | 0.754011183 | 0.863411144 | 105.9909248 | 111.3881161 | ENSG00000228794 |
| NDUFB3 | 0.06949863 | 1.049351946 | 0.564845566 | 0.732937653 | 396.2728459 | 415.785583 | ENSG00000119013 |
| RPL18A | 0.069495063 | 1.049349352 | 0.177246391 | 0.345580119 | 17650.92673 | 18521.563 | ENSG00000105640 |
| CATSPER4 | 0.069488453 | 1.049344544 | 0.936029838 | 0.968425646 | 7.384531763 | 7.786915269 | ENSG00000188782 |
| MBTPS1 | 0.069421287 | 1.049295691 | 0.391050074 | 0.583648646 | 1500.680136 | 1574.754019 | ENSG00000140943 |
| GAL | 0.069402295 | 1.049281879 | 0.536315377 | 0.710664845 | 789.7568996 | 828.4841037 | ENSG00000069482 |
| NEO1 | 0.069332377 | 1.049231028 | 0.379101352 | 0.572276273 | 1399.79549 | 1468.750767 | ENSG00000067141 |
| SLC7A1 | 0.069310123 | 1.049214843 | 0.300311322 | 0.493699667 | 3545.975263 | 3719.61174 | ENSG00000139514 |
| LOC100129215 | 0.069289606 | 1.049199923 | 0.869584049 | 0.932365542 | 26.52843404 | 27.90971516 | ENSG00000226180 |
| UBQLN1 | 0.06922344 | 1.049151804 | 0.301573404 | 0.494940359 | 3124.463799 | 3277.115852 | ENSG00000135018 |
| SNCAIP | 0.069204956 | 1.049138362 | 0.782162531 | 0.8805327 | 83.95283491 | 88.11786444 | ENSG00000064692 |
| SLK | 0.069190033 | 1.04912751 | 0.482493191 | 0.665929783 | 863.6132141 | 905.2894732 | ENSG00000065613 |
| NA | 0.06918587 | 1.049124483 | 0.842056844 | 0.91540158 | 41.38025372 | 43.56254014 | ENSG00000214654 |
| ZMYND15 | 0.069166844 | 1.049110647 | 0.94469975 | 0.972833935 | 5.356758087 | 5.678924688 | ENSG00000141497 |
| THAP9 | 0.068997756 | 1.048987696 | 0.69723092 | 0.827195397 | 163.6138803 | 171.3613372 | ENSG00000168152 |
| PRDM15 | 0.06896851 | 1.048966431 | 0.620541436 | 0.774799082 | 335.7248479 | 352.2499015 | ENSG00000141956 |
| CDS2 | 0.06884626 | 1.048877549 | 0.380409689 | 0.573445561 | 1398.903333 | 1466.851416 | ENSG00000101290 |
| NA | 0.068842242 | 1.048874627 | 0.860346198 | 0.926274214 | 29.76065918 | 31.24779964 | ENSG00000261342 |
| FRMD3 | 0.068814729 | 1.048854625 | 0.856454384 | 0.923894233 | 35.04856654 | 36.72717981 | ENSG00000172159 |
| ID4 | 0.068767563 | 1.048820335 | 0.768802682 | 0.872137894 | 106.6106564 | 111.5140737 | ENSG00000172201 |
| NA | 0.068567305 | 1.048674761 | 0.921952094 | 0.961386103 | 8.889137755 | 9.34707458 | ENSG00000235636 |
| TLE3 | 0.068528725 | 1.048646718 | 0.419429121 | 0.61080375 | 1253.105716 | 1313.567118 | ENSG00000140332 |
| NA | 0.068459879 | 1.048596677 | 0.947118496 | 0.973701697 | 5.071110343 | 5.378915192 | ENSG00000230415 |
| SHMT2 | 0.068430326 | 1.048575197 | 0.234460933 | 0.41940922 | 7589.022172 | 7956.881058 | ENSG00000182199 |
| NA | 0.068347651 | 1.04851511 | 0.89336509 | 0.945501439 | 21.7883483 | 22.83515796 | ENSG00000275557 |
| PLTP | 0.068264733 | 1.048454848 | 0.398416507 | 0.590322714 | 2155.1759 | 2259.326744 | ENSG00000100979 |
| ATP5MC1 | 0.068244722 | 1.048440306 | 0.468709753 | 0.654491566 | 1485.062337 | 1556.381917 | ENSG00000159199 |

| | | | | | | | |
|---|---|---|---|---|---|---|---|
| ATP1B3 | 0.068225632 | 1.048426432 | 0.309506627 | 0.503590188 | 3246.283735 | 3402.399136 | ENSG00000069849 |
| AP2B1 | 0.068214345 | 1.04841823 | 0.316440398 | 0.510080565 | 5261.160101 | 5515.245944 | ENSG00000006125 |
| ATRIP | 0.068096301 | 1.04833245 | 0.589125023 | 0.751417189 | 332.520704 | 348.4143097 | ENSG00000164053 |
| MAP4K5 | 0.068061547 | 1.048307196 | 0.541116753 | 0.714167763 | 480.1597916 | 503.293028 | ENSG00000012983 |
| NA | 0.067992577 | 1.048257082 | 0.946340754 | 0.9733913 | 5.088172683 | 5.277024493 | ENSG00000251654 |
| CAPZB | 0.067875695 | 1.048172159 | 0.272361207 | 0.463454393 | 7547.061144 | 7910.624087 | ENSG00000077549 |
| ELMOD3 | 0.067759585 | 1.048087805 | 0.714620354 | 0.838366554 | 168.8359211 | 177.0460361 | ENSG00000115459 |
| C10orf88 | 0.067740178 | 1.048073705 | 0.719553676 | 0.841294968 | 139.7427153 | 146.2914666 | ENSG00000119965 |
| PCGF3 | 0.067730718 | 1.048066834 | 0.493022545 | 0.675364364 | 631.3025535 | 661.5433422 | ENSG00000185619 |
| FTH1 | 0.067730255 | 1.048066497 | 0.71422613 | 0.838127596 | 3181.407048 | 3334.237811 | ENSG00000167996 |
| NUP58 | 0.067724981 | 1.048062666 | 0.52958687 | 0.705324695 | 1039.469998 | 1089.009617 | ENSG00000139496 |
| NSUN4 | 0.067625721 | 1.04799056 | 0.578310045 | 0.742682814 | 444.5390351 | 465.8244614 | ENSG00000117481 |
| TAX1BP1 | 0.067615332 | 1.047983013 | 0.44344201 | 0.631920207 | 1024.410168 | 1073.072224 | ENSG00000106052 |
| SMYD4 | 0.067491373 | 1.047892972 | 0.569927726 | 0.73651098 | 558.0659258 | 584.7774816 | ENSG00000186532 |
| MORC2 | 0.067480566 | 1.047885123 | 0.34734757 | 0.54072072 | 2173.847965 | 2277.891312 | ENSG00000133422 |
| NA | 0.067458714 | 1.047869251 | 0.846760408 | 0.918413939 | 40.36982448 | 42.44843302 | ENSG00000264204 |
| NA | 0.0674579 | 1.047868659 | 0.894483935 | 0.945983752 | 17.37514557 | 18.25514501 | ENSG00000269119 |
| SLC25A5 | 0.067400749 | 1.04782715 | 0.311335022 | 0.505057619 | 6226.119477 | 6523.320554 | ENSG00000005022 |
| CARMIL1 | 0.067273993 | 1.047735091 | 0.450924235 | 0.63850789 | 910.5178127 | 953.3838607 | ENSG00000079691 |
| GNA12 | 0.067263185 | 1.047727242 | 0.280493838 | 0.471877352 | 3632.478025 | 3805.509403 | ENSG00000146535 |
| USO1 | 0.067212849 | 1.047690688 | 0.363135436 | 0.556043636 | 3288.3444 | 3444.515592 | ENSG00000138768 |
| NEDD1 | 0.067192244 | 1.047675724 | 0.607179351 | 0.765458529 | 391.1052467 | 409.6782778 | ENSG00000139350 |
| SPTBN1 | 0.067163252 | 1.047654671 | 0.393924136 | 0.586514558 | 5613.51698 | 5880.868674 | ENSG00000115306 |
| CEP97 | 0.067063834 | 1.047582478 | 0.726209429 | 0.846194238 | 176.1409287 | 184.0674171 | ENSG00000182504 |
| FAM76A | 0.067036739 | 1.047562803 | 0.682759331 | 0.816825552 | 185.6991285 | 194.4113382 | ENSG00000009780 |
| NA | 0.066966556 | 1.047511844 | 0.922223396 | 0.961440456 | 9.521692366 | 9.984635562 | ENSG00000213763 |
| USP33 | 0.06695731 | 1.04750513 | 0.454102094 | 0.641439541 | 1518.97543 | 1589.87983 | ENSG00000077254 |
| QTRT1 | 0.066900814 | 1.047464111 | 0.417351442 | 0.608543123 | 1317.838259 | 1380.913371 | ENSG00000213339 |
| TPM3P9 | 0.066872681 | 1.047443686 | 0.698422402 | 0.828043791 | 195.6955525 | 205.1957521 | ENSG00000241015 |
| RPRD1B | 0.066827917 | 1.047411186 | 0.589381635 | 0.751692762 | 428.6107841 | 448.3397227 | ENSG00000101413 |
| RING1 | 0.066804836 | 1.047394429 | 0.445197204 | 0.633398709 | 909.7464516 | 952.8968428 | ENSG00000204227 |
| POLG2 | 0.066796707 | 1.047388527 | 0.615830065 | 0.771410815 | 458.6256196 | 481.1315162 | ENSG00000256525 |
| MRPS30 | 0.066661977 | 1.047290719 | 0.435899759 | 0.625355379 | 1140.900353 | 1195.263471 | ENSG00000112996 |
| LCK | 0.066654389 | 1.04728521 | 0.351153579 | 0.544398196 | 1800.872206 | 1885.567569 | ENSG00000182866 |
| PMS2P3 | 0.066628039 | 1.047266082 | 0.771742739 | 0.874456192 | 93.42777494 | 97.96036777 | ENSG00000127957 |
| SMS | 0.066553662 | 1.047212093 | 0.330715342 | 0.525053807 | 3841.732829 | 4022.681588 | ENSG00000102172 |
| DGCR11 | 0.066543649 | 1.047204825 | 0.872488107 | 0.93395696 | 36.14748705 | 37.89759992 | ENSG00000273311 |
| CHD8 | 0.066419984 | 1.047115064 | 0.337877016 | 0.531959964 | 3766.238771 | 3942.960229 | ENSG00000100888 |
| NMNAT3 | 0.066316236 | 1.047039766 | 0.833045574 | 0.910250364 | 46.99663013 | 49.12405233 | ENSG00000163864 |
| GNPDA1 | 0.066301878 | 1.047029346 | 0.355664093 | 0.549185895 | 2155.791655 | 2256.436814 | ENSG00000113552 |
| MCRIP1 | 0.06627764 | 1.047011755 | 0.393571208 | 0.586305751 | 1816.04673 | 1901.355057 | ENSG00000225663 |
| VEZT | 0.066216378 | 1.046967297 | 0.498763181 | 0.679661554 | 797.2745405 | 834.0886057 | ENSG00000028203 |
| RAD54B | 0.066175183 | 1.046937401 | 0.787891379 | 0.884106001 | 86.58475098 | 90.7026244 | ENSG00000197275 |
| NUDT13 | 0.066014293 | 1.046820653 | 0.879827837 | 0.938030898 | 28.93723927 | 30.42329316 | ENSG00000166321 |
| SRSF5 | 0.065974357 | 1.046791676 | 0.733951851 | 0.851336652 | 4270.273824 | 4470.436064 | ENSG00000100650 |
| CDIP1 | 0.065912291 | 1.046746643 | 0.434607947 | 0.624226882 | 1680.909488 | 1759.274791 | ENSG00000089486 |
| FNBP1 | 0.065856399 | 1.046706092 | 0.470269135 | 0.655829056 | 786.7549262 | 823.4150261 | ENSG00000187239 |
| IST1 | 0.065852067 | 1.046702948 | 0.311213063 | 0.504903967 | 3251.699422 | 3402.597885 | ENSG00000182149 |
| ICE1 | 0.065789485 | 1.046657545 | 0.342008141 | 0.536244502 | 1969.530537 | 2061.158251 | ENSG00000164151 |
| NFKBID | 0.065655371 | 1.046560251 | 0.801776472 | 0.893358491 | 84.54988937 | 88.23308677 | ENSG00000167604 |
| TNKS1BP1 | 0.065639078 | 1.046548432 | 0.264066614 | 0.453755853 | 4293.20141 | 4492.842495 | ENSG00000149115 |
| OLIG2 | 0.065631557 | 1.046542976 | 0.751490298 | 0.861772523 | 139.3467937 | 145.9314569 | ENSG00000205927 |
| IFFO1 | 0.06562893 | 1.046541071 | 0.63004902 | 0.781581472 | 393.503033 | 412.429747 | ENSG00000010295 |
| GHET1 | 0.065581251 | 1.046506485 | 0.92376025 | 0.96241072 | 9.802905421 | 10.26207522 | ENSG00000281189 |
| DNPH1 | 0.065557983 | 1.046489607 | 0.481843169 | 0.665443003 | 853.591016 | 893.930439 | ENSG00000112667 |
| TTN-AS1 | 0.06549436 | 1.046443457 | 0.858623462 | 0.925270495 | 34.15492043 | 35.68490748 | ENSG00000237298 |
| CENPBD1 | 0.065340665 | 1.046331982 | 0.735475603 | 0.852171088 | 145.8121309 | 152.6021152 | ENSG00000177946 |
| C6orf223 | 0.065325251 | 1.046320803 | 0.888421767 | 0.942908379 | 20.95112066 | 21.90821114 | ENSG00000181577 |
| NA | 0.065287111 | 1.046293142 | 0.916363216 | 0.958620299 | 12.02597909 | 12.55365048 | ENSG00000268912 |
| NA | 0.065284575 | 1.046291303 | 0.942555758 | 0.971746372 | 5.650598794 | 5.97144811 | ENSG00000242457 |
| UBA3 | 0.065251224 | 1.046267116 | 0.420409156 | 0.611558221 | 1067.057658 | 1116.313825 | ENSG00000144744 |
| RALGDS | 0.06524102 | 1.046259716 | 0.85303809 | 0.922280546 | 45.83015933 | 47.98803664 | ENSG00000160271 |
| ZNF772 | 0.065227405 | 1.046249842 | 0.798193841 | 0.891456743 | 72.74523969 | 75.93050726 | ENSG00000197128 |
| ERCC6L | 0.065153846 | 1.046196498 | 0.495208426 | 0.677066136 | 742.9942795 | 777.4778482 | ENSG00000186871 |
| GFY | 0.065115093 | 1.046168396 | 0.880783059 | 0.938671336 | 26.62321649 | 27.89094417 | ENSG00000261949 |
| ELOVL4 | 0.064994356 | 1.046080847 | 0.658391403 | 0.800802353 | 287.9597053 | 301.1988151 | ENSG00000118402 |
| RITA1 | 0.06490513 | 1.046016153 | 0.53747508 | 0.711203176 | 542.0090381 | 566.653152 | ENSG00000139405 |
| CCDC9 | 0.064857858 | 1.045981879 | 0.45440391 | 0.641621367 | 1008.036778 | 1054.215125 | ENSG00000105321 |
| CRY2 | 0.064833247 | 1.045964035 | 0.553515038 | 0.723938212 | 490.2211353 | 512.7677007 | ENSG00000121671 |
| ZDHHC4 | 0.064813826 | 1.045949956 | 0.637624426 | 0.787271409 | 352.6072672 | 368.3173599 | ENSG00000136247 |
| TMEM205 | 0.064611163 | 1.045803035 | 0.784112212 | 0.881898496 | 85.47272181 | 89.56214859 | ENSG00000105518 |
| PCMT1 | 0.064564636 | 1.045769309 | 0.429079406 | 0.619406318 | 1389.188745 | 1452.974581 | ENSG00000120265 |
| SERGEF | 0.064492239 | 1.045716831 | 0.680530495 | 0.815245229 | 208.475229 | 217.8422783 | ENSG00000129158 |
| NA | 0.064481812 | 1.045709274 | 0.962355048 | 0.981774928 | 3.440152779 | 3.657303005 | ENSG00000224282 |

| | | | | | | | |
|---|---|---|---|---|---|---|---|
| EIF5B | 0.06446055 | 1.045693863 | 0.344470595 | 0.538081221 | 3880.265655 | 4057.476804 | ENSG00000158417 |
| GTF2E2 | 0.064443667 | 1.045681626 | 0.515409081 | 0.693322101 | 804.3480867 | 840.6756264 | ENSG00000197265 |
| PLCG1 | 0.064409386 | 1.045656778 | 0.416684787 | 0.607762324 | 4363.730064 | 4563.733332 | ENSG00000124181 |
| LOC100996419 | 0.064396678 | 1.045647568 | 0.91752524 | 0.959348573 | 11.44333372 | 11.93865934 | ENSG00000245317 |
| UBXN7 | 0.064383215 | 1.04563781 | 0.496566892 | 0.678082046 | 1284.020594 | 1341.809255 | ENSG00000163960 |
| PILRA | 0.064257899 | 1.045546988 | 0.947276797 | 0.973810425 | 6.808200224 | 7.145667061 | ENSG00000085514 |
| MAP1LC3B | 0.064164321 | 1.045479172 | 0.534655249 | 0.70931795 | 841.9438438 | 879.5172718 | ENSG00000140941 |
| SLC25A30 | 0.06411755 | 1.045445279 | 0.756375407 | 0.864335767 | 131.7961634 | 137.4017291 | ENSG00000174032 |
| FBXL6 | 0.064101852 | 1.045433903 | 0.609302162 | 0.766800925 | 499.2472829 | 522.0227438 | ENSG00000182325 |
| TRIM33 | 0.063992691 | 1.045354804 | 0.533417148 | 0.70829298 | 1110.116727 | 1159.455788 | ENSG00000197323 |
| NA | 0.063788136 | 1.045206597 | 0.931992182 | 0.96667654 | 8.898608494 | 9.313219819 | ENSG00000284655 |
| CBSL | 0.063772414 | 1.045195207 | 0.532320805 | 0.707394694 | 854.4084681 | 893.4080375 | ENSG00000274276 |
| NA | 0.06373553 | 1.045168486 | 0.944043865 | 0.972449807 | 5.706821866 | 6.02027502 | ENSG00000249014 |
| MICU2 | 0.06373366 | 1.045167131 | 0.58774594 | 0.750329471 | 803.4479382 | 839.5538982 | ENSG00000165487 |
| EEA1 | 0.063621441 | 1.045085836 | 0.670325705 | 0.808423037 | 302.3068103 | 315.397448 | ENSG00000102189 |
| SLC25A36 | 0.063371043 | 1.044904465 | 0.549071382 | 0.720612448 | 776.5183216 | 810.5590728 | ENSG00000114120 |
| SALL3 | 0.063356442 | 1.04489389 | 0.502778857 | 0.68322482 | 777.4655298 | 812.2655928 | ENSG00000256463 |
| MTSS2 | 0.063180073 | 1.044766159 | 0.655831817 | 0.799052517 | 336.10046 | 351.039526 | ENSG00000132613 |
| NA | 0.063051906 | 1.044673348 | 0.912856381 | 0.956679681 | 14.07269424 | 14.63538399 | ENSG00000189212 |
| DNAJC24 | 0.062900349 | 1.044563609 | 0.679406723 | 0.815010735 | 233.6029042 | 244.2790791 | ENSG00000170946 |
| FOXH1 | 0.062897295 | 1.044561398 | 0.517294512 | 0.694799369 | 1099.219328 | 1148.324602 | ENSG00000160973 |
| DYM | 0.062893538 | 1.044558678 | 0.540976659 | 0.714131985 | 874.7974406 | 913.231964 | ENSG00000141627 |
| TAB3 | 0.062856711 | 1.044532015 | 0.649858317 | 0.795523587 | 276.1004186 | 288.371627 | ENSG00000157625 |
| BARX1 | 0.062793629 | 1.044486343 | 0.866899937 | 0.93034878 | 43.29242434 | 45.05747468 | ENSG00000131668 |
| NF2 | 0.062609303 | 1.044352903 | 0.514557113 | 0.692846128 | 751.6835622 | 784.4681772 | ENSG00000186575 |
| SF1 | 0.062214041 | 1.044066816 | 0.319617843 | 0.51336128 | 5869.329101 | 6127.836811 | ENSG00000168066 |
| TMEM259 | 0.062204239 | 1.044059722 | 0.566521256 | 0.734156409 | 2455.835239 | 2565.184374 | ENSG00000182087 |
| COL9A3 | 0.06211889 | 1.043997958 | 0.508864299 | 0.688013635 | 1764.610048 | 1843.049105 | ENSG00000092758 |
| SRP54 | 0.061992498 | 1.043906499 | 0.510632017 | 0.689850121 | 1096.61384 | 1144.145788 | ENSG00000100883 |
| RALGPS2 | 0.061975794 | 1.043894413 | 0.586273673 | 0.749275703 | 560.6648821 | 584.5562744 | ENSG00000116191 |
| SCOC | 0.061902182 | 1.04384115 | 0.680545978 | 0.815245229 | 456.0654779 | 475.7725737 | ENSG00000153130 |
| COL4A5 | 0.061884722 | 1.043828517 | 0.607028053 | 0.765402948 | 858.3390394 | 896.6952886 | ENSG00000188153 |
| RPL37 | 0.061849648 | 1.043803141 | 0.348314026 | 0.541626995 | 8608.687773 | 8987.070339 | ENSG00000145592 |
| EIF4G3 | 0.061844464 | 1.04379939 | 0.387140678 | 0.580100681 | 2684.656484 | 2802.189846 | ENSG00000075151 |
| PISD | 0.061806086 | 1.043771623 | 0.48299535 | 0.666361675 | 945.8340929 | 987.5316025 | ENSG00000241878 |
| GNB5 | 0.061788736 | 1.043759072 | 0.562882239 | 0.731226453 | 493.0557475 | 514.4020595 | ENSG00000069966 |
| CNIH1 | 0.061774607 | 1.043748849 | 0.520414513 | 0.697473612 | 991.6773683 | 1034.397805 | ENSG00000100528 |
| SNAPIN | 0.06170404 | 1.043697798 | 0.632173207 | 0.783044725 | 337.2459899 | 351.9451055 | ENSG00000143553 |
| PAPSS1 | 0.061691499 | 1.043688724 | 0.374549207 | 0.56747282 | 2547.7423 | 2658.503654 | ENSG00000138801 |
| MTERF4 | 0.061655794 | 1.043662895 | 0.628875394 | 0.780786211 | 491.5936546 | 513.7518684 | ENSG00000122085 |
| FAM219A | 0.06160687 | 1.043627503 | 0.603410686 | 0.762512924 | 418.9200166 | 437.6706033 | ENSG00000164970 |
| GAS7 | 0.061557533 | 1.043591815 | 0.633712214 | 0.783961787 | 314.0769293 | 327.7711381 | ENSG00000007237 |
| NA | 0.061483137 | 1.043538 | 0.909734878 | 0.955016096 | 17.36873405 | 18.01480931 | ENSG00000236698 |
| CBWD2 | 0.061398377 | 1.043476693 | 0.605920481 | 0.764682704 | 508.1953645 | 530.5713944 | ENSG00000136682 |
| KCNQ3 | 0.061349168 | 1.043441102 | 0.806863189 | 0.896600656 | 75.19014642 | 78.2971665 | ENSG00000184156 |
| TRIM26 | 0.061340232 | 1.043434639 | 0.439679988 | 0.628007655 | 1135.423046 | 1184.623322 | ENSG00000234127 |
| DNAJB12 | 0.061314526 | 1.043416047 | 0.560401497 | 0.729387735 | 528.238648 | 551.3212977 | ENSG00000148719 |
| COLGALT2 | 0.061307995 | 1.043411324 | 0.771530982 | 0.874346286 | 104.3258426 | 108.9422476 | ENSG00000198756 |
| ECSIT | 0.061095856 | 1.043257908 | 0.481878903 | 0.665443003 | 1067.598377 | 1114.287123 | ENSG00000130159 |
| TYSND1 | 0.061091375 | 1.043254667 | 0.549152634 | 0.720668054 | 737.8805357 | 769.811921 | ENSG00000156521 |
| TIMM17B | 0.061075993 | 1.043243544 | 0.489138785 | 0.672031004 | 971.8257993 | 1013.971277 | ENSG00000126768 |
| PNRC1 | 0.060958011 | 1.043158233 | 0.7221473 | 0.843051024 | 182.3665812 | 189.9991734 | ENSG00000146278 |
| MRPS6 | 0.060877259 | 1.043099845 | 0.61335607 | 0.769307369 | 446.0907842 | 464.9720002 | ENSG00000243927 |
| RAB7B | 0.060866419 | 1.043092008 | 0.941075255 | 0.970798334 | 9.203197716 | 9.503852543 | ENSG00000276600 |
| KIAA1614 | 0.060816008 | 1.04305556 | 0.855077998 | 0.923059911 | 43.94835512 | 45.86485067 | ENSG00000135835 |
| ZNF408 | 0.060779245 | 1.043028981 | 0.631198796 | 0.782480902 | 335.8668485 | 350.4403831 | ENSG00000175213 |
| NFIC | 0.060774791 | 1.043025761 | 0.806183327 | 0.896221316 | 80.11102537 | 83.70668726 | ENSG00000141905 |
| DENND6B | 0.06075078 | 1.043008402 | 0.884205782 | 0.940480342 | 27.22974168 | 28.40503229 | ENSG00000205593 |
| TMEM9B | 0.060688923 | 1.042963683 | 0.720160046 | 0.841791517 | 234.0702775 | 243.7924187 | ENSG00000175348 |
| ZSWIM7 | 0.060685159 | 1.042960962 | 0.620326377 | 0.774639691 | 406.8899861 | 424.6786099 | ENSG00000214941 |
| TYW1 | 0.060632258 | 1.04292272 | 0.583991041 | 0.747158497 | 479.1006332 | 499.5647663 | ENSG00000198874 |
| REEP4 | 0.060590011 | 1.04289218 | 0.534133682 | 0.708990451 | 808.1742541 | 842.6538371 | ENSG00000168476 |
| FHL1 | 0.060568743 | 1.042876806 | 0.459319717 | 0.646150409 | 1165.226647 | 1214.913363 | ENSG00000022267 |
| NUP93 | 0.060438097 | 1.04278237 | 0.324147489 | 0.518311583 | 3591.836932 | 3745.45762 | ENSG00000102900 |
| EEF2KMT | 0.060398612 | 1.042753831 | 0.645193723 | 0.792814458 | 313.4919238 | 327.0470966 | ENSG00000118894 |
| FAM120A | 0.0603764 | 1.042737776 | 0.453976587 | 0.641311134 | 1414.05663 | 1474.169606 | ENSG00000048828 |
| ZNF7 | 0.060281619 | 1.042669273 | 0.703017663 | 0.831263 | 266.5591083 | 277.8583776 | ENSG00000147789 |
| DNAH11 | 0.060231145 | 1.042632796 | 0.788403857 | 0.884355511 | 107.9870317 | 112.5920645 | ENSG00000105877 |
| CCNG2 | 0.060092467 | 1.042532578 | 0.716694205 | 0.839556455 | 174.6262732 | 181.9882087 | ENSG00000138764 |
| TASOR2 | 0.060035702 | 1.042491558 | 0.437886045 | 0.626459279 | 1490.251605 | 1553.578172 | ENSG00000108021 |
| ATP8 | 0.060001562 | 1.042466889 | 0.853915388 | 0.922557144 | 57.2685547 | 59.65206749 | ENSG00000228253 |
| SCN5A | 0.059910746 | 1.04240127 | 0.623391866 | 0.776590118 | 372.4064239 | 388.3581797 | ENSG00000183873 |
| NNT | 0.059823158 | 1.042337986 | 0.630592802 | 0.781923262 | 381.1166249 | 397.1981026 | ENSG00000112992 |
| NA | 0.059648552 | 1.042211841 | 0.895204255 | 0.946203969 | 24.4518335 | 25.39277634 | ENSG00000231167 |

| | | | | | | | |
|---|---|---|---|---|---|---|---|
| TMEM184B | 0.059513583 | 1.042114344 | 0.507289264 | 0.686735754 | 1699.156753 | 1770.803228 | ENSG00000198792 |
| ZNF720 | 0.059324173 | 1.041977534 | 0.694477531 | 0.825039841 | 279.3852214 | 290.8783802 | ENSG00000197302 |
| KIDINS220 | 0.059293681 | 1.041955512 | 0.484618714 | 0.667900301 | 1261.667324 | 1315.172864 | ENSG00000134313 |
| MFSD4B | 0.059223087 | 1.041904529 | 0.781977913 | 0.88050903 | 108.4111776 | 113.1774378 | ENSG00000173214 |
| NA | 0.059199747 | 1.041887673 | 0.924062041 | 0.96244767 | 12.55861751 | 13.03019662 | ENSG00000225616 |
| ZNF517 | 0.058960307 | 1.041714768 | 0.744284859 | 0.857195413 | 185.8476762 | 193.4736646 | ENSG00000197363 |
| PKN1 | 0.058883658 | 1.041659424 | 0.343535341 | 0.537290635 | 4945.100217 | 5150.529736 | ENSG00000123143 |
| UBA52 | 0.05876593 | 1.041574425 | 0.336073205 | 0.530425 | 9039.864833 | 9416.664283 | ENSG00000221983 |
| COMMD1 | 0.058702085 | 1.041528332 | 0.653119379 | 0.797325732 | 331.6563646 | 345.5491873 | ENSG00000173163 |
| CTDSP2 | 0.058610324 | 1.041462089 | 0.343414062 | 0.537179981 | 3306.648542 | 3443.40186 | ENSG00000175215 |
| NA | 0.058579416 | 1.041439777 | 0.88116439 | 0.939023735 | 31.05723983 | 32.26805179 | ENSG00000259570 |
| INTS8 | 0.058555247 | 1.041422331 | 0.476250705 | 0.660839302 | 1271.884424 | 1324.001493 | ENSG00000164941 |
| NPM1 | 0.058518992 | 1.04139616 | 0.2845627 | 0.475647815 | 44124.86561 | 45950.2805 | ENSG00000181163 |
| EPC2 | 0.058470704 | 1.041361304 | 0.613225891 | 0.769307369 | 414.4098591 | 431.3627743 | ENSG00000135999 |
| CCDC152 | 0.058325441 | 1.041256456 | 0.74619215 | 0.858415077 | 209.468889 | 217.9119229 | ENSG00000198865 |
| NA | 0.058282935 | 1.041225779 | 0.956167649 | 0.97920191 | 4.380453524 | 4.624929397 | ENSG00000259488 |
| SLC25A43 | 0.058272219 | 1.041218044 | 0.761640479 | 0.867730799 | 148.7360312 | 154.9722384 | ENSG00000077713 |
| LOC105369147 | 0.058255024 | 1.041205634 | 0.900536648 | 0.949285116 | 20.89174068 | 21.83736404 | ENSG00000247081 |
| C2CD5 | 0.058226287 | 1.041184895 | 0.50518753 | 0.684740798 | 1109.127885 | 1154.043975 | ENSG00000111731 |
| MNAT1 | 0.058197068 | 1.041163808 | 0.621169116 | 0.775052679 | 384.8583077 | 400.6487099 | ENSG00000020426 |
| HNRNPA1L2 | 0.058180015 | 1.041151501 | 0.801271724 | 0.893214577 | 104.3473396 | 108.6452641 | ENSG00000139675 |
| CPSF4 | 0.05811868 | 1.041107238 | 0.474898076 | 0.659901105 | 1097.53208 | 1143.050573 | ENSG00000160917 |
| WDR53 | 0.058070055 | 1.041072149 | 0.726516445 | 0.846445548 | 202.2403011 | 210.9430476 | ENSG00000185798 |
| NBDY | 0.05805606 | 1.04106205 | 0.569656981 | 0.736315119 | 636.3739343 | 662.0159376 | ENSG00000204272 |
| TMEM164 | 0.057989109 | 1.041013739 | 0.470630193 | 0.656151517 | 1132.23041 | 1178.72938 | ENSG00000157600 |
| ZNF341 | 0.057968076 | 1.040998562 | 0.690307178 | 0.822404913 | 235.7922592 | 245.2941067 | ENSG00000131061 |
| ATP6V1B2 | 0.057921211 | 1.040964746 | 0.440263436 | 0.628529445 | 1745.635454 | 1817.282435 | ENSG00000147416 |
| STK24 | 0.057900685 | 1.040949936 | 0.51649101 | 0.694178403 | 2041.263867 | 2124.273109 | ENSG00000102572 |
| INIP | 0.057894665 | 1.040945593 | 0.682938954 | 0.816931965 | 268.9127729 | 279.7976356 | ENSG00000148153 |
| NARS1 | 0.057840568 | 1.040906561 | 0.309115338 | 0.503262797 | 5465.182906 | 5688.617661 | ENSG00000134440 |
| GAK | 0.057830157 | 1.040899049 | 0.39692411 | 0.589098354 | 2135.976413 | 2223.712063 | ENSG00000178950 |
| LTO1 | 0.057828177 | 1.040897621 | 0.663019141 | 0.804056584 | 324.1764264 | 337.7905045 | ENSG00000149716 |
| NA | 0.057806703 | 1.040882127 | 0.82747473 | 0.907211869 | 68.04441238 | 70.84174236 | ENSG00000227838 |
| TMEM50A | 0.05779408 | 1.040873021 | 0.453792415 | 0.641148698 | 1546.828669 | 1609.556865 | ENSG00000183726 |
| TMEM243 | 0.057757246 | 1.040846446 | 0.641449574 | 0.789732058 | 412.9683688 | 430.1060256 | ENSG00000135185 |
| INTS5 | 0.057706495 | 1.040809831 | 0.5661891 | 0.733828586 | 598.1548902 | 622.7783661 | ENSG00000185085 |
| TANGO2 | 0.057704933 | 1.040808704 | 0.778177448 | 0.878150318 | 129.9134722 | 135.5396252 | ENSG00000183597 |
| PDE6D | 0.057681646 | 1.040791905 | 0.624167233 | 0.77710194 | 484.7291109 | 504.1233631 | ENSG00000156973 |
| TM9SF4 | 0.057619658 | 1.040747186 | 0.493541096 | 0.675874798 | 1780.409145 | 1852.261974 | ENSG00000101337 |
| PTGES | 0.057558666 | 1.040703188 | 0.828958323 | 0.907818102 | 72.56847521 | 75.46144719 | ENSG00000148344 |
| NEIL2 | 0.057552129 | 1.040698472 | 0.604128987 | 0.763097298 | 471.0132229 | 490.0905232 | ENSG00000154328 |
| GPATCH1 | 0.057519386 | 1.040674853 | 0.714095391 | 0.838085358 | 269.4902094 | 280.6892306 | ENSG00000076650 |
| CHST9 | 0.057502101 | 1.040662385 | 0.696103046 | 0.826246173 | 282.2474792 | 293.3433058 | ENSG00000154080 |
| SPIDR | 0.057445796 | 1.040621771 | 0.534313531 | 0.709076829 | 1032.44797 | 1074.123105 | ENSG00000164808 |
| PCYT1A | 0.057269717 | 1.040494773 | 0.606860391 | 0.765295671 | 496.8153871 | 517.2954059 | ENSG00000161217 |
| GOLT1A | 0.057268435 | 1.040493848 | 0.832096097 | 0.909480985 | 66.15887253 | 69.0041434 | ENSG00000174567 |
| NA | 0.057251975 | 1.040481977 | 0.913957308 | 0.957454484 | 17.63613938 | 18.46922364 | ENSG00000272476 |
| CAST | 0.057235429 | 1.040470044 | 0.626295933 | 0.778626832 | 421.5332244 | 438.3354803 | ENSG00000153113 |
| CKAP2 | 0.057219872 | 1.040458824 | 0.571730645 | 0.737943374 | 1358.002846 | 1411.773372 | ENSG00000136108 |
| IQSEC1 | 0.057209147 | 1.040451089 | 0.508656956 | 0.687784872 | 1121.687021 | 1167.275468 | ENSG00000144711 |
| NA | 0.057202639 | 1.040446396 | 0.927493708 | 0.964343189 | 11.85721218 | 12.2836969 | ENSG00000261420 |
| B3GLCT | 0.057152413 | 1.040410174 | 0.79217286 | 0.88692351 | 123.0003198 | 128.0107258 | ENSG00000187676 |
| RAD18 | 0.057130895 | 1.040394657 | 0.599910444 | 0.759476787 | 478.5066379 | 497.4849594 | ENSG00000070950 |
| TRIM15 | 0.05711954 | 1.040386468 | 0.939435249 | 0.970003616 | 8.984521558 | 9.343275734 | ENSG00000204610 |
| RFX5 | 0.057119263 | 1.040386268 | 0.546649983 | 0.718495899 | 1018.732601 | 1060.243628 | ENSG00000143390 |
| LAMTOR5 | 0.057103625 | 1.040374992 | 0.692546652 | 0.823958834 | 280.7496681 | 292.1960454 | ENSG00000134248 |
| PIP4K2B | 0.056917139 | 1.040240519 | 0.374389496 | 0.567389377 | 2392.904563 | 2489.115996 | ENSG00000276293 |
| RPS15 | 0.056900589 | 1.040228586 | 0.295067806 | 0.487936002 | 13684.09401 | 14234.50855 | ENSG00000115268 |
| ZSCAN2 | 0.056852124 | 1.040193642 | 0.685708363 | 0.819137532 | 416.5771309 | 433.0599847 | ENSG00000176371 |
| COPZ1 | 0.056681374 | 1.040070537 | 0.503811416 | 0.683776217 | 1881.043822 | 1955.918979 | ENSG00000111481 |
| PRKAR2A | 0.056640861 | 1.04004133 | 0.537022643 | 0.711203176 | 767.8372917 | 798.2084308 | ENSG00000114302 |
| ABCB6 | 0.056579401 | 1.039997024 | 0.936521355 | 0.968880089 | 9.729522311 | 10.10106037 | ENSG00000115657 |
| ARHGEF25 | 0.056468245 | 1.039916899 | 0.840948642 | 0.914626028 | 61.79675913 | 64.21721147 | ENSG00000240771 |
| HDAC1 | 0.056372064 | 1.039847572 | 0.429409545 | 0.619593344 | 3766.925601 | 3917.154709 | ENSG00000116478 |
| FBXL16 | 0.056354396 | 1.039834837 | 0.702213186 | 0.830927613 | 274.7048087 | 285.2693149 | ENSG00000127585 |
| ZFP36L1 | 0.0563482 | 1.039830372 | 0.491638251 | 0.674215563 | 2269.167931 | 2358.962668 | ENSG00000185650 |
| MAGIX | 0.056315217 | 1.0398066 | 0.91530288 | 0.958169563 | 16.3330495 | 16.99130797 | ENSG00000269313 |
| SMCO4 | 0.056296694 | 1.03979325 | 0.597009726 | 0.757172238 | 712.8489055 | 741.7655045 | ENSG00000166002 |
| SFPQ | 0.056176773 | 1.039706822 | 0.324474743 | 0.518683894 | 10211.02975 | 10615.67598 | ENSG00000116560 |
| MFN1 | 0.056131198 | 1.039673978 | 0.55546756 | 0.725307064 | 656.8436262 | 682.9260771 | ENSG00000171109 |
| ZMYM3 | 0.056129043 | 1.039672425 | 0.451857915 | 0.639387192 | 2530.893983 | 2632.013839 | ENSG00000147130 |
| NOM1 | 0.05612898 | 1.03967238 | 0.537341403 | 0.711203176 | 953.7835312 | 991.2041067 | ENSG00000146909 |
| SHANK2 | 0.056108416 | 1.039657561 | 0.610203674 | 0.767371074 | 883.1635911 | 917.8853724 | ENSG00000162105 |
| TBL1X | 0.055960661 | 1.039551089 | 0.516623141 | 0.694239253 | 1855.719667 | 1928.550244 | ENSG00000101849 |

| | | | | | | | |
|---|---|---|---|---|---|---|---|
| THSD1 | 0.055854163 | 1.039474353 | 0.945762531 | 0.973279442 | 11.21508913 | 11.55607979 | ENSG00000136114 |
| GRPEL2 | 0.055752125 | 1.039400836 | 0.674729292 | 0.811724827 | 389.2466459 | 404.2383747 | ENSG00000164284 |
| FERMT3 | 0.055629905 | 1.039312786 | 0.890358774 | 0.944046876 | 28.321747 | 29.51913941 | ENSG00000149781 |
| CEBPD | 0.055560821 | 1.039263019 | 0.865711414 | 0.930054293 | 45.31082497 | 47.02310979 | ENSG00000221869 |
| MYC | 0.055474658 | 1.039200952 | 0.476239228 | 0.660839302 | 1274.663587 | 1324.674924 | ENSG00000136997 |
| NA | 0.055419488 | 1.039161213 | 0.961932378 | 0.981418842 | 3.903940477 | 4.001902531 | ENSG00000287576 |
| TBCEL | 0.05522052 | 1.039017908 | 0.836380607 | 0.9121504 | 115.6043404 | 119.8316579 | ENSG00000154114 |
| STING1 | 0.055184044 | 1.038991638 | 0.951413829 | 0.976330431 | 5.736511858 | 6.001504025 | ENSG00000184584 |
| WBP1L | 0.055139667 | 1.03895968 | 0.646036261 | 0.793142577 | 657.9736122 | 683.3337573 | ENSG00000166272 |
| NA | 0.055067853 | 1.038907964 | 0.898772196 | 0.948246365 | 30.93394748 | 32.05821004 | ENSG00000244625 |
| TICRR | 0.055040236 | 1.038888077 | 0.567799428 | 0.734939214 | 832.5381604 | 864.6007843 | ENSG00000140534 |
| CCT7 | 0.054972707 | 1.03883945 | 0.307284744 | 0.501471819 | 14388.3756 | 14947.18638 | ENSG00000135624 |
| SNX6 | 0.054920486 | 1.038801848 | 0.604219055 | 0.763159055 | 944.3554186 | 980.0093986 | ENSG00000129515 |
| PRRG4 | 0.054885543 | 1.038776688 | 0.879401678 | 0.937806595 | 41.90096355 | 43.3958627 | ENSG00000135378 |
| ZGRF1 | 0.054861603 | 1.038759451 | 0.643834117 | 0.791682598 | 406.9974188 | 423.1068392 | ENSG00000138658 |
| PPP2R5C | 0.054719817 | 1.038657368 | 0.520115384 | 0.697123122 | 1157.175277 | 1201.776807 | ENSG00000078304 |
| KCNA7 | 0.054664229 | 1.038617349 | 0.938037671 | 0.969503942 | 9.243636223 | 9.579486173 | ENSG00000104848 |
| HEPH | 0.054622648 | 1.038587414 | 0.468329978 | 0.6541584 | 1605.600627 | 1667.385246 | ENSG00000089472 |
| ZNF510 | 0.054516078 | 1.038510698 | 0.73079166 | 0.849344242 | 260.0671328 | 269.5372807 | ENSG00000081386 |
| TRIP12 | 0.054430991 | 1.038449451 | 0.496613976 | 0.678082046 | 2590.755037 | 2690.02283 | ENSG00000153827 |
| RPL8 | 0.054277767 | 1.038339166 | 0.402429981 | 0.594196834 | 48840.68459 | 50712.89454 | ENSG00000161016 |
| GORASP2 | 0.054263396 | 1.038328823 | 0.493431382 | 0.675824507 | 2143.521444 | 2225.211192 | ENSG00000115806 |
| ZNF696 | 0.054101661 | 1.038212427 | 0.612785025 | 0.768999774 | 591.7727151 | 614.5026321 | ENSG00000185730 |
| AFG1L | 0.053969697 | 1.038117465 | 0.812518001 | 0.899366839 | 90.91401748 | 94.34385595 | ENSG00000135537 |
| SCAMP1 | 0.053910703 | 1.038075016 | 0.640555843 | 0.789444284 | 788.1153063 | 817.4887572 | ENSG00000085365 |
| CADPS2 | 0.0538979 | 1.038065804 | 0.656652345 | 0.799445755 | 437.1803448 | 453.4981725 | ENSG00000081803 |
| MED10 | 0.053845046 | 1.038027774 | 0.662880237 | 0.803985331 | 443.865561 | 460.5706109 | ENSG00000133398 |
| SMAGP | 0.05383095 | 1.038017632 | 0.812652231 | 0.899366839 | 95.09789324 | 98.7401726 | ENSG00000170545 |
| NQO1 | 0.053813094 | 1.038004785 | 0.526829009 | 0.702967066 | 953.1410022 | 989.717703 | ENSG00000181019 |
| STK25 | 0.053795774 | 1.037992323 | 0.46687883 | 0.653017299 | 4140.445084 | 4298.919104 | ENSG00000115694 |
| NA | 0.053737716 | 1.037950553 | 0.806309368 | 0.896253917 | 109.1151385 | 113.0628365 | ENSG00000234797 |
| USP31 | 0.053529628 | 1.037800854 | 0.667008329 | 0.806835424 | 382.3343054 | 396.6158669 | ENSG00000103404 |
| ACOT9 | 0.053512071 | 1.037788224 | 0.584997872 | 0.747954646 | 783.5738282 | 812.5856958 | ENSG00000123130 |
| USP46 | 0.053341063 | 1.037665219 | 0.583300353 | 0.746659903 | 1045.570277 | 1083.958989 | ENSG00000109189 |
| DNAH17 | 0.053250202 | 1.037599869 | 0.859406296 | 0.925737318 | 50.22263914 | 52.0644735 | ENSG00000187775 |
| ST7L | 0.053167306 | 1.037540251 | 0.744418082 | 0.857195413 | 226.438131 | 234.8970505 | ENSG00000007341 |
| INPP5E | 0.053102351 | 1.037493538 | 0.657890786 | 0.800351023 | 568.4170285 | 590.5700316 | ENSG00000148384 |
| NUDT4 | 0.052875 | 1.037330055 | 0.567269254 | 0.734612098 | 1349.30086 | 1399.15304 | ENSG00000173598 |
| ADAP1 | 0.052849181 | 1.037311491 | 0.87575384 | 0.935624952 | 41.51173904 | 43.02120726 | ENSG00000105963 |
| ERO1A | 0.052847595 | 1.03731035 | 0.679767831 | 0.815136731 | 501.2681339 | 519.3119944 | ENSG00000197930 |
| SCNN1D | 0.052617328 | 1.0371448 | 0.878275092 | 0.937130141 | 42.18651359 | 43.8399798 | ENSG00000162572 |
| FAM234B | 0.052565466 | 1.037107517 | 0.608717556 | 0.766514251 | 690.5971981 | 715.6999093 | ENSG00000084444 |
| CCDC117 | 0.052509679 | 1.037067414 | 0.618599203 | 0.77331159 | 1042.392005 | 1079.868854 | ENSG00000159873 |
| ATG10 | 0.052498653 | 1.037059488 | 0.771551393 | 0.874346286 | 165.2582396 | 171.4866738 | ENSG00000152348 |
| FNTA | 0.052442901 | 1.037019413 | 0.665380831 | 0.80555022 | 439.8857417 | 456.2271516 | ENSG00000168522 |
| ERICH1 | 0.052429936 | 1.037010093 | 0.557268955 | 0.726688441 | 801.7412904 | 831.4851637 | ENSG00000104714 |
| NA | 0.052381062 | 1.036974963 | 0.92694814 | 0.964343189 | 16.78806552 | 17.5053961 | ENSG00000275734 |
| KCNK12 | 0.052305213 | 1.036920446 | 0.782944108 | 0.881168697 | 199.4514338 | 206.7603313 | ENSG00000184261 |
| LDHA | 0.052229427 | 1.036865977 | 0.547193712 | 0.718911765 | 9102.112648 | 9436.169282 | ENSG00000134333 |
| IPPK | 0.05206283 | 1.036746251 | 0.767733003 | 0.871722555 | 153.9576434 | 159.6489175 | ENSG00000127080 |
| PCCA-DT | 0.051938568 | 1.036656958 | 0.763279975 | 0.868797583 | 199.7871263 | 207.0360591 | ENSG00000274605 |
| SH3BGRL2 | 0.051865038 | 1.036604123 | 0.660181235 | 0.80213708 | 476.0051248 | 493.1255319 | ENSG00000198478 |
| NA | 0.051843615 | 1.036588731 | 0.950609333 | 0.975829043 | 6.331784874 | 6.590238097 | ENSG00000244363 |
| SERTAD3 | 0.051685733 | 1.036475298 | 0.689307682 | 0.821742367 | 358.7037196 | 371.7763696 | ENSG00000167565 |
| PLS3 | 0.051677851 | 1.036469635 | 0.42324096 | 0.614180691 | 6449.063532 | 6683.286907 | ENSG00000102024 |
| LINC01409 | 0.051624916 | 1.036431605 | 0.949049328 | 0.97469657 | 8.000024032 | 8.285593216 | ENSG00000237491 |
| NDUFB6 | 0.051555904 | 1.036382029 | 0.571072648 | 0.737373586 | 738.2049835 | 765.019318 | ENSG00000165264 |
| P2RX3 | 0.051541285 | 1.036371527 | 0.95239941 | 0.976855045 | 6.073947985 | 6.32777059 | ENSG00000109991 |
| NA | 0.051497263 | 1.036339904 | 0.744580025 | 0.857268814 | 215.7624001 | 223.39327 | ENSG00000283041 |
| NA | 0.051492235 | 1.036336292 | 0.854185747 | 0.922728642 | 59.66463457 | 61.8735601 | ENSG00000237638 |
| ZDHHC8 | 0.051478775 | 1.036326623 | 0.59963341 | 0.759385421 | 841.1581939 | 872.5296028 | ENSG00000099904 |
| SLC11A2 | 0.051426254 | 1.036288897 | 0.544664911 | 0.716707097 | 1149.317232 | 1190.37338 | ENSG00000110911 |
| MON2 | 0.051349802 | 1.036233983 | 0.656243863 | 0.799167853 | 478.6257062 | 495.8753606 | ENSG00000061987 |
| NDUFS8 | 0.051300193 | 1.036198351 | 0.519409527 | 0.696529641 | 1414.100345 | 1465.921045 | ENSG00000110717 |
| NKX2-4 | 0.051187194 | 1.036117194 | 0.958489474 | 0.979577828 | 5.088172683 | 5.315116135 | ENSG00000125816 |
| DARS1 | 0.051141481 | 1.036084364 | 0.58907999 | 0.751411462 | 1047.157428 | 1084.261756 | ENSG00000115866 |
| DCUN1D1 | 0.051081665 | 1.036041408 | 0.691972254 | 0.823650512 | 455.4168683 | 471.3423939 | ENSG00000043093 |
| COQ3 | 0.050788917 | 1.035831198 | 0.713968058 | 0.838042051 | 269.505588 | 279.0094686 | ENSG00000132423 |
| PIAS3 | 0.050728388 | 1.035787741 | 0.598035002 | 0.75798268 | 753.0742715 | 779.9933444 | ENSG00000131788 |
| CROCCP2 | 0.050724871 | 1.035785216 | 0.693349674 | 0.82436945 | 549.4490189 | 569.7786642 | ENSG00000215908 |
| NLE1 | 0.050594783 | 1.035691822 | 0.527474238 | 0.703476112 | 1443.301712 | 1494.922936 | ENSG00000073536 |
| PNPLA8 | 0.05058739 | 1.035686515 | 0.757645059 | 0.865096588 | 196.1915107 | 202.8640067 | ENSG00000135241 |
| FAM184A | 0.050561831 | 1.035668168 | 0.74504362 | 0.857529016 | 234.1288607 | 242.2979222 | ENSG00000111879 |
| GLO1 | 0.050520326 | 1.035638372 | 0.524635151 | 0.701203425 | 3190.190964 | 3302.45417 | ENSG00000124767 |

| | | | | | | | |
|---|---|---|---|---|---|---|---|
| NA | 0.050470774 | 1.035602802 | 0.960237547 | 0.980365291 | 4.70082731 | 4.91745282 | ENSG00000226421 |
| DNAAF1 | 0.050427665 | 1.035571858 | 0.951494768 | 0.97635943 | 6.934649492 | 7.19025709 | ENSG00000154099 |
| CDKN1C | 0.050376939 | 1.035535447 | 0.736415404 | 0.852322051 | 295.1864141 | 305.6653175 | ENSG00000129757 |
| GCAT | 0.050307166 | 1.035485367 | 0.632089736 | 0.782993662 | 665.3357146 | 688.7856234 | ENSG00000100116 |
| TECR | 0.050256047 | 1.035448677 | 0.511211727 | 0.690381677 | 1847.00824 | 1912.062011 | ENSG00000099797 |
| RALB | 0.050210915 | 1.035416286 | 0.61915701 | 0.773695794 | 706.4997652 | 731.2657529 | ENSG00000144118 |
| AVP | 0.050155169 | 1.035376277 | 0.954550374 | 0.978185467 | 6.292624143 | 6.533925114 | ENSG00000101200 |
| ZNF140 | 0.050046248 | 1.035298112 | 0.776449795 | 0.876841366 | 168.1945722 | 173.8520189 | ENSG00000196387 |
| KIF23 | 0.050036878 | 1.035291388 | 0.520627659 | 0.697708824 | 1306.707301 | 1352.963747 | ENSG00000137807 |
| GALK1 | 0.049969072 | 1.03524273 | 0.618563938 | 0.77331159 | 583.8277266 | 604.0488248 | ENSG00000108479 |
| NA | 0.04984283 | 1.035152147 | 0.963735352 | 0.982393082 | 4.102397381 | 4.306148907 | ENSG00000232909 |
| CNOT1 | 0.049700382 | 1.035049943 | 0.506447898 | 0.686047752 | 6184.327473 | 6400.358246 | ENSG00000125107 |
| TRAPPC11 | 0.04959 | 1.034970753 | 0.607196034 | 0.765458529 | 686.1748329 | 709.7612654 | ENSG00000168538 |
| RFX1 | 0.049482166 | 1.034893398 | 0.688989431 | 0.821592842 | 502.9094188 | 520.6498479 | ENSG00000132005 |
| SLC6A2 | 0.049462904 | 1.03487958 | 0.959602519 | 0.98010931 | 5.411703382 | 5.633896625 | ENSG00000103546 |
| GTDC1 | 0.049451712 | 1.034871552 | 0.771841832 | 0.874465985 | 247.1459216 | 255.19657 | ENSG00000121964 |
| C11orf49 | 0.049416126 | 1.034846026 | 0.679816057 | 0.815136731 | 384.3533074 | 397.5950014 | ENSG00000149179 |
| PRKRA | 0.049289745 | 1.034755377 | 0.623346657 | 0.776590118 | 800.5469635 | 828.1522551 | ENSG00000180228 |
| PRPSAP2 | 0.049279327 | 1.034747904 | 0.599776944 | 0.759411523 | 825.4686355 | 854.7386113 | ENSG00000141127 |
| AHI1 | 0.04925065 | 1.034727337 | 0.788109521 | 0.884297243 | 147.9967429 | 153.2882798 | ENSG00000135541 |
| RAD17 | 0.04907281 | 1.034599794 | 0.703289825 | 0.831299338 | 446.8025547 | 461.8609995 | ENSG00000152942 |
| CCNB2 | 0.049043079 | 1.034578474 | 0.522244952 | 0.699117944 | 1597.053263 | 1651.90797 | ENSG00000157456 |
| NUMA1 | 0.049018077 | 1.034560544 | 0.528857356 | 0.704661555 | 1802.297559 | 1865.486182 | ENSG00000137497 |
| NA | 0.048963802 | 1.034521624 | 0.957143809 | 0.97933661 | 5.366228826 | 5.59255579 | ENSG00000272189 |
| RAB7A | 0.048930203 | 1.034497532 | 0.434898608 | 0.624499171 | 4157.469407 | 4300.409818 | ENSG00000075785 |
| SNRPD2 | 0.04879756 | 1.034402423 | 0.587874139 | 0.750389758 | 5548.358539 | 5739.523514 | ENSG00000125743 |
| ZNF579 | 0.048715896 | 1.034343873 | 0.722771517 | 0.843567209 | 747.7517653 | 773.9707216 | ENSG00000218891 |
| HACD1 | 0.048703671 | 1.034335108 | 0.771912515 | 0.874465985 | 182.9808934 | 189.1592567 | ENSG00000165996 |
| C12orf76 | 0.048606432 | 1.034265395 | 0.801453183 | 0.893214577 | 155.3021565 | 160.5157525 | ENSG00000174456 |
| SLC37A3 | 0.048594412 | 1.034256778 | 0.603429452 | 0.762512924 | 1026.299151 | 1061.957969 | ENSG00000157800 |
| LOC100506282 | 0.048539362 | 1.034217314 | 0.923572302 | 0.962305433 | 21.2039215 | 22.05524152 | ENSG00000247137 |
| LRRTM4 | 0.048464261 | 1.034163478 | 0.847278183 | 0.918652968 | 98.02849067 | 101.1701929 | ENSG00000176204 |
| SNRK | 0.048450987 | 1.034153963 | 0.81282973 | 0.899402162 | 171.8206293 | 177.4144793 | ENSG00000163788 |
| RAB39A | 0.048346374 | 1.034078976 | 0.811104833 | 0.898809775 | 134.6146078 | 139.0493566 | ENSG00000179331 |
| CDK14 | 0.048300823 | 1.034046327 | 0.662544231 | 0.803690999 | 774.2551025 | 799.5684557 | ENSG00000058091 |
| ZFP91 | 0.048248645 | 1.03400893 | 0.686115911 | 0.819390322 | 434.3886289 | 449.1227138 | ENSG00000186660 |
| SNORD94 | 0.0481593 | 1.033944897 | 0.960889765 | 0.980787031 | 5.478674969 | 5.697695682 | ENSG00000208772 |
| NA | 0.048089287 | 1.033894721 | 0.948605905 | 0.974636169 | 9.385870056 | 9.786078731 | ENSG00000279453 |
| ZNHIT6 | 0.048064315 | 1.033876825 | 0.66057803 | 0.802356197 | 578.8498353 | 598.1128317 | ENSG00000117174 |
| TPRG1L | 0.048014378 | 1.033841039 | 0.614366047 | 0.770149497 | 700.0022051 | 723.529926 | ENSG00000158109 |
| SLC16A1 | 0.047941171 | 1.03378858 | 0.463410043 | 0.649980455 | 9331.644731 | 9645.728761 | ENSG00000155380 |
| CITED2 | 0.047880996 | 1.033745462 | 0.619693979 | 0.774105831 | 820.9547949 | 848.610465 | ENSG00000164442 |
| ZNF33A | 0.047832738 | 1.033710884 | 0.641686032 | 0.789865771 | 567.4954738 | 586.6314466 | ENSG00000189180 |
| BANCR | 0.047825654 | 1.033705808 | 0.947868097 | 0.974202146 | 8.670461597 | 8.938126345 | ENSG00000278910 |
| PUM1 | 0.047800849 | 1.033688035 | 0.531736553 | 0.706922407 | 2727.237326 | 2818.992239 | ENSG00000134644 |
| PRC1-AS1 | 0.047772251 | 1.033667545 | 0.95822175 | 0.979577828 | 5.402232644 | 5.622611705 | ENSG00000258725 |
| TBL2 | 0.047759631 | 1.033658503 | 0.648518757 | 0.794792478 | 625.9818969 | 647.3117306 | ENSG00000106638 |
| TMCO3 | 0.047665367 | 1.033590967 | 0.60800239 | 0.766065623 | 938.60213 | 970.0169513 | ENSG00000150403 |
| ALG5 | 0.047659307 | 1.033586626 | 0.760951206 | 0.867158734 | 235.3511712 | 243.0091415 | ENSG00000120697 |
| LRSAM1 | 0.047618479 | 1.033557376 | 0.686565388 | 0.819791451 | 414.579032 | 428.5060141 | ENSG00000148356 |
| DENND1C | 0.047505969 | 1.033476776 | 0.669382671 | 0.807856359 | 473.610007 | 489.5321799 | ENSG00000205744 |
| WASHC2C | 0.047424133 | 1.033418154 | 0.615089778 | 0.770668544 | 885.418452 | 915.2419835 | ENSG00000172661 |
| FKBP3 | 0.047294108 | 1.03332502 | 0.535022992 | 0.709590766 | 2329.978696 | 2407.24752 | ENSG00000100442 |
| COL27A1 | 0.047288283 | 1.033320848 | 0.897786835 | 0.947692289 | 754.1101515 | 779.4050687 | ENSG00000196739 |
| MEGF8 | 0.047179512 | 1.033242944 | 0.48998488 | 0.672761479 | 3262.885227 | 3371.014324 | ENSG00000105429 |
| SEPTIN9 | 0.047154721 | 1.033225189 | 0.429684712 | 0.619797373 | 10381.28265 | 10725.53075 | ENSG00000184640 |
| HEBP2 | 0.047141647 | 1.033215826 | 0.620304862 | 0.774639691 | 807.2778344 | 833.8560195 | ENSG00000051620 |
| REXO2 | 0.047080116 | 1.03317176 | 0.66943919 | 0.807856359 | 563.0913359 | 582.3320596 | ENSG00000076043 |
| ARL15 | 0.046946686 | 1.03307621 | 0.821869585 | 0.904391889 | 195.8735045 | 202.1981016 | ENSG00000185305 |
| NDUFAF2 | 0.046906002 | 1.033047078 | 0.703510885 | 0.831394807 | 467.2741707 | 482.8084981 | ENSG00000164182 |
| CAPN2 | 0.046741876 | 1.032929561 | 0.564886986 | 0.732937653 | 1160.173252 | 1198.753475 | ENSG00000162909 |
| N4BP3 | 0.046678477 | 1.032884171 | 0.484708984 | 0.66797504 | 2488.441935 | 2569.684724 | ENSG00000145911 |
| ADD1 | 0.046654637 | 1.032867103 | 0.497040678 | 0.678137012 | 3337.340355 | 3447.303522 | ENSG00000087274 |
| NDUFB10 | 0.046583221 | 1.032815975 | 0.560092966 | 0.729190997 | 1602.916148 | 1656.13402 | ENSG00000140990 |
| SPA17 | 0.046533495 | 1.032780377 | 0.949411951 | 0.974818807 | 8.273645486 | 8.521692036 | ENSG00000064199 |
| BIRC5 | 0.046491634 | 1.03275041 | 0.476981077 | 0.661357618 | 3713.895913 | 3834.56716 | ENSG00000089685 |
| MXD3 | 0.046440222 | 1.032713608 | 0.814214407 | 0.89991352 | 154.118721 | 159.3956966 | ENSG00000213347 |
| POLR3K | 0.046365494 | 1.032660117 | 0.638911832 | 0.78834611 | 978.7154931 | 1010.648809 | ENSG00000161980 |
| DIS3 | 0.046099081 | 1.032469441 | 0.556990776 | 0.726427911 | 1160.025208 | 1197.421227 | ENSG00000083520 |
| FLJ16779 | 0.046086166 | 1.032460198 | 0.943940404 | 0.972449807 | 13.2353689 | 13.6832794 | ENSG00000275620 |
| NA | 0.046027544 | 1.032418246 | 0.821104878 | 0.904078763 | 141.9845351 | 146.6045132 | ENSG00000267809 |
| TBCC | 0.04599876 | 1.032397648 | 0.728879415 | 0.848079043 | 435.1291723 | 449.856257 | ENSG00000124659 |
| MFSD9 | 0.045890005 | 1.032319826 | 0.724905916 | 0.84531309 | 369.0627599 | 380.9989837 | ENSG00000135953 |
| TAF13 | 0.045865555 | 1.03230233 | 0.733715246 | 0.851247472 | 415.1319496 | 428.4014952 | ENSG00000197780 |

| | | | | | | | |
|---|---|---|---|---|---|---|---|
| DPY30 | 0.045734888 | 1.032208838 | 0.589611426 | 0.751882356 | 1420.404635 | 1465.510185 | ENSG00000162961 |
| LARP1 | 0.045718124 | 1.032196844 | 0.427361858 | 0.617780027 | 9190.627218 | 9487.060777 | ENSG00000155506 |
| NA | 0.045623293 | 1.032128998 | 0.95831543 | 0.979577828 | 6.27240489 | 6.488785432 | ENSG00000236409 |
| DGKZ | 0.04560896 | 1.032118743 | 0.492297987 | 0.674736395 | 3133.391879 | 3234.36374 | ENSG00000149091 |
| GRASP | 0.045604823 | 1.032115784 | 0.831009743 | 0.908990473 | 123.930977 | 127.7432485 | ENSG00000161835 |
| PROSER1 | 0.045379092 | 1.031954307 | 0.546739356 | 0.718495899 | 2076.10797 | 2141.966997 | ENSG00000120685 |
| LOC730101 | 0.045193695 | 1.031821701 | 0.777041524 | 0.877295785 | 262.669171 | 271.4019499 | ENSG00000216775 |
| RPRD2 | 0.045125096 | 1.03177264 | 0.659965868 | 0.801980553 | 764.0386343 | 787.628076 | ENSG00000163125 |
| MAN1B1-DT | 0.045091155 | 1.031748367 | 0.95211436 | 0.976724832 | 9.541911619 | 9.800612845 | ENSG00000268996 |
| PIGO | 0.045083628 | 1.031742984 | 0.679062391 | 0.814663885 | 565.1968448 | 583.3339788 | ENSG00000165282 |
| NACC2 | 0.045015781 | 1.031694465 | 0.663472058 | 0.804258763 | 620.4987494 | 640.7168091 | ENSG00000148411 |
| ROCK1 | 0.044949694 | 1.031647206 | 0.659127839 | 0.80143484 | 1254.384099 | 1292.90433 | ENSG00000067900 |
| NA | 0.04488694 | 1.031602332 | 0.948800487 | 0.974691536 | 12.16377823 | 12.57242148 | ENSG00000232600 |
| DAPK1 | 0.044864904 | 1.031586575 | 0.49216725 | 0.674641507 | 3128.123447 | 3227.37012 | ENSG00000196730 |
| CUL2 | 0.044787808 | 1.03153145 | 0.679856519 | 0.815136731 | 774.7308489 | 798.4175795 | ENSG00000108094 |
| KMT5C | 0.04476816 | 1.031517402 | 0.663215079 | 0.804083749 | 668.7735072 | 690.4062613 | ENSG00000133247 |
| AKT1 | 0.044710584 | 1.031476237 | 0.586794602 | 0.749424538 | 2044.619835 | 2108.969737 | ENSG00000142208 |
| DHODH | 0.044536276 | 1.03135162 | 0.733633251 | 0.851247472 | 390.4116662 | 403.2515082 | ENSG00000102967 |
| NA | 0.044434192 | 1.031278644 | 0.962741791 | 0.981980854 | 5.088172683 | 5.28506022 | ENSG00000242951 |
| NOL9 | 0.044417419 | 1.031266655 | 0.667525308 | 0.80717496 | 543.5496253 | 560.5110447 | ENSG00000162408 |
| MRPS18A | 0.044408601 | 1.031260351 | 0.696811484 | 0.826911515 | 507.484699 | 522.9913494 | ENSG00000096080 |
| GPR6 | 0.044383445 | 1.03124237 | 0.959468901 | 0.980038391 | 6.286310318 | 6.492584279 | ENSG00000146360 |
| SAR1A | 0.044378363 | 1.031238737 | 0.577646693 | 0.742139457 | 1938.813326 | 1998.444542 | ENSG00000079332 |
| SRA1 | 0.044331068 | 1.031204931 | 0.618937174 | 0.773525393 | 955.7440775 | 985.5588707 | ENSG00000213523 |
| MPLKIP | 0.044213915 | 1.031121196 | 0.658159961 | 0.80059869 | 834.6911354 | 860.3416557 | ENSG00000168303 |
| SUGT1 | 0.044195678 | 1.031108162 | 0.573577008 | 0.739216763 | 1738.225607 | 1792.001829 | ENSG00000165416 |
| YARS1 | 0.044187189 | 1.031102095 | 0.434587935 | 0.624226882 | 6577.906031 | 6782.554662 | ENSG00000134684 |
| SHISA8 | 0.044095213 | 1.031036361 | 0.941099259 | 0.970798334 | 12.64462827 | 13.06025254 | ENSG00000234965 |
| RPS14 | 0.043919112 | 1.030910517 | 0.572289071 | 0.738429799 | 21493.01695 | 22156.91514 | ENSG00000164587 |
| TUBGCP6 | 0.043863609 | 1.030870857 | 0.711121256 | 0.836183312 | 1027.704697 | 1060.498682 | ENSG00000128159 |
| PDCD2 | 0.043569899 | 1.030661009 | 0.615480297 | 0.771024794 | 1324.949094 | 1364.821309 | ENSG00000071994 |
| CD59 | 0.043530304 | 1.030632722 | 0.704116192 | 0.83179868 | 472.80731 | 486.7152944 | ENSG00000085063 |
| CGAS | 0.043478287 | 1.030595564 | 0.893974048 | 0.94574241 | 49.15075538 | 50.51571073 | ENSG00000164430 |
| GATAD2B | 0.043387931 | 1.030531019 | 0.713614839 | 0.837849642 | 416.1346145 | 428.9437041 | ENSG00000143614 |
| SCAF1 | 0.043313238 | 1.030477667 | 0.580480262 | 0.744472349 | 3467.135389 | 3572.51697 | ENSG00000126461 |
| ING1 | 0.043296702 | 1.030465855 | 0.672089722 | 0.809503537 | 717.2982098 | 739.0454138 | ENSG00000153487 |
| UBTD2 | 0.043160854 | 1.030368829 | 0.646559538 | 0.793493211 | 835.617388 | 860.7703139 | ENSG00000168246 |
| WNK2 | 0.043150528 | 1.030361454 | 0.574928129 | 0.740391742 | 2354.297662 | 2426.072293 | ENSG00000165238 |
| ABHD10 | 0.043117818 | 1.030338093 | 0.704478642 | 0.831867939 | 448.1375744 | 461.5344468 | ENSG00000144827 |
| ZNF141 | 0.043080136 | 1.030311182 | 0.760248857 | 0.866678086 | 289.1093243 | 297.8138791 | ENSG00000131127 |
| PRSS8 | 0.043068817 | 1.030303098 | 0.534443852 | 0.709148221 | 1968.197336 | 2027.460855 | ENSG00000052344 |
| PRPF18 | 0.043006421 | 1.030258539 | 0.842162857 | 0.915463132 | 110.5234889 | 113.8446167 | ENSG00000165630 |
| ZNF282 | 0.042922869 | 1.030198875 | 0.592839179 | 0.75444118 | 1546.211613 | 1592.841579 | ENSG00000170265 |
| NA | 0.042838962 | 1.03013896 | 0.964888504 | 0.982987582 | 6.195864867 | 6.395368493 | ENSG00000266903 |
| ADIPOR1 | 0.042766315 | 1.030087088 | 0.570341359 | 0.736994126 | 1421.263578 | 1463.706498 | ENSG00000159346 |
| BLOC1S5 | 0.042745862 | 1.030072485 | 0.79922784 | 0.892074001 | 198.040062 | 204.0618947 | ENSG00000188428 |
| NAF1 | 0.042636033 | 1.029994071 | 0.723045375 | 0.8438337 | 580.1606895 | 597.4695537 | ENSG00000145414 |
| RP2 | 0.042613095 | 1.029977695 | 0.805778376 | 0.895932355 | 238.8600392 | 245.8129411 | ENSG00000102218 |
| TRIM52-AS1 | 0.042594227 | 1.029964225 | 0.834292941 | 0.911076206 | 129.7048682 | 133.5575923 | ENSG00000248275 |
| TMEM191B | 0.042564405 | 1.029942934 | 0.959696976 | 0.98010931 | 8.144137002 | 8.469615933 | ENSG00000278558 |
| CCDC127 | 0.042408613 | 1.02983172 | 0.703721482 | 0.831491391 | 443.3818245 | 456.8555063 | ENSG00000164366 |
| TRAPPC10 | 0.042387193 | 1.02981643 | 0.679522296 | 0.815010735 | 697.7940215 | 719.1214173 | ENSG00000160218 |
| CAD | 0.042377162 | 1.02980927 | 0.513940174 | 0.692401481 | 5950.207301 | 6127.595272 | ENSG00000084774 |
| IL4I1 | 0.042376396 | 1.029808723 | 0.960010696 | 0.980267657 | 7.350407081 | 7.561551717 | ENSG00000104951 |
| TAF15 | 0.042311696 | 1.029762541 | 0.47064197 | 0.656151517 | 9617.071924 | 9903.105777 | ENSG00000270647 |
| GLIPR1L1 | 0.042260088 | 1.029725705 | 0.942410618 | 0.971664352 | 14.40065963 | 14.80823341 | ENSG00000173401 |
| ARHGEF7 | 0.042193052 | 1.029677859 | 0.623219402 | 0.7765209 | 1087.073221 | 1119.656179 | ENSG00000102606 |
| PMS2CL | 0.042123078 | 1.029627919 | 0.807565809 | 0.896736247 | 164.5814132 | 169.5208154 | ENSG00000187953 |
| ZNF100 | 0.042093786 | 1.029607013 | 0.854624535 | 0.922892917 | 140.0155627 | 143.7050661 | ENSG00000197020 |
| BCAT2 | 0.042057686 | 1.029581251 | 0.665058171 | 0.805385316 | 739.2771604 | 761.5136172 | ENSG00000105552 |
| GAS8 | 0.04204092 | 1.029569285 | 0.845211344 | 0.91747413 | 128.074993 | 132.0704702 | ENSG00000141013 |
| RRBP1 | 0.041977599 | 1.029524098 | 0.478639826 | 0.662616562 | 5572.335965 | 5736.816774 | ENSG00000125844 |
| HPS1 | 0.04196866 | 1.029517719 | 0.652256921 | 0.796943154 | 777.6577631 | 800.8688647 | ENSG00000107521 |
| SLC39A10 | 0.041745533 | 1.029358506 | 0.656056534 | 0.79911618 | 2614.529465 | 2690.417254 | ENSG00000196950 |
| ARRDC1-AS1 | 0.04167744 | 1.029309923 | 0.725793875 | 0.845936492 | 613.4586819 | 631.7472583 | ENSG00000203993 |
| TACC3 | 0.041625793 | 1.029273076 | 0.458351401 | 0.645327018 | 4798.071015 | 4938.809562 | ENSG00000013810 |
| RHPN2 | 0.041618185 | 1.029267647 | 0.641144834 | 0.789671608 | 803.1090894 | 826.7354473 | ENSG00000131941 |
| SNHG18 | 0.041520153 | 1.029197711 | 0.851524512 | 0.921524381 | 100.7972212 | 103.7242356 | ENSG00000250786 |
| ANKS6 | 0.041511823 | 1.029191768 | 0.569333664 | 0.736153908 | 1809.301578 | 1862.116155 | ENSG00000165138 |
| ALKAL1 | 0.041484242 | 1.029172093 | 0.958949188 | 0.979831537 | 7.066037113 | 7.265341067 | ENSG00000196711 |
| TCF12 | 0.041393278 | 1.029107204 | 0.590087565 | 0.752075569 | 1685.943084 | 1734.843415 | ENSG00000140262 |
| SCO1 | 0.041370587 | 1.029091018 | 0.634355402 | 0.784489425 | 1565.56999 | 1610.399297 | ENSG00000133028 |
| NA | 0.041353741 | 1.029079001 | 0.742680575 | 0.856105567 | 341.7256832 | 351.4904763 | ENSG00000260834 |
| QSER1 | 0.041339502 | 1.029068845 | 0.624930021 | 0.777404233 | 1780.246218 | 1831.33562 | ENSG00000060749 |

| | | | | | | | |
|---|---|---|---|---|---|---|---|
| LMBRD1 | 0.041187388 | 1.028960348 | 0.788227918 | 0.884323016 | 270.1964176 | 277.6610861 | ENSG00000168216 |
| PAFAH1B1 | 0.041093324 | 1.028893262 | 0.581152366 | 0.745145482 | 1794.311591 | 1845.913578 | ENSG00000007168 |
| NSUN5 | 0.040997776 | 1.028825122 | 0.620330328 | 0.774639691 | 1494.809875 | 1538.82145 | ENSG00000130305 |
| NME6 | 0.040986001 | 1.028816725 | 0.78468016 | 0.882158322 | 248.401079 | 255.8311368 | ENSG00000172113 |
| VAMP2 | 0.040874684 | 1.028737346 | 0.820315464 | 0.903565639 | 152.7518165 | 157.1324167 | ENSG00000220205 |
| RAB2A | 0.040863438 | 1.028729326 | 0.617022972 | 0.772278752 | 1503.114383 | 1546.011309 | ENSG00000104388 |
| CHMP5 | 0.040861517 | 1.028727957 | 0.653902365 | 0.797703095 | 775.684837 | 797.7487206 | ENSG00000086065 |
| UBAP1 | 0.040736626 | 1.028638905 | 0.640689053 | 0.789444284 | 958.8198929 | 986.5597308 | ENSG00000165006 |
| EXOSC2 | 0.040689713 | 1.028605457 | 0.647702312 | 0.794354768 | 1104.879834 | 1136.778222 | ENSG00000130713 |
| ABHD3 | 0.040665499 | 1.028588193 | 0.861989577 | 0.92754963 | 112.5306374 | 116.0214161 | ENSG00000158201 |
| NA | 0.040650043 | 1.028577174 | 0.963053616 | 0.982152407 | 6.056885644 | 6.241401692 | ENSG00000269842 |
| CABIN1 | 0.040574583 | 1.028523376 | 0.532821516 | 0.707857069 | 3197.189879 | 3287.848866 | ENSG00000099991 |
| CMPK1 | 0.040512529 | 1.028479137 | 0.734575016 | 0.851605263 | 637.8357264 | 655.8333421 | ENSG00000162368 |
| AKR7A2 | 0.040507596 | 1.02847562 | 0.683399221 | 0.817274562 | 689.6530262 | 709.2582474 | ENSG00000053371 |
| SOBP | 0.040440016 | 1.028427445 | 0.828153989 | 0.907419807 | 167.4162209 | 171.9080692 | ENSG00000112320 |
| LAMA1 | 0.040402422 | 1.028400646 | 0.652948433 | 0.797321535 | 2637.000887 | 2711.755341 | ENSG00000101680 |
| COG6 | 0.040359296 | 1.028369905 | 0.77930037 | 0.878721973 | 308.4670621 | 316.8264336 | ENSG00000133103 |
| KAT7 | 0.040247449 | 1.028290183 | 0.568768816 | 0.735834234 | 4451.681644 | 4576.773529 | ENSG00000136504 |
| GSDME | 0.040247055 | 1.028289901 | 0.83777409 | 0.912894662 | 149.391556 | 154.0545382 | ENSG00000105928 |
| ZC3H7A | 0.040222722 | 1.028272559 | 0.661760357 | 0.803337329 | 881.1856591 | 905.7459745 | ENSG00000122299 |
| FASTKD5 | 0.040217354 | 1.028268732 | 0.705651047 | 0.832921698 | 545.3950424 | 561.0553003 | ENSG00000215251 |
| ETNK2 | 0.040205612 | 1.028260363 | 0.725190039 | 0.845503613 | 414.3850098 | 426.0606952 | ENSG00000143845 |
| TRAPPC9 | 0.04017849 | 1.028241033 | 0.742059834 | 0.85578601 | 461.8675487 | 474.7901102 | ENSG00000167632 |
| MMS19 | 0.040150814 | 1.028221308 | 0.541226045 | 0.71421029 | 2381.582973 | 2449.141824 | ENSG00000155229 |
| VARS1 | 0.039976707 | 1.028097227 | 0.47986213 | 0.663713784 | 6744.136688 | 6933.63748 | ENSG00000204394 |
| PSPN | 0.039934159 | 1.028066907 | 0.915168962 | 0.958169563 | 37.54210475 | 38.74219149 | ENSG00000125650 |
| FN3KRP | 0.039920362 | 1.028057076 | 0.690830575 | 0.822760165 | 631.7134478 | 649.2624565 | ENSG00000141560 |
| TPD52 | 0.039773589 | 1.027952491 | 0.840282668 | 0.914223597 | 236.4077363 | 242.7958986 | ENSG00000076554 |
| RPS7 | 0.039731712 | 1.027922653 | 0.492309284 | 0.674736395 | 11187.61584 | 11499.08774 | ENSG00000171863 |
| PTPRA | 0.039707321 | 1.027905274 | 0.59442081 | 0.755547815 | 1547.18761 | 1590.021931 | ENSG00000132670 |
| RPL5 | 0.039689347 | 1.027892468 | 0.531526399 | 0.706744406 | 19223.1503 | 19758.18635 | ENSG00000122406 |
| NA | 0.039677222 | 1.02788383 | 0.951710891 | 0.976472701 | 13.11670663 | 13.58562558 | ENSG00000279089 |
| SNAPC2 | 0.039661557 | 1.027872669 | 0.714876307 | 0.838432369 | 465.3794759 | 478.4890889 | ENSG00000104976 |
| TWISTNB | 0.039591126 | 1.02782249 | 0.781511769 | 0.880091088 | 617.2233423 | 634.0227445 | ENSG00000105849 |
| CDK5RAP2 | 0.039583355 | 1.027816954 | 0.61963957 | 0.774090038 | 1424.620471 | 1464.60006 | ENSG00000136861 |
| NA | 0.03958199 | 1.027815982 | 0.944088741 | 0.972449807 | 15.92232796 | 16.30448204 | ENSG00000230091 |
| BCL2L12 | 0.039551385 | 1.027794178 | 0.583078903 | 0.746519084 | 2054.973981 | 2112.909217 | ENSG00000126453 |
| ACBD6 | 0.039536998 | 1.027783928 | 0.642917828 | 0.790856775 | 1084.645452 | 1114.389093 | ENSG00000230124 |
| AMOTL1 | 0.039391711 | 1.02768043 | 0.6019931 | 0.761306908 | 1658.371689 | 1704.497642 | ENSG00000166025 |
| NA | 0.039350337 | 1.027650959 | 0.960709594 | 0.980711148 | 8.79563309 | 9.107176914 | ENSG00000273729 |
| NA | 0.039314436 | 1.027625387 | 0.96187671 | 0.981416072 | 6.728600986 | 6.909018588 | ENSG00000282978 |
| PIN4 | 0.03930875 | 1.027621336 | 0.696764917 | 0.8269092 | 615.3239048 | 632.1256555 | ENSG00000102309 |
| TDRD12 | 0.039216722 | 1.027555788 | 0.933916878 | 0.96733214 | 22.17135669 | 22.85750457 | ENSG00000173809 |
| TMEM39A | 0.039180806 | 1.027530206 | 0.671940184 | 0.809472822 | 761.1384784 | 782.1911763 | ENSG00000176142 |
| NKIRAS2 | 0.039146716 | 1.027505927 | 0.679851918 | 0.815136731 | 843.3362893 | 866.7635448 | ENSG00000168256 |
| BEND4 | 0.03912927 | 1.027493502 | 0.638351129 | 0.787801294 | 1210.188858 | 1243.071727 | ENSG00000188848 |
| SLC25A46 | 0.039101826 | 1.027473956 | 0.76813038 | 0.871908897 | 433.6461611 | 445.2843109 | ENSG00000164209 |
| MTX3 | 0.0390901 | 1.027465605 | 0.746898709 | 0.858676729 | 445.3340503 | 457.4446381 | ENSG00000177034 |
| ELP3 | 0.039019015 | 1.027414981 | 0.701497872 | 0.830452322 | 640.1777858 | 657.4458241 | ENSG00000134014 |
| TDH | 0.039012047 | 1.027410018 | 0.946530121 | 0.9733913 | 15.48625342 | 15.8622287 | ENSG00000154316 |
| TMEM160 | 0.039010449 | 1.027408881 | 0.752863058 | 0.862715058 | 609.6338218 | 626.2569507 | ENSG00000130748 |
| ZNF85 | 0.03896887 | 1.02737927 | 0.816573633 | 0.901347795 | 181.895901 | 186.9581756 | ENSG00000105750 |
| KIAA0930 | 0.038765104 | 1.027234174 | 0.60100867 | 0.7603532 | 2290.911691 | 2353.103941 | ENSG00000100364 |
| DDX55 | 0.038736225 | 1.027213612 | 0.703886085 | 0.831632862 | 724.3455386 | 744.3902828 | ENSG00000111364 |
| SDHC | 0.038722272 | 1.027203677 | 0.577889877 | 0.742308391 | 1832.771056 | 1882.203889 | ENSG00000143252 |
| TBRG1 | 0.038720104 | 1.027202134 | 0.758246854 | 0.86529526 | 377.8899395 | 388.1971649 | ENSG00000154144 |
| CGRRF1 | 0.038607921 | 1.027122262 | 0.813302485 | 0.899577686 | 198.2913897 | 203.438095 | ENSG00000100532 |
| UQCRH | 0.038545705 | 1.027077968 | 0.539570594 | 0.712786985 | 4235.051477 | 4349.555217 | ENSG00000173660 |
| ZNF621 | 0.038518666 | 1.027058719 | 0.775810029 | 0.876322473 | 341.9618277 | 351.380367 | ENSG00000172888 |
| UBE3B | 0.038450919 | 1.027010491 | 0.708937576 | 0.83446266 | 555.5236433 | 570.5449452 | ENSG00000151148 |
| ZNF382 | 0.038439211 | 1.027002157 | 0.879431833 | 0.937806595 | 76.61071849 | 78.67660847 | ENSG00000161298 |
| SPIN3 | 0.038407745 | 1.026979757 | 0.819412137 | 0.90291228 | 257.7746297 | 265.0910152 | ENSG00000204271 |
| AKIRIN2 | 0.038364344 | 1.026948863 | 0.619371315 | 0.773807075 | 1667.021037 | 1711.583947 | ENSG00000135334 |
| HEIH | 0.038362298 | 1.026947406 | 0.963221121 | 0.982247057 | 6.328627961 | 6.500070353 | ENSG00000278970 |
| TDP1 | 0.038358618 | 1.026944787 | 0.640478598 | 0.789444284 | 1092.527152 | 1122.328255 | ENSG00000042088 |
| AGPS | 0.038349134 | 1.026938036 | 0.689093598 | 0.821592842 | 1552.332028 | 1593.597746 | ENSG00000018510 |
| BAG6 | 0.03829731 | 1.026901147 | 0.550119086 | 0.721465527 | 6638.126544 | 6817.243639 | ENSG00000204463 |
| RNASEH2A | 0.038281433 | 1.026889846 | 0.956240927 | 0.97920191 | 10.71983 | 11.0424297 | ENSG00000104889 |
| COPB1 | 0.03827433 | 1.026884791 | 0.655773801 | 0.799049633 | 1364.383098 | 1400.574236 | ENSG00000129083 |
| SLC19A2 | 0.038257763 | 1.026872998 | 0.735662402 | 0.852173027 | 595.1712192 | 610.4632504 | ENSG00000117479 |
| CTF1 | 0.038206196 | 1.026836295 | 0.948080158 | 0.974366067 | 13.42435507 | 13.76921027 | ENSG00000150281 |
| UBXN4 | 0.038191489 | 1.026825827 | 0.604366648 | 0.76323532 | 1737.775612 | 1783.860727 | ENSG00000144224 |
| TM6SF2 | 0.038168044 | 1.026809141 | 0.949414821 | 0.974818807 | 14.64586887 | 15.11996586 | ENSG00000213996 |
| TRIM74 | 0.038120235 | 1.026775114 | 0.892035514 | 0.944983483 | 61.56931129 | 63.39281661 | ENSG00000174353 |

| | | | | | | | |
|---|---|---|---|---|---|---|---|
| NA | 0.038103285 | 1.026763051 | 0.931876934 | 0.966675877 | 24.09733504 | 24.63879055 | ENSG00000280088 |
| KANK1 | 0.038039609 | 1.026717733 | 0.654528853 | 0.798173792 | 1079.482378 | 1108.324331 | ENSG00000107104 |
| TSTD1 | 0.038014356 | 1.026699762 | 0.855624368 | 0.92336477 | 138.1399746 | 141.8564458 | ENSG00000215845 |
| ZNF23 | 0.037974643 | 1.026671501 | 0.908590151 | 0.954583633 | 43.1722889 | 44.21788443 | ENSG00000167377 |
| ZNF710-AS1 | 0.037887916 | 1.026609785 | 0.964090901 | 0.982621686 | 6.702067906 | 6.871476599 | ENSG00000259291 |
| TNPO1 | 0.037867852 | 1.026595508 | 0.620908862 | 0.774962921 | 4065.473456 | 4172.482768 | ENSG00000083312 |
| HECTD3 | 0.037789047 | 1.026539433 | 0.670327249 | 0.808423037 | 998.9668844 | 1025.475344 | ENSG00000126107 |
| RNF4 | 0.037773406 | 1.026528304 | 0.593405569 | 0.75500644 | 1933.972663 | 1984.53328 | ENSG00000063978 |
| AATK | 0.03772878 | 1.026496551 | 0.860328227 | 0.926274214 | 108.2310608 | 111.2118027 | ENSG00000181409 |
| NA | 0.037711035 | 1.026483925 | 0.931202579 | 0.966296058 | 36.50750258 | 37.57666954 | ENSG00000272455 |
| RAB18 | 0.037691164 | 1.026469787 | 0.70801478 | 0.833959068 | 905.4754002 | 928.4761756 | ENSG00000099246 |
| HOOK2 | 0.037635445 | 1.026430144 | 0.609831586 | 0.767202824 | 1591.240466 | 1633.816199 | ENSG00000095066 |
| MGRN1 | 0.037410113 | 1.02626984 | 0.621583762 | 0.775152125 | 1807.66127 | 1855.368764 | ENSG00000102858 |
| ABCD3 | 0.03732262 | 1.026207604 | 0.713121864 | 0.837579253 | 649.2054968 | 665.5862189 | ENSG00000117528 |
| C8orf82 | 0.037305937 | 1.026195737 | 0.743654088 | 0.856878139 | 596.4970913 | 612.2408979 | ENSG00000213563 |
| LMNTD2 | 0.037303236 | 1.026193815 | 0.910694252 | 0.95570436 | 79.99908289 | 82.35488109 | ENSG00000185522 |
| GATAD2A | 0.037266174 | 1.026167454 | 0.54382467 | 0.716160753 | 3841.560576 | 3941.473595 | ENSG00000167491 |
| MAK16 | 0.03714934 | 1.026084355 | 0.786683871 | 0.883178803 | 415.2551669 | 426.0395193 | ENSG00000198042 |
| MIOX | 0.037054938 | 1.026017215 | 0.967390198 | 0.984386182 | 6.091010326 | 6.252686612 | ENSG00000100253 |
| IDH2 | 0.036680609 | 1.025751034 | 0.659305057 | 0.801492846 | 1474.442941 | 1511.411996 | ENSG00000182054 |
| NA | 0.03661762 | 1.02570625 | 0.967662766 | 0.984541474 | 5.402232644 | 5.562499875 | ENSG00000285847 |
| VPS33B | 0.036581035 | 1.02568024 | 0.935983386 | 0.968425646 | 24.08027269 | 24.72191025 | ENSG00000184056 |
| PPP6R1 | 0.036544739 | 1.025654436 | 0.657380512 | 0.799835253 | 2597.669858 | 2664.455663 | ENSG00000105063 |
| PLK4 | 0.036537925 | 1.025649591 | 0.707962542 | 0.833950537 | 719.8345919 | 737.8916777 | ENSG00000142731 |
| MAU2 | 0.036464244 | 1.025597211 | 0.694187574 | 0.824772234 | 898.3995165 | 921.5118635 | ENSG00000129933 |
| SMARCC1 | 0.036350438 | 1.025516311 | 0.543670441 | 0.716110294 | 7947.873386 | 8149.546363 | ENSG00000173473 |
| SFSWAP | 0.036289851 | 1.025473244 | 0.649591583 | 0.795370972 | 1508.434311 | 1547.713498 | ENSG00000061936 |
| ANO10 | 0.036214074 | 1.025419383 | 0.744438187 | 0.857195413 | 934.0076081 | 957.2546524 | ENSG00000160746 |
| STRN4 | 0.036188057 | 1.025400891 | 0.549246509 | 0.720740215 | 3216.480532 | 3298.218234 | ENSG00000090372 |
| CASC4 | 0.036103913 | 1.025341088 | 0.756064272 | 0.864333115 | 511.9426097 | 524.5044021 | ENSG00000166734 |
| SIRT2 | 0.035935468 | 1.025221378 | 0.843568617 | 0.916453737 | 196.4408615 | 201.2604598 | ENSG00000068903 |
| AZI2 | 0.035876777 | 1.025179672 | 0.840017523 | 0.914174887 | 157.932216 | 161.711442 | ENSG00000163512 |
| PYGO2 | 0.035859864 | 1.025167653 | 0.677584612 | 0.813575615 | 997.3306582 | 1022.323369 | ENSG00000163348 |
| STON1 | 0.035785918 | 1.02511511 | 0.840872281 | 0.914596647 | 157.5709001 | 161.5920944 | ENSG00000243244 |
| ASNSD1 | 0.035523654 | 1.024928773 | 0.869308864 | 0.932178344 | 122.2544472 | 124.9897728 | ENSG00000138381 |
| INO80E | 0.035454537 | 1.024879672 | 0.746818953 | 0.858676729 | 1178.49611 | 1208.531201 | ENSG00000169592 |
| ZNF660 | 0.035433597 | 1.024864797 | 0.84273292 | 0.915921653 | 155.9716094 | 159.7870361 | ENSG00000144792 |
| GLRX2 | 0.035419778 | 1.02485498 | 0.866052816 | 0.930085925 | 168.2727808 | 172.5507515 | ENSG00000023572 |
| ATP5PF | 0.035405687 | 1.02484497 | 0.75451528 | 0.863477583 | 492.9670835 | 504.6334376 | ENSG00000154723 |
| HSP90AA1 | 0.03537675 | 1.024824414 | 0.557571227 | 0.726895754 | 41149.31045 | 42169.51189 | ENSG00000080824 |
| NA | 0.03530129 | 1.024770812 | 0.927406923 | 0.964343189 | 38.72180457 | 39.54357847 | ENSG00000242299 |
| PPP2R5E | 0.035261976 | 1.024742887 | 0.735650785 | 0.852173027 | 1174.782907 | 1203.131847 | ENSG00000154001 |
| HGS | 0.035171929 | 1.024678929 | 0.681936224 | 0.816316087 | 1660.562916 | 1702.769621 | ENSG00000185359 |
| SLC35E1 | 0.035083694 | 1.024616262 | 0.754712747 | 0.863520379 | 506.3752707 | 518.586997 | ENSG00000127526 |
| GDAP1L1 | 0.035042086 | 1.024586712 | 0.865301028 | 0.929978419 | 137.3460341 | 140.7596839 | ENSG00000124194 |
| EEF1A2 | 0.034973832 | 1.02453824 | 0.68055166 | 0.815245229 | 1229.559585 | 1259.388482 | ENSG00000101210 |
| EIF3L | 0.034914735 | 1.024496272 | 0.597960038 | 0.757939498 | 3319.362375 | 3400.441864 | ENSG00000100129 |
| CYP2D6 | 0.03483023 | 1.024436265 | 0.945823677 | 0.973288326 | 22.26358358 | 22.95190919 | ENSG00000100197 |
| MMP15 | 0.034737802 | 1.024370635 | 0.645749991 | 0.792972261 | 1956.110157 | 2003.438338 | ENSG00000102996 |
| ZNF436 | 0.034597797 | 1.024271231 | 0.924191327 | 0.962449326 | 39.28984545 | 40.06526429 | ENSG00000125945 |
| DDX59 | 0.034414732 | 1.024141269 | 0.798065662 | 0.89142102 | 318.1787253 | 326.0280554 | ENSG00000118197 |
| KCNT1 | 0.034375184 | 1.024113194 | 0.968664102 | 0.984740121 | 9.329646985 | 9.670091951 | ENSG00000107147 |
| NA | 0.034265772 | 1.02403553 | 0.946387965 | 0.9733913 | 21.97162201 | 22.5388357 | ENSG00000235316 |
| EIF2S3 | 0.034256624 | 1.024029037 | 0.545299375 | 0.717389169 | 11185.37477 | 11453.49348 | ENSG00000130741 |
| LINC01816 | 0.034231255 | 1.02401103 | 0.946145945 | 0.973361629 | 19.68541008 | 20.12279989 | ENSG00000231327 |
| MBTD1 | 0.034182856 | 1.023976678 | 0.690418883 | 0.822485125 | 1285.42714 | 1315.354625 | ENSG00000011258 |
| MMP16 | 0.03417975 | 1.023974473 | 0.795472409 | 0.889328365 | 422.0509274 | 431.6337785 | ENSG00000156103 |
| NA | 0.034159054 | 1.023959783 | 0.938118253 | 0.969503942 | 24.98329185 | 25.56562576 | ENSG00000206228 |
| NA | 0.03415598 | 1.023957602 | 0.959907505 | 0.980219024 | 10.23888226 | 10.5287796 | ENSG00000242588 |
| ENO2 | 0.034153745 | 1.023956016 | 0.629010016 | 0.780901041 | 2342.110297 | 2399.107813 | ENSG00000111674 |
| WDR77 | 0.034143019 | 1.023948403 | 0.698867382 | 0.828237615 | 888.4220265 | 909.3572073 | ENSG00000116455 |
| RPL34 | 0.034136778 | 1.023943973 | 0.577895027 | 0.742308391 | 5876.686791 | 6016.655889 | ENSG00000109475 |
| MEX3C | 0.034100603 | 1.023918298 | 0.645084747 | 0.792759739 | 1758.087488 | 1800.033924 | ENSG00000176624 |
| ISCA2 | 0.034086961 | 1.023908617 | 0.813802683 | 0.89988777 | 286.3376323 | 292.7555594 | ENSG00000165898 |
| TYK2 | 0.033913157 | 1.023785272 | 0.73794297 | 0.853187551 | 657.5233239 | 673.6663507 | ENSG00000105397 |
| KLF16 | 0.033877031 | 1.023759636 | 0.734071058 | 0.851336652 | 809.6098366 | 828.5011713 | ENSG00000129911 |
| SMIM8 | 0.033876591 | 1.023759324 | 0.870345396 | 0.932705584 | 115.6655996 | 118.5104489 | ENSG00000111850 |
| RIMS3 | 0.033831985 | 1.023727671 | 0.731664529 | 0.85014547 | 1009.792975 | 1034.33494 | ENSG00000117016 |
| RAB8B | 0.033748242 | 1.023668249 | 0.772664826 | 0.874752298 | 550.3309842 | 562.9113036 | ENSG00000166128 |
| HTATIP2 | 0.033723879 | 1.023650963 | 0.701035802 | 0.830064368 | 814.1149731 | 833.332287 | ENSG00000109854 |
| PPP1R26 | 0.033679744 | 1.023619647 | 0.645623644 | 0.79292218 | 1605.006038 | 1643.546514 | ENSG00000196422 |
| WDSUB1 | 0.033632594 | 1.023586194 | 0.882108713 | 0.939373634 | 98.36424306 | 100.637995 | ENSG00000196151 |
| POLB | 0.033548666 | 1.023526649 | 0.766431812 | 0.870835943 | 467.5346835 | 478.1272154 | ENSG00000070501 |
| ARHGAP27 | 0.033438385 | 1.023448413 | 0.823569487 | 0.905179201 | 301.6168752 | 309.0150536 | ENSG00000159314 |

| | | | | | | | |
|---|---|---|---|---|---|---|---|
| DENR | 0.033430721 | 1.023442976 | 0.656302496 | 0.799167853 | 2475.649788 | 2533.321742 | ENSG00000139726 |
| SULF2 | 0.033426794 | 1.023440191 | 0.748424462 | 0.859542631 | 704.2392447 | 721.4580207 | ENSG00000196562 |
| EPHA1 | 0.033376952 | 1.023404833 | 0.589573028 | 0.751882356 | 4557.091547 | 4663.590945 | ENSG00000146904 |
| NUP43 | 0.033325277 | 1.023368178 | 0.689061141 | 0.821592842 | 1201.118732 | 1228.514247 | ENSG00000120253 |
| EFCAB7 | 0.033226587 | 1.023298174 | 0.917984809 | 0.959450204 | 53.43148811 | 54.56231491 | ENSG00000203965 |
| CES3 | 0.033030159 | 1.023158858 | 0.918371407 | 0.959665811 | 44.11268734 | 45.21980362 | ENSG00000172828 |
| MCF2L | 0.033018096 | 1.023150303 | 0.764559088 | 0.869725999 | 518.8623616 | 530.801599 | ENSG00000126217 |
| RREB1 | 0.033012911 | 1.023146626 | 0.727138845 | 0.846837128 | 726.5727387 | 743.0376232 | ENSG00000124782 |
| FAM104A | 0.032985322 | 1.023127061 | 0.799969393 | 0.892525635 | 360.0268408 | 368.2965507 | ENSG00000133193 |
| PANX1 | 0.032979283 | 1.023122778 | 0.773469992 | 0.875038864 | 657.3675453 | 671.9496677 | ENSG00000110218 |
| NA | 0.032845691 | 1.023028042 | 0.943598118 | 0.972401864 | 28.49673004 | 29.27599255 | ENSG00000287038 |
| STK40 | 0.032420813 | 1.022726801 | 0.711171788 | 0.83618968 | 922.1145574 | 943.3422282 | ENSG00000196182 |
| CA13 | 0.032381747 | 1.022699108 | 0.933929066 | 0.96733214 | 32.011446 | 32.6844861 | ENSG00000185015 |
| HECTD4 | 0.032360464 | 1.022684021 | 0.770004432 | 0.873233931 | 733.4335014 | 750.3404264 | ENSG00000173064 |
| CCDC77 | 0.032290306 | 1.022634289 | 0.781300566 | 0.879945976 | 503.3456214 | 514.9121742 | ENSG00000120647 |
| ATP5F1E | 0.032270112 | 1.022619975 | 0.743502897 | 0.856819495 | 846.4685377 | 865.660777 | ENSG00000124172 |
| HLA-DRB5 | 0.032260368 | 1.022613069 | 0.921346199 | 0.961120642 | 50.23654457 | 51.31786216 | ENSG00000198502 |
| ZNF90 | 0.032119925 | 1.022513524 | 0.756143271 | 0.864335767 | 663.7726609 | 678.7174789 | ENSG00000213988 |
| DYNLRB1 | 0.032111342 | 1.022507441 | 0.96159472 | 0.981236389 | 11.068616 | 11.33115427 | ENSG00000125971 |
| CCNYL1 | 0.032098539 | 1.022498367 | 0.725851302 | 0.845936492 | 761.5140905 | 778.8831562 | ENSG00000163249 |
| GDF6 | 0.031883052 | 1.022345654 | 0.960251259 | 0.980365291 | 11.80414602 | 12.0700563 | ENSG00000156466 |
| NA | 0.031806277 | 1.02229125 | 0.965762744 | 0.983540123 | 9.2625777 | 9.523611223 | ENSG00000225416 |
| CHGB | 0.031791946 | 1.022281095 | 0.888194757 | 0.942849507 | 92.1386882 | 94.09603418 | ENSG00000089199 |
| MGC32805 | 0.031785023 | 1.022276189 | 0.952659521 | 0.977013699 | 16.30651642 | 16.70258339 | ENSG00000250328 |
| MIDN | 0.031780786 | 1.022273187 | 0.721068557 | 0.842322239 | 3353.699719 | 3428.126151 | ENSG00000167470 |
| SMIM12 | 0.031775627 | 1.022269531 | 0.702412843 | 0.831110804 | 977.2591579 | 999.0776628 | ENSG00000163866 |
| ATPAF2 | 0.031698988 | 1.022215227 | 0.845086739 | 0.917436773 | 203.5673685 | 208.1125126 | ENSG00000171953 |
| GRN | 0.031661603 | 1.022188739 | 0.680352562 | 0.815245229 | 2546.773595 | 2602.65279 | ENSG00000030582 |
| NADSYN1 | 0.031609067 | 1.022151516 | 0.762954033 | 0.868655652 | 669.9003666 | 684.9774601 | ENSG00000172890 |
| LNX2 | 0.031574695 | 1.022127164 | 0.79796235 | 0.891359341 | 404.2386475 | 412.8809432 | ENSG00000139517 |
| ALKBH2 | 0.031545809 | 1.022106699 | 0.805649734 | 0.895843063 | 439.3360561 | 448.5843984 | ENSG00000189046 |
| PDE8A | 0.031522844 | 1.022090429 | 0.883771203 | 0.940072055 | 115.9254133 | 118.7246391 | ENSG00000073417 |
| ARF1 | 0.031387599 | 1.021994618 | 0.57302315 | 0.738883513 | 6272.646396 | 6410.361961 | ENSG00000143761 |
| GLI4 | 0.031385857 | 1.021993384 | 0.916181138 | 0.95853803 | 53.86430804 | 55.01672924 | ENSG00000250571 |
| NA | 0.031248611 | 1.021896165 | 0.950838962 | 0.975956656 | 18.69204318 | 19.20823729 | ENSG00000227939 |
| PROCA1 | 0.031189606 | 1.021854371 | 0.875306774 | 0.935388311 | 228.1606391 | 233.6382635 | ENSG00000167525 |
| HSDL1 | 0.031169218 | 1.02183993 | 0.711833058 | 0.836542636 | 1176.566201 | 1202.305407 | ENSG00000103160 |
| NFKB2 | 0.031149624 | 1.021826052 | 0.748069224 | 0.859335736 | 655.6702402 | 669.7209441 | ENSG00000077150 |
| RPS8 | 0.031115549 | 1.021801918 | 0.596700794 | 0.75696458 | 17248.27076 | 17623.12783 | ENSG00000142937 |
| OPTN | 0.031069282 | 1.02176915 | 0.833656084 | 0.910747762 | 240.5132081 | 245.7228136 | ENSG00000123240 |
| SMIM30 | 0.031065215 | 1.021766269 | 0.834703216 | 0.911277722 | 402.6978272 | 411.2293105 | ENSG00000214194 |
| NA | 0.030998968 | 1.021719352 | 0.954257828 | 0.978111635 | 17.90472478 | 18.39793851 | ENSG00000237087 |
| ADNP | 0.030938292 | 1.021676382 | 0.68847465 | 0.821277542 | 2399.371043 | 2450.246403 | ENSG00000101126 |
| TSPAN10 | 0.030817215 | 1.021590642 | 0.95666541 | 0.979215959 | 17.87946948 | 18.3602849 | ENSG00000182612 |
| QKI | 0.030676324 | 1.021490881 | 0.694844511 | 0.825370003 | 2156.375496 | 2201.767543 | ENSG00000112531 |
| DGKA | 0.030659071 | 1.021478665 | 0.851058232 | 0.92129626 | 198.0524943 | 202.4049577 | ENSG00000065357 |
| GABRA5 | 0.030658031 | 1.021477928 | 0.795314116 | 0.889205032 | 456.6371195 | 465.8123633 | ENSG00000186297 |
| ATG14 | 0.030635369 | 1.021461882 | 0.818137002 | 0.902165959 | 350.9918238 | 358.0872128 | ENSG00000126775 |
| PPP2R3A | 0.030506398 | 1.021370572 | 0.838895954 | 0.913572926 | 286.5123297 | 292.2027268 | ENSG00000073711 |
| NA | 0.030430294 | 1.021316695 | 0.949296897 | 0.974805734 | 21.6777813 | 22.17491552 | ENSG00000265749 |
| HOMER3 | 0.030385018 | 1.021284644 | 0.814167872 | 0.89991352 | 363.7119847 | 371.8489773 | ENSG00000051128 |
| DDX17 | 0.030339865 | 1.02125268 | 0.720845265 | 0.842237882 | 7084.249545 | 7235.579216 | ENSG00000100201 |
| GPBP1 | 0.030335077 | 1.021249291 | 0.688951844 | 0.821592842 | 1670.283111 | 1704.846407 | ENSG00000062194 |
| CENPB | 0.030327606 | 1.021244003 | 0.702044517 | 0.830887169 | 1720.887119 | 1757.135814 | ENSG00000125817 |
| CCDC174 | 0.030296942 | 1.021222297 | 0.831641961 | 0.909313528 | 300.3261047 | 307.0753492 | ENSG00000154781 |
| DCAF7 | 0.030241651 | 1.021183159 | 0.660749633 | 0.802512038 | 2922.372798 | 2983.634555 | ENSG00000136485 |
| NA | 0.030212428 | 1.021162475 | 0.93599852 | 0.968425646 | 33.37263809 | 34.10608879 | ENSG00000264575 |
| SRGAP3 | 0.030188963 | 1.021145866 | 0.877159976 | 0.93639635 | 127.9127353 | 130.7152629 | ENSG00000196220 |
| AP1G1 | 0.030159034 | 1.021124682 | 0.69590706 | 0.826103895 | 1397.120986 | 1426.267174 | ENSG00000166747 |
| SF3A2 | 0.030095162 | 1.021079475 | 0.693951938 | 0.824655819 | 1962.040233 | 2003.308032 | ENSG00000104897 |
| ZMYM5 | 0.030047726 | 1.021045903 | 0.86273335 | 0.928134427 | 159.6941553 | 163.0846558 | ENSG00000132950 |
| RPS20 | 0.030038435 | 1.021039327 | 0.562086928 | 0.730738379 | 13551.30556 | 13836.17334 | ENSG00000008988 |
| SOX2 | 0.029984009 | 1.021000808 | 0.690783953 | 0.822760165 | 5336.154675 | 5449.174985 | ENSG00000181449 |
| BIRC2 | 0.029960241 | 1.020983988 | 0.773500305 | 0.875038864 | 563.0599547 | 574.5714643 | ENSG00000110330 |
| CEP41 | 0.0299328 | 1.020964569 | 0.868810871 | 0.931806069 | 157.6151946 | 160.9730896 | ENSG00000106477 |
| CDC42BPB | 0.029908084 | 1.020947078 | 0.653579159 | 0.797566245 | 3549.331923 | 3623.785821 | ENSG00000198752 |
| COMMD2 | 0.029899184 | 1.020940779 | 0.778489456 | 0.878341972 | 713.520914 | 728.001836 | ENSG00000114744 |
| LOC102724330 | 0.029840024 | 1.020898915 | 0.925571352 | 0.963290836 | 48.33374699 | 49.16115629 | ENSG00000228801 |
| METTL21A | 0.02982583 | 1.020888871 | 0.762663772 | 0.868522909 | 687.1243938 | 701.0387148 | ENSG00000144401 |
| NA | 0.029571898 | 1.020709198 | 0.958209628 | 0.979577828 | 14.788005 | 15.01427631 | ENSG00000265095 |
| SLC4A2 | 0.029562507 | 1.020702554 | 0.678019742 | 0.813965688 | 2344.28504 | 2393.386136 | ENSG00000164889 |
| OSBPL1A | 0.029558825 | 1.020699949 | 0.717200611 | 0.839762217 | 1136.530159 | 1160.121608 | ENSG00000141447 |
| WDYHV1 | 0.029397167 | 1.020585583 | 0.851762518 | 0.921524381 | 243.2227692 | 247.775765 | ENSG00000156795 |
| TYW1B | 0.029382258 | 1.020575036 | 0.928448099 | 0.964674127 | 51.71914987 | 52.99543397 | ENSG00000277149 |

| | | | | | | | |
|---|---|---|---|---|---|---|---|
| ATIC | 0.029360118 | 1.020559374 | 0.63274586 | 0.783596937 | 4205.737934 | 4291.700862 | ENSG00000138363 |
| KAT5 | 0.029245168 | 1.020478062 | 0.85521604 | 0.923101397 | 203.459838 | 207.7875601 | ENSG00000172977 |
| LRRC56 | 0.029060171 | 1.020347215 | 0.929754593 | 0.965487435 | 51.1036576 | 52.25218344 | ENSG00000161328 |
| PSRC1 | 0.029027689 | 1.020324242 | 0.712800863 | 0.837414502 | 1304.380861 | 1331.284523 | ENSG00000134222 |
| ZNF777 | 0.028946716 | 1.020266976 | 0.820789855 | 0.903839639 | 389.7969707 | 398.0881687 | ENSG00000196453 |
| TMEM258 | 0.028842732 | 1.020193442 | 0.812627183 | 0.899366839 | 743.3568431 | 758.3083385 | ENSG00000134825 |
| COX17 | 0.028803368 | 1.020165607 | 0.80722734 | 0.896619594 | 455.5111697 | 464.7255009 | ENSG00000138495 |
| RRP8 | 0.028766549 | 1.020139571 | 0.834231374 | 0.911076206 | 300.8433418 | 306.4642198 | ENSG00000132275 |
| LOC102723996 | 0.028633824 | 1.020045724 | 0.938671767 | 0.969751534 | 33.10976515 | 33.79479437 | ENSG00000277117 |
| NA | 0.02863108 | 1.020043784 | 0.923829131 | 0.96241072 | 56.4557856 | 57.58629381 | ENSG00000255121 |
| TBCCD1 | 0.028585787 | 1.020011761 | 0.846452999 | 0.918323688 | 277.4306581 | 282.6973772 | ENSG00000113838 |
| CCDC92 | 0.028573549 | 1.020003109 | 0.947692628 | 0.974075819 | 25.85977792 | 26.43603637 | ENSG00000119242 |
| RPS6KA3 | 0.028441235 | 1.019909565 | 0.827735389 | 0.907253657 | 356.7275237 | 363.5093955 | ENSG00000177189 |
| NA | 0.028431713 | 1.019902833 | 0.922889086 | 0.961876704 | 60.43045594 | 61.67500327 | ENSG00000272994 |
| OBSL1 | 0.028418905 | 1.019893779 | 0.697600132 | 0.827476534 | 3173.147572 | 3237.419634 | ENSG00000124006 |
| ME2 | 0.028236256 | 1.019764666 | 0.729013871 | 0.848129016 | 1607.757699 | 1638.768055 | ENSG00000082212 |
| NA | 0.028220761 | 1.019753713 | 0.978577633 | 0.990116708 | 4.363391183 | 4.47844867 | ENSG00000236762 |
| SETD3 | 0.0281388 | 1.019695782 | 0.711341683 | 0.836295827 | 1661.345499 | 1693.469778 | ENSG00000183576 |
| PLAAT1 | 0.028124853 | 1.019685925 | 0.914721181 | 0.958005256 | 87.78624092 | 89.33872013 | ENSG00000127252 |
| ZNRF2 | 0.028077164 | 1.019652219 | 0.871087077 | 0.933167325 | 215.1807469 | 219.1756484 | ENSG00000180233 |
| NA | 0.028049291 | 1.019632519 | 0.974867953 | 0.988197476 | 5.716292604 | 5.824967383 | ENSG00000243964 |
| TBRG4 | 0.027963319 | 1.01957176 | 0.652530041 | 0.797031264 | 2880.789652 | 2937.54349 | ENSG00000136270 |
| TOM1L1 | 0.027934949 | 1.01955171 | 0.789587238 | 0.885205049 | 690.9057335 | 704.4427171 | ENSG00000141198 |
| KBTBD8 | 0.02784356 | 1.019487128 | 0.854908798 | 0.923041604 | 335.9368793 | 341.9560896 | ENSG00000163376 |
| ZFYVE9 | 0.027838572 | 1.019483603 | 0.791621432 | 0.886486471 | 718.7696832 | 732.5006015 | ENSG00000157077 |
| PRKAG2 | 0.02782568 | 1.019474493 | 0.874668014 | 0.935130738 | 241.2879739 | 245.6002882 | ENSG00000106617 |
| PDLIM2 | 0.027811017 | 1.019464131 | 0.900581838 | 0.949285116 | 163.6449231 | 166.8810651 | ENSG00000120913 |
| TMEM33 | 0.027800687 | 1.019456832 | 0.809589258 | 0.897492586 | 1056.076631 | 1076.155135 | ENSG00000109133 |
| CHML | 0.027728434 | 1.019405777 | 0.777421094 | 0.877510509 | 1042.714272 | 1062.680352 | ENSG00000203668 |
| DNM1L | 0.027672294 | 1.019366109 | 0.715835448 | 0.8390649 | 1561.234913 | 1591.119426 | ENSG00000087470 |
| IFNLR1 | 0.027568797 | 1.019292984 | 0.853650844 | 0.922378869 | 282.2373094 | 288.01513 | ENSG00000185436 |
| ZDHHC5 | 0.027546267 | 1.019277066 | 0.692774487 | 0.823967294 | 1848.836031 | 1884.534758 | ENSG00000156599 |
| PIK3CD-AS2 | 0.027492513 | 1.019239089 | 0.91771487 | 0.959390744 | 72.63900967 | 73.92429576 | ENSG00000231789 |
| AEBP1 | 0.027437326 | 1.019200101 | 0.700607648 | 0.829664311 | 1849.366077 | 1885.077245 | ENSG00000106624 |
| ARRDC4 | 0.027416813 | 1.01918561 | 0.895192991 | 0.946203969 | 128.8168368 | 130.9498244 | ENSG00000140450 |
| SKA2 | 0.027370828 | 1.019153124 | 0.772266478 | 0.874608633 | 807.6225916 | 822.4514931 | ENSG00000182628 |
| AFDN | 0.027369897 | 1.019152467 | 0.677697929 | 0.813658964 | 3715.361373 | 3786.599049 | ENSG00000130396 |
| FKBP11 | 0.027353404 | 1.019140815 | 0.820967327 | 0.903981014 | 433.1775705 | 441.574972 | ENSG00000134285 |
| OTOG | 0.027328838 | 1.019123462 | 0.935923268 | 0.968425646 | 41.65062056 | 42.38788316 | ENSG00000188162 |
| TOR1B | 0.027294041 | 1.019098882 | 0.814450884 | 0.900013873 | 445.6478922 | 453.8551478 | ENSG00000136816 |
| PCOLCE | 0.027256349 | 1.019072257 | 0.815797551 | 0.900803802 | 435.7153055 | 443.7928403 | ENSG00000106333 |
| C1orf109 | 0.027222033 | 1.019048017 | 0.777173325 | 0.877337703 | 721.6493867 | 735.7412872 | ENSG00000116922 |
| PDIA6 | 0.027207179 | 1.019037525 | 0.616510017 | 0.771897362 | 9749.45441 | 9934.535352 | ENSG00000143870 |
| AKAP1 | 0.027175255 | 1.019014976 | 0.692949291 | 0.824106983 | 2218.038928 | 2259.786509 | ENSG00000121057 |
| RAD52 | 0.027119889 | 1.018975871 | 0.871775641 | 0.933581194 | 175.8994777 | 179.2923601 | ENSG00000002016 |
| NDUFS7 | 0.027106532 | 1.018966437 | 0.920678531 | 0.960748601 | 63.86063432 | 65.14024783 | ENSG00000115286 |
| ERCC2 | 0.027047884 | 1.018925015 | 0.734094846 | 0.851336652 | 2524.84393 | 2573.222769 | ENSG00000104884 |
| TBL3 | 0.027016038 | 1.018902524 | 0.713631363 | 0.837849642 | 1852.960302 | 1888.439454 | ENSG00000183751 |
| FIS1 | 0.026923647 | 1.018837274 | 0.753495375 | 0.863173452 | 1020.76441 | 1039.724908 | ENSG00000214253 |
| CFAP221 | 0.02680446 | 1.018753107 | 0.978208822 | 0.989993298 | 4.753893469 | 4.864827064 | ENSG00000163075 |
| NA | 0.026724421 | 1.01869659 | 0.954632476 | 0.978185467 | 22.10872209 | 22.39180531 | ENSG00000268034 |
| NSG2 | 0.026717604 | 1.018691777 | 0.978169154 | 0.989993298 | 5.153866493 | 5.262490379 | ENSG00000170091 |
| CEP70 | 0.026706071 | 1.018683633 | 0.821812899 | 0.904391889 | 398.9736127 | 406.4011812 | ENSG00000114107 |
| NA | 0.026634659 | 1.01863321 | 0.979100324 | 0.990296129 | 4.439833508 | 4.54984542 | ENSG00000272572 |
| ANAPC11 | 0.026504198 | 1.018541101 | 0.717454173 | 0.839899927 | 2112.366204 | 2151.646016 | ENSG00000141552 |
| CAPN10 | 0.026456585 | 1.018507486 | 0.821539953 | 0.904297073 | 968.4364729 | 987.3410588 | ENSG00000142330 |
| NA | 0.026430678 | 1.018489197 | 0.946545129 | 0.9733913 | 32.03670131 | 32.66235429 | ENSG00000279059 |
| SVIP | 0.026324702 | 1.018414385 | 0.899143773 | 0.948444134 | 120.9212614 | 123.0249022 | ENSG00000198168 |
| RNF115 | 0.026293067 | 1.018392053 | 0.800723873 | 0.8929372 | 521.391693 | 531.006989 | ENSG00000265491 |
| THAP3 | 0.026277222 | 1.018380869 | 0.868111582 | 0.931325542 | 213.4946486 | 217.3353414 | ENSG00000041988 |
| NA | 0.02625936 | 1.01836826 | 0.944984108 | 0.972933398 | 37.1829762 | 37.78541198 | ENSG00000248927 |
| CDK11B | 0.026198579 | 1.018325357 | 0.730772408 | 0.849344242 | 1550.536505 | 1579.674564 | ENSG00000248333 |
| SREBF2 | 0.026153839 | 1.018293778 | 0.669067868 | 0.807850415 | 7624.819881 | 7764.035635 | ENSG00000198911 |
| SLC36A1 | 0.026150357 | 1.01829132 | 0.829254818 | 0.908089146 | 367.7398189 | 374.7131036 | ENSG00000123643 |
| LRP5 | 0.026016089 | 1.018196555 | 0.705043331 | 0.832415373 | 2117.704397 | 2155.796963 | ENSG00000162337 |
| LRWD1 | 0.025994713 | 1.018181469 | 0.775061526 | 0.875992658 | 880.6477741 | 896.9768443 | ENSG00000161036 |
| RFT1 | 0.025820507 | 1.01805853 | 0.816985438 | 0.901565698 | 445.0108052 | 452.7947657 | ENSG00000163933 |
| CIAO1 | 0.025777213 | 1.018027979 | 0.729721097 | 0.848685417 | 1647.740486 | 1677.130121 | ENSG00000144021 |
| SMARCD1 | 0.025685394 | 1.01796319 | 0.67228382 | 0.809571068 | 3992.841783 | 4064.221049 | ENSG00000066117 |
| GTF2IRD2B | 0.025603473 | 1.017905388 | 0.923959663 | 0.96244767 | 67.07451934 | 68.23051054 | ENSG00000174428 |
| KATNAL1 | 0.025441471 | 1.017791093 | 0.88250216 | 0.939477027 | 171.631568 | 174.4938168 | ENSG00000102781 |
| CNOT11 | 0.025374911 | 1.017744137 | 0.782549496 | 0.880885051 | 1063.130853 | 1081.818404 | ENSG00000158435 |
| CCDC32 | 0.025340348 | 1.017719755 | 0.931518524 | 0.966401586 | 53.73164264 | 54.70163597 | ENSG00000128891 |
| DDN | 0.025234714 | 1.017645241 | 0.972248198 | 0.986675141 | 9.676456153 | 9.879933702 | ENSG00000181418 |

| | | | | | | | |
|---|---|---|---|---|---|---|---|
| COPS8 | 0.025180716 | 1.017607153 | 0.767068951 | 0.871187027 | 1143.855364 | 1163.907989 | ENSG00000198612 |
| NA | 0.025164717 | 1.017595867 | 0.981219572 | 0.991683936 | 4.161777366 | 4.268606918 | ENSG00000232946 |
| RAB3GAP2 | 0.025122005 | 1.017565742 | 0.773183246 | 0.874965662 | 1561.968241 | 1588.399868 | ENSG00000118873 |
| SLC30A6 | 0.025062796 | 1.017523981 | 0.834637845 | 0.911277722 | 553.7351558 | 562.9853999 | ENSG00000152683 |
| PCCB | 0.025058763 | 1.017521137 | 0.750601224 | 0.861278241 | 1955.603661 | 1989.516762 | ENSG00000114054 |
| LOC150776 | 0.025038911 | 1.017507135 | 0.869738924 | 0.932423717 | 344.2135089 | 350.7903591 | ENSG00000152117 |
| ZNF114 | 0.024996034 | 1.017476895 | 0.865932307 | 0.930077643 | 232.2430126 | 236.4502734 | ENSG00000178150 |
| CCDC180 | 0.024987622 | 1.017470962 | 0.968324247 | 0.984740121 | 11.50902753 | 11.69496283 | ENSG00000197816 |
| MRPL32 | 0.024840841 | 1.017367449 | 0.75060698 | 0.861278241 | 1544.379673 | 1571.611635 | ENSG00000106591 |
| THSD7B | 0.024815311 | 1.017349446 | 0.909774695 | 0.955016096 | 105.7646571 | 107.6707013 | ENSG00000144229 |
| NA | 0.024775934 | 1.017321679 | 0.976151048 | 0.989065358 | 8.259740058 | 8.345155393 | ENSG00000263535 |
| GOSR2 | 0.024747895 | 1.017301907 | 0.845745416 | 0.917796448 | 423.3648861 | 431.341344 | ENSG00000108433 |
| FOXB1 | 0.024687042 | 1.017258998 | 0.932417567 | 0.966955415 | 65.03351665 | 66.14497818 | ENSG00000171956 |
| NKAIN3-IT1 | 0.024523506 | 1.017143694 | 0.977009247 | 0.989556227 | 6.047414906 | 6.132462953 | ENSG00000274956 |
| PPP1R12A | 0.024497359 | 1.01712526 | 0.809374902 | 0.897454647 | 1253.960291 | 1274.583434 | ENSG00000058272 |
| NTN4 | 0.024454185 | 1.017094822 | 0.952313214 | 0.976820693 | 28.93092544 | 29.51534057 | ENSG00000074527 |
| MAN1B1 | 0.024445139 | 1.017088444 | 0.762967808 | 0.868655652 | 1059.257099 | 1077.502865 | ENSG00000177239 |
| TCAP | 0.024423011 | 1.017072844 | 0.959306472 | 0.980034514 | 21.78519139 | 22.1524573 | ENSG00000173991 |
| NA | 0.024372139 | 1.017036981 | 0.92308408 | 0.961958973 | 84.81797111 | 86.30608451 | ENSG00000269837 |
| OPA3 | 0.024263525 | 1.016960416 | 0.816630634 | 0.901347795 | 755.7544357 | 769.0476211 | ENSG00000125741 |
| PDIK1L | 0.024181234 | 1.01690241 | 0.93913518 | 0.969905908 | 55.41566637 | 56.31073381 | ENSG00000175087 |
| CCDC82 | 0.024143339 | 1.0168757 | 0.866679392 | 0.930273692 | 313.3942775 | 318.8129267 | ENSG00000149231 |
| NA | 0.024098093 | 1.016843809 | 0.97724746 | 0.989678502 | 6.950434057 | 7.028692595 | ENSG00000285752 |
| MTRR | 0.024069984 | 1.016823997 | 0.795958606 | 0.889764584 | 757.9705716 | 770.9968668 | ENSG00000124275 |
| ZRSR2 | 0.024039531 | 1.016802534 | 0.839358305 | 0.913691856 | 482.3428926 | 491.0452644 | ENSG00000169249 |
| MTF2 | 0.024004879 | 1.016778112 | 0.757906577 | 0.865159875 | 1893.78557 | 1924.770696 | ENSG00000143033 |
| BHLHE40-AS1 | 0.023912275 | 1.016712849 | 0.981383752 | 0.991764802 | 6.301997184 | 6.304762716 | ENSG00000235831 |
| NDUFB9 | 0.023870136 | 1.016683153 | 0.804724298 | 0.895189973 | 880.4271363 | 895.4753945 | ENSG00000147684 |
| RALA | 0.023869115 | 1.016682433 | 0.812397363 | 0.899366839 | 1767.962905 | 1796.976957 | ENSG00000006451 |
| WARS1 | 0.02374758 | 1.01659679 | 0.733314138 | 0.851101774 | 2764.470489 | 2809.595487 | ENSG00000140105 |
| ERI1 | 0.023739971 | 1.016591428 | 0.834248225 | 0.911076206 | 433.9114168 | 440.9795792 | ENSG00000104626 |
| NRAS | 0.023683418 | 1.016551579 | 0.79818174 | 0.891456743 | 1683.294173 | 1710.031319 | ENSG00000213281 |
| MRPS5 | 0.023502144 | 1.016423857 | 0.773895403 | 0.875290011 | 1114.656379 | 1132.428511 | ENSG00000144029 |
| SFN | 0.023485421 | 1.016412076 | 0.902495246 | 0.950449757 | 137.5246777 | 139.5510606 | ENSG00000175793 |
| WARS2 | 0.023344776 | 1.016312993 | 0.908380756 | 0.954511501 | 115.8658379 | 117.6507735 | ENSG00000116874 |
| NDC80 | 0.023308661 | 1.016287552 | 0.817128384 | 0.901628712 | 597.8809983 | 607.4886884 | ENSG00000080986 |
| ZBED9 | 0.023280321 | 1.016267588 | 0.927557248 | 0.964343189 | 76.21320332 | 77.28082482 | ENSG00000232040 |
| NA | 0.023270469 | 1.016260649 | 0.975556857 | 0.988571393 | 8.20163785 | 8.398219182 | ENSG00000250731 |
| FAM72D | 0.023218064 | 1.016223734 | 0.920135614 | 0.960668852 | 89.62247288 | 91.09550118 | ENSG00000215784 |
| PPM1L | 0.023192702 | 1.016205869 | 0.875330466 | 0.935388311 | 267.4022438 | 271.6800021 | ENSG00000163590 |
| EZH1 | 0.023126586 | 1.0161593 | 0.940241092 | 0.970453216 | 358.2415405 | 364.175716 | ENSG00000108799 |
| EXT1 | 0.023044417 | 1.016101426 | 0.811178974 | 0.898809775 | 803.2061796 | 815.8136306 | ENSG00000182197 |
| JCAD | 0.0230271 | 1.016089229 | 0.8871192 | 0.942281349 | 212.1523003 | 215.6948107 | ENSG00000165757 |
| PCDHA4 | 0.022658567 | 1.015829705 | 0.919190988 | 0.960223525 | 102.3523152 | 104.0627748 | ENSG00000204967 |
| INPP5F | 0.022481682 | 1.015705165 | 0.754406828 | 0.863474302 | 1797.886303 | 1825.535693 | ENSG00000198825 |
| RAB39B | 0.022401349 | 1.015648609 | 0.878776121 | 0.937550551 | 272.6898807 | 276.7622686 | ENSG00000155961 |
| VASP | 0.022387064 | 1.015638552 | 0.777726681 | 0.877801979 | 2675.553307 | 2717.37684 | ENSG00000125753 |
| DNAJB4 | 0.022334768 | 1.015601737 | 0.89923468 | 0.948444134 | 164.994788 | 167.7660098 | ENSG00000162616 |
| RMND5A | 0.02232881 | 1.015597543 | 0.815296496 | 0.900518787 | 1646.237654 | 1671.136921 | ENSG00000153561 |
| B4GALT2 | 0.022293128 | 1.015572424 | 0.766135499 | 0.870835943 | 1841.347705 | 1869.098519 | ENSG00000117411 |
| ANKRD18A | 0.022261909 | 1.015550449 | 0.937955745 | 0.969503942 | 56.29343021 | 57.21108872 | ENSG00000180071 |
| PCBD2 | 0.022258847 | 1.015548293 | 0.899737008 | 0.948724226 | 161.8298427 | 164.2663608 | ENSG00000132570 |
| NPIPB3 | 0.022244687 | 1.015538326 | 0.962608407 | 0.981946404 | 23.13059891 | 23.44200176 | ENSG00000169246 |
| RALGAPA2 | 0.022197042 | 1.015504788 | 0.833633132 | 0.910747762 | 603.3495035 | 613.0203997 | ENSG00000188559 |
| RBMXL1 | 0.022089187 | 1.015428872 | 0.837639506 | 0.912848275 | 542.2905595 | 550.2702252 | ENSG00000213516 |
| SLAIN2 | 0.022023178 | 1.015382414 | 0.834735895 | 0.911277722 | 621.824945 | 630.9207907 | ENSG00000109171 |
| DZIP1L | 0.022001554 | 1.015367195 | 0.909273691 | 0.954803834 | 147.6813075 | 150.0592457 | ENSG00000158163 |
| BET1 | 0.021987226 | 1.015357111 | 0.859279498 | 0.925654532 | 379.1875873 | 384.7810333 | ENSG00000105829 |
| GPT2 | 0.021864495 | 1.015270737 | 0.747581839 | 0.859000732 | 2667.805002 | 2709.269147 | ENSG00000166123 |
| PHF8 | 0.021829801 | 1.015246322 | 0.811325564 | 0.898880668 | 920.4377944 | 935.0352677 | ENSG00000172943 |
| ING5 | 0.021738121 | 1.015181808 | 0.825816473 | 0.906340611 | 1093.403706 | 1110.735821 | ENSG00000168395 |
| VSTM2B | 0.021701893 | 1.015156315 | 0.962990207 | 0.982152407 | 25.58941108 | 25.87213364 | ENSG00000187135 |
| AMMECR1L | 0.021685691 | 1.015144915 | 0.814949132 | 0.90018876 | 740.0013557 | 751.190872 | ENSG00000144233 |
| NA | 0.021665239 | 1.015130524 | 0.950746501 | 0.975915798 | 42.41278135 | 42.92344182 | ENSG00000213713 |
| PLCXD2 | 0.02163498 | 1.015109233 | 0.946513067 | 0.9733913 | 62.74780839 | 63.59809507 | ENSG00000240891 |
| SLC37A2 | 0.021592477 | 1.015079328 | 0.945425218 | 0.97314845 | 46.10317942 | 46.82499099 | ENSG00000134955 |
| SMAP1 | 0.021579206 | 1.01506999 | 0.888064477 | 0.942849507 | 250.9104769 | 254.8932399 | ENSG00000112305 |
| STAM | 0.021522482 | 1.015030081 | 0.860005027 | 0.926192962 | 374.7053385 | 380.0291671 | ENSG00000136738 |
| DENND4B | 0.021212623 | 1.014812098 | 0.791052698 | 0.886365438 | 1678.620586 | 1704.458437 | ENSG00000198837 |
| CD151 | 0.02113441 | 1.014757083 | 0.801383008 | 0.893214577 | 1091.215147 | 1107.705805 | ENSG00000177697 |
| HNRNPH3 | 0.021113914 | 1.014742666 | 0.768358431 | 0.871924202 | 4597.773483 | 4665.328611 | ENSG00000096746 |
| COX11 | 0.021067981 | 1.01471036 | 0.833889605 | 0.910904105 | 733.7131511 | 745.1703193 | ENSG00000166260 |
| IMP4 | 0.0210669 | 1.014709599 | 0.775564361 | 0.876319163 | 1980.831952 | 2010.01073 | ENSG00000136718 |
| DHFR2 | 0.021046934 | 1.014695556 | 0.946553977 | 0.9733913 | 54.73269885 | 55.46223178 | ENSG00000178700 |

| | | | | | | | |
|---|---|---|---|---|---|---|---|
| OTUD6B-AS1 | 0.020985153 | 1.014652105 | 0.891904694 | 0.944927495 | 271.3454427 | 275.3562993 | ENSG00000253738 |
| CEP76 | 0.02091274 | 1.014601178 | 0.897673141 | 0.947692289 | 214.1333217 | 216.8017178 | ENSG00000101624 |
| ATP5F1D | 0.020878267 | 1.014576934 | 0.849896616 | 0.920406282 | 1490.018745 | 1511.834565 | ENSG00000099624 |
| BMP1 | 0.020865283 | 1.014567803 | 0.893312246 | 0.945499924 | 214.1807731 | 217.1482927 | ENSG00000168487 |
| SBDS | 0.020824721 | 1.014539279 | 0.877299951 | 0.936413113 | 305.9746289 | 310.1893439 | ENSG00000126524 |
| RUVBL1 | 0.020824627 | 1.014539212 | 0.736076291 | 0.852173027 | 3602.822746 | 3654.75083 | ENSG00000175792 |
| NA | 0.020822648 | 1.014537821 | 0.957962207 | 0.979577828 | 35.55527325 | 35.97319402 | ENSG00000218175 |
| NA | 0.020774972 | 1.014504295 | 0.810641611 | 0.898429206 | 961.2261274 | 974.9413088 | ENSG00000250899 |
| GTF2B | 0.020761684 | 1.014494951 | 0.894727252 | 0.946187088 | 221.1178804 | 224.0193313 | ENSG00000137947 |
| SNX17 | 0.020757248 | 1.014491831 | 0.760894466 | 0.867158734 | 1942.547627 | 1970.406362 | ENSG00000115234 |
| ZNF451 | 0.020713277 | 1.014460911 | 0.863627202 | 0.92882647 | 472.6490059 | 479.0249337 | ENSG00000112200 |
| PNMA5 | 0.020694575 | 1.014447761 | 0.983205058 | 0.992498966 | 6.928335667 | 6.994949453 | ENSG00000198883 |
| FUCA1 | 0.020586941 | 1.01437208 | 0.894179001 | 0.94574241 | 214.5851808 | 217.4970175 | ENSG00000179163 |
| FAM86JP | 0.020516512 | 1.014322562 | 0.96587747 | 0.983602914 | 25.8938049 | 26.17301921 | ENSG00000171084 |
| TNFAIP8 | 0.020426259 | 1.014259109 | 0.897256689 | 0.947456442 | 264.0951624 | 267.3837124 | ENSG00000145779 |
| IWS1 | 0.020415683 | 1.014251674 | 0.763077554 | 0.868712073 | 1938.159274 | 1965.681302 | ENSG00000163166 |
| RPL29 | 0.02035535 | 1.014209259 | 0.74330138 | 0.856684769 | 16698.15888 | 16935.6938 | ENSG00000162244 |
| CRIP2 | 0.020319981 | 1.014184395 | 0.894963875 | 0.946203969 | 218.5543114 | 221.7366678 | ENSG00000182809 |
| TM9SF3 | 0.02031584 | 1.014181483 | 0.824467275 | 0.905596573 | 2828.819707 | 2868.255479 | ENSG00000077147 |
| STT3A | 0.020314521 | 1.014180557 | 0.773053106 | 0.874965662 | 3434.245451 | 3482.732385 | ENSG00000134910 |
| SLX4IP | 0.020297855 | 1.014168841 | 0.921890868 | 0.961386103 | 134.1381925 | 136.0306023 | ENSG00000149346 |
| CCDC12 | 0.020245963 | 1.014132363 | 0.807298101 | 0.896619594 | 1153.369585 | 1170.320028 | ENSG00000160799 |
| RNF181 | 0.020189105 | 1.014092396 | 0.841417671 | 0.914973161 | 646.6156053 | 655.515916 | ENSG00000168894 |
| NA | 0.020100477 | 1.0140301 | 0.98223383 | 0.992045368 | 6.781667144 | 6.845107911 | ENSG00000234645 |
| ARHGAP29 | 0.02006481 | 1.014005031 | 0.937267263 | 0.969033102 | 80.39086295 | 81.31223361 | ENSG00000137962 |
| SH3GL1 | 0.019918653 | 1.013902309 | 0.770821978 | 0.87389372 | 3584.024547 | 3634.048185 | ENSG00000141985 |
| ZFP30 | 0.019907054 | 1.013894157 | 0.945294714 | 0.973068161 | 55.08897875 | 55.89831315 | ENSG00000120784 |
| LINC01108 | 0.019822594 | 1.013834803 | 0.902560529 | 0.950449757 | 288.1069903 | 292.1919117 | ENSG00000226673 |
| TM2D2 | 0.019731666 | 1.013770906 | 0.81390318 | 0.899911056 | 1122.651796 | 1137.770863 | ENSG00000169490 |
| CBARP | 0.019535594 | 1.013633137 | 0.878114943 | 0.937013205 | 472.7553563 | 479.3626455 | ENSG00000099625 |
| RBM15-AS1 | 0.019532217 | 1.013630765 | 0.982718401 | 0.992356045 | 5.686602612 | 5.746084558 | ENSG00000227963 |
| PAPOLG | 0.019512699 | 1.013617051 | 0.874936973 | 0.935237434 | 403.3011129 | 408.2267308 | ENSG00000115421 |
| NDUFA12 | 0.019491266 | 1.013601993 | 0.819317357 | 0.90291228 | 975.3364115 | 988.8876371 | ENSG00000184752 |
| BROX | 0.019471432 | 1.013588058 | 0.903749628 | 0.951223557 | 246.6359829 | 249.4921527 | ENSG00000162819 |
| CORO1C | 0.019445624 | 1.013569926 | 0.786579435 | 0.88311505 | 2804.74484 | 2841.720004 | ENSG00000110880 |
| ELOVL2 | 0.019416698 | 1.013549604 | 0.957424032 | 0.979478064 | 41.19570224 | 41.63703499 | ENSG00000197977 |
| NA | 0.019308282 | 1.013473441 | 0.93291972 | 0.967187875 | 106.0656834 | 107.7346119 | ENSG00000224861 |
| LOC100288123 | 0.019301756 | 1.013468857 | 0.964118666 | 0.982621686 | 25.42695799 | 25.76798144 | ENSG00000267244 |
| NPLOC4 | 0.019297459 | 1.013465838 | 0.754776635 | 0.863520379 | 2980.222276 | 3020.270015 | ENSG00000182446 |
| PIGG | 0.019255055 | 1.01343605 | 0.881661478 | 0.939373634 | 573.1050479 | 581.3312715 | ENSG00000174227 |
| FLRT1 | 0.019209491 | 1.013404044 | 0.95353276 | 0.977638777 | 45.81693032 | 46.56676037 | ENSG00000126500 |
| PALD1 | 0.019086638 | 1.013317751 | 0.786520272 | 0.883102122 | 1756.652928 | 1779.542074 | ENSG00000107719 |
| LOC107984875 | 0.019077174 | 1.013311104 | 0.968275969 | 0.984740121 | 21.14957757 | 21.40540793 | ENSG00000235560 |
| CUL3 | 0.018982971 | 1.01324494 | 0.82722682 | 0.907101046 | 1574.842001 | 1594.891687 | ENSG00000036257 |
| SERPINH1 | 0.018981643 | 1.013244008 | 0.727641227 | 0.846909144 | 15650.09023 | 15857.12849 | ENSG00000149257 |
| ZC3HAV1 | 0.01894914 | 1.01322118 | 0.844638837 | 0.917132591 | 941.9599174 | 953.7764026 | ENSG00000105939 |
| FXR2 | 0.018830075 | 1.013137562 | 0.804110759 | 0.894704967 | 1254.687824 | 1271.163364 | ENSG00000129245 |
| MND1 | 0.018775536 | 1.013099263 | 0.888214032 | 0.942849507 | 299.3719736 | 303.3976262 | ENSG00000121211 |
| CDCA2 | 0.018717708 | 1.013058656 | 0.828626282 | 0.907669001 | 950.6616625 | 963.0913285 | ENSG00000184661 |
| CA4 | 0.018696295 | 1.01304362 | 0.954433464 | 0.978183467 | 48.46029396 | 49.14651055 | ENSG00000167434 |
| JAM3 | 0.01863778 | 1.013002532 | 0.826045218 | 0.906395244 | 1336.193181 | 1353.477786 | ENSG00000166086 |
| GRINA | 0.018634106 | 1.012999952 | 0.844308658 | 0.917016342 | 1008.908401 | 1021.526018 | ENSG00000178719 |
| INTS1 | 0.018535683 | 1.012930846 | 0.758103114 | 0.86529526 | 5622.189779 | 5695.695756 | ENSG00000164880 |
| NA | 0.018188055 | 1.012686802 | 0.973879651 | 0.98768183 | 15.15207191 | 15.34054288 | ENSG00000286018 |
| WDR33 | 0.018144155 | 1.012655988 | 0.801304547 | 0.893214577 | 2139.708069 | 2166.019203 | ENSG00000136709 |
| KNSTRN | 0.018102605 | 1.012626823 | 0.82669261 | 0.906810387 | 1126.462577 | 1140.656947 | ENSG00000128944 |
| NDUFAF5 | 0.018066199 | 1.01260127 | 0.910199388 | 0.95534368 | 251.7125353 | 254.7090024 | ENSG00000101247 |
| ALDH9A1 | 0.018054098 | 1.012592776 | 0.796674251 | 0.890296085 | 1799.39139 | 1821.860201 | ENSG00000143149 |
| NRXN3 | 0.018050859 | 1.012590503 | 0.95107192 | 0.976087655 | 56.35418567 | 57.16594903 | ENSG00000021645 |
| NA | 0.018030885 | 1.012576484 | 0.919946101 | 0.960579213 | 229.3855052 | 232.6836423 | ENSG00000183458 |
| PLA2G15 | 0.018028019 | 1.012574473 | 0.920489681 | 0.960700166 | 179.2097161 | 181.3761403 | ENSG00000103066 |
| RPL38 | 0.018022185 | 1.012570378 | 0.811490206 | 0.898880668 | 7127.158959 | 7216.218879 | ENSG00000172809 |
| HRC | 0.017795622 | 1.012411375 | 0.948817562 | 0.974691536 | 73.61591559 | 74.47223865 | ENSG00000130528 |
| PDE9A | 0.017739967 | 1.01237232 | 0.843259761 | 0.91629598 | 918.1114748 | 929.2687229 | ENSG00000160191 |
| KIF26B | 0.017696877 | 1.012342083 | 0.96942264 | 0.985330341 | 23.46793734 | 23.674963 | ENSG00000162849 |
| NA | 0.017684827 | 1.012333628 | 0.988000422 | 0.994484312 | 4.08976973 | 4.159668179 | ENSG00000260532 |
| NAAA | 0.01766565 | 1.012320171 | 0.957559664 | 0.979493552 | 45.2216573 | 45.64653516 | ENSG00000138744 |
| MFSD2B | 0.017531604 | 1.012226117 | 0.979961269 | 0.990886688 | 9.487567685 | 9.617028162 | ENSG00000205639 |
| TEX15 | 0.01749084 | 1.012197517 | 0.905544128 | 0.952625432 | 271.1550811 | 274.2257859 | ENSG00000133863 |
| ASB7 | 0.017467095 | 1.012180858 | 0.915770681 | 0.95827165 | 199.5540184 | 201.7654213 | ENSG00000183475 |
| ANKH | 0.017397054 | 1.012131718 | 0.892614316 | 0.945187103 | 324.690146 | 328.7459242 | ENSG00000154122 |
| ANKRD27 | 0.017350908 | 1.012099345 | 0.818621655 | 0.902535527 | 1596.733836 | 1616.242982 | ENSG00000105186 |
| LMTK2 | 0.01725912 | 1.012034954 | 0.865944739 | 0.930077643 | 869.989098 | 880.435362 | ENSG00000164715 |
| MED8 | 0.017252795 | 1.012030517 | 0.912925464 | 0.956697984 | 204.1669785 | 206.6350349 | ENSG00000159479 |

| | | | | | | | |
|---|---|---|---|---|---|---|---|
| DNAJC15 | 0.017252663 | 1.012030425 | 0.88958812 | 0.943605918 | 418.7645689 | 423.2795368 | ENSG00000120675 |
| RAMAC | 0.017215874 | 1.012004619 | 0.944844534 | 0.972886876 | 92.2570422 | 93.63219868 | ENSG00000169612 |
| COX7C | 0.017215399 | 1.012004286 | 0.784391991 | 0.882048366 | 4813.270569 | 4870.33787 | ENSG00000127184 |
| HP1BP3 | 0.017127789 | 1.011942831 | 0.93332769 | 0.967249799 | 1428.41679 | 1445.803164 | ENSG00000127483 |
| UNC50 | 0.017002915 | 1.011855245 | 0.880122423 | 0.938290997 | 465.6996543 | 471.5029396 | ENSG00000115446 |
| ADAMTS12 | 0.016859793 | 1.01175487 | 0.90189677 | 0.95021014 | 399.7215901 | 403.895304 | ENSG00000151388 |
| CNOT4 | 0.016641234 | 1.011601607 | 0.911145712 | 0.955762377 | 322.5500989 | 326.0269158 | ENSG00000080802 |
| BTN2A3P | 0.016626685 | 1.011591405 | 0.957864477 | 0.979577828 | 58.2287816 | 58.83928396 | ENSG00000124549 |
| FEM1B | 0.016578272 | 1.01155746 | 0.842799563 | 0.915940372 | 1322.115466 | 1336.520747 | ENSG00000169018 |
| SMPD3 | 0.016521708 | 1.011517801 | 0.97074666 | 0.986015158 | 34.63509406 | 35.127217 | ENSG00000103056 |
| PDXDC1 | 0.016453294 | 1.011469834 | 0.808762056 | 0.897311939 | 2740.378611 | 2772.472087 | ENSG00000179889 |
| FKBP14 | 0.016270353 | 1.011341583 | 0.939573773 | 0.970088592 | 168.3005916 | 170.1755069 | ENSG00000106080 |
| TSLP | 0.016259011 | 1.011333632 | 0.986023301 | 0.993685984 | 5.488145708 | 5.536242807 | ENSG00000145777 |
| PPP3CA | 0.016214441 | 1.011302389 | 0.88513485 | 0.940982524 | 1184.38943 | 1196.957222 | ENSG00000138814 |
| CDC42EP3 | 0.016173616 | 1.011273771 | 0.882977092 | 0.939798298 | 535.0794247 | 540.7080843 | ENSG00000163171 |
| PLEKHA1 | 0.01584815 | 1.011045659 | 0.868961226 | 0.931913399 | 646.7406639 | 654.0453747 | ENSG00000107679 |
| UBXN11 | 0.015830569 | 1.011033338 | 0.936924775 | 0.969033102 | 201.5351375 | 204.2572427 | ENSG00000158062 |
| TLCD3A | 0.015763006 | 1.010985991 | 0.886751882 | 0.942266356 | 480.9719593 | 486.697663 | ENSG00000167695 |
| PLEKHG4B | 0.015707315 | 1.010946965 | 0.856498905 | 0.923894233 | 1233.532917 | 1247.607941 | ENSG00000153404 |
| ZNF865 | 0.015684169 | 1.010930746 | 0.911163665 | 0.955762377 | 579.4682361 | 585.9880082 | ENSG00000261221 |
| ATP2A2 | 0.01557899 | 1.010857047 | 0.786345815 | 0.883066732 | 7799.96112 | 7884.153645 | ENSG00000174437 |
| SCN1A | 0.015577895 | 1.010856281 | 0.971601233 | 0.9862886 | 28.76847235 | 29.09533065 | ENSG00000144285 |
| TRAF3 | 0.015570815 | 1.010851319 | 0.885056147 | 0.940967086 | 517.148941 | 522.5287877 | ENSG00000131323 |
| RESF1 | 0.015556712 | 1.010841438 | 0.859025357 | 0.925476317 | 1238.743947 | 1251.937183 | ENSG00000174718 |
| RAP1B | 0.015550647 | 1.010837188 | 0.8450228 | 0.917436773 | 1421.893809 | 1436.826957 | ENSG00000127314 |
| F11R | 0.0154258 | 1.010749717 | 0.839354073 | 0.913691856 | 1608.525617 | 1625.341381 | ENSG00000158769 |
| RALGAPA1P1 | 0.015416523 | 1.010743218 | 0.986422566 | 0.993714308 | 5.481831882 | 5.524957886 | ENSG00000229419 |
| G3BP2 | 0.015402311 | 1.010733261 | 0.812156336 | 0.899301322 | 5336.361822 | 5392.510796 | ENSG00000138757 |
| SRPK1 | 0.015400471 | 1.010731972 | 0.845396258 | 0.91752499 | 1447.669471 | 1462.672101 | ENSG00000096063 |
| BAG2 | 0.015312677 | 1.010670466 | 0.897141441 | 0.947442705 | 438.0255474 | 442.3821877 | ENSG00000112208 |
| ELOVL5 | 0.015300162 | 1.010661699 | 0.838290186 | 0.913181516 | 3633.959495 | 3671.339335 | ENSG00000012660 |
| NA | 0.015283741 | 1.010650196 | 0.978822553 | 0.990148303 | 15.02562264 | 15.17584081 | ENSG00000253931 |
| TUBG1 | 0.015210588 | 1.010598951 | 0.838234948 | 0.913175014 | 1687.656865 | 1705.10302 | ENSG00000131462 |
| ENC1 | 0.01520832 | 1.010597362 | 0.893002574 | 0.945341234 | 546.2057521 | 551.4584286 | ENSG00000171617 |
| CRADD | 0.015174165 | 1.010573438 | 0.956578016 | 0.979215959 | 67.39932781 | 68.26881698 | ENSG00000169372 |
| LARS1 | 0.015054251 | 1.010489444 | 0.785409204 | 0.882568409 | 6051.649088 | 6114.753887 | ENSG00000133706 |
| NA | 0.015049278 | 1.010485961 | 0.985551029 | 0.993485135 | 6.781667144 | 6.800079849 | ENSG00000242798 |
| LPIN3 | 0.014972927 | 1.010432485 | 0.945145866 | 0.972968978 | 126.6518654 | 128.1928134 | ENSG00000132793 |
| COL4A2-AS1 | 0.014883635 | 1.010369949 | 0.990204084 | 0.995467699 | 3.830655064 | 3.893513444 | ENSG00000232814 |
| NIBAN3 | 0.014817632 | 1.010323726 | 0.986863503 | 0.993845887 | 5.42560881 | 5.464846056 | ENSG00000167483 |
| DVL1 | 0.014611841 | 1.01017962 | 0.852969047 | 0.922280546 | 2179.969326 | 2202.911598 | ENSG00000107404 |
| RELT | 0.014548257 | 1.010135099 | 0.943214861 | 0.97211501 | 136.7653656 | 138.4427276 | ENSG00000054967 |
| ZNF566 | 0.01443093 | 1.010052953 | 0.944277153 | 0.972506821 | 160.2113999 | 161.7926979 | ENSG00000186017 |
| SMARCAL1 | 0.014334359 | 1.009985345 | 0.901206627 | 0.949787537 | 515.7629222 | 520.9103803 | ENSG00000138375 |
| PHF20L1 | 0.014181727 | 1.009878497 | 0.905673733 | 0.9527077 | 454.6770013 | 459.0609988 | ENSG00000129292 |
| QPRT | 0.014168948 | 1.009869552 | 0.823411614 | 0.905075215 | 3973.511712 | 4012.735182 | ENSG00000103485 |
| PAIP2B | 0.014113528 | 1.00983076 | 0.88185063 | 0.939373634 | 882.962075 | 891.6565354 | ENSG00000124374 |
| DCAF13 | 0.013991023 | 1.009745015 | 0.845067946 | 0.917436773 | 2179.092186 | 2199.611691 | ENSG00000164934 |
| LEPROTL1 | 0.013879769 | 1.009667151 | 0.891466596 | 0.944698031 | 818.9631528 | 826.7984332 | ENSG00000104660 |
| WNT11 | 0.013835074 | 1.009635872 | 0.990436521 | 0.995539372 | 3.728957436 | 3.792060779 | ENSG00000085741 |
| NA | 0.013822457 | 1.009627041 | 0.981654895 | 0.991822494 | 15.68974638 | 15.81340179 | ENSG00000268649 |
| BRF1 | 0.013747016 | 1.009574248 | 0.901985756 | 0.95021014 | 558.9322117 | 564.7842149 | ENSG00000185024 |
| PREB | 0.013702532 | 1.009543119 | 0.850028418 | 0.920406282 | 1532.190256 | 1546.613407 | ENSG00000138073 |
| NRL | 0.013675076 | 1.009523907 | 0.983569248 | 0.992668425 | 10.46387225 | 10.53582764 | ENSG00000129535 |
| DSCAM | 0.013620267 | 1.009485555 | 0.988309639 | 0.994696218 | 6.117543405 | 6.143747873 | ENSG00000171587 |
| ST3GAL1 | 0.013572 | 1.009451782 | 0.933567823 | 0.96733214 | 214.5904122 | 216.785137 | ENSG00000008513 |
| COP1 | 0.013563007 | 1.00944549 | 0.884754663 | 0.940944853 | 996.6374005 | 1005.157196 | ENSG00000143207 |
| RPL11 | 0.013549899 | 1.009436318 | 0.802571428 | 0.894029261 | 14807.58057 | 14948.11516 | ENSG00000142676 |
| GNPTAB | 0.013424864 | 1.009348837 | 0.871835195 | 0.933591027 | 1120.152545 | 1130.507497 | ENSG00000111670 |
| FOXRED2 | 0.013233725 | 1.009215119 | 0.910791309 | 0.95570436 | 662.2170481 | 668.6524062 | ENSG00000100350 |
| LRRC40 | 0.013164094 | 1.009166411 | 0.905480261 | 0.952612315 | 754.8272961 | 761.0544876 | ENSG00000066557 |
| NA | 0.013110274 | 1.009128764 | 0.98949648 | 0.995302243 | 5.035106525 | 5.070981588 | ENSG00000197320 |
| TESK2 | 0.01304521 | 1.009083255 | 0.949138 | 0.97469657 | 118.902763 | 119.8863622 | ENSG00000070759 |
| PON2 | 0.013042761 | 1.009081542 | 0.931467064 | 0.966401586 | 256.7039109 | 258.960765 | ENSG00000105854 |
| NA | 0.013000945 | 1.009052295 | 0.988598027 | 0.994908619 | 6.289467231 | 6.304762716 | ENSG00000231579 |
| IZUMO1 | 0.012963701 | 1.009026246 | 0.983639289 | 0.992668425 | 14.26927201 | 14.50866195 | ENSG00000182264 |
| FOXJ3 | 0.012924731 | 1.008998991 | 0.897503648 | 0.947625465 | 716.579186 | 722.5413385 | ENSG00000198815 |
| CCDC22 | 0.012756056 | 1.008881029 | 0.927900049 | 0.964537269 | 278.6855073 | 281.1211464 | ENSG00000101997 |
| VAPB | 0.012696283 | 1.00883923 | 0.892642919 | 0.945187103 | 1177.688888 | 1187.131921 | ENSG00000124164 |
| NA | 0.012688778 | 1.008833982 | 0.968664456 | 0.984740121 | 45.90797713 | 46.35972977 | ENSG00000235944 |
| VAPA | 0.012687454 | 1.008833056 | 0.876596353 | 0.936039454 | 1727.205788 | 1741.735995 | ENSG00000101558 |
| ZC2HC1C | 0.012676181 | 1.008825174 | 0.974689682 | 0.988178905 | 30.40584144 | 30.75303216 | ENSG00000119703 |
| SNRNP27 | 0.012656871 | 1.00881167 | 0.947526792 | 0.974013397 | 175.8176911 | 177.5367899 | ENSG00000124380 |
| PLA2G4A | 0.012557099 | 1.008741907 | 0.976728935 | 0.989434526 | 27.5672755 | 27.96277895 | ENSG00000116711 |

| | | | | | | | |
|---|---|---|---|---|---|---|---|
| NA | 0.012543719 | 1.008732552 | 0.99115948 | 0.996103991 | 3.732114349 | 3.792060779 | ENSG00000231991 |
| PPP1R12C | 0.012477439 | 1.00868621 | 0.888255348 | 0.942849507 | 1726.007372 | 1741.937255 | ENSG00000125503 |
| COX6B1 | 0.012434201 | 1.00865598 | 0.871636495 | 0.933486121 | 2608.001388 | 2630.674766 | ENSG00000126267 |
| BRPF1 | 0.01243191 | 1.008654378 | 0.889948156 | 0.943785311 | 794.5257209 | 801.7037716 | ENSG00000156983 |
| EIF2AK2 | 0.01233499 | 1.008586619 | 0.901349998 | 0.949884623 | 881.8230392 | 888.8943938 | ENSG00000055332 |
| ABI2 | 0.012267944 | 1.008539748 | 0.866517962 | 0.930273692 | 1622.775437 | 1635.98031 | ENSG00000138443 |
| USP47 | 0.012257788 | 1.008532649 | 0.908898572 | 0.954691195 | 574.3835664 | 579.2821013 | ENSG00000170242 |
| RSF1 | 0.012207394 | 1.008497421 | 0.879407056 | 0.937806595 | 1355.222127 | 1366.223289 | ENSG00000048649 |
| NA | 0.012186939 | 1.008483122 | 0.988956234 | 0.995026193 | 6.136484882 | 6.151233947 | ENSG00000214719 |
| NA | 0.012172815 | 1.008473249 | 0.979551558 | 0.990615408 | 22.11325448 | 22.34341644 | ENSG00000245205 |
| NA | 0.012009535 | 1.008359119 | 0.988959615 | 0.995026193 | 6.133327969 | 6.143747873 | ENSG00000236439 |
| TRAPPC1 | 0.012000401 | 1.008352735 | 0.897757891 | 0.947692289 | 840.8504177 | 848.0307459 | ENSG00000170043 |
| C3orf33 | 0.011996614 | 1.008350088 | 0.988045436 | 0.994484312 | 8.151630906 | 8.135313641 | ENSG00000174928 |
| NAPEPLD | 0.011972908 | 1.00833352 | 0.954646527 | 0.978185467 | 129.2476799 | 130.1980372 | ENSG00000161048 |
| NA | 0.011972668 | 1.008333352 | 0.989624594 | 0.995302243 | 5.703664953 | 5.723514717 | ENSG00000230896 |
| UGT8 | 0.011960164 | 1.008324612 | 0.915363671 | 0.958169563 | 428.8104211 | 432.335402 | ENSG00000174607 |
| CNOT2 | 0.011930292 | 1.008303734 | 0.887437435 | 0.942496843 | 1146.977059 | 1155.833623 | ENSG00000111596 |
| KCNQ1 | 0.01186405 | 1.008257439 | 0.964432109 | 0.982728532 | 72.04836674 | 72.60381932 | ENSG00000053918 |
| PPM1E | 0.011762926 | 1.008186769 | 0.93474595 | 0.967799644 | 352.1924041 | 354.6220312 | ENSG00000175175 |
| CD1D | 0.011754418 | 1.008180823 | 0.975061905 | 0.988231933 | 41.36890384 | 41.62954887 | ENSG00000158473 |
| NA | 0.011659428 | 1.008114445 | 0.977290166 | 0.989678502 | 29.48260304 | 29.68351507 | ENSG00000257270 |
| ZNF226 | 0.01163219 | 1.008095411 | 0.943748856 | 0.972449807 | 199.3724511 | 200.9832434 | ENSG00000167380 |
| FUBP3 | 0.011625597 | 1.008090805 | 0.873369534 | 0.934693202 | 2664.534961 | 2685.69061 | ENSG00000107164 |
| NA | 0.011538275 | 1.00802979 | 0.986874917 | 0.993845887 | 9.193726977 | 9.27567783 | ENSG00000215210 |
| AHDC1 | 0.011523685 | 1.008019596 | 0.896191038 | 0.946766059 | 878.0602427 | 884.6593555 | ENSG00000126705 |
| NA | 0.0115095 | 1.008009685 | 0.985112045 | 0.993365976 | 13.5336443 | 13.66450841 | ENSG00000272631 |
| HNRNPA1 | 0.011494849 | 1.007999448 | 0.823310574 | 0.905075215 | 59684.66678 | 60161.33824 | ENSG00000135486 |
| UBE2V2 | 0.011348739 | 1.007897367 | 0.920493633 | 0.960700166 | 1209.333854 | 1218.218245 | ENSG00000169139 |
| CHAF1A | 0.011323914 | 1.007880024 | 0.85172205 | 0.921524381 | 4485.444295 | 4521.666893 | ENSG00000167670 |
| ESCO2 | 0.01130031 | 1.007863535 | 0.940705609 | 0.970716428 | 247.0576484 | 248.8186987 | ENSG00000171320 |
| IGLON5 | 0.011292138 | 1.007857826 | 0.885853152 | 0.941584121 | 1454.734749 | 1466.409274 | ENSG00000142549 |
| POU3F1 | 0.01112985 | 1.007744458 | 0.90405852 | 0.95138659 | 1073.661216 | 1082.450168 | ENSG00000185668 |
| ADD2 | 0.011104968 | 1.007727078 | 0.841465281 | 0.914973161 | 6817.680792 | 6870.791047 | ENSG00000075340 |
| BORA | 0.011066224 | 1.007700016 | 0.953014355 | 0.977269451 | 156.622429 | 157.897361 | ENSG00000136122 |
| ASAH2 | 0.01093683 | 1.00760964 | 0.982177851 | 0.992045368 | 21.88067289 | 21.96039886 | ENSG00000188611 |
| BTBD10 | 0.010851326 | 1.007549924 | 0.920804467 | 0.96082592 | 501.9800772 | 505.8504231 | ENSG00000148925 |
| NUP214 | 0.010838232 | 1.007540779 | 0.902731625 | 0.950529936 | 1013.434065 | 1020.531849 | ENSG00000126883 |
| PPP4R4 | 0.010749462 | 1.007478787 | 0.956572108 | 0.979215959 | 130.2702558 | 131.1499186 | ENSG00000119698 |
| LAMB3 | 0.010663793 | 1.007418963 | 0.980982562 | 0.991575633 | 22.88351053 | 23.07114516 | ENSG00000196878 |
| HMGXB4 | 0.010487347 | 1.00729576 | 0.908445935 | 0.954511501 | 974.6395612 | 982.0866728 | ENSG00000100281 |
| ZNF785 | 0.010479009 | 1.007289938 | 0.964532311 | 0.982773054 | 106.2050686 | 107.2295472 | ENSG00000197162 |
| SIK2 | 0.010400709 | 1.007235271 | 0.924042199 | 0.96244767 | 528.2419178 | 531.7940567 | ENSG00000170145 |
| LASP1 | 0.010337953 | 1.007191458 | 0.857734575 | 0.924527631 | 5326.033471 | 5364.297345 | ENSG00000002834 |
| GEMIN4 | 0.010283613 | 1.007153523 | 0.893522822 | 0.945506822 | 1577.05698 | 1588.21884 | ENSG00000179409 |
| CDK16 | 0.010277263 | 1.00714909 | 0.870520518 | 0.932827673 | 4319.91717 | 4350.565822 | ENSG00000102225 |
| FBH1 | 0.010161646 | 1.00706838 | 0.907058305 | 0.95373113 | 1027.457293 | 1035.065105 | ENSG00000134452 |
| CCNF | 0.010049193 | 1.006989886 | 0.903025565 | 0.950731414 | 1146.620381 | 1154.50621 | ENSG00000162063 |
| ZNF813 | 0.010020666 | 1.006969974 | 0.940049753 | 0.970363805 | 431.2541251 | 433.6701575 | ENSG00000198346 |
| VTI1B | 0.010000569 | 1.006955947 | 0.928064716 | 0.964548443 | 465.6075251 | 468.6151039 | ENSG00000100568 |
| RIOX2 | 0.009983525 | 1.006944051 | 0.945014582 | 0.972933398 | 269.6918232 | 271.4788489 | ENSG00000170854 |
| NA | 0.009835797 | 1.006840948 | 0.981495953 | 0.991804865 | 27.22342785 | 27.43371867 | ENSG00000285679 |
| LRRN4 | 0.009813431 | 1.006825339 | 0.988646323 | 0.994908619 | 11.06357995 | 11.09070696 | ENSG00000125872 |
| UBE2M | 0.009773336 | 1.006797358 | 0.877982079 | 0.937000286 | 2729.030263 | 2747.858829 | ENSG00000130725 |
| SFXN1 | 0.009764797 | 1.006791399 | 0.895039896 | 0.946203969 | 2058.629917 | 2071.746066 | ENSG00000164466 |
| NA | 0.009753535 | 1.00678354 | 0.974215538 | 0.987914361 | 53.03339422 | 53.47154207 | ENSG00000228782 |
| PIP4K2A | 0.009724986 | 1.006763618 | 0.921633049 | 0.961257565 | 788.9464156 | 794.1602977 | ENSG00000150867 |
| MTHFSD | 0.009659431 | 1.006717872 | 0.959439615 | 0.980038391 | 140.9256698 | 141.6964815 | ENSG00000103248 |
| LIMD2 | 0.009589007 | 1.006668731 | 0.921585102 | 0.961257565 | 1416.787139 | 1426.889216 | ENSG00000136490 |
| CAPRIN1 | 0.009543271 | 1.006636818 | 0.882331825 | 0.939381593 | 7134.254079 | 7180.472158 | ENSG00000135387 |
| MEAK7 | 0.00953621 | 1.006631891 | 0.945650803 | 0.973218501 | 453.236338 | 455.6023558 | ENSG00000140950 |
| EXO5 | 0.009432356 | 1.00655943 | 0.974757663 | 0.988193771 | 56.70593319 | 56.92561333 | ENSG00000164002 |
| TMEM144 | 0.009427267 | 1.00655588 | 0.978384734 | 0.99001054 | 39.05726386 | 39.25964043 | ENSG00000164124 |
| KDM3B | 0.009380703 | 1.006523393 | 0.882015949 | 0.939373634 | 3174.957497 | 3195.359382 | ENSG00000120733 |
| DNAJC5 | 0.009325067 | 1.006484578 | 0.899789765 | 0.948725879 | 1732.122871 | 1743.580523 | ENSG00000101152 |
| RAD50 | 0.009285273 | 1.006456817 | 0.968267694 | 0.984740121 | 110.3577812 | 110.7980679 | ENSG00000113522 |
| TRAM1 | 0.009230316 | 1.006418478 | 0.917246198 | 0.959219153 | 1641.095167 | 1650.885018 | ENSG00000067167 |
| CTTN | 0.009205648 | 1.00640127 | 0.896603515 | 0.947090479 | 5023.999659 | 5057.402536 | ENSG00000085733 |
| DNASE2 | 0.00906364 | 1.006302213 | 0.943582692 | 0.972401864 | 423.0998636 | 425.3705714 | ENSG00000105612 |
| PDK3 | 0.008997919 | 1.006256372 | 0.92931718 | 0.965198354 | 1052.597927 | 1058.32207 | ENSG00000067992 |
| CNOT3 | 0.008687495 | 1.006039879 | 0.913411894 | 0.957099505 | 1736.915843 | 1747.889397 | ENSG00000088038 |
| LIME1 | 0.008669268 | 1.00602717 | 0.990397292 | 0.995539372 | 10.63707385 | 10.77247612 | ENSG00000203896 |
| PLCG1-AS1 | 0.008659271 | 1.006020199 | 0.990038253 | 0.995445718 | 10.62316842 | 10.76867727 | ENSG00000226648 |
| ETFDH | 0.008591 | 1.005972592 | 0.958146311 | 0.979577828 | 196.7968332 | 197.7564708 | ENSG00000171503 |
| SLC27A6 | 0.008528601 | 1.005929084 | 0.968694393 | 0.984740121 | 111.1536607 | 111.785015 | ENSG00000113396 |

| | | | | | | | |
|---|---|---|---|---|---|---|---|
| KANSL1-AS1 | 0.008521635 | 1.005924227 | 0.990416919 | 0.995539372 | 10.01704919 | 10.05603231 | ENSG00000214401 |
| EOGT | 0.008513358 | 1.005918455 | 0.952930486 | 0.977237516 | 299.8180225 | 301.5588163 | ENSG00000163378 |
| HADHA | 0.008408492 | 1.00584534 | 0.892659709 | 0.945187103 | 4179.935695 | 4203.638993 | ENSG00000084754 |
| AAR2 | 0.008365965 | 1.005815691 | 0.937080402 | 0.969033102 | 792.5515549 | 796.4171422 | ENSG00000131043 |
| CEBPZ | 0.008327717 | 1.005789026 | 0.927772527 | 0.964512901 | 2458.931761 | 2472.388889 | ENSG00000115816 |
| RAB3A | 0.008308467 | 1.005775605 | 0.961554698 | 0.981236389 | 192.7508543 | 193.5201813 | ENSG00000105649 |
| PKN2 | 0.008301555 | 1.005770786 | 0.941652455 | 0.971044712 | 589.6375012 | 592.4422776 | ENSG00000065243 |
| B3GNT7 | 0.008233687 | 1.005723474 | 0.913221253 | 0.956953847 | 2226.140236 | 2237.720564 | ENSG00000156966 |
| TRIM9 | 0.008119978 | 1.005644209 | 0.970102461 | 0.985738981 | 110.974243 | 111.6951736 | ENSG00000100505 |
| XKR6 | 0.008116338 | 1.005641672 | 0.958446111 | 0.979577828 | 228.8839322 | 230.2531353 | ENSG00000171044 |
| DIAPH1 | 0.008022728 | 1.005576422 | 0.9035419 | 0.951166965 | 4014.71987 | 4036.700378 | ENSG00000131504 |
| FBXO31 | 0.007863611 | 1.005465522 | 0.946845969 | 0.973529524 | 509.4004174 | 511.6230509 | ENSG00000103264 |
| TRIM14 | 0.007835966 | 1.005446255 | 0.94394012 | 0.972449807 | 504.261659 | 506.5605431 | ENSG00000106785 |
| CPD | 0.007770002 | 1.005400284 | 0.936799107 | 0.969033102 | 1445.298796 | 1452.202356 | ENSG00000108582 |
| NA | 0.007758016 | 1.005391932 | 0.9584058 | 0.979577828 | 475.9187459 | 478.0319432 | ENSG00000272888 |
| FKBP10 | 0.007734225 | 1.005375352 | 0.919833524 | 0.960515777 | 4399.026906 | 4423.809886 | ENSG00000141756 |
| ATL2 | 0.007602292 | 1.005283415 | 0.933841843 | 0.96733214 | 1306.375615 | 1312.238669 | ENSG00000119787 |
| ZNF444 | 0.007562731 | 1.005255849 | 0.941332242 | 0.970822535 | 860.5018083 | 865.0787815 | ENSG00000167685 |
| MPV17L2 | 0.007523172 | 1.005228285 | 0.937296667 | 0.969033102 | 821.4550749 | 826.0611088 | ENSG00000254858 |
| DEF8 | 0.007484144 | 1.005201092 | 0.926593272 | 0.96415224 | 1366.612928 | 1374.54738 | ENSG00000140995 |
| PLPPR3 | 0.007206316 | 1.005007534 | 0.943076342 | 0.97211501 | 1553.539546 | 1561.061947 | ENSG00000129951 |
| EMC1 | 0.007134445 | 1.004957468 | 0.920336471 | 0.96068883 | 1555.583247 | 1563.410609 | ENSG00000127463 |
| KLHDC10 | 0.007133344 | 1.004956701 | 0.958416452 | 0.979577828 | 431.8123346 | 433.4890491 | ENSG00000128607 |
| NA | 0.007132255 | 1.004955943 | 0.995516932 | 0.998401192 | 3.811713587 | 3.844686535 | ENSG00000279140 |
| EFCAB2 | 0.007130743 | 1.004954889 | 0.972903256 | 0.987015652 | 112.1761389 | 112.6206947 | ENSG00000203666 |
| PRXL2C | 0.006989947 | 1.004856819 | 0.976335648 | 0.989093283 | 89.27872293 | 89.60954978 | ENSG00000158122 |
| PEX7 | 0.006982978 | 1.004851965 | 0.970407868 | 0.985897567 | 154.0435565 | 154.5428786 | ENSG00000112357 |
| FGF12-AS2 | 0.006979094 | 1.004849259 | 0.99164032 | 0.996317108 | 11.28413525 | 11.39875218 | ENSG00000230126 |
| RPL31P11 | 0.006961433 | 1.004836958 | 0.994188477 | 0.997709691 | 5.08501577 | 5.089752583 | ENSG00000213075 |
| ARHGAP21 | 0.006937592 | 1.004820353 | 0.948286332 | 0.97445646 | 746.372324 | 750.1979678 | ENSG00000107863 |
| FAM3A | 0.006915897 | 1.004805243 | 0.953093388 | 0.977296423 | 400.3358871 | 402.2953497 | ENSG00000071889 |
| CACFD1 | 0.00683184 | 1.004746701 | 0.949110278 | 0.97469657 | 639.5817759 | 642.7107565 | ENSG00000160325 |
| PAPOLA | 0.006830057 | 1.004745459 | 0.923351984 | 0.962184058 | 4266.264119 | 4285.212777 | ENSG00000090060 |
| FAM155A | 0.006775594 | 1.00470753 | 0.963541046 | 0.982393082 | 277.9400029 | 279.777814 | ENSG00000204442 |
| NA | 0.006753677 | 1.004692266 | 0.991488458 | 0.996317108 | 13.19620817 | 13.27064394 | ENSG00000282381 |
| PFN2 | 0.006747197 | 1.004687754 | 0.918333185 | 0.959665811 | 2353.247486 | 2363.922665 | ENSG00000070087 |
| SH2D4A | 0.006670044 | 1.004634026 | 0.972462372 | 0.986838459 | 158.9850606 | 160.0258747 | ENSG00000104611 |
| MARCHF6 | 0.0066671 | 1.004631976 | 0.937119102 | 0.969033102 | 1558.845245 | 1566.101091 | ENSG00000145495 |
| NA | 0.006637651 | 1.004611469 | 0.995074096 | 0.998089798 | 4.194624271 | 4.227266083 | ENSG00000232389 |
| NA | 0.006618 | 1.004597786 | 0.979980198 | 0.990886688 | 70.47423361 | 70.84499156 | ENSG00000204253 |
| TTC9 | 0.006546602 | 1.00454807 | 0.94044163 | 0.9705521 | 976.9076057 | 981.406697 | ENSG00000133985 |
| GTF2IRD1 | 0.006503055 | 1.004517749 | 0.94033006 | 0.970490997 | 921.1021361 | 925.4555264 | ENSG00000006704 |
| DCAF15 | 0.006478916 | 1.004500941 | 0.924418191 | 0.962591536 | 1897.101777 | 1905.299204 | ENSG00000132017 |
| ZNF232 | 0.006473516 | 1.004497182 | 0.956737816 | 0.979215959 | 435.3213381 | 437.5000153 | ENSG00000167840 |
| PSMG2 | 0.006441931 | 1.00447519 | 0.952295373 | 0.976820693 | 604.0367928 | 606.4361901 | ENSG00000128789 |
| CBLL1 | 0.006364812 | 1.004421498 | 0.95676001 | 0.979215959 | 557.7272944 | 559.6478651 | ENSG00000105879 |
| DPYD | 0.006241464 | 1.004335625 | 0.993199424 | 0.99717974 | 8.932733175 | 8.975668334 | ENSG00000188641 |
| NPTN | 0.006224811 | 1.004324032 | 0.938577755 | 0.969724993 | 1309.605345 | 1314.866905 | ENSG00000156642 |
| IFNGR1 | 0.006176754 | 1.004290578 | 0.953968361 | 0.977923098 | 485.457072 | 487.536154 | ENSG00000027697 |
| NA | 0.006168354 | 1.00428473 | 0.987100127 | 0.994018677 | 31.95394516 | 32.08066826 | ENSG00000260565 |
| ULK2 | 0.006157235 | 1.00427699 | 0.970092196 | 0.985738981 | 196.5473846 | 197.6643679 | ENSG00000083290 |
| CLTA | 0.006059757 | 1.004209137 | 0.920569256 | 0.960700166 | 3952.740378 | 3968.818412 | ENSG00000122705 |
| DNAJC13 | 0.005952153 | 1.004134241 | 0.953587922 | 0.977641252 | 691.3021512 | 694.3841598 | ENSG00000138246 |
| MRM1 | 0.005940147 | 1.004125884 | 0.956261165 | 0.97920191 | 512.6116491 | 515.0088404 | ENSG00000278619 |
| DCLRE1C | 0.005849034 | 1.004062471 | 0.975002255 | 0.988231933 | 149.2744798 | 150.0893016 | ENSG00000152457 |
| RDH11 | 0.005804335 | 1.004031362 | 0.938593755 | 0.969724993 | 1730.649721 | 1737.113247 | ENSG00000072042 |
| VWA8 | 0.005775346 | 1.004011188 | 0.957053075 | 0.979297835 | 506.8068879 | 508.4973332 | ENSG00000102763 |
| SLC9A1 | 0.005742792 | 1.003988533 | 0.955701057 | 0.978963008 | 829.6345241 | 833.1842689 | ENSG00000090020 |
| PPP1CA | 0.00569646 | 1.00395629 | 0.927056362 | 0.964343189 | 4820.30424 | 4839.352147 | ENSG00000172531 |
| NA | 0.005690739 | 1.003952309 | 0.992845276 | 0.997064737 | 11.81361676 | 11.91284031 | ENSG00000270012 |
| ERP44 | 0.005652486 | 1.00392569 | 0.960264999 | 0.980365291 | 674.5330498 | 676.7729307 | ENSG00000023318 |
| UQCRC2 | 0.00536648 | 1.003726687 | 0.937665946 | 0.969306746 | 3360.329885 | 3371.815951 | ENSG00000140740 |
| TMED9 | 0.005289944 | 1.00367344 | 0.930804072 | 0.9660389 | 3923.963194 | 3938.796367 | ENSG00000184840 |
| NA | 0.005220932 | 1.00362543 | 0.983234786 | 0.992498966 | 83.56814279 | 84.01853133 | ENSG00000174977 |
| AMY2B | 0.005178648 | 1.003596016 | 0.996184942 | 0.998635949 | 4.588283469 | 4.606158403 | ENSG00000240038 |
| PNISR | 0.00508634 | 1.003531804 | 0.985653552 | 0.993533634 | 1922.069713 | 1929.104449 | ENSG00000132424 |
| AK2 | 0.005042575 | 1.003501362 | 0.941797552 | 0.971140304 | 2409.768462 | 2418.557453 | ENSG00000004455 |
| FAM199X | 0.004999148 | 1.003471156 | 0.956316698 | 0.97920191 | 1159.997089 | 1163.351 | ENSG00000123575 |
| RIOK2 | 0.004897252 | 1.003400284 | 0.967316622 | 0.984386182 | 441.5820023 | 442.5497094 | ENSG00000058729 |
| CERS5 | 0.004851797 | 1.003368671 | 0.95811883 | 0.979577828 | 785.3827902 | 787.8847467 | ENSG00000139624 |
| ZNF17 | 0.004814959 | 1.003343051 | 0.98755525 | 0.994259491 | 50.47100529 | 50.57907175 | ENSG00000186272 |
| EGLN1 | 0.004771001 | 1.00331248 | 0.985442126 | 0.993482614 | 86.90266691 | 86.91563629 | ENSG00000135766 |
| PDPK1 | 0.00476835 | 1.003310637 | 0.96635367 | 0.983871618 | 492.4628421 | 494.0678805 | ENSG00000140992 |
| NUB1 | 0.004708723 | 1.00326917 | 0.957808378 | 0.979577828 | 900.4757188 | 902.8551532 | ENSG00000013374 |

| | | | | | | | |
|---|---|---|---|---|---|---|---|
| SLCO2A1 | 0.004708521 | 1.00326903 | 0.989666728 | 0.995302243 | 35.1970165 | 35.31361285 | ENSG00000174640 |
| MIPEP | 0.004683239 | 1.003251448 | 0.964615552 | 0.982803839 | 526.9366632 | 528.6589892 | ENSG00000027001 |
| MATN4 | 0.004514045 | 1.003133798 | 0.993265742 | 0.99717974 | 19.7762615 | 19.68379574 | ENSG00000124159 |
| NA | 0.004494796 | 1.003120414 | 0.985284075 | 0.993482614 | 86.74720407 | 87.07895292 | ENSG00000267886 |
| ULK3 | 0.004460469 | 1.003096546 | 0.967378238 | 0.984386182 | 677.4722008 | 680.1485748 | ENSG00000140474 |
| GMEB1 | 0.004440889 | 1.003082932 | 0.969682571 | 0.985474408 | 488.5968823 | 490.1107912 | ENSG00000162419 |
| PPP1R9B | 0.004418696 | 1.003067502 | 0.966554258 | 0.983967737 | 607.5603782 | 609.3448435 | ENSG00000108819 |
| CELSR3 | 0.004392423 | 1.003049235 | 0.958549337 | 0.979584992 | 1442.820915 | 1447.138432 | ENSG00000008300 |
| PCDHB13 | 0.004380734 | 1.003041108 | 0.989609375 | 0.995302243 | 42.23326593 | 42.29391657 | ENSG00000187372 |
| ZDHHC2 | 0.004352092 | 1.003021195 | 0.975840466 | 0.988804724 | 372.6045725 | 373.5130337 | ENSG00000104219 |
| GPAT3 | 0.004186263 | 1.00290591 | 0.991541551 | 0.996317108 | 33.27094046 | 33.54784867 | ENSG00000138678 |
| APOE | 0.004137664 | 1.002872127 | 0.954586924 | 0.978185467 | 16113.63707 | 16160.09526 | ENSG00000130203 |
| MRTFA | 0.004098773 | 1.002845092 | 0.961819442 | 0.981411668 | 1005.940732 | 1008.461578 | ENSG00000196588 |
| DNAJC19 | 0.004090511 | 1.00283935 | 0.965632769 | 0.983484392 | 800.9762431 | 803.5994441 | ENSG00000205981 |
| IQCD | 0.004084311 | 1.002835039 | 0.992665952 | 0.997023289 | 23.70051892 | 23.71999107 | ENSG00000166578 |
| MIPOL1 | 0.003995035 | 1.002772985 | 0.982333766 | 0.992045368 | 179.6978646 | 179.9060371 | ENSG00000151338 |
| SLC10A3 | 0.003958145 | 1.002747344 | 0.983532359 | 0.992668425 | 183.7072458 | 184.200025 | ENSG00000126903 |
| SAYSD1 | 0.003918591 | 1.002719853 | 0.983698784 | 0.992668425 | 137.8571755 | 138.18241 | ENSG00000112167 |
| SEC14L6 | 0.003872616 | 1.002687898 | 0.992618855 | 0.997023289 | 29.46050465 | 29.70316213 | ENSG00000214491 |
| TUFT1 | 0.003786647 | 1.002628151 | 0.978640266 | 0.990126027 | 326.4197345 | 327.1078697 | ENSG00000143367 |
| ACBD5 | 0.003755564 | 1.00260655 | 0.978951852 | 0.990225052 | 348.7929981 | 349.7014726 | ENSG00000107897 |
| RB1CC1 | 0.003656471 | 1.002537687 | 0.97100238 | 0.986112774 | 865.7546214 | 867.3842925 | ENSG00000023287 |
| WDR43 | 0.003651645 | 1.002534333 | 0.959282014 | 0.980034514 | 2436.705024 | 2441.966816 | ENSG00000163811 |
| GTF3C3 | 0.003634 | 1.002522072 | 0.968405579 | 0.984740121 | 1035.799729 | 1037.529561 | ENSG00000119041 |
| MTMR9 | 0.003549131 | 1.002463099 | 0.977782896 | 0.989847838 | 456.0773315 | 456.8539286 | ENSG00000104643 |
| NA | 0.003512434 | 1.0024376 | 0.994902965 | 0.998026172 | 15.99877029 | 16.09127948 | ENSG00000254851 |
| NA | 0.00350421 | 1.002431886 | 0.995548383 | 0.998401192 | 11.80730293 | 11.88278439 | ENSG00000237758 |
| VAV3 | 0.003369144 | 1.002338042 | 0.981156335 | 0.991683936 | 263.1939247 | 263.5749591 | ENSG00000134215 |
| ZSCAN9 | 0.003246085 | 1.002252548 | 0.985036649 | 0.993365976 | 211.8779798 | 212.2393137 | ENSG00000137185 |
| KIF20B | 0.00321754 | 1.002232717 | 0.973860191 | 0.98768183 | 996.0263278 | 997.5913044 | ENSG00000138182 |
| NA | 0.00318885 | 1.002212787 | 0.99767958 | 0.999412061 | 4.082178128 | 4.099556349 | ENSG00000226986 |
| NA | 0.002839117 | 1.001969864 | 0.993305943 | 0.99717974 | 42.90054658 | 42.87896341 | ENSG00000255310 |
| JPH1 | 0.002833709 | 1.001966107 | 0.978419287 | 0.99001054 | 547.3615343 | 548.77639 | ENSG00000104369 |
| RBFOX2 | 0.002750202 | 1.001908113 | 0.964418026 | 0.982728532 | 3579.522598 | 3585.8689 | ENSG00000100320 |
| COQ7 | 0.002650265 | 1.001838712 | 0.986148543 | 0.993685984 | 215.8504854 | 216.1613457 | ENSG00000167186 |
| AP3B1 | 0.002393196 | 1.001660214 | 0.977831233 | 0.989847838 | 1328.396778 | 1329.669029 | ENSG00000132842 |
| EIF2AK1 | 0.002380806 | 1.001651612 | 0.970679897 | 0.986015158 | 2573.193408 | 2577.409744 | ENSG00000086232 |
| COX5A | 0.002360704 | 1.001637655 | 0.971539165 | 0.986279612 | 2974.953919 | 2978.995944 | ENSG00000178741 |
| TNIK | 0.002250215 | 1.001560947 | 0.977601671 | 0.989847838 | 1510.061541 | 1511.600742 | ENSG00000154310 |
| CIAPIN1 | 0.002175888 | 1.001509348 | 0.978288909 | 0.989993298 | 1914.294371 | 1916.299958 | ENSG00000005194 |
| TMTC1 | 0.002078887 | 1.001442013 | 0.986833837 | 0.993845887 | 360.9212762 | 361.5363146 | ENSG00000133687 |
| HDHD3 | 0.002039202 | 1.001414466 | 0.990177108 | 0.995467699 | 234.2160515 | 234.6667259 | ENSG00000119431 |
| NA | 0.002000672 | 1.001387722 | 0.994431815 | 0.997769557 | 59.71957987 | 59.70490804 | ENSG00000218426 |
| FST | 0.001996189 | 1.001384611 | 0.991526473 | 0.996317108 | 145.3282217 | 145.4589244 | ENSG00000134363 |
| FMR1 | 0.001917149 | 1.001329749 | 0.986122373 | 0.993685984 | 646.2230133 | 646.3837071 | ENSG00000102081 |
| NA | 0.001908045 | 1.001323431 | 0.996848993 | 0.999167121 | 20.4386038 | 20.50592909 | ENSG00000228106 |
| TAOK2 | 0.001671895 | 1.001159541 | 0.981911658 | 0.991973763 | 1535.165426 | 1537.123427 | ENSG00000149930 |
| NA | 0.001630974 | 1.001131144 | 0.997235492 | 0.999318319 | 24.20662427 | 24.17396736 | ENSG00000177337 |
| NA | 0.001614648 | 1.001119815 | 0.998698216 | 0.999580982 | 4.800645802 | 4.789743086 | ENSG00000245768 |
| POR | 0.001573461 | 1.001091235 | 0.980341209 | 0.991143572 | 2921.195081 | 2924.309544 | ENSG00000127948 |
| NA | 0.001477728 | 1.001024808 | 0.99263151 | 0.997023289 | 202.0291188 | 202.0094443 | ENSG00000214182 |
| IFT52 | 0.001452305 | 1.001007168 | 0.989953845 | 0.995432105 | 532.1650702 | 532.1514382 | ENSG00000101052 |
| DNAJA3 | 0.001401073 | 1.000971621 | 0.984313423 | 0.993115724 | 1798.848695 | 1800.567802 | ENSG00000103423 |
| CHD5 | 0.001341564 | 1.000930334 | 0.99619812 | 0.998635949 | 70.35033989 | 70.61345603 | ENSG00000116254 |
| TDRG1 | 0.001294485 | 1.000897671 | 0.9982851 | 0.999580982 | 13.44773123 | 13.4429437 | ENSG00000204091 |
| LRGUK | 0.001258036 | 1.000872384 | 0.997323349 | 0.999318319 | 33.09585972 | 33.17220554 | ENSG00000155530 |
| LOC105375624 | 0.001221364 | 1.000846944 | 0.998994753 | 0.999588012 | 5.058482691 | 5.037126827 | ENSG00000254231 |
| RUVBL2 | 0.001209088 | 1.000838427 | 0.986179993 | 0.993685984 | 3937.885687 | 3940.85804 | ENSG00000183207 |
| LOC283194 | 0.001114152 | 1.000772569 | 0.998876329 | 0.999580982 | 8.448726224 | 8.454094132 | ENSG00000255240 |
| GRB2 | 0.001113839 | 1.000772353 | 0.986279794 | 0.993696159 | 2622.884153 | 2625.186602 | ENSG00000177885 |
| WASH2P | 0.001078213 | 1.000747639 | 0.997167341 | 0.999318319 | 52.80977971 | 52.93119688 | ENSG00000146556 |
| HSPD1 | 0.001046217 | 1.000725446 | 0.986596428 | 0.993835437 | 27342.08524 | 27360.49706 | ENSG00000144381 |
| RNF146 | 0.001021245 | 1.000708124 | 0.992234766 | 0.996860318 | 558.7756817 | 558.8396448 | ENSG00000118518 |
| OARD1 | 0.000994759 | 1.000689752 | 0.992890496 | 0.997064737 | 578.3304783 | 578.3879184 | ENSG00000124596 |
| KRAS | 0.000937371 | 1.000649947 | 0.993367139 | 0.997187161 | 886.9080172 | 887.1032084 | ENSG00000133703 |
| KCNIP2 | 0.000905522 | 1.000627857 | 0.997952292 | 0.999491474 | 42.0614398 | 42.23859127 | ENSG00000120049 |
| CUL4B | 0.000849647 | 1.000589104 | 0.992328902 | 0.996900855 | 1506.066493 | 1506.226636 | ENSG00000158290 |
| CRK | 0.000790847 | 1.000548323 | 0.99161956 | 0.996317108 | 1689.862042 | 1690.597785 | ENSG00000167193 |
| NA | 0.000724205 | 1.000502107 | 0.999473229 | 0.999911419 | 4.055645048 | 4.06201436 | ENSG00000265460 |
| CCDC40 | 0.000705475 | 1.000489117 | 0.996997644 | 0.999262077 | 187.3645039 | 187.9250778 | ENSG00000141519 |
| POLRMT | 0.000674284 | 1.000467487 | 0.992922405 | 0.997064737 | 3707.620418 | 3710.192318 | ENSG00000099821 |
| STK35 | 0.000589641 | 1.000408791 | 0.994264486 | 0.997709691 | 1360.556163 | 1361.629658 | ENSG00000125834 |
| ZNF628 | 0.000541986 | 1.000375747 | 0.997531475 | 0.999412061 | 291.0729599 | 291.2543094 | ENSG00000197483 |
| TIMM29 | 0.000410497 | 1.000284576 | 0.997237771 | 0.999318319 | 474.8230195 | 475.0867673 | ENSG00000142444 |

| | | | | | | | |
|---|---|---|---|---|---|---|---|
| COG1 | 0.000371411 | 1.000257475 | 0.998176252 | 0.999580982 | 267.5766482 | 268.2483177 | ENSG00000166685 |
| YJEFN3 | 0.000340629 | 1.000236134 | 0.998933794 | 0.999580982 | 108.6644821 | 108.9930499 | ENSG00000250067 |
| RHOA | 0.000328189 | 1.000227509 | 0.995814678 | 0.998508399 | 8850.042622 | 8850.824801 | ENSG00000067560 |
| SAMD10 | 0.000205254 | 1.000142281 | 0.999316413 | 0.999855884 | 97.4867723 | 97.72014369 | ENSG00000130590 |
| TMEM170A | 3.45E-05 | 1.000023944 | 0.999782835 | 0.999963694 | 545.11867 | 544.4630009 | ENSG00000166822 |
| MAT2B | 3.15E-05 | 1.000021805 | 0.999727531 | 0.999963694 | 1099.753034 | 1099.105802 | ENSG00000038274 |
| UBE2K | 9.14E-06 | 1.000006334 | 0.999913136 | 0.999963694 | 3225.201271 | 3224.600125 | ENSG00000078140 |
| NA | -1.29E-05 | 0.999991087 | 0.999963694 | 0.999963694 | 71.30849973 | 71.42255233 | ENSG00000261963 |
| APBB3 | -2.38E-05 | 0.999983509 | 0.999900974 | 0.999963694 | 153.3610926 | 153.6944169 | ENSG00000113108 |
| ZNF549 | -3.11E-05 | 0.999978448 | 0.999854172 | 0.999963694 | 179.3364736 | 179.1826569 | ENSG00000121406 |
| CYTH1 | -7.41E-05 | 0.999948621 | 0.999479818 | 0.999911419 | 434.1013877 | 434.0800221 | ENSG00000108669 |
| NA | -0.000158101 | 0.999890419 | 0.999791702 | 0.999963694 | 12.36972904 | 12.38135072 | ENSG00000214535 |
| CLASRP | -0.000190665 | 0.999867849 | 0.998548469 | 0.999580982 | 1389.588929 | 1390.363843 | ENSG00000104859 |
| MMAB | -0.000198506 | 0.999862416 | 0.998749798 | 0.999580982 | 392.8090238 | 393.1989483 | ENSG00000139428 |
| SNX4 | -0.000210782 | 0.999853908 | 0.998514635 | 0.999580982 | 545.6973239 | 545.6948956 | ENSG00000114520 |
| REPS1 | -0.000214631 | 0.99985124 | 0.997811775 | 0.999429498 | 1314.594999 | 1314.173572 | ENSG00000135597 |
| FMNL3 | -0.000262134 | 0.999818319 | 0.997740443 | 0.999412061 | 792.7602188 | 793.1573598 | ENSG00000161791 |
| NA | -0.00045036 | 0.999687883 | 0.998783623 | 0.999580982 | 67.0803295 | 67.27515676 | ENSG00000204860 |
| NA | -0.000541847 | 0.999624491 | 0.998664018 | 0.999580982 | 45.76386416 | 45.77468293 | ENSG00000227081 |
| ZNF570 | -0.000552089 | 0.999617394 | 0.998210325 | 0.999580982 | 77.63189625 | 77.55490366 | ENSG00000171827 |
| NA | -0.000599195 | 0.999584756 | 0.999538458 | 0.999916111 | 4.108711207 | 4.11084127 | ENSG00000275665 |
| ZSCAN10 | -0.00064598 | 0.999552341 | 0.993446263 | 0.997212578 | 4934.204789 | 4933.084939 | ENSG00000130182 |
| KCNAB1 | -0.000696199 | 0.999517548 | 0.998906919 | 0.999580982 | 17.88440783 | 17.84619677 | ENSG00000169282 |
| SYCP2L | -0.000850854 | 0.999410407 | 0.998544295 | 0.999580982 | 24.24272578 | 24.26782234 | ENSG00000153157 |
| USP21 | -0.000883175 | 0.999388017 | 0.993507871 | 0.99722041 | 516.3451616 | 515.8828979 | ENSG00000143258 |
| DDX3X | -0.000967824 | 0.99932938 | 0.986297321 | 0.993696159 | 7216.25029 | 7210.791271 | ENSG00000215301 |
| GLRX3 | -0.000970122 | 0.999327789 | 0.989535754 | 0.995302243 | 2383.346573 | 2382.427312 | ENSG00000108010 |
| ZNF827 | -0.001020188 | 0.999293109 | 0.992899283 | 0.997064737 | 574.3353334 | 573.17392 | ENSG00000151612 |
| DUSP6 | -0.001057594 | 0.999267201 | 0.987946381 | 0.994484312 | 2199.141145 | 2196.954475 | ENSG00000139318 |
| GAR1 | -0.001095194 | 0.999241158 | 0.989361061 | 0.995302243 | 1003.784608 | 1002.905513 | ENSG00000109534 |
| TRIB2 | -0.001132388 | 0.999215397 | 0.996211291 | 0.998635949 | 1848.8742 | 1847.696631 | ENSG00000071575 |
| SNX14 | -0.001186709 | 0.999177774 | 0.99316875 | 0.99717974 | 291.0964489 | 290.9372811 | ENSG00000135317 |
| SDAD1 | -0.001261923 | 0.999125684 | 0.988681717 | 0.994908619 | 1138.259973 | 1136.886763 | ENSG00000198301 |
| NA | -0.001313676 | 0.999089844 | 0.998424584 | 0.999580982 | 13.18555735 | 13.15366947 | ENSG00000236809 |
| PELI2 | -0.001390828 | 0.999036416 | 0.992621728 | 0.997023289 | 244.2384299 | 243.7631272 | ENSG00000139946 |
| NA | -0.001432926 | 0.999007264 | 0.99763117 | 0.999412061 | 23.44268204 | 23.47684421 | ENSG00000244932 |
| USP22 | -0.001498181 | 0.998962079 | 0.980857205 | 0.991502996 | 6005.470241 | 5998.550644 | ENSG00000124422 |
| HSP90AB1 | -0.001545073 | 0.998929611 | 0.977216022 | 0.989678502 | 79234.7167 | 79148.90325 | ENSG00000096384 |
| METTL11B | -0.001632886 | 0.99886881 | 0.998606043 | 0.999580982 | 5.117862675 | 5.078467662 | ENSG00000203740 |
| NA | -0.001699986 | 0.998822354 | 0.998629918 | 0.999580982 | 4.839806533 | 4.816000155 | ENSG00000254577 |
| SCML2 | -0.0017559 | 0.998783643 | 0.985957639 | 0.993685984 | 611.240817 | 610.3099852 | ENSG00000102098 |
| PRRC1 | -0.001773337 | 0.998771572 | 0.984614898 | 0.993135424 | 1214.192091 | 1212.042314 | ENSG00000164244 |
| CBFB | -0.001819927 | 0.998739318 | 0.986142318 | 0.993685984 | 642.0614313 | 640.7778281 | ENSG00000067955 |
| ATP6V0A1 | -0.001858187 | 0.998712832 | 0.984423572 | 0.993115724 | 774.5821809 | 773.8522416 | ENSG00000033627 |
| ZNF546 | -0.002021546 | 0.998599752 | 0.997272846 | 0.999318319 | 16.0898171 | 16.07955653 | ENSG00000187187 |
| FAM50A | -0.002054016 | 0.998577278 | 0.982147157 | 0.992045368 | 1271.947246 | 1270.955252 | ENSG00000071859 |
| TFG | -0.002146835 | 0.998513034 | 0.977710957 | 0.989847838 | 1286.898215 | 1285.168235 | ENSG00000114354 |
| QSOX2 | -0.002148694 | 0.998511748 | 0.981971337 | 0.991979983 | 1405.864057 | 1404.564196 | ENSG00000165661 |
| IGDCC3 | -0.002251492 | 0.998440602 | 0.975160255 | 0.988262147 | 2168.251597 | 2164.477529 | ENSG00000174498 |
| KLHL2 | -0.00229899 | 0.998407731 | 0.987291628 | 0.994103495 | 263.8574471 | 263.3129297 | ENSG00000109466 |
| UBN1 | -0.002302737 | 0.998405137 | 0.976315373 | 0.989093283 | 1613.048989 | 1610.778104 | ENSG00000118900 |
| JAG2 | -0.00233571 | 0.998382319 | 0.984435205 | 0.993115724 | 455.7074544 | 455.5823597 | ENSG00000184916 |
| SOX21-AS1 | -0.002348985 | 0.998373133 | 0.994667317 | 0.997897805 | 42.68146446 | 42.6916915 | ENSG00000227640 |
| YTHDF2 | -0.002351978 | 0.998371061 | 0.97097273 | 0.986112774 | 2587.425978 | 2582.811524 | ENSG00000198492 |
| ZCCHC12 | -0.002352715 | 0.998370551 | 0.99502512 | 0.998089798 | 34.61111653 | 34.52621033 | ENSG00000174460 |
| NA | -0.002381557 | 0.998350592 | 0.993584541 | 0.99724336 | 69.19529401 | 69.22515845 | ENSG00000230074 |
| ECEL1 | -0.002448168 | 0.998304498 | 0.987607491 | 0.994259491 | 249.7425555 | 249.3258419 | ENSG00000171551 |
| NDRG4 | -0.002586201 | 0.998208988 | 0.980439584 | 0.991188962 | 553.1641459 | 552.399615 | ENSG00000103034 |
| LOC101929243 | -0.002616041 | 0.998188342 | 0.996483791 | 0.998855089 | 15.06162646 | 15.0861111 | ENSG00000272288 |
| RNF187 | -0.002737468 | 0.998104331 | 0.971418321 | 0.986210953 | 2310.250932 | 2305.133959 | ENSG00000168159 |
| NA | -0.002770452 | 0.998081512 | 0.997981505 | 0.999491474 | 3.847717405 | 3.852172609 | ENSG00000273091 |
| UNK | -0.002833735 | 0.998037732 | 0.981250227 | 0.991683936 | 488.7887323 | 488.1140327 | ENSG00000132478 |
| NFE2L3 | -0.002886687 | 0.998001101 | 0.968239943 | 0.984740121 | 3665.340371 | 3657.007762 | ENSG00000050344 |
| FAM102A | -0.00302354 | 0.997906437 | 0.972761427 | 0.98701563 | 912.6481034 | 910.6314448 | ENSG00000167106 |
| AQR | -0.003163592 | 0.997809568 | 0.970515735 | 0.9859427 | 1939.705258 | 1934.709379 | ENSG00000021776 |
| LCMT1 | -0.003197867 | 0.997785863 | 0.968661308 | 0.984740121 | 1038.627825 | 1036.328678 | ENSG00000205629 |
| STIL | -0.003258186 | 0.997744146 | 0.977778098 | 0.989847838 | 608.5381711 | 606.4751091 | ENSG00000123473 |
| NKAP | -0.003281038 | 0.997728342 | 0.985400396 | 0.993482614 | 155.4932172 | 155.0572932 | ENSG00000101882 |
| EIF2S2 | -0.003291335 | 0.997721221 | 0.957817552 | 0.979577828 | 5138.079076 | 5125.735608 | ENSG00000125977 |
| RANBP6 | -0.003291889 | 0.997720838 | 0.984491828 | 0.993118791 | 482.8141968 | 482.5234381 | ENSG00000137040 |
| AFTPH | -0.00331722 | 0.99770332 | 0.983156691 | 0.992498966 | 337.6280136 | 336.3058983 | ENSG00000119844 |
| ZNF433-AS1 | -0.00334853 | 0.997681667 | 0.990866725 | 0.995917781 | 72.33629959 | 72.12457364 | ENSG00000219665 |
| N4BP2L2 | -0.003357704 | 0.997675323 | 0.975053759 | 0.988231933 | 711.0969633 | 709.5026539 | ENSG00000244754 |
| NA | -0.003367251 | 0.997668721 | 0.997687513 | 0.999412061 | 3.519752017 | 3.533392118 | ENSG00000248161 |

| | | | | | | | |
|---|---|---|---|---|---|---|---|
| NA | -0.003459618 | 0.997604848 | 0.997735805 | 0.999412061 | 3.810435811 | 3.810831773 | ENSG00000287275 |
| HYKK | -0.003490301 | 0.997583632 | 0.994537637 | 0.997821717 | 20.31225223 | 20.39742839 | ENSG00000188266 |
| FLYWCH2 | -0.003492427 | 0.997582162 | 0.984722655 | 0.993135424 | 216.8133504 | 216.540676 | ENSG00000162076 |
| SPTSSA | -0.003544448 | 0.997546192 | 0.966847007 | 0.9842117 | 1259.187188 | 1255.945976 | ENSG00000165389 |
| DENND4C | -0.003825658 | 0.997351769 | 0.978763225 | 0.990142329 | 288.531715 | 287.248608 | ENSG00000137145 |
| NA | -0.003963303 | 0.997256617 | 0.995914245 | 0.998554213 | 8.03602785 | 8.023125708 | ENSG00000279531 |
| CDH3 | -0.004006891 | 0.997226488 | 0.954088119 | 0.97799177 | 3657.9299 | 3647.402456 | ENSG00000062038 |
| NA | -0.004053652 | 0.997194166 | 0.989717295 | 0.995302243 | 48.12591705 | 47.8935204 | ENSG00000227473 |
| CD9 | -0.004072138 | 0.997181389 | 0.957052506 | 0.979297835 | 2596.353487 | 2588.07621 | ENSG00000010278 |
| ZNF182 | -0.004133981 | 0.997138644 | 0.98321752 | 0.992498966 | 149.8373119 | 149.1609291 | ENSG00000147118 |
| THRAP3 | -0.004190415 | 0.99709964 | 0.944856143 | 0.972886876 | 4560.012779 | 4546.486254 | ENSG00000054118 |
| APPL1 | -0.004312345 | 0.997015373 | 0.968063887 | 0.984740121 | 856.7030532 | 853.5044945 | ENSG00000157500 |
| GNE | -0.004331562 | 0.997002092 | 0.973034709 | 0.987040955 | 370.2688799 | 369.2051645 | ENSG00000159921 |
| DLG5 | -0.004336128 | 0.996998937 | 0.954914859 | 0.978289671 | 2567.90956 | 2561.133384 | ENSG00000151208 |
| LINC02449 | -0.004355871 | 0.996985293 | 0.995600023 | 0.998401192 | 9.17982155 | 9.234886647 | ENSG00000215241 |
| PITHD1 | -0.00437285 | 0.99697356 | 0.954843981 | 0.978279621 | 1552.299775 | 1547.40131 | ENSG00000057757 |
| SPATA20 | -0.004421023 | 0.996940271 | 0.968052743 | 0.984740121 | 552.476571 | 551.1671564 | ENSG00000006282 |
| TOP3B | -0.004429624 | 0.996934328 | 0.994369195 | 0.997760742 | 12.96184514 | 13.02694743 | ENSG00000100038 |
| B3GALNT1 | -0.004461142 | 0.996912548 | 0.983895346 | 0.992787258 | 127.1393223 | 126.5236751 | ENSG00000169255 |
| NA | -0.00448322 | 0.996897292 | 0.995778598 | 0.998508399 | 7.0628802 | 7.081756384 | ENSG00000225774 |
| ATAD2B | -0.004516577 | 0.996874243 | 0.972851546 | 0.987015652 | 329.8204409 | 328.4100845 | ENSG00000119778 |
| SNIP1 | -0.004519179 | 0.996872445 | 0.971403986 | 0.986210953 | 361.2958962 | 359.8390957 | ENSG00000163877 |
| TBC1D7 | -0.004607111 | 0.996811688 | 0.966996226 | 0.9842481 | 519.6211701 | 517.5334398 | ENSG00000145979 |
| RTN4 | -0.004718095 | 0.996735008 | 0.95584049 | 0.978975886 | 6070.426899 | 6050.112706 | ENSG00000115310 |
| NA | -0.004722935 | 0.996731664 | 0.982316807 | 0.992045368 | 104.1507619 | 103.7468055 | ENSG00000197744 |
| ARC | -0.004743237 | 0.996717638 | 0.967702171 | 0.984541474 | 624.2801193 | 622.6723988 | ENSG00000198576 |
| ESCO1 | -0.004861249 | 0.99663611 | 0.971802661 | 0.986331007 | 694.3693903 | 691.4658068 | ENSG00000141446 |
| UBE2J2 | -0.004880473 | 0.996622829 | 0.949005617 | 0.97469657 | 1698.738627 | 1692.912471 | ENSG00000160087 |
| SLC18B1 | -0.004905553 | 0.996605504 | 0.95991018 | 0.980219024 | 661.4210408 | 659.089876 | ENSG00000146409 |
| TTL | -0.004928628 | 0.996589564 | 0.954724808 | 0.978211597 | 1077.873496 | 1073.663546 | ENSG00000114999 |
| TIMELESS | -0.004984289 | 0.996551115 | 0.939364575 | 0.970003616 | 3650.462612 | 3638.461433 | ENSG00000111602 |
| MYL6 | -0.005064142 | 0.996495958 | 0.945058809 | 0.972933398 | 2478.731024 | 2470.144319 | ENSG00000092841 |
| NA | -0.005114849 | 0.996460934 | 0.995167403 | 0.998129371 | 7.486327089 | 7.475182819 | ENSG00000235734 |
| NA | -0.005192109 | 0.996407572 | 0.996022463 | 0.998608695 | 5.077424168 | 5.029640753 | ENSG00000236972 |
| ZNF527 | -0.00521608 | 0.996391017 | 0.986814624 | 0.993845887 | 51.02268289 | 50.76254481 | ENSG00000189164 |
| ASAP1 | -0.005274674 | 0.99635055 | 0.956761725 | 0.979215959 | 769.245567 | 766.6347313 | ENSG00000153317 |
| RNF111 | -0.005313027 | 0.996324063 | 0.963949984 | 0.982557828 | 529.1269802 | 526.8244162 | ENSG00000157450 |
| B4GALT1 | -0.005369531 | 0.996285043 | 0.961459361 | 0.98120631 | 530.8934744 | 528.3223851 | ENSG00000086062 |
| CASC3 | -0.005413662 | 0.996254567 | 0.92836189 | 0.964638631 | 3580.090489 | 3566.717391 | ENSG00000108349 |
| NA | -0.005432023 | 0.996241888 | 0.990074802 | 0.995445718 | 29.35733385 | 29.23790091 | ENSG00000287769 |
| CD83 | -0.005822833 | 0.995972053 | 0.972981946 | 0.987040955 | 228.6917141 | 227.8796513 | ENSG00000112149 |
| NA | -0.006031918 | 0.995827721 | 0.984660485 | 0.993135424 | 62.32506056 | 62.32852409 | ENSG00000268362 |
| C2orf69 | -0.006072256 | 0.995799878 | 0.965515297 | 0.983450237 | 427.7564563 | 425.919184 | ENSG00000178074 |
| GRIK4 | -0.006137612 | 0.995754768 | 0.977363067 | 0.98969825 | 108.2909445 | 107.7454588 | ENSG00000149403 |
| CHST11 | -0.006488329 | 0.995512731 | 0.95661898 | 0.979215959 | 483.8905305 | 481.7406844 | ENSG00000171310 |
| LHX4 | -0.006507206 | 0.995499705 | 0.985773413 | 0.993600415 | 39.48890371 | 39.33483603 | ENSG00000121454 |
| PDS5B | -0.006605295 | 0.995432024 | 0.951660856 | 0.976472701 | 802.3951541 | 798.0299901 | ENSG00000083642 |
| STK16 | -0.006618116 | 0.995423177 | 0.958259873 | 0.979577828 | 352.9616906 | 351.0509225 | ENSG00000115661 |
| ZNF416 | -0.006717535 | 0.995354583 | 0.971151423 | 0.986156064 | 147.8185053 | 146.9139438 | ENSG00000083817 |
| N6AMT1 | -0.006751483 | 0.995331162 | 0.966513799 | 0.983967737 | 217.2578285 | 216.3273703 | ENSG00000156239 |
| WNT4 | -0.006798156 | 0.995298962 | 0.989599745 | 0.995302243 | 18.51884158 | 18.35604802 | ENSG00000162552 |
| AMER1 | -0.006815293 | 0.99528714 | 0.945968976 | 0.973329762 | 740.5770934 | 737.7461048 | ENSG00000184675 |
| CROCCP3 | -0.006819976 | 0.995283908 | 0.979875763 | 0.990886688 | 92.94198655 | 92.83797135 | ENSG00000080947 |
| OXNAD1 | -0.006864984 | 0.995252859 | 0.955722276 | 0.978963008 | 415.7867381 | 413.9756583 | ENSG00000154814 |
| FIP1L1 | -0.006873922 | 0.995246693 | 0.965654957 | 0.983484392 | 201.1924925 | 200.2288195 | ENSG00000145216 |
| RPL36 | -0.006969533 | 0.995180738 | 0.919641289 | 0.960477384 | 5499.616182 | 5473.469557 | ENSG00000130255 |
| SLC35A2 | -0.006996013 | 0.995162472 | 0.947684285 | 0.974075819 | 653.5496609 | 651.0933825 | ENSG00000102100 |
| DCLRE1A | -0.007084199 | 0.995101644 | 0.959214832 | 0.980034514 | 322.3022516 | 320.2190076 | ENSG00000198924 |
| TENT4B | -0.007128287 | 0.995071235 | 0.963639891 | 0.982393082 | 322.3364588 | 320.3853183 | ENSG00000121274 |
| APEH | -0.007228267 | 0.995002277 | 0.899240077 | 0.948444134 | 5204.607638 | 5177.909149 | ENSG00000164062 |
| LUZP2 | -0.007257194 | 0.994982327 | 0.986362498 | 0.993707808 | 28.69971933 | 28.52426827 | ENSG00000187398 |
| GTPBP2 | -0.007277554 | 0.994968286 | 0.93548229 | 0.968291565 | 1254.651527 | 1249.060913 | ENSG00000172432 |
| SLC26A1 | -0.007383495 | 0.994895225 | 0.991143421 | 0.996103991 | 14.86948338 | 14.82744244 | ENSG00000145217 |
| KIFBP | -0.007419394 | 0.994870469 | 0.935468069 | 0.968291565 | 994.7209981 | 989.5087684 | ENSG00000198954 |
| UTP14A | -0.007423714 | 0.99486749 | 0.932134552 | 0.966770106 | 1036.468979 | 1031.687503 | ENSG00000156697 |
| WAKMAR2 | -0.007534126 | 0.994791354 | 0.994815204 | 0.997992151 | 4.147871938 | 4.12212619 | ENSG00000237499 |
| NA | -0.007560681 | 0.994773043 | 0.987427062 | 0.994185852 | 20.85573686 | 20.85096665 | ENSG00000267023 |
| RALGAPB | -0.007627462 | 0.994726998 | 0.934878423 | 0.967882733 | 1228.748845 | 1221.466766 | ENSG00000170471 |
| TXNDC11 | -0.007921783 | 0.994524086 | 0.95966255 | 0.98010931 | 219.0846874 | 217.981273 | ENSG00000153066 |
| TSR2 | -0.007949121 | 0.994505241 | 0.93046415 | 0.965902462 | 1290.858799 | 1284.361544 | ENSG00000158526 |
| PTP4A2 | -0.007979898 | 0.994484025 | 0.919320054 | 0.960250106 | 2652.995688 | 2637.743747 | ENSG00000184007 |
| ZFPL1 | -0.008120971 | 0.994386785 | 0.97519836 | 0.988262147 | 90.9476385 | 90.65043668 | ENSG00000162300 |
| TRAF4 | -0.008162778 | 0.99435797 | 0.896839702 | 0.947229702 | 3161.614349 | 3143.335709 | ENSG00000076604 |
| TRIM13 | -0.008198376 | 0.994333435 | 0.964435593 | 0.982728532 | 235.6166974 | 233.809495 | ENSG00000204977 |

| | | | | | | | |
|---|---|---|---|---|---|---|---|
| ZAN | -0.008332416 | 0.994241056 | 0.993178391 | 0.99717974 | 4.866339612 | 4.808514081 | ENSG00000146839 |
| CTNNBL1 | -0.008515897 | 0.994114617 | 0.912568375 | 0.956487804 | 1457.156941 | 1448.400462 | ENSG00000132792 |
| SSX2IP | -0.008549759 | 0.994091284 | 0.94558764 | 0.973218501 | 466.7939447 | 463.335228 | ENSG00000117155 |
| NA | -0.008572626 | 0.994075528 | 0.972727097 | 0.98701563 | 86.56660626 | 86.29557248 | ENSG00000205746 |
| ZBTB8B | -0.008590923 | 0.99406292 | 0.971269127 | 0.986210953 | 90.16713759 | 89.79879706 | ENSG00000273274 |
| ZNF19 | -0.008596606 | 0.994059005 | 0.984708971 | 0.993135424 | 23.10850052 | 22.98857511 | ENSG00000157429 |
| KMT5A | -0.008774109 | 0.993936707 | 0.946431574 | 0.9733913 | 322.4479354 | 320.6827717 | ENSG00000183955 |
| LARP6 | -0.008825677 | 0.993901181 | 0.939840586 | 0.970206481 | 463.4616905 | 460.901664 | ENSG00000166173 |
| H2AC12 | -0.008857472 | 0.993879277 | 0.994263973 | 0.997709691 | 4.188310445 | 4.155980952 | ENSG00000274997 |
| YY1AP1 | -0.008884372 | 0.993860745 | 0.93711923 | 0.969033102 | 523.2550594 | 520.0206491 | ENSG00000163374 |
| KCTD5 | -0.008951598 | 0.993814435 | 0.922532542 | 0.961675721 | 893.2651702 | 887.7884659 | ENSG00000167977 |
| NA | -0.008989828 | 0.9937881 | 0.984339636 | 0.993115724 | 27.34859935 | 27.29527367 | ENSG00000267519 |
| NA | -0.009036388 | 0.993756028 | 0.983088359 | 0.992498966 | 28.58895693 | 28.39006015 | ENSG00000269352 |
| VANGL2 | -0.009104046 | 0.993709425 | 0.902017762 | 0.95021014 | 1949.824433 | 1937.496106 | ENSG00000162738 |
| HAUS6 | -0.009172537 | 0.993662251 | 0.915901262 | 0.95829931 | 1235.166152 | 1227.036658 | ENSG00000147874 |
| ZDHHC13 | -0.009242189 | 0.993614278 | 0.939844997 | 0.970206481 | 448.0698514 | 444.8832783 | ENSG00000177054 |
| TMEM176B | -0.009272255 | 0.993593572 | 0.986833405 | 0.993845887 | 14.62880653 | 14.57950904 | ENSG00000106565 |
| CLSTN1 | -0.009322681 | 0.993558844 | 0.889786288 | 0.943762108 | 3487.210935 | 3464.457771 | ENSG00000171603 |
| ROGDI | -0.009355156 | 0.993536479 | 0.95352462 | 0.977638777 | 247.6326795 | 246.2220412 | ENSG00000067836 |
| CACUL1 | -0.00942561 | 0.993487961 | 0.926879582 | 0.964343189 | 691.940238 | 686.6657983 | ENSG00000151893 |
| NELFB | -0.009611918 | 0.993359671 | 0.883596512 | 0.939962378 | 3146.833372 | 3126.273012 | ENSG00000188986 |
| CDC37L1 | -0.009629287 | 0.993347712 | 0.94085476 | 0.970762241 | 394.9167875 | 392.0228253 | ENSG00000106993 |
| CALCOCO2 | -0.009657041 | 0.993328602 | 0.941287145 | 0.970822535 | 369.370506 | 366.9335627 | ENSG00000136436 |
| DLGAP5 | -0.009721545 | 0.993284191 | 0.897728 | 0.947692289 | 2723.988437 | 2704.371426 | ENSG00000126787 |
| DEPDC7 | -0.009724687 | 0.993282028 | 0.962387336 | 0.981774928 | 123.8988065 | 123.1049959 | ENSG00000121690 |
| ZNF189 | -0.00973926 | 0.993271995 | 0.94859091 | 0.974636169 | 257.6437458 | 255.5328476 | ENSG00000136870 |
| IFT43 | -0.009833122 | 0.993207374 | 0.949821099 | 0.975127915 | 202.3792803 | 201.0616882 | ENSG00000119650 |
| MOB4 | -0.009874304 | 0.993179023 | 0.982855982 | 0.992440902 | 327.8482964 | 325.5070216 | ENSG00000115540 |
| RHOBTB1 | -0.010010603 | 0.993085197 | 0.943851722 | 0.972449807 | 272.2273482 | 270.2817337 | ENSG00000072422 |
| BAZ2B | -0.010042932 | 0.993062943 | 0.937134886 | 0.969033102 | 387.4640966 | 384.8252256 | ENSG00000123636 |
| DMWD | -0.010142341 | 0.992994518 | 0.925266788 | 0.9631505 | 678.402993 | 674.0350658 | ENSG00000185800 |
| EIF4H | -0.010225436 | 0.992937327 | 0.853124759 | 0.922280546 | 10161.51279 | 10089.59432 | ENSG00000106682 |
| RIT1 | -0.010293307 | 0.992890615 | 0.9338933 | 0.96733214 | 432.4763531 | 428.936218 | ENSG00000143622 |
| PSMG4 | -0.010305915 | 0.992881938 | 0.920946086 | 0.9608655 | 739.5380717 | 734.5595193 | ENSG00000180822 |
| PIGM | -0.010430848 | 0.992795962 | 0.953953508 | 0.977923098 | 227.0133802 | 225.9175204 | ENSG00000143315 |
| NA | -0.01046389 | 0.992773224 | 0.978753089 | 0.990142329 | 34.77238954 | 34.65012121 | ENSG00000204685 |
| SPDEF | -0.010473335 | 0.992766724 | 0.993776762 | 0.997382277 | 3.622727421 | 3.608476096 | ENSG00000124664 |
| AIFM1 | -0.010534709 | 0.992724492 | 0.866249875 | 0.930189756 | 3432.314527 | 3406.972929 | ENSG00000156709 |
| YWHAG | -0.010543569 | 0.992718395 | 0.863131592 | 0.928401191 | 7493.60186 | 7438.04326 | ENSG00000170027 |
| UBALD1 | -0.010584786 | 0.992690034 | 0.914741503 | 0.958005256 | 1083.01714 | 1075.149386 | ENSG00000153443 |
| SLC25A3 | -0.010621513 | 0.992664763 | 0.856171726 | 0.923756361 | 9217.245233 | 9148.385906 | ENSG00000075415 |
| CTAGE7P | -0.010825766 | 0.992524234 | 0.98979372 | 0.995325094 | 7.636054826 | 7.580434331 | ENSG00000233122 |
| TMEM14B | -0.010845156 | 0.992510895 | 0.933133604 | 0.967210907 | 400.5174696 | 397.0050649 | ENSG00000137210 |
| HENMT1 | -0.010891396 | 0.992479084 | 0.9574406 | 0.979478064 | 138.9138761 | 137.9912277 | ENSG00000162639 |
| LTN1 | -0.010931942 | 0.992451191 | 0.927526916 | 0.964343189 | 548.1050619 | 543.2296578 | ENSG00000198862 |
| ZFP92 | -0.011000713 | 0.992403884 | 0.983724018 | 0.992668425 | 17.84712623 | 17.63960422 | ENSG00000189420 |
| ZNF708 | -0.01102859 | 0.992384708 | 0.944220325 | 0.972502334 | 227.8461958 | 226.2441847 | ENSG00000182141 |
| CDH4 | -0.011032055 | 0.992382325 | 0.928285911 | 0.964638631 | 457.6590038 | 454.4197118 | ENSG00000179242 |
| NA | -0.011119816 | 0.992321958 | 0.982357215 | 0.992045368 | 25.13174181 | 25.00358679 | ENSG00000232611 |
| COMMD8 | -0.011123188 | 0.99231964 | 0.974547198 | 0.988144592 | 42.31306056 | 41.98967019 | ENSG00000169019 |
| TUFM | -0.011123923 | 0.992319134 | 0.846620861 | 0.918413939 | 5450.725963 | 5408.118928 | ENSG00000178952 |
| DRG1 | -0.011139395 | 0.992308492 | 0.880436584 | 0.938509763 | 1761.556489 | 1747.849098 | ENSG00000185721 |
| NFXL1 | -0.011161663 | 0.992293176 | 0.944116857 | 0.972449807 | 279.3965787 | 276.9666625 | ENSG00000170448 |
| CEP192 | -0.011187579 | 0.992275351 | 0.907532326 | 0.954013052 | 674.0365804 | 668.4838021 | ENSG00000101639 |
| SCAMP2 | -0.01122064 | 0.992252612 | 0.893461819 | 0.945501439 | 936.7764813 | 929.6614336 | ENSG00000140497 |
| ECHDC3 | -0.011231524 | 0.992245126 | 0.929860368 | 0.965491992 | 376.5853613 | 373.435465 | ENSG00000134463 |
| LINC00888 | -0.011232324 | 0.992244576 | 0.946140791 | 0.973361629 | 186.3614708 | 184.6656127 | ENSG00000240024 |
| PARG | -0.011269569 | 0.992218961 | 0.889256655 | 0.943524322 | 1180.046754 | 1170.701691 | ENSG00000227345 |
| NIFK | -0.011303832 | 0.992195396 | 0.911715251 | 0.956186649 | 662.0480033 | 656.7476017 | ENSG00000155438 |
| PRMT3 | -0.011338668 | 0.992171438 | 0.912586604 | 0.956487804 | 633.9013144 | 628.7867034 | ENSG00000185238 |
| PSPH | -0.011470306 | 0.992080912 | 0.895253739 | 0.946203969 | 1049.297468 | 1040.375179 | ENSG00000146733 |
| SEC31A | -0.011511012 | 0.992052921 | 0.87462258 | 0.935130738 | 1835.227418 | 1820.940342 | ENSG00000138674 |
| LRRC8D | -0.011700055 | 0.991922936 | 0.90765023 | 0.954082882 | 759.6497022 | 753.1673137 | ENSG00000171492 |
| RBM38 | -0.011801349 | 0.991853294 | 0.902221612 | 0.950316853 | 1368.454121 | 1356.615323 | ENSG00000132819 |
| RPAP2 | -0.011812647 | 0.991845526 | 0.934552219 | 0.967761249 | 266.4162731 | 264.2283768 | ENSG00000122484 |
| PKD1P5-LOC105376752 | -0.011896966 | 0.991787559 | 0.9461575 | 0.973361629 | 284.7222714 | 283.0398671 | ENSG00000254681 |
| IREB2 | -0.01190369 | 0.991782937 | 0.89792453 | 0.947703593 | 1841.483955 | 1825.491189 | ENSG00000136381 |
| CERT1 | -0.011948218 | 0.991752327 | 0.902151118 | 0.950296608 | 1276.659994 | 1265.669544 | ENSG00000113163 |
| ICAM5 | -0.011948303 | 0.991752268 | 0.958741049 | 0.979702498 | 102.9683111 | 102.1801478 | ENSG00000105376 |
| C6orf136 | -0.011996011 | 0.991719473 | 0.941092241 | 0.970798334 | 205.4194601 | 203.9168397 | ENSG00000204564 |
| PALM | -0.012096172 | 0.991650624 | 0.922030127 | 0.961386103 | 445.7788965 | 441.7844386 | ENSG00000099864 |
| WSB1 | -0.012229168 | 0.991559212 | 0.973421395 | 0.987379166 | 1543.655824 | 1530.787135 | ENSG00000109046 |
| CALML4 | -0.012267784 | 0.991532671 | 0.987181076 | 0.994046184 | 7.645525565 | 7.584121558 | ENSG00000129007 |
| NA | -0.012283873 | 0.991521614 | 0.962748144 | 0.981980854 | 77.37466072 | 76.59052646 | ENSG00000204745 |

| | | | | | | | |
|---|---|---|---|---|---|---|---|
| OSER1 | -0.012468833 | 0.991394505 | 0.903885224 | 0.951258232 | 584.5581228 | 579.4530156 | ENSG00000132823 |
| SAE1 | -0.012479773 | 0.991386987 | 0.826766047 | 0.906810387 | 7811.675558 | 7744.21811 | ENSG00000142230 |
| RAP1GDS1 | -0.012525937 | 0.991355265 | 0.920098066 | 0.960668852 | 450.8262399 | 446.2994416 | ENSG00000138698 |
| CD58 | -0.012596807 | 0.991306567 | 0.967106138 | 0.984259247 | 53.16280501 | 52.60113147 | ENSG00000116815 |
| PRDM14 | -0.012632227 | 0.99128223 | 0.862132265 | 0.927649312 | 2086.039166 | 2068.653559 | ENSG00000147596 |
| XKR8 | -0.012678781 | 0.991250243 | 0.917940156 | 0.959450204 | 398.4815105 | 395.0702501 | ENSG00000158156 |
| MTMR11 | -0.012679731 | 0.99124959 | 0.971057945 | 0.98611517 | 43.87053732 | 43.57002622 | ENSG00000014914 |
| C2CD2L | -0.01273893 | 0.991208916 | 0.971752846 | 0.986331007 | 49.04472076 | 48.54594191 | ENSG00000172375 |
| IFNGR2 | -0.012778777 | 0.991181539 | 0.922626872 | 0.961698805 | 302.816967 | 300.142335 | ENSG00000159128 |
| KIF22 | -0.012875534 | 0.991115066 | 0.849535835 | 0.920078138 | 2932.384201 | 2906.723116 | ENSG00000079616 |
| ZBED3-AS1 | -0.01304579 | 0.990998109 | 0.978217769 | 0.989993298 | 20.66990761 | 20.42280939 | ENSG00000250802 |
| NA | -0.013080454 | 0.990974298 | 0.988790317 | 0.994963881 | 6.070791072 | 6.069101929 | ENSG00000259287 |
| PRMT9 | -0.013096643 | 0.990963179 | 0.930059267 | 0.965600537 | 240.9486812 | 238.5387285 | ENSG00000164169 |
| FGD1 | -0.013134077 | 0.990937466 | 0.898098364 | 0.947805209 | 667.6421458 | 661.6914232 | ENSG00000102302 |
| HSD17B1 | -0.01320765 | 0.990886933 | 0.976338708 | 0.989093283 | 24.00195123 | 23.75004698 | ENSG00000108786 |
| SNX29 | -0.013331937 | 0.990801572 | 0.931374358 | 0.966360206 | 218.5746284 | 216.5405644 | ENSG00000048471 |
| RNF44 | -0.013542743 | 0.990656807 | 0.823426043 | 0.905075215 | 3816.319303 | 3780.90705 | ENSG00000146083 |
| STXBP2 | -0.013561357 | 0.990644025 | 0.914897914 | 0.958037973 | 437.8176424 | 434.1046705 | ENSG00000076944 |
| RTF2 | -0.013619007 | 0.99060444 | 0.889414173 | 0.943529294 | 941.0718262 | 931.6772726 | ENSG00000022277 |
| USP27X | -0.013652627 | 0.990581356 | 0.935151312 | 0.968111178 | 179.1382121 | 177.4308772 | ENSG00000273820 |
| ACOX3 | -0.013679432 | 0.990562951 | 0.925569505 | 0.963290836 | 241.0726727 | 238.6245478 | ENSG00000087008 |
| TMEM138 | -0.013695304 | 0.990552054 | 0.924072315 | 0.96244767 | 284.9329651 | 282.3913073 | ENSG00000149483 |
| CFAP410 | -0.013695336 | 0.990552032 | 0.970418114 | 0.985897567 | 34.90821185 | 34.55202936 | ENSG00000160226 |
| ZNF707 | -0.013709811 | 0.990542093 | 0.917457122 | 0.959335774 | 299.4186282 | 296.7445768 | ENSG00000181135 |
| TRIM11 | -0.013767028 | 0.990502809 | 0.898957616 | 0.948388004 | 515.2563132 | 510.415902 | ENSG00000154370 |
| INTS13 | -0.01377832 | 0.990495056 | 0.873708319 | 0.934840664 | 1069.593822 | 1058.709047 | ENSG00000064102 |
| TBCD | -0.013792481 | 0.990485334 | 0.83052589 | 0.908658972 | 3998.786304 | 3960.878572 | ENSG00000141556 |
| PYGM | -0.013807976 | 0.990474696 | 0.933793813 | 0.96733214 | 199.7289038 | 197.6705399 | ENSG00000068976 |
| METTL14 | -0.0138128 | 0.990471384 | 0.89346672 | 0.945501439 | 583.0813503 | 577.1177977 | ENSG00000145388 |
| LRRC41 | -0.013826958 | 0.990461664 | 0.875207454 | 0.935364718 | 1157.061712 | 1145.790318 | ENSG00000132128 |
| CAPZA1 | -0.013969964 | 0.99036349 | 0.882198245 | 0.939373634 | 1881.582068 | 1862.187556 | ENSG00000116489 |
| NA | -0.013989155 | 0.990350316 | 0.9850975 | 0.993365976 | 9.574758525 | 9.542382217 | ENSG00000244171 |
| RELA | -0.014008717 | 0.990336888 | 0.908469994 | 0.954511501 | 517.7652526 | 512.8641436 | ENSG00000173039 |
| HEATR3 | -0.014039532 | 0.990315735 | 0.909192381 | 0.954783364 | 450.1209786 | 446.1904718 | ENSG00000155393 |
| ZNF609 | -0.014105122 | 0.990270713 | 0.860046931 | 0.926192962 | 2301.993056 | 2279.878816 | ENSG00000180357 |
| DISP2 | -0.01412327 | 0.990258256 | 0.9651533 | 0.983135546 | 46.24215864 | 45.76730848 | ENSG00000140323 |
| NAALADL1 | -0.014148456 | 0.990240969 | 0.970715685 | 0.986015158 | 32.82727432 | 32.42146894 | ENSG00000168060 |
| PLA2G12A | -0.014165669 | 0.990229155 | 0.920580282 | 0.960700166 | 283.8680564 | 281.298456 | ENSG00000123739 |
| ANAPC15 | -0.01420313 | 0.990203442 | 0.944005545 | 0.972449807 | 143.0059082 | 141.4302151 | ENSG00000110200 |
| NA | -0.0143625 | 0.990094064 | 0.989017453 | 0.99503037 | 4.135244286 | 4.080785355 | ENSG00000230224 |
| TDRD3 | -0.014428504 | 0.990048768 | 0.924681807 | 0.962703775 | 242.2807847 | 239.5353115 | ENSG00000083544 |
| A4GALT | -0.014448064 | 0.990035344 | 0.927261669 | 0.964343189 | 204.5783764 | 202.5316797 | ENSG00000128274 |
| IRF9 | -0.014482752 | 0.99001154 | 0.985387365 | 0.993482614 | 7.291027097 | 7.227799079 | ENSG00000213928 |
| KCTD2 | -0.014528233 | 0.989980331 | 0.870164586 | 0.932664263 | 1101.777238 | 1090.681031 | ENSG00000180901 |
| ARF4 | -0.014539547 | 0.989972567 | 0.86429221 | 0.929307927 | 2029.522257 | 2008.421485 | ENSG00000168374 |
| RPS5 | -0.014586542 | 0.98994032 | 0.816300404 | 0.901144302 | 15313.70479 | 15159.87246 | ENSG00000083845 |
| KCTD9 | -0.014587952 | 0.989939353 | 0.915423154 | 0.958177712 | 307.7136956 | 304.4545443 | ENSG00000104756 |
| HAUS8 | -0.014592094 | 0.98993651 | 0.905308205 | 0.95248537 | 376.6738299 | 372.9377663 | ENSG00000131351 |
| CEP85 | -0.014608594 | 0.989925188 | 0.870332306 | 0.932705584 | 1176.907222 | 1165.044818 | ENSG00000130695 |
| BTN3A1 | -0.014650522 | 0.989896419 | 0.956926867 | 0.979276823 | 73.71583178 | 72.92661345 | ENSG00000026950 |
| MAPK14 | -0.014673957 | 0.98988034 | 0.862237049 | 0.927708201 | 1165.432161 | 1152.942149 | ENSG00000112062 |
| BCOR | -0.014701013 | 0.989861776 | 0.854802558 | 0.92303138 | 1602.637656 | 1586.84467 | ENSG00000183337 |
| PPFIA3 | -0.014854194 | 0.989756681 | 0.882347777 | 0.939381593 | 713.8670467 | 707.1110433 | ENSG00000177380 |
| RPL35A | -0.014868308 | 0.989746999 | 0.812902496 | 0.899428981 | 9516.841723 | 9418.500904 | ENSG00000182899 |
| SNHG7 | -0.014974146 | 0.989674392 | 0.897233595 | 0.947456442 | 740.7501746 | 733.9516165 | ENSG00000233016 |
| DDX41 | -0.01532127 | 0.989436297 | 0.836677423 | 0.912228917 | 1869.358316 | 1850.329885 | ENSG00000183258 |
| HLA-DPB2 | -0.01541658 | 0.989370933 | 0.903846169 | 0.951258232 | 391.2168362 | 386.6159328 | ENSG00000224557 |
| RPL10A | -0.015490036 | 0.98932056 | 0.768490711 | 0.871924202 | 16216.09887 | 16042.34401 | ENSG00000198755 |
| PRMT7 | -0.015491219 | 0.989319748 | 0.846928738 | 0.918489013 | 1112.329947 | 1100.497286 | ENSG00000132600 |
| CAB39 | -0.015511657 | 0.989305733 | 0.876047286 | 0.935668784 | 1197.170159 | 1183.8746 | ENSG00000135932 |
| SDAD1P1 | -0.015530394 | 0.989292884 | 0.981890785 | 0.991973763 | 9.81237616 | 9.763508889 | ENSG00000228451 |
| PEX2 | -0.015540556 | 0.989285917 | 0.891484222 | 0.944698031 | 441.2577725 | 436.3824442 | ENSG00000164751 |
| TRAIP | -0.015565808 | 0.989268601 | 0.90965811 | 0.955001893 | 353.5498531 | 349.9557383 | ENSG00000183763 |
| NA | -0.015609671 | 0.989238524 | 0.97827805 | 0.989993298 | 15.91917105 | 15.84758297 | ENSG00000223803 |
| KIF4A | -0.015658757 | 0.989204867 | 0.853710626 | 0.92238969 | 1137.18264 | 1124.798732 | ENSG00000090889 |
| RNMT | -0.015667065 | 0.989199171 | 0.853976892 | 0.922569813 | 1219.87786 | 1206.154129 | ENSG00000101654 |
| DCAF6 | -0.015707295 | 0.989171587 | 0.908951982 | 0.95469319 | 271.881644 | 268.9958764 | ENSG00000143164 |
| NA | -0.015811065 | 0.98910044 | 0.983667083 | 0.992668425 | 9.971574636 | 9.793126771 | ENSG00000286158 |
| ZBTB11-AS1 | -0.015863087 | 0.989064775 | 0.969609524 | 0.985454207 | 27.65003166 | 27.4374059 | ENSG00000256628 |
| CUL4A | -0.015882227 | 0.989051654 | 0.808757403 | 0.897311939 | 2787.919124 | 2756.787391 | ENSG00000139842 |
| CCDC18 | -0.015924836 | 0.989022443 | 0.929636524 | 0.965421826 | 186.6542292 | 185.0175237 | ENSG00000122483 |
| UBC | -0.016084425 | 0.988913045 | 0.806744418 | 0.896600656 | 9915.936597 | 9806.76761 | ENSG00000150991 |
| RCOR3 | -0.016084885 | 0.988912729 | 0.914979505 | 0.958037973 | 236.3029946 | 233.7688557 | ENSG00000117625 |
| WDR92 | -0.016100901 | 0.988901751 | 0.96125254 | 0.981049261 | 57.98830015 | 57.18340593 | ENSG00000243667 |

| | | | | | | | |
|---|---|---|---|---|---|---|---|
| ATPAF1 | -0.016136337 | 0.988877461 | 0.857383069 | 0.924256255 | 1297.034295 | 1281.975563 | ENSG00000123472 |
| C15orf39 | -0.0162696 | 0.988786122 | 0.853460324 | 0.922280546 | 1442.902777 | 1426.676417 | ENSG00000167173 |
| SNAP47 | -0.01628105 | 0.988778275 | 0.852577212 | 0.92215617 | 954.5543508 | 943.7803076 | ENSG00000143740 |
| MTREX | -0.016331 | 0.988744041 | 0.828016775 | 0.9073231 | 2613.491075 | 2582.823779 | ENSG00000039123 |
| SURF4 | -0.016369349 | 0.988717759 | 0.780836368 | 0.87954426 | 6151.79718 | 6081.898294 | ENSG00000148248 |
| CBLB | -0.016391686 | 0.988702451 | 0.911820851 | 0.956189185 | 404.8435042 | 399.483975 | ENSG00000114423 |
| HES6 | -0.016411825 | 0.988688649 | 0.856030948 | 0.92365824 | 1805.471509 | 1785.941172 | ENSG00000144485 |
| ZNF445 | -0.016426773 | 0.988678406 | 0.887561111 | 0.942534672 | 606.4611797 | 599.6156593 | ENSG00000185219 |
| FOLR1 | -0.016431773 | 0.988674979 | 0.981624891 | 0.991822494 | 9.723306184 | 9.658368997 | ENSG00000110195 |
| NA | -0.016547485 | 0.988595685 | 0.972796726 | 0.98701563 | 20.36216147 | 20.2059196 | ENSG00000232325 |
| TMEM80 | -0.016560982 | 0.988586436 | 0.932321144 | 0.966909523 | 216.9014735 | 214.9615712 | ENSG00000177042 |
| KCNQ2 | -0.016561509 | 0.988586075 | 0.808819953 | 0.89732246 | 2281.477528 | 2255.875319 | ENSG00000075043 |
| RASAL1 | -0.016876956 | 0.988369943 | 0.917720865 | 0.959390744 | 190.7342351 | 188.3648062 | ENSG00000111344 |
| ARAF | -0.016892045 | 0.988359606 | 0.844546151 | 0.917085678 | 899.767308 | 889.1730847 | ENSG00000078061 |
| B3GNT9 | -0.016927672 | 0.988335199 | 0.928118967 | 0.964548443 | 143.461022 | 141.8210537 | ENSG00000237172 |
| QSOX1 | -0.016960329 | 0.988312827 | 0.8040076 | 0.894704967 | 3171.10669 | 3133.329825 | ENSG00000116260 |
| CEP89 | -0.017007445 | 0.988280551 | 0.880432365 | 0.938509763 | 453.144795 | 447.9469978 | ENSG00000121289 |
| MXRA5 | -0.017010816 | 0.988278242 | 0.930614569 | 0.965950404 | 144.9060526 | 143.1034385 | ENSG00000101825 |
| NA | -0.01706299 | 0.988242502 | 0.987835181 | 0.994434699 | 3.761804341 | 3.713177955 | ENSG00000185839 |
| DMAC2 | -0.017081411 | 0.988229884 | 0.826749885 | 0.906810387 | 1377.556951 | 1361.114542 | ENSG00000105341 |
| RPP38 | -0.017200308 | 0.988148444 | 0.933646153 | 0.96733214 | 114.2570764 | 112.9173434 | ENSG00000152464 |
| RHOT1 | -0.017365138 | 0.988035554 | 0.878871441 | 0.937550551 | 651.8974919 | 643.2772698 | ENSG00000126858 |
| USP36 | -0.017373848 | 0.988029588 | 0.823870582 | 0.905349068 | 1293.766667 | 1278.668817 | ENSG00000055483 |
| FAM86DP | -0.017435641 | 0.987987271 | 0.920344389 | 0.96068883 | 184.2874485 | 182.2374561 | ENSG00000244026 |
| NBEA | -0.017459591 | 0.987970869 | 0.881873493 | 0.939373634 | 588.9920485 | 581.6685108 | ENSG00000172915 |
| LPAL2 | -0.01747393 | 0.98796105 | 0.970838615 | 0.986054521 | 20.65284527 | 20.46458826 | ENSG00000213071 |
| COX6C | -0.017526103 | 0.987925322 | 0.859812184 | 0.926120708 | 629.9477578 | 622.6852756 | ENSG00000164919 |
| FGFR3 | -0.017538025 | 0.987917158 | 0.856816708 | 0.924075698 | 670.5693159 | 662.7413419 | ENSG00000068078 |
| JOSD2 | -0.017551705 | 0.987907791 | 0.963650382 | 0.982393082 | 44.56276501 | 44.27898395 | ENSG00000161677 |
| NA | -0.017726554 | 0.987788067 | 0.961099118 | 0.980946697 | 41.08060285 | 40.58034011 | ENSG00000277531 |
| APTX | -0.017755649 | 0.987768147 | 0.857351264 | 0.924256255 | 789.0976468 | 778.733386 | ENSG00000137074 |
| MTPN | -0.017776797 | 0.987753667 | 0.847823428 | 0.918948068 | 1996.468293 | 1971.101182 | ENSG00000105887 |
| OTUD7B | -0.017803401 | 0.987735453 | 0.863505096 | 0.928749039 | 578.2196182 | 571.073241 | ENSG00000264522 |
| AKAP12 | -0.017817287 | 0.987725946 | 0.812310337 | 0.899364384 | 3514.87884 | 3472.216589 | ENSG00000131016 |
| UQCRC1 | -0.01785507 | 0.987700079 | 0.747903903 | 0.859264254 | 5605.549494 | 5536.218691 | ENSG00000010256 |
| KDM4A | -0.01787604 | 0.987685722 | 0.830559601 | 0.908658972 | 985.7592665 | 973.8800825 | ENSG00000066135 |
| RASD1 | -0.01789211 | 0.987674721 | 0.957561437 | 0.979493552 | 43.52057124 | 42.87092769 | ENSG00000108551 |
| WASHC2A | -0.017948906 | 0.987635838 | 0.845576927 | 0.917667336 | 896.2407537 | 885.131585 | ENSG00000099290 |
| CWC25 | -0.018009809 | 0.987594147 | 0.90786061 | 0.954214147 | 361.3674978 | 357.4973484 | ENSG00000273559 |
| CHPT1 | -0.018037664 | 0.987575079 | 0.903708403 | 0.951223557 | 303.3557087 | 299.510508 | ENSG00000111666 |
| PUS7 | -0.018152526 | 0.987496455 | 0.84013925 | 0.914174887 | 1055.531982 | 1042.427192 | ENSG00000091127 |
| ZNF260 | -0.018170506 | 0.987484148 | 0.917972231 | 0.959450204 | 368.397591 | 363.5453054 | ENSG00000254004 |
| UQCRFS1 | -0.018205225 | 0.987460384 | 0.821176077 | 0.904103446 | 1955.999455 | 1930.776239 | ENSG00000169021 |
| SP1 | -0.018250948 | 0.987429089 | 0.831405535 | 0.909306807 | 1334.417848 | 1317.003223 | ENSG00000185591 |
| ACO1 | -0.01830605 | 0.987391376 | 0.806845305 | 0.896600656 | 2080.456456 | 2053.792938 | ENSG00000122729 |
| B3GALT6 | -0.01846067 | 0.987285559 | 0.866443907 | 0.930273692 | 521.3027208 | 514.4566203 | ENSG00000176022 |
| PABPC1 | -0.018517586 | 0.98724661 | 0.801368058 | 0.893214577 | 25758.33979 | 25429.22674 | ENSG00000070756 |
| XPA | -0.018529835 | 0.987238228 | 0.94067242 | 0.970716428 | 81.05310755 | 79.87229796 | ENSG00000136936 |
| G3BP1 | -0.01865825 | 0.987150358 | 0.786439121 | 0.883102122 | 6828.187696 | 6739.119349 | ENSG00000145907 |
| PROM1 | -0.018771341 | 0.987072979 | 0.809371569 | 0.897454647 | 1218.585909 | 1202.744259 | ENSG00000007062 |
| XRN2 | -0.018941672 | 0.986956448 | 0.773645433 | 0.875060689 | 3369.382528 | 3324.672912 | ENSG00000088930 |
| PXK | -0.019176803 | 0.986795606 | 0.912550632 | 0.956487804 | 178.2084645 | 175.543352 | ENSG00000168297 |
| BLMH | -0.019188333 | 0.98678772 | 0.785137637 | 0.882458518 | 1927.932253 | 1902.569389 | ENSG00000108578 |
| FGFR4 | -0.019270035 | 0.986731838 | 0.78585971 | 0.882841791 | 2476.523208 | 2443.975976 | ENSG00000160867 |
| MYO19 | -0.019422131 | 0.986627817 | 0.83467987 | 0.911277722 | 1355.63115 | 1338.166171 | ENSG00000278259 |
| TRMT112 | -0.019422621 | 0.986627483 | 0.770469766 | 0.873601287 | 3881.627968 | 3829.19477 | ENSG00000173113 |
| NA | -0.019456348 | 0.986604417 | 0.959366924 | 0.980038391 | 43.39845896 | 42.64968939 | ENSG00000279369 |
| NA | -0.019498388 | 0.986575669 | 0.968088838 | 0.984740121 | 23.24757744 | 22.99974841 | ENSG00000280273 |
| ZNF547 | -0.019509319 | 0.986568193 | 0.974549206 | 0.988144592 | 12.74001207 | 12.4973375 | ENSG00000152433 |
| SYNGR2 | -0.01959539 | 0.986509336 | 0.828830322 | 0.90773156 | 1059.272146 | 1045.564275 | ENSG00000108639 |
| ZNF117 | -0.019642368 | 0.986477214 | 0.921812734 | 0.961386103 | 144.4085462 | 142.1881114 | ENSG00000152926 |
| FOXA3 | -0.019714112 | 0.986428158 | 0.864826101 | 0.929684296 | 495.7311461 | 488.9448312 | ENSG00000170608 |
| ZNF507 | -0.019744512 | 0.986407373 | 0.843742 | 0.916588373 | 674.5630027 | 664.934227 | ENSG00000168813 |
| NA | -0.019758133 | 0.98639806 | 0.976985962 | 0.989556227 | 10.50303298 | 10.3748128 | ENSG00000286868 |
| NIP7 | -0.019811144 | 0.986361816 | 0.822249784 | 0.904587018 | 1197.304621 | 1180.865291 | ENSG00000132603 |
| RNASEK | -0.019896589 | 0.986303399 | 0.97016306 | 0.985746513 | 18.34691775 | 18.10161624 | ENSG00000219200 |
| CPEB3 | -0.020087043 | 0.986173203 | 0.96501067 | 0.983044287 | 23.54891205 | 23.29282149 | ENSG00000107864 |
| KRTCAP2 | -0.020111745 | 0.986156318 | 0.949078917 | 0.97469657 | 50.10881753 | 49.45086855 | ENSG00000163463 |
| NA | -0.02017524 | 0.986112917 | 0.934433474 | 0.967746424 | 92.16464256 | 90.95325733 | ENSG00000227232 |
| SINHCAF | -0.020197817 | 0.986097485 | 0.803670261 | 0.894661818 | 6368.393891 | 6278.416624 | ENSG00000139146 |
| PMS1 | -0.020210978 | 0.98608849 | 0.834271135 | 0.911076206 | 867.1111683 | 855.704932 | ENSG00000064933 |
| TMEM198B | -0.020231464 | 0.986074487 | 0.923814445 | 0.96241072 | 139.3368345 | 137.5848044 | ENSG00000182796 |
| ZNF800 | -0.020253895 | 0.986059156 | 0.946653388 | 0.973439519 | 53.98000879 | 53.37812514 | ENSG00000048405 |
| SNAI3 | -0.020335143 | 0.986003626 | 0.919297636 | 0.960250106 | 131.4945206 | 129.7245171 | ENSG00000185669 |

| | | | | | | | |
|---|---|---|---|---|---|---|---|
| SPAG8 | -0.020385697 | 0.985969076 | 0.985551834 | 0.993485135 | 3.877407397 | 3.81463062 | ENSG00000137098 |
| NUP37 | -0.020415139 | 0.985948954 | 0.879203459 | 0.937742816 | 488.7665588 | 481.485568 | ENSG00000075188 |
| JADE2 | -0.020465767 | 0.985914355 | 0.846874449 | 0.918483881 | 839.0061956 | 826.9848206 | ENSG00000043143 |
| CSNK1D | -0.020483783 | 0.985902044 | 0.752765429 | 0.862657174 | 2797.219517 | 2758.539457 | ENSG00000141551 |
| PSMD10 | -0.020489415 | 0.985898195 | 0.804289457 | 0.894813662 | 2198.087255 | 2166.11903 | ENSG00000101843 |
| RAB13 | -0.020570532 | 0.985842763 | 0.890732608 | 0.944333494 | 251.0153314 | 247.6354966 | ENSG00000143545 |
| SLC35C2 | -0.020588608 | 0.985830412 | 0.817380854 | 0.901837342 | 945.3328282 | 932.2677044 | ENSG00000080189 |
| TRIM28 | -0.020731432 | 0.985732821 | 0.757122921 | 0.864710144 | 29717.70652 | 29293.25161 | ENSG00000130726 |
| ASPHD1 | -0.0207501 | 0.985720066 | 0.8587867 | 0.925299324 | 428.7992666 | 422.9106555 | ENSG00000174939 |
| EGLN2 | -0.020770366 | 0.985706219 | 0.938109508 | 0.969503942 | 88.06017064 | 86.88474459 | ENSG00000269858 |
| BAIAP2 | -0.020845617 | 0.985654807 | 0.864325567 | 0.929307927 | 395.9454592 | 390.6202487 | ENSG00000175866 |
| KIF18A | -0.020893895 | 0.985621823 | 0.876801132 | 0.93615025 | 330.6021219 | 325.3716831 | ENSG00000121621 |
| GID8 | -0.020930029 | 0.985597137 | 0.755576813 | 0.864169123 | 2391.233866 | 2356.681485 | ENSG00000101193 |
| STAC | -0.020943417 | 0.985587992 | 0.891150351 | 0.944651026 | 244.6234901 | 240.8744558 | ENSG00000144681 |
| TIMM44 | -0.020965832 | 0.985572679 | 0.774778668 | 0.875842184 | 2805.179523 | 2764.938192 | ENSG00000104980 |
| FTX | -0.021200973 | 0.985412056 | 0.911081176 | 0.955762377 | 146.8373601 | 144.9502841 | ENSG00000230590 |
| MRPL40 | -0.021258181 | 0.985372981 | 0.882154765 | 0.939373634 | 248.3074767 | 244.8035761 | ENSG00000185608 |
| NA | -0.021262323 | 0.985370153 | 0.980165752 | 0.991020242 | 7.479915565 | 7.37048096 | ENSG00000260448 |
| TERF2IP | -0.021279981 | 0.985358092 | 0.874189216 | 0.934994283 | 311.895865 | 307.3543891 | ENSG00000166848 |
| SPINDOC | -0.021284063 | 0.985355304 | 0.757694665 | 0.865096588 | 2549.810008 | 2513.065979 | ENSG00000168005 |
| NCBP2 | -0.021400737 | 0.985275619 | 0.768702134 | 0.872077208 | 2133.367851 | 2101.34639 | ENSG00000114503 |
| PPP1R14A | -0.021459264 | 0.98523565 | 0.872406381 | 0.933952288 | 350.194606 | 344.8060484 | ENSG00000167641 |
| MRPS10 | -0.021500058 | 0.985207791 | 0.834772481 | 0.911277722 | 721.1578784 | 709.8145122 | ENSG00000048544 |
| NA | -0.021505641 | 0.985203978 | 0.966984624 | 0.9842481 | 19.12486311 | 18.93079763 | ENSG00000286651 |
| IGBP1 | -0.021610863 | 0.985132126 | 0.837521676 | 0.912773535 | 535.7732158 | 527.7063349 | ENSG00000089289 |
| PSMB4 | -0.021621954 | 0.985124552 | 0.736837186 | 0.852627858 | 5481.035225 | 5400.002267 | ENSG00000159377 |
| EPN2 | -0.021661162 | 0.985097781 | 0.832726029 | 0.910062162 | 579.7873851 | 571.1935763 | ENSG00000072134 |
| NA | -0.021679905 | 0.985084982 | 0.96940602 | 0.985330341 | 15.40034036 | 15.13449997 | ENSG00000279347 |
| RANBP9 | -0.021884239 | 0.984945471 | 0.850827825 | 0.921208255 | 675.6926432 | 665.0794968 | ENSG00000010017 |
| NA | -0.02198286 | 0.984878144 | 0.977632696 | 0.989847838 | 7.284713271 | 7.167687249 | ENSG00000226049 |
| ZNF605 | -0.021991293 | 0.984872387 | 0.889415493 | 0.943529294 | 215.2196145 | 211.9731589 | ENSG00000196458 |
| RWDD4 | -0.022115775 | 0.984787411 | 0.886770784 | 0.942266356 | 237.93523 | 234.025549 | ENSG00000182552 |
| ARL1 | -0.022194233 | 0.984733857 | 0.83662033 | 0.912220332 | 907.4207934 | 892.9546561 | ENSG00000120805 |
| LONRF1 | -0.022211324 | 0.984722192 | 0.788603272 | 0.884433126 | 1087.473006 | 1070.733797 | ENSG00000154359 |
| TRDMT1 | -0.022321565 | 0.984646949 | 0.8926185 | 0.945187103 | 204.0022403 | 200.8279625 | ENSG00000107614 |
| NA | -0.02234305 | 0.984632285 | 0.969434487 | 0.985330341 | 14.25536658 | 14.03167777 | ENSG00000257576 |
| POLE2 | -0.02243962 | 0.984566379 | 0.890360386 | 0.944046876 | 253.3439739 | 249.293971 | ENSG00000100479 |
| C5orf34 | -0.022448522 | 0.984560304 | 0.927447428 | 0.964343189 | 85.60283165 | 84.41977025 | ENSG00000172244 |
| CTSL | -0.022481864 | 0.98453755 | 0.830284634 | 0.90859717 | 669.2756589 | 659.3572332 | ENSG00000135047 |
| PTCHD1 | -0.022565134 | 0.984480726 | 0.939439134 | 0.970003616 | 71.89776719 | 70.77794331 | ENSG00000165186 |
| CCDC134 | -0.022583599 | 0.984468125 | 0.918619515 | 0.959788832 | 103.0961358 | 101.5152304 | ENSG00000100147 |
| TRIT1 | -0.022614718 | 0.984446891 | 0.830307018 | 0.90859717 | 539.2334901 | 530.952763 | ENSG00000043514 |
| LYRM4-AS1 | -0.022727838 | 0.984369704 | 0.974024595 | 0.987774778 | 9.478096946 | 9.377130495 | ENSG00000272142 |
| ATP6V1C1 | -0.022741236 | 0.984360562 | 0.836956719 | 0.912372417 | 560.3508222 | 551.1425625 | ENSG00000155097 |
| C14orf119 | -0.022837866 | 0.984294633 | 0.819255439 | 0.90291228 | 698.6559031 | 687.6179347 | ENSG00000179933 |
| PTMA | -0.022915248 | 0.98424184 | 0.683659081 | 0.817493665 | 39152.14034 | 38534.60164 | ENSG00000187514 |
| NA | -0.022934949 | 0.9842284 | 0.943852765 | 0.972449807 | 48.57895623 | 47.73261718 | ENSG00000233966 |
| ORC4 | -0.022956504 | 0.984213695 | 0.811836987 | 0.899055133 | 859.899808 | 845.8981698 | ENSG00000115947 |
| NA | -0.02297147 | 0.984203485 | 0.939030741 | 0.969852089 | 70.48182521 | 69.68023405 | ENSG00000261505 |
| SFMBT1 | -0.023014893 | 0.984173862 | 0.849403928 | 0.920078138 | 390.0306346 | 384.0027659 | ENSG00000163935 |
| TRIM65 | -0.023026046 | 0.984166254 | 0.784215222 | 0.881903096 | 1367.520074 | 1346.467429 | ENSG00000141569 |
| ZNF263 | -0.023147951 | 0.984083097 | 0.791826989 | 0.886643391 | 876.6984493 | 862.8748723 | ENSG00000006194 |
| ACADSB | -0.023167947 | 0.984069458 | 0.833603461 | 0.910747762 | 475.7944236 | 468.1219854 | ENSG00000196177 |
| GOPC | -0.02320243 | 0.984045937 | 0.844257364 | 0.917016342 | 456.8174091 | 449.0846222 | ENSG00000047932 |
| GRK3 | -0.023322316 | 0.983964167 | 0.808888405 | 0.897335896 | 788.0759729 | 775.165581 | ENSG00000100077 |
| CTNNB1 | -0.02333458 | 0.983955803 | 0.671445026 | 0.809097911 | 10367.91095 | 10200.81187 | ENSG00000168036 |
| FAM162B | -0.023353707 | 0.983942758 | 0.954959357 | 0.978289671 | 27.41557093 | 27.02097159 | ENSG00000183807 |
| MARVELD3 | -0.02338871 | 0.983918886 | 0.812988236 | 0.89947015 | 678.1524773 | 667.2100027 | ENSG00000140832 |
| ANKMY1 | -0.023397272 | 0.983913046 | 0.912621685 | 0.956487804 | 108.0421724 | 106.3763703 | ENSG00000144504 |
| RABGGTA | -0.023429943 | 0.983890765 | 0.882840628 | 0.939729393 | 243.2770002 | 239.887987 | ENSG00000100949 |
| NA | -0.023613845 | 0.983765356 | 0.98294677 | 0.992478505 | 3.768118167 | 3.69440696 | ENSG00000178660 |
| NRDC | -0.02370476 | 0.983703363 | 0.695863145 | 0.826103895 | 3438.619659 | 3382.656226 | ENSG00000078618 |
| DAPK2 | -0.024074016 | 0.983451618 | 0.943155341 | 0.97211501 | 46.59044098 | 46.03813812 | ENSG00000035664 |
| TMBIM6 | -0.024191928 | 0.983371243 | 0.682921241 | 0.816931965 | 11073.42897 | 10888.29146 | ENSG00000139644 |
| POP5 | -0.024262173 | 0.983323364 | 0.836278519 | 0.9121504 | 488.9037113 | 481.1358244 | ENSG00000167272 |
| GCDH | -0.02437337 | 0.983247576 | 0.798485808 | 0.891646271 | 1149.993149 | 1131.439002 | ENSG00000105607 |
| TPST2 | -0.024487061 | 0.983170095 | 0.755972343 | 0.864333115 | 2559.375604 | 2516.68828 | ENSG00000128294 |
| CAPZA2 | -0.024572319 | 0.983111995 | 0.812796457 | 0.899402162 | 1276.21723 | 1253.629615 | ENSG00000198898 |
| NADK2 | -0.024609265 | 0.983086819 | 0.860141675 | 0.926206473 | 335.8897436 | 330.0261188 | ENSG00000152620 |
| NSMCE3 | -0.024670615 | 0.983045014 | 0.830281652 | 0.90859717 | 464.2095551 | 456.0242463 | ENSG00000185115 |
| ADGRA3 | -0.024709305 | 0.983018651 | 0.737474831 | 0.852943558 | 1718.768472 | 1688.977456 | ENSG00000152990 |
| ATF6B | -0.024741367 | 0.982996805 | 0.827842494 | 0.907293051 | 576.7205745 | 567.4845639 | ENSG00000213676 |
| NA | -0.024760719 | 0.98298362 | 0.969749677 | 0.985488568 | 11.11852525 | 10.90343504 | ENSG00000278126 |
| GTF2A2 | -0.024834754 | 0.982933177 | 0.768520221 | 0.871924202 | 1288.087063 | 1265.553302 | ENSG00000140307 |

| | | | | | | | |
|---|---|---|---|---|---|---|---|
| JAKMIP3 | -0.024942626 | 0.982859685 | 0.938589333 | 0.969724993 | 54.0457026 | 53.30716642 | ENSG00000188385 |
| IRF2BP1 | -0.024960308 | 0.982847639 | 0.81863196 | 0.902535527 | 869.6024964 | 854.6926182 | ENSG00000170604 |
| ZNF687 | -0.02498146 | 0.982833229 | 0.727582325 | 0.846909144 | 1754.906646 | 1725.163057 | ENSG00000143373 |
| CD320 | -0.025022551 | 0.982805236 | 0.756255975 | 0.864335767 | 1501.311457 | 1475.25654 | ENSG00000167775 |
| RPS6KA4 | -0.025183711 | 0.982695455 | 0.813158175 | 0.899516491 | 625.6362527 | 614.9579309 | ENSG00000162302 |
| RAD23B | -0.025211218 | 0.982676719 | 0.749587639 | 0.860451957 | 5909.482037 | 5806.297835 | ENSG00000119318 |
| SBSPON | -0.025242637 | 0.982655319 | 0.933202197 | 0.967227912 | 53.17059201 | 52.34994888 | ENSG00000164764 |
| NA | -0.025281458 | 0.982628877 | 0.974811633 | 0.988194432 | 7.784504787 | 7.625462394 | ENSG00000279744 |
| UNC13B | -0.025282744 | 0.982628001 | 0.737749601 | 0.853097773 | 2388.052474 | 2346.634397 | ENSG00000198722 |
| SEC62 | -0.025284209 | 0.982627003 | 0.771931293 | 0.874465985 | 1462.696709 | 1437.190654 | ENSG00000008952 |
| ASAP3 | -0.02539305 | 0.982552874 | 0.746687128 | 0.858611443 | 1746.830135 | 1716.892414 | ENSG00000088280 |
| NA | -0.025420958 | 0.982533867 | 0.963404418 | 0.982379923 | 16.6938618 | 16.44011588 | ENSG00000217094 |
| ITPRIP | -0.025546445 | 0.982448409 | 0.804793095 | 0.895212773 | 622.1052636 | 610.8986161 | ENSG00000148841 |
| DPH1 | -0.02558398 | 0.982422849 | 0.856308422 | 0.923850067 | 265.2876853 | 260.6158786 | ENSG00000108963 |
| HSPB1 | -0.02566261 | 0.982369306 | 0.717021937 | 0.839659101 | 2583.98632 | 2537.801745 | ENSG00000106211 |
| AP3S1 | -0.025713771 | 0.98233447 | 0.786055075 | 0.882900719 | 985.9851283 | 967.9690005 | ENSG00000177879 |
| RAD21 | -0.025740366 | 0.982316362 | 0.74830317 | 0.859456587 | 3602.672575 | 3537.823825 | ENSG00000164754 |
| PCF11 | -0.025851582 | 0.982240639 | 0.802102812 | 0.893630607 | 813.4913105 | 799.7114732 | ENSG00000165494 |
| NA | -0.025852082 | 0.982240298 | 0.889330714 | 0.943529294 | 174.5888939 | 171.8111169 | ENSG00000159247 |
| EGFL8 | -0.025852498 | 0.982240015 | 0.96621715 | 0.983786664 | 16.83156324 | 16.56402677 | ENSG00000241404 |
| THAP5 | -0.025891515 | 0.982213451 | 0.915832714 | 0.958281687 | 136.0228227 | 133.3609705 | ENSG00000177683 |
| PVR | -0.025975822 | 0.982156055 | 0.825191597 | 0.906048641 | 487.5265468 | 479.0620779 | ENSG00000073008 |
| ZFYVE21 | -0.026119893 | 0.982057979 | 0.79320403 | 0.887809861 | 795.5339628 | 781.4283819 | ENSG00000100711 |
| CRABP1 | -0.026158474 | 0.982031717 | 0.804037658 | 0.894704967 | 759.7429138 | 746.1799938 | ENSG00000166426 |
| SLC16A1-AS1 | -0.026201605 | 0.982002359 | 0.900627468 | 0.949285116 | 120.1204435 | 118.0231675 | ENSG00000226419 |
| VHL | -0.026251018 | 0.981968725 | 0.809647545 | 0.897492586 | 755.8634918 | 743.0955625 | ENSG00000134086 |
| PLAG1 | -0.026286211 | 0.981944771 | 0.872529662 | 0.93395696 | 188.7361514 | 185.1727644 | ENSG00000181690 |
| DPYSL2 | -0.026329051 | 0.981915614 | 0.716582103 | 0.839556455 | 2638.177384 | 2590.071901 | ENSG00000092964 |
| PUS1 | -0.026480339 | 0.98181265 | 0.769268783 | 0.87255983 | 1869.423378 | 1836.392575 | ENSG00000177192 |
| SLC35F2 | -0.026506293 | 0.981794988 | 0.796501616 | 0.890210514 | 606.7289008 | 595.2187611 | ENSG00000110660 |
| PGLS | -0.026804027 | 0.981592392 | 0.839110615 | 0.913691856 | 388.2974007 | 381.1058439 | ENSG00000130313 |
| NTMT1 | -0.02683918 | 0.981568475 | 0.792548003 | 0.887182744 | 751.2321689 | 737.5381982 | ENSG00000148335 |
| MEAF6 | -0.026940153 | 0.981499779 | 0.799720292 | 0.892387536 | 653.4595836 | 641.2405014 | ENSG00000163875 |
| SEPTIN6 | -0.026964835 | 0.981482987 | 0.76939563 | 0.872650304 | 885.0613601 | 869.3296285 | ENSG00000125354 |
| NA | -0.026986486 | 0.981468257 | 0.964395 | 0.982728532 | 13.31190892 | 13.0865096 | ENSG00000235454 |
| LAPTM4A | -0.027050215 | 0.981424904 | 0.690268699 | 0.822404913 | 4748.639919 | 4659.394995 | ENSG00000068697 |
| CTU1 | -0.027141538 | 0.981362781 | 0.856677997 | 0.924033637 | 324.1418957 | 318.1494435 | ENSG00000142544 |
| CASP8AP2 | -0.027148048 | 0.981358353 | 0.808686241 | 0.897311939 | 490.5349021 | 481.0269486 | ENSG00000118412 |
| SSR3 | -0.027171198 | 0.981342606 | 0.739972781 | 0.854707161 | 2358.635474 | 2313.754418 | ENSG00000114850 |
| C2orf72 | -0.027180661 | 0.981336168 | 0.932555518 | 0.966994782 | 52.34273741 | 51.58566585 | ENSG00000204128 |
| ZNF680 | -0.027184586 | 0.981333499 | 0.887029377 | 0.942266356 | 151.3226681 | 148.6105099 | ENSG00000173041 |
| NPY1R | -0.027193285 | 0.981327582 | 0.89676544 | 0.947207537 | 114.8251173 | 112.6437026 | ENSG00000164128 |
| CPXM2 | -0.02728842 | 0.981262873 | 0.879149303 | 0.937739018 | 176.7809021 | 173.5880544 | ENSG00000121898 |
| ANKDD1A | -0.027314171 | 0.981245358 | 0.927101201 | 0.964343189 | 55.20595728 | 54.30072347 | ENSG00000166839 |
| EIF2AK3 | -0.027405435 | 0.981183287 | 0.837946731 | 0.912968356 | 307.2480062 | 301.4585746 | ENSG00000172071 |
| PCDHA7 | -0.027469436 | 0.981139761 | 0.954371774 | 0.978174333 | 19.93881228 | 19.5903788 | ENSG00000204963 |
| RNF14 | -0.027490885 | 0.981125174 | 0.840097179 | 0.914174887 | 322.6835762 | 316.2351031 | ENSG00000013561 |
| ATP5MC2 | -0.027508735 | 0.981113034 | 0.681434506 | 0.815977978 | 2892.335104 | 2837.820702 | ENSG00000135390 |
| CMSS1 | -0.027523313 | 0.981103121 | 0.764564891 | 0.869725999 | 854.7607566 | 838.127764 | ENSG00000184220 |
| CWF19L1 | -0.027529045 | 0.981099223 | 0.819687172 | 0.902946927 | 408.5409902 | 400.3580099 | ENSG00000095485 |
| CLP1 | -0.027563546 | 0.981075761 | 0.795163761 | 0.889144202 | 598.675788 | 587.0057756 | ENSG00000172409 |
| CD81 | -0.027578981 | 0.981065265 | 0.712621381 | 0.837256715 | 3260.467831 | 3198.235598 | ENSG00000110651 |
| KYAT3 | -0.027595608 | 0.981053958 | 0.775774715 | 0.876322473 | 720.1369937 | 706.2685613 | ENSG00000137944 |
| MRNIP | -0.02760523 | 0.981047415 | 0.883595661 | 0.939962378 | 291.1434345 | 286.0973743 | ENSG00000161010 |
| SNRPN | -0.02762317 | 0.981035215 | 0.911343166 | 0.955850503 | 82.06088355 | 80.54145219 | ENSG00000128739 |
| NIBAN2 | -0.027670623 | 0.981002948 | 0.68861617 | 0.821393493 | 2801.260257 | 2748.174002 | ENSG00000136830 |
| CNIH4 | -0.027713946 | 0.98097349 | 0.742671701 | 0.856105567 | 1447.634008 | 1420.191176 | ENSG00000143771 |
| MAPK1IP1L | -0.027785836 | 0.980924609 | 0.753920316 | 0.863411144 | 1296.385903 | 1271.383471 | ENSG00000168175 |
| NME4 | -0.027799835 | 0.98091509 | 0.653188575 | 0.797352107 | 4985.204087 | 4890.139222 | ENSG00000103202 |
| SEPTIN5 | -0.027803521 | 0.980912584 | 0.973832964 | 0.98768183 | 12.53909731 | 12.31755166 | ENSG00000184702 |
| TXNDC9 | -0.027864896 | 0.980870855 | 0.849130361 | 0.919961543 | 273.8241811 | 268.3908561 | ENSG00000115514 |
| CLUHP3 | -0.027973335 | 0.980797132 | 0.883617379 | 0.939962378 | 197.4019829 | 193.7836767 | ENSG00000131797 |
| DICER1 | -0.028004096 | 0.980776219 | 0.765892523 | 0.870701835 | 1083.869964 | 1062.38121 | ENSG00000100697 |
| SIAH1 | -0.028025514 | 0.980761659 | 0.922046161 | 0.961386103 | 110.8229292 | 108.6362491 | ENSG00000196470 |
| DDX21 | -0.028060752 | 0.980737704 | 0.705214397 | 0.832512333 | 6779.389334 | 6647.842939 | ENSG00000165732 |
| IRF2BPL | -0.028125786 | 0.980693495 | 0.755311615 | 0.863919123 | 1967.18865 | 1928.292843 | ENSG00000119669 |
| GPHA2 | -0.028290534 | 0.980581512 | 0.971374769 | 0.986210953 | 7.361155596 | 7.197743164 | ENSG00000149735 |
| TINAGL1 | -0.028394399 | 0.980510919 | 0.872424515 | 0.933952288 | 183.8701049 | 180.6282551 | ENSG00000142910 |
| COL24A1 | -0.028406764 | 0.980502515 | 0.952038289 | 0.976700855 | 21.10272754 | 20.67443001 | ENSG00000171502 |
| WDR36 | -0.028432897 | 0.980484754 | 0.783514872 | 0.881593021 | 756.1209605 | 740.663764 | ENSG00000134987 |
| NA | -0.02846083 | 0.980465771 | 0.981530422 | 0.991804865 | 3.782023595 | 3.69440696 | ENSG00000272097 |
| MIIP | -0.02852995 | 0.980418797 | 0.724240962 | 0.844902699 | 1342.723228 | 1317.047584 | ENSG00000116691 |
| RAB11FIP4 | -0.028586372 | 0.980380455 | 0.68086149 | 0.81555528 | 2099.867591 | 2058.265414 | ENSG00000131242 |
| AVPI1 | -0.028686762 | 0.980312238 | 0.871321099 | 0.933202268 | 169.9534522 | 166.6170602 | ENSG00000119986 |

| | | | | | | | |
|---|---|---|---|---|---|---|---|
| DDX20 | -0.028695519 | 0.980306287 | 0.774044628 | 0.875300505 | 832.013091 | 814.7677424 | ENSG00000064703 |
| ZFP69 | -0.028763229 | 0.98026028 | 0.899317977 | 0.948444134 | 102.9647482 | 101.1636945 | ENSG00000187815 |
| XPOT | -0.028793003 | 0.980240049 | 0.649465527 | 0.795321709 | 5212.881028 | 5108.68602 | ENSG00000184575 |
| KDM8 | -0.028855822 | 0.980197368 | 0.875383895 | 0.935391474 | 160.9536497 | 157.9693074 | ENSG00000155666 |
| KCNE5 | -0.028872816 | 0.980185822 | 0.933007015 | 0.967187875 | 50.17017434 | 49.20304678 | ENSG00000176076 |
| PDHA1 | -0.02891085 | 0.980159982 | 0.678354496 | 0.814170153 | 2349.608666 | 2303.284232 | ENSG00000131828 |
| NA | -0.028981878 | 0.980111726 | 0.978295417 | 0.989993298 | 5.717570381 | 5.633896625 | ENSG00000276744 |
| ANAPC1 | -0.029034505 | 0.980075974 | 0.709498352 | 0.834857626 | 2070.987131 | 2028.916909 | ENSG00000153107 |
| RIOX1 | -0.029118914 | 0.980018634 | 0.797195696 | 0.890717689 | 565.9928971 | 554.4934777 | ENSG00000170468 |
| NA | -0.029126335 | 0.980013593 | 0.980512831 | 0.991208946 | 3.791494334 | 3.701893034 | ENSG00000258603 |
| MYO10 | -0.029421461 | 0.979813136 | 0.658753875 | 0.801138078 | 3999.377038 | 3918.149402 | ENSG00000145555 |
| LSMEM1 | -0.029459959 | 0.97978699 | 0.974649227 | 0.988178905 | 5.742825684 | 5.671438614 | ENSG00000181016 |
| IKBKB | -0.029480037 | 0.979773355 | 0.772725679 | 0.874767758 | 655.2378262 | 642.4845487 | ENSG00000104365 |
| ZBED3 | -0.029486871 | 0.979768714 | 0.895858513 | 0.946681207 | 100.0503413 | 98.09100029 | ENSG00000132846 |
| KLHL5 | -0.029501525 | 0.979758762 | 0.835840534 | 0.911983321 | 314.9088353 | 308.2488666 | ENSG00000109790 |
| CDC14A | -0.029585971 | 0.979701415 | 0.876512407 | 0.936003742 | 156.8826261 | 153.3571515 | ENSG00000079335 |
| LGALSL | -0.029600349 | 0.979691651 | 0.815968005 | 0.900938341 | 421.0714885 | 411.9796006 | ENSG00000119862 |
| ZNF585B | -0.029620116 | 0.979678228 | 0.91715046 | 0.959173154 | 63.54076419 | 62.03738611 | ENSG00000245680 |
| CROT | -0.029657843 | 0.979652609 | 0.910780487 | 0.95570436 | 75.09023023 | 73.30842855 | ENSG00000005469 |
| MRAS | -0.029733619 | 0.979601156 | 0.827190895 | 0.907101046 | 312.4634249 | 306.4935515 | ENSG00000158186 |
| RAB22A | -0.030022948 | 0.979404719 | 0.849535319 | 0.920078138 | 216.4773275 | 211.8966492 | ENSG00000124209 |
| CCDC17 | -0.030064098 | 0.979376784 | 0.963468619 | 0.98239134 | 11.71064135 | 11.54143406 | ENSG00000159588 |
| IK | -0.030113753 | 0.979343076 | 0.71135231 | 0.836295827 | 1468.471676 | 1437.844979 | ENSG00000113141 |
| PIK3CD | -0.0301228 | 0.979336934 | 0.763694424 | 0.869002484 | 602.8440972 | 590.4764595 | ENSG00000171608 |
| NA | -0.030184128 | 0.979295304 | 0.963729692 | 0.982393082 | 11.29794298 | 11.02734593 | ENSG00000274104 |
| TRAK2 | -0.030250532 | 0.97925023 | 0.831415948 | 0.909306807 | 310.047434 | 303.3244773 | ENSG00000115993 |
| CNNM4 | -0.030412389 | 0.979140374 | 0.798550101 | 0.891646271 | 436.5536004 | 427.3866908 | ENSG00000158158 |
| C2orf42 | -0.03048532 | 0.979090878 | 0.88880102 | 0.943202869 | 113.1567804 | 110.7311313 | ENSG00000115998 |
| ZNF737 | -0.030500806 | 0.979080368 | 0.878440851 | 0.93725305 | 159.2348023 | 155.5771153 | ENSG00000237440 |
| HSPB11 | -0.030562097 | 0.979038774 | 0.818050583 | 0.902162889 | 383.1021786 | 375.5102939 | ENSG00000081870 |
| NA | -0.030596708 | 0.979015287 | 0.941206052 | 0.970800432 | 35.70490328 | 35.01827827 | ENSG00000224086 |
| KLHL22 | -0.030623052 | 0.97899741 | 0.866020607 | 0.930085925 | 149.7276167 | 146.6638605 | ENSG00000099910 |
| OCLN | -0.030657212 | 0.97897423 | 0.797719979 | 0.891196027 | 421.6519843 | 412.6054789 | ENSG00000197822 |
| PAXBP1 | -0.030834693 | 0.978853803 | 0.790990006 | 0.886348777 | 891.2759937 | 873.2894031 | ENSG00000159086 |
| KHDRBS1 | -0.030843909 | 0.97884755 | 0.719215659 | 0.841017259 | 6818.633592 | 6673.653165 | ENSG00000121774 |
| PTPN12 | -0.03089672 | 0.978811719 | 0.667666054 | 0.80717496 | 2089.266723 | 2045.17372 | ENSG00000127947 |
| TIGD5 | -0.030935248 | 0.97878558 | 0.854340118 | 0.922743843 | 208.4401196 | 204.0012333 | ENSG00000179886 |
| RPA2 | -0.031017598 | 0.978729712 | 0.705915092 | 0.833021159 | 1130.498529 | 1106.133341 | ENSG00000117748 |
| KLK7 | -0.031033139 | 0.978719169 | 0.958053848 | 0.979577828 | 14.45372579 | 14.1105606 | ENSG00000169035 |
| RINT1 | -0.031162431 | 0.978631461 | 0.806151322 | 0.896221316 | 404.0508414 | 394.9290569 | ENSG00000135249 |
| WDR45B | -0.031254314 | 0.978569136 | 0.636376352 | 0.786306621 | 2505.536867 | 2451.621619 | ENSG00000141580 |
| PLPP3 | -0.031277453 | 0.978553441 | 0.793874502 | 0.888278381 | 405.8080529 | 397.0554207 | ENSG00000162407 |
| NA | -0.031277712 | 0.978553265 | 0.965935432 | 0.983607895 | 8.800067779 | 8.66068669 | ENSG00000272944 |
| RPS16 | -0.0313897 | 0.978477309 | 0.645307136 | 0.792859536 | 19607.2615 | 19185.30315 | ENSG00000105193 |
| ASCC1 | -0.031390004 | 0.978477103 | 0.768117996 | 0.871908897 | 545.0788102 | 533.0048384 | ENSG00000138303 |
| HDAC4-AS1 | -0.03139127 | 0.978476244 | 0.979128982 | 0.990296129 | 4.181996619 | 4.065813207 | ENSG00000222020 |
| CSPG5 | -0.031410489 | 0.978463209 | 0.891187706 | 0.944651026 | 105.5070156 | 103.4095804 | ENSG00000114646 |
| CMTM6 | -0.031419108 | 0.978457363 | 0.776882703 | 0.877253417 | 897.719375 | 877.9897563 | ENSG00000091317 |
| NIPAL4 | -0.031455024 | 0.978433005 | 0.979261243 | 0.990375853 | 3.847717405 | 3.747032717 | ENSG00000172548 |
| ZFAND2B | -0.031488174 | 0.978410523 | 0.854249338 | 0.922728642 | 192.5879952 | 188.5901697 | ENSG00000158552 |
| LOC100507642 | -0.031565055 | 0.978358385 | 0.971680649 | 0.986315195 | 6.556774858 | 6.413701455 | ENSG00000240859 |
| NA | -0.031633532 | 0.978311949 | 0.963075313 | 0.982152407 | 9.821846899 | 9.617028162 | ENSG00000271992 |
| POP1 | -0.031731237 | 0.978245696 | 0.789921455 | 0.885419065 | 428.9478143 | 419.8152714 | ENSG00000104356 |
| CUEDC1 | -0.031756954 | 0.978228258 | 0.839802148 | 0.914076405 | 222.6537473 | 218.0874005 | ENSG00000180891 |
| NA | -0.031891872 | 0.978136781 | 0.943205869 | 0.97211501 | 24.59416503 | 24.13642538 | ENSG00000273270 |
| ATG9A | -0.032003236 | 0.978061279 | 0.752004857 | 0.86205208 | 643.1671467 | 629.4188484 | ENSG00000198925 |
| TWNK | -0.032073376 | 0.97801373 | 0.677287104 | 0.813471461 | 1633.796564 | 1598.037929 | ENSG00000107815 |
| RAB9A | -0.032151524 | 0.977960755 | 0.79234054 | 0.88705766 | 402.3645326 | 393.6418142 | ENSG00000123595 |
| BMP2 | -0.032317509 | 0.977848244 | 0.919082259 | 0.96016406 | 61.15936386 | 59.95652866 | ENSG00000125845 |
| ETV6 | -0.03233841 | 0.977834078 | 0.79442532 | 0.8886547 | 403.6526499 | 394.2131627 | ENSG00000139083 |
| TOMM22 | -0.03242503 | 0.97777537 | 0.669988321 | 0.808268285 | 1989.910692 | 1945.013496 | ENSG00000100216 |
| PRX | -0.032426999 | 0.977774036 | 0.802849812 | 0.894151534 | 317.792079 | 310.9419443 | ENSG00000105227 |
| SNX27 | -0.032430445 | 0.9777717 | 0.709101539 | 0.83460265 | 1041.991145 | 1018.303197 | ENSG00000143376 |
| PIN1 | -0.032471199 | 0.97774408 | 0.676022222 | 0.812647287 | 1879.660561 | 1837.942943 | ENSG00000127445 |
| ZNF480 | -0.032644099 | 0.977626909 | 0.767778083 | 0.871722555 | 524.6733464 | 513.0554291 | ENSG00000198464 |
| IBTK | -0.032649918 | 0.977622966 | 0.783687849 | 0.881684509 | 652.5178323 | 637.4987646 | ENSG00000005700 |
| NA | -0.032656092 | 0.977618783 | 0.958160681 | 0.979577828 | 12.31666288 | 12.07765399 | ENSG00000274422 |
| IFT46 | -0.032676574 | 0.977604903 | 0.836147193 | 0.9121504 | 223.7555316 | 218.3922595 | ENSG00000118096 |
| CXorf40A | -0.03268617 | 0.9775984 | 0.879465653 | 0.937806595 | 107.1859056 | 104.8952054 | ENSG00000197620 |
| RNF34 | -0.032686688 | 0.97759805 | 0.679527403 | 0.815010735 | 1185.166789 | 1158.616216 | ENSG00000170633 |
| CSDE1 | -0.032734492 | 0.977565658 | 0.579922162 | 0.744031382 | 12909.11123 | 12618.55515 | ENSG00000009307 |
| ZNF33B | -0.032735216 | 0.977565167 | 0.788805706 | 0.884435871 | 377.9261387 | 369.5103986 | ENSG00000196693 |
| LAMTOR1 | -0.03274694 | 0.977557223 | 0.692158758 | 0.823657812 | 1006.512028 | 983.8773884 | ENSG00000149357 |
| NDUFS5 | -0.032809856 | 0.977514592 | 0.629426906 | 0.781306895 | 5060.258449 | 4946.410717 | ENSG00000168653 |

| | | | | | | | |
|---|---|---|---|---|---|---|---|
| DGUOK-AS1 | -0.032812013 | 0.977513131 | 0.972043667 | 0.986521593 | 6.060042557 | 5.948878269 | ENSG00000237883 |
| TMEM125 | -0.032845694 | 0.97749031 | 0.80373541 | 0.894680606 | 349.7199948 | 341.8370055 | ENSG00000179178 |
| STRBP | -0.032982562 | 0.97739758 | 0.629972174 | 0.781581472 | 2347.1676 | 2293.520722 | ENSG00000165209 |
| TMEM240 | -0.033016717 | 0.977374441 | 0.900184083 | 0.948979683 | 76.0192788 | 74.182638 | ENSG00000205090 |
| MRPS16 | -0.033062785 | 0.977343232 | 0.669692084 | 0.80797247 | 1609.050964 | 1572.387002 | ENSG00000182180 |
| NA | -0.033100662 | 0.977317573 | 0.968502996 | 0.984740121 | 6.689440255 | 6.537612341 | ENSG00000267191 |
| MICA | -0.033116109 | 0.977307109 | 0.800428089 | 0.892875968 | 328.2032911 | 320.8062446 | ENSG00000204520 |
| GK5 | -0.03331662 | 0.977171289 | 0.865501119 | 0.929978419 | 154.3208385 | 150.9673099 | ENSG00000175066 |
| ANXA4 | -0.033379993 | 0.977128366 | 0.831697579 | 0.909313528 | 208.6433945 | 203.8936086 | ENSG00000196975 |
| WDR6 | -0.033412499 | 0.97710635 | 0.607284246 | 0.765517664 | 4711.908562 | 4604.907994 | ENSG00000178252 |
| NAA35 | -0.033443312 | 0.977085481 | 0.718403389 | 0.840477794 | 741.2575955 | 724.2301239 | ENSG00000135040 |
| MITD1 | -0.033528887 | 0.977027526 | 0.79742812 | 0.890923671 | 358.5055557 | 350.132896 | ENSG00000158411 |
| DMPK | -0.033531317 | 0.97702588 | 0.826126538 | 0.906430811 | 399.9322535 | 391.4914725 | ENSG00000104936 |
| SUPT5H | -0.033672102 | 0.976930542 | 0.595695928 | 0.756103843 | 4735.941901 | 4627.858875 | ENSG00000196235 |
| BNIP2 | -0.033739076 | 0.976885191 | 0.761451939 | 0.867676006 | 554.3246187 | 541.2405054 | ENSG00000140299 |
| TMEM63B | -0.033740405 | 0.976884291 | 0.678308097 | 0.814170153 | 1069.451183 | 1044.795483 | ENSG00000137216 |
| NA | -0.033820461 | 0.976830085 | 0.932559876 | 0.966994782 | 29.87122619 | 29.16986498 | ENSG00000263272 |
| RNPEP | -0.033856807 | 0.976805476 | 0.689994988 | 0.822191533 | 1033.293263 | 1008.994334 | ENSG00000176393 |
| INTS9 | -0.033873171 | 0.976794396 | 0.780773769 | 0.879527201 | 422.5910073 | 412.9224359 | ENSG00000104299 |
| ZNF655 | -0.033876711 | 0.976792 | 0.767516405 | 0.871585629 | 492.2654902 | 481.0136254 | ENSG00000197343 |
| SSB | -0.033928668 | 0.976756822 | 0.593789554 | 0.755203695 | 3875.798116 | 3784.979169 | ENSG00000138385 |
| XPO5 | -0.034100809 | 0.976640284 | 0.6355635 | 0.785721312 | 3074.770341 | 3002.209084 | ENSG00000124571 |
| VPS35 | -0.034132802 | 0.976618626 | 0.693859772 | 0.824623085 | 1530.014634 | 1493.221717 | ENSG00000069329 |
| TPST1 | -0.034139155 | 0.976614325 | 0.794227648 | 0.888526301 | 316.6584551 | 309.1170157 | ENSG00000169902 |
| TMEM127 | -0.03415382 | 0.976604398 | 0.728379827 | 0.847604176 | 783.1440145 | 764.5960102 | ENSG00000135956 |
| BAG4 | -0.034236329 | 0.976548547 | 0.760201533 | 0.866677445 | 572.2756672 | 558.7460131 | ENSG00000156735 |
| CPNE2 | -0.034237529 | 0.976547734 | 0.756763652 | 0.864566204 | 669.5612619 | 653.1956702 | ENSG00000140848 |
| TBCA | -0.03425016 | 0.976539184 | 0.606202389 | 0.764778101 | 2545.352286 | 2485.689879 | ENSG00000171530 |
| C17orf58 | -0.034291573 | 0.976511153 | 0.803913608 | 0.894704967 | 379.1213899 | 369.8442226 | ENSG00000186665 |
| NA | -0.034311255 | 0.976497831 | 0.960820054 | 0.98076989 | 10.24963078 | 9.995920482 | ENSG00000248971 |
| TM2D3 | -0.034325235 | 0.976488369 | 0.783558963 | 0.881593021 | 388.3819157 | 379.6804826 | ENSG00000184277 |
| NA | -0.034344033 | 0.976475646 | 0.919756558 | 0.960515777 | 40.98343761 | 40.09630789 | ENSG00000261655 |
| MFF | -0.03442945 | 0.976417834 | 0.723368321 | 0.844046387 | 682.7963301 | 666.6541273 | ENSG00000168958 |
| TCEAL4 | -0.034485325 | 0.976380018 | 0.698022672 | 0.827774653 | 1229.892503 | 1201.268685 | ENSG00000133142 |
| MAD2L1BP | -0.034535647 | 0.976345962 | 0.779914119 | 0.878986213 | 402.480038 | 392.6951171 | ENSG00000124688 |
| ARL2BP | -0.034536093 | 0.97634566 | 0.956553064 | 0.979215959 | 12.82276822 | 12.53542914 | ENSG00000102931 |
| ZNF638 | -0.034544168 | 0.976340195 | 0.617864412 | 0.77281003 | 2240.424647 | 2187.392143 | ENSG00000075292 |
| VWF | -0.034702883 | 0.976232791 | 0.93973109 | 0.970196971 | 23.34917737 | 22.78990666 | ENSG00000110799 |
| ZNF470 | -0.034762074 | 0.976192739 | 0.847151144 | 0.918589821 | 174.1980834 | 170.120508 | ENSG00000197016 |
| CSNK2A1 | -0.034766102 | 0.976190013 | 0.636868087 | 0.786554426 | 4053.2111 | 3955.460688 | ENSG00000101266 |
| ZNF583 | -0.034783239 | 0.976178418 | 0.930069105 | 0.965600537 | 31.64619902 | 30.81628157 | ENSG00000198440 |
| CIB2 | -0.034813193 | 0.97615815 | 0.686317136 | 0.819547825 | 1053.418814 | 1028.188793 | ENSG00000136425 |
| PITPNC1 | -0.034922401 | 0.976084261 | 0.675028018 | 0.811873412 | 1107.316337 | 1080.547837 | ENSG00000154217 |
| IGDCC4 | -0.034936903 | 0.976074449 | 0.780718748 | 0.879518677 | 526.2616932 | 514.37077 | ENSG00000103742 |
| POLK | -0.035009018 | 0.97602566 | 0.864918898 | 0.929688585 | 126.3295147 | 123.1424263 | ENSG00000122008 |
| USP30 | -0.035019817 | 0.976018354 | 0.875827231 | 0.935632495 | 111.1370043 | 108.2890532 | ENSG00000135093 |
| MINDY1 | -0.035026546 | 0.976013801 | 0.844462397 | 0.91704846 | 163.895597 | 159.9870186 | ENSG00000143409 |
| PCBD1 | -0.035075416 | 0.975980741 | 0.735692936 | 0.852173027 | 651.1262437 | 635.4987972 | ENSG00000166228 |
| MXRA7 | -0.035104143 | 0.975961307 | 0.743558486 | 0.856821249 | 858.1198974 | 836.63615 | ENSG00000182534 |
| GRK2 | -0.035142462 | 0.975935385 | 0.609004284 | 0.7667212 | 2139.437446 | 2088.328563 | ENSG00000173020 |
| ACSBG1 | -0.035186806 | 0.975905388 | 0.88830616 | 0.942849507 | 91.20409992 | 89.15363839 | ENSG00000103740 |
| NOX1 | -0.035216578 | 0.97588525 | 0.967521771 | 0.984466017 | 6.580151025 | 6.413701455 | ENSG00000007952 |
| RRAGA | -0.035229181 | 0.975876724 | 0.735770877 | 0.852173027 | 726.1034566 | 709.1816035 | ENSG00000155876 |
| TMED4 | -0.03526063 | 0.975855452 | 0.697113314 | 0.827110823 | 938.3598898 | 916.085699 | ENSG00000158604 |
| LINC00467 | -0.035379953 | 0.975774744 | 0.913701159 | 0.957240251 | 45.63119876 | 44.52538 | ENSG00000153363 |
| C3orf14 | -0.035391215 | 0.975767127 | 0.86875329 | 0.931798233 | 137.0756673 | 133.9299862 | ENSG00000114405 |
| RTKN | -0.035462894 | 0.975718648 | 0.648206347 | 0.794567224 | 1321.776519 | 1290.096249 | ENSG00000114993 |
| CSTB | -0.035469014 | 0.975714509 | 0.654006439 | 0.79772604 | 1382.728827 | 1349.38946 | ENSG00000160213 |
| GLIS3 | -0.035500125 | 0.975693468 | 0.97684035 | 0.989493307 | 3.629041247 | 3.533392118 | ENSG00000107249 |
| SLC45A3 | -0.035517125 | 0.975681971 | 0.779847794 | 0.878986213 | 448.6604791 | 438.275194 | ENSG00000158715 |
| MED15 | -0.035520904 | 0.975679415 | 0.652326188 | 0.796943154 | 1381.532937 | 1348.138986 | ENSG00000099917 |
| SLC31A1 | -0.035687089 | 0.975567033 | 0.628079928 | 0.780425945 | 1883.700618 | 1836.977092 | ENSG00000136868 |
| DMRTA1 | -0.035863294 | 0.975447888 | 0.851555133 | 0.921524381 | 138.4562068 | 134.9828822 | ENSG00000176399 |
| WAPL | -0.035929168 | 0.97540335 | 0.641208279 | 0.789697272 | 1720.041654 | 1676.961841 | ENSG00000062650 |
| UBXN2A | -0.035953192 | 0.975387107 | 0.764198348 | 0.869469133 | 495.2820606 | 483.0359713 | ENSG00000173960 |
| BNIPL | -0.035960784 | 0.975381975 | 0.946924162 | 0.973555911 | 21.65558521 | 21.17398402 | ENSG00000163141 |
| NA | -0.036168587 | 0.975241492 | 0.968222235 | 0.984740121 | 6.467607184 | 6.312798442 | ENSG00000273306 |
| HTT | -0.036176189 | 0.975236353 | 0.672122893 | 0.809503537 | 3046.691187 | 2971.33461 | ENSG00000197386 |
| PECR | -0.036507373 | 0.975012505 | 0.809651237 | 0.897492586 | 256.553191 | 250.1637203 | ENSG00000115425 |
| VASN | -0.036591647 | 0.974955551 | 0.799105292 | 0.892044659 | 325.070967 | 317.3212413 | ENSG00000168140 |
| UTP23 | -0.036676121 | 0.974898467 | 0.86822714 | 0.931362061 | 164.0964966 | 159.9584598 | ENSG00000147679 |
| SCHIP1 | -0.036688188 | 0.974890313 | 0.874205025 | 0.934994283 | 103.1246457 | 100.7812266 | ENSG00000283154 |
| TMEM92 | -0.036790603 | 0.974821109 | 0.793515891 | 0.888105285 | 260.8408389 | 254.2134621 | ENSG00000167105 |
| PDIA5 | -0.0368676 | 0.974769084 | 0.708777832 | 0.834380611 | 691.9594499 | 674.4081755 | ENSG00000065485 |

| | | | | | | | |
|---|---|---|---|---|---|---|---|
| CLEC16A | -0.036882508 | 0.974759011 | 0.746444332 | 0.858598601 | 425.7550581 | 414.9845224 | ENSG00000038532 |
| MLLT10 | -0.036923186 | 0.974731528 | 0.692470314 | 0.823922758 | 902.1739562 | 878.8965785 | ENSG00000078403 |
| STARD4 | -0.036983444 | 0.974690816 | 0.807364002 | 0.896619594 | 597.8384701 | 582.2360455 | ENSG00000164211 |
| LRPPRC | -0.037031756 | 0.974658177 | 0.575553885 | 0.740696842 | 5992.546473 | 5839.449708 | ENSG00000138095 |
| FBXO11 | -0.037047308 | 0.97464767 | 0.734188732 | 0.851336652 | 576.2454143 | 561.6703947 | ENSG00000138081 |
| GDI2 | -0.037076686 | 0.974627824 | 0.577397751 | 0.742025373 | 8729.228899 | 8506.30437 | ENSG00000057608 |
| BLCAP | -0.037086148 | 0.974621432 | 0.724709864 | 0.845161619 | 1266.990837 | 1236.003942 | ENSG00000166619 |
| PPM1B | -0.037117323 | 0.974600371 | 0.819650661 | 0.902946927 | 463.3478311 | 451.1567423 | ENSG00000138032 |
| FAM185A | -0.037143972 | 0.974582369 | 0.866784892 | 0.930279181 | 100.5375052 | 98.11020931 | ENSG00000222011 |
| SRCAP | -0.037186893 | 0.974553375 | 0.909314907 | 0.954803834 | 48.22949381 | 47.02409748 | ENSG00000080603 |
| TM2D1 | -0.037266917 | 0.97449932 | 0.84350688 | 0.916440384 | 149.3006069 | 145.5642875 | ENSG00000162604 |
| MMADHC | -0.037291693 | 0.974482584 | 0.669449233 | 0.807856359 | 1538.402131 | 1498.627869 | ENSG00000168288 |
| NSUN5P1 | -0.037305526 | 0.974473241 | 0.937980322 | 0.969503942 | 588.3081341 | 573.4144111 | ENSG00000223705 |
| ZNF337 | -0.037339165 | 0.974450519 | 0.942824362 | 0.971928735 | 19.98872153 | 19.57215746 | ENSG00000130684 |
| MCL1 | -0.037395376 | 0.974412553 | 0.532745102 | 0.707806288 | 6722.150996 | 6549.257198 | ENSG00000143384 |
| MATN2 | -0.037410917 | 0.974402056 | 0.787426493 | 0.883706479 | 311.032608 | 302.7958551 | ENSG00000132561 |
| VPS37C | -0.037460848 | 0.974368334 | 0.720950901 | 0.842237882 | 621.657824 | 605.9786785 | ENSG00000167987 |
| SLF2 | -0.037475704 | 0.9743583 | 0.747185238 | 0.858811237 | 482.6485868 | 470.0810819 | ENSG00000119906 |
| LRR1 | -0.03756728 | 0.974296454 | 0.857073143 | 0.924108956 | 155.8174469 | 151.4919814 | ENSG00000165501 |
| SERF2 | -0.037656219 | 0.974236393 | 0.734205517 | 0.851336652 | 602.2772289 | 586.3391464 | ENSG00000140264 |
| MED11 | -0.037671605 | 0.974226003 | 0.887054245 | 0.942266356 | 72.83262592 | 71.12666809 | ENSG00000161920 |
| ERLIN2 | -0.037703012 | 0.974204794 | 0.676850656 | 0.813270653 | 1204.158439 | 1172.223375 | ENSG00000147475 |
| FBXO42 | -0.037710064 | 0.974200032 | 0.726973623 | 0.846712067 | 540.540436 | 526.8452741 | ENSG00000037637 |
| MOCS3 | -0.037761447 | 0.974165336 | 0.78742141 | 0.883706479 | 306.0090241 | 298.4027163 | ENSG00000124217 |
| MRPS9 | -0.037875252 | 0.974088493 | 0.708238398 | 0.83401045 | 705.1788389 | 686.6393582 | ENSG00000135972 |
| HBEGF | -0.037905929 | 0.974067781 | 0.892291275 | 0.945066946 | 71.32230746 | 69.32106013 | ENSG00000113070 |
| AP3M1 | -0.03794677 | 0.974040206 | 0.67045761 | 0.808475039 | 945.0205571 | 920.4023366 | ENSG00000185009 |
| ZNF558 | -0.038008643 | 0.973998433 | 0.813733677 | 0.89988777 | 195.8711369 | 190.8034118 | ENSG00000167785 |
| FOXO3B | -0.038069981 | 0.973957023 | 0.785054737 | 0.882418854 | 301.3841732 | 293.5039229 | ENSG00000240445 |
| MYO18A | -0.03807864 | 0.973951178 | 0.628740944 | 0.78073535 | 1719.846534 | 1676.054719 | ENSG00000196535 |
| ZC3H12C | -0.038090021 | 0.973943495 | 0.909619148 | 0.955001893 | 48.27003002 | 46.88780236 | ENSG00000149289 |
| TIPIN | -0.038155095 | 0.973899565 | 0.801454538 | 0.893214577 | 289.2812482 | 281.28811 | ENSG00000075131 |
| RPL27A | -0.038300866 | 0.973801166 | 0.48419205 | 0.667560474 | 13519.84793 | 13164.99744 | ENSG00000166441 |
| DENND2C | -0.038306338 | 0.973797473 | 0.887173114 | 0.942284613 | 70.56576144 | 68.73006455 | ENSG00000175984 |
| TTC23 | -0.038447794 | 0.973701997 | 0.787142959 | 0.883573449 | 277.4462321 | 270.1092107 | ENSG00000103852 |
| ACP6 | -0.038555563 | 0.973629264 | 0.774046362 | 0.875300505 | 325.8176666 | 316.8701878 | ENSG00000162836 |
| RPS13 | -0.038557 | 0.973628294 | 0.496241216 | 0.677862531 | 8604.198129 | 8377.007043 | ENSG00000110700 |
| FUT2 | -0.038718342 | 0.973519416 | 0.854410347 | 0.922743843 | 107.862657 | 105.052533 | ENSG00000176920 |
| DCAF1 | -0.038865846 | 0.973419887 | 0.652805639 | 0.797217744 | 1069.868504 | 1041.239521 | ENSG00000145041 |
| DTD1 | -0.038897569 | 0.973398483 | 0.747131061 | 0.858811237 | 388.1473573 | 377.5763538 | ENSG00000125821 |
| MST1R | -0.038930323 | 0.973376384 | 0.900168296 | 0.948979683 | 53.86421034 | 52.34615003 | ENSG00000164078 |
| TLNRD1 | -0.038993737 | 0.9733336 | 0.699021527 | 0.828314365 | 1258.171639 | 1224.333952 | ENSG00000140406 |
| TAPBP | -0.039070041 | 0.973282122 | 0.725837214 | 0.845936492 | 704.7239733 | 685.841395 | ENSG00000231925 |
| MLYCD | -0.039303569 | 0.97312459 | 0.733682401 | 0.851247472 | 483.9580057 | 471.2185635 | ENSG00000103150 |
| KIFC2 | -0.039426122 | 0.973041929 | 0.85318602 | 0.922280546 | 254.2038861 | 247.8922789 | ENSG00000167702 |
| SLC25A42 | -0.039498814 | 0.972992902 | 0.846042124 | 0.917957191 | 131.2643968 | 127.5635743 | ENSG00000181035 |
| URB1 | -0.039509172 | 0.972985916 | 0.611830911 | 0.768591155 | 1914.886441 | 1862.857029 | ENSG00000142207 |
| BCAR3 | -0.039536396 | 0.972967556 | 0.788447743 | 0.884355511 | 236.9354515 | 230.4977079 | ENSG00000137936 |
| GTPBP4 | -0.039636895 | 0.972899781 | 0.590992992 | 0.752815402 | 2351.160918 | 2286.438376 | ENSG00000107937 |
| SNTB1 | -0.039693827 | 0.972861389 | 0.748749573 | 0.859809454 | 367.4067198 | 357.6359764 | ENSG00000172164 |
| NA | -0.039695132 | 0.972860509 | 0.945615452 | 0.973218501 | 16.39874331 | 15.94848598 | ENSG00000230383 |
| AP5S1 | -0.039765933 | 0.972812767 | 0.896152772 | 0.946766059 | 53.57499972 | 52.26781686 | ENSG00000125843 |
| PPP6R2 | -0.039890599 | 0.972728708 | 0.689955156 | 0.822191533 | 1026.474795 | 999.3964122 | ENSG00000100239 |
| RRP15 | -0.040070175 | 0.972607637 | 0.71470134 | 0.838366554 | 869.5122464 | 845.7710182 | ENSG00000067533 |
| KCNC4 | -0.040197216 | 0.972521995 | 0.752909036 | 0.862715058 | 386.3215656 | 375.7916038 | ENSG00000116396 |
| NAP1L1 | -0.040256582 | 0.972481977 | 0.480489369 | 0.6642839 | 13860.74881 | 13478.28523 | ENSG00000187109 |
| MAPRE2 | -0.040511493 | 0.972310164 | 0.606561591 | 0.765127104 | 1206.874666 | 1173.152847 | ENSG00000166974 |
| SPINT1 | -0.040540913 | 0.972290336 | 0.539440785 | 0.712722396 | 2900.346087 | 2819.731573 | ENSG00000166145 |
| FBXW2 | -0.040562348 | 0.97227589 | 0.626081987 | 0.778465286 | 1042.821945 | 1013.6835 | ENSG00000119402 |
| ZNF671 | -0.040638541 | 0.972224543 | 0.827754687 | 0.907253657 | 158.1288915 | 153.7303184 | ENSG00000083814 |
| CEP170B | -0.04066572 | 0.972206227 | 0.579103983 | 0.743290389 | 1897.771741 | 1845.137182 | ENSG00000099814 |
| ESRRA | -0.040697352 | 0.972184911 | 0.636169671 | 0.786208487 | 1197.356289 | 1164.284525 | ENSG00000173153 |
| TTLL5 | -0.040747453 | 0.97215115 | 0.686754464 | 0.819878795 | 670.7214112 | 652.2777384 | ENSG00000119685 |
| PGAM1 | -0.040777944 | 0.972130604 | 0.608736552 | 0.766514251 | 12902.04817 | 12541.79567 | ENSG00000171314 |
| SAFB | -0.040781746 | 0.972128042 | 0.50291616 | 0.683361299 | 4849.598029 | 4714.865068 | ENSG00000160633 |
| PCCA | -0.040805225 | 0.972112222 | 0.667786284 | 0.80717496 | 925.7425687 | 899.6855612 | ENSG00000175198 |
| FOXD3-AS1 | -0.040838313 | 0.972089927 | 0.617261809 | 0.772316908 | 1503.756619 | 1462.05297 | ENSG00000230798 |
| LSM4 | -0.040907022 | 0.972043632 | 0.51258632 | 0.691410658 | 4909.272661 | 4772.760275 | ENSG00000130520 |
| NMT1 | -0.040927528 | 0.972029816 | 0.495536538 | 0.677275366 | 3493.1957 | 3395.206619 | ENSG00000136448 |
| PTCD3 | -0.040942639 | 0.972019634 | 0.560481925 | 0.729441191 | 2236.716555 | 2173.920345 | ENSG00000132300 |
| TBC1D20 | -0.04100262 | 0.971979223 | 0.704601571 | 0.831947858 | 560.1159781 | 544.2797829 | ENSG00000125875 |
| RNF214 | -0.041043672 | 0.971951565 | 0.737478176 | 0.852943558 | 456.1478811 | 443.2019479 | ENSG00000167257 |
| CPSF1 | -0.041076897 | 0.971929182 | 0.485075292 | 0.668281086 | 4543.535323 | 4416.719778 | ENSG00000071894 |
| EPB41L3 | -0.041085435 | 0.97192343 | 0.726151406 | 0.846179828 | 452.2310047 | 439.5408461 | ENSG00000082397 |

| | | | | | | | |
|---|---|---|---|---|---|---|---|
| NR2C2AP | -0.041094682 | 0.9719172 | 0.741328345 | 0.855338624 | 343.4426515 | 333.8323159 | ENSG00000184162 |
| NA | -0.04121359 | 0.971837098 | 0.95095295 | 0.976019602 | 10.17950228 | 9.856925828 | ENSG00000250616 |
| SIRT7 | -0.041294508 | 0.97178259 | 0.744684974 | 0.85731979 | 432.1307315 | 420.5276387 | ENSG00000187531 |
| TRIM74 | -0.041368674 | 0.971732635 | 0.968559828 | 0.984740121 | 5.488145708 | 5.363943044 | ENSG00000155428 |
| PCDHGA6 | -0.041389366 | 0.971718697 | 0.941206023 | 0.970800432 | 16.80828477 | 16.41754604 | ENSG00000253731 |
| PPIL3 | -0.04143433 | 0.971688413 | 0.718716871 | 0.840687818 | 464.1941387 | 451.2962556 | ENSG00000240344 |
| CHMP6 | -0.041445113 | 0.97168115 | 0.765302351 | 0.870214234 | 307.0430627 | 297.9875558 | ENSG00000176108 |
| SWSAP1 | -0.041486307 | 0.971653405 | 0.924140545 | 0.962449326 | 30.21242059 | 29.40640183 | ENSG00000173928 |
| ATP5F1B | -0.041557166 | 0.971605684 | 0.46705568 | 0.653162956 | 19508.37039 | 18953.15346 | ENSG00000110955 |
| TGS1 | -0.041633702 | 0.97155414 | 0.673605176 | 0.81047769 | 965.2183637 | 936.9555391 | ENSG00000137574 |
| GOLGA8A | -0.041635119 | 0.971553186 | 0.928355539 | 0.964638631 | 312.3112545 | 303.5538806 | ENSG00000175265 |
| DHX15 | -0.041673893 | 0.971527075 | 0.496985601 | 0.678136724 | 5776.608834 | 5611.293073 | ENSG00000109606 |
| NA | -0.041696034 | 0.971512165 | 0.948325843 | 0.97445646 | 11.4698668 | 11.1963965 | ENSG00000280303 |
| NA | -0.041721536 | 0.971494992 | 0.932660147 | 0.967044653 | 18.74826625 | 18.21760302 | ENSG00000216285 |
| CPLANE2 | -0.041796053 | 0.971444814 | 0.875995614 | 0.935667514 | 73.09342433 | 71.11593282 | ENSG00000132881 |
| ZNF280B | -0.041815451 | 0.971431753 | 0.805197018 | 0.89555458 | 218.5633011 | 212.0489671 | ENSG00000275004 |
| RPL7L1 | -0.041848809 | 0.971409292 | 0.551339351 | 0.722322422 | 2789.340756 | 2709.060616 | ENSG00000146223 |
| ZNF12 | -0.04186027 | 0.971401575 | 0.754278786 | 0.863474302 | 353.0016255 | 342.6567254 | ENSG00000164631 |
| CD99 | -0.041900473 | 0.971374505 | 0.565372483 | 0.733122144 | 1587.999582 | 1542.156102 | ENSG00000002586 |
| CHUK | -0.041908974 | 0.971368782 | 0.710906044 | 0.836036334 | 482.7826429 | 468.5098524 | ENSG00000213341 |
| TMEM25 | -0.042021419 | 0.971293075 | 0.806661611 | 0.896591682 | 267.5815639 | 260.4818224 | ENSG00000149582 |
| ZBTB44 | -0.0420979 | 0.971241586 | 0.635637108 | 0.785759898 | 1452.975027 | 1410.277713 | ENSG00000196323 |
| ZMIZ1 | -0.042102056 | 0.971238788 | 0.618331307 | 0.773098296 | 1436.032626 | 1394.610521 | ENSG00000108175 |
| SUN1 | -0.042188509 | 0.971180588 | 0.631504903 | 0.7826627 | 2082.784557 | 2023.674873 | ENSG00000164828 |
| HSPBAP1 | -0.042212439 | 0.97116448 | 0.816601866 | 0.901347795 | 158.8443 | 154.168 | ENSG00000169087 |
| AP3D1 | -0.042248943 | 0.971139907 | 0.437883527 | 0.626459279 | 6983.382059 | 6782.259098 | ENSG00000065000 |
| KCNS3 | -0.042265162 | 0.971128989 | 0.698928042 | 0.828256542 | 499.6065993 | 485.2054994 | ENSG00000170745 |
| GABRQ | -0.042296983 | 0.97110757 | 0.741733501 | 0.855622624 | 360.9899163 | 350.6196193 | ENSG00000268089 |
| KANSL1L | -0.042307542 | 0.971100462 | 0.814584334 | 0.900075694 | 159.3946998 | 154.7799651 | ENSG00000144445 |
| EXOC3 | -0.042324279 | 0.971089196 | 0.641830849 | 0.789939103 | 973.7398717 | 946.2617201 | ENSG00000180104 |
| NA | -0.042390382 | 0.971044703 | 0.370888282 | 0.564139796 | 2190047.35 | 2126633.536 | N_ambiguous |
| PIP5K1A | -0.042501931 | 0.970969625 | 0.557584814 | 0.726895754 | 1839.840506 | 1786.063391 | ENSG00000143398 |
| TTC28-AS1 | -0.042581912 | 0.970915797 | 0.836317301 | 0.9121504 | 142.7828951 | 138.9876361 | ENSG00000235954 |
| SIK3 | -0.042613 | 0.970894875 | 0.653906503 | 0.797703095 | 806.5549925 | 782.7104487 | ENSG00000160584 |
| BAZ2B | -0.042735703 | 0.970812303 | 0.937275456 | 0.969033102 | 16.72364949 | 16.316205 | ENSG00000226266 |
| PTPRF | -0.042821256 | 0.970754735 | 0.449034296 | 0.636658632 | 9733.277642 | 9448.386291 | ENSG00000142949 |
| MARCHF8 | -0.042825565 | 0.970751836 | 0.785460204 | 0.882568409 | 377.1381364 | 365.6406088 | ENSG00000165406 |
| ANK3 | -0.042869507 | 0.970722268 | 0.639909347 | 0.789198819 | 796.7028542 | 773.2152932 | ENSG00000151150 |
| NAA20 | -0.042918607 | 0.970689232 | 0.663590335 | 0.804275652 | 626.7292427 | 608.2577579 | ENSG00000173418 |
| SERHL2 | -0.042947001 | 0.970670128 | 0.89403521 | 0.94574241 | 51.1585052 | 49.86025482 | ENSG00000183569 |
| RAB3D | -0.042972215 | 0.970653164 | 0.794486262 | 0.8886547 | 212.3503513 | 206.2263015 | ENSG00000105514 |
| NFATC2 | -0.042979786 | 0.97064807 | 0.916554909 | 0.958712608 | 29.47628921 | 28.54303926 | ENSG00000101096 |
| NA | -0.043017169 | 0.970622919 | 0.96704208 | 0.9842481 | 5.848958001 | 5.694008455 | ENSG00000245322 |
| WDR61 | -0.04307226 | 0.970585855 | 0.691194364 | 0.823091515 | 581.0401749 | 563.4003372 | ENSG00000140395 |
| NA | -0.043117836 | 0.970555193 | 0.921158448 | 0.961032968 | 26.00634874 | 25.20550443 | ENSG00000260290 |
| SELENOI | -0.043186857 | 0.970508762 | 0.725686591 | 0.845904106 | 1194.474999 | 1158.456966 | ENSG00000138018 |
| ZNF653 | -0.043240872 | 0.970472427 | 0.860364771 | 0.926274214 | 82.97800352 | 80.69118212 | ENSG00000161914 |
| PRKCZ | -0.043285198 | 0.970442609 | 0.641911358 | 0.789985731 | 867.3278601 | 841.9950522 | ENSG00000067606 |
| BRAP | -0.043319952 | 0.970419232 | 0.703215487 | 0.831299338 | 540.7764827 | 524.132661 | ENSG00000089234 |
| SELENBP1 | -0.043324922 | 0.970415889 | 0.727662636 | 0.846909144 | 386.6521243 | 375.1321258 | ENSG00000143416 |
| GPALPP1 | -0.043369461 | 0.970385931 | 0.733307153 | 0.851101774 | 413.4533679 | 401.0822746 | ENSG00000133114 |
| TGFBR1 | -0.043380849 | 0.970378271 | 0.688071768 | 0.820902617 | 819.7341982 | 794.7295593 | ENSG00000106799 |
| SPTLC2 | -0.043408804 | 0.970359469 | 0.607833789 | 0.76605409 | 1392.155504 | 1350.083245 | ENSG00000100596 |
| NA | -0.043477786 | 0.970313072 | 0.966059033 | 0.983679712 | 5.170928834 | 4.939473008 | ENSG00000228434 |
| SCAF11 | -0.043528627 | 0.970278878 | 0.58973324 | 0.751934224 | 1413.623125 | 1371.004231 | ENSG00000139218 |
| SURF2 | -0.043647588 | 0.970198875 | 0.765263808 | 0.870214234 | 291.3398017 | 282.8323582 | ENSG00000148291 |
| ANKS1A | -0.043678011 | 0.970178416 | 0.541547467 | 0.714474773 | 1791.975402 | 1738.547641 | ENSG00000064999 |
| SLC1A5 | -0.043702023 | 0.970162269 | 0.438351186 | 0.626794245 | 7101.121109 | 6888.729752 | ENSG00000105281 |
| TRIM45 | -0.043708745 | 0.970157748 | 0.786039245 | 0.882900719 | 350.3074808 | 340.2848106 | ENSG00000134253 |
| TICAM1 | -0.043789459 | 0.970103473 | 0.912426421 | 0.956487804 | 34.07975588 | 32.93891788 | ENSG00000127666 |
| DESI1 | -0.043828375 | 0.970077305 | 0.659543963 | 0.801678109 | 722.877718 | 701.2776735 | ENSG00000100418 |
| VIRMA | -0.04388331 | 0.970040367 | 0.612489603 | 0.768991559 | 1042.824539 | 1010.90278 | ENSG00000164944 |
| NA | -0.044011028 | 0.969954496 | 0.773233536 | 0.874965662 | 264.0336102 | 256.1207254 | ENSG00000255769 |
| ACSL3 | -0.044066887 | 0.969916941 | 0.595919564 | 0.756284114 | 2115.505263 | 2051.139657 | ENSG00000123983 |
| RFC3 | -0.044082096 | 0.969906717 | 0.621319215 | 0.775143399 | 1019.310834 | 987.8692883 | ENSG00000133119 |
| ALDH7A1 | -0.044115897 | 0.969883993 | 0.467229658 | 0.653162956 | 3604.9033 | 3495.751818 | ENSG00000164904 |
| FAM53C | -0.04411867 | 0.969882129 | 0.553585123 | 0.723938212 | 1552.571961 | 1506.183816 | ENSG00000120709 |
| PPP6C | -0.044211578 | 0.969819672 | 0.661036352 | 0.802702466 | 915.9926392 | 887.8850917 | ENSG00000119414 |
| ZBTB14 | -0.044235925 | 0.969803305 | 0.779050299 | 0.878653821 | 218.9878304 | 212.1738657 | ENSG00000198081 |
| PIAS2 | -0.044250613 | 0.969793431 | 0.650765853 | 0.795936006 | 703.001826 | 681.7700711 | ENSG00000078043 |
| HDDC3 | -0.04433794 | 0.969734731 | 0.828310858 | 0.907514734 | 118.6607107 | 115.0371686 | ENSG00000184508 |
| NA | -0.044351623 | 0.969725534 | 0.949647354 | 0.975003548 | 10.61991381 | 10.2507903 | ENSG00000267655 |
| GOLGA2 | -0.044395327 | 0.969696158 | 0.583436355 | 0.746679284 | 1132.436609 | 1098.203226 | ENSG00000167110 |
| PMPCB | -0.044420018 | 0.969679562 | 0.553075806 | 0.723664454 | 1839.604775 | 1783.144275 | ENSG00000105819 |

| | | | | | | | |
|---|---|---|---|---|---|---|---|
| LAMTOR3 | -0.04454979 | 0.969592342 | 0.740941079 | 0.855134315 | 352.6586496 | 341.6090053 | ENSG00000109270 |
| WDHD1 | -0.044677843 | 0.969506286 | 0.613992538 | 0.769819961 | 1527.057962 | 1479.535404 | ENSG00000198554 |
| NA | -0.04472399 | 0.969475275 | 0.927330344 | 0.964343189 | 20.06072916 | 19.37630017 | ENSG00000249936 |
| METTL2A | -0.044831763 | 0.969402855 | 0.665921989 | 0.80588953 | 733.510177 | 710.2906118 | ENSG00000087995 |
| ISY1 | -0.044833851 | 0.969401452 | 0.856480604 | 0.923894233 | 84.20375661 | 81.59869668 | ENSG00000240682 |
| EHBP1 | -0.044875533 | 0.969373445 | 0.629531721 | 0.781339377 | 725.230413 | 702.8269774 | ENSG00000115504 |
| MED29 | -0.044895818 | 0.969359815 | 0.600963535 | 0.7603532 | 1073.637449 | 1041.278935 | ENSG00000063322 |
| F2RL1 | -0.044935802 | 0.96933295 | 0.709271574 | 0.834749768 | 421.0015328 | 408.2103732 | ENSG00000164251 |
| RABGGTB | -0.045038484 | 0.969263961 | 0.618345157 | 0.773098296 | 922.5758721 | 894.5235929 | ENSG00000137955 |
| MFN2 | -0.04515161 | 0.969187961 | 0.522703608 | 0.699529834 | 2548.82951 | 2470.546992 | ENSG00000116688 |
| SNHG11 | -0.045206608 | 0.969151015 | 0.84066161 | 0.914474838 | 97.2882177 | 94.19693719 | ENSG00000174365 |
| KHDRBS3 | -0.045230328 | 0.969135081 | 0.71304494 | 0.837541981 | 630.1061297 | 610.401212 | ENSG00000131773 |
| CBS | -0.045267512 | 0.969110103 | 0.619886266 | 0.774293844 | 960.172005 | 931.1543701 | ENSG00000160200 |
| TMEM140 | -0.045397699 | 0.969022656 | 0.800701005 | 0.8929372 | 166.3528232 | 161.2724378 | ENSG00000146859 |
| SH3D19 | -0.045401233 | 0.969020282 | 0.562360259 | 0.730808095 | 1403.767898 | 1359.804697 | ENSG00000109686 |
| ELMSAN1 | -0.045435464 | 0.96899729 | 0.747837873 | 0.859241655 | 293.9376833 | 284.5800042 | ENSG00000156030 |
| MAGED1 | -0.045484064 | 0.968964648 | 0.547430807 | 0.719107096 | 2582.804449 | 2502.330845 | ENSG00000179222 |
| TBC1D16 | -0.045491079 | 0.968959937 | 0.432322414 | 0.622246146 | 6219.359938 | 6026.072062 | ENSG00000167291 |
| WDR55 | -0.045638216 | 0.96886112 | 0.686141027 | 0.819390322 | 482.7949623 | 467.742097 | ENSG00000120314 |
| SBDSP1 | -0.045652161 | 0.968851754 | 0.795217935 | 0.889151135 | 182.5759821 | 176.9129902 | ENSG00000225648 |
| RASA1 | -0.045719629 | 0.968806447 | 0.581404014 | 0.745365014 | 993.8371083 | 962.8248389 | ENSG00000145715 |
| PPM1J | -0.045752886 | 0.968784114 | 0.766275488 | 0.870835943 | 244.0805092 | 236.7406469 | ENSG00000155367 |
| NA | -0.045791328 | 0.968758301 | 0.927190265 | 0.964343189 | 20.07147768 | 19.51193401 | ENSG00000233913 |
| TMEM231 | -0.045798512 | 0.968753476 | 0.835344637 | 0.911772818 | 125.8550762 | 121.617722 | ENSG00000205084 |
| GRK5 | -0.045822585 | 0.968737312 | 0.851732548 | 0.921524381 | 86.78193011 | 84.17987259 | ENSG00000198873 |
| MCUR1 | -0.045863916 | 0.96870956 | 0.656015405 | 0.79911618 | 710.5308841 | 688.4045812 | ENSG00000050393 |
| TMEM53 | -0.045890765 | 0.968691532 | 0.813107436 | 0.899516491 | 147.5726947 | 142.716622 | ENSG00000126106 |
| ACTR3 | -0.046032908 | 0.968596096 | 0.530302708 | 0.70592763 | 3275.799891 | 3172.33978 | ENSG00000115091 |
| NA | -0.046058221 | 0.968579101 | 0.935893389 | 0.968425646 | 17.83006389 | 17.17814185 | ENSG00000240731 |
| TNKS2 | -0.04610264 | 0.96854928 | 0.528148829 | 0.704021174 | 2129.254239 | 2061.327436 | ENSG00000107854 |
| BTF3 | -0.04617117 | 0.968503274 | 0.459628687 | 0.646298468 | 10645.6948 | 10309.64239 | ENSG00000145741 |
| NA | -0.046265602 | 0.968439882 | 0.93528533 | 0.96819584 | 14.93655266 | 14.478168 | ENSG00000274565 |
| LIN54 | -0.046366204 | 0.968372353 | 0.787439975 | 0.883706479 | 184.1795347 | 178.0749365 | ENSG00000189308 |
| TLK1 | -0.046493755 | 0.968286741 | 0.612660526 | 0.768999741 | 1110.14214 | 1074.660894 | ENSG00000198586 |
| SLC29A3 | -0.046703555 | 0.968145941 | 0.763553836 | 0.868904241 | 318.3318507 | 308.3866906 | ENSG00000198246 |
| TUT4 | -0.046712227 | 0.968140122 | 0.637980622 | 0.787529817 | 877.4175258 | 849.2738047 | ENSG00000134744 |
| ANKRD24 | -0.046750912 | 0.968114162 | 0.730328994 | 0.84917927 | 423.7638518 | 410.8201164 | ENSG00000089847 |
| USP6NL | -0.046759255 | 0.968108563 | 0.630312999 | 0.781680905 | 691.5931432 | 669.2637186 | ENSG00000148429 |
| GJA1 | -0.046767738 | 0.968102871 | 0.429537551 | 0.619681558 | 11887.60497 | 11507.97706 | ENSG00000152661 |
| FANCG | -0.04680015 | 0.968081122 | 0.558348771 | 0.727526558 | 1126.918022 | 1090.805286 | ENSG00000221829 |
| TEAD4 | -0.046811494 | 0.968073509 | 0.544663577 | 0.716707097 | 1877.534375 | 1817.757635 | ENSG00000197905 |
| PRC1 | -0.046909339 | 0.968007856 | 0.956325336 | 0.97920191 | 7.989275517 | 7.775741968 | ENSG00000198901 |
| LPGAT1 | -0.046954025 | 0.967977873 | 0.762070081 | 0.868113515 | 254.2697526 | 245.9378397 | ENSG00000123684 |
| LSM7 | -0.04700543 | 0.967943384 | 0.593805085 | 0.755203695 | 3302.071074 | 3196.878906 | ENSG00000130332 |
| GNAL | -0.047009971 | 0.967940337 | 0.687620088 | 0.820469369 | 533.3024524 | 515.7988937 | ENSG00000141404 |
| RPL19 | -0.047033321 | 0.967924671 | 0.353353097 | 0.546528107 | 15090.12991 | 14606.7308 | ENSG00000108298 |
| CYCS | -0.047041535 | 0.96791916 | 0.530679933 | 0.70614339 | 5654.113845 | 5471.9731 | ENSG00000172115 |
| NA | -0.047053729 | 0.967910979 | 0.853423336 | 0.922280546 | 81.52458455 | 79.05213998 | ENSG00000272163 |
| STK39 | -0.047073719 | 0.967897568 | 0.665674428 | 0.805800395 | 724.135266 | 700.2293952 | ENSG00000198648 |
| POLR1D | -0.047163532 | 0.967837315 | 0.49795819 | 0.678931121 | 2196.424006 | 2125.260543 | ENSG00000186184 |
| C19orf48 | -0.047230279 | 0.967792538 | 0.514761438 | 0.692846128 | 1882.199552 | 1822.141934 | ENSG00000167747 |
| HERC2 | -0.047240239 | 0.967785857 | 0.801671351 | 0.893310392 | 4170.198949 | 4035.922715 | ENSG00000128731 |
| CWC27 | -0.047272473 | 0.967764234 | 0.587497124 | 0.750166841 | 1116.43635 | 1080.262688 | ENSG00000153015 |
| RPL24 | -0.047275096 | 0.967762474 | 0.470779223 | 0.656213378 | 15202.69837 | 14711.94748 | ENSG00000114391 |
| MCMDC2 | -0.04737291 | 0.967696863 | 0.948964944 | 0.97469657 | 10.93713068 | 10.5208555 | ENSG00000178460 |
| PGAP4 | -0.0473805 | 0.967691772 | 0.651973128 | 0.796865599 | 563.8564733 | 545.3235925 | ENSG00000165152 |
| ALG12 | -0.047440646 | 0.967651429 | 0.737687091 | 0.853078713 | 278.6160552 | 269.5008295 | ENSG00000182858 |
| VWA1 | -0.047445097 | 0.967648444 | 0.656965469 | 0.79954022 | 1171.037579 | 1133.304544 | ENSG00000179403 |
| ARL2 | -0.047461 | 0.967637778 | 0.688704284 | 0.821445729 | 461.2186081 | 446.530508 | ENSG00000213465 |
| MRPS7 | -0.047466358 | 0.967634184 | 0.572271137 | 0.738429799 | 1501.866796 | 1453.075421 | ENSG00000125445 |
| MARK2 | -0.04753761 | 0.967586395 | 0.528653261 | 0.704440257 | 1400.870704 | 1355.321726 | ENSG00000072518 |
| HERPUD2 | -0.047663379 | 0.967502048 | 0.721301061 | 0.842434549 | 344.6485761 | 333.1467638 | ENSG00000122557 |
| CDYL | -0.047740187 | 0.967450541 | 0.503889822 | 0.683823765 | 1699.775658 | 1644.507991 | ENSG00000153046 |
| TBC1D31 | -0.048025811 | 0.967259024 | 0.722478022 | 0.843330877 | 415.7925861 | 402.0479575 | ENSG00000156787 |
| TIAM2 | -0.048094503 | 0.967212971 | 0.780205259 | 0.87926087 | 193.327013 | 186.7812009 | ENSG00000146426 |
| POU5F1 | -0.048099086 | 0.967209898 | 0.345222064 | 0.538671976 | 50384.3021 | 48731.48965 | ENSG00000204531 |
| HTRA3 | -0.048106125 | 0.967205179 | 0.75369518 | 0.863295824 | 224.8349093 | 217.2981343 | ENSG00000170801 |
| PTGES3 | -0.04825905 | 0.967102661 | 0.567538992 | 0.734704734 | 8677.563213 | 8390.792629 | ENSG00000110958 |
| BOP1 | -0.04830398 | 0.967072543 | 0.467332658 | 0.653229513 | 4167.985879 | 4030.942942 | ENSG00000261236 |
| RPF2 | -0.048405803 | 0.967004291 | 0.694054981 | 0.824696436 | 913.2193233 | 882.7481294 | ENSG00000197498 |
| NA | -0.048436359 | 0.96698381 | 0.934697744 | 0.967799644 | 14.19914351 | 13.76172419 | ENSG00000258559 |
| CCT5 | -0.048521133 | 0.966926992 | 0.398536649 | 0.590322714 | 11220.20906 | 10848.47596 | ENSG00000150753 |
| MCRIP2 | -0.048536885 | 0.966916434 | 0.632416355 | 0.783241211 | 789.6080814 | 764.171526 | ENSG00000172366 |
| PIP4K2C | -0.048578353 | 0.966888642 | 0.643660751 | 0.79161303 | 616.7554207 | 596.6979196 | ENSG00000166908 |

| | | | | | | | |
|---|---|---|---|---|---|---|---|
| SPIB | -0.048594608 | 0.966877748 | 0.836829027 | 0.912340541 | 93.51704032 | 90.52564972 | ENSG00000269404 |
| CCT6A | -0.048653066 | 0.966838571 | 0.424694393 | 0.61546308 | 9722.554293 | 9399.267336 | ENSG00000146731 |
| MRPL14 | -0.048727683 | 0.966788566 | 0.569153088 | 0.735971767 | 1599.970668 | 1547.405118 | ENSG00000180992 |
| PPID | -0.048768871 | 0.966760966 | 0.604499846 | 0.763305637 | 1133.702733 | 1095.488208 | ENSG00000171497 |
| PTPRU | -0.048778619 | 0.966754434 | 0.50354876 | 0.683776217 | 1765.490188 | 1706.528318 | ENSG00000060656 |
| NA | -0.04887331 | 0.966690983 | 0.672083579 | 0.809503537 | 896.11686 | 867.0290072 | ENSG00000179899 |
| STEAP3 | -0.04888574 | 0.966682654 | 0.640735727 | 0.789444284 | 580.5765826 | 561.1594526 | ENSG00000115107 |
| PFDN6 | -0.048939244 | 0.966646805 | 0.59390483 | 0.755268722 | 1136.519576 | 1098.988854 | ENSG00000204220 |
| NA | -0.048957727 | 0.96663442 | 0.956913565 | 0.979276823 | 5.991191834 | 5.787425394 | ENSG00000286360 |
| OLMALINC | -0.049007301 | 0.966601206 | 0.666184903 | 0.806102435 | 643.9709639 | 622.7521408 | ENSG00000235823 |
| BSCL2 | -0.049179082 | 0.96648612 | 0.938202451 | 0.969536896 | 12.07145365 | 11.66121969 | ENSG00000168000 |
| C3orf18 | -0.049309705 | 0.966398617 | 0.7771213 | 0.87733241 | 193.6674857 | 187.0376081 | ENSG00000088543 |
| HPCAL1 | -0.049479205 | 0.966285083 | 0.75686811 | 0.864632245 | 222.0943804 | 214.2547231 | ENSG00000115756 |
| ADD3 | -0.049532683 | 0.966249265 | 0.583900836 | 0.747118958 | 1400.171245 | 1352.025657 | ENSG00000148700 |
| LONRF3 | -0.049600065 | 0.966204137 | 0.960654888 | 0.98070932 | 5.327068095 | 5.168635407 | ENSG00000175556 |
| PDCD6IP | -0.049757812 | 0.966098496 | 0.541381647 | 0.714313906 | 1384.664262 | 1337.145415 | ENSG00000170248 |
| CYP2T1P | -0.049917869 | 0.96599132 | 0.933909098 | 0.96733214 | 13.65250196 | 13.2480741 | ENSG00000233622 |
| NKRF | -0.04999126 | 0.965942181 | 0.684266029 | 0.817890607 | 371.320131 | 358.6741234 | ENSG00000186416 |
| DNAJC17 | -0.050024231 | 0.965920106 | 0.740479185 | 0.855027177 | 244.487277 | 236.1132715 | ENSG00000104129 |
| L3MBTL1 | -0.050311857 | 0.965727552 | 0.85540322 | 0.923195905 | 87.83527836 | 84.98714541 | ENSG00000185513 |
| PRPF6 | -0.050344438 | 0.965705743 | 0.42930118 | 0.619564961 | 3251.215528 | 3140.347927 | ENSG00000101161 |
| PPIL4 | -0.050370613 | 0.965688222 | 0.694961105 | 0.825416199 | 555.0800522 | 535.2167177 | ENSG00000131013 |
| UBA2 | -0.050408132 | 0.965663109 | 0.428087122 | 0.618358482 | 3884.681377 | 3750.338907 | ENSG00000126261 |
| SRP68 | -0.050437419 | 0.965643505 | 0.429039066 | 0.619396327 | 3051.716079 | 2947.181319 | ENSG00000167881 |
| SMG6 | -0.050461394 | 0.965627458 | 0.63282655 | 0.783644503 | 763.9826742 | 737.8072043 | ENSG00000070366 |
| MAP2K4 | -0.050530531 | 0.965581184 | 0.600230193 | 0.759725903 | 694.9366118 | 670.7156004 | ENSG00000065559 |
| TBCB | -0.050559304 | 0.965561927 | 0.503174427 | 0.683561888 | 2032.375397 | 1963.129163 | ENSG00000105254 |
| R3HDM1 | -0.050573315 | 0.96555255 | 0.573868435 | 0.739436906 | 1149.51965 | 1109.89435 | ENSG00000048991 |
| RIN3 | -0.050586529 | 0.965543706 | 0.852985895 | 0.922280546 | 71.22504452 | 68.76650724 | ENSG00000100599 |
| C2orf49 | -0.050632938 | 0.965512647 | 0.730003628 | 0.848960736 | 374.1488202 | 361.8700043 | ENSG00000135974 |
| NA | -0.050646966 | 0.965503259 | 0.459130803 | 0.646127338 | 2343324.37 | 2262488.325 | N_unmapped |
| MAPK6 | -0.050654913 | 0.965497941 | 0.444430753 | 0.632891246 | 2696.868029 | 2603.24733 | ENSG00000069956 |
| IARS2 | -0.050707178 | 0.965462964 | 0.501889939 | 0.682417147 | 2465.055499 | 2379.040191 | ENSG00000067704 |
| SHISA5 | -0.050726505 | 0.96545003 | 0.468146322 | 0.654049746 | 2982.18401 | 2878.594511 | ENSG00000164054 |
| CNPY3 | -0.05073561 | 0.965443937 | 0.478724205 | 0.662683876 | 2129.90254 | 2056.605412 | ENSG00000137161 |
| MSH3 | -0.050749636 | 0.965434551 | 0.682029844 | 0.816374395 | 403.4747053 | 389.2474326 | ENSG00000113318 |
| COX7B | -0.050773014 | 0.965418907 | 0.576002298 | 0.740960103 | 1002.005584 | 967.7883478 | ENSG00000131174 |
| ZNF639 | -0.050805676 | 0.96539705 | 0.669508669 | 0.807856359 | 403.3981653 | 389.2522192 | ENSG00000121864 |
| PIK3AP1 | -0.050874273 | 0.965351149 | 0.818163395 | 0.902165959 | 109.2453461 | 105.3664153 | ENSG00000155629 |
| MRPS17 | -0.050920502 | 0.965320216 | 0.814105203 | 0.89991352 | 106.4085616 | 102.764645 | ENSG00000239789 |
| ITGB1BP1 | -0.050943267 | 0.965304984 | 0.685902763 | 0.819285583 | 353.2089518 | 340.9998916 | ENSG00000119185 |
| CEBPA | -0.050958492 | 0.965294797 | 0.908759222 | 0.954598925 | 25.61268954 | 24.6537627 | ENSG00000245848 |
| TMEM50B | -0.051015682 | 0.965256532 | 0.774808916 | 0.875842184 | 198.6930465 | 191.8507971 | ENSG00000142188 |
| ZNF335 | -0.051018338 | 0.965254755 | 0.663435419 | 0.804258763 | 565.2714155 | 545.951964 | ENSG00000198026 |
| NA | -0.05109706 | 0.965202087 | 0.91455382 | 0.957916954 | 41.25705906 | 40.01147634 | ENSG00000271576 |
| GALT | -0.051177245 | 0.965148443 | 0.848231434 | 0.919256309 | 79.13384899 | 76.43429815 | ENSG00000213930 |
| NAA30 | -0.051303104 | 0.965064248 | 0.713894596 | 0.838008895 | 587.0420396 | 566.3273949 | ENSG00000139977 |
| TUBG2 | -0.051340751 | 0.965039065 | 0.725683648 | 0.845904106 | 303.4164042 | 292.814071 | ENSG00000037042 |
| ATP6V1E2 | -0.051360735 | 0.965025697 | 0.839399549 | 0.913691856 | 75.56811876 | 72.99063575 | ENSG00000250565 |
| NA | -0.051511972 | 0.964924539 | 0.934494689 | 0.967755745 | 12.94921749 | 12.57665836 | ENSG00000229994 |
| KIAA1522 | -0.051545285 | 0.964902259 | 0.474206009 | 0.659334897 | 1606.420296 | 1550.25248 | ENSG00000162522 |
| FGD3 | -0.051568397 | 0.964886801 | 0.870810372 | 0.932978752 | 57.20917472 | 55.26768858 | ENSG00000127084 |
| AOPEP | -0.051712819 | 0.964790215 | 0.759288545 | 0.866116069 | 200.5840882 | 193.8991139 | ENSG00000148120 |
| PPME1 | -0.051748228 | 0.964766536 | 0.583026796 | 0.746519084 | 833.0713776 | 803.1697008 | ENSG00000214517 |
| NA | -0.051768992 | 0.964752651 | 0.847550509 | 0.918786988 | 81.02845361 | 78.4643936 | ENSG00000234771 |
| CCDC136 | -0.051811675 | 0.964724108 | 0.686771247 | 0.819878795 | 626.9928072 | 605.4381906 | ENSG00000128596 |
| NA | -0.051813424 | 0.964722939 | 0.807948005 | 0.896899705 | 111.2547796 | 107.3853375 | ENSG00000124224 |
| DRP2 | -0.051936762 | 0.964640467 | 0.910945089 | 0.955762377 | 22.43668748 | 21.57065965 | ENSG00000102385 |
| CCNJ | -0.051976594 | 0.964613834 | 0.62413072 | 0.77710194 | 882.2704183 | 850.406969 | ENSG00000107443 |
| AKAP8 | -0.052015849 | 0.964587588 | 0.555415148 | 0.725307064 | 977.805822 | 943.3716313 | ENSG00000105127 |
| SLC25A17 | -0.052059697 | 0.964558271 | 0.644699629 | 0.792364916 | 672.1107975 | 647.6136978 | ENSG00000100372 |
| TOP1MT | -0.05216359 | 0.964488813 | 0.451079134 | 0.638580864 | 1898.663861 | 1831.280871 | ENSG00000184428 |
| ZNF569 | -0.052213066 | 0.964455737 | 0.785187799 | 0.882461383 | 144.4658364 | 139.4318246 | ENSG00000196437 |
| NA | -0.052258549 | 0.964425332 | 0.964901968 | 0.982987582 | 3.916568128 | 3.74323387 | ENSG00000287760 |
| MTA1 | -0.05230424 | 0.964394789 | 0.388853501 | 0.581598708 | 3405.512817 | 3284.578597 | ENSG00000182979 |
| ZNF277 | -0.052476692 | 0.964279517 | 0.717902218 | 0.84019859 | 293.2975219 | 282.8237728 | ENSG00000198839 |
| FNDC7 | -0.052562516 | 0.964222155 | 0.91353135 | 0.957116452 | 22.0089036 | 21.29745687 | ENSG00000143107 |
| FANCC | -0.052618168 | 0.964184961 | 0.691469437 | 0.823321819 | 302.5982908 | 291.6727191 | ENSG00000158169 |
| DOP1A | -0.052663947 | 0.964154366 | 0.748291066 | 0.859456587 | 213.5311335 | 205.7424925 | ENSG00000083097 |
| TNS3 | -0.052781339 | 0.964075916 | 0.567377259 | 0.73470063 | 853.7960199 | 822.6128344 | ENSG00000136205 |
| ATOX1 | -0.053111646 | 0.963855215 | 0.701282576 | 0.830275647 | 395.1325924 | 381.3243028 | ENSG00000177556 |
| CACTIN | -0.053111883 | 0.963855057 | 0.594019216 | 0.755268722 | 876.7762519 | 845.3849033 | ENSG00000105298 |
| MCTS1 | -0.05324519 | 0.963765999 | 0.596545216 | 0.756870826 | 611.5040807 | 589.4820264 | ENSG00000232119 |
| FNDC3A | -0.053313489 | 0.963720374 | 0.654550743 | 0.798173792 | 532.0834039 | 512.6289837 | ENSG00000102531 |

| | | | | | | | |
|---|---|---|---|---|---|---|---|
| UBXN2B | -0.053413545 | 0.963653539 | 0.606808061 | 0.765281751 | 577.7060416 | 557.0048973 | ENSG00000215114 |
| AHCY | -0.053493782 | 0.963599946 | 0.350127048 | 0.543397647 | 7161.650933 | 6900.326598 | ENSG00000101444 |
| RASGRP1 | -0.053511053 | 0.96358841 | 0.845116976 | 0.917436773 | 66.90131773 | 64.35945532 | ENSG00000172575 |
| MEX3A | -0.053613411 | 0.963520047 | 0.462475021 | 0.649061944 | 2004.701531 | 1931.8899 | ENSG00000254726 |
| LOC112268411 | -0.053644494 | 0.963499288 | 0.94259503 | 0.971746372 | 8.684367025 | 8.379448188 | ENSG00000236106 |
| GAPVD1 | -0.053675247 | 0.96347875 | 0.561808549 | 0.730570307 | 855.7498091 | 824.5181747 | ENSG00000165219 |
| SMG7 | -0.0537122 | 0.963454072 | 0.50030354 | 0.680962061 | 1666.200339 | 1604.960119 | ENSG00000116698 |
| MCFD2 | -0.053728225 | 0.963443371 | 0.455889739 | 0.643266615 | 1684.47821 | 1622.444276 | ENSG00000180398 |
| LOC101927741 | -0.053810528 | 0.963388409 | 0.940977782 | 0.970798334 | 9.285953867 | 8.923154197 | ENSG00000287299 |
| FAM120C | -0.053894172 | 0.963332556 | 0.759857439 | 0.866445046 | 167.7403529 | 161.5347937 | ENSG00000184083 |
| GJD2 | -0.053902549 | 0.963326962 | 0.958770039 | 0.979702498 | 4.66166658 | 4.538560499 | ENSG00000159248 |
| S100PBP | -0.054015547 | 0.963251513 | 0.588618098 | 0.750925649 | 735.6338381 | 708.3752696 | ENSG00000116497 |
| HSPA9 | -0.054056561 | 0.96322413 | 0.345089711 | 0.538556139 | 11825.94156 | 11390.38521 | ENSG00000113013 |
| ZNF529 | -0.054113328 | 0.96318623 | 0.778972684 | 0.878653821 | 137.1923753 | 132.1289415 | ENSG00000186020 |
| C11orf45 | -0.054121034 | 0.963181085 | 0.929803886 | 0.965487435 | 12.37288595 | 11.87854751 | ENSG00000174370 |
| DCUN1D2 | -0.054122519 | 0.963180093 | 0.650697073 | 0.795936006 | 428.1922756 | 411.9667155 | ENSG00000150401 |
| ZNF233 | -0.054169426 | 0.963148778 | 0.901980953 | 0.95021014 | 24.20040814 | 23.26601477 | ENSG00000159915 |
| ZNF420 | -0.05420192 | 0.963127085 | 0.828521947 | 0.907608355 | 78.7705562 | 75.7394365 | ENSG00000197050 |
| MGAT1 | -0.054290565 | 0.963067909 | 0.401872031 | 0.59357069 | 2824.993638 | 2720.788163 | ENSG00000131446 |
| GNPNAT1 | -0.054308393 | 0.963056007 | 0.586585249 | 0.749312221 | 719.307666 | 692.413755 | ENSG00000100522 |
| ABHD12 | -0.054316247 | 0.963050765 | 0.48937015 | 0.67219939 | 1448.655938 | 1395.08968 | ENSG00000100997 |
| CCNT2 | -0.054359457 | 0.963021921 | 0.592425019 | 0.754224846 | 804.1155801 | 773.5299081 | ENSG00000082258 |
| ANKRD40 | -0.054443422 | 0.962965874 | 0.510492909 | 0.689712463 | 1110.357712 | 1069.164646 | ENSG00000154945 |
| PUSL1 | -0.054515828 | 0.962917547 | 0.685482912 | 0.818973783 | 322.1997647 | 310.3863005 | ENSG00000169972 |
| LOC645513 | -0.054608869 | 0.962855449 | 0.790123374 | 0.885591837 | 127.7490044 | 123.1465516 | ENSG00000245958 |
| NA | -0.054659051 | 0.962821958 | 0.937056776 | 0.969033102 | 16.43346935 | 15.78323426 | ENSG00000271043 |
| CAMSAP1 | -0.054758896 | 0.962755326 | 0.493926126 | 0.676002571 | 1726.527007 | 1661.911002 | ENSG00000130559 |
| AAK1 | -0.054794577 | 0.962731515 | 0.698792812 | 0.828207766 | 312.7996809 | 301.4552138 | ENSG00000115977 |
| CCDC30 | -0.054872371 | 0.962679604 | 0.941569407 | 0.971013097 | 9.392183882 | 9.133433982 | ENSG00000186409 |
| KRI1 | -0.054887762 | 0.962669333 | 0.542171263 | 0.714870913 | 1496.609134 | 1441.861219 | ENSG00000129347 |
| LRRC47 | -0.054888408 | 0.962668903 | 0.452312407 | 0.639740396 | 2240.627748 | 2156.399149 | ENSG00000130764 |
| PXDN | -0.054904953 | 0.962657862 | 0.388277285 | 0.581249696 | 6195.531279 | 5963.91234 | ENSG00000130508 |
| CLK3 | -0.054907973 | 0.962655847 | 0.573038467 | 0.738883513 | 798.6638291 | 769.1669687 | ENSG00000179335 |
| CDHR2 | -0.054915794 | 0.962650629 | 0.939020736 | 0.969852089 | 9.776372342 | 9.493555308 | ENSG00000074276 |
| HS3ST1 | -0.054956433 | 0.962623512 | 0.893587107 | 0.945520864 | 30.36214832 | 29.17366382 | ENSG00000002587 |
| SLC35F1 | -0.054960812 | 0.962620591 | 0.726669194 | 0.846463882 | 251.8133233 | 242.0138323 | ENSG00000196376 |
| ZSCAN5A | -0.05499745 | 0.962596145 | 0.857808963 | 0.924554043 | 70.85516746 | 67.89879617 | ENSG00000131848 |
| TFAP4 | -0.05503505 | 0.962571057 | 0.60777115 | 0.766027235 | 591.7583962 | 569.2893184 | ENSG00000090447 |
| CACNB4 | -0.055059318 | 0.962554865 | 0.89162431 | 0.944765273 | 31.33035762 | 30.17872058 | ENSG00000182389 |
| PSMD7 | -0.05512407 | 0.962511665 | 0.49664339 | 0.678082046 | 1920.725199 | 1848.257484 | ENSG00000103035 |
| NA | -0.055136217 | 0.962503561 | 0.938848492 | 0.96979109 | 11.697916 | 11.26779325 | ENSG00000182165 |
| CDC14B | -0.05520219 | 0.962459547 | 0.460451142 | 0.647103538 | 1795.907567 | 1728.030934 | ENSG00000081377 |
| MYMX | -0.05530664 | 0.962389868 | 0.907142994 | 0.953766068 | 27.20586185 | 26.13135196 | ENSG00000262179 |
| PPP1R10 | -0.055317736 | 0.962382467 | 0.515793793 | 0.69358791 | 1025.252297 | 986.5425198 | ENSG00000204569 |
| RRP9 | -0.055454559 | 0.9622912 | 0.441685348 | 0.629949691 | 2020.524521 | 1944.813755 | ENSG00000114767 |
| PI4KAP2 | -0.055462894 | 0.96228564 | 0.741348854 | 0.855338624 | 345.5937402 | 333.0755677 | ENSG00000183506 |
| PIAS4 | -0.05556979 | 0.962214343 | 0.482626164 | 0.666045669 | 1446.887782 | 1392.115189 | ENSG00000105229 |
| C17orf80 | -0.055776713 | 0.962076345 | 0.617621681 | 0.772558564 | 514.3019867 | 494.7380308 | ENSG00000141219 |
| THRA | -0.056111839 | 0.961852888 | 0.650625477 | 0.795936006 | 480.9998452 | 463.1020209 | ENSG00000126351 |
| TBC1D12 | -0.056202622 | 0.961792365 | 0.852598906 | 0.92215617 | 57.70443384 | 55.51562198 | ENSG00000108239 |
| ZNF718 | -0.05623864 | 0.961768353 | 0.741674022 | 0.855607265 | 183.3509811 | 176.2952995 | ENSG00000250312 |
| TP53I13 | -0.056279282 | 0.961741259 | 0.636238179 | 0.786236877 | 465.0995406 | 447.3539237 | ENSG00000167543 |
| GINS1 | -0.056307909 | 0.961722176 | 0.494183521 | 0.676205048 | 1456.190302 | 1399.508526 | ENSG00000101003 |
| PCYT1B | -0.056319558 | 0.96171441 | 0.520832945 | 0.697883011 | 1108.170432 | 1065.560797 | ENSG00000102230 |
| MIR600HG | -0.05636296 | 0.961685479 | 0.706478611 | 0.833271761 | 360.5126816 | 347.0181199 | ENSG00000236901 |
| CDKN2AIP | -0.056389673 | 0.961667673 | 0.651773837 | 0.796727101 | 483.6464861 | 464.7605179 | ENSG00000168564 |
| SUPT7L | -0.056389796 | 0.961667591 | 0.700198033 | 0.829337318 | 436.5823209 | 420.3870438 | ENSG00000119760 |
| ABHD13 | -0.056399932 | 0.961660834 | 0.743817344 | 0.856959697 | 198.9532283 | 191.1804404 | ENSG00000139826 |
| CASP2 | -0.056428985 | 0.961641468 | 0.593951432 | 0.755268722 | 552.9870133 | 531.6264403 | ENSG00000106144 |
| PLXNA3 | -0.056681225 | 0.96147335 | 0.659821665 | 0.801857894 | 484.7675202 | 466.618729 | ENSG00000130827 |
| NA | -0.056756463 | 0.96142321 | 0.893009298 | 0.945341234 | 34.16764578 | 33.07174056 | ENSG00000124097 |
| LARGE2 | -0.056790336 | 0.961400637 | 0.442363479 | 0.630722727 | 2033.497491 | 1955.272188 | ENSG00000165905 |
| ARAP3 | -0.056931263 | 0.961306729 | 0.396206397 | 0.588546073 | 2504.960408 | 2408.061707 | ENSG00000120318 |
| IRS4 | -0.056945377 | 0.961297324 | 0.739709685 | 0.854617835 | 208.1459632 | 200.0870137 | ENSG00000133124 |
| TUBD1 | -0.056956726 | 0.961289762 | 0.721119275 | 0.842328395 | 271.2543209 | 260.588419 | ENSG00000108423 |
| KIAA0100 | -0.056962754 | 0.961285746 | 0.460934542 | 0.647586477 | 2543.711346 | 2445.879451 | ENSG00000007202 |
| SREK1IP1 | -0.05704637 | 0.961230033 | 0.655573397 | 0.798947748 | 519.7324436 | 499.1927701 | ENSG00000153006 |
| BRF2 | -0.05706968 | 0.961214502 | 0.723128623 | 0.843877716 | 207.0029662 | 198.7273463 | ENSG00000104221 |
| ZCCHC10 | -0.057140508 | 0.961167313 | 0.71676329 | 0.839556455 | 288.9424214 | 277.6924646 | ENSG00000155329 |
| ALAD | -0.057221941 | 0.961113062 | 0.64614545 | 0.793142577 | 379.5717758 | 365.1340554 | ENSG00000148218 |
| PDZRN3 | -0.057273983 | 0.961078392 | 0.65681387 | 0.799514587 | 355.7569316 | 341.9143425 | ENSG00000121440 |
| USP10 | -0.057350129 | 0.961027667 | 0.343639947 | 0.53737958 | 4699.609073 | 4516.150103 | ENSG00000103194 |
| GLCCI1 | -0.057407362 | 0.960989543 | 0.670948051 | 0.808859269 | 321.4706462 | 309.2160952 | ENSG00000106415 |
| PAN3 | -0.057420656 | 0.960980688 | 0.597435002 | 0.757636697 | 530.7583285 | 509.6596059 | ENSG00000152520 |

| | | | | | | | |
|---|---|---|---|---|---|---|---|
| TUBGCP3 | -0.057503354 | 0.960925605 | 0.543029332 | 0.715600631 | 745.0122893 | 715.8816077 | ENSG00000126216 |
| PLXNC1 | -0.057539132 | 0.960901774 | 0.846735056 | 0.918413939 | 60.95066211 | 58.51962741 | ENSG00000136040 |
| NA | -0.05753988 | 0.960901276 | 0.952632978 | 0.977013699 | 5.121019588 | 4.954994809 | ENSG00000259005 |
| VSIG10L | -0.05758568 | 0.960870772 | 0.668094246 | 0.807200701 | 333.2705454 | 320.1197535 | ENSG00000186806 |
| LINC00174 | -0.057626949 | 0.960843286 | 0.872001667 | 0.933715344 | 50.12262525 | 48.23998368 | ENSG00000179406 |
| WRNIP1 | -0.057693769 | 0.960798784 | 0.503094596 | 0.683503536 | 1096.391503 | 1053.499892 | ENSG00000124535 |
| PANK2 | -0.057780054 | 0.960741323 | 0.56035505 | 0.729378503 | 686.3243727 | 659.3777245 | ENSG00000125779 |
| TROAP | -0.057854119 | 0.960692002 | 0.557385393 | 0.726781763 | 660.6037468 | 635.112642 | ENSG00000135451 |
| VGF | -0.057892976 | 0.960666127 | 0.731795649 | 0.850244518 | 229.5924481 | 220.8582215 | ENSG00000128564 |
| NA | -0.058079531 | 0.960541911 | 0.760916191 | 0.867158734 | 172.6034379 | 165.9554408 | ENSG00000249673 |
| CD82 | -0.058377797 | 0.960343347 | 0.79399084 | 0.888314996 | 115.1631548 | 110.7955832 | ENSG00000085117 |
| LRRCC1 | -0.058475232 | 0.96027849 | 0.714959777 | 0.838432369 | 237.1563151 | 227.4480216 | ENSG00000133739 |
| SNX12 | -0.058586435 | 0.960204475 | 0.438133887 | 0.626620376 | 1874.308396 | 1800.05881 | ENSG00000147164 |
| TPT1 | -0.058616806 | 0.960184262 | 0.464273364 | 0.650600523 | 5126.102316 | 4920.878415 | ENSG00000133112 |
| C9orf85 | -0.058628908 | 0.960176207 | 0.706737166 | 0.833271761 | 241.7262206 | 232.3212108 | ENSG00000155621 |
| NA | -0.058646326 | 0.960164615 | 0.940118093 | 0.970380303 | 8.254704009 | 7.940444037 | ENSG00000280254 |
| CYB5R4 | -0.058700385 | 0.960128637 | 0.718446621 | 0.840477794 | 242.0229252 | 231.9926424 | ENSG00000065615 |
| COA4 | -0.058732776 | 0.960107081 | 0.453481664 | 0.640808424 | 1252.870609 | 1203.046141 | ENSG00000181924 |
| SMARCD2 | -0.058829983 | 0.960042392 | 0.351303315 | 0.544467173 | 2773.178255 | 2662.462624 | ENSG00000108604 |
| TOR3A | -0.058832043 | 0.960041021 | 0.526865728 | 0.702967066 | 735.2602853 | 705.7412532 | ENSG00000186283 |
| MEA1 | -0.058929778 | 0.959975986 | 0.475054238 | 0.659978814 | 1173.961765 | 1126.678229 | ENSG00000124733 |
| POLR2J | -0.058930882 | 0.959975252 | 0.614341808 | 0.770149497 | 447.1958982 | 429.4183151 | ENSG00000005075 |
| ANGPTL2 | -0.058942002 | 0.959967852 | 0.639243096 | 0.78858703 | 367.7184196 | 353.4016622 | ENSG00000136859 |
| GRSF1 | -0.05896445 | 0.959952916 | 0.513590481 | 0.692196991 | 2616.444485 | 2510.664021 | ENSG00000132463 |
| TMEM214 | -0.058964791 | 0.959952688 | 0.477680937 | 0.661871178 | 1061.962127 | 1019.16128 | ENSG00000119777 |
| DNAJC4 | -0.058981441 | 0.95994161 | 0.739081708 | 0.854158532 | 161.7819103 | 155.3282345 | ENSG00000110011 |
| OGDH | -0.058985122 | 0.959939161 | 0.330619779 | 0.524947056 | 3734.947491 | 3585.008389 | ENSG00000105953 |
| STAT2 | -0.059271687 | 0.959748505 | 0.578939382 | 0.743182055 | 584.4664222 | 560.9960646 | ENSG00000170581 |
| NOSIP | -0.059327277 | 0.959711525 | 0.474996774 | 0.659978814 | 1701.154212 | 1633.031728 | ENSG00000142546 |
| TXNL1 | -0.059359515 | 0.959690079 | 0.42029435 | 0.611558221 | 1509.651218 | 1448.581307 | ENSG00000091164 |
| NEFL | -0.059368206 | 0.959684298 | 0.598251771 | 0.758154903 | 439.9308255 | 422.2048354 | ENSG00000277586 |
| TAF5L | -0.059539436 | 0.959570402 | 0.553318968 | 0.723931509 | 667.7007517 | 640.4377919 | ENSG00000135801 |
| NA | -0.059574828 | 0.959546863 | 0.823750655 | 0.905324635 | 74.05601886 | 71.01685329 | ENSG00000267640 |
| TRPC1 | -0.05958924 | 0.959537277 | 0.745051783 | 0.857529016 | 158.2772438 | 151.7874679 | ENSG00000144935 |
| AMZ1 | -0.059617567 | 0.959518437 | 0.874177614 | 0.934994283 | 39.66526223 | 38.17971448 | ENSG00000174945 |
| FAM131A | -0.059652091 | 0.959495476 | 0.645519196 | 0.792913747 | 368.7164166 | 354.2342044 | ENSG00000175182 |
| ARL16 | -0.059926732 | 0.959312837 | 0.614020097 | 0.769819961 | 416.8292175 | 399.6503663 | ENSG00000214087 |
| NA | -0.059999524 | 0.959264435 | 0.949886881 | 0.975141434 | 5.339695747 | 5.134780645 | ENSG00000279392 |
| SLC30A9 | -0.06004496 | 0.959234226 | 0.503783192 | 0.683776217 | 1557.229365 | 1492.836447 | ENSG00000014824 |
| LDLR | -0.060056649 | 0.959226454 | 0.389903209 | 0.582497668 | 3416.033236 | 3276.249551 | ENSG00000130164 |
| CD63 | -0.060079642 | 0.959211166 | 0.381724452 | 0.574586291 | 6610.067046 | 6340.645014 | ENSG00000135404 |
| DCAF12 | -0.060084174 | 0.959208153 | 0.514830953 | 0.692846128 | 820.0765726 | 786.2308038 | ENSG00000198876 |
| NA | -0.06016063 | 0.95915732 | 0.882822967 | 0.939729393 | 32.84245752 | 31.54835879 | ENSG00000215835 |
| TMEM230 | -0.060168483 | 0.959152099 | 0.47533122 | 0.66010706 | 1240.556982 | 1190.031904 | ENSG00000089063 |
| ZNF133 | -0.060188625 | 0.959138709 | 0.670138547 | 0.80835326 | 268.6113405 | 257.6465008 | ENSG00000125846 |
| ARHGAP10 | -0.060341243 | 0.95903725 | 0.734626276 | 0.851611421 | 163.9342541 | 157.3539814 | ENSG00000071205 |
| DPPA4 | -0.060620844 | 0.958851402 | 0.317219522 | 0.510858486 | 12303.33663 | 11795.63983 | ENSG00000121570 |
| GANC | -0.060668356 | 0.958819825 | 0.753957841 | 0.863411144 | 141.7678358 | 135.6654712 | ENSG00000214013 |
| SPRY3 | -0.06071843 | 0.958786546 | 0.831837767 | 0.909359513 | 61.67356447 | 59.28197534 | ENSG00000168939 |
| NA | -0.060743118 | 0.958770139 | 0.927967303 | 0.964548443 | 11.49008606 | 11.00488771 | ENSG00000236603 |
| RPL14 | -0.060747003 | 0.958767557 | 0.315132477 | 0.508759471 | 17624.99248 | 16898.32559 | ENSG00000188846 |
| USP48 | -0.060947196 | 0.958634525 | 0.394221483 | 0.586631952 | 2000.960389 | 1918.384097 | ENSG00000090686 |
| PPIH | -0.061013323 | 0.958590586 | 0.693491082 | 0.82436945 | 228.4520219 | 219.0955949 | ENSG00000171960 |
| SH3D21 | -0.061078685 | 0.958547158 | 0.718293109 | 0.840457354 | 271.4352269 | 260.5676416 | ENSG00000214193 |
| NBR2 | -0.061089954 | 0.95853967 | 0.84755044 | 0.918786988 | 47.50283317 | 45.55366788 | ENSG00000198496 |
| HYDIN | -0.061209342 | 0.958460351 | 0.902925306 | 0.950679863 | 18.81396006 | 18.08284525 | ENSG00000157423 |
| CREG2 | -0.061371631 | 0.95835254 | 0.929276036 | 0.965198354 | 12.44301445 | 11.87486028 | ENSG00000175874 |
| DIMT1 | -0.061458153 | 0.958295067 | 0.533382581 | 0.70829298 | 1055.044607 | 1010.93141 | ENSG00000086189 |
| SPIRE2 | -0.061560105 | 0.958227349 | 0.696151479 | 0.826246173 | 221.7803204 | 212.7464654 | ENSG00000204991 |
| KIF5C | -0.061675801 | 0.958150508 | 0.395902511 | 0.588262452 | 2113.838587 | 2024.869986 | ENSG00000168280 |
| ATG13 | -0.061728941 | 0.958115215 | 0.557852001 | 0.727192923 | 1056.990738 | 1012.619182 | ENSG00000175224 |
| NOTCH3 | -0.061830317 | 0.958047893 | 0.395343253 | 0.587768127 | 3461.059766 | 3315.55648 | ENSG00000074181 |
| PRDX3 | -0.061950784 | 0.957967898 | 0.308803649 | 0.502976258 | 4101.775813 | 3928.931637 | ENSG00000165672 |
| PTS | -0.062018355 | 0.957923031 | 0.669284666 | 0.807856359 | 326.625377 | 312.9524958 | ENSG00000150787 |
| SPAG7 | -0.062136855 | 0.957844352 | 0.466859944 | 0.653017299 | 1399.216663 | 1341.033928 | ENSG00000091640 |
| PARP8 | -0.062152794 | 0.95783377 | 0.63133111 | 0.782576966 | 382.4744873 | 366.0920056 | ENSG00000151883 |
| ANGEL1 | -0.062246981 | 0.957771239 | 0.49950279 | 0.680219303 | 739.4418759 | 708.3187418 | ENSG00000013523 |
| PIAS1 | -0.062275781 | 0.95775212 | 0.567208301 | 0.734612098 | 499.394237 | 478.4966866 | ENSG00000033800 |
| VPS28 | -0.062403914 | 0.957667061 | 0.537327412 | 0.711203176 | 640.5980532 | 613.1870453 | ENSG00000160948 |
| SMAD4 | -0.062582185 | 0.957548731 | 0.49253754 | 0.674899509 | 988.7828649 | 946.1174203 | ENSG00000141646 |
| PFDN5 | -0.062589802 | 0.957543675 | 0.463868174 | 0.650278552 | 1545.845833 | 1480.644926 | ENSG00000123349 |
| CELSR1 | -0.062646033 | 0.957506355 | 0.572911891 | 0.738883513 | 935.158437 | 896.0457243 | ENSG00000075275 |
| ATG5 | -0.062660351 | 0.957496852 | 0.593609743 | 0.755058882 | 704.2943779 | 673.4822558 | ENSG00000057663 |
| BRCA1 | -0.06267463 | 0.957487375 | 0.515514428 | 0.693363165 | 719.6117742 | 688.8691812 | ENSG00000012048 |

| | | | | | | | |
|---|---|---|---|---|---|---|---|
| PEX19 | -0.062760268 | 0.957430541 | 0.550759125 | 0.721876151 | 616.3534935 | 589.9353414 | ENSG00000162735 |
| NA | -0.062817769 | 0.957392382 | 0.85771278 | 0.924527631 | 51.10534134 | 49.11419313 | ENSG00000239665 |
| SERP1 | -0.06282841 | 0.95738532 | 0.380696836 | 0.573738425 | 3898.364102 | 3731.561901 | ENSG00000120742 |
| PRR14 | -0.062920704 | 0.957324075 | 0.702176055 | 0.830927613 | 218.689149 | 209.1463111 | ENSG00000156858 |
| UBE2D3 | -0.062933518 | 0.957315572 | 0.385103432 | 0.577938906 | 5863.603217 | 5611.868769 | ENSG00000109332 |
| ZNF497 | -0.062971802 | 0.957290169 | 0.893229744 | 0.945466595 | 22.88538967 | 21.87446799 | ENSG00000174586 |
| WASF1 | -0.063025418 | 0.957254592 | 0.495550407 | 0.677275366 | 836.9403814 | 801.0644669 | ENSG00000112290 |
| SHOC2 | -0.063035352 | 0.957248001 | 0.646073972 | 0.793142577 | 376.1797508 | 359.6985722 | ENSG00000108061 |
| IFITM3 | -0.063121705 | 0.957190706 | 0.308036649 | 0.502168287 | 3053.61254 | 2922.630363 | ENSG00000142089 |
| CENPH | -0.063173692 | 0.957156216 | 0.671478756 | 0.809097911 | 652.1742551 | 623.5852948 | ENSG00000153044 |
| ERGIC2 | -0.063267829 | 0.957093762 | 0.687563966 | 0.820455224 | 368.1815082 | 351.7519159 | ENSG00000087502 |
| HCCS | -0.063334459 | 0.957049561 | 0.517717525 | 0.695164609 | 688.0325467 | 658.5788139 | ENSG00000004961 |
| NA | -0.063366594 | 0.957028243 | 0.772101284 | 0.874541661 | 120.4568723 | 115.5591808 | ENSG00000229150 |
| TIMM50 | -0.06337006 | 0.957025944 | 0.386219701 | 0.579235794 | 3245.15426 | 3105.47205 | ENSG00000105197 |
| WASH8P | -0.063395769 | 0.95700889 | 0.680405668 | 0.815245229 | 288.5678014 | 276.6485836 | ENSG00000226210 |
| NCSTN | -0.063408261 | 0.957000603 | 0.432070066 | 0.622027855 | 1411.807993 | 1351.235962 | ENSG00000162736 |
| CLCNKB | -0.063439424 | 0.956979931 | 0.864252844 | 0.929307927 | 35.95968096 | 34.41738321 | ENSG00000184908 |
| SNORD6 | -0.063460677 | 0.956965834 | 0.95582131 | 0.978975886 | 3.850874318 | 3.634295131 | ENSG00000202314 |
| MLST8 | -0.063491475 | 0.956945405 | 0.41750256 | 0.608571958 | 1282.760874 | 1227.344193 | ENSG00000167965 |
| DPM2 | -0.063506712 | 0.956935299 | 0.518501969 | 0.695664941 | 966.6779535 | 925.5364333 | ENSG00000136908 |
| PBDC1 | -0.063553436 | 0.956904307 | 0.531480786 | 0.706734458 | 709.1585149 | 678.8821725 | ENSG00000102390 |
| SLC13A3 | -0.063583155 | 0.956884595 | 0.68464597 | 0.818237581 | 224.0250865 | 214.4815124 | ENSG00000158296 |
| ZNF76 | -0.063585041 | 0.956883345 | 0.617491202 | 0.772447485 | 375.3889074 | 359.4218256 | ENSG00000065029 |
| POLR3G | -0.063667808 | 0.95682845 | 0.499957169 | 0.68058804 | 3511.085541 | 3359.231236 | ENSG00000113356 |
| NA | -0.063675239 | 0.956823522 | 0.908655157 | 0.954597817 | 19.78404849 | 18.90344126 | ENSG00000268262 |
| SBK2 | -0.063790938 | 0.956746791 | 0.793886338 | 0.888278381 | 89.21618603 | 85.58270428 | ENSG00000187550 |
| KRIT1 | -0.063808885 | 0.956734889 | 0.845250393 | 0.91747413 | 48.96827844 | 47.02453551 | ENSG00000001631 |
| CAMK1D | -0.064000522 | 0.956607812 | 0.633288673 | 0.783902509 | 354.3485062 | 339.1503464 | ENSG00000183049 |
| HDAC7 | -0.064076785 | 0.956557246 | 0.409232609 | 0.600769677 | 1961.307492 | 1876.892111 | ENSG00000061273 |
| NA | -0.064102137 | 0.956540437 | 0.837420224 | 0.912716638 | 51.31138985 | 49.14717183 | ENSG00000233230 |
| NA | -0.064209249 | 0.956469421 | 0.760577591 | 0.866999512 | 129.03985 | 123.495826 | ENSG00000255135 |
| TBC1D25 | -0.06424935 | 0.956442836 | 0.732507027 | 0.850804364 | 170.663261 | 162.8621034 | ENSG00000068354 |
| PSMB9 | -0.064254206 | 0.956439617 | 0.852937641 | 0.922280546 | 40.06651303 | 38.39715393 | ENSG00000240065 |
| RPL6 | -0.064268963 | 0.956429833 | 0.268868387 | 0.459112465 | 18412.18426 | 17609.31774 | ENSG00000089009 |
| PCSK4 | -0.064359726 | 0.956369664 | 0.765785704 | 0.870633802 | 130.7075104 | 125.0825772 | ENSG00000115257 |
| ACCS | -0.064433655 | 0.956320658 | 0.817022632 | 0.901565698 | 91.79228499 | 87.95806052 | ENSG00000110455 |
| INTS14 | -0.064457421 | 0.956304904 | 0.500769769 | 0.681443972 | 677.0186806 | 647.3075566 | ENSG00000138614 |
| PGAM5 | -0.064466865 | 0.956298644 | 0.364491332 | 0.557636838 | 2270.210568 | 2170.111815 | ENSG00000247077 |
| HEG1 | -0.064529961 | 0.956256821 | 0.652300112 | 0.796943154 | 276.9558136 | 264.8209013 | ENSG00000173706 |
| PARD3 | -0.064636514 | 0.956186198 | 0.440011367 | 0.628287417 | 1559.970639 | 1491.390242 | ENSG00000148498 |
| MRPS14 | -0.064670398 | 0.95616374 | 0.593547721 | 0.7550489 | 452.3406999 | 432.6678808 | ENSG00000120333 |
| CD44 | -0.064854898 | 0.956041468 | 0.784645734 | 0.882158322 | 95.5304049 | 91.46679581 | ENSG00000026508 |
| SEH1L | -0.064981725 | 0.955957427 | 0.47777744 | 0.661871178 | 1179.910293 | 1127.933593 | ENSG00000085415 |
| FBXW8 | -0.065145437 | 0.955848954 | 0.559976044 | 0.72908999 | 457.4508581 | 437.1918669 | ENSG00000174989 |
| DEXI | -0.065160425 | 0.955839024 | 0.824078188 | 0.905519366 | 70.41347815 | 67.4628667 | ENSG00000182108 |
| KCTD21 | -0.065181321 | 0.95582518 | 0.849520857 | 0.920078138 | 48.98967777 | 46.87045709 | ENSG00000188997 |
| NA | -0.065205437 | 0.955809203 | 0.809717956 | 0.897512864 | 67.38354325 | 64.32571218 | ENSG00000237004 |
| GPRC5A | -0.06520872 | 0.955807027 | 0.916974018 | 0.959042743 | 12.53524134 | 11.90860343 | ENSG00000013588 |
| BAZ2A | -0.065245321 | 0.955782779 | 0.424720664 | 0.61546308 | 3478.451117 | 3323.972641 | ENSG00000076108 |
| TMEM116 | -0.06525745 | 0.955774743 | 0.763111108 | 0.868712073 | 125.4986009 | 119.890934 | ENSG00000198270 |
| TMEM189 | -0.065357774 | 0.955708282 | 0.632353129 | 0.783215242 | 302.2544432 | 288.5320921 | ENSG00000240849 |
| CCDC65 | -0.065368669 | 0.955701065 | 0.891649626 | 0.944765273 | 24.48665724 | 23.53315719 | ENSG00000139537 |
| FBRS | -0.065395455 | 0.95568332 | 0.581925043 | 0.74582662 | 491.4701743 | 469.729211 | ENSG00000156860 |
| MLX | -0.065421106 | 0.955666329 | 0.499315692 | 0.68011443 | 767.4807187 | 733.6429729 | ENSG00000108788 |
| BCORL1 | -0.065438211 | 0.955654998 | 0.609137341 | 0.766741215 | 377.5655218 | 360.8708476 | ENSG00000085185 |
| TSKU | -0.065482443 | 0.955625699 | 0.4887085 | 0.671520682 | 850.8955783 | 813.3057152 | ENSG00000182704 |
| ZNF227 | -0.065574203 | 0.95556492 | 0.693185262 | 0.824297167 | 219.7715859 | 209.750129 | ENSG00000131115 |
| MTMR12 | -0.065602073 | 0.955546461 | 0.464092938 | 0.650446046 | 1231.477333 | 1176.335633 | ENSG00000150712 |
| PTRH2 | -0.065604462 | 0.955544879 | 0.541957694 | 0.714666944 | 616.7578035 | 589.4991887 | ENSG00000141378 |
| PDE5A | -0.065604672 | 0.955544739 | 0.624387991 | 0.777176025 | 326.5482583 | 311.6147455 | ENSG00000138735 |
| MYDGF | -0.065663525 | 0.95550576 | 0.458873174 | 0.645816342 | 1893.866703 | 1809.407717 | ENSG00000074842 |
| RBM15B | -0.065716099 | 0.95547094 | 0.36063659 | 0.553910547 | 2642.244274 | 2524.72656 | ENSG00000259956 |
| MPP6 | -0.065744245 | 0.9554523 | 0.427244042 | 0.617692916 | 1410.044099 | 1347.003448 | ENSG00000105926 |
| LBHD1 | -0.065788887 | 0.955422736 | 0.93598011 | 0.968425646 | 7.334622517 | 7.055499316 | ENSG00000162194 |
| UBAP2L | -0.065836371 | 0.95539129 | 0.260219852 | 0.449102778 | 5334.462805 | 5096.503379 | ENSG00000143569 |
| PCGF1 | -0.065913325 | 0.95534033 | 0.738733366 | 0.853862439 | 170.912499 | 163.5473206 | ENSG00000115289 |
| RAB43 | -0.065926173 | 0.955331822 | 0.738693948 | 0.853862439 | 154.7105289 | 148.1347282 | ENSG00000172780 |
| RPS21 | -0.065950167 | 0.955315934 | 0.369964695 | 0.563197179 | 6451.954291 | 6164.449229 | ENSG00000171858 |
| PHAX | -0.065972097 | 0.955301413 | 0.455545674 | 0.642939652 | 1082.103613 | 1032.997816 | ENSG00000164902 |
| SERPINB8 | -0.066125851 | 0.955199607 | 0.931887228 | 0.966675877 | 7.893891715 | 7.554065643 | ENSG00000166401 |
| PPP1R3G | -0.066145324 | 0.955186715 | 0.939214386 | 0.969933662 | 7.268928707 | 6.980415338 | ENSG00000219607 |
| NA | -0.066279958 | 0.95509758 | 0.94823566 | 0.97445646 | 5.105235024 | 4.894882979 | ENSG00000269680 |
| ERGIC3 | -0.066393162 | 0.955022639 | 0.344682881 | 0.538200859 | 2736.467612 | 2613.890926 | ENSG00000125991 |
| SLC23A1 | -0.06639609 | 0.955020701 | 0.942305291 | 0.971609806 | 5.428765723 | 5.187406401 | ENSG00000170482 |

| | | | | | | | |
|---|---|---|---|---|---|---|---|
| SMARCA4 | -0.066472397 | 0.954970189 | 0.2486887 | 0.436932066 | 10171.63232 | 9713.858715 | ENSG00000127616 |
| EIF2B4 | -0.066496958 | 0.954953932 | 0.415982191 | 0.607215399 | 1149.668243 | 1098.220174 | ENSG00000115211 |
| VPS53 | -0.066606876 | 0.954881177 | 0.506009722 | 0.685654641 | 768.1552302 | 733.7198719 | ENSG00000141252 |
| TBC1D22B | -0.066609837 | 0.954879217 | 0.553693134 | 0.723938212 | 462.9279622 | 442.0166673 | ENSG00000065491 |
| ASPSCR1 | -0.066666331 | 0.954841826 | 0.536438646 | 0.710777371 | 964.065415 | 921.5569177 | ENSG00000169696 |
| NR2C1 | -0.066705976 | 0.954815588 | 0.594453296 | 0.755547815 | 472.6088757 | 451.2367018 | ENSG00000120798 |
| CHM | -0.066881444 | 0.954699465 | 0.536711058 | 0.711036651 | 622.5731778 | 593.88199 | ENSG00000188419 |
| ZNF222 | -0.066941785 | 0.954659535 | 0.869187697 | 0.932102342 | 36.12529096 | 34.34119992 | ENSG00000159885 |
| PORCN | -0.06694591 | 0.954656806 | 0.646109222 | 0.793142577 | 294.1762931 | 280.9378881 | ENSG00000102312 |
| ZFP42 | -0.066963533 | 0.954645144 | 0.354167379 | 0.547467741 | 4331.165451 | 4134.160676 | ENSG00000179059 |
| NRIP3 | -0.06697117 | 0.954640091 | 0.698236641 | 0.827966596 | 177.4040837 | 169.2167923 | ENSG00000175352 |
| ADAL | -0.066988858 | 0.954628387 | 0.72753366 | 0.846909144 | 185.9603404 | 177.6641648 | ENSG00000168803 |
| MYO9B | -0.067028208 | 0.954602349 | 0.383563964 | 0.576294675 | 2256.307918 | 2154.775398 | ENSG00000099331 |
| PPP1CC | -0.067188646 | 0.954496197 | 0.361534819 | 0.554738499 | 6813.393753 | 6501.969842 | ENSG00000186298 |
| COL16A1 | -0.067356321 | 0.954385268 | 0.633733726 | 0.783961787 | 290.7750079 | 277.6692738 | ENSG00000084636 |
| OXA1L | -0.067565291 | 0.954247038 | 0.30997002 | 0.503769234 | 2180.330988 | 2080.612996 | ENSG00000155463 |
| C20orf204 | -0.067576907 | 0.954239355 | 0.924619288 | 0.962692763 | 10.80889998 | 10.3183882 | ENSG00000196421 |
| ATG2B | -0.067592701 | 0.954228908 | 0.627842808 | 0.780183614 | 340.1060529 | 324.2661529 | ENSG00000066739 |
| GBA2 | -0.067768126 | 0.954112886 | 0.28139166 | 0.47271345 | 2943.468452 | 2808.128793 | ENSG00000070610 |
| KIF11 | -0.06793097 | 0.954005196 | 0.471634979 | 0.656794853 | 1590.919919 | 1516.702432 | ENSG00000138160 |
| CXXC5 | -0.067949583 | 0.953992888 | 0.336624662 | 0.530978851 | 2824.68287 | 2694.623846 | ENSG00000171604 |
| MIER3 | -0.068035548 | 0.953936045 | 0.586580521 | 0.749312221 | 362.6717678 | 345.6381526 | ENSG00000155545 |
| SPR | -0.068062365 | 0.953918314 | 0.58972198 | 0.751934224 | 381.6788631 | 364.4768474 | ENSG00000116096 |
| ZDHHC20 | -0.068352811 | 0.953726288 | 0.555721277 | 0.725393397 | 1526.444071 | 1455.024259 | ENSG00000180776 |
| RABL3 | -0.068429666 | 0.953675483 | 0.602716968 | 0.761880879 | 391.310709 | 372.8539938 | ENSG00000144840 |
| RABGEF1 | -0.068535386 | 0.953605601 | 0.860079156 | 0.926192962 | 33.35497438 | 31.89339635 | ENSG00000154710 |
| DHX32 | -0.068609606 | 0.953556543 | 0.608519767 | 0.766449566 | 301.6731733 | 287.4487022 | ENSG00000089876 |
| RTCB | -0.068616657 | 0.953551883 | 0.305968133 | 0.500027633 | 3257.508333 | 3105.455182 | ENSG00000100220 |
| CNOT6L | -0.06862823 | 0.953544233 | 0.698656485 | 0.82815737 | 248.5628331 | 236.5285663 | ENSG00000138767 |
| KIAA1549 | -0.068636902 | 0.953538502 | 0.526880844 | 0.702967066 | 644.5022117 | 614.5408184 | ENSG00000122778 |
| TYMS | -0.068888277 | 0.953372372 | 0.618109843 | 0.772952151 | 292.5249209 | 278.9511174 | ENSG00000176890 |
| PSMC3IP | -0.068903078 | 0.953362591 | 0.683145187 | 0.817076206 | 313.6695374 | 299.565141 | ENSG00000131470 |
| GLRB | -0.069043283 | 0.953269945 | 0.935925736 | 0.968425646 | 6.437917191 | 6.083636043 | ENSG00000109738 |
| AP2S1 | -0.069069296 | 0.953252758 | 0.334587985 | 0.528981192 | 2732.092533 | 2603.934299 | ENSG00000042753 |
| CDCA3 | -0.069114658 | 0.953222785 | 0.493759834 | 0.675874798 | 593.0858696 | 565.4596527 | ENSG00000111665 |
| TTC38 | -0.069137552 | 0.953207659 | 0.442787388 | 0.631223609 | 884.7745475 | 843.4883865 | ENSG00000075234 |
| ACSL1 | -0.069213061 | 0.95315777 | 0.608316783 | 0.766318797 | 300.309621 | 286.1583849 | ENSG00000151726 |
| GRAMD2B | -0.069216259 | 0.953155657 | 0.714997724 | 0.838432369 | 150.7410074 | 143.4711575 | ENSG00000155324 |
| TMEM87A | -0.069219551 | 0.953153482 | 0.604627774 | 0.763311161 | 360.3754314 | 343.41439 | ENSG00000103978 |
| ATP9B | -0.069237092 | 0.953141893 | 0.596443059 | 0.756844835 | 399.0527082 | 380.3024731 | ENSG00000166377 |
| GPR137B | -0.069347147 | 0.953069187 | 0.747147847 | 0.858811237 | 115.96498 | 110.3966058 | ENSG00000077585 |
| SNRPA | -0.069366269 | 0.953056554 | 0.351013353 | 0.544317394 | 1997.300095 | 1903.528885 | ENSG00000077312 |
| TRIM23 | -0.069368914 | 0.953054807 | 0.689467671 | 0.82188023 | 189.9680077 | 181.073208 | ENSG00000113595 |
| PCDHB2 | -0.069507902 | 0.952962995 | 0.720881789 | 0.842237882 | 163.5895195 | 156.2818764 | ENSG00000112852 |
| SWI5 | -0.069859985 | 0.952730457 | 0.594483316 | 0.755547815 | 319.9911752 | 304.9549744 | ENSG00000175854 |
| IDI2-AS1 | -0.069989302 | 0.952645062 | 0.91245695 | 0.956487804 | 11.49639988 | 10.91092112 | ENSG00000232656 |
| FKBP1B | -0.07004361 | 0.952609202 | 0.801685106 | 0.893310392 | 63.95286121 | 60.79980613 | ENSG00000119782 |
| SS18L1 | -0.070124681 | 0.952555673 | 0.558981634 | 0.728050991 | 466.2987606 | 444.4603943 | ENSG00000184402 |
| NA | -0.07025602 | 0.952468959 | 0.695908067 | 0.826103895 | 166.347013 | 158.6592709 | ENSG00000254635 |
| EIF4E2 | -0.070420176 | 0.952360589 | 0.37369237 | 0.566976481 | 2157.614274 | 2055.33193 | ENSG00000135930 |
| LRP10 | -0.070435942 | 0.952350181 | 0.520735737 | 0.697803206 | 591.8470078 | 563.4985406 | ENSG00000197324 |
| MAPK8 | -0.070526392 | 0.952290475 | 0.512601969 | 0.691410658 | 781.1598288 | 743.1889618 | ENSG00000107643 |
| GCA | -0.070538007 | 0.952282809 | 0.616376543 | 0.771888321 | 471.8505104 | 449.1860833 | ENSG00000115271 |
| GTPBP3 | -0.070558792 | 0.952269089 | 0.417298476 | 0.608513765 | 1151.448334 | 1096.770682 | ENSG00000130299 |
| ZCCHC24 | -0.070601376 | 0.952240981 | 0.628792304 | 0.78073535 | 275.8774054 | 262.6334866 | ENSG00000165424 |
| NA | -0.070614221 | 0.952232503 | 0.940845215 | 0.970762241 | 5.035106525 | 4.808514081 | ENSG00000213453 |
| KIF1A | -0.070671491 | 0.952194703 | 0.212718759 | 0.392423238 | 13583.08625 | 12933.76418 | ENSG00000130294 |
| NAXD | -0.070822131 | 0.952095285 | 0.478349233 | 0.662348428 | 755.841416 | 720.1809233 | ENSG00000213995 |
| PFDN1 | -0.070916886 | 0.952032754 | 0.489420198 | 0.672218316 | 587.717258 | 559.4151589 | ENSG00000113068 |
| MRPL15 | -0.070997064 | 0.951979846 | 0.39088742 | 0.5834977 | 1211.319829 | 1152.606859 | ENSG00000137547 |
| TIMM9 | -0.071039806 | 0.951951642 | 0.699681646 | 0.828937588 | 171.9134802 | 163.5976446 | ENSG00000100575 |
| SP2 | -0.07105328 | 0.951942752 | 0.620014108 | 0.77440134 | 319.6541298 | 304.5378788 | ENSG00000167182 |
| FBN3 | -0.071065662 | 0.951934582 | 0.489504199 | 0.67228387 | 1849.519503 | 1761.66326 | ENSG00000142449 |
| R3HDM4 | -0.071140543 | 0.951885174 | 0.327531949 | 0.521697932 | 1757.594319 | 1672.837295 | ENSG00000198858 |
| PEG13 | -0.07127892 | 0.951793878 | 0.938316451 | 0.969600641 | 5.392761905 | 5.127294571 | ENSG00000282164 |
| NA | -0.071282109 | 0.951791774 | 0.874314927 | 0.935034013 | 23.77074512 | 22.62845379 | ENSG00000228506 |
| DNAJC10 | -0.07128243 | 0.951791562 | 0.410577544 | 0.602171734 | 2139.722658 | 2035.836003 | ENSG00000077232 |
| PDP2 | -0.071293543 | 0.951784231 | 0.481013004 | 0.664710335 | 621.8482235 | 592.233567 | ENSG00000172840 |
| ARPC2 | -0.071313452 | 0.951771096 | 0.286992288 | 0.478306614 | 4017.479753 | 3823.996085 | ENSG00000163466 |
| MTMR1 | -0.071326971 | 0.951762178 | 0.54955809 | 0.720961784 | 523.2226185 | 497.4684499 | ENSG00000063601 |
| NSD2 | -0.071343836 | 0.951751052 | 0.272503176 | 0.463610599 | 4935.514741 | 4697.214901 | ENSG00000109685 |
| LCA5 | -0.071478357 | 0.951662312 | 0.802970656 | 0.894151534 | 64.622021 | 61.33591445 | ENSG00000135338 |
| MLLT1 | -0.071627011 | 0.951564259 | 0.357082657 | 0.550616112 | 1712.803204 | 1630.361753 | ENSG00000130382 |
| C16orf70 | -0.071725944 | 0.951499007 | 0.580274522 | 0.74434377 | 357.7848256 | 340.7643424 | ENSG00000125149 |

| | | | | | | | |
|---|---|---|---|---|---|---|---|
| EPCAM | -0.071726279 | 0.951498786 | 0.425918288 | 0.616651967 | 3285.767512 | 3125.930267 | ENSG00000119888 |
| RGS19 | -0.071729526 | 0.951496645 | 0.655757955 | 0.799049633 | 232.0175189 | 220.5826053 | ENSG00000171700 |
| LMTK3 | -0.071735871 | 0.95149246 | 0.610156844 | 0.767371074 | 340.5077171 | 323.9212355 | ENSG00000142235 |
| ARF6 | -0.071745063 | 0.951486398 | 0.425476224 | 0.61626886 | 2276.961726 | 2165.936172 | ENSG00000165527 |
| CDC23 | -0.071761797 | 0.951475361 | 0.434151344 | 0.623909514 | 865.4593978 | 823.3080712 | ENSG00000094880 |
| NDFIP1 | -0.07196323 | 0.951342523 | 0.450004606 | 0.637741655 | 1229.436878 | 1169.163909 | ENSG00000131507 |
| NA | -0.07203464 | 0.951295435 | 0.923434951 | 0.962216415 | 8.287550914 | 7.86536006 | ENSG00000231747 |
| SF3B3 | -0.072057048 | 0.951280659 | 0.223936145 | 0.406387203 | 7564.432719 | 7195.518101 | ENSG00000189091 |
| ZNF136 | -0.072086517 | 0.951261229 | 0.746564782 | 0.858611443 | 112.3568344 | 107.0900029 | ENSG00000196646 |
| ZDHHC7 | -0.072120988 | 0.9512385 | 0.385090193 | 0.577938906 | 1442.386262 | 1371.442463 | ENSG00000153786 |
| ZNF234 | -0.072122667 | 0.951237392 | 0.711571229 | 0.836373711 | 139.5530528 | 132.8125183 | ENSG00000263002 |
| TP73-AS1 | -0.072123335 | 0.951236952 | 0.481389329 | 0.665131193 | 1098.263604 | 1045.486961 | ENSG00000227372 |
| FMNL2 | -0.072143776 | 0.951223475 | 0.358681905 | 0.551935738 | 1276.4796 | 1214.328253 | ENSG00000157827 |
| UBAC2-AS1 | -0.072144815 | 0.951222789 | 0.870576067 | 0.932827673 | 28.05947542 | 26.61918302 | ENSG00000228889 |
| SLC7A7 | -0.072180919 | 0.951198985 | 0.761630753 | 0.867730799 | 96.500997 | 91.96211294 | ENSG00000155465 |
| SPG7 | -0.072207273 | 0.951181609 | 0.334135759 | 0.528670131 | 2162.191456 | 2056.882247 | ENSG00000197912 |
| NA | -0.072238111 | 0.951161278 | 0.945949775 | 0.973329762 | 5.439514238 | 5.168635407 | ENSG00000286408 |
| GSTM1 | -0.072260777 | 0.951146335 | 0.78053242 | 0.879415677 | 76.37122172 | 72.53599818 | ENSG00000134184 |
| AGBL5 | -0.072268691 | 0.951141117 | 0.404601913 | 0.596238518 | 1087.70425 | 1034.58636 | ENSG00000084693 |
| ZBTB49 | -0.07230511 | 0.951117107 | 0.770429971 | 0.873601287 | 103.0625148 | 98.1520998 | ENSG00000168826 |
| SEC23B | -0.072332731 | 0.951098898 | 0.299281824 | 0.492794006 | 2157.742746 | 2052.245059 | ENSG00000101310 |
| CLIC4 | -0.072615875 | 0.950912253 | 0.429231046 | 0.619564961 | 6329.125964 | 6017.259979 | ENSG00000169504 |
| RAB28 | -0.072729923 | 0.950837084 | 0.553716047 | 0.723938212 | 373.7329876 | 355.2894735 | ENSG00000157869 |
| BCAM | -0.072781301 | 0.950803223 | 0.339213152 | 0.533427504 | 1754.336395 | 1668.064889 | ENSG00000187244 |
| RAB1A | -0.072806082 | 0.950786892 | 0.275527124 | 0.46644316 | 3190.918797 | 3033.529597 | ENSG00000138069 |
| BPTF | -0.072822397 | 0.95077614 | 0.298155595 | 0.491376116 | 3442.571953 | 3273.099168 | ENSG00000171634 |
| LSM14A | -0.072844737 | 0.950761417 | 0.282505285 | 0.473456276 | 4537.494537 | 4313.209657 | ENSG00000257103 |
| SPRED3 | -0.072898275 | 0.950726135 | 0.733557753 | 0.851224605 | 124.431445 | 118.0475608 | ENSG00000188766 |
| NA | -0.072941279 | 0.950697796 | 0.537337678 | 0.711203176 | 615.0428796 | 584.4310183 | ENSG00000218283 |
| MRFAP1 | -0.073006089 | 0.950655089 | 0.180697097 | 0.350281351 | 9880.664409 | 9392.981244 | ENSG00000179010 |
| PACRGL | -0.073038094 | 0.950634 | 0.640747295 | 0.789444284 | 213.2209295 | 202.486102 | ENSG00000163138 |
| NA | -0.073045136 | 0.950629359 | 0.902604196 | 0.950449757 | 16.16428259 | 15.37103683 | ENSG00000273702 |
| ZNF253 | -0.073122902 | 0.950578119 | 0.572888127 | 0.738883513 | 341.5589908 | 324.7291611 | ENSG00000256771 |
| H2AC14 | -0.073123268 | 0.950577878 | 0.948809772 | 0.974691536 | 4.04617431 | 3.900999518 | ENSG00000276368 |
| AGTPBP1 | -0.073125834 | 0.950576187 | 0.466467545 | 0.652589785 | 1046.185198 | 993.9530988 | ENSG00000135049 |
| PHTF2 | -0.073144674 | 0.950563774 | 0.592070723 | 0.753877355 | 374.7205217 | 355.6546675 | ENSG00000006576 |
| RASL10A | -0.073155706 | 0.950556505 | 0.862817185 | 0.928170742 | 29.01377929 | 27.56044072 | ENSG00000100276 |
| FBLN5 | -0.07318607 | 0.950536499 | 0.827965936 | 0.9073231 | 43.6614296 | 41.57422357 | ENSG00000140092 |
| NCAN | -0.073190863 | 0.950533341 | 0.66177589 | 0.803337329 | 208.3030946 | 198.3126642 | ENSG00000130287 |
| ACTR3B | -0.073432464 | 0.950374173 | 0.440503222 | 0.628699116 | 796.9326622 | 756.9058098 | ENSG00000133627 |
| NA | -0.073511808 | 0.950321907 | 0.873923134 | 0.934962559 | 23.7896866 | 22.73783056 | ENSG00000235023 |
| SDHAP2 | -0.073564149 | 0.95028743 | 0.758382227 | 0.86529526 | 111.774595 | 106.3177958 | ENSG00000242086 |
| WDR25 | -0.073808718 | 0.950126349 | 0.679983713 | 0.815136731 | 196.1298456 | 186.7877396 | ENSG00000176473 |
| NA | -0.073911396 | 0.95005873 | 0.892125471 | 0.944999342 | 18.38735626 | 17.4866251 | ENSG00000228944 |
| ALKBH4 | -0.073950646 | 0.950032882 | 0.586245474 | 0.749275703 | 354.8990787 | 337.4898422 | ENSG00000160993 |
| RBM41 | -0.073968883 | 0.950020873 | 0.513601652 | 0.692196991 | 488.0303822 | 463.7715729 | ENSG00000089682 |
| SYPL1 | -0.073969966 | 0.95002016 | 0.437428787 | 0.626268718 | 871.1560573 | 827.1557518 | ENSG00000008282 |
| MFSD8 | -0.073991444 | 0.950006017 | 0.716989435 | 0.839659101 | 118.3094669 | 112.3924084 | ENSG00000164073 |
| NA | -0.074063634 | 0.949958481 | 0.924531772 | 0.962655722 | 7.998746256 | 7.595406479 | ENSG00000220157 |
| SNAI1 | -0.074102326 | 0.949933004 | 0.684724363 | 0.818278506 | 157.6088807 | 149.5731265 | ENSG00000124216 |
| DMAC2L | -0.074180504 | 0.94988153 | 0.740825675 | 0.855075943 | 101.4197037 | 96.24481594 | ENSG00000125375 |
| USP27X-AS1 | -0.074184622 | 0.949878819 | 0.899317607 | 0.948444134 | 13.4085705 | 12.72270105 | ENSG00000234390 |
| PSMB6 | -0.074313227 | 0.949794149 | 0.24998682 | 0.43853283 | 2884.313004 | 2739.838005 | ENSG00000142507 |
| QRICH1 | -0.074389839 | 0.949743712 | 0.279881499 | 0.471039716 | 2167.476026 | 2058.383694 | ENSG00000198218 |
| ZNF787 | -0.074463649 | 0.949695124 | 0.400625133 | 0.592545385 | 1599.040673 | 1518.647896 | ENSG00000142409 |
| TBC1D14 | -0.074500042 | 0.949671167 | 0.42777983 | 0.618108004 | 817.2159235 | 776.0827173 | ENSG00000132405 |
| C12orf65 | -0.074562229 | 0.949630233 | 0.573480172 | 0.73914336 | 318.6381987 | 302.6327935 | ENSG00000130921 |
| NA | -0.074586523 | 0.949614242 | 0.829360077 | 0.908150754 | 41.37393989 | 39.32343949 | ENSG00000263266 |
| CRYAB | -0.074753185 | 0.949504547 | 0.899369033 | 0.948444134 | 15.79409726 | 14.98790763 | ENSG00000109846 |
| PLEKHM1 | -0.074766229 | 0.949495963 | 0.596790524 | 0.757026594 | 283.970641 | 269.6001323 | ENSG00000225190 |
| STT3B | -0.074792343 | 0.949478776 | 0.378514042 | 0.571960962 | 2690.823215 | 2553.725341 | ENSG00000163527 |
| PPP4R3A | -0.074825486 | 0.949456964 | 0.362738958 | 0.555712005 | 2001.847353 | 1899.872051 | ENSG00000100796 |
| NCKAP1 | -0.074840326 | 0.949447198 | 0.329073343 | 0.523433345 | 3481.808686 | 3305.023234 | ENSG00000061676 |
| STAG2 | -0.074890859 | 0.949413942 | 0.33808715 | 0.532153481 | 3431.634814 | 3256.850783 | ENSG00000101972 |
| IDH3B | -0.07498913 | 0.949349274 | 0.320133246 | 0.513843386 | 1867.78891 | 1774.031221 | ENSG00000101365 |
| PIK3C2A | -0.075030095 | 0.949322318 | 0.423305501 | 0.614180691 | 1783.63961 | 1692.264091 | ENSG00000011405 |
| LINS1 | -0.075040047 | 0.949315769 | 0.759953079 | 0.866500792 | 81.95169201 | 77.73480112 | ENSG00000140471 |
| ICA1 | -0.075049888 | 0.949309293 | 0.55931517 | 0.728383035 | 359.6180886 | 341.1648169 | ENSG00000003147 |
| CIITA | -0.075060967 | 0.949302003 | 0.91178025 | 0.956189185 | 10.58263222 | 10.04474739 | ENSG00000179583 |
| NA | -0.075111707 | 0.949268617 | 0.926901608 | 0.964343189 | 8.618673215 | 8.150285789 | ENSG00000223529 |
| RAB42 | -0.075211565 | 0.949202914 | 0.68222859 | 0.816506824 | 168.9743967 | 160.4182103 | ENSG00000188060 |
| CASTOR3 | -0.075251528 | 0.949176621 | 0.708328867 | 0.834063991 | 130.1153943 | 123.2804333 | ENSG00000239521 |
| CCNL1 | -0.075251694 | 0.949176512 | 0.414841808 | 0.606314808 | 824.0199821 | 782.403105 | ENSG00000163660 |
| ALG2 | -0.075270621 | 0.94916406 | 0.463815443 | 0.650253815 | 692.9451124 | 658.2635461 | ENSG00000119523 |

| | | | | | | | |
|---|---|---|---|---|---|---|---|
| POLR3A | -0.07531426 | 0.949135349 | 0.396875947 | 0.589098354 | 1421.420093 | 1348.661634 | ENSG00000148606 |
| ZNF773 | -0.075333482 | 0.949122704 | 0.784071958 | 0.881898496 | 82.16465571 | 77.81269625 | ENSG00000152439 |
| TCEAL8 | -0.075345687 | 0.949114674 | 0.388884165 | 0.581598708 | 2031.23552 | 1926.951772 | ENSG00000180964 |
| ZIC5 | -0.075369567 | 0.949098965 | 0.522887638 | 0.699624565 | 841.4891057 | 798.5290517 | ENSG00000139800 |
| NA | -0.075476751 | 0.949028454 | 0.944446951 | 0.972627649 | 4.463209675 | 4.276092992 | ENSG00000235748 |
| PEX14 | -0.075478557 | 0.949027266 | 0.425637864 | 0.616454807 | 792.7064536 | 752.104567 | ENSG00000142655 |
| BICDL1 | -0.075496757 | 0.949015294 | 0.612290369 | 0.768899634 | 624.2253845 | 593.1610409 | ENSG00000135127 |
| ATP5F1C | -0.07554915 | 0.94898083 | 0.302227405 | 0.495706436 | 2720.007084 | 2580.104404 | ENSG00000165629 |
| SNX22 | -0.075713115 | 0.948872983 | 0.768044986 | 0.871908897 | 85.11308959 | 80.9357547 | ENSG00000157734 |
| PNPLA4 | -0.075714215 | 0.94887226 | 0.525246151 | 0.701716439 | 487.3466557 | 462.2414695 | ENSG00000006757 |
| ABL1 | -0.07585483 | 0.94877978 | 0.211777995 | 0.391413379 | 3769.121774 | 3576.169279 | ENSG00000097007 |
| PLD2 | -0.075873638 | 0.948767412 | 0.402405788 | 0.594196834 | 1079.381665 | 1024.786784 | ENSG00000129219 |
| NR1H2 | -0.075954452 | 0.948714267 | 0.383605612 | 0.576294675 | 1169.368167 | 1109.143072 | ENSG00000131408 |
| TFAM | -0.075978963 | 0.948698149 | 0.453047665 | 0.640291705 | 1955.110063 | 1853.880183 | ENSG00000108064 |
| NBPF8 | -0.076110732 | 0.948611503 | 0.817705468 | 0.901961189 | 53.30385876 | 50.73391462 | ENSG00000270231 |
| GSS | -0.076123683 | 0.948602988 | 0.440141462 | 0.628424764 | 865.8579953 | 821.5859496 | ENSG00000100983 |
| MIS12 | -0.076181216 | 0.948565159 | 0.534409889 | 0.709148221 | 402.931198 | 382.0775955 | ENSG00000167842 |
| ECT2 | -0.076206993 | 0.948548211 | 0.438320201 | 0.626794245 | 1535.928497 | 1455.923546 | ENSG00000114346 |
| CDPF1 | -0.076250689 | 0.948519482 | 0.659170859 | 0.80143484 | 184.5852202 | 175.3359236 | ENSG00000205643 |
| PNP | -0.076253449 | 0.948517667 | 0.339730057 | 0.533877978 | 1399.723558 | 1328.046569 | ENSG00000198805 |
| FANCL | -0.076373272 | 0.948438891 | 0.542943964 | 0.715560224 | 534.9879719 | 508.1857929 | ENSG00000115392 |
| SURF6 | -0.076391197 | 0.948427108 | 0.250881326 | 0.439368278 | 2980.359571 | 2826.735276 | ENSG00000148296 |
| WFS1 | -0.076402838 | 0.948419455 | 0.4304656 | 0.620475861 | 1128.841031 | 1071.041946 | ENSG00000109501 |
| RAB9B | -0.076444886 | 0.948391813 | 0.878824485 | 0.937550551 | 19.75298303 | 18.64857145 | ENSG00000123570 |
| ZNF296 | -0.07647672 | 0.948370886 | 0.70844049 | 0.834089445 | 121.1449736 | 115.0446546 | ENSG00000170684 |
| TMEM51 | -0.076476799 | 0.948370835 | 0.571135437 | 0.737403287 | 315.8706331 | 299.9106879 | ENSG00000171729 |
| ADH5 | -0.076575792 | 0.948305763 | 0.211393531 | 0.390897706 | 5844.165888 | 5541.150191 | ENSG00000197894 |
| HSD17B4 | -0.07670108 | 0.948223412 | 0.185492333 | 0.356297533 | 7186.914303 | 6813.801791 | ENSG00000133835 |
| TMEM255B | -0.076709019 | 0.948218195 | 0.768985941 | 0.872292394 | 72.65173502 | 68.75153509 | ENSG00000184497 |
| INKA1 | -0.076903551 | 0.948090346 | 0.796916306 | 0.890505274 | 56.62643165 | 53.56452098 | ENSG00000185614 |
| TACO1 | -0.07694927 | 0.948060302 | 0.441614919 | 0.629897715 | 701.2503496 | 664.5213767 | ENSG00000136463 |
| STAMBP | -0.076965133 | 0.948049877 | 0.393661238 | 0.586309208 | 945.2837907 | 895.7125121 | ENSG00000124356 |
| NA | -0.077000683 | 0.948026516 | 0.946798238 | 0.973529524 | 3.88056431 | 3.615524136 | ENSG00000236861 |
| HNRNPUL1 | -0.077095139 | 0.947964449 | 0.180131986 | 0.349456474 | 6674.431403 | 6327.096548 | ENSG00000105323 |
| TCTE3 | -0.07712712 | 0.947943436 | 0.885069559 | 0.940967086 | 17.45790172 | 16.57867251 | ENSG00000184786 |
| NDUFAF4 | -0.07723096 | 0.947875208 | 0.521239777 | 0.698276691 | 590.7755899 | 559.5982653 | ENSG00000123545 |
| STAT3 | -0.077315138 | 0.947819904 | 0.264951766 | 0.454645173 | 2899.163712 | 2747.082829 | ENSG00000168610 |
| IFT172 | -0.077388191 | 0.947771911 | 0.562583063 | 0.730892646 | 351.3360622 | 333.3834526 | ENSG00000138002 |
| MAP2K5 | -0.077407375 | 0.947759308 | 0.589944417 | 0.75199655 | 266.1672306 | 252.1311789 | ENSG00000137764 |
| RB1 | -0.077444276 | 0.947735066 | 0.627134763 | 0.779408281 | 343.1380623 | 324.5244549 | ENSG00000139687 |
| RUFY1 | -0.077466368 | 0.947720554 | 0.494259165 | 0.676254094 | 542.5269744 | 514.3064842 | ENSG00000176783 |
| MRPL44 | -0.077474898 | 0.947714951 | 0.38169709 | 0.574586291 | 969.384185 | 918.0595126 | ENSG00000135900 |
| BRPF3 | -0.077530917 | 0.947678152 | 0.340514915 | 0.534884603 | 1215.165089 | 1151.359411 | ENSG00000096070 |
| NCBP3 | -0.077537974 | 0.947673516 | 0.361720329 | 0.554809321 | 1552.048883 | 1471.377395 | ENSG00000074356 |
| BFAR | -0.077596617 | 0.947634996 | 0.300414738 | 0.493745352 | 1776.76604 | 1682.981615 | ENSG00000103429 |
| MEMO1 | -0.077598666 | 0.94763365 | 0.730783863 | 0.849344242 | 115.0312635 | 109.0202544 | ENSG00000162959 |
| ARMC4 | -0.077658124 | 0.947594596 | 0.690486571 | 0.822503443 | 130.6210937 | 123.8212165 | ENSG00000169126 |
| SDCCAG8 | -0.07766611 | 0.94758935 | 0.749456067 | 0.860354212 | 89.26717766 | 84.41509534 | ENSG00000054282 |
| NA | -0.077689309 | 0.947574113 | 0.865357841 | 0.929978419 | 25.74545264 | 24.52335342 | ENSG00000243176 |
| CACNG5 | -0.077695973 | 0.947569736 | 0.752355391 | 0.862347236 | 86.55901466 | 82.19166763 | ENSG00000075429 |
| NUDT22 | -0.077698956 | 0.947567777 | 0.558423263 | 0.727526558 | 349.654594 | 331.4278999 | ENSG00000149761 |
| DCX | -0.078061164 | 0.947329907 | 0.738064462 | 0.853248923 | 98.51929994 | 93.53019636 | ENSG00000077279 |
| RPL21 | -0.078073716 | 0.947321665 | 0.194354173 | 0.368356708 | 11536.25945 | 10928.19302 | ENSG00000122026 |
| IRF7 | -0.078099926 | 0.947304455 | 0.650594799 | 0.795936006 | 175.8446301 | 166.3954955 | ENSG00000185507 |
| PELP1 | -0.078150272 | 0.947271397 | 0.175230493 | 0.34255057 | 5589.489848 | 5294.749512 | ENSG00000141456 |
| NBN | -0.07817173 | 0.947257308 | 0.477581945 | 0.661871178 | 668.0839328 | 632.5111175 | ENSG00000104320 |
| NA | -0.078248837 | 0.947206682 | 0.768476467 | 0.871924202 | 142.2171393 | 135.0659305 | ENSG00000185986 |
| CD47 | -0.078250042 | 0.947205891 | 0.59400033 | 0.755268722 | 275.5439382 | 260.9095013 | ENSG00000196776 |
| DUSP11 | -0.078252553 | 0.947204242 | 0.591540696 | 0.753357745 | 307.2936011 | 290.8945145 | ENSG00000144048 |
| EPOR | -0.078422601 | 0.947092603 | 0.615064598 | 0.770668544 | 246.0697008 | 232.7622616 | ENSG00000187266 |
| CARD10 | -0.07848295 | 0.947052986 | 0.398544988 | 0.590322714 | 1099.399633 | 1040.858398 | ENSG00000100065 |
| MTO1 | -0.078494425 | 0.947045454 | 0.515263388 | 0.693226745 | 496.1370648 | 469.2539387 | ENSG00000135297 |
| DYNLL1 | -0.078578653 | 0.946990164 | 0.205462179 | 0.3839133 | 4434.479857 | 4199.404822 | ENSG00000088986 |
| HNRNPL | -0.078593736 | 0.946980264 | 0.140658302 | 0.291930794 | 9603.70102 | 9094.074259 | ENSG00000104824 |
| ZNF202 | -0.078721363 | 0.946896494 | 0.400203612 | 0.59215821 | 942.2866277 | 892.9235588 | ENSG00000166261 |
| CDADC1 | -0.078724554 | 0.9468944 | 0.779412762 | 0.878795238 | 61.88149211 | 58.71157423 | ENSG00000102543 |
| CROCC | -0.078730577 | 0.946890446 | 0.371273451 | 0.564401422 | 2180.196601 | 2065.593395 | ENSG00000058453 |
| ZNRD2 | -0.07880983 | 0.946838431 | 0.435716447 | 0.625189179 | 700.8122006 | 663.9688479 | ENSG00000173465 |
| TCEANC2 | -0.078874258 | 0.946796148 | 0.611196766 | 0.768150869 | 229.8888596 | 217.7421398 | ENSG00000116205 |
| PSMB2 | -0.078904481 | 0.946776314 | 0.235617563 | 0.420544675 | 2698.59967 | 2555.130007 | ENSG00000126067 |
| MUL1 | -0.078931525 | 0.946758567 | 0.456009821 | 0.643266615 | 713.6273772 | 674.9087482 | ENSG00000090432 |
| PACSIN2 | -0.07893938 | 0.946753412 | 0.246429754 | 0.434032981 | 2292.879251 | 2171.249225 | ENSG00000100266 |
| CASC2 | -0.078953763 | 0.946743973 | 0.933880634 | 0.96733214 | 5.058482691 | 4.789743086 | ENSG00000177640 |
| ELOB | -0.078996294 | 0.946716063 | 0.296076282 | 0.488949283 | 2709.935126 | 2565.709923 | ENSG00000103363 |

| | | | | | | | |
|---|---|---|---|---|---|---|---|
| NA | -0.079000526 | 0.946713286 | 0.870137543 | 0.932664263 | 26.22384482 | 25.00738564 | ENSG00000255753 |
| GJC1 | -0.079041329 | 0.946686511 | 0.268667247 | 0.45889584 | 1707.790068 | 1617.172963 | ENSG00000182963 |
| NA | -0.079086912 | 0.9466566 | 0.66760466 | 0.80717496 | 190.9555645 | 181.0335077 | ENSG00000248641 |
| PALM2AKAP2 | -0.079202056 | 0.946581049 | 0.267715814 | 0.45786836 | 2020.758516 | 1912.985972 | ENSG00000157654 |
| KLK5 | -0.079256865 | 0.946545088 | 0.928639564 | 0.964710817 | 6.559931771 | 6.260172686 | ENSG00000167754 |
| NA | -0.079313897 | 0.946507671 | 0.933309348 | 0.967249799 | 6.192707954 | 5.821168536 | ENSG00000249485 |
| STIP1 | -0.079351292 | 0.946483137 | 0.179373308 | 0.348590216 | 11678.19458 | 11053.01605 | ENSG00000168439 |
| PEX11B | -0.079405207 | 0.946447767 | 0.73572317 | 0.852173027 | 104.6741249 | 98.93372811 | ENSG00000131779 |
| ALOXE3 | -0.079405807 | 0.946447373 | 0.931291013 | 0.966327826 | 5.852114914 | 5.506186892 | ENSG00000179148 |
| RAPGEF5 | -0.079539745 | 0.94635951 | 0.637521987 | 0.787197369 | 229.9980359 | 217.5101259 | ENSG00000136237 |
| PTTG1IP | -0.079674292 | 0.946271256 | 0.184872679 | 0.355586368 | 3581.570005 | 3388.906866 | ENSG00000183255 |
| PHF6 | -0.079686091 | 0.946263517 | 0.37265706 | 0.565994096 | 1909.554457 | 1806.108902 | ENSG00000156531 |
| DENND10 | -0.079780291 | 0.946201733 | 0.437550104 | 0.626268718 | 754.2761674 | 713.3388011 | ENSG00000119979 |
| GNPAT | -0.079828489 | 0.946170123 | 0.357261636 | 0.550678478 | 1045.482777 | 988.7398105 | ENSG00000116906 |
| NA | -0.079830971 | 0.946168496 | 0.933659732 | 0.96733214 | 5.144395755 | 4.868625911 | ENSG00000284728 |
| GUSBP1 | -0.079909075 | 0.946117273 | 0.763424028 | 0.868883644 | 80.36423217 | 76.23238051 | ENSG00000183666 |
| HINFP | -0.079916229 | 0.946112582 | 0.459496292 | 0.646251652 | 615.8434119 | 582.5284094 | ENSG00000172273 |
| ZFYVE19 | -0.079942986 | 0.946095035 | 0.587843001 | 0.750389758 | 291.2280471 | 275.7683138 | ENSG00000166140 |
| ZNF26 | -0.08019431 | 0.945930235 | 0.664727397 | 0.805285154 | 175.6476464 | 166.1106814 | ENSG00000198393 |
| NA | -0.080204727 | 0.945923405 | 0.93602529 | 0.968425646 | 4.783583461 | 4.546046573 | ENSG00000287252 |
| FANCF | -0.080217257 | 0.945915189 | 0.583054116 | 0.746519084 | 319.6666598 | 302.2673445 | ENSG00000183161 |
| FILIP1 | -0.080222394 | 0.945911822 | 0.922140868 | 0.961408497 | 6.99718639 | 6.657836001 | ENSG00000118407 |
| RFXANK | -0.08023807 | 0.945901543 | 0.463742521 | 0.650200762 | 573.3907782 | 542.8700635 | ENSG00000064490 |
| MAP2K2 | -0.080316955 | 0.945849824 | 0.27624523 | 0.467322632 | 3099.6324 | 2931.226397 | ENSG00000126934 |
| ZNF672 | -0.080339958 | 0.945834743 | 0.3623074 | 0.555267746 | 955.2407683 | 903.4141662 | ENSG00000171161 |
| MASP2 | -0.08044514 | 0.945765788 | 0.897950944 | 0.947703593 | 13.37760273 | 12.62124838 | ENSG00000009724 |
| GPM6A | -0.0804709 | 0.945748901 | 0.839248683 | 0.913691856 | 31.51669054 | 29.7548002 | ENSG00000150625 |
| PSMB3 | -0.080571303 | 0.945683085 | 0.282013221 | 0.473069106 | 2832.6126 | 2678.098604 | ENSG00000277791 |
| ARMCX4 | -0.080609989 | 0.945657727 | 0.546614524 | 0.718495899 | 385.0243237 | 364.214483 | ENSG00000196440 |
| PELO | -0.080625393 | 0.945647629 | 0.40753728 | 0.599317105 | 847.6230049 | 802.1450775 | ENSG00000152684 |
| ATG12 | -0.080636373 | 0.945640433 | 0.426927943 | 0.617451345 | 577.7166924 | 546.3782491 | ENSG00000145782 |
| CCDC126 | -0.080666319 | 0.945620804 | 0.814676232 | 0.900075694 | 45.79049493 | 43.27238985 | ENSG00000169193 |
| LOC728752 | -0.080723594 | 0.945583264 | 0.878947339 | 0.937577553 | 18.94662546 | 17.86507939 | ENSG00000267309 |
| SELENOF | -0.08073997 | 0.945572531 | 0.28295668 | 0.473955636 | 2075.229161 | 1961.263752 | ENSG00000183291 |
| FAHD2A | -0.080820448 | 0.945519785 | 0.422489393 | 0.613572983 | 678.8748387 | 641.8409987 | ENSG00000115042 |
| METTL26 | -0.080967477 | 0.945423429 | 0.39628159 | 0.588563426 | 1106.795589 | 1047.000747 | ENSG00000130731 |
| NA | -0.080987102 | 0.945410569 | 0.907878077 | 0.954214147 | 10.62307072 | 10.02966362 | ENSG00000286389 |
| EDC3 | -0.081077681 | 0.945351213 | 0.330236737 | 0.524563566 | 981.7028247 | 927.9993836 | ENSG00000179151 |
| MPP7 | -0.081087242 | 0.945344949 | 0.613679134 | 0.769548651 | 228.9349238 | 216.1566708 | ENSG00000150054 |
| DDX31 | -0.081171048 | 0.945290035 | 0.451075283 | 0.638580864 | 603.4774637 | 570.3421515 | ENSG00000125485 |
| PXYLP1 | -0.081200069 | 0.94527102 | 0.500834771 | 0.68147741 | 426.4917682 | 402.6457863 | ENSG00000155893 |
| MRPL52 | -0.08124943 | 0.945238679 | 0.344945664 | 0.538376668 | 948.8340069 | 896.954266 | ENSG00000172590 |
| SMN1 | -0.0813977 | 0.945141539 | 0.811622275 | 0.898880668 | 45.38471175 | 42.89404718 | ENSG00000205571 |
| LOC112268389 | -0.081410587 | 0.945133096 | 0.877636255 | 0.936718131 | 16.78302947 | 15.91474284 | ENSG00000255052 |
| RASL11A | -0.081421652 | 0.945125848 | 0.825735991 | 0.906340611 | 42.44572596 | 40.24290024 | ENSG00000122035 |
| KIF2A | -0.081525779 | 0.945057635 | 0.338176485 | 0.532203699 | 995.6599385 | 940.7657986 | ENSG00000068796 |
| HMGN1 | -0.081548754 | 0.945042585 | 0.250618615 | 0.439032648 | 7489.34914 | 7077.027909 | ENSG00000205581 |
| POU2F3 | -0.08155788 | 0.945036607 | 0.864785814 | 0.929684296 | 23.56973266 | 22.23827655 | ENSG00000137709 |
| SF3B6 | -0.081567771 | 0.945030128 | 0.4310636 | 0.62101304 | 985.6292317 | 930.6275537 | ENSG00000115128 |
| NOMO2 | -0.081683949 | 0.944954029 | 0.311474021 | 0.50519467 | 1438.43187 | 1359.508622 | ENSG00000185164 |
| UBA1 | -0.081694745 | 0.944946958 | 0.123294469 | 0.264882311 | 18443.09054 | 17427.71287 | ENSG00000130985 |
| KBTBD2 | -0.081725187 | 0.944927019 | 0.477482972 | 0.661805968 | 542.5861438 | 512.3522138 | ENSG00000170852 |
| STK11IP | -0.081793413 | 0.944882334 | 0.485117446 | 0.668289485 | 596.5495939 | 564.428594 | ENSG00000144589 |
| CBR1 | -0.081817794 | 0.944866366 | 0.391277147 | 0.583891357 | 807.6300105 | 762.5142767 | ENSG00000159228 |
| MAP2K7 | -0.081830843 | 0.94485782 | 0.295290696 | 0.488173915 | 1375.327353 | 1299.834001 | ENSG00000076984 |
| FAM160A2 | -0.081949372 | 0.944780195 | 0.447691444 | 0.635824769 | 511.1856577 | 482.7857764 | ENSG00000051009 |
| ZFAND5 | -0.081989872 | 0.944753673 | 0.357293057 | 0.550678478 | 2279.888858 | 2153.244019 | ENSG00000107372 |
| DTX3L | -0.082130548 | 0.944661556 | 0.672193195 | 0.809514534 | 154.2391647 | 145.402723 | ENSG00000163840 |
| SNX3 | -0.082187892 | 0.944624008 | 0.272909135 | 0.464130841 | 1539.962742 | 1454.698463 | ENSG00000112335 |
| ZNHIT1 | -0.082209182 | 0.944610068 | 0.345659308 | 0.539172666 | 1158.293673 | 1093.337921 | ENSG00000106400 |
| RBM22 | -0.082237565 | 0.944591485 | 0.405461222 | 0.597125013 | 722.989578 | 683.2856891 | ENSG00000086589 |
| NECAB3 | -0.082244391 | 0.944587016 | 0.537510209 | 0.711203176 | 318.2040783 | 300.80378 | ENSG00000125967 |
| NA | -0.08224759 | 0.944584921 | 0.93695073 | 0.969033102 | 4.449304247 | 4.238551003 | ENSG00000241889 |
| LDHC | -0.08232057 | 0.94453714 | 0.904550929 | 0.951850731 | 11.25128834 | 10.68230837 | ENSG00000166796 |
| UBAC2 | -0.082374862 | 0.944501595 | 0.390331308 | 0.583043235 | 1119.83499 | 1056.722992 | ENSG00000134882 |
| NA | -0.0823792 | 0.944498755 | 0.850037462 | 0.920406282 | 25.07689421 | 23.68624792 | ENSG00000237126 |
| SMIM24 | -0.082379622 | 0.944498479 | 0.77454763 | 0.87565364 | 70.0401359 | 66.28429925 | ENSG00000095932 |
| LRRC4B | -0.08256102 | 0.944379729 | 0.54074654 | 0.713882497 | 606.1271859 | 572.4776898 | ENSG00000131409 |
| LINC01668 | -0.082645156 | 0.944324656 | 0.937402974 | 0.969088951 | 4.671137318 | 4.422135687 | ENSG00000283051 |
| KLF13 | -0.082678933 | 0.944302547 | 0.296486077 | 0.489451585 | 1296.066604 | 1224.107928 | ENSG00000169926 |
| NA | -0.082693196 | 0.944293212 | 0.928564649 | 0.964687063 | 5.484988795 | 5.157350486 | ENSG00000227014 |
| SOD3 | -0.082729991 | 0.944269128 | 0.758331222 | 0.86529526 | 81.48592748 | 76.67233207 | ENSG00000109610 |
| MAPKAPK3 | -0.082731423 | 0.944268191 | 0.235006816 | 0.419939869 | 2048.28279 | 1934.05058 | ENSG00000114738 |
| ABCC5 | -0.08277862 | 0.9442373 | 0.425852175 | 0.616651967 | 1202.091422 | 1135.094201 | ENSG00000114770 |

| | | | | | | | |
|---|---|---|---|---|---|---|---|
| TERT | -0.082782876 | 0.944234515 | 0.472280165 | 0.657495724 | 504.6237717 | 476.6974025 | ENSG00000164362 |
| NUCKS1 | -0.0828618 | 0.944182861 | 0.240852789 | 0.427051143 | 11581.6881 | 10933.92773 | ENSG00000069275 |
| TENM4 | -0.082945581 | 0.944128031 | 0.608505522 | 0.766449566 | 273.9850633 | 258.194395 | ENSG00000149256 |
| CAMK2G | -0.083047263 | 0.944061491 | 0.389440946 | 0.582018075 | 770.0556602 | 727.1916178 | ENSG00000148660 |
| SMG9 | -0.083193788 | 0.943965614 | 0.480747685 | 0.664541885 | 503.6803141 | 475.8433898 | ENSG00000105771 |
| PI4KAP1 | -0.083193829 | 0.943965587 | 0.716644095 | 0.839556455 | 149.7440026 | 141.6926198 | ENSG00000274602 |
| ATP6V0B | -0.083205395 | 0.943958019 | 0.266538343 | 0.456479547 | 1743.52839 | 1645.525982 | ENSG00000117410 |
| MED19 | -0.083251231 | 0.943928029 | 0.555727897 | 0.725393397 | 348.7738687 | 329.0734645 | ENSG00000156603 |
| SARAF | -0.083267605 | 0.943917316 | 0.337724946 | 0.531944773 | 1504.521005 | 1419.685879 | ENSG00000133872 |
| RFLNB | -0.083309991 | 0.943889584 | 0.315516636 | 0.509025309 | 1120.043549 | 1057.175296 | ENSG00000183688 |
| COX3 | -0.083413128 | 0.943822109 | 0.629624205 | 0.781401836 | 47199.94339 | 44548.23508 | ENSG00000198938 |
| CHMP7 | -0.083514071 | 0.943756074 | 0.273208209 | 0.4643838 | 1332.221758 | 1256.975632 | ENSG00000147457 |
| LSM6 | -0.083523401 | 0.94374997 | 0.365705437 | 0.558778613 | 840.5155446 | 793.6883552 | ENSG00000164167 |
| MCM3AP-AS1 | -0.083539741 | 0.943739281 | 0.764088293 | 0.869397288 | 66.4103956 | 62.73235938 | ENSG00000215424 |
| LUC7L3 | -0.083681773 | 0.943646376 | 0.635393757 | 0.785563864 | 2947.366454 | 2781.674671 | ENSG00000108848 |
| MAPKBP1 | -0.083725478 | 0.943617789 | 0.544846036 | 0.716894535 | 302.6835048 | 285.6799036 | ENSG00000137802 |
| SMIM15 | -0.083726643 | 0.943617028 | 0.571056327 | 0.737373586 | 393.4161352 | 370.885284 | ENSG00000188725 |
| FANCD2 | -0.083750436 | 0.943601465 | 0.286614786 | 0.478009399 | 1774.96624 | 1674.530364 | ENSG00000144554 |
| MAST2 | -0.08375709 | 0.943597113 | 0.295347783 | 0.488195003 | 1437.401763 | 1356.691905 | ENSG00000086015 |
| LSM14B | -0.083848051 | 0.943537622 | 0.253350229 | 0.442104617 | 1898.827284 | 1791.402217 | ENSG00000149657 |
| PSME3IP1 | -0.083879588 | 0.943516996 | 0.327461671 | 0.521675658 | 1131.856173 | 1068.403661 | ENSG00000172775 |
| FNBP4 | -0.083949448 | 0.943471309 | 0.375183568 | 0.568118344 | 1040.847058 | 982.4708468 | ENSG00000109920 |
| KIF24 | -0.084053047 | 0.943403562 | 0.65142407 | 0.796404599 | 160.2154665 | 151.1305863 | ENSG00000186638 |
| TGFBRAP1 | -0.084153296 | 0.94333801 | 0.382608591 | 0.575216388 | 804.0996077 | 758.1037523 | ENSG00000135966 |
| PKIA | -0.084158724 | 0.94333446 | 0.547243164 | 0.718925773 | 336.9187986 | 317.5783726 | ENSG00000171033 |
| MED26 | -0.084240002 | 0.943281317 | 0.669424973 | 0.807856359 | 138.1244831 | 130.2418544 | ENSG00000105085 |
| RNGTT | -0.08424652 | 0.943277055 | 0.47635694 | 0.66089522 | 663.0022321 | 624.7825535 | ENSG00000111880 |
| LRRC37A | -0.084307153 | 0.943237412 | 0.883188195 | 0.939883441 | 17.28785704 | 16.3273783 | ENSG00000176681 |
| MYH10 | -0.084353977 | 0.943206799 | 0.193041079 | 0.366655396 | 9853.462855 | 9293.882284 | ENSG00000133026 |
| NAMPT | -0.084474906 | 0.943127741 | 0.43299459 | 0.622850174 | 1592.109458 | 1500.551112 | ENSG00000105835 |
| STK26 | -0.084504379 | 0.943108474 | 0.293229296 | 0.485806004 | 4034.837761 | 3804.297211 | ENSG00000134602 |
| PFN1 | -0.084556873 | 0.943074159 | 0.168361233 | 0.332382519 | 13514.25027 | 12744.91291 | ENSG00000108518 |
| SOCS3 | -0.084648395 | 0.943014334 | 0.467156074 | 0.653162956 | 520.527812 | 490.8783722 | ENSG00000184557 |
| FARSA | -0.084801739 | 0.942914106 | 0.204187303 | 0.382224772 | 3666.502391 | 3458.033137 | ENSG00000179115 |
| PPARD | -0.084822629 | 0.942900453 | 0.37279293 | 0.566107674 | 682.523385 | 643.4442107 | ENSG00000112033 |
| RPL30 | -0.084835733 | 0.942891889 | 0.291447215 | 0.483761659 | 2820.599331 | 2659.847362 | ENSG00000156482 |
| KAT6B | -0.084894504 | 0.942853479 | 0.437134283 | 0.626145331 | 525.4385438 | 495.6153694 | ENSG00000156650 |
| CLUH | -0.084916359 | 0.942839196 | 0.197610725 | 0.373422219 | 4378.841644 | 4129.709851 | ENSG00000132361 |
| SLC39A14 | -0.084944551 | 0.942820772 | 0.158187862 | 0.317610524 | 6870.374999 | 6476.355997 | ENSG00000104635 |
| ZNF431 | -0.085172791 | 0.942671626 | 0.456676644 | 0.643890155 | 533.7089346 | 502.856397 | ENSG00000196705 |
| MFHAS1 | -0.085222894 | 0.942638888 | 0.310877997 | 0.504706837 | 1595.122248 | 1502.906255 | ENSG00000147324 |
| LAP3 | -0.085236032 | 0.942630305 | 0.244313391 | 0.431124858 | 1815.19916 | 1711.365217 | ENSG00000002549 |
| VCAN | -0.085251323 | 0.942620314 | 0.219542838 | 0.40098305 | 5017.093189 | 4729.014162 | ENSG00000038427 |
| PAFAH1B2 | -0.085252749 | 0.942619382 | 0.317206189 | 0.510858486 | 1785.008559 | 1681.60214 | ENSG00000168092 |
| PAQR9 | -0.085285264 | 0.942598138 | 0.846712695 | 0.918413939 | 28.05749858 | 26.31157583 | ENSG00000188582 |
| NA | -0.085328103 | 0.942570149 | 0.927223007 | 0.964343189 | 6.497297176 | 6.057378975 | ENSG00000276007 |
| WDR12 | -0.085372073 | 0.942541422 | 0.281444243 | 0.47271345 | 1826.928562 | 1720.983885 | ENSG00000138442 |
| PPP1R37 | -0.085480761 | 0.942470417 | 0.382328041 | 0.575026598 | 850.5104203 | 802.4237281 | ENSG00000104866 |
| NA | -0.085546875 | 0.942427227 | 0.927880397 | 0.964537269 | 5.733354945 | 5.359706164 | ENSG00000261614 |
| SORD | -0.085660259 | 0.942353163 | 0.268266198 | 0.458464335 | 1410.821195 | 1329.670607 | ENSG00000140263 |
| NA | -0.085669672 | 0.942347015 | 0.871222556 | 0.933190511 | 20.42913306 | 19.34723194 | ENSG00000260276 |
| NA | -0.085715612 | 0.942317008 | 0.888315426 | 0.942849507 | 12.8493013 | 12.09262614 | ENSG00000250397 |
| MMP17 | -0.085800191 | 0.942261765 | 0.405278322 | 0.596903085 | 1152.816502 | 1086.602184 | ENSG00000198598 |
| EML4 | -0.085815946 | 0.942251475 | 0.22070757 | 0.402221642 | 2159.479182 | 2034.3118 | ENSG00000143924 |
| KCNN3 | -0.085853975 | 0.942226638 | 0.748859311 | 0.859882192 | 71.37971061 | 67.30075255 | ENSG00000143603 |
| PUM3 | -0.085875803 | 0.942212382 | 0.239460805 | 0.425361465 | 1662.39533 | 1566.269399 | ENSG00000080608 |
| ARHGEF10L | -0.08602506 | 0.942114909 | 0.275846484 | 0.466822508 | 1292.735335 | 1217.85752 | ENSG00000074964 |
| GLMP | -0.08610978 | 0.942059586 | 0.488526767 | 0.671339445 | 349.9571087 | 329.7777876 | ENSG00000198715 |
| ZFP64 | -0.086165576 | 0.942023153 | 0.405729199 | 0.597424723 | 600.6286902 | 566.0481636 | ENSG00000020256 |
| C5orf15 | -0.086212189 | 0.941992717 | 0.504743144 | 0.684288598 | 599.4237425 | 564.5353736 | ENSG00000113583 |
| HECW2 | -0.086213348 | 0.941991961 | 0.683357467 | 0.817274562 | 114.5741956 | 107.7520688 | ENSG00000138411 |
| EXTL3-AS1 | -0.08626788 | 0.941956355 | 0.927301197 | 0.964343189 | 7.833136256 | 7.374279807 | ENSG00000246339 |
| ABCG4 | -0.0863041 | 0.941932706 | 0.793660051 | 0.888170326 | 48.34657004 | 45.57937529 | ENSG00000172350 |
| TMEM165 | -0.086332648 | 0.941914068 | 0.334210592 | 0.528670131 | 1454.049586 | 1369.094488 | ENSG00000134851 |
| HAP1 | -0.086364433 | 0.941893316 | 0.640502662 | 0.789444284 | 159.7852021 | 150.7869744 | ENSG00000173805 |
| POLR3E | -0.086415288 | 0.941860115 | 0.239065922 | 0.424858357 | 2294.574993 | 2161.503477 | ENSG00000058600 |
| HSD17B12 | -0.08644068 | 0.941843538 | 0.499465235 | 0.68021814 | 501.845728 | 472.3674326 | ENSG00000149084 |
| CTBP1 | -0.086589963 | 0.941746086 | 0.278245426 | 0.469415687 | 1799.665591 | 1694.429243 | ENSG00000159692 |
| MRPL51 | -0.086593041 | 0.941744076 | 0.215662882 | 0.395955923 | 2214.680592 | 2085.113688 | ENSG00000111639 |
| NA | -0.086611212 | 0.941732215 | 0.819376852 | 0.90291228 | 34.85514569 | 32.80036126 | ENSG00000214820 |
| NA | -0.086702115 | 0.941672879 | 0.702965904 | 0.831263 | 111.4631883 | 104.7328765 | ENSG00000226085 |
| CCDC102A | -0.086885927 | 0.94155291 | 0.654054435 | 0.79772604 | 202.9365575 | 191.2221076 | ENSG00000135736 |
| UNC79 | -0.086933658 | 0.941521759 | 0.76638578 | 0.870835943 | 69.57189087 | 65.59411251 | ENSG00000133958 |
| SCRIB | -0.086970338 | 0.941497821 | 0.205588134 | 0.384011032 | 4157.4595 | 3914.880051 | ENSG00000180900 |

| | | | | | | | |
|---|---|---|---|---|---|---|---|
| GATD1 | -0.087012829 | 0.941470093 | 0.714742091 | 0.838366554 | 994.397753 | 936.5036935 | ENSG00000177225 |
| CXorf40B | -0.087098697 | 0.941414059 | 0.382285756 | 0.575026598 | 658.397946 | 619.3549633 | ENSG00000197021 |
| FBXL5 | -0.087248431 | 0.941316357 | 0.309699448 | 0.503645563 | 1051.088894 | 989.3670658 | ENSG00000118564 |
| HNRNPA3 | -0.087385717 | 0.941226786 | 0.105475674 | 0.236040345 | 14834.61023 | 13962.56202 | ENSG00000170144 |
| FKBP5 | -0.087393639 | 0.941221618 | 0.224839383 | 0.407667103 | 1921.896849 | 1808.315048 | ENSG00000096060 |
| PCMTD2 | -0.087484921 | 0.941162067 | 0.335753579 | 0.530010803 | 879.4057405 | 827.4523292 | ENSG00000203880 |
| TEDC1 | -0.087589444 | 0.941093882 | 0.516644459 | 0.694239253 | 361.5587539 | 340.2861557 | ENSG00000185347 |
| NA | -0.087819452 | 0.940943856 | 0.660069825 | 0.802054293 | 137.2276801 | 129.2721897 | ENSG00000182484 |
| DEPDC5 | -0.08782214 | 0.940942102 | 0.63687034 | 0.786554426 | 184.8692069 | 174.176024 | ENSG00000100150 |
| MARS1 | -0.08784694 | 0.940925928 | 0.105166575 | 0.235576179 | 9059.297864 | 8524.094595 | ENSG00000166986 |
| CACNA1I | -0.087938736 | 0.940866061 | 0.746540228 | 0.858611443 | 84.5048056 | 79.73484064 | ENSG00000100346 |
| SERINC2 | -0.087980662 | 0.940838718 | 0.340321011 | 0.534670646 | 934.7448365 | 879.4505902 | ENSG00000168528 |
| MFSD6L | -0.088031959 | 0.940805266 | 0.836565399 | 0.912220332 | 28.45629153 | 26.70175307 | ENSG00000185156 |
| RPL21P28 | -0.088033333 | 0.94080437 | 0.869915552 | 0.932505196 | 16.79881403 | 15.82088786 | ENSG00000220749 |
| COA7 | -0.088049636 | 0.940793739 | 0.250177766 | 0.438674996 | 1382.803255 | 1300.717623 | ENSG00000162377 |
| RPGR | -0.088203101 | 0.940693668 | 0.621461008 | 0.775152125 | 158.5788715 | 149.2139928 | ENSG00000156313 |
| MLEC | -0.088305512 | 0.940626895 | 0.134012056 | 0.28143548 | 6389.186114 | 6009.406248 | ENSG00000110917 |
| C1orf174 | -0.088346471 | 0.94060019 | 0.404844188 | 0.596509035 | 746.6204116 | 702.2379085 | ENSG00000198912 |
| WDR86 | -0.088354554 | 0.94059492 | 0.516280698 | 0.694018523 | 351.7680627 | 331.0470725 | ENSG00000187260 |
| NA | -0.088365181 | 0.940587992 | 0.869854149 | 0.932493308 | 18.04558314 | 17.0514198 | ENSG00000212664 |
| IFI16 | -0.088377519 | 0.940579948 | 0.906101926 | 0.952995863 | 9.640452335 | 9.080919846 | ENSG00000163565 |
| ZNRF1 | -0.088394055 | 0.940569167 | 0.552369761 | 0.723161339 | 551.3348744 | 518.2019093 | ENSG00000186187 |
| UBLCP1 | -0.088398431 | 0.940566314 | 0.499912406 | 0.680577092 | 586.7774231 | 551.4630717 | ENSG00000164332 |
| TMOD2 | -0.088421108 | 0.94055153 | 0.68981797 | 0.822139172 | 99.7311476 | 93.71334301 | ENSG00000128872 |
| FAM193A | -0.088440951 | 0.940538594 | 0.41503849 | 0.606501568 | 718.3191397 | 675.9406838 | ENSG00000125386 |
| NA | -0.088456249 | 0.94052862 | 0.864930413 | 0.929688585 | 17.18931632 | 16.16592542 | ENSG00000259953 |
| CHMP1B | -0.088456422 | 0.940528508 | 0.408849883 | 0.600404094 | 852.8765171 | 802.8876123 | ENSG00000255112 |
| C8orf37 | -0.088494484 | 0.940503694 | 0.79139308 | 0.886482527 | 44.73321566 | 42.16675649 | ENSG00000156172 |
| NPM2 | -0.088502434 | 0.940498512 | 0.840810655 | 0.91458329 | 26.44873711 | 24.76974948 | ENSG00000158806 |
| ZCCHC2 | -0.088566588 | 0.94045669 | 0.523076651 | 0.699675423 | 353.2054866 | 332.0031504 | ENSG00000141664 |
| ARHGEF2 | -0.088641952 | 0.940407564 | 0.234438332 | 0.41940922 | 1408.890774 | 1324.726236 | ENSG00000116584 |
| RPL22L1 | -0.088651351 | 0.940401437 | 0.301372627 | 0.494867229 | 1157.340211 | 1088.219108 | ENSG00000163584 |
| WBP11 | -0.088679273 | 0.940383237 | 0.154363499 | 0.311992704 | 3325.688108 | 3127.406973 | ENSG00000084463 |
| FBXL12 | -0.088772085 | 0.940322741 | 0.353006693 | 0.546129052 | 707.6388386 | 665.5864739 | ENSG00000127452 |
| SAP18 | -0.088825876 | 0.940287682 | 0.224665425 | 0.407446035 | 3950.358737 | 3714.388593 | ENSG00000150459 |
| ADPRHL2 | -0.088879689 | 0.94025261 | 0.361819643 | 0.554809321 | 649.4458654 | 610.8930969 | ENSG00000116863 |
| DHX36 | -0.08894131 | 0.94021245 | 0.30263114 | 0.496102354 | 1601.739124 | 1505.49345 | ENSG00000174953 |
| TIGD3 | -0.088942303 | 0.940211803 | 0.86546176 | 0.929978419 | 23.23357431 | 22.00305379 | ENSG00000173825 |
| ACOT7 | -0.088979898 | 0.940187302 | 0.186753847 | 0.35812683 | 2638.6676 | 2480.035871 | ENSG00000097021 |
| NA | -0.088992255 | 0.94017925 | 0.708155892 | 0.83401045 | 94.87693198 | 89.33678504 | ENSG00000214223 |
| PRUNE1 | -0.089013693 | 0.940165279 | 0.426877042 | 0.617429303 | 662.8887485 | 623.066859 | ENSG00000143363 |
| NA | -0.089031296 | 0.940153808 | 0.928759376 | 0.964781206 | 4.717889651 | 4.440906681 | ENSG00000215861 |
| CSMD2 | -0.089057227 | 0.94013691 | 0.49705899 | 0.678137012 | 451.4380942 | 424.9191604 | ENSG00000121904 |
| ANKRD26 | -0.089076281 | 0.940124493 | 0.623818882 | 0.776902241 | 160.5016929 | 150.6797475 | ENSG00000107890 |
| HMOX2 | -0.089087924 | 0.940116906 | 0.305520534 | 0.499516371 | 1053.727793 | 990.4109786 | ENSG00000103415 |
| FAM98B | -0.089089672 | 0.940115767 | 0.408875669 | 0.600404094 | 1224.022349 | 1149.905817 | ENSG00000171262 |
| IRF3 | -0.089169483 | 0.94006376 | 0.20832621 | 0.387432465 | 1714.895364 | 1612.192659 | ENSG00000126456 |
| LRIG3 | -0.089181924 | 0.940055654 | 0.542558256 | 0.71520446 | 249.3418835 | 234.2437529 | ENSG00000139263 |
| HS3ST5 | -0.089186074 | 0.94005295 | 0.671907898 | 0.809472822 | 169.0589191 | 159.08914 | ENSG00000249853 |
| NUS1 | -0.089197331 | 0.940045615 | 0.256737155 | 0.445373122 | 1901.620323 | 1786.831115 | ENSG00000153989 |
| LLPH | -0.089285717 | 0.939988025 | 0.649114522 | 0.795207122 | 214.1256172 | 200.9489103 | ENSG00000139233 |
| POLG | -0.089328875 | 0.939959906 | 0.254310318 | 0.44298754 | 1798.420324 | 1691.08172 | ENSG00000140521 |
| ABHD15 | -0.08943945 | 0.939887866 | 0.844333866 | 0.917016342 | 23.51163046 | 22.15234568 | ENSG00000168792 |
| PEX6 | -0.089443688 | 0.939885105 | 0.347537228 | 0.540766501 | 814.6076389 | 765.7357703 | ENSG00000124587 |
| MZT2A | -0.089457265 | 0.93987626 | 0.304993868 | 0.498875329 | 1717.410925 | 1614.363254 | ENSG00000173272 |
| AGO1 | -0.089587903 | 0.939791157 | 0.203782874 | 0.381660448 | 2414.696022 | 2268.888239 | ENSG00000092847 |
| RPS17 | -0.089617757 | 0.939771709 | 0.613486399 | 0.769359017 | 174.8847115 | 164.499322 | ENSG00000182774 |
| ADCY1 | -0.089623624 | 0.939767888 | 0.2869843 | 0.478306614 | 1187.798638 | 1116.645093 | ENSG00000164742 |
| GLOD4 | -0.089958614 | 0.939549701 | 0.22360442 | 0.405983965 | 1603.103503 | 1505.984781 | ENSG00000167699 |
| RBM5 | -0.089970014 | 0.939542277 | 0.39392589 | 0.586514558 | 595.5940873 | 559.9964556 | ENSG00000003756 |
| ARL3 | -0.089975515 | 0.939538695 | 0.338347381 | 0.532382236 | 791.1407991 | 742.8960406 | ENSG00000138175 |
| PPP4R3B | -0.089989376 | 0.939529668 | 0.320528957 | 0.514344908 | 1158.312073 | 1087.38944 | ENSG00000275052 |
| MTX1 | -0.089989493 | 0.939529591 | 0.445353886 | 0.633452623 | 580.6818729 | 545.4149712 | ENSG00000173171 |
| NA | -0.089993517 | 0.939526971 | 0.847494464 | 0.918786988 | 22.43678518 | 21.18103206 | ENSG00000231982 |
| SC5D | -0.090059396 | 0.939484069 | 0.317989159 | 0.511824082 | 1658.336888 | 1556.922754 | ENSG00000109929 |
| NA | -0.090140609 | 0.939431185 | 0.93120826 | 0.966296058 | 4.433519683 | 4.189724094 | ENSG00000233155 |
| GSTM2 | -0.090228939 | 0.939373669 | 0.801375574 | 0.893214577 | 39.8617423 | 37.53846628 | ENSG00000213366 |
| ST3GAL2 | -0.090337882 | 0.939302737 | 0.284639735 | 0.475699986 | 1222.017982 | 1147.377044 | ENSG00000157350 |
| TESC | -0.090349463 | 0.939295197 | 0.602489487 | 0.7617532 | 175.2405854 | 164.8217898 | ENSG00000088992 |
| CHORDC1 | -0.090527986 | 0.939178973 | 0.278049458 | 0.46917049 | 1815.451194 | 1704.479205 | ENSG00000110172 |
| KDM4D | -0.090532412 | 0.939176092 | 0.814653801 | 0.900075694 | 36.56264328 | 34.42866813 | ENSG00000186280 |
| ZNF550 | -0.090574918 | 0.939148421 | 0.60614085 | 0.764778101 | 166.9099201 | 156.6592314 | ENSG00000251369 |
| SKOR2 | -0.090744653 | 0.939037936 | 0.92571345 | 0.963290836 | 5.372542652 | 5.010869759 | ENSG00000215474 |
| TRNT1 | -0.090787645 | 0.939009953 | 0.429305478 | 0.619564961 | 499.7977503 | 469.7185161 | ENSG00000072756 |

| | | | | | | | |
|---|---|---|---|---|---|---|---|
| NA | -0.090806745 | 0.938997522 | 0.897519014 | 0.947625465 | 10.16243994 | 9.515575496 | ENSG00000225205 |
| RFC4 | -0.090868604 | 0.93895726 | 0.333642853 | 0.5282536 | 1326.351807 | 1246.177913 | ENSG00000163918 |
| FRG1 | -0.090872315 | 0.938954845 | 0.494292339 | 0.676254094 | 350.8133529 | 329.221371 | ENSG00000109536 |
| NA | -0.090948871 | 0.938905022 | 0.331039967 | 0.525254238 | 778.5748831 | 730.9550166 | ENSG00000281566 |
| SMIM20 | -0.090982786 | 0.93888295 | 0.335614993 | 0.529882297 | 775.0949682 | 727.4423221 | ENSG00000250317 |
| TP53BP2 | -0.091132461 | 0.938785548 | 0.274085664 | 0.465193891 | 1131.299167 | 1061.461584 | ENSG00000143514 |
| APPBP2 | -0.091166909 | 0.938763133 | 0.512352993 | 0.691254284 | 330.4358128 | 309.8749119 | ENSG00000062725 |
| FH | -0.091176817 | 0.938756686 | 0.251774886 | 0.440433764 | 1433.080088 | 1345.055517 | ENSG00000091483 |
| IARS1 | -0.091391546 | 0.938616973 | 0.105894056 | 0.236719386 | 7723.141401 | 7248.405117 | ENSG00000196305 |
| CYP27A1 | -0.091410632 | 0.938604556 | 0.657224065 | 0.799701868 | 124.4845111 | 116.7881754 | ENSG00000135929 |
| TRIP6 | -0.091421777 | 0.938597305 | 0.223111496 | 0.405287509 | 2794.504265 | 2623.26856 | ENSG00000087077 |
| CCDC106 | -0.09143616 | 0.938587948 | 0.486815066 | 0.669985548 | 430.7121136 | 404.8229524 | ENSG00000173581 |
| NSF | -0.09149194 | 0.938551659 | 0.422982542 | 0.614000503 | 503.4595482 | 472.8970828 | ENSG00000073969 |
| CNNM3 | -0.091523666 | 0.93853102 | 0.350767206 | 0.544026727 | 940.24337 | 883.0436964 | ENSG00000168763 |
| PSME4 | -0.091534148 | 0.938524201 | 0.238284688 | 0.42383574 | 2778.389419 | 2606.868729 | ENSG00000068878 |
| SAMD4B | -0.091604709 | 0.9384783 | 0.146585333 | 0.300765256 | 3147.103754 | 2953.480396 | ENSG00000179134 |
| SCAMP3 | -0.091608927 | 0.938475556 | 0.296834389 | 0.489677146 | 928.9276582 | 872.0527847 | ENSG00000116521 |
| LOC339260 | -0.091665049 | 0.938439049 | 0.426279368 | 0.616998189 | 454.1899277 | 426.3653796 | ENSG00000233098 |
| DMAC1 | -0.091708026 | 0.938411094 | 0.289727308 | 0.481726556 | 1022.46125 | 959.3107675 | ENSG00000137038 |
| FAM117A | -0.091719358 | 0.938403723 | 0.435653084 | 0.62514666 | 619.1522595 | 580.8989169 | ENSG00000121104 |
| GNL2 | -0.091732613 | 0.938395101 | 0.257851005 | 0.446392079 | 2653.884087 | 2490.19397 | ENSG00000134697 |
| MIER1 | -0.091823224 | 0.938336165 | 0.504115853 | 0.683938454 | 524.9750492 | 492.1357334 | ENSG00000198160 |
| NA | -0.091913625 | 0.93827737 | 0.791639019 | 0.886486471 | 40.5569315 | 38.06665048 | ENSG00000236565 |
| NA | -0.091967438 | 0.938242373 | 0.934181608 | 0.967539644 | 4.489742754 | 4.231064929 | ENSG00000254428 |
| TFIP11 | -0.092011587 | 0.938213661 | 0.386506703 | 0.579384926 | 622.8920033 | 584.5885607 | ENSG00000100109 |
| NA | -0.092042768 | 0.938193384 | 0.816135506 | 0.901069606 | 42.14805192 | 39.68042324 | ENSG00000230715 |
| FAM171A1 | -0.092042809 | 0.938193357 | 0.270363133 | 0.460858117 | 1478.726298 | 1386.867225 | ENSG00000148468 |
| SKA1 | -0.092141646 | 0.938129085 | 0.552583459 | 0.723326635 | 317.918626 | 297.8018926 | ENSG00000154839 |
| CLBA1 | -0.092172132 | 0.938109261 | 0.774013188 | 0.875300505 | 50.86909918 | 47.70613687 | ENSG00000140104 |
| THOC1 | -0.092284932 | 0.938035916 | 0.320515094 | 0.514344908 | 831.3276882 | 779.6801947 | ENSG00000079134 |
| FAM155B | -0.092350237 | 0.937993456 | 0.337009457 | 0.531367421 | 733.1663892 | 687.6132682 | ENSG00000130054 |
| CDK12 | -0.092384006 | 0.937971501 | 0.227147652 | 0.410699011 | 1857.63198 | 1741.887612 | ENSG00000167258 |
| NA | -0.092454049 | 0.937925964 | 0.909026486 | 0.954717339 | 7.397159414 | 6.935275656 | ENSG00000235370 |
| LINC01572 | -0.092493871 | 0.937900075 | 0.814770345 | 0.900075694 | 36.19404398 | 33.95635887 | ENSG00000261008 |
| PRKCB | -0.092536335 | 0.937872469 | 0.498348018 | 0.679245637 | 310.0345133 | 290.5700085 | ENSG00000166501 |
| ZC3H8 | -0.09254336 | 0.937867902 | 0.664044509 | 0.804615646 | 123.9532482 | 116.5193211 | ENSG00000144161 |
| FIZ1 | -0.092577319 | 0.937845826 | 0.491164651 | 0.673915134 | 431.6156213 | 405.0404632 | ENSG00000179943 |
| UBE2B | -0.092607911 | 0.93782594 | 0.486890868 | 0.669985548 | 334.8096669 | 314.0276437 | ENSG00000119048 |
| ABCC1 | -0.092665731 | 0.937788355 | 0.211081951 | 0.390555345 | 2506.895399 | 2350.634618 | ENSG00000103222 |
| ST6GALNAC4 | -0.092698339 | 0.937767159 | 0.400851621 | 0.592682812 | 485.5008628 | 455.4824041 | ENSG00000136840 |
| DSC2 | -0.092854693 | 0.937665532 | 0.361520533 | 0.554738499 | 657.2007779 | 615.8623623 | ENSG00000134755 |
| ELP1 | -0.092874146 | 0.93765289 | 0.258206408 | 0.446917965 | 1457.985983 | 1367.232502 | ENSG00000070061 |
| SZRD1 | -0.093041064 | 0.93754441 | 0.167887987 | 0.331695551 | 2384.31712 | 2235.185215 | ENSG00000055070 |
| PRUNE2 | -0.09305884 | 0.937532858 | 0.490069749 | 0.672761479 | 418.3591765 | 392.5640784 | ENSG00000106772 |
| GPM6B | -0.093079645 | 0.937519339 | 0.202590885 | 0.379927093 | 1927.636338 | 1806.747912 | ENSG00000046653 |
| GABPA | -0.093117222 | 0.93749492 | 0.42695891 | 0.617451345 | 446.5775421 | 418.3952774 | ENSG00000154727 |
| ZNF281 | -0.093150178 | 0.937473505 | 0.204290224 | 0.382301596 | 1649.12525 | 1545.693405 | ENSG00000162702 |
| NA | -0.093183159 | 0.937452074 | 0.888857662 | 0.943208972 | 11.52924679 | 10.84376125 | ENSG00000146677 |
| ZBED5 | -0.093248051 | 0.937409908 | 0.409328219 | 0.600862443 | 623.4196132 | 584.3036031 | ENSG00000236287 |
| TBP | -0.093258879 | 0.937402873 | 0.42009177 | 0.611384442 | 492.8810576 | 461.9461978 | ENSG00000112592 |
| OTUD5 | -0.093362289 | 0.937335683 | 0.239546184 | 0.425467079 | 1529.671275 | 1434.693165 | ENSG00000068308 |
| NFATC4 | -0.093363793 | 0.937334706 | 0.48990895 | 0.672740052 | 570.2855212 | 535.0350228 | ENSG00000100968 |
| AGPAT4 | -0.093410622 | 0.937304282 | 0.359585045 | 0.552891092 | 671.1757434 | 629.0497036 | ENSG00000026652 |
| PPP1R8 | -0.093447723 | 0.937280178 | 0.269120206 | 0.459415482 | 1007.851024 | 944.3652513 | ENSG00000117751 |
| CHD3 | -0.093613425 | 0.937172532 | 0.231072572 | 0.415473327 | 1477.533692 | 1384.562532 | ENSG00000170004 |
| RPLP1 | -0.093632919 | 0.937159868 | 0.094062305 | 0.216269787 | 22384.71303 | 20977.36926 | ENSG00000137818 |
| MRPS28 | -0.093641497 | 0.937154296 | 0.680558659 | 0.815245229 | 99.24723835 | 93.19275649 | ENSG00000147586 |
| ANKIB1 | -0.093661069 | 0.937141583 | 0.341847629 | 0.536114736 | 853.4396423 | 799.2215862 | ENSG00000001629 |
| CENPW | -0.09379929 | 0.937051802 | 0.342286904 | 0.536395153 | 664.8959423 | 623.2562263 | ENSG00000203760 |
| ARSK | -0.09383609 | 0.9370279 | 0.488182905 | 0.671127481 | 362.0161075 | 339.2406342 | ENSG00000164291 |
| CYP2A13 | -0.093927487 | 0.93696854 | 0.830175192 | 0.90859717 | 26.29387562 | 24.51108082 | ENSG00000197838 |
| WNT5B | -0.093942206 | 0.936958981 | 0.756328048 | 0.864335767 | 65.98812879 | 61.79456566 | ENSG00000111186 |
| EIF3K | -0.094104806 | 0.936853386 | 0.13999248 | 0.290974613 | 3240.64481 | 3035.779108 | ENSG00000178982 |
| LOC101928424 | -0.094125167 | 0.936840164 | 0.919811773 | 0.960515777 | 5.431922636 | 5.048411747 | ENSG00000255337 |
| ARSJ | -0.094242737 | 0.936763821 | 0.669653614 | 0.80797247 | 114.4710248 | 107.1493503 | ENSG00000180801 |
| GTF2H1 | -0.094503237 | 0.93659469 | 0.48312304 | 0.666483247 | 423.7603116 | 397.1255033 | ENSG00000110768 |
| RUNX3 | -0.094519769 | 0.936583957 | 0.824820054 | 0.905753491 | 30.49165681 | 28.48336547 | ENSG00000020633 |
| CMAS | -0.094654902 | 0.936496234 | 0.285781718 | 0.477092269 | 1514.490648 | 1418.145146 | ENSG00000111726 |
| EXOC2 | -0.094706241 | 0.936462909 | 0.243684188 | 0.43036012 | 1214.390684 | 1137.270911 | ENSG00000112685 |
| PEA15 | -0.094833808 | 0.936380108 | 0.18287269 | 0.353293381 | 1675.815536 | 1569.208435 | ENSG00000162734 |
| EXOSC5 | -0.094860904 | 0.936362522 | 0.247901895 | 0.435838903 | 1148.241665 | 1074.954995 | ENSG00000077348 |
| EYA1 | -0.094918564 | 0.936325099 | 0.855294823 | 0.923132672 | 18.66423232 | 17.4898743 | ENSG00000104313 |
| NELFA | -0.095046542 | 0.936242043 | 0.243549946 | 0.430235347 | 1596.107008 | 1494.220857 | ENSG00000185049 |
| ADHFE1 | -0.095073416 | 0.936224604 | 0.93886703 | 0.96979109 | 4.010170492 | 3.780775858 | ENSG00000147576 |

| | | | | | | | |
|---|---|---|---|---|---|---|---|
| NA | -0.095107116 | 0.936202734 | 0.54343704 | 0.71590462 | 288.845369 | 270.8140919 | ENSG00000185596 |
| SNRPB | -0.095145679 | 0.93617771 | 0.160336519 | 0.320707646 | 3142.037935 | 2940.555419 | ENSG00000125835 |
| P4HB | -0.095145732 | 0.936177676 | 0.080307572 | 0.192227888 | 7332.920517 | 6864.457805 | ENSG00000185624 |
| MIR1244-2 | -0.095149962 | 0.936174931 | 0.706331247 | 0.833246982 | 93.38367586 | 87.25753627 | ENSG00000283498 |
| GGA2 | -0.095194679 | 0.936145914 | 0.251480218 | 0.440084446 | 2541.145388 | 2379.631517 | ENSG00000103365 |
| LRFN2 | -0.095309043 | 0.936071708 | 0.781335434 | 0.879945976 | 43.14397438 | 40.3544269 | ENSG00000156564 |
| PSMC2 | -0.095425904 | 0.935995887 | 0.13992306 | 0.290911017 | 2637.366607 | 2468.809213 | ENSG00000161057 |
| PIGB | -0.095448647 | 0.935981133 | 0.841151841 | 0.914793347 | 27.94377466 | 26.19245147 | ENSG00000069943 |
| HNRNPUL2 | -0.09561553 | 0.93587287 | 0.3063156 | 0.500330777 | 790.4460007 | 739.365348 | ENSG00000214753 |
| WDR20 | -0.09566546 | 0.935840481 | 0.398454595 | 0.590322714 | 469.8497735 | 439.8632106 | ENSG00000140153 |
| ARHGEF16 | -0.095767226 | 0.93577447 | 0.35909339 | 0.552318414 | 683.7510021 | 640.4735591 | ENSG00000130762 |
| CCDC71 | -0.095901829 | 0.935687167 | 0.364526174 | 0.557636838 | 563.3029991 | 527.2301236 | ENSG00000177352 |
| LINC02323 | -0.096049654 | 0.935591297 | 0.906717248 | 0.953480708 | 7.648682477 | 7.122659186 | ENSG00000259230 |
| MPHOSPH10 | -0.096120381 | 0.935545431 | 0.332873408 | 0.527395772 | 742.850655 | 694.726466 | ENSG00000124383 |
| TSPAN13 | -0.096253937 | 0.935458828 | 0.425973623 | 0.616651967 | 649.7843461 | 607.5425964 | ENSG00000106537 |
| DOLK | -0.096286869 | 0.935437475 | 0.420330808 | 0.611558221 | 502.8533313 | 470.1789589 | ENSG00000175283 |
| SUCLG1 | -0.096312624 | 0.935420775 | 0.36377914 | 0.556819635 | 684.2525902 | 639.3409184 | ENSG00000163541 |
| STAG3L5P | -0.096332246 | 0.935408053 | 0.752615473 | 0.862591997 | 112.3808119 | 105.4948266 | ENSG00000242294 |
| KIF5A | -0.096336381 | 0.935405372 | 0.381198544 | 0.574121064 | 758.589935 | 709.8732159 | ENSG00000155980 |
| NA | -0.09639251 | 0.93536898 | 0.917651867 | 0.959390744 | 5.769358763 | 5.337136323 | ENSG00000256356 |
| SMARCB1 | -0.096450343 | 0.935331485 | 0.12801154 | 0.272165062 | 3581.588127 | 3349.725684 | ENSG00000099956 |
| SGSH | -0.096473172 | 0.935316685 | 0.471282992 | 0.65650199 | 329.5811395 | 308.6506954 | ENSG00000181523 |
| DGKB | -0.096489069 | 0.935306378 | 0.807163684 | 0.896619594 | 40.68506451 | 37.93536514 | ENSG00000136267 |
| BLOC1S6 | -0.096592019 | 0.935239637 | 0.577268735 | 0.741962465 | 423.3916896 | 395.4537368 | ENSG00000104164 |
| NAT2 | -0.096601293 | 0.935233626 | 0.835530994 | 0.911837216 | 24.94295104 | 23.37115466 | ENSG00000156006 |
| PTPRG-AS1 | -0.096712993 | 0.935161218 | 0.821552768 | 0.904297073 | 28.16688551 | 26.36095239 | ENSG00000241472 |
| NA | -0.096722659 | 0.935154953 | 0.934615066 | 0.967772258 | 3.744742001 | 3.552163113 | ENSG00000273148 |
| NA | -0.096758431 | 0.935131766 | 0.903621727 | 0.951196972 | 7.390845588 | 6.897733667 | ENSG00000256167 |
| KCNJ3 | -0.096809443 | 0.935098701 | 0.924809422 | 0.962782555 | 4.892872691 | 4.576102488 | ENSG00000162989 |
| DERL1 | -0.096887687 | 0.935047988 | 0.375760991 | 0.568849318 | 800.8514026 | 747.8896993 | ENSG00000136986 |
| DDOST | -0.096937099 | 0.935015963 | 0.085921231 | 0.202217662 | 6430.558018 | 6012.040178 | ENSG00000244038 |
| NA | -0.096939777 | 0.935014228 | 0.791571448 | 0.886486471 | 37.63560942 | 35.17461819 | ENSG00000227097 |
| RASGEF1A | -0.097014184 | 0.934966005 | 0.415055912 | 0.606501568 | 448.3406614 | 419.1275695 | ENSG00000198915 |
| ZBTB11 | -0.097047027 | 0.934944721 | 0.512249357 | 0.691228947 | 252.1760147 | 235.6389869 | ENSG00000066422 |
| SRD5A1 | -0.097062436 | 0.934934735 | 0.397879973 | 0.589910045 | 553.8394015 | 517.410087 | ENSG00000145545 |
| EVA1C | -0.097067936 | 0.934931171 | 0.802372294 | 0.893861159 | 31.89772208 | 29.74731413 | ENSG00000166979 |
| MON1B | -0.09710901 | 0.934904554 | 0.295396782 | 0.488218651 | 918.6758551 | 859.1864713 | ENSG00000103111 |
| RER1 | -0.097150988 | 0.934877351 | 0.249276726 | 0.437756392 | 1109.415615 | 1037.667871 | ENSG00000157916 |
| DAAM1 | -0.097232085 | 0.934824801 | 0.451670456 | 0.639173874 | 430.6802741 | 402.1041186 | ENSG00000100592 |
| VSTM4 | -0.097252925 | 0.934811298 | 0.774235395 | 0.875418579 | 45.83409035 | 42.89024833 | ENSG00000165633 |
| NA | -0.097371603 | 0.934734402 | 0.750848867 | 0.861339687 | 60.3179121 | 56.34480337 | ENSG00000223546 |
| NA | -0.097379077 | 0.93472956 | 0.656814994 | 0.799514587 | 113.9927303 | 106.6549092 | ENSG00000242474 |
| SNU13 | -0.097471746 | 0.93466952 | 0.128800897 | 0.27333556 | 2648.082355 | 2475.385873 | ENSG00000100138 |
| KIF21B | -0.097501334 | 0.934650352 | 0.181459527 | 0.351465239 | 1906.703174 | 1782.395282 | ENSG00000116852 |
| LTA4H | -0.097553628 | 0.934616474 | 0.123203904 | 0.26474912 | 2733.275367 | 2554.133392 | ENSG00000111144 |
| H3P47 | -0.09757747 | 0.934601028 | 0.855659324 | 0.92336477 | 16.8221902 | 15.69317813 | ENSG00000179101 |
| CSTF3 | -0.097804858 | 0.934453734 | 0.288489593 | 0.480179616 | 886.3434492 | 827.9659793 | ENSG00000176102 |
| ANKRD39 | -0.097843573 | 0.934428659 | 0.729415731 | 0.848386575 | 60.69913904 | 56.7382298 | ENSG00000213337 |
| UBE2G1 | -0.097854325 | 0.934421694 | 0.198911479 | 0.375152677 | 2738.482345 | 2558.135612 | ENSG00000132388 |
| UPF2 | -0.097908292 | 0.934386741 | 0.2594391 | 0.448305031 | 1164.448994 | 1088.220955 | ENSG00000151461 |
| NKTR | -0.098000339 | 0.934327127 | 0.4208405 | 0.611993552 | 1069.740724 | 1000.327851 | ENSG00000114857 |
| GOLPH3 | -0.09800611 | 0.93432339 | 0.258672467 | 0.447432151 | 1898.805982 | 1773.235263 | ENSG00000113384 |
| STX1A | -0.098031482 | 0.934306958 | 0.689636753 | 0.822028913 | 134.4909095 | 126.0070393 | ENSG00000106089 |
| IGHMBP2 | -0.098118476 | 0.934250622 | 0.396577477 | 0.588906876 | 514.2992357 | 480.5124712 | ENSG00000132740 |
| BBOX1-AS1 | -0.098143295 | 0.93423455 | 0.75377412 | 0.863332686 | 52.89747422 | 49.35266508 | ENSG00000254560 |
| PSMB1 | -0.098144704 | 0.934233637 | 0.113620121 | 0.249388362 | 3579.271594 | 3343.068518 | ENSG00000008018 |
| PLEKHJ1 | -0.098207153 | 0.934193199 | 0.156554169 | 0.315222074 | 1993.53127 | 1862.888943 | ENSG00000104886 |
| MAT2A | -0.098222249 | 0.934183424 | 0.698435771 | 0.828043791 | 3869.996852 | 3615.525837 | ENSG00000168906 |
| USP35 | -0.098226454 | 0.934180701 | 0.496652842 | 0.678082046 | 253.432931 | 236.855646 | ENSG00000118369 |
| NOC2L | -0.098247176 | 0.934167283 | 0.095498156 | 0.218838336 | 5225.198525 | 4881.816462 | ENSG00000188976 |
| ANKRD17 | -0.09833211 | 0.934112289 | 0.225778153 | 0.408815554 | 2706.470899 | 2527.900991 | ENSG00000132466 |
| FRMPD3 | -0.098483641 | 0.934014181 | 0.676933488 | 0.813270653 | 96.62399626 | 90.36864852 | ENSG00000147234 |
| RPL10 | -0.09849578 | 0.934006322 | 0.076896249 | 0.186032511 | 9765.917877 | 9120.951184 | ENSG00000147403 |
| NA | -0.098558694 | 0.933965592 | 0.904961714 | 0.952174874 | 7.820606303 | 7.38176588 | ENSG00000228960 |
| AJAP1 | -0.098614641 | 0.933929374 | 0.493818147 | 0.675904699 | 306.4652428 | 286.059855 | ENSG00000196581 |
| HCFC1 | -0.098694198 | 0.933877874 | 0.174827804 | 0.342015889 | 4719.945649 | 4407.894137 | ENSG00000172534 |
| PAQR8 | -0.098704669 | 0.933871096 | 0.511880903 | 0.690983297 | 290.1578546 | 271.2476482 | ENSG00000170915 |
| PACC1 | -0.098732469 | 0.933853101 | 0.468274091 | 0.654129636 | 323.5467582 | 302.1416017 | ENSG00000065600 |
| NA | -0.098740255 | 0.933848061 | 0.858300583 | 0.925030122 | 15.49897877 | 14.52319606 | ENSG00000229692 |
| ZNF467 | -0.098811278 | 0.93380209 | 0.904680277 | 0.951932795 | 7.036347121 | 6.571467103 | ENSG00000181444 |
| ARMH1 | -0.098845349 | 0.933780037 | 0.925711756 | 0.963290836 | 4.446147334 | 4.163467026 | ENSG00000198520 |
| STC1 | -0.098917407 | 0.933733399 | 0.516951527 | 0.694439342 | 314.584523 | 293.1753545 | ENSG00000159167 |
| FKBP7 | -0.098974644 | 0.933696355 | 0.757905307 | 0.865159875 | 48.24467702 | 44.97187023 | ENSG00000079150 |
| GSE1 | -0.099009306 | 0.933673923 | 0.185625483 | 0.356406749 | 2566.09829 | 2395.486766 | ENSG00000131149 |

| | | | | | | | |
|---|---|---|---|---|---|---|---|
| GNA11 | -0.099009878 | 0.933673552 | 0.332133321 | 0.526583317 | 611.8502885 | 571.4562586 | ENSG00000088256 |
| ZSCAN25 | -0.099063783 | 0.933638667 | 0.435825231 | 0.625296859 | 372.9696467 | 348.1061529 | ENSG00000197037 |
| WDR3 | -0.099094235 | 0.93361896 | 0.21045519 | 0.389867428 | 1482.650976 | 1383.647165 | ENSG00000065183 |
| SOX21 | -0.099173277 | 0.933567811 | 0.735154873 | 0.851957757 | 58.04136631 | 54.18754785 | ENSG00000125285 |
| OAZ1 | -0.099205096 | 0.933547221 | 0.079359258 | 0.190478541 | 4976.975774 | 4645.851741 | ENSG00000104904 |
| EPHA8 | -0.099216485 | 0.933539851 | 0.491517943 | 0.674150341 | 256.9198361 | 239.689828 | ENSG00000070886 |
| IPO11 | -0.099240152 | 0.933524537 | 0.368186441 | 0.561249392 | 861.1866549 | 803.2545005 | ENSG00000086200 |
| TRIM41 | -0.099284684 | 0.933495722 | 0.358948572 | 0.552141492 | 692.728744 | 647.3548236 | ENSG00000146063 |
| F11 | -0.099312659 | 0.933477621 | 0.922961052 | 0.961884848 | 5.605124238 | 5.292546294 | ENSG00000088926 |
| GHDC | -0.099337318 | 0.933461666 | 0.514692725 | 0.692846128 | 219.8169628 | 205.1570076 | ENSG00000167925 |
| THUMPD2 | -0.099357841 | 0.933448387 | 0.507019112 | 0.6864963 | 312.9587817 | 291.96404 | ENSG00000138050 |
| WRN | -0.099364188 | 0.93344428 | 0.292236753 | 0.484724895 | 721.8901765 | 673.8264895 | ENSG00000165392 |
| METTL18 | -0.099460605 | 0.9333819 | 0.718700479 | 0.840687818 | 73.88093812 | 69.09398473 | ENSG00000171806 |
| MELTF-AS1 | -0.099493652 | 0.933360519 | 0.880479538 | 0.938509763 | 14.72734724 | 13.82227406 | ENSG00000228109 |
| C4orf33 | -0.099520008 | 0.933343468 | 0.794459202 | 0.8886547 | 32.11008442 | 30.00273359 | ENSG00000151470 |
| PACSIN1 | -0.099568212 | 0.933312283 | 0.401070502 | 0.592731097 | 400.4275127 | 373.7328782 | ENSG00000124507 |
| ANKRD42 | -0.099622525 | 0.933277147 | 0.679725491 | 0.815136731 | 104.8036334 | 98.129968 | ENSG00000137494 |
| NA | -0.09973024 | 0.933207469 | 0.91530301 | 0.958169563 | 5.511521874 | 5.097238657 | ENSG00000277968 |
| UBE2E1 | -0.099736171 | 0.933203633 | 0.21436374 | 0.394456848 | 1922.833181 | 1793.524017 | ENSG00000170142 |
| FAF2 | -0.099817345 | 0.933151127 | 0.210070113 | 0.389305017 | 1217.254294 | 1136.017595 | ENSG00000113194 |
| NA | -0.099991279 | 0.933038631 | 0.901908994 | 0.95021014 | 7.400316327 | 6.886448747 | ENSG00000228305 |
| RBM42 | -0.099992036 | 0.933038142 | 0.242045396 | 0.428346164 | 2118.049176 | 1975.648154 | ENSG00000126254 |
| ANKRD28 | -0.100069464 | 0.932988068 | 0.277726037 | 0.468710105 | 990.0922686 | 923.0233789 | ENSG00000206560 |
| WDR75 | -0.100074942 | 0.932984525 | 0.179927495 | 0.349227792 | 1692.292285 | 1578.119611 | ENSG00000115368 |
| BNC1 | -0.100188587 | 0.932911035 | 0.930601952 | 0.965950404 | 4.505527319 | 4.215981162 | ENSG00000169594 |
| TMEM223 | -0.100378425 | 0.932788285 | 0.49065327 | 0.673462764 | 334.6313316 | 312.4776071 | ENSG00000168569 |
| SHMT1 | -0.100380364 | 0.932787031 | 0.281854821 | 0.473069106 | 953.5427114 | 889.3066803 | ENSG00000176974 |
| TPRN | -0.100384566 | 0.932784315 | 0.334456529 | 0.528981192 | 705.2150682 | 658.0888414 | ENSG00000176058 |
| KLHL32 | -0.100412878 | 0.932766009 | 0.922067517 | 0.961386103 | 4.49605658 | 4.208495088 | ENSG00000186231 |
| NA | -0.100628548 | 0.93262658 | 0.692744191 | 0.823967294 | 76.21951715 | 70.97226326 | ENSG00000240489 |
| ACTR6 | -0.100738489 | 0.932555511 | 0.463172702 | 0.649713191 | 386.7939522 | 360.1778179 | ENSG00000075089 |
| NA | -0.100782641 | 0.932526972 | 0.916300581 | 0.958608883 | 6.2503065 | 5.855023298 | ENSG00000273783 |
| RUSC2 | -0.100792533 | 0.932520578 | 0.466683376 | 0.652842455 | 295.2389167 | 275.4217389 | ENSG00000198853 |
| ABITRAM | -0.100835127 | 0.932493047 | 0.614630079 | 0.770272086 | 156.2852634 | 145.7323504 | ENSG00000119328 |
| CCDC112 | -0.100901622 | 0.932450068 | 0.498685748 | 0.679606004 | 297.987909 | 277.3870073 | ENSG00000164221 |
| ARL5A | -0.100918925 | 0.932438885 | 0.730463594 | 0.849282495 | 542.1270687 | 505.3886654 | ENSG00000162980 |
| RBPMS2 | -0.101066774 | 0.932343332 | 0.15903477 | 0.318844663 | 3482.022838 | 3246.096371 | ENSG00000166831 |
| MTHFS | -0.10110753 | 0.932316995 | 0.866602245 | 0.930273692 | 17.47190485 | 16.31196812 | ENSG00000136371 |
| RCBTB1 | -0.101115649 | 0.932311748 | 0.257370642 | 0.445975272 | 926.5290227 | 863.7993026 | ENSG00000136144 |
| ATP2B1-AS1 | -0.101185932 | 0.93226633 | 0.894204705 | 0.94574241 | 8.492321644 | 7.955527805 | ENSG00000271614 |
| ATF2 | -0.101267404 | 0.932213684 | 0.391935763 | 0.584591777 | 429.9760049 | 400.7898544 | ENSG00000115966 |
| STAU1 | -0.101390853 | 0.93213392 | 0.125986256 | 0.269105597 | 3155.089016 | 2940.218137 | ENSG00000124214 |
| DNMT3A | -0.101400873 | 0.932127446 | 0.12750827 | 0.27135587 | 3872.244653 | 3609.588216 | ENSG00000119772 |
| CCDC114 | -0.101405065 | 0.932124737 | 0.811178434 | 0.898809775 | 29.71390685 | 27.83506922 | ENSG00000105479 |
| GLB1L2 | -0.101475394 | 0.932079299 | 0.553758816 | 0.723938212 | 201.5261704 | 188.1755991 | ENSG00000149328 |
| SNRPD1 | -0.101496655 | 0.932065563 | 0.143322707 | 0.296004352 | 2859.639007 | 2664.446416 | ENSG00000167088 |
| POC5 | -0.101521052 | 0.932049801 | 0.570467489 | 0.737054335 | 197.3160698 | 183.7326055 | ENSG00000152359 |
| TMSB10 | -0.10166972 | 0.931953759 | 0.165483839 | 0.328310584 | 6968.661056 | 6494.901593 | ENSG00000034510 |
| GON4L | -0.101687999 | 0.931941951 | 0.259878853 | 0.448806807 | 968.8436619 | 902.8019064 | ENSG00000116580 |
| POLR1C | -0.101691874 | 0.931939448 | 0.461276704 | 0.647890508 | 323.07931 | 301.4082591 | ENSG00000171453 |
| USP11 | -0.101691918 | 0.93193942 | 0.097852914 | 0.222911248 | 4834.547212 | 4505.788582 | ENSG00000102226 |
| MTERF2 | -0.101862819 | 0.93182903 | 0.733267523 | 0.851101774 | 53.96738114 | 50.36966803 | ENSG00000120832 |
| MYNN | -0.10189025 | 0.931811312 | 0.539215529 | 0.71257188 | 215.4195446 | 200.4366859 | ENSG00000085274 |
| MBD6 | -0.101912262 | 0.931797095 | 0.403147076 | 0.594898719 | 479.2288487 | 446.6476655 | ENSG00000166987 |
| SLC25A44 | -0.101929792 | 0.931785773 | 0.283755036 | 0.47467075 | 723.5907589 | 674.3823564 | ENSG00000160785 |
| MOAP1 | -0.102069396 | 0.931695612 | 0.334617485 | 0.528981192 | 595.7574274 | 555.0928756 | ENSG00000165943 |
| PRMT5 | -0.102130197 | 0.931656348 | 0.28848461 | 0.480179616 | 1342.301569 | 1251.000131 | ENSG00000100462 |
| NA | -0.102161453 | 0.931636163 | 0.918954271 | 0.960084467 | 4.816430366 | 4.47844867 | ENSG00000274423 |
| NOP58 | -0.102329868 | 0.931527414 | 0.124045472 | 0.266218016 | 2198.681468 | 2048.071303 | ENSG00000055044 |
| PRR35 | -0.102358061 | 0.93150921 | 0.840384772 | 0.91422735 | 24.03173892 | 22.54307258 | ENSG00000161992 |
| OTUD6B | -0.102371486 | 0.931500542 | 0.454656697 | 0.641929399 | 462.7777157 | 430.9789923 | ENSG00000155100 |
| ZNF362 | -0.102446274 | 0.931452255 | 0.340750935 | 0.535164633 | 571.955722 | 532.822353 | ENSG00000160094 |
| YWHAZ | -0.102716371 | 0.931277888 | 0.181386156 | 0.351359847 | 11479.4757 | 10689.24422 | ENSG00000164924 |
| UBR2 | -0.10272165 | 0.93127448 | 0.373248397 | 0.566659875 | 673.1955346 | 626.0670337 | ENSG00000024048 |
| ZIC2 | -0.102734377 | 0.931266265 | 0.181640866 | 0.351779708 | 3789.614999 | 3528.902122 | ENSG00000043355 |
| SLC35E2B | -0.102742828 | 0.93126081 | 0.189868861 | 0.362144541 | 1401.648597 | 1305.641289 | ENSG00000189339 |
| METAP2 | -0.102752573 | 0.931254519 | 0.150381758 | 0.306485101 | 3214.637596 | 2992.63769 | ENSG00000111142 |
| FTO | -0.102756636 | 0.931251897 | 0.277609821 | 0.468699553 | 774.3893387 | 721.1405224 | ENSG00000140718 |
| PSME1 | -0.102835578 | 0.931200942 | 0.310919291 | 0.504706837 | 757.7191614 | 705.6361535 | ENSG00000092010 |
| PRKD3 | -0.102872151 | 0.931177335 | 0.311968239 | 0.505553847 | 1045.072439 | 972.7033929 | ENSG00000115825 |
| CHSY3 | -0.102872765 | 0.931176939 | 0.71772929 | 0.84009 | 60.02238765 | 55.86033312 | ENSG00000198108 |
| HS2ST1 | -0.102948419 | 0.93112811 | 0.502575624 | 0.68301375 | 330.6645686 | 307.7346668 | ENSG00000153936 |
| FRY | -0.102976109 | 0.931110239 | 0.56210193 | 0.730738379 | 172.1808628 | 160.3046998 | ENSG00000073910 |
| RAVER1 | -0.102976136 | 0.931110221 | 0.261648685 | 0.450896953 | 1008.004976 | 938.2575165 | ENSG00000161847 |

| | | | | | | | |
|---|---|---|---|---|---|---|---|
| RPSA | -0.103025159 | 0.931078583 | 0.08625181 | 0.20278968 | 17793.89385 | 16566.51848 | ENSG00000168028 |
| RHD | -0.103267868 | 0.930921958 | 0.925670376 | 0.963290836 | 4.115025033 | 3.859658683 | ENSG00000187010 |
| NA | -0.103420359 | 0.930823565 | 0.771940109 | 0.874465985 | 45.23814092 | 42.12117877 | ENSG00000175741 |
| RCAN2 | -0.103431459 | 0.930816404 | 0.501544818 | 0.682098008 | 237.34933 | 220.6392447 | ENSG00000172348 |
| NA | -0.1034405 | 0.930810571 | 0.824234663 | 0.905588076 | 23.76946735 | 22.20128421 | ENSG00000224152 |
| ZNF334 | -0.103519863 | 0.930759368 | 0.524543928 | 0.701132062 | 241.417911 | 224.526921 | ENSG00000198185 |
| UBE2Z | -0.103550266 | 0.930739754 | 0.124158726 | 0.266399378 | 2350.552943 | 2187.8918 | ENSG00000159202 |
| EMC2 | -0.103610392 | 0.930700965 | 0.314449186 | 0.508019632 | 610.9662109 | 568.6232388 | ENSG00000104412 |
| SNX11 | -0.103630956 | 0.930687699 | 0.38859121 | 0.581422849 | 503.1461572 | 468.3515774 | ENSG00000002919 |
| NFKBIA | -0.103671751 | 0.930661382 | 0.497857754 | 0.678931121 | 278.8306049 | 259.1517241 | ENSG00000100906 |
| NA | -0.103683193 | 0.930654001 | 0.793910186 | 0.888278381 | 34.43741127 | 32.05061235 | ENSG00000233757 |
| MBNL1 | -0.103699945 | 0.930643195 | 0.648421989 | 0.794726435 | 127.2663503 | 118.2536037 | ENSG00000152601 |
| ATP8A1 | -0.10377734 | 0.930593271 | 0.47070118 | 0.656184706 | 312.1293562 | 289.9298109 | ENSG00000124406 |
| STAB2 | -0.103804081 | 0.930576022 | 0.883460674 | 0.939962378 | 14.49929804 | 13.63489053 | ENSG00000136011 |
| GALM | -0.103818144 | 0.930566951 | 0.425159743 | 0.615858592 | 373.2690045 | 346.9233485 | ENSG00000143891 |
| NA | -0.103828134 | 0.930560507 | 0.804143484 | 0.894704967 | 30.08724909 | 27.99239683 | ENSG00000238045 |
| ZNF513 | -0.103951699 | 0.930480809 | 0.337518468 | 0.531935994 | 586.2860351 | 545.8518255 | ENSG00000163795 |
| TMEM161B | -0.104044361 | 0.930421048 | 0.551348619 | 0.722322422 | 183.681427 | 170.6415326 | ENSG00000164180 |
| GLE1 | -0.104061496 | 0.930409997 | 0.447099283 | 0.635261315 | 408.50204 | 379.4506187 | ENSG00000119392 |
| NAA25 | -0.104084391 | 0.930395232 | 0.300699827 | 0.493987821 | 978.9383258 | 910.2693078 | ENSG00000111300 |
| SYK | -0.104087017 | 0.930393539 | 0.630025862 | 0.781581472 | 111.1498274 | 103.3415445 | ENSG00000165025 |
| CASP6 | -0.104140879 | 0.930358804 | 0.337878099 | 0.531959964 | 542.5837762 | 504.6132812 | ENSG00000138794 |
| PHF10 | -0.104178265 | 0.930334695 | 0.344687827 | 0.538200859 | 1004.090964 | 933.6220751 | ENSG00000130024 |
| SP3 | -0.104228642 | 0.930302209 | 0.241210732 | 0.427440453 | 1243.80878 | 1156.62633 | ENSG00000172845 |
| E4F1 | -0.104233972 | 0.930298773 | 0.288036248 | 0.479603253 | 853.9318196 | 794.6425552 | ENSG00000167967 |
| CLVS2 | -0.104392165 | 0.930196769 | 0.858799987 | 0.925299324 | 15.13175496 | 14.00162186 | ENSG00000146352 |
| THAP7 | -0.104436192 | 0.930168383 | 0.488398567 | 0.671262814 | 260.8515123 | 242.8312194 | ENSG00000184436 |
| MAPKAPK5 | -0.104488403 | 0.930134721 | 0.301014325 | 0.494460648 | 823.6045025 | 765.946113 | ENSG00000089022 |
| MGST3 | -0.10466897 | 0.930018313 | 0.228188271 | 0.411810953 | 1147.65021 | 1067.506284 | ENSG00000143198 |
| MUC1 | -0.104841532 | 0.92990708 | 0.828012267 | 0.9073231 | 21.44785296 | 20.01061196 | ENSG00000185499 |
| C11orf80 | -0.10485165 | 0.929900558 | 0.528453446 | 0.704224631 | 202.4922301 | 188.2299368 | ENSG00000173715 |
| THBS4 | -0.10488065 | 0.929881866 | 0.405763074 | 0.597427138 | 403.5387305 | 375.2903295 | ENSG00000113296 |
| TJAP1 | -0.104965702 | 0.929827048 | 0.282208566 | 0.473151553 | 712.8650058 | 663.352311 | ENSG00000137221 |
| C11orf95 | -0.1050834 | 0.929751193 | 0.263897371 | 0.453591196 | 1138.092281 | 1058.676644 | ENSG00000188070 |
| RALY | -0.105214164 | 0.929666926 | 0.081047395 | 0.193523888 | 4896.784269 | 4552.583184 | ENSG00000125970 |
| PHF12 | -0.105284261 | 0.929621757 | 0.200653796 | 0.377248677 | 1252.101382 | 1164.539753 | ENSG00000109118 |
| ZNHIT3 | -0.105338628 | 0.929586725 | 0.316221207 | 0.509815924 | 560.1053046 | 521.0207532 | ENSG00000273611 |
| NA | -0.105340196 | 0.929585715 | 0.894998248 | 0.946203969 | 7.370626335 | 6.818850843 | ENSG00000272140 |
| ATRAID | -0.105357819 | 0.92957436 | 0.361253162 | 0.554489946 | 535.6157237 | 497.9711818 | ENSG00000138085 |
| TSNARE1 | -0.105451236 | 0.92951417 | 0.448024899 | 0.635957225 | 322.9447428 | 300.0517293 | ENSG00000171045 |
| PDF | -0.105476246 | 0.929498057 | 0.543579346 | 0.716041194 | 315.4470507 | 293.258371 | ENSG00000258429 |
| NA | -0.105535118 | 0.929460128 | 0.887474636 | 0.942496843 | 8.634457779 | 8.060667697 | ENSG00000269427 |
| RPRM | -0.105541933 | 0.929455737 | 0.506088862 | 0.685711746 | 305.8164151 | 284.6403796 | ENSG00000177519 |
| SEPTIN7 | -0.105674831 | 0.929370121 | 0.168086143 | 0.33198088 | 2572.777147 | 2390.65711 | ENSG00000122545 |
| SEC24C | -0.105752183 | 0.929320293 | 0.206194599 | 0.384777557 | 1343.627232 | 1248.486902 | ENSG00000176986 |
| BDH2 | -0.105797784 | 0.92929092 | 0.560340565 | 0.729378503 | 209.316914 | 194.4073246 | ENSG00000164039 |
| RNF149 | -0.105838976 | 0.929264387 | 0.43549637 | 0.625018563 | 353.1585388 | 327.9718533 | ENSG00000163162 |
| ARGLU1 | -0.106096764 | 0.929098356 | 0.681937087 | 0.816316087 | 2901.32167 | 2695.857017 | ENSG00000134884 |
| DNAH14 | -0.106107042 | 0.929091737 | 0.474278676 | 0.659368981 | 257.8410976 | 239.5825382 | ENSG00000185842 |
| C5orf22 | -0.106133769 | 0.929074525 | 0.345371849 | 0.53886033 | 567.6353249 | 527.0776539 | ENSG00000082213 |
| LRATD2 | -0.10615978 | 0.929057775 | 0.256380659 | 0.445149075 | 803.6860974 | 746.6657463 | ENSG00000168672 |
| PAIP1 | -0.106234171 | 0.92900987 | 0.185184814 | 0.35592817 | 1994.647816 | 1852.597258 | ENSG00000172239 |
| MRPL49 | -0.106275648 | 0.928983161 | 0.234163123 | 0.419038265 | 1098.482543 | 1020.697086 | ENSG00000149792 |
| BUD23 | -0.106297758 | 0.928968924 | 0.153537569 | 0.310967687 | 2204.332217 | 2048.267622 | ENSG00000071462 |
| NA | -0.106372571 | 0.928920753 | 0.8991728 | 0.948444134 | 7.506448644 | 6.946560577 | ENSG00000253106 |
| NA | -0.106399646 | 0.92890332 | 0.909119342 | 0.95476076 | 5.922341111 | 5.543728881 | ENSG00000270504 |
| DROSHA | -0.106473265 | 0.92885592 | 0.18952514 | 0.361719591 | 2499.443129 | 2322.408706 | ENSG00000113360 |
| TMEM9 | -0.1065368 | 0.928815015 | 0.144531575 | 0.297837248 | 2247.908703 | 2088.837396 | ENSG00000116857 |
| ASCC2 | -0.106633148 | 0.928752988 | 0.25438275 | 0.443072069 | 900.6779939 | 837.0906449 | ENSG00000100325 |
| CACNA2D2 | -0.106646672 | 0.928744282 | 0.236065731 | 0.42106075 | 851.326151 | 791.0285112 | ENSG00000007402 |
| ADIPOR2 | -0.106711851 | 0.928702323 | 0.177300509 | 0.345649273 | 1538.75224 | 1428.352889 | ENSG00000006831 |
| TRMT10C | -0.106814303 | 0.928636374 | 0.313254089 | 0.50683992 | 704.7640356 | 654.7204777 | ENSG00000174173 |
| MAP7 | -0.106823219 | 0.928630635 | 0.112781006 | 0.247957671 | 2264.393878 | 2102.990634 | ENSG00000135525 |
| TMEM178B | -0.106866188 | 0.928602978 | 0.388283829 | 0.581249696 | 414.1792544 | 384.5565459 | ENSG00000261115 |
| TRIM44 | -0.106894822 | 0.928584547 | 0.158907174 | 0.318707138 | 1594.643863 | 1480.748224 | ENSG00000166326 |
| ZFR2 | -0.106967427 | 0.928537816 | 0.462349153 | 0.648934434 | 311.4758833 | 289.4550656 | ENSG00000105278 |
| PRELID3B | -0.106973071 | 0.928534184 | 0.344833712 | 0.538292599 | 722.3064225 | 670.0993581 | ENSG00000101166 |
| ACER3 | -0.107111978 | 0.928444787 | 0.487493338 | 0.670565648 | 249.1494321 | 231.3178659 | ENSG00000078124 |
| BCL7A | -0.1071818 | 0.928399854 | 0.317119035 | 0.510819068 | 511.4700276 | 474.8388339 | ENSG00000110987 |
| VPS16 | -0.107198883 | 0.92838886 | 0.386499414 | 0.579384926 | 407.8030773 | 378.7304247 | ENSG00000215305 |
| UVRAG | -0.107266372 | 0.928345432 | 0.378818172 | 0.572023464 | 409.9359987 | 380.3285153 | ENSG00000198382 |
| ANAPC13 | -0.107373764 | 0.928276329 | 0.383235955 | 0.576019398 | 471.395224 | 437.1177219 | ENSG00000129055 |
| COG8 | -0.107523291 | 0.928180124 | 0.647432264 | 0.794202863 | 92.91357434 | 86.31335579 | ENSG00000213380 |
| KCNK6 | -0.107658071 | 0.928093415 | 0.37600381 | 0.569071911 | 416.7513845 | 386.8071151 | ENSG00000099337 |

| | | | | | | | |
|---|---|---|---|---|---|---|---|
| ZNF30 | -0.107829477 | 0.927983155 | 0.613293817 | 0.769307369 | 118.0452184 | 109.5157863 | ENSG00000168661 |
| RHEB | -0.107853404 | 0.927967765 | 0.135138434 | 0.283207678 | 2534.072781 | 2351.217993 | ENSG00000106615 |
| MEX3D | -0.10787827 | 0.927951771 | 0.233301056 | 0.418222437 | 2155.933565 | 2000.583038 | ENSG00000181588 |
| TRPC4AP | -0.107891142 | 0.927943492 | 0.131554222 | 0.277860508 | 2536.861107 | 2353.407432 | ENSG00000100991 |
| STARD3NL | -0.108028265 | 0.927855298 | 0.31984579 | 0.513518689 | 632.5329593 | 586.5368272 | ENSG00000010270 |
| GCLC | -0.108138427 | 0.927784451 | 0.314894697 | 0.508473455 | 723.1616594 | 670.7060274 | ENSG00000001084 |
| NA | -0.108148285 | 0.927778111 | 0.928539234 | 0.964687063 | 3.800965072 | 3.563448033 | ENSG00000228252 |
| PIH1D1 | -0.10817671 | 0.927759832 | 0.18081917 | 0.350371406 | 1950.39766 | 1809.417791 | ENSG00000104872 |
| PYM1 | -0.108217483 | 0.927733612 | 0.182416932 | 0.352766634 | 1187.686567 | 1101.667371 | ENSG00000170473 |
| MRPL39 | -0.108235215 | 0.92772221 | 0.30536066 | 0.499299026 | 1045.779437 | 969.2757786 | ENSG00000154719 |
| ZNF618 | -0.108290978 | 0.927686352 | 0.209907952 | 0.389121223 | 1223.589537 | 1135.218509 | ENSG00000157657 |
| BCAP31 | -0.108310013 | 0.927674112 | 0.096413082 | 0.220525738 | 3810.493557 | 3535.316255 | ENSG00000185825 |
| HSH2D | -0.108400567 | 0.927615887 | 0.830659926 | 0.908715091 | 23.65880264 | 21.89692621 | ENSG00000196684 |
| NA | -0.108638273 | 0.92746306 | 0.916497494 | 0.958706658 | 4.449304247 | 4.129612264 | ENSG00000260669 |
| ZNF134 | -0.108756917 | 0.927386791 | 0.210494102 | 0.389867428 | 1167.70564 | 1082.737521 | ENSG00000213762 |
| ABCD4 | -0.10884957 | 0.927327234 | 0.279130544 | 0.470138631 | 1108.819215 | 1028.700153 | ENSG00000119688 |
| CAND2 | -0.108920808 | 0.927281445 | 0.469423124 | 0.655190112 | 283.3834481 | 262.8585153 | ENSG00000144712 |
| MIOS | -0.108946035 | 0.927265231 | 0.233430108 | 0.418332395 | 813.5733676 | 754.1598202 | ENSG00000164654 |
| DCLK3 | -0.108968983 | 0.927250482 | 0.695005531 | 0.825416199 | 67.8776223 | 63.04029298 | ENSG00000163673 |
| GTF3C2 | -0.108982191 | 0.927241992 | 0.113700903 | 0.24953612 | 1972.16933 | 1828.820459 | ENSG00000115207 |
| MAL2 | -0.109176389 | 0.927117187 | 0.275106016 | 0.466037706 | 1758.807279 | 1630.12269 | ENSG00000147676 |
| TRERF1 | -0.109178293 | 0.927115964 | 0.233924768 | 0.418856139 | 849.0332571 | 786.8241011 | ENSG00000124496 |
| ECD | -0.109211794 | 0.927094435 | 0.187829497 | 0.3596177 | 1108.328427 | 1027.148791 | ENSG00000122882 |
| SYNGR3 | -0.109224943 | 0.927085985 | 0.310928168 | 0.504706837 | 613.237029 | 568.6607893 | ENSG00000127561 |
| VEGFB | -0.109411019 | 0.926966419 | 0.457569625 | 0.644667079 | 382.5233967 | 354.6921541 | ENSG00000173511 |
| ITPA | -0.109464816 | 0.926931854 | 0.242127077 | 0.428449803 | 813.540205 | 753.8002401 | ENSG00000125877 |
| SHCBP1 | -0.109481091 | 0.926921398 | 0.389776233 | 0.582354914 | 408.622258 | 378.4226656 | ENSG00000171241 |
| SPATA6 | -0.109524721 | 0.926893366 | 0.577249396 | 0.741962465 | 180.5038389 | 167.0053737 | ENSG00000132122 |
| TBCE | -0.109553148 | 0.926875102 | 0.756066083 | 0.864333115 | 39.7366685 | 36.85912643 | ENSG00000285053 |
| PTGES2 | -0.109570952 | 0.926863664 | 0.167654935 | 0.331447101 | 2014.538373 | 1867.837631 | ENSG00000148334 |
| ZNF836 | -0.109595317 | 0.926848011 | 0.713241547 | 0.837666741 | 67.44292323 | 62.25796312 | ENSG00000196267 |
| ATP5PO | -0.109750174 | 0.92674853 | 0.669385 | 0.807856359 | 84.27202862 | 78.25356416 | ENSG00000241837 |
| CHMP2B | -0.109767907 | 0.926737139 | 0.262514108 | 0.452052075 | 742.2794497 | 687.402224 | ENSG00000083937 |
| APOL2 | -0.10979511 | 0.926719665 | 0.3834377 | 0.576275895 | 387.4868941 | 359.2792554 | ENSG00000128335 |
| TMEM131L | -0.109825475 | 0.92670016 | 0.305336962 | 0.499299026 | 627.6409584 | 581.0540375 | ENSG00000121210 |
| ELOC | -0.109887427 | 0.926660366 | 0.200727857 | 0.377311369 | 1404.774216 | 1302.089667 | ENSG00000154582 |
| INPP5K | -0.109894433 | 0.926655866 | 0.404055486 | 0.5957522 | 367.5698719 | 340.2851681 | ENSG00000132376 |
| SPACA6 | -0.109899445 | 0.926652647 | 0.729059646 | 0.848129016 | 69.41131692 | 64.59428032 | ENSG00000182310 |
| BRD7 | -0.109900709 | 0.926651835 | 0.119643045 | 0.259049555 | 2250.291442 | 2084.960726 | ENSG00000166164 |
| MAP3K11 | -0.109987321 | 0.926596205 | 0.179696783 | 0.348999285 | 1524.286112 | 1412.441556 | ENSG00000173327 |
| PABPC1L | -0.110001608 | 0.926587029 | 0.80108595 | 0.893214577 | 846.6281049 | 784.6198549 | ENSG00000101104 |
| CMTM7 | -0.110078436 | 0.926537687 | 0.412126328 | 0.603631213 | 346.2605172 | 320.8463829 | ENSG00000153551 |
| SNORA5C | -0.110083128 | 0.926534674 | 0.909476654 | 0.954919574 | 5.786421104 | 5.288309414 | ENSG00000201772 |
| POMP | -0.110098943 | 0.926524516 | 0.088302262 | 0.206484246 | 2756.97751 | 2554.176605 | ENSG00000132963 |
| HIPK2 | -0.110133515 | 0.926502315 | 0.326456969 | 0.520701675 | 862.3067794 | 798.992782 | ENSG00000064393 |
| DENND11 | -0.110339274 | 0.926370185 | 0.24712201 | 0.434921597 | 899.8604219 | 832.8550393 | ENSG00000257093 |
| PRSS36 | -0.110354802 | 0.926360214 | 0.854273269 | 0.922728642 | 15.41612492 | 14.30575662 | ENSG00000178226 |
| ACADM | -0.110370775 | 0.926349958 | 0.247783364 | 0.435795869 | 1011.329593 | 936.4537554 | ENSG00000117054 |
| DUSP12 | -0.110393033 | 0.926335666 | 0.36130229 | 0.554519431 | 400.6044725 | 371.4202704 | ENSG00000081721 |
| LDHB | -0.110500894 | 0.926266412 | 0.684267387 | 0.817890607 | 66.07611638 | 61.23556109 | ENSG00000111716 |
| VPS52 | -0.110539008 | 0.926241942 | 0.327085837 | 0.521301591 | 627.8718941 | 581.5194421 | ENSG00000223501 |
| DNTTIP2 | -0.11061852 | 0.926190895 | 0.171066184 | 0.336682665 | 1566.212985 | 1450.453932 | ENSG00000067334 |
| MAPK1 | -0.110637253 | 0.926178869 | 0.157780208 | 0.317036092 | 1787.751035 | 1654.887948 | ENSG00000100030 |
| CD3EAP | -0.110641708 | 0.926176009 | 0.532171729 | 0.707348739 | 187.2388287 | 173.4665166 | ENSG00000117877 |
| C1orf35 | -0.110682753 | 0.926149659 | 0.23656313 | 0.421663883 | 835.2181665 | 774.1064501 | ENSG00000143793 |
| SH3BP5L | -0.110857388 | 0.926037558 | 0.164508469 | 0.326900393 | 1199.827473 | 1110.759147 | ENSG00000175137 |
| CCDC93 | -0.110861065 | 0.926035198 | 0.343946733 | 0.537587178 | 579.9563173 | 536.9076932 | ENSG00000125633 |
| PPRC1 | -0.110892851 | 0.926014795 | 0.107103333 | 0.238559449 | 2658.874374 | 2462.240844 | ENSG00000148840 |
| FUZ | -0.110923707 | 0.92599499 | 0.393952721 | 0.586514558 | 321.336485 | 297.5919795 | ENSG00000010361 |
| ALG8 | -0.110991048 | 0.925951768 | 0.206881321 | 0.385515625 | 909.9502755 | 842.4925362 | ENSG00000159063 |
| LITAF | -0.111004715 | 0.925942996 | 0.227044831 | 0.410621477 | 1897.211028 | 1756.956123 | ENSG00000189067 |
| BAZ1B | -0.111018431 | 0.925934193 | 0.069720395 | 0.172453998 | 3442.257855 | 3187.360476 | ENSG00000009954 |
| SLC9A6 | -0.111031497 | 0.925925807 | 0.379109101 | 0.572276273 | 646.9382264 | 598.1913394 | ENSG00000198689 |
| FLII | -0.111035322 | 0.925923353 | 0.050060654 | 0.133885159 | 5741.927213 | 5317.038663 | ENSG00000177731 |
| C19orf12 | -0.111047256 | 0.925915693 | 0.405216411 | 0.596903085 | 342.9856587 | 317.6822387 | ENSG00000131943 |
| EIF2A | -0.111108952 | 0.925876098 | 0.254285168 | 0.44298754 | 1115.332672 | 1031.959749 | ENSG00000144895 |
| YTHDF1 | -0.111109934 | 0.925875468 | 0.130577693 | 0.276585938 | 1700.282192 | 1574.248931 | ENSG00000149658 |
| ZNF438 | -0.111194556 | 0.925821162 | 0.779125194 | 0.878684821 | 33.03204505 | 30.58778044 | ENSG00000183621 |
| NA | -0.111281665 | 0.925765263 | 0.921444613 | 0.961169206 | 4.122616635 | 3.833401615 | ENSG00000261173 |
| RAD9A | -0.111516602 | 0.925614518 | 0.385035533 | 0.577938906 | 365.8175012 | 338.8030558 | ENSG00000172613 |
| CASKIN2 | -0.111569536 | 0.925580557 | 0.279999676 | 0.471174323 | 841.2481357 | 778.3445857 | ENSG00000177303 |
| MSANTD3 | -0.111599737 | 0.925561181 | 0.366341985 | 0.559450102 | 421.1128367 | 389.3911022 | ENSG00000066697 |
| LOC105369760 | -0.111635444 | 0.925538274 | 0.917461239 | 0.959335774 | 4.691356572 | 4.320683021 | ENSG00000258017 |
| ZNF286B | -0.111728682 | 0.92547846 | 0.786757585 | 0.88320806 | 29.85406615 | 27.5834486 | ENSG00000249459 |

| | | | | | | | |
|---|---|---|---|---|---|---|---|
| POLE | -0.111830106 | 0.9254134 | 0.216796785 | 0.397400021 | 2198.668089 | 2035.760228 | ENSG00000177084 |
| KDELR2 | -0.111879747 | 0.925381558 | 0.22933541 | 0.413231819 | 1189.013064 | 1099.812132 | ENSG00000136240 |
| ZC3H7B | -0.111975317 | 0.925320259 | 0.108240315 | 0.240362578 | 3637.314651 | 3366.141807 | ENSG00000100403 |
| NA | -0.1120452 | 0.925275438 | 0.920938442 | 0.9608655 | 3.794651247 | 3.552163113 | ENSG00000274220 |
| TBC1D15 | -0.11205224 | 0.925270923 | 0.301209354 | 0.494693325 | 751.2488783 | 694.4003026 | ENSG00000121749 |
| MEPCE | -0.112145979 | 0.925210806 | 0.14281006 | 0.295307559 | 1455.461988 | 1346.619363 | ENSG00000146834 |
| SPCS3 | -0.112202664 | 0.925174454 | 0.695459816 | 0.825783345 | 1118.404345 | 1034.528325 | ENSG00000129128 |
| TMEM17 | -0.112254353 | 0.925141307 | 0.81751586 | 0.901895175 | 23.46299899 | 21.62273575 | ENSG00000186889 |
| NA | -0.112301533 | 0.925111054 | 0.77132492 | 0.874316298 | 41.17420521 | 38.25534811 | ENSG00000225177 |
| CPO | -0.112345922 | 0.92508259 | 0.835373596 | 0.911772818 | 16.65342329 | 15.33305681 | ENSG00000144410 |
| IRS1 | -0.112480911 | 0.924996036 | 0.193968767 | 0.367889595 | 1226.633542 | 1134.891868 | ENSG00000169047 |
| RBMX | -0.112617448 | 0.924908499 | 0.035403609 | 0.102127704 | 10894.68849 | 10075.91272 | ENSG00000147274 |
| NA | -0.112650571 | 0.924887264 | 0.911207575 | 0.955762377 | 4.839806533 | 4.452191601 | ENSG00000285859 |
| COPE | -0.112659496 | 0.924881543 | 0.22046219 | 0.402011633 | 772.5744159 | 714.5559067 | ENSG00000105669 |
| PARN | -0.112750308 | 0.924823326 | 0.209013112 | 0.388200122 | 1199.993962 | 1109.796576 | ENSG00000140694 |
| ACACA | -0.112837771 | 0.924767261 | 0.104882818 | 0.235111049 | 5749.431167 | 5316.416404 | ENSG00000278540 |
| TBC1D1 | -0.112900536 | 0.92472703 | 0.111753809 | 0.246288841 | 2142.679624 | 1980.795047 | ENSG00000065882 |
| ARFGAP2 | -0.112916731 | 0.924716649 | 0.13315307 | 0.280214117 | 1602.564183 | 1481.494509 | ENSG00000149182 |
| LIN9 | -0.112943345 | 0.924699591 | 0.431994502 | 0.6220157 | 285.1671176 | 263.5860293 | ENSG00000183814 |
| MST1 | -0.113024318 | 0.924647692 | 0.722141107 | 0.843051024 | 58.41461086 | 53.98574182 | ENSG00000173531 |
| C16orf72 | -0.113068639 | 0.924619286 | 0.217759363 | 0.398730689 | 963.1727384 | 890.8125564 | ENSG00000182831 |
| BCL7B | -0.113071793 | 0.924617265 | 0.273939545 | 0.46515382 | 582.9098173 | 538.9359334 | ENSG00000106635 |
| DOCK11 | -0.11309798 | 0.924600482 | 0.346551572 | 0.540064478 | 439.7010925 | 406.3175118 | ENSG00000147251 |
| GFM2 | -0.113102002 | 0.924597904 | 0.299493166 | 0.492966815 | 515.9897687 | 476.7433695 | ENSG00000164347 |
| AGGF1 | -0.113134779 | 0.924576898 | 0.590194171 | 0.752108001 | 159.9871242 | 147.5131984 | ENSG00000164252 |
| RAB23 | -0.113149796 | 0.924567275 | 0.497794517 | 0.678931121 | 278.384556 | 256.7827631 | ENSG00000112210 |
| USP19 | -0.113171394 | 0.924553433 | 0.114931114 | 0.25166981 | 2119.916369 | 1959.592871 | ENSG00000172046 |
| SUMF1 | -0.11319381 | 0.924539069 | 0.426447255 | 0.617131017 | 260.6865788 | 241.0940854 | ENSG00000144455 |
| TOB1-AS1 | -0.113220575 | 0.924521916 | 0.827663648 | 0.907253657 | 23.15840976 | 21.56416126 | ENSG00000229980 |
| GJA4 | -0.113329546 | 0.924452087 | 0.698665622 | 0.82815737 | 65.80417867 | 60.64247852 | ENSG00000187513 |
| MED22 | -0.113352603 | 0.924437313 | 0.129661422 | 0.274896453 | 1818.126901 | 1681.13614 | ENSG00000148297 |
| BPNT1 | -0.113387996 | 0.924414634 | 0.436705286 | 0.625881207 | 291.4266768 | 269.2465094 | ENSG00000162813 |
| DDRGK1 | -0.113418159 | 0.924395307 | 0.194121439 | 0.368050701 | 1243.911756 | 1150.718682 | ENSG00000198171 |
| VPS26B | -0.11345255 | 0.924373272 | 0.217709423 | 0.398718028 | 792.6264711 | 732.8298227 | ENSG00000151502 |
| IGFBP4 | -0.113489473 | 0.924349614 | 0.064151527 | 0.162163721 | 4724.643538 | 4366.204089 | ENSG00000141753 |
| FAM13A | -0.113713349 | 0.924206186 | 0.389241463 | 0.581884278 | 326.7626951 | 302.2666119 | ENSG00000138640 |
| HPRT1 | -0.11373338 | 0.924193354 | 0.21180675 | 0.39142749 | 1257.530185 | 1161.617656 | ENSG00000165704 |
| POLR1B | -0.113845967 | 0.924121233 | 0.133714571 | 0.281046253 | 1598.142795 | 1476.77539 | ENSG00000125630 |
| NA | -0.114045419 | 0.923993483 | 0.889464974 | 0.943529294 | 7.033190208 | 6.470014438 | ENSG00000250896 |
| DHX8 | -0.114118339 | 0.923946781 | 0.142583728 | 0.29500411 | 1386.356035 | 1281.373578 | ENSG00000067596 |
| USPL1 | -0.114128812 | 0.923940074 | 0.348469572 | 0.541777959 | 434.7152188 | 401.3664275 | ENSG00000132952 |
| PCDH9 | -0.114168936 | 0.923914378 | 0.850000302 | 0.920406282 | 13.00859748 | 12.02502824 | ENSG00000184226 |
| GRIP1 | -0.114269415 | 0.923850033 | 0.389338656 | 0.581935697 | 318.6228951 | 294.0876556 | ENSG00000155974 |
| BDH1 | -0.114291119 | 0.923836134 | 0.321704634 | 0.515651101 | 477.1194164 | 440.9605221 | ENSG00000161267 |
| NCKAP5L | -0.114312596 | 0.923822382 | 0.211139825 | 0.390584441 | 1004.624571 | 928.214736 | ENSG00000167566 |
| HS3ST3A1 | -0.11437964 | 0.923779451 | 0.631858595 | 0.782788727 | 154.3604651 | 142.5706994 | ENSG00000153976 |
| NRXN1 | -0.114413576 | 0.923757721 | 0.514907657 | 0.692859017 | 200.8822659 | 185.1631913 | ENSG00000179915 |
| THAP12 | -0.114517178 | 0.923691388 | 0.269326336 | 0.45951754 | 862.4235851 | 795.9194032 | ENSG00000137492 |
| MT1E | -0.114555287 | 0.923666988 | 0.457353862 | 0.644461072 | 275.1681154 | 253.7808531 | ENSG00000169715 |
| NOL6 | -0.114564881 | 0.923660846 | 0.088396157 | 0.206651661 | 2256.562298 | 2084.598215 | ENSG00000165271 |
| ESS2 | -0.11471791 | 0.923562877 | 0.321360898 | 0.515233814 | 524.5661091 | 484.963993 | ENSG00000100056 |
| GTF3C5 | -0.114724072 | 0.923558933 | 0.080510801 | 0.192639708 | 2826.043506 | 2609.740272 | ENSG00000148308 |
| CHPF | -0.114766869 | 0.923531536 | 0.152907289 | 0.310131724 | 2438.051634 | 2251.428079 | ENSG00000123989 |
| GLUD1P3 | -0.114861233 | 0.923471131 | 0.857096688 | 0.924108956 | 17.65182624 | 16.39552586 | ENSG00000250959 |
| MESP2 | -0.11489456 | 0.923449799 | 0.86559705 | 0.930027579 | 10.92006834 | 10.0935743 | ENSG00000188095 |
| CRBN | -0.114930625 | 0.923426714 | 0.408960778 | 0.600465741 | 299.0645357 | 275.9491586 | ENSG00000113851 |
| SDHAP1 | -0.11497144 | 0.92340059 | 0.567520391 | 0.734704734 | 219.6322159 | 203.1507332 | ENSG00000185485 |
| ENGASE | -0.115020048 | 0.923369479 | 0.745771305 | 0.858090723 | 849.6944344 | 784.7732317 | ENSG00000167280 |
| SLC26A11 | -0.115040433 | 0.923356432 | 0.346985041 | 0.540416227 | 394.3919513 | 364.0935034 | ENSG00000181045 |
| LDLRAD3 | -0.115065815 | 0.923340187 | 0.349642774 | 0.543044355 | 396.2070544 | 365.6218462 | ENSG00000179241 |
| ST3GAL5 | -0.115096966 | 0.923320251 | 0.718119206 | 0.840306923 | 61.76629502 | 56.8123261 | ENSG00000115525 |
| TSEN2 | -0.115114102 | 0.923309284 | 0.42984958 | 0.619986935 | 252.600544 | 233.0249439 | ENSG00000154743 |
| PSMD12 | -0.115126202 | 0.92330154 | 0.162578283 | 0.324037628 | 1934.778967 | 1785.630313 | ENSG00000197170 |
| KANSL2 | -0.115169936 | 0.923273551 | 0.30437543 | 0.498171513 | 573.1853462 | 529.3177346 | ENSG00000139620 |
| LOC100506098 | -0.115196454 | 0.923256581 | 0.744410707 | 0.857195413 | 39.10283612 | 36.10893948 | ENSG00000233834 |
| CWC22 | -0.115257229 | 0.923217688 | 0.228145587 | 0.411808562 | 874.4132521 | 807.0547405 | ENSG00000163510 |
| CRB3 | -0.115267885 | 0.92321087 | 0.39556503 | 0.587970348 | 323.7486425 | 298.9302392 | ENSG00000130545 |
| TRPM7 | -0.115271288 | 0.923208692 | 0.234435495 | 0.41940922 | 791.8786967 | 730.798293 | ENSG00000092439 |
| JKAMP | -0.115307438 | 0.923185559 | 0.347209737 | 0.540635576 | 430.1753714 | 396.8056236 | ENSG00000050130 |
| SMG5 | -0.11532634 | 0.923173464 | 0.055700919 | 0.145198428 | 4286.47623 | 3957.953613 | ENSG00000198952 |
| RTEL1-TNFRSF6B | -0.115449215 | 0.92309484 | 0.72157078 | 0.842590274 | 48.23214706 | 44.63855562 | ENSG00000026036 |
| GFER | -0.115610825 | 0.922991441 | 0.321234165 | 0.515075181 | 509.6471975 | 470.4201707 | ENSG00000127554 |
| TMBIM4 | -0.115692251 | 0.922939349 | 0.759751133 | 0.866398174 | 47.39304028 | 43.63955921 | ENSG00000155957 |
| CDC42EP1 | -0.115742794 | 0.922907016 | 0.393497635 | 0.586305751 | 378.5258312 | 349.5393669 | ENSG00000128283 |

| | | | | | | | |
|---|---|---|---|---|---|---|---|
| TRAF7 | -0.115743927 | 0.922906291 | 0.056056654 | 0.146002534 | 3466.225854 | 3198.680983 | ENSG00000131653 |
| NA | -0.115777552 | 0.92288478 | 0.662115032 | 0.803380688 | 74.6246611 | 68.96225291 | ENSG00000264577 |
| TADA3 | -0.115809227 | 0.922864518 | 0.172093802 | 0.338062404 | 1223.353054 | 1128.489969 | ENSG00000171148 |
| SNHG17 | -0.115920364 | 0.922793429 | 0.331034555 | 0.525254238 | 429.5695453 | 396.5340697 | ENSG00000196756 |
| FRMD5 | -0.115985495 | 0.92275177 | 0.374276428 | 0.567389377 | 322.927004 | 297.9251825 | ENSG00000171877 |
| TMED10 | -0.116044773 | 0.922713856 | 0.099024299 | 0.22505412 | 2996.595478 | 2764.161309 | ENSG00000170348 |
| EPS15L1 | -0.116057965 | 0.922705419 | 0.236744194 | 0.421747913 | 888.1595822 | 819.8492537 | ENSG00000127527 |
| NA | -0.116079071 | 0.92269192 | 0.908209239 | 0.954399843 | 5.59121881 | 5.198691322 | ENSG00000261549 |
| MSRA | -0.116174922 | 0.92263062 | 0.702638329 | 0.831165356 | 67.79358837 | 62.52207959 | ENSG00000175806 |
| NRBP1 | -0.116413558 | 0.92247802 | 0.067513014 | 0.168342015 | 2921.908356 | 2694.928754 | ENSG00000115216 |
| NA | -0.116457021 | 0.92245023 | 0.886789579 | 0.942266356 | 7.0628802 | 6.481299358 | ENSG00000272721 |
| PSKH1 | -0.116459555 | 0.92244861 | 0.244596596 | 0.431542439 | 783.7342741 | 723.0502339 | ENSG00000159792 |
| FICD | -0.116591955 | 0.922363958 | 0.600365691 | 0.759741754 | 131.4753837 | 121.3270623 | ENSG00000198855 |
| TMUB2 | -0.116592054 | 0.922363895 | 0.442816951 | 0.631223609 | 340.1041964 | 314.2271169 | ENSG00000168591 |
| FANCE | -0.116688421 | 0.922302286 | 0.302119707 | 0.495573648 | 472.088534 | 435.6370438 | ENSG00000112039 |
| LHFPL4 | -0.116857228 | 0.922194375 | 0.225781591 | 0.408815554 | 755.3698413 | 696.8079049 | ENSG00000156959 |
| PDIA4 | -0.116891046 | 0.922172759 | 0.062651808 | 0.159306983 | 9649.406007 | 8898.164205 | ENSG00000155660 |
| ETV1 | -0.11690239 | 0.922165508 | 0.210459531 | 0.389867428 | 1316.706905 | 1213.461942 | ENSG00000006468 |
| ESRP1 | -0.117034948 | 0.922080781 | 0.10872919 | 0.241166445 | 2714.072424 | 2501.722151 | ENSG00000104413 |
| APBA3 | -0.117120732 | 0.922025955 | 0.388631591 | 0.581422849 | 325.5123632 | 300.496396 | ENSG00000011132 |
| ELMO2 | -0.117153988 | 0.922004701 | 0.343061959 | 0.536974103 | 486.3032893 | 447.8784751 | ENSG00000062598 |
| GYG2 | -0.117314553 | 0.921902092 | 0.063527138 | 0.16106177 | 3117.33415 | 2873.224857 | ENSG00000056998 |
| EIPR1 | -0.117394028 | 0.921851308 | 0.261006536 | 0.450039732 | 550.3408609 | 507.2646514 | ENSG00000032389 |
| RCHY1 | -0.117424531 | 0.921831818 | 0.40111261 | 0.592746062 | 348.922296 | 321.4525739 | ENSG00000163743 |
| MMEL1 | -0.117441605 | 0.921820908 | 0.894942605 | 0.946203969 | 8.08465932 | 7.490704619 | ENSG00000142606 |
| PPP2R5B | -0.117482967 | 0.92179448 | 0.601603299 | 0.760944212 | 126.471944 | 116.8237907 | ENSG00000068971 |
| DCLK1 | -0.117523365 | 0.921768668 | 0.248413365 | 0.436542952 | 691.0679987 | 636.4875274 | ENSG00000133083 |
| OPA1 | -0.117603534 | 0.921717448 | 0.166673509 | 0.33014073 | 1881.718281 | 1733.315707 | ENSG00000198836 |
| RADIL | -0.117689772 | 0.921662353 | 0.447807446 | 0.635877392 | 232.1855117 | 213.9067628 | ENSG00000157927 |
| SNORD62B | -0.117701077 | 0.921655131 | 0.875861864 | 0.935632495 | 8.640771605 | 7.970499952 | ENSG00000231587 |
| PROSER2 | -0.117753455 | 0.921621671 | 0.438356912 | 0.626794245 | 267.7698736 | 246.9569296 | ENSG00000148426 |
| APOO | -0.117808463 | 0.921586531 | 0.312211329 | 0.505808704 | 604.8712847 | 557.686539 | ENSG00000184831 |
| STRCP1 | -0.117869236 | 0.92154771 | 0.867384221 | 0.930814611 | 11.18737597 | 10.29213113 | ENSG00000166763 |
| NA | -0.117975581 | 0.921479783 | 0.772505461 | 0.874732173 | 31.6101952 | 29.05332854 | ENSG00000234614 |
| NA | -0.118037681 | 0.921440119 | 0.6631996 | 0.804083749 | 73.11108803 | 67.40969129 | ENSG00000244313 |
| CARNS1 | -0.118151499 | 0.921367427 | 0.800017717 | 0.892525635 | 24.69260805 | 22.70364938 | ENSG00000172508 |
| NA | -0.118153594 | 0.921366089 | 0.881792938 | 0.939373634 | 7.876829374 | 7.302883056 | ENSG00000267541 |
| MORN4 | -0.118197311 | 0.92133817 | 0.381708067 | 0.574586291 | 318.3934705 | 293.4368746 | ENSG00000171160 |
| PIP5KL1 | -0.118214775 | 0.921327017 | 0.824486704 | 0.905596573 | 158.9284316 | 146.5677843 | ENSG00000167103 |
| STIMATE-MUSTN1 | -0.11824123 | 0.921310123 | 0.878053051 | 0.937001108 | 8.276802399 | 7.651719462 | ENSG00000248592 |
| NDC1 | -0.118310949 | 0.921265602 | 0.213879208 | 0.393989055 | 1738.872067 | 1601.177089 | ENSG00000058804 |
| DESI2 | -0.118377791 | 0.921222919 | 0.175549988 | 0.343080664 | 2635.488057 | 2426.671515 | ENSG00000121644 |
| ZC3H4 | -0.118404523 | 0.921205849 | 0.222155237 | 0.404263639 | 1116.001463 | 1027.767317 | ENSG00000130749 |
| SPSB1 | -0.11842681 | 0.921191619 | 0.621618188 | 0.775152125 | 96.756158 | 89.01332964 | ENSG00000171621 |
| NA | -0.118430568 | 0.921189219 | 0.612571025 | 0.768991559 | 109.8469329 | 100.9434036 | ENSG00000250462 |
| NA | -0.118462971 | 0.92116853 | 0.871151423 | 0.933182319 | 8.978207732 | 8.251738454 | ENSG00000254244 |
| EDF1 | -0.118536754 | 0.921121419 | 0.097829948 | 0.222911248 | 4219.501769 | 3886.908072 | ENSG00000107223 |
| PLEKHM2 | -0.118548369 | 0.921114004 | 0.106120768 | 0.23708321 | 1790.372594 | 1649.307849 | ENSG00000116786 |
| STIM1 | -0.118749393 | 0.920985665 | 0.274086397 | 0.465193891 | 526.5618251 | 484.7893915 | ENSG00000167323 |
| NA | -0.118767429 | 0.920974152 | 0.887004619 | 0.942266356 | 6.994029477 | 6.402416534 | ENSG00000183889 |
| HS6ST1 | -0.11883838 | 0.92092886 | 0.109007343 | 0.241609914 | 2362.318634 | 2175.173942 | ENSG00000136720 |
| NA | -0.118845701 | 0.920924186 | 0.851308792 | 0.921459858 | 11.97922676 | 11.08745776 | ENSG00000279520 |
| SSBP3 | -0.118871945 | 0.920907434 | 0.07816849 | 0.188421744 | 3270.664591 | 3012.33146 | ENSG00000157216 |
| NUP62CL | -0.118881384 | 0.920901409 | 0.750067281 | 0.8608959 | 38.45439925 | 35.45304554 | ENSG00000198088 |
| PURG | -0.118932997 | 0.920868464 | 0.550811205 | 0.721876151 | 137.6965038 | 126.9503353 | ENSG00000172733 |
| TSSC2 | -0.118934049 | 0.920867793 | 0.189207508 | 0.361336626 | 1816.293526 | 1673.31757 | ENSG00000223756 |
| CAAP1 | -0.118935901 | 0.920866611 | 0.302639596 | 0.496102354 | 640.2674193 | 589.1185124 | ENSG00000120159 |
| METTL3 | -0.118971517 | 0.920843877 | 0.532116924 | 0.70732662 | 1919.554768 | 1767.967021 | ENSG00000165819 |
| NA | -0.119006173 | 0.920821757 | 0.892285537 | 0.945066946 | 6.020881826 | 5.573784796 | ENSG00000259209 |
| ZC3H14 | -0.119028478 | 0.920807521 | 0.174888477 | 0.342098472 | 1576.330402 | 1451.800116 | ENSG00000100722 |
| AP1M1 | -0.119045728 | 0.920796511 | 0.202547733 | 0.379923053 | 761.8238287 | 701.0744333 | ENSG00000072958 |
| ZNF324B | -0.11916782 | 0.92071859 | 0.703205593 | 0.831299338 | 67.29360146 | 61.85066385 | ENSG00000249471 |
| DTD2 | -0.119244772 | 0.92066948 | 0.697273858 | 0.827195397 | 54.86398877 | 50.44062675 | ENSG00000129480 |
| PCID2 | -0.119313059 | 0.920625904 | 0.195072137 | 0.369339769 | 1075.686652 | 989.7168983 | ENSG00000126226 |
| NCOA7 | -0.119338832 | 0.920609457 | 0.477699881 | 0.661871178 | 179.3346922 | 165.1476183 | ENSG00000111912 |
| THBS2 | -0.11935087 | 0.920601775 | 0.307214297 | 0.501401002 | 527.2120283 | 485.6564413 | ENSG00000186340 |
| BTNL9 | -0.119384404 | 0.920580377 | 0.790948981 | 0.886348777 | 24.74567421 | 22.72610761 | ENSG00000165810 |
| MOB2 | -0.11941084 | 0.920563509 | 0.353697489 | 0.546969483 | 420.0934776 | 387.0910935 | ENSG00000182208 |
| FPGS | -0.119443006 | 0.920542984 | 0.153094158 | 0.310406434 | 1521.084321 | 1401.026303 | ENSG00000136877 |
| AKAP8L | -0.119533896 | 0.920484992 | 0.189930899 | 0.362158362 | 1181.680501 | 1088.288865 | ENSG00000011243 |
| TNNC2 | -0.119598745 | 0.920443617 | 0.70306785 | 0.831263 | 51.72399052 | 47.46211394 | ENSG00000101470 |
| KCNG3 | -0.119769622 | 0.920334603 | 0.510966042 | 0.690100169 | 156.8746285 | 144.4507301 | ENSG00000171126 |
| AP5Z1 | -0.119846534 | 0.92028554 | 0.272987543 | 0.464179002 | 487.9868094 | 449.1952014 | ENSG00000242802 |
| DCTD | -0.119848812 | 0.920284088 | 0.110234352 | 0.243746537 | 1457.82562 | 1341.332703 | ENSG00000129187 |

| | | | | | | | |
|---|---|---|---|---|---|---|---|
| NINJ2-AS1 | -0.119849566 | 0.920283606 | 0.754491182 | 0.863477583 | 34.45832958 | 31.8219996 | ENSG00000177406 |
| CCR10 | -0.119894839 | 0.920254727 | 0.829998806 | 0.908528105 | 18.95431476 | 17.56550793 | ENSG00000184451 |
| KLF10 | -0.119973026 | 0.920204856 | 0.269988499 | 0.460388889 | 515.6492733 | 474.4301005 | ENSG00000155090 |
| PCGF6 | -0.120303924 | 0.91999382 | 0.349603232 | 0.543039415 | 403.9788338 | 371.1491772 | ENSG00000156374 |
| CDC16 | -0.12035212 | 0.919963087 | 0.128020665 | 0.272165062 | 1386.413769 | 1276.000986 | ENSG00000130177 |
| SLC66A1 | -0.12038401 | 0.919942752 | 0.387348714 | 0.580318573 | 371.8926292 | 342.1146684 | ENSG00000040487 |
| SPINT2 | -0.120543072 | 0.919841331 | 0.12690756 | 0.270525727 | 2583.891763 | 2377.234421 | ENSG00000167642 |
| GTF2F2 | -0.120706176 | 0.919737344 | 0.211569201 | 0.391183517 | 838.2244548 | 770.6776164 | ENSG00000188342 |
| ZKSCAN5 | -0.120711112 | 0.919734197 | 0.310422401 | 0.504327558 | 643.9885898 | 591.5725827 | ENSG00000196652 |
| PRPS1 | -0.120833546 | 0.919656147 | 0.059040824 | 0.152044274 | 6666.790899 | 6130.405068 | ENSG00000147224 |
| FARSB | -0.120839985 | 0.919652043 | 0.163324164 | 0.325094004 | 1241.610744 | 1141.259968 | ENSG00000116120 |
| C1orf131 | -0.120870894 | 0.91963234 | 0.442311898 | 0.630697701 | 211.8807534 | 194.9478846 | ENSG00000143633 |
| ARL6IP1 | -0.120874548 | 0.919630011 | 0.260414533 | 0.449314018 | 1477.938633 | 1358.412426 | ENSG00000170540 |
| MAGED4B | -0.120889344 | 0.919620579 | 0.715069626 | 0.838432369 | 43.84094502 | 40.38123362 | ENSG00000187243 |
| SCCPDH | -0.121010699 | 0.919543227 | 0.263194493 | 0.452803002 | 548.8554895 | 504.7778801 | ENSG00000143653 |
| NA | -0.121017636 | 0.919538805 | 0.800717195 | 0.8929372 | 20.63390379 | 19.02041572 | ENSG00000220842 |
| BYSL | -0.121128314 | 0.919468264 | 0.126168976 | 0.269340683 | 1164.412773 | 1070.569748 | ENSG00000112578 |
| NSFL1C | -0.121212015 | 0.919414921 | 0.091449927 | 0.212131782 | 2569.973142 | 2363.636099 | ENSG00000088833 |
| NA | -0.121228803 | 0.919404222 | 0.915771418 | 0.95827165 | 3.791494334 | 3.514621124 | ENSG00000258611 |
| LXN | -0.12141202 | 0.919287469 | 0.649433979 | 0.795321709 | 68.85708377 | 63.26087 | ENSG00000079257 |
| SUPT3H | -0.121465227 | 0.919253566 | 0.422321773 | 0.613473763 | 244.5677707 | 224.8298449 | ENSG00000196284 |
| POLR2E | -0.121481332 | 0.919243304 | 0.054671521 | 0.143240312 | 5523.575778 | 5078.07048 | ENSG00000099817 |
| TPBG | -0.121495326 | 0.919234387 | 0.301454754 | 0.494877096 | 1023.651142 | 941.5377139 | ENSG00000146242 |
| SPPL3 | -0.121500202 | 0.919231281 | 0.352548463 | 0.545624369 | 628.4121316 | 577.1851406 | ENSG00000157837 |
| HEXIM2 | -0.121539171 | 0.919206452 | 0.772464138 | 0.874732173 | 29.04337158 | 26.75850408 | ENSG00000168517 |
| MINDY3 | -0.121620146 | 0.91915486 | 0.437959562 | 0.626516094 | 237.0536101 | 217.8540013 | ENSG00000148481 |
| LDAH | -0.121762156 | 0.919064389 | 0.300568926 | 0.493816547 | 488.9048164 | 448.8620613 | ENSG00000118961 |
| FKBP4 | -0.121866741 | 0.918997765 | 0.02583767 | 0.079494585 | 9477.260569 | 8709.108387 | ENSG00000004478 |
| AGPAT1 | -0.121877513 | 0.918990904 | 0.25060425 | 0.439032648 | 533.7336112 | 490.5547967 | ENSG00000204310 |
| UMPS | -0.121945794 | 0.91894741 | 0.14506332 | 0.298543277 | 1268.857766 | 1165.923725 | ENSG00000114491 |
| BMP8A | -0.122024947 | 0.918896994 | 0.787747074 | 0.883997594 | 25.74229573 | 23.77305486 | ENSG00000183682 |
| RSL1D1 | -0.12213428 | 0.918827359 | 0.055109818 | 0.144062815 | 4391.049048 | 4033.923897 | ENSG00000171490 |
| METTL1 | -0.122135135 | 0.918826814 | 0.373467726 | 0.566807307 | 302.5585287 | 278.0614893 | ENSG00000037897 |
| MEGF10 | -0.122267631 | 0.918742434 | 0.246457827 | 0.434041178 | 576.5018757 | 529.193895 | ENSG00000145794 |
| RGS17 | -0.122280977 | 0.918733935 | 0.633801847 | 0.783961787 | 89.30653378 | 81.90162896 | ENSG00000091844 |
| ZMYM2 | -0.122338251 | 0.918697463 | 0.061700378 | 0.157449375 | 2450.881122 | 2251.099136 | ENSG00000121741 |
| GPD1L | -0.122354967 | 0.918686818 | 0.522863671 | 0.699624565 | 172.1214829 | 157.9636448 | ENSG00000152642 |
| TCP1 | -0.122360935 | 0.918683017 | 0.083223915 | 0.197463546 | 7084.900468 | 6507.653345 | ENSG00000120438 |
| ZNF367 | -0.122415127 | 0.91864851 | 0.512058583 | 0.69104346 | 175.8042893 | 161.2521295 | ENSG00000165244 |
| DDX24 | -0.122476157 | 0.918609649 | 0.052748066 | 0.139323452 | 3235.720721 | 2972.123328 | ENSG00000089737 |
| PDXP | -0.122662308 | 0.918491129 | 0.504406621 | 0.684032511 | 274.6319368 | 251.9451893 | ENSG00000241360 |
| SIAH2 | -0.122787109 | 0.918411678 | 0.297173018 | 0.490149035 | 497.3524602 | 456.3521131 | ENSG00000181788 |
| ID1 | -0.122787815 | 0.918411228 | 0.128616335 | 0.273107936 | 1566.396852 | 1438.060643 | ENSG00000125968 |
| IDE | -0.122801105 | 0.918402768 | 0.160890572 | 0.321468938 | 1021.704441 | 937.7730378 | ENSG00000119912 |
| HEY1 | -0.122821693 | 0.918389662 | 0.676462363 | 0.812987929 | 71.71321571 | 65.68720304 | ENSG00000164683 |
| TMEM41B | -0.122827299 | 0.918386093 | 0.323550445 | 0.517714232 | 498.5899313 | 457.5334917 | ENSG00000166471 |
| SEZ6L2 | -0.122891158 | 0.918345443 | 0.135739685 | 0.284206882 | 1639.940204 | 1505.324466 | ENSG00000174938 |
| CARM1 | -0.122908904 | 0.918334147 | 0.053370255 | 0.140484408 | 4808.543931 | 4415.499311 | ENSG00000142453 |
| HDAC5 | -0.122958179 | 0.918302782 | 0.087434841 | 0.204921262 | 2303.778108 | 2115.223878 | ENSG00000108840 |
| RELCH | -0.122970186 | 0.918295139 | 0.246085942 | 0.433592002 | 578.7715138 | 531.4490276 | ENSG00000134444 |
| TAF4B | -0.12299821 | 0.918277301 | 0.344495078 | 0.538081221 | 413.7753503 | 379.5135416 | ENSG00000141384 |
| NA | -0.123150462 | 0.918180398 | 0.796117921 | 0.889889002 | 21.0932568 | 19.38053705 | ENSG00000253954 |
| ETNK1 | -0.123151315 | 0.918179855 | 0.293475971 | 0.486046741 | 851.6346411 | 781.1677145 | ENSG00000139163 |
| NA | -0.123160806 | 0.918173815 | 0.751151636 | 0.861500274 | 32.74263903 | 30.1379294 | ENSG00000213430 |
| NDUFAB1 | -0.123200464 | 0.918148576 | 0.15894087 | 0.318707138 | 1455.05174 | 1335.027756 | ENSG00000004779 |
| ULK1 | -0.123287792 | 0.918093001 | 0.123937466 | 0.266017023 | 1810.371876 | 1662.21807 | ENSG00000177169 |
| NA | -0.123308684 | 0.918079706 | 0.865833856 | 0.930066343 | 10.83227614 | 9.932009806 | ENSG00000253570 |
| NGDN | -0.123504018 | 0.917955411 | 0.220416329 | 0.401967552 | 885.9395073 | 813.7995756 | ENSG00000129460 |
| GLYCTK | -0.123547261 | 0.917927896 | 0.662091278 | 0.803380688 | 70.43635065 | 64.83710076 | ENSG00000168237 |
| DHRS4 | -0.123656287 | 0.91785853 | 0.50258668 | 0.68301375 | 181.6870789 | 166.9083098 | ENSG00000157326 |
| WTAP | -0.123687017 | 0.91783898 | 0.105328126 | 0.235795567 | 1351.354701 | 1240.45481 | ENSG00000146457 |
| CD40 | -0.123727334 | 0.91781333 | 0.348841771 | 0.542174707 | 368.0048566 | 337.794415 | ENSG00000101017 |
| CRYZL2P | -0.123878604 | 0.917717101 | 0.83930257 | 0.913691856 | 16.3254579 | 14.9090248 | ENSG00000242193 |
| TSNAX | -0.123944607 | 0.917675117 | 0.559755309 | 0.728853793 | 211.4989099 | 193.9246778 | ENSG00000116918 |
| PLRG1 | -0.12409968 | 0.917576482 | 0.10275204 | 0.231145322 | 1581.108408 | 1450.400462 | ENSG00000171566 |
| CCDC138 | -0.124161046 | 0.917537453 | 0.373383049 | 0.566771616 | 359.6509203 | 329.9378232 | ENSG00000163006 |
| C1orf50 | -0.124193923 | 0.917516544 | 0.735419946 | 0.852171088 | 41.46744456 | 38.09714443 | ENSG00000164008 |
| GPD2 | -0.124212616 | 0.917504656 | 0.275437453 | 0.466419164 | 763.0722815 | 699.1559358 | ENSG00000115159 |
| RBL1 | -0.124299669 | 0.917449295 | 0.493261055 | 0.675641149 | 172.501838 | 158.4833955 | ENSG00000080839 |
| ORAI2 | -0.124305685 | 0.917445469 | 0.222954612 | 0.405081931 | 827.4947103 | 759.1727282 | ENSG00000160991 |
| NA | -0.124364239 | 0.917408234 | 0.827416163 | 0.907201323 | 15.78019183 | 14.53816821 | ENSG00000278784 |
| SLC50A1 | -0.124600051 | 0.917258293 | 0.21580305 | 0.396166177 | 687.2003477 | 629.7562564 | ENSG00000169241 |
| GANAB | -0.124620458 | 0.917245319 | 0.04021631 | 0.112912566 | 11761.8016 | 10788.04469 | ENSG00000089597 |
| PKNOX1 | -0.124683818 | 0.917205036 | 0.241613447 | 0.427785979 | 587.2063498 | 538.6076201 | ENSG00000160199 |

| | | | | | | | |
|---|---|---|---|---|---|---|---|
| H4-16 | -0.124745443 | 0.917165859 | 0.820790167 | 0.903839639 | 16.51760098 | 15.16455589 | ENSG00000197837 |
| BACE2 | -0.124877558 | 0.917081873 | 0.408129101 | 0.599941488 | 317.7491827 | 291.084924 | ENSG00000182240 |
| SPATA2 | -0.124915113 | 0.917058001 | 0.274264221 | 0.465250018 | 484.7065315 | 444.178231 | ENSG00000158480 |
| ENOPH1 | -0.125112388 | 0.91693261 | 0.136428437 | 0.285230079 | 1861.056138 | 1705.516794 | ENSG00000145293 |
| STRN | -0.125153376 | 0.91690656 | 0.276489385 | 0.467559695 | 691.7142633 | 633.4767601 | ENSG00000115808 |
| INPP5A | -0.125189293 | 0.916883733 | 0.268988269 | 0.459272625 | 533.8280255 | 488.9585925 | ENSG00000068383 |
| B4GALT7 | -0.125252914 | 0.916843301 | 0.333960179 | 0.528575402 | 393.1637177 | 360.073554 | ENSG00000027847 |
| ACBD7 | -0.125261433 | 0.916837886 | 0.414775327 | 0.606313267 | 325.641774 | 298.0153099 | ENSG00000176244 |
| GRPEL1 | -0.12526241 | 0.916837265 | 0.147175612 | 0.301709191 | 1034.327068 | 947.8973293 | ENSG00000109519 |
| NME3 | -0.125297176 | 0.916815172 | 0.185323057 | 0.356056516 | 1193.792535 | 1095.252342 | ENSG00000103024 |
| ABLIM1 | -0.125314221 | 0.91680434 | 0.116719924 | 0.254534307 | 1458.558 | 1336.744167 | ENSG00000099204 |
| NMT2 | -0.125366076 | 0.916771388 | 0.334644679 | 0.528981192 | 337.8690964 | 309.6203685 | ENSG00000152465 |
| EPHB4 | -0.125499806 | 0.916686412 | 0.044783918 | 0.122748307 | 4524.68581 | 4147.371225 | ENSG00000196411 |
| GMEB2 | -0.125503876 | 0.916683826 | 0.173179455 | 0.339687587 | 752.4357712 | 689.8278244 | ENSG00000101216 |
| C7orf25 | -0.125557665 | 0.916649649 | 0.897906379 | 0.947703593 | 5.310005754 | 4.924938894 | ENSG00000136197 |
| ALDH18A1 | -0.125636275 | 0.916599704 | 0.061181242 | 0.156404571 | 4320.750031 | 3960.246116 | ENSG00000059573 |
| DOCK4 | -0.125645086 | 0.916594106 | 0.226416478 | 0.40980498 | 666.2218069 | 610.0779714 | ENSG00000128512 |
| SIPA1 | -0.125740786 | 0.916533306 | 0.178467466 | 0.347376183 | 1336.173871 | 1224.926789 | ENSG00000213445 |
| BANF1 | -0.125755048 | 0.916524246 | 0.086659079 | 0.203463311 | 4518.650993 | 4141.511248 | ENSG00000175334 |
| RHBG | -0.125788587 | 0.91650294 | 0.808439734 | 0.897169154 | 19.39976234 | 17.73736966 | ENSG00000132677 |
| UAP1 | -0.125973208 | 0.916385663 | 0.204256285 | 0.382276682 | 843.1718442 | 772.4844813 | ENSG00000117143 |
| TENT4A | -0.125996935 | 0.916370591 | 0.188668714 | 0.360567741 | 705.7495631 | 646.9109926 | ENSG00000112941 |
| EMG1 | -0.12606932 | 0.916324615 | 0.104738933 | 0.234873731 | 1276.401414 | 1169.429921 | ENSG00000126749 |
| NDUFAF6 | -0.12612953 | 0.916286374 | 0.475925361 | 0.66068464 | 185.7544646 | 170.1611876 | ENSG00000156170 |
| ZNF764 | -0.126167892 | 0.916262009 | 0.533722146 | 0.708596437 | 135.793398 | 124.6470371 | ENSG00000169951 |
| CHMP4B | -0.12626502 | 0.916200325 | 0.074770999 | 0.182151116 | 2205.822098 | 2021.101564 | ENSG00000101421 |
| KLF15 | -0.126316886 | 0.916167388 | 0.42055925 | 0.611728547 | 367.454081 | 337.3234198 | ENSG00000163884 |
| NECTIN3 | -0.126317323 | 0.91616711 | 0.248107332 | 0.436158711 | 502.9244445 | 460.4461588 | ENSG00000177707 |
| SLC7A6 | -0.126450062 | 0.91608282 | 0.413489206 | 0.604862584 | 330.2335226 | 302.8805524 | ENSG00000103064 |
| PRADC1 | -0.126488428 | 0.916058458 | 0.546343569 | 0.718354974 | 133.9380518 | 122.5332723 | ENSG00000135617 |
| KLK8 | -0.126497947 | 0.916052414 | 0.88236183 | 0.939381593 | 7.036347121 | 6.39493046 | ENSG00000129455 |
| APCDD1 | -0.126533573 | 0.916029793 | 0.546502382 | 0.718495899 | 114.8364672 | 105.1798047 | ENSG00000154856 |
| DHX16 | -0.126629078 | 0.915969155 | 0.096691677 | 0.221026583 | 1465.703156 | 1342.879531 | ENSG00000204560 |
| SRSF6 | -0.12670152 | 0.915923163 | 0.163336321 | 0.325094004 | 1370.593567 | 1254.776558 | ENSG00000124193 |
| TGFBI | -0.126720582 | 0.915911061 | 0.39162022 | 0.584309222 | 252.3238633 | 231.0941513 | ENSG00000120708 |
| BNC2 | -0.126916961 | 0.915786396 | 0.336331489 | 0.530561564 | 415.8218475 | 380.1809353 | ENSG00000173068 |
| FZD1 | -0.126925695 | 0.915780852 | 0.68994419 | 0.822191533 | 56.44621717 | 51.47150255 | ENSG00000157240 |
| KCTD10 | -0.126926398 | 0.915780406 | 0.122834496 | 0.26423103 | 1297.654425 | 1187.897058 | ENSG00000110906 |
| FRMD4B | -0.127042572 | 0.915706665 | 0.495094217 | 0.677062939 | 146.5155131 | 134.1247442 | ENSG00000114541 |
| STK33 | -0.127237448 | 0.915582982 | 0.447831423 | 0.635877392 | 267.4080918 | 244.8908295 | ENSG00000130413 |
| KCTD6 | -0.127239992 | 0.915581367 | 0.436635846 | 0.62586147 | 196.0425571 | 179.5768478 | ENSG00000168301 |
| YIPF3 | -0.127317804 | 0.915531987 | 0.122867552 | 0.264271463 | 1132.822601 | 1037.279174 | ENSG00000137207 |
| GSTCD | -0.127339911 | 0.915517957 | 0.452353391 | 0.639749542 | 337.4636814 | 308.3964063 | ENSG00000138780 |
| TCEANC | -0.127413055 | 0.915471542 | 0.735608189 | 0.852173027 | 38.92271933 | 35.5533989 | ENSG00000176896 |
| RDH10 | -0.127513242 | 0.91540797 | 0.504719151 | 0.684288598 | 141.9177589 | 130.0106537 | ENSG00000121039 |
| EMC1-AS1 | -0.127546101 | 0.915387121 | 0.921297599 | 0.961120642 | 4.303913501 | 3.946027581 | ENSG00000230424 |
| KDM2B | -0.127564874 | 0.91537521 | 0.046838133 | 0.127100727 | 2753.093541 | 2519.549443 | ENSG00000089094 |
| KLHL3 | -0.127784247 | 0.91523603 | 0.346704508 | 0.540170237 | 352.9185762 | 322.8873168 | ENSG00000146021 |
| ADAMTS7 | -0.127933037 | 0.915141644 | 0.090995695 | 0.211369121 | 2303.505749 | 2108.353114 | ENSG00000136378 |
| DHX30 | -0.128000672 | 0.915098742 | 0.036850291 | 0.105301157 | 3267.657536 | 2990.206732 | ENSG00000132153 |
| SRSF4 | -0.128063962 | 0.915058598 | 0.056383426 | 0.146668128 | 3802.128511 | 3479.332529 | ENSG00000116350 |
| LOC100128770 | -0.128068307 | 0.915055842 | 0.792498647 | 0.887181078 | 36.25164253 | 33.3645904 | ENSG00000205890 |
| CYP2S1 | -0.128106447 | 0.915031652 | 0.076602527 | 0.185394521 | 6928.459621 | 6340.029857 | ENSG00000167600 |
| BNIP1 | -0.12813884 | 0.915011106 | 0.572976467 | 0.738883513 | 104.7800619 | 95.7050204 | ENSG00000113734 |
| CABLES1 | -0.128162215 | 0.914996281 | 0.286256714 | 0.477584115 | 422.8796014 | 387.0967158 | ENSG00000134508 |
| MRPL37 | -0.128165 | 0.914994515 | 0.054782539 | 0.143450068 | 2920.027724 | 2672.059447 | ENSG00000116221 |
| ADPGK | -0.128200904 | 0.914971744 | 0.184731934 | 0.355426313 | 701.1916612 | 641.6122743 | ENSG00000159322 |
| EIF2S1 | -0.128251155 | 0.914939875 | 0.127431303 | 0.271285523 | 2013.609783 | 1841.881149 | ENSG00000134001 |
| DKC1 | -0.12826983 | 0.914928032 | 0.050982708 | 0.13584697 | 4357.646186 | 3987.026197 | ENSG00000130826 |
| STARD10 | -0.128364709 | 0.914867863 | 0.313068434 | 0.506627988 | 342.1832548 | 312.9096261 | ENSG00000214530 |
| THUMPD1 | -0.128409692 | 0.914839338 | 0.20333826 | 0.381020252 | 1055.242614 | 964.591287 | ENSG00000066654 |
| ZNF557 | -0.128667289 | 0.914676006 | 0.600157103 | 0.759725903 | 92.9412875 | 85.03052451 | ENSG00000130544 |
| NA | -0.128698832 | 0.914656007 | 0.671473896 | 0.809097911 | 53.87831117 | 49.40204164 | ENSG00000280385 |
| LOC374443 | -0.128726179 | 0.91463867 | 0.860404493 | 0.926274214 | 11.36245671 | 10.44241071 | ENSG00000256594 |
| NIBAN1 | -0.128917734 | 0.914517236 | 0.278452016 | 0.46955051 | 543.6379212 | 496.4357104 | ENSG00000135842 |
| HHAT | -0.128935417 | 0.914506027 | 0.869413453 | 0.93223656 | 8.700151589 | 7.940444037 | ENSG00000054392 |
| GPATCH11 | -0.12896 | 0.914490444 | 0.599907238 | 0.759476787 | 83.74322353 | 76.62493088 | ENSG00000152133 |
| LYN | -0.129044481 | 0.914436896 | 0.15270751 | 0.309930025 | 982.0619759 | 897.5044301 | ENSG00000254087 |
| LRP2BP | -0.129126975 | 0.914384609 | 0.778367267 | 0.87829679 | 23.13819051 | 21.11668335 | ENSG00000109771 |
| NLN | -0.129129708 | 0.914382877 | 0.102331425 | 0.230590958 | 3674.323967 | 3358.653655 | ENSG00000123213 |
| ADRM1 | -0.129165547 | 0.914360162 | 0.093872789 | 0.215914405 | 1509.523911 | 1380.31077 | ENSG00000130706 |
| SLC35G1 | -0.129192604 | 0.914343014 | 0.46721047 | 0.653162956 | 167.0995076 | 152.939555 | ENSG00000176273 |
| GIN1 | -0.129217181 | 0.914327438 | 0.662917008 | 0.803985331 | 63.91420414 | 58.21895664 | ENSG00000145723 |
| GPANK1 | -0.129247698 | 0.914308097 | 0.276248341 | 0.467322632 | 478.4339312 | 437.1158985 | ENSG00000204438 |

| | | | | | | | |
|---|---|---|---|---|---|---|---|
| FNDC10 | -0.129266402 | 0.914296244 | 0.378860341 | 0.572040534 | 519.5660821 | 474.9505923 | ENSG00000228594 |
| CGNL1 | -0.129369159 | 0.914231125 | 0.102400923 | 0.23068994 | 1887.785427 | 1726.150363 | ENSG00000128849 |
| ZDHHC18 | -0.12941938 | 0.9141993 | 0.155960114 | 0.314340912 | 1029.448334 | 941.3835211 | ENSG00000204160 |
| TTI2 | -0.129420545 | 0.914198562 | 0.336709953 | 0.531068187 | 342.034632 | 312.6288256 | ENSG00000129696 |
| UTP15 | -0.129534287 | 0.914126489 | 0.237765872 | 0.423156584 | 578.6028219 | 528.6154582 | ENSG00000164338 |
| RAD54B | -0.12960143 | 0.914083947 | 0.513320035 | 0.691998545 | 133.3016412 | 121.7541287 | ENSG00000265817 |
| CENPN | -0.129623154 | 0.914070183 | 0.256204455 | 0.444990476 | 465.3111062 | 425.2611317 | ENSG00000166451 |
| HSF1 | -0.129643689 | 0.914057173 | 0.090680297 | 0.210768576 | 2121.860522 | 1939.359955 | ENSG00000185122 |
| AFF2 | -0.129693471 | 0.914025632 | 0.417048734 | 0.608245297 | 235.5712979 | 215.510015 | ENSG00000155966 |
| ZDHHC23 | -0.129743805 | 0.913993744 | 0.331611288 | 0.525800635 | 349.0446188 | 319.4348945 | ENSG00000184307 |
| NA | -0.129823382 | 0.913943331 | 0.55350234 | 0.723938212 | 118.1431578 | 107.9745096 | ENSG00000213621 |
| PUM2 | -0.130034279 | 0.913809738 | 0.115870249 | 0.253096652 | 1818.2328 | 1660.762458 | ENSG00000055917 |
| CENPBD1P1 | -0.130136707 | 0.913744862 | 0.304170155 | 0.497923481 | 407.9828859 | 373.1297131 | ENSG00000213753 |
| MRPL30 | -0.130213735 | 0.913696076 | 0.228223316 | 0.41182854 | 650.4218018 | 593.623433 | ENSG00000185414 |
| PRB3 | -0.130246768 | 0.913675156 | 0.788777878 | 0.884435871 | 21.19317299 | 19.45562103 | ENSG00000197870 |
| NDUFV1 | -0.130343619 | 0.913613821 | 0.068263259 | 0.169710429 | 2543.738525 | 2324.589143 | ENSG00000167792 |
| DIDO1 | -0.130408055 | 0.913573016 | 0.184136636 | 0.35483348 | 2829.038211 | 2585.804727 | ENSG00000101191 |
| ARFIP2 | -0.130491792 | 0.913519993 | 0.118315322 | 0.256955259 | 1112.391476 | 1016.650992 | ENSG00000132254 |
| SRSF3 | -0.130547003 | 0.913485034 | 0.037966876 | 0.107760809 | 12358.5652 | 11288.01965 | ENSG00000112081 |
| EGFL6 | -0.130550704 | 0.91348269 | 0.742619807 | 0.856105567 | 31.72273904 | 28.90316059 | ENSG00000198759 |
| LOC101927702 | -0.130718129 | 0.913376687 | 0.910598218 | 0.955664065 | 4.191467358 | 3.825915541 | ENSG00000254718 |
| PIGU | -0.130852159 | 0.913291835 | 0.19774995 | 0.373471984 | 676.8722 | 617.8456063 | ENSG00000101464 |
| RPLP2 | -0.13088098 | 0.913273591 | 0.046231707 | 0.125720292 | 10406.29903 | 9503.697511 | ENSG00000177600 |
| NA | -0.130953985 | 0.913227377 | 0.618720963 | 0.773359478 | 73.67035723 | 67.16143148 | ENSG00000233328 |
| NA | -0.130969594 | 0.913217496 | 0.880721384 | 0.938659582 | 7.900107843 | 7.137631334 | ENSG00000234337 |
| MED28 | -0.130973144 | 0.913215249 | 0.084470525 | 0.199614524 | 2140.618357 | 1953.818299 | ENSG00000118579 |
| MIR924HG | -0.131023186 | 0.913183574 | 0.654019813 | 0.79772604 | 56.57148636 | 51.71987397 | ENSG00000267374 |
| LAGE3 | -0.131068625 | 0.913154813 | 0.117202366 | 0.25525601 | 1324.424302 | 1209.872812 | ENSG00000196976 |
| GALNT14 | -0.13110168 | 0.913133891 | 0.260657611 | 0.449607088 | 416.3750959 | 380.3085821 | ENSG00000158089 |
| CCNQ | -0.131147105 | 0.91310514 | 0.351541022 | 0.544557714 | 383.6393343 | 350.7796951 | ENSG00000262919 |
| RHBDD3 | -0.131161978 | 0.913095727 | 0.13756653 | 0.287026687 | 1576.663267 | 1440.744157 | ENSG00000100263 |
| SLC25A39 | -0.131238236 | 0.913047464 | 0.019574082 | 0.06337981 | 5513.211755 | 5034.311296 | ENSG00000013306 |
| WASH5P | -0.131268684 | 0.913028194 | 0.745763677 | 0.858090723 | 29.94639073 | 27.40366276 | ENSG00000282458 |
| HCG23 | -0.131349835 | 0.912976838 | 0.737464384 | 0.852943558 | 31.75242904 | 29.05767704 | ENSG00000225914 |
| CPOX | -0.131386867 | 0.912953404 | 0.200586834 | 0.377199309 | 835.0651841 | 762.2388293 | ENSG00000080819 |
| GIPC1 | -0.131453432 | 0.912911281 | 0.091739918 | 0.212485334 | 2341.567894 | 2137.554975 | ENSG00000123159 |
| GFRA3 | -0.131531581 | 0.912861831 | 0.838429113 | 0.913279176 | 13.74157194 | 12.45979551 | ENSG00000146013 |
| MRPL27 | -0.131594446 | 0.912822054 | 0.174301801 | 0.341202955 | 828.1491604 | 756.0679801 | ENSG00000108826 |
| LINC01801 | -0.13160598 | 0.912814757 | 0.784630355 | 0.882158322 | 20.71350303 | 18.89270599 | ENSG00000267767 |
| SLC35B4 | -0.131803482 | 0.912689803 | 0.151871478 | 0.308909787 | 922.9586776 | 841.8032968 | ENSG00000205060 |
| PGP | -0.131815581 | 0.912682149 | 0.201854373 | 0.37889092 | 1615.043629 | 1474.586003 | ENSG00000184207 |
| NOS2 | -0.131891936 | 0.912633846 | 0.633685296 | 0.783961787 | 64.66689419 | 59.07918163 | ENSG00000007171 |
| ARHGAP23 | -0.131970593 | 0.91258409 | 0.078676186 | 0.189227151 | 1977.639203 | 1805.13704 | ENSG00000275832 |
| ZNF586 | -0.13201576 | 0.91255552 | 0.358695692 | 0.551935738 | 269.0487906 | 245.6888472 | ENSG00000083828 |
| NA | -0.132025845 | 0.91254914 | 0.545984525 | 0.718035705 | 114.7296358 | 104.5604651 | ENSG00000213866 |
| NA | -0.132029928 | 0.912546558 | 0.678090526 | 0.813972138 | 62.11014267 | 56.86312838 | ENSG00000279495 |
| C3orf38 | -0.132051754 | 0.912532752 | 0.288563751 | 0.480179616 | 405.7265745 | 369.7659926 | ENSG00000179021 |
| LDB1 | -0.132086438 | 0.912510814 | 0.062643185 | 0.159306914 | 2511.748607 | 2292.937717 | ENSG00000198728 |
| TPMT | -0.132131675 | 0.912482202 | 0.203323928 | 0.381020252 | 605.2882822 | 552.3913645 | ENSG00000137364 |
| KCNB2 | -0.132213887 | 0.912430206 | 0.756372907 | 0.864335767 | 27.24364711 | 24.82661211 | ENSG00000182674 |
| RNPS1 | -0.132219976 | 0.912426355 | 0.017500432 | 0.057786035 | 7017.819516 | 6403.651772 | ENSG00000205937 |
| SLC10A7 | -0.13223236 | 0.912418523 | 0.72267966 | 0.843513119 | 35.51729259 | 32.36939284 | ENSG00000120519 |
| ZFX | -0.132332127 | 0.912355428 | 0.367161702 | 0.560264695 | 401.1058349 | 365.4873746 | ENSG00000005889 |
| DCLRE1B | -0.132350763 | 0.912343643 | 0.365006815 | 0.558171313 | 258.0641334 | 235.388354 | ENSG00000118655 |
| ARID3B | -0.132383973 | 0.912322642 | 0.03289865 | 0.096402146 | 5135.058251 | 4685.36133 | ENSG00000179361 |
| NUP98 | -0.132390934 | 0.91231824 | 0.069251152 | 0.171613966 | 3004.06336 | 2739.99419 | ENSG00000110713 |
| RCE1 | -0.132508952 | 0.912243612 | 0.156417622 | 0.315045012 | 828.0835869 | 755.1834332 | ENSG00000173653 |
| AMDHD2 | -0.132543575 | 0.912221719 | 0.381105628 | 0.574027769 | 311.8105533 | 284.9545968 | ENSG00000162066 |
| EIF2D | -0.132572352 | 0.912203524 | 0.183607651 | 0.35414551 | 783.1322139 | 714.4164026 | ENSG00000143486 |
| LAMB2 | -0.132674989 | 0.91213863 | 0.059183832 | 0.152349047 | 2038.432944 | 1859.978587 | ENSG00000172037 |
| TRA2B | -0.132732057 | 0.912102549 | 0.042995532 | 0.119132784 | 6690.653411 | 6102.582544 | ENSG00000136527 |
| NA | -0.132745566 | 0.912094008 | 0.881908566 | 0.939373634 | 7.287870184 | 6.574716297 | ENSG00000244586 |
| GPATCH3 | -0.132782461 | 0.912070683 | 0.311769413 | 0.505364204 | 356.067955 | 324.9350924 | ENSG00000198746 |
| PPP5C | -0.13281813 | 0.912048134 | 0.092483676 | 0.213700593 | 3329.332348 | 3036.968196 | ENSG00000011485 |
| BAG3 | -0.133002171 | 0.911931794 | 0.259827945 | 0.448762231 | 613.7905107 | 559.6611254 | ENSG00000151929 |
| MAGOH | -0.133153674 | 0.911836033 | 0.151933361 | 0.309001747 | 1488.261083 | 1356.947857 | ENSG00000162385 |
| CCNJL | -0.13315657 | 0.911834202 | 0.122529596 | 0.263666961 | 1080.203379 | 984.4515481 | ENSG00000135083 |
| ABCB10 | -0.133273294 | 0.911760432 | 0.473546301 | 0.658615274 | 258.2585238 | 235.3268249 | ENSG00000135776 |
| SEMA4D | -0.133279188 | 0.911756707 | 0.131021869 | 0.277115066 | 1193.694694 | 1087.982117 | ENSG00000187764 |
| SIPA1L1 | -0.133287515 | 0.911751444 | 0.092660939 | 0.214056817 | 1578.490427 | 1439.082734 | ENSG00000197555 |
| SREBF2-AS1 | -0.133309656 | 0.911737452 | 0.680487487 | 0.815245229 | 51.59082147 | 47.09125735 | ENSG00000184068 |
| PYY | -0.133331068 | 0.91172392 | 0.825635946 | 0.906340611 | 15.75040414 | 14.40352205 | ENSG00000131096 |
| UBFD1 | -0.133347764 | 0.911713369 | 0.033151732 | 0.096990404 | 3065.905271 | 2794.877 | ENSG00000103353 |
| ARVCF | -0.133356899 | 0.911707596 | 0.236508762 | 0.421648075 | 1615.328766 | 1473.682345 | ENSG00000099889 |

| | | | | | | | |
|---|---|---|---|---|---|---|---|
| SNUPN | -0.133378437 | 0.911693985 | 0.368704477 | 0.5616478 | 276.4938824 | 252.113722 | ENSG00000169371 |
| HNRNPU | -0.133411974 | 0.911672792 | 0.043475712 | 0.120104166 | 20025.50893 | 18256.32053 | ENSG00000153187 |
| TASOR | -0.133570507 | 0.911572617 | 0.195731832 | 0.370399609 | 802.9462303 | 731.4073036 | ENSG00000163946 |
| RAB35 | -0.133622781 | 0.911539588 | 0.107318802 | 0.238867141 | 1565.759607 | 1427.06532 | ENSG00000111737 |
| NAPB | -0.133715969 | 0.911480711 | 0.388924916 | 0.581598708 | 237.4833935 | 216.6994294 | ENSG00000125814 |
| NA | -0.133814814 | 0.911418264 | 0.828373508 | 0.907514734 | 14.13029279 | 12.8649449 | ENSG00000273796 |
| TMEM216 | -0.13386369 | 0.911387387 | 0.312604319 | 0.506161696 | 325.8023705 | 297.0590087 | ENSG00000187049 |
| FAM131C | -0.133892061 | 0.911369465 | 0.709424503 | 0.83482373 | 39.25444299 | 35.73753323 | ENSG00000185519 |
| ALKBH7 | -0.133953068 | 0.911330926 | 0.183195933 | 0.35364581 | 1070.570089 | 975.0312214 | ENSG00000125652 |
| FHOD1 | -0.133975338 | 0.911316859 | 0.15805459 | 0.317445143 | 1326.162776 | 1209.084471 | ENSG00000135723 |
| ZNF200 | -0.134002912 | 0.911299441 | 0.517567699 | 0.695065556 | 142.4273142 | 129.9422829 | ENSG00000010539 |
| AHCYL2 | -0.134113933 | 0.911229316 | 0.217553373 | 0.398510992 | 575.5726318 | 524.2148333 | ENSG00000158467 |
| FBXO7 | -0.134159491 | 0.911200541 | 0.080892233 | 0.193228076 | 1341.932459 | 1222.4339 | ENSG00000100225 |
| NBPF10 | -0.134166187 | 0.911196312 | 0.804141015 | 0.894704967 | 16.19091337 | 14.75180881 | ENSG00000271425 |
| LINC00539 | -0.134176671 | 0.911189691 | 0.865451863 | 0.929978419 | 7.973490953 | 7.276625988 | ENSG00000224429 |
| RBM45 | -0.134249084 | 0.911143956 | 0.519468983 | 0.696558973 | 162.4274985 | 147.8377531 | ENSG00000155636 |
| CDKL3 | -0.134503436 | 0.910983333 | 0.884899716 | 0.940949559 | 5.727041119 | 5.243719384 | ENSG00000006837 |
| GFM1 | -0.134556771 | 0.910949655 | 0.137788327 | 0.28736006 | 899.4928898 | 818.9041166 | ENSG00000168827 |
| PFDN4 | -0.134557439 | 0.910949233 | 0.232712825 | 0.417490997 | 550.9232957 | 501.83581 | ENSG00000101132 |
| NA | -0.134756203 | 0.910823738 | 0.675020787 | 0.811873412 | 56.49051165 | 51.59695077 | ENSG00000275131 |
| IL21R | -0.134776636 | 0.910810838 | 0.746169704 | 0.858415077 | 27.88261324 | 25.41534619 | ENSG00000103522 |
| USP39 | -0.134823539 | 0.910781228 | 0.045688171 | 0.124784049 | 2543.39319 | 2316.6422 | ENSG00000168883 |
| P3H3 | -0.13489441 | 0.910736488 | 0.130627515 | 0.276628241 | 1115.979636 | 1016.911739 | ENSG00000110811 |
| A1BG-AS1 | -0.134929136 | 0.910714566 | 0.885066161 | 0.940967086 | 5.948874191 | 5.416019147 | ENSG00000268895 |
| DNAJA4 | -0.134973487 | 0.91068657 | 0.563621998 | 0.731985853 | 121.0301598 | 110.1088689 | ENSG00000140403 |
| ARID1A | -0.135011888 | 0.91066233 | 0.088573266 | 0.207013256 | 3959.022231 | 3605.260345 | ENSG00000117713 |
| FKBP8 | -0.135054584 | 0.910635379 | 0.02244151 | 0.070977976 | 7904.235226 | 7197.764859 | ENSG00000105701 |
| TNNI3 | -0.135101891 | 0.91060552 | 0.707062704 | 0.83342026 | 40.43551829 | 36.81789721 | ENSG00000129991 |
| KPNA4 | -0.135184131 | 0.910553612 | 0.104937718 | 0.235205667 | 1976.698842 | 1799.225669 | ENSG00000186432 |
| FAM117B | -0.135247963 | 0.910513326 | 0.266740062 | 0.456696932 | 520.3405322 | 473.0607569 | ENSG00000138439 |
| CXorf56 | -0.135442651 | 0.910390463 | 0.228335275 | 0.411950354 | 531.5414303 | 483.7308329 | ENSG00000018610 |
| NA | -0.135590313 | 0.910297288 | 0.824123269 | 0.905519366 | 13.34919052 | 12.20905096 | ENSG00000277304 |
| RHOBTB3 | -0.135632999 | 0.910270354 | 0.345690072 | 0.539175276 | 268.1306859 | 244.0261444 | ENSG00000164292 |
| NSMF | -0.135735064 | 0.910205959 | 0.076931549 | 0.186093621 | 2181.464777 | 1985.920472 | ENSG00000165802 |
| IKZF5 | -0.135874103 | 0.910118242 | 0.45677787 | 0.643894208 | 164.5119611 | 149.6069812 | ENSG00000095574 |
| WDR4 | -0.135883536 | 0.910112291 | 0.127749997 | 0.271715728 | 1386.011984 | 1262.188541 | ENSG00000160193 |
| C19orf47 | -0.135893106 | 0.910106254 | 0.328496311 | 0.522919404 | 363.318115 | 330.8230859 | ENSG00000160392 |
| DNPEP | -0.135905506 | 0.910098432 | 0.103455005 | 0.232472729 | 1429.476618 | 1300.847523 | ENSG00000123992 |
| NA | -0.135943649 | 0.910074371 | 0.863680164 | 0.928829533 | 9.646863858 | 8.784159543 | ENSG00000277476 |
| RAB11FIP5 | -0.135995615 | 0.91004159 | 0.174058252 | 0.340834193 | 679.2092005 | 618.4849278 | ENSG00000135631 |
| NUP210 | -0.136067637 | 0.90999616 | 0.044546158 | 0.122313852 | 4251.135935 | 3868.1015 | ENSG00000132182 |
| NA | -0.136069194 | 0.909995178 | 0.861525138 | 0.927265209 | 8.309649304 | 7.546579569 | ENSG00000279307 |
| VEZF1 | -0.13609114 | 0.909981336 | 0.128701116 | 0.273204271 | 897.9120596 | 816.9060524 | ENSG00000136451 |
| TBC1D24 | -0.136161765 | 0.90993679 | 0.348045056 | 0.541345005 | 302.67671 | 275.1265873 | ENSG00000162065 |
| S100A16 | -0.136236055 | 0.909889935 | 0.370430762 | 0.563767758 | 239.5400604 | 217.7444013 | ENSG00000188643 |
| FAM219B | -0.136263323 | 0.909872738 | 0.736046702 | 0.852173027 | 282.2438338 | 256.9099493 | ENSG00000178761 |
| PEX5 | -0.136267185 | 0.909870302 | 0.054770864 | 0.143439761 | 1848.77959 | 1682.391647 | ENSG00000139197 |
| AGAP6 | -0.136272181 | 0.909867151 | 0.803307296 | 0.894365197 | 20.48535613 | 18.7434141 | ENSG00000204149 |
| CSK | -0.136349378 | 0.909818467 | 0.097197902 | 0.221855409 | 1616.326147 | 1471.205238 | ENSG00000103653 |
| SGTA | -0.136387793 | 0.909794241 | 0.033259171 | 0.097243511 | 3141.709737 | 2858.660138 | ENSG00000104969 |
| EIF4G1 | -0.136647502 | 0.909630478 | 0.008188535 | 0.031221209 | 19923.22827 | 18123.2817 | ENSG00000114867 |
| SOD2 | -0.136716326 | 0.909587084 | 0.194336627 | 0.368356708 | 724.6519918 | 658.6012402 | ENSG00000112096 |
| HFE | -0.136754643 | 0.909562926 | 0.798836492 | 0.891886713 | 16.90494635 | 15.31428581 | ENSG00000010704 |
| SNTA1 | -0.136819475 | 0.909522054 | 0.367075923 | 0.560179923 | 236.7219091 | 215.3033193 | ENSG00000101400 |
| TMEM150A | -0.136853083 | 0.909500866 | 0.316145191 | 0.509737709 | 362.5771808 | 329.845394 | ENSG00000168890 |
| LINC01829 | -0.136891875 | 0.909476412 | 0.817819024 | 0.901961189 | 14.20545734 | 12.89500081 | ENSG00000236780 |
| ARHGAP1 | -0.1369323 | 0.909450928 | 0.148512737 | 0.303869815 | 896.1407699 | 815.1113316 | ENSG00000175220 |
| PER3 | -0.136934076 | 0.909449808 | 0.537414613 | 0.711203176 | 152.549218 | 138.7144963 | ENSG00000049246 |
| NA | -0.136982795 | 0.909419097 | 0.437746562 | 0.626390837 | 180.854985 | 164.7176376 | ENSG00000274265 |
| POMGNT2 | -0.137001266 | 0.909407453 | 0.188628113 | 0.360528882 | 583.3067311 | 530.5193812 | ENSG00000144647 |
| PLEKHF2 | -0.137185154 | 0.909291547 | 0.533680812 | 0.708592319 | 107.1827486 | 97.38529175 | ENSG00000175895 |
| GAB1 | -0.137189271 | 0.909288952 | 0.34746864 | 0.54072072 | 269.6506857 | 244.9527564 | ENSG00000109458 |
| RHOT2 | -0.13730812 | 0.909214048 | 0.176734864 | 0.344746767 | 4225.38513 | 3843.075834 | ENSG00000140983 |
| SLC33A1 | -0.137350351 | 0.909187434 | 0.639976744 | 0.789229421 | 91.80725763 | 83.34850103 | ENSG00000169359 |
| NOS1AP | -0.13748532 | 0.90910238 | 0.435446713 | 0.624995692 | 223.3218247 | 202.9046862 | ENSG00000198929 |
| NA | -0.137490117 | 0.909099357 | 0.8941202 | 0.94574241 | 5.53489804 | 4.950757929 | ENSG00000255320 |
| NFKBIB | -0.137594736 | 0.909033435 | 0.15446144 | 0.312122583 | 991.828817 | 901.6563346 | ENSG00000104825 |
| DNTTIP1 | -0.13787346 | 0.908857829 | 0.185625774 | 0.356406749 | 781.922185 | 710.5656239 | ENSG00000101457 |
| ZNF175 | -0.137937571 | 0.908817443 | 0.476825067 | 0.661190767 | 138.4201053 | 125.6245227 | ENSG00000105497 |
| ARHGAP35 | -0.137971371 | 0.908796151 | 0.05691127 | 0.147792278 | 3429.595856 | 3116.415749 | ENSG00000160007 |
| ZMYND11 | -0.138033711 | 0.908756882 | 0.186045841 | 0.35706468 | 595.042312 | 540.6812147 | ENSG00000015171 |
| LSM2 | -0.138038571 | 0.90875382 | 0.10476075 | 0.234894234 | 1131.384584 | 1027.691294 | ENSG00000204392 |
| RARS1 | -0.138080569 | 0.908727366 | 0.077634865 | 0.187501249 | 2151.462058 | 1954.585058 | ENSG00000113643 |
| F2R | -0.138090689 | 0.908720992 | 0.176601097 | 0.344619331 | 617.1431569 | 561.0894815 | ENSG00000181104 |

| | | | | | | | |
|---|---|---|---|---|---|---|---|
| MSH6 | -0.13811004 | 0.908708803 | 0.01774713 | 0.058441072 | 6127.762021 | 5567.850765 | ENSG00000116062 |
| TWIST2 | -0.138177922 | 0.908666047 | 0.62172342 | 0.775152125 | 71.84685062 | 65.37277105 | ENSG00000233608 |
| AP1AR | -0.138288648 | 0.90859631 | 0.237454562 | 0.422736887 | 575.4873799 | 522.338776 | ENSG00000138660 |
| DPCD | -0.138304088 | 0.908586586 | 0.286165547 | 0.477519715 | 345.570845 | 313.7606045 | ENSG00000166171 |
| CPNE3 | -0.138331389 | 0.908569393 | 0.175557193 | 0.343080664 | 862.376923 | 782.7706234 | ENSG00000085719 |
| EIF4EBP1 | -0.138361786 | 0.90855025 | 0.058544885 | 0.151103035 | 2923.490689 | 2655.376893 | ENSG00000187840 |
| GYS1 | -0.138456501 | 0.908490604 | 0.109443564 | 0.242287029 | 1414.246239 | 1284.880218 | ENSG00000104812 |
| NA | -0.13854566 | 0.908434461 | 0.882202302 | 0.939373634 | 6.398756461 | 5.787425394 | ENSG00000254419 |
| GOLGA8B | -0.138577566 | 0.908414371 | 0.607020068 | 0.765402948 | 154.5263833 | 140.7149194 | ENSG00000215252 |
| CADM3 | -0.138649125 | 0.908369314 | 0.668367461 | 0.807372907 | 52.24103978 | 47.66589534 | ENSG00000162706 |
| MYO1E | -0.13867651 | 0.908352071 | 0.098425569 | 0.223913035 | 1273.186251 | 1156.096744 | ENSG00000157483 |
| SMC4 | -0.138714094 | 0.908328408 | 0.14940062 | 0.305176482 | 1454.505219 | 1320.121717 | ENSG00000113810 |
| TRIM29 | -0.138761406 | 0.908298621 | 0.883597691 | 0.939962378 | 5.692916438 | 5.187406401 | ENSG00000137699 |
| WDR82 | -0.138796281 | 0.908276664 | 0.040172508 | 0.112828803 | 3484.054445 | 3163.761066 | ENSG00000164091 |
| KIF3C | -0.138897249 | 0.9082131 | 0.051271132 | 0.136355168 | 2034.179496 | 1847.168105 | ENSG00000084731 |
| ARPC4 | -0.138898788 | 0.908212131 | 0.082354676 | 0.195942672 | 1302.617509 | 1183.19632 | ENSG00000241553 |
| NA | -0.138947805 | 0.908181274 | 0.876246572 | 0.935827704 | 6.127014144 | 5.562499875 | ENSG00000175886 |
| FAM149B1 | -0.139003749 | 0.908146058 | 0.361032007 | 0.554242293 | 268.0797095 | 243.3406326 | ENSG00000138286 |
| TXNDC17 | -0.139023157 | 0.908133841 | 0.122329515 | 0.263332361 | 975.5831916 | 885.7217214 | ENSG00000129235 |
| NA | -0.139073047 | 0.908102437 | 0.705387113 | 0.83266319 | 42.04309968 | 38.29256369 | ENSG00000246465 |
| PDCD10 | -0.139097984 | 0.908086741 | 0.139888722 | 0.29088944 | 1136.523447 | 1031.106939 | ENSG00000114209 |
| PPT2 | -0.139112073 | 0.908077873 | 0.518318423 | 0.695519378 | 113.0991819 | 102.6369352 | ENSG00000221988 |
| ZNF219 | -0.139232591 | 0.908002018 | 0.120548676 | 0.260553839 | 2404.609687 | 2183.18129 | ENSG00000165804 |
| TTC30A | -0.139262029 | 0.907983491 | 0.730612778 | 0.849344242 | 33.43339354 | 30.48307858 | ENSG00000197557 |
| LIPE-AS1 | -0.139319637 | 0.907947235 | 0.825426866 | 0.906253275 | 12.05882599 | 10.93349096 | ENSG00000213904 |
| CDK13 | -0.139371646 | 0.907914504 | 0.121832041 | 0.262562215 | 1214.090228 | 1102.120758 | ENSG00000065883 |
| GMPPA | -0.139527647 | 0.907816335 | 0.118365385 | 0.257033866 | 920.3880957 | 834.9577424 | ENSG00000144591 |
| NA | -0.139595355 | 0.907773731 | 0.911027506 | 0.955762377 | 3.913411215 | 3.563448033 | ENSG00000229207 |
| CCDC47 | -0.139602079 | 0.9077695 | 0.068606663 | 0.170457532 | 1913.180116 | 1737.130776 | ENSG00000108588 |
| NAA50 | -0.139663776 | 0.90773068 | 0.079759078 | 0.191310309 | 4314.342035 | 3915.585246 | ENSG00000121579 |
| NA | -0.139690609 | 0.907713797 | 0.874343071 | 0.935034013 | 6.364631779 | 5.746084558 | ENSG00000260751 |
| ZNF695 | -0.139698256 | 0.907708986 | 0.653561921 | 0.797566245 | 65.91238552 | 59.60246769 | ENSG00000197472 |
| CDIPT | -0.139806857 | 0.907640659 | 0.193343171 | 0.367003516 | 909.6707385 | 825.5070166 | ENSG00000103502 |
| NUP54 | -0.139896851 | 0.907584043 | 0.135184558 | 0.283268581 | 1361.902841 | 1235.118421 | ENSG00000138750 |
| EIF2B5 | -0.139988265 | 0.907526537 | 0.082048172 | 0.195435139 | 1198.674637 | 1087.54966 | ENSG00000145191 |
| NECAB2 | -0.139989916 | 0.907525499 | 0.391051521 | 0.583648646 | 214.8315701 | 194.9935739 | ENSG00000103154 |
| ATP6V1H | -0.140016207 | 0.90750896 | 0.184345593 | 0.354977789 | 597.7205971 | 542.0657763 | ENSG00000047249 |
| NA | -0.140033527 | 0.907498065 | 0.809122558 | 0.897335896 | 15.26008336 | 13.89312115 | ENSG00000269688 |
| MKKS | -0.140129306 | 0.90743782 | 0.376926542 | 0.570049493 | 274.5387177 | 248.6220685 | ENSG00000125863 |
| CEP57 | -0.14023684 | 0.907370185 | 0.140706865 | 0.29194048 | 906.4598977 | 821.9965779 | ENSG00000166037 |
| CACNB2 | -0.140254234 | 0.907359245 | 0.653102098 | 0.797325732 | 52.07858669 | 47.35405127 | ENSG00000165995 |
| POFUT1 | -0.140335641 | 0.907308047 | 0.087769886 | 0.205498618 | 1029.676052 | 934.2515205 | ENSG00000101346 |
| PRPS2 | -0.140437997 | 0.907243678 | 0.175107735 | 0.342419278 | 747.3694864 | 678.0000474 | ENSG00000101911 |
| RASL10B | -0.140497218 | 0.907206437 | 0.109089439 | 0.241705161 | 999.2883406 | 906.4182748 | ENSG00000270885 |
| SMARCC2 | -0.140512247 | 0.907196987 | 0.164413686 | 0.32674708 | 2123.7244 | 1927.604287 | ENSG00000139613 |
| SEPHS2 | -0.140517446 | 0.907193717 | 0.078203704 | 0.18847602 | 1658.985904 | 1504.77271 | ENSG00000179918 |
| PPP2R5D | -0.140539146 | 0.907180072 | 0.069716453 | 0.172453998 | 1376.711456 | 1249.206987 | ENSG00000112640 |
| PDZD7 | -0.140670752 | 0.907097321 | 0.725206158 | 0.845503613 | 39.99826367 | 36.46993687 | ENSG00000186862 |
| NA | -0.140673378 | 0.907095669 | 0.803104862 | 0.894247252 | 18.87649696 | 17.18292838 | ENSG00000260000 |
| LEMD3 | -0.140704787 | 0.907075921 | 0.128453811 | 0.272804279 | 922.9750409 | 836.4988043 | ENSG00000174106 |
| CSTF2 | -0.140705627 | 0.907075393 | 0.075589119 | 0.183560626 | 1476.597224 | 1339.693383 | ENSG00000101811 |
| ACAD8 | -0.140760216 | 0.907041072 | 0.071064974 | 0.175078856 | 1360.122012 | 1233.534395 | ENSG00000151498 |
| SPON2 | -0.1408092 | 0.907010276 | 0.716569609 | 0.839556455 | 33.817582 | 30.77928923 | ENSG00000159674 |
| TIAL1 | -0.140830507 | 0.90699688 | 0.06231866 | 0.158612202 | 1740.955809 | 1579.565848 | ENSG00000151923 |
| DMAP1 | -0.140831586 | 0.906996202 | 0.108314629 | 0.240448356 | 970.8334624 | 881.076341 | ENSG00000178028 |
| ARIH2 | -0.140874592 | 0.906969165 | 0.066911002 | 0.16726848 | 1955.084628 | 1773.473338 | ENSG00000177479 |
| DIAPH3 | -0.140970508 | 0.906908868 | 0.214453544 | 0.394470224 | 640.7747048 | 580.4941484 | ENSG00000139734 |
| ANAPC5 | -0.141006761 | 0.906886079 | 0.042717297 | 0.118521316 | 1886.057544 | 1710.593456 | ENSG00000089053 |
| ACOX2 | -0.141024781 | 0.906874751 | 0.798262823 | 0.891480066 | 17.30176246 | 15.59552431 | ENSG00000168306 |
| ZFHX2 | -0.141036997 | 0.906867073 | 0.27072435 | 0.461304138 | 639.9170699 | 580.6302521 | ENSG00000136367 |
| ARPP19 | -0.141127736 | 0.906810037 | 0.14226664 | 0.294479551 | 2189.167705 | 1984.46013 | ENSG00000128989 |
| LOC101930370 | -0.141240088 | 0.90673942 | 0.67241602 | 0.809585672 | 44.85768809 | 40.71878511 | ENSG00000245213 |
| CADPS | -0.141264496 | 0.90672408 | 0.356599426 | 0.550199771 | 227.9716529 | 206.5598393 | ENSG00000163618 |
| C4A | -0.141284254 | 0.906711662 | 0.872218679 | 0.93389376 | 6.375380294 | 5.746084558 | ENSG00000244731 |
| SPTBN2 | -0.141491692 | 0.9065813 | 0.040180489 | 0.112834119 | 2078.8422 | 1885.067944 | ENSG00000173898 |
| LOC101927151 | -0.141501466 | 0.906575158 | 0.566982051 | 0.734548115 | 86.78952171 | 78.56398251 | ENSG00000267575 |
| CRYBB1 | -0.141684994 | 0.906459838 | 0.72340978 | 0.844046387 | 30.90878987 | 28.07496688 | ENSG00000100122 |
| MOGS | -0.14169425 | 0.906454023 | 0.091185113 | 0.211681085 | 1867.430571 | 1693.828205 | ENSG00000115275 |
| NA | -0.141748865 | 0.906419708 | 0.807057812 | 0.896619594 | 14.67555886 | 13.24003837 | ENSG00000235508 |
| PRKCA | -0.141763181 | 0.906410714 | 0.281983976 | 0.473069106 | 547.1681134 | 495.8602197 | ENSG00000154229 |
| ADAR | -0.141789683 | 0.906394064 | 0.019194492 | 0.062302816 | 7565.979298 | 6857.346085 | ENSG00000160710 |
| CHERP | -0.141817994 | 0.906376276 | 0.084308073 | 0.199306866 | 1663.806566 | 1507.813763 | ENSG00000085872 |
| ST6GALNAC3 | -0.141838984 | 0.90636309 | 0.254057858 | 0.44288077 | 380.4397985 | 344.7469243 | ENSG00000184005 |
| MALT1 | -0.141897841 | 0.906326114 | 0.307664938 | 0.501782996 | 299.9097457 | 272.083328 | ENSG00000172175 |

| | | | | | | | |
|---|---|---|---|---|---|---|---|
| RAF1 | -0.141898927 | 0.906325432 | 0.03198735 | 0.094283326 | 2599.053382 | 2355.379574 | ENSG00000132155 |
| MRPL4 | -0.141904438 | 0.90632197 | 0.119272011 | 0.258397131 | 849.6156692 | 770.3048163 | ENSG00000105364 |
| RNF166 | -0.141968929 | 0.906281456 | 0.292629571 | 0.485159359 | 425.4732512 | 386.0633637 | ENSG00000158717 |
| UBE2T | -0.141974225 | 0.90627813 | 0.167810054 | 0.331576923 | 688.2296433 | 623.2340542 | ENSG00000077152 |
| NA | -0.141997492 | 0.906263514 | 0.047454435 | 0.128439584 | 1873.636568 | 1697.756766 | ENSG00000198618 |
| XRCC1 | -0.142021057 | 0.906248711 | 0.171579884 | 0.33733548 | 626.640962 | 567.7545888 | ENSG00000073050 |
| ZNF35 | -0.142095187 | 0.906202146 | 0.350524512 | 0.543916327 | 254.3259007 | 230.5309014 | ENSG00000169981 |
| NA | -0.14217086 | 0.906154615 | 0.841806497 | 0.915290486 | 9.448406954 | 8.581803865 | ENSG00000235558 |
| NA | -0.142177608 | 0.906150377 | 0.873375316 | 0.934693202 | 6.92705789 | 6.331569437 | ENSG00000226179 |
| PRPF4 | -0.142201226 | 0.906135543 | 0.082538996 | 0.196231601 | 1520.869516 | 1377.821354 | ENSG00000136875 |
| NA | -0.142244871 | 0.90610813 | 0.076243529 | 0.184791104 | 1323.973 | 1198.99807 | ENSG00000188856 |
| ZC3H3 | -0.142262722 | 0.906096919 | 0.20525976 | 0.383573744 | 757.8160857 | 686.5003318 | ENSG00000014164 |
| BSG | -0.142308692 | 0.906068047 | 0.008674885 | 0.03273215 | 20138.30073 | 18246.3429 | ENSG00000172270 |
| CERCAM | -0.142324199 | 0.906058308 | 0.447979165 | 0.635957225 | 147.0204384 | 133.2509727 | ENSG00000167123 |
| FBXL14 | -0.142372354 | 0.906028066 | 0.257070689 | 0.445576373 | 355.4466299 | 321.9260856 | ENSG00000171823 |
| SLC43A2 | -0.142474548 | 0.905963889 | 0.212422273 | 0.392213032 | 521.2964069 | 472.11398 | ENSG00000167703 |
| LRRC20 | -0.142587334 | 0.905893066 | 0.148524693 | 0.303869815 | 682.4983403 | 618.327377 | ENSG00000172731 |
| KARS1 | -0.142707185 | 0.905817812 | 0.017126424 | 0.056814236 | 5032.303415 | 4557.633545 | ENSG00000065427 |
| SPAST | -0.14282456 | 0.90574412 | 0.250025705 | 0.43853283 | 551.5933425 | 498.9251812 | ENSG00000021574 |
| PPP2R2D | -0.142871391 | 0.905714719 | 0.154928469 | 0.312861652 | 715.0901692 | 647.109724 | ENSG00000175470 |
| NA | -0.142952477 | 0.905663815 | 0.884417748 | 0.940651816 | 5.42560881 | 4.947508735 | ENSG00000275198 |
| PPAT | -0.143114435 | 0.905562151 | 0.120812757 | 0.260972453 | 1697.14187 | 1536.170039 | ENSG00000128059 |
| FOXN4 | -0.143221364 | 0.905495035 | 0.835455985 | 0.911809046 | 11.90278443 | 10.87761601 | ENSG00000139445 |
| INTS7 | -0.143239963 | 0.905483362 | 0.183553427 | 0.354114626 | 535.370379 | 484.7063834 | ENSG00000143493 |
| PARP1 | -0.143306583 | 0.90544155 | 0.005878195 | 0.023648027 | 17927.6764 | 16232.57038 | ENSG00000143799 |
| FCF1 | -0.143415394 | 0.905373262 | 0.125775112 | 0.268685561 | 1187.783424 | 1074.556916 | ENSG00000119616 |
| ILF3 | -0.143539974 | 0.905295085 | 0.015554392 | 0.052597171 | 21930.47872 | 19854.2969 | ENSG00000129351 |
| GRIK5 | -0.143544921 | 0.90529198 | 0.068024215 | 0.169206926 | 1401.182734 | 1268.473584 | ENSG00000105737 |
| H19 | -0.143573579 | 0.905273997 | 0.687760544 | 0.820584133 | 40.79445144 | 36.95689186 | ENSG00000130600 |
| ABCD1 | -0.143731904 | 0.905174656 | 0.324834712 | 0.518917992 | 299.0793356 | 271.0257486 | ENSG00000101986 |
| LRRC1 | -0.143763233 | 0.905155 | 0.368822472 | 0.561781369 | 278.508758 | 251.7527331 | ENSG00000137269 |
| BORCS8 | -0.143930527 | 0.905050044 | 0.684180129 | 0.817890607 | 40.28706832 | 36.50660279 | ENSG00000254901 |
| NA | -0.144043451 | 0.904979207 | 0.279711457 | 0.470817559 | 433.0178836 | 391.261345 | ENSG00000225178 |
| FIBP | -0.144093851 | 0.904947592 | 0.092799276 | 0.214216343 | 1268.148401 | 1148.219638 | ENSG00000172500 |
| NPC2 | -0.144122708 | 0.904929491 | 0.118953536 | 0.257888037 | 967.8230628 | 875.1396463 | ENSG00000119655 |
| TEFM | -0.144126779 | 0.904926938 | 0.512607848 | 0.691410658 | 110.238713 | 99.69172754 | ENSG00000172171 |
| LARS2 | -0.144189055 | 0.904887876 | 0.031866498 | 0.094029298 | 2188.422825 | 1980.065057 | ENSG00000011376 |
| SRRM3 | -0.14420399 | 0.904878509 | 0.279023765 | 0.470063197 | 436.7836038 | 394.8769892 | ENSG00000177679 |
| SOX8 | -0.144365409 | 0.90477727 | 0.233635596 | 0.418579231 | 415.6643176 | 376.0466968 | ENSG00000005513 |
| RNF13 | -0.144394543 | 0.904758999 | 0.308799182 | 0.502976258 | 314.5838088 | 284.3011388 | ENSG00000082996 |
| FBXO5 | -0.144619167 | 0.904618141 | 0.141773205 | 0.293589338 | 709.2045083 | 641.2270666 | ENSG00000112029 |
| LIPG | -0.144669786 | 0.904586402 | 0.356556336 | 0.550196914 | 225.1851684 | 203.7427794 | ENSG00000101670 |
| H3-3A | -0.144845732 | 0.904476088 | 0.009942912 | 0.036600187 | 7939.960087 | 7180.981781 | ENSG00000163041 |
| TKFC | -0.144878664 | 0.904455443 | 0.154269735 | 0.311837198 | 620.3945114 | 561.0559616 | ENSG00000149476 |
| EMC8 | -0.1448897 | 0.904448524 | 0.223008233 | 0.405139639 | 657.2346396 | 594.1814499 | ENSG00000131148 |
| MED16 | -0.145015017 | 0.904369964 | 0.050700395 | 0.135264303 | 2197.881875 | 1987.396762 | ENSG00000175221 |
| ZNF589 | -0.145157748 | 0.904280496 | 0.111756186 | 0.246288841 | 2531.481108 | 2290.107128 | ENSG00000164048 |
| LGR4 | -0.145403584 | 0.904126419 | 0.102522384 | 0.230796777 | 1308.972873 | 1183.330058 | ENSG00000205213 |
| MIR589 | -0.145462422 | 0.904089546 | 0.874098566 | 0.934994283 | 5.677131874 | 5.138579492 | ENSG00000207973 |
| XPO4 | -0.145490454 | 0.90407198 | 0.126092432 | 0.269266151 | 1007.853061 | 911.018659 | ENSG00000132953 |
| C19orf44 | -0.145512555 | 0.90405813 | 0.569145966 | 0.735971767 | 93.36610986 | 84.59218163 | ENSG00000105072 |
| COPS5 | -0.145519665 | 0.904053675 | 0.123006188 | 0.264477574 | 950.2123515 | 858.8873923 | ENSG00000121022 |
| RSRC2 | -0.145592523 | 0.90400802 | 0.045169747 | 0.123641425 | 2004.916975 | 1812.435838 | ENSG00000111011 |
| MBD1 | -0.145620185 | 0.903990687 | 0.094811442 | 0.21756039 | 1028.48574 | 929.8802987 | ENSG00000141644 |
| TGFBR3 | -0.145693628 | 0.903944669 | 0.53253659 | 0.707603539 | 104.8938835 | 94.59036362 | ENSG00000069702 |
| MTMR7 | -0.145728772 | 0.903922649 | 0.543400698 | 0.71590462 | 92.22272212 | 83.32779494 | ENSG00000003987 |
| NUDT16 | -0.145813052 | 0.903869845 | 0.124968436 | 0.267743086 | 989.1025775 | 893.9783983 | ENSG00000198585 |
| C22orf39 | -0.145830405 | 0.903858973 | 0.285979527 | 0.477319548 | 330.356627 | 298.6373894 | ENSG00000242259 |
| AGAP3 | -0.145840295 | 0.903852777 | 0.078932252 | 0.189744533 | 1760.466837 | 1592.064325 | ENSG00000133612 |
| HSPA1B | -0.146053226 | 0.903719385 | 0.077484437 | 0.187186726 | 1243.222542 | 1123.505397 | ENSG00000204388 |
| PAICS | -0.146065945 | 0.903711417 | 0.011811588 | 0.042204739 | 10012.72443 | 9047.820824 | ENSG00000128050 |
| RNF130 | -0.146074906 | 0.903705804 | 0.102632395 | 0.230988318 | 1051.824446 | 950.5810572 | ENSG00000113269 |
| BASP1 | -0.146106979 | 0.903685714 | 0.053312528 | 0.140414153 | 4685.72208 | 4234.64489 | ENSG00000176788 |
| NSD3 | -0.146200949 | 0.903626854 | 0.111930673 | 0.246556107 | 1196.100056 | 1080.554813 | ENSG00000147548 |
| API5 | -0.146268538 | 0.903584521 | 0.082380563 | 0.195949866 | 3101.508588 | 2801.473034 | ENSG00000166181 |
| TSEN34 | -0.14627006 | 0.903583568 | 0.212595086 | 0.392385079 | 478.6649268 | 432.3867137 | ENSG00000170892 |
| NUP85 | -0.146410929 | 0.903495344 | 0.074025951 | 0.180668552 | 1377.532426 | 1244.700888 | ENSG00000125450 |
| DDAH2 | -0.146424226 | 0.903487016 | 0.074917613 | 0.182308087 | 1317.762027 | 1190.004516 | ENSG00000213722 |
| FAM49B | -0.14645697 | 0.903466511 | 0.055550441 | 0.144887683 | 1740.038816 | 1571.382343 | ENSG00000153310 |
| FGD4 | -0.146529813 | 0.903420895 | 0.286788711 | 0.478170382 | 458.3599805 | 413.9680204 | ENSG00000139132 |
| ZNF501 | -0.146647569 | 0.903347159 | 0.508296361 | 0.687496516 | 136.5763794 | 123.1853674 | ENSG00000186446 |
| TUBGCP2 | -0.146809989 | 0.903245464 | 0.037000421 | 0.105632441 | 1794.431839 | 1621.067853 | ENSG00000130640 |
| GLYATL1 | -0.146876601 | 0.903203761 | 0.502170256 | 0.682636825 | 111.9466165 | 100.8880783 | ENSG00000166840 |
| SUPT6H | -0.146880918 | 0.903201058 | 0.043598996 | 0.120301293 | 3283.350243 | 2966.032668 | ENSG00000109111 |

| | | | | | | | |
|---|---|---|---|---|---|---|---|
| CRAT | -0.146942385 | 0.903162578 | 0.325797446 | 0.520007739 | 307.8386717 | 277.6501276 | ENSG00000095321 |
| SAP130 | -0.146949724 | 0.903157983 | 0.066363067 | 0.166122783 | 1394.693893 | 1258.915297 | ENSG00000136715 |
| PNPLA2 | -0.14695529 | 0.903154499 | 0.202428137 | 0.379737156 | 511.7153119 | 462.0631807 | ENSG00000177666 |
| TMUB1 | -0.147003848 | 0.903124101 | 0.209333859 | 0.388490944 | 887.2491738 | 801.6189082 | ENSG00000164897 |
| DLK1 | -0.147063915 | 0.9030865 | 0.708918004 | 0.83446266 | 32.62387906 | 29.46607563 | ENSG00000185559 |
| SNRNP70 | -0.147074122 | 0.903080111 | 0.014201037 | 0.048858739 | 9284.047715 | 8384.154655 | ENSG00000104852 |
| ZNF567 | -0.147099616 | 0.903064153 | 0.485675063 | 0.66875218 | 160.8315374 | 144.8535777 | ENSG00000189042 |
| NEK4 | -0.1471003 | 0.903063725 | 0.157491585 | 0.316657522 | 726.2718327 | 655.2559961 | ENSG00000114904 |
| EFNA1 | -0.147193346 | 0.903005484 | 0.63097372 | 0.782290917 | 53.95781271 | 48.75699458 | ENSG00000169242 |
| FGFR2 | -0.14720497 | 0.902998208 | 0.042226897 | 0.117424352 | 1564.498918 | 1412.796811 | ENSG00000066468 |
| ITIH5 | -0.147249894 | 0.90297009 | 0.36547121 | 0.558573214 | 207.1837821 | 187.0328216 | ENSG00000123243 |
| ATP5MPL | -0.147250053 | 0.90296999 | 0.112418701 | 0.24742527 | 772.6821569 | 697.5458277 | ENSG00000156411 |
| TWF2 | -0.147262223 | 0.902962374 | 0.145266731 | 0.298886944 | 712.806881 | 643.7863255 | ENSG00000247596 |
| ELF4 | -0.147466143 | 0.902834752 | 0.416668685 | 0.607762324 | 155.2201971 | 140.2312765 | ENSG00000102034 |
| PUS10 | -0.147567744 | 0.902771173 | 0.688467549 | 0.821277542 | 37.40608705 | 33.73099532 | ENSG00000162927 |
| LRRC32 | -0.147571097 | 0.902769075 | 0.64647765 | 0.793445253 | 56.55946007 | 50.89730259 | ENSG00000137507 |
| CYB5D2 | -0.147578241 | 0.902764604 | 0.519589846 | 0.696633863 | 108.1528371 | 97.85154065 | ENSG00000167740 |
| PELI3 | -0.147588645 | 0.902758094 | 0.516164148 | 0.693936217 | 135.0089661 | 121.8862986 | ENSG00000174516 |
| IQGAP3 | -0.147651692 | 0.902718643 | 0.152003375 | 0.309042404 | 1024.647861 | 925.8886624 | ENSG00000183856 |
| FKBP15 | -0.147975957 | 0.902515768 | 0.291373455 | 0.483761659 | 451.2972132 | 408.016858 | ENSG00000119321 |
| DNAJC11 | -0.148041105 | 0.902475014 | 0.073431169 | 0.179713027 | 1809.54829 | 1633.842153 | ENSG00000007923 |
| RBM7 | -0.14812931 | 0.902419839 | 0.436775971 | 0.625932098 | 305.9150384 | 275.6743782 | ENSG00000076053 |
| NOP2 | -0.148186627 | 0.902383987 | 0.856929173 | 0.924099792 | 7.754814795 | 7.006672407 | ENSG00000111641 |
| KLC1 | -0.148202311 | 0.902374178 | 0.240901935 | 0.427097424 | 522.5234378 | 472.1775417 | ENSG00000126214 |
| DCAF11 | -0.148293266 | 0.902317289 | 0.070274224 | 0.173499728 | 1506.847829 | 1359.431611 | ENSG00000100897 |
| VBP1 | -0.14831496 | 0.902303721 | 0.19375346 | 0.36762982 | 1126.971674 | 1016.139267 | ENSG00000155959 |
| SF3B4 | -0.148402503 | 0.90224897 | 0.065594181 | 0.164665115 | 2048.122081 | 1848.004661 | ENSG00000143368 |
| UBTD1 | -0.148505024 | 0.902184857 | 0.200709018 | 0.377311369 | 596.8125267 | 538.5185114 | ENSG00000165886 |
| WDCP | -0.148533073 | 0.902167317 | 0.200481696 | 0.377154664 | 689.0960421 | 622.4505396 | ENSG00000163026 |
| PIGH | -0.148538401 | 0.902163985 | 0.526113993 | 0.702433961 | 92.05287283 | 83.04980563 | ENSG00000100564 |
| NUP50 | -0.148543921 | 0.902160533 | 0.03217318 | 0.094628066 | 2436.486453 | 2197.654575 | ENSG00000093000 |
| SHISA7 | -0.148561517 | 0.90214953 | 0.709768309 | 0.835055282 | 31.87118901 | 28.67823507 | ENSG00000187902 |
| DCST1 | -0.148563096 | 0.902148543 | 0.895482335 | 0.946391606 | 3.774431993 | 3.405682385 | ENSG00000163357 |
| RARA | -0.14865201 | 0.902092944 | 0.292755188 | 0.485324209 | 343.902974 | 309.8991936 | ENSG00000131759 |
| NKD2 | -0.148662658 | 0.902086286 | 0.273659456 | 0.464871193 | 313.9936921 | 283.1843322 | ENSG00000145506 |
| HNRNPC | -0.148683437 | 0.902073294 | 0.005438536 | 0.022114485 | 21242.02559 | 19160.89161 | ENSG00000092199 |
| PMPCA | -0.14868429 | 0.902072761 | 0.028755684 | 0.08661756 | 2660.651714 | 2400.831248 | ENSG00000165688 |
| LMO2 | -0.148765531 | 0.902021965 | 0.692078939 | 0.823657812 | 38.49355998 | 34.61594004 | ENSG00000135363 |
| SCMH1 | -0.148834867 | 0.901978614 | 0.212726679 | 0.392423238 | 411.3765946 | 371.2962479 | ENSG00000010803 |
| MRPL47 | -0.148880153 | 0.901950302 | 0.112684476 | 0.247804233 | 889.0739361 | 801.5608346 | ENSG00000136522 |
| LINC01618 | -0.148890915 | 0.901943574 | 0.891814838 | 0.944886309 | 4.141558112 | 3.713177955 | ENSG00000250302 |
| CLK1 | -0.148896684 | 0.901939967 | 0.334488384 | 0.528981192 | 297.9795432 | 269.1559439 | ENSG00000013441 |
| NOMO3 | -0.149035789 | 0.901853006 | 0.479796534 | 0.663713784 | 120.5646884 | 108.6604595 | ENSG00000103226 |
| LINC00309 | -0.149150146 | 0.901781522 | 0.829625658 | 0.908334236 | 9.842066152 | 8.851757447 | ENSG00000230923 |
| CLMP | -0.149178176 | 0.901764001 | 0.22747537 | 0.410959013 | 462.9416797 | 417.2777063 | ENSG00000166250 |
| RDH13 | -0.149237592 | 0.901726864 | 0.363736797 | 0.556819635 | 209.813819 | 189.5216798 | ENSG00000160439 |
| CTCF | -0.149272187 | 0.901705241 | 0.023868457 | 0.07461258 | 2613.311356 | 2356.071352 | ENSG00000102974 |
| LOC574538 | -0.149338086 | 0.901664055 | 0.873249758 | 0.934673771 | 8.535819366 | 7.674289303 | ENSG00000249695 |
| NINL | -0.14934562 | 0.901659346 | 0.148306576 | 0.303524082 | 1795.399732 | 1619.922466 | ENSG00000101004 |
| LSM1 | -0.149467505 | 0.901583173 | 0.29342112 | 0.486036914 | 337.3137356 | 303.9527288 | ENSG00000175324 |
| FDFT1 | -0.149477264 | 0.901577074 | 0.010267264 | 0.037562863 | 11991.75447 | 10810.77033 | ENSG00000079459 |
| NCAPG2 | -0.149503295 | 0.901560807 | 0.027406954 | 0.083526452 | 2304.428146 | 2076.955979 | ENSG00000146918 |
| NA | -0.149521877 | 0.901549195 | 0.853203118 | 0.922280546 | 7.556455588 | 6.811364769 | ENSG00000225345 |
| PINLYP | -0.149614851 | 0.901491097 | 0.623747489 | 0.77686553 | 65.7656193 | 59.18388349 | ENSG00000234465 |
| LINC01838 | -0.149635277 | 0.901478333 | 0.891354811 | 0.944688675 | 3.797808159 | 3.424453379 | ENSG00000269037 |
| USP7 | -0.149680638 | 0.90144999 | 0.013873607 | 0.048008483 | 5180.143869 | 4668.986587 | ENSG00000187555 |
| SLC2A4RG | -0.149760463 | 0.901400113 | 0.252084132 | 0.44090874 | 436.4670109 | 393.9071416 | ENSG00000125520 |
| KIAA0513 | -0.149818274 | 0.901363994 | 0.296734132 | 0.489599431 | 308.900769 | 278.0525775 | ENSG00000135709 |
| TSACC | -0.149853724 | 0.901341846 | 0.767071387 | 0.871187027 | 21.16220522 | 19.05003361 | ENSG00000163467 |
| CHD1 | -0.149865439 | 0.901334527 | 0.067176981 | 0.167741783 | 1237.588645 | 1115.092277 | ENSG00000153922 |
| PCDHB8 | -0.149873264 | 0.901329638 | 0.836193932 | 0.9121504 | 9.243636223 | 8.360677193 | ENSG00000120322 |
| HTRA4 | -0.150055353 | 0.901215884 | 0.773629408 | 0.875060689 | 19.50461688 | 17.56495827 | ENSG00000169495 |
| PABPC4 | -0.15007907 | 0.901201069 | 0.015364241 | 0.052096752 | 4445.035243 | 4005.589243 | ENSG00000090621 |
| BUD13 | -0.15013243 | 0.901167738 | 0.131409964 | 0.277650738 | 640.8952688 | 577.6907239 | ENSG00000137656 |
| FAM110A | -0.150266896 | 0.901083748 | 0.320697719 | 0.514531257 | 241.2526917 | 217.5094017 | ENSG00000125898 |
| MRPL36 | -0.150314993 | 0.901053708 | 0.124393829 | 0.266756457 | 711.8051484 | 641.8191933 | ENSG00000171421 |
| COL22A1 | -0.150416741 | 0.900990163 | 0.518247579 | 0.695519378 | 107.0422963 | 96.11666817 | ENSG00000169436 |
| CHST10 | -0.150501339 | 0.900937331 | 0.124290571 | 0.2666514 | 782.436393 | 704.9530862 | ENSG00000115526 |
| HDGFL2 | -0.150669514 | 0.900832315 | 0.019054334 | 0.061913222 | 4218.761541 | 3800.566492 | ENSG00000167674 |
| EIF2AK4 | -0.150682256 | 0.900824358 | 0.018227194 | 0.059675467 | 3567.620566 | 3213.26069 | ENSG00000128829 |
| SSNA1 | -0.15068405 | 0.900823238 | 0.044541833 | 0.122313852 | 2003.917933 | 1805.365246 | ENSG00000176101 |
| MYO7A | -0.150739944 | 0.900788339 | 0.514642711 | 0.692846128 | 102.324602 | 92.22446883 | ENSG00000137474 |
| ETAA1 | -0.150750852 | 0.900781528 | 0.292546772 | 0.485125532 | 445.9956935 | 401.224693 | ENSG00000143971 |
| ABCE1 | -0.150789202 | 0.900757584 | 0.078890341 | 0.189668385 | 1934.985061 | 1742.133833 | ENSG00000164163 |

| | | | | | | | |
|---|---|---|---|---|---|---|---|
| NACC1 | -0.150995207 | 0.900628972 | 0.053594588 | 0.140956732 | 4740.881714 | 4269.488 | ENSG00000160877 |
| LOC728554 | -0.151059637 | 0.900588751 | 0.204435127 | 0.382495523 | 532.7472119 | 479.6727206 | ENSG00000170089 |
| PCBP2 | -0.151290543 | 0.900444622 | 0.035742387 | 0.102944738 | 1992.895792 | 1794.507667 | ENSG00000197111 |
| PHLDB1 | -0.151305133 | 0.900435515 | 0.045072842 | 0.12343086 | 1502.897534 | 1353.804262 | ENSG00000019144 |
| LSM8 | -0.151385524 | 0.900385342 | 0.1633558 | 0.325097862 | 624.4984645 | 562.7099759 | ENSG00000128534 |
| ZSCAN30 | -0.151390748 | 0.900382082 | 0.375189842 | 0.568118344 | 207.0375946 | 186.4945633 | ENSG00000186814 |
| NA | -0.151398318 | 0.900377358 | 0.892456648 | 0.945187103 | 4.730517303 | 4.317433827 | ENSG00000272829 |
| GNAI3 | -0.151553747 | 0.900280361 | 0.050625499 | 0.135103383 | 2061.75183 | 1855.368898 | ENSG00000065135 |
| RPA1 | -0.151607227 | 0.900246988 | 0.015986052 | 0.053816558 | 4450.134901 | 4005.625319 | ENSG00000132383 |
| GDAP2 | -0.151670028 | 0.900207801 | 0.33545406 | 0.52967333 | 222.5961487 | 200.2539773 | ENSG00000196505 |
| AP5M1 | -0.151750558 | 0.900157554 | 0.137828813 | 0.287398473 | 711.0973541 | 639.8300636 | ENSG00000053770 |
| NA | -0.15175345 | 0.900155749 | 0.388376183 | 0.581300136 | 187.3474415 | 168.4241652 | ENSG00000196810 |
| SUZ12 | -0.151814701 | 0.900117533 | 0.128740403 | 0.273256372 | 817.2677944 | 735.0680706 | ENSG00000178691 |
| TRIM8 | -0.151817605 | 0.900115721 | 0.190612145 | 0.363195794 | 555.2092 | 499.3662723 | ENSG00000171206 |
| CLIC1 | -0.151839988 | 0.900101756 | 0.015522322 | 0.052508185 | 4427.177413 | 3983.881077 | ENSG00000213719 |
| NA | -0.15185994 | 0.900089308 | 0.865722276 | 0.930054293 | 6.616154843 | 6.02027502 | ENSG00000275494 |
| CALHM2 | -0.151911512 | 0.900057133 | 0.661035038 | 0.802702466 | 45.92937646 | 41.20182964 | ENSG00000138172 |
| NA | -0.151948503 | 0.900034056 | 0.83097817 | 0.908990473 | 11.07610991 | 9.886981743 | ENSG00000274525 |
| ATP2B4 | -0.151955059 | 0.900029966 | 0.187453985 | 0.35920928 | 501.2894355 | 450.6900243 | ENSG00000058668 |
| CTDSP1 | -0.152026344 | 0.899985495 | 0.216631763 | 0.397254684 | 434.8520258 | 390.8039047 | ENSG00000144579 |
| STEAP1 | -0.152046583 | 0.89997287 | 0.546152023 | 0.718205024 | 90.64808533 | 81.36485937 | ENSG00000164647 |
| RNF10 | -0.152184785 | 0.899886662 | 0.015137595 | 0.051490338 | 3813.042251 | 3430.887352 | ENSG00000022840 |
| ELAC2 | -0.152367862 | 0.899772475 | 0.015875941 | 0.053499033 | 3241.46426 | 2917.107756 | ENSG00000006744 |
| NA | -0.152451246 | 0.899720471 | 0.724326048 | 0.844902699 | 27.17983243 | 24.42514995 | ENSG00000274964 |
| CRY1 | -0.152484833 | 0.899699526 | 0.14307708 | 0.295598758 | 581.2680135 | 522.7995142 | ENSG00000008405 |
| MED17 | -0.152505602 | 0.899686574 | 0.71718103 | 0.839762217 | 29.37015689 | 26.47314032 | ENSG00000042429 |
| GSTP1 | -0.152621988 | 0.899613996 | 0.002789271 | 0.012514243 | 14501.17372 | 13045.21979 | ENSG00000084207 |
| AK6 | -0.152651283 | 0.899595729 | 0.340817425 | 0.535223705 | 228.1464254 | 205.1448466 | ENSG00000085231 |
| ERI2 | -0.152696947 | 0.899567256 | 0.439406228 | 0.627761892 | 138.42454 | 124.398116 | ENSG00000196678 |
| COA6-AS1 | -0.152753556 | 0.899531959 | 0.88996093 | 0.943785311 | 5.455298802 | 4.921251667 | ENSG00000231663 |
| PAGR1 | -0.152763409 | 0.899525816 | 0.367722636 | 0.560766469 | 190.295199 | 171.2581727 | ENSG00000280789 |
| ARMC8 | -0.152766018 | 0.899524189 | 0.127168334 | 0.270819017 | 746.5058385 | 670.9046958 | ENSG00000114098 |
| C19orf73 | -0.152834181 | 0.89948169 | 0.746601228 | 0.858611443 | 27.9027348 | 25.14594226 | ENSG00000221916 |
| SLC1A1 | -0.152889876 | 0.899446966 | 0.277363585 | 0.468482337 | 306.7575126 | 275.7121753 | ENSG00000106688 |
| NA | -0.152969881 | 0.899397089 | 0.853392008 | 0.922280546 | 7.7175332 | 6.939074503 | ENSG00000230170 |
| KCNN4 | -0.153072367 | 0.899333199 | 0.837780582 | 0.912894662 | 9.60760543 | 8.634429622 | ENSG00000104783 |
| DAB2 | -0.153111961 | 0.899308518 | 0.521786364 | 0.698705908 | 109.6865544 | 98.81850577 | ENSG00000153071 |
| ISCA1 | -0.153119928 | 0.899303552 | 0.148499219 | 0.303869815 | 703.7239542 | 632.1727931 | ENSG00000135070 |
| GPRC5B | -0.153160777 | 0.899278089 | 0.020413682 | 0.065742343 | 2978.798765 | 2678.77933 | ENSG00000167191 |
| KIN | -0.15322283 | 0.89923941 | 0.192628868 | 0.366060026 | 437.7629675 | 393.8266098 | ENSG00000151657 |
| PCDHGB6 | -0.153339529 | 0.899166674 | 0.457405284 | 0.644484532 | 134.8271655 | 121.5456011 | ENSG00000253305 |
| AIFM2 | -0.153358552 | 0.899154818 | 0.343285415 | 0.537082521 | 255.6352817 | 230.206945 | ENSG00000042286 |
| RAB5B | -0.15337057 | 0.899147328 | 0.070029318 | 0.173010314 | 1290.755945 | 1160.896546 | ENSG00000111540 |
| CRTAP | -0.153422357 | 0.899115053 | 0.018899986 | 0.061548868 | 2516.50353 | 2262.506362 | ENSG00000170275 |
| SF3B1 | -0.153489765 | 0.899073044 | 0.006212743 | 0.024789444 | 8203.228185 | 7375.603772 | ENSG00000115524 |
| KAT8 | -0.15361484 | 0.898995101 | 0.137112452 | 0.286304887 | 948.3982406 | 853.2531466 | ENSG00000103510 |
| C9orf163 | -0.153659105 | 0.898967519 | 0.858734195 | 0.925299324 | 7.056566374 | 6.38788242 | ENSG00000196366 |
| BRD2 | -0.153673679 | 0.898958438 | 0.034153378 | 0.099325074 | 3470.836951 | 3120.467143 | ENSG00000204256 |
| C16orf87 | -0.153778924 | 0.898892861 | 0.196961957 | 0.37246127 | 409.421505 | 367.8130681 | ENSG00000155330 |
| UQCR11 | -0.15378918 | 0.898886471 | 0.345882329 | 0.539293605 | 260.3689786 | 233.9051021 | ENSG00000127540 |
| MAP7D3 | -0.153844129 | 0.898852235 | 0.075146257 | 0.182756987 | 998.0188119 | 896.5606877 | ENSG00000129680 |
| SLC6A1 | -0.153859316 | 0.898842773 | 0.812648748 | 0.899366839 | 13.25243124 | 11.9316113 | ENSG00000157103 |
| HIBCH | -0.153873953 | 0.898833653 | 0.13956118 | 0.290469052 | 655.4417025 | 589.1012074 | ENSG00000198130 |
| CCDC25 | -0.154077289 | 0.898706979 | 0.182804743 | 0.353293381 | 802.3367588 | 720.2386613 | ENSG00000147419 |
| PSMA3 | -0.154105614 | 0.898689334 | 0.023134657 | 0.072698834 | 2365.999787 | 2126.2171 | ENSG00000100567 |
| SEMA3F-AS1 | -0.154160186 | 0.898655341 | 0.866675372 | 0.930273692 | 5.988034921 | 5.348421244 | ENSG00000235016 |
| MERTK | -0.154231867 | 0.898610692 | 0.36766518 | 0.560755962 | 220.8056019 | 198.5047226 | ENSG00000153208 |
| AVEN | -0.154279655 | 0.898580927 | 0.227737842 | 0.411353002 | 400.1331909 | 359.5695886 | ENSG00000169857 |
| NA | -0.154338845 | 0.898544061 | 0.84327509 | 0.91629598 | 9.045277017 | 8.135313641 | ENSG00000225695 |
| UBA5 | -0.154378439 | 0.898519401 | 0.142604216 | 0.295010085 | 774.0437922 | 695.0821356 | ENSG00000081307 |
| CNDP2 | -0.154403786 | 0.898503615 | 0.011095503 | 0.040141334 | 6229.094452 | 5596.693726 | ENSG00000133313 |
| KIAA2013 | -0.154419531 | 0.89849381 | 0.08639443 | 0.203073476 | 1486.580998 | 1335.609268 | ENSG00000116685 |
| SLC52A2 | -0.154479986 | 0.89845616 | 0.081390372 | 0.194167737 | 932.590125 | 838.0063697 | ENSG00000185803 |
| NBAS | -0.15450225 | 0.898442295 | 0.119242893 | 0.258364249 | 823.8925407 | 739.7186361 | ENSG00000151779 |
| LOXL1-AS1 | -0.15458825 | 0.89838874 | 0.573079233 | 0.738883513 | 79.90191765 | 72.03104508 | ENSG00000261801 |
| MORC4 | -0.154590377 | 0.898387415 | 0.35232195 | 0.545516575 | 190.4670251 | 171.1302397 | ENSG00000133131 |
| TSPYL1 | -0.15465629 | 0.898346371 | 0.062035995 | 0.158044692 | 1013.378811 | 910.3104028 | ENSG00000189241 |
| GSDMD | -0.154848575 | 0.898226646 | 0.208348946 | 0.387432465 | 677.0282264 | 608.7454632 | ENSG00000104518 |
| ZWILCH | -0.154985595 | 0.898141341 | 0.212585967 | 0.392385079 | 731.7178788 | 656.7142732 | ENSG00000174442 |
| STAB1 | -0.155027616 | 0.898115181 | 0.77256261 | 0.874743443 | 20.21864986 | 18.08240721 | ENSG00000010327 |
| WTIP | -0.155185345 | 0.898016996 | 0.201606008 | 0.378693194 | 423.5857497 | 380.3608815 | ENSG00000142279 |
| NR2F6 | -0.155197455 | 0.898009458 | 0.049362528 | 0.132419321 | 2505.758498 | 2250.032009 | ENSG00000160113 |
| CEP131 | -0.155209652 | 0.898001866 | 0.199969696 | 0.376420714 | 858.4231407 | 771.7790306 | ENSG00000141577 |
| PPAN | -0.155213481 | 0.897999483 | 0.451381981 | 0.638863202 | 136.8327431 | 123.0417381 | ENSG00000130810 |

| | | | | | | | |
|---|---|---|---|---|---|---|---|
| CFL1 | -0.155344794 | 0.897917751 | 0.004004865 | 0.017043272 | 9031.146394 | 8109.615345 | ENSG00000172757 |
| DENND2A | -0.155378116 | 0.897897012 | 0.247538949 | 0.435531316 | 364.5533614 | 327.8853014 | ENSG00000146966 |
| LOC107984660 | -0.155398343 | 0.897884424 | 0.885355278 | 0.941108896 | 4.813273453 | 4.253085118 | ENSG00000258457 |
| ATP11C | -0.155429808 | 0.897864841 | 0.084679257 | 0.199958324 | 1818.810793 | 1631.955932 | ENSG00000101974 |
| TUBA4A | -0.155431497 | 0.89786379 | 0.082605168 | 0.196309447 | 934.5228458 | 838.6410793 | ENSG00000127824 |
| FLVCR2 | -0.155458474 | 0.897847001 | 0.578099051 | 0.742463296 | 67.83855926 | 60.86371681 | ENSG00000119686 |
| HSD17B6 | -0.155509195 | 0.897815436 | 0.618719588 | 0.773359478 | 52.14102589 | 46.73113602 | ENSG00000025423 |
| GTF3C4 | -0.1555389 | 0.89779695 | 0.045628863 | 0.124677186 | 1730.634418 | 1553.092252 | ENSG00000125484 |
| ARHGEF15 | -0.155760094 | 0.89765931 | 0.692725267 | 0.823967294 | 32.73948212 | 29.3195949 | ENSG00000198844 |
| BID | -0.155781347 | 0.897646086 | 0.109784853 | 0.242897513 | 706.9391396 | 634.4121178 | ENSG00000015475 |
| PNMA8A | -0.155859838 | 0.89759725 | 0.07178838 | 0.176485719 | 1025.218089 | 920.1186049 | ENSG00000182013 |
| YBX1 | -0.155924439 | 0.897557059 | 0.026907658 | 0.082220027 | 26487.89599 | 23773.98348 | ENSG00000065978 |
| CRNKL1 | -0.15594216 | 0.897546034 | 0.147023999 | 0.301431726 | 765.038097 | 685.7525095 | ENSG00000101343 |
| LIN52 | -0.156019657 | 0.897497822 | 0.338493458 | 0.532566871 | 211.9950787 | 190.0066594 | ENSG00000205659 |
| MCRS1 | -0.156046124 | 0.897481357 | 0.035844003 | 0.103141245 | 1856.838393 | 1666.741364 | ENSG00000187778 |
| PHF21A | -0.156165317 | 0.897407212 | 0.140221237 | 0.291286754 | 578.5376544 | 519.2631993 | ENSG00000135365 |
| ZNF280D | -0.156170996 | 0.897403679 | 0.662092819 | 0.803380688 | 40.11830141 | 35.94368776 | ENSG00000137871 |
| CTSZ | -0.156224531 | 0.897370379 | 0.155055402 | 0.313049762 | 701.2920507 | 629.5812571 | ENSG00000101160 |
| MSMO1 | -0.156478949 | 0.897212143 | 0.151965915 | 0.309034043 | 1741.138834 | 1561.459364 | ENSG00000052802 |
| NRDE2 | -0.156543648 | 0.897171907 | 0.362819662 | 0.555747709 | 189.8294345 | 170.0185057 | ENSG00000119720 |
| POLE3 | -0.156548895 | 0.897168645 | 0.085086989 | 0.200585379 | 954.3255653 | 856.0208526 | ENSG00000148229 |
| ZC4H2 | -0.156620713 | 0.897123984 | 0.161940121 | 0.323043613 | 557.931073 | 500.4119373 | ENSG00000126970 |
| SEPTIN4 | -0.15662767 | 0.897119658 | 0.453545359 | 0.640848494 | 117.9751876 | 105.9754578 | ENSG00000108387 |
| RNF157 | -0.156645882 | 0.897108333 | 0.266447884 | 0.456408973 | 314.8971173 | 282.3744714 | ENSG00000141576 |
| LGALS3 | -0.156664619 | 0.897096682 | 0.621744442 | 0.775152125 | 60.04824431 | 53.85368359 | ENSG00000131981 |
| POLDIP2 | -0.156758797 | 0.897038122 | 0.022483135 | 0.071088411 | 2795.557471 | 2507.140091 | ENSG00000004142 |
| POLD1 | -0.156823333 | 0.896997996 | 0.029477101 | 0.088279137 | 3827.618975 | 3434.305296 | ENSG00000062822 |
| YIPF1 | -0.156835113 | 0.896990672 | 0.15937471 | 0.319266199 | 478.439764 | 429.4110523 | ENSG00000058799 |
| TLCD4 | -0.156873598 | 0.896966744 | 0.734879604 | 0.851745268 | 29.78847004 | 26.60366122 | ENSG00000152078 |
| CHCHD3 | -0.156916944 | 0.896939795 | 0.025609512 | 0.078949883 | 2885.443764 | 2588.011704 | ENSG00000106554 |
| ORC6 | -0.156923069 | 0.896935987 | 0.0456796 | 0.124782882 | 1220.188357 | 1094.241462 | ENSG00000091651 |
| LOC100128568 | -0.156990197 | 0.896894254 | 0.831757966 | 0.909325909 | 10.10937378 | 9.137232829 | ENSG00000266983 |
| CCNA2 | -0.156999758 | 0.89688831 | 0.028443238 | 0.085955809 | 2541.997377 | 2279.421232 | ENSG00000145386 |
| ABHD17A | -0.157046079 | 0.896859514 | 0.120320844 | 0.260122071 | 1533.985794 | 1375.785956 | ENSG00000129968 |
| GPS1 | -0.15711677 | 0.896815569 | 0.009303773 | 0.0346466 | 3434.434986 | 3080.232112 | ENSG00000169727 |
| CYB5R1 | -0.15723177 | 0.896744086 | 0.341526868 | 0.53583425 | 230.9110295 | 207.2802968 | ENSG00000159348 |
| CKAP4 | -0.15724409 | 0.896736427 | 0.031018309 | 0.091968219 | 2579.111653 | 2312.756762 | ENSG00000136026 |
| NELL2 | -0.157413156 | 0.896631347 | 0.055185158 | 0.144178422 | 1234.517242 | 1106.509637 | ENSG00000184613 |
| MET | -0.157454233 | 0.896605819 | 0.219106648 | 0.400546616 | 362.3015825 | 324.584607 | ENSG00000105976 |
| PAK4 | -0.157566605 | 0.896535985 | 0.057999431 | 0.150008577 | 2837.073757 | 2543.691937 | ENSG00000130669 |
| PUF60 | -0.157703617 | 0.896450845 | 0.015392378 | 0.052173066 | 3883.91451 | 3482.218414 | ENSG00000179950 |
| ZNF684 | -0.157716484 | 0.89644285 | 0.509401945 | 0.688589866 | 96.35225394 | 86.18498478 | ENSG00000117010 |
| SART1 | -0.157794489 | 0.896394381 | 0.042289973 | 0.117564475 | 2711.060001 | 2430.619067 | ENSG00000175467 |
| B4GALT3 | -0.157796397 | 0.896393196 | 0.113434247 | 0.249100286 | 667.0860713 | 597.7293047 | ENSG00000158850 |
| BTRC | -0.157845172 | 0.896362891 | 0.093081765 | 0.214654403 | 754.4522177 | 676.2790796 | ENSG00000166167 |
| SFRP2 | -0.157851088 | 0.896359215 | 0.016684896 | 0.055708498 | 6724.209264 | 6026.684999 | ENSG00000145423 |
| UNKL | -0.157864028 | 0.896351175 | 0.135775219 | 0.28424917 | 592.3264371 | 530.8574824 | ENSG00000059145 |
| C7orf26 | -0.157875801 | 0.896343861 | 0.131281602 | 0.277442784 | 742.2233092 | 665.2533339 | ENSG00000146576 |
| RPL26L1 | -0.157983617 | 0.896276878 | 0.128807207 | 0.27333556 | 566.8582665 | 507.849475 | ENSG00000037241 |
| SEM1 | -0.158144981 | 0.896176635 | 0.091635586 | 0.212349831 | 2144.363226 | 1921.950093 | ENSG00000127922 |
| IFT20 | -0.158150211 | 0.896173387 | 0.357142003 | 0.550616112 | 193.8079608 | 173.8013686 | ENSG00000109083 |
| SPIN2A | -0.158217825 | 0.896131387 | 0.641550438 | 0.789751316 | 47.5168363 | 42.61660752 | ENSG00000147059 |
| NEK8 | -0.158248273 | 0.896112475 | 0.275489541 | 0.466422139 | 273.1923255 | 245.1758584 | ENSG00000160602 |
| MRPL45 | -0.158296738 | 0.896082371 | 0.09732761 | 0.222042087 | 1197.689598 | 1072.513617 | ENSG00000278845 |
| WDR11 | -0.158322271 | 0.896066513 | 0.051308642 | 0.136435347 | 1159.609083 | 1038.989398 | ENSG00000120008 |
| GBF1 | -0.158339506 | 0.896055808 | 0.080736974 | 0.19300646 | 1224.27763 | 1096.692681 | ENSG00000107862 |
| PSMC4 | -0.158521935 | 0.895942509 | 0.015281187 | 0.051853079 | 3185.596832 | 2854.084155 | ENSG00000013275 |
| ZCCHC17 | -0.158542099 | 0.895929987 | 0.122462155 | 0.263552436 | 636.5937905 | 570.3577849 | ENSG00000121766 |
| PNN | -0.158544236 | 0.895928659 | 0.460761288 | 0.647441221 | 4359.164608 | 3905.797851 | ENSG00000100941 |
| COPS7A | -0.158590949 | 0.895899651 | 0.097440789 | 0.22224558 | 808.0971505 | 724.4847075 | ENSG00000111652 |
| PBX4 | -0.158637232 | 0.89587091 | 0.714929523 | 0.838432369 | 34.20157507 | 30.63704538 | ENSG00000105717 |
| RTL8B | -0.158811728 | 0.89576256 | 0.492603586 | 0.674940109 | 101.7243906 | 91.1100353 | ENSG00000212747 |
| HMCES | -0.158830936 | 0.895750633 | 0.046617258 | 0.12663114 | 1383.365636 | 1239.090397 | ENSG00000183624 |
| L3HYPDH | -0.158921393 | 0.895694472 | 0.505635095 | 0.685272653 | 103.5371487 | 92.8970955 | ENSG00000126790 |
| RNF26 | -0.159011087 | 0.895638787 | 0.045511249 | 0.124410839 | 1754.935569 | 1572.133455 | ENSG00000173456 |
| MED21 | -0.159091245 | 0.895589026 | 0.176106894 | 0.343937321 | 476.6990136 | 426.5826672 | ENSG00000152944 |
| SUMO2 | -0.159091815 | 0.895588672 | 0.008367701 | 0.031773607 | 4553.057639 | 4077.339479 | ENSG00000188612 |
| PSMF1 | -0.159093367 | 0.895587708 | 0.02510601 | 0.077733845 | 1727.078099 | 1546.674506 | ENSG00000125818 |
| AGMAT | -0.159120075 | 0.895571129 | 0.354106518 | 0.547419318 | 203.4095455 | 182.2543634 | ENSG00000116771 |
| ZNF71 | -0.159202939 | 0.895519691 | 0.276694688 | 0.467693512 | 269.8267359 | 241.4296615 | ENSG00000197951 |
| LYAR | -0.159311477 | 0.895452322 | 0.035095337 | 0.101443697 | 1589.175629 | 1422.93312 | ENSG00000145220 |
| VDAC3 | -0.159322578 | 0.895445431 | 0.020754053 | 0.066618569 | 2657.29697 | 2378.734067 | ENSG00000078668 |
| NA | -0.159338825 | 0.895435347 | 0.835919024 | 0.911992182 | 8.981364645 | 8.071952618 | ENSG00000265511 |
| SRP72 | -0.159379646 | 0.895410012 | 0.02199745 | 0.069943513 | 2284.296101 | 2045.416286 | ENSG00000174780 |

| | | | | | | | |
|---|---|---|---|---|---|---|---|
| TAF1D | -0.159385755 | 0.89540622 | 0.081667914 | 0.194700205 | 1061.101952 | 950.2129404 | ENSG00000166012 |
| KLHDC2 | -0.159551307 | 0.895303476 | 0.112309692 | 0.24724407 | 702.2385828 | 628.3793554 | ENSG00000165516 |
| AIDA | -0.159609312 | 0.89526748 | 0.108848492 | 0.241344413 | 1708.32015 | 1528.86452 | ENSG00000186063 |
| RUNDC3B | -0.159609709 | 0.895267234 | 0.851290166 | 0.921459858 | 6.884740247 | 6.192574783 | ENSG00000105784 |
| ULK4P3 | -0.159685439 | 0.895220241 | 0.871060616 | 0.933167325 | 5.83189566 | 5.213663469 | ENSG00000178081 |
| SEMA3B | -0.159754814 | 0.895177193 | 0.540531329 | 0.713700054 | 85.62866567 | 76.52738868 | ENSG00000012171 |
| YWHAH | -0.159811754 | 0.895141863 | 0.024922947 | 0.077231551 | 2975.452861 | 2663.079676 | ENSG00000128245 |
| NOL4L | -0.159840407 | 0.895124085 | 0.129482989 | 0.274643821 | 611.1822564 | 547.3169189 | ENSG00000197183 |
| TOR1AIP1 | -0.159842752 | 0.89512263 | 0.146664529 | 0.300894442 | 504.8258666 | 451.8133781 | ENSG00000143337 |
| PRKCH | -0.159868585 | 0.895106603 | 0.294795136 | 0.487615622 | 278.6417915 | 249.2845011 | ENSG00000027075 |
| UBBP4 | -0.159925066 | 0.89507156 | 0.379349328 | 0.572545639 | 165.2118933 | 147.8794203 | ENSG00000263563 |
| FAM13B | -0.159953678 | 0.895053809 | 0.338238839 | 0.532256634 | 211.2522275 | 189.3634761 | ENSG00000031003 |
| FBXO27 | -0.16017354 | 0.894917416 | 0.326048959 | 0.520261164 | 209.2652983 | 187.5144806 | ENSG00000161243 |
| TBC1D10B | -0.160206125 | 0.894897204 | 0.094331933 | 0.216835922 | 982.9529536 | 879.4596765 | ENSG00000169221 |
| NUDT5 | -0.160371729 | 0.894794486 | 0.066020002 | 0.165532295 | 1030.557379 | 922.548339 | ENSG00000165609 |
| MAST3 | -0.1605684 | 0.894672514 | 0.261987258 | 0.451186754 | 304.9861027 | 272.67465 | ENSG00000099308 |
| ZDHHC12 | -0.160610525 | 0.894646391 | 0.255416567 | 0.444288188 | 375.1581897 | 335.7825493 | ENSG00000160446 |
| JOSD1 | -0.160840538 | 0.894503766 | 0.11603332 | 0.253275415 | 725.6271315 | 649.2264434 | ENSG00000100221 |
| LPCAT1 | -0.161091684 | 0.894348064 | 0.004324532 | 0.018189324 | 6937.719585 | 6204.685792 | ENSG00000153395 |
| TRMT44 | -0.161113999 | 0.89433423 | 0.247814558 | 0.435800274 | 309.1342451 | 276.7238908 | ENSG00000155275 |
| EIF5 | -0.161135844 | 0.894320689 | 0.019931309 | 0.064367814 | 4226.856213 | 3779.727511 | ENSG00000100664 |
| SPATA13 | -0.161147855 | 0.894313243 | 0.222843206 | 0.404946971 | 375.1919236 | 335.0706201 | ENSG00000182957 |
| SPHK1 | -0.161152748 | 0.89431021 | 0.643808196 | 0.791682598 | 42.65227813 | 38.12307509 | ENSG00000176170 |
| ACTL6A | -0.161213053 | 0.894272829 | 0.036068414 | 0.103678081 | 1677.92838 | 1500.135818 | ENSG00000136518 |
| ZNF767P | -0.161308633 | 0.894213584 | 0.714683756 | 0.838366554 | 188.9473336 | 169.1174669 | ENSG00000133624 |
| FSIP1 | -0.161431785 | 0.894137255 | 0.821630503 | 0.904297073 | 9.229730795 | 8.251738454 | ENSG00000150667 |
| YDJC | -0.161470946 | 0.894112985 | 0.173777888 | 0.340537045 | 765.6118653 | 684.4726582 | ENSG00000161179 |
| IQCC | -0.161529517 | 0.894076686 | 0.590037678 | 0.752063705 | 63.17826816 | 56.65978501 | ENSG00000160051 |
| LMAN2 | -0.161535027 | 0.894073271 | 0.014127465 | 0.048670975 | 3101.24316 | 2772.953952 | ENSG00000169223 |
| CCDC6 | -0.161581883 | 0.894044234 | 0.108906448 | 0.241415153 | 1248.49038 | 1115.682897 | ENSG00000108091 |
| ZSCAN1 | -0.16164299 | 0.894006366 | 0.802926732 | 0.894151534 | 13.35993903 | 11.90860343 | ENSG00000152467 |
| GRK4 | -0.161662133 | 0.893994504 | 0.704094378 | 0.83179868 | 29.69812228 | 26.64219089 | ENSG00000125388 |
| THOP1 | -0.161662597 | 0.893994216 | 0.015632436 | 0.052803276 | 3472.148744 | 3105.006848 | ENSG00000172009 |
| MMP14 | -0.161787162 | 0.893917031 | 0.018553283 | 0.060550545 | 2052.03695 | 1834.298749 | ENSG00000157227 |
| PCDH7 | -0.161802522 | 0.893907513 | 0.469199392 | 0.655027232 | 111.9525244 | 100.3121665 | ENSG00000169851 |
| WASF3 | -0.161803865 | 0.893906681 | 0.157192928 | 0.316194239 | 505.2704197 | 451.1778554 | ENSG00000132970 |
| MAFG-DT | -0.161822518 | 0.893895124 | 0.536537635 | 0.710857712 | 82.45927053 | 73.92864426 | ENSG00000265688 |
| PYGB | -0.162032648 | 0.893764937 | 0.021846694 | 0.069571586 | 2580.97659 | 2306.824074 | ENSG00000100994 |
| SPIN4 | -0.162051892 | 0.893753015 | 0.330957208 | 0.525254238 | 242.1187975 | 216.8502586 | ENSG00000186767 |
| SLC38A1 | -0.162081647 | 0.893734581 | 0.018931347 | 0.061621634 | 6562.254851 | 5863.896949 | ENSG00000111371 |
| NA | -0.162173624 | 0.893677604 | 0.291080806 | 0.483456551 | 272.4796832 | 243.0236756 | ENSG00000132967 |
| NA | -0.162285313 | 0.893608422 | 0.663131076 | 0.804083749 | 36.78743786 | 32.74404828 | ENSG00000213235 |
| NA | -0.162301361 | 0.893598481 | 0.86964247 | 0.932374243 | 5.008573445 | 4.508504585 | ENSG00000202058 |
| AVPR1A | -0.162396663 | 0.893539454 | 0.886158497 | 0.941854661 | 3.910254302 | 3.484565209 | ENSG00000166148 |
| DNAJB9 | -0.162492645 | 0.893480009 | 0.327114358 | 0.521301591 | 214.9980519 | 191.9460375 | ENSG00000128590 |
| COX14 | -0.162497045 | 0.893477284 | 0.592802582 | 0.75444118 | 67.53278996 | 60.53832631 | ENSG00000178449 |
| CLDND1 | -0.162501214 | 0.893474702 | 0.341167537 | 0.535546635 | 380.0913881 | 339.461609 | ENSG00000080822 |
| RAB11A | -0.162619974 | 0.893401156 | 0.027194214 | 0.082952198 | 2544.960167 | 2272.994502 | ENSG00000103769 |
| TMED3 | -0.162655257 | 0.893379307 | 0.057316579 | 0.148636558 | 1085.628476 | 969.7308231 | ENSG00000166557 |
| PIGC | -0.162802059 | 0.893288405 | 0.1688439 | 0.333157972 | 450.7876805 | 402.5044185 | ENSG00000135845 |
| SALL2 | -0.162812138 | 0.893282164 | 0.003036976 | 0.013465864 | 8263.538406 | 7382.042342 | ENSG00000165821 |
| ZNF384 | -0.162843997 | 0.893262438 | 0.15685205 | 0.315748441 | 815.6192407 | 729.0407652 | ENSG00000126746 |
| DLEC1 | -0.162917852 | 0.893216711 | 0.848980306 | 0.919906518 | 7.552020899 | 6.721197025 | ENSG00000008226 |
| NA | -0.163040082 | 0.893141038 | 0.856790751 | 0.924075698 | 7.115946358 | 6.384083573 | ENSG00000276571 |
| RPS10 | -0.16304455 | 0.893138272 | 0.771020269 | 0.87406506 | 15.78019183 | 14.13681767 | ENSG00000124614 |
| TMEM70 | -0.163119181 | 0.893092071 | 0.158286868 | 0.317773918 | 694.0122532 | 619.6808003 | ENSG00000175606 |
| ENPEP | -0.163207862 | 0.893037175 | 0.785799553 | 0.882841791 | 13.38835125 | 11.90860343 | ENSG00000138792 |
| TLCD3B | -0.163291701 | 0.89298528 | 0.205796606 | 0.384267005 | 407.5066659 | 364.3041096 | ENSG00000149926 |
| CORO6 | -0.163313083 | 0.892972045 | 0.667915738 | 0.807195506 | 54.44063958 | 48.71262779 | ENSG00000167549 |
| CELF4 | -0.163321772 | 0.892966667 | 0.648947509 | 0.79510763 | 43.72020822 | 39.0722569 | ENSG00000101489 |
| ABHD2 | -0.163322863 | 0.892965991 | 0.158931249 | 0.318707138 | 628.926986 | 561.6391278 | ENSG00000140526 |
| TP53RK | -0.163357868 | 0.892944326 | 0.192457517 | 0.365809416 | 449.3953851 | 401.1498725 | ENSG00000172315 |
| TRPC5 | -0.163415547 | 0.892908626 | 0.655312284 | 0.798737248 | 49.53879981 | 44.25206561 | ENSG00000072315 |
| OGFOD3 | -0.16347859 | 0.892869609 | 0.127666731 | 0.271599539 | 663.0674898 | 591.5018875 | ENSG00000181396 |
| NDUFC1 | -0.163538805 | 0.892832343 | 0.125334687 | 0.268270364 | 586.8784443 | 523.8980995 | ENSG00000109390 |
| NUP42 | -0.163539116 | 0.892832151 | 0.220215096 | 0.401751821 | 307.5720405 | 274.4789438 | ENSG00000136243 |
| CACNA2D4 | -0.163556656 | 0.892821296 | 0.874685792 | 0.935130738 | 5.138081929 | 4.617443324 | ENSG00000151062 |
| MARK3 | -0.163561518 | 0.892818287 | 0.034005236 | 0.098987443 | 1776.003593 | 1585.677963 | ENSG00000075413 |
| TRIM27 | -0.163565364 | 0.892815907 | 0.016971443 | 0.056411177 | 2112.577665 | 1886.06702 | ENSG00000204713 |
| NA | -0.163578187 | 0.892807971 | 0.8558047 | 0.923467881 | 5.76620185 | 5.127294571 | ENSG00000215895 |
| METTL9 | -0.163636216 | 0.892772061 | 0.064434037 | 0.162700332 | 2367.978192 | 2113.296763 | ENSG00000197006 |
| SLC38A10 | -0.163678896 | 0.89274565 | 0.062258271 | 0.158482504 | 1153.44154 | 1030.332448 | ENSG00000157637 |
| USP14 | -0.163698105 | 0.892733763 | 0.083602181 | 0.198097979 | 2134.968216 | 1904.999368 | ENSG00000101557 |
| PALB2 | -0.163714497 | 0.89272362 | 0.202582711 | 0.379927093 | 412.3215406 | 367.5270431 | ENSG00000083093 |

| | | | | | | | |
|---|---|---|---|---|---|---|---|
| PCSK2 | -0.163819914 | 0.892658392 | 0.817773304 | 0.901961189 | 10.24203918 | 9.061599198 | ENSG00000125851 |
| AMBRA1 | -0.163858324 | 0.892634626 | 0.047412036 | 0.128357388 | 1069.07875 | 953.9680717 | ENSG00000110497 |
| SPAG5 | -0.16388698 | 0.892616896 | 0.049169553 | 0.132016296 | 1676.185473 | 1496.79869 | ENSG00000076382 |
| SAXO2 | -0.163914271 | 0.892600011 | 0.766354424 | 0.870835943 | 21.06988063 | 18.8595125 | ENSG00000188659 |
| GSPT1 | -0.163987983 | 0.892554406 | 0.021588289 | 0.068890727 | 4652.093876 | 4151.653418 | ENSG00000103342 |
| ZC3HC1 | -0.163996566 | 0.892549096 | 0.140247834 | 0.291309354 | 579.9241921 | 517.1647331 | ENSG00000091732 |
| RSPRY1 | -0.164183226 | 0.892433623 | 0.155532648 | 0.313739889 | 552.7339042 | 493.0169111 | ENSG00000159579 |
| ASF1B | -0.164246485 | 0.892394493 | 0.075338349 | 0.183100048 | 931.5561469 | 831.0384015 | ENSG00000105011 |
| RBBP7 | -0.164259053 | 0.892386719 | 0.002499091 | 0.011462529 | 6644.188142 | 5929.10062 | ENSG00000102054 |
| KLF8 | -0.164410049 | 0.892293324 | 0.362999192 | 0.555880941 | 175.2518376 | 156.3518474 | ENSG00000102349 |
| POU5F1P3 | -0.164431882 | 0.892279821 | 0.497955734 | 0.678931121 | 85.33995871 | 76.03564715 | ENSG00000235602 |
| DPYSL4 | -0.164481547 | 0.892249104 | 0.085031051 | 0.200513407 | 847.4205267 | 756.3328208 | ENSG00000151640 |
| GS1-124K5.11 | -0.16477564 | 0.892067238 | 0.246831414 | 0.434492679 | 317.5822722 | 283.7087578 | ENSG00000229180 |
| UNC13D | -0.164787108 | 0.892060147 | 0.40044263 | 0.592464573 | 153.4698782 | 137.1034718 | ENSG00000092929 |
| HSPA14 | -0.164791457 | 0.892057458 | 0.211918966 | 0.391595824 | 409.5435347 | 364.8193599 | ENSG00000284024 |
| PNPT1 | -0.164840851 | 0.892026917 | 0.067569273 | 0.168459632 | 959.8849818 | 855.6946574 | ENSG00000138035 |
| TRIP13 | -0.164873665 | 0.892006628 | 0.054879371 | 0.143642743 | 1133.488198 | 1011.732145 | ENSG00000071539 |
| UQCC3 | -0.164879618 | 0.892002947 | 0.537266968 | 0.711203176 | 91.59677443 | 81.61593033 | ENSG00000204922 |
| PBX3 | -0.164914562 | 0.891981342 | 0.155497226 | 0.313739889 | 541.8525906 | 483.1145192 | ENSG00000167081 |
| SPDYE3 | -0.164915567 | 0.891980721 | 0.662533772 | 0.803690999 | 40.2730652 | 35.79720703 | ENSG00000214300 |
| ST6GAL2 | -0.164980482 | 0.891940586 | 0.208526337 | 0.38769228 | 343.4470862 | 306.6329842 | ENSG00000144057 |
| SNX5 | -0.16508875 | 0.891873652 | 0.03868164 | 0.109571376 | 1836.951219 | 1637.574602 | ENSG00000089006 |
| CDK5RAP1 | -0.165094718 | 0.891869963 | 0.07925617 | 0.190301056 | 842.7554027 | 751.5552302 | ENSG00000101391 |
| NA | -0.165106953 | 0.8918624 | 0.791499854 | 0.886486471 | 13.28981053 | 11.79966469 | ENSG00000276386 |
| NA | -0.165113303 | 0.891858474 | 0.617194375 | 0.772310229 | 55.82332869 | 50.04427753 | ENSG00000242622 |
| MYO18B | -0.165121004 | 0.891853713 | 0.750830203 | 0.861339687 | 18.82974462 | 16.74717342 | ENSG00000133454 |
| TNFSF11 | -0.165167621 | 0.891824896 | 0.38194069 | 0.574745693 | 145.374974 | 129.51055 | ENSG00000120659 |
| CYFIP1 | -0.165219808 | 0.891792636 | 0.024875174 | 0.077135097 | 2260.207079 | 2015.109321 | ENSG00000273749 |
| FAM78B | -0.165223214 | 0.891790531 | 0.562576858 | 0.730892646 | 69.99218085 | 62.57871899 | ENSG00000188859 |
| HSP90B1 | -0.165254278 | 0.891771329 | 0.005484984 | 0.022288685 | 14247.36701 | 12704.39979 | ENSG00000166598 |
| COMMD10 | -0.165343031 | 0.89171647 | 0.266943087 | 0.456919392 | 287.4753076 | 256.0052685 | ENSG00000145781 |
| REXO4 | -0.165363336 | 0.891703919 | 0.030296217 | 0.090274934 | 1535.522691 | 1369.022119 | ENSG00000148300 |
| CTDP1 | -0.165460577 | 0.891643819 | 0.051802853 | 0.137266634 | 1680.033506 | 1497.601676 | ENSG00000060069 |
| SIN3A | -0.165461771 | 0.891643081 | 0.016030283 | 0.053911317 | 2438.136848 | 2173.426262 | ENSG00000169375 |
| SNRPA1 | -0.165478589 | 0.891632687 | 0.020697655 | 0.066483595 | 2276.824122 | 2029.813021 | ENSG00000131876 |
| SLC39A6 | -0.165537548 | 0.891596249 | 0.065231919 | 0.164000054 | 1052.738522 | 938.3597336 | ENSG00000141424 |
| VOPP1 | -0.165588645 | 0.891564671 | 0.04240753 | 0.117820592 | 1566.500391 | 1396.374438 | ENSG00000154978 |
| GYPC | -0.165666769 | 0.891516393 | 0.168403198 | 0.332429958 | 488.5340523 | 435.1150945 | ENSG00000136732 |
| ZHX3 | -0.1657434 | 0.89146904 | 0.415720389 | 0.606967561 | 131.3760688 | 116.9717601 | ENSG00000174306 |
| GNL3 | -0.165760621 | 0.891458399 | 0.008019139 | 0.030688977 | 3189.382126 | 2843.728368 | ENSG00000163938 |
| CNOT8 | -0.165776401 | 0.891448648 | 0.078534848 | 0.18893625 | 951.7291268 | 847.6637192 | ENSG00000155508 |
| SMYD2 | -0.165916262 | 0.891362231 | 0.411649801 | 0.603123907 | 135.0924963 | 120.6187659 | ENSG00000143499 |
| PPIP5K2 | -0.165963399 | 0.891333109 | 0.198857389 | 0.37508883 | 549.5992276 | 489.0266997 | ENSG00000145725 |
| TTC26 | -0.166004932 | 0.891307449 | 0.417163763 | 0.608365189 | 148.7151129 | 132.2629264 | ENSG00000105948 |
| ASRGL1 | -0.166011768 | 0.891303226 | 0.312896561 | 0.506526759 | 240.2287404 | 213.6208493 | ENSG00000162174 |
| TRABD | -0.166035009 | 0.891288867 | 0.035634716 | 0.102714437 | 1795.333948 | 1600.22005 | ENSG00000170638 |
| SOX2-OT | -0.166093632 | 0.891252651 | 0.659612491 | 0.801708828 | 33.98940813 | 30.32900016 | ENSG00000242808 |
| RAX | -0.166162315 | 0.891210222 | 0.881268462 | 0.939080645 | 4.241376604 | 3.74323387 | ENSG00000134438 |
| NA | -0.166196307 | 0.891189224 | 0.86117288 | 0.926993737 | 5.484988795 | 4.894882979 | ENSG00000267364 |
| GULP1 | -0.166257436 | 0.891151463 | 0.031114569 | 0.092209374 | 1855.501426 | 1652.899466 | ENSG00000144366 |
| GJA3 | -0.166302953 | 0.891123348 | 0.775848465 | 0.876322473 | 14.27558583 | 12.73398597 | ENSG00000121743 |
| BRD8 | -0.166314056 | 0.89111649 | 0.041329337 | 0.115482879 | 1490.367171 | 1328.491682 | ENSG00000112983 |
| NA | -0.166470036 | 0.89102015 | 0.388395115 | 0.581300136 | 144.8126456 | 129.283363 | ENSG00000237517 |
| DUSP1 | -0.166677431 | 0.890892071 | 0.339437469 | 0.533734182 | 187.9343262 | 167.4661119 | ENSG00000120129 |
| NKAPD1 | -0.166783085 | 0.89082683 | 0.177858983 | 0.34651933 | 479.8535937 | 427.2097161 | ENSG00000150776 |
| SLC16A2 | -0.166811749 | 0.890809131 | 0.014666703 | 0.05013513 | 2313.891337 | 2061.280401 | ENSG00000147100 |
| CPSF3 | -0.166821271 | 0.890803251 | 0.040148566 | 0.112778649 | 1796.872446 | 1599.730249 | ENSG00000119203 |
| GMPS | -0.166867547 | 0.890774678 | 0.021878199 | 0.069647979 | 3405.877982 | 3032.803978 | ENSG00000163655 |
| ABRACL | -0.166894242 | 0.890758196 | 0.07355289 | 0.179851124 | 1425.125749 | 1268.677898 | ENSG00000146386 |
| PTPN1 | -0.167013481 | 0.890684578 | 0.04496484 | 0.123189704 | 1106.309238 | 985.3018022 | ENSG00000196396 |
| NUBP2 | -0.167139431 | 0.890606823 | 0.040587572 | 0.113770427 | 1846.734746 | 1644.31743 | ENSG00000095906 |
| C8orf58 | -0.167567829 | 0.890342402 | 0.423872772 | 0.61471502 | 140.1034526 | 124.7931914 | ENSG00000241852 |
| MED27 | -0.167624768 | 0.890307264 | 0.166229708 | 0.329402482 | 545.5742573 | 485.1753403 | ENSG00000160563 |
| DSTN | -0.167725976 | 0.89024481 | 0.041969033 | 0.116917789 | 2392.197106 | 2129.13824 | ENSG00000125868 |
| DSN1 | -0.167733528 | 0.89024015 | 0.13687943 | 0.285850519 | 481.5081606 | 428.8298043 | ENSG00000149636 |
| MAML1 | -0.167747475 | 0.890231543 | 0.030934407 | 0.091792874 | 1687.365806 | 1502.303697 | ENSG00000161021 |
| TRIM24 | -0.167764395 | 0.890221102 | 0.023301411 | 0.073123661 | 3148.556496 | 2802.158953 | ENSG00000122779 |
| GALNT7 | -0.167792021 | 0.890204056 | 0.075256947 | 0.18297419 | 1212.734358 | 1078.950327 | ENSG00000109586 |
| THOC6 | -0.167810783 | 0.890192479 | 0.044657808 | 0.122475261 | 1123.042178 | 999.8474968 | ENSG00000131652 |
| TENT5B | -0.167836787 | 0.890176434 | 0.028303927 | 0.085660513 | 2511.994049 | 2235.764837 | ENSG00000158246 |
| NA | -0.167842827 | 0.890172707 | 0.788326035 | 0.884326033 | 15.10028352 | 13.50294391 | ENSG00000269737 |
| CDH13 | -0.167843288 | 0.890172422 | 0.784176522 | 0.881903096 | 14.46329422 | 12.85365998 | ENSG00000140945 |
| USP15 | -0.167947261 | 0.890108271 | 0.193418742 | 0.367109366 | 441.229383 | 392.1344636 | ENSG00000135655 |
| THAP2 | -0.167988651 | 0.890082735 | 0.572880621 | 0.738883513 | 58.90828394 | 52.33822593 | ENSG00000173451 |

| | | | | | | | |
|---|---|---|---|---|---|---|---|
| SLBP | -0.168005877 | 0.890072108 | 0.063405813 | 0.160849075 | 1031.307807 | 917.3982909 | ENSG00000163950 |
| SPOCK2 | -0.16807624 | 0.890028698 | 0.570618612 | 0.737198198 | 65.34294652 | 58.03972045 | ENSG00000107742 |
| MIR4713HG | -0.168347556 | 0.889861333 | 0.851571369 | 0.921524381 | 6.620589532 | 5.930107275 | ENSG00000259240 |
| MADD | -0.168366608 | 0.889849582 | 0.046891505 | 0.127169874 | 1362.734837 | 1213.214719 | ENSG00000110514 |
| SENP6 | -0.16839155 | 0.889834198 | 0.052209865 | 0.138019917 | 1427.733154 | 1270.12657 | ENSG00000112701 |
| MRPS23 | -0.168395525 | 0.889831746 | 0.032376563 | 0.095067684 | 1464.188041 | 1302.566507 | ENSG00000181610 |
| KRT19 | -0.168509561 | 0.889761413 | 0.018611143 | 0.060717994 | 2209.980223 | 1966.078702 | ENSG00000171345 |
| INAVA | -0.168600096 | 0.889705579 | 0.077509297 | 0.18722238 | 847.0361655 | 753.9480433 | ENSG00000163362 |
| FAM110B | -0.168601927 | 0.88970445 | 0.477762144 | 0.661871178 | 104.8207935 | 93.17993424 | ENSG00000169122 |
| AIMP2 | -0.168607741 | 0.889700864 | 0.100042081 | 0.226727797 | 640.310218 | 569.5991242 | ENSG00000106305 |
| CYBC1 | -0.16861012 | 0.889699397 | 0.052860445 | 0.13956061 | 940.5458999 | 836.9676814 | ENSG00000178927 |
| CLASP1 | -0.168666695 | 0.889664508 | 0.056707307 | 0.147386513 | 1364.615844 | 1213.811177 | ENSG00000074054 |
| NA | -0.168760837 | 0.889606456 | 0.808003892 | 0.89690029 | 11.48565137 | 10.17190747 | ENSG00000269906 |
| ATP13A2 | -0.168766162 | 0.889603172 | 0.039920769 | 0.112326025 | 2192.899204 | 1950.579646 | ENSG00000159363 |
| VPS37B | -0.168781682 | 0.889593603 | 0.313414194 | 0.506988374 | 217.1366106 | 193.1159079 | ENSG00000139722 |
| ADAMTSL1 | -0.168842367 | 0.889556184 | 0.665000581 | 0.805385316 | 32.89750051 | 29.34640162 | ENSG00000178031 |
| KHSRP | -0.168842864 | 0.889555877 | 0.004220996 | 0.017828611 | 12544.12004 | 11159.16006 | ENSG00000088247 |
| PGBD1 | -0.168907968 | 0.889515735 | 0.345654037 | 0.539172666 | 192.0725546 | 170.6871103 | ENSG00000137338 |
| SAMD13 | -0.169027847 | 0.889441825 | 0.838525126 | 0.913330082 | 7.939366272 | 7.002435527 | ENSG00000203943 |
| LOC101927551 | -0.169050779 | 0.889427688 | 0.799749486 | 0.892387536 | 12.11189215 | 10.81370533 | ENSG00000230922 |
| ACTR10 | -0.169052637 | 0.889426542 | 0.157592393 | 0.31682584 | 521.4092742 | 463.3384058 | ENSG00000131966 |
| RABEP1 | -0.169166302 | 0.88935647 | 0.042643109 | 0.118350912 | 1056.099128 | 939.3636197 | ENSG00000029725 |
| MCTP1 | -0.169193348 | 0.889339797 | 0.501838352 | 0.682397067 | 79.66735922 | 70.92387438 | ENSG00000175471 |
| TIMM8A | -0.169320488 | 0.889261426 | 0.179247313 | 0.348418427 | 419.454647 | 372.9087065 | ENSG00000126953 |
| HNRNPH1 | -0.169363947 | 0.889234639 | 0.00233251 | 0.010821212 | 10120.62398 | 9000.193535 | ENSG00000169045 |
| ITM2A | -0.169429172 | 0.889194437 | 0.188982582 | 0.361055888 | 504.7475451 | 448.0982877 | ENSG00000078596 |
| PRCP | -0.169488182 | 0.889158068 | 0.070604963 | 0.174107556 | 1459.203483 | 1296.418434 | ENSG00000137509 |
| TULP3 | -0.169619209 | 0.889077317 | 0.17944455 | 0.34865555 | 447.8037012 | 397.5586217 | ENSG00000078246 |
| SPATA2L | -0.169650859 | 0.889057813 | 0.300472312 | 0.493745352 | 311.0095475 | 276.6654908 | ENSG00000158792 |
| OIP5 | -0.169701248 | 0.889026761 | 0.498187354 | 0.679126538 | 84.79657178 | 75.47360818 | ENSG00000104147 |
| RNF145 | -0.169733261 | 0.889007034 | 0.03227869 | 0.094855436 | 1874.576621 | 1665.746587 | ENSG00000145860 |
| C20orf96 | -0.169749408 | 0.888997084 | 0.156916821 | 0.315810226 | 534.9292085 | 475.7118727 | ENSG00000196476 |
| MALSU1 | -0.169777088 | 0.888980027 | 0.065490234 | 0.164493292 | 889.6158871 | 791.1634923 | ENSG00000156928 |
| OS9 | -0.169784375 | 0.888975537 | 0.034362039 | 0.099765879 | 1937.296528 | 1721.819955 | ENSG00000135506 |
| IL17RA | -0.169969893 | 0.88886123 | 0.209716611 | 0.388922119 | 318.6489472 | 283.4750321 | ENSG00000177663 |
| FOXI3 | -0.170037649 | 0.888819486 | 0.37991567 | 0.572945989 | 164.6243322 | 146.3071 | ENSG00000214336 |
| NA | -0.170089757 | 0.888787384 | 0.788199168 | 0.884323016 | 12.78045058 | 11.34612642 | ENSG00000250838 |
| EEF1AKMT1 | -0.170147346 | 0.888751906 | 0.51872214 | 0.695809231 | 73.09224425 | 64.97130888 | ENSG00000150456 |
| NA | -0.170259576 | 0.888682771 | 0.836408549 | 0.9121504 | 8.534639288 | 7.610378626 | ENSG00000286064 |
| MFSD10 | -0.170356693 | 0.88862295 | 0.031575301 | 0.093335986 | 2177.569533 | 1935.372078 | ENSG00000109736 |
| NA | -0.170392535 | 0.888600874 | 0.757686773 | 0.865096588 | 20.53714451 | 18.32329257 | ENSG00000288586 |
| CDH23 | -0.170441883 | 0.888570479 | 0.300354813 | 0.49372737 | 235.1362307 | 209.1651224 | ENSG00000107736 |
| RBM28 | -0.170542227 | 0.888508678 | 0.04423883 | 0.121693796 | 1372.609359 | 1220.030767 | ENSG00000106344 |
| DSCC1 | -0.170554153 | 0.888501334 | 0.077055835 | 0.186336152 | 891.4971277 | 791.3449497 | ENSG00000136982 |
| ENTR1 | -0.170639545 | 0.888448746 | 0.039955397 | 0.112406394 | 1900.959221 | 1689.637877 | ENSG00000165689 |
| SORBS3 | -0.170659497 | 0.888436459 | 0.066877612 | 0.167207562 | 933.5486006 | 829.4564537 | ENSG00000120896 |
| HEXIM1 | -0.170754898 | 0.888377711 | 0.154090174 | 0.311699488 | 1236.064398 | 1099.073377 | ENSG00000186834 |
| RAP2C | -0.170877994 | 0.888301914 | 0.23598278 | 0.421034352 | 739.6010441 | 656.8943939 | ENSG00000123728 |
| DDX42 | -0.170931034 | 0.888269257 | 0.008323918 | 0.031664066 | 3905.813914 | 3469.987078 | ENSG00000198231 |
| TRMT61B | -0.170941509 | 0.888262808 | 0.144342063 | 0.297512877 | 446.7680166 | 396.7817799 | ENSG00000171103 |
| NA | -0.170986416 | 0.888235159 | 0.874398262 | 0.935039083 | 4.697670398 | 4.212293935 | ENSG00000232187 |
| HEBP1 | -0.171027919 | 0.888209607 | 0.1229351 | 0.264355394 | 701.8401732 | 622.6419191 | ENSG00000013583 |
| RRN3 | -0.171213954 | 0.88809508 | 0.07812161 | 0.188382244 | 768.6963321 | 682.4620813 | ENSG00000085721 |
| YKT6 | -0.171230394 | 0.88808496 | 0.011466742 | 0.041232369 | 3381.103391 | 3002.860074 | ENSG00000106636 |
| OCIAD1 | -0.171269178 | 0.888061086 | 0.03511722 | 0.101491121 | 1803.175195 | 1600.399085 | ENSG00000109180 |
| NA | -0.17131257 | 0.888034376 | 0.8771832 | 0.93639635 | 3.827498152 | 3.37562647 | ENSG00000273521 |
| ANAPC10 | -0.171403864 | 0.887978183 | 0.760944833 | 0.867158734 | 20.66053457 | 18.35615964 | ENSG00000164162 |
| NA | -0.171459745 | 0.887943789 | 0.798867558 | 0.891886713 | 11.78392676 | 10.378062 | ENSG00000213881 |
| TMED2 | -0.171476065 | 0.887933745 | 0.039218484 | 0.110747536 | 2944.572348 | 2613.864061 | ENSG00000086598 |
| MIR9-3HG | -0.171509416 | 0.887913218 | 0.823333678 | 0.905075215 | 9.323235461 | 8.255537301 | ENSG00000255571 |
| LINC02289 | -0.17155804 | 0.887883293 | 0.77568258 | 0.876322473 | 15.03637115 | 13.29646297 | ENSG00000258819 |
| ENTPD6 | -0.171569676 | 0.887876131 | 0.051030229 | 0.135922274 | 1120.180145 | 995.3160331 | ENSG00000197586 |
| NA | -0.171597341 | 0.887859106 | 0.853453496 | 0.922280546 | 5.756731112 | 5.078467662 | ENSG00000279833 |
| GIGYF1 | -0.171686687 | 0.887804123 | 0.021960078 | 0.069860641 | 2794.107533 | 2481.545384 | ENSG00000146830 |
| PLXNB1 | -0.171690184 | 0.887801971 | 0.04231956 | 0.117629082 | 3556.928509 | 3159.224104 | ENSG00000164050 |
| LDLRAP1 | -0.171716029 | 0.887786067 | 0.107127773 | 0.238585213 | 558.1281393 | 495.8258725 | ENSG00000157978 |
| LRRC59 | -0.171807393 | 0.887729846 | 0.007839263 | 0.030118761 | 3086.85697 | 2740.681592 | ENSG00000108829 |
| NA | -0.171980755 | 0.887623178 | 0.867891413 | 0.931197143 | 5.383291167 | 4.770972092 | ENSG00000229670 |
| PGS1 | -0.172226635 | 0.887471912 | 0.081220413 | 0.19386209 | 1089.64142 | 967.8400177 | ENSG00000087157 |
| ZCCHC8 | -0.17227046 | 0.887444954 | 0.115701979 | 0.252820146 | 598.8019894 | 531.1943726 | ENSG00000033030 |
| ACIN1 | -0.172274882 | 0.887442234 | 0.01563977 | 0.052818422 | 5812.939768 | 5159.895429 | ENSG00000100813 |
| BAK1 | -0.17230264 | 0.887425159 | 0.049752729 | 0.133253913 | 1123.879578 | 996.736121 | ENSG00000030110 |
| DMKN | -0.172318347 | 0.887415497 | 0.012358533 | 0.043829515 | 2546.567817 | 2259.754456 | ENSG00000161249 |
| ANKRD36C | -0.172386596 | 0.887373518 | 0.704351425 | 0.831811477 | 25.77514263 | 22.80444078 | ENSG00000174501 |

| | | | | | | | |
|---|---|---|---|---|---|---|---|
| BRD1 | -0.172404318 | 0.887362618 | 0.039766243 | 0.111969844 | 1208.540779 | 1072.329332 | ENSG00000100425 |
| SLC29A1 | -0.172463135 | 0.887326441 | 0.007817667 | 0.030066952 | 4483.123425 | 3978.611938 | ENSG00000112759 |
| CYHR1 | -0.172484638 | 0.887313216 | 0.09861702 | 0.224265994 | 993.9602128 | 882.7897514 | ENSG00000187954 |
| METTL22 | -0.172572917 | 0.887258923 | 0.163059268 | 0.324716931 | 638.2277701 | 566.3321503 | ENSG00000067365 |
| WDR81 | -0.172724113 | 0.887165942 | 0.12555224 | 0.268487968 | 585.7727063 | 519.6786459 | ENSG00000167716 |
| SSR2 | -0.172800613 | 0.8871189 | 0.065281194 | 0.164079421 | 1165.998218 | 1033.970994 | ENSG00000163479 |
| DOCK9 | -0.172826718 | 0.887102849 | 0.134304215 | 0.281838124 | 537.9267397 | 476.9609121 | ENSG00000088387 |
| MYLK2 | -0.172860319 | 0.887082188 | 0.857125764 | 0.924108956 | 5.091329596 | 4.538560499 | ENSG00000101306 |
| HK2 | -0.172865812 | 0.88707881 | 0.029591162 | 0.088558468 | 2349.926348 | 2083.848975 | ENSG00000159399 |
| ERAP1 | -0.172982521 | 0.887007052 | 0.272014344 | 0.46303397 | 277.2045857 | 245.5253074 | ENSG00000164307 |
| PCDHA11 | -0.172994304 | 0.886999807 | 0.281398905 | 0.47271345 | 207.2360515 | 183.7515997 | ENSG00000249158 |
| VPS50 | -0.173027268 | 0.886979541 | 0.403809558 | 0.59554439 | 123.6395942 | 109.4436251 | ENSG00000004766 |
| FOXD3 | -0.173088617 | 0.886941824 | 0.236600557 | 0.421690039 | 415.0474346 | 368.1918573 | ENSG00000187140 |
| PPP1CB | -0.173127768 | 0.886917755 | 0.101720365 | 0.229548915 | 1585.294733 | 1405.140367 | ENSG00000213639 |
| MICALL2 | -0.173169143 | 0.886892319 | 0.413278833 | 0.604602564 | 177.5543151 | 157.7575305 | ENSG00000164877 |
| MORF4L1 | -0.17320599 | 0.886869668 | 0.004621557 | 0.019222606 | 10028.30248 | 8892.625082 | ENSG00000185787 |
| NDUFA4 | -0.173234664 | 0.886852041 | 0.007131452 | 0.027832036 | 3156.99945 | 2800.139356 | ENSG00000189043 |
| CCDC59 | -0.173339959 | 0.886787317 | 0.159253812 | 0.319161997 | 565.6250794 | 502.1920611 | ENSG00000133773 |
| TMEM59 | -0.173434577 | 0.886729159 | 0.050448434 | 0.134766688 | 1429.67878 | 1267.419549 | ENSG00000116209 |
| TBXA2R | -0.173541156 | 0.886663654 | 0.775609127 | 0.876319163 | 13.59439976 | 12.09262614 | ENSG00000006638 |
| HDAC2 | -0.173711785 | 0.886558794 | 0.036439122 | 0.104524967 | 7095.394407 | 6289.192996 | ENSG00000196591 |
| FAM217A | -0.173760889 | 0.886528619 | 0.876742715 | 0.936141806 | 4.250847342 | 3.724462875 | ENSG00000145975 |
| UBP1 | -0.173803365 | 0.886502518 | 0.024948672 | 0.077298343 | 1819.802544 | 1613.252826 | ENSG00000153560 |
| CCDC66 | -0.17380766 | 0.88649988 | 0.311503461 | 0.50519821 | 272.8600982 | 241.9640664 | ENSG00000180376 |
| JMJD8 | -0.173834732 | 0.886483244 | 0.106817814 | 0.238111163 | 696.6352022 | 617.8466024 | ENSG00000161999 |
| RBX1 | -0.173985502 | 0.886390607 | 0.038013872 | 0.107877676 | 1912.857067 | 1695.655561 | ENSG00000100387 |
| NCAPH2 | -0.173997724 | 0.886383097 | 0.030966349 | 0.091846387 | 1471.006802 | 1304.233513 | ENSG00000025770 |
| FOXQ1 | -0.174057096 | 0.886346621 | 0.886898308 | 0.942266356 | 3.857188144 | 3.398196311 | ENSG00000164379 |
| MTMR14 | -0.174108943 | 0.886314768 | 0.027033233 | 0.082529064 | 1564.671593 | 1387.370435 | ENSG00000163719 |
| AADAT | -0.174166358 | 0.886279496 | 0.31266113 | 0.506189848 | 239.1372236 | 212.4516804 | ENSG00000109576 |
| CIAO2A | -0.174174258 | 0.886274643 | 0.089684568 | 0.209057198 | 987.0776897 | 874.1459458 | ENSG00000166797 |
| EFNA5 | -0.174178561 | 0.886271999 | 0.434651092 | 0.624240475 | 116.7033586 | 103.4801011 | ENSG00000184349 |
| BCLAF3 | -0.174183211 | 0.886269143 | 0.351232763 | 0.544467173 | 165.7485831 | 146.8808619 | ENSG00000173681 |
| HMGCR | -0.174315051 | 0.886188155 | 0.008781205 | 0.033052571 | 4880.681247 | 4323.950978 | ENSG00000113161 |
| SPIN2B | -0.174365565 | 0.886157127 | 0.439800862 | 0.628035224 | 125.5436695 | 111.4410767 | ENSG00000186787 |
| CSRP2 | -0.17437342 | 0.886152302 | 0.010102443 | 0.037076964 | 2381.505689 | 2110.33657 | ENSG00000175183 |
| PSMA7 | -0.174377283 | 0.88614993 | 0.004448259 | 0.018637396 | 6896.151418 | 6110.88902 | ENSG00000101182 |
| RNFT1 | -0.174424827 | 0.886120727 | 0.680288441 | 0.815245229 | 27.52495786 | 24.43643487 | ENSG00000189050 |
| CTR9 | -0.174424924 | 0.886120668 | 0.03308665 | 0.096845857 | 1544.423998 | 1368.069536 | ENSG00000198730 |
| GABARAPL2 | -0.17447529 | 0.886089733 | 0.061955969 | 0.157908383 | 796.909564 | 706.1009365 | ENSG00000034713 |
| RNF150 | -0.174532935 | 0.886054328 | 0.301395343 | 0.494867229 | 194.1592046 | 171.9440109 | ENSG00000170153 |
| RFNG | -0.174560306 | 0.886037518 | 0.153834954 | 0.311365843 | 375.0731637 | 332.2563084 | ENSG00000169733 |
| DXO | -0.174666495 | 0.885972304 | 0.371263599 | 0.564401422 | 154.296861 | 136.9317217 | ENSG00000204348 |
| JADE3 | -0.174689322 | 0.885958286 | 0.106472235 | 0.237553438 | 589.6319091 | 521.8786277 | ENSG00000102221 |
| LMBR1 | -0.174840783 | 0.885865278 | 0.072327892 | 0.177629234 | 951.6718889 | 842.2748332 | ENSG00000105983 |
| GRIK3 | -0.174878358 | 0.885842207 | 0.436587954 | 0.62586147 | 117.15049 | 103.8635567 | ENSG00000163873 |
| PLEKHG6 | -0.174886414 | 0.88583726 | 0.089497715 | 0.208726644 | 606.7470455 | 537.120243 | ENSG00000008323 |
| DTL | -0.174909445 | 0.885823118 | 0.050554363 | 0.134952407 | 1194.515904 | 1057.390943 | ENSG00000143476 |
| AURKAIP1 | -0.174950667 | 0.885797808 | 0.033494882 | 0.097747464 | 2694.103793 | 2386.570536 | ENSG00000175756 |
| NA | -0.174958803 | 0.885792813 | 0.506649317 | 0.686270441 | 79.29066467 | 70.38071802 | ENSG00000250986 |
| PPIEL | -0.17495892 | 0.885792741 | 0.624670474 | 0.777266072 | 48.83995003 | 43.42409514 | ENSG00000243970 |
| SMU1 | -0.174966597 | 0.885788028 | 0.025007622 | 0.077468038 | 1751.489704 | 1550.658966 | ENSG00000122692 |
| NUP35 | -0.175070254 | 0.885724387 | 0.071467269 | 0.175822244 | 783.7078312 | 693.820712 | ENSG00000163002 |
| MYBL1 | -0.175119942 | 0.885693882 | 0.506296628 | 0.685892968 | 74.25112346 | 65.60832021 | ENSG00000185697 |
| SLC9A7 | -0.175197743 | 0.88564612 | 0.132470513 | 0.279159579 | 461.6648072 | 408.697543 | ENSG00000065923 |
| LIN28A | -0.175221888 | 0.885631298 | 0.001651498 | 0.008063453 | 22323.38759 | 19769.2778 | ENSG00000131914 |
| DDX1 | -0.17525102 | 0.885613414 | 0.006396889 | 0.025409332 | 4692.304261 | 4155.200246 | ENSG00000079785 |
| FAM118A | -0.175282395 | 0.885594155 | 0.213534793 | 0.393483184 | 635.0921762 | 563.3330833 | ENSG00000100376 |
| COL1A1 | -0.175436504 | 0.885499561 | 0.083230022 | 0.197463546 | 1655.071042 | 1465.279255 | ENSG00000108821 |
| MAP9 | -0.175450156 | 0.885491181 | 0.432210704 | 0.622133674 | 131.7743581 | 116.4532203 | ENSG00000164114 |
| ANKHD1 | -0.17550757 | 0.885455942 | 0.72428776 | 0.844902699 | 22.97248281 | 20.24671078 | ENSG00000131503 |
| CWC15 | -0.17576172 | 0.885299972 | 0.091663364 | 0.212361099 | 794.7324232 | 703.1885958 | ENSG00000150316 |
| KBTBD11 | -0.175862994 | 0.885237828 | 0.18210074 | 0.352375769 | 342.6971471 | 303.0999899 | ENSG00000176595 |
| NA | -0.176025322 | 0.885138228 | 0.552538917 | 0.723319415 | 59.52881226 | 52.5705259 | ENSG00000280206 |
| ERCC8 | -0.176090565 | 0.885098201 | 0.402879691 | 0.594693548 | 145.2227884 | 128.5352143 | ENSG00000049167 |
| STXBP5L | -0.176135956 | 0.885070354 | 0.378061154 | 0.571392434 | 127.6890231 | 113.030081 | ENSG00000145087 |
| BICRA | -0.176151826 | 0.885060618 | 0.140344165 | 0.291476776 | 573.8253644 | 507.8888404 | ENSG00000063169 |
| SDF2 | -0.176172585 | 0.885047883 | 0.094500554 | 0.217034802 | 614.1605682 | 543.3062077 | ENSG00000132581 |
| PRKACA | -0.176173585 | 0.885047269 | 0.092806741 | 0.214216343 | 1185.16573 | 1048.403154 | ENSG00000072062 |
| MSI2 | -0.176221822 | 0.885017678 | 0.064452439 | 0.162702466 | 970.7768107 | 858.6117358 | ENSG00000153944 |
| EIF3B | -0.176294839 | 0.884972887 | 0.001550493 | 0.00764481 | 11088.46601 | 9812.494708 | ENSG00000106263 |
| NECAP2 | -0.176342642 | 0.884943564 | 0.069770125 | 0.172534048 | 993.3284551 | 878.8471391 | ENSG00000157191 |
| CFAP157 | -0.176345603 | 0.884941748 | 0.65052934 | 0.795936006 | 35.04728877 | 31.12812563 | ENSG00000160401 |
| NBPF9 | -0.176482861 | 0.884857559 | 0.275073872 | 0.466037706 | 203.2241972 | 179.8430025 | ENSG00000269713 |

| | | | | | | | |
|---|---|---|---|---|---|---|---|
| NT5DC2 | -0.176505338 | 0.884843773 | 0.025200465 | 0.077935162 | 6778.437821 | 5997.083705 | ENSG00000168268 |
| RNF144A | -0.176597722 | 0.884787113 | 0.141469142 | 0.293188984 | 461.7376116 | 408.4305037 | ENSG00000151692 |
| NA | -0.176602696 | 0.884784063 | 0.644278307 | 0.791952125 | 34.84557725 | 30.8731442 | ENSG00000254536 |
| SH3BGRL3 | -0.176623529 | 0.884771286 | 0.094438052 | 0.216959508 | 721.0636071 | 638.2648316 | ENSG00000142669 |
| PHF14 | -0.176704483 | 0.88472164 | 0.029295481 | 0.087829577 | 1291.294423 | 1142.471148 | ENSG00000106443 |
| KIF20A | -0.176725723 | 0.884708615 | 0.030251502 | 0.090212416 | 1650.877869 | 1460.225173 | ENSG00000112984 |
| AK1 | -0.176752736 | 0.88469205 | 0.672803266 | 0.809789244 | 34.28630805 | 30.41580709 | ENSG00000106992 |
| NA | -0.176826484 | 0.884646827 | 0.849075891 | 0.919956306 | 5.733354945 | 5.029640753 | ENSG00000285103 |
| SEC23IP | -0.176887921 | 0.884609155 | 0.032558239 | 0.095525472 | 1452.040792 | 1283.683955 | ENSG00000107651 |
| CDC34 | -0.176932674 | 0.884581715 | 0.009082113 | 0.033957614 | 2664.362068 | 2356.852433 | ENSG00000099804 |
| SRRM1 | -0.176967293 | 0.884560489 | 0.015138171 | 0.051490338 | 3078.578672 | 2724.014441 | ENSG00000133226 |
| UGT3A2 | -0.17702483 | 0.884525212 | 0.106512829 | 0.237615404 | 560.0490065 | 495.6649692 | ENSG00000168671 |
| NACA | -0.177039384 | 0.884516289 | 0.004197999 | 0.017755756 | 4701.285198 | 4158.278626 | ENSG00000196531 |
| NUDCD3 | -0.177040695 | 0.884515485 | 0.025715869 | 0.079185565 | 1589.235911 | 1405.216654 | ENSG00000015676 |
| KNOP1 | -0.177208513 | 0.884412602 | 0.027202575 | 0.082964049 | 1223.269674 | 1082.257154 | ENSG00000103550 |
| HSBP1 | -0.17730034 | 0.884356311 | 0.021738804 | 0.069299447 | 3368.329186 | 2978.567859 | ENSG00000230989 |
| NA | -0.177347665 | 0.884327302 | 0.812796672 | 0.899402162 | 9.421776176 | 8.300565364 | ENSG00000243829 |
| MOCS1 | -0.177360597 | 0.884319375 | 0.06418454 | 0.162193143 | 713.2531405 | 630.5115563 | ENSG00000124615 |
| ABCB4 | -0.177525514 | 0.884218293 | 0.761618304 | 0.867730799 | 18.22312173 | 16.1889333 | ENSG00000005471 |
| TUBB | -0.177561592 | 0.884196181 | 0.001203596 | 0.00616573 | 31521.0438 | 27870.18993 | ENSG00000196230 |
| NA | -0.177597953 | 0.884173896 | 0.704269798 | 0.831811477 | 25.64741559 | 22.60968279 | ENSG00000232472 |
| BCR | -0.177616577 | 0.884162483 | 0.008015976 | 0.03068321 | 4163.292599 | 3680.753524 | ENSG00000186716 |
| SPATA33 | -0.177745369 | 0.884083555 | 0.320700609 | 0.514531257 | 199.6016652 | 176.9190908 | ENSG00000167523 |
| WWOX | -0.177798542 | 0.884050972 | 0.286779614 | 0.478170382 | 204.5365625 | 180.9805555 | ENSG00000186153 |
| MARCHF5 | -0.177901355 | 0.883987972 | 0.171243045 | 0.336816152 | 542.6593315 | 479.1335545 | ENSG00000198060 |
| PARD6G | -0.178033117 | 0.883907241 | 0.160674992 | 0.321211337 | 361.8377948 | 319.6564189 | ENSG00000178184 |
| NA | -0.178188056 | 0.883812318 | 0.872283457 | 0.933909167 | 4.218000437 | 3.675635966 | ENSG00000204434 |
| TRAPPC13 | -0.178189037 | 0.883811717 | 0.335003239 | 0.529337273 | 170.003572 | 150.020278 | ENSG00000113597 |
| TRIB1 | -0.178198102 | 0.883806164 | 0.041858507 | 0.116680037 | 1023.896118 | 904.6550985 | ENSG00000173334 |
| SERBP1 | -0.178201908 | 0.883803832 | 0.005056176 | 0.020787748 | 21344.70272 | 18863.4709 | ENSG00000142864 |
| ASB8 | -0.178283858 | 0.883753631 | 0.480546334 | 0.664313101 | 80.62206906 | 71.08445119 | ENSG00000177981 |
| LEO1 | -0.178439576 | 0.883658247 | 0.050022368 | 0.13386003 | 964.2635335 | 852.4910848 | ENSG00000166477 |
| SIRT6 | -0.178448187 | 0.883652973 | 0.102845523 | 0.231299469 | 619.9211401 | 547.398899 | ENSG00000077463 |
| COPS7B | -0.178507966 | 0.883616359 | 0.031195765 | 0.092376148 | 1535.991838 | 1357.499536 | ENSG00000144524 |
| ZNF607 | -0.17852098 | 0.883608388 | 0.23056327 | 0.414678278 | 278.1997189 | 245.7921234 | ENSG00000198182 |
| DYNLL2 | -0.178759291 | 0.883462442 | 0.0168113 | 0.055979273 | 2980.628825 | 2633.264389 | ENSG00000264364 |
| ATP2A1-AS1 | -0.178818444 | 0.883426219 | 0.546746594 | 0.718495899 | 81.78104596 | 72.54589765 | ENSG00000260442 |
| NCS1 | -0.178824074 | 0.883422772 | 0.037439078 | 0.106687558 | 1353.532962 | 1195.159444 | ENSG00000107130 |
| CSTF1 | -0.178824226 | 0.883422679 | 0.044651256 | 0.122475261 | 962.0986824 | 849.7814892 | ENSG00000101138 |
| FANCA | -0.178837839 | 0.883414343 | 0.125225033 | 0.268128554 | 1202.113918 | 1062.790154 | ENSG00000187741 |
| SNAP29 | -0.178846399 | 0.883409101 | 0.294984512 | 0.487841791 | 181.3965905 | 160.2338527 | ENSG00000099940 |
| EIF4E3 | -0.179018656 | 0.883303629 | 0.428849685 | 0.61917115 | 220.3309528 | 194.1580284 | ENSG00000163412 |
| SRSF10 | -0.179091936 | 0.883258764 | 0.014461728 | 0.049553276 | 3553.156715 | 3137.621829 | ENSG00000188529 |
| NA | -0.179210577 | 0.883186131 | 0.64773301 | 0.794354768 | 31.54134448 | 27.84211726 | ENSG00000226360 |
| ARHGAP28 | -0.179260721 | 0.883155435 | 0.365328889 | 0.558433455 | 217.437637 | 191.8538944 | ENSG00000088756 |
| ALKBH1 | -0.179351664 | 0.883099765 | 0.186649248 | 0.358000328 | 337.8603998 | 298.5715436 | ENSG00000100601 |
| ERP29 | -0.179352195 | 0.88309944 | 0.003168089 | 0.013949657 | 3874.161536 | 3421.383986 | ENSG00000089248 |
| PAXX | -0.179380468 | 0.883082134 | 0.176380221 | 0.344362269 | 533.4015793 | 471.285079 | ENSG00000148362 |
| SLITRK5 | -0.179427685 | 0.883053233 | 0.371796429 | 0.56501107 | 134.956674 | 119.339407 | ENSG00000165300 |
| ALAS1 | -0.179441976 | 0.883044485 | 0.02741345 | 0.083526452 | 1108.208893 | 978.4809131 | ENSG00000023330 |
| NA | -0.179449734 | 0.883039737 | 0.709801495 | 0.835055282 | 24.48350033 | 21.74014825 | ENSG00000282022 |
| MVD | -0.179461551 | 0.883032504 | 0.027115734 | 0.082740043 | 2444.795042 | 2159.296718 | ENSG00000167508 |
| WNT5A | -0.179569077 | 0.882966693 | 0.538321009 | 0.711948165 | 60.48499527 | 53.39970729 | ENSG00000114251 |
| PPM1M | -0.17960684 | 0.882943581 | 0.437623139 | 0.626324885 | 114.5136355 | 100.9628359 | ENSG00000164088 |
| RTL8A | -0.179646653 | 0.882919216 | 0.098844586 | 0.224755926 | 554.7208334 | 489.9782321 | ENSG00000203950 |
| NA | -0.179717317 | 0.882875971 | 0.813763123 | 0.89988777 | 8.240798581 | 7.314167977 | ENSG00000266947 |
| SASH1 | -0.179816162 | 0.882815484 | 0.162222161 | 0.323560058 | 546.7037169 | 482.5553317 | ENSG00000111961 |
| CMTM8 | -0.179833129 | 0.882805101 | 0.403350729 | 0.595101349 | 138.6661113 | 122.364883 | ENSG00000170293 |
| AGPAT2 | -0.179842021 | 0.88279966 | 0.091262065 | 0.211775304 | 956.0100019 | 844.5270917 | ENSG00000169692 |
| PKD2 | -0.179891959 | 0.882769103 | 0.057799753 | 0.149632319 | 811.886886 | 716.9810775 | ENSG00000118762 |
| NDUFS6 | -0.179912864 | 0.882756311 | 0.015719124 | 0.053038093 | 3059.074085 | 2701.210599 | ENSG00000145494 |
| SLC39A7 | -0.180020996 | 0.88269015 | 0.00480658 | 0.019876205 | 3481.137204 | 3072.632514 | ENSG00000112473 |
| DPYS | -0.180089106 | 0.882648479 | 0.819594597 | 0.902946927 | 8.287550914 | 7.351709965 | ENSG00000147647 |
| CTTNBP2 | -0.1800952 | 0.882644751 | 0.167475268 | 0.331233232 | 433.5273412 | 382.8388929 | ENSG00000077063 |
| PARK7 | -0.180213585 | 0.882572325 | 0.002876663 | 0.012850343 | 6038.074952 | 5328.829264 | ENSG00000116288 |
| SAAL1 | -0.180228701 | 0.882563078 | 0.040226701 | 0.112912566 | 906.8579014 | 800.1552949 | ENSG00000166788 |
| PLS1 | -0.180269485 | 0.882538129 | 0.139690226 | 0.290574485 | 532.6044971 | 469.5504842 | ENSG00000120756 |
| HCN4 | -0.180282365 | 0.88253025 | 0.095636792 | 0.219101644 | 615.0813336 | 542.7945959 | ENSG00000138622 |
| SLC27A1 | -0.180295054 | 0.882522488 | 0.250868882 | 0.439368278 | 231.3568604 | 204.1525005 | ENSG00000130304 |
| SAC3D1 | -0.180297565 | 0.882520952 | 0.168962291 | 0.333320606 | 467.9117462 | 412.7961201 | ENSG00000168061 |
| NA | -0.180311151 | 0.882512641 | 0.689175845 | 0.82163805 | 26.29840801 | 23.31159248 | ENSG00000180581 |
| NA | -0.18031503 | 0.882510268 | 0.652538707 | 0.797031264 | 36.18416728 | 32.03184135 | ENSG00000233225 |
| EPHX3 | -0.180362009 | 0.882481532 | 0.120409397 | 0.260283155 | 462.1306921 | 407.8428774 | ENSG00000105131 |
| METTL6 | -0.180561051 | 0.882359788 | 0.263955199 | 0.453648522 | 252.607925 | 223.3613906 | ENSG00000206562 |

| | | | | | | | |
|---|---|---|---|---|---|---|---|
| DENND1A | -0.180613122 | 0.882327942 | 0.051806416 | 0.137266634 | 931.2007766 | 821.1061854 | ENSG00000119522 |
| NA | -0.180699854 | 0.882274899 | 0.767643806 | 0.871676896 | 14.53648194 | 12.7940978 | ENSG00000228501 |
| SLC12A7 | -0.180708339 | 0.88226971 | 0.047979763 | 0.129534876 | 1378.379718 | 1217.066288 | ENSG00000113504 |
| TXLNG | -0.180715768 | 0.882265167 | 0.010492638 | 0.038266538 | 2596.070635 | 2289.80725 | ENSG00000086712 |
| SLC41A1 | -0.180727517 | 0.882257982 | 0.057462971 | 0.1489745 | 904.0212372 | 797.3070798 | ENSG00000133065 |
| KDM4C | -0.180785365 | 0.882222607 | 0.083132173 | 0.197307169 | 689.4326815 | 608.7684139 | ENSG00000107077 |
| TSPYL2 | -0.180799243 | 0.88221412 | 0.080215817 | 0.1921075 | 613.1312952 | 540.8804244 | ENSG00000184205 |
| YRDC | -0.180845414 | 0.882185887 | 0.216542008 | 0.397207995 | 340.4040426 | 300.1023712 | ENSG00000196449 |
| SIGIRR | -0.18088864 | 0.882159455 | 0.058472438 | 0.15093707 | 816.332026 | 720.7144522 | ENSG00000185187 |
| ATP5MC3 | -0.180946135 | 0.8821243 | 0.002282895 | 0.010622941 | 4897.019129 | 4319.759028 | ENSG00000154518 |
| B2M | -0.181021055 | 0.882078492 | 0.063926613 | 0.161661324 | 791.6600584 | 697.7241878 | ENSG00000166710 |
| FBXO44 | -0.181037447 | 0.88206847 | 0.218011233 | 0.399034189 | 667.7477372 | 589.6758228 | ENSG00000132879 |
| MFSD14A | -0.181091744 | 0.882035273 | 0.611608605 | 0.768407937 | 41.81129221 | 36.97555124 | ENSG00000156875 |
| ST8SIA6 | -0.181159908 | 0.8819936 | 0.610168482 | 0.767371074 | 41.24758832 | 36.38269191 | ENSG00000148488 |
| TOX | -0.181215555 | 0.881959581 | 0.249828044 | 0.438338668 | 237.0673428 | 208.7646077 | ENSG00000198846 |
| TMEM74 | -0.181362643 | 0.881869666 | 0.360014538 | 0.553230585 | 154.0169257 | 135.4883849 | ENSG00000164841 |
| PLPP7 | -0.181378815 | 0.881859781 | 0.512028212 | 0.69104346 | 72.77008902 | 64.379214 | ENSG00000160539 |
| SPECC1L | -0.181378986 | 0.881859676 | 0.070145176 | 0.173204196 | 708.4281565 | 625.1655484 | ENSG00000100014 |
| DGCR2 | -0.181428919 | 0.881829155 | 0.009208268 | 0.034373825 | 2912.127488 | 2567.29903 | ENSG00000070413 |
| RPF1 | -0.181448916 | 0.881816932 | 0.041994202 | 0.116952747 | 927.1563233 | 817.5794435 | ENSG00000117133 |
| TSPAN7 | -0.18147468 | 0.881801184 | 0.633505626 | 0.783961787 | 46.2877309 | 40.83086142 | ENSG00000156298 |
| PGGT1B | -0.181481287 | 0.881797146 | 0.246641131 | 0.434240214 | 301.7559294 | 265.5644464 | ENSG00000164219 |
| NA | -0.18159461 | 0.881727884 | 0.847848857 | 0.918948068 | 5.828738747 | 5.097238657 | ENSG00000272311 |
| C11orf58 | -0.181601085 | 0.881723927 | 0.012900345 | 0.045308885 | 5009.728612 | 4416.744878 | ENSG00000110696 |
| MAP3K6 | -0.181640209 | 0.881700016 | 0.21139066 | 0.390897706 | 258.8863506 | 228.3939542 | ENSG00000142733 |
| TCERG1 | -0.181646636 | 0.881696088 | 0.007857837 | 0.03017484 | 2567.918324 | 2263.96131 | ENSG00000113649 |
| TAOK3 | -0.181768785 | 0.881621441 | 0.230638503 | 0.414773338 | 272.132874 | 239.4601159 | ENSG00000135090 |
| TBC1D32 | -0.181862552 | 0.881564142 | 0.513128751 | 0.691911842 | 75.00243803 | 65.96453107 | ENSG00000146350 |
| SETMAR | -0.181964048 | 0.881502125 | 0.166648554 | 0.330126581 | 389.186507 | 342.7020799 | ENSG00000170364 |
| PLD3 | -0.181980831 | 0.88149187 | 0.005660756 | 0.022897525 | 4631.133292 | 4082.463318 | ENSG00000105223 |
| CELSR2 | -0.182016438 | 0.881470114 | 0.040056265 | 0.112587627 | 1511.627842 | 1332.167221 | ENSG00000143126 |
| GPT | -0.182112934 | 0.881411158 | 0.829422532 | 0.908165488 | 6.695754081 | 5.922621201 | ENSG00000167701 |
| NDUFB11 | -0.182246319 | 0.881329671 | 0.015995129 | 0.053827052 | 1560.600555 | 1375.637563 | ENSG00000147123 |
| ACTL10 | -0.182266621 | 0.881317269 | 0.779514645 | 0.878856647 | 12.75263972 | 11.18467355 | ENSG00000182584 |
| WDR74 | -0.182294766 | 0.881300076 | 0.027624077 | 0.084083535 | 1549.853485 | 1366.333742 | ENSG00000133316 |
| SCPEP1 | -0.182322802 | 0.881282949 | 0.058717774 | 0.151380613 | 840.3905011 | 740.2190746 | ENSG00000121064 |
| SLC35D3 | -0.182340957 | 0.88127186 | 0.779003249 | 0.878653821 | 12.83351674 | 11.25975752 | ENSG00000182747 |
| VCP | -0.182387385 | 0.881243499 | 0.000814423 | 0.004408443 | 14429.75118 | 12715.71336 | ENSG00000165280 |
| NA | -0.182467745 | 0.881194414 | 0.366511983 | 0.559549678 | 142.7056787 | 125.9115354 | ENSG00000254615 |
| MAP4K4 | -0.182478192 | 0.881188033 | 0.002337596 | 0.010833958 | 4557.004401 | 4015.560793 | ENSG00000071054 |
| DVL3 | -0.18247893 | 0.881187583 | 0.011236612 | 0.040517385 | 2262.128096 | 1993.949745 | ENSG00000161202 |
| AUNIP | -0.18283079 | 0.880972695 | 0.189710333 | 0.361923964 | 323.3029018 | 284.9718304 | ENSG00000127423 |
| ZNF32 | -0.182831001 | 0.880972567 | 0.351359693 | 0.544467173 | 150.10489 | 132.3173127 | ENSG00000169740 |
| MAK | -0.182962832 | 0.880892068 | 0.827128673 | 0.907100759 | 8.101819358 | 7.156402328 | ENSG00000111837 |
| GLS | -0.182984669 | 0.880878735 | 0.093326292 | 0.215004287 | 879.98698 | 774.8044889 | ENSG00000115419 |
| DAZAP1 | -0.182997434 | 0.880870941 | 0.001345547 | 0.006754706 | 8591.876535 | 7568.043792 | ENSG00000071626 |
| UBXN6 | -0.183011701 | 0.88086223 | 0.043290299 | 0.119752598 | 919.5546111 | 810.3900541 | ENSG00000167671 |
| TNFRSF13C | -0.183059713 | 0.880832916 | 0.710765283 | 0.835923837 | 22.12903904 | 19.48903776 | ENSG00000159958 |
| HMGB1 | -0.183078263 | 0.88082159 | 0.003953035 | 0.016877701 | 14959.31736 | 13175.97907 | ENSG00000189403 |
| PRKCSH | -0.183260871 | 0.880710108 | 0.001801208 | 0.008686856 | 7171.668342 | 6316.554612 | ENSG00000130175 |
| ZC2HC1A | -0.183309396 | 0.880680486 | 0.510163139 | 0.68941769 | 90.39360075 | 79.58379662 | ENSG00000104427 |
| GIT1 | -0.18337453 | 0.880640726 | 0.017573804 | 0.05794572 | 2371.595862 | 2088.959196 | ENSG00000108262 |
| CCDC173 | -0.18343319 | 0.88060492 | 0.749450609 | 0.860354212 | 16.34705263 | 14.43682716 | ENSG00000154479 |
| MAPK9 | -0.183458697 | 0.880589351 | 0.039587489 | 0.111557628 | 987.7221431 | 869.8873819 | ENSG00000050748 |
| EIF5AL1 | -0.183567171 | 0.880523143 | 0.641049446 | 0.789659074 | 36.03867884 | 31.74311678 | ENSG00000253626 |
| NA | -0.183596791 | 0.880505065 | 0.70764403 | 0.833734311 | 23.71442435 | 20.92605063 | ENSG00000220848 |
| TRIM38 | -0.183642696 | 0.880477049 | 0.59533995 | 0.755953244 | 44.70983949 | 39.33808522 | ENSG00000112343 |
| LINC01534 | -0.183718135 | 0.88043101 | 0.760724786 | 0.867113972 | 14.40697346 | 12.64001938 | ENSG00000225975 |
| LMNB1 | -0.183746937 | 0.880413433 | 0.003884977 | 0.016632977 | 4590.247455 | 4041.289457 | ENSG00000113368 |
| YARS2 | -0.183791346 | 0.880386333 | 0.057495353 | 0.149037605 | 721.3682865 | 634.8390641 | ENSG00000139131 |
| ACAA2 | -0.183792734 | 0.880385486 | 0.018861579 | 0.0614485 | 1737.42848 | 1528.870652 | ENSG00000167315 |
| ELF3 | -0.183843415 | 0.880354559 | 0.547909207 | 0.719443801 | 56.09507101 | 49.39411753 | ENSG00000163435 |
| GKAP1 | -0.183930527 | 0.880301403 | 0.448253183 | 0.635989014 | 90.74652835 | 79.79820167 | ENSG00000165113 |
| FGD5-AS1 | -0.184035532 | 0.880237334 | 0.025894087 | 0.079615298 | 1511.238805 | 1329.839649 | ENSG00000225733 |
| HADH | -0.184072396 | 0.880214843 | 0.039101777 | 0.110546156 | 1205.667502 | 1060.78691 | ENSG00000138796 |
| SGMS1 | -0.184137638 | 0.880175038 | 0.18063541 | 0.350198399 | 376.0118033 | 330.8330568 | ENSG00000198964 |
| TPRA1 | -0.184237622 | 0.880114041 | 0.103435347 | 0.232456738 | 490.7111478 | 431.963781 | ENSG00000163870 |
| HLA-L | -0.184238099 | 0.88011375 | 0.564419994 | 0.73261154 | 54.36222042 | 47.91327908 | ENSG00000243753 |
| HSPH1 | -0.184239352 | 0.880112985 | 0.007400741 | 0.028677677 | 4598.78232 | 4046.607152 | ENSG00000120694 |
| LAMB1 | -0.184250468 | 0.880106204 | 0.006411102 | 0.025454877 | 2364.041308 | 2080.730838 | ENSG00000091136 |
| NA | -0.184332215 | 0.880056336 | 0.739590322 | 0.854586473 | 16.52834949 | 14.59459281 | ENSG00000259343 |
| NPR2 | -0.184356416 | 0.880041573 | 0.23732337 | 0.422653151 | 264.9380275 | 233.0021508 | ENSG00000159899 |
| NA | -0.184393805 | 0.880018767 | 0.879658149 | 0.937957895 | 3.900783564 | 3.405682385 | ENSG00000257043 |
| STARD3 | -0.184560713 | 0.879916961 | 0.034322149 | 0.099706382 | 1398.027087 | 1230.960141 | ENSG00000131748 |

| | | | | | | | |
|---|---|---|---|---|---|---|---|
| LCMT2 | -0.184603581 | 0.879890816 | 0.20051358 | 0.377174832 | 295.2976953 | 259.6673581 | ENSG00000168806 |
| DYNC1LI1 | -0.184645553 | 0.879865218 | 0.040036082 | 0.112553432 | 1441.893791 | 1268.227657 | ENSG00000144635 |
| NA | -0.184816591 | 0.879760912 | 0.654628116 | 0.798215625 | 29.76697301 | 26.25949972 | ENSG00000234964 |
| RPL36AL | -0.184852433 | 0.879739056 | 0.007215148 | 0.028058234 | 3539.670778 | 3114.495099 | ENSG00000165502 |
| EIF3J | -0.18487054 | 0.879728014 | 0.026042522 | 0.079992061 | 1612.566026 | 1418.617947 | ENSG00000104131 |
| CANT1 | -0.184918616 | 0.879698699 | 0.044513657 | 0.122280008 | 1038.234361 | 912.841071 | ENSG00000171302 |
| TPRKB | -0.184945317 | 0.879682418 | 0.119510944 | 0.258824005 | 415.9731461 | 365.7693146 | ENSG00000144034 |
| INAFM2 | -0.184946027 | 0.879681985 | 0.069691375 | 0.17245126 | 790.7880895 | 696.0830445 | ENSG00000259330 |
| FBLL1 | -0.184995717 | 0.879651688 | 0.31571459 | 0.509182144 | 177.9483425 | 156.7279286 | ENSG00000188573 |
| AP2A1 | -0.185069372 | 0.879606779 | 0.004276143 | 0.018024571 | 3448.048162 | 3033.024484 | ENSG00000196961 |
| NA | -0.185097782 | 0.879589458 | 0.736360859 | 0.852322051 | 21.44006596 | 18.84062989 | ENSG00000250770 |
| KATNBL1 | -0.185127358 | 0.879571426 | 0.151365931 | 0.308016706 | 343.8630392 | 302.6259687 | ENSG00000134152 |
| TACR3 | -0.18520397 | 0.879524719 | 0.568555512 | 0.735712341 | 56.86029102 | 49.83962007 | ENSG00000169836 |
| CRYBG1 | -0.185235192 | 0.879505685 | 0.098158237 | 0.223442001 | 588.6068753 | 517.5786826 | ENSG00000112297 |
| STX2 | -0.185240061 | 0.879502717 | 0.117260167 | 0.255316058 | 467.4583389 | 411.487072 | ENSG00000111450 |
| MAG | -0.185281158 | 0.879477663 | 0.854436702 | 0.922743843 | 5.108391937 | 4.497219664 | ENSG00000105695 |
| ADCY3 | -0.185320566 | 0.87945364 | 0.050947675 | 0.135787204 | 776.6769336 | 683.1577158 | ENSG00000138031 |
| CDCA5 | -0.185403581 | 0.879403037 | 0.005307666 | 0.021701362 | 2564.309307 | 2254.809615 | ENSG00000146670 |
| ATPSCKMT | -0.185451696 | 0.879373708 | 0.300032231 | 0.493415916 | 173.3499118 | 152.5975434 | ENSG00000150756 |
| NA | -0.18549241 | 0.879348892 | 0.665847318 | 0.805855189 | 28.07653776 | 24.6349917 | ENSG00000269825 |
| PHETA1 | -0.185508174 | 0.879339283 | 0.139612817 | 0.290514 | 466.2958968 | 409.6549435 | ENSG00000198324 |
| PDCD6 | -0.185560211 | 0.879307567 | 0.493758189 | 0.675874798 | 84.11716713 | 74.18070291 | ENSG00000249915 |
| KCND2 | -0.185562514 | 0.879306163 | 0.228425774 | 0.412073514 | 272.815631 | 240.0096993 | ENSG00000184408 |
| PHB | -0.185566108 | 0.879303973 | 0.008903265 | 0.033383192 | 5682.975243 | 4996.948482 | ENSG00000167085 |
| DAGLB | -0.185606835 | 0.879279151 | 0.105286511 | 0.235759356 | 503.8846788 | 443.3555883 | ENSG00000164535 |
| VWA2 | -0.185608977 | 0.879277845 | 0.449522534 | 0.637107184 | 90.62037217 | 79.62111537 | ENSG00000165816 |
| PMVK | -0.185660349 | 0.879246536 | 0.132273061 | 0.278871802 | 521.4887228 | 458.5112408 | ENSG00000163344 |
| MDH1 | -0.185701709 | 0.87922133 | 0.002832425 | 0.01267717 | 4055.405626 | 3564.684699 | ENSG00000014641 |
| ZNF630 | -0.185704759 | 0.879219471 | 0.704325655 | 0.831811477 | 28.86573529 | 25.41198537 | ENSG00000221994 |
| GSTM4 | -0.185726274 | 0.879206359 | 0.19607508 | 0.37097341 | 287.9014077 | 253.238676 | ENSG00000168765 |
| BSDC1 | -0.185745269 | 0.879194783 | 0.051969056 | 0.137520628 | 1011.412259 | 890.0613592 | ENSG00000160058 |
| UMAD1 | -0.185749139 | 0.879192425 | 0.552681082 | 0.723352248 | 73.00445205 | 63.98728462 | ENSG00000219545 |
| IRF2 | -0.185812943 | 0.879153543 | 0.242780626 | 0.429237444 | 219.6095388 | 193.0048193 | ENSG00000168310 |
| RAB38 | -0.185899931 | 0.879100536 | 0.342699874 | 0.536797013 | 162.9351747 | 143.5219598 | ENSG00000123892 |
| MAP3K12 | -0.185925195 | 0.879085141 | 0.185218883 | 0.355956737 | 309.4469296 | 272.346998 | ENSG00000139625 |
| CCDC24 | -0.185939664 | 0.879076325 | 0.396608159 | 0.588906876 | 151.5828652 | 133.5455832 | ENSG00000159214 |
| KLHL20 | -0.185978334 | 0.879052762 | 0.216789039 | 0.397400021 | 242.5392003 | 213.0675074 | ENSG00000076321 |
| NA | -0.185992648 | 0.879044041 | 0.884900767 | 0.940949559 | 4.370982785 | 3.87474245 | ENSG00000287742 |
| PNO1 | -0.186095597 | 0.878981315 | 0.067657666 | 0.168611964 | 646.9777779 | 568.5137589 | ENSG00000115946 |
| CREB3L4 | -0.18615689 | 0.878943972 | 0.233746037 | 0.418655686 | 283.52926 | 249.2759644 | ENSG00000143578 |
| EML6 | -0.18618542 | 0.878926591 | 0.677453464 | 0.81350335 | 29.99442084 | 26.4209526 | ENSG00000214595 |
| TRAPPC4 | -0.186309204 | 0.878851182 | 0.068982142 | 0.17119912 | 649.4674828 | 570.6590682 | ENSG00000196655 |
| POC1A | -0.186310172 | 0.878850592 | 0.035000134 | 0.101200075 | 911.3775447 | 800.6398768 | ENSG00000164087 |
| CLSTN3 | -0.186322729 | 0.878842943 | 0.050836777 | 0.135569615 | 917.8872664 | 807.3837956 | ENSG00000139182 |
| GNB2 | -0.186372243 | 0.878812781 | 0.018261841 | 0.059767784 | 2937.118056 | 2580.773424 | ENSG00000172354 |
| ZFYVE1 | -0.186427667 | 0.87877902 | 0.142298887 | 0.294483259 | 373.2793471 | 328.4134856 | ENSG00000165861 |
| NA | -0.186611556 | 0.878667016 | 0.868246125 | 0.931362061 | 3.850874318 | 3.345570555 | ENSG00000270953 |
| ILVBL | -0.186613785 | 0.878665659 | 0.032028887 | 0.094375738 | 1345.669407 | 1182.619595 | ENSG00000105135 |
| EPHX2 | -0.18662516 | 0.878658731 | 0.28497873 | 0.476180652 | 245.3283607 | 215.8611617 | ENSG00000120915 |
| EFR3B | -0.186656289 | 0.878639773 | 0.12637369 | 0.269746629 | 432.4641389 | 380.2911252 | ENSG00000084710 |
| C8orf33 | -0.186667689 | 0.878632829 | 0.025765354 | 0.079298417 | 1160.943125 | 1020.586164 | ENSG00000182307 |
| PEX12 | -0.186673845 | 0.87862908 | 0.507931034 | 0.687132274 | 66.75867794 | 58.71482342 | ENSG00000108733 |
| POLR2G | -0.186728579 | 0.878595747 | 0.021274804 | 0.068019185 | 1238.761174 | 1088.20373 | ENSG00000168002 |
| UBE2D1 | -0.186797488 | 0.878553783 | 0.157483089 | 0.316657522 | 656.5379394 | 576.4269179 | ENSG00000072401 |
| ZMIZ2 | -0.186824735 | 0.878537191 | 0.056361543 | 0.146668128 | 1618.795888 | 1422.927483 | ENSG00000122515 |
| TMCC2 | -0.186893038 | 0.878495598 | 0.201753097 | 0.378831343 | 287.1639235 | 252.1156571 | ENSG00000133069 |
| NUTM2G | -0.18690512 | 0.878488241 | 0.65667218 | 0.799445755 | 31.46046747 | 27.70356064 | ENSG00000188152 |
| AJUBA | -0.186927652 | 0.878474521 | 0.087868615 | 0.205703791 | 816.2832364 | 717.5359484 | ENSG00000129474 |
| LHFPL2 | -0.187046344 | 0.878402251 | 0.01774926 | 0.058441072 | 1457.095321 | 1279.188488 | ENSG00000145685 |
| CPT1C | -0.187158573 | 0.878333922 | 0.030840499 | 0.091558194 | 1204.998906 | 1058.918691 | ENSG00000169169 |
| CLTB | -0.187318383 | 0.878236632 | 0.069711376 | 0.172453998 | 712.7111064 | 626.0537424 | ENSG00000175416 |
| FASTK | -0.187376385 | 0.878201324 | 0.046571198 | 0.126524565 | 821.8200741 | 722.3048822 | ENSG00000164896 |
| NA | -0.187529498 | 0.878108126 | 0.359707935 | 0.552942429 | 123.4304864 | 108.2698442 | ENSG00000226259 |
| SHQ1 | -0.187593099 | 0.878069415 | 0.079215166 | 0.190227247 | 668.4146643 | 586.4447872 | ENSG00000144736 |
| BTN2A1 | -0.187664925 | 0.878025701 | 0.18785075 | 0.3596177 | 305.3476891 | 268.3595179 | ENSG00000112763 |
| MFSD2A | -0.187674869 | 0.878019649 | 0.091782331 | 0.212492343 | 485.6215996 | 426.638694 | ENSG00000168389 |
| EHMT1 | -0.187721636 | 0.877991187 | 0.002176226 | 0.010226717 | 3279.99895 | 2880.106171 | ENSG00000181090 |
| TPM2 | -0.187758407 | 0.877968809 | 0.008594039 | 0.032473378 | 2786.861739 | 2447.573798 | ENSG00000198467 |
| HLA-C | -0.187783804 | 0.877953354 | 0.003558722 | 0.015392613 | 5671.967225 | 4979.966947 | ENSG00000204525 |
| OFD1 | -0.187862181 | 0.877905659 | 0.210541874 | 0.389867428 | 327.8341881 | 288.0978116 | ENSG00000046651 |
| SLC6A11 | -0.187920195 | 0.877870357 | 0.398428801 | 0.590322714 | 111.9385213 | 98.25927796 | ENSG00000132164 |
| BTBD3 | -0.188292033 | 0.877644126 | 0.047165433 | 0.127839155 | 898.6455901 | 788.1657301 | ENSG00000132640 |
| BHLHA15 | -0.188365024 | 0.877599723 | 0.640386684 | 0.789444284 | 34.47529422 | 30.22374865 | ENSG00000180535 |
| FNTB | -0.188390128 | 0.877584453 | 0.562267186 | 0.730738379 | 63.33915038 | 55.67036167 | ENSG00000257365 |

| | | | | | | | |
|---|---|---|---|---|---|---|---|
| FAM168A | -0.18844184 | 0.877552997 | 0.053309661 | 0.140414153 | 755.8343054 | 663.3763694 | ENSG00000054965 |
| NCK1 | -0.188483827 | 0.877527457 | 0.221036231 | 0.402662227 | 258.5138051 | 227.1107965 | ENSG00000158092 |
| PIPSL | -0.188590363 | 0.877462659 | 0.782717938 | 0.880989463 | 11.41364373 | 10.04843462 | ENSG00000180764 |
| ATG4C | -0.188635409 | 0.877435262 | 0.147890565 | 0.302806422 | 411.845516 | 360.887102 | ENSG00000125703 |
| TCF19 | -0.188644985 | 0.877429438 | 0.164050341 | 0.326199874 | 390.0655561 | 342.7835909 | ENSG00000137310 |
| AP1S3 | -0.188780984 | 0.877346729 | 0.458283852 | 0.645327018 | 88.91721158 | 77.82430759 | ENSG00000152056 |
| PIK3R4 | -0.188837588 | 0.877312307 | 0.062762843 | 0.159558122 | 968.3806936 | 848.7529463 | ENSG00000196455 |
| FAM200A | -0.189017533 | 0.877202888 | 0.328773613 | 0.523226012 | 155.4458861 | 136.299345 | ENSG00000221909 |
| GBA | -0.189069737 | 0.877171147 | 0.329535602 | 0.523943798 | 180.5627152 | 158.2177417 | ENSG00000177628 |
| BLOC1S1 | -0.189118859 | 0.877141281 | 0.741320527 | 0.855338624 | 16.63951786 | 14.62833595 | ENSG00000135441 |
| NA | -0.189122742 | 0.87713892 | 0.830531936 | 0.908658972 | 6.63006027 | 5.821168536 | ENSG00000263503 |
| HEATR5B | -0.189187977 | 0.877099259 | 0.097805853 | 0.222911248 | 455.9087751 | 400.0158238 | ENSG00000008869 |
| ARPC3 | -0.18930264 | 0.877029551 | 0.059573101 | 0.153011065 | 818.2554715 | 717.9024564 | ENSG00000111229 |
| NA | -0.189315838 | 0.877021528 | 0.842655348 | 0.915891052 | 6.011411088 | 5.330088282 | ENSG00000279799 |
| ARHGEF9 | -0.189334021 | 0.877010475 | 0.037773947 | 0.107377658 | 815.5688731 | 715.3527706 | ENSG00000131089 |
| PDZD11 | -0.18936168 | 0.876993661 | 0.126913844 | 0.270525727 | 378.0518889 | 331.4575093 | ENSG00000120509 |
| RIMKLB | -0.189393224 | 0.876974486 | 0.031035398 | 0.092004169 | 1133.077492 | 994.10025 | ENSG00000166532 |
| YME1L1 | -0.189429731 | 0.876952295 | 0.005944469 | 0.023847356 | 3362.827286 | 2948.408346 | ENSG00000136758 |
| RTCA | -0.189441766 | 0.876944979 | 0.061401474 | 0.156893355 | 739.0941723 | 647.6543145 | ENSG00000137996 |
| TRMT1L | -0.189478783 | 0.876922478 | 0.080880117 | 0.193228076 | 582.1876891 | 510.3082371 | ENSG00000121486 |
| GRTP1 | -0.189524115 | 0.876894925 | 0.030206947 | 0.090124848 | 1085.871348 | 952.7851332 | ENSG00000139835 |
| SHFL | -0.189682974 | 0.876798373 | 0.361972389 | 0.554950473 | 143.1863181 | 125.7983598 | ENSG00000130813 |
| SART3 | -0.189703626 | 0.876785822 | 0.008056428 | 0.030818956 | 2603.631127 | 2282.439285 | ENSG00000075856 |
| NDUFA2 | -0.189745959 | 0.876760094 | 0.291588048 | 0.483908747 | 236.0960517 | 207.2490787 | ENSG00000131495 |
| TOX2 | -0.189912588 | 0.876658836 | 0.487949587 | 0.670944257 | 73.3330188 | 64.19789082 | ENSG00000124191 |
| PLPP6 | -0.189921317 | 0.876653532 | 0.263774452 | 0.45342197 | 193.4732756 | 169.8176874 | ENSG00000205808 |
| DLL4 | -0.189943939 | 0.876639786 | 0.802833955 | 0.894151534 | 8.264174747 | 7.257854994 | ENSG00000128917 |
| RHOG | -0.189956533 | 0.876632133 | 0.134624224 | 0.282445706 | 430.9670117 | 377.9674155 | ENSG00000177105 |
| SHPRH | -0.190444949 | 0.876335404 | 0.131935908 | 0.278444559 | 529.8340003 | 464.1462199 | ENSG00000146414 |
| ENTPD7 | -0.190473293 | 0.876318188 | 0.213248995 | 0.393151981 | 306.3186645 | 268.0452605 | ENSG00000198018 |
| SAMD14 | -0.190600895 | 0.876240684 | 0.371229878 | 0.564401422 | 116.9642547 | 102.633248 | ENSG00000167100 |
| NCKIPSD | -0.190619696 | 0.876229265 | 0.035286078 | 0.101867939 | 929.9952427 | 814.6786026 | ENSG00000213672 |
| DHX33 | -0.190666407 | 0.876200895 | 0.004783198 | 0.019797186 | 2196.539076 | 1924.339825 | ENSG00000005100 |
| MAFA-AS1 | -0.190702002 | 0.876179277 | 0.788728438 | 0.884435871 | 9.762466914 | 8.570518945 | ENSG00000254338 |
| LOC148709 | -0.190706902 | 0.876176301 | 0.56821277 | 0.735371516 | 49.39280771 | 43.13383323 | ENSG00000234996 |
| MIS18A | -0.190758957 | 0.876144687 | 0.024853744 | 0.077081543 | 1775.61079 | 1556.131181 | ENSG00000159055 |
| ZNF418 | -0.190795292 | 0.876122622 | 0.329457824 | 0.523943798 | 210.7058942 | 185.0151506 | ENSG00000196724 |
| MRPS21 | -0.190821883 | 0.876106473 | 0.113192284 | 0.248743868 | 1078.197215 | 945.5169499 | ENSG00000266472 |
| SELENOK | -0.190926145 | 0.876043161 | 0.082887193 | 0.196851774 | 599.6581359 | 524.7560143 | ENSG00000113811 |
| NA | -0.19094919 | 0.876029167 | 0.833697342 | 0.910747762 | 6.78798097 | 5.952677116 | ENSG00000260498 |
| LRP11 | -0.190970119 | 0.876016459 | 0.711542253 | 0.836373711 | 20.02472534 | 17.55367335 | ENSG00000120256 |
| RRP7A | -0.190994988 | 0.876001359 | 0.010063583 | 0.036962107 | 1935.62547 | 1696.002757 | ENSG00000189306 |
| SETD4 | -0.191077184 | 0.875951451 | 0.11428193 | 0.250485016 | 564.5461979 | 494.9171957 | ENSG00000185917 |
| NPRL2 | -0.191139518 | 0.875913604 | 0.176162534 | 0.344009736 | 307.5094059 | 269.6090441 | ENSG00000114388 |
| DOC2B | -0.191161539 | 0.875900235 | 0.334508917 | 0.528981192 | 136.4518093 | 119.7146121 | ENSG00000272636 |
| EXOSC8 | -0.191227826 | 0.875859991 | 0.024713042 | 0.07674791 | 1317.728548 | 1154.608811 | ENSG00000120699 |
| STMP1 | -0.191238652 | 0.875853419 | 0.046037636 | 0.125442744 | 868.5548234 | 760.5415935 | ENSG00000243317 |
| TIMM10 | -0.191449441 | 0.875725459 | 0.028739592 | 0.086611315 | 1040.495581 | 911.4188876 | ENSG00000134809 |
| CHD7 | -0.191474819 | 0.875710055 | 0.048176235 | 0.12995173 | 2877.268622 | 2519.508633 | ENSG00000171316 |
| UTP4 | -0.191498445 | 0.875695713 | 0.0026419 | 0.011971879 | 2638.778092 | 2310.578944 | ENSG00000141076 |
| HIRIP3 | -0.191535605 | 0.875673158 | 0.092397748 | 0.213555289 | 526.3782432 | 461.0818963 | ENSG00000149929 |
| LRP5L | -0.191546999 | 0.875666243 | 0.531247204 | 0.706575936 | 73.11612408 | 64.19238012 | ENSG00000100068 |
| RAE1 | -0.191558489 | 0.875659269 | 0.036441474 | 0.104524967 | 806.036124 | 705.6618207 | ENSG00000101146 |
| SEC11A | -0.191635025 | 0.875612815 | 0.006757149 | 0.026629172 | 3410.471519 | 2985.187053 | ENSG00000140612 |
| GMFB | -0.191636603 | 0.875611858 | 0.11851682 | 0.257242152 | 1270.78778 | 1112.127461 | ENSG00000197045 |
| CSDC2 | -0.191641393 | 0.87560895 | 0.626875549 | 0.779190626 | 31.93500368 | 27.96982699 | ENSG00000172346 |
| TONSL | -0.191826376 | 0.875496687 | 0.030273516 | 0.090250821 | 2710.948503 | 2374.483295 | ENSG00000160949 |
| CREG1 | -0.1918272 | 0.875496187 | 0.14059652 | 0.29187006 | 493.366034 | 431.7705201 | ENSG00000143162 |
| NA | -0.191834708 | 0.87549163 | 0.867760856 | 0.931110966 | 3.847717405 | 3.326799561 | ENSG00000272518 |
| NSL1 | -0.191899349 | 0.875452404 | 0.047357845 | 0.128248144 | 906.1230704 | 792.7403985 | ENSG00000117697 |
| CD4 | -0.19209943 | 0.875331 | 0.504202438 | 0.683955792 | 77.53829389 | 68.07840749 | ENSG00000010610 |
| KCTD14 | -0.192267082 | 0.875229286 | 0.534177695 | 0.708998096 | 55.71709868 | 48.71901456 | ENSG00000151364 |
| ARPC1A | -0.192414751 | 0.875139705 | 0.374441442 | 0.567389377 | 107.6769254 | 94.29882789 | ENSG00000241685 |
| UBQLN4 | -0.192485682 | 0.87509668 | 0.011777157 | 0.042122313 | 1725.163823 | 1510.119171 | ENSG00000160803 |
| DDX19A | -0.192508179 | 0.875083034 | 0.160453529 | 0.320907058 | 357.0109992 | 312.4228859 | ENSG00000168872 |
| VMA21 | -0.192583463 | 0.87503737 | 0.189904111 | 0.362144541 | 552.4461516 | 483.4563245 | ENSG00000160131 |
| ARF5 | -0.192614104 | 0.875018786 | 0.010105315 | 0.037080163 | 2058.796737 | 1801.576426 | ENSG00000004059 |
| ARL6IP6 | -0.192694708 | 0.8749699 | 0.037541106 | 0.106863287 | 842.8833326 | 737.8722857 | ENSG00000177917 |
| C22orf46 | -0.192796711 | 0.874908039 | 0.190847093 | 0.363568715 | 327.8182082 | 286.907736 | ENSG00000184208 |
| NA | -0.19280702 | 0.874901787 | 0.771336235 | 0.874316298 | 11.43070607 | 10.02217755 | ENSG00000261143 |
| PPP2R3C | -0.192873257 | 0.874861619 | 0.294218326 | 0.486878791 | 185.0112453 | 162.1744417 | ENSG00000092020 |
| STXBP4 | -0.192941052 | 0.874820509 | 0.289054794 | 0.480780898 | 172.8626503 | 151.1151761 | ENSG00000166263 |
| ALG3 | -0.192992444 | 0.874789346 | 0.02263785 | 0.07155012 | 1000.512643 | 875.5418101 | ENSG00000214160 |
| CNP | -0.193002298 | 0.874783372 | 0.011891285 | 0.042464947 | 1544.028317 | 1350.493333 | ENSG00000173786 |

| | | | | | | | |
|---|---|---|---|---|---|---|---|
| CNOT7 | -0.193059877 | 0.874748459 | 0.068064612 | 0.169284692 | 1782.05436 | 1557.888914 | ENSG00000198791 |
| ZNF138 | -0.193088053 | 0.874731375 | 0.19779447 | 0.373471984 | 296.0012729 | 258.8114816 | ENSG00000197008 |
| PTBP1 | -0.19314042 | 0.874699625 | 0.001266744 | 0.006418214 | 18830.72603 | 16471.22097 | ENSG00000011304 |
| DCBLD2 | -0.193171775 | 0.874680614 | 0.078515584 | 0.188914428 | 1105.420507 | 965.9607649 | ENSG00000057019 |
| C11orf24 | -0.193186159 | 0.874671894 | 0.051120158 | 0.13609035 | 745.9124295 | 652.9207472 | ENSG00000171067 |
| TOP3A | -0.193188847 | 0.874670264 | 0.017824427 | 0.058585823 | 1103.589161 | 965.3682496 | ENSG00000177302 |
| RMDN3 | -0.193255725 | 0.874629718 | 0.039447244 | 0.111230066 | 829.1294186 | 725.5141576 | ENSG00000137824 |
| SIPA1L3 | -0.193284054 | 0.874612544 | 0.028050977 | 0.085079937 | 980.2453618 | 857.3970924 | ENSG00000105738 |
| ATG101 | -0.193308047 | 0.874597999 | 0.06777486 | 0.168730667 | 825.6941664 | 722.4454855 | ENSG00000123395 |
| POLR2K | -0.193379774 | 0.874554518 | 0.103941926 | 0.233368825 | 668.5008553 | 584.7352244 | ENSG00000147669 |
| FAM20C | -0.193389763 | 0.874548462 | 0.037931216 | 0.107678205 | 1148.654859 | 1004.425614 | ENSG00000177706 |
| YPEL5 | -0.193569059 | 0.874439781 | 0.094453057 | 0.216959508 | 467.081742 | 408.1526345 | ENSG00000119801 |
| DHRS1 | -0.193720393 | 0.874348061 | 0.218052349 | 0.399070034 | 282.3251016 | 247.2365594 | ENSG00000157379 |
| MRPL46 | -0.193741717 | 0.874335137 | 0.602430952 | 0.761731154 | 38.91828464 | 34.13278389 | ENSG00000259494 |
| SUB1 | -0.193792723 | 0.874304225 | 0.034570867 | 0.100224692 | 2208.330301 | 1929.854213 | ENSG00000113387 |
| TSC2 | -0.19385336 | 0.874267479 | 0.022958749 | 0.072346473 | 2089.604761 | 1828.127264 | ENSG00000103197 |
| NA | -0.193925688 | 0.87422365 | 0.854923495 | 0.923041604 | 4.74442273 | 4.163467026 | ENSG00000286459 |
| POLR2M | -0.193936437 | 0.874217136 | 0.394174772 | 0.586609541 | 192.6564626 | 168.2162586 | ENSG00000255529 |
| ORMDL1 | -0.193963663 | 0.874200638 | 0.181774219 | 0.351927647 | 349.1186033 | 304.8096643 | ENSG00000128699 |
| RAB1B | -0.19397841 | 0.874191703 | 0.033674496 | 0.098209772 | 860.7637868 | 752.6720369 | ENSG00000174903 |
| NA | -0.193998083 | 0.874179782 | 0.813175721 | 0.899516491 | 7.622149398 | 6.725433905 | ENSG00000251259 |
| NCCRP1 | -0.194065288 | 0.874139061 | 0.448651442 | 0.636263517 | 76.09632249 | 66.56564937 | ENSG00000188505 |
| PIMREG | -0.194117069 | 0.874107687 | 0.006423714 | 0.025494031 | 1900.796196 | 1661.453444 | ENSG00000129195 |
| DOLPP1 | -0.194425348 | 0.873920925 | 0.040110062 | 0.112704652 | 767.1837437 | 670.0403859 | ENSG00000167130 |
| ZNF606 | -0.19453497 | 0.873854524 | 0.296696467 | 0.489580869 | 171.5987211 | 150.0401483 | ENSG00000166704 |
| KCNK5 | -0.194566312 | 0.87383554 | 0.008794367 | 0.033088673 | 1743.604682 | 1523.726101 | ENSG00000164626 |
| QARS1 | -0.19458662 | 0.87382324 | 0.000640021 | 0.003560068 | 5916.958594 | 5170.225743 | ENSG00000172053 |
| PDCD7 | -0.194732944 | 0.873734617 | 0.05189317 | 0.137398288 | 709.0352604 | 619.873775 | ENSG00000090470 |
| ZNF789 | -0.194907926 | 0.87362865 | 0.452411882 | 0.639762821 | 75.38860333 | 65.95054661 | ENSG00000198556 |
| CHCHD1 | -0.195044822 | 0.873545756 | 0.030918565 | 0.091760559 | 849.4594323 | 742.0642713 | ENSG00000172586 |
| NA | -0.195088942 | 0.873519042 | 0.259469351 | 0.44830847 | 181.3706588 | 158.2414109 | ENSG00000286619 |
| SLC45A4 | -0.195090958 | 0.873517821 | 0.02859638 | 0.086306027 | 857.3538276 | 748.9922756 | ENSG00000022567 |
| CYB561 | -0.195112133 | 0.873505001 | 0.04714513 | 0.127802815 | 839.9795015 | 734.1640059 | ENSG00000008283 |
| UIMC1 | -0.195118581 | 0.873501096 | 0.046867678 | 0.127162281 | 805.7474396 | 704.2013937 | ENSG00000087206 |
| HYPK | -0.19512953 | 0.873494467 | 0.83811585 | 0.91309894 | 5.117862675 | 4.459677675 | ENSG00000242028 |
| PPIA | -0.195138739 | 0.873488891 | 0.000283554 | 0.001750632 | 23509.89321 | 20534.58684 | ENSG00000196262 |
| PSMC1 | -0.195142253 | 0.873486764 | 0.039081502 | 0.110518243 | 1574.775277 | 1375.31801 | ENSG00000100764 |
| LAS1L | -0.195250146 | 0.873421442 | 0.003351233 | 0.014624854 | 2466.073985 | 2154.194341 | ENSG00000001497 |
| CCDC51 | -0.195259809 | 0.873415592 | 0.0892073 | 0.208206536 | 549.2097925 | 479.4251305 | ENSG00000164051 |
| RNF128 | -0.195372639 | 0.873347287 | 0.624660384 | 0.777266072 | 31.62094372 | 27.62099058 | ENSG00000133135 |
| RNF139-AS1 | -0.19545272 | 0.873298811 | 0.801681526 | 0.893310392 | 13.41488433 | 11.7931663 | ENSG00000245149 |
| CNTLN | -0.195487449 | 0.873277788 | 0.266221119 | 0.456270818 | 196.2171342 | 171.4598671 | ENSG00000044459 |
| HDDC2 | -0.19573394 | 0.873128598 | 0.019966144 | 0.064457849 | 1235.435061 | 1078.505947 | ENSG00000111906 |
| PDIA3 | -0.195754703 | 0.873116032 | 0.00234276 | 0.010849753 | 8880.891259 | 7753.103649 | ENSG00000167004 |
| NA | -0.195810585 | 0.873082213 | 0.742692125 | 0.856105567 | 13.74916354 | 11.96491641 | ENSG00000237977 |
| PLEKHG3 | -0.195872753 | 0.873044591 | 0.01719102 | 0.056973409 | 1276.196319 | 1114.881966 | ENSG00000126822 |
| FAAP100 | -0.195924709 | 0.87301315 | 0.036156948 | 0.103896571 | 1184.941754 | 1034.352929 | ENSG00000185504 |
| UBE2A | -0.196098276 | 0.872908127 | 0.039164494 | 0.11063477 | 782.7330225 | 683.0814209 | ENSG00000077721 |
| REEP2 | -0.196198606 | 0.872847424 | 0.036511427 | 0.104590848 | 898.7470998 | 784.9571562 | ENSG00000132563 |
| NA | -0.196271102 | 0.872803564 | 0.818199396 | 0.902165959 | 6.715973334 | 5.855023298 | ENSG00000226945 |
| BIN3 | -0.196401189 | 0.872724868 | 0.199721866 | 0.376183442 | 320.0211734 | 279.5906453 | ENSG00000147439 |
| COA6 | -0.196526711 | 0.872648939 | 0.12161483 | 0.262246569 | 357.3790198 | 311.7952958 | ENSG00000168275 |
| SCX | -0.19665815 | 0.872569439 | 0.610101424 | 0.767371074 | 37.98755233 | 33.12662782 | ENSG00000260428 |
| ZNF592 | -0.196673413 | 0.872560207 | 0.012265861 | 0.043550857 | 1427.909723 | 1245.782485 | ENSG00000166716 |
| ZFP41 | -0.196690423 | 0.87254992 | 0.185793269 | 0.356690744 | 261.2113173 | 228.1991962 | ENSG00000181638 |
| ELK1 | -0.196739951 | 0.872519965 | 0.02411486 | 0.075220489 | 1014.982943 | 884.9936578 | ENSG00000126767 |
| EHMT2 | -0.196745996 | 0.872516309 | 0.000618812 | 0.003456617 | 4924.822466 | 4297.289447 | ENSG00000204371 |
| MMP2 | -0.196882288 | 0.872433886 | 0.011485075 | 0.041252789 | 1960.276843 | 1710.102948 | ENSG00000087245 |
| NA | -0.196887855 | 0.87243052 | 0.32985711 | 0.524185173 | 156.5706406 | 136.7963111 | ENSG00000204791 |
| TBK1 | -0.196950841 | 0.872392431 | 0.203551655 | 0.381381558 | 418.2259172 | 364.6220854 | ENSG00000183735 |
| ZHX2 | -0.197071755 | 0.872319318 | 0.054260446 | 0.142365465 | 716.1988687 | 624.5257973 | ENSG00000178764 |
| NSA2 | -0.197162441 | 0.872264487 | 0.014665729 | 0.05013513 | 1437.451394 | 1253.199276 | ENSG00000164346 |
| DPY19L3 | -0.1971659 | 0.872262396 | 0.058061131 | 0.150126256 | 886.8119948 | 773.111928 | ENSG00000178904 |
| YPEL2 | -0.197218522 | 0.872230581 | 0.471376077 | 0.656550632 | 75.73796805 | 65.97267842 | ENSG00000175155 |
| NA | -0.197245114 | 0.872214504 | 0.704243699 | 0.831811477 | 21.52291981 | 18.66354359 | ENSG00000219891 |
| UBE2L6 | -0.197248745 | 0.872212309 | 0.013110167 | 0.045897968 | 1225.846262 | 1069.525429 | ENSG00000156587 |
| BCCIP | -0.197263799 | 0.872203208 | 0.007145829 | 0.027864675 | 2126.288412 | 1853.764197 | ENSG00000107949 |
| GGCT | -0.197307392 | 0.872176853 | 0.11596627 | 0.253218525 | 392.6399487 | 342.8314302 | ENSG00000006625 |
| LOC101928000 | -0.197543467 | 0.872034146 | 0.64134931 | 0.789727124 | 30.55547148 | 26.60833613 | ENSG00000234327 |
| ACTR8 | -0.197547798 | 0.872031528 | 0.071799205 | 0.17648892 | 645.6777246 | 563.2428266 | ENSG00000113812 |
| NA | -0.197553394 | 0.872028146 | 0.63006414 | 0.781581472 | 33.26590441 | 28.96695964 | ENSG00000226221 |
| PPP4R1 | -0.197588112 | 0.872007161 | 0.010571027 | 0.038506962 | 1764.481532 | 1537.747799 | ENSG00000154845 |
| RMI1 | -0.197708292 | 0.871934524 | 0.221146077 | 0.402736883 | 365.3963993 | 318.175437 | ENSG00000178966 |
| TRIM59 | -0.197794618 | 0.871882352 | 0.159206097 | 0.319100876 | 379.2441035 | 330.3370868 | ENSG00000213186 |

| | | | | | | | |
|---|---|---|---|---|---|---|---|
| ZNF274 | -0.197858297 | 0.871843869 | 0.085506351 | 0.201394677 | 535.9563167 | 467.4442861 | ENSG00000171606 |
| EPN1 | -0.197872757 | 0.87183513 | 0.019817957 | 0.064046384 | 4259.32677 | 3713.038586 | ENSG00000063245 |
| NRF1 | -0.197877293 | 0.871832389 | 0.098004039 | 0.223200645 | 414.3410084 | 361.344591 | ENSG00000106459 |
| FAU | -0.197896252 | 0.871820932 | 0.011077855 | 0.040108803 | 1922.868313 | 1676.66845 | ENSG00000149806 |
| PRLR | -0.197909415 | 0.871812978 | 0.616385704 | 0.771888321 | 35.31389734 | 30.86185928 | ENSG00000113494 |
| NA | -0.197926779 | 0.871802485 | 0.819276787 | 0.90291228 | 6.947277144 | 6.01603814 | ENSG00000267742 |
| RTL5 | -0.197961022 | 0.871781793 | 0.359428753 | 0.552696633 | 121.0234627 | 105.310214 | ENSG00000242732 |
| TMED7 | -0.198001538 | 0.87175731 | 0.545827527 | 0.717931119 | 476.6782755 | 415.4147574 | ENSG00000134970 |
| ZNF879 | -0.198072103 | 0.871714672 | 0.556493133 | 0.726110513 | 44.94242108 | 39.24835551 | ENSG00000234284 |
| NA | -0.19809523 | 0.871700698 | 0.091112477 | 0.211587352 | 437.1895975 | 381.0244763 | ENSG00000184809 |
| IQCB1 | -0.198119462 | 0.871686057 | 0.054953771 | 0.143766005 | 796.2893287 | 694.003309 | ENSG00000173226 |
| SYF2 | -0.198192736 | 0.871641785 | 0.065399402 | 0.164287411 | 656.0422669 | 571.9651222 | ENSG00000117614 |
| GTPBP6 | -0.198298108 | 0.871578124 | 0.005943754 | 0.023847356 | 1954.468835 | 1703.629733 | ENSG00000178605 |
| EEF1D | -0.198345794 | 0.871549316 | 0.012152219 | 0.043225528 | 1333.268718 | 1162.260737 | ENSG00000104529 |
| NBPF15 | -0.198478297 | 0.871469273 | 0.212543419 | 0.392385079 | 291.6309817 | 254.1180069 | ENSG00000266338 |
| AP4B1 | -0.198512122 | 0.871448841 | 0.267289161 | 0.45741008 | 170.2881374 | 148.4418973 | ENSG00000134262 |
| LINC01285 | -0.198527401 | 0.871439612 | 0.848662796 | 0.919616246 | 4.730517303 | 4.133411111 | ENSG00000203650 |
| LOC105372401 | -0.198690274 | 0.871341237 | 0.86058171 | 0.926411187 | 4.353920445 | 3.829602768 | ENSG00000286177 |
| MIR4482 | -0.198740748 | 0.871310752 | 0.837052719 | 0.912409457 | 5.958344929 | 5.228747237 | ENSG00000266852 |
| FGF1 | -0.198751169 | 0.871304459 | 0.830488164 | 0.908658972 | 6.073947985 | 5.341373203 | ENSG00000113578 |
| ZNF10 | -0.19875407 | 0.871302707 | 0.475470003 | 0.660200856 | 73.4985311 | 63.94290939 | ENSG00000256223 |
| SNRPC | -0.198818499 | 0.871263796 | 0.005317305 | 0.021731185 | 2329.766515 | 2029.184935 | ENSG00000124562 |
| LMNB2 | -0.19883499 | 0.871253837 | 0.000219019 | 0.001393031 | 12478.98551 | 10872.62558 | ENSG00000176619 |
| MVK | -0.19896988 | 0.87117238 | 0.077067674 | 0.186336152 | 498.9630031 | 434.8836077 | ENSG00000110921 |
| SNED1 | -0.198987616 | 0.87116167 | 0.653062077 | 0.797325732 | 35.41501625 | 30.87270617 | ENSG00000162804 |
| IGF2BP1 | -0.199018607 | 0.871142957 | 0.001270729 | 0.006434889 | 8690.883 | 7570.35575 | ENSG00000159217 |
| VTI1A | -0.199057593 | 0.871119416 | 0.207790318 | 0.386820586 | 242.8326577 | 211.5893685 | ENSG00000151532 |
| G6PD | -0.199118984 | 0.871082348 | 0.018309025 | 0.059879913 | 1293.21511 | 1126.352249 | ENSG00000160211 |
| PPHLN1 | -0.199127812 | 0.871077018 | 0.009640873 | 0.03567673 | 1355.098068 | 1180.683953 | ENSG00000134283 |
| RPN1 | -0.19924655 | 0.871005329 | 0.001568611 | 0.007723869 | 4932.210861 | 4295.440655 | ENSG00000163902 |
| NCOA1 | -0.199366359 | 0.870932999 | 0.09401147 | 0.216179724 | 448.5558125 | 390.2605251 | ENSG00000084676 |
| NOL7 | -0.199442463 | 0.870887057 | 0.023300994 | 0.073123661 | 2033.512726 | 1771.31164 | ENSG00000225921 |
| CXXC4 | -0.199520089 | 0.870840199 | 0.633112222 | 0.783736436 | 28.28574318 | 24.58247757 | ENSG00000168772 |
| UBE2L3 | -0.19957938 | 0.87080441 | 0.001647088 | 0.008047158 | 3634.712538 | 3164.767986 | ENSG00000185651 |
| THOC3 | -0.199592168 | 0.870796692 | 0.005492236 | 0.022308372 | 2210.948364 | 1924.599441 | ENSG00000051596 |
| PLBD2 | -0.19959698 | 0.870793787 | 0.041881667 | 0.11672704 | 1067.508571 | 928.7496016 | ENSG00000151176 |
| NA | -0.199679515 | 0.870743972 | 0.828774714 | 0.907724296 | 6.302094882 | 5.498700818 | ENSG00000239911 |
| RASSF4 | -0.199781144 | 0.870682635 | 0.2119591 | 0.391615515 | 228.8385705 | 199.099517 | ENSG00000107551 |
| MASTL | -0.199900021 | 0.870610894 | 0.027919308 | 0.084794593 | 1001.012841 | 870.8556377 | ENSG00000120539 |
| NDUFB8 | -0.199985017 | 0.870559604 | 0.668077523 | 0.807200701 | 23.4990028 | 20.39275347 | ENSG00000166136 |
| NPHS1 | -0.200086275 | 0.870498505 | 0.668022371 | 0.807200701 | 26.26102872 | 22.73695449 | ENSG00000161270 |
| PPP3R1 | -0.200226181 | 0.870414092 | 0.240584813 | 0.426780143 | 433.5969357 | 377.1514054 | ENSG00000221823 |
| NCL | -0.200332602 | 0.870349888 | 8.70E-05 | 0.000615434 | 35217.98687 | 30651.50707 | ENSG00000115053 |
| NA | -0.200360062 | 0.870333322 | 0.577626267 | 0.742139457 | 39.70244613 | 34.54129409 | ENSG00000181524 |
| LAD1 | -0.200369475 | 0.870327643 | 0.070660393 | 0.174197887 | 483.0369169 | 420.3211091 | ENSG00000159166 |
| RFC1 | -0.200424019 | 0.87029474 | 0.007440303 | 0.028800874 | 1881.536233 | 1636.65961 | ENSG00000035928 |
| CNN3 | -0.200449819 | 0.870279176 | 0.000551331 | 0.003126796 | 4518.030074 | 3931.918566 | ENSG00000117519 |
| ATF1 | -0.200505109 | 0.870245824 | 0.132254298 | 0.278871802 | 432.6683833 | 375.9798231 | ENSG00000123268 |
| AXIN1 | -0.2005714 | 0.870205837 | 0.008158977 | 0.031134133 | 1984.932508 | 1727.035587 | ENSG00000103126 |
| ERF | -0.200676382 | 0.870142517 | 0.018113957 | 0.059376726 | 1362.849087 | 1185.916451 | ENSG00000105722 |
| DTX3 | -0.200726841 | 0.870112083 | 0.110567909 | 0.244367473 | 775.8295513 | 675.7897373 | ENSG00000178498 |
| LOC100505501 | -0.20082925 | 0.870050321 | 0.384021054 | 0.576778462 | 97.30202543 | 84.59911806 | ENSG00000167912 |
| CBX2 | -0.200838772 | 0.870044579 | 0.004660413 | 0.019345597 | 3931.773938 | 3420.718679 | ENSG00000173894 |
| SLC35C1 | -0.200884371 | 0.87001708 | 0.106046752 | 0.236974983 | 463.7664751 | 403.5094837 | ENSG00000181830 |
| CHCHD2 | -0.200912916 | 0.869999866 | 0.028659645 | 0.086440662 | 3288.052175 | 2860.019423 | ENSG00000106153 |
| CLDN10 | -0.200958294 | 0.869972502 | 0.075782588 | 0.183938512 | 516.214736 | 448.8836922 | ENSG00000134873 |
| TIMM22 | -0.201044017 | 0.869920811 | 0.034417021 | 0.099841143 | 745.913527 | 648.579103 | ENSG00000177370 |
| FAM126B | -0.201068755 | 0.869905894 | 0.48039824 | 0.6642839 | 66.79458406 | 57.96777405 | ENSG00000155744 |
| C1D | -0.201072769 | 0.869903474 | 0.153310975 | 0.310712477 | 292.2709173 | 254.0971489 | ENSG00000197223 |
| VSIG10 | -0.201087508 | 0.869894587 | 0.015996515 | 0.053827052 | 1277.471193 | 1111.834653 | ENSG00000176834 |
| ZNF174 | -0.201233434 | 0.869806603 | 0.27171222 | 0.462562165 | 212.1825765 | 184.6057241 | ENSG00000103343 |
| STUB1 | -0.201249357 | 0.869797003 | 0.047978243 | 0.129534876 | 1001.702963 | 871.5526979 | ENSG00000103266 |
| ZNF84 | -0.201276458 | 0.869780664 | 0.074435297 | 0.181486345 | 498.5240648 | 433.7535407 | ENSG00000198040 |
| PIM2 | -0.201323877 | 0.869752076 | 0.001933918 | 0.009245097 | 5135.831372 | 4465.648942 | ENSG00000102096 |
| MTCH2 | -0.201379505 | 0.869718541 | 0.006185373 | 0.024717486 | 1550.083752 | 1348.461351 | ENSG00000109919 |
| RTL6 | -0.201587098 | 0.869593404 | 0.025225369 | 0.077947148 | 1053.743908 | 915.5304128 | ENSG00000188636 |
| PPP2R1B | -0.201656165 | 0.869551774 | 0.007024124 | 0.027500023 | 1925.431792 | 1673.665647 | ENSG00000137713 |
| UBE2I | -0.201751222 | 0.869494483 | 0.002845656 | 0.012727168 | 3050.931849 | 2652.661118 | ENSG00000103275 |
| TSSC4 | -0.201832508 | 0.869445494 | 0.076030862 | 0.184396232 | 691.824582 | 602.1480167 | ENSG00000184281 |
| FAM92A | -0.201865034 | 0.869425892 | 0.190347055 | 0.362765276 | 228.3200329 | 198.442747 | ENSG00000188343 |
| TIMP1 | -0.201901643 | 0.86940383 | 0.09017694 | 0.209809115 | 488.7154242 | 424.4936708 | ENSG00000102265 |
| TTC33 | -0.201951564 | 0.869373748 | 0.309215323 | 0.503332882 | 188.2769938 | 163.2916461 | ENSG00000113638 |
| AUP1 | -0.20222441 | 0.869209345 | 0.021527093 | 0.068719108 | 970.0908444 | 843.4806857 | ENSG00000115307 |
| NA | -0.202274206 | 0.869179344 | 0.391792899 | 0.584519809 | 94.18856026 | 81.76351037 | ENSG00000234742 |

| | | | | | | | |
|---|---|---|---|---|---|---|---|
| APEX1 | -0.202293313 | 0.869167832 | 0.000485052 | 0.002789313 | 8759.846508 | 7613.776379 | ENSG00000100823 |
| FAM118B | -0.202312387 | 0.869156341 | 0.022978492 | 0.072367607 | 866.2315178 | 752.9571858 | ENSG00000197798 |
| UQCR10 | -0.202317982 | 0.86915297 | 0.33462674 | 0.528981192 | 139.0735782 | 120.7636061 | ENSG00000184076 |
| RAB8A | -0.202334981 | 0.869142729 | 0.149849133 | 0.305769441 | 391.0079838 | 340.2852083 | ENSG00000167461 |
| SSPO | -0.20235532 | 0.869130476 | 0.765369711 | 0.870214234 | 13.31950053 | 11.66545657 | ENSG00000197558 |
| CCT3 | -0.202435907 | 0.869081929 | 0.000251111 | 0.001576053 | 12627.34828 | 10973.54578 | ENSG00000163468 |
| LUC7L2 | -0.202513189 | 0.869035376 | 0.074922718 | 0.182308087 | 592.4868005 | 515.0590527 | ENSG00000146963 |
| CLK2 | -0.202573852 | 0.868998835 | 0.036529675 | 0.104626952 | 1200.465552 | 1044.200228 | ENSG00000176444 |
| UTP3 | -0.202708212 | 0.868917908 | 0.042978621 | 0.119103731 | 1380.961762 | 1201.007374 | ENSG00000132467 |
| YY1 | -0.202847002 | 0.86883432 | 0.007183601 | 0.027976647 | 2772.848139 | 2408.601891 | ENSG00000100811 |
| TBC1D17 | -0.202861603 | 0.868825527 | 0.067285648 | 0.167910622 | 606.0358838 | 526.1513683 | ENSG00000104946 |
| HIF1A | -0.202965028 | 0.868763244 | 0.035191175 | 0.101688999 | 2609.97708 | 2266.517373 | ENSG00000100644 |
| TSR3 | -0.203113851 | 0.86867363 | 0.070442001 | 0.173821334 | 1107.117654 | 961.6552715 | ENSG00000007520 |
| CHD4 | -0.203252155 | 0.868590359 | 0.185114432 | 0.355873787 | 258.3340943 | 224.5490444 | ENSG00000111642 |
| ANP32E | -0.203338815 | 0.868538186 | 0.011218459 | 0.040475553 | 3842.849586 | 3336.89242 | ENSG00000143401 |
| DHDDS | -0.203378053 | 0.868514564 | 0.024550519 | 0.076319914 | 833.0299317 | 723.1854381 | ENSG00000117682 |
| SAMD1 | -0.203386286 | 0.868509608 | 0.013161028 | 0.046023866 | 1671.396026 | 1452.0858 | ENSG00000141858 |
| SBK1 | -0.203483374 | 0.868451163 | 0.012843617 | 0.04518671 | 4390.786069 | 3812.903649 | ENSG00000188322 |
| FADS1 | -0.203537917 | 0.86841833 | 0.003317739 | 0.014502589 | 3198.51152 | 2777.184141 | ENSG00000149485 |
| ABHD17C | -0.203546159 | 0.868413369 | 0.090058066 | 0.209653406 | 528.0722639 | 458.2398615 | ENSG00000136379 |
| SLC25A10 | -0.203596927 | 0.86838281 | 0.363792433 | 0.556819635 | 104.760038 | 90.97538914 | ENSG00000183048 |
| THUMPD3-AS1 | -0.203607258 | 0.868376592 | 0.37780947 | 0.571087073 | 181.3688774 | 157.9422145 | ENSG00000206573 |
| CELF5 | -0.203621555 | 0.868367986 | 0.171171973 | 0.336816152 | 279.5290487 | 243.1059595 | ENSG00000161082 |
| POMT1 | -0.203683452 | 0.868330731 | 0.066009634 | 0.165528692 | 548.774124 | 476.7620776 | ENSG00000130714 |
| GPR3 | -0.203949261 | 0.86817076 | 0.206877562 | 0.385515625 | 232.4665294 | 201.7365763 | ENSG00000181773 |
| PPIL1 | -0.204056553 | 0.868106198 | 0.030161517 | 0.090032785 | 1305.840601 | 1132.903702 | ENSG00000137168 |
| CLPP | -0.204063398 | 0.868102079 | 0.03192337 | 0.094154638 | 1097.168795 | 951.9982711 | ENSG00000125656 |
| CCDC3 | -0.204081914 | 0.868090937 | 0.579700323 | 0.74384975 | 39.879406 | 34.53705721 | ENSG00000151468 |
| COG2 | -0.204207252 | 0.868015523 | 0.037473303 | 0.106726004 | 670.1983338 | 581.477122 | ENSG00000135775 |
| VPS25 | -0.204263915 | 0.867981432 | 0.006357013 | 0.025279444 | 1852.620851 | 1607.989045 | ENSG00000131475 |
| YWHAE | -0.204283334 | 0.867969748 | 0.00125302 | 0.006359112 | 21192.90066 | 18393.465 | ENSG00000108953 |
| NUP107 | -0.204354666 | 0.867926834 | 0.00367545 | 0.015823646 | 1854.861092 | 1609.862992 | ENSG00000111581 |
| LINC00599 | -0.204401056 | 0.867898926 | 0.747460552 | 0.858996555 | 532.3799509 | 462.1394707 | ENSG00000253230 |
| KCNMA1 | -0.204495263 | 0.867842255 | 0.346058245 | 0.539522503 | 126.1158746 | 109.2754506 | ENSG00000156113 |
| FABP5 | -0.204498404 | 0.867840366 | 0.018399351 | 0.060132881 | 1027.36666 | 891.7586007 | ENSG00000164687 |
| TAF9 | -0.20454355 | 0.867813209 | 0.126688949 | 0.270295036 | 497.6176183 | 431.2165797 | ENSG00000273841 |
| ZFP14 | -0.20456595 | 0.867799734 | 0.268336005 | 0.458541353 | 159.5040867 | 138.5167123 | ENSG00000142065 |
| SHANK1 | -0.204571296 | 0.867796519 | 0.448159828 | 0.635975449 | 71.26726447 | 61.81680909 | ENSG00000161681 |
| ERAL1 | -0.204677801 | 0.867732457 | 0.015149889 | 0.051520741 | 1267.386548 | 1099.975314 | ENSG00000132591 |
| ULK4 | -0.204735117 | 0.867697984 | 0.28281853 | 0.473767049 | 167.0767328 | 145.0396789 | ENSG00000168038 |
| EBAG9 | -0.204830968 | 0.867640337 | 0.207044761 | 0.38570385 | 255.4635533 | 221.6255305 | ENSG00000147654 |
| EWSR1 | -0.204879763 | 0.867610992 | 0.000199674 | 0.001281227 | 9588.856551 | 8319.639381 | ENSG00000182944 |
| EIF4G2 | -0.204927796 | 0.867582107 | 0.000433627 | 0.002527005 | 17067.44229 | 14806.5729 | ENSG00000110321 |
| CTNND2 | -0.205000023 | 0.867538673 | 0.090142954 | 0.20978271 | 400.6504508 | 347.6540403 | ENSG00000169862 |
| ZBTB12 | -0.205373105 | 0.867314356 | 0.02217812 | 0.070336973 | 900.1902664 | 780.2340863 | ENSG00000204366 |
| HS3ST3B1 | -0.205411558 | 0.867291239 | 0.245051552 | 0.432180557 | 174.2736539 | 151.1079048 | ENSG00000125430 |
| NA | -0.205485286 | 0.867246918 | 0.769847601 | 0.8731095 | 11.09199217 | 9.632000309 | ENSG00000272525 |
| HAUS5 | -0.205506199 | 0.867234347 | 0.06100323 | 0.155992531 | 730.9166097 | 634.6139012 | ENSG00000249115 |
| MRPL55 | -0.205510286 | 0.86723189 | 0.091994294 | 0.212888281 | 409.1505594 | 354.9864296 | ENSG00000162910 |
| MRPL54 | -0.20555575 | 0.867204561 | 0.091258518 | 0.211775304 | 451.9559101 | 391.4273696 | ENSG00000183617 |
| TADA1 | -0.205700563 | 0.867117519 | 0.070625905 | 0.174136029 | 507.5512873 | 440.1108091 | ENSG00000152382 |
| UBL3 | -0.205706399 | 0.867114011 | 0.079649191 | 0.191096194 | 534.4818666 | 462.9944273 | ENSG00000122042 |
| COMTD1 | -0.205782446 | 0.867068305 | 0.113840458 | 0.249694563 | 436.1360613 | 378.4826029 | ENSG00000165644 |
| NA | -0.205806325 | 0.867053954 | 0.751803738 | 0.862034798 | 11.89321599 | 10.26587406 | ENSG00000267390 |
| SYNPO2 | -0.205831545 | 0.867038797 | 0.309688368 | 0.503645563 | 135.3109771 | 117.3591262 | ENSG00000172403 |
| NNAT | -0.20595405 | 0.866965176 | 0.106774917 | 0.238068842 | 387.8669183 | 336.3678025 | ENSG00000053438 |
| HINT1 | -0.205958921 | 0.866962249 | 0.003574914 | 0.015448231 | 3111.910563 | 2697.67773 | ENSG00000169567 |
| MLF2 | -0.205971865 | 0.86695447 | 0.000473486 | 0.002729579 | 5667.369847 | 4913.246898 | ENSG00000089693 |
| FASTKD2 | -0.206003964 | 0.866935182 | 0.044528627 | 0.122301952 | 956.1348431 | 828.424988 | ENSG00000118246 |
| SLC25A38 | -0.206018259 | 0.866926591 | 0.019781168 | 0.06396095 | 1046.99154 | 907.2493199 | ENSG00000144659 |
| RWDD1 | -0.206101889 | 0.866876339 | 0.056986462 | 0.147905333 | 681.7949429 | 590.7632172 | ENSG00000111832 |
| NIF3L1 | -0.206101985 | 0.866876282 | 0.033843463 | 0.098655983 | 810.2722167 | 702.6603403 | ENSG00000196290 |
| SNN | -0.206246095 | 0.866789694 | 0.002639406 | 0.011966425 | 2451.542187 | 2125.414685 | ENSG00000184602 |
| NA | -0.206343266 | 0.866731314 | 0.61134382 | 0.768273474 | 32.31297601 | 28.11630772 | ENSG00000253341 |
| TNIP2 | -0.206367615 | 0.866716686 | 0.017670864 | 0.058224318 | 1373.672328 | 1191.17453 | ENSG00000168884 |
| PDCD11 | -0.20646212 | 0.866659913 | 0.002028159 | 0.009611329 | 4289.793143 | 3717.924748 | ENSG00000148843 |
| GCSH | -0.206526473 | 0.866621255 | 0.678636649 | 0.814350236 | 20.99343831 | 18.18374826 | ENSG00000140905 |
| NA | -0.206642735 | 0.86655142 | 0.597959301 | 0.757939498 | 39.04473391 | 34.03188088 | ENSG00000272655 |
| C12orf43 | -0.206652066 | 0.866545816 | 0.069341879 | 0.1717699 | 515.7208523 | 446.9636264 | ENSG00000157895 |
| ZNF343 | -0.206714324 | 0.866508421 | 0.37872751 | 0.572023464 | 111.6370889 | 96.90994809 | ENSG00000088876 |
| NA | -0.206717876 | 0.866506288 | 0.864548699 | 0.929493914 | 4.138401199 | 3.638532011 | ENSG00000279573 |
| NA | -0.206885889 | 0.866405383 | 0.666812724 | 0.806704114 | 21.7883483 | 18.80633709 | ENSG00000262801 |
| MSL3 | -0.206964809 | 0.866357989 | 0.141497888 | 0.293215772 | 282.8173992 | 245.0058202 | ENSG00000005302 |
| WDR49 | -0.206975573 | 0.866351525 | 0.82267061 | 0.904849165 | 6.100481064 | 5.322602209 | ENSG00000174776 |

| | | | | | | | |
|---|---|---|---|---|---|---|---|
| GLI1 | -0.207028046 | 0.866320015 | 0.181150806 | 0.350940634 | 274.9829852 | 238.6165523 | ENSG00000111087 |
| NA | -0.20707184 | 0.866293717 | 0.57097479 | 0.737373586 | 38.59485164 | 33.54361179 | ENSG00000227615 |
| RNF220 | -0.207212552 | 0.866209228 | 0.008909805 | 0.033400954 | 1459.616414 | 1264.329653 | ENSG00000187147 |
| HGH1 | -0.20726538 | 0.866177511 | 0.036621755 | 0.104825884 | 915.9162946 | 793.1285459 | ENSG00000235173 |
| MED30 | -0.207286456 | 0.866164857 | 0.231688202 | 0.416337903 | 193.2694896 | 167.1082521 | ENSG00000164758 |
| PSMC3 | -0.207302347 | 0.866155317 | 0.001382667 | 0.006903648 | 4257.513351 | 3688.309209 | ENSG00000165916 |
| NA | -0.207417223 | 0.86608635 | 0.725646622 | 0.845904106 | 17.91547329 | 15.47150181 | ENSG00000236166 |
| MMD | -0.207434671 | 0.866075876 | 0.091527538 | 0.212197037 | 590.9948448 | 511.4238412 | ENSG00000108960 |
| TRNP1 | -0.207446336 | 0.866068874 | 0.086590441 | 0.203328678 | 946.7603229 | 819.4560497 | ENSG00000253368 |
| SPATA5L1 | -0.207501257 | 0.866035904 | 0.112663598 | 0.247787721 | 343.1887457 | 297.4051456 | ENSG00000171763 |
| MPST | -0.207686489 | 0.865924719 | 0.018284583 | 0.059831648 | 2095.915853 | 1814.334701 | ENSG00000128309 |
| TRMT61A | -0.207699594 | 0.865916853 | 0.102843113 | 0.231299469 | 620.822265 | 538.1669125 | ENSG00000166166 |
| RRP12 | -0.207785537 | 0.86586527 | 0.00964392 | 0.03567673 | 1981.475789 | 1716.642616 | ENSG00000052749 |
| DNMT1 | -0.20781438 | 0.86584796 | 0.000178644 | 0.001164202 | 6765.39536 | 5858.419137 | ENSG00000130816 |
| NDNF | -0.207826638 | 0.865840603 | 0.572416011 | 0.738490766 | 38.61369542 | 33.42349975 | ENSG00000173376 |
| PGBD5 | -0.207847084 | 0.865828333 | 0.034852658 | 0.100899589 | 790.9863737 | 684.6569989 | ENSG00000177614 |
| DEPDC1B | -0.207877016 | 0.865810369 | 0.014814252 | 0.050546273 | 1016.740259 | 880.1597627 | ENSG00000035499 |
| NA | -0.207885149 | 0.865805488 | 0.719742587 | 0.841462759 | 14.31790348 | 12.40392056 | ENSG00000227885 |
| ETF1 | -0.207896683 | 0.865798567 | 0.004626403 | 0.019234129 | 4657.622054 | 4031.652389 | ENSG00000120705 |
| RAD51AP1 | -0.207902065 | 0.865795337 | 0.049122741 | 0.131909719 | 685.6860604 | 593.0490098 | ENSG00000111247 |
| ICMT | -0.207923103 | 0.865782712 | 0.002031477 | 0.009619671 | 3858.253024 | 3339.899745 | ENSG00000116237 |
| RILPL2 | -0.207924743 | 0.865781727 | 0.358759945 | 0.551942958 | 107.9471945 | 93.29702032 | ENSG00000150977 |
| STOML2 | -0.20797503 | 0.86575155 | 0.001045489 | 0.005474247 | 4125.780297 | 3571.747424 | ENSG00000165283 |
| CAMTA1 | -0.207979998 | 0.865748569 | 0.013685531 | 0.047455123 | 1312.250287 | 1135.818815 | ENSG00000171735 |
| C16orf54 | -0.208038056 | 0.865713729 | 0.672746391 | 0.809789244 | 20.79182449 | 18.01524734 | ENSG00000185905 |
| VPS51 | -0.208088052 | 0.865683729 | 0.006475849 | 0.025673453 | 2036.745703 | 1763.059844 | ENSG00000149823 |
| LRRC71 | -0.208106028 | 0.865672942 | 0.81773438 | 0.901961189 | 6.050571819 | 5.273775299 | ENSG00000160838 |
| MIAT | -0.20815626 | 0.865642802 | 0.582257666 | 0.746040945 | 1019.552804 | 882.7289649 | ENSG00000225783 |
| CALM2 | -0.208195671 | 0.865619155 | 0.005322812 | 0.021748899 | 4258.224836 | 3685.008369 | ENSG00000143933 |
| KDELR1 | -0.208210857 | 0.865610043 | 0.002611289 | 0.011870894 | 3416.198868 | 2956.62205 | ENSG00000105438 |
| LAMA3 | -0.208212739 | 0.865608914 | 0.626059502 | 0.778465286 | 27.52170325 | 23.81015881 | ENSG00000053747 |
| SMC3 | -0.208255522 | 0.865583245 | 0.003766996 | 0.016180185 | 2527.654902 | 2187.176566 | ENSG00000108055 |
| SLC22A31 | -0.208273226 | 0.865572623 | 0.336883043 | 0.531250771 | 166.3677359 | 144.1519315 | ENSG00000259803 |
| KIAA1328 | -0.208328492 | 0.865539465 | 0.489248263 | 0.672081774 | 58.88688461 | 51.08991068 | ENSG00000150477 |
| ATP6V1A | -0.208330181 | 0.865538452 | 0.032333434 | 0.094971135 | 869.7969398 | 752.3296586 | ENSG00000114573 |
| LINC00863 | -0.208454175 | 0.865464066 | 0.446995613 | 0.635225939 | 89.41739388 | 77.6117663 | ENSG00000224914 |
| ZNF256 | -0.208455874 | 0.865463046 | 0.250366948 | 0.438923668 | 217.2704713 | 187.6853547 | ENSG00000152454 |
| PPP1R1A | -0.208483887 | 0.865446242 | 0.154187956 | 0.311780056 | 295.3608109 | 255.2327265 | ENSG00000135447 |
| DNAJC25 | -0.208526819 | 0.865420488 | 0.314874352 | 0.508473455 | 119.0873145 | 102.9778475 | ENSG00000059769 |
| TMEM100 | -0.208552594 | 0.865405027 | 0.757288997 | 0.864846526 | 13.44141741 | 11.58613571 | ENSG00000166292 |
| CMTR1 | -0.208558945 | 0.865401217 | 0.024070445 | 0.075120695 | 1253.030446 | 1084.969217 | ENSG00000137200 |
| CACYBP | -0.208731433 | 0.865297756 | 0.001036949 | 0.005438261 | 4369.521828 | 3780.076322 | ENSG00000116161 |
| NYAP1 | -0.208751165 | 0.865285921 | 0.218594259 | 0.399864391 | 197.0876147 | 170.7486478 | ENSG00000166924 |
| DNAJC1 | -0.208808924 | 0.86525128 | 0.102504127 | 0.230796777 | 380.1057146 | 328.711305 | ENSG00000136770 |
| NA | -0.208832441 | 0.865237176 | 0.668708166 | 0.807626558 | 21.46807222 | 18.63403733 | ENSG00000229212 |
| ARHGEF26 | -0.208950209 | 0.865166549 | 0.104518602 | 0.234521522 | 346.2780605 | 299.7153803 | ENSG00000114790 |
| ZBTB8A | -0.208959617 | 0.865160908 | 0.397666656 | 0.589769848 | 121.6676376 | 105.1752414 | ENSG00000160062 |
| APEX2 | -0.208960467 | 0.865160398 | 0.040976593 | 0.114652804 | 620.3177835 | 536.6253639 | ENSG00000169188 |
| NA | -0.208991544 | 0.865141761 | 0.732749206 | 0.850978995 | 14.91949032 | 12.99309266 | ENSG00000267265 |
| MMS22L | -0.209093639 | 0.86508054 | 0.098174377 | 0.223451297 | 460.4229839 | 398.1140506 | ENSG00000146263 |
| ISOC2 | -0.209112144 | 0.865069444 | 0.006243795 | 0.024897265 | 1744.600551 | 1509.187901 | ENSG00000063241 |
| MFAP1 | -0.209143344 | 0.865050736 | 0.007484049 | 0.028946029 | 1558.469625 | 1347.875706 | ENSG00000140259 |
| KHDC1 | -0.209180787 | 0.865028285 | 0.444519364 | 0.632920173 | 150.0870535 | 130.2355079 | ENSG00000135314 |
| PLA2G3 | -0.209230584 | 0.864998428 | 0.014012232 | 0.048397821 | 1003.334432 | 867.5877793 | ENSG00000100078 |
| NA | -0.20931097 | 0.864950232 | 0.46350643 | 0.649994428 | 78.9791829 | 68.56557728 | ENSG00000206567 |
| NA | -0.209504979 | 0.864833924 | 0.656154868 | 0.799155153 | 27.95146396 | 24.33976874 | ENSG00000256943 |
| SCARB1 | -0.209510187 | 0.864830802 | 0.008960928 | 0.033585812 | 1324.305189 | 1144.76461 | ENSG00000073060 |
| NA | -0.209519395 | 0.864825283 | 0.727223278 | 0.846837128 | 14.81463578 | 12.89175162 | ENSG00000269069 |
| CYP51A1 | -0.209612839 | 0.864769269 | 0.204339059 | 0.382354379 | 253.2484772 | 219.4654009 | ENSG00000001630 |
| SNRPG | -0.209694451 | 0.864720351 | 0.007136031 | 0.027840705 | 1866.081285 | 1613.710298 | ENSG00000143977 |
| SPANXA2-OT1 | -0.20973337 | 0.864697025 | 0.84236118 | 0.915625014 | 4.419614255 | 3.852172609 | ENSG00000277215 |
| SF3B2 | -0.209889273 | 0.864603587 | 0.000101495 | 0.000706384 | 7141.604816 | 6174.424845 | ENSG00000087365 |
| DAP | -0.209941044 | 0.864572562 | 0.019037951 | 0.061881688 | 990.2119981 | 856.5065254 | ENSG00000112977 |
| ZC3H18 | -0.210258441 | 0.864382374 | 0.001247736 | 0.006337499 | 3532.000485 | 3053.874163 | ENSG00000158545 |
| ZNHIT2 | -0.210283933 | 0.864367101 | 0.386127447 | 0.579144299 | 95.8057101 | 82.64410659 | ENSG00000174276 |
| RMC1 | -0.210284286 | 0.864366889 | 0.224466623 | 0.407204268 | 210.3756664 | 182.0866269 | ENSG00000141452 |
| NAA60 | -0.210318072 | 0.864346647 | 0.037107179 | 0.105855696 | 736.546681 | 637.1838635 | ENSG00000122390 |
| RFTN2 | -0.21036421 | 0.864319006 | 0.699176255 | 0.828444746 | 21.69118306 | 18.75751018 | ENSG00000162944 |
| FYN | -0.2104174 | 0.86428714 | 0.006136961 | 0.024555792 | 1656.078766 | 1431.306491 | ENSG00000010810 |
| ZMYM6 | -0.210713795 | 0.864109594 | 0.508155847 | 0.687356625 | 63.04294951 | 54.3630968 | ENSG00000163867 |
| SEC61A1 | -0.210724919 | 0.864102931 | 0.000949124 | 0.005036091 | 4946.898695 | 4274.316646 | ENSG00000058262 |
| PICK1 | -0.210815466 | 0.8640487 | 0.041388388 | 0.115630446 | 602.8179474 | 521.0318149 | ENSG00000100151 |
| MRPS12 | -0.210841286 | 0.864033236 | 0.016163984 | 0.054203597 | 1111.894338 | 960.7761957 | ENSG00000128626 |
| NA | -0.210885503 | 0.864006755 | 0.74598261 | 0.858280569 | 12.75076059 | 11.01606101 | ENSG00000272205 |

| | | | | | | | |
|---|---|---|---|---|---|---|---|
| EIF2B2 | -0.210897507 | 0.863999566 | 0.036838965 | 0.105285025 | 692.2184744 | 598.3381239 | ENSG00000119718 |
| PHKG2 | -0.210943753 | 0.863971871 | 0.097482861 | 0.222286829 | 390.5101092 | 337.2668433 | ENSG00000156873 |
| THEMIS2 | -0.211223086 | 0.863804606 | 0.125037584 | 0.267794977 | 322.1239988 | 278.0334801 | ENSG00000130775 |
| NFYB | -0.211306245 | 0.863754816 | 0.036507453 | 0.104590848 | 864.6861877 | 746.4640201 | ENSG00000120837 |
| ACVR1B | -0.211360607 | 0.86372227 | 0.012168377 | 0.043262747 | 985.4275428 | 851.0922265 | ENSG00000135503 |
| LSM10 | -0.211512436 | 0.863631377 | 0.029733998 | 0.088928499 | 796.0990648 | 688.0356916 | ENSG00000181817 |
| EIF4E | -0.211559333 | 0.863603304 | 0.009739807 | 0.03596222 | 2046.138 | 1766.333668 | ENSG00000151247 |
| CCDC43 | -0.211655945 | 0.863545473 | 0.140879957 | 0.29221444 | 271.6225294 | 234.5898898 | ENSG00000180329 |
| VAC14 | -0.211718691 | 0.863507916 | 0.008696346 | 0.032793098 | 1359.16253 | 1173.690501 | ENSG00000103043 |
| SIMC1 | -0.21174793 | 0.863490416 | 0.092090385 | 0.213071568 | 363.027063 | 313.5061727 | ENSG00000170085 |
| RTN2 | -0.212108859 | 0.863274417 | 0.141734737 | 0.293589338 | 276.5415292 | 238.9186085 | ENSG00000125744 |
| UBB | -0.212150921 | 0.863249249 | 0.003994986 | 0.017009664 | 6428.339378 | 5549.734731 | ENSG00000170315 |
| NA | -0.212178324 | 0.863232853 | 0.69351292 | 0.82436945 | 17.14384176 | 14.81940671 | ENSG00000263968 |
| KDM1A | -0.212254722 | 0.863187141 | 0.000765396 | 0.004173538 | 7929.098331 | 6843.751901 | ENSG00000004487 |
| NA | -0.212305172 | 0.863156956 | 0.685920708 | 0.819285583 | 17.88440783 | 15.39316863 | ENSG00000269044 |
| PLAGL1 | -0.212464044 | 0.863061909 | 0.705755839 | 0.832939297 | 16.63763872 | 14.29403366 | ENSG00000118495 |
| C9orf40 | -0.21264712 | 0.862952395 | 0.161452932 | 0.322210472 | 315.9641377 | 272.4713151 | ENSG00000135045 |
| APBA2 | -0.212656829 | 0.862946587 | 0.02020402 | 0.065157701 | 806.4120218 | 695.8771677 | ENSG00000034053 |
| FBLN1 | -0.212700125 | 0.86292069 | 0.000311915 | 0.001896969 | 5397.701626 | 4657.365062 | ENSG00000077942 |
| PRPF8 | -0.212798375 | 0.862861926 | 0.000980376 | 0.005176721 | 10999.45708 | 9490.893024 | ENSG00000174231 |
| EMC3 | -0.21280065 | 0.862860566 | 0.076497644 | 0.185203324 | 564.5056843 | 487.6523958 | ENSG00000125037 |
| NA | -0.212884509 | 0.862810412 | 0.716843688 | 0.839556455 | 14.34759347 | 12.36637857 | ENSG00000276449 |
| EIF1AD | -0.212891119 | 0.862806459 | 0.043707581 | 0.120511204 | 593.3890833 | 511.5768606 | ENSG00000175376 |
| ZNF142 | -0.212975742 | 0.862755851 | 0.048848946 | 0.131323367 | 644.3487105 | 555.9747457 | ENSG00000115568 |
| CTSA | -0.21304657 | 0.862713496 | 0.034462074 | 0.099940553 | 620.7000928 | 535.5986133 | ENSG00000064601 |
| ITGA4 | -0.213047667 | 0.86271284 | 0.408150546 | 0.599941488 | 86.26457259 | 74.29102709 | ENSG00000115232 |
| HPF1 | -0.213060218 | 0.862705335 | 0.063781311 | 0.161383985 | 453.1264323 | 390.6518017 | ENSG00000056050 |
| NA | -0.213095676 | 0.862684131 | 0.801122575 | 0.893214577 | 7.919147018 | 6.82264969 | ENSG00000273004 |
| PEX13 | -0.213169361 | 0.862640072 | 0.118380605 | 0.257036801 | 450.8464138 | 388.5771162 | ENSG00000162928 |
| FAT2 | -0.213230748 | 0.862603367 | 0.561113017 | 0.730002144 | 49.85551302 | 43.14968144 | ENSG00000086570 |
| VPS36 | -0.213306695 | 0.862557959 | 0.118924637 | 0.257885604 | 332.4980269 | 286.6272301 | ENSG00000136100 |
| NA | -0.213353022 | 0.862530261 | 0.169624309 | 0.334402711 | 375.2814746 | 324.1746485 | ENSG00000264538 |
| ACP1 | -0.213511874 | 0.862435295 | 0.001572693 | 0.007739855 | 2562.506726 | 2209.739532 | ENSG00000143727 |
| SMC2 | -0.213588796 | 0.862389313 | 0.034981591 | 0.101193409 | 1297.478081 | 1118.232153 | ENSG00000136824 |
| MEX3B | -0.213600735 | 0.862382176 | 0.053457854 | 0.140637028 | 588.340928 | 507.7151552 | ENSG00000183496 |
| PRDM10 | -0.213619991 | 0.862370665 | 0.08475134 | 0.200048566 | 460.1407787 | 396.5997954 | ENSG00000170325 |
| PHF3 | -0.21374239 | 0.862297505 | 0.005020084 | 0.02065769 | 1612.580533 | 1389.896158 | ENSG00000118482 |
| SOCS2 | -0.213749797 | 0.862293077 | 0.116840473 | 0.254677328 | 327.2380055 | 282.1607192 | ENSG00000120833 |
| HNRNPAB | -0.213784456 | 0.862272362 | 0.000746259 | 0.004082287 | 16621.20725 | 14331.69807 | ENSG00000197451 |
| SRSF11 | -0.213785092 | 0.862271982 | 0.003322728 | 0.014517547 | 3538.720758 | 3052.105891 | ENSG00000116754 |
| ABCC6P1 | -0.214163475 | 0.862045859 | 0.850937435 | 0.921239019 | 4.102397381 | 3.589705101 | ENSG00000256340 |
| TGFB1 | -0.214177528 | 0.862037462 | 0.144702357 | 0.298056622 | 285.6234114 | 246.2953646 | ENSG00000105329 |
| UTP25 | -0.214272916 | 0.861980468 | 0.062078157 | 0.15813037 | 490.3849638 | 422.642079 | ENSG00000117597 |
| CCDC115 | -0.214292712 | 0.86196864 | 0.121528286 | 0.262120941 | 337.1737118 | 290.4693203 | ENSG00000136710 |
| UBR1 | -0.214325859 | 0.861948836 | 0.057196759 | 0.148388122 | 492.9091767 | 424.8041295 | ENSG00000159459 |
| ASCL2 | -0.214377709 | 0.861917858 | 0.766545564 | 0.870835943 | 11.97922676 | 10.23906734 | ENSG00000183734 |
| NA | -0.214434047 | 0.861884201 | 0.582926015 | 0.746490069 | 42.44572596 | 36.66448006 | ENSG00000213025 |
| TBC1D4 | -0.214549238 | 0.861815387 | 0.009188731 | 0.034307806 | 1190.827272 | 1026.581241 | ENSG00000136111 |
| GABRA2 | -0.214716103 | 0.861715713 | 0.672858226 | 0.809789244 | 22.26792057 | 19.07249183 | ENSG00000151834 |
| PRIMA1 | -0.214732024 | 0.861706204 | 0.325838614 | 0.520028663 | 132.5700649 | 114.3240856 | ENSG00000175785 |
| NA | -0.214732862 | 0.861705703 | 0.698374784 | 0.828043791 | 16.46896951 | 14.26072855 | ENSG00000237214 |
| KIAA1841 | -0.214810926 | 0.861659078 | 0.172671086 | 0.338941528 | 250.7417326 | 216.4788524 | ENSG00000162929 |
| ZKSCAN4 | -0.214899667 | 0.861606078 | 0.235535045 | 0.420544675 | 174.0853442 | 150.0092079 | ENSG00000187626 |
| NAA15 | -0.215009974 | 0.861540203 | 0.014996002 | 0.05107238 | 2778.290833 | 2392.723295 | ENSG00000164134 |
| JPT1 | -0.21508293 | 0.861496637 | 0.001112014 | 0.005779606 | 4321.135707 | 3721.860812 | ENSG00000189159 |
| THG1L | -0.215242044 | 0.861401628 | 0.115930551 | 0.253170359 | 296.7739868 | 255.7218717 | ENSG00000113272 |
| UPF3A | -0.215260531 | 0.86139059 | 0.008672857 | 0.03273215 | 1583.071983 | 1364.253443 | ENSG00000169062 |
| PPP4C | -0.215297818 | 0.861368328 | 0.000801723 | 0.004349514 | 4460.696291 | 3842.555725 | ENSG00000149923 |
| NA | -0.215437368 | 0.861285012 | 0.771416112 | 0.874346286 | 9.299859295 | 8.030611782 | ENSG00000287837 |
| ZNF266 | -0.215439431 | 0.861283781 | 0.136730916 | 0.28570133 | 315.4025235 | 271.8065412 | ENSG00000174652 |
| MEGF11 | -0.215492254 | 0.861252246 | 0.43158743 | 0.621587546 | 74.69539096 | 64.49943766 | ENSG00000157890 |
| NA | -0.215521831 | 0.86123459 | 0.703550024 | 0.831394807 | 16.65026638 | 14.26397775 | ENSG00000232499 |
| SERPINF1 | -0.215541122 | 0.861223074 | 0.08474862 | 0.200048566 | 392.8166154 | 338.2054099 | ENSG00000132386 |
| NA | -0.2155477 | 0.861219147 | 0.730935138 | 0.849457728 | 14.51320347 | 12.58414443 | ENSG00000271966 |
| COQ8B | -0.215602372 | 0.861186512 | 0.016281138 | 0.054556973 | 923.1770228 | 794.8752445 | ENSG00000123815 |
| ADGRL1 | -0.215665898 | 0.861148592 | 0.003614026 | 0.015595426 | 4676.644347 | 4027.801492 | ENSG00000072071 |
| GLDC | -0.215713845 | 0.861119972 | 0.001121284 | 0.00580986 | 4182.212173 | 3600.56946 | ENSG00000178445 |
| SYNCRIP | -0.21581522 | 0.861059465 | 0.006697653 | 0.026417173 | 3798.382574 | 3269.519441 | ENSG00000135316 |
| DDX46 | -0.215818107 | 0.861057742 | 0.000811558 | 0.004395504 | 3365.712898 | 2898.595846 | ENSG00000145833 |
| PCDH19 | -0.216016746 | 0.860939195 | 0.495395129 | 0.677212961 | 59.9943814 | 51.41887679 | ENSG00000165194 |
| TMEM11 | -0.216024869 | 0.860934348 | 0.029232834 | 0.087741109 | 681.5994476 | 586.73614 | ENSG00000178307 |
| DPAGT1 | -0.216076017 | 0.860903825 | 0.013210837 | 0.046145807 | 1005.918732 | 866.3416627 | ENSG00000172269 |
| PTPA | -0.216078766 | 0.860902185 | 0.000184452 | 0.001196581 | 5591.883553 | 4814.318959 | ENSG00000119383 |
| FAM131B | -0.216088189 | 0.860896562 | 0.023075871 | 0.072563318 | 1104.001829 | 950.2368788 | ENSG00000159784 |

| | | | | | | | |
|---|---|---|---|---|---|---|---|
| DRAP1 | -0.216115996 | 0.860879969 | 0.008282739 | 0.031526521 | 2903.543713 | 2499.577161 | ENSG00000175550 |
| GLDN | -0.216270775 | 0.860787614 | 0.714267043 | 0.838127596 | 14.7551581 | 12.6513043 | ENSG00000186417 |
| FAM124A | -0.216346383 | 0.860742504 | 0.009724808 | 0.03592577 | 1422.20908 | 1223.830193 | ENSG00000150510 |
| UTP18 | -0.216348429 | 0.860741283 | 0.004775616 | 0.019771391 | 1961.850247 | 1687.796595 | ENSG00000011260 |
| KCTD1 | -0.216386031 | 0.860718849 | 0.101803265 | 0.229708025 | 326.3538527 | 280.9043682 | ENSG00000134504 |
| DBR1 | -0.216418392 | 0.860699543 | 0.037201759 | 0.106092846 | 616.9367927 | 530.6273323 | ENSG00000138231 |
| SCP2 | -0.216437363 | 0.860688225 | 0.039064902 | 0.110488158 | 652.5182907 | 561.2931195 | ENSG00000116171 |
| CCNB3 | -0.216502693 | 0.860649251 | 0.856938768 | 0.924099792 | 4.327387365 | 3.735747796 | ENSG00000147082 |
| SLC7A11 | -0.216504723 | 0.86064804 | 0.135665623 | 0.284151041 | 448.3732 | 385.5697097 | ENSG00000151012 |
| TNFRSF11B | -0.21652036 | 0.860638712 | 0.740718148 | 0.855075943 | 12.07588833 | 10.43112579 | ENSG00000164761 |
| FBXW11 | -0.216675225 | 0.860546332 | 0.022161325 | 0.070307771 | 1464.48555 | 1259.332526 | ENSG00000072803 |
| VAMP1 | -0.216817108 | 0.860461705 | 0.436808339 | 0.625932098 | 128.1311184 | 110.542912 | ENSG00000139190 |
| NEIL1 | -0.216884632 | 0.860421433 | 0.388650463 | 0.581422849 | 100.756685 | 86.88094574 | ENSG00000140398 |
| C1orf210 | -0.216974461 | 0.860367861 | 0.288557937 | 0.480179616 | 129.7962983 | 111.6876876 | ENSG00000253313 |
| FAM104B | -0.216987258 | 0.86036023 | 0.260181166 | 0.449102778 | 140.0852852 | 120.5165487 | ENSG00000182518 |
| GORASP1 | -0.217128842 | 0.860275799 | 0.020040187 | 0.064674357 | 1010.385534 | 869.3156037 | ENSG00000114745 |
| LOC100129534 | -0.217228463 | 0.860216398 | 0.681137614 | 0.815727872 | 20.66547292 | 17.8730035 | ENSG00000269896 |
| ARHGEF40 | -0.217348403 | 0.860144885 | 0.004775899 | 0.019771391 | 2200.720734 | 1893.797273 | ENSG00000165801 |
| CD276 | -0.217426467 | 0.860098344 | 0.000822846 | 0.004447545 | 4199.17513 | 3611.610985 | ENSG00000103855 |
| NA | -0.217431471 | 0.860095361 | 0.819464266 | 0.90291604 | 5.670818048 | 4.924938894 | ENSG00000204620 |
| RAB36 | -0.217451245 | 0.860083572 | 0.554581017 | 0.724709888 | 40.44814594 | 34.79996275 | ENSG00000100228 |
| RPUSD2 | -0.217566403 | 0.860014922 | 0.036386654 | 0.104443277 | 782.2940389 | 672.945232 | ENSG00000166133 |
| NA | -0.217570215 | 0.86001265 | 0.550791091 | 0.721876151 | 38.14802858 | 32.87979374 | ENSG00000218226 |
| PI4KA | -0.217703675 | 0.859933096 | 0.01719285 | 0.056973409 | 1555.972554 | 1339.167309 | ENSG00000241973 |
| IL4R | -0.217706734 | 0.859931272 | 0.024182507 | 0.075352821 | 852.6846898 | 732.9318879 | ENSG00000077238 |
| NA | -0.217810663 | 0.859869327 | 0.684822203 | 0.818342663 | 18.28753777 | 15.6330663 | ENSG00000279738 |
| LOC101927751 | -0.217968814 | 0.859775071 | 0.826650068 | 0.906810387 | 5.59121881 | 4.84605607 | ENSG00000259583 |
| TANC1 | -0.217971454 | 0.859773498 | 0.130615194 | 0.276628241 | 314.4696038 | 270.0810898 | ENSG00000115183 |
| ATP6V0E1 | -0.217986834 | 0.859764332 | 0.008737107 | 0.032920014 | 1379.253287 | 1186.104304 | ENSG00000113732 |
| PCMTD1 | -0.21813743 | 0.85967459 | 0.109228165 | 0.241918074 | 320.9921563 | 276.220976 | ENSG00000168300 |
| NA | -0.218214012 | 0.859628958 | 0.27548467 | 0.466422139 | 142.6532889 | 122.530796 | ENSG00000188002 |
| GNAI2 | -0.218405286 | 0.859514995 | 0.000327206 | 0.001979256 | 6764.422572 | 5813.926876 | ENSG00000114353 |
| NA | -0.218409347 | 0.859512575 | 0.488479069 | 0.671323674 | 49.7164361 | 42.74420564 | ENSG00000229132 |
| NA | -0.218437814 | 0.859495616 | 0.778766556 | 0.878494178 | 7.956428612 | 6.818850843 | ENSG00000231881 |
| ZMYND19 | -0.218484024 | 0.859468087 | 0.017444212 | 0.057633768 | 1084.698293 | 931.8341791 | ENSG00000165724 |
| NA | -0.218507956 | 0.85945383 | 0.835861696 | 0.911983321 | 4.389924263 | 3.792060779 | ENSG00000272370 |
| ZSCAN18 | -0.218556378 | 0.859424983 | 0.155303995 | 0.313415096 | 325.5728482 | 280.3604877 | ENSG00000121413 |
| UBE2R2 | -0.218557663 | 0.859424218 | 0.001423334 | 0.007079996 | 2170.965006 | 1865.740319 | ENSG00000107341 |
| NUP153 | -0.218619575 | 0.859387337 | 0.005711352 | 0.023071971 | 2249.933914 | 1933.00802 | ENSG00000124789 |
| MTMR4 | -0.218650097 | 0.859369156 | 0.036233322 | 0.104067625 | 773.0518764 | 664.0894609 | ENSG00000108389 |
| NA | -0.21868259 | 0.859349801 | 0.514692746 | 0.692846128 | 48.56948549 | 41.5845208 | ENSG00000179967 |
| KLHL7 | -0.218831779 | 0.85926094 | 0.006538921 | 0.025873688 | 1712.731106 | 1470.791757 | ENSG00000122550 |
| GPRC5C | -0.218836373 | 0.859258204 | 0.13968922 | 0.290574485 | 296.1733694 | 254.5055475 | ENSG00000170412 |
| UNC5CL | -0.219056379 | 0.859127181 | 0.629820413 | 0.781581472 | 32.28073047 | 27.83897968 | ENSG00000124602 |
| RBM11 | -0.219108448 | 0.859096174 | 0.541619217 | 0.714474773 | 45.37070862 | 38.83896924 | ENSG00000185272 |
| MNS1 | -0.219111577 | 0.859094311 | 0.483851036 | 0.667139941 | 55.40294102 | 47.53774757 | ENSG00000138587 |
| NA | -0.219236989 | 0.859019634 | 0.806780988 | 0.896600656 | 5.988034921 | 5.157350486 | ENSG00000217835 |
| UBL4A | -0.219422718 | 0.858909053 | 0.020529468 | 0.06600055 | 952.2057899 | 817.3849632 | ENSG00000102178 |
| NA | -0.219426443 | 0.858906835 | 0.824832322 | 0.905753491 | 4.780426548 | 4.092070275 | ENSG00000230832 |
| HAUS4 | -0.219448024 | 0.858893987 | 0.324323333 | 0.51850329 | 111.3305229 | 95.4006624 | ENSG00000092036 |
| ASIC2 | -0.219704322 | 0.858741416 | 0.604588268 | 0.763311161 | 27.27659171 | 23.47260733 | ENSG00000108684 |
| PAM16 | -0.219720734 | 0.858731647 | 0.448833331 | 0.636422382 | 60.77370223 | 52.1508424 | ENSG00000217930 |
| DUS3L | -0.219722535 | 0.858730575 | 0.011027216 | 0.039962922 | 1079.037374 | 926.9379022 | ENSG00000141994 |
| MRPL20-AS1 | -0.219782159 | 0.858695086 | 0.076513535 | 0.185203324 | 480.3507546 | 413.0711153 | ENSG00000224870 |
| LIN7B | -0.219996558 | 0.858567485 | 0.507109365 | 0.686542364 | 45.87187561 | 39.44746199 | ENSG00000104863 |
| EXOSC4 | -0.22006481 | 0.858526868 | 0.163709604 | 0.325662102 | 229.9118298 | 197.6832505 | ENSG00000178896 |
| POU5F1B | -0.220182224 | 0.858456999 | 0.226711327 | 0.410182097 | 166.7897847 | 143.2548173 | ENSG00000212993 |
| NA | -0.22019453 | 0.858449677 | 0.620862751 | 0.774962921 | 24.46140194 | 21.02326641 | ENSG00000196741 |
| GTF2F1 | -0.220216868 | 0.858436385 | 0.003584988 | 0.015484543 | 2619.466782 | 2248.345947 | ENSG00000125651 |
| MINCR | -0.220218303 | 0.858435532 | 0.470844821 | 0.656230415 | 56.70415175 | 48.73073752 | ENSG00000253716 |
| LAMP1 | -0.220304422 | 0.85838429 | 0.000436122 | 0.002539454 | 4429.995497 | 3802.106852 | ENSG00000185896 |
| SRC | -0.220443052 | 0.858301811 | 0.004964824 | 0.020457546 | 1304.164553 | 1119.801792 | ENSG00000197122 |
| SRRM4 | -0.220501529 | 0.858267022 | 0.410451691 | 0.602082441 | 74.08749029 | 63.73416694 | ENSG00000139767 |
| NA | -0.220574664 | 0.858223515 | 0.742145477 | 0.855831525 | 11.61841446 | 10.02217755 | ENSG00000238058 |
| ZNF419 | -0.220580925 | 0.858219791 | 0.24659282 | 0.434196401 | 165.6528085 | 141.9826182 | ENSG00000105136 |
| PAX9 | -0.220715791 | 0.858139566 | 0.852816968 | 0.922280546 | 4.916248858 | 4.215981162 | ENSG00000198807 |
| HIC1 | -0.220795351 | 0.858092244 | 0.8115375 | 0.898880668 | 6.623746445 | 5.637145819 | ENSG00000177374 |
| HYAL2 | -0.220807986 | 0.858084729 | 0.015987589 | 0.053816558 | 837.8681749 | 719.1061901 | ENSG00000068001 |
| ZNF75A | -0.220908855 | 0.858024736 | 0.219551958 | 0.40098305 | 177.5540069 | 152.1721973 | ENSG00000162086 |
| MFNG | -0.22099535 | 0.857973296 | 0.434126729 | 0.623909514 | 67.17245869 | 57.55568824 | ENSG00000100060 |
| MBD4 | -0.22101087 | 0.857964066 | 0.044388022 | 0.12200617 | 592.0416013 | 507.4209913 | ENSG00000129071 |
| COL2A1 | -0.221050643 | 0.857940414 | 0.076000895 | 0.184347676 | 402.5312476 | 345.7612277 | ENSG00000139219 |
| PRPF3 | -0.221093504 | 0.857914926 | 0.011929743 | 0.042564646 | 1176.975261 | 1010.135002 | ENSG00000117360 |
| MRPS26 | -0.221115589 | 0.857901792 | 0.003070047 | 0.013583254 | 2022.620852 | 1735.153332 | ENSG00000125901 |

| | | | | | | | |
|---|---|---|---|---|---|---|---|
| RIMS2 | -0.221235149 | 0.857830699 | 0.064593474 | 0.162903178 | 412.797144 | 354.1653956 | ENSG00000176406 |
| DHRS11 | -0.221274213 | 0.857807472 | 0.036203809 | 0.104014444 | 691.8020329 | 593.7462135 | ENSG00000278535 |
| NA | -0.221317236 | 0.857781891 | 0.835595926 | 0.911854386 | 5.392761905 | 4.685041227 | ENSG00000272502 |
| RBM39 | -0.221354051 | 0.857760002 | 0.002096746 | 0.009895873 | 4704.700666 | 4036.74312 | ENSG00000131051 |
| KIF3A | -0.221366751 | 0.857752451 | 0.117932333 | 0.256243595 | 292.1535555 | 250.8426221 | ENSG00000131437 |
| RNPEPL1 | -0.221372274 | 0.857749168 | 0.110260434 | 0.243775127 | 344.056385 | 295.0328238 | ENSG00000142327 |
| HYOU1 | -0.221421714 | 0.857719774 | 5.95E-05 | 0.000439159 | 9210.076964 | 7899.848665 | ENSG00000149428 |
| PPP1R7 | -0.221481018 | 0.857684517 | 0.011270561 | 0.040609365 | 980.5317012 | 840.7892485 | ENSG00000115685 |
| PWWP2A | -0.221485759 | 0.857681698 | 0.00612832 | 0.024526515 | 1311.245097 | 1124.944179 | ENSG00000170234 |
| SCAP | -0.221633623 | 0.857593797 | 0.001167797 | 0.006015549 | 3528.919123 | 3026.74597 | ENSG00000114650 |
| ZNF542P | -0.221678787 | 0.857566951 | 0.500867879 | 0.68147741 | 60.12683623 | 51.59921228 | ENSG00000240225 |
| CARMIL2 | -0.221689236 | 0.85756074 | 0.02795087 | 0.084855271 | 986.4022241 | 846.2457642 | ENSG00000159753 |
| DCHS2 | -0.221710588 | 0.857548048 | 0.548427994 | 0.719971981 | 43.73975106 | 37.48160364 | ENSG00000197410 |
| NA | -0.221790719 | 0.857500419 | 0.826036301 | 0.906395244 | 4.757050382 | 4.06201436 | ENSG00000275318 |
| ZNF107 | -0.221848632 | 0.857465997 | 0.136543757 | 0.28539026 | 411.8706585 | 352.8479048 | ENSG00000196247 |
| NA | -0.222020342 | 0.857363948 | 0.807345938 | 0.896619594 | 5.984878009 | 5.138579492 | ENSG00000259032 |
| TMEM150C | -0.222039784 | 0.857352394 | 0.459418991 | 0.646241013 | 60.45332844 | 51.78334662 | ENSG00000249242 |
| IKZF1 | -0.222139532 | 0.857293118 | 0.561467138 | 0.730313141 | 42.62880426 | 36.50269233 | ENSG00000185811 |
| SUDS3 | -0.222181845 | 0.857267975 | 0.003738345 | 0.016068293 | 2146.749775 | 1839.644836 | ENSG00000111707 |
| TBC1D13 | -0.222277982 | 0.857210851 | 0.025735468 | 0.079219593 | 912.8319633 | 782.7754587 | ENSG00000107021 |
| RPS29 | -0.222366313 | 0.857158369 | 0.007531141 | 0.02910387 | 3820.409606 | 3274.299989 | ENSG00000213741 |
| SLC41A2 | -0.222367526 | 0.857157648 | 0.577485789 | 0.742035609 | 35.33609343 | 30.2496793 | ENSG00000136052 |
| NDUFB7 | -0.222392042 | 0.857143083 | 0.012366647 | 0.043841512 | 1396.548675 | 1197.194255 | ENSG00000099795 |
| EXTL2 | -0.222459017 | 0.857103292 | 0.038906858 | 0.11010837 | 705.0149049 | 604.2215711 | ENSG00000162694 |
| H3C2 | -0.222482039 | 0.857089614 | 0.82621048 | 0.90646925 | 5.088172683 | 4.313196947 | ENSG00000286522 |
| TOMM34 | -0.222523313 | 0.857065095 | 0.010970591 | 0.039790399 | 1120.347462 | 960.6448306 | ENSG00000025772 |
| RPUSD4 | -0.222558614 | 0.857044123 | 0.007087143 | 0.027694099 | 1074.37237 | 920.722646 | ENSG00000165526 |
| PTOV1 | -0.222649837 | 0.856989933 | 0.05122506 | 0.136271747 | 952.1751832 | 816.8072138 | ENSG00000104960 |
| LYL1 | -0.222723717 | 0.856946048 | 0.751528504 | 0.861772523 | 12.64650741 | 10.79829515 | ENSG00000104903 |
| PWWP3A | -0.222912073 | 0.856834174 | 0.048790018 | 0.131225395 | 820.5149323 | 703.8396008 | ENSG00000160953 |
| NIPA2 | -0.222972835 | 0.856798088 | 0.005998119 | 0.024031375 | 1861.674388 | 1594.571522 | ENSG00000140157 |
| GMPR2 | -0.222986129 | 0.856790192 | 0.003856861 | 0.016531699 | 1421.146561 | 1217.179008 | ENSG00000100938 |
| SMIM26 | -0.223030643 | 0.856763757 | 0.50768379 | 0.68692604 | 51.44583668 | 43.94424362 | ENSG00000232388 |
| DNAAF5 | -0.223078728 | 0.856735201 | 0.014251821 | 0.048969828 | 1420.139109 | 1216.39581 | ENSG00000164818 |
| ASMT | -0.223269776 | 0.856621756 | 0.742009048 | 0.85578069 | 12.68506678 | 10.82455222 | ENSG00000196433 |
| FGF12 | -0.223344779 | 0.856577223 | 0.159894984 | 0.320170013 | 224.3415293 | 192.0924066 | ENSG00000114279 |
| NA | -0.223374562 | 0.85655954 | 0.374458246 | 0.567389377 | 89.43338902 | 76.61028514 | ENSG00000203761 |
| RSBN1L | -0.223650404 | 0.856395782 | 0.07435974 | 0.181411534 | 471.1675656 | 403.3955352 | ENSG00000187257 |
| MAGOHB | -0.223682547 | 0.856376703 | 0.067119671 | 0.167677044 | 441.5918412 | 378.0712329 | ENSG00000111196 |
| CCAR1 | -0.223867489 | 0.856266929 | 0.00067493 | 0.00373296 | 2387.868742 | 2044.650211 | ENSG00000060339 |
| HMGA1 | -0.223909631 | 0.856241917 | 0.000556245 | 0.003151144 | 51935.05762 | 44468.60265 | ENSG00000137309 |
| DIXDC1 | -0.223971496 | 0.856205201 | 0.113742544 | 0.249538862 | 321.2627185 | 274.8901536 | ENSG00000150764 |
| GNG11 | -0.224038714 | 0.85616531 | 0.510929671 | 0.690100169 | 46.97009706 | 40.28412946 | ENSG00000127920 |
| AARS2 | -0.224235194 | 0.856048716 | 0.01681876 | 0.055994055 | 839.2993978 | 718.7505205 | ENSG00000124608 |
| ZNF598 | -0.224337286 | 0.855988141 | 0.013389696 | 0.046647487 | 1332.00021 | 1140.783065 | ENSG00000167962 |
| NA | -0.224379199 | 0.855963273 | 0.194653481 | 0.368810839 | 235.8849746 | 202.2849488 | ENSG00000277200 |
| POPDC3 | -0.224426018 | 0.855935495 | 0.503487973 | 0.683776217 | 50.85835067 | 43.35452187 | ENSG00000132429 |
| SLC23A2 | -0.22458447 | 0.855841493 | 0.029171981 | 0.087586828 | 633.4263722 | 542.4296252 | ENSG00000089057 |
| IDH3A | -0.224804845 | 0.855710771 | 0.058233868 | 0.150446964 | 418.5051686 | 358.0054072 | ENSG00000166411 |
| MDFIC | -0.224887297 | 0.855661867 | 0.161181716 | 0.32187716 | 251.1399242 | 214.7388669 | ENSG00000135272 |
| ARHGDIA | -0.224890057 | 0.85566023 | 0.001974028 | 0.009388409 | 3621.997359 | 3099.432926 | ENSG00000141522 |
| ANKRD9 | -0.224922285 | 0.855641116 | 0.012959449 | 0.045481997 | 1035.52038 | 886.5387108 | ENSG00000156381 |
| OSTC | -0.22501713 | 0.855584867 | 0.011400862 | 0.041013893 | 1918.769779 | 1640.781052 | ENSG00000198856 |
| IPO8 | -0.225087449 | 0.855543165 | 0.011080037 | 0.04010887 | 1299.184602 | 1110.60674 | ENSG00000133704 |
| TRUB2 | -0.225240689 | 0.855452296 | 0.028360645 | 0.08577614 | 651.7272367 | 557.8232403 | ENSG00000167112 |
| C5orf30 | -0.225302421 | 0.855415693 | 0.083914275 | 0.19862927 | 384.9193562 | 329.2321063 | ENSG00000181751 |
| TNFAIP3 | -0.225336759 | 0.855395333 | 0.484505013 | 0.667842921 | 49.43078836 | 42.23760358 | ENSG00000118503 |
| SMARCAD1 | -0.225363161 | 0.855379679 | 0.01179836 | 0.042173731 | 1339.451534 | 1145.220203 | ENSG00000163104 |
| UBIAD1 | -0.225451396 | 0.855327366 | 0.034716789 | 0.100584799 | 641.7914705 | 548.4450734 | ENSG00000120942 |
| C12orf75 | -0.22549038 | 0.855304254 | 0.124087038 | 0.266276388 | 437.8896122 | 374.5466578 | ENSG00000235162 |
| BAZ1A | -0.225545173 | 0.85527177 | 0.03393406 | 0.098857884 | 588.0586477 | 503.0411757 | ENSG00000198604 |
| ALPL | -0.225708081 | 0.855175199 | 8.00E-05 | 0.000570016 | 5463.091693 | 4671.986502 | ENSG00000162551 |
| OR7E12P | -0.225750387 | 0.855150122 | 0.354334749 | 0.547662123 | 106.1659079 | 90.97659162 | ENSG00000189398 |
| FAXC | -0.225795801 | 0.855123203 | 0.165768877 | 0.328805691 | 209.8808883 | 179.5918199 | ENSG00000146267 |
| EEFSEC | -0.225838822 | 0.855097704 | 0.099873212 | 0.22664979 | 349.2363786 | 298.825649 | ENSG00000132394 |
| AARSD1 | -0.225884961 | 0.855070358 | 0.749226988 | 0.860197794 | 11.32013907 | 9.740939048 | ENSG00000266967 |
| RPGRIP1L | -0.225923419 | 0.855047564 | 0.166368659 | 0.329642583 | 224.1480107 | 191.5189795 | ENSG00000103494 |
| GNRHR2 | -0.225933428 | 0.855041632 | 0.707370062 | 0.833517498 | 13.63987431 | 11.66490691 | ENSG00000211451 |
| GRB10 | -0.226025515 | 0.854987057 | 0.001934953 | 0.009245823 | 2884.024666 | 2465.03073 | ENSG00000106070 |
| SF3A1 | -0.226079529 | 0.854955047 | 0.002479833 | 0.011399114 | 4233.302188 | 3618.809344 | ENSG00000099995 |
| SMN1 | -0.226231506 | 0.854864989 | 0.306527984 | 0.500589448 | 152.7571456 | 130.3039013 | ENSG00000172062 |
| ENTPD1 | -0.226247847 | 0.854855306 | 0.227955569 | 0.411545735 | 173.8218925 | 148.2933296 | ENSG00000138185 |
| RFLNA | -0.22641013 | 0.854759152 | 0.239628112 | 0.425490269 | 208.9526365 | 178.3214442 | ENSG00000178882 |
| RBM10 | -0.226410161 | 0.854759134 | 0.001180078 | 0.006070378 | 2191.242727 | 1873.289088 | ENSG00000182872 |

| | | | | | | | |
|---|---|---|---|---|---|---|---|
| SCAND1 | -0.226456846 | 0.854731474 | 0.023675652 | 0.07413491 | 972.2835889 | 831.0820695 | ENSG00000171222 |
| NUTM2B-AS1 | -0.226600549 | 0.854646341 | 0.255540413 | 0.444420195 | 157.0647197 | 134.3086553 | ENSG00000225484 |
| ITGAE | -0.226618747 | 0.854635561 | 0.1193337 | 0.258470352 | 381.9618501 | 326.7126198 | ENSG00000083457 |
| LIPE | -0.226689713 | 0.854593522 | 0.154640139 | 0.312381508 | 231.2718418 | 197.666303 | ENSG00000079435 |
| ZNF823 | -0.226734043 | 0.854567264 | 0.469850043 | 0.655539918 | 54.98472557 | 46.95594992 | ENSG00000197933 |
| SHROOM2 | -0.226856065 | 0.854494988 | 0.020466924 | 0.065890911 | 857.2560988 | 732.4169555 | ENSG00000146950 |
| DUSP23 | -0.226898051 | 0.85447012 | 0.048336458 | 0.130249192 | 575.4181459 | 491.8229907 | ENSG00000158716 |
| TRAF5 | -0.226908044 | 0.854464202 | 0.100006074 | 0.226727797 | 322.791084 | 276.1391704 | ENSG00000082512 |
| NA | -0.226929264 | 0.854451634 | 0.650783114 | 0.795936006 | 26.50298335 | 22.52711274 | ENSG00000214106 |
| MFSD4A | -0.226986428 | 0.854417779 | 0.82142329 | 0.904268191 | 5.789578017 | 4.981251877 | ENSG00000174514 |
| EIF4EBP2 | -0.226987077 | 0.854417394 | 0.00058198 | 0.003275558 | 6636.456268 | 5669.509902 | ENSG00000148730 |
| NA | -0.226992025 | 0.854414464 | 0.732048047 | 0.850484455 | 13.16967509 | 11.31282131 | ENSG00000287431 |
| CSGALNACT1 | -0.226999069 | 0.854410292 | 0.836888144 | 0.912351324 | 5.570901858 | 4.703374188 | ENSG00000147408 |
| FANCM | -0.227103986 | 0.854348159 | 0.159990216 | 0.320256903 | 219.8964643 | 187.677757 | ENSG00000187790 |
| GRIA1 | -0.227334434 | 0.854211701 | 0.622721029 | 0.776056452 | 25.96087419 | 22.18620044 | ENSG00000155511 |
| PSMD11 | -0.227419094 | 0.854161576 | 0.000480633 | 0.002766478 | 2996.912086 | 2559.141625 | ENSG00000108671 |
| DDX54 | -0.227422647 | 0.854159473 | 0.000345248 | 0.002069475 | 3322.891026 | 2838.808654 | ENSG00000123064 |
| LOC100128398 | -0.227485283 | 0.854122389 | 0.294622798 | 0.487504592 | 128.9180308 | 110.364838 | ENSG00000176593 |
| GMCL1 | -0.227608273 | 0.854049578 | 0.074427199 | 0.181486345 | 681.102001 | 581.1153199 | ENSG00000087338 |
| PARP14 | -0.227621341 | 0.854041842 | 0.076477103 | 0.185203324 | 366.0082688 | 312.5462556 | ENSG00000173193 |
| NEDD8 | -0.227653424 | 0.85402285 | 0.187602819 | 0.359383012 | 204.0064045 | 174.4093117 | ENSG00000129559 |
| NA | -0.227694372 | 0.853998611 | 0.800085543 | 0.892547578 | 6.017724914 | 5.138579492 | ENSG00000279880 |
| IER3IP1 | -0.227727919 | 0.853978753 | 0.473251031 | 0.658405892 | 92.19204745 | 78.53954891 | ENSG00000134049 |
| RANBP10 | -0.22776063 | 0.85395939 | 0.041718422 | 0.116412109 | 558.222343 | 476.9775333 | ENSG00000141084 |
| MMP25 | -0.227765202 | 0.853956684 | 0.042537177 | 0.118092277 | 480.7300047 | 410.6707758 | ENSG00000008516 |
| IDH1 | -0.227801478 | 0.853935212 | 0.000194249 | 0.001252246 | 9888.822144 | 8443.190437 | ENSG00000138413 |
| BLOC1S4 | -0.227992931 | 0.853821898 | 0.072330195 | 0.177629234 | 442.432053 | 378.1239073 | ENSG00000186222 |
| RXRA | -0.228014604 | 0.853809071 | 0.004750771 | 0.019676156 | 1555.62573 | 1328.430814 | ENSG00000186350 |
| RGS12 | -0.228060345 | 0.853782002 | 0.025577024 | 0.078862846 | 589.3830393 | 503.1611846 | ENSG00000159788 |
| POLR2H | -0.22812221 | 0.853745391 | 0.00447101 | 0.018705577 | 1386.481484 | 1183.903343 | ENSG00000163882 |
| NA | -0.228179751 | 0.853711341 | 0.756245112 | 0.864335767 | 9.164036985 | 7.80524823 | ENSG00000259577 |
| TCEAL9 | -0.228469586 | 0.853539848 | 0.005718035 | 0.023085366 | 1319.507994 | 1125.709798 | ENSG00000185222 |
| MAGT1 | -0.228579578 | 0.853474777 | 0.025381494 | 0.078351199 | 1045.675424 | 891.6568442 | ENSG00000102158 |
| HNRNPF | -0.228604532 | 0.853460014 | 4.63E-05 | 0.000351948 | 8912.262008 | 7605.530732 | ENSG00000169813 |
| HCG18 | -0.228827286 | 0.853328249 | 0.08178022 | 0.194922145 | 488.9360173 | 417.2996467 | ENSG00000231074 |
| BABAM1 | -0.228837337 | 0.853322304 | 0.186459353 | 0.357710139 | 223.6305555 | 191.2667379 | ENSG00000105393 |
| ILF2 | -0.22891318 | 0.853277446 | 0.000452028 | 0.002622189 | 8744.474024 | 7460.831173 | ENSG00000143621 |
| ACTR1A | -0.229039394 | 0.8532028 | 0.000507456 | 0.002905528 | 2395.168 | 2043.783751 | ENSG00000138107 |
| TSPAN15 | -0.229084034 | 0.853176401 | 0.082744427 | 0.196613489 | 338.1257306 | 288.6585281 | ENSG00000099282 |
| TMEM238 | -0.22914431 | 0.853140756 | 0.46705786 | 0.653162956 | 50.80084982 | 43.38176662 | ENSG00000233493 |
| ABTB1 | -0.229273236 | 0.853064519 | 0.324282404 | 0.518482581 | 100.8219728 | 86.13714556 | ENSG00000114626 |
| ENO1 | -0.229338139 | 0.853026143 | 5.15E-05 | 0.000386226 | 53845.73517 | 45930.69583 | ENSG00000074800 |
| TYW5 | -0.22934995 | 0.853019159 | 0.197618879 | 0.373422219 | 190.8082195 | 162.604096 | ENSG00000162971 |
| VIM-AS1 | -0.229370948 | 0.853006744 | 0.802811111 | 0.894151534 | 6.027195652 | 5.138579492 | ENSG00000229124 |
| MED20 | -0.229372547 | 0.853005798 | 0.022520444 | 0.071191184 | 951.5672676 | 811.0579571 | ENSG00000124641 |
| PDCL3 | -0.229404436 | 0.852986943 | 0.015598675 | 0.05271806 | 742.0807072 | 633.2854519 | ENSG00000115539 |
| PQBP1 | -0.229499927 | 0.852930487 | 0.014243681 | 0.04896001 | 1180.827638 | 1007.22849 | ENSG00000102103 |
| C14orf132 | -0.229508959 | 0.852925147 | 0.007162975 | 0.027908049 | 1242.33663 | 1060.104688 | ENSG00000227051 |
| RFFL | -0.229514421 | 0.852921918 | 0.143611414 | 0.296435455 | 237.2759469 | 202.4198182 | ENSG00000092871 |
| AKT2 | -0.229563582 | 0.852892854 | 0.001374036 | 0.006871663 | 2930.992959 | 2500.70121 | ENSG00000105221 |
| EXOSC9 | -0.229589004 | 0.852877826 | 0.011395725 | 0.041003371 | 965.0475297 | 823.261729 | ENSG00000123737 |
| ZNF213 | -0.229722674 | 0.852798808 | 0.152741242 | 0.309931737 | 234.1945393 | 199.8193217 | ENSG00000085644 |
| TOP1 | -0.229776758 | 0.852766839 | 0.000613486 | 0.00343204 | 3735.128231 | 3184.524429 | ENSG00000198900 |
| NOP9 | -0.229783621 | 0.852762782 | 0.009612391 | 0.035609845 | 943.5085248 | 804.6143601 | ENSG00000196943 |
| PPA1 | -0.229943439 | 0.85266832 | 0.001220564 | 0.006230221 | 2644.704206 | 2254.687073 | ENSG00000180817 |
| MTERF3 | -0.230070489 | 0.852593233 | 0.05637787 | 0.146668128 | 473.9658059 | 404.5447483 | ENSG00000156469 |
| NA | -0.23008022 | 0.852587483 | 0.840098818 | 0.914174887 | 4.082178128 | 3.514621124 | ENSG00000234134 |
| ZNF711 | -0.230081659 | 0.852586633 | 0.075983307 | 0.184329137 | 387.9389486 | 331.1183576 | ENSG00000147180 |
| ZNF782 | -0.230218307 | 0.852505882 | 0.360702908 | 0.553925441 | 85.33354719 | 72.70191117 | ENSG00000196597 |
| PAK1IP1 | -0.230242613 | 0.852491519 | 0.008511663 | 0.032227816 | 1134.59132 | 966.9924941 | ENSG00000111845 |
| ITPK1 | -0.230260862 | 0.852480736 | 0.000228779 | 0.001446671 | 3871.087713 | 3300.117863 | ENSG00000100605 |
| ANKRD52 | -0.230394793 | 0.8524016 | 0.002334965 | 0.010827181 | 1566.143082 | 1335.496929 | ENSG00000139645 |
| CUTA | -0.230553245 | 0.852307986 | 0.000263167 | 0.001642269 | 3099.113178 | 2641.451009 | ENSG00000112514 |
| KLHDC4 | -0.230576151 | 0.852294454 | 0.016527481 | 0.055262553 | 1235.256049 | 1053.770356 | ENSG00000104731 |
| FBXO43 | -0.230653509 | 0.852248755 | 0.595429099 | 0.755953244 | 27.48442166 | 23.39752335 | ENSG00000156509 |
| GUCD1 | -0.230776755 | 0.852175952 | 0.00305272 | 0.013525967 | 1350.269205 | 1150.871191 | ENSG00000138867 |
| ZNF322 | -0.230842839 | 0.852136918 | 0.126864646 | 0.270483072 | 253.1807088 | 215.5876954 | ENSG00000181315 |
| LZTFL1 | -0.2308523 | 0.85213133 | 0.269987663 | 0.460388889 | 126.8629499 | 107.9398904 | ENSG00000163818 |
| ZNF846 | -0.230917657 | 0.852092728 | 0.380955271 | 0.573941228 | 74.35419656 | 63.42243449 | ENSG00000196605 |
| UQCRQ | -0.231036085 | 0.852022784 | 0.131245812 | 0.277398777 | 242.302184 | 206.4856314 | ENSG00000164405 |
| C9orf135 | -0.231074514 | 0.852000089 | 0.223711503 | 0.406128153 | 170.8141537 | 145.4960284 | ENSG00000204711 |
| SNF8 | -0.231086222 | 0.851993174 | 0.093285107 | 0.215004191 | 382.143147 | 325.1103149 | ENSG00000159210 |
| NA | -0.231104635 | 0.851982301 | 0.036479133 | 0.104579156 | 615.357353 | 524.2388373 | ENSG00000227063 |
| LOC102724250 | -0.231122311 | 0.851971863 | 0.302316191 | 0.495722891 | 137.0007359 | 116.9398807 | ENSG00000271254 |

| | | | | | | | |
|---|---|---|---|---|---|---|---|
| NAP1L4 | -0.23114891 | 0.851956155 | 0.000171936 | 0.001122852 | 4908.108197 | 4182.409335 | ENSG00000205531 |
| SUOX | -0.23117463 | 0.851940966 | 0.055232198 | 0.14424817 | 428.3693105 | 365.0998742 | ENSG00000139531 |
| CYGB | -0.231207409 | 0.85192161 | 0.449168863 | 0.636800711 | 57.03399628 | 48.45968463 | ENSG00000161544 |
| NA | -0.231337106 | 0.851845026 | 0.828387519 | 0.907514734 | 4.472680413 | 3.81463062 | ENSG00000228201 |
| LINC00937 | -0.231346886 | 0.851839251 | 0.593148867 | 0.754731642 | 27.10151097 | 23.08991616 | ENSG00000226091 |
| SLC25A29 | -0.231351746 | 0.851836382 | 0.255070943 | 0.443895292 | 1831.784492 | 1560.729785 | ENSG00000197119 |
| SPECC1 | -0.231519826 | 0.851737145 | 0.001401259 | 0.006979557 | 1615.647156 | 1376.023504 | ENSG00000128487 |
| RNF216P1 | -0.231520022 | 0.85173703 | 0.066064318 | 0.165553823 | 400.0481649 | 340.8284763 | ENSG00000196204 |
| MRPL16 | -0.231575756 | 0.851704126 | 0.008288622 | 0.031537943 | 1475.832012 | 1256.902014 | ENSG00000166902 |
| NA | -0.231596181 | 0.851692068 | 0.530831071 | 0.706211432 | 35.98749181 | 30.65906557 | ENSG00000213058 |
| ERGIC1 | -0.231717826 | 0.851620258 | 0.000322602 | 0.001953954 | 2900.309933 | 2469.93533 | ENSG00000113719 |
| NA | -0.231732533 | 0.851611577 | 0.727670602 | 0.846909144 | 11.44017681 | 9.801050878 | ENSG00000281571 |
| PCDHGA4 | -0.231749431 | 0.851601602 | 0.765362567 | 0.870214234 | 8.581391621 | 7.351709965 | ENSG00000262576 |
| DUSP22 | -0.231760423 | 0.851595114 | 0.029900079 | 0.089367532 | 722.1381292 | 615.0880456 | ENSG00000112679 |
| COBLL1 | -0.231768856 | 0.851590136 | 0.406002243 | 0.597593867 | 81.0987775 | 69.08919819 | ENSG00000082438 |
| TMEM19 | -0.23184479 | 0.851545315 | 0.314326858 | 0.50786627 | 124.5661849 | 105.9095717 | ENSG00000139291 |
| UBE2Q2P1 | -0.23189738 | 0.851514274 | 0.766574318 | 0.870835943 | 9.492002374 | 8.165807589 | ENSG00000189136 |
| ZNF319 | -0.232002072 | 0.851452485 | 0.1816761 | 0.351790627 | 208.1640177 | 177.3221532 | ENSG00000166188 |
| SF3B5 | -0.232146346 | 0.851367341 | 0.002341963 | 0.010848774 | 1639.175555 | 1395.933691 | ENSG00000169976 |
| RPL23A | -0.232229328 | 0.851318373 | 0.006316794 | 0.025139672 | 1552.554071 | 1322.70233 | ENSG00000198242 |
| ELP5 | -0.232237621 | 0.85131348 | 0.01259912 | 0.044563377 | 781.7471495 | 665.2842743 | ENSG00000170291 |
| PFAS | -0.232251985 | 0.851305004 | 7.63E-05 | 0.000545911 | 5523.98977 | 4703.361073 | ENSG00000178921 |
| CHRNA4 | -0.232296163 | 0.851278935 | 0.380070962 | 0.573108755 | 81.56287347 | 69.29567913 | ENSG00000101204 |
| HSF2 | -0.232336197 | 0.851255313 | 0.140200965 | 0.291277288 | 227.177517 | 193.289307 | ENSG00000025156 |
| PSD | -0.232347108 | 0.851248875 | 0.303769376 | 0.497443154 | 138.1725132 | 117.9757663 | ENSG00000059915 |
| DENND2B | -0.232351857 | 0.851246073 | 0.058299337 | 0.150553144 | 391.7138616 | 333.1632733 | ENSG00000166444 |
| PER1 | -0.232416986 | 0.851207645 | 0.032948384 | 0.096517361 | 850.4371948 | 723.9346458 | ENSG00000179094 |
| RCC1L | -0.232442818 | 0.851192404 | 0.004973039 | 0.020478453 | 2200.188516 | 1872.645069 | ENSG00000274523 |
| MT2A | -0.232517442 | 0.851148377 | 0.036129003 | 0.103832369 | 644.4104658 | 547.8831545 | ENSG00000125148 |
| CDC123 | -0.232558713 | 0.851124029 | 0.002182765 | 0.010252245 | 3059.280291 | 2604.106296 | ENSG00000151465 |
| GABBR1 | -0.232680172 | 0.851052377 | 0.341034042 | 0.53542778 | 1491.370723 | 1269.502217 | ENSG00000204681 |
| GEMIN6 | -0.232719237 | 0.851029332 | 0.131443815 | 0.277658955 | 327.8833006 | 278.604216 | ENSG00000152147 |
| MCAT | -0.232733019 | 0.851021203 | 0.058115109 | 0.150244865 | 425.0495112 | 361.6318899 | ENSG00000100294 |
| SLC27A4 | -0.232834912 | 0.8509611 | 0.012907156 | 0.045315633 | 853.5760282 | 726.5207114 | ENSG00000167114 |
| NA | -0.23284631 | 0.850954377 | 0.380955116 | 0.573941228 | 78.47543772 | 66.6058909 | ENSG00000283567 |
| GSN | -0.232906159 | 0.850919077 | 0.004318699 | 0.018175014 | 1198.912931 | 1020.202717 | ENSG00000148180 |
| NDUFA6 | -0.232926385 | 0.850907147 | 0.005907066 | 0.023728122 | 1858.379318 | 1581.288168 | ENSG00000184983 |
| TM9SF2 | -0.232978997 | 0.850876117 | 0.004159136 | 0.017611476 | 2724.768494 | 2317.509152 | ENSG00000125304 |
| COASY | -0.233077155 | 0.850818227 | 0.002713533 | 0.012230443 | 2030.276924 | 1727.050605 | ENSG00000068120 |
| DHX37 | -0.233114823 | 0.850796012 | 0.001223527 | 0.006238473 | 2573.470931 | 2189.598863 | ENSG00000150990 |
| TSPAN4 | -0.233263004 | 0.850708631 | 0.00444868 | 0.018637396 | 1556.271483 | 1324.631706 | ENSG00000214063 |
| SENP3 | -0.233281222 | 0.850697888 | 0.00713994 | 0.02784757 | 1014.808719 | 863.5385872 | ENSG00000161956 |
| CDK7 | -0.233292679 | 0.850691132 | 0.009438021 | 0.035076054 | 1091.64463 | 928.0869147 | ENSG00000134058 |
| IKZF4 | -0.233317266 | 0.850676635 | 0.122694187 | 0.263990487 | 247.0548974 | 210.1308851 | ENSG00000123411 |
| ALG14 | -0.233354687 | 0.85065457 | 0.486481076 | 0.669670251 | 53.41117116 | 45.25255908 | ENSG00000172339 |
| LIMK1 | -0.23336199 | 0.850650264 | 0.001247552 | 0.006337499 | 3081.94935 | 2620.983234 | ENSG00000106683 |
| NA | -0.233396492 | 0.850629921 | 0.691771141 | 0.823566696 | 14.9788703 | 12.74147204 | ENSG00000236801 |
| UBE2N | -0.233496877 | 0.850570735 | 0.001303615 | 0.006579225 | 3686.986845 | 3135.43388 | ENSG00000177889 |
| ZNF532 | -0.233498186 | 0.850569963 | 0.000332948 | 0.002007437 | 4207.639702 | 3578.87623 | ENSG00000074657 |
| LINC01006 | -0.233631749 | 0.850491222 | 0.300030046 | 0.493415916 | 107.3798301 | 91.35103227 | ENSG00000182648 |
| TM4SF1 | -0.233747085 | 0.850423233 | 0.783901756 | 0.881838231 | 7.261337105 | 6.185088709 | ENSG00000169908 |
| KIAA0895L | -0.233755304 | 0.850418388 | 0.380093494 | 0.573108755 | 590.898702 | 502.7590872 | ENSG00000196123 |
| TAF1C | -0.23381397 | 0.850383806 | 0.041824671 | 0.116619922 | 1493.670352 | 1271.137173 | ENSG00000103168 |
| RHOV | -0.233906498 | 0.850329269 | 0.42764823 | 0.618108004 | 63.19583416 | 53.66966087 | ENSG00000104140 |
| RAB5C | -0.233911506 | 0.850326317 | 0.075118559 | 0.182733611 | 371.0801306 | 315.5771709 | ENSG00000108774 |
| GPN2 | -0.234128241 | 0.850198582 | 0.051042356 | 0.135922274 | 513.4972226 | 436.4527501 | ENSG00000142751 |
| CCDC85C | -0.234461412 | 0.850002263 | 0.007667027 | 0.029575197 | 889.5925487 | 755.9568428 | ENSG00000205476 |
| NA | -0.234601559 | 0.849919696 | 0.785473503 | 0.882568409 | 6.884740247 | 5.892565286 | ENSG00000231940 |
| NPHP3 | -0.23461521 | 0.849911654 | 0.58483168 | 0.747954646 | 26.79692175 | 22.78621944 | ENSG00000113971 |
| TNNT2 | -0.234681742 | 0.84987246 | 0.792040561 | 0.886828959 | 6.467607184 | 5.457359983 | ENSG00000118194 |
| MARCHF9 | -0.23479079 | 0.849808224 | 0.356185478 | 0.549793097 | 97.11588791 | 82.43013958 | ENSG00000139266 |
| ZNF771 | -0.234864981 | 0.849764523 | 0.125719768 | 0.268621944 | 255.0633849 | 216.8826877 | ENSG00000179965 |
| ING2 | -0.234877505 | 0.849757147 | 0.105012755 | 0.235316938 | 279.9768565 | 237.5818374 | ENSG00000168556 |
| URB2 | -0.234884221 | 0.849753191 | 0.011216054 | 0.040474756 | 1132.208582 | 961.6083459 | ENSG00000135763 |
| RAB11FIP2 | -0.234938413 | 0.849721272 | 0.235614757 | 0.420544675 | 197.7019495 | 167.7747697 | ENSG00000107560 |
| NA | -0.234963619 | 0.849706426 | 0.848445677 | 0.91943473 | 4.066393563 | 3.477079135 | ENSG00000267676 |
| WASHC1 | -0.2350404 | 0.849661205 | 0.327377931 | 0.521631927 | 107.9063501 | 91.94075403 | ENSG00000181404 |
| YJU2 | -0.235113122 | 0.849618378 | 0.042134913 | 0.117203733 | 506.7555654 | 430.7865276 | ENSG00000105248 |
| ZFP62 | -0.235160498 | 0.849590478 | 0.040320746 | 0.113131029 | 599.6717331 | 509.8111452 | ENSG00000196670 |
| PHF13 | -0.235163448 | 0.849588741 | 0.003425734 | 0.01488927 | 1255.947506 | 1066.596691 | ENSG00000116273 |
| ZUP1 | -0.235196722 | 0.849569146 | 0.125462227 | 0.268373048 | 259.2834598 | 220.3585559 | ENSG00000153975 |
| MTMR2 | -0.235263282 | 0.849529952 | 0.006872859 | 0.027021969 | 1445.493352 | 1227.187773 | ENSG00000087053 |
| NRARP | -0.235290447 | 0.849513956 | 0.10757333 | 0.23920385 | 721.9644168 | 612.8530461 | ENSG00000198435 |
| SNX1 | -0.235392558 | 0.849453831 | 0.002026996 | 0.009608272 | 1385.844518 | 1176.976286 | ENSG00000028528 |

| | | | | | | | |
|---|---|---|---|---|---|---|---|
| TPI1 | -0.235558 | 0.849356425 | 7.36E-05 | 0.000529015 | 12861.6975 | 10922.94735 | ENSG00000111669 |
| MAGI1 | -0.235568501 | 0.849350243 | 0.003109972 | 0.013740219 | 2067.985533 | 1756.673634 | ENSG00000151276 |
| ZNF780B | -0.235599286 | 0.849332119 | 0.302656163 | 0.496102354 | 124.5742801 | 106.1491832 | ENSG00000128000 |
| SLC35D1 | -0.235612589 | 0.849324287 | 0.332677387 | 0.52722041 | 99.77937311 | 84.78661321 | ENSG00000116704 |
| RNF138 | -0.235776449 | 0.849227827 | 0.125418796 | 0.268357419 | 706.0242366 | 599.1643733 | ENSG00000134758 |
| CPSF7 | -0.235841882 | 0.849189311 | 0.001630785 | 0.007981246 | 2071.475467 | 1759.940651 | ENSG00000149532 |
| WDR35 | -0.235844172 | 0.849187964 | 0.017142889 | 0.056848507 | 733.1657279 | 622.1355805 | ENSG00000118965 |
| SPPL2A | -0.235890897 | 0.849160461 | 0.180517369 | 0.350079409 | 274.8194119 | 233.4291056 | ENSG00000138600 |
| APLP1 | -0.236099745 | 0.849037544 | 0.015782876 | 0.053233816 | 790.9732349 | 671.1201917 | ENSG00000105290 |
| OLA1 | -0.23611814 | 0.849026718 | 0.011476901 | 0.041247407 | 2053.45111 | 1742.71845 | ENSG00000138430 |
| HNRNPA0 | -0.236362575 | 0.84888288 | 2.68E-05 | 0.000217115 | 12651.39714 | 10739.77197 | ENSG00000177733 |
| CHMP1A | -0.236402759 | 0.848859236 | 0.001431436 | 0.007112662 | 1991.606314 | 1690.223295 | ENSG00000131165 |
| PSMC5 | -0.236405925 | 0.848857374 | 0.002575265 | 0.011744579 | 1721.431829 | 1461.030654 | ENSG00000087191 |
| SEC13 | -0.236475961 | 0.848816166 | 0.000221183 | 0.001406312 | 2692.181942 | 2284.895683 | ENSG00000157020 |
| CCDC69 | -0.236568477 | 0.848761736 | 0.091397131 | 0.212035853 | 301.474115 | 255.8490318 | ENSG00000198624 |
| TMEM43 | -0.236642419 | 0.848718236 | 0.019495166 | 0.063168428 | 751.9301696 | 637.6877802 | ENSG00000170876 |
| NA | -0.236781902 | 0.848636184 | 0.825756955 | 0.906340611 | 4.49605658 | 3.81463062 | ENSG00000217950 |
| FBXW5 | -0.236831263 | 0.848607149 | 0.002743568 | 0.012339065 | 1750.923474 | 1485.939232 | ENSG00000159069 |
| MCM3AP | -0.237037563 | 0.84848581 | 0.000789668 | 0.004290737 | 1794.904827 | 1523.176844 | ENSG00000160294 |
| JPH3 | -0.237113571 | 0.848441108 | 0.008227995 | 0.031352315 | 1260.099181 | 1068.833235 | ENSG00000154118 |
| ELP6 | -0.237147104 | 0.848421388 | 0.018207843 | 0.059633178 | 796.0779436 | 675.2631759 | ENSG00000163832 |
| TNIP1 | -0.237250006 | 0.848360876 | 0.018462084 | 0.060306002 | 785.9624514 | 666.6432804 | ENSG00000145901 |
| NUBP1 | -0.237253302 | 0.848358937 | 0.127495446 | 0.27135587 | 286.2791241 | 242.6153886 | ENSG00000103274 |
| HNRNPD | -0.237305772 | 0.848328084 | 0.000252922 | 0.001585342 | 18692.20598 | 15857.21707 | ENSG00000138668 |
| BCL2 | -0.23732921 | 0.848314302 | 0.355006509 | 0.548282644 | 79.94098068 | 67.77262378 | ENSG00000171791 |
| RPS19BP1 | -0.237402048 | 0.848271474 | 0.011517289 | 0.041344458 | 1316.498248 | 1117.155176 | ENSG00000187051 |
| ING4 | -0.237406061 | 0.848269115 | 0.133122413 | 0.280214117 | 327.187029 | 277.7719374 | ENSG00000111653 |
| NA | -0.237492626 | 0.848218217 | 0.62441611 | 0.777176025 | 22.81022512 | 19.41439181 | ENSG00000271524 |
| SP6 | -0.237568985 | 0.848173325 | 0.429627396 | 0.619762933 | 59.7663322 | 50.52264715 | ENSG00000189120 |
| NA | -0.237577075 | 0.848168568 | 0.757936813 | 0.865159875 | 10.83858997 | 9.272428636 | ENSG00000233967 |
| RPS6KA1 | -0.237615798 | 0.848145803 | 0.001904012 | 0.009111533 | 1880.718886 | 1594.367446 | ENSG00000117676 |
| SH2D7 | -0.237646937 | 0.848127497 | 0.634077546 | 0.784250466 | 23.69924115 | 20.11206462 | ENSG00000183476 |
| ATP5PB | -0.237839333 | 0.848014399 | 0.002900717 | 0.012929746 | 2973.260499 | 2520.493177 | ENSG00000116459 |
| MCCC1 | -0.237910651 | 0.84797248 | 0.053904775 | 0.141681393 | 507.149142 | 430.3092573 | ENSG00000078070 |
| LSR | -0.238144721 | 0.847834912 | 0.001232936 | 0.006281262 | 3633.756535 | 3080.746405 | ENSG00000105699 |
| ARMC5 | -0.238188795 | 0.847809011 | 0.132926108 | 0.279950464 | 285.6952236 | 242.0343638 | ENSG00000140691 |
| NFYC | -0.23826884 | 0.847761973 | 0.022676065 | 0.071633114 | 758.0797932 | 642.8856357 | ENSG00000066136 |
| TTF2 | -0.238479987 | 0.847637907 | 0.005432839 | 0.022096168 | 1845.996376 | 1565.044992 | ENSG00000116830 |
| RGL2 | -0.238503349 | 0.847624181 | 0.008477558 | 0.032131507 | 981.3794143 | 832.3838067 | ENSG00000237441 |
| MAMDC4 | -0.238524059 | 0.847612014 | 0.229755675 | 0.413627166 | 328.4522434 | 279.0766059 | ENSG00000177943 |
| NUP188 | -0.238528529 | 0.847609388 | 0.000194348 | 0.001252453 | 3088.076108 | 2617.518834 | ENSG00000095319 |
| POP4 | -0.238557904 | 0.847592129 | 0.051181153 | 0.136213596 | 422.1204772 | 357.5015047 | ENSG00000105171 |
| RTRAF | -0.238567171 | 0.847586685 | 0.000294367 | 0.001807154 | 3123.391096 | 2647.616633 | ENSG00000087302 |
| LEPROT | -0.238591188 | 0.847572575 | 0.099899072 | 0.226653127 | 268.1351206 | 227.4977245 | ENSG00000213625 |
| ENSA | -0.2386647 | 0.847529388 | 0.000115388 | 0.000792955 | 4971.771021 | 4214.32774 | ENSG00000143420 |
| DDA1 | -0.238719019 | 0.847497478 | 0.012101306 | 0.043098696 | 1167.374413 | 990.0090474 | ENSG00000130311 |
| HNRNPK | -0.238740645 | 0.847484775 | 3.94E-05 | 0.000305999 | 17613.14248 | 14925.90489 | ENSG00000165119 |
| VCPKMT | -0.238854596 | 0.847417839 | 0.275242793 | 0.466217321 | 126.5427715 | 107.0726174 | ENSG00000100483 |
| NBPF25P | -0.238916045 | 0.847381745 | 0.736022547 | 0.852173027 | 11.36995061 | 9.568201252 | ENSG00000272150 |
| AMMECR1 | -0.239141929 | 0.84724908 | 0.012655983 | 0.044704782 | 1023.897035 | 867.0888463 | ENSG00000101935 |
| EIF3I | -0.23921215 | 0.847207843 | 8.39E-05 | 0.000595333 | 5400.561118 | 4575.285526 | ENSG00000084623 |
| SCAF4 | -0.239247018 | 0.847187368 | 0.005273084 | 0.02157901 | 1509.101351 | 1278.334475 | ENSG00000156304 |
| SEMA4B | -0.239308644 | 0.84715118 | 0.002219811 | 0.010378902 | 1380.518224 | 1168.965719 | ENSG00000185033 |
| NA | -0.239333808 | 0.847136403 | 0.717752517 | 0.84009 | 13.9229665 | 11.85272848 | ENSG00000228302 |
| TOP2B | -0.239388948 | 0.847104026 | 0.00539311 | 0.021977992 | 4134.126438 | 3501.166774 | ENSG00000077097 |
| DDX39A | -0.239476102 | 0.847052854 | 0.000585547 | 0.003291639 | 3231.014414 | 2737.749407 | ENSG00000123136 |
| ST3GAL3 | -0.239489972 | 0.84704471 | 0.633797847 | 0.783961787 | 20.82279226 | 17.63635502 | ENSG00000126091 |
| C17orf75 | -0.23950144 | 0.847037977 | 0.015460091 | 0.052345146 | 709.8745626 | 601.1609178 | ENSG00000108666 |
| TPI1P2 | -0.239511137 | 0.847032284 | 0.825848723 | 0.906340611 | 4.460052762 | 3.773289785 | ENSG00000230359 |
| PSMD13 | -0.23951625 | 0.847029282 | 0.001487559 | 0.007356036 | 3009.718777 | 2548.446869 | ENSG00000185627 |
| KIAA0895 | -0.239533109 | 0.847019384 | 0.799404396 | 0.892109892 | 5.647441881 | 4.79722916 | ENSG00000164542 |
| MRPL18 | -0.239534361 | 0.847018649 | 0.009651854 | 0.035698953 | 1360.374227 | 1151.465164 | ENSG00000112110 |
| AGA | -0.239546823 | 0.847011332 | 0.271209246 | 0.462002957 | 113.4519966 | 96.05005796 | ENSG00000038002 |
| TK1 | -0.239647493 | 0.846952231 | 0.000872024 | 0.004684663 | 2004.559004 | 1697.923779 | ENSG00000167900 |
| CCDC58 | -0.239824701 | 0.846848205 | 0.022736541 | 0.071788596 | 652.5706578 | 552.3287276 | ENSG00000160124 |
| CRAMP1 | -0.239907777 | 0.846799442 | 0.0710824 | 0.175098513 | 352.4886276 | 298.8233875 | ENSG00000007545 |
| INTS10 | -0.239955524 | 0.846771417 | 0.005627322 | 0.02279213 | 1043.278758 | 883.1507714 | ENSG00000104613 |
| ZMYND8 | -0.240013026 | 0.846737667 | 0.000262715 | 0.001639997 | 3136.809491 | 2655.515679 | ENSG00000101040 |
| RNF20 | -0.240132916 | 0.846667305 | 0.001941654 | 0.009265351 | 1480.656477 | 1253.608583 | ENSG00000155827 |
| NA | -0.240138559 | 0.846663993 | 0.667099894 | 0.806893522 | 22.52585515 | 19.23873124 | ENSG00000246889 |
| SCARA3 | -0.240305616 | 0.846565959 | 0.073707033 | 0.180110562 | 377.6642579 | 319.5845841 | ENSG00000168077 |
| ZNF211 | -0.240360511 | 0.846533748 | 0.087408185 | 0.204910359 | 321.4622352 | 272.2559542 | ENSG00000121417 |
| KIF18B | -0.240497694 | 0.846453257 | 0.001776859 | 0.008587304 | 1541.863556 | 1305.539558 | ENSG00000186185 |
| LINC02724 | -0.240553908 | 0.846420275 | 0.825916751 | 0.9063616 | 5.31631958 | 4.534761653 | ENSG00000181908 |

| | | | | | | | |
|---|---|---|---|---|---|---|---|
| NUTM2A-AS1 | -0.240582211 | 0.84640367 | 0.061774292 | 0.157568984 | 372.878194 | 315.6965185 | ENSG00000223482 |
| HIVEP3 | -0.240635987 | 0.846372121 | 0.631835552 | 0.782788727 | 23.84087362 | 20.14525811 | ENSG00000127124 |
| NA | -0.240670215 | 0.846352042 | 0.655786337 | 0.799049633 | 17.14384176 | 14.55325198 | ENSG00000214975 |
| PROM2 | -0.240908907 | 0.846212025 | 0.358936749 | 0.552141492 | 95.18588083 | 80.62749468 | ENSG00000155066 |
| ASAP2 | -0.240979635 | 0.846170541 | 0.035840065 | 0.103141245 | 476.5798628 | 403.1845313 | ENSG00000151693 |
| NA | -0.240997591 | 0.84616001 | 0.424642409 | 0.615445919 | 58.66623162 | 49.50250662 | ENSG00000286132 |
| ARHGAP17 | -0.241153246 | 0.846068721 | 0.005949423 | 0.023862065 | 1311.519199 | 1109.278139 | ENSG00000140750 |
| PAFAH1B3 | -0.241343172 | 0.845957346 | 0.04243715 | 0.117867547 | 537.6996076 | 455.2056257 | ENSG00000079462 |
| FAM98A | -0.241353203 | 0.845951464 | 0.000429308 | 0.002502926 | 2347.962585 | 1986.028375 | ENSG00000119812 |
| TMEM126B | -0.241382647 | 0.845934199 | 0.01056527 | 0.038493556 | 856.5555653 | 724.0233794 | ENSG00000171204 |
| STRADB | -0.241434704 | 0.845903675 | 0.040003634 | 0.112490875 | 702.7637726 | 594.2310412 | ENSG00000082146 |
| DLAT | -0.241476228 | 0.845879329 | 0.000594271 | 0.003333599 | 2259.503975 | 1910.629364 | ENSG00000150768 |
| SCNM1 | -0.241581659 | 0.845817515 | 0.023070238 | 0.072557916 | 634.8594894 | 536.4180069 | ENSG00000163156 |
| ZNF841 | -0.241734633 | 0.845727835 | 0.220071741 | 0.401576219 | 162.2666163 | 137.3696265 | ENSG00000197608 |
| HIP1 | -0.241798894 | 0.845690165 | 0.001724295 | 0.008370375 | 3373.854863 | 2853.343888 | ENSG00000127946 |
| TMC5 | -0.24185359 | 0.845658103 | 0.610745022 | 0.767843457 | 24.82852806 | 20.86126389 | ENSG00000103534 |
| FBXL17 | -0.241951894 | 0.845600483 | 0.352363558 | 0.545516575 | 81.35947821 | 68.59008221 | ENSG00000145743 |
| MZF1-AS1 | -0.242194016 | 0.845458581 | 0.609066377 | 0.7667212 | 26.42348181 | 22.46407813 | ENSG00000267858 |
| WDR5 | -0.242316609 | 0.845386741 | 0.000301331 | 0.001845015 | 2445.315963 | 2067.361888 | ENSG00000196363 |
| RBM27 | -0.242456228 | 0.845304932 | 0.582133166 | 0.745978891 | 329.7615116 | 278.6536152 | ENSG00000091009 |
| CNTN3 | -0.242469449 | 0.845297185 | 0.802117124 | 0.893630607 | 5.372542652 | 4.587387409 | ENSG00000113805 |
| GPR182 | -0.242707563 | 0.845157682 | 0.733775473 | 0.851264059 | 11.02757614 | 9.365845575 | ENSG00000166856 |
| NOP56 | -0.242710794 | 0.84515579 | 0.000672282 | 0.00372169 | 4877.854497 | 4123.196999 | ENSG00000101361 |
| RPIA | -0.242714399 | 0.845153677 | 0.032795008 | 0.096128843 | 569.6698934 | 481.1154674 | ENSG00000153574 |
| UTP6 | -0.242791172 | 0.845108704 | 0.001835652 | 0.008813983 | 1656.488217 | 1399.501845 | ENSG00000108651 |
| NA | -0.243063889 | 0.844948965 | 0.819138607 | 0.90291228 | 4.842963446 | 4.054528287 | ENSG00000272696 |
| CPTP | -0.243115718 | 0.844918611 | 0.149796327 | 0.305729089 | 316.1636618 | 267.1941789 | ENSG00000224051 |
| CIC | -0.243166366 | 0.84488895 | 0.00354166 | 0.01533671 | 1681.940919 | 1421.408782 | ENSG00000079432 |
| NA | -0.243194079 | 0.84487272 | 0.791179843 | 0.886400725 | 5.680288787 | 4.808514081 | ENSG00000250461 |
| AMER2 | -0.243298524 | 0.844811557 | 0.764460076 | 0.869713526 | 9.22341697 | 7.752622474 | ENSG00000165566 |
| ZFYVE27 | -0.243351396 | 0.844780597 | 0.126662261 | 0.270272281 | 564.7994047 | 477.721096 | ENSG00000155256 |
| ZNF700 | -0.243441367 | 0.844727915 | 0.347463106 | 0.54072072 | 96.86664995 | 81.66173129 | ENSG00000196757 |
| COPRS | -0.243477552 | 0.844706728 | 0.023573036 | 0.073888492 | 786.0798359 | 664.4127418 | ENSG00000172301 |
| NUBPL | -0.243659887 | 0.844599977 | 0.156271432 | 0.314818991 | 190.5611312 | 160.7281821 | ENSG00000151413 |
| SMYD5 | -0.243706538 | 0.844572666 | 0.003574049 | 0.015448092 | 1508.88835 | 1274.445971 | ENSG00000135632 |
| NDUFAF7 | -0.243714426 | 0.844568049 | 0.017918382 | 0.058830699 | 950.0398942 | 802.9988041 | ENSG00000003509 |
| HLA-A | -0.243759937 | 0.844541406 | 9.64E-05 | 0.000674002 | 5945.821578 | 5021.134401 | ENSG00000206503 |
| PIGS | -0.243941681 | 0.844435022 | 0.003626741 | 0.015643012 | 1222.52665 | 1031.812974 | ENSG00000087111 |
| EIF4A1 | -0.24404337 | 0.844375503 | 0.232301611 | 0.417039061 | 141.7163405 | 119.932378 | ENSG00000161960 |
| LINC01980 | -0.244053607 | 0.844369512 | 0.380474543 | 0.57349668 | 70.78759451 | 59.84434073 | ENSG00000225548 |
| ZNF337-AS1 | -0.244229142 | 0.844266783 | 0.624057772 | 0.77710194 | 23.1602889 | 19.54914959 | ENSG00000213742 |
| NA | -0.244307491 | 0.844220934 | 0.762818888 | 0.868646183 | 7.734595541 | 6.492584279 | ENSG00000256633 |
| SDR39U1 | -0.244324894 | 0.84421075 | 0.387058349 | 0.580024211 | 87.7899992 | 74.45120614 | ENSG00000100445 |
| AKIRIN1 | -0.244371061 | 0.844183736 | 0.001426509 | 0.007091775 | 4673.491977 | 3944.692952 | ENSG00000174574 |
| NA | -0.244373434 | 0.844182347 | 0.853411595 | 0.922280546 | 3.943101207 | 3.266687731 | ENSG00000261542 |
| NA | -0.244471503 | 0.844124964 | 0.571031658 | 0.737373586 | 34.15038805 | 28.70449214 | ENSG00000224945 |
| POLA1 | -0.244700682 | 0.843990882 | 0.000332903 | 0.002007437 | 2223.02426 | 1875.900065 | ENSG00000101868 |
| R3HCC1 | -0.244757722 | 0.843957513 | 0.029478773 | 0.088279137 | 509.6394331 | 430.3595813 | ENSG00000104679 |
| H3-3B | -0.244808089 | 0.84392805 | 4.03E-05 | 0.000311608 | 13583.18491 | 11462.93111 | ENSG00000132475 |
| NA | -0.2448703 | 0.84389166 | 0.479697585 | 0.663634773 | 45.23438265 | 38.25078482 | ENSG00000178458 |
| DPP3 | -0.244949827 | 0.843845142 | 0.000350747 | 0.002098368 | 2282.621833 | 1926.677018 | ENSG00000254986 |
| FAM174C | -0.245061498 | 0.843779827 | 0.061799982 | 0.157595057 | 580.8860345 | 490.2912383 | ENSG00000228300 |
| SAMM50 | -0.245117794 | 0.843746903 | 0.005745961 | 0.023171376 | 1408.469928 | 1187.688999 | ENSG00000100347 |
| CCT2 | -0.245198409 | 0.843699757 | 3.73E-05 | 0.000291249 | 8435.664579 | 7116.442494 | ENSG00000166226 |
| GSK3A | -0.245206912 | 0.843694784 | 0.011044099 | 0.040010032 | 1097.908489 | 925.7845723 | ENSG00000105723 |
| NA | -0.245311795 | 0.843633451 | 0.727478287 | 0.846909144 | 12.88973981 | 10.78701023 | ENSG00000274943 |
| TMEM266 | -0.245323422 | 0.843626652 | 0.21441394 | 0.394456848 | 136.1982117 | 114.8080062 | ENSG00000169758 |
| SNRPF | -0.245414823 | 0.843573206 | 0.000929121 | 0.004949791 | 2260.321802 | 1906.285378 | ENSG00000139343 |
| NOL10 | -0.245440956 | 0.843557925 | 0.01341659 | 0.046714836 | 749.810387 | 632.5631223 | ENSG00000115761 |
| TPX2 | -0.245496199 | 0.843525625 | 0.000111233 | 0.000766392 | 5925.363342 | 4997.32894 | ENSG00000088325 |
| ND2 | -0.245499362 | 0.843523776 | 0.394907384 | 0.587322536 | 12690.19329 | 10704.38158 | ENSG00000198763 |
| SLC44A2 | -0.245506516 | 0.843519593 | 0.000249856 | 0.001569242 | 3005.904327 | 2536.035647 | ENSG00000129353 |
| TMEM130 | -0.245716677 | 0.843396724 | 0.384645372 | 0.577543422 | 62.7938843 | 52.91991196 | ENSG00000166448 |
| CBX3 | -0.245720808 | 0.843394309 | 0.000252584 | 0.001584227 | 3712.380979 | 3130.44856 | ENSG00000122565 |
| ANXA11 | -0.245757158 | 0.843373059 | 0.000484653 | 0.002787884 | 1805.239191 | 1522.560427 | ENSG00000122359 |
| NA | -0.245793766 | 0.843351659 | 0.570875166 | 0.737373586 | 29.78087843 | 25.1529903 | ENSG00000274021 |
| MARK1 | -0.245857134 | 0.843314617 | 0.0219687 | 0.069876074 | 532.1447759 | 448.8431158 | ENSG00000116141 |
| SAMHD1 | -0.24590249 | 0.843288105 | 0.006647543 | 0.026245288 | 1042.100945 | 878.4549462 | ENSG00000101347 |
| NA | -0.245960818 | 0.843254012 | 0.774245244 | 0.875418579 | 7.870515549 | 6.702864064 | ENSG00000266490 |
| ATN1 | -0.246041357 | 0.843206938 | 0.002650753 | 0.01200906 | 2349.319169 | 1981.455793 | ENSG00000111676 |
| ELOA | -0.246074991 | 0.84318728 | 0.006849023 | 0.026939684 | 1198.093473 | 1009.824892 | ENSG00000011007 |
| ATAD2 | -0.246118877 | 0.843161632 | 0.017548349 | 0.057895091 | 617.7982159 | 520.8951934 | ENSG00000156802 |
| MTRF1 | -0.24613243 | 0.843153711 | 0.183099687 | 0.353550219 | 164.7601319 | 138.9712383 | ENSG00000120662 |
| FAM43A | -0.246143004 | 0.843147531 | 0.268599781 | 0.458822889 | 117.7943944 | 99.40041504 | ENSG00000185112 |

| | | | | | | | |
|---|---|---|---|---|---|---|---|
| UFD1 | -0.246146579 | 0.843145442 | 0.005442898 | 0.022127369 | 1149.634907 | 969.5574643 | ENSG00000070010 |
| CHRNB1 | -0.246151 | 0.843142858 | 0.093218527 | 0.214916306 | 314.4891467 | 265.6108288 | ENSG00000170175 |
| CTDNEP1 | -0.246279192 | 0.843067943 | 0.03579746 | 0.103071326 | 577.8238693 | 487.1447743 | ENSG00000175826 |
| ZNF350 | -0.246416933 | 0.842987455 | 0.322475117 | 0.51652921 | 86.54125326 | 72.8285216 | ENSG00000256683 |
| NA | -0.246675157 | 0.842836585 | 0.616205811 | 0.771777165 | 28.13285853 | 23.8520493 | ENSG00000262152 |
| TACC2 | -0.246819364 | 0.842752342 | 0.004570907 | 0.019048581 | 1078.356518 | 909.3678713 | ENSG00000138162 |
| ATP9A | -0.246867858 | 0.842724014 | 0.109072572 | 0.241704842 | 246.8137169 | 207.8226887 | ENSG00000054793 |
| UPRT | -0.246875084 | 0.842719794 | 0.078211976 | 0.18847602 | 335.6366271 | 282.5952717 | ENSG00000094841 |
| EFNA2 | -0.246972388 | 0.842662958 | 0.090156276 | 0.209787371 | 308.011971 | 259.2528504 | ENSG00000099617 |
| RAB14 | -0.247150904 | 0.842558694 | 0.044623691 | 0.122436047 | 899.0224875 | 756.9726659 | ENSG00000119396 |
| PPIF | -0.247201027 | 0.842529422 | 0.001327922 | 0.006673457 | 2481.506504 | 2090.171544 | ENSG00000108179 |
| MEST | -0.247395343 | 0.84241595 | 0.000399671 | 0.002350097 | 2090.504518 | 1760.738771 | ENSG00000106484 |
| PRPF31 | -0.24744673 | 0.842385945 | 0.012974324 | 0.045516964 | 1074.512905 | 906.055981 | ENSG00000105618 |
| TBKBP1 | -0.247554623 | 0.842322949 | 0.031626938 | 0.093443912 | 639.7839009 | 539.3658512 | ENSG00000198933 |
| CHST15 | -0.24756898 | 0.842314566 | 0.063556641 | 0.16106177 | 340.5922547 | 286.7922586 | ENSG00000182022 |
| TMEM201 | -0.247640148 | 0.842273016 | 0.021201732 | 0.067855794 | 759.6018071 | 639.6647087 | ENSG00000188807 |
| MAPKAPK2 | -0.247700586 | 0.842237732 | 0.006705243 | 0.026441485 | 1114.091878 | 937.7466607 | ENSG00000162889 |
| CDH1 | -0.247704562 | 0.842235411 | 0.000433711 | 0.002527005 | 3954.702834 | 3329.877933 | ENSG00000039068 |
| PRELID1 | -0.247941928 | 0.842096849 | 0.001448555 | 0.007188091 | 3515.494504 | 2960.662927 | ENSG00000169230 |
| TP53 | -0.247966065 | 0.842082761 | 0.000169704 | 0.001109843 | 3421.592803 | 2880.345054 | ENSG00000141510 |
| PKMYT1 | -0.248040561 | 0.84203928 | 0.009410247 | 0.035000907 | 1389.182033 | 1169.96627 | ENSG00000127564 |
| EPS8 | -0.248066368 | 0.842024217 | 0.29466308 | 0.48752772 | 102.3262857 | 85.94899758 | ENSG00000151491 |
| PARP6 | -0.24808596 | 0.842012782 | 0.136551248 | 0.28539026 | 346.1129542 | 291.9442503 | ENSG00000137817 |
| SH2B1 | -0.248144144 | 0.841978825 | 0.031582399 | 0.093338766 | 1180.74168 | 995.0488765 | ENSG00000178188 |
| SYNE4 | -0.24815055 | 0.841975086 | 0.092937737 | 0.214482383 | 258.7826011 | 217.9365713 | ENSG00000181392 |
| IFT122 | -0.248165374 | 0.841966435 | 0.027697647 | 0.084252123 | 605.9849824 | 510.507925 | ENSG00000163913 |
| EMC10 | -0.248326506 | 0.841872402 | 0.002540433 | 0.011605714 | 1516.779919 | 1276.353653 | ENSG00000161671 |
| TOB2 | -0.248346001 | 0.841861026 | 0.000816168 | 0.004416602 | 1965.110824 | 1654.074226 | ENSG00000183864 |
| FCHO1 | -0.248396799 | 0.841831384 | 0.023511393 | 0.073732683 | 687.6090847 | 579.1300067 | ENSG00000130475 |
| PATZ1 | -0.24842994 | 0.841812047 | 0.001662683 | 0.008111653 | 2224.052541 | 1872.305015 | ENSG00000100105 |
| RP9 | -0.248995452 | 0.841482135 | 0.152896428 | 0.310131724 | 174.6717478 | 147.0464401 | ENSG00000164610 |
| NECTIN1 | -0.249113897 | 0.841413052 | 0.000133925 | 0.000900313 | 3437.106341 | 2891.765661 | ENSG00000110400 |
| NA | -0.249118311 | 0.841410478 | 0.763222671 | 0.868785713 | 8.7735347 | 7.479419699 | ENSG00000231073 |
| MTCP1 | -0.249165813 | 0.841382774 | 0.825525594 | 0.906307982 | 4.396238088 | 3.68312204 | ENSG00000214827 |
| GPRASP1 | -0.249178666 | 0.841375278 | 0.434726485 | 0.624300378 | 52.16459745 | 43.94044478 | ENSG00000198932 |
| ASPG | -0.249255246 | 0.841330618 | 0.533849705 | 0.708715023 | 33.72388194 | 28.34871931 | ENSG00000166183 |
| PRAF2 | -0.249332271 | 0.841285701 | 0.560940514 | 0.729870729 | 29.67918081 | 24.98382811 | ENSG00000243279 |
| THAP4 | -0.249366446 | 0.841265773 | 0.000619916 | 0.003460701 | 1676.86484 | 1410.725598 | ENSG00000176946 |
| PPP1R16A | -0.249387967 | 0.841253224 | 0.113778875 | 0.249589025 | 241.0740481 | 203.0197256 | ENSG00000160972 |
| STX10 | -0.249439526 | 0.841223159 | 0.019077216 | 0.061965845 | 907.8762079 | 764.2077942 | ENSG00000104915 |
| GLIS2 | -0.249505677 | 0.841184588 | 0.067199889 | 0.167741783 | 349.3584682 | 294.0130097 | ENSG00000126603 |
| RPN2 | -0.24961252 | 0.841122294 | 7.95E-05 | 0.000567321 | 4906.773907 | 4127.125637 | ENSG00000118705 |
| SUV39H2 | -0.249749531 | 0.841042417 | 0.017221331 | 0.05704739 | 647.1642836 | 544.2506115 | ENSG00000152455 |
| USB1 | -0.249995705 | 0.840898918 | 0.001006657 | 0.005297384 | 1636.077724 | 1376.164147 | ENSG00000103005 |
| RNF217 | -0.250003997 | 0.840894085 | 0.339130894 | 0.533343401 | 80.32013309 | 67.50646904 | ENSG00000146373 |
| SOX13 | -0.250011078 | 0.840889958 | 0.00045118 | 0.002618086 | 2426.269406 | 2040.57978 | ENSG00000143842 |
| GPI | -0.250027806 | 0.840880208 | 7.18E-05 | 0.000518117 | 6236.670261 | 5244.165477 | ENSG00000105220 |
| PDLIM4 | -0.250078 | 0.840850953 | 0.103317655 | 0.232248566 | 246.2455783 | 206.8964031 | ENSG00000131435 |
| NA | -0.250178154 | 0.840792582 | 0.744373514 | 0.857195413 | 9.600013828 | 8.120779527 | ENSG00000271646 |
| TTK | -0.250214576 | 0.840771356 | 0.030648141 | 0.091089262 | 954.0585356 | 801.5099921 | ENSG00000112742 |
| TMX1 | -0.250225127 | 0.840765207 | 0.017814608 | 0.058585823 | 1563.877833 | 1314.182475 | ENSG00000139921 |
| PWWP2B | -0.250237909 | 0.840757758 | 0.054487055 | 0.142857983 | 423.4502955 | 356.4437593 | ENSG00000171813 |
| TRMT1 | -0.250585854 | 0.840555011 | 0.002867543 | 0.012818873 | 2721.767294 | 2288.86039 | ENSG00000104907 |
| NA | -0.250703532 | 0.840486451 | 0.773111552 | 0.874965662 | 8.217324716 | 6.848906758 | ENSG00000248690 |
| SHC4 | -0.250959886 | 0.840337117 | 0.739951432 | 0.854707161 | 8.568763969 | 7.197743164 | ENSG00000185634 |
| PPP4R2 | -0.251154296 | 0.840223885 | 0.0493943 | 0.132485378 | 876.8849624 | 736.1365682 | ENSG00000163605 |
| NA | -0.25117179 | 0.840213697 | 0.784904871 | 0.882357426 | 6.017724914 | 5.018355832 | ENSG00000199476 |
| PSMC6 | -0.251174994 | 0.840211831 | 0.001239754 | 0.006312528 | 1498.516404 | 1258.540753 | ENSG00000100519 |
| MSH2 | -0.25118914 | 0.840203593 | 2.78E-05 | 0.000223596 | 6478.112549 | 5442.35179 | ENSG00000095002 |
| PSMD3 | -0.251191244 | 0.840202368 | 2.61E-05 | 0.000212197 | 7121.083012 | 5982.987353 | ENSG00000108344 |
| MCMBP | -0.251267937 | 0.840157704 | 0.001406137 | 0.007000091 | 2125.845174 | 1785.08523 | ENSG00000197771 |
| C4orf3 | -0.251386939 | 0.840088406 | 0.014603739 | 0.049956755 | 868.6166386 | 729.186567 | ENSG00000164096 |
| NUDT21 | -0.251444543 | 0.840054863 | 0.00012249 | 0.000834947 | 4465.041132 | 3750.116546 | ENSG00000167005 |
| GRM8 | -0.25145436 | 0.840049147 | 0.454235235 | 0.641557992 | 49.03199541 | 41.19533125 | ENSG00000179603 |
| NA | -0.251498337 | 0.84002354 | 0.546639314 | 0.718495899 | 42.67377516 | 36.06142667 | ENSG00000278635 |
| ANTXR1 | -0.251542532 | 0.839997808 | 0.002126534 | 0.010021149 | 1128.717438 | 948.3399493 | ENSG00000169604 |
| TTC37 | -0.251551021 | 0.839992865 | 0.001936933 | 0.009252348 | 1377.556704 | 1157.081772 | ENSG00000198677 |
| KNTC1 | -0.25157443 | 0.839979236 | 0.253925771 | 0.442748437 | 1058.252803 | 889.1818396 | ENSG00000184445 |
| GAPDH | -0.251718805 | 0.839895181 | 4.31E-06 | 4.24E-05 | 36890.29654 | 30983.51665 | ENSG00000111640 |
| PM20D2 | -0.251785379 | 0.839856424 | 0.118486329 | 0.257206094 | 267.1741946 | 224.2394703 | ENSG00000146281 |
| LARP7 | -0.252046696 | 0.839704314 | 0.044383143 | 0.12200617 | 830.6241479 | 698.335621 | ENSG00000174720 |
| GATB | -0.252136551 | 0.839652016 | 0.044790482 | 0.122748307 | 416.3735251 | 349.6045917 | ENSG00000059691 |
| TUBA1B | -0.252206703 | 0.839611189 | 0.000257368 | 0.001609875 | 4223.440218 | 3546.782748 | ENSG00000123416 |
| SYT16 | -0.252350492 | 0.839527511 | 0.517006107 | 0.694462327 | 32.74136125 | 27.48579478 | ENSG00000139973 |

| | | | | | | | |
|---|---|---|---|---|---|---|---|
| VPS26A | -0.252392376 | 0.839503139 | 0.04313074 | 0.11940032 | 852.4611352 | 715.1596439 | ENSG00000122958 |
| NA | -0.252456213 | 0.839465993 | 0.783103058 | 0.88124313 | 5.76620185 | 4.838569996 | ENSG00000275720 |
| NA | -0.25262264 | 0.839369159 | 0.813797836 | 0.89988777 | 4.972569627 | 4.227266083 | ENSG00000276259 |
| APH1A | -0.252650856 | 0.839352743 | 0.000578394 | 0.003257355 | 1801.048994 | 1511.740542 | ENSG00000117362 |
| FZD8 | -0.252712199 | 0.839317054 | 0.040136552 | 0.11276199 | 404.7443469 | 339.9272369 | ENSG00000177283 |
| THOC5 | -0.252753729 | 0.839292894 | 0.001593931 | 0.00783605 | 1400.149124 | 1175.277905 | ENSG00000100296 |
| RIPK2 | -0.252757455 | 0.839290727 | 0.080080893 | 0.191933179 | 326.1515399 | 273.2238667 | ENSG00000104312 |
| NAV1 | -0.252819794 | 0.839254461 | 0.006618132 | 0.026164774 | 1192.341822 | 1000.522766 | ENSG00000134369 |
| PARP11 | -0.253065273 | 0.839111671 | 0.262593899 | 0.452072269 | 121.3041721 | 101.5929108 | ENSG00000111224 |
| NA | -0.253097636 | 0.839092849 | 0.106433103 | 0.237543542 | 363.7398182 | 305.8331346 | ENSG00000166492 |
| AATF | -0.253232715 | 0.839014288 | 0.000384461 | 0.002275093 | 1984.34905 | 1665.081518 | ENSG00000275700 |
| ZNF157 | -0.253269711 | 0.838992773 | 0.667962878 | 0.807199843 | 17.41420861 | 14.48141719 | ENSG00000147117 |
| PSMA1 | -0.253288714 | 0.838981722 | 0.256781035 | 0.445394336 | 116.9364439 | 98.01101816 | ENSG00000129084 |
| RRP1B | -0.253346665 | 0.838948022 | 0.000265469 | 0.001653847 | 3287.722459 | 2758.01425 | ENSG00000160208 |
| FAM122B | -0.253354013 | 0.838943749 | 0.014145368 | 0.048685284 | 1024.726859 | 860.409669 | ENSG00000156504 |
| INSIG2 | -0.253394684 | 0.838920099 | 0.250428296 | 0.438989695 | 131.3282115 | 110.1081044 | ENSG00000125629 |
| GTF2IP4 | -0.253644176 | 0.838775033 | 0.060412361 | 0.154780577 | 559.1946567 | 468.685951 | ENSG00000233369 |
| LOC93622 | -0.253751684 | 0.838712531 | 0.031939747 | 0.094172967 | 425.2029746 | 356.6707231 | ENSG00000170846 |
| SEC22C | -0.253777951 | 0.838697261 | 0.018345616 | 0.05996784 | 664.2689274 | 557.1207725 | ENSG00000093183 |
| NA | -0.253792962 | 0.838688534 | 0.766268711 | 0.870835943 | 6.956747883 | 5.843738377 | ENSG00000235027 |
| SCAMP4 | -0.254038085 | 0.838546048 | 0.009234564 | 0.034423453 | 844.9183524 | 708.490492 | ENSG00000227500 |
| DNA2 | -0.254244255 | 0.838426223 | 0.013392437 | 0.046648268 | 784.106128 | 657.54316 | ENSG00000138346 |
| ENY2 | -0.25426772 | 0.838412586 | 0.000256988 | 0.00160804 | 2448.986374 | 2053.269374 | ENSG00000120533 |
| SEC24D | -0.254289683 | 0.838399823 | 0.000282472 | 0.00174453 | 2038.720276 | 1708.884139 | ENSG00000150961 |
| CC2D1A | -0.254573829 | 0.838234712 | 0.007870977 | 0.030209294 | 1000.261902 | 838.8244979 | ENSG00000132024 |
| ADSL | -0.254586852 | 0.838227145 | 0.337624059 | 0.531944773 | 81.35020287 | 68.34368616 | ENSG00000239900 |
| DNAAF2 | -0.254659585 | 0.838184887 | 0.010687035 | 0.038860803 | 960.1766125 | 804.5203616 | ENSG00000165506 |
| IRAK1BP1 | -0.254662187 | 0.838183376 | 0.220636044 | 0.402130832 | 132.8069835 | 111.2779749 | ENSG00000146243 |
| MYO1B | -0.254667606 | 0.838180227 | 0.000661216 | 0.003668055 | 1974.416163 | 1654.040657 | ENSG00000128641 |
| MAP4K1 | -0.254674162 | 0.838176419 | 0.268032139 | 0.458233343 | 99.48909528 | 83.23718916 | ENSG00000104814 |
| RNF5 | -0.254704752 | 0.838158647 | 0.000932633 | 0.004965648 | 2144.206553 | 1796.516202 | ENSG00000204308 |
| UCK2 | -0.254805392 | 0.83810018 | 0.00194098 | 0.009264518 | 1294.293488 | 1084.920143 | ENSG00000143179 |
| ZBTB33 | -0.254916669 | 0.838035539 | 0.031147316 | 0.092291665 | 798.3414859 | 668.5934819 | ENSG00000177485 |
| NA | -0.254920968 | 0.838033042 | 0.424631108 | 0.615445919 | 50.80834372 | 42.47750125 | ENSG00000261512 |
| SSR4P1 | -0.254963823 | 0.838008148 | 0.704489002 | 0.831867939 | 11.14190141 | 9.358359501 | ENSG00000235374 |
| RBM14 | -0.255080627 | 0.837940304 | 0.001204434 | 0.006165909 | 1859.03458 | 1557.595564 | ENSG00000239306 |
| MYORG | -0.255135078 | 0.837908678 | 0.021206933 | 0.06786072 | 520.4482354 | 435.8368745 | ENSG00000164976 |
| ZRANB2 | -0.255171202 | 0.837887698 | 0.008506381 | 0.0322198 | 1742.85587 | 1461.069267 | ENSG00000132485 |
| NA | -0.255184011 | 0.837880259 | 0.815542241 | 0.900629202 | 5.5178357 | 4.647499239 | ENSG00000230185 |
| ZNF329 | -0.255243954 | 0.837845446 | 0.102131005 | 0.230307341 | 273.7119455 | 228.8418298 | ENSG00000181894 |
| RNF121 | -0.255319697 | 0.837801459 | 0.03051156 | 0.090756098 | 646.5014528 | 541.9739286 | ENSG00000137522 |
| TMEM199 | -0.255422318 | 0.837741868 | 0.344286254 | 0.537891042 | 85.26153955 | 71.56231143 | ENSG00000244045 |
| MSL1 | -0.255862352 | 0.837486389 | 0.004308701 | 0.018145299 | 1788.066222 | 1498.619765 | ENSG00000188895 |
| PSMD8 | -0.255874889 | 0.837479111 | 0.000120824 | 0.000825723 | 3003.182161 | 2515.027089 | ENSG00000099341 |
| DKK3 | -0.255899011 | 0.837465108 | 0.027767812 | 0.084410141 | 540.3047725 | 451.9561229 | ENSG00000050165 |
| NLGN4X | -0.256023319 | 0.837392952 | 0.000104892 | 0.000727029 | 4706.429022 | 3940.424813 | ENSG00000146938 |
| DOHH | -0.256120802 | 0.837336372 | 0.020313114 | 0.065486739 | 675.3733965 | 565.7614857 | ENSG00000129932 |
| ZNF608 | -0.25629192 | 0.837237061 | 0.006635257 | 0.026215701 | 858.0519935 | 718.3159764 | ENSG00000168916 |
| LRRFIP2 | -0.256374069 | 0.837189389 | 0.03029391 | 0.090274934 | 582.2016695 | 487.1048104 | ENSG00000093167 |
| USP1 | -0.256613706 | 0.83705034 | 0.012754249 | 0.044974744 | 1259.151913 | 1053.299383 | ENSG00000162607 |
| ATMIN | -0.256958759 | 0.836850165 | 0.005742795 | 0.023163648 | 1261.230836 | 1054.969509 | ENSG00000166454 |
| AMPD2 | -0.256966991 | 0.83684539 | 0.004852217 | 0.020042564 | 904.901752 | 757.5979719 | ENSG00000116337 |
| CCDC63 | -0.257091135 | 0.836773382 | 0.653123949 | 0.797325732 | 15.24745571 | 12.78281288 | ENSG00000173093 |
| KAT2A | -0.257094856 | 0.836771224 | 0.384655013 | 0.577543422 | 1120.491575 | 937.8507937 | ENSG00000108773 |
| TMEM38B | -0.257124104 | 0.83675426 | 0.098882334 | 0.224786603 | 309.6480397 | 258.6064892 | ENSG00000095209 |
| SSH3 | -0.257132728 | 0.836749258 | 0.020177813 | 0.065095838 | 516.8835875 | 432.4964169 | ENSG00000172830 |
| SYNGAP1 | -0.25715482 | 0.836736445 | 0.18323172 | 0.353678056 | 2238.725206 | 1873.645432 | ENSG00000197283 |
| CBX6 | -0.25721215 | 0.836703195 | 0.003163467 | 0.013933389 | 973.9343749 | 814.8180353 | ENSG00000183741 |
| NEFM | -0.257390957 | 0.836599501 | 0.020876919 | 0.066943395 | 525.4851007 | 439.3856768 | ENSG00000104722 |
| ATXN10 | -0.257520229 | 0.836524542 | 0.000668665 | 0.003703838 | 1936.365638 | 1619.261552 | ENSG00000130638 |
| ERICH6-AS1 | -0.25760926 | 0.83647292 | 0.755039965 | 0.863715017 | 9.15772316 | 7.708144064 | ENSG00000240137 |
| ZNF584 | -0.257651036 | 0.836448698 | 0.274859985 | 0.465866938 | 95.04432342 | 79.55081793 | ENSG00000171574 |
| ZNF626 | -0.257875902 | 0.836318335 | 0.265269137 | 0.454978548 | 127.7560924 | 106.5881471 | ENSG00000188171 |
| GLA | -0.257896317 | 0.836306501 | 0.018979177 | 0.0617302 | 640.5470842 | 535.2011245 | ENSG00000102393 |
| TMEM14A | -0.257909691 | 0.836298748 | 0.190223282 | 0.362566672 | 145.192204 | 121.1723311 | ENSG00000096092 |
| CFAP74 | -0.257971686 | 0.836262812 | 0.757446204 | 0.864972763 | 7.059723287 | 5.911336281 | ENSG00000142609 |
| TFDP2 | -0.257991645 | 0.836251243 | 0.002412822 | 0.011121421 | 1004.187835 | 839.9955708 | ENSG00000114126 |
| TACSTD2 | -0.25799808 | 0.836247513 | 0.569152583 | 0.735971767 | 28.91523857 | 24.31676087 | ENSG00000184292 |
| NA | -0.257998117 | 0.836247491 | 0.591789908 | 0.753623344 | 22.65724277 | 18.9415329 | ENSG00000212769 |
| OSBPL7 | -0.258018935 | 0.836235425 | 0.282049354 | 0.473077169 | 177.1308682 | 148.3810613 | ENSG00000006025 |
| TRIM2 | -0.258100669 | 0.83618805 | 0.017441725 | 0.057633768 | 727.6388874 | 607.8800363 | ENSG00000109654 |
| D21S2088E | -0.258175014 | 0.83614496 | 0.013157308 | 0.046019539 | 721.3728189 | 602.5683841 | ENSG00000228592 |
| PWP1 | -0.258212791 | 0.836123066 | 0.002439277 | 0.011234979 | 1540.706969 | 1287.509106 | ENSG00000136045 |
| FAM8A1 | -0.258292752 | 0.836076726 | 0.145700697 | 0.299513833 | 255.0635803 | 213.0883251 | ENSG00000137414 |

| | | | | | | | |
|---|---|---|---|---|---|---|---|
| NA | -0.258359481 | 0.836038056 | 0.790672735 | 0.886153987 | 6.037944167 | 5.007070912 | ENSG00000270083 |
| UBE2D2 | -0.25841887 | 0.83600364 | 0.001263635 | 0.006407717 | 3257.072392 | 2722.634003 | ENSG00000131508 |
| SGCE | -0.258559812 | 0.835921972 | 0.055376289 | 0.144575874 | 353.3262386 | 294.9008772 | ENSG00000127990 |
| LETM1 | -0.258652933 | 0.835868018 | 0.000215928 | 0.001374318 | 2167.901202 | 1812.293863 | ENSG00000168924 |
| PAXIP1 | -0.258906092 | 0.835721356 | 0.008985479 | 0.033643811 | 863.2337912 | 721.7618374 | ENSG00000157212 |
| PCBP4 | -0.258921803 | 0.835712255 | 0.087454747 | 0.204942001 | 322.9612264 | 269.6355958 | ENSG00000090097 |
| WWP2 | -0.258952493 | 0.835694477 | 0.000966771 | 0.005122392 | 1390.408463 | 1161.829951 | ENSG00000198373 |
| TMEM260 | -0.259322427 | 0.835480217 | 0.083594993 | 0.198097979 | 263.6903487 | 220.3173266 | ENSG00000070269 |
| FUNDC1 | -0.259329033 | 0.835476391 | 0.124580124 | 0.267118003 | 203.8467024 | 170.1875563 | ENSG00000069509 |
| FAM86C1 | -0.259453594 | 0.83540426 | 0.259925942 | 0.448814496 | 102.4774866 | 85.44239553 | ENSG00000158483 |
| POLR2L | -0.259492277 | 0.835381861 | 0.00258986 | 0.011793726 | 1385.549805 | 1157.615341 | ENSG00000177700 |
| BOD1 | -0.259619748 | 0.835308053 | 9.65E-05 | 0.000674098 | 3257.639577 | 2720.223662 | ENSG00000145919 |
| MIS18BP1 | -0.25963772 | 0.835297648 | 0.003220644 | 0.014140825 | 1039.140635 | 867.8301617 | ENSG00000129534 |
| TCF7 | -0.259637848 | 0.835297573 | 0.082314474 | 0.195918384 | 265.365909 | 221.4610516 | ENSG00000081059 |
| PLEKHH2 | -0.259677863 | 0.835274405 | 0.295992989 | 0.488855289 | 95.85364251 | 80.19305382 | ENSG00000152527 |
| MRM3 | -0.259715917 | 0.835252374 | 0.051681813 | 0.137073658 | 332.1648301 | 277.446833 | ENSG00000171861 |
| NLGN2 | -0.259855462 | 0.835171588 | 0.001525413 | 0.007531166 | 2635.417583 | 2202.250257 | ENSG00000169992 |
| NA | -0.259898233 | 0.835146828 | 0.6915818 | 0.823394172 | 12.97131588 | 10.92644292 | ENSG00000270605 |
| HCFC2 | -0.25991436 | 0.835137493 | 0.528079217 | 0.704021174 | 30.09178148 | 25.18673344 | ENSG00000111727 |
| ELAVL1 | -0.25996658 | 0.835107264 | 0.00027093 | 0.001681655 | 3073.996071 | 2566.194344 | ENSG00000066044 |
| NA | -0.260052545 | 0.835057505 | 0.808980721 | 0.897335896 | 4.478994239 | 3.713177955 | ENSG00000213856 |
| ACTR5 | -0.260062447 | 0.835051773 | 0.027921733 | 0.084794593 | 531.1064155 | 443.9098551 | ENSG00000101442 |
| NA | -0.260065977 | 0.83504973 | 0.382060247 | 0.574858306 | 64.83260189 | 54.30452231 | ENSG00000261188 |
| PTPRD-AS1 | -0.260075269 | 0.835044352 | 0.772809398 | 0.874809099 | 7.63477705 | 6.444195403 | ENSG00000225706 |
| ABCF1 | -0.260130438 | 0.83501242 | 1.96E-05 | 0.000163014 | 5506.707355 | 4598.823424 | ENSG00000204574 |
| LURAP1 | -0.260174071 | 0.834987167 | 0.66534189 | 0.80555022 | 13.92424428 | 11.61987885 | ENSG00000171357 |
| NA | -0.260320576 | 0.834902378 | 0.686920379 | 0.819951202 | 16.52066019 | 13.73546712 | ENSG00000268030 |
| NXT2 | -0.260332378 | 0.834895548 | 0.130237512 | 0.275991545 | 372.0677698 | 310.3745373 | ENSG00000101888 |
| PARL | -0.260337519 | 0.834892573 | 0.438833463 | 0.627185333 | 62.40090152 | 52.1364199 | ENSG00000175193 |
| ID3 | -0.260360355 | 0.834879358 | 8.13E-05 | 0.00057875 | 3238.720086 | 2704.055444 | ENSG00000117318 |
| KREMEN1 | -0.260409189 | 0.834851098 | 0.123505967 | 0.265213718 | 209.6208041 | 174.9599541 | ENSG00000183762 |
| HIGD2A | -0.260420929 | 0.834844305 | 0.183176517 | 0.353645164 | 142.2306388 | 118.7310259 | ENSG00000146066 |
| RIBC1 | -0.26051528 | 0.834789709 | 0.659165357 | 0.80143484 | 14.40381654 | 11.98748625 | ENSG00000158423 |
| LBR | -0.260606839 | 0.834736731 | 0.001734476 | 0.008413183 | 1847.3266 | 1541.015872 | ENSG00000143815 |
| NA | -0.26064082 | 0.83471707 | 0.519098685 | 0.696163172 | 37.57919095 | 31.17596486 | ENSG00000264350 |
| SPACA9 | -0.260651014 | 0.834711172 | 0.67242699 | 0.809585672 | 13.58049433 | 11.36121019 | ENSG00000165698 |
| SLC39A13 | -0.260703578 | 0.834680761 | 0.025286569 | 0.078110211 | 517.6973639 | 432.2466684 | ENSG00000165915 |
| PSMA4 | -0.260753175 | 0.834652066 | 0.000144077 | 0.000958826 | 5090.226442 | 4247.488218 | ENSG00000041357 |
| EFEMP1 | -0.260786324 | 0.834632889 | 0.118973592 | 0.257901351 | 221.7297348 | 184.8343368 | ENSG00000115380 |
| PRKACB | -0.26098651 | 0.834517084 | 0.044741771 | 0.122651086 | 498.0428088 | 415.2575733 | ENSG00000142875 |
| NA | -0.261021975 | 0.83449657 | 0.776943281 | 0.877253417 | 5.76620185 | 4.79722916 | ENSG00000278356 |
| FXYD7 | -0.261036843 | 0.83448797 | 0.50674036 | 0.686343601 | 37.13897481 | 31.12432679 | ENSG00000221946 |
| NA | -0.261051293 | 0.834479612 | 0.762450551 | 0.868357323 | 8.029714024 | 6.744204899 | ENSG00000230454 |
| TNPO2 | -0.261127132 | 0.834435747 | 0.000263434 | 0.001642829 | 2694.614657 | 2249.421184 | ENSG00000105576 |
| CCAR2 | -0.261376838 | 0.834291333 | 2.17E-05 | 0.000179727 | 3534.262051 | 2949.022122 | ENSG00000158941 |
| HSPA8 | -0.261438673 | 0.834255574 | 9.70E-07 | 1.08E-05 | 44302.55511 | 36959.56932 | ENSG00000109971 |
| NRM | -0.261493529 | 0.834223854 | 0.033346715 | 0.097418997 | 481.4796355 | 401.3997326 | ENSG00000137404 |
| SLC4A1AP | -0.261563043 | 0.83418366 | 0.02491072 | 0.077206571 | 591.4941704 | 493.1912032 | ENSG00000163798 |
| VKORC1L1 | -0.261591693 | 0.834167094 | 0.002855934 | 0.012770053 | 1481.408168 | 1235.370313 | ENSG00000196715 |
| CMC2 | -0.261620002 | 0.834150726 | 0.261029843 | 0.450039732 | 106.3612079 | 88.56292894 | ENSG00000103121 |
| LY6L | -0.261648776 | 0.834134089 | 0.783441899 | 0.881568341 | 5.402232644 | 4.527275579 | ENSG00000261667 |
| ZNF286A | -0.261731426 | 0.834086304 | 0.064175405 | 0.162193143 | 459.4942058 | 382.8464016 | ENSG00000187607 |
| CACNA2D3 | -0.261810341 | 0.834040681 | 0.31459711 | 0.508170022 | 81.8682368 | 68.33228962 | ENSG00000157445 |
| MESD | -0.261961567 | 0.83395326 | 0.001613409 | 0.007917112 | 1307.409308 | 1090.317111 | ENSG00000117899 |
| CRACR2B | -0.262015316 | 0.83392219 | 0.088075839 | 0.206084788 | 286.9414891 | 239.7908829 | ENSG00000177685 |
| PMM1 | -0.262365386 | 0.833719864 | 0.155722131 | 0.31398542 | 173.8526649 | 144.7612516 | ENSG00000100417 |
| OAZ2 | -0.262435388 | 0.833679412 | 1.89E-05 | 0.000158467 | 5944.219596 | 4954.997681 | ENSG00000180304 |
| USP43 | -0.262464066 | 0.83366284 | 0.096380814 | 0.220506357 | 223.8707287 | 186.4848787 | ENSG00000154914 |
| ZNF214 | -0.262467921 | 0.833660612 | 0.527344064 | 0.703403733 | 31.89772208 | 26.61950943 | ENSG00000149050 |
| ZNF781 | -0.26259251 | 0.833588621 | 0.807166974 | 0.896619594 | 5.588061897 | 4.639463512 | ENSG00000196381 |
| NA | -0.262651699 | 0.833554423 | 0.81392085 | 0.899911056 | 4.45246116 | 3.68312204 | ENSG00000285676 |
| NA | -0.262796361 | 0.833470845 | 0.671498007 | 0.809097911 | 14.69134342 | 12.17107093 | ENSG00000273716 |
| SRD5A3-AS1 | -0.262803934 | 0.83346647 | 0.702568832 | 0.831159372 | 11.01998453 | 9.178024011 | ENSG00000249700 |
| NOC4L | -0.263124128 | 0.833281509 | 0.002668778 | 0.012080645 | 1181.809182 | 985.2690955 | ENSG00000184967 |
| QRSL1 | -0.263233092 | 0.833218576 | 0.021225794 | 0.067909351 | 633.3329426 | 527.460225 | ENSG00000130348 |
| CLN5 | -0.263237532 | 0.833216011 | 0.670024492 | 0.808268285 | 13.68219196 | 11.42500925 | ENSG00000102805 |
| COX6A1 | -0.263298019 | 0.833181078 | 0.264975882 | 0.454645173 | 113.2206928 | 94.4947968 | ENSG00000111775 |
| FAAP20 | -0.263325882 | 0.833164987 | 0.002476924 | 0.011388565 | 974.3248022 | 811.5619797 | ENSG00000162585 |
| CENPO | -0.263331492 | 0.833161748 | 0.026858854 | 0.08212242 | 587.9659923 | 490.3471451 | ENSG00000138092 |
| NA | -0.263702601 | 0.832947458 | 0.664449435 | 0.805001035 | 14.95549414 | 12.49028946 | ENSG00000236969 |
| NA | -0.263719012 | 0.832937983 | 0.604402978 | 0.76323532 | 22.1745136 | 18.59693338 | ENSG00000280649 |
| PDCD5 | -0.263862151 | 0.832855346 | 0.007117318 | 0.027788577 | 1119.995736 | 933.1159832 | ENSG00000105185 |
| EMSY | -0.26398165 | 0.832786363 | 0.023911379 | 0.074721548 | 549.5524979 | 457.4918648 | ENSG00000158636 |
| POLR2B | -0.264081164 | 0.832728922 | 0.001392217 | 0.006941985 | 2878.413739 | 2395.815637 | ENSG00000047315 |

| | | | | | | | |
|---|---|---|---|---|---|---|---|
| INPP4A | -0.264130794 | 0.832700275 | 0.003764397 | 0.01617277 | 864.70888 | 719.7425402 | ENSG00000040933 |
| NUDC | -0.264228529 | 0.832643866 | 9.15E-06 | 8.25E-05 | 6234.186961 | 5191.469403 | ENSG00000090273 |
| ATP5MD | -0.26430882 | 0.832597528 | 0.002912919 | 0.012974775 | 1210.071835 | 1007.530323 | ENSG00000173915 |
| NA | -0.264409264 | 0.832539562 | 0.808409225 | 0.897169154 | 4.11186812 | 3.424453379 | ENSG00000263766 |
| TMEM208 | -0.264495566 | 0.832489761 | 0.014133333 | 0.048670975 | 549.2133554 | 457.4259074 | ENSG00000168701 |
| CLCN6 | -0.26455396 | 0.832456067 | 0.011366997 | 0.040922997 | 712.4386126 | 593.6196031 | ENSG00000011021 |
| TRMT11 | -0.264666564 | 0.832391095 | 0.054942968 | 0.143766005 | 391.8076594 | 326.5505141 | ENSG00000066651 |
| RNASEH1 | -0.264670096 | 0.832389057 | 0.023634481 | 0.07405604 | 580.4382798 | 482.5468261 | ENSG00000171865 |
| UCHL1 | -0.264712457 | 0.832364617 | 6.68E-05 | 0.000486523 | 10954.23538 | 9119.087389 | ENSG00000154277 |
| AAGAB | -0.264730225 | 0.832354365 | 0.004234199 | 0.017880301 | 918.3533544 | 764.1177522 | ENSG00000103591 |
| SUMO3 | -0.264744305 | 0.832346242 | 8.39E-05 | 0.000595333 | 3045.330461 | 2534.305926 | ENSG00000184900 |
| CHRNA3 | -0.264754954 | 0.832340098 | 0.232417671 | 0.417082601 | 107.5846985 | 89.53870268 | ENSG00000080644 |
| CNNM1 | -0.264979186 | 0.832210741 | 0.408444302 | 0.600182883 | 58.43227456 | 48.54280433 | ENSG00000119946 |
| CHCHD7 | -0.265083241 | 0.83215072 | 0.008866558 | 0.033292705 | 693.860391 | 577.7487974 | ENSG00000170791 |
| NA | -0.265171796 | 0.832099642 | 0.74160335 | 0.855578991 | 7.864201723 | 6.578953177 | ENSG00000228589 |
| SRP19 | -0.265198911 | 0.832084003 | 0.231924549 | 0.416637613 | 117.5050861 | 97.60218154 | ENSG00000153037 |
| CLPTM1L | -0.265403271 | 0.831966146 | 0.000185259 | 0.001200973 | 2605.797482 | 2167.615949 | ENSG00000049656 |
| ATP23 | -0.265579958 | 0.831864261 | 0.214320544 | 0.394456848 | 118.327807 | 98.32286223 | ENSG00000166896 |
| GPR139 | -0.265642726 | 0.83182807 | 0.744097853 | 0.857157223 | 8.767123176 | 7.269139914 | ENSG00000180269 |
| ANGEL2 | -0.265650606 | 0.831823526 | 0.020333605 | 0.065518607 | 595.7507151 | 495.6933846 | ENSG00000174606 |
| MOSMO | -0.265706889 | 0.831791075 | 0.252554237 | 0.44125063 | 121.4544035 | 101.0470549 | ENSG00000185716 |
| SAP30BP | -0.265863192 | 0.831700963 | 0.000446938 | 0.002594283 | 1543.217352 | 1283.646047 | ENSG00000161526 |
| MPHOSPH6 | -0.265921205 | 0.83166752 | 0.03384175 | 0.098655983 | 612.5956574 | 509.1511401 | ENSG00000135698 |
| CALM3 | -0.265994006 | 0.831625554 | 1.63E-05 | 0.000139189 | 9279.914981 | 7717.818096 | ENSG00000160014 |
| PRR5 | -0.266121766 | 0.831551911 | 0.099683863 | 0.226331093 | 221.5952879 | 184.2422419 | ENSG00000186654 |
| CACNG8 | -0.266138543 | 0.831542241 | 0.09375253 | 0.215691334 | 228.5107028 | 190.1248364 | ENSG00000142408 |
| GJB3 | -0.266164612 | 0.831527216 | 0.671112386 | 0.808896206 | 12.42595211 | 10.33347197 | ENSG00000188910 |
| CCDC91 | -0.266249426 | 0.831478333 | 0.168171751 | 0.33211457 | 184.8620963 | 153.4396184 | ENSG00000123106 |
| ABHD14A | -0.266252444 | 0.831476593 | 0.754059401 | 0.863411144 | 7.115946358 | 5.911336281 | ENSG00000248487 |
| POLR1A | -0.266316949 | 0.831439417 | 0.000168798 | 0.001105084 | 2730.053891 | 2269.980907 | ENSG00000068654 |
| IL17RC | -0.266336094 | 0.831428384 | 0.153186975 | 0.310529081 | 165.2264978 | 137.2638338 | ENSG00000163702 |
| SLIRP | -0.26638082 | 0.831402609 | 0.000543529 | 0.003088218 | 1415.698779 | 1177.082405 | ENSG00000119705 |
| NA | -0.266523895 | 0.831320161 | 0.592763269 | 0.75444118 | 23.1678805 | 19.1396517 | ENSG00000269973 |
| ZNF184 | -0.2665731 | 0.831291808 | 0.147562174 | 0.302368366 | 161.7939366 | 134.311578 | ENSG00000096654 |
| TIMM21 | -0.266650306 | 0.831247322 | 0.004589523 | 0.019110813 | 815.5688731 | 678.2054608 | ENSG00000075336 |
| DTYMK | -0.266683362 | 0.831228277 | 0.000356879 | 0.002129552 | 1570.304739 | 1305.118623 | ENSG00000168393 |
| RASSF7 | -0.266781085 | 0.831171974 | 0.063688345 | 0.161256803 | 525.900956 | 437.7314251 | ENSG00000099849 |
| ENOSF1 | -0.266826076 | 0.831146054 | 0.100523998 | 0.227513955 | 428.8937786 | 357.1234658 | ENSG00000132199 |
| PLAC8 | -0.266962235 | 0.831067616 | 0.756753537 | 0.864566204 | 6.642687922 | 5.543728881 | ENSG00000145287 |
| MPZL1 | -0.266963405 | 0.831066942 | 0.000722088 | 0.00396069 | 1779.255947 | 1478.940904 | ENSG00000197965 |
| SLC19A1 | -0.267137039 | 0.830966926 | 0.001622623 | 0.007950406 | 1071.143114 | 889.9378058 | ENSG00000173638 |
| NA | -0.267152676 | 0.830957919 | 0.822852423 | 0.904942493 | 4.185153532 | 3.480766362 | ENSG00000251775 |
| ETFB | -0.267152823 | 0.830957835 | 0.364000116 | 0.557091508 | 66.76932875 | 55.54567789 | ENSG00000105379 |
| TXNL4A | -0.267281514 | 0.830883715 | 0.000786435 | 0.004275677 | 1998.376834 | 1660.188968 | ENSG00000141759 |
| PITRM1-AS1 | -0.267404705 | 0.830812769 | 0.809538308 | 0.897492586 | 4.164934279 | 3.461995368 | ENSG00000237399 |
| COPS9 | -0.267415656 | 0.830806463 | 0.002875883 | 0.012849955 | 958.5437159 | 796.5398195 | ENSG00000172428 |
| WIPI2 | -0.267417982 | 0.830805124 | 0.000327748 | 0.001981884 | 1831.622098 | 1521.328867 | ENSG00000157954 |
| FAM53B | -0.267543354 | 0.830732928 | 0.008705911 | 0.032815814 | 616.6085416 | 512.3322721 | ENSG00000189319 |
| ZNF207 | -0.267618675 | 0.830689558 | 0.002627597 | 0.011931552 | 2580.021813 | 2143.028263 | ENSG00000010244 |
| NUDCD2 | -0.267672702 | 0.830658451 | 0.003221245 | 0.014140825 | 944.3565835 | 784.6249954 | ENSG00000170584 |
| TOLLIP | -0.267704797 | 0.830639971 | 0.005871568 | 0.023626497 | 829.872931 | 689.4630282 | ENSG00000078902 |
| NOP10 | -0.267780328 | 0.830596485 | 0.000342713 | 0.002056944 | 1575.668029 | 1309.059913 | ENSG00000182117 |
| ARID4A | -0.267824074 | 0.8305713 | 0.082788087 | 0.196666803 | 237.6570761 | 197.4423652 | ENSG00000032219 |
| GPR161 | -0.267864093 | 0.830548261 | 0.012686862 | 0.044805317 | 563.200332 | 467.5084829 | ENSG00000143147 |
| SYNRG | -0.267958993 | 0.830493629 | 0.00759087 | 0.029298039 | 795.518216 | 660.4531819 | ENSG00000275066 |
| ARL17A | -0.268012685 | 0.830462722 | 0.400516984 | 0.592527282 | 50.20873371 | 41.75780825 | ENSG00000185829 |
| LTBP3 | -0.268172318 | 0.830370837 | 0.060331106 | 0.154687282 | 288.5255966 | 239.4124999 | ENSG00000168056 |
| USP25 | -0.268200379 | 0.830354686 | 0.00943355 | 0.03506647 | 961.0406743 | 797.8067539 | ENSG00000155313 |
| NA | -0.268224523 | 0.83034079 | 0.661104269 | 0.802732346 | 15.09575114 | 12.55420013 | ENSG00000234678 |
| RASSF2 | -0.268296122 | 0.830299582 | 0.084066488 | 0.198887961 | 292.280794 | 242.8995817 | ENSG00000101265 |
| RPS23 | -0.268376432 | 0.830253364 | 0.000330328 | 0.001994232 | 2199.438389 | 1826.201965 | ENSG00000186468 |
| NA | -0.26841784 | 0.830229535 | 0.642740921 | 0.790691638 | 14.94286649 | 12.42269155 | ENSG00000287630 |
| GRM3 | -0.268424185 | 0.830225883 | 0.469457467 | 0.655190112 | 39.88205925 | 33.07031484 | ENSG00000198822 |
| VPS45 | -0.268477601 | 0.830195144 | 0.021624332 | 0.068981992 | 457.3035132 | 379.5872402 | ENSG00000136631 |
| BUB1 | -0.268499932 | 0.830182294 | 2.48E-05 | 0.000202253 | 3499.837809 | 2904.960505 | ENSG00000169679 |
| TRIR | -0.268540951 | 0.83015869 | 0.000184111 | 0.00119479 | 3580.802251 | 2972.811717 | ENSG00000123144 |
| RCN1 | -0.268554032 | 0.830151163 | 0.011538056 | 0.041410987 | 866.97494 | 719.6653776 | ENSG00000049449 |
| CTIF | -0.268849628 | 0.82998109 | 0.032230358 | 0.094728427 | 473.6289711 | 393.6266675 | ENSG00000134030 |
| NA | -0.269177036 | 0.829792754 | 0.635899525 | 0.785927034 | 16.32230099 | 13.57434066 | ENSG00000269982 |
| DNALI1 | -0.269248595 | 0.829751596 | 0.153680614 | 0.311155397 | 238.9468771 | 198.159247 | ENSG00000163879 |
| GDE1 | -0.26925047 | 0.829750518 | 0.001952395 | 0.009302234 | 1064.635955 | 882.8132828 | ENSG00000006007 |
| RAB29 | -0.269496986 | 0.829608749 | 0.043043081 | 0.119246705 | 351.325697 | 291.1732767 | ENSG00000117280 |
| NAPA | -0.269531028 | 0.829589174 | 0.024177584 | 0.075350149 | 445.4398668 | 369.7586584 | ENSG00000105402 |
| TRIM3 | -0.26956435 | 0.829570013 | 0.05390834 | 0.141681393 | 310.6171586 | 257.8721908 | ENSG00000110171 |

| | | | | | | | |
|---|---|---|---|---|---|---|---|
| ODR4 | -0.269589182 | 0.829555734 | 0.068449639 | 0.17012815 | 278.1862945 | 230.5677905 | ENSG00000157181 |
| CLSPN | -0.269617642 | 0.82953937 | 0.003578425 | 0.015459799 | 972.4997399 | 806.7074817 | ENSG00000092853 |
| PSMA5 | -0.269951895 | 0.829347199 | 4.82E-05 | 0.000365386 | 3354.232403 | 2780.858279 | ENSG00000143106 |
| ZNF692 | -0.270006029 | 0.82931608 | 0.052058095 | 0.137657972 | 644.7197229 | 535.2272961 | ENSG00000171163 |
| GIGYF2 | -0.270016571 | 0.82931002 | 0.004421596 | 0.01854069 | 1590.33641 | 1318.317917 | ENSG00000204120 |
| MID1IP1 | -0.270028394 | 0.829303224 | 8.26E-06 | 7.52E-05 | 4317.671193 | 3580.429103 | ENSG00000165175 |
| NAPSB | -0.270066678 | 0.829281218 | 0.81619521 | 0.901081846 | 4.757050382 | 4.006139411 | ENSG00000131401 |
| LRRC8A | -0.27027478 | 0.829161606 | 3.59E-05 | 0.00028199 | 3801.97253 | 3152.747802 | ENSG00000136802 |
| NTM | -0.270325177 | 0.829132642 | 0.068480761 | 0.170182679 | 291.031567 | 241.0027152 | ENSG00000182667 |
| GMPPB | -0.270431726 | 0.829071409 | 0.027960147 | 0.084855634 | 580.4572812 | 481.7580869 | ENSG00000173540 |
| RAPGEFL1 | -0.270527338 | 0.829016466 | 0.068077929 | 0.169295093 | 351.1148305 | 291.4878606 | ENSG00000108352 |
| MDC1 | -0.270551147 | 0.829002785 | 0.000673098 | 0.003723942 | 1481.573132 | 1228.205183 | ENSG00000137337 |
| RELL2 | -0.270582154 | 0.828984968 | 0.156562646 | 0.315222074 | 334.4922546 | 277.717823 | ENSG00000164620 |
| PHACTR1 | -0.270639569 | 0.828951977 | 0.169751291 | 0.334556617 | 137.5297138 | 114.0619445 | ENSG00000112137 |
| CYB5A | -0.27069382 | 0.828920806 | 0.001266313 | 0.006418214 | 1638.892343 | 1358.337978 | ENSG00000166347 |
| CSE1L | -0.270715642 | 0.828908268 | 0.000418551 | 0.002451006 | 6807.792695 | 5642.307131 | ENSG00000124207 |
| JUP | -0.270804809 | 0.828857038 | 0.000330458 | 0.001994367 | 3064.279852 | 2539.740788 | ENSG00000173801 |
| SMPD2 | -0.270867514 | 0.828821014 | 0.205968892 | 0.384511226 | 124.8282611 | 103.4166285 | ENSG00000135587 |
| PLCE1 | -0.271029579 | 0.828727913 | 0.54176087 | 0.714559918 | 28.8513262 | 23.87720706 | ENSG00000138193 |
| LRP4-AS1 | -0.271191172 | 0.828635094 | 0.778726653 | 0.878494178 | 5.342852659 | 4.42962176 | ENSG00000247675 |
| CLDN12 | -0.271244862 | 0.828604257 | 0.012447713 | 0.044078317 | 659.6879046 | 546.1276875 | ENSG00000157224 |
| PCDHGB1 | -0.271251158 | 0.828600641 | 0.550449419 | 0.721705541 | 25.40170269 | 21.09466316 | ENSG00000254221 |
| NAA40 | -0.271322137 | 0.828559876 | 0.012764532 | 0.044993883 | 911.4395181 | 755.4568281 | ENSG00000110583 |
| NA | -0.271441632 | 0.828491251 | 0.700830429 | 0.829874212 | 14.78938048 | 12.35173284 | ENSG00000287001 |
| ZNF143 | -0.271542498 | 0.828433329 | 0.00396462 | 0.016903675 | 796.9940038 | 660.0140746 | ENSG00000166478 |
| ANKRD13D | -0.271546691 | 0.828430922 | 0.034259415 | 0.099582285 | 451.2039039 | 374.3547428 | ENSG00000172932 |
| HSD17B10 | -0.271601993 | 0.828399166 | 0.000470389 | 0.002712564 | 2295.088975 | 1901.106701 | ENSG00000072506 |
| SKP1 | -0.271645701 | 0.828374069 | 0.002125185 | 0.010017341 | 1513.231439 | 1252.897077 | ENSG00000113558 |
| PTPN11 | -0.271701943 | 0.828341777 | 9.74E-06 | 8.74E-05 | 3432.1434 | 2842.731808 | ENSG00000179295 |
| HAUS3 | -0.271766365 | 0.828304789 | 0.092810901 | 0.214216343 | 241.8416509 | 200.2360908 | ENSG00000214367 |
| MAF1 | -0.271832798 | 0.828266648 | 0.000162307 | 0.001067493 | 2733.567321 | 2263.606627 | ENSG00000179632 |
| ATXN7L3 | -0.271846046 | 0.828259042 | 2.80E-05 | 0.000224997 | 2875.591168 | 2382.080858 | ENSG00000087152 |
| AHSA1 | -0.271856398 | 0.828253099 | 8.64E-06 | 7.83E-05 | 3652.534319 | 3025.659826 | ENSG00000100591 |
| MRPL57 | -0.271875655 | 0.828242044 | 0.00323889 | 0.014201464 | 937.9795194 | 777.1686952 | ENSG00000173141 |
| CHST6 | -0.271998125 | 0.828171738 | 0.017549078 | 0.057895091 | 501.7505774 | 415.8477733 | ENSG00000183196 |
| WDR24 | -0.272002254 | 0.828169367 | 0.019831955 | 0.064080449 | 463.786499 | 384.2727194 | ENSG00000127580 |
| RRAGD | -0.272172177 | 0.82807183 | 0.052630253 | 0.139078373 | 371.9718225 | 307.5357836 | ENSG00000025039 |
| NOP14 | -0.272173955 | 0.82807081 | 0.002502811 | 0.011473434 | 1098.492277 | 910.3015313 | ENSG00000087269 |
| KIF1C | -0.272420092 | 0.827929545 | 8.07E-06 | 7.38E-05 | 4537.53342 | 3757.125856 | ENSG00000129250 |
| TXNRD1 | -0.272434379 | 0.827921346 | 1.16E-05 | 0.000102707 | 3444.931562 | 2851.61793 | ENSG00000198431 |
| SET | -0.272510759 | 0.827877515 | 0.000532066 | 0.003030519 | 22068.92114 | 18269.32013 | ENSG00000119335 |
| HNRNPA2B1 | -0.27254954 | 0.827855261 | 3.79E-07 | 4.57E-06 | 41661.38337 | 34490.21797 | ENSG00000122566 |
| PLPPR5 | -0.272655385 | 0.827794527 | 0.391913042 | 0.584591777 | 58.96332694 | 48.85052314 | ENSG00000117598 |
| KIF9 | -0.272793078 | 0.827715524 | 0.283618388 | 0.474563117 | 85.96374164 | 71.21271058 | ENSG00000088727 |
| FSD1 | -0.272898449 | 0.827655072 | 0.00230368 | 0.010706219 | 1944.965791 | 1610.826113 | ENSG00000105255 |
| ARHGAP33 | -0.272945707 | 0.827627962 | 0.024568211 | 0.076362104 | 547.5290461 | 453.7407471 | ENSG00000004777 |
| SYNJ2BP | -0.273079445 | 0.827551244 | 0.034840989 | 0.100897326 | 401.4897303 | 331.9287277 | ENSG00000213463 |
| NAGK | -0.27327824 | 0.82743722 | 0.00516411 | 0.021169942 | 802.0635433 | 664.1040353 | ENSG00000124357 |
| SNRNP40 | -0.2733122 | 0.827417743 | 0.000507646 | 0.002905718 | 2036.618818 | 1684.060075 | ENSG00000060688 |
| TJP2 | -0.273423433 | 0.827353951 | 0.28156018 | 0.472812936 | 81.2236559 | 67.07874981 | ENSG00000119139 |
| TRIM36 | -0.27343496 | 0.82734734 | 0.0671893 | 0.167741783 | 291.3562778 | 241.1964141 | ENSG00000152503 |
| SRSF7 | -0.273459054 | 0.827333523 | 1.81E-06 | 1.92E-05 | 6769.146679 | 5600.601816 | ENSG00000115875 |
| LRRC37B | -0.273614782 | 0.827244224 | 0.160166582 | 0.320540701 | 185.6670784 | 153.8096308 | ENSG00000185158 |
| TPBGL | -0.273735035 | 0.827175274 | 0.63152696 | 0.7826627 | 18.57959703 | 15.51003148 | ENSG00000261594 |
| ZNF530 | -0.273820787 | 0.827126108 | 0.145461404 | 0.299121453 | 151.8457381 | 125.4485357 | ENSG00000183647 |
| BMT2 | -0.273831375 | 0.827120038 | 0.373278861 | 0.566659875 | 64.05465653 | 52.92304953 | ENSG00000164603 |
| NA | -0.273874633 | 0.827095238 | 0.821644692 | 0.904297073 | 3.854031231 | 3.214611628 | ENSG00000255216 |
| MAFA | -0.273977272 | 0.827036397 | 0.281395089 | 0.47271345 | 82.88124424 | 68.49655366 | ENSG00000182759 |
| MED25 | -0.273996566 | 0.827025337 | 0.000658391 | 0.003654573 | 1432.588294 | 1185.071358 | ENSG00000104973 |
| LYPLAL1 | -0.274050047 | 0.82699468 | 0.310557998 | 0.504415208 | 75.63381257 | 62.40879235 | ENSG00000143353 |
| NA | -0.274149167 | 0.826937863 | 0.624395672 | 0.777176025 | 17.18615941 | 14.25704133 | ENSG00000275807 |
| DDX18 | -0.274342634 | 0.826826977 | 0.000128597 | 0.000871131 | 3097.943497 | 2560.786315 | ENSG00000088205 |
| UBE2Q1 | -0.274356188 | 0.826819209 | 0.000845544 | 0.00455694 | 1296.637344 | 1071.783771 | ENSG00000160714 |
| VPS37D | -0.274406908 | 0.826790141 | 0.263681716 | 0.453369079 | 105.8520433 | 87.31698683 | ENSG00000176428 |
| SLC25A19 | -0.27455385 | 0.826705935 | 0.095195908 | 0.218253458 | 256.5669236 | 212.1455619 | ENSG00000125454 |
| UBE2QL1 | -0.274775673 | 0.826578834 | 0.492700111 | 0.675022462 | 35.4951942 | 29.19188516 | ENSG00000215218 |
| ZNF74 | -0.274841983 | 0.826540844 | 0.000163915 | 0.001077303 | 1827.859753 | 1510.43792 | ENSG00000185252 |
| PPP2CA | -0.274940502 | 0.826484402 | 0.001413165 | 0.007031302 | 2197.348049 | 1815.0997 | ENSG00000113575 |
| RASGEF1B | -0.274992402 | 0.82645467 | 0.37662601 | 0.569780955 | 59.06502457 | 48.84650106 | ENSG00000138670 |
| NA | -0.275045975 | 0.826423982 | 0.812528989 | 0.899366839 | 4.118181946 | 3.394397464 | ENSG00000275630 |
| NA | -0.275050329 | 0.826421488 | 0.046099341 | 0.125573953 | 380.2293751 | 314.1609673 | ENSG00000259781 |
| DOCK7 | -0.275145851 | 0.826366772 | 0.008676699 | 0.032732329 | 725.2822014 | 598.6906157 | ENSG00000116641 |
| GADD45GIP1 | -0.275161715 | 0.826357685 | 0.003030869 | 0.013442001 | 1905.408096 | 1574.294271 | ENSG00000179271 |
| NA | -0.275177845 | 0.826348446 | 0.798555976 | 0.891646271 | 4.502370406 | 3.675635966 | ENSG00000234773 |

| | | | | | | | |
|---|---|---|---|---|---|---|---|
| DPP9 | -0.275433772 | 0.826201869 | 0.001702585 | 0.008282341 | 1305.29989 | 1079.16298 | ENSG00000142002 |
| SERAC1 | -0.275496338 | 0.826166039 | 0.073709864 | 0.180110562 | 262.9951595 | 217.1098747 | ENSG00000122335 |
| NA | -0.275684489 | 0.826058301 | 0.608008272 | 0.766065623 | 17.41114939 | 14.38420141 | ENSG00000251495 |
| KCNMB3 | -0.276390087 | 0.825654388 | 0.641816202 | 0.789939103 | 15.96464561 | 13.21053211 | ENSG00000171121 |
| GABPB1 | -0.276436411 | 0.825627877 | 0.025198174 | 0.077935162 | 503.0586581 | 414.9890454 | ENSG00000104064 |
| MKS1 | -0.276482807 | 0.825601326 | 0.053353252 | 0.140462954 | 333.4735574 | 275.3872228 | ENSG00000011143 |
| PCGF2 | -0.276502669 | 0.82558996 | 0.006790762 | 0.026750263 | 673.7526239 | 556.3208826 | ENSG00000277258 |
| XPO7 | -0.276521095 | 0.825579415 | 9.03E-06 | 8.15E-05 | 3231.936111 | 2668.271958 | ENSG00000130227 |
| ZNF622 | -0.276560944 | 0.825556612 | 0.001099882 | 0.005726182 | 1231.34393 | 1016.895676 | ENSG00000173545 |
| CTNS | -0.27656912 | 0.825551934 | 0.065832208 | 0.165217894 | 279.0198842 | 230.727858 | ENSG00000040531 |
| MGST1 | -0.276593883 | 0.825537764 | 0.000101627 | 0.000706778 | 3459.904216 | 2856.082139 | ENSG00000008394 |
| BAP1 | -0.27659856 | 0.825535087 | 1.64E-05 | 0.000139229 | 3003.542395 | 2479.284228 | ENSG00000163930 |
| EPHB1 | -0.276644319 | 0.825508904 | 0.07840262 | 0.18878968 | 271.5923357 | 223.8449445 | ENSG00000154928 |
| TOE1 | -0.276708282 | 0.825472305 | 0.118750788 | 0.257599146 | 229.9088909 | 189.8911107 | ENSG00000132773 |
| LIN28B | -0.276737905 | 0.825455356 | 0.008427293 | 0.031980231 | 2241.81183 | 1849.656659 | ENSG00000187772 |
| TRAPPC2B | -0.27675449 | 0.825445867 | 0.53706061 | 0.711203176 | 25.82693101 | 21.29701884 | ENSG00000256060 |
| NA | -0.276809274 | 0.825414522 | 0.641401052 | 0.789727124 | 17.08120717 | 14.04285108 | ENSG00000273340 |
| SHISA6 | -0.276873438 | 0.825377813 | 0.136383115 | 0.285167492 | 158.7089813 | 131.147219 | ENSG00000188803 |
| MRPS2 | -0.27708187 | 0.825258576 | 0.011911074 | 0.042511043 | 770.4166375 | 636.1987399 | ENSG00000122140 |
| PGAP3 | -0.277111157 | 0.825241822 | 0.022436479 | 0.070974176 | 463.8328453 | 382.9561535 | ENSG00000161395 |
| RIC8A | -0.277142886 | 0.825223674 | 7.96E-05 | 0.000567424 | 1924.801214 | 1588.38183 | ENSG00000177963 |
| ESYT1 | -0.277214918 | 0.825182472 | 0.00012097 | 0.000826414 | 2370.628419 | 1956.074013 | ENSG00000139641 |
| TUSC2 | -0.277216537 | 0.825181546 | 0.008361298 | 0.03176882 | 944.5328291 | 780.2414381 | ENSG00000114383 |
| EMC7 | -0.277227959 | 0.825175013 | 0.00461348 | 0.019197628 | 738.9569745 | 609.5476288 | ENSG00000134153 |
| NA | -0.277277532 | 0.825146659 | 0.640690981 | 0.789444284 | 15.69418107 | 13.01566251 | ENSG00000283638 |
| PPP1R18 | -0.27730042 | 0.825133569 | 0.003398961 | 0.014791346 | 806.3523561 | 665.7294907 | ENSG00000146112 |
| CCL26 | -0.277320057 | 0.825122338 | 0.404104915 | 0.5957522 | 54.2232412 | 44.72437487 | ENSG00000006606 |
| CDH20 | -0.277323961 | 0.825120105 | 0.75696235 | 0.864633317 | 6.606684104 | 5.453561136 | ENSG00000101542 |
| COPS4 | -0.277324529 | 0.82511978 | 0.013184619 | 0.046088972 | 782.1166957 | 645.0385025 | ENSG00000138663 |
| ATG3 | -0.277390505 | 0.825082047 | 0.007732076 | 0.029781024 | 708.770606 | 584.2573642 | ENSG00000144848 |
| ECHS1 | -0.27740213 | 0.825075399 | 2.74E-05 | 0.000220777 | 2875.78988 | 2372.40445 | ENSG00000127884 |
| PHACTR2 | -0.277425252 | 0.825062176 | 0.018816426 | 0.061322953 | 524.4693498 | 432.4826069 | ENSG00000112419 |
| SPCS2 | -0.277520983 | 0.82500743 | 0.00018846 | 0.00121917 | 1896.490682 | 1564.63817 | ENSG00000118363 |
| TNPO3 | -0.277613255 | 0.824954666 | 2.88E-05 | 0.000231443 | 3570.939253 | 2945.492812 | ENSG00000064419 |
| RTN4R | -0.277641932 | 0.824938268 | 0.049213575 | 0.132115352 | 547.2116413 | 451.2696318 | ENSG00000040608 |
| C19orf54 | -0.277649905 | 0.824933709 | 0.015434713 | 0.052287875 | 506.8067902 | 418.1285102 | ENSG00000188493 |
| INSYN2A | -0.277651361 | 0.824932876 | 0.69522791 | 0.825605076 | 12.93027602 | 10.67278052 | ENSG00000188916 |
| RECQL4 | -0.277727258 | 0.824889479 | 0.002742287 | 0.012336297 | 1884.368364 | 1555.468877 | ENSG00000160957 |
| ZNF830 | -0.277759692 | 0.824870935 | 0.068609755 | 0.170457532 | 409.4758641 | 338.0068934 | ENSG00000198783 |
| KLHL9 | -0.277787705 | 0.824854919 | 0.009991354 | 0.036752983 | 671.2052153 | 553.6756215 | ENSG00000198642 |
| CEP78 | -0.277806443 | 0.824844205 | 0.00273537 | 0.012308169 | 820.0777527 | 676.2447952 | ENSG00000148019 |
| HNRNPH2 | -0.277893572 | 0.824794392 | 0.000693346 | 0.003826822 | 1365.442308 | 1125.714808 | ENSG00000126945 |
| NHP2 | -0.27794494 | 0.824765025 | 0.000919362 | 0.004904852 | 1774.547791 | 1464.168327 | ENSG00000145912 |
| NA | -0.278000305 | 0.824733374 | 0.440727996 | 0.628874639 | 45.98579493 | 37.95402452 | ENSG00000260920 |
| ATP1A1 | -0.278005543 | 0.82473038 | 1.25E-06 | 1.37E-05 | 6704.483205 | 5528.828283 | ENSG00000163399 |
| C7orf50 | -0.278164865 | 0.824639307 | 0.00013153 | 0.000886141 | 1984.953869 | 1637.324831 | ENSG00000146540 |
| ARMCX1 | -0.278422719 | 0.824491932 | 0.020704609 | 0.066494408 | 649.1642389 | 534.5937395 | ENSG00000126947 |
| ALKBH5 | -0.278656798 | 0.824358168 | 0.0009381 | 0.004989024 | 1300.187327 | 1071.281692 | ENSG00000091542 |
| NA | -0.278709009 | 0.824328335 | 0.55822099 | 0.727516983 | 25.11280033 | 20.64068687 | ENSG00000260588 |
| NA | -0.278740849 | 0.824310142 | 0.753304025 | 0.863060938 | 7.76112862 | 6.444195403 | ENSG00000227388 |
| NDOR1 | -0.278908071 | 0.824214602 | 0.009168116 | 0.034244633 | 759.2535852 | 625.8917483 | ENSG00000188566 |
| HSPBP1 | -0.279050705 | 0.824133119 | 0.004933798 | 0.02033875 | 1592.410431 | 1312.10298 | ENSG00000133265 |
| MELK | -0.279064017 | 0.824125515 | 0.004167403 | 0.017642452 | 1085.741223 | 894.4558516 | ENSG00000165304 |
| RNF175 | -0.27923028 | 0.824030544 | 0.147588359 | 0.302388087 | 149.336919 | 123.0490094 | ENSG00000145428 |
| PA2G4 | -0.279328789 | 0.82397428 | 1.08E-05 | 9.58E-05 | 2947.300219 | 2428.598966 | ENSG00000170515 |
| C16orf86 | -0.279335642 | 0.823970366 | 0.575927997 | 0.740958594 | 21.02124916 | 17.37020029 | ENSG00000159761 |
| ZPR1 | -0.279338761 | 0.823968585 | 0.000353458 | 0.00211254 | 1524.341163 | 1255.341113 | ENSG00000109917 |
| FBXO28 | -0.279346718 | 0.823964041 | 0.017371051 | 0.057450939 | 697.4806454 | 574.2561075 | ENSG00000143756 |
| DDX51 | -0.27939898 | 0.823934193 | 0.002899826 | 0.012929746 | 916.9675683 | 755.9976827 | ENSG00000185163 |
| DIP2A | -0.279551727 | 0.823846963 | 0.025117624 | 0.077756812 | 772.6559094 | 637.2344828 | ENSG00000160305 |
| NA | -0.279553204 | 0.823846119 | 0.785973155 | 0.882900719 | 5.058482691 | 4.197210168 | ENSG00000271474 |
| CYB561D1 | -0.279613086 | 0.823811924 | 0.149920389 | 0.305881165 | 145.1450457 | 119.6015481 | ENSG00000174151 |
| ABCC8 | -0.279785803 | 0.823713305 | 0.424754203 | 0.61546356 | 42.94867439 | 35.35495368 | ENSG00000006071 |
| NOTCH2NLC | -0.279803217 | 0.823703362 | 0.807105256 | 0.896619594 | 4.096083556 | 3.356855475 | ENSG00000286219 |
| SCN4A | -0.279922992 | 0.82363498 | 0.066485086 | 0.166338362 | 268.6917365 | 221.1719922 | ENSG00000007314 |
| EDRF1 | -0.279952602 | 0.823618076 | 0.017003303 | 0.056476562 | 448.854862 | 369.8290675 | ENSG00000107938 |
| ZNF442 | -0.280111462 | 0.823527389 | 0.499421617 | 0.680208719 | 31.73211209 | 26.05670601 | ENSG00000198342 |
| ADRA2B | -0.28013815 | 0.823512155 | 0.078329835 | 0.188663439 | 245.8886971 | 202.7613918 | ENSG00000274286 |
| MAML3 | -0.280212227 | 0.823469872 | 0.138230844 | 0.28802366 | 171.6246529 | 141.0359126 | ENSG00000196782 |
| TRABD2B | -0.280239992 | 0.823454024 | 0.179683979 | 0.348999285 | 148.9606152 | 122.5332723 | ENSG00000269113 |
| FDX1 | -0.280278528 | 0.823432029 | 0.043431562 | 0.120089598 | 513.6264979 | 422.5566659 | ENSG00000137714 |
| DPYSL3 | -0.280402628 | 0.823361201 | 5.73E-08 | 8.02E-07 | 19145.44379 | 15763.21439 | ENSG00000113657 |
| LOC729732 | -0.280510714 | 0.823299518 | 0.253266668 | 0.442000417 | 99.61120756 | 82.27858618 | ENSG00000283674 |
| JRK | -0.280592658 | 0.823252756 | 0.00826395 | 0.031463443 | 936.0786388 | 771.42215 | ENSG00000234616 |

| | | | | | | | |
|---|---|---|---|---|---|---|---|
| PINX1 | -0.280710783 | 0.823185353 | 0.047456257 | 0.128439584 | 293.2928918 | 241.6795703 | ENSG00000254093 |
| MINPP1 | -0.280806698 | 0.823130627 | 0.016336627 | 0.054723125 | 679.4048686 | 559.0984336 | ENSG00000107789 |
| NA | -0.280868066 | 0.823095614 | 0.381370377 | 0.574329449 | 60.43045594 | 49.49447089 | ENSG00000228222 |
| RHOBTB2 | -0.280873906 | 0.823092282 | 0.138075298 | 0.287764261 | 189.9423691 | 156.4714098 | ENSG00000008853 |
| PRSS16 | -0.280873999 | 0.823092229 | 0.093344764 | 0.215004743 | 235.338115 | 193.8830424 | ENSG00000112812 |
| ZNF354A | -0.280931792 | 0.823059257 | 0.107147304 | 0.23860004 | 207.8961313 | 171.2539358 | ENSG00000169131 |
| GOT2 | -0.281065745 | 0.82298284 | 1.41E-06 | 1.52E-05 | 4583.703309 | 3772.326345 | ENSG00000125166 |
| BMS1 | -0.281116878 | 0.822953672 | 4.58E-06 | 4.48E-05 | 3263.95085 | 2685.911347 | ENSG00000165733 |
| DERA | -0.281128219 | 0.822947203 | 0.090346329 | 0.210124089 | 275.1181085 | 225.8584189 | ENSG00000023697 |
| CCDC154 | -0.28114233 | 0.822939154 | 0.733273344 | 0.851101774 | 9.085715524 | 7.478981666 | ENSG00000197599 |
| BRCC3 | -0.281350141 | 0.822820623 | 0.023715658 | 0.074247638 | 442.9319422 | 364.0421515 | ENSG00000185515 |
| FANCI | -0.281394624 | 0.822795253 | 0.000572396 | 0.003226517 | 1718.694525 | 1413.852948 | ENSG00000140525 |
| RAD1 | -0.28141576 | 0.822783199 | 0.009521515 | 0.035336757 | 711.9563719 | 585.3709216 | ENSG00000113456 |
| NAB1 | -0.281494509 | 0.822738289 | 0.015537571 | 0.052549881 | 449.0196002 | 369.4765438 | ENSG00000138386 |
| TESK1 | -0.281541008 | 0.822711772 | 0.060498682 | 0.154958896 | 311.4566562 | 255.9595792 | ENSG00000107140 |
| RPAP3 | -0.281661029 | 0.822643332 | 0.004565309 | 0.019039927 | 743.2611811 | 611.1669123 | ENSG00000005175 |
| COX8A | -0.281663324 | 0.822642023 | 0.054983964 | 0.143814957 | 268.3536991 | 220.8091713 | ENSG00000176340 |
| RNF7 | -0.281828866 | 0.822547634 | 0.005351135 | 0.021845361 | 638.6590638 | 525.0984639 | ENSG00000114125 |
| NA | -0.281944632 | 0.822481634 | 0.178194945 | 0.34695505 | 140.0981083 | 115.4536029 | ENSG00000286456 |
| PAMR1 | -0.281968847 | 0.822467829 | 0.222649424 | 0.404765514 | 113.932749 | 93.54809129 | ENSG00000149090 |
| STRIP2 | -0.282137381 | 0.822371754 | 0.088069303 | 0.206084788 | 221.6116512 | 182.53389 | ENSG00000128578 |
| ATP6V1F | -0.282153108 | 0.82236279 | 0.035239129 | 0.101779962 | 346.9146665 | 285.4542137 | ENSG00000128524 |
| ADCY10 | -0.282228376 | 0.822319887 | 0.740844367 | 0.855075943 | 6.666064088 | 5.476130977 | ENSG00000143199 |
| NA | -0.282395032 | 0.8222249 | 0.706801651 | 0.833271761 | 9.24047931 | 7.606691399 | ENSG00000123009 |
| SRPRB | -0.282402803 | 0.822220471 | 0.007300987 | 0.028344469 | 604.0197457 | 496.2548655 | ENSG00000144867 |
| LOC100291105 | -0.282482998 | 0.822174768 | 0.762471516 | 0.868357323 | 6.431603366 | 5.322602209 | ENSG00000218018 |
| SDHAF3 | -0.28249846 | 0.822165956 | 0.017420571 | 0.057583888 | 432.2626002 | 355.5237489 | ENSG00000196636 |
| MAPK7 | -0.282872434 | 0.821952863 | 0.004973212 | 0.020478453 | 718.6080269 | 590.8623999 | ENSG00000166484 |
| RBFA | -0.283052808 | 0.821850104 | 0.137132856 | 0.286315235 | 154.5496618 | 127.1415177 | ENSG00000101546 |
| CEMIP2 | -0.283210596 | 0.821760223 | 0.001311128 | 0.006606973 | 1459.308541 | 1198.585816 | ENSG00000135048 |
| IQUB | -0.283384296 | 0.821661289 | 0.759600079 | 0.866364792 | 6.281875628 | 5.187406401 | ENSG00000164675 |
| ZNF876P | -0.283496618 | 0.821597321 | 0.611377224 | 0.768273474 | 17.72964404 | 14.64341972 | ENSG00000198155 |
| ASH1L-AS1 | -0.283515643 | 0.821586486 | 0.452041828 | 0.639455292 | 50.13988299 | 41.40627231 | ENSG00000235919 |
| EMD | -0.283575515 | 0.821552391 | 0.000487453 | 0.002800513 | 1189.608427 | 977.0083965 | ENSG00000102119 |
| ASTE1 | -0.283629391 | 0.821521711 | 0.361820452 | 0.554809321 | 57.8276285 | 47.31183437 | ENSG00000034533 |
| EIF4A3 | -0.283720561 | 0.821469798 | 7.07E-06 | 6.61E-05 | 4988.782821 | 4098.100008 | ENSG00000141543 |
| NA | -0.283903575 | 0.821365596 | 0.497096871 | 0.678138775 | 34.15807735 | 28.10871002 | ENSG00000259917 |
| FZD7 | -0.283910707 | 0.821361536 | 3.16E-05 | 0.000252294 | 6353.978972 | 5219.513655 | ENSG00000155760 |
| PHF23 | -0.283919977 | 0.821356258 | 0.000184996 | 0.001199692 | 1451.253842 | 1192.254559 | ENSG00000040633 |
| PRPF19 | -0.28393451 | 0.821347984 | 2.41E-06 | 2.49E-05 | 4139.691988 | 3399.661366 | ENSG00000110107 |
| FOXN3-AS1 | -0.283971489 | 0.821326932 | 0.560670197 | 0.729583756 | 22.56501589 | 18.52141136 | ENSG00000258920 |
| DEGS1 | -0.284020822 | 0.821298847 | 0.007154068 | 0.027888409 | 736.6994679 | 604.5224079 | ENSG00000143753 |
| NA | -0.284055904 | 0.821278876 | 0.716078451 | 0.839249283 | 8.270488573 | 6.75125294 | ENSG00000228150 |
| C19orf53 | -0.2841151 | 0.821245178 | 0.000618405 | 0.003456428 | 1978.758786 | 1625.685832 | ENSG00000104979 |
| FTSJ3 | -0.284253685 | 0.821166293 | 1.26E-05 | 0.000110347 | 2593.861083 | 2130.390283 | ENSG00000108592 |
| UBE2J1 | -0.284277742 | 0.821152601 | 0.017801016 | 0.058580263 | 553.3871666 | 454.4035371 | ENSG00000198833 |
| MCM10 | -0.284290512 | 0.821145332 | 0.00091272 | 0.004873628 | 1082.389787 | 888.517852 | ENSG00000065328 |
| ZNF385A | -0.284348872 | 0.821112116 | 0.047475741 | 0.128461667 | 311.6956493 | 256.2210993 | ENSG00000161642 |
| CYP2J2 | -0.284371032 | 0.821099503 | 0.547986837 | 0.71949476 | 28.75446923 | 23.46088437 | ENSG00000134716 |
| PODN | -0.284432496 | 0.821064523 | 0.775765541 | 0.876322473 | 5.058482691 | 4.178439173 | ENSG00000174348 |
| RABL6 | -0.284564063 | 0.820989649 | 5.95E-05 | 0.000439356 | 3111.613347 | 2555.286936 | ENSG00000196642 |
| ATP13A1 | -0.284579115 | 0.820981083 | 0.00067248 | 0.00372169 | 2063.378969 | 1694.645855 | ENSG00000105726 |
| NA | -0.284586038 | 0.820977143 | 0.780304119 | 0.879291341 | 5.45214189 | 4.463476522 | ENSG00000242262 |
| GLUD2 | -0.28510559 | 0.820681541 | 0.526946131 | 0.70297413 | 25.70817104 | 21.17723322 | ENSG00000182890 |
| GRM2 | -0.285151323 | 0.820655526 | 0.188498115 | 0.360427532 | 130.4371436 | 106.8326081 | ENSG00000164082 |
| ERN2 | -0.285507374 | 0.820453017 | 0.579604154 | 0.743829344 | 18.84680696 | 15.43071062 | ENSG00000134398 |
| SMPD1 | -0.285522294 | 0.820444532 | 0.369034548 | 0.5619446 | 50.62892599 | 41.49110386 | ENSG00000166311 |
| KPNA3 | -0.285572843 | 0.820415786 | 0.009294638 | 0.034626498 | 1481.21955 | 1214.500933 | ENSG00000102753 |
| RIPPLY3 | -0.285576377 | 0.820413777 | 0.483809293 | 0.667132016 | 39.03142984 | 32.05821004 | ENSG00000183145 |
| KRT10 | -0.285576572 | 0.820413666 | 0.027650052 | 0.084134964 | 551.5928388 | 453.1117715 | ENSG00000186395 |
| ST3GAL4 | -0.285879197 | 0.82024159 | 0.033973451 | 0.098957088 | 397.9955493 | 326.8529999 | ENSG00000110080 |
| TMEM104 | -0.285919862 | 0.820218471 | 0.010398988 | 0.037977311 | 653.3911767 | 535.4230241 | ENSG00000109066 |
| STARD4-AS1 | -0.285965524 | 0.820192511 | 0.333589124 | 0.528213654 | 66.66873615 | 54.6377253 | ENSG00000246859 |
| LINC02649 | -0.286055324 | 0.82014146 | 0.829957988 | 0.908528105 | 3.930473556 | 3.240868696 | ENSG00000215244 |
| GOLGA8H | -0.286159579 | 0.820082195 | 0.628772037 | 0.78073535 | 15.09259422 | 12.33252381 | ENSG00000261794 |
| CCNH | -0.286189512 | 0.82006518 | 0.053400584 | 0.140526257 | 264.6581899 | 217.2055934 | ENSG00000134480 |
| METTL16 | -0.286237128 | 0.820038115 | 0.00019502 | 0.001255907 | 1634.065359 | 1340.293832 | ENSG00000127804 |
| RCC1 | -0.286542235 | 0.819864708 | 3.51E-06 | 3.53E-05 | 3314.926539 | 2718.068454 | ENSG00000180198 |
| PCM1 | -0.2866049 | 0.819829098 | 4.49E-05 | 0.000342071 | 1986.025107 | 1627.728898 | ENSG00000078674 |
| LRRN1 | -0.286794058 | 0.819721613 | 1.74E-05 | 0.000147036 | 2602.544601 | 2133.224416 | ENSG00000175928 |
| VPS4A | -0.286873826 | 0.819676291 | 0.008251371 | 0.031428464 | 645.7433434 | 529.3004522 | ENSG00000132612 |
| SDF4 | -0.287000207 | 0.81960449 | 5.91E-05 | 0.000436538 | 2764.554568 | 2266.165984 | ENSG00000078808 |
| NA | -0.287018483 | 0.819594107 | 0.638457281 | 0.787879835 | 13.43510358 | 10.98980394 | ENSG00000285128 |
| EDC4 | -0.287051397 | 0.819575409 | 0.000603647 | 0.003382603 | 1625.182062 | 1332.93165 | ENSG00000038358 |

| | | | | | | | |
|---|---|---|---|---|---|---|---|
| MAP3K21 | -0.287229843 | 0.819474043 | 0.004458165 | 0.01865659 | 764.1591379 | 625.7187385 | ENSG00000143674 |
| NA | -0.28725066 | 0.819462218 | 0.700535189 | 0.829664311 | 9.143817732 | 7.48646774 | ENSG00000231064 |
| PAM | -0.287327731 | 0.819418443 | 0.006970755 | 0.027325714 | 951.9002613 | 779.3031971 | ENSG00000145730 |
| RRS1 | -0.28737525 | 0.819391453 | 0.002499195 | 0.011462529 | 1257.931632 | 1031.141552 | ENSG00000179041 |
| NA | -0.287414342 | 0.819369251 | 0.624698144 | 0.777266072 | 16.37042879 | 13.46540192 | ENSG00000286536 |
| TMEM69 | -0.287433705 | 0.819358254 | 0.010095344 | 0.037065591 | 529.4674603 | 433.5190421 | ENSG00000159596 |
| LYPLA2 | -0.287509146 | 0.819315409 | 2.64E-06 | 2.71E-05 | 3521.809566 | 2885.316218 | ENSG00000011009 |
| RRP1 | -0.287553936 | 0.819289973 | 8.93E-05 | 0.000629771 | 2049.71036 | 1679.97793 | ENSG00000160214 |
| MAPK3 | -0.287608272 | 0.819259117 | 0.001323916 | 0.006658739 | 868.7283106 | 711.8863151 | ENSG00000102882 |
| SELENOW | -0.287640241 | 0.819240963 | 3.84E-05 | 0.00029877 | 3117.729508 | 2553.467329 | ENSG00000178980 |
| RASGRP2 | -0.288012379 | 0.81902967 | 0.043642035 | 0.120402124 | 539.0985623 | 442.2777981 | ENSG00000068831 |
| KCNK1 | -0.288133967 | 0.818960647 | 0.06991518 | 0.172781452 | 290.3593254 | 237.8313081 | ENSG00000135750 |
| SHLD2P1 | -0.288154769 | 0.818948838 | 0.768286695 | 0.871924202 | 5.464769541 | 4.459677675 | ENSG00000165874 |
| BTG3-AS1 | -0.288183666 | 0.818932435 | 0.697555629 | 0.827476534 | 9.3327062 | 7.644233388 | ENSG00000280594 |
| ZNF365 | -0.288192594 | 0.818927367 | 0.766334036 | 0.870835943 | 5.375699565 | 4.380794851 | ENSG00000138311 |
| LIX1L | -0.28820282 | 0.818921562 | 0.419395512 | 0.610802799 | 46.32441113 | 37.87514169 | ENSG00000271601 |
| INTS11 | -0.288598556 | 0.818696961 | 0.000137082 | 0.000918209 | 3468.745902 | 2840.911335 | ENSG00000127054 |
| PHRF1 | -0.288840161 | 0.818559867 | 3.94E-05 | 0.000305643 | 2609.559451 | 2136.577848 | ENSG00000070047 |
| GHRLOS | -0.288938701 | 0.818503959 | 0.649427683 | 0.795321709 | 12.73054133 | 10.46866777 | ENSG00000240288 |
| ST8SIA2 | -0.289003497 | 0.818467198 | 0.25380888 | 0.442613266 | 85.94795708 | 70.50798972 | ENSG00000140557 |
| KLHL23 | -0.289018579 | 0.818458641 | 0.000952886 | 0.005053033 | 1220.126895 | 998.8057423 | ENSG00000213160 |
| EPB41 | -0.289024973 | 0.818455014 | 2.37E-05 | 0.00019417 | 2097.807782 | 1717.326591 | ENSG00000159023 |
| SLC27A5 | -0.289207962 | 0.818351209 | 0.128874493 | 0.273447034 | 224.3693175 | 183.9562572 | ENSG00000083807 |
| FAM171A2 | -0.289330162 | 0.818281895 | 0.002791058 | 0.012519232 | 924.8650077 | 757.018659 | ENSG00000161682 |
| PAOX | -0.28933233 | 0.818280666 | 0.717451732 | 0.839899927 | 9.959044683 | 8.165369556 | ENSG00000148832 |
| ZNF610 | -0.289340055 | 0.818276284 | 0.342663239 | 0.536797013 | 66.3630419 | 54.19459589 | ENSG00000167554 |
| NA | -0.289496396 | 0.818187615 | 0.653795956 | 0.797703095 | 11.75423677 | 9.617028162 | ENSG00000272716 |
| NA | -0.289736403 | 0.818051512 | 0.01354737 | 0.047090575 | 840.1462462 | 686.4773868 | ENSG00000235655 |
| GTF2I | -0.289773845 | 0.818030282 | 0.001171242 | 0.006029943 | 1553.926997 | 1270.445414 | ENSG00000263001 |
| HMGA2 | -0.28988094 | 0.81796956 | 8.47E-05 | 0.000600963 | 3306.246975 | 2704.369262 | ENSG00000149948 |
| AP1B1 | -0.289920367 | 0.817947206 | 3.08E-05 | 0.000246521 | 2802.52953 | 2292.044241 | ENSG00000100280 |
| HIGD1A | -0.290410176 | 0.817669552 | 0.498268194 | 0.679186784 | 200.087243 | 163.5001737 | ENSG00000181061 |
| GNAO1 | -0.290503571 | 0.817616621 | 0.001991021 | 0.009457094 | 1041.61572 | 852.0873209 | ENSG00000087258 |
| GFAP | -0.290531264 | 0.817600926 | 0.320775956 | 0.514607598 | 60.81286296 | 49.79970495 | ENSG00000131095 |
| SHANK3 | -0.2905485 | 0.817591158 | 0.016134312 | 0.054143282 | 567.9205294 | 464.609228 | ENSG00000251322 |
| EPC1 | -0.290648623 | 0.81753442 | 0.084018317 | 0.198824755 | 242.1941952 | 197.8498877 | ENSG00000120616 |
| CLNS1A | -0.290776832 | 0.817461771 | 6.43E-05 | 0.000470669 | 2307.701779 | 1885.896701 | ENSG00000074201 |
| GABRD | -0.291023896 | 0.817321791 | 0.465947977 | 0.652207522 | 35.91292863 | 29.41768675 | ENSG00000187730 |
| NDUFB2 | -0.291032332 | 0.817317012 | 0.028336448 | 0.085716945 | 362.9833473 | 296.8089255 | ENSG00000090266 |
| SEMA6B | -0.291318992 | 0.817154629 | 0.000570454 | 0.003217525 | 2953.588002 | 2413.484558 | ENSG00000167680 |
| TXN2 | -0.291344346 | 0.817140268 | 0.000998309 | 0.005259424 | 1065.721917 | 870.5533667 | ENSG00000100348 |
| DNAJC8 | -0.291431922 | 0.817090667 | 5.21E-05 | 0.00038987 | 3683.478948 | 3009.181338 | ENSG00000126698 |
| FBXO9 | -0.291520824 | 0.817040317 | 0.012150711 | 0.043225528 | 583.1305454 | 476.6482492 | ENSG00000112146 |
| NA | -0.291666267 | 0.816957953 | 0.636558657 | 0.786374602 | 13.02122513 | 10.63348146 | ENSG00000260066 |
| SLITRK2 | -0.29168214 | 0.816948964 | 0.62120468 | 0.775052679 | 14.70840576 | 11.9836874 | ENSG00000185985 |
| IDI1 | -0.291739893 | 0.816916262 | 0.009976588 | 0.036709564 | 1139.194907 | 929.9363877 | ENSG00000067064 |
| NPDC1 | -0.291740373 | 0.81691599 | 0.308703937 | 0.502946446 | 508.3599374 | 415.5360267 | ENSG00000107281 |
| IMPA2 | -0.291809759 | 0.816876701 | 5.61E-05 | 0.000417408 | 2374.000405 | 1939.102856 | ENSG00000141401 |
| C17orf100 | -0.29182612 | 0.816867438 | 0.459219014 | 0.646127338 | 37.95084946 | 30.99292983 | ENSG00000256806 |
| MSRB1 | -0.291885593 | 0.816833764 | 0.000994757 | 0.005242202 | 922.5718958 | 753.6768788 | ENSG00000198736 |
| SUV39H1 | -0.291978365 | 0.816781239 | 0.037501524 | 0.106767011 | 316.6735406 | 258.9073035 | ENSG00000101945 |
| NA | -0.292012952 | 0.816761658 | 0.4798304 | 0.663713784 | 33.49720822 | 27.50131658 | ENSG00000264070 |
| ARRB2 | -0.292042997 | 0.816744649 | 0.000237773 | 0.001500415 | 1709.57742 | 1396.732776 | ENSG00000141480 |
| NA | -0.2920661 | 0.81673157 | 0.783919645 | 0.881838231 | 5.077424168 | 4.159668179 | ENSG00000259151 |
| FAM122A | -0.292094963 | 0.81671523 | 0.509356817 | 0.688579084 | 26.77985941 | 21.79178632 | ENSG00000187866 |
| NA | -0.292338904 | 0.816577146 | 0.791325629 | 0.886482527 | 4.168091191 | 3.37562647 | ENSG00000273026 |
| KLHL25 | -0.292429261 | 0.816526005 | 0.004344106 | 0.018257068 | 769.6332427 | 628.7352801 | ENSG00000183655 |
| ZNF396 | -0.292437215 | 0.816521503 | 0.692270053 | 0.823737362 | 10.02473849 | 8.202911545 | ENSG00000186496 |
| NOMO1 | -0.292550533 | 0.816457371 | 7.01E-05 | 0.000507426 | 2079.050301 | 1697.114848 | ENSG00000103512 |
| STKLD1 | -0.292699247 | 0.816373214 | 0.669448126 | 0.807856359 | 12.23834142 | 10.04474739 | ENSG00000198870 |
| TTC9C | -0.292810805 | 0.81631009 | 0.024626825 | 0.076505795 | 425.8824921 | 347.6354841 | ENSG00000162222 |
| HTATSF1 | -0.292884207 | 0.816268559 | 6.04E-05 | 0.000444308 | 1991.623917 | 1625.367919 | ENSG00000102241 |
| DRAXIN | -0.293006949 | 0.816199115 | 0.000807111 | 0.004375252 | 991.3444646 | 809.3731712 | ENSG00000162490 |
| NA | -0.293049612 | 0.816174978 | 0.531091316 | 0.706419294 | 32.5377706 | 26.60497532 | ENSG00000286797 |
| MZT2B | -0.293073821 | 0.816161283 | 0.287681447 | 0.4792262 | 79.54575061 | 64.8553221 | ENSG00000152082 |
| SLF1 | -0.293320339 | 0.816021835 | 0.028795473 | 0.086681062 | 391.5001086 | 319.5864563 | ENSG00000133302 |
| HAUS2 | -0.293357954 | 0.816000559 | 0.014050913 | 0.048495275 | 441.8824499 | 360.5557628 | ENSG00000137814 |
| C5orf51 | -0.293386041 | 0.815984673 | 0.019400296 | 0.062914399 | 446.8340035 | 364.1317695 | ENSG00000205765 |
| NUP205 | -0.293434429 | 0.815957305 | 6.71E-06 | 6.30E-05 | 5399.33305 | 4404.855292 | ENSG00000155561 |
| PANK1 | -0.293583174 | 0.815873183 | 0.00500918 | 0.020617399 | 642.1914435 | 523.5069346 | ENSG00000152782 |
| ATP2B1 | -0.293620046 | 0.815852331 | 0.005427905 | 0.022090643 | 842.0471873 | 686.3633754 | ENSG00000070961 |
| DEDD | -0.293664057 | 0.815827443 | 0.03055589 | 0.090844221 | 338.1068117 | 275.8859008 | ENSG00000158796 |
| ARHGAP45 | -0.293751888 | 0.815777777 | 0.000166807 | 0.001093984 | 1795.608757 | 1465.517115 | ENSG00000180448 |
| NA | -0.293900913 | 0.815693515 | 0.354821164 | 0.548158328 | 50.96448298 | 41.5324447 | ENSG00000249565 |

| | | | | | | | |
|---|---|---|---|---|---|---|---|
| CASKIN1 | -0.294016287 | 0.815628285 | 0.087174558 | 0.20446637 | 194.850831 | 158.9396333 | ENSG00000167971 |
| CAND1 | -0.294072656 | 0.815596418 | 6.95E-05 | 0.000503589 | 4722.823428 | 3850.582877 | ENSG00000111530 |
| TEX22 | -0.29410106 | 0.81558036 | 0.65334557 | 0.797491227 | 15.29430574 | 12.57297113 | ENSG00000226174 |
| DNAL1 | -0.294399482 | 0.815411675 | 0.097153687 | 0.221809119 | 171.898372 | 140.1745255 | ENSG00000119661 |
| RARRES2 | -0.294439855 | 0.815388856 | 5.21E-06 | 5.01E-05 | 4462.132655 | 3638.852814 | ENSG00000106538 |
| FBXO15 | -0.294479051 | 0.815366704 | 0.622071327 | 0.775507498 | 15.51792025 | 12.64381823 | ENSG00000141665 |
| DNAAF3 | -0.29453197 | 0.815336796 | 0.424028991 | 0.61489346 | 41.33359909 | 33.79859322 | ENSG00000167646 |
| MBD3 | -0.294543867 | 0.815330072 | 0.003096153 | 0.013689407 | 984.317573 | 803.0391254 | ENSG00000071655 |
| ZDHHC3 | -0.294580436 | 0.815309406 | 0.000108737 | 0.000750306 | 1553.91588 | 1266.606095 | ENSG00000163812 |
| NA | -0.29484481 | 0.815160014 | 0.361368749 | 0.554575507 | 58.69138922 | 47.74367886 | ENSG00000275234 |
| CCDC150 | -0.294878779 | 0.815140821 | 0.072943615 | 0.178780345 | 255.380429 | 208.5687907 | ENSG00000144395 |
| DUS1L | -0.294925535 | 0.815114404 | 0.001055229 | 0.005520074 | 1843.00712 | 1502.95335 | ENSG00000169718 |
| SORL1 | -0.295045421 | 0.815046671 | 0.002760323 | 0.01240473 | 2278.975662 | 1856.583436 | ENSG00000137642 |
| ITSN1 | -0.295060749 | 0.815038012 | 1.51E-05 | 0.000129692 | 3074.722333 | 2506.081253 | ENSG00000205726 |
| C19orf84 | -0.295097536 | 0.81501723 | 0.285995122 | 0.477319548 | 68.91774153 | 56.15709343 | ENSG00000262874 |
| BRK1 | -0.295283749 | 0.81491204 | 0.000212222 | 0.00135352 | 3786.85177 | 3086.504724 | ENSG00000254999 |
| DIPK1B | -0.295358075 | 0.814870058 | 0.024750519 | 0.076838545 | 413.4775786 | 336.7605761 | ENSG00000165716 |
| LPAR4 | -0.295482805 | 0.81479961 | 0.25933993 | 0.44824303 | 98.80810459 | 80.38985852 | ENSG00000147145 |
| PLCXD1 | -0.295503899 | 0.814787697 | 1.21E-05 | 0.000105932 | 3089.105141 | 2517.574722 | ENSG00000182378 |
| ZNF181 | -0.295602061 | 0.81473226 | 0.210056958 | 0.389305017 | 125.1827822 | 101.8707885 | ENSG00000197841 |
| PIGZ | -0.295628849 | 0.814717132 | 0.727188078 | 0.846837128 | 7.231647112 | 5.941392196 | ENSG00000119227 |
| TPM3 | -0.295746709 | 0.814650577 | 1.23E-06 | 1.35E-05 | 13311.27668 | 10843.31445 | ENSG00000143549 |
| EXOC7 | -0.295840543 | 0.814597593 | 0.00012851 | 0.000870861 | 1723.702452 | 1404.248352 | ENSG00000182473 |
| RTN3 | -0.295855561 | 0.814589114 | 6.79E-05 | 0.000493206 | 3555.223567 | 2895.199566 | ENSG00000133318 |
| NME7 | -0.295990818 | 0.814512747 | 0.059228024 | 0.152420465 | 282.7328617 | 229.9368396 | ENSG00000143156 |
| FAM149A | -0.296091173 | 0.814456091 | 0.044005063 | 0.121169194 | 261.7766072 | 213.2782252 | ENSG00000109794 |
| VRK2 | -0.296092605 | 0.814455282 | 0.052056808 | 0.137657972 | 333.1444643 | 271.1471345 | ENSG00000028116 |
| TLK2 | -0.296093213 | 0.814454939 | 0.001619198 | 0.007939212 | 1022.731354 | 832.3522361 | ENSG00000146872 |
| BLM | -0.296115824 | 0.814442175 | 0.002900329 | 0.012929746 | 711.5124127 | 579.1053499 | ENSG00000197299 |
| SNRPB2 | -0.296133013 | 0.814432471 | 0.001243719 | 0.006325766 | 1462.265611 | 1190.092444 | ENSG00000125870 |
| ADAM23 | -0.296161849 | 0.814416193 | 0.000536179 | 0.003051136 | 981.4639293 | 799.2418316 | ENSG00000114948 |
| EEF1AKNMT | -0.296203078 | 0.814392919 | 3.47E-05 | 0.000273076 | 2495.535076 | 2032.598806 | ENSG00000010165 |
| SFTPD | -0.296474164 | 0.814239906 | 0.754410239 | 0.863474302 | 6.027195652 | 4.954994809 | ENSG00000133661 |
| MRGBP | -0.296578894 | 0.8141808 | 0.004455506 | 0.018653607 | 721.3895504 | 587.294715 | ENSG00000101189 |
| NEPRO | -0.296636712 | 0.814148171 | 0.016770171 | 0.055892529 | 674.2352402 | 548.6008636 | ENSG00000163608 |
| TMEM183A | -0.296639959 | 0.814146339 | 1.95E-05 | 0.000163014 | 2310.019244 | 1880.085114 | ENSG00000163444 |
| KCNK17 | -0.296672706 | 0.814127859 | 0.394346278 | 0.586770546 | 51.8206521 | 42.33244624 | ENSG00000124780 |
| LOC100128108 | -0.296888199 | 0.814006264 | 0.592193173 | 0.75398147 | 17.12362251 | 13.8813982 | ENSG00000259658 |
| TAF1A | -0.296891935 | 0.814004156 | 0.092983576 | 0.214535858 | 196.6919786 | 159.8240284 | ENSG00000143498 |
| PUS3 | -0.29695882 | 0.813966418 | 0.030580248 | 0.090902055 | 353.7875684 | 288.1183977 | ENSG00000110060 |
| BRWD1-AS2 | -0.297041594 | 0.813919719 | 0.78048265 | 0.879413061 | 4.703984223 | 3.870943603 | ENSG00000255568 |
| SLC2A11 | -0.297122145 | 0.813874276 | 0.211694572 | 0.391298214 | 112.287886 | 91.52800694 | ENSG00000133460 |
| MVB12B | -0.297209569 | 0.813824959 | 0.043536994 | 0.120148101 | 296.4769891 | 241.5032484 | ENSG00000196814 |
| PRDX6 | -0.297248301 | 0.81380311 | 1.26E-06 | 1.38E-05 | 4582.660844 | 3729.28381 | ENSG00000117592 |
| AK9 | -0.29735412 | 0.813743421 | 0.50372017 | 0.683776217 | 27.86743004 | 22.70733661 | ENSG00000155085 |
| PSMB5 | -0.297384181 | 0.813726466 | 3.54E-06 | 3.55E-05 | 3802.666223 | 3093.90627 | ENSG00000100804 |
| NA | -0.297468738 | 0.813678774 | 0.648775948 | 0.794951114 | 12.70273048 | 10.35224296 | ENSG00000267248 |
| C1orf226 | -0.297533492 | 0.813642254 | 0.0234516 | 0.073582523 | 444.4689292 | 361.3315455 | ENSG00000239887 |
| CACNB3 | -0.297665312 | 0.813567914 | 0.037642999 | 0.107087543 | 300.2938365 | 244.5793119 | ENSG00000167535 |
| RET | -0.297675783 | 0.81356201 | 0.030059065 | 0.089770338 | 400.5239863 | 325.935251 | ENSG00000165731 |
| STIM2 | -0.297718411 | 0.813537971 | 0.016425244 | 0.054950446 | 420.4622124 | 342.212537 | ENSG00000109689 |
| KANK2 | -0.297898554 | 0.813436394 | 0.000339328 | 0.002041253 | 1139.459073 | 927.3190963 | ENSG00000197256 |
| PRIM1 | -0.297918603 | 0.81342509 | 0.118630416 | 0.257398284 | 250.6863813 | 203.7069577 | ENSG00000198056 |
| GIPC2 | -0.298190644 | 0.813271722 | 0.566163883 | 0.733828586 | 21.12738148 | 17.13735066 | ENSG00000137960 |
| CIB1 | -0.298418851 | 0.813143088 | 0.062028115 | 0.158044692 | 219.0687074 | 178.2345659 | ENSG00000185043 |
| PPP1R13B | -0.298440545 | 0.81313086 | 0.045775373 | 0.124966972 | 263.5225891 | 214.2753578 | ENSG00000088808 |
| POMT2 | -0.298500816 | 0.813096891 | 0.002559589 | 0.01168267 | 689.5685265 | 560.6684839 | ENSG00000009830 |
| NA | -0.298548674 | 0.813069919 | 0.603976231 | 0.762992316 | 237.1730469 | 192.8768863 | ENSG00000198744 |
| MRPL43 | -0.298558024 | 0.81306465 | 0.002088832 | 0.009863545 | 855.953084 | 696.2023752 | ENSG00000055950 |
| IFIT5 | -0.298662259 | 0.813005908 | 0.374122438 | 0.567336383 | 51.5522621 | 41.83950224 | ENSG00000152778 |
| TMEM65 | -0.298831816 | 0.812910362 | 0.047212374 | 0.127928968 | 375.7233668 | 305.3311269 | ENSG00000164983 |
| NA | -0.299040345 | 0.812792872 | 0.719031468 | 0.840949657 | 7.979804779 | 6.45504229 | ENSG00000224837 |
| DIS3L | -0.299083191 | 0.812768733 | 0.001107673 | 0.005758657 | 1051.871785 | 854.5389149 | ENSG00000166938 |
| ZNF664 | -0.299099357 | 0.812759626 | 0.000149384 | 0.000989873 | 1648.345418 | 1338.988565 | ENSG00000179195 |
| IPO9 | -0.299167944 | 0.812720988 | 4.63E-06 | 4.52E-05 | 3541.047251 | 2878.212233 | ENSG00000198700 |
| NA | -0.299212309 | 0.812695996 | 0.433881398 | 0.623666654 | 35.7859757 | 29.06841231 | ENSG00000225285 |
| THAP1 | -0.299464781 | 0.812553786 | 0.258424997 | 0.447074219 | 91.13161126 | 73.9610733 | ENSG00000131931 |
| AACS | -0.299496696 | 0.812535811 | 0.000169123 | 0.001106825 | 1459.412959 | 1185.316878 | ENSG00000081760 |
| DLG3 | -0.299513804 | 0.812526176 | 0.000331527 | 0.002000167 | 1182.792996 | 960.878078 | ENSG00000082458 |
| AP2M1 | -0.29961268 | 0.812470491 | 1.06E-07 | 1.41E-06 | 12208.54858 | 9919.501527 | ENSG00000161203 |
| IPO5 | -0.299769298 | 0.812382294 | 2.82E-06 | 2.88E-05 | 11200.3615 | 9097.686845 | ENSG00000065150 |
| NDUFS3 | -0.299815492 | 0.812356283 | 0.001663477 | 0.008112866 | 1584.415744 | 1288.065787 | ENSG00000213619 |
| MAMDC2 | -0.299957205 | 0.812276491 | 0.08250491 | 0.196195279 | 195.5305439 | 158.7448753 | ENSG00000165072 |
| CDC26 | -0.300009064 | 0.812247293 | 0.025260861 | 0.078043808 | 336.9952636 | 273.7139995 | ENSG00000176386 |

| | | | | | | | |
|---|---|---|---|---|---|---|---|
| NA | -0.30003557 | 0.81223237 | 0.65185439 | 0.796773018 | 14.10997584 | 11.34568839 | ENSG00000177855 |
| NEURL4 | -0.300097551 | 0.812197476 | 0.000859856 | 0.004625735 | 1161.9858 | 944.2777689 | ENSG00000215041 |
| RPP25 | -0.300135009 | 0.812176388 | 0.00426516 | 0.017997538 | 737.9536484 | 598.6615475 | ENSG00000178718 |
| DZANK1 | -0.300248183 | 0.812112679 | 0.064576114 | 0.162888012 | 223.450341 | 181.3019324 | ENSG00000089091 |
| NA | -0.300396011 | 0.812029469 | 0.768509193 | 0.871924202 | 5.627222628 | 4.636214318 | ENSG00000272008 |
| SCYL1 | -0.300446271 | 0.81200118 | 0.000247228 | 0.001553786 | 1686.890488 | 1369.234574 | ENSG00000142186 |
| INTS6 | -0.300447514 | 0.812000481 | 0.01096858 | 0.039790399 | 588.8439143 | 477.7287877 | ENSG00000102786 |
| GLT8D2 | -0.30046552 | 0.811990346 | 0.291415594 | 0.483761659 | 70.82863438 | 57.63175991 | ENSG00000120820 |
| SCIN | -0.300501569 | 0.811970057 | 0.708404004 | 0.834089445 | 9.295424606 | 7.520322501 | ENSG00000006747 |
| SETD1A | -0.300554284 | 0.811940389 | 0.000581605 | 0.003274444 | 1705.724599 | 1385.062931 | ENSG00000099381 |
| COL9A1 | -0.30060117 | 0.811914002 | 0.02127316 | 0.068019185 | 476.7725096 | 387.4568858 | ENSG00000112280 |
| LOC105370689 | -0.300627832 | 0.811898997 | 0.702097746 | 0.830897109 | 9.429465476 | 7.723116212 | ENSG00000287346 |
| RANBP3 | -0.300653703 | 0.811884438 | 2.31E-05 | 0.000189571 | 2389.381548 | 1940.338658 | ENSG00000031823 |
| TMEM115 | -0.300697049 | 0.811860045 | 0.014325976 | 0.049206382 | 448.4593237 | 363.9615568 | ENSG00000126062 |
| ZFP69B | -0.300792718 | 0.81180621 | 0.16594641 | 0.329087392 | 133.6877088 | 108.2088563 | ENSG00000187801 |
| CSNK1G2 | -0.30084639 | 0.81177601 | 0.000227053 | 0.001437717 | 4224.514936 | 3429.758684 | ENSG00000133275 |
| DBF4 | -0.3008503 | 0.81177381 | 0.012761663 | 0.044992328 | 811.6594981 | 658.453557 | ENSG00000006634 |
| CSF1 | -0.300941156 | 0.811722688 | 0.122198053 | 0.263167409 | 145.8844317 | 118.4610723 | ENSG00000184371 |
| SLC1A3 | -0.300949274 | 0.811718121 | 0.00746404 | 0.028880692 | 529.7591515 | 429.905167 | ENSG00000079215 |
| ZC3H15 | -0.300981978 | 0.811699721 | 2.30E-05 | 0.000189063 | 2163.001271 | 1755.213598 | ENSG00000065548 |
| C1orf94 | -0.301204605 | 0.811574474 | 0.209384504 | 0.388490944 | 104.9237689 | 85.06730205 | ENSG00000142698 |
| C12orf73 | -0.301217981 | 0.811566949 | 0.16031548 | 0.320707646 | 128.6016107 | 104.3163305 | ENSG00000204954 |
| PDE8B | -0.301274126 | 0.811535367 | 0.533377635 | 0.70829298 | 23.13375582 | 18.73873919 | ENSG00000113231 |
| EEF1E1 | -0.301321444 | 0.81150875 | 0.334796536 | 0.529176105 | 89.17090687 | 72.20461866 | ENSG00000124802 |
| ENPP4 | -0.301448041 | 0.811437543 | 0.212828872 | 0.392533617 | 95.02104495 | 77.11221228 | ENSG00000001561 |
| JMJD6 | -0.30147553 | 0.811422082 | 0.008437901 | 0.032013932 | 528.4045889 | 428.7114041 | ENSG00000070495 |
| CYB5R2 | -0.30153006 | 0.811391413 | 0.007667466 | 0.029575197 | 501.4317671 | 406.841611 | ENSG00000166394 |
| FCHSD1 | -0.301543139 | 0.811384057 | 0.039634588 | 0.111673373 | 628.2793384 | 510.3978728 | ENSG00000197948 |
| MICAL1 | -0.301615587 | 0.811343313 | 0.194107725 | 0.368050701 | 1510.376909 | 1225.751752 | ENSG00000135596 |
| NA | -0.301707975 | 0.811291358 | 0.731303109 | 0.849778798 | 6.411384112 | 5.206177396 | ENSG00000287878 |
| SDHAF1 | -0.302184017 | 0.811023702 | 0.02396377 | 0.074860021 | 327.4642959 | 265.6384713 | ENSG00000205138 |
| RPL41 | -0.30234783 | 0.810931618 | 4.89E-07 | 5.79E-06 | 9312.223103 | 7550.651407 | ENSG00000229117 |
| WRAP73 | -0.302372413 | 0.8109178 | 0.014810913 | 0.050544183 | 615.8087835 | 499.310781 | ENSG00000116213 |
| MIR4292 | -0.302453698 | 0.810872113 | 0.466382039 | 0.652519417 | 31.64935593 | 25.71579371 | ENSG00000265806 |
| NUDT16L1 | -0.302475945 | 0.810859609 | 0.00138375 | 0.006907199 | 909.7488193 | 738.1769701 | ENSG00000168101 |
| ANKLE2 | -0.302615357 | 0.810781257 | 4.57E-05 | 0.000347756 | 1606.590356 | 1302.690141 | ENSG00000176915 |
| FGF22 | -0.302677476 | 0.810746347 | 0.492029349 | 0.674602157 | 32.96762901 | 26.84129738 | ENSG00000070388 |
| DYNLT3 | -0.302769166 | 0.810694822 | 0.341553136 | 0.53583425 | 69.27125532 | 56.12604982 | ENSG00000165169 |
| NA | -0.302777955 | 0.810689884 | 0.23229256 | 0.417039061 | 99.49116981 | 80.95072684 | ENSG00000180385 |
| NA | -0.302837201 | 0.810656592 | 0.367345057 | 0.560452196 | 45.44221259 | 36.87421019 | ENSG00000185275 |
| SHLD1 | -0.303044594 | 0.810540066 | 0.297652166 | 0.490677332 | 80.87289307 | 65.30787268 | ENSG00000171984 |
| TMEM132A | -0.303103512 | 0.810506965 | 0.000108117 | 0.000746586 | 2876.95254 | 2331.912877 | ENSG00000006118 |
| CRYBB2P1 | -0.303103907 | 0.810506743 | 0.087541267 | 0.205092888 | 273.5138794 | 222.1707109 | ENSG00000100058 |
| C1QBP | -0.303199609 | 0.810452979 | 4.75E-05 | 0.000360315 | 6324.16418 | 5124.603089 | ENSG00000108561 |
| HMOX1 | -0.303222095 | 0.810440347 | 0.002086047 | 0.009852906 | 816.0471524 | 661.1314394 | ENSG00000100292 |
| TMX2 | -0.303471593 | 0.810300203 | 0.00103683 | 0.005438261 | 1082.226665 | 876.9875828 | ENSG00000213593 |
| EXT2 | -0.30355657 | 0.810252476 | 1.92E-05 | 0.000160472 | 2025.742947 | 1641.144964 | ENSG00000151348 |
| CNOT10 | -0.303869847 | 0.810076552 | 0.002368764 | 0.010959228 | 794.317485 | 643.1078214 | ENSG00000182973 |
| CREM | -0.303901667 | 0.810058685 | 0.026385573 | 0.080951864 | 314.7078002 | 255.0468803 | ENSG00000095794 |
| GAS6 | -0.303958995 | 0.810026496 | 0.006117723 | 0.024489392 | 562.164858 | 455.3092198 | ENSG00000183087 |
| SLC7A10 | -0.303973363 | 0.810018429 | 0.238853365 | 0.424643479 | 80.10639528 | 64.91912116 | ENSG00000130876 |
| ZNF891 | -0.304081222 | 0.809957873 | 0.267574365 | 0.457746287 | 74.8854595 | 60.57922911 | ENSG00000214029 |
| SNORD100 | -0.304104414 | 0.809944852 | 0.622828139 | 0.776137748 | 14.38803198 | 11.61239278 | ENSG00000221500 |
| TMPO-AS1 | -0.304131882 | 0.809929431 | 0.309212661 | 0.503332882 | 64.17025959 | 52.16680223 | ENSG00000257167 |
| ZNF345 | -0.304137015 | 0.80992655 | 0.331302852 | 0.52552268 | 56.9449263 | 46.28932071 | ENSG00000251247 |
| RFXAP | -0.30422502 | 0.809877145 | 0.112577504 | 0.247686545 | 155.7092401 | 126.3128859 | ENSG00000133111 |
| TLR5 | -0.30434783 | 0.809808208 | 0.319798929 | 0.513518689 | 62.07659671 | 50.34005015 | ENSG00000187554 |
| ZNF395 | -0.304402264 | 0.809777653 | 0.041715253 | 0.116412109 | 341.7260892 | 276.6860224 | ENSG00000186918 |
| DEDD2 | -0.304525457 | 0.809708508 | 0.000146394 | 0.000972846 | 1268.405689 | 1027.079696 | ENSG00000160570 |
| KALRN | -0.304588286 | 0.809673247 | 0.069270562 | 0.171639118 | 222.9043759 | 180.4548561 | ENSG00000160145 |
| NSMAF | -0.304593545 | 0.809670295 | 0.00436036 | 0.018315045 | 582.0583911 | 471.0685701 | ENSG00000035681 |
| NA | -0.304682245 | 0.809620516 | 0.675704434 | 0.812436293 | 10.48724841 | 8.450295285 | ENSG00000275088 |
| SASS6 | -0.304691352 | 0.809615406 | 0.089260249 | 0.208303885 | 218.4080337 | 176.5814991 | ENSG00000156876 |
| LOC442497 | -0.304704438 | 0.809608063 | 0.612963521 | 0.769113472 | 14.80200813 | 12.00625724 | ENSG00000249574 |
| NA | -0.304796541 | 0.809556378 | 0.744415805 | 0.857195413 | 6.339376476 | 5.127294571 | ENSG00000279789 |
| NA | -0.304819307 | 0.809543603 | 0.588429346 | 0.750839896 | 16.48791099 | 13.33769219 | ENSG00000228532 |
| LRRC73 | -0.304834528 | 0.809535062 | 0.734314057 | 0.851409242 | 6.319157223 | 5.108523577 | ENSG00000204052 |
| FBXO41 | -0.30508017 | 0.809397237 | 0.010388449 | 0.037953776 | 487.4213393 | 394.4871787 | ENSG00000163013 |
| SNCB | -0.305087961 | 0.809392866 | 0.07848272 | 0.188899274 | 216.7378177 | 175.3225602 | ENSG00000074317 |
| TMEM88B | -0.305108179 | 0.809381524 | 0.590826014 | 0.752706169 | 16.9549533 | 13.77669634 | ENSG00000205116 |
| ANKLE1 | -0.305161937 | 0.809351365 | 0.020974479 | 0.067221337 | 385.4779642 | 312.1767131 | ENSG00000160117 |
| SMG1P7 | -0.305358058 | 0.809241348 | 0.291343912 | 0.483761659 | 81.23006743 | 66.02672989 | ENSG00000261556 |
| EXOSC6 | -0.305423478 | 0.809204654 | 0.011270884 | 0.040609365 | 520.2387217 | 420.8223834 | ENSG00000223496 |
| NA | -0.305475055 | 0.809175725 | 0.633629263 | 0.783961787 | 12.43226594 | 10.0935743 | ENSG00000225850 |

| | | | | | | | |
|---|---|---|---|---|---|---|---|
| GDF1 | -0.305674664 | 0.809063776 | 0.127067888 | 0.270729535 | 174.6839695 | 141.7896123 | ENSG00000223802 |
| UBR5 | -0.305744067 | 0.809024856 | 0.000820684 | 0.004437154 | 4053.027571 | 3278.646241 | ENSG00000104517 |
| DDIT4L | -0.305853601 | 0.808963435 | 0.264976646 | 0.454645173 | 85.30267712 | 68.94706597 | ENSG00000145358 |
| BRAT1 | -0.305991659 | 0.808886025 | 0.000279959 | 0.001730739 | 1634.874137 | 1322.820435 | ENSG00000106009 |
| SRSF9 | -0.306021778 | 0.808869138 | 4.40E-05 | 0.000335863 | 3683.397791 | 2979.967713 | ENSG00000111786 |
| USP5 | -0.306184046 | 0.808778165 | 1.02E-05 | 9.12E-05 | 4142.071825 | 3350.131599 | ENSG00000111667 |
| GPC1 | -0.306412665 | 0.808650011 | 0.000308742 | 0.001881076 | 1704.039583 | 1377.879842 | ENSG00000063660 |
| PIF1 | -0.306421916 | 0.808644826 | 0.010141041 | 0.037189167 | 1771.603642 | 1433.541032 | ENSG00000140451 |
| NA | -0.306480514 | 0.808611981 | 0.762055273 | 0.868113515 | 5.180399573 | 4.189724094 | ENSG00000232545 |
| TIMP2 | -0.306486187 | 0.808608802 | 0.017126366 | 0.056814236 | 521.7504383 | 421.8426355 | ENSG00000035862 |
| L3MBTL2 | -0.306501896 | 0.808599998 | 0.005847645 | 0.023540454 | 532.4632706 | 430.4027372 | ENSG00000100395 |
| NA | -0.30652253 | 0.808588432 | 0.441082815 | 0.629284035 | 34.29084044 | 27.84635414 | ENSG00000271888 |
| ZNF37BP | -0.306606793 | 0.808541207 | 0.204918772 | 0.383091035 | 134.0537526 | 108.7827701 | ENSG00000234420 |
| GPC3 | -0.306847131 | 0.808406524 | 1.97E-06 | 2.07E-05 | 2925.798473 | 2365.305746 | ENSG00000147257 |
| ST20 | -0.306885611 | 0.808384962 | 0.687420803 | 0.820390029 | 8.921984661 | 7.205229238 | ENSG00000180953 |
| ANXA9 | -0.306893707 | 0.808380425 | 0.29468967 | 0.487528194 | 62.97961586 | 50.8232063 | ENSG00000143412 |
| KLK10 | -0.306979236 | 0.808332503 | 0.337043024 | 0.531367421 | 58.57588387 | 47.30467471 | ENSG00000129451 |
| PCBP1 | -0.306990726 | 0.808326065 | 0.140122814 | 0.29114756 | 184.445354 | 149.1006024 | ENSG00000169564 |
| NFKB1 | -0.307266236 | 0.808171715 | 0.023984517 | 0.074912207 | 316.0225482 | 255.388119 | ENSG00000109320 |
| RAB40C | -0.30732886 | 0.808136634 | 0.00244061 | 0.011238328 | 749.5553913 | 605.2331262 | ENSG00000197562 |
| MGAT4B | -0.307409289 | 0.808091583 | 6.20E-05 | 0.000455469 | 1770.409593 | 1431.41399 | ENSG00000161013 |
| CHRNA5 | -0.307457803 | 0.808064409 | 0.080145696 | 0.192014003 | 208.3773873 | 168.2111859 | ENSG00000169684 |
| KIAA1586 | -0.307462003 | 0.808062056 | 0.107995579 | 0.239901583 | 238.5562241 | 192.7674064 | ENSG00000168116 |
| IFT88 | -0.307535727 | 0.808020765 | 0.204650198 | 0.382704749 | 89.78482827 | 72.52157568 | ENSG00000032742 |
| ITLN2 | -0.307543217 | 0.808016569 | 0.245011719 | 0.432151427 | 86.8564933 | 69.98151737 | ENSG00000158764 |
| C11orf87 | -0.307754655 | 0.807898157 | 0.601209449 | 0.760549856 | 15.1817619 | 12.23866884 | ENSG00000185742 |
| PMF1 | -0.307793297 | 0.807876518 | 0.036880858 | 0.105372255 | 347.9355738 | 281.4505993 | ENSG00000160783 |
| SRM | -0.307948894 | 0.807789392 | 0.000110384 | 0.000761389 | 3219.19967 | 2600.259141 | ENSG00000116649 |
| SMTNL2 | -0.308003453 | 0.807758844 | 0.263751507 | 0.45342197 | 82.42878378 | 66.52755773 | ENSG00000188176 |
| SMNDC1 | -0.30801495 | 0.807752407 | 0.005564592 | 0.022582471 | 647.5762305 | 522.8244572 | ENSG00000119953 |
| PRKCG | -0.308036641 | 0.807740263 | 0.062923165 | 0.159865379 | 210.9839504 | 170.5383681 | ENSG00000126583 |
| ZNF271P | -0.308116724 | 0.807695427 | 0.011901216 | 0.042492225 | 457.7089881 | 370.1144312 | ENSG00000257267 |
| NA | -0.308194244 | 0.807652028 | 0.636531647 | 0.786374602 | 13.31950053 | 10.81370533 | ENSG00000269054 |
| NA | -0.308349154 | 0.807565311 | 0.76498377 | 0.870042292 | 5.035106525 | 4.054528287 | ENSG00000251292 |
| CDK2AP2 | -0.308413847 | 0.807529099 | 0.001283877 | 0.006492597 | 887.7643143 | 716.2710918 | ENSG00000167797 |
| MAP7D1 | -0.308535387 | 0.807461071 | 0.000279935 | 0.001730739 | 3101.099295 | 2504.135713 | ENSG00000116871 |
| NA | -0.308563828 | 0.807445154 | 0.739872094 | 0.854707161 | 8.050030976 | 6.537612341 | ENSG00000233672 |
| TARBP2 | -0.308611077 | 0.80741871 | 0.000245273 | 0.00154307 | 1338.020483 | 1081.03097 | ENSG00000139546 |
| DDX27 | -0.30867689 | 0.807381878 | 3.13E-05 | 0.000249788 | 1785.735934 | 1442.170354 | ENSG00000124228 |
| SMARCA5 | -0.308890318 | 0.807262445 | 1.63E-06 | 1.74E-05 | 4424.950363 | 3571.140259 | ENSG00000153147 |
| NRG2 | -0.308935554 | 0.807237133 | 0.641028399 | 0.789659074 | 11.52293296 | 9.28696275 | ENSG00000158458 |
| EED | -0.308949534 | 0.807229311 | 0.002308212 | 0.010721907 | 701.3853752 | 566.1299776 | ENSG00000074266 |
| CCNE2 | -0.309001136 | 0.807200439 | 0.33419516 | 0.528670131 | 58.21389149 | 47.16677936 | ENSG00000175305 |
| BCAN | -0.309137106 | 0.807124366 | 0.202881683 | 0.380318511 | 104.2343148 | 84.0658209 | ENSG00000132692 |
| C12orf10 | -0.309139685 | 0.807122923 | 0.002580648 | 0.011763338 | 706.3753229 | 570.428632 | ENSG00000139637 |
| SNX33 | -0.309148396 | 0.80711805 | 0.000516832 | 0.002953733 | 943.5459041 | 761.2413533 | ENSG00000173548 |
| UBE2O | -0.309162378 | 0.807110228 | 3.67E-06 | 3.67E-05 | 2583.737879 | 2085.025232 | ENSG00000175931 |
| PRR15 | -0.309316211 | 0.807024171 | 0.098860563 | 0.22476468 | 155.7371486 | 125.6476422 | ENSG00000176532 |
| GET3 | -0.309501785 | 0.80692037 | 3.94E-05 | 0.000305999 | 2275.341262 | 1835.843976 | ENSG00000198356 |
| KPNB1 | -0.309602132 | 0.806864247 | 3.73E-08 | 5.40E-07 | 10590.59402 | 8544.60271 | ENSG00000108424 |
| MTCH1 | -0.309664776 | 0.806829212 | 9.80E-08 | 1.31E-06 | 5522.083553 | 4454.654982 | ENSG00000137409 |
| GMNN | -0.309881518 | 0.806708008 | 1.10E-05 | 9.75E-05 | 1908.898406 | 1539.703177 | ENSG00000112312 |
| GLRX | -0.309911017 | 0.806691513 | 0.026891281 | 0.082190469 | 383.7475337 | 309.5719797 | ENSG00000173221 |
| SYVN1 | -0.310069224 | 0.806603056 | 0.00756035 | 0.029198484 | 699.8376098 | 564.9802954 | ENSG00000162298 |
| E2F3 | -0.310171421 | 0.80654592 | 0.000191425 | 0.00123576 | 1261.977957 | 1017.147757 | ENSG00000112242 |
| PARP10 | -0.310245906 | 0.80650428 | 0.24082968 | 0.427051023 | 82.15764282 | 66.05426078 | ENSG00000178685 |
| DDX5 | -0.310268032 | 0.806491911 | 3.75E-06 | 3.74E-05 | 9243.553465 | 7455.916163 | ENSG00000108654 |
| SEC61B | -0.310277693 | 0.80648651 | 5.21E-05 | 0.00038987 | 1605.428884 | 1294.955748 | ENSG00000106803 |
| TNS2 | -0.310291271 | 0.80647892 | 0.178895964 | 0.347844922 | 119.7525206 | 96.62144675 | ENSG00000111077 |
| TNFRSF8 | -0.310543798 | 0.806337768 | 0.000905032 | 0.00484094 | 1069.462638 | 862.1314784 | ENSG00000120949 |
| ADAM15 | -0.310563696 | 0.806326647 | 0.001141815 | 0.005898104 | 900.6429822 | 726.6126398 | ENSG00000143537 |
| SLCO4C1 | -0.310604869 | 0.806303635 | 0.100247034 | 0.226997987 | 156.8967269 | 126.4161621 | ENSG00000173930 |
| LINC01664 | -0.31062269 | 0.806293675 | 0.754075547 | 0.863411144 | 5.124176501 | 4.12212619 | ENSG00000235478 |
| ABCB1 | -0.310691646 | 0.806255138 | 0.552136762 | 0.723048311 | 20.1150731 | 16.21855118 | ENSG00000085563 |
| LRRC34 | -0.310729399 | 0.80623404 | 0.361571885 | 0.554749447 | 50.13219369 | 40.29822554 | ENSG00000171757 |
| TMEM67 | -0.310762178 | 0.806215722 | 0.105785168 | 0.236533032 | 163.4255932 | 131.9313724 | ENSG00000164953 |
| WDR19 | -0.311119495 | 0.806016068 | 0.058187608 | 0.150348408 | 310.7188563 | 250.9093842 | ENSG00000157796 |
| NLK | -0.311189179 | 0.805977137 | 0.002510449 | 0.011505603 | 635.2432945 | 511.855726 | ENSG00000087095 |
| MTAP | -0.311269553 | 0.805932237 | 0.008539344 | 0.032306405 | 681.0628629 | 548.4590663 | ENSG00000099810 |
| GPR27 | -0.311313285 | 0.805907808 | 0.008254048 | 0.031432202 | 493.9615176 | 397.743194 | ENSG00000170837 |
| LRRC8C | -0.3114396 | 0.805837249 | 0.251383789 | 0.439998786 | 77.78181938 | 62.70923988 | ENSG00000171488 |
| AGPAT5 | -0.311571573 | 0.805763537 | 0.001743797 | 0.00844955 | 1724.07299 | 1388.983339 | ENSG00000155189 |
| BEND7 | -0.311599895 | 0.80574772 | 0.138211823 | 0.288016407 | 125.0539501 | 100.851524 | ENSG00000165626 |
| GPR4 | -0.31162875 | 0.805731604 | 0.730693089 | 0.849344242 | 6.702067906 | 5.348421244 | ENSG00000177464 |

| | | | | | | | |
|---|---|---|---|---|---|---|---|
| MMP11 | -0.311667744 | 0.805709827 | 0.069558 | 0.172190193 | 189.5955751 | 152.5591254 | ENSG00000099953 |
| XYLB | -0.31170297 | 0.805690154 | 0.258274117 | 0.446993416 | 76.46670322 | 61.8326573 | ENSG00000093217 |
| SLC35B1 | -0.311849243 | 0.80560847 | 0.000849701 | 0.004578013 | 1012.587622 | 815.6678027 | ENSG00000121073 |
| OSGEP | -0.312026247 | 0.805509636 | 0.005672274 | 0.022939105 | 520.4251675 | 419.1932407 | ENSG00000092094 |
| ISOC1 | -0.312097721 | 0.805469731 | 0.001782707 | 0.008608432 | 1008.837821 | 812.1585891 | ENSG00000066583 |
| SLC25A35 | -0.312131459 | 0.805450894 | 0.480290235 | 0.66425634 | 39.35869617 | 31.87550142 | ENSG00000125434 |
| TMEM220 | -0.31231994 | 0.805345673 | 0.413760466 | 0.605166455 | 39.16665079 | 31.48780893 | ENSG00000187824 |
| CALR | -0.312333386 | 0.805338167 | 1.51E-09 | 2.74E-08 | 27731.8866 | 22333.29408 | ENSG00000179218 |
| NA | -0.312368015 | 0.805318837 | 0.721653459 | 0.842633731 | 6.911273326 | 5.622611705 | ENSG00000269038 |
| NPY5R | -0.312434377 | 0.805281795 | 0.484984788 | 0.668210541 | 25.56593721 | 20.52794928 | ENSG00000164129 |
| BMERB1 | -0.312552671 | 0.805215768 | 0.03248928 | 0.095368438 | 320.5385158 | 258.6158793 | ENSG00000166780 |
| EPS8L1 | -0.312678432 | 0.805145579 | 0.003369653 | 0.014688193 | 737.2298212 | 593.8572074 | ENSG00000131037 |
| HDAC6 | -0.312688275 | 0.805140087 | 0.019219037 | 0.062360929 | 803.6007857 | 647.8884959 | ENSG00000094631 |
| MRPS11 | -0.312755349 | 0.805102655 | 0.006675367 | 0.026340485 | 529.6853397 | 426.4433864 | ENSG00000181991 |
| CA8 | -0.313025988 | 0.804951638 | 0.1672783 | 0.330949618 | 125.3520301 | 100.6400333 | ENSG00000178538 |
| UROD | -0.313050078 | 0.804938197 | 1.91E-05 | 0.000159822 | 1595.194549 | 1283.923169 | ENSG00000126088 |
| CRYBG2 | -0.313301519 | 0.80479792 | 0.028271373 | 0.085575965 | 287.9531961 | 231.6672519 | ENSG00000176092 |
| SNRNP200 | -0.313313964 | 0.804790977 | 9.35E-09 | 1.51E-07 | 11908.88431 | 9584.28373 | ENSG00000144028 |
| MRPL13 | -0.313599039 | 0.804631967 | 0.002333723 | 0.010824129 | 735.1790774 | 591.1217927 | ENSG00000172172 |
| CHIC2 | -0.313605124 | 0.804628573 | 0.17363164 | 0.340352355 | 128.707743 | 103.5555199 | ENSG00000109220 |
| TMEM39B | -0.313614289 | 0.804623462 | 0.008179627 | 0.031200082 | 469.0331711 | 377.1372464 | ENSG00000121775 |
| CIZ1 | -0.313722474 | 0.804563127 | 1.53E-05 | 0.000131367 | 2291.667771 | 1844.247603 | ENSG00000148337 |
| ZNF793 | -0.313728264 | 0.804559898 | 0.093165523 | 0.214820826 | 158.053727 | 127.1555021 | ENSG00000188227 |
| HARS1 | -0.313916901 | 0.804454706 | 4.23E-05 | 0.000324298 | 1547.39668 | 1245.3826 | ENSG00000170445 |
| JAKMIP2 | -0.313998947 | 0.804408958 | 0.030984935 | 0.091883966 | 272.7454048 | 219.5122524 | ENSG00000176049 |
| PPP3CB | -0.314003014 | 0.80440669 | 0.007077168 | 0.027668854 | 555.0188157 | 445.8607639 | ENSG00000107758 |
| RNF167 | -0.314064434 | 0.804372445 | 4.28E-06 | 4.22E-05 | 2139.96114 | 1721.42676 | ENSG00000108523 |
| PCTP | -0.314111699 | 0.804346093 | 0.006200488 | 0.024756538 | 471.2815977 | 379.2471008 | ENSG00000141179 |
| NCLN | -0.314527849 | 0.80411411 | 0.000124056 | 0.00084376 | 4226.425677 | 3398.731977 | ENSG00000125912 |
| NAP1L5 | -0.314572416 | 0.80408927 | 0.365197102 | 0.558416227 | 44.7831249 | 35.99240305 | ENSG00000177432 |
| MRPL34 | -0.314657536 | 0.80404183 | 0.000134361 | 0.000902265 | 1174.580805 | 944.3865473 | ENSG00000130312 |
| DGKK | -0.314776223 | 0.803975686 | 0.002262658 | 0.010545845 | 654.0339761 | 525.6696379 | ENSG00000274588 |
| RBBP5 | -0.314871106 | 0.803922812 | 0.002191906 | 0.010284754 | 645.8038057 | 518.8313548 | ENSG00000117222 |
| NA | -0.314904871 | 0.803903997 | 0.721492777 | 0.842552273 | 8.089191707 | 6.533925114 | ENSG00000235586 |
| GPATCH4 | -0.314905114 | 0.803903862 | 0.000124872 | 0.000848687 | 1755.04467 | 1411.392259 | ENSG00000160818 |
| PDE7A | -0.314963821 | 0.803871149 | 6.72E-05 | 0.000489289 | 1329.345332 | 1068.208488 | ENSG00000205268 |
| LRIF1 | -0.31518636 | 0.80374716 | 0.133182495 | 0.280214117 | 155.8164396 | 125.3664037 | ENSG00000121931 |
| TOR1A | -0.315332709 | 0.803665631 | 0.00012997 | 0.000877685 | 1131.10337 | 909.1743871 | ENSG00000136827 |
| CENPVL3 | -0.315465377 | 0.803591731 | 0.146399745 | 0.300550827 | 125.3704679 | 100.8671574 | ENSG00000224109 |
| LRRC2 | -0.31558543 | 0.803524863 | 0.177562797 | 0.346124198 | 100.6076337 | 80.9878308 | ENSG00000163827 |
| YWHAQ | -0.315982568 | 0.803303703 | 6.64E-05 | 0.000484626 | 10318.05121 | 8287.445313 | ENSG00000134308 |
| GDPD1 | -0.316019203 | 0.803283304 | 0.241880683 | 0.428095549 | 74.6946919 | 59.99798111 | ENSG00000153982 |
| CISD2 | -0.316096982 | 0.803239999 | 0.132273768 | 0.278871802 | 271.3994784 | 217.7864747 | ENSG00000145354 |
| CAPN15 | -0.316182695 | 0.803192279 | 0.000115638 | 0.000794382 | 2150.874729 | 1727.638074 | ENSG00000103326 |
| PPP2R1A | -0.316317627 | 0.803117161 | 9.15E-09 | 1.48E-07 | 8527.304483 | 6848.31315 | ENSG00000105568 |
| LOC105371414 | -0.316348105 | 0.803100195 | 0.558277682 | 0.727516983 | 21.69612142 | 17.4977984 | ENSG00000260279 |
| METTL17 | -0.31635446 | 0.803096658 | 0.002891881 | 0.012899667 | 742.9789986 | 597.2396989 | ENSG00000165792 |
| NRXN2 | -0.316369432 | 0.803088323 | 0.008579943 | 0.032426725 | 480.4148397 | 385.7832874 | ENSG00000110076 |
| NA | -0.316371312 | 0.803087277 | 0.139488703 | 0.290350812 | 125.4545019 | 100.5730967 | ENSG00000261409 |
| CHEK1 | -0.316460041 | 0.803037887 | 9.33E-05 | 0.000655418 | 1718.269116 | 1379.113846 | ENSG00000149554 |
| TOM1 | -0.316672106 | 0.802919855 | 0.001982629 | 0.009424481 | 628.7078061 | 504.6945371 | ENSG00000100284 |
| RFC5 | -0.316757762 | 0.802872185 | 0.00350173 | 0.015185087 | 558.2784457 | 448.1216707 | ENSG00000111445 |
| TCOF1 | -0.316921919 | 0.802780835 | 3.30E-08 | 4.83E-07 | 8165.533638 | 6555.840304 | ENSG00000070814 |
| TUB | -0.317087924 | 0.802688468 | 0.00032133 | 0.001947523 | 1308.618059 | 1050.845966 | ENSG00000166402 |
| RBM6 | -0.317245182 | 0.802600977 | 0.002307858 | 0.010721907 | 2504.800007 | 2011.429126 | ENSG00000004534 |
| NA | -0.317250991 | 0.802597746 | 0.784586876 | 0.882158322 | 4.442990421 | 3.619761016 | ENSG00000240522 |
| APOB | -0.317325656 | 0.802556209 | 0.500682354 | 0.681425081 | 23.68345658 | 19.0091308 | ENSG00000084674 |
| ACAP3 | -0.317445918 | 0.802489312 | 0.003729145 | 0.016036189 | 822.673883 | 660.8802342 | ENSG00000131584 |
| NA | -0.317487088 | 0.802466411 | 0.684162116 | 0.817890607 | 10.2559446 | 8.199224318 | ENSG00000267500 |
| TMEM132B | -0.317547253 | 0.802432946 | 5.21E-05 | 0.00038987 | 2000.822387 | 1605.273715 | ENSG00000139364 |
| FAM241A | -0.317815479 | 0.802283772 | 0.470138672 | 0.655738783 | 33.35063739 | 26.94632565 | ENSG00000174749 |
| PDE2A | -0.317942633 | 0.802213065 | 0.028229693 | 0.085503133 | 304.3569754 | 244.1732864 | ENSG00000186642 |
| NDUFA5 | -0.317955024 | 0.802206175 | 0.000720769 | 0.00395697 | 1071.925862 | 859.6409082 | ENSG00000128609 |
| NA | -0.318127485 | 0.802110284 | 0.504889426 | 0.68443685 | 23.24747974 | 18.67906539 | ENSG00000234518 |
| DDX39B | -0.318197008 | 0.802071631 | 0.006293852 | 0.025069904 | 1391.363323 | 1116.605642 | ENSG00000198563 |
| GSPT2 | -0.318243668 | 0.802045691 | 0.000779923 | 0.004244008 | 755.3217886 | 606.0583258 | ENSG00000189369 |
| ERCC3 | -0.318266452 | 0.802033025 | 0.000141064 | 0.0009408 | 1606.844224 | 1289.439965 | ENSG00000163161 |
| ITFG2 | -0.318392563 | 0.801962919 | 0.031553303 | 0.093300355 | 299.3978302 | 240.3895878 | ENSG00000111203 |
| EFCAB11 | -0.318439867 | 0.801936625 | 0.283704232 | 0.474650139 | 71.91560365 | 57.49893723 | ENSG00000140025 |
| NA | -0.318599895 | 0.801847676 | 0.483797279 | 0.667132016 | 24.41149269 | 19.5903788 | ENSG00000280120 |
| MPV17L | -0.318679996 | 0.801803158 | 0.190124614 | 0.362415879 | 89.85821138 | 72.02625855 | ENSG00000156968 |
| SRP14 | -0.318744335 | 0.801767401 | 3.78E-07 | 4.56E-06 | 5646.191489 | 4526.993833 | ENSG00000140319 |
| CBX1 | -0.318918696 | 0.801670507 | 9.35E-09 | 1.51E-07 | 7178.891722 | 5754.82667 | ENSG00000108468 |
| NHLRC3 | -0.319069009 | 0.801586986 | 0.049080883 | 0.131835518 | 216.5251623 | 173.5307537 | ENSG00000188811 |

| | | | | | | | |
|---|---|---|---|---|---|---|---|
| WAS | -0.319119443 | 0.801558964 | 0.487852639 | 0.6709105 | 24.15049889 | 19.41807904 | ENSG00000015285 |
| DYNC1I2 | -0.319874219 | 0.801139722 | 1.31E-05 | 0.000113954 | 1769.502733 | 1417.092407 | ENSG00000077380 |
| NCOA2 | -0.319994648 | 0.801072849 | 0.00194078 | 0.009264518 | 709.9095742 | 568.1260579 | ENSG00000140396 |
| EBNA1BP2 | -0.320039747 | 0.801047808 | 5.97E-05 | 0.000440531 | 1369.097319 | 1096.680944 | ENSG00000117395 |
| ZNF326 | -0.32004006 | 0.801047634 | 0.001104512 | 0.005743832 | 850.9618735 | 681.6565775 | ENSG00000162664 |
| CIR1 | -0.320210337 | 0.800953094 | 0.063751892 | 0.161371677 | 281.5114456 | 224.9801958 | ENSG00000138433 |
| RAB11B | -0.320212601 | 0.800951838 | 0.000132604 | 0.000893055 | 1665.748667 | 1334.608153 | ENSG00000185236 |
| TIMM23 | -0.320236895 | 0.80093835 | 7.15E-06 | 6.68E-05 | 2334.470206 | 1869.302771 | ENSG00000265354 |
| DAAM2 | -0.320318994 | 0.800892773 | 0.069629085 | 0.172320131 | 178.1567738 | 142.5219757 | ENSG00000146122 |
| PREX2 | -0.320512864 | 0.800785155 | 0.000395466 | 0.002329063 | 1142.219814 | 914.9504562 | ENSG00000046889 |
| CAPNS1 | -0.320595505 | 0.800739286 | 4.49E-06 | 4.39E-05 | 2836.835336 | 2272.140146 | ENSG00000126247 |
| ZNF740 | -0.320644581 | 0.800712048 | 0.001045583 | 0.005474247 | 685.213501 | 548.8129443 | ENSG00000139651 |
| SPOP | -0.320680945 | 0.800691866 | 0.00335854 | 0.01465329 | 582.0145852 | 465.9410607 | ENSG00000121067 |
| NA | -0.320847539 | 0.800599412 | 0.563479005 | 0.731851428 | 17.7827102 | 14.30955546 | ENSG00000198237 |
| BAHD1 | -0.320911937 | 0.800563676 | 6.80E-05 | 0.000493761 | 1190.621509 | 953.1549618 | ENSG00000140320 |
| LRRC4C | -0.320981833 | 0.800524891 | 0.684188105 | 0.817890607 | 8.868918502 | 7.05874851 | ENSG00000148948 |
| MRFAP1L1 | -0.321154798 | 0.800428922 | 9.58E-05 | 0.000670767 | 1422.949924 | 1138.686851 | ENSG00000178988 |
| LOC101927811 | -0.321357784 | 0.80031631 | 0.684106793 | 0.817890607 | 10.16875376 | 8.10145888 | ENSG00000271133 |
| ZFP36L2 | -0.321499997 | 0.800237423 | 0.003362413 | 0.014663283 | 1456.911062 | 1165.778655 | ENSG00000152518 |
| EFNA4 | -0.321505546 | 0.800234345 | 0.041106281 | 0.114946261 | 223.6912133 | 178.9238766 | ENSG00000243364 |
| AGAP4 | -0.321514689 | 0.800229274 | 0.252408883 | 0.441168072 | 100.8083757 | 80.80859462 | ENSG00000188234 |
| CSKMT | -0.321567305 | 0.800200089 | 0.253055789 | 0.441715576 | 81.09680067 | 65.07297634 | ENSG00000214756 |
| MIR4664 | -0.321785335 | 0.800079167 | 0.239690355 | 0.425560018 | 73.32166892 | 58.80174198 | ENSG00000265660 |
| LMAN2L | -0.32181104 | 0.800064912 | 0.002282836 | 0.010622941 | 665.4265812 | 532.4980534 | ENSG00000114988 |
| XKR7 | -0.321886254 | 0.800023202 | 0.034390631 | 0.09979583 | 270.9827665 | 216.5100705 | ENSG00000260903 |
| PEMT | -0.322127214 | 0.799889593 | 0.136792893 | 0.285798611 | 118.1237353 | 94.67067217 | ENSG00000133027 |
| UBL5 | -0.322184708 | 0.799857716 | 9.47E-06 | 8.52E-05 | 2376.800149 | 1900.877762 | ENSG00000198258 |
| PMEL | -0.322201839 | 0.799848218 | 4.17E-05 | 0.000320974 | 1963.950442 | 1570.398081 | ENSG00000185664 |
| NA | -0.322231943 | 0.799831529 | 0.706310151 | 0.833246982 | 7.636054826 | 6.072351123 | ENSG00000272275 |
| NA | -0.322251568 | 0.799820649 | 0.782137097 | 0.8805327 | 4.369705009 | 3.540878192 | ENSG00000284642 |
| EEF1B2 | -0.322531589 | 0.799665422 | 1.27E-07 | 1.66E-06 | 6857.225528 | 5482.785064 | ENSG00000114942 |
| RNF24 | -0.322583654 | 0.799636563 | 0.024478114 | 0.076145917 | 344.8397497 | 275.9092351 | ENSG00000101236 |
| EXOSC10 | -0.322625318 | 0.799613471 | 0.000549863 | 0.003119423 | 1189.524145 | 951.5909323 | ENSG00000171824 |
| BCLAF1 | -0.322654517 | 0.799597287 | 0.000114056 | 0.000784969 | 2892.477428 | 2311.757087 | ENSG00000029363 |
| AFG3L1P | -0.3227206 | 0.799560663 | 0.079973522 | 0.191725424 | 177.6916106 | 142.3252826 | ENSG00000223959 |
| ATXN7L1 | -0.322790184 | 0.799522099 | 0.001074891 | 0.005611857 | 870.4225193 | 695.8057625 | ENSG00000146776 |
| EML5 | -0.323156099 | 0.79931934 | 0.597027899 | 0.757172238 | 14.86011034 | 11.83720668 | ENSG00000165521 |
| NA | -0.32327381 | 0.799254125 | 0.575102818 | 0.740467702 | 15.62848726 | 12.53163029 | ENSG00000228393 |
| ARMH4 | -0.323374875 | 0.799198137 | 0.021801367 | 0.069474989 | 292.9623709 | 234.3616034 | ENSG00000139971 |
| SLC9A3R1 | -0.323390998 | 0.799189205 | 2.38E-06 | 2.46E-05 | 2747.220086 | 2195.025234 | ENSG00000109062 |
| PEPD | -0.323393007 | 0.799188092 | 4.21E-05 | 0.000323171 | 2208.392635 | 1765.435679 | ENSG00000124299 |
| MB | -0.32348802 | 0.799135462 | 0.657227735 | 0.799701868 | 9.47178312 | 7.595406479 | ENSG00000198125 |
| C10orf82 | -0.323650599 | 0.799045411 | 0.144597465 | 0.297939901 | 143.9152639 | 115.3010618 | ENSG00000165863 |
| NA | -0.32380915 | 0.798957601 | 0.46984025 | 0.655539918 | 30.48671846 | 24.48569981 | ENSG00000279672 |
| RPS15A | -0.32387341 | 0.798922015 | 0.002401786 | 0.01108158 | 636.501203 | 508.3172209 | ENSG00000134419 |
| CCNDBP1 | -0.323912747 | 0.798900232 | 0.014888148 | 0.050742358 | 455.1616847 | 363.7815393 | ENSG00000166946 |
| MAP3K8 | -0.323919773 | 0.798896341 | 0.277658925 | 0.468699553 | 77.23192323 | 61.42509451 | ENSG00000107968 |
| KCNAB3 | -0.323957834 | 0.798875265 | 0.334146114 | 0.528670131 | 54.33322949 | 43.58879721 | ENSG00000170049 |
| ADH6 | -0.32405651 | 0.798820626 | 0.759522773 | 0.866329932 | 5.164615008 | 4.103355196 | ENSG00000172955 |
| TMPO | -0.324082436 | 0.798806271 | 0.000335701 | 0.002023377 | 3034.682819 | 2423.217341 | ENSG00000120802 |
| ZNF668 | -0.324108822 | 0.798791662 | 0.099323498 | 0.225595797 | 160.2832122 | 127.9012374 | ENSG00000167394 |
| MRPL20 | -0.324206712 | 0.798737464 | 0.00020947 | 0.001337807 | 1249.923137 | 998.7197316 | ENSG00000242485 |
| NA | -0.324219979 | 0.798730118 | 0.772301979 | 0.874608633 | 5.925400326 | 4.669519427 | ENSG00000219797 |
| TRMU | -0.3244385 | 0.798609146 | 0.001243482 | 0.006325766 | 1231.996359 | 984.7533189 | ENSG00000100416 |
| WIZ | -0.324459374 | 0.798597592 | 0.001120931 | 0.005809655 | 1240.913014 | 991.1597483 | ENSG00000011451 |
| LINC01252 | -0.324463075 | 0.798595543 | 0.735970673 | 0.852173027 | 6.791137883 | 5.367192238 | ENSG00000247157 |
| SETBP1 | -0.324534671 | 0.798555912 | 0.18013959 | 0.349456474 | 89.10176305 | 71.13371613 | ENSG00000152217 |
| DNAJA1 | -0.324623479 | 0.798506757 | 1.33E-08 | 2.08E-07 | 6689.554514 | 5341.86326 | ENSG00000086061 |
| GNPDA2 | -0.324724199 | 0.798451012 | 0.154191 | 0.311780056 | 134.0561127 | 106.7008847 | ENSG00000163281 |
| DOCK10 | -0.324767018 | 0.798427315 | 0.318240501 | 0.512002556 | 52.32695284 | 41.90797621 | ENSG00000135905 |
| VPS18 | -0.32484559 | 0.798383832 | 0.000298102 | 0.00182706 | 1282.274582 | 1023.196448 | ENSG00000104142 |
| MREG | -0.32487629 | 0.798366843 | 0.009763489 | 0.036025583 | 434.9735594 | 347.0785263 | ENSG00000118242 |
| ZNF112 | -0.324957662 | 0.798321814 | 0.37674162 | 0.569909335 | 38.7685569 | 30.96656114 | ENSG00000062370 |
| ESYT3 | -0.325014946 | 0.798290116 | 0.100034915 | 0.226727797 | 205.7047396 | 164.4029823 | ENSG00000158220 |
| SLC29A4 | -0.325178794 | 0.798199459 | 0.027579037 | 0.083960228 | 317.4984563 | 253.6364509 | ENSG00000164638 |
| MICAL2 | -0.325320529 | 0.798121045 | 0.343164317 | 0.537043608 | 47.59002401 | 37.84877301 | ENSG00000133816 |
| IFRD1 | -0.325349839 | 0.79810483 | 0.000468721 | 0.002703787 | 993.8403629 | 793.7695482 | ENSG00000006652 |
| GUSB | -0.325354974 | 0.79810199 | 0.000771464 | 0.00420415 | 905.4756781 | 723.163355 | ENSG00000169919 |
| SRCIN1 | -0.325536162 | 0.798001763 | 0.158045645 | 0.317445143 | 114.8289733 | 91.91438534 | ENSG00000277363 |
| PPIB | -0.325545574 | 0.797996556 | 3.89E-06 | 3.88E-05 | 2067.490589 | 1650.071814 | ENSG00000166794 |
| NEGR1 | -0.325666717 | 0.797929551 | 0.349824047 | 0.543200309 | 45.36704805 | 36.12716082 | ENSG00000172260 |
| NA | -0.325788847 | 0.797862007 | 0.213987333 | 0.394042844 | 75.83532869 | 60.46269268 | ENSG00000180867 |
| IFRD2 | -0.325909591 | 0.797795234 | 3.93E-05 | 0.000305021 | 1617.896634 | 1291.408466 | ENSG00000214706 |
| SEC11C | -0.325952574 | 0.797771465 | 0.002235167 | 0.010434909 | 691.4296602 | 551.6665501 | ENSG00000166562 |

| | | | | | | | |
|---|---|---|---|---|---|---|---|
| LMF2 | -0.326261694 | 0.797600548 | 0.000479262 | 0.002760301 | 864.8809241 | 689.7139875 | ENSG00000100258 |
| SLC37A1 | -0.326357316 | 0.797547685 | 0.000953135 | 0.005053033 | 730.4799564 | 583.0068277 | ENSG00000160190 |
| SRP9 | -0.326470843 | 0.797484927 | 0.000131515 | 0.000886141 | 4871.97006 | 3884.388892 | ENSG00000143742 |
| PHKG1 | -0.326508835 | 0.797463927 | 0.720455226 | 0.842030342 | 6.944120231 | 5.569985949 | ENSG00000164776 |
| GS1-124K5.4 | -0.326516712 | 0.797459573 | 0.567173369 | 0.734612098 | 18.37660774 | 14.67716286 | ENSG00000237310 |
| NA | -0.326663366 | 0.797378513 | 0.290846471 | 0.483197247 | 67.92437463 | 54.1769242 | ENSG00000223804 |
| HTR6 | -0.326681082 | 0.797368721 | 0.333104119 | 0.527671088 | 46.95303471 | 37.3614916 | ENSG00000158748 |
| RAMP2 | -0.326777671 | 0.797315339 | 0.234096535 | 0.419001939 | 75.46386558 | 60.08716117 | ENSG00000131477 |
| TIMM13 | -0.326950008 | 0.797220101 | 5.76E-06 | 5.50E-05 | 4141.60282 | 3302.443431 | ENSG00000099800 |
| PTPN13 | -0.327010141 | 0.797186873 | 4.73E-07 | 5.61E-06 | 2530.751087 | 2017.468989 | ENSG00000163629 |
| LSAMP | -0.327010855 | 0.797186478 | 0.170269405 | 0.335328142 | 108.3871024 | 86.24531985 | ENSG00000185565 |
| PPP1R14B | -0.327082356 | 0.79714697 | 2.78E-05 | 0.000223539 | 6107.149271 | 4867.282832 | ENSG00000173457 |
| MAP7D2 | -0.327428887 | 0.796955521 | 0.197823266 | 0.373471984 | 94.87733794 | 75.62411099 | ENSG00000184368 |
| ZNF45 | -0.327546509 | 0.796890548 | 0.030396009 | 0.090528625 | 355.563819 | 282.7463786 | ENSG00000124459 |
| NA | -0.327619771 | 0.796850082 | 0.794728007 | 0.888735818 | 4.458774986 | 3.589705101 | ENSG00000275263 |
| IMPAD1 | -0.327713821 | 0.796798137 | 0.01483692 | 0.050614296 | 630.3905369 | 501.8653876 | ENSG00000104331 |
| GHITM | -0.327793805 | 0.796753963 | 2.25E-05 | 0.000185755 | 2666.194121 | 2123.127807 | ENSG00000165678 |
| SF3A3 | -0.327891334 | 0.796700102 | 2.16E-08 | 3.26E-07 | 4872.216465 | 3881.776787 | ENSG00000183431 |
| CNOT9 | -0.327943529 | 0.79667128 | 6.57E-06 | 6.18E-05 | 1707.632959 | 1360.129793 | ENSG00000144580 |
| RBM4 | -0.327948923 | 0.796668301 | 0.208229966 | 0.387327799 | 106.8104137 | 84.97779558 | ENSG00000173933 |
| UNC119 | -0.328059959 | 0.796606988 | 0.000307986 | 0.001877703 | 918.3130887 | 731.7291991 | ENSG00000109103 |
| PYURF | -0.328190355 | 0.796534991 | 0.002195391 | 0.010293291 | 733.8669001 | 584.5208681 | ENSG00000145337 |
| MFSD5 | -0.328229081 | 0.79651361 | 0.009513975 | 0.035315846 | 423.8153697 | 337.3145794 | ENSG00000182544 |
| AUH | -0.32847905 | 0.796375614 | 0.007508707 | 0.029023226 | 406.2567551 | 323.6985001 | ENSG00000148090 |
| BDKRB2 | -0.3285876 | 0.796315696 | 0.610841521 | 0.767912687 | 13.40729273 | 10.67102345 | ENSG00000168398 |
| OTUB2 | -0.329127937 | 0.796017506 | 0.185114505 | 0.355873787 | 103.7872963 | 82.67581146 | ENSG00000089723 |
| MAD2L2 | -0.329142996 | 0.796009196 | 8.28E-07 | 9.37E-06 | 4260.780371 | 3391.36714 | ENSG00000116670 |
| SCUBE1 | -0.329223583 | 0.795964734 | 0.043702777 | 0.120511204 | 222.0145857 | 176.5384464 | ENSG00000159307 |
| NA | -0.329396572 | 0.795869298 | 0.527070299 | 0.703089169 | 21.79032514 | 17.30585158 | ENSG00000269867 |
| CDO1 | -0.329420681 | 0.795855998 | 0.013053403 | 0.045716511 | 344.3166571 | 274.1272963 | ENSG00000129596 |
| NA | -0.329435908 | 0.795847598 | 0.675759031 | 0.812436293 | 8.181418596 | 6.530126268 | ENSG00000223612 |
| NA | -0.329510726 | 0.795806327 | 0.59509617 | 0.755943716 | 15.10965656 | 11.97620133 | ENSG00000161132 |
| PRPF39 | -0.329608828 | 0.795752215 | 0.003812944 | 0.016351018 | 611.5500515 | 486.5355573 | ENSG00000185246 |
| NA | -0.32974448 | 0.795677396 | 0.62675542 | 0.779145815 | 10.82468454 | 8.619345854 | ENSG00000227347 |
| TSPAN33 | -0.329894601 | 0.795594606 | 0.000774277 | 0.004218238 | 928.6194536 | 738.8637473 | ENSG00000158457 |
| CCSER1 | -0.329907782 | 0.795587337 | 0.655382782 | 0.798767968 | 9.623389994 | 7.674289303 | ENSG00000184305 |
| TRIM39 | -0.330000815 | 0.795536034 | 0.03087234 | 0.091638045 | 269.866979 | 214.6428536 | ENSG00000204599 |
| ZNF512 | -0.330108533 | 0.795476638 | 0.005402702 | 0.022002567 | 439.3541857 | 349.7057179 | ENSG00000243943 |
| LRRC37A3 | -0.330131528 | 0.79546396 | 0.465308935 | 0.651608296 | 27.27215702 | 21.73591137 | ENSG00000176809 |
| C2CD3 | -0.330132807 | 0.795463254 | 0.00664841 | 0.026245288 | 523.6850155 | 416.4034884 | ENSG00000168014 |
| LOC100506302 | -0.330150683 | 0.795453398 | 0.544596142 | 0.716707097 | 19.22656073 | 15.32600876 | ENSG00000216895 |
| TTC1 | -0.33021644 | 0.795417142 | 0.000150388 | 0.000995462 | 1437.555331 | 1142.77228 | ENSG00000113312 |
| NA | -0.330245821 | 0.795400943 | 0.663100568 | 0.804083749 | 9.4281877 | 7.505238734 | ENSG00000232630 |
| MTA3 | -0.330267007 | 0.795389263 | 8.57E-07 | 9.65E-06 | 2945.517625 | 2342.39395 | ENSG00000057935 |
| PRMT1 | -0.330326383 | 0.795356529 | 2.66E-07 | 3.30E-06 | 15124.44408 | 12029.39845 | ENSG00000126457 |
| GABRB3 | -0.330337413 | 0.795350448 | 8.22E-07 | 9.32E-06 | 3839.042045 | 3052.289575 | ENSG00000166206 |
| SCFD2 | -0.330461125 | 0.795282249 | 0.014412409 | 0.049429755 | 315.4003361 | 250.9510112 | ENSG00000184178 |
| NA | -0.330509573 | 0.795255543 | 0.706701313 | 0.833271761 | 9.379458533 | 7.50198954 | ENSG00000270933 |
| UBOX5 | -0.330567388 | 0.795223674 | 0.145777784 | 0.29958243 | 133.8580466 | 106.7332424 | ENSG00000185019 |
| SPAG1 | -0.330815856 | 0.795086729 | 0.415561525 | 0.606868713 | 40.01156773 | 31.65196134 | ENSG00000104450 |
| NA | -0.331027674 | 0.794970001 | 0.750731125 | 0.86133779 | 4.813273453 | 3.833401615 | ENSG00000270926 |
| KCNH4 | -0.331044328 | 0.794960825 | 0.557984015 | 0.727313857 | 17.32523633 | 13.8293221 | ENSG00000089558 |
| DOCK8 | -0.331377486 | 0.794777268 | 0.396927821 | 0.589098354 | 41.84454508 | 33.1041696 | ENSG00000107099 |
| DHCR7 | -0.331545239 | 0.794684858 | 3.43E-08 | 5.00E-07 | 4999.568931 | 3972.804275 | ENSG00000172893 |
| CALM1 | -0.331718921 | 0.794589194 | 1.10E-06 | 1.22E-05 | 6012.23519 | 4776.85948 | ENSG00000198668 |
| PPIL2 | -0.331772215 | 0.794559842 | 0.000683349 | 0.003777272 | 815.7428941 | 647.9593253 | ENSG00000100023 |
| C20orf27 | -0.331792815 | 0.794548497 | 8.71E-06 | 7.89E-05 | 2297.173633 | 1825.484868 | ENSG00000101220 |
| UBL7 | -0.331989206 | 0.794440344 | 9.36E-05 | 0.000656938 | 1177.170133 | 934.6497964 | ENSG00000138629 |
| PRPF40A | -0.332017193 | 0.794424932 | 6.85E-07 | 7.88E-06 | 3561.754357 | 2828.806083 | ENSG00000196504 |
| ATP5MF | -0.332022818 | 0.794421835 | 0.024283867 | 0.075605105 | 313.662179 | 249.1770997 | ENSG00000241468 |
| SHKBP1 | -0.332256432 | 0.794293206 | 1.63E-05 | 0.000138839 | 1513.569822 | 1202.466931 | ENSG00000160410 |
| INSIG1 | -0.332434262 | 0.794195305 | 0.000117967 | 0.000808583 | 1285.199294 | 1020.178904 | ENSG00000186480 |
| TCTN2 | -0.332438916 | 0.794192744 | 0.087702547 | 0.205366899 | 147.2132579 | 116.9385666 | ENSG00000168778 |
| TUBB4A | -0.332455266 | 0.794183743 | 0.00122336 | 0.006238473 | 649.6707803 | 515.845794 | ENSG00000104833 |
| C2CD4C | -0.332474048 | 0.794173404 | 0.376168914 | 0.569228825 | 364.0427988 | 289.3102741 | ENSG00000183186 |
| B4GALNT4 | -0.332551139 | 0.794130968 | 0.000166642 | 0.001093293 | 2071.133799 | 1645.735309 | ENSG00000182272 |
| PSMD6 | -0.332713076 | 0.794041835 | 6.81E-07 | 7.83E-06 | 2453.555919 | 1948.279669 | ENSG00000163636 |
| VRTN | -0.332763689 | 0.794013978 | 3.34E-08 | 4.89E-07 | 5372.24285 | 4266.097569 | ENSG00000133980 |
| DDB1 | -0.332861178 | 0.793960325 | 2.39E-09 | 4.19E-08 | 9601.469392 | 7623.225614 | ENSG00000167986 |
| SH3BP1 | -0.332862299 | 0.793959708 | 0.215667154 | 0.395955923 | 90.60015292 | 71.9137442 | ENSG00000100092 |
| CLDN7 | -0.332868335 | 0.793956387 | 2.22E-05 | 0.000183293 | 1406.975732 | 1117.292777 | ENSG00000181885 |
| ZNF844 | -0.332895421 | 0.793941481 | 0.251409158 | 0.440001637 | 66.57275099 | 52.78102892 | ENSG00000223547 |
| RECQL | -0.333146581 | 0.793803275 | 0.004125735 | 0.017482025 | 614.1509021 | 486.8693729 | ENSG00000004700 |
| GALNS | -0.333380535 | 0.793674559 | 0.005217374 | 0.021374628 | 443.9784729 | 352.5901525 | ENSG00000141012 |

| | | | | | | | |
|---|---|---|---|---|---|---|---|
| MRPL50 | -0.33344759 | 0.79363767 | 0.000285023 | 0.001759113 | 904.748454 | 718.3960702 | ENSG00000136897 |
| C11orf96 | -0.333487079 | 0.793615947 | 0.644170403 | 0.791924532 | 9.528006192 | 7.565350564 | ENSG00000187479 |
| MIR1915HG | -0.333628681 | 0.793538057 | 0.06500989 | 0.163689459 | 215.1024254 | 170.8456488 | ENSG00000204682 |
| TRA2A | -0.333799846 | 0.793443915 | 8.25E-06 | 7.52E-05 | 1680.522346 | 1333.463097 | ENSG00000164548 |
| NA | -0.333833503 | 0.793425405 | 0.733173708 | 0.851101774 | 5.667661135 | 4.527275579 | ENSG00000236989 |
| SLC26A6 | -0.33387345 | 0.793403436 | 0.001354025 | 0.006782656 | 1334.396359 | 1059.531461 | ENSG00000225697 |
| EML1 | -0.333975784 | 0.793347159 | 0.001668996 | 0.008136024 | 592.7611362 | 470.252211 | ENSG00000066629 |
| RCC2 | -0.334007995 | 0.793329447 | 2.06E-10 | 4.29E-09 | 14371.43602 | 11400.9275 | ENSG00000179051 |
| STRIP1 | -0.33415964 | 0.793246063 | 0.000945668 | 0.005019187 | 1159.691433 | 920.6104439 | ENSG00000143093 |
| LRPAP1 | -0.33421766 | 0.793214161 | 4.42E-06 | 4.35E-05 | 2110.598793 | 1674.649314 | ENSG00000163956 |
| CALY | -0.334412297 | 0.793107155 | 0.554656834 | 0.724709888 | 19.03767227 | 15.09695799 | ENSG00000130643 |
| RDH5 | -0.334415207 | 0.793105555 | 0.778402021 | 0.87829679 | 4.822744192 | 3.829602768 | ENSG00000135437 |
| SMPDL3A | -0.334445546 | 0.793088876 | 0.435062633 | 0.624541154 | 32.17074218 | 25.53545823 | ENSG00000172594 |
| CRMP1 | -0.334464847 | 0.793078266 | 5.87E-09 | 9.77E-08 | 5908.518331 | 4686.082838 | ENSG00000072832 |
| TMEM109 | -0.334472667 | 0.793073968 | 0.000196339 | 0.001262646 | 996.5775017 | 790.5622623 | ENSG00000110108 |
| NAT16 | -0.334475058 | 0.793072653 | 0.750518151 | 0.861278241 | 4.800645802 | 3.810831773 | ENSG00000167011 |
| CHGA | -0.334555912 | 0.793028208 | 3.20E-05 | 0.000255045 | 1691.252135 | 1341.340532 | ENSG00000100604 |
| COMT | -0.334574832 | 0.793017808 | 0.09339672 | 0.215086334 | 178.2369517 | 141.0471976 | ENSG00000093010 |
| NA | -0.334609124 | 0.792998959 | 0.748592921 | 0.85968283 | 4.74442273 | 3.762004864 | ENSG00000108958 |
| ZNF614 | -0.334678433 | 0.792960863 | 0.014460827 | 0.049553276 | 481.0470035 | 380.9957026 | ENSG00000142556 |
| DHRS4-AS1 | -0.334930596 | 0.792822276 | 0.065503949 | 0.164505446 | 183.090784 | 144.9408226 | ENSG00000215256 |
| KIF13A | -0.334961144 | 0.792805489 | 0.000217262 | 0.001382334 | 959.4895563 | 760.3936157 | ENSG00000137177 |
| PDPN | -0.335064347 | 0.792748778 | 1.30E-06 | 1.41E-05 | 2033.956543 | 1612.327305 | ENSG00000162493 |
| NA | -0.335245974 | 0.792648981 | 0.724314606 | 0.844902699 | 7.967177127 | 6.354027658 | ENSG00000232439 |
| ZNF790 | -0.335308165 | 0.792614813 | 0.327078958 | 0.521301591 | 44.94548029 | 35.53094067 | ENSG00000197863 |
| XRCC6 | -0.335496504 | 0.792511347 | 7.57E-09 | 1.24E-07 | 15759.72813 | 12488.50131 | ENSG00000196419 |
| KEAP1 | -0.335874395 | 0.792303788 | 5.14E-06 | 4.95E-05 | 2082.341883 | 1650.072475 | ENSG00000079999 |
| ISG20L2 | -0.335922414 | 0.792277417 | 2.26E-05 | 0.000186508 | 1372.488923 | 1087.826152 | ENSG00000143319 |
| RBM12B-AS1 | -0.3360005 | 0.792234536 | 0.665023919 | 0.805385316 | 9.299859295 | 7.411821795 | ENSG00000279331 |
| PON1 | -0.336000668 | 0.792234444 | 0.675518646 | 0.812305364 | 8.198480937 | 6.500070353 | ENSG00000005421 |
| TGIF2 | -0.336054037 | 0.792205138 | 0.000419061 | 0.002453212 | 939.9196591 | 744.1087489 | ENSG00000118707 |
| KCTD15 | -0.336102452 | 0.792178553 | 0.00032788 | 0.001982036 | 1173.348723 | 930.1183814 | ENSG00000153885 |
| NUFIP1 | -0.336158553 | 0.792147749 | 0.009511679 | 0.035314394 | 454.870535 | 359.7689728 | ENSG00000083635 |
| NBPF1 | -0.3362214 | 0.792113242 | 0.013772377 | 0.047711633 | 418.5900743 | 332.1513204 | ENSG00000219481 |
| KIAA0319 | -0.336420668 | 0.792003841 | 0.506790875 | 0.686361862 | 22.36971589 | 17.617146 | ENSG00000137261 |
| EIF2S3B | -0.336527254 | 0.79194533 | 0.486955947 | 0.670025354 | 24.20672196 | 19.12880481 | ENSG00000180574 |
| IL17RB | -0.336549533 | 0.7919331 | 0.133256145 | 0.280241619 | 106.7302131 | 84.44146403 | ENSG00000056736 |
| XKRX | -0.336694482 | 0.791853538 | 0.365605265 | 0.558671612 | 46.65475932 | 37.07744194 | ENSG00000182489 |
| LRCH2 | -0.336695351 | 0.791853061 | 0.089795046 | 0.209262087 | 205.3539768 | 162.1185264 | ENSG00000130224 |
| MRPS22 | -0.336743277 | 0.791826756 | 0.002221387 | 0.010381035 | 579.4024749 | 458.3827582 | ENSG00000175110 |
| GPN1 | -0.33683529 | 0.791776256 | 0.003170517 | 0.013954492 | 649.3393876 | 514.1023764 | ENSG00000198522 |
| PTDSS1 | -0.336921025 | 0.791729205 | 4.01E-06 | 3.98E-05 | 1866.964348 | 1478.418932 | ENSG00000156471 |
| TELO2 | -0.337001452 | 0.791685069 | 0.001387946 | 0.006922548 | 2060.892128 | 1632.720062 | ENSG00000100726 |
| C1orf52 | -0.337097489 | 0.79163237 | 0.00124429 | 0.006326934 | 589.3122117 | 466.4448516 | ENSG00000162642 |
| NA | -0.337294061 | 0.791524515 | 0.739073039 | 0.854158532 | 7.105197843 | 5.641382699 | ENSG00000224195 |
| CYP27C1 | -0.337414009 | 0.791458709 | 0.1260679 | 0.269248959 | 115.5991316 | 91.4632202 | ENSG00000186684 |
| TMEM63C | -0.337578358 | 0.791368553 | 0.192842879 | 0.36635403 | 111.0846372 | 87.59811371 | ENSG00000165548 |
| NA | -0.33783564 | 0.791227437 | 0.76839956 | 0.871924202 | 4.380453524 | 3.495850129 | ENSG00000219928 |
| CCDC28B | -0.337885154 | 0.791200282 | 0.036467448 | 0.104578011 | 261.8285684 | 207.2128105 | ENSG00000160050 |
| NA | -0.337886531 | 0.791199527 | 0.388073019 | 0.581174299 | 36.38746484 | 28.7084026 | ENSG00000123965 |
| NA | -0.337945829 | 0.791167008 | 0.407564087 | 0.599317105 | 41.3214751 | 32.82023156 | ENSG00000276791 |
| OGFOD1 | -0.337982021 | 0.791147161 | 4.21E-05 | 0.000323171 | 1112.885863 | 880.6574933 | ENSG00000087263 |
| CITED4 | -0.338079417 | 0.791093752 | 0.09055407 | 0.21052799 | 237.0287683 | 187.391852 | ENSG00000179862 |
| USP2 | -0.338181527 | 0.791037763 | 0.080772211 | 0.193065792 | 153.9178063 | 121.9278542 | ENSG00000036672 |
| TTF1 | -0.338182648 | 0.791037148 | 0.001627299 | 0.007966287 | 622.1521137 | 491.8132976 | ENSG00000125482 |
| PSME2 | -0.338288739 | 0.79097898 | 0.000665371 | 0.003686691 | 1219.411682 | 964.7912703 | ENSG00000100911 |
| CDKL1 | -0.338312222 | 0.790966105 | 0.510391012 | 0.689675344 | 21.44332058 | 16.9762242 | ENSG00000100490 |
| BEX2 | -0.338384058 | 0.790926722 | 0.000228006 | 0.001442275 | 998.1443515 | 789.5920565 | ENSG00000133134 |
| LINC00476 | -0.338424734 | 0.790904422 | 0.432701714 | 0.622647016 | 30.2231691 | 23.82849177 | ENSG00000175611 |
| IGSF1 | -0.338426435 | 0.79090349 | 0.000144824 | 0.000963449 | 1031.36079 | 815.8335412 | ENSG00000147255 |
| PIGQ | -0.338432262 | 0.790900295 | 0.013746293 | 0.047630172 | 446.963437 | 353.94333 | ENSG00000007541 |
| HNRNPR | -0.338457525 | 0.790886446 | 2.78E-09 | 4.83E-08 | 6103.119804 | 4826.276062 | ENSG00000125944 |
| SIX1 | -0.338527281 | 0.790848206 | 0.707183935 | 0.833461357 | 7.205114033 | 5.667201734 | ENSG00000126778 |
| FREM1 | -0.33854223 | 0.790840012 | 0.256120224 | 0.444969276 | 60.62259902 | 47.95786911 | ENSG00000164946 |
| VSTM2L | -0.338606685 | 0.790804681 | 0.273620236 | 0.464871193 | 59.88143159 | 47.29350141 | ENSG00000132821 |
| NA | -0.338735696 | 0.790733967 | 0.551672382 | 0.722593351 | 16.66732872 | 13.19544834 | ENSG00000237398 |
| NA | -0.33888076 | 0.790654462 | 0.500735609 | 0.681443972 | 21.97733447 | 17.45277034 | ENSG00000224094 |
| SHC1 | -0.338895906 | 0.790646161 | 4.04E-05 | 0.000312216 | 6044.403964 | 4780.15272 | ENSG00000160691 |
| LAMTOR2 | -0.339019297 | 0.790578542 | 0.001137835 | 0.005879187 | 683.3246839 | 540.5438606 | ENSG00000116586 |
| TRIM47 | -0.339301178 | 0.79042409 | 0.232377309 | 0.417050552 | 75.77534734 | 59.9003273 | ENSG00000132481 |
| LRRC37A2 | -0.33937435 | 0.790384002 | 0.153974627 | 0.311567392 | 101.0669867 | 79.87317403 | ENSG00000238083 |
| NA | -0.339396533 | 0.790371849 | 0.542426529 | 0.715081677 | 17.07371326 | 13.56685459 | ENSG00000286827 |
| PPIG | -0.339424605 | 0.79035647 | 1.80E-05 | 0.000151874 | 1948.706113 | 1539.292771 | ENSG00000138398 |
| FAM207A | -0.33947888 | 0.790326737 | 0.001599336 | 0.007860537 | 1260.632849 | 996.270207 | ENSG00000160256 |

| | | | | | | | |
|---|---|---|---|---|---|---|---|
| VPS33A | -0.339556702 | 0.790284106 | 0.062220663 | 0.158428049 | 244.8233225 | 193.1177314 | ENSG00000139719 |
| NLRP2 | -0.339623915 | 0.790247288 | 1.41E-07 | 1.83E-06 | 3733.658871 | 2950.906716 | ENSG00000022556 |
| SLC12A9 | -0.339769604 | 0.79016749 | 9.51E-05 | 0.000665904 | 1188.4095 | 939.6107258 | ENSG00000146828 |
| CTDSPL2 | -0.33980066 | 0.790150481 | 0.00212506 | 0.010017341 | 695.7226298 | 549.1431927 | ENSG00000137770 |
| GLRX5 | -0.339814065 | 0.790143139 | 6.60E-07 | 7.60E-06 | 2900.625413 | 2291.693088 | ENSG00000182512 |
| OTUD7A | -0.339817587 | 0.79014121 | 0.105887517 | 0.236719386 | 143.3339713 | 113.4661624 | ENSG00000169918 |
| QTRT2 | -0.339881262 | 0.790106337 | 3.96E-05 | 0.000306745 | 1193.0247 | 942.3626238 | ENSG00000151576 |
| OSMR | -0.339917069 | 0.790086727 | 0.191452745 | 0.364497707 | 80.44698832 | 63.44825353 | ENSG00000145623 |
| GRHL2 | -0.339931515 | 0.790078816 | 0.001029395 | 0.005403543 | 660.7034902 | 522.1669545 | ENSG00000083307 |
| HAUS1 | -0.340028872 | 0.790025501 | 0.000608905 | 0.003408473 | 930.9398805 | 735.0094877 | ENSG00000152240 |
| TIMM8B | -0.340117826 | 0.789976791 | 7.35E-05 | 0.000528504 | 1335.622517 | 1055.596406 | ENSG00000150779 |
| ZNF821 | -0.34020655 | 0.78992821 | 0.236998696 | 0.422156062 | 72.99478591 | 57.49926364 | ENSG00000102984 |
| APBB1 | -0.340286232 | 0.789884582 | 0.000423513 | 0.00247459 | 906.3201141 | 716.0073505 | ENSG00000166313 |
| PHF2 | -0.340286787 | 0.789884279 | 0.00791865 | 0.030354551 | 421.2836781 | 332.7288325 | ENSG00000197724 |
| EID2 | -0.340417159 | 0.789812902 | 0.000982179 | 0.005181809 | 626.9787889 | 495.2138041 | ENSG00000176396 |
| TMEM126A | -0.340486371 | 0.789775013 | 0.009835094 | 0.03623929 | 363.3907391 | 286.6494651 | ENSG00000171202 |
| MED6 | -0.340673408 | 0.78967263 | 0.016214936 | 0.054344959 | 302.7748447 | 239.1805264 | ENSG00000133997 |
| SLFN11 | -0.340755015 | 0.789627962 | 0.522162968 | 0.699117944 | 18.76720773 | 14.8381777 | ENSG00000172716 |
| PHB2 | -0.340824344 | 0.789590018 | 2.39E-10 | 4.91E-09 | 7648.991516 | 6039.514937 | ENSG00000215021 |
| MCM7 | -0.340881216 | 0.789558892 | 6.41E-11 | 1.44E-09 | 11740.47117 | 9269.829947 | ENSG00000166508 |
| NA | -0.340891688 | 0.789553161 | 0.277659861 | 0.468699553 | 54.58415119 | 43.19069586 | ENSG00000235363 |
| GTF3A | -0.341104887 | 0.789436491 | 8.33E-05 | 0.000591774 | 1182.558896 | 933.4359972 | ENSG00000122034 |
| NA | -0.341156205 | 0.78940841 | 0.706518222 | 0.833271761 | 7.522330907 | 6.001504025 | ENSG00000251400 |
| RBM23 | -0.341201627 | 0.789383557 | 0.000162234 | 0.001067396 | 959.8211898 | 757.7312955 | ENSG00000100461 |
| BCAS2 | -0.341326245 | 0.789315374 | 0.000199712 | 0.001281227 | 1148.807654 | 905.8942872 | ENSG00000116752 |
| TMCO1 | -0.341339724 | 0.789308 | 0.000227309 | 0.001438845 | 892.148992 | 703.8566508 | ENSG00000143183 |
| HHIPL1 | -0.341413111 | 0.78926785 | 0.131662971 | 0.278026832 | 117.0627955 | 92.4455955 | ENSG00000182218 |
| SENP1 | -0.341461887 | 0.789241166 | 0.018180588 | 0.059554439 | 391.4333476 | 308.5034015 | ENSG00000079387 |
| NA | -0.341480085 | 0.789231211 | 0.715634647 | 0.838882585 | 5.683445699 | 4.508504585 | ENSG00000233588 |
| UBE4B | -0.34156553 | 0.789184469 | 1.03E-05 | 9.14E-05 | 3229.057137 | 2547.638958 | ENSG00000130939 |
| TM9SF1 | -0.341861344 | 0.789022669 | 0.275274258 | 0.466228009 | 57.5042932 | 45.47478505 | ENSG00000100926 |
| DCUN1D5 | -0.341914006 | 0.788993869 | 2.55E-05 | 0.000206975 | 2031.299605 | 1602.43369 | ENSG00000137692 |
| SLC4A10 | -0.342215837 | 0.788828818 | 0.714015992 | 0.838045241 | 6.153547223 | 4.838569996 | ENSG00000144290 |
| NA | -0.342283373 | 0.788791892 | 0.426661505 | 0.617262067 | 30.16822381 | 23.68949712 | ENSG00000260947 |
| NPHP4 | -0.342446312 | 0.78870281 | 0.007036119 | 0.027541168 | 429.7440997 | 339.3962415 | ENSG00000131697 |
| PITPNM1 | -0.342483593 | 0.788682429 | 4.04E-05 | 0.000312216 | 2104.46999 | 1659.701464 | ENSG00000110697 |
| NA | -0.342691398 | 0.788568836 | 0.51091709 | 0.690100169 | 22.71790054 | 17.76307707 | ENSG00000225159 |
| TPD52L2 | -0.342726568 | 0.788549613 | 4.36E-06 | 4.29E-05 | 2798.42006 | 2207.782661 | ENSG00000101150 |
| ZNF324 | -0.342747241 | 0.788538313 | 0.132928013 | 0.279950464 | 107.7799985 | 84.9737735 | ENSG00000083812 |
| ADAMTS2 | -0.342834974 | 0.788490362 | 0.01795447 | 0.058928307 | 280.2828212 | 220.8763312 | ENSG00000087116 |
| GCFC2 | -0.342884151 | 0.788463485 | 0.109755576 | 0.24286173 | 121.2372005 | 95.52088606 | ENSG00000005436 |
| OTUB1 | -0.342886816 | 0.788462029 | 1.00E-05 | 8.95E-05 | 1989.663814 | 1569.633321 | ENSG00000167770 |
| TRMT6 | -0.342923513 | 0.788441974 | 0.009684025 | 0.035803654 | 558.3843675 | 439.8096685 | ENSG00000089195 |
| WFIKKN1 | -0.343050442 | 0.788372609 | 0.108280164 | 0.240400642 | 194.459825 | 153.0033138 | ENSG00000127578 |
| PRRX2 | -0.343085823 | 0.788353275 | 0.189503529 | 0.361715592 | 115.3217745 | 90.68523885 | ENSG00000167157 |
| PCDHA12 | -0.343121783 | 0.788333625 | 0.069001886 | 0.171225192 | 168.4834745 | 132.8797897 | ENSG00000251664 |
| EPHB3 | -0.343139617 | 0.78832388 | 0.22819135 | 0.411810953 | 76.27395878 | 60.26826111 | ENSG00000182580 |
| NA | -0.34318603 | 0.78829852 | 0.480800188 | 0.664564895 | 23.72261731 | 18.70543408 | ENSG00000256092 |
| CCDC78 | -0.343241088 | 0.788268436 | 0.237899879 | 0.423354427 | 130.3280497 | 103.0556798 | ENSG00000162004 |
| GTF3C6 | -0.343398998 | 0.788182161 | 0.000336942 | 0.002028879 | 1238.471114 | 976.7701712 | ENSG00000155115 |
| PREX1 | -0.343430544 | 0.788164927 | 0.000348035 | 0.002082816 | 804.8582282 | 634.2507765 | ENSG00000124126 |
| PHC2 | -0.343613431 | 0.788065019 | 0.019068706 | 0.061949063 | 323.2866362 | 254.3612166 | ENSG00000134686 |
| IMP3 | -0.343632576 | 0.788054561 | 0.021162401 | 0.067741612 | 333.1239746 | 262.7379024 | ENSG00000177971 |
| ZMYM4 | -0.343667849 | 0.788035294 | 1.56E-05 | 0.000133412 | 1946.619967 | 1532.986136 | ENSG00000146463 |
| NAT14 | -0.343757397 | 0.787986382 | 0.000561481 | 0.003177553 | 1118.622608 | 881.9459956 | ENSG00000090971 |
| GLYR1 | -0.34381148 | 0.787956843 | 3.26E-08 | 4.79E-07 | 3163.811536 | 2492.873437 | ENSG00000140632 |
| SHF | -0.343865655 | 0.787927255 | 0.00123371 | 0.006283479 | 685.7184489 | 540.6637578 | ENSG00000138606 |
| RPL26 | -0.343898226 | 0.787909467 | 0.004503271 | 0.018815063 | 622.1190714 | 490.7548195 | ENSG00000161970 |
| EIF3C | -0.344346385 | 0.787664748 | 4.65E-05 | 0.000352702 | 1159.322135 | 912.9210934 | ENSG00000184110 |
| PCOLCE2 | -0.344444248 | 0.78761132 | 0.025224251 | 0.077947148 | 262.2354566 | 206.4586099 | ENSG00000163710 |
| PLP2 | -0.344579049 | 0.787537731 | 0.213660883 | 0.393637257 | 86.36993079 | 67.8471581 | ENSG00000102007 |
| RNF103-CHMP3 | -0.344597244 | 0.787527799 | 0.727263762 | 0.846837128 | 5.783264191 | 4.576102488 | ENSG00000249884 |
| FAAP24 | -0.344665143 | 0.787490736 | 0.159304921 | 0.319195394 | 92.46094112 | 72.87085012 | ENSG00000131944 |
| TCTN3 | -0.344670718 | 0.787487693 | 0.001306966 | 0.006589583 | 800.2716511 | 629.8454365 | ENSG00000119977 |
| FKBP1A | -0.344682365 | 0.787481335 | 5.41E-05 | 0.000403541 | 2827.557048 | 2226.533757 | ENSG00000088832 |
| EHD1 | -0.344794777 | 0.787419979 | 0.000196665 | 0.001264306 | 1437.691462 | 1131.933425 | ENSG00000110047 |
| NA | -0.344840997 | 0.787394753 | 0.715052103 | 0.838432369 | 5.759888024 | 4.557331494 | ENSG00000267141 |
| NFKBIE | -0.344949576 | 0.787335494 | 0.020503655 | 0.065942947 | 370.3339724 | 291.2014689 | ENSG00000146232 |
| PGRMC2 | -0.345027723 | 0.787292848 | 3.78E-05 | 0.000294787 | 1420.058374 | 1118.138986 | ENSG00000164040 |
| ZCCHC3 | -0.345043184 | 0.78728441 | 2.04E-06 | 2.14E-05 | 2506.967075 | 1974.091963 | ENSG00000247315 |
| SLCO3A1 | -0.345124495 | 0.78724004 | 0.005187715 | 0.021257816 | 413.1928777 | 325.1137789 | ENSG00000176463 |
| PIGF | -0.345173236 | 0.787213444 | 0.16874516 | 0.333034054 | 85.02589875 | 67.01213959 | ENSG00000151665 |
| SSU72 | -0.345270274 | 0.787160497 | 2.72E-07 | 3.37E-06 | 3565.10075 | 2806.092383 | ENSG00000160075 |
| C8orf34-AS1 | -0.345378388 | 0.78710151 | 0.602534795 | 0.76175852 | 12.06513982 | 9.440491519 | ENSG00000248801 |

| | | | | | | | |
|---|---|---|---|---|---|---|---|
| DPM1 | -0.345415116 | 0.787081472 | 0.000942341 | 0.00500583 | 1040.194735 | 818.1019249 | ENSG00000000419 |
| MRAP2 | -0.34550001 | 0.787035158 | 0.1745704 | 0.341618195 | 92.82558674 | 72.91500212 | ENSG00000135324 |
| STARD7 | -0.345600927 | 0.786980107 | 2.31E-07 | 2.90E-06 | 3792.20989 | 2983.654568 | ENSG00000084090 |
| SPDL1 | -0.345964458 | 0.786781828 | 0.000195262 | 0.001257029 | 902.2364628 | 709.7345703 | ENSG00000040275 |
| FOXD2-AS1 | -0.346126414 | 0.786693509 | 0.710378337 | 0.835574798 | 6.083418724 | 4.759687172 | ENSG00000237424 |
| SLC38A9 | -0.346197163 | 0.786654931 | 0.011169185 | 0.040368501 | 416.2846505 | 327.0597754 | ENSG00000177058 |
| NA | -0.346269144 | 0.786615683 | 0.597570427 | 0.757703763 | 13.17410978 | 10.34475689 | ENSG00000256238 |
| CCDC120 | -0.346509233 | 0.786484788 | 0.011837743 | 0.042281889 | 383.1321844 | 301.0967011 | ENSG00000147144 |
| TRMT13 | -0.346615982 | 0.786426596 | 0.08612379 | 0.202591486 | 139.3794452 | 109.5074242 | ENSG00000122435 |
| NA | -0.346624905 | 0.786421732 | 0.203021705 | 0.380542504 | 73.98263575 | 58.25011186 | ENSG00000262539 |
| NA | -0.346639758 | 0.786413636 | 0.71766306 | 0.84009 | 6.077104898 | 4.752201098 | ENSG00000273343 |
| MEIS2 | -0.346724205 | 0.786367605 | 0.720599747 | 0.842146145 | 5.786421104 | 4.568616414 | ENSG00000134138 |
| EXOSC3 | -0.346800616 | 0.786325957 | 0.004144144 | 0.017556014 | 420.5994102 | 330.8207213 | ENSG00000107371 |
| MAOA | -0.346887853 | 0.786278411 | 0.051708664 | 0.137125251 | 189.2132658 | 148.9450269 | ENSG00000189221 |
| LCN12 | -0.347128077 | 0.786147498 | 0.595536389 | 0.755953244 | 12.8789913 | 10.14988728 | ENSG00000184925 |
| GMIP | -0.347151576 | 0.786134693 | 0.002316174 | 0.010756196 | 537.5513304 | 422.638615 | ENSG00000089639 |
| RUFY2 | -0.347169359 | 0.786125003 | 0.093066018 | 0.214644797 | 128.7732414 | 101.0646149 | ENSG00000204130 |
| FAM76B | -0.347220559 | 0.786097105 | 0.018880077 | 0.061497952 | 296.9699858 | 233.5360061 | ENSG00000077458 |
| PCDHB17P | -0.347440268 | 0.785977399 | 0.749120399 | 0.860128699 | 5.071110343 | 3.934304627 | ENSG00000255622 |
| MBNL2 | -0.347486555 | 0.785952182 | 0.522668497 | 0.699529834 | 21.54135763 | 16.85187528 | ENSG00000139793 |
| ZNF48 | -0.34761243 | 0.78588361 | 0.000981785 | 0.005181207 | 996.5117177 | 782.9743101 | ENSG00000180035 |
| MRPL33 | -0.347633903 | 0.785871914 | 0.008568475 | 0.032396595 | 377.3402387 | 296.4475932 | ENSG00000243147 |
| NRP1 | -0.347654949 | 0.785860449 | 0.362607723 | 0.555556877 | 41.38085508 | 32.59475639 | ENSG00000099250 |
| NA | -0.347878396 | 0.785738743 | 0.571752825 | 0.737943374 | 21.46679444 | 16.99185762 | ENSG00000269929 |
| CAMK1G | -0.348085405 | 0.785626008 | 0.491472209 | 0.674137502 | 20.75897758 | 16.35374699 | ENSG00000008118 |
| FHAD1 | -0.348094248 | 0.785621192 | 0.366669044 | 0.559651162 | 35.52498189 | 27.96982699 | ENSG00000142621 |
| PPT1 | -0.348114156 | 0.785610351 | 3.85E-06 | 3.85E-05 | 3488.771907 | 2739.664047 | ENSG00000131238 |
| MEGF9 | -0.348114403 | 0.785610217 | 0.282105465 | 0.473085658 | 55.71384406 | 43.5686005 | ENSG00000106780 |
| DHDH | -0.348127264 | 0.785603213 | 0.447071931 | 0.635261315 | 26.96559096 | 21.05657152 | ENSG00000104808 |
| SHISAL1 | -0.348134707 | 0.78559916 | 0.01995469 | 0.064432094 | 272.8316109 | 214.4327971 | ENSG00000138944 |
| MRPS18C | -0.348272289 | 0.785524246 | 0.123030508 | 0.26449918 | 113.9835527 | 89.59870289 | ENSG00000163319 |
| FASTKD3 | -0.348494442 | 0.785403296 | 0.088390608 | 0.206651661 | 130.1635221 | 102.1073253 | ENSG00000124279 |
| NPW | -0.348505822 | 0.785397101 | 0.233837519 | 0.418779068 | 79.47531384 | 62.43571069 | ENSG00000183971 |
| CACNA1E | -0.348607668 | 0.785341658 | 0.476464217 | 0.66089522 | 23.50215972 | 18.57067631 | ENSG00000198216 |
| DCTN6 | -0.348647155 | 0.785320164 | 0.226324896 | 0.409719245 | 76.21330102 | 59.63263522 | ENSG00000104671 |
| SMAP2 | -0.348706868 | 0.78528766 | 0.066717505 | 0.166897318 | 165.5123409 | 129.9831857 | ENSG00000084070 |
| ECPAS | -0.348791323 | 0.785241691 | 3.29E-08 | 4.82E-07 | 4441.746795 | 3487.275423 | ENSG00000136813 |
| ITGA7 | -0.348821135 | 0.785225465 | 0.154233236 | 0.311797426 | 1584.848069 | 1244.699362 | ENSG00000135424 |
| STAP2 | -0.349343569 | 0.784941167 | 0.016710967 | 0.055745331 | 293.7844153 | 230.4556026 | ENSG00000178078 |
| SLC2A12 | -0.349679494 | 0.784758419 | 0.150326419 | 0.306439711 | 98.93801904 | 77.53233382 | ENSG00000146411 |
| POLD2 | -0.349774848 | 0.784706553 | 2.49E-08 | 3.72E-07 | 3349.768246 | 2628.660996 | ENSG00000106628 |
| NA | -0.349907925 | 0.784634173 | 0.481628974 | 0.665363104 | 20.99787299 | 16.45844885 | ENSG00000285517 |
| NA | -0.350029544 | 0.784568031 | 0.399768593 | 0.591750747 | 45.01630785 | 35.51739425 | ENSG00000231793 |
| MDK | -0.350140107 | 0.784507907 | 2.07E-09 | 3.68E-08 | 6568.577202 | 5153.794492 | ENSG00000110492 |
| NA | -0.350224516 | 0.784462008 | 0.559269516 | 0.72837476 | 16.90188714 | 13.27009429 | ENSG00000229358 |
| GAREM2 | -0.350286761 | 0.784428164 | 0.000567562 | 0.003205117 | 825.5796159 | 648.1437232 | ENSG00000157833 |
| PRDX2 | -0.350291487 | 0.784425594 | 8.71E-07 | 9.79E-06 | 2311.695361 | 1813.297606 | ENSG00000167815 |
| DHX34 | -0.350310081 | 0.784415484 | 5.03E-05 | 0.000378671 | 1520.151567 | 1193.255419 | ENSG00000134815 |
| RETREG2 | -0.350315062 | 0.784412776 | 4.24E-05 | 0.000324999 | 1215.738722 | 953.2165707 | ENSG00000144567 |
| DUSP9 | -0.350345604 | 0.78439617 | 0.202682414 | 0.380052362 | 71.06752979 | 55.72188812 | ENSG00000130829 |
| DAB2IP | -0.350363701 | 0.78438633 | 4.77E-06 | 4.63E-05 | 1899.382652 | 1489.972178 | ENSG00000136848 |
| PTDSS2 | -0.350373588 | 0.784380955 | 4.18E-05 | 0.000321973 | 1426.919979 | 1119.458809 | ENSG00000174915 |
| PJVK | -0.350514053 | 0.784304589 | 0.588505097 | 0.750857513 | 14.81453808 | 11.68042871 | ENSG00000204311 |
| ZBTB26 | -0.350573946 | 0.78427203 | 0.11683715 | 0.254677328 | 144.8396823 | 113.8232577 | ENSG00000171448 |
| RPP21 | -0.350607447 | 0.784253818 | 0.503920631 | 0.683823765 | 19.49514614 | 15.33729369 | ENSG00000241370 |
| ABHD5 | -0.350672601 | 0.784218401 | 0.016133108 | 0.054143282 | 278.6925952 | 218.6637504 | ENSG00000011198 |
| NA | -0.350713647 | 0.78419609 | 0.344633654 | 0.538200859 | 41.53007916 | 32.56794967 | ENSG00000272335 |
| PGLYRP1 | -0.350718806 | 0.784193286 | 0.707315619 | 0.833517498 | 5.713135692 | 4.48973359 | ENSG00000008438 |
| HSPA5 | -0.350816004 | 0.784140454 | 6.24E-10 | 1.19E-08 | 20223.08771 | 15856.96723 | ENSG00000044574 |
| ARTN | -0.350913539 | 0.784087443 | 0.048169959 | 0.12995173 | 215.152237 | 169.0464679 | ENSG00000117407 |
| SIGMAR1 | -0.350947975 | 0.784068728 | 2.09E-06 | 2.18E-05 | 2583.900437 | 2025.702156 | ENSG00000147955 |
| PIGT | -0.350955326 | 0.784064733 | 0.000110615 | 0.000762701 | 992.209834 | 778.1593295 | ENSG00000124155 |
| RBP1 | -0.351004982 | 0.784037746 | 0.011017768 | 0.039945876 | 347.8266679 | 272.5402271 | ENSG00000114115 |
| ODF2L | -0.351029883 | 0.784024214 | 0.016778407 | 0.055909924 | 288.6924844 | 226.6771285 | ENSG00000122417 |
| MTMR8 | -0.351222767 | 0.7839194 | 0.14290226 | 0.29539934 | 105.874044 | 83.07270189 | ENSG00000102043 |
| NCOA6 | -0.351289001 | 0.783883411 | 0.013470455 | 0.04686717 | 304.7475754 | 238.9440927 | ENSG00000198646 |
| MGC12916 | -0.351293224 | 0.783881116 | 0.117357196 | 0.255413136 | 123.0698696 | 96.71855092 | ENSG00000266709 |
| YEATS4 | -0.351404024 | 0.783820916 | 0.007672397 | 0.029588059 | 567.9681309 | 444.649123 | ENSG00000127337 |
| MAP3K4 | -0.351465532 | 0.783787499 | 3.42E-05 | 0.000269842 | 1292.490922 | 1013.553999 | ENSG00000085511 |
| FBXO2 | -0.35151236 | 0.783762059 | 0.000919693 | 0.004905204 | 920.2260858 | 720.7634049 | ENSG00000116661 |
| NKAIN4 | -0.351642328 | 0.783691455 | 0.133116896 | 0.280214117 | 116.7859194 | 91.79091248 | ENSG00000101198 |
| FKBPL | -0.351765539 | 0.783624528 | 0.07445357 | 0.181486345 | 152.5970527 | 119.7677875 | ENSG00000204315 |
| ARHGAP39 | -0.351791102 | 0.783610643 | 0.00505016 | 0.020767621 | 547.5566464 | 429.4747482 | ENSG00000147799 |
| PEF1 | -0.351806118 | 0.783602487 | 0.002811572 | 0.01259905 | 520.4864266 | 407.8578411 | ENSG00000162517 |

| | | | | | | | |
|---|---|---|---|---|---|---|---|
| HSPA12B | -0.35181266 | 0.783598934 | 0.028425419 | 0.085915967 | 234.5750824 | 183.8347194 | ENSG00000132622 |
| EEF1AKMT4 | -0.351841926 | 0.783583039 | 0.341271804 | 0.53566494 | 48.00440613 | 37.52674332 | ENSG00000284753 |
| PTPN9 | -0.351920739 | 0.783540233 | 1.76E-05 | 0.000148673 | 1242.25428 | 973.1730683 | ENSG00000169410 |
| TMEM97 | -0.35202632 | 0.783482893 | 2.01E-06 | 2.11E-05 | 2227.640219 | 1744.995881 | ENSG00000109084 |
| CCDC137 | -0.352040139 | 0.783475389 | 2.73E-05 | 0.000220541 | 1591.098721 | 1247.037619 | ENSG00000185298 |
| PLEKHA3 | -0.352057034 | 0.783466214 | 0.046034293 | 0.125442744 | 225.956875 | 177.0032695 | ENSG00000116095 |
| LZTS2 | -0.352142945 | 0.78341956 | 2.98E-07 | 3.66E-06 | 3124.891823 | 2449.032384 | ENSG00000107816 |
| AP4M1 | -0.352227977 | 0.783373388 | 0.032119058 | 0.094566261 | 225.368149 | 176.354312 | ENSG00000221838 |
| ZNF500 | -0.352308934 | 0.783329429 | 0.037827934 | 0.107498149 | 215.3090753 | 168.7668297 | ENSG00000103199 |
| PHKA1 | -0.352451675 | 0.78325193 | 0.015864009 | 0.053468548 | 361.0822334 | 283.0336548 | ENSG00000067177 |
| FAM161A | -0.352652814 | 0.783142738 | 0.017361744 | 0.057440657 | 289.308578 | 226.3367256 | ENSG00000170264 |
| NA | -0.352668194 | 0.783134389 | 0.259282817 | 0.448195089 | 66.50144241 | 52.3358528 | ENSG00000272106 |
| KRBOX4 | -0.352997127 | 0.782955855 | 0.099885372 | 0.22664979 | 129.6998321 | 101.7105381 | ENSG00000147121 |
| TMEM222 | -0.353161743 | 0.782866523 | 0.009218488 | 0.034384273 | 468.2666959 | 367.153304 | ENSG00000186501 |
| NA | -0.353245941 | 0.782820835 | 0.471224575 | 0.656501394 | 26.09995111 | 20.34772541 | ENSG00000238072 |
| CCZ1B | -0.353299423 | 0.782791815 | 0.008366284 | 0.031773607 | 361.7873819 | 283.0371902 | ENSG00000146574 |
| ZNF544 | -0.353491619 | 0.782687539 | 0.000852803 | 0.004592054 | 586.7718461 | 459.1830862 | ENSG00000198131 |
| PHTF1 | -0.353663122 | 0.782594501 | 0.000869651 | 0.004673273 | 590.9991969 | 462.4433871 | ENSG00000116793 |
| SESN3 | -0.353816231 | 0.782511451 | 0.018040446 | 0.059179048 | 457.3506563 | 357.4042486 | ENSG00000149212 |
| PSMA3-AS1 | -0.353882809 | 0.78247534 | 0.077288452 | 0.186761952 | 202.5708447 | 158.9462836 | ENSG00000257621 |
| AKIP1 | -0.353902821 | 0.782464486 | 0.008635789 | 0.032611187 | 382.3739471 | 299.1754731 | ENSG00000166452 |
| TBC1D9 | -0.353994907 | 0.782414544 | 0.192680393 | 0.366120403 | 73.25785425 | 57.33456157 | ENSG00000109436 |
| C1QTNF4 | -0.354054355 | 0.782382304 | 0.097137281 | 0.221798985 | 138.8090215 | 108.6525353 | ENSG00000172247 |
| BMI1 | -0.354262861 | 0.782269238 | 0.389543174 | 0.582100555 | 33.95212654 | 26.46890344 | ENSG00000168283 |
| COMMD7 | -0.354290196 | 0.782254416 | 0.014109188 | 0.048651104 | 285.8160355 | 223.7995187 | ENSG00000149600 |
| CLK4 | -0.354524824 | 0.782127208 | 0.03583885 | 0.103141245 | 209.8832711 | 164.3757376 | ENSG00000113240 |
| ERICH2 | -0.354567094 | 0.782104292 | 0.569054966 | 0.735971767 | 13.89139738 | 10.83203829 | ENSG00000204334 |
| ZKSCAN7 | -0.355088519 | 0.781821672 | 0.069240747 | 0.171613966 | 158.8509221 | 124.0236838 | ENSG00000196345 |
| RPL13AP20 | -0.355240075 | 0.781739545 | 0.569463595 | 0.736177672 | 14.85369881 | 11.51842618 | ENSG00000234498 |
| NFE2 | -0.355241912 | 0.78173855 | 0.508436499 | 0.687585704 | 22.01018138 | 17.32929749 | ENSG00000123405 |
| NA | -0.355256459 | 0.781730667 | 0.107480413 | 0.23908329 | 140.9773605 | 110.4137765 | ENSG00000280079 |
| GUK1 | -0.355353551 | 0.781678059 | 3.42E-06 | 3.45E-05 | 1431.446703 | 1118.97922 | ENSG00000143774 |
| TARS3 | -0.355416135 | 0.781644151 | 0.012479518 | 0.044174061 | 307.6491819 | 240.523461 | ENSG00000185418 |
| SUMO1 | -0.355521836 | 0.781586885 | 0.000136095 | 0.000913249 | 4164.804472 | 3254.309945 | ENSG00000116030 |
| NOVA2 | -0.355619696 | 0.78153387 | 0.003276032 | 0.014347349 | 554.4816297 | 434.0298097 | ENSG00000104967 |
| SH3BP5 | -0.355743109 | 0.781467018 | 0.161772995 | 0.322744962 | 95.52597021 | 74.80599128 | ENSG00000131370 |
| NA | -0.355755444 | 0.781460337 | 0.760135447 | 0.866655412 | 4.681885833 | 3.634295131 | ENSG00000263235 |
| SMARCA5-AS1 | -0.356050094 | 0.781300751 | 0.208534854 | 0.38769228 | 67.82771305 | 52.99499593 | ENSG00000245112 |
| RFWD3 | -0.356123958 | 0.78126075 | 4.18E-07 | 5.01E-06 | 2585.348873 | 2019.161444 | ENSG00000168411 |
| AKAP5 | -0.356315222 | 0.781157182 | 0.172127682 | 0.338089706 | 78.85972388 | 61.6332244 | ENSG00000179841 |
| RNF144B | -0.356534833 | 0.781038282 | 0.172095542 | 0.338062404 | 101.6015042 | 79.00513655 | ENSG00000137393 |
| FAAH | -0.356575965 | 0.781016014 | 0.019533335 | 0.063269982 | 289.5346504 | 226.4864958 | ENSG00000117480 |
| NA | -0.356625892 | 0.780988986 | 0.654740833 | 0.798300546 | 8.71721393 | 6.739968019 | ENSG00000271869 |
| NXPE3 | -0.356797095 | 0.780896312 | 0.001219186 | 0.006224903 | 584.7040172 | 456.6223218 | ENSG00000144815 |
| PLOD3 | -0.356882977 | 0.780849828 | 3.52E-06 | 3.54E-05 | 1611.21904 | 1258.184574 | ENSG00000106397 |
| WDTC1 | -0.356945106 | 0.780816202 | 0.000379138 | 0.002248619 | 817.3191169 | 638.1639286 | ENSG00000142784 |
| SSRP1 | -0.357222591 | 0.780666036 | 9.91E-10 | 1.85E-08 | 7915.285954 | 6179.62066 | ENSG00000149136 |
| C5orf49 | -0.35742283 | 0.780557691 | 0.595281615 | 0.755953244 | 13.02438204 | 10.25502718 | ENSG00000215217 |
| ASPRV1 | -0.357451437 | 0.780542213 | 0.576553008 | 0.741351009 | 13.95521205 | 10.85460814 | ENSG00000244617 |
| NONO | -0.357485204 | 0.780523945 | 7.75E-10 | 1.46E-08 | 15159.80004 | 11832.8856 | ENSG00000147140 |
| NA | -0.357550448 | 0.780488647 | 0.315275967 | 0.50882621 | 44.67699259 | 34.98029824 | ENSG00000239569 |
| LOC389641 | -0.357624068 | 0.78044882 | 0.446920485 | 0.635167876 | 28.62623852 | 22.39648023 | ENSG00000246582 |
| TSPAN3 | -0.357650119 | 0.780434727 | 3.12E-08 | 4.60E-07 | 2794.416947 | 2180.900331 | ENSG00000140391 |
| KIF15 | -0.357670002 | 0.780423972 | 0.00111869 | 0.005809655 | 664.4518853 | 518.3558995 | ENSG00000163808 |
| SLC6A9 | -0.357692754 | 0.780411664 | 0.001976295 | 0.009396782 | 553.4961627 | 432.5332347 | ENSG00000196517 |
| NA | -0.3579136 | 0.780292209 | 0.541012583 | 0.714131985 | 16.917574 | 13.24058802 | ENSG00000271032 |
| NA | -0.358248558 | 0.780111065 | 0.535264425 | 0.709650959 | 16.66111259 | 13.02271055 | ENSG00000235245 |
| CLTCL1 | -0.358377831 | 0.780041167 | 0.060646153 | 0.155272248 | 181.0702112 | 141.2258212 | ENSG00000070371 |
| NA | -0.358395377 | 0.78003168 | 0.701706098 | 0.83064577 | 6.698910993 | 5.251205458 | ENSG00000279792 |
| NHLRC4 | -0.358439518 | 0.780007814 | 0.682481006 | 0.816696987 | 7.450225573 | 5.757369479 | ENSG00000257108 |
| SCFD1 | -0.358471072 | 0.779990754 | 0.002785405 | 0.012505984 | 756.0422179 | 589.694911 | ENSG00000092108 |
| CALHM6 | -0.358549457 | 0.779948377 | 0.706082212 | 0.833165323 | 6.073947985 | 4.692089268 | ENSG00000188820 |
| SDHD | -0.35867641 | 0.779879746 | 0.311913164 | 0.505553002 | 387.5390581 | 302.1403674 | ENSG00000204370 |
| RRAGB | -0.358732244 | 0.779849565 | 0.100026824 | 0.226727797 | 139.9809343 | 109.4611223 | ENSG00000083750 |
| NA | -0.358798692 | 0.779813647 | 0.483159331 | 0.666483704 | 19.37954308 | 15.11193013 | ENSG00000273373 |
| C4orf46 | -0.358910716 | 0.779753098 | 0.01119422 | 0.040427449 | 484.2059659 | 376.9354001 | ENSG00000205208 |
| PTPRZ1 | -0.359094585 | 0.779653726 | 2.47E-08 | 3.69E-07 | 5592.786791 | 4359.660503 | ENSG00000106278 |
| PRPSAP1 | -0.359178053 | 0.77960862 | 0.000376839 | 0.002235756 | 798.6808762 | 622.804958 | ENSG00000161542 |
| NA | -0.359306603 | 0.779539157 | 0.708205741 | 0.83401045 | 6.589621763 | 5.15355164 | ENSG00000273554 |
| ATP6V0C | -0.359411997 | 0.779482211 | 0.755302019 | 0.863919123 | 4.446147334 | 3.480766362 | ENSG00000185883 |
| LUC7L | -0.359512385 | 0.779427973 | 1.51E-05 | 0.000130183 | 1970.054245 | 1536.41471 | ENSG00000007392 |
| TNFAIP2 | -0.359734723 | 0.779307862 | 0.01116496 | 0.040361102 | 542.0357818 | 422.8171757 | ENSG00000185215 |
| FUNDC2 | -0.359829123 | 0.779256872 | 0.000268099 | 0.001665756 | 762.742437 | 593.9295431 | ENSG00000165775 |
| CDS1 | -0.359914385 | 0.77921082 | 0.008526885 | 0.032272265 | 396.405609 | 308.6584761 | ENSG00000163624 |

| | | | | | | | |
|---|---|---|---|---|---|---|---|
| APIP | -0.359959947 | 0.779186212 | 0.018682969 | 0.060941595 | 307.0707607 | 239.1490447 | ENSG00000149089 |
| NPAS2 | -0.360014315 | 0.779156849 | 0.470875596 | 0.656230415 | 21.14642065 | 16.51476183 | ENSG00000170485 |
| CNFN | -0.360062038 | 0.779131075 | 0.145318446 | 0.298960159 | 88.40855068 | 68.94359353 | ENSG00000105427 |
| NA | -0.360091822 | 0.77911499 | 0.631521354 | 0.7826627 | 10.78424603 | 8.333870472 | ENSG00000236675 |
| KLHL35 | -0.360154064 | 0.779081378 | 0.558386093 | 0.727526558 | 15.33336877 | 11.89351966 | ENSG00000149243 |
| ANO8 | -0.360264547 | 0.779021717 | 0.165633774 | 0.328572875 | 1236.278452 | 963.3760345 | ENSG00000074855 |
| U2SURP | -0.360310864 | 0.778996707 | 1.37E-05 | 0.000118973 | 2301.733738 | 1792.519017 | ENSG00000163714 |
| CELF2 | -0.36036533 | 0.778967299 | 0.001285432 | 0.006496915 | 529.1882468 | 412.3135051 | ENSG00000048740 |
| BANP | -0.360407838 | 0.778944347 | 0.001353013 | 0.006781163 | 657.8251169 | 512.5132689 | ENSG00000172530 |
| SLCO4A1 | -0.360603388 | 0.778838772 | 0.002792054 | 0.012520668 | 556.568663 | 434.048804 | ENSG00000101187 |
| NA | -0.360617402 | 0.778831207 | 0.732473221 | 0.850804364 | 4.439833508 | 3.473280288 | ENSG00000285706 |
| TLCD1 | -0.360677493 | 0.778798768 | 0.045892596 | 0.125139528 | 220.9577723 | 172.2108269 | ENSG00000160606 |
| ZNF730 | -0.36069772 | 0.778787849 | 0.234646421 | 0.419534371 | 75.3922639 | 58.50030676 | ENSG00000183850 |
| FTCD | -0.360874109 | 0.778692637 | 0.329508933 | 0.523943798 | 41.25508223 | 32.12949517 | ENSG00000160282 |
| ZWINT | -0.360889677 | 0.778684235 | 1.23E-07 | 1.61E-06 | 2074.09907 | 1615.145129 | ENSG00000122952 |
| NA | -0.360948065 | 0.778652721 | 0.717935959 | 0.84019859 | 5.617751889 | 4.362023857 | ENSG00000228057 |
| ZNF69 | -0.361111568 | 0.77856448 | 0.133234413 | 0.280227714 | 105.0600948 | 82.0385769 | ENSG00000198429 |
| NA | -0.361146807 | 0.778545463 | 0.345158498 | 0.538618136 | 388.6610768 | 302.7764405 | ENSG00000214826 |
| RABGAP1L | -0.361206313 | 0.778513352 | 3.44E-05 | 0.000271251 | 1182.47617 | 920.3147483 | ENSG00000152061 |
| WDFY3-AS2 | -0.361295802 | 0.778465063 | 0.269011741 | 0.459272625 | 53.34991204 | 41.43847811 | ENSG00000180769 |
| AGFG2 | -0.361432793 | 0.778391147 | 0.122278386 | 0.26327923 | 106.0675626 | 82.71649103 | ENSG00000106351 |
| NA | -0.36149849 | 0.778355702 | 0.525631294 | 0.702129759 | 16.85503711 | 13.14293421 | ENSG00000260806 |
| GABPB1-IT1 | -0.361715897 | 0.778238416 | 0.054385184 | 0.142631244 | 173.067421 | 134.9961425 | ENSG00000285410 |
| DAXX | -0.361899537 | 0.778139361 | 1.00E-05 | 8.96E-05 | 2240.421174 | 1743.827245 | ENSG00000204209 |
| CAPN5 | -0.361906472 | 0.77813562 | 1.26E-05 | 0.000110551 | 1115.023024 | 867.6294466 | ENSG00000149260 |
| NA | -0.3619632 | 0.778105024 | 0.581041784 | 0.745106789 | 12.3198198 | 9.594458321 | ENSG00000260233 |
| NOP16 | -0.362028702 | 0.778069697 | 0.00037499 | 0.002229016 | 680.1467503 | 529.4973056 | ENSG00000048162 |
| PNMA6A | -0.362269573 | 0.777939802 | 0.138053057 | 0.287750266 | 98.78029374 | 77.04180322 | ENSG00000235961 |
| PLCH1 | -0.362380133 | 0.777880188 | 0.002963653 | 0.013184913 | 555.9585151 | 431.9565726 | ENSG00000114805 |
| LTBP2 | -0.362471913 | 0.777830703 | 0.314583428 | 0.508170022 | 42.52089051 | 32.97645987 | ENSG00000119681 |
| GOLGA2P5 | -0.362720303 | 0.777696794 | 0.081676599 | 0.194700205 | 146.8743486 | 114.3780254 | ENSG00000238105 |
| TMEM218 | -0.362805389 | 0.777650929 | 0.008123163 | 0.031016626 | 330.6014455 | 257.0798986 | ENSG00000150433 |
| DNAJC2 | -0.362861212 | 0.77762084 | 0.000308156 | 0.001878121 | 809.6707874 | 629.7046183 | ENSG00000105821 |
| ATP5MG | -0.362934335 | 0.777581427 | 0.191159771 | 0.364052116 | 115.1246705 | 89.20677352 | ENSG00000167283 |
| ECHDC1 | -0.363037134 | 0.777526023 | 0.000564635 | 0.003191502 | 975.5676024 | 758.1292619 | ENSG00000093144 |
| VPS11 | -0.363046695 | 0.77752087 | 0.004305687 | 0.018136728 | 457.4404405 | 355.6578454 | ENSG00000160695 |
| TCTEX1D1 | -0.363078189 | 0.777503897 | 0.49115277 | 0.673915134 | 20.15739074 | 15.73115815 | ENSG00000152760 |
| USF2 | -0.363110419 | 0.777486528 | 6.17E-07 | 7.15E-06 | 2613.493623 | 2032.947333 | ENSG00000105698 |
| UGDH | -0.363228524 | 0.777422882 | 0.000119192 | 0.000816071 | 1767.827082 | 1373.989632 | ENSG00000109814 |
| IL15RA | -0.363487813 | 0.777283172 | 0.476447283 | 0.66089522 | 20.81845527 | 16.24480825 | ENSG00000134470 |
| ATP6AP1 | -0.363498423 | 0.777277455 | 4.34E-06 | 4.28E-05 | 1583.36641 | 1230.278849 | ENSG00000071553 |
| KCTD21-AS1 | -0.36359887 | 0.77722334 | 0.265565248 | 0.455402137 | 75.42145023 | 58.84786934 | ENSG00000246174 |
| FHOD3 | -0.363853052 | 0.777086416 | 0.009750398 | 0.035991611 | 318.3987019 | 247.7716801 | ENSG00000134775 |
| RASA2 | -0.363952428 | 0.777032891 | 0.01126491 | 0.040603625 | 308.8118944 | 239.6216804 | ENSG00000155903 |
| ZGPAT | -0.364103484 | 0.776951537 | 0.069247603 | 0.171613966 | 166.0475349 | 129.2196756 | ENSG00000197114 |
| BRMS1 | -0.364183538 | 0.776908425 | 2.71E-05 | 0.000219204 | 990.4573202 | 769.4271428 | ENSG00000174744 |
| NA | -0.364500393 | 0.776737814 | 0.712384081 | 0.837030972 | 5.366228826 | 4.201009014 | ENSG00000214578 |
| EIF4E1B | -0.364594636 | 0.776687076 | 0.390564956 | 0.583196503 | 36.91892318 | 28.84447448 | ENSG00000175766 |
| LIPT2 | -0.364764605 | 0.776595577 | 0.344335234 | 0.53792222 | 40.55377459 | 31.62333115 | ENSG00000175536 |
| MFSD11 | -0.364796458 | 0.776578431 | 0.042959631 | 0.119068909 | 174.5888939 | 135.5501457 | ENSG00000092931 |
| CABLES2 | -0.364828253 | 0.776561316 | 0.002628692 | 0.011931552 | 428.8143747 | 332.878236 | ENSG00000149679 |
| NA | -0.364932932 | 0.776504972 | 0.243155561 | 0.429654417 | 61.09733063 | 47.52255218 | ENSG00000279041 |
| QRFP | -0.364971902 | 0.776483998 | 0.7293286 | 0.848386575 | 5.484988795 | 4.294863986 | ENSG00000188710 |
| ZNF577 | -0.365094802 | 0.776417853 | 0.094974458 | 0.217799628 | 145.2535608 | 113.0882577 | ENSG00000161551 |
| KIT | -0.365158977 | 0.776383317 | 0.002967652 | 0.013199537 | 532.7392522 | 413.0500022 | ENSG00000157404 |
| C10orf25 | -0.365213411 | 0.776354024 | 0.422278981 | 0.613459684 | 25.1691211 | 19.50021106 | ENSG00000165511 |
| TSTA3 | -0.365239542 | 0.776339963 | 6.51E-07 | 7.50E-06 | 2019.194793 | 1567.928687 | ENSG00000104522 |
| VPS8 | -0.365268672 | 0.776324287 | 0.002212337 | 0.010354409 | 726.7461735 | 563.450996 | ENSG00000156931 |
| NA | -0.365279442 | 0.776318492 | 0.538494377 | 0.712004119 | 15.59751949 | 12.14145305 | ENSG00000279382 |
| ZSWIM3 | -0.365321307 | 0.776295964 | 0.104159709 | 0.233774869 | 118.6463993 | 92.31518623 | ENSG00000132801 |
| NA | -0.365391214 | 0.776258349 | 0.650123999 | 0.795760018 | 8.584548533 | 6.725433905 | ENSG00000279039 |
| NA | -0.365583669 | 0.776154804 | 0.583109786 | 0.746519084 | 12.04807748 | 9.39590149 | ENSG00000262454 |
| CDR2L | -0.365622511 | 0.776133907 | 7.24E-05 | 0.000521527 | 1364.433585 | 1058.864671 | ENSG00000109089 |
| VASH1 | -0.365698579 | 0.776092986 | 0.002635814 | 0.011955947 | 408.5873365 | 316.9602439 | ENSG00000071246 |
| PAF1 | -0.36572811 | 0.7760771 | 1.61E-05 | 0.000137138 | 1079.311491 | 837.624125 | ENSG00000006712 |
| NA | -0.365859214 | 0.776006578 | 0.519832558 | 0.696844831 | 15.9065434 | 12.32503774 | ENSG00000198406 |
| COL14A1 | -0.36591352 | 0.775977368 | 9.96E-05 | 0.00069416 | 949.0711586 | 735.8538645 | ENSG00000187955 |
| ECEL1P2 | -0.365944577 | 0.775960663 | 0.439747672 | 0.628007655 | 24.43161425 | 18.90399091 | ENSG00000244280 |
| TMEM98 | -0.365996616 | 0.775932674 | 1.29E-06 | 1.41E-05 | 1647.891199 | 1278.330613 | ENSG00000006042 |
| HKDC1 | -0.366141017 | 0.775855014 | 0.706770324 | 0.833271761 | 5.783264191 | 4.47844867 | ENSG00000156510 |
| HMGN4 | -0.366246898 | 0.775798075 | 0.000376366 | 0.002234997 | 951.7422203 | 737.7541801 | ENSG00000182952 |
| RAN | -0.36625821 | 0.775791993 | 2.84E-11 | 6.74E-10 | 19237.66572 | 14923.71474 | ENSG00000132341 |
| LBH | -0.366318808 | 0.775759407 | 3.67E-05 | 0.000287811 | 1125.89304 | 873.4213236 | ENSG00000213626 |
| BCL9 | -0.366418082 | 0.775706028 | 2.26E-05 | 0.000186508 | 1513.201381 | 1173.393795 | ENSG00000116128 |

| | | | | | | | |
|---|---|---|---|---|---|---|---|
| ZNF792 | -0.366556992 | 0.775631343 | 0.092110611 | 0.213077641 | 134.1939119 | 104.0691616 | ENSG00000180884 |
| C9orf78 | -0.36662784 | 0.775593254 | 4.94E-05 | 0.000372941 | 1166.021128 | 903.7371348 | ENSG00000136819 |
| NA | -0.36680549 | 0.775497755 | 0.105679423 | 0.236353617 | 108.420047 | 84.17943455 | ENSG00000284648 |
| NA | -0.36692539 | 0.775433307 | 0.241839602 | 0.428063718 | 53.98128657 | 41.84742634 | ENSG00000243199 |
| ATP2B2 | -0.367024435 | 0.775380074 | 0.402037621 | 0.593734284 | 28.57011315 | 22.0805109 | ENSG00000157087 |
| HIRA | -0.367072039 | 0.775354489 | 0.013145484 | 0.045995546 | 312.9138108 | 242.7672055 | ENSG00000100084 |
| DPH2 | -0.367114861 | 0.775331475 | 1.39E-05 | 0.000120764 | 1164.518522 | 902.871113 | ENSG00000132768 |
| PDSS1 | -0.367121025 | 0.775328163 | 0.019777197 | 0.06395927 | 261.5962048 | 202.7480199 | ENSG00000148459 |
| NA | -0.367222573 | 0.775273591 | 0.706598815 | 0.833271761 | 7.906421668 | 6.061066202 | ENSG00000230084 |
| TRMT12 | -0.367357848 | 0.7752009 | 0.004107442 | 0.017416455 | 419.8136778 | 325.4160986 | ENSG00000183665 |
| ERH | -0.367563146 | 0.775090596 | 1.40E-06 | 1.51E-05 | 3158.787659 | 2447.584475 | ENSG00000100632 |
| DNAJC7 | -0.367769833 | 0.774979561 | 2.06E-05 | 0.000171091 | 1330.451175 | 1031.412651 | ENSG00000168259 |
| RAD51 | -0.36778873 | 0.77496941 | 0.000192401 | 0.0012412 | 813.5532008 | 630.4488565 | ENSG00000051180 |
| NA | -0.367797385 | 0.774964761 | 0.445679955 | 0.633793623 | 24.62257725 | 19.05751968 | ENSG00000272079 |
| MEN1 | -0.367953167 | 0.774881085 | 1.61E-06 | 1.72E-05 | 1662.413459 | 1288.129027 | ENSG00000133895 |
| NA | -0.368210327 | 0.774742975 | 0.726120936 | 0.846179828 | 5.412981159 | 4.219780009 | ENSG00000264304 |
| TMEFF2 | -0.368262415 | 0.774715004 | 0.347556413 | 0.540766501 | 36.25795636 | 28.20267662 | ENSG00000144339 |
| SERPING1 | -0.368281929 | 0.774704525 | 0.000744521 | 0.004074091 | 687.7855787 | 532.7234883 | ENSG00000149131 |
| RTL8C | -0.368404863 | 0.774638514 | 0.000121843 | 0.000831457 | 937.4475801 | 726.205077 | ENSG00000134590 |
| PHKA2 | -0.368656614 | 0.774503351 | 0.000149135 | 0.000988582 | 728.442712 | 564.2393064 | ENSG00000044446 |
| TDG | -0.3686775 | 0.774492138 | 0.000714424 | 0.003925626 | 1216.704308 | 941.6160435 | ENSG00000139372 |
| ARG2 | -0.36881243 | 0.774419706 | 0.005829734 | 0.02347855 | 510.9320601 | 395.3284892 | ENSG00000081181 |
| NA | -0.368828267 | 0.774411206 | 0.25873599 | 0.447500265 | 503.7106054 | 390.2635856 | ENSG00000281398 |
| BIN1 | -0.368920686 | 0.774361598 | 1.37E-05 | 0.000119139 | 1353.202764 | 1048.596504 | ENSG00000136717 |
| ZSWIM8 | -0.369107393 | 0.77426139 | 0.001194285 | 0.006128149 | 609.7944109 | 472.6937797 | ENSG00000214655 |
| KLHL17 | -0.369309895 | 0.77415272 | 0.004631924 | 0.019248448 | 665.5057069 | 516.0406409 | ENSG00000187961 |
| SH3BGRL | -0.369361637 | 0.774124956 | 0.003806596 | 0.016335136 | 625.304048 | 483.6037442 | ENSG00000131171 |
| CXXC1 | -0.369425552 | 0.77409066 | 2.54E-06 | 2.62E-05 | 2000.585912 | 1549.37456 | ENSG00000154832 |
| STUM | -0.36957359 | 0.774011234 | 0.368619671 | 0.5616478 | 39.88256291 | 30.93336765 | ENSG00000203685 |
| CHRDL1 | -0.369771105 | 0.773905273 | 0.000383656 | 0.00227106 | 754.7296126 | 583.7654481 | ENSG00000101938 |
| POLR3F | -0.369846987 | 0.773864569 | 0.022674328 | 0.071633114 | 282.4783771 | 218.2637853 | ENSG00000132664 |
| MIR22HG | -0.369887767 | 0.773842695 | 0.397977311 | 0.589934568 | 32.83161131 | 25.43086799 | ENSG00000186594 |
| CRYBB3 | -0.369982718 | 0.773791766 | 0.632967648 | 0.783663094 | 11.45280446 | 8.953648145 | ENSG00000100053 |
| PBRM1 | -0.370285897 | 0.773629172 | 3.12E-05 | 0.000249257 | 1168.456564 | 903.4755434 | ENSG00000163939 |
| AUTS2 | -0.370301946 | 0.773620566 | 8.71E-07 | 9.79E-06 | 1846.655801 | 1428.671598 | ENSG00000158321 |
| C16orf91 | -0.370435484 | 0.773548962 | 0.02706412 | 0.082609751 | 254.8234222 | 197.0525228 | ENSG00000174109 |
| ALOX12 | -0.370565854 | 0.773479063 | 0.217831836 | 0.398784596 | 58.67264314 | 45.43344422 | ENSG00000108839 |
| SPEG | -0.370618696 | 0.773450733 | 0.004841815 | 0.020005773 | 1135.597916 | 879.2412529 | ENSG00000072195 |
| NA | -0.370795546 | 0.773355927 | 0.672834077 | 0.809789244 | 6.973810223 | 5.348421244 | ENSG00000197149 |
| IRX3 | -0.370917896 | 0.773290344 | 0.264699376 | 0.454379756 | 52.0336158 | 40.18972483 | ENSG00000177508 |
| ARSB | -0.370920517 | 0.773288939 | 0.008400929 | 0.031886711 | 334.4082207 | 258.2065644 | ENSG00000113273 |
| LOC728485 | -0.371163055 | 0.773158949 | 0.648622263 | 0.794866769 | 8.03602785 | 6.200060856 | ENSG00000267260 |
| MAEA | -0.371262219 | 0.773105808 | 3.31E-08 | 4.85E-07 | 2360.721854 | 1825.332327 | ENSG00000090316 |
| RMDN2 | -0.371652577 | 0.772896653 | 0.390496151 | 0.58319546 | 30.90435518 | 23.79463701 | ENSG00000115841 |
| PIK3C3 | -0.371819162 | 0.772807413 | 0.000347368 | 0.002080843 | 607.7374509 | 469.667499 | ENSG00000078142 |
| NDST3 | -0.37184447 | 0.772793856 | 0.652337574 | 0.796943154 | 8.618673215 | 6.70666291 | ENSG00000164100 |
| TLE1 | -0.372106324 | 0.772653604 | 6.63E-08 | 9.14E-07 | 3449.724969 | 2664.938922 | ENSG00000196781 |
| LSM3 | -0.372203387 | 0.772601623 | 7.48E-06 | 6.94E-05 | 1555.271134 | 1200.762512 | ENSG00000170860 |
| ZMAT2 | -0.372227596 | 0.772588658 | 9.76E-06 | 8.76E-05 | 1597.605474 | 1234.078865 | ENSG00000146007 |
| TRNAU1AP | -0.372404247 | 0.772494064 | 0.002732674 | 0.012299025 | 472.0084463 | 364.4455805 | ENSG00000180098 |
| MAP2K3 | -0.372420847 | 0.772485176 | 0.000542811 | 0.003085083 | 891.4054271 | 687.8825203 | ENSG00000034152 |
| TRAF3IP1 | -0.372654117 | 0.772360282 | 0.001943456 | 0.009269173 | 513.1683401 | 396.1994326 | ENSG00000204104 |
| MRPL11 | -0.372717288 | 0.772326464 | 2.72E-07 | 3.37E-06 | 1807.627461 | 1395.830661 | ENSG00000174547 |
| H2AZ1 | -0.372743759 | 0.772312293 | 4.15E-10 | 8.13E-09 | 4795.71348 | 3703.257326 | ENSG00000164032 |
| NA | -0.372829795 | 0.772266237 | 0.373915712 | 0.56711847 | 31.63672828 | 24.39520566 | ENSG00000127589 |
| ABCF3 | -0.372957074 | 0.772198109 | 4.38E-06 | 4.31E-05 | 1273.765717 | 983.4467464 | ENSG00000161204 |
| NA | -0.373282803 | 0.772023783 | 0.35597813 | 0.549579231 | 35.15144425 | 27.08433262 | ENSG00000226415 |
| CLPTM1 | -0.373287856 | 0.772021079 | 2.58E-06 | 2.66E-05 | 2467.610484 | 1904.686331 | ENSG00000104853 |
| YAP1 | -0.373298667 | 0.772015294 | 7.17E-05 | 0.000517568 | 2144.062508 | 1654.380447 | ENSG00000137693 |
| L3MBTL3 | -0.373327207 | 0.772000022 | 0.004114036 | 0.017436438 | 407.3170936 | 314.7761103 | ENSG00000198945 |
| FXN | -0.373367067 | 0.771978692 | 0.06192162 | 0.157861802 | 148.6874974 | 114.6610874 | ENSG00000165060 |
| MYL6B | -0.373447563 | 0.771935621 | 0.000148421 | 0.000984553 | 1162.506187 | 897.5014932 | ENSG00000196465 |
| KAT14 | -0.373450966 | 0.7719338 | 0.000830408 | 0.0044858 | 535.8753194 | 413.5403499 | ENSG00000149474 |
| MRPL3 | -0.373527835 | 0.771892671 | 7.47E-05 | 0.000536555 | 2451.234418 | 1891.306341 | ENSG00000114686 |
| POP7 | -0.373642269 | 0.771831447 | 1.43E-06 | 1.55E-05 | 1478.363621 | 1141.515683 | ENSG00000172336 |
| SYT13 | -0.373744248 | 0.771776891 | 0.002532929 | 0.011582853 | 462.9455131 | 357.0208333 | ENSG00000019505 |
| CASZ1 | -0.373787696 | 0.771753649 | 0.1407092 | 0.29194048 | 135.8370161 | 104.5734618 | ENSG00000130940 |
| CCZ1 | -0.373798701 | 0.771747762 | 0.007547381 | 0.029154474 | 330.6754299 | 255.064552 | ENSG00000122674 |
| NA | -0.3738523 | 0.771719091 | 0.670643487 | 0.808646568 | 7.517294857 | 5.813682462 | ENSG00000229413 |
| NA | -0.374070387 | 0.771602441 | 0.4488057 | 0.636422382 | 22.48985134 | 17.45656919 | ENSG00000227486 |
| NA | -0.374106396 | 0.771583183 | 0.030660195 | 0.091096769 | 228.2283097 | 176.4504688 | ENSG00000259203 |
| GTF2IP1 | -0.374208745 | 0.771528446 | 0.038702396 | 0.109613417 | 195.5374591 | 150.9025232 | ENSG00000277053 |
| NA | -0.374271393 | 0.771494944 | 0.556955959 | 0.726427911 | 13.02753895 | 10.0935743 | ENSG00000259891 |
| IP6K1 | -0.374770347 | 0.77122817 | 8.75E-06 | 7.92E-05 | 1196.347679 | 922.2695534 | ENSG00000176095 |

| | | | | | | | |
|---|---|---|---|---|---|---|---|
| THUMPD3 | -0.375067315 | 0.771069434 | 0.005822989 | 0.023456484 | 369.8984015 | 285.0629371 | ENSG00000134077 |
| ZNF487 | -0.375082488 | 0.771061325 | 0.496702516 | 0.678099913 | 18.05001783 | 13.85882836 | ENSG00000243660 |
| CERK | -0.375107111 | 0.771048165 | 7.04E-08 | 9.63E-07 | 2360.624599 | 1819.600728 | ENSG00000100422 |
| ZNF354B | -0.375112513 | 0.771045278 | 0.141335 | 0.293009271 | 184.3001738 | 142.3985515 | ENSG00000178338 |
| ANKS1B | -0.375175562 | 0.771011582 | 0.286329038 | 0.477588268 | 43.05668584 | 33.25488721 | ENSG00000185046 |
| NA | -0.375384005 | 0.770900193 | 0.368608196 | 0.5616478 | 32.9643744 | 25.34026221 | ENSG00000284727 |
| B9D2 | -0.375437465 | 0.770871628 | 0.267860886 | 0.45798281 | 51.43911689 | 39.72490165 | ENSG00000123810 |
| WDR48 | -0.37552894 | 0.770822751 | 3.63E-05 | 0.000285296 | 1091.951602 | 841.2727395 | ENSG00000114742 |
| XRCC5 | -0.375582206 | 0.770794293 | 7.95E-10 | 1.50E-08 | 13616.91513 | 10494.59564 | ENSG00000079246 |
| TMOD4 | -0.375610672 | 0.770779084 | 0.666093697 | 0.806044698 | 7.711219375 | 5.967649264 | ENSG00000163157 |
| CPZ | -0.375651525 | 0.770757258 | 0.612040475 | 0.768689989 | 10.43733917 | 8.120779527 | ENSG00000109625 |
| CENPT | -0.375771974 | 0.770692911 | 0.256925627 | 0.445408246 | 1022.87467 | 788.537833 | ENSG00000102901 |
| NA | -0.376030296 | 0.770554927 | 0.362838555 | 0.555747709 | 35.65952643 | 27.31630618 | ENSG00000179611 |
| TNFRSF1A | -0.376263129 | 0.770430579 | 3.38E-05 | 0.000267236 | 1078.816646 | 830.7933993 | ENSG00000067182 |
| UCHL5 | -0.376357041 | 0.770380429 | 0.000126595 | 0.000859773 | 699.9837446 | 539.1349282 | ENSG00000116750 |
| SLC35E4 | -0.376469754 | 0.770320245 | 0.037896016 | 0.107641395 | 198.6922345 | 152.9114342 | ENSG00000100036 |
| NA | -0.376511878 | 0.770297753 | 0.714175263 | 0.838126026 | 6.427168677 | 4.985050724 | ENSG00000261570 |
| CALN1 | -0.37662177 | 0.770239081 | 0.000696834 | 0.003843784 | 532.8995405 | 410.6899848 | ENSG00000183166 |
| CTSC | -0.376663689 | 0.770216701 | 6.54E-12 | 1.72E-10 | 7579.030167 | 5836.977506 | ENSG00000109861 |
| NA | -0.376786276 | 0.770151258 | 0.631692492 | 0.782745522 | 8.512540898 | 6.590238097 | ENSG00000229582 |
| WASIR2 | -0.376801614 | 0.77014307 | 0.663528081 | 0.804258763 | 6.689440255 | 5.146065566 | ENSG00000231439 |
| BAG5 | -0.376867991 | 0.770107637 | 0.000155699 | 0.001028047 | 884.5999552 | 681.3293719 | ENSG00000166170 |
| NA | -0.376970956 | 0.770052677 | 0.430134262 | 0.620204481 | 26.23006095 | 20.26971865 | ENSG00000215102 |
| OLFML2A | -0.37699284 | 0.770040996 | 0.022401803 | 0.070876582 | 237.5065743 | 182.6216444 | ENSG00000185585 |
| PI4K2A | -0.377079706 | 0.769994633 | 0.000507164 | 0.00290475 | 546.7716056 | 420.8047032 | ENSG00000155252 |
| RAB5IF | -0.377143603 | 0.76996053 | 0.027630254 | 0.084088528 | 245.9651394 | 189.4566783 | ENSG00000101084 |
| GPR160 | -0.377172673 | 0.769945016 | 0.003187811 | 0.014020618 | 549.9079433 | 422.7895155 | ENSG00000173890 |
| FAM171B | -0.377251439 | 0.76990298 | 0.015081953 | 0.051336807 | 292.0535416 | 224.596454 | ENSG00000144369 |
| NA | -0.377253198 | 0.769902042 | 0.56519447 | 0.733099812 | 13.78210814 | 10.64476638 | ENSG00000261499 |
| MISP3 | -0.377265023 | 0.769895731 | 0.257773096 | 0.446392079 | 46.85133709 | 36.02625781 | ENSG00000141854 |
| GNG5 | -0.377397233 | 0.76982518 | 0.103358935 | 0.232313183 | 117.2446937 | 90.35334996 | ENSG00000174021 |
| SNORD12C | -0.377445822 | 0.769799254 | 0.736058031 | 0.852173027 | 4.549122739 | 3.503336203 | ENSG00000209042 |
| ELOVL1 | -0.37760667 | 0.769713433 | 0.000547628 | 0.003108646 | 531.4805545 | 409.0544151 | ENSG00000066322 |
| C1orf56 | -0.377636025 | 0.769697771 | 0.205051343 | 0.38330021 | 64.46222116 | 49.53636138 | ENSG00000143443 |
| CAPN1 | -0.377745756 | 0.769639231 | 1.75E-10 | 3.69E-09 | 4383.161545 | 3373.868084 | ENSG00000014216 |
| NA | -0.377895042 | 0.769559594 | 0.671038099 | 0.808859269 | 6.775353319 | 5.213663469 | ENSG00000272501 |
| PRORP | -0.377908394 | 0.769552472 | 0.13856467 | 0.28858946 | 86.1317118 | 66.1750341 | ENSG00000100890 |
| NA | -0.377921198 | 0.769545643 | 0.733107331 | 0.851101774 | 4.449304247 | 3.416967305 | ENSG00000258682 |
| AMIGO1 | -0.377943586 | 0.769533701 | 0.0777865 | 0.187736319 | 119.6731168 | 92.20635911 | ENSG00000181754 |
| ARAP2 | -0.378066396 | 0.769468196 | 0.423499485 | 0.614269796 | 25.29862958 | 19.35752918 | ENSG00000047365 |
| NA | -0.378117648 | 0.769440861 | 0.758178156 | 0.86529526 | 4.022798143 | 3.124443883 | ENSG00000265566 |
| RASD2 | -0.378290978 | 0.769348424 | 0.048342732 | 0.130249192 | 166.1982699 | 127.5935186 | ENSG00000100302 |
| CIP2A | -0.378334241 | 0.769325353 | 9.85E-05 | 0.00068757 | 751.5317674 | 578.5012459 | ENSG00000163507 |
| RPP40 | -0.378344847 | 0.769319698 | 0.102062152 | 0.230208097 | 101.8873474 | 78.44788411 | ENSG00000124787 |
| JPT2 | -0.378354085 | 0.769314771 | 4.97E-09 | 8.37E-08 | 4538.732866 | 3490.737896 | ENSG00000206053 |
| TPCN2 | -0.378544037 | 0.769213487 | 0.013218758 | 0.046164773 | 276.1772669 | 212.7168475 | ENSG00000162341 |
| TNMD | -0.378547006 | 0.769211904 | 0.261620881 | 0.450890962 | 48.63715614 | 37.39523474 | ENSG00000000005 |
| NIPBL-DT | -0.37869923 | 0.769130746 | 0.102348391 | 0.230601152 | 115.4513581 | 88.97341452 | ENSG00000285967 |
| UFSP1 | -0.378899867 | 0.769023789 | 0.348945435 | 0.542244881 | 39.49027919 | 30.46810643 | ENSG00000176125 |
| SLC25A27 | -0.379013216 | 0.768963371 | 0.246705653 | 0.434312555 | 71.38780588 | 55.08944009 | ENSG00000153291 |
| H3-5 | -0.37912934 | 0.768901479 | 0.562233575 | 0.730738379 | 13.71631664 | 10.49068796 | ENSG00000188375 |
| ARMC6 | -0.379152326 | 0.768889229 | 1.30E-06 | 1.41E-05 | 1661.012197 | 1277.464328 | ENSG00000105676 |
| PSMD9 | -0.379264244 | 0.768829584 | 0.569142335 | 0.735971767 | 13.78022901 | 10.62219654 | ENSG00000110801 |
| RRM1 | -0.379564746 | 0.76866946 | 1.70E-07 | 2.17E-06 | 2026.652768 | 1557.136913 | ENSG00000167325 |
| CENPU | -0.379607937 | 0.768646447 | 4.56E-06 | 4.46E-05 | 1177.299145 | 905.0287034 | ENSG00000151725 |
| SNRPE | -0.37966586 | 0.768615588 | 1.64E-05 | 0.000139367 | 1866.724799 | 1434.064664 | ENSG00000182004 |
| GNAS | -0.379806683 | 0.768540566 | 5.56E-12 | 1.48E-10 | 16874.80617 | 12968.68397 | ENSG00000087460 |
| CACNA1C-AS2 | -0.379857694 | 0.768513392 | 0.650308281 | 0.795928168 | 7.946957874 | 6.072351123 | ENSG00000256271 |
| TRAPPC2 | -0.379992986 | 0.768441326 | 0.043699632 | 0.120511204 | 161.6409542 | 124.1439075 | ENSG00000196459 |
| KCTD8 | -0.380000052 | 0.768437563 | 0.060370709 | 0.154738034 | 137.7826123 | 105.948651 | ENSG00000183783 |
| TULP1 | -0.380010209 | 0.768432153 | 0.136364348 | 0.285160423 | 90.43266378 | 69.71310113 | ENSG00000112041 |
| CNKSR2 | -0.380369295 | 0.768240915 | 0.009757288 | 0.036009873 | 348.6247197 | 267.571972 | ENSG00000149970 |
| RASL12 | -0.380374168 | 0.76823832 | 0.007860372 | 0.03017484 | 285.1038816 | 219.0951569 | ENSG00000103710 |
| SLC16A13 | -0.380380909 | 0.76823473 | 0.135376023 | 0.283605654 | 93.41977738 | 71.84256224 | ENSG00000174327 |
| TCF7L1 | -0.380689338 | 0.768070509 | 4.69E-07 | 5.56E-06 | 3612.202114 | 2774.198181 | ENSG00000152284 |
| PPWD1 | -0.380968405 | 0.767921952 | 6.05E-05 | 0.000445094 | 781.8408496 | 600.4306011 | ENSG00000113593 |
| TSPAN17 | -0.381242501 | 0.767776069 | 1.52E-07 | 1.97E-06 | 1912.853421 | 1468.667553 | ENSG00000048140 |
| NA | -0.381431336 | 0.767675581 | 0.716825104 | 0.839556455 | 4.472680413 | 3.424453379 | ENSG00000233885 |
| SETDB2 | -0.381508074 | 0.767634749 | 0.052815977 | 0.139482945 | 160.2019444 | 122.8575006 | ENSG00000136169 |
| LAPTM4B | -0.381789395 | 0.767485077 | 1.91E-08 | 2.91E-07 | 10672.41807 | 8189.895104 | ENSG00000104341 |
| ZNF432 | -0.381826198 | 0.767465499 | 0.086439937 | 0.20312892 | 117.8121332 | 90.53346221 | ENSG00000256087 |
| TMEM179 | -0.381862353 | 0.767446266 | 0.024671087 | 0.076630457 | 223.8836494 | 172.0116088 | ENSG00000258986 |
| SNAPC4 | -0.38204657 | 0.767348278 | 0.000266419 | 0.001656109 | 1019.563334 | 783.1344665 | ENSG00000165684 |
| RPARP-AS1 | -0.382174073 | 0.767280464 | 0.051965919 | 0.137520628 | 148.2609913 | 113.8141227 | ENSG00000269609 |

| | | | | | | | |
|---|---|---|---|---|---|---|---|
| PES1 | -0.382184774 | 0.767274773 | 6.69E-08 | 9.21E-07 | 4320.858138 | 3316.091868 | ENSG00000100029 |
| SUFU | -0.382231636 | 0.76724985 | 0.001623197 | 0.007950406 | 494.4139554 | 378.9064746 | ENSG00000107882 |
| NA | -0.382239768 | 0.767245525 | 0.718410078 | 0.840477794 | 4.475837326 | 3.424453379 | ENSG00000279253 |
| BRSK2 | -0.382263856 | 0.767232715 | 0.000603078 | 0.00338096 | 872.7451107 | 670.0791875 | ENSG00000174672 |
| PASK | -0.382294321 | 0.767216514 | 7.65E-06 | 7.07E-05 | 1646.470822 | 1263.822023 | ENSG00000115687 |
| NA | -0.382355725 | 0.76718386 | 0.68112499 | 0.815727872 | 6.361474866 | 4.913653973 | ENSG00000229816 |
| NUTM1 | -0.382379814 | 0.76717105 | 0.474653274 | 0.659659852 | 24.20987888 | 18.68341389 | ENSG00000184507 |
| PLAGL2 | -0.382428144 | 0.767145351 | 7.29E-06 | 6.79E-05 | 1087.137524 | 834.2900542 | ENSG00000126003 |
| LINC01950 | -0.382629504 | 0.767038286 | 0.1474603 | 0.302225942 | 86.51522384 | 66.42709275 | ENSG00000251027 |
| ADM | -0.382638876 | 0.767033303 | 8.48E-06 | 7.71E-05 | 1098.593388 | 842.5279827 | ENSG00000148926 |
| FAM32A | -0.382830463 | 0.76693145 | 1.57E-07 | 2.02E-06 | 1729.37566 | 1326.353046 | ENSG00000105058 |
| CENPK | -0.382927986 | 0.766879608 | 0.003346211 | 0.014606378 | 519.9157697 | 398.5204194 | ENSG00000123219 |
| HMG20B | -0.383064336 | 0.766807134 | 2.71E-07 | 3.35E-06 | 1717.840663 | 1317.663506 | ENSG00000064961 |
| PTCD1 | -0.383099199 | 0.766788604 | 0.39201775 | 0.584626937 | 27.9906247 | 21.53355569 | ENSG00000106246 |
| NA | -0.383131368 | 0.766771507 | 0.552672801 | 0.723352248 | 16.50181641 | 12.63264493 | ENSG00000267493 |
| SMARCA1 | -0.383907897 | 0.766358904 | 2.83E-09 | 4.91E-08 | 3170.446211 | 2429.111588 | ENSG00000102038 |
| BIVM | -0.383941256 | 0.766341183 | 0.192181505 | 0.365472196 | 88.6473484 | 67.94244724 | ENSG00000134897 |
| ZNF385D | -0.384153741 | 0.766228322 | 0.133423129 | 0.280560958 | 85.81972637 | 65.90551855 | ENSG00000151789 |
| NUCB1 | -0.384252292 | 0.766175983 | 1.46E-08 | 2.28E-07 | 2334.58293 | 1788.48967 | ENSG00000104805 |
| LRRC3 | -0.38450157 | 0.76604361 | 0.007563957 | 0.02920633 | 288.6111037 | 221.2295161 | ENSG00000160233 |
| ABCC6 | -0.384576515 | 0.766003816 | 0.481009943 | 0.664710335 | 17.4515879 | 13.36026203 | ENSG00000091262 |
| SMIM7 | -0.38461528 | 0.765983234 | 0.021894148 | 0.069686784 | 219.8470587 | 168.3359729 | ENSG00000214046 |
| ABCG1 | -0.384658657 | 0.765960204 | 0.067963222 | 0.169122381 | 122.8023665 | 94.15278519 | ENSG00000160179 |
| LSG1 | -0.384735631 | 0.765919338 | 2.34E-07 | 2.93E-06 | 1679.275983 | 1286.536041 | ENSG00000041802 |
| EPO | -0.384772019 | 0.76590002 | 0.666289223 | 0.806176031 | 7.850296295 | 5.96721123 | ENSG00000130427 |
| NENF | -0.385173231 | 0.765687054 | 0.018119929 | 0.059376726 | 294.8844182 | 225.4815104 | ENSG00000117691 |
| C16orf89 | -0.385358595 | 0.765588681 | 0.286938003 | 0.478306614 | 43.41306344 | 33.29622805 | ENSG00000153446 |
| EIF3CL | -0.385520943 | 0.765502533 | 0.011968028 | 0.042681437 | 343.7396718 | 263.4839556 | ENSG00000205609 |
| CUX2 | -0.385558397 | 0.76548266 | 0.007747784 | 0.029829128 | 283.3483539 | 216.9006858 | ENSG00000111249 |
| C6orf226 | -0.385609434 | 0.765455581 | 0.357047415 | 0.550616112 | 31.86940757 | 24.45900471 | ENSG00000221821 |
| TVP23A | -0.385773641 | 0.765368462 | 0.166175743 | 0.329330755 | 78.98933006 | 60.47788807 | ENSG00000166676 |
| PABPN1 | -0.385795921 | 0.765356642 | 1.58E-05 | 0.000135098 | 2009.033563 | 1538.655094 | ENSG00000100836 |
| MSH4 | -0.385832818 | 0.765337068 | 0.727691169 | 0.846909144 | 4.489742754 | 3.424453379 | ENSG00000057468 |
| NA | -0.386004342 | 0.765246082 | 0.494918911 | 0.676911453 | 17.13309325 | 13.14293421 | ENSG00000206120 |
| PCNA | -0.386017565 | 0.765239068 | 1.39E-08 | 2.17E-07 | 5491.478653 | 4201.418258 | ENSG00000132646 |
| UNC45A | -0.386101489 | 0.765194554 | 1.05E-08 | 1.68E-07 | 3119.798208 | 2387.317695 | ENSG00000140553 |
| ZSWIM1 | -0.386573097 | 0.764944457 | 0.024605676 | 0.076460416 | 222.8406589 | 170.6361335 | ENSG00000168612 |
| MYCBP | -0.386579497 | 0.764941064 | 0.027568879 | 0.083943092 | 234.1564836 | 179.0017717 | ENSG00000214114 |
| KANSL3 | -0.386585715 | 0.764937767 | 3.74E-06 | 3.74E-05 | 1164.177342 | 890.2836078 | ENSG00000114982 |
| ALDH1L1 | -0.386768199 | 0.764841018 | 0.343841217 | 0.537558255 | 36.3779941 | 27.92154973 | ENSG00000144908 |
| C1QL2 | -0.386787553 | 0.764830757 | 0.497958644 | 0.678931121 | 18.46073937 | 14.12921997 | ENSG00000144119 |
| NA | -0.386790701 | 0.764829088 | 0.701842062 | 0.830701395 | 5.386448079 | 4.12212619 | ENSG00000258702 |
| EPDR1 | -0.386831245 | 0.764807595 | 0.038360097 | 0.108767947 | 188.4261201 | 143.9167716 | ENSG00000086289 |
| ZNF775 | -0.386994103 | 0.764721265 | 0.000785355 | 0.004271059 | 594.0515082 | 454.2153807 | ENSG00000196456 |
| PANK4 | -0.387071328 | 0.764680332 | 7.82E-05 | 0.000558846 | 926.8343035 | 709.1657384 | ENSG00000157881 |
| CYP4V2 | -0.38717798 | 0.764623804 | 0.265041717 | 0.454714726 | 59.52712852 | 45.55702869 | ENSG00000145476 |
| PEX10 | -0.387294979 | 0.764561798 | 0.002881657 | 0.012863439 | 445.7828049 | 341.0002264 | ENSG00000157911 |
| CTSD | -0.387337645 | 0.764539187 | 0.000777824 | 0.004235076 | 664.4044641 | 507.7996604 | ENSG00000117984 |
| NA | -0.387557221 | 0.764422834 | 0.03335079 | 0.097418997 | 190.1342191 | 145.6745404 | ENSG00000273015 |
| RNF25 | -0.387578805 | 0.764411398 | 0.001816787 | 0.008732451 | 476.696458 | 364.2529724 | ENSG00000163481 |
| SREK1 | -0.387705836 | 0.764344093 | 0.000844452 | 0.004553552 | 680.5083367 | 520.3419715 | ENSG00000153914 |
| DNAJB1 | -0.387819625 | 0.76428381 | 2.86E-09 | 4.94E-08 | 3298.494957 | 2520.797289 | ENSG00000132002 |
| SNRNP25 | -0.388043754 | 0.764165085 | 1.52E-06 | 1.63E-05 | 1395.59244 | 1066.692847 | ENSG00000161981 |
| SEMA7A | -0.388193905 | 0.764085557 | 4.49E-05 | 0.000342071 | 755.705977 | 577.3631432 | ENSG00000138623 |
| PNRC2 | -0.388193958 | 0.764085529 | 4.99E-07 | 5.90E-06 | 2347.665233 | 1793.498556 | ENSG00000189266 |
| MAZ | -0.388348315 | 0.764003782 | 0.004726758 | 0.019581076 | 502.7520622 | 384.1644504 | ENSG00000103495 |
| NDUFAF8 | -0.38838079 | 0.763986584 | 0.001123054 | 0.005814155 | 586.0323472 | 448.1924549 | ENSG00000224877 |
| DCAF12L2 | -0.388416275 | 0.763967794 | 0.287908605 | 0.479433791 | 39.61031693 | 30.31022916 | ENSG00000198354 |
| RNF207 | -0.388451989 | 0.763948881 | 0.241377359 | 0.427638142 | 59.31329302 | 45.50951588 | ENSG00000158286 |
| RBM17 | -0.388459736 | 0.76394478 | 6.36E-09 | 1.05E-07 | 2565.945811 | 1960.357856 | ENSG00000134453 |
| ARNTL | -0.38853386 | 0.76390553 | 0.054300864 | 0.142430259 | 142.7891112 | 109.1863822 | ENSG00000133794 |
| RARG | -0.388558816 | 0.763892316 | 1.16E-05 | 0.000102707 | 1187.634305 | 907.5432775 | ENSG00000172819 |
| HLA-DPB1 | -0.38861164 | 0.763864346 | 0.010579538 | 0.038515255 | 287.9597053 | 220.3718162 | ENSG00000223865 |
| FUT8 | -0.388631237 | 0.76385397 | 0.001152204 | 0.005946797 | 461.6926407 | 352.4083913 | ENSG00000033170 |
| CD248 | -0.388999952 | 0.763658774 | 0.12838711 | 0.272693869 | 106.7917653 | 81.60205749 | ENSG00000174807 |
| PAK3 | -0.389169151 | 0.763569218 | 0.033998349 | 0.098987443 | 208.2835896 | 158.7969514 | ENSG00000077264 |
| MAP3K3 | -0.389174565 | 0.763566353 | 2.90E-05 | 0.000232979 | 842.14999 | 643.1637367 | ENSG00000198909 |
| TSN | -0.389331827 | 0.763483124 | 1.42E-07 | 1.84E-06 | 2416.930115 | 1844.671761 | ENSG00000211460 |
| TRAM2-AS1 | -0.389377756 | 0.763458818 | 0.188162869 | 0.359970311 | 63.26911957 | 48.39307441 | ENSG00000225791 |
| CSPP1 | -0.389417446 | 0.763437815 | 0.0188076 | 0.061304971 | 257.836956 | 197.0707357 | ENSG00000104218 |
| XRCC2 | -0.389549057 | 0.763368173 | 0.00393817 | 0.016825735 | 545.9527255 | 417.4525714 | ENSG00000196584 |
| PSMB7 | -0.389800489 | 0.763235145 | 1.05E-08 | 1.68E-07 | 3767.774269 | 2875.422226 | ENSG00000136930 |
| GUSBP3 | -0.389825522 | 0.763221902 | 0.450164962 | 0.637871361 | 21.87751598 | 16.80392444 | ENSG00000253203 |
| NDN | -0.389977769 | 0.763141364 | 0.000188997 | 0.001221364 | 701.6770511 | 535.435958 | ENSG00000182636 |

| | | | | | | | |
|---|---|---|---|---|---|---|---|
| PLPPR2 | -0.390134551 | 0.763058436 | 0.008199927 | 0.031251787 | 328.2976902 | 250.3949294 | ENSG00000105520 |
| WWC1 | -0.390195964 | 0.763025954 | 2.08E-06 | 2.18E-05 | 1528.066294 | 1166.032068 | ENSG00000113645 |
| FAM110D | -0.390389804 | 0.762923441 | 0.426476089 | 0.617131017 | 29.78097613 | 22.81258812 | ENSG00000197245 |
| NA | -0.390671589 | 0.762774443 | 0.719812158 | 0.841491014 | 4.416457342 | 3.345570555 | ENSG00000227620 |
| NREP | -0.390732376 | 0.762742304 | 0.013331011 | 0.046477982 | 286.1422346 | 218.1582476 | ENSG00000134986 |
| AKAP17A | -0.390745944 | 0.762735131 | 5.54E-07 | 6.50E-06 | 2202.70392 | 1681.144695 | ENSG00000197976 |
| ADCY6 | -0.390862564 | 0.762673478 | 0.001083853 | 0.005655441 | 439.0605555 | 334.6754818 | ENSG00000174233 |
| PGAM4 | -0.391220655 | 0.762484199 | 0.656314551 | 0.799167853 | 7.718810977 | 5.881280366 | ENSG00000226784 |
| ATP6V0A2 | -0.391331815 | 0.762425451 | 3.53E-05 | 0.000277599 | 769.8658621 | 586.9581512 | ENSG00000185344 |
| GRWD1 | -0.391390758 | 0.762394302 | 1.95E-08 | 2.96E-07 | 2531.950585 | 1930.063506 | ENSG00000105447 |
| PSMD2 | -0.391394829 | 0.762392151 | 1.71E-13 | 5.53E-12 | 10605.72185 | 8085.555931 | ENSG00000175166 |
| IGFBP6 | -0.391590254 | 0.762288885 | 0.043276945 | 0.119733526 | 218.2725196 | 166.35179 | ENSG00000167779 |
| HSPE1 | -0.391689383 | 0.76223651 | 0.01129055 | 0.040672314 | 521.59257 | 397.3336014 | ENSG00000115541 |
| ZNF624 | -0.391710265 | 0.762225477 | 0.378764936 | 0.572023464 | 29.6387423 | 22.61348164 | ENSG00000197566 |
| ABCA3 | -0.391726041 | 0.762217142 | 0.001158963 | 0.00597669 | 457.8620833 | 349.3470223 | ENSG00000167972 |
| NR6A1 | -0.391917246 | 0.762116129 | 1.54E-09 | 2.77E-08 | 5483.345107 | 4178.224425 | ENSG00000148200 |
| ARPC5L | -0.392378993 | 0.761872247 | 1.28E-06 | 1.40E-05 | 1541.376279 | 1174.1359 | ENSG00000136950 |
| WDR46 | -0.392404046 | 0.761859016 | 3.66E-08 | 5.31E-07 | 2214.169405 | 1687.457779 | ENSG00000227057 |
| ANKRD65 | -0.392546693 | 0.761783691 | 0.555158251 | 0.725160549 | 12.81014057 | 9.793564804 | ENSG00000235098 |
| UNC119B | -0.392614418 | 0.761747931 | 1.39E-07 | 1.81E-06 | 2038.323159 | 1552.615332 | ENSG00000175970 |
| HILPDA | -0.392788036 | 0.761656266 | 0.006489357 | 0.025721503 | 289.581305 | 220.4416756 | ENSG00000135245 |
| NA | -0.392898372 | 0.761598017 | 0.199445099 | 0.375776707 | 64.56647434 | 49.18036531 | ENSG00000254612 |
| CDKN2AIPNL | -0.39292022 | 0.761586484 | 3.35E-06 | 3.38E-05 | 1241.287288 | 945.3761939 | ENSG00000237190 |
| MBLAC2 | -0.392970518 | 0.761559932 | 0.133759354 | 0.281076629 | 88.11608545 | 67.06009044 | ENSG00000176055 |
| DEK | -0.392970703 | 0.761559835 | 0.0001405 | 0.000937373 | 4531.63689 | 3450.17151 | ENSG00000124795 |
| PRPF38A | -0.393089695 | 0.761497025 | 1.19E-07 | 1.57E-06 | 1568.733184 | 1194.579551 | ENSG00000134748 |
| ARID1B | -0.393107483 | 0.761487635 | 5.77E-06 | 5.50E-05 | 1909.164865 | 1453.904872 | ENSG00000049618 |
| GATC | -0.393287779 | 0.761392477 | 0.013914517 | 0.048132077 | 232.4917847 | 177.0642574 | ENSG00000257218 |
| MCOLN1 | -0.393354441 | 0.761357297 | 0.003522948 | 0.015262815 | 333.5132968 | 253.7332287 | ENSG00000090674 |
| CHCHD4 | -0.393430113 | 0.761317363 | 0.012347705 | 0.04380788 | 361.1618175 | 274.4905149 | ENSG00000163528 |
| OLFM1 | -0.393500737 | 0.761280096 | 1.35E-05 | 0.000117856 | 1394.338027 | 1061.168527 | ENSG00000130558 |
| PTGER3 | -0.393768337 | 0.761138902 | 0.610989467 | 0.767994492 | 9.801627645 | 7.430154756 | ENSG00000050628 |
| NA | -0.393784174 | 0.761130546 | 0.687010826 | 0.819981438 | 5.458455715 | 4.152182105 | ENSG00000238186 |
| RAVER2 | -0.39385338 | 0.761094036 | 5.01E-05 | 0.000377619 | 791.0118169 | 601.9090665 | ENSG00000162437 |
| H2BU1 | -0.393936057 | 0.761050421 | 0.664112975 | 0.804646001 | 6.378537207 | 4.868625911 | ENSG00000196890 |
| AIF1L | -0.39397191 | 0.761031508 | 8.25E-12 | 2.12E-10 | 5425.243586 | 4128.932322 | ENSG00000126878 |
| PCAT1 | -0.393983881 | 0.761025193 | 0.437770611 | 0.626390837 | 20.61052762 | 15.64055238 | ENSG00000253438 |
| SFXN2 | -0.393996159 | 0.761018717 | 0.007491412 | 0.02896846 | 353.4792435 | 269.2755147 | ENSG00000156398 |
| BCL2L1 | -0.394169524 | 0.760927273 | 0.0002463 | 0.001548479 | 662.6161645 | 504.2074705 | ENSG00000171552 |
| CCDC181 | -0.394293197 | 0.760862046 | 0.161022848 | 0.321655904 | 72.02549424 | 54.82185963 | ENSG00000117477 |
| LKAAEAR1 | -0.394301103 | 0.760857876 | 0.673502433 | 0.81040668 | 6.619311756 | 4.999584838 | ENSG00000171695 |
| MGAT4A | -0.394517383 | 0.760743821 | 0.117918359 | 0.256243272 | 86.55378321 | 65.82663572 | ENSG00000071073 |
| NA | -0.395051278 | 0.760462347 | 0.089817369 | 0.209287795 | 112.8199457 | 85.95422215 | ENSG00000239883 |
| NA | -0.395100048 | 0.76043664 | 0.551291119 | 0.722322422 | 11.77761294 | 8.960696186 | ENSG00000214391 |
| USP16 | -0.395210555 | 0.760378395 | 0.013205055 | 0.046143002 | 359.7998739 | 273.1637633 | ENSG00000156256 |
| TTLL12 | -0.395741809 | 0.760098447 | 3.66E-10 | 7.24E-09 | 6018.360858 | 4574.922859 | ENSG00000100304 |
| CS | -0.395808139 | 0.760063501 | 7.89E-09 | 1.29E-07 | 2907.453758 | 2209.078605 | ENSG00000062485 |
| PDHX | -0.397116944 | 0.759374288 | 6.63E-05 | 0.000483906 | 817.9791893 | 620.710179 | ENSG00000110435 |
| SNX7 | -0.397263113 | 0.759297355 | 0.003225298 | 0.014153686 | 423.7689257 | 321.2602922 | ENSG00000162627 |
| DCTN5 | -0.397285077 | 0.759285795 | 1.73E-07 | 2.21E-06 | 1766.944704 | 1340.976228 | ENSG00000166847 |
| NA | -0.39739176 | 0.75922965 | 0.722450445 | 0.843330877 | 5.028792699 | 3.848373762 | ENSG00000260015 |
| RNF165 | -0.397565875 | 0.759138026 | 0.012583454 | 0.044516461 | 278.3672232 | 211.1922548 | ENSG00000141622 |
| CREBZF | -0.397619921 | 0.759109588 | 0.001899953 | 0.009094454 | 1253.811548 | 952.5095515 | ENSG00000137504 |
| ZNF704 | -0.39765983 | 0.759088589 | 0.065263329 | 0.164056767 | 144.3273609 | 109.4847427 | ENSG00000164684 |
| TMSB15B | -0.397774053 | 0.759028492 | 0.277167506 | 0.468279175 | 48.78432832 | 36.96751551 | ENSG00000158427 |
| AHSP | -0.398098077 | 0.758858036 | 0.639503525 | 0.788803297 | 7.370626335 | 5.622611705 | ENSG00000169877 |
| ZDHHC9 | -0.398649209 | 0.758568196 | 1.86E-07 | 2.36E-06 | 1654.280417 | 1254.646292 | ENSG00000188706 |
| TOMM40 | -0.398811738 | 0.758482743 | 5.89E-08 | 8.19E-07 | 5456.401868 | 4138.508581 | ENSG00000130204 |
| RPP30 | -0.398896614 | 0.758438122 | 9.81E-05 | 0.000685133 | 786.6712831 | 596.354442 | ENSG00000148688 |
| LSM5 | -0.399041931 | 0.758361731 | 0.000302083 | 0.001848399 | 653.9346985 | 496.0126032 | ENSG00000106355 |
| NA | -0.39910102 | 0.758330671 | 0.616096656 | 0.771692601 | 8.554858541 | 6.473813284 | ENSG00000260793 |
| RETREG1 | -0.39910414 | 0.758329031 | 0.335073699 | 0.529337273 | 32.47721054 | 24.5562205 | ENSG00000154153 |
| MARCKS | -0.39917858 | 0.758289904 | 0.024367285 | 0.075852077 | 2879.293201 | 2183.026558 | ENSG00000277443 |
| TIMM17A | -0.399269844 | 0.758241936 | 7.51E-07 | 8.57E-06 | 1445.489993 | 1095.775356 | ENSG00000134375 |
| MACO1 | -0.399304804 | 0.758223563 | 0.005917306 | 0.023758957 | 306.892929 | 232.4952224 | ENSG00000204178 |
| EVA1A | -0.399387632 | 0.758180033 | 0.293840476 | 0.486482833 | 42.10641069 | 31.9492713 | ENSG00000115363 |
| NOL4 | -0.39940238 | 0.758172282 | 0.451998864 | 0.639455292 | 21.5647338 | 16.3006832 | ENSG00000101746 |
| NA | -0.39966034 | 0.75803673 | 0.639334872 | 0.788647755 | 8.875232328 | 6.679856189 | ENSG00000228906 |
| CRTC2 | -0.399827369 | 0.757948973 | 0.000797692 | 0.004331796 | 516.6232177 | 392.0580345 | ENSG00000160741 |
| VEGFC | -0.39982751 | 0.757948899 | 0.360706062 | 0.553925441 | 27.65634548 | 20.96315458 | ENSG00000150630 |
| MAPK15 | -0.399931187 | 0.757894432 | 0.102412692 | 0.23068994 | 189.1274504 | 143.6623886 | ENSG00000181085 |
| ARNTL2 | -0.399938041 | 0.757890831 | 0.006255902 | 0.024934814 | 297.3720107 | 225.3355709 | ENSG00000029153 |
| ACOT8 | -0.400017839 | 0.757848912 | 0.001704615 | 0.008287995 | 376.1996996 | 285.0002374 | ENSG00000101473 |
| PSMB8 | -0.400083529 | 0.757814406 | 0.007186925 | 0.027983712 | 293.3588939 | 222.49098 | ENSG00000204264 |

| | | | | | | | |
|---|---|---|---|---|---|---|---|
| MIR5587 | -0.40008786 | 0.757812131 | 0.678584021 | 0.814350236 | 5.455298802 | 4.12212619 | ENSG00000266124 |
| MAX | -0.40009827 | 0.757806663 | 0.000338744 | 0.002038402 | 551.336806 | 417.8613677 | ENSG00000125952 |
| PPP2R2C | -0.400490019 | 0.757600916 | 0.00615066 | 0.024601795 | 329.8007027 | 249.6521085 | ENSG00000074211 |
| ABAT | -0.400991912 | 0.757337404 | 0.008895444 | 0.033367368 | 278.9202536 | 211.3397232 | ENSG00000183044 |
| TSPAN5 | -0.401072162 | 0.757295278 | 8.77E-05 | 0.000619855 | 665.7961727 | 504.1359706 | ENSG00000168785 |
| NA | -0.401247044 | 0.757203485 | 0.577129023 | 0.74188579 | 10.52009532 | 7.989270947 | ENSG00000281849 |
| TATDN2 | -0.40125294 | 0.75720039 | 0.001113836 | 0.005787452 | 492.9348001 | 373.0784728 | ENSG00000157014 |
| NA | -0.401505717 | 0.757067732 | 0.670308053 | 0.808423037 | 8.331146334 | 6.335256664 | ENSG00000285416 |
| TSEN54 | -0.401617148 | 0.757009259 | 7.34E-06 | 6.83E-05 | 1123.228089 | 850.8448915 | ENSG00000182173 |
| IQCK | -0.401639069 | 0.756997757 | 0.336072911 | 0.530425 | 35.17807503 | 26.72107372 | ENSG00000174628 |
| TARS1 | -0.401671894 | 0.756980534 | 1.99E-11 | 4.81E-10 | 8273.485636 | 6262.060331 | ENSG00000113407 |
| PLA2G7 | -0.401765695 | 0.756931318 | 0.142300208 | 0.294483259 | 76.68024563 | 58.16406937 | ENSG00000146070 |
| PRKAR2B | -0.401823335 | 0.756901077 | 0.000275536 | 0.001707957 | 814.9056665 | 616.2622956 | ENSG00000005249 |
| MYOZ3 | -0.401823875 | 0.756900794 | 0.003892643 | 0.016661949 | 350.0600841 | 265.37473 | ENSG00000164591 |
| STX4 | -0.402171629 | 0.756718369 | 0.000161741 | 0.001064906 | 763.2555022 | 577.5076486 | ENSG00000103496 |
| HDGFL3 | -0.402214438 | 0.756695915 | 5.17E-08 | 7.31E-07 | 2251.833171 | 1703.373397 | ENSG00000166503 |
| NA | -0.402614048 | 0.756486348 | 0.608995491 | 0.7667212 | 9.2625777 | 7.055499316 | ENSG00000273456 |
| ZBTB8OS | -0.402673466 | 0.756455192 | 0.001213929 | 0.00620148 | 433.1577346 | 327.6552629 | ENSG00000176261 |
| BEST2 | -0.402865789 | 0.756354357 | 0.214913289 | 0.395041446 | 55.91230097 | 42.23000589 | ENSG00000039987 |
| NSDHL | -0.40310538 | 0.756228759 | 3.35E-06 | 3.38E-05 | 1281.362919 | 968.9697809 | ENSG00000147383 |
| NAV2 | -0.403272701 | 0.756141058 | 6.01E-06 | 5.70E-05 | 1256.661712 | 950.2454642 | ENSG00000166833 |
| INTS3 | -0.403333318 | 0.756109288 | 3.05E-05 | 0.000244323 | 2002.951776 | 1515.178902 | ENSG00000143624 |
| NA | -0.4034027 | 0.756072926 | 0.652136563 | 0.796943154 | 6.719130247 | 5.029640753 | ENSG00000269934 |
| HDAC8 | -0.403424409 | 0.756061549 | 0.182859437 | 0.353293381 | 62.08163276 | 46.91515874 | ENSG00000147099 |
| ELAVL4 | -0.403462501 | 0.756041587 | 0.643088434 | 0.791014138 | 7.357998683 | 5.581270869 | ENSG00000162374 |
| TTLL4 | -0.403491675 | 0.756026298 | 3.17E-06 | 3.21E-05 | 1966.996823 | 1487.330745 | ENSG00000135912 |
| ANKRD13A | -0.403595501 | 0.755971892 | 0.006924704 | 0.027168178 | 304.4272016 | 229.884317 | ENSG00000076513 |
| LRFN3 | -0.403677195 | 0.755929085 | 0.005748806 | 0.023177805 | 406.9723816 | 307.4623486 | ENSG00000126243 |
| CASP7 | -0.403786867 | 0.755871623 | 0.063815585 | 0.161446635 | 119.9481137 | 90.85235432 | ENSG00000165806 |
| RIOK1 | -0.404053614 | 0.755731879 | 0.00016827 | 0.001102301 | 717.6269871 | 541.7243144 | ENSG00000124784 |
| TDRD6 | -0.404352734 | 0.755575206 | 0.719225399 | 0.841017259 | 4.108711207 | 3.116957809 | ENSG00000180113 |
| E2F1 | -0.404555582 | 0.755468977 | 1.36E-05 | 0.000118349 | 1050.078983 | 793.8259644 | ENSG00000101412 |
| NA | -0.40459059 | 0.755450645 | 0.445135508 | 0.633364921 | 19.07938855 | 14.4029724 | ENSG00000231993 |
| NA | -0.404649541 | 0.755419777 | 0.663863252 | 0.804553817 | 6.705224819 | 5.018355832 | ENSG00000238287 |
| NA | -0.404697287 | 0.755394777 | 0.668794928 | 0.807678712 | 5.991191834 | 4.568616414 | ENSG00000272702 |
| UFC1 | -0.404803586 | 0.755339121 | 0.000786908 | 0.004276995 | 492.9462629 | 371.9949713 | ENSG00000143222 |
| CIAO3 | -0.404954804 | 0.755259953 | 0.004017691 | 0.01708246 | 365.7068138 | 276.5898975 | ENSG00000103245 |
| MAP10 | -0.405010024 | 0.755231045 | 0.325402029 | 0.519510828 | 33.36434743 | 25.1641636 | ENSG00000212916 |
| CDC37 | -0.405067884 | 0.755200757 | 1.96E-08 | 2.98E-07 | 4992.121584 | 3771.268506 | ENSG00000105401 |
| PIGN | -0.405093675 | 0.755187256 | 0.001457498 | 0.007228598 | 416.0431769 | 314.1666299 | ENSG00000197563 |
| BZW1 | -0.405136896 | 0.755164633 | 1.86E-08 | 2.85E-07 | 3399.365591 | 2566.016811 | ENSG00000082153 |
| CXCL6 | -0.405442108 | 0.755004889 | 0.076500145 | 0.185203324 | 105.3318372 | 79.39630147 | ENSG00000124875 |
| NOL8 | -0.405532383 | 0.754957647 | 0.001070476 | 0.005593518 | 556.701682 | 420.2907183 | ENSG00000198000 |
| RELB | -0.405652741 | 0.754894667 | 0.009086545 | 0.033967329 | 277.9280516 | 209.7396085 | ENSG00000104856 |
| EFTUD2 | -0.405709187 | 0.754865132 | 1.02E-09 | 1.90E-08 | 4333.606075 | 3270.837865 | ENSG00000108883 |
| CORO2B | -0.405786821 | 0.754824512 | 0.061512895 | 0.157040749 | 141.0748188 | 106.3493487 | ENSG00000103647 |
| NA | -0.405863196 | 0.754784554 | 0.58281504 | 0.746399526 | 10.62622764 | 8.038097856 | ENSG00000232450 |
| NA | -0.406068091 | 0.754677365 | 0.186684151 | 0.358030221 | 56.55885871 | 42.72543464 | ENSG00000260469 |
| USP44 | -0.406171389 | 0.754623332 | 5.47E-08 | 7.67E-07 | 4024.11848 | 3035.614817 | ENSG00000136014 |
| EPM2AIP1 | -0.406201032 | 0.754607826 | 0.375390596 | 0.568375893 | 224.6501624 | 169.6540133 | ENSG00000178567 |
| NUP62 | -0.40632455 | 0.754543223 | 7.47E-08 | 1.02E-06 | 2277.667822 | 1718.735426 | ENSG00000213024 |
| RCOR2 | -0.406486071 | 0.75445875 | 1.47E-06 | 1.58E-05 | 2653.545991 | 2001.279219 | ENSG00000167771 |
| PSIP1 | -0.406540848 | 0.754430105 | 4.00E-09 | 6.81E-08 | 7825.574245 | 5903.041069 | ENSG00000164985 |
| KIAA1324 | -0.406703088 | 0.754345269 | 0.278088521 | 0.469193687 | 39.85858539 | 29.9496698 | ENSG00000116299 |
| NA | -0.407035124 | 0.754171677 | 0.67326806 | 0.810177265 | 5.455298802 | 4.092070275 | ENSG00000286162 |
| CNPPD1 | -0.407080864 | 0.754147767 | 0.000184095 | 0.00119479 | 690.6845995 | 520.9944877 | ENSG00000115649 |
| TMEM135 | -0.407182842 | 0.754094461 | 0.002358521 | 0.01091729 | 543.2820094 | 409.1053516 | ENSG00000166575 |
| ARL10 | -0.407257686 | 0.754055341 | 0.011235702 | 0.040517385 | 268.4615151 | 202.649307 | ENSG00000175414 |
| RSRP1 | -0.407568553 | 0.753892878 | 0.341429328 | 0.535821437 | 673.7780823 | 508.0864131 | ENSG00000117616 |
| HTR1D | -0.407662129 | 0.75384398 | 0.35469131 | 0.548049082 | 27.69362708 | 20.8204727 | ENSG00000179546 |
| ATP6V1D | -0.407935595 | 0.753701101 | 1.46E-05 | 0.000125935 | 837.0830963 | 631.0382434 | ENSG00000100554 |
| JPH4 | -0.408068337 | 0.753631756 | 2.58E-07 | 3.21E-06 | 3168.437085 | 2388.536189 | ENSG00000092051 |
| EFHC1 | -0.408230555 | 0.753547022 | 0.017888357 | 0.058763346 | 218.4315227 | 164.6344062 | ENSG00000096093 |
| NAA10 | -0.408390532 | 0.753463468 | 0.00396202 | 0.016900279 | 370.0776087 | 279.3838379 | ENSG00000102030 |
| LINC00641 | -0.408511056 | 0.753400525 | 0.055730361 | 0.145254747 | 130.2405658 | 98.15088888 | ENSG00000258441 |
| NA | -0.408672436 | 0.753316254 | 0.523412082 | 0.699924224 | 18.35135244 | 13.91612903 | ENSG00000230805 |
| ZNF616 | -0.40884842 | 0.753224368 | 0.000222013 | 0.001410625 | 546.1192225 | 411.5414096 | ENSG00000204611 |
| NCBP1 | -0.408865107 | 0.753215656 | 9.73E-08 | 1.30E-06 | 1827.646196 | 1376.410503 | ENSG00000136937 |
| OVOS2 | -0.408893112 | 0.753201035 | 0.148080416 | 0.303161651 | 76.264894 | 57.45891049 | ENSG00000177359 |
| RNLS | -0.408920159 | 0.753186915 | 0.05176518 | 0.137216225 | 135.0319363 | 101.7928933 | ENSG00000184719 |
| TMEM250 | -0.409220602 | 0.753030079 | 1.27E-06 | 1.39E-05 | 1387.076359 | 1044.426244 | ENSG00000238227 |
| NA | -0.409525475 | 0.752870964 | 0.651153149 | 0.796243605 | 6.897367898 | 5.236233311 | ENSG00000272619 |
| SLC44A1 | -0.409636138 | 0.752813216 | 6.92E-06 | 6.48E-05 | 1910.944761 | 1437.599418 | ENSG00000070214 |
| EFL1 | -0.409637095 | 0.752812717 | 1.28E-05 | 0.000112128 | 1051.023771 | 790.9505674 | ENSG00000140598 |

| | | | | | | | |
|---|---|---|---|---|---|---|---|
| ZBTB2 | -0.409678549 | 0.752791087 | 0.002061216 | 0.009750531 | 453.9849917 | 342.0546597 | ENSG00000181472 |
| NA | -0.410008803 | 0.752618781 | 0.645417334 | 0.79287896 | 7.553298675 | 5.637145819 | ENSG00000228395 |
| NA | -0.410080984 | 0.752581127 | 0.493740722 | 0.675874798 | 15.95645264 | 11.98000018 | ENSG00000278864 |
| LZTS1 | -0.410311327 | 0.752460978 | 0.001578977 | 0.007766657 | 465.2134599 | 349.7782853 | ENSG00000061337 |
| NA | -0.410354404 | 0.752438512 | 0.617321754 | 0.772320958 | 9.326392374 | 7.048013242 | ENSG00000267108 |
| ZER1 | -0.410428122 | 0.752400065 | 0.00018674 | 0.00120931 | 661.0263219 | 496.8493335 | ENSG00000160445 |
| PRICKLE1 | -0.41043228 | 0.752397896 | 0.008766232 | 0.033016325 | 254.4182026 | 191.6100233 | ENSG00000139174 |
| ILF3-DT | -0.410751317 | 0.75223153 | 0.006901456 | 0.027094173 | 381.2871733 | 287.2320759 | ENSG00000267100 |
| TOR2A | -0.410892582 | 0.752157876 | 0.00237794 | 0.010996192 | 351.2881072 | 264.3978654 | ENSG00000160404 |
| TTLL3 | -0.410948799 | 0.752128568 | 0.306069794 | 0.500105577 | 36.26417248 | 27.23841104 | ENSG00000214021 |
| NA | -0.411065712 | 0.75206762 | 0.365269751 | 0.558431292 | 29.08381009 | 21.93121901 | ENSG00000215859 |
| C4orf48 | -0.411708915 | 0.751732397 | 0.040679702 | 0.113959735 | 222.4000368 | 167.0284932 | ENSG00000243449 |
| SNORD3A | -0.411767628 | 0.751701804 | 0.503248338 | 0.683612189 | 16.75018257 | 12.56124817 | ENSG00000263934 |
| NSUN2 | -0.411998763 | 0.751581384 | 3.50E-10 | 6.98E-09 | 3571.886618 | 2685.216025 | ENSG00000037474 |
| ANP32A | -0.412083862 | 0.751537052 | 1.71E-12 | 4.88E-11 | 7353.269892 | 5525.632451 | ENSG00000140350 |
| CDCP1 | -0.412359376 | 0.751393543 | 0.000618624 | 0.003456611 | 435.949939 | 327.5752808 | ENSG00000163814 |
| ESPN | -0.412473612 | 0.751334049 | 0.001761274 | 0.008518647 | 493.6623552 | 370.6094933 | ENSG00000187017 |
| NA | -0.412938469 | 0.751091997 | 0.709595365 | 0.834918772 | 4.105554294 | 3.086901894 | ENSG00000281501 |
| PBX1 | -0.412983887 | 0.751068352 | 2.03E-10 | 4.22E-09 | 3240.937823 | 2433.3921 | ENSG00000185630 |
| C1S | -0.413087831 | 0.75101424 | 0.019044735 | 0.061892883 | 184.5434062 | 138.505539 | ENSG00000182326 |
| TUG1 | -0.413121364 | 0.750996785 | 2.30E-06 | 2.39E-05 | 1809.708729 | 1359.912698 | ENSG00000253352 |
| DBNL | -0.413360782 | 0.750872165 | 5.23E-06 | 5.03E-05 | 1300.429822 | 976.96395 | ENSG00000136279 |
| CABP1 | -0.413435753 | 0.750833147 | 0.127192703 | 0.270839792 | 96.15587157 | 72.34993719 | ENSG00000157782 |
| LOC101928674 | -0.413464843 | 0.750818008 | 0.256400046 | 0.445149075 | 46.57446102 | 34.9577284 | ENSG00000266970 |
| NA | -0.413579923 | 0.750758119 | 0.490013423 | 0.672761479 | 26.63140945 | 20.09384328 | ENSG00000273253 |
| NA | -0.413965681 | 0.750557403 | 0.481902233 | 0.665443003 | 15.47244569 | 11.60859393 | ENSG00000287016 |
| TRO | -0.414038252 | 0.750519649 | 1.04E-05 | 9.27E-05 | 1165.99001 | 875.759075 | ENSG00000067445 |
| NA | -0.414048333 | 0.750514405 | 0.052632767 | 0.139078373 | 125.4241354 | 94.23166802 | ENSG00000248124 |
| OAF | -0.414104303 | 0.750485289 | 0.264831721 | 0.454522744 | 46.47158332 | 34.92029803 | ENSG00000184232 |
| C6orf132 | -0.414216518 | 0.750426917 | 2.69E-07 | 3.33E-06 | 1625.048081 | 1219.588219 | ENSG00000188112 |
| ZNF230 | -0.414292101 | 0.750387603 | 0.215189111 | 0.395430794 | 55.23682736 | 41.47656975 | ENSG00000159882 |
| HDAC11 | -0.414495491 | 0.750281822 | 0.005680232 | 0.022961268 | 348.4123499 | 261.390396 | ENSG00000163517 |
| CAMSAP3 | -0.414860267 | 0.750092142 | 1.04E-05 | 9.20E-05 | 1247.686075 | 936.2673135 | ENSG00000076826 |
| MRPL21 | -0.415009817 | 0.750014391 | 6.95E-06 | 6.52E-05 | 1047.316311 | 785.7522512 | ENSG00000197345 |
| NA | -0.415302251 | 0.749862379 | 0.463497163 | 0.649994428 | 18.6655101 | 13.98273925 | ENSG00000228218 |
| TAS2R14 | -0.415347419 | 0.749838902 | 0.656505768 | 0.799348173 | 5.991191834 | 4.519789505 | ENSG00000212127 |
| NA | -0.415381471 | 0.749821204 | 0.625996568 | 0.778463526 | 7.942523184 | 6.008990099 | ENSG00000275029 |
| RAP2A | -0.415644027 | 0.749684757 | 2.51E-05 | 0.00020435 | 844.6796821 | 633.0879541 | ENSG00000125249 |
| PTPN2 | -0.415653388 | 0.749679892 | 0.000108192 | 0.000746829 | 716.109483 | 536.3096178 | ENSG00000175354 |
| NA | -0.415699766 | 0.749655793 | 0.33091689 | 0.525238815 | 29.61842535 | 22.12185173 | ENSG00000128692 |
| DHX9 | -0.415909274 | 0.749546936 | 1.96E-13 | 6.30E-12 | 7291.221881 | 5464.445613 | ENSG00000135829 |
| ARHGEF6 | -0.416109385 | 0.749442976 | 0.001780181 | 0.008598873 | 384.0259207 | 287.7691317 | ENSG00000129675 |
| IMPDH1 | -0.416156408 | 0.749418549 | 1.55E-06 | 1.66E-05 | 1394.188359 | 1044.785154 | ENSG00000106348 |
| NABP2 | -0.416211941 | 0.749389703 | 1.02E-06 | 1.14E-05 | 1104.64063 | 828.1924337 | ENSG00000139579 |
| NA | -0.416396165 | 0.749294016 | 0.491956901 | 0.674552726 | 14.26483732 | 10.71985036 | ENSG00000282572 |
| MCM8 | -0.416539991 | 0.749219321 | 0.011187394 | 0.040410671 | 229.0058491 | 171.5909376 | ENSG00000125885 |
| ND1 | -0.416640264 | 0.749167249 | 0.060670583 | 0.155291892 | 17427.72217 | 13056.25071 | ENSG00000198888 |
| IRX6 | -0.416683428 | 0.749144835 | 0.564697114 | 0.73291991 | 10.50303298 | 7.846589066 | ENSG00000159387 |
| GPR75 | -0.416910378 | 0.749026996 | 0.485644049 | 0.66875218 | 16.58960862 | 12.43397647 | ENSG00000119737 |
| PCDHA3 | -0.416933368 | 0.74901506 | 0.401464907 | 0.593077521 | 27.68287856 | 20.76470937 | ENSG00000255408 |
| C11orf68 | -0.417029045 | 0.748965389 | 9.35E-05 | 0.000656021 | 676.0072214 | 506.3953401 | ENSG00000175573 |
| FSTL4 | -0.417088027 | 0.748934769 | 0.093044588 | 0.214622079 | 93.17121583 | 69.69702967 | ENSG00000053108 |
| TFB2M | -0.417104585 | 0.748926173 | 0.000329527 | 0.001990422 | 503.4871637 | 376.9383547 | ENSG00000162851 |
| SPATA3-AS1 | -0.417279639 | 0.748835306 | 0.610927745 | 0.767968993 | 10.65148294 | 7.966812725 | ENSG00000238062 |
| NA | -0.41734666 | 0.748800519 | 0.086850802 | 0.203835984 | 93.32933193 | 69.8168153 | ENSG00000272645 |
| NA | -0.417401526 | 0.748772043 | 0.212685117 | 0.392423238 | 56.26679944 | 41.99358066 | ENSG00000255857 |
| SLC39A3 | -0.417532338 | 0.748704153 | 0.002392906 | 0.011046105 | 460.4202178 | 344.9527524 | ENSG00000141873 |
| SAP30L | -0.417561022 | 0.748689267 | 0.001842257 | 0.008841115 | 525.4718193 | 392.7710369 | ENSG00000164576 |
| NA | -0.417615994 | 0.74866074 | 0.535520715 | 0.709916381 | 12.62440902 | 9.399150683 | ENSG00000265750 |
| ACSS2 | -0.417637255 | 0.748649707 | 2.84E-06 | 2.89E-05 | 917.432243 | 686.9438279 | ENSG00000131069 |
| TMEM269 | -0.417660162 | 0.74863782 | 0.657823756 | 0.80032201 | 6.394321771 | 4.767173245 | ENSG00000274386 |
| THEM5 | -0.41767512 | 0.748630058 | 0.20860154 | 0.38769228 | 48.30731161 | 36.2065933 | ENSG00000196407 |
| EIF1AX | -0.417801392 | 0.748564537 | 4.92E-05 | 0.000371778 | 2873.318943 | 2150.028285 | ENSG00000173674 |
| TFE3 | -0.417833624 | 0.748547813 | 4.81E-10 | 9.36E-09 | 2926.568338 | 2191.317744 | ENSG00000068323 |
| TPGS2 | -0.4179248 | 0.748500508 | 2.23E-08 | 3.36E-07 | 2329.192288 | 1742.874407 | ENSG00000134779 |
| TMEM123 | -0.41796173 | 0.748481348 | 3.35E-05 | 0.000264877 | 1847.598424 | 1382.23496 | ENSG00000152558 |
| LTV1 | -0.418092279 | 0.748413621 | 0.006375679 | 0.025346812 | 273.4793487 | 204.6049395 | ENSG00000135521 |
| SNW1 | -0.418226544 | 0.748343973 | 1.67E-06 | 1.78E-05 | 1066.785721 | 798.0284217 | ENSG00000100603 |
| PDE3A | -0.41857456 | 0.748163475 | 0.022098028 | 0.070167013 | 188.1989806 | 140.4772345 | ENSG00000172572 |
| OXTR | -0.418669745 | 0.748114115 | 0.180812519 | 0.350371406 | 56.70287398 | 42.29347853 | ENSG00000180914 |
| LOC102724064 | -0.418701494 | 0.748097651 | 0.283419376 | 0.474492229 | 45.47445814 | 34.14026997 | ENSG00000255468 |
| NA | -0.418852364 | 0.748019423 | 0.627019284 | 0.77931702 | 8.501792383 | 6.312248789 | ENSG00000272604 |
| TMEM147 | -0.418891472 | 0.747999146 | 5.94E-07 | 6.90E-06 | 1230.73261 | 921.043187 | ENSG00000105677 |
| NA | -0.418975234 | 0.747955719 | 0.59553596 | 0.755953244 | 9.541911619 | 7.100089345 | ENSG00000275202 |

| | | | | | | | |
|---|---|---|---|---|---|---|---|
| IGSF11 | -0.41900514 | 0.747940214 | 0.414764949 | 0.606313267 | 22.65220673 | 16.81802052 | ENSG00000144847 |
| KBTBD4 | -0.41921467 | 0.747831595 | 0.233246491 | 0.418165066 | 47.64012866 | 35.66993533 | ENSG00000123444 |
| CDK20 | -0.419301649 | 0.74778651 | 0.096706227 | 0.221032583 | 88.19243008 | 66.05948535 | ENSG00000156345 |
| NA | -0.419361686 | 0.747755392 | 0.545887562 | 0.717959131 | 11.19181066 | 8.409504103 | ENSG00000213885 |
| CCDC86 | -0.419484164 | 0.747691914 | 2.55E-07 | 3.17E-06 | 1461.553464 | 1093.010595 | ENSG00000110104 |
| IDO1 | -0.419501594 | 0.747682881 | 0.002012231 | 0.009548052 | 384.7019805 | 288.051023 | ENSG00000131203 |
| RFX2 | -0.419728224 | 0.747565438 | 0.053785884 | 0.141399657 | 115.0178617 | 86.03580451 | ENSG00000087903 |
| COX6B2 | -0.419971232 | 0.747439528 | 0.388014513 | 0.581174299 | 26.78489546 | 20.03686903 | ENSG00000160471 |
| DBNDD1 | -0.420270338 | 0.747284582 | 7.10E-05 | 0.000512896 | 1299.96293 | 971.1523453 | ENSG00000003249 |
| GGPS1 | -0.420285623 | 0.747276665 | 0.025200022 | 0.077935162 | 176.5511692 | 131.9567534 | ENSG00000152904 |
| NA | -0.420290135 | 0.747274327 | 0.739671952 | 0.854617835 | 4.088491954 | 3.049359905 | ENSG00000276434 |
| TMIE | -0.420436923 | 0.747198299 | 0.555935918 | 0.725587033 | 10.90744069 | 8.075201812 | ENSG00000181585 |
| SOX9 | -0.420444596 | 0.747194326 | 0.078769633 | 0.189427323 | 115.5417285 | 86.13951869 | ENSG00000125398 |
| MIGA2 | -0.420571632 | 0.747128535 | 0.001467819 | 0.007271959 | 698.9849961 | 522.8750136 | ENSG00000148343 |
| RAB40B | -0.420702491 | 0.74706077 | 0.28084707 | 0.472214424 | 39.80226462 | 29.71769625 | ENSG00000141542 |
| USP41 | -0.420979662 | 0.746917258 | 0.575529551 | 0.740696842 | 9.749839263 | 7.344223892 | ENSG00000161133 |
| TMEM237 | -0.421139258 | 0.746834636 | 0.000132827 | 0.00089423 | 623.9672394 | 465.8618431 | ENSG00000155755 |
| NA | -0.421143401 | 0.746832491 | 0.624802333 | 0.777343522 | 7.450225573 | 5.555013801 | ENSG00000226478 |
| ZDHHC14 | -0.421479607 | 0.74665847 | 0.012015203 | 0.042833192 | 201.9729935 | 150.9024115 | ENSG00000175048 |
| FAM210B | -0.421482196 | 0.74665713 | 0.000528495 | 0.003012955 | 450.57274 | 336.1719537 | ENSG00000124098 |
| SUPT16H | -0.421522774 | 0.746636129 | 1.61E-14 | 5.83E-13 | 9390.23649 | 7010.477755 | ENSG00000092201 |
| PALM3 | -0.421632633 | 0.746579276 | 2.28E-07 | 2.86E-06 | 1657.560882 | 1237.70343 | ENSG00000187867 |
| CCM2L | -0.421689117 | 0.746550047 | 0.069815777 | 0.172574767 | 121.809195 | 90.88835897 | ENSG00000101331 |
| DFFB | -0.422078262 | 0.746348703 | 0.008325186 | 0.031664066 | 224.6239224 | 167.5738397 | ENSG00000169598 |
| TSPY26P | -0.422187186 | 0.746292356 | 0.00768647 | 0.029636164 | 250.2739912 | 187.1060821 | ENSG00000235217 |
| THAP11 | -0.422261637 | 0.746253844 | 1.76E-05 | 0.000149014 | 911.5658015 | 680.5518202 | ENSG00000168286 |
| NAE1 | -0.422329293 | 0.746218849 | 2.46E-08 | 3.67E-07 | 2081.682645 | 1552.928538 | ENSG00000159593 |
| PDAP1 | -0.422399688 | 0.746182439 | 2.12E-11 | 5.11E-10 | 6220.057885 | 4641.495683 | ENSG00000106244 |
| RBBP4 | -0.422495341 | 0.746132967 | 1.19E-08 | 1.88E-07 | 3869.134398 | 2886.592071 | ENSG00000162521 |
| NA | -0.42264651 | 0.74605479 | 0.711674393 | 0.836409207 | 4.783583461 | 3.619761016 | ENSG00000260209 |
| ORC2 | -0.422668478 | 0.74604343 | 4.91E-05 | 0.000371203 | 1118.36965 | 833.5166756 | ENSG00000115942 |
| DDX52 | -0.42269591 | 0.746029244 | 0.000434486 | 0.002530724 | 883.0859086 | 658.4012577 | ENSG00000278053 |
| ZNF563 | -0.422788211 | 0.745981516 | 0.26762567 | 0.457791801 | 41.58049206 | 30.84589945 | ENSG00000188868 |
| ZNF488 | -0.422966716 | 0.745889221 | 0.009230271 | 0.034414372 | 237.2558253 | 176.8460536 | ENSG00000265763 |
| AKR1B1 | -0.42304564 | 0.745848418 | 4.78E-11 | 1.09E-09 | 4100.423828 | 3058.222423 | ENSG00000085662 |
| NCKAP5 | -0.423511999 | 0.745607357 | 0.426068099 | 0.616740562 | 21.04966138 | 15.77998506 | ENSG00000176771 |
| ARRDC3 | -0.423723323 | 0.74549815 | 0.125416741 | 0.268357419 | 77.83666698 | 57.87012023 | ENSG00000113369 |
| BRIX1 | -0.423780448 | 0.745468632 | 0.003304003 | 0.014445954 | 563.1390577 | 419.3710109 | ENSG00000113460 |
| DNAJC3-DT | -0.423878935 | 0.745417743 | 0.20453036 | 0.382541509 | 52.2922268 | 39.08734067 | ENSG00000247400 |
| YPEL3 | -0.4239752 | 0.745368006 | 0.00818661 | 0.031220293 | 272.7950057 | 203.1641278 | ENSG00000090238 |
| EXD2 | -0.424000028 | 0.745355179 | 0.000188543 | 0.001219281 | 651.5407235 | 484.9255347 | ENSG00000081177 |
| LYPD6B | -0.42410468 | 0.745301113 | 0.173912711 | 0.340621175 | 57.0150548 | 42.4061045 | ENSG00000150556 |
| KCNA1 | -0.424301489 | 0.745199448 | 0.693398675 | 0.82436945 | 5.105235024 | 3.810831773 | ENSG00000111262 |
| LOC105374618 | -0.424317502 | 0.745191177 | 0.497437656 | 0.678503791 | 15.80800268 | 11.83775633 | ENSG00000248994 |
| MIR4258 | -0.424320372 | 0.745189695 | 0.69982286 | 0.828998907 | 4.70082731 | 3.552163113 | ENSG00000264349 |
| PSMG1 | -0.424462746 | 0.745116158 | 9.19E-06 | 8.27E-05 | 930.9716748 | 693.5402379 | ENSG00000183527 |
| KIAA0753 | -0.424586133 | 0.745052434 | 0.000153128 | 0.001012149 | 520.8997037 | 387.8804311 | ENSG00000198920 |
| ZBTB46 | -0.424601897 | 0.745044294 | 0.000909649 | 0.004861428 | 513.0276173 | 381.6218586 | ENSG00000130584 |
| NA | -0.424808584 | 0.744937563 | 0.686762315 | 0.819878795 | 5.366228826 | 3.953075621 | ENSG00000280374 |
| CCDC142 | -0.424860993 | 0.744910502 | 0.043459838 | 0.120096115 | 132.4965841 | 98.74353342 | ENSG00000135637 |
| TFPT | -0.424873358 | 0.744904118 | 0.003915165 | 0.016742887 | 320.4710405 | 239.1636191 | ENSG00000105619 |
| RAB27A | -0.424922086 | 0.744878958 | 0.034150391 | 0.099325074 | 136.1408085 | 101.3989172 | ENSG00000069974 |
| USP3 | -0.425005116 | 0.744836091 | 0.007200935 | 0.028016631 | 240.5119303 | 178.9763907 | ENSG00000140455 |
| NA | -0.425203126 | 0.744733869 | 0.593104564 | 0.754727085 | 8.234484755 | 6.113691958 | ENSG00000234692 |
| CENPX | -0.425216754 | 0.744726833 | 3.05E-08 | 4.51E-07 | 1707.128981 | 1271.73629 | ENSG00000169689 |
| JAKMIP2-AS1 | -0.425227777 | 0.744721143 | 0.011376569 | 0.040942395 | 246.6030383 | 183.4766364 | ENSG00000280780 |
| NCKAP5-AS2 | -0.425531383 | 0.744564438 | 0.584150212 | 0.747231695 | 9.720051573 | 7.197743164 | ENSG00000226953 |
| SDSL | -0.425604278 | 0.744526818 | 0.543726849 | 0.716133699 | 11.19181066 | 8.35319112 | ENSG00000139410 |
| MRPS34 | -0.425751569 | 0.74445081 | 5.78E-08 | 8.07E-07 | 1946.383243 | 1449.35334 | ENSG00000074071 |
| RNF8 | -0.425751982 | 0.744450597 | 2.95E-06 | 3.00E-05 | 944.4450747 | 702.9306916 | ENSG00000112130 |
| SLC25A48 | -0.425886904 | 0.744380979 | 0.254679289 | 0.443475905 | 41.48954295 | 30.97459687 | ENSG00000145832 |
| EFNA3 | -0.426098131 | 0.744272001 | 0.16151886 | 0.322307338 | 66.08045337 | 48.98461964 | ENSG00000143590 |
| SPATA6L | -0.426150927 | 0.744244765 | 0.568727979 | 0.735832765 | 9.226573883 | 6.860191679 | ENSG00000106686 |
| GOLM1 | -0.426186651 | 0.744226336 | 2.52E-10 | 5.17E-09 | 3904.880062 | 2905.339746 | ENSG00000135052 |
| COQ10A | -0.426215251 | 0.744211583 | 0.014387604 | 0.049381453 | 202.6767816 | 150.8422997 | ENSG00000135469 |
| LOC100996842 | -0.426516337 | 0.744056284 | 0.046113661 | 0.125594502 | 125.5253294 | 93.52662075 | ENSG00000233223 |
| TMEM107 | -0.426700531 | 0.743961294 | 0.153812444 | 0.311354286 | 60.18789995 | 44.79588324 | ENSG00000179029 |
| NA | -0.426920778 | 0.743847726 | 0.064717731 | 0.163094745 | 104.6789656 | 77.84417789 | ENSG00000235552 |
| SELENOP | -0.426957835 | 0.74382862 | 0.296620457 | 0.489542614 | 36.2706817 | 26.97919272 | ENSG00000250722 |
| TMEM99 | -0.426996409 | 0.743808732 | 0.116778689 | 0.254602543 | 119.6292283 | 88.64451131 | ENSG00000167920 |
| DOK4 | -0.427052932 | 0.743779592 | 9.05E-06 | 8.16E-05 | 827.1042458 | 615.355809 | ENSG00000125170 |
| ELOVL6 | -0.427101252 | 0.743754681 | 9.30E-08 | 1.25E-06 | 2356.915973 | 1752.439608 | ENSG00000170522 |
| RAB11FIP3 | -0.427175915 | 0.743716191 | 3.24E-07 | 3.95E-06 | 1359.093078 | 1011.394164 | ENSG00000090565 |
| LRIG1 | -0.427411481 | 0.743594765 | 1.83E-10 | 3.84E-09 | 2825.330044 | 2100.623625 | ENSG00000144749 |

| | | | | | | | |
|---|---|---|---|---|---|---|---|
| SLC27A3 | -0.427434665 | 0.743582816 | 0.000248743 | 0.001562777 | 602.6246842 | 448.5554905 | ENSG00000143554 |
| DHX57 | -0.427487779 | 0.74355544 | 4.95E-06 | 4.78E-05 | 835.4384513 | 621.3794364 | ENSG00000163214 |
| SLCO4A1-AS1 | -0.427597148 | 0.743499075 | 0.667780674 | 0.80717496 | 6.523927953 | 4.898681826 | ENSG00000232803 |
| COA3 | -0.427614947 | 0.743489902 | 5.28E-06 | 5.07E-05 | 1017.816533 | 757.1486621 | ENSG00000183978 |
| PNPLA3 | -0.427792615 | 0.743398347 | 0.000940605 | 0.004999474 | 567.0739963 | 421.7527144 | ENSG00000100344 |
| TMEM52 | -0.428200631 | 0.743188132 | 0.060713481 | 0.155380235 | 175.5816821 | 130.263883 | ENSG00000178821 |
| SCGB3A2 | -0.428316889 | 0.743128246 | 1.40E-06 | 1.51E-05 | 1035.704601 | 769.7192198 | ENSG00000164265 |
| C2orf50 | -0.428367618 | 0.743102116 | 0.559719673 | 0.728853793 | 10.13590686 | 7.527808575 | ENSG00000150873 |
| LOC107984813 | -0.428427499 | 0.743071273 | 0.674787589 | 0.811742271 | 5.094486509 | 3.784574705 | ENSG00000259804 |
| CNRIP1 | -0.428587414 | 0.742988912 | 0.220539106 | 0.402037429 | 55.86494728 | 41.54807812 | ENSG00000119865 |
| RPS26 | -0.428812365 | 0.742873071 | 4.61E-11 | 1.06E-09 | 3378.623495 | 2510.192813 | ENSG00000197728 |
| ACAD9 | -0.429598233 | 0.742468522 | 0.000529083 | 0.003015382 | 582.8501442 | 433.0306933 | ENSG00000177646 |
| NA | -0.429752834 | 0.742388962 | 0.644237077 | 0.791952125 | 6.923900978 | 5.15355164 | ENSG00000278867 |
| NA | -0.429777622 | 0.742376207 | 0.595326241 | 0.755953244 | 8.971893906 | 6.699176837 | ENSG00000260293 |
| DGKG | -0.429941042 | 0.74229212 | 0.5283721 | 0.704217496 | 11.52924679 | 8.514205962 | ENSG00000058866 |
| CANX | -0.429949088 | 0.74228798 | 2.81E-14 | 9.97E-13 | 11242.38948 | 8344.395584 | ENSG00000127022 |
| DCAF5 | -0.430339216 | 0.74208728 | 8.13E-06 | 7.43E-05 | 817.3631934 | 606.5267246 | ENSG00000139990 |
| HYAL1 | -0.430389516 | 0.742061408 | 0.123121741 | 0.264603241 | 70.88495515 | 52.61610361 | ENSG00000114378 |
| NA | -0.430463442 | 0.742023384 | 0.69879751 | 0.828207766 | 4.138401199 | 3.056845979 | ENSG00000240695 |
| HCK | -0.43054011 | 0.741983953 | 0.438783874 | 0.627185333 | 17.82818476 | 13.30438708 | ENSG00000101336 |
| NA | -0.430825749 | 0.741837062 | 0.711404652 | 0.836304315 | 4.638290413 | 3.473280288 | ENSG00000287574 |
| PLAUR | -0.430997672 | 0.741748664 | 0.002530439 | 0.011577322 | 326.6115919 | 242.2755756 | ENSG00000011422 |
| KHDC4 | -0.431122549 | 0.741684462 | 0.111301675 | 0.245608435 | 1212.01701 | 899.2004818 | ENSG00000132680 |
| DLST | -0.431127871 | 0.741681726 | 1.86E-09 | 3.33E-08 | 2769.468668 | 2053.879905 | ENSG00000119689 |
| KLHL13 | -0.431205373 | 0.741641884 | 0.000603736 | 0.003382603 | 583.2167667 | 431.8240848 | ENSG00000003096 |
| PDE4D | -0.431301703 | 0.741592365 | 0.365480596 | 0.558573214 | 24.76283425 | 18.28096405 | ENSG00000113448 |
| TIMM10B | -0.431308756 | 0.74158874 | 0.001040965 | 0.005454694 | 395.4023805 | 292.8940044 | ENSG00000132286 |
| MSL3P1 | -0.431310826 | 0.741587676 | 0.666471159 | 0.806343525 | 7.073628715 | 5.258803152 | ENSG00000224287 |
| MARVELD1 | -0.431327471 | 0.74157912 | 4.21E-06 | 4.17E-05 | 1286.544274 | 954.3073124 | ENSG00000155254 |
| SLC35A4 | -0.431376525 | 0.741553906 | 5.94E-10 | 1.14E-08 | 2188.715395 | 1623.530351 | ENSG00000176087 |
| DRG2 | -0.431416346 | 0.741533438 | 4.11E-05 | 0.000316944 | 632.6241789 | 469.2398829 | ENSG00000108591 |
| PJA1 | -0.431513801 | 0.741483348 | 1.35E-06 | 1.47E-05 | 1737.296356 | 1288.986728 | ENSG00000181191 |
| ITGA6 | -0.431514829 | 0.74148282 | 4.02E-09 | 6.84E-08 | 2343.090172 | 1736.66311 | ENSG00000091409 |
| FAM83H | -0.431576814 | 0.741450963 | 1.55E-08 | 2.41E-07 | 2185.116345 | 1620.541828 | ENSG00000180921 |
| CENPI | -0.431691409 | 0.741392071 | 0.002955665 | 0.01315569 | 327.3225431 | 242.3261144 | ENSG00000102384 |
| KSR2 | -0.431903992 | 0.741282834 | 0.100916733 | 0.228040811 | 81.72660433 | 60.53777666 | ENSG00000171435 |
| ATP8B1 | -0.431961199 | 0.741253441 | 0.171228941 | 0.336816152 | 60.24472438 | 44.6117489 | ENSG00000081923 |
| USH1G | -0.432068683 | 0.741198218 | 0.14501918 | 0.298543277 | 68.54302381 | 50.87258286 | ENSG00000182040 |
| P2RX2 | -0.432074699 | 0.741195127 | 0.23602851 | 0.42103488 | 47.14141952 | 35.01035416 | ENSG00000187848 |
| BCDIN3D | -0.432130947 | 0.741166229 | 0.411158987 | 0.602690656 | 22.91961205 | 17.11521885 | ENSG00000186666 |
| NA | -0.432194199 | 0.741133735 | 0.616486036 | 0.771897362 | 7.856610121 | 5.836252303 | ENSG00000286311 |
| NA | -0.432319301 | 0.741069471 | 0.539629468 | 0.712813952 | 12.15864449 | 9.020808016 | ENSG00000286753 |
| PSMD14 | -0.432428837 | 0.741013208 | 2.53E-08 | 3.77E-07 | 2317.773833 | 1716.755846 | ENSG00000115233 |
| ADAMTSL3 | -0.432520706 | 0.740966023 | 0.366408866 | 0.559481692 | 28.04053394 | 20.69265135 | ENSG00000156218 |
| CSTF2T | -0.432530079 | 0.740961209 | 4.58E-07 | 5.44E-06 | 1280.734686 | 949.5443761 | ENSG00000177613 |
| CDC25A | -0.432878077 | 0.7407825 | 1.53E-08 | 2.38E-07 | 2057.399481 | 1523.858637 | ENSG00000164045 |
| MPC1 | -0.432885787 | 0.740778541 | 0.011202656 | 0.040450033 | 209.9888997 | 155.6919314 | ENSG00000060762 |
| POM121L9P | -0.432914273 | 0.740763915 | 0.637029575 | 0.786694173 | 6.017724914 | 4.459677675 | ENSG00000128262 |
| NA | -0.43293042 | 0.740755624 | 0.675737773 | 0.812436293 | 5.04142035 | 3.724462875 | ENSG00000248375 |
| SLC49A4 | -0.432932903 | 0.740754349 | 0.014772159 | 0.050430501 | 278.4978966 | 205.8212234 | ENSG00000138463 |
| LTB4R2 | -0.433222488 | 0.740605676 | 0.420321628 | 0.611558221 | 18.53906083 | 13.69032744 | ENSG00000213906 |
| NA | -0.433348649 | 0.740540914 | 0.565335271 | 0.733122144 | 10.53272297 | 7.7451364 | ENSG00000261087 |
| NBL1 | -0.43343358 | 0.74049732 | 0.035444898 | 0.102230895 | 158.1832355 | 116.9340033 | ENSG00000158747 |
| NA | -0.433455737 | 0.740485948 | 0.630783181 | 0.782107003 | 6.923900978 | 5.134780645 | ENSG00000223343 |
| YTHDC2 | -0.433586566 | 0.740418801 | 7.55E-06 | 7.00E-05 | 747.1963976 | 553.1070438 | ENSG00000047188 |
| ZFP37 | -0.433960379 | 0.740226978 | 0.160730346 | 0.32126819 | 82.04606849 | 60.91710701 | ENSG00000136866 |
| LINC01224 | -0.434047145 | 0.740182461 | 0.000120297 | 0.000822724 | 520.8342053 | 385.7744159 | ENSG00000269416 |
| KLHDC3 | -0.434268946 | 0.740068673 | 3.49E-11 | 8.14E-10 | 5095.938488 | 3770.743201 | ENSG00000124702 |
| NA | -0.434387217 | 0.740008005 | 0.645869894 | 0.793066955 | 6.011411088 | 4.448392755 | ENSG00000260908 |
| PRCC | -0.434405152 | 0.739998806 | 8.54E-10 | 1.60E-08 | 2016.601384 | 1491.965335 | ENSG00000143294 |
| MAST1 | -0.434464036 | 0.739968603 | 0.001120695 | 0.005809655 | 361.4821162 | 267.7398519 | ENSG00000105613 |
| FAM111A | -0.434573985 | 0.739912212 | 0.000382216 | 0.002264701 | 430.6680524 | 318.4157014 | ENSG00000166801 |
| MIR573 | -0.434593899 | 0.739901999 | 0.70284307 | 0.831263 | 4.138401199 | 3.045561059 | ENSG00000207697 |
| MRPS15 | -0.434660364 | 0.739867912 | 1.61E-09 | 2.91E-08 | 1883.882208 | 1393.452725 | ENSG00000116898 |
| TFDP1 | -0.434685782 | 0.739854877 | 1.07E-09 | 1.98E-08 | 2534.223238 | 1874.341523 | ENSG00000198176 |
| PGAP2 | -0.434724523 | 0.73983501 | 9.75E-05 | 0.000680988 | 649.656138 | 481.176504 | ENSG00000148985 |
| LOC100507437 | -0.434787747 | 0.739802588 | 0.240328888 | 0.4264728 | 51.48666598 | 38.20184629 | ENSG00000251602 |
| RABIF | -0.434816933 | 0.739787622 | 0.024273808 | 0.075599186 | 169.9374722 | 125.933882 | ENSG00000183155 |
| NA | -0.434981944 | 0.739703012 | 0.613468096 | 0.769359017 | 7.000343303 | 5.187406401 | ENSG00000273382 |
| AAMP | -0.435149401 | 0.739617158 | 2.58E-10 | 5.28E-09 | 3765.110468 | 2785.422224 | ENSG00000127837 |
| MAP3K13 | -0.435245206 | 0.739568044 | 0.053140269 | 0.140099822 | 118.9722151 | 88.04289208 | ENSG00000073803 |
| NA | -0.435327555 | 0.739525831 | 0.583373645 | 0.746670773 | 9.781408392 | 7.257854994 | ENSG00000205940 |
| DOK1 | -0.435338044 | 0.739520454 | 0.003444482 | 0.014957831 | 270.6594312 | 200.2630007 | ENSG00000115325 |
| RAD54L | -0.435375324 | 0.739501345 | 1.41E-05 | 0.000122589 | 945.4324513 | 699.4322854 | ENSG00000085999 |

| | | | | | | | |
|---|---|---|---|---|---|---|---|
| SNAP25 | -0.435526765 | 0.739423723 | 0.360866566 | 0.554034205 | 52.11782248 | 38.54330824 | ENSG00000132639 |
| NA | -0.435537069 | 0.739418441 | 0.319520476 | 0.513348865 | 28.7811977 | 21.31199099 | ENSG00000255836 |
| NA | -0.435626157 | 0.739372783 | 0.638962353 | 0.78834611 | 6.870834819 | 5.078467662 | ENSG00000229525 |
| DBN1 | -0.435851618 | 0.739257245 | 1.68E-12 | 4.79E-11 | 13622.49617 | 10071.094 | ENSG00000113758 |
| TMX4 | -0.435863708 | 0.73925105 | 0.004516055 | 0.018859974 | 275.107653 | 203.1151892 | ENSG00000125827 |
| KIRREL2 | -0.436119975 | 0.739119747 | 0.025040905 | 0.077558177 | 165.7970191 | 122.6554713 | ENSG00000126259 |
| ALPK1 | -0.436740115 | 0.738802106 | 0.719086781 | 0.840961283 | 4.075864302 | 2.989248076 | ENSG00000073331 |
| NA | -0.436876761 | 0.738732134 | 0.582496451 | 0.74614619 | 8.326711644 | 6.113691958 | ENSG00000280300 |
| ERI3 | -0.437312338 | 0.73850913 | 7.19E-08 | 9.83E-07 | 1541.868599 | 1138.336606 | ENSG00000117419 |
| PTK2B | -0.437332986 | 0.73849856 | 0.066081624 | 0.165574804 | 104.3997294 | 76.96573156 | ENSG00000120899 |
| ZBTB17 | -0.437450012 | 0.738438659 | 0.000301915 | 0.001847981 | 550.1009582 | 406.7576556 | ENSG00000116809 |
| HAT1 | -0.437460618 | 0.73843323 | 3.72E-07 | 4.49E-06 | 1760.930744 | 1300.087852 | ENSG00000128708 |
| FAM200B | -0.43750191 | 0.738412095 | 0.014312036 | 0.049167612 | 187.4927346 | 138.5849715 | ENSG00000237765 |
| HSPB8 | -0.437788225 | 0.738265566 | 0.01565222 | 0.052850837 | 181.8324697 | 134.0190546 | ENSG00000152137 |
| LINC02043 | -0.43800329 | 0.738155519 | 0.433028136 | 0.622850174 | 19.61843849 | 14.51950883 | ENSG00000232233 |
| FLJ37453 | -0.438455141 | 0.737924366 | 0.055233506 | 0.14424817 | 107.5284754 | 79.39305228 | ENSG00000179743 |
| NA | -0.438563582 | 0.737868902 | 0.633524478 | 0.783961787 | 6.050571819 | 4.459677675 | ENSG00000249406 |
| ACLY | -0.438681924 | 0.737808378 | 5.08E-17 | 2.41E-15 | 13695.92456 | 10104.9156 | ENSG00000131473 |
| SLC15A3 | -0.438787273 | 0.737754503 | 0.00946909 | 0.035170363 | 414.3524036 | 305.3221035 | ENSG00000110446 |
| HSPA1A | -0.438804712 | 0.737745586 | 4.54E-05 | 0.000345558 | 770.4278596 | 568.0813563 | ENSG00000204389 |
| NA | -0.438980737 | 0.737655578 | 0.703061925 | 0.831263 | 4.185153532 | 3.0681309 | ENSG00000261120 |
| POLR3C | -0.43910386 | 0.737592627 | 4.37E-05 | 0.000333619 | 607.4702636 | 447.9866897 | ENSG00000186141 |
| LOC100129484 | -0.439113366 | 0.737587767 | 0.67326451 | 0.810177265 | 5.108391937 | 3.75451879 | ENSG00000234432 |
| NA | -0.439170776 | 0.737558416 | 0.611517407 | 0.768345443 | 7.40347324 | 5.416019147 | ENSG00000254331 |
| LINC00869 | -0.439337363 | 0.737473256 | 0.186022776 | 0.357057387 | 55.59192718 | 41.11699808 | ENSG00000277147 |
| ALOX12-AS1 | -0.439457446 | 0.737411875 | 0.471743412 | 0.656896499 | 15.65817725 | 11.62031688 | ENSG00000215067 |
| TFAP2E | -0.439527495 | 0.737376071 | 0.575168824 | 0.7404959 | 10.43408456 | 7.685574223 | ENSG00000116819 |
| NA | -0.439530342 | 0.737374616 | 0.495105225 | 0.677062939 | 13.03385278 | 9.605743241 | ENSG00000225093 |
| LRRC14 | -0.439642217 | 0.737317438 | 0.000590053 | 0.003313951 | 546.1734536 | 403.2381363 | ENSG00000160959 |
| FAM181B | -0.439995699 | 0.737136806 | 0.311144837 | 0.504837468 | 29.39787005 | 21.63457033 | ENSG00000182103 |
| AP4S1 | -0.44029481 | 0.736983993 | 0.028237674 | 0.085503133 | 167.9640425 | 123.9229956 | ENSG00000100478 |
| SMIM4 | -0.440388263 | 0.736936255 | 0.005717129 | 0.023085366 | 248.6739037 | 183.4751477 | ENSG00000168273 |
| CACNB1 | -0.440414789 | 0.736922706 | 0.015153309 | 0.051522031 | 224.1526182 | 165.471735 | ENSG00000067191 |
| CYB561D2 | -0.44042054 | 0.736919768 | 0.097186142 | 0.221855409 | 82.64362661 | 61.09375527 | ENSG00000114395 |
| ITGB1BP2 | -0.440459168 | 0.736900037 | 0.321850837 | 0.515796231 | 26.65794253 | 19.68803262 | ENSG00000147166 |
| UCKL1 | -0.440560207 | 0.736848431 | 0.000476218 | 0.002743619 | 616.8883792 | 455.3067838 | ENSG00000198276 |
| CPE | -0.44064067 | 0.736807336 | 0.000116131 | 0.000797178 | 510.3963624 | 375.7981022 | ENSG00000109472 |
| RCN2 | -0.440696033 | 0.736779061 | 7.25E-06 | 6.77E-05 | 2634.713532 | 1940.373874 | ENSG00000117906 |
| NTHL1 | -0.44080756 | 0.736722107 | 1.06E-07 | 1.42E-06 | 1211.079926 | 892.1627708 | ENSG00000065057 |
| MLLT3 | -0.440845137 | 0.736702919 | 0.044883691 | 0.122985559 | 145.3219079 | 106.7422255 | ENSG00000171843 |
| NA | -0.440897162 | 0.736676353 | 0.421406478 | 0.612624336 | 19.44523689 | 14.26397775 | ENSG00000180211 |
| COPS3 | -0.440917076 | 0.736666184 | 1.28E-07 | 1.68E-06 | 1607.945986 | 1183.654238 | ENSG00000141030 |
| PKD1L1 | -0.440926951 | 0.736661142 | 0.624302419 | 0.777176025 | 7.522330907 | 5.585069716 | ENSG00000158683 |
| KCTD3 | -0.441065299 | 0.736590503 | 2.32E-08 | 3.49E-07 | 1803.046386 | 1327.28305 | ENSG00000136636 |
| NA | -0.44128175 | 0.736479999 | 0.424263946 | 0.61515336 | 20.58261907 | 15.11941621 | ENSG00000266651 |
| HDHD2 | -0.44154781 | 0.73634419 | 0.554602405 | 0.724709888 | 9.931136129 | 7.336737818 | ENSG00000167220 |
| NA | -0.441552492 | 0.736341801 | 0.667811634 | 0.80717496 | 5.06795343 | 3.713177955 | ENSG00000231006 |
| GASK1A | -0.441612722 | 0.73631106 | 0.06772432 | 0.16871002 | 106.890013 | 78.75592933 | ENSG00000144649 |
| FLT4 | -0.4417366 | 0.736247839 | 6.49E-05 | 0.000474556 | 549.8983748 | 404.9328386 | ENSG00000037280 |
| TRMT2A | -0.441823666 | 0.736203408 | 0.000122847 | 0.000837075 | 645.7304074 | 475.9115373 | ENSG00000099899 |
| ZNF223 | -0.442179547 | 0.736021826 | 0.579677548 | 0.74384975 | 8.868918502 | 6.537612341 | ENSG00000178386 |
| LMF1 | -0.442181869 | 0.736020641 | 0.024807844 | 0.076986674 | 152.983218 | 112.8324002 | ENSG00000260807 |
| ST8SIA3 | -0.442492215 | 0.735862329 | 0.061716336 | 0.157468417 | 129.5919184 | 95.43517844 | ENSG00000177511 |
| NUP160 | -0.442564492 | 0.735825464 | 2.63E-10 | 5.37E-09 | 3398.687886 | 2500.217054 | ENSG00000030066 |
| LBP | -0.442709 | 0.735751763 | 0.678669135 | 0.814350236 | 5.570901858 | 4.05072944 | ENSG00000129988 |
| METTL25 | -0.442713279 | 0.735749581 | 0.148640967 | 0.304040579 | 62.17839203 | 45.56016627 | ENSG00000127720 |
| CCP110 | -0.44275742 | 0.73572707 | 0.001828884 | 0.008786038 | 351.5689521 | 258.5039775 | ENSG00000103540 |
| B3GAT1 | -0.442762011 | 0.735724729 | 4.56E-07 | 5.42E-06 | 1239.53473 | 912.6618037 | ENSG00000109956 |
| SLC4A8 | -0.44294787 | 0.735629953 | 0.002363645 | 0.010938278 | 323.3937532 | 238.0907412 | ENSG00000050438 |
| DNAJB5 | -0.443041899 | 0.73558201 | 5.85E-07 | 6.82E-06 | 1333.828227 | 981.233115 | ENSG00000137094 |
| PDK2 | -0.443086625 | 0.735559206 | 0.012788371 | 0.045052207 | 182.2668605 | 134.1397163 | ENSG00000005882 |
| NA | -0.443207445 | 0.735497608 | 0.423348107 | 0.614194428 | 18.05633165 | 13.24003837 | ENSG00000282012 |
| NOLC1 | -0.443258284 | 0.735471691 | 6.45E-14 | 2.19E-12 | 6451.446931 | 4745.291452 | ENSG00000166197 |
| NA | -0.443321748 | 0.735439338 | 0.477793999 | 0.661871178 | 16.72039488 | 12.35933053 | ENSG00000255201 |
| NA | -0.443458986 | 0.735369382 | 0.661849556 | 0.803355337 | 6.534676468 | 4.831083922 | ENSG00000274225 |
| LOC105371267 | -0.443752362 | 0.735219857 | 0.199932491 | 0.376388907 | 45.53887417 | 33.53232687 | ENSG00000277639 |
| MDM1 | -0.443862777 | 0.73516359 | 0.013974497 | 0.048285481 | 193.5111586 | 142.2143685 | ENSG00000111554 |
| UCMA | -0.444062838 | 0.735061651 | 0.436647345 | 0.62586147 | 16.45506408 | 12.15273797 | ENSG00000165623 |
| NA | -0.444407439 | 0.734886096 | 0.569471204 | 0.736177672 | 8.820287032 | 6.481299358 | ENSG00000213104 |
| IKZF2 | -0.444906592 | 0.734631879 | 0.333534264 | 0.528171911 | 27.306184 | 20.04066787 | ENSG00000030419 |
| TTC14 | -0.445004708 | 0.734581919 | 0.005953677 | 0.023873962 | 317.0151032 | 233.414835 | ENSG00000163728 |
| SVBP | -0.445023994 | 0.7345721 | 0.110042722 | 0.243409932 | 79.85863049 | 58.6702334 | ENSG00000177868 |
| ATP6V1G1 | -0.445270431 | 0.734446633 | 6.93E-07 | 7.95E-06 | 1932.246832 | 1418.713196 | ENSG00000136888 |
| ATP6V0E2-AS1 | -0.445337574 | 0.734412453 | 0.416575307 | 0.607751088 | 18.79374081 | 13.8217244 | ENSG00000204934 |

| | | | | | | | |
|---|---|---|---|---|---|---|---|
| CDC27 | -0.445809569 | 0.73417222 | 8.59E-05 | 0.000608515 | 757.2615368 | 555.5223781 | ENSG00000004897 |
| MORN2 | -0.445952323 | 0.734099578 | 0.102478491 | 0.230796777 | 76.15580017 | 56.05239157 | ENSG00000188010 |
| B3GNT5 | -0.4461065 | 0.73402113 | 0.063284385 | 0.160628977 | 101.6666944 | 74.42253567 | ENSG00000176597 |
| EIF3E | -0.4461096 | 0.734019553 | 1.83E-07 | 2.33E-06 | 5931.416311 | 4352.479568 | ENSG00000104408 |
| PEBP1 | -0.446211773 | 0.733967571 | 2.26E-12 | 6.29E-11 | 7976.442334 | 5853.498957 | ENSG00000089220 |
| TMED1 | -0.446399673 | 0.733871984 | 0.008826823 | 0.033170384 | 220.8082551 | 161.7934624 | ENSG00000099203 |
| LOC283335 | -0.446671238 | 0.733733857 | 0.214191376 | 0.394340217 | 46.82470631 | 34.39905025 | ENSG00000257337 |
| PIK3C2B | -0.446787943 | 0.733674505 | 2.05E-05 | 0.000170569 | 776.8859285 | 570.2403724 | ENSG00000133056 |
| CCDC85B | -0.446788707 | 0.733674116 | 1.18E-05 | 0.000103983 | 1559.297337 | 1143.951963 | ENSG00000175602 |
| LIMK2 | -0.446878386 | 0.733628512 | 7.53E-07 | 8.58E-06 | 1137.328309 | 834.4827252 | ENSG00000182541 |
| POLR2I | -0.446883001 | 0.733626165 | 0.206961714 | 0.385626663 | 46.3028164 | 33.88452408 | ENSG00000105258 |
| DCLK2 | -0.447036251 | 0.73354824 | 0.517423169 | 0.694921813 | 11.82624441 | 8.698228678 | ENSG00000170390 |
| OXCT1 | -0.447057941 | 0.733537212 | 1.74E-06 | 1.85E-05 | 842.7478185 | 618.0507731 | ENSG00000083720 |
| TAP2 | -0.447177569 | 0.733476389 | 0.046162637 | 0.125638294 | 134.8453103 | 98.91649445 | ENSG00000204267 |
| PPP6R3 | -0.44718241 | 0.733473928 | 1.83E-07 | 2.33E-06 | 1958.505215 | 1435.681439 | ENSG00000110075 |
| SLC35F3 | -0.447248501 | 0.733440328 | 0.177964985 | 0.346656777 | 51.33674285 | 37.62395911 | ENSG00000183780 |
| CAVIN4 | -0.447316175 | 0.733405924 | 0.026015628 | 0.07993595 | 152.2803395 | 111.6653409 | ENSG00000170681 |
| GPR162 | -0.447437832 | 0.733344082 | 0.016365942 | 0.054792132 | 207.0790026 | 151.9903732 | ENSG00000250510 |
| E2F5 | -0.447517233 | 0.733303722 | 0.002672469 | 0.012083812 | 501.0934288 | 367.1494246 | ENSG00000133740 |
| GON7 | -0.447582377 | 0.733270611 | 0.039988829 | 0.112466307 | 133.7813112 | 98.02296435 | ENSG00000170270 |
| NA | -0.447601778 | 0.73326075 | 0.5070383 | 0.6864963 | 12.33372522 | 9.114662988 | ENSG00000236683 |
| SRRT | -0.447787581 | 0.73316632 | 2.99E-13 | 9.38E-12 | 6449.149385 | 4729.355017 | ENSG00000087087 |
| GVINP1 | -0.448105783 | 0.733004631 | 0.693976303 | 0.824655819 | 4.191467358 | 3.045561059 | ENSG00000254838 |
| LOC106699570 | -0.44826481 | 0.732923836 | 0.650347264 | 0.795928168 | 5.697351127 | 4.208495088 | ENSG00000246777 |
| NA | -0.448378808 | 0.732865925 | 0.079173257 | 0.190151243 | 106.8686887 | 78.04510785 | ENSG00000288066 |
| MBIP | -0.448443319 | 0.732833155 | 0.000445612 | 0.002587397 | 411.3667179 | 301.4133317 | ENSG00000151332 |
| CHRNB4 | -0.448503603 | 0.732802534 | 0.072829541 | 0.178595225 | 100.6198554 | 73.92484541 | ENSG00000117971 |
| CSNK2B | -0.448527209 | 0.732790544 | 0.206319658 | 0.38491693 | 46.64854319 | 34.27513936 | ENSG00000204435 |
| TARDBP | -0.448570626 | 0.732768491 | 6.14E-12 | 1.61E-10 | 2758.030287 | 2021.239579 | ENSG00000120948 |
| SLC25A21 | -0.448806498 | 0.732648698 | 0.117499941 | 0.255633748 | 75.90249567 | 55.53351691 | ENSG00000183032 |
| ALYREF | -0.448971457 | 0.73256493 | 3.85E-11 | 8.93E-10 | 2826.17975 | 2070.159635 | ENSG00000183684 |
| NISCH | -0.449104786 | 0.732497233 | 1.80E-09 | 3.23E-08 | 3276.514195 | 2400.979576 | ENSG00000010322 |
| ZNF689 | -0.449286327 | 0.732405065 | 0.000140354 | 0.000936738 | 571.837368 | 418.6703524 | ENSG00000156853 |
| LOC729870 | -0.44955769 | 0.732267316 | 0.443147957 | 0.631553963 | 16.86766476 | 12.33632266 | ENSG00000287642 |
| NA | -0.449955014 | 0.732065675 | 0.413075832 | 0.604535668 | 24.5068765 | 18.07210998 | ENSG00000223509 |
| NA | -0.450087971 | 0.731998211 | 0.550188506 | 0.721465527 | 10.75455604 | 7.869158907 | ENSG00000286548 |
| EIF5A | -0.450164367 | 0.731959451 | 4.65E-11 | 1.06E-09 | 18911.25207 | 13842.62139 | ENSG00000132507 |
| S1PR5 | -0.450171653 | 0.731955754 | 0.004005638 | 0.017043272 | 265.181628 | 194.1333489 | ENSG00000180739 |
| PRKCD | -0.45019555 | 0.73194363 | 4.45E-05 | 0.000339845 | 683.8614639 | 500.1328571 | ENSG00000163932 |
| AREL1 | -0.450320155 | 0.731880415 | 5.11E-07 | 6.02E-06 | 1060.111501 | 775.70564 | ENSG00000119682 |
| GLI2 | -0.450560726 | 0.731758383 | 1.76E-05 | 0.000148977 | 1091.189058 | 798.6887758 | ENSG00000074047 |
| PHF5A | -0.450600858 | 0.731738028 | 7.12E-09 | 1.17E-07 | 1528.662507 | 1118.135027 | ENSG00000100410 |
| ABT1 | -0.450663812 | 0.731706098 | 8.75E-07 | 9.83E-06 | 1310.030355 | 957.7818976 | ENSG00000146109 |
| NCR1 | -0.450923474 | 0.731574415 | 0.640962345 | 0.789656747 | 6.526987168 | 4.718346336 | ENSG00000189430 |
| MRPL48 | -0.451096605 | 0.731486627 | 0.001130863 | 0.005851313 | 358.3511002 | 261.823291 | ENSG00000175581 |
| KLHDC9 | -0.4511817 | 0.731443483 | 0.091273396 | 0.211775304 | 93.17386908 | 68.31775551 | ENSG00000162755 |
| IFI6 | -0.451244165 | 0.731411814 | 0.136520181 | 0.285389695 | 62.19555207 | 45.57623772 | ENSG00000126709 |
| C13orf42 | -0.451245522 | 0.731411126 | 8.37E-07 | 9.45E-06 | 1278.285398 | 934.7624308 | ENSG00000226792 |
| ITFG1 | -0.451306527 | 0.731380199 | 0.001424047 | 0.007081644 | 338.6519575 | 247.2922825 | ENSG00000129636 |
| POSTN | -0.451315938 | 0.731375428 | 0.40291225 | 0.594694246 | 19.27015615 | 14.03916385 | ENSG00000133110 |
| SEC63 | -0.45133292 | 0.731366819 | 1.29E-08 | 2.04E-07 | 2114.362672 | 1545.731287 | ENSG00000025796 |
| LHFPL5 | -0.451761234 | 0.731149719 | 0.40189482 | 0.59357069 | 23.07112123 | 16.97297501 | ENSG00000197753 |
| SLC29A2 | -0.451809389 | 0.731125315 | 2.71E-05 | 0.000218949 | 775.6831009 | 567.5847724 | ENSG00000174669 |
| NA | -0.451995018 | 0.731031249 | 0.68051628 | 0.815245229 | 5.074267255 | 3.68312204 | ENSG00000284959 |
| SERF1A | -0.45211049 | 0.73097274 | 0.482506262 | 0.665929783 | 13.73525811 | 10.05971954 | ENSG00000205572 |
| URGCP | -0.452181492 | 0.730936766 | 0.000238731 | 0.001505495 | 445.2546389 | 325.5470491 | ENSG00000106608 |
| SHISA3 | -0.452300822 | 0.730876311 | 0.084173592 | 0.199054384 | 97.87677092 | 71.56479618 | ENSG00000178343 |
| ALDH4A1 | -0.452429198 | 0.730811277 | 9.63E-08 | 1.29E-06 | 1438.278956 | 1050.42836 | ENSG00000159423 |
| ME1 | -0.452540983 | 0.730754654 | 0.012827222 | 0.045154745 | 200.6936705 | 146.8134788 | ENSG00000065833 |
| RABEP2 | -0.452573513 | 0.730738177 | 0.000148333 | 0.000984319 | 507.4427722 | 371.0752328 | ENSG00000177548 |
| TMEM45A | -0.452704634 | 0.730671766 | 0.002197606 | 0.010299906 | 271.4658864 | 198.5377013 | ENSG00000181458 |
| CPNE9 | -0.452761194 | 0.730643121 | 0.061406302 | 0.156893355 | 103.9212395 | 75.9165228 | ENSG00000144550 |
| SPC25 | -0.452813149 | 0.730616809 | 0.000273569 | 0.001697464 | 567.4057572 | 414.5830602 | ENSG00000152253 |
| TRAPPC2L | -0.452823254 | 0.730611692 | 1.71E-06 | 1.82E-05 | 981.6906781 | 717.2962824 | ENSG00000167515 |
| NEXN | -0.453446627 | 0.73029607 | 0.064044892 | 0.16193834 | 103.2403465 | 75.54142932 | ENSG00000162614 |
| SULT1A2 | -0.453472053 | 0.7302832 | 0.648776872 | 0.794951114 | 6.169331787 | 4.48973359 | ENSG00000197165 |
| TTC7A | -0.453484473 | 0.730276913 | 9.07E-05 | 0.000638396 | 543.4398324 | 396.935197 | ENSG00000068724 |
| NA | -0.453513479 | 0.730262231 | 0.59777873 | 0.757813336 | 8.723527756 | 6.274706801 | ENSG00000273893 |
| NA | -0.453516061 | 0.730260924 | 0.662685579 | 0.803809851 | 6.037944167 | 4.384593698 | ENSG00000246100 |
| COL5A3 | -0.453739248 | 0.73014796 | 0.123878179 | 0.265920567 | 66.40733638 | 48.59543009 | ENSG00000080573 |
| MKRN2OS | -0.453881968 | 0.730075733 | 0.672140387 | 0.809503537 | 4.813273453 | 3.540878192 | ENSG00000225526 |
| HMGN3 | -0.453890839 | 0.730071244 | 0.000210442 | 0.001343087 | 828.6040338 | 604.7608254 | ENSG00000118418 |
| NA | -0.453892464 | 0.730070421 | 0.055484499 | 0.144797198 | 98.71568231 | 71.97363279 | ENSG00000277013 |
| MYBBP1A | -0.454032677 | 0.729999471 | 5.79E-13 | 1.77E-11 | 4691.790805 | 3425.679337 | ENSG00000132382 |

| | | | | | | | |
|---|---|---|---|---|---|---|---|
| C9orf116 | -0.454161584 | 0.729934247 | 0.0593634 | 0.152641683 | 96.75685705 | 70.60085701 | ENSG00000160345 |
| NA | -0.454274817 | 0.729876959 | 0.360355785 | 0.553616887 | 29.78205851 | 21.84528814 | ENSG00000277103 |
| SOX10 | -0.454487224 | 0.729769507 | 0.040775201 | 0.114192744 | 119.4166554 | 87.10444554 | ENSG00000100146 |
| CNKSR3 | -0.45452804 | 0.729748861 | 0.11567827 | 0.252820146 | 77.01156333 | 56.26168367 | ENSG00000153721 |
| NACAD | -0.454531176 | 0.729747275 | 0.032193944 | 0.09465142 | 140.6092496 | 102.670352 | ENSG00000136274 |
| ATCAY | -0.454718807 | 0.729652373 | 7.80E-07 | 8.87E-06 | 1303.052366 | 950.3424164 | ENSG00000167654 |
| ST8SIA1 | -0.45507384 | 0.729472835 | 0.132558777 | 0.279313822 | 70.71134758 | 51.7012146 | ENSG00000111728 |
| RFC2 | -0.45508523 | 0.729467076 | 1.14E-09 | 2.10E-08 | 1690.822217 | 1233.539396 | ENSG00000049541 |
| GUCY1A2 | -0.455103859 | 0.729457657 | 0.321100448 | 0.514905321 | 30.24348605 | 22.01291299 | ENSG00000152402 |
| PRMT6 | -0.455124829 | 0.729447054 | 4.59E-07 | 5.45E-06 | 1215.960059 | 887.4776667 | ENSG00000198890 |
| NA | -0.455138618 | 0.729440082 | 0.413105583 | 0.604535668 | 18.66235318 | 13.57758986 | ENSG00000240376 |
| SLC7A8 | -0.455180016 | 0.729419151 | 2.00E-13 | 6.40E-12 | 6036.165969 | 4402.23021 | ENSG00000092068 |
| NDST1 | -0.455395421 | 0.729310252 | 4.28E-10 | 8.36E-09 | 2606.419558 | 1901.244373 | ENSG00000070614 |
| QPCTL | -0.455809142 | 0.729101138 | 9.31E-05 | 0.000654296 | 535.0202326 | 390.0763279 | ENSG00000011478 |
| ANKRD54 | -0.45599067 | 0.729009404 | 0.013684818 | 0.047455123 | 211.7279665 | 154.7896414 | ENSG00000100124 |
| GPHN | -0.456008348 | 0.729000471 | 0.002029407 | 0.009614782 | 329.4069532 | 240.1552008 | ENSG00000171723 |
| SETD9 | -0.456124 | 0.728942034 | 0.047883445 | 0.129387925 | 117.1736707 | 85.29310364 | ENSG00000155542 |
| RRM2 | -0.456216537 | 0.72889528 | 1.83E-09 | 3.27E-08 | 3703.804496 | 2698.653431 | ENSG00000171848 |
| LPAR2 | -0.456369243 | 0.728818132 | 2.40E-05 | 0.000196171 | 810.8963904 | 591.5290694 | ENSG00000064547 |
| ARV1 | -0.456387746 | 0.728808785 | 0.02670869 | 0.081740234 | 132.7601334 | 96.71387601 | ENSG00000173409 |
| FASN | -0.456503457 | 0.728750333 | 9.70E-12 | 2.46E-10 | 33598.67425 | 24484.37853 | ENSG00000169710 |
| NA | -0.456798079 | 0.728601526 | 0.504038069 | 0.683882985 | 14.04931807 | 10.19079009 | ENSG00000260805 |
| STRA6 | -0.456798927 | 0.728601097 | 0.002828855 | 0.012664249 | 254.4299584 | 185.3379045 | ENSG00000137868 |
| NXT1 | -0.45683058 | 0.728585112 | 0.001118799 | 0.005809655 | 366.6975275 | 266.6454632 | ENSG00000132661 |
| NA | -0.456870561 | 0.728564921 | 0.211993011 | 0.391615515 | 52.50203358 | 38.46161426 | ENSG00000272686 |
| RHOH | -0.45703229 | 0.728483252 | 0.006340368 | 0.025222662 | 238.3973339 | 173.661163 | ENSG00000168421 |
| TCTA | -0.457046335 | 0.72847616 | 0.064488456 | 0.162768849 | 106.6329502 | 77.9240484 | ENSG00000145022 |
| BAD | -0.457183716 | 0.728406794 | 0.281700534 | 0.472981414 | 31.39477366 | 22.81203847 | ENSG00000002330 |
| APOM | -0.457803519 | 0.728093927 | 0.451662745 | 0.639173874 | 16.13143568 | 11.71373382 | ENSG00000204444 |
| METTL8 | -0.457851721 | 0.728069601 | 4.21E-05 | 0.000323238 | 615.2273635 | 447.7844456 | ENSG00000123600 |
| HNRNPLL | -0.458575411 | 0.727704476 | 4.35E-08 | 6.24E-07 | 1825.19538 | 1328.307802 | ENSG00000143889 |
| SYS1 | -0.458701049 | 0.727641106 | 0.004623043 | 0.019224473 | 242.7762392 | 176.4701872 | ENSG00000204070 |
| MTDH | -0.458793619 | 0.727594419 | 2.30E-06 | 2.39E-05 | 1569.251752 | 1140.914229 | ENSG00000147649 |
| NA | -0.458826066 | 0.727578055 | 0.416605419 | 0.607751088 | 17.56591318 | 12.79029895 | ENSG00000261094 |
| NA | -0.458893017 | 0.727544291 | 0.656937185 | 0.79954022 | 5.091329596 | 3.664351046 | ENSG00000269846 |
| NA | -0.458921929 | 0.727529711 | 0.429998795 | 0.620105639 | 16.21428953 | 11.85272848 | ENSG00000272221 |
| MEI4 | -0.459080314 | 0.727449844 | 0.595161903 | 0.755943716 | 8.481573129 | 6.185088709 | ENSG00000269964 |
| IPO13 | -0.459290865 | 0.727343685 | 3.47E-05 | 0.000273182 | 634.0388053 | 460.9176951 | ENSG00000117408 |
| ARFRP1 | -0.459448032 | 0.727264453 | 5.89E-06 | 5.61E-05 | 701.6569221 | 510.5560991 | ENSG00000101246 |
| CLDN3 | -0.459623118 | 0.727176197 | 0.005297247 | 0.021668322 | 325.8326166 | 236.661827 | ENSG00000165215 |
| SPATC1L | -0.459640183 | 0.727167596 | 2.53E-05 | 0.000205964 | 900.4038914 | 654.2089457 | ENSG00000160284 |
| FDXACB1 | -0.459887322 | 0.72704304 | 0.404882036 | 0.596509035 | 17.96726168 | 13.10907944 | ENSG00000255561 |
| THYN1 | -0.459927698 | 0.727022693 | 9.49E-08 | 1.28E-06 | 1126.810799 | 818.7064358 | ENSG00000151500 |
| POLR2F | -0.459988704 | 0.726991951 | 4.00E-08 | 5.76E-07 | 1980.488991 | 1440.786254 | ENSG00000100142 |
| NA | -0.460239964 | 0.726865349 | 0.393198025 | 0.585908683 | 23.21916522 | 16.95745321 | ENSG00000234825 |
| FBRSL1 | -0.460303513 | 0.726833332 | 6.37E-06 | 6.01E-05 | 2986.007095 | 2170.454504 | ENSG00000112787 |
| CLSTN2 | -0.460449146 | 0.726759966 | 0.209281212 | 0.388490944 | 52.11092993 | 37.84497416 | ENSG00000158258 |
| RSPO4 | -0.460449749 | 0.726759662 | 0.252466739 | 0.441227585 | 286.5876896 | 208.4524146 | ENSG00000101282 |
| GPR68 | -0.460567318 | 0.726700439 | 0.535219334 | 0.709650959 | 14.18514038 | 10.27704736 | ENSG00000119714 |
| LOC100287042 | -0.460624301 | 0.726671736 | 0.306063313 | 0.500105577 | 27.56086398 | 20.01061196 | ENSG00000263843 |
| ZNF440 | -0.460643052 | 0.726662291 | 0.093743161 | 0.215691334 | 110.6540798 | 80.10459793 | ENSG00000171295 |
| ASPHD2 | -0.460684972 | 0.726641178 | 0.005305634 | 0.021697842 | 223.3043715 | 162.1692171 | ENSG00000128203 |
| NR3C1 | -0.460879356 | 0.726543279 | 0.005896509 | 0.023695988 | 240.6948957 | 174.7934914 | ENSG00000113580 |
| MCM3 | -0.461046545 | 0.726459087 | 1.44E-17 | 7.33E-16 | 10873.18836 | 7899.25393 | ENSG00000112118 |
| ARMCX5 | -0.461055676 | 0.726454489 | 0.011368971 | 0.040922997 | 179.0421292 | 129.9645263 | ENSG00000125962 |
| SHE | -0.461109723 | 0.726427275 | 0.297514381 | 0.490493821 | 34.14870431 | 24.8753274 | ENSG00000169291 |
| CTPS1 | -0.461333327 | 0.726314694 | 8.22E-12 | 2.12E-10 | 2786.27036 | 2023.839655 | ENSG00000171793 |
| ASF1A | -0.461602335 | 0.726179277 | 0.001082737 | 0.005651212 | 445.463461 | 322.9848272 | ENSG00000111875 |
| PARP12 | -0.461699805 | 0.726130217 | 0.04781936 | 0.129233599 | 172.6495138 | 125.7330637 | ENSG00000059378 |
| CENPM | -0.46189194 | 0.726033519 | 3.30E-06 | 3.34E-05 | 694.2086734 | 503.8471344 | ENSG00000100162 |
| RTN1 | -0.462099369 | 0.725929138 | 0.103683005 | 0.23287213 | 85.52747171 | 61.78196664 | ENSG00000139970 |
| TDP2 | -0.462293399 | 0.725831514 | 2.21E-07 | 2.78E-06 | 1147.264263 | 833.015602 | ENSG00000111802 |
| PLPBP | -0.462344158 | 0.725805977 | 2.00E-06 | 2.10E-05 | 808.3899839 | 587.0310853 | ENSG00000147471 |
| NA | -0.462781639 | 0.725585917 | 0.272754126 | 0.463952366 | 33.43782823 | 24.26369708 | ENSG00000180015 |
| HYAL3 | -0.46282886 | 0.725562169 | 0.228818331 | 0.412701338 | 36.02033872 | 26.113019 | ENSG00000186792 |
| HLA-DMB | -0.463025594 | 0.725463234 | 0.586785262 | 0.749424538 | 7.579831754 | 5.513672966 | ENSG00000242574 |
| MANF | -0.463165955 | 0.725392656 | 9.35E-09 | 1.51E-07 | 2285.162515 | 1657.580826 | ENSG00000145050 |
| C3orf80 | -0.463195725 | 0.725377688 | 0.229279174 | 0.41317435 | 41.00997069 | 29.676029 | ENSG00000180044 |
| PTPRN2 | -0.463197984 | 0.725376552 | 0.010254509 | 0.037531016 | 199.4116869 | 144.6046969 | ENSG00000155093 |
| P4HTM | -0.46342361 | 0.725263118 | 0.011807571 | 0.042198518 | 242.8371826 | 176.2141552 | ENSG00000178467 |
| HLA-F-AS1 | -0.463515306 | 0.725217023 | 0.323766734 | 0.517881475 | 29.12928465 | 21.13980285 | ENSG00000214922 |
| SRSF8 | -0.463805469 | 0.725071177 | 0.000785168 | 0.004271059 | 499.940668 | 362.3470196 | ENSG00000263465 |
| NVL | -0.463835421 | 0.725056124 | 9.13E-06 | 8.23E-05 | 897.3179666 | 650.7777941 | ENSG00000143748 |
| PNKP | -0.46386776 | 0.725039872 | 2.97E-07 | 3.65E-06 | 1260.730804 | 914.7457359 | ENSG00000039650 |

| | | | | | | | |
|---|---|---|---|---|---|---|---|
| CPNE5 | -0.463933891 | 0.725006638 | 0.470169024 | 0.655738783 | 16.23323101 | 11.79272827 | ENSG00000124772 |
| CLN6 | -0.464056367 | 0.724945092 | 2.80E-06 | 2.86E-05 | 910.3174615 | 660.5324399 | ENSG00000128973 |
| SCAMP5 | -0.464060408 | 0.724943061 | 2.20E-10 | 4.57E-09 | 1925.171069 | 1395.742477 | ENSG00000198794 |
| ARIH1 | -0.464182958 | 0.724881483 | 4.92E-06 | 4.76E-05 | 1239.808268 | 897.7778159 | ENSG00000166233 |
| CDK5RAP3 | -0.464414633 | 0.724765088 | 0.105766366 | 0.236519525 | 84.97786864 | 61.84449187 | ENSG00000108465 |
| DCUN1D4 | -0.464524369 | 0.724709962 | 2.73E-05 | 0.000220493 | 683.9815091 | 495.2892858 | ENSG00000109184 |
| LOC101928415 | -0.464595957 | 0.724674002 | 0.184452637 | 0.355110126 | 51.14399841 | 37.05388441 | ENSG00000286724 |
| FRAT2 | -0.464597707 | 0.724673123 | 2.83E-07 | 3.49E-06 | 1332.5846 | 965.9383243 | ENSG00000181274 |
| BEND3 | -0.46459983 | 0.724672056 | 1.08E-05 | 9.58E-05 | 1109.080975 | 803.7893584 | ENSG00000178409 |
| SRI | -0.464774919 | 0.724584114 | 2.17E-06 | 2.26E-05 | 863.4354801 | 624.965414 | ENSG00000075142 |
| IL17RE | -0.464849187 | 0.724546814 | 0.120094445 | 0.259723505 | 66.13421859 | 47.97675172 | ENSG00000163701 |
| FCRL5 | -0.46505087 | 0.724445533 | 0.362448464 | 0.555404698 | 21.68852981 | 15.64803845 | ENSG00000143297 |
| LRRC6 | -0.465085657 | 0.724428065 | 0.165063318 | 0.327721856 | 54.13279575 | 39.13561792 | ENSG00000129295 |
| CTBP2 | -0.465131254 | 0.724405169 | 1.57E-14 | 5.69E-13 | 3585.315822 | 2597.278997 | ENSG00000175029 |
| LINC00458 | -0.465132801 | 0.724404392 | 0.011624107 | 0.041655299 | 199.4191057 | 144.1497329 | ENSG00000234787 |
| EFEMP2 | -0.46513887 | 0.724401345 | 0.136340539 | 0.285142807 | 85.56415194 | 62.10695938 | ENSG00000172638 |
| SSBP2 | -0.465149837 | 0.724395838 | 0.000526911 | 0.0030069 | 364.5736936 | 264.2333781 | ENSG00000145687 |
| TAFA5 | -0.465536441 | 0.724201746 | 4.38E-07 | 5.23E-06 | 1070.602688 | 774.9226658 | ENSG00000219438 |
| GTF3C1 | -0.465610416 | 0.724164613 | 9.63E-14 | 3.17E-12 | 5135.376641 | 3718.409807 | ENSG00000077235 |
| TMEM203 | -0.465822877 | 0.724057975 | 1.65E-07 | 2.11E-06 | 952.882158 | 689.9171161 | ENSG00000187713 |
| EPHA4 | -0.466237108 | 0.723850111 | 0.016688175 | 0.055709408 | 169.7078522 | 122.5601906 | ENSG00000116106 |
| NA | -0.466254483 | 0.723841394 | 0.017140401 | 0.056848507 | 244.7064569 | 177.4269038 | ENSG00000262001 |
| PDXDC2P-NPIPB14P | -0.466336502 | 0.723800243 | 0.302003885 | 0.49542751 | 27.77194854 | 20.15329384 | ENSG00000196696 |
| MLPH | -0.466783564 | 0.723575987 | 0.027341388 | 0.08337369 | 148.1779421 | 107.2129261 | ENSG00000115648 |
| TRIM25 | -0.466834017 | 0.723550683 | 2.26E-06 | 2.35E-05 | 1014.145024 | 733.1289158 | ENSG00000121060 |
| MFAP2 | -0.467119094 | 0.723407724 | 3.23E-09 | 5.54E-08 | 1788.760382 | 1294.537155 | ENSG00000117122 |
| HCG11 | -0.46743599 | 0.72324884 | 0.002707953 | 0.012220405 | 291.6233522 | 210.9824131 | ENSG00000228223 |
| PPM1G | -0.467543103 | 0.723195145 | 1.81E-16 | 8.08E-15 | 8265.793297 | 5977.735487 | ENSG00000115241 |
| COL17A1 | -0.467607498 | 0.723162866 | 0.149703343 | 0.305673874 | 51.22547679 | 36.9938842 | ENSG00000065618 |
| DPH7 | -0.467673261 | 0.723129902 | 1.26E-05 | 0.000110551 | 837.7865237 | 606.4701255 | ENSG00000148399 |
| TBC1D10A | -0.467757395 | 0.723087732 | 0.067186589 | 0.167741783 | 89.44876762 | 64.60683906 | ENSG00000099992 |
| NA | -0.467780394 | 0.723076205 | 0.473322051 | 0.658451617 | 14.75703723 | 10.64845361 | ENSG00000226239 |
| CEP152 | -0.467781057 | 0.723075873 | 0.003599182 | 0.015538606 | 258.8048949 | 187.3944802 | ENSG00000103995 |
| NT5C3B | -0.468214748 | 0.72285854 | 6.94E-09 | 1.14E-07 | 1446.785476 | 1045.422603 | ENSG00000141698 |
| PSMD1 | -0.468222488 | 0.722854662 | 8.66E-14 | 2.89E-12 | 5396.834077 | 3900.344697 | ENSG00000173692 |
| CCNL2 | -0.468270183 | 0.722830765 | 0.048322237 | 0.130231836 | 2054.801803 | 1485.582579 | ENSG00000221978 |
| TTC4 | -0.468372353 | 0.722779577 | 0.554970229 | 0.724966044 | 8.634457779 | 6.252686612 | ENSG00000243725 |
| FAM111A-DT | -0.46853802 | 0.722696584 | 0.23372166 | 0.418652483 | 38.26678855 | 27.70356064 | ENSG00000245571 |
| HSD11B2 | -0.468637367 | 0.722646819 | 3.67E-05 | 0.000287435 | 636.7283425 | 460.0525175 | ENSG00000176387 |
| NA | -0.468662013 | 0.722634474 | 0.425136327 | 0.615858592 | 16.71102183 | 12.13396698 | ENSG00000249572 |
| TAX1BP3 | -0.468675351 | 0.722627793 | 0.626217545 | 0.778581605 | 5.706821866 | 4.129612264 | ENSG00000213977 |
| NA | -0.468862256 | 0.722534181 | 0.318334213 | 0.512037619 | 24.79124647 | 17.93636452 | ENSG00000249129 |
| NA | -0.468981594 | 0.722474416 | 0.370757373 | 0.564125864 | 19.51220848 | 14.06542092 | ENSG00000220472 |
| TMEM159 | -0.46934622 | 0.722291842 | 0.000438099 | 0.002547763 | 451.0126478 | 326.2434481 | ENSG00000011638 |
| HMGCS1 | -0.469377718 | 0.722276072 | 8.05E-15 | 3.01E-13 | 5624.258569 | 4061.474295 | ENSG00000112972 |
| ATXN7L2 | -0.469453894 | 0.722237936 | 0.013567856 | 0.047108777 | 213.1644208 | 154.0791548 | ENSG00000162650 |
| PRSS46P | -0.469856828 | 0.722036249 | 0.648993506 | 0.795111426 | 5.752296422 | 4.159668179 | ENSG00000261603 |
| UBR7 | -0.469900765 | 0.722014259 | 1.74E-05 | 0.000147535 | 688.1491571 | 496.5467277 | ENSG00000012963 |
| TCF24 | -0.470096361 | 0.721916377 | 0.201779929 | 0.378831343 | 40.36982448 | 29.10975315 | ENSG00000261787 |
| NA | -0.470138976 | 0.721895054 | 0.651263121 | 0.796260352 | 6.127014144 | 4.388280925 | ENSG00000226329 |
| DTX4 | -0.470146277 | 0.7218914 | 2.14E-06 | 2.23E-05 | 856.4557695 | 617.8238722 | ENSG00000110042 |
| ACOT4 | -0.470262144 | 0.721833425 | 0.37576466 | 0.568849318 | 20.6743423 | 14.98845728 | ENSG00000177465 |
| MAN1C1 | -0.470273795 | 0.721827596 | 0.001303847 | 0.006579225 | 330.7729633 | 238.899391 | ENSG00000117643 |
| NA | -0.470358093 | 0.72178542 | 0.324589646 | 0.518767319 | 25.43327181 | 18.26219305 | ENSG00000223611 |
| TMEM167A | -0.470433917 | 0.721747486 | 1.78E-07 | 2.27E-06 | 3179.326179 | 2293.865209 | ENSG00000174695 |
| NA | -0.470577323 | 0.721675747 | 0.592584211 | 0.754323885 | 7.54255016 | 5.438588988 | ENSG00000285646 |
| INHBA | -0.470615335 | 0.721656733 | 0.524118764 | 0.7006341 | 10.37164536 | 7.478981666 | ENSG00000122641 |
| NA | -0.471091669 | 0.721418503 | 0.654274051 | 0.797941387 | 4.724203477 | 3.405682385 | ENSG00000286281 |
| VPS29 | -0.471542078 | 0.721193311 | 0.00046784 | 0.002701232 | 388.2973781 | 279.6794586 | ENSG00000111237 |
| NA | -0.471593403 | 0.721167655 | 0.477802607 | 0.661871178 | 12.41204669 | 8.956897339 | ENSG00000228204 |
| ARMH3 | -0.472352276 | 0.720788413 | 3.38E-05 | 0.000267236 | 527.2810895 | 379.7398929 | ENSG00000120029 |
| MTM1 | -0.472358021 | 0.720785542 | 0.000527801 | 0.003010854 | 332.4352946 | 239.5105918 | ENSG00000171100 |
| TAF11 | -0.472405797 | 0.720761673 | 7.25E-05 | 0.000521666 | 460.5483433 | 331.8133938 | ENSG00000064995 |
| FAM184B | -0.472624538 | 0.7206524 | 0.311683271 | 0.505327089 | 25.45980489 | 18.40542458 | ENSG00000047662 |
| NA | -0.47276021 | 0.720584633 | 0.526275924 | 0.702433961 | 12.74316899 | 9.264942562 | ENSG00000259529 |
| RIDA | -0.472792373 | 0.720568568 | 0.000166531 | 0.001092945 | 546.8451615 | 393.5416438 | ENSG00000132541 |
| CASD1 | -0.472815236 | 0.720557149 | 0.000975793 | 0.005155457 | 365.1102857 | 263.0397353 | ENSG00000127995 |
| SCNN1G | -0.472980457 | 0.720474634 | 0.578716644 | 0.74299961 | 7.605087057 | 5.476130977 | ENSG00000166828 |
| ADRA2C | -0.473380923 | 0.720274671 | 3.91E-06 | 3.90E-05 | 681.825813 | 491.4118841 | ENSG00000184160 |
| GNAZ | -0.473418766 | 0.720255778 | 0.00253574 | 0.01158999 | 269.6638769 | 194.2305647 | ENSG00000128266 |
| SLC25A22 | -0.473659031 | 0.720135837 | 7.57E-06 | 7.01E-05 | 1157.530017 | 833.4506638 | ENSG00000177542 |
| NA | -0.473908837 | 0.720011155 | 0.603762924 | 0.762842868 | 7.331465604 | 5.330088282 | ENSG00000279412 |
| NA | -0.473924952 | 0.720003113 | 0.609759133 | 0.767176415 | 7.098884018 | 5.067182742 | ENSG00000237950 |
| NA | -0.473932086 | 0.719999552 | 0.348432489 | 0.541765751 | 23.15397508 | 16.76269522 | ENSG00000243547 |

| | | | | | | | |
|---|---|---|---|---|---|---|---|
| RNF40 | -0.474028089 | 0.719951642 | 5.44E-09 | 9.10E-08 | 1773.743576 | 1277.881893 | ENSG00000103549 |
| CDC42EP4 | -0.474453385 | 0.719739436 | 0.000107938 | 0.000745904 | 438.1744858 | 315.4377382 | ENSG00000179604 |
| SLC2A8 | -0.474544667 | 0.719693899 | 9.96E-06 | 8.91E-05 | 802.252612 | 577.0679916 | ENSG00000136856 |
| LANCL3 | -0.474817357 | 0.719557879 | 0.513342398 | 0.691998545 | 10.47334299 | 7.527808575 | ENSG00000147036 |
| NA | -0.475030101 | 0.719451779 | 0.269731825 | 0.460054717 | 211.2039043 | 152.1221369 | ENSG00000263146 |
| PRPF38B | -0.47503696 | 0.719448358 | 3.94E-11 | 9.12E-10 | 2351.215307 | 1691.855768 | ENSG00000134186 |
| PCDHA2 | -0.475119578 | 0.719407159 | 0.318699222 | 0.512518124 | 26.00328953 | 18.79549021 | ENSG00000204969 |
| HERC2P9 | -0.475625363 | 0.719154992 | 0.002713557 | 0.012230443 | 423.4010779 | 305.019475 | ENSG00000206149 |
| OGFR | -0.475880363 | 0.71902789 | 1.55E-08 | 2.42E-07 | 1269.979033 | 913.4692511 | ENSG00000060491 |
| TP53I3 | -0.475931703 | 0.719002303 | 0.000149952 | 0.000993283 | 497.5071037 | 357.5630507 | ENSG00000115129 |
| TTC13 | -0.476509575 | 0.718714364 | 0.029576633 | 0.088529283 | 135.4108933 | 97.0016129 | ENSG00000143643 |
| XXYLT1 | -0.476731043 | 0.718604043 | 2.76E-06 | 2.82E-05 | 704.7165318 | 506.7293075 | ENSG00000173950 |
| OPTC | -0.476885826 | 0.71852695 | 0.456961912 | 0.644055671 | 18.13099254 | 13.07522468 | ENSG00000188770 |
| 1-Mar | -0.477167325 | 0.718386764 | 0.001404336 | 0.006993004 | 317.7681241 | 227.9638215 | ENSG00000186205 |
| NA | -0.477694288 | 0.718124412 | 0.530632234 | 0.70614339 | 9.492002374 | 6.800079849 | ENSG00000257718 |
| LHX1-DT | -0.47770081 | 0.718121166 | 0.405968721 | 0.597593867 | 16.90178944 | 12.17530781 | ENSG00000277268 |
| TXNDC15 | -0.477804858 | 0.718069376 | 1.83E-05 | 0.000153715 | 592.5889267 | 425.2440726 | ENSG00000113621 |
| NA | -0.477903535 | 0.718020264 | 0.592478046 | 0.754240546 | 6.72228716 | 4.819799001 | ENSG00000251593 |
| GNB3 | -0.478115103 | 0.717914976 | 0.02888951 | 0.086921781 | 128.9103415 | 92.61910619 | ENSG00000111664 |
| NRTN | -0.478238718 | 0.717853465 | 0.086197246 | 0.202712823 | 87.99033524 | 63.21215471 | ENSG00000171119 |
| B4GAT1 | -0.478385559 | 0.717780404 | 2.08E-06 | 2.17E-05 | 733.2767834 | 526.1608298 | ENSG00000174684 |
| SETSIP | -0.478674224 | 0.717636799 | 0.444868766 | 0.633174452 | 15.75809344 | 11.3236682 | ENSG00000230667 |
| POLR3H | -0.478706342 | 0.717620823 | 3.14E-07 | 3.84E-06 | 909.6336448 | 652.9605507 | ENSG00000100413 |
| ZNF568 | -0.47872676 | 0.717610667 | 0.197838995 | 0.373471984 | 41.30055678 | 29.60127143 | ENSG00000198453 |
| CTPS2 | -0.478739781 | 0.71760419 | 1.06E-10 | 2.32E-09 | 2069.826801 | 1484.739806 | ENSG00000047230 |
| ZIC3 | -0.478846802 | 0.717550959 | 1.17E-07 | 1.55E-06 | 1596.126145 | 1144.954487 | ENSG00000156925 |
| FGFR1 | -0.47917738 | 0.717386559 | 5.42E-19 | 3.17E-17 | 9482.28686 | 6802.387864 | ENSG00000077782 |
| TIAF1 | -0.47954362 | 0.717204467 | 0.397888501 | 0.589910045 | 18.14667941 | 12.98885578 | ENSG00000221995 |
| SSTR2 | -0.479575823 | 0.717188459 | 0.01002242 | 0.036841637 | 184.9201008 | 132.8503867 | ENSG00000180616 |
| UBAC1 | -0.479688479 | 0.717132458 | 9.98E-11 | 2.19E-09 | 1682.678155 | 1206.766032 | ENSG00000130560 |
| CHAF1B | -0.479706782 | 0.71712336 | 2.72E-06 | 2.78E-05 | 782.4359197 | 560.7468171 | ENSG00000159259 |
| ATXN7L3B | -0.47980174 | 0.71707616 | 3.64E-10 | 7.20E-09 | 1690.786814 | 1212.864362 | ENSG00000253719 |
| UBXN8 | -0.479860703 | 0.717046854 | 0.03217158 | 0.094628066 | 124.3626919 | 89.14863706 | ENSG00000104691 |
| POLE4 | -0.480050029 | 0.716952762 | 9.72E-05 | 0.000679039 | 469.2700146 | 336.630708 | ENSG00000115350 |
| CHAC2 | -0.480051083 | 0.716952238 | 0.012183417 | 0.043299606 | 192.9095491 | 138.1255474 | ENSG00000143942 |
| TASP1 | -0.480204173 | 0.716876163 | 0.01362449 | 0.047287696 | 168.2135137 | 120.6223499 | ENSG00000089123 |
| TMEM62 | -0.480232661 | 0.716862007 | 0.210720076 | 0.390119896 | 39.06179625 | 28.09005065 | ENSG00000137842 |
| NA | -0.480425405 | 0.716766241 | 0.423831214 | 0.614702849 | 15.8111596 | 11.34992527 | ENSG00000280187 |
| MFSD13A | -0.480549429 | 0.716704626 | 0.002524771 | 0.011556954 | 252.2633032 | 180.833422 | ENSG00000138111 |
| CD53 | -0.480731806 | 0.71661403 | 0.555544356 | 0.725307064 | 8.535917064 | 6.094920964 | ENSG00000143119 |
| UNG | -0.480733837 | 0.716613021 | 1.21E-12 | 3.52E-11 | 4470.55539 | 3202.720355 | ENSG00000076248 |
| LAT2 | -0.480839156 | 0.716560709 | 0.460863208 | 0.647535342 | 15.16154265 | 10.82835107 | ENSG00000086730 |
| CCDC163 | -0.480859113 | 0.716550797 | 0.188621179 | 0.360528882 | 41.2335852 | 29.47736055 | ENSG00000280670 |
| APOC1 | -0.480941083 | 0.716510086 | 0.062432248 | 0.158842186 | 96.15893079 | 69.07922737 | ENSG00000130208 |
| HDAC3 | -0.481066757 | 0.716447673 | 3.61E-10 | 7.16E-09 | 1556.883857 | 1115.565167 | ENSG00000171720 |
| NA | -0.481068707 | 0.716446704 | 0.076868281 | 0.185989129 | 89.34381538 | 63.76604633 | ENSG00000213707 |
| CTNNBIP1 | -0.481085891 | 0.71643817 | 1.57E-06 | 1.67E-05 | 958.0477878 | 686.016626 | ENSG00000178585 |
| FTSJ1 | -0.48132972 | 0.716317096 | 3.76E-07 | 4.53E-06 | 969.697982 | 694.9831366 | ENSG00000068438 |
| CDCA7 | -0.481628171 | 0.716168927 | 1.11E-09 | 2.05E-08 | 2088.606689 | 1496.334313 | ENSG00000144354 |
| TERF1 | -0.481760819 | 0.716103082 | 2.21E-11 | 5.30E-10 | 10479.75966 | 7503.545483 | ENSG00000147601 |
| LHFPL6 | -0.481906649 | 0.716030701 | 0.065152998 | 0.163890562 | 95.75035884 | 68.61309009 | ENSG00000183722 |
| RCBTB2 | -0.482084421 | 0.715942475 | 0.000831867 | 0.004492372 | 300.8693939 | 215.4909176 | ENSG00000136161 |
| DNAJC28 | -0.482193165 | 0.715888513 | 0.256080436 | 0.444941858 | 31.49774906 | 22.64397559 | ENSG00000177692 |
| GALNT18 | -0.482382239 | 0.715794697 | 0.003670174 | 0.015808275 | 241.7420278 | 173.3440314 | ENSG00000110328 |
| PRTFDC1 | -0.482386619 | 0.715792524 | 0.004696991 | 0.019479531 | 202.3231549 | 144.7726481 | ENSG00000099256 |
| NA | -0.482416274 | 0.715777811 | 0.02871984 | 0.086565867 | 135.4102919 | 96.72461127 | ENSG00000230330 |
| TUBB1 | -0.482428709 | 0.715771642 | 0.66827456 | 0.807313302 | 5.372542652 | 3.87474245 | ENSG00000101162 |
| H2AC16 | -0.482436332 | 0.715767859 | 0.60661408 | 0.765141239 | 6.679969516 | 4.767173245 | ENSG00000276903 |
| NAGS | -0.482549952 | 0.715711491 | 0.009740496 | 0.03596222 | 168.1257215 | 120.2729639 | ENSG00000161653 |
| COPS2 | -0.482606187 | 0.715683594 | 0.000138543 | 0.000926455 | 1091.303549 | 780.2896659 | ENSG00000166200 |
| ATG2A | -0.482974806 | 0.715500755 | 1.63E-06 | 1.74E-05 | 868.046914 | 621.0592386 | ENSG00000110046 |
| DERL2 | -0.483087115 | 0.715445058 | 3.39E-05 | 0.000267464 | 504.4532984 | 360.7994593 | ENSG00000072849 |
| ARID5A | -0.483369206 | 0.71530518 | 0.002486433 | 0.011423787 | 240.5930027 | 172.0071487 | ENSG00000196843 |
| MAGI2-AS3 | -0.483582621 | 0.715199374 | 0.028676549 | 0.086477572 | 123.0241996 | 87.98245384 | ENSG00000234456 |
| SHISAL2B | -0.483585304 | 0.715198044 | 0.025216033 | 0.077947148 | 134.8990755 | 96.71150288 | ENSG00000145642 |
| BOLA2B | -0.483752145 | 0.715115339 | 0.415956713 | 0.607215399 | 18.51390322 | 13.22875345 | ENSG00000169627 |
| BAIAP2-DT | -0.483774721 | 0.715104149 | 0.000591945 | 0.003322564 | 350.320672 | 250.959923 | ENSG00000226137 |
| TAF6L | -0.483812661 | 0.715085343 | 0.000715365 | 0.003929632 | 343.1020585 | 245.7617813 | ENSG00000162227 |
| UPF3B | -0.484488264 | 0.714750553 | 1.22E-07 | 1.60E-06 | 1051.775883 | 752.2893944 | ENSG00000125351 |
| CASP9 | -0.484621105 | 0.714684743 | 6.74E-05 | 0.000489811 | 453.7632488 | 324.1584251 | ENSG00000132906 |
| BUD31 | -0.484632604 | 0.714679047 | 7.94E-07 | 9.01E-06 | 775.368492 | 553.8936735 | ENSG00000106245 |
| NEIL3 | -0.484770975 | 0.714610504 | 0.004045437 | 0.017177121 | 212.0291057 | 151.4949445 | ENSG00000109674 |
| EMC9 | -0.484776799 | 0.714607619 | 0.000442949 | 0.002575163 | 430.9417337 | 308.3995128 | ENSG00000100908 |
| NA | -0.484853923 | 0.714569418 | 0.002601476 | 0.011837897 | 255.9653368 | 182.9836295 | ENSG00000282048 |

| | | | | | | | |
|---|---|---|---|---|---|---|---|
| LINC00862 | -0.485313118 | 0.714342015 | 0.407146964 | 0.598941331 | 17.18743718 | 12.22738392 | ENSG00000203721 |
| TMEM94 | -0.486054888 | 0.713974826 | 3.76E-08 | 5.43E-07 | 1394.802514 | 996.7271153 | ENSG00000177728 |
| LOC107985164 | -0.486071677 | 0.713966517 | 0.153366032 | 0.310756099 | 49.72600454 | 35.59908823 | ENSG00000267327 |
| KPTN | -0.486168449 | 0.713918628 | 0.003121026 | 0.013775924 | 279.0036111 | 199.2624041 | ENSG00000118162 |
| MAF | -0.486339371 | 0.713834052 | 0.036744429 | 0.105063446 | 128.4883678 | 91.94075403 | ENSG00000178573 |
| CDC45 | -0.486466945 | 0.713770932 | 9.11E-05 | 0.000641014 | 570.0308185 | 407.2385502 | ENSG00000093009 |
| NA | -0.486480831 | 0.713764062 | 0.537374919 | 0.711203176 | 9.52041459 | 6.76253786 | ENSG00000286416 |
| LGI2 | -0.486716818 | 0.713647319 | 0.004837397 | 0.019994718 | 225.6573218 | 161.3301765 | ENSG00000153012 |
| STX1B | -0.486882874 | 0.713565182 | 0.001349178 | 0.006765603 | 284.4864352 | 203.2415135 | ENSG00000099365 |
| GRAMD1B | -0.487334135 | 0.71334202 | 0.000318186 | 0.00193036 | 438.5461595 | 313.2201879 | ENSG00000023171 |
| STK38 | -0.487346168 | 0.713336071 | 1.81E-05 | 0.000152241 | 530.8933615 | 378.8226307 | ENSG00000112079 |
| LZTR1 | -0.48785004 | 0.713086976 | 0.014954831 | 0.050950891 | 161.8692965 | 115.7506896 | ENSG00000099949 |
| NA | -0.487897283 | 0.713063625 | 0.667723685 | 0.80717496 | 5.088172683 | 3.566697227 | ENSG00000272918 |
| NA | -0.487922808 | 0.71305101 | 0.381954392 | 0.574745693 | 16.87209945 | 12.02502824 | ENSG00000207425 |
| SPATA31C2 | -0.487937492 | 0.713043752 | 0.648129009 | 0.794567224 | 4.777269635 | 3.38691139 | ENSG00000177910 |
| NA | -0.488064633 | 0.712980916 | 0.286336508 | 0.477588268 | 25.53309031 | 18.16877611 | ENSG00000203279 |
| SERPINE3 | -0.488150612 | 0.712938427 | 0.640742661 | 0.789444284 | 5.967815668 | 4.324919901 | ENSG00000253309 |
| ZNF426-DT | -0.48818257 | 0.712922634 | 0.377782894 | 0.571087073 | 17.76564786 | 12.6513043 | ENSG00000278611 |
| WFDC2 | -0.48820986 | 0.712909149 | 0.000190462 | 0.001229972 | 427.8125218 | 305.1060268 | ENSG00000101443 |
| DHRSX | -0.48829275 | 0.71286819 | 0.001346439 | 0.006757352 | 320.2268386 | 227.9448273 | ENSG00000169084 |
| TSPAN14 | -0.488465321 | 0.712782924 | 6.48E-11 | 1.45E-09 | 1779.821591 | 1268.716388 | ENSG00000108219 |
| NA | -0.488638536 | 0.71269735 | 0.485733681 | 0.66875218 | 11.50587062 | 8.16157071 | ENSG00000286835 |
| CDC25C | -0.488699282 | 0.712667342 | 0.000170374 | 0.001113833 | 365.1842475 | 260.3168568 | ENSG00000158402 |
| MRPL1 | -0.488827095 | 0.712604206 | 0.002270887 | 0.010577689 | 388.5498632 | 276.3857494 | ENSG00000169288 |
| SCLT1 | -0.489619891 | 0.712212721 | 0.006170246 | 0.024667674 | 184.297791 | 131.4321448 | ENSG00000151466 |
| LINC00665 | -0.489661554 | 0.712192154 | 3.31E-10 | 6.63E-09 | 1843.749046 | 1313.69678 | ENSG00000232677 |
| MOK | -0.489963818 | 0.712042955 | 0.349482697 | 0.542897687 | 22.81653895 | 16.15420247 | ENSG00000080823 |
| EXO1 | -0.490116522 | 0.711967592 | 8.77E-06 | 7.93E-05 | 785.4373295 | 558.4620263 | ENSG00000174371 |
| LHX1 | -0.490214497 | 0.711919243 | 0.599549765 | 0.759331376 | 6.689440255 | 4.73711733 | ENSG00000273706 |
| GDF11 | -0.490377058 | 0.71183903 | 1.15E-05 | 0.000101169 | 530.5038439 | 377.5470707 | ENSG00000135414 |
| YAE1 | -0.490387718 | 0.71183377 | 0.002162131 | 0.010173377 | 266.9165683 | 190.1251713 | ENSG00000241127 |
| DACT2 | -0.490481876 | 0.711787313 | 0.046183734 | 0.125638294 | 126.616538 | 90.29802466 | ENSG00000164488 |
| NA | -0.490518672 | 0.711769159 | 0.676508769 | 0.812987929 | 4.43036277 | 3.184555713 | ENSG00000250327 |
| KLK1 | -0.490521624 | 0.711767703 | 0.386771051 | 0.579687421 | 24.3818027 | 17.41913882 | ENSG00000167748 |
| BTBD6 | -0.490553819 | 0.711751819 | 0.00458028 | 0.019076609 | 233.3380771 | 166.4923135 | ENSG00000184887 |
| DCHS1 | -0.490636388 | 0.711711085 | 2.00E-06 | 2.10E-05 | 808.1994495 | 575.3311042 | ENSG00000166341 |
| CHURC1 | -0.490652965 | 0.711702908 | 0.056928487 | 0.147816277 | 84.79784955 | 60.47821448 | ENSG00000258289 |
| SFXN3 | -0.490869754 | 0.71159597 | 0.210516279 | 0.389867428 | 38.60422468 | 27.42937017 | ENSG00000107819 |
| EIF6 | -0.491036464 | 0.711513747 | 3.30E-09 | 5.65E-08 | 1453.578568 | 1034.549694 | ENSG00000242372 |
| NA | -0.491073758 | 0.711495354 | 0.049293491 | 0.132291566 | 110.5622437 | 78.91804348 | ENSG00000229320 |
| MRPL9 | -0.491184953 | 0.711440518 | 1.10E-06 | 1.22E-05 | 813.8719361 | 578.7448937 | ENSG00000143436 |
| GET4 | -0.491809716 | 0.711132494 | 0.005355487 | 0.021856227 | 202.0664981 | 143.9605259 | ENSG00000239857 |
| HPCA | -0.492418011 | 0.710832717 | 0.229546263 | 0.4134467 | 45.60278654 | 32.47475597 | ENSG00000121905 |
| CASTOR2 | -0.492485628 | 0.710799402 | 0.000182928 | 0.001188778 | 389.3095888 | 276.4431532 | ENSG00000274070 |
| ANP32AP1 | -0.492545217 | 0.710770043 | 0.479398278 | 0.663418777 | 13.22589816 | 9.410873637 | ENSG00000259516 |
| UBTF | -0.492580339 | 0.71075274 | 2.26E-12 | 6.29E-11 | 3335.604873 | 2371.762733 | ENSG00000108312 |
| E2F4 | -0.492916972 | 0.710586915 | 1.23E-08 | 1.95E-07 | 2215.259035 | 1575.165272 | ENSG00000205250 |
| MYPOP | -0.492972068 | 0.710559779 | 0.008742347 | 0.032933062 | 209.9585333 | 149.5873745 | ENSG00000176182 |
| NAT10 | -0.49300197 | 0.710545051 | 2.67E-15 | 1.06E-13 | 3297.486008 | 2342.884315 | ENSG00000135372 |
| TPPP | -0.493222533 | 0.71043643 | 0.102172064 | 0.230358848 | 62.07857354 | 44.13206519 | ENSG00000171368 |
| NA | -0.493264694 | 0.710415668 | 0.670970121 | 0.808859269 | 4.393081176 | 3.147013724 | ENSG00000240449 |
| SLITRK1 | -0.493375009 | 0.710361349 | 0.622635299 | 0.776001791 | 5.405389557 | 3.870943603 | ENSG00000178235 |
| COMMD5 | -0.493456351 | 0.710321298 | 6.48E-06 | 6.11E-05 | 741.0065157 | 526.722909 | ENSG00000170619 |
| MROH6 | -0.493642323 | 0.71022974 | 0.097733596 | 0.222776346 | 68.5948122 | 48.6175619 | ENSG00000204839 |
| NA | -0.493763331 | 0.710170171 | 0.503773052 | 0.683776217 | 10.72298691 | 7.663004382 | ENSG00000257511 |
| DPH3 | -0.493851278 | 0.71012688 | 5.45E-06 | 5.22E-05 | 627.2380989 | 445.7227972 | ENSG00000154813 |
| FAM43B | -0.493913166 | 0.710096418 | 0.063316544 | 0.160688598 | 79.25526221 | 56.21285676 | ENSG00000183114 |
| FAM66C | -0.493918114 | 0.710093982 | 0.256608665 | 0.445275256 | 28.60296005 | 20.32939245 | ENSG00000226711 |
| TCEAL1 | -0.493994998 | 0.710056141 | 0.11156332 | 0.245991187 | 64.52160114 | 46.01188106 | ENSG00000172465 |
| CCDC107 | -0.494577291 | 0.70976961 | 0.236731809 | 0.421747913 | 39.94755766 | 28.3074901 | ENSG00000159884 |
| TTLL10 | -0.494834412 | 0.709643124 | 0.483563284 | 0.666941651 | 12.37919978 | 8.874765321 | ENSG00000162571 |
| FAM83E | -0.49497525 | 0.709573851 | 0.090078222 | 0.209658392 | 74.47365558 | 53.00683051 | ENSG00000105523 |
| ZBTB40 | -0.495359052 | 0.709385107 | 5.93E-06 | 5.63E-05 | 699.623436 | 496.6029693 | ENSG00000184677 |
| ORMDL3 | -0.495606973 | 0.709263213 | 8.76E-11 | 1.93E-09 | 1529.393023 | 1085.103282 | ENSG00000172057 |
| HMX2 | -0.495815851 | 0.709160531 | 0.01285295 | 0.045202385 | 152.8801449 | 108.5787655 | ENSG00000188816 |
| NA | -0.495833551 | 0.70915183 | 0.3825212 | 0.575178316 | 19.67772078 | 13.97949005 | ENSG00000238000 |
| USF1 | -0.4960717 | 0.709034779 | 9.77E-09 | 1.57E-07 | 1300.111764 | 921.9139241 | ENSG00000158773 |
| UCK1 | -0.496135236 | 0.709003554 | 2.61E-08 | 3.88E-07 | 1202.889804 | 852.8261113 | ENSG00000130717 |
| PCSK7 | -0.49630154 | 0.70892183 | 2.37E-07 | 2.96E-06 | 883.0464095 | 626.5427843 | ENSG00000160613 |
| MYT1 | -0.496531948 | 0.708808619 | 0.070112565 | 0.173169813 | 76.43248084 | 54.24809771 | ENSG00000196132 |
| ADCK2 | -0.496882332 | 0.708636493 | 0.011955463 | 0.042644831 | 148.1617516 | 104.9055026 | ENSG00000133597 |
| DBP | -0.496890848 | 0.70863231 | 0.027001123 | 0.082458199 | 156.2556711 | 110.711261 | ENSG00000105516 |
| CFAP20 | -0.497046589 | 0.708555816 | 7.88E-06 | 7.25E-05 | 537.7001863 | 381.107158 | ENSG00000070761 |
| MRPL58 | -0.497140847 | 0.708509525 | 8.05E-06 | 7.37E-05 | 568.7885893 | 403.3124558 | ENSG00000167862 |

| | | | | | | | |
|---|---|---|---|---|---|---|---|
| TVP23C | -0.497446465 | 0.708359451 | 0.240342527 | 0.4264728 | 30.50250302 | 21.58574342 | ENSG00000175106 |
| ANKHD1-EIF4EBP3 | -0.497487928 | 0.708339093 | 0.617132054 | 0.772310229 | 6.219338731 | 4.410850766 | ENSG00000254996 |
| LPAR3 | -0.497579854 | 0.708293961 | 6.63E-05 | 0.000483906 | 399.9174763 | 283.2185133 | ENSG00000171517 |
| NA | -0.497584746 | 0.708291559 | 0.014859443 | 0.050681803 | 151.4563182 | 107.2619763 | ENSG00000276710 |
| SPNS2 | -0.497658684 | 0.70825526 | 0.163989073 | 0.326113034 | 49.46165843 | 34.90835184 | ENSG00000183018 |
| NA | -0.497892792 | 0.70814034 | 0.667518014 | 0.80717496 | 4.403829691 | 3.143214877 | ENSG00000228952 |
| LOC105371730 | -0.497990023 | 0.708092616 | 0.656174766 | 0.799155153 | 4.393081176 | 3.135728803 | ENSG00000214708 |
| CFAP36 | -0.498020575 | 0.708077621 | 0.001971192 | 0.009381952 | 248.3961407 | 175.8251401 | ENSG00000163001 |
| WDR62 | -0.498193884 | 0.707992565 | 3.48E-11 | 8.14E-10 | 1820.815604 | 1288.890284 | ENSG00000075702 |
| NUDT11 | -0.498238108 | 0.707970863 | 2.51E-07 | 3.14E-06 | 782.6291526 | 553.8037289 | ENSG00000196368 |
| VPS9D1 | -0.498262331 | 0.707958977 | 0.111568032 | 0.245991187 | 81.4847474 | 57.91734691 | ENSG00000075399 |
| GALNTL6 | -0.498418999 | 0.707882101 | 0.521743654 | 0.698699197 | 10.33248463 | 7.393050801 | ENSG00000174473 |
| S100A1 | -0.498481906 | 0.707851235 | 0.456603988 | 0.643844972 | 12.7791728 | 9.084607073 | ENSG00000160678 |
| NA | -0.498623871 | 0.707781584 | 0.092339347 | 0.213500183 | 69.03462237 | 48.79409854 | ENSG00000274317 |
| MSC | -0.498948273 | 0.707622451 | 0.443651179 | 0.632021449 | 17.18172472 | 12.20525211 | ENSG00000178860 |
| TP53INP2 | -0.499165688 | 0.70751582 | 0.017254604 | 0.057126979 | 139.3579482 | 98.6271086 | ENSG00000078804 |
| TBCK | -0.499176935 | 0.707510304 | 0.00548291 | 0.022285144 | 261.6693175 | 185.0555667 | ENSG00000145348 |
| NA | -0.499481531 | 0.707360943 | 0.0012306 | 0.006271088 | 275.2643937 | 194.5864492 | ENSG00000284691 |
| NA | -0.499530595 | 0.707336888 | 0.589790821 | 0.751955911 | 6.375380294 | 4.527275579 | ENSG00000230295 |
| L3MBTL4 | -0.499752468 | 0.707228114 | 0.222794389 | 0.404909904 | 34.50054952 | 24.35755205 | ENSG00000154655 |
| KCNN1 | -0.499929364 | 0.707141403 | 0.006443165 | 0.025554807 | 226.3713548 | 159.75913 | ENSG00000105642 |
| SV2A | -0.499995598 | 0.707108939 | 8.70E-12 | 2.22E-10 | 2520.327954 | 1781.337249 | ENSG00000159164 |
| TPGS1 | -0.500132318 | 0.707041931 | 0.125602449 | 0.268544655 | 51.53322292 | 36.42771997 | ENSG00000141933 |
| NA | -0.500537127 | 0.706843569 | 0.687034194 | 0.819981438 | 4.458774986 | 3.173270792 | ENSG00000274356 |
| ABHD4 | -0.50057179 | 0.706826586 | 1.54E-07 | 1.99E-06 | 839.2375373 | 593.4437906 | ENSG00000100439 |
| KIF14 | -0.500603742 | 0.706810932 | 8.85E-06 | 7.99E-05 | 619.641145 | 437.5022853 | ENSG00000118193 |
| ZNF774 | -0.501147855 | 0.706544408 | 0.255052913 | 0.443895292 | 27.69047016 | 19.54155189 | ENSG00000196391 |
| ZMAT1 | -0.501470993 | 0.706386172 | 0.388611608 | 0.581422849 | 17.77827551 | 12.48985142 | ENSG00000166432 |
| MAGED4 | -0.501486352 | 0.706378652 | 0.238997121 | 0.424776816 | 32.94740976 | 23.41673238 | ENSG00000154545 |
| ZNF302 | -0.502212281 | 0.706023309 | 5.83E-05 | 0.000431729 | 439.354787 | 310.3223899 | ENSG00000089335 |
| CRYM | -0.502260785 | 0.705999573 | 0.001218611 | 0.006223679 | 259.8504335 | 183.5496821 | ENSG00000103316 |
| TSC22D4 | -0.503047402 | 0.705614737 | 6.69E-06 | 6.28E-05 | 695.8409912 | 491.3853006 | ENSG00000166925 |
| NA | -0.503199785 | 0.705540211 | 0.418667759 | 0.609982568 | 16.71733566 | 11.72826794 | ENSG00000227542 |
| MAIP1 | -0.503362008 | 0.705460882 | 0.001929495 | 0.00922633 | 225.6925516 | 159.1043354 | ENSG00000162972 |
| RANBP1 | -0.503405593 | 0.70543957 | 2.25E-18 | 1.23E-16 | 7680.161021 | 5418.089566 | ENSG00000099901 |
| SHROOM1 | -0.503732306 | 0.705279834 | 0.011671834 | 0.04178717 | 148.0202168 | 104.5538551 | ENSG00000164403 |
| MAN2A2 | -0.503846563 | 0.70522398 | 0.006928707 | 0.027172376 | 398.8032749 | 281.8109841 | ENSG00000196547 |
| SLC25A1 | -0.504125082 | 0.705087846 | 4.74E-13 | 1.46E-11 | 4936.137749 | 3480.210942 | ENSG00000100075 |
| NA | -0.504248747 | 0.70502741 | 0.515662125 | 0.693461169 | 10.31542229 | 7.336737818 | ENSG00000227543 |
| CCDC153 | -0.504293579 | 0.705005502 | 0.235230832 | 0.420178126 | 32.42088977 | 22.86130341 | ENSG00000248712 |
| CHDH | -0.504326971 | 0.704989184 | 0.000276522 | 0.001713495 | 344.6807466 | 243.0625317 | ENSG00000016391 |
| UBE2E3 | -0.504452627 | 0.704927784 | 7.28E-09 | 1.20E-07 | 1211.453006 | 854.0659528 | ENSG00000170035 |
| NA | -0.504504129 | 0.704902619 | 0.36894064 | 0.561915179 | 19.47304775 | 13.69412629 | ENSG00000235381 |
| ARL6IP5 | -0.504813469 | 0.704751492 | 1.17E-05 | 0.000103265 | 592.4792467 | 416.9502288 | ENSG00000144746 |
| RBMX2 | -0.504903752 | 0.70470739 | 6.18E-05 | 0.000454002 | 455.0657147 | 320.3654883 | ENSG00000134597 |
| ANKRD36 | -0.504918487 | 0.704700192 | 0.115883144 | 0.253096652 | 69.2021115 | 48.96033795 | ENSG00000135976 |
| NR1I3 | -0.505021058 | 0.704650092 | 0.326721838 | 0.520989637 | 22.91823658 | 16.14335558 | ENSG00000143257 |
| CDC7 | -0.505216687 | 0.704554548 | 5.17E-08 | 7.31E-07 | 1034.529885 | 728.465059 | ENSG00000097046 |
| NA | -0.505301249 | 0.704513253 | 0.025220871 | 0.077947148 | 114.9174419 | 81.09633151 | ENSG00000276529 |
| LINC02762 | -0.505459422 | 0.704436016 | 0.537135705 | 0.711203176 | 8.382934716 | 5.903850207 | ENSG00000250303 |
| NA | -0.505625222 | 0.704355064 | 0.558461224 | 0.727526558 | 7.394002501 | 5.213663469 | ENSG00000249631 |
| KLHL18 | -0.505627491 | 0.704353957 | 8.61E-08 | 1.17E-06 | 822.6313473 | 579.3096809 | ENSG00000114648 |
| NA | -0.505757714 | 0.704290382 | 0.551505181 | 0.722473927 | 9.627824683 | 6.728683099 | ENSG00000273014 |
| TBXAS1 | -0.505784607 | 0.704277253 | 0.306781589 | 0.500827092 | 24.18018888 | 17.11153163 | ENSG00000059377 |
| FBXL19 | -0.505819612 | 0.704260165 | 3.99E-11 | 9.20E-10 | 3243.859092 | 2284.61282 | ENSG00000099364 |
| CBLN3 | -0.50582108 | 0.704259449 | 0.476469296 | 0.66089522 | 11.41048682 | 8.109494606 | ENSG00000139899 |
| ATP7B | -0.505872691 | 0.704234255 | 0.00036734 | 0.002187752 | 302.6652624 | 213.0900772 | ENSG00000123191 |
| RTKN2 | -0.506012699 | 0.704165915 | 7.20E-05 | 0.000519089 | 498.9818091 | 351.1965272 | ENSG00000182010 |
| CCND3 | -0.506326515 | 0.704012761 | 8.55E-07 | 9.63E-06 | 843.4154447 | 593.8180164 | ENSG00000112576 |
| CLINT1 | -0.506332886 | 0.704009652 | 9.35E-12 | 2.38E-10 | 2140.08532 | 1505.896396 | ENSG00000113282 |
| SDC1 | -0.506516959 | 0.703919834 | 2.67E-10 | 5.42E-09 | 1985.833805 | 1397.677474 | ENSG00000115884 |
| MCF2 | -0.506566284 | 0.703895767 | 0.254790875 | 0.44358835 | 29.21657319 | 20.54672027 | ENSG00000101977 |
| ZNF268 | -0.506902053 | 0.703731963 | 0.001863561 | 0.008931792 | 280.2122867 | 196.8358075 | ENSG00000090612 |
| USP49 | -0.507052956 | 0.703658358 | 1.49E-05 | 0.000128234 | 670.9178088 | 472.5700836 | ENSG00000164663 |
| PSENEN | -0.507066315 | 0.703651843 | 0.221976006 | 0.40397715 | 32.51005744 | 22.95147116 | ENSG00000205155 |
| NA | -0.507109144 | 0.703630954 | 0.647557721 | 0.794244908 | 4.469523501 | 3.165784718 | ENSG00000287817 |
| PIGL | -0.507563767 | 0.70340926 | 0.002441453 | 0.011239415 | 254.3642646 | 179.3125568 | ENSG00000108474 |
| PAXBP1-AS1 | -0.507590064 | 0.703396439 | 0.294080503 | 0.486777407 | 24.16628346 | 17.07767687 | ENSG00000238197 |
| SLC66A2 | -0.507651923 | 0.70336628 | 0.000235158 | 0.001484474 | 445.1275505 | 313.2669363 | ENSG00000122490 |
| SNPH | -0.507777256 | 0.703305178 | 0.005646997 | 0.022856832 | 173.6480895 | 122.0015125 | ENSG00000101298 |
| ARMC7 | -0.507796283 | 0.703295903 | 0.000289622 | 0.001783342 | 313.6990547 | 220.6296087 | ENSG00000125449 |
| SEMA4G | -0.507809056 | 0.703289676 | 0.025624831 | 0.078983972 | 117.5220508 | 82.83715272 | ENSG00000095539 |
| MBD2 | -0.507959616 | 0.703216284 | 0.000287607 | 0.001771519 | 407.8095115 | 286.6047719 | ENSG00000134046 |
| NA | -0.508175119 | 0.703111249 | 0.283087822 | 0.474132451 | 33.72891799 | 23.76936763 | ENSG00000273162 |

| | | | | | | | |
|---|---|---|---|---|---|---|---|
| MIR25 | -0.50884945 | 0.702782684 | 0.337786345 | 0.531951068 | 23.31505269 | 16.45888688 | ENSG00000207547 |
| LRFN1 | -0.509017792 | 0.702700684 | 0.001102215 | 0.005735109 | 261.9847303 | 183.9605654 | ENSG00000128011 |
| PPM1H | -0.509066593 | 0.702676915 | 2.65E-11 | 6.28E-10 | 1932.590943 | 1357.27234 | ENSG00000111110 |
| USP18 | -0.509089619 | 0.7026657 | 0.017804418 | 0.058581054 | 127.1315353 | 89.24986649 | ENSG00000184979 |
| XPO6 | -0.509140023 | 0.702641151 | 5.07E-15 | 1.96E-13 | 4066.065084 | 2856.703317 | ENSG00000169180 |
| RUSC1 | -0.509205027 | 0.702609493 | 4.88E-09 | 8.23E-08 | 1260.843355 | 886.1929632 | ENSG00000160753 |
| TXNL4B | -0.50952799 | 0.702452223 | 0.012736078 | 0.044927765 | 147.9603331 | 104.1186498 | ENSG00000140830 |
| NIPAL2 | -0.509591842 | 0.702421134 | 0.200182058 | 0.376705682 | 38.39008091 | 26.89662267 | ENSG00000104361 |
| TYRO3 | -0.5096148 | 0.702409957 | 2.18E-15 | 8.77E-14 | 3381.017719 | 2374.325343 | ENSG00000092445 |
| STRAP | -0.509700481 | 0.702368242 | 6.70E-10 | 1.28E-08 | 4114.579564 | 2888.92654 | ENSG00000023734 |
| GDF10 | -0.509769416 | 0.702334682 | 0.543051844 | 0.715600631 | 7.843982469 | 5.543728881 | ENSG00000266524 |
| CLDND2 | -0.510032991 | 0.70220638 | 0.432695641 | 0.622647016 | 12.73054133 | 8.938126345 | ENSG00000160318 |
| TMEM200C | -0.510033783 | 0.702205995 | 0.004075653 | 0.017297492 | 220.1760913 | 154.6678888 | ENSG00000206432 |
| KPNA1 | -0.510175016 | 0.702137255 | 1.95E-06 | 2.06E-05 | 613.6360099 | 430.6686771 | ENSG00000114030 |
| CCDC96 | -0.510568438 | 0.701945809 | 0.642068616 | 0.790074341 | 5.723884207 | 3.975645462 | ENSG00000173013 |
| NDUFA11 | -0.510701156 | 0.701881238 | 0.057192611 | 0.148388122 | 85.54610492 | 60.21585859 | ENSG00000174886 |
| TCTN1 | -0.510880832 | 0.70179383 | 0.045249548 | 0.12381868 | 88.46161684 | 62.16608353 | ENSG00000204852 |
| ZNF613 | -0.510994263 | 0.701738653 | 0.01213454 | 0.043193438 | 154.5465049 | 108.2963245 | ENSG00000176024 |
| ATF7IP2 | -0.511038326 | 0.701717221 | 0.001545607 | 0.007628021 | 252.3213077 | 176.8868447 | ENSG00000166669 |
| CCK | -0.51124148 | 0.701618416 | 0.576009127 | 0.740960103 | 7.867260938 | 5.476130977 | ENSG00000187094 |
| FLAD1 | -0.511444995 | 0.701519448 | 3.04E-11 | 7.15E-10 | 1852.929898 | 1300.29583 | ENSG00000160688 |
| NA | -0.511584159 | 0.701451782 | 0.499864986 | 0.680562524 | 13.45720197 | 9.471097087 | ENSG00000260526 |
| RBM8A | -0.511872163 | 0.701311766 | 1.71E-06 | 1.82E-05 | 820.8496774 | 575.1271794 | ENSG00000265241 |
| ZNF514 | -0.511915237 | 0.701290827 | 0.009643437 | 0.03567673 | 210.4666155 | 148.012243 | ENSG00000144026 |
| SEPT5-GP1BB | -0.51243084 | 0.701040238 | 0.598252695 | 0.758154903 | 6.437917191 | 4.512303431 | ENSG00000284874 |
| PRR36 | -0.512693552 | 0.700912592 | 1.68E-05 | 0.000142246 | 710.978098 | 498.4981635 | ENSG00000183248 |
| DNAJC9 | -0.512810689 | 0.700855685 | 9.55E-06 | 8.57E-05 | 531.8224253 | 372.7321699 | ENSG00000213551 |
| FBXL19-AS1 | -0.512841853 | 0.700840546 | 0.281512532 | 0.472775758 | 26.96115628 | 18.83259416 | ENSG00000260852 |
| LLGL1 | -0.513088883 | 0.700720552 | 8.82E-13 | 2.65E-11 | 3179.272166 | 2228.561703 | ENSG00000131899 |
| NA | -0.513114 | 0.700708353 | 0.514271633 | 0.692702668 | 9.763744691 | 6.848906758 | ENSG00000285077 |
| ATG9B | -0.51332151 | 0.700607574 | 0.250246547 | 0.438754092 | 34.96009793 | 24.59051329 | ENSG00000181652 |
| C9orf129 | -0.513353902 | 0.700591844 | 0.00896413 | 0.033591021 | 159.9065176 | 112.171943 | ENSG00000204352 |
| C18orf32 | -0.513360745 | 0.700588521 | 0.302890433 | 0.496398557 | 29.36945784 | 20.65989589 | ENSG00000177576 |
| NCDN | -0.513365916 | 0.70058601 | 1.67E-08 | 2.59E-07 | 1155.107381 | 809.5680408 | ENSG00000020129 |
| TAP1 | -0.51358869 | 0.700477837 | 8.92E-05 | 0.000629771 | 385.496402 | 269.9632796 | ENSG00000168394 |
| NA | -0.51389755 | 0.700327891 | 0.601707069 | 0.761023531 | 5.42560881 | 3.8033457 | ENSG00000286129 |
| RHPN1 | -0.513914916 | 0.700319461 | 0.059210216 | 0.152395797 | 882.4998883 | 618.2812841 | ENSG00000158106 |
| ASIC1 | -0.51401159 | 0.700272535 | 0.007121436 | 0.027798798 | 163.8364124 | 114.7516932 | ENSG00000110881 |
| PDK1 | -0.51418348 | 0.700189106 | 0.007339381 | 0.028469669 | 208.0279023 | 145.2097885 | ENSG00000152256 |
| NA | -0.514713899 | 0.699931722 | 0.549960573 | 0.721319741 | 7.287870184 | 5.078467662 | ENSG00000217261 |
| FEN1 | -0.514867678 | 0.69985712 | 1.99E-12 | 5.61E-11 | 2272.312612 | 1590.962393 | ENSG00000168496 |
| CARHSP1 | -0.514905635 | 0.699838707 | 2.66E-10 | 5.42E-09 | 1516.047952 | 1061.124519 | ENSG00000153048 |
| HEATR1 | -0.515183164 | 0.699704093 | 2.98E-11 | 7.04E-10 | 2771.707548 | 1938.444506 | ENSG00000119285 |
| KATNB1 | -0.515416823 | 0.699590778 | 2.79E-10 | 5.65E-09 | 1535.590677 | 1075.01537 | ENSG00000140854 |
| DUSP2 | -0.515455395 | 0.699572074 | 0.012870161 | 0.045229925 | 136.1142755 | 95.38711597 | ENSG00000158050 |
| ETHE1 | -0.515552318 | 0.699525077 | 0.10631397 | 0.237428982 | 73.046672 | 51.24822599 | ENSG00000105755 |
| LRRC46 | -0.515578156 | 0.699512549 | 0.58028613 | 0.74434377 | 6.464450271 | 4.519789505 | ENSG00000141294 |
| NA | -0.515658636 | 0.699473528 | 0.567457934 | 0.734702435 | 7.298618699 | 5.085953736 | ENSG00000232811 |
| LINC02805 | -0.515664984 | 0.69947045 | 0.638046171 | 0.787529817 | 4.442990421 | 3.116957809 | ENSG00000232265 |
| NA | -0.515689175 | 0.699458722 | 0.438624221 | 0.627079706 | 12.36972904 | 8.690742605 | ENSG00000271119 |
| DNAJB11 | -0.51584983 | 0.699380836 | 3.38E-10 | 6.76E-09 | 1273.569319 | 890.8208154 | ENSG00000090520 |
| CYB5B | -0.515870591 | 0.699370772 | 9.16E-13 | 2.74E-11 | 2647.974569 | 1851.74498 | ENSG00000103018 |
| THAP7-AS1 | -0.516185354 | 0.699218202 | 0.491434649 | 0.674135874 | 10.39817844 | 7.317966823 | ENSG00000230513 |
| PRXL2A | -0.516416015 | 0.699106418 | 1.21E-09 | 2.21E-08 | 1351.498025 | 944.9172078 | ENSG00000122378 |
| DYRK3 | -0.516566054 | 0.699033715 | 0.002712078 | 0.012230085 | 203.4030363 | 142.1511191 | ENSG00000143479 |
| MPZL2 | -0.516958079 | 0.698843792 | 0.235934873 | 0.421016555 | 28.87470237 | 20.14157089 | ENSG00000149573 |
| NA | -0.516963238 | 0.698841293 | 0.608333251 | 0.766318797 | 5.448984977 | 3.810831773 | ENSG00000256591 |
| SLC8A1 | -0.517172575 | 0.698739898 | 0.022373126 | 0.070825037 | 119.2761053 | 83.25639819 | ENSG00000183023 |
| ARHGEF28 | -0.517199286 | 0.698726961 | 0.310261432 | 0.50411023 | 21.31824678 | 14.95091529 | ENSG00000214944 |
| ERBB3 | -0.51729536 | 0.698680432 | 4.03E-09 | 6.86E-08 | 1205.135389 | 842.2527332 | ENSG00000065361 |
| DDR1 | -0.517350087 | 0.698653929 | 2.74E-17 | 1.34E-15 | 6246.460878 | 4364.337783 | ENSG00000204580 |
| TSPAN6 | -0.51745864 | 0.698601362 | 1.13E-09 | 2.08E-08 | 1658.835319 | 1158.02177 | ENSG00000000003 |
| PTENP1 | -0.517498811 | 0.69858191 | 0.642241516 | 0.790182173 | 5.058482691 | 3.582219028 | ENSG00000237984 |
| HACD2 | -0.517604228 | 0.698530867 | 1.68E-05 | 0.000142425 | 844.2882556 | 589.0068254 | ENSG00000206527 |
| MYOM2 | -0.518037704 | 0.698321016 | 0.296432242 | 0.489406304 | 30.02608767 | 20.88372211 | ENSG00000036448 |
| C19orf57 | -0.518094877 | 0.698293342 | 0.043364414 | 0.119939719 | 86.59441712 | 60.50783236 | ENSG00000132016 |
| SERTM1 | -0.518343326 | 0.698173098 | 0.278355774 | 0.469483037 | 25.41433034 | 17.68843113 | ENSG00000180440 |
| NA | -0.518382513 | 0.698154135 | 0.511396936 | 0.690531168 | 8.83922851 | 6.181289862 | ENSG00000219507 |
| UBXN10 | -0.518714721 | 0.69799339 | 0.211366752 | 0.390897706 | 33.69419195 | 23.51014932 | ENSG00000162543 |
| PCNX3 | -0.518929286 | 0.697889589 | 3.73E-11 | 8.68E-10 | 1794.327593 | 1252.789524 | ENSG00000197136 |
| CHRNA7 | -0.51896559 | 0.697872027 | 0.045803744 | 0.125007598 | 90.81557447 | 63.31663333 | ENSG00000175344 |
| RAB34 | -0.518981718 | 0.697864226 | 2.28E-15 | 9.13E-14 | 3277.757424 | 2287.075365 | ENSG00000109113 |
| B3GALT1 | -0.51902231 | 0.697844591 | 0.003096255 | 0.013689407 | 303.0924449 | 211.1406796 | ENSG00000172318 |
| FOXP4-AS1 | -0.51908406 | 0.697814722 | 0.554644808 | 0.724709888 | 7.370626335 | 5.119808498 | ENSG00000234753 |

| | | | | | | | |
|---|---|---|---|---|---|---|---|
| DBI | -0.51939883 | 0.697662488 | 5.01E-13 | 1.53E-11 | 1917.610216 | 1337.823091 | ENSG00000155368 |
| NELFCD | -0.519566101 | 0.697581603 | 5.07E-11 | 1.15E-09 | 1358.196929 | 947.5671925 | ENSG00000101158 |
| NRSN2 | -0.519858625 | 0.697440174 | 2.43E-05 | 0.000198946 | 615.713798 | 429.5815999 | ENSG00000125841 |
| ARF3 | -0.519906753 | 0.697416909 | 1.62E-15 | 6.63E-14 | 4208.400254 | 2934.084686 | ENSG00000134287 |
| SLC15A1 | -0.52000528 | 0.697369281 | 0.171446037 | 0.337108089 | 43.12110187 | 30.10038741 | ENSG00000088386 |
| DDX23 | -0.520739653 | 0.69701439 | 4.48E-16 | 1.92E-14 | 4662.667698 | 3250.09927 | ENSG00000174243 |
| ARHGAP22 | -0.520913503 | 0.696930403 | 0.00013703 | 0.000918209 | 405.9738583 | 283.3033936 | ENSG00000128805 |
| NA | -0.520932238 | 0.696921352 | 0.59906191 | 0.758920946 | 5.484988795 | 3.822116694 | ENSG00000231381 |
| MACROH2A2 | -0.520942825 | 0.696916238 | 4.15E-10 | 8.13E-09 | 1773.381975 | 1235.21363 | ENSG00000099284 |
| NUDT10 | -0.520989216 | 0.696893829 | 0.000104256 | 0.000722887 | 557.3759076 | 388.8410582 | ENSG00000122824 |
| RNF38 | -0.521213399 | 0.696785545 | 4.70E-06 | 4.58E-05 | 633.6954839 | 441.1316823 | ENSG00000137075 |
| ALKBH8 | -0.52129744 | 0.696744957 | 0.000989743 | 0.005220234 | 300.6607672 | 208.9692248 | ENSG00000137760 |
| AXIN2 | -0.521502973 | 0.696645702 | 0.000569914 | 0.003215459 | 261.4177718 | 182.0782648 | ENSG00000168646 |
| MTG1 | -0.521528565 | 0.696633345 | 0.058253265 | 0.150455131 | 74.77311106 | 52.17385027 | ENSG00000148824 |
| SNHG26 | -0.521530652 | 0.696632337 | 0.019403189 | 0.062914399 | 134.7196577 | 94.00696574 | ENSG00000228649 |
| NA | -0.521613254 | 0.696592452 | 0.603881864 | 0.762941136 | 5.511521874 | 3.840887688 | ENSG00000287601 |
| NA | -0.521908663 | 0.696449831 | 0.055617348 | 0.145000975 | 85.700771 | 59.89371729 | ENSG00000213612 |
| KREMEN2 | -0.522191546 | 0.696313285 | 0.00907128 | 0.03392395 | 195.0964462 | 135.6014574 | ENSG00000131650 |
| PCDHB11 | -0.522294582 | 0.696263557 | 0.541162502 | 0.714177287 | 7.364312509 | 5.097238657 | ENSG00000197479 |
| GPR37L1 | -0.522745804 | 0.696045825 | 0.607137405 | 0.765458529 | 5.40854647 | 3.75451879 | ENSG00000170075 |
| TM7SF2 | -0.5229759 | 0.695934821 | 8.36E-08 | 1.14E-06 | 1041.59275 | 724.6804843 | ENSG00000149809 |
| C2CD4D | -0.523128737 | 0.695861099 | 0.250163247 | 0.438674996 | 30.70539461 | 21.4172425 | ENSG00000225556 |
| KCNA5 | -0.523719165 | 0.695576374 | 0.476675487 | 0.661082271 | 11.19181066 | 7.715080485 | ENSG00000130037 |
| DHX38 | -0.523956232 | 0.695462084 | 2.67E-12 | 7.34E-11 | 2267.679267 | 1577.842731 | ENSG00000140829 |
| ZNF350-AS1 | -0.524205282 | 0.695342038 | 0.487707132 | 0.670809943 | 9.785843081 | 6.788794928 | ENSG00000269235 |
| IRF8 | -0.524586988 | 0.69515809 | 0.191640664 | 0.36467997 | 34.11448193 | 23.68993515 | ENSG00000140968 |
| TAF1B | -0.525127193 | 0.694897843 | 0.000124817 | 0.000848625 | 349.981184 | 243.2919173 | ENSG00000115750 |
| S100A13 | -0.525158051 | 0.694882979 | 0.239579872 | 0.425477961 | 32.57702903 | 22.73403171 | ENSG00000189171 |
| RAMP2-AS1 | -0.525209714 | 0.694858096 | 0.204542266 | 0.382541509 | 37.21226023 | 25.96317745 | ENSG00000197291 |
| HMSD | -0.525240378 | 0.694843328 | 0.295928651 | 0.488836151 | 23.59132739 | 16.40582309 | ENSG00000221887 |
| LYPD6 | -0.525305013 | 0.694812198 | 0.003910495 | 0.016726773 | 177.5551869 | 123.4879019 | ENSG00000187123 |
| NCAPG | -0.525350829 | 0.694790133 | 1.08E-07 | 1.43E-06 | 1160.723103 | 806.0808474 | ENSG00000109805 |
| PNKD | -0.525834705 | 0.694557142 | 1.52E-05 | 0.000130692 | 669.6752638 | 465.8613824 | ENSG00000127838 |
| PWWP3B | -0.525847049 | 0.694551199 | 0.356965702 | 0.550599395 | 16.75649639 | 11.63864984 | ENSG00000157502 |
| NA | -0.525910457 | 0.694520673 | 0.034773403 | 0.10073308 | 153.6414339 | 107.0126574 | ENSG00000280128 |
| NA | -0.526095574 | 0.694431562 | 0.379769236 | 0.572904271 | 15.32389803 | 10.61471047 | ENSG00000233426 |
| NA | -0.526199222 | 0.694381674 | 0.528344864 | 0.704217496 | 8.317240906 | 5.727313564 | ENSG00000253170 |
| SIX5 | -0.526365265 | 0.694301761 | 0.0096982 | 0.03584048 | 164.8277048 | 114.1959378 | ENSG00000177045 |
| NA | -0.526508732 | 0.69423272 | 0.603341287 | 0.762512924 | 6.352004128 | 4.384593698 | ENSG00000214784 |
| NA | -0.526517869 | 0.694228324 | 0.050687529 | 0.135249448 | 80.93306981 | 56.29608808 | ENSG00000269958 |
| PCDHB7 | -0.526593472 | 0.694191944 | 0.458712244 | 0.645687912 | 13.14314201 | 9.039029357 | ENSG00000113212 |
| ALG1 | -0.526641185 | 0.694168986 | 7.97E-06 | 7.33E-05 | 525.2576529 | 364.7659068 | ENSG00000033011 |
| NA | -0.526680035 | 0.694150293 | 0.552036023 | 0.722967471 | 7.328308691 | 5.048411747 | ENSG00000225137 |
| TMEM26 | -0.526741005 | 0.694120958 | 0.252206317 | 0.440938768 | 29.93879913 | 20.6852769 | ENSG00000196932 |
| NA | -0.526754154 | 0.694114632 | 0.225208328 | 0.408093511 | 31.84918831 | 22.22374243 | ENSG00000241506 |
| GFOD2 | -0.526855923 | 0.69406567 | 6.66E-08 | 9.17E-07 | 887.907495 | 616.3911365 | ENSG00000141098 |
| MAPK4 | -0.526906773 | 0.694041207 | 0.002251963 | 0.010508029 | 199.5174359 | 138.486768 | ENSG00000141639 |
| LNP1 | -0.527415661 | 0.693796438 | 0.004604169 | 0.019163187 | 180.9673335 | 125.2869712 | ENSG00000206535 |
| NA | -0.527557574 | 0.693728195 | 0.679483797 | 0.815010735 | 4.397515865 | 3.045561059 | ENSG00000226009 |
| LRP3 | -0.527950125 | 0.69353946 | 5.18E-09 | 8.69E-08 | 1745.673179 | 1210.567427 | ENSG00000130881 |
| ZNF880 | -0.527988012 | 0.693521247 | 0.005606498 | 0.022731462 | 166.4209748 | 115.4304834 | ENSG00000221923 |
| NUP88 | -0.528203867 | 0.693417491 | 1.29E-11 | 3.23E-10 | 1675.842181 | 1161.381048 | ENSG00000108559 |
| TEAD2 | -0.528504322 | 0.693273095 | 1.62E-12 | 4.64E-11 | 2654.619444 | 1840.620004 | ENSG00000074219 |
| FJX1 | -0.528592906 | 0.693230528 | 1.58E-05 | 0.000135309 | 506.4591691 | 351.1304666 | ENSG00000179431 |
| NA | -0.528679416 | 0.693188961 | 0.213894349 | 0.393989055 | 35.29249801 | 24.413427 | ENSG00000275764 |
| RTBDN | -0.528726042 | 0.693166558 | 0.519916583 | 0.696907062 | 8.346930898 | 5.734799638 | ENSG00000132026 |
| NA | -0.529318417 | 0.692882 | 0.353213833 | 0.546358303 | 20.1328345 | 13.93771118 | ENSG00000280433 |
| ELOF1 | -0.52954552 | 0.692772938 | 1.59E-09 | 2.87E-08 | 1406.817225 | 975.0693936 | ENSG00000130165 |
| ACE | -0.529607595 | 0.692743131 | 0.140673029 | 0.291930794 | 53.17237345 | 36.98347535 | ENSG00000159640 |
| CBY1 | -0.529654087 | 0.692720807 | 0.016121279 | 0.054138754 | 151.6591347 | 104.905391 | ENSG00000100211 |
| ORC1 | -0.529748269 | 0.692675586 | 2.87E-11 | 6.80E-10 | 1397.994661 | 968.0629989 | ENSG00000085840 |
| CETN2 | -0.52975627 | 0.692671745 | 2.17E-05 | 0.000180042 | 508.0024473 | 351.4922914 | ENSG00000147400 |
| SEPTIN2 | -0.529981894 | 0.692563426 | 1.50E-23 | 1.28E-21 | 8885.323071 | 6154.008887 | ENSG00000168385 |
| PANO1 | -0.530025689 | 0.692542402 | 0.578848906 | 0.743117381 | 6.070791072 | 4.238551003 | ENSG00000274897 |
| GNL1 | -0.530451766 | 0.692337901 | 6.23E-13 | 1.90E-11 | 2020.180681 | 1398.525244 | ENSG00000204590 |
| MXD4 | -0.530457516 | 0.692335142 | 1.06E-07 | 1.42E-06 | 1011.554268 | 700.2912761 | ENSG00000123933 |
| NA | -0.530634349 | 0.692250287 | 0.597608241 | 0.757703763 | 5.511521874 | 3.8033457 | ENSG00000275329 |
| MED24 | -0.530690054 | 0.692223558 | 9.70E-16 | 4.03E-14 | 2683.471455 | 1857.844943 | ENSG00000008838 |
| MLH1 | -0.530834993 | 0.692154018 | 2.58E-09 | 4.49E-08 | 1159.510226 | 802.1176893 | ENSG00000076242 |
| NAA38 | -0.531105098 | 0.692024443 | 0.000136636 | 0.000916213 | 368.1948123 | 255.029097 | ENSG00000183011 |
| RBCK1 | -0.53113465 | 0.692010268 | 5.28E-08 | 7.44E-07 | 1247.080271 | 863.7175373 | ENSG00000125826 |
| ADAMTS3 | -0.531777043 | 0.691702203 | 0.0523686 | 0.1384198 | 100.2746322 | 69.06943951 | ENSG00000156140 |
| FADS6 | -0.532092219 | 0.691551108 | 0.313424053 | 0.506988374 | 20.07463459 | 13.94194806 | ENSG00000172782 |
| KCNH2 | -0.532114233 | 0.691540556 | 0.000210542 | 0.001343265 | 436.2960566 | 301.7017299 | ENSG00000055118 |

| | | | | | | | |
|---|---|---|---|---|---|---|---|
| VILL | -0.532312973 | 0.691445299 | 0.041830649 | 0.116619922 | 96.17609083 | 66.75011012 | ENSG00000136059 |
| NA | -0.532752922 | 0.691234475 | 0.529829383 | 0.705507653 | 8.548544716 | 5.945191042 | ENSG00000280828 |
| C1orf198 | -0.532952015 | 0.69113909 | 3.34E-08 | 4.88E-07 | 776.6937029 | 536.8439026 | ENSG00000119280 |
| IFT22 | -0.533002788 | 0.691114768 | 0.00102371 | 0.005379486 | 232.9521072 | 161.1606879 | ENSG00000128581 |
| MYD88 | -0.533112248 | 0.691062334 | 3.40E-06 | 3.42E-05 | 651.0615495 | 449.9740186 | ENSG00000172936 |
| SOWAHD | -0.533137326 | 0.691050321 | 0.051947038 | 0.137501628 | 81.83905047 | 56.37398322 | ENSG00000187808 |
| SUSD5 | -0.533215657 | 0.691012801 | 0.041697526 | 0.116388848 | 92.41043051 | 63.62380248 | ENSG00000173705 |
| TMEM191A | -0.53348763 | 0.690882546 | 0.074371983 | 0.181417523 | 72.89516282 | 50.33548686 | ENSG00000226287 |
| CCDC88A | -0.533611885 | 0.690823045 | 1.46E-09 | 2.64E-08 | 1111.171954 | 767.8043954 | ENSG00000115355 |
| NA | -0.533750288 | 0.690756775 | 0.329818985 | 0.524185173 | 21.01040295 | 14.4705703 | ENSG00000247373 |
| NA | -0.533870745 | 0.690699103 | 0.428369447 | 0.618622357 | 13.04144438 | 9.043266237 | ENSG00000239377 |
| RBPMS-AS1 | -0.534465225 | 0.690414551 | 0.659775296 | 0.80185412 | 4.433519683 | 3.049359905 | ENSG00000254109 |
| H2BC4 | -0.534467669 | 0.690413381 | 0.445790592 | 0.633889072 | 11.48692914 | 7.932957964 | ENSG00000180596 |
| OSBPL11 | -0.534644843 | 0.690328598 | 4.75E-08 | 6.76E-07 | 817.8768525 | 564.6142883 | ENSG00000144909 |
| NA | -0.534699479 | 0.690302455 | 0.621735091 | 0.775152125 | 4.469523501 | 3.075616974 | ENSG00000237991 |
| HDGF | -0.534890255 | 0.690211178 | 4.25E-17 | 2.04E-15 | 9700.670406 | 6694.818797 | ENSG00000143321 |
| GTPBP8 | -0.535477127 | 0.689930466 | 0.00578804 | 0.023320767 | 165.3497901 | 113.8996155 | ENSG00000163607 |
| LTBP1 | -0.536178013 | 0.689595367 | 1.53E-10 | 3.25E-09 | 1253.50417 | 864.2080191 | ENSG00000049323 |
| HM13-AS1 | -0.536287327 | 0.689543118 | 0.623400657 | 0.776590118 | 4.475837326 | 3.075616974 | ENSG00000230613 |
| TMEM74B | -0.536304583 | 0.68953487 | 0.284125311 | 0.475123329 | 31.13811685 | 21.59041833 | ENSG00000125895 |
| GLCE | -0.536461478 | 0.689459887 | 0.000285479 | 0.001760755 | 382.4930752 | 263.5411044 | ENSG00000138604 |
| CA14 | -0.536474255 | 0.689453781 | 6.45E-08 | 8.93E-07 | 864.8311051 | 596.1485594 | ENSG00000118298 |
| ARL6 | -0.536530887 | 0.689426717 | 0.047909434 | 0.129401553 | 85.16823028 | 58.6204188 | ENSG00000113966 |
| CTAGE8 | -0.536532206 | 0.689426087 | 0.595877839 | 0.756282946 | 6.056885644 | 4.197210168 | ENSG00000244693 |
| GJD3 | -0.536561247 | 0.689412209 | 0.234107028 | 0.419001939 | 29.09012392 | 20.07072379 | ENSG00000183153 |
| COMMD4 | -0.53657686 | 0.689404748 | 1.45E-10 | 3.10E-09 | 1614.162641 | 1113.634312 | ENSG00000140365 |
| KCTD17 | -0.536932065 | 0.689235031 | 7.14E-08 | 9.77E-07 | 862.3779524 | 594.5345634 | ENSG00000100379 |
| PI4KB | -0.537173933 | 0.68911949 | 1.11E-08 | 1.76E-07 | 916.1455616 | 631.6243896 | ENSG00000143393 |
| GALR2 | -0.537605402 | 0.688913425 | 0.514757491 | 0.692846128 | 7.965899351 | 5.476130977 | ENSG00000182687 |
| NA | -0.537722147 | 0.68885768 | 0.492465255 | 0.674850354 | 9.67329924 | 6.69537799 | ENSG00000285689 |
| EDARADD | -0.537867538 | 0.688788262 | 0.037499839 | 0.106767011 | 87.45690006 | 60.10593217 | ENSG00000186197 |
| SLAIN1 | -0.537943819 | 0.688751844 | 4.66E-06 | 4.54E-05 | 541.7857093 | 372.7710889 | ENSG00000139737 |
| SLC24A4 | -0.538051911 | 0.688700242 | 0.296324093 | 0.489314926 | 20.86205069 | 14.36543041 | ENSG00000140090 |
| ATP10A | -0.539043756 | 0.688226927 | 7.86E-05 | 0.000561666 | 340.7132771 | 234.3306715 | ENSG00000206190 |
| ANPEP | -0.539266476 | 0.688120688 | 0.069812262 | 0.172574767 | 69.07901454 | 47.64277584 | ENSG00000166825 |
| AP2A2 | -0.539296529 | 0.688106354 | 8.25E-12 | 2.12E-10 | 1572.077855 | 1081.664572 | ENSG00000183020 |
| NA | -0.539439751 | 0.688038047 | 0.444494044 | 0.632920173 | 11.61831676 | 7.993069793 | ENSG00000250539 |
| WRAP53 | -0.53948684 | 0.68801559 | 7.90E-08 | 1.08E-06 | 727.1414036 | 500.4772822 | ENSG00000141499 |
| NA | -0.539942924 | 0.687798119 | 0.288740004 | 0.480429772 | 24.83672102 | 17.03209915 | ENSG00000287678 |
| VPS9D1-AS1 | -0.5400579 | 0.687743307 | 0.058017802 | 0.150035152 | 85.17958016 | 58.84720807 | ENSG00000261373 |
| NIPSNAP3B | -0.540228525 | 0.687661974 | 0.506239819 | 0.685866141 | 10.43102534 | 7.164000022 | ENSG00000165028 |
| MED12 | -0.540239965 | 0.687656521 | 1.81E-10 | 3.80E-09 | 1196.781085 | 823.1490946 | ENSG00000184634 |
| LRFN4 | -0.540289467 | 0.687632926 | 5.61E-06 | 5.36E-05 | 846.5527448 | 582.6396668 | ENSG00000173621 |
| PROKR1 | -0.540385336 | 0.687587234 | 0.087406745 | 0.204910359 | 58.83509623 | 40.47844941 | ENSG00000169618 |
| NA | -0.540522508 | 0.687521861 | 0.476032225 | 0.660734012 | 9.90460305 | 6.76253786 | ENSG00000214776 |
| RNF135 | -0.540589912 | 0.68748974 | 0.003908013 | 0.016720017 | 179.4473337 | 123.3179753 | ENSG00000181481 |
| AGAP9 | -0.54073234 | 0.687421872 | 0.113494746 | 0.24917219 | 53.51296649 | 36.92347514 | ENSG00000204172 |
| NAT8L | -0.540798612 | 0.687390295 | 1.17E-07 | 1.55E-06 | 966.9745979 | 664.8691625 | ENSG00000185818 |
| BRD3OS | -0.541018176 | 0.687285689 | 0.000182723 | 0.001187857 | 329.2046404 | 226.1155989 | ENSG00000235106 |
| KLHDC8B | -0.541244934 | 0.687177672 | 0.000143221 | 0.000954156 | 353.5075202 | 243.2703351 | ENSG00000185909 |
| PTPRT | -0.541383552 | 0.687111649 | 0.027401131 | 0.083526452 | 110.9212745 | 76.26089908 | ENSG00000196090 |
| NA | -0.541552702 | 0.687031093 | 0.100039018 | 0.226727797 | 63.9126181 | 44.00424383 | ENSG00000259959 |
| NA | -0.541761892 | 0.686931481 | 0.516495099 | 0.694178403 | 8.840506286 | 6.034809134 | ENSG00000184441 |
| DUSP16 | -0.541765163 | 0.686929924 | 1.95E-14 | 6.98E-13 | 2470.790989 | 1696.765368 | ENSG00000111266 |
| FPGT | -0.542299747 | 0.686675432 | 0.1909583 | 0.363743179 | 36.33449638 | 24.93511282 | ENSG00000254685 |
| MIR4442 | -0.542336055 | 0.686658151 | 0.275048466 | 0.466037706 | 29.03074393 | 19.92033259 | ENSG00000283971 |
| NA | -0.542341082 | 0.686655758 | 0.028407141 | 0.085874726 | 96.67716012 | 66.4077721 | ENSG00000287086 |
| CCDC34 | -0.542413648 | 0.686621221 | 1.36E-06 | 1.47E-05 | 564.3787314 | 387.1780834 | ENSG00000109881 |
| NA | -0.542495047 | 0.686582482 | 0.002202591 | 0.010316607 | 227.2904668 | 156.3274541 | ENSG00000214796 |
| NTN5 | -0.542513835 | 0.686573541 | 0.573953326 | 0.739436906 | 6.619311756 | 4.594873483 | ENSG00000142233 |
| EEF1AKMT3 | -0.54262186 | 0.686522134 | 0.183482739 | 0.354015104 | 36.16888638 | 24.72472141 | ENSG00000123427 |
| VAT1L | -0.542700333 | 0.686484793 | 2.06E-13 | 6.56E-12 | 2576.490418 | 1768.133877 | ENSG00000171724 |
| MRTO4 | -0.542728244 | 0.686471512 | 1.29E-12 | 3.72E-11 | 1745.309691 | 1198.110744 | ENSG00000053372 |
| CNPY4 | -0.542969452 | 0.686356749 | 0.003672925 | 0.015816449 | 176.1946939 | 121.0610192 | ENSG00000166997 |
| NA | -0.543193649 | 0.686250096 | 0.065071386 | 0.163796424 | 73.45503338 | 50.57994782 | ENSG00000229321 |
| DHRS3 | -0.543313453 | 0.686193111 | 0.000686963 | 0.003793851 | 239.9821405 | 164.7836981 | ENSG00000162496 |
| GDF7 | -0.543417164 | 0.686143784 | 0.273208011 | 0.4643838 | 25.34726105 | 17.43031212 | ENSG00000143869 |
| NA | -0.543504712 | 0.686102148 | 0.278728476 | 0.469845709 | 22.62755278 | 15.53964936 | ENSG00000260362 |
| GAS1 | -0.543828086 | 0.685948378 | 0.012066678 | 0.042994176 | 131.7862867 | 90.20576993 | ENSG00000180447 |
| CHRAC1 | -0.544046999 | 0.685844301 | 3.61E-07 | 4.37E-06 | 652.1494809 | 447.2854412 | ENSG00000104472 |
| MON1A | -0.544404699 | 0.685674274 | 0.001271259 | 0.006435813 | 297.4246936 | 204.3900251 | ENSG00000164077 |
| ZNF443 | -0.544601849 | 0.685580581 | 0.048837666 | 0.131315437 | 78.07368326 | 53.5161321 | ENSG00000180855 |
| RIT2 | -0.544742892 | 0.685513559 | 0.434505051 | 0.624209861 | 13.24937203 | 9.167288744 | ENSG00000152214 |
| LOC100130950 | -0.545581598 | 0.685115154 | 0.596535928 | 0.756870826 | 5.428765723 | 3.68312204 | ENSG00000261879 |

| | | | | | | | |
|---|---|---|---|---|---|---|---|
| NA | -0.545794454 | 0.68501408 | 0.546795754 | 0.718495899 | 7.109632533 | 4.868625911 | ENSG00000275325 |
| LOC102724159 | -0.545827621 | 0.684998332 | 2.20E-07 | 2.77E-06 | 896.6490698 | 614.4127278 | ENSG00000275464 |
| LINC01842 | -0.545879291 | 0.684973799 | 0.652808925 | 0.797217744 | 4.486585841 | 3.056845979 | ENSG00000267147 |
| CSAD | -0.545899431 | 0.684964237 | 0.160329532 | 0.320707646 | 59.97110293 | 41.29028553 | ENSG00000139631 |
| LINC00852 | -0.546574242 | 0.684643924 | 0.418181096 | 0.609369324 | 13.29296745 | 9.031543283 | ENSG00000231177 |
| NRIP2 | -0.546691612 | 0.684588227 | 0.167978879 | 0.331804386 | 41.354322 | 28.36369146 | ENSG00000053702 |
| RNF208 | -0.546761858 | 0.684554895 | 0.039363669 | 0.111061994 | 109.7675291 | 75.12400733 | ENSG00000212864 |
| LINC00632 | -0.546829844 | 0.684522637 | 0.030542691 | 0.090834119 | 101.5074959 | 69.49836122 | ENSG00000203930 |
| MMP24 | -0.547159247 | 0.684366361 | 0.000170792 | 0.00111617 | 324.0057803 | 221.6185941 | ENSG00000125966 |
| SLC1A4 | -0.54733979 | 0.684280723 | 0.075597187 | 0.183560626 | 73.10940429 | 50.1599379 | ENSG00000115902 |
| L1TD1 | -0.547515125 | 0.684197565 | 2.87E-20 | 1.88E-18 | 14400.65761 | 9852.906151 | ENSG00000240563 |
| UNQ6494 | -0.547533531 | 0.684188836 | 0.515835245 | 0.693593328 | 8.06256093 | 5.494901971 | ENSG00000237372 |
| WDR76 | -0.54755547 | 0.684178432 | 4.35E-05 | 0.000332828 | 382.4870923 | 261.3431693 | ENSG00000092470 |
| SCD | -0.54777339 | 0.684075094 | 1.04E-19 | 6.47E-18 | 28811.44875 | 19708.10872 | ENSG00000099194 |
| RIC8B | -0.547929167 | 0.684001234 | 0.000112633 | 0.000775458 | 318.5644096 | 217.81536 | ENSG00000111785 |
| PDE3B | -0.547996239 | 0.683969435 | 0.000389302 | 0.002298606 | 362.3058066 | 247.6445913 | ENSG00000152270 |
| PTBP2 | -0.54807333 | 0.683932888 | 3.38E-07 | 4.11E-06 | 784.9993008 | 536.1342433 | ENSG00000117569 |
| NA | -0.548436409 | 0.683760786 | 0.237496019 | 0.422736887 | 25.19881109 | 17.23120563 | ENSG00000283544 |
| PCDHGB7 | -0.548591232 | 0.683687412 | 0.001578936 | 0.007766657 | 198.2662321 | 135.6440007 | ENSG00000254122 |
| PCGF5 | -0.548609255 | 0.683678871 | 0.020574291 | 0.066133178 | 124.9479155 | 85.35222778 | ENSG00000180628 |
| SHLD3 | -0.548713776 | 0.683629342 | 0.252329668 | 0.441112815 | 24.47402959 | 16.67577667 | ENSG00000253251 |
| PNMA3 | -0.548742754 | 0.68361561 | 0.0003413 | 0.002050543 | 327.3192134 | 223.7280188 | ENSG00000183837 |
| INTS6L | -0.548975344 | 0.683505407 | 0.000497241 | 0.002850562 | 285.4298928 | 195.4480999 | ENSG00000165359 |
| ARHGAP18 | -0.54911855 | 0.683437564 | 0.012365727 | 0.043841512 | 123.7394127 | 84.64849462 | ENSG00000146376 |
| EMILIN3 | -0.549147648 | 0.683423779 | 0.000981708 | 0.005181207 | 256.6372475 | 174.968714 | ENSG00000183798 |
| TCF3 | -0.54920815 | 0.68339512 | 8.15E-17 | 3.77E-15 | 6997.996922 | 4782.244656 | ENSG00000071564 |
| NA | -0.549396204 | 0.683306046 | 0.453482426 | 0.640808424 | 11.56219139 | 7.944242884 | ENSG00000240668 |
| PNPLA6 | -0.549500086 | 0.683256845 | 4.98E-14 | 1.71E-12 | 2678.141914 | 1829.31482 | ENSG00000032444 |
| URM1 | -0.549534253 | 0.683240664 | 1.39E-09 | 2.53E-08 | 1244.790947 | 850.8841369 | ENSG00000167118 |
| SFXN4 | -0.549664636 | 0.683178919 | 0.000117641 | 0.000806945 | 319.4577626 | 218.3268602 | ENSG00000183605 |
| POU4F1 | -0.549686126 | 0.683168743 | 0.617994562 | 0.772868503 | 5.40854647 | 3.656864972 | ENSG00000152192 |
| ZNF580 | -0.549961861 | 0.683038185 | 0.001779453 | 0.008597597 | 225.6990608 | 154.2175998 | ENSG00000213015 |
| WIF1 | -0.550342448 | 0.682858021 | 0.088758088 | 0.207288609 | 58.04008853 | 39.55910027 | ENSG00000156076 |
| CCT6P3 | -0.550467623 | 0.682798776 | 0.249377889 | 0.437852387 | 24.59406734 | 16.7358885 | ENSG00000234585 |
| NA | -0.550767305 | 0.682656957 | 0.47828611 | 0.662348428 | 10.00126463 | 6.770023934 | ENSG00000225022 |
| MGAT4C | -0.550914957 | 0.682587095 | 0.098042441 | 0.223216685 | 60.10564746 | 40.86438977 | ENSG00000182050 |
| DNMT3B | -0.550942513 | 0.682574057 | 2.46E-28 | 3.18E-26 | 38489.18972 | 26271.17079 | ENSG00000088305 |
| YTHDF3-AS1 | -0.550989588 | 0.682551785 | 0.350087982 | 0.543382519 | 17.15646942 | 11.72133152 | ENSG00000270673 |
| NA | -0.551122351 | 0.682488976 | 0.352408445 | 0.545516575 | 15.62848726 | 10.65973853 | ENSG00000260005 |
| GEMIN5 | -0.551384711 | 0.682364874 | 1.31E-12 | 3.77E-11 | 1528.61144 | 1043.484245 | ENSG00000082516 |
| ITGA1 | -0.551602223 | 0.682262003 | 0.006928485 | 0.027172376 | 142.2330966 | 96.89746069 | ENSG00000213949 |
| NCAM2 | -0.551612945 | 0.682256933 | 0.232338005 | 0.417039061 | 29.44650152 | 19.95754817 | ENSG00000154654 |
| TUT1 | -0.551900158 | 0.682121122 | 0.357172987 | 0.550616112 | 16.91126018 | 11.61283081 | ENSG00000255508 |
| CA9 | -0.551955626 | 0.682094897 | 0.529976124 | 0.705644935 | 7.056566374 | 4.79722916 | ENSG00000107159 |
| ZNF25 | -0.552084163 | 0.682034128 | 0.504657305 | 0.684288598 | 9.67329924 | 6.594036944 | ENSG00000175395 |
| ZFYVE26 | -0.552956802 | 0.681621713 | 1.38E-08 | 2.17E-07 | 940.5842412 | 641.6897717 | ENSG00000072121 |
| NA | -0.55305362 | 0.681575971 | 0.314289104 | 0.50786627 | 21.34793677 | 14.59448119 | ENSG00000250132 |
| TMEM221 | -0.553066549 | 0.681569863 | 0.146778174 | 0.301094265 | 44.73567351 | 30.55056486 | ENSG00000188051 |
| EDNRA | -0.553678714 | 0.68128072 | 0.029082271 | 0.087359936 | 89.85948916 | 61.265617 | ENSG00000151617 |
| SRSF1 | -0.553721229 | 0.681260644 | 1.30E-05 | 0.000113402 | 1399.907508 | 953.30419 | ENSG00000136450 |
| NA | -0.553753388 | 0.681245458 | 0.00987956 | 0.036388666 | 126.8408515 | 86.39972469 | ENSG00000228709 |
| CXXC5-AS1 | -0.554271342 | 0.681000923 | 0.590904712 | 0.752754687 | 5.055325778 | 3.454509294 | ENSG00000250635 |
| FAM167A | -0.554469868 | 0.680907218 | 0.191399106 | 0.364433021 | 34.44570193 | 23.536518 | ENSG00000154319 |
| COQ2 | -0.554524272 | 0.680881541 | 0.000205505 | 0.001315205 | 376.197903 | 256.2804467 | ENSG00000173085 |
| EXOC6 | -0.554888018 | 0.680709893 | 0.007344608 | 0.028483985 | 136.6266946 | 92.87298832 | ENSG00000138190 |
| ZDHHC6 | -0.555379338 | 0.680478112 | 9.53E-08 | 1.28E-06 | 661.7640015 | 450.4290942 | ENSG00000023041 |
| ZNF615 | -0.555773412 | 0.680292264 | 0.004720534 | 0.019559666 | 152.6595896 | 103.7918335 | ENSG00000197619 |
| NA | -0.555850151 | 0.680256079 | 0.40565115 | 0.597357255 | 13.28221893 | 9.103378067 | ENSG00000260167 |
| RCAN1 | -0.556152365 | 0.680113595 | 0.003263042 | 0.014297215 | 172.6470333 | 117.2905406 | ENSG00000159200 |
| C1QL1 | -0.556157969 | 0.680110953 | 0.000486941 | 0.002798968 | 275.0872384 | 186.9091339 | ENSG00000131094 |
| YIF1A | -0.556538118 | 0.679931768 | 7.71E-12 | 2.00E-10 | 1299.303535 | 883.6095258 | ENSG00000174851 |
| ZNF423 | -0.556819368 | 0.67979923 | 1.75E-11 | 4.27E-10 | 1449.020485 | 984.8095195 | ENSG00000102935 |
| SFRP5 | -0.556877177 | 0.679771991 | 0.057972542 | 0.149959958 | 72.01286658 | 48.81991757 | ENSG00000120057 |
| DM1-AS | -0.557243669 | 0.679599328 | 0.382162409 | 0.574965346 | 14.35834199 | 9.812335799 | ENSG00000267395 |
| NA | -0.557344757 | 0.679551711 | 0.489223022 | 0.672081774 | 9.13119008 | 6.233915618 | ENSG00000215006 |
| CD24 | -0.557345649 | 0.679551291 | 1.06E-12 | 3.14E-11 | 30869.16534 | 20976.57747 | ENSG00000272398 |
| FAM136A | -0.557726818 | 0.679371773 | 4.35E-13 | 1.35E-11 | 2218.726435 | 1507.067916 | ENSG00000035141 |
| NA | -0.557751088 | 0.679360344 | 0.26286362 | 0.452401739 | 22.70083819 | 15.36311272 | ENSG00000233325 |
| MGAT5B | -0.55811159 | 0.679190606 | 0.000226105 | 0.001432213 | 292.1036462 | 198.3713503 | ENSG00000167889 |
| H4C3 | -0.558429693 | 0.679040866 | 0.273734457 | 0.464894568 | 21.23864754 | 14.43302832 | ENSG00000197061 |
| DISP3 | -0.559280958 | 0.678640315 | 0.001559547 | 0.007681275 | 203.5061094 | 138.2924481 | ENSG00000204624 |
| RGMA | -0.559459148 | 0.6785565 | 3.53E-08 | 5.13E-07 | 874.7540406 | 593.5583601 | ENSG00000182175 |
| NA | -0.559599664 | 0.678490413 | 0.404573214 | 0.596238518 | 13.17736439 | 8.986953254 | ENSG00000286511 |
| MTHFD1 | -0.559741832 | 0.678423556 | 1.90E-17 | 9.46E-16 | 5165.196041 | 3503.96383 | ENSG00000100714 |

| | | | | | | | |
|---|---|---|---|---|---|---|---|
| NA | -0.560018319 | 0.678293551 | 0.306304927 | 0.500330777 | 18.80005463 | 12.76024304 | ENSG00000272070 |
| SMO | -0.560276869 | 0.678172002 | 1.37E-14 | 5.02E-13 | 2629.337712 | 1782.75037 | ENSG00000128602 |
| KLKB1 | -0.560311504 | 0.678155722 | 0.012868729 | 0.045229925 | 142.0362106 | 96.22929414 | ENSG00000164344 |
| MRO | -0.560611789 | 0.678014584 | 0.207653304 | 0.386604354 | 29.46672077 | 19.94301405 | ENSG00000134042 |
| NA | -0.560830428 | 0.67791184 | 0.464349685 | 0.650601375 | 11.35298597 | 7.771943121 | ENSG00000279452 |
| NOL11 | -0.560863212 | 0.677896435 | 4.63E-12 | 1.25E-10 | 1986.103729 | 1345.759656 | ENSG00000130935 |
| TMEM121 | -0.560940554 | 0.677860094 | 0.063552771 | 0.16106177 | 65.87440487 | 44.54426262 | ENSG00000184986 |
| GNB1L | -0.561017364 | 0.677824005 | 0.000266225 | 0.001655998 | 375.8965609 | 255.266948 | ENSG00000185838 |
| HOMER2 | -0.561154106 | 0.677759763 | 6.19E-06 | 5.87E-05 | 478.3045431 | 323.9098389 | ENSG00000103942 |
| PRKD1 | -0.561291854 | 0.677695053 | 0.002294477 | 0.010668799 | 178.7361645 | 120.9437183 | ENSG00000184304 |
| FBXO6 | -0.561376915 | 0.677655098 | 0.274983499 | 0.466033665 | 24.6175412 | 16.80392444 | ENSG00000116663 |
| NA | -0.562144693 | 0.677294557 | 0.131885245 | 0.278369335 | 41.41941445 | 28.13887756 | ENSG00000261663 |
| SDCBP2 | -0.562267842 | 0.677236745 | 0.030507638 | 0.090756098 | 83.75131879 | 56.68571567 | ENSG00000125775 |
| IRAK3 | -0.562348675 | 0.677198802 | 0.125622611 | 0.268545499 | 46.60612785 | 31.65338706 | ENSG00000090376 |
| SSBP1 | -0.562504895 | 0.677125476 | 4.28E-15 | 1.67E-13 | 3326.96365 | 2252.994961 | ENSG00000106028 |
| NA | -0.56307922 | 0.676855972 | 0.538516982 | 0.712004119 | 7.519173994 | 5.085953736 | ENSG00000267890 |
| NA | -0.563119314 | 0.676837161 | 0.344045756 | 0.537651269 | 19.48705087 | 13.1461834 | ENSG00000286606 |
| CDK2 | -0.563329156 | 0.676738722 | 1.99E-05 | 0.000165841 | 378.1838779 | 255.7097107 | ENSG00000123374 |
| KIAA1143 | -0.563667578 | 0.676579993 | 1.29E-07 | 1.68E-06 | 875.814973 | 591.7611772 | ENSG00000163807 |
| DHPS | -0.563710521 | 0.676559855 | 0.144785717 | 0.298162053 | 46.92018781 | 31.89752161 | ENSG00000095059 |
| ATP6V0A4 | -0.563776474 | 0.676528926 | 0.404243908 | 0.595900469 | 13.90836202 | 9.414672484 | ENSG00000105929 |
| LOC102723582 | -0.563839937 | 0.676499167 | 0.640554224 | 0.789444284 | 4.707141136 | 3.233382622 | ENSG00000272758 |
| LOC100505715 | -0.56402895 | 0.676410542 | 0.464340993 | 0.650601375 | 10.34185767 | 7.01415848 | ENSG00000267058 |
| NA | -0.56434348 | 0.67626309 | 0.461583637 | 0.648154491 | 9.649923073 | 6.492584279 | ENSG00000262227 |
| ATP8B2 | -0.564834556 | 0.676032937 | 4.14E-14 | 1.43E-12 | 1715.485638 | 1159.878118 | ENSG00000143515 |
| ZIK1 | -0.564875493 | 0.676013755 | 9.79E-08 | 1.31E-06 | 682.9966661 | 461.734103 | ENSG00000171649 |
| NA | -0.565234459 | 0.675845573 | 0.635800875 | 0.785909925 | 5.239779557 | 3.552163113 | ENSG00000260288 |
| GLIPR2 | -0.565353173 | 0.675789962 | 0.00206922 | 0.009785895 | 209.2536554 | 141.4092943 | ENSG00000122694 |
| TCEA2 | -0.565443224 | 0.675747782 | 1.89E-08 | 2.88E-07 | 826.2607794 | 558.9235777 | ENSG00000171703 |
| CPXM1 | -0.565458736 | 0.675740516 | 6.14E-12 | 1.61E-10 | 1788.343136 | 1209.296271 | ENSG00000088882 |
| POLA2 | -0.565536595 | 0.675704049 | 0.014644688 | 0.050087591 | 120.6424311 | 81.51970224 | ENSG00000014138 |
| EML3 | -0.565903814 | 0.675532079 | 3.16E-07 | 3.86E-06 | 620.6750481 | 419.5283074 | ENSG00000149499 |
| N4BP2L2-IT2 | -0.566268846 | 0.675361177 | 0.552906931 | 0.723596742 | 7.531801645 | 5.078467662 | ENSG00000281026 |
| MFSD1 | -0.56629732 | 0.675347848 | 0.007955486 | 0.030483144 | 147.5136078 | 99.72624358 | ENSG00000118855 |
| DCTN3 | -0.566453445 | 0.675274767 | 5.05E-05 | 0.000379567 | 338.2027743 | 228.6235546 | ENSG00000137100 |
| DGCR6L | -0.566895928 | 0.675067688 | 2.87E-09 | 4.95E-08 | 1277.870902 | 862.9431315 | ENSG00000128185 |
| POGLUT2 | -0.566918223 | 0.675057256 | 9.84E-07 | 1.10E-05 | 605.1510919 | 408.7921708 | ENSG00000134901 |
| GPR153 | -0.567218428 | 0.6749168 | 7.39E-05 | 0.000531282 | 343.0983228 | 231.5063487 | ENSG00000158292 |
| AZGP1 | -0.567680351 | 0.67470074 | 0.482366569 | 0.66583615 | 8.545387803 | 5.734799638 | ENSG00000160862 |
| NA | -0.567872965 | 0.674610667 | 0.034364162 | 0.099765879 | 100.9690473 | 68.36333322 | ENSG00000186301 |
| PCP4L1 | -0.568204483 | 0.674455665 | 0.328007061 | 0.522275161 | 22.90117424 | 15.55885839 | ENSG00000248485 |
| SEZ6 | -0.568221099 | 0.674447897 | 0.049661463 | 0.133070443 | 70.85466379 | 47.80758953 | ENSG00000063015 |
| PRPF4B | -0.568293721 | 0.674413948 | 4.07E-10 | 8.01E-09 | 1816.27599 | 1225.613919 | ENSG00000112739 |
| DERL3 | -0.568433406 | 0.674348653 | 0.013566443 | 0.047108777 | 114.5242864 | 77.33421502 | ENSG00000099958 |
| MTURN | -0.569474587 | 0.673862156 | 0.00572256 | 0.023097111 | 157.3452337 | 106.1784747 | ENSG00000180354 |
| THADA | -0.569547483 | 0.673828109 | 3.36E-08 | 4.90E-07 | 695.0950135 | 468.3471257 | ENSG00000115970 |
| PRSS22 | -0.569984469 | 0.67362404 | 0.315154175 | 0.508759471 | 17.95769324 | 12.09262614 | ENSG00000005001 |
| ZDHHC22 | -0.570000761 | 0.673616433 | 2.21E-11 | 5.30E-10 | 1459.59624 | 983.9928029 | ENSG00000177108 |
| ADGRB1 | -0.570135576 | 0.673553489 | 2.86E-07 | 3.52E-06 | 567.8348641 | 382.6983609 | ENSG00000181790 |
| LINC01547 | -0.570251699 | 0.673499276 | 0.328938052 | 0.523307971 | 18.00948162 | 12.1264809 | ENSG00000183250 |
| PHOSPHO2 | -0.570309354 | 0.673472362 | 0.283464216 | 0.474492229 | 22.73506057 | 15.42746143 | ENSG00000144362 |
| DDX49 | -0.570512725 | 0.673377432 | 7.33E-09 | 1.20E-07 | 996.5139277 | 671.3173716 | ENSG00000105671 |
| SEPHS1 | -0.570656899 | 0.673310142 | 1.31E-21 | 9.50E-20 | 15131.14254 | 10187.06818 | ENSG00000086475 |
| NA | -0.570835559 | 0.673226766 | 0.45802103 | 0.645204968 | 9.226573883 | 6.222630697 | ENSG00000213790 |
| PCDHGA12 | -0.571229156 | 0.673043121 | 0.578440964 | 0.742799475 | 5.121019588 | 3.443224373 | ENSG00000253159 |
| TIMM23B | -0.57132653 | 0.672997695 | 0.099586538 | 0.22616553 | 48.2668731 | 32.41398287 | ENSG00000204152 |
| AMPH | -0.571394311 | 0.672966077 | 5.34E-05 | 0.00039926 | 313.36685 | 210.8429804 | ENSG00000078053 |
| SPINT1-AS1 | -0.571439189 | 0.672945144 | 0.187817008 | 0.3596177 | 33.35171977 | 22.42609811 | ENSG00000261183 |
| C7orf31 | -0.571661239 | 0.672841577 | 0.17173214 | 0.337462611 | 34.70650033 | 23.25472985 | ENSG00000153790 |
| NA | -0.571840667 | 0.6727579 | 0.522947199 | 0.699653749 | 7.781347874 | 5.236233311 | ENSG00000267799 |
| H2BC10 | -0.571855248 | 0.672751101 | 0.639233773 | 0.78858703 | 5.105235024 | 3.428252226 | ENSG00000278588 |
| HID1 | -0.572004673 | 0.672681425 | 0.000628979 | 0.003508124 | 255.71213 | 172.1432206 | ENSG00000167861 |
| NA | -0.572042041 | 0.672664002 | 0.197369041 | 0.373154932 | 31.99004667 | 21.48060353 | ENSG00000261762 |
| DUSP26 | -0.572121061 | 0.672627159 | 0.104109268 | 0.233716228 | 52.11586828 | 35.05125696 | ENSG00000133878 |
| BMP6 | -0.572254067 | 0.672565151 | 0.026978026 | 0.082414824 | 93.60926721 | 63.02108395 | ENSG00000153162 |
| NA | -0.572347853 | 0.672521431 | 0.539332332 | 0.71267542 | 6.709659508 | 4.527275579 | ENSG00000280434 |
| SYBU | -0.572624496 | 0.672392484 | 0.404546251 | 0.596238518 | 11.86983983 | 7.993069793 | ENSG00000147642 |
| WASHC4 | -0.572736058 | 0.672340491 | 1.17E-08 | 1.85E-07 | 844.1737349 | 567.1927243 | ENSG00000136051 |
| NA | -0.572742811 | 0.672337344 | 0.351956288 | 0.545049953 | 14.30843274 | 9.617028162 | ENSG00000258741 |
| ANAPC2 | -0.572825379 | 0.672298866 | 2.39E-10 | 4.92E-09 | 1314.799033 | 884.6623589 | ENSG00000176248 |
| DPY19L2P3 | -0.573037796 | 0.672199887 | 0.145356498 | 0.299005253 | 38.59791085 | 25.99334499 | ENSG00000227855 |
| TMEM102 | -0.57313644 | 0.672153926 | 0.066140539 | 0.165672491 | 82.51183302 | 55.50780949 | ENSG00000181284 |
| SEMA6C | -0.573366819 | 0.672046601 | 4.84E-08 | 6.87E-07 | 807.0422388 | 543.0397669 | ENSG00000143434 |
| NA | -0.573464716 | 0.672001 | 0.375875406 | 0.568924032 | 15.07237497 | 10.17614435 | ENSG00000264107 |

| | | | | | | | |
|---|---|---|---|---|---|---|---|
| NA | -0.573629082 | 0.671924443 | 0.493615011 | 0.675874798 | 8.326711644 | 5.630097779 | ENSG00000275120 |
| CAMK4 | -0.573673723 | 0.671903652 | 0.005046227 | 0.020756053 | 153.4861664 | 102.9210965 | ENSG00000152495 |
| SCLY | -0.574018556 | 0.671743073 | 0.014097124 | 0.048627599 | 108.5325909 | 72.97555198 | ENSG00000132330 |
| TCN2 | -0.57406507 | 0.671721415 | 1.64E-11 | 4.02E-10 | 1288.049014 | 864.5879705 | ENSG00000185339 |
| CPNE1 | -0.574162437 | 0.671676083 | 3.60E-10 | 7.15E-09 | 1309.016709 | 879.787999 | ENSG00000214078 |
| FASTKD1 | -0.574351995 | 0.671587836 | 1.39E-07 | 1.81E-06 | 680.5676416 | 457.1585501 | ENSG00000138399 |
| NME1 | -0.574381806 | 0.671573959 | 7.33E-07 | 8.38E-06 | 531.2542867 | 356.7234605 | ENSG00000239672 |
| ABCA1 | -0.574415065 | 0.671558477 | 0.001476466 | 0.00730898 | 208.8328844 | 139.9973275 | ENSG00000165029 |
| NA | -0.574527905 | 0.671505953 | 0.373631929 | 0.566963662 | 14.44297727 | 9.70339706 | ENSG00000256338 |
| NUTM2A | -0.57453609 | 0.671502144 | 0.447764414 | 0.635877392 | 10.13718464 | 6.781308855 | ENSG00000184923 |
| SLC38A7 | -0.574986255 | 0.671292647 | 4.29E-08 | 6.17E-07 | 695.1346854 | 467.0467344 | ENSG00000103042 |
| SREBF1 | -0.575037035 | 0.671269019 | 2.42E-13 | 7.64E-12 | 2145.526848 | 1440.511227 | ENSG00000072310 |
| GPRIN1 | -0.575160969 | 0.671211357 | 4.12E-06 | 4.07E-05 | 496.1399361 | 333.2548668 | ENSG00000169258 |
| RWDD2B | -0.575161042 | 0.671211323 | 0.039136271 | 0.110605618 | 80.05974065 | 53.63971658 | ENSG00000156253 |
| NRIP1 | -0.575203054 | 0.671191777 | 0.00336204 | 0.014663283 | 190.6457438 | 127.8493761 | ENSG00000180530 |
| NA | -0.575249256 | 0.671170282 | 0.058435006 | 0.150861457 | 67.59916019 | 45.47098621 | ENSG00000213888 |
| ZBED5-AS1 | -0.575305441 | 0.671144145 | 0.386610721 | 0.579493983 | 12.7791728 | 8.551747951 | ENSG00000247271 |
| MX1 | -0.575376734 | 0.67111098 | 0.094941123 | 0.217799628 | 59.94910224 | 40.12548774 | ENSG00000157601 |
| PPOX | -0.575561064 | 0.671025239 | 3.43E-06 | 3.45E-05 | 551.5879383 | 370.5668073 | ENSG00000143224 |
| NA | -0.576134051 | 0.670758785 | 0.605388472 | 0.764063326 | 5.061639604 | 3.383112544 | ENSG00000275601 |
| IKBKE | -0.576160271 | 0.670746594 | 0.00056886 | 0.003211466 | 219.0391151 | 146.8392979 | ENSG00000263528 |
| NA | -0.576859638 | 0.670421519 | 0.379923719 | 0.572945989 | 14.55679889 | 9.83490564 | ENSG00000272927 |
| NA | -0.577044545 | 0.670335598 | 0.613000119 | 0.769113472 | 4.671137318 | 3.173270792 | ENSG00000266648 |
| DHX40 | -0.577379098 | 0.670180169 | 1.15E-07 | 1.53E-06 | 895.3640722 | 599.207077 | ENSG00000108406 |
| POLR2C | -0.577425293 | 0.67015871 | 1.48E-12 | 4.23E-11 | 1467.290706 | 983.3079665 | ENSG00000102978 |
| NA | -0.57752953 | 0.670110292 | 0.45712353 | 0.644203135 | 9.3327062 | 6.260172686 | ENSG00000279861 |
| NA | -0.5775346 | 0.670107937 | 0.11902424 | 0.257980969 | 46.0343287 | 30.98577017 | ENSG00000238123 |
| FAM81A | -0.577573886 | 0.670089689 | 0.245133297 | 0.432220409 | 25.1438658 | 16.84851447 | ENSG00000157470 |
| RDH14 | -0.577829887 | 0.669970795 | 0.000556304 | 0.003151144 | 233.1049919 | 155.9831407 | ENSG00000240857 |
| ACAN | -0.578041072 | 0.669872731 | 0.235366421 | 0.420379807 | 28.97146165 | 19.53070501 | ENSG00000157766 |
| PDRG1 | -0.578043786 | 0.66987147 | 3.46E-08 | 5.04E-07 | 948.4462329 | 635.5957897 | ENSG00000088356 |
| CCDC189 | -0.578241147 | 0.669779838 | 0.536032205 | 0.710391197 | 8.518854723 | 5.795349501 | ENSG00000196118 |
| SNHG9 | -0.578420484 | 0.669696585 | 0.373701559 | 0.566976481 | 14.60670814 | 9.861162708 | ENSG00000255198 |
| KIF21A | -0.57893773 | 0.669456523 | 1.13E-11 | 2.86E-10 | 1166.733305 | 781.1512771 | ENSG00000139116 |
| DLEU2 | -0.578984297 | 0.669434915 | 0.470624302 | 0.656151517 | 9.732776922 | 6.533925114 | ENSG00000231607 |
| MYCN | -0.579031468 | 0.669413027 | 2.35E-12 | 6.49E-11 | 1739.258954 | 1164.353389 | ENSG00000134323 |
| HS3ST2 | -0.579207421 | 0.66933139 | 0.113435115 | 0.249100286 | 43.97233264 | 29.34216474 | ENSG00000122254 |
| DUSP7 | -0.579314874 | 0.669281539 | 4.76E-08 | 6.77E-07 | 646.4475223 | 432.9252354 | ENSG00000164086 |
| ZMYND12 | -0.5795404 | 0.669176924 | 0.566488856 | 0.734156409 | 5.79273493 | 3.908485592 | ENSG00000066185 |
| LIN7A | -0.57968967 | 0.66910769 | 0.001122674 | 0.005814155 | 211.0846633 | 141.1455127 | ENSG00000111052 |
| SUPV3L1 | -0.579829628 | 0.669042782 | 8.73E-09 | 1.42E-07 | 779.2334147 | 521.2055001 | ENSG00000156502 |
| ZNF385C | -0.580030592 | 0.668949593 | 0.013731609 | 0.04758819 | 116.377077 | 77.91568626 | ENSG00000187595 |
| ADPRH | -0.580062613 | 0.668934745 | 0.01488342 | 0.050735573 | 116.5722793 | 78.21277298 | ENSG00000144843 |
| CDC6 | -0.580127339 | 0.668904734 | 1.39E-17 | 7.08E-16 | 2949.626696 | 1972.578558 | ENSG00000094804 |
| NA | -0.580405645 | 0.66877571 | 0.59899922 | 0.758920946 | 4.721046564 | 3.203326707 | ENSG00000214646 |
| SNHG3 | -0.580441113 | 0.668759269 | 0.059354638 | 0.15264033 | 1156.862534 | 773.854123 | ENSG00000242125 |
| TSC22D1 | -0.580781312 | 0.668601589 | 2.74E-15 | 1.09E-13 | 1949.686973 | 1304.08019 | ENSG00000102804 |
| NA | -0.581062464 | 0.668471305 | 0.020950907 | 0.067157403 | 103.0859887 | 68.82282022 | ENSG00000261061 |
| NAP1L2 | -0.581284413 | 0.668368472 | 2.14E-05 | 0.000177732 | 507.4334218 | 339.3283257 | ENSG00000186462 |
| NA | -0.581388185 | 0.668320399 | 0.410269424 | 0.601910354 | 11.33720141 | 7.614177473 | ENSG00000231856 |
| ZNF571-AS1 | -0.581735222 | 0.668159655 | 0.575561335 | 0.740696842 | 5.124176501 | 3.405682385 | ENSG00000267470 |
| CDK5 | -0.581749013 | 0.668153268 | 7.26E-06 | 6.77E-05 | 400.46166 | 267.8113603 | ENSG00000164885 |
| FAHD1 | -0.581971578 | 0.6680502 | 1.68E-05 | 0.000142675 | 358.9064006 | 239.7091486 | ENSG00000180185 |
| BEX3 | -0.581997319 | 0.668038281 | 4.81E-26 | 4.96E-24 | 7860.143766 | 5251.23983 | ENSG00000166681 |
| ZNF22 | -0.58201383 | 0.668030635 | 0.000231072 | 0.001460171 | 316.3646144 | 211.3048892 | ENSG00000165512 |
| CPLX1 | -0.582339845 | 0.667879693 | 0.25629427 | 0.445063056 | 25.42833346 | 16.96450125 | ENSG00000168993 |
| KANK4 | -0.58255153 | 0.667781703 | 0.429356445 | 0.619564961 | 10.75771295 | 7.178972169 | ENSG00000132854 |
| RPE | -0.582612769 | 0.667753358 | 2.28E-10 | 4.71E-09 | 1046.811002 | 698.5492439 | ENSG00000197713 |
| RUFY4 | -0.582708592 | 0.667709008 | 0.570746485 | 0.737312006 | 6.361474866 | 4.298662833 | ENSG00000188282 |
| SOX12 | -0.582960452 | 0.667592452 | 2.15E-12 | 6.02E-11 | 3963.303587 | 2645.992271 | ENSG00000177732 |
| TEDC2 | -0.582968533 | 0.667588712 | 1.83E-08 | 2.80E-07 | 1005.017674 | 671.7339578 | ENSG00000162062 |
| NA | -0.583171062 | 0.667495001 | 0.474649076 | 0.659659852 | 8.684367025 | 5.746084558 | ENSG00000227008 |
| STK36 | -0.583653064 | 0.66727203 | 5.65E-07 | 6.62E-06 | 674.8699822 | 450.9612291 | ENSG00000163482 |
| EMP2 | -0.583703402 | 0.667248748 | 3.29E-06 | 3.33E-05 | 446.032201 | 297.53088 | ENSG00000213853 |
| BICC1 | -0.583973786 | 0.667123706 | 0.152432555 | 0.309550819 | 35.81556799 | 23.9340697 | ENSG00000122870 |
| SLC43A3 | -0.584038696 | 0.667093692 | 0.000221909 | 0.001410444 | 288.7871917 | 192.6824632 | ENSG00000134802 |
| ZNF276 | -0.584196481 | 0.667020737 | 0.022188346 | 0.070357365 | 145.1641826 | 97.2754056 | ENSG00000158805 |
| ST3GAL6 | -0.584639772 | 0.666815816 | 0.082493856 | 0.196194164 | 54.667411 | 36.39017798 | ENSG00000064225 |
| KCNJ11 | -0.584846442 | 0.666720299 | 0.00051158 | 0.002925523 | 230.8186072 | 154.151316 | ENSG00000187486 |
| NA | -0.584985859 | 0.666655873 | 0.34310859 | 0.53700174 | 16.01277341 | 10.73113528 | ENSG00000288060 |
| NA | -0.585636683 | 0.666355201 | 0.256858874 | 0.445408246 | 25.12986267 | 16.78146622 | ENSG00000234743 |
| RASSF5 | -0.585637325 | 0.666354904 | 0.117170366 | 0.255216307 | 42.68777828 | 28.46090724 | ENSG00000266094 |
| NA | -0.585740687 | 0.666307165 | 0.002201176 | 0.010312587 | 175.2024094 | 116.6874956 | ENSG00000254682 |
| NGEF | -0.585822779 | 0.666269252 | 0.024078879 | 0.075130966 | 86.02114479 | 57.40259751 | ENSG00000066248 |

| | | | | | | | |
|---|---|---|---|---|---|---|---|
| CENPQ | -0.58597217 | 0.666200263 | 0.001623165 | 0.007950406 | 186.9789399 | 124.3114207 | ENSG00000031691 |
| LDOC1 | -0.586239301 | 0.666076921 | 2.27E-09 | 4.00E-08 | 845.1531586 | 563.3860666 | ENSG00000182195 |
| NA | -0.586267006 | 0.66606413 | 0.332598894 | 0.527186172 | 2396.113519 | 1595.974296 | ENSG00000248527 |
| NA | -0.586608835 | 0.665906333 | 0.513811788 | 0.692278836 | 7.58614558 | 5.010869759 | ENSG00000260643 |
| CTAGE4 | -0.58677818 | 0.665828172 | 0.459206196 | 0.646127338 | 9.371866931 | 6.230116771 | ENSG00000225932 |
| USP6 | -0.587333061 | 0.665572135 | 0.341746546 | 0.536046927 | 16.097311 | 10.69359329 | ENSG00000129204 |
| DLGAP3 | -0.587376728 | 0.66555199 | 1.28E-05 | 0.000112018 | 572.8315689 | 381.2631715 | ENSG00000116544 |
| P2RY2 | -0.587433475 | 0.665525811 | 0.064069692 | 0.161978949 | 60.33822905 | 40.06581395 | ENSG00000175591 |
| SSC5D | -0.587601773 | 0.665448179 | 0.221221281 | 0.40276181 | 26.02213331 | 17.31757453 | ENSG00000179954 |
| ATAD3A | -0.587739545 | 0.665384634 | 7.69E-13 | 2.32E-11 | 1820.839115 | 1212.075869 | ENSG00000197785 |
| C1orf112 | -0.587811269 | 0.665351555 | 5.48E-05 | 0.000408261 | 365.989425 | 243.0826881 | ENSG00000000460 |
| TIGD1 | -0.58798949 | 0.665269367 | 0.066478773 | 0.166338362 | 112.393635 | 75.07445625 | ENSG00000221944 |
| COL8A2 | -0.5880054 | 0.665262031 | 0.005728058 | 0.023114266 | 130.8106812 | 87.10108472 | ENSG00000171812 |
| NA | -0.588200429 | 0.665172104 | 0.163803545 | 0.325779042 | 35.72709937 | 23.7613319 | ENSG00000271976 |
| TOMM5 | -0.588489846 | 0.665038678 | 0.236282038 | 0.421324927 | 22.72105745 | 15.06310322 | ENSG00000175768 |
| GLB1L3 | -0.588726241 | 0.664929716 | 3.70E-10 | 7.31E-09 | 1462.322443 | 973.3271156 | ENSG00000166105 |
| BORCS7 | -0.589200122 | 0.664711343 | 0.200176453 | 0.376705682 | 31.22926135 | 20.86605042 | ENSG00000166275 |
| GNG4 | -0.589748558 | 0.664458703 | 1.78E-08 | 2.73E-07 | 1257.928964 | 835.1648528 | ENSG00000168243 |
| PEX3 | -0.589847332 | 0.664413212 | 0.001841788 | 0.008841115 | 190.9684852 | 126.9039931 | ENSG00000034693 |
| ECI2 | -0.590082679 | 0.664304835 | 0.000205176 | 0.001313553 | 266.2687328 | 176.6619192 | ENSG00000198721 |
| NA | -0.590207138 | 0.664247529 | 0.564919667 | 0.732937653 | 6.95359097 | 4.553094614 | ENSG00000243485 |
| HTR1B | -0.590214986 | 0.664243916 | 0.291509332 | 0.483821435 | 18.81080315 | 12.54660244 | ENSG00000135312 |
| LRRC42 | -0.590331388 | 0.664190324 | 1.59E-11 | 3.92E-10 | 1110.059497 | 737.0630375 | ENSG00000116212 |
| CYBRD1 | -0.590348677 | 0.664182365 | 0.009225286 | 0.034402705 | 123.0254774 | 81.5113401 | ENSG00000071967 |
| SEMA3D | -0.590608019 | 0.664062981 | 0.023652969 | 0.074088919 | 88.35982151 | 58.55337055 | ENSG00000153993 |
| NA | -0.591090744 | 0.663840823 | 0.075435305 | 0.183263593 | 54.16258344 | 35.86469331 | ENSG00000235859 |
| MBOAT2 | -0.591095131 | 0.663838804 | 3.89E-05 | 0.000302462 | 380.7349995 | 252.7764492 | ENSG00000143797 |
| ZNF829 | -0.59113039 | 0.663822581 | 0.121166611 | 0.261493009 | 39.88078147 | 26.52565446 | ENSG00000185869 |
| PNMA1 | -0.591146668 | 0.663815091 | 4.98E-15 | 1.93E-13 | 2062.230989 | 1369.393296 | ENSG00000176903 |
| SYNDIG1L | -0.591231989 | 0.663775834 | 0.300020989 | 0.493415916 | 21.88057519 | 14.37247845 | ENSG00000183379 |
| BRINP1 | -0.591244177 | 0.663770226 | 1.88E-09 | 3.35E-08 | 884.5051576 | 587.4539063 | ENSG00000078725 |
| PERP | -0.591566309 | 0.663622033 | 1.45E-08 | 2.26E-07 | 934.5517087 | 619.9391341 | ENSG00000112378 |
| NA | -0.591913885 | 0.663462171 | 0.571001357 | 0.737373586 | 6.034787254 | 3.953075621 | ENSG00000285454 |
| NA | -0.591992937 | 0.663425818 | 0.08648507 | 0.203209214 | 55.47750421 | 36.72013177 | ENSG00000264164 |
| DUSP4 | -0.592043196 | 0.663402707 | 1.05E-05 | 9.29E-05 | 382.047748 | 253.3770178 | ENSG00000120875 |
| LCA5L | -0.592155158 | 0.663351225 | 0.608693825 | 0.766514251 | 4.738108905 | 3.173270792 | ENSG00000157578 |
| SLC5A2 | -0.59219612 | 0.663332391 | 0.421635779 | 0.612861542 | 11.55262295 | 7.68926145 | ENSG00000140675 |
| RHOF | -0.592241233 | 0.663311649 | 0.063740464 | 0.161366721 | 80.22544834 | 53.49889845 | ENSG00000139725 |
| PTRHD1 | -0.592338765 | 0.663266807 | 0.005839514 | 0.023512831 | 127.4987591 | 84.57286099 | ENSG00000184924 |
| LETM2 | -0.59241281 | 0.663232767 | 0.249841971 | 0.438338668 | 23.60701426 | 15.56915562 | ENSG00000165046 |
| CCDC87 | -0.592473961 | 0.663204655 | 0.575564687 | 0.740696842 | 5.117862675 | 3.364341549 | ENSG00000182791 |
| SLTM | -0.592659159 | 0.663119525 | 2.41E-20 | 1.59E-18 | 3613.583022 | 2396.733922 | ENSG00000137776 |
| PRR7 | -0.592714199 | 0.663094227 | 0.005894539 | 0.023693209 | 193.2267886 | 128.142346 | ENSG00000131188 |
| SCN4B | -0.592714221 | 0.663094217 | 0.003543677 | 0.01533828 | 163.8031747 | 108.5558692 | ENSG00000177098 |
| NRN1 | -0.593082286 | 0.662925068 | 0.413172174 | 0.604535668 | 10.88090761 | 7.19025709 | ENSG00000124785 |
| ERG28 | -0.593187756 | 0.662876606 | 9.60E-11 | 2.11E-09 | 1185.192864 | 785.1159322 | ENSG00000133935 |
| SMPD4 | -0.593223633 | 0.662860122 | 4.24E-17 | 2.04E-15 | 2737.247362 | 1814.576647 | ENSG00000136699 |
| NA | -0.594257034 | 0.662385486 | 0.167738786 | 0.331542141 | 36.91210569 | 24.38337108 | ENSG00000284610 |
| TMEM145 | -0.594421921 | 0.662309785 | 0.005891776 | 0.023687241 | 134.378561 | 89.08147719 | ENSG00000167619 |
| MED31 | -0.594497031 | 0.662275305 | 0.105620253 | 0.236249791 | 43.46109355 | 28.68203392 | ENSG00000108590 |
| SLC25A23 | -0.595059841 | 0.662016995 | 6.07E-10 | 1.16E-08 | 1456.224374 | 964.4305675 | ENSG00000125648 |
| TTC27 | -0.59513418 | 0.661982884 | 1.52E-09 | 2.75E-08 | 797.31891 | 527.532283 | ENSG00000018699 |
| METTL7B | -0.595134416 | 0.661982776 | 0.201887984 | 0.378915635 | 26.02350878 | 17.26126155 | ENSG00000170439 |
| SCRT1 | -0.595155168 | 0.661973253 | 0.520976433 | 0.697974353 | 6.739349501 | 4.440906681 | ENSG00000261678 |
| FMN2 | -0.595331998 | 0.661892121 | 0.040894769 | 0.114458937 | 77.63642864 | 51.35942623 | ENSG00000155816 |
| KLK13 | -0.596021551 | 0.661575837 | 0.063440678 | 0.160876442 | 55.68740868 | 36.8666125 | ENSG00000167759 |
| TBPL1 | -0.596510057 | 0.661351861 | 0.000390575 | 0.002305387 | 292.3462999 | 193.0285514 | ENSG00000028839 |
| ALDH16A1 | -0.596563434 | 0.661327393 | 8.12E-09 | 1.32E-07 | 1069.372869 | 706.7654871 | ENSG00000161618 |
| RFESD | -0.597274771 | 0.661001399 | 0.481958519 | 0.665447848 | 9.557696184 | 6.225879891 | ENSG00000175449 |
| MPP3 | -0.597463598 | 0.660914889 | 0.006264623 | 0.024964207 | 138.5305746 | 91.64399372 | ENSG00000161647 |
| NA | -0.597484159 | 0.66090547 | 0.283268194 | 0.474305963 | 20.78107597 | 13.70529959 | ENSG00000254461 |
| MKX | -0.597725034 | 0.660795133 | 0.178070003 | 0.346784643 | 36.82491485 | 24.36894859 | ENSG00000150051 |
| CYP2U1 | -0.597737235 | 0.660789545 | 0.010528296 | 0.038373931 | 173.2964397 | 114.2619984 | ENSG00000155016 |
| CDC25B | -0.598973237 | 0.660223668 | 3.79E-09 | 6.48E-08 | 862.2855004 | 569.467942 | ENSG00000101224 |
| MCC | -0.599231692 | 0.660105401 | 8.34E-07 | 9.42E-06 | 571.4131092 | 377.0954591 | ENSG00000171444 |
| HNRNPDL | -0.599266809 | 0.660089334 | 6.94E-26 | 7.07E-24 | 7487.143416 | 4942.240659 | ENSG00000152795 |
| CKB | -0.599332218 | 0.660059407 | 1.83E-19 | 1.12E-17 | 7094.98394 | 4683.102232 | ENSG00000166165 |
| TEX10 | -0.599663513 | 0.659907851 | 8.41E-14 | 2.81E-12 | 1989.166729 | 1312.474865 | ENSG00000136891 |
| EEPD1 | -0.599859918 | 0.659818019 | 2.68E-05 | 0.000217047 | 335.1259143 | 220.8910886 | ENSG00000122547 |
| NA | -0.600282366 | 0.65962484 | 0.337559918 | 0.531944773 | 15.14260117 | 10.0259764 | ENSG00000262877 |
| TIMP4 | -0.60033529 | 0.659600643 | 0.045098456 | 0.123464517 | 70.11342131 | 46.3677655 | ENSG00000157150 |
| ARPIN | -0.600558024 | 0.659498817 | 0.002872273 | 0.01283692 | 153.8285409 | 101.3462915 | ENSG00000242498 |
| NA | -0.60142974 | 0.65910045 | 0.281329017 | 0.47271345 | 19.39217073 | 12.81655602 | ENSG00000284428 |
| ZNF205 | -0.601974858 | 0.658851458 | 7.60E-05 | 0.000544421 | 341.7231125 | 225.2348911 | ENSG00000122386 |

| | | | | | | | |
|---|---|---|---|---|---|---|---|
| CFAP43 | -0.602246509 | 0.658727411 | 0.487755152 | 0.670826209 | 7.281556358 | 4.79722916 | ENSG00000197748 |
| NA | -0.602338942 | 0.658685208 | 0.590137028 | 0.752086892 | 6.32537335 | 4.133411111 | ENSG00000263011 |
| ASGR1 | -0.60265031 | 0.658543064 | 0.0010143 | 0.005333062 | 194.0757494 | 127.6129509 | ENSG00000141505 |
| FNDC11 | -0.602652548 | 0.658542042 | 0.059560577 | 0.153000101 | 64.80459564 | 42.7935822 | ENSG00000125531 |
| NA | -0.6029327 | 0.658414175 | 0.486860536 | 0.669985548 | 8.234484755 | 5.389762079 | ENSG00000184100 |
| ZCWPW1 | -0.602950322 | 0.658406132 | 0.243976781 | 0.430735917 | 24.13649577 | 15.8622287 | ENSG00000078487 |
| SLC27A2 | -0.603317184 | 0.658238728 | 0.044346041 | 0.121945033 | 77.51511312 | 50.87853159 | ENSG00000140284 |
| NA | -0.603995656 | 0.657929243 | 0.358432383 | 0.551667978 | 13.12735745 | 8.638116849 | ENSG00000231713 |
| TSR1 | -0.604231963 | 0.657821487 | 8.26E-18 | 4.30E-16 | 3419.736359 | 2249.982155 | ENSG00000167721 |
| PRR5L | -0.604311711 | 0.657785125 | 0.008563822 | 0.032385609 | 127.9705444 | 84.00245988 | ENSG00000135362 |
| HSPA1L | -0.604415814 | 0.657737662 | 0.067745658 | 0.168713486 | 58.15578929 | 38.21638041 | ENSG00000204390 |
| DNAH9 | -0.604604843 | 0.657651487 | 0.359849278 | 0.553113826 | 14.40381654 | 9.542382217 | ENSG00000007174 |
| NA | -0.60462089 | 0.657644172 | 0.308966635 | 0.503153292 | 19.34975539 | 12.67756137 | ENSG00000263301 |
| NA | -0.604652899 | 0.657629582 | 0.547491185 | 0.719107096 | 6.715973334 | 4.384593698 | ENSG00000228169 |
| FBXL15 | -0.605298972 | 0.657335145 | 4.90E-05 | 0.00037032 | 307.0718431 | 201.6714547 | ENSG00000107872 |
| SNRPD3 | -0.605357771 | 0.657308356 | 4.92E-08 | 6.98E-07 | 727.7529497 | 478.058295 | ENSG00000100028 |
| SS18L2 | -0.606066182 | 0.656985674 | 8.30E-07 | 9.38E-06 | 840.6298321 | 552.0386974 | ENSG00000008324 |
| ZNF180 | -0.606228407 | 0.656911803 | 0.001690956 | 0.008227929 | 173.109235 | 113.6408353 | ENSG00000167384 |
| RASGRF1 | -0.60627205 | 0.656891931 | 0.254798833 | 0.44358835 | 19.40291925 | 12.76024304 | ENSG00000058335 |
| FADS2 | -0.606274954 | 0.656890609 | 1.84E-21 | 1.32E-19 | 10644.12906 | 6991.998946 | ENSG00000134824 |
| SERPINF2 | -0.606418536 | 0.656825236 | 0.115063636 | 0.251840981 | 44.94369885 | 29.39424084 | ENSG00000167711 |
| TMEM151A | -0.60667572 | 0.656708157 | 0.158816687 | 0.318561522 | 30.43237452 | 20.03686903 | ENSG00000179292 |
| NA | -0.606883946 | 0.65661338 | 0.50377004 | 0.683776217 | 7.99243243 | 5.277574146 | ENSG00000248578 |
| LOC105372482 | -0.607151433 | 0.65649165 | 0.14462621 | 0.297966004 | 32.94109593 | 21.6420564 | ENSG00000268516 |
| MPC2 | -0.607333742 | 0.656408697 | 0.006979228 | 0.027353141 | 117.4595139 | 76.94696056 | ENSG00000143158 |
| WIPF3 | -0.607343297 | 0.656404349 | 9.51E-06 | 8.55E-05 | 454.4782513 | 297.7532805 | ENSG00000122574 |
| NA | -0.607343509 | 0.656404253 | 0.13353028 | 0.28075442 | 46.70033157 | 30.8134704 | ENSG00000236397 |
| CNTF | -0.607894289 | 0.656153704 | 0.491341308 | 0.674098864 | 7.344093255 | 4.816000155 | ENSG00000242689 |
| MTHFD2L | -0.608699566 | 0.655787557 | 0.023145533 | 0.072720682 | 94.06062265 | 61.74161349 | ENSG00000163738 |
| NA | -0.608905406 | 0.655693998 | 0.047321359 | 0.128186797 | 63.99192424 | 41.87748226 | ENSG00000273247 |
| TIFA | -0.609201589 | 0.655559399 | 0.107895019 | 0.239775333 | 42.4179151 | 27.83831841 | ENSG00000145365 |
| KIF3B | -0.609801271 | 0.655286961 | 4.39E-12 | 1.18E-10 | 1179.464831 | 772.3412582 | ENSG00000101350 |
| SIRPA | -0.610334497 | 0.655044809 | 0.000138727 | 0.000927214 | 256.9147023 | 168.0081689 | ENSG00000198053 |
| NA | -0.611268324 | 0.654620949 | 0.270258549 | 0.460764599 | 23.31760824 | 15.16031901 | ENSG00000230562 |
| ZNF778 | -0.611279489 | 0.654615882 | 3.10E-06 | 3.15E-05 | 500.5706072 | 328.0349197 | ENSG00000170100 |
| PLEKHA6 | -0.611521463 | 0.654506097 | 3.45E-06 | 3.47E-05 | 388.2731301 | 254.2483045 | ENSG00000143850 |
| ABCF2 | -0.611631402 | 0.654456223 | 9.81E-06 | 8.79E-05 | 385.5181921 | 251.9461685 | ENSG00000033050 |
| MOB3C | -0.611725486 | 0.654413545 | 0.017016119 | 0.056504518 | 89.32222065 | 58.38855686 | ENSG00000142961 |
| NA | -0.611763107 | 0.65439648 | 0.213527363 | 0.393483184 | 29.63430761 | 19.50064909 | ENSG00000279413 |
| NA | -0.611898136 | 0.654335235 | 0.466321294 | 0.65250214 | 7.929895533 | 5.213663469 | ENSG00000213938 |
| GPR45 | -0.612027493 | 0.654276567 | 0.560295711 | 0.729378503 | 5.31631958 | 3.522107198 | ENSG00000135973 |
| SFXN5 | -0.612123112 | 0.654233205 | 1.44E-10 | 3.07E-09 | 910.2470399 | 595.792675 | ENSG00000144040 |
| POLR3B | -0.612277713 | 0.6541631 | 1.31E-05 | 0.000114687 | 516.2179906 | 337.1210867 | ENSG00000013503 |
| NA | -0.612519474 | 0.654053487 | 0.586834916 | 0.749424538 | 4.667980405 | 3.056845979 | ENSG00000213261 |
| TANGO6 | -0.612566827 | 0.65403202 | 2.21E-06 | 2.29E-05 | 417.4417709 | 272.9333186 | ENSG00000103047 |
| SMOC1 | -0.612746853 | 0.653950412 | 0.153109724 | 0.310406434 | 31.9085683 | 20.86550077 | ENSG00000198732 |
| CA5BP1 | -0.613108173 | 0.653786652 | 0.00119177 | 0.00612033 | 165.9354947 | 108.5295005 | ENSG00000186312 |
| RPUSD1 | -0.613162333 | 0.653762109 | 3.98E-08 | 5.73E-07 | 878.5657164 | 574.3719198 | ENSG00000007376 |
| MTA2 | -0.613353508 | 0.653675482 | 1.78E-24 | 1.60E-22 | 4206.467383 | 2749.279953 | ENSG00000149480 |
| ADCY5 | -0.61368783 | 0.653524021 | 9.23E-08 | 1.25E-06 | 528.4875178 | 345.6892813 | ENSG00000173175 |
| TMED6 | -0.614183882 | 0.653299354 | 0.461166059 | 0.647813532 | 9.826281588 | 6.376159466 | ENSG00000157315 |
| EFS | -0.614189715 | 0.653296713 | 0.011994781 | 0.042768616 | 1052.206448 | 687.7319093 | ENSG00000100842 |
| CAMK2N1 | -0.614415849 | 0.65319432 | 0.011712717 | 0.041908011 | 103.1196097 | 67.40545441 | ENSG00000162545 |
| NA | -0.614557898 | 0.653130009 | 0.546799647 | 0.718495899 | 5.775672589 | 3.75451879 | ENSG00000248996 |
| LOC100287015 | -0.614689269 | 0.653070538 | 0.000700186 | 0.003856539 | 188.3849075 | 123.1540376 | ENSG00000246089 |
| LOC112267871 | -0.615240308 | 0.652821145 | 0.266385174 | 0.45634373 | 18.83417931 | 12.23486999 | ENSG00000228549 |
| NA | -0.615415845 | 0.652741719 | 0.052856258 | 0.13956061 | 57.21666862 | 37.29009485 | ENSG00000287222 |
| NA | -0.615597687 | 0.65265945 | 0.328850392 | 0.523240833 | 15.52866876 | 10.22877011 | ENSG00000277159 |
| NA | -0.615909697 | 0.652518316 | 0.395064571 | 0.587509168 | 11.03704687 | 7.216514158 | ENSG00000253549 |
| SRSF2 | -0.616252387 | 0.652363339 | 1.23E-13 | 4.00E-12 | 2517.417193 | 1641.747945 | ENSG00000161547 |
| MICOS10 | -0.616459958 | 0.652269485 | 0.034975165 | 0.101191025 | 69.77400835 | 45.55355626 | ENSG00000173436 |
| RNF182 | -0.617385127 | 0.651851333 | 0.01607923 | 0.054026912 | 101.500002 | 65.98297565 | ENSG00000180537 |
| ZNF512B | -0.617569853 | 0.651767874 | 2.96E-11 | 7.01E-10 | 1147.975861 | 748.8571286 | ENSG00000196700 |
| NA | -0.61761695 | 0.651746597 | 0.26111494 | 0.450133564 | 22.92267127 | 14.99605497 | ENSG00000203497 |
| ABHD16A | -0.618700417 | 0.651257318 | 0.407203221 | 0.598955168 | 10.54978531 | 6.856392832 | ENSG00000204427 |
| HECTD2 | -0.618997653 | 0.651123154 | 0.000444787 | 0.002583416 | 206.3502901 | 134.1763822 | ENSG00000165338 |
| ILKAP | -0.619101595 | 0.651076244 | 1.18E-07 | 1.55E-06 | 580.8642072 | 378.588284 | ENSG00000132323 |
| PARP16 | -0.619574655 | 0.650862791 | 0.005060329 | 0.020800207 | 129.481156 | 84.18984341 | ENSG00000138617 |
| HFM1 | -0.619824896 | 0.650749906 | 0.31545029 | 0.509015816 | 15.32705495 | 9.958378493 | ENSG00000162669 |
| IGSF8 | -0.620231437 | 0.650566556 | 2.09E-16 | 9.22E-15 | 2565.77742 | 1668.484094 | ENSG00000162729 |
| NA | -0.620428504 | 0.650477697 | 0.565573164 | 0.733286675 | 4.783583461 | 3.116957809 | ENSG00000273449 |
| RSPH9 | -0.620924133 | 0.650254267 | 0.334644489 | 0.528981192 | 15.90960261 | 10.29213113 | ENSG00000172426 |
| DDX56 | -0.621157668 | 0.650149016 | 1.44E-14 | 5.25E-13 | 1507.511013 | 980.1894169 | ENSG00000136271 |
| NA | -0.621504638 | 0.649992674 | 0.446617099 | 0.634882752 | 8.929576263 | 5.776140473 | ENSG00000258317 |

| | | | | | | | |
|---|---|---|---|---|---|---|---|
| NA | -0.621900113 | 0.64981452 | 0.347010549 | 0.540416227 | 14.60039431 | 9.429206598 | ENSG00000229644 |
| NA | -0.622420469 | 0.649580185 | 0.569413107 | 0.736177672 | 4.800645802 | 3.124443883 | ENSG00000227775 |
| ANAPC4 | -0.622571924 | 0.649511996 | 3.94E-08 | 5.68E-07 | 590.6117539 | 383.7563698 | ENSG00000053900 |
| SNHG21 | -0.622614131 | 0.649492994 | 0.459835084 | 0.646482852 | 9.190570065 | 6.03155994 | ENSG00000250988 |
| GEMIN7 | -0.62297621 | 0.649330009 | 0.576350713 | 0.74114231 | 4.83664962 | 3.147013724 | ENSG00000142252 |
| LOC105377623 | -0.623522555 | 0.649084155 | 0.497364584 | 0.678454052 | 7.953271699 | 5.161149333 | ENSG00000250326 |
| GINS3 | -0.623593398 | 0.649052284 | 0.000285173 | 0.00175945 | 233.9680762 | 151.9937256 | ENSG00000181938 |
| NA | -0.624113077 | 0.648818528 | 0.114688842 | 0.251198653 | 43.95211339 | 28.49107478 | ENSG00000232640 |
| NA | -0.624186216 | 0.648785636 | 0.00103971 | 0.005451198 | 190.0782066 | 123.4094571 | ENSG00000282936 |
| PIK3R3 | -0.624220022 | 0.648770433 | 9.35E-05 | 0.000656021 | 265.0552764 | 171.9396624 | ENSG00000117461 |
| FMNL1 | -0.624363423 | 0.64870595 | 0.000391808 | 0.002311195 | 198.5929197 | 128.8246002 | ENSG00000184922 |
| CFAP46 | -0.625144156 | 0.64835499 | 0.00688433 | 0.027055593 | 114.6924519 | 74.43480828 | ENSG00000171811 |
| PGD | -0.62550936 | 0.648190886 | 1.60E-29 | 2.17E-27 | 10227.98378 | 6629.424131 | ENSG00000142657 |
| REPIN1 | -0.625901466 | 0.648014739 | 1.01E-15 | 4.20E-14 | 2515.378686 | 1630.773395 | ENSG00000214022 |
| ESR2 | -0.626050304 | 0.64794789 | 0.477386645 | 0.661721941 | 9.084437747 | 5.926420048 | ENSG00000140009 |
| FARS2 | -0.626203853 | 0.647878931 | 4.91E-06 | 4.75E-05 | 345.2331006 | 223.7046761 | ENSG00000145982 |
| DENND1B | -0.62621976 | 0.647871788 | 0.089091401 | 0.207988414 | 51.46101989 | 33.22808049 | ENSG00000213047 |
| TUBB8 | -0.626529227 | 0.64773283 | 0.540081019 | 0.713207097 | 7.434441008 | 4.79722916 | ENSG00000261456 |
| PRXL2B | -0.626552615 | 0.64772233 | 9.81E-14 | 3.23E-12 | 1956.908495 | 1267.444317 | ENSG00000157870 |
| ANKRD18B | -0.626554137 | 0.647721646 | 0.011570812 | 0.041512473 | 109.6887191 | 70.88198389 | ENSG00000230453 |
| STXBP6 | -0.626624072 | 0.647690248 | 0.00263645 | 0.011955947 | 147.5204253 | 95.45328816 | ENSG00000168952 |
| ACVR2B-AS1 | -0.626645294 | 0.647680721 | 0.099922936 | 0.226679522 | 39.81055528 | 25.77166866 | ENSG00000229589 |
| NA | -0.626802459 | 0.647610167 | 0.178189632 | 0.34695505 | 29.59317004 | 19.28332127 | ENSG00000240429 |
| FUCA2 | -0.627067449 | 0.647491227 | 1.17E-10 | 2.53E-09 | 870.6382566 | 563.6276761 | ENSG00000001036 |
| CISH | -0.627266442 | 0.647401924 | 0.060405058 | 0.154780577 | 59.18831692 | 38.28788878 | ENSG00000114737 |
| RAC3 | -0.62737432 | 0.647353516 | 8.30E-12 | 2.13E-10 | 1698.25547 | 1099.238865 | ENSG00000169750 |
| SQLE | -0.627596752 | 0.647253716 | 8.33E-24 | 7.15E-22 | 4135.859424 | 2676.365237 | ENSG00000104549 |
| SEPTIN8 | -0.627709076 | 0.647203325 | 8.87E-14 | 2.95E-12 | 1540.408836 | 997.2675234 | ENSG00000164402 |
| ZMPSTE24 | -0.627881727 | 0.647125877 | 4.97E-09 | 8.37E-08 | 1042.40509 | 674.1927425 | ENSG00000084073 |
| TRH | -0.628031644 | 0.647058635 | 0.568999671 | 0.735971767 | 5.875588778 | 3.840887688 | ENSG00000170893 |
| SKP2 | -0.628760143 | 0.64673198 | 1.16E-12 | 3.39E-11 | 1654.627617 | 1069.63873 | ENSG00000145604 |
| PLCB3 | -0.629346212 | 0.64646931 | 1.56E-22 | 1.22E-20 | 4697.415735 | 3037.171994 | ENSG00000149782 |
| NA | -0.629467619 | 0.64641491 | 0.148805346 | 0.304276068 | 32.81968271 | 21.15422534 | ENSG00000285728 |
| ATP1B2 | -0.629476209 | 0.646411061 | 3.80E-05 | 0.000295945 | 298.7638024 | 193.2149875 | ENSG00000129244 |
| PDE4DIP | -0.629767637 | 0.646280498 | 1.83E-09 | 3.27E-08 | 800.5426339 | 517.4399112 | ENSG00000178104 |
| NA | -0.629770268 | 0.64627932 | 0.546721326 | 0.718495899 | 5.395918818 | 3.514621124 | ENSG00000185065 |
| PCDHB14 | -0.63002691 | 0.646164362 | 0.069053584 | 0.171325291 | 48.67443773 | 31.3525015 | ENSG00000120327 |
| NA | -0.630047238 | 0.646155258 | 0.555441596 | 0.725307064 | 6.008254175 | 3.931055433 | ENSG00000236184 |
| FBXL13 | -0.630728713 | 0.645850111 | 0.381065586 | 0.574027769 | 12.27938129 | 7.970499952 | ENSG00000161040 |
| CCDC15 | -0.630871814 | 0.645786052 | 0.042734887 | 0.118534629 | 64.00327412 | 41.28126211 | ENSG00000149548 |
| NTRK3 | -0.631235108 | 0.645623453 | 0.000969837 | 0.005134236 | 171.0782067 | 110.5700048 | ENSG00000140538 |
| TNFRSF25 | -0.631444268 | 0.645529858 | 0.010265328 | 0.037562863 | 139.6973385 | 90.48171252 | ENSG00000215788 |
| LANCL2 | -0.631781066 | 0.645379176 | 1.63E-07 | 2.09E-06 | 499.3526184 | 322.4936756 | ENSG00000132434 |
| NA | -0.632353188 | 0.645123293 | 0.400593475 | 0.592545385 | 10.66232915 | 6.916504662 | ENSG00000256542 |
| LY6G5C | -0.632358744 | 0.645120808 | 0.329763 | 0.52417045 | 13.8977112 | 9.005724249 | ENSG00000204428 |
| NA | -0.632381822 | 0.645110488 | 0.086087489 | 0.2025318 | 45.56234804 | 29.53042433 | ENSG00000259642 |
| ACPP | -0.632715105 | 0.644961476 | 0.277048852 | 0.468121381 | 18.02338705 | 11.54099603 | ENSG00000014257 |
| NA | -0.632897777 | 0.644879817 | 0.213370868 | 0.393298425 | 22.83044438 | 14.71046797 | ENSG00000231113 |
| STARD5 | -0.63291083 | 0.644873982 | 0.298320199 | 0.491603678 | 16.11131413 | 10.45738285 | ENSG00000172345 |
| FEM1A | -0.633041981 | 0.644815361 | 0.077194309 | 0.186583114 | 51.72842521 | 33.16373178 | ENSG00000141965 |
| PCSK9 | -0.633184995 | 0.644751445 | 1.20E-16 | 5.44E-15 | 2171.113268 | 1399.643771 | ENSG00000169174 |
| NFATC2IP | -0.63409733 | 0.644343844 | 5.01E-10 | 9.73E-09 | 1094.704594 | 705.9950011 | ENSG00000176953 |
| NA | -0.634369152 | 0.644222453 | 0.567136121 | 0.734612098 | 4.763364208 | 3.056845979 | ENSG00000226054 |
| ZNF677 | -0.634900947 | 0.643985028 | 6.37E-06 | 6.01E-05 | 384.9775487 | 247.7472465 | ENSG00000197928 |
| LRRC37A11P | -0.635238782 | 0.643834244 | 0.411919465 | 0.603423598 | 9.649923073 | 6.200060856 | ENSG00000214553 |
| ARPC1B | -0.635759543 | 0.643601885 | 0.169323352 | 0.333890733 | 29.57482993 | 18.92980995 | ENSG00000130429 |
| TXNDC16 | -0.635782495 | 0.643591646 | 5.19E-05 | 0.000389158 | 355.1013166 | 228.5044931 | ENSG00000087301 |
| CCDC184 | -0.635798927 | 0.643584316 | 0.096403008 | 0.220525738 | 41.19264303 | 26.56699529 | ENSG00000177875 |
| PSME3 | -0.636014465 | 0.643488172 | 7.94E-27 | 8.66E-25 | 6010.335144 | 3866.94795 | ENSG00000131467 |
| FOXRED1 | -0.63610152 | 0.643449343 | 2.61E-10 | 5.35E-09 | 1273.350538 | 819.8509111 | ENSG00000110074 |
| PARPBP | -0.636420414 | 0.643307131 | 1.38E-06 | 1.50E-05 | 396.8157292 | 255.3463402 | ENSG00000185480 |
| SCOC-AS1 | -0.636519502 | 0.643262948 | 0.552203743 | 0.723084935 | 4.839806533 | 3.105672888 | ENSG00000196951 |
| NA | -0.636666158 | 0.643197561 | 0.465644436 | 0.651911536 | 7.867358636 | 5.029640753 | ENSG00000212789 |
| SLC51A | -0.637039578 | 0.643031101 | 0.026084739 | 0.080068658 | 77.04934858 | 49.44619364 | ENSG00000163959 |
| DIRAS1 | -0.637192732 | 0.642962841 | 5.27E-10 | 1.02E-08 | 894.0005874 | 575.1208329 | ENSG00000176490 |
| SEC61A2 | -0.63735303 | 0.642891406 | 3.68E-07 | 4.45E-06 | 437.4032603 | 281.475757 | ENSG00000065665 |
| MDH1B | -0.638136068 | 0.642542564 | 0.135738926 | 0.284206882 | 34.24596724 | 22.02474757 | ENSG00000138400 |
| ST7 | -0.638162274 | 0.642530893 | 0.001183286 | 0.0060835 | 202.4626378 | 129.8698985 | ENSG00000004866 |
| PEX26 | -0.638304089 | 0.642467736 | 7.10E-15 | 2.67E-13 | 1477.587149 | 948.9439417 | ENSG00000215193 |
| NAP1L3 | -0.638380979 | 0.642433496 | 2.26E-12 | 6.29E-11 | 1449.588038 | 932.1051924 | ENSG00000186310 |
| SLC35A3 | -0.638441451 | 0.642406569 | 0.012153244 | 0.043225528 | 128.4247485 | 82.36265329 | ENSG00000117620 |
| ROPN1L | -0.639156001 | 0.642088471 | 0.56096949 | 0.729870729 | 4.74442273 | 3.026790064 | ENSG00000145491 |
| TEX14 | -0.639615516 | 0.641883991 | 0.212353392 | 0.392124915 | 25.62847411 | 16.54525578 | ENSG00000121101 |
| H2AJ | -0.639709662 | 0.641842105 | 2.27E-05 | 0.000187056 | 314.5125979 | 202.2294087 | ENSG00000246705 |

| | | | | | | | |
|---|---|---|---|---|---|---|---|
| PLEKHB1 | -0.640266165 | 0.641594569 | 0.000336242 | 0.002025321 | 201.2387412 | 128.8950092 | ENSG00000021300 |
| NA | -0.640470606 | 0.641503657 | 0.510939585 | 0.690100169 | 6.342533389 | 4.043243366 | ENSG00000279722 |
| KCNV1 | -0.641140551 | 0.64120583 | 0.556765026 | 0.726235695 | 4.860025786 | 3.105672888 | ENSG00000164794 |
| ANOS1 | -0.641558553 | 0.641020077 | 1.11E-16 | 5.05E-15 | 4323.097726 | 2770.432632 | ENSG00000011201 |
| PROKR2 | -0.641589476 | 0.641006337 | 0.145613697 | 0.299368194 | 30.04946384 | 19.3091403 | ENSG00000101292 |
| ACP2 | -0.64210288 | 0.640778266 | 2.49E-05 | 0.00020285 | 311.0192062 | 199.52476 | ENSG00000134575 |
| NA | -0.64221377 | 0.640729016 | 0.476431512 | 0.66089522 | 7.020562556 | 4.508504585 | ENSG00000287620 |
| PHLDB2 | -0.642556707 | 0.640576729 | 2.56E-10 | 5.26E-09 | 744.3264806 | 476.683919 | ENSG00000144824 |
| NA | -0.642679625 | 0.640522154 | 0.051789113 | 0.137260035 | 60.1854421 | 38.62576668 | ENSG00000271109 |
| SMARCA2 | -0.6429125 | 0.640418771 | 3.42E-05 | 0.000269454 | 325.3431078 | 207.9086196 | ENSG00000080503 |
| ZNF460-AS1 | -0.644016671 | 0.639928812 | 0.527599127 | 0.703541422 | 5.366228826 | 3.443224373 | ENSG00000267871 |
| NA | -0.644175192 | 0.639858502 | 0.585546262 | 0.74855252 | 4.416457342 | 2.873261296 | ENSG00000226855 |
| ALKBH3 | -0.644195635 | 0.639849435 | 0.000564412 | 0.003191216 | 185.0476324 | 118.5034008 | ENSG00000166199 |
| LYPD3 | -0.644204898 | 0.639845327 | 0.023035712 | 0.072498536 | 78.26495452 | 49.945198 | ENSG00000124466 |
| UHRF2 | -0.644478255 | 0.639724103 | 1.63E-10 | 3.44E-09 | 1028.727116 | 657.4801883 | ENSG00000147854 |
| ZDHHC16 | -0.644736769 | 0.639609482 | 3.47E-14 | 1.21E-12 | 1171.408275 | 749.2395562 | ENSG00000171307 |
| PDGFRL | -0.6449837 | 0.639500016 | 0.031166781 | 0.092330451 | 77.01828312 | 49.43622281 | ENSG00000104213 |
| MAGEH1 | -0.64644023 | 0.638854709 | 0.000526946 | 0.0030069 | 255.8404584 | 163.7770411 | ENSG00000187601 |
| ATXN7 | -0.646792406 | 0.638698778 | 0.458333925 | 0.645327018 | 8.131509351 | 5.243719384 | ENSG00000163635 |
| POLR2J3 | -0.647087835 | 0.638568001 | 0.544249967 | 0.716466289 | 5.458455715 | 3.495850129 | ENSG00000285437 |
| CENPV | -0.647336784 | 0.63845782 | 6.20E-15 | 2.36E-13 | 2184.435038 | 1394.204918 | ENSG00000166582 |
| BTNL2 | -0.648306457 | 0.63802884 | 0.57307887 | 0.738883513 | 4.797488889 | 3.038074985 | ENSG00000204290 |
| NPEPPS | -0.648317233 | 0.638024074 | 4.66E-15 | 1.81E-13 | 1631.111935 | 1039.962495 | ENSG00000141279 |
| GPRIN2 | -0.648642196 | 0.637880378 | 6.79E-05 | 0.000493206 | 285.1514081 | 181.9663001 | ENSG00000204175 |
| NA | -0.648796924 | 0.637811969 | 0.44358117 | 0.631972743 | 8.595297048 | 5.468644903 | ENSG00000261559 |
| UTP20 | -0.648801952 | 0.637809746 | 7.77E-13 | 2.35E-11 | 1690.375386 | 1077.2811 | ENSG00000120800 |
| WDR73 | -0.648901141 | 0.637765897 | 0.004920084 | 0.02029124 | 115.2769764 | 73.53791736 | ENSG00000177082 |
| NA | -0.649090442 | 0.637682218 | 0.417884986 | 0.608985715 | 8.6079247 | 5.476130977 | ENSG00000233859 |
| NA | -0.649999953 | 0.637280335 | 0.542241416 | 0.714888491 | 5.994348747 | 3.848373762 | ENSG00000260018 |
| HRAS | -0.650380193 | 0.637112393 | 7.65E-10 | 1.45E-08 | 761.1528723 | 484.713105 | ENSG00000174775 |
| PODNL1 | -0.650548881 | 0.637037903 | 0.202157038 | 0.379382193 | 23.04280671 | 14.66543991 | ENSG00000132000 |
| NA | -0.650686364 | 0.636977199 | 2.39E-05 | 0.000195638 | 493.3317214 | 314.3386117 | ENSG00000284976 |
| FOXP4 | -0.651681344 | 0.636538048 | 1.77E-07 | 2.26E-06 | 817.7049814 | 520.1629643 | ENSG00000137166 |
| NA | -0.652377838 | 0.636230819 | 0.43717517 | 0.626145331 | 8.207951676 | 5.273775299 | ENSG00000267053 |
| RBM19 | -0.653014287 | 0.635950206 | 2.88E-15 | 1.13E-13 | 1727.817642 | 1099.663734 | ENSG00000122965 |
| NA | -0.653324437 | 0.635813504 | 0.44547284 | 0.633547699 | 7.648682477 | 4.887396905 | ENSG00000278987 |
| LINC01529 | -0.653547289 | 0.635715298 | 0.174928803 | 0.342141244 | 29.6638999 | 18.96833962 | ENSG00000225872 |
| NA | -0.653549772 | 0.635714204 | 0.504375986 | 0.684032511 | 6.037944167 | 3.870943603 | ENSG00000232546 |
| HDX | -0.653562316 | 0.635708676 | 0.017481009 | 0.057732186 | 101.6642365 | 64.39221077 | ENSG00000165259 |
| NA | -0.653770415 | 0.635616986 | 0.234666711 | 0.419534371 | 22.29001896 | 14.13301882 | ENSG00000269918 |
| SSTR5 | -0.654004208 | 0.635513991 | 0.188628929 | 0.360528882 | 24.39433266 | 15.46825261 | ENSG00000162009 |
| TOMM40L | -0.654302026 | 0.635382814 | 3.39E-06 | 3.41E-05 | 340.3910242 | 216.3087025 | ENSG00000158882 |
| NA | -0.654522539 | 0.635285705 | 0.001878319 | 0.008995548 | 164.6249109 | 104.8474778 | ENSG00000220785 |
| NA | -0.654986777 | 0.635081312 | 0.447372062 | 0.635565981 | 8.240798581 | 5.281261373 | ENSG00000273391 |
| CHRNB2 | -0.654991143 | 0.63507939 | 0.003492365 | 0.01514802 | 127.5372208 | 80.86661945 | ENSG00000160716 |
| AQP3 | -0.655193228 | 0.634990438 | 0.02394329 | 0.074808653 | 72.20816658 | 45.73670291 | ENSG00000165272 |
| DPP9-AS1 | -0.65527478 | 0.634954544 | 0.513028505 | 0.691827001 | 5.399075731 | 3.424453379 | ENSG00000205790 |
| FNDC4 | -0.655660075 | 0.634784992 | 0.000232 | 0.001465536 | 201.258284 | 127.703007 | ENSG00000115226 |
| NA | -0.655674392 | 0.634778693 | 0.554854234 | 0.724865607 | 4.750736556 | 2.977963155 | ENSG00000261351 |
| MAP6D1 | -0.656068169 | 0.634605456 | 0.114341795 | 0.250586595 | 38.81412916 | 24.63879055 | ENSG00000180834 |
| DNAJB6 | -0.656370053 | 0.634472679 | 4.16E-26 | 4.30E-24 | 8297.875646 | 5264.09848 | ENSG00000105993 |
| PGRMC1 | -0.656814869 | 0.634277087 | 4.74E-16 | 2.02E-14 | 6261.246845 | 3970.79343 | ENSG00000101856 |
| RNASEH2B | -0.657082986 | 0.634159221 | 8.54E-10 | 1.60E-08 | 687.4974278 | 435.7355334 | ENSG00000136104 |
| ANKRD34B | -0.657181096 | 0.634116096 | 0.024020033 | 0.074997859 | 68.7130685 | 43.51652439 | ENSG00000189127 |
| INSRR | -0.657192741 | 0.634110978 | 0.563722606 | 0.732061945 | 4.413300429 | 2.835719307 | ENSG00000027644 |
| DPP10-AS1 | -0.657860899 | 0.633817369 | 0.056381194 | 0.146668128 | 53.53456121 | 34.09155468 | ENSG00000235026 |
| NA | -0.658317262 | 0.633616907 | 0.493749815 | 0.675874798 | 6.020881826 | 3.840887688 | ENSG00000259408 |
| NA | -0.65834993 | 0.63360256 | 0.374296182 | 0.567389377 | 12.0916729 | 7.708032445 | ENSG00000231566 |
| HOGA1 | -0.658360238 | 0.633598033 | 0.574433754 | 0.739857901 | 4.472680413 | 2.873261296 | ENSG00000241935 |
| POLD3 | -0.65878241 | 0.633412652 | 1.37E-10 | 2.96E-09 | 749.6478136 | 474.5868869 | ENSG00000077514 |
| NA | -0.659511165 | 0.633092774 | 0.448126857 | 0.635975449 | 7.569083239 | 4.79722916 | ENSG00000286379 |
| TFEB | -0.659686371 | 0.633015894 | 0.034004123 | 0.098987443 | 67.59156859 | 42.79259451 | ENSG00000112561 |
| HSP90B2P | -0.660497531 | 0.632660079 | 0.487400822 | 0.67048815 | 8.072129366 | 5.119808498 | ENSG00000259706 |
| TRIM46 | -0.660551879 | 0.632636246 | 3.25E-05 | 0.000258458 | 457.6386869 | 290.0791918 | ENSG00000163462 |
| SNORD14A | -0.660569631 | 0.632628462 | 0.370886515 | 0.564139796 | 11.92931751 | 7.56914941 | ENSG00000272034 |
| OGG1 | -0.660762204 | 0.632544023 | 1.19E-08 | 1.88E-07 | 585.3795507 | 370.5193658 | ENSG00000114026 |
| ZNF358 | -0.660984687 | 0.632446484 | 0.001456296 | 0.007224567 | 187.8253301 | 118.5987529 | ENSG00000198816 |
| NA | -0.661304344 | 0.632306369 | 0.307637827 | 0.501782934 | 14.14913657 | 8.911869277 | ENSG00000270157 |
| EIF2B1 | -0.662013503 | 0.631995634 | 7.11E-15 | 2.67E-13 | 1298.568974 | 821.1486573 | ENSG00000111361 |
| BRICD5 | -0.662356529 | 0.631845384 | 0.028242699 | 0.085503133 | 85.52588566 | 54.23747406 | ENSG00000182685 |
| LSS | -0.663547217 | 0.631324123 | 4.94E-22 | 3.67E-20 | 5262.769402 | 3323.472695 | ENSG00000160285 |
| BMP3 | -0.663746997 | 0.631236705 | 0.498054318 | 0.678995125 | 6.954868746 | 4.362023857 | ENSG00000152785 |
| ADRA1D | -0.663765123 | 0.631228775 | 0.352877399 | 0.54602018 | 12.36341522 | 7.763907395 | ENSG00000171873 |
| UNC5B-AS1 | -0.664077888 | 0.631091944 | 0.358104244 | 0.551437692 | 13.70231351 | 8.608060934 | ENSG00000237512 |

| | | | | | | | |
|---|---|---|---|---|---|---|---|
| ANAPC7 | -0.664252962 | 0.631015364 | 9.82E-13 | 2.92E-11 | 983.3570834 | 620.8224072 | ENSG00000196510 |
| MRPL12 | -0.664863347 | 0.630748446 | 0.009992354 | 0.036752983 | 94.39036948 | 59.64914471 | ENSG00000262814 |
| CCNE1 | -0.665610649 | 0.630421809 | 7.66E-14 | 2.58E-12 | 1170.994435 | 738.3483853 | ENSG00000105173 |
| SAP30-DT | -0.665746121 | 0.630362614 | 0.410082643 | 0.601779233 | 10.24203918 | 6.406215381 | ENSG00000272870 |
| CNTFR | -0.666099184 | 0.630208368 | 0.009029345 | 0.033787578 | 108.4680998 | 68.24493304 | ENSG00000122756 |
| TMEM198 | -0.666615894 | 0.629982696 | 3.72E-05 | 0.000290897 | 249.3948519 | 157.2075007 | ENSG00000188760 |
| RUNDC3A | -0.666783064 | 0.629909701 | 0.013018381 | 0.045619716 | 87.73358073 | 55.32846169 | ENSG00000108309 |
| STX18 | -0.667039698 | 0.62979766 | 6.87E-09 | 1.13E-07 | 570.6321198 | 358.960323 | ENSG00000168818 |
| NA | -0.667376919 | 0.629650466 | 0.347467818 | 0.54072072 | 15.43634418 | 9.722279674 | ENSG00000214439 |
| LINC01018 | -0.668674124 | 0.629084567 | 0.248742929 | 0.436985919 | 21.68231369 | 13.78093322 | ENSG00000250056 |
| ESPL1 | -0.669149822 | 0.628877174 | 3.71E-17 | 1.80E-15 | 1860.104735 | 1170.434469 | ENSG00000135476 |
| TNFSF12 | -0.669277386 | 0.628821571 | 0.155527658 | 0.313739889 | 28.69528464 | 18.01101046 | ENSG00000239697 |
| TEX261 | -0.669374989 | 0.628779031 | 1.01E-06 | 1.13E-05 | 400.355024 | 252.1451005 | ENSG00000144043 |
| CDK4 | -0.669383997 | 0.628775105 | 1.11E-21 | 8.10E-20 | 2320.901452 | 1458.947861 | ENSG00000135446 |
| ZNF599 | -0.669506083 | 0.628721898 | 0.049472031 | 0.132674666 | 59.68692836 | 37.62450876 | ENSG00000153896 |
| CLDN5 | -0.669748974 | 0.628616056 | 0.211655913 | 0.391265777 | 20.71350303 | 12.98516856 | ENSG00000184113 |
| U2AF2 | -0.66975587 | 0.628613051 | 1.20E-24 | 1.10E-22 | 8646.229674 | 5435.990094 | ENSG00000063244 |
| DHX29 | -0.67057722 | 0.628255273 | 3.51E-12 | 9.52E-11 | 1057.652118 | 664.079235 | ENSG00000067248 |
| SKIV2L | -0.67066075 | 0.628218899 | 5.29E-14 | 1.81E-12 | 1279.977509 | 804.5488492 | ENSG00000204351 |
| CARMIL3 | -0.670664048 | 0.628217463 | 0.000636106 | 0.003540418 | 327.0375717 | 205.9530358 | ENSG00000186648 |
| NA | -0.671262026 | 0.627957129 | 0.553667363 | 0.723938212 | 4.446147334 | 2.816948313 | ENSG00000262766 |
| C10orf143 | -0.672566966 | 0.627389389 | 0.071643352 | 0.176175918 | 46.24027951 | 29.04270489 | ENSG00000237489 |
| SNORD17 | -0.672730372 | 0.627318332 | 0.259791856 | 0.448762231 | 20.22308455 | 12.78325091 | ENSG00000212232 |
| PWP2 | -0.673166125 | 0.627128885 | 0.083964656 | 0.198723142 | 43.13826191 | 27.19002216 | ENSG00000241945 |
| ZEB2 | -0.673285067 | 0.627077184 | 0.38812325 | 0.581197085 | 11.47430149 | 7.178972169 | ENSG00000169554 |
| GPX3 | -0.673389468 | 0.627031807 | 1.31E-09 | 2.39E-08 | 687.4823423 | 430.7461342 | ENSG00000211445 |
| TBC1D30 | -0.673512791 | 0.62697821 | 0.005165128 | 0.021169942 | 102.902715 | 64.41208107 | ENSG00000111490 |
| NDUFA9 | -0.673828258 | 0.626841126 | 0.02912541 | 0.087474554 | 66.99166549 | 41.96776162 | ENSG00000139180 |
| EPB41L1 | -0.673828719 | 0.626840926 | 1.25E-11 | 3.12E-10 | 883.0542942 | 553.7860488 | ENSG00000088367 |
| NCF1B | -0.674190298 | 0.626683843 | 0.553977129 | 0.724128085 | 5.114705763 | 3.252153616 | ENSG00000182487 |
| DHRS13 | -0.674776823 | 0.626429117 | 2.20E-07 | 2.77E-06 | 529.3371626 | 331.3958686 | ENSG00000167536 |
| LOC101929798 | -0.675152356 | 0.626266079 | 0.415118353 | 0.606527675 | 8.664147771 | 5.367192238 | ENSG00000275585 |
| NA | -0.676050801 | 0.62587619 | 0.420713575 | 0.611904999 | 8.776593914 | 5.446075062 | ENSG00000228060 |
| NA | -0.676053025 | 0.625875226 | 0.052917387 | 0.139691048 | 50.96448298 | 32.01730724 | ENSG00000279673 |
| TSPOAP1 | -0.676064459 | 0.625870265 | 0.00131425 | 0.006619104 | 264.0349857 | 165.7064483 | ENSG00000005379 |
| NA | -0.67628707 | 0.625773699 | 0.015605396 | 0.052731159 | 89.20553521 | 55.97339712 | ENSG00000182057 |
| NA | -0.676439923 | 0.625707402 | 0.002378848 | 0.010996578 | 154.0453379 | 96.21878211 | ENSG00000226067 |
| TCF15 | -0.677010568 | 0.625459958 | 8.04E-06 | 7.36E-05 | 323.5005695 | 202.6147994 | ENSG00000125878 |
| GNG13 | -0.67776948 | 0.625131029 | 0.411508021 | 0.603059197 | 9.546947669 | 6.027761094 | ENSG00000127588 |
| STBD1 | -0.678412938 | 0.624852276 | 0.257831084 | 0.446392079 | 16.9953918 | 10.64096754 | ENSG00000118804 |
| LOC100379224 | -0.678686651 | 0.624733738 | 0.456010188 | 0.643266615 | 7.559612501 | 4.692089268 | ENSG00000186019 |
| NA | -0.678686955 | 0.624733607 | 0.477095428 | 0.661466686 | 6.583307938 | 4.129612264 | ENSG00000225335 |
| REC8 | -0.679307016 | 0.624465158 | 1.23E-07 | 1.61E-06 | 568.2193835 | 354.6560463 | ENSG00000100918 |
| UBE2F | -0.679374661 | 0.624435879 | 0.002557181 | 0.011676471 | 130.3839797 | 81.16619092 | ENSG00000184182 |
| NA | -0.679752919 | 0.62427218 | 0.03657173 | 0.104715039 | 55.22795798 | 34.48118226 | ENSG00000234009 |
| FERMT1 | -0.679874549 | 0.624219552 | 1.72E-08 | 2.65E-07 | 579.2520707 | 361.0111245 | ENSG00000101311 |
| GLT1D1 | -0.680151787 | 0.624099609 | 1.93E-06 | 2.04E-05 | 325.1304598 | 202.774715 | ENSG00000151948 |
| ACTL6B | -0.680697881 | 0.623863417 | 0.223921605 | 0.406387203 | 24.77979889 | 15.5400874 | ENSG00000077080 |
| NAB2 | -0.681839884 | 0.623369778 | 7.02E-08 | 9.61E-07 | 661.9415174 | 412.5648078 | ENSG00000166886 |
| RAPGEF4 | -0.682030565 | 0.623287392 | 0.059326719 | 0.152589703 | 52.42865047 | 32.84648863 | ENSG00000091428 |
| DZIP1 | -0.682338017 | 0.623154578 | 8.82E-13 | 2.65E-11 | 1025.209415 | 639.296091 | ENSG00000134874 |
| PCDHGC4 | -0.682800719 | 0.622954751 | 0.025329966 | 0.078205164 | 69.26021371 | 43.18309817 | ENSG00000242419 |
| SP110 | -0.683444327 | 0.622676904 | 0.014415012 | 0.049429755 | 82.17342739 | 51.27773225 | ENSG00000135899 |
| MGC16275 | -0.6837129 | 0.622560997 | 0.397662947 | 0.589769848 | 8.895451581 | 5.562499875 | ENSG00000246731 |
| PRDM16 | -0.683776118 | 0.622533717 | 0.272887182 | 0.464130841 | 14.953615 | 9.324504739 | ENSG00000142611 |
| TBC1D19 | -0.684159139 | 0.622368463 | 0.079098049 | 0.190069135 | 38.46524546 | 23.87351984 | ENSG00000109680 |
| GCLM | -0.685394521 | 0.621835756 | 1.99E-06 | 2.10E-05 | 443.2943254 | 275.3285049 | ENSG00000023909 |
| ACSS1 | -0.68543183 | 0.621819675 | 0.057050786 | 0.148050863 | 47.15404717 | 29.35388769 | ENSG00000154930 |
| EBP | -0.685687525 | 0.621709476 | 8.79E-07 | 9.87E-06 | 336.1202507 | 209.0129791 | ENSG00000147155 |
| ATAD3B | -0.685854809 | 0.621637392 | 1.64E-15 | 6.69E-14 | 2560.001995 | 1592.00176 | ENSG00000160072 |
| LOC101928069 | -0.686107499 | 0.621528521 | 0.421061469 | 0.612218817 | 8.859447763 | 5.52115904 | ENSG00000255517 |
| C21orf59-TCP10L | -0.686213088 | 0.621483034 | 0.411133702 | 0.602690656 | 8.783005438 | 5.464846056 | ENSG00000265590 |
| NA | -0.686748468 | 0.621252446 | 0.57479716 | 0.740274517 | 4.757050382 | 2.899080331 | ENSG00000280010 |
| NA | -0.686801566 | 0.621229582 | 0.490729445 | 0.673517442 | 6.183237215 | 3.840887688 | ENSG00000260160 |
| PROS1 | -0.68702228 | 0.621134549 | 0.252189342 | 0.440938768 | 16.49550259 | 10.21704715 | ENSG00000184500 |
| NDUFC2 | -0.687199497 | 0.621058255 | 0.032138033 | 0.094592076 | 63.80766586 | 39.75875641 | ENSG00000151366 |
| LMNTD2-AS1 | -0.687611094 | 0.620881094 | 0.41629919 | 0.607486748 | 16.13587037 | 10.00351818 | ENSG00000254815 |
| PDE10A | -0.687626658 | 0.620874396 | 0.07159391 | 0.176101066 | 45.22471651 | 27.94681912 | ENSG00000112541 |
| PCDHA6 | -0.68763009 | 0.620872918 | 0.00336971 | 0.014688193 | 110.4560137 | 68.63989681 | ENSG00000081842 |
| CEP162 | -0.687763285 | 0.6208156 | 0.001196014 | 0.006135326 | 163.5242316 | 101.3191583 | ENSG00000135315 |
| C5 | -0.687939979 | 0.62073957 | 0.433078137 | 0.622850174 | 7.7876617 | 4.808514081 | ENSG00000106804 |
| SMIM3 | -0.688347369 | 0.620564309 | 8.26E-10 | 1.55E-08 | 655.0239681 | 406.0559288 | ENSG00000256235 |
| FBXO3 | -0.689443657 | 0.620092929 | 2.34E-07 | 2.93E-06 | 401.9683179 | 249.2248357 | ENSG00000110429 |
| NA | -0.689466202 | 0.620083239 | 0.27363021 | 0.464871193 | 14.88980033 | 9.219364847 | ENSG00000257740 |

| | | | | | | | |
|---|---|---|---|---|---|---|---|
| NA | -0.689557108 | 0.620044168 | 0.5454182 | 0.717494564 | 4.505527319 | 2.805663392 | ENSG00000233747 |
| NA | -0.689671434 | 0.619995034 | 0.233925752 | 0.418856139 | 17.93431707 | 11.15505567 | ENSG00000277182 |
| NA | -0.690060613 | 0.619827808 | 0.388816998 | 0.581598708 | 9.968417723 | 6.151233947 | ENSG00000229932 |
| SMOC2 | -0.69024133 | 0.619750171 | 0.008824319 | 0.0331677 | 96.19067267 | 59.70925654 | ENSG00000112562 |
| NA | -0.690291318 | 0.619728698 | 0.152057225 | 0.309117978 | 26.02350878 | 16.12089736 | ENSG00000258568 |
| LINGO1 | -0.690300352 | 0.619724817 | 3.21E-13 | 1.00E-11 | 2075.536201 | 1285.861479 | ENSG00000169783 |
| GPR143 | -0.690358873 | 0.61969968 | 4.57E-07 | 5.44E-06 | 388.231782 | 240.8389923 | ENSG00000101850 |
| C11orf71 | -0.690701738 | 0.619552422 | 0.268128387 | 0.458313339 | 16.19407028 | 10.06731723 | ENSG00000180425 |
| ADPRHL1 | -0.691738915 | 0.619107175 | 0.105613807 | 0.236249791 | 32.03364209 | 19.79273448 | ENSG00000153531 |
| CR1 | -0.691760784 | 0.619097791 | 0.398994493 | 0.59074644 | 8.954831566 | 5.555013801 | ENSG00000203710 |
| NA | -0.691941954 | 0.619020051 | 0.089870928 | 0.209342168 | 56.28899553 | 35.09011305 | ENSG00000272849 |
| NA | -0.691969763 | 0.619008119 | 0.004177944 | 0.017679 | 125.4122819 | 77.77344241 | ENSG00000274712 |
| CYP27B1 | -0.692127718 | 0.61894035 | 0.068954272 | 0.17115287 | 43.47509668 | 26.97963076 | ENSG00000111012 |
| STS | -0.692234716 | 0.618894447 | 0.004806429 | 0.019876205 | 113.4666988 | 70.24453453 | ENSG00000101846 |
| GRIA3 | -0.693195366 | 0.61848248 | 0.00060053 | 0.003367691 | 197.4571009 | 122.2621565 | ENSG00000125675 |
| HNRNPM | -0.693304255 | 0.618435802 | 1.17E-27 | 1.37E-25 | 14475.76401 | 8952.038608 | ENSG00000099783 |
| KLRG1 | -0.693404476 | 0.618392842 | 0.056852735 | 0.14772306 | 50.97385602 | 31.55573324 | ENSG00000139187 |
| NA | -0.693510792 | 0.618347272 | 0.26852701 | 0.458781521 | 14.9460234 | 9.219364847 | ENSG00000235459 |
| PROC | -0.693857575 | 0.618198657 | 0.229739033 | 0.413627166 | 17.14699868 | 10.57336963 | ENSG00000115718 |
| PIANP | -0.694050952 | 0.6181158 | 0.000114688 | 0.000788946 | 226.4011425 | 140.2018818 | ENSG00000139200 |
| NA | -0.694088193 | 0.618099845 | 0.300448652 | 0.493745352 | 13.46479357 | 8.338107352 | ENSG00000283692 |
| HMGN2 | -0.694177554 | 0.61806156 | 9.64E-16 | 4.02E-14 | 6303.993973 | 3895.67335 | ENSG00000198830 |
| CHMP4A | -0.694485789 | 0.617929524 | 0.22056273 | 0.402037429 | 19.69488082 | 12.24290572 | ENSG00000254505 |
| CNOT2-DT | -0.694504209 | 0.617921635 | 0.317977315 | 0.511824082 | 12.91499511 | 7.974298799 | ENSG00000257815 |
| LZTS3 | -0.69451035 | 0.617919004 | 1.49E-11 | 3.67E-10 | 1231.19533 | 761.5871412 | ENSG00000088899 |
| SYNDIG1 | -0.695564259 | 0.617467771 | 0.002531033 | 0.011577322 | 119.8640798 | 73.96151133 | ENSG00000101463 |
| CTB-178M22.2 | -0.695639708 | 0.61743548 | 0.276298978 | 0.467365634 | 21.7480075 | 13.57109147 | ENSG00000253978 |
| PAK5 | -0.695691034 | 0.617413514 | 0.227467148 | 0.410959013 | 17.17668867 | 10.57336963 | ENSG00000101349 |
| RPGRIP1 | -0.695934798 | 0.617309202 | 0.319780514 | 0.513518689 | 12.37604287 | 7.685574223 | ENSG00000092200 |
| SLC12A8 | -0.696357048 | 0.617128553 | 0.000143706 | 0.000956697 | 223.4486799 | 137.9954645 | ENSG00000221955 |
| AKT1S1 | -0.696963387 | 0.61686924 | 1.06E-12 | 3.13E-11 | 1425.54198 | 879.1628046 | ENSG00000204673 |
| NA | -0.697038027 | 0.616837326 | 0.35296508 | 0.546110259 | 15.32083882 | 9.546069444 | ENSG00000214222 |
| RIPOR3 | -0.697649656 | 0.616575874 | 0.034950859 | 0.101136491 | 66.64683315 | 41.24426978 | ENSG00000042062 |
| NA | -0.697833159 | 0.616497454 | 0.516843181 | 0.694373435 | 5.061639604 | 3.143214877 | ENSG00000232842 |
| FAM151B | -0.69799185 | 0.616429645 | 0.439647166 | 0.628007655 | 7.275242532 | 4.501018511 | ENSG00000152380 |
| SETD6 | -0.69833738 | 0.616282026 | 1.70E-10 | 3.58E-09 | 668.3700842 | 412.2924726 | ENSG00000103037 |
| NA | -0.698381469 | 0.616263193 | 0.42646645 | 0.617131017 | 7.843982469 | 4.868625911 | ENSG00000205267 |
| SNHG20 | -0.698615972 | 0.61616303 | 0.045870816 | 0.125119921 | 66.81087228 | 41.33476393 | ENSG00000234912 |
| RFX8 | -0.698730357 | 0.616114179 | 0.467529722 | 0.653434572 | 7.214584772 | 4.452191601 | ENSG00000196460 |
| SPNS1 | -0.698836519 | 0.616068844 | 0.307877177 | 0.501996621 | 12.39626212 | 7.681775377 | ENSG00000169682 |
| NA | -0.699256046 | 0.615889721 | 0.083556196 | 0.198085257 | 40.80519996 | 25.12293438 | ENSG00000196656 |
| PCBP1-AS1 | -0.69926172 | 0.615887298 | 0.000456834 | 0.00264675 | 221.5169665 | 136.9252233 | ENSG00000179818 |
| LMO3 | -0.699338944 | 0.615854332 | 0.003062829 | 0.01355618 | 134.145303 | 82.58454441 | ENSG00000048540 |
| HM13 | -0.699492884 | 0.615788622 | 8.35E-19 | 4.81E-17 | 2330.123824 | 1434.565446 | ENSG00000101294 |
| LOC102724474 | -0.700273775 | 0.615455403 | 0.261592117 | 0.450883315 | 15.16154265 | 9.384616569 | ENSG00000283828 |
| SLC30A2 | -0.701018423 | 0.615137817 | 0.25817191 | 0.446899988 | 15.18807572 | 9.39590149 | ENSG00000158014 |
| NA | -0.701299207 | 0.615018108 | 0.208848674 | 0.387970471 | 18.9966324 | 11.69876167 | ENSG00000257851 |
| NA | -0.702158789 | 0.614651779 | 0.531365369 | 0.706652587 | 5.153866493 | 3.192041786 | ENSG00000215630 |
| C8G | -0.702538564 | 0.614489999 | 0.488137808 | 0.671127481 | 6.083418724 | 3.705691881 | ENSG00000176919 |
| UCP1 | -0.702759399 | 0.614395946 | 0.41515412 | 0.606532118 | 8.429784747 | 5.22494839 | ENSG00000109424 |
| KIAA1549L | -0.703983002 | 0.613875075 | 0.002683593 | 0.01212819 | 109.3140991 | 67.09047277 | ENSG00000110427 |
| ATP6V0D1 | -0.704035941 | 0.613852549 | 6.63E-16 | 2.81E-14 | 1281.647056 | 786.6920916 | ENSG00000159720 |
| MCM4 | -0.704088745 | 0.613830082 | 1.17E-31 | 1.82E-29 | 8426.954086 | 5172.758363 | ENSG00000104738 |
| CHD1L | -0.704420957 | 0.613688751 | 0.543412727 | 0.71590462 | 4.406986603 | 2.689238579 | ENSG00000131778 |
| ZW10 | -0.704466441 | 0.613669403 | 1.12E-09 | 2.05E-08 | 661.6947075 | 405.7176044 | ENSG00000086827 |
| OTULINL | -0.704612375 | 0.613607331 | 0.000346405 | 0.00207574 | 207.1340682 | 126.9404358 | ENSG00000145569 |
| BOLA3-AS1 | -0.705005243 | 0.613440259 | 0.000175002 | 0.001140865 | 204.9431424 | 126.05053 | ENSG00000225439 |
| NA | -0.705124527 | 0.613389541 | 0.229582816 | 0.4134467 | 21.41245051 | 13.19924719 | ENSG00000262691 |
| SPSB3 | -0.705720526 | 0.613136193 | 0.261132841 | 0.450133564 | 16.37230793 | 10.10485922 | ENSG00000162032 |
| NT5C3A | -0.705770682 | 0.613114878 | 1.03E-06 | 1.15E-05 | 392.7612642 | 240.4123321 | ENSG00000122643 |
| NA | -0.705776927 | 0.613112224 | 0.424359246 | 0.615227963 | 8.587705446 | 5.315116135 | ENSG00000283355 |
| CHST1 | -0.705865202 | 0.61307471 | 0.040322843 | 0.113131029 | 56.68215106 | 34.64924515 | ENSG00000175264 |
| LOC728673 | -0.706352832 | 0.612867526 | 0.33954829 | 0.533776924 | 11.2341283 | 6.867677753 | ENSG00000170161 |
| NA | -0.706558527 | 0.612780152 | 0.258340817 | 0.447067105 | 16.60982787 | 10.09682349 | ENSG00000272078 |
| ARMC2 | -0.706849349 | 0.612656638 | 0.042782082 | 0.118630028 | 55.55464559 | 34.15166651 | ENSG00000118690 |
| FOXI2 | -0.706940107 | 0.612618098 | 0.274663881 | 0.465619717 | 13.97415353 | 8.544261877 | ENSG00000186766 |
| KCP | -0.706980076 | 0.612601126 | 0.000398642 | 0.002345534 | 159.7636074 | 97.96372859 | ENSG00000135253 |
| NA | -0.7079985 | 0.612168833 | 0.299676126 | 0.49318037 | 13.31950053 | 8.1127438 | ENSG00000276564 |
| RNF224 | -0.708071917 | 0.612137681 | 0.514794068 | 0.692846128 | 5.170928834 | 3.184555713 | ENSG00000233198 |
| LRP2 | -0.708606527 | 0.611910887 | 0.190585908 | 0.36318314 | 26.65606339 | 16.38369128 | ENSG00000081479 |
| ALOX15 | -0.70882646 | 0.611817611 | 0.288533664 | 0.480179616 | 15.73461957 | 9.601944394 | ENSG00000161905 |
| CCDC157 | -0.708899041 | 0.611786831 | 0.015447967 | 0.052323211 | 72.6859574 | 44.56715887 | ENSG00000187860 |
| CHN1 | -0.709435276 | 0.611559479 | 5.09E-12 | 1.36E-10 | 893.7728616 | 545.9428919 | ENSG00000128656 |
| LINC02387 | -0.709467249 | 0.611545926 | 0.230390015 | 0.414487336 | 18.99525692 | 11.53719718 | ENSG00000256232 |

| | | | | | | | |
|---|---|---|---|---|---|---|---|
| NA | -0.709786527 | 0.611410602 | 0.468395356 | 0.654200416 | 6.652158661 | 4.043243366 | ENSG00000267342 |
| WDR86-AS1 | -0.710014679 | 0.611313919 | 0.194260181 | 0.368253882 | 25.66901031 | 15.6630106 | ENSG00000243836 |
| GPR62 | -0.710631274 | 0.611052704 | 0.419556896 | 0.610893827 | 8.959266255 | 5.449873909 | ENSG00000180929 |
| ZNF511 | -0.710769775 | 0.610994045 | 0.003812614 | 0.016351018 | 102.0263266 | 62.22520767 | ENSG00000198546 |
| LOC105374338 | -0.711175403 | 0.610822283 | 0.526609932 | 0.702829156 | 4.528903485 | 2.756836483 | ENSG00000283183 |
| BMF | -0.711175696 | 0.610822158 | 1.00E-07 | 1.34E-06 | 508.1660054 | 310.7251577 | ENSG00000104081 |
| NKAIN3 | -0.711280842 | 0.610777643 | 0.275938239 | 0.466925895 | 13.98678118 | 8.514205962 | ENSG00000185942 |
| NXF1 | -0.711329613 | 0.610756995 | 4.00E-14 | 1.39E-12 | 1025.520033 | 626.2768041 | ENSG00000162231 |
| TMEM163 | -0.711484517 | 0.610691421 | 0.081640551 | 0.194664348 | 35.03140651 | 21.4284158 | ENSG00000152128 |
| AMD1 | -0.711688772 | 0.610604966 | 0.000461064 | 0.002667924 | 3930.515804 | 2399.722927 | ENSG00000123505 |
| DLX4 | -0.711740082 | 0.61058325 | 0.350814194 | 0.544054076 | 10.11065156 | 6.181289862 | ENSG00000108813 |
| SERP2 | -0.711933157 | 0.610501542 | 0.182095195 | 0.352375769 | 21.64621217 | 13.29310216 | ENSG00000151778 |
| SH3GL2 | -0.711942035 | 0.610497785 | 0.003933255 | 0.016812488 | 109.4685546 | 67.00834075 | ENSG00000107295 |
| LOC653160 | -0.711944262 | 0.610496842 | 0.504316748 | 0.684010728 | 5.094486509 | 3.116957809 | ENSG00000241014 |
| PRRT3 | -0.712288886 | 0.610351027 | 0.000239102 | 0.001507316 | 209.4716399 | 127.7395613 | ENSG00000163704 |
| NA | -0.712386805 | 0.610309603 | 0.420385439 | 0.611558221 | 7.834511731 | 4.789743086 | ENSG00000269425 |
| ADORA1 | -0.71241136 | 0.610299215 | 0.002408005 | 0.011104745 | 135.5789611 | 82.77649124 | ENSG00000163485 |
| SLC20A1 | -0.712570622 | 0.610231846 | 9.07E-23 | 7.25E-21 | 2448.411328 | 1493.425754 | ENSG00000144136 |
| ANKRD36B | -0.712618934 | 0.610211412 | 0.022960071 | 0.072346473 | 61.9989743 | 37.84183659 | ENSG00000196912 |
| MCM6 | -0.713021211 | 0.610041286 | 8.54E-30 | 1.18E-27 | 5133.474836 | 3130.8564 | ENSG00000076003 |
| SYT2 | -0.713112214 | 0.610002807 | 7.66E-13 | 2.32E-11 | 803.3529982 | 490.1087445 | ENSG00000143858 |
| DHFR | -0.71438382 | 0.609465381 | 3.68E-15 | 1.45E-13 | 1363.956102 | 830.8491627 | ENSG00000228716 |
| FOS | -0.714394172 | 0.609461008 | 5.29E-07 | 6.23E-06 | 358.4630427 | 218.3526792 | ENSG00000170345 |
| EVL | -0.714884604 | 0.609253862 | 2.41E-12 | 6.67E-11 | 1173.792682 | 715.6776433 | ENSG00000196405 |
| NA | -0.715055289 | 0.609181785 | 0.48044487 | 0.6642839 | 6.411384112 | 3.957312501 | ENSG00000263345 |
| NA | -0.715183557 | 0.609127626 | 0.323661219 | 0.517846775 | 11.39342448 | 6.987901412 | ENSG00000226608 |
| ETV2 | -0.715186222 | 0.609126501 | 0.256487073 | 0.445189306 | 15.83316029 | 9.598257167 | ENSG00000105672 |
| LOC100996756 | -0.7156274 | 0.608940258 | 0.383850308 | 0.576568734 | 8.522011636 | 5.22494839 | ENSG00000287978 |
| SRPK3 | -0.715829083 | 0.608855136 | 0.09986868 | 0.22664979 | 35.37841107 | 21.69512019 | ENSG00000184343 |
| MFSD6 | -0.715908833 | 0.608821481 | 0.016890933 | 0.056193959 | 73.77273127 | 44.79164636 | ENSG00000151690 |
| TNFSF13 | -0.716261512 | 0.608672668 | 0.041891993 | 0.116738265 | 52.1113359 | 31.70188756 | ENSG00000161955 |
| N4BP2L1 | -0.716902043 | 0.608402488 | 0.147886718 | 0.302806422 | 27.09707628 | 16.46962215 | ENSG00000139597 |
| NA | -0.71753875 | 0.60813404 | 0.277293271 | 0.468448955 | 13.02438204 | 7.932957964 | ENSG00000234618 |
| ENKD1 | -0.717693198 | 0.608068939 | 5.98E-05 | 0.000441262 | 214.5384284 | 130.2646475 | ENSG00000124074 |
| NA | -0.718032405 | 0.607925986 | 0.498873382 | 0.679761746 | 5.174085747 | 3.154499798 | ENSG00000229539 |
| SLC1A6 | -0.71816997 | 0.607868022 | 2.07E-05 | 0.000171879 | 282.4116161 | 171.6028838 | ENSG00000105143 |
| IL17C | -0.718453125 | 0.607748728 | 0.218643928 | 0.399915776 | 19.7138223 | 11.96111756 | ENSG00000124391 |
| IRF1 | -0.718498461 | 0.60772963 | 0.017415055 | 0.057575922 | 67.81330396 | 41.24740735 | ENSG00000125347 |
| FAIM2 | -0.718536331 | 0.607713678 | 0.221522983 | 0.403231877 | 17.52803022 | 10.64096754 | ENSG00000135472 |
| NSG1 | -0.718744895 | 0.60762583 | 1.54E-14 | 5.61E-13 | 1037.585173 | 630.5046199 | ENSG00000168824 |
| NA | -0.719021808 | 0.607509213 | 0.375081359 | 0.568093316 | 9.712459971 | 5.892565286 | ENSG00000214433 |
| NA | -0.719505088 | 0.607305741 | 0.352434201 | 0.545516575 | 9.584229263 | 5.794911468 | ENSG00000215007 |
| TAF7 | -0.719895456 | 0.607141437 | 0.000297049 | 0.001821811 | 2994.820859 | 1818.666245 | ENSG00000178913 |
| NA | -0.720329056 | 0.606958989 | 0.53468658 | 0.70931795 | 5.683445699 | 3.458308141 | ENSG00000226330 |
| DPP4 | -0.720410893 | 0.60692456 | 0.011300977 | 0.040701966 | 82.05416376 | 49.83680891 | ENSG00000197635 |
| ARSI | -0.72045024 | 0.606908007 | 0.00035944 | 0.002143456 | 172.364813 | 104.7148699 | ENSG00000183876 |
| PPP1R1B | -0.721029304 | 0.606664457 | 8.59E-07 | 9.67E-06 | 299.2238319 | 181.4745586 | ENSG00000131771 |
| ZNF670-ZNF695 | -0.721068316 | 0.606648053 | 0.371159857 | 0.564401422 | 9.187413152 | 5.622611705 | ENSG00000135747 |
| NAA16 | -0.721188563 | 0.606597492 | 3.24E-08 | 4.76E-07 | 464.4385889 | 281.5158954 | ENSG00000172766 |
| FEZF1 | -0.721622581 | 0.606415031 | 3.37E-05 | 0.000266456 | 215.4233779 | 130.6702349 | ENSG00000128610 |
| NA | -0.721788 | 0.606345504 | 0.333809918 | 0.528382698 | 10.79183764 | 6.552696109 | ENSG00000270878 |
| LINC01574 | -0.721816835 | 0.606333385 | 0.422565025 | 0.613587969 | 7.350407081 | 4.440906681 | ENSG00000248859 |
| GPAM | -0.722112401 | 0.606209178 | 4.07E-11 | 9.39E-10 | 723.3321479 | 437.9811961 | ENSG00000119927 |
| ZNF506 | -0.722339251 | 0.606113864 | 0.520880524 | 0.697896309 | 5.074267255 | 3.0681309 | ENSG00000081665 |
| PXMP2 | -0.722400324 | 0.606088207 | 0.322220944 | 0.516255444 | 10.76214764 | 6.530126268 | ENSG00000176894 |
| RHBDL2 | -0.722409807 | 0.606084223 | 0.567172258 | 0.734612098 | 4.516275834 | 2.719294494 | ENSG00000158315 |
| IFT57 | -0.722656229 | 0.605980709 | 2.84E-09 | 4.92E-08 | 561.8653647 | 340.1383525 | ENSG00000114446 |
| CDK2AP1 | -0.722783442 | 0.605927277 | 3.21E-13 | 1.00E-11 | 2951.363066 | 1787.621775 | ENSG00000111328 |
| MPZ | -0.722819594 | 0.605912093 | 0.034330229 | 0.099707731 | 66.52768239 | 40.22737844 | ENSG00000158887 |
| NA | -0.723231635 | 0.605739067 | 0.254117209 | 0.442914535 | 15.82378725 | 9.5831734 | ENSG00000260401 |
| MYH14 | -0.723630688 | 0.605571541 | 3.16E-18 | 1.70E-16 | 1406.153094 | 851.1987518 | ENSG00000105357 |
| ZNF385B | -0.723831763 | 0.605487146 | 0.056472897 | 0.146839039 | 42.53657737 | 25.68529977 | ENSG00000144331 |
| ST14 | -0.724194447 | 0.605334949 | 8.84E-22 | 6.53E-20 | 1869.553525 | 1131.556388 | ENSG00000149418 |
| DTX1 | -0.724423292 | 0.605238936 | 7.01E-07 | 8.04E-06 | 329.6916088 | 199.5797504 | ENSG00000135144 |
| LIG1 | -0.724439431 | 0.605232166 | 3.17E-19 | 1.88E-17 | 2125.183831 | 1286.575455 | ENSG00000105486 |
| GEMIN7-AS1 | -0.72449539 | 0.605208691 | 0.43724204 | 0.626166278 | 6.748820239 | 4.043243366 | ENSG00000267348 |
| ARL9 | -0.724689662 | 0.6051272 | 0.276453418 | 0.467541533 | 15.05974732 | 9.174774817 | ENSG00000196503 |
| CCDC88B | -0.724970681 | 0.60500934 | 3.17E-07 | 3.87E-06 | 474.6621751 | 287.6968991 | ENSG00000168071 |
| ZNF236-DT | -0.725065096 | 0.604969747 | 0.254936258 | 0.443744234 | 19.43586385 | 11.69821202 | ENSG00000264278 |
| STAMBPL1 | -0.725813487 | 0.604656003 | 0.000972264 | 0.005139743 | 128.7954375 | 77.7911141 | ENSG00000138134 |
| SYT10 | -0.726008949 | 0.604574088 | 0.185584463 | 0.356406749 | 25.10460737 | 15.14947212 | ENSG00000110975 |
| NPIPB15 | -0.726078822 | 0.604544808 | 0.249510037 | 0.438009765 | 16.17700794 | 9.868648782 | ENSG00000196436 |
| NA | -0.726180357 | 0.604502262 | 0.447613643 | 0.635762991 | 7.516017081 | 4.598672329 | ENSG00000287430 |
| BAG1 | -0.726727118 | 0.604273207 | 2.14E-12 | 6.02E-11 | 1176.210619 | 710.4196441 | ENSG00000107262 |

| | | | | | | | |
|---|---|---|---|---|---|---|---|
| AMER3 | -0.726866497 | 0.604214832 | 0.461659032 | 0.64821125 | 7.572240152 | 4.628728244 | ENSG00000178171 |
| LINC01431 | -0.727658431 | 0.603883253 | 0.495139262 | 0.677062939 | 5.17724266 | 3.124443883 | ENSG00000232645 |
| H2AC20 | -0.72777858 | 0.603832963 | 0.15564089 | 0.313924067 | 27.56223946 | 16.71766716 | ENSG00000184260 |
| DHCR24 | -0.728181151 | 0.603664492 | 1.08E-38 | 2.67E-36 | 13545.5691 | 8176.710401 | ENSG00000116133 |
| HES1 | -0.729014209 | 0.603316018 | 0.000197572 | 0.001269254 | 212.0236863 | 127.6039274 | ENSG00000114315 |
| NA | -0.729290899 | 0.603200321 | 0.216366296 | 0.396924966 | 17.10971708 | 10.34095804 | ENSG00000198225 |
| GUSBP1 | -0.729488239 | 0.603117817 | 0.154100702 | 0.311699488 | 25.19693195 | 15.2133828 | ENSG00000215190 |
| NPPC | -0.73023916 | 0.602803977 | 0.143901762 | 0.296770363 | 23.44277973 | 14.17435966 | ENSG00000163273 |
| DLK2 | -0.730442437 | 0.602719047 | 0.376778647 | 0.569918832 | 10.31532459 | 6.20754693 | ENSG00000171462 |
| NA | -0.730820298 | 0.602561208 | 0.130889096 | 0.277023924 | 25.53309031 | 15.31428581 | ENSG00000268573 |
| NA | -0.731005179 | 0.602483995 | 0.455330288 | 0.642684606 | 6.289467231 | 3.810831773 | ENSG00000232150 |
| AARD | -0.731418266 | 0.60231151 | 0.310627564 | 0.504483988 | 14.3691882 | 8.72079852 | ENSG00000205002 |
| NA | -0.731449814 | 0.602298339 | 0.318383083 | 0.512054153 | 11.59178368 | 7.006672407 | ENSG00000230373 |
| THAP10 | -0.731485202 | 0.602283566 | 0.007830607 | 0.030097983 | 95.20489737 | 57.48907802 | ENSG00000129028 |
| ARHGEF19 | -0.732768316 | 0.60174814 | 1.63E-15 | 6.68E-14 | 1410.487419 | 848.9827558 | ENSG00000142632 |
| GTF2IRD2 | -0.732818311 | 0.601727288 | 0.135140213 | 0.283207678 | 29.89016766 | 17.8796135 | ENSG00000196275 |
| PCYT2 | -0.733237553 | 0.601552454 | 9.78E-17 | 4.46E-15 | 1193.523582 | 718.4554091 | ENSG00000185813 |
| OAT | -0.733722663 | 0.601350214 | 2.17E-25 | 2.13E-23 | 2426.821745 | 1458.796602 | ENSG00000065154 |
| ZNF703 | -0.733777555 | 0.601327334 | 0.06418942 | 0.162193143 | 44.04383662 | 26.3599647 | ENSG00000183779 |
| DHX35 | -0.734390409 | 0.601071946 | 1.00E-09 | 1.86E-08 | 527.6294848 | 317.2773126 | ENSG00000101452 |
| HTR3A | -0.735332306 | 0.60067965 | 7.20E-05 | 0.000519089 | 197.4851298 | 118.792237 | ENSG00000166736 |
| SCNN1A | -0.735379472 | 0.600660012 | 1.40E-18 | 7.81E-17 | 1609.664149 | 967.2706353 | ENSG00000111319 |
| NA | -0.73600508 | 0.6003996 | 0.543993648 | 0.716179733 | 4.469523501 | 2.647897744 | ENSG00000285090 |
| COL9A2 | -0.736482294 | 0.600201033 | 0.000383096 | 0.002268921 | 149.1701289 | 89.53914072 | ENSG00000049089 |
| PRRT1 | -0.73677985 | 0.600077254 | 0.10518806 | 0.23559583 | 31.10093295 | 18.55091763 | ENSG00000204314 |
| CA7 | -0.736998265 | 0.599986413 | 0.135082583 | 0.283150938 | 27.15142022 | 16.16168854 | ENSG00000168748 |
| CAMK1 | -0.737207211 | 0.599899523 | 0.143100901 | 0.295612139 | 24.49868353 | 14.64666891 | ENSG00000134072 |
| NA | -0.737519849 | 0.599769536 | 0.554049081 | 0.724171063 | 4.74442273 | 2.89203229 | ENSG00000240950 |
| PAQR4 | -0.737587475 | 0.599741423 | 2.27E-06 | 2.36E-05 | 349.7586367 | 210.003725 | ENSG00000162073 |
| ACAA1 | -0.737607118 | 0.599733257 | 1.84E-09 | 3.28E-08 | 564.8285008 | 338.8998335 | ENSG00000060971 |
| PARD6A | -0.738320054 | 0.59943696 | 0.000110796 | 0.000763661 | 204.9732157 | 122.5559537 | ENSG00000102981 |
| LINC01134 | -0.738908507 | 0.599192509 | 0.422678116 | 0.613702797 | 8.865761589 | 5.382714038 | ENSG00000236423 |
| CDH26 | -0.739873656 | 0.598791789 | 0.146509533 | 0.300677168 | 24.52078192 | 14.62041185 | ENSG00000124215 |
| DNAJC3 | -0.74026217 | 0.598630558 | 6.47E-10 | 1.24E-08 | 727.5272833 | 435.1342003 | ENSG00000102580 |
| NA | -0.74055295 | 0.598509914 | 0.29261992 | 0.485159359 | 12.1542098 | 7.287910909 | ENSG00000226243 |
| TCF25 | -0.740579724 | 0.598498807 | 1.19E-18 | 6.70E-17 | 1286.333324 | 770.2054506 | ENSG00000141002 |
| PYGO1 | -0.740685669 | 0.598454857 | 0.16968966 | 0.334470717 | 26.16318706 | 15.77249899 | ENSG00000171016 |
| POMC | -0.741189161 | 0.598246037 | 0.015247418 | 0.051757443 | 76.87880023 | 46.02228991 | ENSG00000115138 |
| DPM3 | -0.741424028 | 0.598148652 | 1.45E-06 | 1.56E-05 | 338.8378844 | 203.0738887 | ENSG00000179085 |
| LEMD1-DT | -0.741494129 | 0.598119588 | 0.067197759 | 0.167741783 | 39.44974298 | 23.70545695 | ENSG00000224717 |
| LOC105374042 | -0.74174775 | 0.59801445 | 0.43719363 | 0.626145331 | 6.411384112 | 3.852172609 | ENSG00000239482 |
| NA | -0.742051639 | 0.597888497 | 0.537632076 | 0.711290805 | 4.694513485 | 2.847004228 | ENSG00000254094 |
| STAC2 | -0.742898569 | 0.597537612 | 1.81E-05 | 0.000152142 | 229.5219288 | 136.9858445 | ENSG00000141750 |
| MCM5 | -0.743228561 | 0.597400951 | 2.42E-35 | 4.82E-33 | 8473.265906 | 5061.752723 | ENSG00000100297 |
| NA | -0.744023477 | 0.597071877 | 0.408299182 | 0.600105067 | 8.009494771 | 4.752201098 | ENSG00000279601 |
| ACOT2 | -0.744230664 | 0.596986137 | 0.000685431 | 0.003786966 | 140.9940168 | 84.34479789 | ENSG00000119673 |
| C6orf118 | -0.744259575 | 0.596974174 | 0.263039947 | 0.452632683 | 15.30367878 | 9.223601727 | ENSG00000112539 |
| CA12 | -0.744555271 | 0.59685183 | 0.002633108 | 0.011946639 | 115.8331864 | 69.24371465 | ENSG00000074410 |
| NA | -0.74487101 | 0.596721222 | 0.473481216 | 0.658574174 | 6.969375534 | 4.178439173 | ENSG00000279179 |
| ANKRD35 | -0.74526011 | 0.596560305 | 0.000209578 | 0.001338036 | 171.2825866 | 102.2247378 | ENSG00000198483 |
| SLC2A6 | -0.745704081 | 0.59637675 | 3.97E-09 | 6.77E-08 | 611.9675828 | 364.9062869 | ENSG00000160326 |
| DUT | -0.745728299 | 0.596366739 | 2.77E-20 | 1.82E-18 | 1963.056051 | 1170.350999 | ENSG00000128951 |
| NA | -0.745940647 | 0.596278967 | 0.345744822 | 0.539215291 | 9.285953867 | 5.543728881 | ENSG00000257303 |
| P2RY11 | -0.746596345 | 0.596008022 | 0.031949363 | 0.094186335 | 50.08809461 | 29.85625287 | ENSG00000244165 |
| NA | -0.74735599 | 0.59569428 | 0.082596736 | 0.196309447 | 32.17893514 | 19.10634659 | ENSG00000287271 |
| CEP290 | -0.747920718 | 0.595461147 | 7.54E-05 | 0.000540707 | 200.2628426 | 119.4377221 | ENSG00000198707 |
| HEXA | -0.748862572 | 0.595072531 | 0.013796929 | 0.047778827 | 72.06149805 | 42.96956919 | ENSG00000213614 |
| OPRL1 | -0.749044651 | 0.594997433 | 0.472074807 | 0.657299702 | 5.74913951 | 3.405682385 | ENSG00000125510 |
| NA | -0.74906388 | 0.594989503 | 0.526169271 | 0.702433961 | 4.685042746 | 2.824434387 | ENSG00000233971 |
| PRTG | -0.749133512 | 0.594960786 | 0.004882694 | 0.020159469 | 100.2449422 | 59.39273756 | ENSG00000166450 |
| NA | -0.749747869 | 0.594707482 | 0.437508043 | 0.626268718 | 7.020562556 | 4.197210168 | ENSG00000205740 |
| ZNF576 | -0.750271206 | 0.594491791 | 0.001326601 | 0.006668624 | 121.153843 | 72.18066338 | ENSG00000124444 |
| SMG7-AS1 | -0.750568112 | 0.594369457 | 0.295786147 | 0.488644305 | 13.56974581 | 8.158321516 | ENSG00000232860 |
| SPSB4 | -0.75061764 | 0.594349053 | 5.91E-12 | 1.56E-10 | 683.1032568 | 405.6016177 | ENSG00000175093 |
| CYP46A1 | -0.750698602 | 0.5943157 | 0.285133024 | 0.476309641 | 14.3943458 | 8.518004809 | ENSG00000036530 |
| SLC6A8 | -0.750778618 | 0.594282738 | 1.58E-23 | 1.34E-21 | 2976.925402 | 1769.077858 | ENSG00000130821 |
| GINS2 | -0.751278391 | 0.594076905 | 2.53E-19 | 1.53E-17 | 1525.921527 | 906.4975327 | ENSG00000131153 |
| COQ4 | -0.75165012 | 0.593923853 | 3.69E-10 | 7.29E-09 | 538.5659989 | 319.9882449 | ENSG00000167113 |
| ADAMTS19-AS1 | -0.752227805 | 0.593686081 | 0.03047866 | 0.090716473 | 53.8043267 | 31.91541654 | ENSG00000249421 |
| NA | -0.752362959 | 0.593630466 | 0.436927814 | 0.625978002 | 6.431603366 | 3.822116694 | ENSG00000188801 |
| PAQR5 | -0.752711607 | 0.593487024 | 8.05E-05 | 0.000573309 | 210.5138489 | 124.5245116 | ENSG00000137819 |
| PCDH1 | -0.753682213 | 0.593087877 | 7.32E-31 | 1.09E-28 | 5315.140244 | 3152.016743 | ENSG00000156453 |
| SLC25A41 | -0.754149876 | 0.592895653 | 0.326237636 | 0.520441414 | 10.44365299 | 6.203859703 | ENSG00000181240 |
| GLYATL2 | -0.75418404 | 0.592881613 | 0.311689249 | 0.505327089 | 11.06545909 | 6.590238097 | ENSG00000156689 |

| | | | | | | | |
|---|---|---|---|---|---|---|---|
| ZNF32-AS2 | -0.754674935 | 0.592679912 | 0.225929481 | 0.409043373 | 15.63164417 | 9.245621915 | ENSG00000230565 |
| CPT1B | -0.754677857 | 0.592678711 | 0.212973532 | 0.392722261 | 16.93779326 | 10.05603231 | ENSG00000205560 |
| NA | -0.754812069 | 0.592623578 | 0.316995853 | 0.510709417 | 11.13558759 | 6.631578933 | ENSG00000275582 |
| CCBE1 | -0.754974173 | 0.592556993 | 0.001461619 | 0.007247097 | 123.4973603 | 72.92193854 | ENSG00000183287 |
| FLVCR1 | -0.755246776 | 0.592445038 | 2.87E-19 | 1.71E-17 | 1423.730748 | 843.6323507 | ENSG00000162769 |
| TSPEAR-AS1 | -0.755544573 | 0.59232276 | 0.014411888 | 0.049429755 | 67.94647302 | 40.21229467 | ENSG00000235890 |
| CDH24 | -0.75593843 | 0.592161077 | 5.81E-11 | 1.31E-09 | 637.3977129 | 377.7745213 | ENSG00000139880 |
| CLDN6 | -0.756505774 | 0.591928254 | 1.49E-38 | 3.64E-36 | 13898.98977 | 8226.727945 | ENSG00000184697 |
| CYB561A3 | -0.756982935 | 0.59173251 | 0.00015059 | 0.00099644 | 246.3914501 | 145.9675103 | ENSG00000162144 |
| GGT7 | -0.757564027 | 0.591494219 | 2.36E-11 | 5.63E-10 | 1509.040754 | 893.4357605 | ENSG00000131067 |
| EFNB3 | -0.757694553 | 0.591440707 | 3.54E-10 | 7.03E-09 | 564.4611793 | 334.0461713 | ENSG00000108947 |
| NA | -0.757856884 | 0.591374162 | 0.507952913 | 0.687132274 | 5.336538834 | 3.195840633 | ENSG00000236015 |
| PLXNA4 | -0.758439256 | 0.59113549 | 0.044470122 | 0.122195589 | 48.40397319 | 28.47175413 | ENSG00000221866 |
| NA | -0.759269181 | 0.590795531 | 2.17E-05 | 0.000179867 | 209.1590683 | 123.76969 | ENSG00000234664 |
| NA | -0.759642933 | 0.590642497 | 0.222581442 | 0.404721323 | 18.41704625 | 10.92644292 | ENSG00000279069 |
| CRPPA | -0.759782557 | 0.590585337 | 0.367803563 | 0.560828567 | 9.226573883 | 5.419817994 | ENSG00000214960 |
| ENOX1 | -0.760219013 | 0.590406695 | 0.121961007 | 0.26277904 | 24.96444807 | 14.75180881 | ENSG00000120658 |
| UBQLN2 | -0.760743034 | 0.590192285 | 1.17E-14 | 4.33E-13 | 1035.20968 | 611.0755908 | ENSG00000188021 |
| NXPH3 | -0.761113265 | 0.590040846 | 0.000406891 | 0.002388759 | 162.5087437 | 96.00981643 | ENSG00000182575 |
| PCDHB9 | -0.761167327 | 0.590018736 | 0.111673563 | 0.246165297 | 27.72331707 | 16.45140081 | ENSG00000177839 |
| KCND1 | -0.761229284 | 0.589993398 | 1.41E-06 | 1.52E-05 | 291.0491179 | 171.9481362 | ENSG00000102057 |
| NA | -0.761436972 | 0.58990847 | 0.531381089 | 0.706652587 | 4.713454962 | 2.805663392 | ENSG00000285803 |
| CH17-340M24.3 | -0.761508959 | 0.589879035 | 0.337657472 | 0.531944773 | 10.7450853 | 6.425424408 | ENSG00000197180 |
| SPATA41 | -0.761960108 | 0.589694601 | 0.347161591 | 0.540606027 | 10.02789541 | 5.978934184 | ENSG00000189419 |
| CES4A | -0.761996517 | 0.58967972 | 0.46313996 | 0.649713191 | 5.958344929 | 3.570934107 | ENSG00000172824 |
| ULK4P2 | -0.762002814 | 0.589677146 | 0.399875788 | 0.591862152 | 8.339339296 | 4.954994809 | ENSG00000260128 |
| FOXO4 | -0.762021892 | 0.589669348 | 1.09E-11 | 2.74E-10 | 595.8816142 | 351.0540601 | ENSG00000184481 |
| TTYH2 | -0.762622367 | 0.589423969 | 0.001341551 | 0.006738296 | 121.2207168 | 71.57326994 | ENSG00000141540 |
| NA | -0.763301631 | 0.589146515 | 0.120829804 | 0.260978858 | 27.12676628 | 16.04625142 | ENSG00000275437 |
| ENHO | -0.763600804 | 0.589024356 | 0.000381449 | 0.002260883 | 178.7113 | 105.1638449 | ENSG00000168913 |
| HUNK | -0.763689428 | 0.588988174 | 8.21E-08 | 1.12E-06 | 384.0238536 | 226.1683279 | ENSG00000142149 |
| AFAP1L2 | -0.763939092 | 0.588886256 | 0.002607639 | 0.011860118 | 105.2807478 | 61.94028194 | ENSG00000169129 |
| TRPA1 | -0.764404417 | 0.588696348 | 0.31813308 | 0.511874164 | 11.71507604 | 6.912817435 | ENSG00000104321 |
| CXCL12 | -0.764807798 | 0.58853177 | 1.00E-13 | 3.29E-12 | 783.8272903 | 460.8239518 | ENSG00000107562 |
| GATD3A | -0.764883049 | 0.588501073 | 0.100648666 | 0.2276849 | 28.11391705 | 16.61241565 | ENSG00000160221 |
| CLCN2 | -0.765213095 | 0.588366457 | 1.34E-05 | 0.000116932 | 250.6424178 | 147.4964027 | ENSG00000114859 |
| NA | -0.765373774 | 0.588300932 | 0.043838411 | 0.120782088 | 42.12723131 | 24.83789703 | ENSG00000230555 |
| BRD9 | -0.766153034 | 0.587983251 | 4.13E-13 | 1.28E-11 | 1065.466907 | 627.0366044 | ENSG00000028310 |
| MBNL3 | -0.766218935 | 0.587956393 | 0.006209002 | 0.024779852 | 87.00829557 | 50.97998426 | ENSG00000076770 |
| NA | -0.766462817 | 0.58785701 | 0.085034821 | 0.200513407 | 32.31485515 | 19.06555541 | ENSG00000227896 |
| CHST7 | -0.766738423 | 0.587744719 | 2.20E-09 | 3.90E-08 | 656.6656665 | 386.2503092 | ENSG00000147119 |
| NASP | -0.766770746 | 0.587731551 | 1.16E-44 | 4.05E-42 | 17308.41126 | 10173.57458 | ENSG00000132780 |
| MPP2 | -0.767188066 | 0.587561566 | 5.18E-28 | 6.41E-26 | 2335.566435 | 1372.254562 | ENSG00000108852 |
| SEPTIN3 | -0.767491307 | 0.587438079 | 4.55E-22 | 3.40E-20 | 1632.101273 | 958.5600309 | ENSG00000100167 |
| NA | -0.768680559 | 0.586954038 | 0.137664303 | 0.287166032 | 24.17071815 | 14.23815871 | ENSG00000277283 |
| FBXW4 | -0.768687091 | 0.58695138 | 1.31E-11 | 3.26E-10 | 689.7691103 | 404.4985807 | ENSG00000107829 |
| SH2B2 | -0.768761587 | 0.586921073 | 6.58E-08 | 9.07E-07 | 429.1458878 | 251.8916161 | ENSG00000160999 |
| GLTPD2 | -0.768794423 | 0.586907715 | 0.333785035 | 0.528382698 | 10.62632533 | 6.297714675 | ENSG00000182327 |
| RNF31 | -0.768916167 | 0.586858189 | 0.375862433 | 0.568924032 | 7.595616319 | 4.459677675 | ENSG00000092098 |
| PDS5A | -0.769201821 | 0.586742003 | 1.74E-20 | 1.16E-18 | 2491.26743 | 1460.649522 | ENSG00000121892 |
| LOC100631378 | -0.769286694 | 0.586707487 | 0.328262443 | 0.522636897 | 9.862285406 | 5.753570632 | ENSG00000225868 |
| CD2BP2 | -0.770476889 | 0.586223664 | 9.47E-17 | 4.36E-15 | 2274.278465 | 1334.402773 | ENSG00000169217 |
| NA | -0.770882247 | 0.586058974 | 0.224618881 | 0.407426731 | 15.30811347 | 8.97946718 | ENSG00000271870 |
| EAPP | -0.771196985 | 0.585931134 | 5.69E-08 | 7.97E-07 | 367.5020059 | 215.0850038 | ENSG00000129518 |
| TOLLIP-AS1 | -0.771486111 | 0.585813721 | 0.273045342 | 0.464192108 | 13.70872503 | 8.083237538 | ENSG00000255153 |
| NA | -0.771654041 | 0.585745536 | 0.485164734 | 0.668304955 | 4.747579643 | 2.805663392 | ENSG00000235554 |
| DYRK1B | -0.771956688 | 0.585622672 | 1.42E-05 | 0.000122657 | 255.4713781 | 149.6081922 | ENSG00000105204 |
| CCNC | -0.772222581 | 0.58551475 | 9.20E-08 | 1.24E-06 | 673.3365581 | 393.6739882 | ENSG00000112237 |
| JTB | -0.772587398 | 0.585366708 | 0.221925525 | 0.403924941 | 14.73493884 | 8.619345854 | ENSG00000143543 |
| MPDU1 | -0.772765457 | 0.585294466 | 7.14E-12 | 1.85E-10 | 663.6546 | 388.3975452 | ENSG00000129255 |
| NA | -0.772783454 | 0.585287165 | 0.353413913 | 0.546576558 | 8.270488573 | 4.85734099 | ENSG00000231731 |
| GHRHR | -0.773151015 | 0.585138068 | 0.106617688 | 0.237792085 | 30.2679446 | 17.72652277 | ENSG00000106128 |
| PCDHA9 | -0.773399853 | 0.585037151 | 0.126852681 | 0.270483072 | 26.75658094 | 15.62926745 | ENSG00000204961 |
| BMS1P1 | -0.773774615 | 0.584885199 | 0.108643678 | 0.241063321 | 29.4983876 | 17.29511631 | ENSG00000204177 |
| TLR2 | -0.774224765 | 0.584702731 | 0.159858072 | 0.320130687 | 21.9210137 | 12.80906995 | ENSG00000137462 |
| NA | -0.775551812 | 0.584165146 | 0.493701662 | 0.675874798 | 4.714732738 | 2.768121403 | ENSG00000232470 |
| KIF6 | -0.775881828 | 0.584031534 | 0.093895861 | 0.215940674 | 31.80687067 | 18.63783618 | ENSG00000164627 |
| NA | -0.775995019 | 0.583985714 | 0.275111608 | 0.466037706 | 13.85529586 | 8.147036595 | ENSG00000279259 |
| SOAT1 | -0.775997141 | 0.583984855 | 0.00131833 | 0.006634244 | 119.4797936 | 69.77613574 | ENSG00000057252 |
| PDSS2 | -0.776077968 | 0.583952138 | 4.67E-06 | 4.56E-05 | 275.8085774 | 161.2509673 | ENSG00000164494 |
| NA | -0.776176161 | 0.583912394 | 0.175784384 | 0.343452222 | 20.3760669 | 11.9992092 | ENSG00000271533 |
| NA | -0.776537681 | 0.583766092 | 0.214704892 | 0.394854184 | 16.27682643 | 9.421720525 | ENSG00000275457 |
| SCRT2 | -0.776797527 | 0.583660958 | 0.268167767 | 0.45833838 | 12.53839826 | 7.344223892 | ENSG00000215397 |
| SLC22A5 | -0.776966542 | 0.583592585 | 0.001549102 | 0.007639982 | 109.5549713 | 63.96971611 | ENSG00000197375 |

| | | | | | | | |
|---|---|---|---|---|---|---|---|
| GPR137 | -0.776976363 | 0.583588612 | 4.14E-10 | 8.13E-09 | 640.177485 | 373.3702318 | ENSG00000173264 |
| RAB15 | -0.777536934 | 0.583361898 | 7.44E-23 | 6.02E-21 | 2481.982792 | 1448.031996 | ENSG00000139998 |
| TAF2 | -0.777655983 | 0.583313762 | 1.16E-13 | 3.78E-12 | 726.9153086 | 423.6240247 | ENSG00000064313 |
| SHISAL2A | -0.777733914 | 0.583282254 | 0.274061707 | 0.465193891 | 14.1662966 | 8.296766517 | ENSG00000182183 |
| DACT3 | -0.777824211 | 0.583245748 | 0.0230557 | 0.072524499 | 58.16388455 | 34.04272777 | ENSG00000197380 |
| FAM189B | -0.778767037 | 0.582864711 | 2.96E-14 | 1.04E-12 | 1284.215992 | 748.4463879 | ENSG00000160767 |
| HCN3 | -0.778833474 | 0.58283787 | 0.000893783 | 0.004786299 | 224.915884 | 131.5852758 | ENSG00000143630 |
| ATP1A2 | -0.778899057 | 0.582811376 | 9.43E-12 | 2.39E-10 | 1925.034105 | 1122.743442 | ENSG00000018625 |
| C4orf19 | -0.779027428 | 0.58275952 | 0.290259362 | 0.482351567 | 12.8746543 | 7.505238734 | ENSG00000154274 |
| LOC105371067 | -0.779348324 | 0.582629912 | 0.20588754 | 0.384398072 | 15.8692618 | 9.230649767 | ENSG00000285567 |
| ERG | -0.779972116 | 0.582378049 | 0.1257195 | 0.268621944 | 26.77482336 | 15.47530065 | ENSG00000157554 |
| SUSD4 | -0.780642619 | 0.582107447 | 0.008507807 | 0.0322198 | 76.56632631 | 44.40146912 | ENSG00000143502 |
| ATAD1 | -0.780781659 | 0.58205135 | 3.87E-11 | 8.98E-10 | 770.4505442 | 448.1986898 | ENSG00000138138 |
| FIG4 | -0.78135181 | 0.581821369 | 1.88E-12 | 5.31E-11 | 670.8879234 | 390.0898743 | ENSG00000112367 |
| NA | -0.781706082 | 0.581678513 | 0.230420865 | 0.414498987 | 14.82854121 | 8.668172763 | ENSG00000229097 |
| NA | -0.781867773 | 0.581613325 | 0.004567329 | 0.019044065 | 134.2697755 | 78.43120011 | ENSG00000260708 |
| SRRM2-AS1 | -0.78216915 | 0.581491839 | 0.218316113 | 0.399513314 | 15.54257419 | 9.103378067 | ENSG00000205913 |
| SIVA1 | -0.782190675 | 0.581483163 | 1.23E-18 | 6.89E-17 | 1575.465393 | 916.0557313 | ENSG00000184990 |
| NA | -0.782263783 | 0.581453698 | 0.087166408 | 0.20446637 | 33.61606588 | 19.59417765 | ENSG00000261754 |
| VIPR1 | -0.783596314 | 0.580916892 | 0.009465296 | 0.035163319 | 77.5042669 | 44.8774656 | ENSG00000114812 |
| SDF2L1 | -0.783875185 | 0.580804612 | 2.66E-15 | 1.06E-13 | 1602.434328 | 930.8590582 | ENSG00000128228 |
| SLC25A13 | -0.784454426 | 0.580571466 | 6.29E-21 | 4.32E-19 | 2131.171167 | 1236.403562 | ENSG00000004864 |
| FLRT2 | -0.784602825 | 0.58051175 | 0.001393038 | 0.006944211 | 105.3129934 | 61.15667826 | ENSG00000185070 |
| NA | -0.784757957 | 0.580449332 | 0.002115682 | 0.009980159 | 98.35665145 | 57.01578108 | ENSG00000235501 |
| RUSC1-AS1 | -0.785461946 | 0.58016616 | 0.017350026 | 0.057412137 | 58.38068157 | 33.99390086 | ENSG00000225855 |
| RTN4RL2 | -0.785645988 | 0.580092154 | 1.98E-05 | 0.000165094 | 335.2222902 | 194.3770539 | ENSG00000186907 |
| SAP30L-AS1 | -0.786036244 | 0.579935258 | 0.174587652 | 0.341618195 | 18.22184396 | 10.49448681 | ENSG00000245275 |
| CPLX2 | -0.786115858 | 0.579903255 | 0.080630891 | 0.192832468 | 33.00363283 | 19.26834912 | ENSG00000145920 |
| CIPC | -0.786625502 | 0.579698436 | 7.93E-08 | 1.08E-06 | 418.7590518 | 242.9935568 | ENSG00000198894 |
| MBLAC1 | -0.788002679 | 0.579145328 | 0.086713748 | 0.203565877 | 29.81047073 | 17.25377548 | ENSG00000214309 |
| CDH22 | -0.788349864 | 0.579005973 | 0.271682854 | 0.462554658 | 12.33372522 | 7.111374266 | ENSG00000149654 |
| KCNA2 | -0.788678939 | 0.578873918 | 0.185398122 | 0.356153482 | 21.69602372 | 12.57297113 | ENSG00000177301 |
| PRELID3A | -0.789833778 | 0.57841073 | 0.000126821 | 0.000860987 | 167.2803985 | 97.04905436 | ENSG00000141391 |
| BSND | -0.789929076 | 0.578372524 | 0.428119998 | 0.618358482 | 5.961501842 | 3.473280288 | ENSG00000162399 |
| NA | -0.789975439 | 0.578353938 | 0.427772426 | 0.618108004 | 5.988034921 | 3.492051283 | ENSG00000226666 |
| GLIDR | -0.790705843 | 0.578061205 | 0.062498029 | 0.158981398 | 35.08329259 | 20.3255936 | ENSG00000278175 |
| LINC00652 | -0.790708451 | 0.57806016 | 0.482025727 | 0.665464792 | 5.855271827 | 3.338084481 | ENSG00000179935 |
| LRRC69 | -0.790778459 | 0.578032109 | 0.342973467 | 0.536880932 | 8.297021652 | 4.778458166 | ENSG00000214954 |
| FGFR1OP | -0.792104817 | 0.577500933 | 4.50E-05 | 0.000342589 | 198.9961473 | 114.8677916 | ENSG00000213066 |
| GK | -0.792370671 | 0.577394524 | 1.94E-05 | 0.000161822 | 202.7773742 | 116.8135564 | ENSG00000198814 |
| CDHR1 | -0.792453305 | 0.577361453 | 3.68E-08 | 5.34E-07 | 503.1793425 | 291.0832524 | ENSG00000148600 |
| NA | -0.792770692 | 0.57723445 | 0.323340705 | 0.517557356 | 10.19844376 | 5.903850207 | ENSG00000214772 |
| STAG1 | -0.79329349 | 0.577025312 | 1.65E-11 | 4.04E-10 | 546.8635242 | 315.7159423 | ENSG00000118007 |
| KCNN2 | -0.79341554 | 0.576976498 | 1.35E-06 | 1.46E-05 | 309.7362227 | 178.6004212 | ENSG00000080709 |
| SPRYD4 | -0.793876907 | 0.576792013 | 0.000130763 | 0.000882579 | 182.4106803 | 104.8668613 | ENSG00000176422 |
| ARNT2 | -0.794669516 | 0.576475214 | 8.89E-08 | 1.20E-06 | 332.104751 | 191.6253219 | ENSG00000172379 |
| RORB | -0.794749742 | 0.576443158 | 0.000194769 | 0.001254725 | 148.7053339 | 85.85981752 | ENSG00000198963 |
| CFAP69 | -0.794780583 | 0.576430835 | 0.04540953 | 0.124151089 | 42.78504122 | 24.81956407 | ENSG00000105792 |
| CA11 | -0.795255619 | 0.576241065 | 2.41E-17 | 1.19E-15 | 1008.866857 | 581.3239281 | ENSG00000063180 |
| NA | -0.795766492 | 0.576037048 | 0.16121314 | 0.321905229 | 18.56559391 | 10.68979445 | ENSG00000264016 |
| TMEM200A | -0.795944226 | 0.575966087 | 4.79E-07 | 5.67E-06 | 418.7370359 | 240.8660936 | ENSG00000164484 |
| NA | -0.796961743 | 0.575560008 | 0.413762238 | 0.605166455 | 6.686283342 | 3.882228524 | ENSG00000272277 |
| LINC01783 | -0.798127773 | 0.575095011 | 0.338069583 | 0.532153481 | 8.873353191 | 5.078467662 | ENSG00000233421 |
| NA | -0.798160836 | 0.575081831 | 0.319241005 | 0.513129982 | 9.679613065 | 5.573784796 | ENSG00000269318 |
| TAF1A-AS1 | -0.798212415 | 0.575061272 | 0.318837381 | 0.512695819 | 10.12327921 | 5.817481309 | ENSG00000225265 |
| FOXM1 | -0.798333757 | 0.575012906 | 7.11E-28 | 8.67E-26 | 2953.789496 | 1698.923428 | ENSG00000111206 |
| LEAP2 | -0.798901404 | 0.574786704 | 0.334219701 | 0.528670131 | 10.85309676 | 6.252686612 | ENSG00000164406 |
| WDR27 | -0.799174087 | 0.574678074 | 0.031558213 | 0.093300355 | 370.1566444 | 212.9121954 | ENSG00000184465 |
| PAK6 | -0.79930762 | 0.574624886 | 7.64E-06 | 7.06E-05 | 200.9953134 | 115.4193101 | ENSG00000137843 |
| NOG | -0.799847274 | 0.574409982 | 0.266307291 | 0.456336064 | 12.57755899 | 7.19025709 | ENSG00000183691 |
| ACP7 | -0.799883691 | 0.574395483 | 0.2533988 | 0.442142686 | 14.80191043 | 8.431524291 | ENSG00000183760 |
| LOC145783 | -0.801065368 | 0.573925202 | 0.342178711 | 0.536395153 | 9.687204668 | 5.562499875 | ENSG00000285331 |
| DONSON | -0.801357559 | 0.573808976 | 8.10E-10 | 1.52E-08 | 488.4407204 | 279.9914143 | ENSG00000159147 |
| AVIL | -0.801726368 | 0.573662307 | 0.155833771 | 0.314142171 | 19.29363002 | 11.1287986 | ENSG00000135407 |
| HS6ST3 | -0.802708976 | 0.573271723 | 0.028190308 | 0.08542825 | 46.33132632 | 26.6345932 | ENSG00000185352 |
| LOC101929295 | -0.80308258 | 0.573123286 | 0.372017937 | 0.565162332 | 7.979804779 | 4.624929397 | ENSG00000245552 |
| CTBP1-AS | -0.80310165 | 0.573115711 | 0.199760798 | 0.376218538 | 16.85503711 | 9.680827219 | ENSG00000280927 |
| DDX58 | -0.803594581 | 0.572919926 | 0.001200924 | 0.0061554 | 112.237398 | 64.14168946 | ENSG00000107201 |
| BEND5 | -0.803812432 | 0.57283342 | 0.000879964 | 0.004722876 | 109.5776484 | 62.8411865 | ENSG00000162373 |
| PHYHIP | -0.803856726 | 0.572815832 | 1.50E-06 | 1.61E-05 | 245.4470078 | 140.7186695 | ENSG00000168490 |
| MAP1LC3B2 | -0.804364564 | 0.572614233 | 0.327908996 | 0.522208741 | 8.951674653 | 5.097238657 | ENSG00000258102 |
| KCNQ4 | -0.804523487 | 0.572551159 | 0.000347799 | 0.002082747 | 127.3830584 | 72.9304123 | ENSG00000117013 |
| IRF4 | -0.804649589 | 0.572501116 | 0.03478289 | 0.100744816 | 45.27797807 | 26.03088698 | ENSG00000137265 |
| BEX1 | -0.805761387 | 0.572060094 | 7.32E-22 | 5.43E-20 | 1261.80247 | 721.6333145 | ENSG00000133169 |

| | | | | | | | |
|---|---|---|---|---|---|---|---|
| SLC16A14 | -0.806184211 | 0.57189246 | 0.006045112 | 0.024214418 | 93.17890513 | 53.01234121 | ENSG00000163053 |
| EID1 | -0.806654038 | 0.571706248 | 3.22E-23 | 2.64E-21 | 2775.267986 | 1587.115471 | ENSG00000255302 |
| DIABLO | -0.807028233 | 0.571557982 | 0.158451992 | 0.31803652 | 23.1932335 | 13.24383722 | ENSG00000184047 |
| RORC | -0.807255389 | 0.571467996 | 0.382835556 | 0.575510924 | 8.20163785 | 4.639463512 | ENSG00000143365 |
| LINC00511 | -0.807893745 | 0.571215192 | 0.212624663 | 0.392391268 | 16.31420572 | 9.377130495 | ENSG00000227036 |
| GPX2 | -0.807920193 | 0.571204721 | 0.381921394 | 0.574745693 | 7.242395627 | 4.159668179 | ENSG00000176153 |
| NA | -0.808427563 | 0.571003873 | 0.453952004 | 0.641311134 | 5.538054953 | 3.173270792 | ENSG00000268295 |
| CRELD2 | -0.808479298 | 0.570983398 | 4.36E-17 | 2.08E-15 | 1006.659861 | 574.7628614 | ENSG00000184164 |
| GNG2 | -0.808529691 | 0.570963454 | 1.99E-07 | 2.53E-06 | 314.7925332 | 179.4591088 | ENSG00000186469 |
| TRAM1L1 | -0.808594002 | 0.570938002 | 0.100820957 | 0.22793004 | 24.77230499 | 14.18184573 | ENSG00000174599 |
| NEK9 | -0.80895545 | 0.570794979 | 2.52E-17 | 1.24E-15 | 1116.149943 | 637.665411 | ENSG00000119638 |
| NA | -0.809145546 | 0.570719774 | 0.046469649 | 0.126267186 | 38.82793689 | 22.24251343 | ENSG00000213222 |
| LOC112694756 | -0.810066167 | 0.570355699 | 0.330377545 | 0.524726562 | 9.229730795 | 5.322602209 | ENSG00000285043 |
| PLEKHO1 | -0.810121839 | 0.57033369 | 4.13E-15 | 1.61E-13 | 792.1475904 | 451.2932925 | ENSG00000023902 |
| NA | -0.810919741 | 0.570018346 | 0.440316516 | 0.628529445 | 5.478674969 | 3.124443883 | ENSG00000254554 |
| NA | -0.81196227 | 0.569606585 | 3.66E-06 | 3.66E-05 | 242.5239194 | 138.2964617 | ENSG00000159712 |
| RAB33A | -0.812197916 | 0.569513555 | 0.004371828 | 0.018356924 | 98.43160543 | 56.32278318 | ENSG00000134594 |
| LOC100288748 | -0.812333858 | 0.569459893 | 0.110820018 | 0.244720389 | 24.75198804 | 14.03916385 | ENSG00000261437 |
| KISS1R | -0.81273546 | 0.569301395 | 0.071231327 | 0.17534884 | 31.24750377 | 17.72228589 | ENSG00000116014 |
| HAUS7 | -0.812769858 | 0.569287821 | 0.407345866 | 0.599091277 | 6.044257993 | 3.443224373 | ENSG00000213397 |
| CYP3A5 | -0.813228809 | 0.569106748 | 0.354735471 | 0.548071626 | 8.522011636 | 4.879910831 | ENSG00000106258 |
| MCM2 | -0.813556286 | 0.568977581 | 4.22E-37 | 9.10E-35 | 4696.760796 | 2672.626991 | ENSG00000073111 |
| NCBP2AS2 | -0.813698932 | 0.568921327 | 4.26E-14 | 1.47E-12 | 835.3830774 | 475.5913311 | ENSG00000270170 |
| EYA2 | -0.813731946 | 0.568908308 | 0.083824678 | 0.198543973 | 30.30592525 | 17.2462894 | ENSG00000064655 |
| IL12RB2 | -0.814020738 | 0.568794438 | 0.034412788 | 0.099841143 | 44.38442966 | 25.35567239 | ENSG00000081985 |
| CXCL2 | -0.814024377 | 0.568793003 | 0.391118115 | 0.583701034 | 6.626903358 | 3.773289785 | ENSG00000081041 |
| HAS3 | -0.814887649 | 0.568452754 | 3.95E-30 | 5.63E-28 | 2496.190128 | 1419.208809 | ENSG00000103044 |
| ENOX2 | -0.815221143 | 0.568321365 | 2.07E-11 | 4.99E-10 | 515.9840336 | 292.9209227 | ENSG00000165675 |
| GSG1L | -0.815243059 | 0.568312732 | 0.301887612 | 0.495324452 | 10.23572535 | 5.787425394 | ENSG00000169181 |
| FAM47E | -0.815747595 | 0.568114018 | 0.484963788 | 0.668210541 | 4.916248858 | 2.786892398 | ENSG00000189157 |
| NA | -0.815790851 | 0.568096984 | 0.46182997 | 0.648316025 | 4.803802715 | 2.708009574 | ENSG00000286181 |
| DGCR5 | -0.816029921 | 0.568002852 | 0.417876257 | 0.608985715 | 6.017724914 | 3.413168459 | ENSG00000273032 |
| IGSF9 | -0.816160376 | 0.567951493 | 3.28E-17 | 1.60E-15 | 1026.169748 | 583.1375718 | ENSG00000085552 |
| TMA7 | -0.816589887 | 0.567782431 | 0.000342642 | 0.002056944 | 133.0159935 | 75.28122332 | ENSG00000232112 |
| ASTN1 | -0.81681549 | 0.56769365 | 4.24E-07 | 5.08E-06 | 254.9323055 | 144.6154321 | ENSG00000152092 |
| NA | -0.816870359 | 0.56767206 | 0.180540755 | 0.350088131 | 20.35900456 | 11.64668557 | ENSG00000262049 |
| NPEPL1 | -0.818465346 | 0.567044811 | 0.00060514 | 0.003389442 | 146.7271013 | 83.40397822 | ENSG00000215440 |
| NA | -0.819941204 | 0.566465028 | 0.001486701 | 0.007353752 | 118.952364 | 67.48053839 | ENSG00000279207 |
| NA | -0.820128146 | 0.566391631 | 0.222346758 | 0.404413623 | 14.37412655 | 8.067715738 | ENSG00000282386 |
| NA | -0.820272219 | 0.566335072 | 0.230044782 | 0.413986795 | 12.70716517 | 7.186458243 | ENSG00000279329 |
| 2-Mar | -0.820506289 | 0.566243195 | 2.74E-05 | 0.000220806 | 204.9750198 | 116.2194232 | ENSG00000117791 |
| ADCYAP1R1 | -0.821217115 | 0.565964271 | 2.80E-13 | 8.81E-12 | 817.5428664 | 462.8855204 | ENSG00000078549 |
| MITF | -0.821304122 | 0.56593014 | 0.162376519 | 0.32377476 | 18.88734317 | 10.7424202 | ENSG00000187098 |
| CLDN9 | -0.821703368 | 0.565773548 | 0.342525996 | 0.536679077 | 8.47653708 | 4.808514081 | ENSG00000213937 |
| TIGD7 | -0.822142562 | 0.565601338 | 0.034096383 | 0.099206022 | 42.07859984 | 23.88904164 | ENSG00000140993 |
| NA | -0.822588312 | 0.565426611 | 0.121157619 | 0.261493009 | 23.42877661 | 13.30438708 | ENSG00000234062 |
| NA | -0.822607405 | 0.565419128 | 0.43204988 | 0.622027855 | 5.395918818 | 3.026790064 | ENSG00000263745 |
| LINC01315 | -0.823266802 | 0.565160757 | 2.65E-10 | 5.41E-09 | 429.2349503 | 242.5341327 | ENSG00000229891 |
| SEC14L4 | -0.823497242 | 0.565070492 | 0.075892167 | 0.184180362 | 32.81780358 | 18.52878582 | ENSG00000133488 |
| KTN1 | -0.823547903 | 0.565050649 | 5.89E-31 | 8.88E-29 | 2576.66592 | 1455.379635 | ENSG00000126777 |
| SLC18A2 | -0.823645754 | 0.565012326 | 0.017509467 | 0.057805567 | 69.01252398 | 39.14409168 | ENSG00000165646 |
| TUBA1A | -0.824503182 | 0.564676625 | 1.93E-39 | 5.17E-37 | 5707.072196 | 3222.234486 | ENSG00000167552 |
| NA | -0.824544566 | 0.564660428 | 0.035756802 | 0.102970256 | 39.6336931 | 22.42609811 | ENSG00000240253 |
| TRIM17 | -0.82468321 | 0.564606166 | 0.073574294 | 0.179874154 | 33.85989965 | 19.24209205 | ENSG00000162931 |
| MAP2K6 | -0.824837587 | 0.564545753 | 1.57E-13 | 5.06E-12 | 926.8120848 | 522.4545395 | ENSG00000108984 |
| LINC02751 | -0.825651435 | 0.564227374 | 0.270554218 | 0.46113237 | 11.11221142 | 6.320284516 | ENSG00000254946 |
| HTRA1 | -0.825840295 | 0.564153517 | 8.80E-14 | 2.93E-12 | 618.1863353 | 348.5189 | ENSG00000166033 |
| COL11A2 | -0.826407039 | 0.56393194 | 0.004893473 | 0.020194974 | 77.01956089 | 43.49059374 | ENSG00000204248 |
| CDH15 | -0.826506311 | 0.563893137 | 0.177753723 | 0.346387078 | 16.38937027 | 9.249420762 | ENSG00000129910 |
| NA | -0.827089348 | 0.563665296 | 0.420400728 | 0.611558221 | 5.99750566 | 3.356855475 | ENSG00000273674 |
| C5orf38 | -0.827918485 | 0.563341443 | 0.07936092 | 0.190478541 | 31.33982836 | 17.56831909 | ENSG00000186493 |
| NA | -0.828318368 | 0.563185319 | 0.077358853 | 0.186907702 | 27.58749476 | 15.59596235 | ENSG00000285825 |
| KCNH7 | -0.828848594 | 0.562978373 | 0.317031297 | 0.510722126 | 9.150131557 | 5.16483656 | ENSG00000184611 |
| TMEM158 | -0.829712638 | 0.562641301 | 4.11E-05 | 0.000316944 | 318.7689174 | 179.3318371 | ENSG00000249992 |
| GTSE1-DT | -0.830174643 | 0.562461151 | 0.210737125 | 0.390119896 | 15.13944426 | 8.495434967 | ENSG00000277232 |
| NA | -0.830695823 | 0.562257996 | 0.319638997 | 0.51336128 | 9.679710764 | 5.446075062 | ENSG00000257557 |
| NA | -0.830912206 | 0.562173672 | 0.074889165 | 0.182295218 | 32.9195989 | 18.51001482 | ENSG00000255397 |
| NA | -0.832483934 | 0.561561551 | 0.24070278 | 0.426907687 | 12.18389979 | 6.800079849 | ENSG00000233270 |
| TRAPPC5 | -0.832508082 | 0.561552152 | 0.028373665 | 0.085801518 | 48.99796843 | 27.68533929 | ENSG00000181029 |
| RNF144A-AS1 | -0.832599154 | 0.561516704 | 0.114515262 | 0.250877762 | 21.88067289 | 12.24615491 | ENSG00000228203 |
| MLLT11 | -0.832790746 | 0.561442139 | 5.05E-08 | 7.15E-07 | 406.5490324 | 227.9903018 | ENSG00000213190 |
| EXOC3-AS1 | -0.833528233 | 0.56115521 | 0.199822224 | 0.376257757 | 21.43266976 | 12.04391085 | ENSG00000221990 |
| GPRASP2 | -0.833730207 | 0.561076655 | 2.53E-06 | 2.61E-05 | 247.4476696 | 138.9520292 | ENSG00000158301 |
| TAF5 | -0.833934177 | 0.560997335 | 1.84E-12 | 5.22E-11 | 585.5750613 | 328.0990134 | ENSG00000148835 |

| | | | | | | | |
|---|---|---|---|---|---|---|---|
| NA | -0.834034079 | 0.560958489 | 0.210405839 | 0.389867428 | 14.0303766 | 7.842790219 | ENSG00000248015 |
| NCAPH | -0.83539325 | 0.560430256 | 8.30E-18 | 4.31E-16 | 1303.857003 | 730.5832755 | ENSG00000121152 |
| HES3 | -0.835573682 | 0.56036017 | 1.18E-07 | 1.56E-06 | 295.8226292 | 165.7772552 | ENSG00000173673 |
| INPPL1 | -0.835905422 | 0.560231333 | 7.45E-28 | 8.94E-26 | 2309.894223 | 1294.700369 | ENSG00000165458 |
| DYNC2H1 | -0.836016486 | 0.560188206 | 3.57E-08 | 5.19E-07 | 377.6285171 | 211.2283711 | ENSG00000187240 |
| SCUBE3 | -0.836083799 | 0.560162069 | 0.003570208 | 0.01543509 | 81.34241587 | 45.62115416 | ENSG00000146197 |
| PLEKHD1 | -0.836278056 | 0.560086649 | 0.109806372 | 0.242916124 | 26.79248706 | 15.10108325 | ENSG00000175985 |
| IL20RA | -0.837670397 | 0.559546372 | 0.032532158 | 0.095479179 | 43.67839424 | 24.37263582 | ENSG00000016402 |
| NA | -0.837788593 | 0.559500532 | 0.22747418 | 0.410959013 | 13.48816974 | 7.509037581 | ENSG00000287707 |
| NA | -0.838668389 | 0.559159437 | 0.278773591 | 0.469879022 | 12.65725592 | 7.179410203 | ENSG00000279798 |
| ZNF44 | -0.838844863 | 0.559091043 | 0.004052731 | 0.017204149 | 75.98713096 | 42.47001518 | ENSG00000197857 |
| NA | -0.838957498 | 0.559047395 | 0.003199643 | 0.014059315 | 80.85840892 | 45.27991545 | ENSG00000278133 |
| NA | -0.838992378 | 0.559033879 | 0.351572232 | 0.544557714 | 7.308089437 | 4.06201436 | ENSG00000279822 |
| BISPR | -0.83912283 | 0.558983332 | 0.185626111 | 0.356406749 | 15.39274876 | 8.581803865 | ENSG00000282851 |
| NA | -0.839332201 | 0.558902215 | 0.095668857 | 0.219148016 | 25.87190191 | 14.51191114 | ENSG00000260285 |
| SLC25A53 | -0.840641877 | 0.558395075 | 9.76E-07 | 1.09E-05 | 241.2333443 | 134.9278833 | ENSG00000269743 |
| NA | -0.840922724 | 0.558286384 | 0.276568477 | 0.467615547 | 9.868599232 | 5.513672966 | ENSG00000220323 |
| GRHL1 | -0.841111394 | 0.558213378 | 0.00264082 | 0.011969909 | 82.44714654 | 46.04843536 | ENSG00000134317 |
| C8orf76 | -0.841702421 | 0.557984743 | 0.237971161 | 0.423399971 | 15.08825723 | 8.367725234 | ENSG00000189376 |
| CFTR | -0.841968016 | 0.557882029 | 0.295111956 | 0.487965474 | 11.18233992 | 6.256373839 | ENSG00000001626 |
| H1-10 | -0.842186967 | 0.557797368 | 2.47E-28 | 3.18E-26 | 2238.724642 | 1248.75192 | ENSG00000184897 |
| NA | -0.842483914 | 0.55768257 | 0.175316013 | 0.342681606 | 16.03161719 | 8.911869277 | ENSG00000232000 |
| LYSMD2 | -0.842905018 | 0.557519813 | 0.03155361 | 0.093300355 | 41.39476051 | 23.09371501 | ENSG00000140280 |
| PDZK1 | -0.843414859 | 0.557322824 | 0.417428668 | 0.608571958 | 7.559514803 | 4.212293935 | ENSG00000174827 |
| ADGRF3 | -0.843961375 | 0.557111741 | 0.00707919 | 0.027668854 | 81.85187352 | 45.71154515 | ENSG00000173567 |
| NA | -0.845372473 | 0.556567097 | 0.430316452 | 0.620322401 | 6.886619383 | 3.870943603 | ENSG00000225883 |
| PLXDC1 | -0.846038133 | 0.556310356 | 0.229541742 | 0.4134467 | 13.83507661 | 7.734401133 | ENSG00000161381 |
| MARCKSL1 | -0.846548363 | 0.556113644 | 1.86E-25 | 1.86E-23 | 11657.21784 | 6482.587608 | ENSG00000175130 |
| HSD17B7 | -0.846575905 | 0.556103027 | 1.25E-14 | 4.60E-13 | 735.6028099 | 409.344677 | ENSG00000132196 |
| NA | -0.846997013 | 0.55594073 | 0.455323733 | 0.642684606 | 5.088172683 | 2.873261296 | ENSG00000276116 |
| SAP30 | -0.847227339 | 0.555851981 | 1.40E-07 | 1.82E-06 | 317.7745357 | 176.5343211 | ENSG00000164105 |
| NA | -0.847711968 | 0.555665291 | 0.302317677 | 0.495722891 | 10.34185767 | 5.667201734 | ENSG00000285602 |
| NA | -0.847730409 | 0.555658188 | 0.260078094 | 0.449025374 | 11.04651761 | 6.124976879 | ENSG00000259404 |
| NA | -0.847978178 | 0.555562768 | 0.459634192 | 0.646298468 | 4.992788881 | 2.805663392 | ENSG00000261305 |
| NA | -0.84823579 | 0.555463573 | 0.148907319 | 0.304417411 | 18.67498084 | 10.42732694 | ENSG00000269883 |
| SGSM2 | -0.848280607 | 0.555446318 | 6.33E-09 | 1.05E-07 | 763.8621781 | 424.9910439 | ENSG00000141258 |
| TMPRSS2 | -0.849233346 | 0.555079629 | 1.19E-06 | 1.31E-05 | 225.2160385 | 125.1230336 | ENSG00000184012 |
| ANGPTL4 | -0.849749814 | 0.554880953 | 0.058601014 | 0.151163701 | 38.96621705 | 21.53311766 | ENSG00000167772 |
| NA | -0.849944218 | 0.554806187 | 0.023539855 | 0.073796969 | 50.94436143 | 28.33754601 | ENSG00000277534 |
| TSC22D1-AS1 | -0.850217454 | 0.554701121 | 0.13571284 | 0.284206882 | 23.03393733 | 12.83532702 | ENSG00000278156 |
| ADAMTS8 | -0.850450438 | 0.554611548 | 1.23E-12 | 3.57E-11 | 586.9324201 | 325.8672638 | ENSG00000134917 |
| GALNT16 | -0.850698291 | 0.554516275 | 2.20E-06 | 2.29E-05 | 241.982607 | 134.5313193 | ENSG00000100626 |
| ANKRD61 | -0.851384719 | 0.554252501 | 0.332842074 | 0.527391211 | 8.743844707 | 4.894882979 | ENSG00000157999 |
| LOC100130111 | -0.851644943 | 0.554152538 | 0.077969731 | 0.188074715 | 28.59852536 | 15.873402 | ENSG00000256802 |
| HMGN5 | -0.851750169 | 0.554112121 | 9.64E-13 | 2.87E-11 | 703.1251031 | 389.4016227 | ENSG00000198157 |
| SLC9A3 | -0.852046873 | 0.553998174 | 0.061493957 | 0.157030723 | 33.93190728 | 18.93448486 | ENSG00000066230 |
| DNLZ | -0.852386416 | 0.553867804 | 0.385724634 | 0.578680618 | 6.961182572 | 3.900999518 | ENSG00000213221 |
| NA | -0.853553578 | 0.553419898 | 0.164563417 | 0.326939476 | 18.50493615 | 10.19447731 | ENSG00000285531 |
| MARCHF4 | -0.854102624 | 0.553209323 | 0.002213139 | 0.010355547 | 82.14511287 | 45.52729919 | ENSG00000144583 |
| RBM25 | -0.854323587 | 0.553124601 | 1.55E-30 | 2.28E-28 | 3281.614973 | 1815.726664 | ENSG00000119707 |
| RIMS4 | -0.854846412 | 0.552924187 | 3.56E-30 | 5.12E-28 | 1980.03057 | 1094.366209 | ENSG00000101098 |
| SMIM10L1 | -0.855767803 | 0.55257117 | 9.35E-10 | 1.75E-08 | 347.8150098 | 192.220666 | ENSG00000256537 |
| RTN4RL1 | -0.855957881 | 0.552498372 | 0.33742533 | 0.53183443 | 7.953271699 | 4.350738936 | ENSG00000185924 |
| NA | -0.856365022 | 0.552342474 | 0.048830332 | 0.131314768 | 37.08265404 | 20.42660823 | ENSG00000224043 |
| MAPK8IP1 | -0.856682223 | 0.552221046 | 2.37E-21 | 1.68E-19 | 1207.805774 | 667.2971361 | ENSG00000121653 |
| CUL1 | -0.857176694 | 0.552031809 | 7.13E-35 | 1.35E-32 | 2584.103758 | 1426.693293 | ENSG00000055130 |
| FABP3 | -0.857211864 | 0.552018352 | 2.04E-06 | 2.14E-05 | 267.85962 | 147.5241484 | ENSG00000121769 |
| GPR1 | -0.857288297 | 0.551989107 | 0.298655456 | 0.492019081 | 11.12483907 | 6.117490805 | ENSG00000183671 |
| ADAP2 | -0.858389747 | 0.551567843 | 0.183346873 | 0.353826629 | 16.54541183 | 9.042828204 | ENSG00000184060 |
| KCNJ8 | -0.859664167 | 0.551080825 | 0.236746805 | 0.421747913 | 14.25664436 | 7.839102992 | ENSG00000121361 |
| STK32B | -0.859911674 | 0.55098629 | 0.000274163 | 0.00170001 | 122.7867774 | 67.72434652 | ENSG00000152953 |
| LINC01122 | -0.859963699 | 0.550966421 | 0.115879508 | 0.253096652 | 23.15387738 | 12.7264999 | ENSG00000233723 |
| GRIA2 | -0.860203066 | 0.550875014 | 0.134645406 | 0.282458173 | 24.49996131 | 13.5293126 | ENSG00000120251 |
| ANKRD13B | -0.860238749 | 0.550861389 | 1.73E-14 | 6.23E-13 | 971.2537226 | 535.0054276 | ENSG00000198720 |
| NA | -0.860408203 | 0.550796691 | 0.419484389 | 0.61083624 | 5.65691262 | 3.143214877 | ENSG00000228110 |
| NA | -0.860517589 | 0.550754931 | 0.127102225 | 0.270771568 | 22.22885753 | 12.35553166 | ENSG00000261189 |
| MFGE8 | -0.860741862 | 0.55066932 | 2.70E-61 | 2.00E-58 | 19038.13264 | 10484.36887 | ENSG00000140545 |
| PSMA2 | -0.861462761 | 0.550394225 | 0.009014407 | 0.033738493 | 64.44525652 | 35.40323094 | ENSG00000106588 |
| SLC24A3 | -0.861838885 | 0.550250751 | 0.000393671 | 0.002319233 | 121.9758874 | 67.35381634 | ENSG00000185052 |
| NA | -0.862404179 | 0.550035188 | 0.359629638 | 0.552913788 | 6.99718639 | 3.889714598 | ENSG00000250308 |
| ADAM11 | -0.86249451 | 0.550000749 | 7.05E-07 | 8.08E-06 | 245.6286129 | 135.34118 | ENSG00000073670 |
| NA | -0.862623811 | 0.549951458 | 0.121508971 | 0.262109784 | 20.66990761 | 11.38366841 | ENSG00000272578 |
| STX16-NPEPL1 | -0.86304855 | 0.549789573 | 0.313908439 | 0.507632756 | 10.71845453 | 5.877593139 | ENSG00000254995 |
| MANEAL | -0.863432587 | 0.549643242 | 0.002407002 | 0.011102879 | 85.84794319 | 46.99685272 | ENSG00000185090 |

| | | | | | | | |
|---|---|---|---|---|---|---|---|
| ATP1A3 | -0.863498244 | 0.549618228 | 0.051358993 | 0.136530061 | 35.40178724 | 19.42881431 | ENSG00000105409 |
| LPCAT3 | -0.864180752 | 0.549358277 | 2.23E-12 | 6.22E-11 | 573.4121775 | 315.2322365 | ENSG00000111684 |
| NA | -0.864622913 | 0.549189934 | 0.139930491 | 0.290911017 | 18.02032784 | 9.875696822 | ENSG00000215302 |
| ADI1 | -0.864905182 | 0.549082493 | 3.54E-18 | 1.90E-16 | 825.6483463 | 453.2375687 | ENSG00000182551 |
| DRD4 | -0.864948617 | 0.549065962 | 0.012099328 | 0.043098696 | 51.98951671 | 28.58861698 | ENSG00000069696 |
| SH2D5 | -0.865076148 | 0.549017428 | 4.11E-15 | 1.61E-13 | 699.4167861 | 384.4287732 | ENSG00000189410 |
| GDAP1 | -0.865168643 | 0.54898223 | 7.09E-17 | 3.29E-15 | 796.3146743 | 437.4679212 | ENSG00000104381 |
| CLIP3 | -0.865334353 | 0.548919177 | 5.56E-08 | 7.79E-07 | 330.687148 | 181.2063168 | ENSG00000105270 |
| H4C5 | -0.865887817 | 0.548708634 | 0.130997899 | 0.277115066 | 18.69204318 | 10.23581815 | ENSG00000276966 |
| NOL12 | -0.866260574 | 0.54856688 | 0.021921958 | 0.069751342 | 63.89851727 | 35.29626757 | ENSG00000273899 |
| KHK | -0.867051536 | 0.548266209 | 1.98E-07 | 2.52E-06 | 264.1447634 | 144.9909636 | ENSG00000138030 |
| IQGAP1 | -0.867137903 | 0.548233388 | 2.73E-28 | 3.47E-26 | 3237.864714 | 1774.592349 | ENSG00000140575 |
| C1orf21 | -0.867739991 | 0.548004638 | 6.31E-17 | 2.97E-15 | 705.2224418 | 386.5080933 | ENSG00000116667 |
| NAGPA | -0.869247024 | 0.547432494 | 6.62E-07 | 7.63E-06 | 223.2153993 | 122.0991663 | ENSG00000103174 |
| NA | -0.86939387 | 0.547376776 | 0.231746631 | 0.416402527 | 12.08535907 | 6.665322075 | ENSG00000273284 |
| NA | -0.869594298 | 0.547300736 | 0.090361559 | 0.210133142 | 25.40801651 | 13.81000145 | ENSG00000249637 |
| NA | -0.870136382 | 0.54709513 | 0.254267087 | 0.44298754 | 11.17918301 | 6.064865049 | ENSG00000255921 |
| CAMKV | -0.871288232 | 0.546658503 | 4.89E-31 | 7.42E-29 | 2172.235482 | 1187.981549 | ENSG00000164076 |
| NA | -0.87129774 | 0.5466549 | 0.434985376 | 0.624534268 | 5.022478873 | 2.756836483 | ENSG00000235084 |
| EXOSC1 | -0.871852575 | 0.546444706 | 1.88E-12 | 5.31E-11 | 676.4868089 | 370.0298145 | ENSG00000171311 |
| GJC3 | -0.872690554 | 0.5461274 | 0.214826101 | 0.394998707 | 16.18332177 | 8.949849299 | ENSG00000176402 |
| MSI1 | -0.872930491 | 0.54603658 | 5.28E-25 | 4.99E-23 | 1439.663569 | 785.9098649 | ENSG00000135097 |
| HAPLN3 | -0.873200099 | 0.545934547 | 2.33E-23 | 1.94E-21 | 1905.898981 | 1041.00794 | ENSG00000140511 |
| NA | -0.873222719 | 0.545925988 | 0.292871198 | 0.485432453 | 8.826600858 | 4.838569996 | ENSG00000228519 |
| LOC105376114 | -0.874014448 | 0.545626474 | 0.21052577 | 0.389867428 | 13.7321012 | 7.445126904 | ENSG00000254473 |
| LOC399900 | -0.874239208 | 0.545541477 | 0.333349901 | 0.527970182 | 7.654996303 | 4.201009014 | ENSG00000279491 |
| TTLL11 | -0.874499545 | 0.545443042 | 0.117742066 | 0.255980228 | 22.36981359 | 12.14850109 | ENSG00000175764 |
| LOC730183 | -0.874630752 | 0.545393438 | 0.189384356 | 0.361599841 | 16.61120334 | 9.156003823 | ENSG00000261840 |
| NA | -0.874758855 | 0.545345013 | 0.036366656 | 0.104402044 | 38.43565316 | 21.02326641 | ENSG00000272667 |
| SLITRK4 | -0.874776069 | 0.545338506 | 4.76E-06 | 4.63E-05 | 190.8109705 | 103.8773264 | ENSG00000179542 |
| MYBL2 | -0.875711845 | 0.544984897 | 4.64E-44 | 1.54E-41 | 6258.44021 | 3410.40628 | ENSG00000101057 |
| NA | -0.875923915 | 0.544904793 | 0.327504132 | 0.521697932 | 7.592459406 | 4.152182105 | ENSG00000272989 |
| NKAIN1 | -0.876061291 | 0.544852908 | 2.41E-20 | 1.59E-18 | 2011.946302 | 1096.016767 | ENSG00000084628 |
| RND2 | -0.876690942 | 0.544615164 | 2.53E-35 | 4.99E-33 | 3200.174136 | 1743.014814 | ENSG00000108830 |
| NA | -0.876929352 | 0.544525172 | 0.386300365 | 0.579263023 | 7.644247788 | 4.178439173 | ENSG00000181227 |
| TPPP3 | -0.877103963 | 0.544459272 | 0.342059289 | 0.536265171 | 7.592459406 | 4.148383258 | ENSG00000159713 |
| NA | -0.877685743 | 0.544239758 | 0.089669788 | 0.209049038 | 26.21565186 | 14.38843829 | ENSG00000228158 |
| NPEPPSP1 | -0.877720931 | 0.544226484 | 0.001591558 | 0.007826465 | 87.43154706 | 47.5978594 | ENSG00000274487 |
| CACHD1 | -0.877743757 | 0.544217873 | 7.48E-28 | 8.94E-26 | 1654.230259 | 900.4190725 | ENSG00000158966 |
| NA | -0.87864695 | 0.543877275 | 0.055799218 | 0.145413767 | 38.06852704 | 20.79096644 | ENSG00000259865 |
| GAL3ST3 | -0.878721902 | 0.543849019 | 2.43E-08 | 3.64E-07 | 280.3240564 | 152.3794511 | ENSG00000175229 |
| NA | -0.879123501 | 0.543697651 | 0.400195819 | 0.59215821 | 5.716292604 | 3.116957809 | ENSG00000260577 |
| TNKS2-AS1 | -0.880314925 | 0.543248833 | 0.418806245 | 0.610090886 | 5.855271827 | 3.203326707 | ENSG00000228701 |
| CRELD1 | -0.880325746 | 0.543244758 | 0.099831152 | 0.226609991 | 24.08480508 | 13.10539222 | ENSG00000163703 |
| UCKL1-AS1 | -0.8805375 | 0.543165028 | 0.213275681 | 0.393162071 | 12.09482981 | 6.590238097 | ENSG00000280213 |
| HINT2 | -0.880585043 | 0.543147129 | 0.438975061 | 0.62733933 | 5.822424922 | 3.177069639 | ENSG00000137133 |
| EPN3 | -0.880741442 | 0.543088251 | 0.001375056 | 0.006874912 | 84.69161954 | 45.92832332 | ENSG00000049283 |
| ADAT3 | -0.88178201 | 0.542696681 | 7.09E-05 | 0.000512896 | 139.0527576 | 75.40948271 | ENSG00000213638 |
| SNHG19 | -0.883119398 | 0.54219383 | 0.062885479 | 0.159791537 | 405.1133447 | 219.7737096 | ENSG00000260260 |
| NA | -0.883455971 | 0.542067354 | 0.319291901 | 0.513144724 | 9.825003812 | 5.408971107 | ENSG00000272267 |
| PSORS1C2 | -0.883475216 | 0.542060123 | 0.362188669 | 0.555144293 | 6.361474866 | 3.454509294 | ENSG00000204538 |
| RSPH1 | -0.883524073 | 0.542041767 | 0.416026827 | 0.60723273 | 5.124176501 | 2.786892398 | ENSG00000160188 |
| C9orf24 | -0.884035677 | 0.541849584 | 0.067971977 | 0.169122381 | 30.71802227 | 16.56315071 | ENSG00000164972 |
| OGFR-AS1 | -0.884203571 | 0.541786529 | 0.261929097 | 0.451186754 | 11.89459147 | 6.432472449 | ENSG00000229873 |
| NA | -0.884492505 | 0.541678035 | 0.306743284 | 0.500827092 | 9.402932397 | 5.085953736 | ENSG00000223461 |
| NPR1 | -0.884680352 | 0.54160751 | 1.18E-14 | 4.37E-13 | 571.4946027 | 309.4743258 | ENSG00000169418 |
| NA | -0.884873261 | 0.541535094 | 0.31010379 | 0.503942449 | 8.214265501 | 4.448392755 | ENSG00000283213 |
| TCEAL3 | -0.884991785 | 0.541490606 | 4.68E-10 | 9.14E-09 | 349.4225912 | 189.1033818 | ENSG00000196507 |
| SEMA4A | -0.885117681 | 0.541443355 | 2.97E-09 | 5.12E-08 | 316.8170522 | 171.6895791 | ENSG00000196189 |
| PPP2R2B | -0.885504356 | 0.541298255 | 3.30E-25 | 3.14E-23 | 1423.232016 | 770.2877572 | ENSG00000156475 |
| HLF | -0.885798845 | 0.541187775 | 0.408822207 | 0.600404094 | 6.864423295 | 3.656864972 | ENSG00000108924 |
| NA | -0.885893054 | 0.541152436 | 0.450834984 | 0.638430286 | 5.194305001 | 2.824434387 | ENSG00000230747 |
| RMST | -0.886026571 | 0.541102357 | 0.00063336 | 0.00352725 | 115.2931669 | 62.5379278 | ENSG00000255794 |
| GPLD1 | -0.886386077 | 0.540967536 | 0.000186912 | 0.001209999 | 130.9949396 | 70.6228772 | ENSG00000112293 |
| SLC4A4 | -0.886476707 | 0.540933553 | 0.036935964 | 0.105513169 | 40.06651303 | 21.51010979 | ENSG00000080493 |
| DPY19L2 | -0.88648284 | 0.540931254 | 0.029392892 | 0.088093121 | 42.86582054 | 23.21773751 | ENSG00000177990 |
| TMEM121B | -0.887161081 | 0.54067701 | 9.38E-07 | 1.05E-05 | 232.8325731 | 125.8084338 | ENSG00000183307 |
| LOC101928718 | -0.887338192 | 0.540610639 | 0.38589095 | 0.578883276 | 5.713135692 | 3.086901894 | ENSG00000284830 |
| NPY | -0.887342167 | 0.540609149 | 0.35615204 | 0.549793097 | 7.778190961 | 4.219780009 | ENSG00000122585 |
| FOXP1 | -0.887388613 | 0.540591745 | 0.001938087 | 0.009255476 | 82.69984968 | 44.52494197 | ENSG00000114861 |
| NGRN | -0.887501122 | 0.540549589 | 2.22E-12 | 6.20E-11 | 539.4211833 | 291.6090317 | ENSG00000182768 |
| PNCK | -0.887558134 | 0.540528228 | 0.002732016 | 0.012299025 | 79.98763531 | 43.34834989 | ENSG00000130822 |
| SAFB2 | -0.888179636 | 0.540295422 | 5.42E-26 | 5.55E-24 | 2390.201781 | 1292.088029 | ENSG00000130254 |
| MAN2A1 | -0.888671 | 0.540111436 | 1.53E-24 | 1.39E-22 | 1558.440033 | 841.0396179 | ENSG00000112893 |

| | | | | | | | |
|---|---|---|---|---|---|---|---|
| TIA1 | -0.888988898 | 0.539992435 | 2.52E-19 | 1.53E-17 | 1578.165371 | 851.9617779 | ENSG00000116001 |
| FAM133B | -0.889738847 | 0.539711807 | 3.02E-12 | 8.27E-11 | 759.626942 | 409.4433411 | ENSG00000234545 |
| IFT81 | -0.88984503 | 0.539672085 | 5.77E-09 | 9.61E-08 | 300.7185611 | 162.1987234 | ENSG00000122970 |
| DKKL1 | -0.889940651 | 0.539636317 | 0.048623493 | 0.130910492 | 36.25479944 | 19.71852657 | ENSG00000104901 |
| LRRC17 | -0.890438585 | 0.539450098 | 0.208184191 | 0.387281521 | 15.53626037 | 8.443358864 | ENSG00000128606 |
| APCDD1L | -0.891075104 | 0.539212145 | 5.05E-05 | 0.000379408 | 137.145623 | 74.01457512 | ENSG00000198768 |
| LRRC45 | -0.891103073 | 0.539201691 | 1.87E-14 | 6.72E-13 | 874.8631943 | 472.1406207 | ENSG00000169683 |
| LINC01089 | -0.891828797 | 0.538930523 | 0.090640878 | 0.210703378 | 182.5182108 | 98.49575192 | ENSG00000212694 |
| ZNF674-AS1 | -0.891993662 | 0.53886894 | 0.018985755 | 0.061733682 | 70.92086127 | 38.41702423 | ENSG00000230844 |
| EFHD2 | -0.892104765 | 0.538827443 | 4.29E-21 | 2.99E-19 | 1287.538031 | 693.3877682 | ENSG00000142634 |
| ABCB9 | -0.892403735 | 0.538715793 | 0.000479679 | 0.002761841 | 107.9405725 | 58.25759793 | ENSG00000150967 |
| NA | -0.892764485 | 0.538581103 | 0.294864113 | 0.487686192 | 9.060963883 | 4.887396905 | ENSG00000260774 |
| NDRG2 | -0.893058437 | 0.538471376 | 9.60E-10 | 1.79E-08 | 385.5462136 | 207.8818531 | ENSG00000165795 |
| NA | -0.894539402 | 0.537918905 | 0.427077748 | 0.617568648 | 5.720727294 | 3.0681309 | ENSG00000259051 |
| FBXW4P1 | -0.894973873 | 0.537756934 | 0.17071754 | 0.336067859 | 14.66481034 | 7.86536006 | ENSG00000230701 |
| HSBP1L1 | -0.896639333 | 0.537136501 | 0.172759587 | 0.339043434 | 14.93339575 | 8.090723612 | ENSG00000226742 |
| NA | -0.896863635 | 0.537052996 | 0.112701933 | 0.24781322 | 20.71981685 | 11.19259765 | ENSG00000280604 |
| NA | -0.898453768 | 0.536461384 | 0.20789216 | 0.386932445 | 13.5994358 | 7.351709965 | ENSG00000253333 |
| MYO5C | -0.899655613 | 0.536014669 | 4.61E-13 | 1.42E-11 | 498.643133 | 267.2665549 | ENSG00000128833 |
| ETNPPL | -0.900273222 | 0.535785253 | 0.030292072 | 0.090274934 | 37.41061943 | 20.12703677 | ENSG00000164089 |
| NA | -0.90088287 | 0.535558891 | 0.036209194 | 0.104014444 | 33.49267583 | 17.95513551 | ENSG00000257122 |
| SRSF12 | -0.902175756 | 0.535079159 | 6.28E-05 | 0.000461269 | 148.333074 | 79.54365827 | ENSG00000154548 |
| ZNF878 | -0.903286691 | 0.534667285 | 0.008343792 | 0.031721813 | 55.7563571 | 29.79614104 | ENSG00000257446 |
| CNPY2 | -0.90331446 | 0.534656994 | 0.057541837 | 0.149095542 | 29.33721229 | 15.59921154 | ENSG00000257727 |
| NKX3-2 | -0.904785139 | 0.534112244 | 0.021450482 | 0.068521758 | 41.82697908 | 22.30587445 | ENSG00000109705 |
| LINC02696 | -0.904836831 | 0.534093107 | 0.307514595 | 0.501731365 | 7.704905549 | 4.092070275 | ENSG00000254427 |
| TRABD2A | -0.904969797 | 0.534043884 | 1.91E-05 | 0.000160143 | 163.6759362 | 87.52500511 | ENSG00000186854 |
| TMEM229A | -0.905232911 | 0.533946496 | 0.317529377 | 0.511257882 | 9.012430111 | 4.84605607 | ENSG00000234224 |
| TRIL | -0.90573565 | 0.533760463 | 1.03E-06 | 1.14E-05 | 193.1558633 | 103.0492443 | ENSG00000255690 |
| FAIM | -0.90604376 | 0.533646482 | 0.000912717 | 0.004873628 | 106.909924 | 57.1357815 | ENSG00000158234 |
| NA | -0.906326318 | 0.533541975 | 0.157782503 | 0.317036092 | 19.60000068 | 10.49492484 | ENSG00000267904 |
| GH2 | -0.90683598 | 0.533353524 | 0.360059643 | 0.553253641 | 7.189329469 | 3.882228524 | ENSG00000136487 |
| SCYL2 | -0.90691117 | 0.533325727 | 6.57E-13 | 2.00E-11 | 673.2636786 | 358.5988876 | ENSG00000136021 |
| ME3 | -0.907354761 | 0.533161769 | 0.021691929 | 0.069161915 | 44.80196868 | 23.92658362 | ENSG00000151376 |
| RAB6B | -0.9073776 | 0.533153329 | 1.29E-11 | 3.23E-10 | 421.25604 | 224.5068275 | ENSG00000154917 |
| CORO1A | -0.907640987 | 0.533056002 | 1.37E-07 | 1.79E-06 | 236.1483285 | 125.9819445 | ENSG00000102879 |
| BBS5 | -0.907729526 | 0.533023289 | 0.166706805 | 0.330171397 | 14.8582312 | 7.944242884 | ENSG00000163093 |
| OVOL2 | -0.907925556 | 0.532950868 | 7.38E-06 | 6.86E-05 | 212.9678204 | 113.4237307 | ENSG00000125850 |
| NA | -0.90926184 | 0.532457455 | 0.489802983 | 0.672644375 | 5.102078111 | 2.7005235 | ENSG00000257195 |
| MYL5 | -0.909917439 | 0.532215547 | 0.001545853 | 0.007628021 | 81.93590745 | 43.79115289 | ENSG00000215375 |
| PANX2 | -0.910269707 | 0.53208561 | 4.86E-06 | 4.71E-05 | 210.8258191 | 112.0577797 | ENSG00000073150 |
| NOS1 | -0.912487465 | 0.531268299 | 2.62E-05 | 0.000212614 | 172.1560135 | 91.11273484 | ENSG00000089250 |
| NA | -0.913190222 | 0.531009574 | 0.123447181 | 0.265118198 | 21.10272754 | 11.28656425 | ENSG00000257489 |
| MCOLN3 | -0.913214036 | 0.531000809 | 0.08514602 | 0.200699013 | 26.69206721 | 14.04989912 | ENSG00000055732 |
| NA | -0.914425561 | 0.53055508 | 0.080044752 | 0.191871371 | 23.96920203 | 12.75275697 | ENSG00000213740 |
| NA | -0.915984276 | 0.529982168 | 0.234120216 | 0.419001939 | 10.43418226 | 5.562499875 | ENSG00000253200 |
| ATP6V1FNB | -0.916782968 | 0.529688845 | 0.227853477 | 0.411481667 | 11.16212067 | 5.948878269 | ENSG00000272899 |
| HSPA14 | -0.917136541 | 0.529559046 | 4.65E-09 | 7.86E-08 | 392.3322252 | 207.394205 | ENSG00000187522 |
| HSD3B7 | -0.918084822 | 0.529211082 | 0.199319028 | 0.375658161 | 12.97575057 | 6.871476599 | ENSG00000099377 |
| NA | -0.918313011 | 0.529127384 | 0.143819413 | 0.296699577 | 15.69102415 | 8.311850284 | ENSG00000279692 |
| NRCAM | -0.918471098 | 0.529069407 | 0.000185742 | 0.001203265 | 118.4003183 | 62.61571132 | ENSG00000091129 |
| ADGRG5 | -0.918530809 | 0.52904751 | 2.89E-10 | 5.84E-09 | 359.1784738 | 190.0755715 | ENSG00000159618 |
| NA | -0.918557621 | 0.529037678 | 0.411620641 | 0.603123907 | 5.914749509 | 3.165784718 | ENSG00000267898 |
| NA | -0.919424778 | 0.528719786 | 0.427686771 | 0.618108004 | 5.147552668 | 2.696724653 | ENSG00000260060 |
| NTNG2 | -0.919617024 | 0.528649336 | 0.026789449 | 0.081933275 | 37.99188932 | 20.13832169 | ENSG00000196358 |
| TLL1 | -0.91970826 | 0.528615905 | 0.091764308 | 0.212492343 | 23.66451511 | 12.55420013 | ENSG00000038295 |
| C1QTNF2 | -0.920619427 | 0.528282151 | 0.09116841 | 0.211681085 | 25.38779726 | 13.41668663 | ENSG00000145861 |
| FAM228B | -0.920679875 | 0.528260017 | 0.223354212 | 0.405609148 | 11.89637291 | 6.316485669 | ENSG00000219626 |
| CRTC3 | -0.921159666 | 0.528084365 | 5.38E-08 | 7.56E-07 | 250.0642071 | 132.1142958 | ENSG00000140577 |
| NA | -0.921289182 | 0.528036959 | 0.271265434 | 0.462013743 | 9.206354629 | 4.894882979 | ENSG00000283959 |
| EMX1 | -0.921311891 | 0.528028648 | 1.12E-05 | 9.90E-05 | 155.5601888 | 82.11234677 | ENSG00000135638 |
| F13A1 | -0.922317252 | 0.527660812 | 0.083465169 | 0.197920093 | 24.07080196 | 12.64761707 | ENSG00000124491 |
| LINC02570 | -0.922751236 | 0.527502108 | 9.91E-07 | 1.10E-05 | 189.8004435 | 99.98056374 | ENSG00000237923 |
| NUCB2 | -0.923278068 | 0.527309514 | 2.22E-08 | 3.34E-07 | 301.3297915 | 158.8794099 | ENSG00000070081 |
| ZNF8-ERVK3-1 | -0.92353784 | 0.527214575 | 0.375169603 | 0.568118344 | 5.984878009 | 3.203326707 | ENSG00000267216 |
| GRIN2B | -0.923957446 | 0.527061258 | 0.002628797 | 0.011931552 | 71.56298431 | 37.66518832 | ENSG00000273079 |
| PNMT | -0.924013797 | 0.527040671 | 0.001797118 | 0.008673904 | 87.03788786 | 46.07199289 | ENSG00000141744 |
| NA | -0.924545024 | 0.526846641 | 0.288918463 | 0.480638465 | 9.117186954 | 4.759687172 | ENSG00000226976 |
| NAALAD2 | -0.925114717 | 0.52663864 | 0.019538893 | 0.063276924 | 49.32909073 | 26.05725567 | ENSG00000077616 |
| SCD5 | -0.925542991 | 0.526482327 | 0.0207519 | 0.066618569 | 41.82382216 | 21.99469165 | ENSG00000145284 |
| FDPS | -0.925623879 | 0.526452809 | 1.76E-24 | 1.59E-22 | 1745.645812 | 919.4628628 | ENSG00000160752 |
| NA | -0.926088905 | 0.526283144 | 0.158736459 | 0.318435061 | 15.68786724 | 8.236766307 | ENSG00000226074 |
| DMC1 | -0.927366827 | 0.525817176 | 0.000631874 | 0.003521095 | 86.72580474 | 45.54563215 | ENSG00000100206 |
| ANGPT1 | -0.927788679 | 0.525663446 | 0.008068265 | 0.030845138 | 53.4612758 | 28.10458476 | ENSG00000154188 |

| | | | | | | | |
|---|---|---|---|---|---|---|---|
| SMARCE1 | -0.92915451 | 0.525166025 | 0.027955449 | 0.084855272 | 38.38436844 | 20.1040289 | ENSG00000073584 |
| ATP6V0E2 | -0.929745882 | 0.524950799 | 2.77E-16 | 1.21E-14 | 619.3971983 | 325.1676156 | ENSG00000171130 |
| NA | -0.930209767 | 0.524782033 | 0.231799899 | 0.416457863 | 10.4215546 | 5.464846056 | ENSG00000218690 |
| ANAPC1P1 | -0.930427249 | 0.52470293 | 0.229281223 | 0.41317435 | 11.64494754 | 6.091122117 | ENSG00000233673 |
| ZDHHC15 | -0.930984164 | 0.524500421 | 0.000263865 | 0.00164441 | 109.8651753 | 57.5219451 | ENSG00000102383 |
| NA | -0.93179563 | 0.524205491 | 0.085970602 | 0.202308169 | 23.16778281 | 12.16022405 | ENSG00000224831 |
| DZIP3 | -0.932422474 | 0.523977775 | 3.41E-14 | 1.20E-12 | 570.4262666 | 299.3756789 | ENSG00000198919 |
| NA | -0.932685297 | 0.523882328 | 0.065926554 | 0.165383781 | 24.18018888 | 12.63253331 | ENSG00000253683 |
| EFNB1 | -0.93280163 | 0.523840087 | 5.31E-11 | 1.20E-09 | 427.5895912 | 224.1398814 | ENSG00000090776 |
| PPFIA2 | -0.933293058 | 0.52366168 | 0.09251794 | 0.213753115 | 21.08062915 | 11.04611693 | ENSG00000139220 |
| NA | -0.933798002 | 0.52347843 | 0.020341591 | 0.065532948 | 40.1853707 | 21.11343416 | ENSG00000286707 |
| NA | -0.933801067 | 0.523477318 | 0.285543456 | 0.476780398 | 8.479693993 | 4.440906681 | ENSG00000286873 |
| LRRC38 | -0.933963244 | 0.523418476 | 0.338661337 | 0.532785776 | 6.606684104 | 3.484565209 | ENSG00000162494 |
| C6orf141 | -0.934750286 | 0.52313301 | 0.004455569 | 0.018653607 | 83.3103059 | 43.81427238 | ENSG00000197261 |
| NA | -0.935742099 | 0.522773495 | 0.287652203 | 0.4792262 | 7.923581707 | 4.159668179 | ENSG00000272092 |
| HSPA6 | -0.936155182 | 0.522623832 | 0.406636745 | 0.598333236 | 6.027195652 | 3.177069639 | ENSG00000173110 |
| H2AX | -0.936394615 | 0.522537103 | 5.87E-23 | 4.79E-21 | 3409.572092 | 1781.261472 | ENSG00000188486 |
| NR0B1 | -0.937282544 | 0.522215598 | 0.2884969 | 0.480179616 | 9.359336977 | 4.954994809 | ENSG00000169297 |
| TTYH1 | -0.937624907 | 0.522091687 | 3.14E-12 | 8.58E-11 | 424.9925891 | 221.6446363 | ENSG00000167614 |
| RBP4 | -0.938492244 | 0.521777904 | 0.161389814 | 0.322172101 | 14.63827726 | 7.681775377 | ENSG00000138207 |
| NA | -0.939176741 | 0.521530401 | 0.369001509 | 0.5619446 | 6.431603366 | 3.296743646 | ENSG00000233514 |
| NA | -0.93963002 | 0.521366568 | 0.36164219 | 0.554809321 | 6.070791072 | 3.203326707 | ENSG00000280422 |
| STOX1 | -0.940279034 | 0.521132078 | 0.001775781 | 0.008584333 | 72.60951507 | 37.8906635 | ENSG00000165730 |
| NA | -0.940330951 | 0.521113325 | 0.407911861 | 0.599685764 | 5.435079549 | 2.873261296 | ENSG00000287787 |
| SCG3 | -0.940449416 | 0.521070536 | 9.29E-16 | 3.89E-14 | 1045.987034 | 544.5055526 | ENSG00000104112 |
| PGAM1P5 | -0.940841987 | 0.520928767 | 0.015311329 | 0.051945847 | 45.76060955 | 23.85518687 | ENSG00000257150 |
| NA | -0.941066356 | 0.520847758 | 0.530749654 | 0.706167559 | 4.690078795 | 2.460625833 | ENSG00000234241 |
| NA | -0.941606986 | 0.520652614 | 0.259358922 | 0.44824303 | 10.07327226 | 5.255004305 | ENSG00000233045 |
| RTP1 | -0.941720547 | 0.520611633 | 0.001999207 | 0.009493542 | 74.71628663 | 38.92957502 | ENSG00000175077 |
| ADGRG2 | -0.941946566 | 0.520530078 | 1.45E-07 | 1.87E-06 | 248.3115206 | 129.0831572 | ENSG00000173698 |
| STAM-AS1 | -0.942069571 | 0.520485699 | 0.334914814 | 0.529296293 | 6.712816421 | 3.522107198 | ENSG00000260589 |
| P2RX5 | -0.942396708 | 0.52036769 | 0.026879612 | 0.082168353 | 35.63930717 | 18.53258467 | ENSG00000083454 |
| NA | -0.943251523 | 0.520059457 | 0.324771536 | 0.518861792 | 6.689440255 | 3.503336203 | ENSG00000280020 |
| SLC49A3 | -0.943994716 | 0.519791621 | 6.95E-05 | 0.000503316 | 120.1074099 | 62.42800172 | ENSG00000169026 |
| NA | -0.944230862 | 0.519706547 | 0.356449568 | 0.55007796 | 7.184894779 | 3.731948949 | ENSG00000264456 |
| BNC2-AS1 | -0.944403665 | 0.519644301 | 0.11607233 | 0.253300901 | 19.67338379 | 10.17950517 | ENSG00000234779 |
| CELF3 | -0.944447078 | 0.519628665 | 0.0949481 | 0.217799628 | 20.52145765 | 10.69359329 | ENSG00000159409 |
| ZNF428 | -0.944724017 | 0.519528926 | 3.59E-06 | 3.60E-05 | 216.1997372 | 112.3092887 | ENSG00000131116 |
| NYNRIN | -0.94483364 | 0.519489451 | 9.23E-29 | 1.21E-26 | 1298.69734 | 674.5875233 | ENSG00000205978 |
| CYP7B1 | -0.944892232 | 0.519468354 | 0.309826767 | 0.503757285 | 9.771839955 | 5.010869759 | ENSG00000172817 |
| C12orf57 | -0.945041302 | 0.519414681 | 0.000536174 | 0.003051136 | 96.91340228 | 50.20006781 | ENSG00000111678 |
| NA | -0.945142305 | 0.519378318 | 0.102734684 | 0.231145322 | 19.67466157 | 10.16822024 | ENSG00000285693 |
| RHCG | -0.945254072 | 0.519338083 | 0.281189335 | 0.472704139 | 8.012651684 | 4.1709531 | ENSG00000140519 |
| LAYN | -0.945377823 | 0.519293538 | 0.000215603 | 0.001372716 | 110.4763306 | 57.20250334 | ENSG00000204381 |
| ADGRA1 | -0.945647848 | 0.519196352 | 0.284370773 | 0.475379085 | 9.407968447 | 4.943709888 | ENSG00000197177 |
| NA | -0.94609516 | 0.519035399 | 0.001135625 | 0.005870873 | 166.3638799 | 86.72515546 | ENSG00000247092 |
| TAS1R3 | -0.946192938 | 0.519000222 | 0.295356133 | 0.488195003 | 8.690680851 | 4.512303431 | ENSG00000169962 |
| NA | -0.946698009 | 0.518818558 | 0.184238326 | 0.354918733 | 12.72738442 | 6.64655108 | ENSG00000237742 |
| CCKBR | -0.946867685 | 0.518757543 | 2.06E-13 | 6.56E-12 | 501.4182524 | 260.3927137 | ENSG00000110148 |
| SPHK2 | -0.946867828 | 0.518757492 | 3.14E-19 | 1.87E-17 | 812.312753 | 421.5218083 | ENSG00000063176 |
| LOC102723313 | -0.947058158 | 0.518689059 | 0.281082811 | 0.472567925 | 7.903362454 | 4.092070275 | ENSG00000248896 |
| NA | -0.947721385 | 0.518450665 | 0.157154586 | 0.316151426 | 18.15812698 | 9.512326302 | ENSG00000235043 |
| NA | -0.948774563 | 0.51807233 | 0.236028259 | 0.42103488 | 9.957669208 | 5.138579492 | ENSG00000273188 |
| ARL4C | -0.950416284 | 0.517483123 | 2.75E-16 | 1.20E-14 | 591.1818919 | 305.8058495 | ENSG00000188042 |
| NA | -0.951122175 | 0.517229987 | 0.138757139 | 0.288957845 | 16.04424484 | 8.300565364 | ENSG00000239218 |
| GPR61 | -0.951858308 | 0.516966139 | 0.40342677 | 0.595121916 | 5.481831882 | 2.865775222 | ENSG00000156097 |
| GCHFR | -0.952436607 | 0.516758956 | 0.325640101 | 0.519846135 | 7.383253986 | 3.822116694 | ENSG00000137880 |
| NA | -0.952624837 | 0.516691539 | 0.279679407 | 0.47080637 | 8.643928518 | 4.452191601 | ENSG00000269086 |
| WNT3A | -0.9533782 | 0.516421797 | 0.398131681 | 0.590012783 | 5.283472675 | 2.738065489 | ENSG00000154342 |
| UNC5A | -0.95356916 | 0.516353446 | 1.54E-08 | 2.40E-07 | 300.8125921 | 155.4823129 | ENSG00000113763 |
| NA | -0.954779522 | 0.515920429 | 0.423178467 | 0.614140609 | 4.780426548 | 2.498167822 | ENSG00000271366 |
| ADM5 | -0.954951479 | 0.515858939 | 0.001632247 | 0.00798629 | 73.04549192 | 37.61647303 | ENSG00000224420 |
| NA | -0.955693065 | 0.515593841 | 0.028147799 | 0.085327331 | 39.03210626 | 20.22848944 | ENSG00000260924 |
| EMC6 | -0.956511156 | 0.515301552 | 0.106464841 | 0.237553438 | 22.3848991 | 11.60903196 | ENSG00000127774 |
| MATR3 | -0.957433952 | 0.514972054 | 0.100679418 | 0.227698882 | 21.53188689 | 11.14756959 | ENSG00000015479 |
| NA | -0.957597273 | 0.514913759 | 0.350226577 | 0.543506605 | 6.772196406 | 3.495850129 | ENSG00000262692 |
| ANKRD44 | -0.957658342 | 0.514891964 | 0.107208323 | 0.238678565 | 18.13593089 | 9.373331648 | ENSG00000065413 |
| NA | -0.95798647 | 0.51477487 | 0.148281985 | 0.303507267 | 16.32989259 | 8.469615933 | ENSG00000273723 |
| BAIAP2L2 | -0.958254963 | 0.514679076 | 0.000262217 | 0.001637445 | 115.692734 | 59.57417235 | ENSG00000128298 |
| NA | -0.959092375 | 0.514380417 | 0.017888198 | 0.058763346 | 45.95137715 | 23.6904848 | ENSG00000275964 |
| MID2 | -0.959954676 | 0.514073063 | 0.001632766 | 0.007986722 | 69.62230378 | 35.75630423 | ENSG00000080561 |
| DCTPP1 | -0.960063756 | 0.514034196 | 4.11E-22 | 3.09E-20 | 1013.976926 | 520.9127535 | ENSG00000179958 |
| OLIG1 | -0.960107859 | 0.514018483 | 0.025682527 | 0.079118723 | 35.05793958 | 18.09413017 | ENSG00000184221 |
| TMEM35A | -0.961433233 | 0.513546482 | 0.000102313 | 0.000711281 | 112.1621358 | 57.4766938 | ENSG00000126950 |

| | | | | | | | |
|---|---|---|---|---|---|---|---|
| NA | -0.961943703 | 0.513364805 | 0.005112697 | 0.020982887 | 61.7929258 | 31.75071447 | ENSG00000227946 |
| ARHGAP30 | -0.962860144 | 0.513038805 | 0.124893414 | 0.267634933 | 16.14416103 | 8.319336358 | ENSG00000186517 |
| COMMD9 | -0.963644587 | 0.512759924 | 1.80E-09 | 3.23E-08 | 350.3003777 | 179.7236633 | ENSG00000110442 |
| NA | -0.963740778 | 0.512725737 | 3.87E-06 | 3.86E-05 | 154.0265918 | 78.95773536 | ENSG00000278922 |
| TP53TG5 | -0.964023745 | 0.512625182 | 0.283767773 | 0.47467075 | 8.025279335 | 4.092070275 | ENSG00000124251 |
| TMPRSS6 | -0.964202714 | 0.512561593 | 0.239598224 | 0.425477961 | 10.6421099 | 5.457359983 | ENSG00000187045 |
| NGB | -0.965241767 | 0.512192571 | 0.17868928 | 0.347714418 | 13.21652512 | 6.732481945 | ENSG00000165553 |
| MACROH2A1 | -0.965265747 | 0.512184058 | 9.84E-38 | 2.22E-35 | 2634.686419 | 1349.619172 | ENSG00000113648 |
| RNU6-8 | -0.966519735 | 0.511739062 | 0.427714067 | 0.618108004 | 4.803802715 | 2.479396828 | ENSG00000202337 |
| PIGW | -0.966613153 | 0.511705926 | 1.06E-09 | 1.97E-08 | 310.9872311 | 159.0368491 | ENSG00000277161 |
| DDX47 | -0.966649085 | 0.511693182 | 0.364637578 | 0.557744728 | 6.930214803 | 3.608476096 | ENSG00000213782 |
| MIR1244-3 | -0.967157154 | 0.511513012 | 0.065675725 | 0.164847493 | 29.24300857 | 14.83773967 | ENSG00000283429 |
| TREX2 | -0.967562742 | 0.51136923 | 0.240611305 | 0.426786289 | 10.07968379 | 5.168635407 | ENSG00000183479 |
| NA | -0.967637556 | 0.511342712 | 0.24259386 | 0.429094759 | 9.492002374 | 4.887396905 | ENSG00000267419 |
| MIR4453HG | -0.967705914 | 0.511318484 | 0.021861829 | 0.069607823 | 36.89494566 | 18.83639301 | ENSG00000268471 |
| SLC4A3 | -0.968139908 | 0.511164692 | 1.96E-08 | 2.98E-07 | 350.8567756 | 179.559949 | ENSG00000114923 |
| SHISA2 | -0.968680443 | 0.510973209 | 1.09E-10 | 2.37E-09 | 419.2726359 | 213.7928227 | ENSG00000180730 |
| H2AC8 | -0.968904904 | 0.510893716 | 0.072248371 | 0.177475323 | 25.81440106 | 13.28941493 | ENSG00000277075 |
| B3GAT2 | -0.969440846 | 0.510703961 | 0.06937729 | 0.17183465 | 25.75186416 | 13.2480741 | ENSG00000112309 |
| NA | -0.970106036 | 0.510468543 | 0.392875389 | 0.585526245 | 5.372542652 | 2.745551562 | ENSG00000217275 |
| ANKRD45 | -0.970317275 | 0.510393806 | 0.452848737 | 0.640205835 | 5.117862675 | 2.550243925 | ENSG00000183831 |
| NA | -0.972942015 | 0.509466074 | 0.183908194 | 0.354503845 | 15.31768191 | 7.790276083 | ENSG00000269388 |
| ARHGAP8 | -0.973313896 | 0.509334767 | 0.033198012 | 0.097110474 | 33.79598728 | 17.21612187 | ENSG00000241484 |
| EPHB2 | -0.97437509 | 0.508960257 | 3.96E-20 | 2.55E-18 | 856.8319604 | 436.3161689 | ENSG00000133216 |
| GFRA2 | -0.974548524 | 0.508899076 | 1.82E-05 | 0.000153221 | 134.9345756 | 68.85386382 | ENSG00000168546 |
| NA | -0.975271045 | 0.508644276 | 0.109076254 | 0.241704842 | 17.57666169 | 8.926841424 | ENSG00000255389 |
| SIT1 | -0.97551102 | 0.508559676 | 0.000753986 | 0.004119805 | 83.52336729 | 42.4625291 | ENSG00000137078 |
| NA | -0.97582154 | 0.508450228 | 0.281347328 | 0.47271345 | 9.401654621 | 4.782257013 | ENSG00000286964 |
| CARD9 | -0.976514226 | 0.508206162 | 0.110196069 | 0.243690961 | 24.2237843 | 12.41195629 | ENSG00000187796 |
| WDR90 | -0.977045035 | 0.508019213 | 6.75E-14 | 2.29E-12 | 763.6843161 | 388.6203695 | ENSG00000161996 |
| STMN3 | -0.977710758 | 0.507784845 | 6.24E-27 | 6.88E-25 | 2022.246518 | 1027.385814 | ENSG00000197457 |
| NA | -0.979261879 | 0.507239191 | 0.237445397 | 0.422736887 | 9.474940033 | 4.816000155 | ENSG00000287225 |
| H2BC15 | -0.979325062 | 0.507216976 | 0.109371741 | 0.242185883 | 17.8376555 | 9.107176914 | ENSG00000233822 |
| NA | -0.979409673 | 0.50718723 | 0.21690982 | 0.397456783 | 10.16687463 | 5.157350486 | ENSG00000213467 |
| NA | -0.980765308 | 0.506710873 | 0.000923866 | 0.004926047 | 150.6991783 | 76.78499831 | ENSG00000225138 |
| NA | -0.981573904 | 0.506426954 | 0.412657205 | 0.604265516 | 4.846120358 | 2.468111907 | ENSG00000249593 |
| NA | -0.981790076 | 0.506351076 | 0.437425309 | 0.626268718 | 4.760207295 | 2.411798924 | ENSG00000244265 |
| NA | -0.981804099 | 0.506346155 | 0.004246291 | 0.017923199 | 58.78193237 | 29.88305959 | ENSG00000253636 |
| NA | -0.98254545 | 0.506086028 | 0.206394385 | 0.38495658 | 10.17634537 | 5.146065566 | ENSG00000228038 |
| GCGR | -0.982623307 | 0.506058717 | 0.328857799 | 0.523240833 | 7.72196789 | 3.957312501 | ENSG00000215644 |
| LINC01003 | -0.982748855 | 0.50601468 | 0.116915379 | 0.254810634 | 21.90848375 | 10.97808099 | ENSG00000261455 |
| NA | -0.983038158 | 0.505913219 | 0.203604423 | 0.381441162 | 12.93521437 | 6.51515412 | ENSG00000270938 |
| MAGEF1 | -0.983693572 | 0.505683436 | 3.13E-25 | 3.01E-23 | 1708.959064 | 864.6322199 | ENSG00000177383 |
| NA | -0.984034916 | 0.505563805 | 0.108750383 | 0.241184588 | 20.26677767 | 10.220846 | ENSG00000286482 |
| EXOG | -0.984515845 | 0.505395301 | 2.58E-07 | 3.21E-06 | 211.3667634 | 106.9961479 | ENSG00000157036 |
| NA | -0.986220879 | 0.504798357 | 0.049639731 | 0.133066643 | 26.57646415 | 13.44663093 | ENSG00000232931 |
| OBI1 | -0.987653395 | 0.50429737 | 6.60E-06 | 6.21E-05 | 169.2760017 | 85.28138069 | ENSG00000152193 |
| LRRC10B | -0.987655931 | 0.504296483 | 0.031280456 | 0.092597345 | 31.77264829 | 15.94804795 | ENSG00000204950 |
| NA | -0.987955146 | 0.504191903 | 0.394391119 | 0.586790159 | 4.717889651 | 2.370458089 | ENSG00000285872 |
| NA | -0.989925354 | 0.503503826 | 0.260414977 | 0.449314018 | 8.320397819 | 4.219780009 | ENSG00000234297 |
| GPR137C | -0.990066242 | 0.503454658 | 0.001040965 | 0.005454694 | 74.0578003 | 37.17746888 | ENSG00000180998 |
| NA | -0.990244444 | 0.503392475 | 0.410781097 | 0.602279634 | 5.306848842 | 2.647897744 | ENSG00000281332 |
| TGFB3 | -0.990313581 | 0.503368352 | 0.19979776 | 0.376249916 | 11.51977605 | 5.757369479 | ENSG00000119699 |
| CYLD | -0.990814614 | 0.503193568 | 1.66E-08 | 2.57E-07 | 301.9881277 | 151.6500735 | ENSG00000083799 |
| NA | -0.990904709 | 0.503162145 | 0.228917107 | 0.412798988 | 11.89321599 | 6.035247167 | ENSG00000276488 |
| PKDCC | -0.991086842 | 0.503098627 | 2.07E-15 | 8.34E-14 | 881.5789198 | 443.9995932 | ENSG00000162878 |
| BAALC | -0.991462378 | 0.502967687 | 0.041521939 | 0.115968597 | 30.54482066 | 15.49126049 | ENSG00000164929 |
| IL1RAPL2 | -0.992498644 | 0.502606542 | 0.210502133 | 0.389867428 | 10.73877148 | 5.348421244 | ENSG00000189108 |
| NA | -0.993833803 | 0.502141614 | 0.327700158 | 0.521921005 | 7.462950922 | 3.75451879 | ENSG00000270761 |
| NA | -0.993909994 | 0.502115096 | 0.054135994 | 0.142136022 | 25.45476884 | 12.81286879 | ENSG00000268297 |
| ERVK3-1 | -0.995111287 | 0.501697173 | 0.098321983 | 0.223732307 | 17.78774625 | 8.919355351 | ENSG00000142396 |
| BEGAIN | -0.995198529 | 0.501666835 | 0.001188082 | 0.006106467 | 74.87420733 | 37.58316793 | ENSG00000183092 |
| REEP1 | -0.995265352 | 0.5016436 | 0.050395522 | 0.134683577 | 28.00009543 | 14.01290678 | ENSG00000068615 |
| GFI1 | -0.995291312 | 0.501634573 | 0.365237387 | 0.558431292 | 6.089732549 | 3.026790064 | ENSG00000162676 |
| ZNF559 | -0.99540603 | 0.501594686 | 6.20E-07 | 7.17E-06 | 181.0074637 | 90.66354507 | ENSG00000188321 |
| ISM1 | -0.99598166 | 0.501394592 | 0.031496185 | 0.093191299 | 29.51988463 | 14.82320556 | ENSG00000101230 |
| CYP2C8 | -0.997727632 | 0.500788163 | 0.339652641 | 0.533877978 | 7.112789446 | 3.608476096 | ENSG00000138115 |
| NA | -0.997782437 | 0.50076914 | 0.232348405 | 0.417039061 | 11.84330675 | 5.956475963 | ENSG00000287710 |
| NA | -0.998478358 | 0.500527639 | 0.279472461 | 0.470500734 | 8.259740058 | 4.140897185 | ENSG00000278192 |
| LOC101928058 | -0.998719676 | 0.500443924 | 0.340432668 | 0.534800735 | 7.109632533 | 3.600990022 | ENSG00000253764 |
| SNORC | -0.999870161 | 0.500045001 | 0.108905971 | 0.241415153 | 20.70265681 | 10.31849982 | ENSG00000182600 |
| PRRG2 | -1.000177425 | 0.499938513 | 0.017228058 | 0.057057358 | 47.54454946 | 23.92333443 | ENSG00000126460 |
| EBLN2 | -1.001013004 | 0.499649043 | 0.096990178 | 0.221572287 | 18.55937778 | 9.305733745 | ENSG00000255423 |
| NA | -1.001829248 | 0.499366433 | 0.112887741 | 0.248162897 | 16.63763872 | 8.323135205 | ENSG00000266865 |

| | | | | | | | |
|---|---|---|---|---|---|---|---|
| NA | -1.003639745 | 0.49874015 | 0.274010095 | 0.465192187 | 8.228170929 | 4.099556349 | ENSG00000236814 |
| CNIH2 | -1.004093162 | 0.498583428 | 7.49E-07 | 8.55E-06 | 210.1884617 | 104.6164516 | ENSG00000174871 |
| ANO1 | -1.004298814 | 0.498512362 | 3.02E-06 | 3.07E-05 | 158.5992861 | 79.17561283 | ENSG00000131620 |
| CFAP73 | -1.004640568 | 0.498394285 | 0.281995152 | 0.473069106 | 7.648682477 | 3.829602768 | ENSG00000186710 |
| GABRA1 | -1.005076414 | 0.498243741 | 0.335362408 | 0.52957373 | 6.073947985 | 2.977963155 | ENSG00000022355 |
| ZNF649 | -1.006618346 | 0.497711509 | 5.14E-15 | 1.98E-13 | 498.9410024 | 248.208494 | ENSG00000198093 |
| ALKBH6 | -1.007556945 | 0.49738781 | 0.020859499 | 0.066922269 | 34.59405419 | 17.1710938 | ENSG00000239382 |
| FNDC5 | -1.008573738 | 0.497037381 | 0.042536712 | 0.118092277 | 29.96345308 | 14.93594314 | ENSG00000160097 |
| ACOXL | -1.008708341 | 0.49699101 | 2.89E-13 | 9.09E-12 | 412.9116647 | 205.5562083 | ENSG00000153093 |
| KCND3 | -1.009293087 | 0.496789613 | 0.224474264 | 0.407204268 | 9.739090748 | 4.876111984 | ENSG00000171385 |
| ANO4 | -1.01040897 | 0.496405508 | 0.003042342 | 0.013486432 | 62.2347128 | 31.07225069 | ENSG00000151572 |
| NKX1-2 | -1.012038224 | 0.495845228 | 2.38E-13 | 7.53E-12 | 486.7943842 | 241.001281 | ENSG00000229544 |
| NA | -1.012631752 | 0.495641278 | 0.122919349 | 0.264352195 | 19.71391999 | 9.721730021 | ENSG00000262519 |
| PCDHB4 | -1.014056786 | 0.495151946 | 0.239334753 | 0.42525456 | 9.081280834 | 4.538560499 | ENSG00000081818 |
| ADGRB2 | -1.01424557 | 0.495087157 | 3.84E-28 | 4.81E-26 | 1178.683518 | 584.1476299 | ENSG00000121753 |
| NA | -1.014869875 | 0.494872962 | 0.059403977 | 0.152682471 | 23.16472359 | 11.54523291 | ENSG00000229036 |
| NA | -1.015325253 | 0.494716783 | 0.379431299 | 0.572622726 | 5.772515676 | 2.903317211 | ENSG00000279637 |
| NA | -1.01566344 | 0.494600828 | 0.009053723 | 0.033871963 | 46.21058951 | 22.73315565 | ENSG00000272841 |
| PKNOX2 | -1.016754991 | 0.494226752 | 0.005913344 | 0.023748195 | 51.26345744 | 25.31769237 | ENSG00000165495 |
| E2F8 | -1.017147745 | 0.494092224 | 2.45E-09 | 4.30E-08 | 243.5529819 | 120.3554223 | ENSG00000129173 |
| ITPRIPL1 | -1.017498593 | 0.49397208 | 8.43E-12 | 2.16E-10 | 384.4311025 | 190.1891851 | ENSG00000198885 |
| NA | -1.018139184 | 0.493752794 | 0.413887575 | 0.605206495 | 4.77095581 | 2.329117253 | ENSG00000234026 |
| TMEM81 | -1.018666601 | 0.493572322 | 0.002585155 | 0.011780741 | 57.86806701 | 28.52101907 | ENSG00000174529 |
| NOX4 | -1.019230667 | 0.493379382 | 0.04869027 | 0.131052203 | 25.72523338 | 12.67767299 | ENSG00000086991 |
| NA | -1.020586447 | 0.492915944 | 0.394538267 | 0.586961972 | 4.777269635 | 2.321631179 | ENSG00000228818 |
| TMSB15A | -1.020696106 | 0.492878479 | 3.27E-17 | 1.60E-15 | 618.2125753 | 304.2209218 | ENSG00000158164 |
| LRRC61 | -1.021073336 | 0.49274962 | 0.331485337 | 0.525735858 | 7.072350938 | 3.495850129 | ENSG00000127399 |
| ACAT2 | -1.021088361 | 0.492744488 | 2.63E-38 | 6.26E-36 | 2464.503498 | 1213.766237 | ENSG00000120437 |
| NA | -1.02131426 | 0.49266734 | 0.426793124 | 0.61740429 | 4.90234343 | 2.400514004 | ENSG00000237840 |
| NA | -1.021397678 | 0.492638854 | 0.306101058 | 0.500112571 | 6.322314136 | 3.135728803 | ENSG00000287966 |
| CHRM4 | -1.022343639 | 0.492315942 | 0.100499229 | 0.227485676 | 17.88440783 | 8.712762793 | ENSG00000180720 |
| NA | -1.022453347 | 0.492278505 | 0.319245809 | 0.513129982 | 6.967496398 | 3.431939453 | ENSG00000270964 |
| ANO2 | -1.022767917 | 0.492171179 | 0.000630421 | 0.003515108 | 74.27449962 | 36.59297169 | ENSG00000047617 |
| NLGN3 | -1.022902465 | 0.492125281 | 7.42E-11 | 1.65E-09 | 359.7109994 | 177.1310825 | ENSG00000196338 |
| SYTL4 | -1.025662179 | 0.4911848 | 0.343305095 | 0.537082521 | 5.620908802 | 2.786892398 | ENSG00000102362 |
| RRP36 | -1.026527994 | 0.49089011 | 2.78E-38 | 6.45E-36 | 1569.137111 | 770.1423757 | ENSG00000124541 |
| NA | -1.027119333 | 0.490688943 | 0.389281955 | 0.581897875 | 5.816111096 | 2.89203229 | ENSG00000250012 |
| LINC01347 | -1.027442886 | 0.490578908 | 0.346503115 | 0.540053587 | 6.497297176 | 3.214611628 | ENSG00000214837 |
| NUDT8 | -1.027902114 | 0.490422776 | 0.000173304 | 0.00113099 | 98.90358609 | 48.61799993 | ENSG00000167799 |
| CNTN1 | -1.029259653 | 0.489961518 | 3.26E-11 | 7.64E-10 | 386.6123622 | 189.0645172 | ENSG00000018236 |
| RND1 | -1.029954978 | 0.489725431 | 1.58E-07 | 2.04E-06 | 215.9403521 | 105.9378041 | ENSG00000172602 |
| CALML6 | -1.031975256 | 0.489040124 | 0.374061991 | 0.567291133 | 5.00225962 | 2.486882902 | ENSG00000169885 |
| PARP2 | -1.032477953 | 0.488869751 | 2.87E-19 | 1.71E-17 | 727.1495739 | 355.9375868 | ENSG00000129484 |
| SLC30A10 | -1.032954945 | 0.488708145 | 0.263456764 | 0.453128031 | 7.728281715 | 3.75451879 | ENSG00000196660 |
| PRR29 | -1.03401898 | 0.48834784 | 0.238978368 | 0.424776816 | 8.539073977 | 4.208495088 | ENSG00000224383 |
| MYCNOS | -1.035703083 | 0.487778109 | 0.002975117 | 0.013229562 | 62.58349881 | 30.68207344 | ENSG00000233718 |
| NA | -1.035944182 | 0.487696599 | 0.408886396 | 0.600404094 | 4.975628842 | 2.408000077 | ENSG00000286299 |
| RAMP3 | -1.036809949 | 0.487404019 | 0.097302779 | 0.222012766 | 17.03455253 | 8.360677193 | ENSG00000122679 |
| RHPN1-AS1 | -1.037460229 | 0.487184376 | 0.056945989 | 0.147841008 | 23.12871977 | 11.23350046 | ENSG00000254389 |
| FBXL22 | -1.039005186 | 0.486662938 | 0.300118427 | 0.493470097 | 6.405070286 | 3.124443883 | ENSG00000197361 |
| AIFM3 | -1.039653822 | 0.486444183 | 0.249727459 | 0.438258568 | 9.204475492 | 4.482247516 | ENSG00000183773 |
| NA | -1.040553371 | 0.48614097 | 0.254125022 | 0.442914535 | 8.382934716 | 4.024472372 | ENSG00000255639 |
| DQX1 | -1.041786724 | 0.485725548 | 0.153491158 | 0.310907665 | 13.55711816 | 6.631578933 | ENSG00000144045 |
| ZNF799 | -1.042118486 | 0.485613864 | 0.016037933 | 0.053927258 | 38.69645157 | 18.69359951 | ENSG00000196466 |
| IFIH1 | -1.042478906 | 0.485492561 | 0.058585475 | 0.151156538 | 23.68789127 | 11.5640039 | ENSG00000115267 |
| VWA7 | -1.042537647 | 0.485472794 | 0.200933687 | 0.377621675 | 11.9337522 | 5.791112621 | ENSG00000204396 |
| PRR19 | -1.043016474 | 0.485311693 | 0.00479413 | 0.019838002 | 61.55600722 | 29.87502386 | ENSG00000188368 |
| PRKAR1B | -1.043139332 | 0.485270366 | 2.28E-33 | 3.78E-31 | 1947.5789 | 945.2722649 | ENSG00000188191 |
| CLCNKA | -1.043396577 | 0.485183846 | 0.075945271 | 0.184285107 | 20.89479989 | 10.19871419 | ENSG00000186510 |
| CDK8 | -1.045334384 | 0.484532592 | 3.47E-17 | 1.69E-15 | 680.2518526 | 329.1069929 | ENSG00000132964 |
| WDR45 | -1.045406628 | 0.484508329 | 4.20E-05 | 0.000323106 | 117.8594869 | 57.0834906 | ENSG00000196998 |
| NA | -1.046187086 | 0.484246294 | 7.19E-06 | 6.71E-05 | 129.2056554 | 62.60822525 | ENSG00000211772 |
| LINC01273 | -1.046293501 | 0.484210577 | 0.140874157 | 0.29221444 | 14.44109814 | 7.036728322 | ENSG00000231742 |
| LAG3 | -1.046379093 | 0.484181851 | 0.002783614 | 0.012500969 | 58.52291541 | 28.4726302 | ENSG00000089692 |
| JAM2 | -1.046422379 | 0.484167324 | 8.38E-06 | 7.63E-05 | 139.7059374 | 67.54389941 | ENSG00000154721 |
| SLC5A6 | -1.046971411 | 0.483983104 | 7.99E-52 | 3.70E-49 | 3823.677843 | 1851.336439 | ENSG00000138074 |
| NA | -1.047835871 | 0.483693189 | 0.101552733 | 0.229282293 | 17.95967007 | 8.765826582 | ENSG00000213144 |
| CAMK2B | -1.051990466 | 0.48230228 | 0.127500423 | 0.27135587 | 15.02058659 | 7.246570073 | ENSG00000058404 |
| CACNG4 | -1.05271113 | 0.482061417 | 5.45E-06 | 5.22E-05 | 130.7385759 | 62.87405357 | ENSG00000075461 |
| CNTN2 | -1.05301371 | 0.481960324 | 7.43E-05 | 0.00053355 | 126.2497049 | 60.76237576 | ENSG00000184144 |
| YAF2 | -1.053389813 | 0.481834696 | 0.070809695 | 0.174519533 | 19.98872153 | 9.639598003 | ENSG00000015153 |
| SHBG | -1.053552892 | 0.481780233 | 0.132415382 | 0.279106868 | 14.1958889 | 6.781308855 | ENSG00000129214 |
| SPX | -1.053692798 | 0.481733514 | 0.371625406 | 0.564843798 | 5.581748071 | 2.677953659 | ENSG00000134548 |
| NA | -1.053985112 | 0.481635917 | 0.055178134 | 0.144178422 | 22.13978756 | 10.68979445 | ENSG00000214189 |

| | | | | | | | |
|---|---|---|---|---|---|---|---|
| SPINK2 | -1.054793813 | 0.481366012 | 0.152098326 | 0.309147341 | 12.07904525 | 5.794911468 | ENSG00000128040 |
| C1orf54 | -1.054982093 | 0.481303196 | 1.01E-05 | 9.03E-05 | 121.1139081 | 58.41905081 | ENSG00000118292 |
| NA | -1.055066305 | 0.481275102 | 0.304803114 | 0.49869535 | 7.045817859 | 3.356855475 | ENSG00000285763 |
| PCDHB16 | -1.056077528 | 0.480937882 | 0.000427077 | 0.002493053 | 92.66658366 | 44.40851716 | ENSG00000272674 |
| DLD | -1.057616109 | 0.480425253 | 3.70E-39 | 9.52E-37 | 1770.558893 | 850.1368324 | ENSG00000091140 |
| CEP83-DT | -1.057815318 | 0.48035892 | 0.154636715 | 0.312381508 | 12.94468511 | 6.200060856 | ENSG00000278916 |
| NA | -1.058380766 | 0.480170685 | 0.001442142 | 0.007162018 | 67.14474553 | 32.43374155 | ENSG00000273117 |
| FEZF1-AS1 | -1.058501726 | 0.480130428 | 3.61E-10 | 7.16E-09 | 277.0013858 | 133.3195583 | ENSG00000230316 |
| GGN | -1.058980655 | 0.479971066 | 0.023471886 | 0.073633705 | 32.81790127 | 15.75328996 | ENSG00000179168 |
| NA | -1.059711191 | 0.479728085 | 0.173266646 | 0.339786692 | 13.02122513 | 6.275256453 | ENSG00000258674 |
| NA | -1.059778133 | 0.479705826 | 0.319442644 | 0.513268311 | 5.745982597 | 2.756836483 | ENSG00000286529 |
| ST8SIA5 | -1.060451741 | 0.479481899 | 0.000569742 | 0.003215459 | 79.80072368 | 38.37458409 | ENSG00000101638 |
| NA | -1.060645351 | 0.479417557 | 0.299997437 | 0.493415916 | 6.391164859 | 3.038074985 | ENSG00000259969 |
| LINC02728 | -1.061887973 | 0.479004803 | 0.35135989 | 0.544467173 | 5.653755707 | 2.696724653 | ENSG00000251323 |
| RWDD3 | -1.062027164 | 0.478958591 | 0.059711896 | 0.153303821 | 22.38234354 | 10.73862136 | ENSG00000122481 |
| NA | -1.063214567 | 0.478564548 | 0.21100053 | 0.390521652 | 11.77445603 | 5.694008455 | ENSG00000279342 |
| LINC02249 | -1.063821643 | 0.478363214 | 0.304958263 | 0.498861116 | 6.497297176 | 3.094387968 | ENSG00000225930 |
| ZNF521 | -1.064464054 | 0.478150254 | 1.13E-09 | 2.07E-08 | 261.6327123 | 124.9607046 | ENSG00000198795 |
| SEC14L2 | -1.06542534 | 0.477831762 | 0.004717413 | 0.019555477 | 49.15410769 | 23.57405999 | ENSG00000100003 |
| NA | -1.067395004 | 0.477179839 | 0.109237745 | 0.241918074 | 17.712484 | 8.525928916 | ENSG00000272449 |
| KLHL14 | -1.067877696 | 0.477020213 | 0.009429468 | 0.035058329 | 40.36478843 | 19.25662616 | ENSG00000197705 |
| ABCB7 | -1.06833171 | 0.476870119 | 1.09E-23 | 9.30E-22 | 820.0568344 | 391.5061811 | ENSG00000131269 |
| TRPM3 | -1.068703323 | 0.476747301 | 5.85E-07 | 6.82E-06 | 184.9994069 | 88.12893457 | ENSG00000083067 |
| NA | -1.068867676 | 0.476692993 | 0.020367719 | 0.065605719 | 34.34262882 | 16.49642887 | ENSG00000257354 |
| NA | -1.069176295 | 0.47659103 | 0.308596072 | 0.502897789 | 6.517516429 | 3.086901894 | ENSG00000228477 |
| PCSK1N | -1.069268928 | 0.47656043 | 5.06E-05 | 0.000380221 | 144.0437877 | 68.56382515 | ENSG00000102109 |
| TMEM132E | -1.069781668 | 0.476391089 | 0.009679102 | 0.035792592 | 38.32932545 | 18.25514501 | ENSG00000181291 |
| C15orf61 | -1.069790154 | 0.476388287 | 0.058813118 | 0.1515533 | 21.88373211 | 10.44989678 | ENSG00000189227 |
| CCT6B | -1.070100309 | 0.476285882 | 0.117517404 | 0.255634331 | 14.65849652 | 7.044214395 | ENSG00000132141 |
| RTEL1 | -1.070299598 | 0.476220094 | 0.032810679 | 0.09615957 | 26.75460411 | 12.74895812 | ENSG00000258366 |
| C1orf216 | -1.070421152 | 0.476179972 | 1.09E-10 | 2.37E-09 | 329.1483196 | 157.1903786 | ENSG00000142686 |
| NMU | -1.071133035 | 0.475945064 | 9.10E-08 | 1.23E-06 | 206.7892284 | 98.04063604 | ENSG00000109255 |
| LOC729603 | -1.07190663 | 0.475689923 | 0.118288417 | 0.256926932 | 15.28977335 | 7.302883056 | ENSG00000213073 |
| FAM222A | -1.072608111 | 0.475458685 | 0.00216678 | 0.010192663 | 61.64763275 | 29.36148539 | ENSG00000139438 |
| CAPN10-DT | -1.073048683 | 0.475313511 | 0.001093194 | 0.005696457 | 65.5124125 | 31.25583537 | ENSG00000260942 |
| ARHGAP6 | -1.073211729 | 0.475259797 | 0.036720857 | 0.105028451 | 26.61246797 | 12.61756116 | ENSG00000047648 |
| MRGPRF | -1.073244465 | 0.475249013 | 2.49E-05 | 0.0002031 | 108.2096615 | 51.37439838 | ENSG00000172935 |
| LOXL1 | -1.074613795 | 0.474798146 | 2.74E-10 | 5.56E-09 | 311.4127299 | 147.9954071 | ENSG00000129038 |
| CTXN1 | -1.074717022 | 0.474764174 | 1.83E-14 | 6.56E-13 | 775.0381134 | 367.7668376 | ENSG00000178531 |
| PRR22 | -1.075381451 | 0.474545573 | 0.035862438 | 0.103178272 | 33.74419889 | 16.0913911 | ENSG00000212123 |
| NOVA1 | -1.075539915 | 0.474493452 | 4.61E-08 | 6.59E-07 | 195.2148979 | 92.62549296 | ENSG00000139910 |
| PLAC9 | -1.075614253 | 0.474469004 | 0.000886098 | 0.004750632 | 62.34400203 | 29.58629928 | ENSG00000189129 |
| SLITRK6 | -1.077550349 | 0.473832694 | 0.225005131 | 0.407847931 | 10.59347843 | 4.980813844 | ENSG00000184564 |
| GPR50 | -1.07906271 | 0.47333624 | 3.32E-07 | 4.04E-06 | 187.6309397 | 88.83755744 | ENSG00000102195 |
| PDIA2 | -1.081589022 | 0.472508104 | 0.106038704 | 0.236974983 | 15.30811347 | 7.235285152 | ENSG00000185615 |
| KLHL4 | -1.082983393 | 0.472051643 | 2.96E-23 | 2.44E-21 | 774.3790787 | 365.1342384 | ENSG00000102271 |
| NSMCE4A | -1.084341507 | 0.471607476 | 1.89E-24 | 1.69E-22 | 944.5349566 | 445.4450396 | ENSG00000107672 |
| NA | -1.084344823 | 0.471606392 | 0.041095275 | 0.114932824 | 23.99130042 | 11.32735543 | ENSG00000278493 |
| BCL6B | -1.084778867 | 0.471464527 | 2.03E-14 | 7.24E-13 | 364.7652353 | 171.9176422 | ENSG00000161940 |
| LINC02139 | -1.085502473 | 0.471228116 | 0.350739345 | 0.544026727 | 5.00225962 | 2.340402174 | ENSG00000278214 |
| STAR | -1.085568189 | 0.471206652 | 0.000925185 | 0.004931658 | 65.73542564 | 30.96276229 | ENSG00000147465 |
| OTX2 | -1.085957586 | 0.471079486 | 1.47E-20 | 9.93E-19 | 696.4976589 | 328.3714835 | ENSG00000165588 |
| NA | -1.085971056 | 0.471075088 | 0.270896628 | 0.461555258 | 7.456539398 | 3.533392118 | ENSG00000280195 |
| LOC728392 | -1.086445052 | 0.470920342 | 2.22E-09 | 3.92E-08 | 215.7485924 | 101.6392529 | ENSG00000286190 |
| NA | -1.087483973 | 0.470581343 | 0.346829064 | 0.540224378 | 5.04142035 | 2.359173168 | ENSG00000260855 |
| KCNJ12 | -1.088305244 | 0.470313535 | 1.92E-05 | 0.00016038 | 107.0848093 | 50.27146456 | ENSG00000184185 |
| LOC101928837 | -1.088354963 | 0.470297327 | 0.221297721 | 0.402861405 | 8.915670835 | 4.231064929 | ENSG00000255100 |
| DUSP28 | -1.090152228 | 0.46971181 | 0.045008795 | 0.1232919 | 29.1089677 | 13.73959239 | ENSG00000188542 |
| NA | -1.090220706 | 0.469689515 | 0.314153899 | 0.507759937 | 5.742825684 | 2.659182664 | ENSG00000268157 |
| NA | -1.091398227 | 0.469306313 | 0.373855519 | 0.56711722 | 4.972569627 | 2.310346259 | ENSG00000272155 |
| GATA5 | -1.091755346 | 0.469190157 | 0.186134858 | 0.357198535 | 9.474940033 | 4.440906681 | ENSG00000130700 |
| ADAMTS17 | -1.092143998 | 0.469063778 | 0.003168666 | 0.013949657 | 51.38221741 | 24.1477103 | ENSG00000140470 |
| U2AF1L4 | -1.093049486 | 0.468769469 | 0.042143884 | 0.117211095 | 27.12233159 | 12.70013121 | ENSG00000161265 |
| FLT3 | -1.093702864 | 0.468557217 | 0.111448313 | 0.24587302 | 14.78484809 | 6.935275656 | ENSG00000122025 |
| SCN9A | -1.093910887 | 0.46848966 | 0.000560122 | 0.0031718 | 74.25240123 | 34.80364998 | ENSG00000169432 |
| NA | -1.093993755 | 0.468462751 | 0.201146021 | 0.377982394 | 9.523571502 | 4.459677675 | ENSG00000279605 |
| ACSM1 | -1.094692555 | 0.468235896 | 0.288347074 | 0.480077674 | 6.58646485 | 3.094387968 | ENSG00000166743 |
| RTF1 | -1.096128762 | 0.467769998 | 1.35E-51 | 5.96E-49 | 2220.576821 | 1038.507413 | ENSG00000137815 |
| SYT9 | -1.096227015 | 0.467738142 | 0.000148112 | 0.000983206 | 84.23670122 | 39.32300145 | ENSG00000170743 |
| GPR85 | -1.09679742 | 0.467553247 | 0.374396683 | 0.567389377 | 5.174085747 | 2.411798924 | ENSG00000164604 |
| NA | -1.098024537 | 0.467155728 | 0.043489321 | 0.12011937 | 27.74975245 | 13.0344335 | ENSG00000279233 |
| FAM78A | -1.098351147 | 0.467049981 | 7.96E-09 | 1.30E-07 | 194.2465909 | 90.71281001 | ENSG00000126882 |
| MCUB | -1.099623643 | 0.466638212 | 3.83E-14 | 1.33E-12 | 389.3175712 | 181.7257411 | ENSG00000005059 |
| NTF3 | -1.100670408 | 0.46629976 | 0.101719284 | 0.229548915 | 16.40190023 | 7.666803229 | ENSG00000185652 |

| | | | | | | | |
|---|---|---|---|---|---|---|---|
| FEZ1 | -1.100804055 | 0.466256565 | 7.19E-28 | 8.71E-26 | 928.3800471 | 432.5173377 | ENSG00000149557 |
| NA | -1.102616403 | 0.46567121 | 0.35620565 | 0.549793097 | 5.097643422 | 2.351687094 | ENSG00000248476 |
| RBM3 | -1.104282081 | 0.465133875 | 2.32E-32 | 3.64E-30 | 1626.00231 | 756.0075016 | ENSG00000102317 |
| NA | -1.10501734 | 0.464896884 | 0.01079287 | 0.0391995 | 44.66446263 | 20.89599471 | ENSG00000260257 |
| CASQ1 | -1.105213428 | 0.4648337 | 0.082821489 | 0.196720937 | 18.93272003 | 8.701477872 | ENSG00000143318 |
| E2F2 | -1.105344865 | 0.464791353 | 6.17E-10 | 1.18E-08 | 221.5275196 | 103.0420846 | ENSG00000007968 |
| NA | -1.106342664 | 0.464470005 | 0.054601265 | 0.143096697 | 27.85608016 | 13.04203119 | ENSG00000174403 |
| LOX | -1.106917969 | 0.464284825 | 0.017824833 | 0.058585823 | 31.02892531 | 14.40677125 | ENSG00000113083 |
| CALCB | -1.107259001 | 0.464175087 | 4.98E-06 | 4.81E-05 | 121.4223307 | 56.35565026 | ENSG00000175868 |
| NA | -1.109179486 | 0.463557598 | 0.257143016 | 0.445660058 | 6.662907176 | 3.086901894 | ENSG00000255031 |
| VWA5B1 | -1.109333141 | 0.46350823 | 0.028583714 | 0.086281849 | 31.01166758 | 14.27526267 | ENSG00000158816 |
| NA | -1.109383505 | 0.463492049 | 0.125484837 | 0.268374795 | 13.34919052 | 6.185088709 | ENSG00000261324 |
| LINC01918 | -1.110709744 | 0.463066166 | 0.002205821 | 0.010329126 | 54.40907045 | 25.24684527 | ENSG00000226508 |
| FAM27E3 | -1.113659673 | 0.462120286 | 0.256978222 | 0.44545776 | 6.669221001 | 3.075616974 | ENSG00000274026 |
| LINC02523 | -1.114426804 | 0.461874626 | 0.063190526 | 0.160456667 | 19.85595843 | 9.080370193 | ENSG00000224506 |
| KCNC1 | -1.114914636 | 0.461718474 | 0.075291544 | 0.183010291 | 17.72333022 | 8.154084636 | ENSG00000129159 |
| CARD6 | -1.115145226 | 0.461644682 | 0.281960133 | 0.473069106 | 6.676812603 | 3.075616974 | ENSG00000132357 |
| H3C12 | -1.11515857 | 0.461640413 | 0.082366966 | 0.195942672 | 16.89863253 | 7.726365406 | ENSG00000197153 |
| DPH5 | -1.115386104 | 0.461567611 | 5.93E-24 | 5.18E-22 | 699.3748744 | 322.8123445 | ENSG00000117543 |
| POLR2J3 | -1.115460248 | 0.46154389 | 0.273668563 | 0.464871193 | 6.67869174 | 3.075616974 | ENSG00000168255 |
| KCNC3 | -1.116465183 | 0.461222506 | 4.72E-08 | 6.73E-07 | 179.1450296 | 82.67894904 | ENSG00000131398 |
| SARM1 | -1.117086908 | 0.461023786 | 8.20E-05 | 0.000583394 | 91.21248827 | 41.9708992 | ENSG00000004139 |
| DLG1 | -1.11709457 | 0.461021338 | 4.09E-35 | 7.90E-33 | 1165.899016 | 537.8054687 | ENSG00000075711 |
| MSANTD2 | -1.117795124 | 0.460797526 | 4.83E-11 | 1.10E-09 | 265.8445717 | 122.651346 | ENSG00000120458 |
| NA | -1.121378334 | 0.459654467 | 0.307724849 | 0.501836546 | 5.925498024 | 2.726780568 | ENSG00000227440 |
| SLC16A9 | -1.12252768 | 0.459288422 | 2.60E-11 | 6.17E-10 | 281.7263634 | 129.1387058 | ENSG00000165449 |
| ARHGDIG | -1.122589114 | 0.459268864 | 0.104234993 | 0.233913471 | 14.24461807 | 6.560182182 | ENSG00000242173 |
| NA | -1.125976885 | 0.458191663 | 0.311719252 | 0.505327089 | 6.087853413 | 2.805663392 | ENSG00000275956 |
| ADAMTS19 | -1.126411897 | 0.458053527 | 1.84E-19 | 1.13E-17 | 547.9738697 | 250.7092498 | ENSG00000145808 |
| ASXL1 | -1.127182971 | 0.457808777 | 1.18E-24 | 1.08E-22 | 963.0894414 | 440.5436095 | ENSG00000171456 |
| NA | -1.127873539 | 0.457589693 | 0.331074246 | 0.525263659 | 5.144395755 | 2.310346259 | ENSG00000222881 |
| NA | -1.129368162 | 0.457115878 | 0.357314092 | 0.550678478 | 5.108391937 | 2.291575264 | ENSG00000284526 |
| MAP2 | -1.129719072 | 0.457004707 | 2.82E-07 | 3.48E-06 | 176.2409426 | 80.28080816 | ENSG00000078018 |
| NA | -1.131533613 | 0.456430273 | 0.231413038 | 0.415924093 | 8.370307064 | 3.829602768 | ENSG00000231831 |
| HERC2P3 | -1.131747864 | 0.456362495 | 0.000307485 | 0.00187574 | 104.1631941 | 47.76398719 | ENSG00000180229 |
| NA | -1.131795873 | 0.456347308 | 0.214424989 | 0.394456848 | 8.382934716 | 3.833401615 | ENSG00000255139 |
| HGSNAT | -1.131976039 | 0.456290323 | 4.85E-06 | 4.70E-05 | 118.4130436 | 54.045304 | ENSG00000165102 |
| SLC22A4 | -1.132862088 | 0.456010172 | 0.208601702 | 0.38769228 | 8.204794763 | 3.724462875 | ENSG00000197208 |
| NA | -1.135091349 | 0.455306087 | 5.22E-05 | 0.000390156 | 89.44994769 | 40.8127517 | ENSG00000263823 |
| NA | -1.135392009 | 0.45521121 | 0.047490989 | 0.128477301 | 20.38869455 | 9.27567783 | ENSG00000224825 |
| C15orf62 | -1.138308134 | 0.454292019 | 0.072524771 | 0.17805989 | 17.24553939 | 7.775192315 | ENSG00000188277 |
| BBS1 | -1.139066731 | 0.454053206 | 0.267274689 | 0.45741008 | 7.447166358 | 3.37562647 | ENSG00000174483 |
| THEM6 | -1.142015994 | 0.453125946 | 2.26E-08 | 3.41E-07 | 188.041353 | 85.05601713 | ENSG00000130193 |
| LOXL3 | -1.142367265 | 0.453015631 | 6.82E-17 | 3.19E-15 | 419.9050928 | 190.307983 | ENSG00000115318 |
| NA | -1.142836917 | 0.452868181 | 0.232025868 | 0.41670227 | 7.661310129 | 3.484565209 | ENSG00000285722 |
| H2AW | -1.143713188 | 0.452593199 | 2.10E-13 | 6.68E-12 | 353.590081 | 159.790062 | ENSG00000181218 |
| MIR3176 | -1.143998473 | 0.45250371 | 0.055896476 | 0.145646742 | 19.84520992 | 8.986953254 | ENSG00000266235 |
| DAZL | -1.144120185 | 0.452465537 | 0.22256828 | 0.404721323 | 7.58614558 | 3.4357383 | ENSG00000092345 |
| PTGS1 | -1.14419599 | 0.452441763 | 0.001811445 | 0.00871808 | 50.08356222 | 22.60968279 | ENSG00000095303 |
| RN7SL3 | -1.145511384 | 0.452029432 | 0.179063319 | 0.348133804 | 10.79371677 | 4.91745282 | ENSG00000278771 |
| HERC2P2 | -1.146030401 | 0.451866841 | 0.007910092 | 0.030334294 | 1772.969765 | 801.3036673 | ENSG00000276550 |
| CNTNAP3B | -1.146143347 | 0.451831467 | 0.005156306 | 0.021143136 | 43.92873722 | 19.78893563 | ENSG00000154529 |
| SMTNL1 | -1.146441514 | 0.451738095 | 0.229901743 | 0.413809742 | 9.272048439 | 4.212293935 | ENSG00000214872 |
| PINK1-AS | -1.146879892 | 0.45160085 | 0.025976271 | 0.079828256 | 26.67047248 | 12.00625724 | ENSG00000117242 |
| LY6H | -1.1485124 | 0.451090122 | 0.282577317 | 0.473534176 | 5.988034921 | 2.677953659 | ENSG00000176956 |
| LEF1 | -1.150079365 | 0.450600442 | 0.323508434 | 0.517691704 | 5.484988795 | 2.498167822 | ENSG00000138795 |
| MYB | -1.150929463 | 0.450335008 | 0.000253018 | 0.001585342 | 80.91285055 | 36.27689074 | ENSG00000118513 |
| NA | -1.152127335 | 0.449961249 | 0.211622472 | 0.391242982 | 7.672058644 | 3.454509294 | ENSG00000272087 |
| CBLN1 | -1.152453539 | 0.449859521 | 1.99E-09 | 3.55E-08 | 236.2963725 | 106.6097695 | ENSG00000102924 |
| RUBCNL | -1.152858183 | 0.449733363 | 0.00011929 | 0.000816443 | 79.13146619 | 35.7117142 | ENSG00000102445 |
| CYP39A1 | -1.155235692 | 0.448992829 | 0.002711162 | 0.012228931 | 48.58399228 | 21.87490603 | ENSG00000146233 |
| DANCR | -1.155786259 | 0.448821515 | 5.41E-37 | 1.15E-34 | 1431.357776 | 642.5573393 | ENSG00000226950 |
| NA | -1.155815734 | 0.448812345 | 0.067626785 | 0.16858034 | 19.14636014 | 8.581803865 | ENSG00000214077 |
| COX16 | -1.156426398 | 0.448622412 | 0.036815566 | 0.10525061 | 29.25317836 | 13.14673305 | ENSG00000133983 |
| NA | -1.156487212 | 0.448603502 | 0.284007546 | 0.47498609 | 6.156704136 | 2.749350409 | ENSG00000274818 |
| PVALB | -1.156665336 | 0.448548118 | 0.030386194 | 0.090513937 | 26.66543643 | 11.87854751 | ENSG00000100362 |
| PCDHB12 | -1.157885173 | 0.448169019 | 5.82E-05 | 0.000430985 | 86.20066022 | 38.49013283 | ENSG00000120328 |
| SNORD3B-2 | -1.158338427 | 0.448028239 | 0.004107262 | 0.017416455 | 41.30381139 | 18.46878561 | ENSG00000262074 |
| RGS11 | -1.15882082 | 0.447878457 | 0.034986909 | 0.101193409 | 22.85066363 | 10.18699124 | ENSG00000076344 |
| ADNP-AS1 | -1.159310196 | 0.447726558 | 0.235950089 | 0.421016555 | 8.912513922 | 3.923019706 | ENSG00000259456 |
| NA | -1.159414898 | 0.447694066 | 0.152182549 | 0.309237075 | 10.67307767 | 4.778458166 | ENSG00000235912 |
| NA | -1.159976729 | 0.447519754 | 0.223604025 | 0.405983965 | 9.431344613 | 4.279891839 | ENSG00000248489 |
| NA | -1.161137813 | 0.447159734 | 0.299923501 | 0.493415916 | 5.366228826 | 2.400514004 | ENSG00000273387 |
| SCN3B | -1.161230118 | 0.447131125 | 0.119675739 | 0.259059816 | 12.33688214 | 5.536242807 | ENSG00000166257 |

| | | | | | | | |
|---|---|---|---|---|---|---|---|
| PSMG3 | -1.161895486 | 0.446924957 | 5.58E-16 | 2.37E-14 | 736.2430764 | 329.7074273 | ENSG00000157778 |
| LOC728975 | -1.162385592 | 0.446773155 | 0.039269808 | 0.110838089 | 23.00099273 | 10.23581815 | ENSG00000254614 |
| FIGNL2 | -1.164255369 | 0.446194499 | 4.91E-06 | 4.75E-05 | 152.9486873 | 68.22313609 | ENSG00000261308 |
| NA | -1.164848328 | 0.446011147 | 0.146266791 | 0.300377696 | 10.94660142 | 4.906167899 | ENSG00000248873 |
| OR7E14P | -1.165088301 | 0.445936965 | 0.057225279 | 0.148420559 | 18.37354853 | 8.165369556 | ENSG00000184669 |
| SNORA11F | -1.165219374 | 0.445896452 | 0.071127529 | 0.175163118 | 20.44481992 | 9.156003823 | ENSG00000221164 |
| NA | -1.165540791 | 0.445797122 | 0.124683585 | 0.267297543 | 11.50271371 | 5.097238657 | ENSG00000286808 |
| RAMP1 | -1.16818679 | 0.44498025 | 0.092427485 | 0.213597381 | 14.05690968 | 6.290228601 | ENSG00000132329 |
| NA | -1.168300177 | 0.444945279 | 0.121671775 | 0.262308327 | 12.99912674 | 5.753570632 | ENSG00000235082 |
| H2BC20P | -1.169173879 | 0.444675899 | 0.005431979 | 0.022096168 | 42.62524139 | 19.02421457 | ENSG00000261716 |
| SLC6A12 | -1.169792848 | 0.444485158 | 0.178427668 | 0.347335196 | 9.2625777 | 4.103355196 | ENSG00000111181 |
| H2BC8 | -1.170238805 | 0.444347783 | 0.012870542 | 0.045229925 | 30.21813305 | 13.43914486 | ENSG00000273802 |
| LRRN3 | -1.172360365 | 0.443694826 | 0.284179638 | 0.475145279 | 5.45214189 | 2.419284998 | ENSG00000173114 |
| TNFRSF18 | -1.172623978 | 0.44361376 | 0.057787677 | 0.149627942 | 22.08040757 | 9.902503544 | ENSG00000186891 |
| TMX3 | -1.172784975 | 0.443564258 | 1.18E-12 | 3.44E-11 | 368.4445015 | 163.3656795 | ENSG00000166479 |
| RSPO2 | -1.173153425 | 0.443450991 | 0.12016168 | 0.259838592 | 13.77126193 | 6.046094055 | ENSG00000147655 |
| TTI1 | -1.173285703 | 0.443410333 | 8.46E-34 | 1.48E-31 | 1311.267053 | 581.4984181 | ENSG00000101407 |
| PRRT2 | -1.174210631 | 0.443126149 | 6.97E-06 | 6.52E-05 | 164.2570107 | 72.94670699 | ENSG00000167371 |
| YY2 | -1.17463236 | 0.442996633 | 0.165450871 | 0.328280318 | 10.94481998 | 4.894882979 | ENSG00000230797 |
| NA | -1.175294192 | 0.442793457 | 0.329035808 | 0.523418562 | 5.455298802 | 2.411798924 | ENSG00000262410 |
| SFRP1 | -1.176997963 | 0.442270842 | 1.37E-56 | 7.91E-54 | 13909.63302 | 6150.892453 | ENSG00000104332 |
| NA | -1.17709022 | 0.442242561 | 0.11146613 | 0.24587302 | 13.1475767 | 5.776140473 | ENSG00000280046 |
| NA | -1.17772695 | 0.442047421 | 0.019808478 | 0.064026915 | 29.82890854 | 13.26684509 | ENSG00000273619 |
| NA | -1.178779838 | 0.44172493 | 0.184473551 | 0.355113503 | 9.303016208 | 4.084584202 | ENSG00000226900 |
| RNF39 | -1.179112634 | 0.441623046 | 0.000880156 | 0.004722876 | 57.53082628 | 25.44160325 | ENSG00000204618 |
| PFKFB4 | -1.179913608 | 0.441377928 | 1.78E-12 | 5.06E-11 | 351.2842664 | 154.5835582 | ENSG00000114268 |
| NA | -1.180272965 | 0.441268 | 0.248233856 | 0.436339746 | 7.112789446 | 3.147013724 | ENSG00000286084 |
| NA | -1.180295115 | 0.441261225 | 0.309967198 | 0.503769234 | 5.372542652 | 2.351687094 | ENSG00000279812 |
| SLITRK3 | -1.182673133 | 0.440534486 | 0.000898741 | 0.00481007 | 58.89447621 | 25.87257168 | ENSG00000121871 |
| PP2D1 | -1.182878531 | 0.440471771 | 0.253615063 | 0.442400148 | 6.325471048 | 2.816948313 | ENSG00000183977 |
| H2BC6 | -1.187009314 | 0.439212399 | 0.045401094 | 0.124146336 | 21.01997138 | 9.193107779 | ENSG00000274290 |
| LOC101928521 | -1.188317891 | 0.438814198 | 0.117849432 | 0.256127738 | 13.6367174 | 6.069101929 | ENSG00000242686 |
| NA | -1.188434571 | 0.438778709 | 0.004629275 | 0.019241754 | 41.4769153 | 18.16129004 | ENSG00000279348 |
| ABHD8 | -1.188540709 | 0.43874643 | 3.43E-10 | 6.85E-09 | 226.2765724 | 99.11004151 | ENSG00000127220 |
| CAMK2N2 | -1.188833194 | 0.438657489 | 1.67E-06 | 1.77E-05 | 139.5956635 | 61.15244138 | ENSG00000163888 |
| EPHB6 | -1.18897752 | 0.438613609 | 1.03E-26 | 1.10E-24 | 738.8195662 | 323.872735 | ENSG00000106123 |
| HTR7 | -1.188987226 | 0.438610658 | 0.000709659 | 0.003901754 | 59.8770946 | 26.1650951 | ENSG00000148680 |
| ITPKA | -1.190717716 | 0.438084867 | 0.122133483 | 0.263058916 | 12.54471208 | 5.468644903 | ENSG00000137825 |
| ODC1 | -1.190797023 | 0.438060785 | 5.84E-60 | 4.16E-57 | 4256.00613 | 1863.638365 | ENSG00000115758 |
| NA | -1.192918373 | 0.437417131 | 0.002214588 | 0.010359711 | 50.88991979 | 22.411564 | ENSG00000257086 |
| PRSS35 | -1.197310423 | 0.43608751 | 0.000133673 | 0.000899599 | 81.03228694 | 35.30601516 | ENSG00000146250 |
| H2AC11 | -1.198059256 | 0.435861217 | 0.00241714 | 0.011137305 | 58.8135992 | 25.60630532 | ENSG00000196787 |
| ZNF439 | -1.198674987 | 0.435675235 | 0.178521096 | 0.347444082 | 9.249950049 | 3.964360542 | ENSG00000171291 |
| CNMD | -1.200574448 | 0.435102 | 1.47E-49 | 5.81E-47 | 3178.378722 | 1381.899814 | ENSG00000136110 |
| NA | -1.200857785 | 0.435016556 | 0.289927509 | 0.482016186 | 5.620908802 | 2.486882902 | ENSG00000235772 |
| KIF26A | -1.203759182 | 0.434142576 | 1.79E-45 | 6.50E-43 | 3907.371217 | 1696.18963 | ENSG00000066735 |
| LOC105373383 | -1.206382198 | 0.433353963 | 0.159156503 | 0.319035975 | 10.47018607 | 4.579901335 | ENSG00000260081 |
| NA | -1.206421461 | 0.433342169 | 0.155710489 | 0.31398542 | 9.626546907 | 4.189724094 | ENSG00000275888 |
| LINC01963 | -1.206475126 | 0.43332605 | 0.002076378 | 0.009814739 | 49.66978147 | 21.48049191 | ENSG00000260804 |
| NA | -1.207696689 | 0.432959298 | 0.088584339 | 0.207013256 | 15.41296801 | 6.736718825 | ENSG00000279656 |
| ACOT1 | -1.208150878 | 0.432823015 | 0.008708405 | 0.032818539 | 33.29106201 | 14.35414549 | ENSG00000184227 |
| CCDC183-AS1 | -1.208648005 | 0.432673898 | 0.317991491 | 0.511824082 | 5.369385739 | 2.280290344 | ENSG00000228544 |
| WNT2B | -1.208701722 | 0.432657788 | 0.049077457 | 0.131835518 | 20.70916604 | 9.009523095 | ENSG00000134245 |
| SUGT1P4-STRA6LP | -1.21015066 | 0.432223476 | 0.231963972 | 0.416637613 | 7.975370089 | 3.44702322 | ENSG00000254876 |
| FGFBP3 | -1.210908814 | 0.431996397 | 1.40E-39 | 3.81E-37 | 2067.901416 | 893.3415699 | ENSG00000174721 |
| NA | -1.211991161 | 0.431672424 | 0.243880543 | 0.430644458 | 6.322314136 | 2.719294494 | ENSG00000231563 |
| RGPD6 | -1.212130699 | 0.431630675 | 0.324607145 | 0.518767319 | 5.349166485 | 2.261519349 | ENSG00000183054 |
| ZNF596 | -1.212143189 | 0.431626938 | 0.10742503 | 0.239017467 | 14.34453426 | 6.230116771 | ENSG00000172748 |
| NA | -1.215105455 | 0.430741593 | 0.022096602 | 0.070167013 | 27.74225854 | 11.89351966 | ENSG00000267480 |
| TMEM114 | -1.21529737 | 0.430684297 | 0.053771461 | 0.141381792 | 20.73748055 | 8.919355351 | ENSG00000232258 |
| YWHAB | -1.216267368 | 0.430394824 | 1.77E-78 | 2.52E-75 | 6249.440994 | 2688.889676 | ENSG00000166913 |
| PARTICL | -1.216508044 | 0.43032303 | 0.002026685 | 0.009608272 | 46.6340364 | 20.17206483 | ENSG00000286532 |
| TEX26-AS1 | -1.216785244 | 0.430240355 | 0.234926786 | 0.419877826 | 6.328627961 | 2.708009574 | ENSG00000224743 |
| DUSP15 | -1.217408433 | 0.430054548 | 0.000487033 | 0.002798968 | 63.8872651 | 27.50087854 | ENSG00000149599 |
| NCAM1 | -1.220722757 | 0.429067711 | 1.34E-11 | 3.32E-10 | 239.7096991 | 102.6885733 | ENSG00000149294 |
| CFAP298 | -1.221151712 | 0.428940156 | 3.14E-05 | 0.00025013 | 86.97229175 | 37.21880972 | ENSG00000159079 |
| ACD | -1.222083605 | 0.428663176 | 7.02E-29 | 9.43E-27 | 770.025256 | 330.1001522 | ENSG00000102977 |
| SEMA5B | -1.22258134 | 0.428515311 | 2.93E-25 | 2.83E-23 | 767.3204605 | 329.0890351 | ENSG00000082684 |
| AFP | -1.222585398 | 0.428514106 | 0.031090287 | 0.092152146 | 24.82419107 | 10.66353738 | ENSG00000081051 |
| NA | -1.223698226 | 0.428183698 | 0.177088387 | 0.345308382 | 8.995270073 | 3.882228524 | ENSG00000287234 |
| NA | -1.226083643 | 0.427476304 | 4.81E-11 | 1.10E-09 | 256.6220643 | 110.0198004 | ENSG00000267321 |
| PTX3 | -1.226482813 | 0.427358045 | 0.037687563 | 0.107181416 | 23.59250747 | 10.13491514 | ENSG00000163661 |
| FRMPD1 | -1.229782042 | 0.426381858 | 0.124397111 | 0.266756457 | 11.89962752 | 5.048411747 | ENSG00000070601 |
| NA | -1.232908754 | 0.425458773 | 0.009831986 | 0.036235041 | 31.75370681 | 13.58562558 | ENSG00000235494 |

| | | | | | | | |
|---|---|---|---|---|---|---|---|
| MAP3K7 | -1.235061981 | 0.424824248 | 2.74E-36 | 5.72E-34 | 1247.50913 | 530.6548405 | ENSG00000135341 |
| NA | -1.236104198 | 0.424517462 | 0.237460957 | 0.422736887 | 6.352004128 | 2.659182664 | ENSG00000270175 |
| LINC00242 | -1.236833737 | 0.424302847 | 0.214357045 | 0.394456848 | 7.450225573 | 3.184555713 | ENSG00000229214 |
| NA | -1.237165494 | 0.424205287 | 0.19006989 | 0.362386107 | 8.122038612 | 3.473280288 | ENSG00000259319 |
| NA | -1.23771774 | 0.424042937 | 0.012385207 | 0.043895177 | 28.46072622 | 12.06636907 | ENSG00000277459 |
| TP53BP1 | -1.239436152 | 0.423538155 | 2.42E-51 | 1.04E-48 | 2579.395181 | 1092.836678 | ENSG00000067369 |
| PCLAF | -1.240919712 | 0.423102844 | 1.14E-05 | 0.000100503 | 112.3588112 | 47.26625665 | ENSG00000166803 |
| SLC15A2 | -1.241565786 | 0.42291341 | 0.000748948 | 0.004093484 | 56.78898244 | 24.05005648 | ENSG00000163406 |
| RANGRF | -1.242020891 | 0.422780022 | 0.142803234 | 0.295307559 | 10.37480227 | 4.350738936 | ENSG00000108961 |
| NA | -1.24231824 | 0.422692893 | 0.162656874 | 0.324089712 | 9.524849279 | 3.964360542 | ENSG00000260296 |
| NA | -1.242966211 | 0.422503087 | 0.073506124 | 0.179849834 | 14.61617887 | 6.170004941 | ENSG00000282886 |
| NA | -1.245640254 | 0.421720701 | 0.235578515 | 0.420544675 | 7.598773232 | 3.270924611 | ENSG00000178162 |
| TSPYL4 | -1.24599395 | 0.421617323 | 7.29E-25 | 6.76E-23 | 841.9186635 | 355.6994497 | ENSG00000187189 |
| LRRC49 | -1.246237676 | 0.421546102 | 1.12E-10 | 2.43E-09 | 194.7774479 | 82.08977693 | ENSG00000137821 |
| NA | -1.246864483 | 0.421362993 | 0.191711873 | 0.364766231 | 7.340936343 | 3.086901894 | ENSG00000270055 |
| INSYN1 | -1.247829769 | 0.421081159 | 1.40E-10 | 3.02E-09 | 211.7714642 | 89.4537595 | ENSG00000205363 |
| NA | -1.248179846 | 0.420978994 | 0.20746176 | 0.386325354 | 8.137725478 | 3.394397464 | ENSG00000258417 |
| PCDHGA8 | -1.249127472 | 0.420702568 | 0.085562174 | 0.20147495 | 15.54573111 | 6.545210035 | ENSG00000253767 |
| FXYD6 | -1.251157354 | 0.420111052 | 3.09E-07 | 3.78E-06 | 122.1755244 | 51.38980856 | ENSG00000137726 |
| SYPL2 | -1.251758078 | 0.419936159 | 3.74E-05 | 0.000291498 | 77.71662923 | 32.65063134 | ENSG00000143028 |
| SLC25A12 | -1.251916453 | 0.419890062 | 2.02E-12 | 5.67E-11 | 260.7985439 | 109.4515492 | ENSG00000115840 |
| NA | -1.252263664 | 0.41978902 | 0.219601421 | 0.401023051 | 8.251547096 | 3.477079135 | ENSG00000270587 |
| NA | -1.252503414 | 0.419719264 | 0.141017937 | 0.292417369 | 9.838909239 | 4.133411111 | ENSG00000271781 |
| TAGLN3 | -1.253314501 | 0.419483363 | 4.47E-06 | 4.38E-05 | 103.7006842 | 43.52076127 | ENSG00000144834 |
| COL21A1 | -1.253515757 | 0.419424849 | 6.05E-05 | 0.000445251 | 73.47643271 | 30.72622544 | ENSG00000124749 |
| GPR135 | -1.254588028 | 0.419113231 | 0.076168213 | 0.184681012 | 14.03984734 | 5.903850207 | ENSG00000181619 |
| CYP4X1 | -1.255195804 | 0.418936705 | 0.000761837 | 0.004156915 | 52.703452 | 22.09234547 | ENSG00000186377 |
| LINC01126 | -1.25630453 | 0.418614871 | 0.154986129 | 0.312943994 | 9.173507724 | 3.889714598 | ENSG00000279873 |
| NA | -1.258264493 | 0.418046551 | 0.143940615 | 0.296784442 | 10.60926299 | 4.440906681 | ENSG00000230042 |
| HDAC9 | -1.258638677 | 0.417938139 | 2.77E-09 | 4.81E-08 | 204.7674603 | 85.58953753 | ENSG00000048052 |
| P2RX5-TAX1BP3 | -1.258928458 | 0.4178542 | 0.00597939 | 0.023971881 | 36.45000174 | 15.27338301 | ENSG00000257950 |
| VIPR2 | -1.26083966 | 0.417301017 | 0.002808386 | 0.012587817 | 50.83695133 | 21.38002693 | ENSG00000106018 |
| SYP | -1.261009987 | 0.417251752 | 5.54E-09 | 9.26E-08 | 199.8013999 | 83.70756333 | ENSG00000102003 |
| KEL | -1.261491676 | 0.417112463 | 0.012229636 | 0.043437209 | 31.58178299 | 13.22181703 | ENSG00000197993 |
| GRIN1 | -1.261771068 | 0.417031693 | 7.83E-09 | 1.28E-07 | 150.8397436 | 62.87829045 | ENSG00000176884 |
| ERBB2 | -1.262119516 | 0.416930981 | 3.23E-53 | 1.58E-50 | 12945.20586 | 5397.128691 | ENSG00000141736 |
| LOC145694 | -1.262126543 | 0.41692895 | 0.22116413 | 0.402736883 | 7.631620137 | 3.233382622 | ENSG00000259426 |
| HACL1 | -1.262342929 | 0.416866421 | 2.60E-28 | 3.32E-26 | 688.1921285 | 286.5480125 | ENSG00000131373 |
| NA | -1.26272439 | 0.416756212 | 0.267719835 | 0.45786836 | 5.733354945 | 2.377944162 | ENSG00000252061 |
| FAM133A | -1.264553874 | 0.416228058 | 0.03330799 | 0.097370708 | 20.66675069 | 8.604373707 | ENSG00000179083 |
| DBX1 | -1.266316577 | 0.415719816 | 0.249277187 | 0.437756392 | 7.533079422 | 3.158298644 | ENSG00000109851 |
| MAPRE3 | -1.267628114 | 0.415342062 | 1.09E-08 | 1.73E-07 | 148.5914146 | 61.82473319 | ENSG00000084764 |
| FZD2 | -1.268616637 | 0.41505757 | 8.52E-30 | 1.18E-27 | 1397.861619 | 580.2393273 | ENSG00000180340 |
| DRC3 | -1.269832874 | 0.414707811 | 0.059056285 | 0.152052806 | 15.7409334 | 6.500070353 | ENSG00000171962 |
| GAS6-DT | -1.271160579 | 0.414326333 | 0.087574582 | 0.205145008 | 13.39909976 | 5.581270869 | ENSG00000272695 |
| NA | -1.271617428 | 0.414195152 | 0.008693601 | 0.032789419 | 37.1103672 | 15.28802874 | ENSG00000215105 |
| NA | -1.272443214 | 0.413958138 | 0.199927785 | 0.376388907 | 7.340936343 | 2.996734149 | ENSG00000229020 |
| DOK3 | -1.274858309 | 0.413265745 | 4.26E-07 | 5.09E-06 | 117.6580685 | 48.77196673 | ENSG00000146094 |
| C1QTNF1 | -1.276495102 | 0.412797145 | 2.34E-05 | 0.000192133 | 87.77864932 | 36.15341789 | ENSG00000173918 |
| GPR26 | -1.27779219 | 0.412426177 | 0.057448585 | 0.148958041 | 15.80672491 | 6.522640194 | ENSG00000154478 |
| PSMB10 | -1.279424828 | 0.411959716 | 0.209804485 | 0.389007236 | 6.719130247 | 2.775607477 | ENSG00000205220 |
| LOC100129129 | -1.279469127 | 0.411947066 | 0.274270082 | 0.465250018 | 6.014568001 | 2.52442489 | ENSG00000255020 |
| NA | -1.27985173 | 0.411837832 | 0.291738673 | 0.484115369 | 5.114705763 | 2.130560422 | ENSG00000278291 |
| DBNDD2 | -1.281213598 | 0.411449251 | 0.01219222 | 0.043322586 | 28.25793232 | 11.58982294 | ENSG00000244274 |
| CLIP4 | -1.281676854 | 0.411317154 | 0.137487688 | 0.286959099 | 9.203197716 | 3.792060779 | ENSG00000115295 |
| NCR3 | -1.281852276 | 0.411267144 | 0.269279202 | 0.45951754 | 5.094486509 | 2.119275502 | ENSG00000204475 |
| NA | -1.28188147 | 0.411258821 | 0.282009931 | 0.473069106 | 5.031949612 | 2.081733513 | ENSG00000197332 |
| ELP2 | -1.28272698 | 0.411017868 | 8.24E-60 | 5.46E-57 | 1921.34869 | 789.7339655 | ENSG00000134759 |
| CASR | -1.28272853 | 0.411017427 | 0.121402696 | 0.261941509 | 12.88983751 | 5.390200112 | ENSG00000036828 |
| NA | -1.284869145 | 0.410408027 | 0.137287984 | 0.286574558 | 12.95741046 | 5.401485033 | ENSG00000272853 |
| NA | -1.285211151 | 0.410310748 | 7.02E-05 | 0.000508177 | 89.14417839 | 36.83385704 | ENSG00000272086 |
| LRRC55 | -1.285468563 | 0.410237545 | 1.78E-05 | 0.000150503 | 93.30793259 | 38.20520711 | ENSG00000183908 |
| SMPDL3B | -1.28754597 | 0.409647248 | 3.22E-25 | 3.08E-23 | 578.6975293 | 237.3925188 | ENSG00000130768 |
| NA | -1.288310374 | 0.409430257 | 0.264601306 | 0.454253484 | 5.091329596 | 2.100504507 | ENSG00000260176 |
| CEP135 | -1.290231078 | 0.408885532 | 4.59E-13 | 1.41E-11 | 287.849642 | 117.4588267 | ENSG00000174799 |
| SLC10A4 | -1.295059374 | 0.407519394 | 5.74E-05 | 0.00042608 | 80.58626064 | 32.80047288 | ENSG00000145248 |
| NA | -1.29588271 | 0.407286892 | 0.06627784 | 0.165954266 | 14.32737422 | 5.824967383 | ENSG00000255182 |
| NA | -1.296530501 | 0.407104056 | 0.18057696 | 0.350121704 | 7.67837247 | 3.143214877 | ENSG00000269935 |
| NXPH2 | -1.297122523 | 0.406937031 | 0.000183305 | 0.001190809 | 59.69757918 | 24.22235624 | ENSG00000144227 |
| NRSN1 | -1.298038691 | 0.406678692 | 0.282091327 | 0.473085658 | 5.049011952 | 2.051677598 | ENSG00000152954 |
| LOC642361 | -1.29961652 | 0.406234164 | 0.000552056 | 0.003129948 | 52.68333045 | 21.47355548 | ENSG00000272447 |
| CCDC148 | -1.302121742 | 0.405529356 | 0.136042924 | 0.284648855 | 9.19688389 | 3.69440696 | ENSG00000153237 |
| EME2 | -1.302396222 | 0.405452209 | 0.000701816 | 0.003863218 | 543.7489993 | 220.6579931 | ENSG00000197774 |
| MIF | -1.303185612 | 0.405230421 | 0.256439877 | 0.445149075 | 6.053728732 | 2.479396828 | ENSG00000240972 |

| | | | | | | | |
|---|---|---|---|---|---|---|---|
| FAM135B | -1.304286674 | 0.404921268 | 0.097472903 | 0.222286829 | 11.07936452 | 4.508504585 | ENSG00000147724 |
| TSGA10IP | -1.304714629 | 0.404801172 | 0.000973352 | 0.005144026 | 46.1297125 | 18.75046214 | ENSG00000175513 |
| AQP5 | -1.304877181 | 0.404755564 | 0.000951219 | 0.005045764 | 48.45250696 | 19.56032289 | ENSG00000161798 |
| NA | -1.304977732 | 0.404727355 | 0.015594404 | 0.052713238 | 25.70817104 | 10.47246662 | ENSG00000288559 |
| PCDHGA2 | -1.305243972 | 0.404652672 | 0.052967825 | 0.139764474 | 15.28789422 | 6.181289862 | ENSG00000081853 |
| TPR | -1.311073432 | 0.403020902 | 8.82E-67 | 7.78E-64 | 3374.809483 | 1360.69615 | ENSG00000047410 |
| NFIX | -1.311394648 | 0.402931179 | 0.092171934 | 0.213185038 | 11.81045984 | 4.710860262 | ENSG00000008441 |
| ASTL | -1.311746582 | 0.402832899 | 0.218906281 | 0.400247568 | 6.639531009 | 2.629126749 | ENSG00000188886 |
| TRIM54 | -1.312042309 | 0.402750334 | 8.05E-06 | 7.37E-05 | 97.21127171 | 39.38030212 | ENSG00000138100 |
| ZNF321P | -1.312540666 | 0.402611234 | 0.039103302 | 0.110546156 | 17.99369706 | 7.235285152 | ENSG00000213801 |
| NA | -1.313064532 | 0.402465066 | 0.255969594 | 0.444941858 | 5.15070958 | 2.070448592 | ENSG00000218418 |
| NA | -1.31312454 | 0.402448326 | 0.037843917 | 0.1075106 | 17.99369706 | 7.227799079 | ENSG00000286342 |
| ADAM33 | -1.313490857 | 0.402346153 | 0.003788414 | 0.016264642 | 323.4946616 | 130.3219884 | ENSG00000149451 |
| PRR18 | -1.313609091 | 0.40231318 | 0.159503783 | 0.319490232 | 8.532760151 | 3.4357383 | ENSG00000176381 |
| MT1H | -1.313980858 | 0.402209522 | 0.001284596 | 0.006494464 | 46.59034328 | 18.60723061 | ENSG00000205358 |
| CHST13 | -1.314793361 | 0.401983068 | 6.00E-05 | 0.000441693 | 75.56378177 | 30.50184957 | ENSG00000180767 |
| PCDHB15 | -1.318583781 | 0.400928316 | 1.59E-06 | 1.70E-05 | 103.8702479 | 41.69389757 | ENSG00000113248 |
| PTN | -1.3204807 | 0.400401504 | 1.64E-33 | 2.79E-31 | 784.6113389 | 314.0343569 | ENSG00000105894 |
| NA | -1.322095922 | 0.399953471 | 0.065185236 | 0.16392716 | 14.4568827 | 5.716028643 | ENSG00000276728 |
| NA | -1.32229304 | 0.399898829 | 6.30E-07 | 7.28E-06 | 125.7896755 | 50.25280519 | ENSG00000273209 |
| ATL1 | -1.322361431 | 0.399879872 | 1.22E-06 | 1.34E-05 | 97.25221388 | 38.92957502 | ENSG00000198513 |
| MIR762HG | -1.323909619 | 0.399450982 | 0.189899618 | 0.362144541 | 6.976967136 | 2.816948313 | ENSG00000260083 |
| SPTB | -1.324048358 | 0.39941257 | 0.000281315 | 0.001737964 | 62.98474961 | 25.18304621 | ENSG00000070182 |
| FRMD6-AS1 | -1.324077057 | 0.399404625 | 0.173845496 | 0.340561507 | 7.929895533 | 3.222097701 | ENSG00000273888 |
| INCA1 | -1.324625084 | 0.399252934 | 0.014117696 | 0.048662614 | 28.32056692 | 11.31986935 | ENSG00000196388 |
| NA | -1.329076735 | 0.398022879 | 0.103732545 | 0.232926942 | 12.36213744 | 4.992536797 | ENSG00000255224 |
| NA | -1.330454936 | 0.39764283 | 0.053245818 | 0.140318212 | 15.53310345 | 6.173803788 | ENSG00000234373 |
| NA | -1.332214385 | 0.397158178 | 0.140888542 | 0.29221444 | 11.62157138 | 4.703812222 | ENSG00000271933 |
| NA | -1.333564923 | 0.396786564 | 0.270664438 | 0.461244454 | 5.346009572 | 2.168102411 | ENSG00000254783 |
| NA | -1.33380413 | 0.396720779 | 0.194990635 | 0.369260901 | 7.929895533 | 3.177069639 | ENSG00000232320 |
| NA | -1.333865682 | 0.396703854 | 0.21691353 | 0.397456783 | 7.142479438 | 2.865775222 | ENSG00000274292 |
| FAM111B | -1.333916869 | 0.396689779 | 6.24E-20 | 3.99E-18 | 617.2576624 | 244.5908829 | ENSG00000189057 |
| NA | -1.334793358 | 0.396448849 | 0.207267419 | 0.386002245 | 6.047414906 | 2.389229083 | ENSG00000260992 |
| TEKT2 | -1.335134881 | 0.39635501 | 0.025394377 | 0.078377915 | 19.96218845 | 7.92547189 | ENSG00000092850 |
| GPD1 | -1.340593343 | 0.394858228 | 0.092282614 | 0.21339563 | 11.50271371 | 4.587387409 | ENSG00000167588 |
| DPY19L2P2 | -1.340673989 | 0.394836156 | 0.032993195 | 0.096602825 | 18.34701545 | 7.276625988 | ENSG00000170629 |
| REM2 | -1.34436565 | 0.393827115 | 0.000150647 | 0.000996463 | 71.2734806 | 28.14647525 | ENSG00000139890 |
| LRRN2 | -1.347760206 | 0.392901558 | 1.97E-15 | 7.99E-14 | 289.1328959 | 113.461161 | ENSG00000170382 |
| NA | -1.352943627 | 0.391492445 | 0.215624292 | 0.395955923 | 6.053728732 | 2.340402174 | ENSG00000238082 |
| ZNF20 | -1.353226342 | 0.391415735 | 0.18751886 | 0.359259308 | 7.036347121 | 2.749350409 | ENSG00000132010 |
| MAGEE1 | -1.353637516 | 0.391304195 | 1.23E-16 | 5.58E-15 | 321.0476803 | 125.2827343 | ENSG00000198934 |
| NA | -1.354957315 | 0.390946388 | 0.232747751 | 0.417513243 | 6.127014144 | 2.377944162 | ENSG00000286681 |
| NA | -1.356699875 | 0.390474469 | 0.168955774 | 0.333320606 | 8.022122422 | 3.147013724 | ENSG00000260144 |
| CP | -1.35733588 | 0.390302368 | 0.043104302 | 0.119344957 | 17.80786781 | 6.957845497 | ENSG00000047457 |
| URAHP | -1.359014533 | 0.389848494 | 0.117271263 | 0.255316058 | 9.640452335 | 3.724462875 | ENSG00000222019 |
| PHACTR3 | -1.359800678 | 0.389636118 | 0.076268062 | 0.184826393 | 13.1476744 | 5.127294571 | ENSG00000087495 |
| FOXO6 | -1.362954013 | 0.388785411 | 4.83E-08 | 6.86E-07 | 125.9677178 | 48.88740386 | ENSG00000204060 |
| DPYSL5 | -1.363160551 | 0.388729755 | 6.98E-40 | 2.02E-37 | 1059.469153 | 411.9844359 | ENSG00000157851 |
| CCNI2 | -1.363226155 | 0.388712079 | 0.028052417 | 0.085079937 | 20.34184452 | 7.86536006 | ENSG00000205089 |
| PLA2R1 | -1.365662262 | 0.388056261 | 3.07E-05 | 0.000245893 | 67.53199321 | 26.2482148 | ENSG00000153246 |
| MYLPF | -1.366269742 | 0.387892896 | 0.230441219 | 0.414498987 | 5.395918818 | 2.119275502 | ENSG00000180209 |
| ZSWIM5 | -1.366660403 | 0.387787874 | 4.22E-10 | 8.25E-09 | 197.0015288 | 76.57318119 | ENSG00000162415 |
| TTLL7 | -1.369731668 | 0.386963214 | 4.54E-05 | 0.00034515 | 92.86110954 | 36.19629606 | ENSG00000137941 |
| NA | -1.371455215 | 0.386501196 | 0.064325052 | 0.162447269 | 17.389051 | 6.830573797 | ENSG00000283662 |
| UBE2E1-AS1 | -1.371957742 | 0.386366592 | 0.206100428 | 0.384640554 | 6.934649492 | 2.64041167 | ENSG00000223791 |
| PSMG3-AS1 | -1.373091606 | 0.386063052 | 1.01E-07 | 1.35E-06 | 119.9184237 | 46.48787754 | ENSG00000230487 |
| KIF19 | -1.374277934 | 0.385745723 | 0.017175146 | 0.056935104 | 26.28124797 | 10.0104546 | ENSG00000196169 |
| WDR34 | -1.37453471 | 0.385677073 | 1.22E-39 | 3.44E-37 | 1306.100956 | 504.089892 | ENSG00000119333 |
| NA | -1.375140678 | 0.385515113 | 0.294209057 | 0.486878791 | 5.391484129 | 2.089219587 | ENSG00000284624 |
| IL18R1 | -1.377422209 | 0.384905927 | 0.23853704 | 0.424203176 | 6.265993366 | 2.389229083 | ENSG00000115604 |
| RGS7BP | -1.380270417 | 0.384146784 | 0.000417424 | 0.002445173 | 50.84769985 | 19.53406582 | ENSG00000186479 |
| PPP1R14B-AS1 | -1.380641375 | 0.384048022 | 0.197517913 | 0.37332215 | 6.322314136 | 2.449340913 | ENSG00000256940 |
| NA | -1.381276338 | 0.383879031 | 0.196351899 | 0.371459232 | 6.994029477 | 2.64041167 | ENSG00000267394 |
| ALOX12B | -1.381387565 | 0.383849436 | 0.240518169 | 0.426702761 | 5.438236462 | 2.100504507 | ENSG00000179477 |
| NA | -1.382687048 | 0.383503846 | 0.216685906 | 0.397275357 | 6.434760279 | 2.498167822 | ENSG00000243230 |
| TSNAXIP1 | -1.383323191 | 0.383334781 | 0.060843216 | 0.155669265 | 15.45844257 | 5.952677116 | ENSG00000102904 |
| NA | -1.384142393 | 0.383117175 | 0.043468519 | 0.120102197 | 15.21776572 | 5.824967383 | ENSG00000226361 |
| NA | -1.387805802 | 0.382145566 | 0.188038922 | 0.359787707 | 7.10647562 | 2.677953659 | ENSG00000244471 |
| MAP1A | -1.390265271 | 0.38149465 | 3.38E-07 | 4.11E-06 | 146.9102395 | 56.28133073 | ENSG00000166963 |
| CDYL2 | -1.391524487 | 0.381161818 | 9.39E-08 | 1.27E-06 | 110.6873175 | 42.11737992 | ENSG00000166446 |
| TMEM255A | -1.392687044 | 0.380854793 | 0.001443082 | 0.007164197 | 44.95504873 | 17.10718313 | ENSG00000125355 |
| KRT7 | -1.393231712 | 0.380711034 | 1.68E-14 | 6.08E-13 | 268.17368 | 102.1005005 | ENSG00000135480 |
| H4C1 | -1.393395261 | 0.380667878 | 0.22893929 | 0.412798988 | 5.378856477 | 2.040392677 | ENSG00000278637 |
| PTPRO | -1.393485877 | 0.380643969 | 0.007408736 | 0.028702656 | 26.97831631 | 10.24330422 | ENSG00000151490 |

| | | | | | | | |
|---|---|---|---|---|---|---|---|
| NA | -1.395900367 | 0.380007457 | 0.147829962 | 0.302782686 | 8.065717842 | 3.026790064 | ENSG00000259172 |
| NA | -1.39856455 | 0.379306355 | 0.087696726 | 0.205366899 | 14.83357726 | 5.52820708 | ENSG00000238207 |
| SCN2B | -1.400290287 | 0.378852904 | 0.043143697 | 0.119414916 | 17.59244626 | 6.736718825 | ENSG00000149575 |
| PRRT4 | -1.400473603 | 0.378804768 | 0.002881311 | 0.012863439 | 40.14117392 | 15.27349463 | ENSG00000224940 |
| NA | -1.401687719 | 0.378486115 | 0.16778093 | 0.331554724 | 7.255023279 | 2.749350409 | ENSG00000213683 |
| MYO16 | -1.401836279 | 0.378447143 | 0.246112663 | 0.433592002 | 5.322633406 | 1.991565768 | ENSG00000041515 |
| SHD | -1.402425316 | 0.378292659 | 0.000112594 | 0.000775458 | 65.33880493 | 24.74404206 | ENSG00000105251 |
| NTN3 | -1.402826982 | 0.378187351 | 0.22434432 | 0.407048226 | 5.448984977 | 2.051677598 | ENSG00000162068 |
| NA | -1.403083347 | 0.378120154 | 0.015128315 | 0.051475711 | 26.84880783 | 10.27759702 | ENSG00000256667 |
| NA | -1.403641983 | 0.377973768 | 0.125469541 | 0.268373048 | 9.095186262 | 3.454509294 | ENSG00000277383 |
| CHN2 | -1.404226457 | 0.377820672 | 0.217543347 | 0.398510992 | 5.399075731 | 2.021621683 | ENSG00000106069 |
| HTR7P1 | -1.406060682 | 0.37734062 | 0.000308996 | 0.001882001 | 80.11152903 | 30.48775349 | ENSG00000183935 |
| FRRS1L | -1.406294203 | 0.377279547 | 0.000319753 | 0.001938602 | 49.41953618 | 18.73169115 | ENSG00000260230 |
| ROM1 | -1.407665136 | 0.376921204 | 0.064578672 | 0.162888012 | 12.81014057 | 4.827285075 | ENSG00000149489 |
| CCDC74A | -1.408210438 | 0.376778765 | 5.12E-17 | 2.42E-15 | 290.6413579 | 109.6661775 | ENSG00000163040 |
| AQP4-AS1 | -1.408280902 | 0.376760363 | 0.243694263 | 0.43036012 | 5.472361143 | 2.051677598 | ENSG00000260372 |
| NA | -1.409709455 | 0.37638748 | 0.074731041 | 0.182077706 | 11.82752218 | 4.42962176 | ENSG00000280087 |
| NA | -1.413535686 | 0.37539057 | 0.148545186 | 0.303878198 | 7.308089437 | 2.738065489 | ENSG00000270184 |
| NA | -1.414034681 | 0.375260753 | 0.099154445 | 0.225294654 | 10.75899073 | 3.983131536 | ENSG00000278012 |
| PNMA6F | -1.414446318 | 0.375153697 | 0.107476474 | 0.23908329 | 10.06380153 | 3.74323387 | ENSG00000225110 |
| CCDC160 | -1.415687281 | 0.37483114 | 2.30E-06 | 2.39E-05 | 91.3591568 | 34.21827672 | ENSG00000203952 |
| PSKH2 | -1.415739342 | 0.374817614 | 0.221144854 | 0.402736883 | 5.511521874 | 2.051677598 | ENSG00000147613 |
| EZH2 | -1.42042739 | 0.373601618 | 1.26E-51 | 5.72E-49 | 1537.381607 | 574.7288406 | ENSG00000106462 |
| NA | -1.420594163 | 0.373558433 | 0.243202287 | 0.429696014 | 6.222495644 | 2.291575264 | ENSG00000286004 |
| LIG3 | -1.426889405 | 0.371931951 | 6.32E-75 | 7.32E-72 | 2204.612017 | 820.3119899 | ENSG00000005156 |
| TMEM117 | -1.427953552 | 0.371657712 | 0.009409624 | 0.035000907 | 27.51992181 | 10.18699124 | ENSG00000139173 |
| CD300A | -1.429422708 | 0.37127943 | 0.084431419 | 0.199573004 | 11.06673687 | 4.103355196 | ENSG00000167851 |
| HSF2BP | -1.429809761 | 0.371179835 | 0.014532629 | 0.049744639 | 24.25357199 | 9.069634925 | ENSG00000160207 |
| BHMT | -1.430622593 | 0.370970766 | 0.025891626 | 0.079615298 | 24.35714876 | 9.092204766 | ENSG00000145692 |
| SOX7 | -1.435236103 | 0.369786355 | 0.112378631 | 0.247366455 | 9.326392374 | 3.443224373 | ENSG00000171056 |
| TNFRSF19 | -1.437682446 | 0.369159849 | 0.117556985 | 0.255652641 | 9.154566247 | 3.345570555 | ENSG00000127863 |
| CD247 | -1.438806405 | 0.36887236 | 0.004539827 | 0.018946445 | 30.46522143 | 11.16958978 | ENSG00000198821 |
| NA | -1.440602217 | 0.368413487 | 0.179914473 | 0.349227792 | 7.291027097 | 2.647897744 | ENSG00000268230 |
| PCDHA5 | -1.442601542 | 0.367903284 | 0.132666173 | 0.279476568 | 9.123598478 | 3.31551464 | ENSG00000204965 |
| CTNNAL1 | -1.442923582 | 0.367821169 | 4.13E-39 | 1.05E-36 | 1358.195846 | 498.8285235 | ENSG00000119326 |
| NA | -1.443462283 | 0.367683851 | 0.028706614 | 0.086540076 | 17.6095086 | 6.413701455 | ENSG00000269993 |
| NA | -1.445554013 | 0.36715114 | 0.013306479 | 0.046427388 | 21.71506289 | 8.011840788 | ENSG00000271147 |
| C17orf49 | -1.447137424 | 0.3667484 | 0.076287074 | 0.184848298 | 12.16683745 | 4.448392755 | ENSG00000258315 |
| RADX | -1.448137607 | 0.366494231 | 1.58E-10 | 3.34E-09 | 220.899302 | 80.53155272 | ENSG00000147231 |
| H2BC13 | -1.449034982 | 0.366266337 | 0.069161499 | 0.171486103 | 12.39626212 | 4.594873483 | ENSG00000185130 |
| MAP6 | -1.450205052 | 0.365969405 | 1.84E-06 | 1.95E-05 | 99.93316738 | 36.37834341 | ENSG00000171533 |
| LOC105375304 | -1.452019742 | 0.365509361 | 0.240079164 | 0.42612787 | 5.5178357 | 1.972794774 | ENSG00000225792 |
| NA | -1.45245151 | 0.365399988 | 0.117291411 | 0.255329929 | 8.51569781 | 3.135728803 | ENSG00000259994 |
| NA | -1.452691992 | 0.365339084 | 0.083450691 | 0.197914715 | 10.4840915 | 3.870943603 | ENSG00000261526 |
| KCNK15 | -1.45369565 | 0.365085013 | 0.145394004 | 0.299021169 | 7.639211739 | 2.816948313 | ENSG00000124249 |
| DDX11-AS1 | -1.454658912 | 0.364841333 | 0.003254234 | 0.014265366 | 32.8165258 | 11.91988835 | ENSG00000245614 |
| ZNF763 | -1.454743974 | 0.364819823 | 0.166031456 | 0.329185602 | 7.528644732 | 2.756836483 | ENSG00000197054 |
| LY6G5B | -1.454979863 | 0.364760177 | 0.007888155 | 0.030268959 | 30.84625297 | 11.39919021 | ENSG00000240053 |
| NA | -1.456167114 | 0.364460125 | 0.133000894 | 0.28005437 | 8.317240906 | 2.977963155 | ENSG00000280219 |
| NA | -1.460084844 | 0.363471753 | 0.07326991 | 0.179413992 | 14.34759347 | 5.240032157 | ENSG00000275180 |
| CNTD2 | -1.461529514 | 0.363107966 | 0.011052686 | 0.040028073 | 23.57101044 | 8.529178109 | ENSG00000105219 |
| LRRC37A6P | -1.465518269 | 0.362105434 | 0.014674329 | 0.050151949 | 21.99439681 | 8.000555867 | ENSG00000230445 |
| ALDH3A1 | -1.466111288 | 0.361956621 | 0.070591833 | 0.174098342 | 12.99784896 | 4.639463512 | ENSG00000108602 |
| DLX3 | -1.466373084 | 0.361890945 | 0.199580148 | 0.375954718 | 5.733354945 | 2.081733513 | ENSG00000064195 |
| LCMT1-AS2 | -1.468694042 | 0.361309216 | 0.206564192 | 0.385152757 | 5.624065715 | 2.021621683 | ENSG00000260034 |
| LINC00115 | -1.469693182 | 0.361059077 | 0.034873182 | 0.100927477 | 16.36273949 | 5.933906122 | ENSG00000225880 |
| NA | -1.474179055 | 0.359938154 | 0.158082757 | 0.317467311 | 6.689440255 | 2.408000077 | ENSG00000276436 |
| VN1R1 | -1.474347309 | 0.359896179 | 0.215831278 | 0.396178756 | 6.576896414 | 2.321631179 | ENSG00000178201 |
| CD34 | -1.474357372 | 0.359893669 | 0.023112017 | 0.072640009 | 18.9340955 | 6.788794928 | ENSG00000174059 |
| FABP7 | -1.474518226 | 0.359853545 | 5.15E-12 | 1.38E-10 | 172.7066087 | 62.12799188 | ENSG00000164434 |
| MAPK8IP2 | -1.475422667 | 0.359628019 | 5.41E-12 | 1.44E-10 | 243.5393922 | 87.64836634 | ENSG00000008735 |
| PLCL2 | -1.475937482 | 0.359499712 | 9.41E-14 | 3.11E-12 | 201.7727551 | 72.5516316 | ENSG00000154822 |
| IL12A | -1.475988203 | 0.359487073 | 0.220347141 | 0.401880921 | 5.720727294 | 2.051677598 | ENSG00000168811 |
| LRRTM1 | -1.476420046 | 0.359379483 | 0.006406033 | 0.025440202 | 26.02350878 | 9.414672484 | ENSG00000162951 |
| TBXT | -1.477083424 | 0.359214272 | 0.183274378 | 0.353723557 | 5.763044937 | 2.070448592 | ENSG00000164458 |
| H2BC5 | -1.482166825 | 0.357950792 | 1.33E-07 | 1.73E-06 | 105.5001004 | 37.77043984 | ENSG00000158373 |
| EXOSC10-AS1 | -1.483988016 | 0.357499216 | 0.132621383 | 0.279413972 | 8.522011636 | 3.01930399 | ENSG00000230337 |
| SCML1 | -1.485115597 | 0.357219911 | 0.000196979 | 0.001265886 | 51.14765899 | 18.27022878 | ENSG00000047634 |
| LOC102724532 | -1.489012661 | 0.356256277 | 0.007362918 | 0.028546854 | 26.99034261 | 9.609542088 | ENSG00000264920 |
| TUBB3 | -1.489711386 | 0.356083777 | 0.054957491 | 0.143766005 | 12.61809519 | 4.527275579 | ENSG00000258947 |
| LINC00545 | -1.490187427 | 0.355966301 | 0.023830974 | 0.074507973 | 17.53434405 | 6.278943681 | ENSG00000236094 |
| CXCL16 | -1.490495856 | 0.355890208 | 0.004381911 | 0.018386764 | 37.68236175 | 13.24708641 | ENSG00000161921 |
| PROZ | -1.492208866 | 0.355467886 | 0.141766356 | 0.293589338 | 7.820606303 | 2.816948313 | ENSG00000126231 |
| NA | -1.49891517 | 0.353819344 | 0.033885991 | 0.098764421 | 16.26923483 | 5.685972729 | ENSG00000263731 |

| | | | | | | | |
|---|---|---|---|---|---|---|---|
| NA | -1.500864579 | 0.353341576 | 0.018221177 | 0.059666306 | 23.67408354 | 8.443358864 | ENSG00000287737 |
| NA | -1.501578784 | 0.353166698 | 0.162839074 | 0.324348134 | 6.930214803 | 2.486882902 | ENSG00000235489 |
| ACSL6 | -1.501823028 | 0.353106913 | 0.152426525 | 0.309550819 | 7.575397065 | 2.629126749 | ENSG00000164398 |
| FRG1-DT | -1.502924613 | 0.352837398 | 0.026420266 | 0.080978721 | 17.81743624 | 6.357826505 | ENSG00000245685 |
| TRPC4 | -1.503224078 | 0.352764166 | 1.77E-16 | 7.92E-15 | 243.9351184 | 86.08463142 | ENSG00000133107 |
| DLEU7 | -1.503241139 | 0.352759994 | 2.03E-07 | 2.58E-06 | 107.1216849 | 37.83803774 | ENSG00000186047 |
| UHRF1 | -1.506225238 | 0.352031092 | 5.37E-81 | 8.30E-78 | 2415.711555 | 850.8774237 | ENSG00000276043 |
| ZNF441 | -1.512376826 | 0.350533243 | 0.028760996 | 0.086619484 | 16.58329479 | 5.783626547 | ENSG00000197044 |
| LINC00648 | -1.513224236 | 0.350327407 | 0.004653214 | 0.019323923 | 30.19613236 | 10.56208471 | ENSG00000259129 |
| NA | -1.513466176 | 0.350268662 | 0.030722563 | 0.091266546 | 17.44083938 | 6.057378975 | ENSG00000215014 |
| C1R | -1.513785036 | 0.350191255 | 2.74E-08 | 4.06E-07 | 117.6338955 | 41.12718369 | ENSG00000159403 |
| NUTM2B | -1.515146573 | 0.34986092 | 0.184306084 | 0.354938587 | 6.822105651 | 2.359173168 | ENSG00000188199 |
| NA | -1.515641432 | 0.349740935 | 0.015571251 | 0.052644577 | 20.3760669 | 7.051262436 | ENSG00000217648 |
| NA | -1.517558394 | 0.34927653 | 0.178217033 | 0.346961605 | 6.017724914 | 2.138046496 | ENSG00000232186 |
| LOC101927278 | -1.517744259 | 0.349231535 | 0.044602827 | 0.122415045 | 16.94981955 | 5.952677116 | ENSG00000287261 |
| NA | -1.520048227 | 0.348674261 | 0.143525907 | 0.296291955 | 6.973810223 | 2.456826987 | ENSG00000248126 |
| S1PR1 | -1.520411845 | 0.348586392 | 0.003660977 | 0.015775993 | 30.90553526 | 10.81001811 | ENSG00000170989 |
| NA | -1.521069698 | 0.348427477 | 0.024204983 | 0.07541018 | 18.02338705 | 6.320284516 | ENSG00000266777 |
| EGR1 | -1.52166789 | 0.348283036 | 1.00E-22 | 7.94E-21 | 620.4802216 | 215.6201244 | ENSG00000120738 |
| CHRD | -1.52240874 | 0.348104232 | 0.012618306 | 0.044614208 | 19.77004537 | 6.878962673 | ENSG00000090539 |
| NA | -1.527781065 | 0.346810368 | 0.184198787 | 0.35487945 | 5.964658755 | 2.081733513 | ENSG00000233461 |
| WNT7B | -1.5283096 | 0.346683337 | 0.088730815 | 0.207266124 | 10.06695844 | 3.503336203 | ENSG00000188064 |
| NA | -1.531093108 | 0.346015097 | 0.084932868 | 0.200400481 | 11.25434756 | 3.957312501 | ENSG00000231043 |
| RAI2 | -1.531803456 | 0.34584477 | 0.00092705 | 0.004940182 | 37.57495166 | 12.97388364 | ENSG00000131831 |
| NA | -1.533286751 | 0.345489375 | 0.043451476 | 0.120096115 | 13.02122513 | 4.527275579 | ENSG00000223969 |
| PCDHGB4 | -1.533683973 | 0.345394263 | 1.68E-08 | 2.60E-07 | 118.0928652 | 40.76337514 | ENSG00000253953 |
| CAMK2A | -1.533871111 | 0.345349464 | 0.036399889 | 0.10446509 | 23.30941528 | 8.102008533 | ENSG00000070808 |
| NA | -1.535487233 | 0.344962816 | 0.000941907 | 0.005004959 | 38.61369542 | 13.23635114 | ENSG00000256663 |
| NA | -1.536462021 | 0.344729814 | 0.001950906 | 0.009297529 | 32.72241977 | 11.23718768 | ENSG00000254252 |
| EFCAB6 | -1.538266781 | 0.344298839 | 0.045383669 | 0.124117001 | 13.87749195 | 4.767173245 | ENSG00000186976 |
| H4C12 | -1.538519776 | 0.344238467 | 0.084179792 | 0.199054384 | 9.832595414 | 3.345570555 | ENSG00000273542 |
| CCDC177 | -1.538812147 | 0.344168712 | 3.27E-14 | 1.15E-12 | 241.1873887 | 83.18566271 | ENSG00000267909 |
| CTAGE15 | -1.540766499 | 0.343702798 | 0.130566765 | 0.276585938 | 7.030033295 | 2.419284998 | ENSG00000271079 |
| NA | -1.543591761 | 0.343030376 | 0.014431546 | 0.049477298 | 20.173273 | 6.976616492 | ENSG00000276672 |
| NA | -1.547838968 | 0.342022 | 0.03369121 | 0.09824306 | 16.15678869 | 5.547527727 | ENSG00000283236 |
| AMBP | -1.550375826 | 0.341421111 | 0.00093318 | 0.004967133 | 37.39345939 | 12.77532681 | ENSG00000106927 |
| NA | -1.55048405 | 0.341395501 | 0.120576412 | 0.260583399 | 8.059404017 | 2.738065489 | ENSG00000276408 |
| RPRML | -1.552686171 | 0.340874794 | 0.006215096 | 0.024793499 | 30.05833322 | 10.21704715 | ENSG00000179673 |
| DNAJC9-AS1 | -1.556827355 | 0.339897733 | 0.088661391 | 0.207167198 | 12.30581667 | 4.193522941 | ENSG00000236756 |
| CD8B | -1.557279389 | 0.339791251 | 0.18306292 | 0.353536331 | 6.120700318 | 2.081733513 | ENSG00000172116 |
| NA | -1.557814027 | 0.339665353 | 1.48E-05 | 0.000127523 | 61.7532614 | 21.01198149 | ENSG00000274372 |
| NA | -1.557902877 | 0.339644435 | 0.183945561 | 0.354539001 | 5.93181185 | 1.991565768 | ENSG00000280332 |
| MMUT | -1.558896091 | 0.33941069 | 5.02E-28 | 6.25E-26 | 477.9577113 | 162.6095354 | ENSG00000146085 |
| LOC644656 | -1.559387619 | 0.339295072 | 0.084826321 | 0.200200056 | 10.14665537 | 3.44702322 | ENSG00000268403 |
| LRRC4 | -1.562912682 | 0.338467055 | 4.15E-17 | 2.00E-15 | 284.9719153 | 96.20314869 | ENSG00000128594 |
| NA | -1.563214172 | 0.33839633 | 0.167968864 | 0.331804386 | 6.163017962 | 2.089219587 | ENSG00000263603 |
| KCNK2 | -1.564656434 | 0.338058205 | 0.005859325 | 0.023582352 | 25.32072797 | 8.510407115 | ENSG00000082482 |
| CTU2 | -1.567385532 | 0.337419315 | 1.02E-29 | 1.40E-27 | 506.8573833 | 171.2280051 | ENSG00000174177 |
| NA | -1.570697213 | 0.336645664 | 0.060334216 | 0.154687282 | 12.36647443 | 4.159668179 | ENSG00000237436 |
| GDPD2 | -1.57338803 | 0.33601836 | 6.02E-24 | 5.24E-22 | 354.2270929 | 119.1104679 | ENSG00000130055 |
| CACNA1G | -1.574311533 | 0.335803336 | 5.29E-21 | 3.64E-19 | 327.2791657 | 110.1051816 | ENSG00000006283 |
| CEND1 | -1.577199825 | 0.335131726 | 1.89E-10 | 3.93E-09 | 144.6838135 | 48.52414496 | ENSG00000184524 |
| TMEM202-AS1 | -1.578069093 | 0.33492986 | 0.150821752 | 0.307060718 | 7.0628802 | 2.329117253 | ENSG00000261423 |
| CMPK2 | -1.584344226 | 0.333476216 | 0.021602553 | 0.068924378 | 18.57644012 | 6.215144624 | ENSG00000134326 |
| CD177 | -1.585490275 | 0.333211414 | 0.001254907 | 0.006366121 | 33.87874343 | 11.22221554 | ENSG00000204936 |
| H2BC11 | -1.586315643 | 0.333020838 | 2.36E-07 | 2.96E-06 | 84.69803106 | 28.24726665 | ENSG00000124635 |
| CFL1P1 | -1.58680844 | 0.332907103 | 0.135667021 | 0.284151041 | 7.208270946 | 2.408000077 | ENSG00000223820 |
| VARS2 | -1.58696419 | 0.332871166 | 0.076623541 | 0.185421167 | 10.54347149 | 3.559649186 | ENSG00000137411 |
| ATOH7 | -1.598497261 | 0.330220763 | 0.152453862 | 0.309550819 | 6.269247977 | 2.089219587 | ENSG00000179774 |
| CCDC74B | -1.599236141 | 0.330051683 | 7.62E-13 | 2.31E-11 | 191.3544551 | 63.33661525 | ENSG00000152076 |
| FAM3B | -1.601446357 | 0.329546429 | 0.004007776 | 0.017048454 | 28.14163021 | 9.249420762 | ENSG00000183844 |
| METTL7A | -1.602357775 | 0.329338305 | 1.28E-39 | 3.53E-37 | 899.5345462 | 296.329027 | ENSG00000185432 |
| ZAR1 | -1.602800448 | 0.329237267 | 0.009631948 | 0.035660913 | 22.28695974 | 7.423106716 | ENSG00000182223 |
| GABRA3 | -1.605752356 | 0.328564301 | 6.18E-15 | 2.35E-13 | 201.7322189 | 66.47591966 | ENSG00000011677 |
| SLC26A8 | -1.60810729 | 0.328028418 | 0.14052135 | 0.291779375 | 7.553298675 | 2.535709811 | ENSG00000112053 |
| LINC02701 | -1.614990226 | 0.326467159 | 0.065349963 | 0.164229995 | 10.44680991 | 3.394397464 | ENSG00000250508 |
| FBP1 | -1.619593551 | 0.325427133 | 0.000744373 | 0.004074091 | 35.55339411 | 11.4808842 | ENSG00000165140 |
| NA | -1.6199227 | 0.325352896 | 0.174295555 | 0.341202955 | 9.251325523 | 2.989248076 | ENSG00000280056 |
| KLK4 | -1.624468462 | 0.324329361 | 0.039180689 | 0.110663653 | 12.67747518 | 4.103355196 | ENSG00000167749 |
| NA | -1.626349515 | 0.32390676 | 0.160773725 | 0.321304746 | 7.617714709 | 2.505653896 | ENSG00000280233 |
| ZBED6CL | -1.626580531 | 0.323854898 | 0.085220105 | 0.200848095 | 8.54223089 | 2.786892398 | ENSG00000188707 |
| STIMATE | -1.628595809 | 0.323402826 | 0.059060546 | 0.152052806 | 10.79815146 | 3.540878192 | ENSG00000213533 |
| TMEM233 | -1.630073725 | 0.323071698 | 0.132430783 | 0.279107588 | 6.408227199 | 2.070448592 | ENSG00000224982 |
| NA | -1.630877169 | 0.322891827 | 5.26E-06 | 5.05E-05 | 63.92623043 | 20.61431818 | ENSG00000206791 |

| | | | | | | | |
|---|---|---|---|---|---|---|---|
| NA | -1.631329901 | 0.322790517 | 0.082344469 | 0.195942672 | 10.53272297 | 3.368140396 | ENSG00000274825 |
| SLC30A3 | -1.631554188 | 0.322740338 | 4.14E-22 | 3.10E-20 | 297.9391798 | 96.04246026 | ENSG00000115194 |
| NA | -1.631777315 | 0.322690427 | 0.134773176 | 0.282630238 | 6.417697938 | 2.070448592 | ENSG00000228559 |
| NA | -1.634131327 | 0.32216433 | 0.15329153 | 0.310707044 | 7.215862548 | 2.280290344 | ENSG00000236264 |
| GALR3 | -1.634505608 | 0.322080761 | 0.04977387 | 0.133272018 | 12.65715822 | 4.013187451 | ENSG00000128310 |
| NA | -1.635153206 | 0.321936218 | 0.11496523 | 0.251685044 | 7.539393247 | 2.456826987 | ENSG00000267868 |
| TPP2 | -1.635649374 | 0.321825517 | 2.91E-78 | 3.85E-75 | 1791.457195 | 577.0964791 | ENSG00000134900 |
| SLC7A4 | -1.640651122 | 0.320711697 | 0.194487433 | 0.368571583 | 5.448984977 | 1.762953022 | ENSG00000099960 |
| PHGDH | -1.641659667 | 0.320487575 | 1.33E-177 | 6.16E-174 | 14736.58269 | 4723.511788 | ENSG00000092621 |
| EPHA10 | -1.64195942 | 0.320420994 | 0.001146273 | 0.005917832 | 36.68258332 | 11.78144335 | ENSG00000183317 |
| TSFM | -1.642413724 | 0.320320109 | 0.179907123 | 0.349227792 | 5.399075731 | 1.740383181 | ENSG00000123297 |
| LRRC8E | -1.643522362 | 0.320074054 | 0.025450991 | 0.078513428 | 15.98308342 | 5.108523577 | ENSG00000171017 |
| GSG1 | -1.650662928 | 0.318493773 | 0.161449325 | 0.322210472 | 5.399075731 | 1.721612187 | ENSG00000111305 |
| C2orf15 | -1.652014195 | 0.318195603 | 0.003062549 | 0.01355618 | 26.90187399 | 8.544261877 | ENSG00000273045 |
| WFDC1 | -1.652156172 | 0.318164291 | 0.054206219 | 0.142262541 | 10.91691143 | 3.492051283 | ENSG00000103175 |
| NA | -1.652388284 | 0.318113106 | 0.171666654 | 0.337442447 | 5.511521874 | 1.770439096 | ENSG00000224550 |
| NA | -1.654757213 | 0.317591188 | 0.042337412 | 0.117643417 | 12.01207366 | 3.852172609 | ENSG00000242375 |
| MPPED2 | -1.658594277 | 0.316747629 | 1.35E-06 | 1.46E-05 | 81.77680667 | 25.80128655 | ENSG00000066382 |
| KLF1 | -1.660172876 | 0.316401232 | 0.043455153 | 0.120096115 | 11.9665991 | 3.8033457 | ENSG00000105610 |
| PAX3 | -1.662081807 | 0.315982856 | 0.15851374 | 0.318126004 | 6.367788692 | 1.972794774 | ENSG00000135903 |
| NA | -1.665533262 | 0.315227813 | 0.074853581 | 0.182295218 | 9.696675406 | 3.00801907 | ENSG00000261338 |
| NA | -1.674977161 | 0.313171067 | 0.189434655 | 0.361658621 | 6.275561803 | 1.912682944 | ENSG00000273486 |
| TM4SF18 | -1.676321961 | 0.312879283 | 4.36E-07 | 5.21E-06 | 76.52726328 | 23.81720685 | ENSG00000163762 |
| H3C8 | -1.677245557 | 0.312679045 | 9.28E-05 | 0.000652281 | 50.21632532 | 15.87015281 | ENSG00000273983 |
| NA | -1.677319474 | 0.312663025 | 0.070483385 | 0.173860287 | 11.26066138 | 3.563448033 | ENSG00000286747 |
| NA | -1.679860335 | 0.312112851 | 0.155680724 | 0.313970244 | 6.764604804 | 2.138046496 | ENSG00000260949 |
| FAM163A | -1.681441346 | 0.311771002 | 0.001810143 | 0.008715203 | 27.68603547 | 8.66068669 | ENSG00000143340 |
| NA | -1.683591961 | 0.311306594 | 0.107391015 | 0.238970474 | 7.923581707 | 2.516938816 | ENSG00000278341 |
| SLC16A11 | -1.684993587 | 0.311004296 | 0.043688818 | 0.120511204 | 13.61640045 | 4.287377912 | ENSG00000174326 |
| CRB2 | -1.686009818 | 0.310785302 | 1.68E-15 | 6.86E-14 | 311.3046434 | 96.99307624 | ENSG00000148204 |
| TMEM176A | -1.686921714 | 0.310588924 | 0.113891572 | 0.249777115 | 7.870515549 | 2.479396828 | ENSG00000002933 |
| LINC00237 | -1.689283667 | 0.310080849 | 0.068853481 | 0.170971386 | 11.31500532 | 3.540878192 | ENSG00000225127 |
| STEAP3-AS1 | -1.691574657 | 0.309588834 | 0.072903141 | 0.178728416 | 10.02336302 | 3.094387968 | ENSG00000229867 |
| NA | -1.693815847 | 0.309108269 | 0.17049849 | 0.335743626 | 5.594375723 | 1.751668102 | ENSG00000237080 |
| NA | -1.694485792 | 0.308964761 | 0.084578517 | 0.1997933 | 9.739090748 | 2.959192161 | ENSG00000279694 |
| TMEM151B | -1.694593964 | 0.308941596 | 3.07E-26 | 3.20E-24 | 394.8473581 | 122.2466347 | ENSG00000178233 |
| NA | -1.701050085 | 0.307562158 | 0.148872874 | 0.304380567 | 7.859669336 | 2.408000077 | ENSG00000226912 |
| HSPB6 | -1.703572181 | 0.307024953 | 0.093731648 | 0.215691334 | 7.837668644 | 2.419284998 | ENSG00000004776 |
| QRFPR | -1.706642377 | 0.306372269 | 0.002670764 | 0.012081998 | 27.91989484 | 8.525490882 | ENSG00000186867 |
| C1orf229 | -1.707836376 | 0.306118815 | 0.094516205 | 0.217034802 | 7.946957874 | 2.456826987 | ENSG00000221953 |
| LILRB3 | -1.709017521 | 0.305868296 | 0.160738099 | 0.32126819 | 5.666383359 | 1.751668102 | ENSG00000204577 |
| SLC39A1 | -1.716769777 | 0.304229134 | 2.28E-05 | 0.000187649 | 56.80910399 | 17.35522814 | ENSG00000143570 |
| NA | -1.717846462 | 0.304002172 | 0.121819459 | 0.262562215 | 6.65659335 | 1.991565768 | ENSG00000260095 |
| LHX2 | -1.719921962 | 0.303565141 | 0.050601051 | 0.135057587 | 10.15612611 | 3.075616974 | ENSG00000106689 |
| H2BC17 | -1.721127677 | 0.303311546 | 0.072649494 | 0.178248044 | 10.17950228 | 3.07941582 | ENSG00000274641 |
| LAMP5 | -1.723082385 | 0.302900867 | 0.000436835 | 0.002542805 | 34.22061424 | 10.38978495 | ENSG00000125869 |
| NA | -1.72335171 | 0.302844327 | 0.089473846 | 0.208720233 | 7.946957874 | 2.408000077 | ENSG00000272121 |
| PCDHGA1 | -1.723805442 | 0.302749096 | 0.009795133 | 0.03612796 | 19.35922613 | 5.843738377 | ENSG00000204956 |
| SIAH3 | -1.724554467 | 0.302591954 | 0.088194244 | 0.206309745 | 7.876829374 | 2.370458089 | ENSG00000215475 |
| ERICH5 | -1.72509557 | 0.302478484 | 0.000373724 | 0.002222198 | 37.5470431 | 11.36121019 | ENSG00000177459 |
| NA | -1.727857769 | 0.30189991 | 0.059297543 | 0.152535831 | 10.40764918 | 3.192041786 | ENSG00000235100 |
| UPK3B | -1.727907821 | 0.301889436 | 2.52E-07 | 3.15E-06 | 76.28352722 | 23.06365909 | ENSG00000243566 |
| NA | -1.729065076 | 0.301647373 | 0.039497818 | 0.111338792 | 12.55743743 | 3.810831773 | ENSG00000267980 |
| NA | -1.72947331 | 0.301562029 | 0.065934 | 0.165383781 | 9.134346993 | 2.768121403 | ENSG00000271862 |
| BTN1A1 | -1.733571743 | 0.300706562 | 0.05510516 | 0.144062815 | 10.12959303 | 3.00801907 | ENSG00000124557 |
| SEC14L1P1 | -1.736378443 | 0.30012212 | 0.060864231 | 0.155701539 | 10.5466284 | 3.214611628 | ENSG00000213693 |
| H4C11 | -1.740898323 | 0.299183326 | 0.03913342 | 0.110605618 | 13.67903504 | 4.065813207 | ENSG00000197238 |
| ECRG4 | -1.743133031 | 0.298720255 | 0.001610439 | 0.00790463 | 29.75109074 | 8.825500379 | ENSG00000119147 |
| H2BC7 | -1.743560341 | 0.29863179 | 0.010249256 | 0.0375192 | 21.18360455 | 6.406653414 | ENSG00000277224 |
| CABP7 | -1.744417351 | 0.298454446 | 0.09197975 | 0.212881204 | 7.943800961 | 2.340402174 | ENSG00000100314 |
| RSPH4A | -1.745923849 | 0.298142955 | 0.029271031 | 0.087798882 | 12.69641665 | 3.792060779 | ENSG00000111834 |
| SMAD9 | -1.74849946 | 0.297611162 | 0.001163698 | 0.005997771 | 29.09515997 | 8.698228678 | ENSG00000120693 |
| S100A6 | -1.751606592 | 0.296970886 | 0.000465084 | 0.00268867 | 33.46308354 | 9.935808652 | ENSG00000197956 |
| PARP3 | -1.753847423 | 0.296509982 | 0.000222842 | 0.001414923 | 38.58084851 | 11.51137814 | ENSG00000041880 |
| NA | -1.755707005 | 0.296128037 | 0.15762337 | 0.316842236 | 5.941282589 | 1.800495011 | ENSG00000234791 |
| PRSS27 | -1.756118454 | 0.296043595 | 0.002048549 | 0.009693084 | 29.28740074 | 8.754541661 | ENSG00000172382 |
| PLPPR1 | -1.760508392 | 0.295144141 | 8.43E-06 | 7.67E-05 | 67.75460039 | 20.05564002 | ENSG00000148123 |
| PPP2R5A | -1.763049335 | 0.294624777 | 1.90E-55 | 1.07E-52 | 847.3358089 | 249.5929928 | ENSG00000066027 |
| HRH3 | -1.766435176 | 0.293934137 | 6.86E-14 | 2.32E-12 | 168.9095747 | 49.54753468 | ENSG00000101180 |
| SAMD15 | -1.766565331 | 0.29390762 | 0.036959179 | 0.105547214 | 14.27558583 | 4.24234985 | ENSG00000100583 |
| NA | -1.767251325 | 0.293767902 | 0.183701487 | 0.354252768 | 5.691037301 | 1.642729363 | ENSG00000274248 |
| HLA-V | -1.771914242 | 0.292819951 | 0.143078475 | 0.295598758 | 5.763044937 | 1.654014283 | ENSG00000181126 |
| VWA5B2 | -1.774496992 | 0.292296206 | 0.031237521 | 0.092485019 | 11.73086061 | 3.424453379 | ENSG00000145198 |
| MTSS1 | -1.776118412 | 0.291967884 | 4.18E-10 | 8.19E-09 | 104.572931 | 30.47504285 | ENSG00000170873 |

| | | | | | | | |
|---|---|---|---|---|---|---|---|
| NA | -1.778305471 | 0.291525609 | 0.032613449 | 0.095657172 | 12.97121819 | 3.75451879 | ENSG00000274849 |
| NA | -1.779200467 | 0.291344813 | 2.92E-10 | 5.90E-09 | 113.8721889 | 33.30751297 | ENSG00000206417 |
| SMKR1 | -1.781836046 | 0.290813058 | 0.00270862 | 0.012220437 | 23.60070043 | 6.841420684 | ENSG00000240204 |
| NA | -1.784357212 | 0.290305294 | 0.150601409 | 0.306797814 | 5.789578017 | 1.642729363 | ENSG00000236060 |
| ZHX1 | -1.790352762 | 0.289101347 | 1.05E-08 | 1.68E-07 | 92.13878589 | 26.7773867 | ENSG00000165156 |
| BRINP2 | -1.790859556 | 0.288999809 | 0.001009748 | 0.005312138 | 28.53401163 | 8.214196466 | ENSG00000198797 |
| ADAM20P1 | -1.796772448 | 0.287817766 | 0.171231353 | 0.336816152 | 5.938027978 | 1.684070198 | ENSG00000259158 |
| NA | -1.797253619 | 0.287721788 | 0.131433151 | 0.277658955 | 6.103637977 | 1.781724017 | ENSG00000268279 |
| CLEC4G | -1.797346191 | 0.287703327 | 0.004217537 | 0.017823251 | 23.62860899 | 6.758739013 | ENSG00000182566 |
| NA | -1.798302622 | 0.287512658 | 0.052029494 | 0.137641258 | 9.478096946 | 2.708009574 | ENSG00000241627 |
| NA | -1.803650195 | 0.28644892 | 0.11985346 | 0.259383936 | 5.994348747 | 1.721612187 | ENSG00000226445 |
| CDK5R2 | -1.806275164 | 0.285928203 | 0.033608313 | 0.098047611 | 14.23632741 | 4.013187451 | ENSG00000171450 |
| TRIM31 | -1.808643911 | 0.285459126 | 0.133991937 | 0.28143548 | 7.01928478 | 1.961509853 | ENSG00000204616 |
| NA | -1.81001553 | 0.285187859 | 0.117052493 | 0.255049483 | 6.097324152 | 1.751668102 | ENSG00000286650 |
| TNFRSF11A | -1.812996349 | 0.284599227 | 5.42E-19 | 3.17E-17 | 213.2805049 | 60.57119339 | ENSG00000141655 |
| WDR97 | -1.817969877 | 0.283619793 | 0.019887529 | 0.064248818 | 15.69239963 | 4.452191601 | ENSG00000179698 |
| NA | -1.828230684 | 0.281609775 | 0.005072407 | 0.020835991 | 19.69360304 | 5.59255579 | ENSG00000270354 |
| NA | -1.83073251 | 0.281121849 | 0.004024194 | 0.017098671 | 26.25165568 | 7.268590261 | ENSG00000268798 |
| H2BC21 | -1.835377546 | 0.280218178 | 1.31E-07 | 1.71E-06 | 77.30087165 | 21.69457054 | ENSG00000184678 |
| ZGLP1 | -1.839530635 | 0.279412674 | 0.032063358 | 0.094443194 | 14.78998184 | 4.144696031 | ENSG00000220201 |
| UGT3A1 | -1.840496585 | 0.279225657 | 2.72E-06 | 2.78E-05 | 64.88409718 | 17.98844062 | ENSG00000145626 |
| NPTX1 | -1.844092137 | 0.278530624 | 7.54E-09 | 1.24E-07 | 107.5729578 | 29.927538 | ENSG00000171246 |
| WASHC5 | -1.852266833 | 0.276956858 | 5.08E-66 | 4.28E-63 | 899.4196271 | 248.9772288 | ENSG00000164961 |
| SLC17A6 | -1.862004438 | 0.275093806 | 0.037446242 | 0.106691568 | 11.4622752 | 3.203326707 | ENSG00000091664 |
| KCNK13 | -1.86253337 | 0.274992968 | 0.021262744 | 0.068015826 | 15.98614264 | 4.339454016 | ENSG00000152315 |
| PCDHB10 | -1.873215341 | 0.27296439 | 1.99E-06 | 2.10E-05 | 62.0887207 | 17.02136388 | ENSG00000120324 |
| NA | -1.877755589 | 0.272106705 | 0.028605259 | 0.08631528 | 11.37763991 | 3.105672888 | ENSG00000273008 |
| SCT | -1.878617913 | 0.271944111 | 0.007649537 | 0.029518324 | 17.76564786 | 4.849854916 | ENSG00000070031 |
| GPR155 | -1.882751242 | 0.271166104 | 0.000172179 | 0.001124046 | 41.16601225 | 11.01562298 | ENSG00000163328 |
| NA | -1.884243503 | 0.270885767 | 0.090506946 | 0.210444829 | 7.54255016 | 2.032906604 | ENSG00000224080 |
| H2AC13 | -1.886103929 | 0.270536671 | 0.016925719 | 0.056279382 | 13.98678118 | 3.784574705 | ENSG00000196747 |
| C3orf70 | -1.890646971 | 0.269686092 | 5.89E-07 | 6.85E-06 | 59.91625533 | 16.17721034 | ENSG00000187068 |
| AMH | -1.895879956 | 0.268709651 | 1.41E-22 | 1.11E-20 | 290.7242117 | 78.19692476 | ENSG00000104899 |
| HESX1 | -1.90560513 | 0.266904377 | 2.24E-11 | 5.36E-10 | 132.1193859 | 35.40433025 | ENSG00000163666 |
| RDX | -1.914207585 | 0.265317624 | 1.60E-103 | 4.24E-100 | 1630.292521 | 432.1162104 | ENSG00000137710 |
| NA | -1.917277853 | 0.264753589 | 0.102581259 | 0.23090127 | 6.603527191 | 1.781724017 | ENSG00000267199 |
| B4GALNT1 | -1.918425962 | 0.26454298 | 4.84E-05 | 0.000366548 | 41.32275287 | 10.86589306 | ENSG00000135454 |
| ACKR3 | -1.919368189 | 0.264370263 | 0.00278915 | 0.012514243 | 20.89174068 | 5.555013801 | ENSG00000144476 |
| NA | -1.921997697 | 0.263888851 | 0.023043356 | 0.072510283 | 11.81361676 | 3.135728803 | ENSG00000258748 |
| GNA15 | -1.924237204 | 0.263479532 | 0.080189637 | 0.192094447 | 7.929895533 | 2.111789428 | ENSG00000060558 |
| PTENP1-AS | -1.925496866 | 0.26324958 | 0.112596084 | 0.24769802 | 6.334941787 | 1.623958368 | ENSG00000281128 |
| RHBDL3 | -1.935956003 | 0.261347995 | 3.98E-34 | 7.16E-32 | 525.0728158 | 137.2719811 | ENSG00000141314 |
| CRHR2 | -1.941316678 | 0.260378697 | 0.001622733 | 0.007950406 | 28.09181866 | 7.396849648 | ENSG00000106113 |
| LHFPL3 | -1.946874705 | 0.25937751 | 0.09496916 | 0.217799628 | 6.620589532 | 1.721612187 | ENSG00000187416 |
| H4C8 | -1.967345847 | 0.255723057 | 0.000123118 | 0.000838613 | 38.36227005 | 9.687875259 | ENSG00000158406 |
| NA | -1.969534339 | 0.255335433 | 0.012647333 | 0.044691263 | 18.9890408 | 4.872424757 | ENSG00000272931 |
| LINC00574 | -1.970544199 | 0.255156766 | 0.0902164 | 0.209874578 | 6.669221001 | 1.684070198 | ENSG00000231690 |
| NA | -1.973463054 | 0.254641055 | 0.051658738 | 0.137032066 | 9.636017645 | 2.486882902 | ENSG00000270344 |
| KCNIP3 | -1.974050804 | 0.254537336 | 2.64E-08 | 3.92E-07 | 80.58596754 | 20.66738197 | ENSG00000115041 |
| SLC9A2 | -1.988646807 | 0.25197512 | 0.156135563 | 0.314602042 | 5.664504222 | 1.4704296 | ENSG00000115616 |
| H2BC12 | -1.989298288 | 0.251861361 | 1.01E-26 | 1.09E-24 | 342.22794 | 85.97090615 | ENSG00000197903 |
| CLEC14A | -1.995388506 | 0.25080039 | 0.076210242 | 0.184758747 | 6.940963318 | 1.770439096 | ENSG00000176435 |
| TMSB15B | -1.996062558 | 0.250683239 | 0.038362796 | 0.108767947 | 9.498316199 | 2.359173168 | ENSG00000269226 |
| SCUBE2 | -2.003493432 | 0.249395367 | 0.012981021 | 0.045523225 | 13.81170044 | 3.4357383 | ENSG00000175356 |
| NA | -2.004095377 | 0.249291332 | 0.074923961 | 0.182308087 | 6.994029477 | 1.770439096 | ENSG00000236643 |
| CDHR4 | -2.005364451 | 0.249072138 | 0.05155262 | 0.136868109 | 8.379777803 | 2.100504507 | ENSG00000187492 |
| PDYN | -2.011420208 | 0.248028841 | 0.000179844 | 0.001170378 | 37.02780645 | 9.140482023 | ENSG00000101327 |
| NA | -2.012949631 | 0.247766041 | 0.077072207 | 0.186336152 | 6.884740247 | 1.702841192 | ENSG00000272599 |
| NA | -2.017453919 | 0.246993688 | 0.085354897 | 0.201114627 | 6.944120231 | 1.714126113 | ENSG00000271918 |
| SMIM27 | -2.023270286 | 0.245999914 | 0.011090222 | 0.040137897 | 14.08975658 | 3.484565209 | ENSG00000235453 |
| CLIC2 | -2.028786104 | 0.245061184 | 0.078155322 | 0.188414509 | 6.947277144 | 1.691556272 | ENSG00000155962 |
| MT1M | -2.033358173 | 0.244285786 | 0.002224802 | 0.010394373 | 21.17601295 | 5.168635407 | ENSG00000205364 |
| LINC01465 | -2.040093921 | 0.243147907 | 0.037635581 | 0.107082877 | 9.739090748 | 2.340402174 | ENSG00000221949 |
| HAS1 | -2.042493657 | 0.242743798 | 3.68E-05 | 0.000288085 | 36.71552793 | 8.900584356 | ENSG00000105509 |
| H2BC9 | -2.044616318 | 0.242386908 | 3.48E-09 | 5.94E-08 | 85.29595733 | 20.76079891 | ENSG00000275713 |
| NA | -2.046320018 | 0.242100839 | 0.121109953 | 0.26144373 | 5.872334167 | 1.451658605 | ENSG00000271855 |
| ABCA5 | -2.051698047 | 0.241200023 | 2.24E-06 | 2.33E-05 | 48.52589007 | 11.69876167 | ENSG00000154265 |
| DGKI | -2.061686767 | 0.239535806 | 0.003203644 | 0.014073558 | 22.42721674 | 5.258253499 | ENSG00000157680 |
| NA | -2.068238202 | 0.238450514 | 0.073519553 | 0.179851124 | 7.304932525 | 1.762953022 | ENSG00000237410 |
| DRD5 | -2.069635523 | 0.238219675 | 0.072061994 | 0.177087908 | 7.24555254 | 1.740383181 | ENSG00000169676 |
| ZBTB16 | -2.073133804 | 0.237642734 | 0.042428369 | 0.11786082 | 8.723527756 | 2.070448592 | ENSG00000109906 |
| LHB | -2.082516168 | 0.236102272 | 0.105080339 | 0.235411458 | 5.928654937 | 1.432887611 | ENSG00000104826 |
| NA | -2.082573165 | 0.236092945 | 0.002842384 | 0.012715603 | 18.74826625 | 4.380794851 | ENSG00000271851 |
| CHRDL2 | -2.083222691 | 0.235986676 | 0.011499612 | 0.041296998 | 14.25852349 | 3.296743646 | ENSG00000054938 |

| | | | | | | | |
|---|---|---|---|---|---|---|---|
| ELMOD1 | -2.084923696 | 0.2357086 | 0.000129245 | 0.000873604 | 30.52460141 | 7.167687249 | ENSG00000110675 |
| HPDL | -2.097937675 | 0.233591928 | 1.66E-33 | 2.79E-31 | 538.9619054 | 125.7826232 | ENSG00000186603 |
| TTBK1 | -2.098888032 | 0.233438103 | 3.36E-08 | 4.89E-07 | 75.7135095 | 17.51949218 | ENSG00000146216 |
| LOC101927124 | -2.104980057 | 0.232454449 | 0.099986447 | 0.226727797 | 5.905181073 | 1.354004787 | ENSG00000250365 |
| ACTL8 | -2.105087232 | 0.232437181 | 4.11E-05 | 0.000316944 | 38.65611076 | 9.043266237 | ENSG00000117148 |
| NA | -2.106536269 | 0.232203839 | 0.028847385 | 0.08680913 | 10.44996682 | 2.449340913 | ENSG00000284634 |
| DPF1 | -2.123001251 | 0.229568842 | 1.87E-34 | 3.47E-32 | 351.5176295 | 80.66068817 | ENSG00000011332 |
| FOSB | -2.125103117 | 0.229234626 | 7.60E-05 | 0.000544408 | 33.19577591 | 7.674289303 | ENSG00000125740 |
| ENTPD3 | -2.126320868 | 0.229041215 | 0.079067094 | 0.190044032 | 6.017724914 | 1.384060702 | ENSG00000168032 |
| NA | -2.127181671 | 0.228904595 | 0.067417166 | 0.168193533 | 7.347250168 | 1.642729363 | ENSG00000281641 |
| C1QTNF3 | -2.127546368 | 0.228846738 | 0.056218019 | 0.146340557 | 7.648682477 | 1.781724017 | ENSG00000082196 |
| SLC38A5 | -2.12913807 | 0.228594394 | 1.47E-85 | 2.48E-82 | 1110.689398 | 254.440801 | ENSG00000017483 |
| NA | -2.130507887 | 0.22837745 | 0.002193023 | 0.010287394 | 19.71825699 | 4.538560499 | ENSG00000239881 |
| NA | -2.141115431 | 0.226704443 | 0.052987528 | 0.139796562 | 7.529922509 | 1.702841192 | ENSG00000261572 |
| ANKRD63 | -2.144361318 | 0.226194959 | 0.044675898 | 0.122506745 | 8.889137755 | 1.961509853 | ENSG00000230778 |
| NA | -2.153135542 | 0.224823455 | 0.050126433 | 0.134041742 | 7.609521747 | 1.702841192 | ENSG00000281468 |
| LOC105376748 | -2.15533115 | 0.224481561 | 0.037107086 | 0.105855696 | 9.237322397 | 2.081733513 | ENSG00000233654 |
| FLJ38576 | -2.155412953 | 0.224468833 | 0.004896533 | 0.020203106 | 16.72996331 | 3.735747796 | ENSG00000279943 |
| ZNF594 | -2.175267132 | 0.221400881 | 2.25E-18 | 1.23E-16 | 200.3170737 | 44.6615635 | ENSG00000180626 |
| NA | -2.176664434 | 0.22118655 | 0.036630349 | 0.104834295 | 9.52041459 | 2.149331417 | ENSG00000272468 |
| NA | -2.182015775 | 0.22036763 | 0.031929844 | 0.094158747 | 9.554539271 | 2.149331417 | ENSG00000240093 |
| RGS8 | -2.182119473 | 0.220351791 | 0.044461801 | 0.122190841 | 9.101500088 | 1.961509853 | ENSG00000135824 |
| BOC | -2.191230379 | 0.21896461 | 2.01E-21 | 1.44E-19 | 182.3141164 | 39.94982717 | ENSG00000144857 |
| BTBD17 | -2.19697269 | 0.218094805 | 1.62E-37 | 3.58E-35 | 472.5324933 | 103.0343838 | ENSG00000204347 |
| NA | -2.201766613 | 0.217371302 | 0.063782114 | 0.161383985 | 8.187732422 | 1.838037 | ENSG00000248774 |
| H4C9 | -2.211955488 | 0.215841549 | 8.93E-13 | 2.68E-11 | 106.9408917 | 23.17259783 | ENSG00000276180 |
| DRD2 | -2.216279741 | 0.215195566 | 0.008975621 | 0.033618803 | 12.76654515 | 2.738065489 | ENSG00000149295 |
| NA | -2.226379058 | 0.21369439 | 0.044177927 | 0.121609045 | 7.989275517 | 1.691556272 | ENSG00000241168 |
| MAP3K2-DT | -2.227997346 | 0.213454821 | 0.028745431 | 0.086614831 | 11.32447606 | 2.411798924 | ENSG00000236682 |
| NA | -2.239111764 | 0.211816699 | 1.24E-20 | 8.40E-19 | 204.6297589 | 43.09954043 | ENSG00000278630 |
| LRRC3B | -2.273888805 | 0.206771778 | 6.10E-07 | 7.07E-06 | 49.60724457 | 10.21704715 | ENSG00000179796 |
| NA | -2.284740106 | 0.205222369 | 0.036743346 | 0.105063446 | 8.558015454 | 1.800495011 | ENSG00000282787 |
| NA | -2.28502885 | 0.205181299 | 0.000130918 | 0.000883137 | 28.61233309 | 5.873794292 | ENSG00000206341 |
| ABALON | -2.29215979 | 0.204169633 | 0.000931655 | 0.004961867 | 23.55650365 | 4.801028007 | ENSG00000281376 |
| TCEAL2 | -2.297640214 | 0.203395517 | 3.82E-37 | 8.34E-35 | 376.8386431 | 76.29387778 | ENSG00000184905 |
| GSAP | -2.307534954 | 0.202005299 | 0.00635737 | 0.025279444 | 13.66640739 | 2.775607477 | ENSG00000186088 |
| CDH9 | -2.318682723 | 0.20045041 | 0.082362692 | 0.195942672 | 6.906838637 | 1.395345622 | ENSG00000113100 |
| NA | -2.325567678 | 0.199496083 | 0.064239005 | 0.162252074 | 7.098884018 | 1.451658605 | ENSG00000256897 |
| COL15A1 | -2.328697057 | 0.199063821 | 0.047903251 | 0.129401553 | 8.409565493 | 1.642729363 | ENSG00000204291 |
| ARHGAP36 | -2.33376912 | 0.198365203 | 0.000234891 | 0.001483294 | 24.46771577 | 4.906167899 | ENSG00000147256 |
| ATOH8 | -2.338057041 | 0.197776506 | 0.000884943 | 0.004745814 | 23.97985284 | 4.669519427 | ENSG00000168874 |
| SLC1A2 | -2.339733992 | 0.197546749 | 5.13E-06 | 4.94E-05 | 38.18846709 | 7.516523654 | ENSG00000110436 |
| PDE1B | -2.348003977 | 0.196417588 | 1.49E-08 | 2.32E-07 | 62.90454901 | 12.34000988 | ENSG00000123360 |
| TMEM215 | -2.361164276 | 0.194634009 | 0.004998891 | 0.02057962 | 14.24273893 | 2.805663392 | ENSG00000188133 |
| NA | -2.362662208 | 0.194432028 | 0.03337357 | 0.097470178 | 8.820287032 | 1.714126113 | ENSG00000277342 |
| NUP210L | -2.367986009 | 0.193715861 | 0.04906341 | 0.131826796 | 7.219019461 | 1.432887611 | ENSG00000143552 |
| BEX5 | -2.378990197 | 0.19224391 | 8.47E-05 | 0.000600971 | 26.77354558 | 5.127294571 | ENSG00000184515 |
| NOTO | -2.382504511 | 0.191776186 | 0.025723604 | 0.079196224 | 8.846820112 | 1.672785277 | ENSG00000214513 |
| PHETA2 | -2.383148004 | 0.191690666 | 2.43E-11 | 5.80E-10 | 89.44816626 | 17.11846805 | ENSG00000177096 |
| NA | -2.397965505 | 0.189731943 | 0.001647292 | 0.008047158 | 18.29394929 | 3.522107198 | ENSG00000284636 |
| PITX3 | -2.409296365 | 0.188247634 | 0.011470492 | 0.041232369 | 12.82276822 | 2.411798924 | ENSG00000107859 |
| NA | -2.424353865 | 0.186293099 | 0.004325043 | 0.018189324 | 14.74135037 | 2.756836483 | ENSG00000250046 |
| TCEAL5 | -2.432130055 | 0.185291672 | 5.67E-11 | 1.28E-09 | 81.56817997 | 15.10444406 | ENSG00000204065 |
| ORM2 | -2.446318364 | 0.183478336 | 0.000868349 | 0.004667627 | 20.46839149 | 3.735747796 | ENSG00000228278 |
| PCDHGA3 | -2.453063751 | 0.182622477 | 0.01717077 | 0.05693078 | 9.474940033 | 1.751668102 | ENSG00000254245 |
| NRGN | -2.455176017 | 0.182355293 | 0.032175789 | 0.094628066 | 7.595616319 | 1.402831696 | ENSG00000154146 |
| TCHH | -2.471156634 | 0.180346504 | 0.024817601 | 0.076986674 | 9.670240025 | 1.781724017 | ENSG00000159450 |
| NA | -2.475609581 | 0.179790715 | 0.039426943 | 0.11120666 | 7.790818613 | 1.42160269 | ENSG00000279161 |
| NA | -2.501422667 | 0.176602459 | 0.078485176 | 0.188899274 | 6.031630341 | 1.1028222 | ENSG00000269752 |
| NA | -2.513221577 | 0.175164026 | 0.00638146 | 0.025363815 | 13.68219196 | 2.381743009 | ENSG00000261485 |
| CFAP65 | -2.519751263 | 0.17437302 | 0.030660497 | 0.091096769 | 8.048655502 | 1.432887611 | ENSG00000181378 |
| KCNJ10 | -2.526985946 | 0.17350078 | 0.000156817 | 0.00103506 | 25.34420184 | 4.313196947 | ENSG00000177807 |
| SHPK | -2.537000193 | 0.172300621 | 6.51E-17 | 3.05E-15 | 117.7957699 | 20.3255936 | ENSG00000197417 |
| NME5 | -2.565933883 | 0.1688795 | 0.080127719 | 0.191995753 | 6.186491826 | 1.065280211 | ENSG00000112981 |
| SLC2A9 | -2.594120868 | 0.165612002 | 0.046170459 | 0.125638294 | 6.355161041 | 1.084051205 | ENSG00000109667 |
| NA | -2.594844738 | 0.165528927 | 0.05102217 | 0.135922274 | 6.269247977 | 1.053995291 | ENSG00000279696 |
| THBD | -2.597233818 | 0.165255041 | 0.000716514 | 0.003934776 | 18.71098465 | 3.094387968 | ENSG00000178726 |
| CDH7 | -2.600870311 | 0.164839019 | 3.71E-05 | 0.000289954 | 27.012441 | 4.42962176 | ENSG00000081138 |
| NA | -2.644278215 | 0.159953203 | 0.004571495 | 0.019048581 | 12.90552438 | 2.051677598 | ENSG00000213642 |
| ANKS4B | -2.654815021 | 0.158789231 | 0.043790533 | 0.120686057 | 6.623746445 | 1.084051205 | ENSG00000175311 |
| NA | -2.666590053 | 0.157498495 | 0.022120488 | 0.070226298 | 8.845542335 | 1.42160269 | ENSG00000272764 |
| SLC34A3 | -2.673215839 | 0.156776819 | 0.037722859 | 0.107248883 | 6.719130247 | 1.084051205 | ENSG00000198569 |
| NA | -2.679760067 | 0.156067272 | 0.018336134 | 0.059947418 | 8.935890088 | 1.42160269 | ENSG00000235790 |
| PTGER2 | -2.681745898 | 0.155852597 | 5.09E-08 | 7.20E-07 | 50.8678214 | 7.948041731 | ENSG00000125384 |

| | | | | | | | |
|---|---|---|---|---|---|---|---|
| LOC105371050 | -2.713674923 | 0.152441232 | 0.035699467 | 0.102853088 | 6.798729485 | 1.035224296 | ENSG00000260874 |
| IGFLR1 | -2.722669956 | 0.151493736 | 0.043755147 | 0.12060647 | 7.017405643 | 1.1028222 | ENSG00000126246 |
| H2AC6 | -2.766972356 | 0.146912356 | 6.21E-54 | 3.26E-51 | 512.5062984 | 75.64332002 | ENSG00000180573 |
| GRIK1 | -2.777462683 | 0.145847981 | 0.013796671 | 0.047778827 | 9.246793136 | 1.305177878 | ENSG00000171189 |
| PLEK2 | -2.810489407 | 0.1425471 | 0.0121349 | 0.043193438 | 9.643609248 | 1.365289707 | ENSG00000100558 |
| TRPC3 | -2.871361542 | 0.136657681 | 0.003525296 | 0.015269419 | 12.82908205 | 1.800495011 | ENSG00000138741 |
| SEMA6D | -2.91055014 | 0.132995548 | 2.34E-08 | 3.52E-07 | 50.85519375 | 6.612258286 | ENSG00000137872 |
| SYT11 | -2.91160448 | 0.132898389 | 8.98E-35 | 1.68E-32 | 273.9300051 | 36.37520583 | ENSG00000132718 |
| ZNF229 | -2.969960428 | 0.127630016 | 9.13E-12 | 2.32E-10 | 72.45287215 | 9.215566 | ENSG00000278318 |
| EDN3 | -2.994135352 | 0.125509167 | 0.021413407 | 0.068426912 | 8.426627834 | 1.1028222 | ENSG00000124205 |
| PCDHB5 | -3.046345914 | 0.121048247 | 5.09E-39 | 1.28E-36 | 249.8048744 | 30.16374844 | ENSG00000113209 |
| LUARIS | -3.071086727 | 0.118990085 | 0.013139372 | 0.045982841 | 8.88282393 | 1.1028222 | ENSG00000231638 |
| FRZB | -3.110533167 | 0.115780712 | 1.59E-27 | 1.84E-25 | 163.5777788 | 18.96410274 | ENSG00000162998 |
| GSC2 | -3.210236522 | 0.108049438 | 0.029262148 | 0.087786445 | 6.408227199 | 0.697672811 | ENSG00000063515 |
| H3C10 | -3.221319231 | 0.107222588 | 1.92E-09 | 3.42E-08 | 48.02294164 | 5.134780645 | ENSG00000278828 |
| NA | -3.23849761 | 0.105953444 | 0.00798774 | 0.030587765 | 9.779529255 | 1.046509217 | ENSG00000286478 |
| CRHR1 | -3.306516424 | 0.101073982 | 0.024282065 | 0.075605105 | 6.636374096 | 0.637560981 | ENSG00000120088 |
| LRRTM3 | -3.34394387 | 0.098485567 | 7.66E-05 | 0.000547826 | 21.23361149 | 2.149331417 | ENSG00000198739 |
| MTTP | -3.3847875 | 0.095736475 | 0.016990185 | 0.056453226 | 7.33777943 | 0.7352148 | ENSG00000138823 |
| SSTR3 | -3.405688325 | 0.094359506 | 0.001023676 | 0.005379486 | 14.4644743 | 1.335233792 | ENSG00000278195 |
| MYZAP | -3.41575661 | 0.093703282 | 0.016696122 | 0.055725901 | 7.26765093 | 0.667616896 | ENSG00000263155 |
| GABRA4 | -3.423256757 | 0.09321741 | 0.01807056 | 0.059256859 | 7.38137485 | 0.686387891 | ENSG00000109158 |
| SPEF1 | -3.455666967 | 0.091146624 | 0.00046015 | 0.002664301 | 15.15522882 | 1.402831696 | ENSG00000101222 |
| GNG8 | -3.461671498 | 0.090768058 | 0.013566651 | 0.047108777 | 7.556455588 | 0.686387891 | ENSG00000167414 |
| NA | -3.506532403 | 0.087989037 | 0.000398937 | 0.002346524 | 15.62533034 | 1.372775781 | ENSG00000286340 |
| PAPLN | -3.6631207 | 0.078938849 | 5.89E-30 | 8.34E-28 | 166.1684822 | 13.20673326 | ENSG00000100767 |
| POU3F4 | -3.788802617 | 0.072353037 | 1.26E-05 | 0.000110697 | 23.46172121 | 1.642729363 | ENSG00000196767 |
| TRIM4 | -3.913991952 | 0.066339321 | 4.93E-08 | 6.98E-07 | 36.18910563 | 2.381743009 | ENSG00000146833 |
| NA | -3.944991333 | 0.064929084 | 0.002730955 | 0.01229726 | 10.67623458 | 0.716443806 | ENSG00000277511 |
| DMRT2 | -4.133813496 | 0.056963694 | 0.000105607 | 0.000731432 | 18.0360147 | 1.005168381 | ENSG00000173253 |
| SLC12A1 | -4.286022191 | 0.051260018 | 0.006537028 | 0.025873688 | 7.205114033 | 0.348836406 | ENSG00000074803 |
| ORM1 | -4.293187271 | 0.051006069 | 0.000792238 | 0.004303438 | 13.26643437 | 0.637560981 | ENSG00000229314 |
| HTR1A | -4.306845795 | 0.050525455 | 6.17E-08 | 8.56E-07 | 34.09869736 | 1.732897107 | ENSG00000178394 |
| TAF9B | -4.710876231 | 0.03818431 | 2.03E-73 | 1.98E-70 | 422.0349927 | 15.9143048 | ENSG00000187325 |
| NA | -6.111981425 | 0.014458068 | 1.65E-05 | 0.000140164 | 13.16651818 | 0 | ENSG00000270325 |
| TNFAIP8L3 | -6.297546956 | 0.012713042 | 1.94E-05 | 0.000161822 | 14.91327419 | 0 | ENSG00000183578 |
| ZFP28 | -6.319457209 | 0.012521427 | 5.91E-06 | 5.63E-05 | 15.16597733 | 0 | ENSG00000196867 |
