## Supplementary material for "Chemokine Receptor 1 and its associated immune pathway are downregulated in SF3B1^MT^ blood and non-blood cancers": Table_S4

| Table S4. Significantly dysregulated genes expressed in 3 biological replicates of SF3B1$^{K700E}$ compared to SF3B1$^{WT}$ ES cells (K700E ES line 1). Genes with fold change (FC) ≥1.5 and p-value < 0.05 are shown. Up- and downregulated genes begin in rows 4 and 1619, respectively. | | | | | | | |
|---|---|---|---|---|---|---|---|
| **Gene Symbol** | **log2 FC** | **FC** | **p-value** | **padj** | **WT Basemean** | **MT Basemean** | **Ensembl ID** |
| HORMAD1 | 7.709351052 | 209.2887643 | 2.44E-10 | 3.22E-09 | 0.323808629 | 75.24437789 | ENSG00000143452 |
| KHDC3L | 7.238165959 | 150.97501 | 1.96E-08 | 1.99E-07 | 0 | 27.85557538 | ENSG00000203908 |
| PRAM1 | 6.507107875 | 90.95669163 | 1.44E-06 | 1.10E-05 | 0 | 16.79604861 | ENSG00000133246 |
| MAGEA4 | 6.464545728 | 88.31249863 | 1.23E-67 | 3.48E-65 | 5.655543462 | 501.7202351 | ENSG00000147381 |
| MEG3 | 6.200776765 | 73.5562877 | 1.64E-06 | 1.24E-05 | 0.326599532 | 26.41062234 | ENSG00000214548 |
| CRYGD | 6.002812812 | 64.12490206 | 2.24E-05 | 1.38E-04 | 0 | 11.83819395 | ENSG00000118231 |
| NPAP1 | 5.970279957 | 62.69506392 | 5.17E-06 | 3.58E-05 | 0.353832839 | 22.51295481 | ENSG00000185823 |
| TDRD1 | 5.888482557 | 59.23929461 | 1.33E-10 | 1.81E-09 | 0.979798595 | 59.25309434 | ENSG00000095627 |
| CD48 | 5.71803415 | 52.63805037 | 2.07E-04 | 1.04E-03 | 0 | 9.745504575 | ENSG00000117091 |
| DMRTB1 | 5.68093887 | 51.30184727 | 2.00E-05 | 1.25E-04 | 0.353832839 | 18.45368483 | ENSG00000143006 |
| IL6 | 5.639623035 | 49.85350505 | 1.38E-04 | 7.23E-04 | 0 | 9.202643856 | ENSG00000136244 |
| CYTOR | 5.567756627 | 47.43094235 | 8.13E-07 | 6.50E-06 | 0.677641468 | 31.78737952 | ENSG00000222041 |
| MOV10L1 | 5.457889884 | 43.95300455 | 4.21E-04 | 1.97E-03 | 0 | 8.094059608 | ENSG00000073146 |
| LCN1 | 5.350583359 | 40.80243534 | 5.21E-04 | 2.38E-03 | 0 | 7.526185506 | ENSG00000160349 |
| CPA4 | 5.332015031 | 40.28064923 | 1.37E-03 | 5.67E-03 | 0 | 7.465145408 | ENSG00000128510 |
| MAS1 | 5.291909685 | 39.17631199 | 3.63E-06 | 2.58E-05 | 0.677641468 | 26.2108426 | ENSG00000130368 |
| TH | 5.282015601 | 38.90855781 | 7.44E-04 | 3.26E-03 | 0 | 7.172612117 | ENSG00000180176 |
| LGALS12 | 5.25966229 | 38.31034971 | 1.88E-08 | 1.93E-07 | 1.007031902 | 38.43003436 | ENSG00000133317 |
| KRT80 | 5.250022216 | 38.05521369 | 1.48E-03 | 6.07E-03 | 0 | 7.044352774 | ENSG00000167767 |
| DPEP3 | 5.23372785 | 37.62782122 | 7.70E-108 | 7.82E-105 | 16.04062268 | 603.5524554 | ENSG00000141096 |
| C1GALT1C1L | 5.212732984 | 37.08420623 | 9.79E-04 | 4.18E-03 | 0 | 6.834092744 | ENSG00000223658 |
| TCL1A | 5.159910435 | 35.75096886 | 1.21E-03 | 5.07E-03 | 0 | 6.604738015 | ENSG00000100721 |
| FOXC2 | 5.092767788 | 34.12525185 | 1.60E-03 | 6.48E-03 | 0 | 6.310314614 | ENSG00000176692 |
| C11orf16 | 5.03551052 | 32.79742217 | 3.54E-04 | 1.69E-03 | 0.323808629 | 11.78334511 | ENSG00000176029 |
| OR2W3 | 4.668332813 | 25.42776602 | 1.47E-03 | 6.03E-03 | 0.323808629 | 9.12897289 | ENSG00000238243 |
| PLA2G2C | 4.603287774 | 24.30679503 | 2.79E-03 | 1.05E-02 | 0.353832839 | 8.773509391 | ENSG00000187980 |
| NECTIN4 | 4.571522879 | 23.77746289 | 6.40E-11 | 8.96E-10 | 2.011272902 | 47.72087016 | ENSG00000143217 |
| TBX3 | 4.518753338 | 22.92346678 | 2.44E-04 | 1.21E-03 | 0.65040816 | 15.28575431 | ENSG00000135111 |
| PRAME | 4.502780945 | 22.67107577 | 3.11E-04 | 1.50E-03 | 0.707665678 | 15.35942528 | ENSG00000185686 |
| TCL1B | 4.402024718 | 21.14177672 | 1.21E-09 | 1.45E-08 | 2.032924404 | 42.46241295 | ENSG00000213231 |
| PRSS21 | 4.384836363 | 20.89138686 | 2.46E-08 | 2.46E-07 | 1.739139985 | 35.16961075 | ENSG00000007038 |
| C5orf63 | 4.37132788 | 20.6966861 | 5.32E-03 | 1.85E-02 | 0.323808629 | 7.418638398 | ENSG00000164241 |
| ICAM2 | 4.329727043 | 20.10840914 | 4.51E-03 | 1.60E-02 | 0.353832839 | 7.229611531 | ENSG00000108622 |
| CALML5 | 4.229934169 | 18.76450293 | 9.44E-03 | 3.00E-02 | 0.353832839 | 6.777353794 | ENSG00000178372 |
| EBF2 | 4.22617039 | 18.7156129 | 9.30E-04 | 3.99E-03 | 0.647617257 | 12.4227396 | ENSG00000221818 |
| PF4 | 4.221940043 | 18.66081438 | 3.19E-06 | 2.29E-05 | 4.000894302 | 74.7345877 | ENSG00000163737 |
| SERPINB5 | 4.18711038 | 18.21569809 | 7.78E-03 | 2.55E-02 | 0.323808629 | 6.543970491 | ENSG00000206075 |
| DPPA5 | 4.17031804 | 18.00490447 | 3.54E-34 | 2.77E-32 | 10.27056419 | 186.0106071 | ENSG00000203909 |
| PCDHGB5 | 4.138211679 | 17.60864118 | 8.90E-22 | 3.43E-20 | 6.074330806 | 106.0931174 | ENSG00000276547 |
| CH25H | 4.131105927 | 17.52212601 | 8.70E-03 | 2.80E-02 | 0.323808629 | 6.312465188 | ENSG00000138135 |
| F7 | 4.086801334 | 16.99220691 | 8.92E-05 | 4.87E-04 | 1.007031902 | 17.04637604 | ENSG00000057593 |
| GDF15 | 4.063999448 | 16.72575521 | 4.51E-133 | 7.91E-130 | 43.62260166 | 731.6650691 | ENSG00000130513 |
| SDR42E1 | 4.051002407 | 16.57575188 | 1.09E-02 | 3.39E-02 | 0.326599532 | 5.969644667 | ENSG00000184860 |
| CHI3L1 | 4.035098312 | 16.3940263 | 1.16E-02 | 3.56E-02 | 0.326599532 | 5.88952198 | ENSG00000133048 |
| AGBL4 | 4.016627476 | 16.18547136 | 1.76E-04 | 8.95E-04 | 1.004240999 | 16.26366818 | ENSG00000186094 |
| ANXA1 | 3.95626295 | 15.52221943 | 4.33E-09 | 4.87E-08 | 2.628865949 | 41.38285855 | ENSG00000135046 |
| NPIPA5 | 3.952946233 | 15.48657526 | 2.15E-04 | 1.07E-03 | 0.974216789 | 15.43068521 | ENSG00000183793 |
| IZUMO2 | 3.92758043 | 15.21666641 | 3.67E-07 | 3.11E-06 | 1.986830497 | 30.42820738 | ENSG00000161652 |
| TRIM58 | 3.881638408 | 14.7397322 | 9.51E-37 | 8.74E-35 | 12.41309754 | 182.3874626 | ENSG00000162722 |
| AADACL3 | 3.808707343 | 14.01313013 | 9.48E-06 | 6.28E-05 | 1.709115775 | 23.60082682 | ENSG00000188984 |
| SLC8A3 | 3.761170692 | 13.55892309 | 6.49E-11 | 9.07E-10 | 3.774855292 | 50.16713292 | ENSG00000100678 |
| CYP2B6 | 3.761143052 | 13.55866332 | 6.81E-150 | 1.88E-146 | 61.10925564 | 828.9639954 | ENSG00000197408 |
| BCO1 | 3.681035694 | 12.82632261 | 6.13E-03 | 2.09E-02 | 0.680432371 | 8.628318033 | ENSG00000135697 |
| SIX3 | 3.670681853 | 12.734601 | 8.55E-04 | 3.70E-03 | 1.001450097 | 12.75318972 | ENSG00000138083 |
| TMEM105 | 3.646491493 | 12.52285403 | 7.34E-03 | 2.43E-02 | 0.707665678 | 8.489578395 | ENSG00000185332 |
| MYBPHL | 3.646094208 | 12.519406 | 8.80E-03 | 2.83E-02 | 0.647617257 | 8.315072506 | ENSG00000221986 |
| COL8A1 | 3.645819131 | 12.51701916 | 1.55E-08 | 1.61E-07 | 3.015513901 | 37.72744919 | ENSG00000144810 |
| ADTRP | 3.644992323 | 12.50984772 | 8.25E-03 | 2.68E-02 | 0.647617257 | 8.310771359 | ENSG00000111863 |
| CER1 | 3.641423747 | 12.47894225 | 3.34E-17 | 8.95E-16 | 6.887938652 | 87.08284441 | ENSG00000147869 |
| KRT81 | 3.611101273 | 12.21939772 | 1.25E-04 | 6.58E-04 | 2.008481999 | 24.58061869 | ENSG00000205426 |
| SCG2 | 3.608959547 | 12.2012711 | 1.35E-10 | 1.83E-09 | 4.079803322 | 49.13652079 | ENSG00000171951 |
| SPDYE2 | 3.560266724 | 11.79633443 | 9.23E-03 | 2.94E-02 | 0.65040816 | 7.818458333 | ENSG00000205238 |
| VAV1 | 3.549641049 | 11.70977174 | 1.06E-57 | 2.27E-55 | 28.08122695 | 326.0017978 | ENSG00000141968 |
| PRPH2 | 3.539320095 | 11.62629968 | 7.42E-06 | 5.01E-05 | 2.035715307 | 23.42606047 | ENSG00000112619 |
| MPO | 3.524388994 | 11.50659437 | 1.21E-02 | 3.68E-02 | 0.707665678 | 7.809856038 | ENSG00000005381 |
| CELP | 3.49696534 | 11.28993551 | 1.15E-02 | 3.55E-02 | 0.677641468 | 7.578883773 | ENSG00000170827 |
| FOXL1 | 3.496275548 | 11.28453877 | 2.05E-03 | 8.06E-03 | 0.974216789 | 11.24289543 | ENSG00000176678 |
| PITX2 | 3.476115717 | 11.1279483 | 6.00E-10 | 7.50E-09 | 4.324702931 | 48.28444311 | ENSG00000164093 |
| SP5 | 3.469673202 | 11.078366 | 2.57E-03 | 9.81E-03 | 1.007031902 | 11.11222505 | ENSG00000204335 |

| | | | | | | | |
|---|---|---|---|---|---|---|---|
| LCP2 | 3.459119213 | 10.99761829 | 1.53E-02 | 4.51E-02 | 0.65040816 | 7.307583645 | ENSG00000043462 |
| LEFTY2 | 3.44596579 | 10.89780586 | 2.79E-63 | 6.90E-61 | 135.7222926 | 1476.540506 | ENSG00000143768 |
| PKP1 | 3.376722505 | 10.38711072 | 7.80E-21 | 2.80E-19 | 10.04240999 | 104.2090706 | ENSG00000081277 |
| KIF25 | 3.368397745 | 10.32734674 | 3.44E-05 | 2.05E-04 | 2.038506209 | 20.83460635 | ENSG00000125337 |
| TEX11 | 3.351548526 | 10.20743536 | 1.81E-02 | 5.16E-02 | 0.707665678 | 6.951859682 | ENSG00000120498 |
| CTCFL | 3.351414341 | 10.20648601 | 1.52E-16 | 3.84E-15 | 9.621031907 | 96.51052974 | ENSG00000124092 |
| TRDN | 3.334204556 | 10.08545702 | 4.82E-07 | 3.98E-06 | 2.985489691 | 30.28757763 | ENSG00000186439 |
| HCG20 | 3.304152622 | 9.877545741 | 2.29E-02 | 6.28E-02 | 0.653199063 | 6.554190321 | ENSG00000228022 |
| PTPRB | 3.303855818 | 9.875513855 | 6.16E-15 | 1.32E-13 | 7.800191998 | 76.26261962 | ENSG00000127329 |
| RGS4 | 3.280316748 | 9.715691949 | 6.32E-05 | 3.56E-04 | 1.978457788 | 19.41492715 | ENSG00000117152 |
| TMEM82 | 3.26650214 | 9.623102793 | 4.47E-07 | 3.72E-06 | 3.336531627 | 32.13477376 | ENSG00000162460 |
| PLAAT4 | 3.249189433 | 9.508313245 | 9.57E-05 | 5.20E-04 | 1.984039594 | 18.99628509 | ENSG00000133321 |
| FGF17 | 3.237723815 | 9.433046734 | 1.23E-04 | 6.53E-04 | 2.062948614 | 19.09738047 | ENSG00000158815 |
| GIP | 3.233672871 | 9.406596828 | 2.31E-02 | 6.32E-02 | 0.65040816 | 6.238794221 | ENSG00000159224 |
| MCCC2 | 3.230398919 | 9.385274353 | 1.73E-06 | 1.30E-05 | 3.015513901 | 28.25324475 | ENSG00000131844 |
| KRT17 | 3.228281938 | 9.371512697 | 7.49E-10 | 9.23E-09 | 5.32894393 | 50.0872707 | ENSG00000128422 |
| CBY2 | 3.213209356 | 9.274113323 | 8.17E-03 | 2.66E-02 | 0.971425886 | 9.20479443 | ENSG00000174015 |
| ST6GALNAC6 | 3.154415713 | 8.903766217 | 3.00E-02 | 7.84E-02 | 0.65040816 | 5.900274849 | ENSG00000160408 |
| EVPL | 3.15145577 | 8.885517298 | 6.64E-22 | 2.60E-20 | 15.56178692 | 138.9842591 | ENSG00000167880 |
| PIWIL2 | 3.099197064 | 8.56941704 | 1.17E-13 | 2.20E-12 | 8.643148334 | 74.34873729 | ENSG00000197181 |
| LAPTM5 | 3.099163248 | 8.569216181 | 7.97E-05 | 4.40E-04 | 2.30505732 | 19.93440413 | ENSG00000162511 |
| MYCT1 | 3.088252613 | 8.504654411 | 1.02E-02 | 3.21E-02 | 1.03426521 | 8.63261918 | ENSG00000120279 |
| CCDC60 | 3.086961143 | 8.497044632 | 1.37E-02 | 4.10E-02 | 0.977007692 | 8.42235915 | ENSG00000183273 |
| ZNF648 | 3.074770211 | 8.425546177 | 8.25E-04 | 3.58E-03 | 1.681882467 | 14.1034991 | ENSG00000179930 |
| OR52N2 | 3.068403113 | 8.388443349 | 3.67E-02 | 9.27E-02 | 0.677641468 | 5.643495699 | ENSG00000180988 |
| S100A6 | 3.062310864 | 8.353095107 | 3.22E-46 | 4.78E-44 | 35.15872271 | 295.2482229 | ENSG00000197956 |
| MYL7 | 3.043729029 | 8.246197582 | 4.61E-05 | 2.67E-04 | 2.634447755 | 21.92625858 | ENSG00000106631 |
| CECR3 | 3.035189724 | 8.197532541 | 4.25E-02 | 1.05E-01 | 0.680432371 | 5.496153767 | ENSG00000241832 |
| LEFTY1 | 3.030830244 | 8.172798945 | 1.55E-154 | 4.99E-151 | 136.1337463 | 1110.347549 | ENSG00000243709 |
| FAM90A1 | 3.017583944 | 8.098102728 | 4.59E-05 | 2.66E-04 | 2.691705273 | 21.70550615 | ENSG00000171847 |
| ODF3L2 | 3.004576951 | 8.025420309 | 4.40E-03 | 1.57E-02 | 1.360864742 | 10.78230797 | ENSG00000181781 |
| GZMM | 3.003289259 | 8.018260332 | 5.68E-04 | 2.57E-03 | 2.014063805 | 16.10610642 | ENSG00000197540 |
| GSTT2 | 2.991995276 | 7.955735299 | 1.41E-02 | 4.20E-02 | 0.977007692 | 7.917403145 | ENSG00000099984 |
| ALOX15B | 2.989806011 | 7.943671763 | 2.09E-04 | 1.05E-03 | 2.307848223 | 18.52520522 | ENSG00000179593 |
| SLC16A3 | 2.983863439 | 7.911018463 | 1.27E-100 | 1.07E-97 | 117.5315276 | 926.4134256 | ENSG00000141526 |
| SMC1B | 2.981177075 | 7.896301491 | 5.23E-03 | 1.82E-02 | 1.333631434 | 10.55725439 | ENSG00000077935 |
| C1orf87 | 2.965078506 | 7.808678994 | 1.78E-02 | 5.09E-02 | 1.03426521 | 7.92600544 | ENSG00000162598 |
| DUSP27 | 2.940705907 | 7.677868797 | 2.13E-07 | 1.86E-06 | 4.667372158 | 35.91870293 | ENSG00000198842 |
| SYT4 | 2.937606972 | 7.661394296 | 7.35E-34 | 5.72E-32 | 27.97699964 | 214.8768637 | ENSG00000132872 |
| PRDM1 | 2.927750325 | 7.609229233 | 6.36E-05 | 3.59E-04 | 3.100004727 | 23.18218476 | ENSG00000057657 |
| PLA2G4C | 2.925155979 | 7.595558128 | 1.00E-24 | 4.87E-23 | 19.42113304 | 147.3044263 | ENSG00000105499 |
| EPHA3 | 2.89781722 | 7.452979131 | 9.86E-05 | 5.34E-04 | 3.012722998 | 22.38228443 | ENSG00000044524 |
| CDKN1A | 2.881340427 | 7.36834405 | 7.52E-180 | 3.62E-176 | 238.6018974 | 1761.026169 | ENSG00000124762 |
| GALR1 | 2.865794114 | 7.289369911 | 2.65E-04 | 1.30E-03 | 2.716147677 | 19.65691274 | ENSG00000166573 |
| H1-4 | 2.861344684 | 7.266923324 | 2.21E-02 | 6.11E-02 | 1.001450097 | 7.296830776 | ENSG00000168298 |
| SCARA5 | 2.856648154 | 7.243305143 | 2.57E-04 | 1.27E-03 | 2.694496175 | 19.43374927 | ENSG00000168079 |
| SRRM5 | 2.85337481 | 7.226889371 | 6.68E-04 | 2.96E-03 | 2.367896644 | 16.96410277 | ENSG00000226763 |
| SPAAR | 2.832474973 | 7.122950544 | 2.33E-02 | 6.37E-02 | 1.03426521 | 7.221009236 | ENSG00000235387 |
| NTSR2 | 2.812820666 | 7.026570253 | 3.31E-02 | 8.51E-02 | 0.979798595 | 7.000256802 | ENSG00000169006 |
| ITPRID1 | 2.79875885 | 6.958415608 | 4.89E-07 | 4.03E-06 | 5.336640917 | 37.19534134 | ENSG00000180347 |
| MOGAT1 | 2.798452594 | 6.956938629 | 2.61E-02 | 7.01E-02 | 0.974216789 | 6.90776371 | ENSG00000124003 |
| CYTH4 | 2.792264881 | 6.927164244 | 3.01E-02 | 7.87E-02 | 0.974216789 | 6.865818312 | ENSG00000100055 |
| USH2A | 2.78148893 | 6.875615792 | 8.11E-03 | 2.64E-02 | 1.621834046 | 11.3184565 | ENSG00000042781 |
| HCLS1 | 2.739709869 | 6.679359978 | 1.57E-16 | 3.95E-15 | 15.76079278 | 105.0520129 | ENSG00000180353 |
| C14orf39 | 2.734259354 | 6.654172903 | 8.38E-09 | 9.05E-08 | 7.386310738 | 49.02761661 | ENSG00000179008 |
| GRAMD1C | 2.7293319 | 6.631484671 | 7.66E-08 | 7.16E-07 | 6.768717691 | 44.59541808 | ENSG00000178075 |
| ANGPT4 | 2.724800429 | 6.61068801 | 6.49E-07 | 5.26E-06 | 5.698846465 | 37.56370828 | ENSG00000101280 |
| CD163 | 2.721424315 | 6.59523614 | 4.52E-02 | 1.10E-01 | 0.979798595 | 6.546121064 | ENSG00000177575 |
| MEIOB | 2.717584023 | 6.577703705 | 2.59E-34 | 2.04E-32 | 37.02932331 | 242.5791785 | ENSG00000162039 |
| SP140L | 2.680839568 | 6.412289532 | 1.09E-10 | 1.50E-09 | 9.628528734 | 61.97413434 | ENSG00000185404 |
| FAM215A | 2.653274373 | 6.290934637 | 4.55E-02 | 1.11E-01 | 1.001450097 | 6.318916909 | ENSG00000267496 |
| NR1I2 | 2.651859895 | 6.284769767 | 3.40E-20 | 1.18E-18 | 20.05758668 | 126.1017135 | ENSG00000144852 |
| DNAJC22 | 2.645157162 | 6.255638576 | 4.28E-02 | 1.05E-01 | 1.007031902 | 6.293643062 | ENSG00000178401 |
| GDA | 2.628490033 | 6.183784457 | 1.83E-02 | 5.21E-02 | 1.328049628 | 8.255922518 | ENSG00000119125 |
| MUC4 | 2.613954767 | 6.12179514 | 7.13E-16 | 1.69E-14 | 18.63388262 | 114.3595018 | ENSG00000145113 |
| IL1A | 2.612756185 | 6.116711302 | 3.99E-04 | 1.88E-03 | 3.300925611 | 20.33207349 | ENSG00000115008 |
| PTPN7 | 2.60171894 | 6.070094347 | 2.15E-02 | 5.96E-02 | 1.385307146 | 8.234949819 | ENSG00000143851 |
| ASB2 | 2.599200165 | 6.059505926 | 5.48E-04 | 2.49E-03 | 2.985489691 | 18.2033574 | ENSG00000100628 |
| CTSLP2 | 2.599137942 | 6.059244586 | 1.51E-03 | 6.17E-03 | 2.748962791 | 16.38170769 | ENSG00000280913 |
| S100A2 | 2.580854862 | 5.982941109 | 3.82E-03 | 1.39E-02 | 2.800638503 | 16.35805138 | ENSG00000196754 |
| PTPRH | 2.580458169 | 5.981296225 | 2.51E-03 | 9.63E-03 | 2.392339049 | 14.11425197 | ENSG00000080031 |
| ATP1A4 | 2.571304311 | 5.943465218 | 9.75E-05 | 5.29E-04 | 3.95970648 | 23.73553788 | ENSG00000132681 |
| TTC22 | 2.550623473 | 5.8588742 | 1.59E-11 | 2.38E-10 | 13.0662966 | 76.50298768 | ENSG00000006555 |

| | | | | | | | |
|---|---|---|---|---|---|---|---|
| TNFRSF10C | 2.548111886 | 5.84868336 | 4.04E-17 | 1.07E-15 | 18.69393104 | 109.4237044 | ENSG00000173535 |
| DLGAP2 | 2.544437737 | 5.833807318 | 1.13E-11 | 1.72E-10 | 12.70130015 | 74.05943907 | ENSG00000198010 |
| GJC2 | 2.542920056 | 5.827673519 | 9.69E-25 | 4.71E-23 | 28.46141721 | 165.7094414 | ENSG00000198835 |
| GPAT2 | 2.542774811 | 5.827086839 | 4.71E-05 | 2.72E-04 | 4.340772627 | 25.31305142 | ENSG00000186281 |
| CCDC80 | 2.539990112 | 5.81585021 | 2.80E-37 | 2.67E-35 | 52.2524712 | 304.714116 | ENSG00000091986 |
| ARSL | 2.534581524 | 5.794087708 | 1.02E-65 | 2.68E-63 | 89.72549983 | 520.6957959 | ENSG00000157399 |
| STMN2 | 2.534260157 | 5.792797195 | 5.52E-24 | 2.52E-22 | 30.05410293 | 173.0651617 | ENSG00000104435 |
| IQSEC3 | 2.531613022 | 5.782178001 | 5.03E-27 | 2.74E-25 | 31.79025185 | 183.6135534 | ENSG00000120645 |
| CMKLR1 | 2.520336874 | 5.737160482 | 1.36E-15 | 3.13E-14 | 21.47359376 | 122.9101387 | ENSG00000174600 |
| OTOF | 2.509136082 | 5.692790791 | 7.95E-05 | 4.39E-04 | 4.319121125 | 24.68817791 | ENSG00000115155 |
| HAO2 | 2.50721825 | 5.685228167 | 5.20E-03 | 1.81E-02 | 2.367896644 | 13.34848824 | ENSG00000116882 |
| UNC93A | 2.507014364 | 5.684424769 | 1.22E-04 | 6.46E-04 | 4.422472549 | 24.86483437 | ENSG00000112494 |
| NODAL | 2.496970809 | 5.644989154 | 1.78E-36 | 1.60E-34 | 46.58731573 | 262.5781183 | ENSG00000156574 |
| RGN | 2.495114013 | 5.637728539 | 2.65E-05 | 1.62E-04 | 5.456737759 | 30.38196083 | ENSG00000130988 |
| NLRP1 | 2.490439673 | 5.619491828 | 1.79E-07 | 1.58E-06 | 8.082812805 | 45.19045614 | ENSG00000091592 |
| GSTT2B | 2.483989281 | 5.594422764 | 8.47E-17 | 2.19E-15 | 21.21473964 | 118.158243 | ENSG00000133433 |
| CHL1 | 2.46748711 | 5.530795922 | 3.73E-02 | 9.40E-02 | 1.388098049 | 7.50736338 | ENSG00000134121 |
| KLHDC8A | 2.464306491 | 5.518615956 | 1.88E-38 | 1.94E-36 | 51.90422017 | 287.2848457 | ENSG00000162873 |
| PTGDR2 | 2.454461828 | 5.481086215 | 1.73E-02 | 4.97E-02 | 1.65464916 | 9.103699043 | ENSG00000183134 |
| FOXA2 | 2.45427706 | 5.480384292 | 1.60E-05 | 1.01E-04 | 5.677194963 | 31.10954727 | ENSG00000125798 |
| SOX15 | 2.453598122 | 5.477805808 | 3.46E-24 | 1.60E-22 | 31.39802209 | 171.9525489 | ENSG00000129194 |
| HLA-DQA1 | 2.450904366 | 5.467587357 | 2.74E-07 | 2.36E-06 | 8.346573013 | 45.663414 | ENSG00000196735 |
| CHADL | 2.448644217 | 5.459028455 | 3.02E-03 | 1.13E-02 | 2.626075046 | 14.49901789 | ENSG00000100399 |
| SIGLEC15 | 2.440188021 | 5.427124562 | 3.32E-04 | 1.60E-03 | 3.986939787 | 21.72056016 | ENSG00000197046 |
| C6orf223 | 2.439542251 | 5.424695849 | 2.32E-17 | 6.30E-16 | 22.00881116 | 119.6314139 | ENSG00000181577 |
| ADGRD1 | 2.428978891 | 5.385121484 | 7.34E-05 | 4.09E-04 | 5.276592496 | 28.63424255 | ENSG00000111452 |
| MMP9 | 2.418413645 | 5.345828833 | 9.70E-44 | 1.29E-41 | 67.13335019 | 359.0082833 | ENSG00000100985 |
| HHLA1 | 2.415433285 | 5.334796669 | 2.25E-57 | 4.77E-55 | 88.49330479 | 471.337735 | ENSG00000132297 |
| FAT4 | 2.40733059 | 5.304918511 | 2.77E-33 | 2.12E-31 | 48.81278831 | 258.9880381 | ENSG00000196159 |
| TDRD5 | 2.406803219 | 5.302979675 | 7.74E-08 | 7.22E-07 | 10.47515185 | 55.30431554 | ENSG00000162782 |
| VSIR | 2.388513001 | 5.236173862 | 8.91E-04 | 3.84E-03 | 3.663131159 | 19.18101081 | ENSG00000107738 |
| HR | 2.381113158 | 5.209385343 | 2.58E-02 | 6.96E-02 | 1.956806287 | 10.34672178 | ENSG00000168453 |
| PADI1 | 2.378028378 | 5.198258499 | 8.54E-07 | 6.80E-06 | 8.02834619 | 41.71089763 | ENSG00000142623 |
| VSIG1 | 2.377742716 | 5.197229314 | 7.62E-19 | 2.32E-17 | 25.6363363 | 133.583803 | ENSG00000101842 |
| TRPM2 | 2.367066925 | 5.158912312 | 1.40E-04 | 7.28E-04 | 4.936714172 | 25.71018774 | ENSG00000142185 |
| PCDHGB3 | 2.364034967 | 5.148081762 | 2.25E-04 | 1.12E-03 | 4.340772627 | 22.36587334 | ENSG00000262209 |
| IFIT3 | 2.35951169 | 5.131966277 | 1.01E-03 | 4.32E-03 | 3.630316045 | 18.77768323 | ENSG00000119917 |
| SLC52A1 | 2.352794301 | 5.108126668 | 3.64E-04 | 1.73E-03 | 4.400821048 | 22.31155754 | ENSG00000132517 |
| LPAR5 | 2.35086389 | 5.101296265 | 4.02E-03 | 1.45E-02 | 2.979907885 | 15.31963046 | ENSG00000184574 |
| PCDHGA11 | 2.348611914 | 5.093339606 | 1.34E-24 | 6.39E-23 | 36.6424752 | 187.0054408 | ENSG00000253873 |
| SEMA3C | 2.346941466 | 5.087445613 | 1.59E-04 | 8.19E-04 | 5.018414094 | 25.57170857 | ENSG00000075223 |
| PAPPA2 | 2.324513596 | 5.008968697 | 1.38E-08 | 1.44E-07 | 12.7904969 | 64.69787638 | ENSG00000116183 |
| BIRC7 | 2.322897938 | 5.003362349 | 3.57E-02 | 9.06E-02 | 1.95959719 | 9.924311612 | ENSG00000101197 |
| DENND2D | 2.318582795 | 4.988419505 | 7.16E-08 | 6.72E-07 | 10.00680398 | 49.98832588 | ENSG00000162777 |
| ARID5B | 2.31634748 | 4.980696422 | 6.68E-06 | 4.55E-05 | 7.465219758 | 36.8880145 | ENSG00000150347 |
| MLKL | 2.307481132 | 4.950180503 | 5.07E-03 | 1.78E-02 | 3.591919126 | 17.95922123 | ENSG00000168404 |
| RNASEL | 2.30635339 | 4.946312501 | 4.95E-09 | 5.49E-08 | 12.0809162 | 59.66015949 | ENSG00000135828 |
| BTG2 | 2.304645055 | 4.940458901 | 1.16E-34 | 9.22E-33 | 63.3019131 | 313.2811087 | ENSG00000159388 |
| KCNE3 | 2.304344979 | 4.93943141 | 2.12E-15 | 4.79E-14 | 26.75038642 | 131.1765537 | ENSG00000175538 |
| PCDH11Y | 2.303711888 | 4.937264339 | 2.53E-04 | 1.25E-03 | 5.012832289 | 24.77852042 | ENSG00000099715 |
| TBX1 | 2.300080091 | 4.92485105 | 2.65E-02 | 7.10E-02 | 1.956806287 | 9.743093538 | ENSG00000184058 |
| IL32 | 2.283034679 | 4.867006426 | 3.14E-05 | 1.89E-04 | 7.939149439 | 38.032365 | ENSG00000008517 |
| GJB5 | 2.270608747 | 4.825266909 | 3.07E-06 | 2.21E-05 | 8.972538768 | 43.43896975 | ENSG00000189280 |
| FLI1 | 2.265810307 | 4.80924461 | 8.53E-06 | 5.69E-05 | 7.389101641 | 35.4263899 | ENSG00000151702 |
| AGMO | 2.265165099 | 4.807094279 | 2.79E-02 | 7.41E-02 | 2.014063805 | 9.690655735 | ENSG00000187546 |
| TFF3 | 2.2584256 | 4.78469048 | 2.81E-08 | 2.79E-07 | 11.71871066 | 56.05906594 | ENSG00000160180 |
| GATA4 | 2.257106249 | 4.780316859 | 4.80E-02 | 1.16E-01 | 1.649067354 | 7.949128713 | ENSG00000136574 |
| KCNMB1 | 2.252521526 | 4.765149659 | 1.35E-07 | 1.22E-06 | 10.60832733 | 50.749528 | ENSG00000145936 |
| CFHR3 | 2.248227006 | 4.750986168 | 1.09E-02 | 3.39E-02 | 2.721729483 | 12.81018913 | ENSG00000116785 |
| TMC8 | 2.246112035 | 4.744026389 | 5.03E-16 | 1.21E-14 | 25.71245442 | 121.9994441 | ENSG00000167895 |
| TRIM4 | 2.243651556 | 4.735942469 | 1.21E-23 | 5.38E-22 | 38.01661873 | 180.2587465 | ENSG00000146833 |
| RAB17 | 2.240501204 | 4.725612076 | 6.39E-190 | 4.43E-186 | 412.6543296 | 1948.649129 | ENSG00000124839 |
| SERPINB9 | 2.236615779 | 4.712902314 | 3.70E-104 | 3.40E-101 | 231.1099568 | 1088.956536 | ENSG00000170542 |
| ANGPTL6 | 2.2358425 | 4.710376895 | 2.70E-02 | 7.22E-02 | 2.06853042 | 9.598162643 | ENSG00000130812 |
| CD93 | 2.231201355 | 4.695247978 | 2.63E-02 | 7.05E-02 | 2.367896644 | 11.03210236 | ENSG00000125810 |
| MAGEB17 | 2.223474101 | 4.670166873 | 5.57E-05 | 3.18E-04 | 6.327603124 | 29.62505986 | ENSG00000182798 |
| LGR5 | 2.222795024 | 4.667969142 | 1.78E-02 | 5.09E-02 | 2.277824012 | 10.77800682 | ENSG00000139292 |
| IP6K3 | 2.220739567 | 4.66132326 | 4.12E-03 | 1.48E-02 | 3.284855915 | 15.46671192 | ENSG00000161896 |
| KCNJ9 | 2.210973816 | 4.629876839 | 2.11E-02 | 5.86E-02 | 2.299475514 | 10.72100741 | ENSG00000162728 |
| TDRD10 | 2.196822963 | 4.584686109 | 6.72E-03 | 2.26E-02 | 3.037165403 | 13.86392454 | ENSG00000163239 |
| ACTA1 | 2.191793667 | 4.568731524 | 3.97E-100 | 3.19E-97 | 201.4160322 | 920.4551158 | ENSG00000143632 |
| TLE6 | 2.181237782 | 4.53542511 | 3.90E-10 | 5.00E-09 | 17.40986013 | 78.82262075 | ENSG00000104953 |
| PPFIA4 | 2.171477652 | 4.504845577 | 5.23E-58 | 1.15E-55 | 140.8864852 | 632.7567348 | ENSG00000143847 |

| | | | | | | | |
|---|---|---|---|---|---|---|---|
| TGFB2 | 2.165140453 | 4.485100936 | 3.51E-04 | 1.68E-03 | 5.88090675 | 26.70960735 | ENSG00000092969 |
| INMT | 2.164448939 | 4.482951649 | 8.33E-03 | 2.70E-02 | 3.045538112 | 13.58617269 | ENSG00000241644 |
| PTGS2 | 2.156071346 | 4.456995012 | 2.96E-02 | 7.76E-02 | 1.981248691 | 8.897467587 | ENSG00000073756 |
| PLD6 | 2.1557813 | 4.456099047 | 2.26E-04 | 1.13E-03 | 5.970979382 | 26.74563406 | ENSG00000179598 |
| NRK | 2.151601574 | 4.443207674 | 1.38E-49 | 2.25E-47 | 102.6935512 | 456.2834802 | ENSG00000123572 |
| TSPYL5 | 2.146913713 | 4.428793446 | 9.85E-32 | 6.86E-30 | 59.16572814 | 261.9820263 | ENSG00000180543 |
| HAPLN1 | 2.139376617 | 4.405716359 | 1.05E-14 | 2.21E-13 | 27.30426426 | 120.5294777 | ENSG00000145681 |
| CREB5 | 2.135680276 | 4.394442886 | 1.68E-06 | 1.27E-05 | 10.92376325 | 48.312128 | ENSG00000146592 |
| CFAP300 | 2.13037924 | 4.378325582 | 2.00E-04 | 1.01E-03 | 6.754763177 | 29.45862323 | ENSG00000137691 |
| BCAS1 | 2.129166669 | 4.374647185 | 2.49E-03 | 9.57E-03 | 3.98973069 | 17.53223734 | ENSG00000064787 |
| SLC7A14 | 2.122891904 | 4.355661694 | 3.90E-05 | 2.30E-04 | 8.055579497 | 35.05587239 | ENSG00000013293 |
| CD8A | 2.122691179 | 4.355055726 | 5.95E-03 | 2.04E-02 | 3.336531627 | 14.57053828 | ENSG00000153563 |
| LCN15 | 2.102856117 | 4.295589457 | 2.65E-02 | 7.10E-02 | 2.974326079 | 12.84406527 | ENSG00000177984 |
| CSMD1 | 2.099966564 | 4.286994495 | 4.54E-04 | 2.10E-03 | 5.666031352 | 24.33245394 | ENSG00000183117 |
| LAMA2 | 2.099604821 | 4.285919703 | 9.78E-26 | 5.03E-24 | 71.49887752 | 305.3650506 | ENSG00000196569 |
| RASSF9 | 2.09074801 | 4.259688724 | 3.36E-04 | 1.61E-03 | 7.155365644 | 30.19723511 | ENSG00000198774 |
| ARHGDIB | 2.085162509 | 4.243228906 | 3.71E-02 | 9.36E-02 | 2.011272902 | 8.514852242 | ENSG00000111348 |
| ERVH48-1 | 2.082459598 | 4.235286596 | 1.25E-27 | 7.06E-26 | 534.3344308 | 2262.065906 | ENSG00000233056 |
| DPPA2 | 2.080326485 | 4.229029093 | 1.03E-39 | 1.19E-37 | 87.8121967 | 370.8104321 | ENSG00000163530 |
| COL4A4 | 2.080273953 | 4.228875105 | 7.54E-07 | 6.05E-06 | 11.66982585 | 49.36372494 | ENSG00000081052 |
| CR1L | 2.077466878 | 4.220654916 | 3.98E-07 | 3.33E-06 | 12.2930007 | 52.09417916 | ENSG00000197721 |
| CFP | 2.072115624 | 4.205028632 | 8.42E-03 | 2.73E-02 | 3.698737175 | 15.5231783 | ENSG00000126759 |
| RASAL3 | 2.071026427 | 4.201855145 | 1.30E-05 | 8.37E-05 | 9.122659821 | 38.10603596 | ENSG00000105122 |
| PCDHGA7 | 2.064316057 | 4.182356545 | 9.31E-03 | 2.96E-02 | 3.421022453 | 14.13522467 | ENSG00000253537 |
| MS4A15 | 2.062359793 | 4.176689201 | 2.11E-03 | 8.29E-03 | 5.135720033 | 21.23442628 | ENSG00000166961 |
| PADI3 | 2.060368897 | 4.170929413 | 3.45E-13 | 6.20E-12 | 29.6127882 | 123.9167915 | ENSG00000142619 |
| GOLGA6C | 2.057742284 | 4.163342606 | 4.81E-02 | 1.16E-01 | 1.984039594 | 8.331744058 | ENSG00000167195 |
| ADGRG3 | 2.052631405 | 4.148619677 | 2.06E-02 | 5.77E-02 | 3.130028937 | 12.72791587 | ENSG00000182885 |
| ABCC3 | 2.051235106 | 4.144606416 | 6.58E-03 | 2.22E-02 | 3.660340256 | 15.23305605 | ENSG00000108846 |
| LCP1 | 2.049931734 | 4.140863753 | 1.15E-14 | 2.42E-13 | 30.27926606 | 125.6435492 | ENSG00000136167 |
| COL6A3 | 2.043003258 | 4.12102512 | 2.46E-42 | 3.08E-40 | 106.4860278 | 437.8849167 | ENSG00000163359 |
| TMEM232 | 2.042609546 | 4.119900645 | 1.14E-04 | 6.09E-04 | 7.326262317 | 30.24993338 | ENSG00000186952 |
| EBI3 | 2.039200334 | 4.110176461 | 1.67E-04 | 8.55E-04 | 7.620046736 | 31.52253111 | ENSG00000105246 |
| PCDH17 | 2.039034674 | 4.109704529 | 2.27E-08 | 2.29E-07 | 17.97318673 | 73.36652227 | ENSG00000118946 |
| CHRNE | 2.035681111 | 4.100162565 | 9.85E-03 | 3.10E-02 | 3.336531627 | 13.66388434 | ENSG00000108556 |
| SCARF1 | 2.034385777 | 4.096482856 | 1.44E-36 | 1.30E-34 | 88.02842354 | 360.2712063 | ENSG00000074660 |
| SERPINA11 | 2.033485311 | 4.093926812 | 4.27E-02 | 1.05E-01 | 2.005691096 | 8.209675972 | ENSG00000186910 |
| TNN | 2.032396307 | 4.090837718 | 4.94E-10 | 6.26E-09 | 20.45539825 | 83.56375155 | ENSG00000120332 |
| MYOM3 | 2.031051696 | 4.087026777 | 3.06E-02 | 7.99E-02 | 2.329499725 | 9.54761495 | ENSG00000142661 |
| HSD52 | 2.025316585 | 4.070811969 | 4.07E-02 | 1.01E-01 | 2.335081531 | 9.512121272 | ENSG00000224609 |
| CFAP44 | 2.024696489 | 4.069062639 | 4.95E-03 | 1.74E-02 | 4.92555056 | 20.23769029 | ENSG00000206530 |
| COL13A1 | 2.024048425 | 4.067235211 | 1.50E-47 | 2.31E-45 | 120.8365586 | 492.2968631 | ENSG00000197467 |
| DNAH2 | 2.023637823 | 4.06607781 | 3.15E-04 | 1.52E-03 | 7.726189063 | 31.29533907 | ENSG00000183914 |
| C20orf144 | 2.022991442 | 4.064256462 | 3.23E-02 | 8.34E-02 | 2.332290628 | 9.486314388 | ENSG00000149609 |
| NHLRC1 | 2.017830941 | 4.049744641 | 4.57E-06 | 3.20E-05 | 10.66000304 | 43.21527324 | ENSG00000187566 |
| HTR2C | 2.015909087 | 4.044353455 | 1.32E-14 | 2.72E-13 | 33.80643084 | 136.6178281 | ENSG00000147246 |
| APOBEC3H | 2.014267107 | 4.039753058 | 2.55E-02 | 6.87E-02 | 2.683332564 | 10.81403353 | ENSG00000100298 |
| TAFA2 | 1.999934391 | 3.999818097 | 3.14E-04 | 1.52E-03 | 6.645829947 | 26.6843335 | ENSG00000198673 |
| GABRP | 1.997592233 | 3.993329819 | 4.64E-16 | 1.12E-14 | 49.55027804 | 198.4642119 | ENSG00000094755 |
| PDE4B | 1.997299131 | 3.992518607 | 9.42E-59 | 2.14E-56 | 153.9363998 | 614.357844 | ENSG00000184588 |
| PDCD1 | 1.994301426 | 3.984231362 | 7.15E-10 | 8.84E-09 | 21.67383892 | 86.38968558 | ENSG00000188389 |
| OTOGL | 1.994269749 | 3.98414388 | 2.90E-03 | 1.09E-02 | 4.751862984 | 18.7922042 | ENSG00000165899 |
| TOGARAM2 | 1.987961872 | 3.966762088 | 6.53E-05 | 3.68E-04 | 8.387760835 | 33.2110994 | ENSG00000189350 |
| CXCL1 | 1.987379005 | 3.965159788 | 8.07E-03 | 2.63E-02 | 3.968079188 | 15.83480629 | ENSG00000163739 |
| TXNIP | 1.983673506 | 3.954988517 | 3.38E-27 | 1.85E-25 | 113.2875606 | 447.4201002 | ENSG00000265972 |
| SLC25A45 | 1.982812779 | 3.952629633 | 5.44E-03 | 1.89E-02 | 4.291887817 | 17.07164988 | ENSG00000162241 |
| LPO | 1.98179893 | 3.949852913 | 2.32E-03 | 8.98E-03 | 5.345013626 | 21.14140015 | ENSG00000167419 |
| SPTY2D1OS | 1.977231367 | 3.949852913 | 7.78E-04 | 3.39E-03 | 6.812020695 | 26.56441599 | ENSG00000247595 |
| CEACAM1 | 1.973881455 | 3.928235587 | 2.79E-06 | 2.02E-05 | 12.48921566 | 48.77487814 | ENSG00000079385 |
| SLC39A4 | 1.97311471 | 3.926148415 | 5.93E-12 | 9.31E-11 | 26.55417146 | 103.8977151 | ENSG00000147804 |
| FALEC | 1.968174978 | 3.912728422 | 2.07E-02 | 5.78E-02 | 3.306507417 | 13.05002416 | ENSG00000228126 |
| KCNJ2 | 1.966055314 | 3.906983908 | 3.53E-05 | 2.10E-04 | 9.721392269 | 37.8742702 | ENSG00000123700 |
| WNT3 | 1.964395036 | 3.902490273 | 3.43E-35 | 2.84E-33 | 88.05988721 | 343.2635406 | ENSG00000108379 |
| HSPA2 | 1.961860178 | 3.895639502 | 1.56E-140 | 3.34E-137 | 659.0975074 | 2566.760206 | ENSG00000126803 |
| ATP8B3 | 1.960292232 | 3.891407952 | 6.77E-20 | 2.26E-18 | 54.53483788 | 211.5739617 | ENSG00000130270 |
| GDF3 | 1.959273262 | 3.888660435 | 3.46E-39 | 3.77E-37 | 115.430933 | 449.4332489 | ENSG00000184344 |
| C4orf51 | 1.957662574 | 3.884321389 | 1.56E-02 | 4.59E-02 | 3.948542868 | 15.44735676 | ENSG00000237136 |
| ADGRE5 | 1.954700748 | 3.876355128 | 1.42E-14 | 2.91E-13 | 33.91324889 | 131.1663217 | ENSG00000123146 |
| PLA2G2A | 1.95215738 | 3.869527413 | 1.38E-13 | 2.56E-12 | 34.14419398 | 132.0933969 | ENSG00000188257 |
| DCC | 1.950151667 | 3.864151523 | 1.47E-04 | 7.61E-04 | 8.752757286 | 33.66765829 | ENSG00000187323 |
| BGN | 1.949702689 | 3.862949157 | 2.01E-27 | 1.11E-25 | 76.65517302 | 295.9645171 | ENSG00000182492 |
| UNC5C | 1.947176283 | 3.85619039 | 7.16E-07 | 5.77E-06 | 14.47883706 | 55.65064371 | ENSG00000182168 |
| SNTG1 | 1.944799367 | 3.849842344 | 1.56E-02 | 4.56E-02 | 3.36934674 | 12.9467782 | ENSG00000147481 |

| | | | | | | | |
|---|---|---|---|---|---|---|---|
| COL5A2 | 1.935541441 | 3.825216599 | 9.58E-141 | 2.31E-137 | 469.9839871 | 1798.740606 | ENSG00000204262 |
| BCL2L14 | 1.934870779 | 3.823438794 | 5.99E-04 | 2.69E-03 | 6.91517196 | 26.65475851 | ENSG00000121380 |
| CFD | 1.927637999 | 3.804318405 | 4.87E-11 | 6.91E-10 | 26.12633568 | 99.21492873 | ENSG00000197766 |
| GDNF | 1.927345444 | 3.80354703 | 2.50E-02 | 6.76E-02 | 3.744831081 | 14.13145656 | ENSG00000168621 |
| CAV1 | 1.923699906 | 3.793948001 | 1.55E-30 | 1.01E-28 | 79.12766039 | 300.4091044 | ENSG00000105974 |
| SERPINE1 | 1.922231384 | 3.790088101 | 4.62E-18 | 1.32E-16 | 44.62492764 | 168.7778941 | ENSG00000106366 |
| ABHD1 | 1.920668354 | 3.785984104 | 4.63E-03 | 1.64E-02 | 4.664581255 | 17.67501766 | ENSG00000143994 |
| FRMPD4 | 1.920132619 | 3.784578466 | 1.69E-08 | 1.75E-07 | 21.15748212 | 79.82849207 | ENSG00000169933 |
| RTL4 | 1.912075275 | 3.763500792 | 2.17E-02 | 6.02E-02 | 3.300925611 | 12.50501286 | ENSG00000187823 |
| PLK2 | 1.909319633 | 3.756319121 | 9.22E-66 | 2.47E-63 | 206.0195259 | 773.3135758 | ENSG00000145632 |
| ENPP6 | 1.907986517 | 3.752849714 | 2.99E-02 | 7.83E-02 | 3.069980516 | 11.38352517 | ENSG00000164303 |
| BANK1 | 1.907544906 | 3.751701136 | 8.37E-05 | 4.60E-04 | 9.326371608 | 35.02441939 | ENSG00000153064 |
| MOGAT3 | 1.905130687 | 3.745428255 | 2.13E-02 | 5.92E-02 | 3.344904336 | 12.53458786 | ENSG00000106384 |
| ANKRD33 | 1.898447396 | 3.728117668 | 1.46E-03 | 5.97E-03 | 6.78757829 | 25.07483393 | ENSG00000167612 |
| ITGA11 | 1.894814694 | 3.718742089 | 2.82E-02 | 7.47E-02 | 3.018304804 | 11.21547101 | ENSG00000137809 |
| IER3 | 1.894700614 | 3.718448045 | 1.47E-76 | 6.05E-74 | 249.1957066 | 927.3018331 | ENSG00000137331 |
| GP1BA | 1.893598073 | 3.715607404 | 3.24E-03 | 1.20E-02 | 5.660449546 | 21.0827832 | ENSG00000185245 |
| CD79B | 1.892022899 | 3.711552816 | 1.04E-04 | 5.62E-04 | 10.48910637 | 38.69434973 | ENSG00000007312 |
| TUSC8 | 1.887956329 | 3.701105676 | 2.16E-02 | 6.00E-02 | 3.804879502 | 13.87629494 | ENSG00000237361 |
| EID3 | 1.880905498 | 3.683061526 | 3.55E-04 | 1.69E-03 | 7.998321979 | 29.48416965 | ENSG00000255150 |
| KCNAB2 | 1.879482874 | 3.679431495 | 1.20E-55 | 2.42E-53 | 189.9218459 | 697.4349834 | ENSG00000069424 |
| RBPJL | 1.879328965 | 3.679038987 | 5.93E-03 | 2.04E-02 | 4.637347948 | 17.15984183 | ENSG00000124232 |
| DPPA3 | 1.875426948 | 3.669101839 | 1.91E-12 | 3.15E-11 | 35.27814383 | 129.6826584 | ENSG00000187569 |
| UPK1A | 1.871347664 | 3.658741943 | 2.81E-03 | 1.06E-02 | 7.612549908 | 27.4390839 | ENSG00000105668 |
| MYOM1 | 1.853163964 | 3.612916629 | 3.72E-04 | 1.77E-03 | 8.662684654 | 31.38379148 | ENSG00000101605 |
| OR2H2 | 1.845568617 | 3.593945713 | 1.72E-02 | 4.97E-02 | 4.223466687 | 15.36561654 | ENSG00000204657 |
| RIN1 | 1.839088189 | 3.577838307 | 3.43E-13 | 6.16E-12 | 39.22756258 | 140.1912246 | ENSG00000174791 |
| C4orf47 | 1.836683834 | 3.571880549 | 2.59E-05 | 1.58E-04 | 13.45226884 | 47.90021023 | ENSG00000205129 |
| ADGRL4 | 1.836680343 | 3.571871905 | 1.27E-04 | 6.67E-04 | 9.963500975 | 35.67456676 | ENSG00000162618 |
| NTN1 | 1.836427659 | 3.571246357 | 1.51E-27 | 8.46E-26 | 88.68796815 | 316.5759784 | ENSG00000065320 |
| ADAMTS4 | 1.8348418 | 3.567322878 | 7.32E-56 | 1.50E-53 | 187.9045909 | 670.4607333 | ENSG00000158859 |
| ESPNL | 1.82831362 | 3.551217245 | 4.72E-10 | 5.99E-09 | 25.64191811 | 91.20744353 | ENSG00000144488 |
| RORC | 1.824104365 | 3.540871196 | 4.93E-04 | 2.26E-03 | 8.618705929 | 30.66589183 | ENSG00000143365 |
| FOXJ1 | 1.821028446 | 3.533329874 | 5.92E-36 | 5.14E-34 | 121.7486118 | 430.9959994 | ENSG00000129654 |
| DNM3 | 1.818659892 | 3.527533765 | 6.09E-15 | 1.31E-13 | 43.50529572 | 153.9193841 | ENSG00000197959 |
| DDX60L | 1.818170914 | 3.526338367 | 3.54E-04 | 1.69E-03 | 8.662684654 | 30.61157603 | ENSG00000181381 |
| BHLHE41 | 1.814373594 | 3.51706889 | 3.39E-06 | 2.42E-05 | 16.97173663 | 59.22458542 | ENSG00000123095 |
| PTAFR | 1.807181296 | 3.499578805 | 2.30E-12 | 3.78E-11 | 43.43910192 | 152.0934212 | ENSG00000169403 |
| PTPRVP | 1.806094525 | 3.496943593 | 9.66E-03 | 3.05E-02 | 4.982808078 | 17.49244252 | ENSG00000243323 |
| SNCG | 1.804395521 | 3.492827807 | 1.08E-28 | 6.43E-27 | 93.31012228 | 325.1860194 | ENSG00000173267 |
| PMFBP1 | 1.803962786 | 3.491780295 | 4.22E-04 | 1.97E-03 | 9.027005384 | 31.50936721 | ENSG00000118557 |
| RBM43 | 1.803580929 | 3.4908562 | 9.37E-07 | 7.40E-06 | 18.23204085 | 63.32444372 | ENSG00000184898 |
| PTPRR | 1.802710408 | 3.488750457 | 7.05E-03 | 2.35E-02 | 5.366665128 | 18.68088899 | ENSG00000153233 |
| OR52A1 | 1.796280061 | 3.473235082 | 3.41E-04 | 1.63E-03 | 9.010935688 | 31.35421648 | ENSG00000182070 |
| TUBB8B | 1.793959423 | 3.467652722 | 2.00E-02 | 5.62E-02 | 3.720388677 | 12.82471011 | ENSG00000173213 |
| CYSRT1 | 1.792466593 | 3.46406642 | 8.35E-05 | 4.59E-04 | 10.71726056 | 37.07542383 | ENSG00000197191 |
| GPA33 | 1.79236502 | 3.463822542 | 1.30E-02 | 3.92E-02 | 4.395239242 | 15.14271353 | ENSG00000143167 |
| COLEC11 | 1.788567837 | 3.454717726 | 2.58E-03 | 9.85E-03 | 6.906799251 | 24.06330439 | ENSG00000118004 |
| NOS3 | 1.784608786 | 3.445250267 | 3.93E-16 | 9.54E-15 | 53.40057573 | 184.0526351 | ENSG00000164867 |
| LAMP3 | 1.783811377 | 3.443346529 | 1.96E-02 | 5.53E-02 | 4.049779111 | 13.90156879 | ENSG00000078081 |
| C2orf91 | 1.782736765 | 3.440782659 | 4.08E-02 | 1.01E-01 | 3.276483206 | 11.35018207 | ENSG00000205086 |
| PDE6G | 1.782447352 | 3.440092485 | 7.69E-18 | 2.18E-16 | 53.83678421 | 185.1267917 | ENSG00000185527 |
| DNAH3 | 1.776752295 | 3.426539445 | 5.99E-05 | 3.40E-04 | 12.41588844 | 42.48579669 | ENSG00000158486 |
| TNFSF9 | 1.775849313 | 3.424395446 | 9.93E-14 | 1.88E-12 | 40.68878769 | 139.4477907 | ENSG00000125657 |
| OAZ3 | 1.774955585 | 3.422274743 | 3.59E-02 | 9.11E-02 | 3.578640333 | 12.3807942 | ENSG00000143450 |
| ZNF843 | 1.771360038 | 3.41375622 | 1.54E-02 | 4.53E-02 | 4.307957513 | 14.76601687 | ENSG00000176723 |
| C12orf56 | 1.769727398 | 3.409895195 | 2.20E-03 | 8.56E-03 | 7.492453066 | 25.32972297 | ENSG00000185306 |
| TBCE | 1.769170805 | 3.408579908 | 8.41E-03 | 2.72E-02 | 5.426713549 | 18.35904117 | ENSG00000284770 |
| AMDHD1 | 1.767862446 | 3.405490118 | 4.88E-03 | 1.72E-02 | 5.698846465 | 19.37674986 | ENSG00000139344 |
| DMTN | 1.76569032 | 3.400366661 | 7.47E-08 | 6.99E-07 | 23.78846325 | 80.71284677 | ENSG00000158856 |
| LIPA | 1.765349768 | 3.399564089 | 8.05E-90 | 4.85E-87 | 454.174761 | 1545.823216 | ENSG00000107798 |
| KCNK7 | 1.7624809 | 3.392810611 | 1.36E-02 | 4.07E-02 | 4.343563529 | 14.72622204 | ENSG00000173338 |
| HAPLN2 | 1.760704729 | 3.388636129 | 3.43E-03 | 1.26E-02 | 6.637457238 | 22.5425298 | ENSG00000132702 |
| SATB2 | 1.760066163 | 3.387136583 | 3.38E-15 | 7.41E-14 | 48.33048592 | 163.9195173 | ENSG00000119042 |
| SLC22A2 | 1.758155395 | 3.382653481 | 3.95E-02 | 9.87E-02 | 3.012722998 | 10.1827083 | ENSG00000112499 |
| ADCY8 | 1.757625116 | 3.381410376 | 2.14E-11 | 3.15E-10 | 34.56365705 | 116.6544004 | ENSG00000155897 |
| ZIC1 | 1.753852056 | 3.372578575 | 2.85E-02 | 7.53E-02 | 5.130138228 | 17.10741613 | ENSG00000152977 |
| GREM2 | 1.753837422 | 3.372544365 | 2.74E-02 | 7.29E-02 | 4.077012419 | 13.6090355 | ENSG00000180875 |
| MYO1F | 1.753772857 | 3.372393439 | 7.81E-22 | 3.04E-20 | 88.58481689 | 298.1220452 | ENSG00000142347 |
| PLAT | 1.752983149 | 3.37054795 | 1.96E-34 | 1.56E-32 | 130.5815543 | 439.1449078 | ENSG00000104368 |
| RAC2 | 1.752949536 | 3.370469421 | 9.29E-11 | 1.29E-09 | 44.23508847 | 149.7993466 | ENSG00000128340 |
| TNS4 | 1.744898987 | 3.351713853 | 1.25E-04 | 6.59E-04 | 12.57928829 | 41.86307375 | ENSG00000131746 |
| GJA5 | 1.744398849 | 3.35055212 | 1.88E-09 | 2.22E-08 | 32.86570488 | 110.08621 | ENSG00000265107 |

| | | | | | | | |
|---|---|---|---|---|---|---|---|
| SYTL2 | 1.740359107 | 3.341183242 | 4.35E-15 | 9.44E-14 | 49.95040495 | 166.6809036 | ENSG00000137501 |
| POLN | 1.739612552 | 3.339454718 | 3.10E-02 | 8.06E-02 | 3.747621984 | 12.37649306 | ENSG00000130997 |
| CD19 | 1.737484842 | 3.334533266 | 4.62E-03 | 1.64E-02 | 6.022655094 | 20.08174606 | ENSG00000177455 |
| MT1G | 1.733137548 | 3.324500393 | 5.24E-90 | 3.26E-87 | 455.2158842 | 1514.065127 | ENSG00000125144 |
| EMP1 | 1.732541347 | 3.32312681 | 8.42E-03 | 2.72E-02 | 5.312198513 | 17.70459266 | ENSG00000134531 |
| DBH | 1.730774643 | 3.319059847 | 3.20E-04 | 1.54E-03 | 10.10245842 | 33.37538546 | ENSG00000123454 |
| APOBEC3D | 1.724074214 | 3.303680602 | 7.99E-03 | 2.61E-02 | 5.619937445 | 18.65131399 | ENSG00000243811 |
| CLC | 1.718199099 | 3.290254314 | 9.33E-04 | 4.00E-03 | 8.689917962 | 28.5965983 | ENSG00000105205 |
| PPP1R3B | 1.715559222 | 3.284239235 | 8.28E-77 | 3.47E-74 | 405.2136403 | 1330.689519 | ENSG00000173281 |
| S100A4 | 1.714713898 | 3.282315451 | 6.26E-26 | 3.23E-24 | 94.42867816 | 309.8233349 | ENSG00000196154 |
| DLK1 | 1.712532728 | 3.277356762 | 1.44E-10 | 1.96E-09 | 34.27669374 | 112.5427047 | ENSG00000185559 |
| FMO4 | 1.708324694 | 3.267811339 | 4.31E-05 | 2.51E-04 | 14.25259789 | 46.79135341 | ENSG00000076258 |
| FMN1 | 1.70645732 | 3.263584334 | 4.97E-11 | 7.04E-10 | 37.05655662 | 120.4665596 | ENSG00000248905 |
| GTSF1 | 1.706431792 | 3.263526585 | 1.39E-02 | 4.16E-02 | 5.345013626 | 17.45694884 | ENSG00000170627 |
| BTBD19 | 1.704857033 | 3.259966261 | 9.97E-03 | 3.14E-02 | 5.369456031 | 17.48599079 | ENSG00000222009 |
| MARCHF11 | 1.701767111 | 3.252991632 | 7.48E-03 | 2.47E-02 | 6.019864191 | 19.53888534 | ENSG00000183654 |
| LOXHD1 | 1.701021728 | 3.251311374 | 1.33E-04 | 6.94E-04 | 12.14096462 | 39.26705802 | ENSG00000167210 |
| LYST | 1.69564422 | 3.23921498 | 3.93E-02 | 9.83E-02 | 3.366555838 | 10.87265049 | ENSG00000143669 |
| PRTN3 | 1.692419081 | 3.231981817 | 3.30E-02 | 8.49E-02 | 3.693155369 | 11.93498819 | ENSG00000196415 |
| NKAIN2 | 1.691632712 | 3.23022064 | 3.46E-02 | 8.83E-02 | 3.687573563 | 11.91401549 | ENSG00000188580 |
| TEX19 | 1.690021377 | 3.226614847 | 1.83E-02 | 5.20E-02 | 5.227707687 | 17.02917145 | ENSG00000182459 |
| MECOM | 1.688559867 | 3.223347806 | 3.76E-03 | 1.37E-02 | 7.375147127 | 23.76242927 | ENSG00000085276 |
| RASGRP4 | 1.684464624 | 3.214210967 | 1.72E-13 | 3.17E-12 | 52.57907074 | 169.2019155 | ENSG00000171777 |
| HLA-DQB1 | 1.681758915 | 3.208188508 | 1.28E-50 | 2.20E-48 | 223.2996772 | 715.4947243 | ENSG00000179344 |
| IL6R | 1.680494407 | 3.205377795 | 1.37E-26 | 7.36E-25 | 104.4805 | 335.55478 | ENSG00000160712 |
| GRIN2D | 1.676831546 | 3.197249978 | 1.54E-111 | 1.65E-108 | 568.8535598 | 1816.977319 | ENSG00000105464 |
| CHRNA9 | 1.673972707 | 3.190920592 | 6.93E-07 | 5.60E-06 | 21.32000608 | 68.08602629 | ENSG00000174343 |
| LGALS1 | 1.670876218 | 3.184079192 | 1.01E-33 | 7.78E-32 | 149.5161546 | 475.6467751 | ENSG00000100097 |
| SLC44A5 | 1.667062624 | 3.175673569 | 6.44E-10 | 8.00E-09 | 33.04708945 | 105.0775472 | ENSG00000137968 |
| MYO15B | 1.663037665 | 3.16682614 | 2.70E-27 | 1.48E-25 | 126.2146996 | 399.3605418 | ENSG00000266714 |
| CABYR | 1.663007505 | 3.166759936 | 4.35E-19 | 1.35E-17 | 82.27164384 | 260.685457 | ENSG00000154040 |
| TBC1D10C | 1.662270122 | 3.165141773 | 1.03E-03 | 4.36E-03 | 9.010935688 | 28.53099659 | ENSG00000175463 |
| ADAMTS14 | 1.660485987 | 3.161229961 | 2.17E-15 | 4.89E-14 | 61.15901633 | 192.8032938 | ENSG00000138316 |
| ETV3L | 1.660320372 | 3.160867088 | 3.54E-03 | 1.30E-02 | 8.124000627 | 25.50799697 | ENSG00000253831 |
| ARSD | 1.655565741 | 3.150467094 | 8.92E-23 | 3.71E-21 | 88.42276848 | 278.2249943 | ENSG00000006756 |
| IL15 | 1.652540948 | 3.14386866 | 1.20E-03 | 5.03E-03 | 10.95187244 | 34.06747822 | ENSG00000164136 |
| ATXN1 | 1.652281226 | 3.143302734 | 3.41E-07 | 2.89E-06 | 23.50584244 | 73.75989048 | ENSG00000124788 |
| MX2 | 1.649852702 | 3.138015987 | 1.29E-13 | 2.41E-12 | 57.47423367 | 179.8333923 | ENSG00000183486 |
| CAPN11 | 1.641225814 | 3.119307574 | 6.84E-08 | 6.44E-07 | 28.89204389 | 89.80847656 | ENSG00000137225 |
| PCDH10 | 1.64092468 | 3.118656547 | 3.50E-14 | 6.97E-13 | 72.88004232 | 226.5948676 | ENSG00000138650 |
| CD7 | 1.64069396 | 3.118157843 | 2.87E-04 | 1.40E-03 | 13.83034392 | 43.40939475 | ENSG00000173762 |
| ZNF37A | 1.636097879 | 3.108239946 | 2.27E-16 | 5.64E-15 | 61.97107257 | 192.9401433 | ENSG00000075407 |
| CATSPER2 | 1.633686399 | 3.10304883 | 3.34E-03 | 1.23E-02 | 8.542587812 | 26.74133292 | ENSG00000166762 |
| PSTPIP1 | 1.631881244 | 3.099168605 | 5.09E-03 | 1.78E-02 | 7.296238107 | 22.70492575 | ENSG00000140368 |
| CEMIP | 1.626906865 | 3.088501144 | 2.21E-08 | 2.23E-07 | 30.41052651 | 94.01586986 | ENSG00000103888 |
| PLCB2 | 1.619234559 | 3.072119975 | 4.53E-55 | 8.83E-53 | 257.6741406 | 790.5855932 | ENSG00000137841 |
| DNAH1 | 1.612370664 | 3.057538497 | 5.28E-05 | 3.02E-04 | 16.84952461 | 51.35153871 | ENSG00000114841 |
| EMP3 | 1.605230575 | 3.042443718 | 1.66E-10 | 2.24E-09 | 46.28603449 | 140.1737596 | ENSG00000142227 |
| POLR2J3 | 1.60269126 | 3.037093366 | 1.41E-02 | 4.20E-02 | 5.731661579 | 17.31122444 | ENSG00000285437 |
| CDA | 1.597998772 | 3.027231009 | 4.84E-20 | 1.65E-18 | 91.11870412 | 275.7155288 | ENSG00000158825 |
| CCDC9B | 1.597233138 | 3.025624893 | 6.55E-05 | 3.69E-04 | 15.30014189 | 46.42702715 | ENSG00000188549 |
| PTK6 | 1.595532992 | 3.022061441 | 1.75E-02 | 5.02E-02 | 6.017073288 | 18.2119597 | ENSG00000101213 |
| SLFN13 | 1.593572932 | 3.017958425 | 1.26E-38 | 1.33E-36 | 189.1059227 | 569.9458251 | ENSG00000154760 |
| ZYG11A | 1.592586047 | 3.015894678 | 2.30E-08 | 2.32E-07 | 29.69169722 | 89.61325844 | ENSG00000203995 |
| FAM163B | 1.587501936 | 3.005285258 | 8.30E-12 | 1.28E-10 | 51.80365965 | 156.2271556 | ENSG00000196990 |
| DEGS2 | 1.586761865 | 3.003744007 | 5.08E-04 | 2.33E-03 | 13.01115427 | 39.11944351 | ENSG00000168350 |
| RGS7 | 1.576260602 | 2.981959372 | 9.73E-08 | 8.99E-07 | 28.86481058 | 85.95327536 | ENSG00000182901 |
| CTHRC1 | 1.573791806 | 2.976860892 | 1.34E-05 | 8.62E-05 | 20.1309139 | 59.83198177 | ENSG00000164932 |
| ABI3 | 1.573787932 | 2.976852897 | 1.16E-08 | 1.23E-07 | 37.49380424 | 111.8686406 | ENSG00000108798 |
| GABRG2 | 1.573347021 | 2.975943262 | 7.17E-03 | 2.38E-02 | 9.96003435 | 29.67749767 | ENSG00000113327 |
| DUSP10 | 1.571941122 | 2.973044633 | 2.49E-04 | 1.23E-03 | 15.27378447 | 45.13586776 | ENSG00000143507 |
| MSH5 | 1.571939236 | 2.973040747 | 8.35E-03 | 2.71E-02 | 7.347913819 | 21.80849164 | ENSG00000204410 |
| SPARCL1 | 1.571690121 | 2.972527425 | 2.90E-03 | 1.09E-02 | 10.14085533 | 30.0248798 | ENSG00000152583 |
| HPSE2 | 1.566778032 | 2.962423773 | 2.24E-03 | 8.71E-03 | 10.92463913 | 32.04389821 | ENSG00000172987 |
| PLAC8L1 | 1.565212 | 2.959209834 | 3.83E-02 | 9.63E-02 | 4.628975239 | 13.75637743 | ENSG00000173261 |
| GGT1 | 1.564716322 | 2.95819329 | 1.08E-05 | 7.05E-05 | 21.1568064 | 62.46161318 | ENSG00000100031 |
| SNORD15B | 1.560679091 | 2.949926668 | 4.02E-02 | 1.00E-01 | 4.316330222 | 12.76179201 | ENSG00000207445 |
| KCNH3 | 1.559977606 | 2.948492668 | 9.28E-03 | 2.96E-02 | 9.680880168 | 28.61514786 | ENSG00000135519 |
| CAPN9 | 1.55750817 | 2.943450101 | 4.83E-02 | 1.16E-01 | 3.984148884 | 11.74570086 | ENSG00000135773 |
| DCST2 | 1.556186555 | 2.940754918 | 4.97E-02 | 1.19E-01 | 4.041406403 | 11.84679624 | ENSG00000163354 |
| PWWP3B | 1.553579325 | 2.935445201 | 3.39E-05 | 2.02E-04 | 17.60519921 | 51.8118657 | ENSG00000157502 |
| TGM2 | 1.552682931 | 2.93362188 | 4.34E-12 | 6.89E-11 | 51.22956977 | 150.0784556 | ENSG00000198959 |
| NMI | 1.550728928 | 2.929651237 | 3.72E-08 | 3.64E-07 | 37.09878358 | 109.1779386 | ENSG00000123609 |

| | | | | | | | |
|---|---|---|---|---|---|---|---|
| MMRN2 | 1.546230191 | 2.920529973 | 1.97E-09 | 2.32E-08 | 39.01527793 | 113.9954666 | ENSG00000173269 |
| ALOX5 | 1.544253057 | 2.916530289 | 1.28E-07 | 1.16E-06 | 31.95086079 | 92.83094315 | ENSG00000012779 |
| DPEP1 | 1.543741933 | 2.915497191 | 1.61E-13 | 2.98E-12 | 62.57646286 | 181.8937925 | ENSG00000015413 |
| AIPL1 | 1.537485954 | 2.902882052 | 6.12E-04 | 2.74E-03 | 13.60218973 | 39.58083658 | ENSG00000129221 |
| WDR66 | 1.536808166 | 2.901518578 | 5.06E-04 | 2.32E-03 | 15.65888082 | 45.45315397 | ENSG00000158023 |
| KCTD19 | 1.536158279 | 2.900211832 | 2.19E-02 | 6.05E-02 | 6.412093949 | 18.55209661 | ENSG00000168676 |
| TSPAN32 | 1.53578564 | 2.899462824 | 3.31E-02 | 8.51E-02 | 5.43787716 | 15.62454626 | ENSG00000064201 |
| ARRB1 | 1.535483099 | 2.898854853 | 3.57E-58 | 7.92E-56 | 472.4525181 | 1367.80266 | ENSG00000137486 |
| ENPP2 | 1.535254414 | 2.898395386 | 1.00E-06 | 7.88E-06 | 34.59260522 | 100.7846335 | ENSG00000136960 |
| C5AR2 | 1.534736748 | 2.897355573 | 2.54E-02 | 6.85E-02 | 5.284965205 | 15.38658924 | ENSG00000134830 |
| CRYZL2P | 1.534299951 | 2.896478489 | 8.77E-05 | 4.81E-04 | 17.14330902 | 49.50461515 | ENSG00000242193 |
| SMIM10 | 1.530880372 | 2.88962118 | 2.89E-07 | 2.48E-06 | 32.82750812 | 94.38585434 | ENSG00000184785 |
| FGF8 | 1.530198507 | 2.888255773 | 1.21E-05 | 7.84E-05 | 23.84013896 | 68.65820154 | ENSG00000107831 |
| PPARGC1A | 1.522216466 | 2.87231996 | 1.31E-05 | 8.39E-05 | 22.8693888 | 65.54914844 | ENSG00000109819 |
| ITGB2 | 1.518632603 | 2.865193556 | 4.70E-03 | 1.66E-02 | 9.019308396 | 25.81962496 | ENSG00000160255 |
| LDHD | 1.516836206 | 2.861628131 | 3.17E-22 | 1.26E-20 | 106.4390581 | 304.4487282 | ENSG00000166816 |
| MOG | 1.516361503 | 2.860686699 | 1.71E-02 | 4.94E-02 | 6.313648609 | 18.12323472 | ENSG00000204655 |
| MT1H | 1.510583472 | 2.849252487 | 8.01E-11 | 1.11E-09 | 48.93567606 | 139.3369964 | ENSG00000205358 |
| C1orf74 | 1.510514247 | 2.849115774 | 2.85E-22 | 1.13E-20 | 112.7827309 | 321.0243091 | ENSG00000162757 |
| CERS3 | 1.504925209 | 2.838099578 | 3.39E-02 | 8.68E-02 | 5.489552873 | 15.39089038 | ENSG00000154227 |
| LCN2 | 1.503974271 | 2.836229489 | 7.32E-03 | 2.43E-02 | 8.292106398 | 23.58604538 | ENSG00000148346 |
| CCDC169 | 1.502183996 | 2.832712127 | 8.57E-14 | 1.63E-12 | 64.17086344 | 181.8182315 | ENSG00000242715 |
| PPFIBP2 | 1.499522052 | 2.827490256 | 1.08E-30 | 7.06E-29 | 170.240019 | 481.3782023 | ENSG00000166387 |
| SULT1C2 | 1.496579809 | 2.821729727 | 5.86E-04 | 2.64E-03 | 14.0677467 | 39.63138427 | ENSG00000198203 |
| BNIP3 | 1.49535659 | 2.819338278 | 5.16E-61 | 1.22E-58 | 815.7135219 | 2298.686115 | ENSG00000176171 |
| MUC19 | 1.490571172 | 2.810002029 | 4.30E-03 | 1.54E-02 | 9.873428343 | 27.94214979 | ENSG00000205592 |
| TBX10 | 1.489157607 | 2.807250113 | 2.09E-03 | 8.21E-03 | 11.48218376 | 32.05196746 | ENSG00000167800 |
| C22orf15 | 1.482872688 | 2.79504728 | 5.62E-07 | 4.59E-06 | 31.27301917 | 87.55741981 | ENSG00000169314 |
| C1orf220 | 1.480800045 | 2.791034669 | 7.95E-03 | 2.60E-02 | 9.054238691 | 25.21787472 | ENSG00000213057 |
| ANXA3 | 1.477332684 | 2.784334773 | 5.94E-37 | 5.56E-35 | 198.1940157 | 551.9373827 | ENSG00000138772 |
| TMPRSS9 | 1.477241629 | 2.784159047 | 5.98E-04 | 2.69E-03 | 13.62105033 | 38.00924173 | ENSG00000178297 |
| NEK7 | 1.474628581 | 2.779120869 | 1.31E-23 | 5.75E-22 | 129.3734014 | 359.1287154 | ENSG00000151414 |
| RNASE1 | 1.473860277 | 2.777641247 | 3.38E-02 | 8.65E-02 | 6.134379227 | 16.93908939 | ENSG00000129538 |
| IFI35 | 1.473537687 | 2.777020229 | 3.61E-05 | 2.14E-04 | 23.089846 | 64.01574298 | ENSG00000068079 |
| ADGRG1 | 1.473316717 | 2.776594921 | 1.12E-06 | 8.77E-06 | 32.04496363 | 89.02226106 | ENSG00000205336 |
| FMO5 | 1.472834298 | 2.775666618 | 2.21E-03 | 8.62E-03 | 11.97967996 | 33.35252265 | ENSG00000131781 |
| PYY | 1.471964249 | 2.773993196 | 4.57E-04 | 2.11E-03 | 16.53899477 | 45.61313888 | ENSG00000131096 |
| ALG1L | 1.46975744 | 2.769753219 | 1.38E-06 | 1.06E-05 | 27.84661507 | 77.03053392 | ENSG00000189366 |
| LRFN5 | 1.469674451 | 2.769593897 | 7.38E-03 | 2.44E-02 | 8.632660444 | 23.98371474 | ENSG00000165379 |
| SLC44A4 | 1.465579165 | 2.761743177 | 5.89E-07 | 4.80E-06 | 33.27591936 | 92.18587832 | ENSG00000204385 |
| CR2 | 1.465021699 | 2.760676229 | 5.47E-26 | 2.83E-24 | 135.4580199 | 373.841501 | ENSG00000117322 |
| EXOC8 | 1.464473183 | 2.759626812 | 1.99E-19 | 6.37E-18 | 104.2336854 | 287.6873614 | ENSG00000116903 |
| GABRE | 1.463385038 | 2.757546163 | 1.04E-13 | 1.97E-12 | 74.36359465 | 204.4298706 | ENSG00000102287 |
| APOBEC3G | 1.461673011 | 2.754275761 | 5.95E-05 | 3.38E-04 | 20.48474673 | 56.33843532 | ENSG00000239713 |
| AIF1 | 1.46093036 | 2.752858317 | 2.51E-07 | 2.17E-06 | 35.82799147 | 98.96217815 | ENSG00000204472 |
| MYOZ1 | 1.45869673 | 2.748599546 | 6.01E-04 | 2.70E-03 | 16.73154294 | 45.95890348 | ENSG00000177791 |
| B3GNT3 | 1.456435023 | 2.744293953 | 1.27E-04 | 6.71E-04 | 17.92621694 | 49.37420524 | ENSG00000179913 |
| MYH2 | 1.455923239 | 2.74332061 | 2.28E-08 | 2.29E-07 | 40.34141253 | 110.3881818 | ENSG00000125414 |
| PEX5L | 1.455922847 | 2.743319864 | 2.47E-07 | 2.14E-06 | 35.22088631 | 96.85900848 | ENSG00000114757 |
| NR4A2 | 1.455425518 | 2.742374345 | 3.54E-02 | 9.01E-02 | 6.335300111 | 17.38489541 | ENSG00000153234 |
| NAALADL1 | 1.453913834 | 2.739502336 | 2.43E-07 | 2.11E-06 | 34.47916622 | 94.42511612 | ENSG00000168060 |
| ISLR | 1.453001774 | 2.737770992 | 7.97E-03 | 2.60E-02 | 11.49546255 | 31.36658689 | ENSG00000129009 |
| OCA2 | 1.452307761 | 2.736454295 | 4.36E-03 | 1.56E-02 | 9.718601366 | 26.54559387 | ENSG00000104044 |
| NKPD1 | 1.452060534 | 2.735985404 | 2.21E-03 | 8.60E-03 | 11.8443893 | 32.23318554 | ENSG00000179846 |
| PCED1B | 1.445802507 | 2.724143132 | 1.19E-14 | 2.48E-13 | 78.18047675 | 213.3859552 | ENSG00000179715 |
| COLCA1 | 1.445251793 | 2.723103453 | 5.00E-02 | 1.19E-01 | 5.056811014 | 13.69560991 | ENSG00000196167 |
| HPGD | 1.444188143 | 2.721096542 | 1.11E-04 | 5.95E-04 | 18.99608816 | 51.7156045 | ENSG00000164120 |
| FAM157A | 1.443986127 | 2.720715541 | 8.94E-13 | 1.53E-11 | 78.51740091 | 212.889874 | ENSG00000236438 |
| AZU1 | 1.442199627 | 2.717348544 | 1.33E-02 | 4.01E-02 | 8.423366851 | 22.76113166 | ENSG00000172232 |
| NRROS | 1.439551962 | 2.712366182 | 1.01E-14 | 2.14E-13 | 81.17541519 | 219.7648461 | ENSG00000174004 |
| MYBPC2 | 1.438400488 | 2.710202195 | 2.77E-17 | 7.48E-16 | 102.1332157 | 276.5001573 | ENSG00000086967 |
| ABHD12B | 1.437289874 | 2.708116633 | 3.33E-10 | 4.32E-09 | 52.35390761 | 141.6017685 | ENSG00000131969 |
| WDR31 | 1.436912955 | 2.707409202 | 2.06E-02 | 5.77E-02 | 7.068083915 | 19.08447703 | ENSG00000148225 |
| HCN1 | 1.436584501 | 2.706792884 | 1.63E-07 | 1.45E-06 | 42.4004942 | 115.2922415 | ENSG00000164588 |
| OXCT2 | 1.435348639 | 2.704475146 | 5.28E-04 | 2.40E-03 | 17.76301725 | 47.9101696 | ENSG00000198754 |
| PHLDA3 | 1.434118063 | 2.702169293 | 1.17E-74 | 4.51E-72 | 914.1972132 | 2472.52064 | ENSG00000174307 |
| TAGLN | 1.4335246 | 2.701057963 | 1.48E-70 | 4.77E-68 | 541.4533288 | 1461.936618 | ENSG00000149591 |
| PRCD | 1.432490876 | 2.699123287 | 2.27E-02 | 6.24E-02 | 6.684226866 | 18.03504277 | ENSG00000214140 |
| FOXB1 | 1.432101357 | 2.698394639 | 7.76E-10 | 9.57E-09 | 68.32734846 | 184.4116062 | ENSG00000171956 |
| SH2D2A | 1.431753899 | 2.697744836 | 8.03E-18 | 2.27E-16 | 95.38614953 | 257.1543089 | ENSG00000027869 |
| PYROXD2 | 1.43163507 | 2.697522643 | 5.21E-07 | 4.28E-06 | 32.55796595 | 87.64184365 | ENSG00000119943 |
| CAV2 | 1.431210702 | 2.696729285 | 1.95E-05 | 1.22E-04 | 26.57370778 | 71.52362729 | ENSG00000105971 |
| FAM124B | 1.430280491 | 2.694991068 | 3.13E-04 | 1.51E-03 | 16.64214603 | 44.91700544 | ENSG00000124019 |

| | | | | | | | |
|---|---|---|---|---|---|---|---|
| FAM107A | 1.426176389 | 2.687335395 | 3.52E-02 | 8.95E-02 | 6.801532805 | 18.17378241 | ENSG00000168309 |
| F10 | 1.42607742 | 2.687151049 | 5.29E-08 | 5.07E-07 | 37.42067719 | 100.5891549 | ENSG00000126218 |
| MISP | 1.424706429 | 2.684598667 | 1.74E-02 | 5.00E-02 | 7.595604331 | 20.54018295 | ENSG00000099812 |
| KCNJ6 | 1.420909314 | 2.677542201 | 1.05E-04 | 5.66E-04 | 19.95423526 | 53.55177519 | ENSG00000157542 |
| MASP1 | 1.415547943 | 2.667610333 | 2.79E-07 | 2.40E-06 | 42.94707538 | 114.8321871 | ENSG00000127241 |
| TNFAIP6 | 1.415382939 | 2.667305252 | 4.40E-02 | 1.08E-01 | 5.701637368 | 15.20616466 | ENSG00000123610 |
| FGF4 | 1.414521394 | 2.665712872 | 2.30E-05 | 1.42E-04 | 25.54626367 | 68.31295787 | ENSG00000075388 |
| LGR6 | 1.413540472 | 2.663901009 | 6.74E-05 | 3.78E-04 | 24.42905924 | 65.04714862 | ENSG00000133067 |
| SBK3 | 1.412503078 | 2.661986175 | 3.20E-03 | 1.18E-02 | 11.37045962 | 30.27252362 | ENSG00000231274 |
| PCDH11X | 1.41187231 | 2.660822567 | 9.28E-18 | 2.61E-16 | 112.8869582 | 299.8635296 | ENSG00000102290 |
| CAVIN3 | 1.410319681 | 2.657960531 | 1.60E-21 | 6.01E-20 | 171.4677593 | 456.5908312 | ENSG00000170955 |
| WDR72 | 1.407879135 | 2.653467971 | 4.10E-06 | 2.90E-05 | 30.37982658 | 80.69937976 | ENSG00000166415 |
| GBGT1 | 1.406776838 | 2.651441354 | 8.60E-17 | 2.22E-15 | 110.1589712 | 291.6382665 | ENSG00000148288 |
| MAFB | 1.40540266 | 2.648917038 | 6.49E-07 | 5.26E-06 | 32.38061159 | 85.85217997 | ENSG00000204103 |
| WFDC1 | 1.401372417 | 2.641527477 | 6.24E-03 | 2.12E-02 | 11.46332316 | 30.16067536 | ENSG00000103175 |
| PRDM16 | 1.400927715 | 2.640713367 | 5.70E-04 | 2.58E-03 | 15.70844135 | 41.51165093 | ENSG00000142611 |
| ALDH1A1 | 1.400536442 | 2.639997278 | 1.55E-04 | 7.97E-04 | 20.81971898 | 54.94429043 | ENSG00000165092 |
| GJB7 | 1.399997021 | 2.639010371 | 4.76E-03 | 1.68E-02 | 11.89048321 | 31.15202571 | ENSG00000164411 |
| NDRG1 | 1.399244203 | 2.63763366 | 1.31E-08 | 1.37E-07 | 46.35637064 | 122.0438006 | ENSG00000104419 |
| GNGT1 | 1.397808431 | 2.63500999 | 3.95E-04 | 1.87E-03 | 20.09809878 | 52.91721488 | ENSG00000127928 |
| SCTR | 1.395910946 | 2.631546606 | 9.29E-03 | 2.96E-02 | 9.699740767 | 25.48755731 | ENSG00000080293 |
| INPP5J | 1.393449283 | 2.62706024 | 5.73E-20 | 1.92E-18 | 129.7992852 | 341.490288 | ENSG00000185133 |
| FAM20A | 1.392095491 | 2.624596224 | 1.20E-02 | 3.66E-02 | 8.684336156 | 22.84151481 | ENSG00000108950 |
| SORBS2 | 1.391878268 | 2.624201075 | 4.00E-47 | 6.07E-45 | 403.797196 | 1061.096175 | ENSG00000154556 |
| TFAP2C | 1.390171078 | 2.621097605 | 2.75E-15 | 6.10E-14 | 102.6008879 | 268.7206392 | ENSG00000087510 |
| NPFFR2 | 1.389329453 | 2.619568982 | 3.56E-14 | 7.06E-13 | 90.08252389 | 235.4915417 | ENSG00000056291 |
| CA4 | 1.388400946 | 2.61788359 | 2.12E-09 | 2.47E-08 | 50.91325796 | 133.5012693 | ENSG00000167434 |
| PTPN14 | 1.388183461 | 2.617488975 | 7.22E-87 | 3.98E-84 | 1146.724876 | 3000.574179 | ENSG00000152104 |
| NTSR1 | 1.387250692 | 2.615797196 | 3.81E-04 | 1.81E-03 | 19.07778809 | 49.85927304 | ENSG00000101188 |
| QPCT | 1.385753182 | 2.613083421 | 5.06E-12 | 7.97E-11 | 70.38136079 | 183.7138247 | ENSG00000115828 |
| POLH | 1.383964319 | 2.609845348 | 3.62E-19 | 1.13E-17 | 134.0761794 | 350.1129236 | ENSG00000170734 |
| TENM1 | 1.383692864 | 2.60935433 | 9.19E-06 | 6.10E-05 | 38.99816909 | 101.2223702 | ENSG00000009694 |
| NPIPB6 | 1.382713907 | 2.607584324 | 1.16E-03 | 4.88E-03 | 16.11115899 | 41.99536166 | ENSG00000198156 |
| ZMAT3 | 1.381376969 | 2.605169007 | 6.68E-23 | 2.79E-21 | 201.7079016 | 525.6525355 | ENSG00000172667 |
| SMIM10L2A | 1.380057691 | 2.602787791 | 1.03E-25 | 5.31E-24 | 170.4625913 | 443.4684932 | ENSG00000178947 |
| FGF16 | 1.376470963 | 2.596324957 | 2.48E-06 | 1.81E-05 | 30.96528023 | 80.55580594 | ENSG00000196468 |
| PPP1R3C | 1.376435275 | 2.596260733 | 1.35E-04 | 7.04E-04 | 22.26419866 | 57.98208361 | ENSG00000119938 |
| TOR4A | 1.376426019 | 2.596244075 | 4.94E-28 | 2.84E-26 | 222.4400635 | 576.6895569 | ENSG00000198113 |
| TRPM6 | 1.374627554 | 2.593009612 | 2.12E-03 | 8.32E-03 | 15.19208454 | 39.18478475 | ENSG00000119121 |
| NPNT | 1.371013217 | 2.58652156 | 3.25E-02 | 8.39E-02 | 6.975220381 | 18.09634333 | ENSG00000168743 |
| SPOCD1 | 1.369414154 | 2.583656283 | 6.34E-04 | 2.83E-03 | 17.13214541 | 44.10929631 | ENSG00000134668 |
| PROX1 | 1.36841155 | 2.58186139 | 4.08E-03 | 1.47E-02 | 11.78434088 | 30.35694745 | ENSG00000117707 |
| PCDHGA5 | 1.364206117 | 2.574346264 | 3.29E-02 | 8.46E-02 | 7.260632091 | 18.80725822 | ENSG00000253485 |
| NABP1 | 1.362734085 | 2.571720909 | 7.38E-06 | 4.98E-05 | 31.43920992 | 80.92846192 | ENSG00000173559 |
| PYGM | 1.36005674 | 2.566952749 | 1.73E-27 | 9.68E-26 | 209.7902389 | 537.79358 | ENSG00000068976 |
| INPP5D | 1.359273671 | 2.565559831 | 2.97E-35 | 2.48E-33 | 256.1992364 | 657.1493621 | ENSG00000168918 |
| IRF2BP2 | 1.356690585 | 2.560970412 | 1.54E-51 | 2.69E-49 | 462.4178945 | 1185.747265 | ENSG00000168264 |
| MVP | 1.356127474 | 2.559971012 | 7.79E-05 | 4.31E-04 | 28.63318977 | 72.97263313 | ENSG00000013364 |
| PKP3 | 1.352041533 | 2.552731026 | 2.03E-39 | 2.29E-37 | 353.2691465 | 902.7075175 | ENSG00000184363 |
| KLK8 | 1.351886769 | 2.552457199 | 2.73E-02 | 7.28E-02 | 7.389101641 | 18.84920362 | ENSG00000129455 |
| ACTG2 | 1.351847801 | 2.552388256 | 3.88E-06 | 2.75E-05 | 32.05122116 | 81.76281407 | ENSG00000163017 |
| ERVMER34-1 | 1.348886792 | 2.547155068 | 1.55E-31 | 1.07E-29 | 219.7716366 | 559.5619921 | ENSG00000226887 |
| KIAA1257 | 1.34661322 | 2.543144116 | 1.53E-06 | 1.16E-05 | 35.26207414 | 89.92031271 | ENSG00000114656 |
| SERPINB1 | 1.345842043 | 2.54178507 | 1.85E-24 | 8.72E-23 | 194.0243399 | 493.2182558 | ENSG00000021355 |
| POU6F2 | 1.343880749 | 2.538331949 | 4.43E-03 | 1.58E-02 | 15.66332332 | 39.50904361 | ENSG00000106536 |
| TRAF1 | 1.342416317 | 2.53575668 | 1.45E-05 | 9.28E-05 | 29.95912421 | 75.69663563 | ENSG00000056558 |
| MAMLD1 | 1.338609386 | 2.529074237 | 1.68E-02 | 4.87E-02 | 9.321465523 | 23.6048675 | ENSG00000013619 |
| HES4 | 1.338543185 | 2.528958188 | 4.91E-04 | 2.25E-03 | 20.06528367 | 50.68015818 | ENSG00000188290 |
| MAP1LC3C | 1.338170883 | 2.528305649 | 1.54E-02 | 4.53E-02 | 11.55618669 | 29.07223978 | ENSG00000197769 |
| SOX3 | 1.33772018 | 2.527515921 | 8.02E-52 | 1.42E-49 | 385.999961 | 975.9422606 | ENSG00000134595 |
| HACD4 | 1.335638538 | 2.523871638 | 4.80E-05 | 2.77E-04 | 25.47871842 | 64.26874191 | ENSG00000188921 |
| LMCD1 | 1.334593869 | 2.522044741 | 1.23E-23 | 5.45E-22 | 172.0584314 | 434.2943461 | ENSG00000071282 |
| FZD10 | 1.332710898 | 2.518755175 | 2.18E-02 | 6.04E-02 | 10.91626642 | 27.24172731 | ENSG00000111432 |
| ZNF124 | 1.329126517 | 2.512505087 | 3.77E-19 | 1.17E-17 | 161.4944462 | 406.6476553 | ENSG00000196418 |
| S100A14 | 1.327519885 | 2.509708636 | 2.58E-04 | 1.27E-03 | 24.01382654 | 60.28477255 | ENSG00000189334 |
| MEIS3 | 1.327428827 | 2.509550237 | 3.04E-72 | 1.03E-69 | 640.1458977 | 1605.749073 | ENSG00000105419 |
| ARHGAP9 | 1.324586215 | 2.504610417 | 4.90E-06 | 3.41E-05 | 32.30660866 | 81.01398236 | ENSG00000123329 |
| LIF | 1.324229717 | 2.50399159 | 1.66E-13 | 3.06E-12 | 85.05812767 | 213.173805 | ENSG00000128342 |
| PLIN4 | 1.323638207 | 2.502965156 | 1.18E-04 | 6.28E-04 | 27.57918808 | 69.07443256 | ENSG00000167676 |
| FAM183A | 1.321366822 | 2.499027578 | 4.49E-02 | 1.09E-01 | 6.300369816 | 15.81168302 | ENSG00000186973 |
| COLQ | 1.320657826 | 2.497799761 | 4.80E-02 | 1.16E-01 | 6.585781526 | 16.58981715 | ENSG00000206561 |
| WNT9B | 1.320620694 | 2.497735473 | 2.81E-02 | 7.44E-02 | 7.590022525 | 19.06995605 | ENSG00000158955 |
| BBC3 | 1.319123465 | 2.495144668 | 1.09E-37 | 1.06E-35 | 282.3703145 | 704.3539904 | ENSG00000105327 |

| | | | | | | | |
|---|---|---|---|---|---|---|---|
| GRHL3 | 1.318474823 | 2.494023093 | 1.14E-02 | 3.52E-02 | 10.97332378 | 27.38046695 | ENSG00000158055 |
| PTH1R | 1.31846018 | 2.493997779 | 1.41E-02 | 4.21E-02 | 10.34177622 | 25.77391145 | ENSG00000160801 |
| PDGFB | 1.318443446 | 2.49396885 | 1.69E-18 | 5.03E-17 | 127.8840671 | 318.838842 | ENSG00000100311 |
| VIM | 1.318303998 | 2.4937278 | 1.47E-23 | 6.44E-22 | 212.9997276 | 531.8287147 | ENSG00000026025 |
| OVGP1 | 1.318215708 | 2.493575193 | 1.74E-02 | 5.01E-02 | 9.005353882 | 22.51966699 | ENSG00000085465 |
| S100A11 | 1.31730013 | 2.491993197 | 3.02E-82 | 1.36E-79 | 1194.75104 | 2976.996805 | ENSG00000163191 |
| KIAA1107 | 1.317027864 | 2.491522952 | 7.11E-06 | 4.82E-05 | 37.63276168 | 94.04302171 | ENSG00000189195 |
| CEBPZOS | 1.315692924 | 2.489218586 | 7.81E-03 | 2.56E-02 | 11.35159902 | 28.27690106 | ENSG00000218739 |
| RAB37 | 1.315101252 | 2.488197928 | 2.07E-09 | 2.42E-08 | 69.63653749 | 173.4784366 | ENSG00000172794 |
| SAMSN1 | 1.312595852 | 2.483880647 | 3.94E-02 | 9.86E-02 | 6.790369193 | 16.74765149 | ENSG00000155307 |
| STAT6 | 1.311854995 | 2.482605444 | 1.70E-88 | 9.92E-86 | 1011.431859 | 2509.530845 | ENSG00000166888 |
| KCNN4 | 1.311240761 | 2.481548689 | 1.31E-02 | 3.96E-02 | 10.0912948 | 24.97399901 | ENSG00000104783 |
| WASIR2 | 1.307502206 | 2.475126406 | 3.57E-02 | 9.06E-02 | 7.026896093 | 17.37467558 | ENSG00000231439 |
| C11orf86 | 1.307220857 | 2.474643762 | 8.27E-03 | 2.69E-02 | 11.62094104 | 28.82111885 | ENSG00000173237 |
| TTC39C | 1.304119939 | 2.469330493 | 4.39E-43 | 5.69E-41 | 339.3529972 | 838.017383 | ENSG00000168234 |
| KLK7 | 1.298335458 | 2.459449543 | 2.50E-03 | 9.59E-03 | 15.17813003 | 37.19964249 | ENSG00000169035 |
| FOSL1 | 1.298004316 | 2.45888509 | 3.21E-20 | 1.12E-18 | 162.026346 | 397.8451039 | ENSG00000175592 |
| CSRP1 | 1.297785502 | 2.458512179 | 3.31E-02 | 8.50E-02 | 7.389101641 | 18.14635799 | ENSG00000159176 |
| CRYBB1 | 1.297557507 | 2.458123681 | 6.38E-06 | 4.36E-05 | 32.46510242 | 79.73089222 | ENSG00000100122 |
| ARMCX3 | 1.296523586 | 2.456362675 | 4.71E-45 | 6.54E-43 | 380.1253118 | 934.1334785 | ENSG00000102401 |
| KLHDC7A | 1.295650809 | 2.454877116 | 4.47E-53 | 8.13E-51 | 536.6664986 | 1317.669773 | ENSG00000179023 |
| C16orf89 | 1.293815293 | 2.451755805 | 5.21E-07 | 4.29E-06 | 45.59423835 | 111.6373958 | ENSG00000153446 |
| KANK3 | 1.293011336 | 2.450389918 | 2.31E-07 | 2.01E-06 | 47.04083321 | 115.468898 | ENSG00000186994 |
| CPZ | 1.289978195 | 2.445243597 | 1.27E-02 | 3.86E-02 | 10.96495107 | 26.8626075 | ENSG00000109625 |
| TAPBPL | 1.288520601 | 2.442774349 | 2.63E-02 | 7.06E-02 | 10.3033793 | 25.27890271 | ENSG00000139192 |
| SLC30A1 | 1.287751938 | 2.441473194 | 3.94E-32 | 2.84E-30 | 508.027578 | 1242.075147 | ENSG00000170385 |
| NFASC | 1.287345687 | 2.440785792 | 1.48E-39 | 1.67E-37 | 329.2546811 | 803.8162782 | ENSG00000163531 |
| TLR3 | 1.285160653 | 2.437091898 | 6.29E-03 | 2.14E-02 | 12.71246377 | 30.99849252 | ENSG00000164342 |
| FAXDC2 | 1.285130895 | 2.437041629 | 4.01E-06 | 2.84E-05 | 38.6781905 | 94.4700056 | ENSG00000170271 |
| ICAM3 | 1.28434309 | 2.43571121 | 6.42E-10 | 7.98E-09 | 65.85207019 | 160.3122446 | ENSG00000076662 |
| ATP2A3 | 1.277635513 | 2.424413049 | 9.87E-83 | 4.65E-80 | 1039.182006 | 2519.846883 | ENSG00000074370 |
| UHMK1 | 1.27552723 | 2.420872721 | 8.89E-32 | 6.26E-30 | 422.6941247 | 1023.021264 | ENSG00000152332 |
| C4orf50 | 1.273718569 | 2.417839649 | 1.92E-02 | 5.44E-02 | 11.0089298 | 26.64615621 | ENSG00000181215 |
| CD300A | 1.270350053 | 2.412200876 | 1.16E-02 | 3.56E-02 | 11.62863802 | 28.08277954 | ENSG00000167851 |
| NMNAT2 | 1.270140106 | 2.411849869 | 4.31E-42 | 5.37E-40 | 370.2857746 | 892.4884672 | ENSG00000157064 |
| CBX7 | 1.269570994 | 2.410898634 | 1.23E-11 | 1.87E-10 | 87.27630358 | 210.0833136 | ENSG00000100307 |
| MYMX | 1.268654218 | 2.409367089 | 3.21E-04 | 1.54E-03 | 28.58833516 | 68.94455568 | ENSG00000262179 |
| CRABP2 | 1.268542459 | 2.409180454 | 6.50E-102 | 5.70E-99 | 1498.208663 | 3607.715721 | ENSG00000143320 |
| RASEF | 1.267849219 | 2.408023079 | 1.70E-08 | 1.75E-07 | 86.69220137 | 208.6821413 | ENSG00000165105 |
| RYR2 | 1.263580113 | 2.400907985 | 2.78E-12 | 4.51E-11 | 109.384436 | 262.0841756 | ENSG00000198626 |
| ITPR2 | 1.261686582 | 2.39775887 | 8.89E-54 | 1.67E-51 | 850.575506 | 2038.371152 | ENSG00000123104 |
| LRRC7 | 1.260417155 | 2.395650011 | 2.24E-02 | 6.17E-02 | 9.032587189 | 21.61301305 | ENSG00000033122 |
| RASIP1 | 1.25802544 | 2.391681767 | 1.32E-20 | 4.69E-19 | 172.3618647 | 411.6318804 | ENSG00000105538 |
| MXRA8 | 1.257036922 | 2.390043575 | 1.17E-21 | 4.45E-20 | 170.478661 | 407.3666515 | ENSG00000162576 |
| STARD13 | 1.255639555 | 2.387729746 | 2.32E-08 | 2.33E-07 | 57.48105477 | 137.3419253 | ENSG00000133121 |
| IFIT1 | 1.254078082 | 2.38514683 | 1.40E-04 | 7.28E-04 | 28.96325592 | 68.84372076 | ENSG00000185745 |
| PADI2 | 1.252335467 | 2.382267578 | 1.44E-15 | 3.28E-14 | 146.2643139 | 347.5930262 | ENSG00000117115 |
| DOC2A | 1.252053743 | 2.381802424 | 1.40E-17 | 3.84E-16 | 130.6649691 | 311.1857663 | ENSG00000149927 |
| SULF1 | 1.24916211 | 2.377033292 | 4.70E-19 | 1.45E-17 | 303.1521859 | 719.3065988 | ENSG00000137573 |
| LPL | 1.247309763 | 2.373983261 | 1.52E-14 | 3.12E-13 | 113.898696 | 270.2701953 | ENSG00000175445 |
| KIFAP3 | 1.246887709 | 2.373288865 | 2.40E-08 | 2.41E-07 | 57.2492338 | 136.1521338 | ENSG00000075945 |
| OLFML3 | 1.246816651 | 2.373171974 | 5.87E-76 | 2.36E-73 | 811.3709095 | 1924.605689 | ENSG00000116774 |
| LRRC4C | 1.246178459 | 2.372122409 | 2.84E-02 | 7.52E-02 | 9.315883717 | 22.17899705 | ENSG00000148948 |
| SRMS | 1.244202893 | 2.368876347 | 4.23E-02 | 1.04E-01 | 8.384969932 | 19.86959592 | ENSG00000125508 |
| ELK4 | 1.243729157 | 2.368098609 | 1.22E-28 | 7.21E-27 | 318.6530116 | 754.1438446 | ENSG00000158711 |
| NEB | 1.242703119 | 2.366415027 | 2.24E-04 | 1.12E-03 | 27.87943018 | 65.79436279 | ENSG00000183091 |
| MSS51 | 1.241709337 | 2.364785515 | 2.03E-02 | 5.69E-02 | 10.34177622 | 24.52604242 | ENSG00000166343 |
| EFCAB13 | 1.241566896 | 2.364552044 | 4.12E-02 | 1.02E-01 | 8.003903785 | 18.91024372 | ENSG00000178852 |
| CALCR | 1.238333997 | 2.359259311 | 1.69E-04 | 8.64E-04 | 26.64336821 | 63.02167849 | ENSG00000004948 |
| LEPR | 1.237568478 | 2.358007779 | 4.04E-05 | 2.37E-04 | 32.20046633 | 76.13033171 | ENSG00000116678 |
| PAX5 | 1.235645503 | 2.354866873 | 8.86E-03 | 2.84E-02 | 12.40193393 | 29.14511724 | ENSG00000196092 |
| TLR5 | 1.234075895 | 2.352306244 | 2.32E-07 | 2.02E-06 | 65.1969561 | 153.2184044 | ENSG00000187554 |
| PPP1R15B | 1.233691423 | 2.351679447 | 3.52E-68 | 1.03E-65 | 1020.432819 | 2398.773568 | ENSG00000158615 |
| METTL27 | 1.231792524 | 2.348586164 | 2.19E-09 | 2.55E-08 | 77.7669958 | 182.3016938 | ENSG00000165171 |
| TTN | 1.225919081 | 2.33904413 | 1.48E-10 | 2.00E-09 | 90.67076844 | 211.6562471 | ENSG00000155657 |
| RNF212 | 1.225379896 | 2.338170112 | 4.32E-08 | 4.19E-07 | 62.90131063 | 147.3686893 | ENSG00000178222 |
| CRLF1 | 1.223937997 | 2.335834399 | 4.96E-15 | 1.07E-13 | 131.3579165 | 306.835841 | ENSG00000006016 |
| GABRB2 | 1.223483154 | 2.335098089 | 2.76E-02 | 7.36E-02 | 8.953678169 | 20.96716684 | ENSG00000145864 |
| RHBDL1 | 1.222602154 | 2.333672568 | 1.47E-07 | 1.32E-06 | 63.65662182 | 148.2495606 | ENSG00000103269 |
| MYCBPAP | 1.220297843 | 2.329948139 | 8.83E-03 | 2.84E-02 | 13.11239051 | 30.44756254 | ENSG00000136449 |
| LRRN4 | 1.218366592 | 2.32683126 | 1.78E-02 | 5.09E-02 | 11.62584712 | 27.07744125 | ENSG00000125872 |
| ADAMTSL2 | 1.218007371 | 2.326251968 | 3.52E-05 | 2.10E-04 | 38.1618337 | 88.81629007 | ENSG00000197859 |
| PDZD2 | 1.216297204 | 2.323496068 | 4.28E-29 | 2.60E-27 | 278.3173811 | 645.8774432 | ENSG00000133401 |

| | | | | | | | |
|---|---|---|---|---|---|---|---|
| ACAP1 | 1.215664668 | 2.322477577 | 4.90E-49 | 7.68E-47 | 557.5110092 | 1293.649977 | ENSG00000072818 |
| ACER2 | 1.215348848 | 2.321969219 | 4.44E-11 | 6.33E-10 | 83.49076024 | 193.973439 | ENSG00000177076 |
| HMCN1 | 1.215303325 | 2.321895953 | 3.05E-15 | 6.72E-14 | 243.4954704 | 564.4251542 | ENSG00000143341 |
| YPEL4 | 1.211968895 | 2.316535666 | 2.28E-03 | 8.85E-03 | 20.64583124 | 47.88032203 | ENSG00000166793 |
| TIGD6 | 1.2117913 | 2.316250519 | 6.64E-05 | 3.73E-04 | 30.49992342 | 70.53685066 | ENSG00000164296 |
| TNNC1 | 1.21124532 | 2.315374113 | 1.34E-04 | 6.99E-04 | 29.02242846 | 67.2264124 | ENSG00000114854 |
| CPEB4 | 1.21052822 | 2.314223529 | 1.08E-17 | 3.00E-16 | 152.6060348 | 353.1542487 | ENSG00000113742 |
| EDA2R | 1.210025184 | 2.313416751 | 1.32E-14 | 2.73E-13 | 121.7962573 | 281.9941299 | ENSG00000131080 |
| ADARB2 | 1.209896576 | 2.313210532 | 1.73E-06 | 1.30E-05 | 63.5425071 | 146.3913206 | ENSG00000185736 |
| TRIM22 | 1.208649247 | 2.311211436 | 3.09E-32 | 2.24E-30 | 457.3160514 | 1055.805974 | ENSG00000132274 |
| PLD1 | 1.207094471 | 2.308722011 | 8.24E-04 | 3.57E-03 | 22.74650105 | 52.58004047 | ENSG00000075651 |
| ELAVL2 | 1.206581875 | 2.307901857 | 9.19E-25 | 4.49E-23 | 274.7126218 | 634.6095159 | ENSG00000107105 |
| SUCLG2 | 1.203418048 | 2.30284618 | 4.66E-50 | 7.75E-48 | 675.7860838 | 1555.482946 | ENSG00000172340 |
| HTR1D | 1.199860164 | 2.297174041 | 1.32E-04 | 6.90E-04 | 29.09017387 | 66.74862035 | ENSG00000179546 |
| SPSB2 | 1.199518456 | 2.29663001 | 2.63E-12 | 4.29E-11 | 114.998079 | 264.638016 | ENSG00000111671 |
| NEFH | 1.197184873 | 2.292918177 | 3.80E-13 | 6.78E-12 | 119.6164031 | 274.1114148 | ENSG00000100285 |
| C1QTNF12 | 1.195954051 | 2.29096283 | 1.28E-03 | 5.31E-03 | 23.95935992 | 54.92709795 | ENSG00000184163 |
| CST1 | 1.194113951 | 2.28804266 | 1.20E-04 | 6.38E-04 | 30.81815024 | 70.49301515 | ENSG00000170373 |
| RRNAD1 | 1.193850373 | 2.287624676 | 1.40E-24 | 6.63E-23 | 245.4255942 | 561.1593817 | ENSG00000143303 |
| YOD1 | 1.193123493 | 2.286472382 | 3.57E-17 | 9.55E-16 | 154.1517508 | 352.780484 | ENSG00000180667 |
| PHYHD1 | 1.188990119 | 2.27993093 | 2.25E-11 | 3.31E-10 | 95.27530128 | 216.9120207 | ENSG00000175287 |
| LACTB2 | 1.187558063 | 2.277668935 | 2.19E-06 | 1.62E-05 | 47.27803583 | 107.7932321 | ENSG00000147592 |
| GRIK2 | 1.18668559 | 2.276291926 | 2.53E-04 | 1.25E-03 | 29.33997956 | 66.80561976 | ENSG00000164418 |
| PAH | 1.185849037 | 2.274972391 | 6.12E-04 | 2.74E-03 | 24.79617087 | 56.30483176 | ENSG00000171759 |
| CDK10 | 1.18511802 | 2.273819949 | 1.11E-25 | 5.68E-24 | 265.9576741 | 604.2297788 | ENSG00000185324 |
| PLAAT5 | 1.18486438 | 2.273420225 | 3.89E-09 | 4.39E-08 | 73.06748425 | 165.8229193 | ENSG00000168004 |
| POLD4 | 1.183163033 | 2.270740797 | 2.64E-04 | 1.30E-03 | 28.4069506 | 64.44352037 | ENSG00000175482 |
| ANXA2 | 1.181122453 | 2.26753128 | 3.81E-72 | 1.27E-69 | 859.2970382 | 1947.686833 | ENSG00000182718 |
| LMO7 | 1.181076883 | 2.267459658 | 9.25E-25 | 4.51E-23 | 236.0315652 | 534.9399368 | ENSG00000136153 |
| GPR20 | 1.179484589 | 2.264958456 | 2.90E-03 | 1.09E-02 | 19.95164451 | 44.9697037 | ENSG00000204882 |
| ACHE | 1.177076661 | 2.261181284 | 4.85E-08 | 4.67E-07 | 81.31955411 | 184.1558505 | ENSG00000087085 |
| DHRS12 | 1.175656337 | 2.258956261 | 1.57E-04 | 8.07E-04 | 29.04128906 | 65.6209414 | ENSG00000102796 |
| FZD5 | 1.173878101 | 2.256173634 | 2.20E-112 | 2.50E-109 | 1651.262559 | 3725.686592 | ENSG00000163251 |
| CLTRN | 1.169407158 | 2.249192526 | 4.71E-02 | 1.14E-01 | 8.564239314 | 19.38105101 | ENSG00000147003 |
| KRBA2 | 1.16869716 | 2.248085896 | 2.19E-02 | 6.05E-02 | 10.95936927 | 24.7301233 | ENSG00000184619 |
| SFR1 | 1.16855174 | 2.247859306 | 3.20E-06 | 2.29E-05 | 47.77361702 | 107.5095495 | ENSG00000156384 |
| EXOC3L2 | 1.167690103 | 2.246517193 | 8.51E-07 | 6.78E-06 | 51.52682081 | 115.7291848 | ENSG00000283632 |
| RBMS2 | 1.167072843 | 2.245556221 | 7.23E-17 | 1.88E-15 | 165.4810226 | 372.046458 | ENSG00000076067 |
| OPCML | 1.166631887 | 2.244869978 | 1.39E-04 | 7.26E-04 | 37.93280363 | 85.34398889 | ENSG00000183715 |
| MIXL1 | 1.166329381 | 2.24439932 | 1.03E-02 | 3.23E-02 | 22.02535642 | 49.68368265 | ENSG00000185155 |
| TNNI1 | 1.166048314 | 2.243962107 | 4.50E-04 | 2.09E-03 | 28.13740842 | 63.38306067 | ENSG00000159173 |
| P4HA2 | 1.163436587 | 2.239903512 | 1.64E-14 | 3.35E-13 | 146.1570956 | 327.5050647 | ENSG00000072682 |
| ACP5 | 1.156766885 | 2.229572147 | 1.65E-09 | 1.96E-08 | 96.45478146 | 214.4977866 | ENSG00000102575 |
| DACT1 | 1.15409348 | 2.225444435 | 2.90E-35 | 2.43E-33 | 401.3378241 | 893.4943754 | ENSG00000165617 |
| SNORA33 | 1.153124514 | 2.223950248 | 3.83E-02 | 9.63E-02 | 11.59390789 | 25.59510442 | ENSG00000200534 |
| SYT5 | 1.15088033 | 2.220493472 | 1.52E-04 | 7.87E-04 | 36.62784496 | 81.52752855 | ENSG00000129990 |
| TENT5C | 1.149776072 | 2.218794527 | 1.06E-03 | 4.49E-03 | 23.9197237 | 52.89732668 | ENSG00000183508 |
| IL2RB | 1.148248938 | 2.216447112 | 7.93E-05 | 4.38E-04 | 38.64346037 | 85.4937724 | ENSG00000100385 |
| PCSK1 | 1.147354314 | 2.215073106 | 6.86E-04 | 3.03E-03 | 26.88826782 | 59.70882919 | ENSG00000175426 |
| POMK | 1.145991117 | 2.212981081 | 3.70E-05 | 2.19E-04 | 38.32435767 | 84.97106034 | ENSG00000185900 |
| ASH1L | 1.145649388 | 2.212456957 | 1.74E-30 | 1.13E-28 | 448.4632972 | 991.6971499 | ENSG00000116539 |
| TRPV2 | 1.145319347 | 2.211950877 | 4.93E-04 | 2.26E-03 | 28.48932624 | 63.00877505 | ENSG00000187688 |
| ZFHX3 | 1.142878111 | 2.208211121 | 1.06E-05 | 6.92E-05 | 45.81334411 | 101.1339057 | ENSG00000140836 |
| DDR2 | 1.141600011 | 2.206255709 | 1.66E-02 | 4.80E-02 | 15.09922101 | 33.20115215 | ENSG00000162733 |
| SEMA3G | 1.138178293 | 2.201029213 | 2.49E-09 | 2.88E-08 | 87.78408751 | 193.2719505 | ENSG00000010319 |
| RASAL2 | 1.137661246 | 2.200240527 | 1.02E-43 | 1.35E-41 | 603.0068902 | 1325.921812 | ENSG00000075391 |
| MYOF | 1.136896328 | 2.199074267 | 1.26E-04 | 6.67E-04 | 34.09251827 | 74.89107708 | ENSG00000138119 |
| DISC1 | 1.136465902 | 2.198418275 | 1.72E-05 | 1.08E-04 | 41.63365598 | 91.67233213 | ENSG00000162946 |
| HES2 | 1.135719647 | 2.197281405 | 1.86E-02 | 5.29E-02 | 13.90155595 | 30.69546682 | ENSG00000069812 |
| FAM50B | 1.134404294 | 2.195278984 | 3.80E-08 | 3.71E-07 | 69.54590128 | 152.6190894 | ENSG00000145945 |
| PLXDC2 | 1.133307336 | 2.193610431 | 7.70E-28 | 4.38E-26 | 394.0820981 | 865.0165185 | ENSG00000120594 |
| MMP23B | 1.131799335 | 2.191318721 | 6.65E-04 | 2.95E-03 | 28.26886904 | 61.78080633 | ENSG00000189409 |
| PLA2G4A | 1.131784102 | 2.191295584 | 4.77E-04 | 2.20E-03 | 28.95958914 | 63.5172387 | ENSG00000116711 |
| ZBTB41 | 1.131570415 | 2.19097104 | 2.75E-07 | 2.37E-06 | 65.25401346 | 143.0015411 | ENSG00000177888 |
| GAS5 | 1.130806065 | 2.189810556 | 8.56E-100 | 6.61E-97 | 3880.315331 | 8494.623767 | ENSG00000234741 |
| PPP1R17 | 1.130148256 | 2.18881232 | 2.05E-03 | 8.05E-03 | 21.99764755 | 48.17904658 | ENSG00000106341 |
| ELAVL3 | 1.129863692 | 2.188380632 | 4.07E-03 | 1.47E-02 | 20.80644018 | 45.49804344 | ENSG00000196361 |
| PCDHGA4 | 1.129827214 | 2.1883253 | 4.87E-02 | 1.17E-01 | 9.01372659 | 19.74322669 | ENSG00000262576 |
| CEACAM19 | 1.126313337 | 2.183001826 | 1.65E-02 | 4.78E-02 | 14.88067883 | 32.64590891 | ENSG00000186567 |
| PLOD2 | 1.126200459 | 2.182831033 | 4.16E-35 | 3.41E-33 | 422.3638825 | 921.697817 | ENSG00000152952 |
| VASH2 | 1.124014656 | 2.179526366 | 1.01E-82 | 4.65E-80 | 1463.45764 | 3190.432561 | ENSG00000143494 |
| ACOT11 | 1.122182473 | 2.176760185 | 8.64E-10 | 1.06E-08 | 98.63627531 | 214.3671041 | ENSG00000162390 |
| RAG1 | 1.120450584 | 2.174148651 | 1.27E-02 | 3.86E-02 | 15.48865986 | 33.59639836 | ENSG00000166349 |

| | | | | | | | |
|---|---|---|---|---|---|---|---|
| LAMC3 | 1.116825784 | 2.168692917 | 8.53E-67 | 2.35E-64 | 1040.687386 | 2257.145066 | ENSG00000050555 |
| UCMA | 1.115743107 | 2.167066021 | 6.76E-03 | 2.27E-02 | 17.28697239 | 37.58225783 | ENSG00000165623 |
| HNMT | 1.115559543 | 2.166790308 | 3.87E-03 | 1.40E-02 | 21.14631851 | 45.66745468 | ENSG00000150540 |
| TIRAP | 1.114989045 | 2.165933644 | 7.26E-03 | 2.41E-02 | 17.54323577 | 37.87453066 | ENSG00000150455 |
| CORO2A | 1.113694516 | 2.163991026 | 8.07E-73 | 2.83E-70 | 1397.217769 | 3022.351036 | ENSG00000106789 |
| MTMR11 | 1.113332417 | 2.163447958 | 1.53E-05 | 9.68E-05 | 46.07307411 | 99.38969508 | ENSG00000014914 |
| KC6 | 1.111440623 | 2.160612905 | 1.69E-06 | 1.28E-05 | 55.40136074 | 119.8419465 | ENSG00000267313 |
| MAML2 | 1.111239208 | 2.160311284 | 9.24E-10 | 1.13E-08 | 98.81554469 | 213.414985 | ENSG00000184384 |
| RNF152 | 1.111023836 | 2.159988806 | 4.49E-03 | 1.60E-02 | 23.24611244 | 50.34404985 | ENSG00000176641 |
| ELOVL2 | 1.11024105 | 2.158817144 | 7.21E-05 | 4.02E-04 | 43.27175989 | 93.19741998 | ENSG00000197977 |
| ANO9 | 1.109320607 | 2.157440253 | 3.23E-20 | 1.13E-18 | 278.3511473 | 600.176155 | ENSG00000185101 |
| HSH2D | 1.105470649 | 2.15169061 | 4.95E-03 | 1.74E-02 | 24.8429405 | 53.41329602 | ENSG00000196684 |
| RBM15 | 1.104682574 | 2.150515566 | 2.13E-28 | 1.24E-26 | 380.3111638 | 816.8375572 | ENSG00000162775 |
| SERTM2 | 1.10417198 | 2.149754597 | 4.67E-03 | 1.66E-02 | 20.61022522 | 44.4228023 | ENSG00000260802 |
| RGS2 | 1.103169577 | 2.14826144 | 1.59E-15 | 3.62E-14 | 190.8803564 | 410.2984788 | ENSG00000116741 |
| FHIT | 1.1007358 | 2.144640451 | 7.62E-03 | 2.51E-02 | 18.27725887 | 39.24985343 | ENSG00000189283 |
| CBLC | 1.094541 | 2.135451304 | 9.80E-18 | 2.74E-16 | 202.5007339 | 432.3565044 | ENSG00000142273 |
| TAGLN2 | 1.091969678 | 2.131648669 | 6.97E-93 | 4.80E-90 | 1656.738334 | 3531.330022 | ENSG00000158710 |
| SYTL1 | 1.09108878 | 2.130347499 | 2.52E-11 | 3.69E-10 | 118.5833772 | 252.604616 | ENSG00000142765 |
| ADAMTSL4 | 1.089994914 | 2.12873286 | 2.98E-10 | 3.89E-09 | 137.1453209 | 291.2284993 | ENSG00000143382 |
| LGALS8 | 1.08903559 | 2.127317824 | 4.90E-13 | 8.59E-12 | 151.9685421 | 322.7400292 | ENSG00000116977 |
| VAMP4 | 1.08827901 | 2.126202506 | 6.69E-10 | 8.29E-09 | 134.9989206 | 287.0533099 | ENSG00000117533 |
| CNTD2 | 1.087633987 | 2.125252103 | 1.44E-03 | 5.90E-03 | 24.75777395 | 52.60720443 | ENSG00000105219 |
| PRODH | 1.087337817 | 2.124815855 | 5.38E-04 | 2.45E-03 | 29.41956431 | 62.53743472 | ENSG00000100033 |
| GAL3ST1 | 1.087190014 | 2.12459818 | 2.30E-19 | 7.31E-18 | 220.2141906 | 467.4852784 | ENSG00000128242 |
| PTPN6 | 1.083969857 | 2.119861275 | 3.95E-28 | 2.29E-26 | 345.2056315 | 731.4139545 | ENSG00000111679 |
| GCSAM | 1.083494154 | 2.119162404 | 1.34E-02 | 4.02E-02 | 16.18707695 | 34.45869586 | ENSG00000174500 |
| PHF11 | 1.082476312 | 2.117667832 | 7.73E-07 | 6.19E-06 | 77.34071163 | 163.4812291 | ENSG00000136147 |
| CAPN6 | 1.082332033 | 2.117456061 | 1.68E-13 | 3.10E-12 | 173.6889504 | 367.470431 | ENSG00000077274 |
| TRIML2 | 1.081828807 | 2.116717601 | 3.75E-32 | 2.71E-30 | 520.3906493 | 1101.243498 | ENSG00000179046 |
| TP73 | 1.081719734 | 2.116557575 | 3.62E-09 | 4.11E-08 | 93.52758843 | 197.8818715 | ENSG00000078900 |
| VWA5B1 | 1.081272146 | 2.115901027 | 2.06E-03 | 8.08E-03 | 32.56095701 | 68.55038186 | ENSG00000158816 |
| ESRRG | 1.080515321 | 2.114791336 | 4.01E-04 | 1.89E-03 | 31.88802147 | 67.30600205 | ENSG00000196482 |
| STAC3 | 1.080429148 | 2.114665021 | 1.33E-03 | 5.51E-03 | 28.48865052 | 60.19792556 | ENSG00000185482 |
| EMILIN1 | 1.079804547 | 2.113749696 | 1.92E-13 | 3.53E-12 | 150.565977 | 318.6912396 | ENSG00000138080 |
| ILDR1 | 1.076861914 | 2.109442721 | 4.89E-04 | 2.25E-03 | 31.64571261 | 66.904292 | ENSG00000145103 |
| A2ML1 | 1.076111295 | 2.108345486 | 9.92E-13 | 1.68E-11 | 159.8306975 | 336.8429527 | ENSG00000166535 |
| BOLA1 | 1.074123504 | 2.105442541 | 3.54E-11 | 5.09E-10 | 126.7773505 | 266.7148152 | ENSG00000178096 |
| TP53INP1 | 1.073504452 | 2.104539302 | 3.66E-10 | 4.71E-09 | 110.8748094 | 233.2122486 | ENSG00000164938 |
| SPAG4 | 1.072281484 | 2.102756047 | 1.48E-04 | 7.64E-04 | 40.60305756 | 85.29720931 | ENSG00000061656 |
| ADA2 | 1.071020842 | 2.100919439 | 7.96E-43 | 1.02E-40 | 948.7988753 | 1992.396946 | ENSG00000093072 |
| APOBEC3B | 1.071009334 | 2.100902681 | 3.17E-08 | 3.12E-07 | 86.69112533 | 182.3998209 | ENSG00000179750 |
| S100A16 | 1.067917938 | 2.096405702 | 6.67E-20 | 2.23E-18 | 251.5970189 | 527.1118101 | ENSG00000188643 |
| ZMAT1 | 1.067428422 | 2.095694499 | 7.98E-03 | 2.61E-02 | 18.67507044 | 39.20224981 | ENSG00000166432 |
| VIL1 | 1.066286983 | 2.094037071 | 9.30E-13 | 1.58E-11 | 144.5862615 | 302.428668 | ENSG00000127831 |
| MC1R | 1.065576732 | 2.093006413 | 3.99E-02 | 9.94E-02 | 14.17109813 | 29.48605976 | ENSG00000258839 |
| TLE2 | 1.065517226 | 2.092920086 | 3.47E-26 | 1.82E-24 | 379.7976362 | 795.2678587 | ENSG00000065717 |
| C1orf167 | 1.06354132 | 2.090055597 | 2.69E-02 | 7.19E-02 | 12.73132436 | 26.57301829 | ENSG00000215910 |
| MINAR1 | 1.063127353 | 2.089455963 | 2.10E-02 | 5.84E-02 | 16.30525877 | 33.87415021 | ENSG00000169330 |
| HCP5 | 1.062161995 | 2.088058303 | 8.29E-05 | 4.57E-04 | 43.51925024 | 91.12381319 | ENSG00000206337 |
| CEP350 | 1.059273243 | 2.083881503 | 4.79E-32 | 3.42E-30 | 557.9010996 | 1162.154198 | ENSG00000135837 |
| DUOX1 | 1.056761416 | 2.080256485 | 5.97E-03 | 2.05E-02 | 21.22993345 | 44.28352962 | ENSG00000137857 |
| BAIAP3 | 1.056600242 | 2.080024097 | 9.66E-18 | 2.70E-16 | 222.0262575 | 461.7726855 | ENSG00000007516 |
| ZFP92 | 1.056527571 | 2.079919326 | 1.33E-02 | 4.00E-02 | 18.74349157 | 39.02290973 | ENSG00000189420 |
| GCNT4 | 1.056480306 | 2.079851184 | 3.88E-04 | 1.84E-03 | 34.98224423 | 72.83119777 | ENSG00000176928 |
| KCNS1 | 1.055975333 | 2.079123322 | 8.58E-06 | 5.73E-05 | 60.01079966 | 124.9049373 | ENSG00000124134 |
| TRPV4 | 1.055438386 | 2.07834965 | 1.20E-04 | 6.37E-04 | 43.49588387 | 90.11794187 | ENSG00000111199 |
| SNHG8 | 1.055158809 | 2.07794693 | 1.04E-34 | 8.39E-33 | 568.3507714 | 1182.055865 | ENSG00000269893 |
| NUDT18 | 1.054693557 | 2.077276926 | 5.55E-04 | 2.52E-03 | 32.71490811 | 67.99002555 | ENSG00000275074 |
| FGF18 | 1.054629662 | 2.077184927 | 3.34E-02 | 8.58E-02 | 17.0572666 | 35.67214361 | ENSG00000156427 |
| ATF6 | 1.054445155 | 2.076919292 | 1.75E-36 | 1.58E-34 | 480.1202245 | 997.4119482 | ENSG00000118217 |
| RGS16 | 1.054407639 | 2.076865285 | 2.21E-23 | 9.63E-22 | 327.7983252 | 681.6646701 | ENSG00000143333 |
| HBQ1 | 1.054275458 | 2.076675008 | 7.42E-04 | 3.25E-03 | 30.6791928 | 63.78718175 | ENSG00000086506 |
| PYCR2 | 1.046840474 | 2.066000315 | 7.18E-31 | 4.75E-29 | 444.3984506 | 918.61571 | ENSG00000143811 |
| RNF213 | 1.04554523 | 2.064146303 | 1.41E-31 | 9.73E-30 | 1053.382352 | 2173.365728 | ENSG00000173821 |
| LAMC2 | 1.045273251 | 2.063757203 | 5.41E-22 | 2.12E-20 | 277.1621431 | 571.5892129 | ENSG00000058085 |
| NEURL1 | 1.04504612 | 2.063432321 | 1.15E-02 | 3.55E-02 | 19.3254786 | 39.88064563 | ENSG00000107954 |
| IRAK1 | 1.044249064 | 2.062292636 | 1.00E-54 | 1.93E-52 | 1083.033414 | 2234.123158 | ENSG00000184216 |
| SH2D3A | 1.043830244 | 2.061694032 | 6.10E-38 | 6.09E-36 | 539.7738369 | 1112.581916 | ENSG00000125731 |
| CTSK | 1.041765732 | 2.058745835 | 1.01E-09 | 1.22E-08 | 113.4545904 | 233.8718102 | ENSG00000143387 |
| CCDC172 | 1.041754209 | 2.058729393 | 2.25E-03 | 8.73E-03 | 25.61747571 | 52.82795686 | ENSG00000182645 |
| LENG9 | 1.040784422 | 2.057345969 | 1.67E-07 | 1.49E-06 | 74.85251799 | 153.9952057 | ENSG00000275183 |
| ELN | 1.040552091 | 2.057014681 | 1.34E-14 | 2.77E-13 | 168.3144762 | 346.2752544 | ENSG00000049540 |

| | | | | | | | |
|---|---|---|---|---|---|---|---|
| OPN3 | 1.0401977 | 2.056509448 | 1.13E-02 | 3.49E-02 | 17.70364455 | 36.40215319 | ENSG00000054277 |
| GIPC3 | 1.039227282 | 2.055126617 | 1.91E-06 | 1.43E-05 | 73.66111046 | 151.4776951 | ENSG00000179855 |
| CARF | 1.039024344 | 2.054837551 | 6.51E-03 | 2.20E-02 | 25.99575095 | 53.53159599 | ENSG00000138380 |
| AKT3 | 1.037559229 | 2.052751841 | 9.40E-19 | 2.84E-17 | 284.4122873 | 583.9000311 | ENSG00000117020 |
| OBSCN | 1.035520124 | 2.04985253 | 1.55E-68 | 4.60E-66 | 1252.834707 | 2567.052249 | ENSG00000154358 |
| LTBR | 1.03405134 | 2.047766672 | 2.08E-04 | 1.05E-03 | 51.02740958 | 104.1144269 | ENSG00000111321 |
| PYROXD1 | 1.03305802 | 2.046357235 | 1.16E-05 | 7.49E-05 | 58.03389347 | 118.6911443 | ENSG00000121350 |
| LY75 | 1.032987993 | 2.046257909 | 2.49E-03 | 9.58E-03 | 29.26262215 | 59.7249677 | ENSG00000054219 |
| ADGRA2 | 1.032048672 | 2.04492605 | 3.86E-07 | 3.25E-06 | 100.699187 | 206.4064106 | ENSG00000020181 |
| SPIB | 1.031036237 | 2.043491492 | 1.80E-08 | 1.85E-07 | 98.24085433 | 200.9414493 | ENSG00000269404 |
| TUBB6 | 1.030399242 | 2.042589426 | 1.43E-84 | 7.28E-82 | 3084.244985 | 6300.897145 | ENSG00000176014 |
| MCHR1 | 1.03002673 | 2.042062087 | 1.99E-04 | 1.01E-03 | 47.50706591 | 96.82729502 | ENSG00000128285 |
| MEF2D | 1.029611582 | 2.041474549 | 9.05E-30 | 5.71E-28 | 492.5829706 | 1005.306022 | ENSG00000116604 |
| GRIP2 | 1.027039858 | 2.037838693 | 2.46E-15 | 5.50E-14 | 200.7152998 | 409.1097961 | ENSG00000144596 |
| NAXE | 1.025477607 | 2.035633173 | 1.95E-74 | 7.24E-72 | 1427.384374 | 2905.21086 | ENSG00000163382 |
| TRPV1 | 1.023939987 | 2.033464757 | 1.99E-04 | 1.00E-03 | 48.9747487 | 99.36254323 | ENSG00000196689 |
| SLC17A9 | 1.023542121 | 2.032904046 | 2.29E-39 | 2.55E-37 | 749.4157448 | 1523.805144 | ENSG00000101194 |
| SIT1 | 1.022540472 | 2.031493111 | 6.72E-06 | 4.57E-05 | 87.74895706 | 178.3271508 | ENSG00000137078 |
| ISLR2 | 1.022513917 | 2.031455718 | 1.90E-03 | 7.56E-03 | 29.30504927 | 59.5668729 | ENSG00000167178 |
| ADCY7 | 1.021801777 | 2.030453203 | 5.61E-08 | 5.34E-07 | 92.20021452 | 187.2525516 | ENSG00000121281 |
| RENBP | 1.021426057 | 2.029924482 | 7.06E-17 | 1.84E-15 | 233.8223255 | 474.8135743 | ENSG00000102032 |
| ZCCHC12 | 1.019708455 | 2.027509192 | 7.42E-04 | 3.25E-03 | 36.34889094 | 73.47595949 | ENSG00000174460 |
| APOBEC3C | 1.019453233 | 2.027150545 | 1.62E-25 | 8.12E-24 | 375.7254547 | 761.8458994 | ENSG00000244509 |
| SPN | 1.01885032 | 2.02630356 | 6.62E-04 | 2.94E-03 | 35.96503389 | 72.64674462 | ENSG00000197471 |
| CDK18 | 1.018203122 | 2.025394757 | 3.99E-70 | 1.26E-67 | 1265.421731 | 2563.304371 | ENSG00000117266 |
| MYO7A | 1.018061122 | 2.025195414 | 4.12E-09 | 4.64E-08 | 107.4885171 | 217.6928689 | ENSG00000137474 |
| MRVI1 | 1.017365779 | 2.024219555 | 2.22E-06 | 1.64E-05 | 72.43294563 | 146.8500059 | ENSG00000072952 |
| CHRM3 | 1.015161661 | 2.021129357 | 1.57E-12 | 2.62E-11 | 155.4752946 | 313.992042 | ENSG00000133019 |
| RASGRF2 | 1.015107059 | 2.021052865 | 5.26E-19 | 1.62E-17 | 361.0345331 | 729.5804426 | ENSG00000113319 |
| PIFO | 1.012333212 | 2.017170752 | 3.76E-05 | 2.22E-04 | 52.37065302 | 105.456667 | ENSG00000173947 |
| BATF2 | 1.011940866 | 2.016622249 | 3.26E-03 | 1.21E-02 | 30.65965648 | 61.92923276 | ENSG00000168062 |
| CDH11 | 1.011347514 | 2.015793024 | 2.85E-15 | 6.31E-14 | 197.9265887 | 399.3268956 | ENSG00000140937 |
| IRF6 | 1.011034877 | 2.015356241 | 5.18E-24 | 2.38E-22 | 407.1729493 | 820.8540399 | ENSG00000117595 |
| STAB1 | 1.010533149 | 2.01465548 | 1.53E-02 | 4.51E-02 | 21.22869415 | 42.63287816 | ENSG00000010327 |
| DNAJA4 | 1.007664672 | 2.010653767 | 5.53E-08 | 5.28E-07 | 127.1015595 | 254.8900699 | ENSG00000140403 |
| NPL | 1.006888046 | 2.009571691 | 2.12E-05 | 1.32E-04 | 60.40996267 | 121.5409951 | ENSG00000135838 |
| GADD45G | 1.005671817 | 2.007878284 | 6.88E-13 | 1.19E-11 | 162.9331808 | 327.2198193 | ENSG00000130222 |
| AMIGO2 | 1.005492859 | 2.007629233 | 2.71E-10 | 3.56E-09 | 132.9250085 | 266.4376085 | ENSG00000139211 |
| TRHDE | 1.005201814 | 2.007224262 | 2.74E-03 | 1.04E-02 | 31.45998554 | 63.44032055 | ENSG00000072657 |
| ZDHHC21 | 1.005175526 | 2.007187687 | 2.86E-06 | 2.07E-05 | 103.1396471 | 207.5085126 | ENSG00000175893 |
| SPINK5 | 1.005149387 | 2.007151322 | 9.05E-06 | 6.01E-05 | 63.83656692 | 128.140087 | ENSG00000133710 |
| SARDH | 1.00286203 | 2.003971555 | 1.35E-21 | 5.10E-20 | 311.4941794 | 623.8121724 | ENSG00000123453 |
| SULT1C4 | 1.002569714 | 2.003565554 | 1.47E-02 | 4.36E-02 | 21.29767886 | 42.74634395 | ENSG00000198075 |
| RRM2B | 1.002540463 | 2.003524933 | 4.94E-46 | 7.23E-44 | 1218.219113 | 2441.336512 | ENSG00000048392 |
| LAMC1 | 1.001004419 | 2.001392905 | 5.69E-116 | 7.31E-113 | 4192.538256 | 8389.687545 | ENSG00000135862 |
| PLD5 | 0.999949149 | 1.999929507 | 2.19E-05 | 1.35E-04 | 55.10267024 | 110.1951263 | ENSG00000180287 |
| STEAP2 | 0.998330856 | 1.997687413 | 1.22E-12 | 2.06E-11 | 198.417427 | 397.1121503 | ENSG00000157214 |
| RAB20 | 0.997981609 | 1.997203873 | 1.39E-22 | 5.67E-21 | 326.8603149 | 652.8567273 | ENSG00000139832 |
| TIGD4 | 0.997261614 | 1.996207391 | 1.85E-02 | 5.25E-02 | 25.27624594 | 50.45054299 | ENSG00000169989 |
| KCNQ5 | 0.995312158 | 1.993511818 | 5.50E-03 | 1.91E-02 | 26.78790745 | 53.41007937 | ENSG00000185760 |
| MARCHF1 | 0.994585701 | 1.992508254 | 3.45E-03 | 1.27E-02 | 29.23471312 | 58.22783732 | ENSG00000145416 |
| RNF180 | 0.994466144 | 1.992343141 | 7.12E-03 | 2.37E-02 | 24.23515962 | 48.21318319 | ENSG00000164197 |
| PDK1 | 0.993149532 | 1.990525746 | 2.67E-14 | 5.40E-13 | 218.5127718 | 434.6513903 | ENSG00000152256 |
| TMEM63A | 0.992824765 | 1.990077706 | 7.56E-47 | 1.14E-44 | 1673.066554 | 3327.767314 | ENSG00000196187 |
| CSPG5 | 0.992438545 | 1.98954502 | 9.12E-09 | 9.79E-08 | 110.8409182 | 220.816964 | ENSG00000114646 |
| ABO | 0.991793575 | 1.988655774 | 2.30E-02 | 6.31E-02 | 15.71123225 | 31.25527167 | ENSG00000175164 |
| REXO5 | 0.991647934 | 1.988455028 | 1.57E-06 | 1.20E-05 | 83.71313246 | 166.9089012 | ENSG00000005189 |
| SESN1 | 0.991371197 | 1.98807364 | 1.37E-21 | 5.19E-20 | 322.8172689 | 642.2252383 | ENSG00000080546 |
| SLC5A3 | 0.991215985 | 1.987859765 | 6.12E-22 | 2.40E-20 | 533.7151371 | 1060.564224 | ENSG00000198743 |
| FGF11 | 0.990160704 | 1.986406247 | 1.93E-02 | 5.46E-02 | 17.71413244 | 35.20080328 | ENSG00000161958 |
| USH1C | 0.99012207 | 1.986353053 | 3.61E-03 | 1.32E-02 | 32.6010688 | 64.98636898 | ENSG00000006611 |
| NAV3 | 0.989693505 | 1.985763078 | 2.32E-03 | 8.98E-03 | 31.64782779 | 62.97973309 | ENSG00000067798 |
| ASB16 | 0.988512484 | 1.984138156 | 3.65E-02 | 9.24E-02 | 13.43899004 | 26.60636139 | ENSG00000161664 |
| RORA | 0.988162377 | 1.983656712 | 2.89E-04 | 1.40E-03 | 47.5825083 | 94.39848521 | ENSG00000069667 |
| NFAM1 | 0.986837944 | 1.981836497 | 1.03E-02 | 3.23E-02 | 24.23862625 | 47.90129473 | ENSG00000235568 |
| CRYAB | 0.983399717 | 1.977119013 | 2.65E-02 | 7.10E-02 | 16.59537641 | 32.91720904 | ENSG00000109846 |
| ZNF91 | 0.982758822 | 1.976240903 | 1.95E-18 | 5.74E-17 | 283.0559284 | 559.8168506 | ENSG00000167232 |
| ZBTB18 | 0.982741905 | 1.976217731 | 9.85E-16 | 2.30E-14 | 271.0105429 | 535.4327708 | ENSG00000179456 |
| LHFPL1 | 0.982617532 | 1.976047371 | 2.96E-04 | 1.44E-03 | 46.79680949 | 92.29531554 | ENSG00000182508 |
| LPAR6 | 0.980539547 | 1.973203221 | 1.32E-03 | 5.47E-03 | 34.91469898 | 68.7071317 | ENSG00000139679 |
| C3 | 0.979249344 | 1.971439373 | 9.14E-17 | 2.35E-15 | 270.6338193 | 533.0351898 | ENSG00000125730 |
| PPL | 0.97689428 | 1.968223809 | 1.36E-44 | 1.84E-42 | 1015.712095 | 1998.206231 | ENSG00000118898 |
| SVOPL | 0.975932215 | 1.966911731 | 1.89E-03 | 7.53E-03 | 33.14553479 | 65.18642129 | ENSG00000157703 |

| | | | | | | | |
|---|---|---|---|---|---|---|---|
| RCN3 | 0.975828815 | 1.966770764 | 1.91E-17 | 5.20E-16 | 292.8537511 | 575.8509036 | ENSG00000142552 |
| PURPL | 0.975093099 | 1.965768046 | 1.14E-07 | 1.04E-06 | 139.34336 | 273.7448891 | ENSG00000250337 |
| COQ8A | 0.974500253 | 1.96496042 | 1.14E-46 | 1.71E-44 | 972.1954496 | 1911.34784 | ENSG00000163050 |
| IPP | 0.974139788 | 1.964469526 | 1.39E-06 | 1.07E-05 | 84.64377085 | 166.2192316 | ENSG00000197429 |
| IER5 | 0.974023624 | 1.964311356 | 1.81E-30 | 1.16E-28 | 523.884011 | 1029.279971 | ENSG00000162783 |
| ADAMTS16 | 0.973902331 | 1.964146215 | 5.78E-13 | 1.01E-11 | 215.7143237 | 424.3880205 | ENSG00000145536 |
| ATP1B1 | 0.973328632 | 1.963365313 | 6.33E-38 | 6.30E-36 | 801.3925043 | 1574.690328 | ENSG00000143153 |
| C1orf94 | 0.972237678 | 1.961881193 | 2.04E-07 | 1.79E-06 | 110.2331374 | 216.6243279 | ENSG00000142698 |
| ZMAT4 | 0.970451117 | 1.959453201 | 1.68E-13 | 3.10E-12 | 198.4059 | 389.1616645 | ENSG00000165061 |
| MSRB3 | 0.969990319 | 1.95882745 | 2.37E-02 | 6.47E-02 | 19.48030558 | 38.29774644 | ENSG00000174099 |
| HPCAL4 | 0.969836589 | 1.958618734 | 5.19E-04 | 2.37E-03 | 38.99083552 | 76.43497494 | ENSG00000116983 |
| HLA-DPA1 | 0.969708719 | 1.958445145 | 1.58E-03 | 6.42E-03 | 33.27611952 | 65.01327248 | ENSG00000231389 |
| ELFN1 | 0.969566803 | 1.958252503 | 6.52E-08 | 6.15E-07 | 112.4415011 | 220.2751451 | ENSG00000225968 |
| EBF4 | 0.968590801 | 1.956928168 | 1.82E-03 | 7.26E-03 | 37.90876154 | 73.83921967 | ENSG00000088881 |
| TOMM20 | 0.967109871 | 1.954920407 | 9.97E-84 | 4.81E-81 | 4486.619348 | 8770.783287 | ENSG00000173726 |
| BAHCC1 | 0.966758853 | 1.95444482 | 4.12E-03 | 1.48E-02 | 29.3706795 | 57.52284111 | ENSG00000266074 |
| ACTN3 | 0.965297064 | 1.952465511 | 2.13E-20 | 7.47E-19 | 359.1393268 | 700.8475342 | ENSG00000248746 |
| KDELR3 | 0.964627263 | 1.951559248 | 5.63E-09 | 6.20E-08 | 122.2396872 | 238.445723 | ENSG00000100196 |
| HLA-DRA | 0.961885693 | 1.9478542 | 5.24E-04 | 2.39E-03 | 44.27456143 | 86.14660337 | ENSG00000204287 |
| SCX | 0.961873184 | 1.94783731 | 8.59E-04 | 3.71E-03 | 39.90567961 | 77.89283143 | ENSG00000260428 |
| CALB1 | 0.961414732 | 1.947218436 | 3.90E-22 | 1.54E-20 | 356.4032425 | 694.367833 | ENSG00000104327 |
| MDM2 | 0.960301614 | 1.945716629 | 5.93E-19 | 1.81E-17 | 404.1945689 | 786.4490957 | ENSG00000135679 |
| PCDHGA6 | 0.959744212 | 1.944965025 | 4.56E-02 | 1.11E-01 | 17.65475974 | 34.34791368 | ENSG00000253731 |
| LACC1 | 0.958598138 | 1.943420562 | 2.58E-04 | 1.27E-03 | 45.53139902 | 88.45008581 | ENSG00000179630 |
| FAM71E1 | 0.958454922 | 1.943227648 | 1.15E-07 | 1.05E-06 | 98.65984183 | 191.7253385 | ENSG00000142530 |
| ZNF808 | 0.958334714 | 1.943065742 | 1.51E-04 | 7.80E-04 | 57.9195786 | 112.3098424 | ENSG00000198482 |
| IAH1 | 0.958152784 | 1.942820728 | 2.88E-13 | 5.22E-12 | 190.6340173 | 370.4068319 | ENSG00000134330 |
| DARS2 | 0.956113081 | 1.940075881 | 7.80E-34 | 6.05E-32 | 764.6372531 | 1484.624042 | ENSG00000117593 |
| PLEKHA2 | 0.955948395 | 1.93985443 | 1.30E-07 | 1.17E-06 | 148.0537782 | 286.8441344 | ENSG00000169499 |
| FAAH2 | 0.954938296 | 1.938496723 | 1.13E-05 | 7.31E-05 | 71.19564436 | 138.0262214 | ENSG00000165591 |
| MUC12 | 0.954537127 | 1.937957762 | 3.22E-02 | 8.33E-02 | 16.16351043 | 31.24021765 | ENSG00000205277 |
| PSD4 | 0.954261065 | 1.937586965 | 2.19E-03 | 8.53E-03 | 40.36266372 | 78.46500668 | ENSG00000125637 |
| PLAAT3 | 0.953741829 | 1.93688974 | 4.31E-40 | 5.07E-38 | 1089.4194 | 2109.787281 | ENSG00000176485 |
| IL23A | 0.950405189 | 1.932415311 | 2.29E-02 | 6.29E-02 | 20.28382585 | 39.01726362 | ENSG00000110944 |
| ZBTB7B | 0.950393157 | 1.932399195 | 5.96E-41 | 7.33E-39 | 750.4902467 | 1450.431564 | ENSG00000160685 |
| S100A10 | 0.949367739 | 1.9310262 | 1.98E-35 | 1.68E-33 | 621.1083094 | 1199.124832 | ENSG00000197747 |
| PKD1L2 | 0.948745479 | 1.930193494 | 2.94E-03 | 1.10E-02 | 31.32190398 | 60.56278486 | ENSG00000166473 |
| DNAJC6 | 0.948627016 | 1.930035007 | 1.84E-07 | 1.63E-06 | 95.61218854 | 184.4873972 | ENSG00000116675 |
| SLFN5 | 0.948486676 | 1.929847271 | 4.10E-07 | 3.43E-06 | 93.53057949 | 180.272183 | ENSG00000166750 |
| DGKH | 0.947671429 | 1.928757049 | 7.66E-07 | 6.14E-06 | 87.82894212 | 169.2121232 | ENSG00000102780 |
| ARMCX6 | 0.947627772 | 1.928698684 | 3.18E-17 | 8.56E-16 | 279.4059127 | 538.3442374 | ENSG00000198960 |
| ALS2CL | 0.946042739 | 1.926580862 | 1.60E-10 | 2.16E-09 | 165.0451264 | 317.7388905 | ENSG00000178038 |
| DIAPH2 | 0.943940045 | 1.92377496 | 5.07E-44 | 6.84E-42 | 965.0770399 | 1855.629947 | ENSG00000147202 |
| MT1X | 0.943899907 | 1.923721439 | 2.08E-04 | 1.04E-03 | 58.4142839 | 112.162773 | ENSG00000187193 |
| COL25A1 | 0.942425558 | 1.921756513 | 2.91E-08 | 2.88E-07 | 117.6231149 | 226.1842948 | ENSG00000188517 |
| IGSF21 | 0.94152898 | 1.920562589 | 1.09E-17 | 3.03E-16 | 407.7119087 | 783.6586801 | ENSG00000117154 |
| TCIRG1 | 0.940614972 | 1.919346216 | 2.19E-17 | 5.96E-16 | 287.6543158 | 552.2008862 | ENSG00000110719 |
| BFSP1 | 0.940002465 | 1.918531517 | 2.39E-04 | 1.19E-03 | 61.83319117 | 118.4626136 | ENSG00000125864 |
| TRANK1 | 0.939344378 | 1.917656576 | 2.72E-14 | 5.47E-13 | 259.5908351 | 497.0339532 | ENSG00000168016 |
| ITLN2 | 0.938757639 | 1.91687683 | 1.16E-06 | 9.06E-06 | 91.23042825 | 174.6109497 | ENSG00000158764 |
| TENM2 | 0.938150027 | 1.91606968 | 1.38E-03 | 5.69E-03 | 42.62760925 | 81.67325295 | ENSG00000145934 |
| MDM4 | 0.937692371 | 1.915461955 | 3.54E-20 | 1.22E-18 | 371.9105997 | 711.9546403 | ENSG00000198625 |
| AKAP7 | 0.937273311 | 1.914905651 | 1.41E-14 | 2.91E-13 | 261.3051323 | 500.2524072 | ENSG00000118507 |
| KLHL12 | 0.937082983 | 1.914653044 | 6.43E-41 | 7.80E-39 | 933.5606728 | 1786.241512 | ENSG00000117153 |
| SCYL3 | 0.936386765 | 1.91372929 | 5.89E-08 | 5.59E-07 | 109.309357 | 209.0462618 | ENSG00000000457 |
| DLG2 | 0.936182017 | 1.913457712 | 4.54E-08 | 4.39E-07 | 119.7714302 | 228.9292821 | ENSG00000150672 |
| ALPG | 0.936162628 | 1.913431995 | 2.50E-02 | 6.76E-02 | 21.25784248 | 40.77384466 | ENSG00000163286 |
| SELENBP1 | 0.935767315 | 1.912907768 | 4.38E-20 | 1.49E-18 | 406.1244541 | 776.0661783 | ENSG00000143416 |
| SORCS2 | 0.934990621 | 1.911878206 | 8.24E-05 | 4.54E-04 | 65.09687114 | 124.5174877 | ENSG00000184985 |
| CCL26 | 0.934792214 | 1.911615293 | 1.04E-04 | 5.63E-04 | 56.95564954 | 108.7907616 | ENSG00000006606 |
| B3GALT4 | 0.934349121 | 1.911028271 | 8.90E-03 | 2.85E-02 | 24.41721991 | 46.64831263 | ENSG00000235863 |
| IFITM1 | 0.933862816 | 1.910384208 | 2.87E-08 | 2.85E-07 | 832.045351 | 1589.161133 | ENSG00000185885 |
| CD46 | 0.932629649 | 1.908751974 | 5.90E-38 | 5.93E-36 | 1558.452396 | 2973.30696 | ENSG00000117335 |
| TUBB2A | 0.93145384 | 1.907196958 | 1.22E-92 | 8.09E-90 | 2824.085074 | 5385.091255 | ENSG00000137267 |
| ADAMTS5 | 0.929906253 | 1.905152195 | 2.09E-03 | 8.22E-03 | 34.72618101 | 66.12589741 | ENSG00000154736 |
| CGN | 0.927812567 | 1.90238938 | 1.44E-68 | 4.33E-66 | 1907.074123 | 3628.497243 | ENSG00000143375 |
| RIMBP2 | 0.926908254 | 1.901197295 | 1.39E-02 | 4.15E-02 | 24.49333802 | 46.45472415 | ENSG00000060709 |
| P4HA1 | 0.926742757 | 1.900979214 | 1.03E-23 | 4.61E-22 | 529.6917907 | 1007.2023 | ENSG00000122884 |
| BRCA2 | 0.926699504 | 1.900922222 | 2.65E-10 | 3.48E-09 | 219.9736846 | 418.6092438 | ENSG00000139618 |
| RNF227 | 0.926055212 | 1.900073481 | 3.10E-02 | 8.07E-02 | 18.79516728 | 35.61164866 | ENSG00000179859 |
| OSCAR | 0.925721145 | 1.899633555 | 4.37E-04 | 2.04E-03 | 51.48264192 | 97.96518169 | ENSG00000170909 |
| QRICH2 | 0.923491207 | 1.896699606 | 1.48E-03 | 6.05E-03 | 38.74661163 | 73.48617932 | ENSG00000129646 |
| TRIM6 | 0.923426551 | 1.896614605 | 1.22E-13 | 2.29E-12 | 223.0530259 | 423.1258846 | ENSG00000121236 |

| | | | | | | | |
|---|---|---|---|---|---|---|---|
| CACNG6 | 0.922437563 | 1.895314895 | 1.89E-02 | 5.36E-02 | 21.1030155 | 39.92527464 | ENSG00000130433 |
| APOBEC3F | 0.920171658 | 1.892340439 | 1.72E-05 | 1.08E-04 | 78.21887367 | 148.3646013 | ENSG00000128394 |
| LMO3 | 0.919706569 | 1.891730493 | 3.21E-07 | 2.74E-06 | 140.910127 | 266.9842252 | ENSG00000048540 |
| TPO | 0.919138908 | 1.890986295 | 4.34E-02 | 1.07E-01 | 18.78467939 | 35.4807057 | ENSG00000115705 |
| CASC9 | 0.918750878 | 1.890477761 | 4.39E-03 | 1.57E-02 | 35.7557403 | 67.38880835 | ENSG00000249395 |
| DYNC1I1 | 0.918290592 | 1.889874708 | 2.79E-02 | 7.41E-02 | 18.05189559 | 34.09652018 | ENSG00000158560 |
| OXSM | 0.918187197 | 1.889739269 | 4.36E-02 | 1.07E-01 | 19.56567228 | 36.80117963 | ENSG00000151093 |
| ZBTB37 | 0.917950119 | 1.889428754 | 2.23E-07 | 1.95E-06 | 165.7443441 | 312.6118667 | ENSG00000185278 |
| HMGB3 | 0.916453269 | 1.887469418 | 6.43E-124 | 9.53E-121 | 6576.241435 | 12413.56698 | ENSG00000029993 |
| XPR1 | 0.915457562 | 1.886167189 | 3.25E-55 | 6.46E-53 | 1443.101576 | 2722.263768 | ENSG00000143324 |
| SAMD4A | 0.91508264 | 1.885677084 | 5.10E-06 | 3.54E-05 | 87.04707335 | 164.3836003 | ENSG00000020577 |
| NPIPA1 | 0.913377871 | 1.883450178 | 1.85E-07 | 1.63E-06 | 115.1184881 | 216.6020223 | ENSG00000183426 |
| PLBD1 | 0.912701765 | 1.882567723 | 7.46E-11 | 1.04E-09 | 179.888084 | 338.5874664 | ENSG00000121316 |
| PURA | 0.912579574 | 1.882408283 | 4.95E-06 | 3.44E-05 | 92.43172319 | 174.2842677 | ENSG00000185129 |
| ASPM | 0.911447007 | 1.880931105 | 3.00E-29 | 1.86E-27 | 1074.123801 | 2019.579232 | ENSG00000066279 |
| ENG | 0.909023951 | 1.877774668 | 1.20E-05 | 7.75E-05 | 82.96024876 | 155.9424311 | ENSG00000106991 |
| UEVLD | 0.908103499 | 1.876577013 | 3.03E-05 | 1.82E-04 | 80.12006205 | 150.3126445 | ENSG00000151116 |
| LRRK1 | 0.907646999 | 1.875983318 | 1.98E-05 | 1.24E-04 | 107.5403049 | 201.7321838 | ENSG00000154237 |
| CYTL1 | 0.907646483 | 1.875982648 | 1.39E-07 | 1.25E-06 | 115.7668691 | 217.0773912 | ENSG00000170891 |
| PIPOX | 0.907606154 | 1.875930206 | 2.17E-27 | 1.19E-25 | 826.5124824 | 1549.959938 | ENSG00000179761 |
| MAOB | 0.906484657 | 1.874472495 | 5.69E-04 | 2.57E-03 | 47.26986329 | 88.54688005 | ENSG00000069535 |
| SWT1 | 0.905298158 | 1.872931529 | 3.66E-07 | 3.10E-06 | 107.3699718 | 201.1484622 | ENSG00000116668 |
| KITLG | 0.904909667 | 1.872427251 | 5.83E-06 | 4.00E-05 | 87.94046609 | 164.7575949 | ENSG00000049130 |
| TMEM38A | 0.904826661 | 1.872319524 | 4.54E-11 | 6.46E-10 | 189.1979104 | 353.9649019 | ENSG00000072954 |
| DPH6 | 0.904459354 | 1.871842895 | 1.96E-08 | 1.99E-07 | 129.4081315 | 242.3280455 | ENSG00000134146 |
| RHBG | 0.904053375 | 1.871316227 | 1.98E-02 | 5.56E-02 | 20.37860441 | 38.15658366 | ENSG00000132677 |
| GPSM3 | 0.903866407 | 1.871073727 | 3.92E-05 | 2.30E-04 | 84.43303779 | 157.6584359 | ENSG00000213654 |
| ADGRE2 | 0.903451644 | 1.870535886 | 9.08E-03 | 2.90E-02 | 28.3475779 | 53.12909245 | ENSG00000127507 |
| INKA2 | 0.90290072 | 1.869821718 | 1.58E-12 | 2.63E-11 | 308.1555695 | 575.2725676 | ENSG00000197852 |
| MME | 0.901864335 | 1.868478981 | 2.69E-11 | 3.92E-10 | 238.6110707 | 445.7152003 | ENSG00000196549 |
| PTAR1 | 0.90014645 | 1.86625542 | 3.92E-26 | 2.05E-24 | 709.5969963 | 1324.161432 | ENSG00000188647 |
| CAPG | 0.899067142 | 1.86485976 | 4.30E-31 | 2.89E-29 | 693.1475106 | 1291.77724 | ENSG00000042493 |
| NDUFS2 | 0.897524562 | 1.862866853 | 4.46E-74 | 1.62E-71 | 2181.738259 | 4063.36283 | ENSG00000158864 |
| UBE2D4 | 0.8965725 | 1.861637916 | 1.03E-02 | 3.21E-02 | 27.383849 | 50.91381406 | ENSG00000078967 |
| CSF2RA | 0.896173842 | 1.861123563 | 1.49E-02 | 4.39E-02 | 24.57571367 | 45.59457722 | ENSG00000198223 |
| PDCD4 | 0.895383403 | 1.860104151 | 4.94E-14 | 9.61E-13 | 255.8938128 | 475.8185974 | ENSG00000150593 |
| FCSK | 0.893866398 | 1.858149266 | 6.54E-08 | 6.17E-07 | 151.1645462 | 280.9048766 | ENSG00000157353 |
| C1QTNF6 | 0.890324848 | 1.853593446 | 1.19E-04 | 6.30E-04 | 61.96078484 | 114.6875593 | ENSG00000133466 |
| CDC42BPA | 0.890220101 | 1.85345887 | 4.30E-33 | 3.28E-31 | 824.489921 | 1528.198201 | ENSG00000143776 |
| ALDH1A3 | 0.889381318 | 1.852381582 | 2.51E-03 | 9.63E-03 | 39.26663522 | 72.63356861 | ENSG00000184254 |
| JAK1 | 0.889374553 | 1.852372896 | 1.15E-28 | 6.84E-27 | 664.2869545 | 1230.074339 | ENSG00000162434 |
| MGST2 | 0.888256953 | 1.850938491 | 6.57E-05 | 3.69E-04 | 69.97027043 | 129.4344815 | ENSG00000085871 |
| HLA-G | 0.887758123 | 1.850298616 | 7.20E-04 | 3.16E-03 | 48.35859511 | 89.30163045 | ENSG00000204632 |
| TSEN15 | 0.886385286 | 1.84853875 | 4.58E-26 | 2.37E-24 | 666.1274698 | 1231.605848 | ENSG00000198860 |
| CHODL | 0.886247187 | 1.848361811 | 2.36E-02 | 6.44E-02 | 23.71880282 | 43.60864145 | ENSG00000154645 |
| DYDC2 | 0.886056927 | 1.848118068 | 4.67E-05 | 2.70E-04 | 67.88499459 | 125.415812 | ENSG00000133665 |
| MGMT | 0.884969914 | 1.84672611 | 1.78E-15 | 4.04E-14 | 274.5315254 | 507.1166506 | ENSG00000170430 |
| ZDHHC1 | 0.883584183 | 1.844953152 | 4.85E-04 | 2.23E-03 | 53.60304822 | 98.7874118 | ENSG00000159714 |
| KRCC1 | 0.88326298 | 1.844542435 | 6.42E-09 | 7.02E-08 | 138.0596525 | 254.6661008 | ENSG00000172086 |
| CD74 | 0.881993438 | 1.84291999 | 1.29E-29 | 8.01E-28 | 613.1882207 | 1129.909236 | ENSG00000019582 |
| GMPR | 0.880492751 | 1.841003987 | 4.93E-02 | 1.18E-01 | 16.53811889 | 30.57769989 | ENSG00000137198 |
| RNF112 | 0.879930545 | 1.840286703 | 1.43E-03 | 5.88E-03 | 45.59268674 | 84.12543439 | ENSG00000128482 |
| DNER | 0.879598887 | 1.839863692 | 1.78E-04 | 9.07E-04 | 60.63062003 | 111.4913804 | ENSG00000187957 |
| SCAPER | 0.878986807 | 1.839083274 | 1.05E-05 | 6.85E-05 | 107.6555837 | 198.0213805 | ENSG00000140386 |
| PAQR7 | 0.87852852 | 1.838499164 | 3.84E-07 | 3.23E-06 | 127.8414398 | 235.1358056 | ENSG00000182749 |
| C16orf74 | 0.878483695 | 1.838442042 | 2.07E-02 | 5.79E-02 | 26.79839534 | 49.21717651 | ENSG00000154102 |
| SDCCAG8 | 0.877943652 | 1.837753988 | 1.14E-05 | 7.41E-05 | 93.75402775 | 171.9323881 | ENSG00000054282 |
| LAMA3 | 0.877893409 | 1.837689988 | 1.19E-02 | 3.64E-02 | 28.90532268 | 53.04843672 | ENSG00000053747 |
| ADAMTS10 | 0.876278499 | 1.835634082 | 2.60E-10 | 3.42E-09 | 179.9242536 | 330.321615 | ENSG00000142303 |
| BTN3A2 | 0.87565733 | 1.8348439 | 1.53E-04 | 7.91E-04 | 74.78776364 | 137.2671577 | ENSG00000186470 |
| GORAB | 0.875430109 | 1.834554938 | 9.91E-09 | 1.06E-07 | 207.6936745 | 381.4185436 | ENSG00000120370 |
| SPART | 0.87535391 | 1.834458045 | 2.24E-40 | 2.67E-38 | 931.3623597 | 1708.764602 | ENSG00000133104 |
| FAM167B | 0.875291346 | 1.834378494 | 3.40E-02 | 8.70E-02 | 21.69337524 | 39.73975541 | ENSG00000183615 |
| ATRNL1 | 0.875029152 | 1.834045145 | 5.24E-09 | 5.80E-08 | 156.7981259 | 287.7766378 | ENSG00000107518 |
| DOCK2 | 0.874710838 | 1.833640529 | 2.59E-07 | 2.24E-06 | 119.5987818 | 219.3125883 | ENSG00000134516 |
| JMJD4 | 0.873501206 | 1.832103753 | 5.05E-03 | 1.77E-02 | 34.62562049 | 63.46884158 | ENSG00000081692 |
| ARRDC2 | 0.872890869 | 1.831328839 | 7.75E-06 | 5.21E-05 | 96.80793857 | 176.8966945 | ENSG00000105643 |
| SPATA18 | 0.872715551 | 1.831106307 | 2.54E-02 | 6.86E-02 | 20.84974319 | 38.07646097 | ENSG00000163071 |
| MRPL24 | 0.872455842 | 1.830776708 | 1.12E-21 | 4.28E-20 | 436.7229862 | 799.4529344 | ENSG00000143314 |
| SPATS2L | 0.871624941 | 1.829722599 | 1.84E-26 | 9.82E-25 | 943.9901055 | 1726.814748 | ENSG00000196141 |
| RO60 | 0.870072983 | 1.82775536 | 5.27E-11 | 7.45E-10 | 319.3112432 | 584.4547229 | ENSG00000116747 |
| MEIOC | 0.870008051 | 1.827673099 | 7.60E-06 | 5.12E-05 | 104.3833309 | 190.5971265 | ENSG00000180336 |
| DCAF6 | 0.869734297 | 1.827326328 | 1.40E-14 | 2.88E-13 | 285.5928067 | 521.8003879 | ENSG00000143164 |

| | | | | | | | |
|---|---|---|---|---|---|---|---|
| MATN4 | 0.869082058 | 1.826500386 | 4.51E-02 | 1.10E-01 | 20.76054644 | 37.66642121 | ENSG00000124159 |
| PRKG2 | 0.868321877 | 1.825538225 | 3.14E-02 | 8.14E-02 | 30.12878159 | 54.80313976 | ENSG00000138669 |
| SDE2 | 0.867913227 | 1.825021205 | 1.01E-17 | 2.82E-16 | 404.5167748 | 737.3304041 | ENSG00000143751 |
| AP3B2 | 0.865543069 | 1.822025397 | 1.75E-12 | 2.90E-11 | 264.0785375 | 480.607647 | ENSG00000103723 |
| PRELID2 | 0.865427066 | 1.8218789 | 9.42E-04 | 4.04E-03 | 50.28450131 | 91.76750884 | ENSG00000186314 |
| COL6A2 | 0.865323205 | 1.821747746 | 9.28E-68 | 2.67E-65 | 3325.788053 | 6059.551733 | ENSG00000142173 |
| LYSMD1 | 0.864979383 | 1.82131364 | 8.41E-07 | 6.71E-06 | 106.1089919 | 193.1442242 | ENSG00000163155 |
| GDPD5 | 0.86482037 | 1.821112906 | 1.22E-06 | 9.52E-06 | 115.048152 | 209.1581101 | ENSG00000158555 |
| RPH3AL | 0.86453701 | 1.820755256 | 1.67E-02 | 4.84E-02 | 27.86894229 | 50.58336394 | ENSG00000181031 |
| GNG12 | 0.864493199 | 1.820699965 | 5.60E-20 | 1.89E-18 | 524.9044849 | 956.461386 | ENSG00000172380 |
| KTI12 | 0.864122611 | 1.820232338 | 2.89E-02 | 7.62E-02 | 21.66267531 | 39.49237206 | ENSG00000198841 |
| GRIN2A | 0.863664108 | 1.819653943 | 2.13E-04 | 1.07E-03 | 75.87948653 | 137.6078519 | ENSG00000183454 |
| ZADH2 | 0.863577774 | 1.819545054 | 3.87E-05 | 2.28E-04 | 80.58226454 | 146.4434737 | ENSG00000180011 |
| KLHL11 | 0.863430665 | 1.819359528 | 1.88E-04 | 9.54E-04 | 68.71363299 | 124.7239797 | ENSG00000178502 |
| PGPEP1 | 0.863408639 | 1.819331751 | 1.41E-28 | 8.24E-27 | 615.593275 | 1119.619241 | ENSG00000130517 |
| PSMB9 | 0.862343695 | 1.817989283 | 2.06E-03 | 8.11E-03 | 42.08804935 | 76.53230221 | ENSG00000240065 |
| SENP8 | 0.861364862 | 1.81675624 | 1.11E-03 | 4.67E-03 | 46.52168551 | 84.49165076 | ENSG00000166192 |
| CACNB4 | 0.860458029 | 1.815614643 | 9.37E-03 | 2.98E-02 | 32.9186199 | 59.90671881 | ENSG00000182389 |
| C1orf116 | 0.860040666 | 1.815089473 | 9.01E-06 | 5.99E-05 | 104.7203303 | 189.7093068 | ENSG00000182795 |
| DES | 0.857927673 | 1.812433011 | 1.24E-03 | 5.17E-03 | 47.33740853 | 85.82072698 | ENSG00000175084 |
| ZNF814 | 0.857682317 | 1.812124801 | 1.75E-14 | 3.58E-13 | 273.0596123 | 494.9574144 | ENSG00000204514 |
| EFNA1 | 0.857569772 | 1.811983442 | 4.29E-04 | 2.00E-03 | 56.67302873 | 102.4732018 | ENSG00000169242 |
| ACAD10 | 0.857134338 | 1.811436632 | 1.34E-11 | 2.03E-10 | 303.6894688 | 550.0452787 | ENSG00000111271 |
| EDIL3 | 0.856478868 | 1.810613816 | 1.76E-18 | 5.21E-17 | 772.6093411 | 1398.034497 | ENSG00000164176 |
| TMCO4 | 0.856403116 | 1.810518748 | 7.18E-06 | 4.86E-05 | 109.1888967 | 197.4220655 | ENSG00000162542 |
| PKLR | 0.856013187 | 1.81002947 | 1.15E-04 | 6.13E-04 | 66.97341697 | 121.4275293 | ENSG00000143627 |
| TNRC18 | 0.854586746 | 1.808240717 | 8.43E-09 | 9.08E-08 | 158.8505866 | 287.5029146 | ENSG00000182095 |
| COX6B2 | 0.85400429 | 1.807510828 | 2.62E-02 | 7.04E-02 | 28.12779641 | 50.78717225 | ENSG00000160471 |
| ARNT | 0.853618986 | 1.807028156 | 2.63E-32 | 1.92E-30 | 807.2431725 | 1458.817297 | ENSG00000143437 |
| CD70 | 0.853351335 | 1.806692945 | 2.03E-02 | 5.70E-02 | 26.30069898 | 47.67488408 | ENSG00000125726 |
| KLF5 | 0.852879849 | 1.806102597 | 2.64E-02 | 7.07E-02 | 23.01026126 | 41.66329401 | ENSG00000102554 |
| IL13RA1 | 0.852522656 | 1.805655484 | 1.56E-24 | 7.39E-23 | 555.9691233 | 1003.547224 | ENSG00000131724 |
| TRIM56 | 0.85185592 | 1.8048212 | 1.14E-30 | 7.44E-29 | 745.759784 | 1345.789821 | ENSG00000169871 |
| CCDC121 | 0.851801478 | 1.804753095 | 4.32E-02 | 1.06E-01 | 18.45461323 | 33.2826198 | ENSG00000176714 |
| MELTF | 0.850653284 | 1.803317322 | 7.75E-05 | 4.30E-04 | 86.5457471 | 155.7692518 | ENSG00000163975 |
| PCAT6 | 0.850383372 | 1.802979973 | 6.36E-04 | 2.84E-03 | 65.94318196 | 119.1348119 | ENSG00000228288 |
| RASSF10 | 0.850328113 | 1.802910916 | 1.99E-04 | 1.00E-03 | 63.54489769 | 114.6165599 | ENSG00000189431 |
| MPPE1 | 0.850266472 | 1.802833886 | 2.19E-05 | 1.36E-04 | 102.0581736 | 183.6000863 | ENSG00000154889 |
| METAP1D | 0.850034514 | 1.802544047 | 1.37E-04 | 7.15E-04 | 67.04203826 | 120.6496678 | ENSG00000172878 |
| GJA4 | 0.849795109 | 1.802244953 | 3.47E-04 | 1.66E-03 | 69.10307185 | 124.0991183 | ENSG00000187513 |
| CEL | 0.849652078 | 1.802066285 | 2.64E-03 | 1.00E-02 | 39.79136473 | 71.7255576 | ENSG00000170835 |
| DAGLA | 0.846400332 | 1.798009111 | 3.30E-11 | 4.78E-10 | 260.3080759 | 468.1375948 | ENSG00000134780 |
| NLRP7 | 0.846346123 | 1.797941553 | 2.48E-02 | 6.71E-02 | 27.77627892 | 49.66029892 | ENSG00000167634 |
| FCHO2 | 0.845374299 | 1.796730836 | 1.25E-06 | 9.71E-06 | 129.5551493 | 233.0928826 | ENSG00000157107 |
| NMRK1 | 0.845247343 | 1.796572733 | 8.38E-04 | 3.63E-03 | 53.2322698 | 95.87490336 | ENSG00000106733 |
| NUAK2 | 0.844675474 | 1.795860731 | 1.21E-09 | 1.46E-08 | 222.1308482 | 399.1932996 | ENSG00000163545 |
| HLA-F | 0.841362449 | 1.791741424 | 2.35E-03 | 9.07E-03 | 43.85509836 | 78.53652707 | ENSG00000204642 |
| SEMG1 | 0.841055383 | 1.791360106 | 1.98E-02 | 5.57E-02 | 24.36275329 | 43.66133972 | ENSG00000124233 |
| IL17C | 0.840785209 | 1.79102467 | 3.04E-02 | 7.95E-02 | 20.70799484 | 37.09209538 | ENSG00000124391 |
| EXD3 | 0.839546949 | 1.789488099 | 1.48E-06 | 1.13E-05 | 126.4023417 | 226.113871 | ENSG00000187609 |
| NEBL | 0.838635956 | 1.788358479 | 6.04E-05 | 3.42E-04 | 78.89651513 | 141.3302079 | ENSG00000078114 |
| VENTX | 0.838412 | 1.788080886 | 1.73E-09 | 2.05E-08 | 194.1843852 | 346.6882503 | ENSG00000151650 |
| CASP8 | 0.838369546 | 1.788028269 | 1.30E-03 | 5.41E-03 | 49.63775993 | 88.61760694 | ENSG00000064012 |
| ITPRIPL2 | 0.83753482 | 1.786994036 | 3.26E-06 | 2.34E-05 | 107.4042264 | 191.860601 | ENSG00000205730 |
| TRIM72 | 0.836999486 | 1.786331068 | 7.62E-03 | 2.51E-02 | 37.29375924 | 66.43967597 | ENSG00000177238 |
| USP53 | 0.836961793 | 1.786284398 | 1.02E-13 | 1.93E-12 | 330.7624005 | 590.8427555 | ENSG00000145390 |
| KIRREL1 | 0.836829347 | 1.786120416 | 4.25E-46 | 6.26E-44 | 1562.48134 | 2790.156667 | ENSG00000183853 |
| GUCY2C | 0.836758826 | 1.78603311 | 6.95E-03 | 2.33E-02 | 34.8705201 | 62.44279105 | ENSG00000070019 |
| HIVEP2 | 0.835768234 | 1.784807194 | 2.56E-08 | 2.56E-07 | 194.8302876 | 347.0638688 | ENSG00000010818 |
| EFR3A | 0.834998395 | 1.783855055 | 3.16E-26 | 1.66E-24 | 689.0264825 | 1229.690918 | ENSG00000132294 |
| PDP1 | 0.834692445 | 1.783476795 | 3.64E-13 | 6.52E-12 | 282.7099927 | 504.3407128 | ENSG00000164951 |
| WNT9A | 0.833961025 | 1.782572834 | 2.58E-04 | 1.27E-03 | 62.79325328 | 111.9457887 | ENSG00000143816 |
| DAP3 | 0.831991716 | 1.780141245 | 1.98E-38 | 2.04E-36 | 1487.769993 | 2647.273565 | ENSG00000132676 |
| PLK3 | 0.831564961 | 1.779614749 | 5.52E-07 | 4.51E-06 | 121.8509241 | 216.804201 | ENSG00000173846 |
| MST1L | 0.831476995 | 1.779506244 | 1.25E-03 | 5.23E-03 | 50.30981959 | 89.42584911 | ENSG00000186715 |
| KBTBD3 | 0.831449615 | 1.779472472 | 3.31E-02 | 8.51E-02 | 25.93311179 | 46.0406437 | ENSG00000182359 |
| C17orf113 | 0.831271882 | 1.779253264 | 2.17E-03 | 8.47E-03 | 47.45559035 | 84.25799488 | ENSG00000267221 |
| CPNE7 | 0.83012157 | 1.777835167 | 1.27E-13 | 2.38E-12 | 366.7215275 | 652.2550197 | ENSG00000178773 |
| ABCG2 | 0.829453884 | 1.777012567 | 6.08E-03 | 2.08E-02 | 40.43810611 | 72.11085655 | ENSG00000118777 |
| MYH11 | 0.828182445 | 1.775447186 | 3.79E-02 | 9.54E-02 | 20.78132206 | 36.87779467 | ENSG00000133392 |
| SMYD3 | 0.826397852 | 1.773252341 | 9.12E-13 | 1.56E-11 | 311.950651 | 553.6415075 | ENSG00000185420 |
| MGAT3 | 0.825636033 | 1.772316218 | 6.30E-20 | 2.11E-18 | 446.495854 | 791.4470667 | ENSG00000128268 |
| CEP170 | 0.825084285 | 1.771638537 | 1.86E-31 | 1.27E-29 | 1011.13228 | 1791.613616 | ENSG00000143702 |

| | | | | | | | |
|---|---|---|---|---|---|---|---|
| PDGFRB | 0.824794625 | 1.771282869 | 4.62E-13 | 8.15E-12 | 300.5109313 | 532.1387557 | ENSG00000113721 |
| SUGP1 | 0.824728076 | 1.771201165 | 1.79E-30 | 1.15E-28 | 885.337807 | 1567.982708 | ENSG00000105705 |
| TATDN3 | 0.82465695 | 1.771113846 | 5.30E-11 | 7.48E-10 | 217.1375522 | 384.8418778 | ENSG00000203705 |
| MPI | 0.824594963 | 1.771037749 | 5.49E-21 | 2.00E-19 | 545.0428204 | 965.5731849 | ENSG00000178802 |
| FZD9 | 0.824369621 | 1.770761143 | 1.07E-06 | 8.40E-06 | 131.5370737 | 233.3149615 | ENSG00000188763 |
| KIAA1614 | 0.823832122 | 1.770101541 | 2.62E-03 | 9.99E-03 | 46.16717695 | 81.657678 | ENSG00000135835 |
| WDR26 | 0.823731238 | 1.769977767 | 5.97E-46 | 8.67E-44 | 1561.224727 | 2762.559924 | ENSG00000162923 |
| LCTL | 0.822066113 | 1.767936077 | 1.27E-06 | 9.85E-06 | 125.9195638 | 222.5880539 | ENSG00000188501 |
| TRPS1 | 0.82164124 | 1.767415498 | 6.54E-05 | 3.68E-04 | 78.16739811 | 138.2639058 | ENSG00000104447 |
| P3H2 | 0.821446529 | 1.767176978 | 2.21E-15 | 4.97E-14 | 415.6259159 | 734.8644647 | ENSG00000090530 |
| ZNF654 | 0.820214122 | 1.76566803 | 2.79E-05 | 1.69E-04 | 93.55482173 | 165.0958659 | ENSG00000175105 |
| ALG10 | 0.818969872 | 1.764145889 | 3.26E-02 | 8.41E-02 | 24.24054127 | 42.91895973 | ENSG00000139133 |
| UAP1L1 | 0.81822981 | 1.763241163 | 3.21E-10 | 4.17E-09 | 205.0152849 | 361.1364479 | ENSG00000197355 |
| HLA-DRB1 | 0.817151596 | 1.761923878 | 3.08E-11 | 4.47E-10 | 281.2283923 | 494.8382904 | ENSG00000196126 |
| TDRKH | 0.817138508 | 1.761907895 | 6.48E-21 | 2.34E-19 | 499.6741696 | 880.2340423 | ENSG00000182134 |
| LMNA | 0.817105557 | 1.761867654 | 8.99E-33 | 6.67E-31 | 1249.406828 | 2202.659964 | ENSG00000160789 |
| F8A1 | 0.816972655 | 1.761705357 | 1.09E-04 | 5.83E-04 | 80.79817908 | 142.5232281 | ENSG00000277203 |
| ZNF79 | 0.816434635 | 1.761048492 | 7.08E-06 | 4.80E-05 | 107.3789081 | 189.3796381 | ENSG00000196152 |
| ORAI3 | 0.816100583 | 1.760640772 | 1.19E-02 | 3.65E-02 | 36.63756911 | 64.64113743 | ENSG00000175938 |
| ENO3 | 0.815877842 | 1.760368964 | 2.63E-12 | 4.29E-11 | 272.3378288 | 478.8736378 | ENSG00000108515 |
| KLF11 | 0.81521141 | 1.759555976 | 4.26E-10 | 5.43E-09 | 215.3279128 | 378.4119061 | ENSG00000172059 |
| EPS8L2 | 0.814490671 | 1.758677159 | 2.53E-23 | 1.09E-21 | 591.5311272 | 1040.3844 | ENSG00000177106 |
| TUSC1 | 0.81436617 | 1.758525396 | 5.38E-04 | 2.45E-03 | 64.52940221 | 113.5795081 | ENSG00000198680 |
| NOS2 | 0.81419674 | 1.758318887 | 2.71E-04 | 1.33E-03 | 67.92618242 | 119.3214156 | ENSG00000007171 |
| TMEM59L | 0.814086152 | 1.758184111 | 6.47E-16 | 1.54E-14 | 375.9090665 | 660.9346547 | ENSG00000105696 |
| PPIC | 0.813847033 | 1.757892725 | 1.84E-04 | 9.33E-04 | 72.13146423 | 126.8591475 | ENSG00000168938 |
| ZC3HAV1L | 0.8128569 | 1.756686683 | 2.52E-04 | 1.25E-03 | 70.23961244 | 123.3051245 | ENSG00000146858 |
| FTX | 0.81238821 | 1.756116079 | 6.01E-08 | 5.70E-07 | 154.2570172 | 271.1825195 | ENSG00000230590 |
| ARMCX2 | 0.809620641 | 1.752750493 | 4.41E-52 | 7.88E-50 | 2813.533928 | 4930.499149 | ENSG00000184867 |
| GEMIN8 | 0.809367117 | 1.752442511 | 1.31E-08 | 1.37E-07 | 171.918635 | 301.0076196 | ENSG00000046647 |
| BLZF1 | 0.809247263 | 1.75229693 | 1.41E-08 | 1.48E-07 | 184.2482808 | 323.0424914 | ENSG00000117475 |
| GPRIN3 | 0.80916055 | 1.752191612 | 1.55E-03 | 6.30E-03 | 50.17365305 | 87.83355413 | ENSG00000185477 |
| CENPF | 0.80858814 | 1.751496542 | 6.39E-41 | 7.80E-39 | 2956.018067 | 5176.503737 | ENSG00000117724 |
| CREBRF | 0.808049397 | 1.750842606 | 1.38E-02 | 4.12E-02 | 32.9201715 | 57.54785449 | ENSG00000164463 |
| TMEM256 | 0.807302103 | 1.749935931 | 4.12E-02 | 1.02E-01 | 23.99564166 | 42.12818262 | ENSG00000205544 |
| NEDD9 | 0.806114927 | 1.748496523 | 6.30E-11 | 8.84E-10 | 229.7863377 | 401.4698722 | ENSG00000111859 |
| TNFRSF1B | 0.805010656 | 1.747158697 | 1.37E-05 | 8.75E-05 | 97.36440716 | 170.3099546 | ENSG00000028137 |
| ELFN2 | 0.803908485 | 1.745824435 | 1.40E-04 | 7.28E-04 | 79.26749371 | 138.194003 | ENSG00000166897 |
| GSTM3 | 0.802158857 | 1.74370847 | 3.93E-03 | 1.42E-02 | 46.08403756 | 80.35361517 | ENSG00000134202 |
| IL1RAP | 0.801925447 | 1.743426383 | 1.77E-05 | 1.11E-04 | 114.6035708 | 199.7908771 | ENSG00000196083 |
| HHLA3 | 0.801300153 | 1.74267091 | 1.18E-04 | 6.26E-04 | 77.59589897 | 135.2532766 | ENSG00000197568 |
| SMIM10L2B | 0.801138106 | 1.74247518 | 9.57E-07 | 7.55E-06 | 128.7221173 | 224.1260573 | ENSG00000196972 |
| RDM1 | 0.800383143 | 1.741563581 | 1.21E-03 | 5.07E-03 | 73.5428534 | 128.0056364 | ENSG00000278023 |
| TMEM184A | 0.800152173 | 1.741284785 | 4.19E-02 | 1.04E-01 | 21.06182768 | 36.62720678 | ENSG00000164855 |
| MBTPS2 | 0.796984904 | 1.73746619 | 2.60E-06 | 1.90E-05 | 156.1086451 | 271.4067491 | ENSG00000012174 |
| LRRC75A | 0.796525124 | 1.736912556 | 4.59E-14 | 8.95E-13 | 334.0048292 | 579.6329853 | ENSG00000181350 |
| RBMS3 | 0.79616755 | 1.736482113 | 1.51E-02 | 4.45E-02 | 31.52003396 | 54.899934 | ENSG00000144642 |
| ZNF552 | 0.796164681 | 1.73647866 | 8.20E-06 | 5.48E-05 | 115.5275513 | 200.4053008 | ENSG00000178935 |
| PRRC2C | 0.79458857 | 1.734582633 | 3.76E-37 | 3.57E-35 | 4575.798097 | 7935.076245 | ENSG00000117523 |
| ECHDC3 | 0.793879042 | 1.733729762 | 8.21E-16 | 1.93E-14 | 395.5770142 | 685.8363519 | ENSG00000134463 |
| SLC39A8 | 0.793594233 | 1.733387533 | 7.10E-20 | 2.37E-18 | 535.3890855 | 928.3673695 | ENSG00000138821 |
| XK | 0.792841518 | 1.732483387 | 2.61E-02 | 7.02E-02 | 28.94005282 | 50.31473532 | ENSG00000047597 |
| P2RY1 | 0.792119979 | 1.731617131 | 7.35E-18 | 2.09E-16 | 703.741757 | 1217.298323 | ENSG00000169860 |
| COL11A1 | 0.791940881 | 1.73140218 | 5.02E-12 | 7.91E-11 | 319.0357174 | 551.7021214 | ENSG00000060718 |
| YY1AP1 | 0.791886384 | 1.731336779 | 1.13E-19 | 3.73E-18 | 549.6276409 | 951.8044491 | ENSG00000163374 |
| PARVB | 0.791811809 | 1.731247286 | 3.11E-10 | 4.06E-09 | 247.3440916 | 428.1066205 | ENSG00000188677 |
| ARID4B | 0.791505679 | 1.730879966 | 5.92E-19 | 1.81E-17 | 534.309439 | 924.2140316 | ENSG00000054267 |
| SUSD2 | 0.79099981 | 1.730273154 | 5.56E-10 | 6.97E-09 | 217.3211641 | 376.115148 | ENSG00000099994 |
| NID1 | 0.790257737 | 1.729383389 | 1.17E-22 | 4.80E-21 | 1308.242462 | 2262.379648 | ENSG00000116962 |
| PPP1R12B | 0.790058723 | 1.729144843 | 1.18E-11 | 1.79E-10 | 261.66068 | 452.4240141 | ENSG00000077157 |
| GRID2IP | 0.789115291 | 1.728014461 | 1.93E-02 | 5.46E-02 | 31.1265649 | 53.7241305 | ENSG00000215045 |
| CIART | 0.788826722 | 1.727668857 | 2.39E-05 | 1.47E-04 | 122.77518 | 212.218233 | ENSG00000159208 |
| WIPF1 | 0.788784795 | 1.727618649 | 1.07E-06 | 8.41E-06 | 155.0552311 | 268.3329292 | ENSG00000115935 |
| EPRS1 | 0.788036917 | 1.726723302 | 2.29E-78 | 9.84E-76 | 4536.270925 | 7833.361612 | ENSG00000136628 |
| ABRAXAS1 | 0.787985423 | 1.726661671 | 1.42E-09 | 1.69E-08 | 299.4324872 | 517.3702731 | ENSG00000163322 |
| P2RX4 | 0.787326758 | 1.725873541 | 1.72E-05 | 1.08E-04 | 103.9247071 | 179.0953135 | ENSG00000135124 |
| IFI27L2 | 0.784280027 | 1.72223263 | 2.66E-06 | 1.94E-05 | 129.4966525 | 223.2769115 | ENSG00000119632 |
| ETV3 | 0.783912602 | 1.721794068 | 3.12E-20 | 1.09E-18 | 728.8940534 | 1254.996519 | ENSG00000117036 |
| DCAF8 | 0.783888045 | 1.721764761 | 1.56E-11 | 2.34E-10 | 303.4764332 | 521.9670913 | ENSG00000132716 |
| ANXA9 | 0.783705612 | 1.721547052 | 9.33E-04 | 4.00E-03 | 66.14673049 | 113.8031925 | ENSG00000143412 |
| CLDN4 | 0.783702859 | 1.721543767 | 3.72E-12 | 5.96E-11 | 306.9492946 | 528.3094225 | ENSG00000189143 |
| FOXL2NB | 0.783477248 | 1.72127457 | 1.89E-02 | 5.37E-02 | 31.26763752 | 53.7555835 | ENSG00000206262 |
| JUNB | 0.783268851 | 1.72102595 | 4.35E-05 | 2.53E-04 | 95.22641647 | 163.6167642 | ENSG00000171223 |

| | | | | | | | |
|---|---|---|---|---|---|---|---|
| WWC3 | 0.783186911 | 1.720928205 | 2.19E-33 | 1.68E-31 | 1220.013369 | 2099.038414 | ENSG00000047644 |
| RIMKLA | 0.782013394 | 1.719528937 | 4.86E-09 | 5.41E-08 | 247.977591 | 425.7270377 | ENSG00000177181 |
| IVNS1ABP | 0.781678799 | 1.719130184 | 3.80E-67 | 1.06E-64 | 5769.322553 | 9917.285426 | ENSG00000116679 |
| CREB3L4 | 0.78146192 | 1.718871767 | 1.28E-10 | 1.74E-09 | 297.7799021 | 511.0116466 | ENSG00000143578 |
| SPRY3 | 0.781206025 | 1.718566914 | 6.19E-04 | 2.77E-03 | 64.78778077 | 111.3854508 | ENSG00000168939 |
| TOB1 | 0.781088128 | 1.718426478 | 7.56E-14 | 1.44E-12 | 347.0041073 | 596.6978925 | ENSG00000141232 |
| EML2 | 0.780905359 | 1.718208791 | 3.70E-14 | 7.33E-13 | 375.3267161 | 644.2905338 | ENSG00000125746 |
| ANKRD29 | 0.780314814 | 1.717505613 | 1.55E-03 | 6.28E-03 | 59.84962714 | 102.7684187 | ENSG00000154065 |
| MDN1 | 0.779341068 | 1.716346775 | 4.92E-09 | 5.46E-08 | 5207.823077 | 8938.105752 | ENSG00000112159 |
| CGAS | 0.778918313 | 1.715843905 | 3.86E-03 | 1.40E-02 | 51.6143027 | 88.24789504 | ENSG00000164430 |
| BDNF | 0.778853548 | 1.71576688 | 4.71E-04 | 2.17E-03 | 67.48506785 | 115.8749092 | ENSG00000176697 |
| HECW1 | 0.778762548 | 1.715658659 | 1.46E-05 | 9.28E-05 | 124.6238168 | 214.2248568 | ENSG00000002746 |
| ONECUT2 | 0.778489327 | 1.715333774 | 3.08E-02 | 8.02E-02 | 28.43119284 | 48.96416547 | ENSG00000119547 |
| COL23A1 | 0.778350894 | 1.715169188 | 9.15E-13 | 1.56E-11 | 333.488948 | 571.6824995 | ENSG00000050767 |
| NOTCH4 | 0.778175896 | 1.714961152 | 9.45E-03 | 3.00E-02 | 36.76747812 | 63.04830941 | ENSG00000204301 |
| GAL | 0.77812174 | 1.714896777 | 7.21E-20 | 2.39E-18 | 829.6928001 | 1423.669543 | ENSG00000069482 |
| MTR | 0.777943625 | 1.714685069 | 3.87E-19 | 1.20E-17 | 683.8645541 | 1172.464869 | ENSG00000116984 |
| CDC42SE1 | 0.777640092 | 1.714324349 | 9.68E-39 | 1.03E-36 | 1434.582116 | 2458.872585 | ENSG00000197622 |
| KCNT2 | 0.777254823 | 1.713866602 | 1.68E-06 | 1.27E-05 | 147.5901362 | 252.6532736 | ENSG00000162687 |
| PHETA2 | 0.775792427 | 1.712130212 | 1.02E-04 | 5.49E-04 | 93.97484838 | 161.0965579 | ENSG00000177096 |
| STC1 | 0.775742594 | 1.712071074 | 1.48E-10 | 2.00E-09 | 330.386953 | 564.8010757 | ENSG00000159167 |
| CA7 | 0.775416866 | 1.71168457 | 2.78E-02 | 7.40E-02 | 28.51867473 | 48.67970144 | ENSG00000168748 |
| ACTN2 | 0.775156119 | 1.711375235 | 3.85E-03 | 1.40E-02 | 48.10300745 | 82.30299118 | ENSG00000077522 |
| DCP1B | 0.774916348 | 1.711090834 | 4.84E-13 | 8.50E-12 | 323.512932 | 553.1755834 | ENSG00000151065 |
| FES | 0.774868887 | 1.711034545 | 2.35E-05 | 1.44E-04 | 106.0187191 | 181.4345621 | ENSG00000182511 |
| TM7SF3 | 0.774616895 | 1.710735709 | 1.61E-19 | 5.19E-18 | 1058.603866 | 1812.207757 | ENSG00000064115 |
| RAB4A | 0.774445751 | 1.710532779 | 1.28E-19 | 4.19E-18 | 654.3049322 | 1119.244132 | ENSG00000168118 |
| POU2F2 | 0.77378584 | 1.709750534 | 4.40E-05 | 2.55E-04 | 100.0207066 | 171.004731 | ENSG00000028277 |
| KIAA1217 | 0.77360292 | 1.709533768 | 4.44E-14 | 8.72E-13 | 368.4099556 | 630.2821809 | ENSG00000120549 |
| NTPCR | 0.772537048 | 1.70827122 | 3.77E-29 | 2.30E-27 | 1017.599366 | 1737.286724 | ENSG00000135778 |
| ADSS2 | 0.772519223 | 1.708250114 | 1.54E-40 | 1.85E-38 | 1526.261645 | 2607.887631 | ENSG00000035687 |
| OSGEPL1 | 0.772187915 | 1.707857867 | 2.30E-04 | 1.15E-03 | 82.82918846 | 141.4724368 | ENSG00000128694 |
| CLMN | 0.772084487 | 1.707735434 | 2.69E-05 | 1.64E-04 | 114.8898584 | 195.9870597 | ENSG00000165959 |
| ZNF280D | 0.771911714 | 1.707530933 | 6.14E-03 | 2.09E-02 | 42.13760988 | 71.93555716 | ENSG00000137871 |
| BATF3 | 0.771850928 | 1.70745899 | 3.37E-03 | 1.24E-02 | 48.64101576 | 83.17415144 | ENSG00000123685 |
| DYRK3 | 0.771463432 | 1.707000442 | 3.82E-09 | 4.31E-08 | 213.6608239 | 364.9550467 | ENSG00000143479 |
| MAPK13 | 0.771306011 | 1.706814191 | 2.49E-11 | 3.65E-10 | 306.553435 | 522.4704419 | ENSG00000156711 |
| TIMP4 | 0.770655362 | 1.7060446 | 6.91E-04 | 3.05E-03 | 73.64994685 | 125.7704698 | ENSG00000157150 |
| KRTCAP2 | 0.769443271 | 1.704611855 | 2.83E-03 | 1.07E-02 | 52.62671624 | 89.59013516 | ENSG00000163463 |
| ZSCAN31 | 0.769102207 | 1.704208919 | 7.69E-04 | 3.35E-03 | 72.44881517 | 123.674818 | ENSG00000235109 |
| CDC42EP5 | 0.768987357 | 1.704073257 | 9.50E-05 | 5.17E-04 | 117.6236273 | 200.0097699 | ENSG00000167617 |
| SEC14L5 | 0.767798588 | 1.702669692 | 1.43E-10 | 1.95E-09 | 243.5703875 | 414.7441564 | ENSG00000103184 |
| CNIH3 | 0.767246296 | 1.702018002 | 8.84E-05 | 4.84E-04 | 100.9426841 | 171.5567575 | ENSG00000143786 |
| NIT1 | 0.766243898 | 1.700835835 | 8.67E-17 | 2.23E-15 | 501.3119101 | 852.0168426 | ENSG00000158793 |
| ABL2 | 0.766011463 | 1.700561832 | 2.75E-18 | 7.98E-17 | 543.0093324 | 923.2890517 | ENSG00000143322 |
| DDX60 | 0.765182716 | 1.699585236 | 5.90E-03 | 2.03E-02 | 49.18615747 | 83.70490222 | ENSG00000137628 |
| PLEKHN1 | 0.764783121 | 1.699114553 | 2.44E-02 | 6.62E-02 | 34.52505997 | 58.70054682 | ENSG00000187583 |
| DENND3 | 0.764091113 | 1.698299745 | 2.32E-04 | 1.15E-03 | 118.8635946 | 202.0172235 | ENSG00000105339 |
| REL | 0.76384723 | 1.698012677 | 7.06E-03 | 2.35E-02 | 43.81044391 | 74.42025768 | ENSG00000162924 |
| CDH8 | 0.763155725 | 1.69719899 | 8.95E-04 | 3.85E-03 | 67.7879007 | 115.1285006 | ENSG00000150394 |
| TRIM5 | 0.763023588 | 1.69704355 | 2.38E-08 | 2.39E-07 | 252.8512657 | 429.001134 | ENSG00000132256 |
| LGALS3BP | 0.762851438 | 1.696841061 | 5.03E-36 | 4.39E-34 | 1259.122274 | 2136.221324 | ENSG00000108679 |
| OGFRL1 | 0.762716502 | 1.696682363 | 2.06E-06 | 1.54E-05 | 140.0374346 | 237.5301878 | ENSG00000119900 |
| OMA1 | 0.762289404 | 1.696180148 | 1.78E-05 | 1.12E-04 | 113.9908838 | 193.1270196 | ENSG00000162600 |
| SLFN12 | 0.761119572 | 1.694805332 | 3.39E-04 | 1.63E-03 | 99.52456181 | 168.6348228 | ENSG00000172123 |
| MFSD6L | 0.760916089 | 1.694566307 | 2.99E-02 | 7.83E-02 | 29.8830061 | 50.54759769 | ENSG00000185156 |
| HPN | 0.760395824 | 1.693955322 | 4.13E-08 | 4.03E-07 | 224.4964664 | 379.9791814 | ENSG00000105707 |
| FN1 | 0.76036573 | 1.693919987 | 1.97E-26 | 1.05E-24 | 2693.31661 | 4560.816455 | ENSG00000115414 |
| ZNF292 | 0.760033955 | 1.693530483 | 1.20E-11 | 1.82E-10 | 372.1999295 | 630.5096518 | ENSG00000188994 |
| TMEM185A | 0.759634057 | 1.693061122 | 5.93E-13 | 1.03E-11 | 340.223387 | 575.8390968 | ENSG00000269556 |
| OSBPL5 | 0.758620143 | 1.69187167 | 5.98E-12 | 9.37E-11 | 317.5348192 | 537.0162457 | ENSG00000021762 |
| ARHGAP4 | 0.758430361 | 1.691649124 | 8.51E-22 | 3.28E-20 | 730.8979657 | 1235.752875 | ENSG00000089820 |
| TTC32 | 0.758395551 | 1.691608308 | 1.16E-03 | 4.87E-03 | 65.77615223 | 111.1270541 | ENSG00000183891 |
| ZFHX4 | 0.758252873 | 1.691441022 | 2.00E-02 | 5.62E-02 | 32.71211721 | 55.41752087 | ENSG00000091656 |
| RALGPS2 | 0.757135372 | 1.690131351 | 2.36E-15 | 5.28E-14 | 588.9175672 | 995.5071401 | ENSG00000116191 |
| BROX | 0.756843742 | 1.689789739 | 4.24E-10 | 5.42E-09 | 259.0667444 | 437.7117616 | ENSG00000162819 |
| ELF3 | 0.756407453 | 1.689278802 | 2.14E-03 | 8.38E-03 | 58.93111626 | 99.6373389 | ENSG00000163435 |
| VEPH1 | 0.754620928 | 1.687188221 | 5.70E-03 | 1.97E-02 | 58.4321806 | 98.63709526 | ENSG00000197415 |
| PLEKHH1 | 0.754615508 | 1.687181882 | 6.40E-18 | 1.82E-16 | 681.4840303 | 1149.686836 | ENSG00000054690 |
| TP53I11 | 0.754417196 | 1.68694998 | 1.74E-27 | 9.72E-26 | 1184.476572 | 1998.834018 | ENSG00000175274 |
| C2orf72 | 0.753596798 | 1.685990957 | 3.19E-03 | 1.18E-02 | 54.9853643 | 92.75484903 | ENSG00000204128 |
| CLDN19 | 0.753490996 | 1.685867317 | 5.20E-06 | 3.60E-05 | 153.972206 | 259.3937767 | ENSG00000164007 |
| ZC3H6 | 0.751768469 | 1.683855652 | 4.18E-04 | 1.96E-03 | 81.27422395 | 137.140528 | ENSG00000188177 |

| | | | | | | | |
|---|---|---|---|---|---|---|---|
| FAM162A | 0.75125509 | 1.683256563 | 2.63E-21 | 9.78E-20 | 760.0103916 | 1278.239446 | ENSG00000114023 |
| STEAP1B | 0.750492557 | 1.682367117 | 1.18E-03 | 4.96E-03 | 76.61733968 | 129.1128941 | ENSG00000105889 |
| NPAS1 | 0.749781159 | 1.68153774 | 2.82E-05 | 1.71E-04 | 121.4934245 | 204.2177326 | ENSG00000130751 |
| HMGCL | 0.749132007 | 1.68078129 | 9.82E-06 | 6.48E-05 | 215.9937519 | 362.313578 | ENSG00000117305 |
| TNXB | 0.748547697 | 1.68010069 | 2.98E-07 | 2.55E-06 | 162.3255121 | 272.5640214 | ENSG00000168477 |
| EMBP1 | 0.747476882 | 1.678854127 | 1.64E-02 | 4.77E-02 | 40.67060281 | 68.44173814 | ENSG00000231752 |
| KLF7 | 0.747400759 | 1.678765545 | 1.23E-10 | 1.69E-09 | 275.4845663 | 462.7220783 | ENSG00000118263 |
| RGS5 | 0.747273792 | 1.678617809 | 1.96E-08 | 1.99E-07 | 221.911467 | 372.4685893 | ENSG00000143248 |
| VWDE | 0.747133451 | 1.678454526 | 1.55E-07 | 1.39E-06 | 209.1909429 | 350.72974 | ENSG00000146530 |
| NPTXR | 0.747113327 | 1.678431114 | 4.16E-15 | 9.05E-14 | 448.4546874 | 752.7653784 | ENSG00000221890 |
| EDEM3 | 0.745178089 | 1.676181167 | 4.49E-10 | 5.72E-09 | 301.495636 | 505.1688737 | ENSG00000116406 |
| RINL | 0.745114619 | 1.676107427 | 1.19E-07 | 1.08E-06 | 191.7171674 | 321.006844 | ENSG00000187994 |
| SLC52A3 | 0.744991259 | 1.675964115 | 6.04E-08 | 5.72E-07 | 187.2213676 | 313.8745356 | ENSG00000101276 |
| RYR1 | 0.744595071 | 1.67550393 | 1.22E-25 | 6.16E-24 | 1055.920383 | 1768.812606 | ENSG00000196218 |
| SMIM14 | 0.744334646 | 1.675201508 | 4.54E-07 | 3.77E-06 | 204.5015189 | 342.3969236 | ENSG00000163683 |
| SUMF2 | 0.743970904 | 1.674779198 | 2.79E-22 | 1.11E-20 | 1112.575901 | 1862.350786 | ENSG00000129103 |
| GADD45B | 0.743771606 | 1.674547855 | 4.55E-05 | 2.64E-04 | 111.4335933 | 186.6591432 | ENSG00000099860 |
| JUN | 0.743114934 | 1.673785824 | 9.26E-10 | 1.13E-08 | 367.6645815 | 615.1174712 | ENSG00000177606 |
| HSPG2 | 0.742276376 | 1.672813229 | 3.18E-39 | 3.49E-37 | 2831.48077 | 4735.312908 | ENSG00000142798 |
| C1orf61 | 0.741546778 | 1.671967469 | 1.10E-03 | 4.63E-03 | 67.67761602 | 113.2390986 | ENSG00000125462 |
| CCDC113 | 0.741416339 | 1.671816308 | 1.88E-06 | 1.41E-05 | 148.2989532 | 247.8244718 | ENSG00000103021 |
| CCDC114 | 0.741209019 | 1.67157608 | 4.00E-02 | 9.96E-02 | 31.21576165 | 52.21519328 | ENSG00000105479 |
| PLA2G6 | 0.740347307 | 1.670577957 | 8.42E-06 | 5.63E-05 | 128.080282 | 213.6760774 | ENSG00000184381 |
| CHGB | 0.740311867 | 1.670536919 | 1.19E-04 | 6.31E-04 | 96.77982939 | 161.4307882 | ENSG00000089199 |
| DPP6 | 0.739768814 | 1.669908221 | 1.31E-06 | 1.01E-05 | 148.0287353 | 247.2899408 | ENSG00000130226 |
| ZNF678 | 0.738762377 | 1.668743685 | 7.67E-08 | 7.16E-07 | 190.0830937 | 317.4019703 | ENSG00000181450 |
| TTC17 | 0.738292025 | 1.668199724 | 1.04E-24 | 5.03E-23 | 985.8975468 | 1644.134442 | ENSG00000052841 |
| CST3 | 0.737597624 | 1.667396976 | 1.88E-12 | 3.11E-11 | 358.6778753 | 598.6279133 | ENSG00000101439 |
| DECR2 | 0.736995562 | 1.666701287 | 2.41E-04 | 1.19E-03 | 99.81834623 | 166.1668243 | ENSG00000242612 |
| RPL39L | 0.736585672 | 1.666227822 | 6.27E-16 | 1.50E-14 | 511.0911235 | 851.2981069 | ENSG00000163923 |
| PFKFB2 | 0.736032117 | 1.66558862 | 3.34E-14 | 6.65E-13 | 478.4594669 | 796.1586409 | ENSG00000123836 |
| TSTD1 | 0.734864731 | 1.66424142 | 7.24E-06 | 4.90E-05 | 145.0848338 | 241.0498201 | ENSG00000215845 |
| TMC4 | 0.73432563 | 1.663619649 | 2.27E-03 | 8.80E-03 | 68.29975173 | 113.4851128 | ENSG00000167608 |
| LHX4 | 0.733886031 | 1.66311281 | 1.37E-02 | 4.11E-02 | 41.47392292 | 68.9528854 | ENSG00000121454 |
| ASAH2B | 0.733780964 | 1.662991695 | 4.88E-03 | 1.72E-02 | 63.0261134 | 104.9560243 | ENSG00000204147 |
| HYLS1 | 0.733761977 | 1.662969809 | 5.12E-07 | 4.22E-06 | 170.7814939 | 283.9733535 | ENSG00000198331 |
| NUFIP2 | 0.733026483 | 1.662122234 | 6.71E-33 | 5.02E-31 | 2150.516601 | 3573.191666 | ENSG00000108256 |
| GUSBP11 | 0.732688271 | 1.661732628 | 4.61E-02 | 1.12E-01 | 25.02990687 | 41.58076029 | ENSG00000228315 |
| CSRNP3 | 0.732612065 | 1.661644853 | 2.14E-04 | 1.07E-03 | 99.13568654 | 164.9732406 | ENSG00000178662 |
| THSD4 | 0.730994813 | 1.659783204 | 1.43E-04 | 7.43E-04 | 96.99538051 | 160.8376402 | ENSG00000187720 |
| ARHGEF11 | 0.730842103 | 1.659607524 | 1.10E-25 | 5.65E-24 | 993.5222994 | 1647.97843 | ENSG00000132694 |
| PHLDA2 | 0.730809531 | 1.659570055 | 1.28E-04 | 6.71E-04 | 99.51272248 | 165.0350863 | ENSG00000181649 |
| CNTNAP2 | 0.730765822 | 1.659519776 | 1.28E-36 | 1.17E-34 | 1507.404715 | 2501.958024 | ENSG00000174469 |
| NIT2 | 0.730625173 | 1.659357997 | 7.29E-09 | 7.91E-08 | 298.6477764 | 494.8996152 | ENSG00000114021 |
| PTPDC1 | 0.730386652 | 1.659083678 | 6.41E-06 | 4.38E-05 | 134.3289761 | 222.6649536 | ENSG00000158079 |
| RGL3 | 0.730359067 | 1.659051955 | 3.69E-18 | 1.06E-16 | 613.8580403 | 1018.758811 | ENSG00000205517 |
| PCNX2 | 0.73013639 | 1.658795904 | 3.15E-31 | 2.13E-29 | 1365.144572 | 2264.251973 | ENSG00000135749 |
| SCG5 | 0.729909739 | 1.658535324 | 5.77E-10 | 7.22E-09 | 344.0884271 | 570.5351439 | ENSG00000166922 |
| C12orf66 | 0.72830975 | 1.656696982 | 2.00E-04 | 1.01E-03 | 96.20657848 | 159.6032077 | ENSG00000174206 |
| ARL8A | 0.727267079 | 1.655500078 | 6.06E-21 | 2.21E-19 | 775.2086726 | 1283.356165 | ENSG00000143862 |
| SLC8A2 | 0.725611978 | 1.65360193 | 4.04E-06 | 2.86E-05 | 208.6990796 | 345.0422094 | ENSG00000118160 |
| ITGB1 | 0.723943677 | 1.651690846 | 4.44E-55 | 8.73E-53 | 5979.88533 | 9877.58802 | ENSG00000150093 |
| CILP2 | 0.723536142 | 1.651224339 | 1.57E-12 | 2.62E-11 | 362.0922768 | 598.1861359 | ENSG00000160161 |
| MCEE | 0.723439297 | 1.6511135 | 1.45E-04 | 7.52E-04 | 97.27541057 | 160.4340401 | ENSG00000124370 |
| RAB26 | 0.722820621 | 1.650405599 | 1.35E-03 | 5.56E-03 | 74.92373002 | 123.6605759 | ENSG00000167964 |
| TRMT2B | 0.722161296 | 1.649651521 | 3.63E-15 | 7.94E-14 | 460.9970183 | 760.381646 | ENSG00000188917 |
| THEM4 | 0.72172622 | 1.649154108 | 3.44E-07 | 2.92E-06 | 188.4709085 | 310.3608526 | ENSG00000159445 |
| RCOR3 | 0.721429012 | 1.648814403 | 7.00E-09 | 7.62E-08 | 248.2231295 | 409.556644 | ENSG00000117625 |
| MDGA2 | 0.720195979 | 1.647405806 | 2.05E-02 | 5.75E-02 | 35.61957376 | 58.59352064 | ENSG00000139915 |
| GPATCH2 | 0.720071297 | 1.647263439 | 8.39E-09 | 9.05E-08 | 278.9691918 | 459.6447445 | ENSG00000092978 |
| SNAPC5 | 0.719919505 | 1.647090133 | 2.89E-08 | 2.87E-07 | 205.3729478 | 338.3979065 | ENSG00000174446 |
| KCNA6 | 0.718620476 | 1.64560773 | 8.09E-03 | 2.64E-02 | 51.18259998 | 84.48922761 | ENSG00000130035 |
| TGFB1I1 | 0.718594208 | 1.645577768 | 1.01E-04 | 5.45E-04 | 101.959328 | 167.7429808 | ENSG00000140682 |
| IBA57 | 0.718452423 | 1.645416053 | 4.87E-09 | 5.42E-08 | 245.24167 | 403.7921768 | ENSG00000181873 |
| CRISPLD2 | 0.717975001 | 1.644871634 | 6.71E-05 | 3.77E-04 | 106.7969211 | 175.7760003 | ENSG00000103196 |
| EXPH5 | 0.716865498 | 1.643607135 | 1.68E-02 | 4.86E-02 | 38.2051367 | 62.72699462 | ENSG00000110723 |
| ZNF160 | 0.716134132 | 1.642774128 | 5.19E-12 | 8.17E-11 | 401.4702728 | 659.1345044 | ENSG00000170949 |
| SIX4 | 0.715909075 | 1.642517879 | 3.02E-14 | 6.06E-13 | 485.7882831 | 798.3117642 | ENSG00000100625 |
| ZFP36 | 0.715724223 | 1.642307438 | 4.41E-04 | 2.05E-03 | 96.11295121 | 158.0464064 | ENSG00000128016 |
| C15orf40 | 0.715397479 | 1.641935527 | 4.46E-06 | 3.13E-05 | 163.6541253 | 268.6023819 | ENSG00000169609 |
| DEPDC4 | 0.715228645 | 1.641743388 | 4.83E-02 | 1.16E-01 | 28.16272671 | 46.25090373 | ENSG00000166153 |
| FAM241A | 0.714454919 | 1.640863147 | 3.15E-02 | 8.16E-02 | 35.04420768 | 57.68901728 | ENSG00000174749 |
| ADAMTS20 | 0.713727196 | 1.640035673 | 1.53E-05 | 9.71E-05 | 141.751093 | 232.1343238 | ENSG00000173157 |

| | | | | | | | |
|---|---|---|---|---|---|---|---|
| AR | 0.713494681 | 1.639771375 | 2.35E-09 | 2.73E-08 | 266.8210426 | 437.0917528 | ENSG00000169083 |
| TFPI | 0.713013795 | 1.63922489 | 7.54E-04 | 3.29E-03 | 78.01727706 | 128.0892668 | ENSG00000003436 |
| SP8 | 0.71240578 | 1.638534194 | 9.60E-05 | 5.21E-04 | 113.0257155 | 184.880505 | ENSG00000164651 |
| ZNF8 | 0.712310939 | 1.638426482 | 1.10E-05 | 7.14E-05 | 171.9998224 | 282.2473893 | ENSG00000278129 |
| CLEC11A | 0.711892153 | 1.637950948 | 5.72E-06 | 3.93E-05 | 149.4348181 | 244.9092919 | ENSG00000105472 |
| TMC7 | 0.711731951 | 1.637769074 | 2.20E-03 | 8.57E-03 | 69.62548602 | 113.9683026 | ENSG00000170537 |
| TMEM179B | 0.71040185 | 1.636259819 | 1.75E-02 | 5.02E-02 | 45.08729339 | 73.99301332 | ENSG00000185475 |
| TFR2 | 0.709969973 | 1.635770071 | 4.09E-05 | 2.40E-04 | 122.5369383 | 200.1934232 | ENSG00000106327 |
| ZNF669 | 0.709856208 | 1.635641086 | 2.60E-08 | 2.59E-07 | 225.10585 | 368.5760469 | ENSG00000188295 |
| STX6 | 0.70950516 | 1.635243138 | 4.04E-23 | 1.70E-21 | 977.0283368 | 1598.778094 | ENSG00000135823 |
| NDUFAF1 | 0.709499518 | 1.635236743 | 1.62E-10 | 2.19E-09 | 339.6071454 | 555.4311591 | ENSG00000137806 |
| TMEM37 | 0.706882469 | 1.632273111 | 4.83E-08 | 4.65E-07 | 234.0031465 | 381.9501121 | ENSG00000171227 |
| DDHD2 | 0.705840569 | 1.631094725 | 2.73E-09 | 3.13E-08 | 287.6793587 | 468.846589 | ENSG00000085788 |
| SFT2D2 | 0.704714369 | 1.629821954 | 8.15E-07 | 6.51E-06 | 187.001386 | 304.9958901 | ENSG00000213064 |
| CCDC122 | 0.70460255 | 1.629695636 | 1.17E-02 | 3.60E-02 | 43.7371167 | 71.31766841 | ENSG00000151773 |
| IRAK4 | 0.704164573 | 1.629200964 | 2.88E-02 | 7.60E-02 | 35.57539487 | 58.15416635 | ENSG00000198001 |
| SEC14L6 | 0.704061242 | 1.629084278 | 3.54E-02 | 9.01E-02 | 30.95132572 | 50.47392673 | ENSG00000214491 |
| PBXIP1 | 0.703887147 | 1.628887704 | 1.28E-18 | 3.83E-17 | 667.8073493 | 1087.524293 | ENSG00000163346 |
| NANOS1 | 0.703726602 | 1.628706448 | 6.87E-03 | 2.31E-02 | 75.50427759 | 123.0206484 | ENSG00000188613 |
| SLC5A12 | 0.702949487 | 1.627829374 | 8.92E-03 | 2.86E-02 | 49.61331752 | 80.62759891 | ENSG00000148942 |
| ATP7A | 0.701997414 | 1.62675548 | 9.01E-03 | 2.88E-02 | 58.08365416 | 94.24522459 | ENSG00000165240 |
| PEAK1 | 0.701564616 | 1.626267539 | 7.57E-15 | 1.61E-13 | 877.3205108 | 1425.927475 | ENSG00000173517 |
| CXCL5 | 0.701093902 | 1.625737016 | 4.05E-03 | 1.46E-02 | 73.19843241 | 119.1458252 | ENSG00000163735 |
| DRAM1 | 0.700517993 | 1.625088168 | 3.07E-05 | 1.85E-04 | 129.0429349 | 209.9937646 | ENSG00000136048 |
| ZNF687 | 0.700139232 | 1.624661578 | 1.68E-32 | 1.24E-30 | 1843.470891 | 2995.318437 | ENSG00000143373 |
| ARHGAP31 | 0.699179756 | 1.623581444 | 1.45E-06 | 1.11E-05 | 195.6435832 | 317.243082 | ENSG00000031081 |
| TNMD | 0.699170226 | 1.623570718 | 7.52E-03 | 2.48E-02 | 51.08849714 | 82.93001527 | ENSG00000000005 |
| CUL9 | 0.698971927 | 1.623347573 | 1.92E-05 | 1.20E-04 | 186.4291231 | 302.7072012 | ENSG00000112659 |
| ACBD3 | 0.698674052 | 1.623012433 | 5.83E-20 | 1.95E-18 | 748.1075197 | 1214.703622 | ENSG00000182827 |
| PAQR6 | 0.698655694 | 1.622991781 | 4.33E-04 | 2.02E-03 | 90.02295103 | 146.107638 | ENSG00000160781 |
| CERS2 | 0.698006123 | 1.622261196 | 1.39E-38 | 1.45E-36 | 2206.766077 | 3580.016393 | ENSG00000143418 |
| RICTOR | 0.696734252 | 1.620831651 | 1.26E-06 | 9.77E-06 | 190.9091413 | 309.2500872 | ENSG00000164327 |
| NUF2 | 0.696692997 | 1.620785303 | 1.43E-12 | 2.41E-11 | 421.1395477 | 682.8592652 | ENSG00000143228 |
| TRIM32 | 0.696398474 | 1.620454456 | 1.72E-11 | 2.56E-10 | 437.647279 | 709.313953 | ENSG00000119401 |
| MDFI | 0.695156306 | 1.619059838 | 1.61E-25 | 8.11E-24 | 2129.106865 | 3447.820176 | ENSG00000112559 |
| HMBOX1 | 0.695020962 | 1.618907956 | 6.01E-05 | 3.41E-04 | 125.3648101 | 202.88192 | ENSG00000147421 |
| ZNF34 | 0.695020368 | 1.618907289 | 6.40E-04 | 2.85E-03 | 88.4163108 | 143.1217617 | ENSG00000196378 |
| SLCO1A2 | 0.69267436 | 1.616276876 | 1.80E-02 | 5.15E-02 | 44.07854663 | 71.11788867 | ENSG00000084453 |
| C1orf43 | 0.692372422 | 1.615938645 | 2.52E-37 | 2.42E-35 | 2154.112301 | 3480.599473 | ENSG00000143612 |
| HELZ | 0.692080625 | 1.615611841 | 2.20E-12 | 3.62E-11 | 420.7383817 | 679.7935082 | ENSG00000198265 |
| INO80D | 0.691964685 | 1.61548201 | 5.24E-07 | 4.30E-06 | 229.1860536 | 370.1960205 | ENSG00000114933 |
| PABPC5 | 0.691818603 | 1.61531844 | 1.15E-02 | 3.56E-02 | 45.55584143 | 73.50500144 | ENSG00000174740 |
| LOXL2 | 0.691445218 | 1.614900433 | 1.16E-23 | 5.15E-22 | 952.3070305 | 1537.221845 | ENSG00000134013 |
| GLRX2 | 0.691321324 | 1.614761756 | 6.83E-05 | 3.83E-04 | 176.8027606 | 286.0302582 | ENSG00000023572 |
| MORN1 | 0.689993721 | 1.613276497 | 1.24E-03 | 5.17E-03 | 95.59141292 | 153.8715322 | ENSG00000116151 |
| TARBP1 | 0.689912339 | 1.613185495 | 2.11E-13 | 3.86E-12 | 542.1372278 | 874.5340375 | ENSG00000059588 |
| AGPAT3 | 0.688705919 | 1.61183707 | 7.99E-06 | 5.37E-05 | 160.154306 | 258.0902774 | ENSG00000160216 |
| HTR6 | 0.687316713 | 1.61028574 | 1.20E-02 | 3.67E-02 | 49.31183612 | 79.23911224 | ENSG00000158748 |
| SUCO | 0.686449569 | 1.609318156 | 1.19E-20 | 4.26E-19 | 868.0248775 | 1396.799048 | ENSG00000094975 |
| CEP85L | 0.68632548 | 1.609179741 | 1.45E-03 | 5.94E-03 | 84.68371937 | 136.0327494 | ENSG00000111860 |
| HLTF | 0.685897396 | 1.608702328 | 2.70E-19 | 8.48E-18 | 890.5555904 | 1433.276992 | ENSG00000071794 |
| STX17 | 0.685654881 | 1.60843193 | 3.20E-09 | 3.65E-08 | 297.5618355 | 478.1809815 | ENSG00000136874 |
| TOR1AIP2 | 0.68450202 | 1.607147141 | 3.77E-13 | 6.74E-12 | 526.4592621 | 845.9719216 | ENSG00000169905 |
| ACTRT3 | 0.684371833 | 1.60700212 | 6.33E-03 | 2.15E-02 | 55.39577893 | 89.09378145 | ENSG00000184378 |
| PTP4A3 | 0.684326054 | 1.606951129 | 1.77E-16 | 4.46E-15 | 849.8022343 | 1364.291976 | ENSG00000184489 |
| GLP1R | 0.683796465 | 1.606361352 | 1.52E-03 | 6.18E-03 | 86.89244653 | 139.3477614 | ENSG00000112164 |
| PHYH | 0.683725728 | 1.606282593 | 1.03E-07 | 9.50E-07 | 267.8268352 | 430.0521979 | ENSG00000107537 |
| ZBTB5 | 0.683136073 | 1.605626211 | 1.60E-10 | 2.16E-09 | 339.9429935 | 545.4915451 | ENSG00000168795 |
| MYEF2 | 0.682291284 | 1.604686291 | 1.93E-19 | 6.19E-18 | 1002.397781 | 1607.743865 | ENSG00000104177 |
| DHTKD1 | 0.681907701 | 1.604259694 | 7.79E-29 | 4.65E-27 | 1503.336261 | 2411.199307 | ENSG00000181192 |
| RPS27L | 0.681853397 | 1.60419931 | 9.43E-18 | 2.64E-16 | 690.8720402 | 1108.64979 | ENSG00000185088 |
| ZNF670 | 0.681793925 | 1.604133182 | 3.15E-06 | 2.26E-05 | 216.5467922 | 347.6895784 | ENSG00000277462 |
| STUM | 0.681121589 | 1.603385785 | 2.72E-02 | 7.26E-02 | 41.88501328 | 67.03014031 | ENSG00000203685 |
| ZNF81 | 0.680271315 | 1.602441084 | 2.74E-04 | 1.34E-03 | 114.7381857 | 183.9657576 | ENSG00000197779 |
| TRAF3IP2 | 0.679221376 | 1.601275312 | 6.67E-14 | 1.29E-12 | 470.0752238 | 752.7766401 | ENSG00000056972 |
| SMIM29 | 0.678948982 | 1.600973005 | 1.98E-03 | 7.84E-03 | 82.45669519 | 132.0361249 | ENSG00000186577 |
| TIE1 | 0.678855231 | 1.600868972 | 2.65E-03 | 1.01E-02 | 83.09150921 | 133.1944634 | ENSG00000066056 |
| SDHC | 0.67862727 | 1.600616038 | 1.05E-37 | 1.03E-35 | 1925.120147 | 3080.787087 | ENSG00000143252 |
| CYP11A1 | 0.678532952 | 1.600511399 | 4.17E-02 | 1.03E-01 | 34.52872675 | 55.2153301 | ENSG00000140459 |
| SRGAP2 | 0.678344595 | 1.600302452 | 2.08E-19 | 6.63E-18 | 887.3757865 | 1419.43301 | ENSG00000266028 |
| NES | 0.677866096 | 1.599771767 | 2.75E-52 | 4.96E-50 | 6401.071849 | 10240.50488 | ENSG00000132688 |
| SIPA1L2 | 0.676956927 | 1.598763927 | 1.63E-11 | 2.43E-10 | 413.9202156 | 661.4167537 | ENSG00000116991 |
| ALPK3 | 0.676316791 | 1.5980547 | 4.81E-32 | 3.42E-30 | 1746.329656 | 2789.750843 | ENSG00000136383 |

| | | | | | | | |
|---|---|---|---|---|---|---|---|
| COLEC12 | 0.676268251 | 1.598000933 | 1.62E-02 | 4.72E-02 | 50.55519476 | 80.95509284 | ENSG00000158270 |
| ATG16L2 | 0.676191353 | 1.597915759 | 4.59E-08 | 4.43E-07 | 243.8708298 | 389.3074194 | ENSG00000168010 |
| GAP43 | 0.67590347 | 1.597596935 | 3.32E-11 | 4.80E-10 | 403.1391646 | 643.4279574 | ENSG00000172020 |
| ITPKB | 0.675566165 | 1.597223457 | 8.62E-09 | 9.28E-08 | 297.4544169 | 475.374421 | ENSG00000143772 |
| ZCCHC4 | 0.675440131 | 1.59708393 | 1.65E-04 | 8.47E-04 | 131.2583199 | 209.6791804 | ENSG00000168228 |
| ZNF486 | 0.675385681 | 1.597023654 | 1.82E-10 | 2.44E-09 | 366.0226348 | 584.8994812 | ENSG00000256229 |
| PTPN3 | 0.675128108 | 1.596738553 | 1.67E-05 | 1.05E-04 | 155.653488 | 248.9236482 | ENSG00000070159 |
| IGFBP2 | 0.674466591 | 1.596006571 | 1.81E-08 | 1.86E-07 | 5889.046162 | 9398.797282 | ENSG00000115457 |
| IKBIP | 0.674308761 | 1.595831978 | 1.95E-07 | 1.72E-06 | 230.7146238 | 368.4762902 | ENSG00000166130 |
| SOCS1 | 0.674274371 | 1.595793938 | 3.91E-07 | 3.29E-06 | 395.766922 | 632.1345448 | ENSG00000185338 |
| IGSF9B | 0.67266794 | 1.594018022 | 2.80E-21 | 1.04E-19 | 1505.846819 | 2399.633442 | ENSG00000080854 |
| AK5 | 0.672591217 | 1.593933253 | 2.77E-03 | 1.05E-02 | 79.39999346 | 126.5569153 | ENSG00000154027 |
| VSNL1 | 0.672194017 | 1.593494475 | 1.12E-22 | 4.64E-21 | 1314.357458 | 2094.902982 | ENSG00000163032 |
| STARD8 | 0.672055216 | 1.593341173 | 4.59E-05 | 2.66E-04 | 137.0430086 | 218.0706317 | ENSG00000130052 |
| PGF | 0.6718585 | 1.59312393 | 2.95E-04 | 1.43E-03 | 103.5935649 | 165.2526221 | ENSG00000119630 |
| NUP133 | 0.671766805 | 1.593022678 | 6.49E-30 | 4.13E-28 | 1856.210813 | 2958.070433 | ENSG00000069248 |
| MCAM | 0.670835437 | 1.591994594 | 1.39E-14 | 2.87E-13 | 529.4616093 | 842.6531599 | ENSG00000076706 |
| KDM3A | 0.669079553 | 1.590058179 | 4.52E-14 | 8.85E-13 | 595.6100035 | 946.3034854 | ENSG00000115548 |
| PSMD4 | 0.668691977 | 1.589631071 | 2.66E-49 | 4.27E-47 | 4373.307834 | 6951.244747 | ENSG00000159352 |
| SLC16A5 | 0.668308348 | 1.589208426 | 9.25E-08 | 8.57E-07 | 278.9842976 | 443.9099617 | ENSG00000170190 |
| PRR13 | 0.667987152 | 1.58885465 | 6.31E-11 | 8.84E-10 | 442.540402 | 703.0935026 | ENSG00000205352 |
| DDX28 | 0.667655662 | 1.588489619 | 1.11E-08 | 1.18E-07 | 345.2446672 | 548.3683421 | ENSG00000182810 |
| RUNX1T1 | 0.667623809 | 1.588454547 | 1.08E-05 | 7.02E-05 | 188.4506085 | 299.1741388 | ENSG00000079102 |
| FBLN2 | 0.666930294 | 1.587691148 | 5.66E-27 | 3.08E-25 | 1178.977857 | 1871.312583 | ENSG00000163520 |
| MACROD1 | 0.666843758 | 1.587595918 | 1.35E-09 | 1.61E-08 | 372.8960681 | 591.6671603 | ENSG00000133315 |
| KLHL28 | 0.666813169 | 1.587562257 | 1.16E-03 | 4.86E-03 | 110.6073824 | 175.4092567 | ENSG00000179454 |
| ZNF619 | 0.666588908 | 1.587315496 | 7.24E-04 | 3.18E-03 | 94.03856358 | 149.2244878 | ENSG00000177873 |
| KCTD13 | 0.666328262 | 1.587028748 | 4.80E-06 | 3.34E-05 | 172.7395395 | 274.3991076 | ENSG00000174943 |
| CD1D | 0.666172224 | 1.586857109 | 3.88E-02 | 9.72E-02 | 43.46151716 | 69.04133781 | ENSG00000158473 |
| SH3PXD2A | 0.665634811 | 1.586266105 | 2.00E-15 | 4.52E-14 | 573.2393502 | 909.0591897 | ENSG00000107957 |
| SLC19A2 | 0.665023012 | 1.585593565 | 3.30E-14 | 6.58E-13 | 625.1813796 | 991.0663941 | ENSG00000117479 |
| KDM5B | 0.664850228 | 1.585403677 | 8.66E-44 | 1.16E-41 | 5058.809699 | 8018.454128 | ENSG00000117139 |
| C18orf25 | 0.663896476 | 1.584355929 | 1.10E-10 | 1.52E-09 | 379.2589891 | 600.6213853 | ENSG00000152242 |
| DCX | 0.662503865 | 1.582827312 | 5.04E-03 | 1.77E-02 | 103.4985862 | 164.2523542 | ENSG00000077279 |
| ARL4D | 0.660903887 | 1.581072898 | 4.15E-06 | 2.93E-05 | 230.8915395 | 364.5337211 | ENSG00000175906 |
| ARL4A | 0.660800233 | 1.580959306 | 1.57E-08 | 1.63E-07 | 326.3155607 | 515.9337471 | ENSG00000122644 |
| SPTSSB | 0.660575773 | 1.580713354 | 5.23E-05 | 3.00E-04 | 157.1005953 | 248.0118933 | ENSG00000196542 |
| POGK | 0.660458508 | 1.580584875 | 7.93E-30 | 5.03E-28 | 1360.126133 | 2149.393731 | ENSG00000143157 |
| SMIM24 | 0.659706614 | 1.579761331 | 5.19E-03 | 1.81E-02 | 73.57382873 | 116.4094402 | ENSG00000095932 |
| ZNF425 | 0.659601935 | 1.579646711 | 4.80E-06 | 3.34E-05 | 186.5058928 | 294.8376622 | ENSG00000204947 |
| SERPINB6 | 0.658475457 | 1.57841378 | 2.02E-11 | 2.99E-10 | 449.9555104 | 709.5301864 | ENSG00000124570 |
| PRNP | 0.658444491 | 1.578379902 | 7.61E-12 | 1.18E-10 | 472.8897047 | 746.3641761 | ENSG00000171867 |
| MIAT | 0.657842939 | 1.577721912 | 4.07E-13 | 7.22E-12 | 1070.831109 | 1688.028839 | ENSG00000225783 |
| SPRTN | 0.656926707 | 1.576720244 | 4.73E-07 | 3.91E-06 | 209.0877547 | 329.7426971 | ENSG00000010072 |
| THBS3 | 0.656713878 | 1.57648766 | 7.12E-03 | 2.37E-02 | 60.92631947 | 96.16691572 | ENSG00000169231 |
| METTL23 | 0.655825548 | 1.575517248 | 6.88E-05 | 3.85E-04 | 152.8256162 | 241.2632846 | ENSG00000181038 |
| IARS2 | 0.65578194 | 1.575469625 | 4.90E-40 | 5.73E-38 | 2589.382161 | 4080.326842 | ENSG00000067704 |
| HLA-DRB5 | 0.655022149 | 1.574640128 | 1.60E-02 | 4.67E-02 | 52.77249479 | 83.01579618 | ENSG00000198502 |
| NEDD4 | 0.654700868 | 1.574289502 | 2.59E-06 | 1.89E-05 | 193.4121285 | 304.451424 | ENSG00000069869 |
| TNFRSF10B | 0.654268211 | 1.573817451 | 4.46E-22 | 1.76E-20 | 1923.223399 | 3025.83357 | ENSG00000120889 |
| PGM1 | 0.653954043 | 1.573474766 | 9.07E-15 | 1.92E-13 | 1344.153479 | 2114.115615 | ENSG00000079739 |
| RAB24 | 0.653747128 | 1.573249111 | 6.40E-03 | 2.17E-02 | 63.91460006 | 100.7655388 | ENSG00000169228 |
| RASGRP2 | 0.653365472 | 1.572832973 | 1.20E-14 | 2.50E-13 | 566.292002 | 890.9230821 | ENSG00000068831 |
| SLC50A1 | 0.653136629 | 1.572583507 | 6.35E-16 | 1.52E-14 | 721.8099608 | 1134.405468 | ENSG00000169241 |
| TSHZ3 | 0.652481017 | 1.571869031 | 5.05E-06 | 3.50E-05 | 202.7869846 | 319.1948385 | ENSG00000121297 |
| LPP | 0.650989435 | 1.570244739 | 6.03E-08 | 5.71E-07 | 407.06135 | 639.0470453 | ENSG00000145012 |
| NLGN1 | 0.650781089 | 1.570017989 | 9.95E-04 | 4.24E-03 | 116.7820736 | 183.5653862 | ENSG00000169760 |
| HASPIN | 0.650594839 | 1.569815314 | 3.47E-09 | 3.94E-08 | 316.1154576 | 495.9971498 | ENSG00000177602 |
| DST | 0.650545014 | 1.5697611 | 3.07E-13 | 5.55E-12 | 1327.706084 | 2083.383018 | ENSG00000151914 |
| C5orf24 | 0.650483697 | 1.569694384 | 1.44E-04 | 7.46E-04 | 172.2149732 | 270.8176418 | ENSG00000181904 |
| FGF19 | 0.650103951 | 1.569281264 | 5.11E-10 | 6.46E-09 | 357.4923747 | 560.6186837 | ENSG00000162344 |
| TTPAL | 0.650088705 | 1.569264681 | 8.64E-09 | 9.29E-08 | 304.7456368 | 478.5310777 | ENSG00000124120 |
| MRTFB | 0.649561061 | 1.56869085 | 1.51E-07 | 1.36E-06 | 290.5069182 | 455.5072298 | ENSG00000186260 |
| GADD45A | 0.649261597 | 1.568365266 | 5.91E-10 | 7.39E-09 | 408.3888872 | 640.6703029 | ENSG00000116717 |
| SYNPO | 0.648680585 | 1.567733771 | 1.42E-03 | 5.85E-03 | 94.06523331 | 147.2304827 | ENSG00000171992 |
| RNF2 | 0.648419297 | 1.567449863 | 1.22E-11 | 1.85E-10 | 640.8377833 | 1005.276659 | ENSG00000121481 |
| POGZ | 0.647933519 | 1.566922167 | 2.05E-28 | 1.20E-26 | 1341.570559 | 2102.2176 | ENSG00000143442 |
| LAMA5 | 0.647835548 | 1.566815763 | 2.03E-32 | 1.48E-30 | 3617.258085 | 5667.219763 | ENSG00000130702 |
| PLEKHA4 | 0.647689599 | 1.566657266 | 3.77E-08 | 3.69E-07 | 358.9640365 | 561.5810105 | ENSG00000105559 |
| KIZ | 0.646970061 | 1.565876097 | 9.38E-13 | 1.59E-11 | 470.5503048 | 736.8781222 | ENSG00000088970 |
| JAG2 | 0.64642809 | 1.565287962 | 1.12E-11 | 1.71E-10 | 478.7068082 | 749.3245607 | ENSG00000184916 |
| MFNG | 0.645670288 | 1.564465981 | 4.33E-03 | 1.55E-02 | 70.56274535 | 110.3698927 | ENSG00000100060 |
| MAP3K15 | 0.644979645 | 1.563717224 | 2.38E-02 | 6.48E-02 | 48.91314867 | 76.51752077 | ENSG00000180815 |

| | | | | | | | |
|---|---|---|---|---|---|---|---|
| GAL3ST4 | 0.644877414 | 1.563606421 | 8.93E-04 | 3.84E-03 | 95.14663157 | 148.7281462 | ENSG00000197093 |
| SNHG5 | 0.644833003 | 1.563558289 | 1.62E-38 | 1.68E-36 | 3786.568189 | 5918.649787 | ENSG00000203875 |
| AK7 | 0.644431116 | 1.563122794 | 1.06E-02 | 3.31E-02 | 67.71708898 | 105.6292828 | ENSG00000140057 |
| RIPOR2 | 0.644000595 | 1.562656405 | 1.25E-14 | 2.61E-13 | 623.1256397 | 973.5369198 | ENSG00000111913 |
| PLXNB3 | 0.643862552 | 1.56250689 | 1.97E-02 | 5.56E-02 | 48.14563473 | 75.21695347 | ENSG00000198753 |
| REPS2 | 0.643672422 | 1.562300984 | 1.95E-03 | 7.70E-03 | 104.1369918 | 162.409514 | ENSG00000169891 |
| SYT14 | 0.641519317 | 1.559971117 | 1.01E-04 | 5.45E-04 | 207.4739811 | 324.0093614 | ENSG00000143469 |
| EIF4E1B | 0.641206848 | 1.559633283 | 4.44E-02 | 1.08E-01 | 38.77663584 | 60.38130632 | ENSG00000175766 |
| VWA5A | 0.640967078 | 1.5593741 | 6.31E-14 | 1.22E-12 | 540.7852113 | 843.087389 | ENSG00000110002 |
| THNSL1 | 0.639778702 | 1.558090142 | 4.95E-05 | 2.85E-04 | 151.679827 | 236.6754144 | ENSG00000185875 |
| H19 | 0.639497526 | 1.557786506 | 3.41E-02 | 8.71E-02 | 42.85018164 | 66.61956751 | ENSG00000130600 |
| SLC45A3 | 0.639235141 | 1.557503214 | 2.51E-09 | 2.90E-08 | 471.3060546 | 734.2458435 | ENSG00000158715 |
| DTNA | 0.638653577 | 1.556875497 | 2.26E-05 | 1.39E-04 | 181.243455 | 282.1153861 | ENSG00000134769 |
| PDLIM3 | 0.637947217 | 1.55611342 | 5.33E-03 | 1.85E-02 | 72.09518249 | 112.3205952 | ENSG00000154553 |
| FZD6 | 0.637908107 | 1.556071236 | 1.53E-04 | 7.91E-04 | 148.2193685 | 230.6509025 | ENSG00000164930 |
| TPCN1 | 0.637674756 | 1.555819567 | 4.27E-10 | 5.45E-09 | 524.343935 | 815.2138034 | ENSG00000186815 |
| RAB42 | 0.637082218 | 1.555180698 | 8.06E-06 | 5.40E-05 | 177.4894874 | 276.0599789 | ENSG00000188060 |
| EFCAB2 | 0.636837889 | 1.554917341 | 3.01E-04 | 1.46E-03 | 117.8358751 | 183.4331167 | ENSG00000203666 |
| DSEL | 0.636708173 | 1.554777541 | 5.03E-05 | 2.89E-04 | 182.9741342 | 284.5791018 | ENSG00000171451 |
| SNAPIN | 0.636692242 | 1.554760373 | 4.21E-09 | 4.73E-08 | 354.237543 | 550.2227713 | ENSG00000143553 |
| MYL9 | 0.636436 | 1.554484251 | 2.58E-27 | 1.42E-25 | 1444.761458 | 2246.499038 | ENSG00000101335 |
| PTCD2 | 0.636386046 | 1.554430427 | 2.18E-05 | 1.35E-04 | 157.9240153 | 245.390319 | ENSG00000049883 |
| PLEKHF2 | 0.635523854 | 1.553501736 | 2.07E-03 | 8.15E-03 | 112.5935372 | 175.0569856 | ENSG00000175895 |
| AHCTF1 | 0.635204429 | 1.553157816 | 2.96E-24 | 1.38E-22 | 1539.404885 | 2390.919628 | ENSG00000153207 |
| ZNF18 | 0.634825386 | 1.552749804 | 1.56E-05 | 9.88E-05 | 166.7415269 | 258.8450278 | ENSG00000154957 |
| FGF2 | 0.634599876 | 1.552507111 | 6.50E-16 | 1.55E-14 | 920.9376954 | 1430.360231 | ENSG00000138685 |
| CKS1B | 0.634337104 | 1.552224363 | 7.46E-25 | 3.66E-23 | 3390.992942 | 5262.788057 | ENSG00000173207 |
| SBK2 | 0.633802052 | 1.551648797 | 1.13E-03 | 4.74E-03 | 93.71802142 | 145.541654 | ENSG00000187550 |
| NSUN7 | 0.63326354 | 1.551069724 | 2.98E-05 | 1.80E-04 | 165.8775948 | 257.2494856 | ENSG00000179299 |
| PLEKHG4 | 0.633164895 | 1.550963672 | 5.32E-10 | 6.69E-09 | 407.7610064 | 632.1242824 | ENSG00000196155 |
| USP51 | 0.632878503 | 1.550655818 | 1.40E-03 | 5.77E-03 | 91.55403672 | 142.0126139 | ENSG00000247746 |
| STPG1 | 0.63271901 | 1.550484399 | 3.51E-05 | 2.09E-04 | 158.2506148 | 245.3599384 | ENSG00000001460 |
| FAM78B | 0.632319864 | 1.550055491 | 7.24E-03 | 2.40E-02 | 73.53898645 | 114.1056973 | ENSG00000188859 |
| APELA | 0.632106803 | 1.549826591 | 8.78E-06 | 5.85E-05 | 230.8755082 | 358.283362 | ENSG00000248329 |
| CENPL | 0.632024339 | 1.549738007 | 1.43E-05 | 9.14E-05 | 193.9316765 | 300.3666076 | ENSG00000120334 |
| VPS72 | 0.631505475 | 1.549180745 | 1.55E-20 | 5.49E-19 | 1755.252395 | 2718.333302 | ENSG00000163159 |
| RPS27 | 0.631253825 | 1.548910544 | 3.45E-34 | 2.71E-32 | 10243.62012 | 15865.26323 | ENSG00000177954 |
| ZNF888 | 0.631184192 | 1.548835786 | 4.06E-04 | 1.91E-03 | 119.8864208 | 185.4397768 | ENSG00000213793 |
| EFHC2 | 0.63113359 | 1.548781463 | 3.27E-02 | 8.42E-02 | 40.07398554 | 62.08706709 | ENSG00000183690 |
| NCSTN | 0.630534282 | 1.548138219 | 8.64E-24 | 3.90E-22 | 1482.951462 | 2295.838296 | ENSG00000162736 |
| NXPH4 | 0.629351727 | 1.546869754 | 1.34E-02 | 4.04E-02 | 65.07578322 | 100.7892072 | ENSG00000182379 |
| ANKRA2 | 0.629137569 | 1.546640148 | 2.26E-03 | 8.79E-03 | 84.06225938 | 130.1211886 | ENSG00000164331 |
| POU2F1 | 0.62907032 | 1.546568056 | 2.85E-14 | 5.74E-13 | 1028.461343 | 1591.150758 | ENSG00000143190 |
| ZNF548 | 0.628767924 | 1.546243922 | 2.17E-03 | 8.48E-03 | 88.82181935 | 137.4147843 | ENSG00000188785 |
| STK17A | 0.628530648 | 1.545989637 | 1.02E-07 | 9.36E-07 | 282.8052468 | 437.6499159 | ENSG00000164543 |
| KCTD18 | 0.627736832 | 1.545139219 | 4.17E-03 | 1.50E-02 | 81.42609676 | 125.5459493 | ENSG00000155729 |
| CALB2 | 0.627726803 | 1.545128477 | 4.59E-02 | 1.12E-01 | 35.12120167 | 54.27129237 | ENSG00000172137 |
| ANP32E | 0.62770118 | 1.545101035 | 7.56E-38 | 7.48E-36 | 4036.870044 | 6238.77006 | ENSG00000143401 |
| NEK2 | 0.627272656 | 1.544642161 | 2.61E-12 | 4.27E-11 | 589.9687028 | 910.5315793 | ENSG00000117650 |
| C11orf54 | 0.627238597 | 1.544605696 | 2.44E-02 | 6.63E-02 | 55.43138495 | 85.61771217 | ENSG00000182919 |
| FAM160B2 | 0.626815258 | 1.544152519 | 3.44E-09 | 3.91E-08 | 347.0100014 | 535.5992622 | ENSG00000158863 |
| FAM20B | 0.624386609 | 1.541555262 | 2.85E-15 | 6.31E-14 | 945.9205302 | 1458.46986 | ENSG00000116199 |
| DUSP5 | 0.624152489 | 1.54130512 | 1.49E-17 | 4.08E-16 | 913.1787891 | 1406.906268 | ENSG00000138166 |
| GAN | 0.624146441 | 1.541298659 | 4.90E-04 | 2.25E-03 | 123.9897906 | 191.3023832 | ENSG00000261609 |
| RILP | 0.623510881 | 1.54061981 | 1.46E-04 | 7.57E-04 | 155.0434287 | 238.5761145 | ENSG00000167705 |
| DDX25 | 0.623333699 | 1.540430612 | 4.70E-06 | 3.28E-05 | 204.0316194 | 314.4114897 | ENSG00000109832 |
| IPMK | 0.623268836 | 1.540361357 | 1.62E-05 | 1.03E-04 | 187.5455396 | 289.2662199 | ENSG00000151151 |
| SIL1 | 0.623054832 | 1.540132883 | 8.34E-03 | 2.70E-02 | 73.85298291 | 113.997605 | ENSG00000120725 |
| BCAN | 0.621842853 | 1.538839592 | 1.39E-03 | 5.72E-03 | 109.4974746 | 168.7991273 | ENSG00000132692 |
| TNFRSF10D | 0.621777396 | 1.538769774 | 4.24E-08 | 4.13E-07 | 337.3979058 | 519.3533011 | ENSG00000173530 |
| DCAF12L1 | 0.621033295 | 1.537976324 | 1.61E-04 | 8.29E-04 | 136.8487087 | 210.4643235 | ENSG00000198889 |
| ZBTB4 | 0.62030184 | 1.537196759 | 2.82E-10 | 3.70E-09 | 473.6887575 | 728.5872696 | ENSG00000174282 |
| GATAD2B | 0.619668779 | 1.536522379 | 5.35E-10 | 6.72E-09 | 437.1537761 | 672.4017796 | ENSG00000143614 |
| SYDE2 | 0.618345045 | 1.5351132 | 2.21E-03 | 8.60E-03 | 115.5160243 | 177.0870358 | ENSG00000097096 |
| ZBTB34 | 0.618282265 | 1.5350464 | 8.36E-08 | 7.78E-07 | 370.5466687 | 568.2806221 | ENSG00000177125 |
| MACF1 | 0.618277346 | 1.535041166 | 5.84E-24 | 2.67E-22 | 4749.831584 | 7289.978958 | ENSG00000127603 |
| MUC3A | 0.618044916 | 1.534793879 | 3.93E-07 | 3.30E-06 | 271.5823175 | 416.7101977 | ENSG00000169894 |
| INAVA | 0.618030302 | 1.534778331 | 1.12E-15 | 2.62E-14 | 889.773909 | 1366.058504 | ENSG00000163362 |
| PRKAA1 | 0.617630318 | 1.534352876 | 1.87E-07 | 1.65E-06 | 361.5592626 | 555.0162667 | ENSG00000132356 |
| TADA2A | 0.617182595 | 1.533876782 | 1.51E-06 | 1.15E-05 | 291.947768 | 447.3693531 | ENSG00000276234 |
| PIP5K1A | 0.616823728 | 1.533495283 | 1.09E-27 | 6.17E-26 | 1932.650335 | 2964.397777 | ENSG00000143398 |
| SERPINF2 | 0.615407709 | 1.53199088 | 2.88E-02 | 7.60E-02 | 47.2201026 | 72.42814276 | ENSG00000167711 |
| GALNT2 | 0.614882129 | 1.531432871 | 1.04E-31 | 7.22E-30 | 3307.197896 | 5063.496649 | ENSG00000143641 |

| | | | | | | | |
|---|---|---|---|---|---|---|---|
| MARVELD2 | 0.614707241 | 1.531247237 | 4.61E-08 | 4.45E-07 | 305.2641457 | 467.2349205 | ENSG00000152939 |
| UFSP1 | 0.614540775 | 1.531070564 | 4.81E-02 | 1.16E-01 | 41.48911674 | 63.75786722 | ENSG00000176125 |
| TTC39A | 0.614037456 | 1.530536506 | 1.27E-03 | 5.30E-03 | 111.2979392 | 169.9625725 | ENSG00000085831 |
| TMEM54 | 0.613902359 | 1.530393189 | 3.50E-02 | 8.92E-02 | 51.72726613 | 79.00760693 | ENSG00000121900 |
| CHROMR | 0.613576229 | 1.530047274 | 9.51E-04 | 4.07E-03 | 105.3384869 | 161.2105567 | ENSG00000223960 |
| EPPK1 | 0.612600755 | 1.529013087 | 4.87E-07 | 4.02E-06 | 283.2357102 | 432.7149362 | ENSG00000261150 |
| ICAM1 | 0.612041609 | 1.528420601 | 9.04E-03 | 2.89E-02 | 66.65818121 | 101.7905048 | ENSG00000090339 |
| TTLL6 | 0.612037266 | 1.528416001 | 5.93E-03 | 2.04E-02 | 80.13277727 | 122.3962895 | ENSG00000170703 |
| PIGM | 0.611306546 | 1.527642059 | 2.73E-06 | 1.98E-05 | 238.4862311 | 364.5539003 | ENSG00000143315 |
| RGL1 | 0.610396458 | 1.526678688 | 4.20E-03 | 1.51E-02 | 91.91480281 | 140.2455525 | ENSG00000143344 |
| NAT9 | 0.610263386 | 1.526537876 | 1.13E-15 | 2.62E-14 | 756.3987871 | 1154.578313 | ENSG00000109065 |
| CRIM1 | 0.609005425 | 1.525207388 | 8.72E-09 | 9.38E-08 | 371.9937022 | 567.6103077 | ENSG00000150938 |
| TRMT1L | 0.608938272 | 1.525136397 | 3.72E-13 | 6.66E-12 | 611.5760739 | 933.356683 | ENSG00000121486 |
| RELL1 | 0.608444809 | 1.524614824 | 5.63E-03 | 1.94E-02 | 75.06181158 | 114.5961202 | ENSG00000181826 |
| COQ5 | 0.608321783 | 1.524484818 | 6.43E-10 | 8.00E-09 | 513.8095865 | 783.001548 | ENSG00000110871 |
| PHF7 | 0.608211837 | 1.524368643 | 1.29E-03 | 5.37E-03 | 117.6958785 | 179.3889007 | ENSG00000010318 |
| TRIM68 | 0.608188995 | 1.524344509 | 2.50E-04 | 1.23E-03 | 168.7721729 | 256.7354064 | ENSG00000167333 |
| BCL11B | 0.608050441 | 1.52419812 | 1.02E-02 | 3.20E-02 | 74.74637566 | 114.0900981 | ENSG00000127152 |
| ANKRD50 | 0.607641857 | 1.523766515 | 4.51E-11 | 6.42E-10 | 639.2549083 | 974.0498293 | ENSG00000151458 |
| COL16A1 | 0.606796777 | 1.522874207 | 6.60E-07 | 5.34E-06 | 305.4149425 | 464.5875941 | ENSG00000084636 |
| ANK1 | 0.606480022 | 1.522539885 | 6.03E-07 | 4.90E-06 | 280.5465957 | 427.0034156 | ENSG00000029534 |
| ZBED8 | 0.605960489 | 1.521991698 | 2.17E-03 | 8.48E-03 | 104.9427025 | 159.7532332 | ENSG00000221886 |
| SYT1 | 0.605921333 | 1.52195039 | 1.06E-12 | 1.80E-11 | 611.6028317 | 931.3193755 | ENSG00000067715 |
| RNPEP | 0.605825971 | 1.521849793 | 1.85E-19 | 5.95E-18 | 1085.414226 | 1652.26514 | ENSG00000176393 |
| ZSWIM6 | 0.605509295 | 1.521515779 | 1.95E-06 | 1.46E-05 | 280.9347711 | 427.101579 | ENSG00000130449 |
| FGD6 | 0.605403259 | 1.521403953 | 3.81E-12 | 6.09E-11 | 561.9891507 | 855.3197142 | ENSG00000180263 |
| PIGC | 0.605253955 | 1.521246512 | 4.44E-10 | 5.65E-09 | 473.4940431 | 719.9801968 | ENSG00000135845 |
| LRRC27 | 0.605195319 | 1.521184685 | 3.76E-02 | 9.47E-02 | 48.0110198 | 72.82340109 | ENSG00000148814 |
| JAZF1 | 0.604434496 | 1.520382681 | 1.01E-03 | 4.30E-03 | 131.8017098 | 200.5211592 | ENSG00000153814 |
| FLNC | 0.603288011 | 1.519174939 | 3.19E-16 | 7.83E-15 | 763.354183 | 1159.950515 | ENSG00000128591 |
| SFI1 | 0.603219574 | 1.519102875 | 2.05E-07 | 1.80E-06 | 299.2707511 | 454.3190743 | ENSG00000198089 |
| PARS2 | 0.603078008 | 1.518953819 | 1.60E-06 | 1.21E-05 | 281.0698616 | 427.2972633 | ENSG00000162396 |
| PRKG1 | 0.603031043 | 1.518904372 | 1.16E-04 | 6.20E-04 | 155.5445547 | 236.5600707 | ENSG00000185532 |
| GALNT12 | 0.602955958 | 1.518825323 | 2.36E-06 | 1.73E-05 | 332.9766966 | 505.9117867 | ENSG00000119514 |
| SETDB1 | 0.602902899 | 1.518769466 | 2.48E-13 | 4.51E-12 | 648.1626799 | 984.0726501 | ENSG00000143379 |
| H6PD | 0.602361768 | 1.518199907 | 1.49E-07 | 1.33E-06 | 293.8292071 | 446.1852078 | ENSG00000049239 |
| SCN8A | 0.601928517 | 1.51774405 | 1.49E-10 | 2.02E-09 | 507.8960279 | 770.8874923 | ENSG00000196876 |
| PARP1 | 0.601318042 | 1.517101953 | 6.38E-62 | 1.54E-59 | 18831.54173 | 28568.52438 | ENSG00000143799 |
| IMPDH2 | 0.600875982 | 1.516637166 | 2.44E-39 | 2.70E-37 | 4184.24869 | 6345.868291 | ENSG00000178035 |
| FHDC1 | 0.600496037 | 1.5162378 | 1.56E-17 | 4.27E-16 | 883.5607825 | 1339.320878 | ENSG00000137460 |
| CPT1A | 0.600315158 | 1.516047712 | 6.47E-04 | 2.88E-03 | 138.3558644 | 209.5057527 | ENSG00000110090 |
| GAS2L1 | 0.600040053 | 1.515758647 | 2.25E-06 | 1.66E-05 | 294.939078 | 447.2658404 | ENSG00000185340 |
| THBS2 | 0.599889312 | 1.51560028 | 2.59E-10 | 3.41E-09 | 553.85818 | 840.145324 | ENSG00000186340 |
| DGAT2 | 0.599387257 | 1.515072947 | 5.90E-03 | 2.03E-02 | 89.2449492 | 135.16211 | ENSG00000062282 |
| NSMCE1 | 0.598863714 | 1.514523238 | 9.58E-06 | 6.34E-05 | 203.8281078 | 308.4727771 | ENSG00000169189 |
| KDM6B | 0.598038496 | 1.513657182 | 2.77E-05 | 1.68E-04 | 188.0014846 | 284.5017054 | ENSG00000132510 |
| PCLO | 0.597361778 | 1.512947345 | 3.58E-03 | 1.31E-02 | 93.72727001 | 141.6759846 | ENSG00000186472 |
| CORO7 | 0.596769756 | 1.512326621 | 2.28E-02 | 6.27E-02 | 52.30223189 | 78.94845694 | ENSG00000262246 |
| NBEAL2 | 0.595845353 | 1.511357913 | 1.21E-23 | 5.36E-22 | 1929.003987 | 2914.630635 | ENSG00000160796 |
| CYBA | 0.595434972 | 1.510928061 | 1.79E-27 | 9.93E-26 | 3063.985223 | 4628.599753 | ENSG00000051523 |
| CEP112 | 0.595421791 | 1.510914257 | 3.55E-02 | 9.02E-02 | 45.87319237 | 69.4978967 | ENSG00000154240 |
| IKBKE | 0.59532288 | 1.510810672 | 3.67E-06 | 2.61E-05 | 230.0874188 | 347.6793465 | ENSG00000263528 |
| MYSM1 | 0.594675385 | 1.510132758 | 3.21E-14 | 6.42E-13 | 1042.464545 | 1573.757557 | ENSG00000162601 |
| ADAM19 | 0.594622903 | 1.510077825 | 3.29E-15 | 7.22E-14 | 826.5124327 | 1248.108922 | ENSG00000135074 |
| SPTBN5 | 0.593631525 | 1.509040499 | 9.54E-03 | 3.02E-02 | 73.87618602 | 111.4083136 | ENSG00000137877 |
| CSRNP1 | 0.593599027 | 1.509006507 | 2.43E-04 | 1.20E-03 | 142.1979527 | 214.6244163 | ENSG00000144655 |
| GNPAT | 0.593312502 | 1.508706842 | 7.06E-21 | 2.55E-19 | 1098.210705 | 1656.948193 | ENSG00000116906 |
| PYCARD | 0.593251345 | 1.508642888 | 2.38E-03 | 9.17E-03 | 107.4831354 | 162.1984178 | ENSG00000103490 |
| SMAD3 | 0.592542366 | 1.507901682 | 2.52E-09 | 2.91E-08 | 456.8337121 | 688.8125414 | ENSG00000166949 |
| FLNB | 0.59253191 | 1.507890754 | 3.86E-36 | 3.40E-34 | 4207.811939 | 6343.296404 | ENSG00000136068 |
| XIAP | 0.592487472 | 1.507844309 | 1.15E-11 | 1.75E-10 | 738.4677152 | 1113.935381 | ENSG00000101966 |
| ENAH | 0.591993019 | 1.507327615 | 3.85E-50 | 6.52E-48 | 6554.125418 | 9879.978272 | ENSG00000154380 |
| FRMD6 | 0.591477382 | 1.506788975 | 3.01E-06 | 2.17E-05 | 261.9292342 | 394.8995434 | ENSG00000139926 |
| XPC | 0.5910042 | 1.506294852 | 7.52E-13 | 1.30E-11 | 695.9329553 | 1047.72188 | ENSG00000154767 |
| COCH | 0.590838828 | 1.506122199 | 4.99E-05 | 2.87E-04 | 232.3077099 | 349.8787653 | ENSG00000100473 |
| MAGI3 | 0.590737618 | 1.506016544 | 1.64E-04 | 8.41E-04 | 213.0134948 | 321.0199716 | ENSG00000081026 |
| ADCK5 | 0.590550378 | 1.505821098 | 1.37E-04 | 7.16E-04 | 153.228534 | 230.9249047 | ENSG00000173137 |
| CCDC141 | 0.590544366 | 1.505814822 | 2.52E-03 | 9.64E-03 | 105.0462541 | 158.0598308 | ENSG00000163492 |
| PRDM5 | 0.59000671 | 1.505253748 | 2.32E-07 | 2.01E-06 | 313.248313 | 471.7292194 | ENSG00000138738 |
| ARPC5 | 0.589650284 | 1.504881913 | 3.76E-23 | 1.59E-21 | 2358.653722 | 3550.472394 | ENSG00000162704 |
| QSOX1 | 0.589380018 | 1.504600023 | 6.21E-35 | 5.05E-33 | 3331.081328 | 5012.421645 | ENSG00000116260 |
| NUCKS1 | 0.589283207 | 1.504499062 | 4.43E-39 | 4.80E-37 | 12165.91395 | 18304.8125 | ENSG00000069275 |
| LRATD1 | 0.588976571 | 1.504179324 | 9.26E-04 | 3.97E-03 | 116.0703777 | 174.4641408 | ENSG00000162981 |

| | | | | | | | |
|---|---|---|---|---|---|---|---|
| PUDP | 0.588785432 | 1.503980052 | 3.06E-13 | 5.53E-12 | 812.0742951 | 1221.81872 | ENSG00000130021 |
| SLC12A4 | 0.588750131 | 1.503943252 | 3.85E-20 | 1.32E-18 | 1201.360713 | 1806.561984 | ENSG00000124067 |
| CTC1 | 0.58871333 | 1.503904889 | 1.97E-10 | 2.63E-09 | 503.4705643 | 757.0147476 | ENSG00000178971 |
| HIPK3 | 0.588328621 | 1.503503912 | 2.91E-05 | 1.76E-04 | 207.5954662 | 311.893434 | ENSG00000110422 |
| FNIP1 | 0.588295126 | 1.503469005 | 6.46E-04 | 2.88E-03 | 166.4532122 | 250.492541 | ENSG00000217128 |
| ABCB10 | 0.588120878 | 1.503287427 | 4.76E-06 | 3.32E-05 | 271.3208357 | 408.1937463 | ENSG00000135776 |
| VANGL2 | 0.587631837 | 1.502777933 | 2.15E-24 | 1.01E-22 | 2048.115685 | 3077.406783 | ENSG00000162738 |
| RGS9 | 0.586941933 | 1.502059469 | 5.89E-05 | 3.35E-04 | 171.376084 | 257.5629916 | ENSG00000108370 |
| MRPS14 | 0.586167392 | 1.501253273 | 5.23E-10 | 6.59E-09 | 475.11022 | 712.6766112 | ENSG00000120333 |
| ZNF860 | 0.586072849 | 1.501154896 | 1.32E-02 | 3.98E-02 | 69.83390373 | 104.9810255 | ENSG00000197385 |
| CX3CL1 | 0.585582105 | 1.500644354 | 3.23E-02 | 8.34E-02 | 48.45963119 | 72.77284128 | ENSG00000006210 |
| HIF3A | 0.585202022 | 1.500249056 | 3.22E-06 | 2.31E-05 | 270.6454585 | 405.9845924 | ENSG00000124440 |
| INA | 0.585082241 | 1.500124501 | 5.84E-08 | 5.55E-07 | 409.0446401 | 614.2833611 | ENSG00000148798 |
| ASPHD2 | -0.514929405 | 0.699827176 | 9.30E-04 | 3.99E-03 | 234.5470489 | 164.1932285 | ENSG00000128203 |
| CDK20 | -0.515695511 | 0.69945565 | 3.18E-02 | 8.23E-02 | 92.63997764 | 64.82854676 | ENSG00000156345 |
| TACC2 | -0.515731723 | 0.699438093 | 1.07E-10 | 1.47E-09 | 1132.767979 | 792.1593508 | ENSG00000138162 |
| GNB3 | -0.516504809 | 0.699063392 | 1.43E-02 | 4.25E-02 | 135.4009625 | 94.43373053 | ENSG00000111664 |
| JAM3 | -0.516798059 | 0.698921311 | 1.16E-13 | 2.18E-12 | 1403.604434 | 981.1585304 | ENSG00000166086 |
| PASK | -0.517148426 | 0.698751594 | 1.38E-15 | 3.16E-14 | 1729.433749 | 1207.869311 | ENSG00000115687 |
| INSYN1 | -0.517541368 | 0.698561303 | 1.31E-03 | 5.45E-03 | 222.4534175 | 155.4541467 | ENSG00000205363 |
| CRNDE | -0.517555692 | 0.698554367 | 3.57E-02 | 9.06E-02 | 96.6385566 | 67.50954989 | ENSG00000245694 |
| ZMAT2 | -0.517579534 | 0.698542823 | 2.44E-12 | 4.00E-11 | 1678.120465 | 1171.675558 | ENSG00000146007 |
| AAMP | -0.517751857 | 0.698459391 | 4.53E-17 | 1.20E-15 | 3955.169461 | 2762.10568 | ENSG00000127837 |
| RPIA | -0.51886759 | 0.697919434 | 2.51E-07 | 2.17E-06 | 598.4119325 | 417.768501 | ENSG00000153574 |
| STS | -0.518974985 | 0.697867482 | 2.11E-02 | 5.86E-02 | 119.2117335 | 83.28143809 | ENSG00000101846 |
| SMIM7 | -0.519208074 | 0.697754741 | 9.91E-04 | 4.23E-03 | 230.9036543 | 161.0293386 | ENSG00000214046 |
| CCDC86 | -0.519598131 | 0.697566116 | 3.88E-12 | 6.19E-11 | 1535.355195 | 1070.985048 | ENSG00000110104 |
| SMAP2 | -0.519880813 | 0.697429448 | 4.30E-03 | 1.54E-02 | 173.8762802 | 121.2965985 | ENSG00000084070 |
| ZNF584 | -0.520512144 | 0.697124316 | 3.57E-02 | 9.06E-02 | 99.84335222 | 69.75923746 | ENSG00000171574 |
| ELOF1 | -0.52066512 | 0.697050401 | 3.61E-10 | 4.66E-09 | 1477.866719 | 1030.01816 | ENSG00000130165 |
| PSME3 | -0.520768403 | 0.6970005 | 4.05E-26 | 2.10E-24 | 6313.428082 | 4400.000577 | ENSG00000131467 |
| CCND3 | -0.521143439 | 0.696819335 | 1.06E-08 | 1.12E-07 | 886.0162633 | 617.3768577 | ENSG00000112576 |
| GLDC | -0.521534587 | 0.696630437 | 2.36E-25 | 1.18E-23 | 4392.912576 | 3060.043623 | ENSG00000178445 |
| TEDC2 | -0.522034193 | 0.696389235 | 2.05E-09 | 2.40E-08 | 1055.781264 | 735.3796895 | ENSG00000162062 |
| LRRC42 | -0.522235948 | 0.696291855 | 1.58E-12 | 2.64E-11 | 1166.071145 | 812.0789371 | ENSG00000116212 |
| NPEPPS | -0.522434402 | 0.696196081 | 4.39E-15 | 9.50E-14 | 1713.328523 | 1193.195206 | ENSG00000141279 |
| CHRNA3 | -0.522490133 | 0.696169188 | 1.96E-02 | 5.53E-02 | 113.0183819 | 78.681991 | ENSG00000080644 |
| UBAC1 | -0.523204603 | 0.695824507 | 5.01E-17 | 1.32E-15 | 1767.532594 | 1229.916548 | ENSG00000130560 |
| PNMT | -0.523565662 | 0.695650387 | 4.09E-02 | 1.01E-01 | 91.42365215 | 63.64978708 | ENSG00000141744 |
| WARS1 | -0.52392252 | 0.695478336 | 1.11E-17 | 3.06E-16 | 2903.72849 | 2018.467305 | ENSG00000140105 |
| RPE | -0.524177797 | 0.695355285 | 8.04E-12 | 1.25E-10 | 1099.608139 | 764.7382408 | ENSG00000197713 |
| LTV1 | -0.524311139 | 0.69529102 | 2.55E-04 | 1.26E-03 | 287.299495 | 199.9909478 | ENSG00000135521 |
| CDH4 | -0.524538951 | 0.695181237 | 1.24E-05 | 7.98E-05 | 480.7656017 | 334.350486 | ENSG00000179242 |
| CARHSP1 | -0.525109123 | 0.694906546 | 1.28E-14 | 2.67E-13 | 1592.487522 | 1106.242268 | ENSG00000153048 |
| CDH23 | -0.525365768 | 0.694782938 | 5.46E-04 | 2.48E-03 | 246.9680436 | 171.4806634 | ENSG00000107736 |
| DCTN5 | -0.525469426 | 0.69473302 | 8.94E-17 | 2.30E-15 | 1855.97859 | 1289.52453 | ENSG00000166847 |
| FTSJ1 | -0.525477834 | 0.694728971 | 8.27E-10 | 1.02E-08 | 1018.590642 | 707.3861801 | ENSG00000068438 |
| C1QTNF4 | -0.525575515 | 0.694681935 | 1.07E-02 | 3.35E-02 | 145.8383563 | 101.4444676 | ENSG00000172247 |
| UBR7 | -0.526058202 | 0.694449551 | 2.04E-08 | 2.07E-07 | 722.8259162 | 502.0273015 | ENSG00000012963 |
| NOL8 | -0.526341409 | 0.694313242 | 3.66E-06 | 2.60E-05 | 584.7604817 | 406.1037101 | ENSG00000198000 |
| FBRS | -0.526819335 | 0.694083272 | 3.86E-06 | 2.74E-05 | 516.2811483 | 358.5186539 | ENSG00000156860 |
| H1-2 | -0.527715545 | 0.693652238 | 6.38E-03 | 2.16E-02 | 155.3832701 | 107.9908492 | ENSG00000187837 |
| RAB34 | -0.528118269 | 0.693458634 | 8.65E-23 | 3.60E-21 | 3443.106345 | 2387.42707 | ENSG00000109113 |
| SYT11 | -0.528577561 | 0.693237902 | 4.39E-04 | 2.05E-03 | 287.7900579 | 199.6440987 | ENSG00000132718 |
| TOR2A | -0.529315531 | 0.692883386 | 2.06E-05 | 1.28E-04 | 369.0219669 | 255.9365479 | ENSG00000160404 |
| SMPD4 | -0.529945158 | 0.692581061 | 6.51E-16 | 1.55E-14 | 2875.418474 | 1991.23209 | ENSG00000136699 |
| PAK6 | -0.529949062 | 0.692579187 | 9.71E-04 | 4.15E-03 | 211.1194398 | 146.179431 | ENSG00000137843 |
| CYFIP1 | -0.530328805 | 0.692396912 | 4.59E-21 | 1.69E-19 | 2374.102079 | 1643.683565 | ENSG00000273749 |
| GNG5 | -0.530426926 | 0.692349822 | 1.53E-02 | 4.49E-02 | 123.152216 | 85.24640115 | ENSG00000174021 |
| PSMD1 | -0.530627903 | 0.69225338 | 9.36E-32 | 6.56E-30 | 5668.888369 | 3924.209276 | ENSG00000173692 |
| MAP3K13 | -0.531106274 | 0.692023879 | 1.77E-02 | 5.08E-02 | 124.9543954 | 86.29610799 | ENSG00000073803 |
| CCDC34 | -0.531446745 | 0.691860583 | 8.32E-07 | 6.63E-06 | 592.8368496 | 410.3275144 | ENSG00000109881 |
| NCLN | -0.531671058 | 0.69175302 | 4.61E-13 | 8.14E-12 | 4440.003292 | 3071.406388 | ENSG00000125912 |
| ERI3 | -0.531794982 | 0.691693603 | 5.87E-16 | 1.41E-14 | 1619.659037 | 1120.623514 | ENSG00000117419 |
| NT5C3B | -0.531844289 | 0.691669963 | 4.81E-14 | 9.38E-13 | 1519.767013 | 1051.381318 | ENSG00000141698 |
| STAMBPL1 | -0.531848961 | 0.691667723 | 7.23E-03 | 2.40E-02 | 135.2869231 | 93.57599464 | ENSG00000138134 |
| ORMDL3 | -0.532195284 | 0.691501706 | 5.12E-13 | 8.98E-12 | 1606.525795 | 1110.624247 | ENSG00000172057 |
| CDKN2AIPNL | -0.533333196 | 0.690956506 | 5.47E-13 | 9.58E-12 | 1303.937461 | 901.0127458 | ENSG00000237190 |
| SCNN1A | -0.533373137 | 0.690937377 | 2.27E-14 | 4.63E-13 | 1690.939974 | 1168.767872 | ENSG00000111319 |
| PHF3 | -0.533473271 | 0.690889422 | 2.57E-15 | 5.73E-14 | 1693.827854 | 1170.26118 | ENSG00000118482 |
| BAK1 | -0.53355747 | 0.690849101 | 4.04E-12 | 6.43E-11 | 1180.53838 | 815.2345094 | ENSG00000030110 |
| ZNF793 | -0.534385713 | 0.690452603 | 6.33E-03 | 2.15E-02 | 166.0303946 | 114.5122415 | ENSG00000188227 |
| GK | -0.534451226 | 0.69042125 | 1.12E-03 | 4.73E-03 | 213.0015674 | 147.1543886 | ENSG00000198814 |

| | | | | | | | |
|---|---|---|---|---|---|---|---|
| STOX1 | -0.535222111 | 0.690052431 | 4.50E-02 | 1.10E-01 | 76.27813707 | 52.75669693 | ENSG00000165730 |
| MYB | -0.535320654 | 0.690005299 | 3.39E-02 | 8.66E-02 | 84.99807925 | 58.63788919 | ENSG00000118513 |
| KCNJ11 | -0.536247975 | 0.689561927 | 5.73E-04 | 2.59E-03 | 242.4572133 | 167.0812687 | ENSG00000187486 |
| SAP30L | -0.536332834 | 0.689521368 | 2.78E-07 | 2.39E-06 | 551.9571406 | 380.5874018 | ENSG00000164576 |
| PRRT3 | -0.536951764 | 0.68922562 | 1.63E-03 | 6.59E-03 | 220.0773114 | 151.9256396 | ENSG00000163704 |
| ANKRD42 | -0.537881877 | 0.688781416 | 1.58E-02 | 4.63E-02 | 110.0978467 | 75.86871837 | ENSG00000137494 |
| KIF26A | -0.537942896 | 0.688752284 | 7.92E-05 | 4.38E-04 | 4104.889243 | 2827.253483 | ENSG00000066735 |
| ENOX2 | -0.538022778 | 0.688714149 | 2.12E-07 | 1.86E-06 | 541.9789342 | 373.1069298 | ENSG00000165675 |
| PAQR5 | -0.538781856 | 0.688351876 | 6.94E-04 | 3.06E-03 | 221.120662 | 152.0498703 | ENSG00000137819 |
| FAM53B | -0.539077444 | 0.688210857 | 3.72E-08 | 3.64E-07 | 647.716347 | 445.6555536 | ENSG00000189319 |
| SIX5 | -0.539283313 | 0.688112658 | 2.56E-03 | 9.80E-03 | 173.1293787 | 118.9850468 | ENSG00000177045 |
| ZNF613 | -0.539297589 | 0.688105848 | 4.34E-03 | 1.55E-02 | 162.3353243 | 111.8172568 | ENSG00000176024 |
| CA12 | -0.53937098 | 0.688070845 | 1.59E-02 | 4.65E-02 | 121.6962604 | 83.72939468 | ENSG00000074410 |
| NCOA7 | -0.539878698 | 0.68782874 | 1.94E-03 | 7.67E-03 | 188.3825508 | 129.5223946 | ENSG00000111912 |
| SLC35B1 | -0.539954783 | 0.687792465 | 2.39E-10 | 3.16E-09 | 1063.705908 | 731.3578154 | ENSG00000121073 |
| NDRG2 | -0.540024672 | 0.687759148 | 8.81E-06 | 5.87E-05 | 404.9548983 | 278.4338973 | ENSG00000165795 |
| DONSON | -0.54043794 | 0.687562163 | 4.60E-07 | 3.82E-06 | 513.1055138 | 353.0464169 | ENSG00000159147 |
| STAG1 | -0.540916689 | 0.687334038 | 1.69E-07 | 1.50E-06 | 574.4551633 | 394.9603293 | ENSG00000118007 |
| SETBP1 | -0.540917405 | 0.687333697 | 2.77E-02 | 7.37E-02 | 93.59533383 | 64.30289063 | ENSG00000152217 |
| PGD | -0.541701808 | 0.68696009 | 5.55E-37 | 5.22E-35 | 10743.77204 | 7380.363062 | ENSG00000142657 |
| CCDC69 | -0.541777881 | 0.686923868 | 4.06E-05 | 2.38E-04 | 316.6770325 | 217.5342106 | ENSG00000198624 |
| C9orf116 | -0.541873541 | 0.686878322 | 1.79E-02 | 5.13E-02 | 101.6451314 | 69.82593578 | ENSG00000160345 |
| UNC45A | -0.542345869 | 0.68665348 | 1.02E-24 | 4.92E-23 | 3277.123946 | 2250.270827 | ENSG00000140553 |
| ASF1A | -0.542488215 | 0.686585734 | 1.84E-06 | 1.38E-05 | 467.9343173 | 321.3060895 | ENSG00000111875 |
| PCDH7 | -0.542588555 | 0.686537983 | 1.45E-02 | 4.31E-02 | 117.6117511 | 80.81582016 | ENSG00000169851 |
| POLR2F | -0.543148466 | 0.686271588 | 2.36E-18 | 6.89E-17 | 2080.422695 | 1427.638737 | ENSG00000100142 |
| ZNF205 | -0.5434411 | 0.686132401 | 6.29E-05 | 3.55E-04 | 359.0027231 | 246.5341365 | ENSG00000122386 |
| BEND3 | -0.544102222 | 0.685818049 | 3.62E-11 | 5.20E-10 | 1165.048981 | 799.2760238 | ENSG00000178409 |
| ATP6V1G1 | -0.544440521 | 0.68565725 | 4.92E-15 | 1.06E-13 | 2029.837891 | 1391.799345 | ENSG00000136888 |
| VPS29 | -0.544644234 | 0.685560439 | 3.47E-06 | 2.48E-05 | 407.8935814 | 279.8444411 | ENSG00000111237 |
| WFDC2 | -0.544681811 | 0.685542584 | 2.85E-06 | 2.06E-05 | 449.3638745 | 308.2243156 | ENSG00000101443 |
| RAB27A | -0.544794991 | 0.685488805 | 6.43E-03 | 2.18E-02 | 143.0047394 | 97.94742564 | ENSG00000069974 |
| C1orf52 | -0.544807792 | 0.685482722 | 7.61E-08 | 7.12E-07 | 619.0314063 | 424.0218405 | ENSG00000162642 |
| MINDY3 | -0.54490428 | 0.685436878 | 4.75E-04 | 2.19E-03 | 249.0064377 | 170.6898863 | ENSG00000148481 |
| SOCS2 | -0.545044881 | 0.685370081 | 6.32E-05 | 3.56E-04 | 343.7571727 | 235.5362012 | ENSG00000120833 |
| AUH | -0.54531844 | 0.685240135 | 2.87E-06 | 2.08E-05 | 426.7310126 | 292.5511117 | ENSG00000148090 |
| AKAP17A | -0.545353331 | 0.685223563 | 1.43E-16 | 3.62E-15 | 2313.886157 | 1585.556109 | ENSG00000197976 |
| AMD1 | -0.54553855 | 0.685135597 | 6.58E-15 | 1.41E-13 | 4129.266992 | 2829.920435 | ENSG00000123505 |
| BRINP1 | -0.545567641 | 0.685121782 | 2.16E-08 | 2.18E-07 | 929.1720559 | 636.8584774 | ENSG00000078725 |
| USF2 | -0.545820515 | 0.685001705 | 1.27E-24 | 6.06E-23 | 2745.338369 | 1880.785461 | ENSG00000105698 |
| TUBD1 | -0.546617122 | 0.684623575 | 3.14E-04 | 1.52E-03 | 284.94758 | 195.114294 | ENSG00000108423 |
| HCN4 | -0.546815226 | 0.684529573 | 1.18E-08 | 1.25E-07 | 646.1327977 | 442.4677044 | ENSG00000138622 |
| ALDH4A1 | -0.547227595 | 0.68433394 | 1.45E-15 | 3.31E-14 | 1510.796426 | 1033.824612 | ENSG00000159423 |
| CENPX | -0.547389546 | 0.684257124 | 1.70E-15 | 3.86E-14 | 1793.244937 | 1226.950045 | ENSG00000169689 |
| SLC6A9 | -0.547525646 | 0.684192576 | 4.37E-07 | 3.64E-06 | 581.4231622 | 397.5953522 | ENSG00000196517 |
| NEXN | -0.547698203 | 0.684110746 | 1.72E-02 | 4.96E-02 | 108.4597428 | 74.31647868 | ENSG00000162614 |
| SETDB2 | -0.547811132 | 0.684057199 | 3.63E-03 | 1.33E-02 | 168.2535888 | 114.9537343 | ENSG00000136169 |
| CDR2L | -0.547900082 | 0.684015024 | 2.05E-11 | 3.04E-10 | 1433.380449 | 980.6562885 | ENSG00000109089 |
| EPHB3 | -0.548199117 | 0.683873259 | 3.75E-02 | 9.45E-02 | 80.12843476 | 54.79238689 | ENSG00000182580 |
| U2AF2 | -0.548784698 | 0.683595736 | 4.83E-19 | 1.49E-17 | 9082.902399 | 6208.741572 | ENSG00000063244 |
| B3GNT7 | -0.548816925 | 0.683580465 | 3.42E-20 | 1.19E-18 | 2338.283061 | 1597.752383 | ENSG00000156966 |
| MPDU1 | -0.549101876 | 0.683445463 | 2.53E-08 | 2.53E-07 | 697.1346008 | 476.3577448 | ENSG00000129255 |
| NAPA | -0.549739382 | 0.683143525 | 4.29E-06 | 3.02E-05 | 467.8979603 | 319.5955009 | ENSG00000105402 |
| ARHGAP29 | -0.549885297 | 0.683074435 | 2.80E-02 | 7.43E-02 | 84.44823161 | 57.64707188 | ENSG00000137962 |
| KIF3A | -0.550220495 | 0.682915747 | 6.07E-05 | 3.43E-04 | 306.8965429 | 209.7560801 | ENSG00000131437 |
| HTR7P1 | -0.550357327 | 0.682850979 | 3.04E-02 | 7.94E-02 | 84.14954111 | 57.47418353 | ENSG00000183935 |
| SLC25A13 | -0.551841735 | 0.682148746 | 1.48E-21 | 5.60E-20 | 2238.652834 | 1527.487348 | ENSG00000004864 |
| ALG1 | -0.551853928 | 0.682142981 | 2.60E-07 | 2.24E-06 | 551.7500502 | 376.222453 | ENSG00000033011 |
| OGG1 | -0.552349704 | 0.681908605 | 6.11E-08 | 5.78E-07 | 614.9455087 | 419.3820534 | ENSG00000114026 |
| PPM1H | -0.552852267 | 0.681671104 | 5.08E-19 | 1.56E-17 | 2030.013265 | 1384.024092 | ENSG00000111110 |
| WASHC4 | -0.553582126 | 0.681326334 | 9.18E-11 | 1.27E-09 | 886.7153291 | 604.04475 | ENSG00000136051 |
| MLLT6 | -0.55381116 | 0.681218179 | 6.18E-06 | 4.23E-05 | 389.9290797 | 265.6100112 | ENSG00000275023 |
| MRPL12 | -0.554155837 | 0.681055447 | 2.43E-02 | 6.61E-02 | 99.1606045 | 67.58213636 | ENSG00000262814 |
| DAP | -0.554328193 | 0.680974088 | 7.28E-13 | 1.26E-11 | 1040.180166 | 708.4888214 | ENSG00000112977 |
| RFNG | -0.554394318 | 0.680942876 | 2.33E-06 | 1.71E-05 | 393.9852044 | 268.2627659 | ENSG00000169733 |
| CYB5B | -0.555103615 | 0.680608175 | 1.43E-19 | 4.66E-18 | 2781.646975 | 1893.147215 | ENSG00000103018 |
| ZBTB40 | -0.555281147 | 0.680524427 | 9.24E-10 | 1.13E-08 | 734.8903752 | 499.8268167 | ENSG00000184677 |
| ZDHHC16 | -0.555619865 | 0.680364672 | 4.69E-11 | 6.67E-10 | 1230.507595 | 837.0610665 | ENSG00000171307 |
| ZNF616 | -0.555634059 | 0.680357978 | 3.89E-08 | 3.81E-07 | 573.6774369 | 390.4143861 | ENSG00000204611 |
| CYB561D2 | -0.556685371 | 0.679862372 | 2.37E-02 | 6.47E-02 | 86.81333735 | 58.98877897 | ENSG00000114395 |
| GPC1 | -0.556897544 | 0.679762394 | 8.20E-13 | 1.41E-11 | 1790.127788 | 1217.37257 | ENSG00000063660 |
| SCAMP4 | -0.557265039 | 0.679589261 | 8.33E-10 | 1.02E-08 | 887.5152579 | 603.1047714 | ENSG00000227500 |
| PTPRT | -0.557671163 | 0.679397981 | 1.16E-02 | 3.58E-02 | 116.5335072 | 79.30014022 | ENSG00000196090 |

| | | | | | | | |
|---|---|---|---|---|---|---|---|
| WFIKKN1 | -0.557694029 | 0.679387214 | 1.35E-03 | 5.58E-03 | 204.30344 | 138.9557381 | ENSG00000127578 |
| SGSM2 | -0.55782413 | 0.67932595 | 6.44E-09 | 7.03E-08 | 802.3179342 | 544.6260736 | ENSG00000141258 |
| RANBP1 | -0.557860917 | 0.679308628 | 3.95E-35 | 3.26E-33 | 8067.364831 | 5479.584276 | ENSG00000099901 |
| TOM1 | -0.558242613 | 0.679128926 | 3.53E-09 | 4.00E-08 | 660.428135 | 448.6113219 | ENSG00000100284 |
| ILKAP | -0.558446159 | 0.679033116 | 1.57E-07 | 1.41E-06 | 610.1484518 | 414.1988663 | ENSG00000132323 |
| KRT10 | -0.558518351 | 0.678999138 | 5.45E-07 | 4.46E-06 | 579.4387351 | 393.7492805 | ENSG00000186395 |
| TSPAN14 | -0.559151721 | 0.67870111 | 6.48E-19 | 1.98E-17 | 1869.621106 | 1269.441166 | ENSG00000108219 |
| EIF6 | -0.559158756 | 0.678697801 | 4.51E-13 | 7.98E-12 | 1526.911978 | 1035.790708 | ENSG00000242372 |
| SH3BP5 | -0.559197455 | 0.678679596 | 2.07E-02 | 5.78E-02 | 100.3601846 | 68.15969726 | ENSG00000131370 |
| RNF24 | -0.559477643 | 0.678547801 | 7.06E-06 | 4.79E-05 | 362.2104586 | 245.7578803 | ENSG00000101236 |
| C19orf54 | -0.559889181 | 0.678354269 | 8.07E-08 | 7.52E-07 | 532.3376022 | 361.0275437 | ENSG00000188493 |
| OIP5 | -0.560114087 | 0.678248526 | 2.11E-02 | 5.86E-02 | 89.07509167 | 60.45308718 | ENSG00000104147 |
| TRAPPC2L | -0.56011932 | 0.678246066 | 2.86E-12 | 4.63E-11 | 1031.169667 | 699.0536183 | ENSG00000167515 |
| UBXN8 | -0.560630587 | 0.67800575 | 6.62E-03 | 2.23E-02 | 130.6501387 | 88.62889285 | ENSG00000104691 |
| CCDC88A | -0.560752817 | 0.677948309 | 1.21E-11 | 1.84E-10 | 1167.189449 | 791.7250243 | ENSG00000115355 |
| FARS2 | -0.56088159 | 0.677887799 | 1.59E-05 | 1.01E-04 | 362.6594335 | 245.7654287 | ENSG00000145982 |
| ATP6V1D | -0.561163275 | 0.677755455 | 1.91E-11 | 2.84E-10 | 879.2944643 | 595.7813097 | ENSG00000100554 |
| SAP30 | -0.56129359 | 0.677694238 | 3.67E-05 | 2.17E-04 | 333.8099288 | 226.2104048 | ENSG00000164105 |
| CYB561A3 | -0.562659211 | 0.677053052 | 2.06E-04 | 1.04E-03 | 258.8177024 | 175.1212913 | ENSG00000162144 |
| MAF | -0.5627 | 0.67703391 | 5.46E-03 | 1.89E-02 | 134.9750418 | 91.43490815 | ENSG00000178573 |
| ZWINT | -0.562767682 | 0.677002149 | 2.69E-22 | 1.08E-20 | 2178.70122 | 1475.143628 | ENSG00000122952 |
| HIRA | -0.56287034 | 0.676953977 | 9.41E-05 | 5.12E-04 | 328.7037574 | 222.5531238 | ENSG00000100084 |
| PIP5K1B | -0.563569693 | 0.6766259 | 1.75E-02 | 5.02E-02 | 103.9643434 | 70.20237197 | ENSG00000107242 |
| PHKG2 | -0.563980293 | 0.676433355 | 3.57E-06 | 2.55E-05 | 410.1877874 | 277.24682 | ENSG00000156873 |
| WIPF3 | -0.564354397 | 0.676257973 | 3.59E-07 | 3.04E-06 | 477.3991708 | 323.0486889 | ENSG00000122574 |
| PCSK7 | -0.56480656 | 0.676046056 | 1.10E-11 | 1.68E-10 | 927.6157769 | 627.1304315 | ENSG00000160613 |
| SLC6A8 | -0.565152559 | 0.675883941 | 3.36E-18 | 9.69E-17 | 3127.07818 | 2113.376441 | ENSG00000130821 |
| TMA7 | -0.565449932 | 0.67574464 | 4.22E-03 | 1.51E-02 | 139.7247896 | 94.49555202 | ENSG00000232112 |
| CCDC120 | -0.565485348 | 0.675728051 | 5.43E-06 | 3.75E-05 | 402.4680192 | 272.0302961 | ENSG00000147144 |
| DDX58 | -0.566097245 | 0.675441513 | 1.03E-02 | 3.22E-02 | 117.8938083 | 79.55880948 | ENSG00000107201 |
| DZANK1 | -0.56618105 | 0.675402278 | 2.20E-04 | 1.10E-03 | 234.7063064 | 158.513706 | ENSG00000089091 |
| POLR3H | -0.566285211 | 0.675353516 | 2.16E-10 | 2.88E-09 | 955.545043 | 645.4854504 | ENSG00000100413 |
| CYLD | -0.566562263 | 0.675223835 | 1.07E-04 | 5.74E-04 | 317.235328 | 214.2912826 | ENSG00000083799 |
| PLEKHA3 | -0.567105991 | 0.674969402 | 7.75E-04 | 3.38E-03 | 237.3856599 | 160.1896435 | ENSG00000116095 |
| CDC25B | -0.567455032 | 0.674806123 | 1.05E-08 | 1.12E-07 | 905.7879741 | 611.268164 | ENSG00000101224 |
| C9orf78 | -0.567542973 | 0.67476499 | 4.20E-15 | 9.12E-14 | 1224.843879 | 826.3988996 | ENSG00000136819 |
| CEND1 | -0.567709928 | 0.674686908 | 9.82E-03 | 3.10E-02 | 152.0022842 | 102.4143243 | ENSG00000184524 |
| LLGL1 | -0.568377893 | 0.674374601 | 2.77E-17 | 7.48E-16 | 3339.755366 | 2252.30882 | ENSG00000131899 |
| ZNF488 | -0.568464089 | 0.674334311 | 1.54E-04 | 7.96E-04 | 249.2142918 | 167.9320077 | ENSG00000265763 |
| MCM6 | -0.571252681 | 0.673032146 | 7.33E-37 | 6.83E-35 | 5392.290391 | 3628.952172 | ENSG00000076003 |
| SUSD5 | -0.572164366 | 0.67260697 | 1.85E-02 | 5.27E-02 | 97.068032 | 65.38162731 | ENSG00000173705 |
| FAM155A | -0.572442115 | 0.672477491 | 2.10E-04 | 1.05E-03 | 291.9787064 | 196.7439486 | ENSG00000204442 |
| FOXP4 | -0.572477319 | 0.672461082 | 5.52E-07 | 4.52E-06 | 858.9954702 | 577.5040451 | ENSG00000137166 |
| MDM1 | -0.572781513 | 0.672319308 | 8.92E-04 | 3.84E-03 | 203.2804506 | 136.8318562 | ENSG00000111554 |
| PRKAR1B | -0.573347944 | 0.672055394 | 5.37E-14 | 1.04E-12 | 2045.945108 | 1375.016497 | ENSG00000188191 |
| NR6A1 | -0.57351232 | 0.671978826 | 4.13E-32 | 2.96E-30 | 5759.745803 | 3871.125466 | ENSG00000148200 |
| MAP3K14 | -0.573574118 | 0.671950043 | 4.02E-03 | 1.45E-02 | 155.6117366 | 104.6492184 | ENSG00000006062 |
| SLC35E4 | -0.573578685 | 0.671947915 | 5.48E-04 | 2.49E-03 | 208.7302551 | 140.2197457 | ENSG00000100036 |
| VPS18 | -0.574159956 | 0.671677238 | 1.94E-13 | 3.56E-12 | 1346.997412 | 904.6902429 | ENSG00000104142 |
| BAG1 | -0.574179604 | 0.67166809 | 1.83E-10 | 2.45E-09 | 1235.632779 | 829.8717644 | ENSG00000107262 |
| TXNL4B | -0.574928814 | 0.671319375 | 5.11E-03 | 1.79E-02 | 155.4454337 | 104.5924794 | ENSG00000140830 |
| STK36 | -0.575322345 | 0.671136281 | 1.14E-07 | 1.04E-06 | 708.9339354 | 475.588225 | ENSG00000163482 |
| ASAP2 | -0.575354962 | 0.671121108 | 2.91E-07 | 2.50E-06 | 500.5976746 | 336.0987131 | ENSG00000151693 |
| PANX2 | -0.575989012 | 0.670826222 | 1.04E-03 | 4.40E-03 | 221.5003398 | 148.8647231 | ENSG00000073150 |
| FZD8 | -0.577008586 | 0.670352306 | 1.29E-06 | 9.95E-06 | 425.1873749 | 285.0644727 | ENSG00000177283 |
| WDR45 | -0.577412719 | 0.670164551 | 5.93E-03 | 2.04E-02 | 123.7965668 | 83.03352169 | ENSG00000196998 |
| TOLLIP | -0.577961863 | 0.66990951 | 1.67E-09 | 1.99E-08 | 871.7913743 | 583.8455401 | ENSG00000078902 |
| LSG1 | -0.57842127 | 0.66969622 | 3.47E-20 | 1.20E-18 | 1763.962127 | 1181.537982 | ENSG00000041802 |
| HDX | -0.578977635 | 0.669438006 | 1.10E-02 | 3.41E-02 | 106.7898998 | 71.47281913 | ENSG00000165259 |
| TMEM200A | -0.579178821 | 0.669344659 | 3.06E-06 | 2.20E-05 | 439.911475 | 294.712359 | ENSG00000164484 |
| PCTP | -0.580008373 | 0.668959895 | 8.25E-08 | 7.69E-07 | 495.0635425 | 331.211046 | ENSG00000141179 |
| LTBP1 | -0.580127384 | 0.668904713 | 6.64E-16 | 1.58E-14 | 1316.697319 | 880.8350053 | ENSG00000049323 |
| PIGQ | -0.580734821 | 0.668623135 | 3.23E-07 | 2.76E-06 | 469.4873668 | 313.8901532 | ENSG00000007541 |
| KCNN2 | -0.580838631 | 0.668575025 | 1.26E-04 | 6.64E-04 | 325.4002042 | 217.6234807 | ENSG00000080709 |
| TEX261 | -0.580846994 | 0.66857115 | 8.60E-07 | 6.84E-06 | 420.5674481 | 281.2533796 | ENSG00000144043 |
| DHRS13 | -0.580885979 | 0.668553083 | 5.35E-07 | 4.39E-06 | 556.0877949 | 371.9564186 | ENSG00000167536 |
| THAP10 | -0.581411232 | 0.668309722 | 1.83E-02 | 5.21E-02 | 100.0219459 | 66.82821 | ENSG00000129028 |
| RHOF | -0.581540436 | 0.668249873 | 2.98E-02 | 7.80E-02 | 84.28953769 | 56.39355674 | ENSG00000139725 |
| TCTN3 | -0.581673842 | 0.668188083 | 7.91E-10 | 9.73E-09 | 840.6056851 | 561.8082026 | ENSG00000119977 |
| RCN1 | -0.58180677 | 0.66812652 | 1.21E-09 | 1.46E-08 | 910.7986301 | 609.0133639 | ENSG00000049449 |
| SLIT3 | -0.581806936 | 0.668126443 | 4.90E-02 | 1.18E-01 | 68.31454523 | 45.77096111 | ENSG00000184347 |
| CXCL12 | -0.582074156 | 0.668002702 | 2.18E-11 | 3.22E-10 | 823.3493133 | 550.191282 | ENSG00000107562 |
| KHDC1 | -0.582105591 | 0.667988147 | 2.13E-03 | 8.33E-03 | 157.6420702 | 105.3472298 | ENSG00000135314 |

| | | | | | | | |
|---|---|---|---|---|---|---|---|
| HEG1 | -0.583139457 | 0.667509624 | 2.13E-05 | 1.32E-04 | 290.9114629 | 194.1855892 | ENSG00000173706 |
| PDGFA | -0.583248188 | 0.667459318 | 6.30E-11 | 8.84E-10 | 777.753272 | 519.1836357 | ENSG00000197461 |
| EDNRA | -0.583280696 | 0.667444279 | 1.44E-02 | 4.27E-02 | 94.39777807 | 63.02733671 | ENSG00000151617 |
| PRXL2A | -0.583494767 | 0.667345249 | 2.57E-15 | 5.73E-14 | 1419.741238 | 948.0519914 | ENSG00000122378 |
| MED24 | -0.584215665 | 0.667011867 | 5.03E-28 | 2.88E-26 | 2818.792526 | 1880.151464 | ENSG00000008838 |
| QPCTL | -0.584268021 | 0.666987662 | 3.59E-08 | 3.53E-07 | 561.9488757 | 374.4967735 | ENSG00000011478 |
| ZNF302 | -0.584668262 | 0.666802648 | 5.18E-06 | 3.58E-05 | 461.5243385 | 307.8626666 | ENSG00000089335 |
| HMGN2 | -0.585390183 | 0.666469065 | 1.53E-26 | 8.19E-25 | 6622.39424 | 4414.017082 | ENSG00000198830 |
| MLH1 | -0.585497723 | 0.666419387 | 8.66E-14 | 1.65E-12 | 1217.909445 | 811.4634778 | ENSG00000076242 |
| SEMA4G | -0.585541983 | 0.666398943 | 5.77E-03 | 1.99E-02 | 123.4427339 | 82.13090844 | ENSG00000095539 |
| CIPC | -0.585786565 | 0.666285977 | 6.98E-07 | 5.62E-06 | 439.851976 | 292.943426 | ENSG00000198894 |
| PLAGL2 | -0.586029884 | 0.666173613 | 2.35E-13 | 4.28E-12 | 1141.99707 | 761.1127037 | ENSG00000126003 |
| ALKBH8 | -0.58606397 | 0.666157874 | 1.22E-05 | 7.85E-05 | 315.8103094 | 210.2113124 | ENSG00000137760 |
| CNPY4 | -0.58632164 | 0.666038907 | 5.93E-04 | 2.67E-03 | 185.0816251 | 123.3521645 | ENSG00000166997 |
| NACAD | -0.586327402 | 0.666036247 | 3.89E-03 | 1.41E-02 | 147.7150143 | 98.47523235 | ENSG00000136274 |
| ITPRIPL1 | -0.587720081 | 0.665393611 | 1.97E-06 | 1.48E-05 | 403.8164938 | 268.6072039 | ENSG00000198885 |
| PEF1 | -0.587989989 | 0.665269137 | 3.23E-08 | 3.18E-07 | 546.6976938 | 363.4563414 | ENSG00000162517 |
| SERAC1 | -0.588629516 | 0.664974297 | 4.15E-05 | 2.43E-04 | 276.2753713 | 183.8977448 | ENSG00000122335 |
| KCTD17 | -0.588835451 | 0.664879384 | 3.78E-11 | 5.42E-10 | 905.9376834 | 602.4731552 | ENSG00000100379 |
| CTSD | -0.588869167 | 0.664863845 | 4.76E-08 | 4.59E-07 | 697.8870331 | 463.8532649 | ENSG00000117984 |
| SLC43A3 | -0.589359884 | 0.664637738 | 2.52E-04 | 1.25E-03 | 303.3411057 | 201.39023 | ENSG00000134802 |
| LARP6 | -0.589962902 | 0.66435999 | 8.29E-08 | 7.72E-07 | 486.8480298 | 323.5985465 | ENSG00000166173 |
| IFRD1 | -0.59031315 | 0.664198721 | 4.01E-14 | 7.89E-13 | 1043.975909 | 693.3079427 | ENSG00000006652 |
| RIMS3 | -0.590502457 | 0.664111572 | 1.99E-13 | 3.64E-12 | 1060.662671 | 704.4370755 | ENSG00000117016 |
| GDAP2 | -0.590749501 | 0.663997861 | 1.32E-04 | 6.94E-04 | 233.8238771 | 155.4194465 | ENSG00000196505 |
| KATNB1 | -0.591005459 | 0.663880067 | 3.94E-17 | 1.05E-15 | 1613.075646 | 1070.968091 | ENSG00000140854 |
| TMED1 | -0.59116677 | 0.663805842 | 4.58E-04 | 2.12E-03 | 231.9275196 | 153.9185785 | ENSG00000099203 |
| CISH | -0.592291129 | 0.663288709 | 4.77E-02 | 1.15E-01 | 62.18315707 | 41.25676189 | ENSG00000114737 |
| BCL2L1 | -0.592727019 | 0.663088335 | 1.82E-09 | 2.15E-08 | 696.0182723 | 461.644123 | ENSG00000171552 |
| SLITRK3 | -0.592950425 | 0.662985662 | 4.55E-02 | 1.11E-01 | 61.87541813 | 41.08655715 | ENSG00000121871 |
| MIGA2 | -0.593597513 | 0.662688361 | 2.54E-09 | 2.94E-08 | 734.1820579 | 486.2148371 | ENSG00000148343 |
| GDAP1 | -0.594220154 | 0.662402419 | 9.10E-10 | 1.11E-08 | 836.4888249 | 553.8738979 | ENSG00000104381 |
| EMC9 | -0.594412179 | 0.662314258 | 1.52E-06 | 1.16E-05 | 452.7289298 | 299.9305005 | ENSG00000100908 |
| PTRHD1 | -0.594916105 | 0.662082955 | 5.57E-03 | 1.93E-02 | 133.9260583 | 88.53130511 | ENSG00000184924 |
| PCDHGB7 | -0.595393994 | 0.661863678 | 2.89E-04 | 1.41E-03 | 208.2690406 | 137.8675935 | ENSG00000254122 |
| SIVA1 | -0.59547697 | 0.661825612 | 8.97E-12 | 1.38E-10 | 1655.015665 | 1094.865292 | ENSG00000184990 |
| GPRASP2 | -0.595584638 | 0.661776223 | 2.17E-04 | 1.08E-03 | 259.9474972 | 172.2190027 | ENSG00000158301 |
| EFCAB11 | -0.5960658 | 0.661555547 | 2.59E-02 | 6.98E-02 | 75.54343825 | 50.00284686 | ENSG00000140025 |
| C19orf47 | -0.596429605 | 0.661388743 | 9.59E-06 | 6.34E-05 | 381.6929305 | 252.7906562 | ENSG00000160392 |
| ATAD3A | -0.596787915 | 0.661224499 | 8.85E-14 | 1.68E-12 | 1912.790129 | 1264.582384 | ENSG00000197785 |
| CHD7 | -0.596862882 | 0.661190141 | 2.88E-23 | 1.24E-21 | 3022.178052 | 1997.763587 | ENSG00000171316 |
| ARL6IP5 | -0.596965357 | 0.661143178 | 5.50E-10 | 6.91E-09 | 622.3422562 | 411.3116501 | ENSG00000144746 |
| DHRSX | -0.597008092 | 0.661123594 | 3.91E-06 | 2.77E-05 | 336.3714497 | 222.4458187 | ENSG00000169084 |
| WDR76 | -0.597060185 | 0.661099723 | 1.25E-06 | 9.67E-06 | 401.7405787 | 265.2819842 | ENSG00000092470 |
| SLC27A4 | -0.597124245 | 0.661070369 | 3.20E-10 | 4.16E-09 | 896.6836752 | 592.6889708 | ENSG00000167114 |
| CACNA2D2 | -0.597809491 | 0.660756451 | 1.44E-12 | 2.41E-11 | 894.2818362 | 591.0079082 | ENSG00000007402 |
| SRCIN1 | -0.598128257 | 0.660610471 | 1.30E-02 | 3.93E-02 | 120.6279407 | 79.57575361 | ENSG00000277363 |
| LRP3 | -0.598284498 | 0.660538933 | 1.72E-20 | 6.10E-19 | 1833.772302 | 1211.505886 | ENSG00000130881 |
| TAFA5 | -0.599412535 | 0.660022662 | 1.33E-15 | 3.05E-14 | 1124.599161 | 742.4060015 | ENSG00000219438 |
| PEX12 | -0.599878623 | 0.659809464 | 4.80E-02 | 1.16E-01 | 70.14029122 | 46.34797054 | ENSG00000108733 |
| ZNF138 | -0.60052021 | 0.659516103 | 9.53E-06 | 6.30E-05 | 310.9379493 | 205.1026082 | ENSG00000197008 |
| ASRGL1 | -0.600986394 | 0.659303025 | 1.98E-04 | 9.99E-04 | 252.3104665 | 166.2047291 | ENSG00000162174 |
| TBC1D10A | -0.601049828 | 0.659274037 | 1.23E-02 | 3.75E-02 | 93.96782711 | 61.95908032 | ENSG00000099992 |
| XYLB | -0.601093921 | 0.659253888 | 2.36E-02 | 6.45E-02 | 80.32791619 | 53.05703902 | ENSG00000093217 |
| KLHL18 | -0.60158743 | 0.659028412 | 8.03E-12 | 1.25E-10 | 864.1245068 | 569.6317371 | ENSG00000114648 |
| OSBPL11 | -0.601904 | 0.658883818 | 3.45E-12 | 5.55E-11 | 859.1375564 | 566.1686081 | ENSG00000144909 |
| NUCB1 | -0.602045836 | 0.658819044 | 1.46E-25 | 7.38E-24 | 2452.336086 | 1615.281967 | ENSG00000104805 |
| RBCK1 | -0.603514286 | 0.658148805 | 2.55E-11 | 3.73E-10 | 1310.083505 | 862.2665645 | ENSG00000125826 |
| RPUSD1 | -0.603564674 | 0.658125819 | 9.19E-10 | 1.12E-08 | 923.004237 | 607.8335318 | ENSG00000007376 |
| CTSA | -0.603662039 | 0.658081404 | 5.17E-10 | 6.53E-09 | 652.0035289 | 429.2176095 | ENSG00000064601 |
| TMEM158 | -0.604008486 | 0.657923393 | 1.29E-03 | 5.37E-03 | 334.8801507 | 220.4757425 | ENSG00000249992 |
| TMEM200C | -0.604156051 | 0.657856101 | 1.56E-04 | 8.03E-04 | 231.3021927 | 152.3733236 | ENSG00000206432 |
| GNB1L | -0.604922658 | 0.657506628 | 1.50E-06 | 1.14E-05 | 394.8619397 | 259.6572985 | ENSG00000185838 |
| GNG7 | -0.606575508 | 0.656753774 | 1.01E-07 | 9.31E-07 | 464.0080264 | 304.7587387 | ENSG00000176533 |
| SEMA4F | -0.607046258 | 0.656539512 | 1.89E-04 | 9.58E-04 | 242.3251139 | 159.0003914 | ENSG00000135622 |
| GNAZ | -0.60726166 | 0.656441494 | 4.17E-05 | 2.43E-04 | 283.2979249 | 186.1824051 | ENSG00000128266 |
| ICAM5 | -0.608729762 | 0.655773832 | 8.65E-03 | 2.79E-02 | 108.140677 | 70.76405482 | ENSG00000105376 |
| PDAP1 | -0.609128011 | 0.655592833 | 3.47E-29 | 2.12E-27 | 6533.853133 | 4283.397221 | ENSG00000106244 |
| MCM4 | -0.609639021 | 0.65536066 | 9.23E-35 | 7.45E-33 | 8852.130384 | 5801.404537 | ENSG00000104738 |
| HMG20B | -0.610014148 | 0.655190276 | 2.12E-22 | 8.54E-21 | 1804.490581 | 1182.673439 | ENSG00000064961 |
| SLC22A31 | -0.610599978 | 0.65492428 | 1.31E-03 | 5.45E-03 | 174.744667 | 114.3705457 | ENSG00000259803 |
| HEY2 | -0.610784578 | 0.654840485 | 3.65E-04 | 1.73E-03 | 184.5993228 | 120.9545715 | ENSG00000135547 |
| FUCA1 | -0.610836188 | 0.654817059 | 1.87E-04 | 9.50E-04 | 225.3861924 | 147.3654664 | ENSG00000179163 |

| | | | | | | | |
|---|---|---|---|---|---|---|---|
| ZNF689 | -0.611149753 | 0.654674752 | 3.80E-09 | 4.29E-08 | 600.6740743 | 393.1187851 | ENSG00000156853 |
| DPH7 | -0.611186675 | 0.654657998 | 1.18E-11 | 1.79E-10 | 880.020214 | 576.3709321 | ENSG00000148399 |
| MDK | -0.611501955 | 0.654514947 | 1.54E-40 | 1.85E-38 | 6900.051246 | 4517.068051 | ENSG00000110492 |
| EMILIN3 | -0.61229771 | 0.654154032 | 4.28E-05 | 2.49E-04 | 269.5691296 | 176.1102307 | ENSG00000183798 |
| ARG2 | -0.612963974 | 0.653852002 | 4.35E-08 | 4.22E-07 | 536.690102 | 350.7746905 | ENSG00000081181 |
| SLCO3A1 | -0.614035788 | 0.653366419 | 1.02E-07 | 9.36E-07 | 434.0443595 | 283.6025755 | ENSG00000176463 |
| EIF2B1 | -0.614273924 | 0.653258581 | 9.57E-16 | 2.24E-14 | 1364.095503 | 891.0263585 | ENSG00000111361 |
| EIF2B2 | -0.615273017 | 0.652806344 | 2.08E-11 | 3.08E-10 | 727.1606826 | 474.836134 | ENSG00000119718 |
| FAHD1 | -0.616040343 | 0.652459229 | 8.13E-07 | 6.50E-06 | 377.0226042 | 246.060373 | ENSG00000180185 |
| FLRT2 | -0.616395702 | 0.652298537 | 1.21E-02 | 3.69E-02 | 110.6365307 | 72.33832117 | ENSG00000185070 |
| ADGRB2 | -0.617008473 | 0.652021538 | 4.31E-17 | 1.14E-15 | 1238.152085 | 807.238068 | ENSG00000121753 |
| TRAF3IP1 | -0.617120915 | 0.651970722 | 7.65E-09 | 8.29E-08 | 539.0379868 | 351.2858073 | ENSG00000204104 |
| SKP2 | -0.617990474 | 0.651577877 | 3.18E-19 | 9.96E-18 | 1738.052881 | 1132.174857 | ENSG00000145604 |
| OXCT1 | -0.618088257 | 0.651533716 | 3.14E-13 | 5.66E-12 | 885.2606128 | 577.1071208 | ENSG00000083720 |
| STX18 | -0.618361031 | 0.65141054 | 4.37E-09 | 4.90E-08 | 599.3925189 | 390.0982086 | ENSG00000168818 |
| SPSB4 | -0.618383485 | 0.651400402 | 1.73E-10 | 2.32E-09 | 717.5394122 | 467.1376237 | ENSG00000175093 |
| SCG3 | -0.619201625 | 0.651031103 | 6.54E-13 | 1.14E-11 | 1098.789851 | 715.6727194 | ENSG00000104112 |
| SPHK2 | -0.620609728 | 0.650395992 | 1.93E-07 | 1.70E-06 | 853.3484002 | 554.6362025 | ENSG00000063176 |
| MGAT5B | -0.620829104 | 0.6502971 | 6.78E-06 | 4.61E-05 | 306.8448672 | 199.6209754 | ENSG00000167889 |
| EME2 | -0.62113955 | 0.650157181 | 1.30E-06 | 1.01E-05 | 571.1884795 | 371.1358302 | ENSG00000197774 |
| FAM32A | -0.622276691 | 0.649644925 | 2.17E-21 | 8.10E-20 | 1816.533925 | 1179.972609 | ENSG00000105058 |
| TFPT | -0.622298646 | 0.649635039 | 2.88E-05 | 1.74E-04 | 336.667037 | 218.7331679 | ENSG00000105619 |
| RNF208 | -0.622522226 | 0.649534371 | 1.18E-02 | 3.63E-02 | 115.3307359 | 75.11019987 | ENSG00000212864 |
| EFR3B | -0.622646929 | 0.649478229 | 7.03E-08 | 6.60E-07 | 454.2549703 | 294.8919901 | ENSG00000084710 |
| SLC22A5 | -0.622710694 | 0.649449524 | 5.72E-03 | 1.97E-02 | 115.0820062 | 74.89321554 | ENSG00000197375 |
| TMEM191A | -0.62307312 | 0.649286393 | 2.56E-02 | 6.90E-02 | 76.58240938 | 49.69202448 | ENSG00000226287 |
| PCDH1 | -0.623353793 | 0.649160088 | 5.55E-32 | 3.93E-30 | 5583.427717 | 3624.468953 | ENSG00000156453 |
| ZNF100 | -0.623483424 | 0.649101761 | 2.04E-03 | 8.03E-03 | 147.0560701 | 95.47561646 | ENSG00000197020 |
| CHRNB4 | -0.623501264 | 0.649093735 | 1.26E-02 | 3.83E-02 | 105.7178013 | 68.72944936 | ENSG00000117971 |
| DUSP26 | -0.62366615 | 0.649019554 | 4.72E-02 | 1.14E-01 | 54.74816168 | 35.58610224 | ENSG00000133878 |
| TRIM39 | -0.623896024 | 0.648916149 | 1.21E-05 | 7.80E-05 | 283.4695342 | 183.8533884 | ENSG00000204599 |
| TMEM198 | -0.623915745 | 0.648907279 | 2.63E-05 | 1.60E-04 | 261.9574555 | 169.8356702 | ENSG00000188760 |
| URM1 | -0.623935796 | 0.648898261 | 2.26E-15 | 5.08E-14 | 1307.657337 | 848.8709509 | ENSG00000167118 |
| EPN3 | -0.624704754 | 0.648552489 | 1.15E-02 | 3.56E-02 | 88.96077679 | 57.61319574 | ENSG00000049283 |
| MT1E | -0.624786228 | 0.648515864 | 2.24E-05 | 1.39E-04 | 289.0356808 | 187.6464408 | ENSG00000169715 |
| UNG | -0.625321657 | 0.648275224 | 9.64E-36 | 8.26E-34 | 4695.992278 | 3044.10886 | ENSG00000076248 |
| RPS6KA2 | -0.625591311 | 0.648154067 | 6.29E-12 | 9.83E-11 | 779.3624278 | 504.9298506 | ENSG00000071242 |
| RPL26L1 | -0.625617268 | 0.648142405 | 4.08E-10 | 5.22E-09 | 595.4288703 | 385.9566412 | ENSG00000037241 |
| ABCA1 | -0.625925701 | 0.648003854 | 1.23E-04 | 6.51E-04 | 219.3467919 | 142.1556547 | ENSG00000165029 |
| C7orf50 | -0.626283213 | 0.647843293 | 1.16E-21 | 4.45E-20 | 2085.157776 | 1350.716829 | ENSG00000146540 |
| ABITRAM | -0.627682642 | 0.647215183 | 2.23E-03 | 8.69E-03 | 164.2022212 | 106.529485 | ENSG00000119328 |
| MCTP1 | -0.628144003 | 0.647008243 | 1.37E-02 | 4.10E-02 | 83.6937963 | 54.18041682 | ENSG00000175471 |
| NXF1 | -0.628989029 | 0.646629383 | 7.34E-13 | 1.27E-11 | 1077.228649 | 696.3659335 | ENSG00000162231 |
| CYP2U1 | -0.629507506 | 0.646397039 | 1.51E-03 | 6.18E-03 | 182.0767624 | 117.9388171 | ENSG00000155016 |
| MGAT4C | -0.63078702 | 0.645824009 | 4.90E-02 | 1.17E-01 | 63.12140442 | 40.71899582 | ENSG00000182050 |
| MYCN | -0.630788921 | 0.645823158 | 2.03E-18 | 5.96E-17 | 1827.058602 | 1180.167015 | ENSG00000134323 |
| HTRA3 | -0.63093361 | 0.645758391 | 9.25E-05 | 5.04E-04 | 236.1626255 | 152.3426762 | ENSG00000170801 |
| MAOA | -0.63125467 | 0.645614699 | 2.09E-04 | 1.05E-03 | 198.7398331 | 128.3691692 | ENSG00000189221 |
| NUDT8 | -0.631373979 | 0.64556131 | 1.90E-02 | 5.38E-02 | 103.9115916 | 67.1091785 | ENSG00000167799 |
| MTHFD1 | -0.631556471 | 0.645479655 | 2.07E-42 | 2.61E-40 | 5425.486862 | 3501.800667 | ENSG00000100714 |
| MARCKSL1 | -0.63170713 | 0.645412252 | 1.97E-09 | 2.31E-08 | 12246.62068 | 7904.226673 | ENSG00000175130 |
| CDC27 | -0.632445188 | 0.645082155 | 2.10E-09 | 2.45E-08 | 795.5153524 | 513.2855051 | ENSG00000004897 |
| ATP6V0E1 | -0.632575669 | 0.645023814 | 1.67E-18 | 4.96E-17 | 1448.810323 | 934.7357802 | ENSG00000113732 |
| ZSCAN18 | -0.632862857 | 0.644895426 | 2.15E-06 | 1.59E-05 | 341.9885962 | 220.5220075 | ENSG00000121413 |
| DNAL1 | -0.633541889 | 0.644591966 | 5.39E-04 | 2.45E-03 | 180.576777 | 116.5196893 | ENSG00000119661 |
| FJX1 | -0.634272233 | 0.644265732 | 4.70E-08 | 4.53E-07 | 532.0408778 | 342.6559022 | ENSG00000179431 |
| TUBA1A | -0.634452673 | 0.644185158 | 1.68E-41 | 2.08E-39 | 5994.689562 | 3861.585386 | ENSG00000167552 |
| DUSP16 | -0.634790457 | 0.64403435 | 1.80E-22 | 7.30E-21 | 2595.337207 | 1671.580498 | ENSG00000111266 |
| DNAJB11 | -0.636129125 | 0.643437032 | 2.27E-16 | 5.63E-15 | 1337.843964 | 861.4732668 | ENSG00000090520 |
| SLC4A8 | -0.63634341 | 0.643341468 | 1.40E-06 | 1.07E-05 | 339.6806728 | 218.3900564 | ENSG00000050438 |
| UPF3B | -0.637066439 | 0.643019129 | 1.63E-15 | 3.71E-14 | 1104.815071 | 710.7670058 | ENSG00000125351 |
| NISCH | -0.637297927 | 0.642915961 | 2.62E-23 | 1.13E-21 | 3441.781011 | 2212.181924 | ENSG00000010322 |
| P2RY11 | -0.637924438 | 0.642636826 | 4.76E-02 | 1.15E-01 | 52.62237374 | 33.86124677 | ENSG00000244165 |
| TAF7 | -0.638102802 | 0.642557381 | 4.63E-33 | 3.50E-31 | 3145.762336 | 2020.998718 | ENSG00000178913 |
| SLC20A1 | -0.638208278 | 0.642510404 | 5.00E-31 | 3.34E-29 | 2571.88985 | 1652.575666 | ENSG00000144136 |
| SLC35D1 | -0.638372386 | 0.642437322 | 7.54E-03 | 2.49E-02 | 104.8308662 | 67.46409685 | ENSG00000116704 |
| UBE2F | -0.639076598 | 0.642123811 | 2.00E-03 | 7.88E-03 | 136.9473542 | 88.00159618 | ENSG00000184182 |
| NAGPA | -0.639205721 | 0.642066342 | 6.43E-05 | 3.63E-04 | 234.467264 | 150.4712907 | ENSG00000103174 |
| MYL6B | -0.639343208 | 0.642005157 | 4.55E-14 | 8.91E-13 | 1221.076607 | 783.7566015 | ENSG00000196465 |
| GET4 | -0.63960173 | 0.641890125 | 1.24E-04 | 6.57E-04 | 212.2582957 | 136.2051047 | ENSG00000239857 |
| SFXN5 | -0.64021073 | 0.641619223 | 1.32E-10 | 1.80E-09 | 956.1828608 | 613.2108952 | ENSG00000144040 |
| SLC25A27 | -0.640264324 | 0.641595388 | 3.75E-02 | 9.45E-02 | 74.97069981 | 47.94297967 | ENSG00000153291 |
| VPS8 | -0.640303186 | 0.641578105 | 1.11E-11 | 1.70E-10 | 763.3444077 | 489.8392902 | ENSG00000156931 |

| | | | | | | | |
|---|---|---|---|---|---|---|---|
| COMMD4 | -0.640449751 | 0.64151293 | 8.43E-19 | 2.56E-17 | 1695.669998 | 1087.720862 | ENSG00000140365 |
| SCN4A | -0.640657704 | 0.641420468 | 8.89E-06 | 5.91E-05 | 282.2402053 | 180.9355248 | ENSG00000007314 |
| DRG2 | -0.640693451 | 0.641404575 | 3.98E-11 | 5.69E-10 | 664.5173871 | 426.4712835 | ENSG00000108591 |
| ZDHHC6 | -0.64070409 | 0.641399845 | 1.62E-11 | 2.42E-10 | 695.171625 | 445.8776326 | ENSG00000023041 |
| NBPF1 | -0.641191155 | 0.64118334 | 2.83E-08 | 2.81E-07 | 439.7180879 | 282.1204992 | ENSG00000219481 |
| CUL1 | -0.642012092 | 0.640818591 | 1.23E-21 | 4.68E-20 | 2714.503149 | 1739.207192 | ENSG00000055130 |
| RIPOR3 | -0.64228618 | 0.640696858 | 4.58E-02 | 1.11E-01 | 70.01875492 | 44.89871635 | ENSG00000042062 |
| LPCAT3 | -0.642616565 | 0.640550152 | 1.41E-10 | 1.92E-09 | 602.3476721 | 385.7566131 | ENSG00000111684 |
| CFAP36 | -0.643687348 | 0.640074905 | 4.01E-05 | 2.36E-04 | 260.9499112 | 167.1175558 | ENSG00000163001 |
| GPLD1 | -0.644018707 | 0.639927909 | 3.93E-03 | 1.42E-02 | 137.5769867 | 88.07633323 | ENSG00000112293 |
| ADAMTS8 | -0.644105909 | 0.639889231 | 1.57E-10 | 2.12E-09 | 616.5237884 | 394.3903168 | ENSG00000134917 |
| BUD31 | -0.644233674 | 0.639832564 | 8.96E-13 | 1.53E-11 | 814.4704018 | 521.1445702 | ENSG00000106245 |
| ZFP69B | -0.644620572 | 0.639660999 | 3.49E-03 | 1.28E-02 | 140.4296644 | 90.0259697 | ENSG00000187801 |
| DCTN3 | -0.645618738 | 0.639218586 | 6.54E-07 | 5.29E-06 | 355.2450504 | 226.8955128 | ENSG00000137100 |
| RIDA | -0.646682119 | 0.638747605 | 9.51E-10 | 1.15E-08 | 574.4400575 | 366.957945 | ENSG00000132541 |
| CCM2L | -0.647434482 | 0.638414586 | 3.08E-03 | 1.15E-02 | 127.9700726 | 81.82169149 | ENSG00000101331 |
| MRTO4 | -0.647500551 | 0.63838535 | 7.59E-22 | 2.96E-20 | 1833.382588 | 1170.336553 | ENSG00000053372 |
| PIGH | -0.647936998 | 0.638192254 | 9.00E-03 | 2.88E-02 | 96.70283539 | 61.75231583 | ENSG00000100564 |
| PCDHA12 | -0.648179889 | 0.638084817 | 2.14E-04 | 1.07E-03 | 176.9848578 | 112.9403741 | ENSG00000251664 |
| BEX1 | -0.648305571 | 0.638029232 | 2.51E-17 | 6.81E-16 | 1325.416586 | 845.3984382 | ENSG00000133169 |
| FBN2 | -0.648574689 | 0.637910226 | 1.03E-10 | 1.42E-09 | 939.8937825 | 599.5848118 | ENSG00000138829 |
| TMEM199 | -0.648734586 | 0.637839529 | 1.59E-02 | 4.65E-02 | 89.55412759 | 57.03833688 | ENSG00000244045 |
| KSR2 | -0.649302856 | 0.637588337 | 1.27E-02 | 3.84E-02 | 85.83720554 | 54.59072915 | ENSG00000171435 |
| DHX38 | -0.649964485 | 0.637296002 | 1.42E-27 | 8.00E-26 | 2382.01058 | 1517.707216 | ENSG00000140829 |
| ARRDC3 | -0.650079313 | 0.63724528 | 1.17E-02 | 3.60E-02 | 81.76489904 | 52.06649428 | ENSG00000113369 |
| COL9A2 | -0.650569177 | 0.637028941 | 1.02E-03 | 4.33E-03 | 156.6762262 | 99.89033783 | ENSG00000049089 |
| TNFSF13 | -0.650798187 | 0.636927829 | 4.49E-02 | 1.09E-01 | 54.73017696 | 34.8165704 | ENSG00000161955 |
| GORASP1 | -0.651087707 | 0.636800023 | 2.89E-14 | 5.79E-13 | 1061.354606 | 675.8646787 | ENSG00000114745 |
| OBI1 | -0.651532332 | 0.636603797 | 9.88E-04 | 4.22E-03 | 177.8369506 | 113.2256622 | ENSG00000152193 |
| SH2B2 | -0.651588157 | 0.636579165 | 1.71E-07 | 1.52E-06 | 450.8323593 | 287.1509708 | ENSG00000160999 |
| CPNE1 | -0.653495859 | 0.63573796 | 4.93E-20 | 1.67E-18 | 1374.955173 | 873.7648393 | ENSG00000214078 |
| ZMPSTE24 | -0.654282912 | 0.635391232 | 2.86E-13 | 5.19E-12 | 1095.089538 | 696.037585 | ENSG00000084073 |
| IQGAP1 | -0.654668816 | 0.635221296 | 2.31E-31 | 1.57E-29 | 3401.01211 | 2160.714774 | ENSG00000140575 |
| PCNX3 | -0.654835051 | 0.635148106 | 1.15E-19 | 3.80E-18 | 1884.900235 | 1197.066885 | ENSG00000197136 |
| CNIH2 | -0.654905209 | 0.63511722 | 2.82E-04 | 1.38E-03 | 220.7763531 | 140.1009126 | ENSG00000174871 |
| POU6F1 | -0.656171597 | 0.634559963 | 5.33E-03 | 1.85E-02 | 102.9948325 | 65.42599585 | ENSG00000184271 |
| ANKRD18B | -0.656795816 | 0.634285464 | 3.57E-03 | 1.31E-02 | 115.2025034 | 73.01079831 | ENSG00000230453 |
| RAPGEF4 | -0.65685058 | 0.634261387 | 4.00E-02 | 9.95E-02 | 55.07264603 | 35.01607756 | ENSG00000091428 |
| SLC25A53 | -0.657325083 | 0.634052812 | 1.51E-05 | 9.61E-05 | 253.4199371 | 160.8672031 | ENSG00000269743 |
| PTDSS2 | -0.658071555 | 0.633724829 | 1.31E-20 | 4.69E-19 | 1498.947244 | 950.2812819 | ENSG00000174915 |
| CENPV | -0.658111548 | 0.633707261 | 3.21E-24 | 1.49E-22 | 2294.728659 | 1454.713676 | ENSG00000166582 |
| PJA1 | -0.658147841 | 0.63369132 | 1.97E-25 | 9.84E-24 | 1824.895579 | 1156.25293 | ENSG00000181191 |
| PRRT2 | -0.658150801 | 0.633690019 | 1.40E-03 | 5.77E-03 | 172.5138128 | 109.0970345 | ENSG00000167371 |
| MX1 | -0.65822519 | 0.633657346 | 3.16E-02 | 8.19E-02 | 62.97810447 | 39.86612465 | ENSG00000157601 |
| TNFRSF11A | -0.65868291 | 0.633456339 | 2.75E-05 | 1.67E-04 | 224.0424365 | 141.9991774 | ENSG00000141655 |
| NRTN | -0.659132739 | 0.63325886 | 1.53E-02 | 4.51E-02 | 92.43375036 | 58.51797168 | ENSG00000171119 |
| SCHIP1 | -0.659143147 | 0.633254291 | 5.56E-03 | 1.92E-02 | 108.3321491 | 68.64662464 | ENSG00000283154 |
| HOMER2 | -0.660190801 | 0.632794602 | 2.19E-09 | 2.55E-08 | 502.4192655 | 317.8117496 | ENSG00000103942 |
| INPPL1 | -0.66083239 | 0.632513251 | 5.37E-25 | 2.66E-23 | 2426.393825 | 1534.272329 | ENSG00000165458 |
| POMC | -0.661266569 | 0.632322925 | 1.48E-02 | 4.38E-02 | 80.77306096 | 51.09664967 | ENSG00000115138 |
| NUBPL | -0.6613265 | 0.632296658 | 1.73E-04 | 8.83E-04 | 200.1572796 | 126.7074923 | ENSG00000151413 |
| EXOSC1 | -0.661440526 | 0.632246686 | 3.55E-11 | 5.12E-10 | 710.557771 | 448.895265 | ENSG00000171311 |
| CRELD2 | -0.661574677 | 0.632187898 | 3.52E-12 | 5.65E-11 | 1057.497801 | 668.4014845 | ENSG00000184164 |
| WDR90 | -0.661904273 | 0.632043486 | 3.00E-11 | 4.36E-10 | 802.2373998 | 507.0919403 | ENSG00000161996 |
| MAGEF1 | -0.663662693 | 0.631273593 | 1.75E-19 | 5.64E-18 | 1795.30969 | 1133.560096 | ENSG00000177383 |
| EFS | -0.663801099 | 0.631213034 | 2.33E-14 | 4.75E-13 | 1105.345696 | 697.732363 | ENSG00000100842 |
| USP49 | -0.665411273 | 0.630508938 | 1.50E-11 | 2.25E-10 | 704.7008183 | 443.9097255 | ENSG00000164663 |
| CHST1 | -0.665767686 | 0.630353192 | 4.05E-02 | 1.01E-01 | 59.5536154 | 37.64785954 | ENSG00000175264 |
| ATP7B | -0.665851434 | 0.630316601 | 5.70E-06 | 3.92E-05 | 317.9219427 | 200.2482905 | ENSG00000123191 |
| MRPS18C | -0.665990323 | 0.630255923 | 2.79E-03 | 1.05E-02 | 119.7506177 | 75.57644554 | ENSG00000163319 |
| ZFYVE26 | -0.666117152 | 0.630200519 | 3.61E-16 | 8.82E-15 | 988.0085554 | 622.5129742 | ENSG00000072121 |
| DTX4 | -0.666528923 | 0.630020674 | 3.11E-13 | 5.61E-12 | 899.6355349 | 566.4337533 | ENSG00000110042 |
| FGFR1 | -0.666700255 | 0.629945859 | 7.24E-65 | 1.86E-62 | 9960.326261 | 6274.20026 | ENSG00000077782 |
| MAMDC4 | -0.667182913 | 0.629735143 | 2.87E-06 | 2.08E-05 | 345.0557489 | 217.3887588 | ENSG00000177943 |
| ST3GAL5 | -0.667593875 | 0.629555785 | 3.13E-02 | 8.13E-02 | 64.86333531 | 40.80665473 | ENSG00000115525 |
| TSPAN33 | -0.667960921 | 0.629395635 | 1.16E-13 | 2.19E-12 | 975.4719569 | 613.920695 | ENSG00000158457 |
| THEM6 | -0.66822289 | 0.629281358 | 3.90E-04 | 1.85E-03 | 197.5425315 | 124.2220104 | ENSG00000130193 |
| KREMEN1 | -0.669050498 | 0.628920472 | 3.72E-04 | 1.77E-03 | 220.1619513 | 138.3166346 | ENSG00000183762 |
| CDK4 | -0.670132 | 0.628449185 | 5.76E-30 | 3.68E-28 | 2437.900701 | 1531.668207 | ENSG00000135446 |
| AREL1 | -0.670171677 | 0.628431901 | 1.38E-17 | 3.78E-16 | 1113.497441 | 699.448313 | ENSG00000119682 |
| FBXW4 | -0.670307053 | 0.628372934 | 3.23E-12 | 5.21E-11 | 724.5900381 | 455.4833194 | ENSG00000107829 |
| REC8 | -0.671125671 | 0.628016483 | 1.56E-10 | 2.11E-09 | 596.8417982 | 374.6481561 | ENSG00000100918 |
| UCK1 | -0.672896836 | 0.627245954 | 1.01E-16 | 2.59E-15 | 1263.555298 | 792.3131508 | ENSG00000130717 |

| | | | | | | | |
|---|---|---|---|---|---|---|---|
| FHOD3 | -0.672965246 | 0.627216212 | 3.07E-07 | 2.62E-06 | 334.4523646 | 209.9238617 | ENSG00000134775 |
| SEMA7A | -0.67351167 | 0.626978697 | 1.72E-13 | 3.17E-12 | 793.8206156 | 497.8164069 | ENSG00000138623 |
| PRXL2B | -0.673606442 | 0.626937512 | 2.01E-22 | 8.13E-21 | 2055.725296 | 1288.66176 | ENSG00000157870 |
| JPH4 | -0.673985038 | 0.62677301 | 1.17E-28 | 6.95E-27 | 3328.475031 | 2086.466429 | ENSG00000092051 |
| CRB2 | -0.673999969 | 0.626766524 | 1.45E-05 | 9.24E-05 | 327.0492715 | 204.9942492 | ENSG00000148204 |
| PNMA1 | -0.674646247 | 0.626485817 | 3.33E-29 | 2.05E-27 | 2166.232704 | 1357.309681 | ENSG00000176903 |
| TTLL4 | -0.674883127 | 0.626382961 | 3.61E-26 | 1.89E-24 | 2066.273311 | 1294.754763 | ENSG00000135912 |
| PARD6A | -0.675157753 | 0.626263736 | 8.83E-05 | 4.83E-04 | 215.3061492 | 134.6725112 | ENSG00000102981 |
| BEND5 | -0.676964228 | 0.625480049 | 3.14E-03 | 1.17E-02 | 115.1031822 | 71.99524019 | ENSG00000162373 |
| DLEU7 | -0.677104278 | 0.625419333 | 6.55E-03 | 2.21E-02 | 112.5561794 | 70.58335767 | ENSG00000186047 |
| MVB12B | -0.677479437 | 0.62525672 | 4.17E-07 | 3.48E-06 | 311.430264 | 194.7115179 | ENSG00000196814 |
| FMNL1 | -0.678291863 | 0.624904718 | 1.07E-04 | 5.76E-04 | 208.6095947 | 130.2465039 | ENSG00000184922 |
| DPP4 | -0.678318737 | 0.624893077 | 1.20E-02 | 3.66E-02 | 86.2102624 | 53.9537457 | ENSG00000197635 |
| BIVM | -0.67882142 | 0.624675382 | 5.15E-03 | 1.80E-02 | 93.10900124 | 58.12433089 | ENSG00000134897 |
| MPP2 | -0.680321035 | 0.624026398 | 4.53E-28 | 2.61E-26 | 2453.331315 | 1530.554256 | ENSG00000108852 |
| KIF9 | -0.680686885 | 0.623868173 | 7.85E-03 | 2.57E-02 | 90.30258077 | 56.31935273 | ENSG00000088727 |
| WDR62 | -0.680984094 | 0.623739663 | 6.30E-21 | 2.29E-19 | 1912.659042 | 1192.690359 | ENSG00000075702 |
| PTBP2 | -0.681290703 | 0.623607117 | 5.34E-14 | 1.04E-12 | 824.5569026 | 514.4298498 | ENSG00000117569 |
| MAP3K8 | -0.683390931 | 0.62269995 | 1.90E-02 | 5.38E-02 | 81.13056058 | 50.65378773 | ENSG00000107968 |
| EID1 | -0.683752968 | 0.622543707 | 4.61E-29 | 2.79E-27 | 2915.413815 | 1815.030117 | ENSG00000255302 |
| ARSB | -0.684537041 | 0.622205459 | 1.45E-07 | 1.30E-06 | 351.2683984 | 218.4586327 | ENSG00000113273 |
| C12orf10 | -0.684727685 | 0.622123244 | 5.25E-11 | 7.43E-10 | 742.0299096 | 461.8278011 | ENSG00000139637 |
| SIRT2 | -0.6848621 | 0.622065284 | 2.40E-04 | 1.19E-03 | 206.3115955 | 128.3737308 | ENSG00000068903 |
| HTRA1 | -0.685167785 | 0.621933492 | 5.93E-11 | 8.34E-10 | 649.3634993 | 404.2874218 | ENSG00000166033 |
| ANAPC2 | -0.685897601 | 0.621618954 | 3.03E-14 | 6.07E-13 | 1381.211905 | 858.2444115 | ENSG00000176248 |
| OTUD7A | -0.686281988 | 0.621453354 | 8.62E-04 | 3.72E-03 | 150.5766651 | 93.59428373 | ENSG00000169918 |
| SLC25A10 | -0.686673205 | 0.621284857 | 3.20E-03 | 1.19E-02 | 110.0517528 | 68.46245038 | ENSG00000183048 |
| EPHB6 | -0.687336891 | 0.620999111 | 8.55E-13 | 1.47E-11 | 776.1096375 | 481.7775439 | ENSG00000106123 |
| CLDN6 | -0.687783518 | 0.620806893 | 7.27E-59 | 1.67E-56 | 14600.46626 | 9064.501592 | ENSG00000184697 |
| ATG2A | -0.688583086 | 0.620462926 | 1.92E-13 | 3.53E-12 | 911.8909535 | 565.6838555 | ENSG00000110046 |
| DGCR6L | -0.688742943 | 0.62039418 | 2.68E-14 | 5.40E-13 | 1342.406334 | 832.6640827 | ENSG00000128185 |
| SNHG19 | -0.688893022 | 0.620329646 | 1.18E-07 | 1.08E-06 | 425.50903 | 263.6490888 | ENSG00000260260 |
| SUFU | -0.690378572 | 0.619691218 | 3.48E-10 | 4.50E-09 | 519.3575993 | 321.8215258 | ENSG00000107882 |
| ZNF664 | -0.69105435 | 0.619401014 | 2.14E-22 | 8.59E-21 | 1731.370856 | 1072.561125 | ENSG00000179195 |
| AKT1S1 | -0.691955455 | 0.619014258 | 1.72E-16 | 4.33E-15 | 1497.604778 | 927.2029188 | ENSG00000204673 |
| ATF5 | -0.69323323 | 0.618466248 | 1.78E-05 | 1.12E-04 | 220.1446934 | 136.1766142 | ENSG00000169136 |
| KCNQ1 | -0.693274986 | 0.618448348 | 1.47E-02 | 4.35E-02 | 75.68989252 | 46.94084592 | ENSG00000053918 |
| TMX4 | -0.693329806 | 0.618424849 | 5.30E-06 | 3.66E-05 | 288.9913017 | 178.981563 | ENSG00000125827 |
| LZTR1 | -0.694215746 | 0.618045199 | 6.54E-04 | 2.91E-03 | 170.0422524 | 105.0237766 | ENSG00000099949 |
| COL21A1 | -0.695129892 | 0.617653706 | 8.99E-03 | 2.88E-02 | 77.18201771 | 47.70634918 | ENSG00000124749 |
| PCYT2 | -0.695679495 | 0.617418452 | 4.36E-15 | 9.45E-14 | 1253.770341 | 773.9934019 | ENSG00000185813 |
| EXT2 | -0.695899996 | 0.617324093 | 4.35E-28 | 2.51E-26 | 2127.819975 | 1313.705328 | ENSG00000151348 |
| SMIM10L1 | -0.696867144 | 0.616910393 | 2.70E-08 | 2.69E-07 | 365.3646177 | 225.5123812 | ENSG00000256537 |
| DDX49 | -0.697756409 | 0.616530252 | 5.12E-14 | 9.96E-13 | 1046.832404 | 645.4902967 | ENSG00000105671 |
| SPC25 | -0.698040171 | 0.616408999 | 2.04E-09 | 2.40E-08 | 596.0492301 | 367.3392154 | ENSG00000152253 |
| MFSD1 | -0.698848792 | 0.616063603 | 4.45E-04 | 2.07E-03 | 154.9798767 | 95.60172523 | ENSG00000118855 |
| HES1 | -0.699594661 | 0.615745182 | 8.30E-05 | 4.57E-04 | 222.7232351 | 137.3204195 | ENSG00000114315 |
| ARMCX5 | -0.699770681 | 0.615670061 | 7.21E-05 | 4.02E-04 | 188.0797179 | 115.7316079 | ENSG00000125962 |
| PIM2 | -0.700071644 | 0.615541638 | 1.35E-45 | 1.94E-43 | 5394.586272 | 3319.983865 | ENSG00000102096 |
| RTN1 | -0.700111329 | 0.615524706 | 6.88E-03 | 2.31E-02 | 89.8194394 | 55.20295969 | ENSG00000139970 |
| CPXM1 | -0.700819934 | 0.615222455 | 3.43E-28 | 2.00E-26 | 1878.514685 | 1155.535285 | ENSG00000088882 |
| ANKRD36B | -0.702340186 | 0.614574501 | 1.60E-02 | 4.67E-02 | 65.12342873 | 40.0075479 | ENSG00000196912 |
| DEXI | -0.703412278 | 0.614117969 | 1.64E-02 | 4.77E-02 | 73.95297985 | 45.44104403 | ENSG00000182108 |
| EAPP | -0.703483308 | 0.614087734 | 1.73E-08 | 1.78E-07 | 386.014516 | 236.8348784 | ENSG00000129518 |
| NAT8L | -0.703893029 | 0.61391336 | 6.57E-15 | 1.41E-13 | 1015.821593 | 624.0727196 | ENSG00000185818 |
| RORB | -0.705621237 | 0.613178392 | 7.19E-04 | 3.16E-03 | 156.2264266 | 95.69584797 | ENSG00000198963 |
| SEC61A2 | -0.708803293 | 0.611827435 | 2.95E-10 | 3.86E-09 | 459.4568841 | 281.1046806 | ENSG00000065665 |
| ARMH4 | -0.709212041 | 0.611654116 | 1.72E-06 | 1.30E-05 | 307.7452812 | 188.1753378 | ENSG00000139971 |
| DCHS1 | -0.709694528 | 0.611449592 | 3.04E-15 | 6.70E-14 | 848.926389 | 518.9158738 | ENSG00000166341 |
| ZNF253 | -0.710403894 | 0.611149019 | 5.27E-08 | 5.05E-07 | 358.7801876 | 219.4090916 | ENSG00000256771 |
| MEGF11 | -0.711772738 | 0.610569429 | 8.36E-03 | 2.71E-02 | 78.45906735 | 47.89752662 | ENSG00000157890 |
| FAXC | -0.712882134 | 0.610100097 | 1.03E-05 | 6.74E-05 | 220.4817808 | 134.5404837 | ENSG00000146267 |
| ZNF772 | -0.713201737 | 0.609964956 | 1.33E-02 | 4.00E-02 | 76.40680678 | 46.70934061 | ENSG00000197128 |
| CYB5D2 | -0.713290782 | 0.609927309 | 1.87E-03 | 7.43E-03 | 113.6241356 | 69.38443091 | ENSG00000167740 |
| ANOS1 | -0.713436324 | 0.609865781 | 3.85E-36 | 3.40E-34 | 4541.34789 | 2770.347092 | ENSG00000011201 |
| NT5C3A | -0.713907645 | 0.609666574 | 5.90E-09 | 6.47E-08 | 412.5945324 | 251.6312638 | ENSG00000122643 |
| PCDHA6 | -0.713997908 | 0.609628431 | 1.11E-03 | 4.70E-03 | 116.0236081 | 70.78743856 | ENSG00000081842 |
| CCDC74B | -0.714013532 | 0.609621829 | 2.78E-05 | 1.69E-04 | 201.0229266 | 122.6960985 | ENSG00000152076 |
| TMEM123 | -0.715255066 | 0.609097435 | 1.66E-22 | 6.76E-21 | 1940.942144 | 1182.890944 | ENSG00000152558 |
| P2RX2 | -0.715610125 | 0.60894755 | 4.30E-02 | 1.06E-01 | 49.52256917 | 30.11066071 | ENSG00000187848 |
| SH3RF3 | -0.71598407 | 0.608789731 | 4.89E-02 | 1.17E-01 | 42.01962822 | 25.58220098 | ENSG00000172985 |
| RNF150 | -0.716453868 | 0.608591518 | 2.44E-05 | 1.50E-04 | 203.9300197 | 123.9875305 | ENSG00000170153 |
| NBPF15 | -0.717020219 | 0.608352653 | 2.03E-06 | 1.51E-05 | 306.2849689 | 186.2367209 | ENSG00000266338 |

| | | | | | | | |
|---|---|---|---|---|---|---|---|
| FGFR1OP | -0.717306724 | 0.608231852 | 4.35E-05 | 2.53E-04 | 209.0393454 | 127.2718708 | ENSG00000213066 |
| NASP | -0.717343942 | 0.608216162 | 4.27E-96 | 3.17E-93 | 18181.00777 | 11058.0248 | ENSG00000132780 |
| PMM1 | -0.717371331 | 0.608204615 | 5.59E-05 | 3.19E-04 | 182.6058713 | 110.9711099 | ENSG00000100417 |
| ABCG1 | -0.717458705 | 0.608167782 | 1.54E-03 | 6.27E-03 | 128.9942502 | 78.42789546 | ENSG00000160179 |
| AJUBA | -0.718870611 | 0.607572884 | 3.93E-11 | 5.62E-10 | 857.5538581 | 521.3749789 | ENSG00000129474 |
| KTN1 | -0.71888902 | 0.607565131 | 2.85E-35 | 2.40E-33 | 2706.515449 | 1644.716855 | ENSG00000126777 |
| REM2 | -0.719200786 | 0.607433851 | 9.81E-03 | 3.09E-02 | 74.85829995 | 45.47223656 | ENSG00000139890 |
| NOL12 | -0.719647032 | 0.607245992 | 1.17E-02 | 3.58E-02 | 67.12074712 | 40.79428432 | ENSG00000273899 |
| MCM2 | -0.719825603 | 0.607170834 | 2.88E-39 | 3.18E-37 | 4933.799769 | 2995.175663 | ENSG00000073111 |
| MACROH2A1 | -0.719898729 | 0.60714006 | 8.05E-31 | 5.28E-29 | 2767.669226 | 1680.280174 | ENSG00000113648 |
| PHKA2 | -0.721853956 | 0.606317784 | 2.19E-15 | 4.93E-14 | 765.1804045 | 463.9747879 | ENSG00000044446 |
| DHX35 | -0.72191884 | 0.606290516 | 1.03E-11 | 1.58E-10 | 554.1973683 | 335.7620959 | ENSG00000101452 |
| SEPTIN8 | -0.722394778 | 0.606090537 | 1.99E-23 | 8.68E-22 | 1618.077289 | 980.2889998 | ENSG00000164402 |
| RNF38 | -0.722874221 | 0.605889151 | 3.50E-13 | 6.28E-12 | 665.6578699 | 403.3044375 | ENSG00000137075 |
| SLC49A4 | -0.723731485 | 0.605529233 | 9.20E-07 | 7.28E-06 | 292.5538724 | 177.1636935 | ENSG00000138463 |
| SLC13A4 | -0.72376164 | 0.605516576 | 2.37E-02 | 6.46E-02 | 56.56621068 | 34.28418997 | ENSG00000164707 |
| GFOD2 | -0.725935146 | 0.604605016 | 1.29E-15 | 2.97E-14 | 932.7116733 | 563.6632196 | ENSG00000141098 |
| NOS1 | -0.726607622 | 0.604323261 | 6.70E-05 | 3.76E-04 | 180.8283344 | 109.3228694 | ENSG00000089250 |
| APCDD1 | -0.726797495 | 0.604243731 | 6.47E-04 | 2.88E-03 | 120.6281409 | 72.87797735 | ENSG00000154856 |
| ALDH16A1 | -0.727101767 | 0.604116306 | 2.74E-17 | 7.41E-16 | 1123.367292 | 678.5355047 | ENSG00000161618 |
| CHN1 | -0.728206343 | 0.603653951 | 5.76E-19 | 1.77E-17 | 938.8499067 | 566.7189988 | ENSG00000128656 |
| ENKD1 | -0.728743972 | 0.603429038 | 1.92E-05 | 1.20E-04 | 225.3373076 | 136.0502328 | ENSG00000124074 |
| SLC27A1 | -0.729349811 | 0.60317569 | 8.02E-06 | 5.38E-05 | 243.0123674 | 146.5459262 | ENSG00000130304 |
| POLR2C | -0.729411006 | 0.603150105 | 6.77E-28 | 3.86E-26 | 1541.246412 | 929.5434698 | ENSG00000102978 |
| TMEM45A | -0.729953673 | 0.602923274 | 3.29E-07 | 2.80E-06 | 285.16218 | 172.0832072 | ENSG00000181458 |
| MCM5 | -0.730590593 | 0.602657155 | 2.29E-38 | 2.34E-36 | 8900.855631 | 5363.42806 | ENSG00000100297 |
| ARNT2 | -0.730761549 | 0.602585746 | 2.99E-08 | 2.96E-07 | 348.8380627 | 210.064231 | ENSG00000172379 |
| GSTM4 | -0.731028969 | 0.60247406 | 3.86E-07 | 3.25E-06 | 302.4394282 | 182.2495165 | ENSG00000168765 |
| EFNB3 | -0.731165485 | 0.602417053 | 3.74E-12 | 5.99E-11 | 592.9592249 | 357.2629882 | ENSG00000108947 |
| SLC4A3 | -0.731830759 | 0.602139323 | 1.14E-08 | 1.21E-07 | 368.5292135 | 221.7045353 | ENSG00000114923 |
| SAFB2 | -0.732392542 | 0.601904896 | 1.37E-24 | 6.51E-23 | 2510.981118 | 1511.801816 | ENSG00000130254 |
| POLR3B | -0.733475441 | 0.601453271 | 8.23E-12 | 1.27E-10 | 542.2439337 | 326.0152343 | ENSG00000013503 |
| ULK4 | -0.733953433 | 0.601254031 | 1.72E-04 | 8.78E-04 | 175.4900539 | 105.5475426 | ENSG00000168038 |
| KLHL14 | -0.734163793 | 0.601166369 | 4.89E-02 | 1.17E-01 | 42.40348527 | 25.42840732 | ENSG00000197705 |
| MYO5C | -0.734624134 | 0.600974576 | 3.64E-10 | 4.69E-09 | 523.7626748 | 314.8927957 | ENSG00000128833 |
| CCDC88B | -0.735505156 | 0.600607686 | 5.62E-09 | 6.20E-08 | 498.597165 | 299.1873575 | ENSG00000168071 |
| ZNF599 | -0.735673151 | 0.600537753 | 3.01E-02 | 7.87E-02 | 62.7085623 | 37.8349963 | ENSG00000153896 |
| TCF25 | -0.738514211 | 0.599356294 | 1.80E-24 | 8.51E-23 | 1351.17437 | 809.5195117 | ENSG00000141002 |
| FAM78A | -0.738838762 | 0.599221477 | 1.40E-05 | 8.96E-05 | 204.042783 | 122.1658808 | ENSG00000126882 |
| C12orf57 | -0.740427011 | 0.598562163 | 2.68E-03 | 1.02E-02 | 101.7884313 | 60.79564723 | ENSG00000111678 |
| KIF3B | -0.741264344 | 0.598214861 | 8.19E-24 | 3.71E-22 | 1238.930799 | 741.3828951 | ENSG00000101350 |
| RBM25 | -0.741488301 | 0.598122004 | 3.07E-38 | 3.10E-36 | 3446.956928 | 2060.820269 | ENSG00000119707 |
| LANCL2 | -0.742722212 | 0.59761066 | 4.71E-09 | 5.26E-08 | 524.5595004 | 313.2335783 | ENSG00000132434 |
| BCL6B | -0.746448547 | 0.596069084 | 5.05E-09 | 5.59E-08 | 383.1433412 | 228.5076839 | ENSG00000161940 |
| BEND7 | -0.746532439 | 0.596034424 | 5.04E-04 | 2.31E-03 | 131.3647376 | 78.4437614 | ENSG00000165626 |
| EML3 | -0.746547341 | 0.596028267 | 3.72E-13 | 6.66E-12 | 651.9330295 | 388.254508 | ENSG00000149499 |
| TSPAN18 | -0.747606877 | 0.595590696 | 6.88E-13 | 1.19E-11 | 591.0980219 | 352.0082685 | ENSG00000157570 |
| MYO5B | -0.748036606 | 0.595413317 | 2.09E-09 | 2.45E-08 | 406.8559617 | 242.2167424 | ENSG00000167306 |
| PCDHB10 | -0.748756647 | 0.595116223 | 1.85E-02 | 5.27E-02 | 65.22542871 | 38.83415544 | ENSG00000120324 |
| ITFG1 | -0.749877748 | 0.594653945 | 2.10E-08 | 2.12E-07 | 355.7101318 | 211.5271821 | ENSG00000129636 |
| SMO | -0.749991198 | 0.594607185 | 6.43E-36 | 5.57E-34 | 2762.064457 | 1642.244325 | ENSG00000128602 |
| MMP25 | -0.750885136 | 0.594238862 | 7.90E-12 | 1.23E-10 | 504.9914375 | 300.1767872 | ENSG00000008516 |
| MSANTD2 | -0.75089046 | 0.59423667 | 7.06E-07 | 5.69E-06 | 279.2816367 | 166.0073725 | ENSG00000120458 |
| PLPBP | -0.750959349 | 0.594208295 | 9.44E-18 | 2.64E-16 | 849.174531 | 504.5480593 | ENSG00000147471 |
| CCNC | -0.751309612 | 0.594064049 | 1.66E-12 | 2.76E-11 | 707.355092 | 420.3475784 | ENSG00000112237 |
| CYP27A1 | -0.753451402 | 0.59318277 | 3.13E-04 | 1.51E-03 | 130.7627387 | 77.57528475 | ENSG00000135929 |
| HSPA1B | -0.75424087 | 0.592858259 | 1.37E-22 | 5.63E-21 | 1305.939522 | 774.2409847 | ENSG00000204388 |
| TENM4 | -0.754828034 | 0.59261702 | 4.36E-07 | 3.63E-06 | 287.7839494 | 170.4365964 | ENSG00000149256 |
| SHF | -0.755001306 | 0.592545849 | 1.42E-13 | 2.64E-12 | 720.3012407 | 427.0345776 | ENSG00000138606 |
| ADI1 | -0.756416768 | 0.591964774 | 2.78E-18 | 8.04E-17 | 867.318214 | 513.4858547 | ENSG00000182551 |
| PLEKHB1 | -0.756483254 | 0.591937494 | 9.80E-06 | 6.47E-05 | 211.392812 | 124.984539 | ENSG00000021300 |
| NAA16 | -0.760969581 | 0.590099614 | 6.39E-10 | 7.96E-09 | 487.8068143 | 287.645416 | ENSG00000172766 |
| C1orf216 | -0.761672852 | 0.589812028 | 1.50E-08 | 1.56E-07 | 345.759109 | 203.846149 | ENSG00000142686 |
| RIN3 | -0.761699123 | 0.589801288 | 1.62E-02 | 4.72E-02 | 74.83210579 | 44.11253138 | ENSG00000100599 |
| H1-10 | -0.762238503 | 0.58958082 | 1.88E-31 | 1.28E-29 | 2351.701779 | 1386.415536 | ENSG00000184897 |
| LOXL1 | -0.762977321 | 0.589278967 | 1.08E-06 | 8.45E-06 | 327.1750382 | 192.8764133 | ENSG00000129038 |
| ATP6V0D1 | -0.763086685 | 0.589234299 | 6.67E-19 | 2.03E-17 | 1346.271523 | 792.8767542 | ENSG00000159720 |
| SLC35A2 | -0.764762341 | 0.588550314 | 1.99E-13 | 3.64E-12 | 686.5228069 | 403.9645626 | ENSG00000102100 |
| SHISA6 | -0.764814073 | 0.58852921 | 3.04E-04 | 1.47E-03 | 166.7093876 | 98.36363245 | ENSG00000188803 |
| ZFP30 | -0.766037978 | 0.588030145 | 1.64E-02 | 4.76E-02 | 57.86491182 | 33.93680784 | ENSG00000120784 |
| DNAJC3 | -0.766827063 | 0.587708609 | 1.20E-14 | 2.51E-13 | 764.2313811 | 449.1800016 | ENSG00000102580 |
| P2RY2 | -0.7671735 | 0.587567498 | 1.20E-02 | 3.66E-02 | 63.37121011 | 37.25610887 | ENSG00000175591 |
| LYPD3 | -0.767272423 | 0.587527211 | 4.44E-03 | 1.58E-02 | 82.1964016 | 48.22367559 | ENSG00000124466 |

| | | | | | | | |
|---|---|---|---|---|---|---|---|
| MARCHF4 | -0.767868711 | 0.587284428 | 4.13E-03 | 1.48E-02 | 86.29821985 | 50.61508951 | ENSG00000144583 |
| GCLM | -0.767986444 | 0.587236504 | 2.07E-10 | 2.76E-09 | 465.6862052 | 273.6403287 | ENSG00000023909 |
| RCAN1 | -0.768272237 | 0.587120185 | 1.40E-05 | 8.94E-05 | 181.3432518 | 106.4512524 | ENSG00000159200 |
| NCAPH | -0.768717069 | 0.586939184 | 1.71E-19 | 5.52E-18 | 1369.485368 | 803.5592628 | ENSG00000121152 |
| SLC66A2 | -0.769749502 | 0.586519304 | 4.02E-09 | 4.53E-08 | 467.6335755 | 274.2977518 | ENSG00000122490 |
| YPEL2 | -0.769980805 | 0.586425277 | 8.09E-03 | 2.64E-02 | 79.5522297 | 46.5875451 | ENSG00000175155 |
| B4GAT1 | -0.770355877 | 0.586272838 | 5.03E-17 | 1.32E-15 | 770.2577158 | 451.7539665 | ENSG00000174684 |
| ZNF576 | -0.771644268 | 0.585749504 | 3.79E-04 | 1.80E-03 | 127.2807919 | 74.64477822 | ENSG00000124444 |
| CCBE1 | -0.77178384 | 0.585692839 | 2.45E-04 | 1.21E-03 | 129.7132797 | 75.93862123 | ENSG00000183287 |
| GFAP | -0.771939138 | 0.585629795 | 9.56E-03 | 3.03E-02 | 63.87919421 | 37.37629895 | ENSG00000131095 |
| DFFB | -0.772792573 | 0.585283465 | 8.65E-07 | 6.87E-06 | 235.9496651 | 137.9899342 | ENSG00000169598 |
| NDUFC2 | -0.772969611 | 0.585211648 | 1.60E-02 | 4.67E-02 | 67.0430774 | 39.39368771 | ENSG00000151366 |
| PHC2 | -0.773539857 | 0.58498038 | 8.54E-09 | 9.20E-08 | 339.5976966 | 198.6968443 | ENSG00000134686 |
| GLYCTK | -0.773948677 | 0.584814636 | 5.46E-03 | 1.89E-02 | 73.99473126 | 43.27389019 | ENSG00000168237 |
| PDS5A | -0.774019819 | 0.584785798 | 1.49E-45 | 2.12E-43 | 2616.855356 | 1530.353058 | ENSG00000121892 |
| MYBL2 | -0.774370024 | 0.584643863 | 1.84E-35 | 1.57E-33 | 6574.235971 | 3843.109229 | ENSG00000101057 |
| NMNAT1 | -0.774533255 | 0.584577718 | 1.16E-02 | 3.56E-02 | 60.57739272 | 35.4387603 | ENSG00000173614 |
| C1QL1 | -0.774800505 | 0.584469439 | 2.41E-07 | 2.09E-06 | 288.9490748 | 169.0421789 | ENSG00000131094 |
| GINS2 | -0.77508957 | 0.584352344 | 7.61E-24 | 3.46E-22 | 1602.76672 | 935.8721283 | ENSG00000131153 |
| CHST7 | -0.775421643 | 0.584217855 | 1.46E-12 | 2.45E-11 | 689.875059 | 403.3568632 | ENSG00000147119 |
| CAVIN4 | -0.7781816 | 0.583101282 | 1.11E-04 | 5.92E-04 | 159.9368398 | 93.3114067 | ENSG00000170681 |
| ADGRG2 | -0.778322524 | 0.583044327 | 1.12E-06 | 8.73E-06 | 260.7987879 | 151.9890907 | ENSG00000173698 |
| COL26A1 | -0.779866468 | 0.582420698 | 1.34E-39 | 1.52E-37 | 2889.785824 | 1683.3954 | ENSG00000160963 |
| PEX26 | -0.77992277 | 0.582397969 | 9.50E-30 | 5.97E-28 | 1552.149139 | 904.0782423 | ENSG00000215193 |
| HRAS | -0.780133409 | 0.582312943 | 5.76E-17 | 1.51E-15 | 799.5941918 | 465.6832444 | ENSG00000174775 |
| AQP5 | -0.781190567 | 0.5818864 | 2.41E-02 | 6.56E-02 | 50.88219461 | 29.53848546 | ENSG00000161798 |
| ARFRP1 | -0.781563304 | 0.581736082 | 4.91E-17 | 1.29E-15 | 737.0287662 | 428.5583389 | ENSG00000101246 |
| FAM222A | -0.782609277 | 0.581314469 | 1.09E-02 | 3.40E-02 | 64.7535262 | 37.58844909 | ENSG00000139438 |
| ND1 | -0.783008481 | 0.581153637 | 7.76E-04 | 3.38E-03 | 18308.1132 | 10639.81274 | ENSG00000198888 |
| C10orf143 | -0.783092748 | 0.581119694 | 3.29E-02 | 8.47E-02 | 48.57067961 | 28.19085969 | ENSG00000237489 |
| CCDC160 | -0.784358336 | 0.580610136 | 2.18E-03 | 8.52E-03 | 95.98871202 | 55.80632747 | ENSG00000203952 |
| TM7SF2 | -0.784443862 | 0.580575718 | 1.12E-19 | 3.72E-18 | 1094.188197 | 635.2984347 | ENSG00000149809 |
| SLC38A7 | -0.785382228 | 0.580198219 | 5.47E-16 | 1.32E-14 | 730.2082479 | 423.6935288 | ENSG00000103042 |
| SYS1 | -0.786787715 | 0.57963326 | 3.43E-07 | 2.91E-06 | 255.0071531 | 147.8155614 | ENSG00000204070 |
| KPNA1 | -0.787883854 | 0.57919303 | 6.52E-14 | 1.26E-12 | 644.6012365 | 373.68257 | ENSG00000114030 |
| NYNRIN | -0.788292576 | 0.579028965 | 3.78E-29 | 2.30E-27 | 1364.156852 | 789.7574324 | ENSG00000205978 |
| MAPK8IP2 | -0.788656762 | 0.578882817 | 2.88E-05 | 1.74E-04 | 255.851424 | 148.0718197 | ENSG00000008735 |
| GGT7 | -0.78869298 | 0.578868284 | 3.49E-22 | 1.39E-20 | 1585.049396 | 916.7995007 | ENSG00000131067 |
| GFRA2 | -0.790557562 | 0.578120621 | 1.97E-04 | 9.92E-04 | 141.7491411 | 82.09917076 | ENSG00000168546 |
| EPHA4 | -0.791424544 | 0.577773306 | 8.35E-05 | 4.59E-04 | 178.2436473 | 103.0773509 | ENSG00000116106 |
| UBQLN2 | -0.794176266 | 0.576672341 | 1.82E-20 | 6.45E-19 | 1087.469916 | 627.1312371 | ENSG00000188021 |
| HAPLN3 | -0.795017919 | 0.576336015 | 5.19E-39 | 5.60E-37 | 2002.041685 | 1153.916347 | ENSG00000140511 |
| ACAT2 | -0.796665183 | 0.575678332 | 6.30E-29 | 3.77E-27 | 2588.908921 | 1490.282975 | ENSG00000120437 |
| TAF5 | -0.797152667 | 0.575483844 | 5.78E-13 | 1.01E-11 | 615.1066301 | 354.064677 | ENSG00000148835 |
| MORN2 | -0.797758385 | 0.575242277 | 2.88E-03 | 1.08E-02 | 80.00131665 | 46.09711008 | ENSG00000188010 |
| KLRG1 | -0.798504696 | 0.574944779 | 2.31E-02 | 6.32E-02 | 53.53271206 | 30.64895981 | ENSG00000139187 |
| MMP2 | -0.798711334 | 0.574862436 | 1.37E-28 | 8.03E-27 | 2059.175227 | 1184.447491 | ENSG00000087245 |
| C1R | -0.799066897 | 0.574720774 | 6.63E-04 | 2.94E-03 | 123.5642702 | 70.83961589 | ENSG00000159403 |
| VASH1 | -0.802181899 | 0.573481201 | 4.10E-11 | 5.85E-10 | 429.1902212 | 246.1883598 | ENSG00000071246 |
| ABCF2 | -0.803618647 | 0.572910369 | 4.06E-11 | 5.80E-10 | 404.9709311 | 231.9931732 | ENSG00000033050 |
| CCDC15 | -0.804034986 | 0.57274506 | 7.57E-03 | 2.49E-02 | 67.21448653 | 38.45100706 | ENSG00000149548 |
| SUSD4 | -0.804076743 | 0.572728482 | 4.46E-03 | 1.59E-02 | 80.41069215 | 45.9788101 | ENSG00000143502 |
| MTA2 | -0.804588456 | 0.572525376 | 8.83E-56 | 1.79E-53 | 4418.582343 | 2529.381625 | ENSG00000149480 |
| TP53BP1 | -0.804667735 | 0.572493916 | 2.35E-37 | 2.27E-35 | 2709.455172 | 1550.675821 | ENSG00000067369 |
| TMEM99 | -0.805070178 | 0.572334239 | 6.52E-04 | 2.90E-03 | 125.6296463 | 71.86644781 | ENSG00000167920 |
| BRD9 | -0.805185077 | 0.572288659 | 1.61E-21 | 6.04E-20 | 1119.104012 | 640.0301513 | ENSG00000028310 |
| SOAT1 | -0.805361155 | 0.572218817 | 1.99E-04 | 1.00E-03 | 125.5054824 | 71.78606466 | ENSG00000057252 |
| RTL5 | -0.805366749 | 0.572216598 | 5.32E-04 | 2.42E-03 | 127.1149135 | 72.629007 | ENSG00000242732 |
| LTBP2 | -0.806699454 | 0.571688251 | 4.30E-02 | 1.06E-01 | 44.66803049 | 25.67280396 | ENSG00000119681 |
| DNAJB6 | -0.806742908 | 0.571671032 | 1.63E-63 | 4.08E-61 | 8716.27997 | 4982.263466 | ENSG00000105993 |
| WWOX | -0.808145579 | 0.57111549 | 1.54E-06 | 1.17E-05 | 214.8419437 | 122.7450287 | ENSG00000186153 |
| SLC24A3 | -0.80840361 | 0.571013354 | 1.33E-04 | 6.96E-04 | 128.1336726 | 73.16809961 | ENSG00000185052 |
| KCNQ4 | -0.808831011 | 0.570844215 | 1.30E-04 | 6.81E-04 | 133.7984646 | 76.45835867 | ENSG00000117013 |
| UVRAG | -0.808920405 | 0.570808845 | 8.70E-12 | 1.34E-10 | 430.6212219 | 245.8000983 | ENSG00000198382 |
| HIPK2 | -0.809883702 | 0.570427839 | 8.05E-22 | 3.13E-20 | 905.8026171 | 516.6645381 | ENSG00000064393 |
| ACAA1 | -0.811334607 | 0.569854454 | 2.32E-12 | 3.80E-11 | 593.3636574 | 337.97795 | ENSG00000060971 |
| ZNF594 | -0.812186117 | 0.569518212 | 1.90E-05 | 1.19E-04 | 210.4571554 | 120.1699735 | ENSG00000180626 |
| DLD | -0.812574311 | 0.569364989 | 1.83E-32 | 1.34E-30 | 1859.892505 | 1059.623743 | ENSG00000091140 |
| RND1 | -0.812602762 | 0.569353761 | 2.17E-06 | 1.61E-05 | 226.8433504 | 129.1742068 | ENSG00000172602 |
| SLC4A4 | -0.813117149 | 0.569150797 | 2.54E-02 | 6.84E-02 | 42.08804935 | 23.94553745 | ENSG00000080493 |
| FOXM1 | -0.813406887 | 0.569036505 | 1.34E-38 | 1.40E-36 | 3102.913947 | 1765.751622 | ENSG00000111206 |
| LBH | -0.814163059 | 0.568738329 | 1.13E-22 | 4.68E-21 | 1182.675694 | 672.4080135 | ENSG00000213626 |
| ATAD3B | -0.81422116 | 0.568715425 | 3.01E-23 | 1.29E-21 | 2689.424273 | 1529.436312 | ENSG00000160072 |

| | | | | | | | |
|---|---|---|---|---|---|---|---|
| EPHB2 | -0.814297388 | 0.568685377 | 7.19E-20 | 2.39E-18 | 900.0553614 | 512.2368891 | ENSG00000133216 |
| RET | -0.816937639 | 0.567645587 | 1.34E-09 | 1.60E-08 | 420.7700839 | 238.8662672 | ENSG00000165731 |
| FRY | -0.817400647 | 0.56746344 | 4.06E-05 | 2.38E-04 | 180.8782584 | 102.764911 | ENSG00000073910 |
| DUSP4 | -0.820655936 | 0.566184463 | 1.10E-10 | 1.51E-09 | 401.3336818 | 227.4544815 | ENSG00000120875 |
| CFAP298 | -0.820800535 | 0.566127718 | 1.47E-03 | 6.01E-03 | 91.36830965 | 51.67312607 | ENSG00000159079 |
| SDF2L1 | -0.822653862 | 0.565400921 | 3.71E-19 | 1.15E-17 | 1683.48562 | 951.9637137 | ENSG00000128228 |
| SKIV2L | -0.822696928 | 0.565384043 | 6.13E-27 | 3.32E-25 | 1344.477928 | 760.4267717 | ENSG00000204351 |
| BRICD5 | -0.823043412 | 0.565248275 | 1.38E-03 | 5.71E-03 | 89.85241777 | 50.80652742 | ENSG00000182685 |
| SCYL2 | -0.823135032 | 0.565212379 | 6.88E-16 | 1.63E-14 | 707.2513772 | 400.034315 | ENSG00000136021 |
| PM20D2 | -0.824338465 | 0.5647411 | 1.70E-07 | 1.52E-06 | 280.6856042 | 158.8083899 | ENSG00000146281 |
| RUBCNL | -0.825576984 | 0.564256492 | 2.42E-03 | 9.33E-03 | 83.12787897 | 46.88950472 | ENSG00000102445 |
| KHK | -0.826111362 | 0.564047528 | 2.24E-07 | 1.95E-06 | 277.5013088 | 156.4578795 | ENSG00000138030 |
| SEPTIN3 | -0.826961588 | 0.563715215 | 5.35E-35 | 4.37E-33 | 1714.337982 | 966.3104823 | ENSG00000100167 |
| PIANP | -0.827083739 | 0.563667488 | 7.71E-07 | 6.18E-06 | 237.8516925 | 134.0836644 | ENSG00000139200 |
| ZNF181 | -0.827477519 | 0.563513657 | 5.58E-04 | 2.53E-03 | 131.4861859 | 74.06695686 | ENSG00000197841 |
| TCEAL2 | -0.827564067 | 0.563479853 | 1.25E-10 | 1.71E-09 | 395.8398233 | 222.9674462 | ENSG00000184905 |
| ARMC7 | -0.828215603 | 0.563225437 | 3.70E-09 | 4.19E-08 | 329.5074779 | 185.3515848 | ENSG00000125449 |
| COMMD9 | -0.828282367 | 0.563199373 | 9.32E-10 | 1.13E-08 | 367.9708682 | 207.1974482 | ENSG00000110442 |
| FZD2 | -0.829698022 | 0.562647001 | 8.66E-18 | 2.44E-16 | 1468.615104 | 826.5750231 | ENSG00000180340 |
| EPB41L1 | -0.829921349 | 0.56255991 | 6.44E-16 | 1.54E-14 | 927.6571279 | 521.5710758 | ENSG00000088367 |
| FOXI3 | -0.830397634 | 0.56237422 | 2.22E-05 | 1.37E-04 | 172.9182453 | 97.29915628 | ENSG00000214336 |
| C1QTNF1 | -0.830844408 | 0.562200091 | 1.36E-03 | 5.61E-03 | 92.21752352 | 51.8957565 | ENSG00000173918 |
| EFHD2 | -0.831302031 | 0.562021789 | 4.51E-23 | 1.89E-21 | 1352.550643 | 759.9038476 | ENSG00000142634 |
| ETHE1 | -0.83154625 | 0.561926658 | 2.74E-03 | 1.04E-02 | 76.72503361 | 43.12250757 | ENSG00000105755 |
| ERBB2 | -0.832013255 | 0.56174479 | 1.64E-12 | 2.74E-11 | 13599.06045 | 7639.18773 | ENSG00000141736 |
| SLC30A3 | -0.836920156 | 0.559837427 | 2.33E-09 | 2.71E-08 | 312.9601104 | 175.25974 | ENSG00000115194 |
| DZIP3 | -0.837022376 | 0.559797762 | 5.09E-16 | 1.23E-14 | 599.190522 | 335.3687277 | ENSG00000198919 |
| RADX | -0.838488926 | 0.559228997 | 4.47E-07 | 3.72E-06 | 232.0257648 | 129.8609018 | ENSG00000147231 |
| SCN9A | -0.839142256 | 0.558975805 | 3.53E-03 | 1.30E-02 | 77.9979409 | 43.52475066 | ENSG00000169432 |
| CLCN2 | -0.839387183 | 0.558880916 | 1.04E-07 | 9.52E-07 | 263.2801988 | 147.2261816 | ENSG00000114859 |
| FKBP7 | -0.839772895 | 0.558731516 | 1.40E-02 | 4.18E-02 | 50.67202514 | 28.30379244 | ENSG00000079150 |
| PCSK1N | -0.839818266 | 0.558713945 | 4.05E-04 | 1.91E-03 | 151.3269453 | 84.74815733 | ENSG00000102109 |
| RGS7BP | -0.840027457 | 0.558632937 | 1.95E-02 | 5.50E-02 | 53.42270279 | 29.89367638 | ENSG00000186479 |
| SLC25A48 | -0.840153523 | 0.558584125 | 2.21E-02 | 6.09E-02 | 43.57949882 | 24.33460452 | ENSG00000145832 |
| FOXO6 | -0.840587747 | 0.558416027 | 8.88E-05 | 4.86E-04 | 132.3277908 | 73.97150759 | ENSG00000204060 |
| HPCA | -0.840686951 | 0.55837763 | 2.82E-02 | 7.47E-02 | 47.91381376 | 26.86555158 | ENSG00000121905 |
| ZNF69 | -0.840771932 | 0.55834474 | 2.84E-04 | 1.38E-03 | 110.3547858 | 61.51731499 | ENSG00000198429 |
| MAPK4 | -0.84404572 | 0.557079171 | 4.44E-06 | 3.12E-05 | 209.5859266 | 116.5662206 | ENSG00000141639 |
| ATP10A | -0.848228709 | 0.5554663 | 1.88E-10 | 2.51E-09 | 357.8720014 | 198.7011455 | ENSG00000206190 |
| XKR7 | -0.848233765 | 0.555464353 | 1.59E-07 | 1.42E-06 | 284.6198661 | 158.0031223 | ENSG00000260903 |
| SLC2A8 | -0.848840301 | 0.555230875 | 2.47E-19 | 7.80E-18 | 842.7686576 | 467.8789192 | ENSG00000136856 |
| TAGLN3 | -0.849276037 | 0.555063204 | 4.13E-04 | 1.94E-03 | 108.9105063 | 60.32671795 | ENSG00000144834 |
| EVL | -0.850378061 | 0.554639373 | 1.70E-23 | 7.42E-22 | 1232.938468 | 683.6436453 | ENSG00000196405 |
| HTR7 | -0.851229272 | 0.554312224 | 6.40E-03 | 2.17E-02 | 62.90689244 | 34.96122872 | ENSG00000148680 |
| POGLUT2 | -0.851235963 | 0.554309653 | 1.64E-15 | 3.72E-14 | 635.7275817 | 352.7100481 | ENSG00000134901 |
| ZDHHC15 | -0.852593469 | 0.55378832 | 7.51E-04 | 3.28E-03 | 115.4053393 | 63.82778219 | ENSG00000102383 |
| MT2A | -0.853404326 | 0.553477154 | 2.10E-17 | 5.72E-16 | 676.8841153 | 374.7704726 | ENSG00000125148 |
| NOVA1 | -0.853441557 | 0.553462871 | 1.06E-05 | 6.92E-05 | 205.0857844 | 113.7155336 | ENSG00000139910 |
| TPR | -0.854039885 | 0.553233382 | 4.48E-50 | 7.51E-48 | 3544.78047 | 1960.445644 | ENSG00000047410 |
| EFNB1 | -0.855428088 | 0.5527013 | 5.80E-11 | 8.17E-10 | 449.2117646 | 248.5827177 | ENSG00000090776 |
| GLT1D1 | -0.857534353 | 0.551894972 | 2.71E-10 | 3.56E-09 | 341.5204485 | 188.6579703 | ENSG00000151948 |
| TRIL | -0.858941552 | 0.551356918 | 4.79E-07 | 3.95E-06 | 202.8942029 | 111.7747904 | ENSG00000255690 |
| CCNE2 | -0.863497425 | 0.54961854 | 1.06E-02 | 3.31E-02 | 61.15514938 | 33.56441233 | ENSG00000175305 |
| CEP290 | -0.863864094 | 0.549478869 | 1.40E-06 | 1.07E-05 | 210.3393228 | 115.6439369 | ENSG00000198707 |
| TFE3 | -0.864821767 | 0.549114241 | 4.44E-53 | 8.13E-51 | 3074.174175 | 1687.666838 | ENSG00000068323 |
| SLC16A14 | -0.865080362 | 0.549015825 | 1.75E-03 | 7.03E-03 | 97.88375503 | 53.68701929 | ENSG00000163053 |
| SHD | -0.866318646 | 0.548544799 | 4.74E-03 | 1.68E-02 | 68.65163265 | 37.70406546 | ENSG00000105251 |
| DCTPP1 | -0.866377079 | 0.548522582 | 9.84E-24 | 4.41E-22 | 1065.071953 | 584.3520468 | ENSG00000179958 |
| E2F8 | -0.866380502 | 0.548521281 | 4.61E-08 | 4.45E-07 | 255.8452914 | 140.4211308 | ENSG00000129173 |
| CDH24 | -0.866494925 | 0.548477778 | 1.85E-16 | 4.64E-15 | 669.5313707 | 367.2722751 | ENSG00000139880 |
| TCEA2 | -0.866559138 | 0.548453366 | 6.73E-22 | 2.63E-20 | 867.9733281 | 476.0108835 | ENSG00000171703 |
| TXNDC16 | -0.867243684 | 0.548193192 | 3.14E-09 | 3.58E-08 | 373.0445879 | 204.8275279 | ENSG00000087301 |
| EZH2 | -0.867586761 | 0.548062845 | 1.54E-31 | 1.06E-29 | 1614.973167 | 884.8937786 | ENSG00000106462 |
| GRHL1 | -0.869449209 | 0.54735578 | 1.68E-03 | 6.79E-03 | 86.6094254 | 47.34391303 | ENSG00000134317 |
| MYL5 | -0.869967177 | 0.547159299 | 1.64E-03 | 6.63E-03 | 86.07285656 | 47.01561349 | ENSG00000215375 |
| ZNF578 | -0.870069204 | 0.547120605 | 4.20E-02 | 1.04E-01 | 38.9917114 | 21.28524655 | ENSG00000258405 |
| CUX2 | -0.870372744 | 0.547005505 | 1.20E-09 | 1.44E-08 | 297.6381168 | 162.7797405 | ENSG00000111249 |
| TMEM121B | -0.871509999 | 0.546574478 | 1.76E-06 | 1.32E-05 | 244.5772193 | 133.4668722 | ENSG00000183307 |
| UBE2QL1 | -0.87329974 | 0.545896843 | 3.08E-02 | 8.03E-02 | 37.28191991 | 20.3024985 | ENSG00000215218 |
| GAS1 | -0.873736498 | 0.545731605 | 2.14E-04 | 1.07E-03 | 138.4344612 | 75.53988579 | ENSG00000180447 |
| FIG4 | -0.875072589 | 0.545226433 | 8.88E-18 | 2.50E-16 | 704.6502187 | 384.1589082 | ENSG00000112367 |
| PDRG1 | -0.875816679 | 0.544945297 | 8.39E-22 | 3.25E-20 | 996.3220891 | 542.9869564 | ENSG00000088356 |
| VIPR1 | -0.876326817 | 0.544752638 | 1.28E-03 | 5.31E-03 | 81.41174193 | 44.38677558 | ENSG00000114812 |

| | | | | | | | |
|---|---|---|---|---|---|---|---|
| PIK3R3 | -0.877736813 | 0.544220492 | 5.10E-08 | 4.90E-07 | 278.4512834 | 151.7059716 | ENSG00000117461 |
| MACROH2A2 | -0.878747477 | 0.543839378 | 1.10E-34 | 8.79E-33 | 1862.763278 | 1012.41408 | ENSG00000099284 |
| SIRPA | -0.88007407 | 0.543339535 | 1.86E-08 | 1.91E-07 | 269.8699352 | 146.6787472 | ENSG00000198053 |
| ST8SIA1 | -0.8807146 | 0.543098356 | 3.16E-03 | 1.17E-02 | 74.2960125 | 40.43479225 | ENSG00000111728 |
| ACADSB | -0.880943175 | 0.543012316 | 1.22E-14 | 2.54E-13 | 499.7733787 | 271.5231774 | ENSG00000196177 |
| PROKR1 | -0.881536152 | 0.542789173 | 5.79E-03 | 1.99E-02 | 61.81536971 | 33.65716588 | ENSG00000169618 |
| HM13 | -0.884056922 | 0.541841604 | 9.64E-39 | 1.03E-36 | 2447.71646 | 1326.227092 | ENSG00000101294 |
| H2AW | -0.884142311 | 0.541809536 | 3.21E-11 | 4.65E-10 | 371.4354819 | 201.2377507 | ENSG00000181218 |
| SYPL2 | -0.884335888 | 0.541736842 | 1.27E-03 | 5.28E-03 | 81.63989612 | 44.21200923 | ENSG00000143028 |
| SH2D5 | -0.885662019 | 0.541239104 | 4.38E-20 | 1.49E-18 | 734.6748241 | 397.3210959 | ENSG00000189410 |
| ABHD8 | -0.886276779 | 0.54100852 | 2.91E-07 | 2.50E-06 | 237.7087048 | 128.465182 | ENSG00000127220 |
| CDK5 | -0.886864916 | 0.540788015 | 4.94E-12 | 7.81E-11 | 420.6590724 | 227.3321651 | ENSG00000164885 |
| HDAC9 | -0.88715241 | 0.54068026 | 2.02E-07 | 1.78E-06 | 215.11444 | 116.4277293 | ENSG00000048052 |
| CD2BP2 | -0.88805298 | 0.540342858 | 2.65E-47 | 4.06E-45 | 2389.012872 | 1290.677277 | ENSG00000169217 |
| TMEM109 | -0.888222752 | 0.540279275 | 2.43E-26 | 1.29E-24 | 1046.896895 | 565.7276484 | ENSG00000110108 |
| MAPK8IP1 | -0.892498905 | 0.538680257 | 8.54E-28 | 4.85E-26 | 1268.804771 | 683.5513943 | ENSG00000121653 |
| HEXA | -0.892938345 | 0.538516202 | 3.22E-03 | 1.19E-02 | 75.68374713 | 40.74426967 | ENSG00000213614 |
| JAM2 | -0.897250325 | 0.536909068 | 1.14E-05 | 7.41E-05 | 146.7343767 | 78.67715682 | ENSG00000154721 |
| PRR36 | -0.897531931 | 0.536804276 | 5.43E-15 | 1.17E-13 | 746.922408 | 401.0770491 | ENSG00000183248 |
| WTIP | -0.898615089 | 0.536401401 | 1.19E-12 | 2.01E-11 | 444.9307139 | 238.4180502 | ENSG00000142279 |
| TRAPPC5 | -0.898651683 | 0.536387796 | 7.39E-03 | 2.45E-02 | 51.46677239 | 27.6181514 | ENSG00000181029 |
| NLRP2 | -0.901376036 | 0.535375849 | 2.12E-66 | 5.76E-64 | 3921.792349 | 2099.188997 | ENSG00000022556 |
| SEMA5B | -0.902506645 | 0.534956451 | 1.18E-16 | 3.01E-15 | 806.0041195 | 430.7118384 | ENSG00000082684 |
| DZIP1 | -0.903588705 | 0.534555369 | 7.58E-25 | 3.71E-23 | 1076.872826 | 575.9527925 | ENSG00000134874 |
| COL12A1 | -0.906513035 | 0.533472928 | 9.99E-03 | 3.14E-02 | 50.62784626 | 27.09222269 | ENSG00000111799 |
| CAMK2N2 | -0.908777476 | 0.53263625 | 5.53E-05 | 3.16E-04 | 146.6486097 | 78.20339334 | ENSG00000163888 |
| ATAD1 | -0.909196208 | 0.532481679 | 3.32E-19 | 1.04E-17 | 809.4016776 | 431.2140561 | ENSG00000138138 |
| TP53INP2 | -0.909219959 | 0.532472913 | 2.69E-05 | 1.64E-04 | 146.3878405 | 77.88880285 | ENSG00000078804 |
| HTR1A | -0.909543115 | 0.532353655 | 2.35E-02 | 6.43E-02 | 35.81770374 | 19.06565491 | ENSG00000178394 |
| ZNF585B | -0.909979092 | 0.532192804 | 2.28E-03 | 8.85E-03 | 66.73497505 | 35.49145857 | ENSG00000245680 |
| GPR137C | -0.910235126 | 0.532098364 | 1.23E-03 | 5.15E-03 | 77.79191377 | 41.41647423 | ENSG00000180998 |
| MPPED2 | -0.9109333 | 0.531840925 | 3.97E-03 | 1.43E-02 | 85.91974444 | 45.66692165 | ENSG00000066382 |
| FAAP24 | -0.911607403 | 0.531592479 | 2.47E-04 | 1.22E-03 | 97.11268644 | 51.61853769 | ENSG00000131944 |
| NUCB2 | -0.912543882 | 0.531247524 | 3.12E-09 | 3.56E-08 | 316.5724928 | 168.1764164 | ENSG00000070081 |
| LYPD6B | -0.915018933 | 0.530336911 | 9.05E-03 | 2.89E-02 | 59.88946352 | 31.70161072 | ENSG00000150556 |
| BBS9 | -0.917066697 | 0.529584684 | 6.75E-11 | 9.43E-10 | 455.6435808 | 241.4966679 | ENSG00000122507 |
| SLC16A9 | -0.919567126 | 0.528667621 | 3.39E-10 | 4.40E-09 | 295.9518919 | 156.4355498 | ENSG00000165449 |
| FAM133B | -0.919597088 | 0.528656641 | 4.26E-20 | 1.46E-18 | 798.0155486 | 422.0517401 | ENSG00000234545 |
| ZNF443 | -0.920814837 | 0.528210602 | 3.66E-03 | 1.34E-02 | 82.02240172 | 43.19995876 | ENSG00000180855 |
| NXPH2 | -0.920986276 | 0.528147837 | 2.73E-03 | 1.04E-02 | 62.71155336 | 33.11430516 | ENSG00000144227 |
| GPAM | -0.922454439 | 0.527610639 | 1.64E-21 | 6.14E-20 | 759.7849914 | 400.8740222 | ENSG00000119927 |
| NDUFA9 | -0.923122708 | 0.527366302 | 1.95E-03 | 7.73E-03 | 70.36250019 | 37.06305343 | ENSG00000139180 |
| PDSS2 | -0.923556516 | 0.52720775 | 1.27E-08 | 1.33E-07 | 289.7348488 | 152.9239931 | ENSG00000164494 |
| TMX3 | -0.924208091 | 0.526969697 | 2.38E-10 | 3.15E-09 | 387.0248896 | 204.0356784 | ENSG00000166479 |
| H2BC21 | -0.92579799 | 0.526389278 | 1.30E-03 | 5.42E-03 | 81.20926944 | 42.84045458 | ENSG00000184678 |
| RIMS4 | -0.926197305 | 0.526243603 | 1.90E-49 | 3.07E-47 | 2079.892408 | 1094.579932 | ENSG00000101098 |
| COBLL1 | -0.926357224 | 0.526185273 | 1.95E-03 | 7.73E-03 | 85.20928786 | 45.0221294 | ENSG00000082438 |
| BICC1 | -0.926823415 | 0.52601527 | 2.16E-02 | 6.00E-02 | 37.61689215 | 19.79754249 | ENSG00000122870 |
| PIGL | -0.928259132 | 0.525492059 | 1.14E-08 | 1.20E-07 | 267.1823723 | 140.6305668 | ENSG00000108474 |
| TCEAL3 | -0.931805533 | 0.524201892 | 3.81E-12 | 6.09E-11 | 367.052082 | 192.2313911 | ENSG00000196507 |
| GPX3 | -0.931808887 | 0.524200674 | 3.73E-21 | 1.38E-19 | 722.188838 | 378.5737811 | ENSG00000211445 |
| KCNMA1 | -0.932705441 | 0.523875014 | 3.02E-05 | 1.82E-04 | 132.4470487 | 69.28333552 | ENSG00000156113 |
| OLIG1 | -0.932755769 | 0.523856739 | 2.45E-02 | 6.65E-02 | 36.82473564 | 19.22348924 | ENSG00000184221 |
| NTRK3 | -0.93611986 | 0.522636627 | 2.67E-06 | 1.94E-05 | 179.7217811 | 94.16913048 | ENSG00000140538 |
| ZNF501 | -0.93657721 | 0.522470972 | 2.71E-05 | 1.65E-04 | 143.4846143 | 75.10428118 | ENSG00000186446 |
| VAT1L | -0.936869569 | 0.522365105 | 7.51E-50 | 1.24E-47 | 2706.424152 | 1414.006833 | ENSG00000171724 |
| SMPDL3B | -0.937348136 | 0.522191856 | 9.16E-20 | 3.04E-18 | 607.8998641 | 317.4810269 | ENSG00000130768 |
| FUT8 | -0.941129369 | 0.520825009 | 4.43E-16 | 1.07E-14 | 484.9581299 | 252.7056993 | ENSG00000033170 |
| FIGNL2 | -0.941783667 | 0.520588855 | 2.85E-06 | 2.07E-05 | 160.6870959 | 83.73825743 | ENSG00000261308 |
| SNAP25 | -0.94196628 | 0.520522965 | 4.56E-02 | 1.11E-01 | 54.76499511 | 28.49120177 | ENSG00000132639 |
| MEGF9 | -0.94252898 | 0.520319983 | 5.97E-03 | 2.05E-02 | 58.51061406 | 30.50428938 | ENSG00000106780 |
| PCDHGB4 | -0.942739373 | 0.520244109 | 5.67E-05 | 3.23E-04 | 124.0599266 | 64.56290485 | ENSG00000253953 |
| GLIPR2 | -0.945474692 | 0.519258671 | 1.38E-07 | 1.24E-06 | 219.7867551 | 114.0532595 | ENSG00000122694 |
| TMEM221 | -0.94870306 | 0.518098007 | 1.13E-02 | 3.49E-02 | 46.98935766 | 24.32868583 | ENSG00000188051 |
| ANKRD36 | -0.948796943 | 0.518064293 | 1.37E-03 | 5.66E-03 | 72.68641811 | 37.55080484 | ENSG00000135976 |
| TRPC4 | -0.950712442 | 0.517376904 | 2.11E-09 | 2.46E-08 | 256.2208879 | 132.4394646 | ENSG00000133107 |
| CBLN1 | -0.950984604 | 0.517279311 | 3.90E-08 | 3.82E-07 | 248.2554321 | 128.587238 | ENSG00000102924 |
| SPTB | -0.951161178 | 0.517216004 | 1.71E-03 | 6.87E-03 | 66.15769394 | 34.2610667 | ENSG00000070182 |
| ATF7IP2 | -0.95148435 | 0.517100158 | 6.08E-10 | 7.58E-09 | 265.0356086 | 137.1047739 | ENSG00000166669 |
| RRP36 | -0.952945603 | 0.516576671 | 4.10E-36 | 3.60E-34 | 1648.251751 | 850.9037515 | ENSG00000124541 |
| PINLYP | -0.955510648 | 0.515659038 | 3.20E-03 | 1.18E-02 | 69.05765366 | 35.59040338 | ENSG00000234465 |
| SLC1A2 | -0.956293008 | 0.515379476 | 1.48E-02 | 4.39E-02 | 40.11728854 | 20.66199057 | ENSG00000110436 |
| LNP1 | -0.956315059 | 0.515371599 | 1.30E-07 | 1.18E-06 | 190.0717668 | 97.88585251 | ENSG00000206535 |

| | | | | | | | |
|---|---|---|---|---|---|---|---|
| DANCR | -0.957576277 | 0.514921253 | 6.83E-40 | 7.94E-38 | 1503.515956 | 774.4283635 | ENSG00000226950 |
| WDR73 | -0.959640415 | 0.514185056 | 2.00E-05 | 1.25E-04 | 121.0984038 | 62.22392873 | ENSG00000177082 |
| ZNF704 | -0.960547704 | 0.513861794 | 8.03E-06 | 5.38E-05 | 151.6127573 | 77.84579138 | ENSG00000164684 |
| NCAM1 | -0.961056288 | 0.513680677 | 5.99E-09 | 6.57E-08 | 251.811582 | 129.3500392 | ENSG00000149294 |
| FAM163A | -0.962976299 | 0.5129975 | 4.69E-02 | 1.14E-01 | 29.07968598 | 14.99563206 | ENSG00000143340 |
| CAMK2N1 | -0.963146153 | 0.512937107 | 1.28E-04 | 6.73E-04 | 108.3314734 | 55.46754764 | ENSG00000162545 |
| VEGFC | -0.963803751 | 0.512703357 | 4.22E-02 | 1.04E-01 | 29.04966177 | 14.96820764 | ENSG00000150630 |
| TBC1D9 | -0.96413452 | 0.512585822 | 6.79E-04 | 3.00E-03 | 76.95856945 | 39.52356459 | ENSG00000109436 |
| ZNF823 | -0.966798768 | 0.511640095 | 3.66E-03 | 1.34E-02 | 57.75386341 | 29.55569005 | ENSG00000197933 |
| ZNF649 | -0.968205941 | 0.511141296 | 5.71E-16 | 1.37E-14 | 524.1057075 | 268.0374213 | ENSG00000198093 |
| DHDH | -0.968686226 | 0.510971161 | 3.75E-02 | 9.45E-02 | 28.31775385 | 14.42749749 | ENSG00000104808 |
| PARP16 | -0.97027585 | 0.510408461 | 9.45E-06 | 6.26E-05 | 136.0181553 | 69.36696585 | ENSG00000138617 |
| NAA38 | -0.970311518 | 0.510395842 | 2.52E-13 | 4.59E-12 | 386.7735451 | 197.3954651 | ENSG00000183011 |
| GATD3A | -0.970915859 | 0.510182084 | 3.27E-02 | 8.43E-02 | 29.53878527 | 15.09001526 | ENSG00000160221 |
| C3orf80 | -0.971294673 | 0.510048141 | 9.57E-03 | 3.03E-02 | 43.08112673 | 21.95798415 | ENSG00000180044 |
| EXOG | -0.971668201 | 0.509916101 | 1.46E-08 | 1.52E-07 | 221.9981852 | 113.1823597 | ENSG00000157036 |
| CTXN1 | -0.97355953 | 0.509248055 | 1.42E-14 | 2.93E-13 | 814.2818611 | 414.5855528 | ENSG00000178531 |
| MSI1 | -0.976504541 | 0.508209574 | 9.52E-37 | 8.74E-35 | 1512.235423 | 768.2694925 | ENSG00000135097 |
| TRABD2A | -0.977910608 | 0.507714508 | 5.70E-06 | 3.92E-05 | 171.9347799 | 87.33613434 | ENSG00000186854 |
| DOK3 | -0.978264267 | 0.507590064 | 5.27E-05 | 3.02E-04 | 123.6022668 | 62.63370803 | ENSG00000146094 |
| SOX21 | -0.97872586 | 0.507427685 | 2.31E-03 | 8.95E-03 | 60.97731946 | 31.02376637 | ENSG00000125285 |
| RNF39 | -0.981962333 | 0.506290622 | 2.98E-03 | 1.11E-02 | 60.44401708 | 30.71052084 | ENSG00000204618 |
| LRRC45 | -0.983370486 | 0.505796695 | 1.28E-19 | 4.20E-18 | 919.099873 | 464.8126477 | ENSG00000169683 |
| TSSC2 | -0.983445081 | 0.505770543 | 3.80E-48 | 5.92E-46 | 1907.763826 | 964.7778767 | ENSG00000223756 |
| MAN2A1 | -0.983914488 | 0.505606008 | 1.38E-43 | 1.82E-41 | 1637.025443 | 828.0742372 | ENSG00000112893 |
| TAF2 | -0.986401134 | 0.50473529 | 1.10E-24 | 5.31E-23 | 763.5762798 | 385.6759147 | ENSG00000064313 |
| H2AC11 | -0.989399708 | 0.503687311 | 2.10E-03 | 8.23E-03 | 61.78881213 | 31.14772456 | ENSG00000196787 |
| RAB11FIP3 | -0.990921751 | 0.503156201 | 8.32E-38 | 8.19E-36 | 1427.650312 | 718.3870841 | ENSG00000090565 |
| MAP2 | -0.99333164 | 0.502316427 | 1.34E-07 | 1.21E-06 | 185.147819 | 93.10061363 | ENSG00000078018 |
| ADGRG5 | -0.993719136 | 0.502181527 | 1.43E-11 | 2.16E-10 | 377.2642615 | 189.542325 | ENSG00000159618 |
| RHBDL3 | -0.993955275 | 0.502099337 | 9.96E-18 | 2.78E-16 | 551.583259 | 276.8362472 | ENSG00000141314 |
| ZSWIM5 | -0.994454454 | 0.501925639 | 3.76E-07 | 3.18E-06 | 206.9084882 | 103.986428 | ENSG00000162415 |
| KCNIP3 | -0.994541268 | 0.501895436 | 5.90E-04 | 2.65E-03 | 84.63694974 | 42.46805906 | ENSG00000115041 |
| CNTNAP3B | -0.997301194 | 0.50093621 | 8.98E-03 | 2.87E-02 | 46.13850418 | 23.18218476 | ENSG00000154529 |
| ZHX1 | -0.998033129 | 0.50068213 | 1.39E-04 | 7.24E-04 | 96.79011712 | 48.39549787 | ENSG00000165156 |
| IFT57 | -0.999656285 | 0.500119137 | 5.09E-18 | 1.46E-16 | 590.237971 | 294.9742754 | ENSG00000114446 |
| ZNF778 | -1.000064462 | 0.49997766 | 7.61E-16 | 1.79E-14 | 525.7937594 | 262.6757487 | ENSG00000170100 |
| FAM111B | -1.003014545 | 0.498956329 | 8.67E-17 | 2.23E-15 | 648.4790812 | 323.8166153 | ENSG00000189057 |
| ZNF558 | -1.004549432 | 0.49842577 | 1.06E-08 | 1.13E-07 | 205.7388202 | 102.5509013 | ENSG00000167785 |
| YWHAB | -1.009007662 | 0.496887908 | 7.33E-115 | 8.84E-112 | 6564.644323 | 3261.835308 | ENSG00000166913 |
| PLXNA4 | -1.010408779 | 0.496405574 | 4.10E-03 | 1.48E-02 | 50.84571272 | 25.19690202 | ENSG00000221866 |
| ZNF846 | -1.010701955 | 0.496304708 | 4.71E-04 | 2.17E-03 | 78.10946487 | 38.79005947 | ENSG00000196605 |
| E2F2 | -1.012828048 | 0.495573845 | 9.18E-09 | 9.85E-08 | 232.6948334 | 115.3223801 | ENSG00000007968 |
| PCDHB15 | -1.014811783 | 0.494892889 | 8.65E-05 | 4.74E-04 | 109.1061208 | 53.91907605 | ENSG00000113248 |
| PVALB | -1.016938668 | 0.494163833 | 3.81E-02 | 9.58E-02 | 28.00443311 | 13.81552742 | ENSG00000100362 |
| HID1 | -1.018481213 | 0.49363575 | 2.76E-11 | 4.02E-10 | 268.5998189 | 132.4728077 | ENSG00000167861 |
| IGFBP6 | -1.02030128 | 0.493013385 | 5.82E-09 | 6.39E-08 | 229.2695806 | 112.8723613 | ENSG00000167779 |
| ERICH5 | -1.020671944 | 0.492886734 | 1.39E-02 | 4.17E-02 | 39.43541671 | 19.4735441 | ENSG00000177459 |
| PPP2R5A | -1.023526165 | 0.491912573 | 3.32E-31 | 2.24E-29 | 890.0444519 | 437.7596441 | ENSG00000066027 |
| PLPPR1 | -1.026152758 | 0.491017804 | 5.64E-03 | 1.95E-02 | 71.19485649 | 35.04403502 | ENSG00000148123 |
| MXRA5 | -1.030009256 | 0.489707007 | 3.76E-07 | 3.18E-06 | 152.2112143 | 74.49528572 | ENSG00000101825 |
| KCTD8 | -1.030297264 | 0.489609256 | 6.24E-07 | 5.07E-06 | 144.7402126 | 70.87752061 | ENSG00000183783 |
| ADAMTS17 | -1.034606562 | 0.488148986 | 8.35E-03 | 2.71E-02 | 53.97342631 | 26.25091 | ENSG00000140470 |
| MFGE8 | -1.036818745 | 0.487401047 | 6.89E-190 | 4.43E-186 | 19998.02296 | 9747.035294 | ENSG00000140545 |
| EPHA10 | -1.037187757 | 0.487276396 | 9.42E-03 | 2.99E-02 | 38.53106051 | 18.7922042 | ENSG00000183317 |
| UHRF1 | -1.03725233 | 0.487254586 | 4.18E-40 | 4.95E-38 | 2537.673382 | 1236.097374 | ENSG00000276043 |
| GPR155 | -1.038023491 | 0.486994205 | 7.30E-03 | 2.42E-02 | 43.24173568 | 21.05750936 | ENSG00000163328 |
| EMX1 | -1.039135668 | 0.486618924 | 8.85E-07 | 7.03E-06 | 163.4120166 | 79.35876928 | ENSG00000135638 |
| C9orf24 | -1.040265557 | 0.486237964 | 2.27E-02 | 6.24E-02 | 32.27379354 | 15.67456092 | ENSG00000164972 |
| HUNK | -1.042525522 | 0.485476874 | 7.77E-15 | 1.65E-13 | 403.411536 | 196.0615547 | ENSG00000142149 |
| NGRN | -1.043660481 | 0.485095102 | 1.55E-19 | 5.00E-18 | 566.6746709 | 274.9360802 | ENSG00000182768 |
| ECRG4 | -1.04394317 | 0.48500006 | 2.06E-02 | 5.75E-02 | 31.24598602 | 15.15131582 | ENSG00000119147 |
| EPHB1 | -1.051080182 | 0.482606689 | 5.00E-12 | 7.89E-11 | 285.3030524 | 137.6073188 | ENSG00000154928 |
| TMEM121 | -1.05524221 | 0.481216425 | 7.07E-04 | 3.11E-03 | 69.18668679 | 33.30627611 | ENSG00000184986 |
| TRPM3 | -1.055936272 | 0.480984973 | 9.57E-08 | 8.85E-07 | 194.2886125 | 93.37138072 | ENSG00000083067 |
| RTN4RL2 | -1.057393534 | 0.480499377 | 7.02E-11 | 9.80E-10 | 352.129727 | 169.0395195 | ENSG00000186907 |
| AMBP | -1.058358443 | 0.480178115 | 1.02E-02 | 3.20E-02 | 39.27433221 | 18.91427229 | ENSG00000106927 |
| ITGB1BP2 | -1.06065491 | 0.47941438 | 2.39E-02 | 6.51E-02 | 28.00423295 | 13.41140634 | ENSG00000147166 |
| LIMCH1 | -1.063202754 | 0.478568467 | 2.46E-02 | 6.68E-02 | 28.1920752 | 13.4850773 | ENSG00000064042 |
| SLC25A12 | -1.063981259 | 0.478310292 | 3.30E-11 | 4.78E-10 | 273.9485745 | 131.0974849 | ENSG00000115840 |
| ANGPT1 | -1.064183668 | 0.47824319 | 1.74E-03 | 6.98E-03 | 56.16608378 | 26.87551094 | ENSG00000154188 |
| ALKBH3 | -1.064186693 | 0.478242188 | 5.30E-09 | 5.86E-08 | 194.3629789 | 92.85002573 | ENSG00000166199 |
| MAN2A2 | -1.06600175 | 0.477640889 | 1.23E-16 | 3.13E-15 | 418.8960904 | 199.8350157 | ENSG00000196547 |

| | | | | | | | |
|---|---|---|---|---|---|---|---|
| ABAT | -1.06655774 | 0.47745685 | 3.03E-12 | 4.90E-11 | 292.9608573 | 139.6938169 | ENSG00000183044 |
| NAP1L3 | -1.066743078 | 0.477395516 | 8.64E-46 | 1.24E-43 | 1522.714105 | 726.5604666 | ENSG00000186310 |
| N4BP2L1 | -1.066794256 | 0.477378581 | 3.52E-02 | 8.95E-02 | 28.45930203 | 13.64532268 | ENSG00000139597 |
| PRSS27 | -1.068314523 | 0.4768758 | 2.88E-02 | 7.60E-02 | 30.77185618 | 14.66492148 | ENSG00000172382 |
| EPOP | -1.071851232 | 0.47570819 | 2.19E-06 | 1.62E-05 | 151.7718146 | 72.08882989 | ENSG00000273604 |
| C2orf15 | -1.075730383 | 0.474430813 | 2.50E-02 | 6.77E-02 | 28.26029617 | 13.39688536 | ENSG00000273045 |
| C19orf57 | -1.076745479 | 0.474097116 | 6.93E-05 | 3.88E-04 | 90.97117381 | 43.24672623 | ENSG00000132016 |
| BTBD17 | -1.078033144 | 0.473674153 | 2.11E-16 | 5.26E-15 | 496.418313 | 235.0210375 | ENSG00000204347 |
| ENHO | -1.078984473 | 0.47336191 | 1.18E-07 | 1.08E-06 | 187.7623301 | 88.96579469 | ENSG00000168913 |
| SLC39A1 | -1.079712886 | 0.473122971 | 8.04E-04 | 3.49E-03 | 59.67717886 | 28.25754589 | ENSG00000143570 |
| TIMM23B | -1.07992502 | 0.473053408 | 2.34E-03 | 9.05E-03 | 50.70184919 | 24.01060612 | ENSG00000204152 |
| KIAA0319 | -1.080238945 | 0.472950485 | 4.31E-02 | 1.06E-01 | 23.49256365 | 11.05575867 | ENSG00000137261 |
| ADCYAP1R1 | -1.081470566 | 0.472546902 | 5.37E-31 | 3.57E-29 | 858.8173776 | 406.1120582 | ENSG00000078549 |
| RBM19 | -1.083183873 | 0.471986051 | 2.84E-49 | 4.53E-47 | 1815.013429 | 856.4878126 | ENSG00000122965 |
| H2AC8 | -1.087066838 | 0.470717424 | 2.95E-02 | 7.74E-02 | 27.12479472 | 12.79298454 | ENSG00000277075 |
| HESX1 | -1.087365892 | 0.47061986 | 6.98E-07 | 5.62E-06 | 138.7908847 | 65.33350276 | ENSG00000163666 |
| RTEL1 | -1.091893096 | 0.469145361 | 3.03E-02 | 7.91E-02 | 28.10479347 | 13.20785849 | ENSG00000258366 |
| METTL7A | -1.09385443 | 0.468507994 | 7.31E-33 | 5.45E-31 | 944.8186445 | 442.6637039 | ENSG00000185432 |
| PCLAF | -1.093868743 | 0.468503346 | 4.66E-06 | 3.26E-05 | 118.0340051 | 55.42827374 | ENSG00000166803 |
| MYCNOS | -1.094613148 | 0.468261669 | 8.27E-04 | 3.58E-03 | 65.73611569 | 30.74360348 | ENSG00000233718 |
| PRSS35 | -1.096968149 | 0.46749792 | 1.43E-04 | 7.45E-04 | 85.13010344 | 39.84756299 | ENSG00000146250 |
| ARL4C | -1.097197883 | 0.467423482 | 3.33E-23 | 1.42E-21 | 620.9694641 | 290.5159548 | ENSG00000188042 |
| FXYD6 | -1.09886168 | 0.466884734 | 1.44E-06 | 1.10E-05 | 128.3403754 | 60.01373288 | ENSG00000137726 |
| MAP1A | -1.099968005 | 0.466526842 | 1.55E-06 | 1.18E-05 | 154.3607321 | 72.03209094 | ENSG00000166963 |
| CDK8 | -1.100698849 | 0.466290567 | 3.81E-22 | 1.51E-20 | 714.593722 | 333.6225843 | ENSG00000132964 |
| RBM3 | -1.105297554 | 0.464806596 | 3.75E-50 | 6.41E-48 | 1708.113395 | 794.1079091 | ENSG00000102317 |
| GPR137 | -1.105568701 | 0.464719246 | 3.84E-21 | 1.41E-19 | 672.5157474 | 312.4387605 | ENSG00000173264 |
| PDGFRL | -1.108274919 | 0.463848339 | 8.94E-05 | 4.88E-04 | 80.8975003 | 37.57795669 | ENSG00000104213 |
| CCDC177 | -1.109707635 | 0.463387928 | 1.45E-10 | 1.97E-09 | 253.3846064 | 117.5150804 | ENSG00000267909 |
| EGR1 | -1.113188326 | 0.462271291 | 1.13E-15 | 2.62E-14 | 651.655001 | 300.7782891 | ENSG00000120738 |
| CFAP69 | -1.115615002 | 0.461494385 | 4.64E-03 | 1.65E-02 | 44.94574521 | 20.80341382 | ENSG00000105792 |
| CEP135 | -1.116194071 | 0.461309187 | 1.13E-13 | 2.13E-12 | 302.3812068 | 139.5612565 | ENSG00000174799 |
| ADGRF3 | -1.116387267 | 0.461247416 | 5.31E-05 | 3.04E-04 | 85.98345965 | 39.64967336 | ENSG00000173567 |
| ADAM33 | -1.118188839 | 0.46067179 | 1.24E-14 | 2.59E-13 | 339.8284416 | 156.5984972 | ENSG00000149451 |
| NCAM2 | -1.121626673 | 0.459575351 | 1.76E-02 | 5.05E-02 | 30.92496829 | 14.23847063 | ENSG00000154654 |
| CLIP3 | -1.122488398 | 0.459300928 | 1.42E-15 | 3.24E-14 | 347.3884399 | 159.6623577 | ENSG00000105270 |
| ADPRHL1 | -1.123041784 | 0.459124783 | 9.70E-03 | 3.07E-02 | 33.6514037 | 15.4812329 | ENSG00000153531 |
| ABCA5 | -1.129509552 | 0.457071081 | 2.01E-03 | 7.91E-03 | 50.96860046 | 23.24967658 | ENSG00000154265 |
| MAGEE1 | -1.129525236 | 0.457066112 | 2.58E-14 | 5.22E-13 | 337.2383729 | 154.0003124 | ENSG00000198934 |
| PTN | -1.130338729 | 0.456808459 | 9.44E-32 | 6.60E-30 | 824.227163 | 376.7636657 | ENSG00000105894 |
| FNDC4 | -1.131432723 | 0.456462193 | 4.78E-10 | 6.06E-09 | 211.4025362 | 96.48231182 | ENSG00000115226 |
| DLL1 | -1.133465188 | 0.455819583 | 4.60E-05 | 2.67E-04 | 96.91388074 | 44.21925445 | ENSG00000198719 |
| KRT7 | -1.135940293 | 0.455038244 | 2.12E-11 | 3.14E-10 | 281.7153637 | 128.4613897 | ENSG00000135480 |
| CRELD1 | -1.138356492 | 0.454276792 | 2.28E-02 | 6.26E-02 | 25.30415497 | 11.51419555 | ENSG00000163703 |
| ZNF493 | -1.138577652 | 0.454207158 | 4.07E-08 | 3.97E-07 | 156.9124407 | 71.3596138 | ENSG00000196268 |
| LRP11 | -1.139980309 | 0.453765771 | 3.68E-02 | 9.29E-02 | 21.03459437 | 9.564286502 | ENSG00000120256 |
| TMEM163 | -1.141547743 | 0.453273039 | 6.66E-03 | 2.24E-02 | 36.79750234 | 16.62128225 | ENSG00000152128 |
| LTB4R | -1.143006284 | 0.452815019 | 3.34E-06 | 2.38E-05 | 112.6879523 | 51.03560957 | ENSG00000213903 |
| NSMCE4A | -1.145401763 | 0.45206378 | 1.27E-36 | 1.16E-34 | 992.1474433 | 448.240023 | ENSG00000107672 |
| OVOL2 | -1.149107528 | 0.450904081 | 1.24E-10 | 1.70E-09 | 223.7282398 | 100.7886742 | ENSG00000125850 |
| CRTC3 | -1.150079175 | 0.450600502 | 6.26E-10 | 7.80E-09 | 262.6730936 | 118.6042973 | ENSG00000140577 |
| ARSI | -1.153294526 | 0.449597361 | 7.33E-08 | 6.87E-07 | 181.0812944 | 81.27480219 | ENSG00000183876 |
| SHISA2 | -1.153638563 | 0.44949016 | 3.86E-20 | 1.32E-18 | 440.4465292 | 198.047763 | ENSG00000180730 |
| ZNF428 | -1.153710012 | 0.449467899 | 3.67E-09 | 4.16E-08 | 227.1599134 | 102.2510923 | ENSG00000131116 |
| SEM1 | -1.155305989 | 0.448970952 | 1.74E-54 | 3.32E-52 | 2252.45642 | 1011.127737 | ENSG00000127922 |
| RTF1 | -1.156809793 | 0.448503207 | 1.69E-74 | 6.40E-72 | 2332.565919 | 1046.573749 | ENSG00000137815 |
| PARTICL | -1.156852489 | 0.448489934 | 1.30E-03 | 5.40E-03 | 48.99205769 | 21.9746557 | ENSG00000286532 |
| MTSS1 | -1.159790575 | 0.447577502 | 2.00E-06 | 1.50E-05 | 109.8459258 | 49.11124694 | ENSG00000170873 |
| C18orf32 | -1.16012481 | 0.447473822 | 2.21E-02 | 6.11E-02 | 30.85308054 | 13.79670529 | ENSG00000177576 |
| TSPYL4 | -1.162442992 | 0.44675538 | 1.25E-26 | 6.73E-25 | 884.3885962 | 394.6234638 | ENSG00000187189 |
| SMKR1 | -1.162849779 | 0.446629429 | 3.08E-02 | 8.03E-02 | 24.78779816 | 11.00090983 | ENSG00000240204 |
| CCDC74A | -1.164248415 | 0.446196649 | 9.11E-15 | 1.93E-13 | 305.3047331 | 136.2594205 | ENSG00000163040 |
| KIAA1324 | -1.164402055 | 0.446149134 | 3.93E-03 | 1.42E-02 | 41.86759214 | 18.6341094 | ENSG00000116299 |
| AMOTL1 | -1.164785947 | 0.446030433 | 3.74E-63 | 9.14E-61 | 1742.021149 | 776.9198188 | ENSG00000166025 |
| SCML1 | -1.166653021 | 0.445453572 | 1.05E-03 | 4.46E-03 | 53.71544807 | 23.85089378 | ENSG00000047634 |
| MAPRE3 | -1.167351558 | 0.445237941 | 2.73E-08 | 2.71E-07 | 156.08643 | 69.40137503 | ENSG00000084764 |
| KCNJ12 | -1.172929175 | 0.443519925 | 7.58E-07 | 6.08E-06 | 112.4880706 | 49.89368222 | ENSG00000184185 |
| ARAP2 | -1.173244157 | 0.443423103 | 2.48E-02 | 6.72E-02 | 26.57024115 | 11.72849627 | ENSG00000047365 |
| SFRP1 | -1.173885523 | 0.443226018 | 1.90E-176 | 7.32E-173 | 14611.66082 | 6477.0009 | ENSG00000104332 |
| SLC5A6 | -1.174444284 | 0.443054388 | 7.19E-88 | 4.08E-85 | 4016.68699 | 1779.39431 | ENSG00000138074 |
| KCNAB3 | -1.174654699 | 0.442989774 | 6.54E-04 | 2.90E-03 | 57.07064014 | 25.28992815 | ENSG00000170049 |
| H2BC7 | -1.174659012 | 0.44298845 | 4.28E-02 | 1.05E-01 | 22.24832912 | 9.827517372 | ENSG00000277224 |
| PCGF5 | -1.177824808 | 0.442017438 | 2.88E-06 | 2.08E-05 | 131.2660921 | 58.22998789 | ENSG00000180628 |

| | | | | | | | |
|---|---|---|---|---|---|---|---|
| SEMA6D | -1.178057711 | 0.441946086 | 7.93E-04 | 3.45E-03 | 53.42290295 | 23.59196406 | ENSG00000137872 |
| H3C8 | -1.178346107 | 0.44185775 | 6.34E-04 | 2.83E-03 | 52.75084329 | 23.3104441 | ENSG00000273983 |
| CNTFR | -1.179044314 | 0.44164396 | 1.07E-05 | 6.98E-05 | 113.9394835 | 50.22979055 | ENSG00000122756 |
| ST8SIA2 | -1.181279424 | 0.440960268 | 1.75E-05 | 1.10E-04 | 90.28862626 | 39.9131647 | ENSG00000140557 |
| SMAD9 | -1.181387815 | 0.44092714 | 1.28E-02 | 3.88E-02 | 30.55506576 | 13.4517342 | ENSG00000120693 |
| ATP6V0E2 | -1.183519901 | 0.440275997 | 5.03E-29 | 3.04E-27 | 650.6473701 | 286.4104809 | ENSG00000171130 |
| NOX4 | -1.18378086 | 0.440196366 | 1.75E-02 | 5.04E-02 | 27.02443435 | 11.95596089 | ENSG00000086991 |
| ABCB7 | -1.185879825 | 0.439556393 | 2.54E-38 | 2.58E-36 | 861.409235 | 378.7840291 | ENSG00000131269 |
| H2AX | -1.187971917 | 0.438919443 | 3.20E-57 | 6.71E-55 | 3581.892341 | 1572.475067 | ENSG00000188486 |
| PARP2 | -1.195481511 | 0.436640691 | 9.46E-36 | 8.15E-34 | 763.8033949 | 333.5180239 | ENSG00000129484 |
| SYT9 | -1.201667286 | 0.434772536 | 1.20E-05 | 7.77E-05 | 88.49175319 | 38.52037688 | ENSG00000170743 |
| ZNF442 | -1.203717449 | 0.434155134 | 7.76E-03 | 2.55E-02 | 33.32288915 | 14.43609979 | ENSG00000198342 |
| THBD | -1.203738421 | 0.434148823 | 3.80E-02 | 9.55E-02 | 19.65486903 | 8.540126088 | ENSG00000178726 |
| HSF2BP | -1.204925616 | 0.433791709 | 2.59E-02 | 6.98E-02 | 25.48621525 | 11.0240331 | ENSG00000160207 |
| WDR34 | -1.207738083 | 0.432946876 | 3.79E-45 | 5.34E-43 | 1372.096542 | 594.1941278 | ENSG00000119333 |
| CYP26A1 | -1.208489339 | 0.432721486 | 3.74E-05 | 2.20E-04 | 76.06413756 | 32.92957945 | ENSG00000095596 |
| GRPR | -1.209572722 | 0.432396658 | 2.79E-12 | 4.52E-11 | 232.7315154 | 100.5848537 | ENSG00000126010 |
| FBXO43 | -1.210268718 | 0.432188108 | 1.17E-02 | 3.58E-02 | 28.86481058 | 12.44801345 | ENSG00000156509 |
| ITGB8 | -1.212726079 | 0.431452583 | 2.97E-03 | 1.11E-02 | 43.73241077 | 18.91857344 | ENSG00000105855 |
| SCUBE3 | -1.215222172 | 0.430706747 | 1.18E-05 | 7.62E-05 | 85.43448789 | 36.76594641 | ENSG00000146197 |
| ADAM11 | -1.215966173 | 0.430484688 | 7.32E-14 | 1.40E-12 | 257.9956339 | 111.0410127 | ENSG00000073670 |
| DPH5 | -1.220074291 | 0.429260613 | 2.14E-34 | 1.69E-32 | 734.6060395 | 315.1963666 | ENSG00000117543 |
| CDHR1 | -1.221340673 | 0.428883978 | 9.39E-24 | 4.21E-22 | 528.5570402 | 226.8438685 | ENSG00000148600 |
| DLG1 | -1.225440543 | 0.4276669 | 2.25E-54 | 4.26E-52 | 1224.700416 | 523.922628 | ENSG00000075711 |
| SLC1A4 | -1.227477421 | 0.427063522 | 4.40E-05 | 2.55E-04 | 76.80844839 | 32.83923693 | ENSG00000115902 |
| SLC27A2 | -1.229071148 | 0.426592011 | 3.62E-05 | 2.14E-04 | 81.43530845 | 34.76118852 | ENSG00000140284 |
| PRR22 | -1.229194576 | 0.426555516 | 3.84E-03 | 1.40E-02 | 35.4450103 | 15.1238914 | ENSG00000212123 |
| MYOM2 | -1.231242948 | 0.425950312 | 8.48E-03 | 2.74E-02 | 31.54726727 | 13.4517342 | ENSG00000036448 |
| TM4SF18 | -1.232172176 | 0.425676049 | 3.73E-05 | 2.20E-04 | 80.38258296 | 34.2277236 | ENSG00000163762 |
| CCDC157 | -1.234839608 | 0.424889734 | 3.17E-05 | 1.90E-04 | 76.35704609 | 32.42059487 | ENSG00000187860 |
| JAKMIP2 | -1.238842283 | 0.423712536 | 1.22E-15 | 2.83E-14 | 286.4938964 | 121.5184049 | ENSG00000176049 |
| BMI1 | -1.242253302 | 0.422711919 | 3.57E-03 | 1.31E-02 | 35.66546751 | 15.06904256 | ENSG00000168283 |
| GRM3 | -1.243650413 | 0.422302762 | 1.96E-03 | 7.75E-03 | 41.90232228 | 17.70244208 | ENSG00000198822 |
| C1QTNF2 | -1.25154481 | 0.419998241 | 1.20E-02 | 3.66E-02 | 26.67060151 | 11.24074485 | ENSG00000145861 |
| PCDHB12 | -1.252817626 | 0.419627861 | 5.04E-06 | 3.50E-05 | 90.54979572 | 37.92481789 | ENSG00000120328 |
| CDH7 | -1.255942396 | 0.418719962 | 1.04E-02 | 3.24E-02 | 28.37692639 | 11.93068704 | ENSG00000081138 |
| L3MBTL4 | -1.257467115 | 0.41827767 | 3.81E-03 | 1.38E-02 | 36.23226072 | 15.1658368 | ENSG00000154655 |
| PLAC9 | -1.260499579 | 0.417399397 | 7.47E-05 | 4.16E-04 | 65.48774946 | 27.39524839 | ENSG00000189129 |
| ASXL1 | -1.263587191 | 0.416507047 | 1.17E-39 | 1.35E-37 | 1011.641691 | 421.2857159 | ENSG00000171456 |
| PYGO1 | -1.263940168 | 0.416405154 | 1.83E-02 | 5.20E-02 | 27.48488508 | 11.44912688 | ENSG00000171016 |
| ADAMTS19 | -1.265848679 | 0.415854665 | 5.13E-29 | 3.08E-27 | 575.6324161 | 239.4239094 | ENSG00000145808 |
| SFXN3 | -1.266541045 | 0.415655139 | 2.46E-03 | 9.45E-03 | 40.54791522 | 16.9095144 | ENSG00000107819 |
| TMEFF2 | -1.267603947 | 0.415349019 | 2.53E-03 | 9.70E-03 | 38.08503986 | 15.86223071 | ENSG00000144339 |
| STK32B | -1.271558232 | 0.414212147 | 5.67E-07 | 4.63E-06 | 129.0008712 | 53.64452874 | ENSG00000152953 |
| ISM1 | -1.274289833 | 0.413428619 | 5.67E-03 | 1.96E-02 | 31.01137414 | 12.84353224 | ENSG00000101230 |
| FGFBP3 | -1.27629252 | 0.412855114 | 2.61E-72 | 8.98E-70 | 2172.34029 | 897.3538167 | ENSG00000174721 |
| TTI1 | -1.27827127 | 0.412289244 | 4.19E-45 | 5.86E-43 | 1377.243346 | 567.4143203 | ENSG00000101407 |
| GDPD2 | -1.281849599 | 0.411267907 | 2.54E-19 | 8.01E-18 | 372.1009958 | 152.8825929 | ENSG00000130055 |
| GRIA2 | -1.283245802 | 0.410870085 | 2.87E-02 | 7.58E-02 | 25.74180291 | 10.60995265 | ENSG00000120251 |
| KCND1 | -1.285371387 | 0.410265178 | 3.60E-15 | 7.88E-14 | 305.7524686 | 125.3655368 | ENSG00000102057 |
| ZNF732 | -1.285723188 | 0.410165147 | 8.04E-11 | 1.12E-09 | 180.3235046 | 74.03119061 | ENSG00000186777 |
| NKAIN1 | -1.288443503 | 0.409392477 | 5.20E-61 | 1.22E-58 | 2113.480034 | 865.6952359 | ENSG00000084628 |
| HACL1 | -1.288913778 | 0.409259049 | 3.39E-35 | 2.82E-33 | 722.8876923 | 295.5445606 | ENSG00000131373 |
| KCNC1 | -1.303399519 | 0.405170342 | 3.49E-02 | 8.90E-02 | 18.622719 | 7.553609926 | ENSG00000129159 |
| S1PR1 | -1.304859847 | 0.404760428 | 5.76E-03 | 1.98E-02 | 32.45202378 | 13.07503754 | ENSG00000170989 |
| GALNT16 | -1.304983145 | 0.404725837 | 3.81E-13 | 6.79E-12 | 254.1793906 | 102.7396677 | ENSG00000100626 |
| TEK | -1.308436588 | 0.403758185 | 5.84E-04 | 2.63E-03 | 48.65217937 | 19.62062557 | ENSG00000120156 |
| LTB4R2 | -1.30923759 | 0.403534076 | 2.61E-02 | 7.01E-02 | 19.47001785 | 7.820608906 | ENSG00000213906 |
| ALDH1L1 | -1.309264952 | 0.403526423 | 1.83E-03 | 7.31E-03 | 38.21004279 | 15.38443866 | ENSG00000144908 |
| ADM | -1.309697025 | 0.403405588 | 9.22E-58 | 2.00E-55 | 1154.018813 | 465.7154909 | ENSG00000148926 |
| TTLL7 | -1.312161171 | 0.402717153 | 2.06E-06 | 1.54E-05 | 97.54993407 | 39.25253704 | ENSG00000137941 |
| H2BC15 | -1.313387198 | 0.402375063 | 2.67E-02 | 7.13E-02 | 18.73511886 | 7.521884358 | ENSG00000233822 |
| PDE1B | -1.319586604 | 0.400649726 | 2.42E-04 | 1.20E-03 | 66.08301528 | 26.41439045 | ENSG00000123360 |
| SOX9 | -1.322782857 | 0.39976308 | 9.46E-08 | 8.76E-07 | 121.3690972 | 48.49874383 | ENSG00000125398 |
| ZNF626 | -1.326565804 | 0.398716219 | 6.05E-09 | 6.63E-08 | 134.2155002 | 53.57489846 | ENSG00000188171 |
| CXCL16 | -1.32962311 | 0.397872169 | 2.59E-03 | 9.89E-03 | 39.59168315 | 15.76920458 | ENSG00000161921 |
| DRC3 | -1.331458082 | 0.397366434 | 4.46E-02 | 1.09E-01 | 16.53062206 | 6.543970491 | ENSG00000171962 |
| UGT3A1 | -1.337811165 | 0.39562043 | 1.12E-04 | 5.98E-04 | 68.13443687 | 26.89003192 | ENSG00000145626 |
| ODC1 | -1.34420041 | 0.393872225 | 1.95E-135 | 3.76E-132 | 4470.751577 | 1760.72819 | ENSG00000115758 |
| ELP2 | -1.348747329 | 0.392632818 | 1.50E-91 | 9.62E-89 | 2018.188311 | 792.4878866 | ENSG00000134759 |
| CYP39A1 | -1.350285319 | 0.392214474 | 4.29E-04 | 2.01E-03 | 51.0237428 | 19.96344609 | ENSG00000146233 |
| HGSNAT | -1.371781143 | 0.386413889 | 2.15E-08 | 2.17E-07 | 124.3743234 | 48.13090992 | ENSG00000165102 |
| CPNE9 | -1.383992787 | 0.383156906 | 9.81E-08 | 9.06E-07 | 109.1421271 | 41.81844474 | ENSG00000144550 |

| | | | | | | | |
|---|---|---|---|---|---|---|---|
| ST8SIA5 | -1.396849702 | 0.379757483 | 2.68E-06 | 1.95E-05 | 83.8332293 | 31.87746157 | ENSG00000101638 |
| DYNC2H1 | -1.398415764 | 0.379345475 | 1.19E-21 | 4.51E-20 | 396.6718162 | 150.4279883 | ENSG00000187240 |
| PCDHGB6 | -1.400030613 | 0.378921101 | 2.07E-09 | 2.42E-08 | 141.6353018 | 53.58539087 | ENSG00000253305 |
| PLCL2 | -1.400206049 | 0.378875026 | 3.11E-11 | 4.51E-10 | 211.9608445 | 80.42217937 | ENSG00000154822 |
| LOXL3 | -1.402659762 | 0.378231189 | 1.15E-25 | 5.85E-24 | 441.1027193 | 166.8823008 | ENSG00000115318 |
| ACD | -1.406416582 | 0.377247545 | 8.23E-45 | 1.13E-42 | 808.874982 | 305.0128463 | ENSG00000102977 |
| DGKI | -1.407603898 | 0.376937204 | 1.17E-02 | 3.60E-02 | 23.55472725 | 8.861973909 | ENSG00000157680 |
| SLC7A11 | -1.4108399 | 0.376092672 | 1.14E-22 | 4.70E-21 | 471.0520824 | 177.02791 | ENSG00000151012 |
| TEX13B | -1.412710135 | 0.37560544 | 7.58E-03 | 2.49E-02 | 28.90619856 | 10.80543124 | ENSG00000170925 |
| INCA1 | -1.414188622 | 0.375220714 | 7.09E-03 | 2.36E-02 | 29.75433639 | 11.2031006 | ENSG00000196388 |
| LRRC49 | -1.414651294 | 0.3751004 | 9.88E-14 | 1.87E-12 | 204.6080246 | 76.70574203 | ENSG00000137821 |
| MAP3K7 | -1.416239721 | 0.374687636 | 8.00E-71 | 2.62E-68 | 1310.456561 | 490.9484317 | ENSG00000135341 |
| TTLL11 | -1.416898516 | 0.374516577 | 1.32E-02 | 3.98E-02 | 23.50285138 | 8.786152369 | ENSG00000175764 |
| SLC38A5 | -1.422932604 | 0.372953429 | 1.49E-42 | 1.89E-40 | 1166.717523 | 434.7657291 | ENSG00000017483 |
| RAI2 | -1.424429414 | 0.372566687 | 9.73E-04 | 4.16E-03 | 39.47784384 | 14.69664705 | ENSG00000131831 |
| ZNF521 | -1.430630601 | 0.370968707 | 6.72E-17 | 1.76E-15 | 274.8299278 | 102.1179987 | ENSG00000198795 |
| PDYN | -1.468996357 | 0.361233512 | 2.04E-03 | 8.03E-03 | 38.91595671 | 14.07392411 | ENSG00000101327 |
| SARM1 | -1.47341295 | 0.360129341 | 1.62E-07 | 1.44E-06 | 95.82618806 | 34.58883321 | ENSG00000004139 |
| FEZF1 | -1.47931698 | 0.358658573 | 1.15E-14 | 2.41E-13 | 226.3028393 | 81.2280105 | ENSG00000128610 |
| ARHGAP36 | -1.483384185 | 0.357648877 | 4.47E-03 | 1.59E-02 | 25.70196653 | 9.194041561 | ENSG00000147256 |
| CDYL2 | -1.488955057 | 0.356270502 | 1.44E-09 | 1.72E-08 | 116.2685077 | 41.46325381 | ENSG00000166446 |
| CTNNAL1 | -1.489000575 | 0.356259261 | 1.78E-81 | 7.82E-79 | 1426.73209 | 508.5527104 | ENSG00000119326 |
| SPEF1 | -1.494111208 | 0.354999474 | 3.66E-02 | 9.25E-02 | 15.92331674 | 5.717166665 | ENSG00000101222 |
| DBNDD2 | -1.497039427 | 0.354279667 | 2.50E-03 | 9.61E-03 | 29.68120933 | 10.52982996 | ENSG00000244274 |
| LONRF2 | -1.497533262 | 0.354158418 | 3.71E-02 | 9.36E-02 | 13.88760144 | 4.934731385 | ENSG00000170500 |
| GPR50 | -1.499922863 | 0.353572295 | 3.50E-14 | 6.97E-13 | 197.074696 | 69.6874566 | ENSG00000102195 |
| PAPLN | -1.501110327 | 0.353281393 | 1.46E-11 | 2.20E-10 | 174.5190283 | 61.51033023 | ENSG00000100767 |
| ZNF878 | -1.502684797 | 0.352896054 | 2.07E-05 | 1.29E-04 | 58.57237734 | 20.68941499 | ENSG00000257446 |
| CDK5R2 | -1.503746741 | 0.352636388 | 4.69E-02 | 1.14E-01 | 14.94930012 | 5.254428633 | ENSG00000171450 |
| SPAG1 | -1.510047162 | 0.351099741 | 3.04E-04 | 1.47E-03 | 42.03569791 | 14.75149589 | ENSG00000104450 |
| CACNA1G | -1.514860419 | 0.34993032 | 1.00E-22 | 4.15E-21 | 343.7950813 | 120.2304927 | ENSG00000006283 |
| B4GALNT1 | -1.52263443 | 0.34804978 | 1.73E-04 | 8.83E-04 | 43.40561108 | 15.11313854 | ENSG00000135454 |
| TPP2 | -1.524759584 | 0.347537465 | 1.61E-107 | 1.55E-104 | 1881.721269 | 653.8349137 | ENSG00000134900 |
| LIG3 | -1.525617942 | 0.347330753 | 2.17E-123 | 2.98E-120 | 2315.702901 | 804.1623396 | ENSG00000005156 |
| GRIN2B | -1.526161754 | 0.347199854 | 3.16E-06 | 2.27E-05 | 75.16862963 | 26.21540421 | ENSG00000273079 |
| MID2 | -1.527402032 | 0.346901496 | 2.62E-06 | 1.91E-05 | 73.12474177 | 25.39102354 | ENSG00000080561 |
| CRHR2 | -1.537713701 | 0.344430856 | 2.43E-03 | 9.36E-03 | 29.51924895 | 10.18700944 | ENSG00000106113 |
| PRTG | -1.541394883 | 0.343553126 | 1.22E-08 | 1.28E-07 | 105.3028809 | 36.21259329 | ENSG00000166450 |
| MMUT | -1.541429712 | 0.343544833 | 7.20E-32 | 5.09E-30 | 502.0760085 | 172.7712593 | ENSG00000146085 |
| SRSF12 | -1.543479958 | 0.34305696 | 1.71E-11 | 2.54E-10 | 155.8316061 | 53.43132464 | ENSG00000154548 |
| ADAP2 | -1.548562712 | 0.341850464 | 1.77E-02 | 5.08E-02 | 17.3819511 | 5.967494094 | ENSG00000184060 |
| YAF2 | -1.549917972 | 0.341529482 | 1.05E-02 | 3.28E-02 | 20.99898836 | 7.134967866 | ENSG00000015153 |
| HS6ST3 | -1.554170882 | 0.340524172 | 2.31E-04 | 1.15E-03 | 48.66892479 | 16.69091254 | ENSG00000185352 |
| LRRC4 | -1.554784006 | 0.340379486 | 5.42E-20 | 1.83E-18 | 299.3601111 | 101.8080004 | ENSG00000128594 |
| HRH3 | -1.565106387 | 0.337952786 | 5.86E-13 | 1.02E-11 | 177.45636 | 59.95270489 | ENSG00000101180 |
| FAIM2 | -1.566658177 | 0.337589473 | 1.43E-02 | 4.25E-02 | 18.41342541 | 6.190397102 | ENSG00000135472 |
| ETV2 | -1.573446775 | 0.336004678 | 2.13E-02 | 5.91E-02 | 16.6234856 | 5.553153181 | ENSG00000105672 |
| TMSB15A | -1.579830304 | 0.334521234 | 3.80E-43 | 4.95E-41 | 649.3598694 | 217.0007762 | ENSG00000158164 |
| PCDHB5 | -1.580405174 | 0.334387964 | 1.45E-18 | 4.33E-17 | 262.38414 | 87.6870057 | ENSG00000113209 |
| DPF1 | -1.583232176 | 0.333733363 | 1.91E-25 | 9.57E-24 | 369.2792694 | 123.1728366 | ENSG00000011332 |
| SLC9A3 | -1.595231479 | 0.33096912 | 4.56E-04 | 2.11E-03 | 35.643816 | 11.78979683 | ENSG00000066230 |
| CALCB | -1.604095071 | 0.328941954 | 3.42E-10 | 4.43E-09 | 127.5559159 | 41.90852679 | ENSG00000175868 |
| LRRTM3 | -1.613174721 | 0.326878247 | 9.08E-03 | 2.90E-02 | 22.31029257 | 7.351679617 | ENSG00000198739 |
| BOC | -1.616202274 | 0.326193 | 2.19E-16 | 5.45E-15 | 191.5048827 | 62.52506432 | ENSG00000144857 |
| ABHD16A | -1.617878113 | 0.325814313 | 4.48E-02 | 1.09E-01 | 11.07946611 | 3.598942982 | ENSG00000204427 |
| SCN2B | -1.628687025 | 0.323382379 | 1.60E-02 | 4.67E-02 | 18.47414955 | 6.011590066 | ENSG00000149575 |
| ZNF799 | -1.629455509 | 0.323210168 | 1.68E-04 | 8.63E-04 | 40.64077876 | 13.10461253 | ENSG00000196466 |
| DCLK2 | -1.639300542 | 0.321012072 | 4.22E-02 | 1.04E-01 | 12.41867935 | 4.017585042 | ENSG00000170390 |
| CTU2 | -1.64451489 | 0.319853929 | 2.07E-36 | 1.85E-34 | 532.461292 | 170.3656275 | ENSG00000174177 |
| ZGLP1 | -1.645228567 | 0.319695742 | 3.43E-02 | 8.75E-02 | 15.53734451 | 5.010552925 | ENSG00000220201 |
| NMU | -1.650284826 | 0.318577255 | 7.83E-17 | 2.03E-15 | 217.1967617 | 69.29219828 | ENSG00000109255 |
| H3C10 | -1.650986151 | 0.318422426 | 4.14E-05 | 2.42E-04 | 50.44578597 | 16.04050471 | ENSG00000278828 |
| ZNF433 | -1.655665998 | 0.317391193 | 1.71E-02 | 4.94E-02 | 16.45729485 | 5.185058814 | ENSG00000197647 |
| ABALON | -1.657864239 | 0.316907951 | 5.60E-03 | 1.94E-02 | 24.74872552 | 7.812006611 | ENSG00000281376 |
| DUSP15 | -1.660929563 | 0.316235325 | 5.04E-06 | 3.50E-05 | 67.12477732 | 21.17769944 | ENSG00000149599 |
| HMX2 | -1.667757526 | 0.314742188 | 2.39E-13 | 4.35E-12 | 160.5807903 | 50.48951378 | ENSG00000188816 |
| ZNF880 | -1.680274188 | 0.312023331 | 2.69E-15 | 5.99E-14 | 174.82837 | 54.61411289 | ENSG00000221923 |
| C8orf76 | -1.68890815 | 0.31016157 | 1.74E-02 | 5.00E-02 | 15.85278043 | 4.932580812 | ENSG00000189376 |
| SAMD15 | -1.701158233 | 0.307539104 | 2.86E-02 | 7.56E-02 | 14.99798477 | 4.627404542 | ENSG00000100583 |
| KCNA5 | -1.706901305 | 0.306317288 | 3.37E-02 | 8.64E-02 | 11.75431667 | 3.580120856 | ENSG00000130037 |
| SHISA3 | -1.708814981 | 0.305911239 | 7.15E-09 | 7.78E-08 | 102.8410447 | 31.4846264 | ENSG00000178343 |
| CCDC163 | -1.714976474 | 0.304607533 | 4.97E-05 | 2.86E-04 | 43.30525072 | 13.21592775 | ENSG00000280670 |
| SMPD1 | -1.7187171 | 0.303818768 | 8.69E-06 | 5.79E-05 | 53.17867906 | 16.20935238 | ENSG00000166311 |

| | | | | | | | |
|---|---|---|---|---|---|---|---|
| ZNF559 | -1.722949279 | 0.302928815 | 3.83E-16 | 9.32E-15 | 190.1529543 | 57.74574412 | ENSG00000188321 |
| H2BC9 | -1.737118078 | 0.299968294 | 1.97E-08 | 2.00E-07 | 89.62271198 | 26.91153766 | ENSG00000275713 |
| CREG2 | -1.741756839 | 0.299005341 | 2.87E-02 | 7.58E-02 | 13.07120269 | 3.874544257 | ENSG00000175874 |
| H2BC5 | -1.743004855 | 0.298746796 | 3.16E-10 | 4.11E-09 | 110.8423577 | 33.11430516 | ENSG00000158373 |
| H2BC11 | -1.74977209 | 0.297348749 | 1.82E-08 | 1.87E-07 | 88.97664633 | 26.36222522 | ENSG00000124635 |
| GRIN1 | -1.751049688 | 0.297085544 | 6.92E-14 | 1.33E-12 | 158.4438387 | 47.1535291 | ENSG00000176884 |
| DPYSL5 | -1.761041031 | 0.295035195 | 7.08E-86 | 3.80E-83 | 1112.929223 | 328.3805688 | ENSG00000157851 |
| NFIX | -1.776385692 | 0.291913798 | 2.21E-02 | 6.11E-02 | 12.40472483 | 3.624216828 | ENSG00000008441 |
| TNFAIP8L3 | -1.776943211 | 0.291801011 | 2.76E-02 | 7.36E-02 | 15.67542607 | 4.587609718 | ENSG00000183578 |
| ZNF677 | -1.778031989 | 0.291580877 | 1.10E-29 | 6.89E-28 | 404.3751144 | 118.0479817 | ENSG00000197928 |
| PCDHGA1 | -1.778927585 | 0.291399926 | 8.72E-03 | 2.81E-02 | 20.32501367 | 5.904575996 | ENSG00000204956 |
| CCDC173 | -1.77920322 | 0.291344258 | 2.00E-02 | 5.63E-02 | 17.18015434 | 5.075621597 | ENSG00000154479 |
| THSD7A | -1.786261415 | 0.289922375 | 4.36E-05 | 2.54E-04 | 43.26876882 | 12.62493038 | ENSG00000005108 |
| PDE10A | -1.787139715 | 0.289745927 | 2.72E-05 | 1.66E-04 | 47.49610246 | 13.69130876 | ENSG00000112541 |
| SLC1A6 | -1.79250447 | 0.288670489 | 2.18E-24 | 1.02E-22 | 296.6230245 | 85.75941431 | ENSG00000105143 |
| CLEC4G | -1.794677643 | 0.288235983 | 2.74E-03 | 1.04E-02 | 24.83022529 | 7.124214997 | ENSG00000182566 |
| RDX | -1.813992801 | 0.284402726 | 9.63E-131 | 1.55E-127 | 1712.504403 | 487.00101 | ENSG00000137710 |
| ADM5 | -1.827817246 | 0.281690488 | 2.86E-08 | 2.84E-07 | 76.73041525 | 21.57053462 | ENSG00000224420 |
| LHX2 | -1.846287389 | 0.278107125 | 3.02E-02 | 7.90E-02 | 10.66837575 | 2.98267176 | ENSG00000106689 |
| AMH | -1.84871305 | 0.277639925 | 1.14E-25 | 5.79E-24 | 305.3995116 | 84.63847176 | ENSG00000104899 |
| H4C8 | -1.853134744 | 0.276790293 | 8.57E-05 | 4.70E-04 | 40.30002455 | 11.15847159 | ENSG00000158406 |
| ZNF440 | -1.862705264 | 0.274960205 | 1.69E-08 | 1.75E-07 | 116.1945416 | 31.9296268 | ENSG00000171295 |
| FOSB | -1.880070851 | 0.271670374 | 1.25E-04 | 6.61E-04 | 34.87821709 | 9.450820711 | ENSG00000125740 |
| CBLN3 | -1.883114754 | 0.271097788 | 3.13E-02 | 8.12E-02 | 11.98805267 | 3.224396894 | ENSG00000139899 |
| EFCAB6 | -1.893631213 | 0.269128818 | 1.18E-02 | 3.62E-02 | 14.57919742 | 3.920790803 | ENSG00000186976 |
| WASHC5 | -1.895381518 | 0.268802504 | 5.74E-69 | 1.76E-66 | 944.7733527 | 254.2506817 | ENSG00000164961 |
| BAALC | -1.908429197 | 0.266382424 | 1.25E-04 | 6.59E-04 | 32.08403627 | 8.554647066 | ENSG00000164929 |
| CHRDL2 | -1.926774105 | 0.263016624 | 7.84E-03 | 2.57E-02 | 14.97912417 | 3.946064649 | ENSG00000054938 |
| PTGER2 | -1.955429189 | 0.257844077 | 4.50E-07 | 3.73E-06 | 53.43406656 | 13.76282915 | ENSG00000125384 |
| ZNF596 | -1.95630218 | 0.2576881 | 2.19E-02 | 6.04E-02 | 15.07669363 | 3.865941962 | ENSG00000172748 |
| TTBK1 | -1.973276827 | 0.254673927 | 7.17E-09 | 7.78E-08 | 79.54345667 | 20.19065024 | ENSG00000146216 |
| PHGDH | -2.011864276 | 0.247952508 | 0.00E+00 | 0.00E+00 | 15479.36997 | 3837.192421 | ENSG00000092621 |
| ATOH8 | -2.027799159 | 0.245228887 | 1.98E-03 | 7.80E-03 | 25.18984009 | 6.118876709 | ENSG00000168874 |
| NKX2-5 | -2.043504096 | 0.242573844 | 4.72E-02 | 1.14E-01 | 8.091185514 | 1.945607904 | ENSG00000183072 |
| TMEM215 | -2.050239531 | 0.241443992 | 6.88E-03 | 2.31E-02 | 14.96516965 | 3.622066255 | ENSG00000188133 |
| ALDH3A1 | -2.061612621 | 0.239548117 | 9.72E-03 | 3.07E-02 | 13.65386544 | 3.260423609 | ENSG00000108602 |
| FABP7 | -2.083014704 | 0.236020699 | 1.26E-18 | 3.78E-17 | 181.4238756 | 42.66381022 | ENSG00000164434 |
| BEX5 | -2.102996595 | 0.232774254 | 1.38E-04 | 7.21E-04 | 28.12153889 | 6.512244923 | ENSG00000184515 |
| SLITRK6 | -2.105917018 | 0.23230353 | 3.07E-02 | 8.00E-02 | 11.13584774 | 2.568330847 | ENSG00000184564 |
| ZNF737 | -2.123946876 | 0.229418418 | 2.00E-19 | 6.39E-18 | 167.2739534 | 38.33485766 | ENSG00000237440 |
| ADRA1D | -2.128651433 | 0.228671515 | 1.30E-02 | 3.93E-02 | 12.98950277 | 2.989123481 | ENSG00000171873 |
| H4C9 | -2.153463353 | 0.224772376 | 1.00E-13 | 1.89E-12 | 112.3559342 | 25.27272356 | ENSG00000276180 |
| EDN3 | -2.173058692 | 0.221740055 | 3.31E-02 | 8.50E-02 | 8.858023732 | 1.973032325 | ENSG00000124205 |
| LAMP5 | -2.247141394 | 0.210641062 | 6.77E-06 | 4.60E-05 | 35.94059149 | 7.572432051 | ENSG00000125869 |
| DGKK | -2.247417989 | 0.210600682 | 7.81E-76 | 3.07E-73 | 686.9909915 | 144.4914262 | ENSG00000274588 |
| DAZL | -2.268480988 | 0.207548299 | 3.70E-02 | 9.35E-02 | 7.971088672 | 1.674307777 | ENSG00000092345 |
| H4C11 | -2.290091266 | 0.20446258 | 5.37E-03 | 1.87E-02 | 14.36711293 | 2.915452515 | ENSG00000197238 |
| SLC30A10 | -2.307600084 | 0.20199618 | 3.13E-02 | 8.14E-02 | 8.115627918 | 1.625910657 | ENSG00000196660 |
| FAM135B | -2.330336032 | 0.198837802 | 9.17E-03 | 2.92E-02 | 11.63980164 | 2.341126692 | ENSG00000147724 |
| RSPH4A | -2.336367624 | 0.19800824 | 7.55E-03 | 2.49E-02 | 13.33563862 | 2.658673365 | ENSG00000111834 |
| FAM133A | -2.340078312 | 0.197499607 | 3.25E-04 | 1.56E-03 | 21.70944494 | 4.314159016 | ENSG00000179083 |
| PCDHGA3 | -2.353599774 | 0.195657216 | 1.71E-02 | 4.93E-02 | 9.955128266 | 1.924635205 | ENSG00000254245 |
| FOXB2 | -2.357413323 | 0.195140709 | 1.32E-03 | 5.48E-03 | 18.3645406 | 3.548395288 | ENSG00000204612 |
| FRZB | -2.374626534 | 0.192826263 | 3.38E-23 | 1.44E-21 | 171.8173987 | 33.04278477 | ENSG00000162998 |
| ANKS4B | -2.387977377 | 0.191050061 | 4.86E-02 | 1.17E-01 | 6.961265867 | 1.354610529 | ENSG00000175311 |
| DLK2 | -2.43896646 | 0.184415719 | 1.17E-02 | 3.60E-02 | 10.8317756 | 2.021429444 | ENSG00000171462 |
| FBP1 | -2.440643761 | 0.184201439 | 1.70E-06 | 1.28E-05 | 37.34734997 | 6.861517164 | ENSG00000165140 |
| UNC45B | -2.478887251 | 0.17938271 | 3.48E-02 | 8.87E-02 | 7.443568256 | 1.356761103 | ENSG00000141161 |
| ZNF439 | -2.531089133 | 0.173008025 | 2.02E-02 | 5.67E-02 | 9.715810463 | 1.72485547 | ENSG00000171291 |
| POU3F4 | -2.537650793 | 0.172222938 | 8.01E-05 | 4.42E-04 | 24.64604982 | 4.215214203 | ENSG00000196767 |
| HAS1 | -2.551964067 | 0.170522727 | 5.66E-07 | 4.62E-06 | 38.57416336 | 6.609039162 | ENSG00000105509 |
| TMEM114 | -2.59262 | 0.165784381 | 3.94E-04 | 1.86E-03 | 21.77575089 | 3.603244129 | ENSG00000232258 |
| H2AC6 | -2.623929908 | 0.162225227 | 1.86E-65 | 4.84E-63 | 538.4221469 | 87.44474752 | ENSG00000180573 |
| ZNF229 | -2.652417923 | 0.159053285 | 1.13E-11 | 1.72E-10 | 76.1245494 | 12.07561793 | ENSG00000278318 |
| WDR97 | -2.667434499 | 0.157406334 | 2.94E-03 | 1.10E-02 | 16.49414016 | 2.572631995 | ENSG00000179698 |
| H2BC12 | -2.694606297 | 0.154469478 | 3.87E-49 | 6.12E-47 | 359.5234224 | 55.659246 | ENSG00000197903 |
| SHPK | -2.720038553 | 0.151770305 | 2.22E-19 | 7.07E-18 | 123.7391091 | 18.88684787 | ENSG00000197417 |
| ZNF568 | -2.723909565 | 0.151363623 | 5.40E-07 | 4.43E-06 | 43.37578703 | 6.5477386 | ENSG00000198453 |
| ZNF44 | -2.924483703 | 0.131717259 | 2.72E-14 | 5.48E-13 | 79.8295441 | 10.55080266 | ENSG00000197857 |
| TMED6 | -2.958869802 | 0.128614945 | 6.79E-03 | 2.28E-02 | 10.3250308 | 1.354610529 | ENSG00000157315 |
| GPR85 | -3.053427429 | 0.120455533 | 4.42E-02 | 1.08E-01 | 5.432295355 | 0.643695642 | ENSG00000164604 |
| MT1M | -3.077049882 | 0.118499273 | 3.00E-05 | 1.81E-04 | 22.23784123 | 2.637700666 | ENSG00000205364 |
| SIAH3 | -3.083499196 | 0.117970726 | 1.12E-02 | 3.48E-02 | 8.276036702 | 0.963392889 | ENSG00000215475 |

| | | | | | | | |
|---|---|---|---|---|---|---|---|
| CFAP65 | -3.085457994 | 0.117810661 | 1.10E-02 | 3.40E-02 | 8.450600159 | 1.009639435 | ENSG00000181378 |
| ZNF491 | -3.132989181 | 0.113992501 | 3.16E-02 | 8.19E-02 | 5.731661579 | 0.641545068 | ENSG00000177599 |
| ZNF829 | -3.185737216 | 0.109899961 | 1.05E-08 | 1.11E-07 | 41.89741619 | 4.63170569 | ENSG00000185869 |
| TBXT | -3.214318303 | 0.107744169 | 2.84E-02 | 7.52E-02 | 6.052679304 | 0.639394495 | ENSG00000164458 |
| MAP6 | -3.371370854 | 0.096630949 | 7.58E-17 | 1.97E-15 | 104.9437785 | 10.05472153 | ENSG00000171533 |
| NUTM2B | -3.434925975 | 0.092466464 | 1.88E-02 | 5.34E-02 | 7.163062631 | 0.666818915 | ENSG00000188199 |
| ORM2 | -3.444997815 | 0.091823179 | 2.34E-05 | 1.44E-04 | 21.50764817 | 1.998306171 | ENSG00000228278 |
| MTTP | -3.560234162 | 0.08477401 | 9.10E-03 | 2.90E-02 | 7.707328464 | 0.641545068 | ENSG00000138823 |
| ZNF441 | -3.730096107 | 0.075357969 | 9.79E-05 | 5.31E-04 | 17.41544194 | 1.30621341 | ENSG00000197044 |
| PITX3 | -3.77006356 | 0.073298955 | 4.89E-04 | 2.25E-03 | 13.46622335 | 0.986516162 | ENSG00000107859 |
| ORM1 | -3.833869398 | 0.070127816 | 5.38E-04 | 2.45E-03 | 13.93927715 | 0.959091742 | ENSG00000229314 |
| ZFP28 | -4.009342624 | 0.062096569 | 1.76E-04 | 8.96E-04 | 15.93659554 | 1.009639435 | ENSG00000196867 |
| ZNF563 | -4.052224135 | 0.060278021 | 3.74E-10 | 4.80E-09 | 43.66745627 | 2.637700666 | ENSG00000188868 |
| PKIB | -4.663384566 | 0.039462206 | 1.54E-73 | 5.50E-71 | 383.1093379 | 15.1868095 | ENSG00000135549 |
| GDF7 | -4.753744957 | 0.03706638 | 1.37E-06 | 1.05E-05 | 26.61701078 | 0.984365589 | ENSG00000143869 |
| NKX3-2 | -5.47425816 | 0.022494901 | 3.41E-09 | 3.87E-08 | 43.93333166 | 1.009639435 | ENSG00000109705 |
| ZNF354C | -5.52761245 | 0.021678182 | 1.24E-06 | 9.60E-06 | 30.25223291 | 0.639394495 | ENSG00000177932 |
| ZNF502 | -5.649082319 | 0.019927682 | 2.35E-05 | 1.45E-04 | 17.72808696 | 0.344971094 | ENSG00000196653 |
| TCEAL5 | -5.707550587 | 0.019136216 | 1.70E-15 | 3.86E-14 | 85.69602077 | 1.623760083 | ENSG00000204065 |
| DMRT2 | -5.741603223 | 0.018689824 | 1.81E-05 | 1.14E-04 | 18.93691562 | 0.319697247 | ENSG00000173253 |
| ZNF595 | -6.122191821 | 0.014356105 | 1.15E-11 | 1.75E-10 | 68.80418119 | 1.009639435 | ENSG00000272602 |
| ZNF736 | -6.303595468 | 0.012659854 | 5.13E-20 | 1.74E-18 | 129.6285887 | 1.623760083 | ENSG00000234444 |
| ZNF572 | -6.761092111 | 0.009219523 | 3.97E-07 | 3.32E-06 | 19.66045084 | 0 | ENSG00000180938 |

| **Table S4. Significantly dysregulated genes expressed in 3 biological replicates of SF3B1$^{K700E}$ compared to SF3B1$^{WT}$ ES cells (K700E ES line 2).** Genes with fold change (FC) ≥1.5 and p-value < 0.05 are shown. Up- and downregulated genes begin in rows 4 and 1363, respectively. | | | | | | | |
|---|---|---|---|---|---|---|---|
| **Gene Symbol** | **log2 FC** | **FC** | **p-value** | **padj** | **WT Basemean** | **MT Basemean** | **Ensembl ID** |
| CRYZ | 8.242771432 | 302.9154665 | 4.02E-29 | 4.57E-27 | 1.279874314 | 400.6412587 | ENSG00000116791 |
| ZXDA | 8.190694607 | 292.1761499 | 3.54E-11 | 8.44E-10 | 0 | 53.38454532 | ENSG00000198205 |
| HORMAD1 | 8.146699087 | 283.4006165 | 1.66E-11 | 4.11E-10 | 0.319968578 | 100.866641 | ENSG00000143452 |
| ZNF717 | 7.745476067 | 214.5955081 | 7.43E-10 | 1.45E-08 | 0 | 39.22583136 | ENSG00000227124 |
| CPA4 | 7.287506785 | 156.2277348 | 2.12E-03 | 1.07E-02 | 0 | 28.55344195 | ENSG00000128510 |
| KHDC3L | 7.092427378 | 136.4688032 | 5.14E-08 | 7.50E-07 | 0 | 24.94765513 | ENSG00000203908 |
| TYW3 | 6.463206412 | 88.23055235 | 5.68E-25 | 5.06E-23 | 2.050290591 | 175.4681275 | ENSG00000162623 |
| ZNF248 | 6.129825244 | 70.02631396 | 4.86E-15 | 1.84E-13 | 1.283433786 | 92.51505542 | ENSG00000198105 |
| PRAME | 6.044934624 | 66.0247315 | 3.27E-04 | 2.12E-03 | 0.701617244 | 44.91995124 | ENSG00000185686 |
| LTF | 5.942188469 | 61.48610311 | 3.92E-05 | 3.24E-04 | 0 | 11.22692653 | ENSG00000012223 |
| CRYGD | 5.757325535 | 54.09133258 | 1.22E-04 | 8.89E-04 | 0 | 9.872160761 | ENSG00000118231 |
| REG1A | 5.66857426 | 50.86404385 | 1.63E-04 | 1.15E-03 | 0 | 9.295417497 | ENSG00000115386 |
| PRAM1 | 5.664084035 | 50.70598138 | 1.44E-04 | 1.03E-03 | 0 | 9.260893626 | ENSG00000133246 |
| MOV10L1 | 5.394077911 | 42.05128292 | 6.20E-04 | 3.69E-03 | 0 | 7.693519787 | ENSG00000073146 |
| MAGEA6 | 5.393743784 | 42.04154502 | 6.34E-04 | 3.77E-03 | 0 | 7.691344323 | ENSG00000197172 |
| ZNF454 | 5.352430224 | 40.85470201 | 3.84E-06 | 3.98E-05 | 0.670777201 | 27.05318098 | ENSG00000178187 |
| COL3A1 | 5.202709248 | 36.827441 | 3.81E-02 | 1.14E-01 | 0.319968578 | 13.11163692 | ENSG00000168542 |
| KRT81 | 5.13476368 | 35.13321458 | 9.76E-05 | 7.26E-04 | 1.988610503 | 69.83792788 | ENSG00000205426 |
| TDRD1 | 5.132118094 | 35.06884703 | 1.08E-07 | 1.50E-06 | 0.970584153 | 34.68635488 | ENSG00000095627 |
| DPEP3 | 5.10978467 | 34.53014938 | 4.41E-100 | 1.19E-96 | 15.88160346 | 548.4444865 | ENSG00000141096 |
| F7 | 5.093349157 | 34.13900622 | 9.48E-08 | 1.33E-06 | 0.997864724 | 33.92282557 | ENSG00000057593 |
| MAGEA4 | 5.073056547 | 33.66217632 | 1.10E-35 | 1.73E-33 | 5.601980206 | 189.2650926 | ENSG00000147381 |
| GAB4 | 5.070949133 | 33.61304025 | 3.24E-04 | 2.10E-03 | 0.319968578 | 11.96730375 | ENSG00000215568 |
| CYTOR | 5.06928834 | 33.57436805 | 1.36E-05 | 1.25E-04 | 0.670777201 | 22.25741921 | ENSG00000222041 |
| ANKRD1 | 4.960190706 | 31.12907292 | 4.83E-02 | 1.37E-01 | 0.323528051 | 11.08308266 | ENSG00000148677 |
| NPAP1 | 4.852613072 | 28.89229842 | 7.57E-04 | 4.39E-03 | 0.350808622 | 10.28264311 | ENSG00000185823 |
| TBX3 | 4.820399335 | 28.25431553 | 5.67E-05 | 4.51E-04 | 0.643496629 | 18.63377717 | ENSG00000135111 |
| DMRTB1 | 4.763140466 | 27.15489662 | 1.16E-03 | 6.39E-03 | 0.350808622 | 9.673551435 | ENSG00000143006 |
| ICAM2 | 4.420757578 | 21.41808485 | 4.32E-03 | 1.96E-02 | 0.350808622 | 7.621350622 | ENSG00000108622 |
| OR2W3 | 4.358664362 | 20.51581215 | 4.63E-03 | 2.07E-02 | 0.319968578 | 7.302333068 | ENSG00000238243 |
| SIX3 | 4.328593149 | 20.09261103 | 2.35E-05 | 2.06E-04 | 0.990745779 | 19.9421958 | ENSG00000138083 |
| GIP | 4.327452113 | 20.07672595 | 7.03E-04 | 4.12E-03 | 0.643496629 | 13.2016613 | ENSG00000159224 |
| MEG3 | 4.303574019 | 19.74717015 | 8.68E-03 | 3.50E-02 | 0.323528051 | 7.038080968 | ENSG00000214548 |
| IZUMO2 | 4.246100974 | 18.97596007 | 1.69E-08 | 2.70E-07 | 1.968448877 | 37.6242172 | ENSG00000161652 |
| SCG2 | 4.218302549 | 18.61382377 | 2.27E-15 | 8.98E-14 | 4.042460567 | 74.24162961 | ENSG00000171951 |
| TEX101 | 4.155477169 | 17.82063906 | 9.10E-03 | 3.62E-02 | 0.350808622 | 6.350577289 | ENSG00000131126 |
| C11orf16 | 4.154057553 | 17.80311213 | 9.10E-03 | 3.62E-02 | 0.319968578 | 6.34310494 | ENSG00000176029 |
| LGALS12 | 4.149642016 | 17.74870692 | 7.99E-05 | 6.09E-04 | 0.997864724 | 17.64844348 | ENSG00000133317 |
| ANXA1 | 4.100254174 | 17.15139687 | 2.80E-10 | 5.83E-09 | 2.601267089 | 45.25304017 | ENSG00000135046 |
| NPIPA5 | 4.041305562 | 16.46471417 | 1.43E-04 | 1.02E-03 | 0.963465208 | 16.23673679 | ENSG00000183793 |
| KRT17 | 4.038527269 | 16.4330375 | 1.06E-13 | 3.39E-12 | 5.278452155 | 86.97004662 | ENSG00000128422 |
| MLKL | 4.006410435 | 16.07125219 | 5.81E-04 | 3.50E-03 | 3.550494407 | 57.42425013 | ENSG00000168404 |
| DNAJC22 | 3.966491219 | 15.63265827 | 2.48E-04 | 1.66E-03 | 0.997864724 | 15.52530301 | ENSG00000178401 |
| MPO | 3.888265851 | 14.80759925 | 3.52E-03 | 1.65E-02 | 0.701617244 | 9.939978998 | ENSG00000005381 |
| CYP2B6 | 3.851905541 | 14.43906622 | 4.67E-156 | 2.94E-152 | 60.50673429 | 874.1315679 | ENSG00000197408 |
| TCL1B | 3.845138652 | 14.37149909 | 1.08E-06 | 1.24E-05 | 2.012331602 | 28.64876321 | ENSG00000213231 |
| CNTN5 | 3.827175716 | 14.19366952 | 4.73E-03 | 2.11E-02 | 0.639937157 | 9.320010009 | ENSG00000149972 |
| ASB2 | 3.814918256 | 14.07358777 | 1.11E-09 | 2.13E-08 | 2.955635184 | 41.84295216 | ENSG00000100628 |
| SLC22A1 | 3.753544777 | 13.48744131 | 4.93E-03 | 2.18E-02 | 0.674336673 | 8.967698088 | ENSG00000175003 |
| TEX11 | 3.739480999 | 13.35660088 | 6.33E-03 | 2.69E-02 | 0.701617244 | 8.986047756 | ENSG00000120498 |
| DPPA5 | 3.684131461 | 12.85387516 | 4.43E-25 | 3.98E-23 | 10.16338202 | 131.5190824 | ENSG00000203909 |
| EPHA3 | 3.61663793 | 12.26638242 | 2.14E-08 | 3.34E-07 | 2.982915755 | 36.55299915 | ENSG00000044524 |
| EBF2 | 3.600286359 | 12.12813959 | 9.73E-03 | 3.82E-02 | 0.639937157 | 7.951245495 | ENSG00000221818 |
| CER1 | 3.572584253 | 11.89748106 | 3.20E-15 | 1.25E-13 | 6.81661496 | 82.14234499 | ENSG00000147869 |
| GDF15 | 3.552840496 | 11.73576917 | 6.94E-85 | 9.36E-82 | 43.18274214 | 508.061081 | ENSG00000130513 |
| PRSS3 | 3.453886554 | 10.95780214 | 1.45E-02 | 5.29E-02 | 0.674336673 | 7.289563833 | ENSG00000010438 |
| MAS1 | 3.397113581 | 10.53496474 | 1.72E-02 | 6.08E-02 | 0.670777201 | 7.006016107 | ENSG00000130368 |
| S100A2 | 3.391054704 | 10.49081389 | 4.49E-02 | 1.30E-01 | 2.775628934 | 28.90074053 | ENSG00000196754 |
| OR52N2 | 3.385145892 | 10.44793478 | 1.63E-02 | 5.82E-02 | 0.670777201 | 6.933846943 | ENSG00000180988 |
| PKP1 | 3.380436192 | 10.41388295 | 5.36E-20 | 3.20E-18 | 9.943052517 | 103.3975939 | ENSG00000081277 |
| TRDN | 3.376301009 | 10.38407648 | 9.34E-07 | 1.09E-05 | 2.955635184 | 30.84764577 | ENSG00000186439 |
| SCARA5 | 3.370952592 | 10.34565148 | 1.54E-05 | 1.40E-04 | 2.670066121 | 27.4665012 | ENSG00000168079 |
| AGBL4 | 3.368642321 | 10.32909765 | 3.79E-03 | 1.75E-02 | 0.994305252 | 10.2515242 | ENSG00000186094 |
| TRIM58 | 3.343907492 | 10.153516 | 1.35E-23 | 1.09E-21 | 12.29315006 | 124.4638813 | ENSG00000162722 |
| F5 | 3.337622578 | 10.10937974 | 2.32E-02 | 7.75E-02 | 0.639937157 | 6.615775347 | ENSG00000198734 |
| GPAT2 | 3.337487023 | 10.10842991 | 2.84E-09 | 5.06E-08 | 4.297189585 | 43.46603741 | ENSG00000186281 |
| GSTT2 | 3.298223689 | 9.837036047 | 6.26E-03 | 2.67E-02 | 0.96702468 | 9.696252029 | ENSG00000099984 |
| COL8A1 | 3.287789748 | 9.766148723 | 1.46E-06 | 1.63E-05 | 2.986475227 | 29.08797808 | ENSG00000144810 |
| PCDHGB5 | 3.282925969 | 9.733279396 | 6.31E-11 | 1.46E-09 | 6.013273708 | 58.17616708 | ENSG00000276547 |
| PAX7 | 3.28245932 | 9.730131619 | 1.40E-05 | 1.29E-04 | 2.751907835 | 26.09802023 | ENSG00000009709 |
| RASSF6 | 3.269151569 | 9.640791327 | 2.58E-02 | 8.44E-02 | 0.639937157 | 6.31510746 | ENSG00000169435 |

| | | | | | | | |
|---|---|---|---|---|---|---|---|
| STMN2 | 3.267997783 | 9.633084248 | 2.90E-47 | 7.60E-45 | 29.76183306 | 284.7198593 | ENSG00000104435 |
| C1orf87 | 3.267568508 | 9.630218343 | 6.13E-03 | 2.62E-02 | 1.025145295 | 9.667025044 | ENSG00000162598 |
| NECTIN4 | 3.249851878 | 9.5126802 | 1.37E-04 | 9.83E-04 | 1.992169976 | 18.90521808 | ENSG00000143217 |
| GPRC5D | 3.246688378 | 9.491843923 | 2.38E-02 | 7.91E-02 | 0.674336673 | 6.320404345 | ENSG00000111291 |
| MYL7 | 3.206346012 | 9.230098297 | 2.03E-05 | 1.80E-04 | 2.608386034 | 24.33014515 | ENSG00000106631 |
| VAV1 | 3.188480854 | 9.116505065 | 7.47E-39 | 1.33E-36 | 27.81118154 | 251.6480035 | ENSG00000141968 |
| NCF1 | 3.16888659 | 8.993524381 | 2.83E-02 | 9.08E-02 | 0.674336673 | 5.989563514 | ENSG00000158517 |
| TMEM82 | 3.142154648 | 8.828416223 | 1.75E-06 | 1.93E-05 | 3.302884333 | 29.19221211 | ENSG00000162460 |
| TBX1 | 3.137191388 | 8.798096273 | 4.53E-04 | 2.82E-03 | 1.937608833 | 17.32725046 | ENSG00000184058 |
| EVPL | 3.135624518 | 8.788546097 | 1.11E-21 | 7.59E-20 | 15.40862987 | 136.2147931 | ENSG00000167880 |
| FMOD | 3.126126086 | 8.730874029 | 1.08E-02 | 4.17E-02 | 0.994305252 | 8.680745395 | ENSG00000122176 |
| ZNF648 | 3.103317351 | 8.593925982 | 7.98E-04 | 4.61E-03 | 1.665082452 | 14.24053094 | ENSG00000179930 |
| LEFTY2 | 3.089445072 | 8.511686838 | 1.91E-120 | 6.01E-117 | 134.4144698 | 1139.740463 | ENSG00000143768 |
| MCCC2 | 3.08255541 | 8.471135758 | 9.39E-06 | 9.03E-05 | 2.986475227 | 25.25702487 | ENSG00000131844 |
| SRRM5 | 3.073941731 | 8.420709129 | 1.62E-04 | 1.14E-03 | 2.346538071 | 19.59830219 | ENSG00000226763 |
| TTC22 | 3.067111079 | 8.380934272 | 1.37E-18 | 7.21E-17 | 12.94020616 | 108.2753835 | ENSG00000006555 |
| BTBD16 | 3.065053299 | 8.368988698 | 1.53E-02 | 5.55E-02 | 0.997864724 | 8.329379432 | ENSG00000138152 |
| PRPH2 | 3.026024805 | 8.145621647 | 4.57E-04 | 2.85E-03 | 2.015891075 | 16.25291099 | ENSG00000112619 |
| GDA | 3.010693712 | 8.059518846 | 4.21E-03 | 1.92E-02 | 1.31427383 | 10.64800781 | ENSG00000119125 |
| SYT4 | 2.989073274 | 7.939638236 | 4.92E-35 | 7.50E-33 | 27.69702525 | 220.3241896 | ENSG00000132872 |
| CTCFL | 2.939237674 | 7.67005899 | 3.09E-11 | 7.45E-10 | 9.532038347 | 71.92250678 | ENSG00000124092 |
| KIF25 | 2.90523159 | 7.491380444 | 1.00E-03 | 5.61E-03 | 2.019450547 | 14.97003084 | ENSG00000125337 |
| ACTA1 | 2.892002181 | 7.422999034 | 1.41E-163 | 1.33E-159 | 199.4105523 | 1480.560833 | ENSG00000143632 |
| ZIC1 | 2.891364041 | 7.419716382 | 7.17E-07 | 8.50E-06 | 5.080648543 | 37.1673877 | ENSG00000152977 |
| CTSLP2 | 2.87643089 | 7.34331196 | 4.23E-04 | 2.66E-03 | 2.724627264 | 19.69194244 | ENSG00000280913 |
| NINJ2 | 2.870919562 | 7.315312838 | 3.78E-03 | 1.75E-02 | 1.61052131 | 11.99312577 | ENSG00000171840 |
| FRMD1 | 2.820234895 | 7.062773817 | 2.99E-02 | 9.46E-02 | 0.963465208 | 6.92419913 | ENSG00000153303 |
| PLA2G4C | 2.818707936 | 7.055302479 | 1.47E-21 | 9.99E-20 | 19.23312519 | 135.5407211 | ENSG00000105499 |
| DUSP27 | 2.810987537 | 7.017647766 | 1.54E-06 | 1.72E-05 | 4.620717636 | 32.46789315 | ENSG00000198842 |
| TCEA3 | 2.792552031 | 6.928543146 | 2.70E-02 | 8.75E-02 | 1.025145295 | 6.957493496 | ENSG00000204219 |
| UNC5C | 2.78576822 | 6.896040368 | 7.68E-16 | 3.21E-14 | 14.33988118 | 98.38929668 | ENSG00000182168 |
| PPP1R36 | 2.771587809 | 6.828590448 | 2.56E-02 | 8.39E-02 | 1.279874314 | 8.889002534 | ENSG00000165807 |
| RGN | 2.766233541 | 6.803294499 | 3.24E-06 | 3.39E-05 | 5.404176594 | 36.37368545 | ENSG00000130988 |
| UNC93A | 2.751908731 | 6.736077472 | 1.50E-05 | 1.37E-04 | 4.379031299 | 29.18038883 | ENSG00000112494 |
| HLA-DQA1 | 2.746997971 | 6.713187656 | 2.41E-09 | 4.39E-08 | 8.260172702 | 55.51326622 | ENSG00000196735 |
| LAIR1 | 2.689104159 | 6.449128245 | 1.79E-02 | 6.30E-02 | 1.372394445 | 8.667976159 | ENSG00000167613 |
| NTSR2 | 2.687206168 | 6.440649433 | 4.77E-02 | 1.36E-01 | 0.970584153 | 6.340929476 | ENSG00000169006 |
| ANKDD1B | 2.6865147 | 6.43756324 | 7.35E-03 | 3.05E-02 | 1.634242409 | 10.59636377 | ENSG00000189045 |
| RGS4 | 2.664732755 | 6.341098356 | 5.03E-03 | 2.22E-02 | 1.95777046 | 12.49675391 | ENSG00000117152 |
| PRSS21 | 2.658266998 | 6.312742912 | 9.87E-03 | 3.86E-02 | 1.723203067 | 10.62218579 | ENSG00000007038 |
| CFAP300 | 2.643451691 | 6.248247887 | 3.10E-07 | 3.96E-06 | 6.687610381 | 41.53452838 | ENSG00000137691 |
| FAM90A1 | 2.642486171 | 6.244067665 | 1.03E-03 | 5.76E-03 | 2.666506649 | 16.57287451 | ENSG00000171847 |
| SOX15 | 2.563814105 | 5.912687812 | 2.77E-27 | 2.86E-25 | 31.08531076 | 183.8609742 | ENSG00000129194 |
| PLAAT4 | 2.557108771 | 5.885270684 | 7.90E-03 | 3.23E-02 | 1.964889405 | 11.63863838 | ENSG00000133321 |
| SP140L | 2.552631183 | 5.867033312 | 3.78E-09 | 6.62E-08 | 9.530563807 | 56.15319325 | ENSG00000185404 |
| KCNJ9 | 2.5353758 | 5.797278524 | 6.23E-03 | 2.66E-02 | 2.274179566 | 13.32547449 | ENSG00000162728 |
| SLC22A2 | 2.533241911 | 5.788710111 | 8.39E-04 | 4.81E-03 | 2.982915755 | 17.24760895 | ENSG00000112499 |
| AADACL3 | 2.523797682 | 5.750939585 | 1.57E-02 | 5.66E-02 | 1.692363024 | 9.600436312 | ENSG00000188984 |
| VSIG1 | 2.512826304 | 5.707370832 | 1.18E-20 | 7.40E-19 | 25.37776775 | 145.1254347 | ENSG00000101842 |
| CGB2 | 2.510910978 | 5.699798737 | 3.47E-03 | 1.63E-02 | 2.62854766 | 15.06272513 | ENSG00000104818 |
| PLD6 | 2.506477712 | 5.682310676 | 7.29E-06 | 7.19E-05 | 5.911270367 | 33.70972304 | ENSG00000179598 |
| SERPINB9 | 2.50271778 | 5.667520796 | 2.47E-126 | 9.32E-123 | 228.8358408 | 1296.265401 | ENSG00000170542 |
| SEMA3C | 2.493416157 | 5.631097603 | 7.45E-05 | 5.72E-04 | 4.967966786 | 27.97715019 | ENSG00000075223 |
| PIWIL2 | 2.481779171 | 5.585859054 | 5.83E-07 | 7.02E-06 | 8.552860709 | 47.94299254 | ENSG00000197181 |
| LEFTY1 | 2.476284883 | 5.564626602 | 4.82E-89 | 7.00E-86 | 134.7943384 | 748.6697517 | ENSG00000243709 |
| MUC4 | 2.468176593 | 5.533439795 | 4.73E-13 | 1.41E-11 | 18.44491156 | 102.3188309 | ENSG00000145113 |
| C14orf39 | 2.463351171 | 5.514962862 | 7.85E-07 | 9.23E-06 | 7.314504856 | 40.24322991 | ENSG00000179008 |
| GJC2 | 2.45494283 | 5.482913945 | 1.84E-19 | 1.06E-17 | 28.17711778 | 154.2784058 | ENSG00000198835 |
| PTPN7 | 2.44252535 | 5.435924241 | 4.46E-02 | 1.29E-01 | 1.372394445 | 7.317561313 | ENSG00000143851 |
| ATP1A4 | 2.434260738 | 5.404873073 | 5.83E-04 | 3.50E-03 | 3.919100392 | 21.35607796 | ENSG00000132681 |
| AGMO | 2.419654573 | 5.350429003 | 1.56E-02 | 5.64E-02 | 1.995729448 | 10.6545342 | ENSG00000187546 |
| VSIR | 2.408002877 | 5.307391148 | 8.16E-04 | 4.69E-03 | 3.626412384 | 19.29640479 | ENSG00000107738 |
| MEIOB | 2.393883022 | 5.255700344 | 8.99E-25 | 7.79E-23 | 36.66740868 | 191.955045 | ENSG00000162039 |
| CCDC80 | 2.386078927 | 5.227346984 | 5.76E-33 | 8.00E-31 | 51.72611964 | 270.9814435 | ENSG00000091986 |
| IFIT3 | 2.382905685 | 5.215861946 | 1.05E-03 | 5.83E-03 | 3.592012868 | 18.86568087 | ENSG00000119917 |
| TREH | 2.382337974 | 5.213809873 | 4.16E-02 | 1.22E-01 | 1.345113874 | 6.941319293 | ENSG00000118094 |
| NHLRC1 | 2.366367284 | 5.156411085 | 2.03E-08 | 3.18E-07 | 10.55214963 | 54.53975581 | ENSG00000187566 |
| TSKS | 2.364431712 | 5.149497693 | 1.22E-02 | 4.63E-02 | 2.683108803 | 13.74778012 | ENSG00000126467 |
| COLEC11 | 2.360083957 | 5.134002353 | 1.69E-05 | 1.53E-04 | 6.833217114 | 35.37073714 | ENSG00000118004 |
| CHADL | 2.348131329 | 5.091643216 | 8.23E-03 | 3.34E-02 | 2.597707617 | 13.37806449 | ENSG00000100399 |
| MS4A15 | 2.34449367 | 5.078821149 | 2.40E-04 | 1.62E-03 | 5.087767488 | 25.4879826 | ENSG00000166961 |
| TNN | 2.330604741 | 5.030161571 | 4.28E-13 | 1.29E-11 | 20.25827049 | 101.7398168 | ENSG00000120332 |
| HR | 2.30487987 | 4.941263085 | 3.15E-02 | 9.88E-02 | 1.937608833 | 9.705899842 | ENSG00000168453 |
| USH2A | 2.297441709 | 4.915852775 | 4.38E-02 | 1.27E-01 | 1.603402365 | 8.023414659 | ENSG00000042781 |

| | | | | | | | |
|---|---|---|---|---|---|---|---|
| NLRP10 | 2.29560836 | 4.909609772 | 2.73E-02 | 8.84E-02 | 1.964889405 | 9.715547655 | ENSG00000182261 |
| CSRP1 | 2.289339564 | 4.888322825 | 2.38E-05 | 2.08E-04 | 7.318064329 | 35.71077428 | ENSG00000159176 |
| FAT4 | 2.289142662 | 4.887655703 | 5.98E-27 | 6.10E-25 | 48.32507078 | 236.0571827 | ENSG00000196159 |
| GPER1 | 2.27806816 | 4.850280423 | 4.80E-02 | 1.37E-01 | 1.644920826 | 8.010361878 | ENSG00000164850 |
| RERG | 2.277956012 | 4.8499034 | 8.06E-03 | 3.29E-02 | 2.69378722 | 12.93835516 | ENSG00000134533 |
| CFHR3 | 2.276988012 | 4.84665037 | 1.39E-02 | 5.11E-02 | 2.697346693 | 12.95547532 | ENSG00000116785 |
| ADGRD1 | 2.260727234 | 4.792329927 | 9.54E-04 | 5.39E-03 | 5.219136332 | 25.16962746 | ENSG00000111452 |
| CCDC87 | 2.24860945 | 4.752245773 | 1.75E-04 | 1.22E-03 | 5.322334881 | 25.21502865 | ENSG00000182791 |
| CALHM3 | 2.235517461 | 4.709315768 | 2.73E-02 | 8.84E-02 | 1.985051031 | 9.362006229 | ENSG00000183128 |
| ARID5B | 2.233408059 | 4.702435185 | 2.35E-05 | 2.06E-04 | 7.392787098 | 34.46627446 | ENSG00000150347 |
| DENND2D | 2.231034553 | 4.694705153 | 8.13E-07 | 9.55E-06 | 9.905093528 | 46.57791654 | ENSG00000162777 |
| CCDC146 | 2.229280089 | 4.688999381 | 8.54E-03 | 3.45E-02 | 3.251882663 | 15.39886285 | ENSG00000135205 |
| DCC | 2.224090075 | 4.672161278 | 1.11E-05 | 1.05E-04 | 8.670297148 | 40.32143101 | ENSG00000187323 |
| EID3 | 2.218297671 | 4.65344021 | 1.14E-05 | 1.07E-04 | 7.916483025 | 36.91590484 | ENSG00000255150 |
| C2orf80 | 2.215395359 | 4.644088159 | 4.00E-02 | 1.19E-01 | 2.046731118 | 9.381301855 | ENSG00000188674 |
| TSPYL5 | 2.214899656 | 4.642492746 | 9.53E-33 | 1.30E-30 | 58.57860866 | 271.9482596 | ENSG00000180543 |
| IQSEC3 | 2.207068205 | 4.617359954 | 1.17E-17 | 5.64E-16 | 31.47763785 | 145.2867252 | ENSG00000120645 |
| BCAS1 | 2.206151747 | 4.614427753 | 1.80E-03 | 9.28E-03 | 3.949940435 | 18.28676214 | ENSG00000064787 |
| RCSD1 | 2.200358822 | 4.595936363 | 4.94E-05 | 3.97E-04 | 7.052656893 | 32.14792963 | ENSG00000198771 |
| PTH1R | 2.195925489 | 4.581834946 | 2.64E-06 | 2.82E-05 | 10.2393 | 46.99123676 | ENSG00000160801 |
| NR1I2 | 2.193049496 | 4.572710219 | 6.18E-12 | 1.60E-10 | 19.85882446 | 90.75896066 | ENSG00000144852 |
| TLE6 | 2.183425356 | 4.54230744 | 1.96E-09 | 3.62E-08 | 17.24095522 | 78.28757726 | ENSG00000104953 |
| ANGPT4 | 2.17876949 | 4.527672131 | 2.30E-04 | 1.56E-03 | 5.642303459 | 25.48892856 | ENSG00000101280 |
| PF4 | 2.176211243 | 4.519650589 | 5.17E-03 | 2.28E-02 | 3.964178325 | 17.93416667 | ENSG00000163737 |
| DPPA3 | 2.158507305 | 4.464526907 | 1.07E-16 | 4.71E-15 | 34.91781476 | 156.2724791 | ENSG00000187569 |
| GRAMD1C | 2.151527826 | 4.442980551 | 1.43E-04 | 1.03E-03 | 6.705407743 | 29.60080254 | ENSG00000178075 |
| CAV1 | 2.145609117 | 4.424790399 | 1.39E-39 | 2.65E-37 | 78.33779405 | 346.8233624 | ENSG00000105974 |
| OTOF | 2.143141341 | 4.417228124 | 2.15E-03 | 1.08E-02 | 4.277027959 | 18.95932112 | ENSG00000115155 |
| TGFB2 | 2.13689539 | 4.398145677 | 9.75E-04 | 5.49E-03 | 5.818750236 | 25.87018392 | ENSG00000092969 |
| PRDM1 | 2.134544502 | 4.390984683 | 1.18E-02 | 4.50E-02 | 3.071876414 | 13.27572238 | ENSG00000057657 |
| SPAG17 | 2.13089513 | 4.379891496 | 5.25E-03 | 2.30E-02 | 3.629971857 | 15.91961115 | ENSG00000155761 |
| RORC | 2.128186513 | 4.371676095 | 2.03E-05 | 1.80E-04 | 8.52913961 | 37.44346308 | ENSG00000143365 |
| LCN15 | 2.126618439 | 4.366927076 | 1.68E-02 | 5.98E-02 | 2.941397294 | 12.91688407 | ENSG00000177984 |
| IL18 | 2.123688224 | 4.35806654 | 9.10E-03 | 3.62E-02 | 3.571851242 | 15.73093319 | ENSG00000150782 |
| EMP1 | 2.122921725 | 4.355751729 | 9.29E-04 | 5.26E-03 | 5.257095321 | 23.00091785 | ENSG00000134531 |
| SERPINA11 | 2.121971085 | 4.352882524 | 3.46E-02 | 1.06E-01 | 1.985051031 | 8.644329606 | ENSG00000186910 |
| IL1A | 2.114055141 | 4.329064037 | 1.13E-02 | 4.34E-02 | 3.264925345 | 14.21783034 | ENSG00000115008 |
| MECOM | 2.107373181 | 4.309059982 | 2.71E-04 | 1.80E-03 | 7.300266966 | 31.39109467 | ENSG00000085276 |
| RGS7 | 2.102618574 | 4.294882237 | 1.25E-14 | 4.47E-13 | 28.58368275 | 122.4788363 | ENSG00000182901 |
| ANGPTL6 | 2.096425854 | 4.27648611 | 4.37E-02 | 1.27E-01 | 2.050290591 | 8.623804476 | ENSG00000130812 |
| ADGRE5 | 2.094905842 | 4.27198281 | 8.90E-16 | 3.70E-14 | 33.58248958 | 143.1337907 | ENSG00000123146 |
| CDKN1A | 2.088734977 | 4.253749207 | 1.28E-81 | 1.51E-78 | 236.2146949 | 1006.171369 | ENSG00000124762 |
| ARSL | 2.085370671 | 4.243841195 | 1.54E-38 | 2.68E-36 | 88.83301827 | 377.441112 | ENSG00000157399 |
| LMX1B | 2.080108259 | 4.228389446 | 8.27E-03 | 3.36E-02 | 3.302884333 | 13.99027758 | ENSG00000136944 |
| ESPNL | 2.077405769 | 4.220476143 | 1.12E-12 | 3.19E-11 | 25.38488669 | 107.2944734 | ENSG00000144488 |
| PENK | 2.069672156 | 4.197912675 | 4.35E-02 | 1.27E-01 | 1.988610503 | 8.352080027 | ENSG00000181195 |
| TRIM4 | 2.067367409 | 4.191211743 | 1.62E-18 | 8.48E-17 | 37.63651829 | 157.9205131 | ENSG00000146833 |
| RBM43 | 2.066531087 | 4.188782826 | 2.72E-08 | 4.16E-07 | 18.05881369 | 75.26099271 | ENSG00000184898 |
| BEX5 | 2.050392851 | 4.142187475 | 8.38E-14 | 2.71E-12 | 27.84410652 | 115.2430919 | ENSG00000184515 |
| HJV | 2.049417897 | 4.139389186 | 3.46E-02 | 1.06E-01 | 2.748348363 | 11.16563469 | ENSG00000168509 |
| FCGBP | 2.049155144 | 4.138635361 | 2.45E-06 | 2.63E-05 | 12.04790425 | 49.60958707 | ENSG00000275395 |
| ITGA11 | 2.039404556 | 4.110758322 | 1.84E-02 | 6.43E-02 | 2.9900347 | 12.26362071 | ENSG00000137809 |
| TNS4 | 2.037611362 | 4.105652041 | 2.67E-06 | 2.85E-05 | 12.46039296 | 50.70613268 | ENSG00000131746 |
| SLC7A14 | 2.033263819 | 4.093298333 | 1.10E-04 | 8.06E-04 | 7.97460364 | 32.58328804 | ENSG00000013293 |
| TOGARAM2 | 2.028036904 | 4.078495059 | 5.64E-05 | 4.48E-04 | 8.305250636 | 33.78718909 | ENSG00000189350 |
| LGR5 | 2.027574066 | 4.077186825 | 4.35E-02 | 1.27E-01 | 2.254017939 | 9.321239514 | ENSG00000139292 |
| ENPP6 | 2.022244134 | 4.062151745 | 1.80E-02 | 6.31E-02 | 3.04103637 | 12.20667979 | ENSG00000164303 |
| CFAP44 | 2.020261681 | 4.056573649 | 3.48E-03 | 1.63E-02 | 4.871887182 | 19.94966815 | ENSG00000206530 |
| COL13A1 | 2.016506031 | 4.046027236 | 9.27E-45 | 2.22E-42 | 119.6194118 | 484.8510327 | ENSG00000197467 |
| SLC39A4 | 2.013133264 | 4.036579382 | 2.35E-12 | 6.44E-11 | 26.2952653 | 105.8023902 | ENSG00000147804 |
| LOXL4 | 2.012825126 | 4.035717322 | 1.25E-02 | 4.69E-02 | 4.508035878 | 18.41899364 | ENSG00000138131 |
| ABCC3 | 2.012709723 | 4.035394512 | 8.58E-03 | 3.46E-02 | 3.622852912 | 14.66936295 | ENSG00000108846 |
| PRDM16 | 2.011234125 | 4.031269197 | 7.58E-08 | 1.08E-06 | 15.55095646 | 62.68185474 | ENSG00000142611 |
| HCLS1 | 2.010098986 | 4.028098566 | 1.91E-07 | 2.51E-06 | 15.61027228 | 62.77454903 | ENSG00000180353 |
| ADGRG3 | 2.009954151 | 4.027694198 | 3.09E-02 | 9.73E-02 | 3.102716458 | 12.27355207 | ENSG00000182885 |
| PKDREJ | 2.007018056 | 4.019505586 | 2.40E-02 | 7.98E-02 | 2.944956766 | 11.89078366 | ENSG00000130943 |
| C8orf31 | 2.006267454 | 4.017414873 | 1.29E-04 | 9.32E-04 | 8.305250636 | 33.32584072 | ENSG00000177335 |
| PRELP | 1.993122262 | 3.980976243 | 2.06E-02 | 7.04E-02 | 3.044595843 | 11.99530123 | ENSG00000188783 |
| GALR1 | 1.991711383 | 3.977084962 | 2.46E-02 | 8.14E-02 | 2.690227748 | 10.60383612 | ENSG00000166573 |
| FLI1 | 1.984359542 | 3.956869655 | 2.43E-04 | 1.64E-03 | 7.318064329 | 28.87754548 | ENSG00000151702 |
| KCNE3 | 1.981421539 | 3.948819814 | 9.42E-11 | 2.12E-09 | 26.49334824 | 103.9794661 | ENSG00000175538 |
| TRPM2 | 1.976071443 | 3.934203114 | 5.22E-03 | 2.29E-02 | 4.886125072 | 19.39127455 | ENSG00000142185 |
| TUSC8 | 1.974589062 | 3.930162765 | 1.55E-02 | 5.61E-02 | 3.769934186 | 14.57042581 | ENSG00000237361 |
| NLRP1 | 1.972547427 | 3.924604914 | 2.74E-04 | 1.82E-03 | 8.001884211 | 31.30788023 | ENSG00000091592 |

| | | | | | | | |
|---|---|---|---|---|---|---|---|
| HSPA2 | 1.971958662 | 3.923003606 | 3.04E-129 | 1.44E-125 | 652.5103022 | 2558.942635 | ENSG00000126803 |
| DNM3 | 1.961173492 | 3.893785717 | 2.89E-17 | 1.34E-15 | 43.06294144 | 168.2096826 | ENSG00000197959 |
| BTG2 | 1.960814907 | 3.892818028 | 1.45E-23 | 1.16E-21 | 62.66703689 | 244.3399744 | ENSG00000159388 |
| PCDHGA11 | 1.949896276 | 3.863467539 | 5.53E-15 | 2.08E-13 | 36.27836173 | 140.3958828 | ENSG00000253873 |
| SH3TC2 | 1.949651906 | 3.862813183 | 2.33E-02 | 7.78E-02 | 3.368123893 | 12.89446702 | ENSG00000169247 |
| PADI1 | 1.949126924 | 3.861407799 | 4.17E-04 | 2.63E-03 | 7.947323068 | 30.70871992 | ENSG00000142623 |
| COL4A4 | 1.944107823 | 3.847997396 | 1.16E-05 | 1.09E-04 | 11.55357383 | 44.52108126 | ENSG00000081052 |
| TRADD | 1.935112654 | 3.824079864 | 9.53E-03 | 3.76E-02 | 4.031782149 | 15.3064521 | ENSG00000102871 |
| BMPER | 1.928352 | 3.806201658 | 3.27E-02 | 1.02E-01 | 2.917676195 | 11.19892905 | ENSG00000164619 |
| PSTPIP1 | 1.926350058 | 3.800923683 | 4.33E-04 | 2.71E-03 | 7.221984725 | 27.56912685 | ENSG00000140368 |
| PAPPA2 | 1.9236261 | 3.793753914 | 1.95E-05 | 1.75E-04 | 12.65552682 | 48.43508095 | ENSG00000116183 |
| CHODL | 1.92327492 | 3.792830553 | 7.49E-09 | 1.26E-07 | 23.49383033 | 88.52366231 | ENSG00000154645 |
| PDCD1 | 1.921951557 | 3.789353042 | 2.57E-09 | 4.63E-08 | 21.45510788 | 81.42544637 | ENSG00000188389 |
| KCNH3 | 1.917701205 | 3.778205582 | 3.44E-04 | 2.21E-03 | 9.58987963 | 36.2573446 | ENSG00000135519 |
| TDRD10 | 1.917027261 | 3.776441034 | 2.59E-02 | 8.47E-02 | 3.006636854 | 11.30221712 | ENSG00000163239 |
| INMT | 1.916369106 | 3.774718623 | 2.83E-02 | 9.07E-02 | 3.017315271 | 11.32368821 | ENSG00000241644 |
| SIGLEC15 | 1.914226255 | 3.769116153 | 9.74E-03 | 3.82E-02 | 3.946380963 | 14.9183868 | ENSG00000197046 |
| TNFAIP6 | 1.913698405 | 3.76773737 | 1.89E-03 | 9.69E-03 | 5.645862931 | 21.22819738 | ENSG00000123610 |
| ASCL5 | 1.906562899 | 3.749148313 | 2.36E-02 | 7.87E-02 | 4.155142324 | 15.29775025 | ENSG00000232237 |
| DDX60L | 1.900335115 | 3.732998981 | 3.50E-04 | 2.25E-03 | 8.577777016 | 32.03688567 | ENSG00000181381 |
| FOXS1 | 1.893307338 | 3.714858702 | 5.00E-02 | 1.41E-01 | 2.312138554 | 8.588334646 | ENSG00000179772 |
| SNTG1 | 1.890816932 | 3.708451579 | 2.21E-02 | 7.46E-02 | 3.33728385 | 12.33484392 | ENSG00000147481 |
| LGALS1 | 1.890747691 | 3.7082736 | 2.14E-41 | 4.55E-39 | 148.0521489 | 548.4196534 | ENSG00000100097 |
| TPRG1 | 1.890213025 | 3.706899559 | 3.42E-02 | 1.05E-01 | 2.955635184 | 11.00779208 | ENSG00000188001 |
| TBX10 | 1.884805995 | 3.693032592 | 2.92E-05 | 2.50E-04 | 11.3700081 | 41.7558383 | ENSG00000167800 |
| PCDHGC5 | 1.876214551 | 3.671105438 | 1.77E-02 | 6.23E-02 | 3.622852912 | 13.34571608 | ENSG00000240764 |
| SLC25A45 | 1.874888073 | 3.667731612 | 1.61E-02 | 5.77E-02 | 4.249747387 | 15.68988292 | ENSG00000162241 |
| TNFRSF10C | 1.874608587 | 3.667021151 | 1.23E-07 | 1.67E-06 | 18.50659164 | 67.87192465 | ENSG00000173535 |
| KLHDC8A | 1.874218763 | 3.666030434 | 1.14E-18 | 6.14E-17 | 51.38242997 | 188.7865085 | ENSG00000162873 |
| KCNAB2 | 1.861075133 | 3.632782856 | 6.98E-53 | 2.40E-50 | 188.0508915 | 682.0030005 | ENSG00000069424 |
| ZP3 | 1.857380217 | 3.623490765 | 3.63E-03 | 1.69E-02 | 6.767087831 | 24.2987427 | ENSG00000188372 |
| ETV3L | 1.854716857 | 3.616807606 | 5.80E-04 | 3.49E-03 | 8.046962145 | 28.92606809 | ENSG00000253831 |
| C4orf50 | 1.852462135 | 3.611159478 | 1.98E-04 | 1.37E-03 | 10.90415346 | 39.38339042 | ENSG00000181215 |
| ZCCHC12 | 1.85075994 | 3.606901287 | 2.49E-12 | 6.80E-11 | 35.993072 | 129.3797677 | ENSG00000174460 |
| ADGRE2 | 1.848384392 | 3.600967034 | 2.84E-10 | 5.92E-09 | 28.0691907 | 101.163525 | ENSG00000127507 |
| EMP3 | 1.8455264 | 3.593840546 | 2.22E-12 | 6.11E-11 | 45.84271467 | 164.1180931 | ENSG00000142227 |
| LRRC4C | 1.843700397 | 3.589294739 | 2.36E-04 | 1.59E-03 | 9.224833118 | 33.18150239 | ENSG00000148948 |
| DLGAP2 | 1.83122699 | 3.558395798 | 2.38E-05 | 2.08E-04 | 12.57515965 | 44.7293384 | ENSG00000198010 |
| IP6K3 | 1.821405209 | 3.53425273 | 3.19E-02 | 9.99E-02 | 3.251882663 | 11.58576485 | ENSG00000161896 |
| KCNMB1 | 1.821316 | 3.534034197 | 1.48E-04 | 1.05E-03 | 10.50114796 | 37.28155309 | ENSG00000145936 |
| NTN1 | 1.820386654 | 3.531758398 | 2.46E-26 | 2.41E-24 | 87.79983815 | 309.8810685 | ENSG00000065320 |
| RBPJL | 1.818200645 | 3.526411039 | 9.07E-03 | 3.62E-02 | 4.589877592 | 16.26255881 | ENSG00000124232 |
| HLA-DQB1 | 1.818053557 | 3.526051527 | 2.54E-49 | 7.74E-47 | 221.1106011 | 778.7121781 | ENSG00000179344 |
| PDE4B | 1.816713077 | 3.522776816 | 9.44E-42 | 2.03E-39 | 152.4023989 | 536.4951106 | ENSG00000184588 |
| FAM20A | 1.813493858 | 3.514924884 | 3.64E-04 | 2.32E-03 | 8.597938643 | 30.23883765 | ENSG00000108950 |
| CAVIN3 | 1.809808845 | 3.505958321 | 1.53E-34 | 2.29E-32 | 169.7169073 | 596.1848547 | ENSG00000170955 |
| CD79B | 1.809004331 | 3.50400378 | 1.75E-04 | 1.22E-03 | 10.38994074 | 36.18422948 | ENSG00000007312 |
| TNFSF9 | 1.807999747 | 3.501564705 | 1.62E-13 | 5.13E-12 | 40.27810863 | 141.2211984 | ENSG00000125657 |
| NMI | 1.807306059 | 3.499881459 | 5.83E-12 | 1.52E-10 | 36.71902074 | 129.0513859 | ENSG00000123609 |
| IER3 | 1.803241059 | 3.490033922 | 1.70E-53 | 5.93E-51 | 246.7232673 | 861.6564757 | ENSG00000137331 |
| ISLR | 1.793103981 | 3.465597197 | 6.49E-04 | 3.85E-03 | 11.37949131 | 39.2787049 | ENSG00000129009 |
| ABCC2 | 1.789407324 | 3.456728569 | 1.77E-03 | 9.14E-03 | 8.601498115 | 29.77625977 | ENSG00000023839 |
| C6orf223 | 1.789043028 | 3.455855819 | 3.36E-08 | 5.09E-07 | 21.78931435 | 75.40041302 | ENSG00000181577 |
| DEGS2 | 1.784374835 | 3.444691623 | 4.79E-05 | 3.87E-04 | 12.87733087 | 44.38355287 | ENSG00000168350 |
| RASSF9 | 1.784349726 | 3.444631671 | 4.39E-03 | 1.98E-02 | 7.090615882 | 24.21162884 | ENSG00000198774 |
| HAPLN1 | 1.779866217 | 3.433943298 | 3.86E-09 | 6.75E-08 | 27.02505284 | 92.98756466 | ENSG00000145681 |
| PLAT | 1.773315567 | 3.418386601 | 5.24E-34 | 7.67E-32 | 129.2943616 | 441.0563299 | ENSG00000104368 |
| TFF3 | 1.772105455 | 3.415520508 | 2.07E-04 | 1.42E-03 | 11.60101602 | 39.61455907 | ENSG00000160180 |
| PTPRB | 1.772076947 | 3.415453018 | 1.64E-03 | 8.59E-03 | 7.726993566 | 26.19601141 | ENSG00000127329 |
| DPPA2 | 1.770192266 | 3.410994115 | 1.12E-22 | 8.33E-21 | 86.93957149 | 296.291952 | ENSG00000163530 |
| HTR2C | 1.768856825 | 3.407838165 | 6.89E-11 | 1.57E-09 | 33.46861262 | 114.0106263 | ENSG00000147246 |
| MX2 | 1.761473059 | 3.390441281 | 6.55E-15 | 2.44E-13 | 56.91975543 | 192.4488144 | ENSG00000183486 |
| ADAMTSL2 | 1.758137607 | 3.382611774 | 2.25E-11 | 5.47E-10 | 37.79191372 | 127.7190372 | ENSG00000197859 |
| SORBS2 | 1.755580592 | 3.376621784 | 1.02E-72 | 8.34E-70 | 399.728839 | 1351.561604 | ENSG00000154556 |
| BANK1 | 1.751299747 | 3.366617334 | 2.51E-03 | 1.23E-02 | 9.230756855 | 31.10631744 | ENSG00000153064 |
| GPA33 | 1.749985936 | 3.363552872 | 2.16E-02 | 7.31E-02 | 4.351750728 | 14.54677926 | ENSG00000143167 |
| DCST2 | 1.744963456 | 3.351863633 | 3.21E-02 | 1.00E-01 | 4.000942105 | 13.38553684 | ENSG00000163354 |
| FAM124B | 1.742535605 | 3.346227667 | 1.39E-05 | 1.27E-04 | 16.47765789 | 55.27237713 | ENSG00000124019 |
| PADI3 | 1.737388514 | 3.334310628 | 1.62E-09 | 3.02E-08 | 29.31822497 | 97.98243022 | ENSG00000142619 |
| GABRG2 | 1.737320551 | 3.334153558 | 2.35E-03 | 1.17E-02 | 9.852896649 | 32.95707105 | ENSG00000113327 |
| HLA-DMB | 1.736072783 | 3.33127114 | 3.11E-03 | 1.49E-02 | 7.882083508 | 26.37023914 | ENSG00000242574 |
| ALPK1 | 1.735617747 | 3.330220601 | 2.11E-02 | 7.19E-02 | 4.23906897 | 14.20288565 | ENSG00000073331 |
| SYTL2 | 1.735394863 | 3.32970615 | 3.51E-14 | 1.19E-12 | 49.45221941 | 164.4677351 | ENSG00000137501 |
| RIN1 | 1.72876524 | 3.314440239 | 1.95E-11 | 4.78E-10 | 38.83483022 | 128.5521087 | ENSG00000174791 |

| | | | | | | | |
|---|---|---|---|---|---|---|---|
| TEX19 | 1.728474956 | 3.313773407 | 1.57E-02 | 5.65E-02 | 5.171694134 | 17.33160139 | ENSG00000182459 |
| SLC8A3 | 1.727197221 | 3.310839835 | 4.00E-02 | 1.19E-01 | 3.739094142 | 12.21850307 | ENSG00000100678 |
| THPO | 1.726324688 | 3.308838065 | 9.39E-03 | 3.71E-02 | 5.625701305 | 18.6219539 | ENSG00000090534 |
| MYOM1 | 1.724500356 | 3.304656582 | 1.92E-03 | 9.78E-03 | 8.577777016 | 28.34760087 | ENSG00000101605 |
| SERPINE1 | 1.720213905 | 3.294852553 | 7.01E-12 | 1.80E-10 | 44.18564088 | 145.399094 | ENSG00000106366 |
| MSRB3 | 1.716720728 | 3.286884423 | 1.09E-05 | 1.03E-04 | 19.28265232 | 63.69887442 | ENSG00000174099 |
| GABRB2 | 1.715699735 | 3.284559124 | 9.75E-04 | 5.49E-03 | 8.863346079 | 29.23014095 | ENSG00000145864 |
| HCN1 | 1.706084619 | 3.262741339 | 6.52E-11 | 1.50E-09 | 41.97019232 | 137.404201 | ENSG00000164588 |
| IFI35 | 1.704590646 | 3.259364379 | 6.07E-07 | 7.30E-06 | 22.86665652 | 74.51430002 | ENSG00000068079 |
| GDNF | 1.698460854 | 3.245545213 | 3.90E-02 | 1.17E-01 | 3.708254098 | 11.92530753 | ENSG00000168621 |
| ZNF37A | 1.69768527 | 3.243800896 | 3.39E-17 | 1.56E-15 | 61.34920358 | 199.2624641 | ENSG00000075407 |
| DNAH3 | 1.69754156 | 3.24347779 | 1.47E-04 | 1.05E-03 | 12.29670953 | 39.79510227 | ENSG00000158486 |
| PPP1R17 | 1.69483038 | 3.237388218 | 4.65E-07 | 5.74E-06 | 21.77507646 | 70.56461958 | ENSG00000106341 |
| SATB2 | 1.694046079 | 3.235628736 | 4.46E-12 | 1.18E-10 | 47.84022356 | 154.9054683 | ENSG00000119042 |
| MXRA8 | 1.693286071 | 3.233924664 | 3.59E-42 | 7.80E-40 | 168.7899271 | 545.8466614 | ENSG00000162576 |
| PLK2 | 1.692290251 | 3.231693218 | 1.57E-47 | 4.19E-45 | 203.9867768 | 658.4850753 | ENSG00000145632 |
| JSRP1 | 1.688764389 | 3.223804793 | 9.35E-03 | 3.70E-02 | 5.700424074 | 18.25971062 | ENSG00000167476 |
| PLA2G2A | 1.688561664 | 3.223351821 | 3.84E-10 | 7.85E-09 | 33.80637856 | 108.8910016 | ENSG00000188257 |
| OVGP1 | 1.684142869 | 3.213494202 | 3.22E-03 | 1.53E-02 | 8.914347749 | 28.69917774 | ENSG00000085465 |
| DNAH2 | 1.678146958 | 3.200166478 | 5.95E-03 | 2.55E-02 | 7.647516116 | 24.3918885 | ENSG00000183914 |
| KLK8 | 1.673728478 | 3.190380458 | 3.92E-03 | 1.80E-02 | 7.318064329 | 23.3049907 | ENSG00000129455 |
| MAP1LC3C | 1.670287331 | 3.182779762 | 8.16E-04 | 4.69E-03 | 11.44948555 | 36.1724062 | ENSG00000197769 |
| ADAD2 | 1.66783796 | 3.177380706 | 2.25E-02 | 7.56E-02 | 4.589877592 | 14.6606611 | ENSG00000140955 |
| LAMP3 | 1.664023338 | 3.168990508 | 3.88E-02 | 1.16E-01 | 4.011620523 | 12.66445524 | ENSG00000078081 |
| COL6A3 | 1.653688337 | 3.146370004 | 9.44E-23 | 7.02E-21 | 105.4464686 | 331.1029504 | ENSG00000163359 |
| GCNT4 | 1.649575508 | 3.137413117 | 1.22E-09 | 2.31E-08 | 34.63344091 | 108.8091846 | ENSG00000176928 |
| IL32 | 1.644724079 | 3.126880488 | 1.01E-02 | 3.93E-02 | 7.866955895 | 24.30214767 | ENSG00000008517 |
| VIL1 | 1.644276513 | 3.125910588 | 4.97E-34 | 7.40E-32 | 143.1562351 | 447.229601 | ENSG00000127831 |
| LAMA2 | 1.631559543 | 3.098477609 | 3.35E-14 | 1.14E-12 | 70.81422292 | 218.6300921 | ENSG00000196569 |
| SPSB2 | 1.628010855 | 3.090865455 | 6.95E-23 | 5.26E-21 | 113.8288319 | 352.6557458 | ENSG00000111671 |
| DLK1 | 1.622467761 | 3.079012578 | 6.85E-09 | 1.16E-07 | 33.92706898 | 104.7058446 | ENSG00000185559 |
| SLC52A1 | 1.616021655 | 3.06528593 | 3.33E-02 | 1.03E-01 | 4.358869673 | 13.27912735 | ENSG00000132517 |
| OXCT2 | 1.612599292 | 3.058023074 | 1.06E-04 | 7.84E-04 | 17.58344969 | 53.70110387 | ENSG00000198754 |
| CCDC9B | 1.605672156 | 3.043375094 | 1.56E-04 | 1.11E-03 | 15.14558669 | 46.20886333 | ENSG00000188549 |
| EBI3 | 1.597658102 | 3.026516261 | 5.08E-03 | 2.24E-02 | 7.541953304 | 22.93215365 | ENSG00000105246 |
| FOXB1 | 1.596609207 | 3.024316665 | 8.30E-14 | 2.69E-12 | 67.62905378 | 204.768641 | ENSG00000171956 |
| TAFA2 | 1.594130812 | 3.019125675 | 8.41E-03 | 3.41E-02 | 6.578488096 | 19.92072471 | ENSG00000198673 |
| IL6R | 1.589139741 | 3.008698914 | 2.90E-22 | 2.09E-20 | 103.4285182 | 311.7549742 | ENSG00000160712 |
| ADAMTS4 | 1.588677423 | 3.007734919 | 8.87E-40 | 1.71E-37 | 186.043925 | 559.434664 | ENSG00000158859 |
| PCDHGB3 | 1.588048894 | 3.006424844 | 4.19E-02 | 1.23E-01 | 4.297189585 | 12.92000549 | ENSG00000262209 |
| PRTN3 | 1.587465936 | 3.005210266 | 5.00E-02 | 1.41E-01 | 3.657252428 | 10.96920083 | ENSG00000196415 |
| C1orf74 | 1.586555057 | 3.003313456 | 3.73E-24 | 3.14E-22 | 111.6691141 | 335.242443 | ENSG00000162757 |
| NPFFR2 | 1.5844375 | 2.99890849 | 1.38E-18 | 7.26E-17 | 89.19506397 | 266.8363907 | ENSG00000056291 |
| HLA-DRA | 1.582489802 | 2.994862572 | 3.56E-10 | 7.32E-09 | 43.84670589 | 131.0992358 | ENSG00000204287 |
| LCP1 | 1.580921497 | 2.991608727 | 5.11E-08 | 7.48E-07 | 29.97476428 | 89.84172875 | ENSG00000136167 |
| FAM163B | 1.579289796 | 2.988227102 | 1.27E-11 | 3.18E-10 | 51.2839861 | 153.6840909 | ENSG00000196990 |
| TM4SF1 | 1.578284617 | 2.98614582 | 6.96E-03 | 2.91E-02 | 7.552631721 | 22.65078139 | ENSG00000169908 |
| B3GNT3 | 1.577702718 | 2.984941625 | 3.19E-05 | 2.70E-04 | 17.74329431 | 53.14422332 | ENSG00000179913 |
| SLFN13 | 1.575605963 | 2.980606583 | 9.66E-30 | 1.13E-27 | 187.2484662 | 557.6235335 | ENSG00000154760 |
| RNF212 | 1.574079279 | 2.977454122 | 1.34E-13 | 4.29E-12 | 62.26403138 | 185.7867349 | ENSG00000178222 |
| SH2D2A | 1.573718104 | 2.976708818 | 1.45E-20 | 8.99E-19 | 94.43764207 | 280.97602 | ENSG00000027869 |
| TGM2 | 1.570136138 | 2.969327324 | 6.71E-12 | 1.73E-10 | 50.71905104 | 150.5107519 | ENSG00000198959 |
| SCX | 1.56997474 | 2.968995157 | 2.79E-09 | 4.98E-08 | 39.51008277 | 117.5797864 | ENSG00000260428 |
| LPL | 1.56982112 | 2.968679031 | 7.68E-24 | 6.33E-22 | 112.7761011 | 334.4148363 | ENSG00000175445 |
| ADAMTS14 | 1.567287147 | 2.963469372 | 1.90E-10 | 4.06E-09 | 60.56632944 | 179.2383263 | ENSG00000138316 |
| ALOX5 | 1.566774248 | 2.962416002 | 1.38E-07 | 1.87E-06 | 31.6377618 | 93.51438785 | ENSG00000012779 |
| EHBP1L1 | 1.562772636 | 2.954210517 | 2.95E-26 | 2.86E-24 | 149.5523527 | 441.5914464 | ENSG00000173442 |
| RAC2 | 1.559509686 | 2.947536515 | 7.46E-08 | 1.07E-06 | 43.78919566 | 129.5967267 | ENSG00000128340 |
| NRK | 1.55229647 | 2.932836142 | 1.63E-20 | 1.00E-18 | 101.6738647 | 298.0099366 | ENSG00000123572 |
| C16orf89 | 1.54868928 | 2.925512292 | 6.54E-10 | 1.29E-08 | 45.1503013 | 131.9382666 | ENSG00000153446 |
| PDE6G | 1.54821088 | 2.924542349 | 2.43E-12 | 6.66E-11 | 53.30015651 | 155.80144 | ENSG00000185527 |
| TMEM232 | 1.544103647 | 2.91622826 | 8.50E-03 | 3.43E-02 | 7.252824769 | 21.17220242 | ENSG00000186952 |
| FAM66B | 1.540615951 | 2.909186831 | 7.93E-04 | 4.58E-03 | 13.04073496 | 38.18913718 | ENSG00000215374 |
| CMKLR1 | 1.53568115 | 2.89925283 | 1.13E-04 | 8.30E-04 | 21.2703731 | 61.60394234 | ENSG00000174600 |
| FOXA2 | 1.534093661 | 2.896064354 | 2.23E-02 | 7.51E-02 | 5.622141833 | 16.29367771 | ENSG00000125798 |
| LGR6 | 1.533966351 | 2.895808803 | 1.35E-05 | 1.25E-04 | 24.19516824 | 70.01776572 | ENSG00000133067 |
| C12orf56 | 1.527728361 | 2.883314804 | 1.56E-02 | 5.61E-02 | 7.420067669 | 21.22261695 | ENSG00000185306 |
| FAM183A | 1.526664389 | 2.881189175 | 2.62E-02 | 8.55E-02 | 6.238357891 | 18.05797987 | ENSG00000186973 |
| PLD5 | 1.526142424 | 2.880146954 | 1.45E-10 | 3.17E-09 | 54.55750493 | 157.0772993 | ENSG00000180287 |
| GJA5 | 1.525891824 | 2.879646707 | 1.45E-07 | 1.94E-06 | 32.54786108 | 93.57083431 | ENSG00000265107 |
| F10 | 1.52515838 | 2.878183112 | 7.50E-09 | 1.26E-07 | 37.0475828 | 106.6771744 | ENSG00000126218 |
| CFD | 1.524615471 | 2.87710021 | 1.42E-06 | 1.60E-05 | 25.86497922 | 74.34101825 | ENSG00000197766 |
| HPGD | 1.524014624 | 2.875902216 | 3.38E-05 | 2.84E-04 | 18.8063986 | 54.17656657 | ENSG00000164120 |
| VIM | 1.521404914 | 2.870704662 | 1.08E-32 | 1.45E-30 | 210.887955 | 606.0254654 | ENSG00000026025 |

| | | | | | | | |
|---|---|---|---|---|---|---|---|
| INPP5J | 1.520500166 | 2.868904941 | 1.46E-22 | 1.08E-20 | 128.5017505 | 369.324436 | ENSG00000185133 |
| NODAL | 1.516031604 | 2.860032625 | 1.73E-10 | 3.74E-09 | 46.13036866 | 131.7546746 | ENSG00000156574 |
| OR2A7 | 1.515672303 | 2.859320427 | 1.74E-02 | 6.15E-02 | 6.667448754 | 19.02373439 | ENSG00000243896 |
| CYSRT1 | 1.515603657 | 2.859184378 | 3.67E-03 | 1.71E-02 | 10.61027024 | 30.33654528 | ENSG00000197191 |
| ABI3 | 1.511892236 | 2.851838404 | 2.20E-08 | 3.41E-07 | 37.11490729 | 106.1914014 | ENSG00000108798 |
| CABYR | 1.508235967 | 2.844620049 | 2.07E-15 | 8.27E-14 | 81.45830787 | 231.7266163 | ENSG00000154040 |
| CD19 | 1.507901054 | 2.843959763 | 1.99E-02 | 6.84E-02 | 5.962272037 | 16.95441341 | ENSG00000177455 |
| TAGLN | 1.506999555 | 2.842183209 | 9.46E-73 | 8.12E-70 | 536.0808659 | 1522.956167 | ENSG00000149591 |
| KCTD19 | 1.505850647 | 2.839920704 | 2.17E-02 | 7.34E-02 | 6.351039648 | 17.96528557 | ENSG00000168676 |
| MYOCD | 1.499373834 | 2.827199782 | 3.08E-04 | 2.01E-03 | 16.6837495 | 47.37909115 | ENSG00000141052 |
| MN1 | 1.499250315 | 2.826957738 | 2.26E-36 | 3.69E-34 | 201.9053668 | 570.6705759 | ENSG00000169184 |
| LHX4 | 1.498914467 | 2.826299721 | 2.86E-08 | 4.36E-07 | 41.07107695 | 116.0051082 | ENSG00000121454 |
| LRFN5 | 1.49870454 | 2.825888494 | 7.79E-03 | 3.20E-02 | 8.546936973 | 24.22222261 | ENSG00000165379 |
| COL5A2 | 1.498125333 | 2.824754197 | 1.27E-65 | 7.51E-63 | 465.3195026 | 1314.873304 | ENSG00000204262 |
| PHOX2A | 1.494900475 | 2.818447071 | 3.09E-02 | 9.71E-02 | 5.662465085 | 15.90684192 | ENSG00000165462 |
| NRROS | 1.488367241 | 2.80571261 | 6.02E-15 | 2.25E-13 | 80.38007597 | 225.2049327 | ENSG00000174004 |
| PCDH11Y | 1.488269344 | 2.805522229 | 4.27E-02 | 1.25E-01 | 4.960847841 | 13.91498699 | ENSG00000099715 |
| PTAFR | 1.483628892 | 2.796512719 | 2.06E-07 | 2.69E-06 | 43.0193642 | 120.2711363 | ENSG00000169403 |
| GSTT2B | 1.482557831 | 2.794437351 | 1.15E-04 | 8.43E-04 | 21.0108894 | 58.55865194 | ENSG00000133433 |
| KCNJ2 | 1.481301941 | 2.792005804 | 4.13E-03 | 1.89E-02 | 9.626643411 | 26.82439871 | ENSG00000123700 |
| MYOZ1 | 1.477389736 | 2.784444884 | 2.65E-04 | 1.77E-03 | 16.56186386 | 46.08221225 | ENSG00000177791 |
| SLC44A4 | 1.47375146 | 2.777431748 | 3.32E-07 | 4.23E-06 | 32.94463736 | 91.69938762 | ENSG00000204385 |
| DMBX1 | 1.469725037 | 2.769691011 | 1.03E-02 | 4.01E-02 | 8.870465024 | 24.6628779 | ENSG00000197587 |
| LPO | 1.468477621 | 2.767297253 | 3.55E-02 | 1.08E-01 | 5.291494837 | 14.65971514 | ENSG00000167419 |
| FLRT3 | 1.46835546 | 2.767062939 | 2.14E-02 | 7.27E-02 | 6.547648052 | 18.19094642 | ENSG00000125848 |
| TBC1D10C | 1.467706515 | 2.765818555 | 1.10E-02 | 4.22E-02 | 8.921466694 | 24.72104832 | ENSG00000175463 |
| AMDHD1 | 1.465486909 | 2.761566576 | 3.30E-02 | 1.02E-01 | 5.642303459 | 15.54894956 | ENSG00000139344 |
| C1orf94 | 1.459286847 | 2.749724056 | 9.13E-18 | 4.44E-16 | 109.1170686 | 300.5032872 | ENSG00000142698 |
| MMP9 | 1.459261107 | 2.749674997 | 1.69E-11 | 4.18E-10 | 66.45593749 | 182.8435031 | ENSG00000100985 |
| CATSPER2 | 1.452043691 | 2.735953464 | 1.50E-02 | 5.46E-02 | 8.454416841 | 23.31057113 | ENSG00000166762 |
| RGS16 | 1.451368078 | 2.734672518 | 1.46E-42 | 3.20E-40 | 324.5106691 | 888.3832394 | ENSG00000143333 |
| PNLIPRP2 | 1.447717179 | 2.727760876 | 2.51E-02 | 8.27E-02 | 6.756409413 | 18.26935843 | ENSG00000266200 |
| GJB5 | 1.446500286 | 2.725461019 | 8.28E-03 | 3.36E-02 | 8.879948232 | 24.27944708 | ENSG00000189280 |
| AIPL1 | 1.445002531 | 2.72263301 | 1.60E-03 | 8.41E-03 | 13.46746156 | 36.76872863 | ENSG00000129221 |
| SLC30A1 | 1.437409832 | 2.708341818 | 1.63E-36 | 2.67E-34 | 502.8794913 | 1363.251526 | ENSG00000170385 |
| DANT2 | 1.436811005 | 2.707217887 | 4.87E-02 | 1.38E-01 | 6.302402242 | 17.07075426 | ENSG00000235244 |
| ATXN1 | 1.436540648 | 2.706710608 | 3.35E-05 | 2.81E-04 | 23.27439055 | 62.90271315 | ENSG00000124788 |
| C22orf15 | 1.430780212 | 2.69592472 | 1.46E-05 | 1.33E-04 | 30.9631458 | 83.67987238 | ENSG00000169314 |
| ATP8B3 | 1.428743248 | 2.692120988 | 7.07E-09 | 1.20E-07 | 54.00800297 | 145.0480116 | ENSG00000130270 |
| FAM107A | 1.427245741 | 2.689328035 | 4.52E-02 | 1.30E-01 | 6.739807259 | 18.023456 | ENSG00000168309 |
| GABRP | 1.425452442 | 2.685987223 | 2.77E-09 | 4.95E-08 | 49.05368926 | 132.0335879 | ENSG00000094755 |
| PPP1R3C | 1.420975069 | 2.677664242 | 6.03E-05 | 4.75E-04 | 22.03692442 | 59.18637683 | ENSG00000119938 |
| ATOH1 | 1.416610865 | 2.66957645 | 3.94E-02 | 1.17E-01 | 7.130939134 | 18.84175077 | ENSG00000172238 |
| S100A4 | 1.415595026 | 2.667697395 | 1.29E-15 | 5.29E-14 | 93.4955337 | 249.2895543 | ENSG00000196154 |
| HAPLN2 | 1.414900584 | 2.666413606 | 3.25E-02 | 1.01E-01 | 6.567809678 | 17.56255913 | ENSG00000132702 |
| PTGER3 | 1.411438788 | 2.660023126 | 7.60E-03 | 3.14E-02 | 10.19541727 | 27.18541249 | ENSG00000050628 |
| ZFP57 | 1.410389131 | 2.658088484 | 3.05E-02 | 9.63E-02 | 9.413432853 | 24.89818656 | ENSG00000204644 |
| FMO5 | 1.407425011 | 2.652632859 | 3.64E-03 | 1.69E-02 | 11.85574504 | 31.50866502 | ENSG00000131781 |
| CSPG5 | 1.407202089 | 2.652223012 | 3.00E-17 | 1.39E-15 | 109.7285562 | 291.5431568 | ENSG00000114646 |
| NOS3 | 1.406317713 | 2.650597689 | 3.57E-09 | 6.27E-08 | 52.85919202 | 140.2315138 | ENSG00000164867 |
| KLHDC7A | 1.403658065 | 2.645715742 | 6.14E-60 | 2.70E-57 | 531.308364 | 1406.13398 | ENSG00000179023 |
| TLR3 | 1.403296965 | 2.645053614 | 3.22E-03 | 1.53E-02 | 12.58939754 | 33.30843701 | ENSG00000164342 |
| IL15 | 1.400571722 | 2.640061838 | 1.34E-02 | 4.96E-02 | 10.84987165 | 28.36377507 | ENSG00000164136 |
| KCNK3 | 1.394836247 | 2.629587033 | 1.53E-02 | 5.54E-02 | 8.935704584 | 23.47872397 | ENSG00000171303 |
| PCSK2 | 1.394154715 | 2.628345104 | 5.66E-03 | 2.45E-02 | 10.65178871 | 27.93543752 | ENSG00000125851 |
| GPR20 | 1.393507949 | 2.62716707 | 5.10E-04 | 3.13E-03 | 19.75710045 | 51.63736332 | ENSG00000204882 |
| OCA2 | 1.391970527 | 2.624368894 | 8.92E-03 | 3.57E-02 | 9.623083938 | 25.22278455 | ENSG00000104044 |
| FOXJ1 | 1.389820937 | 2.620461543 | 8.65E-18 | 4.24E-16 | 120.5259516 | 316.4618685 | ENSG00000129654 |
| NFASC | 1.38862247 | 2.618285593 | 3.59E-43 | 8.07E-41 | 325.9918809 | 853.4478607 | ENSG00000163531 |
| SPARCL1 | 1.388333654 | 2.617761486 | 1.28E-02 | 4.78E-02 | 10.04625107 | 26.16460896 | ENSG00000152583 |
| LIPA | 1.38818088 | 2.617484293 | 9.51E-49 | 2.81E-46 | 449.6030303 | 1177.887509 | ENSG00000107798 |
| RAB17 | 1.386564423 | 2.614553195 | 2.44E-48 | 7.08E-46 | 408.5835666 | 1067.8712 | ENSG00000124839 |
| TRPM6 | 1.385827718 | 2.613218427 | 1.88E-03 | 9.63E-03 | 15.04861737 | 39.1536622 | ENSG00000119121 |
| LMCD1 | 1.385623491 | 2.612848528 | 5.35E-25 | 4.79E-23 | 170.3360986 | 445.5076739 | ENSG00000071282 |
| LYNX1 | 1.385354834 | 2.612362011 | 1.57E-13 | 4.98E-12 | 89.66035995 | 234.2180117 | ENSG00000180155 |
| BHLHE41 | 1.383609123 | 2.609202875 | 1.71E-03 | 8.91E-03 | 16.80858422 | 43.6203071 | ENSG00000123095 |
| CEACAM1 | 1.382874309 | 2.607874256 | 3.78E-03 | 1.75E-02 | 12.36787283 | 32.08975921 | ENSG00000079385 |
| KCNN4 | 1.381750263 | 2.605843176 | 1.46E-02 | 5.34E-02 | 9.990494714 | 25.98747073 | ENSG00000104783 |
| FGF16 | 1.379349526 | 2.601510492 | 4.36E-06 | 4.47E-05 | 30.6562199 | 79.85757807 | ENSG00000196468 |
| WDR66 | 1.377876062 | 2.598854855 | 4.12E-03 | 1.89E-02 | 15.49520011 | 40.35209841 | ENSG00000158023 |
| GGT1 | 1.377046348 | 2.597360647 | 3.23E-04 | 2.10E-03 | 20.94445464 | 54.26318598 | ENSG00000100031 |
| SLC6A17 | 1.376883493 | 2.597067468 | 1.70E-02 | 6.03E-02 | 8.897745595 | 23.12546609 | ENSG00000197106 |
| ALG1L | 1.376828894 | 2.596969182 | 3.20E-05 | 2.70E-04 | 27.57158014 | 71.52188316 | ENSG00000189366 |
| SMOC2 | 1.376626809 | 2.596605438 | 1.79E-14 | 6.19E-13 | 100.0321978 | 259.9906036 | ENSG00000112562 |

| | | | | | | | |
|---|---|---|---|---|---|---|---|
| NPNT | 1.372773098 | 2.589678669 | 4.15E-02 | 1.22E-01 | 6.905575619 | 17.92858624 | ENSG00000168743 |
| ZNF354C | 1.372731015 | 2.589603132 | 1.51E-05 | 1.38E-04 | 29.95132252 | 77.39255149 | ENSG00000177932 |
| ERVMER34-1 | 1.369469153 | 2.58375478 | 1.36E-28 | 1.49E-26 | 217.6045998 | 562.1451845 | ENSG00000226887 |
| GBGT1 | 1.36740704 | 2.580064335 | 4.63E-15 | 1.76E-13 | 109.0704906 | 280.9091477 | ENSG00000148288 |
| POLR3GL | 1.365708774 | 2.577029004 | 1.82E-03 | 9.34E-03 | 14.54597279 | 37.49822853 | ENSG00000121851 |
| WDR72 | 1.360060017 | 2.56695858 | 8.24E-06 | 8.04E-05 | 30.07320815 | 77.21151383 | ENSG00000166415 |
| MINAR1 | 1.354488721 | 2.557064799 | 2.57E-03 | 1.26E-02 | 16.1520449 | 41.01111015 | ENSG00000169330 |
| LEPR | 1.354017916 | 2.55623047 | 7.24E-06 | 7.16E-05 | 31.87441413 | 81.73727512 | ENSG00000116678 |
| TXNIP | 1.353760278 | 2.555774017 | 2.51E-15 | 9.87E-14 | 112.1862759 | 286.0640313 | ENSG00000265972 |
| PPP1R3B | 1.351685767 | 2.552101605 | 1.38E-47 | 3.72E-45 | 401.1848035 | 1023.329224 | ENSG00000173281 |
| KLK7 | 1.351403148 | 2.551601706 | 2.31E-03 | 1.15E-02 | 15.03082001 | 38.21250019 | ENSG00000169035 |
| ALPG | 1.348979785 | 2.547319257 | 2.88E-04 | 1.90E-03 | 21.04737385 | 53.75113953 | ENSG00000163286 |
| MCHR1 | 1.34884779 | 2.547086208 | 2.53E-08 | 3.87E-07 | 47.03927273 | 119.7379023 | ENSG00000128285 |
| TNNC1 | 1.345674389 | 2.541489709 | 1.77E-05 | 1.59E-04 | 28.73640843 | 73.11224114 | ENSG00000114854 |
| SEMA3G | 1.342835948 | 2.536494352 | 5.42E-13 | 1.61E-11 | 86.90013797 | 220.594444 | ENSG00000010319 |
| CREB5 | 1.342744063 | 2.536332809 | 8.11E-03 | 3.30E-02 | 10.81043812 | 27.53677845 | ENSG00000146592 |
| PYY | 1.337808936 | 2.527671421 | 1.45E-03 | 7.75E-03 | 16.37949335 | 41.2435083 | ENSG00000131096 |
| ENPP2 | 1.335273308 | 2.52323278 | 9.66E-05 | 7.20E-04 | 34.24083449 | 86.77241293 | ENSG00000136960 |
| DUSP10 | 1.335212198 | 2.523125903 | 4.24E-03 | 1.93E-02 | 15.13045908 | 37.97217819 | ENSG00000143507 |
| NOD2 | 1.334416742 | 2.521735116 | 7.97E-03 | 3.26E-02 | 11.56661651 | 29.1760379 | ENSG00000167207 |
| APOBEC3B | 1.332474782 | 2.518342981 | 6.28E-12 | 1.62E-10 | 85.82994089 | 216.4149627 | ENSG00000179750 |
| ARL11 | 1.332129788 | 2.517740839 | 4.11E-03 | 1.88E-02 | 13.55998169 | 34.13855505 | ENSG00000152213 |
| MVP | 1.33190881 | 2.517355224 | 9.29E-05 | 6.96E-04 | 28.35147962 | 71.18451593 | ENSG00000013364 |
| HNMT | 1.327658592 | 2.509949942 | 2.39E-04 | 1.61E-03 | 20.9385309 | 52.45549014 | ENSG00000150540 |
| NPL | 1.32578425 | 2.506691146 | 9.22E-08 | 1.30E-06 | 59.81579546 | 150.0712535 | ENSG00000135838 |
| PLCB2 | 1.325368289 | 2.505968516 | 6.08E-29 | 6.83E-27 | 255.1325545 | 638.8708527 | ENSG00000137841 |
| ELF3 | 1.317039752 | 2.491543483 | 2.58E-09 | 4.64E-08 | 58.33900726 | 145.569495 | ENSG00000163435 |
| SCARF1 | 1.314993185 | 2.488011553 | 1.59E-10 | 3.45E-09 | 87.16704609 | 216.7587407 | ENSG00000074660 |
| PRODH | 1.314899761 | 2.487850443 | 2.43E-05 | 2.12E-04 | 29.12754031 | 72.4860293 | ENSG00000100033 |
| TMPRSS9 | 1.314610922 | 2.487352405 | 6.02E-03 | 2.58E-02 | 13.48406371 | 33.64769615 | ENSG00000178297 |
| ADARB2 | 1.313547212 | 2.485519138 | 1.95E-07 | 2.55E-06 | 62.92143484 | 155.854808 | ENSG00000185736 |
| TENM2 | 1.311317572 | 2.481680813 | 1.09E-05 | 1.03E-04 | 42.21126827 | 104.6737071 | ENSG00000145934 |
| MYO18B | 1.308740579 | 2.477251903 | 7.02E-04 | 4.12E-03 | 19.58393382 | 48.4503092 | ENSG00000133454 |
| PTCH2 | 1.306683253 | 2.473721786 | 4.33E-02 | 1.26E-01 | 7.238586879 | 17.9534623 | ENSG00000117425 |
| HPSE2 | 1.305518138 | 2.471724825 | 1.59E-02 | 5.72E-02 | 10.82259108 | 26.51502897 | ENSG00000172987 |
| WFDC1 | 1.304287378 | 2.469617101 | 9.63E-03 | 3.79E-02 | 11.35340595 | 27.94508533 | ENSG00000103175 |
| TPCN2 | 1.304154949 | 2.469390418 | 4.29E-02 | 1.25E-01 | 7.452992645 | 18.5519602 | ENSG00000287725 |
| MICAL2 | 1.303161344 | 2.467690298 | 4.59E-07 | 5.67E-06 | 49.4901784 | 121.9852348 | ENSG00000133816 |
| CR2 | 1.300814535 | 2.463679409 | 2.30E-18 | 1.20E-16 | 134.1149677 | 330.2412916 | ENSG00000117322 |
| TNNI1 | 1.299760905 | 2.461880789 | 1.21E-04 | 8.79E-04 | 27.8533104 | 68.86645465 | ENSG00000159173 |
| PWWP3B | 1.29817678 | 2.459179051 | 2.58E-03 | 1.26E-02 | 17.42688521 | 42.91490525 | ENSG00000157502 |
| CTHRC1 | 1.293055828 | 2.450465487 | 6.14E-04 | 3.67E-03 | 19.92998776 | 48.74662616 | ENSG00000164932 |
| ACTG2 | 1.291423649 | 2.447694745 | 1.28E-05 | 1.19E-04 | 31.73236687 | 77.72121686 | ENSG00000163017 |
| RNASEL | 1.290887449 | 2.446785191 | 6.77E-03 | 2.85E-02 | 11.96250306 | 29.21708817 | ENSG00000135828 |
| RGS2 | 1.290375039 | 2.445916307 | 2.68E-21 | 1.77E-19 | 188.9722535 | 462.0744509 | ENSG00000116741 |
| NABP1 | 1.289742363 | 2.444843915 | 3.04E-05 | 2.59E-04 | 31.1303887 | 76.10503686 | ENSG00000173559 |
| CD7 | 1.283010277 | 2.433462051 | 7.08E-03 | 2.95E-02 | 13.68779106 | 33.5092218 | ENSG00000173762 |
| SOX3 | 1.278587364 | 2.426013139 | 1.02E-35 | 1.62E-33 | 382.1607968 | 927.3548454 | ENSG00000134595 |
| DDR2 | 1.276489872 | 2.422488593 | 3.89E-03 | 1.79E-02 | 14.95253776 | 36.15434008 | ENSG00000162733 |
| ANXA2 | 1.274738966 | 2.419550358 | 1.45E-69 | 1.01E-66 | 850.7786567 | 2058.097923 | ENSG00000182718 |
| OPTC | 1.272153059 | 2.415217405 | 4.77E-03 | 2.12E-02 | 18.85473052 | 45.80613688 | ENSG00000188770 |
| ITGB2 | 1.270969833 | 2.413237378 | 2.51E-02 | 8.27E-02 | 8.932145111 | 21.55374132 | ENSG00000160255 |
| ADAMTS5 | 1.270600575 | 2.412619788 | 8.55E-06 | 8.34E-05 | 34.37751668 | 82.97787551 | ENSG00000154736 |
| HHLA1 | 1.268080409 | 2.408408992 | 2.47E-11 | 5.98E-10 | 87.62222232 | 210.6988772 | ENSG00000132297 |
| CCDC110 | 1.267587051 | 2.40758553 | 7.77E-03 | 3.19E-02 | 13.22813949 | 31.85255863 | ENSG00000168491 |
| SPIB | 1.263949315 | 2.401522482 | 6.71E-11 | 1.54E-09 | 97.26015454 | 233.8437545 | ENSG00000269404 |
| GDF3 | 1.262939525 | 2.399842166 | 1.08E-13 | 3.45E-12 | 114.2914587 | 274.4618087 | ENSG00000184344 |
| PHF24 | 1.261362428 | 2.397220187 | 2.48E-02 | 8.20E-02 | 9.285317997 | 22.23812358 | ENSG00000122733 |
| PNMA6F | 1.260818258 | 2.396316151 | 2.21E-02 | 7.47E-02 | 10.46466351 | 24.87766143 | ENSG00000225110 |
| LRRC15 | 1.255719454 | 2.387861985 | 3.74E-02 | 1.13E-01 | 7.906999816 | 18.91609539 | ENSG00000172061 |
| SYNPO | 1.252401162 | 2.382376061 | 4.86E-11 | 1.14E-09 | 93.1509543 | 221.50392 | ENSG00000171992 |
| IL2RB | 1.251266158 | 2.380502523 | 7.81E-06 | 7.67E-05 | 38.26844678 | 90.87218008 | ENSG00000100385 |
| SSTR1 | 1.248268991 | 2.375562215 | 6.91E-03 | 2.89E-02 | 13.81115124 | 32.90665652 | ENSG00000139874 |
| PEX5L | 1.247719187 | 2.374657073 | 4.23E-05 | 3.46E-04 | 34.85969415 | 82.97598359 | ENSG00000114757 |
| POLD4 | 1.247155248 | 2.373729017 | 8.44E-05 | 6.39E-04 | 28.12255664 | 66.77348712 | ENSG00000175482 |
| TAPBPL | 1.247108952 | 2.373652847 | 2.18E-02 | 7.38E-02 | 10.19778154 | 24.27386664 | ENSG00000139192 |
| OAS3 | 1.245146609 | 2.370426417 | 1.79E-03 | 9.24E-03 | 22.33701071 | 52.81309894 | ENSG00000111331 |
| MUC19 | 1.244775922 | 2.369817436 | 2.84E-02 | 9.09E-02 | 9.772250144 | 23.32116491 | ENSG00000205592 |
| CD1D | 1.244626539 | 2.369572067 | 1.40E-05 | 1.28E-04 | 43.01814284 | 101.9390885 | ENSG00000158473 |
| SULT2B1 | 1.243317961 | 2.367423751 | 4.31E-02 | 1.26E-01 | 7.552631721 | 17.94504399 | ENSG00000088002 |
| NEURL1 | 1.243040527 | 2.366968534 | 1.60E-03 | 8.41E-03 | 19.13348612 | 45.31071613 | ENSG00000107954 |
| BAHCC1 | 1.240150983 | 2.362232525 | 7.62E-05 | 5.84E-04 | 29.08009811 | 68.74387096 | ENSG00000266074 |
| SBK3 | 1.238768898 | 2.359970618 | 1.23E-02 | 4.65E-02 | 11.25732635 | 26.56232207 | ENSG00000231274 |
| F8 | 1.23822829 | 2.359086453 | 1.14E-04 | 8.36E-04 | 28.08935233 | 66.2799583 | ENSG00000185010 |

| | | | | | | | |
|---|---|---|---|---|---|---|---|
| NOS2 | 1.237669598 | 2.358173061 | 4.86E-09 | 8.43E-08 | 67.25127007 | 158.4742722 | ENSG00000007171 |
| MYO1F | 1.233656577 | 2.351622647 | 1.06E-09 | 2.03E-08 | 87.70167362 | 205.9173978 | ENSG00000142347 |
| IRX1 | 1.232198177 | 2.349246626 | 1.24E-02 | 4.68E-02 | 11.80830284 | 27.83432491 | ENSG00000170549 |
| PTCHD1 | 1.229709399 | 2.34519746 | 1.58E-07 | 2.11E-06 | 74.77812192 | 175.114019 | ENSG00000165186 |
| OPN3 | 1.229168816 | 2.34431887 | 2.39E-03 | 1.19E-02 | 17.53008375 | 41.0885762 | ENSG00000054277 |
| UCMA | 1.229136617 | 2.344266548 | 2.96E-03 | 1.42E-02 | 17.11403557 | 40.22705571 | ENSG00000165623 |
| BGN | 1.227151539 | 2.341043171 | 2.13E-09 | 3.91E-08 | 75.90616028 | 177.58635 | ENSG00000182492 |
| RHBDL1 | 1.226611726 | 2.340167386 | 1.04E-08 | 1.72E-07 | 63.02999847 | 147.3359726 | ENSG00000103269 |
| SULT1C2 | 1.225066816 | 2.337662759 | 8.43E-03 | 3.41E-02 | 13.93095194 | 32.53665734 | ENSG00000198203 |
| RHBG | 1.224824802 | 2.337270646 | 1.22E-03 | 6.66E-03 | 20.17523357 | 47.17518494 | ENSG00000132677 |
| SIT1 | 1.22196952 | 2.332649457 | 1.85E-10 | 3.98E-09 | 86.86665433 | 203.0646847 | ENSG00000137078 |
| NEK7 | 1.220454307 | 2.330200841 | 1.74E-14 | 6.01E-13 | 128.0996091 | 298.1067936 | ENSG00000151414 |
| FGF4 | 1.218619909 | 2.327239855 | 4.74E-04 | 2.94E-03 | 25.28524761 | 59.01186556 | ENSG00000075388 |
| SPOCD1 | 1.217004394 | 2.324635295 | 7.15E-03 | 2.98E-02 | 16.96486937 | 39.40876094 | ENSG00000134668 |
| CDA | 1.216674919 | 2.324104467 | 6.14E-11 | 1.42E-09 | 90.21161578 | 209.4780669 | ENSG00000158825 |
| NAV3 | 1.21570375 | 2.322540494 | 1.26E-04 | 9.13E-04 | 31.32580189 | 72.81079525 | ENSG00000067798 |
| IRAK1 | 1.215029984 | 2.321456075 | 6.83E-73 | 6.15E-70 | 1072.231524 | 2490.168321 | ENSG00000184216 |
| NUTM2D | 1.214089269 | 2.319942855 | 9.72E-03 | 3.81E-02 | 13.55522701 | 31.43649586 | ENSG00000214562 |
| ADGRG1 | 1.212813192 | 2.317891751 | 3.33E-04 | 2.15E-03 | 31.71693377 | 73.60018953 | ENSG00000205336 |
| GLP1R | 1.212122065 | 2.316781625 | 6.59E-10 | 1.30E-08 | 86.03066743 | 199.0795774 | ENSG00000112164 |
| S100A11 | 1.211319923 | 2.315493846 | 3.97E-71 | 3.00E-68 | 1182.932471 | 2739.218025 | ENSG00000163191 |
| CAPN13 | 1.210247304 | 2.313772957 | 1.46E-06 | 1.63E-05 | 47.17211611 | 108.9989241 | ENSG00000162949 |
| METTL27 | 1.209337029 | 2.312313532 | 1.77E-08 | 2.81E-07 | 76.99295951 | 177.9089834 | ENSG00000165171 |
| MYOF | 1.207705979 | 2.309700805 | 3.64E-05 | 3.03E-04 | 33.75537689 | 77.95066154 | ENSG00000138119 |
| OXSM | 1.204863596 | 2.305154736 | 3.23E-03 | 1.53E-02 | 19.38020647 | 44.51606792 | ENSG00000151093 |
| SYT5 | 1.203138779 | 2.302400449 | 7.48E-05 | 5.74E-04 | 36.25225022 | 83.58015724 | ENSG00000129990 |
| AMIGO2 | 1.202441043 | 2.3012872 | 1.94E-12 | 5.37E-11 | 131.6171785 | 302.3219127 | ENSG00000139211 |
| PHLDA3 | 1.201636555 | 2.300004295 | 2.03E-40 | 4.03E-38 | 904.9827338 | 2082.95405 | ENSG00000174307 |
| CA3 | 1.201339036 | 2.299530026 | 4.94E-02 | 1.40E-01 | 8.911983485 | 20.49746797 | ENSG00000164879 |
| CRABP2 | 1.201125907 | 2.299190342 | 3.68E-76 | 3.66E-73 | 1483.429603 | 3409.046588 | ENSG00000143320 |
| FAM50B | 1.198091538 | 2.29435962 | 4.37E-08 | 6.46E-07 | 68.85942712 | 158.0404698 | ENSG00000145945 |
| ILDR1 | 1.191489066 | 2.283883507 | 9.10E-05 | 6.84E-04 | 31.33055657 | 71.64766091 | ENSG00000145103 |
| B3GALT4 | 1.1896617 | 2.280992495 | 6.10E-04 | 3.65E-03 | 24.1726162 | 55.12832235 | ENSG00000235863 |
| WNT3 | 1.188641473 | 2.279380022 | 4.47E-10 | 9.06E-09 | 87.1848173 | 198.5218033 | ENSG00000108379 |
| ELN | 1.188186697 | 2.278661613 | 4.12E-18 | 2.09E-16 | 166.6399974 | 379.7855195 | ENSG00000049540 |
| DHRS12 | 1.186097517 | 2.275364251 | 1.94E-04 | 1.34E-03 | 28.75301058 | 65.48095916 | ENSG00000102796 |
| CAPN11 | 1.183801014 | 2.271745173 | 3.49E-04 | 2.24E-03 | 28.61096332 | 64.89674354 | ENSG00000137225 |
| CAV2 | 1.179320066 | 2.264700178 | 9.94E-04 | 5.59E-03 | 26.3201816 | 59.46869505 | ENSG00000105971 |
| APOBEC3G | 1.177858335 | 2.262406758 | 2.22E-03 | 1.11E-02 | 20.28079638 | 45.78996267 | ENSG00000239713 |
| FZD10 | 1.177314222 | 2.261553652 | 3.92E-02 | 1.17E-01 | 10.81191266 | 24.27850112 | ENSG00000111432 |
| ZNF124 | 1.176433133 | 2.260172887 | 2.08E-14 | 7.17E-13 | 159.8545274 | 362.0125277 | ENSG00000196418 |
| STARD13 | 1.176081237 | 2.259621664 | 3.19E-07 | 4.06E-06 | 56.90996674 | 128.6457489 | ENSG00000133121 |
| MESP2 | 1.174103576 | 2.256526272 | 2.40E-02 | 7.96E-02 | 11.35696542 | 25.52222293 | ENSG00000188095 |
| WNT9A | 1.173020222 | 2.25483243 | 1.88E-07 | 2.47E-06 | 62.16706206 | 140.2566734 | ENSG00000143816 |
| EXOC8 | 1.168718534 | 2.248119203 | 2.12E-11 | 5.18E-10 | 103.1954253 | 231.9526131 | ENSG00000116903 |
| MAP10 | 1.168257035 | 2.247400175 | 1.32E-04 | 9.53E-04 | 34.69540033 | 77.74486341 | ENSG00000212916 |
| CHRNA9 | 1.166261488 | 2.244293702 | 2.14E-03 | 1.08E-02 | 21.10429926 | 47.42174978 | ENSG00000174343 |
| SNCG | 1.166171146 | 2.244153167 | 9.91E-10 | 1.91E-08 | 92.38882602 | 207.055699 | ENSG00000173267 |
| PSMB9 | 1.16409142 | 2.240920425 | 1.49E-05 | 1.36E-04 | 41.66830044 | 93.43909727 | ENSG00000240065 |
| GCNA | 1.163063244 | 2.239323941 | 4.91E-02 | 1.39E-01 | 9.758012254 | 22.01151677 | ENSG00000147174 |
| SPTBN5 | 1.159463044 | 2.233742745 | 3.31E-07 | 4.20E-06 | 73.15069297 | 163.4778826 | ENSG00000137877 |
| CDK10 | 1.155179836 | 2.227120836 | 1.29E-15 | 5.28E-14 | 263.3236488 | 586.520134 | ENSG00000185324 |
| RASGRP4 | 1.154819503 | 2.226564652 | 1.60E-06 | 1.78E-05 | 52.0496477 | 116.0872087 | ENSG00000171777 |
| GSG1L | 1.147044812 | 2.214597957 | 2.96E-02 | 9.38E-02 | 10.64466976 | 23.50795096 | ENSG00000169181 |
| STAT6 | 1.146578207 | 2.213881813 | 1.25E-52 | 4.20E-50 | 1001.400291 | 2216.433807 | ENSG00000166888 |
| TIGD6 | 1.145437792 | 2.212132486 | 5.14E-04 | 3.15E-03 | 30.19656833 | 66.72997785 | ENSG00000164296 |
| HCP5 | 1.144103553 | 2.210087598 | 2.54E-05 | 2.20E-04 | 43.0807388 | 95.40463572 | ENSG00000206337 |
| FAXDC2 | 1.142618693 | 2.207814088 | 4.19E-05 | 3.43E-04 | 38.29425281 | 84.62895824 | ENSG00000170271 |
| DAGLA | 1.141804835 | 2.206568959 | 3.35E-22 | 2.38E-20 | 257.6997269 | 569.0404402 | ENSG00000134780 |
| DNAH1 | 1.141325462 | 2.205835892 | 1.94E-02 | 6.70E-02 | 16.68997872 | 36.75522435 | ENSG00000114841 |
| KIFAP3 | 1.140525174 | 2.204612615 | 3.52E-06 | 3.67E-05 | 56.6739248 | 125.0584797 | ENSG00000075945 |
| CPZ | 1.140112097 | 2.203981475 | 3.91E-02 | 1.17E-01 | 10.85551605 | 24.00147978 | ENSG00000109625 |
| NUDT7 | 1.140058722 | 2.203899936 | 2.57E-02 | 8.43E-02 | 11.6282966 | 25.59344613 | ENSG00000140876 |
| ANXA3 | 1.139122059 | 2.202469528 | 2.22E-19 | 1.27E-17 | 196.2239092 | 432.1892929 | ENSG00000138772 |
| PHYHD1 | 1.134698908 | 2.195727329 | 1.35E-09 | 2.55E-08 | 94.33711327 | 206.8516772 | ENSG00000175287 |
| MEF2C | 1.132892712 | 2.192980087 | 1.94E-02 | 6.70E-02 | 14.50209007 | 31.84631579 | ENSG00000081189 |
| C11orf86 | 1.131227567 | 2.19045043 | 2.88E-02 | 9.19E-02 | 11.50613163 | 25.28502235 | ENSG00000173237 |
| NEK11 | 1.130655383 | 2.189581852 | 2.50E-02 | 8.26E-02 | 12.32635437 | 26.92551131 | ENSG00000114670 |
| TUSC1 | 1.129105881 | 2.187231432 | 4.55E-07 | 5.63E-06 | 63.88286685 | 139.9751855 | ENSG00000198680 |
| RNF227 | 1.128757508 | 2.186703337 | 8.02E-03 | 3.27E-02 | 18.61334966 | 40.67206192 | ENSG00000179859 |
| MASP1 | 1.125498779 | 2.181769632 | 5.78E-05 | 4.58E-04 | 42.50721026 | 92.87462878 | ENSG00000127241 |
| INHBA | 1.125108242 | 2.181179107 | 4.22E-02 | 1.24E-01 | 10.78671702 | 23.65730262 | ENSG00000122641 |
| TIRAP | 1.1247658 | 2.180661438 | 8.41E-03 | 3.41E-02 | 17.3737986 | 37.71395802 | ENSG00000150455 |
| NTSR1 | 1.124465608 | 2.180207741 | 4.46E-03 | 2.01E-02 | 18.88824031 | 41.15544848 | ENSG00000101188 |

| | | | | | | | |
|---|---|---|---|---|---|---|---|
| FMN1 | 1.12363662 | 2.17895533 | 1.92E-04 | 1.33E-03 | 36.69468925 | 79.72612457 | ENSG00000248905 |
| FHIT | 1.123350806 | 2.178523697 | 7.31E-03 | 3.03E-02 | 18.09054346 | 39.50597412 | ENSG00000189283 |
| HPCAL4 | 1.120235928 | 2.173825188 | 5.40E-05 | 4.31E-04 | 38.5999835 | 84.02856843 | ENSG00000116983 |
| LAMB3 | 1.116599682 | 2.168353061 | 3.27E-03 | 1.55E-02 | 23.79808648 | 51.67020619 | ENSG00000196878 |
| ARSD | 1.115774344 | 2.167112942 | 5.58E-09 | 9.61E-08 | 87.5546185 | 189.5918661 | ENSG00000006756 |
| PCSK1 | 1.112371494 | 2.16200745 | 1.26E-03 | 6.87E-03 | 26.61731881 | 57.69256961 | ENSG00000175426 |
| CALB1 | 1.109732405 | 2.158056154 | 2.72E-24 | 2.30E-22 | 352.8556146 | 761.4100943 | ENSG00000104327 |
| LAMC2 | 1.108439526 | 2.156123066 | 2.68E-23 | 2.07E-21 | 274.417571 | 591.3006543 | ENSG00000058085 |
| KIAA1257 | 1.10625338 | 2.152858321 | 6.32E-04 | 3.75E-03 | 34.90477208 | 75.33526471 | ENSG00000114656 |
| SFR1 | 1.10619342 | 2.152768848 | 1.20E-05 | 1.12E-04 | 47.30112069 | 101.8295576 | ENSG00000156384 |
| FGF8 | 1.106065866 | 2.152578524 | 3.62E-03 | 1.69E-02 | 23.60028287 | 50.69771437 | ENSG00000107831 |
| SLC44A5 | 1.104394423 | 2.150086084 | 2.55E-04 | 1.70E-03 | 32.7159937 | 70.38653539 | ENSG00000137968 |
| S100A16 | 1.102862549 | 2.147804304 | 1.29E-20 | 8.02E-19 | 249.1157213 | 534.6318417 | ENSG00000188643 |
| MYBPC2 | 1.100935479 | 2.144937304 | 4.36E-10 | 8.84E-09 | 101.1228882 | 216.7325398 | ENSG00000086967 |
| XK | 1.100548515 | 2.14436206 | 1.79E-03 | 9.24E-03 | 28.64625256 | 61.48682349 | ENSG00000047597 |
| SERPINB1 | 1.100378953 | 2.144110045 | 1.10E-16 | 4.84E-15 | 192.0927673 | 411.7755339 | ENSG00000021355 |
| HECW1 | 1.094853909 | 2.135914517 | 2.30E-08 | 3.56E-07 | 123.3558362 | 263.7994757 | ENSG00000002746 |
| HACD4 | 1.094205866 | 2.134955303 | 1.71E-03 | 8.91E-03 | 25.22504207 | 53.7928522 | ENSG00000188921 |
| ANO9 | 1.093159076 | 2.133406784 | 7.53E-04 | 4.38E-03 | 275.6215797 | 588.0464482 | ENSG00000185101 |
| NMUR1 | 1.093034058 | 2.133221921 | 2.73E-02 | 8.83E-02 | 12.90817091 | 27.52807659 | ENSG00000171596 |
| NDRG1 | 1.091150677 | 2.130438901 | 4.61E-05 | 3.75E-04 | 45.90647969 | 97.61082267 | ENSG00000104419 |
| ZBTB7B | 1.090184973 | 2.129013316 | 7.35E-49 | 2.20E-46 | 743.0087863 | 1582.236844 | ENSG00000160685 |
| MAFB | 1.08847447 | 2.126490588 | 5.84E-04 | 3.51E-03 | 32.05945439 | 68.15793138 | ENSG00000204103 |
| TRPV3 | 1.088264759 | 2.126181503 | 2.28E-02 | 7.63E-02 | 13.87283133 | 29.51122966 | ENSG00000167723 |
| IRF2BP2 | 1.087515874 | 2.125078116 | 9.98E-29 | 1.10E-26 | 457.7908968 | 973.6573989 | ENSG00000168264 |
| TMEM38A | 1.087178824 | 2.124581701 | 3.18E-15 | 1.24E-13 | 187.3323928 | 397.661611 | ENSG00000072954 |
| CPEB4 | 1.086321901 | 2.12332013 | 3.47E-13 | 1.05E-11 | 151.089041 | 320.8179935 | ENSG00000113742 |
| DYRK3 | 1.084935338 | 2.121280405 | 2.22E-17 | 1.04E-15 | 211.5412141 | 448.9740405 | ENSG00000143479 |
| MEF2D | 1.082406904 | 2.117565954 | 1.06E-32 | 1.43E-30 | 487.6602998 | 1032.599103 | ENSG00000116604 |
| MYO15B | 1.079483364 | 2.11327917 | 1.59E-06 | 1.77E-05 | 124.9826809 | 264.1621314 | ENSG00000266714 |
| DOC2A | 1.078220799 | 2.111430558 | 1.27E-06 | 1.45E-05 | 129.363771 | 273.3086917 | ENSG00000149927 |
| CD247 | 1.078125 | 2.111290357 | 5.29E-04 | 3.22E-03 | 31.68492467 | 66.94286947 | ENSG00000198821 |
| MYH3 | 1.077461602 | 2.11031974 | 1.58E-03 | 8.34E-03 | 28.28596073 | 59.85704389 | ENSG00000109063 |
| GNGT1 | 1.076778827 | 2.109321239 | 8.96E-03 | 3.58E-02 | 19.89558824 | 41.96336039 | ENSG00000127928 |
| C16orf74 | 1.076134837 | 2.108379891 | 2.58E-03 | 1.26E-02 | 26.52863748 | 55.92157311 | ENSG00000154102 |
| RIMBP3 | 1.074402717 | 2.105850059 | 4.86E-02 | 1.38E-01 | 10.20845995 | 21.53009477 | ENSG00000275793 |
| RNF152 | 1.074203792 | 2.105559715 | 8.90E-03 | 3.57E-02 | 23.0027539 | 48.56296153 | ENSG00000176641 |
| GJB7 | 1.073537 | 2.104586782 | 4.94E-02 | 1.40E-01 | 11.77537787 | 24.65228413 | ENSG00000164411 |
| SUCLG2 | 1.07190914 | 2.102213418 | 9.74E-34 | 1.40E-31 | 669.1324506 | 1405.528366 | ENSG00000172340 |
| ALS2CL | 1.07135674 | 2.101408645 | 1.78E-07 | 2.35E-06 | 163.4263793 | 343.4677696 | ENSG00000178038 |
| NAALADL1 | 1.07062164 | 2.100338183 | 3.30E-04 | 2.13E-03 | 34.14058502 | 71.6464314 | ENSG00000168060 |
| CBX7 | 1.067800527 | 2.096235096 | 4.14E-07 | 5.15E-06 | 86.4123161 | 180.9474639 | ENSG00000100307 |
| TMEM88B | 1.066099354 | 2.093764751 | 1.49E-02 | 5.43E-02 | 17.63417203 | 37.08651668 | ENSG00000205116 |
| LGALS3BP | 1.064638153 | 2.091645202 | 2.95E-56 | 1.12E-53 | 1246.661339 | 2606.71907 | ENSG00000108679 |
| COX6B2 | 1.064550166 | 2.09151764 | 5.31E-03 | 2.32E-02 | 27.85953962 | 58.25921356 | ENSG00000160471 |
| GAS5 | 1.060886762 | 2.086213432 | 7.46E-58 | 2.94E-55 | 3842.104162 | 8014.312912 | ENSG00000234741 |
| GCSAM | 1.060085545 | 2.085055152 | 1.85E-02 | 6.44E-02 | 16.02009124 | 33.57051365 | ENSG00000174500 |
| KIRREL1 | 1.056045548 | 2.079224514 | 1.30E-60 | 5.86E-58 | 1546.945058 | 3215.959807 | ENSG00000183853 |
| FZD5 | 1.055013281 | 2.077737333 | 3.24E-64 | 1.75E-61 | 1634.875148 | 3396.381371 | ENSG00000163251 |
| FBXO48 | 1.054875476 | 2.077538879 | 1.81E-02 | 6.34E-02 | 18.2162679 | 37.83001532 | ENSG00000204923 |
| LMOD1 | 1.054499255 | 2.076997177 | 2.71E-02 | 8.78E-02 | 14.28176056 | 29.64497422 | ENSG00000163431 |
| LRRTM1 | 1.050729343 | 2.071576852 | 2.30E-03 | 1.15E-02 | 27.06301182 | 56.12888429 | ENSG00000162951 |
| EPSTI1 | 1.044471965 | 2.062611293 | 2.81E-02 | 9.03E-02 | 15.35879725 | 31.81368383 | ENSG00000133106 |
| HES4 | 1.043988215 | 2.061919794 | 1.58E-02 | 5.67E-02 | 19.86118873 | 40.97641833 | ENSG00000188290 |
| NAXE | 1.043800064 | 2.061650904 | 2.59E-59 | 1.06E-56 | 1413.178788 | 2913.268657 | ENSG00000163382 |
| RYR2 | 1.043446111 | 2.061145157 | 1.89E-08 | 2.98E-07 | 108.316982 | 222.7071586 | ENSG00000198626 |
| MUC12 | 1.04162462 | 2.058544477 | 2.07E-02 | 7.07E-02 | 16.0085231 | 32.85255348 | ENSG00000205277 |
| MT1H | 1.037782095 | 2.053068971 | 5.21E-05 | 4.17E-04 | 48.45199042 | 99.39848784 | ENSG00000205358 |
| ALDH1A1 | 1.035502808 | 2.049827927 | 7.04E-03 | 2.94E-02 | 20.61500285 | 42.18031938 | ENSG00000165092 |
| SH3RF2 | 1.035179041 | 2.049367961 | 3.62E-02 | 1.10E-01 | 14.50000513 | 29.50716228 | ENSG00000156463 |
| NEDD9 | 1.032371458 | 2.045383628 | 5.88E-17 | 2.65E-15 | 227.5158964 | 464.9600294 | ENSG00000111859 |
| KCNN3 | 1.029958735 | 2.041965845 | 1.12E-06 | 1.29E-05 | 74.23156846 | 151.4377043 | ENSG00000143603 |
| TUBB6 | 1.029096307 | 2.040745544 | 4.92E-63 | 2.58E-60 | 3053.364317 | 6231.726987 | ENSG00000176014 |
| CDH11 | 1.028171957 | 2.039438436 | 1.44E-14 | 5.07E-13 | 195.9499083 | 399.8023434 | ENSG00000140937 |
| UHMK1 | 1.027399718 | 2.038347066 | 7.13E-15 | 2.63E-13 | 418.4636654 | 852.5295313 | ENSG00000152332 |
| GRIN2D | 1.026660433 | 2.037302817 | 4.30E-27 | 4.42E-25 | 563.2429326 | 1146.728153 | ENSG00000105464 |
| PTPN14 | 1.023670267 | 2.033084624 | 4.00E-39 | 7.33E-37 | 1135.397543 | 2307.082461 | ENSG00000152104 |
| CBLC | 1.023093608 | 2.032272144 | 4.49E-14 | 1.50E-12 | 200.5036069 | 407.3494877 | ENSG00000142273 |
| C1orf116 | 1.021847229 | 2.030517173 | 9.59E-08 | 1.34E-06 | 103.7042991 | 210.2167202 | ENSG00000182795 |
| FAM78B | 1.020196329 | 2.028194948 | 6.69E-06 | 6.65E-05 | 72.78770523 | 147.8224806 | ENSG00000188859 |
| RCN3 | 1.019865038 | 2.027729259 | 5.04E-18 | 2.54E-16 | 289.9768678 | 587.6938489 | ENSG00000142552 |
| PSEN2 | 1.018686068 | 2.026072877 | 9.31E-25 | 8.02E-23 | 427.9877735 | 866.3271721 | ENSG00000143801 |
| TOR4A | 1.018026028 | 2.02514615 | 1.88E-13 | 5.86E-12 | 220.23368 | 445.4159982 | ENSG00000198113 |
| POU6F2 | 1.017987734 | 2.025092397 | 4.66E-02 | 1.33E-01 | 15.51922669 | 31.24346696 | ENSG00000106536 |

| | | | | | | | |
|---|---|---|---|---|---|---|---|
| IER5 | 1.017845902 | 2.02489332 | 4.46E-30 | 5.27E-28 | 518.6925762 | 1050.274001 | ENSG00000162783 |
| LIF | 1.017543655 | 2.024469145 | 9.89E-07 | 1.15E-05 | 84.21230063 | 170.6931018 | ENSG00000128342 |
| TGM1 | 1.016000503 | 2.022304866 | 2.10E-02 | 7.17E-02 | 19.68565783 | 39.95003437 | ENSG00000092295 |
| ACTN3 | 1.015952161 | 2.022237105 | 5.73E-19 | 3.18E-17 | 355.5784362 | 718.92557 | ENSG00000248746 |
| BMP8B | 1.015124891 | 2.021077846 | 1.02E-03 | 5.72E-03 | 33.59553226 | 67.77565743 | ENSG00000116985 |
| SUGP1 | 1.015037792 | 2.020955832 | 1.73E-43 | 3.98E-41 | 876.5422429 | 1771.359919 | ENSG00000105705 |
| MGAT3 | 1.011348263 | 2.01579407 | 7.99E-28 | 8.47E-26 | 442.0489247 | 891.210414 | ENSG00000128268 |
| ZBTB41 | 1.009316928 | 2.0129578 | 5.05E-06 | 5.13E-05 | 64.60967973 | 129.9527272 | ENSG00000177888 |
| PCLO | 1.00808954 | 2.011245984 | 1.21E-07 | 1.65E-06 | 92.80934955 | 186.5499807 | ENSG00000186472 |
| CGAS | 1.007206441 | 2.010015241 | 2.33E-04 | 1.58E-03 | 51.11285267 | 102.3720606 | ENSG00000164430 |
| FRMPD4 | 1.006627146 | 2.009208308 | 1.59E-02 | 5.72E-02 | 20.95276879 | 41.99114696 | ENSG00000169933 |
| TTC39C | 1.006451494 | 2.008963696 | 1.62E-21 | 1.10E-19 | 335.9764257 | 674.8354103 | ENSG00000168234 |
| DARS2 | 1.005499372 | 2.007638297 | 1.08E-30 | 1.31E-28 | 757.0154547 | 1520.633215 | ENSG00000117593 |
| ZFHX3 | 1.0047545 | 2.006602009 | 2.13E-04 | 1.45E-03 | 45.35163823 | 91.00129017 | ENSG00000140836 |
| HLA-G | 1.004518498 | 2.006273787 | 1.27E-04 | 9.22E-04 | 47.87965709 | 95.86893756 | ENSG00000204632 |
| AIF1 | 1.004393174 | 2.006099514 | 9.50E-04 | 5.37E-03 | 35.46286752 | 71.35323586 | ENSG00000204472 |
| ANKRD65 | 1.00224006 | 2.003107794 | 4.73E-02 | 1.35E-01 | 13.32065962 | 26.63761266 | ENSG00000235098 |
| FGD3 | 1.001648332 | 2.00228638 | 7.15E-05 | 5.51E-04 | 59.50027608 | 119.4666723 | ENSG00000127084 |
| C3orf62 | 1.001227908 | 2.001702966 | 5.52E-03 | 2.40E-02 | 24.63819151 | 49.46118136 | ENSG00000188315 |
| NMNAT2 | 1.000487087 | 2.00067536 | 2.01E-22 | 1.46E-20 | 366.6356208 | 732.9801523 | ENSG00000157064 |
| PYCR2 | 1.000288961 | 2.000400626 | 1.32E-24 | 1.13E-22 | 439.9805835 | 880.6435084 | ENSG00000143811 |
| NIT1 | 0.999521329 | 1.999336532 | 3.51E-25 | 3.20E-23 | 496.3816253 | 991.7845124 | ENSG00000158793 |
| ERN2 | 0.998390903 | 1.997770562 | 1.27E-02 | 4.74E-02 | 19.60053597 | 39.07307472 | ENSG00000134398 |
| ERVH48-1 | 0.997748084 | 1.996880617 | 5.92E-16 | 2.51E-14 | 529.1435355 | 1055.462769 | ENSG00000233056 |
| TENM1 | 0.997159163 | 1.996065638 | 3.04E-03 | 1.46E-02 | 38.63140836 | 76.78658125 | ENSG00000009694 |
| LIX1L | 0.996596688 | 1.995287568 | 2.48E-04 | 1.67E-03 | 48.17323482 | 96.20725074 | ENSG00000271601 |
| CGN | 0.995887522 | 1.994307013 | 3.46E-68 | 2.18E-65 | 1888.079628 | 3765.971964 | ENSG00000143375 |
| OLFML3 | 0.993418298 | 1.990896605 | 3.86E-39 | 7.15E-37 | 803.3274539 | 1598.691035 | ENSG00000116774 |
| ARHGAP9 | 0.992496823 | 1.98962539 | 1.51E-03 | 8.00E-03 | 31.97997694 | 63.72100792 | ENSG00000123329 |
| GPRIN3 | 0.992428693 | 1.989531433 | 9.31E-05 | 6.97E-04 | 49.67374412 | 98.76213373 | ENSG00000185477 |
| CYTL1 | 0.991999196 | 1.988939228 | 1.50E-08 | 2.42E-07 | 114.608121 | 227.8197625 | ENSG00000170891 |
| HMCN1 | 0.991325531 | 1.988010713 | 7.19E-10 | 1.40E-08 | 241.104276 | 478.4212654 | ENSG00000143341 |
| BRCA2 | 0.989999779 | 1.986184686 | 8.40E-12 | 2.15E-10 | 217.7472051 | 432.7310905 | ENSG00000139618 |
| RAB26 | 0.988986971 | 1.984790824 | 4.08E-05 | 3.35E-04 | 74.18532147 | 147.3111395 | ENSG00000167964 |
| STEAP2 | 0.987622332 | 1.982914307 | 8.51E-12 | 2.17E-10 | 196.4125345 | 390.14875 | ENSG00000157214 |
| TAGLN2 | 0.986894678 | 1.981914434 | 5.06E-56 | 1.87E-53 | 1640.236533 | 3250.556078 | ENSG00000158710 |
| KLF7 | 0.985280396 | 1.97969804 | 1.35E-15 | 5.50E-14 | 272.739777 | 540.1525939 | ENSG00000118263 |
| PADI2 | 0.984503867 | 1.978632758 | 6.97E-09 | 1.18E-07 | 144.8385046 | 285.9428451 | ENSG00000117115 |
| PDLIM3 | 0.981691882 | 1.974779921 | 3.07E-06 | 3.23E-05 | 71.37409781 | 141.0177437 | ENSG00000154553 |
| GADD45G | 0.981006098 | 1.973841435 | 1.06E-11 | 2.67E-10 | 161.3022295 | 318.4846314 | ENSG00000130222 |
| TRPV1 | 0.97999637 | 1.972460446 | 8.40E-04 | 4.81E-03 | 48.50182304 | 95.6182327 | ENSG00000196689 |
| HFE | 0.976821538 | 1.968124571 | 2.29E-02 | 7.67E-02 | 17.58108542 | 34.58450723 | ENSG00000010704 |
| RBM15 | 0.976675032 | 1.967924718 | 4.87E-20 | 2.91E-18 | 376.55668 | 740.2666377 | ENSG00000162775 |
| HLA-DRB1 | 0.974991662 | 1.965629837 | 3.75E-15 | 1.44E-13 | 278.4499119 | 546.6335537 | ENSG00000196126 |
| EFCAB2 | 0.973280875 | 1.963300322 | 9.81E-09 | 1.62E-07 | 116.6628869 | 229.0958327 | ENSG00000203666 |
| IFIT1 | 0.972394959 | 1.962095087 | 6.05E-03 | 2.59E-02 | 28.6868813 | 56.13607309 | ENSG00000185745 |
| SNHG8 | 0.970426232 | 1.959419404 | 3.14E-22 | 2.25E-20 | 562.6475397 | 1103.487977 | ENSG00000269893 |
| KCNH5 | 0.970127262 | 1.959013394 | 2.50E-03 | 1.23E-02 | 33.62845724 | 65.94448299 | ENSG00000140015 |
| EXPH5 | 0.9679234 | 1.95602309 | 1.01E-03 | 5.65E-03 | 37.83223697 | 73.88853969 | ENSG00000110723 |
| RYR1 | 0.967913386 | 1.956009512 | 1.53E-32 | 2.02E-30 | 1045.408952 | 2045.240304 | ENSG00000196218 |
| RRNAD1 | 0.967668728 | 1.955677832 | 5.03E-14 | 1.67E-12 | 243.0158773 | 475.0523136 | ENSG00000143303 |
| MEIS3 | 0.967558759 | 1.955528767 | 2.94E-30 | 3.52E-28 | 633.8202494 | 1238.80523 | ENSG00000105419 |
| ACTRT3 | 0.967127372 | 1.954944122 | 2.51E-04 | 1.68E-03 | 54.83831931 | 107.1971447 | ENSG00000184378 |
| ASH1L | 0.966664211 | 1.95431661 | 2.79E-21 | 1.84E-19 | 444.0627314 | 867.0190267 | ENSG00000116539 |
| ZFHX4 | 0.964710756 | 1.951672194 | 3.71E-03 | 1.72E-02 | 32.38178723 | 63.18548288 | ENSG00000091656 |
| PYROXD2 | 0.963556147 | 1.950110868 | 2.67E-03 | 1.30E-02 | 32.24093518 | 62.77767045 | ENSG00000119943 |
| CADM3 | 0.963383606 | 1.949877657 | 1.03E-04 | 7.64E-04 | 54.33094621 | 106.1158273 | ENSG00000162706 |
| ZMAT4 | 0.961032112 | 1.946702078 | 4.09E-11 | 9.68E-10 | 196.4273572 | 382.5867791 | ENSG00000165061 |
| RNF128 | 0.960708336 | 1.946265239 | 3.08E-03 | 1.47E-02 | 32.8832366 | 63.86674371 | ENSG00000133135 |
| HLA-F | 0.960554173 | 1.946057278 | 7.74E-04 | 4.48E-03 | 43.42709823 | 84.50034262 | ENSG00000204642 |
| NFAM1 | 0.958007181 | 1.942624661 | 1.33E-02 | 4.94E-02 | 24.00804305 | 46.52504301 | ENSG00000235568 |
| RIN2 | 0.957336695 | 1.941722044 | 2.04E-02 | 6.98E-02 | 20.15031726 | 39.14618985 | ENSG00000132669 |
| ACER2 | 0.956552426 | 1.940666784 | 2.52E-05 | 2.19E-04 | 82.66345941 | 160.347967 | ENSG00000177076 |
| DPP6 | 0.955958684 | 1.939868266 | 2.02E-10 | 4.30E-09 | 146.5608435 | 284.2975537 | ENSG00000130226 |
| EVA1C | 0.951175223 | 1.933447007 | 2.41E-03 | 1.19E-02 | 33.17356034 | 64.08296766 | ENSG00000166979 |
| YOD1 | 0.951160302 | 1.933427011 | 1.50E-10 | 3.25E-09 | 152.6177206 | 295.2281633 | ENSG00000180667 |
| KBTBD3 | 0.950740817 | 1.93286492 | 1.84E-02 | 6.43E-02 | 25.67429456 | 49.51783873 | ENSG00000182359 |
| PROS1 | 0.950221924 | 1.932169853 | 2.73E-02 | 8.83E-02 | 17.15435882 | 33.18367785 | ENSG00000184500 |
| CD70 | 0.948802858 | 1.930270264 | 7.97E-03 | 3.25E-02 | 26.03906174 | 50.30435219 | ENSG00000125726 |
| FAM167A | 0.94522559 | 1.925489946 | 3.32E-03 | 1.57E-02 | 35.82435456 | 69.08263564 | ENSG00000154319 |
| SLC45A1 | 0.944676365 | 1.924757063 | 1.85E-05 | 1.67E-04 | 74.35940399 | 143.1369847 | ENSG00000162426 |
| KIAA1614 | 0.943315883 | 1.922942845 | 1.49E-03 | 7.94E-03 | 45.70244685 | 87.83530799 | ENSG00000135835 |
| ZMAT3 | 0.943187027 | 1.922771103 | 3.92E-12 | 1.04E-10 | 199.7082743 | 383.7513311 | ENSG00000172667 |
| PRDX4 | 0.942125684 | 1.921357104 | 1.43E-62 | 6.91E-60 | 2196.625528 | 4219.848584 | ENSG00000123131 |

| | | | | | | | |
|---|---|---|---|---|---|---|---|
| NEB | 0.941759268 | 1.92086918 | 8.44E-03 | 3.41E-02 | 27.60597965 | 52.87824726 | ENSG00000183091 |
| SELENBP1 | 0.94160231 | 1.92066021 | 1.25E-18 | 6.64E-17 | 402.1082509 | 772.0979791 | ENSG00000143416 |
| KCNS1 | 0.937527925 | 1.915243632 | 1.44E-04 | 1.03E-03 | 59.39972112 | 113.8827457 | ENSG00000124134 |
| PLA2G4A | 0.937362598 | 1.915024167 | 5.89E-03 | 2.53E-02 | 28.67116887 | 54.96062102 | ENSG00000116711 |
| MYMX | 0.93567544 | 1.912785952 | 1.33E-02 | 4.94E-02 | 28.29068926 | 54.24220934 | ENSG00000262179 |
| RNF112 | 0.934494206 | 1.911220463 | 1.90E-03 | 9.72E-03 | 45.12983419 | 86.39486877 | ENSG00000128482 |
| PLIN4 | 0.933063959 | 1.909326673 | 9.27E-03 | 3.67E-02 | 27.29757922 | 52.27305501 | ENSG00000167676 |
| ZDHHC21 | 0.932731117 | 1.908886225 | 5.94E-05 | 4.69E-04 | 102.1092365 | 195.2001313 | ENSG00000175893 |
| TP73 | 0.932072805 | 1.908015386 | 1.05E-06 | 1.21E-05 | 92.59939298 | 176.7658797 | ENSG00000078900 |
| CILP2 | 0.93202803 | 1.90795617 | 1.11E-18 | 5.99E-17 | 358.5003084 | 683.9853326 | ENSG00000160161 |
| FAM71E1 | 0.930219165 | 1.905565456 | 5.58E-07 | 6.75E-06 | 97.67528685 | 186.04775 | ENSG00000142530 |
| DUSP2 | 0.928904105 | 1.903829267 | 2.02E-09 | 3.72E-08 | 141.5584772 | 269.5679605 | ENSG00000158050 |
| CALB2 | 0.926902512 | 1.901189728 | 2.16E-03 | 1.08E-02 | 34.77340324 | 66.10310361 | ENSG00000172137 |
| GAL3ST1 | 0.925799661 | 1.899736942 | 3.57E-13 | 1.08E-11 | 218.0313002 | 414.0294654 | ENSG00000128242 |
| UBE2D4 | 0.923574075 | 1.896808555 | 6.75E-03 | 2.84E-02 | 27.11164923 | 51.44215897 | ENSG00000078967 |
| BBC3 | 0.923461396 | 1.896660414 | 7.57E-16 | 3.17E-14 | 279.5643489 | 529.9581262 | ENSG00000105327 |
| BDNF | 0.923128088 | 1.896222276 | 2.77E-05 | 2.38E-04 | 66.81150079 | 126.6876399 | ENSG00000176697 |
| COL1A1 | 0.920830004 | 1.893204169 | 2.30E-29 | 2.64E-27 | 1721.290925 | 3260.386711 | ENSG00000108821 |
| PLEKHN1 | 0.920266729 | 1.892465145 | 5.83E-03 | 2.51E-02 | 34.18062895 | 64.86628705 | ENSG00000187583 |
| LAMC3 | 0.920050497 | 1.892181522 | 5.81E-39 | 1.05E-36 | 1030.315893 | 1950.232191 | ENSG00000050555 |
| RPL39L | 0.919702138 | 1.891724683 | 8.23E-22 | 5.73E-20 | 506.0411675 | 956.8095762 | ENSG00000163923 |
| CDH6 | 0.919495296 | 1.891453483 | 2.28E-03 | 1.14E-02 | 35.5625066 | 67.40187442 | ENSG00000113361 |
| PURA | 0.918697164 | 1.890407376 | 1.46E-05 | 1.33E-04 | 91.49210049 | 173.1764351 | ENSG00000185129 |
| NDUFS2 | 0.918080177 | 1.889599093 | 4.51E-62 | 2.13E-59 | 2160.102531 | 4081.00057 | ENSG00000158864 |
| WIPF1 | 0.917603736 | 1.888975167 | 1.64E-08 | 2.63E-07 | 153.5097432 | 290.3931479 | ENSG00000115935 |
| PKNOX2 | 0.917342838 | 1.888633595 | 3.75E-04 | 2.39E-03 | 53.31767454 | 100.895868 | ENSG00000165495 |
| AKT3 | 0.913880346 | 1.884106278 | 1.48E-15 | 6.02E-14 | 281.595673 | 530.2382689 | ENSG00000117020 |
| RASSF10 | 0.91212128 | 1.881810403 | 7.51E-05 | 5.76E-04 | 62.91731671 | 118.326858 | ENSG00000189431 |
| SLC22A3 | 0.911377266 | 1.880840182 | 7.72E-03 | 3.18E-02 | 39.81550797 | 75.21268101 | ENSG00000146477 |
| ACAP1 | 0.910626662 | 1.879861874 | 4.36E-24 | 3.64E-22 | 552.0212009 | 1036.97215 | ENSG00000072818 |
| ACOT11 | 0.910591837 | 1.879816498 | 1.46E-06 | 1.64E-05 | 97.66371871 | 183.362643 | ENSG00000162390 |
| RPH3AL | 0.90836901 | 1.876922408 | 1.02E-02 | 3.98E-02 | 27.60005592 | 51.6892909 | ENSG00000181031 |
| SMS | 0.907969992 | 1.876403364 | 1.10E-60 | 5.06E-58 | 3995.328304 | 7497.70089 | ENSG00000102172 |
| NPAS1 | 0.907951666 | 1.876379528 | 1.89E-07 | 2.48E-06 | 120.2821803 | 225.8365368 | ENSG00000130751 |
| ITPR2 | 0.907442739 | 1.875717731 | 8.70E-21 | 5.46E-19 | 842.1622324 | 1578.511519 | ENSG00000123104 |
| TRPV4 | 0.906949509 | 1.875076569 | 2.17E-03 | 1.09E-02 | 43.07300947 | 80.47877656 | ENSG00000111199 |
| PYGM | 0.906282135 | 1.874209381 | 3.30E-11 | 7.92E-10 | 207.7056832 | 388.914109 | ENSG00000068976 |
| LAMC1 | 0.905763544 | 1.873535798 | 9.61E-46 | 2.39E-43 | 4151.019431 | 7776.220394 | ENSG00000135862 |
| TMEM47 | 0.905292342 | 1.872923978 | 3.12E-18 | 1.60E-16 | 577.0920416 | 1081.149305 | ENSG00000147027 |
| PAQR7 | 0.905008337 | 1.872555316 | 1.65E-07 | 2.19E-06 | 126.5848881 | 236.9976526 | ENSG00000182749 |
| MST1L | 0.903777422 | 1.870958323 | 7.16E-04 | 4.19E-03 | 49.81014698 | 93.11449928 | ENSG00000186715 |
| ESRRG | 0.903767971 | 1.870946067 | 5.95E-03 | 2.55E-02 | 31.57252224 | 58.9409259 | ENSG00000196482 |
| THBS3 | 0.90243407 | 1.869217009 | 1.37E-04 | 9.83E-04 | 60.31813455 | 112.9384623 | ENSG00000169231 |
| CAPN6 | 0.902407451 | 1.869182521 | 3.59E-09 | 6.30E-08 | 171.9875541 | 321.2586488 | ENSG00000077274 |
| ZNF502 | 0.90238785 | 1.869157126 | 4.65E-02 | 1.33E-01 | 17.55380485 | 32.78407283 | ENSG00000196653 |
| TMEM151A | 0.902112589 | 1.86880053 | 7.08E-03 | 2.95E-02 | 31.65052515 | 59.17405909 | ENSG00000179292 |
| HMGB3 | 0.90097227 | 1.867323998 | 5.60E-90 | 9.61E-87 | 6510.825532 | 12158.04375 | ENSG00000029993 |
| NLRC5 | 0.900709192 | 1.866983519 | 1.36E-02 | 5.04E-02 | 29.66308371 | 55.45727126 | ENSG00000140853 |
| TMC8 | 0.899970002 | 1.866027182 | 1.27E-02 | 4.75E-02 | 25.45249051 | 47.53061828 | ENSG00000167895 |
| PLXDC2 | 0.899480467 | 1.865394109 | 2.33E-19 | 1.32E-17 | 390.1543669 | 727.8455829 | ENSG00000120594 |
| OPCML | 0.896922519 | 1.862089632 | 9.08E-03 | 3.62E-02 | 37.55943126 | 69.95815488 | ENSG00000183715 |
| TMEM184A | 0.896775024 | 1.86189927 | 2.39E-02 | 7.96E-02 | 20.85312971 | 38.81875398 | ENSG00000164855 |
| CYP2A13 | 0.89673163 | 1.861843268 | 1.01E-02 | 3.94E-02 | 27.34413169 | 50.75247982 | ENSG00000197838 |
| RAB24 | 0.896470101 | 1.861505787 | 1.83E-04 | 1.28E-03 | 63.27732921 | 117.9996331 | ENSG00000169228 |
| ABHD12B | 0.896143441 | 1.861084346 | 4.45E-04 | 2.78E-03 | 51.83671647 | 96.34071175 | ENSG00000131969 |
| JMJD4 | 0.895019587 | 1.859635133 | 6.32E-03 | 2.69E-02 | 34.27907281 | 63.8656298 | ENSG00000081692 |
| CSF2RA | 0.893414082 | 1.857566787 | 1.61E-02 | 5.77E-02 | 24.33749484 | 45.06046277 | ENSG00000198223 |
| FAM71F1 | 0.891719811 | 1.855386579 | 1.07E-02 | 4.11E-02 | 28.02202784 | 51.85169535 | ENSG00000135248 |
| SLC17A9 | 0.891418061 | 1.854998553 | 4.12E-18 | 2.09E-16 | 741.9475137 | 1377.003245 | ENSG00000101194 |
| TCIRG1 | 0.890322302 | 1.853590174 | 1.97E-12 | 5.45E-11 | 284.8117078 | 528.1023498 | ENSG00000110719 |
| BOLA1 | 0.889916311 | 1.853068627 | 1.05E-07 | 1.46E-06 | 125.5327415 | 232.3097704 | ENSG00000178096 |
| TMEM220 | 0.889813909 | 1.852937101 | 2.32E-03 | 1.15E-02 | 40.7321158 | 75.48629738 | ENSG00000187824 |
| METTL23 | 0.889482821 | 1.852511913 | 1.62E-08 | 2.60E-07 | 151.2948533 | 280.6154577 | ENSG00000181038 |
| GAP43 | 0.88938459 | 1.852385784 | 1.06E-17 | 5.15E-16 | 399.1564545 | 738.8489943 | ENSG00000172020 |
| MRPL24 | 0.888800681 | 1.85163621 | 2.01E-21 | 1.35E-19 | 432.3751184 | 800.7085596 | ENSG00000143314 |
| PTPN6 | 0.888239936 | 1.850916658 | 9.35E-17 | 4.14E-15 | 341.7886418 | 632.311145 | ENSG00000111679 |
| BAIAP3 | 0.887900628 | 1.850481392 | 1.79E-10 | 3.85E-09 | 219.817989 | 407.0530028 | ENSG00000007516 |
| KCNIP1 | 0.887397106 | 1.849835659 | 2.63E-02 | 8.58E-02 | 22.04404337 | 40.87473863 | ENSG00000182132 |
| CACNG6 | 0.887332502 | 1.849752825 | 3.30E-02 | 1.02E-01 | 20.89820765 | 38.65873591 | ENSG00000130433 |
| HTR1D | 0.886157468 | 1.848246868 | 9.78E-03 | 3.83E-02 | 28.80045278 | 53.25933466 | ENSG00000179546 |
| SVOPL | 0.88523238 | 1.84706211 | 6.36E-03 | 2.70E-02 | 32.81919225 | 60.52882487 | ENSG00000157703 |
| ALDH1A3 | 0.884181923 | 1.845717715 | 5.43E-03 | 2.37E-02 | 38.88466284 | 71.66109255 | ENSG00000184254 |
| FCRL5 | 0.882013127 | 1.842945141 | 2.79E-02 | 8.98E-02 | 22.55617115 | 41.5478647 | ENSG00000143297 |
| EPHA8 | 0.881961169 | 1.84287877 | 1.51E-14 | 5.30E-13 | 267.1926372 | 492.5176166 | ENSG00000070886 |

| | | | | | | | |
|---|---|---|---|---|---|---|---|
| GABRD | 0.879066629 | 1.839185031 | 4.47E-03 | 2.01E-02 | 37.35214444 | 68.73176414 | ENSG00000187730 |
| IFI27L2 | 0.878925108 | 1.839004625 | 1.80E-07 | 2.38E-06 | 128.1974682 | 235.9472432 | ENSG00000119632 |
| LACTB2 | 0.87693033 | 1.836463637 | 1.27E-03 | 6.91E-03 | 46.80679026 | 86.12058999 | ENSG00000147592 |
| CDK18 | 0.876289348 | 1.835647886 | 8.45E-41 | 1.75E-38 | 1252.850806 | 2300.184437 | ENSG00000117266 |
| ADCY7 | 0.8745255 | 1.833404984 | 2.67E-05 | 2.31E-04 | 91.29343331 | 167.4325334 | ENSG00000121281 |
| PDP1 | 0.8744559 | 1.833316536 | 1.80E-12 | 5.03E-11 | 279.8935213 | 513.0928139 | ENSG00000164951 |
| KCNA6 | 0.873655397 | 1.832299573 | 9.17E-04 | 5.20E-03 | 50.66301536 | 93.0566124 | ENSG00000130035 |
| KCNJ6 | 0.873508331 | 1.8321128 | 4.12E-02 | 1.21E-01 | 19.75682112 | 36.28562562 | ENSG00000157542 |
| SEC14L5 | 0.87264633 | 1.831018452 | 5.81E-12 | 1.52E-10 | 241.1559659 | 441.5668836 | ENSG00000103184 |
| CNTNAP2 | 0.872622536 | 1.830988254 | 5.79E-25 | 5.13E-23 | 1492.365306 | 2732.225839 | ENSG00000174469 |
| GLRX2 | 0.870859519 | 1.828752097 | 6.14E-07 | 7.37E-06 | 175.0069537 | 320.5573144 | ENSG00000023572 |
| DYDC2 | 0.870358413 | 1.828117009 | 1.25E-04 | 9.07E-04 | 67.20619214 | 122.8479848 | ENSG00000133665 |
| CA7 | 0.868447626 | 1.82569735 | 1.14E-02 | 4.34E-02 | 28.23523839 | 51.48103376 | ENSG00000168748 |
| HEYL | 0.868422038 | 1.825664969 | 1.12E-07 | 1.54E-06 | 135.8802997 | 248.2991643 | ENSG00000163909 |
| EDEM3 | 0.868195856 | 1.825378768 | 1.70E-11 | 4.19E-10 | 298.4834031 | 544.5537842 | ENSG00000116406 |
| MSTO1 | 0.867875808 | 1.824973871 | 7.72E-18 | 3.81E-16 | 400.8817425 | 731.185931 | ENSG00000125459 |
| BHLHA15 | 0.867200319 | 1.824119593 | 4.22E-03 | 1.92E-02 | 35.85310967 | 65.29585411 | ENSG00000180535 |
| NKX3-1 | 0.867027779 | 1.82390145 | 4.31E-03 | 1.95E-02 | 35.1313308 | 64.02943171 | ENSG00000167034 |
| STEAP1B | 0.864824416 | 1.821118014 | 8.38E-05 | 6.36E-04 | 75.83942002 | 138.2488122 | ENSG00000105889 |
| MAOB | 0.863638755 | 1.819621965 | 1.47E-03 | 7.80E-03 | 46.79995065 | 85.09013866 | ENSG00000069535 |
| USH1C | 0.863392208 | 1.81931103 | 9.93E-03 | 3.88E-02 | 32.27741963 | 58.84342917 | ENSG00000006611 |
| SYN1 | 0.862844806 | 1.81862086 | 1.00E-05 | 9.56E-05 | 88.81166144 | 161.65955 | ENSG00000008056 |
| HBQ1 | 0.859598794 | 1.814533629 | 1.03E-02 | 4.00E-02 | 30.36945563 | 55.17561546 | ENSG00000086506 |
| BFSP1 | 0.859411187 | 1.814297683 | 3.19E-04 | 2.07E-03 | 61.22523302 | 110.8520212 | ENSG00000125864 |
| SARDH | 0.859191607 | 1.814021565 | 4.59E-14 | 1.52E-12 | 308.4113541 | 559.0180341 | ENSG00000123453 |
| CSDC2 | 0.857992634 | 1.812514622 | 7.60E-03 | 3.14E-02 | 33.21032412 | 60.12441741 | ENSG00000172346 |
| PRRG1 | 0.856964429 | 1.811223308 | 7.73E-06 | 7.60E-05 | 96.05973152 | 174.1056058 | ENSG00000130962 |
| LMNA | 0.856612405 | 1.810781415 | 4.57E-30 | 5.36E-28 | 1236.936075 | 2240.968886 | ENSG00000160789 |
| TMCO4 | 0.856386005 | 1.810497274 | 5.63E-06 | 5.68E-05 | 108.1272132 | 195.3851637 | ENSG00000162542 |
| KANK3 | 0.856222898 | 1.810292597 | 1.51E-03 | 8.00E-03 | 46.56746819 | 84.38088035 | ENSG00000186994 |
| LENG9 | 0.856007633 | 1.810022502 | 6.14E-05 | 4.82E-04 | 74.10940349 | 134.1685516 | ENSG00000275183 |
| PKDCC | 0.855661857 | 1.80958874 | 2.14E-22 | 1.55E-20 | 916.8593299 | 1660.323301 | ENSG00000162878 |
| ADAMTSL4 | 0.855066178 | 1.808841727 | 1.95E-05 | 1.75E-04 | 135.8200685 | 245.2677477 | ENSG00000143382 |
| CCSAP | 0.85458692 | 1.808240936 | 6.65E-12 | 1.72E-10 | 412.5897692 | 745.9248026 | ENSG00000154429 |
| DPEP1 | 0.854474074 | 1.808099502 | 1.45E-03 | 7.74E-03 | 61.96480925 | 111.8823046 | ENSG00000015413 |
| TRHDE | 0.853624976 | 1.80703566 | 1.15E-02 | 4.41E-02 | 31.13839737 | 56.47466982 | ENSG00000072657 |
| FES | 0.85359276 | 1.806995308 | 1.06E-05 | 1.00E-04 | 104.9586723 | 189.725663 | ENSG00000182511 |
| LTBR | 0.850491225 | 1.803114765 | 4.27E-03 | 1.94E-02 | 50.54290975 | 90.8040783 | ENSG00000111321 |
| TFAP2C | 0.84829248 | 1.800368814 | 1.00E-05 | 9.54E-05 | 101.5816239 | 182.7177979 | ENSG00000087510 |
| VAMP4 | 0.847516716 | 1.799400982 | 1.15E-06 | 1.32E-05 | 133.6707493 | 240.2717776 | ENSG00000117533 |
| TMEM169 | 0.847030791 | 1.798795015 | 2.26E-03 | 1.13E-02 | 50.53075679 | 90.70391166 | ENSG00000163449 |
| CASP8 | 0.846644315 | 1.79831321 | 1.57E-03 | 8.28E-03 | 49.14648872 | 88.26007262 | ENSG00000064012 |
| PLD1 | 0.846467996 | 1.798093442 | 2.56E-02 | 8.39E-02 | 22.52177164 | 40.47853857 | ENSG00000075651 |
| ACTC1 | 0.846320295 | 1.797909366 | 4.61E-09 | 8.01E-08 | 253.0671874 | 455.7263553 | ENSG00000159251 |
| THEM4 | 0.846272699 | 1.797850052 | 2.58E-09 | 4.65E-08 | 186.6096981 | 335.0621404 | ENSG00000159445 |
| KLKB1 | 0.846108072 | 1.797644909 | 5.12E-07 | 6.27E-06 | 147.7155776 | 265.9811857 | ENSG00000164344 |
| HTR6 | 0.845918849 | 1.797409146 | 1.54E-03 | 8.15E-03 | 48.83127483 | 87.65221046 | ENSG00000158748 |
| MST1R | 0.84515586 | 1.796458814 | 1.53E-03 | 8.09E-03 | 56.0167751 | 100.4199538 | ENSG00000164078 |
| INPP5D | 0.844702861 | 1.795894823 | 8.87E-11 | 2.00E-09 | 253.6721152 | 455.3259723 | ENSG00000168918 |
| ATP1B1 | 0.844490014 | 1.795629886 | 2.30E-26 | 2.26E-24 | 793.3592569 | 1425.351469 | ENSG00000143153 |
| DGKH | 0.839551579 | 1.789493842 | 3.71E-05 | 3.08E-04 | 86.96092833 | 155.3803639 | ENSG00000102780 |
| CST3 | 0.83843108 | 1.788104533 | 9.86E-15 | 3.58E-13 | 355.0983954 | 635.5482348 | ENSG00000101439 |
| FN1 | 0.838237297 | 1.787864371 | 2.59E-25 | 2.40E-23 | 2666.960164 | 4766.577664 | ENSG00000115414 |
| PAH | 0.837760452 | 1.787273538 | 2.50E-02 | 8.25E-02 | 24.55546008 | 43.85192725 | ENSG00000171759 |
| TSEN15 | 0.837672601 | 1.787164707 | 2.21E-21 | 1.47E-19 | 659.4710755 | 1178.399723 | ENSG00000198860 |
| REXO5 | 0.83574391 | 1.784777103 | 2.10E-04 | 1.44E-03 | 82.87283116 | 148.2371459 | ENSG00000005189 |
| FAM241A | 0.835469807 | 1.784438038 | 1.23E-02 | 4.64E-02 | 34.68652751 | 62.14739124 | ENSG00000174749 |
| CHCHD10 | 0.835327312 | 1.784261797 | 1.41E-10 | 3.08E-09 | 551.3870628 | 984.8397749 | ENSG00000250479 |
| PCED1B | 0.834185891 | 1.782850697 | 1.28E-04 | 9.27E-04 | 77.39777061 | 138.2623595 | ENSG00000179715 |
| AHNAK2 | 0.833913162 | 1.782513696 | 9.58E-04 | 5.41E-03 | 60.29469278 | 107.4046667 | ENSG00000185567 |
| KLHL12 | 0.832159227 | 1.780347948 | 3.45E-26 | 3.31E-24 | 924.329913 | 1644.558357 | ENSG00000117153 |
| KDELR3 | 0.831340373 | 1.779337734 | 1.07E-06 | 1.23E-05 | 121.0291548 | 215.2248494 | ENSG00000100196 |
| FCSK | 0.830956319 | 1.778864126 | 1.14E-07 | 1.56E-06 | 149.6745177 | 265.9140299 | ENSG00000157353 |
| KLK10 | 0.82873196 | 1.776123573 | 3.78E-04 | 2.40E-03 | 60.91802779 | 108.1680281 | ENSG00000129451 |
| GABRE | 0.826508346 | 1.773388156 | 3.05E-03 | 1.46E-02 | 73.63818378 | 130.4780803 | ENSG00000102287 |
| ARL4D | 0.826306066 | 1.773139528 | 1.30E-08 | 2.11E-07 | 228.6163231 | 404.7707473 | ENSG00000175906 |
| QPCT | 0.826076549 | 1.772857464 | 2.01E-04 | 1.38E-03 | 69.6867688 | 123.3957846 | ENSG00000115828 |
| ANXA9 | 0.825927465 | 1.772674271 | 4.06E-04 | 2.56E-03 | 65.49514203 | 115.9250152 | ENSG00000143412 |
| GEMIN8 | 0.825696145 | 1.772390065 | 2.75E-08 | 4.20E-07 | 170.2207215 | 301.4227899 | ENSG00000046647 |
| PKP3 | 0.825495886 | 1.772144058 | 6.91E-13 | 2.03E-11 | 349.7295081 | 620.3454793 | ENSG00000184363 |
| TNFRSF18 | 0.825165325 | 1.771738059 | 3.66E-02 | 1.11E-01 | 22.96243064 | 40.80720394 | ENSG00000186891 |
| ACKR3 | 0.82461452 | 1.771061757 | 3.86E-02 | 1.16E-01 | 21.72763426 | 38.57285155 | ENSG00000144476 |
| FZD9 | 0.824379054 | 1.770772722 | 2.06E-06 | 2.25E-05 | 130.2083252 | 230.9122734 | ENSG00000188763 |
| TMEM37 | 0.82370258 | 1.769942607 | 3.34E-11 | 8.01E-10 | 231.6701746 | 410.2318792 | ENSG00000171227 |

| | | | | | | | |
|---|---|---|---|---|---|---|---|
| DDX28 | 0.82347744 | 1.769666421 | 2.74E-13 | 8.46E-12 | 341.8017362 | 605.3652241 | ENSG00000182810 |
| XPR1 | 0.82275464 | 1.768780028 | 1.66E-33 | 2.36E-31 | 1428.774595 | 2526.723258 | ENSG00000143324 |
| RASIP1 | 0.82209979 | 1.767977347 | 8.06E-08 | 1.14E-06 | 170.6557361 | 301.388266 | ENSG00000105538 |
| UFSP1 | 0.821881779 | 1.767710202 | 5.90E-03 | 2.54E-02 | 41.07196667 | 72.81564063 | ENSG00000176125 |
| ZNF132 | 0.820963715 | 1.766585671 | 2.86E-02 | 9.13E-02 | 25.5616128 | 45.09281118 | ENSG00000131849 |
| ZNF79 | 0.819010477 | 1.764195542 | 9.25E-06 | 8.91E-05 | 106.2990315 | 187.740661 | ENSG00000196152 |
| CEP350 | 0.818812842 | 1.763953881 | 3.94E-12 | 1.05E-10 | 552.3550496 | 973.7026712 | ENSG00000135837 |
| RPS27 | 0.81770328 | 1.762597764 | 5.09E-34 | 7.52E-32 | 10141.68507 | 17876.1448 | ENSG00000177954 |
| CPNE7 | 0.817023177 | 1.761767051 | 3.72E-02 | 1.12E-01 | 363.0801442 | 639.756897 | ENSG00000178773 |
| DMTN | 0.816098361 | 1.760638061 | 3.92E-02 | 1.17E-01 | 23.5492812 | 41.40853971 | ENSG00000158856 |
| STPG1 | 0.815310278 | 1.759676562 | 1.14E-07 | 1.56E-06 | 156.6806221 | 275.6437872 | ENSG00000001460 |
| GEM | 0.815281779 | 1.759641801 | 5.82E-05 | 4.60E-04 | 84.63902723 | 148.9699352 | ENSG00000164949 |
| AMZ1 | 0.814758849 | 1.759004105 | 4.50E-03 | 2.03E-02 | 41.25344746 | 72.62306324 | ENSG00000174945 |
| AP3B2 | 0.814298784 | 1.758443259 | 4.76E-11 | 1.12E-09 | 261.4812298 | 459.3020418 | ENSG00000103723 |
| PKD1L2 | 0.813563349 | 1.757547096 | 1.78E-02 | 6.25E-02 | 31.010588 | 54.6013312 | ENSG00000166473 |
| CLHC1 | 0.812731054 | 1.756533454 | 9.15E-03 | 3.64E-02 | 40.14051057 | 70.62714093 | ENSG00000162994 |
| OSBPL5 | 0.811969755 | 1.75560679 | 2.14E-11 | 5.21E-10 | 314.3780754 | 551.8464625 | ENSG00000021762 |
| POMK | 0.811217992 | 1.754692212 | 7.20E-03 | 3.00E-02 | 37.94344419 | 66.63227022 | ENSG00000185900 |
| SPATA18 | 0.811029498 | 1.75446297 | 4.77E-02 | 1.36E-01 | 20.64584289 | 36.12416713 | ENSG00000163071 |
| SYT14 | 0.809464149 | 1.752560379 | 4.89E-07 | 6.02E-06 | 205.3734086 | 360.1150052 | ENSG00000143469 |
| SLC46A1 | 0.807709389 | 1.750430024 | 4.98E-03 | 2.20E-02 | 41.36968869 | 72.43003434 | ENSG00000076351 |
| TP53I11 | 0.807579399 | 1.750272313 | 4.25E-28 | 4.56E-26 | 1172.7128 | 2053.292636 | ENSG00000175274 |
| GRIK2 | 0.807419388 | 1.750078199 | 2.25E-02 | 7.57E-02 | 29.04094391 | 50.89993957 | ENSG00000164418 |
| RNF180 | 0.807078065 | 1.749664202 | 3.55E-02 | 1.08E-01 | 23.99616942 | 41.89242073 | ENSG00000164197 |
| PPP1R15B | 0.806951417 | 1.749510614 | 2.10E-27 | 2.19E-25 | 1010.357047 | 1766.65491 | ENSG00000158615 |
| MGMT | 0.806820964 | 1.749352425 | 1.65E-10 | 3.57E-09 | 271.791998 | 475.7200701 | ENSG00000170430 |
| FOSL1 | 0.8060357 | 1.748400505 | 2.63E-07 | 3.40E-06 | 160.4318692 | 280.1525963 | ENSG00000175592 |
| HLA-DPA1 | 0.805696099 | 1.747988991 | 1.31E-02 | 4.87E-02 | 32.94847616 | 57.48677148 | ENSG00000231389 |
| NIT2 | 0.804432069 | 1.746458147 | 2.13E-11 | 5.19E-10 | 295.714335 | 516.0319259 | ENSG00000114021 |
| OBSCN | 0.803631593 | 1.745489396 | 1.81E-31 | 2.26E-29 | 1240.427989 | 2164.634164 | ENSG00000154358 |
| MTMR11 | 0.803367842 | 1.745170319 | 4.50E-03 | 2.03E-02 | 45.62327489 | 79.47558766 | ENSG00000014914 |
| VEPH1 | 0.80308003 | 1.744822198 | 6.11E-03 | 2.61E-02 | 57.86633915 | 100.9398287 | ENSG00000197415 |
| BTN3A2 | 0.803006686 | 1.744733498 | 2.30E-04 | 1.56E-03 | 74.05275742 | 129.0103356 | ENSG00000186470 |
| DYRK4 | 0.801115541 | 1.742447926 | 1.76E-02 | 6.19E-02 | 32.30617474 | 56.1858252 | ENSG00000010219 |
| SMYD3 | 0.800007423 | 1.741110085 | 6.16E-12 | 1.60E-10 | 308.8191137 | 538.0817504 | ENSG00000185420 |
| LYSMD1 | 0.799665748 | 1.740697785 | 1.34E-05 | 1.24E-04 | 105.0550313 | 182.7322482 | ENSG00000163155 |
| TSNAX | 0.799510881 | 1.740510938 | 6.51E-07 | 7.77E-06 | 219.9543913 | 383.3109594 | ENSG00000116918 |
| SNHG5 | 0.799458064 | 1.740447219 | 2.73E-41 | 5.73E-39 | 3749.169288 | 6524.201423 | ENSG00000203875 |
| ATP2B2 | 0.799443166 | 1.740429248 | 1.66E-02 | 5.91E-02 | 29.71291632 | 51.75370417 | ENSG00000157087 |
| ARL8A | 0.792148734 | 1.731651645 | 1.98E-23 | 1.57E-21 | 767.4722587 | 1328.870035 | ENSG00000143862 |
| LAD1 | 0.792063542 | 1.731549394 | 7.49E-16 | 3.14E-14 | 502.3474562 | 869.8010337 | ENSG00000159166 |
| SLFN5 | 0.791751815 | 1.731175294 | 7.08E-05 | 5.47E-04 | 92.60679126 | 160.0938572 | ENSG00000166750 |
| CDC42BPA | 0.791402152 | 1.730755763 | 1.93E-21 | 1.30E-19 | 816.2979153 | 1412.265852 | ENSG00000143776 |
| S100A10 | 0.791110677 | 1.730406125 | 3.58E-21 | 2.34E-19 | 614.9313981 | 1064.006793 | ENSG00000197747 |
| COQ8A | 0.790494407 | 1.729667112 | 1.53E-25 | 1.44E-23 | 962.4736279 | 1665.731634 | ENSG00000163050 |
| TNRC18 | 0.790463725 | 1.729630328 | 3.25E-06 | 3.39E-05 | 157.2562094 | 272.1885089 | ENSG00000182095 |
| RASAL2 | 0.790430124 | 1.729590045 | 8.99E-18 | 4.39E-16 | 597.0386066 | 1031.809709 | ENSG00000075391 |
| ZNF469 | 0.789964218 | 1.729031578 | 3.11E-03 | 1.49E-02 | 55.72586654 | 96.42922307 | ENSG00000225614 |
| ADGRD2 | 0.789418435 | 1.728377595 | 4.61E-02 | 1.32E-01 | 24.99019534 | 43.32276063 | ENSG00000180264 |
| PSMD4 | 0.789213918 | 1.728132597 | 1.76E-52 | 5.84E-50 | 4329.793616 | 7481.70548 | ENSG00000159352 |
| TMEM238 | 0.788707151 | 1.727525673 | 2.85E-03 | 1.38E-02 | 52.83430186 | 91.36079089 | ENSG00000233493 |
| DNER | 0.787698349 | 1.726318126 | 1.10E-03 | 6.09E-03 | 60.0375995 | 103.5528095 | ENSG00000187957 |
| TOMM20 | 0.787008005 | 1.725492263 | 3.05E-51 | 9.59E-49 | 4442.025632 | 7663.766886 | ENSG00000173726 |
| TTC39A | 0.786265113 | 1.724603978 | 3.90E-05 | 3.23E-04 | 110.2142675 | 189.648975 | ENSG00000085831 |
| SPATS2L | 0.78443814 | 1.722421389 | 1.37E-18 | 7.21E-17 | 934.718841 | 1609.617039 | ENSG00000196141 |
| SLC8A2 | 0.783985149 | 1.721880651 | 5.07E-07 | 6.22E-06 | 206.5933567 | 355.6983233 | ENSG00000118160 |
| ASPM | 0.783567781 | 1.721382588 | 5.61E-19 | 3.12E-17 | 1063.463797 | 1829.671363 | ENSG00000066279 |
| ZBTB18 | 0.783334951 | 1.721104804 | 8.68E-10 | 1.68E-08 | 268.3167851 | 461.4325593 | ENSG00000179456 |
| ELK4 | 0.783280502 | 1.721039849 | 3.85E-12 | 1.03E-10 | 315.5138436 | 542.5197878 | ENSG00000158711 |
| DECR2 | 0.782970143 | 1.720669651 | 6.27E-05 | 4.91E-04 | 98.8439801 | 169.9425823 | ENSG00000242612 |
| ADAMTS10 | 0.782914146 | 1.720602866 | 1.02E-05 | 9.72E-05 | 178.1318912 | 306.7456287 | ENSG00000142303 |
| CDKN2D | 0.781878708 | 1.719368414 | 1.39E-02 | 5.12E-02 | 34.9154505 | 60.1530773 | ENSG00000129355 |
| RAB42 | 0.781062753 | 1.718396254 | 4.24E-08 | 6.28E-07 | 175.7334617 | 301.9462808 | ENSG00000188060 |
| C1orf56 | 0.779992579 | 1.717122041 | 6.18E-04 | 3.68E-03 | 67.03894924 | 115.2097976 | ENSG00000143443 |
| SCG5 | 0.779454599 | 1.716481846 | 1.35E-12 | 3.80E-11 | 340.6985624 | 584.3818676 | ENSG00000166922 |
| DNAJC6 | 0.779238847 | 1.716225169 | 5.92E-05 | 4.68E-04 | 94.66272626 | 162.3885554 | ENSG00000116675 |
| ZYG11A | 0.77890066 | 1.71582291 | 2.15E-02 | 7.30E-02 | 29.39650721 | 50.4838768 | ENSG00000203995 |
| SOD3 | 0.778104124 | 1.714875837 | 1.16E-04 | 8.45E-04 | 84.74339484 | 145.2803668 | ENSG00000109610 |
| CREB3L4 | 0.77762369 | 1.714304859 | 3.20E-10 | 6.63E-09 | 294.8602714 | 504.7971279 | ENSG00000143578 |
| CHGB | 0.776162795 | 1.712569806 | 7.28E-05 | 5.60E-04 | 95.82071494 | 163.9921771 | ENSG00000089199 |
| IFITM1 | 0.7745795 | 1.710691366 | 6.71E-05 | 5.22E-04 | 823.7637687 | 1409.023651 | ENSG00000185885 |
| DUSP23 | 0.773058992 | 1.708889357 | 8.27E-18 | 4.07E-16 | 598.4297654 | 1023.443927 | ENSG00000158716 |
| CD74 | 0.772184211 | 1.707853483 | 7.02E-18 | 3.49E-16 | 607.0848832 | 1036.892763 | ENSG00000019582 |
| DEGS1 | 0.770754926 | 1.706162341 | 3.33E-23 | 2.54E-21 | 766.1547826 | 1307.521051 | ENSG00000143753 |

| | | | | | | | |
|---|---|---|---|---|---|---|---|
| C21orf91 | 0.770302973 | 1.705627936 | 2.02E-08 | 3.17E-07 | 335.5043157 | 572.1476286 | ENSG00000154642 |
| MRPL55 | 0.770216723 | 1.70552597 | 1.63E-13 | 5.15E-12 | 425.5176215 | 726.1115193 | ENSG00000162910 |
| COL1A2 | 0.770042962 | 1.705320566 | 4.83E-35 | 7.41E-33 | 2327.271441 | 3968.274787 | ENSG00000164692 |
| HS3ST2 | 0.76992581 | 1.705182093 | 5.32E-03 | 2.33E-02 | 45.73211784 | 77.92077215 | ENSG00000122254 |
| PPL | 0.769441798 | 1.704610115 | 4.17E-25 | 3.78E-23 | 1005.65782 | 1713.734349 | ENSG00000118898 |
| DHX58 | 0.768428576 | 1.703413367 | 1.22E-03 | 6.67E-03 | 65.10637442 | 110.7197897 | ENSG00000108771 |
| RNF213 | 0.767420839 | 1.702223931 | 1.84E-15 | 7.39E-14 | 1042.9555 | 1774.239086 | ENSG00000173821 |
| MYL9 | 0.76618255 | 1.700763511 | 5.49E-32 | 7.00E-30 | 1430.345403 | 2433.443397 | ENSG00000101335 |
| ILDR2 | 0.766146886 | 1.700721468 | 4.99E-02 | 1.41E-01 | 24.87423345 | 42.3060245 | ENSG00000143195 |
| SUSD2 | 0.766133078 | 1.700705191 | 1.08E-07 | 1.49E-06 | 215.164067 | 365.896028 | ENSG00000099994 |
| ABL2 | 0.765707371 | 1.700203424 | 1.49E-14 | 5.22E-13 | 537.6116304 | 913.6811333 | ENSG00000143322 |
| RAB4A | 0.765400589 | 1.699841923 | 1.26E-17 | 6.05E-16 | 647.7635808 | 1100.961362 | ENSG00000168118 |
| PARVB | 0.764968487 | 1.69933288 | 2.48E-09 | 4.50E-08 | 244.8822706 | 415.8729873 | ENSG00000188677 |
| MME | 0.764964244 | 1.699327881 | 1.01E-08 | 1.67E-07 | 236.2407285 | 401.0679152 | ENSG00000196549 |
| SNHG12 | 0.764743774 | 1.699068213 | 2.28E-06 | 2.46E-05 | 320.1046894 | 543.9176973 | ENSG00000197989 |
| GDPD5 | 0.764664977 | 1.698975416 | 8.11E-05 | 6.17E-04 | 113.9258018 | 193.3782914 | ENSG00000158555 |
| ATP2C2 | 0.764427369 | 1.698695623 | 1.38E-03 | 7.40E-03 | 67.22607443 | 114.3139212 | ENSG00000064270 |
| ZNF562 | 0.763267066 | 1.697329978 | 1.61E-09 | 3.00E-08 | 308.7067119 | 523.3529016 | ENSG00000171466 |
| F8A1 | 0.762939992 | 1.696945218 | 7.63E-04 | 4.43E-03 | 79.98479981 | 136.0174133 | ENSG00000277203 |
| ADAMTS1 | 0.762198362 | 1.696073113 | 2.33E-15 | 9.19E-14 | 496.736857 | 841.9055824 | ENSG00000154734 |
| PCDHGC4 | 0.762046571 | 1.695894673 | 2.25E-03 | 1.13E-02 | 72.03094261 | 121.9437627 | ENSG00000242419 |
| CEP112 | 0.761164526 | 1.694858142 | 7.42E-03 | 3.07E-02 | 45.40947951 | 77.15194595 | ENSG00000154240 |
| ADGRA2 | 0.760926891 | 1.694578994 | 4.15E-04 | 2.61E-03 | 99.67015215 | 169.36338 | ENSG00000020181 |
| C6orf141 | 0.760906048 | 1.694554512 | 1.12E-03 | 6.18E-03 | 86.63829 | 146.9448288 | ENSG00000197261 |
| SNAPC5 | 0.759376391 | 1.692758767 | 2.56E-08 | 3.92E-07 | 203.3260932 | 344.2191788 | ENSG00000174446 |
| ZNF687 | 0.758544144 | 1.691782547 | 1.15E-32 | 1.52E-30 | 1825.088199 | 3088.096168 | ENSG00000143373 |
| PCDH11X | 0.757619927 | 1.690699107 | 5.79E-05 | 4.59E-04 | 111.7832704 | 188.601687 | ENSG00000102290 |
| TDRKH | 0.756589279 | 1.689491719 | 1.16E-16 | 5.09E-15 | 494.7322809 | 835.6340069 | ENSG00000182134 |
| ZNF286B | 0.753200168 | 1.685527502 | 3.43E-02 | 1.06E-01 | 31.04526684 | 52.19312995 | ENSG00000249459 |
| FIGNL1 | 0.753039318 | 1.685339589 | 2.08E-06 | 2.27E-05 | 169.8699646 | 286.7412248 | ENSG00000132436 |
| SCAPER | 0.752877449 | 1.685150506 | 6.16E-05 | 4.83E-04 | 106.5958638 | 179.466047 | ENSG00000140386 |
| ELFN2 | 0.752548802 | 1.684766671 | 6.02E-04 | 3.61E-03 | 78.48990933 | 131.9775929 | ENSG00000166897 |
| SLC45A3 | 0.75213381 | 1.684282117 | 5.44E-13 | 1.61E-11 | 466.6189798 | 786.4645081 | ENSG00000158715 |
| GSTM3 | 0.751588024 | 1.683645057 | 1.03E-02 | 4.01E-02 | 45.63367397 | 76.80632837 | ENSG00000134202 |
| PTER | 0.751414627 | 1.683442712 | 5.16E-05 | 4.13E-04 | 118.2179573 | 199.0123263 | ENSG00000165983 |
| ALKAL2 | 0.751185458 | 1.683175322 | 3.08E-02 | 9.70E-02 | 35.51414853 | 59.71393507 | ENSG00000189292 |
| DISP1 | 0.751087849 | 1.683061447 | 3.58E-04 | 2.29E-03 | 88.55576336 | 149.1453924 | ENSG00000154309 |
| CD55 | 0.749662432 | 1.681399363 | 8.55E-11 | 1.93E-09 | 312.6916878 | 526.2664979 | ENSG00000196352 |
| IGSF21 | 0.74930823 | 1.680986607 | 5.64E-10 | 1.12E-08 | 403.6832815 | 678.7589423 | ENSG00000117154 |
| RGL1 | 0.748967279 | 1.680589387 | 3.26E-04 | 2.11E-03 | 91.01498319 | 152.9362413 | ENSG00000143344 |
| CGREF1 | 0.748908628 | 1.680521066 | 2.11E-03 | 1.07E-02 | 58.73636835 | 98.61393893 | ENSG00000138028 |
| TULP1 | 0.747064148 | 1.6783739 | 1.48E-04 | 1.06E-03 | 94.04739992 | 158.0073434 | ENSG00000112041 |
| PFKFB2 | 0.746533312 | 1.67775646 | 4.39E-13 | 1.32E-11 | 473.7581297 | 794.1415701 | ENSG00000123836 |
| TMEM79 | 0.746047141 | 1.677191171 | 3.43E-02 | 1.06E-01 | 32.14721984 | 53.92290824 | ENSG00000163472 |
| ERO1B | 0.745336353 | 1.676365055 | 1.71E-03 | 8.91E-03 | 65.17517345 | 109.0273731 | ENSG00000086619 |
| HMCN2 | 0.745088693 | 1.676077306 | 3.79E-12 | 1.01E-10 | 399.3717762 | 669.2188424 | ENSG00000148357 |
| CREG1 | 0.744484718 | 1.675375775 | 1.51E-13 | 4.80E-12 | 513.0914852 | 860.1071024 | ENSG00000143162 |
| POGK | 0.744127704 | 1.674961232 | 2.22E-31 | 2.76E-29 | 1346.634221 | 2255.079223 | ENSG00000143157 |
| SP8 | 0.74250192 | 1.673074769 | 8.78E-05 | 6.61E-04 | 111.9193939 | 186.9821021 | ENSG00000164651 |
| NPIPA1 | 0.741017343 | 1.671354009 | 9.21E-05 | 6.91E-04 | 113.9895668 | 190.5027825 | ENSG00000183426 |
| ZFP3 | 0.739316693 | 1.669384977 | 4.24E-02 | 1.24E-01 | 27.74090797 | 46.30323863 | ENSG00000180787 |
| CHRNB2 | 0.739216859 | 1.66926946 | 1.32E-05 | 1.22E-04 | 132.6337303 | 221.2046495 | ENSG00000160716 |
| PDGFRB | 0.73890072 | 1.668903712 | 5.17E-10 | 1.04E-08 | 297.5140141 | 496.2785761 | ENSG00000113721 |
| EIF4E1B | 0.737989776 | 1.667850268 | 1.55E-02 | 5.60E-02 | 38.39745136 | 64.02725624 | ENSG00000175766 |
| SORCS2 | 0.737659801 | 1.667468839 | 3.83E-03 | 1.77E-02 | 64.4614294 | 107.5200616 | ENSG00000184985 |
| RBMS2 | 0.737552628 | 1.667344973 | 1.87E-06 | 2.05E-05 | 163.829969 | 273.3780324 | ENSG00000076067 |
| BLZF1 | 0.73720699 | 1.666945561 | 8.62E-07 | 1.01E-05 | 182.4077239 | 304.15519 | ENSG00000117475 |
| ZBTB37 | 0.736931921 | 1.666627767 | 3.38E-05 | 2.84E-04 | 164.1205203 | 272.9203296 | ENSG00000185278 |
| SLC50A1 | 0.736330502 | 1.665933141 | 5.47E-20 | 3.25E-18 | 714.6709086 | 1189.954244 | ENSG00000169241 |
| FGF2 | 0.734729943 | 1.664085941 | 2.90E-22 | 2.09E-20 | 911.7183456 | 1517.791613 | ENSG00000138685 |
| C1QTNF6 | 0.734449857 | 1.663762905 | 2.42E-03 | 1.20E-02 | 61.34711865 | 102.019676 | ENSG00000133466 |
| KCNT2 | 0.734438059 | 1.6637493 | 2.00E-05 | 1.78E-04 | 146.1239971 | 242.9016153 | ENSG00000162687 |
| ETV3 | 0.733618087 | 1.662803958 | 1.40E-15 | 5.67E-14 | 721.5856352 | 1199.881316 | ENSG00000117036 |
| GIPC3 | 0.733556744 | 1.662733258 | 2.17E-03 | 1.09E-02 | 72.93600787 | 121.3506773 | ENSG00000179855 |
| SDE2 | 0.732981823 | 1.662070782 | 3.13E-12 | 8.43E-11 | 400.5264847 | 665.0652855 | ENSG00000143751 |
| CDH2 | 0.731322034 | 1.660159706 | 8.71E-15 | 3.20E-13 | 574.568778 | 953.7110928 | ENSG00000170558 |
| CCDC177 | 0.731230251 | 1.660054091 | 4.00E-07 | 5.00E-06 | 250.8419507 | 416.9213165 | ENSG00000267909 |
| GAS2L1 | 0.729356178 | 1.657899067 | 7.30E-09 | 1.23E-07 | 291.9820799 | 484.6221254 | ENSG00000185340 |
| CNTN2 | 0.729107652 | 1.657613494 | 2.10E-04 | 1.44E-03 | 131.2960403 | 217.8659527 | ENSG00000184144 |
| DIAPH2 | 0.72879405 | 1.657253213 | 2.01E-23 | 1.59E-21 | 955.514278 | 1582.875258 | ENSG00000147202 |
| NEFH | 0.728209008 | 1.656581299 | 4.58E-05 | 3.73E-04 | 118.4350067 | 195.9839451 | ENSG00000100285 |
| RPL36A | 0.72770972 | 1.65600809 | 4.25E-02 | 1.24E-01 | 28.04191013 | 46.49704553 | ENSG00000241343 |
| CHP2 | 0.727014154 | 1.655209871 | 1.99E-02 | 6.84E-02 | 37.88323864 | 62.57326978 | ENSG00000166869 |
| HYAL1 | 0.726641095 | 1.654781914 | 9.63E-04 | 5.43E-03 | 73.72183109 | 122.0024276 | ENSG00000114378 |

| | | | | | | | |
|---|---|---|---|---|---|---|---|
| KCTD13 | 0.726149561 | 1.654218217 | 2.74E-06 | 2.92E-05 | 171.0187494 | 283.2015026 | ENSG00000174943 |
| UAP1L1 | 0.726056851 | 1.654111918 | 1.31E-06 | 1.49E-05 | 202.9924716 | 335.6184968 | ENSG00000197355 |
| EPM2A | 0.725399331 | 1.653358215 | 6.08E-04 | 3.64E-03 | 79.33921824 | 131.2650452 | ENSG00000112425 |
| P3H2 | 0.725146756 | 1.653068783 | 4.98E-12 | 1.31E-10 | 411.4549691 | 680.5356551 | ENSG00000090530 |
| PRNP | 0.72358464 | 1.651279849 | 1.17E-12 | 3.33E-11 | 468.1582077 | 773.034516 | ENSG00000171867 |
| GCK | 0.723345612 | 1.651006284 | 9.56E-07 | 1.11E-05 | 167.2458144 | 276.007995 | ENSG00000106633 |
| DCAF6 | 0.72235996 | 1.649878699 | 2.09E-09 | 3.84E-08 | 282.7554673 | 466.5888833 | ENSG00000143164 |
| ICAM3 | 0.722254742 | 1.649758375 | 2.49E-03 | 1.23E-02 | 65.19386054 | 107.5551526 | ENSG00000076662 |
| SH3PXD2A | 0.721380932 | 1.648759454 | 6.64E-16 | 2.80E-14 | 567.5680131 | 935.5448677 | ENSG00000107957 |
| BTN3A1 | 0.721228108 | 1.648584811 | 2.70E-03 | 1.31E-02 | 76.66589814 | 126.3192694 | ENSG00000026950 |
| STC1 | 0.720337349 | 1.647567244 | 5.01E-08 | 7.36E-07 | 327.1524876 | 538.1460778 | ENSG00000159167 |
| LDHD | 0.720249001 | 1.647466354 | 3.55E-04 | 2.28E-03 | 105.3904329 | 173.6212007 | ENSG00000166816 |
| COL12A1 | 0.717660471 | 1.644513067 | 4.30E-02 | 1.26E-01 | 50.11915781 | 82.56469356 | ENSG00000111799 |
| EBF4 | 0.717336588 | 1.644143917 | 3.33E-02 | 1.03E-01 | 37.54338777 | 61.50037073 | ENSG00000088881 |
| OMA1 | 0.71729371 | 1.644095053 | 9.67E-05 | 7.20E-04 | 112.8638665 | 185.405196 | ENSG00000162600 |
| ZNF34 | 0.716820775 | 1.643556184 | 6.53E-04 | 3.87E-03 | 87.53534659 | 144.0287211 | ENSG00000196378 |
| PTAR1 | 0.716647313 | 1.643358583 | 1.33E-16 | 5.81E-15 | 702.5289573 | 1154.0591 | ENSG00000188647 |
| GPATCH2 | 0.716439359 | 1.643121722 | 6.13E-09 | 1.05E-07 | 276.1719191 | 453.9286634 | ENSG00000092978 |
| TOR1AIP2 | 0.716045161 | 1.642672822 | 8.36E-13 | 2.43E-11 | 521.1882021 | 856.028543 | ENSG00000169905 |
| PAQR6 | 0.715548113 | 1.642106974 | 4.98E-03 | 2.20E-02 | 89.13785924 | 146.4375122 | ENSG00000160781 |
| PMVK | 0.715539084 | 1.642096696 | 2.98E-13 | 9.12E-12 | 542.3328763 | 891.1851294 | ENSG00000163344 |
| SCYL3 | 0.714368669 | 1.640765053 | 9.82E-05 | 7.30E-04 | 108.2215127 | 177.5536227 | ENSG00000000457 |
| GUCY2C | 0.714091259 | 1.640449587 | 2.77E-02 | 8.92E-02 | 34.52075915 | 56.76474393 | ENSG00000070019 |
| TTC32 | 0.713970591 | 1.640312384 | 3.51E-03 | 1.64E-02 | 65.12297657 | 106.7138737 | ENSG00000183891 |
| PIFO | 0.713514253 | 1.63979362 | 6.90E-03 | 2.89E-02 | 51.85807331 | 84.87268521 | ENSG00000173947 |
| TNFRSF1B | 0.713289017 | 1.639537633 | 2.95E-04 | 1.94E-03 | 96.39157373 | 158.1250817 | ENSG00000028137 |
| ENO3 | 0.713124368 | 1.63935053 | 9.16E-08 | 1.29E-06 | 269.6524419 | 441.9220037 | ENSG00000108515 |
| RENBP | 0.711452137 | 1.637451456 | 8.15E-08 | 1.15E-06 | 231.4768201 | 379.3620273 | ENSG00000102032 |
| PPFIBP2 | 0.711326581 | 1.637308957 | 5.89E-06 | 5.92E-05 | 168.5636739 | 275.9541029 | ENSG00000166387 |
| RNF2 | 0.710940452 | 1.6368708 | 3.32E-11 | 7.96E-10 | 634.406287 | 1039.058766 | ENSG00000121481 |
| TRIM72 | 0.710683083 | 1.636578816 | 3.32E-02 | 1.03E-01 | 36.93401132 | 60.32028418 | ENSG00000177238 |
| TICAM1 | 0.710607759 | 1.636493372 | 4.69E-02 | 1.34E-01 | 35.43942576 | 57.9543627 | ENSG00000127666 |
| TGFB1I1 | 0.70900627 | 1.634677762 | 2.50E-04 | 1.67E-03 | 100.953281 | 165.0292502 | ENSG00000140682 |
| SWT1 | 0.708764554 | 1.634403902 | 1.85E-04 | 1.29E-03 | 106.3135749 | 173.6444254 | ENSG00000116668 |
| COLEC12 | 0.707950048 | 1.633481424 | 1.28E-02 | 4.78E-02 | 50.05630866 | 81.87213359 | ENSG00000158270 |
| PLCD4 | 0.70725961 | 1.632699867 | 9.63E-03 | 3.79E-02 | 52.64954093 | 85.93482254 | ENSG00000115556 |
| RGS9 | 0.706443677 | 1.631776735 | 1.85E-06 | 2.03E-05 | 169.6703554 | 276.8405721 | ENSG00000108370 |
| GNG12 | 0.706026194 | 1.631304605 | 9.64E-13 | 2.77E-11 | 519.6670248 | 847.9071169 | ENSG00000172380 |
| NDUFAF1 | 0.705674697 | 1.630907204 | 8.53E-11 | 1.93E-09 | 336.2409434 | 548.2374589 | ENSG00000137806 |
| PSD4 | 0.70560706 | 1.630830744 | 3.64E-02 | 1.10E-01 | 39.95905593 | 65.35809192 | ENSG00000125637 |
| P2RX4 | 0.705408373 | 1.630606163 | 2.50E-04 | 1.67E-03 | 102.9054071 | 167.6183735 | ENSG00000135124 |
| EFR3A | 0.704053899 | 1.629075988 | 9.02E-15 | 3.30E-13 | 682.1429292 | 1111.323034 | ENSG00000132294 |
| SURF1 | 0.704048104 | 1.629069443 | 1.38E-10 | 3.01E-09 | 452.1161492 | 736.2928818 | ENSG00000148290 |
| NTPCR | 0.703928088 | 1.628933929 | 4.26E-21 | 2.76E-19 | 1007.547019 | 1640.362409 | ENSG00000135778 |
| ANKRD29 | 0.703131839 | 1.628035139 | 8.28E-03 | 3.36E-02 | 59.26481896 | 96.52922176 | ENSG00000154065 |
| JOSD2 | 0.702748405 | 1.627602504 | 2.78E-02 | 8.96E-02 | 46.34774908 | 75.74090167 | ENSG00000161677 |
| SLC9B2 | 0.702576928 | 1.62740906 | 4.61E-03 | 2.06E-02 | 58.58216813 | 95.35537806 | ENSG00000164038 |
| SYT11 | 0.702024443 | 1.626785958 | 7.41E-08 | 1.06E-06 | 284.8772 | 463.864786 | ENSG00000132718 |
| TRIM56 | 0.701883189 | 1.626626688 | 6.37E-18 | 3.18E-16 | 738.3854256 | 1200.959525 | ENSG00000169871 |
| C1orf21 | 0.701660817 | 1.626375984 | 3.89E-18 | 1.98E-16 | 733.421297 | 1192.952314 | ENSG00000116667 |
| IRF6 | 0.699287968 | 1.623703227 | 1.29E-10 | 2.83E-09 | 403.1518301 | 654.4091226 | ENSG00000117595 |
| RINL | 0.698973956 | 1.623349856 | 2.51E-06 | 2.69E-05 | 189.8162235 | 308.1295785 | ENSG00000187994 |
| NCAN | 0.698908033 | 1.62327568 | 8.70E-07 | 1.02E-05 | 216.6331253 | 351.9190141 | ENSG00000130287 |
| GPR156 | 0.698059863 | 1.622321626 | 1.06E-02 | 4.09E-02 | 51.33882657 | 83.24591144 | ENSG00000175697 |
| TFR2 | 0.697852703 | 1.62208869 | 6.23E-05 | 4.88E-04 | 121.330157 | 196.6388169 | ENSG00000106327 |
| NRP2 | 0.697163185 | 1.621313618 | 5.88E-14 | 1.93E-12 | 689.5079294 | 1118.047048 | ENSG00000118257 |
| CEP170 | 0.696949044 | 1.621072983 | 6.43E-17 | 2.89E-15 | 1001.126699 | 1622.552639 | ENSG00000143702 |
| ANKRD50 | 0.696674122 | 1.620764098 | 3.39E-13 | 1.03E-11 | 632.8958599 | 1025.241866 | ENSG00000151458 |
| IBA57 | 0.696420269 | 1.620478937 | 7.48E-08 | 1.07E-06 | 242.7815887 | 393.55806 | ENSG00000181873 |
| NPTXR | 0.695876611 | 1.6198684 | 4.71E-11 | 1.11E-09 | 444.011476 | 719.30965 | ENSG00000221890 |
| RGS5 | 0.695297701 | 1.619218526 | 2.61E-07 | 3.38E-06 | 219.7011112 | 355.6086254 | ENSG00000143248 |
| LMO7 | 0.695258122 | 1.619174104 | 7.91E-08 | 1.12E-06 | 233.691379 | 378.2062497 | ENSG00000136153 |
| COL11A1 | 0.695018933 | 1.618905679 | 8.93E-09 | 1.48E-07 | 315.8827028 | 510.8320926 | ENSG00000060718 |
| NUP133 | 0.694753837 | 1.618608231 | 7.63E-24 | 6.32E-22 | 1837.771949 | 2974.735282 | ENSG00000069248 |
| SDCCAG8 | 0.693683894 | 1.61740827 | 9.50E-04 | 5.37E-03 | 92.83215478 | 149.9352382 | ENSG00000054282 |
| GALNT12 | 0.693175722 | 1.616838659 | 7.35E-08 | 1.05E-06 | 329.6936787 | 533.3334497 | ENSG00000119514 |
| PDE4DIP | 0.691434056 | 1.614887939 | 9.70E-17 | 4.29E-15 | 832.5484552 | 1344.897599 | ENSG00000178104 |
| COL25A1 | 0.691208458 | 1.614635435 | 1.24E-04 | 8.97E-04 | 116.4472859 | 188.0942024 | ENSG00000188517 |
| WDR26 | 0.690846643 | 1.614230549 | 2.31E-23 | 1.79E-21 | 1545.773774 | 2494.912034 | ENSG00000162923 |
| CACNA1B | 0.689970082 | 1.613250063 | 2.04E-11 | 4.98E-10 | 376.9149386 | 607.8249109 | ENSG00000148408 |
| TUBB4A | 0.689607985 | 1.61284521 | 1.26E-16 | 5.53E-15 | 675.6594039 | 1089.899016 | ENSG00000104833 |
| SPIRE2 | 0.689301578 | 1.612502702 | 9.67E-08 | 1.35E-06 | 230.6453086 | 371.9698378 | ENSG00000204991 |
| RO60 | 0.689154658 | 1.612338498 | 2.09E-06 | 2.27E-05 | 316.0900669 | 509.9630865 | ENSG00000116747 |
| PLAAT5 | 0.688959916 | 1.612120871 | 1.85E-03 | 9.46E-03 | 72.34615651 | 116.5095144 | ENSG00000168004 |

| | | | | | | | |
|---|---|---|---|---|---|---|---|
| STX6 | 0.688905112 | 1.612059633 | 2.82E-20 | 1.71E-18 | 967.2532743 | 1559.95025 | ENSG00000135823 |
| PLPPR4 | 0.687988674 | 1.611035934 | 1.79E-02 | 6.29E-02 | 41.88062127 | 67.35458132 | ENSG00000117600 |
| HLA-DRB5 | 0.687428439 | 1.61041045 | 9.32E-03 | 3.69E-02 | 52.24417117 | 84.10771548 | ENSG00000198502 |
| TRANK1 | 0.687386378 | 1.6103635 | 6.84E-07 | 8.15E-06 | 257.0410771 | 413.2697496 | ENSG00000168016 |
| BCAN | 0.687007923 | 1.609941117 | 4.53E-04 | 2.82E-03 | 108.4095538 | 174.7296422 | ENSG00000132692 |
| JPH2 | 0.686937287 | 1.609862294 | 2.59E-02 | 8.47E-02 | 38.13440819 | 61.3249135 | ENSG00000149596 |
| ELAVL2 | 0.686003908 | 1.6088211 | 2.57E-07 | 3.33E-06 | 271.985989 | 437.754616 | ENSG00000107105 |
| VASH2 | 0.685418139 | 1.608168012 | 8.01E-21 | 5.07E-19 | 1448.944009 | 2330.267905 | ENSG00000143494 |
| ST8SIA3 | 0.684778228 | 1.607454864 | 4.48E-04 | 2.79E-03 | 134.7777619 | 216.9455041 | ENSG00000177511 |
| DGAT2 | 0.684496023 | 1.607140461 | 1.09E-03 | 6.03E-03 | 88.37247697 | 141.8707733 | ENSG00000062282 |
| CHRNA4 | 0.684121072 | 1.606722825 | 1.77E-03 | 9.15E-03 | 84.82554204 | 136.053997 | ENSG00000101204 |
| RASGRF2 | 0.683802717 | 1.606368313 | 2.70E-10 | 5.65E-09 | 357.4338979 | 574.0347683 | ENSG00000113319 |
| SCCPDH | 0.682645712 | 1.605080564 | 7.28E-14 | 2.37E-12 | 570.8030137 | 916.461931 | ENSG00000143653 |
| SH2D3A | 0.681972528 | 1.604331782 | 3.19E-14 | 1.09E-12 | 534.4116653 | 857.3049195 | ENSG00000125731 |
| HYLS1 | 0.681735991 | 1.604068766 | 1.88E-05 | 1.69E-04 | 169.0980482 | 271.2654857 | ENSG00000198331 |
| ADA2 | 0.680947777 | 1.603192625 | 4.30E-16 | 1.83E-14 | 939.4856462 | 1505.309636 | ENSG00000093072 |
| ZNF91 | 0.680203333 | 1.602365576 | 9.89E-09 | 1.63E-07 | 280.2381269 | 449.1975985 | ENSG00000167232 |
| ATRNL1 | 0.679870888 | 1.601996381 | 1.99E-05 | 1.77E-04 | 155.2189615 | 248.7914637 | ENSG00000107518 |
| NUDT6 | 0.67965016 | 1.601751299 | 1.47E-02 | 5.37E-02 | 45.9939658 | 73.66968878 | ENSG00000170917 |
| TMEM63A | 0.678498453 | 1.600473127 | 3.99E-04 | 2.52E-03 | 1656.658209 | 2651.347259 | ENSG00000196187 |
| HS1BP3 | 0.677686278 | 1.599572383 | 1.47E-10 | 3.19E-09 | 386.3918571 | 617.9692476 | ENSG00000118960 |
| APELA | 0.676096862 | 1.597811106 | 2.93E-06 | 3.10E-05 | 228.56334 | 365.669843 | ENSG00000248329 |
| RCOR3 | 0.675976165 | 1.597677437 | 4.55E-08 | 6.70E-07 | 245.750572 | 392.7100868 | ENSG00000117625 |
| PTGES | 0.675513787 | 1.59716547 | 3.88E-03 | 1.79E-02 | 75.46844977 | 120.7526739 | ENSG00000148344 |
| ADAM19 | 0.675338559 | 1.596971492 | 1.22E-17 | 5.85E-16 | 818.3011209 | 1306.726527 | ENSG00000135074 |
| MAML2 | 0.675204279 | 1.59682286 | 6.27E-04 | 3.73E-03 | 97.83660601 | 156.0367486 | ENSG00000184384 |
| ISG15 | 0.674704823 | 1.59627014 | 1.85E-09 | 3.43E-08 | 329.3505216 | 525.7640267 | ENSG00000187608 |
| SH2D3C | 0.674397464 | 1.5959301 | 1.33E-03 | 7.19E-03 | 90.7065566 | 144.8761273 | ENSG00000095370 |
| TCEAL2 | 0.674291734 | 1.595813144 | 2.38E-09 | 4.34E-08 | 391.8935356 | 625.1775116 | ENSG00000184905 |
| IAH1 | 0.67408739 | 1.595587128 | 7.28E-06 | 7.19E-05 | 188.743636 | 301.2000825 | ENSG00000134330 |
| SLC5A5 | 0.673936937 | 1.595420739 | 3.99E-02 | 1.18E-01 | 38.97123308 | 62.03975225 | ENSG00000105641 |
| ZFP36 | 0.673735599 | 1.595198103 | 1.48E-03 | 7.88E-03 | 95.15649802 | 152.0349 | ENSG00000128016 |
| EHHADH | 0.67367358 | 1.59512953 | 4.22E-02 | 1.24E-01 | 32.63771146 | 52.14082351 | ENSG00000113790 |
| LGALS8 | 0.673302011 | 1.594718754 | 6.63E-05 | 5.16E-04 | 150.4840882 | 239.6190814 | ENSG00000116977 |
| CLU | 0.673104617 | 1.594500574 | 1.39E-40 | 2.80E-38 | 3877.674218 | 6184.062362 | ENSG00000120885 |
| CD47 | 0.672656392 | 1.594005262 | 8.73E-08 | 1.23E-06 | 286.5651405 | 456.5072359 | ENSG00000196776 |
| TNNC2 | 0.672651685 | 1.594000062 | 1.30E-02 | 4.84E-02 | 53.79212267 | 85.65042417 | ENSG00000101470 |
| GPR89A | 0.672118368 | 1.593410921 | 3.38E-04 | 2.17E-03 | 129.0360731 | 205.3532558 | ENSG00000117262 |
| DES | 0.671976394 | 1.593254123 | 1.53E-02 | 5.52E-02 | 46.8601562 | 74.75367606 | ENSG00000175084 |
| TTPAL | 0.671826579 | 1.593088681 | 1.18E-08 | 1.92E-07 | 301.7024386 | 480.947815 | ENSG00000124120 |
| MRPS21 | 0.671742664 | 1.592996021 | 5.06E-14 | 1.68E-12 | 1121.302933 | 1786.354424 | ENSG00000266472 |
| SDHC | 0.671740494 | 1.592993626 | 2.27E-30 | 2.73E-28 | 1906.04882 | 3035.850307 | ENSG00000143252 |
| TMEM130 | 0.67089188 | 1.59205688 | 4.21E-03 | 1.92E-02 | 65.30417803 | 103.9060674 | ENSG00000166448 |
| ATF6 | 0.670764008 | 1.591915776 | 5.16E-13 | 1.53E-11 | 475.3410905 | 756.71057 | ENSG00000118217 |
| PLOD2 | 0.670450506 | 1.591569885 | 1.04E-07 | 1.45E-06 | 418.187658 | 665.1145525 | ENSG00000152952 |
| GLMP | 0.66999082 | 1.591062843 | 5.53E-10 | 1.11E-08 | 363.950732 | 579.1624754 | ENSG00000198715 |
| SPART | 0.669866109 | 1.590925314 | 1.01E-18 | 5.50E-17 | 922.089015 | 1466.92566 | ENSG00000133104 |
| RAB13 | 0.669765606 | 1.590814487 | 2.40E-06 | 2.58E-05 | 261.0452988 | 415.5357786 | ENSG00000143545 |
| MAPK11 | 0.669453835 | 1.590470745 | 1.49E-02 | 5.43E-02 | 50.60253048 | 80.68506915 | ENSG00000185386 |
| CNIH3 | 0.668632648 | 1.5895657 | 1.45E-03 | 7.74E-03 | 99.96164552 | 158.6582204 | ENSG00000143786 |
| UEVLD | 0.667983618 | 1.588850758 | 2.74E-03 | 1.33E-02 | 79.30954726 | 126.0496048 | ENSG00000151116 |
| DAP3 | 0.66762379 | 1.588454526 | 3.54E-17 | 1.62E-15 | 1473.107572 | 2339.393845 | ENSG00000132676 |
| CX3CL1 | 0.667106619 | 1.587885207 | 1.75E-02 | 6.17E-02 | 47.98257631 | 76.1434172 | ENSG00000006210 |
| NMB | 0.666805451 | 1.587553764 | 8.91E-06 | 8.65E-05 | 178.1766636 | 283.1372573 | ENSG00000197696 |
| SAMD4A | 0.6665174 | 1.587236822 | 1.70E-03 | 8.86E-03 | 86.17599483 | 136.9275517 | ENSG00000020577 |
| TAF10 | 0.666394907 | 1.587102062 | 5.39E-04 | 3.28E-03 | 98.6695921 | 156.5321694 | ENSG00000166337 |
| ANP32E | 0.666012704 | 1.586681658 | 6.96E-23 | 5.26E-21 | 3996.489267 | 6341.507058 | ENSG00000143401 |
| LCTL | 0.665101506 | 1.585679835 | 1.30E-04 | 9.40E-04 | 124.6769241 | 197.5801898 | ENSG00000188501 |
| GALNT6 | 0.664847213 | 1.585400365 | 1.38E-06 | 1.55E-05 | 215.0652921 | 340.5712278 | ENSG00000139629 |
| CLDN11 | 0.664637539 | 1.585169967 | 1.59E-03 | 8.38E-03 | 85.9117826 | 136.3090958 | ENSG00000013297 |
| GALE | 0.664072486 | 1.584549233 | 1.89E-10 | 4.05E-09 | 516.7949073 | 818.46067 | ENSG00000117308 |
| ZDHHC1 | 0.663762992 | 1.584209345 | 1.01E-02 | 3.94E-02 | 53.07270806 | 84.01456969 | ENSG00000159714 |
| RORA | 0.663728668 | 1.584171653 | 3.30E-02 | 1.02E-01 | 47.10568134 | 74.54114063 | ENSG00000069667 |
| ORMDL2 | 0.663371938 | 1.58377999 | 7.89E-06 | 7.74E-05 | 184.9241485 | 292.7329638 | ENSG00000123353 |
| DISC1 | 0.662483609 | 1.582805089 | 2.62E-02 | 8.55E-02 | 41.21904795 | 65.30361001 | ENSG00000162946 |
| INAVA | 0.662151163 | 1.582440399 | 7.33E-17 | 3.27E-15 | 880.9281917 | 1394.631253 | ENSG00000163362 |
| RAD51C | 0.66204245 | 1.58232116 | 4.75E-06 | 4.85E-05 | 189.74923 | 300.5642955 | ENSG00000108384 |
| RBFOX1 | 0.66161899 | 1.581856785 | 4.14E-02 | 1.22E-01 | 35.03405599 | 55.44639395 | ENSG00000078328 |
| MPPE1 | 0.661226575 | 1.581426576 | 1.82E-03 | 9.33E-03 | 101.0641572 | 159.4890438 | ENSG00000154889 |
| TMEM177 | 0.660762789 | 1.580918274 | 6.74E-16 | 2.83E-14 | 954.9826501 | 1510.294507 | ENSG00000144120 |
| DCXR | 0.660228343 | 1.580332731 | 9.04E-19 | 4.95E-17 | 953.5648467 | 1507.429655 | ENSG00000169738 |
| SNAP47 | 0.660003201 | 1.580086129 | 1.07E-18 | 5.77E-17 | 992.7220621 | 1568.975407 | ENSG00000143740 |
| SCN1B | 0.659562518 | 1.579603553 | 2.09E-02 | 7.13E-02 | 46.25965257 | 72.89573365 | ENSG00000105711 |
| SPINK5 | 0.659025023 | 1.579015161 | 1.26E-02 | 4.74E-02 | 63.21119992 | 99.87177508 | ENSG00000133710 |

| | | | | | | | |
|---|---|---|---|---|---|---|---|
| APOBEC3F | 0.658159574 | 1.578068219 | 4.60E-03 | 2.06E-02 | 77.43928907 | 122.4317541 | ENSG00000128394 |
| KRTCAP2 | 0.657917778 | 1.577803757 | 1.86E-02 | 6.47E-02 | 52.11399753 | 82.19192915 | ENSG00000163463 |
| DDIT4L | 0.657760906 | 1.577632204 | 1.16E-03 | 6.41E-03 | 88.71469211 | 139.9001785 | ENSG00000145358 |
| ZNFX1 | 0.655715767 | 1.575397364 | 8.34E-09 | 1.39E-07 | 349.1705229 | 550.4017746 | ENSG00000124201 |
| EMILIN1 | 0.655639849 | 1.575314465 | 7.94E-05 | 6.06E-04 | 149.0491233 | 234.9879721 | ENSG00000138080 |
| TARBP1 | 0.655530779 | 1.575195373 | 3.59E-13 | 1.08E-11 | 536.7670762 | 845.2187635 | ENSG00000059588 |
| ENDOG | 0.655300033 | 1.574943455 | 4.13E-05 | 3.39E-04 | 176.5231755 | 278.4764695 | ENSG00000167136 |
| CENPL | 0.655223058 | 1.574859426 | 2.84E-05 | 2.43E-04 | 192.0005265 | 302.067592 | ENSG00000120334 |
| CAPG | 0.654647256 | 1.574231002 | 1.10E-14 | 3.97E-13 | 686.2672059 | 1079.986583 | ENSG00000042493 |
| ANKRD33B | 0.654347619 | 1.573904079 | 3.05E-10 | 6.33E-09 | 384.1698483 | 604.5369121 | ENSG00000164236 |
| HHIPL1 | 0.65412249 | 1.573658494 | 2.53E-04 | 1.70E-03 | 121.7456204 | 191.7586838 | ENSG00000182218 |
| ARHGAP4 | 0.65381224 | 1.573320117 | 5.99E-14 | 1.97E-12 | 723.6760487 | 1138.506024 | ENSG00000089820 |
| PLEKHG4 | 0.653487268 | 1.572965762 | 1.68E-09 | 3.12E-08 | 403.7080676 | 634.831715 | ENSG00000196155 |
| LIN9 | 0.652865529 | 1.572288027 | 3.77E-07 | 4.74E-06 | 296.5758224 | 466.3755496 | ENSG00000183814 |
| NUDT19 | 0.652499098 | 1.571888731 | 2.82E-02 | 9.05E-02 | 45.1559457 | 71.08775426 | ENSG00000213965 |
| MMRN2 | 0.652427626 | 1.571810861 | 3.07E-02 | 9.67E-02 | 38.6237046 | 60.77210033 | ENSG00000173269 |
| C3 | 0.652113571 | 1.571468736 | 2.77E-07 | 3.58E-06 | 267.9627225 | 420.5921563 | ENSG00000125730 |
| THSD4 | 0.651355814 | 1.57064356 | 1.16E-03 | 6.39E-03 | 96.03987538 | 150.7368643 | ENSG00000187720 |
| GRIP2 | 0.651240026 | 1.570517508 | 7.01E-04 | 4.12E-03 | 198.7326823 | 312.318678 | ENSG00000144596 |
| CHST13 | 0.651090048 | 1.57035425 | 4.51E-03 | 2.03E-02 | 78.58868426 | 123.6233374 | ENSG00000180767 |
| GAL | 0.650454799 | 1.569662942 | 2.97E-12 | 8.04E-11 | 821.3350372 | 1290.243566 | ENSG00000069482 |
| CXCL5 | 0.650207592 | 1.569394002 | 1.60E-02 | 5.75E-02 | 72.44637982 | 113.765248 | ENSG00000163735 |
| PRKAB2 | 0.649922461 | 1.569083862 | 2.21E-12 | 6.10E-11 | 721.8848329 | 1131.656535 | ENSG00000131791 |
| CLDN19 | 0.6492561 | 1.56835929 | 1.08E-04 | 7.96E-04 | 152.4441967 | 238.7706136 | ENSG00000164007 |
| PIAS3 | 0.647613358 | 1.566574476 | 6.40E-15 | 2.39E-13 | 783.1875619 | 1226.667044 | ENSG00000131788 |
| LACTB | 0.647311316 | 1.566246533 | 4.39E-08 | 6.48E-07 | 290.4145516 | 455.0095293 | ENSG00000103642 |
| SLC52A3 | 0.646981421 | 1.565888427 | 6.30E-06 | 6.27E-05 | 185.3660289 | 290.2801393 | ENSG00000101276 |
| VGLL3 | 0.646601102 | 1.565475687 | 3.94E-03 | 1.81E-02 | 75.30031715 | 117.8961341 | ENSG00000206538 |
| KLC3 | 0.646532634 | 1.565401393 | 3.94E-03 | 1.81E-02 | 85.66623131 | 134.2399428 | ENSG00000104892 |
| PLK3 | 0.646469267 | 1.565332638 | 4.98E-04 | 3.06E-03 | 120.6487014 | 188.8243647 | ENSG00000173846 |
| SLC34A2 | 0.64626918 | 1.565115557 | 2.28E-02 | 7.64E-02 | 46.86519021 | 73.19763104 | ENSG00000157765 |
| SPRTN | 0.646092658 | 1.564924069 | 1.83E-05 | 1.64E-04 | 207.0035073 | 323.8635472 | ENSG00000010072 |
| PBXIP1 | 0.645921691 | 1.564738628 | 3.90E-15 | 1.50E-13 | 661.1744485 | 1034.449332 | ENSG00000163346 |
| RRP15 | 0.645124802 | 1.563874565 | 2.92E-10 | 6.07E-09 | 904.284676 | 1414.801133 | ENSG00000067533 |
| ARL4A | 0.644878967 | 1.563608104 | 3.79E-08 | 5.66E-07 | 323.080765 | 505.4158674 | ENSG00000122644 |
| PSMB4 | 0.643893061 | 1.562539933 | 8.56E-39 | 1.51E-36 | 5700.126335 | 8906.72536 | ENSG00000159377 |
| RRM2B | 0.643774434 | 1.562411458 | 1.84E-18 | 9.59E-17 | 1206.047191 | 1884.324592 | ENSG00000048392 |
| METAP1D | 0.643749096 | 1.562384017 | 7.32E-03 | 3.04E-02 | 66.380325 | 103.6209113 | ENSG00000172878 |
| CAND2 | 0.643470952 | 1.562082827 | 7.10E-07 | 8.43E-06 | 294.7123254 | 460.7178885 | ENSG00000144712 |
| MELTF | 0.643432137 | 1.5620408 | 7.19E-03 | 3.00E-02 | 85.70744486 | 133.634467 | ENSG00000163975 |
| NES | 0.642382982 | 1.560905268 | 1.08E-43 | 2.51E-41 | 6337.300145 | 9892.377601 | ENSG00000132688 |
| IPP | 0.641423968 | 1.55986802 | 5.61E-03 | 2.43E-02 | 83.79533658 | 130.7349326 | ENSG00000197429 |
| CTC1 | 0.640691836 | 1.559076626 | 6.21E-12 | 1.61E-10 | 498.4580007 | 777.10536 | ENSG00000178971 |
| MRVI1 | 0.639333374 | 1.557609269 | 5.53E-03 | 2.40E-02 | 71.71186375 | 111.7173255 | ENSG00000072952 |
| ACBD3 | 0.639132898 | 1.557392839 | 2.65E-15 | 1.04E-13 | 740.6322979 | 1153.73528 | ENSG00000182827 |
| KLHL11 | 0.638624163 | 1.556843755 | 1.13E-02 | 4.33E-02 | 68.04332252 | 105.6182311 | ENSG00000178502 |
| RALGPS2 | 0.637617166 | 1.555757462 | 4.81E-08 | 7.07E-07 | 583.0842907 | 906.823972 | ENSG00000116191 |
| TMEM267 | 0.63698 | 1.555070514 | 3.26E-03 | 1.54E-02 | 75.75076485 | 117.8095147 | ENSG00000151881 |
| FRAT1 | 0.636208597 | 1.554239246 | 2.34E-03 | 1.16E-02 | 91.4651254 | 142.3232518 | ENSG00000165879 |
| PLEKHA4 | 0.635787454 | 1.553785609 | 3.52E-07 | 4.43E-06 | 355.4586361 | 551.7300263 | ENSG00000105559 |
| ABRAXAS1 | 0.635480312 | 1.553454851 | 3.62E-07 | 4.55E-06 | 296.4363926 | 460.4842085 | ENSG00000163322 |
| FBXL6 | 0.635036617 | 1.552977165 | 1.12E-08 | 1.83E-07 | 519.2323674 | 807.1325785 | ENSG00000182325 |
| ARHGEF11 | 0.634104735 | 1.551974373 | 2.27E-15 | 8.98E-14 | 983.7080503 | 1525.893143 | ENSG00000132694 |
| NRAV | 0.633972824 | 1.551832476 | 5.45E-04 | 3.31E-03 | 123.2751641 | 191.1096987 | ENSG00000248008 |
| PIGC | 0.633359903 | 1.551173329 | 1.48E-10 | 3.22E-09 | 468.8210547 | 727.0872778 | ENSG00000135845 |
| WIF1 | 0.633156538 | 1.550954688 | 1.33E-02 | 4.94E-02 | 60.36201728 | 93.75783127 | ENSG00000156076 |
| DPH6 | 0.632953507 | 1.550736437 | 2.02E-04 | 1.39E-03 | 128.1254151 | 198.6004989 | ENSG00000134146 |
| SQSTM1 | 0.632443836 | 1.550188695 | 4.89E-22 | 3.44E-20 | 1428.728399 | 2215.492566 | ENSG00000161011 |
| ETNPPL | 0.632124755 | 1.549845877 | 3.90E-02 | 1.17E-01 | 38.90571419 | 60.36568537 | ENSG00000164089 |
| TRIM11 | 0.631570089 | 1.54925013 | 1.39E-11 | 3.46E-10 | 535.8692078 | 830.3365089 | ENSG00000154370 |
| H19 | 0.631542049 | 1.549220019 | 2.92E-02 | 9.30E-02 | 42.42447883 | 65.6742716 | ENSG00000130600 |
| GORAB | 0.631328378 | 1.548990588 | 1.26E-05 | 1.17E-04 | 205.6127568 | 318.8982352 | ENSG00000120370 |
| TNS1 | 0.631222438 | 1.548876846 | 2.67E-05 | 2.31E-04 | 165.0511123 | 255.4803541 | ENSG00000079308 |
| ZNF669 | 0.631144067 | 1.548792709 | 4.94E-06 | 5.03E-05 | 222.8527961 | 345.3680512 | ENSG00000188295 |
| NANOS1 | 0.630675041 | 1.548289273 | 9.79E-03 | 3.83E-02 | 74.76952843 | 115.7585434 | ENSG00000188613 |
| CERS2 | 0.630538872 | 1.548143145 | 1.34E-25 | 1.26E-23 | 2184.760526 | 3382.177691 | ENSG00000143418 |
| MPI | 0.630439532 | 1.548036548 | 1.97E-11 | 4.83E-10 | 539.5987402 | 835.4357968 | ENSG00000178802 |
| ACTN2 | 0.630311854 | 1.547899552 | 2.51E-02 | 8.27E-02 | 47.62820821 | 73.75434363 | ENSG00000077522 |
| SV2A | 0.630141442 | 1.547716724 | 1.42E-22 | 1.05E-20 | 2621.080884 | 4055.78361 | ENSG00000159164 |
| TATDN3 | 0.629905617 | 1.547463754 | 2.66E-06 | 2.83E-05 | 214.9671532 | 332.8919177 | ENSG00000203705 |
| SRD5A3 | 0.629622481 | 1.547160086 | 1.90E-08 | 3.00E-07 | 358.1690766 | 554.4377909 | ENSG00000128039 |
| LIFR | 0.629443209 | 1.546967845 | 1.95E-05 | 1.75E-04 | 205.2260224 | 317.3111439 | ENSG00000113594 |
| PYROXD1 | 0.629012477 | 1.546506049 | 1.63E-02 | 5.82E-02 | 57.46241778 | 88.69181515 | ENSG00000121350 |
| SERPINF2 | 0.628708741 | 1.546180492 | 2.80E-02 | 9.01E-02 | 46.7403555 | 72.41575206 | ENSG00000167711 |

| | | | | | | | |
|---|---|---|---|---|---|---|---|
| RPS27L | 0.628564634 | 1.546026056 | 1.33E-09 | 2.50E-08 | 683.9723316 | 1058.002813 | ENSG00000185088 |
| IARS2 | 0.628099279 | 1.545527451 | 1.95E-32 | 2.55E-30 | 2563.617958 | 3962.282442 | ENSG00000067704 |
| NUMBL | 0.627279169 | 1.544649135 | 1.98E-11 | 4.85E-10 | 508.8411541 | 785.749406 | ENSG00000105245 |
| SUMF2 | 0.62716371 | 1.544525521 | 6.93E-15 | 2.57E-13 | 1101.599401 | 1700.422528 | ENSG00000129103 |
| C5orf24 | 0.626627895 | 1.543951993 | 3.28E-04 | 2.12E-03 | 170.4728325 | 263.6758937 | ENSG00000181904 |
| ATP2A3 | 0.626503126 | 1.543818473 | 9.23E-16 | 3.82E-14 | 1028.883371 | 1588.525572 | ENSG00000074370 |
| ELFN1 | 0.626126834 | 1.543415857 | 1.92E-03 | 9.79E-03 | 111.3194745 | 171.7900262 | ENSG00000225968 |
| LRRN2 | 0.625717048 | 1.542977525 | 3.79E-08 | 5.66E-07 | 300.6920899 | 464.0194345 | ENSG00000170382 |
| DNAH14 | 0.625245838 | 1.542473643 | 4.44E-07 | 5.50E-06 | 268.1457034 | 413.5508813 | ENSG00000185842 |
| NAGLU | 0.624438527 | 1.541610739 | 5.77E-07 | 6.96E-06 | 298.7539485 | 460.9332095 | ENSG00000108784 |
| ZNF552 | 0.624171668 | 1.541325609 | 8.57E-04 | 4.89E-03 | 114.3735536 | 176.1587526 | ENSG00000178935 |
| QRICH2 | 0.623857993 | 1.540990527 | 4.67E-02 | 1.34E-01 | 38.36661131 | 59.15185296 | ENSG00000129646 |
| ZNF18 | 0.623310803 | 1.540406166 | 3.49E-05 | 2.92E-04 | 165.0840373 | 254.3449337 | ENSG00000154957 |
| ZBTB42 | 0.623284544 | 1.540378128 | 2.51E-03 | 1.23E-02 | 89.17342781 | 137.2304677 | ENSG00000179627 |
| NBPF26 | 0.622752549 | 1.539810218 | 7.82E-03 | 3.21E-02 | 70.03521316 | 107.6822552 | ENSG00000273136 |
| PIGM | 0.622366886 | 1.539398649 | 8.79E-06 | 8.54E-05 | 236.0856641 | 363.62729 | ENSG00000143315 |
| FAM157A | 0.622307459 | 1.53933524 | 1.42E-02 | 5.23E-02 | 77.7601218 | 119.4900353 | ENSG00000236438 |
| THEM5 | 0.622223458 | 1.539245615 | 1.90E-02 | 6.59E-02 | 50.23895851 | 77.37514779 | ENSG00000196407 |
| RNF187 | 0.621217091 | 1.538172272 | 2.30E-26 | 2.26E-24 | 2402.666389 | 3695.854524 | ENSG00000168159 |
| RDM1 | 0.621213771 | 1.538168733 | 1.62E-02 | 5.80E-02 | 72.80725647 | 111.8420847 | ENSG00000278023 |
| EPRS1 | 0.621021815 | 1.537964086 | 1.89E-33 | 2.67E-31 | 4491.209372 | 6906.850064 | ENSG00000136628 |
| INKA1 | 0.62083061 | 1.537760269 | 1.65E-02 | 5.90E-02 | 58.8926535 | 90.5547709 | ENSG00000185614 |
| GPSM1 | 0.620492796 | 1.537400237 | 5.26E-11 | 1.23E-09 | 593.2372992 | 912.2451107 | ENSG00000160360 |
| ZMYM1 | 0.620207656 | 1.537096409 | 9.90E-06 | 9.47E-05 | 218.0164775 | 335.2202665 | ENSG00000197056 |
| COCH | 0.61984655 | 1.536711722 | 1.26E-05 | 1.17E-04 | 229.9949724 | 353.3149684 | ENSG00000100473 |
| ZNF808 | 0.619260077 | 1.536087157 | 3.59E-02 | 1.09E-01 | 57.35001535 | 87.98193738 | ENSG00000198482 |
| PRKAG1 | 0.619169507 | 1.535990728 | 3.39E-09 | 5.98E-08 | 418.4186659 | 642.5345764 | ENSG00000181929 |
| CRABP1 | 0.618528494 | 1.535308413 | 2.00E-13 | 6.23E-12 | 790.1332076 | 1212.907606 | ENSG00000166426 |
| TP53BP2 | 0.616509552 | 1.533161369 | 3.04E-18 | 1.56E-16 | 1176.526156 | 1802.971074 | ENSG00000143514 |
| BPHL | 0.616371333 | 1.53301449 | 7.88E-08 | 1.12E-06 | 363.1446693 | 556.2041006 | ENSG00000137274 |
| LGALS3 | 0.616075415 | 1.532700078 | 3.07E-02 | 9.68E-02 | 62.45621675 | 95.80312683 | ENSG00000131981 |
| APOBEC3C | 0.615619157 | 1.532215433 | 2.14E-08 | 3.34E-07 | 371.9810659 | 570.037976 | ENSG00000244509 |
| TMEFF1 | 0.615521853 | 1.532112094 | 2.43E-05 | 2.12E-04 | 251.7902889 | 385.5862894 | ENSG00000241697 |
| YY1AP1 | 0.615304605 | 1.531881399 | 2.05E-11 | 5.00E-10 | 544.1886963 | 833.6962551 | ENSG00000163374 |
| HPDL | 0.615227581 | 1.531799615 | 2.56E-09 | 4.62E-08 | 560.5025917 | 859.2022311 | ENSG00000186603 |
| GBAP1 | 0.615141911 | 1.531708657 | 4.48E-02 | 1.29E-01 | 49.50177269 | 75.93565452 | ENSG00000160766 |
| ADORA1 | 0.615106 | 1.53167053 | 9.60E-04 | 5.42E-03 | 140.9911517 | 215.7369685 | ENSG00000163485 |
| PROCR | 0.614118233 | 1.530622204 | 8.55E-04 | 4.89E-03 | 132.6061966 | 202.9828678 | ENSG00000101000 |
| EDA2R | 0.613723047 | 1.530202989 | 1.80E-03 | 9.26E-03 | 120.5903014 | 184.7160113 | ENSG00000131080 |
| PGBD2 | 0.612774335 | 1.529197063 | 3.79E-04 | 2.41E-03 | 135.074568 | 206.6618424 | ENSG00000185220 |
| OCEL1 | 0.612595168 | 1.529007166 | 1.10E-05 | 1.04E-04 | 229.0516949 | 350.00862 | ENSG00000099330 |
| FAM227B | 0.612054319 | 1.528434067 | 3.36E-02 | 1.04E-01 | 48.33841894 | 73.70828003 | ENSG00000166262 |
| NUAK2 | 0.611675543 | 1.528032833 | 5.25E-06 | 5.32E-05 | 219.9030847 | 336.2786427 | ENSG00000163545 |
| ACP5 | 0.611138335 | 1.527463955 | 3.30E-03 | 1.56E-02 | 95.517654 | 145.6093157 | ENSG00000102575 |
| JAG2 | 0.610818328 | 1.527125181 | 5.20E-10 | 1.04E-08 | 473.9292625 | 723.9601704 | ENSG00000184916 |
| ABCB10 | 0.61059249 | 1.526886146 | 2.26E-06 | 2.44E-05 | 268.5866934 | 410.6121886 | ENSG00000135776 |
| DDHD2 | 0.610332722 | 1.526611244 | 1.54E-06 | 1.72E-05 | 284.8469453 | 434.3454013 | ENSG00000085788 |
| ADCK5 | 0.609544342 | 1.525777235 | 1.68E-04 | 1.18E-03 | 151.6969429 | 231.6193562 | ENSG00000173137 |
| AGMAT | 0.609369734 | 1.525592583 | 3.07E-05 | 2.61E-04 | 211.5525029 | 323.1116732 | ENSG00000116771 |
| RAB37 | 0.608523359 | 1.524697837 | 1.28E-02 | 4.78E-02 | 68.92672546 | 105.2029463 | ENSG00000172794 |
| CPT1C | 0.607523043 | 1.52364103 | 2.00E-15 | 8.03E-14 | 1253.174562 | 1909.384306 | ENSG00000169169 |
| ZSCAN31 | 0.607475963 | 1.523591308 | 9.38E-03 | 3.71E-02 | 71.72106763 | 109.4755008 | ENSG00000235109 |
| TPCN1 | 0.607074305 | 1.523167188 | 1.47E-07 | 1.97E-06 | 519.215383 | 790.4169537 | ENSG00000186815 |
| AGPAT3 | 0.606978229 | 1.523065756 | 4.10E-04 | 2.58E-03 | 158.5802981 | 241.3807136 | ENSG00000160216 |
| ARID4B | 0.606951478 | 1.523037514 | 1.84E-10 | 3.97E-09 | 529.0418627 | 805.2273953 | ENSG00000054267 |
| ETS1 | 0.606889793 | 1.522972396 | 1.70E-18 | 8.88E-17 | 1541.090382 | 2346.323469 | ENSG00000134954 |
| MAN2B2 | 0.606599309 | 1.522665779 | 9.64E-06 | 9.24E-05 | 205.8517478 | 313.4681268 | ENSG00000013288 |
| TLE2 | 0.606478367 | 1.522538138 | 6.97E-08 | 9.99E-07 | 376.0179599 | 572.8060865 | ENSG00000065717 |
| FZD7 | 0.604817924 | 1.520786809 | 1.58E-24 | 1.35E-22 | 6608.104681 | 10050.82822 | ENSG00000155760 |
| ZNF814 | 0.604222869 | 1.520159674 | 5.10E-07 | 6.25E-06 | 270.3389571 | 411.1058854 | ENSG00000204514 |
| MTX1 | 0.603793847 | 1.519707684 | 3.50E-09 | 6.15E-08 | 603.8717714 | 917.2627924 | ENSG00000173171 |
| MX1 | 0.603304537 | 1.519192341 | 1.33E-02 | 4.94E-02 | 62.34706831 | 94.78684219 | ENSG00000157601 |
| KCNF1 | 0.603243526 | 1.519128096 | 3.02E-02 | 9.55E-02 | 46.80114586 | 71.01492268 | ENSG00000162975 |
| ARNT | 0.602961308 | 1.518830956 | 6.59E-14 | 2.16E-12 | 799.2465018 | 1213.596371 | ENSG00000143437 |
| TSTD1 | 0.602685944 | 1.518541087 | 1.29E-03 | 6.97E-03 | 143.6579638 | 217.9712912 | ENSG00000215845 |
| PLEKHF2 | 0.602570337 | 1.518419408 | 6.65E-03 | 2.81E-02 | 111.4650812 | 169.1309389 | ENSG00000175895 |
| CKS1B | 0.602279736 | 1.518113584 | 1.09E-21 | 7.54E-20 | 3357.33 | 5095.685991 | ENSG00000173207 |
| TRIM6 | 0.602033912 | 1.517854931 | 1.20E-05 | 1.12E-04 | 220.8446088 | 335.0283515 | ENSG00000121236 |
| SHANK1 | 0.601926316 | 1.517741734 | 7.87E-03 | 3.22E-02 | 74.11532723 | 112.4835247 | ENSG00000161681 |
| AGRN | 0.601714437 | 1.51751885 | 1.13E-34 | 1.70E-32 | 8341.292866 | 12658.95489 | ENSG00000188157 |
| NBEAL2 | 0.600513679 | 1.516256341 | 1.29E-18 | 6.86E-17 | 1909.830601 | 2895.999937 | ENSG00000160796 |
| PLAAT1 | 0.600298138 | 1.516029827 | 4.23E-03 | 1.92E-02 | 91.29549209 | 138.5429537 | ENSG00000127252 |
| PRRC2C | 0.59891644 | 1.51457859 | 2.50E-05 | 2.18E-04 | 4530.777469 | 6861.688364 | ENSG00000117523 |
| NDUFB1 | 0.598491122 | 1.514132146 | 1.69E-07 | 2.24E-06 | 442.986837 | 670.8257737 | ENSG00000183648 |

| | | | | | | | |
|---|---|---|---|---|---|---|---|
| SIAE | 0.596681927 | 1.512234556 | 4.65E-03 | 2.08E-02 | 97.02881556 | 146.3904596 | ENSG00000110013 |
| SSBP4 | 0.596625034 | 1.512174922 | 2.02E-10 | 4.31E-09 | 798.1004647 | 1207.133659 | ENSG00000130511 |
| DCAF8 | 0.595863071 | 1.511376473 | 5.91E-06 | 5.93E-05 | 300.5034907 | 453.858073 | ENSG00000132716 |
| RHOB | 0.595241856 | 1.510725825 | 1.08E-07 | 1.50E-06 | 362.2008326 | 547.427991 | ENSG00000143878 |
| PLXNA2 | 0.594578476 | 1.510031322 | 2.48E-02 | 8.21E-02 | 64.57975556 | 97.66029124 | ENSG00000076356 |
| VPS72 | 0.594020825 | 1.509447756 | 4.06E-17 | 1.85E-15 | 1737.858325 | 2622.123431 | ENSG00000163159 |
| KRCC1 | 0.593954768 | 1.509378645 | 5.39E-04 | 3.27E-03 | 136.6889543 | 206.2592112 | ENSG00000172086 |
| CXCL3 | 0.593355487 | 1.508751794 | 3.81E-02 | 1.14E-01 | 43.30966179 | 65.35837546 | ENSG00000163734 |
| GSTO2 | 0.592885418 | 1.508260283 | 5.43E-04 | 3.30E-03 | 140.6249623 | 212.0029879 | ENSG00000065621 |
| CCS | 0.592576869 | 1.507937746 | 6.43E-09 | 1.10E-07 | 653.8646978 | 986.5312291 | ENSG00000173992 |
| GADD45B | 0.591697268 | 1.507018647 | 1.90E-03 | 9.69E-03 | 110.3094568 | 166.3628284 | ENSG00000099860 |
| DCAF12L1 | 0.591693684 | 1.507014903 | 6.20E-04 | 3.69E-03 | 135.4906423 | 204.2230893 | ENSG00000198889 |
| TADA2A | 0.591321691 | 1.506626375 | 1.16E-05 | 1.09E-04 | 289.0554274 | 435.1001037 | ENSG00000276234 |
| C15orf40 | 0.591231118 | 1.506531792 | 2.01E-04 | 1.39E-03 | 162.0133299 | 244.2003861 | ENSG00000169609 |
| RTN4IP1 | 0.59118447 | 1.50648308 | 7.73E-07 | 9.11E-06 | 300.3922829 | 452.4311677 | ENSG00000130347 |
| JUN | 0.591004212 | 1.506294864 | 5.91E-06 | 5.93E-05 | 363.9734849 | 548.2357779 | ENSG00000177606 |
| JAKMIP3 | 0.590707254 | 1.505984847 | 2.40E-02 | 7.98E-02 | 56.2095447 | 84.68807462 | ENSG00000188385 |
| COQ5 | 0.590446557 | 1.505712738 | 2.53E-09 | 4.57E-08 | 508.7304794 | 765.3445894 | ENSG00000110871 |
| PARS2 | 0.590332584 | 1.505593791 | 2.43E-06 | 2.62E-05 | 278.248295 | 419.0730215 | ENSG00000162396 |
| ZNF425 | 0.589958295 | 1.505203234 | 9.67E-05 | 7.20E-04 | 184.6285114 | 278.0468595 | ENSG00000204947 |
| SCN8A | 0.58917908 | 1.504390478 | 1.04E-09 | 2.00E-08 | 502.8444303 | 756.2886862 | ENSG00000196876 |
| MBTPS2 | 0.58831047 | 1.503484995 | 8.24E-04 | 4.74E-03 | 154.5256323 | 232.4178609 | ENSG00000012174 |
| PDE12 | 0.588079696 | 1.503244516 | 3.75E-07 | 4.71E-06 | 377.9128028 | 568.3906115 | ENSG00000174840 |
| C1orf35 | 0.5877365 | 1.502886959 | 1.20E-12 | 3.40E-11 | 868.6115214 | 1306.042297 | ENSG00000143793 |
| NFKBIZ | 0.587483394 | 1.502623317 | 1.10E-03 | 6.10E-03 | 124.2217478 | 186.615907 | ENSG00000144802 |
| MYO7A | 0.586982996 | 1.502102222 | 2.92E-03 | 1.41E-02 | 106.416468 | 159.9959089 | ENSG00000137474 |
| RPS6KC1 | 0.586199065 | 1.501286233 | 6.56E-08 | 9.46E-07 | 470.6195653 | 705.9819774 | ENSG00000136643 |
| IMPDH2 | 0.585699974 | 1.500766963 | 2.53E-27 | 2.63E-25 | 4142.803297 | 6217.597914 | ENSG00000178035 |
| GAMT | 0.585434873 | 1.500491215 | 1.13E-08 | 1.85E-07 | 429.8696254 | 645.1330437 | ENSG00000130005 |
| F12 | 0.585084755 | 1.500127116 | 4.07E-02 | 1.20E-01 | 47.81085806 | 71.62278485 | ENSG00000131187 |
| ZER1 | -0.514744221 | 0.699917012 | 1.19E-07 | 1.63E-06 | 687.4665116 | 481.2546531 | ENSG00000160445 |
| ARMC5 | -0.514969041 | 0.69980795 | 2.97E-04 | 1.95E-03 | 297.1238243 | 208.0149271 | ENSG00000140691 |
| TET1 | -0.515060045 | 0.699763808 | 3.35E-07 | 4.25E-06 | 1361.038478 | 951.9589271 | ENSG00000138336 |
| FOXI3 | -0.515190561 | 0.699700505 | 7.24E-03 | 3.01E-02 | 171.2168585 | 119.8433658 | ENSG00000214336 |
| TCN2 | -0.515213777 | 0.699689246 | 2.33E-12 | 6.41E-11 | 1339.550445 | 937.0926921 | ENSG00000185339 |
| CFAP20 | -0.515585685 | 0.699508898 | 1.91E-06 | 2.10E-05 | 559.2058626 | 391.1404945 | ENSG00000070761 |
| EID1 | -0.515838297 | 0.699386427 | 3.39E-15 | 1.31E-13 | 2886.237595 | 2019.024311 | ENSG00000255302 |
| CACFD1 | -0.515887278 | 0.699362682 | 8.64E-07 | 1.01E-05 | 665.1379898 | 465.3003501 | ENSG00000160325 |
| ATP7B | -0.516023149 | 0.699296821 | 2.84E-04 | 1.88E-03 | 314.7677588 | 219.9924981 | ENSG00000123191 |
| ZNF512 | -0.516070176 | 0.699274026 | 2.02E-05 | 1.79E-04 | 456.9240699 | 319.6353956 | ENSG00000243943 |
| ATP6V0E1 | -0.516368485 | 0.699129451 | 1.16E-12 | 3.29E-11 | 1434.384811 | 1002.843243 | ENSG00000113732 |
| C1orf216 | -0.516541361 | 0.69904568 | 7.65E-05 | 5.85E-04 | 342.3126688 | 239.4268305 | ENSG00000142686 |
| CUL7 | -0.516861436 | 0.698890608 | 1.22E-05 | 1.13E-04 | 590.9624581 | 412.7433685 | ENSG00000044090 |
| HSPA14 | -0.51698374 | 0.698831362 | 5.23E-05 | 4.18E-04 | 408.0335316 | 285.2498766 | ENSG00000187522 |
| MED24 | -0.517006388 | 0.698820391 | 1.54E-19 | 8.95E-18 | 2790.784904 | 1950.353931 | ENSG00000008838 |
| TKFC | -0.517168512 | 0.698741866 | 5.06E-07 | 6.21E-06 | 645.1937908 | 450.8061609 | ENSG00000149476 |
| BRINP1 | -0.517177325 | 0.698737597 | 4.44E-08 | 6.55E-07 | 919.8878561 | 642.9140648 | ENSG00000078725 |
| GSTM4 | -0.517226128 | 0.698713961 | 5.54E-04 | 3.35E-03 | 299.4083506 | 209.3271296 | ENSG00000168765 |
| FBXL14 | -0.517370674 | 0.698643959 | 5.12E-05 | 4.11E-04 | 369.6591648 | 258.2086868 | ENSG00000171823 |
| ANKRD42 | -0.517613971 | 0.698526149 | 3.23E-02 | 1.01E-01 | 108.9928187 | 76.2368895 | ENSG00000137494 |
| NAA10 | -0.517641937 | 0.698512609 | 9.97E-05 | 7.40E-04 | 384.88456 | 269.1234487 | ENSG00000102030 |
| FAM8A1 | -0.517804432 | 0.698433938 | 8.67E-04 | 4.94E-03 | 265.2624523 | 185.4162413 | ENSG00000137414 |
| ARHGAP18 | -0.517816565 | 0.698428064 | 1.91E-02 | 6.61E-02 | 128.6837899 | 89.78072045 | ENSG00000146376 |
| MIF4GD | -0.518159513 | 0.698262058 | 1.01E-03 | 5.64E-03 | 251.590348 | 175.5416215 | ENSG00000125457 |
| PNMA1 | -0.519477162 | 0.697624609 | 2.26E-15 | 8.95E-14 | 2144.669595 | 1496.260994 | ENSG00000176903 |
| RHBDL3 | -0.520018263 | 0.697363005 | 1.59E-06 | 1.77E-05 | 546.0767774 | 381.019142 | ENSG00000141314 |
| TES | -0.520437247 | 0.697160508 | 1.63E-06 | 1.81E-05 | 534.2844146 | 372.3265733 | ENSG00000135269 |
| NCAPG | -0.520946592 | 0.696914418 | 1.15E-08 | 1.88E-07 | 1207.098954 | 841.0741919 | ENSG00000109805 |
| MPP2 | -0.521025426 | 0.696876338 | 8.77E-18 | 4.29E-16 | 2428.928494 | 1692.481056 | ENSG00000108852 |
| PCYT1B | -0.52123344 | 0.696775866 | 3.59E-11 | 8.54E-10 | 1152.4799 | 802.8656865 | ENSG00000102230 |
| TTLL4 | -0.521506151 | 0.696644168 | 1.04E-14 | 3.79E-13 | 2045.676962 | 1425.439529 | ENSG00000135912 |
| POU6F1 | -0.522514251 | 0.69615755 | 2.38E-02 | 7.92E-02 | 101.9615187 | 71.04292016 | ENSG00000184271 |
| MXD3 | -0.523228145 | 0.695813153 | 4.89E-03 | 2.17E-02 | 160.2797539 | 111.4962991 | ENSG00000213347 |
| CENPM | -0.52330562 | 0.695775787 | 1.55E-08 | 2.49E-07 | 721.9652774 | 502.2799629 | ENSG00000100162 |
| RFLNB | -0.524869447 | 0.695022001 | 1.00E-10 | 2.25E-09 | 1164.844876 | 809.5280925 | ENSG00000183688 |
| ULK4 | -0.525181344 | 0.69487176 | 4.54E-03 | 2.04E-02 | 173.7638438 | 120.7067782 | ENSG00000168038 |
| RELB | -0.525908712 | 0.694521513 | 6.73E-04 | 3.97E-03 | 289.037097 | 200.7692085 | ENSG00000104856 |
| ELMSAN1 | -0.526197024 | 0.694382732 | 2.59E-04 | 1.73E-03 | 305.6798872 | 212.1247936 | ENSG00000156030 |
| PIGQ | -0.526526651 | 0.694224097 | 4.47E-05 | 3.65E-04 | 464.8459186 | 322.8688399 | ENSG00000007541 |
| PLPP6 | -0.526794647 | 0.69409515 | 2.19E-03 | 1.10E-02 | 201.2138432 | 139.7949259 | ENSG00000205808 |
| PRPF38A | -0.527372249 | 0.693817315 | 3.38E-14 | 1.15E-12 | 1631.453912 | 1131.822816 | ENSG00000134748 |
| ADCK2 | -0.527527929 | 0.69374245 | 6.28E-03 | 2.68E-02 | 154.0838042 | 106.9395345 | ENSG00000133597 |
| UTF1 | -0.527671973 | 0.693673188 | 1.36E-04 | 9.78E-04 | 482.1041381 | 334.5861106 | ENSG00000171794 |
| MSI1 | -0.527698854 | 0.693660263 | 2.03E-12 | 5.61E-11 | 1497.242637 | 1038.632123 | ENSG00000135097 |

| | | | | | | | |
|---|---|---|---|---|---|---|---|
| SYS1 | -0.528000586 | 0.693515203 | 7.22E-04 | 4.22E-03 | 252.4904054 | 175.0483762 | ENSG00000204070 |
| TOR2A | -0.528780156 | 0.693140559 | 1.54E-04 | 1.09E-03 | 365.3367795 | 253.2359548 | ENSG00000160404 |
| ZNF484 | -0.528790941 | 0.693135377 | 1.68E-02 | 5.97E-02 | 120.5751738 | 83.51574397 | ENSG00000127081 |
| GALNT18 | -0.529260757 | 0.692909693 | 8.13E-04 | 4.68E-03 | 251.4172331 | 174.3688393 | ENSG00000110328 |
| PCDH1 | -0.53014874 | 0.692483336 | 4.44E-17 | 2.02E-15 | 5527.714632 | 3827.855528 | ENSG00000156453 |
| ZDHHC6 | -0.530377737 | 0.692373428 | 6.80E-07 | 8.11E-06 | 688.2244433 | 476.4508816 | ENSG00000023041 |
| ZNF84 | -0.530541473 | 0.692294853 | 4.70E-06 | 4.80E-05 | 518.4630684 | 359.0147187 | ENSG00000198040 |
| ADAMTS7 | -0.5305629 | 0.692284571 | 2.54E-16 | 1.09E-14 | 2395.640376 | 1658.692439 | ENSG00000136378 |
| CNPY4 | -0.530827484 | 0.69215762 | 2.44E-03 | 1.20E-02 | 183.2398203 | 126.8795549 | ENSG00000166997 |
| TNIP1 | -0.531405886 | 0.691880178 | 1.60E-07 | 2.14E-06 | 817.3645267 | 565.5022833 | ENSG00000145901 |
| PARPBP | -0.531451013 | 0.691858537 | 1.76E-05 | 1.59E-04 | 412.6814775 | 285.6367124 | ENSG00000185480 |
| ZNF266 | -0.531746391 | 0.6917169 | 1.07E-04 | 7.90E-04 | 328.0154757 | 226.8115602 | ENSG00000174652 |
| RBBP4 | -0.532295739 | 0.691453558 | 6.89E-18 | 3.43E-16 | 4023.835132 | 2781.628962 | ENSG00000162521 |
| SCD | -0.532313766 | 0.691444919 | 6.39E-23 | 4.86E-21 | 29963.57974 | 20717.88981 | ENSG00000099194 |
| LRRC37B | -0.532758009 | 0.691232037 | 2.05E-03 | 1.04E-02 | 193.0963026 | 133.5435693 | ENSG00000185158 |
| NUCB1 | -0.533544113 | 0.690855498 | 8.64E-19 | 4.74E-17 | 2427.916444 | 1677.427454 | ENSG00000104805 |
| SFXN5 | -0.53376575 | 0.690749372 | 3.96E-10 | 8.09E-09 | 946.6346603 | 653.9425929 | ENSG00000144040 |
| ENKD1 | -0.533996829 | 0.690638742 | 1.45E-03 | 7.72E-03 | 223.1178987 | 154.0395242 | ENSG00000124074 |
| LRRC45 | -0.534114201 | 0.690582557 | 6.60E-08 | 9.50E-07 | 909.851367 | 628.7542596 | ENSG00000169683 |
| SOAT1 | -0.534320957 | 0.690483595 | 1.33E-02 | 4.94E-02 | 124.2605966 | 85.76655411 | ENSG00000057252 |
| FGFR1 | -0.534624965 | 0.69033811 | 1.58E-32 | 2.07E-30 | 9861.498191 | 6807.876783 | ENSG00000077782 |
| MGAT5B | -0.53593561 | 0.689711243 | 1.48E-04 | 1.06E-03 | 303.7918834 | 209.5871034 | ENSG00000167889 |
| NLK | -0.536633152 | 0.689377849 | 1.08E-07 | 1.50E-06 | 660.6496097 | 455.2420954 | ENSG00000087095 |
| OCRL | -0.536758368 | 0.689318018 | 6.46E-10 | 1.28E-08 | 1258.519196 | 867.2437619 | ENSG00000122126 |
| MINDY3 | -0.53722747 | 0.689093918 | 7.64E-04 | 4.43E-03 | 246.5224367 | 169.7792319 | ENSG00000148481 |
| REC8 | -0.537430012 | 0.688997182 | 7.76E-07 | 9.13E-06 | 590.9509934 | 407.2125264 | ENSG00000100918 |
| HEMK1 | -0.537674956 | 0.688880213 | 6.17E-03 | 2.64E-02 | 170.8983122 | 117.7926054 | ENSG00000114735 |
| MCM5 | -0.538011247 | 0.688719654 | 7.04E-25 | 6.16E-23 | 8812.009594 | 6069.05396 | ENSG00000100297 |
| ZNF780B | -0.539263122 | 0.688122288 | 1.16E-02 | 4.42E-02 | 129.5592103 | 89.13492945 | ENSG00000128000 |
| PIP5K1B | -0.541179175 | 0.687208995 | 2.23E-02 | 7.52E-02 | 102.9300179 | 70.75095413 | ENSG00000107242 |
| ACADSB | -0.541435137 | 0.687087082 | 1.60E-06 | 1.78E-05 | 494.8140964 | 339.7919758 | ENSG00000196177 |
| SOX9 | -0.5419632 | 0.686835636 | 1.48E-02 | 5.41E-02 | 120.1683295 | 82.55963727 | ENSG00000125398 |
| USP43 | -0.54276264 | 0.686455146 | 5.41E-04 | 3.29E-03 | 232.8210443 | 159.7854334 | ENSG00000154914 |
| COL26A1 | -0.542797 | 0.686438797 | 6.00E-20 | 3.54E-18 | 2861.000175 | 1964.310179 | ENSG00000160963 |
| RADX | -0.542850928 | 0.686413138 | 5.48E-04 | 3.32E-03 | 229.7283959 | 157.5993201 | ENSG00000147231 |
| MCM4 | -0.54290265 | 0.68638853 | 2.54E-25 | 2.36E-23 | 8764.033061 | 6015.790676 | ENSG00000104738 |
| TMEM109 | -0.543749305 | 0.685985837 | 2.03E-10 | 4.32E-09 | 1036.419881 | 711.2847855 | ENSG00000110108 |
| NISCH | -0.54377175 | 0.685975165 | 2.36E-23 | 1.83E-21 | 3407.537975 | 2337.598209 | ENSG00000010322 |
| UBXN8 | -0.543792665 | 0.68596522 | 1.38E-02 | 5.08E-02 | 129.3338207 | 88.85159263 | ENSG00000104691 |
| PTBP2 | -0.544219784 | 0.685762166 | 8.19E-09 | 1.37E-07 | 816.3833426 | 559.5804724 | ENSG00000117569 |
| PTDSS2 | -0.544284271 | 0.685731513 | 1.59E-12 | 4.45E-11 | 1483.994361 | 1018.045767 | ENSG00000174915 |
| PSMD1 | -0.544378847 | 0.685686561 | 8.05E-28 | 8.50E-26 | 5612.641109 | 3848.234504 | ENSG00000173692 |
| ZNF302 | -0.544604237 | 0.685579446 | 4.21E-06 | 4.33E-05 | 456.9300198 | 313.2685285 | ENSG00000089335 |
| ABHD8 | -0.54654199 | 0.68465923 | 1.46E-03 | 7.76E-03 | 235.3247572 | 161.1716969 | ENSG00000127220 |
| NCAPH | -0.547033921 | 0.684425814 | 9.60E-10 | 1.85E-08 | 1356.025844 | 927.8840096 | ENSG00000121152 |
| ACSS2 | -0.547178342 | 0.684357303 | 5.87E-10 | 1.17E-08 | 954.1087048 | 652.7483921 | ENSG00000131069 |
| DPYSL4 | -0.547193728 | 0.684350004 | 9.72E-10 | 1.87E-08 | 881.3154325 | 603.4094585 | ENSG00000151640 |
| SLC16A9 | -0.548335285 | 0.683808715 | 3.23E-04 | 2.09E-03 | 292.9909024 | 200.2858945 | ENSG00000165449 |
| SLC22A5 | -0.54868234 | 0.683644238 | 1.36E-02 | 5.02E-02 | 113.9394548 | 77.9055439 | ENSG00000197375 |
| NMT2 | -0.549052175 | 0.683469008 | 3.35E-05 | 2.82E-04 | 351.3829968 | 240.1843278 | ENSG00000152465 |
| CHD7 | -0.550004924 | 0.683017797 | 1.89E-13 | 5.88E-12 | 2992.257894 | 2043.049147 | ENSG00000171316 |
| ARSB | -0.550272879 | 0.682890951 | 2.03E-05 | 1.80E-04 | 347.78 | 237.4320221 | ENSG00000113273 |
| SLC25A13 | -0.550603701 | 0.682734376 | 5.38E-17 | 2.43E-15 | 2216.402946 | 1513.094324 | ENSG00000004864 |
| GORASP1 | -0.550714417 | 0.682681983 | 6.97E-10 | 1.37E-08 | 1050.754319 | 717.1971236 | ENSG00000114745 |
| IFRD1 | -0.551883104 | 0.682129185 | 1.48E-11 | 3.68E-10 | 1033.583488 | 705.1241883 | ENSG00000006652 |
| ESPN | -0.552024633 | 0.682062272 | 1.10E-06 | 1.26E-05 | 513.4091156 | 350.3782403 | ENSG00000187017 |
| NR6A1 | -0.55336684 | 0.681428012 | 1.61E-20 | 9.96E-19 | 5702.650349 | 3885.523242 | ENSG00000148200 |
| NOMO1 | -0.553717962 | 0.681262187 | 7.06E-17 | 3.16E-15 | 2162.20487 | 1472.844043 | ENSG00000103512 |
| KCNQ1 | -0.554101843 | 0.681080937 | 4.14E-02 | 1.22E-01 | 74.92815169 | 51.0434999 | ENSG00000053918 |
| KIF3B | -0.554229687 | 0.681020586 | 7.75E-11 | 1.76E-09 | 1226.628458 | 835.1853622 | ENSG00000101350 |
| ZNF501 | -0.554731272 | 0.680783855 | 1.04E-02 | 4.05E-02 | 142.0379849 | 96.73424189 | ENSG00000186446 |
| DUSP16 | -0.555396993 | 0.680469785 | 1.02E-13 | 3.28E-12 | 2569.612955 | 1748.171404 | ENSG00000111266 |
| UNG | -0.555737576 | 0.680309162 | 7.52E-23 | 5.66E-21 | 4649.267012 | 3162.545297 | ENSG00000076248 |
| SCAMP4 | -0.556532529 | 0.679934402 | 3.53E-09 | 6.20E-08 | 878.7281507 | 597.6501177 | ENSG00000227500 |
| SYT9 | -0.55681456 | 0.679801495 | 3.92E-02 | 1.17E-01 | 87.60175521 | 59.49090118 | ENSG00000170743 |
| TMTC2 | -0.557390719 | 0.679530062 | 1.42E-02 | 5.20E-02 | 117.7852809 | 80.06784272 | ENSG00000179104 |
| TAFA5 | -0.557506732 | 0.67947542 | 3.88E-12 | 1.03E-10 | 1113.419427 | 756.7501502 | ENSG00000219438 |
| WTIP | -0.557743171 | 0.679364072 | 2.98E-06 | 3.15E-05 | 440.5288647 | 299.2252095 | ENSG00000142279 |
| PHLDB2 | -0.557760877 | 0.679355735 | 5.96E-08 | 8.63E-07 | 774.1012683 | 525.7297004 | ENSG00000144824 |
| KCNIP3 | -0.558573883 | 0.678973003 | 3.50E-02 | 1.07E-01 | 83.80512527 | 56.86680249 | ENSG00000115041 |
| RBCK1 | -0.558983938 | 0.678780047 | 2.20E-09 | 4.02E-08 | 1296.980819 | 880.8818885 | ENSG00000125826 |
| CCP110 | -0.56013296 | 0.678239654 | 4.81E-05 | 3.88E-04 | 365.6363077 | 247.9563323 | ENSG00000103540 |
| NPTX1 | -0.56027763 | 0.678171645 | 2.82E-02 | 9.05E-02 | 111.8641962 | 75.74307713 | ENSG00000171246 |
| MAP2K6 | -0.561833123 | 0.677440844 | 2.04E-09 | 3.74E-08 | 963.8476728 | 652.6012159 | ENSG00000108984 |

| | | | | | | | |
|---|---|---|---|---|---|---|---|
| PM20D2 | -0.561969139 | 0.677376978 | 1.03E-03 | 5.77E-03 | 277.8589687 | 188.1947776 | ENSG00000146281 |
| ZBTB26 | -0.562228418 | 0.677255252 | 5.20E-03 | 2.28E-02 | 150.6353393 | 102.0616722 | ENSG00000171448 |
| WASHC4 | -0.56250393 | 0.677125929 | 8.52E-09 | 1.42E-07 | 877.9419703 | 594.2951083 | ENSG00000136051 |
| ARRDC3 | -0.563271278 | 0.676765871 | 2.96E-02 | 9.39E-02 | 80.94973955 | 54.83200539 | ENSG00000113369 |
| EML3 | -0.563436653 | 0.676688299 | 3.62E-08 | 5.43E-07 | 645.4936239 | 436.7635768 | ENSG00000149499 |
| SIRT2 | -0.563486437 | 0.676664948 | 2.73E-03 | 1.32E-02 | 204.2883376 | 138.0863351 | ENSG00000068903 |
| TTC26 | -0.565247132 | 0.675839636 | 3.71E-03 | 1.72E-02 | 154.6602818 | 104.4003313 | ENSG00000105948 |
| DHX35 | -0.565261582 | 0.675832867 | 5.43E-07 | 6.59E-06 | 548.7215706 | 370.5552503 | ENSG00000101452 |
| RORB | -0.565724857 | 0.67561588 | 4.11E-03 | 1.88E-02 | 154.6499083 | 104.5897873 | ENSG00000198963 |
| ADGRG2 | -0.566360873 | 0.675318098 | 3.15E-04 | 2.05E-03 | 258.2319908 | 174.201138 | ENSG00000173698 |
| FAM32A | -0.567163685 | 0.674942411 | 7.98E-16 | 3.33E-14 | 1798.53045 | 1213.864942 | ENSG00000105058 |
| AHCYL2 | -0.567699223 | 0.674691914 | 1.06E-07 | 1.47E-06 | 598.5782701 | 403.6372914 | ENSG00000158467 |
| EAPP | -0.568360976 | 0.674382509 | 1.00E-05 | 9.57E-05 | 382.1901106 | 257.5351819 | ENSG00000129518 |
| SLC35A3 | -0.568400337 | 0.67436411 | 8.91E-03 | 3.57E-02 | 133.5619319 | 90.15980035 | ENSG00000117620 |
| DHRSX | -0.568554615 | 0.674291999 | 4.09E-05 | 3.36E-04 | 333.0371395 | 224.5555259 | ENSG00000169084 |
| ASAP2 | -0.568766392 | 0.674193026 | 5.27E-07 | 6.42E-06 | 495.6272002 | 334.1332961 | ENSG00000151693 |
| WDR27 | -0.568946236 | 0.674108987 | 1.29E-03 | 6.99E-03 | 384.969072 | 259.6593542 | ENSG00000184465 |
| HDAC9 | -0.569758499 | 0.673729559 | 7.99E-04 | 4.61E-03 | 212.9524311 | 143.4796489 | ENSG00000048052 |
| CHN1 | -0.569864771 | 0.673679932 | 4.66E-10 | 9.42E-09 | 929.5102779 | 626.004937 | ENSG00000128656 |
| DTX4 | -0.569880921 | 0.67367239 | 1.40E-10 | 3.05E-09 | 890.714628 | 600.0091863 | ENSG00000110042 |
| UHRF1 | -0.570312746 | 0.673470778 | 4.25E-15 | 1.63E-13 | 2512.288588 | 1691.975259 | ENSG00000276043 |
| PPM1L | -0.570657623 | 0.673309804 | 8.28E-04 | 4.75E-03 | 278.0950107 | 187.1910942 | ENSG00000163590 |
| GPX3 | -0.570762536 | 0.673260843 | 5.02E-09 | 8.68E-08 | 714.980276 | 481.5853986 | ENSG00000211445 |
| ZNF778 | -0.571077989 | 0.673113646 | 2.07E-06 | 2.26E-05 | 520.6038465 | 350.3410465 | ENSG00000170100 |
| HIPK2 | -0.571311265 | 0.673004816 | 2.48E-09 | 4.50E-08 | 896.7904454 | 603.3911385 | ENSG00000064393 |
| MRPL48 | -0.571331508 | 0.672995373 | 9.54E-05 | 7.12E-04 | 372.6776497 | 250.8203444 | ENSG00000175581 |
| COL2A1 | -0.573619127 | 0.671929079 | 3.41E-06 | 3.56E-05 | 418.6322075 | 281.3686042 | ENSG00000139219 |
| LBH | -0.574090708 | 0.671709478 | 1.42E-12 | 3.99E-11 | 1170.887359 | 786.3897846 | ENSG00000213626 |
| ATP6V0D1 | -0.574209691 | 0.671654083 | 7.21E-13 | 2.12E-11 | 1332.863315 | 895.116889 | ENSG00000159720 |
| FMNL1 | -0.574364961 | 0.6715818 | 2.73E-03 | 1.33E-02 | 206.5388478 | 138.8651357 | ENSG00000184922 |
| CLIP3 | -0.574600949 | 0.671471956 | 4.14E-05 | 3.40E-04 | 343.9145966 | 231.0878985 | ENSG00000105270 |
| TIGD1 | -0.575095046 | 0.671242028 | 2.00E-02 | 6.88E-02 | 116.8936155 | 78.62711995 | ENSG00000221944 |
| WFDC2 | -0.576075077 | 0.670786204 | 6.76E-07 | 8.07E-06 | 444.9176585 | 298.3581085 | ENSG00000101443 |
| C1QL1 | -0.576710456 | 0.670490848 | 2.12E-04 | 1.45E-03 | 286.0871068 | 191.9079628 | ENSG00000131094 |
| HDAC11 | -0.577616574 | 0.670069863 | 2.94E-05 | 2.51E-04 | 362.3282854 | 242.592697 | ENSG00000163517 |
| UBR7 | -0.578288894 | 0.669757671 | 8.80E-09 | 1.46E-07 | 715.6807248 | 479.2609789 | ENSG00000012963 |
| EPOP | -0.578317336 | 0.669744468 | 4.86E-03 | 2.16E-02 | 150.2447917 | 100.654768 | ENSG00000273604 |
| ADI1 | -0.578393279 | 0.669709214 | 2.32E-09 | 4.23E-08 | 858.6616555 | 574.9757194 | ENSG00000182551 |
| ARMH3 | -0.578834755 | 0.669504309 | 6.60E-08 | 9.50E-07 | 548.3636686 | 367.2261716 | ENSG00000120029 |
| SFRP1 | -0.579254395 | 0.669309597 | 1.33E-19 | 7.72E-18 | 14465.82615 | 9681.878026 | ENSG00000104332 |
| KALRN | -0.57963907 | 0.669131158 | 4.81E-04 | 2.98E-03 | 231.8059409 | 155.0466125 | ENSG00000160145 |
| TGFB1 | -0.580101479 | 0.668916724 | 9.39E-05 | 7.02E-04 | 297.0520762 | 198.8282625 | ENSG00000105329 |
| KCNC4 | -0.5806473 | 0.668663698 | 7.06E-06 | 6.99E-05 | 401.7618909 | 268.7657243 | ENSG00000116396 |
| PIK3R3 | -0.580815948 | 0.668585537 | 1.71E-04 | 1.20E-03 | 275.6491129 | 184.3378344 | ENSG00000117461 |
| ABAT | -0.58158733 | 0.668228152 | 1.09E-04 | 8.02E-04 | 290.0648864 | 193.6485755 | ENSG00000183044 |
| UBE2R2 | -0.582260576 | 0.667916391 | 5.66E-20 | 3.35E-18 | 2257.768412 | 1508.344165 | ENSG00000107341 |
| CACNA2D3 | -0.583211978 | 0.667476071 | 2.74E-02 | 8.84E-02 | 85.13689098 | 56.83880501 | ENSG00000157445 |
| FBXW4 | -0.583281752 | 0.66744379 | 1.51E-08 | 2.44E-07 | 717.3391941 | 478.9806909 | ENSG00000107829 |
| WDTC1 | -0.583507313 | 0.667339446 | 5.76E-10 | 1.15E-08 | 850.0130474 | 567.0523987 | ENSG00000142784 |
| MPP1 | -0.583615137 | 0.667289572 | 2.48E-04 | 1.67E-03 | 278.9522771 | 186.0609184 | ENSG00000130830 |
| GSTO1 | -0.585372677 | 0.666477152 | 5.90E-09 | 1.01E-07 | 977.4832681 | 651.1017985 | ENSG00000148834 |
| MMD | -0.585496485 | 0.666419959 | 1.41E-07 | 1.90E-06 | 614.6265827 | 409.6523504 | ENSG00000108960 |
| CPXM1 | -0.586523001 | 0.665945952 | 1.72E-14 | 5.97E-13 | 1859.870223 | 1238.88436 | ENSG00000088882 |
| PLA2R1 | -0.58661071 | 0.665905467 | 4.37E-02 | 1.27E-01 | 70.23062636 | 46.7330166 | ENSG00000153246 |
| RCN1 | -0.587683551 | 0.66541046 | 4.52E-09 | 7.88E-08 | 901.65646 | 600.2311586 | ENSG00000049449 |
| ZNF615 | -0.58812877 | 0.665205144 | 2.90E-03 | 1.40E-02 | 158.7617527 | 105.5816473 | ENSG00000197619 |
| CDC42EP5 | -0.588204249 | 0.665170343 | 1.53E-02 | 5.54E-02 | 116.4884995 | 77.34497484 | ENSG00000167617 |
| IGDCC3 | -0.588705118 | 0.664939451 | 2.38E-17 | 1.11E-15 | 2254.985795 | 1499.293597 | ENSG00000174498 |
| BRD9 | -0.589083455 | 0.664765098 | 8.32E-12 | 2.13E-10 | 1108.076906 | 736.4795726 | ENSG00000028310 |
| CHST7 | -0.58916507 | 0.664727493 | 1.36E-07 | 1.84E-06 | 682.9359748 | 454.3041704 | ENSG00000147119 |
| NTRK3 | -0.589306889 | 0.664662153 | 1.64E-03 | 8.60E-03 | 177.9183751 | 118.4051021 | ENSG00000140538 |
| ZFYVE26 | -0.589327189 | 0.6646528 | 8.84E-12 | 2.25E-10 | 978.1831565 | 649.9617936 | ENSG00000072121 |
| FASTKD1 | -0.589828497 | 0.664421887 | 3.22E-09 | 5.70E-08 | 707.7980565 | 470.4098326 | ENSG00000138399 |
| CBY1 | -0.590576941 | 0.664077286 | 5.11E-03 | 2.25E-02 | 157.7286249 | 104.7164383 | ENSG00000100211 |
| KCND2 | -0.592393545 | 0.663241624 | 3.51E-05 | 2.93E-04 | 283.7248563 | 188.2283985 | ENSG00000184408 |
| CHRAC1 | -0.592418823 | 0.663230003 | 1.61E-07 | 2.14E-06 | 678.2208797 | 450.0158409 | ENSG00000104472 |
| RNF7 | -0.593077712 | 0.66292717 | 1.60E-09 | 3.00E-08 | 664.1971289 | 440.3322425 | ENSG00000114125 |
| CD2BP2 | -0.593226458 | 0.662858824 | 2.75E-13 | 8.48E-12 | 2365.226679 | 1568.30387 | ENSG00000169217 |
| SERAC1 | -0.593686697 | 0.662647397 | 6.67E-05 | 5.19E-04 | 273.5101671 | 181.3720176 | ENSG00000122335 |
| GLYCTK | -0.593874671 | 0.662561064 | 3.79E-02 | 1.14E-01 | 73.25358603 | 48.62453692 | ENSG00000168237 |
| KRT7 | -0.594100015 | 0.662457582 | 1.53E-04 | 1.09E-03 | 278.8929868 | 184.7904083 | ENSG00000135480 |
| MLH1 | -0.595088572 | 0.662003811 | 3.28E-13 | 1.00E-11 | 1205.852032 | 798.0422537 | ENSG00000076242 |
| TMEM35A | -0.595235572 | 0.661936361 | 8.72E-03 | 3.50E-02 | 116.6477593 | 77.11692762 | ENSG00000126950 |
| HID1 | -0.596197717 | 0.661495058 | 4.70E-05 | 3.80E-04 | 265.9343991 | 175.8652735 | ENSG00000167861 |

| | | | | | | | |
|---|---|---|---|---|---|---|---|
| EOGT | -0.596460588 | 0.661374539 | 2.32E-05 | 2.04E-04 | 311.8151245 | 206.2641292 | ENSG00000163378 |
| TRIM61 | -0.596823319 | 0.661208273 | 2.51E-03 | 1.23E-02 | 169.4944929 | 112.1510755 | ENSG00000183439 |
| ATXN7L3B | -0.59689671 | 0.661174638 | 2.98E-19 | 1.69E-17 | 1758.399728 | 1162.625934 | ENSG00000253719 |
| FAM78A | -0.597049838 | 0.661104464 | 6.35E-04 | 3.77E-03 | 202.0165741 | 133.6574511 | ENSG00000126882 |
| ULK2 | -0.597257152 | 0.661009471 | 4.57E-04 | 2.85E-03 | 204.4057996 | 135.1557772 | ENSG00000083290 |
| NUBPL | -0.597678142 | 0.660816611 | 6.48E-04 | 3.84E-03 | 198.1807899 | 130.8270169 | ENSG00000151413 |
| TCTN3 | -0.598858029 | 0.660276393 | 1.59E-08 | 2.55E-07 | 832.2982806 | 549.5254977 | ENSG00000119977 |
| MAPRE3 | -0.598961727 | 0.660228935 | 2.93E-03 | 1.41E-02 | 154.5306924 | 102.0884402 | ENSG00000084764 |
| DRG2 | -0.598979507 | 0.660220799 | 3.43E-09 | 6.05E-08 | 657.9199223 | 434.3306675 | ENSG00000108591 |
| POLR2C | -0.599056359 | 0.66018563 | 3.48E-15 | 1.35E-13 | 1525.964171 | 1007.495844 | ENSG00000102978 |
| FUNDC2 | -0.599483731 | 0.659990091 | 1.43E-10 | 3.12E-09 | 793.2364565 | 523.3163609 | ENSG00000165775 |
| EXT2 | -0.600162911 | 0.659679459 | 3.61E-17 | 1.65E-15 | 2106.724372 | 1389.25616 | ENSG00000151348 |
| NUCB2 | -0.600721218 | 0.65942422 | 7.66E-05 | 5.86E-04 | 313.3707268 | 206.7825341 | ENSG00000070081 |
| AASS | -0.601014642 | 0.659290116 | 5.35E-26 | 5.10E-24 | 4244.507132 | 2798.150131 | ENSG00000008311 |
| TRAPPC2 | -0.601265646 | 0.659175421 | 1.17E-03 | 6.42E-03 | 168.1028271 | 110.702386 | ENSG00000196459 |
| SEC61A2 | -0.602801713 | 0.658473957 | 5.09E-07 | 6.24E-06 | 454.895721 | 299.4926783 | ENSG00000065665 |
| NHLRC3 | -0.602980431 | 0.658392391 | 3.84E-04 | 2.44E-03 | 225.1800367 | 148.1076367 | ENSG00000188811 |
| TRPM3 | -0.603353422 | 0.658222194 | 1.15E-03 | 6.35E-03 | 192.3922693 | 126.4796663 | ENSG00000083067 |
| AFF2 | -0.604001795 | 0.657926444 | 1.17E-04 | 8.53E-04 | 244.9848843 | 161.1787904 | ENSG00000155966 |
| FOXM1 | -0.60407747 | 0.657891934 | 5.81E-21 | 3.73E-19 | 3071.948692 | 2021.531542 | ENSG00000111206 |
| TCEAL3 | -0.604431871 | 0.657730342 | 2.24E-06 | 2.42E-05 | 363.3935263 | 239.1118803 | ENSG00000196507 |
| ARL4C | -0.604803078 | 0.657561129 | 8.66E-09 | 1.44E-07 | 614.8249717 | 404.1037257 | ENSG00000188042 |
| RECQL | -0.605602326 | 0.657196943 | 3.73E-08 | 5.59E-07 | 638.6884992 | 419.485584 | ENSG00000004700 |
| TIA1 | -0.605980271 | 0.657024798 | 3.74E-14 | 1.26E-12 | 1641.3046 | 1078.671681 | ENSG00000116001 |
| ATG4A | -0.606216856 | 0.656917063 | 1.34E-02 | 4.96E-02 | 94.53344234 | 62.1495667 | ENSG00000101844 |
| RNF150 | -0.606687341 | 0.656702867 | 7.08E-04 | 4.15E-03 | 201.9246387 | 132.4525611 | ENSG00000170153 |
| TEX261 | -0.606810608 | 0.65664676 | 7.14E-07 | 8.48E-06 | 416.3695961 | 273.5868566 | ENSG00000144043 |
| PDSS2 | -0.60789231 | 0.656154604 | 1.08E-04 | 7.96E-04 | 286.851651 | 188.3060325 | ENSG00000164494 |
| INSIG2 | -0.609113577 | 0.655599393 | 4.26E-03 | 1.94E-02 | 136.5726869 | 89.52365715 | ENSG00000125629 |
| SEMA5B | -0.609452725 | 0.655445293 | 1.16E-10 | 2.57E-09 | 798.0087564 | 523.2087648 | ENSG00000082684 |
| BEND3 | -0.611803047 | 0.654378364 | 1.74E-11 | 4.28E-10 | 1153.460403 | 754.8688745 | ENSG00000178409 |
| NDUFAF7 | -0.61267785 | 0.65398169 | 2.64E-11 | 6.38E-10 | 988.0379116 | 646.2225591 | ENSG00000003509 |
| ZNF184 | -0.612696651 | 0.653973168 | 1.24E-03 | 6.74E-03 | 168.2614765 | 109.9287144 | ENSG00000096654 |
| CDC27 | -0.614482951 | 0.65316394 | 5.14E-08 | 7.50E-07 | 787.5265219 | 514.284658 | ENSG00000004897 |
| CCDC120 | -0.614605433 | 0.65310849 | 4.41E-06 | 4.52E-05 | 398.4650088 | 260.4374064 | ENSG00000147144 |
| ADAM11 | -0.61568021 | 0.652622119 | 1.50E-04 | 1.07E-03 | 255.4534389 | 166.6943316 | ENSG00000073670 |
| HES1 | -0.615785557 | 0.652574465 | 2.91E-04 | 1.92E-03 | 220.5098437 | 143.957287 | ENSG00000114315 |
| ZNF165 | -0.616913828 | 0.652064314 | 2.37E-02 | 7.88E-02 | 79.32022567 | 51.71946385 | ENSG00000197279 |
| ATP6V1D | -0.617273759 | 0.651901654 | 3.27E-12 | 8.76E-11 | 870.5422651 | 567.4508899 | ENSG00000100554 |
| KCTD8 | -0.617281897 | 0.651897977 | 7.17E-03 | 2.99E-02 | 143.2932484 | 93.40351184 | ENSG00000183783 |
| PRR5L | -0.61733954 | 0.651871931 | 5.35E-03 | 2.34E-02 | 133.088043 | 86.66091747 | ENSG00000135362 |
| SLC49A4 | -0.617869569 | 0.651632485 | 1.73E-05 | 1.56E-04 | 289.6394068 | 188.8266128 | ENSG00000138463 |
| CCNC | -0.618822293 | 0.651202303 | 2.02E-08 | 3.17E-07 | 700.2735945 | 456.2536502 | ENSG00000112237 |
| PCSK1N | -0.619317529 | 0.650978802 | 1.14E-02 | 4.34E-02 | 149.8190071 | 97.5689944 | ENSG00000102109 |
| NBPF1 | -0.619705249 | 0.650803877 | 6.89E-07 | 8.20E-06 | 435.3245243 | 283.4974407 | ENSG00000219481 |
| SLC35A2 | -0.619976741 | 0.650681418 | 6.51E-10 | 1.28E-08 | 679.6941608 | 442.4060097 | ENSG00000102100 |
| CIPC | -0.619983735 | 0.650678264 | 2.97E-07 | 3.80E-06 | 435.4985813 | 283.3412791 | ENSG00000198894 |
| MYCBP | -0.620379979 | 0.650499576 | 2.89E-04 | 1.90E-03 | 243.5339806 | 158.461503 | ENSG00000214114 |
| WARS1 | -0.621088214 | 0.650180316 | 2.68E-20 | 1.63E-18 | 2874.965521 | 1868.610495 | ENSG00000140105 |
| ZNF107 | -0.621239802 | 0.650112004 | 8.52E-06 | 8.31E-05 | 428.3525918 | 278.5521687 | ENSG00000196247 |
| PTPRZ1 | -0.621253296 | 0.650105923 | 1.54E-22 | 1.13E-20 | 5816.463418 | 3780.778731 | ENSG00000106278 |
| UPF3B | -0.623144015 | 0.649254487 | 7.42E-15 | 2.74E-13 | 1093.819703 | 710.108839 | ENSG00000125351 |
| NGRN | -0.624804906 | 0.648507468 | 1.40E-08 | 2.27E-07 | 560.993415 | 364.00845 | ENSG00000182768 |
| FBN2 | -0.624809667 | 0.648505328 | 3.31E-09 | 5.84E-08 | 930.6058727 | 603.0236937 | ENSG00000138829 |
| LTBP1 | -0.626304499 | 0.647833735 | 1.09E-14 | 3.92E-13 | 1303.616942 | 844.2121564 | ENSG00000049323 |
| ZNF253 | -0.626396117 | 0.647792596 | 3.52E-06 | 3.66E-05 | 355.2211452 | 230.0515605 | ENSG00000256771 |
| DFFB | -0.626411325 | 0.647785767 | 5.73E-05 | 4.55E-04 | 233.6056985 | 151.3240334 | ENSG00000169598 |
| PERP | -0.626796141 | 0.647613003 | 4.06E-11 | 9.62E-10 | 971.9276889 | 629.6029679 | ENSG00000112378 |
| NBAS | -0.626913689 | 0.647560239 | 3.48E-10 | 7.18E-09 | 856.8263043 | 554.4182844 | ENSG00000151779 |
| ARIH1 | -0.628194827 | 0.64698545 | 2.99E-13 | 9.15E-12 | 1289.37731 | 833.9212629 | ENSG00000166233 |
| ZNF844 | -0.628288488 | 0.646943449 | 2.75E-02 | 8.88E-02 | 69.23276163 | 44.76197035 | ENSG00000223547 |
| MTHFD2L | -0.628771007 | 0.64672711 | 1.51E-02 | 5.48E-02 | 97.82625865 | 63.37928977 | ENSG00000163738 |
| KLHL3 | -0.629449726 | 0.646422928 | 3.06E-06 | 3.22E-05 | 367.0222774 | 237.0174723 | ENSG00000146021 |
| PDRG1 | -0.630450771 | 0.645974548 | 1.15E-10 | 2.55E-09 | 986.3418578 | 637.0939063 | ENSG00000088356 |
| DYNC1I2 | -0.630532048 | 0.645938157 | 4.49E-20 | 2.70E-18 | 1840.257151 | 1188.405744 | ENSG00000077380 |
| DHRS1 | -0.630546673 | 0.64593161 | 9.78E-06 | 9.35E-05 | 293.6139325 | 189.6248769 | ENSG00000157379 |
| OAT | -0.631043647 | 0.64570914 | 1.19E-23 | 9.66E-22 | 2523.838857 | 1629.533245 | ENSG00000065154 |
| SEPTIN8 | -0.631537043 | 0.645488348 | 5.61E-18 | 2.82E-16 | 1601.971314 | 1033.870938 | ENSG00000164402 |
| RRP36 | -0.632037673 | 0.645264395 | 1.12E-19 | 6.57E-18 | 1631.883639 | 1053.113849 | ENSG00000124541 |
| RCBTB2 | -0.63253255 | 0.645043093 | 8.76E-06 | 8.52E-05 | 312.9069839 | 201.8943616 | ENSG00000136161 |
| CCDC88A | -0.633566133 | 0.644581133 | 5.43E-15 | 2.04E-13 | 1155.605169 | 744.7991659 | ENSG00000115355 |
| ANKRD36B | -0.633984364 | 0.644394299 | 3.50E-02 | 1.07E-01 | 64.48039582 | 41.59704973 | ENSG00000196912 |
| BRICD5 | -0.634352844 | 0.644229735 | 3.71E-02 | 1.12E-01 | 88.94658975 | 57.4343924 | ENSG00000182685 |
| CBLN1 | -0.636130668 | 0.643436343 | 4.21E-04 | 2.64E-03 | 245.743758 | 158.346151 | ENSG00000102924 |

| | | | | | | | |
|---|---|---|---|---|---|---|---|
| PLPBP | -0.636550191 | 0.643249265 | 9.19E-13 | 2.64E-11 | 840.7077942 | 540.7987114 | ENSG00000147471 |
| AREL1 | -0.636822996 | 0.643127642 | 1.92E-11 | 4.73E-10 | 1102.49267 | 708.5946316 | ENSG00000119682 |
| PEX12 | -0.637207709 | 0.642956167 | 3.43E-02 | 1.06E-01 | 69.42789549 | 44.64998043 | ENSG00000108733 |
| CYP27A1 | -0.637490525 | 0.642830138 | 2.74E-03 | 1.33E-02 | 129.4586554 | 83.15813517 | ENSG00000135929 |
| ARMC7 | -0.638848749 | 0.642225231 | 3.16E-06 | 3.32E-05 | 326.2397703 | 209.4989709 | ENSG00000125449 |
| RPP40 | -0.640142617 | 0.641649516 | 7.78E-03 | 3.20E-02 | 105.963656 | 68.02997817 | ENSG00000124787 |
| PHF3 | -0.640271654 | 0.641592128 | 3.04E-18 | 1.56E-16 | 1677.035896 | 1075.566031 | ENSG00000118482 |
| SCG3 | -0.640549968 | 0.641468369 | 2.75E-09 | 4.91E-08 | 1087.844265 | 697.8504495 | ENSG00000104112 |
| SLC20A1 | -0.640675965 | 0.641412349 | 1.90E-26 | 1.90E-24 | 2546.290049 | 1633.234943 | ENSG00000144136 |
| IFT57 | -0.640733364 | 0.64138683 | 2.52E-09 | 4.56E-08 | 584.3297649 | 374.8876991 | ENSG00000114446 |
| ZNF268 | -0.64078328 | 0.641364639 | 1.79E-05 | 1.61E-04 | 291.4135854 | 186.8719087 | ENSG00000090612 |
| ALDH16A1 | -0.642070222 | 0.640792771 | 7.72E-14 | 2.51E-12 | 1112.142832 | 712.8302664 | ENSG00000161618 |
| POLR3B | -0.642614182 | 0.64055121 | 1.97E-08 | 3.09E-07 | 536.8474172 | 343.6767485 | ENSG00000013503 |
| ARFRP1 | -0.642917842 | 0.6404164 | 5.56E-11 | 1.29E-09 | 729.7055929 | 467.3855618 | ENSG00000101246 |
| GDAP2 | -0.6435186 | 0.640149777 | 5.87E-05 | 4.64E-04 | 231.4972873 | 148.1455655 | ENSG00000196505 |
| SLC44A3 | -0.643714318 | 0.64006294 | 2.37E-02 | 7.91E-02 | 67.88172402 | 43.41997381 | ENSG00000143036 |
| PMM1 | -0.644226521 | 0.639835737 | 6.18E-04 | 3.69E-03 | 180.8001517 | 115.6682354 | ENSG00000100417 |
| PHKA2 | -0.645744219 | 0.639162991 | 1.28E-11 | 3.21E-10 | 757.5773372 | 484.3210357 | ENSG00000044446 |
| GPR160 | -0.64581753 | 0.639130513 | 8.66E-10 | 1.68E-08 | 571.889508 | 365.5439699 | ENSG00000173890 |
| DOCK4 | -0.645994529 | 0.639052105 | 6.66E-11 | 1.53E-09 | 692.8638486 | 442.6654164 | ENSG00000128512 |
| ETHE1 | -0.646306868 | 0.638913767 | 2.37E-02 | 7.89E-02 | 75.96544995 | 48.53061312 | ENSG00000105755 |
| MTHFD1 | -0.646724887 | 0.638728669 | 3.18E-35 | 4.92E-33 | 5371.672593 | 3430.544173 | ENSG00000100714 |
| KMT5A | -0.647545704 | 0.63836537 | 1.25E-06 | 1.42E-05 | 335.3424384 | 214.042871 | ENSG00000183955 |
| GLDC | -0.647604889 | 0.638339182 | 3.56E-33 | 4.97E-31 | 4349.416556 | 2776.012206 | ENSG00000178445 |
| TMEM123 | -0.648111651 | 0.638114998 | 1.18E-15 | 4.84E-14 | 1921.463981 | 1226.204881 | ENSG00000152558 |
| E2F8 | -0.649063635 | 0.637694068 | 3.39E-05 | 2.84E-04 | 253.2877707 | 161.5111443 | ENSG00000129173 |
| FXYD6 | -0.649781025 | 0.637377049 | 2.16E-03 | 1.08E-02 | 127.0575562 | 80.98819605 | ENSG00000137726 |
| ZNF576 | -0.650273616 | 0.637159461 | 2.91E-03 | 1.40E-02 | 125.9965368 | 80.3595252 | ENSG00000124444 |
| CDK5 | -0.652196721 | 0.636310697 | 1.13E-07 | 1.56E-06 | 416.4825833 | 265.0397402 | ENSG00000164885 |
| EXOC3L1 | -0.653310007 | 0.635819863 | 3.70E-02 | 1.12E-01 | 68.43509094 | 43.541328 | ENSG00000179044 |
| FEM1A | -0.653311714 | 0.635819111 | 4.97E-02 | 1.41E-01 | 53.79448693 | 34.16749849 | ENSG00000141965 |
| GLIPR2 | -0.654098165 | 0.635472604 | 3.19E-04 | 2.07E-03 | 217.6086918 | 138.2113778 | ENSG00000122694 |
| FZD2 | -0.654141999 | 0.635453296 | 1.98E-10 | 4.23E-09 | 1453.752661 | 924.083377 | ENSG00000180340 |
| SLC25A53 | -0.657512911 | 0.633970269 | 5.47E-05 | 4.36E-04 | 250.8792732 | 159.0628387 | ENSG00000269743 |
| C1QTNF4 | -0.65878634 | 0.633410927 | 1.87E-03 | 9.60E-03 | 144.3628868 | 91.56185923 | ENSG00000172247 |
| TMEM266 | -0.659177337 | 0.633239284 | 1.08E-03 | 6.01E-03 | 141.6405982 | 89.64964582 | ENSG00000169758 |
| C12orf57 | -0.65997478 | 0.632889361 | 6.74E-03 | 2.84E-02 | 100.7910721 | 63.77823239 | ENSG00000111678 |
| SHC3 | -0.660484651 | 0.632665727 | 3.06E-05 | 2.60E-04 | 250.3893925 | 158.4134748 | ENSG00000148082 |
| ZNF521 | -0.662977254 | 0.631573588 | 1.92E-05 | 1.72E-04 | 272.1057897 | 171.94749 | ENSG00000198795 |
| CUL1 | -0.663392666 | 0.631391758 | 2.20E-26 | 2.18E-24 | 2687.448652 | 1696.953238 | ENSG00000055130 |
| ARHGAP29 | -0.66365954 | 0.631274972 | 1.06E-02 | 4.08E-02 | 83.60556779 | 52.81309894 | ENSG00000137962 |
| RBM25 | -0.663757218 | 0.631232233 | 3.44E-26 | 3.31E-24 | 3412.778488 | 2153.994266 | ENSG00000119707 |
| POU3F1 | -0.665011942 | 0.630683484 | 5.83E-11 | 1.35E-09 | 1116.619496 | 704.4661976 | ENSG00000185668 |
| CRIP2 | -0.665241866 | 0.630582979 | 3.73E-05 | 3.09E-04 | 227.2893377 | 143.2284495 | ENSG00000182809 |
| KIF20A | -0.666545197 | 0.630013568 | 5.96E-18 | 2.99E-16 | 1716.896587 | 1081.502144 | ENSG00000112984 |
| DUSP26 | -0.667355631 | 0.629659757 | 3.99E-02 | 1.18E-01 | 54.19838215 | 34.11273304 | ENSG00000133878 |
| HSPA1B | -0.668275113 | 0.62925858 | 4.14E-14 | 1.39E-12 | 1292.906675 | 813.7708662 | ENSG00000204388 |
| RPE | -0.668683605 | 0.629080433 | 4.23E-13 | 1.28E-11 | 1088.656017 | 684.7158714 | ENSG00000197713 |
| COMMD9 | -0.670067962 | 0.62847708 | 1.07E-06 | 1.23E-05 | 364.3199745 | 229.0863005 | ENSG00000110442 |
| CCNE2 | -0.671976329 | 0.627646294 | 4.83E-02 | 1.38E-01 | 60.54677821 | 37.99873525 | ENSG00000175305 |
| FNDC11 | -0.674570115 | 0.626518878 | 3.22E-02 | 1.00E-01 | 67.39598708 | 42.2133302 | ENSG00000125531 |
| TFPT | -0.675149492 | 0.626267322 | 1.33E-05 | 1.23E-04 | 333.2841386 | 208.9771087 | ENSG00000105619 |
| ITFG1 | -0.675429435 | 0.626145812 | 5.38E-07 | 6.54E-06 | 352.1833111 | 220.4221082 | ENSG00000129636 |
| PPM1H | -0.675803348 | 0.625983551 | 6.39E-21 | 4.09E-19 | 2009.843501 | 1257.731613 | ENSG00000111110 |
| ZNF613 | -0.679777412 | 0.624261582 | 5.25E-04 | 3.21E-03 | 160.7218875 | 100.3312315 | ENSG00000176024 |
| RNF38 | -0.679813432 | 0.624245996 | 4.22E-09 | 7.38E-08 | 659.0474281 | 411.3751281 | ENSG00000137075 |
| SMIM10L1 | -0.679946191 | 0.624188555 | 5.31E-07 | 6.46E-06 | 361.7213249 | 225.8872349 | ENSG00000256537 |
| CCDC15 | -0.680813892 | 0.623813253 | 2.57E-02 | 8.41E-02 | 66.563001 | 41.45488686 | ENSG00000149548 |
| TTBK1 | -0.68091553 | 0.623769307 | 1.49E-02 | 5.42E-02 | 78.74168927 | 49.19503734 | ENSG00000146216 |
| ZNF891 | -0.681877437 | 0.623353551 | 1.36E-02 | 5.03E-02 | 77.87725275 | 48.48454952 | ENSG00000214029 |
| HM13 | -0.682243617 | 0.623195354 | 5.63E-24 | 4.68E-22 | 2423.343428 | 1510.5623 | ENSG00000101294 |
| KREMEN1 | -0.682390831 | 0.623131766 | 7.05E-05 | 5.44E-04 | 218.0055204 | 135.7682739 | ENSG00000183762 |
| FAM111B | -0.682488119 | 0.623089746 | 4.14E-08 | 6.15E-07 | 641.9607243 | 400.231627 | ENSG00000189057 |
| NCAM1 | -0.682620725 | 0.623032477 | 1.27E-05 | 1.18E-04 | 249.2891928 | 155.3615627 | ENSG00000149294 |
| TMEM102 | -0.682841054 | 0.622937335 | 2.67E-02 | 8.69E-02 | 85.80916829 | 53.59547238 | ENSG00000181284 |
| FBRS | -0.683469441 | 0.622666064 | 1.76E-08 | 2.79E-07 | 511.1397169 | 318.5146364 | ENSG00000156860 |
| PCDHA12 | -0.687495207 | 0.620930969 | 1.43E-04 | 1.03E-03 | 175.2201387 | 108.8085222 | ENSG00000251664 |
| ASGR1 | -0.687700525 | 0.620842607 | 9.59E-05 | 7.15E-04 | 201.8312027 | 125.2537271 | ENSG00000141505 |
| MAGEE1 | -0.688358185 | 0.620559657 | 3.30E-07 | 4.20E-06 | 333.8822524 | 207.2453229 | ENSG00000198934 |
| ZNF616 | -0.688515455 | 0.620492013 | 6.42E-10 | 1.27E-08 | 567.9636459 | 352.4237038 | ENSG00000204611 |
| CYFIP1 | -0.688801821 | 0.620368861 | 5.87E-28 | 6.26E-26 | 2350.576236 | 1458.074105 | ENSG00000273749 |
| ARSI | -0.689343 | 0.620136194 | 3.12E-04 | 2.03E-03 | 179.2504202 | 111.238857 | ENSG00000183876 |
| MMP25 | -0.690512086 | 0.619633871 | 7.76E-10 | 1.51E-08 | 499.9552816 | 309.8533546 | ENSG00000008516 |
| SLC9A7 | -0.691501825 | 0.619208927 | 1.57E-08 | 2.52E-07 | 480.1118123 | 297.1431193 | ENSG00000065923 |

| | | | | | | | |
|---|---|---|---|---|---|---|---|
| CYB5D2 | -0.691838256 | 0.619064547 | 4.29E-03 | 1.95E-02 | 112.4745141 | 69.6726426 | ENSG00000167740 |
| ACAT2 | -0.692625435 | 0.618726858 | 1.87E-26 | 1.88E-24 | 2563.04893 | 1586.004571 | ENSG00000120437 |
| CAVIN4 | -0.693812251 | 0.618218078 | 4.50E-04 | 2.81E-03 | 158.3697311 | 97.78862332 | ENSG00000170681 |
| ZNF737 | -0.694544546 | 0.617904358 | 4.65E-04 | 2.89E-03 | 165.5994191 | 102.2566657 | ENSG00000237440 |
| SLC2A8 | -0.694635552 | 0.617865381 | 6.05E-13 | 1.79E-11 | 834.3354494 | 515.7232913 | ENSG00000136856 |
| HAS3 | -0.695483948 | 0.617502144 | 5.14E-29 | 5.82E-27 | 2595.979226 | 1603.003818 | ENSG00000103044 |
| TCF25 | -0.69565203 | 0.617430206 | 4.87E-19 | 2.72E-17 | 1337.762282 | 825.9762735 | ENSG00000141002 |
| SCYL2 | -0.69722311 | 0.616758197 | 3.62E-10 | 7.42E-09 | 700.2006517 | 431.8981346 | ENSG00000136021 |
| NAA16 | -0.697232352 | 0.616754246 | 5.84E-07 | 7.02E-06 | 483.000734 | 297.7588529 | ENSG00000172766 |
| BCL6B | -0.697792536 | 0.616514813 | 1.82E-07 | 2.40E-06 | 379.3468779 | 233.736778 | ENSG00000161940 |
| PCLAF | -0.700057637 | 0.615547615 | 2.24E-03 | 1.12E-02 | 116.8523764 | 71.94861234 | ENSG00000166803 |
| TMEM121 | -0.700494272 | 0.615361346 | 1.75E-02 | 6.16E-02 | 68.50533836 | 42.07703131 | ENSG00000184986 |
| KCNJ11 | -0.701635033 | 0.614874962 | 4.15E-05 | 3.40E-04 | 240.0525384 | 147.742882 | ENSG00000187486 |
| ADGRG5 | -0.702012471 | 0.61471412 | 1.63E-07 | 2.16E-06 | 373.5492313 | 229.5541752 | ENSG00000159618 |
| RAB11FIP3 | -0.702239789 | 0.61461727 | 8.36E-21 | 5.26E-19 | 1413.455614 | 868.9240943 | ENSG00000090565 |
| ING4 | -0.703671146 | 0.614007786 | 5.88E-06 | 5.91E-05 | 340.2614368 | 208.9345454 | ENSG00000111653 |
| ZNF181 | -0.703693362 | 0.613998331 | 2.71E-03 | 1.32E-02 | 130.1991736 | 79.95869067 | ENSG00000197841 |
| PTN | -0.703734057 | 0.613981011 | 2.40E-13 | 7.43E-12 | 815.9821421 | 501.042273 | ENSG00000105894 |
| NAA38 | -0.703808808 | 0.6139492 | 1.26E-07 | 1.71E-06 | 382.9053548 | 235.1241031 | ENSG00000183011 |
| SEPTIN6 | -0.704481484 | 0.613663004 | 1.39E-14 | 4.92E-13 | 920.4464585 | 565.0378136 | ENSG00000125354 |
| ZNF138 | -0.70570947 | 0.613140892 | 5.98E-07 | 7.19E-06 | 307.8438272 | 188.8350311 | ENSG00000197008 |
| SLC2A1 | -0.705755978 | 0.613121126 | 9.07E-29 | 1.01E-26 | 4904.647224 | 3006.449432 | ENSG00000117394 |
| PFKFB4 | -0.70620338 | 0.612931018 | 1.31E-06 | 1.49E-05 | 365.3186255 | 223.7328802 | ENSG00000114268 |
| MXRA5 | -0.706350449 | 0.612868538 | 1.25E-03 | 6.78E-03 | 150.6969932 | 92.23515325 | ENSG00000101825 |
| CCDC74B | -0.708294649 | 0.612043183 | 8.75E-05 | 6.60E-04 | 199.0060722 | 121.9111737 | ENSG00000152076 |
| BARX1 | -0.708502015 | 0.611955217 | 4.93E-02 | 1.40E-01 | 45.02218644 | 27.54983123 | ENSG00000131668 |
| CHRNB4 | -0.708902419 | 0.611785399 | 5.52E-03 | 2.40E-02 | 104.6419577 | 64.11626202 | ENSG00000117971 |
| SLC30A3 | -0.709971065 | 0.6113324 | 1.73E-06 | 1.91E-05 | 309.8495985 | 189.507801 | ENSG00000115194 |
| ATP6V0E2 | -0.710212914 | 0.611229926 | 3.64E-12 | 9.73E-11 | 644.1707822 | 393.7963222 | ENSG00000171130 |
| TMEM191A | -0.710547215 | 0.611088309 | 1.26E-02 | 4.73E-02 | 75.80977519 | 46.32376376 | ENSG00000226287 |
| NASP | -0.711013116 | 0.610890997 | 4.73E-67 | 2.88E-64 | 18000.47091 | 10996.27933 | ENSG00000132780 |
| FAM210B | -0.711569077 | 0.610655628 | 1.20E-09 | 2.29E-08 | 468.5879356 | 286.0310205 | ENSG00000124098 |
| SEPHS1 | -0.712407183 | 0.610300982 | 1.82E-51 | 5.83E-49 | 15736.26242 | 9603.894965 | ENSG00000086475 |
| SLITRK4 | -0.71250814 | 0.610258276 | 3.69E-05 | 3.07E-04 | 198.4420531 | 121.1863083 | ENSG00000179542 |
| PRKCH | -0.714392602 | 0.609461671 | 1.98E-05 | 1.77E-04 | 289.7739779 | 176.5855045 | ENSG00000027075 |
| DLD | -0.714421934 | 0.60944928 | 2.64E-24 | 2.24E-22 | 1841.351094 | 1122.130967 | ENSG00000091140 |
| PEX26 | -0.717798999 | 0.608024348 | 1.89E-19 | 1.08E-17 | 1536.662791 | 934.2348275 | ENSG00000215193 |
| ASF1A | -0.718154319 | 0.607874616 | 1.72E-09 | 3.20E-08 | 463.2858141 | 281.6841214 | ENSG00000111875 |
| CFAP36 | -0.719860527 | 0.607156136 | 7.45E-06 | 7.34E-05 | 258.3287063 | 156.9388679 | ENSG00000163001 |
| POGLUT2 | -0.720360029 | 0.606945958 | 6.39E-11 | 1.47E-09 | 629.3582323 | 382.1869108 | ENSG00000134901 |
| TRAPPC5 | -0.722499919 | 0.606046368 | 3.78E-02 | 1.14E-01 | 50.95837312 | 30.8933305 | ENSG00000181029 |
| UTP14A | -0.723182779 | 0.60575958 | 2.42E-16 | 1.04E-14 | 1077.934975 | 653.108859 | ENSG00000156697 |
| NOVA1 | -0.723223106 | 0.605742648 | 5.50E-05 | 4.38E-04 | 203.0233372 | 123.0782801 | ENSG00000139910 |
| H2AW | -0.723521202 | 0.6056175 | 3.81E-08 | 5.69E-07 | 367.722725 | 222.7544517 | ENSG00000181218 |
| EFNB1 | -0.723932783 | 0.60544475 | 3.38E-08 | 5.12E-07 | 444.6985754 | 269.4783876 | ENSG00000090776 |
| POMC | -0.725203547 | 0.604911693 | 1.21E-02 | 4.60E-02 | 79.95276455 | 48.41332632 | ENSG00000115138 |
| RAB33A | -0.725524795 | 0.604777011 | 5.26E-03 | 2.31E-02 | 102.3710583 | 62.01052526 | ENSG00000134594 |
| USP4 | -0.727088867 | 0.604121708 | 3.04E-16 | 1.29E-14 | 954.0840684 | 576.1866843 | ENSG00000114316 |
| CAB39L | -0.727518133 | 0.603941982 | 1.05E-02 | 4.05E-02 | 75.95477153 | 45.86988773 | ENSG00000102547 |
| TTLL7 | -0.72753407 | 0.60393531 | 3.91E-03 | 1.80E-02 | 96.57989413 | 58.37070902 | ENSG00000137941 |
| ZNF823 | -0.72900305 | 0.603320684 | 2.66E-02 | 8.64E-02 | 57.1872478 | 34.50921664 | ENSG00000197933 |
| EXOG | -0.729363605 | 0.603169923 | 7.43E-05 | 5.71E-04 | 219.8152937 | 132.3903233 | ENSG00000157036 |
| RBM3 | -0.730003809 | 0.602902322 | 4.53E-21 | 2.93E-19 | 1691.034937 | 1019.779169 | ENSG00000102317 |
| ZNF440 | -0.733356475 | 0.60150287 | 4.49E-03 | 2.02E-02 | 115.0713064 | 69.0843596 | ENSG00000171295 |
| KTN1 | -0.733849198 | 0.601297473 | 1.84E-29 | 2.13E-27 | 2679.71154 | 1611.136006 | ENSG00000126777 |
| CLCN2 | -0.735271037 | 0.60070516 | 3.73E-06 | 3.87E-05 | 260.675576 | 156.6463348 | ENSG00000114859 |
| DUSP4 | -0.73542755 | 0.600639996 | 2.04E-08 | 3.19E-07 | 397.3194001 | 238.5879676 | ENSG00000120875 |
| LNP1 | -0.735926437 | 0.600432329 | 9.37E-05 | 7.01E-04 | 188.201253 | 112.9636219 | ENSG00000206535 |
| REM2 | -0.736408383 | 0.600231782 | 9.37E-03 | 3.70E-02 | 74.12036124 | 44.4202522 | ENSG00000139890 |
| MYO5B | -0.737474589 | 0.599788352 | 6.19E-09 | 1.06E-07 | 402.8220728 | 241.6543729 | ENSG00000167306 |
| C19orf54 | -0.739548327 | 0.598926832 | 3.13E-10 | 6.50E-09 | 527.0704647 | 315.7350385 | ENSG00000188493 |
| NTS | -0.739549079 | 0.59892652 | 5.06E-05 | 4.07E-04 | 184.7145236 | 110.6008219 | ENSG00000133636 |
| DNAJB6 | -0.739931707 | 0.598767696 | 3.87E-51 | 1.20E-48 | 8629.556256 | 5166.799878 | ENSG00000105993 |
| MMP2 | -0.740097418 | 0.598698924 | 7.55E-25 | 6.57E-23 | 2038.707699 | 1220.818064 | ENSG00000087245 |
| CFAP298 | -0.740325523 | 0.598604271 | 7.85E-03 | 3.22E-02 | 90.44826811 | 54.20390164 | ENSG00000159079 |
| MYL5 | -0.741110629 | 0.598278602 | 5.66E-03 | 2.45E-02 | 85.21252962 | 50.94891369 | ENSG00000215375 |
| ATF7IP2 | -0.741588162 | 0.598080605 | 2.78E-06 | 2.95E-05 | 262.4156606 | 156.9283695 | ENSG00000166669 |
| IGFBP6 | -0.74416248 | 0.597014352 | 1.12E-05 | 1.05E-04 | 227.0058797 | 135.5836633 | ENSG00000167779 |
| MAP3K13 | -0.747559012 | 0.595610457 | 8.64E-04 | 4.93E-03 | 123.7324514 | 73.63951583 | ENSG00000073803 |
| ZNF577 | -0.748473197 | 0.595233159 | 1.77E-04 | 1.23E-03 | 151.0644302 | 89.86755077 | ENSG00000161551 |
| ALKBH3 | -0.748503586 | 0.595220621 | 5.87E-05 | 4.64E-04 | 192.4426862 | 114.465353 | ENSG00000166199 |
| SFRP2 | -0.74923803 | 0.594917685 | 5.63E-46 | 1.42E-43 | 6993.180566 | 4160.630961 | ENSG00000145423 |
| WWOX | -0.749298683 | 0.594892674 | 3.53E-05 | 2.95E-04 | 212.7125248 | 126.3843021 | ENSG00000186153 |
| DNAJC3 | -0.749431672 | 0.594837839 | 2.81E-13 | 8.64E-12 | 756.642705 | 450.2016607 | ENSG00000102580 |

| | | | | | | | |
|---|---|---|---|---|---|---|---|
| GDPD2 | -0.75003262 | 0.594590113 | 1.84E-08 | 2.91E-07 | 368.3968085 | 219.1628746 | ENSG00000130055 |
| ZFP69B | -0.750394571 | 0.594440958 | 2.72E-04 | 1.80E-03 | 139.0307377 | 82.63738686 | ENSG00000187801 |
| SLIT2 | -0.750446813 | 0.594419433 | 3.21E-05 | 2.71E-04 | 197.4136538 | 117.3066216 | ENSG00000145147 |
| UBQLN2 | -0.75090125 | 0.594232225 | 2.53E-16 | 1.09E-14 | 1076.62515 | 640.1589345 | ENSG00000188021 |
| LDOC1 | -0.752615433 | 0.593526589 | 6.92E-15 | 2.57E-13 | 878.9513764 | 522.0223628 | ENSG00000182195 |
| EVL | -0.75326261 | 0.593260399 | 3.16E-17 | 1.46E-15 | 1220.760895 | 724.3827072 | ENSG00000196405 |
| RAD54B | -0.754724173 | 0.592659685 | 1.06E-03 | 5.91E-03 | 138.6277322 | 82.1011994 | ENSG00000265817 |
| DANCR | -0.755032649 | 0.592532976 | 8.98E-22 | 6.24E-20 | 1488.611215 | 882.2925539 | ENSG00000226950 |
| MAP7D3 | -0.755351046 | 0.592402221 | 1.79E-19 | 1.04E-17 | 1037.91978 | 614.7276389 | ENSG00000129680 |
| RTN4RL2 | -0.75644622 | 0.591952689 | 5.39E-07 | 6.55E-06 | 348.6444889 | 206.4863125 | ENSG00000186907 |
| GINS2 | -0.758616474 | 0.591062881 | 3.24E-23 | 2.49E-21 | 1586.919775 | 937.5325694 | ENSG00000131153 |
| NDUFC2 | -0.759663686 | 0.590634 | 1.80E-02 | 6.32E-02 | 66.35957855 | 39.30112195 | ENSG00000151366 |
| NAP1L3 | -0.762230141 | 0.589584237 | 2.03E-25 | 1.90E-23 | 1507.533724 | 888.8936252 | ENSG00000186310 |
| VPS8 | -0.763639504 | 0.589008556 | 3.31E-14 | 1.13E-12 | 755.8081409 | 444.9329679 | ENSG00000156931 |
| TAF5 | -0.763792193 | 0.588946221 | 3.12E-12 | 8.41E-11 | 608.9809736 | 358.5907118 | ENSG00000148835 |
| TGIF2 | -0.76469739 | 0.588576812 | 1.71E-17 | 8.09E-16 | 977.5175173 | 575.3940006 | ENSG00000118707 |
| TMX3 | -0.765041104 | 0.588436603 | 3.51E-08 | 5.29E-07 | 383.1928079 | 225.5178754 | ENSG00000166479 |
| CUX2 | -0.765806855 | 0.588124357 | 3.66E-07 | 4.61E-06 | 294.6800114 | 173.2121884 | ENSG00000111249 |
| ELF4 | -0.768308862 | 0.587105281 | 1.52E-04 | 1.08E-03 | 161.4267849 | 94.71618608 | ENSG00000102034 |
| TPR | -0.77094159 | 0.586034868 | 1.56E-37 | 2.69E-35 | 3509.767242 | 2056.548833 | ENSG00000047410 |
| ZSCAN18 | -0.772643253 | 0.585344046 | 2.04E-08 | 3.19E-07 | 338.5976012 | 198.2819758 | ENSG00000121413 |
| BMP7 | -0.777297497 | 0.583458724 | 2.20E-08 | 3.41E-07 | 390.1632141 | 227.4925375 | ENSG00000101144 |
| XKR7 | -0.778512105 | 0.582967716 | 3.95E-07 | 4.95E-06 | 281.8118327 | 164.1158747 | ENSG00000260903 |
| ZNF610 | -0.778857446 | 0.582828186 | 1.40E-02 | 5.16E-02 | 69.01212664 | 40.27898329 | ENSG00000167554 |
| DUSP6 | -0.779843985 | 0.582429775 | 4.56E-31 | 5.63E-29 | 2287.081299 | 1331.772784 | ENSG00000139318 |
| ZDHHC15 | -0.781429243 | 0.581790142 | 9.28E-04 | 5.25E-03 | 114.2549481 | 66.50299218 | ENSG00000102383 |
| CAMK2N2 | -0.782805419 | 0.581235441 | 5.69E-04 | 3.43E-03 | 145.1878904 | 84.41178834 | ENSG00000163888 |
| NLRP12 | -0.782929848 | 0.581185314 | 4.45E-05 | 3.63E-04 | 185.2162778 | 107.7810244 | ENSG00000142405 |
| ZNF841 | -0.783919903 | 0.58078661 | 2.40E-04 | 1.62E-03 | 168.7614775 | 98.05932917 | ENSG00000197608 |
| KCNMA1 | -0.784004384 | 0.580752601 | 7.50E-04 | 4.36E-03 | 131.1563579 | 76.10560396 | ENSG00000156113 |
| TAF7 | -0.784591002 | 0.580516508 | 2.34E-36 | 3.78E-34 | 3114.542162 | 1808.133981 | ENSG00000178913 |
| MCTP1 | -0.786661484 | 0.579683978 | 4.02E-03 | 1.85E-02 | 82.85175366 | 48.02147719 | ENSG00000175471 |
| TCEAL1 | -0.788991273 | 0.578748609 | 7.03E-03 | 2.94E-02 | 67.10062933 | 38.85516977 | ENSG00000172465 |
| TRMT10A | -0.791110246 | 0.57789919 | 3.27E-02 | 1.02E-01 | 45.41303898 | 26.31329822 | ENSG00000145331 |
| SLC1A6 | -0.791616356 | 0.577696494 | 1.36E-07 | 1.84E-06 | 293.708843 | 169.6126874 | ENSG00000105143 |
| ENHO | -0.791796064 | 0.577624538 | 4.56E-05 | 3.71E-04 | 185.8591897 | 107.478349 | ENSG00000168913 |
| PARP11 | -0.793188704 | 0.577067224 | 9.21E-04 | 5.23E-03 | 126.1554917 | 72.74978695 | ENSG00000111224 |
| PDGFRL | -0.795769478 | 0.576035856 | 3.43E-03 | 1.61E-02 | 80.10015131 | 46.14132864 | ENSG00000104213 |
| PEF1 | -0.799252599 | 0.574646801 | 2.34E-12 | 6.43E-11 | 541.303614 | 310.9899318 | ENSG00000162517 |
| TEK | -0.804891534 | 0.572405114 | 2.20E-02 | 7.44E-02 | 48.16494682 | 27.61859542 | ENSG00000120156 |
| B3GNT7 | -0.805226728 | 0.572272137 | 2.95E-35 | 4.61E-33 | 2315.135563 | 1324.644606 | ENSG00000156966 |
| SMPDL3B | -0.806028471 | 0.571954199 | 1.66E-13 | 5.22E-12 | 601.8399926 | 344.2352577 | ENSG00000130768 |
| FAM229B | -0.80898916 | 0.570781642 | 6.31E-04 | 3.75E-03 | 108.8424835 | 62.16763282 | ENSG00000203778 |
| SHF | -0.810354136 | 0.570241865 | 1.42E-14 | 5.01E-13 | 713.1568758 | 406.7327854 | ENSG00000138606 |
| MFGE8 | -0.811466462 | 0.569802374 | 3.77E-89 | 5.94E-86 | 19799.31405 | 11281.70147 | ENSG00000140545 |
| BMP6 | -0.812716142 | 0.569309018 | 1.16E-03 | 6.41E-03 | 97.35623414 | 55.4862147 | ENSG00000153162 |
| SHISA6 | -0.812978106 | 0.569205653 | 3.08E-05 | 2.62E-04 | 165.0579519 | 93.98226262 | ENSG00000188803 |
| BEND5 | -0.813060995 | 0.56917295 | 7.97E-04 | 4.60E-03 | 113.9551411 | 64.90733731 | ENSG00000162373 |
| AIFM1 | -0.813062375 | 0.569172406 | 5.15E-46 | 1.33E-43 | 3569.522673 | 2031.45358 | ENSG00000156709 |
| SMARCA2 | -0.813164283 | 0.569132202 | 1.90E-08 | 2.99E-07 | 338.3392349 | 192.385558 | ENSG00000080503 |
| DEXI | -0.814525033 | 0.56859565 | 1.32E-02 | 4.91E-02 | 73.23372989 | 41.68728501 | ENSG00000182108 |
| CXCL12 | -0.815429486 | 0.568239298 | 2.19E-15 | 8.70E-14 | 815.1633684 | 463.1554323 | ENSG00000107562 |
| WDR34 | -0.81626968 | 0.567908465 | 3.35E-22 | 2.38E-20 | 1358.327223 | 771.6591298 | ENSG00000119333 |
| XKRX | -0.816936998 | 0.567645839 | 2.31E-02 | 7.73E-02 | 48.52198466 | 27.49695769 | ENSG00000182489 |
| SKIV2L | -0.819030802 | 0.566822604 | 5.83E-25 | 5.15E-23 | 1331.175279 | 754.5710148 | ENSG00000204351 |
| HEXA | -0.819515615 | 0.566632158 | 4.91E-03 | 2.18E-02 | 74.94625454 | 42.42622181 | ENSG00000213614 |
| HPCA | -0.820345013 | 0.566306497 | 2.64E-02 | 8.59E-02 | 47.42417538 | 26.88909552 | ENSG00000121905 |
| LRAT | -0.820436939 | 0.566270414 | 1.67E-11 | 4.13E-10 | 503.4624775 | 285.2035294 | ENSG00000121207 |
| MYO5C | -0.824657382 | 0.564616274 | 4.79E-12 | 1.26E-10 | 518.5861237 | 292.6291813 | ENSG00000128833 |
| ZNF594 | -0.826161396 | 0.564027967 | 1.19E-05 | 1.11E-04 | 208.3299596 | 117.7001947 | ENSG00000180626 |
| MACROH2A2 | -0.829066972 | 0.562893162 | 2.80E-32 | 3.63E-30 | 1844.263328 | 1038.013858 | ENSG00000099284 |
| ZSWIM5 | -0.830521101 | 0.562326094 | 1.99E-05 | 1.78E-04 | 204.8862238 | 115.1747066 | ENSG00000162415 |
| RIN3 | -0.833300868 | 0.561243654 | 7.50E-03 | 3.10E-02 | 74.07233423 | 41.6788667 | ENSG00000100599 |
| KSR2 | -0.835083607 | 0.560550553 | 4.30E-03 | 1.95E-02 | 84.99367466 | 47.52794836 | ENSG00000171435 |
| CRTC3 | -0.836818431 | 0.559876903 | 6.10E-07 | 7.33E-06 | 260.0724027 | 145.6292738 | ENSG00000140577 |
| C3orf80 | -0.840274833 | 0.558537158 | 3.11E-02 | 9.76E-02 | 42.6483678 | 23.78809371 | ENSG00000180044 |
| PDE1B | -0.840380713 | 0.558496168 | 7.71E-03 | 3.17E-02 | 65.41774951 | 36.57134881 | ENSG00000123360 |
| LMF1 | -0.84102492 | 0.558246838 | 7.25E-05 | 5.58E-04 | 159.1016036 | 88.92593726 | ENSG00000260807 |
| BBS9 | -0.841532831 | 0.558050338 | 5.39E-10 | 1.08E-08 | 451.1815437 | 251.5776833 | ENSG00000122507 |
| SLC5A6 | -0.841593049 | 0.558027046 | 6.31E-48 | 1.75E-45 | 3976.59353 | 2219.488069 | ENSG00000138074 |
| DPF1 | -0.842966778 | 0.557495947 | 1.14E-09 | 2.18E-08 | 365.5757961 | 203.8552656 | ENSG00000011332 |
| STS | -0.843775909 | 0.557183365 | 1.34E-03 | 7.23E-03 | 118.0086112 | 65.75095964 | ENSG00000101846 |
| THSD7A | -0.844659238 | 0.556842319 | 2.56E-02 | 8.40E-02 | 42.83193352 | 23.86555976 | ENSG00000005108 |
| SLC39A1 | -0.845447509 | 0.55653815 | 1.06E-02 | 4.10E-02 | 59.08214296 | 32.9256686 | ENSG00000143570 |

| | | | | | | | |
|---|---|---|---|---|---|---|---|
| TSPAN18 | -0.846814945 | 0.556010894 | 1.35E-14 | 4.78E-13 | 585.2511019 | 325.3293999 | ENSG00000157570 |
| RIPOR3 | -0.848516323 | 0.555355574 | 7.91E-03 | 3.24E-02 | 69.31760415 | 38.58439128 | ENSG00000042062 |
| GPR137 | -0.848795241 | 0.555248216 | 7.38E-14 | 2.40E-12 | 665.7964648 | 369.9801583 | ENSG00000173264 |
| GGT7 | -0.849061255 | 0.555145845 | 8.03E-06 | 7.86E-05 | 1569.372531 | 871.3197934 | ENSG00000131067 |
| COL9A2 | -0.849596925 | 0.554939759 | 4.55E-05 | 3.70E-04 | 155.1377308 | 86.10999622 | ENSG00000049089 |
| CAMK2N1 | -0.85403127 | 0.553236685 | 3.39E-04 | 2.18E-03 | 107.2423351 | 59.35263775 | ENSG00000162545 |
| PDS5A | -0.854708484 | 0.552977052 | 1.01E-35 | 1.62E-33 | 2590.880249 | 1432.326848 | ENSG00000121892 |
| ZNF329 | -0.855067278 | 0.552839545 | 3.93E-08 | 5.86E-07 | 284.6491417 | 157.1963101 | ENSG00000181894 |
| PLXNA4 | -0.85533593 | 0.552736608 | 1.47E-02 | 5.38E-02 | 50.33740238 | 27.78268088 | ENSG00000221866 |
| SELENOP | -0.855552873 | 0.552653497 | 4.16E-02 | 1.22E-01 | 37.72519962 | 20.8821283 | ENSG00000250722 |
| TRIM9 | -0.856873658 | 0.552147775 | 3.90E-04 | 2.47E-03 | 115.4079027 | 63.73689858 | ENSG00000100505 |
| ADCYAP1R1 | -0.856989133 | 0.552103582 | 1.10E-17 | 5.30E-16 | 850.2270948 | 469.3744203 | ENSG00000078549 |
| CYP26A1 | -0.859271387 | 0.551230879 | 2.50E-03 | 1.23E-02 | 75.31602958 | 41.49064024 | ENSG00000095596 |
| PCDH19 | -0.859304703 | 0.55121815 | 2.56E-02 | 8.39E-02 | 62.39153585 | 34.28741226 | ENSG00000165194 |
| DCTPP1 | -0.861069028 | 0.550544457 | 7.92E-23 | 5.93E-21 | 1054.497889 | 580.357559 | ENSG00000179958 |
| SCUBE3 | -0.861160796 | 0.550509438 | 1.69E-03 | 8.82E-03 | 84.59542384 | 46.58633485 | ENSG00000146197 |
| TRABD2A | -0.86168083 | 0.550311038 | 4.17E-05 | 3.42E-04 | 170.2305619 | 93.65650776 | ENSG00000186854 |
| RIPK4 | -0.869760462 | 0.547237704 | 1.74E-04 | 1.21E-03 | 126.2666989 | 69.22094204 | ENSG00000183421 |
| TIMM23B | -0.871384583 | 0.546621995 | 1.88E-02 | 6.52E-02 | 50.19863526 | 27.40454694 | ENSG00000204152 |
| ANKRD36 | -0.8716145 | 0.546534889 | 6.53E-03 | 2.76E-02 | 71.96806731 | 39.38511439 | ENSG00000135976 |
| ASXL1 | -0.87214895 | 0.546332461 | 2.03E-17 | 9.55E-16 | 1001.631786 | 547.1219106 | ENSG00000171456 |
| ZNF558 | -0.872174324 | 0.546322852 | 6.91E-07 | 8.22E-06 | 203.7032927 | 111.3088507 | ENSG00000167785 |
| ZNF704 | -0.873161936 | 0.545948989 | 2.66E-05 | 2.30E-04 | 150.1051348 | 82.02924115 | ENSG00000164684 |
| ST8SIA2 | -0.874791445 | 0.545332694 | 1.04E-03 | 5.77E-03 | 89.38755424 | 48.78304195 | ENSG00000140557 |
| ZHX1 | -0.878904323 | 0.543780257 | 4.90E-04 | 3.03E-03 | 95.82279987 | 52.1383645 | ENSG00000165156 |
| GALNT16 | -0.881401371 | 0.542839884 | 2.78E-07 | 3.58E-06 | 251.665428 | 136.7379278 | ENSG00000100626 |
| IQGAP1 | -0.881870187 | 0.542663512 | 2.40E-43 | 5.46E-41 | 3367.311908 | 1826.798095 | ENSG00000140575 |
| TFE3 | -0.882636676 | 0.542375277 | 2.92E-48 | 8.36E-46 | 3043.600591 | 1651.030016 | ENSG00000068323 |
| ADGRF3 | -0.883771895 | 0.541948664 | 1.22E-03 | 6.68E-03 | 85.12832364 | 46.18862174 | ENSG00000173567 |
| ZNF438 | -0.889004799 | 0.539986484 | 3.99E-02 | 1.18E-01 | 34.35499079 | 18.54448785 | ENSG00000183621 |
| STK32B | -0.889136545 | 0.539937175 | 1.24E-04 | 9.03E-04 | 127.6992989 | 69.03460749 | ENSG00000152953 |
| EPHB1 | -0.889372914 | 0.539848719 | 5.87E-09 | 1.01E-07 | 282.4529906 | 152.4741149 | ENSG00000154928 |
| DLL1 | -0.889850005 | 0.539670224 | 7.39E-04 | 4.31E-03 | 95.96187247 | 51.73251663 | ENSG00000198719 |
| PIGL | -0.892936048 | 0.538517059 | 9.10E-09 | 1.51E-07 | 264.5359193 | 142.4748516 | ENSG00000108474 |
| SMAD9 | -0.895886017 | 0.537417044 | 4.77E-02 | 1.36E-01 | 30.25944362 | 16.2302104 | ENSG00000120693 |
| MANEAL | -0.897963399 | 0.536643758 | 1.02E-03 | 5.68E-03 | 89.28138104 | 47.92710189 | ENSG00000185090 |
| CDHR1 | -0.900220326 | 0.535804898 | 3.50E-11 | 8.36E-10 | 523.3172106 | 280.5859901 | ENSG00000148600 |
| RBM19 | -0.902467065 | 0.534971127 | 8.29E-37 | 1.40E-34 | 1796.900935 | 961.5794207 | ENSG00000122965 |
| PSPN | -0.902662623 | 0.534898616 | 2.56E-02 | 8.39E-02 | 39.04539718 | 20.94029873 | ENSG00000125650 |
| PRRT2 | -0.907019427 | 0.533285709 | 7.11E-04 | 4.17E-03 | 170.83104 | 91.19313251 | ENSG00000167371 |
| MAN2A2 | -0.910851985 | 0.531870902 | 1.03E-09 | 1.97E-08 | 414.7471756 | 220.7358718 | ENSG00000196547 |
| SLC38A5 | -0.91195052 | 0.531466065 | 9.20E-27 | 9.34E-25 | 1155.092709 | 614.0496378 | ENSG00000017483 |
| YWHAB | -0.912298941 | 0.531337727 | 2.38E-59 | 1.00E-56 | 6499.358099 | 3453.135235 | ENSG00000166913 |
| TBC1D19 | -0.913886414 | 0.53075339 | 1.94E-02 | 6.69E-02 | 40.00321799 | 21.23472377 | ENSG00000109680 |
| UBE2QL1 | -0.914523236 | 0.530519161 | 2.96E-02 | 9.40E-02 | 36.91145928 | 19.54665815 | ENSG00000215218 |
| MMGT1 | -0.915513619 | 0.530155095 | 4.07E-15 | 1.56E-13 | 650.5755188 | 344.8942695 | ENSG00000169446 |
| PIM2 | -0.91950486 | 0.528690438 | 6.03E-63 | 3.08E-60 | 5341.16595 | 2823.53961 | ENSG00000102096 |
| LOXL3 | -0.922249653 | 0.527685537 | 6.33E-13 | 1.87E-11 | 436.6886046 | 230.3973461 | ENSG00000115318 |
| CNTFR | -0.925615228 | 0.526455966 | 2.03E-04 | 1.40E-03 | 112.8135014 | 59.44722396 | ENSG00000122756 |
| ADPRHL1 | -0.92609374 | 0.526281381 | 3.20E-02 | 1.00E-01 | 33.31560761 | 17.52150886 | ENSG00000153531 |
| SCN9A | -0.928690574 | 0.525334933 | 2.14E-03 | 1.08E-02 | 77.21684848 | 40.58305614 | ENSG00000169432 |
| PCDHGB6 | -0.929878255 | 0.524902635 | 1.37E-05 | 1.26E-04 | 140.2225672 | 73.64386676 | ENSG00000253305 |
| ANKRD18B | -0.932143175 | 0.524079225 | 1.03E-04 | 7.61E-04 | 114.0704926 | 59.69530185 | ENSG00000230453 |
| GPAM | -0.932186537 | 0.524063473 | 2.03E-21 | 1.36E-19 | 752.2546713 | 394.1358649 | ENSG00000119927 |
| TCEA2 | -0.932951341 | 0.523785729 | 8.07E-17 | 3.59E-15 | 859.3001181 | 450.4358258 | ENSG00000171703 |
| ATAD1 | -0.933553212 | 0.523567259 | 3.06E-18 | 1.57E-16 | 801.252603 | 419.6894902 | ENSG00000138138 |
| JAM2 | -0.933709949 | 0.523510381 | 1.37E-05 | 1.26E-04 | 145.2931221 | 76.02974628 | ENSG00000154721 |
| TTLL3 | -0.937007745 | 0.522315077 | 3.74E-02 | 1.13E-01 | 37.71391081 | 19.70376572 | ENSG00000214021 |
| ADAM33 | -0.939042526 | 0.521578922 | 5.41E-07 | 6.57E-06 | 336.4343234 | 175.6944211 | ENSG00000149451 |
| ATF5 | -0.940986759 | 0.520876495 | 8.02E-08 | 1.14E-06 | 217.94295 | 113.6172641 | ENSG00000169136 |
| GPR137C | -0.940999631 | 0.520871848 | 8.56E-04 | 4.89E-03 | 77.02115595 | 40.11099841 | ENSG00000180998 |
| RTL5 | -0.944229094 | 0.519707184 | 3.10E-05 | 2.63E-04 | 125.8666687 | 65.4100195 | ENSG00000242732 |
| CCDC74A | -0.944561475 | 0.519587463 | 2.21E-09 | 4.04E-08 | 302.2703228 | 157.1725909 | ENSG00000163040 |
| YPEL2 | -0.945609098 | 0.519210297 | 1.60E-03 | 8.41E-03 | 78.7600453 | 40.87161721 | ENSG00000175155 |
| SLC9A6 | -0.950066334 | 0.517608662 | 3.47E-14 | 1.18E-12 | 672.7991857 | 348.0370566 | ENSG00000198689 |
| SHISA2 | -0.953058305 | 0.516536318 | 2.69E-11 | 6.50E-10 | 436.0341503 | 225.1479489 | ENSG00000180730 |
| TMEM132E | -0.953213731 | 0.516480673 | 1.47E-02 | 5.37E-02 | 39.86117073 | 20.56093528 | ENSG00000181291 |
| EPHA10 | -0.956005487 | 0.515482199 | 2.41E-02 | 7.99E-02 | 38.14745087 | 19.69317195 | ENSG00000183317 |
| NOL12 | -0.956298877 | 0.51537738 | 2.18E-03 | 1.09E-02 | 66.45476843 | 34.30001354 | ENSG00000273899 |
| HUNK | -0.960636001 | 0.513830345 | 4.36E-10 | 8.84E-09 | 399.3893198 | 205.1367053 | ENSG00000142149 |
| TXNDC16 | -0.962749501 | 0.513078152 | 1.02E-09 | 1.96E-08 | 369.3179172 | 189.5507002 | ENSG00000087301 |
| DPYSL5 | -0.963910812 | 0.512665311 | 3.05E-30 | 3.62E-28 | 1101.832317 | 564.8836891 | ENSG00000157851 |
| SMARCA1 | -0.965991692 | 0.511926398 | 8.52E-58 | 3.28E-55 | 3297.183538 | 1687.541859 | ENSG00000102038 |
| SHROOM2 | -0.96651012 | 0.511742472 | 2.07E-23 | 1.62E-21 | 891.5333245 | 456.0967333 | ENSG00000146950 |

| | | | | | | | |
|---|---|---|---|---|---|---|---|
| NKAIN1 | -0.968612044 | 0.510997436 | 1.11E-33 | 1.59E-31 | 2092.464026 | 1069.412666 | ENSG00000084628 |
| C1GALT1C1 | -0.970532886 | 0.510317533 | 3.29E-13 | 1.00E-11 | 369.9008511 | 188.7986153 | ENSG00000171155 |
| FAAP24 | -0.970899889 | 0.510187731 | 2.19E-04 | 1.49E-03 | 96.15967608 | 48.98817766 | ENSG00000131944 |
| SMIM3 | -0.971653173 | 0.509921413 | 2.21E-20 | 1.35E-18 | 681.2022698 | 347.2777329 | ENSG00000256235 |
| H2AX | -0.972691733 | 0.509554465 | 2.89E-32 | 3.72E-30 | 3545.992291 | 1807.163418 | ENSG00000188486 |
| ATP8B1 | -0.972960096 | 0.509459689 | 3.55E-03 | 1.66E-02 | 62.65666395 | 31.93409206 | ENSG00000081923 |
| GOPC | -0.973692559 | 0.5092011 | 1.90E-12 | 5.29E-11 | 475.0726823 | 241.7037032 | ENSG00000047932 |
| EDNRB | -0.973756201 | 0.509178638 | 1.09E-08 | 1.78E-07 | 252.263795 | 128.2724174 | ENSG00000136160 |
| IRAK1BP1 | -0.973915708 | 0.509122345 | 7.00E-06 | 6.94E-05 | 138.1200797 | 70.40148008 | ENSG00000146243 |
| DUSP15 | -0.975358637 | 0.508613395 | 1.89E-03 | 9.66E-03 | 66.44142027 | 33.8357117 | ENSG00000149599 |
| SLC16A2 | -0.976075636 | 0.508360684 | 1.34E-45 | 3.29E-43 | 2406.418421 | 1223.243606 | ENSG00000147100 |
| EPHB3 | -0.980866366 | 0.50667538 | 8.12E-04 | 4.67E-03 | 79.32022567 | 40.22488024 | ENSG00000182580 |
| AMOTL1 | -0.983209287 | 0.505853213 | 5.55E-40 | 1.08E-37 | 1724.657289 | 872.2746441 | ENSG00000166025 |
| FAM133B | -0.983449113 | 0.505769129 | 1.81E-20 | 1.11E-18 | 789.9899116 | 399.635021 | ENSG00000234545 |
| GDF7 | -0.984658161 | 0.505345448 | 4.66E-02 | 1.34E-01 | 26.36050486 | 13.30589532 | ENSG00000143869 |
| H2AC11 | -0.986174246 | 0.504814675 | 4.57E-03 | 2.05E-02 | 61.168918 | 30.94715 | ENSG00000196787 |
| COL21A1 | -0.986950679 | 0.504543065 | 6.76E-04 | 3.98E-03 | 76.4144231 | 38.59432264 | ENSG00000124749 |
| HRH3 | -0.987551048 | 0.504333146 | 1.02E-06 | 1.18E-05 | 175.6616875 | 88.69978195 | ENSG00000101180 |
| ZNF677 | -0.988279448 | 0.504078579 | 6.58E-11 | 1.51E-09 | 400.3532658 | 201.5824899 | ENSG00000197928 |
| ITGB1BP2 | -0.991022944 | 0.50312091 | 3.46E-02 | 1.06E-01 | 27.72430582 | 13.97098195 | ENSG00000147166 |
| MCTS1 | -0.991260556 | 0.503038053 | 1.08E-19 | 6.35E-18 | 635.956551 | 319.8961272 | ENSG00000232119 |
| YPEL1 | -0.993846819 | 0.502137084 | 2.88E-05 | 2.47E-04 | 144.284579 | 72.44876288 | ENSG00000100027 |
| CSTF2 | -0.994136201 | 0.502036374 | 5.69E-39 | 1.03E-36 | 1535.619796 | 770.8113971 | ENSG00000101811 |
| RNF39 | -0.995328266 | 0.501621724 | 7.09E-03 | 2.96E-02 | 59.83328734 | 29.9837389 | ENSG00000204618 |
| KPNA1 | -0.997839536 | 0.500749321 | 8.44E-19 | 4.65E-17 | 638.1672449 | 319.7613117 | ENSG00000114030 |
| TAF2 | -0.999385242 | 0.500213104 | 1.93E-23 | 1.54E-21 | 755.9777475 | 378.0763616 | ENSG00000064313 |
| ST8SIA5 | -1.001727366 | 0.499401699 | 8.44E-04 | 4.84E-03 | 82.99619134 | 41.49130265 | ENSG00000101638 |
| SMPD1 | -1.003060202 | 0.498940539 | 3.94E-03 | 1.81E-02 | 52.65429561 | 26.29712402 | ENSG00000166311 |
| ME3 | -1.003615549 | 0.498748515 | 9.11E-03 | 3.62E-02 | 46.59505425 | 23.19640575 | ENSG00000151376 |
| CPNE9 | -1.005569277 | 0.498073557 | 5.48E-05 | 4.37E-04 | 108.0750163 | 53.76920565 | ENSG00000144550 |
| LRRC6 | -1.014786265 | 0.494901642 | 3.48E-03 | 1.63E-02 | 56.29522522 | 27.78268088 | ENSG00000129295 |
| ZFP42 | -1.018510768 | 0.493625638 | 9.90E-48 | 2.71E-45 | 4504.218823 | 2223.221721 | ENSG00000179059 |
| APLN | -1.019733056 | 0.493207603 | 6.77E-05 | 5.26E-04 | 115.3859617 | 56.90321828 | ENSG00000171388 |
| ZNF599 | -1.022390181 | 0.49230006 | 3.52E-03 | 1.65E-02 | 62.07782207 | 30.64525261 | ENSG00000153896 |
| TMEM99 | -1.023721919 | 0.491845832 | 3.67E-05 | 3.05E-04 | 124.4073468 | 61.10417067 | ENSG00000167920 |
| CRELD1 | -1.025573421 | 0.49121502 | 4.47E-02 | 1.29E-01 | 25.04712075 | 12.33796534 | ENSG00000163703 |
| CCBE1 | -1.025780766 | 0.491144427 | 4.76E-06 | 4.85E-05 | 128.4326204 | 63.0195055 | ENSG00000183287 |
| H2BC9 | -1.026975233 | 0.490737956 | 6.05E-04 | 3.62E-03 | 88.70579314 | 43.64707508 | ENSG00000275713 |
| SLC25A27 | -1.027431547 | 0.490582764 | 4.76E-03 | 2.12E-02 | 74.24135716 | 36.46420428 | ENSG00000153291 |
| CCDC157 | -1.028108142 | 0.490352745 | 7.11E-04 | 4.17E-03 | 75.59300516 | 37.04016954 | ENSG00000187860 |
| DZIP3 | -1.028366431 | 0.490264963 | 1.34E-16 | 5.83E-15 | 593.2435546 | 291.0719926 | ENSG00000198919 |
| E2F2 | -1.033223584 | 0.488617153 | 7.15E-10 | 1.40E-08 | 230.3846558 | 112.5957982 | ENSG00000007968 |
| CRHR2 | -1.036456375 | 0.487523486 | 3.05E-02 | 9.63E-02 | 29.21383122 | 14.26852842 | ENSG00000106113 |
| DYNC2H1 | -1.038630479 | 0.486789354 | 1.14E-12 | 3.26E-11 | 392.7090036 | 191.0914217 | ENSG00000187240 |
| FGFBP3 | -1.039759517 | 0.486408546 | 1.74E-45 | 4.22E-43 | 2150.608327 | 1046.307346 | ENSG00000174721 |
| ACD | -1.0405475 | 0.486142948 | 1.62E-27 | 1.70E-25 | 800.8146144 | 389.4216365 | ENSG00000102977 |
| AMBP | -1.041544618 | 0.485807067 | 1.34E-02 | 4.95E-02 | 38.8870271 | 18.84610169 | ENSG00000106927 |
| FIG4 | -1.041735128 | 0.48574292 | 6.00E-22 | 4.21E-20 | 697.7078192 | 338.6904169 | ENSG00000112367 |
| MAN2A1 | -1.04218714 | 0.485590755 | 9.08E-33 | 1.25E-30 | 1620.739221 | 786.7594346 | ENSG00000112893 |
| TMEM221 | -1.042400731 | 0.485518869 | 6.52E-03 | 2.76E-02 | 46.52030533 | 22.56243802 | ENSG00000188051 |
| PLCL2 | -1.045897195 | 0.484343607 | 4.98E-08 | 7.32E-07 | 209.8381465 | 101.5486369 | ENSG00000154822 |
| LRCH2 | -1.049310556 | 0.483199024 | 4.64E-08 | 6.84E-07 | 213.5630034 | 103.0238109 | ENSG00000130224 |
| RAB27A | -1.049697545 | 0.483069427 | 7.43E-07 | 8.78E-06 | 141.5857578 | 68.4236239 | ENSG00000069974 |
| C10orf143 | -1.050044983 | 0.482953106 | 4.07E-03 | 1.86E-02 | 48.08694391 | 23.19735171 | ENSG00000237489 |
| RASL10A | -1.052665405 | 0.482076696 | 2.47E-02 | 8.18E-02 | 30.17493216 | 14.56172396 | ENSG00000100276 |
| PVALB | -1.053589432 | 0.481768031 | 4.43E-02 | 1.28E-01 | 27.72814462 | 13.36189028 | ENSG00000100362 |
| HS6ST3 | -1.054389417 | 0.481500961 | 4.22E-03 | 1.92E-02 | 48.18630366 | 23.20699952 | ENSG00000185352 |
| TM4SF18 | -1.057770285 | 0.480373914 | 7.02E-04 | 4.12E-03 | 79.58474339 | 38.2388893 | ENSG00000163762 |
| DBNDD2 | -1.070359238 | 0.476200408 | 3.28E-02 | 1.02E-01 | 29.39058348 | 14.03662473 | ENSG00000244274 |
| AKR1C3 | -1.071732685 | 0.47574728 | 5.45E-03 | 2.37E-02 | 43.9579131 | 20.92629999 | ENSG00000196139 |
| PARP2 | -1.072089267 | 0.475629708 | 1.26E-23 | 1.02E-21 | 756.2188235 | 359.7415916 | ENSG00000129484 |
| RAI2 | -1.077685727 | 0.473788233 | 1.11E-02 | 4.27E-02 | 39.0797967 | 18.52490868 | ENSG00000131831 |
| CYP39A1 | -1.078648649 | 0.47347211 | 2.74E-03 | 1.33E-02 | 50.52719732 | 23.89790817 | ENSG00000146233 |
| ZNF428 | -1.081233549 | 0.472624542 | 5.50E-08 | 7.98E-07 | 224.8440758 | 106.3274894 | ENSG00000131116 |
| SYPL2 | -1.085967797 | 0.471076152 | 2.42E-04 | 1.63E-03 | 80.82757459 | 38.06437803 | ENSG00000143028 |
| ODC1 | -1.089577722 | 0.469898894 | 1.64E-73 | 1.55E-70 | 4426.181025 | 2079.718362 | ENSG00000115758 |
| ICAM5 | -1.091213373 | 0.46936645 | 4.03E-05 | 3.31E-04 | 107.0842705 | 50.26162092 | ENSG00000105376 |
| ZNF585B | -1.091677753 | 0.469215393 | 1.48E-03 | 7.86E-03 | 66.08171325 | 31.0470331 | ENSG00000245680 |
| H2AC8 | -1.092849429 | 0.468834477 | 3.41E-02 | 1.05E-01 | 26.84832673 | 12.62056711 | ENSG00000277075 |
| ZNF559 | -1.092907369 | 0.468815649 | 1.31E-08 | 2.13E-07 | 188.2418811 | 88.29677196 | ENSG00000188321 |
| DPH5 | -1.0966812 | 0.467590914 | 1.48E-26 | 1.50E-24 | 727.3335543 | 339.9070872 | ENSG00000117543 |
| TBL1X | -1.096893777 | 0.46752202 | 2.73E-55 | 9.72E-53 | 1929.931495 | 902.1796148 | ENSG00000101849 |
| MROH8 | -1.096964171 | 0.467499209 | 7.72E-03 | 3.18E-02 | 38.3654161 | 17.94939492 | ENSG00000101353 |
| H4C8 | -1.098607274 | 0.466967071 | 8.55E-03 | 3.45E-02 | 39.89765517 | 18.68324574 | ENSG00000158406 |

| | | | | | | | |
|---|---|---|---|---|---|---|---|
| ZNF443 | -1.102914471 | 0.46557501 | 1.85E-04 | 1.29E-03 | 81.19259495 | 37.81601658 | ENSG00000180855 |
| ISM1 | -1.104087784 | 0.465196522 | 1.51E-02 | 5.49E-02 | 30.70010263 | 14.30399825 | ENSG00000101230 |
| LRRC49 | -1.106716444 | 0.464349683 | 3.94E-08 | 5.87E-07 | 202.5663553 | 94.16698879 | ENSG00000137821 |
| PPP2R5A | -1.108985664 | 0.46361988 | 3.64E-29 | 4.17E-27 | 881.216632 | 408.3598225 | ENSG00000066027 |
| SLC7A11 | -1.116406371 | 0.461241308 | 1.19E-12 | 3.39E-11 | 466.2950678 | 214.9979888 | ENSG00000151012 |
| ABCB7 | -1.118017496 | 0.460726506 | 1.58E-31 | 1.99E-29 | 852.8402099 | 392.9028321 | ENSG00000131269 |
| LRRC4 | -1.121649491 | 0.459568082 | 1.90E-12 | 5.28E-11 | 296.37418 | 136.2115561 | ENSG00000128594 |
| CEP135 | -1.123004325 | 0.459136705 | 1.15E-10 | 2.56E-09 | 299.3677742 | 137.4175373 | ENSG00000174799 |
| TMSB15B | -1.126352275 | 0.458072457 | 5.68E-03 | 2.46E-02 | 50.73923882 | 23.2337675 | ENSG00000158427 |
| ZNF649 | -1.127405943 | 0.457738028 | 9.38E-22 | 6.49E-20 | 518.8936338 | 237.5022993 | ENSG00000198093 |
| EMC6 | -1.129483449 | 0.457079351 | 3.99E-02 | 1.19E-01 | 23.27675482 | 10.6318336 | ENSG00000127774 |
| LRRC37A3 | -1.131087863 | 0.456571318 | 2.03E-02 | 6.95E-02 | 28.36543818 | 12.97287903 | ENSG00000176809 |
| TSPYL4 | -1.131891999 | 0.456316903 | 7.76E-29 | 8.68E-27 | 875.577277 | 399.7051529 | ENSG00000187189 |
| BCORL1 | -1.13535891 | 0.455221654 | 1.30E-16 | 5.67E-15 | 392.670259 | 178.74693 | ENSG00000085185 |
| SRSF12 | -1.137887381 | 0.45442453 | 4.50E-07 | 5.57E-06 | 154.2578606 | 70.17515683 | ENSG00000154548 |
| DLG1 | -1.139355164 | 0.453962438 | 3.23E-40 | 6.35E-38 | 1212.512439 | 550.3990524 | ENSG00000075711 |
| MAMDC4 | -1.148483828 | 0.451099056 | 1.01E-03 | 5.64E-03 | 341.5983399 | 154.1798381 | ENSG00000177943 |
| GPR155 | -1.153650856 | 0.44948633 | 4.08E-03 | 1.87E-02 | 42.80849176 | 19.21581732 | ENSG00000163328 |
| SHD | -1.154793928 | 0.449130335 | 4.44E-04 | 2.78E-03 | 67.95349771 | 30.59673 | ENSG00000105251 |
| RTF1 | -1.161287797 | 0.447113249 | 9.75E-63 | 4.85E-60 | 2309.374757 | 1032.778314 | ENSG00000137815 |
| OLIG1 | -1.164137996 | 0.446230801 | 5.67E-03 | 2.45E-02 | 36.45864732 | 16.26255881 | ENSG00000184221 |
| ADAMTS19 | -1.16462347 | 0.446080667 | 2.07E-21 | 1.38E-19 | 569.8801511 | 254.1214054 | ENSG00000145808 |
| FOSB | -1.177959267 | 0.441976244 | 7.71E-03 | 3.17E-02 | 34.52312341 | 15.29245336 | ENSG00000125740 |
| COBLL1 | -1.180306868 | 0.441257631 | 2.37E-04 | 1.60E-03 | 84.34575441 | 37.32354931 | ENSG00000082438 |
| NLGN4X | -1.183125964 | 0.440396234 | 1.81E-72 | 1.42E-69 | 4894.629026 | 2155.186244 | ENSG00000146938 |
| DOK3 | -1.1875093 | 0.43906021 | 5.37E-07 | 6.53E-06 | 122.3653959 | 53.76920565 | ENSG00000146094 |
| NSMCE4A | -1.189346517 | 0.438501439 | 1.59E-37 | 2.71E-35 | 982.3048413 | 430.8704914 | ENSG00000107672 |
| TTI1 | -1.194102554 | 0.437058241 | 1.04E-40 | 2.10E-38 | 1363.702011 | 595.726038 | ENSG00000101407 |
| HGSNAT | -1.197576563 | 0.436007071 | 1.33E-05 | 1.23E-04 | 123.1527198 | 53.65910765 | ENSG00000165102 |
| MEGF9 | -1.202593895 | 0.434493381 | 8.58E-04 | 4.90E-03 | 57.94014605 | 25.09244496 | ENSG00000106780 |
| PCDHGB4 | -1.205034731 | 0.433758901 | 2.61E-06 | 2.79E-05 | 122.8226832 | 53.27066348 | ENSG00000253953 |
| TMEM255A | -1.210195621 | 0.432210007 | 1.63E-03 | 8.55E-03 | 46.75578859 | 20.22356806 | ENSG00000125355 |
| CEP290 | -1.216370959 | 0.430363921 | 4.69E-11 | 1.10E-09 | 208.2721189 | 89.57530119 | ENSG00000198707 |
| CDK8 | -1.224990837 | 0.42780023 | 2.62E-25 | 2.42E-23 | 707.4400243 | 302.5576205 | ENSG00000132964 |
| C19orf57 | -1.233722992 | 0.425218717 | 6.13E-05 | 4.82E-04 | 90.05713623 | 38.39353785 | ENSG00000132016 |
| SLFN11 | -1.233745711 | 0.425212021 | 3.34E-02 | 1.03E-01 | 19.51869426 | 8.302611457 | ENSG00000172716 |
| TMEM45A | -1.236283544 | 0.424464692 | 3.93E-15 | 1.51E-13 | 282.3216218 | 119.9003067 | ENSG00000181458 |
| ELP2 | -1.237009217 | 0.42425124 | 1.56E-68 | 1.02E-65 | 1998.193114 | 847.6341106 | ENSG00000134759 |
| BIVM | -1.237740167 | 0.424036345 | 1.68E-05 | 1.52E-04 | 92.19193829 | 39.0226602 | ENSG00000134897 |
| DOCK2 | -1.239299566 | 0.423578255 | 3.07E-07 | 3.93E-06 | 118.4014969 | 50.1897353 | ENSG00000134516 |
| BOC | -1.2406452 | 0.423183358 | 1.22E-10 | 2.70E-09 | 189.6050978 | 80.18652698 | ENSG00000144857 |
| ZNF583 | -1.242370244 | 0.422677656 | 7.47E-03 | 3.09E-02 | 32.91171238 | 13.86769389 | ENSG00000198440 |
| OVOL2 | -1.243255252 | 0.422418448 | 4.02E-12 | 1.07E-10 | 221.482766 | 93.62682927 | ENSG00000125850 |
| CUL4B | -1.244274522 | 0.422120113 | 5.90E-32 | 7.48E-30 | 1566.319497 | 660.8771843 | ENSG00000158290 |
| LAMA4 | -1.255464389 | 0.418858719 | 1.25E-02 | 4.71E-02 | 27.65314252 | 11.58358938 | ENSG00000112769 |
| CACNA1G | -1.255668869 | 0.418799356 | 2.07E-16 | 8.98E-15 | 340.3771711 | 142.5643089 | ENSG00000006283 |
| FAM133A | -1.256566909 | 0.418538746 | 3.27E-02 | 1.02E-01 | 21.49306687 | 9.02028808 | ENSG00000179083 |
| H2BC21 | -1.26490314 | 0.416127304 | 1.02E-04 | 7.57E-04 | 80.39372904 | 33.46098273 | ENSG00000184678 |
| LTB4R | -1.266723492 | 0.415602578 | 2.82E-05 | 2.42E-04 | 111.581628 | 46.43936956 | ENSG00000213903 |
| JAKMIP2 | -1.268836717 | 0.414994259 | 1.29E-14 | 4.58E-13 | 283.651608 | 117.6412462 | ENSG00000176049 |
| GRIN2B | -1.272783111 | 0.413860622 | 6.57E-05 | 5.12E-04 | 74.42700781 | 30.7670583 | ENSG00000273079 |
| PCDHGA1 | -1.281217245 | 0.411448211 | 3.93E-02 | 1.17E-01 | 20.13282538 | 8.236968684 | ENSG00000204956 |
| ADM | -1.289925587 | 0.408972123 | 1.32E-51 | 4.29E-49 | 1142.51727 | 467.3380577 | ENSG00000148926 |
| IGSF11 | -1.291284908 | 0.408586967 | 1.68E-02 | 5.98E-02 | 23.55640014 | 9.614435052 | ENSG00000144847 |
| SPAG1 | -1.2990538 | 0.406392646 | 1.79E-03 | 9.24E-03 | 41.60898461 | 16.91147123 | ENSG00000104450 |
| CAPS2 | -1.300555217 | 0.405969932 | 2.68E-02 | 8.69E-02 | 20.48452373 | 8.334959865 | ENSG00000180881 |
| FRZB | -1.321041892 | 0.400245783 | 1.10E-10 | 2.46E-09 | 170.1139635 | 68.06790701 | ENSG00000162998 |
| KCND1 | -1.324232453 | 0.399361606 | 6.94E-15 | 2.57E-13 | 302.6964646 | 121.0514498 | ENSG00000102057 |
| S100A6 | -1.324260354 | 0.399353883 | 3.50E-03 | 1.64E-02 | 34.80276874 | 13.91281153 | ENSG00000197956 |
| PAPLN | -1.335146431 | 0.396351837 | 4.10E-10 | 8.36E-09 | 172.8116157 | 68.47621389 | ENSG00000100767 |
| MID2 | -1.337937875 | 0.395585685 | 1.66E-04 | 1.17E-03 | 72.40427712 | 28.66645047 | ENSG00000080561 |
| CP | -1.338761171 | 0.395360002 | 3.82E-02 | 1.15E-01 | 18.5187446 | 7.290509791 | ENSG00000047457 |
| CHRM4 | -1.341664583 | 0.394565143 | 3.48E-02 | 1.07E-01 | 18.59911177 | 7.353977101 | ENSG00000180720 |
| ALDH1L1 | -1.343127377 | 0.394165284 | 1.97E-03 | 1.00E-02 | 37.83104176 | 14.87762009 | ENSG00000144908 |
| LETM2 | -1.351361785 | 0.391921932 | 1.08E-02 | 4.17E-02 | 24.5519006 | 9.610084125 | ENSG00000165046 |
| MT2A | -1.352850394 | 0.391517746 | 5.76E-34 | 8.36E-32 | 670.1685531 | 262.4112906 | ENSG00000125148 |
| B4GALNT1 | -1.367095268 | 0.387671003 | 1.58E-03 | 8.32E-03 | 42.97665053 | 16.70851098 | ENSG00000135454 |
| HACL1 | -1.369513293 | 0.387021792 | 1.14E-36 | 1.88E-34 | 715.7034777 | 277.082829 | ENSG00000131373 |
| SPOCK2 | -1.373457399 | 0.385965179 | 6.97E-05 | 5.39E-04 | 67.94960718 | 26.17454032 | ENSG00000107742 |
| LIG3 | -1.375888477 | 0.385315339 | 1.78E-91 | 3.36E-88 | 2292.77463 | 883.3355206 | ENSG00000005156 |
| PARP16 | -1.377382864 | 0.384916424 | 1.18E-08 | 1.92E-07 | 134.6621054 | 51.92074309 | ENSG00000138617 |
| PRTG | -1.378045233 | 0.384739742 | 1.79E-07 | 2.36E-06 | 104.2502155 | 40.07117765 | ENSG00000166450 |
| PTGER2 | -1.385879587 | 0.382656129 | 1.28E-04 | 9.24E-04 | 52.90190569 | 20.24939008 | ENSG00000125384 |
| PDE10A | -1.387467671 | 0.382235142 | 6.13E-04 | 3.66E-03 | 47.02887364 | 17.96216415 | ENSG00000112541 |

| | | | | | | | |
|---|---|---|---|---|---|---|---|
| ABCA5 | -1.389215252 | 0.381772409 | 2.14E-04 | 1.46E-03 | 50.46432203 | 19.25251666 | ENSG00000154265 |
| CTNNAL1 | -1.392649616 | 0.380864673 | 3.67E-65 | 2.10E-62 | 1412.503944 | 537.9463545 | ENSG00000119326 |
| GPR50 | -1.393006573 | 0.38077045 | 2.75E-12 | 7.47E-11 | 195.1385845 | 74.24643204 | ENSG00000102195 |
| PDGFA | -1.399284054 | 0.379117234 | 8.96E-41 | 1.84E-38 | 770.0551965 | 291.7934828 | ENSG00000197461 |
| KCNJ12 | -1.400235719 | 0.378867234 | 5.37E-08 | 7.83E-07 | 111.3678326 | 42.24908358 | ENSG00000184185 |
| CALCB | -1.402086035 | 0.378381633 | 7.01E-09 | 1.19E-07 | 126.272902 | 47.79051945 | ENSG00000175868 |
| PNPLA4 | -1.402988367 | 0.378145048 | 2.74E-26 | 2.67E-24 | 506.8192614 | 191.5404953 | ENSG00000006757 |
| MTSS1 | -1.412679022 | 0.375613541 | 5.49E-08 | 7.97E-07 | 108.7570823 | 40.88996688 | ENSG00000170873 |
| SCML1 | -1.418803547 | 0.374022367 | 1.75E-04 | 1.22E-03 | 53.19370397 | 19.84855555 | ENSG00000047634 |
| LAMP5 | -1.419637106 | 0.373806327 | 1.45E-03 | 7.75E-03 | 35.58770224 | 13.29624751 | ENSG00000125869 |
| CFAP69 | -1.423192589 | 0.372886226 | 1.08E-03 | 5.99E-03 | 44.49821118 | 16.61297881 | ENSG00000105792 |
| TPP2 | -1.432811033 | 0.370408463 | 1.94E-81 | 2.15E-78 | 1863.092078 | 689.9000047 | ENSG00000134900 |
| MAP3K7 | -1.4370619 | 0.36931867 | 9.35E-65 | 5.19E-62 | 1297.395848 | 479.253055 | ENSG00000135341 |
| PRR22 | -1.439570516 | 0.368677042 | 1.91E-03 | 9.77E-03 | 35.09337182 | 12.93211231 | ENSG00000212123 |
| ADAP2 | -1.445432647 | 0.367182028 | 2.65E-02 | 8.61E-02 | 17.20536049 | 6.326930736 | ENSG00000184060 |
| FOXB2 | -1.44898717 | 0.366278476 | 2.28E-02 | 7.63E-02 | 18.17950412 | 6.678296698 | ENSG00000204612 |
| INTS6L | -1.450549709 | 0.365881986 | 1.29E-17 | 6.16E-16 | 296.8510186 | 108.7038367 | ENSG00000165359 |
| CTU2 | -1.450996166 | 0.365768777 | 2.81E-28 | 3.06E-26 | 527.1360609 | 192.9733895 | ENSG00000174177 |
| TMEM121B | -1.452191036 | 0.365465965 | 1.04E-15 | 4.30E-14 | 242.1499918 | 88.47173473 | ENSG00000183307 |
| ZDHHC9 | -1.45291389 | 0.365282897 | 2.30E-80 | 2.41E-77 | 1720.445658 | 628.3858461 | ENSG00000188706 |
| SHISA3 | -1.467916323 | 0.36150404 | 4.02E-07 | 5.03E-06 | 101.791606 | 36.86236889 | ENSG00000178343 |
| ZBTB33 | -1.474508592 | 0.359855948 | 5.36E-46 | 1.37E-43 | 830.272243 | 298.7715014 | ENSG00000177485 |
| GRIN1 | -1.483087695 | 0.357722385 | 5.87E-12 | 1.53E-10 | 156.8713068 | 56.13730259 | ENSG00000176884 |
| THBD | -1.484920431 | 0.357268239 | 2.16E-02 | 7.31E-02 | 19.46057364 | 6.948791642 | ENSG00000178726 |
| ZNF568 | -1.490993038 | 0.355767582 | 3.37E-04 | 2.17E-03 | 42.94964929 | 15.2526326 | ENSG00000198453 |
| ARHGAP36 | -1.504821524 | 0.352373778 | 4.37E-03 | 1.97E-02 | 25.44656678 | 8.977345901 | ENSG00000147256 |
| NMU | -1.518010134 | 0.34916718 | 3.22E-15 | 1.25E-13 | 215.0534185 | 75.07297716 | ENSG00000109255 |
| FOXL2 | -1.529495492 | 0.346398481 | 3.11E-03 | 1.49E-02 | 29.80068177 | 10.34611042 | ENSG00000183770 |
| CCT6B | -1.529946794 | 0.346290138 | 3.45E-02 | 1.06E-01 | 15.24522577 | 5.253537224 | ENSG00000132141 |
| ZNF878 | -1.541884191 | 0.343436626 | 2.51E-05 | 2.18E-04 | 57.98939384 | 19.95184361 | ENSG00000257446 |
| ZNF880 | -1.544512279 | 0.342811573 | 8.29E-13 | 2.41E-11 | 173.0725733 | 59.31622196 | ENSG00000221923 |
| CMPK2 | -1.546812848 | 0.34226535 | 1.38E-02 | 5.08E-02 | 19.3194161 | 6.620126274 | ENSG00000134326 |
| CHRD | -1.553648511 | 0.340647492 | 1.15E-02 | 4.40E-02 | 20.56044171 | 7.025311733 | ENSG00000090539 |
| MMUT | -1.556148267 | 0.340057763 | 6.29E-31 | 7.71E-29 | 497.0627749 | 168.9147579 | ENSG00000146085 |
| SHBG | -1.557943454 | 0.339634883 | 3.67E-02 | 1.11E-01 | 14.7618531 | 5.015107141 | ENSG00000129214 |
| TMEM114 | -1.559083982 | 0.339366489 | 1.82E-02 | 6.36E-02 | 21.57018006 | 7.325979621 | ENSG00000232258 |
| SLC17A6 | -1.581437376 | 0.334148806 | 4.78E-02 | 1.36E-01 | 11.92217981 | 3.983709849 | ENSG00000091664 |
| WASHC5 | -1.587880736 | 0.332659759 | 2.15E-59 | 9.23E-57 | 935.3907344 | 311.1234655 | ENSG00000164961 |
| GPD1 | -1.589729333 | 0.332233779 | 4.43E-02 | 1.28E-01 | 11.96250306 | 3.960063296 | ENSG00000167588 |
| C2orf15 | -1.589875919 | 0.332200024 | 2.58E-03 | 1.26E-02 | 27.98023004 | 9.316888587 | ENSG00000273045 |
| DGKK | -1.593455832 | 0.331376722 | 3.12E-39 | 5.83E-37 | 680.19411 | 225.2316578 | ENSG00000274588 |
| NKX3-2 | -1.603834257 | 0.329001426 | 1.95E-04 | 1.35E-03 | 43.49706632 | 14.32329387 | ENSG00000109705 |
| ZNF829 | -1.606469129 | 0.328401101 | 2.19E-04 | 1.49E-03 | 41.47733644 | 13.66161221 | ENSG00000185869 |
| MT1E | -1.619796814 | 0.325381287 | 6.84E-19 | 3.79E-17 | 286.1677533 | 93.20691003 | ENSG00000169715 |
| ANOS1 | -1.632241945 | 0.322586519 | 1.40E-94 | 2.94E-91 | 4495.998759 | 1450.089654 | ENSG00000011201 |
| ZGLP1 | -1.635399628 | 0.321881234 | 2.93E-02 | 9.32E-02 | 15.38490877 | 4.951639832 | ENSG00000220201 |
| METTL7A | -1.644336181 | 0.319893552 | 1.83E-55 | 6.64E-53 | 935.4959406 | 299.2365812 | ENSG00000185432 |
| WNT5B | -1.64767198 | 0.319154749 | 4.38E-06 | 4.49E-05 | 68.62038438 | 21.85005828 | ENSG00000111186 |
| H2BC5 | -1.669884208 | 0.314278567 | 2.79E-09 | 4.98E-08 | 109.7154873 | 34.50732473 | ENSG00000158373 |
| RTP1 | -1.680985005 | 0.311869634 | 3.33E-07 | 4.23E-06 | 77.69902595 | 24.24274774 | ENSG00000175077 |
| PCDHA9 | -1.691125802 | 0.309685169 | 2.10E-03 | 1.06E-02 | 27.8295893 | 8.644329606 | ENSG00000204961 |
| H3C6 | -1.69771219 | 0.308274574 | 3.77E-02 | 1.13E-01 | 12.8026081 | 3.929890353 | ENSG00000274750 |
| EMX1 | -1.700789857 | 0.30761764 | 6.66E-14 | 2.18E-12 | 161.7752031 | 49.75532286 | ENSG00000135638 |
| PCDHB15 | -1.717156633 | 0.304147567 | 4.62E-09 | 8.03E-08 | 108.0293797 | 32.87553762 | ENSG00000113248 |
| KAZALD1 | -1.727194206 | 0.3020388 | 4.00E-03 | 1.84E-02 | 23.13679249 | 7.006962066 | ENSG00000107821 |
| GLTPD2 | -1.742829809 | 0.298783046 | 4.43E-02 | 1.28E-01 | 11.05212446 | 3.311150869 | ENSG00000182327 |
| H3C8 | -1.745856715 | 0.298156829 | 7.82E-06 | 7.68E-05 | 52.22400954 | 15.55330049 | ENSG00000273983 |
| H3C10 | -1.772893665 | 0.292621228 | 2.42E-05 | 2.11E-04 | 49.94271103 | 14.62613722 | ENSG00000278828 |
| SEC14L4 | -1.781237526 | 0.29093373 | 5.70E-04 | 3.44E-03 | 34.12990661 | 9.904509168 | ENSG00000133488 |
| ZNF799 | -1.796181982 | 0.287935588 | 1.55E-04 | 1.10E-03 | 40.24281939 | 11.54499813 | ENSG00000196466 |
| RDX | -1.797115635 | 0.287749308 | 7.40E-99 | 1.75E-95 | 1695.472876 | 487.7164196 | ENSG00000137710 |
| FAM135B | -1.801708948 | 0.286834617 | 2.94E-02 | 9.35E-02 | 11.52273378 | 3.299327592 | ENSG00000147724 |
| LRRTM3 | -1.811270891 | 0.284939811 | 3.42E-03 | 1.61E-02 | 22.08080715 | 6.289285443 | ENSG00000198739 |
| PCDHB12 | -1.815535799 | 0.284098713 | 1.14E-08 | 1.87E-07 | 89.64612206 | 25.4018147 | ENSG00000120328 |
| ETV2 | -1.817089301 | 0.283792958 | 1.16E-02 | 4.41E-02 | 16.46489454 | 4.668092106 | ENSG00000105672 |
| PHGDH | -1.834298137 | 0.280427913 | 0.00E+00 | 0.00E+00 | 15325.66663 | 4297.344873 | ENSG00000092621 |
| POU3F4 | -1.872617443 | 0.273077538 | 2.90E-03 | 1.40E-02 | 24.40125986 | 6.691065933 | ENSG00000196767 |
| CXCL16 | -1.884885053 | 0.270765334 | 9.23E-05 | 6.92E-04 | 39.18772378 | 10.62095629 | ENSG00000161921 |
| ABALON | -1.886620094 | 0.270439897 | 2.24E-03 | 1.12E-02 | 24.49494904 | 6.612653925 | ENSG00000281376 |
| GRPR | -1.897422963 | 0.268422411 | 3.76E-20 | 2.28E-18 | 230.4386066 | 61.88654411 | ENSG00000126010 |
| TCHH | -1.920246673 | 0.264209332 | 4.50E-02 | 1.30E-01 | 10.05781921 | 2.678412645 | ENSG00000159450 |
| ZNF442 | -1.92892091 | 0.262625533 | 2.05E-04 | 1.40E-03 | 33.00067304 | 8.677623972 | ENSG00000198342 |
| RSPH9 | -1.934394665 | 0.261630989 | 9.94E-03 | 3.88E-02 | 16.54317678 | 4.309253795 | ENSG00000172426 |
| AMH | -1.968164211 | 0.255578041 | 2.31E-19 | 1.32E-17 | 302.3578089 | 77.44440644 | ENSG00000104899 |

| | | | | | | | |
|---|---|---|---|---|---|---|---|
| H2BC17 | -1.983064619 | 0.25295197 | 4.21E-02 | 1.23E-01 | 10.58654915 | 2.696762312 | ENSG00000274641 |
| NKAIN3 | -1.984290779 | 0.252737075 | 1.51E-02 | 5.48E-02 | 14.547168 | 3.700162124 | ENSG00000185942 |
| ZNF441 | -1.993032856 | 0.251210234 | 5.99E-03 | 2.57E-02 | 17.24807416 | 4.346899088 | ENSG00000197044 |
| ADM5 | -2.029954084 | 0.244862867 | 2.53E-09 | 4.58E-08 | 75.96873009 | 18.63472313 | ENSG00000224420 |
| GLIPR1L1 | -2.032825861 | 0.244375937 | 6.69E-03 | 2.82E-02 | 14.97625886 | 3.660341367 | ENSG00000173401 |
| FABP7 | -2.034917786 | 0.244021846 | 1.73E-19 | 1.00E-17 | 179.6166881 | 43.89297751 | ENSG00000164434 |
| SLC9A3 | -2.085583987 | 0.235600746 | 4.38E-05 | 3.58E-04 | 35.29117543 | 8.344607677 | ENSG00000066230 |
| CNTD2 | -2.109863385 | 0.231668952 | 1.13E-03 | 6.25E-03 | 24.51394161 | 5.710366716 | ENSG00000105219 |
| SPEF1 | -2.114426539 | 0.230937355 | 8.48E-03 | 3.43E-02 | 15.76180275 | 3.639816237 | ENSG00000101222 |
| ZBTB16 | -2.176859819 | 0.221156597 | 4.26E-02 | 1.25E-01 | 9.07210744 | 2.010204592 | ENSG00000109906 |
| ZNF44 | -2.193906553 | 0.218558811 | 8.89E-10 | 1.72E-08 | 79.02845354 | 17.31883216 | ENSG00000197857 |
| MAP6 | -2.243392181 | 0.211189179 | 1.91E-12 | 5.29E-11 | 103.9296627 | 21.94275258 | ENSG00000171533 |
| ANKRD53 | -2.251944733 | 0.209940916 | 3.54E-02 | 1.08E-01 | 7.92360197 | 1.670661906 | ENSG00000144031 |
| TCEAL5 | -2.297785315 | 0.203375061 | 3.65E-11 | 8.67E-10 | 84.82493107 | 17.26160769 | ENSG00000204065 |
| H2BC11 | -2.351179681 | 0.195985703 | 1.35E-12 | 3.81E-11 | 88.08632309 | 17.29613156 | ENSG00000124635 |
| H4C12 | -2.357450825 | 0.195135636 | 1.46E-02 | 5.32E-02 | 10.22506211 | 2.003678201 | ENSG00000273542 |
| HMX2 | -2.411564076 | 0.187951968 | 3.37E-21 | 2.21E-19 | 158.9939559 | 29.88841764 | ENSG00000188816 |
| ORM2 | -2.414162552 | 0.187613747 | 6.43E-04 | 3.82E-03 | 21.28786498 | 4.023530606 | ENSG00000228278 |
| H4C11 | -2.419782869 | 0.186884281 | 3.37E-03 | 1.59E-02 | 14.22719942 | 2.664413905 | ENSG00000197238 |
| RASAL3 | -2.432791697 | 0.185206714 | 2.25E-02 | 7.57E-02 | 9.034148451 | 1.682485183 | ENSG00000105122 |
| HRAT92 | -2.470717217 | 0.180401443 | 1.28E-03 | 6.93E-03 | 18.50303217 | 3.362794902 | ENSG00000223855 |
| FBP1 | -2.555446752 | 0.170111579 | 1.38E-06 | 1.56E-05 | 36.97641951 | 6.275286703 | ENSG00000165140 |
| TBXT | -2.594511074 | 0.165567215 | 4.85E-02 | 1.38E-01 | 5.993112081 | 0.984104186 | ENSG00000164458 |
| H2BC12 | -2.597873266 | 0.165181811 | 2.65E-39 | 5.01E-37 | 355.9239832 | 58.90037009 | ENSG00000197903 |
| MT1F | -2.63517211 | 0.160966 | 5.45E-48 | 1.54E-45 | 387.6805787 | 62.319895 | ENSG00000198417 |
| DLX4 | -2.658469259 | 0.158387539 | 1.51E-02 | 5.49E-02 | 10.51419064 | 1.675958792 | ENSG00000108813 |
| ZNF563 | -2.701954146 | 0.153684743 | 1.75E-07 | 2.31E-06 | 43.24353251 | 6.642826869 | ENSG00000188868 |
| SHPK | -2.722041926 | 0.151559698 | 1.16E-18 | 6.21E-17 | 122.5101129 | 18.5603785 | ENSG00000197417 |
| PCDHB5 | -2.728983885 | 0.150832175 | 1.03E-36 | 1.72E-34 | 259.7852812 | 39.17295782 | ENSG00000113209 |
| WDR97 | -2.80695799 | 0.142896453 | 1.60E-03 | 8.39E-03 | 16.31870299 | 2.355044163 | ENSG00000179698 |
| H2AC6 | -2.81722805 | 0.141882833 | 8.98E-69 | 6.06E-66 | 533.0123469 | 75.68613621 | ENSG00000180573 |
| H4C9 | -2.833906864 | 0.140251989 | 7.23E-18 | 3.59E-16 | 111.2180816 | 15.6394684 | ENSG00000276180 |
| MT1M | -2.877259897 | 0.136100107 | 7.08E-05 | 5.46E-04 | 22.02179681 | 3.001781127 | ENSG00000205364 |
| CFAP65 | -3.061286146 | 0.119801166 | 1.25E-02 | 4.70E-02 | 8.370490196 | 1.009926202 | ENSG00000181378 |
| PCDHGC3 | -3.206846377 | 0.108303639 | 1.73E-21 | 1.17E-19 | 129.0345985 | 14.02168003 | ENSG00000240184 |
| ZNF439 | -3.266182829 | 0.103939589 | 5.35E-03 | 2.34E-02 | 9.619524466 | 1.007750739 | ENSG00000171291 |
| TMED6 | -3.361194093 | 0.097314993 | 4.38E-03 | 1.98E-02 | 10.21794316 | 0.993751999 | ENSG00000157315 |
| HAS1 | -3.384080347 | 0.095783413 | 4.92E-08 | 7.23E-07 | 38.18393531 | 3.681812457 | ENSG00000105509 |
| DNAJA4 | -3.431462611 | 0.092688707 | 9.36E-19 | 5.11E-17 | 125.8603877 | 11.70617307 | ENSG00000140403 |
| ZNF229 | -3.659489283 | 0.079137797 | 8.77E-15 | 3.21E-13 | 75.34864907 | 5.956269148 | ENSG00000278318 |
| ZNF736 | -3.685067783 | 0.077747075 | 3.88E-24 | 3.26E-22 | 128.3433804 | 10.03361925 | ENSG00000234444 |
| DMRT2 | -3.819823798 | 0.070813891 | 6.07E-05 | 4.78E-04 | 18.75687147 | 1.319295944 | ENSG00000173253 |
| PKLR | -4.182780684 | 0.055062706 | 1.42E-14 | 5.01E-13 | 66.30412769 | 3.662516831 | ENSG00000143627 |
| NCR1 | -5.20614079 | 0.027089155 | 1.40E-03 | 7.52E-03 | 6.787249457 | 0 | ENSG00000189430 |
| BMP3 | -5.304442673 | 0.025304843 | 1.10E-03 | 6.07E-03 | 7.231467934 | 0 | ENSG00000152785 |
| ZNF280D | -5.974772384 | 0.015900629 | 6.31E-08 | 9.11E-07 | 41.72405679 | 0.65856024 | ENSG00000137871 |
| PSMD5 | -6.595887355 | 0.010338084 | 4.59E-58 | 1.84E-55 | 449.1866511 | 4.618623537 | ENSG00000095261 |
| ZNF572 | -6.732502711 | 0.009404046 | 5.04E-07 | 6.20E-06 | 19.46769259 | 0 | ENSG00000180938 |
| NLRP7 | -7.227593881 | 0.006672329 | 3.03E-08 | 4.60E-07 | 27.50781512 | 0 | ENSG00000167634 |
| NLRP2 | -8.104963942 | 0.003632141 | 1.04E-30 | 1.27E-28 | 3882.973088 | 14.06724917 | ENSG00000022556 |

**Table S4. Significantly dysregulated genes expressed in 3 biological replicates of SF3B1$^{K700E}$ compared to SF3B1$^{WT}$ ES cells (K700E ES line 3).**
Genes with a fold change (FC) ≥1.5 and p-values < 0.05 are shown. Up- and downregulated genes begin in rows 4 and 1346, respectively.

| Gene Symbol | log2 FC | FC | p-value | padj | WT Basemean | MT Basemean | Ensembl ID |
|---|---|---|---|---|---|---|---|
| FOXC2 | 6.16985815 | 71.99666373 | 1.57E-05 | 1.34E-04 | 0 | 12.63633215 | ENSG00000176692 |
| PRAC1 | 6.119885244 | 69.54549912 | 2.51E-05 | 2.05E-04 | 0 | 12.19352916 | ENSG00000159182 |
| NPAP1 | 6.007325678 | 64.32580395 | 5.27E-06 | 5.06E-05 | 0.337436127 | 22.01714987 | ENSG00000185823 |
| C7orf57 | 6.000294458 | 64.01306391 | 5.10E-05 | 3.83E-04 | 0 | 11.29025147 | ENSG00000164746 |
| MOV10L1 | 5.97977569 | 63.10908014 | 5.03E-05 | 3.79E-04 | 0 | 11.10254153 | ENSG00000073146 |
| KRT80 | 5.857995762 | 58.00059368 | 8.58E-05 | 6.08E-04 | 0 | 10.16822024 | ENSG00000167767 |
| KHDC3L | 5.836282162 | 57.13417936 | 8.12E-05 | 5.78E-04 | 0 | 10.05223347 | ENSG00000203908 |
| DDX43 | 5.829745098 | 56.87588159 | 1.67E-04 | 1.10E-03 | 0 | 9.94654392 | ENSG00000080007 |
| TBX3 | 5.707734363 | 52.2635915 | 4.33E-07 | 5.17E-06 | 0.618649183 | 33.2060603 | ENSG00000135111 |
| CYTOR | 5.700794743 | 52.01279798 | 4.51E-07 | 5.37E-06 | 0.645182262 | 33.16103224 | ENSG00000222041 |
| GDF15 | 5.541707786 | 46.5822295 | 2.69E-279 | 4.99E-275 | 41.52248755 | 1939.484302 | ENSG00000130513 |
| REG1A | 5.503257001 | 45.35711588 | 5.84E-04 | 3.29E-03 | 0 | 7.978097646 | ENSG00000115386 |
| HORMAD1 | 5.437236471 | 43.32826226 | 8.67E-05 | 6.13E-04 | 0.307746135 | 14.8381777 | ENSG00000143452 |
| SIX3 | 5.3836572 | 41.74863722 | 4.64E-08 | 6.64E-07 | 0.952928397 | 39.73833646 | ENSG00000138083 |
| KLF18 | 5.361259529 | 41.10549977 | 8.57E-04 | 4.61E-03 | 0 | 7.224000232 | ENSG00000283039 |
| ANXA1 | 5.218125142 | 37.22306998 | 1.43E-18 | 7.95E-17 | 2.50112981 | 94.38756991 | ENSG00000135046 |
| CH25H | 5.190377514 | 36.51399237 | 2.46E-04 | 1.55E-03 | 0.307746135 | 12.52034537 | ENSG00000138135 |
| LGALS12 | 4.89793122 | 29.81427232 | 1.23E-06 | 1.35E-05 | 0.959242223 | 28.41901676 | ENSG00000133317 |
| ANKRD1 | 4.884622445 | 29.54050215 | 1.09E-03 | 5.70E-03 | 0.310903048 | 10.07425365 | ENSG00000148677 |
| GDA | 4.880899918 | 29.46437825 | 9.47E-09 | 1.53E-07 | 1.263831445 | 37.45534657 | ENSG00000119125 |
| DELEC1 | 4.833458176 | 28.51122608 | 1.21E-03 | 6.17E-03 | 0.310903048 | 9.717931174 | ENSG00000173077 |
| PITX2 | 4.806506655 | 27.98354158 | 2.97E-22 | 2.28E-20 | 4.118181946 | 115.6707075 | ENSG00000164093 |
| CCR7 | 4.553232871 | 23.47792286 | 2.94E-03 | 1.31E-02 | 0.310903048 | 8.060667697 | ENSG00000126353 |
| MYL7 | 4.531944526 | 23.13402716 | 1.26E-12 | 3.66E-11 | 2.507443636 | 58.81346493 | ENSG00000106631 |
| MAB21L4 | 4.51351268 | 22.84034725 | 3.39E-03 | 1.48E-02 | 0.310903048 | 7.797762156 | ENSG00000172478 |
| MAGEA4 | 4.508638073 | 22.76330407 | 1.57E-21 | 1.13E-19 | 5.385170303 | 122.8021265 | ENSG00000147381 |
| PAX7 | 4.387670267 | 20.93246441 | 1.21E-11 | 3.04E-10 | 2.646422858 | 53.69591794 | ENSG00000009709 |
| DMRTB1 | 4.317361038 | 19.93678725 | 6.15E-03 | 2.46E-02 | 0.337436127 | 6.811364769 | ENSG00000143006 |
| SCG2 | 4.175350343 | 18.06781751 | 1.56E-13 | 5.04E-12 | 3.886878136 | 69.45878097 | ENSG00000171951 |
| TDRD1 | 4.141973122 | 17.65461092 | 2.39E-04 | 1.51E-03 | 0.932709143 | 16.67902587 | ENSG00000095627 |
| CER1 | 4.132084733 | 17.53401805 | 1.59E-23 | 1.34E-21 | 6.553617945 | 116.6196745 | ENSG00000147869 |
| AADACL3 | 3.980605373 | 15.78634604 | 2.67E-06 | 2.74E-05 | 1.627800651 | 25.28438726 | ENSG00000188984 |
| TBX1 | 3.91322193 | 15.06597286 | 8.80E-07 | 9.87E-06 | 1.862261374 | 28.5811309 | ENSG00000184058 |
| COL8A1 | 3.835740133 | 14.27817944 | 3.05E-09 | 5.25E-08 | 2.871412842 | 40.84550716 | ENSG00000144810 |
| VGLL2 | 3.772595074 | 13.66671943 | 7.80E-04 | 4.24E-03 | 0.92955223 | 12.9325428 | ENSG00000170162 |
| TCL1B | 3.756926121 | 13.51908983 | 3.61E-06 | 3.62E-05 | 1.935546786 | 25.93312154 | ENSG00000213231 |
| OTUD6A | 3.751165934 | 13.46522035 | 5.76E-03 | 2.32E-02 | 0.648339175 | 8.61185978 | ENSG00000189401 |
| PANCR | 3.731322375 | 13.28128078 | 3.03E-05 | 2.43E-04 | 1.657490643 | 21.33781002 | ENSG00000250103 |
| NKX2-5 | 3.703223862 | 13.0251119 | 6.38E-20 | 4.06E-18 | 7.704905549 | 99.64290063 | ENSG00000183072 |
| GALR1 | 3.661327536 | 12.652298 | 1.19E-07 | 1.56E-06 | 2.587042873 | 32.35062184 | ENSG00000166573 |
| PTPRB | 3.620449993 | 12.298837 | 3.50E-17 | 1.70E-15 | 7.430006319 | 90.44274481 | ENSG00000127329 |
| NECTIN4 | 3.618729926 | 12.28418233 | 6.66E-06 | 6.26E-05 | 1.915327532 | 23.49137832 | ENSG00000143217 |
| SGCG | 3.554851356 | 11.75213685 | 1.27E-06 | 1.39E-05 | 2.824660509 | 33.50595818 | ENSG00000102683 |
| TEX11 | 3.480067374 | 11.15847042 | 1.52E-02 | 5.16E-02 | 0.674872254 | 7.235285152 | ENSG00000120498 |
| RGS4 | 3.429717613 | 10.77575921 | 4.05E-05 | 3.13E-04 | 1.882480627 | 20.52470009 | ENSG00000117152 |
| CNTN5 | 3.413679458 | 10.65663062 | 2.38E-02 | 7.43E-02 | 0.61549227 | 6.736718825 | ENSG00000149972 |
| EBF2 | 3.411512531 | 10.64063638 | 2.37E-02 | 7.43E-02 | 0.61549227 | 6.661085195 | ENSG00000221818 |
| HCLS1 | 3.383283939 | 10.4344593 | 2.41E-27 | 2.78E-25 | 15.0085603 | 156.3188687 | ENSG00000180353 |
| SERPINE1 | 3.366546271 | 10.3141017 | 3.98E-69 | 3.69E-66 | 42.4879459 | 436.8225074 | ENSG00000106366 |
| TRPC6 | 3.35563858 | 10.23641459 | 3.01E-09 | 5.18E-08 | 4.90234343 | 49.3794718 | ENSG00000137672 |
| DPPA5 | 3.350992619 | 10.20350293 | 2.55E-18 | 1.39E-16 | 9.773215429 | 100.3034695 | ENSG00000203909 |
| TRDN | 3.320077854 | 9.987183325 | 1.51E-06 | 1.62E-05 | 2.84172285 | 28.52481792 | ENSG00000186439 |
| PLA2G2A | 3.238396603 | 9.437446775 | 7.84E-47 | 2.91E-44 | 32.50690053 | 306.6625937 | ENSG00000188257 |
| SP5 | 3.195835317 | 9.163097163 | 1.01E-02 | 3.70E-02 | 0.959242223 | 8.724047713 | ENSG00000204335 |
| PRDM1 | 3.146191508 | 8.853153945 | 1.46E-05 | 1.27E-04 | 2.954168993 | 25.61390302 | ENSG00000057657 |
| MYCT1 | 3.144792264 | 8.844571589 | 1.14E-02 | 4.10E-02 | 0.985775302 | 8.529178109 | ENSG00000120279 |
| LCP1 | 3.141183664 | 8.822476391 | 2.18E-39 | 5.77E-37 | 28.82163621 | 255.0657545 | ENSG00000136167 |
| PADI3 | 3.133743585 | 8.77709538 | 3.40E-39 | 8.89E-37 | 28.1890816 | 248.3195113 | ENSG00000142619 |
| AGMO | 3.123282119 | 8.713679918 | 4.90E-04 | 2.81E-03 | 1.918484445 | 16.69878455 | ENSG00000187546 |
| TEX37 | 3.113486281 | 8.654714813 | 1.29E-04 | 8.74E-04 | 2.196540588 | 19.16645842 | ENSG00000172073 |
| LAPTM5 | 3.111222535 | 8.641145268 | 1.28E-04 | 8.67E-04 | 2.193383675 | 19.11763151 | ENSG00000162511 |
| CCDC60 | 3.061517111 | 8.348500601 | 1.55E-02 | 5.25E-02 | 0.92955223 | 7.891617128 | ENSG00000183273 |
| NLRP10 | 3.054597776 | 8.308556076 | 5.74E-04 | 3.23E-03 | 1.888794453 | 15.79831802 | ENSG00000182261 |
| CDKN1A | 3.04537355 | 8.255602741 | 1.12E-201 | 1.04E-197 | 227.1284045 | 1877.927819 | ENSG00000124762 |
| IZUMO2 | 3.042124282 | 8.23703022 | 1.80E-03 | 8.69E-03 | 1.891951366 | 15.69730339 | ENSG00000161652 |
| C1orf87 | 3.025641904 | 8.143460035 | 1.77E-02 | 5.84E-02 | 0.985775302 | 7.861561213 | ENSG00000162598 |
| UNC93A | 3.001564942 | 8.008682589 | 1.29E-06 | 1.41E-05 | 4.211686612 | 33.35579022 | ENSG00000112494 |
| NPIPA5 | 3.000147677 | 8.000818936 | 1.79E-02 | 5.88E-02 | 0.926395317 | 7.554065643 | ENSG00000183793 |
| NTSR2 | 2.996283236 | 7.979416409 | 2.15E-02 | 6.88E-02 | 0.932709143 | 7.546579569 | ENSG00000169006 |
| FOXL1 | 2.991469284 | 7.952835246 | 2.76E-02 | 8.39E-02 | 0.926395317 | 7.482780512 | ENSG00000176678 |
| FLI1 | 2.980427004 | 7.892197198 | 2.67E-10 | 5.42E-09 | 7.036347121 | 55.42231666 | ENSG00000151702 |
| PCDHGB5 | 2.976868111 | 7.872752428 | 1.88E-08 | 2.88E-07 | 5.783264191 | 45.22685166 | ENSG00000276547 |

| | | | | | | | |
|---|---|---|---|---|---|---|---|
| LAMP3 | 2.975022855 | 7.862689348 | 2.70E-06 | 2.76E-05 | 3.857188144 | 30.10363661 | ENSG00000078081 |
| POTEF | 2.969451909 | 7.832386231 | 3.30E-05 | 2.62E-04 | 3.454058207 | 27.33639127 | ENSG00000196604 |
| PTPRH | 2.965579132 | 7.811389132 | 6.72E-04 | 3.72E-03 | 2.279296739 | 17.63679306 | ENSG00000080031 |
| TCEA3 | 2.964839269 | 7.807384213 | 1.85E-02 | 6.04E-02 | 0.985775302 | 7.535294649 | ENSG00000204219 |
| SNTG1 | 2.957759732 | 7.769165974 | 1.85E-05 | 1.55E-04 | 3.208848969 | 24.80404227 | ENSG00000147481 |
| CYP2B6 | 2.948660555 | 7.720319505 | 7.38E-65 | 5.95E-62 | 58.17778998 | 449.148198 | ENSG00000197408 |
| SRRM5 | 2.912327775 | 7.528319074 | 6.37E-04 | 3.54E-03 | 2.255920572 | 16.85231331 | ENSG00000226763 |
| GRAMD1C | 2.867947676 | 7.300259137 | 3.15E-07 | 3.85E-06 | 6.44738793 | 46.79190912 | ENSG00000178075 |
| ODF3L2 | 2.846766558 | 7.193862367 | 1.12E-02 | 4.04E-02 | 1.29667835 | 9.208079926 | ENSG00000181781 |
| F7 | 2.843432575 | 7.17725697 | 2.77E-02 | 8.42E-02 | 0.959242223 | 6.860191679 | ENSG00000057593 |
| LRFN5 | 2.829689567 | 7.109211554 | 2.33E-10 | 4.81E-09 | 8.217422414 | 58.62096845 | ENSG00000165379 |
| NTRK2 | 2.815238862 | 7.038357826 | 1.45E-02 | 4.97E-02 | 1.320054516 | 9.125947908 | ENSG00000148053 |
| PTGDR2 | 2.811500914 | 7.020145411 | 5.02E-03 | 2.07E-02 | 1.574734492 | 11.15505567 | ENSG00000183134 |
| CBY2 | 2.800287916 | 6.965794517 | 4.56E-02 | 1.25E-01 | 0.923238405 | 6.556383336 | ENSG00000174015 |
| CREB5 | 2.798450493 | 6.956928498 | 6.56E-12 | 1.72E-10 | 10.39502152 | 72.70147314 | ENSG00000146592 |
| NODAL | 2.762858729 | 6.78739855 | 1.28E-40 | 3.95E-38 | 44.3565211 | 300.5240301 | ENSG00000156574 |
| PRPH2 | 2.760445342 | 6.776053857 | 2.71E-03 | 1.22E-02 | 1.938703699 | 13.00014071 | ENSG00000112619 |
| KRT81 | 2.760117966 | 6.774516413 | 6.52E-03 | 2.58E-02 | 1.912170619 | 12.86070802 | ENSG00000205426 |
| SERPINB9 | 2.736618 | 6.665060635 | 4.40E-134 | 1.63E-130 | 220.0317079 | 1465.935419 | ENSG00000170542 |
| KRT17 | 2.734469756 | 6.655143416 | 3.55E-05 | 2.79E-04 | 5.074267255 | 33.69191599 | ENSG00000128422 |
| DPEP3 | 2.721935093 | 6.597571555 | 2.85E-16 | 1.24E-14 | 15.27083188 | 100.8295038 | ENSG00000141096 |
| SPAAR | 2.710058917 | 6.543483689 | 4.75E-02 | 1.29E-01 | 0.985775302 | 6.293477795 | ENSG00000235387 |
| ARHGDIB | 2.695098465 | 6.475979709 | 4.27E-03 | 1.80E-02 | 1.915327532 | 12.37766349 | ENSG00000111348 |
| SPDYE6 | 2.694582016 | 6.473661883 | 3.72E-02 | 1.06E-01 | 0.95608531 | 6.181289862 | ENSG00000260097 |
| MCCC2 | 2.679649415 | 6.407001884 | 4.66E-04 | 2.69E-03 | 2.871412842 | 18.40162574 | ENSG00000131844 |
| CTCFL | 2.675499634 | 6.388599206 | 4.38E-08 | 6.27E-07 | 9.1670962 | 57.68653555 | ENSG00000124092 |
| ARHGAP24 | 2.67496857 | 6.38624796 | 7.16E-06 | 6.69E-05 | 4.747579643 | 30.44922382 | ENSG00000138639 |
| SGCD | 2.664247416 | 6.3389655 | 1.11E-02 | 4.03E-02 | 1.548201413 | 9.973350641 | ENSG00000170624 |
| GOLGA6C | 2.635923424 | 6.215728221 | 9.10E-03 | 3.40E-02 | 1.888794453 | 11.76580993 | ENSG00000167195 |
| SEMA3C | 2.613419367 | 6.119523695 | 1.33E-05 | 1.16E-04 | 4.777269635 | 29.21120581 | ENSG00000075223 |
| MEIOB | 2.607077686 | 6.092683026 | 1.64E-29 | 2.22E-27 | 35.2594557 | 213.9454041 | ENSG00000162039 |
| TMEM232 | 2.603635939 | 6.078165422 | 4.25E-07 | 5.09E-06 | 6.973810223 | 42.58742768 | ENSG00000186952 |
| HAO2 | 2.603346462 | 6.076945956 | 4.32E-03 | 1.82E-02 | 2.255920572 | 13.5813887 | ENSG00000116882 |
| GCNT4 | 2.602828011 | 6.074762521 | 9.61E-28 | 1.14E-25 | 33.30063045 | 202.4513712 | ENSG00000176928 |
| KCNMB1 | 2.598115599 | 6.054952319 | 2.51E-09 | 4.39E-08 | 10.09674613 | 61.38886662 | ENSG00000145936 |
| GPER1 | 2.57417421 | 5.955300098 | 1.95E-02 | 6.31E-02 | 1.581048318 | 9.482270388 | ENSG00000164850 |
| ERVH48-1 | 2.562599536 | 5.907712161 | 6.25E-38 | 1.43E-35 | 508.809632 | 3004.655753 | ENSG00000233056 |
| CALHM3 | 2.555653183 | 5.87933581 | 7.86E-03 | 3.02E-02 | 1.909013706 | 11.19215962 | ENSG00000183128 |
| PLAAT4 | 2.540414991 | 5.81756325 | 8.89E-03 | 3.34E-02 | 1.888794453 | 11.06868677 | ENSG00000133321 |
| ITPRID1 | 2.516180959 | 5.720657451 | 1.03E-04 | 7.17E-04 | 5.078701945 | 29.18582481 | ENSG00000180347 |
| FGF17 | 2.512064085 | 5.704356252 | 1.10E-02 | 3.99E-02 | 1.965236778 | 11.03863085 | ENSG00000158815 |
| CHL1 | 2.499828773 | 5.656182902 | 4.72E-02 | 1.28E-01 | 1.323211429 | 7.291160102 | ENSG00000134121 |
| PADI1 | 2.493447885 | 5.631221445 | 7.26E-07 | 8.32E-06 | 7.642368652 | 43.0935917 | ENSG00000142623 |
| ACTA1 | 2.487007762 | 5.60613996 | 9.79E-74 | 1.07E-70 | 191.7435819 | 1075.235985 | ENSG00000143632 |
| MECOM | 2.485036949 | 5.598486844 | 2.03E-06 | 2.13E-05 | 7.020562556 | 39.16903465 | ENSG00000085276 |
| TMEM82 | 2.481021205 | 5.582925114 | 1.01E-03 | 5.33E-03 | 3.176002064 | 17.71479981 | ENSG00000162460 |
| ABHD12B | 2.478971962 | 5.575000608 | 2.77E-30 | 4.01E-28 | 49.84594459 | 276.9625373 | ENSG00000131969 |
| BAHCC1 | 2.474290075 | 5.556937735 | 2.63E-21 | 1.85E-19 | 27.9609347 | 155.5079087 | ENSG00000266074 |
| CFAP100 | 2.474062269 | 5.556060347 | 1.89E-02 | 6.14E-02 | 1.604424484 | 8.851757447 | ENSG00000163885 |
| TAGLN | 2.473045198 | 5.552144817 | 8.28E-187 | 5.11E-183 | 515.4779282 | 2861.259272 | ENSG00000149591 |
| FAT4 | 2.460795075 | 5.505200373 | 3.44E-27 | 3.89E-25 | 46.4684264 | 256.0427307 | ENSG00000196159 |
| CMKLR1 | 2.460206288 | 5.502954072 | 3.18E-13 | 9.95E-12 | 20.44995367 | 112.2909557 | ENSG00000174600 |
| PCDH11Y | 2.459540151 | 5.500413776 | 1.63E-04 | 1.07E-03 | 4.77095581 | 26.19883824 | ENSG00000099715 |
| MUC4 | 2.446431355 | 5.450661601 | 6.79E-12 | 1.77E-10 | 17.73595787 | 97.11384111 | ENSG00000145113 |
| PLK2 | 2.444013775 | 5.441535367 | 3.91E-105 | 1.21E-101 | 196.1459384 | 1065.924064 | ENSG00000145632 |
| SLC52A1 | 2.441223187 | 5.431020041 | 2.41E-04 | 1.52E-03 | 4.191467358 | 22.56834196 | ENSG00000132517 |
| KCNJ2 | 2.436187919 | 5.412097833 | 6.12E-08 | 8.50E-07 | 9.256263875 | 50.04328985 | ENSG00000123700 |
| CCDC146 | 2.424730327 | 5.369286274 | 2.16E-03 | 1.02E-02 | 3.126092818 | 16.9650509 | ENSG00000135205 |
| VAV1 | 2.407988461 | 5.307338113 | 3.13E-16 | 1.36E-14 | 26.74375789 | 140.9233983 | ENSG00000141968 |
| PIWIL2 | 2.40676904 | 5.302854044 | 1.72E-06 | 1.83E-05 | 8.225014016 | 43.68871254 | ENSG00000197181 |
| LEFTY1 | 2.399592028 | 5.276539308 | 1.17E-73 | 1.20E-70 | 129.613626 | 682.6245303 | ENSG00000243709 |
| RERG | 2.392401517 | 5.250306028 | 7.14E-03 | 2.78E-02 | 2.590199786 | 13.49165899 | ENSG00000134533 |
| ZNF648 | 2.385140583 | 5.22394817 | 2.79E-02 | 8.48E-02 | 1.601267572 | 8.35319112 | ENSG00000179930 |
| ADGRL4 | 2.375969844 | 5.190846563 | 4.67E-07 | 5.54E-06 | 9.484410772 | 49.29222683 | ENSG00000162618 |
| ANXA3 | 2.366644329 | 5.157401378 | 8.79E-91 | 1.63E-87 | 188.680026 | 972.9615607 | ENSG00000138772 |
| ZIC1 | 2.361949551 | 5.140645584 | 3.68E-04 | 2.19E-03 | 4.886558866 | 24.83409819 | ENSG00000152977 |
| FAM20A | 2.351150491 | 5.102309774 | 2.92E-06 | 2.97E-05 | 8.26733166 | 42.17738013 | ENSG00000108950 |
| PTP4A1 | 2.34308018 | 5.073847577 | 1.91E-02 | 6.20E-02 | 1.941860612 | 9.793564804 | ENSG00000112245 |
| MYOCD | 2.338064208 | 5.056237421 | 1.35E-10 | 2.91E-09 | 16.04246341 | 81.67053146 | ENSG00000141052 |
| MAFB | 2.329757553 | 5.027208596 | 8.22E-19 | 4.75E-17 | 30.82603372 | 155.2732356 | ENSG00000204103 |
| GATA4 | 2.328384567 | 5.022426573 | 4.39E-02 | 1.21E-01 | 1.568420667 | 7.940444037 | ENSG00000136574 |
| ADGRG3 | 2.323344677 | 5.00491191 | 8.45E-03 | 3.21E-02 | 2.983858985 | 14.616613 | ENSG00000182885 |
| TNFRSF10C | 2.304115879 | 4.938647089 | 5.06E-12 | 1.36E-10 | 17.79533785 | 87.92211877 | ENSG00000173535 |
| LMX1B | 2.299241207 | 4.921988227 | 3.18E-03 | 1.40E-02 | 3.176002064 | 15.60301039 | ENSG00000136944 |
| BCAS1 | 2.294721693 | 4.906593297 | 1.22E-03 | 6.24E-03 | 3.797808159 | 18.72040623 | ENSG00000064787 |

| | | | | | | | |
|---|---|---|---|---|---|---|---|
| SMIM38 | 2.292711119 | 4.899760117 | 1.55E-02 | 5.25E-02 | 2.530819802 | 12.51329733 | ENSG00000284713 |
| PARVG | 2.291335537 | 4.895090514 | 7.58E-04 | 4.14E-03 | 4.135244286 | 20.23173863 | ENSG00000138964 |
| ITGAX | 2.27851717 | 4.851790208 | 4.41E-02 | 1.21E-01 | 1.5450445 | 7.625462394 | ENSG00000140678 |
| SERHL | 2.275790812 | 4.842630114 | 1.43E-03 | 7.13E-03 | 4.218000437 | 20.28480242 | ENSG00000172250 |
| DCC | 2.268760165 | 4.819088065 | 6.30E-06 | 5.95E-05 | 8.336182383 | 40.04368214 | ENSG00000187323 |
| SCARA5 | 2.263471444 | 4.801454294 | 1.80E-02 | 5.90E-02 | 2.56682362 | 12.32178854 | ENSG00000168079 |
| PTCHD4 | 2.250503127 | 4.758487652 | 1.50E-10 | 3.20E-09 | 18.68878857 | 88.32683012 | ENSG00000244694 |
| INMT | 2.243095542 | 4.734117591 | 9.95E-03 | 3.66E-02 | 2.901102834 | 13.69825155 | ENSG00000241644 |
| BTG2 | 2.242048334 | 4.73068249 | 1.44E-31 | 2.23E-29 | 60.25684837 | 285.2989328 | ENSG00000159388 |
| PRKN | 2.231954213 | 4.697698791 | 1.34E-04 | 9.00E-04 | 6.811357136 | 31.78869449 | ENSG00000185345 |
| CDKN2A | 2.224114169 | 4.672239307 | 1.92E-02 | 6.23E-02 | 2.252763659 | 10.49872369 | ENSG00000147889 |
| SCNN1B | 2.21250335 | 4.634787999 | 8.17E-04 | 4.42E-03 | 4.70082731 | 21.94955197 | ENSG00000168447 |
| VSIR | 2.205881308 | 4.613562841 | 4.50E-03 | 1.88E-02 | 3.486905112 | 16.1889333 | ENSG00000107738 |
| LAMA2 | 2.192408707 | 4.57067965 | 1.88E-25 | 1.88E-23 | 68.09166837 | 310.1420459 | ENSG00000196569 |
| TWIST1 | 2.191467574 | 4.567698968 | 9.73E-03 | 3.59E-02 | 2.924479001 | 13.2480741 | ENSG00000122691 |
| PKD1L2 | 2.189137151 | 4.560326601 | 7.35E-13 | 2.23E-11 | 29.81688225 | 136.4043103 | ENSG00000166473 |
| GABRP | 2.186132905 | 4.550840127 | 4.21E-24 | 3.71E-22 | 47.16549475 | 215.0885794 | ENSG00000094755 |
| ARSL | 2.176701295 | 4.521186075 | 2.81E-40 | 8.39E-38 | 85.41522096 | 386.4848622 | ENSG00000157399 |
| CATSPER2 | 2.169967147 | 4.50013146 | 5.13E-05 | 3.85E-04 | 8.128352438 | 36.97230205 | ENSG00000166762 |
| DLGAP2 | 2.160786571 | 4.471585838 | 1.45E-07 | 1.88E-06 | 12.0916729 | 54.08653321 | ENSG00000198010 |
| PF4 | 2.159440212 | 4.46741479 | 8.98E-03 | 3.36E-02 | 3.810435811 | 17.00964093 | ENSG00000163737 |
| SPAG17 | 2.153073958 | 4.447744633 | 1.01E-02 | 3.69E-02 | 3.490062025 | 15.62276907 | ENSG00000155761 |
| MMP9 | 2.146441827 | 4.427345084 | 1.03E-25 | 1.04E-23 | 63.90422974 | 282.8378605 | ENSG00000100985 |
| COLEC11 | 2.145243913 | 4.423670449 | 3.15E-04 | 1.91E-03 | 6.570680286 | 29.23702485 | ENSG00000118004 |
| SYTL2 | 2.141757568 | 4.412993334 | 1.86E-22 | 1.44E-20 | 47.5540202 | 209.6327568 | ENSG00000137501 |
| CSMD1 | 2.141417871 | 4.411954372 | 5.90E-04 | 3.31E-03 | 5.392761905 | 23.85518687 | ENSG00000183117 |
| TMPRSS4 | 2.132601658 | 4.385075424 | 4.53E-02 | 1.24E-01 | 1.968393691 | 8.4803512 | ENSG00000137648 |
| SOX1 | 2.115386545 | 4.333060994 | 2.99E-02 | 8.95E-02 | 2.279296739 | 9.793564804 | ENSG00000182968 |
| SYT4 | 2.114613806 | 4.330740733 | 3.32E-13 | 1.03E-11 | 26.63140945 | 115.4452407 | ENSG00000132872 |
| PURPL | 2.104499194 | 4.300484464 | 2.94E-36 | 6.06E-34 | 132.6789633 | 570.2258157 | ENSG00000250337 |
| CEACAM1 | 2.101441383 | 4.291379184 | 9.21E-07 | 1.03E-05 | 11.89321599 | 50.8311304 | ENSG00000079385 |
| GBX2 | 2.086103646 | 4.245997865 | 1.43E-19 | 8.88E-18 | 44.57825648 | 188.5220221 | ENSG00000168505 |
| COL6A3 | 2.082282739 | 4.234767427 | 8.80E-36 | 1.79E-33 | 101.3935766 | 428.2823624 | ENSG00000163359 |
| STARD13 | 2.059590125 | 4.168678539 | 7.05E-25 | 6.56E-23 | 54.72185264 | 228.1498196 | ENSG00000133121 |
| DDIT3 | 2.054637692 | 4.154392978 | 2.05E-11 | 4.95E-10 | 24.98782423 | 104.1560801 | ENSG00000175197 |
| SMIM10L2A | 2.05410746 | 4.1528664 | 7.79E-49 | 3.01E-46 | 162.2849564 | 674.4967833 | ENSG00000178947 |
| MT1G | 2.053717528 | 4.151744113 | 1.61E-95 | 3.31E-92 | 433.3383172 | 1799.29288 | ENSG00000125144 |
| TNFSF9 | 2.051215125 | 4.144549016 | 7.01E-17 | 3.26E-15 | 38.73067395 | 160.8616661 | ENSG00000125657 |
| PADI4 | 2.047624349 | 4.134246324 | 3.05E-02 | 9.07E-02 | 2.507443636 | 10.47246662 | ENSG00000159339 |
| BANK1 | 2.043024641 | 4.121086201 | 3.66E-05 | 2.87E-04 | 8.876510104 | 36.60750581 | ENSG00000153064 |
| LEFTY2 | 2.035376606 | 4.099297248 | 6.00E-34 | 1.07E-31 | 129.2469058 | 528.2066216 | ENSG00000143768 |
| TAC1 | 2.031603046 | 4.088589001 | 2.55E-16 | 1.12E-14 | 36.1499449 | 148.0854632 | ENSG00000006128 |
| PDGFB | 2.030012054 | 4.084082625 | 1.38E-40 | 4.20E-38 | 121.7507997 | 497.3027291 | ENSG00000100311 |
| RTL9 | 2.026618021 | 4.074485851 | 6.65E-03 | 2.62E-02 | 4.132087374 | 16.9241481 | ENSG00000243978 |
| SP140L | 2.017176842 | 4.047908957 | 9.86E-05 | 6.88E-04 | 9.164036985 | 37.14752459 | ENSG00000185404 |
| ALDH1A1 | 1.997190156 | 3.99221704 | 2.71E-09 | 4.71E-08 | 19.82311153 | 78.99213977 | ENSG00000165092 |
| EBI3 | 1.991304059 | 3.975962251 | 2.66E-04 | 1.66E-03 | 7.251866366 | 28.98628029 | ENSG00000105246 |
| PPP1R3B | 1.984841825 | 3.958192633 | 4.09E-96 | 9.47E-93 | 385.7688207 | 1526.37829 | ENSG00000173281 |
| TAFA2 | 1.984515412 | 3.957297181 | 6.51E-04 | 3.62E-03 | 6.325471048 | 25.10785062 | ENSG00000198673 |
| HAPLN1 | 1.983031286 | 3.953228333 | 4.68E-11 | 1.07E-09 | 25.98750496 | 102.86511 | ENSG00000145681 |
| DPPA2 | 1.974725296 | 3.930533908 | 5.74E-28 | 7.05E-26 | 83.600892 | 328.1563056 | ENSG00000163530 |
| PI15 | 1.972420506 | 3.924259665 | 4.46E-02 | 1.22E-01 | 2.61041904 | 10.1611722 | ENSG00000137558 |
| BMPER | 1.971358833 | 3.92137288 | 4.09E-02 | 1.14E-01 | 2.805719032 | 11.1212009 | ENSG00000164619 |
| PCDH10 | 1.955209342 | 3.877721904 | 7.78E-20 | 4.92E-18 | 69.39849387 | 268.224219 | ENSG00000138650 |
| CAPN13 | 1.952283591 | 3.869865945 | 3.54E-14 | 1.24E-12 | 45.35935875 | 175.596456 | ENSG00000162949 |
| PLD6 | 1.949525534 | 3.862474836 | 2.25E-03 | 1.05E-02 | 5.683445699 | 22.03603249 | ENSG00000179598 |
| GADD45B | 1.948753769 | 3.86040917 | 6.76E-35 | 1.29E-32 | 106.0718996 | 409.9139788 | ENSG00000099860 |
| DMBX1 | 1.941826983 | 3.841918682 | 2.53E-04 | 1.59E-03 | 8.528325462 | 32.92850903 | ENSG00000197587 |
| CGB2 | 1.941195999 | 3.840238729 | 3.81E-02 | 1.08E-01 | 2.527662889 | 9.770994963 | ENSG00000104818 |
| ACTG2 | 1.935680195 | 3.825584514 | 1.18E-11 | 2.97E-10 | 30.51197376 | 116.8336415 | ENSG00000163017 |
| IER5L | 1.935336201 | 3.824672456 | 1.22E-07 | 1.61E-06 | 17.07687018 | 65.54919607 | ENSG00000188483 |
| IP6K3 | 1.934181563 | 3.821612664 | 2.60E-02 | 7.98E-02 | 3.126092818 | 12.05128531 | ENSG00000161896 |
| DUSP19 | 1.933917716 | 3.820913815 | 4.31E-05 | 3.30E-04 | 10.12327921 | 38.85872792 | ENSG00000162999 |
| C2orf78 | 1.916954171 | 3.776249719 | 2.76E-03 | 1.24E-02 | 6.553520247 | 24.44392095 | ENSG00000187833 |
| CEMIP | 1.916696799 | 3.775576107 | 1.53E-10 | 3.26E-09 | 28.95302383 | 109.312881 | ENSG00000103888 |
| FIBIN | 1.909312884 | 3.756301546 | 7.11E-03 | 2.78E-02 | 5.074267255 | 19.05003361 | ENSG00000176971 |
| TENM1 | 1.908046845 | 3.753006647 | 1.75E-08 | 2.69E-07 | 37.14687468 | 139.0334286 | ENSG00000009694 |
| ZFHX3 | 1.905883813 | 3.747383981 | 1.24E-13 | 4.02E-12 | 43.61091899 | 163.3425515 | ENSG00000140836 |
| CHRNA9 | 1.883931276 | 3.690794151 | 2.57E-08 | 3.82E-07 | 20.29331075 | 74.94410987 | ENSG00000174343 |
| ZYG11A | 1.882187388 | 3.686335521 | 5.99E-11 | 1.35E-09 | 28.26552393 | 104.222802 | ENSG00000203995 |
| NLRC3 | 1.881782715 | 3.685301657 | 1.94E-02 | 6.28E-02 | 3.774431993 | 14.03960188 | ENSG00000167984 |
| EMP1 | 1.877682367 | 3.674842369 | 6.43E-03 | 2.55E-02 | 5.055325778 | 18.56968862 | ENSG00000134531 |
| OVGP1 | 1.876310707 | 3.671350126 | 3.07E-04 | 1.87E-03 | 8.571920882 | 31.55584486 | ENSG00000085465 |
| IFIT3 | 1.876140215 | 3.670916288 | 2.03E-02 | 6.55E-02 | 3.454058207 | 12.77152796 | ENSG00000119917 |
| ISLR2 | 1.869331918 | 3.653633488 | 7.90E-11 | 1.76E-09 | 27.89524089 | 102.0927912 | ENSG00000167178 |

| | | | | | | | |
|---|---|---|---|---|---|---|---|
| HES4 | 1.868771039 | 3.652213333 | 9.31E-07 | 1.04E-05 | 19.0996078 | 69.85969348 | ENSG00000188290 |
| GRIK2 | 1.857588952 | 3.624015064 | 5.55E-10 | 1.07E-08 | 27.92620866 | 101.2936657 | ENSG00000164418 |
| PAX5 | 1.852689511 | 3.611728661 | 8.04E-05 | 5.73E-04 | 11.80730293 | 42.50320867 | ENSG00000196092 |
| ALDH1A3 | 1.849451483 | 3.603631479 | 2.55E-12 | 7.02E-11 | 37.38902471 | 134.5639631 | ENSG00000184254 |
| ALPK2 | 1.849227209 | 3.603071321 | 4.35E-02 | 1.20E-01 | 2.921322088 | 10.45738285 | ENSG00000198796 |
| SULT2B1 | 1.847950114 | 3.599883238 | 1.29E-03 | 6.54E-03 | 7.261337105 | 26.26318695 | ENSG00000088002 |
| GP1BA | 1.843722604 | 3.589349989 | 7.47E-03 | 2.89E-02 | 5.386448079 | 19.3955092 | ENSG00000185245 |
| NUPR1 | 1.843438292 | 3.588642707 | 9.10E-03 | 3.40E-02 | 4.724203477 | 17.08516294 | ENSG00000176046 |
| TRPM2 | 1.839368883 | 3.578534487 | 9.22E-03 | 3.44E-02 | 4.697670398 | 16.96125205 | ENSG00000142185 |
| KCNK7 | 1.832623722 | 3.561842496 | 1.27E-02 | 4.49E-02 | 4.135244286 | 14.71426682 | ENSG00000173338 |
| ZNF843 | 1.830312562 | 3.556141085 | 1.69E-02 | 5.61E-02 | 4.099240469 | 14.68844778 | ENSG00000176723 |
| DNAH1 | 1.83001316 | 3.555403156 | 8.60E-06 | 7.80E-05 | 16.04740176 | 56.94157317 | ENSG00000114841 |
| STK32A | 1.829017594 | 3.552950513 | 2.09E-04 | 1.33E-03 | 9.474940033 | 33.80607929 | ENSG00000169302 |
| ZBBX | 1.82487794 | 3.542770327 | 5.70E-03 | 2.31E-02 | 5.769358763 | 20.30638457 | ENSG00000169064 |
| SLC9A9 | 1.818380413 | 3.526850476 | 4.61E-02 | 1.26E-01 | 3.208848969 | 11.33539115 | ENSG00000181804 |
| STMN2 | 1.816737144 | 3.522835586 | 2.43E-09 | 4.25E-08 | 28.61992469 | 100.3412347 | ENSG00000104435 |
| SLC7A14 | 1.815606293 | 3.520075306 | 1.24E-03 | 6.32E-03 | 7.668901731 | 26.88153891 | ENSG00000013293 |
| MYOM1 | 1.815556339 | 3.519953424 | 7.26E-04 | 3.98E-03 | 8.247112406 | 29.1214761 | ENSG00000101605 |
| INPP4B | 1.811242805 | 3.509444783 | 1.36E-02 | 4.71E-02 | 5.461612628 | 18.9968582 | ENSG00000109452 |
| ADGRD1 | 1.807629781 | 3.500666875 | 1.30E-02 | 4.55E-02 | 5.01932196 | 17.70775177 | ENSG00000111452 |
| WNT3 | 1.806913594 | 3.498929494 | 5.67E-25 | 5.31E-23 | 83.83535272 | 293.1096204 | ENSG00000108379 |
| ASB2 | 1.794817211 | 3.469715109 | 4.53E-02 | 1.24E-01 | 2.84172285 | 9.875696822 | ENSG00000100628 |
| CTNNA3 | 1.79211989 | 3.463234049 | 1.08E-02 | 3.92E-02 | 5.805362581 | 19.93497833 | ENSG00000183230 |
| BHLHE41 | 1.790681664 | 3.459783263 | 1.20E-05 | 1.06E-04 | 16.16428259 | 55.50769787 | ENSG00000123095 |
| ULBP1 | 1.789602508 | 3.457196265 | 4.12E-04 | 2.41E-03 | 9.170350811 | 31.84077059 | ENSG00000111981 |
| ITGAM | 1.786801513 | 3.450490626 | 8.12E-03 | 3.10E-02 | 5.763044937 | 19.88702749 | ENSG00000169896 |
| FMN1 | 1.786451528 | 3.449653668 | 1.89E-10 | 3.93E-09 | 35.28598878 | 121.1381499 | ENSG00000248905 |
| FAM163B | 1.785816552 | 3.448135699 | 1.15E-14 | 4.26E-13 | 49.30896917 | 170.4960395 | ENSG00000196990 |
| RBM20 | 1.780186442 | 3.43470559 | 2.14E-04 | 1.36E-03 | 12.64137366 | 43.11929911 | ENSG00000203867 |
| ZMAT3 | 1.775482463 | 3.423524798 | 1.30E-35 | 2.61E-33 | 192.0310111 | 656.7387528 | ENSG00000172667 |
| PLA2G4C | 1.771083272 | 3.41310139 | 2.24E-06 | 2.33E-05 | 18.4923085 | 63.00913776 | ENSG00000105499 |
| CYP26B1 | 1.767186778 | 3.403895574 | 3.98E-03 | 1.70E-02 | 7.870417851 | 26.46934148 | ENSG00000003137 |
| IER3 | 1.766556267 | 3.402408272 | 2.41E-45 | 8.60E-43 | 237.2288112 | 807.196778 | ENSG00000137331 |
| AATBC | 1.762894806 | 3.393784141 | 3.86E-03 | 1.65E-02 | 6.411384112 | 21.68708446 | ENSG00000215458 |
| TSPYL5 | 1.762450381 | 3.392738839 | 2.23E-15 | 8.95E-14 | 56.32627712 | 191.3527804 | ENSG00000180543 |
| PRTN3 | 1.762136647 | 3.392001122 | 3.79E-02 | 1.08E-01 | 3.516595104 | 11.95418114 | ENSG00000196415 |
| GPR179 | 1.754939512 | 3.375121672 | 4.62E-02 | 1.26E-01 | 3.19937823 | 10.79873319 | ENSG00000277399 |
| SEZ6L | 1.75473722 | 3.374648453 | 3.88E-03 | 1.66E-02 | 6.328627961 | 21.43970072 | ENSG00000100095 |
| PAPPA2 | 1.751085729 | 3.366117948 | 3.54E-04 | 2.12E-03 | 12.16821292 | 41.28494934 | ENSG00000116183 |
| INSL3 | 1.738849583 | 3.337689116 | 3.75E-02 | 1.07E-01 | 3.493218937 | 11.66490691 | ENSG00000248099 |
| INPP5D | 1.737384241 | 3.334300753 | 4.86E-51 | 2.05E-48 | 243.9148992 | 813.3663054 | ENSG00000168918 |
| ABCC3 | 1.731545341 | 3.320833385 | 3.64E-02 | 1.05E-01 | 3.483748199 | 11.56725309 | ENSG00000108846 |
| PCDH11X | 1.729736142 | 3.316671534 | 1.72E-23 | 1.45E-21 | 107.4877438 | 355.5163257 | ENSG00000102290 |
| IL32 | 1.726412013 | 3.309038353 | 6.93E-03 | 2.72E-02 | 7.565828629 | 24.7101873 | ENSG00000008517 |
| KCNJ6 | 1.726363871 | 3.308927933 | 4.76E-06 | 4.63E-05 | 18.99535462 | 63.08520942 | ENSG00000157542 |
| TENM2 | 1.72274697 | 3.300642694 | 5.56E-12 | 1.48E-10 | 40.58840294 | 133.6781423 | ENSG00000145934 |
| GOLGA6A | 1.720702963 | 3.295969662 | 8.79E-03 | 3.31E-02 | 5.673974961 | 18.74622526 | ENSG00000159289 |
| HAPLN2 | 1.71488183 | 3.282697541 | 7.20E-03 | 2.80E-02 | 6.31600031 | 20.84348058 | ENSG00000132702 |
| CD8A | 1.710281766 | 3.27224726 | 4.95E-02 | 1.33E-01 | 3.176002064 | 10.42363971 | ENSG00000153563 |
| COL4A4 | 1.703257248 | 3.25635333 | 2.79E-04 | 1.72E-03 | 11.10905451 | 36.14973066 | ENSG00000081052 |
| ALOX5 | 1.701135077 | 3.251566832 | 1.24E-08 | 1.96E-07 | 30.42280608 | 98.72128999 | ENSG00000012779 |
| GSTT2B | 1.700863041 | 3.250953771 | 3.26E-05 | 2.59E-04 | 20.2028653 | 65.70458859 | ENSG00000133433 |
| CCDC80 | 1.696881986 | 3.24199527 | 8.94E-14 | 2.96E-12 | 49.73675305 | 161.5390306 | ENSG00000091986 |
| MVP | 1.696279316 | 3.240641245 | 1.52E-07 | 1.96E-06 | 27.26386636 | 87.90995778 | ENSG00000013364 |
| SLC16A3 | 1.694854391 | 3.2374421 | 6.85E-19 | 3.99E-17 | 111.9079368 | 361.2593843 | ENSG00000141526 |
| C14orf39 | 1.69463354 | 3.236946543 | 3.80E-03 | 1.63E-02 | 7.033190208 | 22.71862153 | ENSG00000179008 |
| TNRC18 | 1.693389343 | 3.234156166 | 1.08E-33 | 1.87E-31 | 151.2126798 | 489.1132205 | ENSG00000182095 |
| HTR2C | 1.687099243 | 3.220086052 | 2.53E-09 | 4.42E-08 | 32.18336983 | 103.6651115 | ENSG00000147246 |
| TRADD | 1.682210214 | 3.209192241 | 3.78E-02 | 1.07E-01 | 3.877407397 | 12.3512948 | ENSG00000102871 |
| NLRP1 | 1.680139095 | 3.20458846 | 4.04E-03 | 1.71E-02 | 7.69543481 | 24.52985181 | ENSG00000091592 |
| SLFN12 | 1.671693562 | 3.185883611 | 4.99E-21 | 3.47E-19 | 94.76233625 | 301.6356609 | ENSG00000172123 |
| TTC22 | 1.668918159 | 3.179760618 | 6.97E-04 | 3.84E-03 | 12.44173668 | 39.43909985 | ENSG00000006555 |
| PMFBP1 | 1.665741618 | 3.172767089 | 8.97E-03 | 3.36E-02 | 8.592140136 | 27.39325391 | ENSG00000118557 |
| GUCA1B | 1.663671408 | 3.168217559 | 1.51E-05 | 1.30E-04 | 17.2486963 | 54.60821904 | ENSG00000112599 |
| TUBB8B | 1.662724118 | 3.166137956 | 4.83E-02 | 1.30E-01 | 3.543128183 | 11.1963965 | ENSG00000173213 |
| RGN | 1.660812026 | 3.161944459 | 2.56E-02 | 7.90E-02 | 5.197461913 | 16.3273783 | ENSG00000130988 |
| AZU1 | 1.657057199 | 3.15372573 | 5.66E-03 | 2.29E-02 | 8.022122422 | 25.31488121 | ENSG00000172232 |
| TEX41 | 1.656120195 | 3.151678109 | 1.71E-02 | 5.67E-02 | 5.114705763 | 16.06078553 | ENSG00000226674 |
| NPFFR2 | 1.654246399 | 3.147587314 | 1.91E-17 | 9.49E-16 | 85.77080181 | 269.2870773 | ENSG00000056291 |
| ARID5B | 1.651416305 | 3.141418833 | 7.08E-03 | 2.77E-02 | 7.109632533 | 22.14062273 | ENSG00000150347 |
| CFAP44 | 1.648632685 | 3.13536344 | 4.18E-02 | 1.17E-01 | 4.685042746 | 14.86498443 | ENSG00000206530 |
| ARHGAP25 | 1.648411179 | 3.134882086 | 3.93E-05 | 3.05E-04 | 15.83325799 | 49.80339218 | ENSG00000163219 |
| KIAA1211L | 1.643709029 | 3.124681253 | 6.89E-03 | 2.71E-02 | 7.056566374 | 21.95714966 | ENSG00000196872 |
| PPARGC1A | 1.643144758 | 3.123459359 | 4.10E-06 | 4.06E-05 | 21.77759979 | 67.776311 | ENSG00000109819 |
| PPP1R3C | 1.632190426 | 3.099832854 | 7.15E-06 | 6.68E-05 | 21.19001607 | 65.96618003 | ENSG00000119938 |

| | | | | | | | |
|---|---|---|---|---|---|---|---|
| PTPRR | 1.63103106 | 3.097342794 | 2.32E-02 | 7.30E-02 | 5.108391937 | 15.79083195 | ENSG00000153233 |
| DDR2 | 1.630971569 | 3.097215075 | 1.46E-04 | 9.73E-04 | 14.37728346 | 44.37901089 | ENSG00000162733 |
| GDF3 | 1.630271425 | 3.095712351 | 8.05E-20 | 5.06E-18 | 109.8911297 | 340.2211373 | ENSG00000184344 |
| MYCBPAP | 1.629794617 | 3.094689395 | 9.77E-04 | 5.16E-03 | 12.4853321 | 38.69815111 | ENSG00000136449 |
| ATP8B3 | 1.622697432 | 3.079502783 | 3.69E-11 | 8.60E-10 | 51.93182047 | 159.6616281 | ENSG00000130270 |
| PADI2 | 1.618925568 | 3.071462072 | 1.96E-25 | 1.95E-23 | 139.2729221 | 426.7610676 | ENSG00000117115 |
| TMEM154 | 1.617892519 | 3.069263523 | 5.31E-04 | 3.03E-03 | 14.8252866 | 45.26439365 | ENSG00000170006 |
| OTOF | 1.614897965 | 3.062899363 | 4.10E-02 | 1.15E-01 | 4.11186812 | 12.60247739 | ENSG00000115155 |
| PTH1R | 1.614487798 | 3.062028684 | 1.27E-03 | 6.43E-03 | 9.845223065 | 30.14497744 | ENSG00000160801 |
| LPO | 1.608711149 | 3.049792628 | 2.08E-02 | 6.68E-02 | 5.088172683 | 15.53585052 | ENSG00000167419 |
| TMC8 | 1.603057628 | 3.037864724 | 4.72E-06 | 4.59E-05 | 24.47530737 | 74.55579638 | ENSG00000167895 |
| MAGEB17 | 1.599226943 | 3.029809199 | 1.33E-02 | 4.65E-02 | 6.024038739 | 18.29648585 | ENSG00000182798 |
| KCTD19 | 1.598888837 | 3.029099224 | 1.60E-02 | 5.39E-02 | 6.10679489 | 18.46553641 | ENSG00000168676 |
| HLA-DQA1 | 1.595063585 | 3.021078318 | 7.01E-03 | 2.75E-02 | 7.943800961 | 23.99743072 | ENSG00000196735 |
| ITPR2 | 1.594785965 | 3.020497024 | 1.12E-54 | 6.12E-52 | 809.8102554 | 2444.801801 | ENSG00000123104 |
| RYR2 | 1.591760015 | 3.014168386 | 4.50E-16 | 1.93E-14 | 104.1568803 | 313.3606712 | ENSG00000198626 |
| EID3 | 1.589123062 | 3.008664131 | 6.15E-03 | 2.46E-02 | 7.612678659 | 22.90589344 | ENSG00000255150 |
| PTH2R | 1.585610767 | 3.001348335 | 1.91E-02 | 6.21E-02 | 5.650598794 | 17.02461308 | ENSG00000144407 |
| TEX19 | 1.585185432 | 3.000463609 | 3.42E-02 | 9.94E-02 | 4.972569627 | 15.0518183 | ENSG00000182459 |
| ZNF354C | 1.583879975 | 2.997749794 | 4.22E-06 | 4.17E-05 | 28.80259704 | 85.9436614 | ENSG00000177932 |
| RGS6 | 1.580081997 | 2.989868425 | 1.02E-05 | 9.07E-05 | 22.65724277 | 67.7174101 | ENSG00000182732 |
| TFAP2C | 1.571493111 | 2.972121535 | 1.81E-18 | 9.99E-17 | 97.68117784 | 289.7471023 | ENSG00000087510 |
| FLNC | 1.568934808 | 2.966855799 | 5.33E-62 | 4.11E-59 | 726.7068022 | 2156.208944 | ENSG00000128591 |
| GNGT1 | 1.56464192 | 2.958040735 | 5.39E-05 | 4.02E-04 | 19.13245471 | 56.66033467 | ENSG00000127928 |
| PHF24 | 1.56413608 | 2.957003764 | 1.04E-02 | 3.81E-02 | 8.929576263 | 26.50027346 | ENSG00000122733 |
| MYO15B | 1.55788832 | 2.944225802 | 8.47E-12 | 2.17E-10 | 120.1722319 | 354.1887073 | ENSG00000266714 |
| TPCN2 | 1.553069319 | 2.934407678 | 1.59E-02 | 5.37E-02 | 7.165953302 | 21.25611604 | ENSG00000287725 |
| RAB17 | 1.551361344 | 2.930935752 | 8.37E-54 | 4.19E-51 | 392.8720644 | 1151.39302 | ENSG00000124839 |
| SATB2 | 1.550713468 | 2.929619843 | 7.72E-10 | 1.46E-08 | 46.00275957 | 134.7838873 | ENSG00000119042 |
| GJC2 | 1.54705725 | 2.922204715 | 1.34E-06 | 1.45E-05 | 27.09519715 | 79.02500685 | ENSG00000198835 |
| SSTR1 | 1.545370629 | 2.91879043 | 9.71E-04 | 5.14E-03 | 13.27906202 | 38.86186549 | ENSG00000139874 |
| PSD2 | 1.543918559 | 2.915854152 | 2.87E-02 | 8.65E-02 | 5.114705763 | 14.90957446 | ENSG00000146005 |
| MUC19 | 1.540982393 | 2.909925854 | 1.28E-02 | 4.52E-02 | 9.395340795 | 27.60320728 | ENSG00000205592 |
| TEX13B | 1.539599857 | 2.907138605 | 1.59E-05 | 1.35E-04 | 27.5342332 | 79.63762486 | ENSG00000170925 |
| PIFO | 1.539489108 | 2.906915445 | 1.04E-10 | 2.27E-09 | 49.86488606 | 144.5622567 | ENSG00000173947 |
| HSPA2 | 1.536499265 | 2.900897389 | 6.40E-57 | 3.82E-54 | 627.4487701 | 1819.932689 | ENSG00000126803 |
| BBC3 | 1.534528403 | 2.896937186 | 3.73E-43 | 1.21E-40 | 268.820546 | 778.1448667 | ENSG00000105327 |
| UAP1L1 | 1.531057805 | 2.889976587 | 6.21E-34 | 1.10E-31 | 195.1898531 | 563.6285521 | ENSG00000197355 |
| DDIT4 | 1.526571526 | 2.881003725 | 1.39E-08 | 2.18E-07 | 690.8145741 | 1989.75534 | ENSG00000168209 |
| KLF5 | 1.522377578 | 2.872640743 | 2.26E-05 | 1.87E-04 | 21.90404906 | 62.90042226 | ENSG00000102554 |
| CFD | 1.51884336 | 2.865612152 | 7.81E-06 | 7.20E-05 | 24.87212348 | 71.1221048 | ENSG00000197766 |
| IRX1 | 1.514343679 | 2.856688398 | 1.81E-03 | 8.72E-03 | 11.35426375 | 32.52672045 | ENSG00000170549 |
| ATXN1 | 1.51339595 | 2.854812412 | 1.19E-05 | 1.05E-04 | 22.38046441 | 63.69563727 | ENSG00000124788 |
| CRLF1 | 1.509807249 | 2.847719896 | 6.63E-17 | 3.10E-15 | 125.0620454 | 355.898682 | ENSG00000006016 |
| COL5A2 | 1.508172658 | 2.844495223 | 1.51E-57 | 9.32E-55 | 447.4179493 | 1273.017862 | ENSG00000204262 |
| FAM124B | 1.506737139 | 2.841666284 | 3.01E-04 | 1.84E-03 | 15.84272873 | 45.0202591 | ENSG00000124019 |
| ANGPT4 | 1.496463541 | 2.82150233 | 3.30E-02 | 9.67E-02 | 5.42560881 | 15.33349484 | ENSG00000101280 |
| DPPA3 | 1.494150142 | 2.816981602 | 5.82E-07 | 6.78E-06 | 33.57680746 | 94.62421839 | ENSG00000187569 |
| SPSB2 | 1.493004239 | 2.814745021 | 3.32E-16 | 1.44E-14 | 109.45277 | 308.9554915 | ENSG00000111671 |
| LEPR | 1.488021697 | 2.805040687 | 1.19E-06 | 1.31E-05 | 30.64789377 | 86.0987275 | ENSG00000116678 |
| DACT1 | 1.48751749 | 2.804060525 | 2.78E-52 | 1.32E-49 | 382.060977 | 1071.416498 | ENSG00000165617 |
| ONECUT1 | 1.483813308 | 2.796870213 | 4.51E-05 | 3.44E-04 | 30.80373993 | 85.6150217 | ENSG00000169856 |
| ACRBP | 1.479186296 | 2.78791446 | 3.02E-03 | 1.34E-02 | 11.38079683 | 31.76524858 | ENSG00000111644 |
| EXPH5 | 1.478425478 | 2.786444615 | 1.17E-06 | 1.29E-05 | 36.37671632 | 101.5022337 | ENSG00000110723 |
| DANT2 | 1.476848114 | 2.783399734 | 3.21E-02 | 9.46E-02 | 6.061320334 | 16.85231331 | ENSG00000235244 |
| DGKH | 1.47559027 | 2.780974026 | 8.36E-13 | 2.52E-11 | 83.61983347 | 232.2747485 | ENSG00000102780 |
| WDR66 | 1.475570984 | 2.78093685 | 1.81E-03 | 8.72E-03 | 14.90182662 | 41.54036881 | ENSG00000158023 |
| LOXHD1 | 1.470582026 | 2.77133675 | 4.51E-03 | 1.88E-02 | 11.56209369 | 31.99842462 | ENSG00000167210 |
| SYNPR | 1.469882848 | 2.769993994 | 3.11E-03 | 1.38E-02 | 11.95575289 | 32.97364871 | ENSG00000163630 |
| FAM71E1 | 1.469159539 | 2.768605579 | 8.09E-17 | 3.75E-15 | 93.92262811 | 259.8620044 | ENSG00000142530 |
| TNFAIP6 | 1.468996201 | 2.768292142 | 3.51E-02 | 1.01E-01 | 5.428765723 | 14.96544941 | ENSG00000123610 |
| PLAC1 | 1.467987807 | 2.766357879 | 1.35E-02 | 4.71E-02 | 8.045498589 | 22.09928189 | ENSG00000170965 |
| DNAH10 | 1.465817586 | 2.762199623 | 2.37E-02 | 7.41E-02 | 6.626903358 | 18.40542458 | ENSG00000197653 |
| HAS2 | 1.46344597 | 2.757662631 | 1.14E-15 | 4.72E-14 | 90.81547677 | 250.1090076 | ENSG00000170961 |
| RBMS2 | 1.462963717 | 2.756740974 | 1.86E-22 | 1.44E-20 | 157.5292815 | 434.782473 | ENSG00000076067 |
| ANKRD33 | 1.456800638 | 2.744989513 | 2.83E-02 | 8.57E-02 | 6.464450271 | 17.65143879 | ENSG00000167612 |
| ZP3 | 1.456294559 | 2.744026775 | 4.12E-02 | 1.15E-01 | 6.506767914 | 17.67345898 | ENSG00000188372 |
| VIM | 1.456079054 | 2.743616913 | 1.35E-27 | 1.58E-25 | 202.7693016 | 556.7636856 | ENSG00000026025 |
| UPK1A | 1.45597696 | 2.743422763 | 3.61E-02 | 1.04E-01 | 7.254925581 | 19.66926163 | ENSG00000105668 |
| UNC5C | 1.449417376 | 2.730977402 | 1.24E-03 | 6.31E-03 | 13.78832427 | 37.54551432 | ENSG00000182168 |
| NDRG1 | 1.442825249 | 2.718527173 | 6.49E-08 | 8.96E-07 | 44.14100186 | 119.9367265 | ENSG00000104419 |
| SCARF1 | 1.440411288 | 2.713982255 | 1.08E-13 | 3.54E-12 | 83.81257791 | 227.4906765 | ENSG00000074660 |
| DUSP10 | 1.438375077 | 2.710154459 | 2.27E-03 | 1.06E-02 | 14.54910959 | 39.1164089 | ENSG00000143507 |
| ALX1 | 1.43576394 | 2.705253781 | 9.47E-05 | 6.64E-04 | 23.92748574 | 64.98248218 | ENSG00000180318 |
| HMCN1 | 1.433854585 | 2.701675843 | 2.44E-06 | 2.52E-05 | 231.8525028 | 626.1803762 | ENSG00000143341 |

| | | | | | | | |
|---|---|---|---|---|---|---|---|
| AHNAK2 | 1.431910745 | 2.698038145 | 1.01E-09 | 1.87E-08 | 57.97745394 | 156.2533091 | ENSG00000185567 |
| CAV1 | 1.430141767 | 2.69473194 | 4.58E-13 | 1.41E-11 | 75.32606643 | 203.0153855 | ENSG00000105974 |
| CDH5 | 1.427410796 | 2.689635732 | 4.44E-02 | 1.22E-01 | 8.994089994 | 24.35386482 | ENSG00000179776 |
| SLC39A4 | 1.426758794 | 2.688420469 | 1.95E-05 | 1.63E-04 | 25.28472416 | 67.72759571 | ENSG00000147804 |
| KCNK3 | 1.423821832 | 2.682953094 | 1.40E-02 | 4.82E-02 | 8.590862359 | 23.05237417 | ENSG00000171303 |
| ISLR | 1.423311082 | 2.68200343 | 1.30E-02 | 4.57E-02 | 10.94472229 | 29.23746288 | ENSG00000129009 |
| FOSL1 | 1.422731838 | 2.680926818 | 1.85E-22 | 1.44E-20 | 154.2670733 | 412.9922953 | ENSG00000175592 |
| B3GNT3 | 1.416807862 | 2.669941 | 4.23E-04 | 2.47E-03 | 17.06108561 | 45.67334188 | ENSG00000179913 |
| KITLG | 1.414304203 | 2.665311592 | 1.18E-12 | 3.46E-11 | 83.7216288 | 222.7722899 | ENSG00000049130 |
| MASP1 | 1.412981311 | 2.662868731 | 1.03E-06 | 1.14E-05 | 40.87542615 | 109.0053854 | ENSG00000127241 |
| C22orf23 | 1.411386077 | 2.659925939 | 1.59E-02 | 5.37E-02 | 8.700151589 | 23.02231826 | ENSG00000128346 |
| EMP3 | 1.407175758 | 2.652174604 | 3.49E-07 | 4.23E-06 | 44.08152417 | 116.4850283 | ENSG00000142227 |
| PLIN4 | 1.405783401 | 2.649616204 | 2.53E-05 | 2.06E-04 | 26.24849876 | 69.69702967 | ENSG00000167676 |
| RNASEL | 1.40460596 | 2.647454629 | 4.44E-03 | 1.86E-02 | 11.50271371 | 30.43425167 | ENSG00000135828 |
| PEX5L | 1.402644819 | 2.643858232 | 3.27E-05 | 2.59E-04 | 33.52058438 | 88.55412877 | ENSG00000114757 |
| GABRG2 | 1.400628902 | 2.640166476 | 2.52E-02 | 7.79E-02 | 9.476217809 | 25.07030863 | ENSG00000113327 |
| TGM1 | 1.396163356 | 2.632007055 | 8.88E-04 | 4.76E-03 | 18.92522612 | 49.99446294 | ENSG00000092295 |
| PRODH | 1.393104705 | 2.626432861 | 1.93E-05 | 1.61E-04 | 28.00768704 | 73.44373598 | ENSG00000100033 |
| THBS1 | 1.387668295 | 2.616554474 | 4.60E-33 | 7.55E-31 | 411.2225299 | 1076.045813 | ENSG00000137801 |
| RDM1 | 1.387409693 | 2.616085501 | 1.43E-09 | 2.59E-08 | 70.01536162 | 182.8511332 | ENSG00000278023 |
| MAML2 | 1.379301284 | 2.601423502 | 7.28E-15 | 2.73E-13 | 94.07739189 | 244.503017 | ENSG00000184384 |
| SOX3 | 1.379082231 | 2.601028542 | 1.48E-44 | 4.98E-42 | 367.4637396 | 956.1607194 | ENSG00000134595 |
| JUN | 1.377275911 | 2.597773974 | 8.33E-29 | 1.10E-26 | 350.0000801 | 909.2672947 | ENSG00000177606 |
| FGF18 | 1.374503083 | 2.592785906 | 4.35E-03 | 1.83E-02 | 16.22513575 | 42.34274348 | ENSG00000156427 |
| DENND2D | 1.373991427 | 2.591866529 | 1.11E-02 | 4.01E-02 | 9.524849279 | 24.69510353 | ENSG00000162777 |
| GRPR | 1.371217634 | 2.586888073 | 2.45E-23 | 2.04E-21 | 221.5835472 | 573.0350059 | ENSG00000126010 |
| DISP1 | 1.370435728 | 2.585486421 | 6.79E-12 | 1.77E-10 | 85.14535778 | 220.2236865 | ENSG00000154309 |
| CAPS2 | 1.369988147 | 2.584684426 | 6.32E-04 | 3.52E-03 | 19.69803773 | 51.03034851 | ENSG00000180881 |
| LACC1 | 1.364648741 | 2.575136203 | 1.22E-07 | 1.60E-06 | 43.35052655 | 111.5826593 | ENSG00000179630 |
| DHRS2 | 1.364525965 | 2.574917064 | 2.69E-05 | 2.18E-04 | 28.54348237 | 73.26438817 | ENSG00000100867 |
| LGALS1 | 1.364172191 | 2.574285728 | 6.20E-18 | 3.28E-16 | 142.3586741 | 365.8202914 | ENSG00000100097 |
| TRIM22 | 1.361846401 | 2.570139027 | 2.35E-25 | 2.29E-23 | 435.4405192 | 1118.986286 | ENSG00000132274 |
| COL23A1 | 1.361285435 | 2.569139868 | 2.56E-34 | 4.70E-32 | 317.5118733 | 815.3300852 | ENSG00000050767 |
| POU6F2 | 1.360832072 | 2.568332649 | 6.42E-03 | 2.55E-02 | 14.92254954 | 38.06296326 | ENSG00000106536 |
| NAV3 | 1.36003138 | 2.566907627 | 9.61E-05 | 6.72E-04 | 30.12274925 | 77.42372149 | ENSG00000067798 |
| CREBRF | 1.358784402 | 2.564689907 | 1.54E-05 | 1.32E-04 | 31.34792363 | 80.17274549 | ENSG00000164463 |
| ARSD | 1.357989262 | 2.563276769 | 8.90E-12 | 2.27E-10 | 84.18729563 | 215.704773 | ENSG00000006756 |
| POMK | 1.355434758 | 2.558742127 | 2.75E-06 | 2.81E-05 | 36.48294634 | 93.61711491 | ENSG00000185900 |
| GUCY2C | 1.354470918 | 2.557033246 | 5.92E-06 | 5.63E-05 | 33.19134122 | 85.13577602 | ENSG00000070019 |
| MIXL1 | 1.35211046 | 2.55285299 | 4.22E-03 | 1.78E-02 | 20.96256824 | 53.67291007 | ENSG00000185155 |
| IL15 | 1.349486873 | 2.548214763 | 1.71E-02 | 5.68E-02 | 10.43408456 | 26.32286075 | ENSG00000164136 |
| DSEL | 1.345132375 | 2.540535062 | 1.39E-17 | 7.08E-16 | 174.2034277 | 442.0415615 | ENSG00000171451 |
| MRVI1 | 1.345028407 | 2.540351984 | 7.62E-11 | 1.70E-09 | 68.95384305 | 175.1274673 | ENSG00000072952 |
| MT1X | 1.343384217 | 2.537458482 | 3.10E-08 | 4.57E-07 | 55.61342421 | 140.8459412 | ENSG00000187193 |
| SP8 | 1.342048362 | 2.535110025 | 7.48E-14 | 2.52E-12 | 107.6160722 | 272.2684013 | ENSG00000164651 |
| PLXDC2 | 1.341252168 | 2.533711335 | 7.92E-42 | 2.49E-39 | 375.1541384 | 950.9591052 | ENSG00000120594 |
| PYROXD2 | 1.340237755 | 2.531930413 | 6.74E-05 | 4.90E-04 | 31.00101676 | 78.58830448 | ENSG00000119943 |
| LHFPL1 | 1.337417968 | 2.526986519 | 7.45E-07 | 8.51E-06 | 44.559315 | 112.162808 | ENSG00000182508 |
| KANK3 | 1.335947703 | 2.524412554 | 2.16E-07 | 2.72E-06 | 44.77808886 | 113.0183581 | ENSG00000186994 |
| PDP1 | 1.335565711 | 2.523744237 | 5.15E-33 | 8.37E-31 | 269.1347036 | 679.2653017 | ENSG00000164951 |
| CSRNP3 | 1.331162997 | 2.516054193 | 2.28E-13 | 7.24E-12 | 94.36945116 | 237.5450598 | ENSG00000178662 |
| KCNAB2 | 1.329076164 | 2.512417397 | 2.29E-21 | 1.63E-19 | 180.8263774 | 453.5299805 | ENSG00000069424 |
| PTAFR | 1.325969157 | 2.507012444 | 8.00E-06 | 7.35E-05 | 41.36063583 | 103.5886018 | ENSG00000169403 |
| MYO1F | 1.324108602 | 2.503781388 | 1.58E-10 | 3.35E-09 | 84.33642201 | 211.0247416 | ENSG00000142347 |
| HPCAL4 | 1.322074007 | 2.500252859 | 4.09E-06 | 4.05E-05 | 37.11550095 | 92.73724286 | ENSG00000116983 |
| NOS3 | 1.315334386 | 2.488600045 | 1.30E-07 | 1.70E-06 | 50.82993845 | 126.5668797 | ENSG00000164867 |
| VWC2 | 1.315311936 | 2.488561319 | 1.58E-02 | 5.34E-02 | 9.945041557 | 24.63879055 | ENSG00000188730 |
| BGN | 1.313777175 | 2.485915357 | 6.82E-10 | 1.30E-08 | 72.98473646 | 181.173673 | ENSG00000182492 |
| CPQ | 1.30889386 | 2.477515117 | 2.29E-03 | 1.07E-02 | 15.8692618 | 39.40232231 | ENSG00000104324 |
| HPGD | 1.303113313 | 2.467608144 | 1.31E-03 | 6.59E-03 | 18.08286473 | 44.5662828 | ENSG00000164120 |
| ADGRG1 | 1.302955904 | 2.467338925 | 9.03E-05 | 6.36E-04 | 30.50062388 | 75.28958546 | ENSG00000205336 |
| BCHE | 1.302413507 | 2.466411476 | 6.73E-07 | 7.74E-06 | 48.93029778 | 120.3168926 | ENSG00000114200 |
| CCDC169 | 1.295807658 | 2.455144023 | 2.72E-08 | 4.03E-07 | 61.09279824 | 149.7450998 | ENSG00000242715 |
| DDX60L | 1.292523775 | 2.449561945 | 2.78E-02 | 8.44E-02 | 8.247112406 | 20.20916879 | ENSG00000181381 |
| THRB | 1.291989435 | 2.448654853 | 2.71E-08 | 4.03E-07 | 54.39704416 | 133.1452832 | ENSG00000151090 |
| FN1 | 1.291873111 | 2.448457426 | 5.72E-37 | 1.21E-34 | 2564.409727 | 6277.590035 | ENSG00000115414 |
| WNT8B | 1.289505165 | 2.444441984 | 2.55E-02 | 7.87E-02 | 9.05780697 | 21.99089281 | ENSG00000075290 |
| ZNF808 | 1.282919037 | 2.433308158 | 5.03E-07 | 5.94E-06 | 55.14953882 | 134.0059462 | ENSG00000198482 |
| CR1L | 1.281529858 | 2.430966241 | 1.24E-02 | 4.40E-02 | 11.70117061 | 28.5966527 | ENSG00000197721 |
| FOXB2 | 1.280095668 | 2.428550806 | 1.62E-03 | 7.95E-03 | 17.48127789 | 42.52307897 | ENSG00000204612 |
| ARRB1 | 1.279814591 | 2.428077702 | 7.68E-23 | 6.16E-21 | 449.8543852 | 1091.294142 | ENSG00000137486 |
| SULF1 | 1.27916241 | 2.42698032 | 1.41E-16 | 6.33E-15 | 288.6858926 | 699.4362588 | ENSG00000137573 |
| LMCD1 | 1.273374234 | 2.417262642 | 1.66E-18 | 9.24E-17 | 163.7859018 | 396.3506998 | ENSG00000071282 |
| FRMPD4 | 1.272975352 | 2.416594399 | 1.19E-03 | 6.11E-03 | 20.14664223 | 48.52728253 | ENSG00000169933 |
| SAMD4A | 1.271564795 | 2.414232793 | 2.27E-09 | 4.00E-08 | 82.86230277 | 200.425218 | ENSG00000020577 |

| | | | | | | | |
|---|---|---|---|---|---|---|---|
| HEATR4 | 1.271560622 | 2.41422581 | 2.85E-02 | 8.60E-02 | 8.293864739 | 20.01809803 | ENSG00000187105 |
| FGF8 | 1.271237045 | 2.413684393 | 8.08E-04 | 4.38E-03 | 22.69580215 | 54.68655221 | ENSG00000107831 |
| TGM2 | 1.268238943 | 2.408673661 | 3.01E-07 | 3.69E-06 | 48.77109931 | 117.4151127 | ENSG00000198959 |
| MUC12 | 1.265767732 | 2.404551344 | 8.67E-03 | 3.27E-02 | 15.39274876 | 37.02101733 | ENSG00000205277 |
| ITGB2 | 1.265622965 | 2.404310072 | 3.02E-02 | 9.01E-02 | 8.587705446 | 20.65565901 | ENSG00000160255 |
| NLGN1 | 1.264573323 | 2.402561436 | 3.80E-13 | 1.18E-11 | 111.1682652 | 267.0171958 | ENSG00000169760 |
| IQSEC3 | 1.263774919 | 2.401232199 | 6.37E-05 | 4.67E-04 | 30.26676452 | 72.75866219 | ENSG00000120645 |
| POLD4 | 1.259958035 | 2.394887745 | 1.34E-04 | 9.02E-04 | 27.04213099 | 64.69331958 | ENSG00000175482 |
| ATRNL1 | 1.257855103 | 2.391399401 | 7.86E-16 | 3.31E-14 | 149.2550347 | 357.2032473 | ENSG00000107518 |
| ALG10 | 1.256911575 | 2.389835928 | 6.57E-04 | 3.65E-03 | 23.07437584 | 55.26400136 | ENSG00000139133 |
| GADD45G | 1.25512201 | 2.386873338 | 1.83E-16 | 8.12E-15 | 155.1027149 | 370.4967875 | ENSG00000130222 |
| ELFN1 | 1.252743146 | 2.382940858 | 1.77E-10 | 3.72E-09 | 107.0436717 | 254.5550985 | ENSG00000225968 |
| PTPN14 | 1.250242914 | 2.378814729 | 2.20E-38 | 5.29E-36 | 1091.755724 | 2596.455928 | ENSG00000152104 |
| ARL11 | 1.248632063 | 2.37616013 | 1.01E-02 | 3.70E-02 | 13.03700969 | 30.99661705 | ENSG00000152213 |
| POLR3GL | 1.245712094 | 2.371355721 | 1.28E-02 | 4.52E-02 | 13.98805895 | 33.14159997 | ENSG00000121851 |
| ZNF91 | 1.244876108 | 2.36998201 | 3.88E-27 | 4.31E-25 | 269.4602112 | 638.6458822 | ENSG00000167232 |
| PDE6G | 1.243838217 | 2.368277631 | 2.15E-07 | 2.71E-06 | 51.25200986 | 121.3381324 | ENSG00000185527 |
| ACHE | 1.24297525 | 2.366861439 | 1.41E-08 | 2.21E-07 | 77.41707606 | 183.2852392 | ENSG00000087085 |
| MEF2C | 1.240534194 | 2.36286007 | 6.09E-03 | 2.44E-02 | 13.94446353 | 32.99946774 | ENSG00000081189 |
| TRPV4 | 1.238681959 | 2.359828407 | 1.44E-05 | 1.25E-04 | 41.41931675 | 97.36261029 | ENSG00000111199 |
| KIAA1107 | 1.238328885 | 2.359250951 | 3.73E-05 | 2.91E-04 | 35.82197951 | 84.62939721 | ENSG00000189195 |
| TRPV2 | 1.23656729 | 2.356371959 | 2.58E-04 | 1.61E-03 | 27.12676628 | 63.91015393 | ENSG00000187688 |
| POLH | 1.233708956 | 2.351708029 | 1.75E-11 | 4.26E-10 | 127.6472091 | 300.5165843 | ENSG00000170734 |
| PDZD2 | 1.233682685 | 2.351665205 | 7.20E-18 | 3.78E-16 | 264.98714 | 622.5240919 | ENSG00000133401 |
| LPP | 1.230614062 | 2.346668511 | 9.52E-17 | 4.36E-15 | 387.5593828 | 908.7313581 | ENSG00000145012 |
| TOGARAM2 | 1.229712006 | 2.345201697 | 4.31E-02 | 1.19E-01 | 7.986118604 | 18.73549 | ENSG00000189350 |
| KCNE3 | 1.228844407 | 2.343791778 | 8.73E-04 | 4.69E-03 | 25.47676954 | 59.40880901 | ENSG00000175538 |
| NOTUM | 1.227921064 | 2.342292202 | 4.84E-02 | 1.30E-01 | 7.340936343 | 17.16360773 | ENSG00000185269 |
| CCDC121 | 1.227683038 | 2.341905786 | 2.70E-03 | 1.22E-02 | 17.57034787 | 41.12773334 | ENSG00000176714 |
| HRK | 1.222755947 | 2.333921354 | 3.17E-07 | 3.86E-06 | 53.30947353 | 124.1883859 | ENSG00000135116 |
| CABYR | 1.220428922 | 2.330159841 | 7.19E-10 | 1.37E-08 | 78.32570998 | 182.517707 | ENSG00000154040 |
| CXCR2 | 1.220289621 | 2.329934861 | 1.44E-02 | 4.95E-02 | 12.35582361 | 28.8247158 | ENSG00000180871 |
| VLDLR | 1.21979052 | 2.329128958 | 1.62E-32 | 2.59E-30 | 466.8461164 | 1086.33844 | ENSG00000147852 |
| KIAA1257 | 1.219095273 | 2.328006801 | 6.02E-05 | 4.43E-04 | 33.56290203 | 78.24877762 | ENSG00000114656 |
| CLC | 1.219024241 | 2.327892183 | 3.95E-02 | 1.11E-01 | 8.273645486 | 19.26031339 | ENSG00000105205 |
| NRK | 1.217660624 | 2.325692928 | 2.26E-11 | 5.41E-10 | 97.76591082 | 227.1092591 | ENSG00000123572 |
| MORN3 | 1.216265459 | 2.323444944 | 2.59E-02 | 7.96E-02 | 10.46702916 | 24.4256996 | ENSG00000139714 |
| RBM15 | 1.215098548 | 2.321566405 | 2.45E-20 | 1.62E-18 | 362.0901071 | 840.6628218 | ENSG00000162775 |
| HNMT | 1.213578555 | 2.319121741 | 1.56E-03 | 7.68E-03 | 20.13401457 | 46.66397615 | ENSG00000150540 |
| SPATA18 | 1.212678137 | 2.317674777 | 1.93E-03 | 9.21E-03 | 19.85280152 | 45.83815557 | ENSG00000163071 |
| GRIN2D | 1.21259954 | 2.317548515 | 2.77E-38 | 6.45E-36 | 541.5927921 | 1253.973236 | ENSG00000105464 |
| LTBR | 1.212009946 | 2.316601584 | 2.27E-05 | 1.87E-04 | 48.59967914 | 112.1976504 | ENSG00000111321 |
| GADD45A | 1.211746405 | 2.316178442 | 1.74E-28 | 2.27E-26 | 388.7837452 | 899.9897213 | ENSG00000116717 |
| VENTX | 1.21011003 | 2.313552809 | 8.40E-18 | 4.35E-16 | 184.8870585 | 427.4044028 | ENSG00000151650 |
| WDR72 | 1.204727751 | 2.304937692 | 6.63E-04 | 3.68E-03 | 28.91829779 | 66.51897235 | ENSG00000166415 |
| RENBP | 1.204532521 | 2.304625801 | 9.12E-21 | 6.22E-19 | 222.5785827 | 513.5151796 | ENSG00000102032 |
| KIRREL3 | 1.203808452 | 2.30346943 | 1.36E-02 | 4.73E-02 | 13.76179119 | 31.53608618 | ENSG00000149571 |
| RNF213 | 1.19628427 | 2.291487268 | 1.85E-07 | 2.36E-06 | 1002.897651 | 2298.042022 | ENSG00000173821 |
| WNK4 | 1.195068438 | 2.289556929 | 1.25E-02 | 4.44E-02 | 13.02122513 | 29.81111318 | ENSG00000126562 |
| DNAJC6 | 1.194769774 | 2.289083 | 1.08E-10 | 2.36E-09 | 91.02340441 | 208.2171028 | ENSG00000116675 |
| NEB | 1.193325471 | 2.286792512 | 1.82E-03 | 8.74E-03 | 26.54539869 | 60.77150235 | ENSG00000183091 |
| SNHG8 | 1.192762659 | 2.285900581 | 1.18E-33 | 2.02E-31 | 541.0086432 | 1238.059346 | ENSG00000269893 |
| FAXDC2 | 1.192594191 | 2.285633665 | 7.25E-05 | 5.22E-04 | 36.82038247 | 84.23139904 | ENSG00000170271 |
| SMIM29 | 1.190328022 | 2.282046236 | 6.00E-09 | 9.97E-08 | 78.50512771 | 178.7957288 | ENSG00000186577 |
| PHLDA2 | 1.190079311 | 2.28165286 | 2.05E-10 | 4.27E-09 | 94.74467255 | 215.7783281 | ENSG00000181649 |
| EFCAB10 | 1.189706773 | 2.28106376 | 3.35E-02 | 9.77E-02 | 9.94819847 | 22.59459903 | ENSG00000185055 |
| KCNS1 | 1.189652611 | 2.280978125 | 1.37E-06 | 1.48E-05 | 57.11940568 | 130.3396198 | ENSG00000124134 |
| AMIGO2 | 1.187724025 | 2.277930964 | 9.17E-13 | 2.74E-11 | 126.5608186 | 287.7529486 | ENSG00000139211 |
| SLC66A1L | 1.187684674 | 2.277868833 | 4.01E-02 | 1.13E-01 | 9.828160724 | 22.34297841 | ENSG00000174899 |
| MINAR1 | 1.184795871 | 2.273312269 | 1.15E-02 | 4.13E-02 | 15.53172798 | 35.09997225 | ENSG00000169330 |
| RIPOR2 | 1.182662432 | 2.269953008 | 1.45E-44 | 4.98E-42 | 593.2124165 | 1346.730939 | ENSG00000111913 |
| TRANK1 | 1.17973326 | 2.265348891 | 3.28E-15 | 1.29E-13 | 247.1664491 | 559.310337 | ENSG00000168016 |
| PAH | 1.177928854 | 2.262517348 | 1.48E-03 | 7.33E-03 | 23.61017117 | 53.2869697 | ENSG00000171759 |
| TAPBPL | 1.176910869 | 2.260921448 | 3.57E-02 | 1.03E-01 | 9.806062334 | 22.28385426 | ENSG00000139192 |
| SLIT2 | 1.176368204 | 2.26007117 | 9.12E-17 | 4.20E-15 | 189.819385 | 429.3929341 | ENSG00000145147 |
| IPP | 1.174555369 | 2.257233035 | 1.25E-08 | 1.97E-07 | 80.57787228 | 181.7026216 | ENSG00000197429 |
| KIAA1217 | 1.174300597 | 2.256834455 | 2.65E-27 | 3.03E-25 | 350.7086187 | 791.6441149 | ENSG00000120549 |
| IL6R | 1.173945549 | 2.256279116 | 1.89E-10 | 3.93E-09 | 99.45012994 | 224.7210177 | ENSG00000160712 |
| IGSF21 | 1.173354034 | 2.255354216 | 7.61E-23 | 6.13E-21 | 388.1431406 | 875.4010145 | ENSG00000117154 |
| IL20RB | 1.172546498 | 2.254092154 | 9.71E-04 | 5.14E-03 | 26.05507791 | 58.75225379 | ENSG00000174564 |
| HCP5 | 1.171728282 | 2.252814121 | 2.22E-05 | 1.83E-04 | 41.42266906 | 93.54104325 | ENSG00000206337 |
| OPRK1 | 1.170955598 | 2.251607873 | 4.74E-02 | 1.28E-01 | 8.604767787 | 19.3955092 | ENSG00000082556 |
| TMEM38A | 1.167703793 | 2.24653851 | 1.31E-16 | 5.92E-15 | 180.1301058 | 404.1311881 | ENSG00000072954 |
| INSM1 | 1.16656685 | 2.244768781 | 5.37E-03 | 2.19E-02 | 19.05285547 | 42.93246523 | ENSG00000173404 |
| NLRP7 | 1.163799096 | 2.240466407 | 2.49E-03 | 1.14E-02 | 26.4530741 | 58.98086654 | ENSG00000167634 |

| | | | | | | | |
|---|---|---|---|---|---|---|---|
| ADGRE5 | 1.16276057 | 2.238854186 | 3.22E-04 | 1.95E-03 | 32.29138128 | 72.05967528 | ENSG00000123146 |
| KDM6B | 1.161043758 | 2.236191526 | 2.11E-16 | 9.29E-15 | 178.9707229 | 400.0049366 | ENSG00000132510 |
| MYOF | 1.160983441 | 2.236098036 | 1.42E-04 | 9.47E-04 | 32.45699128 | 72.57420144 | ENSG00000138119 |
| STAC3 | 1.158888638 | 2.232853563 | 7.03E-04 | 3.87E-03 | 27.12173023 | 60.57966715 | ENSG00000185482 |
| F10 | 1.157005566 | 2.229941041 | 6.48E-05 | 4.74E-04 | 35.62352261 | 79.44936526 | ENSG00000126218 |
| IFI35 | 1.154862859 | 2.226631565 | 3.81E-03 | 1.64E-02 | 21.98552743 | 49.04235835 | ENSG00000068079 |
| TLE6 | 1.154231113 | 2.225656753 | 8.12E-03 | 3.10E-02 | 16.57698096 | 36.88549511 | ENSG00000104953 |
| CCN1 | 1.152387785 | 2.222814852 | 2.06E-50 | 8.47E-48 | 1605.64571 | 3569.336539 | ENSG00000142871 |
| DPH6 | 1.14960458 | 2.218530797 | 5.23E-12 | 1.40E-10 | 123.1991826 | 273.0452918 | ENSG00000134146 |
| CCN2 | 1.14936526 | 2.218162808 | 1.53E-37 | 3.41E-35 | 842.8899094 | 1869.250869 | ENSG00000118523 |
| KLHL11 | 1.14647124 | 2.213717674 | 3.93E-06 | 3.91E-05 | 65.43073872 | 144.7075752 | ENSG00000178502 |
| NEBL | 1.146467796 | 2.213712389 | 1.95E-07 | 2.48E-06 | 75.10305328 | 166.2447779 | ENSG00000078114 |
| METTL27 | 1.145330991 | 2.211968731 | 1.89E-07 | 2.40E-06 | 74.03688199 | 163.647794 | ENSG00000165171 |
| TGFB1I1 | 1.141282438 | 2.20577011 | 4.74E-10 | 9.24E-09 | 97.06954154 | 214.1443989 | ENSG00000140682 |
| OPCML | 1.140724005 | 2.204916474 | 2.31E-04 | 1.46E-03 | 36.11138553 | 79.74177707 | ENSG00000183715 |
| FAM71F1 | 1.137651737 | 2.200226025 | 1.00E-03 | 5.27E-03 | 26.94537171 | 59.08622967 | ENSG00000135248 |
| MFAP3L | 1.134715848 | 2.195753112 | 5.86E-08 | 8.16E-07 | 85.59326321 | 187.4552046 | ENSG00000198948 |
| GMPR | 1.129434747 | 2.187730074 | 1.87E-02 | 6.10E-02 | 15.73787418 | 34.68072678 | ENSG00000137198 |
| MDM2 | 1.126304604 | 2.182988612 | 7.30E-18 | 3.82E-16 | 384.7844209 | 839.1398999 | ENSG00000135679 |
| KLHDC8A | 1.123884851 | 2.179330274 | 1.65E-05 | 1.40E-04 | 49.40563075 | 108.0014279 | ENSG00000162873 |
| DNAH3 | 1.122054548 | 2.176567177 | 3.04E-02 | 9.07E-02 | 11.82308749 | 25.66272993 | ENSG00000158486 |
| EXD3 | 1.121723893 | 2.176068382 | 4.11E-09 | 6.97E-08 | 120.3473126 | 262.1194388 | ENSG00000187609 |
| NBEAL1 | 1.120510098 | 2.174238341 | 2.30E-05 | 1.89E-04 | 51.21778749 | 111.2303504 | ENSG00000144426 |
| EFCC1 | 1.120086739 | 2.173600405 | 3.15E-02 | 9.32E-02 | 12.91321368 | 28.15721052 | ENSG00000114654 |
| DYNC1I1 | 1.119298732 | 2.172413498 | 8.20E-03 | 3.12E-02 | 17.18615941 | 37.39198555 | ENSG00000158560 |
| ZFHX4 | 1.112395911 | 2.162044041 | 5.44E-04 | 3.09E-03 | 31.13821454 | 67.44354605 | ENSG00000091656 |
| CALD1 | 1.108551043 | 2.156289736 | 3.32E-78 | 4.10E-75 | 3601.274319 | 7765.555509 | ENSG00000122786 |
| ANKRD37 | 1.108227282 | 2.155805888 | 2.59E-02 | 7.96E-02 | 13.41488433 | 28.9754334 | ENSG00000186352 |
| FOXL2NB | 1.105644209 | 2.15194948 | 2.98E-03 | 1.32E-02 | 29.77131 | 64.15907501 | ENSG00000206262 |
| ACER2 | 1.10409007 | 2.149632548 | 1.27E-06 | 1.39E-05 | 79.48340911 | 170.5989893 | ENSG00000177076 |
| PPFIBP2 | 1.103013912 | 2.148029657 | 1.24E-12 | 3.60E-11 | 162.0789079 | 348.0809867 | ENSG00000166387 |
| KMT2C | 1.102544872 | 2.147331417 | 2.45E-14 | 8.69E-13 | 467.7672803 | 1004.19489 | ENSG00000055609 |
| SFR1 | 1.100999942 | 2.145033148 | 5.00E-05 | 3.77E-04 | 45.48137333 | 97.44496555 | ENSG00000156384 |
| GJA5 | 1.100488372 | 2.144272668 | 1.35E-03 | 6.77E-03 | 31.29485747 | 67.1816282 | ENSG00000265107 |
| DNM3 | 1.099778234 | 2.143217451 | 1.06E-04 | 7.34E-04 | 41.4068845 | 89.02549062 | ENSG00000197959 |
| CDH8 | 1.098571271 | 2.14142518 | 9.47E-06 | 8.52E-05 | 64.53610793 | 138.1452029 | ENSG00000150394 |
| OXCT2 | 1.098411656 | 2.141188272 | 1.78E-02 | 5.85E-02 | 16.90938104 | 36.27094201 | ENSG00000198754 |
| ACSM3 | 1.098382958 | 2.14114568 | 1.40E-02 | 4.83E-02 | 18.35323158 | 39.20756433 | ENSG00000005187 |
| BDNF | 1.097457203 | 2.139772182 | 1.47E-06 | 1.58E-05 | 64.243545 | 137.6626996 | ENSG00000176697 |
| SERPINB1 | 1.096347104 | 2.138126341 | 1.21E-14 | 4.47E-13 | 184.7115718 | 394.8234076 | ENSG00000021355 |
| NHLRC1 | 1.095610641 | 2.137035155 | 4.79E-02 | 1.29E-01 | 10.14665537 | 21.72473807 | ENSG00000187566 |
| ONECUT2 | 1.095338164 | 2.136631578 | 2.49E-03 | 1.14E-02 | 27.05801325 | 58.10775639 | ENSG00000119547 |
| MDN1 | 1.09502249 | 2.136164116 | 4.66E-06 | 4.55E-05 | 4958.188881 | 10591.49696 | ENSG00000112159 |
| CBX7 | 1.095001495 | 2.13613303 | 1.30E-07 | 1.69E-06 | 83.09468896 | 177.2589038 | ENSG00000100307 |
| MYL9 | 1.09472311 | 2.135720877 | 6.26E-50 | 2.52E-47 | 1375.349256 | 2938.323605 | ENSG00000101335 |
| CALB1 | 1.093218788 | 2.133495086 | 8.99E-19 | 5.16E-17 | 339.2885634 | 723.2700146 | ENSG00000104327 |
| TULP4 | 1.089458162 | 2.127941017 | 1.72E-13 | 5.54E-12 | 228.9238822 | 486.3275935 | ENSG00000130338 |
| TFPI2 | 1.086790066 | 2.124009275 | 1.25E-08 | 1.97E-07 | 93.12949955 | 197.7047211 | ENSG00000105825 |
| PURA | 1.085421086 | 2.121994748 | 3.71E-07 | 4.48E-06 | 87.97522709 | 186.7464701 | ENSG00000185129 |
| RNF180 | 1.085220263 | 2.121699388 | 4.80E-03 | 1.98E-02 | 23.07555592 | 48.92593353 | ENSG00000164197 |
| PCDHAC1 | 1.085077718 | 2.121489765 | 5.44E-06 | 5.21E-05 | 63.00634433 | 133.9296513 | ENSG00000248383 |
| DLG2 | 1.084507631 | 2.120651615 | 3.29E-09 | 5.63E-08 | 114.0331688 | 241.7782118 | ENSG00000150672 |
| TXNRD2 | 1.081694291 | 2.116520249 | 9.63E-11 | 2.11E-09 | 125.5410163 | 265.7825789 | ENSG00000184470 |
| PDCD1 | 1.081586882 | 2.116362679 | 5.12E-03 | 2.10E-02 | 20.63074688 | 43.71183203 | ENSG00000188389 |
| SYTL1 | 1.081287384 | 2.115923376 | 1.26E-08 | 1.98E-07 | 112.8832567 | 239.2396908 | ENSG00000142765 |
| GPR20 | 1.079453516 | 2.113235449 | 1.12E-02 | 4.03E-02 | 18.99969161 | 39.99865408 | ENSG00000204882 |
| FAAH2 | 1.079221859 | 2.112896149 | 4.75E-06 | 4.63E-05 | 67.78411763 | 143.0012213 | ENSG00000165591 |
| IRX4 | 1.078370575 | 2.11164977 | 4.68E-04 | 2.70E-03 | 44.70164653 | 94.32768132 | ENSG00000113430 |
| C4orf47 | 1.078366689 | 2.111644083 | 3.64E-02 | 1.05E-01 | 12.80698366 | 27.02477044 | ENSG00000205129 |
| PNMA2 | 1.078217237 | 2.111425344 | 7.90E-10 | 1.49E-08 | 113.7913119 | 240.5244487 | ENSG00000240694 |
| RNF227 | 1.077626681 | 2.110561225 | 1.21E-02 | 4.32E-02 | 17.89703548 | 37.77836394 | ENSG00000179859 |
| EBF4 | 1.076098682 | 2.108327054 | 1.21E-03 | 6.17E-03 | 36.10299717 | 75.83395275 | ENSG00000088881 |
| SULT1C2 | 1.075816587 | 2.107914845 | 3.21E-02 | 9.46E-02 | 13.39466507 | 28.20983628 | ENSG00000198203 |
| MED12L | 1.075301662 | 2.107162625 | 2.81E-07 | 3.48E-06 | 157.2020077 | 330.5940267 | ENSG00000144893 |
| MYH3 | 1.07353077 | 2.104577694 | 2.60E-03 | 1.18E-02 | 27.19699247 | 57.43276504 | ENSG00000109063 |
| CPNE4 | 1.069133529 | 2.098172843 | 1.72E-03 | 8.35E-03 | 32.74569824 | 68.35343374 | ENSG00000196353 |
| HIVEP2 | 1.066413516 | 2.094220739 | 1.06E-10 | 2.32E-09 | 185.5194177 | 387.9983848 | ENSG00000010818 |
| FAT3 | 1.064827903 | 2.091920322 | 2.96E-07 | 3.64E-06 | 862.6612855 | 1804.443138 | ENSG00000165323 |
| SPOCD1 | 1.062555297 | 2.088627619 | 1.83E-02 | 5.98E-02 | 16.31283025 | 34.09155468 | ENSG00000134668 |
| SLC44A5 | 1.060515131 | 2.085676104 | 2.58E-03 | 1.17E-02 | 31.45858833 | 65.78270697 | ENSG00000137968 |
| CPEB4 | 1.060477424 | 2.085621592 | 6.06E-12 | 1.60E-10 | 145.2783124 | 303.0299072 | ENSG00000113742 |
| KLF7 | 1.060368999 | 2.085464854 | 9.53E-17 | 4.36E-15 | 262.2476032 | 547.1457729 | ENSG00000118263 |
| HES2 | 1.060132731 | 2.085123348 | 3.87E-02 | 1.10E-01 | 13.23230969 | 27.7374154 | ENSG00000069812 |
| SOX15 | 1.059105429 | 2.083639121 | 1.31E-03 | 6.62E-03 | 29.89016766 | 62.4139053 | ENSG00000129194 |
| UBE2D4 | 1.057739034 | 2.081666613 | 2.70E-03 | 1.22E-02 | 26.06898334 | 54.26274344 | ENSG00000078967 |

| | | | | | | | |
|---|---|---|---|---|---|---|---|
| NMI | 1.054747115 | 2.077354042 | 8.00E-04 | 4.34E-03 | 35.30640343 | 73.57936982 | ENSG00000123609 |
| DHRS12 | 1.053976434 | 2.076244626 | 1.86E-03 | 8.90E-03 | 27.64687474 | 57.42472931 | ENSG00000102796 |
| WNT9A | 1.053353924 | 2.075348938 | 6.47E-05 | 4.74E-04 | 59.77717841 | 124.3599615 | ENSG00000143816 |
| RUNX2 | 1.050519414 | 2.071275436 | 3.43E-02 | 9.97E-02 | 14.71166038 | 30.6515795 | ENSG00000124813 |
| HSPG2 | 1.049144803 | 2.069302848 | 3.63E-24 | 3.22E-22 | 2695.702641 | 5577.676893 | ENSG00000142798 |
| NEFH | 1.048870872 | 2.068909978 | 8.75E-09 | 1.42E-07 | 113.8834411 | 235.5505713 | ENSG00000100285 |
| RAG1 | 1.048658499 | 2.068605445 | 2.94E-02 | 8.82E-02 | 14.75072341 | 30.47548089 | ENSG00000166349 |
| QPCT | 1.047678086 | 2.067200159 | 2.71E-05 | 2.19E-04 | 67.00872783 | 138.1432191 | ENSG00000115828 |
| PLK3 | 1.046802426 | 2.065945829 | 1.19E-09 | 2.17E-08 | 116.0098532 | 239.3748463 | ENSG00000173846 |
| MYH2 | 1.046630442 | 2.065699562 | 1.07E-03 | 5.61E-03 | 38.4120816 | 79.14489565 | ENSG00000125414 |
| MSRB3 | 1.045862787 | 2.064600701 | 1.59E-02 | 5.34E-02 | 18.53915853 | 38.42677181 | ENSG00000174099 |
| FBXO48 | 1.045077981 | 2.063477892 | 2.30E-02 | 7.23E-02 | 17.51855949 | 36.13899539 | ENSG00000204923 |
| EXOC3L2 | 1.044753044 | 2.063013189 | 9.44E-05 | 6.62E-04 | 49.05734841 | 101.0907604 | ENSG00000283632 |
| CORO2A | 1.044528777 | 2.062692517 | 1.70E-42 | 5.43E-40 | 1330.248073 | 2743.089042 | ENSG00000106789 |
| AOC3 | 1.042687368 | 2.060061444 | 2.71E-03 | 1.22E-02 | 33.23789816 | 68.26524137 | ENSG00000131471 |
| SUCLG2 | 1.042288803 | 2.059492401 | 8.53E-27 | 9.25E-25 | 643.4085378 | 1323.761054 | ENSG00000172340 |
| MGMT | 1.042040659 | 2.059138199 | 1.13E-16 | 5.15E-15 | 261.343885 | 537.815614 | ENSG00000170430 |
| TIE1 | 1.039550642 | 2.055587297 | 2.43E-06 | 2.51E-05 | 79.10741361 | 162.6151577 | ENSG00000066056 |
| SLC2A14 | 1.038652139 | 2.054307487 | 1.69E-08 | 2.62E-07 | 103.6868765 | 212.7860456 | ENSG00000173262 |
| CPEB2 | 1.038582061 | 2.054207703 | 1.01E-02 | 3.69E-02 | 24.08923977 | 49.46496463 | ENSG00000137449 |
| RND3 | 1.037986403 | 2.053359739 | 4.46E-31 | 6.83E-29 | 648.7214899 | 1331.41028 | ENSG00000115963 |
| SLC5A3 | 1.034531225 | 2.048447935 | 2.78E-19 | 1.67E-17 | 508.1398557 | 1040.959462 | ENSG00000198743 |
| HIVEP1 | 1.034022407 | 2.047725605 | 9.47E-18 | 4.88E-16 | 316.9196443 | 649.6724956 | ENSG00000095951 |
| COL25A1 | 1.033982759 | 2.04766933 | 5.30E-09 | 8.89E-08 | 111.9726459 | 229.459784 | ENSG00000188517 |
| DUOX1 | 1.033432975 | 2.04688915 | 1.30E-02 | 4.56E-02 | 20.20424077 | 41.52908389 | ENSG00000137857 |
| PLA2G6 | 1.031021436 | 2.043470528 | 6.61E-09 | 1.09E-07 | 121.9428451 | 249.1593164 | ENSG00000184381 |
| SUGCT | 1.030811885 | 2.043173735 | 1.31E-04 | 8.83E-04 | 50.6845477 | 103.484338 | ENSG00000175600 |
| ROBO2 | 1.03039337 | 2.042581112 | 5.33E-07 | 6.28E-06 | 89.34549912 | 182.2370181 | ENSG00000185008 |
| FZD5 | 1.030063762 | 2.042114503 | 6.64E-47 | 2.51E-44 | 1572.01474 | 3209.252065 | ENSG00000163251 |
| NTS | 1.028163559 | 2.039426564 | 9.96E-13 | 2.95E-11 | 177.6119137 | 361.9065184 | ENSG00000133636 |
| IFITM1 | 1.027946216 | 2.039119346 | 2.03E-18 | 1.12E-16 | 792.1476353 | 1614.767591 | ENSG00000185885 |
| HLA-DQB1 | 1.027044385 | 2.037845087 | 7.94E-14 | 2.67E-12 | 212.6090978 | 432.8606635 | ENSG00000179344 |
| PPP1R15A | 1.026004548 | 2.036376819 | 1.02E-21 | 7.54E-20 | 354.8841061 | 722.8510645 | ENSG00000087074 |
| IRX2 | 1.025983147 | 2.036346611 | 1.13E-13 | 3.70E-12 | 271.8485866 | 553.9514835 | ENSG00000170561 |
| TMEM64 | 1.024483399 | 2.034230835 | 3.84E-21 | 2.69E-19 | 490.3653459 | 997.1309413 | ENSG00000180694 |
| HHLA3 | 1.024456509 | 2.03419292 | 9.62E-07 | 1.07E-05 | 73.87324882 | 150.2079879 | ENSG00000197568 |
| REPS2 | 1.02345686 | 2.032783909 | 3.84E-07 | 4.62E-06 | 99.15471837 | 201.0069754 | ENSG00000169891 |
| ADGRB3 | 1.022910308 | 2.032013953 | 6.33E-03 | 2.52E-02 | 24.95487963 | 50.53013322 | ENSG00000135298 |
| DIAPH2 | 1.022409645 | 2.031308898 | 3.42E-35 | 6.67E-33 | 918.7910674 | 1865.275447 | ENSG00000147202 |
| SORCS2 | 1.019856287 | 2.027716959 | 5.77E-05 | 4.28E-04 | 61.97747727 | 125.8401386 | ENSG00000184985 |
| CLMN | 1.019641723 | 2.027415412 | 1.18E-07 | 1.56E-06 | 109.3936006 | 221.5753181 | ENSG00000165959 |
| EDA2R | 1.018217111 | 2.025414396 | 1.58E-07 | 2.04E-06 | 115.9492931 | 235.3204699 | ENSG00000131080 |
| GAN | 1.016477259 | 2.022973272 | 3.64E-08 | 5.29E-07 | 118.0408815 | 238.9942337 | ENSG00000261609 |
| SLFN13 | 1.016413664 | 2.022884101 | 4.62E-11 | 1.06E-09 | 180.0504089 | 364.2350146 | ENSG00000154760 |
| SORBS2 | 1.016213875 | 2.022603984 | 9.41E-19 | 5.38E-17 | 384.3571255 | 778.2032582 | ENSG00000154556 |
| GRID2IP | 1.015343035 | 2.021383467 | 2.79E-03 | 1.25E-02 | 29.63736682 | 59.88524353 | ENSG00000215045 |
| FAM86EP | 1.014417611 | 2.020087257 | 8.78E-04 | 4.71E-03 | 43.09071282 | 86.78219261 | ENSG00000251669 |
| TENT5C | 1.012810951 | 2.017838836 | 7.90E-03 | 3.03E-02 | 22.77728052 | 45.84239245 | ENSG00000183508 |
| CASC9 | 1.010964791 | 2.015258338 | 1.82E-03 | 8.73E-03 | 34.04869042 | 68.47128427 | ENSG00000249395 |
| SHC2 | 1.008989184 | 2.012500559 | 3.93E-08 | 5.67E-07 | 107.6174477 | 216.7638897 | ENSG00000129946 |
| HSD17B14 | 1.008434191 | 2.011726515 | 1.21E-10 | 2.61E-09 | 178.5780484 | 358.7966001 | ENSG00000087076 |
| ZDHHC21 | 1.007789359 | 2.010827548 | 6.07E-06 | 5.76E-05 | 98.17309213 | 197.7035186 | ENSG00000175893 |
| NPIPB6 | 1.007769302 | 2.010799592 | 3.27E-02 | 9.58E-02 | 15.33780346 | 30.74499643 | ENSG00000198156 |
| CST1 | 1.007166016 | 2.00995892 | 3.70E-03 | 1.59E-02 | 29.33849007 | 59.04956375 | ENSG00000170373 |
| EVPL | 1.006214486 | 2.008633689 | 4.46E-02 | 1.22E-01 | 14.813358 | 29.8299958 | ENSG00000167880 |
| ZNF490 | 1.006123738 | 2.008507346 | 7.04E-03 | 2.76E-02 | 23.45540738 | 47.09070769 | ENSG00000188033 |
| RCN3 | 1.005745049 | 2.007980208 | 9.63E-16 | 4.02E-14 | 278.8217355 | 559.4330538 | ENSG00000142552 |
| M1AP | 1.003556919 | 2.004937021 | 1.04E-05 | 9.23E-05 | 86.89239942 | 173.7356973 | ENSG00000159374 |
| FAM157A | 1.000776631 | 2.00107693 | 2.32E-05 | 1.90E-04 | 74.77163789 | 149.3878299 | ENSG00000236438 |
| CARF | 1.00042858 | 2.000594227 | 5.08E-03 | 2.08E-02 | 24.74577191 | 49.5743414 | ENSG00000138380 |
| SH2D3A | 1.000344428 | 2.000477535 | 7.37E-27 | 8.08E-25 | 513.8648297 | 1028.313525 | ENSG00000125731 |
| OGFRL1 | 1.000084395 | 2.000117 | 1.01E-08 | 1.62E-07 | 133.311112 | 266.2624456 | ENSG00000119900 |
| LPL | 1.00004294 | 2.000059528 | 2.08E-08 | 3.14E-07 | 108.4407699 | 216.7134138 | ENSG00000175445 |
| PLAAT3 | 0.999749293 | 1.999652476 | 3.84E-27 | 4.29E-25 | 1037.142326 | 2073.386772 | ENSG00000176485 |
| CPT1A | 0.998619986 | 1.99808781 | 9.38E-08 | 1.27E-06 | 131.7418195 | 262.7719232 | ENSG00000110090 |
| CCDC113 | 0.996966437 | 1.995799007 | 5.57E-10 | 1.07E-08 | 141.1916019 | 281.7151051 | ENSG00000103021 |
| CAV2 | 0.995913821 | 1.994343368 | 9.24E-03 | 3.44E-02 | 25.30682255 | 50.39223787 | ENSG00000105971 |
| ELK4 | 0.992719437 | 1.989932421 | 5.01E-17 | 2.39E-15 | 303.3854138 | 603.3880814 | ENSG00000158711 |
| TUBB2A | 0.989678461 | 1.985742371 | 3.69E-59 | 2.36E-56 | 2688.5815 | 5338.280746 | ENSG00000137267 |
| CDK10 | 0.989436813 | 1.985409793 | 1.08E-08 | 1.72E-07 | 253.2002517 | 503.265028 | ENSG00000185324 |
| PLD1 | 0.988229267 | 1.983748686 | 1.00E-02 | 3.68E-02 | 21.65568291 | 42.93147755 | ENSG00000075651 |
| EPHA7 | 0.985626617 | 1.980173191 | 8.08E-10 | 1.52E-08 | 155.8437846 | 308.1027123 | ENSG00000135333 |
| NABP1 | 0.983341781 | 1.977039617 | 2.99E-03 | 1.33E-02 | 29.93248531 | 59.16500088 | ENSG00000173559 |
| DOC2A | 0.982523742 | 1.975918911 | 2.70E-05 | 2.19E-04 | 124.3910064 | 246.2382646 | ENSG00000149927 |
| SDK2 | 0.981965759 | 1.975154844 | 9.61E-14 | 3.17E-12 | 288.9115665 | 570.3664247 | ENSG00000069188 |

| | | | | | | | |
|---|---|---|---|---|---|---|---|
| PTPRM | 0.981528481 | 1.97455627 | 2.79E-10 | 5.65E-09 | 158.7460675 | 312.9089648 | ENSG00000173482 |
| ADARB2 | 0.981462343 | 1.974465751 | 3.71E-04 | 2.21E-03 | 60.50935612 | 119.0460075 | ENSG00000185736 |
| EDIL3 | 0.980843935 | 1.973619582 | 2.09E-16 | 9.22E-15 | 735.612604 | 1450.232006 | ENSG00000164176 |
| SCAPER | 0.978456145 | 1.970355766 | 6.20E-07 | 7.17E-06 | 102.4994873 | 201.4455415 | ENSG00000140386 |
| TIGD6 | 0.978354123 | 1.970216434 | 3.52E-03 | 1.52E-02 | 29.03705776 | 57.16594903 | ENSG00000164296 |
| RAD51B | 0.977721642 | 1.969352876 | 1.54E-04 | 1.02E-03 | 49.41816071 | 97.22416529 | ENSG00000182185 |
| REXO5 | 0.976336803 | 1.967463407 | 1.04E-05 | 9.20E-05 | 79.68196371 | 157.0717552 | ENSG00000005189 |
| TTC32 | 0.97380402 | 1.964012375 | 8.59E-05 | 6.09E-04 | 62.62323825 | 123.0123434 | ENSG00000183891 |
| PRKG1 | 0.972975316 | 1.962884544 | 2.21E-09 | 3.90E-08 | 148.0639099 | 290.7073542 | ENSG00000185532 |
| CCDC172 | 0.97293373 | 1.962827963 | 8.11E-03 | 3.10E-02 | 24.38495961 | 47.99172387 | ENSG00000182645 |
| PLAU | 0.971940438 | 1.961477026 | 9.69E-12 | 2.46E-10 | 199.0869011 | 390.4546706 | ENSG00000122861 |
| PLBD1 | 0.969048542 | 1.957549165 | 2.24E-10 | 4.63E-09 | 171.2650433 | 335.0158444 | ENSG00000121316 |
| DUSP8 | 0.967605325 | 1.955591886 | 5.86E-04 | 3.29E-03 | 42.84442121 | 83.85261834 | ENSG00000184545 |
| PDGFRB | 0.966861149 | 1.954583406 | 5.63E-15 | 2.16E-13 | 286.0857033 | 559.1325434 | ENSG00000113721 |
| PGBD4 | 0.965901191 | 1.953283275 | 3.65E-02 | 1.05E-01 | 19.842053 | 38.80296459 | ENSG00000182405 |
| SESN1 | 0.965142494 | 1.952256335 | 4.09E-17 | 1.98E-15 | 307.3131213 | 599.9306091 | ENSG00000080546 |
| WIPI1 | 0.964773268 | 1.951756761 | 1.70E-14 | 6.14E-13 | 332.0842613 | 647.4794099 | ENSG00000070540 |
| VPS13C | 0.964231053 | 1.951023361 | 6.31E-09 | 1.05E-07 | 208.0852077 | 405.5348874 | ENSG00000129003 |
| CHMP4C | 0.963988492 | 1.950695363 | 3.60E-02 | 1.03E-01 | 21.10904136 | 40.94217329 | ENSG00000164695 |
| LDHD | 0.963699737 | 1.950304971 | 2.53E-07 | 3.15E-06 | 101.3374512 | 197.3695427 | ENSG00000166816 |
| CLHC1 | 0.96346726 | 1.949990723 | 2.36E-03 | 1.09E-02 | 38.59928633 | 75.51181144 | ENSG00000162994 |
| STRADA | 0.962646223 | 1.948881299 | 8.47E-03 | 3.21E-02 | 26.90177629 | 52.32358019 | ENSG00000266173 |
| NRROS | 0.961224746 | 1.946962027 | 6.30E-06 | 5.96E-05 | 77.28992774 | 150.2718986 | ENSG00000174004 |
| RASGRF2 | 0.959506137 | 1.944644091 | 6.61E-18 | 3.48E-16 | 343.6980079 | 668.4371654 | ENSG00000113319 |
| RASGEF1C | 0.958458993 | 1.943233131 | 1.47E-02 | 5.02E-02 | 21.88067289 | 42.54521077 | ENSG00000146090 |
| NRP2 | 0.958191735 | 1.942873183 | 5.31E-25 | 5.00E-23 | 662.9841624 | 1288.23642 | ENSG00000118257 |
| NTN1 | 0.955080577 | 1.938687909 | 7.61E-06 | 7.04E-05 | 84.4287466 | 163.4999908 | ENSG00000065320 |
| STK17A | 0.954157099 | 1.937447341 | 8.18E-14 | 2.74E-12 | 269.2150996 | 521.5869888 | ENSG00000164543 |
| PRNP | 0.953977831 | 1.93720661 | 2.64E-19 | 1.59E-17 | 450.1711583 | 871.7110677 | ENSG00000171867 |
| KDELR3 | 0.953662296 | 1.936782966 | 1.24E-07 | 1.63E-06 | 116.3782571 | 225.0612688 | ENSG00000100196 |
| PCED1B | 0.952168894 | 1.934779148 | 1.50E-05 | 1.30E-04 | 74.41871029 | 144.1539698 | ENSG00000179715 |
| HS1BP3 | 0.952030767 | 1.934593917 | 4.28E-19 | 2.52E-17 | 371.5319146 | 718.5186182 | ENSG00000118960 |
| DDX60 | 0.951622332 | 1.934046299 | 3.52E-04 | 2.10E-03 | 46.82796092 | 90.4424184 | ENSG00000137628 |
| PHLDA3 | 0.951542119 | 1.93393877 | 2.35E-21 | 1.67E-19 | 870.1839617 | 1683.822258 | ENSG00000174307 |
| SENP8 | 0.951300656 | 1.933615116 | 1.66E-03 | 8.11E-03 | 44.29082729 | 85.84247225 | ENSG00000166192 |
| RNF152 | 0.950674692 | 1.932776331 | 2.71E-02 | 8.27E-02 | 22.12084608 | 42.81125389 | ENSG00000176641 |
| PRKG2 | 0.949905221 | 1.931745747 | 2.41E-02 | 7.51E-02 | 28.70012529 | 55.23405706 | ENSG00000138669 |
| LIF | 0.949466819 | 1.931158823 | 1.34E-05 | 1.17E-04 | 80.97223054 | 156.7399779 | ENSG00000128342 |
| TRPS1 | 0.947451243 | 1.928462701 | 7.73E-06 | 7.13E-05 | 74.41545568 | 143.5731598 | ENSG00000104447 |
| ACAD10 | 0.947039702 | 1.927912669 | 4.64E-11 | 1.06E-09 | 289.1427952 | 557.6866506 | ENSG00000111271 |
| IGSF9B | 0.945945716 | 1.926451301 | 3.10E-22 | 2.36E-20 | 1433.686365 | 2761.532342 | ENSG00000080854 |
| RPH3AL | 0.945795127 | 1.926250227 | 9.30E-03 | 3.46E-02 | 26.53780708 | 51.07102807 | ENSG00000181031 |
| ANKRD50 | 0.945478526 | 1.925827555 | 7.53E-18 | 3.93E-16 | 608.579692 | 1171.746964 | ENSG00000151458 |
| ILDR1 | 0.945074353 | 1.925288108 | 5.06E-03 | 2.08E-02 | 30.12462839 | 58.05837983 | ENSG00000145103 |
| JAK1 | 0.945031922 | 1.925231484 | 1.31E-22 | 1.03E-20 | 632.4481286 | 1216.972948 | ENSG00000162434 |
| KC6 | 0.94491842 | 1.925080026 | 6.86E-04 | 3.79E-03 | 52.73767438 | 101.7568887 | ENSG00000267313 |
| ADAMTSL2 | 0.943991869 | 1.923844066 | 3.50E-03 | 1.52E-02 | 36.33627782 | 69.96808256 | ENSG00000197859 |
| RHOB | 0.943740841 | 1.923509348 | 8.98E-15 | 3.34E-13 | 348.2771137 | 669.7182043 | ENSG00000143878 |
| EXOC8 | 0.94198769 | 1.921173335 | 1.23E-06 | 1.35E-05 | 99.22819917 | 190.5020882 | ENSG00000116903 |
| NDUFA13 | 0.940678092 | 1.919430192 | 8.50E-03 | 3.22E-02 | 31.69610827 | 60.85743321 | ENSG00000186010 |
| STPG1 | 0.940644087 | 1.919384951 | 9.94E-10 | 1.85E-08 | 150.658349 | 289.1079411 | ENSG00000001460 |
| HLA-E | 0.940026187 | 1.918563062 | 2.04E-32 | 3.24E-30 | 956.3658463 | 1834.197949 | ENSG00000204592 |
| NEDD9 | 0.939170522 | 1.917425498 | 3.63E-12 | 9.84E-11 | 218.7716121 | 418.8981923 | ENSG00000111859 |
| MACROD1 | 0.935993943 | 1.913208284 | 3.96E-16 | 1.70E-14 | 355.0128178 | 678.9291759 | ENSG00000133315 |
| TNN | 0.935695343 | 1.912812341 | 2.63E-02 | 8.08E-02 | 19.4780838 | 37.23053267 | ENSG00000120332 |
| CPS1 | 0.935333763 | 1.912332996 | 1.46E-17 | 7.40E-16 | 377.2865712 | 720.9151102 | ENSG00000021826 |
| RAC2 | 0.935160591 | 1.912103466 | 7.00E-03 | 2.74E-02 | 42.09951815 | 80.80403132 | ENSG00000128340 |
| METAP1D | 0.934702486 | 1.911496403 | 1.19E-04 | 8.17E-04 | 63.82956885 | 122.1086277 | ENSG00000172878 |
| RAI1 | 0.93424466 | 1.910889905 | 5.89E-07 | 6.85E-06 | 115.7943339 | 221.3288189 | ENSG00000108557 |
| BNIP3 | 0.933743892 | 1.910226739 | 5.41E-18 | 2.87E-16 | 776.593952 | 1482.124583 | ENSG00000176171 |
| GOLGA6L10 | 0.933240256 | 1.909560006 | 1.21E-03 | 6.18E-03 | 43.67345589 | 83.47102647 | ENSG00000278662 |
| FABP6 | 0.932719495 | 1.908870848 | 2.01E-03 | 9.55E-03 | 40.70222455 | 77.89767971 | ENSG00000170231 |
| RGS9 | 0.930235566 | 1.905587119 | 3.27E-10 | 6.57E-09 | 163.1469357 | 311.0541322 | ENSG00000108370 |
| DST | 0.929629154 | 1.904786306 | 3.40E-04 | 2.04E-03 | 1264.122716 | 2407.863839 | ENSG00000151914 |
| CRIM1 | 0.929133406 | 1.904131884 | 5.08E-15 | 1.96E-13 | 354.1215393 | 674.5429248 | ENSG00000150938 |
| RIMBP2 | 0.927219231 | 1.901607148 | 2.16E-02 | 6.90E-02 | 23.31760824 | 44.34994267 | ENSG00000060709 |
| CSF2RA | 0.926511467 | 1.900674477 | 1.79E-02 | 5.88E-02 | 23.40224353 | 44.46256863 | ENSG00000198223 |
| SH2D2A | 0.926333913 | 1.900440573 | 2.16E-06 | 2.25E-05 | 90.80916294 | 172.4424656 | ENSG00000027869 |
| ABI3 | 0.926179029 | 1.900236559 | 3.33E-03 | 1.46E-02 | 35.6861572 | 68.06233604 | ENSG00000108798 |
| AHNAK | 0.925880261 | 1.899843078 | 4.83E-05 | 3.66E-04 | 83.73415875 | 158.5661401 | ENSG00000124942 |
| RGS2 | 0.925875991 | 1.899837455 | 1.22E-09 | 2.23E-08 | 181.7119282 | 344.8641849 | ENSG00000116741 |
| KCNQ5 | 0.924432348 | 1.897937321 | 1.04E-02 | 3.80E-02 | 25.50340031 | 48.44190132 | ENSG00000185760 |
| USP53 | 0.923610517 | 1.896856469 | 7.33E-14 | 2.48E-12 | 314.9002515 | 596.4993938 | ENSG00000145390 |
| UPP1 | 0.921889812 | 1.894595433 | 3.23E-28 | 4.07E-26 | 716.3181696 | 1357.248219 | ENSG00000183696 |
| SMG1 | 0.917556375 | 1.888913157 | 4.72E-05 | 3.58E-04 | 1692.385263 | 3196.722745 | ENSG00000157106 |

| | | | | | | | |
|---|---|---|---|---|---|---|---|
| C8orf88 | 0.917282725 | 1.888554903 | 5.28E-08 | 7.44E-07 | 125.9537146 | 237.8752369 | ENSG00000253250 |
| RASIP1 | 0.91458751 | 1.885030034 | 3.00E-08 | 4.43E-07 | 164.1009465 | 308.6954684 | ENSG00000105538 |
| RRM2B | 0.914526021 | 1.884949694 | 2.77E-15 | 1.09E-13 | 1159.693853 | 2184.960263 | ENSG00000048392 |
| TP73 | 0.914206564 | 1.884532354 | 5.79E-06 | 5.52E-05 | 89.03794838 | 167.6255894 | ENSG00000078900 |
| USHBP1 | 0.912137757 | 1.881831895 | 5.68E-07 | 6.65E-06 | 110.8401044 | 208.3720573 | ENSG00000130307 |
| FAM167B | 0.911780978 | 1.881366575 | 4.83E-02 | 1.30E-01 | 20.65284527 | 39.0245293 | ENSG00000183615 |
| ZADH2 | 0.910520441 | 1.879723472 | 3.29E-05 | 2.61E-04 | 76.72750162 | 143.9664746 | ENSG00000180011 |
| BRCA2 | 0.909040172 | 1.877795781 | 9.73E-10 | 1.82E-08 | 209.3817732 | 393.5070564 | ENSG00000139618 |
| EFHC2 | 0.908954761 | 1.877684615 | 2.89E-03 | 1.29E-02 | 38.14802858 | 71.78734857 | ENSG00000183690 |
| MEIS3 | 0.906378091 | 1.874334041 | 3.16E-22 | 2.40E-20 | 609.4503905 | 1141.668742 | ENSG00000105419 |
| TM7SF3 | 0.905910312 | 1.873726406 | 2.02E-19 | 1.23E-17 | 1007.739668 | 1887.807131 | ENSG00000064115 |
| TENT5A | 0.904301859 | 1.871638563 | 3.10E-04 | 1.89E-03 | 63.27473434 | 118.0773934 | ENSG00000112773 |
| CBLC | 0.90382496 | 1.871019973 | 2.22E-10 | 4.60E-09 | 192.7936756 | 360.2997937 | ENSG00000142273 |
| DMTN | 0.902984448 | 1.869930238 | 3.22E-02 | 9.47E-02 | 22.6458929 | 42.23325508 | ENSG00000158856 |
| THSD4 | 0.902460233 | 1.869250907 | 1.50E-05 | 1.29E-04 | 92.34533806 | 172.6530718 | ENSG00000187720 |
| ALPG | 0.901037465 | 1.867408383 | 4.00E-02 | 1.13E-01 | 20.2358099 | 37.78128672 | ENSG00000163286 |
| GBGT1 | 0.900792674 | 1.867091556 | 5.54E-05 | 4.13E-04 | 104.8830373 | 195.8269837 | ENSG00000148288 |
| TP53INP1 | 0.898932508 | 1.864685738 | 1.11E-05 | 9.86E-05 | 105.5598863 | 196.7639098 | ENSG00000164938 |
| P4HA2 | 0.898695346 | 1.86437923 | 4.32E-08 | 6.20E-07 | 139.1499228 | 259.284332 | ENSG00000072682 |
| CEP85L | 0.898685074 | 1.864365956 | 1.89E-04 | 1.22E-03 | 80.63459901 | 149.8386284 | ENSG00000111860 |
| SPN | 0.898307906 | 1.863878613 | 6.39E-03 | 2.54E-02 | 34.24399041 | 63.81684861 | ENSG00000197471 |
| KIAA0825 | 0.897886779 | 1.86333462 | 1.03E-02 | 3.77E-02 | 30.5695723 | 57.15846296 | ENSG00000185261 |
| PILRB | 0.897337753 | 1.862625653 | 1.29E-02 | 4.52E-02 | 35.13438191 | 65.5994487 | ENSG00000121716 |
| CRTAC1 | 0.896712191 | 1.861818181 | 3.00E-04 | 1.84E-03 | 57.58567388 | 107.2994066 | ENSG00000095713 |
| FUT1 | 0.895555419 | 1.860325948 | 1.10E-12 | 3.24E-11 | 238.224636 | 443.2124684 | ENSG00000174951 |
| SIX4 | 0.894954867 | 1.85955171 | 7.82E-20 | 4.93E-18 | 462.474442 | 860.1367037 | ENSG00000100625 |
| ADCY8 | 0.894470454 | 1.858927435 | 7.05E-03 | 2.76E-02 | 32.91003047 | 61.15700468 | ENSG00000155897 |
| CHRM3 | 0.892678732 | 1.856620216 | 1.96E-08 | 2.98E-07 | 148.0244561 | 274.71658 | ENSG00000133019 |
| PDLIM3 | 0.892430739 | 1.856301097 | 6.67E-05 | 4.86E-04 | 68.63031235 | 127.4180813 | ENSG00000154553 |
| KMT2A | 0.891557661 | 1.855178057 | 9.85E-14 | 3.24E-12 | 721.9007069 | 1339.636894 | ENSG00000118058 |
| C1QTNF6 | 0.88973059 | 1.852830092 | 3.06E-04 | 1.87E-03 | 58.98976232 | 109.3778909 | ENSG00000133466 |
| OBSCN | 0.888416573 | 1.851143289 | 3.73E-16 | 1.61E-14 | 1192.744682 | 2208.663232 | ENSG00000154358 |
| PLAT | 0.888073661 | 1.850703347 | 5.72E-07 | 6.68E-06 | 124.3282742 | 229.7626047 | ENSG00000104368 |
| HPN | 0.887941865 | 1.850534286 | 1.16E-10 | 2.51E-09 | 213.7415416 | 395.0439931 | ENSG00000105707 |
| MAP1B | 0.886987789 | 1.849310903 | 1.08E-17 | 5.55E-16 | 1316.847342 | 2435.116697 | ENSG00000131711 |
| SIK1 | 0.884403956 | 1.846001795 | 3.94E-02 | 1.11E-01 | 20.99018369 | 38.57325254 | ENSG00000142178 |
| VWA5A | 0.883856617 | 1.84530158 | 1.45E-19 | 8.98E-18 | 514.8447949 | 950.2214057 | ENSG00000110002 |
| HLA-DRB1 | 0.88362838 | 1.845009673 | 2.99E-11 | 7.05E-10 | 267.7561561 | 493.5074905 | ENSG00000196126 |
| SEMA3G | 0.883218463 | 1.844485519 | 4.00E-05 | 3.09E-04 | 83.56182896 | 154.1458682 | ENSG00000010319 |
| FBXL2 | 0.882982563 | 1.844183946 | 1.52E-03 | 7.53E-03 | 66.64022623 | 122.9911993 | ENSG00000153558 |
| SAR1B | 0.882511502 | 1.843581891 | 1.07E-11 | 2.71E-10 | 284.8837323 | 524.461962 | ENSG00000152700 |
| NR5A2 | 0.882310969 | 1.843325653 | 2.93E-07 | 3.60E-06 | 159.1223561 | 293.2008387 | ENSG00000116833 |
| RIN1 | 0.882012645 | 1.842944526 | 6.71E-03 | 2.65E-02 | 37.34482793 | 68.83151722 | ENSG00000174791 |
| P2RX4 | 0.880796435 | 1.841391555 | 7.54E-06 | 6.99E-05 | 98.94876756 | 182.1245038 | ENSG00000135124 |
| RBFOX1 | 0.880043388 | 1.84043065 | 1.41E-02 | 4.86E-02 | 33.6892536 | 61.89448098 | ENSG00000078328 |
| ZNF578 | 0.879690332 | 1.839980315 | 7.32E-03 | 2.84E-02 | 37.1280309 | 68.30635897 | ENSG00000258405 |
| CAMK2D | 0.878852899 | 1.838912583 | 7.61E-09 | 1.25E-07 | 175.3693949 | 322.5197094 | ENSG00000145349 |
| ENPP2 | 0.878439407 | 1.838385606 | 2.53E-02 | 7.81E-02 | 32.9196966 | 60.65343702 | ENSG00000136960 |
| KCTD18 | 0.876966017 | 1.836509064 | 1.02E-04 | 7.12E-04 | 77.53503927 | 142.0007279 | ENSG00000155729 |
| RAB37 | 0.876812973 | 1.836314255 | 3.76E-04 | 2.23E-03 | 66.28028575 | 121.9466252 | ENSG00000172794 |
| NXPH4 | 0.876141181 | 1.835459372 | 5.15E-04 | 2.95E-03 | 61.93781288 | 113.8735733 | ENSG00000182379 |
| MAOB | 0.876121049 | 1.83543376 | 3.15E-03 | 1.39E-02 | 45.00298114 | 82.42666715 | ENSG00000069535 |
| PHYH | 0.875520003 | 1.834669253 | 2.22E-11 | 5.31E-10 | 255.003539 | 467.3598523 | ENSG00000107537 |
| KIFAP3 | 0.874510542 | 1.833385974 | 1.17E-03 | 6.02E-03 | 54.49380344 | 99.84872874 | ENSG00000075945 |
| FAM189A2 | 0.872874835 | 1.831308486 | 3.39E-11 | 7.94E-10 | 275.9523972 | 505.3849387 | ENSG00000135063 |
| TRAF1 | 0.871862768 | 1.830024252 | 3.93E-02 | 1.11E-01 | 28.52957694 | 52.20467908 | ENSG00000056558 |
| RAB20 | 0.870661842 | 1.82850154 | 5.34E-14 | 1.82E-12 | 311.1614174 | 569.0826629 | ENSG00000139832 |
| HIPK3 | 0.870572855 | 1.82838876 | 2.47E-09 | 4.31E-08 | 197.6517245 | 360.8061725 | ENSG00000110422 |
| GNG12 | 0.869120028 | 1.826548457 | 2.77E-12 | 7.58E-11 | 499.6830417 | 912.0882071 | ENSG00000172380 |
| LIPA | 0.866299624 | 1.822981125 | 3.71E-14 | 1.29E-12 | 432.3187104 | 788.214119 | ENSG00000107798 |
| B4GALT6 | 0.865485667 | 1.821952904 | 5.45E-12 | 1.45E-10 | 393.338603 | 715.8690397 | ENSG00000118276 |
| ADCY2 | 0.86491333 | 1.821230254 | 3.73E-27 | 4.19E-25 | 1905.702328 | 3469.846274 | ENSG00000078295 |
| SPP1 | 0.864371339 | 1.820546184 | 9.87E-09 | 1.59E-07 | 277.0911549 | 503.9952867 | ENSG00000118785 |
| SSC4D | 0.86403113 | 1.820116922 | 8.28E-12 | 2.13E-10 | 274.7403315 | 499.6518509 | ENSG00000146700 |
| ICA1L | 0.862215398 | 1.817827619 | 1.83E-02 | 5.99E-02 | 31.37908679 | 57.24439383 | ENSG00000163596 |
| STAT6 | 0.860883509 | 1.816150183 | 2.58E-19 | 1.56E-17 | 962.9145033 | 1749.340005 | ENSG00000166888 |
| PIEZO2 | 0.860444917 | 1.815598142 | 5.67E-12 | 1.50E-10 | 479.6803322 | 870.757163 | ENSG00000154864 |
| QRICH2 | 0.859917766 | 1.814934856 | 8.47E-03 | 3.21E-02 | 36.88923319 | 67.12520359 | ENSG00000129646 |
| HMGCL | 0.859616354 | 1.814555714 | 1.14E-06 | 1.26E-05 | 205.6873616 | 372.3740868 | ENSG00000117305 |
| ZSCAN20 | 0.859121559 | 1.81393349 | 7.75E-04 | 4.22E-03 | 61.06882071 | 110.8818489 | ENSG00000121903 |
| FBXO36 | 0.857700895 | 1.812148136 | 2.32E-03 | 1.08E-02 | 53.72147285 | 97.26214531 | ENSG00000153832 |
| DMXL2 | 0.857298982 | 1.811643369 | 4.02E-07 | 4.84E-06 | 173.003486 | 313.5800944 | ENSG00000104093 |
| HLA-F | 0.856417347 | 1.810536607 | 3.64E-03 | 1.57E-02 | 41.7555728 | 75.62399937 | ENSG00000204642 |
| WEE1 | 0.856174016 | 1.810231261 | 8.30E-26 | 8.40E-24 | 1356.017668 | 2454.155845 | ENSG00000166483 |
| IGF1R | 0.855961865 | 1.809965082 | 7.79E-21 | 5.33E-19 | 1929.434266 | 3491.501897 | ENSG00000140443 |

| | | | | | | | |
|---|---|---|---|---|---|---|---|
| GIPC3 | 0.854843577 | 1.808562652 | 3.93E-04 | 2.32E-03 | 70.12604897 | 126.9493476 | ENSG00000179855 |
| TTC39B | 0.853004463 | 1.806258608 | 1.61E-08 | 2.49E-07 | 175.2586325 | 316.0935206 | ENSG00000155158 |
| ATP7A | 0.85264995 | 1.805814812 | 2.49E-03 | 1.14E-02 | 55.31377334 | 99.54568485 | ENSG00000165240 |
| B3GALT4 | 0.851737692 | 1.804673302 | 2.85E-02 | 8.62E-02 | 23.24432283 | 41.92642079 | ENSG00000235863 |
| NCALD | 0.851678261 | 1.804598962 | 1.36E-02 | 4.71E-02 | 29.19823307 | 52.78135534 | ENSG00000104490 |
| NFAM1 | 0.851676459 | 1.804596708 | 3.80E-02 | 1.08E-01 | 23.08374888 | 41.53624355 | ENSG00000235568 |
| ZNF486 | 0.851375006 | 1.804219674 | 7.34E-12 | 1.90E-10 | 348.4550809 | 628.2076681 | ENSG00000256229 |
| BLVRB | 0.850034494 | 1.802544022 | 6.02E-15 | 2.31E-13 | 368.9593785 | 665.0528504 | ENSG00000090013 |
| FGF16 | 0.849781259 | 1.802227651 | 1.33E-02 | 4.63E-02 | 29.47628921 | 53.27298524 | ENSG00000196468 |
| NLRC5 | 0.849167095 | 1.801460596 | 2.67E-02 | 8.17E-02 | 28.52591637 | 51.4987473 | ENSG00000140853 |
| FOXO3 | 0.848912253 | 1.801142408 | 2.33E-14 | 8.29E-13 | 350.3271964 | 630.5451878 | ENSG00000118689 |
| HMBOX1 | 0.848141852 | 1.800180853 | 2.20E-06 | 2.29E-05 | 119.3608383 | 214.6767084 | ENSG00000147421 |
| PXN | 0.846371213 | 1.797972822 | 2.76E-23 | 2.28E-21 | 975.6986371 | 1753.802857 | ENSG00000089159 |
| REV3L | 0.845238645 | 1.796561901 | 4.51E-05 | 3.44E-04 | 89.48970979 | 160.5667293 | ENSG00000009413 |
| PLOD2 | 0.844781115 | 1.795992238 | 7.31E-12 | 1.90E-10 | 402.1106646 | 721.2327595 | ENSG00000152952 |
| UHMK1 | 0.844709537 | 1.795903134 | 4.98E-12 | 1.34E-10 | 402.395786 | 722.205874 | ENSG00000152332 |
| DDHD2 | 0.844126349 | 1.795177314 | 5.21E-11 | 1.18E-09 | 273.8996539 | 491.5060655 | ENSG00000085788 |
| SARDH | 0.843925612 | 1.79492755 | 1.37E-12 | 3.93E-11 | 296.5595499 | 531.9262266 | ENSG00000123453 |
| BRWD1 | 0.843782479 | 1.794749479 | 1.35E-15 | 5.55E-14 | 614.4721703 | 1102.105442 | ENSG00000185658 |
| G0S2 | 0.843693962 | 1.794639365 | 4.31E-02 | 1.19E-01 | 20.82288995 | 37.47422919 | ENSG00000123689 |
| STC2 | 0.843148086 | 1.793960451 | 7.10E-31 | 1.06E-28 | 1865.337908 | 3344.880298 | ENSG00000113739 |
| TPSG1 | 0.840725038 | 1.790949972 | 6.58E-03 | 2.60E-02 | 38.52274631 | 69.03376128 | ENSG00000116176 |
| MPI | 0.840201355 | 1.790299995 | 1.55E-17 | 7.83E-16 | 518.8627297 | 929.1746934 | ENSG00000178802 |
| CEP19 | 0.838941977 | 1.788737862 | 3.53E-02 | 1.02E-01 | 27.5792041 | 49.22919222 | ENSG00000174007 |
| NCR3LG1 | 0.838505556 | 1.788196843 | 4.89E-03 | 2.02E-02 | 40.35394221 | 72.03049543 | ENSG00000188211 |
| ATF3 | 0.837806444 | 1.787330516 | 1.51E-14 | 5.50E-13 | 416.0262122 | 743.5725608 | ENSG00000162772 |
| PGPEP1 | 0.837199197 | 1.786578365 | 3.12E-20 | 2.03E-18 | 586.0614511 | 1047.01395 | ENSG00000130517 |
| JPH2 | 0.836964912 | 1.786288259 | 7.15E-03 | 2.79E-02 | 36.66867789 | 65.37959585 | ENSG00000149596 |
| GPRIN3 | 0.836760293 | 1.786034927 | 1.75E-03 | 8.49E-03 | 47.76510475 | 85.28181872 | ENSG00000185477 |
| MX2 | 0.836125045 | 1.785248672 | 2.62E-03 | 1.19E-02 | 54.7299479 | 97.67533042 | ENSG00000183486 |
| INHBE | 0.835508606 | 1.784486028 | 4.26E-13 | 1.32E-11 | 305.4191931 | 544.827949 | ENSG00000139269 |
| BHLHE40 | 0.835394778 | 1.78434524 | 2.59E-03 | 1.18E-02 | 67.7118169 | 120.3736436 | ENSG00000134107 |
| SNAPC5 | 0.835042119 | 1.783909119 | 5.31E-09 | 8.89E-08 | 195.5085432 | 348.8195623 | ENSG00000174446 |
| ALG10B | 0.83444061 | 1.783165501 | 3.06E-04 | 1.87E-03 | 86.02370034 | 153.0869832 | ENSG00000175548 |
| DEPDC4 | 0.832798563 | 1.781137092 | 2.44E-02 | 7.59E-02 | 26.81586323 | 47.78188212 | ENSG00000166153 |
| ARSG | 0.832152721 | 1.78033992 | 6.14E-04 | 3.44E-03 | 61.13451452 | 108.8557671 | ENSG00000141337 |
| LACTB2 | 0.830626459 | 1.778457452 | 3.43E-03 | 1.49E-02 | 45.00495798 | 80.0488346 | ENSG00000147592 |
| GRIN2A | 0.83057827 | 1.778398048 | 9.64E-04 | 5.11E-03 | 72.25600129 | 127.986945 | ENSG00000183454 |
| ZC3H6 | 0.829834716 | 1.777481711 | 1.40E-04 | 9.36E-04 | 77.36152941 | 137.6012737 | ENSG00000188177 |
| PTAR1 | 0.828749156 | 1.776144743 | 1.02E-19 | 6.39E-18 | 675.5367291 | 1199.510876 | ENSG00000188647 |
| BDP1 | 0.828128289 | 1.77538054 | 4.79E-11 | 1.09E-09 | 499.9158264 | 887.1611711 | ENSG00000145734 |
| TNXB | 0.827644531 | 1.774785327 | 3.69E-07 | 4.47E-06 | 154.5439494 | 274.182948 | ENSG00000168477 |
| UTF1 | 0.827185142 | 1.774220283 | 8.06E-03 | 3.08E-02 | 463.5428832 | 822.3157469 | ENSG00000171794 |
| TTPAL | 0.827141192 | 1.774166233 | 1.84E-11 | 4.46E-10 | 290.0973696 | 514.7757272 | ENSG00000124120 |
| TRIM5 | 0.825912692 | 1.772656119 | 7.42E-10 | 1.41E-08 | 240.7362212 | 426.183292 | ENSG00000132256 |
| RLN1 | 0.825661637 | 1.772347671 | 3.75E-02 | 1.07E-01 | 22.28508061 | 39.46579495 | ENSG00000107014 |
| ADAMTS5 | 0.825597785 | 1.772269231 | 1.09E-02 | 3.97E-02 | 33.05669899 | 58.65514963 | ENSG00000154736 |
| TRIML2 | 0.824983403 | 1.771514659 | 4.17E-12 | 1.13E-10 | 495.4556083 | 878.0985297 | ENSG00000179046 |
| FNIP1 | 0.824411148 | 1.770812114 | 3.94E-06 | 3.92E-05 | 158.4405687 | 280.4632771 | ENSG00000217128 |
| EMBP1 | 0.824381093 | 1.770775224 | 9.60E-03 | 3.56E-02 | 38.71864766 | 68.57912371 | ENSG00000231752 |
| APOBEC3F | 0.824138869 | 1.770477941 | 3.93E-04 | 2.32E-03 | 74.45787102 | 131.8015757 | ENSG00000128394 |
| IFI27L2 | 0.823472637 | 1.769660529 | 8.21E-06 | 7.49E-05 | 123.2706866 | 218.5572251 | ENSG00000119632 |
| NOCT | 0.823078436 | 1.769177054 | 3.04E-13 | 9.53E-12 | 384.1734837 | 679.4607923 | ENSG00000151014 |
| ZNF850 | 0.820944927 | 1.766562666 | 2.44E-05 | 1.99E-04 | 102.3885144 | 180.9525463 | ENSG00000267041 |
| NT5C2 | 0.820837016 | 1.766430535 | 2.59E-18 | 1.41E-16 | 616.894475 | 1089.765745 | ENSG00000076685 |
| EPPK1 | 0.820500607 | 1.766018684 | 3.36E-08 | 4.89E-07 | 269.6652902 | 475.9098312 | ENSG00000261150 |
| NHSL2 | 0.819632867 | 1.764956794 | 9.42E-09 | 1.52E-07 | 203.9740387 | 359.7202978 | ENSG00000204131 |
| KCNB1 | 0.818007186 | 1.762969097 | 4.42E-02 | 1.22E-01 | 25.39844807 | 44.70615352 | ENSG00000158445 |
| MAN2B2 | 0.817832228 | 1.762755311 | 5.09E-09 | 8.55E-08 | 197.9350121 | 349.0423379 | ENSG00000013288 |
| PLCB1 | 0.816834147 | 1.761536229 | 1.23E-05 | 1.08E-04 | 132.134276 | 232.3047013 | ENSG00000182621 |
| TRPV6 | 0.816184393 | 1.760743056 | 4.68E-02 | 1.27E-01 | 27.89386542 | 49.13241447 | ENSG00000165125 |
| GPR37 | 0.81615633 | 1.760708806 | 2.90E-04 | 1.79E-03 | 112.0676389 | 196.846145 | ENSG00000170775 |
| SIPA1L2 | 0.814534303 | 1.758730349 | 2.68E-13 | 8.47E-12 | 394.078395 | 692.6157674 | ENSG00000116991 |
| FDXR | 0.813584276 | 1.757572591 | 9.22E-23 | 7.34E-21 | 813.0745608 | 1428.908957 | ENSG00000161513 |
| IDS | 0.813418583 | 1.757370745 | 1.48E-06 | 1.59E-05 | 145.7969477 | 256.0555127 | ENSG00000010404 |
| RPL39L | 0.812511366 | 1.756265996 | 2.33E-16 | 1.02E-14 | 486.5911019 | 854.1881518 | ENSG00000163923 |
| LGI4 | 0.811917259 | 1.755542909 | 4.90E-03 | 2.02E-02 | 61.28168671 | 107.8153585 | ENSG00000153902 |
| SUGP1 | 0.811635562 | 1.755200159 | 3.03E-24 | 2.70E-22 | 842.8411278 | 1479.078147 | ENSG00000105705 |
| ACP5 | 0.808036418 | 1.750826855 | 1.22E-04 | 8.34E-04 | 91.84535115 | 160.5832388 | ENSG00000102575 |
| PIM1 | 0.804737485 | 1.746827907 | 2.66E-27 | 3.03E-25 | 1173.975911 | 2050.172103 | ENSG00000137193 |
| ARL4D | 0.80410365 | 1.746060623 | 2.86E-07 | 3.53E-06 | 219.8393468 | 383.4187151 | ENSG00000175906 |
| RPS27L | 0.80337176 | 1.745175059 | 1.00E-17 | 5.13E-16 | 657.6742317 | 1147.550635 | ENSG00000185088 |
| VCL | 0.802670873 | 1.744327426 | 9.96E-40 | 2.84E-37 | 10195.56786 | 17784.01797 | ENSG00000035403 |
| ZSCAN31 | 0.801887548 | 1.743380585 | 7.12E-04 | 3.91E-03 | 68.96025457 | 120.3812413 | ENSG00000235109 |
| RLIM | 0.800789034 | 1.742053624 | 2.38E-21 | 1.68E-19 | 736.3836417 | 1282.229526 | ENSG00000131263 |

| | | | | | | | |
|---|---|---|---|---|---|---|---|
| ZBTB34 | 0.80065719 | 1.74189443 | 3.21E-10 | 6.46E-09 | 352.8008986 | 614.2975055 | ENSG00000177125 |
| KIFC3 | 0.800363497 | 1.741539864 | 2.03E-07 | 2.57E-06 | 169.2392238 | 294.9413822 | ENSG00000140859 |
| VSNL1 | 0.800185353 | 1.741324833 | 1.78E-17 | 8.91E-16 | 1251.203459 | 2178.036616 | ENSG00000163032 |
| EPB41L4A | 0.799446602 | 1.740433393 | 1.24E-02 | 4.40E-02 | 36.42484413 | 63.45562798 | ENSG00000129595 |
| OLFML3 | 0.79897493 | 1.739864471 | 3.85E-19 | 2.27E-17 | 772.453792 | 1343.844775 | ENSG00000116774 |
| IFIT1 | 0.798780196 | 1.739629641 | 4.08E-02 | 1.14E-01 | 27.58296237 | 47.79955381 | ENSG00000185745 |
| TRIM56 | 0.797985073 | 1.738671131 | 1.15E-18 | 6.51E-17 | 709.9898725 | 1234.735787 | ENSG00000169871 |
| PLXND1 | 0.79763614 | 1.738250664 | 8.88E-27 | 9.57E-25 | 1407.322271 | 2445.328481 | ENSG00000004399 |
| TMEM8B | 0.797269063 | 1.737808442 | 8.24E-09 | 1.34E-07 | 222.4545163 | 386.4810634 | ENSG00000137103 |
| BATF2 | 0.797156967 | 1.737673421 | 3.16E-02 | 9.34E-02 | 29.18056937 | 50.80867218 | ENSG00000168062 |
| ESPNL | 0.796156553 | 1.736468876 | 3.53E-02 | 1.02E-01 | 24.40833578 | 42.37236136 | ENSG00000144488 |
| DLL3 | 0.795264381 | 1.735395365 | 3.16E-09 | 5.43E-08 | 289.8130898 | 502.1959313 | ENSG00000090932 |
| CCDC167 | 0.794923532 | 1.734985412 | 3.06E-06 | 3.10E-05 | 172.2425279 | 298.6905564 | ENSG00000198937 |
| ADA2 | 0.794373493 | 1.73432406 | 9.91E-19 | 5.65E-17 | 903.3617739 | 1565.877622 | ENSG00000093072 |
| EML2 | 0.79433298 | 1.734275359 | 8.61E-12 | 2.20E-10 | 357.3482626 | 619.5584585 | ENSG00000125746 |
| C1orf74 | 0.794171905 | 1.734081741 | 1.61E-05 | 1.37E-04 | 107.3753954 | 186.125745 | ENSG00000162757 |
| GABRE | 0.792260417 | 1.731785703 | 4.32E-03 | 1.82E-02 | 70.8095952 | 122.7438156 | ENSG00000102287 |
| TNNC1 | 0.792079305 | 1.731568313 | 4.86E-02 | 1.31E-01 | 27.6298124 | 48.04902454 | ENSG00000114854 |
| PLCB2 | 0.791960272 | 1.731425452 | 9.12E-08 | 1.23E-06 | 245.3264815 | 424.8903238 | ENSG00000137841 |
| KATNAL2 | 0.791822424 | 1.731260023 | 1.50E-05 | 1.29E-04 | 129.3778875 | 223.9463259 | ENSG00000167216 |
| MAP3K15 | 0.791258481 | 1.730583414 | 4.27E-03 | 1.80E-02 | 46.55759408 | 80.6643754 | ENSG00000180815 |
| LHPP | 0.791093242 | 1.730385213 | 1.72E-16 | 7.73E-15 | 794.7853917 | 1374.959051 | ENSG00000107902 |
| CAPN6 | 0.79108805 | 1.730378986 | 1.07E-06 | 1.19E-05 | 165.3692353 | 285.9259733 | ENSG00000077274 |
| AKAP13 | 0.788859285 | 1.727707853 | 1.41E-10 | 3.02E-09 | 694.2815075 | 1199.582359 | ENSG00000170776 |
| SERINC5 | 0.788345266 | 1.727092396 | 5.80E-20 | 3.72E-18 | 1679.431859 | 2899.634045 | ENSG00000164300 |
| FAM214A | 0.787935228 | 1.726601597 | 7.46E-04 | 4.08E-03 | 67.07136242 | 115.8318339 | ENSG00000047346 |
| ACOT11 | 0.786743232 | 1.725175619 | 9.63E-05 | 6.74E-04 | 93.9117819 | 161.8471874 | ENSG00000162390 |
| SYNM | 0.784972426 | 1.723059387 | 8.28E-04 | 4.47E-03 | 83.77262043 | 143.9739607 | ENSG00000182253 |
| AMOTL2 | 0.784691973 | 1.722724465 | 1.13E-21 | 8.26E-20 | 1054.71444 | 1816.731369 | ENSG00000114019 |
| MPP1 | 0.783327155 | 1.721095503 | 6.79E-09 | 1.12E-07 | 268.2332705 | 461.5262282 | ENSG00000130830 |
| ZNF292 | 0.78193453 | 1.719434943 | 1.14E-09 | 2.09E-08 | 354.3236342 | 608.64756 | ENSG00000188994 |
| AASS | 0.78030313 | 1.717491703 | 2.34E-26 | 2.46E-24 | 4081.332894 | 7010.317738 | ENSG00000008311 |
| C12orf60 | 0.779764707 | 1.716850844 | 4.89E-02 | 1.31E-01 | 24.1833458 | 41.48372941 | ENSG00000182993 |
| C18orf25 | 0.779512539 | 1.716550783 | 2.47E-11 | 5.88E-10 | 361.0823538 | 619.8742131 | ENSG00000152242 |
| TUBB6 | 0.779492076 | 1.716526436 | 1.09E-27 | 1.29E-25 | 2936.01691 | 5040.449442 | ENSG00000176014 |
| PTPDC1 | 0.779469212 | 1.716499233 | 2.83E-05 | 2.27E-04 | 127.8904415 | 219.1193269 | ENSG00000158079 |
| CLYBL | 0.77917738 | 1.716152049 | 2.61E-05 | 2.12E-04 | 109.6591268 | 188.1665757 | ENSG00000125246 |
| MACF1 | 0.778343105 | 1.715159928 | 3.78E-05 | 2.94E-04 | 4522.165416 | 7756.245084 | ENSG00000127603 |
| CGREF1 | 0.77830303 | 1.715112285 | 2.67E-03 | 1.21E-02 | 56.47788399 | 96.80839225 | ENSG00000138028 |
| PPP1R15B | 0.777786511 | 1.714498344 | 1.43E-14 | 5.24E-13 | 971.5246154 | 1665.979025 | ENSG00000158615 |
| ATP1B1 | 0.777771753 | 1.714480806 | 4.56E-16 | 1.95E-14 | 762.857213 | 1307.766655 | ENSG00000143153 |
| SLC48A1 | 0.776704905 | 1.713213446 | 2.39E-08 | 3.59E-07 | 241.5597388 | 413.6633762 | ENSG00000211584 |
| RAB24 | 0.776522234 | 1.712996535 | 2.28E-03 | 1.06E-02 | 60.84137288 | 104.5054662 | ENSG00000169228 |
| AGPAT3 | 0.775653972 | 1.711965907 | 5.49E-06 | 5.25E-05 | 152.4776163 | 261.0062706 | ENSG00000160216 |
| CDH2 | 0.775519408 | 1.711806235 | 1.08E-15 | 4.47E-14 | 552.4811784 | 945.2782765 | ENSG00000170558 |
| MBTPS2 | 0.774430914 | 1.710515188 | 1.23E-05 | 1.08E-04 | 148.5921887 | 254.3528947 | ENSG00000012174 |
| OMA1 | 0.773601233 | 1.709531769 | 9.02E-05 | 6.36E-04 | 108.5279608 | 185.0736848 | ENSG00000162600 |
| RILPL1 | 0.773191138 | 1.709045893 | 9.65E-07 | 1.08E-05 | 165.4602594 | 282.8603674 | ENSG00000188026 |
| CEP350 | 0.772479153 | 1.708202668 | 3.04E-10 | 6.11E-09 | 531.1366842 | 907.2707022 | ENSG00000135837 |
| SPATS2L | 0.772387158 | 1.708093747 | 3.48E-16 | 1.50E-14 | 898.7524069 | 1534.881117 | ENSG00000196141 |
| KIF17 | 0.772373003 | 1.708076988 | 5.85E-10 | 1.13E-08 | 283.4078316 | 484.3099226 | ENSG00000117245 |
| EHBP1L1 | 0.770676491 | 1.706069585 | 7.44E-06 | 6.91E-05 | 143.802021 | 245.4351884 | ENSG00000173442 |
| LHX6 | 0.76975169 | 1.704976306 | 1.40E-02 | 4.84E-02 | 41.90549593 | 71.70411724 | ENSG00000106852 |
| TTC39C | 0.769456493 | 1.704627478 | 6.41E-11 | 1.44E-09 | 323.0628263 | 550.9831337 | ENSG00000168234 |
| TRIM68 | 0.769090818 | 1.704195466 | 1.52E-05 | 1.30E-04 | 160.7086511 | 273.6102854 | ENSG00000167333 |
| LINGO2 | 0.769058879 | 1.704157738 | 2.56E-03 | 1.17E-02 | 57.42893325 | 97.70396062 | ENSG00000174482 |
| SNHG32 | 0.768858105 | 1.703920594 | 1.27E-12 | 3.68E-11 | 585.2847835 | 997.7681053 | ENSG00000204387 |
| GAS5 | 0.768068678 | 1.702988483 | 2.48E-20 | 1.63E-18 | 3694.378444 | 6291.818238 | ENSG00000234741 |
| ZBTB42 | 0.767835342 | 1.70271307 | 2.03E-04 | 1.30E-03 | 85.74752334 | 145.939819 | ENSG00000179627 |
| STK17B | 0.766813008 | 1.701506907 | 3.97E-05 | 3.07E-04 | 147.9158659 | 251.2659526 | ENSG00000081320 |
| DPEP1 | 0.764595678 | 1.698893808 | 3.12E-03 | 1.38E-02 | 59.58493764 | 101.0732003 | ENSG00000015413 |
| AKAP7 | 0.764172807 | 1.698395915 | 9.94E-09 | 1.60E-07 | 248.7855758 | 421.9550382 | ENSG00000118507 |
| PAQR7 | 0.763962055 | 1.698147829 | 3.24E-05 | 2.58E-04 | 121.7153973 | 206.7709952 | ENSG00000182749 |
| APLN | 0.762041001 | 1.695888125 | 6.35E-05 | 4.66E-04 | 110.9512728 | 187.794405 | ENSG00000171388 |
| DHTKD1 | 0.761968014 | 1.695802331 | 2.02E-25 | 2.00E-23 | 1431.237009 | 2426.357706 | ENSG00000181192 |
| ASNS | 0.761922895 | 1.695749298 | 2.74E-02 | 8.35E-02 | 29.90595223 | 50.76733135 | ENSG00000070669 |
| PLEKHA2 | 0.760221238 | 1.693750342 | 1.24E-04 | 8.43E-04 | 140.9702499 | 238.2678904 | ENSG00000169499 |
| XIAP | 0.7601615 | 1.69368021 | 6.83E-12 | 1.78E-10 | 702.9585987 | 1190.085451 | ENSG00000101966 |
| CNN1 | 0.760121932 | 1.693633759 | 2.97E-10 | 5.99E-09 | 321.0194635 | 543.9885735 | ENSG00000130176 |
| WLS | 0.760120191 | 1.693631715 | 1.32E-02 | 4.61E-02 | 42.30339443 | 71.56198501 | ENSG00000116729 |
| UTRN | 0.759908079 | 1.693382727 | 9.28E-11 | 2.05E-09 | 862.7632763 | 1460.462735 | ENSG00000152818 |
| SERPINA5 | 0.759754579 | 1.693202565 | 1.05E-02 | 3.82E-02 | 44.14415877 | 74.71268597 | ENSG00000188488 |
| C17orf113 | 0.758727851 | 1.691997986 | 8.61E-03 | 3.25E-02 | 45.18743492 | 76.24603856 | ENSG00000267221 |
| FOSL2 | 0.758492299 | 1.691721752 | 3.48E-04 | 2.08E-03 | 117.6332191 | 199.0856358 | ENSG00000075426 |
| SYDE2 | 0.758479368 | 1.691706589 | 1.96E-04 | 1.26E-03 | 109.9963675 | 185.5773641 | ENSG00000097096 |

| | | | | | | | |
|---|---|---|---|---|---|---|---|
| ADAMTS16 | 0.757612666 | 1.690690598 | 1.06E-06 | 1.18E-05 | 205.3387784 | 347.3631171 | ENSG00000145536 |
| TRIB3 | 0.757221165 | 1.690231861 | 8.61E-11 | 1.91E-09 | 336.4565067 | 568.7986846 | ENSG00000101255 |
| SIL1 | 0.757135974 | 1.690132057 | 1.97E-03 | 9.38E-03 | 70.29737143 | 118.7940605 | ENSG00000120725 |
| NFATC1 | 0.756851453 | 1.68979877 | 3.17E-05 | 2.53E-04 | 150.5746082 | 254.5302672 | ENSG00000131196 |
| NEK7 | 0.756327538 | 1.689185231 | 1.12E-03 | 5.81E-03 | 123.1757088 | 207.508328 | ENSG00000151414 |
| PIK3R1 | 0.756065804 | 1.688878806 | 2.68E-07 | 3.33E-06 | 238.3628184 | 401.9773965 | ENSG00000145675 |
| ZNF285 | 0.755974604 | 1.688772047 | 2.23E-02 | 7.05E-02 | 37.23732013 | 62.84162453 | ENSG00000267508 |
| FLNB | 0.754916537 | 1.687533962 | 1.38E-21 | 9.98E-20 | 4006.035308 | 6759.520988 | ENSG00000136068 |
| ZNF8 | 0.754682283 | 1.687259975 | 6.55E-06 | 6.17E-05 | 163.7196067 | 276.4482662 | ENSG00000278129 |
| PCLO | 0.754586671 | 1.687148159 | 4.94E-04 | 2.83E-03 | 89.23818672 | 150.542067 | ENSG00000186472 |
| UNC5D | 0.753791882 | 1.686218956 | 5.17E-21 | 3.58E-19 | 1246.535975 | 2101.213997 | ENSG00000156687 |
| RGS7 | 0.75329018 | 1.685632668 | 4.06E-02 | 1.14E-01 | 27.48442166 | 46.31145252 | ENSG00000182901 |
| SESN2 | 0.753216978 | 1.685547142 | 6.96E-19 | 4.03E-17 | 931.2259866 | 1568.60206 | ENSG00000130766 |
| TTLL1 | 0.752291351 | 1.684466049 | 6.41E-05 | 4.69E-04 | 121.504215 | 204.3995494 | ENSG00000100271 |
| IKBIP | 0.751384179 | 1.683407183 | 5.15E-06 | 4.96E-05 | 219.619979 | 369.4645658 | ENSG00000166130 |
| GIHCG | 0.750876067 | 1.682814398 | 3.97E-02 | 1.12E-01 | 27.97168322 | 46.96679681 | ENSG00000257698 |
| SNCG | 0.750804495 | 1.682730916 | 3.62E-04 | 2.16E-03 | 88.84067155 | 149.3910791 | ENSG00000173267 |
| SQSTM1 | 0.750330366 | 1.682177992 | 2.78E-25 | 2.70E-23 | 1373.784789 | 2311.511616 | ENSG00000161011 |
| SLC9B2 | 0.748933853 | 1.68055045 | 3.48E-03 | 1.51E-02 | 56.32943403 | 94.67783183 | ENSG00000164038 |
| RASAL2 | 0.74594245 | 1.677069468 | 1.95E-13 | 6.25E-12 | 574.0972121 | 962.5457822 | ENSG00000075391 |
| ZNF805 | 0.74593445 | 1.677060169 | 3.11E-04 | 1.89E-03 | 92.29039277 | 154.6744904 | ENSG00000204524 |
| TSGA10 | 0.745437236 | 1.676482282 | 1.67E-02 | 5.57E-02 | 37.915447 | 63.50087928 | ENSG00000135951 |
| KLHL29 | 0.744782987 | 1.675722185 | 1.89E-08 | 2.88E-07 | 252.4409395 | 423.5944471 | ENSG00000119771 |
| SOCS1 | 0.743014146 | 1.673668895 | 2.18E-06 | 2.27E-05 | 376.7236715 | 630.6420453 | ENSG00000185338 |
| ZBTB4 | 0.742809897 | 1.673431963 | 3.38E-12 | 9.20E-11 | 450.9032457 | 754.8983874 | ENSG00000174282 |
| PTPN6 | 0.741667393 | 1.672107259 | 1.61E-10 | 3.41E-09 | 328.652572 | 549.0775617 | ENSG00000111679 |
| TRPV1 | 0.741344887 | 1.67173351 | 2.88E-02 | 8.67E-02 | 46.63454007 | 78.12359292 | ENSG00000196689 |
| USH1C | 0.739929301 | 1.670093994 | 3.23E-02 | 9.49E-02 | 31.03396136 | 51.9072575 | ENSG00000006611 |
| TMEM54 | 0.739839786 | 1.669990373 | 1.44E-02 | 4.94E-02 | 49.24563552 | 82.18786878 | ENSG00000121900 |
| CACNA1C | 0.73915526 | 1.669198188 | 3.88E-03 | 1.66E-02 | 60.36919682 | 100.7128558 | ENSG00000151067 |
| MTHFD2 | 0.738930487 | 1.668938147 | 1.97E-30 | 2.88E-28 | 2355.312616 | 3929.984501 | ENSG00000065911 |
| CEP112 | 0.736911203 | 1.666603833 | 1.67E-02 | 5.57E-02 | 43.66280507 | 73.08167956 | ENSG00000154240 |
| ZNF814 | 0.736660589 | 1.666314348 | 9.11E-09 | 1.48E-07 | 259.9429535 | 433.5823318 | ENSG00000204514 |
| ZMAT4 | 0.735780063 | 1.66529765 | 6.33E-07 | 7.31E-06 | 188.870989 | 314.5892362 | ENSG00000165061 |
| PCDHGC3 | 0.73557442 | 1.665060293 | 5.63E-05 | 4.19E-04 | 124.0768488 | 206.1757224 | ENSG00000240184 |
| SH3RF3 | 0.734547692 | 1.663875736 | 2.24E-02 | 7.08E-02 | 39.99766231 | 66.47438232 | ENSG00000172985 |
| TNFRSF10B | 0.734225463 | 1.663504147 | 1.50E-17 | 7.55E-16 | 1831.110484 | 3045.949321 | ENSG00000120889 |
| MRTFB | 0.733837892 | 1.663057317 | 1.73E-08 | 2.66E-07 | 276.5640109 | 459.9990475 | ENSG00000186260 |
| TRIM72 | 0.732997238 | 1.662088541 | 2.64E-02 | 8.10E-02 | 35.5128579 | 58.9514719 | ENSG00000177238 |
| FGF13 | 0.732388683 | 1.66138759 | 9.95E-19 | 5.66E-17 | 1149.636298 | 1909.136222 | ENSG00000129682 |
| CR2 | 0.732018118 | 1.660960907 | 2.12E-05 | 1.76E-04 | 128.9590707 | 214.0990444 | ENSG00000117322 |
| TOR4A | 0.731641704 | 1.660527602 | 3.51E-06 | 3.53E-05 | 211.7781613 | 351.0726163 | ENSG00000198113 |
| PABPC5 | 0.731478095 | 1.660339301 | 1.15E-02 | 4.13E-02 | 43.37390271 | 71.98535574 | ENSG00000174740 |
| SLFN5 | 0.730646615 | 1.659382659 | 1.03E-03 | 5.39E-03 | 89.0485992 | 147.4874825 | ENSG00000166750 |
| SOX4 | 0.730640389 | 1.659375498 | 2.73E-34 | 4.97E-32 | 5295.344605 | 8785.773259 | ENSG00000124766 |
| ITPR1 | 0.730020847 | 1.658663059 | 1.40E-04 | 9.35E-04 | 169.442326 | 280.8988491 | ENSG00000150995 |
| RHBDF1 | 0.72880527 | 1.657266102 | 1.31E-11 | 3.25E-10 | 548.906406 | 910.3922974 | ENSG00000007384 |
| CASP10 | 0.728671054 | 1.657111931 | 1.55E-02 | 5.24E-02 | 43.25386497 | 71.63228246 | ENSG00000003400 |
| CRISPLD2 | 0.728425305 | 1.656829683 | 1.66E-04 | 1.09E-03 | 101.6636352 | 168.4212424 | ENSG00000103196 |
| SLC25A14 | 0.728324017 | 1.656713365 | 4.08E-06 | 4.04E-05 | 188.3259335 | 311.4376592 | ENSG00000102078 |
| VEPH1 | 0.727036158 | 1.655235116 | 1.45E-02 | 4.97E-02 | 55.63494389 | 92.01766148 | ENSG00000197415 |
| MYO5A | 0.726775424 | 1.654935998 | 3.04E-14 | 1.07E-12 | 685.6798594 | 1134.16034 | ENSG00000197535 |
| SUSD2 | 0.726471186 | 1.654587039 | 2.45E-07 | 3.06E-06 | 206.8867845 | 342.3191655 | ENSG00000099994 |
| GALNT10 | 0.726303213 | 1.654394406 | 3.30E-12 | 9.01E-11 | 593.3359945 | 980.6834341 | ENSG00000164574 |
| TMC7 | 0.725978601 | 1.654022203 | 3.33E-03 | 1.45E-02 | 66.29213929 | 109.5531135 | ENSG00000170537 |
| TXNIP | 0.725674746 | 1.653673876 | 1.28E-03 | 6.48E-03 | 107.8738114 | 177.7923839 | ENSG00000265972 |
| ESYT2 | 0.725303171 | 1.653248017 | 4.25E-18 | 2.27E-16 | 966.025706 | 1596.217492 | ENSG00000117868 |
| APC2 | 0.724782043 | 1.652650941 | 9.09E-03 | 3.40E-02 | 50.14431768 | 82.84712354 | ENSG00000115266 |
| TRAPPC6A | 0.724238595 | 1.652028522 | 6.80E-14 | 2.30E-12 | 654.6915528 | 1082.124975 | ENSG00000007255 |
| ZCCHC4 | 0.724173048 | 1.651953466 | 4.31E-05 | 3.30E-04 | 124.9609265 | 206.3917764 | ENSG00000168228 |
| MAN1A1 | 0.722828138 | 1.650414199 | 8.39E-06 | 7.64E-05 | 142.9566004 | 235.8723862 | ENSG00000111885 |
| VPS13D | 0.722307698 | 1.649818933 | 1.77E-08 | 2.71E-07 | 722.9989057 | 1192.456952 | ENSG00000048707 |
| ADAMTS20 | 0.721959798 | 1.649421134 | 5.05E-05 | 3.79E-04 | 134.9671294 | 222.294908 | ENSG00000173157 |
| DMD | 0.721587837 | 1.648995929 | 1.92E-04 | 1.24E-03 | 131.165082 | 215.9455065 | ENSG00000198947 |
| NFASC | 0.721359909 | 1.648735428 | 7.29E-09 | 1.20E-07 | 313.4558223 | 517.158275 | ENSG00000163531 |
| SMTN | 0.721042218 | 1.648372406 | 2.03E-08 | 3.07E-07 | 260.9381018 | 430.0843947 | ENSG00000183963 |
| ENG | 0.719973308 | 1.647151559 | 1.75E-03 | 8.49E-03 | 78.97602599 | 130.1712305 | ENSG00000106991 |
| AK5 | 0.719327994 | 1.646414958 | 2.22E-03 | 1.04E-02 | 75.59139722 | 124.1347724 | ENSG00000154027 |
| GRIP2 | 0.719308117 | 1.646392274 | 5.77E-04 | 3.25E-03 | 191.0850956 | 315.1564771 | ENSG00000144596 |
| MAP3K20 | 0.719033211 | 1.646078583 | 1.65E-07 | 2.12E-06 | 269.9710595 | 443.6367865 | ENSG00000091436 |
| ASH1L | 0.718542749 | 1.645519073 | 5.70E-09 | 9.51E-08 | 426.9882301 | 702.2610196 | ENSG00000116539 |
| TSSK6 | 0.717057404 | 1.64382578 | 2.07E-02 | 6.65E-02 | 40.00081922 | 65.94729742 | ENSG00000178093 |
| FAM160A1 | 0.716565383 | 1.643265261 | 1.45E-12 | 4.16E-11 | 600.950575 | 987.4039388 | ENSG00000164142 |
| C16orf96 | 0.715222233 | 1.641736091 | 4.82E-02 | 1.30E-01 | 28.47147473 | 46.61839844 | ENSG00000205832 |
| HELB | 0.714895841 | 1.641364711 | 1.30E-05 | 1.13E-04 | 154.3404564 | 253.26917 | ENSG00000127311 |

| | | | | | | | |
|---|---|---|---|---|---|---|---|
| RASEF | 0.714782569 | 1.641235846 | 1.15E-03 | 5.96E-03 | 82.54676963 | 135.1594188 | ENSG00000165105 |
| RORA | 0.71466572 | 1.641102921 | 1.61E-02 | 5.40E-02 | 45.29809962 | 74.37513448 | ENSG00000069667 |
| DOCK3 | 0.713447691 | 1.639717966 | 1.39E-05 | 1.21E-04 | 250.4949299 | 411.0208549 | ENSG00000088538 |
| RTL10 | 0.713285664 | 1.639533823 | 5.08E-14 | 1.74E-12 | 511.6821946 | 838.9179064 | ENSG00000215012 |
| DLGAP1 | 0.713063332 | 1.639281175 | 1.10E-08 | 1.75E-07 | 273.4063716 | 447.8693888 | ENSG00000170579 |
| PMEPA1 | 0.712712302 | 1.638882362 | 5.36E-10 | 1.04E-08 | 499.5876805 | 818.7157538 | ENSG00000124225 |
| PARD6B | 0.712406596 | 1.63853512 | 2.59E-03 | 1.18E-02 | 120.3280629 | 197.0360762 | ENSG00000124171 |
| LTBP4 | 0.712323151 | 1.638440351 | 2.33E-22 | 1.79E-20 | 1712.942078 | 2807.572237 | ENSG00000090006 |
| GAL3ST4 | 0.711555187 | 1.637568422 | 4.10E-04 | 2.40E-03 | 90.57667905 | 148.4379869 | ENSG00000197093 |
| IL13RA1 | 0.709602196 | 1.635353128 | 2.50E-12 | 6.90E-11 | 529.3123284 | 864.8202705 | ENSG00000131724 |
| PRR14L | 0.709414565 | 1.635140455 | 1.16E-12 | 3.39E-11 | 869.6918073 | 1422.036284 | ENSG00000183530 |
| STARD8 | 0.708881904 | 1.634536852 | 4.94E-05 | 3.73E-04 | 130.4793635 | 213.0487364 | ENSG00000130052 |
| SLC12A4 | 0.708869843 | 1.634523187 | 6.57E-20 | 4.17E-18 | 1143.739821 | 1869.615142 | ENSG00000124067 |
| RASGRP4 | 0.707885186 | 1.633407986 | 9.06E-03 | 3.39E-02 | 50.0476561 | 81.88362241 | ENSG00000171777 |
| WIPF1 | 0.706945673 | 1.632344622 | 5.81E-05 | 4.31E-04 | 147.5988444 | 241.1114307 | ENSG00000115935 |
| ERN1 | 0.706684888 | 1.632049583 | 3.88E-04 | 2.29E-03 | 103.9890078 | 169.7044001 | ENSG00000178607 |
| FGD6 | 0.705983805 | 1.631256675 | 3.36E-12 | 9.15E-11 | 535.0080935 | 872.1856385 | ENSG00000180263 |
| AZIN1 | 0.705827316 | 1.631079742 | 9.44E-18 | 4.87E-16 | 4567.199771 | 7448.074494 | ENSG00000155096 |
| VPS13B | 0.70546575 | 1.630671015 | 1.57E-08 | 2.44E-07 | 323.0259507 | 526.6728628 | ENSG00000132549 |
| VCPIP1 | 0.705060217 | 1.630212707 | 1.10E-08 | 1.75E-07 | 306.3908675 | 499.3145714 | ENSG00000175073 |
| MUC3A | 0.704887242 | 1.630017262 | 7.88E-08 | 1.07E-06 | 258.543202 | 421.6462604 | ENSG00000169894 |
| FAM50B | 0.704502326 | 1.629582425 | 2.88E-03 | 1.29E-02 | 66.21066092 | 107.9558502 | ENSG00000145945 |
| DTNA | 0.703381536 | 1.628316939 | 2.12E-05 | 1.76E-04 | 172.5384205 | 281.2254505 | ENSG00000134769 |
| PSD3 | 0.703127362 | 1.628030087 | 8.35E-09 | 1.36E-07 | 343.5661393 | 559.8069767 | ENSG00000156011 |
| SYT12 | 0.701888117 | 1.626632244 | 2.62E-05 | 2.13E-04 | 169.0184731 | 274.6523429 | ENSG00000173227 |
| RRAS | 0.701334772 | 1.62600847 | 1.10E-07 | 1.46E-06 | 264.0402171 | 429.7348341 | ENSG00000126458 |
| NFAT5 | 0.701150622 | 1.625800934 | 5.41E-07 | 6.36E-06 | 218.745079 | 355.633014 | ENSG00000102908 |
| LMO7 | 0.699818299 | 1.624300206 | 4.32E-07 | 5.17E-06 | 224.7110155 | 364.6963251 | ENSG00000136153 |
| CRYBG3 | 0.699685224 | 1.624150387 | 9.99E-05 | 6.96E-04 | 146.4698431 | 237.7906201 | ENSG00000080200 |
| CRTC1 | 0.699136623 | 1.623532903 | 2.34E-09 | 4.13E-08 | 350.9214621 | 569.4663015 | ENSG00000105662 |
| TMEM59L | 0.699022602 | 1.623404595 | 2.35E-09 | 4.13E-08 | 357.871122 | 581.0035474 | ENSG00000105696 |
| ZNF460 | 0.698066699 | 1.622329314 | 3.70E-02 | 1.06E-01 | 32.96309662 | 53.39265925 | ENSG00000197714 |
| KIAA2026 | 0.697845066 | 1.622080103 | 4.48E-06 | 4.39E-05 | 191.4585355 | 310.7959016 | ENSG00000183354 |
| XKR5 | 0.697830479 | 1.622063702 | 2.21E-03 | 1.03E-02 | 93.98328587 | 152.3339766 | ENSG00000275591 |
| IL18BP | 0.696906595 | 1.621025285 | 4.69E-03 | 1.95E-02 | 111.2326812 | 180.8003719 | ENSG00000137496 |
| PHYHD1 | 0.696401082 | 1.620457386 | 8.20E-04 | 4.43E-03 | 90.71240366 | 146.8021939 | ENSG00000175287 |
| SEMA6A | 0.69638387 | 1.620438053 | 1.49E-26 | 1.58E-24 | 4622.351362 | 7489.517945 | ENSG00000092421 |
| KLHL24 | 0.696364532 | 1.620416333 | 2.87E-07 | 3.54E-06 | 319.4169333 | 516.7580463 | ENSG00000114796 |
| SLC44A4 | 0.696102093 | 1.620121592 | 4.31E-02 | 1.19E-01 | 31.67598671 | 51.33674477 | ENSG00000204385 |
| ITGB1 | 0.694636936 | 1.618477081 | 7.89E-24 | 6.80E-22 | 5692.774782 | 9212.569901 | ENSG00000150093 |
| TNC | 0.694106633 | 1.617882274 | 7.94E-05 | 5.67E-04 | 131.3160875 | 212.2832424 | ENSG00000041982 |
| ZBTB37 | 0.694082001 | 1.617854651 | 3.14E-04 | 1.91E-03 | 157.8220324 | 254.9456425 | ENSG00000185278 |
| P2RY1 | 0.69401375 | 1.617778115 | 5.59E-09 | 9.33E-08 | 670.0521614 | 1083.921546 | ENSG00000169860 |
| GEM | 0.693134229 | 1.616792158 | 2.67E-03 | 1.21E-02 | 81.3816743 | 131.4038007 | ENSG00000164949 |
| BCL2L13 | 0.692826084 | 1.616446865 | 2.82E-14 | 9.97E-13 | 748.047914 | 1208.357083 | ENSG00000099968 |
| ADGRL3 | 0.69263758 | 1.616235672 | 5.38E-07 | 6.33E-06 | 240.0714211 | 387.4933914 | ENSG00000150471 |
| ZNF185 | 0.69251383 | 1.616097042 | 5.75E-08 | 8.03E-07 | 282.6516915 | 456.3960822 | ENSG00000147394 |
| TPCN1 | 0.692272921 | 1.6158272 | 7.73E-10 | 1.46E-08 | 499.2468095 | 806.279436 | ENSG00000186815 |
| OTUD1 | 0.692130568 | 1.615667772 | 3.25E-04 | 1.97E-03 | 111.2843719 | 179.6178622 | ENSG00000165312 |
| KDM6A | 0.691933203 | 1.615446758 | 2.84E-14 | 1.00E-12 | 991.2385354 | 1600.857568 | ENSG00000147050 |
| SLC30A1 | 0.691389215 | 1.614837747 | 1.52E-06 | 1.63E-05 | 483.5426988 | 780.824612 | ENSG00000170385 |
| FRMD6 | 0.69120841 | 1.614635381 | 3.46E-07 | 4.19E-06 | 249.3313077 | 402.7072123 | ENSG00000139926 |
| SMAD7 | 0.69113156 | 1.614549374 | 5.99E-17 | 2.82E-15 | 1022.003776 | 1650.453486 | ENSG00000101665 |
| KMT2D | 0.68981057 | 1.613071704 | 1.39E-03 | 6.91E-03 | 1917.450356 | 3093.072537 | ENSG00000167548 |
| TES | 0.689458988 | 1.612678649 | 5.22E-12 | 1.40E-10 | 513.7502114 | 827.793854 | ENSG00000135269 |
| ZSCAN23 | 0.689359899 | 1.612567889 | 4.26E-02 | 1.18E-01 | 36.68574024 | 59.29001107 | ENSG00000187987 |
| UBR4 | 0.688910945 | 1.61206615 | 2.82E-03 | 1.26E-02 | 4228.930687 | 6817.24014 | ENSG00000127481 |
| TGFBR2 | 0.688805747 | 1.611948606 | 1.25E-06 | 1.37E-05 | 217.5850197 | 350.4724857 | ENSG00000163513 |
| DCAKD | 0.688032675 | 1.61108507 | 1.38E-14 | 5.07E-13 | 691.1098877 | 1113.270741 | ENSG00000172992 |
| MBOAT1 | 0.687986471 | 1.611033474 | 2.15E-07 | 2.71E-06 | 248.4304759 | 400.0380672 | ENSG00000172197 |
| NHS | 0.687789086 | 1.610813073 | 3.44E-07 | 4.18E-06 | 306.8806847 | 493.7432142 | ENSG00000188158 |
| ZNF516 | 0.687209427 | 1.610165996 | 5.33E-11 | 1.20E-09 | 516.6036749 | 832.401908 | ENSG00000101493 |
| SERINC3 | 0.686329463 | 1.609184183 | 5.73E-14 | 1.95E-12 | 745.1834164 | 1198.126999 | ENSG00000132824 |
| SLC2A3 | 0.685817657 | 1.608613416 | 6.87E-23 | 5.59E-21 | 12379.75373 | 19912.25716 | ENSG00000059804 |
| SPART | 0.685441149 | 1.608193662 | 1.74E-16 | 7.79E-15 | 886.6379738 | 1425.472683 | ENSG00000133104 |
| CTH | 0.684417809 | 1.607053334 | 4.74E-08 | 6.75E-07 | 332.5379769 | 533.6768038 | ENSG00000116761 |
| MED13 | 0.684252416 | 1.606869109 | 2.18E-09 | 3.86E-08 | 502.2571412 | 806.5403175 | ENSG00000108510 |
| ARHGEF39 | 0.683745083 | 1.606304142 | 4.10E-06 | 4.06E-05 | 205.4029014 | 329.6031861 | ENSG00000137135 |
| FZD6 | 0.683608287 | 1.606151841 | 3.92E-05 | 3.05E-04 | 141.1101235 | 226.430907 | ENSG00000164930 |
| ZBTB10 | 0.683446662 | 1.605971913 | 6.29E-06 | 5.95E-05 | 243.1078274 | 390.6013258 | ENSG00000205189 |
| MYEF2 | 0.682209723 | 1.604595574 | 3.98E-15 | 1.56E-13 | 954.3818786 | 1530.57183 | ENSG00000104177 |
| NMNAT1 | 0.68190742 | 1.604259381 | 2.09E-02 | 6.69E-02 | 57.66033476 | 92.78825993 | ENSG00000173614 |
| FRYL | 0.68078314 | 1.603009683 | 1.91E-08 | 2.91E-07 | 513.7895827 | 822.7895774 | ENSG00000075539 |
| TTN | 0.680493245 | 1.602687607 | 8.90E-03 | 3.34E-02 | 86.33195014 | 138.469964 | ENSG00000155657 |
| ID2 | 0.680092813 | 1.602242829 | 2.74E-07 | 3.39E-06 | 270.2452596 | 432.5870226 | ENSG00000115738 |

| | | | | | | | |
|---|---|---|---|---|---|---|---|
| FGF2 | 0.678337313 | 1.600294373 | 1.89E-15 | 7.67E-14 | 876.6686464 | 1403.500221 | ENSG00000138685 |
| CDHR3 | 0.678170675 | 1.600109543 | 2.23E-04 | 1.42E-03 | 127.4330653 | 204.2777255 | ENSG00000128536 |
| MED13L | 0.677656817 | 1.599539719 | 2.14E-09 | 3.79E-08 | 663.0491724 | 1060.292014 | ENSG00000123066 |
| SCN1B | 0.674953057 | 1.596544822 | 2.22E-02 | 7.04E-02 | 44.48287267 | 70.78444169 | ENSG00000105711 |
| TMEM169 | 0.674752979 | 1.596323424 | 1.91E-02 | 6.20E-02 | 48.58714919 | 77.35970764 | ENSG00000163449 |
| PRELID2 | 0.674712193 | 1.596278295 | 2.12E-02 | 6.77E-02 | 47.86186403 | 76.33141977 | ENSG00000186314 |
| GNA14 | 0.674449273 | 1.595987413 | 6.41E-07 | 7.39E-06 | 291.0801008 | 463.8279572 | ENSG00000156049 |
| FES | 0.673887877 | 1.595366487 | 5.27E-04 | 3.01E-03 | 100.9217913 | 161.2095148 | ENSG00000182511 |
| CDH6 | 0.673340138 | 1.594760899 | 4.47E-02 | 1.23E-01 | 34.19417886 | 54.7096717 | ENSG00000113361 |
| KNL1 | 0.672327333 | 1.593641733 | 5.83E-08 | 8.13E-07 | 351.8509166 | 560.4304013 | ENSG00000137812 |
| NUAK2 | 0.671540752 | 1.59277309 | 2.64E-06 | 2.71E-05 | 211.449309 | 337.0846928 | ENSG00000163545 |
| FYTTD1 | 0.671170281 | 1.592364133 | 8.90E-11 | 1.96E-09 | 519.8862903 | 827.1694015 | ENSG00000122068 |
| LCAT | 0.670998496 | 1.592174538 | 6.21E-04 | 3.46E-03 | 104.602621 | 166.6329084 | ENSG00000213398 |
| ZNF124 | 0.670867058 | 1.592029488 | 1.35E-04 | 9.05E-04 | 153.7076912 | 245.1636974 | ENSG00000196418 |
| COQ5 | 0.669821768 | 1.590876417 | 1.52E-10 | 3.24E-09 | 489.1837595 | 777.6292815 | ENSG00000110871 |
| AKAP9 | 0.6697631 | 1.590811725 | 1.08E-09 | 2.01E-08 | 486.4932376 | 773.6804343 | ENSG00000127914 |
| CUX1 | 0.669751773 | 1.590799235 | 6.14E-15 | 2.34E-13 | 1065.633794 | 1694.742558 | ENSG00000257923 |
| ZNF697 | 0.669561381 | 1.590589311 | 1.56E-10 | 3.31E-09 | 648.1315234 | 1030.176926 | ENSG00000143067 |
| KCTD13 | 0.669410796 | 1.590423298 | 7.51E-05 | 5.39E-04 | 164.4392771 | 261.9312908 | ENSG00000174943 |
| ZNF652 | 0.669363163 | 1.590370789 | 8.82E-06 | 7.97E-05 | 212.8377105 | 338.5299159 | ENSG00000198740 |
| CENPF | 0.668816534 | 1.58976832 | 1.26E-18 | 7.07E-17 | 2814.319726 | 4473.568281 | ENSG00000117724 |
| ZNF81 | 0.668364782 | 1.589270592 | 8.94E-04 | 4.79E-03 | 109.2275847 | 173.3339087 | ENSG00000197779 |
| CDC42BPA | 0.6682704 | 1.589166625 | 2.74E-12 | 7.52E-11 | 784.9235349 | 1247.243477 | ENSG00000143776 |
| PEAK1 | 0.66772682 | 1.588567971 | 1.37E-10 | 2.94E-09 | 835.288205 | 1326.167692 | ENSG00000173517 |
| SIRT5 | 0.667593528 | 1.588421208 | 4.24E-08 | 6.10E-07 | 313.2319147 | 497.0050525 | ENSG00000124523 |
| ICAM1 | 0.666979823 | 1.587745657 | 7.08E-03 | 2.77E-02 | 63.46244273 | 100.5468396 | ENSG00000090339 |
| NPAS1 | 0.666685555 | 1.587421835 | 6.59E-04 | 3.65E-03 | 115.6567302 | 183.4804436 | ENSG00000130751 |
| DUSP5 | 0.665839651 | 1.586491346 | 6.25E-15 | 2.37E-13 | 869.4062799 | 1378.568085 | ENSG00000138166 |
| NUFIP2 | 0.665764823 | 1.586409062 | 4.75E-18 | 2.53E-16 | 2047.416166 | 3247.653259 | ENSG00000108256 |
| SNX30 | 0.665422742 | 1.586032949 | 3.93E-08 | 5.67E-07 | 386.1528365 | 611.679025 | ENSG00000148158 |
| ZNF641 | 0.664513982 | 1.585034214 | 1.07E-04 | 7.39E-04 | 128.8434676 | 204.2119511 | ENSG00000167528 |
| MYO9A | 0.663526665 | 1.583949857 | 6.81E-06 | 6.39E-05 | 302.4120556 | 478.9736136 | ENSG00000066933 |
| TMEM200B | 0.663524155 | 1.583947101 | 1.56E-05 | 1.33E-04 | 190.6932929 | 302.2748708 | ENSG00000253304 |
| NMRK1 | 0.663307365 | 1.583709104 | 1.95E-02 | 6.33E-02 | 50.6694622 | 80.53809138 | ENSG00000106733 |
| CA5B | 0.663001309 | 1.583373169 | 1.38E-03 | 6.90E-03 | 93.97431879 | 148.6859203 | ENSG00000169239 |
| CHAC1 | 0.662625029 | 1.58296025 | 5.35E-12 | 1.43E-10 | 735.2207412 | 1164.176909 | ENSG00000128965 |
| BAIAP3 | 0.662497746 | 1.582820599 | 9.00E-06 | 8.13E-05 | 211.3638996 | 334.9861636 | ENSG00000007516 |
| ADPRM | 0.662090022 | 1.582373337 | 2.26E-03 | 1.05E-02 | 81.29062749 | 128.6414535 | ENSG00000170222 |
| ADCY7 | 0.662001706 | 1.582276474 | 2.26E-03 | 1.05E-02 | 87.77915298 | 138.9067779 | ENSG00000121281 |
| NPTX2 | 0.661580542 | 1.581814629 | 1.31E-11 | 3.26E-10 | 741.2225461 | 1172.905368 | ENSG00000106236 |
| MGA | 0.66024904 | 1.580355403 | 1.88E-11 | 4.57E-10 | 953.7160259 | 1506.49264 | ENSG00000174197 |
| EVI5 | 0.65984408 | 1.579911865 | 2.23E-03 | 1.04E-02 | 97.1149184 | 153.0422815 | ENSG00000067208 |
| PRKAB2 | 0.659444057 | 1.579473856 | 9.45E-11 | 2.08E-09 | 694.1484141 | 1095.220548 | ENSG00000131791 |
| AFAP1L1 | 0.659390952 | 1.579415717 | 2.38E-04 | 1.50E-03 | 154.5001585 | 243.5755205 | ENSG00000157510 |
| PTPRG | 0.658938515 | 1.578920481 | 1.98E-13 | 6.32E-12 | 1695.327984 | 2675.823232 | ENSG00000144724 |
| RHBDD1 | 0.65841146 | 1.578343765 | 5.31E-08 | 7.47E-07 | 332.525447 | 524.1716287 | ENSG00000144468 |
| HRH2 | 0.658277172 | 1.578196858 | 9.79E-06 | 8.78E-05 | 299.4221911 | 472.67269 | ENSG00000113749 |
| ZNF425 | 0.657668808 | 1.577531494 | 6.83E-05 | 4.95E-04 | 177.534464 | 279.8517584 | ENSG00000204947 |
| TTC3 | 0.657550914 | 1.577402588 | 4.23E-12 | 1.14E-10 | 729.8583609 | 1150.944944 | ENSG00000182670 |
| PCDHAC2 | 0.657325573 | 1.577156225 | 6.31E-04 | 3.52E-03 | 124.6487456 | 196.1532185 | ENSG00000243232 |
| CBX4 | 0.657081301 | 1.576889209 | 1.25E-03 | 6.35E-03 | 103.1342142 | 162.8855493 | ENSG00000141582 |
| STEAP1B | 0.656375842 | 1.576118319 | 7.27E-03 | 2.82E-02 | 72.92613059 | 114.9160688 | ENSG00000105889 |
| GFPT1 | 0.656009526 | 1.575718176 | 1.13E-13 | 3.69E-12 | 1811.21223 | 2852.751011 | ENSG00000198380 |
| ACSF2 | 0.655765826 | 1.575452029 | 1.96E-06 | 2.07E-05 | 244.1125593 | 384.5710171 | ENSG00000167107 |
| ITGA3 | 0.654971763 | 1.574585135 | 2.74E-15 | 1.09E-13 | 782.4428197 | 1232.09567 | ENSG00000005884 |
| MT1F | 0.654444075 | 1.574009311 | 3.31E-07 | 4.03E-06 | 372.7812545 | 585.9249736 | ENSG00000198417 |
| CDK6 | 0.654133135 | 1.573670106 | 3.38E-06 | 3.41E-05 | 342.1419066 | 538.275419 | ENSG00000105810 |
| MAVS | 0.65407474 | 1.573606411 | 1.76E-17 | 8.79E-16 | 1036.429037 | 1630.842098 | ENSG00000088888 |
| IQCG | 0.653596087 | 1.57308441 | 1.68E-02 | 5.60E-02 | 67.94509755 | 106.7820693 | ENSG00000114473 |
| USP32 | 0.653441295 | 1.572915638 | 1.83E-15 | 7.42E-14 | 1582.214364 | 2488.033439 | ENSG00000170832 |
| GCNT1 | 0.653105304 | 1.572549363 | 1.71E-05 | 1.45E-04 | 180.8051735 | 284.354529 | ENSG00000187210 |
| FAT1 | 0.652892201 | 1.572317096 | 1.32E-10 | 2.85E-09 | 2422.489502 | 3808.550866 | ENSG00000083857 |
| ICAM3 | 0.651910318 | 1.571247357 | 1.07E-02 | 3.90E-02 | 62.68902976 | 98.33348587 | ENSG00000076662 |
| SHC3 | 0.649874656 | 1.56903187 | 1.55E-06 | 1.66E-05 | 240.7591164 | 377.7730871 | ENSG00000148082 |
| PDLIM5 | 0.649143033 | 1.56823638 | 2.29E-12 | 6.35E-11 | 783.7524566 | 1228.421148 | ENSG00000163110 |
| ATF7 | 0.648859146 | 1.56792782 | 1.14E-03 | 5.90E-03 | 100.2469191 | 157.1648457 | ENSG00000170653 |
| MPPE1 | 0.64879533 | 1.567858465 | 7.50E-03 | 2.90E-02 | 97.18307006 | 152.317619 | ENSG00000154889 |
| MEIOC | 0.648489356 | 1.567525982 | 2.69E-03 | 1.21E-02 | 99.37910699 | 155.4544068 | ENSG00000180336 |
| FCSK | 0.648449527 | 1.567482707 | 1.04E-04 | 7.21E-04 | 143.9220587 | 225.4192402 | ENSG00000157353 |
| FAM49A | 0.648154293 | 1.56716197 | 1.65E-04 | 1.08E-03 | 143.3956364 | 224.6361862 | ENSG00000197872 |
| MTMR10 | 0.647536743 | 1.566491285 | 1.06E-05 | 9.42E-05 | 195.6583686 | 306.4272274 | ENSG00000166912 |
| SLC35F6 | 0.646452874 | 1.565314851 | 1.24E-04 | 8.43E-04 | 176.9409951 | 276.8217191 | ENSG00000213699 |
| RAPH1 | 0.646400322 | 1.565257834 | 2.07E-03 | 9.80E-03 | 99.66357466 | 156.0181979 | ENSG00000173166 |
| KLHL15 | 0.645675237 | 1.564471348 | 3.27E-07 | 3.99E-06 | 312.3661771 | 487.9986754 | ENSG00000174010 |
| IL1RAP | 0.645446579 | 1.564223409 | 8.05E-04 | 4.37E-03 | 109.1124853 | 170.4017465 | ENSG00000196083 |

| | | | | | | | |
|---|---|---|---|---|---|---|---|
| FUOM | 0.645176586 | 1.5639307 | 1.01E-05 | 9.05E-05 | 257.5957907 | 402.6408337 | ENSG00000148803 |
| DFFA | 0.642642281 | 1.561185838 | 4.79E-16 | 2.04E-14 | 1493.557492 | 2331.702324 | ENSG00000160049 |
| ZDHHC1 | 0.642170886 | 1.56067581 | 1.89E-02 | 6.17E-02 | 51.03333371 | 79.62221468 | ENSG00000159714 |
| ABRAXAS1 | 0.641315972 | 1.559751257 | 1.45E-06 | 1.56E-05 | 285.0469595 | 444.5356444 | ENSG00000163322 |
| RICTOR | 0.641308495 | 1.559743173 | 6.73E-05 | 4.90E-04 | 181.7560273 | 282.9648057 | ENSG00000164327 |
| UEVLD | 0.641163141 | 1.559586034 | 8.12E-03 | 3.10E-02 | 76.26448804 | 118.7976361 | ENSG00000151116 |
| PPL | 0.640700979 | 1.559086507 | 9.09E-16 | 3.81E-14 | 966.9892101 | 1507.592158 | ENSG00000118898 |
| ATF6 | 0.640597248 | 1.558974411 | 2.66E-10 | 5.42E-09 | 457.066572 | 712.3716212 | ENSG00000118217 |
| GPC6 | 0.640465758 | 1.558832329 | 1.58E-07 | 2.03E-06 | 340.619577 | 530.2552648 | ENSG00000183098 |
| PCYOX1L | 0.640366778 | 1.558725385 | 5.72E-10 | 1.10E-08 | 459.6476093 | 715.89384 | ENSG00000145882 |
| TOB1 | 0.640023211 | 1.558354231 | 1.10E-07 | 1.46E-06 | 330.3224797 | 514.8854304 | ENSG00000141232 |
| SLC16A10 | 0.640004975 | 1.558334533 | 4.93E-06 | 4.76E-05 | 207.0756729 | 322.7636292 | ENSG00000112394 |
| KIF13B | 0.639720425 | 1.558027206 | 8.00E-06 | 7.35E-05 | 269.921323 | 420.070977 | ENSG00000197892 |
| THSD7A | 0.63941935 | 1.557702096 | 4.69E-02 | 1.27E-01 | 41.18505143 | 64.41719402 | ENSG00000005108 |
| ZBTB5 | 0.638862772 | 1.557101264 | 3.03E-05 | 2.43E-04 | 323.6490346 | 504.425875 | ENSG00000168795 |
| RAB12 | 0.638551307 | 1.556765137 | 4.44E-06 | 4.36E-05 | 313.5292054 | 487.6632822 | ENSG00000206418 |
| TMEM192 | 0.638361226 | 1.55656004 | 8.02E-06 | 7.35E-05 | 239.8332021 | 372.6543376 | ENSG00000170088 |
| TCAF1 | 0.63832832 | 1.556524538 | 2.06E-07 | 2.60E-06 | 348.1955528 | 541.3857123 | ENSG00000198420 |
| GRID2 | 0.638243792 | 1.556433342 | 1.83E-09 | 3.27E-08 | 473.5081139 | 736.590943 | ENSG00000152208 |
| B3GALT5 | 0.637595361 | 1.555733949 | 3.20E-12 | 8.74E-11 | 788.2439582 | 1225.913049 | ENSG00000183778 |
| FAM160B2 | 0.637294417 | 1.555409458 | 6.29E-08 | 8.72E-07 | 330.3607686 | 514.0818935 | ENSG00000158863 |
| H6PD | 0.636512484 | 1.554566663 | 8.45E-07 | 9.53E-06 | 279.7337218 | 435.1466164 | ENSG00000049239 |
| AKR1E2 | 0.63590381 | 1.553910929 | 3.27E-02 | 9.59E-02 | 44.17078955 | 68.73755063 | ENSG00000165568 |
| AOC2 | 0.635651027 | 1.553638684 | 1.52E-02 | 5.15E-02 | 60.66175975 | 94.22786917 | ENSG00000131480 |
| MFSD14C | 0.635509458 | 1.553486235 | 4.68E-08 | 6.68E-07 | 508.946311 | 790.490708 | ENSG00000196312 |
| A2ML1 | 0.635445969 | 1.553417872 | 1.74E-04 | 1.14E-03 | 152.1773641 | 236.1004493 | ENSG00000166535 |
| SMG1P3 | 0.635405698 | 1.55337451 | 4.52E-02 | 1.24E-01 | 58.62056167 | 90.94307172 | ENSG00000180747 |
| RAB25 | 0.633776585 | 1.551621406 | 9.45E-09 | 1.52E-07 | 420.6787163 | 652.1754096 | ENSG00000132698 |
| CACNA1B | 0.633536021 | 1.551362701 | 1.10E-07 | 1.47E-06 | 362.4265586 | 562.6194104 | ENSG00000148408 |
| ZNF79 | 0.63312887 | 1.550924945 | 2.76E-03 | 1.24E-02 | 102.2134336 | 158.9268513 | ENSG00000196152 |
| MARVELD2 | 0.632797839 | 1.550569121 | 5.71E-07 | 6.67E-06 | 290.6215219 | 450.1532547 | ENSG00000152939 |
| CHD9 | 0.631719792 | 1.549410898 | 6.04E-09 | 1.00E-07 | 472.2371946 | 731.2273264 | ENSG00000177200 |
| CYB5RL | 0.631646459 | 1.549332143 | 1.54E-05 | 1.32E-04 | 203.1983632 | 315.2180373 | ENSG00000215883 |
| APP | 0.630647683 | 1.548259912 | 7.72E-29 | 1.03E-26 | 19600.40333 | 30345.7315 | ENSG00000142192 |
| EPHA2 | 0.630415259 | 1.548010502 | 6.77E-16 | 2.86E-14 | 1180.218197 | 1827.203979 | ENSG00000142627 |
| SLC16A5 | 0.630098303 | 1.547670446 | 4.64E-06 | 4.53E-05 | 265.5753099 | 411.64819 | ENSG00000170190 |
| KLHL28 | 0.629874264 | 1.547430123 | 2.05E-03 | 9.69E-03 | 105.2958333 | 162.6436763 | ENSG00000179454 |
| SLC2A1 | 0.629738507 | 1.547284518 | 1.48E-20 | 9.96E-19 | 4716.173946 | 7295.666243 | ENSG00000117394 |
| TRAF3IP2 | 0.629716437 | 1.547260848 | 1.11E-09 | 2.05E-08 | 447.5100559 | 692.1098266 | ENSG00000056972 |
| PKP3 | 0.629676097 | 1.547217585 | 7.23E-07 | 8.28E-06 | 336.2740298 | 520.6396708 | ENSG00000184363 |
| XPC | 0.629440629 | 1.546965078 | 4.56E-10 | 8.91E-09 | 662.5632636 | 1025.059317 | ENSG00000154767 |
| SMAD3 | 0.629255428 | 1.546766505 | 6.81E-08 | 9.35E-07 | 434.9329255 | 672.6444496 | ENSG00000166949 |
| INKA2 | 0.628414311 | 1.545864975 | 3.34E-05 | 2.65E-04 | 293.4273538 | 453.2443222 | ENSG00000197852 |
| TFRC | 0.628264008 | 1.545703932 | 5.86E-17 | 2.77E-15 | 3046.886472 | 4708.384741 | ENSG00000072274 |
| SNHG29 | 0.62800417 | 1.545425566 | 2.02E-18 | 1.12E-16 | 3273.871328 | 5059.022858 | ENSG00000175061 |
| CEL | 0.627937831 | 1.545354505 | 4.78E-02 | 1.29E-01 | 37.88260009 | 58.55012136 | ENSG00000170835 |
| MELTF | 0.627220031 | 1.544585819 | 8.15E-03 | 3.11E-02 | 82.41400654 | 126.8659015 | ENSG00000163975 |
| ZNF300 | 0.625689489 | 1.54294805 | 7.17E-05 | 5.18E-04 | 189.3164366 | 292.3833258 | ENSG00000145908 |
| FBLN2 | 0.625478249 | 1.542722148 | 6.52E-15 | 2.47E-13 | 1122.42413 | 1730.57988 | ENSG00000163520 |
| EPS8L2 | 0.625205382 | 1.54243039 | 4.92E-10 | 9.58E-09 | 563.1401553 | 868.8527644 | ENSG00000177106 |
| MORN1 | 0.62514165 | 1.542362254 | 4.15E-03 | 1.76E-02 | 91.01571511 | 140.3897034 | ENSG00000116151 |
| LOXL2 | 0.624590663 | 1.541773314 | 3.76E-14 | 1.31E-12 | 906.6400372 | 1397.258762 | ENSG00000134013 |
| JUND | 0.623513842 | 1.540622971 | 2.02E-11 | 4.88E-10 | 1011.726477 | 1558.159778 | ENSG00000130522 |
| CTSB | 0.622111837 | 1.539126528 | 5.20E-14 | 1.78E-12 | 1030.791351 | 1585.560931 | ENSG00000164733 |
| GOLGA4 | 0.621583517 | 1.538562998 | 2.30E-12 | 6.37E-11 | 875.1381465 | 1346.869192 | ENSG00000144674 |
| RIPOR1 | 0.61932432 | 1.536155561 | 1.47E-04 | 9.73E-04 | 187.6701981 | 288.3535088 | ENSG00000039523 |
| AFAP1 | 0.619043396 | 1.535856467 | 1.36E-03 | 6.83E-03 | 117.3085084 | 180.4143913 | ENSG00000196526 |
| GOLGA3 | 0.618728124 | 1.535520874 | 5.46E-09 | 9.13E-08 | 826.6825351 | 1270.049647 | ENSG00000090615 |
| LRRK1 | 0.618570331 | 1.535352937 | 8.24E-03 | 3.14E-02 | 102.3989924 | 157.2642517 | ENSG00000154237 |
| MAFF | 0.618413677 | 1.535186231 | 1.12E-03 | 5.81E-03 | 126.0078632 | 193.3096153 | ENSG00000185022 |
| RBM33 | 0.616798039 | 1.533467976 | 1.55E-15 | 6.36E-14 | 1321.13048 | 2025.71696 | ENSG00000184863 |
| CAP2 | 0.61661806 | 1.533276685 | 5.73E-07 | 6.68E-06 | 294.7213638 | 451.836824 | ENSG00000112186 |
| DHRS7B | 0.615958011 | 1.532575355 | 9.87E-06 | 8.83E-05 | 232.6068841 | 356.8151571 | ENSG00000109016 |
| TEP1 | 0.615707901 | 1.532309686 | 7.91E-06 | 7.28E-05 | 341.5419436 | 523.118773 | ENSG00000129566 |
| DENND3 | 0.615284062 | 1.531859585 | 1.16E-02 | 4.16E-02 | 113.1600577 | 173.7316123 | ENSG00000105339 |
| ZNF106 | 0.615227013 | 1.531799012 | 1.42E-10 | 3.04E-09 | 1538.832666 | 2356.70336 | ENSG00000103994 |
| SAT1 | 0.614323415 | 1.530839907 | 7.69E-09 | 1.26E-07 | 666.6552958 | 1019.919048 | ENSG00000130066 |
| SPINK5 | 0.614214932 | 1.5307248 | 2.65E-02 | 8.12E-02 | 60.7774605 | 93.25754324 | ENSG00000133710 |
| NID2 | 0.613707271 | 1.530186257 | 5.69E-05 | 4.22E-04 | 206.3350092 | 315.5833344 | ENSG00000087303 |
| LIFR | 0.613547312 | 1.530016607 | 4.21E-05 | 3.23E-04 | 197.3391377 | 301.5993927 | ENSG00000113594 |
| NCEH1 | 0.612940817 | 1.529373537 | 1.67E-02 | 5.57E-02 | 71.30948442 | 108.6930001 | ENSG00000144959 |
| ATF7IP | 0.612548328 | 1.528957524 | 2.86E-18 | 1.54E-16 | 1847.555108 | 2823.706342 | ENSG00000171681 |
| GALNT12 | 0.61151421 | 1.527861967 | 1.52E-05 | 1.31E-04 | 316.9990708 | 484.1836067 | ENSG00000119514 |
| BCL6 | 0.611050715 | 1.527371189 | 4.28E-02 | 1.19E-01 | 49.8042283 | 76.1835536 | ENSG00000113916 |
| MALAT1 | 0.610324736 | 1.526602793 | 3.90E-11 | 9.04E-10 | 1022.211673 | 1560.911748 | ENSG00000251562 |

| | | | | | | | |
|---|---|---|---|---|---|---|---|
| KLHL8 | 0.610311859 | 1.526589167 | 3.98E-05 | 3.08E-04 | 209.4072391 | 319.1688111 | ENSG00000145332 |
| NSUN7 | 0.610267275 | 1.526541992 | 2.05E-04 | 1.31E-03 | 157.9266013 | 241.0184518 | ENSG00000179299 |
| ACTG1 | 0.609527056 | 1.525758953 | 8.15E-30 | 1.15E-27 | 53058.21798 | 80953.41052 | ENSG00000184009 |
| PLGRKT | 0.609499058 | 1.525729343 | 2.56E-04 | 1.60E-03 | 172.6279942 | 263.2731346 | ENSG00000107020 |
| SIAE | 0.608363859 | 1.52452928 | 5.51E-03 | 2.24E-02 | 93.30250575 | 141.8961376 | ENSG00000110013 |
| NIN | 0.608354819 | 1.524519727 | 9.14E-07 | 1.03E-05 | 416.5699074 | 634.6259647 | ENSG00000100503 |
| NIPA1 | 0.608348044 | 1.524512568 | 1.04E-08 | 1.67E-07 | 426.5099204 | 650.5224947 | ENSG00000170113 |
| GDPD5 | 0.608313772 | 1.524476353 | 2.50E-03 | 1.15E-02 | 109.5440274 | 166.7186245 | ENSG00000158555 |
| TMEM241 | 0.607772913 | 1.523904942 | 2.14E-03 | 1.01E-02 | 107.5101353 | 163.8827936 | ENSG00000134490 |
| KLF11 | 0.606390473 | 1.522445383 | 6.91E-05 | 5.01E-04 | 205.0179162 | 311.5240281 | ENSG00000172059 |
| NEDD4 | 0.606119634 | 1.522159599 | 1.37E-04 | 9.19E-04 | 184.1245894 | 279.9458366 | ENSG00000069869 |
| IRAK1 | 0.605458953 | 1.521462687 | 6.98E-15 | 2.63E-13 | 1030.999655 | 1569.191098 | ENSG00000184216 |
| DENND5B | 0.605152888 | 1.521139946 | 8.36E-05 | 5.94E-04 | 222.7249581 | 338.5150469 | ENSG00000170456 |
| AFF4 | 0.605046321 | 1.521027589 | 5.19E-10 | 1.01E-08 | 960.2713949 | 1460.121879 | ENSG00000072364 |
| BLOC1S3 | 0.604908784 | 1.520882591 | 1.07E-03 | 5.58E-03 | 116.7689547 | 177.7276375 | ENSG00000189114 |
| GALNT17 | 0.604452417 | 1.520401567 | 1.95E-16 | 8.66E-15 | 1880.118005 | 2857.867484 | ENSG00000185274 |
| ARMCX3 | 0.604174877 | 1.520109106 | 1.11E-07 | 1.47E-06 | 361.8807138 | 550.0685308 | ENSG00000102401 |
| OCEL1 | 0.603783039 | 1.519696298 | 1.29E-04 | 8.71E-04 | 220.2465281 | 334.1743989 | ENSG00000099330 |
| RGP1 | 0.603357114 | 1.519247707 | 4.79E-09 | 8.09E-08 | 514.2791142 | 781.288225 | ENSG00000107185 |
| SDK1 | 0.602791685 | 1.518652391 | 4.25E-06 | 4.19E-05 | 397.8822239 | 603.6049141 | ENSG00000146555 |
| CDA | 0.601741925 | 1.517547764 | 5.41E-03 | 2.20E-02 | 86.74720407 | 131.6006457 | ENSG00000158825 |
| CLGN | 0.600644044 | 1.516393359 | 3.73E-04 | 2.22E-03 | 162.2622567 | 245.7348227 | ENSG00000153132 |
| SCRN1 | 0.600181887 | 1.515907671 | 1.49E-19 | 9.17E-18 | 2903.275369 | 4400.093692 | ENSG00000136193 |
| AEBP2 | 0.599870762 | 1.515580794 | 3.48E-10 | 6.94E-09 | 785.0897084 | 1189.672363 | ENSG00000139154 |
| KIAA0556 | 0.599823671 | 1.515531324 | 3.25E-06 | 3.29E-05 | 304.8207631 | 462.2420107 | ENSG00000047578 |
| SLC22A18 | 0.599812559 | 1.515519651 | 1.05E-02 | 3.82E-02 | 73.84228105 | 112.0483585 | ENSG00000110628 |
| ZSWIM9 | 0.599556937 | 1.51525115 | 1.16E-06 | 1.28E-05 | 368.341714 | 557.632224 | ENSG00000185453 |
| ZNF18 | 0.598830511 | 1.514488382 | 1.62E-04 | 1.06E-03 | 158.7367922 | 240.5428933 | ENSG00000154957 |
| SLC45A1 | 0.598004544 | 1.513621561 | 1.28E-02 | 4.51E-02 | 71.49601273 | 108.2440251 | ENSG00000162426 |
| ITGB5 | 0.597748595 | 1.513353052 | 6.36E-24 | 5.51E-22 | 6817.6943 | 10316.76943 | ENSG00000082781 |
| CCND2 | 0.597744243 | 1.513348487 | 6.96E-10 | 1.32E-08 | 1291.909029 | 1954.914692 | ENSG00000118971 |
| CLDN11 | 0.596886945 | 1.512449472 | 6.11E-03 | 2.45E-02 | 82.60634501 | 124.9385728 | ENSG00000013297 |
| NBPF26 | 0.596137438 | 1.51166393 | 1.86E-02 | 6.07E-02 | 67.34172927 | 101.785622 | ENSG00000273136 |
| GEMIN8 | 0.596088589 | 1.511612746 | 6.08E-04 | 3.40E-03 | 163.6794764 | 247.4939056 | ENSG00000046647 |
| TNFRSF12A | 0.59598102 | 1.511500042 | 1.58E-11 | 3.89E-10 | 910.9801871 | 1377.153476 | ENSG00000006327 |
| CAVIN3 | 0.595957743 | 1.511475656 | 1.71E-03 | 8.30E-03 | 163.1946575 | 246.7327283 | ENSG00000170955 |
| ZNF391 | 0.595196931 | 1.510678783 | 9.72E-04 | 5.14E-03 | 127.5877314 | 192.4901815 | ENSG00000124613 |
| KCNMB4 | 0.594850114 | 1.510315667 | 7.26E-08 | 9.91E-07 | 363.4750662 | 549.0034654 | ENSG00000135643 |
| EFR3A | 0.594414091 | 1.509859275 | 5.19E-11 | 1.18E-09 | 655.9237175 | 990.3850564 | ENSG00000132294 |
| ASAH2B | 0.593983165 | 1.509408355 | 3.27E-02 | 9.58E-02 | 59.98332462 | 90.6261147 | ENSG00000204147 |
| ANXA2 | 0.593201327 | 1.508590584 | 1.18E-11 | 2.97E-10 | 818.0717843 | 1233.357157 | ENSG00000182718 |
| EMILIN1 | 0.592930684 | 1.508307607 | 6.20E-04 | 3.46E-03 | 143.3188858 | 216.5190939 | ENSG00000138080 |
| FOXK1 | 0.591991947 | 1.507326496 | 2.70E-06 | 2.76E-05 | 370.9545909 | 558.6608383 | ENSG00000164916 |
| CCDC8 | 0.591984508 | 1.507318723 | 9.48E-11 | 2.08E-09 | 681.2615078 | 1026.559069 | ENSG00000169515 |
| P4HA1 | 0.591791021 | 1.507116583 | 3.73E-08 | 5.40E-07 | 504.279225 | 759.2642814 | ENSG00000122884 |
| CD99L2 | 0.591717806 | 1.507040101 | 2.12E-07 | 2.68E-06 | 483.3056152 | 728.3296089 | ENSG00000102181 |
| CDKL5 | 0.591518474 | 1.506831893 | 4.40E-05 | 3.36E-04 | 279.0247022 | 420.0295246 | ENSG00000008086 |
| CNNM2 | 0.591506225 | 1.5068191 | 5.45E-08 | 7.66E-07 | 525.7042805 | 791.5213662 | ENSG00000148842 |
| HECTD1 | 0.591258388 | 1.506560269 | 3.61E-11 | 8.42E-10 | 1828.926759 | 2754.917152 | ENSG00000092148 |
| GABARAPL1 | 0.591233319 | 1.50653409 | 5.67E-12 | 1.50E-10 | 1149.412202 | 1731.107244 | ENSG00000139112 |
| LRIG2 | 0.59111247 | 1.506407899 | 3.89E-07 | 4.69E-06 | 400.6820199 | 603.1920638 | ENSG00000198799 |
| TSC22D3 | 0.590813989 | 1.506096268 | 1.78E-09 | 3.20E-08 | 604.2772894 | 909.5941738 | ENSG00000157514 |
| OGDHL | 0.590784024 | 1.506064987 | 2.05E-07 | 2.60E-06 | 594.559523 | 895.4446229 | ENSG00000197444 |
| NDUFAF3 | 0.590783149 | 1.506064074 | 7.28E-04 | 3.99E-03 | 134.4144672 | 202.5126855 | ENSG00000178057 |
| KLC3 | 0.590639615 | 1.505914243 | 1.29E-02 | 4.53E-02 | 82.37453754 | 124.0190089 | ENSG00000104892 |
| ZFYVE28 | 0.59059319 | 1.505865784 | 1.76E-03 | 8.51E-03 | 117.6938769 | 177.3050312 | ENSG00000159733 |
| RUNX1T1 | 0.590173852 | 1.505428148 | 3.25E-04 | 1.97E-03 | 179.4304819 | 269.7674223 | ENSG00000079102 |
| ANK1 | 0.590091104 | 1.505341805 | 1.96E-05 | 1.64E-04 | 267.0972865 | 402.1873902 | ENSG00000029534 |
| CTNNA2 | 0.589145959 | 1.504355941 | 6.37E-05 | 4.67E-04 | 192.5996533 | 289.534999 | ENSG00000066032 |
| NIPSNAP2 | 0.587935819 | 1.503094609 | 6.84E-10 | 1.30E-08 | 549.9464276 | 826.5932035 | ENSG00000146729 |
| CLIP2 | 0.587915142 | 1.503073066 | 2.05E-17 | 1.01E-15 | 1838.730618 | 2763.595378 | ENSG00000106665 |
| TFPI | 0.58789797 | 1.503055175 | 1.28E-02 | 4.50E-02 | 74.26700572 | 111.9136008 | ENSG00000003436 |
| USP34 | 0.587864053 | 1.50301984 | 3.30E-11 | 7.72E-10 | 1287.88024 | 1935.400483 | ENSG00000115464 |
| ZNF77 | 0.587252519 | 1.50238287 | 2.83E-02 | 8.57E-02 | 70.36168977 | 105.9730846 | ENSG00000175691 |
| ZNF264 | 0.587245806 | 1.50237588 | 1.20E-03 | 6.15E-03 | 185.0775929 | 277.9360411 | ENSG00000083844 |
| PJA2 | 0.587199499 | 1.502327658 | 1.04E-12 | 3.06E-11 | 1228.110247 | 1843.89053 | ENSG00000198961 |
| SFMBT2 | 0.586940645 | 1.502058129 | 1.94E-03 | 9.27E-03 | 137.6944293 | 206.4245319 | ENSG00000198879 |
| ZNF311 | 0.585952191 | 1.501029355 | 1.41E-02 | 4.87E-02 | 67.94459388 | 101.8186007 | ENSG00000197935 |
| KAZN | 0.585842634 | 1.500915372 | 1.36E-07 | 1.77E-06 | 384.5828297 | 577.3910492 | ENSG00000189337 |
| HELZ | 0.585127955 | 1.500172036 | 5.55E-07 | 6.51E-06 | 400.5530901 | 600.6617934 | ENSG00000198265 |
| FEN1 | -0.514867678 | 0.69985712 | 1.99E-12 | 5.61E-11 | 2272.312612 | 1590.962393 | ENSG00000168496 |
| CARHSP1 | -0.514905635 | 0.699838707 | 2.66E-10 | 5.42E-09 | 1516.047952 | 1061.124519 | ENSG00000153048 |
| HEATR1 | -0.515183164 | 0.699704093 | 2.98E-11 | 7.04E-10 | 2771.707548 | 1938.444506 | ENSG00000119285 |
| KATNB1 | -0.515416823 | 0.699590778 | 2.79E-10 | 5.65E-09 | 1535.590677 | 1075.01537 | ENSG00000140854 |
| DUSP2 | -0.515455395 | 0.699572074 | 1.29E-02 | 4.52E-02 | 136.1142755 | 95.38711597 | ENSG00000158050 |

| | | | | | | | |
|---|---|---|---|---|---|---|---|
| DNAJB11 | -0.51584983 | 0.699380836 | 3.38E-10 | 6.76E-09 | 1273.569319 | 890.8208154 | ENSG00000090520 |
| CYB5B | -0.515870591 | 0.699370772 | 9.16E-13 | 2.74E-11 | 2647.974569 | 1851.74498 | ENSG00000103018 |
| PRXL2A | -0.516416015 | 0.699106418 | 1.21E-09 | 2.21E-08 | 1351.498025 | 944.9172078 | ENSG00000122378 |
| DYRK3 | -0.516566054 | 0.699033715 | 2.71E-03 | 1.22E-02 | 203.4030363 | 142.1511191 | ENSG00000143479 |
| SLC8A1 | -0.517172575 | 0.698739898 | 2.24E-02 | 7.08E-02 | 119.2761053 | 83.25639819 | ENSG00000183023 |
| ERBB3 | -0.51729536 | 0.698680432 | 4.03E-09 | 6.86E-08 | 1205.135389 | 842.2527332 | ENSG00000065361 |
| DDR1 | -0.517350087 | 0.698653929 | 2.74E-17 | 1.34E-15 | 6246.460878 | 4364.337783 | ENSG00000204580 |
| TSPAN6 | -0.51745864 | 0.698601362 | 1.13E-09 | 2.08E-08 | 1658.835319 | 1158.02177 | ENSG00000000003 |
| HACD2 | -0.517604228 | 0.698530867 | 1.68E-05 | 1.42E-04 | 844.2882556 | 589.0068254 | ENSG00000206527 |
| C19orf57 | -0.518094877 | 0.698293342 | 4.34E-02 | 1.20E-01 | 86.59441712 | 60.50783236 | ENSG00000132016 |
| PCNX3 | -0.518929286 | 0.697889589 | 3.73E-11 | 8.68E-10 | 1794.327593 | 1252.789524 | ENSG00000197136 |
| CHRNA7 | -0.51896559 | 0.697872027 | 4.58E-02 | 1.25E-01 | 90.81557447 | 63.31663333 | ENSG00000175344 |
| RAB34 | -0.518981718 | 0.697864226 | 2.28E-15 | 9.13E-14 | 3277.757424 | 2287.075365 | ENSG00000109113 |
| B3GALT1 | -0.51902231 | 0.697844591 | 3.10E-03 | 1.37E-02 | 303.0924449 | 211.1406796 | ENSG00000172318 |
| DBI | -0.51939883 | 0.697662488 | 5.01E-13 | 1.53E-11 | 1917.610216 | 1337.823091 | ENSG00000155368 |
| NELFCD | -0.519566101 | 0.697581603 | 5.07E-11 | 1.15E-09 | 1358.196929 | 947.5671925 | ENSG00000101158 |
| NRSN2 | -0.519858625 | 0.697440174 | 2.43E-05 | 1.99E-04 | 615.713798 | 429.5815999 | ENSG00000125841 |
| ARF3 | -0.519906753 | 0.697416909 | 1.62E-15 | 6.63E-14 | 4208.400254 | 2934.084686 | ENSG00000134287 |
| DDX23 | -0.520739653 | 0.69701439 | 4.48E-16 | 1.92E-14 | 4662.667698 | 3250.09927 | ENSG00000174243 |
| ARHGAP22 | -0.520913503 | 0.696930403 | 1.37E-04 | 9.18E-04 | 405.9738583 | 283.3033936 | ENSG00000128805 |
| MACROH2A2 | -0.520942825 | 0.696916238 | 4.15E-10 | 8.13E-09 | 1773.381975 | 1235.21363 | ENSG00000099284 |
| NUDT10 | -0.520989216 | 0.696893829 | 1.04E-04 | 7.23E-04 | 557.3759076 | 388.8410582 | ENSG00000122824 |
| RNF38 | -0.521213399 | 0.696785545 | 4.70E-06 | 4.58E-05 | 633.6954839 | 441.1316823 | ENSG00000137075 |
| ALKBH8 | -0.52129744 | 0.696744957 | 9.90E-04 | 5.22E-03 | 300.6607672 | 208.9692248 | ENSG00000137760 |
| AXIN2 | -0.521502973 | 0.696645702 | 5.70E-04 | 3.22E-03 | 261.4177718 | 182.0782648 | ENSG00000168646 |
| SNHG26 | -0.521530652 | 0.696632337 | 1.94E-02 | 6.29E-02 | 134.7196577 | 94.00696574 | ENSG00000228649 |
| KREMEN2 | -0.522191546 | 0.696313285 | 9.07E-03 | 3.39E-02 | 195.0964462 | 135.6014574 | ENSG00000131650 |
| TM7SF2 | -0.5229759 | 0.695934821 | 8.36E-08 | 1.14E-06 | 1041.59275 | 724.6804843 | ENSG00000149809 |
| DHX38 | -0.523956232 | 0.695462084 | 2.67E-12 | 7.34E-11 | 2267.679267 | 1577.842731 | ENSG00000140829 |
| TAF1B | -0.525127193 | 0.694897843 | 1.25E-04 | 8.49E-04 | 349.981184 | 243.2919173 | ENSG00000115750 |
| LYPD6 | -0.525305013 | 0.694812198 | 3.91E-03 | 1.67E-02 | 177.5551869 | 123.4879019 | ENSG00000187123 |
| NCAPG | -0.525350829 | 0.694790133 | 1.08E-07 | 1.43E-06 | 1160.723103 | 806.0808474 | ENSG00000109805 |
| PNKD | -0.525834705 | 0.694557142 | 1.52E-05 | 1.31E-04 | 669.6752638 | 465.8613824 | ENSG00000127838 |
| SIX5 | -0.526365265 | 0.694301761 | 9.70E-03 | 3.58E-02 | 164.8277048 | 114.1959378 | ENSG00000177045 |
| ALG1 | -0.526641185 | 0.694168986 | 7.97E-06 | 7.33E-05 | 525.2576529 | 364.7659068 | ENSG00000033011 |
| GFOD2 | -0.526855923 | 0.69406567 | 6.66E-08 | 9.17E-07 | 887.907495 | 616.3911365 | ENSG00000141098 |
| MAPK4 | -0.526906773 | 0.694041207 | 2.25E-03 | 1.05E-02 | 199.5174359 | 138.486768 | ENSG00000141639 |
| LNP1 | -0.527415661 | 0.693796438 | 4.60E-03 | 1.92E-02 | 180.9673335 | 125.2869712 | ENSG00000206535 |
| LRP3 | -0.527950125 | 0.69353946 | 5.18E-09 | 8.69E-08 | 1745.673179 | 1210.567427 | ENSG00000130881 |
| ZNF880 | -0.527988012 | 0.693521247 | 5.61E-03 | 2.27E-02 | 166.4209748 | 115.4304834 | ENSG00000221923 |
| NUP88 | -0.528203867 | 0.693417491 | 1.29E-11 | 3.23E-10 | 1675.842181 | 1161.381048 | ENSG00000108559 |
| TEAD2 | -0.528504322 | 0.693273095 | 1.62E-12 | 4.64E-11 | 2654.619444 | 1840.620004 | ENSG00000074219 |
| FJX1 | -0.528592906 | 0.693230528 | 1.58E-05 | 1.35E-04 | 506.4591691 | 351.1304666 | ENSG00000179431 |
| ELOF1 | -0.52954552 | 0.692772938 | 1.59E-09 | 2.87E-08 | 1406.817225 | 975.0693936 | ENSG00000130165 |
| CBY1 | -0.529654087 | 0.692720807 | 1.61E-02 | 5.41E-02 | 151.6591347 | 104.905391 | ENSG00000100211 |
| ORC1 | -0.529748269 | 0.692675586 | 2.87E-11 | 6.80E-10 | 1397.994661 | 968.0629989 | ENSG00000085840 |
| CETN2 | -0.52975627 | 0.692671745 | 2.17E-05 | 1.80E-04 | 508.0024473 | 351.4922914 | ENSG00000147400 |
| SEPTIN2 | -0.529981894 | 0.692563426 | 1.50E-23 | 1.28E-21 | 8885.323071 | 6154.008887 | ENSG00000168385 |
| GNL1 | -0.530451766 | 0.692337901 | 6.23E-13 | 1.90E-11 | 2020.180681 | 1398.525244 | ENSG00000204590 |
| MXD4 | -0.530457516 | 0.692335142 | 1.06E-07 | 1.42E-06 | 1011.554268 | 700.2912761 | ENSG00000123933 |
| MED24 | -0.530690054 | 0.692223558 | 9.70E-16 | 4.03E-14 | 2683.471455 | 1857.844943 | ENSG00000008838 |
| MLH1 | -0.530834993 | 0.692154018 | 2.58E-09 | 4.49E-08 | 1159.510226 | 802.1176893 | ENSG00000076242 |
| NAA38 | -0.531105098 | 0.692024443 | 1.37E-04 | 9.16E-04 | 368.1948123 | 255.029097 | ENSG00000183011 |
| RBCK1 | -0.53113465 | 0.692010268 | 5.28E-08 | 7.44E-07 | 1247.080271 | 863.7175373 | ENSG00000125826 |
| KCNH2 | -0.532114233 | 0.691540556 | 2.11E-04 | 1.34E-03 | 436.2960566 | 301.7017299 | ENSG00000055118 |
| VILL | -0.532312973 | 0.691445299 | 4.18E-02 | 1.17E-01 | 96.17609083 | 66.75011012 | ENSG00000136059 |
| C1orf198 | -0.532952015 | 0.69113909 | 3.34E-08 | 4.88E-07 | 776.6937029 | 536.8439026 | ENSG00000119280 |
| IFT22 | -0.533002788 | 0.691114768 | 1.02E-03 | 5.38E-03 | 232.9521072 | 161.1606879 | ENSG00000128581 |
| MYD88 | -0.533112248 | 0.691062334 | 3.40E-06 | 3.42E-05 | 651.0615495 | 449.9740186 | ENSG00000172936 |
| SUSD5 | -0.533215657 | 0.691012801 | 4.17E-02 | 1.16E-01 | 92.41043051 | 63.62380248 | ENSG00000173705 |
| CCDC88A | -0.533611885 | 0.690823045 | 1.46E-09 | 2.64E-08 | 1111.171954 | 767.8043954 | ENSG00000115355 |
| OSBPL11 | -0.534644843 | 0.690328598 | 4.75E-08 | 6.76E-07 | 817.8768525 | 564.6142883 | ENSG00000144909 |
| HDGF | -0.534890255 | 0.690211178 | 4.25E-17 | 2.04E-15 | 9700.670406 | 6694.818797 | ENSG00000143321 |
| GTPBP8 | -0.535477127 | 0.689930466 | 5.79E-03 | 2.33E-02 | 165.3497901 | 113.8996155 | ENSG00000163607 |
| LTBP1 | -0.536178013 | 0.689595367 | 1.53E-10 | 3.25E-09 | 1253.50417 | 864.2080191 | ENSG00000049323 |
| GLCE | -0.536461478 | 0.689459887 | 2.85E-04 | 1.76E-03 | 382.4930752 | 263.5411044 | ENSG00000138604 |
| CA14 | -0.536474255 | 0.689453781 | 6.45E-08 | 8.93E-07 | 864.8311051 | 596.1485594 | ENSG00000118298 |
| ARL6 | -0.536530887 | 0.689426717 | 4.79E-02 | 1.29E-01 | 85.16823028 | 58.6204188 | ENSG00000113966 |
| COMMD4 | -0.53657686 | 0.689404748 | 1.45E-10 | 3.10E-09 | 1614.162641 | 1113.634312 | ENSG00000140365 |
| KCTD17 | -0.536932065 | 0.689235031 | 7.14E-08 | 9.77E-07 | 862.3779524 | 594.5345634 | ENSG00000100379 |
| PI4KB | -0.537173933 | 0.68911949 | 1.11E-08 | 1.76E-07 | 916.1455616 | 631.6243896 | ENSG00000143393 |
| EDARADD | -0.537867538 | 0.688788262 | 3.75E-02 | 1.07E-01 | 87.45690006 | 60.10593217 | ENSG00000186197 |
| SLAIN1 | -0.537943819 | 0.688751844 | 4.66E-06 | 4.54E-05 | 541.7857093 | 372.7710889 | ENSG00000139737 |
| ATP10A | -0.539043756 | 0.688226927 | 7.86E-05 | 5.62E-04 | 340.7132771 | 234.3306715 | ENSG00000206190 |
| AP2A2 | -0.539296529 | 0.688106354 | 8.25E-12 | 2.12E-10 | 1572.077855 | 1081.664572 | ENSG00000183020 |

| | | | | | | | |
|---|---|---|---|---|---|---|---|
| WRAP53 | -0.53948684 | 0.68801559 | 7.90E-08 | 1.08E-06 | 727.1414036 | 500.4772822 | ENSG00000141499 |
| MED12 | -0.540239965 | 0.687656521 | 1.81E-10 | 3.80E-09 | 1196.781085 | 823.1490946 | ENSG00000184634 |
| LRFN4 | -0.540289467 | 0.687632926 | 5.61E-06 | 5.36E-05 | 846.5527448 | 582.6396668 | ENSG00000173621 |
| RNF135 | -0.540589912 | 0.68748974 | 3.91E-03 | 1.67E-02 | 179.4473337 | 123.3179753 | ENSG00000181481 |
| NAT8L | -0.540798612 | 0.687390295 | 1.17E-07 | 1.55E-06 | 966.9745979 | 664.8691625 | ENSG00000185818 |
| BRD3OS | -0.541018176 | 0.687285689 | 1.83E-04 | 1.19E-03 | 329.2046404 | 226.1155989 | ENSG00000235106 |
| KLHDC8B | -0.541244934 | 0.687177672 | 1.43E-04 | 9.54E-04 | 353.5075202 | 243.2703351 | ENSG00000185909 |
| PTPRT | -0.541383552 | 0.687111649 | 2.74E-02 | 8.35E-02 | 110.9212745 | 76.26089908 | ENSG00000196090 |
| DUSP16 | -0.541765163 | 0.686929924 | 1.95E-14 | 6.98E-13 | 2470.790989 | 1696.765368 | ENSG00000111266 |
| CCDC34 | -0.542413648 | 0.686621221 | 1.36E-06 | 1.47E-05 | 564.3787314 | 387.1780834 | ENSG00000109881 |
| VAT1L | -0.542700333 | 0.686484793 | 2.06E-13 | 6.56E-12 | 2576.490418 | 1768.133877 | ENSG00000171724 |
| MRTO4 | -0.542728244 | 0.686471512 | 1.29E-12 | 3.72E-11 | 1745.309691 | 1198.110744 | ENSG00000053372 |
| CNPY4 | -0.542969452 | 0.686356749 | 3.67E-03 | 1.58E-02 | 176.1946939 | 121.0610192 | ENSG00000166997 |
| DHRS3 | -0.543313453 | 0.686193111 | 6.87E-04 | 3.79E-03 | 239.9821405 | 164.7836981 | ENSG00000162496 |
| GAS1 | -0.543828086 | 0.685948378 | 1.21E-02 | 4.30E-02 | 131.7862867 | 90.20576993 | ENSG00000180447 |
| CHRAC1 | -0.544046999 | 0.685844301 | 3.61E-07 | 4.37E-06 | 652.1494809 | 447.2854412 | ENSG00000104472 |
| MON1A | -0.544404699 | 0.685674274 | 1.27E-03 | 6.44E-03 | 297.4246936 | 204.3900251 | ENSG00000164077 |
| ZNF443 | -0.544601849 | 0.685580581 | 4.88E-02 | 1.31E-01 | 78.07368326 | 53.5161321 | ENSG00000180855 |
| RNF208 | -0.546761858 | 0.684554895 | 3.94E-02 | 1.11E-01 | 109.7675291 | 75.12400733 | ENSG00000212864 |
| MMP24 | -0.547159247 | 0.684366361 | 1.71E-04 | 1.12E-03 | 324.0057803 | 221.6185941 | ENSG00000125966 |
| L1TD1 | -0.547515125 | 0.684197565 | 2.87E-20 | 1.88E-18 | 14400.65761 | 9852.906151 | ENSG00000240563 |
| WDR76 | -0.54755547 | 0.684178432 | 4.35E-05 | 3.33E-04 | 382.4870923 | 261.3431693 | ENSG00000092470 |
| SCD | -0.54777339 | 0.684075094 | 1.04E-19 | 6.47E-18 | 28811.44875 | 19708.10872 | ENSG00000099194 |
| RIC8B | -0.547929167 | 0.684001234 | 1.13E-04 | 7.75E-04 | 318.5644096 | 217.81536 | ENSG00000111785 |
| PDE3B | -0.547996239 | 0.683969435 | 3.89E-04 | 2.30E-03 | 362.3058066 | 247.6445913 | ENSG00000152270 |
| PTBP2 | -0.54807333 | 0.683932888 | 3.38E-07 | 4.11E-06 | 784.9993008 | 536.1342433 | ENSG00000117569 |
| PCDHGB7 | -0.548591232 | 0.683687412 | 1.58E-03 | 7.77E-03 | 198.2662321 | 135.6440007 | ENSG00000254122 |
| PCGF5 | -0.548609255 | 0.683678871 | 2.06E-02 | 6.61E-02 | 124.9479155 | 85.35222778 | ENSG00000180628 |
| PNMA3 | -0.548742754 | 0.68361561 | 3.41E-04 | 2.05E-03 | 327.3192134 | 223.7280188 | ENSG00000183837 |
| INTS6L | -0.548975344 | 0.683505407 | 4.97E-04 | 2.85E-03 | 285.4298928 | 195.4480999 | ENSG00000165359 |
| ARHGAP18 | -0.54911855 | 0.683437564 | 1.24E-02 | 4.38E-02 | 123.7394127 | 84.64849462 | ENSG00000146376 |
| EMILIN3 | -0.549147648 | 0.683423779 | 9.82E-04 | 5.18E-03 | 256.6372475 | 174.968714 | ENSG00000183798 |
| TCF3 | -0.54920815 | 0.68339512 | 8.15E-17 | 3.77E-15 | 6997.996922 | 4782.244656 | ENSG00000071564 |
| PNPLA6 | -0.549500086 | 0.683256845 | 4.98E-14 | 1.71E-12 | 2678.141914 | 1829.31482 | ENSG00000032444 |
| URM1 | -0.549534253 | 0.683240664 | 1.39E-09 | 2.53E-08 | 1244.790947 | 850.8841369 | ENSG00000167118 |
| SFXN4 | -0.549664636 | 0.683178919 | 1.18E-04 | 8.07E-04 | 319.4577626 | 218.3268602 | ENSG00000183605 |
| ZNF580 | -0.549961861 | 0.683038185 | 1.78E-03 | 8.60E-03 | 225.6990608 | 154.2175998 | ENSG00000213015 |
| DNMT3B | -0.550942513 | 0.682574057 | 2.46E-28 | 3.18E-26 | 38489.18972 | 26271.17079 | ENSG00000088305 |
| GEMIN5 | -0.551384711 | 0.682364874 | 1.31E-12 | 3.77E-11 | 1528.61144 | 1043.484245 | ENSG00000082516 |
| ITGA1 | -0.551602223 | 0.682262003 | 6.93E-03 | 2.72E-02 | 142.2330966 | 96.89746069 | ENSG00000213949 |
| ZFYVE26 | -0.552956802 | 0.681621713 | 1.38E-08 | 2.17E-07 | 940.5842412 | 641.6897717 | ENSG00000072121 |
| EDNRA | -0.553678714 | 0.68128072 | 2.91E-02 | 8.74E-02 | 89.85948916 | 61.265617 | ENSG00000151617 |
| SRSF1 | -0.553721229 | 0.681260644 | 1.30E-05 | 1.13E-04 | 1399.907508 | 953.30419 | ENSG00000136450 |
| COQ2 | -0.554524272 | 0.680881541 | 2.06E-04 | 1.32E-03 | 376.197903 | 256.2804467 | ENSG00000173085 |
| EXOC6 | -0.554888018 | 0.680709893 | 7.34E-03 | 2.85E-02 | 136.6266946 | 92.87298832 | ENSG00000138190 |
| ZDHHC6 | -0.555379338 | 0.680478112 | 9.53E-08 | 1.28E-06 | 661.7640015 | 450.4290942 | ENSG00000023041 |
| ZNF615 | -0.555773412 | 0.680292264 | 4.72E-03 | 1.96E-02 | 152.6595896 | 103.7918335 | ENSG00000197619 |
| RCAN1 | -0.556152365 | 0.680113595 | 3.26E-03 | 1.43E-02 | 172.6470333 | 117.2905406 | ENSG00000159200 |
| C1QL1 | -0.556157969 | 0.680110953 | 4.87E-04 | 2.80E-03 | 275.0872384 | 186.9091339 | ENSG00000131094 |
| YIF1A | -0.556538118 | 0.679931768 | 7.71E-12 | 2.00E-10 | 1299.303535 | 883.6095258 | ENSG00000174851 |
| ZNF423 | -0.556819368 | 0.67979923 | 1.75E-11 | 4.27E-10 | 1449.020485 | 984.8095195 | ENSG00000102935 |
| CD24 | -0.557345649 | 0.679551291 | 1.06E-12 | 3.14E-11 | 30869.16534 | 20976.57747 | ENSG00000272398 |
| FAM136A | -0.557726818 | 0.679371773 | 4.35E-13 | 1.35E-11 | 2218.726435 | 1507.067916 | ENSG00000035141 |
| MGAT5B | -0.55811159 | 0.679190606 | 2.26E-04 | 1.43E-03 | 292.1036462 | 198.3713503 | ENSG00000167889 |
| DISP3 | -0.559280958 | 0.678640315 | 1.56E-03 | 7.68E-03 | 203.5061094 | 138.2924481 | ENSG00000204624 |
| RGMA | -0.559459148 | 0.6785565 | 3.53E-08 | 5.13E-07 | 874.7540406 | 593.5583601 | ENSG00000182175 |
| MTHFD1 | -0.559741832 | 0.678423556 | 1.90E-17 | 9.46E-16 | 5165.196041 | 3503.96383 | ENSG00000100714 |
| SMO | -0.560276869 | 0.678172002 | 1.37E-14 | 5.02E-13 | 2629.337712 | 1782.75037 | ENSG00000128602 |
| KLKB1 | -0.560311504 | 0.678155722 | 1.29E-02 | 4.52E-02 | 142.0362106 | 96.22929414 | ENSG00000164344 |
| NOL11 | -0.560863212 | 0.677896435 | 4.63E-12 | 1.25E-10 | 1986.103729 | 1345.759656 | ENSG00000130935 |
| GNB1L | -0.561017364 | 0.677824005 | 2.66E-04 | 1.66E-03 | 375.8965609 | 255.266948 | ENSG00000185838 |
| HOMER2 | -0.561154106 | 0.677759763 | 6.19E-06 | 5.87E-05 | 478.3045431 | 323.9098389 | ENSG00000103942 |
| PRKD1 | -0.561291854 | 0.677695053 | 2.29E-03 | 1.07E-02 | 178.7361645 | 120.9437183 | ENSG00000184304 |
| SDCBP2 | -0.562267842 | 0.677236745 | 3.05E-02 | 9.08E-02 | 83.75131879 | 56.68571567 | ENSG00000125775 |
| SSBP1 | -0.562504895 | 0.677125476 | 4.28E-15 | 1.67E-13 | 3326.96365 | 2252.994961 | ENSG00000106028 |
| CDK2 | -0.563329156 | 0.676738722 | 1.99E-05 | 1.66E-04 | 378.1838779 | 255.7097107 | ENSG00000123374 |
| KIAA1143 | -0.563667578 | 0.676579993 | 1.29E-07 | 1.68E-06 | 875.814973 | 591.7611772 | ENSG00000163807 |
| ATP8B2 | -0.564834556 | 0.676032937 | 4.14E-14 | 1.43E-12 | 1715.485638 | 1159.878118 | ENSG00000143515 |
| ZIK1 | -0.564875493 | 0.676013755 | 9.79E-08 | 1.31E-06 | 682.9966661 | 461.734103 | ENSG00000171649 |
| GLIPR2 | -0.565353173 | 0.675789962 | 2.07E-03 | 9.79E-03 | 209.2536554 | 141.4092943 | ENSG00000122694 |
| TCEA2 | -0.565443224 | 0.675747782 | 1.89E-08 | 2.88E-07 | 826.2607794 | 558.9235777 | ENSG00000171703 |
| CPXM1 | -0.565458736 | 0.675740516 | 6.14E-12 | 1.61E-10 | 1788.343136 | 1209.296271 | ENSG00000088882 |
| POLA2 | -0.565536595 | 0.675704049 | 1.46E-02 | 5.01E-02 | 120.6424311 | 81.51970224 | ENSG00000014138 |
| EML3 | -0.565903814 | 0.675532079 | 3.16E-07 | 3.86E-06 | 620.6750481 | 419.5283074 | ENSG00000149499 |
| MFSD1 | -0.56629732 | 0.675347848 | 7.96E-03 | 3.05E-02 | 147.5136078 | 99.72624358 | ENSG00000118855 |

| | | | | | | | |
|---|---|---|---|---|---|---|---|
| DCTN3 | -0.566453445 | 0.675274767 | 5.05E-05 | 3.80E-04 | 338.2027743 | 228.6235546 | ENSG00000137100 |
| DGCR6L | -0.566895928 | 0.675067688 | 2.87E-09 | 4.95E-08 | 1277.870902 | 862.9431315 | ENSG00000128185 |
| POGLUT2 | -0.566918223 | 0.675057256 | 9.84E-07 | 1.10E-05 | 605.1510919 | 408.7921708 | ENSG00000134901 |
| GPR153 | -0.567218428 | 0.6749168 | 7.39E-05 | 5.31E-04 | 343.0983228 | 231.5063487 | ENSG00000158292 |
| SEZ6 | -0.568221099 | 0.674447897 | 4.97E-02 | 1.33E-01 | 70.85466379 | 47.80758953 | ENSG00000063015 |
| PRPF4B | -0.568293721 | 0.674413948 | 4.07E-10 | 8.01E-09 | 1816.27599 | 1225.613919 | ENSG00000112739 |
| DERL3 | -0.568433406 | 0.674348653 | 1.36E-02 | 4.71E-02 | 114.5242864 | 77.33421502 | ENSG00000099958 |
| MTURN | -0.569474587 | 0.673862156 | 5.72E-03 | 2.31E-02 | 157.3452337 | 106.1784747 | ENSG00000180354 |
| THADA | -0.569547483 | 0.673828109 | 3.36E-08 | 4.90E-07 | 695.0950135 | 468.3471257 | ENSG00000115970 |
| ZDHHC22 | -0.570000761 | 0.673616433 | 2.21E-11 | 5.30E-10 | 1459.59624 | 983.9928029 | ENSG00000177108 |
| ADGRB1 | -0.570135576 | 0.673553489 | 2.86E-07 | 3.52E-06 | 567.8348641 | 382.6983609 | ENSG00000181790 |
| DDX49 | -0.570512725 | 0.673377432 | 7.33E-09 | 1.20E-07 | 996.5139277 | 671.3173716 | ENSG00000105671 |
| SEPHS1 | -0.570656899 | 0.673310142 | 1.31E-21 | 9.50E-20 | 15131.14254 | 10187.06818 | ENSG00000086475 |
| AMPH | -0.571394311 | 0.672966077 | 5.34E-05 | 3.99E-04 | 313.36685 | 210.8429804 | ENSG00000078053 |
| HID1 | -0.572004673 | 0.672681425 | 6.29E-04 | 3.51E-03 | 255.71213 | 172.1432206 | ENSG00000167861 |
| BMP6 | -0.572254067 | 0.672565151 | 2.70E-02 | 8.24E-02 | 93.60926721 | 63.02108395 | ENSG00000153162 |
| WASHC4 | -0.572736058 | 0.672340491 | 1.17E-08 | 1.85E-07 | 844.1737349 | 567.1927243 | ENSG00000136051 |
| ANAPC2 | -0.572825379 | 0.672298866 | 2.39E-10 | 4.92E-09 | 1314.799033 | 884.6623589 | ENSG00000176248 |
| SEMA6C | -0.573366819 | 0.672046601 | 4.84E-08 | 6.87E-07 | 807.0422388 | 543.0397669 | ENSG00000143434 |
| CAMK4 | -0.573673723 | 0.671903652 | 5.05E-03 | 2.08E-02 | 153.4861664 | 102.9210965 | ENSG00000152495 |
| SCLY | -0.574018556 | 0.671743073 | 1.41E-02 | 4.86E-02 | 108.5325909 | 72.97555198 | ENSG00000132330 |
| TCN2 | -0.57406507 | 0.671721415 | 1.64E-11 | 4.02E-10 | 1288.049014 | 864.5879705 | ENSG00000185339 |
| CPNE1 | -0.574162437 | 0.671676083 | 3.60E-10 | 7.15E-09 | 1309.016709 | 879.787999 | ENSG00000214078 |
| FASTKD1 | -0.574351995 | 0.671587836 | 1.39E-07 | 1.81E-06 | 680.5676416 | 457.1585501 | ENSG00000138399 |
| NME1 | -0.574381806 | 0.671573959 | 7.33E-07 | 8.38E-06 | 531.2542867 | 356.7234605 | ENSG00000239672 |
| ABCA1 | -0.574415065 | 0.671558477 | 1.48E-03 | 7.31E-03 | 208.8328844 | 139.9973275 | ENSG00000165029 |
| SLC38A7 | -0.574986255 | 0.671292647 | 4.29E-08 | 6.17E-07 | 695.1346854 | 467.0467344 | ENSG00000103042 |
| SREBF1 | -0.575037035 | 0.671269019 | 2.42E-13 | 7.64E-12 | 2145.526848 | 1440.511227 | ENSG00000072310 |
| GPRIN1 | -0.575160969 | 0.671211357 | 4.12E-06 | 4.07E-05 | 496.1399361 | 333.2548668 | ENSG00000169258 |
| RWDD2B | -0.575161042 | 0.671211323 | 3.91E-02 | 1.11E-01 | 80.05974065 | 53.63971658 | ENSG00000156253 |
| NRIP1 | -0.575203054 | 0.671191777 | 3.36E-03 | 1.47E-02 | 190.6457438 | 127.8493761 | ENSG00000180530 |
| PPOX | -0.575561064 | 0.671025239 | 3.43E-06 | 3.45E-05 | 551.5879383 | 370.5668073 | ENSG00000143224 |
| IKBKE | -0.576160271 | 0.670746594 | 5.69E-04 | 3.21E-03 | 219.0391151 | 146.8392979 | ENSG00000263528 |
| DHX40 | -0.577379098 | 0.670180169 | 1.15E-07 | 1.53E-06 | 895.3640722 | 599.207077 | ENSG00000108406 |
| POLR2C | -0.577425293 | 0.67015871 | 1.48E-12 | 4.23E-11 | 1467.290706 | 983.3079665 | ENSG00000102978 |
| RDH14 | -0.577829887 | 0.669970795 | 5.56E-04 | 3.15E-03 | 233.1049919 | 155.9831407 | ENSG00000240857 |
| PDRG1 | -0.578043786 | 0.66987147 | 3.46E-08 | 5.04E-07 | 948.4462329 | 635.5957897 | ENSG00000088356 |
| KIF21A | -0.57893773 | 0.669456523 | 1.13E-11 | 2.86E-10 | 1166.733305 | 781.1512771 | ENSG00000139116 |
| MYCN | -0.579031468 | 0.669413027 | 2.35E-12 | 6.49E-11 | 1739.258954 | 1164.353389 | ENSG00000134323 |
| DUSP7 | -0.579314874 | 0.669281539 | 4.76E-08 | 6.77E-07 | 646.4475223 | 432.9252354 | ENSG00000164086 |
| LIN7A | -0.57968967 | 0.66910769 | 1.12E-03 | 5.81E-03 | 211.0846633 | 141.1455127 | ENSG00000111052 |
| SUPV3L1 | -0.579829628 | 0.669042782 | 8.73E-09 | 1.42E-07 | 779.2334147 | 521.2055001 | ENSG00000156502 |
| ZNF385C | -0.580030592 | 0.668949593 | 1.37E-02 | 4.76E-02 | 116.377077 | 77.91568626 | ENSG00000187595 |
| ADPRH | -0.580062613 | 0.668934745 | 1.49E-02 | 5.07E-02 | 116.5722793 | 78.21277298 | ENSG00000144843 |
| CDC6 | -0.580127339 | 0.668904734 | 1.39E-17 | 7.08E-16 | 2949.626696 | 1972.578558 | ENSG00000094804 |
| TSC22D1 | -0.580781312 | 0.668601589 | 2.74E-15 | 1.09E-13 | 1949.686973 | 1304.08019 | ENSG00000102804 |
| NAP1L2 | -0.581284413 | 0.668368472 | 2.14E-05 | 1.78E-04 | 507.4334218 | 339.3283257 | ENSG00000186462 |
| CDK5 | -0.581749013 | 0.668153268 | 7.26E-06 | 6.77E-05 | 400.46166 | 267.8113603 | ENSG00000164885 |
| FAHD1 | -0.581971578 | 0.6680502 | 1.68E-05 | 1.43E-04 | 358.9064006 | 239.7091486 | ENSG00000180185 |
| BEX3 | -0.581997319 | 0.668038281 | 4.81E-26 | 4.96E-24 | 7860.143766 | 5251.23983 | ENSG00000166681 |
| ZNF22 | -0.58201383 | 0.668030635 | 2.31E-04 | 1.46E-03 | 316.3646144 | 211.3048892 | ENSG00000165512 |
| RPE | -0.582612769 | 0.667753358 | 2.28E-10 | 4.71E-09 | 1046.811002 | 698.5492439 | ENSG00000197713 |
| SOX12 | -0.582960452 | 0.667592452 | 2.15E-12 | 6.02E-11 | 3963.303587 | 2645.992271 | ENSG00000177732 |
| TEDC2 | -0.582968533 | 0.667588712 | 1.83E-08 | 2.80E-07 | 1005.017674 | 671.7339578 | ENSG00000162062 |
| STK36 | -0.583653064 | 0.66727203 | 5.65E-07 | 6.62E-06 | 674.8699822 | 450.9612291 | ENSG00000163482 |
| EMP2 | -0.583703402 | 0.667248748 | 3.29E-06 | 3.33E-05 | 446.032201 | 297.53088 | ENSG00000213853 |
| SLC43A3 | -0.584038696 | 0.667093692 | 2.22E-04 | 1.41E-03 | 288.7871917 | 192.6824632 | ENSG00000134802 |
| ZNF276 | -0.584196481 | 0.667020737 | 2.22E-02 | 7.04E-02 | 145.1641826 | 97.2754056 | ENSG00000158805 |
| KCNJ11 | -0.584846442 | 0.666720299 | 5.12E-04 | 2.93E-03 | 230.8186072 | 154.151316 | ENSG00000187486 |
| NGEF | -0.585822779 | 0.666269252 | 2.41E-02 | 7.51E-02 | 86.02114479 | 57.40259751 | ENSG00000066248 |
| CENPQ | -0.58597217 | 0.666200263 | 1.62E-03 | 7.95E-03 | 186.9789399 | 124.3114207 | ENSG00000031691 |
| LDOC1 | -0.586239301 | 0.666076921 | 2.27E-09 | 4.00E-08 | 845.1531586 | 563.3860666 | ENSG00000182195 |
| DLGAP3 | -0.587376728 | 0.66555199 | 1.28E-05 | 1.12E-04 | 572.8315689 | 381.2631715 | ENSG00000116544 |
| ATAD3A | -0.587739545 | 0.665384634 | 7.69E-13 | 2.32E-11 | 1820.839115 | 1212.075869 | ENSG00000197785 |
| C1orf112 | -0.587811269 | 0.665351555 | 5.48E-05 | 4.08E-04 | 365.989425 | 243.0826881 | ENSG00000000460 |
| COL8A2 | -0.5880054 | 0.665262031 | 5.73E-03 | 2.31E-02 | 130.8106812 | 87.10108472 | ENSG00000171812 |
| GLB1L3 | -0.588726241 | 0.664929716 | 3.70E-10 | 7.31E-09 | 1462.322443 | 973.3271156 | ENSG00000166105 |
| GNG4 | -0.589748558 | 0.664458703 | 1.78E-08 | 2.73E-07 | 1257.928964 | 835.1648528 | ENSG00000168243 |
| PEX3 | -0.589847332 | 0.664413212 | 1.84E-03 | 8.84E-03 | 190.9684852 | 126.9039931 | ENSG00000034693 |
| ECI2 | -0.590082679 | 0.664304835 | 2.05E-04 | 1.31E-03 | 266.2687328 | 176.6619192 | ENSG00000198721 |
| LRRC42 | -0.590331388 | 0.664190324 | 1.59E-11 | 3.92E-10 | 1110.059497 | 737.0630375 | ENSG00000116212 |
| CYBRD1 | -0.590348677 | 0.664182365 | 9.23E-03 | 3.44E-02 | 123.0254774 | 81.5113401 | ENSG00000071967 |
| SEMA3D | -0.590608019 | 0.664062981 | 2.37E-02 | 7.41E-02 | 88.35982151 | 58.55337055 | ENSG00000153993 |
| MBOAT2 | -0.591095131 | 0.663838804 | 3.89E-05 | 3.02E-04 | 380.7349995 | 252.7764492 | ENSG00000143797 |
| PNMA1 | -0.591146668 | 0.663815091 | 4.98E-15 | 1.93E-13 | 2062.230989 | 1369.393296 | ENSG00000176903 |

| | | | | | | | |
|---|---|---|---|---|---|---|---|
| BRINP1 | -0.591244177 | 0.663770226 | 1.88E-09 | 3.35E-08 | 884.5051576 | 587.4539063 | ENSG00000078725 |
| PERP | -0.591566309 | 0.663622033 | 1.45E-08 | 2.26E-07 | 934.5517087 | 619.9391341 | ENSG00000112378 |
| DUSP4 | -0.592043196 | 0.663402707 | 1.05E-05 | 9.29E-05 | 382.047748 | 253.3770178 | ENSG00000120875 |
| PTRHD1 | -0.592338765 | 0.663266807 | 5.84E-03 | 2.35E-02 | 127.4987591 | 84.57286099 | ENSG00000184924 |
| SLTM | -0.592659159 | 0.663119525 | 2.41E-20 | 1.59E-18 | 3613.583022 | 2396.733922 | ENSG00000137776 |
| PRR7 | -0.592714199 | 0.663094227 | 5.89E-03 | 2.37E-02 | 193.2267886 | 128.142346 | ENSG00000131188 |
| SCN4B | -0.592714221 | 0.663094217 | 3.54E-03 | 1.53E-02 | 163.8031747 | 108.5558692 | ENSG00000177098 |
| ERG28 | -0.593187756 | 0.662876606 | 9.60E-11 | 2.11E-09 | 1185.192864 | 785.1159322 | ENSG00000133935 |
| SMPD4 | -0.593223633 | 0.662860122 | 4.24E-17 | 2.04E-15 | 2737.247362 | 1814.576647 | ENSG00000136699 |
| TMEM145 | -0.594421921 | 0.662309785 | 5.89E-03 | 2.37E-02 | 134.378561 | 89.08147719 | ENSG00000167619 |
| SLC25A23 | -0.595059841 | 0.662016995 | 6.07E-10 | 1.16E-08 | 1456.224374 | 964.4305675 | ENSG00000125648 |
| TTC27 | -0.59513418 | 0.661982884 | 1.52E-09 | 2.75E-08 | 797.31891 | 527.532283 | ENSG00000018699 |
| FMN2 | -0.595331998 | 0.661892121 | 4.09E-02 | 1.14E-01 | 77.63642864 | 51.35942623 | ENSG00000155816 |
| TBPL1 | -0.596510057 | 0.661351861 | 3.91E-04 | 2.31E-03 | 292.3462999 | 193.0285514 | ENSG00000028839 |
| ALDH16A1 | -0.596563434 | 0.661327393 | 8.12E-09 | 1.32E-07 | 1069.372869 | 706.7654871 | ENSG00000161618 |
| MPP3 | -0.597463598 | 0.660914889 | 6.26E-03 | 2.50E-02 | 138.5305746 | 91.64399372 | ENSG00000161647 |
| CYP2U1 | -0.597737235 | 0.660789545 | 1.05E-02 | 3.84E-02 | 173.2964397 | 114.2619984 | ENSG00000155016 |
| CDC25B | -0.598973237 | 0.660223668 | 3.79E-09 | 6.48E-08 | 862.2855004 | 569.467942 | ENSG00000101224 |
| MCC | -0.599231692 | 0.660105401 | 8.34E-07 | 9.42E-06 | 571.4131092 | 377.0954591 | ENSG00000171444 |
| HNRNPDL | -0.599266809 | 0.660089334 | 6.94E-26 | 7.07E-24 | 7487.143416 | 4942.240659 | ENSG00000152795 |
| CKB | -0.599332218 | 0.660059407 | 1.83E-19 | 1.12E-17 | 7094.98394 | 4683.102232 | ENSG00000166165 |
| TEX10 | -0.599663513 | 0.659907851 | 8.41E-14 | 2.81E-12 | 1989.166729 | 1312.474865 | ENSG00000136891 |
| EEPD1 | -0.599859918 | 0.659818019 | 2.68E-05 | 2.17E-04 | 335.1259143 | 220.8910886 | ENSG00000122547 |
| TIMP4 | -0.60033529 | 0.659600643 | 4.51E-02 | 1.23E-01 | 70.11342131 | 46.3677655 | ENSG00000157150 |
| ARPIN | -0.600558024 | 0.659498817 | 2.87E-03 | 1.28E-02 | 153.8285409 | 101.3462915 | ENSG00000242498 |
| ZNF205 | -0.601974858 | 0.658851458 | 7.60E-05 | 5.44E-04 | 341.7231125 | 225.2348911 | ENSG00000122386 |
| ASGR1 | -0.60265031 | 0.658543064 | 1.01E-03 | 5.33E-03 | 194.0757494 | 127.6129509 | ENSG00000141505 |
| SLC27A2 | -0.603317184 | 0.658238728 | 4.43E-02 | 1.22E-01 | 77.51511312 | 50.87853159 | ENSG00000140284 |
| TSR1 | -0.604231963 | 0.657821487 | 8.26E-18 | 4.30E-16 | 3419.736359 | 2249.982155 | ENSG00000167721 |
| PRR5L | -0.604311711 | 0.657785125 | 8.56E-03 | 3.24E-02 | 127.9705444 | 84.00245988 | ENSG00000135362 |
| FBXL15 | -0.605298972 | 0.657335145 | 4.90E-05 | 3.70E-04 | 307.0718431 | 201.6714547 | ENSG00000107872 |
| SNRPD3 | -0.605357771 | 0.657308356 | 4.92E-08 | 6.98E-07 | 727.7529497 | 478.058295 | ENSG00000100028 |
| SS18L2 | -0.606066182 | 0.656985674 | 8.30E-07 | 9.38E-06 | 840.6298321 | 552.0386974 | ENSG00000008324 |
| ZNF180 | -0.606228407 | 0.656911803 | 1.69E-03 | 8.23E-03 | 173.109235 | 113.6408353 | ENSG00000167384 |
| FADS2 | -0.606274954 | 0.656890609 | 1.84E-21 | 1.32E-19 | 10644.12906 | 6991.998946 | ENSG00000134824 |
| MPC2 | -0.607333742 | 0.656408697 | 6.98E-03 | 2.74E-02 | 117.4595139 | 76.94696056 | ENSG00000143158 |
| WIPF3 | -0.607343297 | 0.656404349 | 9.51E-06 | 8.55E-05 | 454.4782513 | 297.7532805 | ENSG00000122574 |
| MTHFD2L | -0.608699566 | 0.655787557 | 2.31E-02 | 7.27E-02 | 94.06062265 | 61.74161349 | ENSG00000163738 |
| KIF3B | -0.609801271 | 0.655286961 | 4.39E-12 | 1.18E-10 | 1179.464831 | 772.3412582 | ENSG00000101350 |
| SIRPA | -0.610334497 | 0.655044809 | 1.39E-04 | 9.27E-04 | 256.9147023 | 168.0081689 | ENSG00000198053 |
| ZNF778 | -0.611279489 | 0.654615882 | 3.10E-06 | 3.15E-05 | 500.5706072 | 328.0349197 | ENSG00000170100 |
| PLEKHA6 | -0.611521463 | 0.654506097 | 3.45E-06 | 3.47E-05 | 388.2731301 | 254.2483045 | ENSG00000143850 |
| ABCF2 | -0.611631402 | 0.654456223 | 9.81E-06 | 8.79E-05 | 385.5181921 | 251.9461685 | ENSG00000033050 |
| MOB3C | -0.611725486 | 0.654413545 | 1.70E-02 | 5.65E-02 | 89.32222065 | 58.38855686 | ENSG00000142961 |
| SFXN5 | -0.612123112 | 0.654233205 | 1.44E-10 | 3.07E-09 | 910.2470399 | 595.792675 | ENSG00000144040 |
| POLR3B | -0.612277713 | 0.6541631 | 1.31E-05 | 1.15E-04 | 516.2179906 | 337.1210867 | ENSG00000013503 |
| TANGO6 | -0.612566827 | 0.65403202 | 2.21E-06 | 2.29E-05 | 417.4417709 | 272.9333186 | ENSG00000103047 |
| RPUSD1 | -0.613162333 | 0.653762109 | 3.98E-08 | 5.73E-07 | 878.5657164 | 574.3719198 | ENSG00000007376 |
| MTA2 | -0.613353508 | 0.653675482 | 1.78E-24 | 1.60E-22 | 4206.467383 | 2749.279953 | ENSG00000149480 |
| ADCY5 | -0.61368783 | 0.653524021 | 9.23E-08 | 1.25E-06 | 528.4875178 | 345.6892813 | ENSG00000173175 |
| EFS | -0.614189715 | 0.653296713 | 1.20E-02 | 4.28E-02 | 1052.206448 | 687.7319093 | ENSG00000100842 |
| CAMK2N1 | -0.614415849 | 0.65319432 | 1.17E-02 | 4.19E-02 | 103.1196097 | 67.40545441 | ENSG00000162545 |
| SRSF2 | -0.616252387 | 0.652363339 | 1.23E-13 | 4.00E-12 | 2517.417193 | 1641.747945 | ENSG00000161547 |
| MICOS10 | -0.616459958 | 0.652269485 | 3.50E-02 | 1.01E-01 | 69.77400835 | 45.55355626 | ENSG00000173436 |
| RNF182 | -0.617385127 | 0.651851333 | 1.61E-02 | 5.40E-02 | 101.500002 | 65.98297565 | ENSG00000180537 |
| ZNF512B | -0.617569853 | 0.651767874 | 2.96E-11 | 7.01E-10 | 1147.975861 | 748.8571286 | ENSG00000196700 |
| HECTD2 | -0.618997653 | 0.651123154 | 4.45E-04 | 2.58E-03 | 206.3502901 | 134.1763822 | ENSG00000165338 |
| ILKAP | -0.619101595 | 0.651076244 | 1.18E-07 | 1.55E-06 | 580.8642072 | 378.588284 | ENSG00000132323 |
| PARP16 | -0.619574655 | 0.650862791 | 5.06E-03 | 2.08E-02 | 129.481156 | 84.18984341 | ENSG00000138617 |
| IGSF8 | -0.620231437 | 0.650566556 | 2.09E-16 | 9.22E-15 | 2565.77742 | 1668.484094 | ENSG00000162729 |
| DDX56 | -0.621157668 | 0.650149016 | 1.44E-14 | 5.25E-13 | 1507.511013 | 980.1894169 | ENSG00000136271 |
| ANAPC4 | -0.622571924 | 0.649511996 | 3.94E-08 | 5.68E-07 | 590.6117539 | 383.7563698 | ENSG00000053900 |
| GINS3 | -0.623593398 | 0.649052284 | 2.85E-04 | 1.76E-03 | 233.9680762 | 151.9937256 | ENSG00000181938 |
| PIK3R3 | -0.624220022 | 0.648770433 | 9.35E-05 | 6.56E-04 | 265.0552764 | 171.9396624 | ENSG00000117461 |
| FMNL1 | -0.624363423 | 0.64870595 | 3.92E-04 | 2.31E-03 | 198.5929197 | 128.8246002 | ENSG00000184922 |
| CFAP46 | -0.625144156 | 0.64835499 | 6.88E-03 | 2.71E-02 | 114.6924519 | 74.43480828 | ENSG00000171811 |
| PGD | -0.62550936 | 0.648190886 | 1.60E-29 | 2.17E-27 | 10227.98378 | 6629.424131 | ENSG00000142657 |
| REPIN1 | -0.625901466 | 0.648014739 | 1.01E-15 | 4.20E-14 | 2515.378686 | 1630.773395 | ENSG00000214022 |
| FARS2 | -0.626203853 | 0.647878931 | 4.91E-06 | 4.75E-05 | 345.2331006 | 223.7046761 | ENSG00000145982 |
| PRXL2B | -0.626552615 | 0.64772233 | 9.81E-14 | 3.23E-12 | 1956.908495 | 1267.444317 | ENSG00000157870 |
| ANKRD18B | -0.626554137 | 0.647721646 | 1.16E-02 | 4.15E-02 | 109.6887191 | 70.88198389 | ENSG00000230453 |
| STXBP6 | -0.626624072 | 0.647690248 | 2.64E-03 | 1.20E-02 | 147.5204253 | 95.45328816 | ENSG00000168952 |
| FUCA2 | -0.627067449 | 0.647491227 | 1.17E-10 | 2.53E-09 | 870.6382566 | 563.6276761 | ENSG00000001036 |
| RAC3 | -0.62737432 | 0.647353516 | 8.30E-12 | 2.13E-10 | 1698.25547 | 1099.238865 | ENSG00000169750 |
| SQLE | -0.627596752 | 0.647253716 | 8.33E-24 | 7.15E-22 | 4135.859424 | 2676.365237 | ENSG00000104549 |

| | | | | | | | |
|---|---|---|---|---|---|---|---|
| SEPTIN8 | -0.627709076 | 0.647203325 | 8.87E-14 | 2.95E-12 | 1540.408836 | 997.2675234 | ENSG00000164402 |
| ZMPSTE24 | -0.627881727 | 0.647125877 | 4.97E-09 | 8.37E-08 | 1042.40509 | 674.1927425 | ENSG00000084073 |
| SKP2 | -0.628760143 | 0.64673198 | 1.16E-12 | 3.39E-11 | 1654.627617 | 1069.63873 | ENSG00000145604 |
| PLCB3 | -0.629346212 | 0.64646931 | 1.56E-22 | 1.22E-20 | 4697.415735 | 3037.171994 | ENSG00000149782 |
| ATP1B2 | -0.629476209 | 0.646411061 | 3.80E-05 | 2.96E-04 | 298.7638024 | 193.2149875 | ENSG00000129244 |
| PDE4DIP | -0.629767637 | 0.646280498 | 1.83E-09 | 3.27E-08 | 800.5426339 | 517.4399112 | ENSG00000178104 |
| CCDC15 | -0.630871814 | 0.645786052 | 4.27E-02 | 1.19E-01 | 64.00327412 | 41.28126211 | ENSG00000149548 |
| NTRK3 | -0.631235108 | 0.645623453 | 9.70E-04 | 5.13E-03 | 171.0782067 | 110.5700048 | ENSG00000140538 |
| TNFRSF25 | -0.631444268 | 0.645529858 | 1.03E-02 | 3.76E-02 | 139.6973385 | 90.48171252 | ENSG00000215788 |
| LANCL2 | -0.631781066 | 0.645379176 | 1.63E-07 | 2.09E-06 | 499.3526184 | 322.4936756 | ENSG00000132434 |
| PCSK9 | -0.633184995 | 0.644751445 | 1.20E-16 | 5.44E-15 | 2171.113268 | 1399.643771 | ENSG00000169174 |
| NFATC2IP | -0.63409733 | 0.644343844 | 5.01E-10 | 9.73E-09 | 1094.704594 | 705.9950011 | ENSG00000176953 |
| ZNF677 | -0.634900947 | 0.643985028 | 6.37E-06 | 6.01E-05 | 384.9775487 | 247.7472465 | ENSG00000197928 |
| TXNDC16 | -0.635782495 | 0.643591646 | 5.19E-05 | 3.89E-04 | 355.1013166 | 228.5044931 | ENSG00000087301 |
| PSME3 | -0.636014465 | 0.643488172 | 7.94E-27 | 8.66E-25 | 6010.335144 | 3866.94795 | ENSG00000131467 |
| FOXRED1 | -0.63610152 | 0.643449343 | 2.61E-10 | 5.35E-09 | 1273.350538 | 819.8509111 | ENSG00000110074 |
| PARPBP | -0.636420414 | 0.643307131 | 1.38E-06 | 1.50E-05 | 396.8157292 | 255.3463402 | ENSG00000185480 |
| SLC51A | -0.637039578 | 0.643031101 | 2.61E-02 | 8.01E-02 | 77.04934858 | 49.44619364 | ENSG00000163959 |
| DIRAS1 | -0.637192732 | 0.642962841 | 5.27E-10 | 1.02E-08 | 894.0005874 | 575.1208329 | ENSG00000176490 |
| SEC61A2 | -0.63735303 | 0.642891406 | 3.68E-07 | 4.45E-06 | 437.4032603 | 281.475757 | ENSG00000065665 |
| ST7 | -0.638162274 | 0.642530893 | 1.18E-03 | 6.08E-03 | 202.4626378 | 129.8698985 | ENSG00000004866 |
| PEX26 | -0.638304089 | 0.642467736 | 7.10E-15 | 2.67E-13 | 1477.587149 | 948.9439417 | ENSG00000215193 |
| NAP1L3 | -0.638380979 | 0.642433496 | 2.26E-12 | 6.29E-11 | 1449.588038 | 932.1051924 | ENSG00000186310 |
| SLC35A3 | -0.638441451 | 0.642406569 | 1.22E-02 | 4.32E-02 | 128.4247485 | 82.36265329 | ENSG00000117620 |
| H2AJ | -0.639709662 | 0.641842105 | 2.27E-05 | 1.87E-04 | 314.5125979 | 202.2294087 | ENSG00000246705 |
| PLEKHB1 | -0.640266165 | 0.641594569 | 3.36E-04 | 2.03E-03 | 201.2387412 | 128.8950092 | ENSG00000021300 |
| ANOS1 | -0.641558553 | 0.641020077 | 1.11E-16 | 5.05E-15 | 4323.097726 | 2770.432632 | ENSG00000011201 |
| ACP2 | -0.64210288 | 0.640778266 | 2.49E-05 | 2.03E-04 | 311.0192062 | 199.52476 | ENSG00000134575 |
| PHLDB2 | -0.642556707 | 0.640576729 | 2.56E-10 | 5.26E-09 | 744.3264806 | 476.683919 | ENSG00000144824 |
| SMARCA2 | -0.6429125 | 0.640418771 | 3.42E-05 | 2.69E-04 | 325.3431078 | 207.9086196 | ENSG00000080503 |
| ALKBH3 | -0.644195635 | 0.639849435 | 5.64E-04 | 3.19E-03 | 185.0476324 | 118.5034008 | ENSG00000166199 |
| LYPD3 | -0.644204898 | 0.639845327 | 2.30E-02 | 7.25E-02 | 78.26495452 | 49.945198 | ENSG00000124466 |
| UHRF2 | -0.644478255 | 0.639724103 | 1.63E-10 | 3.44E-09 | 1028.727116 | 657.4801883 | ENSG00000147854 |
| ZDHHC16 | -0.644736769 | 0.639609482 | 3.47E-14 | 1.21E-12 | 1171.408275 | 749.2395562 | ENSG00000171307 |
| PDGFRL | -0.6449837 | 0.639500016 | 3.12E-02 | 9.23E-02 | 77.01828312 | 49.43622281 | ENSG00000104213 |
| MAGEH1 | -0.64644023 | 0.638854709 | 5.27E-04 | 3.01E-03 | 255.8404584 | 163.7770411 | ENSG00000187601 |
| CENPV | -0.647336784 | 0.63845782 | 6.20E-15 | 2.36E-13 | 2184.435038 | 1394.204918 | ENSG00000166582 |
| NPEPPS | -0.648317233 | 0.638024074 | 4.66E-15 | 1.81E-13 | 1631.111935 | 1039.962495 | ENSG00000141279 |
| GPRIN2 | -0.648642196 | 0.637880378 | 6.79E-05 | 4.93E-04 | 285.1514081 | 181.9663001 | ENSG00000204175 |
| UTP20 | -0.648801952 | 0.637809746 | 7.77E-13 | 2.35E-11 | 1690.375386 | 1077.2811 | ENSG00000120800 |
| WDR73 | -0.648901141 | 0.637765897 | 4.92E-03 | 2.03E-02 | 115.2769764 | 73.53791736 | ENSG00000177082 |
| HRAS | -0.650380193 | 0.637112393 | 7.65E-10 | 1.45E-08 | 761.1528723 | 484.713105 | ENSG00000174775 |
| FOXP4 | -0.651681344 | 0.636538048 | 1.77E-07 | 2.26E-06 | 817.7049814 | 520.1629643 | ENSG00000137166 |
| RBM19 | -0.653014287 | 0.635950206 | 2.88E-15 | 1.13E-13 | 1727.817642 | 1099.663734 | ENSG00000122965 |
| HDX | -0.653562316 | 0.635708676 | 1.75E-02 | 5.77E-02 | 101.6642365 | 64.39221077 | ENSG00000165259 |
| TOMM40L | -0.654302026 | 0.635382814 | 3.39E-06 | 3.41E-05 | 340.3910242 | 216.3087025 | ENSG00000158882 |
| CHRNB2 | -0.654991143 | 0.63507939 | 3.49E-03 | 1.51E-02 | 127.5372208 | 80.86661945 | ENSG00000160716 |
| AQP3 | -0.655193228 | 0.634990438 | 2.39E-02 | 7.48E-02 | 72.20816658 | 45.73670291 | ENSG00000165272 |
| FNDC4 | -0.655660075 | 0.634784992 | 2.32E-04 | 1.47E-03 | 201.258284 | 127.703007 | ENSG00000115226 |
| DNAJB6 | -0.656370053 | 0.634472679 | 4.16E-26 | 4.30E-24 | 8297.875646 | 5264.09848 | ENSG00000105993 |
| PGRMC1 | -0.656814869 | 0.634277087 | 4.74E-16 | 2.02E-14 | 6261.246845 | 3970.79343 | ENSG00000101856 |
| RNASEH2B | -0.657082986 | 0.634159221 | 8.54E-10 | 1.60E-08 | 687.4974278 | 435.7355334 | ENSG00000136104 |
| ANKRD34B | -0.657181096 | 0.634116096 | 2.40E-02 | 7.50E-02 | 68.7130685 | 43.51652439 | ENSG00000189127 |
| POLD3 | -0.65878241 | 0.633412652 | 1.37E-10 | 2.96E-09 | 749.6478136 | 474.5868869 | ENSG00000077514 |
| TFEB | -0.659686371 | 0.633015894 | 3.40E-02 | 9.90E-02 | 67.59156859 | 42.79259451 | ENSG00000112561 |
| TRIM46 | -0.660551879 | 0.632636246 | 3.25E-05 | 2.58E-04 | 457.6386869 | 290.0791918 | ENSG00000163462 |
| OGG1 | -0.660762204 | 0.632544023 | 1.19E-08 | 1.88E-07 | 585.3795507 | 370.5193658 | ENSG00000114026 |
| ZNF358 | -0.660984687 | 0.632446484 | 1.46E-03 | 7.22E-03 | 187.8253301 | 118.5987529 | ENSG00000198816 |
| EIF2B1 | -0.662013503 | 0.631995634 | 7.11E-15 | 2.67E-13 | 1298.568974 | 821.1486573 | ENSG00000111361 |
| BRICD5 | -0.662356529 | 0.631845384 | 2.82E-02 | 8.55E-02 | 85.52588566 | 54.23747406 | ENSG00000182685 |
| LSS | -0.663547217 | 0.631324123 | 4.94E-22 | 3.67E-20 | 5262.769402 | 3323.472695 | ENSG00000160285 |
| ANAPC7 | -0.664252962 | 0.631015364 | 9.82E-13 | 2.92E-11 | 983.3570834 | 620.8224072 | ENSG00000196510 |
| MRPL12 | -0.664863347 | 0.630748446 | 9.99E-03 | 3.68E-02 | 94.39036948 | 59.64914471 | ENSG00000262814 |
| CCNE1 | -0.665610649 | 0.630421809 | 7.66E-14 | 2.58E-12 | 1170.994435 | 738.3483853 | ENSG00000105173 |
| CNTFR | -0.666099184 | 0.630208368 | 9.03E-03 | 3.38E-02 | 108.4680998 | 68.24493304 | ENSG00000122756 |
| TMEM198 | -0.666615894 | 0.629982696 | 3.72E-05 | 2.91E-04 | 249.3948519 | 157.2075007 | ENSG00000188760 |
| RUNDC3A | -0.666783064 | 0.629909701 | 1.30E-02 | 4.56E-02 | 87.73358073 | 55.32846169 | ENSG00000108309 |
| STX18 | -0.667039698 | 0.62979766 | 6.87E-09 | 1.13E-07 | 570.6321198 | 358.960323 | ENSG00000168818 |
| ESPL1 | -0.669149822 | 0.628877174 | 3.71E-17 | 1.80E-15 | 1860.104735 | 1170.434469 | ENSG00000135476 |
| TEX261 | -0.669374989 | 0.628779031 | 1.01E-06 | 1.13E-05 | 400.355024 | 252.1451005 | ENSG00000144043 |
| CDK4 | -0.669383997 | 0.628775105 | 1.11E-21 | 8.10E-20 | 2320.901452 | 1458.947861 | ENSG00000135446 |
| ZNF599 | -0.669506083 | 0.628721898 | 4.95E-02 | 1.33E-01 | 59.68692836 | 37.62450876 | ENSG00000153896 |
| U2AF2 | -0.66975587 | 0.628613051 | 1.20E-24 | 1.10E-22 | 8646.229674 | 5435.990094 | ENSG00000063244 |
| DHX29 | -0.67057722 | 0.628255273 | 3.51E-12 | 9.52E-11 | 1057.652118 | 664.079235 | ENSG00000067248 |
| SKIV2L | -0.67066075 | 0.628218899 | 5.29E-14 | 1.81E-12 | 1279.977509 | 804.5488492 | ENSG00000204351 |

| | | | | | | | |
|---|---|---|---|---|---|---|---|
| CARMIL3 | -0.670664048 | 0.628217463 | 6.36E-04 | 3.54E-03 | 327.0375717 | 205.9530358 | ENSG00000186648 |
| GPX3 | -0.673389468 | 0.627031807 | 1.31E-09 | 2.39E-08 | 687.4823423 | 430.7461342 | ENSG00000211445 |
| TBC1D30 | -0.673512791 | 0.62697821 | 5.17E-03 | 2.12E-02 | 102.902715 | 64.41208107 | ENSG00000111490 |
| NDUFA9 | -0.673828258 | 0.626841126 | 2.91E-02 | 8.75E-02 | 66.99166549 | 41.96776162 | ENSG00000139180 |
| EPB41L1 | -0.673828719 | 0.626840926 | 1.25E-11 | 3.12E-10 | 883.0542942 | 553.7860488 | ENSG00000088367 |
| DHRS13 | -0.674776823 | 0.626429117 | 2.20E-07 | 2.77E-06 | 529.3371626 | 331.3958686 | ENSG00000167536 |
| TSPOAP1 | -0.676064459 | 0.625870265 | 1.31E-03 | 6.62E-03 | 264.0349857 | 165.7064483 | ENSG00000005379 |
| TCF15 | -0.677010568 | 0.625459958 | 8.04E-06 | 7.36E-05 | 323.5005695 | 202.6147994 | ENSG00000125878 |
| REC8 | -0.679307016 | 0.624465158 | 1.23E-07 | 1.61E-06 | 568.2193835 | 354.6560463 | ENSG00000100918 |
| UBE2F | -0.679374661 | 0.624435879 | 2.56E-03 | 1.17E-02 | 130.3839797 | 81.16619092 | ENSG00000184182 |
| FERMT1 | -0.679874549 | 0.624219552 | 1.72E-08 | 2.65E-07 | 579.2520707 | 361.0111245 | ENSG00000101311 |
| GLT1D1 | -0.680151787 | 0.624099609 | 1.93E-06 | 2.04E-05 | 325.1304598 | 202.774715 | ENSG00000151948 |
| NAB2 | -0.681839884 | 0.623369778 | 7.02E-08 | 9.61E-07 | 661.9415174 | 412.5648078 | ENSG00000166886 |
| DZIP1 | -0.682338017 | 0.623154578 | 8.82E-13 | 2.65E-11 | 1025.209415 | 639.296091 | ENSG00000134874 |
| PCDHGC4 | -0.682800719 | 0.622954751 | 2.53E-02 | 7.82E-02 | 69.26021371 | 43.18309817 | ENSG00000242419 |
| SP110 | -0.683444327 | 0.622676904 | 1.44E-02 | 4.94E-02 | 82.17342739 | 51.27773225 | ENSG00000135899 |
| GCLM | -0.685394521 | 0.621835756 | 1.99E-06 | 2.10E-05 | 443.2943254 | 275.3285049 | ENSG00000023909 |
| EBP | -0.685687525 | 0.621709476 | 8.79E-07 | 9.87E-06 | 336.1202507 | 209.0129791 | ENSG00000147155 |
| ATAD3B | -0.685854809 | 0.621637392 | 1.64E-15 | 6.69E-14 | 2560.001995 | 1592.00176 | ENSG00000160072 |
| NDUFC2 | -0.687199497 | 0.621058255 | 3.21E-02 | 9.46E-02 | 63.80766586 | 39.75875641 | ENSG00000151366 |
| PCDHA6 | -0.68763009 | 0.620872918 | 3.37E-03 | 1.47E-02 | 110.4560137 | 68.63989681 | ENSG00000081842 |
| CEP162 | -0.687763285 | 0.6208156 | 1.20E-03 | 6.14E-03 | 163.5242316 | 101.3191583 | ENSG00000135315 |
| SMIM3 | -0.688347369 | 0.620564309 | 8.26E-10 | 1.55E-08 | 655.0239681 | 406.0559288 | ENSG00000256235 |
| FBXO3 | -0.689443657 | 0.620092929 | 2.34E-07 | 2.93E-06 | 401.9683179 | 249.2248357 | ENSG00000110429 |
| SMOC2 | -0.69024133 | 0.619750171 | 8.82E-03 | 3.32E-02 | 96.19067267 | 59.70925654 | ENSG00000112562 |
| LINGO1 | -0.690300352 | 0.619724817 | 3.21E-13 | 1.00E-11 | 2075.536201 | 1285.861479 | ENSG00000169783 |
| GPR143 | -0.690358873 | 0.61969968 | 4.57E-07 | 5.44E-06 | 388.231782 | 240.8389923 | ENSG00000101850 |
| STS | -0.692234716 | 0.618894447 | 4.81E-03 | 1.99E-02 | 113.4666988 | 70.24453453 | ENSG00000101846 |
| GRIA3 | -0.693195366 | 0.61848248 | 6.01E-04 | 3.37E-03 | 197.4571009 | 122.2621565 | ENSG00000125675 |
| HNRNPM | -0.693304255 | 0.618435802 | 1.17E-27 | 1.37E-25 | 14475.76401 | 8952.038608 | ENSG00000099783 |
| PIANP | -0.694050952 | 0.6181158 | 1.15E-04 | 7.89E-04 | 226.4011425 | 140.2018818 | ENSG00000139200 |
| HMGN2 | -0.694177554 | 0.61806156 | 9.64E-16 | 4.02E-14 | 6303.993973 | 3895.67335 | ENSG00000198830 |
| LZTS3 | -0.69451035 | 0.617919004 | 1.49E-11 | 3.67E-10 | 1231.19533 | 761.5871412 | ENSG00000088899 |
| SYNDIG1 | -0.695564259 | 0.617467771 | 2.53E-03 | 1.16E-02 | 119.8640798 | 73.96151133 | ENSG00000101463 |
| SLC12A8 | -0.696357048 | 0.617128553 | 1.44E-04 | 9.57E-04 | 223.4486799 | 137.9954645 | ENSG00000221955 |
| AKT1S1 | -0.696963387 | 0.61686924 | 1.06E-12 | 3.13E-11 | 1425.54198 | 879.1628046 | ENSG00000204673 |
| RIPOR3 | -0.697649656 | 0.616575874 | 3.50E-02 | 1.01E-01 | 66.64683315 | 41.24426978 | ENSG00000042062 |
| SETD6 | -0.69833738 | 0.616282026 | 1.70E-10 | 3.58E-09 | 668.3700842 | 412.2924726 | ENSG00000103037 |
| SNHG20 | -0.698615972 | 0.61616303 | 4.59E-02 | 1.25E-01 | 66.81087228 | 41.33476393 | ENSG00000234912 |
| LMO3 | -0.699338944 | 0.615854332 | 3.06E-03 | 1.36E-02 | 134.145303 | 82.58454441 | ENSG00000048540 |
| HM13 | -0.699492884 | 0.615788622 | 8.35E-19 | 4.81E-17 | 2330.123824 | 1434.565446 | ENSG00000101294 |
| KIAA1549L | -0.703983002 | 0.613875075 | 2.68E-03 | 1.21E-02 | 109.3140991 | 67.09047277 | ENSG00000110427 |
| ATP6V0D1 | -0.704035941 | 0.613852549 | 6.63E-16 | 2.81E-14 | 1281.647056 | 786.6920916 | ENSG00000159720 |
| MCM4 | -0.704088745 | 0.613830082 | 1.17E-31 | 1.82E-29 | 8426.954086 | 5172.758363 | ENSG00000104738 |
| ZW10 | -0.704466441 | 0.613669403 | 1.12E-09 | 2.05E-08 | 661.6947075 | 405.7176044 | ENSG00000086827 |
| OTULINL | -0.704612375 | 0.613607331 | 3.46E-04 | 2.08E-03 | 207.1340682 | 126.9404358 | ENSG00000145569 |
| NT5C3A | -0.705770682 | 0.613114878 | 1.03E-06 | 1.15E-05 | 392.7612642 | 240.4123321 | ENSG00000122643 |
| CHST1 | -0.705865202 | 0.61307471 | 4.03E-02 | 1.13E-01 | 56.68215106 | 34.64924515 | ENSG00000175264 |
| ARMC2 | -0.706849349 | 0.612656638 | 4.28E-02 | 1.19E-01 | 55.55464559 | 34.15166651 | ENSG00000118690 |
| KCP | -0.706980076 | 0.612601136 | 3.99E-04 | 2.35E-03 | 159.7636074 | 97.96372859 | ENSG00000135253 |
| CCDC157 | -0.708899041 | 0.611786831 | 1.54E-02 | 5.23E-02 | 72.6859574 | 44.56715887 | ENSG00000187860 |
| CHN1 | -0.709435276 | 0.611559479 | 5.09E-12 | 1.36E-10 | 893.7728616 | 545.9428919 | ENSG00000128656 |
| ZNF511 | -0.710769775 | 0.610994045 | 3.81E-03 | 1.64E-02 | 102.0263266 | 62.22520767 | ENSG00000198546 |
| BMF | -0.711175696 | 0.610822158 | 1.00E-07 | 1.34E-06 | 508.1660054 | 310.7251577 | ENSG00000104081 |
| NXF1 | -0.711329613 | 0.610756995 | 4.00E-14 | 1.39E-12 | 1025.520033 | 626.2768041 | ENSG00000162231 |
| AMD1 | -0.711688772 | 0.610604966 | 4.61E-04 | 2.67E-03 | 3930.515804 | 2399.722927 | ENSG00000123505 |
| SH3GL2 | -0.711942035 | 0.610497785 | 3.93E-03 | 1.68E-02 | 109.4685546 | 67.00834075 | ENSG00000107295 |
| PRRT3 | -0.712288886 | 0.610351027 | 2.39E-04 | 1.51E-03 | 209.4716399 | 127.7395613 | ENSG00000163704 |
| ADORA1 | -0.71241136 | 0.610299215 | 2.41E-03 | 1.11E-02 | 135.5789611 | 82.77649124 | ENSG00000163485 |
| SLC20A1 | -0.712570622 | 0.610231846 | 9.07E-23 | 7.25E-21 | 2448.411328 | 1493.425754 | ENSG00000144136 |
| ANKRD36B | -0.712618934 | 0.610211412 | 2.30E-02 | 7.23E-02 | 61.9989743 | 37.84183659 | ENSG00000196912 |
| MCM6 | -0.713021211 | 0.610041286 | 8.54E-30 | 1.18E-27 | 5133.474836 | 3130.8564 | ENSG00000076003 |
| SYT2 | -0.713112214 | 0.610002807 | 7.66E-13 | 2.32E-11 | 803.3529982 | 490.1087445 | ENSG00000143858 |
| DHFR | -0.71438382 | 0.609465381 | 3.68E-15 | 1.45E-13 | 1363.956102 | 830.8491627 | ENSG00000228716 |
| FOS | -0.714394172 | 0.609461008 | 5.29E-07 | 6.23E-06 | 358.4630427 | 218.3526792 | ENSG00000170345 |
| EVL | -0.714884604 | 0.609253862 | 2.41E-12 | 6.67E-11 | 1173.792682 | 715.6776433 | ENSG00000196405 |
| MFSD6 | -0.715908833 | 0.608821481 | 1.69E-02 | 5.62E-02 | 73.77273127 | 44.79164636 | ENSG00000151690 |
| TNFSF13 | -0.716261512 | 0.608672668 | 4.19E-02 | 1.17E-01 | 52.1113359 | 31.70188756 | ENSG00000161955 |
| ENKD1 | -0.717693198 | 0.608068939 | 5.98E-05 | 4.41E-04 | 214.5384284 | 130.2646475 | ENSG00000124074 |
| SLC1A6 | -0.71816997 | 0.607868022 | 2.07E-05 | 1.72E-04 | 282.4116161 | 171.6028838 | ENSG00000105143 |
| IRF1 | -0.718498461 | 0.60772963 | 1.74E-02 | 5.76E-02 | 67.81330396 | 41.24740735 | ENSG00000125347 |
| NSG1 | -0.718744895 | 0.60762583 | 1.54E-14 | 5.61E-13 | 1037.585173 | 630.5046199 | ENSG00000168824 |
| TAF7 | -0.719895456 | 0.607141437 | 2.97E-04 | 1.82E-03 | 2994.820859 | 1818.666245 | ENSG00000178913 |
| DPP4 | -0.720410893 | 0.60692456 | 1.13E-02 | 4.07E-02 | 82.05416376 | 49.83680891 | ENSG00000197635 |
| ARSI | -0.72045024 | 0.606908007 | 3.59E-04 | 2.14E-03 | 172.364813 | 104.7148699 | ENSG00000183876 |

| | | | | | | | |
|---|---|---|---|---|---|---|---|
| PPP1R1B | -0.721029304 | 0.606664457 | 8.59E-07 | 9.67E-06 | 299.2238319 | 181.4745586 | ENSG00000131771 |
| NAA16 | -0.721188563 | 0.606597492 | 3.24E-08 | 4.76E-07 | 464.4385889 | 281.5158954 | ENSG00000172766 |
| FEZF1 | -0.721622581 | 0.606415031 | 3.37E-05 | 2.66E-04 | 215.4233779 | 130.6702349 | ENSG00000128610 |
| GPAM | -0.722112401 | 0.606209178 | 4.07E-11 | 9.39E-10 | 723.3321479 | 437.9811961 | ENSG00000119927 |
| IFT57 | -0.722656229 | 0.605980709 | 2.84E-09 | 4.92E-08 | 561.8653647 | 340.1383525 | ENSG00000114446 |
| CDK2AP1 | -0.722783442 | 0.605927277 | 3.21E-13 | 1.00E-11 | 2951.363066 | 1787.621775 | ENSG00000111328 |
| MPZ | -0.722819594 | 0.605912093 | 3.43E-02 | 9.97E-02 | 66.52768239 | 40.22737844 | ENSG00000158887 |
| MYH14 | -0.723630688 | 0.605571541 | 3.16E-18 | 1.70E-16 | 1406.153094 | 851.1987518 | ENSG00000105357 |
| ST14 | -0.724194447 | 0.605334949 | 8.84E-22 | 6.53E-20 | 1869.553525 | 1131.556388 | ENSG00000149418 |
| DTX1 | -0.724423292 | 0.605238936 | 7.01E-07 | 8.04E-06 | 329.6916088 | 199.5797504 | ENSG00000135144 |
| LIG1 | -0.724439431 | 0.605232166 | 3.17E-19 | 1.88E-17 | 2125.183831 | 1286.575455 | ENSG00000105486 |
| CCDC88B | -0.724970681 | 0.60500934 | 3.17E-07 | 3.87E-06 | 474.6621751 | 287.6968991 | ENSG00000168071 |
| STAMBPL1 | -0.725813487 | 0.604656003 | 9.72E-04 | 5.14E-03 | 128.7954375 | 77.7911141 | ENSG00000138134 |
| BAG1 | -0.726727118 | 0.604273207 | 2.14E-12 | 6.02E-11 | 1176.210619 | 710.4196441 | ENSG00000107262 |
| DHCR24 | -0.728181151 | 0.603664492 | 1.08E-38 | 2.67E-36 | 13545.5691 | 8176.710401 | ENSG00000116133 |
| HES1 | -0.729014209 | 0.603316018 | 1.98E-04 | 1.27E-03 | 212.0236863 | 127.6039274 | ENSG00000114315 |
| THAP10 | -0.731485202 | 0.602283566 | 7.83E-03 | 3.01E-02 | 95.20489737 | 57.48907802 | ENSG00000129028 |
| ARHGEF19 | -0.732768316 | 0.60174814 | 1.63E-15 | 6.68E-14 | 1410.487419 | 848.9827558 | ENSG00000142632 |
| PCYT2 | -0.733237553 | 0.601552454 | 9.78E-17 | 4.46E-15 | 1193.523582 | 718.4554091 | ENSG00000185813 |
| OAT | -0.733722663 | 0.601350214 | 2.17E-25 | 2.13E-23 | 2426.821745 | 1458.796602 | ENSG00000065154 |
| DHX35 | -0.734390409 | 0.601071946 | 1.00E-09 | 1.86E-08 | 527.6294848 | 317.2773126 | ENSG00000101452 |
| HTR3A | -0.735332306 | 0.60067965 | 7.20E-05 | 5.19E-04 | 197.4851298 | 118.792237 | ENSG00000166736 |
| SCNN1A | -0.735379472 | 0.600660012 | 1.40E-18 | 7.81E-17 | 1609.664149 | 967.2706353 | ENSG00000111319 |
| COL9A2 | -0.736482294 | 0.600201033 | 3.83E-04 | 2.27E-03 | 149.1701289 | 89.53914072 | ENSG00000049089 |
| PAQR4 | -0.737587475 | 0.599741423 | 2.27E-06 | 2.36E-05 | 349.7586367 | 210.003725 | ENSG00000162073 |
| ACAA1 | -0.737607118 | 0.599733257 | 1.84E-09 | 3.28E-08 | 564.8285008 | 338.8998335 | ENSG00000060971 |
| PARD6A | -0.738320054 | 0.59943696 | 1.11E-04 | 7.64E-04 | 204.9732157 | 122.5559537 | ENSG00000102981 |
| DNAJC3 | -0.74026217 | 0.598630558 | 6.47E-10 | 1.24E-08 | 727.5272833 | 435.1342003 | ENSG00000102580 |
| TCF25 | -0.740579724 | 0.598498807 | 1.19E-18 | 6.70E-17 | 1286.333324 | 770.2054506 | ENSG00000141002 |
| POMC | -0.741189161 | 0.598246037 | 1.52E-02 | 5.18E-02 | 76.87880023 | 46.02228991 | ENSG00000115138 |
| DPM3 | -0.741424028 | 0.598148652 | 1.45E-06 | 1.56E-05 | 338.8378844 | 203.0738887 | ENSG00000179085 |
| STAC2 | -0.742898569 | 0.597537612 | 1.81E-05 | 1.52E-04 | 229.5219288 | 136.9858445 | ENSG00000141750 |
| MCM5 | -0.743228561 | 0.597400951 | 2.42E-35 | 4.82E-33 | 8473.265906 | 5061.752723 | ENSG00000100297 |
| ACOT2 | -0.744230664 | 0.596986137 | 6.85E-04 | 3.79E-03 | 140.9940168 | 84.34479789 | ENSG00000119673 |
| CA12 | -0.744555271 | 0.59685183 | 2.63E-03 | 1.19E-02 | 115.8331864 | 69.24371465 | ENSG00000074410 |
| ANKRD35 | -0.74526011 | 0.596560305 | 2.10E-04 | 1.34E-03 | 171.2825866 | 102.2247378 | ENSG00000198483 |
| SLC2A6 | -0.745704081 | 0.59637675 | 3.97E-09 | 6.77E-08 | 611.9675828 | 364.9062869 | ENSG00000160326 |
| DUT | -0.745728299 | 0.596366739 | 2.77E-20 | 1.82E-18 | 1963.056051 | 1170.350999 | ENSG00000128951 |
| P2RY11 | -0.746596345 | 0.596008022 | 3.19E-02 | 9.42E-02 | 50.08809461 | 29.85625287 | ENSG00000244165 |
| CEP290 | -0.747920718 | 0.595461147 | 7.54E-05 | 5.41E-04 | 200.2628426 | 119.4377221 | ENSG00000198707 |
| HEXA | -0.748862572 | 0.595072531 | 1.38E-02 | 4.78E-02 | 72.06149805 | 42.96956919 | ENSG00000213614 |
| PRTG | -0.749133512 | 0.594960786 | 4.88E-03 | 2.02E-02 | 100.2449422 | 59.39273756 | ENSG00000166450 |
| ZNF576 | -0.750271206 | 0.594491791 | 1.33E-03 | 6.67E-03 | 121.153843 | 72.18066338 | ENSG00000124444 |
| SPSB4 | -0.75061764 | 0.594349053 | 5.91E-12 | 1.56E-10 | 683.1032568 | 405.6016177 | ENSG00000175093 |
| SLC6A8 | -0.750778618 | 0.594282738 | 1.58E-23 | 1.34E-21 | 2976.925402 | 1769.077858 | ENSG00000130821 |
| GINS2 | -0.751278391 | 0.594076905 | 2.53E-19 | 1.53E-17 | 1525.921527 | 906.4975327 | ENSG00000131153 |
| COQ4 | -0.75165012 | 0.593923853 | 3.69E-10 | 7.29E-09 | 538.5659989 | 319.9882449 | ENSG00000167113 |
| PAQR5 | -0.752711607 | 0.593487024 | 8.05E-05 | 5.73E-04 | 210.5138489 | 124.5245116 | ENSG00000137819 |
| PCDH1 | -0.753682213 | 0.593087877 | 7.32E-31 | 1.09E-28 | 5315.140244 | 3152.016743 | ENSG00000156453 |
| CCBE1 | -0.754974173 | 0.592556993 | 1.46E-03 | 7.25E-03 | 123.4973603 | 72.92193854 | ENSG00000183287 |
| FLVCR1 | -0.755246776 | 0.592445038 | 2.87E-19 | 1.71E-17 | 1423.730748 | 843.6323507 | ENSG00000162769 |
| CDH24 | -0.75593843 | 0.592161077 | 5.81E-11 | 1.31E-09 | 637.3977129 | 377.7745213 | ENSG00000139880 |
| CLDN6 | -0.756505774 | 0.591928254 | 1.49E-38 | 3.64E-36 | 13898.98977 | 8226.727945 | ENSG00000184697 |
| CYB561A3 | -0.756982935 | 0.59173251 | 1.51E-04 | 9.96E-04 | 246.3914501 | 145.9675103 | ENSG00000162144 |
| GGT7 | -0.757564027 | 0.591494219 | 2.36E-11 | 5.63E-10 | 1509.040754 | 893.4357605 | ENSG00000131067 |
| EFNB3 | -0.757694553 | 0.591440707 | 3.54E-10 | 7.03E-09 | 564.4611793 | 334.0461713 | ENSG00000108947 |
| PLXNA4 | -0.758439256 | 0.59113549 | 4.45E-02 | 1.22E-01 | 48.40397319 | 28.47175413 | ENSG00000221866 |
| UBQLN2 | -0.760743034 | 0.590192285 | 1.17E-14 | 4.33E-13 | 1035.20968 | 611.0755908 | ENSG00000188021 |
| NXPH3 | -0.761113265 | 0.590040846 | 4.07E-04 | 2.39E-03 | 162.5087437 | 96.00981643 | ENSG00000182575 |
| KCND1 | -0.761229284 | 0.589993398 | 1.41E-06 | 1.52E-05 | 291.0491179 | 171.9481362 | ENSG00000102057 |
| FOXO4 | -0.762021892 | 0.589669348 | 1.09E-11 | 2.74E-10 | 595.8816142 | 351.0540601 | ENSG00000184481 |
| TTYH2 | -0.762622367 | 0.589423969 | 1.34E-03 | 6.74E-03 | 121.2207168 | 71.57326994 | ENSG00000141540 |
| ENHO | -0.763600804 | 0.589024356 | 3.81E-04 | 2.26E-03 | 178.7113 | 105.1638449 | ENSG00000168913 |
| HUNK | -0.763689428 | 0.588988174 | 8.21E-08 | 1.12E-06 | 384.0238536 | 226.1683279 | ENSG00000142149 |
| AFAP1L2 | -0.763939092 | 0.588886256 | 2.61E-03 | 1.19E-02 | 105.2807478 | 61.94028194 | ENSG00000169129 |
| CXCL12 | -0.764807798 | 0.58853177 | 1.00E-13 | 3.29E-12 | 783.8272903 | 460.8239518 | ENSG00000107562 |
| CLCN2 | -0.765213095 | 0.588366457 | 1.34E-05 | 1.17E-04 | 250.6424178 | 147.4964027 | ENSG00000114859 |
| BRD9 | -0.766153034 | 0.587983251 | 4.13E-13 | 1.28E-11 | 1065.466907 | 627.0366044 | ENSG00000028310 |
| MBNL3 | -0.766218935 | 0.587956393 | 6.21E-03 | 2.48E-02 | 87.00829557 | 50.97998426 | ENSG00000076770 |
| CHST7 | -0.766738423 | 0.587744719 | 2.20E-09 | 3.90E-08 | 656.6656665 | 386.2503092 | ENSG00000147119 |
| NASP | -0.766770746 | 0.587731551 | 1.16E-44 | 4.05E-42 | 17308.41126 | 10173.57458 | ENSG00000132780 |
| MPP2 | -0.767188066 | 0.587561566 | 5.18E-28 | 6.41E-26 | 2335.566435 | 1372.254562 | ENSG00000108852 |
| SEPTIN3 | -0.767491307 | 0.587438079 | 4.55E-22 | 3.40E-20 | 1632.101273 | 958.5600309 | ENSG00000100167 |
| FBXW4 | -0.768687091 | 0.58695138 | 1.31E-11 | 3.26E-10 | 689.7691103 | 404.4985807 | ENSG00000107829 |
| SH2B2 | -0.768761587 | 0.586921073 | 6.58E-08 | 9.07E-07 | 429.1458878 | 251.8916161 | ENSG00000160999 |

| | | | | | | | |
|---|---|---|---|---|---|---|---|
| PDS5A | -0.769201821 | 0.586742003 | 1.74E-20 | 1.16E-18 | 2491.26743 | 1460.649522 | ENSG00000121892 |
| CD2BP2 | -0.770476889 | 0.586223664 | 9.47E-17 | 4.36E-15 | 2274.278465 | 1334.402773 | ENSG00000169217 |
| EAPP | -0.771196985 | 0.585931134 | 5.69E-08 | 7.97E-07 | 367.5020059 | 215.0850038 | ENSG00000129518 |
| DYRK1B | -0.771956688 | 0.585622672 | 1.42E-05 | 1.23E-04 | 255.4713781 | 149.6081922 | ENSG00000105204 |
| CCNC | -0.772222581 | 0.58551475 | 9.20E-08 | 1.24E-06 | 673.3365581 | 393.6739882 | ENSG00000112237 |
| MPDU1 | -0.772765457 | 0.585294466 | 7.14E-12 | 1.85E-10 | 663.6546 | 388.3975452 | ENSG00000129255 |
| SOAT1 | -0.775997141 | 0.583984855 | 1.32E-03 | 6.63E-03 | 119.4797936 | 69.77613574 | ENSG00000057252 |
| PDSS2 | -0.776077968 | 0.583952138 | 4.67E-06 | 4.56E-05 | 275.8085774 | 161.2509673 | ENSG00000164494 |
| SLC22A5 | -0.776966542 | 0.583592585 | 1.55E-03 | 7.64E-03 | 109.5549713 | 63.96971611 | ENSG00000197375 |
| GPR137 | -0.776976363 | 0.583588612 | 4.14E-10 | 8.13E-09 | 640.177485 | 373.3702318 | ENSG00000173264 |
| RAB15 | -0.777536934 | 0.583361898 | 7.44E-23 | 6.02E-21 | 2481.982792 | 1448.031996 | ENSG00000139998 |
| TAF2 | -0.777655983 | 0.583313762 | 1.16E-13 | 3.78E-12 | 726.9153086 | 423.6240247 | ENSG00000064313 |
| DACT3 | -0.777824211 | 0.583245748 | 2.31E-02 | 7.25E-02 | 58.16388455 | 34.04272777 | ENSG00000197380 |
| FAM189B | -0.778767037 | 0.582864711 | 2.96E-14 | 1.04E-12 | 1284.215992 | 748.4463879 | ENSG00000160767 |
| HCN3 | -0.778833474 | 0.58283787 | 8.94E-04 | 4.79E-03 | 224.915884 | 131.5852758 | ENSG00000143630 |
| ATP1A2 | -0.778899057 | 0.582811376 | 9.43E-12 | 2.39E-10 | 1925.034105 | 1122.743442 | ENSG00000018625 |
| SUSD4 | -0.780642619 | 0.582107447 | 8.51E-03 | 3.22E-02 | 76.56632631 | 44.40146912 | ENSG00000143502 |
| ATAD1 | -0.780781659 | 0.58205135 | 3.87E-11 | 8.98E-10 | 770.4505442 | 448.1986898 | ENSG00000138138 |
| FIG4 | -0.78135181 | 0.581821369 | 1.88E-12 | 5.31E-11 | 670.8879234 | 390.0898743 | ENSG00000112367 |
| SIVA1 | -0.782190675 | 0.581483163 | 1.23E-18 | 6.89E-17 | 1575.465393 | 916.0557313 | ENSG00000184990 |
| VIPR1 | -0.783596314 | 0.580916892 | 9.47E-03 | 3.52E-02 | 77.5042669 | 44.8774656 | ENSG00000114812 |
| SDF2L1 | -0.783875185 | 0.580804612 | 2.66E-15 | 1.06E-13 | 1602.434328 | 930.8590582 | ENSG00000128228 |
| SLC25A13 | -0.784454426 | 0.580571466 | 6.29E-21 | 4.32E-19 | 2131.171167 | 1236.403562 | ENSG00000004864 |
| FLRT2 | -0.784602825 | 0.58051175 | 1.39E-03 | 6.94E-03 | 105.3129934 | 61.15667826 | ENSG00000185070 |
| RTN4RL2 | -0.785645988 | 0.580092154 | 1.98E-05 | 1.65E-04 | 335.2222902 | 194.3770539 | ENSG00000186907 |
| CIPC | -0.786625502 | 0.579698436 | 7.93E-08 | 1.08E-06 | 418.7590518 | 242.9935568 | ENSG00000198894 |
| PRELID3A | -0.789833778 | 0.57841073 | 1.27E-04 | 8.61E-04 | 167.2803985 | 97.04905436 | ENSG00000141391 |
| FGFR1OP | -0.792104817 | 0.577500933 | 4.50E-05 | 3.43E-04 | 198.9961473 | 114.8677916 | ENSG00000213066 |
| GK | -0.792370671 | 0.577394524 | 1.94E-05 | 1.62E-04 | 202.7773742 | 116.8135564 | ENSG00000198814 |
| CDHR1 | -0.792453305 | 0.577361453 | 3.68E-08 | 5.34E-07 | 503.1793425 | 291.0832524 | ENSG00000148600 |
| STAG1 | -0.79329349 | 0.577025312 | 1.65E-11 | 4.04E-10 | 546.8635242 | 315.7159423 | ENSG00000118007 |
| KCNN2 | -0.79341554 | 0.576976498 | 1.35E-06 | 1.46E-05 | 309.7362227 | 178.6004212 | ENSG00000080709 |
| SPRYD4 | -0.793876907 | 0.576792013 | 1.31E-04 | 8.83E-04 | 182.4106803 | 104.8668613 | ENSG00000176422 |
| ARNT2 | -0.794669516 | 0.576475214 | 8.89E-08 | 1.20E-06 | 332.104751 | 191.6253219 | ENSG00000172379 |
| RORB | -0.794749742 | 0.576443158 | 1.95E-04 | 1.25E-03 | 148.7053339 | 85.85981752 | ENSG00000198963 |
| CFAP69 | -0.794780583 | 0.576430835 | 4.54E-02 | 1.24E-01 | 42.78504122 | 24.81956407 | ENSG00000105792 |
| CA11 | -0.795255619 | 0.576241065 | 2.41E-17 | 1.19E-15 | 1008.866857 | 581.3239281 | ENSG00000063180 |
| TMEM200A | -0.795944226 | 0.575966087 | 4.79E-07 | 5.67E-06 | 418.7370359 | 240.8660936 | ENSG00000164484 |
| FOXM1 | -0.798333757 | 0.575012906 | 7.11E-28 | 8.67E-26 | 2953.789496 | 1698.923428 | ENSG00000111206 |
| WDR27 | -0.799174087 | 0.574678074 | 3.16E-02 | 9.33E-02 | 370.1566444 | 212.9121954 | ENSG00000184465 |
| PAK6 | -0.79930762 | 0.574624886 | 7.64E-06 | 7.06E-05 | 200.9953134 | 115.4193101 | ENSG00000137843 |
| DONSON | -0.801357559 | 0.573808976 | 8.10E-10 | 1.52E-08 | 488.4407204 | 279.9914143 | ENSG00000159147 |
| HS6ST3 | -0.802708976 | 0.573271723 | 2.82E-02 | 8.54E-02 | 46.33132632 | 26.6345932 | ENSG00000185352 |
| DDX58 | -0.803594581 | 0.572919926 | 1.20E-03 | 6.16E-03 | 112.237398 | 64.14168946 | ENSG00000107201 |
| BEND5 | -0.803812432 | 0.57283342 | 8.80E-04 | 4.72E-03 | 109.5776484 | 62.8411865 | ENSG00000162373 |
| PHYHIP | -0.803856726 | 0.572815832 | 1.50E-06 | 1.61E-05 | 245.4470078 | 140.7186695 | ENSG00000168490 |
| KCNQ4 | -0.804523487 | 0.572551159 | 3.48E-04 | 2.08E-03 | 127.3830584 | 72.9304123 | ENSG00000117013 |
| IRF4 | -0.804649589 | 0.572501116 | 3.48E-02 | 1.01E-01 | 45.27797807 | 26.03088698 | ENSG00000137265 |
| BEX1 | -0.805761387 | 0.572060094 | 7.32E-22 | 5.43E-20 | 1261.80247 | 721.6333145 | ENSG00000133169 |
| SLC16A14 | -0.806184211 | 0.57189246 | 6.05E-03 | 2.42E-02 | 93.17890513 | 53.01234121 | ENSG00000163053 |
| EID1 | -0.806654038 | 0.571706248 | 3.22E-23 | 2.64E-21 | 2775.267986 | 1587.115471 | ENSG00000255302 |
| CRELD2 | -0.808479298 | 0.570983398 | 4.36E-17 | 2.08E-15 | 1006.659861 | 574.7628614 | ENSG00000184164 |
| GNG2 | -0.808529691 | 0.570963454 | 1.99E-07 | 2.53E-06 | 314.7925332 | 179.4591088 | ENSG00000186469 |
| NEK9 | -0.80895545 | 0.570794979 | 2.52E-17 | 1.24E-15 | 1116.149943 | 637.665411 | ENSG00000119638 |
| PLEKHO1 | -0.810121839 | 0.57033369 | 4.13E-15 | 1.61E-13 | 792.1475904 | 451.2932925 | ENSG00000023902 |
| RAB33A | -0.812197916 | 0.569513555 | 4.37E-03 | 1.84E-02 | 98.43160543 | 56.32278318 | ENSG00000134594 |
| MCM2 | -0.813556286 | 0.568977581 | 4.22E-37 | 9.10E-35 | 4696.760796 | 2672.626991 | ENSG00000073111 |
| NCBP2AS2 | -0.813698932 | 0.568921327 | 4.26E-14 | 1.47E-12 | 835.3830774 | 475.5913311 | ENSG00000270170 |
| IL12RB2 | -0.814020738 | 0.568794438 | 3.44E-02 | 9.98E-02 | 44.38442966 | 25.35567239 | ENSG00000081985 |
| HAS3 | -0.814887649 | 0.568452754 | 3.95E-30 | 5.63E-28 | 2496.190128 | 1419.208809 | ENSG00000103044 |
| ENOX2 | -0.815221143 | 0.568321365 | 2.07E-11 | 4.99E-10 | 515.9840336 | 292.9209227 | ENSG00000165675 |
| IGSF9 | -0.816160376 | 0.567951493 | 3.28E-17 | 1.60E-15 | 1026.169748 | 583.1375718 | ENSG00000085552 |
| TMA7 | -0.816589887 | 0.567782431 | 3.43E-04 | 2.06E-03 | 133.0159935 | 75.28122332 | ENSG00000232112 |
| ASTN1 | -0.81681549 | 0.56769365 | 4.24E-07 | 5.08E-06 | 254.9323055 | 144.6154321 | ENSG00000152092 |
| NPEPL1 | -0.818465346 | 0.567044811 | 6.05E-04 | 3.39E-03 | 146.7271013 | 83.40397822 | ENSG00000215440 |
| MTARC1 | -0.820506289 | 0.566243195 | 2.74E-05 | 2.21E-04 | 204.9750198 | 116.2194232 | ENSG00000117791 |
| ADCYAP1R1 | -0.821217115 | 0.565964271 | 2.80E-13 | 8.81E-12 | 817.5428664 | 462.8855204 | ENSG00000078549 |
| TIGD7 | -0.822142562 | 0.565601338 | 3.41E-02 | 9.92E-02 | 42.07859984 | 23.88904164 | ENSG00000140993 |
| KTN1 | -0.823547903 | 0.565050649 | 5.89E-31 | 8.88E-29 | 2576.66592 | 1455.379635 | ENSG00000126777 |
| SLC18A2 | -0.823645754 | 0.565012326 | 1.75E-02 | 5.78E-02 | 69.01252398 | 39.14409168 | ENSG00000165646 |
| TUBA1A | -0.824503182 | 0.564676625 | 1.93E-39 | 5.17E-37 | 5707.072196 | 3222.234486 | ENSG00000167552 |
| MAP2K6 | -0.824837587 | 0.564545753 | 1.57E-13 | 5.06E-12 | 926.8120848 | 522.4545395 | ENSG00000108984 |
| HTRA1 | -0.825840295 | 0.564153517 | 8.80E-14 | 2.93E-12 | 618.1863353 | 348.5189 | ENSG00000166033 |
| COL11A2 | -0.826407039 | 0.56393194 | 4.89E-03 | 2.02E-02 | 77.01956089 | 43.49059374 | ENSG00000204248 |
| TMEM158 | -0.829712638 | 0.562641301 | 4.11E-05 | 3.17E-04 | 318.7689174 | 179.3318371 | ENSG00000249992 |

| | | | | | | | |
|---|---|---|---|---|---|---|---|
| TRAPPC5 | -0.832508082 | 0.561552152 | 2.84E-02 | 8.58E-02 | 48.99796843 | 27.68533929 | ENSG00000181029 |
| MLLT11 | -0.832790746 | 0.561442139 | 5.05E-08 | 7.15E-07 | 406.5490324 | 227.9903018 | ENSG00000213190 |
| GPRASP2 | -0.833730207 | 0.561076655 | 2.53E-06 | 2.61E-05 | 247.4476696 | 138.9520292 | ENSG00000158301 |
| TAF5 | -0.833934177 | 0.560997335 | 1.84E-12 | 5.22E-11 | 585.5750613 | 328.0990134 | ENSG00000148835 |
| NCAPH | -0.83539325 | 0.560430256 | 8.30E-18 | 4.31E-16 | 1303.857003 | 730.5832755 | ENSG00000121152 |
| HES3 | -0.835573682 | 0.56036017 | 1.18E-07 | 1.56E-06 | 295.8226292 | 165.7772552 | ENSG00000173673 |
| INPPL1 | -0.835905422 | 0.560231333 | 7.45E-28 | 8.94E-26 | 2309.894223 | 1294.700369 | ENSG00000165458 |
| DYNC2H1 | -0.836016486 | 0.560188206 | 3.57E-08 | 5.19E-07 | 377.6285171 | 211.2283711 | ENSG00000187240 |
| SCUBE3 | -0.836083799 | 0.560162069 | 3.57E-03 | 1.54E-02 | 81.34241587 | 45.62115416 | ENSG00000146197 |
| IL20RA | -0.837670397 | 0.559546372 | 3.25E-02 | 9.55E-02 | 43.67839424 | 24.37263582 | ENSG00000016402 |
| ZNF44 | -0.838844863 | 0.559091043 | 4.05E-03 | 1.72E-02 | 75.98713096 | 42.47001518 | ENSG00000197857 |
| SLC25A53 | -0.840641877 | 0.558395075 | 9.76E-07 | 1.09E-05 | 241.2333443 | 134.9278833 | ENSG00000269743 |
| GRHL1 | -0.841111394 | 0.558213378 | 2.64E-03 | 1.20E-02 | 82.44714654 | 46.04843536 | ENSG00000134317 |
| H1-10 | -0.842186967 | 0.557797368 | 2.47E-28 | 3.18E-26 | 2238.724642 | 1248.75192 | ENSG00000184897 |
| LYSMD2 | -0.842905018 | 0.557519813 | 3.16E-02 | 9.33E-02 | 41.39476051 | 23.09371501 | ENSG00000140280 |
| ADGRF3 | -0.843961375 | 0.557111741 | 7.08E-03 | 2.77E-02 | 81.85187352 | 45.71154515 | ENSG00000173567 |
| MARCKSL1 | -0.846548363 | 0.556113644 | 1.86E-25 | 1.86E-23 | 11657.21784 | 6482.587608 | ENSG00000175130 |
| HSD17B7 | -0.846575905 | 0.556103027 | 1.25E-14 | 4.60E-13 | 735.6028099 | 409.344677 | ENSG00000132196 |
| SAP30 | -0.847227339 | 0.555851981 | 1.40E-07 | 1.82E-06 | 317.7745357 | 176.5343211 | ENSG00000164105 |
| SGSM2 | -0.848280607 | 0.555446318 | 6.33E-09 | 1.05E-07 | 763.8621781 | 424.9910439 | ENSG00000141258 |
| TMPRSS2 | -0.849233346 | 0.555079629 | 1.19E-06 | 1.31E-05 | 225.2160385 | 125.1230336 | ENSG00000184012 |
| ADAMTS8 | -0.850450438 | 0.554611548 | 1.23E-12 | 3.57E-11 | 586.9324201 | 325.8672638 | ENSG00000134917 |
| GALNT16 | -0.850698291 | 0.554516275 | 2.20E-06 | 2.29E-05 | 241.982607 | 134.5313193 | ENSG00000100626 |
| HMGN5 | -0.851750169 | 0.554112121 | 9.64E-13 | 2.87E-11 | 703.1251031 | 389.4016227 | ENSG00000198157 |
| MARCHF4 | -0.854102624 | 0.553209323 | 2.21E-03 | 1.04E-02 | 82.14511287 | 45.52729919 | ENSG00000144583 |
| RBM25 | -0.854323587 | 0.553124601 | 1.55E-30 | 2.28E-28 | 3281.614973 | 1815.726664 | ENSG00000119707 |
| RIMS4 | -0.854846412 | 0.552924187 | 3.56E-30 | 5.12E-28 | 1980.03057 | 1094.366209 | ENSG00000101098 |
| SMIM10L1 | -0.855767803 | 0.55257117 | 9.35E-10 | 1.75E-08 | 347.8150098 | 192.220666 | ENSG00000256537 |
| MAPK8IP1 | -0.856682223 | 0.552221046 | 2.37E-21 | 1.68E-19 | 1207.805774 | 667.2971361 | ENSG00000121653 |
| CUL1 | -0.857176694 | 0.552031809 | 7.13E-35 | 1.35E-32 | 2584.103758 | 1426.693293 | ENSG00000055130 |
| FABP3 | -0.857211864 | 0.552018352 | 2.04E-06 | 2.14E-05 | 267.85962 | 147.5241484 | ENSG00000121769 |
| STK32B | -0.859911674 | 0.55098629 | 2.74E-04 | 1.70E-03 | 122.7867774 | 67.72434652 | ENSG00000152953 |
| ANKRD13B | -0.860238749 | 0.550861389 | 1.73E-14 | 6.23E-13 | 971.2537226 | 535.0054276 | ENSG00000198720 |
| MFGE8 | -0.860741862 | 0.55066932 | 2.70E-61 | 2.00E-58 | 19038.13264 | 10484.36887 | ENSG00000140545 |
| PSMA2 | -0.861462761 | 0.550394225 | 9.01E-03 | 3.37E-02 | 64.44525652 | 35.40323094 | ENSG00000106588 |
| SLC24A3 | -0.861838885 | 0.550250751 | 3.94E-04 | 2.32E-03 | 121.9758874 | 67.35381634 | ENSG00000185052 |
| ADAM11 | -0.86249451 | 0.550000749 | 7.05E-07 | 8.08E-06 | 245.6286129 | 135.34118 | ENSG00000073670 |
| MANEAL | -0.863432587 | 0.549643242 | 2.41E-03 | 1.11E-02 | 85.84794319 | 46.99685272 | ENSG00000185090 |
| LPCAT3 | -0.864180752 | 0.549358277 | 2.23E-12 | 6.22E-11 | 573.4121775 | 315.2322365 | ENSG00000111684 |
| ADI1 | -0.864905182 | 0.549082493 | 3.54E-18 | 1.90E-16 | 825.6483463 | 453.2375687 | ENSG00000182551 |
| DRD4 | -0.864948617 | 0.549065962 | 1.21E-02 | 4.31E-02 | 51.98951671 | 28.58861698 | ENSG00000069696 |
| SH2D5 | -0.865076148 | 0.549017428 | 4.11E-15 | 1.61E-13 | 699.4167861 | 384.4287732 | ENSG00000189410 |
| GDAP1 | -0.865168643 | 0.54898223 | 7.09E-17 | 3.29E-15 | 796.3146743 | 437.4679212 | ENSG00000104381 |
| CLIP3 | -0.865334353 | 0.548919177 | 5.56E-08 | 7.79E-07 | 330.687148 | 181.2063168 | ENSG00000105270 |
| NOL12 | -0.866260574 | 0.54856688 | 2.19E-02 | 6.98E-02 | 63.89851727 | 35.29626757 | ENSG00000273899 |
| KHK | -0.867051536 | 0.548266209 | 1.98E-07 | 2.52E-06 | 264.1447634 | 144.9909636 | ENSG00000138030 |
| IQGAP1 | -0.867137903 | 0.548233388 | 2.73E-28 | 3.47E-26 | 3237.864714 | 1774.592349 | ENSG00000140575 |
| C1orf21 | -0.867739991 | 0.548004638 | 6.31E-17 | 2.97E-15 | 705.2224418 | 386.5080933 | ENSG00000116667 |
| NAGPA | -0.869247024 | 0.547432494 | 6.62E-07 | 7.63E-06 | 223.2153993 | 122.0991663 | ENSG00000103174 |
| CAMKV | -0.871288232 | 0.546658503 | 4.89E-31 | 7.42E-29 | 2172.235482 | 1187.981549 | ENSG00000164076 |
| EXOSC1 | -0.871852575 | 0.546444706 | 1.88E-12 | 5.31E-11 | 676.4868089 | 370.0298145 | ENSG00000171311 |
| MSI1 | -0.872930491 | 0.54603658 | 5.28E-25 | 4.99E-23 | 1439.663569 | 785.9098649 | ENSG00000135097 |
| HAPLN3 | -0.873200099 | 0.545934547 | 2.33E-23 | 1.94E-21 | 1905.898981 | 1041.00794 | ENSG00000140511 |
| SLITRK4 | -0.874776069 | 0.545338506 | 4.76E-06 | 4.63E-05 | 190.8109705 | 103.8773264 | ENSG00000179542 |
| MYBL2 | -0.875711845 | 0.544984897 | 4.64E-44 | 1.54E-41 | 6258.44021 | 3410.40628 | ENSG00000101057 |
| NKAIN1 | -0.876061291 | 0.544852908 | 2.41E-20 | 1.59E-18 | 2011.946302 | 1096.016767 | ENSG00000084628 |
| RND2 | -0.876690942 | 0.544615164 | 2.53E-35 | 4.99E-33 | 3200.174136 | 1743.014814 | ENSG00000108830 |
| CACHD1 | -0.877743757 | 0.544217873 | 7.48E-28 | 8.94E-26 | 1654.230259 | 900.4190725 | ENSG00000158966 |
| GAL3ST3 | -0.878721902 | 0.543849019 | 2.43E-08 | 3.64E-07 | 280.3240564 | 152.3794511 | ENSG00000175229 |
| EPN3 | -0.880741442 | 0.543088251 | 1.38E-03 | 6.87E-03 | 84.69161954 | 45.92832332 | ENSG00000049283 |
| ADAT3 | -0.88178201 | 0.542696681 | 7.09E-05 | 5.13E-04 | 139.0527576 | 75.40948271 | ENSG00000213638 |
| NPR1 | -0.884680352 | 0.54160751 | 1.18E-14 | 4.37E-13 | 571.4946027 | 309.4743258 | ENSG00000169418 |
| TCEAL3 | -0.884991785 | 0.541490606 | 4.68E-10 | 9.14E-09 | 349.4225912 | 189.1033818 | ENSG00000196507 |
| SEMA4A | -0.885117681 | 0.541443355 | 2.97E-09 | 5.12E-08 | 316.8170522 | 171.6895791 | ENSG00000196189 |
| PPP2R2B | -0.885504356 | 0.541298255 | 3.30E-25 | 3.14E-23 | 1423.232016 | 770.2877572 | ENSG00000156475 |
| RMST | -0.886026571 | 0.541102357 | 6.33E-04 | 3.53E-03 | 115.2931669 | 62.5379278 | ENSG00000255794 |
| GPLD1 | -0.886386077 | 0.540967536 | 1.87E-04 | 1.21E-03 | 130.9949396 | 70.6228772 | ENSG00000112293 |
| SLC4A4 | -0.886476707 | 0.540933553 | 3.69E-02 | 1.06E-01 | 40.06651303 | 21.51010979 | ENSG00000080493 |
| DPY19L2 | -0.88648284 | 0.540931254 | 2.94E-02 | 8.81E-02 | 42.86582054 | 23.21773751 | ENSG00000177990 |
| TMEM121B | -0.887161081 | 0.54067701 | 9.38E-07 | 1.05E-05 | 232.8325731 | 125.8084338 | ENSG00000183307 |
| FOXP1 | -0.887388613 | 0.540591745 | 1.94E-03 | 9.26E-03 | 82.69984968 | 44.52494197 | ENSG00000114861 |
| NGRN | -0.887501122 | 0.540549589 | 2.22E-12 | 6.20E-11 | 539.4211833 | 291.6090317 | ENSG00000182768 |
| PNCK | -0.887558134 | 0.540528228 | 2.73E-03 | 1.23E-02 | 79.98763531 | 43.34834989 | ENSG00000130822 |
| SAFB2 | -0.888179636 | 0.540295422 | 5.42E-26 | 5.55E-24 | 2390.201781 | 1292.088029 | ENSG00000130254 |
| MAN2A1 | -0.888671 | 0.540111436 | 1.53E-24 | 1.39E-22 | 1558.440033 | 841.0396179 | ENSG00000112893 |

| | | | | | | | |
|---|---|---|---|---|---|---|---|
| TIA1 | -0.888988898 | 0.539992435 | 2.52E-19 | 1.53E-17 | 1578.165371 | 851.9617779 | ENSG00000116001 |
| FAM133B | -0.889738847 | 0.539711807 | 3.02E-12 | 8.27E-11 | 759.626942 | 409.4433411 | ENSG00000234545 |
| IFT81 | -0.88984503 | 0.539672085 | 5.77E-09 | 9.61E-08 | 300.7185611 | 162.1987234 | ENSG00000122970 |
| DKKL1 | -0.889940651 | 0.539636317 | 4.86E-02 | 1.31E-01 | 36.25479944 | 19.71852657 | ENSG00000104901 |
| APCDD1L | -0.891075104 | 0.539212145 | 5.05E-05 | 3.79E-04 | 137.145623 | 74.01457512 | ENSG00000198768 |
| LRRC45 | -0.891103073 | 0.539201691 | 1.87E-14 | 6.72E-13 | 874.8631943 | 472.1406207 | ENSG00000169683 |
| EFHD2 | -0.892104765 | 0.538827443 | 4.29E-21 | 2.99E-19 | 1287.538031 | 693.3877682 | ENSG00000142634 |
| ABCB9 | -0.892403735 | 0.538715793 | 4.80E-04 | 2.76E-03 | 107.9405725 | 58.25759793 | ENSG00000150967 |
| NDRG2 | -0.893058437 | 0.538471376 | 9.60E-10 | 1.79E-08 | 385.5462136 | 207.8818531 | ENSG00000165795 |
| MYO5C | -0.899655613 | 0.536014669 | 4.61E-13 | 1.42E-11 | 498.643133 | 267.2665549 | ENSG00000128833 |
| ETNPPL | -0.900273222 | 0.535785253 | 3.03E-02 | 9.03E-02 | 37.41061943 | 20.12703677 | ENSG00000164089 |
| SRSF12 | -0.902175756 | 0.535079159 | 6.28E-05 | 4.61E-04 | 148.333074 | 79.54365827 | ENSG00000154548 |
| ZNF878 | -0.903286691 | 0.534667285 | 8.34E-03 | 3.17E-02 | 55.7563571 | 29.79614104 | ENSG00000257446 |
| NKX3-2 | -0.904785139 | 0.534112244 | 2.15E-02 | 6.85E-02 | 41.82697908 | 22.30587445 | ENSG00000109705 |
| TRABD2A | -0.904969797 | 0.534043884 | 1.91E-05 | 1.60E-04 | 163.6759362 | 87.52500511 | ENSG00000186854 |
| TRIL | -0.90573565 | 0.533760463 | 1.03E-06 | 1.14E-05 | 193.1558633 | 103.0492443 | ENSG00000255690 |
| FAIM | -0.90604376 | 0.533646482 | 9.13E-04 | 4.87E-03 | 106.909924 | 57.1357815 | ENSG00000158234 |
| SCYL2 | -0.90691117 | 0.533325727 | 6.57E-13 | 2.00E-11 | 673.2636786 | 358.5988876 | ENSG00000136021 |
| ME3 | -0.907354761 | 0.533161769 | 2.17E-02 | 6.92E-02 | 44.80196868 | 23.92658362 | ENSG00000151376 |
| RAB6B | -0.9073776 | 0.533153329 | 1.29E-11 | 3.23E-10 | 421.25604 | 224.5068275 | ENSG00000154917 |
| CORO1A | -0.907640987 | 0.533056002 | 1.37E-07 | 1.79E-06 | 236.1483285 | 125.9819445 | ENSG00000102879 |
| OVOL2 | -0.907925556 | 0.532950868 | 7.38E-06 | 6.86E-05 | 212.9678204 | 113.4237307 | ENSG00000125850 |
| MYL5 | -0.909917439 | 0.532215547 | 1.55E-03 | 7.63E-03 | 81.93590745 | 43.79115289 | ENSG00000215375 |
| PANX2 | -0.910269707 | 0.53208561 | 4.86E-06 | 4.71E-05 | 210.8258191 | 112.0577797 | ENSG00000073150 |
| NOS1 | -0.912487465 | 0.531268299 | 2.62E-05 | 2.13E-04 | 172.1560135 | 91.11273484 | ENSG00000089250 |
| HSPA14 | -0.917136541 | 0.529559046 | 4.65E-09 | 7.86E-08 | 392.3322252 | 207.394205 | ENSG00000187522 |
| NRCAM | -0.918471098 | 0.529069407 | 1.86E-04 | 1.20E-03 | 118.4003183 | 62.61571132 | ENSG00000091129 |
| ADGRG5 | -0.918530809 | 0.52904751 | 2.89E-10 | 5.84E-09 | 359.1784738 | 190.0755715 | ENSG00000159618 |
| NTNG2 | -0.919617024 | 0.528649336 | 2.68E-02 | 8.19E-02 | 37.99188932 | 20.13832169 | ENSG00000196358 |
| CRTC3 | -0.921159666 | 0.528084365 | 5.38E-08 | 7.56E-07 | 250.0642071 | 132.1142958 | ENSG00000140577 |
| EMX1 | -0.921311891 | 0.528028648 | 1.12E-05 | 9.90E-05 | 155.5601888 | 82.11234677 | ENSG00000135638 |
| NUCB2 | -0.923278068 | 0.527309514 | 2.22E-08 | 3.34E-07 | 301.3297915 | 158.8794099 | ENSG00000070081 |
| GRIN2B | -0.923957446 | 0.527061258 | 2.63E-03 | 1.19E-02 | 71.56298431 | 37.66518832 | ENSG00000273079 |
| PNMT | -0.924013797 | 0.527040671 | 1.80E-03 | 8.67E-03 | 87.03788786 | 46.07199289 | ENSG00000141744 |
| NAALAD2 | -0.925114717 | 0.52663864 | 1.95E-02 | 6.33E-02 | 49.32909073 | 26.05725567 | ENSG00000077616 |
| SCD5 | -0.925542991 | 0.526482327 | 2.08E-02 | 6.66E-02 | 41.82382216 | 21.99469165 | ENSG00000145284 |
| FDPS | -0.925623879 | 0.526452809 | 1.76E-24 | 1.59E-22 | 1745.645812 | 919.4628628 | ENSG00000160752 |
| DMC1 | -0.927366827 | 0.525817176 | 6.32E-04 | 3.52E-03 | 86.72580474 | 45.54563215 | ENSG00000100206 |
| ANGPT1 | -0.927788679 | 0.525663446 | 8.07E-03 | 3.08E-02 | 53.4612758 | 28.10458476 | ENSG00000154188 |
| SMARCE1 | -0.92915451 | 0.525166025 | 2.80E-02 | 8.49E-02 | 38.38436844 | 20.1040289 | ENSG00000073584 |
| ATP6V0E2 | -0.929745882 | 0.524950799 | 2.77E-16 | 1.21E-14 | 619.3971983 | 325.1676156 | ENSG00000171130 |
| ZDHHC15 | -0.930984164 | 0.524500421 | 2.64E-04 | 1.64E-03 | 109.8651753 | 57.5219451 | ENSG00000102383 |
| DZIP3 | -0.932422474 | 0.523977775 | 3.41E-14 | 1.20E-12 | 570.4262666 | 299.3756789 | ENSG00000198919 |
| EFNB1 | -0.93280163 | 0.523840087 | 5.31E-11 | 1.20E-09 | 427.5895912 | 224.1398814 | ENSG00000090776 |
| C6orf141 | -0.934750286 | 0.52313301 | 4.46E-03 | 1.87E-02 | 83.3103059 | 43.81427238 | ENSG00000197261 |
| H2AX | -0.936394615 | 0.522537103 | 5.87E-23 | 4.79E-21 | 3409.572092 | 1781.261472 | ENSG00000188486 |
| TTYH1 | -0.937624907 | 0.522091687 | 3.14E-12 | 8.58E-11 | 424.9925891 | 221.6446363 | ENSG00000167614 |
| STOX1 | -0.940279034 | 0.521132078 | 1.78E-03 | 8.58E-03 | 72.60951507 | 37.8906635 | ENSG00000165730 |
| SCG3 | -0.940449416 | 0.521070536 | 9.29E-16 | 3.89E-14 | 1045.987034 | 544.5055526 | ENSG00000104112 |
| RTP1 | -0.941720547 | 0.520611633 | 2.00E-03 | 9.49E-03 | 74.71628663 | 38.92957502 | ENSG00000175077 |
| ADGRG2 | -0.941946566 | 0.520530078 | 1.45E-07 | 1.87E-06 | 248.3115206 | 129.0831572 | ENSG00000173698 |
| P2RX5 | -0.942396708 | 0.52036769 | 2.69E-02 | 8.22E-02 | 35.63930717 | 18.53258467 | ENSG00000083454 |
| SLC49A3 | -0.943994716 | 0.519791621 | 6.95E-05 | 5.03E-04 | 120.1074099 | 62.42800138 | ENSG00000169026 |
| ZNF428 | -0.944724017 | 0.519528926 | 3.59E-06 | 3.60E-05 | 216.1997372 | 112.3092887 | ENSG00000131116 |
| NYNRIN | -0.94483364 | 0.519489451 | 9.23E-29 | 1.21E-26 | 1298.69734 | 674.5875233 | ENSG00000205978 |
| C12orf57 | -0.945041302 | 0.519414681 | 5.36E-04 | 3.05E-03 | 96.91340228 | 50.20006781 | ENSG00000111678 |
| LAYN | -0.945377823 | 0.519293538 | 2.16E-04 | 1.37E-03 | 110.4763306 | 57.20250334 | ENSG00000204381 |
| CCKBR | -0.946867685 | 0.518757543 | 2.06E-13 | 6.56E-12 | 501.4182524 | 260.3927137 | ENSG00000110148 |
| SPHK2 | -0.946867828 | 0.518757492 | 3.14E-19 | 1.87E-17 | 812.312753 | 421.5218083 | ENSG00000063176 |
| ARL4C | -0.950416284 | 0.517483123 | 2.75E-16 | 1.20E-14 | 591.1818919 | 305.8058495 | ENSG00000188042 |
| UNC5A | -0.95356916 | 0.516353446 | 1.54E-08 | 2.40E-07 | 300.8125921 | 155.4823129 | ENSG00000113763 |
| ADM5 | -0.954951479 | 0.515858939 | 1.63E-03 | 7.99E-03 | 73.04549192 | 37.61647303 | ENSG00000224420 |
| BAIAP2L2 | -0.958254963 | 0.514679076 | 2.62E-04 | 1.64E-03 | 115.692734 | 59.57417235 | ENSG00000128298 |
| MID2 | -0.959954676 | 0.514073063 | 1.63E-03 | 7.99E-03 | 69.62230378 | 35.75630423 | ENSG00000080561 |
| DCTPP1 | -0.960063756 | 0.514034196 | 4.11E-22 | 3.09E-20 | 1013.976926 | 520.9127535 | ENSG00000179958 |
| OLIG1 | -0.960107859 | 0.514018483 | 2.57E-02 | 7.91E-02 | 35.05793958 | 18.09413017 | ENSG00000184221 |
| TMEM35A | -0.961433233 | 0.513546482 | 1.02E-04 | 7.11E-04 | 112.1621358 | 57.4766938 | ENSG00000126950 |
| COMMD9 | -0.963644587 | 0.512759924 | 1.80E-09 | 3.23E-08 | 350.3003777 | 179.7236633 | ENSG00000110442 |
| MACROH2A1 | -0.965265747 | 0.512184058 | 9.84E-38 | 2.22E-35 | 2634.686419 | 1349.619172 | ENSG00000113648 |
| PIGW | -0.966613153 | 0.511705926 | 1.06E-09 | 1.97E-08 | 310.9872311 | 159.0368491 | ENSG00000277161 |
| SLC4A3 | -0.968139908 | 0.511164692 | 1.96E-08 | 2.98E-07 | 350.8567756 | 179.559949 | ENSG00000114923 |
| SHISA2 | -0.968680443 | 0.510973209 | 1.09E-10 | 2.37E-09 | 419.2726359 | 213.7928227 | ENSG00000180730 |
| ARHGAP8 | -0.973313896 | 0.509334767 | 3.32E-02 | 9.71E-02 | 33.79598728 | 17.21612187 | ENSG00000241484 |
| EPHB2 | -0.97437509 | 0.508960257 | 3.96E-20 | 2.55E-18 | 856.8319604 | 436.3161689 | ENSG00000133216 |
| GFRA2 | -0.974548524 | 0.508899076 | 1.82E-05 | 1.53E-04 | 134.9345756 | 68.85386382 | ENSG00000168546 |

| | | | | | | | |
|---|---|---|---|---|---|---|---|
| SIT1 | -0.97551102 | 0.508559676 | 7.54E-04 | 4.12E-03 | 83.52336729 | 42.4625291 | ENSG00000137078 |
| WDR90 | -0.977045035 | 0.508019213 | 6.75E-14 | 2.29E-12 | 763.6843161 | 388.6203695 | ENSG00000161996 |
| STMN3 | -0.977710758 | 0.507784845 | 6.24E-27 | 6.88E-25 | 2022.246518 | 1027.385814 | ENSG00000197457 |
| MAGEF1 | -0.983693572 | 0.505683436 | 3.13E-25 | 3.01E-23 | 1708.959064 | 864.6322199 | ENSG00000177383 |
| EXOG | -0.984515845 | 0.505395301 | 2.58E-07 | 3.21E-06 | 211.3667634 | 106.9961479 | ENSG00000157036 |
| OBI1 | -0.987653395 | 0.50429737 | 6.60E-06 | 6.21E-05 | 169.2760017 | 85.28138069 | ENSG00000152193 |
| LRRC10B | -0.987655931 | 0.504296483 | 3.13E-02 | 9.26E-02 | 31.77264829 | 15.94804795 | ENSG00000204950 |
| GPR137C | -0.990066242 | 0.503454658 | 1.04E-03 | 5.45E-03 | 74.0578003 | 37.17746888 | ENSG00000180998 |
| CYLD | -0.990814614 | 0.503193568 | 1.66E-08 | 2.57E-07 | 301.9881277 | 151.6500735 | ENSG00000083799 |
| PKDCC | -0.991086842 | 0.503098627 | 2.07E-15 | 8.34E-14 | 881.5789198 | 443.9995932 | ENSG00000162878 |
| BAALC | -0.991462378 | 0.502967687 | 4.15E-02 | 1.16E-01 | 30.54482066 | 15.49126049 | ENSG00000164929 |
| BEGAIN | -0.995198529 | 0.501666835 | 1.19E-03 | 6.11E-03 | 74.87420733 | 37.58316793 | ENSG00000183092 |
| ZNF559 | -0.99540603 | 0.501594686 | 6.20E-07 | 7.17E-06 | 181.0074637 | 90.66354507 | ENSG00000188321 |
| ISM1 | -0.99598166 | 0.501394592 | 3.15E-02 | 9.32E-02 | 29.51988463 | 14.82320556 | ENSG00000101230 |
| PRRG2 | -1.000177425 | 0.499938513 | 1.72E-02 | 5.71E-02 | 47.54454946 | 23.92333443 | ENSG00000126460 |
| CNIH2 | -1.004093162 | 0.498583428 | 7.49E-07 | 8.55E-06 | 210.1884617 | 104.6164516 | ENSG00000174871 |
| ANO1 | -1.004298814 | 0.498512362 | 3.02E-06 | 3.07E-05 | 158.5992861 | 79.17561283 | ENSG00000131620 |
| ZNF649 | -1.006618346 | 0.497711509 | 5.14E-15 | 1.98E-13 | 498.9410024 | 248.208494 | ENSG00000198093 |
| ALKBH6 | -1.007556945 | 0.49738781 | 2.09E-02 | 6.69E-02 | 34.59405419 | 17.1710938 | ENSG00000239382 |
| FNDC5 | -1.008573738 | 0.497037381 | 4.25E-02 | 1.18E-01 | 29.96345308 | 14.93594314 | ENSG00000160097 |
| ACOXL | -1.008708341 | 0.49699101 | 2.89E-13 | 9.09E-12 | 412.9116647 | 205.5562083 | ENSG00000153093 |
| ANO4 | -1.01040897 | 0.496405508 | 3.04E-03 | 1.35E-02 | 62.2347128 | 31.07225069 | ENSG00000151572 |
| NKX1-2 | -1.012038224 | 0.495845228 | 2.38E-13 | 7.53E-12 | 486.7943842 | 241.001281 | ENSG00000229544 |
| ADGRB2 | -1.01424557 | 0.495087157 | 3.84E-28 | 4.81E-26 | 1178.683518 | 584.1476299 | ENSG00000121753 |
| PKNOX2 | -1.016754991 | 0.494226752 | 5.91E-03 | 2.37E-02 | 51.26345744 | 25.31769237 | ENSG00000165495 |
| E2F8 | -1.017147745 | 0.494092224 | 2.45E-09 | 4.30E-08 | 243.5529819 | 120.3554223 | ENSG00000129173 |
| ITPRIPL1 | -1.017498593 | 0.49397208 | 8.43E-12 | 2.16E-10 | 384.4311025 | 190.1891851 | ENSG00000198885 |
| TMEM81 | -1.018666601 | 0.493572322 | 2.59E-03 | 1.18E-02 | 57.86806701 | 28.52101907 | ENSG00000174529 |
| NOX4 | -1.019230667 | 0.493379382 | 4.87E-02 | 1.31E-01 | 25.72523338 | 12.67767299 | ENSG00000086991 |
| TMSB15A | -1.020696106 | 0.492878479 | 3.27E-17 | 1.60E-15 | 618.2125753 | 304.2209218 | ENSG00000158164 |
| ACAT2 | -1.021088361 | 0.492744488 | 2.63E-38 | 6.26E-36 | 2464.503498 | 1213.766237 | ENSG00000120437 |
| ANO2 | -1.022767917 | 0.492171179 | 6.30E-04 | 3.52E-03 | 74.27449962 | 36.59297169 | ENSG00000047617 |
| NLGN3 | -1.022902465 | 0.492125281 | 7.42E-11 | 1.65E-09 | 359.7109994 | 177.1310825 | ENSG00000196338 |
| RRP36 | -1.026527994 | 0.49089011 | 2.78E-38 | 6.45E-36 | 1569.137111 | 770.1423757 | ENSG00000124541 |
| NUDT8 | -1.027902114 | 0.490422776 | 1.73E-04 | 1.13E-03 | 98.90358609 | 48.61799993 | ENSG00000167799 |
| CNTN1 | -1.029259653 | 0.489961518 | 3.26E-11 | 7.64E-10 | 386.6123622 | 189.0645172 | ENSG00000018236 |
| RND1 | -1.029954978 | 0.489725431 | 1.58E-07 | 2.04E-06 | 215.9403521 | 105.9378041 | ENSG00000172602 |
| PARP2 | -1.032477953 | 0.488869751 | 2.87E-19 | 1.71E-17 | 727.1495739 | 355.9375868 | ENSG00000129484 |
| MYCNOS | -1.035703083 | 0.487778109 | 2.98E-03 | 1.32E-02 | 62.58349881 | 30.68207344 | ENSG00000233718 |
| ZNF799 | -1.042118486 | 0.485613864 | 1.60E-02 | 5.39E-02 | 38.69645157 | 18.69359951 | ENSG00000196466 |
| PRR19 | -1.043016474 | 0.485311693 | 4.79E-03 | 1.98E-02 | 61.55600722 | 29.87502386 | ENSG00000188368 |
| PRKAR1B | -1.043139332 | 0.485270366 | 2.28E-33 | 3.78E-31 | 1947.5789 | 945.2722649 | ENSG00000188191 |
| CDK8 | -1.045334384 | 0.484532592 | 3.47E-17 | 1.69E-15 | 680.2518526 | 329.1069929 | ENSG00000132964 |
| WDR45 | -1.045406628 | 0.484508329 | 4.20E-05 | 3.23E-04 | 117.8594869 | 57.0834906 | ENSG00000196998 |
| LAG3 | -1.046379093 | 0.484181851 | 2.78E-03 | 1.25E-02 | 58.52291541 | 28.4726302 | ENSG00000089692 |
| JAM2 | -1.046422379 | 0.484167324 | 8.38E-06 | 7.63E-05 | 139.7059374 | 67.54389941 | ENSG00000154721 |
| SLC5A6 | -1.046971411 | 0.483983104 | 7.99E-52 | 3.70E-49 | 3823.677843 | 1851.336439 | ENSG00000138074 |
| CACNG4 | -1.05271113 | 0.482061417 | 5.45E-06 | 5.22E-05 | 130.7385759 | 62.87405357 | ENSG00000075461 |
| CNTN2 | -1.05301371 | 0.481960324 | 7.43E-05 | 5.34E-04 | 126.2497049 | 60.76237576 | ENSG00000184144 |
| C1orf54 | -1.054982093 | 0.481303196 | 1.01E-05 | 9.03E-05 | 121.1139081 | 58.41905081 | ENSG00000118292 |
| PCDHB16 | -1.056077528 | 0.480937882 | 4.27E-04 | 2.49E-03 | 92.66658366 | 44.40851716 | ENSG00000272674 |
| DLD | -1.057616109 | 0.480425253 | 3.70E-39 | 9.52E-37 | 1770.558893 | 850.1368324 | ENSG00000091140 |
| GGN | -1.058980655 | 0.479971066 | 2.35E-02 | 7.36E-02 | 32.81790127 | 15.75328996 | ENSG00000179168 |
| ST8SIA5 | -1.060451741 | 0.479481899 | 5.70E-04 | 3.22E-03 | 79.80072368 | 38.37458409 | ENSG00000101638 |
| ZNF521 | -1.064464054 | 0.478150254 | 1.13E-09 | 2.07E-08 | 261.6327123 | 124.9607046 | ENSG00000198795 |
| SEC14L2 | -1.06542534 | 0.477831762 | 4.72E-03 | 1.96E-02 | 49.15410769 | 23.57405999 | ENSG00000100003 |
| KLHL14 | -1.067877696 | 0.477020213 | 9.43E-03 | 3.51E-02 | 40.36478843 | 19.25662616 | ENSG00000197705 |
| ABCB7 | -1.06833171 | 0.476870119 | 1.09E-23 | 9.30E-22 | 820.0568344 | 391.5061811 | ENSG00000131269 |
| TRPM3 | -1.068703323 | 0.476747301 | 5.85E-07 | 6.82E-06 | 184.9994069 | 88.12893457 | ENSG00000083067 |
| PCSK1N | -1.069268928 | 0.47656043 | 5.06E-05 | 3.80E-04 | 144.0437877 | 68.56382515 | ENSG00000102109 |
| TMEM132E | -1.069781668 | 0.476391089 | 9.68E-03 | 3.58E-02 | 38.32932545 | 18.25514501 | ENSG00000181291 |
| RTEL1 | -1.070299598 | 0.476220094 | 3.28E-02 | 9.62E-02 | 26.75460411 | 12.74895812 | ENSG00000258366 |
| C1orf216 | -1.070421152 | 0.476179972 | 1.09E-10 | 2.37E-09 | 329.1483196 | 157.1903786 | ENSG00000142686 |
| NMU | -1.071133035 | 0.475945064 | 9.10E-08 | 1.23E-06 | 206.7892284 | 98.04063604 | ENSG00000109255 |
| FAM222A | -1.072608111 | 0.475458685 | 2.17E-03 | 1.02E-02 | 61.64763275 | 29.36148539 | ENSG00000139438 |
| ARHGAP6 | -1.073211729 | 0.475259797 | 3.67E-02 | 1.05E-01 | 26.61246797 | 12.61756116 | ENSG00000047648 |
| MRGPRF | -1.073244465 | 0.475249013 | 2.49E-05 | 2.03E-04 | 108.2096615 | 51.37439838 | ENSG00000172935 |
| LOXL1 | -1.074613795 | 0.474798146 | 2.74E-10 | 5.56E-09 | 311.4127299 | 147.9954071 | ENSG00000129038 |
| CTXN1 | -1.074717022 | 0.474764174 | 1.83E-14 | 6.56E-13 | 775.0381134 | 367.7668376 | ENSG00000178531 |
| PRR22 | -1.075381451 | 0.474545573 | 3.59E-02 | 1.03E-01 | 33.74419889 | 16.0913911 | ENSG00000212123 |
| NOVA1 | -1.075539915 | 0.474493452 | 4.61E-08 | 6.59E-07 | 195.2148979 | 92.62549296 | ENSG00000139910 |
| PLAC9 | -1.075614253 | 0.474469004 | 8.86E-04 | 4.75E-03 | 62.34400203 | 29.58629928 | ENSG00000189129 |
| GPR50 | -1.07906271 | 0.47333624 | 3.32E-07 | 4.04E-06 | 187.6309397 | 88.83755744 | ENSG00000102195 |
| KLHL4 | -1.082983393 | 0.472051643 | 2.96E-23 | 2.44E-21 | 774.3790787 | 365.1342384 | ENSG00000102271 |
| NSMCE4A | -1.084341507 | 0.471607476 | 1.89E-24 | 1.69E-22 | 944.5349566 | 445.4450396 | ENSG00000107672 |

| | | | | | | | |
|---|---|---|---|---|---|---|---|
| BCL6B | -1.084778867 | 0.471464527 | 2.03E-14 | 7.24E-13 | 364.7652353 | 171.9176422 | ENSG00000161940 |
| STAR | -1.085568189 | 0.471206652 | 9.25E-04 | 4.93E-03 | 65.73542564 | 30.96276229 | ENSG00000147465 |
| OTX2 | -1.085957586 | 0.471079486 | 1.47E-20 | 9.93E-19 | 696.4976589 | 328.3714835 | ENSG00000165588 |
| KCNJ12 | -1.088305244 | 0.470313535 | 1.92E-05 | 1.60E-04 | 107.0848093 | 50.27146456 | ENSG00000184185 |
| DUSP28 | -1.090152228 | 0.46971181 | 4.50E-02 | 1.23E-01 | 29.1089677 | 13.73959239 | ENSG00000188542 |
| ADAMTS17 | -1.092143998 | 0.469063778 | 3.17E-03 | 1.39E-02 | 51.38221741 | 24.1477103 | ENSG00000140470 |
| U2AF1L4 | -1.093049486 | 0.468769469 | 4.21E-02 | 1.17E-01 | 27.12233159 | 12.70013121 | ENSG00000161265 |
| SCN9A | -1.093910887 | 0.46848966 | 5.60E-04 | 3.17E-03 | 74.25240123 | 34.80364998 | ENSG00000169432 |
| RTF1 | -1.096128762 | 0.467769998 | 1.35E-51 | 5.96E-49 | 2220.576821 | 1038.507413 | ENSG00000137815 |
| SYT9 | -1.096227015 | 0.467738142 | 1.48E-04 | 9.83E-04 | 84.23670122 | 39.32300145 | ENSG00000170743 |
| FAM78A | -1.098351147 | 0.467049981 | 7.96E-09 | 1.30E-07 | 194.2465909 | 90.71281001 | ENSG00000126882 |
| MCUB | -1.099623643 | 0.466638212 | 3.83E-14 | 1.33E-12 | 389.3175712 | 181.7257411 | ENSG00000005059 |
| FEZ1 | -1.100804055 | 0.466256565 | 7.19E-28 | 8.71E-26 | 928.3800471 | 432.5173377 | ENSG00000149557 |
| RBM3 | -1.104282081 | 0.465133875 | 2.32E-32 | 3.64E-30 | 1626.00231 | 756.0075016 | ENSG00000102317 |
| E2F2 | -1.105344865 | 0.464791353 | 6.17E-10 | 1.18E-08 | 221.5275196 | 103.0420846 | ENSG00000007968 |
| LOX | -1.106917969 | 0.464284825 | 1.78E-02 | 5.86E-02 | 31.02892531 | 14.40677125 | ENSG00000113083 |
| CALCB | -1.107259001 | 0.464175087 | 4.98E-06 | 4.81E-05 | 121.4223307 | 56.35565026 | ENSG00000175868 |
| VWA5B1 | -1.109333141 | 0.46350823 | 2.86E-02 | 8.63E-02 | 31.01166758 | 14.27526267 | ENSG00000158816 |
| DPH5 | -1.115386104 | 0.461567611 | 5.93E-24 | 5.18E-22 | 699.3748744 | 322.8123445 | ENSG00000117543 |
| KCNC3 | -1.116465183 | 0.461222506 | 4.72E-08 | 6.73E-07 | 179.1450296 | 82.67894904 | ENSG00000131398 |
| SARM1 | -1.117086908 | 0.461023786 | 8.20E-05 | 5.83E-04 | 91.21248827 | 41.9708992 | ENSG00000004139 |
| DLG1 | -1.11709457 | 0.461021338 | 4.09E-35 | 7.90E-33 | 1165.899016 | 537.8054687 | ENSG00000075711 |
| MSANTD2 | -1.117795124 | 0.460797526 | 4.83E-11 | 1.10E-09 | 265.8445717 | 122.651346 | ENSG00000120458 |
| SLC16A9 | -1.12252768 | 0.459288422 | 2.60E-11 | 6.17E-10 | 281.7263634 | 129.1387058 | ENSG00000165449 |
| ADAMTS19 | -1.126411897 | 0.458053527 | 1.84E-19 | 1.13E-17 | 547.9738697 | 250.7092498 | ENSG00000145808 |
| ASXL1 | -1.127182971 | 0.457808777 | 1.18E-24 | 1.08E-22 | 963.0894414 | 440.5436095 | ENSG00000171456 |
| MAP2 | -1.129719072 | 0.457004707 | 2.82E-07 | 3.48E-06 | 176.2409426 | 80.28080816 | ENSG00000078018 |
| HGSNAT | -1.131976039 | 0.456290323 | 4.85E-06 | 4.70E-05 | 118.4130436 | 54.045304 | ENSG00000165102 |
| THEM6 | -1.142015994 | 0.453125946 | 2.26E-08 | 3.41E-07 | 188.041353 | 85.05601713 | ENSG00000130193 |
| LOXL3 | -1.142367265 | 0.453015631 | 6.82E-17 | 3.19E-15 | 419.9050928 | 190.307983 | ENSG00000115318 |
| H2AW | -1.143713188 | 0.452593199 | 2.10E-13 | 6.68E-12 | 353.590081 | 159.790062 | ENSG00000181218 |
| PTGS1 | -1.14419599 | 0.452441763 | 1.81E-03 | 8.72E-03 | 50.08356222 | 22.60968279 | ENSG00000095303 |
| CNTNAP3B | -1.146143347 | 0.451831467 | 5.16E-03 | 2.11E-02 | 43.92873722 | 19.78893563 | ENSG00000154529 |
| MYB | -1.150929463 | 0.450335008 | 2.53E-04 | 1.59E-03 | 80.91285055 | 36.27689074 | ENSG00000118513 |
| CBLN1 | -1.152453539 | 0.449859521 | 1.99E-09 | 3.55E-08 | 236.2963725 | 106.6097695 | ENSG00000102924 |
| RUBCNL | -1.152858183 | 0.449733363 | 1.19E-04 | 8.16E-04 | 79.13146619 | 35.7117142 | ENSG00000102445 |
| CYP39A1 | -1.155235692 | 0.448992829 | 2.71E-03 | 1.22E-02 | 48.58399228 | 21.87490603 | ENSG00000146233 |
| DANCR | -1.155786259 | 0.448821515 | 5.41E-37 | 1.15E-34 | 1431.357776 | 642.5573393 | ENSG00000226950 |
| COX16 | -1.156426398 | 0.448622412 | 3.68E-02 | 1.05E-01 | 29.25317836 | 13.14673305 | ENSG00000133983 |
| PVALB | -1.156665336 | 0.448548118 | 3.04E-02 | 9.05E-02 | 26.66543643 | 11.87854751 | ENSG00000100362 |
| PCDHB12 | -1.157885173 | 0.448169019 | 5.82E-05 | 4.31E-04 | 86.20066022 | 38.49013283 | ENSG00000120328 |
| RGS11 | -1.15882082 | 0.447878457 | 3.50E-02 | 1.01E-01 | 22.85066363 | 10.18699124 | ENSG00000076344 |
| PSMG3 | -1.161895486 | 0.446924957 | 5.58E-16 | 2.37E-14 | 736.2430764 | 329.7074273 | ENSG00000157778 |
| FIGNL2 | -1.164255369 | 0.446194499 | 4.91E-06 | 4.75E-05 | 152.9486873 | 68.22313609 | ENSG00000261308 |
| H2BC8 | -1.170238805 | 0.444347783 | 1.29E-02 | 4.52E-02 | 30.21813305 | 13.43914486 | ENSG00000273802 |
| TMX3 | -1.172784975 | 0.443564258 | 1.18E-12 | 3.44E-11 | 368.4445015 | 163.3656795 | ENSG00000166479 |
| TTI1 | -1.173285703 | 0.443410333 | 8.46E-34 | 1.48E-31 | 1311.267053 | 581.4984181 | ENSG00000101407 |
| PRRT2 | -1.174210631 | 0.443126149 | 6.97E-06 | 6.52E-05 | 164.2570107 | 72.94670699 | ENSG00000167371 |
| SFRP1 | -1.176997963 | 0.442270842 | 1.37E-56 | 7.91E-54 | 13909.63302 | 6150.892453 | ENSG00000104332 |
| RNF39 | -1.179112634 | 0.441623046 | 8.80E-04 | 4.72E-03 | 57.53082628 | 25.44160325 | ENSG00000204618 |
| PFKFB4 | -1.179913608 | 0.441377928 | 1.78E-12 | 5.06E-11 | 351.2842664 | 154.5835582 | ENSG00000114268 |
| SLITRK3 | -1.182673133 | 0.440534486 | 8.99E-04 | 4.81E-03 | 58.89447621 | 25.87257168 | ENSG00000121871 |
| H2BC6 | -1.187009314 | 0.439212399 | 4.54E-02 | 1.24E-01 | 21.01997138 | 9.193107779 | ENSG00000274290 |
| ABHD8 | -1.188540709 | 0.43874643 | 3.43E-10 | 6.85E-09 | 226.2765724 | 99.11004151 | ENSG00000127220 |
| CAMK2N2 | -1.188833194 | 0.438657489 | 1.67E-06 | 1.77E-05 | 139.5956635 | 61.15244138 | ENSG00000163888 |
| EPHB6 | -1.18897752 | 0.438613609 | 1.03E-26 | 1.10E-24 | 738.8195662 | 323.872735 | ENSG00000106123 |
| HTR7 | -1.188987226 | 0.438610658 | 7.10E-04 | 3.90E-03 | 59.8770946 | 26.1650951 | ENSG00000148680 |
| ODC1 | -1.190797023 | 0.438060785 | 5.84E-60 | 4.16E-57 | 4256.00613 | 1863.638365 | ENSG00000115758 |
| PRSS35 | -1.197310423 | 0.43608751 | 1.34E-04 | 9.00E-04 | 81.03228694 | 35.30601516 | ENSG00000146250 |
| H2AC11 | -1.198059256 | 0.435861217 | 2.42E-03 | 1.11E-02 | 58.8135992 | 25.60630532 | ENSG00000196787 |
| CNMD | -1.200574448 | 0.435102 | 1.47E-49 | 5.81E-47 | 3178.378722 | 1381.899814 | ENSG00000136110 |
| KIF26A | -1.203759182 | 0.434142576 | 1.79E-45 | 6.50E-43 | 3907.371217 | 1696.18963 | ENSG00000066735 |
| ACOT1 | -1.208150878 | 0.432823015 | 8.71E-03 | 3.28E-02 | 33.29106201 | 14.35414549 | ENSG00000184227 |
| WNT2B | -1.208701722 | 0.432657788 | 4.91E-02 | 1.32E-01 | 20.70916604 | 9.009523095 | ENSG00000134245 |
| FGFBP3 | -1.210908814 | 0.431996397 | 1.40E-39 | 3.81E-37 | 2067.901416 | 893.3415699 | ENSG00000174721 |
| YWHAB | -1.216267368 | 0.430394824 | 1.77E-78 | 2.52E-75 | 6249.440994 | 2688.889676 | ENSG00000166913 |
| PARTICL | -1.216508044 | 0.43032303 | 2.03E-03 | 9.61E-03 | 46.6340364 | 20.17206483 | ENSG00000286532 |
| DUSP15 | -1.217408433 | 0.430054548 | 4.87E-04 | 2.80E-03 | 63.8872651 | 27.50087854 | ENSG00000149599 |
| NCAM1 | -1.220722757 | 0.429067711 | 1.34E-11 | 3.32E-10 | 239.7096991 | 102.6885733 | ENSG00000149294 |
| CFAP298 | -1.221151712 | 0.428940156 | 3.14E-05 | 2.50E-04 | 86.97229175 | 37.21880972 | ENSG00000159079 |
| ACD | -1.222083605 | 0.428663176 | 7.02E-29 | 9.43E-27 | 770.025256 | 330.1001522 | ENSG00000102977 |
| SEMA5B | -1.22258134 | 0.428515311 | 2.93E-25 | 2.83E-23 | 767.3204605 | 329.0890351 | ENSG00000082684 |
| AFP | -1.222585398 | 0.428514106 | 3.11E-02 | 9.22E-02 | 24.82419107 | 10.66353738 | ENSG00000081051 |
| PTX3 | -1.226482813 | 0.427358045 | 3.77E-02 | 1.07E-01 | 23.59250747 | 10.13491514 | ENSG00000163661 |
| MAP3K7 | -1.235061981 | 0.424824248 | 2.74E-36 | 5.72E-34 | 1247.50913 | 530.6548405 | ENSG00000135341 |

| | | | | | | | |
|---|---|---|---|---|---|---|---|
| TP53BP1 | -1.239436152 | 0.423538155 | 2.42E-51 | 1.04E-48 | 2579.395181 | 1092.836678 | ENSG00000067369 |
| PCLAF | -1.240919712 | 0.423102844 | 1.14E-05 | 1.01E-04 | 112.3588112 | 47.26625665 | ENSG00000166803 |
| SLC15A2 | -1.241565786 | 0.42291341 | 7.49E-04 | 4.09E-03 | 56.78898244 | 24.05005648 | ENSG00000163406 |
| TSPYL4 | -1.24599395 | 0.421617323 | 7.29E-25 | 6.76E-23 | 841.9186635 | 355.6994497 | ENSG00000187189 |
| LRRC49 | -1.246237676 | 0.421546102 | 1.12E-10 | 2.43E-09 | 194.7774479 | 82.08977693 | ENSG00000137821 |
| INSYN1 | -1.247829769 | 0.421081159 | 1.40E-10 | 3.02E-09 | 211.7714642 | 89.4537595 | ENSG00000205363 |
| FXYD6 | -1.251157354 | 0.420111052 | 3.09E-07 | 3.78E-06 | 122.1755244 | 51.38980856 | ENSG00000137726 |
| SYPL2 | -1.251758078 | 0.419936159 | 3.74E-05 | 2.91E-04 | 77.71662923 | 32.65063134 | ENSG00000143028 |
| SLC25A12 | -1.251916453 | 0.419890062 | 2.02E-12 | 5.67E-11 | 260.7985439 | 109.4515492 | ENSG00000115840 |
| TAGLN3 | -1.253314501 | 0.419483363 | 4.47E-06 | 4.38E-05 | 103.7006842 | 43.52076127 | ENSG00000144834 |
| COL21A1 | -1.253515757 | 0.419424849 | 6.05E-05 | 4.45E-04 | 73.47643271 | 30.72622544 | ENSG00000124749 |
| CYP4X1 | -1.255195804 | 0.418936705 | 7.62E-04 | 4.16E-03 | 52.703452 | 22.09234547 | ENSG00000186377 |
| HDAC9 | -1.258638677 | 0.417938139 | 2.77E-09 | 4.81E-08 | 204.7674603 | 85.58953753 | ENSG00000048052 |
| VIPR2 | -1.26083966 | 0.417301017 | 2.81E-03 | 1.26E-02 | 50.83695133 | 21.38002693 | ENSG00000106018 |
| SYP | -1.261009987 | 0.417251752 | 5.54E-09 | 9.26E-08 | 199.8013999 | 83.70756333 | ENSG00000102003 |
| KEL | -1.261491676 | 0.417112463 | 1.22E-02 | 4.34E-02 | 31.58178299 | 13.22181703 | ENSG00000197993 |
| GRIN1 | -1.261771068 | 0.417031693 | 7.83E-09 | 1.28E-07 | 150.8397436 | 62.87829045 | ENSG00000176884 |
| ERBB2 | -1.262119516 | 0.416930981 | 3.23E-53 | 1.58E-50 | 12945.20586 | 5397.128691 | ENSG00000141736 |
| HACL1 | -1.262342929 | 0.416866421 | 2.60E-28 | 3.32E-26 | 688.1921285 | 286.5480125 | ENSG00000131373 |
| FAM133A | -1.264553874 | 0.416228058 | 3.33E-02 | 9.74E-02 | 20.66675069 | 8.604373707 | ENSG00000179083 |
| MAPRE3 | -1.267628114 | 0.415342062 | 1.09E-08 | 1.73E-07 | 148.5914146 | 61.82473319 | ENSG00000084764 |
| FZD2 | -1.268616637 | 0.41505757 | 8.52E-30 | 1.18E-27 | 1397.861619 | 580.2393273 | ENSG00000180340 |
| DOK3 | -1.274858309 | 0.413265745 | 4.26E-07 | 5.09E-06 | 117.6580685 | 48.77196673 | ENSG00000146094 |
| C1QTNF1 | -1.276495102 | 0.412797145 | 2.34E-05 | 1.92E-04 | 87.77864932 | 36.15341789 | ENSG00000173918 |
| DBNDD2 | -1.281213598 | 0.411449251 | 1.22E-02 | 4.33E-02 | 28.25793232 | 11.58982294 | ENSG00000244274 |
| ELP2 | -1.28272698 | 0.411017868 | 8.24E-60 | 5.46E-57 | 1921.34869 | 789.7339655 | ENSG00000134759 |
| LRRC55 | -1.285468563 | 0.410237545 | 1.78E-05 | 1.51E-04 | 93.30793259 | 38.20520711 | ENSG00000183908 |
| SMPDL3B | -1.28754597 | 0.409647248 | 3.22E-25 | 3.08E-23 | 578.6975293 | 237.3925188 | ENSG00000130768 |
| CEP135 | -1.290231078 | 0.408885532 | 4.59E-13 | 1.41E-11 | 287.849642 | 117.4588267 | ENSG00000174799 |
| SLC10A4 | -1.295059374 | 0.407519394 | 5.74E-05 | 4.26E-04 | 80.58626064 | 32.80047288 | ENSG00000145248 |
| NXPH2 | -1.297122523 | 0.406937031 | 1.83E-04 | 1.19E-03 | 59.69757918 | 24.22235624 | ENSG00000144227 |
| EME2 | -1.302396222 | 0.405452209 | 7.02E-04 | 3.86E-03 | 543.7489993 | 220.6579931 | ENSG00000197774 |
| TSGA10IP | -1.304714629 | 0.404801172 | 9.73E-04 | 5.14E-03 | 46.1297125 | 18.75046214 | ENSG00000175513 |
| AQP5 | -1.304877181 | 0.404755564 | 9.51E-04 | 5.05E-03 | 48.45250696 | 19.56032289 | ENSG00000161798 |
| TPR | -1.311073432 | 0.403020902 | 8.82E-67 | 7.78E-64 | 3374.809483 | 1360.69615 | ENSG00000047410 |
| TRIM54 | -1.312042309 | 0.402750334 | 8.05E-06 | 7.37E-05 | 97.21127171 | 39.38030212 | ENSG00000138100 |
| ADAM33 | -1.313490857 | 0.402346153 | 3.79E-03 | 1.63E-02 | 323.4946616 | 130.3219884 | ENSG00000149451 |
| MT1H | -1.313980858 | 0.402209522 | 1.28E-03 | 6.49E-03 | 46.59034328 | 18.60723061 | ENSG00000205358 |
| CHST13 | -1.314793361 | 0.401983068 | 6.00E-05 | 4.42E-04 | 75.56378177 | 30.50184957 | ENSG00000180767 |
| PCDHB15 | -1.318583781 | 0.400928316 | 1.59E-06 | 1.70E-05 | 103.8702479 | 41.69389757 | ENSG00000113248 |
| PTN | -1.3204807 | 0.400401504 | 1.64E-33 | 2.79E-31 | 784.6113389 | 314.0343569 | ENSG00000105894 |
| ATL1 | -1.322361431 | 0.399879872 | 1.22E-06 | 1.34E-05 | 97.25221388 | 38.92957502 | ENSG00000198513 |
| SPTB | -1.324048358 | 0.39941257 | 2.81E-04 | 1.74E-03 | 62.98474961 | 25.18304621 | ENSG00000070182 |
| INCA1 | -1.324625084 | 0.399252934 | 1.41E-02 | 4.87E-02 | 28.32056692 | 11.31986935 | ENSG00000196388 |
| FAM111B | -1.333916869 | 0.396689779 | 6.24E-20 | 3.99E-18 | 617.2576624 | 244.5908829 | ENSG00000189057 |
| TEKT2 | -1.335134881 | 0.39635501 | 2.54E-02 | 7.84E-02 | 19.96218845 | 7.92547189 | ENSG00000092850 |
| REM2 | -1.34436565 | 0.393827115 | 1.51E-04 | 9.96E-04 | 71.2734806 | 28.14647525 | ENSG00000139890 |
| LRRN2 | -1.347760206 | 0.392901558 | 1.97E-15 | 7.99E-14 | 289.1328959 | 113.461161 | ENSG00000170382 |
| MAGEE1 | -1.353637516 | 0.391304195 | 1.23E-16 | 5.58E-15 | 321.0476803 | 125.2827343 | ENSG00000198934 |
| CP | -1.35733588 | 0.390302368 | 4.31E-02 | 1.19E-01 | 17.80786781 | 6.957845497 | ENSG00000047457 |
| FOXO6 | -1.362954013 | 0.388785411 | 4.83E-08 | 6.86E-07 | 125.9677178 | 48.88740386 | ENSG00000204060 |
| DPYSL5 | -1.363160551 | 0.388729755 | 6.98E-40 | 2.02E-37 | 1059.469153 | 411.9844359 | ENSG00000157851 |
| CCNI2 | -1.363226155 | 0.388712079 | 2.81E-02 | 8.51E-02 | 20.34184452 | 7.86536006 | ENSG00000205089 |
| PLA2R1 | -1.365662262 | 0.388056261 | 3.07E-05 | 2.46E-04 | 67.53199321 | 26.2482148 | ENSG00000153246 |
| ZSWIM5 | -1.366660403 | 0.387787874 | 4.22E-10 | 8.25E-09 | 197.0015288 | 76.57318119 | ENSG00000162415 |
| TTLL7 | -1.369731668 | 0.386963214 | 4.54E-05 | 3.45E-04 | 92.86110954 | 36.19629606 | ENSG00000137941 |
| KIF19 | -1.374277934 | 0.385745723 | 1.72E-02 | 5.69E-02 | 26.28124797 | 10.0104546 | ENSG00000196169 |
| WDR34 | -1.37453471 | 0.385677073 | 1.22E-39 | 3.44E-37 | 1306.100956 | 504.089892 | ENSG00000119333 |
| RGS7BP | -1.380270417 | 0.384146784 | 4.17E-04 | 2.45E-03 | 50.84769985 | 19.53406582 | ENSG00000186479 |
| MAP1A | -1.390265271 | 0.38149465 | 3.38E-07 | 4.11E-06 | 146.9102395 | 56.28133073 | ENSG00000166963 |
| CDYL2 | -1.391524487 | 0.381161818 | 9.39E-08 | 1.27E-06 | 110.6873175 | 42.11737992 | ENSG00000166446 |
| TMEM255A | -1.392687044 | 0.380854793 | 1.44E-03 | 7.16E-03 | 44.95504873 | 17.10718313 | ENSG00000125355 |
| KRT7 | -1.393231712 | 0.380711034 | 1.68E-14 | 6.08E-13 | 268.17368 | 102.1005005 | ENSG00000135480 |
| PTPRO | -1.393485877 | 0.380643969 | 7.41E-03 | 2.87E-02 | 26.97831631 | 10.24330422 | ENSG00000151490 |
| SCN2B | -1.400290287 | 0.378852904 | 4.31E-02 | 1.19E-01 | 17.59244626 | 6.736718825 | ENSG00000149575 |
| PRRT4 | -1.400473603 | 0.378804768 | 2.88E-03 | 1.29E-02 | 40.14117392 | 15.27349463 | ENSG00000224940 |
| SHD | -1.402425316 | 0.378292659 | 1.13E-04 | 7.75E-04 | 65.33880493 | 24.74404206 | ENSG00000105251 |
| HTR7P1 | -1.406060682 | 0.37734062 | 3.09E-04 | 1.88E-03 | 80.11152903 | 30.48775349 | ENSG00000183935 |
| FRRS1L | -1.406294203 | 0.377279547 | 3.20E-04 | 1.94E-03 | 49.41953618 | 18.73169115 | ENSG00000260230 |
| CCDC74A | -1.408210438 | 0.376778765 | 5.12E-17 | 2.42E-15 | 290.6413579 | 109.6661775 | ENSG00000163040 |
| CCDC160 | -1.415687281 | 0.37483114 | 2.30E-06 | 2.39E-05 | 91.3591568 | 34.21827672 | ENSG00000203952 |
| EZH2 | -1.42042739 | 0.373601618 | 1.26E-51 | 5.72E-49 | 1537.381607 | 574.7288406 | ENSG00000106462 |
| LIG3 | -1.426889405 | 0.371931951 | 6.32E-75 | 7.32E-72 | 2204.612017 | 820.3119899 | ENSG00000005156 |
| TMEM117 | -1.427953552 | 0.371657712 | 9.41E-03 | 3.50E-02 | 27.51992181 | 10.18699124 | ENSG00000139173 |
| HSF2BP | -1.429809761 | 0.371179835 | 1.45E-02 | 4.97E-02 | 24.25357199 | 9.069634925 | ENSG00000160207 |

| | | | | | | | |
|---|---|---|---|---|---|---|---|
| BHMT | -1.430622593 | 0.370970766 | 2.59E-02 | 7.96E-02 | 24.35714876 | 9.092204766 | ENSG00000145692 |
| CD247 | -1.438806405 | 0.36887236 | 4.54E-03 | 1.89E-02 | 30.46522143 | 11.16958978 | ENSG00000198821 |
| CTNNAL1 | -1.442923582 | 0.367821169 | 4.13E-39 | 1.05E-36 | 1358.195846 | 498.8285235 | ENSG00000119326 |
| RADX | -1.448137607 | 0.366494231 | 1.58E-10 | 3.34E-09 | 220.899302 | 80.53155272 | ENSG00000147231 |
| MAP6 | -1.450205052 | 0.365969405 | 1.84E-06 | 1.95E-05 | 99.93316738 | 36.37834341 | ENSG00000171533 |
| LY6G5B | -1.454979863 | 0.364760177 | 7.89E-03 | 3.03E-02 | 30.84625297 | 11.39919021 | ENSG00000240053 |
| CNTD2 | -1.461529514 | 0.363107966 | 1.11E-02 | 4.00E-02 | 23.57101044 | 8.529178109 | ENSG00000105219 |
| CD34 | -1.474357372 | 0.359893669 | 2.31E-02 | 7.26E-02 | 18.9340955 | 6.788794928 | ENSG00000174059 |
| FABP7 | -1.474518226 | 0.359853545 | 5.15E-12 | 1.38E-10 | 172.7066087 | 62.12799188 | ENSG00000164434 |
| MAPK8IP2 | -1.475422667 | 0.359628019 | 5.41E-12 | 1.44E-10 | 243.5393922 | 87.64836634 | ENSG00000008735 |
| PLCL2 | -1.475937482 | 0.359499712 | 9.41E-14 | 3.11E-12 | 201.7727551 | 72.5516316 | ENSG00000154822 |
| LRRTM1 | -1.476420046 | 0.359379483 | 6.41E-03 | 2.54E-02 | 26.02350878 | 9.414672484 | ENSG00000162951 |
| H2BC5 | -1.482166825 | 0.357950792 | 1.33E-07 | 1.73E-06 | 105.5001004 | 37.77043984 | ENSG00000158373 |
| SCML1 | -1.485115597 | 0.357219911 | 1.97E-04 | 1.27E-03 | 51.14765899 | 18.27022878 | ENSG00000047634 |
| CXCL16 | -1.490495856 | 0.355890208 | 4.38E-03 | 1.84E-02 | 37.68236175 | 13.24708641 | ENSG00000161921 |
| TRPC4 | -1.503224078 | 0.352764166 | 1.77E-16 | 7.92E-15 | 243.9351184 | 86.08463142 | ENSG00000133107 |
| DLEU7 | -1.503241139 | 0.352759994 | 2.03E-07 | 2.58E-06 | 107.1216849 | 37.83803774 | ENSG00000186047 |
| UHRF1 | -1.506225238 | 0.352031092 | 5.37E-81 | 8.30E-78 | 2415.711555 | 850.8774237 | ENSG00000276043 |
| ZNF441 | -1.512376826 | 0.350533243 | 2.88E-02 | 8.66E-02 | 16.58329479 | 5.783626547 | ENSG00000197044 |
| C1R | -1.513785036 | 0.350191255 | 2.74E-08 | 4.06E-07 | 117.6338955 | 41.12718369 | ENSG00000159403 |
| S1PR1 | -1.520411845 | 0.348586392 | 3.66E-03 | 1.58E-02 | 30.90553526 | 10.81001811 | ENSG00000170989 |
| EGR1 | -1.52166789 | 0.348283036 | 1.00E-22 | 7.94E-21 | 620.4802216 | 215.6201244 | ENSG00000120738 |
| CHRD | -1.52240874 | 0.348104232 | 1.26E-02 | 4.46E-02 | 19.77004537 | 6.878962673 | ENSG00000090539 |
| RAI2 | -1.531803456 | 0.34584477 | 9.27E-04 | 4.94E-03 | 37.57495166 | 12.97388364 | ENSG00000131831 |
| PCDHGB4 | -1.533683973 | 0.345394263 | 1.68E-08 | 2.60E-07 | 118.0928652 | 40.76337514 | ENSG00000253953 |
| CAMK2A | -1.533871111 | 0.345349464 | 3.64E-02 | 1.04E-01 | 23.30941528 | 8.102008533 | ENSG00000070808 |
| EFCAB6 | -1.538266781 | 0.344298839 | 4.54E-02 | 1.24E-01 | 13.87749195 | 4.767173245 | ENSG00000186976 |
| CCDC177 | -1.538812147 | 0.344168712 | 3.27E-14 | 1.15E-12 | 241.1873887 | 83.18566271 | ENSG00000267909 |
| AMBP | -1.550375826 | 0.341421111 | 9.33E-04 | 4.97E-03 | 37.39345939 | 12.77532681 | ENSG00000106927 |
| RPRML | -1.552686171 | 0.340874794 | 6.22E-03 | 2.48E-02 | 30.05833322 | 10.21704715 | ENSG00000179673 |
| MMUT | -1.558896091 | 0.33941069 | 5.02E-28 | 6.25E-26 | 477.9577113 | 162.6095354 | ENSG00000146085 |
| LRRC4 | -1.562912682 | 0.338467055 | 4.15E-17 | 2.00E-15 | 284.9719153 | 96.20314869 | ENSG00000128594 |
| KCNK2 | -1.564656434 | 0.338058205 | 5.86E-03 | 2.36E-02 | 25.32072797 | 8.510407115 | ENSG00000082482 |
| CTU2 | -1.567385532 | 0.337419315 | 1.02E-29 | 1.40E-27 | 506.8573833 | 171.2280051 | ENSG00000174177 |
| GDPD2 | -1.57338803 | 0.33601836 | 6.02E-24 | 5.24E-22 | 354.2270929 | 119.1104679 | ENSG00000130055 |
| CACNA1G | -1.574311533 | 0.335803336 | 5.29E-21 | 3.64E-19 | 327.2791657 | 110.1051816 | ENSG00000006283 |
| CEND1 | -1.577199825 | 0.335131726 | 1.89E-10 | 3.93E-09 | 144.6838135 | 48.52414496 | ENSG00000184524 |
| CMPK2 | -1.584344226 | 0.333476216 | 2.16E-02 | 6.89E-02 | 18.57644012 | 6.215144624 | ENSG00000134326 |
| CD177 | -1.585490275 | 0.333211414 | 1.25E-03 | 6.37E-03 | 33.87874343 | 11.22221554 | ENSG00000204936 |
| H2BC11 | -1.586315643 | 0.333020838 | 2.36E-07 | 2.96E-06 | 84.69803106 | 28.24726665 | ENSG00000124635 |
| CCDC74B | -1.599236141 | 0.330051683 | 7.62E-13 | 2.31E-11 | 191.3544551 | 63.33661525 | ENSG00000152076 |
| FAM3B | -1.601446357 | 0.329546429 | 4.01E-03 | 1.70E-02 | 28.14163021 | 9.249420762 | ENSG00000183844 |
| METTL7A | -1.602357775 | 0.329338305 | 1.28E-39 | 3.53E-37 | 899.5345462 | 296.329027 | ENSG00000185432 |
| ZAR1 | -1.602800448 | 0.329237267 | 9.63E-03 | 3.57E-02 | 22.28695974 | 7.423106716 | ENSG00000182223 |
| GABRA3 | -1.605752356 | 0.328564301 | 6.18E-15 | 2.35E-13 | 201.7322189 | 66.47591966 | ENSG00000011677 |
| FBP1 | -1.619593551 | 0.325427133 | 7.44E-04 | 4.07E-03 | 35.55339411 | 11.4808842 | ENSG00000165140 |
| KLK4 | -1.624468462 | 0.324329361 | 3.92E-02 | 1.11E-01 | 12.67747518 | 4.103355196 | ENSG00000167749 |
| SLC30A3 | -1.631554188 | 0.322740338 | 4.14E-22 | 3.10E-20 | 297.9391798 | 96.04246026 | ENSG00000115194 |
| GALR3 | -1.634505608 | 0.322080761 | 4.98E-02 | 1.33E-01 | 12.65715822 | 4.013187451 | ENSG00000128310 |
| TPP2 | -1.635649374 | 0.321825517 | 2.91E-78 | 3.85E-75 | 1791.457195 | 577.0964791 | ENSG00000134900 |
| PHGDH | -1.641659667 | 0.320487575 | 1.33E-177 | 6.16E-174 | 14736.58269 | 4723.511788 | ENSG00000092621 |
| EPHA10 | -1.64195942 | 0.320420994 | 1.15E-03 | 5.92E-03 | 36.68258332 | 11.78144335 | ENSG00000183317 |
| LRRC8E | -1.643522362 | 0.320074054 | 2.55E-02 | 7.85E-02 | 15.98308342 | 5.108523577 | ENSG00000171017 |
| C2orf15 | -1.652014195 | 0.318195603 | 3.06E-03 | 1.36E-02 | 26.90187399 | 8.544261877 | ENSG00000273045 |
| MPPED2 | -1.658594277 | 0.316747629 | 1.35E-06 | 1.46E-05 | 81.77680667 | 25.80128655 | ENSG00000066382 |
| KLF1 | -1.660172876 | 0.316401232 | 4.35E-02 | 1.20E-01 | 11.9665991 | 3.8033457 | ENSG00000105610 |
| TM4SF18 | -1.676321961 | 0.312879283 | 4.36E-07 | 5.21E-06 | 76.52726328 | 23.81720685 | ENSG00000163762 |
| H3C8 | -1.677245557 | 0.312679045 | 9.28E-05 | 6.52E-04 | 50.21632532 | 15.87015281 | ENSG00000273983 |
| FAM163A | -1.681441346 | 0.311771002 | 1.81E-03 | 8.72E-03 | 27.68603547 | 8.66068669 | ENSG00000143340 |
| SLC16A11 | -1.684993587 | 0.311004296 | 4.37E-02 | 1.21E-01 | 13.61640045 | 4.287377912 | ENSG00000174326 |
| CRB2 | -1.686009818 | 0.310785302 | 1.68E-15 | 6.86E-14 | 311.3046434 | 96.99307624 | ENSG00000148204 |
| TMEM151B | -1.694593964 | 0.308941596 | 3.07E-26 | 3.20E-24 | 394.8473581 | 122.2466347 | ENSG00000178233 |
| QRFPR | -1.706642377 | 0.306372269 | 2.67E-03 | 1.21E-02 | 27.91989484 | 8.525490882 | ENSG00000186867 |
| SLC39A1 | -1.716769777 | 0.304229134 | 2.28E-05 | 1.88E-04 | 56.80910399 | 17.35522814 | ENSG00000143570 |
| LAMP5 | -1.723082385 | 0.302900867 | 4.37E-04 | 2.54E-03 | 34.22061424 | 10.38978495 | ENSG00000125869 |
| PCDHGA1 | -1.723805442 | 0.302749096 | 9.80E-03 | 3.61E-02 | 19.35922613 | 5.843738377 | ENSG00000204956 |
| ERICH5 | -1.72509557 | 0.302478484 | 3.74E-04 | 2.22E-03 | 37.5470431 | 11.36121019 | ENSG00000177459 |
| UPK3B | -1.727907821 | 0.301889436 | 2.52E-07 | 3.15E-06 | 76.28352722 | 23.06365909 | ENSG00000243566 |
| H4C11 | -1.740898323 | 0.299183326 | 3.91E-02 | 1.11E-01 | 13.67903504 | 4.065813207 | ENSG00000197238 |
| ECRG4 | -1.743133031 | 0.298720255 | 1.61E-03 | 7.90E-03 | 29.75109074 | 8.825500379 | ENSG00000119147 |
| H2BC7 | -1.743560341 | 0.29863179 | 1.02E-02 | 3.75E-02 | 21.18360455 | 6.406653414 | ENSG00000277224 |
| RSPH4A | -1.745923849 | 0.298142955 | 2.93E-02 | 8.78E-02 | 12.69641665 | 3.792060779 | ENSG00000111834 |
| SMAD9 | -1.74849946 | 0.297611162 | 1.16E-03 | 6.00E-03 | 29.09515997 | 8.698228678 | ENSG00000120693 |
| S100A6 | -1.751606592 | 0.296970886 | 4.65E-04 | 2.69E-03 | 33.46308354 | 9.935808652 | ENSG00000197956 |
| PARP3 | -1.753847423 | 0.296509982 | 2.23E-04 | 1.41E-03 | 38.58084851 | 11.51137814 | ENSG00000041880 |

| | | | | | | | |
|---|---|---|---|---|---|---|---|
| PRSS27 | -1.756118454 | 0.296043595 | 2.05E-03 | 9.69E-03 | 29.28740074 | 8.754541661 | ENSG00000172382 |
| PLPPR1 | -1.760508392 | 0.295144141 | 8.43E-06 | 7.67E-05 | 67.75460039 | 20.05564002 | ENSG00000148123 |
| PPP2R5A | -1.763049335 | 0.294624777 | 1.90E-55 | 1.07E-52 | 847.3358089 | 249.5929928 | ENSG00000066027 |
| HRH3 | -1.766435176 | 0.293934137 | 6.86E-14 | 2.32E-12 | 168.9095747 | 49.54753468 | ENSG00000101180 |
| SAMD15 | -1.766565331 | 0.29390762 | 3.70E-02 | 1.06E-01 | 14.27558583 | 4.24234985 | ENSG00000100583 |
| VWA5B2 | -1.774496992 | 0.292296206 | 3.12E-02 | 9.25E-02 | 11.73086061 | 3.424453379 | ENSG00000145198 |
| MTSS1 | -1.776118412 | 0.291967884 | 4.18E-10 | 8.19E-09 | 104.572931 | 30.47504285 | ENSG00000170873 |
| SMKR1 | -1.781836046 | 0.290813058 | 2.71E-03 | 1.22E-02 | 23.60070043 | 6.841420684 | ENSG00000240204 |
| ZHX1 | -1.790352762 | 0.289101347 | 1.05E-08 | 1.68E-07 | 92.13878589 | 26.7773867 | ENSG00000165156 |
| BRINP2 | -1.790859556 | 0.288999809 | 1.01E-03 | 5.31E-03 | 28.53401163 | 8.214196466 | ENSG00000198797 |
| CLEC4G | -1.797346191 | 0.287703327 | 4.22E-03 | 1.78E-02 | 23.62860899 | 6.758739013 | ENSG00000182566 |
| CDK5R2 | -1.806275164 | 0.285928203 | 3.36E-02 | 9.80E-02 | 14.23632741 | 4.013187451 | ENSG00000171450 |
| TNFRSF11A | -1.812996349 | 0.284599227 | 5.42E-19 | 3.17E-17 | 213.2805049 | 60.57119339 | ENSG00000141655 |
| WDR97 | -1.817969877 | 0.283619793 | 1.99E-02 | 6.42E-02 | 15.69239963 | 4.452191601 | ENSG00000179698 |
| H2BC21 | -1.835377546 | 0.280218178 | 1.31E-07 | 1.71E-06 | 77.30087165 | 21.69457054 | ENSG00000184678 |
| ZGLP1 | -1.839530635 | 0.279412674 | 3.21E-02 | 9.44E-02 | 14.78998184 | 4.144696031 | ENSG00000220201 |
| UGT3A1 | -1.840496585 | 0.279225657 | 2.72E-06 | 2.78E-05 | 64.88409718 | 17.98844062 | ENSG00000145626 |
| NPTX1 | -1.844092137 | 0.278530624 | 7.54E-09 | 1.24E-07 | 107.5729578 | 29.927538 | ENSG00000171246 |
| WASHC5 | -1.852266833 | 0.276956858 | 5.08E-66 | 4.28E-63 | 899.4196271 | 248.9772288 | ENSG00000164961 |
| SLC17A6 | -1.862004438 | 0.275093806 | 3.74E-02 | 1.07E-01 | 11.4622752 | 3.203326707 | ENSG00000091664 |
| KCNK13 | -1.86253337 | 0.274992968 | 2.13E-02 | 6.80E-02 | 15.98614264 | 4.339454016 | ENSG00000152315 |
| PCDHB10 | -1.873215341 | 0.27296439 | 1.99E-06 | 2.10E-05 | 62.0887207 | 17.02136388 | ENSG00000120324 |
| SCT | -1.878617913 | 0.271944111 | 7.65E-03 | 2.95E-02 | 17.76564786 | 4.849854916 | ENSG00000070031 |
| GPR155 | -1.882751242 | 0.271166104 | 1.72E-04 | 1.12E-03 | 41.16601225 | 11.01562298 | ENSG00000163328 |
| H2AC13 | -1.886103929 | 0.270536671 | 1.69E-02 | 5.63E-02 | 13.98678118 | 3.784574705 | ENSG00000196747 |
| C3orf70 | -1.890646971 | 0.269686092 | 5.89E-07 | 6.85E-06 | 59.91625533 | 16.17721034 | ENSG00000187068 |
| AMH | -1.895879956 | 0.268709651 | 1.41E-22 | 1.11E-20 | 290.7242117 | 78.19692476 | ENSG00000104899 |
| HESX1 | -1.90560513 | 0.266904377 | 2.24E-11 | 5.36E-10 | 132.1193859 | 35.40433025 | ENSG00000163666 |
| RDX | -1.914207585 | 0.265317624 | 1.60E-103 | 4.24E-100 | 1630.292521 | 432.1162104 | ENSG00000137710 |
| B4GALNT1 | -1.918425962 | 0.26454298 | 4.84E-05 | 3.67E-04 | 41.32275287 | 10.86589306 | ENSG00000135454 |
| ACKR3 | -1.919368189 | 0.264370263 | 2.79E-03 | 1.25E-02 | 20.89174068 | 5.555013801 | ENSG00000144476 |
| RHBDL3 | -1.935956003 | 0.261347995 | 3.98E-34 | 7.16E-32 | 525.0728158 | 137.2719811 | ENSG00000141314 |
| CRHR2 | -1.941316678 | 0.260378697 | 1.62E-03 | 7.95E-03 | 28.09181866 | 7.396849648 | ENSG00000106113 |
| H4C8 | -1.967345847 | 0.255723057 | 1.23E-04 | 8.39E-04 | 38.36227005 | 9.687875259 | ENSG00000158406 |
| KCNIP3 | -1.974050804 | 0.254537336 | 2.64E-08 | 3.92E-07 | 80.58596754 | 20.66738197 | ENSG00000115041 |
| H2BC12 | -1.989298288 | 0.251861361 | 1.01E-26 | 1.09E-24 | 342.22794 | 85.97090615 | ENSG00000197903 |
| TMSB15B | -1.996062558 | 0.250683239 | 3.84E-02 | 1.09E-01 | 9.498316199 | 2.359173168 | ENSG00000269226 |
| SCUBE2 | -2.003493432 | 0.249395367 | 1.30E-02 | 4.55E-02 | 13.81170044 | 3.4357383 | ENSG00000175356 |
| PDYN | -2.011420208 | 0.248028841 | 1.80E-04 | 1.17E-03 | 37.02780645 | 9.140482023 | ENSG00000101327 |
| SMIM27 | -2.023270286 | 0.245999914 | 1.11E-02 | 4.01E-02 | 14.08975658 | 3.484565209 | ENSG00000235453 |
| MT1M | -2.033358173 | 0.244285786 | 2.22E-03 | 1.04E-02 | 21.17601295 | 5.168635407 | ENSG00000205364 |
| HAS1 | -2.042493657 | 0.242743798 | 3.68E-05 | 2.88E-04 | 36.71552793 | 8.900584356 | ENSG00000105509 |
| H2BC9 | -2.044616318 | 0.242386908 | 3.48E-09 | 5.94E-08 | 85.29595733 | 20.76079891 | ENSG00000275713 |
| ABCA5 | -2.051698047 | 0.241200023 | 2.24E-06 | 2.33E-05 | 48.52589007 | 11.69876167 | ENSG00000154265 |
| DGKI | -2.061686767 | 0.239535806 | 3.20E-03 | 1.41E-02 | 22.42721674 | 5.258253499 | ENSG00000157680 |
| ZBTB16 | -2.073133804 | 0.237642734 | 4.24E-02 | 1.18E-01 | 8.723527756 | 2.070448592 | ENSG00000109906 |
| CHRDL2 | -2.083222691 | 0.235986676 | 1.15E-02 | 4.13E-02 | 14.25852349 | 3.296743646 | ENSG00000054938 |
| ELMOD1 | -2.084923696 | 0.2357086 | 1.29E-04 | 8.74E-04 | 30.52460141 | 7.167687249 | ENSG00000110675 |
| HPDL | -2.097937675 | 0.233591928 | 1.66E-33 | 2.79E-31 | 538.9619054 | 125.7826232 | ENSG00000186603 |
| TTBK1 | -2.098888032 | 0.233438103 | 3.36E-08 | 4.89E-07 | 75.7135095 | 17.51949218 | ENSG00000146216 |
| ACTL8 | -2.105087232 | 0.232437181 | 4.11E-05 | 3.17E-04 | 38.65611076 | 9.043266237 | ENSG00000117148 |
| DPF1 | -2.123001251 | 0.229568842 | 1.87E-34 | 3.47E-32 | 351.5176295 | 80.66068817 | ENSG00000011332 |
| FOSB | -2.125103117 | 0.229234626 | 7.60E-05 | 5.44E-04 | 33.19577591 | 7.674289303 | ENSG00000125740 |
| SLC38A5 | -2.12913807 | 0.228594394 | 1.47E-85 | 2.48E-82 | 1110.689398 | 254.440801 | ENSG00000017483 |
| ANKRD63 | -2.144361318 | 0.226194959 | 4.47E-02 | 1.23E-01 | 8.889137755 | 1.961509853 | ENSG00000230778 |
| ZNF594 | -2.175267132 | 0.221400881 | 2.25E-18 | 1.23E-16 | 200.3170737 | 44.6615635 | ENSG00000180626 |
| RGS8 | -2.182119473 | 0.220351791 | 4.45E-02 | 1.22E-01 | 9.101500088 | 1.961509853 | ENSG00000135824 |
| BOC | -2.191230379 | 0.21896461 | 2.01E-21 | 1.44E-19 | 182.3141164 | 39.94982717 | ENSG00000144857 |
| BTBD17 | -2.19697269 | 0.218094805 | 1.62E-37 | 3.58E-35 | 472.5324933 | 103.0343838 | ENSG00000204347 |
| H4C9 | -2.211955488 | 0.215841549 | 8.93E-13 | 2.68E-11 | 106.9408917 | 23.17259783 | ENSG00000276180 |
| DRD2 | -2.216279741 | 0.215195566 | 8.98E-03 | 3.36E-02 | 12.76654515 | 2.738065489 | ENSG00000149295 |
| LRRC3B | -2.273888805 | 0.206771778 | 6.10E-07 | 7.07E-06 | 49.60724457 | 10.21704715 | ENSG00000179796 |
| ABALON | -2.29215979 | 0.204169633 | 9.32E-04 | 4.96E-03 | 23.55650365 | 4.801028007 | ENSG00000281376 |
| TCEAL2 | -2.297640214 | 0.203395517 | 3.82E-37 | 8.34E-35 | 376.8386431 | 76.29387778 | ENSG00000184905 |
| GSAP | -2.307534954 | 0.202005299 | 6.36E-03 | 2.53E-02 | 13.66640739 | 2.775607477 | ENSG00000186088 |
| COL15A1 | -2.328697057 | 0.199063821 | 4.79E-02 | 1.29E-01 | 8.409565493 | 1.642729363 | ENSG00000204291 |
| ARHGAP36 | -2.33376912 | 0.198365203 | 2.35E-04 | 1.48E-03 | 24.46771577 | 4.906167899 | ENSG00000147256 |
| ATOH8 | -2.338057041 | 0.197776506 | 8.85E-04 | 4.75E-03 | 23.97985284 | 4.669519427 | ENSG00000168874 |
| SLC1A2 | -2.339733992 | 0.197546749 | 5.13E-06 | 4.94E-05 | 38.18846709 | 7.516523654 | ENSG00000110436 |
| PDE1B | -2.348003977 | 0.196417588 | 1.49E-08 | 2.32E-07 | 62.90454901 | 12.34000988 | ENSG00000123360 |
| TMEM215 | -2.361164276 | 0.194634009 | 5.00E-03 | 2.06E-02 | 14.24273893 | 2.805663392 | ENSG00000188133 |
| NUP210L | -2.367986009 | 0.193715861 | 4.91E-02 | 1.32E-01 | 7.219019461 | 1.432887611 | ENSG00000143552 |
| BEX5 | -2.378990197 | 0.19224391 | 8.47E-05 | 6.01E-04 | 26.77354558 | 5.127294571 | ENSG00000184515 |
| NOTO | -2.382504511 | 0.191776186 | 2.57E-02 | 7.92E-02 | 8.846820112 | 1.672785277 | ENSG00000214513 |
| PHETA2 | -2.383148004 | 0.191690666 | 2.43E-11 | 5.80E-10 | 89.44816626 | 17.11846805 | ENSG00000177096 |

| | | | | | | | |
|---|---|---|---|---|---|---|---|
| PITX3 | -2.409296365 | 0.188247634 | 1.15E-02 | 4.12E-02 | 12.82276822 | 2.411798924 | ENSG00000107859 |
| TCEAL5 | -2.432130055 | 0.185291672 | 5.67E-11 | 1.28E-09 | 81.56817997 | 15.10444406 | ENSG00000204065 |
| ORM2 | -2.446318364 | 0.183478336 | 8.68E-04 | 4.67E-03 | 20.46839149 | 3.735747796 | ENSG00000228278 |
| PCDHGA3 | -2.453063751 | 0.182622477 | 1.72E-02 | 5.69E-02 | 9.474940033 | 1.751668102 | ENSG00000254245 |
| NRGN | -2.455176017 | 0.182355293 | 3.22E-02 | 9.46E-02 | 7.595616319 | 1.402831696 | ENSG00000154146 |
| TCHH | -2.471156634 | 0.180346504 | 2.48E-02 | 7.70E-02 | 9.670240025 | 1.781724017 | ENSG00000159450 |
| CFAP65 | -2.519751263 | 0.17437302 | 3.07E-02 | 9.11E-02 | 8.048655502 | 1.432887611 | ENSG00000181378 |
| KCNJ10 | -2.526985946 | 0.17350078 | 1.57E-04 | 1.04E-03 | 25.34420184 | 4.313196947 | ENSG00000177807 |
| SHPK | -2.537000193 | 0.172300621 | 6.51E-17 | 3.05E-15 | 117.7957699 | 20.3255936 | ENSG00000197417 |
| SLC2A9 | -2.594120868 | 0.165612002 | 4.62E-02 | 1.26E-01 | 6.355161041 | 1.084051205 | ENSG00000109667 |
| THBD | -2.597233818 | 0.165255041 | 7.17E-04 | 3.93E-03 | 18.71098465 | 3.094387968 | ENSG00000178726 |
| CDH7 | -2.600870311 | 0.164839019 | 3.71E-05 | 2.90E-04 | 27.012441 | 4.42962176 | ENSG00000081138 |
| ANKS4B | -2.654815021 | 0.158789231 | 4.38E-02 | 1.21E-01 | 6.623746445 | 1.084051205 | ENSG00000175311 |
| SLC34A3 | -2.673215839 | 0.156776819 | 3.77E-02 | 1.07E-01 | 6.719130247 | 1.084051205 | ENSG00000198569 |
| PTGER2 | -2.681745898 | 0.155852597 | 5.09E-08 | 7.20E-07 | 50.8678214 | 7.948041731 | ENSG00000125384 |
| IGFLR1 | -2.722669956 | 0.151493736 | 4.38E-02 | 1.21E-01 | 7.017405643 | 1.1028222 | ENSG00000126246 |
| H2AC6 | -2.766972356 | 0.146912356 | 6.21E-54 | 3.26E-51 | 512.5062984 | 75.64332002 | ENSG00000180573 |
| GRIK1 | -2.777462683 | 0.145847981 | 1.38E-02 | 4.78E-02 | 9.246793136 | 1.305177878 | ENSG00000171189 |
| PLEK2 | -2.810489407 | 0.1425471 | 1.21E-02 | 4.32E-02 | 9.643609248 | 1.365289707 | ENSG00000100558 |
| TRPC3 | -2.871361542 | 0.136657681 | 3.53E-03 | 1.53E-02 | 12.82908205 | 1.800495011 | ENSG00000138741 |
| SEMA6D | -2.91055014 | 0.132995548 | 2.34E-08 | 3.52E-07 | 50.85519375 | 6.612258286 | ENSG00000137872 |
| SYT11 | -2.91160448 | 0.132898389 | 8.98E-35 | 1.68E-32 | 273.9300051 | 36.37520583 | ENSG00000132718 |
| ZNF229 | -2.969960428 | 0.127630016 | 9.13E-12 | 2.32E-10 | 72.45287215 | 9.215566 | ENSG00000278318 |
| EDN3 | -2.994135352 | 0.125509167 | 2.14E-02 | 6.84E-02 | 8.426627834 | 1.1028222 | ENSG00000124205 |
| PCDHB5 | -3.046345914 | 0.121048247 | 5.09E-39 | 1.28E-36 | 249.8048744 | 30.16374844 | ENSG00000113209 |
| LUARIS | -3.071086727 | 0.118990085 | 1.31E-02 | 4.60E-02 | 8.88282393 | 1.1028222 | ENSG00000231638 |
| FRZB | -3.110533167 | 0.115780712 | 1.59E-27 | 1.84E-25 | 163.5777788 | 18.96410274 | ENSG00000162998 |
| GSC2 | -3.210236522 | 0.108049438 | 2.93E-02 | 8.78E-02 | 6.408227199 | 0.697672811 | ENSG00000063515 |
| H3C10 | -3.221319231 | 0.107222588 | 1.92E-09 | 3.42E-08 | 48.02294164 | 5.134780645 | ENSG00000278828 |
| CRHR1 | -3.306516424 | 0.101073982 | 2.43E-02 | 7.56E-02 | 6.636374096 | 0.637560981 | ENSG00000120088 |
| LRRTM3 | -3.34394387 | 0.098485567 | 7.66E-05 | 5.48E-04 | 21.23361149 | 2.149331417 | ENSG00000198739 |
| MTTP | -3.3847875 | 0.095736475 | 1.70E-02 | 5.65E-02 | 7.33777943 | 0.7352148 | ENSG00000138823 |
| SSTR3 | -3.405688325 | 0.094359506 | 1.02E-03 | 5.38E-03 | 14.4644743 | 1.335233792 | ENSG00000278195 |
| MYZAP | -3.41575661 | 0.093703282 | 1.67E-02 | 5.57E-02 | 7.26765093 | 0.667616896 | ENSG00000263155 |
| GABRA4 | -3.423256757 | 0.09321741 | 1.81E-02 | 5.93E-02 | 7.38137485 | 0.686387891 | ENSG00000109158 |
| SPEF1 | -3.455666967 | 0.091146624 | 4.60E-04 | 2.66E-03 | 15.15522882 | 1.402831696 | ENSG00000101222 |
| GNG8 | -3.461671498 | 0.090768058 | 1.36E-02 | 4.71E-02 | 7.556455588 | 0.686387891 | ENSG00000167414 |
| PAPLN | -3.6631207 | 0.078938849 | 5.89E-30 | 8.34E-28 | 166.1684822 | 13.20673326 | ENSG00000100767 |
| POU3F4 | -3.788802617 | 0.072353037 | 1.26E-05 | 1.11E-04 | 23.46172121 | 1.642729363 | ENSG00000196767 |
| TRIM4 | -3.913991952 | 0.066339321 | 4.93E-08 | 6.98E-07 | 36.18910563 | 2.381743009 | ENSG00000146833 |
| DMRT2 | -4.133813496 | 0.056963694 | 1.06E-04 | 7.31E-04 | 18.0360147 | 1.005168381 | ENSG00000173253 |
| SLC12A1 | -4.286022191 | 0.051260018 | 6.54E-03 | 2.59E-02 | 7.205114033 | 0.348836406 | ENSG00000074803 |
| ORM1 | -4.293187271 | 0.051006069 | 7.92E-04 | 4.30E-03 | 13.26643437 | 0.637560981 | ENSG00000229314 |
| HTR1A | -4.306845795 | 0.050525455 | 6.17E-08 | 8.56E-07 | 34.09869736 | 1.732897107 | ENSG00000178394 |
| TAF9B | -4.710876231 | 0.03818431 | 2.03E-73 | 1.98E-70 | 422.0349927 | 15.9143048 | ENSG00000187325 |
| TNFAIP8L3 | -6.297546956 | 0.012713042 | 1.94E-05 | 1.62E-04 | 14.91327419 | 0 | ENSG00000183578 |
| ZFP28 | -6.319457209 | 0.012521427 | 5.91E-06 | 5.63E-05 | 15.16597733 | 0 | ENSG00000196867 |
