## Supplementary material for "Chemokine Receptor 1 and its associated immune pathway are downregulated in SF3B1^MT^ blood and non-blood cancers": Table_S5

**Table S5. GSEA of dysreguated genes in SF3B1$^{K700E}$ ES cells (K700E ES line 1).**

**GSEA Positive** Hematopoietic/Immune pathways are highlighted in red.

| Gene sets | SIZE | ES | NES | NOM p-val | FDR q-val |
|---|---|---|---|---|---|
| GOBP_PIRNA_METABOLIC_PROCESS | 17 | 0.815325 | 1.93974 | 0 | 0.065123 |
| GOBP_GRANULOCYTE_ACTIVATION | 33 | 0.716879 | 1.935354 | 0 | 0.034953 |
| GOBP_POSITIVE_REGULATION_OF_VASCULAR_ENDOTHELIAL_GROWTH_FACTOR_PRODU( | 24 | 0.752959 | 1.906069 | 0 | 0.049779 |
| GOBP_REGULATION_OF_TRANSPOSITION | 20 | 0.796145 | 1.899141 | 0 | 0.045724 |
| GOBP_VASCULAR_ENDOTHELIAL_GROWTH_FACTOR_PRODUCTION | 30 | 0.717025 | 1.837482 | 0 | 0.13503 |
| GOBP_DNA_METHYLATION_OR_DEMETHYLATION | 82 | 0.600672 | 1.836242 | 0 | 0.115724 |
| GOBP_TRANSPOSITION | 24 | 0.731077 | 1.815935 | 0 | 0.147445 |
| GOBP_NEGATIVE_REGULATION_OF_REPRODUCTIVE_PROCESS | 46 | 0.637312 | 1.8123 | 0.001304 | 0.138697 |
| GOBP_ANTERIOR_POSTERIOR_AXIS_SPECIFICATION | 45 | 0.646102 | 1.805805 | 0 | 0.138705 |
| GOBP_NEUTROPHIL_MEDIATED_IMMUNITY | 30 | 0.688511 | 1.802606 | 0.001335 | 0.132777 |
| GOBP_GRANULOCYTE_MIGRATION | 90 | 0.587611 | 1.801027 | 0 | 0.123926 |
| GOBP_GRANULOCYTE_CHEMOTAXIS | 72 | 0.594912 | 1.800437 | 0 | 0.114881 |
| GOBP_MONOCYTE_CHEMOTAXIS | 32 | 0.674603 | 1.797871 | 0.001353 | 0.110314 |
| GOBP_HOMOLOGOUS_CHROMOSOME_PAIRING_AT_MEIOSIS | 39 | 0.64612 | 1.79278 | 0 | 0.111543 |
| GOBP_DNA_METHYLATION | 61 | 0.605069 | 1.787543 | 0 | 0.114635 |
| GOBP_REGULATION_OF_LEUKOCYTE_DEGRANULATION | 38 | 0.663877 | 1.787469 | 0 | 0.107651 |
| GOBP_POSITIVE_REGULATION_OF_FATTY_ACID_METABOLIC_PROCESS | 26 | 0.69752 | 1.778942 | 0 | 0.117929 |
| GOBP_RESPONSE_TO_TUMOR_CELL | 29 | 0.681938 | 1.771694 | 0.001377 | 0.124451 |
| GOBP_POSITIVE_REGULATION_OF_T_CELL_PROLIFERATION | 64 | 0.599759 | 1.76385 | 0 | 0.134415 |
| GOBP_REGULATION_OF_HUMORAL_IMMUNE_RESPONSE | 22 | 0.699881 | 1.763676 | 0.001414 | 0.128412 |
| GOBP_MYELOID_LEUKOCYTE_MIGRATION | 145 | 0.543261 | 1.755485 | 0 | 0.139143 |
| GOBP_LEUKOCYTE_CHEMOTAXIS | 147 | 0.546422 | 1.754518 | 0 | 0.134825 |
| GOBP_LEUKOCYTE_MIGRATION | 256 | 0.516032 | 1.744428 | 0 | 0.150862 |
| GOBP_NUCLEOSIDE_CATABOLIC_PROCESS | 21 | 0.701743 | 1.729614 | 0.001471 | 0.181524 |
| GOBP_EMBRYONIC_EYE_MORPHOGENESIS | 27 | 0.671075 | 1.713149 | 0 | 0.220823 |
| GOBP_REGULATION_OF_LEUKOCYTE_ADHESION_TO_VASCULAR_ENDOTHELIAL_CELL | 21 | 0.698189 | 1.711391 | 0.007174 | 0.217784 |
| GOBP_BLOOD_VESSEL_REMODELING | 33 | 0.655472 | 1.710488 | 0.002639 | 0.212552 |
| GOBP_REGULATION_OF_WOUND_HEALING | 101 | 0.547387 | 1.708583 | 0 | 0.210636 |
| GOBP_NEUTROPHIL_ACTIVATION_INVOLVED_IN_IMMUNE_RESPONSE | 17 | 0.734699 | 1.706978 | 0.001435 | 0.208025 |
| GOBP_RIBONUCLEOSIDE_CATABOLIC_PROCESS | 16 | 0.738626 | 1.705213 | 0.007564 | 0.206484 |
| GOBP_EXTRACELLULAR_MATRIX_DISASSEMBLY | 45 | 0.614234 | 1.704062 | 0.003947 | 0.203096 |
| GOBP_EMBRYONIC_CAMERA_TYPE_EYE_MORPHOGENESIS | 19 | 0.717701 | 1.70367 | 0.004298 | 0.197797 |
| GOBP_POSITIVE_REGULATION_OF_LEUKOCYTE_PROLIFERATION | 102 | 0.548487 | 1.703348 | 0 | 0.1925 |
| GOBP_GENE_SILENCING_BY_RNA | 123 | 0.532486 | 1.702829 | 0.001098 | 0.188133 |
| GOBP_INTERLEUKIN_17_PRODUCTION | 25 | 0.661059 | 1.699049 | 0.006878 | 0.193012 |
| GOBP_INTERLEUKIN_8_PRODUCTION | 55 | 0.590131 | 1.696291 | 0.001238 | 0.194356 |
| GOBP_T_CELL_SELECTION | 32 | 0.627413 | 1.68544 | 0.008276 | 0.221368 |
| GOBP_SYNAPTONEMAL_COMPLEX_ORGANIZATION | 21 | 0.685847 | 1.68513 | 0.00551 | 0.216273 |
| GOBP_DETECTION_OF_STIMULUS_INVOLVED_IN_SENSORY_PERCEPTION | 70 | 0.565976 | 1.68425 | 0.001198 | 0.213501 |
| GOBP_EMBRYONIC_AXIS_SPECIFICATION | 33 | 0.631341 | 1.684061 | 0.006693 | 0.208785 |
| GOBP_POSITIVE_REGULATION_OF_LEUKOCYTE_ADHESION_TO_VASCULAR_ENDOTHELIAL | 16 | 0.736251 | 1.683884 | 0.001479 | 0.20444 |
| GOBP_KERATINIZATION | 16 | 0.734521 | 1.683796 | 0.004412 | 0.199869 |
| GOBP_HOMOLOGOUS_CHROMOSOME_SEGREGATION | 48 | 0.593348 | 1.6824 | 0.001261 | 0.198518 |
| GOBP_ALPHA_BETA_T_CELL_PROLIFERATION | 30 | 0.633873 | 1.679391 | 0.002681 | 0.202075 |
| GOBP_INOSITOL_PHOSPHATE_BIOSYNTHETIC_PROCESS | 22 | 0.674092 | 1.677708 | 0.00289 | 0.202308 |
| GOBP_MYELOID_LEUKOCYTE_MEDIATED_IMMUNITY | 77 | 0.548463 | 1.677399 | 0 | 0.198782 |
| GOBP_NEGATIVE_REGULATION_OF_INTERFERON_GAMMA_PRODUCTION | 24 | 0.670479 | 1.675977 | 0.001385 | 0.198358 |
| GOBP_NUCLEUS_LOCALIZATION | 24 | 0.668082 | 1.674045 | 0.01122 | 0.198692 |
| GOBP_TISSUE_REGENERATION | 51 | 0.592519 | 1.673575 | 0.001258 | 0.195497 |
| GOBP_POSITIVE_REGULATION_OF_LEUKOCYTE_CELL_CELL_ADHESION | 168 | 0.506064 | 1.669941 | 0 | 0.200793 |
| GOBP_PEPTIDE_CROSS_LINKING | 16 | 0.726117 | 1.664774 | 0.005797 | 0.210652 |
| GOBP_CATECHOL_CONTAINING_COMPOUND_BIOSYNTHETIC_PROCESS | 16 | 0.708541 | 1.662523 | 0.00761 | 0.212897 |
| GOBP_BASE_CONVERSION_OR_SUBSTITUTION_EDITING | 15 | 0.735878 | 1.662504 | 0.007299 | 0.208897 |
| GOBP_NEUTROPHIL_MIGRATION | 73 | 0.551262 | 1.66159 | 0.002375 | 0.207314 |
| GOBP_PEPTIDE_ANTIGEN_ASSEMBLY_WITH_MHC_PROTEIN_COMPLEX | 15 | 0.715925 | 1.661487 | 0.007396 | 0.203753 |
| GOBP_POSITIVE_REGULATION_OF_CELL_CELL_ADHESION | 203 | 0.501361 | 1.661313 | 0 | 0.200508 |
| GOBP_MYELOID_CELL_ACTIVATION_INVOLVED_IN_IMMUNE_RESPONSE | 67 | 0.566932 | 1.65919 | 0.004796 | 0.202446 |
| GOBP_REGULATION_OF_LEUKOCYTE_MIGRATION | 150 | 0.508139 | 1.658017 | 0.002179 | 0.201909 |
| GOBP_ICOSANOID_TRANSPORT | 37 | 0.612793 | 1.656855 | 0.002597 | 0.201209 |
| GOBP_NUCLEOBASE_CONTAINING_SMALL_MOLECULE_CATABOLIC_PROCESS | 26 | 0.647609 | 1.654469 | 0.008357 | 0.204236 |
| GOBP_TISSUE_REMODELING | 127 | 0.518083 | 1.65368 | 0.001111 | 0.202705 |
| GOBP_MYELOID_LEUKOCYTE_ACTIVATION | 157 | 0.511586 | 1.653651 | 0 | 0.199482 |
| GOBP_ARACHIDONIC_ACID_SECRETION | 20 | 0.683418 | 1.653507 | 0.008721 | 0.19665 |
| GOBP_ANTIGEN_PROCESSING_AND_PRESENTATION_OF_PEPTIDE_OR_POLYSACCHARIDE | 25 | 0.641863 | 1.651624 | 0.007194 | 0.197451 |
| GOBP_POSITIVE_REGULATION_OF_CELL_ADHESION_MEDIATED_BY_INTEGRIN | 15 | 0.708866 | 1.651162 | 0.013595 | 0.195401 |
| GOBP_AXIS_SPECIFICATION | 76 | 0.551072 | 1.649389 | 0.002361 | 0.197098 |
| GOBP_ANTIGEN_PROCESSING_AND_PRESENTATION_OF_EXOGENOUS_PEPTIDE_ANTIGEN | 20 | 0.674368 | 1.649368 | 0.007257 | 0.194241 |
| GOBP_REGULATION_OF_T_CELL_PROLIFERATION | 114 | 0.51916 | 1.649048 | 0.001115 | 0.192186 |
| GOBP_LEUKOCYTE_ADHESION_TO_VASCULAR_ENDOTHELIAL_CELL | 32 | 0.619664 | 1.646669 | 0.009346 | 0.194918 |
| GOBP_REGULATION_OF_MYELOID_LEUKOCYTE_MEDIATED_IMMUNITY | 46 | 0.575062 | 1.646621 | 0.003708 | 0.192243 |
| GOBP_POSITIVE_REGULATION_OF_IMMUNE_RESPONSE | 355 | 0.483277 | 1.643489 | 0 | 0.197125 |
| GOBP_ACTIVATION_OF_NF_KAPPAB_INDUCING_KINASE_ACTIVITY | 17 | 0.702525 | 1.641275 | 0.019374 | 0.200288 |
| GOBP_CELLULAR_DEFENSE_RESPONSE | 22 | 0.665926 | 1.637804 | 0.002809 | 0.206106 |
| GOBP_DNA_MODIFICATION | 101 | 0.521007 | 1.635385 | 0.004614 | 0.209916 |
| GOBP_LEUKOCYTE_MEDIATED_IMMUNITY | 256 | 0.485716 | 1.63165 | 0 | 0.21685 |
| GOBP_POSITIVE_REGULATION_OF_INTERLEUKIN_6_PRODUCTION | 67 | 0.552161 | 1.629237 | 0.006002 | 0.220091 |
| GOBP_EMBRYONIC_CAMERA_TYPE_EYE_DEVELOPMENT | 29 | 0.631547 | 1.628675 | 0.006667 | 0.218524 |
| GOBP_REGULATION_OF_LEUKOCYTE_PROLIFERATION | 165 | 0.502726 | 1.627283 | 0 | 0.219646 |
| GOBP_CELLULAR_RESPONSE_TO_ALKALOID | 30 | 0.617979 | 1.627145 | 0.012195 | 0.217157 |
| GOBP_POSITIVE_T_CELL_SELECTION | 22 | 0.657303 | 1.626575 | 0.011173 | 0.216008 |
| GOBP_POSITIVE_REGULATION_OF_LEUKOCYTE_MIGRATION | 98 | 0.525738 | 1.62582 | 0.001163 | 0.215231 |
| GOBP_CELL_CHEMOTAXIS | 197 | 0.490392 | 1.625729 | 0 | 0.212885 |
| GOBP_IMMUNE_RESPONSE_REGULATING_CELL_SURFACE_RECEPTOR_SIGNALING_PATH\ | 180 | 0.493341 | 1.624252 | 0 | 0.213941 |
| GOBP_POSITIVE_REGULATION_OF_CALCIUM_ION_TRANSMEMBRANE_TRANSPORT | 57 | 0.564194 | 1.623891 | 0.005019 | 0.21217 |
| GOBP_SINGLE_STRANDED_VIRAL_RNA_REPLICATION_VIA_DOUBLE_STRANDED_DNA_INT | 17 | 0.692506 | 1.622896 | 0.008889 | 0.211914 |
| GOBP_T_CELL_MEDIATED_IMMUNITY | 77 | 0.534433 | 1.61987 | 0.003525 | 0.217089 |
| GOBP_MACROPHAGE_DIFFERENTIATION | 39 | 0.592688 | 1.619454 | 0.011873 | 0.215739 |
| GOBP_POSITIVE_REGULATION_OF_TYROSINE_PHOSPHORYLATION_OF_STAT_PROTEIN | 37 | 0.604079 | 1.619412 | 0.005249 | 0.213385 |
| GOBP_PROTEIN_AUTOPROCESSING | 21 | 0.664524 | 1.616701 | 0.013755 | 0.217427 |
| GOBP_LEUKOCYTE_DEGRANULATION | 57 | 0.550523 | 1.616281 | 0.007308 | 0.216116 |
| GOBP_ACTIVATION_OF_IMMUNE_RESPONSE | 220 | 0.481999 | 1.615652 | 0 | 0.215257 |
| GOBP_T_CELL_LINEAGE_COMMITMENT | 17 | 0.685179 | 1.615353 | 0.013433 | 0.213634 |
| GOBP_POSITIVE_REGULATION_OF_CHEMOKINE_PRODUCTION | 47 | 0.57659 | 1.615336 | 0.002513 | 0.211399 |
| GOBP_TYROSINE_PHOSPHORYLATION_OF_STAT_PROTEIN | 51 | 0.57331 | 1.614146 | 0.006266 | 0.212511 |
| GOBP_POSITIVE_REGULATION_OF_RELEASE_OF_SEQUESTERED_CALCIUM_ION_INTO_CY | 29 | 0.622169 | 1.611113 | 0.01071 | 0.217991 |
| GOBP_ARACHIDONIC_ACID_METABOLIC_PROCESS | 39 | 0.586501 | 1.610826 | 0.011873 | 0.216389 |
| GOBP_POSITIVE_REGULATION_OF_LEUKOCYTE_APOPTOTIC_PROCESS | 19 | 0.667965 | 1.610055 | 0.012876 | 0.215876 |
| GOBP_POSITIVE_REGULATION_OF_FATTY_ACID_BIOSYNTHETIC_PROCESS | 15 | 0.706996 | 1.60985 | 0.010836 | 0.214269 |
| GOBP_POSITIVE_REGULATION_OF_WOUND_HEALING | 44 | 0.578505 | 1.609606 | 0.007772 | 0.212752 |
| GOBP_REGULATION_OF_ADAPTIVE_IMMUNE_RESPONSE | 134 | 0.504378 | 1.608796 | 0 | 0.212777 |
| GOBP_SKELETAL_MUSCLE_TISSUE_REGENERATION | 29 | 0.620076 | 1.607924 | 0.014765 | 0.212888 |
| GOBP_RESPONSE_TO_ACTIVITY | 45 | 0.562656 | 1.607706 | 0.01023 | 0.211419 |
| GOBP_HUMORAL_IMMUNE_RESPONSE | 100 | 0.513143 | 1.606379 | 0.001136 | 0.212843 |
| GOBP_IMMUNE_EFFECTOR_PROCESS | 395 | 0.469483 | 1.605233 | 0 | 0.21344 |
| GOBP_POSITIVE_REGULATION_OF_CELL_ADHESION | 337 | 0.467677 | 1.603288 | 0 | 0.215855 |
| GOBP_POSITIVE_REGULATION_OF_INFLAMMATORY_RESPONSE | 92 | 0.509142 | 1.602694 | 0.003425 | 0.215302 |
| GOBP_POSITIVE_REGULATION_OF_CELL_MIGRATION_INVOLVED_IN_SPROUTING_ANGIOGE | 18 | 0.68568 | 1.602459 | 0.013373 | 0.213818 |
| GOBP_POSITIVE_REGULATION_OF_BLOOD_VESSEL_ENDOTHELIAL_CELL_MIGRATION | 50 | 0.562254 | 1.602071 | 0.003827 | 0.212971 |
| GOBP_LEUKOCYTE_CELL_CELL_ADHESION | 257 | 0.47446 | 1.600182 | 0 | 0.215841 |
| GOBP_NEUTROPHIL_CHEMOTAXIS | 57 | 0.550082 | 1.599372 | 0.008631 | 0.215925 |
| GOBP_BROWN_FAT_CELL_DIFFERENTIATION | 42 | 0.570615 | 1.597285 | 0.012739 | 0.218945 |
| GOBP_ANTIGEN_RECEPTOR_MEDIATED_SIGNALING_PATHWAY | 133 | 0.499314 | 1.597211 | 0.001111 | 0.217205 |
| GOBP_POSITIVE_REGULATION_OF_ALPHA_BETA_T_CELL_PROLIFERATION | 17 | 0.685157 | 1.596582 | 0.020802 | 0.216741 |
| GOBP_REGULATION_OF_COAGULATION | 53 | 0.565281 | 1.596368 | 0.003851 | 0.215335 |
| GOBP_T_CELL_RECEPTOR_SIGNALING_PATHWAY | 103 | 0.506246 | 1.595786 | 0.003421 | 0.21496 |
| GOBP_REGULATION_OF_CELL_MIGRATION_INVOLVED_IN_SPROUTING_ANGIOGENESIS | 33 | 0.60248 | 1.595252 | 0.010738 | 0.214402 |
| GOBP_REGULATION_OF_FATTY_ACID_METABOLIC_PROCESS | 64 | 0.53584 | 1.594468 | 0.004854 | 0.214321 |
| GOBP_DNA_DEALKYLATION | 27 | 0.614364 | 1.592893 | 0.018182 | 0.216163 |
| GOBP_REGULATION_OF_FATTY_ACID_BIOSYNTHETIC_PROCESS | 31 | 0.617556 | 1.590551 | 0.014865 | 0.219959 |
| GOBP_REGULATION_OF_MACROPHAGE_DERIVED_FOAM_CELL_DIFFERENTIATION | 21 | 0.656784 | 1.590543 | 0.017699 | 0.218182 |
| GOBP_POSITIVE_REGULATION_OF_MACROPHAGE_DIFFERENTIATION | 15 | 0.69301 | 1.588807 | 0.017442 | 0.220713 |
| GOBP_ANTIGEN_PROCESSING_AND_PRESENTATION_OF_EXOGENOUS_ANTIGEN | 31 | 0.605304 | 1.588498 | 0.021651 | 0.219563 |
| GOBP_NEUROTRANSMITTER_METABOLIC_PROCESS | 24 | 0.626012 | 1.588193 | 0.012397 | 0.218665 |
| GOBP_REGENERATION | 143 | 0.491415 | 1.587685 | 0 | 0.218081 |
| GOBP_T_HELPER_17_CELL_DIFFERENTIATION | 18 | 0.670692 | 1.586509 | 0.017143 | 0.219278 |
| GOBP_TRIPARTITE_REGIONAL_SUBDIVISION | 16 | 0.686802 | 1.586415 | 0.014663 | 0.21785 |
| GOBP_ADAPTIVE_IMMUNE_RESPONSE | 282 | 0.476413 | 1.585662 | 0 | 0.217853 |
| GOBP_POSITIVE_REGULATION_OF_CD4_POSITIVE_ALPHA_BETA_T_CELL_ACTIVATION | 26 | 0.621483 | 1.58542 | 0.021337 | 0.216735 |
| GOBP_MITOCHONDRIAL_CALCIUM_ION_HOMEOSTASIS | 24 | 0.62119 | 1.58404 | 0.01844 | 0.218374 |
| GOBP_GLYCOSYL_COMPOUND_CATABOLIC_PROCESS | 30 | 0.606225 | 1.58352 | 0.022504 | 0.218057 |
| GOBP_REGULATION_OF_IMMUNE_EFFECTOR_PROCESS | 234 | 0.470435 | 1.582645 | 0 | 0.218271 |
| GOBP_LEUKOCYTE_MEDIATED_CYTOTOXICITY | 75 | 0.53146 | 1.58257 | 0.003521 | 0.216755 |
| GOBP_REGULATION_OF_RESPONSE_TO_WOUNDING | 129 | 0.490832 | 1.580937 | 0 | 0.218876 |
| GOBP_REGULATION_OF_T_HELPER_1_TYPE_IMMUNE_RESPONSE | 15 | 0.688013 | 1.580166 | 0.016272 | 0.218979 |
| GOBP_NEGATIVE_REGULATION_OF_CATION_CHANNEL_ACTIVITY | 33 | 0.596436 | 1.580128 | 0.014706 | 0.217449 |
| GOBP_CHROMOSOME_ORGANIZATION_INVOLVED_IN_MEIOTIC_CELL_CYCLE | 56 | 0.545468 | 1.579996 | 0.00625 | 0.216188 |
| GOBP_ANTIGEN_PROCESSING_AND_PRESENTATION_OF_EXOGENOUS_PEPTIDE_ANTIGEN | 27 | 0.620102 | 1.579629 | 0.017615 | 0.215378 |
| GOBP_NEGATIVE_REGULATION_OF_POTASSIUM_ION_TRANSPORT | 20 | 0.652682 | 1.579575 | 0.022472 | 0.213957 |
| GOBP_POSITIVE_REGULATION_OF_LEUKOCYTE_CHEMOTAXIS | 63 | 0.535069 | 1.578578 | 0.009804 | 0.214653 |
| GOBP_T_CELL_PROLIFERATION | 141 | 0.493907 | 1.577558 | 0.001096 | 0.215227 |
| GOBP_DETECTION_OF_STIMULUS | 152 | 0.487836 | 1.577117 | 0.001093 | 0.21476 |
| GOBP_LEUKOCYTE_PROLIFERATION | 214 | 0.475389 | 1.57605 | 0 | 0.215977 |
| GOBP_REGULATION_OF_EXTRACELLULAR_MATRIX_ORGANIZATION | 44 | 0.558986 | 1.575136 | 0.01269 | 0.216756 |
| GOBP_POSITIVE_REGULATION_OF_INTERLEUKIN_17_PRODUCTION | 15 | 0.693848 | 1.574576 | 0.017442 | 0.216586 |
| GOBP_DEFENSE_RESPONSE_TO_BACTERIUM | 128 | 0.492235 | 1.574116 | 0.001089 | 0.216056 |
| GOBP_AXONAL_TRANSPORT_OF_MITOCHONDRION | 18 | 0.660926 | 1.571364 | 0.01958 | 0.221279 |
| GOBP_SYNAPTIC_TRANSMISSION_CHOLINERGIC | 22 | 0.631595 | 1.571315 | 0.014265 | 0.219858 |
| GOBP_NEGATIVE_REGULATION_OF_B_CELL_ACTIVATION | 27 | 0.615473 | 1.570875 | 0.015493 | 0.219485 |
| GOBP_POSITIVE_REGULATION_OF_INTERLEUKIN_8_PRODUCTION | 42 | 0.56585 | 1.570675 | 0.012469 | 0.218494 |
| GOBP_REGULATION_OF_LEUKOCYTE_MEDIATED_IMMUNITY | 162 | 0.482163 | 1.570424 | 0 | 0.217625 |
| GOBP_MONONUCLEAR_CELL_MIGRATION | 126 | 0.49406 | 1.568881 | 0.001122 | 0.219816 |
| GOBP_PLASMINOGEN_ACTIVATION | 19 | 0.66162 | 1.566624 | 0.02886 | 0.223666 |
| GOBP_POSITIVE_REGULATION_OF_DEFENSE_RESPONSE | 191 | 0.476963 | 1.566191 | 0.002153 | 0.223337 |

**GSEA Negative** Hematopoietic/Immune pathways are highlighted in green.

| Gene sets | SIZE | ES | NES | NOM p-val | FDR q-val |
|---|---|---|---|---|---|
| GOBP_DNA_REPLICATION_DEPENDENT_CHROMATIN_ORGANIZATION | 23 | -0.807189 | -2.380612 | 0 | 0 |
| GOBP_NUCLEOSOME_ASSEMBLY | 93 | -0.5637 | -2.244686 | 0 | 0.001189 |
| GOBP_PROTEIN_DNA_COMPLEX_ASSEMBLY | 162 | -0.487446 | -2.136381 | 0 | 0.017526 |
| GOBP_DNA_REPLICATION_INDEPENDENT_CHROMATIN_ORGANIZATION | 29 | -0.650943 | -2.022136 | 0 | 0.066358 |
| GOBP_NUCLEOSOME_ORGANIZATION | 138 | -0.48578 | -2.020401 | 0 | 0.054721 |
| GOBP_PROTEIN_DNA_COMPLEX_SUBUNIT_ORGANIZATION | 208 | -0.436068 | -1.952347 | 0 | 0.110601 |
| GOBP_CHROMATIN_ASSEMBLY_OR_DISASSEMBLY | 193 | -0.43737 | -1.930282 | 0 | 0.124389 |
| GOBP_N_GLYCAN_PROCESSING | 19 | -0.6777 | -1.92352 | 0.003356 | 0.117608 |
| GOBP_L_GLUTAMATE_TRANSMEMBRANE_TRANSPORT | 24 | -0.641265 | -1.921984 | 0 | 0.106364 |
| GOBP_C4_DICARBOXYLATE_TRANSPORT | 25 | -0.610228 | -1.901026 | 0 | 0.11963 |
| GOBP_TRACHEA_DEVELOPMENT | 15 | -0.692317 | -1.850454 | 0.003125 | 0.188199 |
| GOBP_PROTEIN_LIPID_COMPLEX_ASSEMBLY | 22 | -0.610195 | -1.818824 | 0.003268 | 0.2317 |
| GOBP_DNA_PACKAGING | 207 | -0.405649 | -1.804416 | 0 | 0.246635 |
| GOBP_POSTTRANSLATIONAL_PROTEIN_TARGETING_TO_ENDOPLASMIC_RETI( | 15 | -0.660176 | -1.75154 | 0.003067 | 0.374439 |
| GOBP_TRABECULA_FORMATION | 21 | -0.600965 | -1.749182 | 0.00692 | 0.356713 |
| GOBP_VIRAL_PROTEIN_PROCESSING | 30 | -0.548021 | -1.739428 | 0 | 0.362289 |
| GOBP_OOCYTE_MATURATION | 25 | -0.563651 | -1.719802 | 0.007018 | 0.400503 |
| GOBP_TYPE_B_PANCREATIC_CELL_PROLIFERATION | 19 | -0.608728 | -1.719399 | 0.006734 | 0.379854 |
| GOBP_CHROMATIN_REMODELING | 222 | -0.391314 | -1.710375 | 0 | 0.387374 |
| GOBP_DNA_STRAND_ELONGATION_INVOLVED_IN_DNA_REPLICATION | 15 | -0.659385 | -1.70768 | 0.008876 | 0.376577 |
| GOBP_MATURATION_OF_5_8S_RRNA | 35 | -0.511399 | -1.67357 | 0.019608 | 0.472849 |
| GOBP_C21_STEROID_HORMONE_BIOSYNTHETIC_PROCESS | 16 | -0.602751 | -1.659317 | 0.028481 | 0.505563 |
| GOBP_CERAMIDE_TRANSPORT | 16 | -0.596813 | -1.64888 | 0.02454 | 0.520129 |
| GOBP_TRANSCRIPTION_BY_RNA_POLYMERASE_I | 53 | -0.466661 | -1.642886 | 0.005495 | 0.521343 |
| GOBP_MATURATION_OF_5_8S_RRNA_FROM_TRICISTRONIC_RRNA_TRANSCR | 24 | -0.558119 | -1.642072 | 0.018382 | 0.50383 |
| GOBP_PROTEIN_LOCALIZATION_TO_MICROTUBULE | 17 | -0.607253 | -1.63646 | 0.032468 | 0.505266 |
| GOBP_DNA_UNWINDING_INVOLVED_IN_DNA_REPLICATION | 22 | -0.557134 | -1.630556 | 0.030303 | 0.509641 |
| GOBP_NADPH_REGENERATION | 17 | -0.59016 | -1.626654 | 0.03413 | 0.505972 |
| GOBP_DNA_CONFORMATION_CHANGE | 303 | -0.357022 | -1.621367 | 0 | 0.50872 |
| GOBP_POSITIVE_REGULATION_OF_DENDRITIC_SPINE_MORPHOGENESIS | 19 | -0.550425 | -1.619096 | 0.013289 | 0.499452 |
| GOBP_MATURATION_OF_SSU_RRNA_FROM_TRICISTRONIC_RRNA_TRANSCRI | 37 | -0.48049 | -1.604758 | 0 | 0.536942 |
| GOBP_MATURATION_OF_SSU_RRNA | 50 | -0.457381 | -1.603933 | 0.015306 | 0.522777 |
| GOBP_WALKING_BEHAVIOR | 28 | -0.516194 | -1.596717 | 0.017794 | 0.532887 |
| GOBP_PYRIMIDINE_NUCLEOSIDE_TRIPHOSPHATE_METABOLIC_PROCESS | 24 | -0.527167 | -1.595521 | 0.02649 | 0.521119 |
| GOBP_ACIDIC_AMINO_ACID_TRANSPORT | 54 | -0.429702 | -1.58639 | 0 | 0.539674 |
| GOBP_DNA_REPLICATION_INITIATION | 35 | -0.483781 | -1.586152 | 0.023622 | 0.525693 |
| GOBP_TELOMERE_ORGANIZATION | 149 | -0.380233 | -1.578852 | 0 | 0.538705 |
| GOBP_TRABECULA_MORPHOGENESIS | 41 | -0.466541 | -1.578031 | 0.022026 | 0.52722 |
| GOBP_OLEFINIC_COMPOUND_BIOSYNTHETIC_PROCESS | 17 | -0.578392 | -1.575502 | 0.02454 | 0.522557 |
| GOBP_PROTEIN_INSERTION_INTO_ER_MEMBRANE | 22 | -0.532189 | -1.573402 | 0.034247 | 0.517273 |
| GOBP_OLFACTORY_LOBE_DEVELOPMENT | 27 | -0.505882 | -1.572 | 0.010204 | 0.509517 |
| GOBP_CLEAVAGE_INVOLVED_IN_RRNA_PROCESSING | 27 | -0.511899 | -1.568437 | 0.011111 | 0.509667 |
| GOBP_GLUCOSE_6_PHOSPHATE_METABOLIC_PROCESS | 20 | -0.544122 | -1.565051 | 0.038961 | 0.509303 |
| GOBP_NEGATIVE_REGULATION_OF_CARTILAGE_DEVELOPMENT | 23 | -0.526052 | -1.551044 | 0.029316 | 0.545931 |
| GOBP_REGULATION_OF_TRANSCRIPTION_BY_RNA_POLYMERASE_I | 33 | -0.478686 | -1.539196 | 0.021097 | 0.576275 |
| GOBP_MAINTENANCE_OF_SYNAPSE_STRUCTURE | 19 | -0.545472 | -1.530807 | 0.062284 | 0.595601 |
| GOBP_MATURATION_OF_LSU_RRNA_FROM_TRICISTRONIC_RRNA_TRANSCRI | 15 | -0.573829 | -1.529432 | 0.04717 | 0.587968 |
| GOBP_INTRACILIARY_TRANSPORT | 39 | -0.45344 | -1.527299 | 0.012245 | 0.584481 |
| GOBP_HEART_TRABECULA_MORPHOGENESIS | 28 | -0.489317 | -1.522149 | 0.029412 | 0.591909 |
| GOBP_RNA_SPLICING_VIA_TRANSESTERIFICATION_REACTIONS | 284 | -0.332182 | -1.512005 | 0 | 0.621313 |
| GOBP_STEROID_HORMONE_BIOSYNTHETIC_PROCESS | 27 | -0.493709 | -1.501809 | 0.045283 | 0.649511 |
| GOBP_DICARBOXYLIC_ACID_TRANSPORT | 72 | -0.395386 | -1.490299 | 0.005848 | 0.685194 |
| GOBP_COCHLEA_DEVELOPMENT | 40 | -0.433737 | -1.483746 | 0.033473 | 0.70039 |
| GOBP_BRAIN_MORPHOGENESIS | 32 | -0.444328 | -1.483247 | 0.040323 | 0.689471 |
| GOBP_POSITIVE_REGULATION_OF_DOUBLE_STRAND_BREAK_REPAIR_VIA_N( | 16 | -0.537187 | -1.483101 | 0.039877 | 0.677545 |
| GOBP_RRNA_TRANSCRIPTION | 34 | -0.465135 | -1.477996 | 0.054688 | 0.686829 |
| GOBP_POSITIVE_REGULATION_OF_TORC1_SIGNALING | 20 | -0.517448 | -1.477865 | 0.057229 | 0.675407 |
| GOBP_RETINAL_GANGLION_CELL_AXON_GUIDANCE | 18 | -0.521319 | -1.472415 | 0.072327 | 0.686425 |
| GOBP_NEGATIVE_REGULATION_OF_RNA_SPLICING | 25 | -0.493525 | -1.465754 | 0.045455 | 0.702483 |
| GOBP_DNA_TEMPLATED_TRANSCRIPTION_INITIATION | 118 | -0.357703 | -1.457838 | 0 | 0.72404 |
| GOBP_POSITIVE_REGULATION_OF_SYNAPSE_ASSEMBLY | 52 | -0.418439 | -1.456906 | 0.034483 | 0.716016 |
| GOBP_GTP_METABOLIC_PROCESS | 23 | -0.494294 | -1.453072 | 0.048701 | 0.719983 |
| GOBP_PYRIMIDINE_NUCLEOSIDE_TRIPHOSPHATE_BIOSYNTHETIC_PROCESS | 19 | -0.504515 | -1.445482 | 0.092527 | 0.741798 |
| GOBP_RIBOSOMAL_SMALL_SUBUNIT_BIOGENESIS | 73 | -0.385802 | -1.438086 | 0.006944 | 0.762617 |
| GOBP_TAIL_ANCHORED_MEMBRANE_PROTEIN_INSERTION_INTO_ER_MEMBR | 17 | -0.536457 | -1.435433 | 0.087948 | 0.762258 |
| GOBP_HINDBRAIN_DEVELOPMENT | 126 | -0.339486 | -1.434041 | 0.021053 | 0.756851 |
| GOBP_POSITIVE_REGULATION_OF_NEURON_DIFFERENTIATION | 70 | -0.382751 | -1.433032 | 0.035503 | 0.749527 |
| GOBP_L_AMINO_ACID_TRANSPORT | 56 | -0.390677 | -1.43292 | 0.022099 | 0.739038 |
| GOBP_CHAPERONE_COFACTOR_DEPENDENT_PROTEIN_REFOLDING | 32 | -0.434012 | -1.432158 | 0.044355 | 0.73159 |
| GOBP_NEGATIVE_REGULATION_OF_DNA_TEMPLATED_TRANSCRIPTION_ELON | 16 | -0.509461 | -1.425634 | 0.095808 | 0.748785 |
| GOBP_CELL_CYCLE_DNA_REPLICATION | 41 | -0.435767 | -1.424866 | 0.040486 | 0.741535 |
| GOBP_EMBRYONIC_PLACENTA_MORPHOGENESIS | 22 | -0.4958 | -1.424776 | 0.075812 | 0.731562 |
| GOBP_REGULATION_OF_MRNA_SPLICING_VIA_SPLICEOSOME | 98 | -0.363522 | -1.424252 | 0 | 0.724088 |
| GOBP_BASE_EXCISION_REPAIR | 43 | -0.411696 | -1.422182 | 0.04717 | 0.722864 |
| GOBP_NEGATIVE_REGULATION_OF_MRNA_SPLICING_VIA_SPLICEOSOME | 20 | -0.503246 | -1.41852 | 0.069686 | 0.727964 |
| GOBP_AMINO_ACID_TRANSMEMBRANE_TRANSPORT | 78 | -0.373487 | -1.411539 | 0.034722 | 0.746749 |
| GOBP_EPITHELIAL_TUBE_BRANCHING_INVOLVED_IN_LUNG_MORPHOGENESI | 20 | -0.489493 | -1.406356 | 0.093333 | 0.75845 |
| GOBP_SPHINGOMYELIN_METABOLIC_PROCESS | 16 | -0.513739 | -1.406265 | 0.089337 | 0.749129 |
| GOBP_POSITIVE_REGULATION_OF_TRANSCRIPTION_BY_RNA_POLYMERASE_ | 23 | -0.470072 | -1.405895 | 0.09589 | 0.741032 |
| GOBP_PROTEIN_LOCALIZATION_TO_CILIUM | 69 | -0.365928 | -1.404897 | 0.012903 | 0.736162 |
| GOBP_DNA_TEMPLATED_TRANSCRIPTION_ELONGATION | 73 | -0.360968 | -1.39928 | 0.023669 | 0.750933 |
| GOBP_NEGATIVE_REGULATION_OF_MRNA_PROCESSING | 29 | -0.448522 | -1.39639 | 0.076364 | 0.754172 |
| GOBP_L_GLUTAMATE_IMPORT_ACROSS_PLASMA_MEMBRANE | 15 | -0.525763 | -1.396075 | 0.103858 | 0.746417 |
| GOBP_CYTOPLASMIC_SEQUESTERING_OF_PROTEIN | 21 | -0.47271 | -1.394224 | 0.083916 | 0.744949 |
| GOBP_VIRAL_RELEASE_FROM_HOST_CELL | 28 | -0.439605 | -1.388806 | 0.061818 | 0.75884 |
| GOBP_PYRIMIDINE_NUCLEOTIDE_CATABOLIC_PROCESS | 18 | -0.514714 | -1.38827 | 0.090909 | 0.752559 |
| GOBP_REGULATION_OF_MRNA_PROCESSING | 134 | -0.333396 | -1.379014 | 0 | 0.783974 |
| GOBP_POSITIVE_REGULATION_OF_TELOMERE_CAPPING | 17 | -0.506105 | -1.377598 | 0.126183 | 0.780979 |
| GOBP_PYRIMIDINE_DEOXYRIBONUCLEOTIDE_METABOLIC_PROCESS | 22 | -0.469931 | -1.377301 | 0.062016 | 0.773261 |
| GOBP_NEURON_FATE_COMMITMENT | 37 | -0.411922 | -1.376726 | 0.072961 | 0.767156 |
| GOBP_ENDONUCLEOLYTIC_CLEAVAGE_INVOLVED_IN_RRNA_PROCESSING | 15 | -0.529242 | -1.37293 | 0.117647 | 0.774861 |
| GOBP_POSITIVE_REGULATION_OF_MRNA_SPLICING_VIA_SPLICEOSOME | 21 | -0.473564 | -1.36727 | 0.111486 | 0.790434 |
| GOBP_CELL_DIFFERENTIATION_IN_SPINAL_CORD | 35 | -0.418208 | -1.364395 | 0.073643 | 0.794073 |
| GOBP_RESPONSE_TO_COCAINE | 42 | -0.397544 | -1.361571 | 0.080508 | 0.797325 |
| GOBP_COLUMNAR_CUBOIDAL_EPITHELIAL_CELL_DEVELOPMENT | 35 | -0.419942 | -1.357663 | 0.102273 | 0.806916 |
| GOBP_RETINA_MORPHOGENESIS_IN_CAMERA_TYPE_EYE | 43 | -0.395577 | -1.353838 | 0.089552 | 0.815873 |
| GOBP_NEGATIVE_REGULATION_OF_CHONDROCYTE_DIFFERENTIATION | 18 | -0.474855 | -1.353787 | 0.139241 | 0.807669 |
| GOBP_PEPTIDYL_ASPARAGINE_MODIFICATION | 23 | -0.449467 | -1.353182 | 0.086667 | 0.801983 |
| GOBP_SNO_S_RNA_METABOLIC_PROCESS | 16 | -0.493127 | -1.350392 | 0.124638 | 0.805212 |
| GOBP_POSITIVE_REGULATION_OF_DENDRITE_MORPHOGENESIS | 34 | -0.430147 | -1.3476 | 0.068548 | 0.808467 |
| GOBP_REGULATION_OF_CAMP_DEPENDENT_PROTEIN_KINASE_ACTIVITY | 18 | -0.488792 | -1.346134 | 0.144231 | 0.80676 |
| GOBP_RRNA_METABOLIC_PROCESS | 258 | -0.291335 | -1.346003 | 0 | 0.799432 |
| GOBP_PYRIMIDINE_RIBONUCLEOSIDE_TRIPHOSPHATE_METABOLIC_PROCES | 18 | -0.47924 | -1.343286 | 0.10625 | 0.803243 |
| GOBP_RESPONSE_TO_PROSTAGLANDIN_E | 15 | -0.505758 | -1.341493 | 0.146032 | 0.802994 |
| GOBP_RESPONSE_TO_SALT | 17 | -0.489366 | -1.340731 | 0.115502 | 0.798242 |
| GOBP_POSITIVE_REGULATION_OF_SMOOTHENED_SIGNALING_PATHWAY | 29 | -0.426621 | -1.340665 | 0.100346 | 0.79102 |
| GOBP_POSITIVE_REGULATION_OF_MRNA_PROCESSING | 30 | -0.41416 | -1.340054 | 0.09215 | 0.786243 |
| GOBP_REGULATION_OF_PROTEIN_SUMOYLATION | 21 | -0.467113 | -1.337243 | 0.130282 | 0.790697 |
| GOBP_NUCLEAR_MEMBRANE_REASSEMBLY | 23 | -0.449109 | -1.33511 | 0.107527 | 0.79184 |
| GOBP_ALTERNATIVE_MRNA_SPLICING_VIA_SPLICEOSOME | 72 | -0.35413 | -1.334603 | 0.045198 | 0.786966 |
| GOBP_SULFUR_COMPOUND_TRANSPORT | 43 | -0.388109 | -1.33023 | 0.073529 | 0.798033 |
| GOBP_REGULATION_OF_TELOMERASE_ACTIVITY | 49 | -0.368359 | -1.328949 | 0.07 | 0.796237 |
| GOBP_KETONE_BIOSYNTHETIC_PROCESS | 36 | -0.40166 | -1.328865 | 0.094262 | 0.789591 |
| GOBP_ENTEROENDOCRINE_CELL_DIFFERENTIATION | 21 | -0.45595 | -1.328211 | 0.112245 | 0.785368 |
| GOBP_MODIFIED_AMINO_ACID_TRANSPORT | 25 | -0.436256 | -1.328025 | 0.12782 | 0.779168 |
| GOBP_ANAPHASE_PROMOTING_COMPLEX_DEPENDENT_CATABOLIC_PROCE | 16 | -0.486002 | -1.327886 | 0.137195 | 0.773013 |
| GOBP_MATURATION_OF_LSU_RRNA | 28 | -0.41732 | -1.326347 | 0.103806 | 0.773041 |
| GOBP_POSITIVE_REGULATION_OF_TELOMERASE_ACTIVITY | 34 | -0.396453 | -1.324465 | 0.077922 | 0.774249 |
| GOBP_REGULATION_OF_PRESYNAPSE_ORGANIZATION | 30 | -0.402163 | -1.318456 | 0.087121 | 0.792406 |
| GOBP_DE_NOVO_PROTEIN_FOLDING | 41 | -0.389763 | -1.317528 | 0.060606 | 0.789461 |
| GOBP_CHAPERONE_MEDIATED_PROTEIN_FOLDING | 67 | -0.344167 | -1.315338 | 0.078788 | 0.79172 |
| GOBP_POSITIVE_REGULATION_OF_TELOMERE_MAINTENANCE_VIA_TELOMER | 37 | -0.393053 | -1.310566 | 0.096 | 0.804915 |
| GOBP_G1_TO_G0_TRANSITION | 17 | -0.479601 | -1.307615 | 0.128205 | 0.810323 |
| GOBP_RNA_SPLICING | 391 | -0.280504 | -1.307375 | 0 | 0.804869 |
| GOBP_METENCEPHALON_DEVELOPMENT | 95 | -0.334362 | -1.303679 | 0.029197 | 0.813762 |
| GOBP_POSITIVE_REGULATION_OF_HEART_RATE | 22 | -0.439947 | -1.297575 | 0.156463 | 0.833206 |
| GOBP_NUCLEOLUS_ORGANIZATION | 16 | -0.473459 | -1.297491 | 0.143302 | 0.827068 |
| GOBP_REGULATION_OF_HORMONE_BIOSYNTHETIC_PROCESS | 20 | -0.448767 | -1.297386 | 0.151515 | 0.820989 |
| GOBP_CELLULAR_RESPONSE_TO_PROSTAGLANDIN_STIMULUS | 16 | -0.471104 | -1.296158 | 0.167722 | 0.820029 |
| GOBP_TELOMERE_CAPPING | 39 | -0.385462 | -1.294491 | 0.123967 | 0.820576 |
| GOBP_ADHERENS_JUNCTION_ORGANIZATION | 46 | -0.375516 | -1.290534 | 0.070755 | 0.831241 |
| GOBP_RIBOSOME_BIOGENESIS | 297 | -0.283025 | -1.287675 | 0 | 0.837169 |
| GOBP_NUCLEAR_TRANSCRIBED_MRNA_CATABOLIC_PROCESS_EXONUCLEO | 19 | -0.444199 | -1.286884 | 0.154799 | 0.833888 |
| GOBP_CALCIUM_DEPENDENT_CELL_CELL_ADHESION_VIA_PLASMA_MEMBRA | 33 | -0.402282 | -1.285389 | 0.129707 | 0.834125 |
| GOBP_PYRIMIDINE_NUCLEOTIDE_METABOLIC_PROCESS | 48 | -0.374772 | -1.284956 | 0.107692 | 0.829638 |
| GOBP_RNA_SURVEILLANCE | 15 | -0.477978 | -1.282963 | 0.153846 | 0.832352 |
| GOBP_NEGATIVE_REGULATION_OF_NUCLEOCYTOPLASMIC_TRANSPORT | 20 | -0.449051 | -1.282176 | 0.144876 | 0.829598 |
| GOBP_STEROL_HOMEOSTASIS | 61 | -0.345667 | -1.278961 | 0.086207 | 0.837072 |
| GOBP_RIBONUCLEOPROTEIN_COMPLEX_BIOGENESIS | 420 | -0.275042 | -1.276557 | 0 | 0.841062 |
| GOBP_LIPOPROTEIN_BIOSYNTHETIC_PROCESS | 94 | -0.313916 | -1.276051 | 0.051282 | 0.837002 |
| GOBP_PYRIMIDINE_RIBONUCLEOSIDE_TRIPHOSPHATE_BIOSYNTHETIC_PROC | 15 | -0.489944 | -1.274617 | 0.162791 | 0.836796 |
| GOBP_HISTONE_DEUBIQUITINATION | 40 | -0.367232 | -1.271281 | 0.089686 | 0.845239 |
| GOBP_GLIAL_CELL_APOPTOTIC_PROCESS | 15 | -0.464934 | -1.269662 | 0.188623 | 0.846143 |
| GOBP_REGULATION_OF_KIDNEY_DEVELOPMENT | 23 | -0.432429 | -1.269145 | 0.153846 | 0.842443 |
| GOBP_POSITIVE_REGULATION_OF_RNA_SPLICING | 35 | -0.395446 | -1.268537 | 0.129032 | 0.839247 |
| GOBP_POSITIVE_REGULATION_OF_DENDRITIC_SPINE_DEVELOPMENT | 35 | -0.380061 | -1.267322 | 0.144681 | 0.838647 |
| GOBP_DENDRITIC_SPINE_MORPHOGENESIS | 56 | -0.341689 | -1.265315 | 0.131707 | 0.841318 |
| GOBP_SOMITE_DEVELOPMENT | 70 | -0.334126 | -1.264798 | 0.071429 | 0.837769 |
| GOBP_NUCLEOSIDE_TRIPHOSPHATE_METABOLIC_PROCESS | 101 | -0.313318 | -1.26328 | 0.052632 | 0.838864 |
| GOBP_POSITIVE_REGULATION_OF_GLIAL_CELL_DIFFERENTIATION | 36 | -0.379753 | -1.262809 | 0.109804 | 0.835315 |
| GOBP_VIRAL_GENE_EXPRESSION | 95 | -0.320035 | -1.258844 | 0.0625 | 0.845985 |
| GOBP_ORGANIC_ACID_TRANSMEMBRANE_TRANSPORT | 122 | -0.31175 | -1.258169 | 0.03 | 0.843099 |
| GOBP_NEGATIVE_REGULATION_OF_MYELOID_CELL_DIFFERENTIATION | 71 | -0.325044 | -1.258043 | 0.104046 | 0.83816 |

| | | | | | |
|---|---|---|---|---|---|
| GOBP_POSITIVE_REGULATION_OF_CHEMOTAXIS | 103 | 0.501808 | 1.565289 | 0.002283 | 0.224068 |
| GOBP_REGULATION_OF_CELL_KILLING | 55 | 0.542492 | 1.564644 | 0.009913 | 0.224215 |
| GOBP_NEGATIVE_REGULATION_OF_ALPHA_BETA_T_CELL_ACTIVATION | 30 | 0.603707 | 1.561291 | 0.026455 | 0.230973 |
| GOBP_REGULATION_OF_INFLAMMATORY_RESPONSE | 246 | 0.461747 | 1.559879 | 0 | 0.232922 |
| GOBP_NEGATIVE_REGULATION_OF_LYMPHOCYTE_ACTIVATION | 108 | 0.491409 | 1.559259 | 0.002328 | 0.232996 |
| GOBP_MYELOID_CELL_APOPTOTIC_PROCESS | 26 | 0.60501 | 1.55845 | 0.030899 | 0.233421 |
| GOBP_ALPHA_BETA_T_CELL_ACTIVATION | 108 | 0.503059 | 1.557398 | 0.004425 | 0.234211 |
| GOBP_NEGATIVE_REGULATION_OF_IMMUNE_EFFECTOR_PROCESS | 71 | 0.517506 | 1.55573 | 0.009547 | 0.236995 |
| GOBP_SMOOTH_MUSCLE_CELL_PROLIFERATION | 116 | 0.489075 | 1.553409 | 0.003363 | 0.241331 |
| GOBP_ADAPTIVE_IMMUNE_RESPONSE_BASED_ON_SOMATIC_RECOMBINATION_OF_IMMUN | 204 | 0.47056 | 1.552604 | 0 | 0.241934 |
| GOBP_NEGATIVE_REGULATION_OF_T_CELL_RECEPTOR_SIGNALING_PATHWAY | 18 | 0.646967 | 1.552267 | 0.015291 | 0.241306 |
| GOBP_T_CELL_ACTIVATION_INVOLVED_IN_IMMUNE_RESPONSE | 67 | 0.524325 | 1.551814 | 0.009744 | 0.240846 |
| GOBP_INFLAMMATORY_CELL_APOPTOTIC_PROCESS | 18 | 0.655881 | 1.550934 | 0.028653 | 0.241603 |
| GOBP_REGULATION_OF_ALPHA_BETA_T_CELL_ACTIVATION | 72 | 0.512583 | 1.55081 | 0.009581 | 0.240517 |
| GOBP_REGULATION_OF_T_CELL_ACTIVATION | 234 | 0.460003 | 1.550694 | 0 | 0.239354 |
| GOBP_RESPONSE_TO_GROWTH_HORMONE | 28 | 0.589611 | 1.55065 | 0.021828 | 0.237988 |
| GOBP_RESPONSE_TO_HYDROPEROXIDE | 15 | 0.668973 | 1.549606 | 0.041481 | 0.239195 |
| GOBP_REGULATION_OF_CELL_ACTIVATION | 424 | 0.450972 | 1.547446 | 0 | 0.243159 |
| GOBP_SKELETAL_MUSCLE_ADAPTATION | 16 | 0.678808 | 1.546931 | 0.02071 | 0.242937 |
| GOBP_NEGATIVE_REGULATION_OF_ION_TRANSMEMBRANE_TRANSPORT | 69 | 0.519582 | 1.545974 | 0.004825 | 0.243983 |
| GOBP_TYPE_2_IMMUNE_RESPONSE | 25 | 0.617463 | 1.545917 | 0.027739 | 0.24268 |
| GOBP_INTERMEDIATE_FILAMENT_BASED_PROCESS | 38 | 0.571328 | 1.545607 | 0.007782 | 0.241955 |
| GOBP_REGULATION_OF_ANION_TRANSMEMBRANE_TRANSPORT | 20 | 0.63805 | 1.544686 | 0.03592 | 0.242882 |
| GOBP_RESPONSE_TO_CHEMOKINE | 45 | 0.549971 | 1.544511 | 0.016539 | 0.241974 |
| GOBP_POSITIVE_REGULATION_OF_CD4_POSITIVE_ALPHA_BETA_T_CELL_DIFFERENTIATIO | 21 | 0.634055 | 1.543441 | 0.023564 | 0.243271 |
| GOBP_REGULATION_OF_COLLAGEN_METABOLIC_PROCESS | 31 | 0.589656 | 1.5429 | 0.02027 | 0.243297 |
| GOBP_MEIOSIS_I_CELL_CYCLE_PROCESS | 100 | 0.497058 | 1.540628 | 0.005574 | 0.24722 |
| GOBP_COLLAGEN_METABOLIC_PROCESS | 73 | 0.51352 | 1.54032 | 0.012034 | 0.246574 |
| GOBP_REGULATION_OF_HAIR_CYCLE | 23 | 0.603263 | 1.540184 | 0.028694 | 0.245535 |
| GOBP_REGULATION_OF_T_CELL_RECEPTOR_SIGNALING_PATHWAY | 33 | 0.577066 | 1.539944 | 0.022546 | 0.24481 |
| GOBP_NEGATIVE_REGULATION_OF_PEPTIDASE_ACTIVITY | 157 | 0.470801 | 1.539518 | 0.002198 | 0.244541 |
| GOBP_HYDROGEN_PEROXIDE_CATABOLIC_PROCESS | 15 | 0.679748 | 1.539055 | 0.03096 | 0.244389 |
| GOBP_POSITIVE_REGULATION_OF_ALPHA_BETA_T_CELL_ACTIVATION | 48 | 0.538713 | 1.537725 | 0.02439 | 0.246229 |
| GOBP_RIBONUCLEOSIDE_METABOLIC_PROCESS | 29 | 0.592441 | 1.536727 | 0.018006 | 0.247408 |
| GOBP_VIRAL_RNA_GENOME_REPLICATION | 32 | 0.573971 | 1.535003 | 0.01455 | 0.250482 |
| GOBP_INTERFERON_GAMMA_PRODUCTION | 66 | 0.524974 | 1.534856 | 0.006143 | 0.249491 |
| GOBP_REGULATION_OF_LYMPHOCYTE_ACTIVATION | 330 | 0.452891 | 1.534173 | 0 | 0.249824 |
| GOBP_LYMPHOCYTE_MEDIATED_IMMUNITY | 186 | 0.465528 | 1.532602 | 0.002153 | 0.252408 |
| GOBP_FOAM_CELL_DIFFERENTIATION | 27 | 0.606508 | 1.53216 | 0.023004 | 0.252139 |
| GOBP_REGULATION_OF_LEUKOCYTE_CHEMOTAXIS | 82 | 0.501564 | 1.531924 | 0.010702 | 0.251344 |
| GOBP_T_CELL_CYTOKINE_PRODUCTION | 26 | 0.595997 | 1.531562 | 0.027894 | 0.250843 |
| GOBP_POSITIVE_REGULATION_OF_HEMOPOIESIS | 117 | 0.481503 | 1.531227 | 0.003352 | 0.250338 |
| GOBP_T_CELL_ACTIVATION | 348 | 0.449843 | 1.530798 | 0 | 0.250124 |
| GOBP_REGULATION_OF_MACROPHAGE_DIFFERENTIATION | 20 | 0.643496 | 1.529904 | 0.033237 | 0.250949 |
| GOBP_T_HELPER_17_TYPE_IMMUNE_RESPONSE | 29 | 0.59048 | 1.52666 | 0.02449 | 0.257706 |
| GOBP_SENSORY_PERCEPTION | 384 | 0.449196 | 1.526505 | 0 | 0.256772 |
| GOBP_RESPONSE_TO_LITHIUM_ION | 17 | 0.661662 | 1.526134 | 0.030166 | 0.256411 |
| GOBP_CELLULAR_RESPONSE_TO_INTERFERON_GAMMA | 76 | 0.502968 | 1.525903 | 0.008363 | 0.255731 |
| GOBP_REGULATION_OF_CELL_CELL_ADHESION | 324 | 0.450725 | 1.525006 | 0 | 0.256603 |
| GOBP_RESPONSE_TO_AMPHETAMINE | 28 | 0.581024 | 1.524159 | 0.029024 | 0.257551 |
| GOBP_REGULATION_OF_VASCULATURE_DEVELOPMENT | 234 | 0.453361 | 1.524015 | 0 | 0.25664 |
| GOBP_REGULATION_OF_B_CELL_ACTIVATION | 99 | 0.484409 | 1.523675 | 0.004499 | 0.256205 |
| GOBP_POSITIVE_REGULATION_OF_RESPONSE_TO_WOUNDING | 55 | 0.534568 | 1.521116 | 0.01107 | 0.261748 |
| GOBP_BONE_CELL_DEVELOPMENT | 30 | 0.578966 | 1.520539 | 0.035135 | 0.261949 |
| GOBP_EMBRYONIC_PATTERN_SPECIFICATION | 57 | 0.53417 | 1.519776 | 0.016129 | 0.262756 |
| GOBP_POSITIVE_REGULATION_OF_TRANSFORMING_GROWTH_FACTOR_BETA_PRODUCTIC | 17 | 0.655866 | 1.519077 | 0.039648 | 0.263353 |
| GOBP_REGULATION_OF_GRANULOCYTE_CHEMOTAXIS | 36 | 0.560681 | 1.5183 | 0.031373 | 0.264004 |
| GOBP_BLASTODERM_SEGMENTATION | 20 | 0.63557 | 1.517701 | 0.032258 | 0.264201 |
| GOBP_B_CELL_MEDIATED_IMMUNITY | 97 | 0.48958 | 1.517543 | 0.009112 | 0.263362 |
| GOBP_VASCULAR_ENDOTHELIAL_GROWTH_FACTOR_RECEPTOR_SIGNALING_PATHWAY | 47 | 0.53691 | 1.516881 | 0.026958 | 0.263815 |
| GOBP_LYMPH_VESSEL_DEVELOPMENT | 29 | 0.576652 | 1.516338 | 0.024032 | 0.263938 |
| GOBP_POSITIVE_REGULATION_OF_CYTOKINE_PRODUCTION_INVOLVED_IN_INFLAMMATOR | 15 | 0.661826 | 1.516278 | 0.035008 | 0.262822 |
| GOBP_CALCIUM_ION_IMPORT_INTO_CYTOSOL | 20 | 0.633785 | 1.516129 | 0.044715 | 0.261986 |
| GOBP_CELL_KILLING | 95 | 0.485996 | 1.515999 | 0.004695 | 0.261096 |
| GOBP_NEGATIVE_REGULATION_OF_IMMUNE_RESPONSE | 126 | 0.473329 | 1.51485 | 0.005447 | 0.262924 |
| GOBP_POLYOL_BIOSYNTHETIC_PROCESS | 53 | 0.524669 | 1.513534 | 0.020434 | 0.264898 |
| GOBP_TRIGLYCERIDE_BIOSYNTHETIC_PROCESS | 34 | 0.568287 | 1.510601 | 0.02656 | 0.27071 |
| GOBP_INTERLEUKIN_6_PRODUCTION | 105 | 0.477984 | 1.51044 | 0.002301 | 0.269857 |
| GOBP_POSITIVE_REGULATION_OF_PEPTIDYL_SERINE_PHOSPHORYLATION | 83 | 0.490654 | 1.508957 | 0.019296 | 0.272504 |
| GOBP_NEGATIVE_REGULATION_OF_POTASSIUM_ION_TRANSMEMBRANE_TRANSPORT | 16 | 0.654806 | 1.508629 | 0.033973 | 0.27202 |
| GOBP_POSITIVE_REGULATION_OF_IMMUNE_EFFECTOR_PROCESS | 165 | 0.463514 | 1.508595 | 0.002148 | 0.270934 |
| GOBP_POSITIVE_REGULATION_OF_REGULATORY_T_CELL_DIFFERENTIATION | 16 | 0.667282 | 1.508508 | 0.040724 | 0.269964 |
| GOBP_RESPONSE_TO_BACTERIUM | 378 | 0.446343 | 1.508074 | 0 | 0.269883 |
| GOBP_NEGATIVE_REGULATION_OF_CELL_ACTIVATION | 143 | 0.470498 | 1.507158 | 0.004391 | 0.270899 |
| GOBP_POSITIVE_REGULATION_OF_SMOOTH_MUSCLE_CELL_PROLIFERATION | 67 | 0.504174 | 1.505718 | 0.014581 | 0.273489 |
| GOBP_STARTLE_RESPONSE | 23 | 0.587782 | 1.505247 | 0.028409 | 0.273528 |
| GOBP_REGULATION_OF_CALCIUM_ION_TRANSMEMBRANE_TRANSPORT | 119 | 0.476365 | 1.505095 | 0.003308 | 0.272664 |
| GOBP_EXTERNAL_ENCAPSULATING_STRUCTURE_ORGANIZATION | 250 | 0.452499 | 1.504573 | 0.001046 | 0.272851 |
| GOBP_MAMMARY_GLAND_MORPHOGENESIS | 33 | 0.565311 | 1.501508 | 0.02038 | 0.279494 |
| GOBP_NEGATIVE_REGULATION_OF_EXOCYTOSIS | 27 | 0.593514 | 1.501401 | 0.030812 | 0.278528 |
| GOBP_HUMORAL_IMMUNE_RESPONSE_MEDIATED_BY_CIRCULATING_IMMUNOGLOBULIN | 26 | 0.579156 | 1.498756 | 0.03125 | 0.284258 |
| GOBP_HEMOSTASIS | 171 | 0.459973 | 1.498387 | 0.002128 | 0.283936 |
| GOBP_POSITIVE_REGULATION_OF_ACUTE_INFLAMMATORY_RESPONSE | 16 | 0.64375 | 1.497442 | 0.05772 | 0.285048 |
| GOBP_NEGATIVE_REGULATION_OF_ANTIGEN_RECEPTOR_MEDIATED_SIGNALING_PATHWA | 23 | 0.600433 | 1.496012 | 0.034578 | 0.287721 |
| GOBP_ZYMOGEN_ACTIVATION | 48 | 0.520575 | 1.495812 | 0.0246 | 0.287011 |
| GOBP_POSITIVE_REGULATION_OF_EMBRYONIC_DEVELOPMENT | 20 | 0.60883 | 1.495626 | 0.025937 | 0.286279 |
| GOBP_CARDIAC_CELL_DEVELOPMENT | 63 | 0.513514 | 1.495533 | 0.012469 | 0.285357 |
| GOBP_REGULATION_OF_HAIR_FOLLICLE_DEVELOPMENT | 15 | 0.650675 | 1.495266 | 0.051095 | 0.284931 |
| GOBP_PHENOL_CONTAINING_COMPOUND_BIOSYNTHETIC_PROCESS | 32 | 0.569452 | 1.494745 | 0.030303 | 0.285188 |
| GOBP_NEGATIVE_REGULATION_OF_LEUKOCYTE_CELL_CELL_ADHESION | 89 | 0.490228 | 1.494317 | 0.012717 | 0.28505 |
| GOBP_REGULATION_OF_ANTIGEN_RECEPTOR_MEDIATED_SIGNALING_PATHWAY | 46 | 0.540659 | 1.494164 | 0.026752 | 0.284243 |
| GOBP_AROMATIC_AMINO_ACID_FAMILY_METABOLIC_PROCESS | 18 | 0.626883 | 1.494142 | 0.039074 | 0.283126 |
| GOBP_CALCIUM_ION_TRANSMEMBRANE_TRANSPORT | 249 | 0.44408 | 1.493712 | 0 | 0.283111 |
| GOBP_POSITIVE_REGULATION_OF_BEHAVIOR | 16 | 0.640181 | 1.492906 | 0.051051 | 0.284159 |
| GOBP_ARTERY_MORPHOGENESIS | 59 | 0.51709 | 1.492637 | 0.018657 | 0.283619 |
| GOBP_KERATINOCYTE_DIFFERENTIATION | 79 | 0.494873 | 1.492251 | 0.009445 | 0.28343 |
| GOBP_REPLICATIVE_SENESCENCE | 15 | 0.646482 | 1.492072 | 0.039764 | 0.282672 |
| GOBP_POSITIVE_REGULATION_OF_RESPONSE_TO_EXTERNAL_STIMULUS | 297 | 0.43675 | 1.491693 | 0.001025 | 0.282595 |
| GOBP_POSITIVE_REGULATION_OF_CELL_ACTIVATION | 249 | 0.446346 | 1.491347 | 0 | 0.282377 |
| GOBP_SENSORY_PERCEPTION_OF_PAIN | 77 | 0.495208 | 1.490616 | 0.010613 | 0.283206 |
| GOBP_MALE_MEIOTIC_NUCLEAR_DIVISION | 36 | 0.555033 | 1.490607 | 0.028075 | 0.28211 |
| GOBP_AMINOGLYCAN_CATABOLIC_PROCESS | 22 | 0.601105 | 1.490238 | 0.038298 | 0.281964 |
| GOBP_MAMMARY_GLAND_DUCT_MORPHOGENESIS | 23 | 0.595693 | 1.490111 | 0.042213 | 0.281146 |
| GOBP_SENSORY_PERCEPTION_OF_LIGHT_STIMULUS | 141 | 0.456877 | 1.483476 | 0.003268 | 0.296887 |
| GOBP_POSITIVE_REGULATION_OF_VASCULATURE_DEVELOPMENT | 133 | 0.462126 | 1.483407 | 0.008782 | 0.295881 |
| GOBP_3_PHOSPHOADENOSINE_5_PHOSPHOSULFATE_METABOLIC_PROCESS | 15 | 0.649271 | 1.483394 | 0.04271 | 0.294768 |
| GOBP_REGULATION_OF_T_CELL_MEDIATED_IMMUNITY | 57 | 0.501064 | 1.48295 | 0.013871 | 0.294885 |
| GOBP_CELL_ACTIVATION_INVOLVED_IN_IMMUNE_RESPONSE | 196 | 0.44587 | 1.482639 | 0.004251 | 0.294559 |
| GOBP_REGULATION_OF_ACUTE_INFLAMMATORY_RESPONSE | 28 | 0.582804 | 1.481938 | 0.036536 | 0.295362 |
| GOBP_ESTABLISHMENT_OF_MITOCHONDRION_LOCALIZATION | 29 | 0.577675 | 1.481743 | 0.038095 | 0.294672 |
| GOBP_POSITIVE_REGULATION_OF_RECEPTOR_SIGNALING_PATHWAY_VIA_STAT | 25 | 0.583217 | 1.481584 | 0.040872 | 0.293922 |
| GOBP_EMBRYONIC_FORELIMB_MORPHOGENESIS | 17 | 0.620576 | 1.481483 | 0.045858 | 0.29309 |
| GOBP_EPIDERMAL_CELL_DIFFERENTIATION | 132 | 0.462789 | 1.480925 | 0.006565 | 0.293381 |
| GOBP_LEUKOCYTE_APOPTOTIC_PROCESS | 84 | 0.482134 | 1.480612 | 0.020047 | 0.293109 |
| GOBP_NEGATIVE_REGULATION_OF_IMMUNE_SYSTEM_PROCESS | 290 | 0.437286 | 1.478665 | 0 | 0.297248 |
| GOBP_REGULATION_OF_MEIOTIC_NUCLEAR_DIVISION | 22 | 0.598665 | 1.478485 | 0.054852 | 0.296605 |
| GOBP_EPIDERMIS_DEVELOPMENT | 225 | 0.445969 | 1.477348 | 0.00209 | 0.298638 |
| GOBP_GLYCEROLIPID_CATABOLIC_PROCESS | 51 | 0.519876 | 1.477164 | 0.031328 | 0.297953 |
| GOBP_SENSORY_PERCEPTION_OF_CHEMICAL_STIMULUS | 57 | 0.511344 | 1.476928 | 0.02396 | 0.297474 |
| GOBP_RESPONSE_TO_INTERLEUKIN_6 | 31 | 0.560861 | 1.476861 | 0.042384 | 0.296559 |
| GOBP_POSITIVE_REGULATION_OF_INTERLEUKIN_10_PRODUCTION | 27 | 0.577268 | 1.476525 | 0.052126 | 0.296295 |
| GOBP_POSITIVE_REGULATION_OF_RELEASE_OF_CYTOCHROME_C_FROM_MITOCHONDRIA | 24 | 0.581373 | 1.476101 | 0.049716 | 0.296377 |
| GOBP_T_CELL_DIFFERENTIATION | 185 | 0.445554 | 1.475559 | 0.003195 | 0.296593 |
| GOBP_WOUND_HEALING | 341 | 0.43171 | 1.474749 | 0.001016 | 0.297809 |
| GOBP_NEGATIVE_REGULATION_OF_WOUND_HEALING | 53 | 0.518648 | 1.473171 | 0.023661 | 0.300853 |
| GOBP_REGULATION_OF_NATURAL_KILLER_CELL_ACTIVATION | 27 | 0.578784 | 1.472922 | 0.047493 | 0.300426 |
| GOBP_CD4_POSITIVE_ALPHA_BETA_T_CELL_ACTIVATION | 70 | 0.488895 | 1.471551 | 0.027879 | 0.302959 |
| GOBP_CELL_MIGRATION_INVOLVED_IN_SPROUTING_ANGIOGENESIS | 48 | 0.514958 | 1.470781 | 0.025707 | 0.303864 |
| GOBP_REGULATION_OF_SENSORY_PERCEPTION | 24 | 0.590314 | 1.470592 | 0.054441 | 0.30335 |
| GOBP_NEUTRAL_LIPID_BIOSYNTHETIC_PROCESS | 40 | 0.535042 | 1.470091 | 0.036785 | 0.30363 |
| GOBP_INOSITOL_PHOSPHATE_METABOLIC_PROCESS | 48 | 0.518402 | 1.470065 | 0.028571 | 0.302612 |
| GOBP_PRODUCTION_OF_MOLECULAR_MEDIATOR_INVOLVED_IN_INFLAMMATORY_RESPOI | 54 | 0.511912 | 1.469393 | 0.035194 | 0.303293 |
| GOBP_LEUKOCYTE_DIFFERENTIATION | 404 | 0.428576 | 1.468612 | 0 | 0.304288 |
| GOBP_SUPEROXIDE_METABOLIC_PROCESS | 51 | 0.508602 | 1.468125 | 0.031888 | 0.304578 |
| GOBP_POSITIVE_REGULATION_OF_MYELOID_LEUKOCYTE_DIFFERENTIATION | 40 | 0.524317 | 1.468113 | 0.035387 | 0.303554 |
| GOBP_ICOSANOID_METABOLIC_PROCESS | 75 | 0.487387 | 1.465143 | 0.02439 | 0.310534 |
| GOBP_MEIOTIC_CHROMOSOME_SEGREGATION | 74 | 0.483759 | 1.464606 | 0.02561 | 0.310943 |
| GOBP_NEGATIVE_REGULATION_OF_LEUKOCYTE_MEDIATED_IMMUNITY | 42 | 0.537179 | 1.464486 | 0.032342 | 0.310177 |
| GOBP_FEMALE_GAMETE_GENERATION | 128 | 0.456206 | 1.46444 | 0.011765 | 0.309239 |
| GOBP_COLLAGEN_CATABOLIC_PROCESS | 27 | 0.568737 | 1.464038 | 0.037868 | 0.309223 |
| GOBP_PHAGOCYTOSIS | 180 | 0.451059 | 1.46391 | 0.004251 | 0.308496 |
| GOBP_NEGATIVE_REGULATION_OF_LEUKOCYTE_MIGRATION | 34 | 0.544863 | 1.463627 | 0.046729 | 0.308229 |
| GOBP_CELL_DIFFERENTIATION_INVOLVED_IN_EMBRYONIC_PLACENTA_DEVELOPMENT | 23 | 0.583347 | 1.46334 | 0.051893 | 0.307924 |
| GOBP_ENDOTHELIAL_CELL_MIGRATION | 194 | 0.440282 | 1.462837 | 0.00639 | 0.308192 |
| GOBP_CELLULAR_RESPONSE_TO_REACTIVE_OXYGEN_SPECIES | 122 | 0.462619 | 1.46277 | 0.007883 | 0.307329 |
| GOBP_MONONUCLEAR_CELL_DIFFERENTIATION | 309 | 0.43083 | 1.462728 | 0 | 0.3064 |
| GOBP_POSITIVE_REGULATION_OF_PROTEIN_DEPOLYMERIZATION | 17 | 0.622843 | 1.461933 | 0.0701 | 0.307492 |
| GOBP_REGULATION_OF_PLATELET_AGGREGATION | 21 | 0.60088 | 1.461861 | 0.066396 | 0.306658 |
| GOBP_POSITIVE_REGULATION_OF_COLLAGEN_METABOLIC_PROCESS | 21 | 0.589306 | 1.461339 | 0.051724 | 0.306962 |
| GOBP_CATECHOL_CONTAINING_COMPOUND_METABOLIC_PROCESS | 49 | 0.517981 | 1.460748 | 0.026854 | 0.307446 |
| GOBP_ANIMAL_ORGAN_REGENERATION | 49 | 0.512119 | 1.460561 | 0.029077 | 0.306951 |
| GOBP_NEGATIVE_REGULATION_OF_TISSUE_REMODELING | 18 | 0.602945 | 1.460169 | 0.046828 | 0.306983 |
| GOBP_POSITIVE_REGULATION_OF_COLD_INDUCED_THERMOGENESIS | 79 | 0.478711 | 1.459942 | 0.020833 | 0.306568 |
| GOBP_REGULATION_OF_TISSUE_REMODELING | 56 | 0.510689 | 1.45988 | 0.03317 | 0.305725 |
| GOBP_MYELOID_CELL_DEVELOPMENT | 60 | 0.504926 | 1.458385 | 0.021066 | 0.308471 |
| GOBP_POSITIVE_REGULATION_OF_EXTRACELLULAR_MATRIX_ORGANIZATION | 25 | 0.578727 | 1.457903 | 0.05146 | 0.308796 |
| GOBP_POSITIVE_REGULATION_OF_CYTOKINE_PRODUCTION | 321 | 0.427995 | 1.457555 | 0 | 0.30864 |

| | | | | | |
|---|---|---|---|---|---|
| GOBP_SPINAL_CORD_DEVELOPMENT | 76 | -0.324357 | -1.258035 | 0.045161 | 0.832769 |
| GOBP_PROTEIN_LIPID_COMPLEX_SUBUNIT_ORGANIZATION | 33 | -0.395557 | -1.25583 | 0.149194 | 0.836411 |
| GOBP_REGULATION_OF_DENDRITIC_SPINE_MORPHOGENESIS | 43 | -0.365075 | -1.255498 | 0.107296 | 0.832477 |
| GOBP_MIDDLE_EAR_MORPHOGENESIS | 15 | -0.461833 | -1.253687 | 0.202279 | 0.8349 |
| GOBP_ANION_HOMEOSTASIS | 30 | -0.391983 | -1.251318 | 0.152263 | 0.839849 |
| GOBP_AMINO_ACID_TRANSPORT | 123 | -0.302163 | -1.250842 | 0.030928 | 0.836619 |
| GOBP_REGULATION_OF_REACTIVE_OXYGEN_SPECIES_BIOSYNTHETIC_PROC | 34 | -0.391567 | -1.25052 | 0.17623 | 0.832587 |
| GOBP_INTERMEMBRANE_LIPID_TRANSFER | 44 | -0.361141 | -1.249028 | 0.118182 | 0.833434 |
| GOBP_AUTOPHAGY_OF_NUCLEUS | 16 | -0.455739 | -1.248689 | 0.180685 | 0.829883 |
| GOBP_MITOCHONDRION_MORPHOGENESIS | 20 | -0.440172 | -1.248682 | 0.167155 | 0.824828 |
| GOBP_HISTONE_H2A_UBIQUITINATION | 26 | -0.405586 | -1.247014 | 0.174377 | 0.826191 |
| GOBP_REGULATION_OF_RNA_SPLICING | 147 | -0.302987 | -1.246715 | 0.084337 | 0.822393 |
| GOBP_MIDBRAIN_DEVELOPMENT | 76 | -0.331022 | -1.240017 | 0.125714 | 0.844752 |
| GOBP_L_ALPHA_AMINO_ACID_TRANSMEMBRANE_TRANSPORT | 55 | -0.34814 | -1.239521 | 0.124352 | 0.841679 |
| GOBP_POSITIVE_REGULATION_OF_CELL_JUNCTION_ASSEMBLY | 88 | -0.31159 | -1.237916 | 0.049296 | 0.843373 |
| GOBP_TYPE_B_PANCREATIC_CELL_DEVELOPMENT | 19 | -0.44213 | -1.23577 | 0.19281 | 0.84725 |
| GOBP_NEURAL_NUCLEUS_DEVELOPMENT | 54 | -0.346051 | -1.233481 | 0.145631 | 0.851759 |
| GOBP_MONOUBIQUITINATED_PROTEIN_DEUBIQUITINATION | 34 | -0.382062 | -1.233232 | 0.126582 | 0.848009 |
| GOBP_DNA_REPLICATION_CHECKPOINT_SIGNALING | 16 | -0.462126 | -1.232921 | 0.198813 | 0.844318 |
| GOBP_SMOOTHENED_SIGNALING_PATHWAY | 121 | -0.300685 | -1.231367 | 0.05 | 0.845679 |
| GOBP_TRANSCRIPTION_PREINITIATION_COMPLEX_ASSEMBLY | 36 | -0.368495 | -1.230799 | 0.162698 | 0.843377 |
| GOBP_NEURON_PROJECTION_GUIDANCE | 207 | -0.286509 | -1.230739 | 0.073171 | 0.83875 |
| GOBP_NEURON_FATE_SPECIFICATION | 16 | -0.464035 | -1.230652 | 0.203175 | 0.834382 |
| GOBP_MODULATION_BY_HOST_OF_VIRAL_PROCESS | 25 | -0.3999 | -1.229574 | 0.188755 | 0.833941 |
| GOBP_RESPONSE_TO_PROSTAGLANDIN | 20 | -0.425678 | -1.226994 | 0.209726 | 0.84025 |
| GOBP_NEGATIVE_REGULATION_OF_GLIOGENESIS | 30 | -0.394331 | -1.226889 | 0.2 | 0.835994 |
| GOBP_DNA_LIGATION | 15 | -0.443919 | -1.224499 | 0.203448 | 0.840857 |
| GOBP_NUCLEOLAR_LARGE_RRNA_TRANSCRIPTION_BY_RNA_POLYMERASE_ | 18 | -0.438554 | -1.223906 | 0.165049 | 0.838829 |
| GOBP_POSITIVE_REGULATION_OF_CILIUM_ASSEMBLY | 23 | -0.403083 | -1.219431 | 0.202658 | 0.852157 |
| GOBP_REGULATION_OF_KETONE_BIOSYNTHETIC_PROCESS | 16 | -0.451674 | -1.218692 | 0.228916 | 0.850587 |
| GOBP_CELL_MIGRATION_IN_HINDBRAIN | 16 | -0.441638 | -1.21767 | 0.217949 | 0.850277 |
| GOBP_CENTRAL_NERVOUS_SYSTEM_NEURON_DIFFERENTIATION | 138 | -0.296628 | -1.216787 | 0.105263 | 0.849343 |
| GOBP_TRNA_5_END_PROCESSING | 16 | -0.436704 | -1.213671 | 0.222571 | 0.857146 |
| GOBP_NEURON_PROJECTION_ORGANIZATION | 81 | -0.309602 | -1.209681 | 0.098765 | 0.868992 |
| GOBP_MODULATION_BY_HOST_OF_SYMBIONT_PROCESS | 53 | -0.335135 | -1.209106 | 0.154229 | 0.866622 |
| GOBP_GLANDULAR_EPITHELIAL_CELL_DEVELOPMENT | 24 | -0.400497 | -1.20693 | 0.202952 | 0.870994 |
| GOBP_NEGATIVE_REGULATION_OF_GENE_EXPRESSION_EPIGENETIC | 75 | -0.316575 | -1.206801 | 0.104575 | 0.866923 |
| GOBP_SECONDARY_PALATE_DEVELOPMENT | 22 | -0.410008 | -1.206536 | 0.205776 | 0.863514 |
| GOBP_NUCLEAR_TRANSCRIBED_MRNA_CATABOLIC_PROCESS_NONSENSE_I | 37 | -0.36291 | -1.204646 | 0.158537 | 0.866582 |
| GOBP_REGULATION_OF_TELOMERE_CAPPING | 26 | -0.400466 | -1.203491 | 0.213333 | 0.866964 |
| GOBP_HISTONE_H2A_MONOUBIQUITINATION | 17 | -0.444343 | -1.203362 | 0.217252 | 0.863105 |
| GOBP_PROTEIN_NEDDYLATION | 17 | -0.43289 | -1.203211 | 0.225166 | 0.859288 |
| GOBP_SODIUM_ION_TRANSPORT | 193 | -0.280648 | -1.202521 | 0.125 | 0.857574 |
| GOBP_TELOMERASE_RNA_LOCALIZATION | 19 | -0.425858 | -1.201986 | 0.198142 | 0.855258 |
| GOBP_MORPHOGENESIS_OF_AN_EPITHELIAL_FOLD | 17 | -0.443751 | -1.200721 | 0.235849 | 0.855894 |
| GOBP_SPLEEN_DEVELOPMENT | 25 | -0.402592 | -1.200491 | 0.171617 | 0.852541 |
| GOBP_POSITIVE_REGULATION_OF_AMYLOID_BETA_FORMATION | 16 | -0.439877 | -1.200451 | 0.225309 | 0.848435 |
| GOBP_ENDOPLASMIC_RETICULUM_TO_CYTOSOL_TRANSPORT | 28 | -0.374887 | -1.197272 | 0.184502 | 0.857656 |
| GOBP_OLIGOPEPTIDE_TRANSPORT | 15 | -0.441795 | -1.197225 | 0.23125 | 0.853623 |
| GOBP_REGULATION_OF_DNA_DEPENDENT_DNA_REPLICATION | 49 | -0.335284 | -1.195651 | 0.139665 | 0.85577 |
| GOBP_ANION_TRANSMEMBRANE_TRANSPORT | 203 | -0.27747 | -1.194803 | 0.022727 | 0.854951 |
| GOBP_FACULTATIVE_HETEROCHROMATIN_ASSEMBLY | 30 | -0.371069 | -1.194547 | 0.204461 | 0.85174 |
| GOBP_TRANSCRIPTION_ELONGATION_FROM_RNA_POLYMERASE_II_PROMOT | 55 | -0.327405 | -1.193337 | 0.147208 | 0.85259 |
| GOBP_AXIS_ELONGATION | 21 | -0.407187 | -1.191859 | 0.200658 | 0.854505 |
| GOBP_DNA_STRAND_ELONGATION | 36 | -0.363216 | -1.189784 | 0.16055 | 0.858674 |
| GOBP_HISTONE_UBIQUITINATION | 47 | -0.332172 | -1.189671 | 0.133971 | 0.854988 |
| GOBP_REGULATION_OF_CARTILAGE_DEVELOPMENT | 59 | -0.326415 | -1.189303 | 0.161111 | 0.852452 |
| GOBP_REGULATION_OF_ALTERNATIVE_MRNA_SPLICING_VIA_SPLICEOSOME | 57 | -0.334404 | -1.187479 | 0.162562 | 0.855999 |
| GOBP_POSITIVE_REGULATION_OF_RECEPTOR_INTERNALIZATION | 25 | -0.385427 | -1.184208 | 0.207018 | 0.864718 |
| GOBP_REGULATION_OF_CHONDROCYTE_DIFFERENTIATION | 44 | -0.341832 | -1.183979 | 0.144186 | 0.861656 |
| GOBP_NONRIBOSOMAL_PEPTIDE_BIOSYNTHETIC_PROCESS | 15 | -0.442295 | -1.182501 | 0.259868 | 0.863945 |
| GOBP_REGULATION_OF_BRANCHING_INVOLVED_IN_URETERIC_BUD_MORPH | 16 | -0.430934 | -1.180635 | 0.249249 | 0.867442 |
| GOBP_REGULATION_OF_SPINDLE_ORGANIZATION | 44 | -0.342955 | -1.18019 | 0.199153 | 0.865132 |
| GOBP_ADENYLATE_CYCLASE_ACTIVATING_ADRENERGIC_RECEPTOR_SIGNA | 21 | -0.401751 | -1.179224 | 0.20922 | 0.865218 |
| GOBP_REGULATION_OF_TRANSLATIONAL_ELONGATION | 19 | -0.426517 | -1.179051 | 0.216718 | 0.862037 |
| GOBP_LIGAND_GATED_ION_CHANNEL_SIGNALING_PATHWAY | 20 | -0.411731 | -1.178007 | 0.223242 | 0.862083 |
| GOBP_ATP_SYNTHESIS_COUPLED_PROTON_TRANSPORT | 24 | -0.390018 | -1.174856 | 0.235075 | 0.870764 |
| GOBP_REGULATION_OF_MORPHOGENESIS_OF_A_BRANCHING_STRUCTURE | 35 | -0.358854 | -1.173655 | 0.190083 | 0.871604 |
| GOBP_RNA_POLYMERASE_II_PREINITIATION_COMPLEX_ASSEMBLY | 24 | -0.395451 | -1.168559 | 0.215054 | 0.888221 |
| GOBP_ANTIBACTERIAL_HUMORAL_RESPONSE | 21 | -0.39459 | -1.16799 | 0.242105 | 0.886621 |
| GOBP_REGULATION_OF_GENE_EXPRESSION_EPIGENETIC | 126 | -0.28433 | -1.167924 | 0.157895 | 0.883007 |
| GOBP_REGULATION_OF_MEGAKARYOCYTE_DIFFERENTIATION | 32 | -0.370501 | -1.16753 | 0.22963 | 0.880663 |
| GOBP_ANIMAL_ORGAN_MATURATION | 26 | -0.374515 | -1.167225 | 0.256809 | 0.877967 |
| GOBP_STRIATUM_DEVELOPMENT | 17 | -0.423015 | -1.165987 | 0.248366 | 0.879081 |
| GOBP_REGULATION_OF_NEURON_DIFFERENTIATION | 158 | -0.278297 | -1.165045 | 0.083333 | 0.879091 |
| GOBP_RESPONSE_TO_ISCHEMIA | 44 | -0.329291 | -1.164621 | 0.199153 | 0.876924 |
| GOBP_SODIUM_ION_IMPORT_ACROSS_PLASMA_MEMBRANE | 18 | -0.425515 | -1.162863 | 0.250729 | 0.880131 |
| GOBP_SERINE_FAMILY_AMINO_ACID_METABOLIC_PROCESS | 33 | -0.357532 | -1.162214 | 0.210526 | 0.879078 |
| GOBP_POSITIVE_REGULATION_OF_NEURON_DEATH | 71 | -0.307472 | -1.160756 | 0.168675 | 0.88138 |
| GOBP_BONE_MATURATION | 23 | -0.39339 | -1.160266 | 0.243421 | 0.879727 |
| GOBP_GAMMA_AMINOBUTYRIC_ACID_TRANSPORT | 16 | -0.432648 | -1.157873 | 0.253333 | 0.885626 |
| GOBP_POSITIVE_REGULATION_OF_HISTONE_H3_K4_METHYLATION | 25 | -0.390598 | -1.157609 | 0.242215 | 0.882806 |
| GOBP_TRNA_WOBBLE_BASE_MODIFICATION | 17 | -0.407872 | -1.157473 | 0.251592 | 0.879633 |
| GOBP_NEGATIVE_REGULATION_OF_INTRACELLULAR_PROTEIN_TRANSPORT | 37 | -0.348993 | -1.156112 | 0.223077 | 0.881093 |
| GOBP_INTRACELLULAR_PROTEIN_TRANSMEMBRANE_TRANSPORT | 52 | -0.323382 | -1.155024 | 0.198157 | 0.881855 |
| GOBP_REGULATION_OF_HEART_RATE | 81 | -0.299293 | -1.154429 | 0.18 | 0.880526 |
| GOBP_NEGATIVE_REGULATION_OF_ATP_METABOLIC_PROCESS | 27 | -0.378597 | -1.151715 | 0.26936 | 0.887914 |
| GOBP_RIBOSOMAL_LARGE_SUBUNIT_BIOGENESIS | 71 | -0.305714 | -1.151161 | 0.184211 | 0.886563 |
| GOBP_POSITIVE_REGULATION_OF_AMYLOID_PRECURSOR_PROTEIN_CATABC | 21 | -0.399926 | -1.151025 | 0.24911 | 0.88346 |
| GOBP_NUCLEOSIDE_TRIPHOSPHATE_BIOSYNTHETIC_PROCESS | 79 | -0.297609 | -1.150649 | 0.1625 | 0.881241 |
| GOBP_REGULATION_OF_POSTSYNAPSE_ORGANIZATION | 73 | -0.302828 | -1.149325 | 0.175 | 0.882944 |
| GOBP_MONOVALENT_INORGANIC_CATION_HOMEOSTASIS | 123 | -0.278552 | -1.148881 | 0.144144 | 0.881236 |
| GOBP_POSITIVE_REGULATION_OF_METAPHASE_ANAPHASE_TRANSITION_OF_ | 15 | -0.430112 | -1.146266 | 0.31085 | 0.888597 |
| GOBP_NEGATIVE_REGULATION_OF_DNA_DEPENDENT_DNA_REPLICATION | 15 | -0.429528 | -1.144401 | 0.299685 | 0.892579 |
| GOBP_REGULATION_OF_DENDRITE_MORPHOGENESIS | 61 | -0.305976 | -1.143811 | 0.170455 | 0.891387 |
| GOBP_PROTEIN_TRANSMEMBRANE_TRANSPORT | 63 | -0.314248 | -1.143053 | 0.185629 | 0.890762 |
| GOBP_MRNA_PROCESSING | 445 | -0.239102 | -1.142106 | 0.125 | 0.890841 |
| GOBP_REGULATION_OF_WNT_SIGNALING_PATHWAY_PLANAR_CELL_POLARI | 15 | -0.443336 | -1.14113 | 0.294118 | 0.891103 |
| GOBP_PTERIDINE_CONTAINING_COMPOUND_BIOSYNTHETIC_PROCESS | 15 | -0.425905 | -1.139424 | 0.285266 | 0.894337 |
| GOBP_REGULATION_OF_CYTOKINESIS | 79 | -0.293275 | -1.139302 | 0.206897 | 0.891336 |
| GOBP_GPI_ANCHOR_METABOLIC_PROCESS | 30 | -0.353758 | -1.138749 | 0.261993 | 0.890082 |
| GOBP_RIBONUCLEOSIDE_TRIPHOSPHATE_METABOLIC_PROCESS | 82 | -0.292525 | -1.138572 | 0.196078 | 0.887372 |
| GOBP_SPLICEOSOMAL_SNRNP_ASSEMBLY | 38 | -0.346893 | -1.135692 | 0.229437 | 0.895411 |
| GOBP_DOUBLE_STRAND_BREAK_REPAIR_VIA_NONHOMOLOGOUS_END_JOIN | 68 | -0.298161 | -1.134232 | 0.172775 | 0.897904 |
| GOBP_NEGATIVE_REGULATION_OF_ACTIN_FILAMENT_BUNDLE_ASSEMBLY | 29 | -0.355898 | -1.133044 | 0.25 | 0.899123 |
| GOBP_SOMITOGENESIS | 54 | -0.309302 | -1.130495 | 0.253521 | 0.905932 |
| GOBP_NEGATIVE_REGULATION_OF_DOUBLE_STRAND_BREAK_REPAIR_VIA_H | 19 | -0.394872 | -1.129845 | 0.278146 | 0.905 |
| GOBP_POSITIVE_REGULATION_OF_TELOMERE_MAINTENANCE | 50 | -0.326457 | -1.129178 | 0.232877 | 0.904583 |
| GOBP_REGULATION_OF_LONG_TERM_NEURONAL_SYNAPTIC_PLASTICITY | 22 | -0.379883 | -1.12863 | 0.25188 | 0.903182 |
| GOBP_REGULATION_OF_DNA_TEMPLATED_TRANSCRIPTION_ELONGATION | 51 | -0.317786 | -1.127944 | 0.219048 | 0.902643 |
| GOBP_NEURAL_TUBE_FORMATION | 102 | -0.283065 | -1.127188 | 0.175573 | 0.902481 |
| GOBP_VENTRICULAR_SYSTEM_DEVELOPMENT | 28 | -0.371878 | -1.123665 | 0.301754 | 0.913475 |
| GOBP_EPHRIN_RECEPTOR_SIGNALING_PATHWAY | 50 | -0.321725 | -1.12296 | 0.25 | 0.912879 |
| GOBP_NEGATIVE_REGULATION_OF_DNA_RECOMBINATION | 40 | -0.33277 | -1.122418 | 0.246637 | 0.911791 |
| GOBP_NEUROTRANSMITTER_UPTAKE | 37 | -0.342735 | -1.121538 | 0.265873 | 0.912142 |
| GOBP_POSITIVE_REGULATION_OF_CHROMOSOME_ORGANIZATION | 85 | -0.289562 | -1.121152 | 0.253333 | 0.910351 |
| GOBP_V_D_J_RECOMBINATION | 18 | -0.410991 | -1.120619 | 0.282051 | 0.909201 |
| GOBP_REGULATION_OF_TRANSCRIPTION_OF_NUCLEOLAR_LARGE_RRNA_BY | 15 | -0.416388 | -1.120236 | 0.317784 | 0.9076 |
| GOBP_POSITIVE_REGULATION_OF_BLOOD_CIRCULATION | 34 | -0.341126 | -1.118545 | 0.275304 | 0.91117 |
| GOBP_DENDRITE_MORPHOGENESIS | 133 | -0.267703 | -1.117569 | 0.197368 | 0.911873 |
| GOBP_SIALYLATION | 18 | -0.408253 | -1.112441 | 0.306962 | 0.929079 |
| GOBP_DOSAGE_COMPENSATION | 20 | -0.386117 | -1.112379 | 0.316456 | 0.925948 |
| GOBP_CARDIOBLAST_DIFFERENTIATION | 15 | -0.434646 | -1.111418 | 0.303303 | 0.926618 |
| GOBP_HORMONE_BIOSYNTHETIC_PROCESS | 46 | -0.318077 | -1.11005 | 0.258883 | 0.928671 |
| GOBP_ANIMAL_ORGAN_FORMATION | 47 | -0.309744 | -1.10725 | 0.275701 | 0.936746 |
| GOBP_PYRIMIDINE_CONTAINING_COMPOUND_BIOSYNTHETIC_PROCESS | 35 | -0.341631 | -1.103301 | 0.2875 | 0.950301 |
| GOBP_LEFT_RIGHT_PATTERN_FORMATION | 22 | -0.370489 | -1.102639 | 0.315603 | 0.949792 |
| GOBP_TUBE_FORMATION | 139 | -0.262957 | -1.102312 | 0.195652 | 0.947775 |
| GOBP_PROTEIN_N_LINKED_GLYCOSYLATION | 65 | -0.285883 | -1.10171 | 0.275862 | 0.947057 |
| GOBP_C21_STEROID_HORMONE_METABOLIC_PROCESS | 28 | -0.360027 | -1.1005 | 0.296992 | 0.948856 |
| GOBP_HISTONE_EXCHANGE | 17 | -0.391697 | -1.100252 | 0.328947 | 0.946469 |
| GOBP_REGULATION_OF_MYELINATION | 41 | -0.320255 | -1.099933 | 0.264317 | 0.944571 |
| GOBP_NEUTRAL_AMINO_ACID_TRANSPORT | 38 | -0.328692 | -1.099819 | 0.306452 | 0.941781 |
| GOBP_LABYRINTHINE_LAYER_MORPHOGENESIS | 18 | -0.403175 | -1.099259 | 0.336283 | 0.940959 |
| GOBP_VESICLE_FUSION_TO_PLASMA_MEMBRANE | 18 | -0.392648 | -1.098055 | 0.337423 | 0.942722 |
| GOBP_EPITHELIAL_TUBE_FORMATION | 127 | -0.265912 | -1.096981 | 0.23301 | 0.944019 |
| GOBP_SMOOTH_MUSCLE_TISSUE_DEVELOPMENT | 16 | -0.412313 | -1.094773 | 0.340426 | 0.950067 |
| GOBP_LIPID_HOMEOSTASIS | 122 | -0.265862 | -1.092814 | 0.234234 | 0.954829 |
| GOBP_GLYCOSPHINGOLIPID_METABOLIC_PROCESS | 54 | -0.302618 | -1.091614 | 0.268293 | 0.956753 |
| GOBP_REGULATION_OF_SYSTEMIC_ARTERIAL_BLOOD_PRESSURE_MEDIATEI | 31 | -0.355097 | -1.09063 | 0.316176 | 0.957866 |
| GOBP_POSITIVE_REGULATION_OF_CHROMATIN_ORGANIZATION | 15 | -0.417309 | -1.090342 | 0.335463 | 0.955789 |
| GOBP_POSITIVE_REGULATION_OF_STEM_CELL_DIFFERENTIATION | 16 | -0.404483 | -1.088231 | 0.318328 | 0.961378 |
| GOBP_REGULATION_OF_SYNAPSE_STRUCTURE_OR_ACTIVITY | 187 | -0.242017 | -1.08775 | 0.233333 | 0.960232 |
| GOBP_CYTOKINESIS | 157 | -0.253041 | -1.087376 | 0.28169 | 0.958652 |
| GOBP_RIBONUCLEOSIDE_TRIPHOSPHATE_BIOSYNTHETIC_PROCESS | 69 | -0.291583 | -1.085616 | 0.290323 | 0.962774 |
| GOBP_RESPONSE_TO_INTERLEUKIN_4 | 29 | -0.342621 | -1.084642 | 0.315603 | 0.963527 |
| GOBP_COCHLEA_MORPHOGENESIS | 17 | -0.388928 | -1.083464 | 0.330128 | 0.965081 |
| GOBP_AMINO_ACID_IMPORT | 46 | -0.317229 | -1.083321 | 0.309179 | 0.962401 |
| GOBP_TRANSCRIPTION_INITIATION_FROM_RNA_POLYMERASE_II_PROMOTER | 71 | -0.297267 | -1.082965 | 0.315068 | 0.96081 |
| GOBP_MORPHOGENESIS_OF_EMBRYONIC_EPITHELIUM | 134 | -0.260811 | -1.082567 | 0.191011 | 0.959412 |
| GOBP_MONOSACCHARIDE_CATABOLIC_PROCESS | 34 | -0.335285 | -1.082279 | 0.353175 | 0.957488 |
| GOBP_LIPID_IMPORT_INTO_CELL | 16 | -0.39693 | -1.081469 | 0.346939 | 0.957714 |
| GOBP_LIPOPROTEIN_METABOLIC_PROCESS | 122 | -0.25972 | -1.08063 | 0.185567 | 0.957947 |
| GOBP_PROTEIN_DNA_COMPLEX_DISASSEMBLY | 19 | -0.376604 | -1.079159 | 0.334426 | 0.960815 |
| GOBP_RESPONSE_TO_AUDITORY_STIMULUS | 22 | -0.368117 | -1.078479 | 0.343396 | 0.960506 |
| GOBP_NEGATIVE_REGULATION_OF_DNA_REPLICATION | 27 | -0.34571 | -1.077126 | 0.341912 | 0.963458 |
| GOBP_DIACYLGLYCEROL_METABOLIC_PROCESS | 24 | -0.349729 | -1.076665 | 0.327068 | 0.962251 |

| | | | | | |
|---|---|---|---|---|---|
| GOBP_POSITIVE_REGULATION_OF_CELLULAR_CARBOHYDRATE_METABOLIC_PROCESS | 43 | 0.526332 | 1.457523 | 0.027569 | 0.307722 |
| GOBP_BLOOD_VESSEL_ENDOTHELIAL_CELL_MIGRATION | 104 | 0.463737 | 1.457152 | 0.012543 | 0.307625 |
| GOBP_INTERLEUKIN_10_PRODUCTION | 37 | 0.543237 | 1.456701 | 0.045213 | 0.307798 |
| GOBP_MATERNAL_PROCESS_INVOLVED_IN_FEMALE_PREGNANCY | 45 | 0.516476 | 1.456134 | 0.031332 | 0.308229 |
| GOBP_T_HELPER_1_TYPE_IMMUNE_RESPONSE | 31 | 0.551699 | 1.455186 | 0.037992 | 0.30984 |
| GOBP_LYMPHOCYTE_MIGRATION | 68 | 0.486589 | 1.454424 | 0.027545 | 0.310893 |
| GOBP_POSITIVE_REGULATION_OF_CALCIUM_ION_TRANSPORT | 93 | 0.473239 | 1.454391 | 0.017523 | 0.310021 |
| GOBP_POSITIVE_REGULATION_OF_CELL_KILLING | 35 | 0.532634 | 1.45435 | 0.037185 | 0.309142 |
| GOBP_RESPIRATORY_BURST | 23 | 0.587825 | 1.453924 | 0.054054 | 0.309349 |
| GOBP_TRIGLYCERIDE_CATABOLIC_PROCESS | 21 | 0.598398 | 1.45348 | 0.0625 | 0.309547 |
| GOBP_NEGATIVE_REGULATION_OF_SMOOTH_MUSCLE_CELL_PROLIFERATION | 42 | 0.521936 | 1.453465 | 0.043767 | 0.308636 |
| GOBP_REGULATION_OF_LEUKOCYTE_MEDIATED_CYTOTOXICITY | 45 | 0.511869 | 1.452611 | 0.033248 | 0.310012 |
| GOBP_NEGATIVE_REGULATION_OF_LYMPHOCYTE_MEDIATED_IMMUNITY | 35 | 0.54012 | 1.451902 | 0.036269 | 0.310841 |
| GOBP_RECEPTOR_SIGNALING_PATHWAY_VIA_STAT | 106 | 0.464152 | 1.451834 | 0.018265 | 0.310042 |
| GOBP_T_CELL_DIFFERENTIATION_INVOLVED_IN_IMMUNE_RESPONSE | 52 | 0.512935 | 1.450702 | 0.036524 | 0.312065 |
| GOBP_INFLAMMATORY_RESPONSE | 496 | 0.41908 | 1.449561 | 0 | 0.314193 |
| GOBP_REGULATION_OF_DEFENSE_RESPONSE | 437 | 0.42069 | 1.448939 | 0 | 0.314835 |
| GOBP_MESENCHYME_MORPHOGENESIS | 50 | 0.511788 | 1.448471 | 0.033854 | 0.315065 |
| GOBP_LYMPH_VESSEL_MORPHOGENESIS | 22 | 0.585538 | 1.44782 | 0.04878 | 0.315878 |
| GOBP_PROTEIN_KINASE_C_SIGNALING | 26 | 0.576866 | 1.447802 | 0.06069 | 0.314967 |
| GOBP_RESPONSE_TO_DIETARY_EXCESS | 24 | 0.575446 | 1.447339 | 0.0625 | 0.315368 |
| GOBP_T_CELL_MEDIATED_CYTOTOXICITY | 32 | 0.54562 | 1.447142 | 0.053191 | 0.3149 |
| GOBP_REGULATION_OF_CELL_ADHESION_MEDIATED_BY_INTEGRIN | 40 | 0.52179 | 1.446925 | 0.052151 | 0.314477 |
| GOBP_POSITIVE_REGULATION_OF_TYPE_I_INTERFERON_MEDIATED_SIGNALING_PATHWAY | 16 | 0.627026 | 1.44684 | 0.066568 | 0.313739 |
| GOBP_DNA_DAMAGE_RESPONSE_SIGNAL_TRANSDUCTION_BY_P53_CLASS_MEDIATOR_R | 16 | 0.628467 | 1.446322 | 0.071111 | 0.314323 |
| GOBP_POSITIVE_REGULATION_OF_ALCOHOL_BIOSYNTHETIC_PROCESS | 19 | 0.603838 | 1.445025 | 0.064607 | 0.316771 |
| GOBP_OOGENESIS | 82 | 0.470351 | 1.444794 | 0.02217 | 0.316496 |
| GOBP_RESPONSE_TO_REACTIVE_OXYGEN_SPECIES | 170 | 0.446899 | 1.444703 | 0.006431 | 0.31582 |
| GOBP_TISSUE_MIGRATION | 273 | 0.4262 | 1.444417 | 0.003125 | 0.315603 |
| GOBP_CYTOKINE_PRODUCTION_INVOLVED_IN_IMMUNE_RESPONSE | 71 | 0.484847 | 1.444226 | 0.026347 | 0.315179 |
| GOBP_NEGATIVE_REGULATION_OF_ION_TRANSPORT | 107 | 0.461159 | 1.444128 | 0.019231 | 0.314507 |
| GOBP_NEGATIVE_REGULATION_OF_ADAPTIVE_IMMUNE_RESPONSE | 39 | 0.52216 | 1.444041 | 0.040951 | 0.313839 |
| GOBP_POSITIVE_REGULATION_OF_RESPONSE_TO_BIOTIC_STIMULUS | 117 | 0.456694 | 1.443807 | 0.010953 | 0.313603 |
| GOBP_REGULATION_OF_PLATELET_ACTIVATION | 32 | 0.551997 | 1.442278 | 0.050754 | 0.316707 |
| GOBP_PYRIMIDINE_NUCLEOSIDE_METABOLIC_PROCESS | 19 | 0.604235 | 1.442141 | 0.050725 | 0.316197 |
| GOBP_PEPTIDYL_ARGININE_MODIFICATION | 24 | 0.586774 | 1.441715 | 0.064972 | 0.316378 |
| GOBP_TOLERANCE_INDUCTION | 19 | 0.60649 | 1.441539 | 0.07478 | 0.315916 |
| GOBP_REGULATION_OF_CALCIUM_ION_TRANSPORT | 192 | 0.435295 | 1.440649 | 0.008475 | 0.317358 |
| GOBP_REGULATION_OF_ACTIN_FILAMENT_BASED_MOVEMENT | 36 | 0.535741 | 1.43997 | 0.06469 | 0.318197 |
| GOBP_ANTERIOR_POSTERIOR_PATTERN_SPECIFICATION | 149 | 0.443838 | 1.439879 | 0.010989 | 0.317523 |
| GOBP_POSITIVE_REGULATION_OF_T_CELL_MEDIATED_IMMUNITY | 41 | 0.521781 | 1.438845 | 0.047244 | 0.319403 |
| GOBP_MUSCLE_ADAPTATION | 82 | 0.470775 | 1.437921 | 0.021505 | 0.320933 |
| GOBP_GLYCEROL_ETHER_METABOLIC_PROCESS | 21 | 0.583821 | 1.437456 | 0.075177 | 0.321279 |
| GOBP_NEGATIVE_REGULATION_OF_VIRAL_GENOME_REPLICATION | 44 | 0.515554 | 1.436837 | 0.051833 | 0.321835 |
| GOBP_MALE_SEX_DIFFERENTIATION | 129 | 0.445115 | 1.436559 | 0.010965 | 0.321651 |
| GOBP_CYTOKINE_PRODUCTION_INVOLVED_IN_INFLAMMATORY_RESPONSE | 36 | 0.536068 | 1.436169 | 0.050453 | 0.321753 |
| GOBP_RESPONSE_TO_ESTRADIOL | 96 | 0.463165 | 1.436094 | 0.016393 | 0.321066 |
| GOBP_CELL_MIGRATION_INVOLVED_IN_HEART_DEVELOPMENT | 17 | 0.614837 | 1.436004 | 0.063798 | 0.320418 |
| GOBP_RESPONSE_TO_ELECTRICAL_STIMULUS | 33 | 0.54451 | 1.435458 | 0.047556 | 0.320936 |
| GOBP_HYDROGEN_PEROXIDE_METABOLIC_PROCESS | 33 | 0.54022 | 1.435207 | 0.050734 | 0.320691 |
| GOBP_DOPAMINE_METABOLIC_PROCESS | 36 | 0.53434 | 1.433179 | 0.05148 | 0.324998 |
| GOBP_VASCULAR_ASSOCIATED_SMOOTH_MUSCLE_CELL_PROLIFERATION | 46 | 0.50804 | 1.43283 | 0.050649 | 0.32501 |
| GOBP_ARF_PROTEIN_SIGNAL_TRANSDUCTION | 20 | 0.58676 | 1.432234 | 0.076479 | 0.32569 |
| GOBP_CELL_ADHESION_MEDIATED_BY_INTEGRIN | 70 | 0.480863 | 1.431814 | 0.034856 | 0.325831 |
| GOBP_CELL_COMMUNICATION_BY_ELECTRICAL_COUPLING | 30 | 0.548312 | 1.4314 | 0.065217 | 0.326116 |
| GOBP_MATERNAL_PLACENTA_DEVELOPMENT | 29 | 0.538621 | 1.431316 | 0.062587 | 0.325445 |
| GOBP_REGULATION_OF_NEUTROPHIL_CHEMOTAXIS | 25 | 0.559447 | 1.430666 | 0.05791 | 0.32628 |
| GOBP_REGULATION_OF_LYMPHOCYTE_MIGRATION | 42 | 0.517804 | 1.430027 | 0.055627 | 0.327175 |
| GOBP_LIPID_DROPLET_ORGANIZATION | 22 | 0.582975 | 1.429478 | 0.0767 | 0.327774 |
| GOBP_TRANSFORMING_GROWTH_FACTOR_BETA_PRODUCTION | 33 | 0.527409 | 1.428712 | 0.061058 | 0.32889 |
| GOBP_REGULATION_OF_NEUTROPHIL_MIGRATION | 35 | 0.533093 | 1.427902 | 0.052989 | 0.330171 |
| GOBP_LEUKOCYTE_HOMEOSTASIS | 69 | 0.472741 | 1.427784 | 0.027027 | 0.329603 |
| GOBP_POSITIVE_REGULATION_OF_G_PROTEIN_COUPLED_RECEPTOR_SIGNALING_PATHW | 21 | 0.578048 | 1.42445 | 0.054913 | 0.337729 |
| GOBP_RESPONSE_TO_INTERFERON_ALPHA | 17 | 0.593009 | 1.422514 | 0.076135 | 0.342088 |
| GOBP_INNATE_IMMUNE_RESPONSE | 500 | 0.412652 | 1.42161 | 0 | 0.343571 |
| GOBP_O_GLYCAN_PROCESSING | 36 | 0.530442 | 1.421508 | 0.057796 | 0.342953 |
| GOBP_CALCIUM_ION_TRANSPORT | 332 | 0.421781 | 1.421399 | 0.00203 | 0.342383 |
| GOBP_POSITIVE_REGULATION_OF_MYELOID_LEUKOCYTE_MEDIATED_IMMUNITY | 17 | 0.596139 | 1.421202 | 0.073855 | 0.341999 |
| GOBP_VASCULAR_WOUND_HEALING | 17 | 0.613822 | 1.420785 | 0.073314 | 0.342137 |
| GOBP_POSITIVE_REGULATION_OF_RHO_PROTEIN_SIGNAL_TRANSDUCTION | 23 | 0.566042 | 1.41985 | 0.07972 | 0.343847 |
| GOBP_SENSORY_PERCEPTION_OF_SMELL | 30 | 0.539254 | 1.418666 | 0.063087 | 0.346186 |
| GOBP_SECONDARY_METABOLIC_PROCESS | 39 | 0.513486 | 1.417795 | 0.061697 | 0.347595 |
| GOBP_ETHER_METABOLIC_PROCESS | 26 | 0.56579 | 1.417742 | 0.063536 | 0.3468 |
| GOBP_PHENOL_CONTAINING_COMPOUND_METABOLIC_PROCESS | 83 | 0.468381 | 1.416934 | 0.029762 | 0.348178 |
| GOBP_POSITIVE_REGULATION_OF_MUSCLE_CONTRACTION | 38 | 0.519188 | 1.416598 | 0.060027 | 0.348198 |
| GOBP_REGULATION_OF_LEUKOCYTE_APOPTOTIC_PROCESS | 62 | 0.481374 | 1.415311 | 0.04896 | 0.350784 |
| GOBP_CRANIAL_SKELETAL_SYSTEM_DEVELOPMENT | 57 | 0.488347 | 1.41523 | 0.046798 | 0.350105 |
| GOBP_MALE_MEIOSIS_I | 19 | 0.584051 | 1.413384 | 0.09389 | 0.354273 |
| GOBP_POSITIVE_REGULATION_OF_ERBB_SIGNALING_PATHWAY | 28 | 0.541145 | 1.413191 | 0.076923 | 0.353973 |
| GOBP_FERTILIZATION | 116 | 0.446467 | 1.412472 | 0.023385 | 0.35504 |
| GOBP_ENDODERM_DEVELOPMENT | 70 | 0.466251 | 1.41223 | 0.029586 | 0.354786 |
| GOBP_NEUROINFLAMMATORY_RESPONSE | 23 | 0.566351 | 1.41201 | 0.079167 | 0.354477 |
| GOBP_POSITIVE_REGULATION_OF_SMALL_MOLECULE_METABOLIC_PROCESS | 105 | 0.450532 | 1.411978 | 0.022805 | 0.353663 |
| GOBP_CD4_POSITIVE_ALPHA_BETA_T_CELL_DIFFERENTIATION | 58 | 0.480918 | 1.411286 | 0.040915 | 0.35466 |
| GOBP_CARDIAC_EPITHELIAL_TO_MESENCHYMAL_TRANSITION | 34 | 0.528784 | 1.4108 | 0.055172 | 0.355078 |
| GOBP_INTERLEUKIN_12_PRODUCTION | 41 | 0.503823 | 1.410695 | 0.070556 | 0.354474 |
| GOBP_MUSCLE_CELL_DEVELOPMENT | 141 | 0.433653 | 1.410164 | 0.013086 | 0.355096 |
| GOBP_MITOTIC_G1_S_TRANSITION_CHECKPOINT_SIGNALING | 27 | 0.542378 | 1.409539 | 0.076923 | 0.355886 |
| GOBP_REGULATION_OF_VIRAL_ENTRY_INTO_HOST_CELL | 38 | 0.523382 | 1.409122 | 0.063745 | 0.356148 |
| GOBP_RESPONSE_TO_OXYGEN_RADICAL | 24 | 0.564181 | 1.408266 | 0.074965 | 0.357548 |
| GOBP_CELLULAR_RESPONSE_TO_HYDROGEN_PEROXIDE | 72 | 0.4691 | 1.408024 | 0.031063 | 0.35727 |
| GOBP_REGULATION_OF_BIOLOGICAL_PROCESS_INVOLVED_IN_SYMBIOTIC_INTERACTION | 46 | 0.502595 | 1.407685 | 0.045 | 0.357376 |
| GOBP_COMPLEMENT_ACTIVATION | 28 | 0.542911 | 1.40749 | 0.074689 | 0.357023 |
| GOBP_ALPHA_BETA_T_CELL_DIFFERENTIATION | 79 | 0.459124 | 1.407397 | 0.032105 | 0.356395 |
| GOBP_REGULATION_OF_LYMPHOCYTE_MEDIATED_IMMUNITY | 112 | 0.448762 | 1.406181 | 0.024719 | 0.358746 |
| GOBP_POLYOL_METABOLIC_PROCESS | 100 | 0.452469 | 1.405804 | 0.026196 | 0.358858 |
| GOBP_SKELETAL_MUSCLE_CONTRACTION | 33 | 0.529591 | 1.405384 | 0.062827 | 0.359196 |
| GOBP_NEGATIVE_REGULATION_OF_CALCIUM_ION_TRANSMEMBRANE_TRANSPORT | 31 | 0.527347 | 1.404991 | 0.063428 | 0.359412 |
| GOBP_INTERLEUKIN_6_MEDIATED_SIGNALING_PATHWAY | 17 | 0.588019 | 1.404162 | 0.1 | 0.360977 |
| GOBP_MEMBRANE_RAFT_ORGANIZATION | 20 | 0.569191 | 1.403987 | 0.071848 | 0.360571 |
| GOBP_MONOCARBOXYLIC_ACID_TRANSPORT | 77 | 0.459703 | 1.403938 | 0.041032 | 0.359831 |
| GOBP_NEGATIVE_REGULATION_OF_COAGULATION | 36 | 0.514197 | 1.40383 | 0.069645 | 0.359256 |
| GOBP_POSITIVE_REGULATION_OF_NEUTROPHIL_MIGRATION | 25 | 0.555665 | 1.403395 | 0.086957 | 0.359651 |
| GOBP_HINDLIMB_MORPHOGENESIS | 22 | 0.578357 | 1.403296 | 0.087075 | 0.359039 |
| GOBP_B_CELL_RECEPTOR_SIGNALING_PATHWAY | 41 | 0.501477 | 1.403098 | 0.068702 | 0.358751 |
| GOBP_REGULATION_OF_MUSCLE_ADAPTATION | 66 | 0.470973 | 1.403032 | 0.03995 | 0.358049 |
| GOBP_RESPONSE_TO_LEAD_ION | 17 | 0.596854 | 1.402973 | 0.083578 | 0.357354 |
| GOBP_THYROID_HORMONE_METABOLIC_PROCESS | 16 | 0.602234 | 1.402766 | 0.097242 | 0.357046 |
| GOBP_ANTIGEN_PROCESSING_AND_PRESENTATION_OF_PEPTIDE_ANTIGEN | 46 | 0.495403 | 1.402504 | 0.052767 | 0.356916 |
| GOBP_REGULATION_OF_CD4_POSITIVE_ALPHA_BETA_T_CELL_ACTIVATION | 44 | 0.497991 | 1.402476 | 0.05283 | 0.356155 |
| GOBP_POSITIVE_REGULATION_OF_INNATE_IMMUNE_RESPONSE | 95 | 0.449256 | 1.401506 | 0.031953 | 0.357978 |
| GOBP_POSITIVE_REGULATION_OF_LEUKOCYTE_DEGRANULATION | 21 | 0.563093 | 1.400875 | 0.097561 | 0.358797 |
| GOBP_NATURAL_KILLER_CELL_MEDIATED_IMMUNITY | 42 | 0.495074 | 1.399996 | 0.060209 | 0.360356 |
| GOBP_MUSCLE_CELL_PROLIFERATION | 156 | 0.43054 | 1.399981 | 0.013072 | 0.35954 |
| GOBP_ADAPTIVE_THERMOGENESIS | 133 | 0.4357 | 1.399672 | 0.026696 | 0.359588 |
| GOBP_AMINE_BIOSYNTHETIC_PROCESS | 32 | 0.533495 | 1.399362 | 0.069057 | 0.359603 |
| GOBP_LIVER_REGENERATION | 24 | 0.551652 | 1.398518 | 0.088953 | 0.361116 |
| GOBP_INTERFERON_GAMMA_MEDIATED_SIGNALING_PATHWAY | 20 | 0.580023 | 1.39803 | 0.083807 | 0.361552 |
| GOBP_GLYCEROPHOSPHOLIPID_CATABOLIC_PROCESS | 27 | 0.536791 | 1.39786 | 0.075949 | 0.36118 |
| GOBP_SPROUTING_ANGIOGENESIS | 113 | 0.442222 | 1.397412 | 0.026048 | 0.361564 |
| GOBP_POSITIVE_REGULATION_OF_CELL_SUBSTRATE_ADHESION | 104 | 0.446579 | 1.397354 | 0.038288 | 0.360881 |
| GOBP_STRIATED_MUSCLE_CELL_DEVELOPMENT | 52 | 0.485519 | 1.396401 | 0.045802 | 0.362644 |
| GOBP_B_CELL_DIFFERENTIATION | 99 | 0.447714 | 1.395135 | 0.034483 | 0.365258 |
| GOBP_REGULATION_OF_STRIATED_MUSCLE_CONTRACTION | 75 | 0.456764 | 1.394858 | 0.033735 | 0.365219 |
| GOBP_CALCIUM_ION_TRANSPORT_INTO_CYTOSOL | 126 | 0.438917 | 1.394376 | 0.025612 | 0.365785 |
| GOBP_BONE_RESORPTION | 49 | 0.493743 | 1.393799 | 0.055276 | 0.3664 |
| GOBP_REGULATION_OF_B_CELL_PROLIFERATION | 40 | 0.511117 | 1.393272 | 0.056136 | 0.367077 |
| GOBP_POSITIVE_REGULATION_OF_GRANULOCYTE_CHEMOTAXIS | 19 | 0.581132 | 1.393152 | 0.07958 | 0.366556 |
| GOBP_NEGATIVE_REGULATION_OF_VASCULAR_ASSOCIATED_SMOOTH_MUSCLE_CELL_PI | 21 | 0.570293 | 1.393 | 0.095041 | 0.3661 |
| GOBP_REGULATION_OF_MONONUCLEAR_CELL_MIGRATION | 79 | 0.463142 | 1.392743 | 0.036276 | 0.365982 |
| GOBP_BRANCHING_INVOLVED_IN_MAMMARY_GLAND_DUCT_MORPHOGENESIS | 18 | 0.579167 | 1.392581 | 0.092375 | 0.365653 |
| GOBP_POSITIVE_REGULATION_OF_INTERLEUKIN_12_PRODUCTION | 28 | 0.539453 | 1.391456 | 0.077975 | 0.368035 |
| GOBP_CELL_MATRIX_ADHESION | 198 | 0.418725 | 1.390745 | 0.013514 | 0.369103 |
| GOBP_MICROGLIAL_CELL_ACTIVATION | 28 | 0.530738 | 1.390534 | 0.078947 | 0.368873 |
| GOBP_NEGATIVE_REGULATION_OF_TRANSMEMBRANE_TRANSPORT | 91 | 0.451826 | 1.389645 | 0.025671 | 0.370441 |
| GOBP_REGULATION_OF_PEPTIDYL_SERINE_PHOSPHORYLATION | 114 | 0.437129 | 1.38948 | 0.032366 | 0.370059 |
| GOBP_RESPONSE_TO_MOLECULE_OF_BACTERIAL_ORIGIN | 220 | 0.417408 | 1.389296 | 0.012422 | 0.36973 |
| GOBP_POSITIVE_REGULATION_OF_LOCOMOTION | 466 | 0.404704 | 1.388369 | 0.001005 | 0.371458 |
| GOBP_POSITIVE_REGULATION_OF_TRANSMEMBRANE_TRANSPORT | 172 | 0.424241 | 1.387851 | 0.012931 | 0.37205 |
| GOBP_DENDRITIC_CELL_MIGRATION | 20 | 0.579219 | 1.387296 | 0.083212 | 0.372891 |
| GOBP_MITOCHONDRION_LOCALIZATION | 48 | 0.486178 | 1.387195 | 0.06879 | 0.372372 |
| GOBP_SEQUESTERING_OF_CALCIUM_ION | 96 | 0.445093 | 1.386997 | 0.039638 | 0.372082 |
| GOBP_REGULATION_OF_ACTION_POTENTIAL | 45 | 0.504448 | 1.386329 | 0.070423 | 0.373276 |
| GOBP_DIGESTION | 76 | 0.464479 | 1.38599 | 0.039627 | 0.373421 |
| GOBP_ACROSOME_REACTION | 26 | 0.545028 | 1.385417 | 0.093315 | 0.374207 |
| GOBP_MUSCLE_CONTRACTION | 277 | 0.411303 | 1.385103 | 0.006122 | 0.374234 |
| GOBP_B_CELL_ACTIVATION | 197 | 0.419851 | 1.384626 | 0.01046 | 0.374795 |
| GOBP_ICOSANOID_BIOSYNTHETIC_PROCESS | 32 | 0.529448 | 1.383783 | 0.079576 | 0.376309 |
| GOBP_MUSCLE_SYSTEM_PROCESS | 344 | 0.408454 | 1.383731 | 0.004049 | 0.375642 |
| GOBP_REGULATORY_T_CELL_DIFFERENTIATION | 25 | 0.548694 | 1.383466 | 0.085399 | 0.375593 |
| GOBP_MAMMARY_GLAND_EPITHELIUM_DEVELOPMENT | 57 | 0.476551 | 1.383184 | 0.065823 | 0.375644 |
| GOBP_NEGATIVE_REGULATION_OF_HYDROLASE_ACTIVITY | 244 | 0.412604 | 1.383096 | 0.013444 | 0.375059 |
| GOBP_POSITIVE_REGULATION_OF_ALPHA_BETA_T_CELL_DIFFERENTIATION | 36 | 0.506247 | 1.383024 | 0.077025 | 0.374455 |
| GOBP_POSITIVE_REGULATION_OF_ADAPTIVE_IMMUNE_RESPONSE | 83 | 0.44626 | 1.383023 | 0.046099 | 0.373648 |
| GOBP_NEGATIVE_REGULATION_OF_LIPID_STORAGE | 16 | 0.599732 | 1.382703 | 0.088698 | 0.373772 |
| GOBP_MACROPHAGE_ACTIVATION | 66 | 0.466539 | 1.38231 | 0.058894 | 0.374059 |
| GOBP_CYTOSOLIC_CALCIUM_ION_TRANSPORT | 145 | 0.428105 | 1.380976 | 0.021858 | 0.377255 |

| | | | | | |
|---|---|---|---|---|---|
| GOBP_NEGATIVE_REGULATION_OF_OSSIFICATION | 27 | -0.346083 | -1.07622 | 0.316176 | 0.961126 |
| GOBP_PROTEIN_INSERTION_INTO_MEMBRANE | 53 | -0.302288 | -1.076069 | 0.32093 | 0.958641 |
| GOBP_NUCLEOSOME_MOBILIZATION | 20 | -0.368547 | -1.074775 | 0.348592 | 0.960963 |
| GOBP_ASPARTATE_FAMILY_AMINO_ACID_METABOLIC_PROCESS | 45 | -0.308968 | -1.074174 | 0.317972 | 0.960167 |
| GOBP_FEMALE_MEIOTIC_NUCLEAR_DIVISION | 29 | -0.336745 | -1.073625 | 0.32971 | 0.959327 |
| GOBP_PORE_COMPLEX_ASSEMBLY | 21 | -0.367183 | -1.073396 | 0.338462 | 0.957208 |
| GOBP_POSTSYNAPTIC_SIGNAL_TRANSDUCTION | 31 | -0.337562 | -1.072209 | 0.343348 | 0.959036 |
| GOBP_PSEUDOURIDINE_SYNTHESIS | 18 | -0.385311 | -1.070212 | 0.376997 | 0.964392 |
| GOBP_SPINAL_CORD_MOTOR_NEURON_DIFFERENTIATION | 21 | -0.375071 | -1.068111 | 0.35514 | 0.970431 |
| GOBP_RETROGRADE_VESICLE_MEDIATED_TRANSPORT_GOLGI_TO_ENDOPLA | 48 | -0.312221 | -1.06277 | 0.364078 | 0.990302 |
| GOBP_DORSAL_VENTRAL_NEURAL_TUBE_PATTERNING | 19 | -0.367534 | -1.062748 | 0.379661 | 0.987303 |
| GOBP_MOTOR_NEURON_AXON_GUIDANCE | 28 | -0.341283 | -1.062364 | 0.329365 | 0.985829 |
| GOBP_CAMERA_TYPE_EYE_PHOTORECEPTOR_CELL_DIFFERENTIATION | 20 | -0.37714 | -1.06061 | 0.36014 | 0.99015 |
| GOBP_ADRENERGIC_RECEPTOR_SIGNALING_PATHWAY | 26 | -0.349111 | -1.060561 | 0.374545 | 0.987291 |
| GOBP_MITOCHONDRIAL_TRANSMEMBRANE_TRANSPORT | 99 | -0.268788 | -1.059793 | 0.367647 | 0.987749 |
| GOBP_NEURAL_TUBE_DEVELOPMENT | 143 | -0.250874 | -1.059665 | 0.273973 | 0.985213 |
| GOBP_REGULATION_OF_SYNAPSE_ASSEMBLY | 90 | -0.265109 | -1.05905 | 0.333333 | 0.984768 |
| GOBP_REGULATION_OF_EXECUTION_PHASE_OF_APOPTOSIS | 16 | -0.385992 | -1.058993 | 0.367742 | 0.981974 |
| GOBP_REGULATION_OF_TELOMERE_MAINTENANCE_VIA_TELOMERE_LENGTH | 58 | -0.294976 | -1.057971 | 0.339713 | 0.983296 |
| GOBP_REGULATION_OF_UBIQUITIN_PROTEIN_TRANSFERASE_ACTIVITY | 52 | -0.301798 | -1.057762 | 0.333333 | 0.981208 |
| GOBP_REGULATION_OF_HORMONE_METABOLIC_PROCESS | 31 | -0.340094 | -1.056083 | 0.349091 | 0.985106 |
| GOBP_SPLICEOSOMAL_COMPLEX_ASSEMBLY | 59 | -0.2955 | -1.055796 | 0.331633 | 0.983343 |
| GOBP_OLIGODENDROCYTE_DEVELOPMENT | 33 | -0.325935 | -1.055583 | 0.330961 | 0.981255 |
| GOBP_HEPARAN_SULFATE_PROTEOGLYCAN_BIOSYNTHETIC_PROCESS | 24 | -0.357212 | -1.0538 | 0.382637 | 0.985556 |
| GOBP_KILLING_OF_CELLS_OF_ANOTHER_ORGANISM | 15 | -0.392558 | -1.053465 | 0.384848 | 0.983984 |
| GOBP_DEOXYRIBONUCLEOSIDE_TRIPHOSPHATE_METABOLIC_PROCESS | 15 | -0.396742 | -1.051901 | 0.381703 | 0.987386 |
| GOBP_REGULATION_OF_EARLY_ENDOSOME_TO_LATE_ENDOSOME_TRANSP( | 18 | -0.381745 | -1.051135 | 0.37299 | 0.987645 |
| GOBP_CELLULAR_COMPONENT_MAINTENANCE | 57 | -0.290978 | -1.050648 | 0.364486 | 0.986652 |
| GOBP_CYTOSKELETON_DEPENDENT_CYTOKINESIS | 94 | -0.264482 | -1.050565 | 0.355372 | 0.984141 |
| GOBP_STEM_CELL_DIVISION | 24 | -0.352068 | -1.050426 | 0.375 | 0.981785 |
| GOBP_NEGATIVE_REGULATION_OF_INTRACELLULAR_TRANSPORT | 49 | -0.306198 | -1.04823 | 0.333333 | 0.988196 |
| GOBP_STEM_CELL_DIFFERENTIATION | 180 | -0.240088 | -1.04644 | 0.319444 | 0.992769 |
| GOBP_NEGATIVE_REGULATION_OF_UBIQUITIN_PROTEIN_TRANSFERASE_ACT | 22 | -0.356169 | -1.043928 | 0.368243 | 1 |
| GOBP_SODIUM_ION_TRANSMEMBRANE_TRANSPORT | 139 | -0.246271 | -1.043835 | 0.315789 | 0.997557 |
| GOBP_REGULATION_OF_NERVOUS_SYSTEM_DEVELOPMENT | 382 | -0.238057 | -1.042853 | 0.333333 | 0.998612 |
| GOBP_MODIFICATION_OF_SYNAPTIC_STRUCTURE | 20 | -0.36689 | -1.040366 | 0.361204 | 1 |
| GOBP_UBIQUITIN_DEPENDENT_ERAD_PATHWAY | 84 | -0.265329 | -1.038814 | 0.394366 | 1 |
| GOBP_POSITIVE_REGULATION_OF_DNA_TEMPLATED_TRANSCRIPTION_INITIAT | 42 | -0.295219 | -1.036576 | 0.391892 | 1 |
| GOBP_PYRIMIDINE_RIBONUCLEOTIDE_BIOSYNTHETIC_PROCESS | 22 | -0.35223 | -1.036529 | 0.389892 | 1 |
| GOBP_POSITIVE_REGULATION_OF_CALCIUM_ION_DEPENDENT_EXOCYTOSIS | 17 | -0.375338 | -1.036507 | 0.415625 | 1 |
| GOBP_POSITIVE_REGULATION_OF_TELOMERASE_RNA_LOCALIZATION_TO_CA | 15 | -0.384136 | -1.036439 | 0.385093 | 1 |
| GOBP_ESTABLISHMENT_OF_PROTEIN_LOCALIZATION_TO_ENDOPLASMIC_RET | 42 | -0.301507 | -1.036023 | 0.380567 | 1 |
| GOBP_ESTABLISHMENT_OF_PROTEIN_LOCALIZATION_TO_CHROMOSOME | 26 | -0.333905 | -1.035806 | 0.413559 | 1 |
| GOBP_PROTEIN_IMPORT_INTO_MITOCHONDRIAL_MATRIX | 20 | -0.355957 | -1.033992 | 0.383901 | 1 |
| GOBP_NEGATIVE_REGULATION_OF_OSTEOBLAST_DIFFERENTIATION | 40 | -0.303268 | -1.031398 | 0.392857 | 1 |
| GOBP_REGULATION_OF_INTRACELLULAR_ESTROGEN_RECEPTOR_SIGNALIN( | 29 | -0.325344 | -1.031244 | 0.384921 | 1 |
| GOBP_PROTEIN_LOCALIZATION_TO_NUCLEOLUS | 15 | -0.388876 | -1.031089 | 0.406154 | 1 |
| GOBP_RIBOSOME_ASSEMBLY | 60 | -0.28302 | -1.030273 | 0.419689 | 1 |
| GOBP_SRP_DEPENDENT_COTRANSLATIONAL_PROTEIN_TARGETING_TO_MEM | 18 | -0.363951 | -1.029963 | 0.373802 | 1 |
| GOBP_RESPONSE_TO_PLATELET_DERIVED_GROWTH_FACTOR | 23 | -0.353803 | -1.028992 | 0.393939 | 1 |
| GOBP_POSITIVE_REGULATION_OF_ISOTYPE_SWITCHING | 23 | -0.336984 | -1.028974 | 0.385455 | 1 |
| GOBP_CELL_JUNCTION_MAINTENANCE | 38 | -0.314332 | -1.028911 | 0.38565 | 1 |
| GOBP_SEX_DETERMINATION | 15 | -0.376568 | -1.028906 | 0.38024 | 1 |
| GOBP_RESPONSE_TO_ACETYLCHOLINE | 25 | -0.336474 | -1.028742 | 0.406143 | 1 |
| GOBP_REGULATION_OF_DNA_TEMPLATED_TRANSCRIPTION_INITIATION | 51 | -0.294007 | -1.027883 | 0.375546 | 1 |
| GOBP_POTASSIUM_ION_HOMEOSTASIS | 25 | -0.34084 | -1.027882 | 0.395683 | 1 |
| GOBP_PROTEIN_KINASE_C_ACTIVATING_G_PROTEIN_COUPLED_RECEPTOR_ | 22 | -0.352426 | -1.025355 | 0.386207 | 1 |
| GOBP_NUCLEOSIDE_DIPHOSPHATE_METABOLIC_PROCESS | 117 | -0.24757 | -1.024829 | 0.443182 | 1 |
| GOBP_SULFUR_AMINO_ACID_METABOLIC_PROCESS | 31 | -0.322801 | -1.021729 | 0.413919 | 1 |
| GOBP_POSITIVE_REGULATION_OF_NEURON_APOPTOTIC_PROCESS | 41 | -0.309733 | -1.02168 | 0.377049 | 1 |
| GOBP_AXON_DEVELOPMENT | 433 | -0.212516 | -1.019453 | 0.3 | 1 |
| GOBP_NEGATIVE_REGULATION_OF_OXIDATIVE_STRESS_INDUCED_NEURON_ | 17 | -0.370095 | -1.019396 | 0.413793 | 1 |
| GOBP_PHOSPHATIDYLINOSITOL_3_PHOSPHATE_BIOSYNTHETIC_PROCESS | 19 | -0.366011 | -1.018827 | 0.425249 | 1 |
| GOBP_OLIGOSACCHARIDE_METABOLIC_PROCESS | 46 | -0.295739 | -1.018662 | 0.387755 | 1 |
| GOBP_TRANSLATIONAL_ELONGATION | 53 | -0.28822 | -1.016607 | 0.38835 | 1 |
| GOBP_CRANIAL_NERVE_DEVELOPMENT | 38 | -0.303935 | -1.016262 | 0.426667 | 1 |
| GOBP_GLUTAMINE_FAMILY_AMINO_ACID_BIOSYNTHETIC_PROCESS | 16 | -0.360472 | -1.01395 | 0.418462 | 1 |
| GOBP_MOTILE_CILIUM_ASSEMBLY | 44 | -0.301676 | -1.013475 | 0.419643 | 1 |
| GOBP_REGULATION_OF_CHROMOSOME_ORGANIZATION | 201 | -0.231245 | -1.01274 | 0.425532 | 1 |
| GOBP_NADH_DEHYDROGENASE_COMPLEX_ASSEMBLY | 58 | -0.271558 | -1.012674 | 0.409357 | 1 |
| GOBP_NUCLEOTIDE_PHOSPHORYLATION | 87 | -0.260855 | -1.012648 | 0.379845 | 1 |
| GOBP_NEGATIVE_REGULATION_OF_CANONICAL_WNT_SIGNALING_PATHWAY | 123 | -0.242222 | -1.012388 | 0.486487 | 1 |
| GOBP_AXON_EXTENSION | 115 | -0.24917 | -1.01224 | 0.401709 | 1 |
| GOBP_OLIGODENDROCYTE_DIFFERENTIATION | 77 | -0.266142 | -1.011797 | 0.413408 | 1 |
| GOBP_REGULATION_OF_RECEPTOR_INTERNALIZATION | 53 | -0.280836 | -1.011329 | 0.40796 | 1 |
| GOBP_MITOTIC_CHROMOSOME_CONDENSATION | 19 | -0.356911 | -1.011113 | 0.424242 | 1 |
| GOBP_LENS_MORPHOGENESIS_IN_CAMERA_TYPE_EYE | 21 | -0.350986 | -1.010881 | 0.430976 | 1 |
| GOBP_SOMATIC_STEM_CELL_POPULATION_MAINTENANCE | 34 | -0.303603 | -1.008845 | 0.428 | 1 |
| GOBP_REGULATION_OF_MITOTIC_SPINDLE_ASSEMBLY | 23 | -0.341804 | -1.008305 | 0.425676 | 1 |
| GOBP_HISTONE_MONOUBIQUITINATION | 29 | -0.315508 | -1.008195 | 0.422053 | 1 |
| GOBP_REGULATION_OF_SYSTEMIC_ARTERIAL_BLOOD_PRESSURE | 68 | -0.269709 | -1.008153 | 0.434524 | 1 |
| GOBP_LATE_ENDOSOME_TO_VACUOLE_TRANSPORT_VIA_MULTIVESICULAR_ | 17 | -0.365088 | -1.007914 | 0.426866 | 1 |
| GOBP_STRIATED_MUSCLE_CELL_APOPTOTIC_PROCESS | 33 | -0.314392 | -1.006743 | 0.445256 | 1 |
| GOBP_LABYRINTHINE_LAYER_DEVELOPMENT | 40 | -0.301281 | -1.006377 | 0.454545 | 1 |
| GOBP_PROTEIN_MANNOSYLATION | 22 | -0.340579 | -1.005109 | 0.428571 | 1 |
| GOBP_POSITIVE_REGULATION_OF_UBIQUITIN_PROTEIN_TRANSFERASE_ACTI | 28 | -0.323021 | -1.004069 | 0.444828 | 1 |
| GOBP_PROTEIN_FOLDING | 185 | -0.233232 | -1.003533 | 0.485294 | 1 |
| GOBP_POSTSYNAPSE_ORGANIZATION | 147 | -0.234531 | -1.001874 | 0.43038 | 1 |
| GOBP_PYRIMIDINE_RIBONUCLEOTIDE_METABOLIC_PROCESS | 28 | -0.31625 | -1.000726 | 0.477444 | 1 |
| GOBP_POSITIVE_REGULATION_OF_TRANSLATIONAL_INITIATION | 23 | -0.333112 | -0.999092 | 0.45 | 1 |
| GOBP_ASPARTATE_FAMILY_AMINO_ACID_CATABOLIC_PROCESS | 16 | -0.382426 | -0.998403 | 0.441718 | 1 |
| GOBP_PROTEIN_LOCALIZATION_TO_ENDOPLASMIC_RETICULUM | 66 | -0.263574 | -0.997581 | 0.439306 | 1 |
| GOBP_REGULATION_OF_ADENYLATE_CYCLASE_ACTIVITY | 28 | -0.331082 | -0.99695 | 0.4375 | 1 |
| GOBP_REGULATION_OF_CARBOHYDRATE_CATABOLIC_PROCESS | 48 | -0.278733 | -0.996376 | 0.42381 | 1 |
| GOBP_POSTSYNAPTIC_SPECIALIZATION_ORGANIZATION | 31 | -0.30675 | -0.9963 | 0.459144 | 1 |
| GOBP_NEGATIVE_REGULATION_OF_STEM_CELL_DIFFERENTIATION | 22 | -0.336547 | -0.99369 | 0.445902 | 1 |
| GOBP_ERAD_PATHWAY | 106 | -0.242788 | -0.992881 | 0.504673 | 1 |
| GOBP_NEURON_PROJECTION_EXTENSION_INVOLVED_IN_NEURON_PROJECT | 37 | -0.290181 | -0.992608 | 0.443137 | 1 |
| GOBP_PRESYNAPSE_ORGANIZATION | 47 | -0.280323 | -0.991963 | 0.506224 | 1 |
| GOBP_NEGATIVE_REGULATION_OF_WNT_SIGNALING_PATHWAY | 152 | -0.234092 | -0.991681 | 0.467391 | 1 |
| GOBP_3_UTR_MEDIATED_MRNA_STABILIZATION | 19 | -0.34189 | -0.990492 | 0.456081 | 1 |
| GOBP_GANGLIOSIDE_METABOLIC_PROCESS | 25 | -0.326448 | -0.990332 | 0.44403 | 1 |
| GOBP_MRNA_CLEAVAGE | 18 | -0.357001 | -0.987809 | 0.45 | 1 |
| GOBP_RESPONSE_TO_HEPATOCYTE_GROWTH_FACTOR | 17 | -0.360621 | -0.987282 | 0.481848 | 1 |
| GOBP_NEURON_PROJECTION_EXTENSION | 161 | -0.228394 | -0.984345 | 0.517241 | 1 |
| GOBP_REGULATION_OF_CYTOPLASMIC_TRANSPORT | 27 | -0.322645 | -0.984216 | 0.478599 | 1 |
| GOBP_REGULATION_OF_GLYCOLYTIC_PROCESS | 39 | -0.297864 | -0.983173 | 0.461538 | 1 |
| GOBP_MEMBRANE_DEPOLARIZATION_DURING_ACTION_POTENTIAL | 32 | -0.308587 | -0.980423 | 0.487705 | 1 |
| GOBP_LIPOSACCHARIDE_METABOLIC_PROCESS | 95 | -0.252075 | -0.9802 | 0.528169 | 1 |
| GOBP_GLUTAMATE_RECEPTOR_SIGNALING_PATHWAY | 48 | -0.281319 | -0.979522 | 0.527778 | 1 |
| GOBP_PROTEIN_TARGETING_TO_MITOCHONDRION | 99 | -0.24773 | -0.978839 | 0.457627 | 1 |
| GOBP_MITOTIC_METAPHASE_PLATE_CONGRESSION | 52 | -0.276838 | -0.978561 | 0.51 | 1 |
| GOBP_POSITIVE_REGULATION_OF_DNA_BIOSYNTHETIC_PROCESS | 71 | -0.260082 | -0.978366 | 0.462963 | 1 |
| GOBP_SOMATIC_DIVERSIFICATION_OF_IMMUNE_RECEPTORS | 69 | -0.261964 | -0.976468 | 0.468927 | 1 |
| GOBP_CELLULAR_MODIFIED_AMINO_ACID_CATABOLIC_PROCESS | 23 | -0.32804 | -0.975981 | 0.475884 | 1 |
| GOBP_SYNAPTIC_VESICLE_CYTOSKELETAL_TRANSPORT | 19 | -0.334191 | -0.97528 | 0.473333 | 1 |
| GOBP_NCRNA_TRANSCRIPTION | 54 | -0.27036 | -0.974933 | 0.512195 | 1 |
| GOBP_POSITIVE_REGULATION_OF_OLIGODENDROCYTE_DIFFERENTIATION | 18 | -0.356206 | -0.9732 | 0.513678 | 1 |
| GOBP_PHOSPHATIDYLGLYCEROL_METABOLIC_PROCESS | 27 | -0.310731 | -0.972298 | 0.485185 | 1 |
| GOBP_PROTEIN_CONTAINING_COMPLEX_LOCALIZATION | 150 | -0.230302 | -0.969598 | 0.6 | 1 |
| GOBP_REGULATION_OF_ACTIN_FILAMENT_BUNDLE_ASSEMBLY | 96 | -0.244089 | -0.969327 | 0.543307 | 1 |
| GOBP_AXONEME_ASSEMBLY | 68 | -0.262518 | -0.968676 | 0.484472 | 1 |
| GOBP_RIBONUCLEOPROTEIN_COMPLEX_SUBUNIT_ORGANIZATION | 184 | -0.224518 | -0.96658 | 0.55 | 1 |
| GOBP_HEMATOPOIETIC_STEM_CELL_DIFFERENTIATION | 24 | -0.323362 | -0.965528 | 0.498361 | 1 |
| GOBP_REGULATION_OF_NMDA_RECEPTOR_ACTIVITY | 17 | -0.356241 | -0.965459 | 0.482972 | 1 |
| GOBP_PROTEIN_TRANSMEMBRANE_IMPORT_INTO_INTRACELLULAR_ORGANE | 36 | -0.296946 | -0.964683 | 0.468254 | 1 |
| GOBP_PROTON_TRANSMEMBRANE_TRANSPORT | 131 | -0.23364 | -0.963313 | 0.571429 | 1 |
| GOBP_CILIUM_ORGANIZATION | 360 | -0.206282 | -0.961363 | 0.785714 | 1 |
| GOBP_NCRNA_PROCESSING | 373 | -0.203772 | -0.961081 | 0.4375 | 1 |
| GOBP_TELOMERE_MAINTENANCE_VIA_TELOMERE_LENGTHENING | 76 | -0.252723 | -0.958333 | 0.514451 | 1 |
| GOBP_OUTER_DYNEIN_ARM_ASSEMBLY | 15 | -0.366193 | -0.958309 | 0.51895 | 1 |
| GOBP_DENDRITIC_SPINE_DEVELOPMENT | 88 | -0.247662 | -0.95828 | 0.57047 | 1 |
| GOBP_MICROTUBULE_DEPOLYMERIZATION | 45 | -0.277146 | -0.957417 | 0.561321 | 1 |
| GOBP_NON_MOTILE_CILIUM_ASSEMBLY | 59 | -0.259299 | -0.955713 | 0.538462 | 1 |
| GOBP_CELLULAR_MODIFIED_AMINO_ACID_BIOSYNTHETIC_PROCESS | 42 | -0.273723 | -0.954913 | 0.487437 | 1 |
| GOBP_CYTOPLASMIC_TRANSLATION | 146 | -0.224017 | -0.952796 | 0.631579 | 1 |
| GOBP_MULTIVESICULAR_BODY_ORGANIZATION | 31 | -0.298734 | -0.952773 | 0.546219 | 1 |
| GOBP_PROTEIN_LOCALIZATION_TO_ENDOSOME | 24 | -0.320981 | -0.952588 | 0.540541 | 1 |
| GOBP_MICROTUBULE_BUNDLE_FORMATION | 98 | -0.240459 | -0.952485 | 0.577236 | 1 |
| GOBP_NEGATIVE_REGULATION_OF_ADENYLATE_CYCLASE_ACTIVITY | 17 | -0.339021 | -0.94858 | 0.515528 | 1 |
| GOBP_REGULATION_OF_RECEPTOR_LOCALIZATION_TO_SYNAPSE | 16 | -0.358935 | -0.948255 | 0.490683 | 1 |
| GOBP_REGULATION_OF_CELLULAR_PH | 71 | -0.253292 | -0.948053 | 0.584416 | 1 |
| GOBP_REGULATION_OF_AMYLOID_PRECURSOR_PROTEIN_CATABOLIC_PROC | 37 | -0.287671 | -0.948043 | 0.557312 | 1 |
| GOBP_NEURON_PROJECTION_REGENERATION | 49 | -0.273441 | -0.945964 | 0.527094 | 1 |
| GOBP_POSITIVE_REGULATION_OF_ENDOCYTOSIS | 80 | -0.246408 | -0.944649 | 0.596027 | 1 |
| GOBP_NEGATIVE_REGULATION_OF_GLIAL_CELL_DIFFERENTIATION | 18 | -0.340315 | -0.944425 | 0.508834 | 1 |
| GOBP_PEPTIDYL_CYSTEINE_MODIFICATION | 44 | -0.274599 | -0.94422 | 0.584746 | 1 |
| GOBP_CARBOHYDRATE_CATABOLIC_PROCESS | 125 | -0.22689 | -0.94336 | 0.625 | 1 |
| GOBP_IRE1_MEDIATED_UNFOLDED_PROTEIN_RESPONSE | 18 | -0.340118 | -0.941744 | 0.527331 | 1 |
| GOBP_SPERM_FLAGELLUM_ASSEMBLY | 24 | -0.310439 | -0.939227 | 0.543919 | 1 |
| GOBP_GLYCOSPHINGOLIPID_BIOSYNTHETIC_PROCESS | 30 | -0.293434 | -0.937519 | 0.536232 | 1 |
| GOBP_REGULATION_OF_SYSTEMIC_ARTERIAL_BLOOD_PRESSURE_BY_HORM | 24 | -0.310313 | -0.937452 | 0.561069 | 1 |
| GOBP_REGULATION_OF_HIPPO_SIGNALING | 20 | -0.326565 | -0.936135 | 0.509868 | 1 |
| GOBP_CEREBELLAR_CORTEX_DEVELOPMENT | 47 | -0.269581 | -0.935604 | 0.542601 | 1 |
| GOBP_PYRUVATE_METABOLIC_PROCESS | 89 | -0.234509 | -0.934646 | 0.638298 | 1 |
| GOBP_SYNAPTIC_VESICLE_MEMBRANE_ORGANIZATION | 23 | -0.314272 | -0.934073 | 0.551495 | 1 |
| GOBP_RNA_DEPENDENT_DNA_BIOSYNTHETIC_PROCESS | 68 | -0.249676 | -0.933345 | 0.576923 | 1 |

| | | | | | |
|---|---|---|---|---|---|
| GOBP_TROPHECTODERMAL_CELL_DIFFERENTIATION | 16 | 0.593514 | 1.379893 | 0.095442 | 0.379506 |
| GOBP_MULTI_MULTICELLULAR_ORGANISM_PROCESS | 155 | 0.425939 | 1.379816 | 0.012807 | 0.378891 |
| GOBP_POSITIVE_REGULATION_OF_MYELOID_CELL_DIFFERENTIATION | 77 | 0.458164 | 1.379807 | 0.060355 | 0.378114 |
| GOBP_DETECTION_OF_VISIBLE_LIGHT | 23 | 0.56191 | 1.378183 | 0.107955 | 0.381904 |
| GOBP_POSITIVE_REGULATION_OF_LYMPHOCYTE_DIFFERENTIATION | 77 | 0.458607 | 1.37709 | 0.03494 | 0.384135 |
| GOBP_POSITIVE_REGULATION_OF_ION_TRANSMEMBRANE_TRANSPORT | 132 | 0.425691 | 1.376649 | 0.040087 | 0.38446 |
| GOBP_NEGATIVE_REGULATION_OF_ENDOTHELIAL_CELL_PROLIFERATION | 35 | 0.516761 | 1.376197 | 0.078844 | 0.384925 |
| GOBP_REGULATION_OF_ENDOTHELIAL_CELL_CHEMOTAXIS | 19 | 0.577567 | 1.374721 | 0.105409 | 0.388212 |
| GOBP_RESPONSE_TO_INTERFERON_GAMMA | 92 | 0.448145 | 1.374579 | 0.054374 | 0.387793 |
| GOBP_NUCLEAR_MIGRATION | 21 | 0.577017 | 1.374208 | 0.089489 | 0.388059 |
| GOBP_COLLAGEN_BIOSYNTHETIC_PROCESS | 35 | 0.514142 | 1.372902 | 0.063802 | 0.390987 |
| GOBP_INTERMEDIATE_FILAMENT_ORGANIZATION | 18 | 0.576042 | 1.372896 | 0.120766 | 0.390191 |
| GOBP_REGULATION_OF_CHEMOTAXIS | 165 | 0.423924 | 1.372445 | 0.01594 | 0.390632 |
| GOBP_POSITIVE_REGULATION_OF_CYTOKINE_PRODUCTION_INVOLVED_IN_IMMUNE_RES | 50 | 0.484905 | 1.372426 | 0.069913 | 0.389862 |
| GOBP_REGULATION_OF_LEUKOCYTE_DIFFERENTIATION | 207 | 0.414785 | 1.372379 | 0.018848 | 0.389206 |
| GOBP_POSITIVE_REGULATION_OF_COAGULATION | 18 | 0.575276 | 1.37111 | 0.109012 | 0.391966 |
| GOBP_ENDODERMAL_CELL_DIFFERENTIATION | 40 | 0.498386 | 1.371005 | 0.063554 | 0.391443 |
| GOBP_NEGATIVE_REGULATION_OF_BONE_REMODELING | 15 | 0.594 | 1.370714 | 0.113095 | 0.391509 |
| GOBP_TEMPERATURE_HOMEOSTASIS | 147 | 0.421271 | 1.370203 | 0.022247 | 0.392142 |
| GOBP_REGULATION_OF_CARBOHYDRATE_BIOSYNTHETIC_PROCESS | 82 | 0.453459 | 1.370166 | 0.058754 | 0.391419 |
| GOBP_SMOOTH_MUSCLE_CONTRACTION | 93 | 0.441131 | 1.369212 | 0.043182 | 0.393217 |
| GOBP_RESPONSE_TO_AMINE | 39 | 0.500556 | 1.367822 | 0.085938 | 0.396367 |
| GOBP_RESPONSE_TO_WOUNDING | 446 | 0.397833 | 1.367642 | 0.001009 | 0.396148 |
| GOBP_ENDOCRINE_PANCREAS_DEVELOPMENT | 33 | 0.521484 | 1.366947 | 0.085938 | 0.397352 |
| GOBP_CIRCADIAN_SLEEP_WAKE_CYCLE | 16 | 0.590737 | 1.366386 | 0.099851 | 0.398118 |
| GOBP_NEGATIVE_REGULATION_OF_CD4_POSITIVE_ALPHA_BETA_T_CELL_ACTIVATION | 22 | 0.559331 | 1.366343 | 0.110957 | 0.397452 |
| GOBP_POSITIVE_REGULATION_OF_T_CELL_CYTOKINE_PRODUCTION | 19 | 0.55864 | 1.366113 | 0.117479 | 0.397343 |
| GOBP_NEGATIVE_REGULATION_OF_EPITHELIAL_CELL_DIFFERENTIATION | 34 | 0.512286 | 1.366051 | 0.104478 | 0.396728 |
| GOBP_BLOOD_VESSEL_ENDOTHELIAL_CELL_PROLIFERATION_INVOLVED_IN_SPROUTING_ | 21 | 0.55467 | 1.365898 | 0.108538 | 0.396388 |
| GOBP_XENOBIOTIC_CATABOLIC_PROCESS | 17 | 0.585975 | 1.365636 | 0.104956 | 0.396318 |
| GOBP_POSITIVE_REGULATION_OF_PEPTIDYL_THREONINE_PHOSPHORYLATION | 26 | 0.53623 | 1.365541 | 0.088643 | 0.395765 |
| GOBP_ACTIVATION_OF_CYSTEINE_TYPE_ENDOPEPTIDASE_ACTIVITY_INVOLVED_IN_APOP | 68 | 0.46335 | 1.365487 | 0.06057 | 0.395108 |
| GOBP_POSITIVE_REGULATION_OF_CARBOHYDRATE_METABOLIC_PROCESS | 61 | 0.461264 | 1.365257 | 0.067402 | 0.394952 |
| GOBP_POSITIVE_REGULATION_OF_EPITHELIAL_CELL_APOPTOTIC_PROCESS | 24 | 0.536793 | 1.365222 | 0.109397 | 0.39425 |
| GOBP_NEGATIVE_REGULATION_OF_INTERLEUKIN_2_PRODUCTION | 20 | 0.563192 | 1.365101 | 0.113235 | 0.393815 |
| GOBP_GERM_CELL_DEVELOPMENT | 213 | 0.40735 | 1.364982 | 0.017617 | 0.393377 |
| GOBP_INTEGRIN_MEDIATED_SIGNALING_PATHWAY | 93 | 0.43896 | 1.364765 | 0.042334 | 0.39319 |
| GOBP_VASODILATION | 37 | 0.510061 | 1.364661 | 0.086735 | 0.392709 |
| GOBP_BONE_MORPHOGENESIS | 81 | 0.450193 | 1.364427 | 0.048894 | 0.39262 |
| GOBP_REGULATION_OF_LYMPHOCYTE_DIFFERENTIATION | 130 | 0.426335 | 1.363822 | 0.035516 | 0.39352 |
| GOBP_GLYCOSYL_COMPOUND_METABOLIC_PROCESS | 76 | 0.449353 | 1.363672 | 0.061466 | 0.393101 |
| GOBP_POSITIVE_REGULATION_OF_MITOCHONDRION_ORGANIZATION | 66 | 0.457633 | 1.363412 | 0.088095 | 0.393074 |
| GOBP_OSTEOCLAST_DEVELOPMENT | 15 | 0.596862 | 1.362919 | 0.117647 | 0.393684 |
| GOBP_POSITIVE_REGULATION_OF_CALCIUM_ION_TRANSPORT_INTO_CYTOSOL | 41 | 0.496687 | 1.362902 | 0.084321 | 0.392963 |
| GOBP_PLATELET_ACTIVATION | 100 | 0.431127 | 1.362612 | 0.05079 | 0.393085 |
| GOBP_REGULATION_OF_BONE_RESORPTION | 31 | 0.51781 | 1.362262 | 0.094911 | 0.393332 |
| GOBP_REGULATION_OF_T_CELL_MEDIATED_CYTOTOXICITY | 23 | 0.549865 | 1.361886 | 0.082504 | 0.393611 |
| GOBP_ANTIGEN_PROCESSING_AND_PRESENTATION | 77 | 0.448753 | 1.361877 | 0.055952 | 0.392869 |
| GOBP_MALE_GAMETE_GENERATION | 401 | 0.397865 | 1.361767 | 0.004032 | 0.392443 |
| GOBP_B_CELL_PROLIFERATION | 52 | 0.466941 | 1.360926 | 0.072152 | 0.394121 |
| GOBP_EMBRYONIC_HINDLIMB_MORPHOGENESIS | 19 | 0.566682 | 1.360283 | 0.099707 | 0.395177 |
| GOBP_POSITIVE_REGULATION_OF_LEUKOCYTE_MEDIATED_IMMUNITY | 92 | 0.442713 | 1.359715 | 0.052326 | 0.395996 |
| GOBP_UNSATURATED_FATTY_ACID_METABOLIC_PROCESS | 81 | 0.449862 | 1.359664 | 0.050648 | 0.395369 |
| GOBP_HEMATOPOIETIC_STEM_CELL_HOMEOSTASIS | 19 | 0.559469 | 1.358892 | 0.115385 | 0.396893 |
| GOBP_REGULATION_OF_EPITHELIAL_CELL_MIGRATION | 205 | 0.410927 | 1.358752 | 0.020942 | 0.396514 |
| GOBP_EYE_PHOTORECEPTOR_CELL_DEVELOPMENT | 26 | 0.538767 | 1.358698 | 0.101813 | 0.395891 |
| GOBP_DETECTION_OF_ABIOTIC_STIMULUS | 91 | 0.439381 | 1.358343 | 0.058411 | 0.396115 |
| GOBP_MYELOID_LEUKOCYTE_DIFFERENTIATION | 164 | 0.413601 | 1.358011 | 0.031385 | 0.396275 |
| GOBP_REGULATION_OF_MAST_CELL_ACTIVATION | 30 | 0.513037 | 1.357722 | 0.097394 | 0.39635 |
| GOBP_INTERLEUKIN_2_PRODUCTION | 46 | 0.48166 | 1.356741 | 0.0725 | 0.398333 |
| GOBP_CYTOKINE_MEDIATED_SIGNALING_PATHWAY | 311 | 0.398894 | 1.356099 | 0.00715 | 0.399308 |
| GOBP_GLAND_MORPHOGENESIS | 97 | 0.433298 | 1.354914 | 0.051843 | 0.401868 |
| GOBP_SYNAPTIC_TRANSMISSION_DOPAMINERGIC | 23 | 0.539804 | 1.354309 | 0.099719 | 0.402898 |
| GOBP_POSITIVE_REGULATION_OF_DEFENSE_RESPONSE_TO_VIRUS_BY_HOST | 24 | 0.533304 | 1.354208 | 0.106849 | 0.402429 |
| GOBP_NEGATIVE_REGULATION_OF_CALCIUM_ION_TRANSPORT | 47 | 0.483977 | 1.353285 | 0.068921 | 0.404279 |
| GOBP_NEGATIVE_REGULATION_OF_REGULATED_SECRETORY_PATHWAY | 19 | 0.560804 | 1.35299 | 0.120815 | 0.404363 |
| GOBP_RESPONSE_TO_TYPE_I_INTERFERON | 54 | 0.471253 | 1.352419 | 0.076142 | 0.405265 |
| GOBP_DETECTION_OF_EXTERNAL_BIOTIC_STIMULUS | 15 | 0.596113 | 1.352314 | 0.10503 | 0.404773 |
| GOBP_ACTIVATED_T_CELL_PROLIFERATION | 32 | 0.511654 | 1.351794 | 0.094595 | 0.405464 |
| GOBP_NEGATIVE_REGULATION_OF_CELL_CELL_ADHESION | 133 | 0.42409 | 1.351246 | 0.046002 | 0.406364 |
| GOBP_BONE_DEVELOPMENT | 182 | 0.41566 | 1.351023 | 0.020364 | 0.406305 |
| GOBP_NEGATIVE_REGULATION_OF_RESPONSE_TO_WOUNDING | 68 | 0.45406 | 1.350439 | 0.055419 | 0.407249 |
| GOBP_RESPONSE_TO_INTERFERON_BETA | 21 | 0.562214 | 1.350365 | 0.11731 | 0.406736 |
| GOBP_POSITIVE_REGULATION_OF_VASOCONSTRICTION | 24 | 0.535414 | 1.350257 | 0.122535 | 0.406308 |
| GOBP_NEGATIVE_REGULATION_OF_SODIUM_ION_TRANSPORT | 17 | 0.576477 | 1.350194 | 0.113043 | 0.40575 |
| GOBP_MEMORY | 110 | 0.426934 | 1.349974 | 0.047458 | 0.405691 |
| GOBP_MYOTUBE_DIFFERENTIATION | 93 | 0.433657 | 1.349879 | 0.064327 | 0.405231 |
| GOBP_RESPONSE_TO_CORTICOSTEROID | 120 | 0.427265 | 1.349717 | 0.048619 | 0.404958 |
| GOBP_POSITIVE_REGULATION_OF_DOUBLE_STRAND_BREAK_REPAIR_VIA_HOMOLOGOUS | 36 | 0.506817 | 1.349675 | 0.094805 | 0.404347 |
| GOBP_OUTFLOW_TRACT_MORPHOGENESIS | 73 | 0.455543 | 1.348748 | 0.058824 | 0.406221 |
| GOBP_CARDIAC_NEURAL_CREST_CELL_DIFFERENTIATION_INVOLVED_IN_HEART_DEVEL | 15 | 0.576862 | 1.348512 | 0.135095 | 0.406172 |
| GOBP_STRIATED_MUSCLE_CELL_DIFFERENTIATION | 214 | 0.405113 | 1.34835 | 0.018104 | 0.4059 |
| GOBP_AMEBOIDAL_TYPE_CELL_MIGRATION | 377 | 0.396656 | 1.348249 | 0.002033 | 0.405447 |
| GOBP_REGULATION_OF_FILOPODIUM_ASSEMBLY | 47 | 0.479225 | 1.347975 | 0.094578 | 0.405547 |
| GOBP_POSITIVE_REGULATION_OF_ERK1_AND_ERK2_CASCADE | 148 | 0.416058 | 1.347849 | 0.03433 | 0.405165 |
| GOBP_CELLULAR_RESPONSE_TO_BIOTIC_STIMULUS | 159 | 0.413379 | 1.347819 | 0.030205 | 0.404522 |
| GOBP_POSITIVE_REGULATION_OF_EPITHELIAL_CELL_MIGRATION | 138 | 0.414857 | 1.346798 | 0.048098 | 0.406593 |
| GOBP_DEFENSE_RESPONSE_TO_GRAM_NEGATIVE_BACTERIUM | 31 | 0.50663 | 1.346741 | 0.101351 | 0.405995 |
| GOBP_NEGATIVE_REGULATION_OF_CYSTEINE_TYPE_ENDOPEPTIDASE_ACTIVITY | 66 | 0.452758 | 1.346506 | 0.074341 | 0.40593 |
| GOBP_COLLAGEN_FIBRIL_ORGANIZATION | 55 | 0.470024 | 1.345623 | 0.070197 | 0.407605 |
| GOBP_NEGATIVE_REGULATION_OF_ION_TRANSMEMBRANE_TRANSPORTER_ACTIVITY | 52 | 0.468646 | 1.345383 | 0.088832 | 0.407617 |
| GOBP_NEGATIVE_REGULATION_OF_VIRAL_PROCESS | 73 | 0.450392 | 1.344925 | 0.07177 | 0.408151 |
| GOBP_IMMUNE_RESPONSE_REGULATING_SIGNALING_PATHWAY | 299 | 0.400243 | 1.344642 | 0.008138 | 0.408202 |
| GOBP_REGULATION_OF_SPROUTING_ANGIOGENESIS | 35 | 0.495774 | 1.344569 | 0.090674 | 0.407679 |
| GOBP_CELL_COMMUNICATION_BY_ELECTRICAL_COUPLING_INVOLVED_IN_CARDIAC_CON | 26 | 0.52145 | 1.34425 | 0.106901 | 0.407928 |
| GOBP_REGULATION_OF_REPRODUCTIVE_PROCESS | 126 | 0.424369 | 1.343431 | 0.038932 | 0.409722 |
| GOBP_ANGIOGENESIS_INVOLVED_IN_WOUND_HEALING | 26 | 0.521624 | 1.34268 | 0.095833 | 0.411175 |
| GOBP_CELLULAR_PROCESS_INVOLVED_IN_REPRODUCTION_IN_MULTICELLULAR_ORGAN | 287 | 0.39993 | 1.342536 | 0.00818 | 0.410839 |
| GOBP_POSITIVE_REGULATION_OF_FIBROBLAST_PROLIFERATION | 37 | 0.490317 | 1.341912 | 0.102632 | 0.411863 |
| GOBP_APOPTOTIC_PROCESS_INVOLVED_IN_DEVELOPMENT | 37 | 0.492005 | 1.341706 | 0.098318 | 0.411769 |
| GOBP_ENDOTHELIAL_CELL_PROLIFERATION | 130 | 0.416625 | 1.341577 | 0.048246 | 0.411381 |
| GOBP_CARDIAC_SEPTUM_MORPHOGENESIS | 63 | 0.458805 | 1.341381 | 0.087782 | 0.411297 |
| GOBP_RESPONSE_TO_SALT_STRESS | 21 | 0.541321 | 1.340935 | 0.110465 | 0.411789 |
| GOBP_RESPONSE_TO_PAIN | 27 | 0.517965 | 1.340069 | 0.109397 | 0.413509 |
| GOBP_CELLULAR_RESPONSE_TO_OXYGEN_RADICAL | 20 | 0.558573 | 1.339892 | 0.127956 | 0.41337 |
| GOBP_PHOTORECEPTOR_CELL_DEVELOPMENT | 38 | 0.49409 | 1.339703 | 0.101469 | 0.413177 |
| GOBP_REGULATION_OF_ACTIVIN_RECEPTOR_SIGNALING_PATHWAY | 22 | 0.537404 | 1.339322 | 0.12028 | 0.413551 |
| GOBP_SENSORY_PERCEPTION_OF_MECHANICAL_STIMULUS | 140 | 0.412966 | 1.33857 | 0.04185 | 0.415028 |
| GOBP_NEGATIVE_REGULATION_OF_SIGNAL_TRANSDUCTION_IN_ABSENCE_OF_LIGAND | 25 | 0.52715 | 1.338401 | 0.140647 | 0.414806 |
| GOBP_VITAMIN_BIOSYNTHETIC_PROCESS | 17 | 0.56489 | 1.338242 | 0.123839 | 0.414494 |
| GOBP_REGULATION_OF_EPITHELIAL_CELL_DIFFERENTIATION | 115 | 0.423617 | 1.33762 | 0.046067 | 0.415555 |
| GOBP_REGULATION_OF_NITRIC_OXIDE_METABOLIC_PROCESS | 42 | 0.484084 | 1.337593 | 0.106021 | 0.41491 |
| GOBP_MULTICELLULAR_ORGANISMAL_MOVEMENT | 44 | 0.478844 | 1.337238 | 0.100645 | 0.415189 |
| GOBP_RESPONSE_TO_ETHANOL | 91 | 0.438663 | 1.337106 | 0.053864 | 0.414873 |
| GOBP_REGULATION_OF_T_HELPER_CELL_DIFFERENTIATION | 26 | 0.531414 | 1.336318 | 0.127956 | 0.416395 |
| GOBP_DEFENSE_RESPONSE_TO_FUNGUS | 17 | 0.572181 | 1.336093 | 0.13985 | 0.416281 |
| GOBP_FORELIMB_MORPHOGENESIS | 21 | 0.547786 | 1.335421 | 0.132022 | 0.417496 |
| GOBP_REGULATION_OF_NATURAL_KILLER_CELL_MEDIATED_IMMUNITY | 24 | 0.524294 | 1.333489 | 0.125356 | 0.422378 |
| GOBP_DENDRITIC_CELL_DIFFERENTIATION | 34 | 0.502752 | 1.332989 | 0.096774 | 0.423179 |
| GOBP_AMINO_SUGAR_METABOLIC_PROCESS | 36 | 0.489531 | 1.332467 | 0.096306 | 0.423959 |
| GOBP_DEMETHYLATION | 60 | 0.456085 | 1.332311 | 0.079853 | 0.423671 |
| GOBP_CARDIOCYTE_DIFFERENTIATION | 115 | 0.424185 | 1.332087 | 0.045506 | 0.423565 |
| GOBP_REGULATION_OF_VESICLE_FUSION | 20 | 0.556559 | 1.332084 | 0.120739 | 0.422853 |
| GOBP_NEGATIVE_REGULATION_OF_ALPHA_BETA_T_CELL_DIFFERENTIATION | 19 | 0.555318 | 1.332052 | 0.145954 | 0.422233 |
| GOBP_MAINTENANCE_OF_BLOOD_BRAIN_BARRIER | 32 | 0.498695 | 1.331765 | 0.105263 | 0.422308 |
| GOBP_CELL_SUBSTRATE_ADHESION | 314 | 0.393149 | 1.331324 | 0.013252 | 0.422838 |
| GOBP_NEGATIVE_REGULATION_OF_CYTOKINE_PRODUCTION | 193 | 0.403879 | 1.331021 | 0.033862 | 0.422992 |
| GOBP_CELLULAR_RESPONSE_TO_MOLECULE_OF_BACTERIAL_ORIGIN | 133 | 0.419188 | 1.330767 | 0.046205 | 0.423041 |
| GOBP_EATING_BEHAVIOR | 26 | 0.51855 | 1.330684 | 0.113233 | 0.42255 |
| GOBP_REGULATION_OF_ERBB_SIGNALING_PATHWAY | 67 | 0.45435 | 1.330436 | 0.086747 | 0.422579 |
| GOBP_EPITHELIAL_CELL_DIFFERENTIATION_INVOLVED_IN_KIDNEY_DEVELOPMENT | 42 | 0.477822 | 1.330059 | 0.109114 | 0.422993 |
| GOBP_PRODUCTION_OF_MOLECULAR_MEDIATOR_OF_IMMUNE_RESPONSE | 157 | 0.417261 | 1.329549 | 0.041439 | 0.42379 |
| GOBP_PLATELET_AGGREGATION | 55 | 0.457945 | 1.328668 | 0.093674 | 0.425531 |
| GOBP_CELL_PROLIFERATION_INVOLVED_IN_HEART_MORPHOGENESIS | 16 | 0.569778 | 1.328089 | 0.141176 | 0.426561 |
| GOBP_MEIOTIC_CELL_CYCLE | 209 | 0.40141 | 1.327905 | 0.025237 | 0.426377 |
| GOBP_ENDOCARDIAL_CUSHION_MORPHOGENESIS | 36 | 0.483722 | 1.325974 | 0.108401 | 0.431256 |
| GOBP_CHEMOKINE_PRODUCTION | 63 | 0.455144 | 1.324926 | 0.089242 | 0.433599 |
| GOBP_REGULATION_OF_MAST_CELL_ACTIVATION_INVOLVED_IN_IMMUNE_RESPONSE | 26 | 0.52388 | 1.324806 | 0.112925 | 0.433209 |
| GOBP_MEGAKARYOCYTE_DEVELOPMENT | 15 | 0.575334 | 1.322853 | 0.147761 | 0.438226 |
| GOBP_STRIATED_MUSCLE_ADAPTATION | 34 | 0.483409 | 1.322506 | 0.124157 | 0.438529 |
| GOBP_REGULATION_OF_CELL_SUBSTRATE_ADHESION | 183 | 0.40167 | 1.321888 | 0.04492 | 0.439729 |
| GOBP_RESPONSE_TO_MECHANICAL_STIMULUS | 167 | 0.404304 | 1.32156 | 0.032189 | 0.440038 |
| GOBP_REGULATION_OF_MEIOTIC_CELL_CYCLE | 38 | 0.482925 | 1.321401 | 0.122503 | 0.439816 |
| GOBP_REGULATION_OF_PEPTIDASE_ACTIVITY | 327 | 0.394153 | 1.321128 | 0.010256 | 0.439934 |
| GOBP_NEUTRAL_LIPID_CATABOLIC_PROCESS | 29 | 0.518535 | 1.321053 | 0.122283 | 0.439456 |
| GOBP_AMINE_TRANSPORT | 79 | 0.427343 | 1.318631 | 0.07907 | 0.445772 |
| GOBP_NEGATIVE_REGULATION_OF_VASCULATURE_DEVELOPMENT | 82 | 0.4366 | 1.317547 | 0.08024 | 0.448439 |
| GOBP_TRIGLYCERIDE_METABOLIC_PROCESS | 75 | 0.432316 | 1.31697 | 0.070755 | 0.449468 |
| GOBP_REGULATION_OF_ENDOTHELIAL_CELL_MIGRATION | 150 | 0.408352 | 1.316725 | 0.039911 | 0.449452 |
| GOBP_NEGATIVE_REGULATION_OF_TRANSPORTER_ACTIVITY | 59 | 0.451233 | 1.316516 | 0.085242 | 0.449338 |
| GOBP_FATTY_ACID_BIOSYNTHETIC_PROCESS | 118 | 0.418966 | 1.316233 | 0.065022 | 0.449496 |
| GOBP_DEFENSE_RESPONSE_TO_SYMBIONT | 216 | 0.399393 | 1.315994 | 0.034737 | 0.44948 |
| GOBP_REGULATION_OF_CALCIUM_ION_IMPORT | 32 | 0.498031 | 1.31574 | 0.12786 | 0.449513 |
| GOBP_MYELOID_CELL_HOMEOSTASIS | 143 | 0.413038 | 1.315509 | 0.050756 | 0.449438 |
| GOBP_DNA_DAMAGE_RESPONSE_SIGNAL_TRANSDUCTION_BY_P53_CLASS_MEDIATOR | 67 | 0.441449 | 1.315472 | 0.083028 | 0.448808 |
| GOBP_PHYSIOLOGICAL_CARDIAC_MUSCLE_HYPERTROPHY | 16 | 0.575085 | 1.315427 | 0.143472 | 0.448221 |
| GOBP_OVULATION | 19 | 0.549266 | 1.314862 | 0.136827 | 0.449173 |

| | | | | | |
|---|---|---|---|---|---|
| GOBP_NEGATIVE_REGULATION_OF_LIPID_CATABOLIC_PROCESS | 16 | -0.345461 | -0.931932 | 0.548485 | 1 |
| GOBP_NEUROMUSCULAR_PROCESS_CONTROLLING_POSTURE | 15 | -0.355956 | -0.930862 | 0.546547 | 1 |
| GOBP_DICARBOXYLIC_ACID_METABOLIC_PROCESS | 83 | -0.237563 | -0.930389 | 0.613793 | 1 |
| GOBP_VACUOLAR_ACIDIFICATION | 22 | -0.31921 | -0.930207 | 0.55 | 1 |
| GOBP_SYNAPTIC_VESICLE_TRANSPORT | 39 | -0.272872 | -0.930027 | 0.590164 | 1 |
| GOBP_SULFUR_COMPOUND_BIOSYNTHETIC_PROCESS | 129 | -0.225995 | -0.929717 | 0.680412 | 1 |
| GOBP_REGULATION_OF_SISTER_CHROMATID_COHESION | 21 | -0.314442 | -0.929208 | 0.580858 | 1 |
| GOBP_PROTEIN_PALMITOYLATION | 31 | -0.29556 | -0.92896 | 0.538732 | 1 |
| GOBP_SUBPALLIUM_DEVELOPMENT | 19 | -0.348681 | -0.928813 | 0.555921 | 1 |
| GOBP_VENTRAL_SPINAL_CORD_DEVELOPMENT | 33 | -0.283782 | -0.925904 | 0.554656 | 1 |
| GOBP_NUCLEAR_MEMBRANE_ORGANIZATION | 42 | -0.268942 | -0.924876 | 0.578723 | 1 |
| GOBP_REGULATION_OF_SMOOTHENED_SIGNALING_PATHWAY | 71 | -0.2468 | -0.922154 | 0.658824 | 1 |
| GOBP_HISTONE_H2A_ACETYLATION | 23 | -0.31232 | -0.921883 | 0.581315 | 1 |
| GOBP_POSITIVE_REGULATION_OF_ACTIN_CYTOSKELETON_REORGANIZATION | 16 | -0.342121 | -0.921593 | 0.5625 | 1 |
| GOBP_RIBOSOMAL_SMALL_SUBUNIT_ASSEMBLY | 19 | -0.334317 | -0.921518 | 0.582781 | 1 |
| GOBP_REGULATION_OF_TELOMERE_MAINTENANCE | 77 | -0.239974 | -0.920775 | 0.64375 | 1 |
| GOBP_REGULATION_OF_PH | 77 | -0.236641 | -0.920612 | 0.651934 | 1 |
| GOBP_POSITIVE_REGULATION_OF_NEUROBLAST_PROLIFERATION | 18 | -0.328774 | -0.92003 | 0.551601 | 1 |
| GOBP_PROTEIN_TARGETING_TO_PEROXISOME | 18 | -0.332758 | -0.919921 | 0.545752 | 1 |
| GOBP_DEVELOPMENTAL_GROWTH_INVOLVED_IN_MORPHOGENESIS | 213 | -0.208548 | -0.918949 | 0.763636 | 1 |
| GOBP_NEGATIVE_REGULATION_OF_INTRACELLULAR_STEROID_HORMONE_R | 33 | -0.283325 | -0.918721 | 0.636364 | 1 |
| GOBP_MITOCHONDRIAL_TRANSLATION | 70 | -0.250198 | -0.918094 | 0.625731 | 1 |
| GOBP_NADH_METABOLIC_PROCESS | 28 | -0.296205 | -0.917883 | 0.611111 | 1 |
| GOBP_PHOSPHOLIPID_TRANSPORT | 83 | -0.241382 | -0.916418 | 0.682759 | 1 |
| GOBP_AUDITORY_RECEPTOR_CELL_MORPHOGENESIS | 15 | -0.340799 | -0.916032 | 0.565916 | 1 |
| GOBP_REGULATION_OF_TRANSLATIONAL_FIDELITY | 21 | -0.316684 | -0.914508 | 0.59322 | 1 |
| GOBP_CENTRAL_NERVOUS_SYSTEM_PROJECTION_NEURON_AXONOGENESI | 25 | -0.312006 | -0.913433 | 0.603774 | 1 |
| GOBP_PROTEIN_O_LINKED_MANNOSYLATION | 17 | -0.338501 | -0.913059 | 0.561905 | 1 |
| GOBP_IRON_SULFUR_CLUSTER_ASSEMBLY | 25 | -0.298201 | -0.912347 | 0.60687 | 1 |
| GOBP_NUCLEUS_ORGANIZATION | 126 | -0.216947 | -0.911999 | 0.744898 | 1 |
| GOBP_REGULATION_OF_MICROTUBULE_CYTOSKELETON_ORGANIZATION | 149 | -0.219609 | -0.911862 | 0.736111 | 1 |
| GOBP_NEGATIVE_REGULATION_OF_LYASE_ACTIVITY | 22 | -0.305621 | -0.911435 | 0.608108 | 1 |
| GOBP_MRNA_EXPORT_FROM_NUCLEUS | 54 | -0.252747 | -0.911193 | 0.632124 | 1 |
| GOBP_NEGATIVE_REGULATION_OF_TELOMERE_MAINTENANCE_VIA_TELOME | 19 | -0.311326 | -0.910686 | 0.592105 | 1 |
| GOBP_PROTEIN_SUMOYLATION | 47 | -0.258954 | -0.908172 | 0.627706 | 1 |
| GOBP_OVARIAN_FOLLICLE_DEVELOPMENT | 47 | -0.260474 | -0.907852 | 0.604061 | 1 |
| GOBP_NEUROTRANSMITTER_RECEPTOR_TRANSPORT | 22 | -0.30966 | -0.907027 | 0.587248 | 1 |
| GOBP_REGULATION_OF_MICROTUBULE_DEPOLYMERIZATION | 31 | -0.277812 | -0.907022 | 0.609023 | 1 |
| GOBP_HEART_FORMATION | 26 | -0.289488 | -0.906539 | 0.616438 | 1 |
| GOBP_MITOCHONDRIAL_ATP_SYNTHESIS_COUPLED_PROTON_TRANSPORT | 18 | -0.321535 | -0.906519 | 0.540193 | 1 |
| GOBP_MITOTIC_CYTOKINESIS | 69 | -0.240965 | -0.905448 | 0.696429 | 1 |
| GOBP_MITOCHONDRIAL_ELECTRON_TRANSPORT_NADH_TO_UBIQUINONE | 48 | -0.253874 | -0.905227 | 0.62 | 1 |
| GOBP_INNER_EAR_AUDITORY_RECEPTOR_CELL_DIFFERENTIATION | 32 | -0.279673 | -0.904959 | 0.596774 | 1 |
| GOBP_VASCULAR_TRANSPORT | 77 | -0.227224 | -0.904508 | 0.683544 | 1 |
| GOBP_AMINOGLYCAN_BIOSYNTHETIC_PROCESS | 74 | -0.234891 | -0.903983 | 0.7125 | 1 |
| GOBP_TRNA_PROCESSING | 121 | -0.2208 | -0.903305 | 0.75 | 1 |
| GOBP_MITOTIC_CYTOKINETIC_PROCESS | 23 | -0.304347 | -0.902784 | 0.605072 | 1 |
| GOBP_REGULATION_OF_OLIGODENDROCYTE_DIFFERENTIATION | 31 | -0.291268 | -0.902284 | 0.605948 | 1 |
| GOBP_MIDBODY_ABSCISSION | 17 | -0.330926 | -0.902177 | 0.56314 | 1 |
| GOBP_METAPHASE_PLATE_CONGRESSION | 65 | -0.239018 | -0.901203 | 0.716867 | 1 |
| GOBP_DICARBOXYLIC_ACID_CATABOLIC_PROCESS | 17 | -0.330465 | -0.900469 | 0.560127 | 1 |
| GOBP_REGULATION_OF_CARDIAC_MUSCLE_CELL_DIFFERENTIATION | 16 | -0.338496 | -0.900219 | 0.606154 | 1 |
| GOBP_TRNA_METABOLIC_PROCESS | 181 | -0.204793 | -0.89901 | 0.791045 | 1 |
| GOBP_PROTEIN_LOCALIZATION_TO_SYNAPSE | 59 | -0.246561 | -0.898961 | 0.7 | 1 |
| GOBP_PROTEIN_LOCALIZATION_TO_POSTSYNAPSE | 31 | -0.286611 | -0.898609 | 0.645757 | 1 |
| GOBP_CHONDROCYTE_PROLIFERATION | 18 | -0.319757 | -0.898227 | 0.599398 | 1 |
| GOBP_SOMATIC_DIVERSIFICATION_OF_IMMUNOGLOBULINS_INVOLVED_IN_IM | 42 | -0.260264 | -0.898008 | 0.671429 | 1 |
| GOBP_POSITIVE_REGULATION_OF_ATP_METABOLIC_PROCESS | 29 | -0.297597 | -0.89772 | 0.609375 | 1 |
| GOBP_EXIT_FROM_MITOSIS | 28 | -0.297463 | -0.897553 | 0.625483 | 1 |
| GOBP_REGULATION_OF_MACROAUTOPHAGY | 151 | -0.211463 | -0.894927 | 0.747126 | 1 |
| GOBP_POSITIVE_REGULATION_OF_MESENCHYMAL_CELL_PROLIFERATION | 16 | -0.327083 | -0.894594 | 0.626298 | 1 |
| GOBP_NEGATIVE_REGULATION_OF_UBIQUITIN_DEPENDENT_PROTEIN_CATA | 44 | -0.256314 | -0.892579 | 0.647343 | 1 |
| GOBP_PROTEIN_LOCALIZATION_TO_GOLGI_APPARATUS | 26 | -0.290498 | -0.892449 | 0.603279 | 1 |
| GOBP_PYRIMIDINE_CONTAINING_COMPOUND_METABOLIC_PROCESS | 79 | -0.226112 | -0.892352 | 0.730496 | 1 |
| GOBP_ESTABLISHMENT_OF_PLANAR_POLARITY_OF_EMBRYONIC_EPITHELIU | 15 | -0.346229 | -0.89223 | 0.610922 | 1 |
| GOBP_REGULATION_OF_PROTEIN_EXIT_FROM_ENDOPLASMIC_RETICULUM | 24 | -0.292223 | -0.89188 | 0.613383 | 1 |
| GOBP_REGULATION_OF_STEM_CELL_DIFFERENTIATION | 55 | -0.249927 | -0.89165 | 0.688525 | 1 |
| GOBP_MEMBRANE_DEPOLARIZATION_DURING_CARDIAC_MUSCLE_CELL_AC | 20 | -0.309471 | -0.891443 | 0.627063 | 1 |
| GOBP_REGULATION_OF_ANIMAL_ORGAN_MORPHOGENESIS | 109 | -0.218564 | -0.890374 | 0.827273 | 1 |
| GOBP_REGULATION_OF_PROTEIN_LOCALIZATION_TO_SYNAPSE | 18 | -0.318626 | -0.890139 | 0.596215 | 1 |
| GOBP_POSITIVE_REGULATION_OF_MAP_KINASE_ACTIVITY | 100 | -0.23293 | -0.890086 | 0.70073 | 1 |
| GOBP_CELLULAR_RESPONSE_TO_PH | 21 | -0.313048 | -0.888932 | 0.618421 | 1 |
| GOBP_NEGATIVE_REGULATION_OF_LIPID_METABOLIC_PROCESS | 74 | -0.233573 | -0.888777 | 0.736842 | 1 |
| GOBP_CEREBELLAR_CORTEX_FORMATION | 24 | -0.304618 | -0.888665 | 0.630769 | 1 |
| GOBP_POLARIZED_EPITHELIAL_CELL_DIFFERENTIATION | 24 | -0.290158 | -0.888493 | 0.595156 | 1 |
| GOBP_PEROXISOMAL_TRANSPORT | 21 | -0.313057 | -0.888312 | 0.593548 | 1 |
| GOBP_REGULATION_OF_DNA_METHYLATION_DEPENDENT_HETEROCHROMA | 16 | -0.325068 | -0.886991 | 0.617089 | 1 |
| GOBP_AUDITORY_RECEPTOR_CELL_DEVELOPMENT | 19 | -0.320085 | -0.886141 | 0.59396 | 1 |
| GOBP_PHOSPHATE_ION_TRANSPORT | 22 | -0.306842 | -0.885935 | 0.629371 | 1 |
| GOBP_PROTEIN_LOCALIZATION_TO_CHROMATIN | 31 | -0.280045 | -0.884802 | 0.642553 | 1 |
| GOBP_MELANOCYTE_DIFFERENTIATION | 22 | -0.300917 | -0.884764 | 0.623762 | 1 |
| GOBP_REGULATION_OF_ENDOCYTIC_RECYCLING | 17 | -0.321565 | -0.881322 | 0.618462 | 1 |
| GOBP_POSITIVE_REGULATION_OF_CELL_MORPHOGENESIS_INVOLVED_IN_DI | 72 | -0.230936 | -0.881289 | 0.705882 | 1 |
| GOBP_REGULATION_OF_NON_CANONICAL_WNT_SIGNALING_PATHWAY | 24 | -0.297073 | -0.879041 | 0.646388 | 1 |
| GOBP_NEGATIVE_REGULATION_OF_ORGANELLE_ASSEMBLY | 41 | -0.254888 | -0.878338 | 0.714286 | 1 |
| GOBP_REGULATION_OF_ISOTYPE_SWITCHING | 32 | -0.273755 | -0.878256 | 0.645161 | 1 |
| GOBP_REGULATION_OF_VACUOLE_ORGANIZATION | 49 | -0.251791 | -0.878207 | 0.696262 | 1 |
| GOBP_LATE_ENDOSOME_TO_VACUOLE_TRANSPORT | 24 | -0.291581 | -0.87796 | 0.642066 | 1 |
| GOBP_HISTONE_H4_K16_ACETYLATION | 27 | -0.286622 | -0.877289 | 0.662116 | 1 |
| GOBP_PEPTIDYL_L_CYSTEINE_S_PALMITOYLATION | 23 | -0.306127 | -0.876161 | 0.6609 | 1 |
| GOBP_REGULATION_OF_NEURON_PROJECTION_REGENERATION | 26 | -0.282983 | -0.876061 | 0.659091 | 1 |
| GOBP_PYRIMIDINE_NUCLEOBASE_METABOLIC_PROCESS | 20 | -0.314329 | -0.875651 | 0.66443 | 1 |
| GOBP_CANONICAL_WNT_SIGNALING_PATHWAY | 270 | -0.192279 | -0.875524 | 0.970588 | 1 |
| GOBP_CHONDROCYTE_DIFFERENTIATION | 92 | -0.221331 | -0.875248 | 0.754098 | 1 |
| GOBP_CENTRAL_NERVOUS_SYSTEM_NEURON_AXONOGENESIS | 33 | -0.270362 | -0.875232 | 0.661818 | 1 |
| GOBP_INNER_EAR_RECEPTOR_CELL_DEVELOPMENT | 38 | -0.262042 | -0.874304 | 0.680769 | 1 |
| GOBP_NEGATIVE_REGULATION_OF_PROTEIN_LOCALIZATION_TO_NUCLEUS | 30 | -0.273645 | -0.874096 | 0.693227 | 1 |
| GOBP_REGULATION_OF_URINE_VOLUME | 16 | -0.32861 | -0.873909 | 0.629139 | 1 |
| GOBP_RESPONSE_TO_EPIDERMAL_GROWTH_FACTOR | 44 | -0.252726 | -0.872503 | 0.684647 | 1 |
| GOBP_NEGATIVE_REGULATION_OF_DNA_REPAIR | 35 | -0.269349 | -0.872357 | 0.669683 | 1 |
| GOBP_PHOSPHATIDYLSERINE_METABOLIC_PROCESS | 21 | -0.299766 | -0.871518 | 0.622837 | 1 |
| GOBP_TELOMERE_MAINTENANCE | 127 | -0.210724 | -0.870257 | 0.842105 | 1 |
| GOBP_POSITIVE_REGULATION_OF_MITOCHONDRIAL_TRANSLATION | 15 | -0.334216 | -0.869907 | 0.641745 | 1 |
| GOBP_SOMATIC_DIVERSIFICATION_OF_IMMUNOGLOBULINS | 57 | -0.235701 | -0.869813 | 0.709677 | 1 |
| GOBP_LUNG_ALVEOLUS_DEVELOPMENT | 32 | -0.269777 | -0.867662 | 0.669145 | 1 |
| GOBP_REGULATION_OF_MRNA_POLYADENYLATION | 20 | -0.306832 | -0.866996 | 0.662207 | 1 |
| GOBP_RECEPTOR_LOCALIZATION_TO_SYNAPSE | 51 | -0.2479 | -0.863932 | 0.782178 | 1 |
| GOBP_BILE_ACID_METABOLIC_PROCESS | 36 | -0.265126 | -0.863082 | 0.693227 | 1 |
| GOBP_GLANDULAR_EPITHELIAL_CELL_DIFFERENTIATION | 34 | -0.258694 | -0.861226 | 0.696356 | 1 |
| GOBP_RESPONSE_TO_INCREASED_OXYGEN_LEVELS | 24 | -0.290455 | -0.861105 | 0.658273 | 1 |
| GOBP_NEGATIVE_REGULATION_OF_ENDOPLASMIC_RETICULUM_STRESS_IND | 17 | -0.319763 | -0.860703 | 0.620805 | 1 |
| GOBP_CELLULAR_CARBOHYDRATE_BIOSYNTHETIC_PROCESS | 63 | -0.234454 | -0.860294 | 0.725807 | 1 |
| GOBP_CEREBELLAR_CORTEX_MORPHOGENESIS | 33 | -0.259602 | -0.859506 | 0.697417 | 1 |
| GOBP_NEGATIVE_REGULATION_OF_STRIATED_MUSCLE_CELL_APOPTOTIC_P | 19 | -0.294589 | -0.858035 | 0.65411 | 1 |
| GOBP_PROTEIN_DEGLYCOSYLATION | 25 | -0.290444 | -0.857852 | 0.678445 | 1 |
| GOBP_CELL_CELL_SIGNALING_BY_WNT | 398 | -0.18063 | -0.85774 | 1 | 1 |
| GOBP_CEREBRAL_CORTEX_NEURON_DIFFERENTIATION | 21 | -0.303041 | -0.856991 | 0.653125 | 1 |
| GOBP_VIRAL_TRANSCRIPTION | 48 | -0.250121 | -0.855939 | 0.757848 | 1 |
| GOBP_CYTOPLASMIC_TRANSLATIONAL_INITIATION | 27 | -0.277431 | -0.855288 | 0.692015 | 1 |
| GOBP_MRNA_MODIFICATION | 26 | -0.279711 | -0.855077 | 0.659091 | 1 |
| GOBP_PROTEIN_LOCALIZATION_TO_CHROMOSOME | 91 | -0.212514 | -0.853272 | 0.837037 | 1 |
| GOBP_POSITIVE_REGULATION_OF_CYTOKINESIS | 31 | -0.271656 | -0.852918 | 0.692015 | 1 |
| GOBP_NLS_BEARING_PROTEIN_IMPORT_INTO_NUCLEUS | 18 | -0.308273 | -0.851726 | 0.660256 | 1 |
| GOBP_CENTROMERE_COMPLEX_ASSEMBLY | 30 | -0.263652 | -0.850367 | 0.732203 | 1 |
| GOBP_HISTONE_H3_DEACETYLATION | 17 | -0.306384 | -0.849469 | 0.65109 | 1 |
| GOBP_MAINTENANCE_OF_CELL_POLARITY | 18 | -0.303824 | -0.849374 | 0.654237 | 1 |
| GOBP_INTRACELLULAR_ESTROGEN_RECEPTOR_SIGNALING_PATHWAY | 43 | -0.243941 | -0.849192 | 0.736607 | 1 |
| GOBP_NUCLEAR_BODY_ORGANIZATION | 15 | -0.312327 | -0.848701 | 0.664557 | 1 |
| GOBP_NERVE_DEVELOPMENT | 64 | -0.231746 | -0.848022 | 0.788462 | 1 |
| GOBP_CELL_CELL_SIGNALING_INVOLVED_IN_CARDIAC_CONDUCTION | 27 | -0.278376 | -0.847992 | 0.690909 | 1 |
| GOBP_NEGATIVE_REGULATION_OF_CHROMOSOME_ORGANIZATION | 82 | -0.226127 | -0.847653 | 0.832168 | 1 |
| GOBP_VESICLE_MEDIATED_TRANSPORT_BETWEEN_ENDOSOMAL_COMPART | 45 | -0.239314 | -0.846386 | 0.75814 | 1 |
| GOBP_HEART_FIELD_SPECIFICATION | 16 | -0.313046 | -0.845754 | 0.673981 | 1 |
| GOBP_TETRAHYDROFOLATE_METABOLIC_PROCESS | 20 | -0.291525 | -0.845592 | 0.707071 | 1 |
| GOBP_NCRNA_3_END_PROCESSING | 39 | -0.250664 | -0.844261 | 0.739837 | 1 |
| GOBP_DNA_DEPENDENT_DNA_REPLICATION | 152 | -0.197283 | -0.843422 | 0.898734 | 1 |
| GOBP_MITOCHONDRIAL_TRANSPORT | 190 | -0.191504 | -0.842964 | 0.955224 | 1 |
| GOBP_POSITIVE_REGULATION_OF_BLOOD_PRESSURE | 28 | -0.2669 | -0.842891 | 0.724 | 1 |
| GOBP_ESTABLISHMENT_OF_EPITHELIAL_CELL_POLARITY | 31 | -0.262899 | -0.841988 | 0.726141 | 1 |
| GOBP_POSITIVE_REGULATION_OF_PROTEIN_AUTOPHOSPHORYLATION | 25 | -0.274377 | -0.841426 | 0.711744 | 1 |
| GOBP_MRNA_SPLICE_SITE_SELECTION | 33 | -0.26267 | -0.840945 | 0.751938 | 1 |
| GOBP_PROTEIN_LOCALIZATION_TO_MITOCHONDRION | 121 | -0.202916 | -0.840656 | 0.920354 | 1 |
| GOBP_CORTICAL_ACTIN_CYTOSKELETON_ORGANIZATION | 37 | -0.249508 | -0.84038 | 0.690678 | 1 |
| GOBP_NEGATIVE_REGULATION_OF_CYTOSKELETON_ORGANIZATION | 149 | -0.199112 | -0.840326 | 0.957895 | 1 |
| GOBP_PROTEIN_LOCALIZATION_TO_CYTOSKELETON | 56 | -0.236023 | -0.83895 | 0.789474 | 1 |
| GOBP_ESTABLISHMENT_OF_EPITHELIAL_CELL_APICAL_BASAL_POLARITY | 18 | -0.290058 | -0.838488 | 0.672783 | 1 |
| GOBP_SUBSTANTIA_NIGRA_DEVELOPMENT | 40 | -0.252899 | -0.836539 | 0.742358 | 1 |
| GOBP_HINDBRAIN_MORPHOGENESIS | 40 | -0.251517 | -0.835083 | 0.710901 | 1 |
| GOBP_POSITIVE_REGULATION_OF_CHROMOSOME_SEPARATION | 17 | -0.305828 | -0.831749 | 0.680912 | 1 |
| GOBP_REGULATION_OF_CELL_MORPHOGENESIS_INVOLVED_IN_DIFFERENTIA | 88 | -0.210201 | -0.831497 | 0.847682 | 1 |
| GOBP_ENDOPLASMIC_RETICULUM_TO_GOLGI_VESICLE_MEDIATED_TRANSP | 122 | -0.204151 | -0.831183 | 0.90625 | 1 |
| GOBP_HEPARAN_SULFATE_PROTEOGLYCAN_METABOLIC_PROCESS | 30 | -0.272917 | -0.830463 | 0.72695 | 1 |
| GOBP_AXONAL_FASCICULATION | 21 | -0.295567 | -0.829983 | 0.695513 | 1 |
| GOBP_HISTONE_H4_ACETYLATION | 72 | -0.214265 | -0.829659 | 0.819876 | 1 |
| GOBP_NEGATIVE_REGULATION_OF_ACTIN_FILAMENT_POLYMERIZATION | 58 | -0.224294 | -0.829369 | 0.848341 | 1 |
| GOBP_REGULATION_OF_SPINDLE_ASSEMBLY | 30 | -0.264416 | -0.829188 | 0.714829 | 1 |
| GOBP_REGULATION_OF_ORGANELLE_ASSEMBLY | 211 | -0.177663 | -0.828826 | 0.956522 | 1 |

| | | | | | |
|---|---|---|---|---|---|
| GOBP_REGULATION_OF_MUSCLE_SYSTEM_PROCESS | 188 | 0.39841 | 1.314739 | 0.03072 | 0.448862 |
| GOBP_POSITIVE_REGULATION_OF_INTERLEUKIN_2_PRODUCTION | 25 | 0.524587 | 1.314712 | 0.131469 | 0.448227 |
| GOBP_NEGATIVE_REGULATION_OF_BLOOD_VESSEL_ENDOTHELIAL_CELL_MIGRATION | 29 | 0.507316 | 1.314495 | 0.133238 | 0.448149 |
| GOBP_POSITIVE_REGULATION_OF_PHOSPHATASE_ACTIVITY | 28 | 0.507052 | 1.314294 | 0.121984 | 0.448056 |
| GOBP_REGULATION_OF_RELEASE_OF_SEQUESTERED_CALCIUM_ION_INTO_CYTOSOL | 62 | 0.447734 | 1.313973 | 0.102091 | 0.448321 |
| GOBP_CELLULAR_COMPONENT_DISASSEMBLY_INVOLVED_IN_EXECUTION_PHASE_OF_AP | 25 | 0.513531 | 1.313785 | 0.135402 | 0.448124 |
| GOBP_REGULATION_OF_BROWN_FAT_CELL_DIFFERENTIATION | 19 | 0.546458 | 1.313509 | 0.143895 | 0.448202 |
| GOBP_POSITIVE_REGULATION_OF_PEPTIDYL_TYROSINE_PHOSPHORYLATION | 131 | 0.415425 | 1.313084 | 0.061269 | 0.448852 |
| GOBP_ALDITOL_METABOLIC_PROCESS | 16 | 0.570746 | 1.312781 | 0.139319 | 0.449082 |
| GOBP_REGULATION_OF_ANION_TRANSPORT | 70 | 0.441624 | 1.312618 | 0.09842 | 0.448821 |
| GOBP_SPERM_MOTILITY | 75 | 0.439635 | 1.312583 | 0.078916 | 0.44822 |
| GOBP_AMELOGENESIS | 18 | 0.545001 | 1.311787 | 0.126939 | 0.449834 |
| GOBP_ENDODERM_FORMATION | 48 | 0.460699 | 1.311597 | 0.108557 | 0.449726 |
| GOBP_LONG_CHAIN_FATTY_ACID_TRANSPORT | 49 | 0.461064 | 1.311338 | 0.115906 | 0.449781 |
| GOBP_POSITIVE_REGULATION_OF_MONONUCLEAR_CELL_MIGRATION | 42 | 0.476083 | 1.311077 | 0.112125 | 0.449831 |
| GOBP_RESPONSE_TO_ANGIOTENSIN | 22 | 0.526904 | 1.309995 | 0.140376 | 0.452331 |
| GOBP_CELL_PROLIFERATION_IN_FOREBRAIN | 19 | 0.5568 | 1.309808 | 0.139368 | 0.452171 |
| GOBP_REGULATION_OF_BODY_FLUID_LEVELS | 293 | 0.388252 | 1.309442 | 0.014403 | 0.452581 |
| GOBP_REGULATION_OF_CD4_POSITIVE_ALPHA_BETA_T_CELL_DIFFERENTIATION | 35 | 0.48355 | 1.308474 | 0.120155 | 0.454856 |
| GOBP_FATTY_ACID_METABOLIC_PROCESS | 300 | 0.386848 | 1.308433 | 0.015353 | 0.454277 |
| GOBP_FEAR_RESPONSE | 35 | 0.49111 | 1.307495 | 0.117048 | 0.456363 |
| GOBP_CARDIAC_MUSCLE_TISSUE_DEVELOPMENT | 181 | 0.398891 | 1.306194 | 0.051173 | 0.459579 |
| GOBP_REGULATION_OF_PEPTIDYL_TYROSINE_PHOSPHORYLATION | 191 | 0.395737 | 1.305741 | 0.04661 | 0.460187 |
| GOBP_PLATELET_DERIVED_GROWTH_FACTOR_RECEPTOR_SIGNALING_PATHWAY | 48 | 0.46366 | 1.305275 | 0.127742 | 0.460929 |
| GOBP_REGULATION_OF_CATION_TRANSMEMBRANE_TRANSPORT | 275 | 0.386472 | 1.30492 | 0.027027 | 0.461266 |
| GOBP_REGULATION_OF_INNATE_IMMUNE_RESPONSE | 162 | 0.401593 | 1.304139 | 0.04086 | 0.463006 |
| GOBP_CALCIUM_MEDIATED_SIGNALING_USING_INTRACELLULAR_CALCIUM_SOURCE | 16 | 0.565425 | 1.303304 | 0.174803 | 0.464659 |
| GOBP_CELLULAR_RESPONSE_TO_MECHANICAL_STIMULUS | 59 | 0.438113 | 1.303261 | 0.110974 | 0.464075 |
| GOBP_RETROGRADE_AXONAL_TRANSPORT | 21 | 0.534885 | 1.302807 | 0.152327 | 0.464706 |
| GOBP_POSITIVE_REGULATION_OF_CALCIUM_ION_TRANSMEMBRANE_TRANSPORTER_ACT | 35 | 0.478209 | 1.302675 | 0.125 | 0.464379 |
| GOBP_VENTRICULAR_CARDIAC_MUSCLE_TISSUE_DEVELOPMENT | 49 | 0.456143 | 1.302364 | 0.121982 | 0.464627 |
| GOBP_POSITIVE_REGULATION_OF_PROTEIN_SECRETION | 105 | 0.418437 | 1.301092 | 0.067873 | 0.467771 |
| GOBP_REGIONALIZATION | 261 | 0.382776 | 1.300811 | 0.029774 | 0.467924 |
| GOBP_DIGESTIVE_SYSTEM_PROCESS | 62 | 0.442715 | 1.30064 | 0.09291 | 0.467724 |
| GOBP_FC_GAMMA_RECEPTOR_SIGNALING_PATHWAY | 23 | 0.52302 | 1.300121 | 0.156522 | 0.46855 |
| GOBP_POSITIVE_REGULATION_OF_LIPID_METABOLIC_PROCESS | 115 | 0.410337 | 1.300065 | 0.05708 | 0.46804 |
| GOBP_ADIPOSE_TISSUE_DEVELOPMENT | 41 | 0.465446 | 1.30005 | 0.109626 | 0.46739 |
| GOBP_UTERUS_DEVELOPMENT | 15 | 0.568109 | 1.299925 | 0.16293 | 0.467053 |
| GOBP_ENDOCRINE_SYSTEM_DEVELOPMENT | 96 | 0.419115 | 1.299816 | 0.076212 | 0.466679 |
| GOBP_MAST_CELL_ACTIVATION | 47 | 0.465696 | 1.297829 | 0.114068 | 0.472014 |
| GOBP_REGULATION_OF_PRODUCTION_OF_MOLECULAR_MEDIATOR_OF_IMMUNE_RESPON | 119 | 0.405373 | 1.297741 | 0.068462 | 0.471559 |
| GOBP_FILOPODIUM_ASSEMBLY | 63 | 0.44134 | 1.297611 | 0.112624 | 0.471273 |
| GOBP_NEGATIVE_REGULATION_OF_SYNAPTIC_TRANSMISSION | 62 | 0.441131 | 1.297406 | 0.095465 | 0.471185 |
| GOBP_POSITIVE_REGULATION_OF_INTERFERON_GAMMA_PRODUCTION | 43 | 0.471332 | 1.297379 | 0.112658 | 0.470571 |
| GOBP_CARDIAC_VENTRICLE_DEVELOPMENT | 111 | 0.416006 | 1.297246 | 0.080627 | 0.470268 |
| GOBP_REGULATION_OF_SUPEROXIDE_ANION_GENERATION | 16 | 0.552444 | 1.296872 | 0.166667 | 0.470753 |
| GOBP_NEGATIVE_REGULATION_OF_INFLAMMATORY_RESPONSE | 106 | 0.412762 | 1.296743 | 0.077803 | 0.47044 |
| GOBP_POSITIVE_REGULATION_OF_SMOOTH_MUSCLE_CONTRACTION | 27 | 0.502169 | 1.296131 | 0.148299 | 0.471606 |
| GOBP_REGULATION_OF_T_CELL_DIFFERENTIATION | 105 | 0.415456 | 1.29523 | 0.07545 | 0.47352 |
| GOBP_POSITIVE_REGULATION_OF_PATHWAY_RESTRICTED_SMAD_PROTEIN_PHOSPHORY | 42 | 0.462327 | 1.294491 | 0.133598 | 0.475058 |
| GOBP_REGULATION_OF_PHAGOCYTOSIS | 68 | 0.436058 | 1.294167 | 0.105968 | 0.475315 |
| GOBP_REGULATION_OF_CYTOSOLIC_CALCIUM_ION_CONCENTRATION | 244 | 0.386743 | 1.293906 | 0.0375 | 0.47538 |
| GOBP_REGULATION_OF_CELLULAR_CARBOHYDRATE_METABOLIC_PROCESS | 126 | 0.405272 | 1.293319 | 0.068233 | 0.476413 |
| GOBP_PROTEIN_KINASE_B_SIGNALING | 161 | 0.394836 | 1.292613 | 0.050921 | 0.477912 |
| GOBP_NEGATIVE_REGULATION_OF_CELL_MATRIX_ADHESION | 31 | 0.489829 | 1.292477 | 0.134487 | 0.477639 |
| GOBP_NEGATIVE_REGULATION_OF_TORC1_SIGNALING | 19 | 0.531518 | 1.292269 | 0.15767 | 0.477545 |
| GOBP_POSITIVE_REGULATION_OF_TOLL_LIKE_RECEPTOR_SIGNALING_PATHWAY | 20 | 0.532265 | 1.291874 | 0.172316 | 0.478097 |
| GOBP_NEGATIVE_REGULATION_OF_HEMOPOIESIS | 77 | 0.427195 | 1.29166 | 0.090584 | 0.478053 |
| GOBP_INTERLEUKIN_1_PRODUCTION | 70 | 0.433094 | 1.291531 | 0.106173 | 0.477778 |
| GOBP_POSITIVE_REGULATION_OF_ENDOTHELIAL_CELL_MIGRATION | 99 | 0.415467 | 1.29096 | 0.083529 | 0.478851 |
| GOBP_ACUTE_PHASE_RESPONSE | 27 | 0.498466 | 1.290932 | 0.155772 | 0.478222 |
| GOBP_NEGATIVE_REGULATION_OF_PRODUCTION_OF_MOLECULAR_MEDIATOR_OF_IMMUN | 23 | 0.513164 | 1.290676 | 0.172316 | 0.478316 |
| GOBP_ATRIOVENTRICULAR_VALVE_DEVELOPMENT | 25 | 0.505357 | 1.290393 | 0.155462 | 0.478528 |
| GOBP_ANATOMICAL_STRUCTURE_HOMEOSTASIS | 242 | 0.381322 | 1.289215 | 0.037815 | 0.481378 |
| GOBP_PODOSOME_ASSEMBLY | 16 | 0.54885 | 1.289147 | 0.161616 | 0.480905 |
| GOBP_SEGMENTATION | 83 | 0.418564 | 1.289026 | 0.071514 | 0.480559 |
| GOBP_POSITIVE_REGULATION_OF_ENDOTHELIAL_CELL_CHEMOTAXIS | 15 | 0.563764 | 1.288854 | 0.154848 | 0.480393 |
| GOBP_RESPONSE_TO_HEAT | 95 | 0.419721 | 1.288731 | 0.084091 | 0.480072 |
| GOBP_PHOSPHATIDYLCHOLINE_METABOLIC_PROCESS | 61 | 0.439835 | 1.288562 | 0.115055 | 0.479934 |
| GOBP_NEGATIVE_REGULATION_OF_CD4_POSITIVE_ALPHA_BETA_T_CELL_DIFFERENTIATIO | 17 | 0.552355 | 1.288552 | 0.179104 | 0.479277 |
| GOBP_LIPID_CATABOLIC_PROCESS | 251 | 0.383882 | 1.288255 | 0.048354 | 0.47947 |
| GOBP_REGULATION_OF_ALPHA_BETA_T_CELL_DIFFERENTIATION | 48 | 0.459018 | 1.287948 | 0.125161 | 0.479733 |
| GOBP_POSITIVE_REGULATION_OF_LYMPHOCYTE_MEDIATED_IMMUNITY | 75 | 0.431619 | 1.28771 | 0.103695 | 0.479804 |
| GOBP_NEGATIVE_REGULATION_OF_EPITHELIAL_CELL_MIGRATION | 57 | 0.437147 | 1.287503 | 0.123472 | 0.479766 |
| GOBP_UNSATURATED_FATTY_ACID_BIOSYNTHETIC_PROCESS | 39 | 0.475612 | 1.287406 | 0.1469 | 0.479381 |
| GOBP_EMBRYONIC_CRANIAL_SKELETON_MORPHOGENESIS | 39 | 0.47247 | 1.287349 | 0.126467 | 0.478864 |
| GOBP_REGULATION_OF_VIRAL_PROCESS | 134 | 0.399546 | 1.287343 | 0.069845 | 0.478195 |
| GOBP_INORGANIC_ION_IMPORT_ACROSS_PLASMA_MEMBRANE | 83 | 0.420482 | 1.286457 | 0.098062 | 0.480108 |
| GOBP_NEGATIVE_REGULATION_OF_PROTEOLYSIS | 228 | 0.388593 | 1.286109 | 0.042842 | 0.480474 |
| GOBP_GLIAL_CELL_PROLIFERATION | 32 | 0.488407 | 1.285975 | 0.160265 | 0.480243 |
| GOBP_TROPHOBLAST_CELL_MIGRATION | 15 | 0.573891 | 1.285808 | 0.155488 | 0.480054 |
| GOBP_RESPONSE_TO_VITAMIN | 62 | 0.436765 | 1.285268 | 0.115713 | 0.481135 |
| GOBP_ENDOCARDIAL_CUSHION_FORMATION | 28 | 0.491632 | 1.285265 | 0.169444 | 0.480462 |
| GOBP_POSITIVE_REGULATION_OF_HISTONE_ACETYLATION | 29 | 0.488366 | 1.284948 | 0.164675 | 0.480739 |
| GOBP_NEGATIVE_REGULATION_OF_MONONUCLEAR_CELL_MIGRATION | 18 | 0.549235 | 1.284764 | 0.167139 | 0.480635 |
| GOBP_REGULATION_OF_MYELOID_CELL_APOPTOTIC_PROCESS | 21 | 0.515499 | 1.284667 | 0.160767 | 0.480232 |
| GOBP_REGULATION_OF_PEPTIDYL_THREONINE_PHOSPHORYLATION | 38 | 0.468079 | 1.284172 | 0.152091 | 0.481041 |
| GOBP_REGULATION_OF_FAT_CELL_DIFFERENTIATION | 119 | 0.404899 | 1.284088 | 0.079775 | 0.480624 |
| GOBP_REGULATION_OF_RESPIRATORY_GASEOUS_EXCHANGE | 17 | 0.546134 | 1.283243 | 0.167916 | 0.482469 |
| GOBP_REGULATION_OF_DEFENSE_RESPONSE_TO_VIRUS_BY_HOST | 32 | 0.487869 | 1.282956 | 0.150538 | 0.482708 |
| GOBP_CARDIAC_MUSCLE_CELL_DIFFERENTIATION | 88 | 0.41013 | 1.282892 | 0.087457 | 0.482215 |
| GOBP_NEURON_CELLULAR_HOMEOSTASIS | 29 | 0.495124 | 1.282047 | 0.167568 | 0.483991 |
| GOBP_RESPONSE_TO_MANGANESE_ION | 18 | 0.537515 | 1.281531 | 0.171014 | 0.484935 |
| GOBP_NEGATIVE_REGULATION_OF_MRNA_CATABOLIC_PROCESS | 55 | 0.445609 | 1.281327 | 0.136986 | 0.484915 |
| GOBP_GLUCOSAMINE_CONTAINING_COMPOUND_METABOLIC_PROCESS | 21 | 0.519785 | 1.280832 | 0.167622 | 0.485762 |
| GOBP_LYMPHOCYTE_CHEMOTAXIS | 24 | 0.51283 | 1.280674 | 0.15491 | 0.485567 |
| GOBP_NIK_NF_KAPPAB_SIGNALING | 103 | 0.415932 | 1.280123 | 0.08099 | 0.486531 |
| GOBP_MUSCLE_TISSUE_DEVELOPMENT | 316 | 0.377674 | 1.279487 | 0.027579 | 0.487928 |
| GOBP_ORGANIC_HYDROXY_COMPOUND_BIOSYNTHETIC_PROCESS | 192 | 0.38713 | 1.279334 | 0.055437 | 0.487759 |
| GOBP_MEIOTIC_CELL_CYCLE_PROCESS | 159 | 0.3898 | 1.278958 | 0.070815 | 0.488243 |
| GOBP_RESPONSE_TO_HYDROGEN_PEROXIDE | 105 | 0.41466 | 1.278085 | 0.102593 | 0.490255 |
| GOBP_REGULATION_OF_MUSCLE_CONTRACTION | 136 | 0.399699 | 1.278008 | 0.074033 | 0.489842 |
| GOBP_REGULATION_OF_INFLAMMATORY_RESPONSE_TO_ANTIGENIC_STIMULUS | 31 | 0.484188 | 1.277749 | 0.143631 | 0.489943 |
| GOBP_CILIUM_OR_FLAGELLUM_DEPENDENT_CELL_MOTILITY | 95 | 0.409627 | 1.277687 | 0.100114 | 0.489445 |
| GOBP_PROTEIN_HOMOOLIGOMERIZATION | 160 | 0.389842 | 1.277558 | 0.065946 | 0.489147 |
| GOBP_REGULATION_OF_CAMP_MEDIATED_SIGNALING | 26 | 0.499816 | 1.277059 | 0.178025 | 0.490065 |
| GOBP_REGULATION_OF_VIRAL_GENOME_REPLICATION | 71 | 0.430217 | 1.276722 | 0.116364 | 0.490432 |
| GOBP_RESPONSE_TO_VIRUS | 294 | 0.377513 | 1.276302 | 0.029652 | 0.491008 |
| GOBP_PROSTANOID_BIOSYNTHETIC_PROCESS | 22 | 0.52502 | 1.275821 | 0.169188 | 0.491761 |
| GOBP_REGULATION_OF_EMBRYONIC_DEVELOPMENT | 57 | 0.436482 | 1.275587 | 0.128109 | 0.491812 |
| GOBP_RESPONSE_TO_STEROL | 23 | 0.514087 | 1.275015 | 0.160689 | 0.49282 |
| GOBP_AORTA_MORPHOGENESIS | 29 | 0.485716 | 1.274658 | 0.170699 | 0.493226 |
| GOBP_POSITIVE_REGULATION_OF_CALCIUM_ION_IMPORT | 16 | 0.552051 | 1.274035 | 0.184615 | 0.494509 |
| GOBP_POSITIVE_REGULATION_OF_PROTEIN_KINASE_B_SIGNALING | 90 | 0.414749 | 1.273939 | 0.107351 | 0.494139 |
| GOBP_SENSORY_PERCEPTION_OF_TASTE | 22 | 0.508112 | 1.273473 | 0.17316 | 0.49489 |
| GOBP_SYNCYTIUM_FORMATION | 43 | 0.457545 | 1.273248 | 0.142672 | 0.494957 |
| GOBP_TOLL_LIKE_RECEPTOR_3_SIGNALING_PATHWAY | 15 | 0.555403 | 1.273199 | 0.179331 | 0.49446 |
| GOBP_REGULATION_OF_CELL_MATRIX_ADHESION | 105 | 0.404093 | 1.273164 | 0.101754 | 0.493894 |
| GOBP_REGULATION_OF_LIPID_STORAGE | 40 | 0.46152 | 1.272683 | 0.160315 | 0.494718 |
| GOBP_DNA_CATABOLIC_PROCESS | 26 | 0.502942 | 1.27188 | 0.175824 | 0.496588 |
| GOBP_NODAL_SIGNALING_PATHWAY | 15 | 0.559744 | 1.271848 | 0.185075 | 0.496013 |
| GOBP_POSITIVE_REGULATION_OF_NITRIC_OXIDE_METABOLIC_PROCESS | 27 | 0.496685 | 1.271712 | 0.171468 | 0.495808 |
| GOBP_MUSCLE_ORGAN_DEVELOPMENT | 272 | 0.379202 | 1.269828 | 0.032922 | 0.500904 |
| GOBP_CELLULAR_RESPONSE_TO_GROWTH_HORMONE_STIMULUS | 20 | 0.518704 | 1.269774 | 0.173853 | 0.500404 |
| GOBP_ENDOTHELIAL_CELL_CHEMOTAXIS | 25 | 0.495827 | 1.269773 | 0.178522 | 0.499741 |
| GOBP_AMINE_METABOLIC_PROCESS | 98 | 0.404564 | 1.269728 | 0.101382 | 0.499186 |
| GOBP_PULMONARY_VALVE_MORPHOGENESIS | 16 | 0.560085 | 1.269564 | 0.184087 | 0.499004 |
| GOBP_NUCLEOSIDE_METABOLIC_PROCESS | 54 | 0.441702 | 1.268879 | 0.138821 | 0.500453 |
| GOBP_GLOMERULAR_EPITHELIAL_CELL_DIFFERENTIATION | 20 | 0.509279 | 1.268707 | 0.181435 | 0.500337 |
| GOBP_SALIVARY_GLAND_DEVELOPMENT | 29 | 0.482667 | 1.268407 | 0.14779 | 0.500563 |
| GOBP_XENOBIOTIC_METABOLIC_PROCESS | 64 | 0.425204 | 1.267198 | 0.127404 | 0.503734 |
| GOBP_POSITIVE_REGULATION_OF_LIPID_BIOSYNTHETIC_PROCESS | 64 | 0.432746 | 1.267196 | 0.117216 | 0.503074 |
| GOBP_BLASTOCYST_DEVELOPMENT | 101 | 0.404994 | 1.266953 | 0.106897 | 0.503145 |
| GOBP_NEGATIVE_REGULATION_OF_BLOOD_PRESSURE | 36 | 0.469176 | 1.266514 | 0.181333 | 0.503775 |
| GOBP_CARBOHYDRATE_DERIVATIVE_CATABOLIC_PROCESS | 144 | 0.394182 | 1.265941 | 0.08 | 0.504899 |
| GOBP_CD8_POSITIVE_ALPHA_BETA_T_CELL_ACTIVATION | 16 | 0.557358 | 1.265793 | 0.16692 | 0.504672 |
| GOBP_REGULATION_OF_LAMELLIPODIUM_ORGANIZATION | 46 | 0.451202 | 1.264239 | 0.143774 | 0.508957 |
| GOBP_CELL_FATE_DETERMINATION | 25 | 0.49957 | 1.264149 | 0.174468 | 0.508559 |
| GOBP_NEGATIVE_REGULATION_OF_LEUKOCYTE_PROLIFERATION | 55 | 0.433826 | 1.263565 | 0.143742 | 0.509691 |
| GOBP_ERYTHROCYTE_DEVELOPMENT | 28 | 0.491134 | 1.263505 | 0.15775 | 0.509214 |
| GOBP_SKIN_DEVELOPMENT | 184 | 0.379815 | 1.262905 | 0.085745 | 0.510409 |
| GOBP_BRANCHING_INVOLVED_IN_SALIVARY_GLAND_MORPHOGENESIS | 17 | 0.535788 | 1.262325 | 0.201504 | 0.511557 |
| GOBP_ENTRY_INTO_HOST | 128 | 0.399748 | 1.262282 | 0.098343 | 0.511004 |
| GOBP_IRON_ION_TRANSPORT | 45 | 0.455187 | 1.262263 | 0.159383 | 0.510411 |
| GOBP_TOOTH_MINERALIZATION | 17 | 0.535476 | 1.262226 | 0.205318 | 0.509869 |
| GOBP_CARDIAC_SEPTUM_DEVELOPMENT | 95 | 0.40491 | 1.26157 | 0.111494 | 0.511316 |
| GOBP_HOMEOSTASIS_OF_NUMBER_OF_CELLS | 237 | 0.377224 | 1.26044 | 0.053125 | 0.514243 |
| GOBP_RESPONSE_TO_OSMOTIC_STRESS | 71 | 0.42223 | 1.260284 | 0.12561 | 0.514044 |
| GOBP_NEGATIVE_REGULATION_OF_T_CELL_DIFFERENTIATION | 31 | 0.478341 | 1.259838 | 0.187416 | 0.514834 |
| GOBP_REGULATION_OF_DEFENSE_RESPONSE_TO_BACTERIUM | 15 | 0.544697 | 1.25979 | 0.184962 | 0.514347 |
| GOBP_POSITIVE_REGULATION_OF_PROTEIN_CONTAINING_COMPLEX_DISASSEMBLY | 31 | 0.476929 | 1.259458 | 0.172414 | 0.514652 |
| GOBP_INOSITOL_PHOSPHATE_CATABOLIC_PROCESS | 19 | 0.529553 | 1.257657 | 0.172767 | 0.519574 |
| GOBP_ENDOCARDIAL_CUSHION_DEVELOPMENT | 43 | 0.450638 | 1.257485 | 0.167095 | 0.519445 |
| GOBP_MULTICELLULAR_ORGANISMAL_HOMEOSTASIS | 404 | 0.368439 | 1.256115 | 0.028426 | 0.523077 |
| GOBP_NEGATIVE_REGULATION_OF_LYMPHOCYTE_DIFFERENTIATION | 39 | 0.454764 | 1.256051 | 0.162197 | 0.522585 |
| GOBP_RESPONSE_TO_OXYGEN_LEVELS | 283 | 0.37233 | 1.255301 | 0.04918 | 0.524278 |
| GOBP_NEGATIVE_REGULATION_OF_DEVELOPMENTAL_GROWTH | 97 | 0.402532 | 1.254706 | 0.122685 | 0.525464 |

| | | | | | |
|---|---|---|---|---|---|
| GOBP_DNA_GEOMETRIC_CHANGE | 94 | -0.210137 | -0.828473 | 0.885714 | 1 |
| GOBP_NCRNA_CATABOLIC_PROCESS | 43 | -0.240668 | -0.828252 | 0.781893 | 1 |
| GOBP_METAPHASE_ANAPHASE_TRANSITION_OF_CELL_CYCLE | 61 | -0.22586 | -0.827435 | 0.85 | 1 |
| GOBP_NUCLEOBASE_BIOSYNTHETIC_PROCESS | 19 | -0.289794 | -0.826255 | 0.711111 | 1 |
| GOBP_OLIGOSACCHARIDE_BIOSYNTHETIC_PROCESS | 19 | -0.284271 | -0.823102 | 0.727891 | 1 |
| GOBP_KERATAN_SULFATE_METABOLIC_PROCESS | 17 | -0.299384 | -0.821027 | 0.7 | 1 |
| GOBP_LABYRINTHINE_LAYER_BLOOD_VESSEL_DEVELOPMENT | 18 | -0.293476 | -0.82039 | 0.71134 | 1 |
| GOBP_REGULATION_OF_NEURONAL_SYNAPTIC_PLASTICITY | 49 | -0.232733 | -0.818995 | 0.783019 | 1 |
| GOBP_CHROMOSOME_LOCALIZATION | 78 | -0.215077 | -0.818668 | 0.88961 | 1 |
| GOBP_NUCLEOSIDE_PHOSPHATE_CATABOLIC_PROCESS | 74 | -0.217009 | -0.818631 | 0.843137 | 1 |
| GOBP_TRANSLATIONAL_INITIATION | 112 | -0.202354 | -0.818042 | 0.91 | 1 |
| GOBP_POSITIVE_REGULATION_OF_CHONDROCYTE_DIFFERENTIATION | 17 | -0.297758 | -0.81793 | 0.689759 | 1 |
| GOBP_PROTEIN_LOCALIZATION_TO_CHROMOSOME_TELOMERIC_REGION | 30 | -0.261707 | -0.817586 | 0.751073 | 1 |
| GOBP_NEUROBLAST_PROLIFERATION | 49 | -0.234889 | -0.816695 | 0.793296 | 1 |
| GOBP_NEGATIVE_REGULATION_OF_PROTEASOMAL_UBIQUITIN_DEPENDENT_ | 32 | -0.250295 | -0.815884 | 0.769231 | 1 |
| GOBP_TRNA_MODIFICATION | 85 | -0.20825 | -0.815381 | 0.897436 | 1 |
| GOBP_REGULATION_OF_AMPA_RECEPTOR_ACTIVITY | 23 | -0.268809 | -0.812209 | 0.773163 | 1 |
| GOBP_FATTY_ACID_TRANSMEMBRANE_TRANSPORT | 15 | -0.308362 | -0.812065 | 0.710692 | 1 |
| GOBP_PIGMENT_CELL_DIFFERENTIATION | 32 | -0.256652 | -0.811265 | 0.765385 | 1 |
| GOBP_AMYLOID_BETA_FORMATION | 39 | -0.241483 | -0.810314 | 0.822511 | 1 |
| GOBP_POSITIVE_REGULATION_OF_APOPTOTIC_SIGNALING_PATHWAY | 112 | -0.200314 | -0.810128 | 0.907407 | 1 |
| GOBP_CELL_VOLUME_HOMEOSTASIS | 29 | -0.26093 | -0.809584 | 0.802974 | 1 |
| GOBP_RESPONSE_TO_IRON_ION | 24 | -0.267163 | -0.809277 | 0.789286 | 1 |
| GOBP_PROTEIN_EXIT_FROM_ENDOPLASMIC_RETICULUM | 48 | -0.237433 | -0.808847 | 0.836449 | 1 |
| GOBP_RETINA_LAYER_FORMATION | 19 | -0.281602 | -0.80866 | 0.759494 | 1 |
| GOBP_INNERVATION | 24 | -0.271859 | -0.808517 | 0.768421 | 1 |
| GOBP_GLYCOLIPID_BIOSYNTHETIC_PROCESS | 66 | -0.222051 | -0.807635 | 0.877095 | 1 |
| GOBP_ESTABLISHMENT_OF_PROTEIN_LOCALIZATION_TO_MEMBRANE | 236 | -0.180432 | -0.806977 | 1 | 1 |
| GOBP_EXCITATORY_SYNAPSE_ASSEMBLY | 23 | -0.270497 | -0.806262 | 0.712329 | 1 |
| GOBP_NEURAL_TUBE_PATTERNING | 29 | -0.255588 | -0.806133 | 0.763838 | 1 |
| GOBP_NEGATIVE_REGULATION_OF_RECEPTOR_INTERNALIZATION | 15 | -0.299674 | -0.805297 | 0.733333 | 1 |
| GOBP_MITOCHONDRIAL_RNA_PROCESSING | 20 | -0.284121 | -0.805015 | 0.748175 | 1 |
| GOBP_SEMAPHORIN_PLEXIN_SIGNALING_PATHWAY | 42 | -0.237174 | -0.804541 | 0.843318 | 1 |
| GOBP_MRNA_CIS_SPLICING_VIA_SPLICEOSOME | 26 | -0.264073 | -0.804303 | 0.766323 | 1 |
| GOBP_REGULATION_OF_CANONICAL_WNT_SIGNALING_PATHWAY | 226 | -0.182939 | -0.801027 | 0.969697 | 1 |
| GOBP_BILE_ACID_BIOSYNTHETIC_PROCESS | 27 | -0.256672 | -0.800848 | 0.781818 | 1 |
| GOBP_SERINE_FAMILY_AMINO_ACID_BIOSYNTHETIC_PROCESS | 18 | -0.286292 | -0.800455 | 0.75641 | 1 |
| GOBP_REGULATION_OF_PROTEIN_KINASE_C_SIGNALING | 17 | -0.299207 | -0.798873 | 0.732087 | 1 |
| GOBP_REGULATION_OF_ACTOMYOSIN_STRUCTURE_ORGANIZATION | 90 | -0.206975 | -0.79742 | 0.90566 | 1 |
| GOBP_POSITIVE_REGULATION_OF_HISTONE_METHYLATION | 45 | -0.227541 | -0.795881 | 0.826271 | 1 |
| GOBP_REGULATION_OF_MICROTUBULE_POLYMERIZATION_OR_DEPOLYMERI | 84 | -0.202899 | -0.795611 | 0.912752 | 1 |
| GOBP_DEOXYRIBONUCLEOTIDE_BIOSYNTHETIC_PROCESS | 15 | -0.311737 | -0.795427 | 0.766026 | 1 |
| GOBP_REGULATION_OF_JUN_KINASE_ACTIVITY | 51 | -0.228453 | -0.795329 | 0.805 | 1 |
| GOBP_CELLULAR_RESPONSE_TO_AMYLOID_BETA | 33 | -0.244776 | -0.795142 | 0.81746 | 1 |
| GOBP_REGULATION_OF_LIPOPROTEIN_PARTICLE_CLEARANCE | 15 | -0.291385 | -0.793793 | 0.768519 | 1 |
| GOBP_PROTEIN_MODIFICATION_BY_SMALL_PROTEIN_REMOVAL | 146 | -0.191727 | -0.793465 | 0.966292 | 1 |
| GOBP_PROTEIN_TARGETING | 294 | -0.175599 | -0.79294 | 1 | 1 |
| GOBP_REGULATION_OF_ATP_METABOLIC_PROCESS | 77 | -0.20712 | -0.792552 | 0.865169 | 1 |
| GOBP_NEGATIVE_REGULATION_OF_NOTCH_SIGNALING_PATHWAY | 32 | -0.248282 | -0.792543 | 0.818841 | 1 |
| GOBP_BARBED_END_ACTIN_FILAMENT_CAPPING | 25 | -0.259678 | -0.792201 | 0.776596 | 1 |
| GOBP_PEROXISOME_ORGANIZATION | 33 | -0.243687 | -0.791602 | 0.788288 | 1 |
| GOBP_REGULATION_OF_EXIT_FROM_MITOSIS | 16 | -0.29853 | -0.791129 | 0.723404 | 1 |
| GOBP_VESICLE_TRANSPORT_ALONG_MICROTUBULE | 45 | -0.228605 | -0.790842 | 0.84434 | 1 |
| GOBP_GLUTAMINE_METABOLIC_PROCESS | 22 | -0.274297 | -0.790264 | 0.774086 | 1 |
| GOBP_NEGATIVE_REGULATION_OF_PEPTIDE_SECRETION | 32 | -0.243429 | -0.789623 | 0.836 | 1 |
| GOBP_PLASMA_MEMBRANE_TUBULATION | 17 | -0.288273 | -0.788116 | 0.726984 | 1 |
| GOBP_REGULATION_OF_LIPOPROTEIN_METABOLIC_PROCESS | 15 | -0.291309 | -0.787711 | 0.747692 | 1 |
| GOBP_POSITIVE_REGULATION_OF_PROTEIN_EXPORT_FROM_NUCLEUS | 16 | -0.289338 | -0.786356 | 0.760563 | 1 |
| GOBP_REGULATION_OF_TRANSCRIPTION_INVOLVED_IN_G1_S_TRANSITION_( | 20 | -0.277078 | -0.784127 | 0.738351 | 1 |
| GOBP_REGULATION_OF_ESTABLISHMENT_OF_PROTEIN_LOCALIZATION_TO_M | 47 | -0.221482 | -0.780668 | 0.876712 | 1 |
| GOBP_PROTEIN_QUALITY_CONTROL_FOR_MISFOLDED_OR_INCOMPLETELY_S | 27 | -0.252381 | -0.780038 | 0.778182 | 1 |
| GOBP_NEGATIVE_REGULATION_OF_TELOMERE_MAINTENANCE | 32 | -0.239299 | -0.778746 | 0.805785 | 1 |
| GOBP_POSITIVE_REGULATION_OF_SIGNALING_RECEPTOR_ACTIVITY | 35 | -0.238756 | -0.778186 | 0.857143 | 1 |
| GOBP_CELL_SURFACE_RECEPTOR_SIGNALING_PATHWAY_INVOLVED_IN_HE | 25 | -0.254726 | -0.777072 | 0.814286 | 1 |
| GOBP_NEGATIVE_REGULATION_OF_MUSCLE_CELL_APOPTOTIC_PROCESS | 31 | -0.247797 | -0.775461 | 0.823322 | 1 |
| GOBP_FOLIC_ACID_CONTAINING_COMPOUND_METABOLIC_PROCESS | 27 | -0.256912 | -0.775003 | 0.814545 | 1 |
| GOBP_UBIQUINONE_METABOLIC_PROCESS | 18 | -0.27822 | -0.774959 | 0.750789 | 1 |
| GOBP_NEGATIVE_REGULATION_OF_TOLL_LIKE_RECEPTOR_SIGNALING_PATH | 27 | -0.249631 | -0.774841 | 0.843416 | 1 |
| GOBP_FOREBRAIN_CELL_MIGRATION | 56 | -0.211609 | -0.773515 | 0.918129 | 1 |
| GOBP_REGULATION_OF_DOUBLE_STRAND_BREAK_REPAIR_VIA_NONHOMOLO | 31 | -0.243325 | -0.77181 | 0.843478 | 1 |
| GOBP_DNA_METHYLATION_DEPENDENT_HETEROCHROMATIN_ASSEMBLY | 22 | -0.266364 | -0.770879 | 0.801282 | 1 |
| GOBP_MITOCHONDRIAL_RESPIRATORY_CHAIN_COMPLEX_ASSEMBLY | 94 | -0.19413 | -0.769508 | 0.935185 | 1 |
| GOBP_REGULATION_OF_DENDRITIC_SPINE_DEVELOPMENT | 54 | -0.216122 | -0.76851 | 0.868132 | 1 |
| GOBP_POSITIVE_REGULATION_OF_MUSCLE_HYPERTROPHY | 20 | -0.264153 | -0.767693 | 0.815069 | 1 |
| GOBP_NUCLEOTIDE_BINDING_DOMAIN_LEUCINE_RICH_REPEAT_CONTAINING | 22 | -0.264464 | -0.766846 | 0.798701 | 1 |
| GOBP_POSITIVE_REGULATION_OF_LYASE_ACTIVITY | 16 | -0.279897 | -0.765981 | 0.778816 | 1 |
| GOBP_CEREBELLAR_PURKINJE_CELL_LAYER_MORPHOGENESIS | 18 | -0.270836 | -0.764536 | 0.810897 | 1 |
| GOBP_VIRAL_TRANSLATION | 18 | -0.277288 | -0.764371 | 0.798137 | 1 |
| GOBP_DNA_REPLICATION | 267 | -0.171279 | -0.763785 | 1 | 1 |
| GOBP_MACROAUTOPHAGY | 298 | -0.162988 | -0.760469 | 1 | 1 |
| GOBP_REGULATION_OF_RESPONSE_TO_EXTRACELLULAR_STIMULUS | 20 | -0.268494 | -0.760237 | 0.808581 | 1 |
| GOBP_NEGATIVE_REGULATION_OF_DEFENSE_RESPONSE_TO_VIRUS | 25 | -0.251808 | -0.759697 | 0.844068 | 1 |
| GOBP_GANGLIOSIDE_BIOSYNTHETIC_PROCESS | 16 | -0.286695 | -0.75713 | 0.820261 | 1 |
| GOBP_POSITIVE_REGULATION_OF_VACUOLE_ORGANIZATION | 21 | -0.258043 | -0.757036 | 0.791391 | 1 |
| GOBP_POSITIVE_REGULATION_OF_GLYCOLYTIC_PROCESS | 15 | -0.2906 | -0.756521 | 0.792899 | 1 |
| GOBP_POSITIVE_REGULATION_OF_ORGANELLE_ASSEMBLY | 75 | -0.196929 | -0.756514 | 0.968153 | 1 |
| GOBP_RESPIRATORY_ELECTRON_TRANSPORT_CHAIN | 106 | -0.18824 | -0.756379 | 0.95082 | 1 |
| GOBP_DNA_TEMPLATED_TRANSCRIPTION_TERMINATION | 21 | -0.257929 | -0.756328 | 0.841216 | 1 |
| GOBP_REGULATION_OF_FEEDING_BEHAVIOR | 17 | -0.282653 | -0.755111 | 0.801223 | 1 |
| GOBP_PROTEIN_TARGETING_TO_MEMBRANE | 113 | -0.18304 | -0.754563 | 0.97 | 1 |
| GOBP_ATP_BIOSYNTHETIC_PROCESS | 52 | -0.213033 | -0.754075 | 0.905473 | 1 |
| GOBP_VIRAL_BUDDING | 27 | -0.247739 | -0.753827 | 0.848185 | 1 |
| GOBP_POSITIVE_REGULATION_OF_TOR_SIGNALING | 43 | -0.223234 | -0.753529 | 0.896861 | 1 |
| GOBP_POSITIVE_REGULATION_OF_MULTICELLULAR_ORGANISM_GROWTH | 25 | -0.251315 | -0.752995 | 0.847273 | 1 |
| GOBP_REGULATION_OF_CYTOPLASMIC_TRANSLATION | 27 | -0.243444 | -0.752693 | 0.834615 | 1 |
| GOBP_REGULATION_OF_FIBROBLAST_MIGRATION | 34 | -0.231233 | -0.751116 | 0.863281 | 1 |
| GOBP_I_KAPPAB_PHOSPHORYLATION | 16 | -0.282845 | -0.749448 | 0.785714 | 1 |
| GOBP_INDOLE_CONTAINING_COMPOUND_METABOLIC_PROCESS | 15 | -0.276725 | -0.748799 | 0.825 | 1 |
| GOBP_POSITIVE_REGULATION_OF_CYCLASE_ACTIVITY | 16 | -0.281779 | -0.748301 | 0.792793 | 1 |
| GOBP_HISTONE_H4_K5_ACETYLATION | 16 | -0.273126 | -0.747307 | 0.795918 | 1 |
| GOBP_MYELIN_ASSEMBLY | 22 | -0.258895 | -0.747252 | 0.833333 | 1 |
| GOBP_CYTOPLASMIC_PATTERN_RECOGNITION_RECEPTOR_SIGNALING_PATH | 54 | -0.209474 | -0.746805 | 0.887641 | 1 |
| GOBP_CARDIOLIPIN_METABOLIC_PROCESS | 15 | -0.287375 | -0.745514 | 0.81686 | 1 |
| GOBP_NEGATIVE_REGULATION_OF_DNA_BINDING | 45 | -0.219285 | -0.744594 | 0.892019 | 1 |
| GOBP_GLUCOSE_CATABOLIC_PROCESS | 17 | -0.270238 | -0.74386 | 0.814286 | 1 |
| GOBP_MITOCHONDRIAL_DNA_REPLICATION | 15 | -0.27712 | -0.74264 | 0.772358 | 1 |
| GOBP_REGULATION_OF_INTRACELLULAR_STEROID_HORMONE_RECEPTOR_S | 64 | -0.202817 | -0.741292 | 0.957895 | 1 |
| GOBP_RESPIRATORY_SYSTEM_PROCESS | 33 | -0.231286 | -0.740845 | 0.893773 | 1 |
| GOBP_AMYLOID_PRECURSOR_PROTEIN_METABOLIC_PROCESS | 67 | -0.198403 | -0.739355 | 0.953488 | 1 |
| GOBP_REGULATION_OF_CLATHRIN_DEPENDENT_ENDOCYTOSIS | 18 | -0.261907 | -0.739172 | 0.83125 | 1 |
| GOBP_AUTOPHAGOSOME_ORGANIZATION | 105 | -0.182091 | -0.736303 | 0.976563 | 1 |
| GOBP_REGULATION_OF_MESENCHYMAL_CELL_PROLIFERATION | 22 | -0.248282 | -0.735506 | 0.872852 | 1 |
| GOBP_DEVELOPMENTAL_PIGMENTATION | 39 | -0.216632 | -0.735459 | 0.880658 | 1 |
| GOBP_CENTRIOLE_ASSEMBLY | 43 | -0.21048 | -0.735065 | 0.918182 | 1 |
| GOBP_CYTOPLASMIC_MICROTUBULE_ORGANIZATION | 55 | -0.206773 | -0.733909 | 0.91623 | 1 |
| GOBP_DENDRITIC_SPINE_MAINTENANCE | 15 | -0.275007 | -0.731813 | 0.816619 | 1 |
| GOBP_NEGATIVE_REGULATION_OF_SIGNALING_RECEPTOR_ACTIVITY | 39 | -0.218988 | -0.731525 | 0.913793 | 1 |
| GOBP_REGULATION_OF_FIBROBLAST_GROWTH_FACTOR_RECEPTOR_SIGNAL | 30 | -0.235965 | -0.731228 | 0.889299 | 1 |
| GOBP_FOREBRAIN_NEURON_DEVELOPMENT | 21 | -0.25921 | -0.730311 | 0.835052 | 1 |
| GOBP_AUTOPHAGOSOME_MATURATION | 58 | -0.202723 | -0.728951 | 0.946809 | 1 |
| GOBP_P38MAPK_CASCADE | 46 | -0.21157 | -0.728275 | 0.922727 | 1 |
| GOBP_PROTEIN_ACYLATION | 238 | -0.159453 | -0.7259 | 1 | 1 |
| GOBP_REGULATION_OF_DNA_BIOSYNTHETIC_PROCESS | 117 | -0.173567 | -0.725852 | 1 | 1 |
| GOBP_SEMAPHORIN_PLEXIN_SIGNALING_PATHWAY_INVOLVED_IN_NEURON_ | 16 | -0.260825 | -0.725715 | 0.820755 | 1 |
| GOBP_POSITIVE_REGULATION_OF_JUN_KINASE_ACTIVITY | 38 | -0.216123 | -0.725616 | 0.936759 | 1 |
| GOBP_NEUROTROPHIN_TRK_RECEPTOR_SIGNALING_PATHWAY | 22 | -0.239221 | -0.723495 | 0.859589 | 1 |
| GOBP_HIPPO_SIGNALING | 39 | -0.216223 | -0.723184 | 0.938389 | 1 |
| GOBP_POSITIVE_REGULATION_OF_NOTCH_SIGNALING_PATHWAY | 36 | -0.224573 | -0.723147 | 0.941964 | 1 |
| GOBP_REGULATION_OF_CHROMATIN_ASSEMBLY | 29 | -0.22715 | -0.723103 | 0.882353 | 1 |
| GOBP_CYTOKINETIC_PROCESS | 39 | -0.211873 | -0.720232 | 0.92511 | 1 |
| GOBP_CELLULAR_PIGMENTATION | 50 | -0.208508 | -0.720148 | 0.960591 | 1 |
| GOBP_MICROTUBULE_POLYMERIZATION_OR_DEPOLYMERIZATION | 118 | -0.174345 | -0.7199 | 0.978723 | 1 |
| GOBP_NEGATIVE_REGULATION_OF_RESPONSE_TO_DNA_DAMAGE_STIMULUS | 80 | -0.185751 | -0.719576 | 0.95092 | 1 |
| GOBP_SCF_DEPENDENT_PROTEASOMAL_UBIQUITIN_DEPENDENT_PROTEIN_( | 47 | -0.20898 | -0.717911 | 0.905759 | 1 |
| GOBP_NUCLEAR_ENVELOPE_ORGANIZATION | 55 | -0.206009 | -0.715795 | 0.945946 | 1 |
| GOBP_N_TERMINAL_PROTEIN_AMINO_ACID_ACETYLATION | 15 | -0.268338 | -0.715504 | 0.863787 | 1 |
| GOBP_PROTEIN_MONOUBIQUITINATION | 78 | -0.188317 | -0.713401 | 0.955224 | 1 |
| GOBP_AMYLOID_PRECURSOR_PROTEIN_CATABOLIC_PROCESS | 50 | -0.198079 | -0.711746 | 0.949239 | 1 |
| GOBP_FLUID_TRANSPORT | 20 | -0.245479 | -0.709227 | 0.849673 | 1 |
| GOBP_REGULATION_OF_GLYCOGEN_METABOLIC_PROCESS | 29 | -0.230895 | -0.709028 | 0.908046 | 1 |
| GOBP_REGULATION_OF_MITOCHONDRIAL_GENE_EXPRESSION | 28 | -0.235245 | -0.705904 | 0.909836 | 1 |
| GOBP_MITOCHONDRIAL_GENE_EXPRESSION | 102 | -0.171531 | -0.705709 | 0.970588 | 1 |
| GOBP_REGULATION_OF_CARDIAC_MUSCLE_CONTRACTION_BY_REGULATION | 18 | -0.253906 | -0.705604 | 0.888158 | 1 |
| GOBP_PROTEIN_LOCALIZATION_TO_CELL_JUNCTION | 87 | -0.179162 | -0.70453 | 1 | 1 |
| GOBP_ENDOSOME_TO_LYSOSOME_TRANSPORT | 58 | -0.19423 | -0.70332 | 0.934673 | 1 |
| GOBP_ELECTRON_TRANSPORT_CHAIN | 149 | -0.169497 | -0.702498 | 0.989796 | 1 |
| GOBP_MRNA_TRANSCRIPTION | 39 | -0.211301 | -0.701642 | 0.945607 | 1 |
| GOBP_NEGATIVE_REGULATION_OF_STEROID_METABOLIC_PROCESS | 22 | -0.233783 | -0.701449 | 0.884354 | 1 |
| GOBP_POSITIVE_REGULATION_OF_CATION_CHANNEL_ACTIVITY | 57 | -0.195877 | -0.697781 | 0.95858 | 1 |
| GOBP_NEURON_PROJECTION_ARBORIZATION | 29 | -0.222847 | -0.697487 | 0.928826 | 1 |
| GOBP_ESTABLISHMENT_OF_PROTEIN_LOCALIZATION_TO_TELOMERE | 17 | -0.25397 | -0.696352 | 0.880734 | 1 |
| GOBP_NEGATIVE_REGULATION_OF_MACROAUTOPHAGY | 34 | -0.211358 | -0.694623 | 0.925651 | 1 |
| GOBP_3_UTR_MEDIATED_MRNA_DESTABILIZATION | 15 | -0.265627 | -0.694292 | 0.882353 | 1 |
| GOBP_NEGATIVE_REGULATION_OF_CILIUM_ASSEMBLY | 17 | -0.250534 | -0.693913 | 0.889241 | 1 |
| GOBP_RNA_3_END_PROCESSING | 104 | -0.172019 | -0.69295 | 0.992754 | 1 |
| GOBP_OLIGOSACCHARIDE_LIPID_INTERMEDIATE_BIOSYNTHETIC_PROCESS | 19 | -0.243565 | -0.691704 | 0.851266 | 1 |
| GOBP_ESTABLISHMENT_OR_MAINTENANCE_OF_BIPOLAR_CELL_POLARITY | 48 | -0.194443 | -0.690669 | 0.977478 | 1 |

| | | | | | |
|---|---|---|---|---|---|
| GOBP_CELLULAR_OXIDANT_DETOXIFICATION | 73 | 0.414515 | 1.254681 | 0.135613 | 0.524878 |
| GOBP_LYMPHOCYTE_ACTIVATION_INVOLVED_IN_IMMUNE_RESPONSE | 134 | 0.394244 | 1.25418 | 0.096739 | 0.525751 |
| GOBP_REGULATION_OF_EPITHELIAL_CELL_APOPTOTIC_PROCESS | 66 | 0.418821 | 1.253985 | 0.135922 | 0.5257 |
| GOBP_REGULATION_OF_ION_TRANSMEMBRANE_TRANSPORT | 380 | 0.366134 | 1.253522 | 0.037449 | 0.526435 |
| GOBP_MUSCLE_ORGAN_MORPHOGENESIS | 65 | 0.419146 | 1.25314 | 0.139706 | 0.526951 |
| GOBP_REGULATION_OF_T_CELL_MIGRATION | 29 | 0.479404 | 1.252282 | 0.165545 | 0.528942 |
| GOBP_REGULATION_OF_PRODUCTION_OF_SMALL_RNA_INVOLVED_IN_GENE_SILENCING_ | 20 | 0.517765 | 1.251273 | 0.188489 | 0.531499 |
| GOBP_CELL_DIFFERENTIATION_INVOLVED_IN_KIDNEY_DEVELOPMENT | 49 | 0.444683 | 1.251216 | 0.155666 | 0.531008 |
| GOBP_REGULATION_OF_CELLULAR_KETONE_METABOLIC_PROCESS | 98 | 0.402257 | 1.251127 | 0.111738 | 0.530659 |
| GOBP_POSITIVE_REGULATION_OF_ANION_TRANSPORT | 37 | 0.464035 | 1.249621 | 0.167336 | 0.534751 |
| GOBP_LIVER_MORPHOGENESIS | 20 | 0.521995 | 1.249393 | 0.193833 | 0.534774 |
| GOBP_CELLULAR_RESPONSE_TO_LIPOPROTEIN_PARTICLE_STIMULUS | 26 | 0.49823 | 1.249088 | 0.180791 | 0.535028 |
| GOBP_EMBRYO_IMPLANTATION | 48 | 0.431905 | 1.248784 | 0.136364 | 0.535342 |
| GOBP_POSITIVE_CHEMOTAXIS | 41 | 0.45468 | 1.248705 | 0.165385 | 0.534935 |
| GOBP_RESPONSE_TO_NUTRIENT | 113 | 0.391774 | 1.248274 | 0.113839 | 0.535625 |
| GOBP_PHOSPHATIDYLETHANOLAMINE_METABOLIC_PROCESS | 24 | 0.491205 | 1.247655 | 0.19323 | 0.536941 |
| GOBP_BLASTOCYST_FORMATION | 37 | 0.465342 | 1.247345 | 0.161917 | 0.537246 |
| GOBP_ACUTE_INFLAMMATORY_RESPONSE | 63 | 0.427176 | 1.24713 | 0.150485 | 0.537279 |
| GOBP_CELLULAR_RESPONSE_TO_XENOBIOTIC_STIMULUS | 114 | 0.398229 | 1.24706 | 0.12051 | 0.536856 |
| GOBP_SUPEROXIDE_ANION_GENERATION | 29 | 0.471917 | 1.246018 | 0.180263 | 0.539499 |
| GOBP_RESPONSE_TO_TEMPERATURE_STIMULUS | 153 | 0.386293 | 1.245889 | 0.098501 | 0.539286 |
| GOBP_REGULATION_OF_RELEASE_OF_CYTOCHROME_C_FROM_MITOCHONDRIA | 41 | 0.450622 | 1.245332 | 0.169713 | 0.540491 |
| GOBP_NEGATIVE_REGULATION_OF_RESPONSE_TO_CYTOKINE_STIMULUS | 58 | 0.425707 | 1.244901 | 0.15621 | 0.541171 |
| GOBP_HEART_VALVE_DEVELOPMENT | 61 | 0.422053 | 1.244885 | 0.148649 | 0.54056 |
| GOBP_DIVALENT_INORGANIC_CATION_HOMEOSTASIS | 367 | 0.366688 | 1.244809 | 0.031536 | 0.540127 |
| GOBP_RELAXATION_OF_CARDIAC_MUSCLE | 16 | 0.539989 | 1.244672 | 0.206687 | 0.539886 |
| GOBP_GENITALIA_DEVELOPMENT | 28 | 0.479384 | 1.244268 | 0.182944 | 0.540468 |
| GOBP_CELLULAR_RESPONSE_TO_DSRNA | 21 | 0.504633 | 1.244013 | 0.193501 | 0.540567 |
| GOBP_RESPONSE_TO_EXTRACELLULAR_STIMULUS | 393 | 0.363791 | 1.243285 | 0.031377 | 0.54225 |
| GOBP_POSITIVE_REGULATION_OF_KERATINOCYTE_DIFFERENTIATION | 16 | 0.530198 | 1.242719 | 0.218289 | 0.54341 |
| GOBP_EPITHELIAL_CELL_APOPTOTIC_PROCESS | 87 | 0.403286 | 1.242247 | 0.148912 | 0.54427 |
| GOBP_POSITIVE_REGULATION_OF_NIK_NF_KAPPAB_SIGNALING | 51 | 0.43625 | 1.24199 | 0.166256 | 0.544457 |
| GOBP_POSITIVE_REGULATION_OF_ION_TRANSPORT | 217 | 0.370019 | 1.241972 | 0.08805 | 0.543843 |
| GOBP_IMMUNOGLOBULIN_PRODUCTION | 94 | 0.403687 | 1.241132 | 0.130533 | 0.545942 |
| GOBP_T_CELL_DIFFERENTIATION_IN_THYMUS | 57 | 0.433069 | 1.240914 | 0.161006 | 0.545982 |
| GOBP_PROSTATE_GLAND_DEVELOPMENT | 36 | 0.469522 | 1.240722 | 0.183246 | 0.545967 |
| GOBP_NECROTIC_CELL_DEATH | 50 | 0.43346 | 1.240697 | 0.15444 | 0.545383 |
| GOBP_CELLULAR_RESPONSE_TO_INTERFERON_BETA | 15 | 0.535402 | 1.240434 | 0.207849 | 0.54555 |
| GOBP_CELLULAR_RESPONSE_TO_OXYGEN_LEVELS | 141 | 0.385909 | 1.240146 | 0.102371 | 0.545771 |
| GOBP_NEGATIVE_REGULATION_OF_MUSCLE_HYPERTROPHY | 24 | 0.493854 | 1.239861 | 0.202797 | 0.546066 |
| GOBP_DETECTION_OF_BIOTIC_STIMULUS | 24 | 0.489282 | 1.239569 | 0.202817 | 0.546296 |
| GOBP_REGULATION_OF_PLATELET_DERIVED_GROWTH_FACTOR_RECEPTOR_SIGNALING_ | 19 | 0.514641 | 1.239507 | 0.213139 | 0.545823 |
| GOBP_REGULATION_OF_RHO_PROTEIN_SIGNAL_TRANSDUCTION | 74 | 0.413435 | 1.239385 | 0.131673 | 0.545549 |
| GOBP_REGULATION_OF_G_PROTEIN_COUPLED_RECEPTOR_SIGNALING_PATHWAY | 113 | 0.393797 | 1.239006 | 0.122886 | 0.546091 |
| GOBP_EPITHELIAL_STRUCTURE_MAINTENANCE | 22 | 0.49794 | 1.238819 | 0.19225 | 0.546041 |
| GOBP_DNA_DOUBLE_STRAND_BREAK_PROCESSING | 16 | 0.538718 | 1.238516 | 0.218563 | 0.546364 |
| GOBP_CELLULAR_RESPONSE_TO_CORTICOSTEROID_STIMULUS | 48 | 0.435876 | 1.237861 | 0.164808 | 0.54791 |
| GOBP_NEGATIVE_REGULATION_OF_MYOBLAST_DIFFERENTIATION | 19 | 0.51209 | 1.236655 | 0.220117 | 0.551113 |
| GOBP_DETECTION_OF_LIGHT_STIMULUS | 34 | 0.460246 | 1.236377 | 0.205645 | 0.551353 |
| GOBP_POSITIVE_REGULATION_OF_FILOPODIUM_ASSEMBLY | 27 | 0.477693 | 1.235901 | 0.201669 | 0.552225 |
| GOBP_MUSCLE_CELL_DIFFERENTIATION | 293 | 0.366192 | 1.235301 | 0.059244 | 0.553462 |
| GOBP_CELLULAR_RESPONSE_TO_TOXIC_SUBSTANCE | 92 | 0.401674 | 1.234684 | 0.136259 | 0.55477 |
| GOBP_HOMOLOGOUS_RECOMBINATION | 50 | 0.435054 | 1.234665 | 0.164331 | 0.554141 |
| GOBP_LIPID_STORAGE | 67 | 0.415766 | 1.234138 | 0.172209 | 0.555149 |
| GOBP_NATURAL_KILLER_CELL_DIFFERENTIATION | 20 | 0.506099 | 1.233663 | 0.193732 | 0.555995 |
| GOBP_APOPTOTIC_PROCESS_INVOLVED_IN_MORPHOGENESIS | 23 | 0.494329 | 1.23358 | 0.234352 | 0.555597 |
| GOBP_MACROPHAGE_MIGRATION | 33 | 0.461068 | 1.233326 | 0.191099 | 0.555731 |
| GOBP_CARDIAC_CHAMBER_MORPHOGENESIS | 109 | 0.393295 | 1.232616 | 0.137413 | 0.557311 |
| GOBP_LYMPHOCYTE_COSTIMULATION | 27 | 0.47489 | 1.23247 | 0.193906 | 0.557155 |
| GOBP_RENAL_SYSTEM_PROCESS_INVOLVED_IN_REGULATION_OF_SYSTEMIC_ARTERIAL_ | 15 | 0.543055 | 1.232276 | 0.221739 | 0.557091 |
| GOBP_RESPONSE_TO_CHOLESTEROL | 19 | 0.511622 | 1.231611 | 0.212339 | 0.558538 |
| GOBP_MACROPHAGE_CYTOKINE_PRODUCTION | 15 | 0.547283 | 1.231488 | 0.22406 | 0.558239 |
| GOBP_REGULATION_OF_MEMBRANE_REPOLARIZATION | 27 | 0.476357 | 1.231391 | 0.210526 | 0.557892 |
| GOBP_ACTIN_CYTOSKELETON_REORGANIZATION | 97 | 0.399182 | 1.231243 | 0.144009 | 0.557712 |
| GOBP_PRESYNAPTIC_ENDOCYTOSIS | 57 | 0.423202 | 1.231099 | 0.170343 | 0.55751 |
| GOBP_POSITIVE_REGULATION_OF_T_CELL_MEDIATED_CYTOTOXICITY | 20 | 0.518252 | 1.230826 | 0.217765 | 0.557685 |
| GOBP_FEEDING_BEHAVIOR | 70 | 0.410768 | 1.230439 | 0.157647 | 0.558282 |
| GOBP_CARDIAC_VENTRICLE_MORPHOGENESIS | 63 | 0.416689 | 1.228998 | 0.148325 | 0.56216 |
| GOBP_ARTERY_DEVELOPMENT | 84 | 0.402118 | 1.22893 | 0.153216 | 0.561728 |
| GOBP_REGULATION_OF_SMALL_MOLECULE_METABOLIC_PROCESS | 274 | 0.361914 | 1.228473 | 0.085655 | 0.562539 |
| GOBP_POSITIVE_REGULATION_OF_TUMOR_NECROSIS_FACTOR_SUPERFAMILY_CYTOKINE | 73 | 0.404622 | 1.228433 | 0.160714 | 0.561992 |
| GOBP_CATECHOLAMINE_UPTAKE | 17 | 0.528782 | 1.228385 | 0.21203 | 0.561507 |
| GOBP_EPITHELIAL_CELL_DIFFERENTIATION | 489 | 0.3566 | 1.228357 | 0.03144 | 0.560962 |
| GOBP_NEGATIVE_REGULATION_OF_GROWTH | 214 | 0.368529 | 1.227736 | 0.091675 | 0.562259 |
| GOBP_REGULATION_OF_POSITIVE_CHEMOTAXIS | 20 | 0.501095 | 1.227543 | 0.229985 | 0.562221 |
| GOBP_OUTFLOW_TRACT_SEPTUM_MORPHOGENESIS | 22 | 0.503637 | 1.227258 | 0.21262 | 0.562562 |
| GOBP_RESPONSE_TO_STEROID_HORMONE | 266 | 0.365907 | 1.226694 | 0.06893 | 0.563818 |
| GOBP_NEGATIVE_REGULATION_OF_INTERLEUKIN_1_PRODUCTION | 20 | 0.509581 | 1.226686 | 0.228448 | 0.563177 |
| GOBP_SKELETAL_MUSCLE_CELL_DIFFERENTIATION | 52 | 0.427138 | 1.226672 | 0.180285 | 0.562568 |
| GOBP_IRON_ION_TRANSMEMBRANE_TRANSPORT | 17 | 0.522373 | 1.226653 | 0.210756 | 0.561965 |
| GOBP_POSITIVE_REGULATION_OF_PEPTIDE_SECRETION | 72 | 0.409732 | 1.226064 | 0.150179 | 0.563192 |
| GOBP_FAT_CELL_DIFFERENTIATION | 205 | 0.370202 | 1.225724 | 0.09428 | 0.563651 |
| GOBP_REGULATION_OF_RESPONSE_TO_BIOTIC_STIMULUS | 251 | 0.364383 | 1.224483 | 0.079958 | 0.567049 |
| GOBP_REGULATION_OF_HOMOTYPIC_CELL_CELL_ADHESION | 27 | 0.482915 | 1.224044 | 0.200855 | 0.567807 |
| GOBP_REGULATION_OF_TRANSMEMBRANE_TRANSPORT | 450 | 0.355293 | 1.223966 | 0.029087 | 0.567387 |
| GOBP_EXECUTION_PHASE_OF_APOPTOSIS | 64 | 0.407114 | 1.223691 | 0.171951 | 0.567611 |
| GOBP_PURINE_DEOXYRIBONUCLEOTIDE_METABOLIC_PROCESS | 15 | 0.524904 | 1.222812 | 0.233786 | 0.569848 |
| GOBP_NEGATIVE_REGULATION_OF_CELL_ADHESION | 218 | 0.370447 | 1.222565 | 0.088727 | 0.569968 |
| GOBP_PHOTORECEPTOR_CELL_DIFFERENTIATION | 48 | 0.430262 | 1.222562 | 0.193256 | 0.569327 |
| GOBP_CELLULAR_RESPONSE_TO_EXOGENOUS_DSRNA | 16 | 0.529568 | 1.22248 | 0.211731 | 0.568948 |
| GOBP_POSITIVE_REGULATION_OF_REACTIVE_OXYGEN_SPECIES_METABOLIC_PROCESS | 53 | 0.429644 | 1.22243 | 0.18 | 0.568459 |
| GOBP_CALCIUM_ION_REGULATED_EXOCYTOSIS_OF_NEUROTRANSMITTER | 15 | 0.526316 | 1.222419 | 0.220211 | 0.567847 |
| GOBP_REGULATION_OF_B_CELL_RECEPTOR_SIGNALING_PATHWAY | 15 | 0.54513 | 1.222188 | 0.233577 | 0.567948 |
| GOBP_MUSCLE_HYPERTROPHY | 65 | 0.413712 | 1.221838 | 0.173445 | 0.56841 |
| GOBP_CELLULAR_RESPONSE_TO_VIRUS | 63 | 0.412295 | 1.220758 | 0.187576 | 0.571174 |
| GOBP_INFLAMMASOME_COMPLEX_ASSEMBLY | 16 | 0.524064 | 1.220629 | 0.252924 | 0.570949 |
| GOBP_REGULATION_OF_TUBE_SIZE | 108 | 0.38794 | 1.220264 | 0.138636 | 0.571429 |
| GOBP_MUSCLE_CELL_MIGRATION | 71 | 0.403107 | 1.219129 | 0.166467 | 0.574399 |
| GOBP_REGULATION_OF_REGULATED_SECRETORY_PATHWAY | 121 | 0.386455 | 1.21875 | 0.138165 | 0.575 |
| GOBP_REGULATION_OF_CARBOHYDRATE_METABOLIC_PROCESS | 155 | 0.374551 | 1.218698 | 0.117838 | 0.574546 |
| GOBP_POSITIVE_REGULATION_OF_SMALL_GTPASE_MEDIATED_SIGNAL_TRANSDUCTION | 53 | 0.42726 | 1.21795 | 0.185606 | 0.576384 |
| GOBP_MEIOTIC_CHROMOSOME_SEPARATION | 21 | 0.505508 | 1.217949 | 0.219686 | 0.575734 |
| GOBP_REGULATION_OF_PROTEIN_SECRETION | 199 | 0.366512 | 1.217789 | 0.101064 | 0.575586 |
| GOBP_MONOCARBOXYLIC_ACID_BIOSYNTHETIC_PROCESS | 159 | 0.372943 | 1.217167 | 0.134698 | 0.577019 |
| GOBP_OOCYTE_DIFFERENTIATION | 50 | 0.427611 | 1.216947 | 0.186548 | 0.57709 |
| GOBP_EPIDERMIS_MORPHOGENESIS | 22 | 0.490616 | 1.216932 | 0.217631 | 0.576479 |
| GOBP_RHYTHMIC_BEHAVIOR | 36 | 0.445323 | 1.216168 | 0.21875 | 0.578302 |
| GOBP_EMBRYONIC_HEART_TUBE_DEVELOPMENT | 73 | 0.405535 | 1.215837 | 0.182473 | 0.578738 |
| GOBP_CELL_CELL_ADHESION_MEDIATED_BY_CADHERIN | 26 | 0.468793 | 1.2158 | 0.207736 | 0.5782 |
| GOBP_POSITIVE_REGULATION_OF_PHAGOCYTOSIS | 42 | 0.441023 | 1.215286 | 0.188608 | 0.579234 |
| GOBP_REACTIVE_NITROGEN_SPECIES_METABOLIC_PROCESS | 60 | 0.416216 | 1.215122 | 0.185819 | 0.579151 |
| GOBP_EPITHELIAL_CELL_PROLIFERATION | 323 | 0.35809 | 1.214426 | 0.07996 | 0.580744 |
| GOBP_PROXIMAL_DISTAL_PATTERN_FORMATION | 18 | 0.512433 | 1.214336 | 0.218978 | 0.580375 |
| GOBP_ACTIVATION_OF_PROTEIN_KINASE_ACTIVITY | 122 | 0.380646 | 1.214335 | 0.141899 | 0.579729 |
| GOBP_REGULATION_OF_B_CELL_MEDIATED_IMMUNITY | 47 | 0.437642 | 1.212888 | 0.209451 | 0.583824 |
| GOBP_REGULATION_OF_BONE_REMODELING | 38 | 0.444194 | 1.212763 | 0.208279 | 0.583589 |
| GOBP_REGULATION_OF_CILIUM_DEPENDENT_CELL_MOTILITY | 19 | 0.50982 | 1.212547 | 0.215976 | 0.583646 |
| GOBP_RETINA_HOMEOSTASIS | 51 | 0.415392 | 1.21253 | 0.195624 | 0.583062 |
| GOBP_PEROXISOME_PROLIFERATOR_ACTIVATED_RECEPTOR_SIGNALING_PATHWAY | 20 | 0.509867 | 1.212465 | 0.232323 | 0.582622 |
| GOBP_POSITIVE_REGULATION_OF_FATTY_ACID_TRANSPORT | 18 | 0.509664 | 1.212379 | 0.231905 | 0.582252 |
| GOBP_POSITIVE_REGULATION_OF_ACTIVATED_T_CELL_PROLIFERATION | 17 | 0.523418 | 1.212296 | 0.232384 | 0.581877 |
| GOBP_RESPONSE_TO_ISOQUINOLINE_ALKALOID | 27 | 0.470054 | 1.212276 | 0.218968 | 0.581297 |
| GOBP_POSITIVE_REGULATION_OF_STEM_CELL_PROLIFERATION | 21 | 0.494685 | 1.211897 | 0.234072 | 0.581855 |
| GOBP_NEGATIVE_REGULATION_OF_T_CELL_PROLIFERATION | 44 | 0.431612 | 1.211805 | 0.205772 | 0.581505 |
| GOBP_REGULATION_OF_MACROPHAGE_MIGRATION | 24 | 0.479489 | 1.211649 | 0.227763 | 0.581372 |
| GOBP_SKELETAL_SYSTEM_DEVELOPMENT | 407 | 0.35593 | 1.21127 | 0.054545 | 0.581995 |
| GOBP_REGULATION_OF_CALCIUM_ION_TRANSMEMBRANE_TRANSPORTER_ACTIVITY | 73 | 0.396877 | 1.210613 | 0.198824 | 0.583474 |
| GOBP_TUMOR_NECROSIS_FACTOR_SUPERFAMILY_CYTOKINE_PRODUCTION | 114 | 0.383572 | 1.210468 | 0.167416 | 0.583282 |
| GOBP_NEGATIVE_REGULATION_OF_POSTTRANSCRIPTIONAL_GENE_SILENCING | 15 | 0.517903 | 1.210157 | 0.245083 | 0.583636 |
| GOBP_NEGATIVE_REGULATION_OF_PEPTIDYL_TYROSINE_PHOSPHORYLATION | 51 | 0.421072 | 1.208802 | 0.19603 | 0.587552 |
| GOBP_EPIBOLY | 33 | 0.443468 | 1.208711 | 0.21123 | 0.587213 |
| GOBP_CORONARY_VASCULATURE_DEVELOPMENT | 41 | 0.436947 | 1.208694 | 0.195822 | 0.586624 |
| GOBP_DNA_CATABOLIC_PROCESS_ENDONUCLEOLYTIC | 21 | 0.486918 | 1.20841 | 0.25641 | 0.586889 |
| GOBP_NEGATIVE_REGULATION_OF_CELLULAR_RESPONSE_TO_GROWTH_FACTOR_STIMU | 88 | 0.391342 | 1.207377 | 0.171129 | 0.589634 |
| GOBP_REGULATION_OF_MYELOID_LEUKOCYTE_DIFFERENTIATION | 86 | 0.391604 | 1.207183 | 0.169683 | 0.589645 |
| GOBP_RESPONSE_TO_ACID_CHEMICAL | 109 | 0.38015 | 1.206722 | 0.15162 | 0.590439 |
| GOBP_CELLULAR_RESPONSE_TO_CHEMICAL_STRESS | 291 | 0.358045 | 1.20645 | 0.094608 | 0.590747 |
| GOBP_SISTER_CHROMATID_COHESION | 59 | 0.41108 | 1.205862 | 0.189591 | 0.59204 |
| GOBP_CALCIUM_ION_IMPORT | 73 | 0.39781 | 1.205805 | 0.186962 | 0.591591 |
| GOBP_T_CELL_MIGRATION | 44 | 0.434655 | 1.205591 | 0.191677 | 0.59166 |
| GOBP_POSITIVE_REGULATION_OF_PEPTIDYL_LYSINE_ACETYLATION | 35 | 0.455846 | 1.20554 | 0.196501 | 0.591195 |
| GOBP_MYELIN_MAINTENANCE | 16 | 0.528584 | 1.20528 | 0.245614 | 0.591355 |
| GOBP_REGULATION_OF_VASCULAR_ENDOTHELIAL_GROWTH_FACTOR_RECEPTOR_SIGNA | 22 | 0.489727 | 1.20504 | 0.219178 | 0.591511 |
| GOBP_POSITIVE_REGULATION_OF_ORGANIC_ACID_TRANSPORT | 34 | 0.456454 | 1.204751 | 0.228947 | 0.591817 |
| GOBP_BEHAVIORAL_RESPONSE_TO_PAIN | 15 | 0.520483 | 1.204735 | 0.244573 | 0.591234 |
| GOBP_POSITIVE_REGULATION_OF_SPROUTING_ANGIOGENESIS | 18 | 0.509116 | 1.204456 | 0.246418 | 0.591481 |
| GOBP_REACTIVE_OXYGEN_SPECIES_METABOLIC_PROCESS | 171 | 0.363392 | 1.203277 | 0.126461 | 0.594729 |
| GOBP_GLYCOSAMINOGLYCAN_CATABOLIC_PROCESS | 19 | 0.494092 | 1.20327 | 0.23839 | 0.594113 |
| GOBP_BILE_ACID_AND_BILE_SALT_TRANSPORT | 17 | 0.509683 | 1.202547 | 0.253602 | 0.595886 |
| GOBP_RESPONSE_TO_MONOSACCHARIDE | 155 | 0.369648 | 1.201884 | 0.144708 | 0.597485 |
| GOBP_INTEGRIN_ACTIVATION | 20 | 0.500859 | 1.201772 | 0.218475 | 0.597207 |
| GOBP_GLYCOSIDE_METABOLIC_PROCESS | 18 | 0.502453 | 1.201652 | 0.251445 | 0.596941 |
| GOBP_NEGATIVE_REGULATION_OF_TRANSMEMBRANE_RECEPTOR_PROTEIN_SERINE_THI | 112 | 0.377796 | 1.201582 | 0.158014 | 0.596544 |
| GOBP_GLOMERULAR_EPITHELIUM_DEVELOPMENT | 22 | 0.48753 | 1.201359 | 0.24 | 0.596641 |
| GOBP_REGULATION_OF_ORGANIC_ACID_TRANSPORT | 57 | 0.417909 | 1.200987 | 0.189055 | 0.597299 |
| GOBP_HETEROPHILIC_CELL_CELL_ADHESION_VIA_PLASMA_MEMBRANE_CELL_ADHESION | 32 | 0.451246 | 1.200766 | 0.225503 | 0.597395 |
| GOBP_GLAND_DEVELOPMENT | 337 | 0.352864 | 1.200747 | 0.087133 | 0.596813 |

| | | | | | |
|---|---|---|---|---|---|
| GOBP_POSITIVE_REGULATION_BY_HOST_OF_VIRAL_TRANSCRIPTION | 17 | -0.255787 | -0.689896 | 0.892537 | 1 |
| GOBP_REGULATION_OF_PROTEIN_STABILITY | 272 | -0.155156 | -0.689858 | 1 | 1 |
| GOBP_PROTEIN_K11_LINKED_UBIQUITINATION | 29 | -0.223413 | -0.689259 | 0.941177 | 1 |
| GOBP_MITOTIC_SPINDLE_ASSEMBLY | 70 | -0.18426 | -0.689039 | 0.987654 | 1 |
| GOBP_PROTEIN_K63_LINKED_DEUBIQUITINATION | 33 | -0.21031 | -0.687605 | 0.928571 | 1 |
| GOBP_REGULATION_OF_CHROMOSOME_SEPARATION | 66 | -0.187085 | -0.68728 | 0.973404 | 1 |
| GOBP_POSITIVE_REGULATION_OF_STRESS_FIBER_ASSEMBLY | 49 | -0.19974 | -0.686857 | 0.961538 | 1 |
| GOBP_PROTEIN_LOCALIZATION_TO_CILIARY_MEMBRANE | 17 | -0.251741 | -0.68627 | 0.889213 | 1 |
| GOBP_VITAMIN_TRANSMEMBRANE_TRANSPORT | 15 | -0.257253 | -0.684297 | 0.870057 | 1 |
| GOBP_ANDROGEN_METABOLIC_PROCESS | 17 | -0.248633 | -0.683849 | 0.898413 | 1 |
| GOBP_SUPPRESSION_OF_VIRAL_RELEASE_BY_HOST | 16 | -0.255499 | -0.680337 | 0.877493 | 1 |
| GOBP_PHAGOCYTOSIS_RECOGNITION | 16 | -0.244773 | -0.675766 | 0.874214 | 1 |
| GOBP_VACUOLE_ORGANIZATION | 176 | -0.158541 | -0.67549 | 1 | 1 |
| GOBP_ENDOCYTIC_RECYCLING | 74 | -0.17628 | -0.674548 | 0.96732 | 1 |
| GOBP_NEGATIVE_REGULATION_OF_OXIDATIVE_STRESS_INDUCED_INTRINSIC | 18 | -0.244838 | -0.673541 | 0.901639 | 1 |
| GOBP_REGULATION_OF_MRNA_3_END_PROCESSING | 28 | -0.223918 | -0.672071 | 0.916667 | 1 |
| GOBP_REGULATION_OF_CILIUM_ASSEMBLY | 82 | -0.173593 | -0.672071 | 0.993103 | 1 |
| GOBP_AMINO_ACID_ACTIVATION | 45 | -0.195583 | -0.672062 | 0.933014 | 1 |
| GOBP_POSITIVE_REGULATION_OF_MACROAUTOPHAGY | 69 | -0.177856 | -0.671762 | 0.988506 | 1 |
| GOBP_NEGATIVE_REGULATION_OF_VIRAL_TRANSCRIPTION | 15 | -0.249938 | -0.671094 | 0.86076 | 1 |
| GOBP_CYTOPLASMIC_PATTERN_RECOGNITION_RECEPTOR_SIGNALING_PATH | 31 | -0.215607 | -0.670917 | 0.916335 | 1 |
| GOBP_NEUROTROPHIN_SIGNALING_PATHWAY | 34 | -0.205877 | -0.670903 | 0.92766 | 1 |
| GOBP_MICROTUBULE_ANCHORING | 24 | -0.220674 | -0.669546 | 0.9375 | 1 |
| GOBP_TRNA_METHYLATION | 40 | -0.196433 | -0.668149 | 0.956757 | 1 |
| GOBP_PROTEIN_MATURATION_BY_IRON_SULFUR_CLUSTER_TRANSFER | 15 | -0.247411 | -0.668023 | 0.892857 | 1 |
| GOBP_VESICLE_DOCKING_INVOLVED_IN_EXOCYTOSIS | 34 | -0.203532 | -0.667145 | 0.957447 | 1 |
| GOBP_CEREBELLAR_PURKINJE_CELL_LAYER_DEVELOPMENT | 26 | -0.220677 | -0.666918 | 0.932384 | 1 |
| GOBP_PIGMENT_GRANULE_ORGANIZATION | 27 | -0.212737 | -0.66553 | 0.910394 | 1 |
| GOBP_MICROTUBULE_CYTOSKELETON_ORGANIZATION_INVOLVED_IN_MITOS | 149 | -0.155007 | -0.663865 | 1 | 1 |
| GOBP_VACUOLAR_LOCALIZATION | 68 | -0.175353 | -0.661469 | 1 | 1 |
| GOBP_MRNA_METHYLATION | 16 | -0.249377 | -0.660198 | 0.892308 | 1 |
| GOBP_MISMATCH_REPAIR | 30 | -0.209348 | -0.657863 | 0.933071 | 1 |
| GOBP_CHONDROITIN_SULFATE_PROTEOGLYCAN_BIOSYNTHETIC_PROCESS | 23 | -0.223632 | -0.656597 | 0.92963 | 1 |
| GOBP_NEGATIVE_REGULATION_OF_RESPONSE_TO_OXIDATIVE_STRESS | 16 | -0.242505 | -0.655533 | 0.919732 | 1 |
| GOBP_ATP_SYNTHESIS_COUPLED_ELECTRON_TRANSPORT | 85 | -0.168484 | -0.65208 | 0.992806 | 1 |
| GOBP_RIBOSE_PHOSPHATE_BIOSYNTHETIC_PROCESS | 173 | -0.150759 | -0.651525 | 1 | 1 |
| GOBP_PROTEIN_STABILIZATION | 166 | -0.156174 | -0.649932 | 1 | 1 |
| GOBP_UBIQUITIN_DEPENDENT_PROTEIN_CATABOLIC_PROCESS_VIA_THE_MU | 34 | -0.199449 | -0.649567 | 0.972763 | 1 |
| GOBP_NEGATIVE_REGULATION_OF_HORMONE_SECRETION | 48 | -0.187703 | -0.64843 | 0.986175 | 1 |
| GOBP_CELLULAR_RESPONSE_TO_ESTRADIOL_STIMULUS | 33 | -0.205596 | -0.648181 | 0.954373 | 1 |
| GOBP_DNA_REPAIR | 500 | -0.135984 | -0.647797 | 1 | 1 |
| GOBP_REGULATION_OF_POSTSYNAPTIC_MEMBRANE_NEUROTRANSMITTER_ | 52 | -0.182695 | -0.647605 | 0.985149 | 1 |
| GOBP_NEGATIVE_REGULATION_OF_METAPHASE_ANAPHASE_TRANSITION_OF | 39 | -0.198299 | -0.647349 | 0.982533 | 1 |
| GOBP_SPINDLE_ORGANIZATION | 192 | -0.153223 | -0.646379 | 1 | 1 |
| GOBP_HISTONE_H3_ACETYLATION | 74 | -0.174094 | -0.646025 | 0.99359 | 1 |
| GOBP_NEGATIVE_REGULATION_OF_PROTEOLYSIS_INVOLVED_IN_CELLULAR_ | 57 | -0.181624 | -0.645918 | 0.99005 | 1 |
| GOBP_FIBROBLAST_APOPTOTIC_PROCESS | 18 | -0.227553 | -0.645818 | 0.915825 | 1 |
| GOBP_REGULATION_OF_TRANSCRIPTION_ELONGATION_FROM_RNA_POLYME | 36 | -0.19671 | -0.644678 | 0.944882 | 1 |
| GOBP_VERY_LONG_CHAIN_FATTY_ACID_METABOLIC_PROCESS | 30 | -0.20628 | -0.6443 | 0.947791 | 1 |
| GOBP_CHROMATIN_DISASSEMBLY | 19 | -0.226358 | -0.644044 | 0.916955 | 1 |
| GOBP_REGULATION_OF_AUTOPHAGOSOME_MATURATION | 15 | -0.24814 | -0.643622 | 0.89759 | 1 |
| GOBP_MAINTENANCE_OF_PROTEIN_LOCATION_IN_CELL | 60 | -0.176256 | -0.643482 | 0.989744 | 1 |
| GOBP_AMYLOID_BETA_METABOLIC_PROCESS | 46 | -0.184499 | -0.64328 | 0.985222 | 1 |
| GOBP_POSITIVE_REGULATION_OF_PROTEIN_TYROSINE_KINASE_ACTIVITY | 43 | -0.186964 | -0.642168 | 0.976526 | 1 |
| GOBP_PROTEASOMAL_PROTEIN_CATABOLIC_PROCESS | 464 | -0.133238 | -0.641016 | 1 | 1 |
| GOBP_MANNOSYLATION | 35 | -0.198557 | -0.640349 | 0.988462 | 1 |
| GOBP_NEUROTRANSMITTER_RECEPTOR_TRANSPORT_TO_PLASMA_MEMBRA | 16 | -0.238673 | -0.639546 | 0.90566 | 1 |
| GOBP_CARBOHYDRATE_DERIVATIVE_TRANSPORT | 71 | -0.169644 | -0.636776 | 1 | 1 |
| GOBP_COTRANSLATIONAL_PROTEIN_TARGETING_TO_MEMBRANE | 24 | -0.214312 | -0.636321 | 0.968532 | 1 |
| GOBP_REGULATION_OF_MITOTIC_SISTER_CHROMATID_SEGREGATION | 42 | -0.184783 | -0.634838 | 0.970874 | 1 |
| GOBP_REGULATION_OF_TRANSLATIONAL_INITIATION | 74 | -0.167367 | -0.633557 | 0.993976 | 1 |
| GOBP_SNRNA_PROCESSING | 25 | -0.206524 | -0.625057 | 0.97417 | 1 |
| GOBP_ASPARTATE_FAMILY_AMINO_ACID_BIOSYNTHETIC_PROCESS | 19 | -0.226739 | -0.624822 | 0.945017 | 1 |
| GOBP_NEGATIVE_REGULATION_OF_TELOMERE_MAINTENANCE_VIA_TELOME | 24 | -0.208342 | -0.624702 | 0.952206 | 1 |
| GOBP_ARP2_3_COMPLEX_MEDIATED_ACTIN_NUCLEATION | 39 | -0.187121 | -0.62412 | 0.982143 | 1 |
| GOBP_MRNA_3_END_PROCESSING | 61 | -0.171594 | -0.623943 | 0.994595 | 1 |
| GOBP_MITOTIC_SISTER_CHROMATID_SEGREGATION | 166 | -0.143408 | -0.623851 | 1 | 1 |
| GOBP_MITOTIC_SPINDLE_ORGANIZATION | 125 | -0.153184 | -0.619393 | 1 | 1 |
| GOBP_INTRA_GOLGI_VESICLE_MEDIATED_TRANSPORT | 29 | -0.204516 | -0.619171 | 0.967972 | 1 |
| GOBP_REGULATION_OF_DNA_DAMAGE_CHECKPOINT | 16 | -0.228188 | -0.619048 | 0.940789 | 1 |
| GOBP_RNA_PHOSPHODIESTER_BOND_HYDROLYSIS_EXONUCLEOLYTIC | 41 | -0.187898 | -0.617672 | 0.975369 | 1 |
| GOBP_REGULATION_OF_CHOLESTEROL_BIOSYNTHETIC_PROCESS | 18 | -0.224443 | -0.617615 | 0.941538 | 1 |
| GOBP_PROTEIN_LOCALIZATION_TO_CELL_SURFACE | 58 | -0.169979 | -0.616478 | 1 | 1 |
| GOBP_FOLIC_ACID_METABOLIC_PROCESS | 15 | -0.235389 | -0.611183 | 0.927928 | 1 |
| GOBP_RESPONSE_TO_ENDOPLASMIC_RETICULUM_STRESS | 240 | -0.138412 | -0.610355 | 1 | 1 |
| GOBP_REGULATION_OF_VIRAL_TRANSCRIPTION | 19 | -0.219212 | -0.60873 | 0.954839 | 1 |
| GOBP_QUINONE_METABOLIC_PROCESS | 31 | -0.192382 | -0.607981 | 0.962199 | 1 |
| GOBP_PROTEIN_LOCALIZATION_TO_PHAGOPHORE_ASSEMBLY_SITE | 17 | -0.213167 | -0.601898 | 0.948718 | 1 |
| GOBP_NEGATIVE_REGULATION_OF_BLOOD_CIRCULATION | 19 | -0.215187 | -0.601712 | 0.94702 | 1 |
| GOBP_RNA_POLYADENYLATION | 47 | -0.171456 | -0.600636 | 0.975124 | 1 |
| GOBP_PROTEIN_LOCALIZATION_TO_CHROMOSOME_CENTROMERIC_REGION | 25 | -0.196949 | -0.598508 | 0.973881 | 1 |
| GOBP_GOLGI_VESICLE_TRANSPORT | 281 | -0.134306 | -0.59757 | 1 | 1 |
| GOBP_POSITIVE_REGULATION_OF_MRNA_METABOLIC_PROCESS | 116 | -0.145718 | -0.59536 | 1 | 1 |
| GOBP_STORE_OPERATED_CALCIUM_ENTRY | 18 | -0.213465 | -0.590346 | 0.961078 | 1 |
| GOBP_CHONDROITIN_SULFATE_BIOSYNTHETIC_PROCESS | 18 | -0.210472 | -0.589796 | 0.968641 | 1 |
| GOBP_RNA_INTERFERENCE | 17 | -0.217918 | -0.586064 | 0.951456 | 1 |
| GOBP_INTRINSIC_APOPTOTIC_SIGNALING_PATHWAY_IN_RESPONSE_TO_OXID | 42 | -0.172272 | -0.584152 | 0.991525 | 1 |
| GOBP_EXTRACELLULAR_VESICLE_BIOGENESIS | 24 | -0.194453 | -0.580717 | 0.981191 | 1 |
| GOBP_PROTEASOME_MEDIATED_UBIQUITIN_DEPENDENT_PROTEIN_CATABOL | 394 | -0.127635 | -0.575688 | 1 | 1 |
| GOBP_TETRAPYRROLE_BIOSYNTHETIC_PROCESS | 31 | -0.176937 | -0.574881 | 0.989209 | 1 |
| GOBP_VIRION_ASSEMBLY | 36 | -0.173751 | -0.571705 | 1 | 1 |
| GOBP_REGULATION_OF_PROTEASOMAL_PROTEIN_CATABOLIC_PROCESS | 173 | -0.131137 | -0.571399 | 1 | 1 |
| GOBP_LYSOSOMAL_TRANSPORT | 117 | -0.13924 | -0.570709 | 1 | 1 |
| GOBP_RETROGRADE_TRANSPORT_ENDOSOME_TO_GOLGI | 91 | -0.141286 | -0.569978 | 1 | 1 |
| GOBP_MULTIVESICULAR_BODY_SORTING_PATHWAY | 37 | -0.173574 | -0.569609 | 0.988462 | 1 |
| GOBP_POSITIVE_REGULATION_OF_SIGNAL_TRANSDUCTION_BY_P53_CLASS_ | 25 | -0.189401 | -0.569591 | 0.986577 | 1 |
| GOBP_BLASTOCYST_GROWTH | 19 | -0.20147 | -0.567061 | 0.975352 | 1 |
| GOBP_OXIDATIVE_PHOSPHORYLATION | 129 | -0.138469 | -0.566387 | 1 | 1 |
| GOBP_PIGMENT_BIOSYNTHETIC_PROCESS | 57 | -0.154191 | -0.565476 | 1 | 1 |
| GOBP_NEGATIVE_REGULATION_OF_RECEPTOR_MEDIATED_ENDOCYTOSIS | 28 | -0.183812 | -0.561453 | 0.98855 | 1 |
| GOBP_POSITIVE_REGULATION_OF_NEURON_MIGRATION | 16 | -0.205738 | -0.55574 | 0.984756 | 1 |
| GOBP_CLATHRIN_DEPENDENT_ENDOCYTOSIS | 45 | -0.161837 | -0.553844 | 0.991031 | 1 |
| GOBP_GOLGI_TO_ENDOSOME_TRANSPORT | 17 | -0.200933 | -0.552152 | 0.966867 | 1 |
| GOBP_REGULATION_OF_OXIDATIVE_PHOSPHORYLATION | 21 | -0.187267 | -0.547086 | 0.975352 | 1 |
| GOBP_REGULATION_OF_OXIDATIVE_STRESS_INDUCED_INTRINSIC_APOPTOT | 26 | -0.180958 | -0.546703 | 0.989474 | 1 |
| GOBP_REGULATION_OF_MITOTIC_CELL_CYCLE_SPINDLE_ASSEMBLY_CHECK | 15 | -0.207902 | -0.544503 | 0.961877 | 1 |
| GOBP_LYSOSOMAL_PROTEIN_CATABOLIC_PROCESS | 19 | -0.192331 | -0.539743 | 0.986622 | 1 |
| GOBP_REGULATION_OF_ARP2_3_COMPLEX_MEDIATED_ACTIN_NUCLEATION | 21 | -0.183516 | -0.537832 | 0.986577 | 1 |
| GOBP_PROTEIN_K48_LINKED_DEUBIQUITINATION | 31 | -0.172285 | -0.536448 | 0.983264 | 1 |
| GOBP_2_OXOGLUTARATE_METABOLIC_PROCESS | 16 | -0.192765 | -0.535699 | 0.978328 | 1 |
| GOBP_FATTY_ACID_BETA_OXIDATION | 68 | -0.140307 | -0.53523 | 0.994444 | 1 |
| GOBP_MIRNA_METABOLIC_PROCESS | 27 | -0.175672 | -0.534522 | 0.986885 | 1 |
| GOBP_POSITIVE_REGULATION_OF_NITRIC_OXIDE_SYNTHASE_ACTIVITY | 16 | -0.198809 | -0.534373 | 0.993399 | 1 |
| GOBP_INNER_MITOCHONDRIAL_MEMBRANE_ORGANIZATION | 40 | -0.160486 | -0.533212 | 0.995652 | 1 |
| GOBP_RESPONSE_TO_MISFOLDED_PROTEIN | 24 | -0.179824 | -0.530927 | 0.989209 | 1 |
| GOBP_REGULATION_OF_GOLGI_ORGANIZATION | 18 | -0.188066 | -0.525744 | 0.980769 | 1 |
| GOBP_POLY_A_PLUS_MRNA_EXPORT_FROM_NUCLEUS | 15 | -0.1944 | -0.51833 | 0.987768 | 1 |
| GOBP_MRNA_TRANSPORT | 116 | -0.124735 | -0.514369 | 1 | 1 |
| GOBP_MICROTUBULE_NUCLEATION | 35 | -0.158371 | -0.512661 | 1 | 1 |
| GOBP_NUCLEOTIDE_EXCISION_REPAIR | 56 | -0.143343 | -0.512618 | 1 | 1 |
| GOBP_VESICLE_CARGO_LOADING | 21 | -0.178361 | -0.511913 | 0.992647 | 1 |
| GOBP_ENDOPLASMIC_RETICULUM_CALCIUM_ION_HOMEOSTASIS | 25 | -0.168078 | -0.510458 | 0.988848 | 1 |
| GOBP_POSITIVE_REGULATION_OF_DNA_TEMPLATED_TRANSCRIPTION_ELON | 27 | -0.165372 | -0.508689 | 1 | 1 |
| GOBP_SELECTIVE_AUTOPHAGY | 77 | -0.1302 | -0.508487 | 1 | 1 |
| GOBP_VACUOLAR_TRANSPORT | 149 | -0.124067 | -0.5037 | 1 | 1 |
| GOBP_REGULATION_OF_ENDOPLASMIC_RETICULUM_STRESS_INDUCED_INT | 28 | -0.159947 | -0.500794 | 0.992481 | 1 |
| GOBP_NEGATIVE_REGULATION_OF_LIPID_TRANSPORT | 19 | -0.177023 | -0.498648 | 0.990228 | 1 |
| GOBP_NEGATIVE_REGULATION_OF_AMYLOID_PRECURSOR_PROTEIN_CATAB | 19 | -0.175073 | -0.4923 | 0.993197 | 1 |
| GOBP_PROTEIN_LOCALIZATION_TO_CONDENSED_CHROMOSOME | 19 | -0.17152 | -0.490927 | 1 | 1 |
| GOBP_CHAPERONE_MEDIATED_AUTOPHAGY | 16 | -0.17575 | -0.471083 | 0.993485 | 1 |
| GOBP_TRANSLATIONAL_TERMINATION | 18 | -0.166546 | -0.460664 | 1 | 1 |
| GOBP_EMBRYONIC_BRAIN_DEVELOPMENT | 15 | -0.163592 | -0.446538 | 0.993399 | 1 |
| GOBP_LATE_ENDOSOME_TO_LYSOSOME_TRANSPORT | 18 | -0.152503 | -0.423622 | 0.997076 | 1 |
| GOBP_VESICLE_TARGETING | 40 | -0.10172 | -0.342259 | 1 | 0.999983 |

| | | | | | |
|---|---|---|---|---|---|
| GOBP_CELL_DIFFERENTIATION_INVOLVED_IN_METANEPHROS_DEVELOPMENT | 17 | 0.515475 | 1.200641 | 0.244573 | 0.59653 |
| GOBP_VENTRICULAR_SEPTUM_DEVELOPMENT | 68 | 0.407271 | 1.200264 | 0.187345 | 0.597086 |
| GOBP_RESPONSE_TO_INORGANIC_SUBSTANCE | 431 | 0.349067 | 1.200141 | 0.070636 | 0.59686 |
| GOBP_REGULATION_OF_METAL_ION_TRANSPORT | 311 | 0.352494 | 1.199841 | 0.087629 | 0.597186 |
| GOBP_EYE_PHOTORECEPTOR_CELL_DIFFERENTIATION | 35 | 0.450122 | 1.199747 | 0.222513 | 0.59687 |
| GOBP_ECTODERM_DEVELOPMENT | 18 | 0.507266 | 1.199373 | 0.245763 | 0.59743 |
| GOBP_MITOCHONDRIAL_CALCIUM_ION_TRANSMEMBRANE_TRANSPORT | 18 | 0.506813 | 1.199128 | 0.254731 | 0.597599 |
| GOBP_PATTERN_SPECIFICATION_PROCESS | 347 | 0.353627 | 1.198902 | 0.068298 | 0.597708 |
| GOBP_VENTRICULAR_CARDIAC_MUSCLE_TISSUE_MORPHOGENESIS | 42 | 0.433638 | 1.198381 | 0.206536 | 0.598772 |
| GOBP_POSITIVE_REGULATION_OF_T_CELL_MIGRATION | 21 | 0.486196 | 1.198285 | 0.226629 | 0.598491 |
| GOBP_CELL_MATURATION | 139 | 0.372506 | 1.198039 | 0.146697 | 0.598697 |
| GOBP_FATTY_ACID_TRANSPORT | 71 | 0.401081 | 1.197802 | 0.190024 | 0.598889 |
| GOBP_POSITIVE_REGULATION_OF_INSULIN_SECRETION | 57 | 0.414641 | 1.197761 | 0.222363 | 0.5984 |
| GOBP_PATHWAY_RESTRICTED_SMAD_PROTEIN_PHOSPHORYLATION | 57 | 0.410892 | 1.196642 | 0.215539 | 0.601376 |
| GOBP_REGULATION_OF_TYPE_I_INTERFERON_MEDIATED_SIGNALING_PATHWAY | 36 | 0.444636 | 1.196476 | 0.242503 | 0.601258 |
| GOBP_BONE_REMODELING | 70 | 0.404115 | 1.196256 | 0.195065 | 0.601376 |
| GOBP_NEGATIVE_REGULATION_OF_TOR_SIGNALING | 43 | 0.42999 | 1.196123 | 0.215938 | 0.601162 |
| GOBP_RESPONSE_TO_OXIDATIVE_STRESS | 373 | 0.34915 | 1.196009 | 0.081218 | 0.60089 |
| GOBP_POSITIVE_REGULATION_OF_PHOSPHATIDYLINOSITOL_3_KINASE_SIGNALING | 53 | 0.415701 | 1.195658 | 0.203222 | 0.601512 |
| GOBP_CELL_FATE_COMMITMENT | 180 | 0.365439 | 1.194877 | 0.139037 | 0.603582 |
| GOBP_GOLGI_TO_VACUOLE_TRANSPORT | 20 | 0.493324 | 1.194451 | 0.262626 | 0.604426 |
| GOBP_EXTRINSIC_APOPTOTIC_SIGNALING_PATHWAY | 189 | 0.361698 | 1.194446 | 0.135394 | 0.603812 |
| GOBP_REGULATION_OF_CYSTEINE_TYPE_ENDOPEPTIDASE_ACTIVITY | 190 | 0.363916 | 1.194298 | 0.128828 | 0.603655 |
| GOBP_PROTEIN_HOMOTETRAMERIZATION | 48 | 0.420612 | 1.193763 | 0.216602 | 0.604755 |
| GOBP_POSITIVE_REGULATION_OF_VASCULAR_ASSOCIATED_SMOOTH_MUSCLE_CELL_PR | 24 | 0.476657 | 1.193547 | 0.246844 | 0.60485 |
| GOBP_REGULATION_OF_EXTRINSIC_APOPTOTIC_SIGNALING_PATHWAY | 128 | 0.375721 | 1.19312 | 0.160131 | 0.605601 |
| GOBP_RESPONSE_TO_TUMOR_NECROSIS_FACTOR | 180 | 0.36514 | 1.192482 | 0.128068 | 0.607055 |
| GOBP_REGULATION_OF_SECRETION | 483 | 0.347838 | 1.192275 | 0.055332 | 0.607131 |
| GOBP_NEGATIVE_REGULATION_OF_FAT_CELL_DIFFERENTIATION | 47 | 0.418445 | 1.191793 | 0.220551 | 0.608106 |
| GOBP_EYE_MORPHOGENESIS | 127 | 0.37234 | 1.191616 | 0.181617 | 0.608045 |
| GOBP_REGULATION_OF_CELLULAR_RESPIRATION | 43 | 0.425121 | 1.190728 | 0.22846 | 0.610358 |
| GOBP_REGULATION_OF_BMP_SIGNALING_PATHWAY | 87 | 0.383959 | 1.190614 | 0.196283 | 0.610123 |
| GOBP_RETINOIC_ACID_RECEPTOR_SIGNALING_PATHWAY | 29 | 0.448698 | 1.189594 | 0.238558 | 0.61285 |
| GOBP_ADULT_LOCOMOTORY_BEHAVIOR | 69 | 0.394825 | 1.189429 | 0.212515 | 0.612773 |
| GOBP_HOMOTYPIC_CELL_CELL_ADHESION | 75 | 0.399252 | 1.189104 | 0.204057 | 0.613278 |
| GOBP_RESPONSE_TO_FOOD | 26 | 0.46699 | 1.189087 | 0.25 | 0.612717 |
| GOBP_REGULATION_OF_ICOSANOID_SECRETION | 16 | 0.510437 | 1.188905 | 0.251863 | 0.612711 |
| GOBP_POSITIVE_REGULATION_OF_CELL_MATRIX_ADHESION | 47 | 0.421792 | 1.188598 | 0.220608 | 0.613138 |
| GOBP_POSITIVE_REGULATION_OF_ATP_DEPENDENT_ACTIVITY | 40 | 0.434132 | 1.187848 | 0.227506 | 0.614895 |
| GOBP_REGULATION_OF_GLIAL_CELL_PROLIFERATION | 21 | 0.486669 | 1.187617 | 0.227536 | 0.615005 |
| GOBP_REGULATION_OF_RECEPTOR_BINDING | 18 | 0.512936 | 1.18755 | 0.275281 | 0.614601 |
| GOBP_NEGATIVE_REGULATION_OF_CYTOKINE_PRODUCTION_INVOLVED_IN_IMMUNE_RES | 16 | 0.518072 | 1.187041 | 0.25656 | 0.615646 |
| GOBP_POSITIVE_REGULATION_OF_MIRNA_TRANSCRIPTION | 41 | 0.426573 | 1.186674 | 0.234375 | 0.616244 |
| GOBP_MULTICELLULAR_ORGANISMAL_RESPONSE_TO_STRESS | 68 | 0.400735 | 1.186617 | 0.219162 | 0.615798 |
| GOBP_REGULATION_OF_PEPTIDE_TRANSPORT | 141 | 0.36739 | 1.186125 | 0.167031 | 0.616806 |
| GOBP_REGULATION_OF_POTASSIUM_ION_TRANSPORT | 83 | 0.388922 | 1.185984 | 0.200231 | 0.616636 |
| GOBP_METAL_ION_HOMEOSTASIS | 489 | 0.344835 | 1.185325 | 0.070707 | 0.618226 |
| GOBP_NEGATIVE_REGULATION_OF_INNATE_IMMUNE_RESPONSE | 53 | 0.42383 | 1.185242 | 0.225527 | 0.617864 |
| GOBP_CELLULAR_RESPONSE_TO_REACTIVE_NITROGEN_SPECIES | 15 | 0.511596 | 1.184716 | 0.256923 | 0.618945 |
| GOBP_RESPONSE_TO_NITRIC_OXIDE | 15 | 0.520494 | 1.184313 | 0.253731 | 0.619689 |
| GOBP_POSITIVE_REGULATION_OF_SECRETION | 244 | 0.35097 | 1.184153 | 0.129938 | 0.61957 |
| GOBP_LEARNING | 134 | 0.370882 | 1.183972 | 0.184152 | 0.619523 |
| GOBP_NEUROMUSCULAR_PROCESS | 125 | 0.374551 | 1.183621 | 0.182628 | 0.620032 |
| GOBP_HISTONE_H3_K9_METHYLATION | 36 | 0.4422 | 1.183482 | 0.255007 | 0.619869 |
| GOBP_TAXIS | 449 | 0.345722 | 1.183333 | 0.072508 | 0.619743 |
| GOBP_CELLULAR_COMPONENT_ASSEMBLY_INVOLVED_IN_MORPHOGENESIS | 89 | 0.381772 | 1.183294 | 0.207569 | 0.619251 |
| GOBP_SMOOTH_MUSCLE_CELL_DIFFERENTIATION | 47 | 0.419298 | 1.183047 | 0.238693 | 0.619449 |
| GOBP_RESPONSE_TO_MINERALOCORTICOID | 25 | 0.467543 | 1.18304 | 0.257703 | 0.618848 |
| GOBP_PULMONARY_VALVE_DEVELOPMENT | 20 | 0.487126 | 1.182747 | 0.27983 | 0.619162 |
| GOBP_POSITIVE_REGULATION_OF_CREB_TRANSCRIPTION_FACTOR_ACTIVITY | 17 | 0.501463 | 1.182662 | 0.244344 | 0.618798 |
| GOBP_ORGANIC_HYDROXY_COMPOUND_METABOLIC_PROCESS | 428 | 0.344089 | 1.182635 | 0.072654 | 0.61827 |
| GOBP_REGULATION_OF_CARDIAC_CONDUCTION | 20 | 0.491312 | 1.182626 | 0.229577 | 0.617681 |
| GOBP_POSITIVE_REGULATION_OF_SYNAPTIC_TRANSMISSION | 127 | 0.367118 | 1.182402 | 0.180804 | 0.6178 |
| GOBP_POSITIVE_REGULATION_OF_NATURAL_KILLER_CELL_ACTIVATION | 20 | 0.48344 | 1.181919 | 0.261803 | 0.61873 |
| GOBP_MESODERM_DEVELOPMENT | 108 | 0.374127 | 1.181757 | 0.176339 | 0.618628 |
| GOBP_REGULATION_OF_LIPID_BIOSYNTHETIC_PROCESS | 134 | 0.371135 | 1.18168 | 0.165017 | 0.61828 |
| GOBP_POSITIVE_REGULATION_OF_DENDRITE_DEVELOPMENT | 15 | 0.519552 | 1.181548 | 0.281818 | 0.618092 |
| GOBP_REGULATION_OF_EXTRINSIC_APOPTOTIC_SIGNALING_PATHWAY_IN_ABSENCE_OF_ | 35 | 0.435288 | 1.181318 | 0.24124 | 0.618293 |
| GOBP_REGULATION_OF_FATTY_ACID_TRANSPORT | 26 | 0.460592 | 1.181105 | 0.252825 | 0.618378 |
| GOBP_TRANSPORT_OF_VIRUS | 17 | 0.506737 | 1.180849 | 0.262195 | 0.618577 |
| GOBP_POSITIVE_REGULATION_OF_MAPK_CASCADE | 374 | 0.343862 | 1.180679 | 0.097363 | 0.618501 |
| GOBP_REGULATION_OF_COLLATERAL_SPROUTING | 19 | 0.489583 | 1.180354 | 0.272464 | 0.618947 |
| GOBP_REGULATION_OF_GASTRULATION | 15 | 0.515394 | 1.180123 | 0.266867 | 0.619103 |
| GOBP_SKELETAL_SYSTEM_MORPHOGENESIS | 168 | 0.361373 | 1.180093 | 0.17679 | 0.618589 |
| GOBP_NEGATIVE_REGULATION_OF_CELL_SUBSTRATE_ADHESION | 51 | 0.416971 | 1.179632 | 0.227735 | 0.619468 |
| GOBP_RHO_PROTEIN_SIGNAL_TRANSDUCTION | 120 | 0.373272 | 1.179103 | 0.19265 | 0.62058 |
| GOBP_REGULATION_OF_HORMONE_SECRETION | 183 | 0.362514 | 1.179028 | 0.147346 | 0.62021 |
| GOBP_REGULATION_OF_LIPID_LOCALIZATION | 122 | 0.367439 | 1.178926 | 0.19539 | 0.619943 |
| GOBP_POSITIVE_REGULATION_OF_HORMONE_SECRETION | 98 | 0.380072 | 1.178835 | 0.187643 | 0.619618 |
| GOBP_PROSTANOID_METABOLIC_PROCESS | 36 | 0.441793 | 1.178538 | 0.239843 | 0.619971 |
| GOBP_MATURE_B_CELL_DIFFERENTIATION | 23 | 0.478113 | 1.177858 | 0.248945 | 0.62163 |
| GOBP_REGULATION_OF_CALCIUM_MEDIATED_SIGNALING | 58 | 0.408281 | 1.177229 | 0.247246 | 0.623081 |
| GOBP_REGULATION_OF_RENAL_SYSTEM_PROCESS | 25 | 0.467927 | 1.177107 | 0.268908 | 0.622873 |
| GOBP_HISTONE_H3_K9_MODIFICATION | 50 | 0.413899 | 1.176638 | 0.241206 | 0.623802 |
| GOBP_CELLULAR_RESPONSE_TO_LOW_DENSITY_LIPOPROTEIN_PARTICLE_STIMULUS | 15 | 0.519131 | 1.176419 | 0.268657 | 0.623877 |
| GOBP_POSITIVE_REGULATION_OF_PRODUCTION_OF_MOLECULAR_MEDIATOR_OF_IMMUN | 86 | 0.385398 | 1.175948 | 0.207018 | 0.624905 |
| GOBP_RELAXATION_OF_MUSCLE | 27 | 0.464062 | 1.175853 | 0.263448 | 0.624584 |
| GOBP_PROTEIN_KINASE_A_SIGNALING | 25 | 0.457252 | 1.175337 | 0.267394 | 0.625639 |
| GOBP_POSITIVE_REGULATION_OF_CELLULAR_EXTRAVASATION | 15 | 0.516228 | 1.175239 | 0.269747 | 0.625349 |
| GOBP_ENDOTHELIAL_CELL_APOPTOTIC_PROCESS | 43 | 0.41901 | 1.17523 | 0.235741 | 0.624775 |
| GOBP_REGULATION_OF_THE_FORCE_OF_HEART_CONTRACTION | 21 | 0.485257 | 1.175115 | 0.24758 | 0.624574 |
| GOBP_NEGATIVE_REGULATION_OF_PEPTIDYL_SERINE_PHOSPHORYLATION | 26 | 0.46037 | 1.173944 | 0.264993 | 0.627747 |
| GOBP_REGULATION_OF_MITOCHONDRIAL_FISSION | 28 | 0.453484 | 1.17342 | 0.234483 | 0.628863 |
| GOBP_POSITIVE_REGULATION_OF_PROTEIN_MATURATION | 25 | 0.456952 | 1.172536 | 0.258741 | 0.631263 |
| GOBP_POSITIVE_REGULATION_OF_EPIDERMAL_CELL_DIFFERENTIATION | 23 | 0.465272 | 1.171564 | 0.261411 | 0.633871 |
| GOBP_SECRETION_BY_TISSUE | 31 | 0.447964 | 1.171228 | 0.26162 | 0.634282 |
| GOBP_CARDIAC_CHAMBER_DEVELOPMENT | 146 | 0.357808 | 1.170613 | 0.190683 | 0.63571 |
| GOBP_REGULATION_OF_CALCIUM_ION_TRANSPORT_INTO_CYTOSOL | 80 | 0.386345 | 1.170482 | 0.209677 | 0.635547 |
| GOBP_PROTEIN_LOCALIZATION_TO_EXTRACELLULAR_REGION | 284 | 0.347928 | 1.169427 | 0.141249 | 0.63843 |
| GOBP_HAIR_CELL_DIFFERENTIATION | 40 | 0.423827 | 1.169291 | 0.243207 | 0.638233 |
| GOBP_SPINDLE_LOCALIZATION | 48 | 0.413256 | 1.168948 | 0.251588 | 0.638801 |
| GOBP_REGULATION_OF_EXOCYTOSIS | 179 | 0.354149 | 1.168841 | 0.169346 | 0.638516 |
| GOBP_RESPONSE_TO_CATECHOLAMINE | 75 | 0.390854 | 1.16847 | 0.228091 | 0.639173 |
| GOBP_POSITIVE_REGULATION_OF_ENDOTHELIAL_CELL_PROLIFERATION | 78 | 0.380373 | 1.168027 | 0.210714 | 0.63999 |
| GOBP_NEGATIVE_REGULATION_OF_BMP_SIGNALING_PATHWAY | 49 | 0.412287 | 1.167807 | 0.23385 | 0.640111 |
| GOBP_ERK1_AND_ERK2_CASCADE | 247 | 0.351555 | 1.16745 | 0.164241 | 0.640713 |
| GOBP_POSITIVE_REGULATION_OF_TRANSMEMBRANE_RECEPTOR_PROTEIN_SERINE_THR | 103 | 0.368266 | 1.167244 | 0.199079 | 0.640854 |
| GOBP_NEGATIVE_REGULATION_OF_PHOSPHATASE_ACTIVITY | 30 | 0.445476 | 1.167025 | 0.259947 | 0.640966 |
| GOBP_REGULATION_OF_LYMPHOCYTE_APOPTOTIC_PROCESS | 41 | 0.419816 | 1.166819 | 0.251276 | 0.64103 |
| GOBP_REGULATION_OF_TRANSMEMBRANE_RECEPTOR_PROTEIN_SERINE_THREONINE_K | 223 | 0.349031 | 1.166519 | 0.157233 | 0.641382 |
| GOBP_POSITIVE_REGULATION_OF_GLUCOSE_TRANSMEMBRANE_TRANSPORT | 35 | 0.435971 | 1.166483 | 0.273779 | 0.640904 |
| GOBP_NEGATIVE_REGULATION_OF_LOCOMOTION | 280 | 0.346771 | 1.165997 | 0.121677 | 0.641913 |
| GOBP_REGULATION_OF_HEMOPOIESIS | 280 | 0.344248 | 1.165863 | 0.144319 | 0.641727 |
| GOBP_POSITIVE_REGULATION_OF_B_CELL_PROLIFERATION | 27 | 0.459002 | 1.165247 | 0.246217 | 0.643134 |
| GOBP_POSITIVE_REGULATION_OF_I_KAPPAB_KINASE_NF_KAPPAB_SIGNALING | 163 | 0.357628 | 1.164962 | 0.199784 | 0.643482 |
| GOBP_ADENYLATE_CYCLASE_ACTIVATING_G_PROTEIN_COUPLED_RECEPTOR_SIGNALING | 111 | 0.368609 | 1.164491 | 0.220377 | 0.644397 |
| GOBP_TUMOR_NECROSIS_FACTOR_MEDIATED_SIGNALING_PATHWAY | 87 | 0.374944 | 1.164117 | 0.228637 | 0.645047 |
| GOBP_BIOLOGICAL_PROCESS_INVOLVED_IN_INTRASPECIES_INTERACTION_BETWEEN_OR | 44 | 0.414788 | 1.162634 | 0.239848 | 0.649305 |
| GOBP_BODY_FLUID_SECRETION | 74 | 0.384063 | 1.162194 | 0.214634 | 0.650141 |
| GOBP_LENS_FIBER_CELL_DIFFERENTIATION | 33 | 0.432001 | 1.161943 | 0.263158 | 0.650415 |
| GOBP_PHOSPHATIDIC_ACID_METABOLIC_PROCESS | 34 | 0.434618 | 1.161726 | 0.277854 | 0.65052 |
| GOBP_REGULATION_OF_LAMELLIPODIUM_ASSEMBLY | 34 | 0.43112 | 1.161684 | 0.275538 | 0.650043 |
| GOBP_PROGRAMMED_NECROTIC_CELL_DEATH | 37 | 0.427701 | 1.161449 | 0.250321 | 0.650223 |
| GOBP_MATING_BEHAVIOR | 17 | 0.489673 | 1.161377 | 0.299712 | 0.649856 |
| GOBP_REGULATION_OF_ALCOHOL_BIOSYNTHETIC_PROCESS | 40 | 0.42712 | 1.161328 | 0.250328 | 0.649382 |
| GOBP_INNER_EAR_MORPHOGENESIS | 80 | 0.383525 | 1.160621 | 0.250875 | 0.651129 |
| GOBP_CORONARY_VASCULATURE_MORPHOGENESIS | 16 | 0.495121 | 1.160402 | 0.294461 | 0.651243 |
| GOBP_COLLATERAL_SPROUTING | 25 | 0.45164 | 1.160221 | 0.279167 | 0.651247 |
| GOBP_NEGATIVE_REGULATION_OF_RESPONSE_TO_EXTERNAL_STIMULUS | 297 | 0.342362 | 1.160067 | 0.143296 | 0.651144 |
| GOBP_POSITIVE_REGULATION_OF_B_CELL_ACTIVATION | 60 | 0.39738 | 1.160026 | 0.260976 | 0.650657 |
| GOBP_CALCIUM_MEDIATED_SIGNALING | 145 | 0.359169 | 1.159862 | 0.180239 | 0.650577 |
| GOBP_POSITIVE_REGULATION_OF_REPRODUCTIVE_PROCESS | 64 | 0.394601 | 1.159195 | 0.250888 | 0.652222 |
| GOBP_PHOSPHOLIPASE_C_ACTIVATING_G_PROTEIN_COUPLED_RECEPTOR_SIGNALING_P | 60 | 0.400316 | 1.158552 | 0.234017 | 0.653748 |
| GOBP_POSITIVE_REGULATION_OF_INTERLEUKIN_4_PRODUCTION | 16 | 0.502478 | 1.158466 | 0.283159 | 0.65341 |
| GOBP_NEGATIVE_REGULATION_OF_PROTEIN_KINASE_B_SIGNALING | 39 | 0.423266 | 1.158408 | 0.263158 | 0.652993 |
| GOBP_HISTONE_LYSINE_DEMETHYLATION | 24 | 0.452533 | 1.158099 | 0.280166 | 0.653408 |
| GOBP_ISOPRENOID_BIOSYNTHETIC_PROCESS | 24 | 0.457831 | 1.157296 | 0.269394 | 0.655508 |
| GOBP_POSITIVE_REGULATION_OF_PATTERN_RECOGNITION_RECEPTOR_SIGNALING_PATH | 33 | 0.438136 | 1.157123 | 0.271505 | 0.655481 |
| GOBP_REGULATION_OF_BLOOD_CIRCULATION | 202 | 0.350345 | 1.156936 | 0.199153 | 0.655509 |
| GOBP_NATURAL_KILLER_CELL_ACTIVATION | 54 | 0.400966 | 1.156501 | 0.240786 | 0.656281 |
| GOBP_NEGATIVE_REGULATION_OF_MAPK_CASCADE | 147 | 0.356156 | 1.155927 | 0.206174 | 0.657512 |
| GOBP_CARDIAC_MUSCLE_TISSUE_MORPHOGENESIS | 51 | 0.400328 | 1.155669 | 0.23871 | 0.657741 |
| GOBP_REPLACEMENT_OSSIFICATION | 26 | 0.447806 | 1.155152 | 0.293878 | 0.658951 |
| GOBP_MYELOID_DENDRITIC_CELL_ACTIVATION | 18 | 0.481569 | 1.154409 | 0.300719 | 0.660801 |
| GOBP_HEART_MORPHOGENESIS | 223 | 0.345059 | 1.154345 | 0.188421 | 0.66042 |
| GOBP_HEART_VALVE_MORPHOGENESIS | 52 | 0.40322 | 1.154341 | 0.248455 | 0.659819 |
| GOBP_BASEMENT_MEMBRANE_ORGANIZATION | 29 | 0.445693 | 1.15434 | 0.277624 | 0.659215 |
| GOBP_LINOLEIC_ACID_METABOLIC_PROCESS | 16 | 0.48927 | 1.153519 | 0.28264 | 0.66126 |
| GOBP_CELLULAR_SODIUM_ION_HOMEOSTASIS | 20 | 0.472863 | 1.153474 | 0.306452 | 0.660815 |
| GOBP_NEGATIVE_REGULATION_OF_PROTEIN_MATURATION | 20 | 0.463844 | 1.153335 | 0.311047 | 0.660658 |
| GOBP_NEGATIVE_REGULATION_OF_CHEMOKINE_PRODUCTION | 15 | 0.498198 | 1.153139 | 0.287481 | 0.660708 |
| GOBP_VIRAL_GENOME_REPLICATION | 111 | 0.365201 | 1.153099 | 0.218286 | 0.660235 |
| GOBP_RESPONSE_TO_PURINE_CONTAINING_COMPOUND | 105 | 0.365095 | 1.152497 | 0.228861 | 0.661625 |
| GOBP_LAMELLIPODIUM_MORPHOGENESIS | 17 | 0.495587 | 1.152446 | 0.299703 | 0.661186 |
| GOBP_MESENCHYME_DEVELOPMENT | 250 | 0.343407 | 1.150533 | 0.164241 | 0.667033 |
| GOBP_REGULATION_OF_SMOOTH_MUSCLE_CONTRACTION | 54 | 0.397923 | 1.149895 | 0.247177 | 0.668557 |
| GOBP_IMMUNOGLOBULIN_PRODUCTION_INVOLVED_IN_IMMUNOGLOBULIN_MEDIATED_IMM | 56 | 0.397107 | 1.148287 | 0.254068 | 0.673332 |

| | | | | | |
|---|---|---|---|---|---|
| GOBP_REGULATION_OF_PHOSPHATIDYLINOSITOL_3_KINASE_SIGNALING | 78 | 0.378763 | 1.147963 | 0.24911 | 0.673807 |
| GOBP_SECOND_MESSENGER_MEDIATED_SIGNALING | 221 | 0.345338 | 1.147786 | 0.186916 | 0.673794 |
| GOBP_CARBOHYDRATE_HOMEOSTASIS | 186 | 0.345054 | 1.147205 | 0.212474 | 0.675098 |
| GOBP_LYMPHOCYTE_APOPTOTIC_PROCESS | 57 | 0.38888 | 1.146732 | 0.256627 | 0.676058 |
| GOBP_SENSORY_SYSTEM_DEVELOPMENT | 310 | 0.33731 | 1.145638 | 0.172131 | 0.679113 |
| GOBP_HEART_PROCESS | 206 | 0.344523 | 1.145612 | 0.197228 | 0.678596 |
| GOBP_ESTABLISHMENT_OF_SPINDLE_ORIENTATION | 32 | 0.427106 | 1.145495 | 0.284197 | 0.678347 |
| GOBP_CARDIAC_RIGHT_VENTRICLE_MORPHOGENESIS | 17 | 0.479416 | 1.145248 | 0.311844 | 0.678572 |
| GOBP_REGULATION_OF_MONOCYTE_CHEMOTAXIS | 16 | 0.492634 | 1.145217 | 0.32782 | 0.678065 |
| GOBP_REGULATION_OF_RAS_PROTEIN_SIGNAL_TRANSDUCTION | 166 | 0.350949 | 1.144382 | 0.215897 | 0.680328 |
| GOBP_RESPONSE_TO_ALCOHOL | 180 | 0.346663 | 1.1441 | 0.212089 | 0.680654 |
| GOBP_REGULATION_OF_B_CELL_DIFFERENTIATION | 23 | 0.460445 | 1.144041 | 0.289898 | 0.68025 |
| GOBP_MEMBRANE_FUSION | 135 | 0.354703 | 1.144009 | 0.214751 | 0.679738 |
| GOBP_RAS_PROTEIN_SIGNAL_TRANSDUCTION | 316 | 0.334677 | 1.143932 | 0.174002 | 0.679384 |
| GOBP_NEGATIVE_REGULATION_OF_RHO_PROTEIN_SIGNAL_TRANSDUCTION | 21 | 0.469526 | 1.143566 | 0.315642 | 0.67998 |
| GOBP_SLEEP | 18 | 0.484464 | 1.143504 | 0.30848 | 0.679549 |
| GOBP_REGULATION_OF_ACTIN_CYTOSKELETON_REORGANIZATION | 34 | 0.428499 | 1.14305 | 0.288022 | 0.680525 |
| GOBP_CELL_CELL_RECOGNITION | 36 | 0.420521 | 1.143042 | 0.278732 | 0.679938 |
| GOBP_CELLULAR_RESPONSE_TO_INORGANIC_SUBSTANCE | 177 | 0.350127 | 1.142946 | 0.214671 | 0.67962 |
| GOBP_TONGUE_DEVELOPMENT | 15 | 0.488557 | 1.142913 | 0.306184 | 0.679154 |
| GOBP_CELL_AGING | 105 | 0.362267 | 1.142666 | 0.255079 | 0.679403 |
| GOBP_NEGATIVE_REGULATION_OF_CALCIUM_MEDIATED_SIGNALING | 18 | 0.476702 | 1.141869 | 0.324324 | 0.681416 |
| GOBP_NEGATIVE_REGULATION_OF_RNA_CATABOLIC_PROCESS | 67 | 0.384978 | 1.141249 | 0.272509 | 0.682834 |
| GOBP_STRIATED_MUSCLE_CONTRACTION | 139 | 0.358305 | 1.140741 | 0.225 | 0.68394 |
| GOBP_APOPTOTIC_CELL_CLEARANCE | 41 | 0.414038 | 1.140617 | 0.30343 | 0.683746 |
| GOBP_REPRODUCTIVE_SYSTEM_DEVELOPMENT | 357 | 0.333451 | 1.140093 | 0.15 | 0.684906 |
| GOBP_CELLULAR_RESPONSE_TO_ORGANIC_CYCLIC_COMPOUND | 461 | 0.333292 | 1.139627 | 0.131048 | 0.685785 |
| GOBP_MACROPHAGE_CHEMOTAXIS | 23 | 0.459713 | 1.139572 | 0.307582 | 0.685358 |
| GOBP_MYELOID_LEUKOCYTE_CYTOKINE_PRODUCTION | 27 | 0.440153 | 1.139249 | 0.288435 | 0.685814 |
| GOBP_MONOCYTE_DIFFERENTIATION | 29 | 0.442744 | 1.13884 | 0.296044 | 0.686556 |
| GOBP_PROTEIN_COMPLEX_OLIGOMERIZATION | 213 | 0.340866 | 1.138477 | 0.202717 | 0.687107 |
| GOBP_SENSORY_ORGAN_MORPHOGENESIS | 213 | 0.34335 | 1.137856 | 0.210913 | 0.688598 |
| GOBP_POSITIVE_REGULATION_OF_LYMPHOCYTE_MIGRATION | 24 | 0.44869 | 1.137654 | 0.324859 | 0.688672 |
| GOBP_POSITIVE_REGULATION_OF_INTERLEUKIN_1_PRODUCTION | 46 | 0.401255 | 1.137593 | 0.27451 | 0.688268 |
| GOBP_EXPLORATION_BEHAVIOR | 23 | 0.462174 | 1.137379 | 0.322535 | 0.688402 |
| GOBP_REPLICATION_FORK_PROCESSING | 38 | 0.417833 | 1.137357 | 0.293578 | 0.687854 |
| GOBP_REGULATION_OF_CHROMATIN_BINDING | 23 | 0.468046 | 1.137292 | 0.323529 | 0.687453 |
| GOBP_MYOBLAST_FUSION | 32 | 0.429178 | 1.136983 | 0.292165 | 0.687849 |
| GOBP_MEMBRANE_DEPOLARIZATION | 71 | 0.375733 | 1.136898 | 0.27262 | 0.687495 |
| GOBP_LYMPHANGIOGENESIS | 17 | 0.483941 | 1.136256 | 0.316964 | 0.689053 |
| GOBP_RENAL_FILTRATION | 22 | 0.457754 | 1.136149 | 0.29972 | 0.68879 |
| GOBP_LONG_TERM_SYNAPTIC_POTENTIATION | 75 | 0.37791 | 1.136065 | 0.268116 | 0.688446 |
| GOBP_RESPONSE_TO_GONADOTROPIN | 21 | 0.475907 | 1.135909 | 0.316619 | 0.688345 |
| GOBP_ANTIMICROBIAL_HUMORAL_RESPONSE | 42 | 0.409302 | 1.135213 | 0.301546 | 0.690067 |
| GOBP_REGULATION_OF_VOLTAGE_GATED_CALCIUM_CHANNEL_ACTIVITY | 29 | 0.437116 | 1.1352 | 0.302098 | 0.689509 |
| GOBP_GRANULOCYTE_DIFFERENTIATION | 26 | 0.439862 | 1.135103 | 0.30274 | 0.689247 |
| GOBP_IMPORT_ACROSS_PLASMA_MEMBRANE | 142 | 0.353537 | 1.133928 | 0.245614 | 0.692693 |
| GOBP_MAINTENANCE_OF_LOCATION | 263 | 0.337386 | 1.133838 | 0.212435 | 0.692361 |
| GOBP_ANATOMICAL_STRUCTURE_MATURATION | 191 | 0.341933 | 1.133577 | 0.220697 | 0.692651 |
| GOBP_CELLULAR_RESPONSE_TO_LIPID | 432 | 0.330814 | 1.132964 | 0.140688 | 0.69412 |
| GOBP_RESPONSE_TO_FUNGUS | 26 | 0.447514 | 1.132879 | 0.290366 | 0.693794 |
| GOBP_REGULATION_OF_NERVOUS_SYSTEM_PROCESS | 114 | 0.359195 | 1.132745 | 0.260773 | 0.693623 |
| GOBP_POST_EMBRYONIC_ANIMAL_ORGAN_DEVELOPMENT | 16 | 0.483349 | 1.132692 | 0.326379 | 0.693187 |
| GOBP_CILIUM_MOVEMENT | 126 | 0.353113 | 1.13254 | 0.235028 | 0.693046 |
| GOBP_REGULATION_OF_AMINO_ACID_TRANSPORT | 34 | 0.426282 | 1.132166 | 0.30343 | 0.693691 |
| GOBP_FC_RECEPTOR_MEDIATED_STIMULATORY_SIGNALING_PATHWAY | 26 | 0.449673 | 1.132134 | 0.3083 | 0.693217 |
| GOBP_POSITIVE_REGULATION_OF_AMINO_ACID_TRANSPORT | 19 | 0.466078 | 1.131943 | 0.301136 | 0.693273 |
| GOBP_BONE_GROWTH | 26 | 0.436545 | 1.131773 | 0.301676 | 0.693287 |
| GOBP_REGULATION_OF_RESPONSE_TO_CYTOKINE_STIMULUS | 133 | 0.351096 | 1.131557 | 0.241455 | 0.693407 |
| GOBP_FIBROBLAST_PROLIFERATION | 66 | 0.379094 | 1.131495 | 0.269976 | 0.693 |
| GOBP_NEGATIVE_REGULATION_OF_DEFENSE_RESPONSE | 174 | 0.345821 | 1.131357 | 0.224622 | 0.692836 |
| GOBP_NEGATIVE_REGULATION_OF_ANION_TRANSPORT | 18 | 0.472177 | 1.1311 | 0.313584 | 0.693103 |
| GOBP_REGULATION_OF_TRANSPORTER_ACTIVITY | 231 | 0.334774 | 1.130969 | 0.213542 | 0.692939 |
| GOBP_REGULATION_OF_PROTEIN_MATURATION | 55 | 0.396956 | 1.130775 | 0.267427 | 0.692979 |
| GOBP_REGULATION_OF_POTASSIUM_ION_TRANSMEMBRANE_TRANSPORTER_ACTIVITY | 45 | 0.39951 | 1.130137 | 0.281445 | 0.694511 |
| GOBP_PEPTIDYL_TYROSINE_MODIFICATION | 296 | 0.335005 | 1.129904 | 0.181448 | 0.694679 |
| GOBP_NEGATIVE_REGULATION_OF_TRANSFORMING_GROWTH_FACTOR_BETA_RECEPTOR | 67 | 0.378846 | 1.129703 | 0.268765 | 0.694744 |
| GOBP_GLYCOSYLCERAMIDE_METABOLIC_PROCESS | 15 | 0.499126 | 1.129262 | 0.326895 | 0.6956 |
| GOBP_LEUKOCYTE_TETHERING_OR_ROLLING | 19 | 0.478781 | 1.129045 | 0.333826 | 0.695704 |
| GOBP_SINGLE_FERTILIZATION | 85 | 0.365563 | 1.129043 | 0.260369 | 0.695115 |
| GOBP_NEGATIVE_REGULATION_OF_CARDIAC_MUSCLE_TISSUE_GROWTH | 21 | 0.457863 | 1.128746 | 0.329004 | 0.695496 |
| GOBP_CELLULAR_RESPONSE_TO_UV | 81 | 0.371082 | 1.128659 | 0.261877 | 0.695174 |
| GOBP_POSITIVE_REGULATION_OF_SYNAPTIC_TRANSMISSION_GLUTAMATERGIC | 23 | 0.454462 | 1.128543 | 0.337989 | 0.694955 |
| GOBP_EMBRYONIC_ORGAN_DEVELOPMENT | 355 | 0.330329 | 1.128169 | 0.182004 | 0.69559 |
| GOBP_SKELETAL_MUSCLE_ORGAN_DEVELOPMENT | 133 | 0.352512 | 1.127088 | 0.246137 | 0.698573 |
| GOBP_SPECIFICATION_OF_SYMMETRY | 115 | 0.355828 | 1.126996 | 0.276715 | 0.698286 |
| GOBP_CIRCULATORY_SYSTEM_PROCESS | 473 | 0.32701 | 1.126944 | 0.158317 | 0.69786 |
| GOBP_METANEPHROS_DEVELOPMENT | 68 | 0.370591 | 1.126824 | 0.276191 | 0.697652 |
| GOBP_CELL_CELL_JUNCTION_ASSEMBLY | 117 | 0.35433 | 1.12678 | 0.249723 | 0.697176 |
| GOBP_INSULIN_SECRETION | 146 | 0.347409 | 1.126216 | 0.246456 | 0.698427 |
| GOBP_MESODERMAL_CELL_DIFFERENTIATION | 24 | 0.445749 | 1.125229 | 0.319061 | 0.701121 |
| GOBP_NEGATIVE_REGULATION_OF_VIRAL_ENTRY_INTO_HOST_CELL | 19 | 0.465119 | 1.124717 | 0.306241 | 0.702181 |
| GOBP_PROTEIN_LOCALIZATION_TO_CELL_CELL_JUNCTION | 20 | 0.469247 | 1.124475 | 0.329446 | 0.702381 |
| GOBP_REGULATION_OF_GENE_SILENCING_BY_RNA | 45 | 0.404511 | 1.123984 | 0.298357 | 0.703393 |
| GOBP_REGULATION_OF_MITOCHONDRION_ORGANIZATION | 132 | 0.35323 | 1.123921 | 0.251101 | 0.702993 |
| GOBP_ORGAN_OR_TISSUE_SPECIFIC_IMMUNE_RESPONSE | 17 | 0.480098 | 1.123849 | 0.313522 | 0.702646 |
| GOBP_NEGATIVE_REGULATION_OF_EXTRINSIC_APOPTOTIC_SIGNALING_PATHWAY | 79 | 0.374469 | 1.123834 | 0.289256 | 0.702097 |
| GOBP_MYELOID_CELL_DIFFERENTIATION | 322 | 0.332522 | 1.123293 | 0.204733 | 0.703312 |
| GOBP_RESPONSE_TO_XENOBIOTIC_STIMULUS | 310 | 0.330471 | 1.123098 | 0.21058 | 0.703369 |
| GOBP_MESONEPHROS_DEVELOPMENT | 80 | 0.370472 | 1.122643 | 0.285063 | 0.704256 |
| GOBP_POSITIVE_REGULATION_OF_PROTEIN_ACETYLATION | 44 | 0.39432 | 1.122237 | 0.293035 | 0.704969 |
| GOBP_ASTROCYTE_ACTIVATION | 15 | 0.492149 | 1.122229 | 0.318253 | 0.704404 |
| GOBP_EMBRYONIC_SKELETAL_SYSTEM_MORPHOGENESIS | 58 | 0.391278 | 1.122156 | 0.306197 | 0.704064 |
| GOBP_MEMBRANE_INVAGINATION | 60 | 0.388011 | 1.12211 | 0.298137 | 0.703649 |
| GOBP_NEGATIVE_REGULATION_OF_INTERLEUKIN_1_BETA_PRODUCTION | 17 | 0.475468 | 1.121622 | 0.301471 | 0.704744 |
| GOBP_VENTRICULAR_CARDIAC_MUSCLE_CELL_ACTION_POTENTIAL | 27 | 0.430406 | 1.121392 | 0.331962 | 0.704948 |
| GOBP_CELLULAR_RESPONSE_TO_ATP | 16 | 0.487335 | 1.121384 | 0.342944 | 0.704379 |
| GOBP_T_CELL_HOMEOSTASIS | 29 | 0.42779 | 1.121002 | 0.326503 | 0.705036 |
| GOBP_REGULATION_OF_CARDIAC_MUSCLE_CONTRACTION | 62 | 0.385555 | 1.120875 | 0.306733 | 0.704857 |
| GOBP_POSITIVE_REGULATION_OF_MITOCHONDRIAL_FISSION | 20 | 0.465119 | 1.12082 | 0.328889 | 0.704434 |
| GOBP_REGULATION_OF_SYSTEM_PROCESS | 439 | 0.324487 | 1.120813 | 0.187751 | 0.703867 |
| GOBP_GASTRULATION | 161 | 0.343841 | 1.119968 | 0.24866 | 0.706091 |
| GOBP_ACTIN_FILAMENT_BASED_TRANSPORT | 18 | 0.471905 | 1.119913 | 0.325037 | 0.705679 |
| GOBP_POSITIVE_REGULATION_OF_EPIDERMIS_DEVELOPMENT | 27 | 0.434865 | 1.119886 | 0.316556 | 0.705183 |
| GOBP_NEGATIVE_REGULATION_OF_INFLAMMATORY_RESPONSE_TO_ANTIGENIC_STIMULU | 23 | 0.454794 | 1.119762 | 0.320621 | 0.704987 |
| GOBP_POSITIVE_REGULATION_OF_NUCLEAR_DIVISION | 42 | 0.405953 | 1.118775 | 0.314286 | 0.707629 |
| GOBP_REGULATION_OF_SUPEROXIDE_METABOLIC_PROCESS | 23 | 0.450384 | 1.118582 | 0.335684 | 0.707697 |
| GOBP_SYNAPTIC_MEMBRANE_ADHESION | 25 | 0.445895 | 1.118045 | 0.300423 | 0.708866 |
| GOBP_POSITIVE_REGULATION_OF_NEUROTRANSMITTER_TRANSPORT | 21 | 0.458224 | 1.118033 | 0.349711 | 0.708328 |
| GOBP_SMOOTH_MUSCLE_CELL_MIGRATION | 59 | 0.38348 | 1.116526 | 0.303704 | 0.712761 |
| GOBP_PYRIMIDINE_CONTAINING_COMPOUND_CATABOLIC_PROCESS | 36 | 0.414727 | 1.116369 | 0.320479 | 0.712672 |
| GOBP_MONOCARBOXYLIC_ACID_METABOLIC_PROCESS | 455 | 0.324563 | 1.11633 | 0.190332 | 0.712202 |
| GOBP_NEUTRAL_LIPID_METABOLIC_PROCESS | 101 | 0.359372 | 1.115698 | 0.290509 | 0.713701 |
| GOBP_RESPONSE_TO_CARBOHYDRATE | 178 | 0.341412 | 1.115648 | 0.264454 | 0.713271 |
| GOBP_MAINTENANCE_OF_LOCATION_IN_CELL | 174 | 0.340515 | 1.115429 | 0.262084 | 0.713396 |
| GOBP_RESPONSE_TO_AXON_INJURY | 71 | 0.366468 | 1.115295 | 0.301549 | 0.713235 |
| GOBP_LONG_CHAIN_FATTY_ACID_METABOLIC_PROCESS | 85 | 0.360525 | 1.11488 | 0.295954 | 0.713983 |
| GOBP_ACTIN_FILAMENT_BASED_MOVEMENT | 115 | 0.351961 | 1.114668 | 0.289655 | 0.714144 |
| GOBP_SUBSTRATE_ADHESION_DEPENDENT_CELL_SPREADING | 99 | 0.358266 | 1.114106 | 0.286511 | 0.715373 |
| GOBP_RELEASE_OF_CYTOCHROME_C_FROM_MITOCHONDRIA | 51 | 0.391886 | 1.113606 | 0.305627 | 0.716448 |
| GOBP_DETECTION_OF_MECHANICAL_STIMULUS | 41 | 0.398999 | 1.113291 | 0.324538 | 0.716927 |
| GOBP_CELLULAR_GLUCOSE_HOMEOSTASIS | 123 | 0.3519 | 1.112168 | 0.28491 | 0.720038 |
| GOBP_HORMONE_TRANSPORT | 231 | 0.332953 | 1.111741 | 0.252874 | 0.720907 |
| GOBP_POSITIVE_REGULATION_OF_INTERLEUKIN_1_BETA_PRODUCTION | 40 | 0.400899 | 1.111514 | 0.317073 | 0.721089 |
| GOBP_PEPTIDE_CATABOLIC_PROCESS | 22 | 0.45287 | 1.11144 | 0.363242 | 0.720737 |
| GOBP_REGULATION_OF_LIPID_METABOLIC_PROCESS | 261 | 0.329269 | 1.111435 | 0.235537 | 0.720161 |
| GOBP_PROTEIN_PROCESSING | 191 | 0.33602 | 1.111272 | 0.275716 | 0.720089 |
| GOBP_CELL_CELL_ADHESION_VIA_PLASMA_MEMBRANE_ADHESION_MOLECULES | 225 | 0.332798 | 1.111227 | 0.244514 | 0.719661 |
| GOBP_POSITIVE_REGULATION_OF_DNA_BINDING_TRANSCRIPTION_FACTOR_ACTIVITY | 218 | 0.337152 | 1.111142 | 0.247368 | 0.71935 |
| GOBP_SEX_DIFFERENTIATION | 216 | 0.333756 | 1.111086 | 0.253919 | 0.718946 |
| GOBP_NEGATIVE_REGULATION_OF_LEUKOCYTE_APOPTOTIC_PROCESS | 37 | 0.40604 | 1.110877 | 0.319205 | 0.719073 |
| GOBP_HOMOPHILIC_CELL_ADHESION_VIA_PLASMA_MEMBRANE_ADHESION_MOLECULES | 144 | 0.343208 | 1.110425 | 0.281628 | 0.71998 |
| GOBP_POSITIVE_REGULATION_OF_POSTTRANSCRIPTIONAL_GENE_SILENCING | 31 | 0.420132 | 1.110196 | 0.344043 | 0.720138 |
| GOBP_REPRODUCTIVE_BEHAVIOR | 23 | 0.446677 | 1.110187 | 0.344491 | 0.719577 |
| GOBP_SARCOMERE_ORGANIZATION | 32 | 0.417604 | 1.109809 | 0.328534 | 0.720184 |
| GOBP_RESPONSE_TO_BMP | 145 | 0.341766 | 1.109746 | 0.27927 | 0.719815 |
| GOBP_RESPONSE_TO_METAL_ION | 279 | 0.330009 | 1.109723 | 0.230448 | 0.7193 |
| GOBP_VIRAL_LIFE_CYCLE | 271 | 0.325111 | 1.109129 | 0.233846 | 0.720703 |
| GOBP_POSITIVE_REGULATION_OF_NF_KAPPAB_TRANSCRIPTION_FACTOR_ACTIVITY | 127 | 0.346896 | 1.108838 | 0.27088 | 0.721093 |
| GOBP_CELLULAR_SENESCENCE | 74 | 0.366551 | 1.108679 | 0.297773 | 0.721029 |
| GOBP_POSITIVE_REGULATION_OF_NEUROTRANSMITTER_SECRETION | 16 | 0.481489 | 1.108295 | 0.352484 | 0.72172 |
| GOBP_EMBRYONIC_HEMOPOIESIS | 20 | 0.453054 | 1.108204 | 0.335312 | 0.721436 |
| GOBP_REGULATION_OF_CIRCADIAN_RHYTHM | 104 | 0.355839 | 1.106967 | 0.299087 | 0.724956 |
| GOBP_CELLULAR_CARBOHYDRATE_METABOLIC_PROCESS | 242 | 0.329234 | 1.106458 | 0.252862 | 0.726008 |
| GOBP_UROGENITAL_SYSTEM_DEVELOPMENT | 293 | 0.327759 | 1.106327 | 0.244151 | 0.725863 |
| GOBP_POSITIVE_REGULATION_OF_SODIUM_ION_TRANSMEMBRANE_TRANSPORTER_ACTI | 16 | 0.482123 | 1.105933 | 0.347507 | 0.72663 |
| GOBP_NEGATIVE_REGULATION_OF_ENDOTHELIAL_CELL_APOPTOTIC_PROCESS | 25 | 0.438636 | 1.105289 | 0.349866 | 0.728143 |
| GOBP_PINOCYTOSIS | 23 | 0.443136 | 1.105212 | 0.337111 | 0.727809 |
| GOBP_POSITIVE_REGULATION_OF_TRANSCRIPTION_FROM_RNA_POLYMERASE_II_PROMO | 22 | 0.451243 | 1.105026 | 0.336088 | 0.72788 |
| GOBP_MAMMARY_GLAND_EPITHELIAL_CELL_PROLIFERATION | 23 | 0.447284 | 1.105019 | 0.344398 | 0.727323 |
| GOBP_RAB_PROTEIN_SIGNAL_TRANSDUCTION | 21 | 0.451819 | 1.104958 | 0.344118 | 0.726932 |
| GOBP_BINDING_OF_SPERM_TO_ZONA_PELLUCIDA | 21 | 0.453259 | 1.104708 | 0.355396 | 0.727222 |
| GOBP_RESPONSE_TO_PROGESTERONE | 33 | 0.413723 | 1.104645 | 0.317204 | 0.726847 |
| GOBP_REGULATION_OF_NUCLEAR_DIVISION | 117 | 0.350169 | 1.104494 | 0.300786 | 0.72677 |
| GOBP_ATRIAL_SEPTUM_DEVELOPMENT | 19 | 0.46401 | 1.104219 | 0.367799 | 0.727101 |
| GOBP_REGULATION_OF_INSULIN_SECRETION | 120 | 0.34874 | 1.10392 | 0.280277 | 0.727501 |
| GOBP_DECIDUALIZATION | 19 | 0.464038 | 1.103673 | 0.34438 | 0.727739 |

| | | | | | |
|---|---|---|---|---|---|
| GOBP_FORMATION_OF_PRIMARY_GERM_LAYER | 105 | 0.35442 | 1.10285 | 0.309091 | 0.72993 |
| GOBP_CELLULAR_RESPONSE_TO_CADMIUM_ION | 28 | 0.424857 | 1.102247 | 0.350416 | 0.731324 |
| GOBP_POSITIVE_REGULATION_OF_LAMELLIPODIUM_ORGANIZATION | 32 | 0.413557 | 1.10206 | 0.337255 | 0.731405 |
| GOBP_REGULATION_OF_CELLULAR_EXTRAVASATION | 27 | 0.420647 | 1.101501 | 0.337047 | 0.73266 |
| GOBP_POSITIVE_REGULATION_OF_MACROPHAGE_MIGRATION | 15 | 0.484647 | 1.101359 | 0.349345 | 0.732554 |
| GOBP_REGULATION_OF_DNA_BINDING_TRANSCRIPTION_FACTOR_ACTIVITY | 366 | 0.320985 | 1.101118 | 0.235474 | 0.732757 |
| GOBP_CRANIOFACIAL_SUTURE_MORPHOGENESIS | 15 | 0.486947 | 1.101086 | 0.358006 | 0.732277 |
| GOBP_DEVELOPMENTAL_MATURATION | 239 | 0.326987 | 1.10108 | 0.246106 | 0.731717 |
| GOBP_REGULATION_OF_NUCLEOTIDE_BIOSYNTHETIC_PROCESS | 32 | 0.418009 | 1.101062 | 0.327078 | 0.731189 |
| GOBP_SULFATION | 16 | 0.48137 | 1.100891 | 0.369501 | 0.731165 |
| GOBP_PHOSPHOLIPID_CATABOLIC_PROCESS | 42 | 0.397365 | 1.100528 | 0.325786 | 0.731775 |
| GOBP_REGULATION_OF_RECEPTOR_SIGNALING_PATHWAY_VIA_STAT | 59 | 0.375302 | 1.100067 | 0.321739 | 0.73268 |
| GOBP_CELLULAR_RESPONSE_TO_LIGHT_STIMULUS | 95 | 0.356115 | 1.099423 | 0.312862 | 0.73421 |
| GOBP_LIMBIC_SYSTEM_DEVELOPMENT | 90 | 0.355314 | 1.099091 | 0.32243 | 0.734774 |
| GOBP_CELLULAR_RESPONSE_TO_ACID_CHEMICAL | 72 | 0.363836 | 1.098702 | 0.303722 | 0.7355 |
| GOBP_REGULATION_OF_NIK_NF_KAPPAB_SIGNALING | 80 | 0.361237 | 1.098422 | 0.320887 | 0.735819 |
| GOBP_MESODERM_MORPHOGENESIS | 62 | 0.374121 | 1.097879 | 0.318239 | 0.737034 |
| GOBP_POSITIVE_REGULATION_OF_LIPID_LOCALIZATION | 81 | 0.356963 | 1.097497 | 0.322468 | 0.737729 |
| GOBP_REGULATED_EXOCYTOSIS | 194 | 0.333556 | 1.096479 | 0.274655 | 0.740557 |
| GOBP_NEGATIVE_REGULATION_OF_STRIATED_MUSCLE_CELL_DIFFERENTIATION | 33 | 0.411857 | 1.095516 | 0.34912 | 0.743173 |
| GOBP_ORGANIC_HYDROXY_COMPOUND_CATABOLIC_PROCESS | 56 | 0.377567 | 1.095507 | 0.332915 | 0.742627 |
| GOBP_RESPONSE_TO_KETONE | 151 | 0.337111 | 1.095301 | 0.298491 | 0.742724 |
| GOBP_TRANSFORMING_GROWTH_FACTOR_BETA_RECEPTOR_SIGNALING_PATHWAY | 171 | 0.335822 | 1.095289 | 0.299363 | 0.742178 |
| GOBP_BRANCHING_INVOLVED_IN_BLOOD_VESSEL_MORPHOGENESIS | 29 | 0.41655 | 1.095086 | 0.351902 | 0.742256 |
| GOBP_SENSORY_ORGAN_DEVELOPMENT | 446 | 0.320028 | 1.095029 | 0.239719 | 0.741881 |
| GOBP_NEGATIVE_REGULATION_OF_VIRAL_LIFE_CYCLE | 22 | 0.442511 | 1.094895 | 0.359375 | 0.741729 |
| GOBP_RHYTHMIC_PROCESS | 256 | 0.323004 | 1.094485 | 0.273292 | 0.742499 |
| GOBP_RESPONSE_TO_INTERLEUKIN_1 | 90 | 0.355899 | 1.09416 | 0.31682 | 0.742976 |
| GOBP_HEART_GROWTH | 64 | 0.368774 | 1.094142 | 0.322503 | 0.742462 |
| GOBP_REGULATION_OF_HEART_CONTRACTION | 165 | 0.334277 | 1.0941 | 0.306034 | 0.74203 |
| GOBP_VENTRICULAR_SEPTUM_MORPHOGENESIS | 41 | 0.389736 | 1.093376 | 0.332464 | 0.743885 |
| GOBP_REGULATION_OF_DEVELOPMENTAL_GROWTH | 270 | 0.325402 | 1.093163 | 0.267081 | 0.743985 |
| GOBP_NEGATIVE_REGULATION_OF_LIPID_LOCALIZATION | 31 | 0.414153 | 1.093044 | 0.374168 | 0.743808 |
| GOBP_OLEFINIC_COMPOUND_METABOLIC_PROCESS | 109 | 0.350539 | 1.092976 | 0.324263 | 0.743467 |
| GOBP_REGULATION_OF_SMOOTH_MUSCLE_CELL_DIFFERENTIATION | 25 | 0.431858 | 1.09297 | 0.356546 | 0.74291 |
| GOBP_PROTEIN_HYDROXYLATION | 25 | 0.43728 | 1.09289 | 0.358025 | 0.742594 |
| GOBP_POSITIVE_REGULATION_OF_FAT_CELL_DIFFERENTIATION | 59 | 0.376678 | 1.092436 | 0.335784 | 0.743517 |
| GOBP_MOLTING_CYCLE | 82 | 0.361524 | 1.091726 | 0.337587 | 0.745282 |
| GOBP_REGULATION_OF_ACTIN_FILAMENT_BASED_PROCESS | 351 | 0.320588 | 1.091276 | 0.253049 | 0.746215 |
| GOBP_COGNITION | 267 | 0.32565 | 1.091044 | 0.286742 | 0.746394 |
| GOBP_EMBRYONIC_ORGAN_MORPHOGENESIS | 217 | 0.323904 | 1.090237 | 0.296842 | 0.748486 |
| GOBP_EPITHELIAL_TO_MESENCHYMAL_TRANSITION_INVOLVED_IN_ENDOCARDIAL_CUSHI | 20 | 0.450947 | 1.089925 | 0.372607 | 0.748962 |
| GOBP_ACTIN_MEDIATED_CELL_CONTRACTION | 88 | 0.353562 | 1.089493 | 0.318287 | 0.749771 |
| GOBP_NEGATIVE_REGULATION_OF_INTERLEUKIN_6_PRODUCTION | 32 | 0.410214 | 1.088412 | 0.3397 | 0.752813 |
| GOBP_ORGANELLE_TRANSPORT_ALONG_MICROTUBULE | 84 | 0.35679 | 1.088053 | 0.326389 | 0.753475 |
| GOBP_NEGATIVE_REGULATION_OF_PROTEIN_TYROSINE_KINASE_ACTIVITY | 29 | 0.412045 | 1.087461 | 0.358799 | 0.754839 |
| GOBP_TOLL_LIKE_RECEPTOR_4_SIGNALING_PATHWAY | 28 | 0.427666 | 1.087251 | 0.355281 | 0.754978 |
| GOBP_AGING | 251 | 0.323119 | 1.086799 | 0.296066 | 0.755837 |
| GOBP_RESPONSE_TO_ANTIBIOTIC | 41 | 0.392583 | 1.086707 | 0.339719 | 0.755572 |
| GOBP_REGULATION_OF_MONOCYTE_DIFFERENTIATION | 16 | 0.478313 | 1.086594 | 0.38864 | 0.755373 |
| GOBP_OSTEOBLAST_PROLIFERATION | 27 | 0.421363 | 1.086434 | 0.352859 | 0.755333 |
| GOBP_REGULATION_OF_SKELETAL_MUSCLE_CELL_DIFFERENTIATION | 17 | 0.472942 | 1.085925 | 0.365527 | 0.75639 |
| GOBP_DEVELOPMENT_OF_PRIMARY_SEXUAL_CHARACTERISTICS | 185 | 0.330783 | 1.085051 | 0.329424 | 0.758682 |
| GOBP_NEGATIVE_REGULATION_OF_EPITHELIAL_CELL_PROLIFERATION | 107 | 0.345338 | 1.084866 | 0.31573 | 0.758717 |
| GOBP_VASCULAR_ASSOCIATED_SMOOTH_MUSCLE_CELL_MIGRATION | 18 | 0.458512 | 1.084859 | 0.373563 | 0.758167 |
| GOBP_METHYLATION | 340 | 0.319298 | 1.084675 | 0.272358 | 0.758189 |
| GOBP_CHLORIDE_TRANSPORT | 85 | 0.351173 | 1.084518 | 0.343023 | 0.758117 |
| GOBP_ALCOHOL_METABOLIC_PROCESS | 285 | 0.322984 | 1.084245 | 0.292608 | 0.758463 |
| GOBP_NEGATIVE_REGULATION_OF_COLD_INDUCED_THERMOGENESIS | 41 | 0.390446 | 1.083922 | 0.367589 | 0.758937 |
| GOBP_RETINAL_METABOLIC_PROCESS | 15 | 0.470776 | 1.081815 | 0.375918 | 0.765193 |
| GOBP_RESPONSE_TO_DEXAMETHASONE | 33 | 0.407623 | 1.081734 | 0.375335 | 0.764882 |
| GOBP_NEGATIVE_REGULATION_OF_TRANSPORT | 341 | 0.31778 | 1.08115 | 0.298264 | 0.766223 |
| GOBP_GLUTAMATE_SECRETION | 22 | 0.428295 | 1.080995 | 0.380674 | 0.766143 |
| GOBP_PANCREAS_DEVELOPMENT | 55 | 0.372523 | 1.080857 | 0.34901 | 0.765978 |
| GOBP_EXOCRINE_SYSTEM_DEVELOPMENT | 37 | 0.399085 | 1.080399 | 0.363878 | 0.766883 |
| GOBP_METANEPHROS_MORPHOGENESIS | 26 | 0.418263 | 1.080139 | 0.372355 | 0.767189 |
| GOBP_CELLULAR_RESPONSE_TO_CARBOHYDRATE_STIMULUS | 117 | 0.345769 | 1.080098 | 0.334463 | 0.766748 |
| GOBP_MATING | 31 | 0.409677 | 1.080066 | 0.369418 | 0.766274 |
| GOBP_CAMERA_TYPE_EYE_MORPHOGENESIS | 100 | 0.348127 | 1.079746 | 0.335593 | 0.766771 |
| GOBP_POSITIVE_REGULATION_OF_IMMUNOGLOBULIN_PRODUCTION | 38 | 0.38939 | 1.078927 | 0.350129 | 0.768976 |
| GOBP_REGULATION_OF_SMALL_GTPASE_MEDIATED_SIGNAL_TRANSDUCTION | 274 | 0.31854 | 1.07845 | 0.290589 | 0.769992 |
| GOBP_RESPONSE_TO_DOPAMINE | 61 | 0.366166 | 1.078082 | 0.353155 | 0.770594 |
| GOBP_NEGATIVE_REGULATION_OF_ERK1_AND_ERK2_CASCADE | 62 | 0.365422 | 1.077551 | 0.359649 | 0.771725 |
| GOBP_NEGATIVE_REGULATION_OF_SIGNAL_TRANSDUCTION_BY_P53_CLASS_MEDIATOR | 32 | 0.402525 | 1.077466 | 0.364384 | 0.771436 |
| GOBP_POSITIVE_REGULATION_OF_BMP_SIGNALING_PATHWAY | 33 | 0.400067 | 1.077079 | 0.374659 | 0.772116 |
| GOBP_REGULATION_OF_CELLULAR_RESPONSE_TO_GROWTH_FACTOR_STIMULUS | 251 | 0.323329 | 1.076912 | 0.310094 | 0.772103 |
| GOBP_PROSTAGLANDIN_TRANSPORT | 15 | 0.47628 | 1.076356 | 0.382055 | 0.77342 |
| GOBP_LIPOPOLYSACCHARIDE_MEDIATED_SIGNALING_PATHWAY | 43 | 0.383479 | 1.076185 | 0.357692 | 0.773419 |
| GOBP_REGULATION_OF_NECROTIC_CELL_DEATH | 32 | 0.413178 | 1.075453 | 0.378162 | 0.775241 |
| GOBP_REGULATION_OF_CARDIAC_MUSCLE_CELL_ACTION_POTENTIAL | 25 | 0.419729 | 1.075431 | 0.34887 | 0.774721 |
| GOBP_POSITIVE_REGULATION_OF_PROTEIN_IMPORT | 38 | 0.394928 | 1.07527 | 0.371968 | 0.774685 |
| GOBP_MAMMARY_GLAND_DEVELOPMENT | 113 | 0.341797 | 1.074695 | 0.32809 | 0.776053 |
| GOBP_MAST_CELL_ACTIVATION_INVOLVED_IN_IMMUNE_RESPONSE | 40 | 0.394029 | 1.074308 | 0.361979 | 0.776759 |
| GOBP_NEGATIVE_REGULATION_OF_DNA_DAMAGE_RESPONSE_SIGNAL_TRANSDUCTION_ | 15 | 0.46779 | 1.074002 | 0.386167 | 0.777177 |
| GOBP_LIPID_EXPORT_FROM_CELL | 33 | 0.401379 | 1.073674 | 0.364754 | 0.777701 |
| GOBP_POSITIVE_REGULATION_OF_JNK_CASCADE | 81 | 0.356414 | 1.07251 | 0.34126 | 0.781054 |
| GOBP_REGULATION_OF_MICROTUBULE_BASED_MOVEMENT | 42 | 0.384501 | 1.072317 | 0.379665 | 0.781121 |
| GOBP_CELLULAR_RESPONSE_TO_DEXAMETHASONE_STIMULUS | 20 | 0.441722 | 1.071747 | 0.392351 | 0.782415 |
| GOBP_REGULATION_OF_HORMONE_LEVELS | 372 | 0.315591 | 1.071626 | 0.299492 | 0.782223 |
| GOBP_CELLULAR_RESPONSE_TO_RETINOIC_ACID | 53 | 0.378268 | 1.071088 | 0.367758 | 0.783464 |
| GOBP_DOPAMINERGIC_NEURON_DIFFERENTIATION | 27 | 0.41676 | 1.070939 | 0.371389 | 0.78336 |
| GOBP_IMPORT_INTO_CELL | 185 | 0.324397 | 1.070418 | 0.342887 | 0.784509 |
| GOBP_CENTROSOME_SEPARATION | 15 | 0.464234 | 1.070343 | 0.385561 | 0.784172 |
| GOBP_REGULATION_OF_T_HELPER_17_TYPE_IMMUNE_RESPONSE | 18 | 0.455774 | 1.070331 | 0.374817 | 0.783624 |
| GOBP_POSITIVE_REGULATION_OF_OXIDATIVE_STRESS_INDUCED_CELL_DEATH | 15 | 0.469861 | 1.069661 | 0.390649 | 0.785242 |
| GOBP_ENDOCHONDRAL_BONE_MORPHOGENESIS | 49 | 0.378688 | 1.06887 | 0.390525 | 0.7873 |
| GOBP_PARAXIAL_MESODERM_DEVELOPMENT | 17 | 0.450173 | 1.06792 | 0.40416 | 0.789808 |
| GOBP_POSITIVE_REGULATION_OF_MEMBRANE_PERMEABILITY | 41 | 0.394134 | 1.067631 | 0.383226 | 0.790208 |
| GOBP_POSITIVE_REGULATION_OF_AMINE_TRANSPORT | 33 | 0.400687 | 1.067319 | 0.383065 | 0.790633 |
| GOBP_POSITIVE_REGULATION_OF_EPITHELIAL_CELL_DIFFERENTIATION | 44 | 0.381343 | 1.067215 | 0.369171 | 0.790361 |
| GOBP_ADENYLATE_CYCLASE_MODULATING_G_PROTEIN_COUPLED_RECEPTOR_SIGNALIN | 177 | 0.323831 | 1.066505 | 0.352878 | 0.792123 |
| GOBP_PITUITARY_GLAND_DEVELOPMENT | 31 | 0.404885 | 1.066181 | 0.377807 | 0.792572 |
| GOBP_CELLULAR_EXTRAVASATION | 51 | 0.371379 | 1.065975 | 0.381071 | 0.792689 |
| GOBP_MYOTUBE_CELL_DEVELOPMENT | 29 | 0.41453 | 1.06588 | 0.377384 | 0.792427 |
| GOBP_HETEROTYPIC_CELL_CELL_ADHESION | 46 | 0.381287 | 1.065402 | 0.382542 | 0.793448 |
| GOBP_SEMI_LUNAR_VALVE_DEVELOPMENT | 39 | 0.394762 | 1.064664 | 0.388674 | 0.795337 |
| GOBP_RESPONSE_TO_PEPTIDE_HORMONE | 325 | 0.31394 | 1.063772 | 0.3286 | 0.797668 |
| GOBP_MYD88_DEPENDENT_TOLL_LIKE_RECEPTOR_SIGNALING_PATHWAY | 17 | 0.456472 | 1.063657 | 0.400593 | 0.797419 |
| GOBP_CIRCADIAN_RHYTHM | 175 | 0.326214 | 1.0636 | 0.34298 | 0.797023 |
| GOBP_NEGATIVE_REGULATION_OF_MUSCLE_CELL_DIFFERENTIATION | 50 | 0.373151 | 1.063543 | 0.380832 | 0.796628 |
| GOBP_RESPONSE_TO_MURAMYL_DIPEPTIDE | 19 | 0.445754 | 1.063116 | 0.410587 | 0.797395 |
| GOBP_MESENCHYMAL_CELL_DIFFERENTIATION | 209 | 0.320929 | 1.063052 | 0.351464 | 0.797003 |
| GOBP_POSITIVE_REGULATION_OF_CELL_DEATH | 478 | 0.30958 | 1.06134 | 0.308853 | 0.801996 |
| GOBP_RESPONSE_TO_TOXIC_SUBSTANCE | 186 | 0.317175 | 1.060278 | 0.369427 | 0.804821 |
| GOBP_LYMPHOCYTE_HOMEOSTASIS | 49 | 0.370231 | 1.059848 | 0.381935 | 0.805675 |
| GOBP_SYNAPTIC_VESICLE_RECYCLING | 68 | 0.357965 | 1.059481 | 0.373026 | 0.806241 |
| GOBP_ASTROCYTE_DEVELOPMENT | 31 | 0.401006 | 1.059405 | 0.383399 | 0.80589 |
| GOBP_ACYLGLYCEROL_HOMEOSTASIS | 21 | 0.431266 | 1.059196 | 0.393557 | 0.805958 |
| GOBP_INFLAMMATORY_RESPONSE_TO_ANTIGENIC_STIMULUS | 40 | 0.381718 | 1.059094 | 0.394567 | 0.805698 |
| GOBP_CELLULAR_LIPID_CATABOLIC_PROCESS | 184 | 0.322549 | 1.058872 | 0.374603 | 0.80586 |
| GOBP_NEGATIVE_REGULATION_OF_CELL_GROWTH | 164 | 0.324469 | 1.058617 | 0.369936 | 0.806101 |
| GOBP_NUCLEOSIDE_MONOPHOSPHATE_CATABOLIC_PROCESS | 16 | 0.450758 | 1.058555 | 0.391367 | 0.805711 |
| GOBP_PEPTIDYL_TYROSINE_AUTOPHOSPHORYLATION | 18 | 0.448484 | 1.05837 | 0.411679 | 0.805703 |
| GOBP_SIGNAL_RELEASE | 371 | 0.312542 | 1.058189 | 0.339757 | 0.805709 |
| GOBP_REGULATION_OF_CYSTEINE_TYPE_ENDOPEPTIDASE_ACTIVITY_INVOLVED_IN_APOP | 15 | 0.467341 | 1.058175 | 0.399096 | 0.805174 |
| GOBP_CIRCADIAN_REGULATION_OF_GENE_EXPRESSION | 63 | 0.357065 | 1.057676 | 0.384161 | 0.80631 |
| GOBP_APICAL_JUNCTION_ASSEMBLY | 58 | 0.361898 | 1.057223 | 0.392991 | 0.807215 |
| GOBP_INTERFERON_ALPHA_PRODUCTION | 20 | 0.436759 | 1.056928 | 0.396825 | 0.807558 |
| GOBP_RESPONSE_TO_CADMIUM_ION | 46 | 0.372465 | 1.056525 | 0.392768 | 0.808311 |
| GOBP_POSITIVE_REGULATION_OF_PHOSPHOPROTEIN_PHOSPHATASE_ACTIVITY | 17 | 0.454352 | 1.055144 | 0.407953 | 0.812208 |
| GOBP_BIOMINERALIZATION | 124 | 0.331207 | 1.054842 | 0.358744 | 0.812642 |
| GOBP_RENAL_TUBULAR_SECRETION | 18 | 0.448897 | 1.054418 | 0.424069 | 0.813432 |
| GOBP_ALCOHOL_BIOSYNTHETIC_PROCESS | 118 | 0.332289 | 1.053963 | 0.363128 | 0.814323 |
| GOBP_TRANSCYTOSIS | 18 | 0.445087 | 1.053466 | 0.414051 | 0.815366 |
| GOBP_METHYLGUANOSINE_CAP_DECAPPING | 15 | 0.459538 | 1.053236 | 0.420732 | 0.81553 |
| GOBP_ADULT_BEHAVIOR | 117 | 0.333404 | 1.052462 | 0.388764 | 0.817439 |
| GOBP_ENDOCRINE_HORMONE_SECRETION | 37 | 0.39055 | 1.052406 | 0.401316 | 0.817044 |
| GOBP_CARDIAC_CONDUCTION_SYSTEM_DEVELOPMENT | 27 | 0.410422 | 1.051741 | 0.397241 | 0.818671 |
| GOBP_DETOXIFICATION | 106 | 0.334416 | 1.051102 | 0.391892 | 0.820239 |
| GOBP_REGULATION_OF_CARDIOCYTE_DIFFERENTIATION | 24 | 0.410047 | 1.050283 | 0.396333 | 0.822314 |
| GOBP_EXOCYTOSIS | 304 | 0.310776 | 1.048958 | 0.367722 | 0.826064 |
| GOBP_SPERMATID_DIFFERENTIATION | 120 | 0.326617 | 1.048827 | 0.400662 | 0.825884 |
| GOBP_CERAMIDE_CATABOLIC_PROCESS | 18 | 0.444805 | 1.048745 | 0.441558 | 0.825589 |
| GOBP_RESPONSE_TO_ESTROGEN | 53 | 0.36834 | 1.048303 | 0.383562 | 0.826442 |
| GOBP_REGULATION_OF_GTPASE_ACTIVITY | 310 | 0.306387 | 1.048205 | 0.388211 | 0.826163 |
| GOBP_ASSOCIATIVE_LEARNING | 72 | 0.350315 | 1.048127 | 0.407801 | 0.825828 |
| GOBP_FATTY_ACID_DERIVATIVE_BIOSYNTHETIC_PROCESS | 34 | 0.393537 | 1.047722 | 0.412385 | 0.826528 |
| GOBP_FATTY_ACID_DERIVATIVE_METABOLIC_PROCESS | 51 | 0.366664 | 1.047661 | 0.408039 | 0.826138 |
| GOBP_POSITIVE_REGULATION_OF_DEVELOPMENTAL_GROWTH | 132 | 0.326136 | 1.047631 | 0.378468 | 0.825644 |
| GOBP_REGULATION_OF_TRIGLYCERIDE_BIOSYNTHETIC_PROCESS | 18 | 0.442798 | 1.046673 | 0.417997 | 0.828215 |
| GOBP_ERBB_SIGNALING_PATHWAY | 107 | 0.33088 | 1.04664 | 0.393629 | 0.827725 |
| GOBP_POSITIVE_REGULATION_OF_ESTABLISHMENT_OF_PROTEIN_LOCALIZATION | 273 | 0.311976 | 1.046585 | 0.393595 | 0.827312 |
| GOBP_ACTIN_FILAMENT_ORGANIZATION | 403 | 0.307011 | 1.046361 | 0.353955 | 0.827501 |
| GOBP_RESPONSE_TO_ALKALOID | 87 | 0.342439 | 1.046143 | 0.408019 | 0.827631 |
| GOBP_POSITIVE_REGULATION_OF_OSTEOBLAST_DIFFERENTIATION | 59 | 0.355655 | 1.045669 | 0.41118 | 0.828636 |
| GOBP_CARDIAC_MUSCLE_CONTRACTION | 112 | 0.334187 | 1.045522 | 0.407029 | 0.828495 |
| GOBP_REGULATION_OF_CELL_CELL_ADHESION_MEDIATED_BY_CADHERIN | 19 | 0.434526 | 1.045354 | 0.420455 | 0.828463 |

| | | | | | |
|---|---|---|---|---|---|
| GOBP_POSITIVE_REGULATION_OF_MUSCLE_CELL_DIFFERENTIATION | 45 | 0.378015 | 1.045288 | 0.410354 | 0.828093 |
| GOBP_POSITIVE_REGULATION_OF_LIPID_TRANSPORT | 64 | 0.350492 | 1.045143 | 0.406948 | 0.827978 |
| GOBP_POSITIVE_REGULATION_OF_MYELOID_LEUKOCYTE_CYTOKINE_PRODUCTION_INVOL | 20 | 0.429055 | 1.044956 | 0.442771 | 0.828002 |
| GOBP_NEGATIVE_REGULATION_OF_ORGAN_GROWTH | 27 | 0.400449 | 1.044892 | 0.389518 | 0.827616 |
| GOBP_REGULATION_OF_GLUCOSE_METABOLIC_PROCESS | 104 | 0.335766 | 1.044666 | 0.386183 | 0.827727 |
| GOBP_REGULATION_OF_INSULIN_SECRETION_INVOLVED_IN_CELLULAR_RESPONSE_TO_( | 43 | 0.369734 | 1.043857 | 0.406045 | 0.829854 |
| GOBP_POSITIVE_REGULATION_OF_CARDIAC_MUSCLE_CELL_PROLIFERATION | 15 | 0.456277 | 1.043783 | 0.426426 | 0.829493 |
| GOBP_LOCOMOTORY_BEHAVIOR | 170 | 0.324627 | 1.042966 | 0.392553 | 0.831596 |
| GOBP_RESPONSE_TO_PEPTIDE | 395 | 0.306537 | 1.041903 | 0.378296 | 0.834479 |
| GOBP_CARDIAC_MUSCLE_CELL_PROLIFERATION | 38 | 0.378829 | 1.041864 | 0.430968 | 0.834011 |
| GOBP_CARDIAC_MUSCLE_CELL_CONTRACTION | 64 | 0.349709 | 1.041856 | 0.394958 | 0.833458 |
| GOBP_NEGATIVE_REGULATION_OF_NEURON_APOPTOTIC_PROCESS | 128 | 0.325937 | 1.041508 | 0.406321 | 0.834009 |
| GOBP_POTASSIUM_ION_TRANSPORT | 185 | 0.315961 | 1.040479 | 0.40812 | 0.836785 |
| GOBP_REGULATION_OF_NUCLEAR_TRANSCRIBED_MRNA_POLY_A_TAIL_SHORTENING | 15 | 0.458856 | 1.039827 | 0.433483 | 0.838301 |
| GOBP_REGULATION_OF_MUSCLE_CELL_DIFFERENTIATION | 106 | 0.33165 | 1.039463 | 0.405436 | 0.838891 |
| GOBP_POSITIVE_REGULATION_OF_PEPTIDASE_ACTIVITY | 165 | 0.320345 | 1.039111 | 0.408942 | 0.839484 |
| GOBP_POTASSIUM_ION_IMPORT_ACROSS_PLASMA_MEMBRANE | 38 | 0.383681 | 1.038812 | 0.423529 | 0.839874 |
| GOBP_REGULATION_OF_ANIMAL_ORGAN_FORMATION | 22 | 0.413746 | 1.038559 | 0.409029 | 0.840118 |
| GOBP_PEPTIDE_TRANSPORT | 197 | 0.31309 | 1.038543 | 0.402938 | 0.839578 |
| GOBP_REGULATION_OF_IMMUNOGLOBULIN_PRODUCTION | 54 | 0.357682 | 1.038481 | 0.418043 | 0.839205 |
| GOBP_NEGATIVE_REGULATION_OF_SECRETION | 125 | 0.32258 | 1.038395 | 0.410456 | 0.838891 |
| GOBP_REGULATION_OF_BEHAVIOR | 49 | 0.372424 | 1.038137 | 0.419437 | 0.839149 |
| GOBP_SMALL_GTPASE_MEDIATED_SIGNAL_TRANSDUCTION | 464 | 0.301395 | 1.038045 | 0.38446 | 0.83887 |
| GOBP_PROTEIN_DEMETHYLATION | 30 | 0.399506 | 1.037388 | 0.413333 | 0.840396 |
| GOBP_POSITIVE_REGULATION_OF_LIPID_STORAGE | 18 | 0.437423 | 1.037309 | 0.41568 | 0.840076 |
| GOBP_POSITIVE_REGULATION_OF_HYDROLASE_ACTIVITY | 467 | 0.301271 | 1.036877 | 0.378894 | 0.840831 |
| GOBP_REGULATION_OF_EXCRETION | 17 | 0.438419 | 1.03673 | 0.42547 | 0.840725 |
| GOBP_NEGATIVE_REGULATION_OF_CALCIUM_ION_TRANSPORT_INTO_CYTOSOL | 16 | 0.450104 | 1.0364 | 0.438395 | 0.841188 |
| GOBP_NEGATIVE_REGULATION_OF_NUCLEAR_DIVISION | 51 | 0.369131 | 1.036383 | 0.424433 | 0.840657 |
| GOBP_T_HELPER_1_CELL_DIFFERENTIATION | 16 | 0.451256 | 1.036185 | 0.432061 | 0.840691 |
| GOBP_BIOLOGICAL_PROCESS_INVOLVED_IN_INTERACTION_WITH_HOST | 165 | 0.318207 | 1.035859 | 0.414347 | 0.841147 |
| GOBP_CELLULAR_KETONE_METABOLIC_PROCESS | 156 | 0.321055 | 1.035634 | 0.421222 | 0.8413 |
| GOBP_POSITIVE_REGULATION_OF_CYSTEINE_TYPE_ENDOPEPTIDASE_ACTIVITY | 123 | 0.325261 | 1.035005 | 0.418242 | 0.842802 |
| GOBP_CAMERA_TYPE_EYE_DEVELOPMENT | 262 | 0.307075 | 1.034359 | 0.405992 | 0.844385 |
| GOBP_REGULATION_OF_NEUROTRANSMITTER_UPTAKE | 17 | 0.437405 | 1.034023 | 0.439462 | 0.844906 |
| GOBP_PROTEIN_K63_LINKED_UBIQUITINATION | 54 | 0.359275 | 1.033996 | 0.411914 | 0.844413 |
| GOBP_POSITIVE_REGULATION_OF_POTASSIUM_ION_TRANSPORT | 43 | 0.367876 | 1.033721 | 0.427296 | 0.844723 |
| GOBP_REGULATION_OF_MYOTUBE_DIFFERENTIATION | 34 | 0.391577 | 1.03354 | 0.418478 | 0.84475 |
| GOBP_BODY_MORPHOGENESIS | 42 | 0.372911 | 1.033114 | 0.421671 | 0.845514 |
| GOBP_PALLIUM_DEVELOPMENT | 145 | 0.320685 | 1.033053 | 0.413068 | 0.84512 |
| GOBP_OVULATION_CYCLE | 58 | 0.353391 | 1.032685 | 0.422705 | 0.845734 |
| GOBP_ORGANIC_ACID_BIOSYNTHETIC_PROCESS | 239 | 0.308649 | 1.032571 | 0.398543 | 0.845513 |
| GOBP_REGULATION_OF_ASTROCYTE_DIFFERENTIATION | 24 | 0.417803 | 1.032426 | 0.439227 | 0.84539 |
| GOBP_SUBSTRATE_DEPENDENT_CELL_MIGRATION | 23 | 0.419892 | 1.032181 | 0.432545 | 0.845595 |
| GOBP_DOPAMINE_SECRETION | 33 | 0.384493 | 1.031754 | 0.440568 | 0.846401 |
| GOBP_ERYTHROCYTE_HOMEOSTASIS | 119 | 0.323308 | 1.031548 | 0.410515 | 0.846492 |
| GOBP_REGULATION_OF_B_CELL_APOPTOTIC_PROCESS | 18 | 0.433191 | 1.031233 | 0.428788 | 0.846908 |
| GOBP_CELLULAR_RESPONSE_TO_STEROL | 16 | 0.440975 | 1.031212 | 0.425816 | 0.846392 |
| GOBP_DOPAMINE_TRANSPORT | 45 | 0.368067 | 1.031102 | 0.420499 | 0.846167 |
| GOBP_SMAD_PROTEIN_SIGNAL_TRANSDUCTION | 72 | 0.343232 | 1.030966 | 0.427873 | 0.846011 |
| GOBP_GLYCEROLIPID_BIOSYNTHETIC_PROCESS | 229 | 0.307573 | 1.03095 | 0.406904 | 0.845489 |
| GOBP_NEGATIVE_REGULATION_OF_INTRACELLULAR_SIGNAL_TRANSDUCTION | 456 | 0.300443 | 1.030532 | 0.40101 | 0.846241 |
| GOBP_NEGATIVE_REGULATION_OF_NECROTIC_CELL_DEATH | 16 | 0.446488 | 1.030461 | 0.457447 | 0.845894 |
| GOBP_LEUKOTRIENE_METABOLIC_PROCESS | 15 | 0.457491 | 1.030423 | 0.43299 | 0.845446 |
| GOBP_GLOMERULUS_DEVELOPMENT | 55 | 0.351434 | 1.03036 | 0.425505 | 0.84507 |
| GOBP_EAR_MORPHOGENESIS | 95 | 0.323896 | 1.030203 | 0.413873 | 0.845021 |
| GOBP_REGULATION_OF_ATP_DEPENDENT_ACTIVITY | 63 | 0.352645 | 1.030171 | 0.42029 | 0.844549 |
| GOBP_HEPATICOBILIARY_SYSTEM_DEVELOPMENT | 110 | 0.329812 | 1.030082 | 0.430508 | 0.844264 |
| GOBP_REGULATION_OF_LIPASE_ACTIVITY | 61 | 0.350702 | 1.029765 | 0.429963 | 0.84471 |
| GOBP_CELL_FATE_COMMITMENT_INVOLVED_IN_FORMATION_OF_PRIMARY_GERM_LAYER | 26 | 0.403191 | 1.029598 | 0.439024 | 0.844696 |
| GOBP_REGULATION_OF_MRNA_CATABOLIC_PROCESS | 156 | 0.316176 | 1.028464 | 0.415682 | 0.847611 |
| GOBP_FOREBRAIN_NEURON_DIFFERENTIATION | 31 | 0.390523 | 1.028304 | 0.431607 | 0.847562 |
| GOBP_CELLULAR_RESPONSE_TO_HEAT | 61 | 0.349432 | 1.027869 | 0.421557 | 0.848383 |
| GOBP_REGULATION_OF_MUSCLE_HYPERTROPHY | 45 | 0.368566 | 1.027649 | 0.428755 | 0.848513 |
| GOBP_RENAL_VESICLE_DEVELOPMENT | 16 | 0.448926 | 1.02673 | 0.453353 | 0.850858 |
| GOBP_ISOPRENOID_METABOLIC_PROCESS | 82 | 0.336311 | 1.026512 | 0.425329 | 0.850981 |
| GOBP_CARDIAC_ATRIUM_DEVELOPMENT | 29 | 0.402213 | 1.026403 | 0.432692 | 0.850771 |
| GOBP_OSSIFICATION | 340 | 0.299121 | 1.025947 | 0.421699 | 0.851607 |
| GOBP_BEHAVIOR | 499 | 0.299476 | 1.02569 | 0.413065 | 0.851853 |
| GOBP_RESPONSE_TO_MUSCLE_STRETCH | 20 | 0.41778 | 1.024961 | 0.466476 | 0.853564 |
| GOBP_POSITIVE_REGULATION_OF_TRANSPORTER_ACTIVITY | 101 | 0.326763 | 1.02483 | 0.437923 | 0.853411 |
| GOBP_REGULATION_OF_POSTSYNAPTIC_MEMBRANE_POTENTIAL | 110 | 0.325044 | 1.024187 | 0.431743 | 0.85492 |
| GOBP_PERIPHERAL_NERVOUS_SYSTEM_DEVELOPMENT | 69 | 0.344718 | 1.023949 | 0.443385 | 0.855096 |
| GOBP_SMOOTH_MUSCLE_CELL_APOPTOTIC_PROCESS | 18 | 0.424954 | 1.023541 | 0.460317 | 0.855849 |
| GOBP_EPITHELIAL_TO_MESENCHYMAL_TRANSITION | 139 | 0.314093 | 1.023199 | 0.427948 | 0.856336 |
| GOBP_NEGATIVE_REGULATION_OF_G_PROTEIN_COUPLED_RECEPTOR_SIGNALING_PATHW | 43 | 0.370776 | 1.022668 | 0.443022 | 0.857472 |
| GOBP_REGULATION_OF_PHOSPHOLIPASE_ACTIVITY | 46 | 0.365059 | 1.022578 | 0.430303 | 0.857186 |
| GOBP_POSITIVE_REGULATION_OF_DNA_REPLICATION | 39 | 0.372569 | 1.022495 | 0.442513 | 0.856881 |
| GOBP_TRANSITION_METAL_ION_TRANSPORT | 82 | 0.341327 | 1.022396 | 0.437939 | 0.85663 |
| GOBP_NEGATIVE_REGULATION_OF_CELL_DEVELOPMENT | 151 | 0.312697 | 1.022202 | 0.439103 | 0.85668 |
| GOBP_OSTEOCLAST_DIFFERENTIATION | 71 | 0.342313 | 1.021759 | 0.415663 | 0.857528 |
| GOBP_REGULATION_OF_NEUROTRANSMITTER_TRANSPORT | 88 | 0.334738 | 1.021405 | 0.448399 | 0.858049 |
| GOBP_CELLULAR_RESPONSE_TO_EXTERNAL_STIMULUS | 268 | 0.301477 | 1.021375 | 0.43763 | 0.857577 |
| GOBP_POSITIVE_REGULATION_OF_EPITHELIAL_TO_MESENCHYMAL_TRANSITION | 45 | 0.366275 | 1.021111 | 0.444301 | 0.857818 |
| GOBP_NEGATIVE_REGULATION_OF_MYELOID_LEUKOCYTE_DIFFERENTIATION | 37 | 0.377328 | 1.021104 | 0.440055 | 0.857263 |
| GOBP_CHEMICAL_SYNAPTIC_TRANSMISSION_POSTSYNAPTIC | 84 | 0.330889 | 1.020618 | 0.44289 | 0.85825 |
| GOBP_REGULATION_OF_OSTEOBLAST_PROLIFERATION | 22 | 0.41141 | 1.020337 | 0.439416 | 0.858548 |
| GOBP_CELL_SUBSTRATE_JUNCTION_ORGANIZATION | 95 | 0.32688 | 1.019887 | 0.455172 | 0.859392 |
| GOBP_MESENCHYMAL_CELL_DEVELOPMENT | 82 | 0.332382 | 1.019321 | 0.446759 | 0.860637 |
| GOBP_EPITHELIAL_CELL_DEVELOPMENT | 175 | 0.314178 | 1.018902 | 0.454248 | 0.861371 |
| GOBP_RESPONSE_TO_CALCIUM_ION | 122 | 0.317515 | 1.018866 | 0.44581 | 0.860896 |
| GOBP_NEGATIVE_REGULATION_OF_PROTEIN_BINDING | 79 | 0.334968 | 1.018643 | 0.461894 | 0.861062 |
| GOBP_NEGATIVE_REGULATION_OF_MYOTUBE_DIFFERENTIATION | 15 | 0.453517 | 1.018482 | 0.468148 | 0.861027 |
| GOBP_SMALL_MOLECULE_BIOSYNTHETIC_PROCESS | 452 | 0.295131 | 1.018039 | 0.440763 | 0.861859 |
| GOBP_DEFENSE_RESPONSE_TO_GRAM_POSITIVE_BACTERIUM | 46 | 0.366749 | 1.017446 | 0.444583 | 0.863219 |
| GOBP_LENS_DEVELOPMENT_IN_CAMERA_TYPE_EYE | 70 | 0.339759 | 1.017301 | 0.455504 | 0.863094 |
| GOBP_NEGATIVE_REGULATION_OF_INTRINSIC_APOPTOTIC_SIGNALING_PATHWAY_BY_P53 | 21 | 0.414228 | 1.017251 | 0.481322 | 0.862692 |
| GOBP_BIOLOGICAL_PROCESS_INVOLVED_IN_SYMBIOTIC_INTERACTION | 228 | 0.303337 | 1.017039 | 0.452256 | 0.862787 |
| GOBP_REGULATION_OF_KERATINOCYTE_DIFFERENTIATION | 28 | 0.390021 | 1.016928 | 0.465517 | 0.862555 |
| GOBP_GLIOGENESIS | 235 | 0.303247 | 1.01649 | 0.463235 | 0.86335 |
| GOBP_DIENCEPHALON_DEVELOPMENT | 59 | 0.348557 | 1.016328 | 0.447304 | 0.863296 |
| GOBP_NEUROMUSCULAR_JUNCTION_DEVELOPMENT | 38 | 0.374243 | 1.015922 | 0.457474 | 0.863969 |
| GOBP_CARBOHYDRATE_BIOSYNTHETIC_PROCESS | 171 | 0.311515 | 1.015745 | 0.453674 | 0.863956 |
| GOBP_REGULATION_OF_DEFENSE_RESPONSE_TO_VIRUS | 61 | 0.346107 | 1.015728 | 0.440741 | 0.863443 |
| GOBP_REGULATION_OF_FATTY_ACID_OXIDATION | 25 | 0.401568 | 1.015369 | 0.452617 | 0.864002 |
| GOBP_RETINOIC_ACID_METABOLIC_PROCESS | 19 | 0.423567 | 1.015057 | 0.474601 | 0.864398 |
| GOBP_ORGAN_GROWTH | 123 | 0.319528 | 1.014825 | 0.456954 | 0.864545 |
| GOBP_REGULATION_OF_JNK_CASCADE | 123 | 0.32047 | 1.014502 | 0.463442 | 0.865043 |
| GOBP_MORPHOGENESIS_OF_A_BRANCHING_STRUCTURE | 154 | 0.312967 | 1.014322 | 0.463283 | 0.865035 |
| GOBP_SPHINGOID_METABOLIC_PROCESS | 21 | 0.421802 | 1.014135 | 0.466472 | 0.865043 |
| GOBP_PURINE_NUCLEOSIDE_METABOLIC_PROCESS | 24 | 0.404606 | 1.01411 | 0.446429 | 0.86456 |
| GOBP_POSITIVE_REGULATION_OF_NUCLEOTIDE_BIOSYNTHETIC_PROCESS | 17 | 0.432945 | 1.013828 | 0.466102 | 0.864888 |
| GOBP_DNA_DEPENDENT_DNA_REPLICATION_MAINTENANCE_OF_FIDELITY | 46 | 0.36136 | 1.013755 | 0.456962 | 0.864555 |
| GOBP_VESICLE_MEDIATED_TRANSPORT_IN_SYNAPSE | 180 | 0.307401 | 1.012877 | 0.443386 | 0.866664 |
| GOBP_POSTTRANSCRIPTIONAL_REGULATION_OF_GENE_EXPRESSION | 497 | 0.295031 | 1.012239 | 0.461307 | 0.868129 |
| GOBP_NEGATIVE_REGULATION_OF_SMOOTH_MUSCLE_CELL_MIGRATION | 23 | 0.404098 | 1.01175 | 0.460581 | 0.869073 |
| GOBP_GLYCEROLIPID_METABOLIC_PROCESS | 337 | 0.298549 | 1.011684 | 0.469325 | 0.868707 |
| GOBP_RESPIRATORY_GASEOUS_EXCHANGE_BY_RESPIRATORY_SYSTEM | 56 | 0.348663 | 1.011643 | 0.457862 | 0.868267 |
| GOBP_REGULATION_OF_MACROPHAGE_CHEMOTAXIS | 16 | 0.434467 | 1.011499 | 0.457227 | 0.868141 |
| GOBP_DNA_DAMAGE_RESPONSE_SIGNAL_TRANSDUCTION_RESULTING_IN_TRANSCRIPTI( | 17 | 0.433153 | 1.011075 | 0.448476 | 0.868887 |
| GOBP_ZINC_ION_HOMEOSTASIS | 30 | 0.390063 | 1.011044 | 0.469496 | 0.868431 |
| GOBP_DETECTION_OF_MECHANICAL_STIMULUS_INVOLVED_IN_SENSORY_PERCEPTION | 29 | 0.393877 | 1.011019 | 0.462277 | 0.867946 |
| GOBP_RESPONSE_TO_LIGHT_STIMULUS | 255 | 0.303751 | 1.010883 | 0.465357 | 0.867783 |
| GOBP_RESPONSE_TO_RADIATION | 366 | 0.29718 | 1.010762 | 0.470588 | 0.867614 |
| GOBP_HISTONE_H3_K27_METHYLATION | 18 | 0.421552 | 1.010701 | 0.473761 | 0.867253 |
| GOBP_RETINOL_METABOLIC_PROCESS | 32 | 0.380648 | 1.010654 | 0.452855 | 0.866833 |
| GOBP_MEMBRANE_REPOLARIZATION_DURING_ACTION_POTENTIAL | 22 | 0.41261 | 1.010377 | 0.453258 | 0.867082 |
| GOBP_NEGATIVE_REGULATION_OF_HISTONE_ACETYLATION | 15 | 0.429389 | 1.010186 | 0.460177 | 0.867082 |
| GOBP_POSITIVE_REGULATION_OF_CYTOSOLIC_CALCIUM_ION_CONCENTRATION_INVOLVE | 20 | 0.420517 | 1.010016 | 0.478448 | 0.86704 |
| GOBP_NEGATIVE_REGULATION_OF_NIK_NF_KAPPAB_SIGNALING | 24 | 0.407675 | 1.009546 | 0.460383 | 0.867941 |
| GOBP_CELLULAR_RESPONSE_TO_CALCIUM_ION | 72 | 0.337774 | 1.008881 | 0.469734 | 0.869471 |
| GOBP_RESPONSE_TO_TRANSFORMING_GROWTH_FACTOR_BETA | 217 | 0.302312 | 1.008769 | 0.457113 | 0.869239 |
| GOBP_DIOL_METABOLIC_PROCESS | 29 | 0.386872 | 1.008313 | 0.452957 | 0.870081 |
| GOBP_MULTICELLULAR_ORGANISM_AGING | 34 | 0.38125 | 1.008168 | 0.470748 | 0.86997 |
| GOBP_CAMP_MEDIATED_SIGNALING | 43 | 0.360035 | 1.007915 | 0.456186 | 0.870225 |
| GOBP_MAGNESIUM_ION_TRANSPORT | 17 | 0.430332 | 1.007729 | 0.47654 | 0.870199 |
| GOBP_CYCLIC_NUCLEOTIDE_MEDIATED_SIGNALING | 62 | 0.342165 | 1.007255 | 0.469136 | 0.871081 |
| GOBP_POSITIVE_REGULATION_OF_DOUBLE_STRAND_BREAK_REPAIR | 63 | 0.342652 | 1.005905 | 0.459492 | 0.874657 |
| GOBP_VASCULAR_PROCESS_IN_CIRCULATORY_SYSTEM | 217 | 0.300978 | 1.00535 | 0.477941 | 0.875855 |
| GOBP_ATP_TRANSPORT | 15 | 0.442017 | 1.005348 | 0.465857 | 0.875294 |
| GOBP_REGULATION_OF_VESICLE_MEDIATED_TRANSPORT | 452 | 0.293323 | 1.005249 | 0.478958 | 0.87504 |
| GOBP_ENGULFMENT_OF_APOPTOTIC_CELL | 18 | 0.425271 | 1.004891 | 0.460432 | 0.875595 |
| GOBP_NOTCH_SIGNALING_PATHWAY | 152 | 0.306089 | 1.004526 | 0.501082 | 0.876154 |
| GOBP_DETECTION_OF_TEMPERATURE_STIMULUS_INVOLVED_IN_SENSORY_PERCEPTION | 15 | 0.439112 | 1.003982 | 0.476331 | 0.877305 |
| GOBP_NEGATIVE_REGULATION_OF_ATP_DEPENDENT_ACTIVITY | 17 | 0.42022 | 1.003787 | 0.468481 | 0.877332 |
| GOBP_REGULATION_OF_OSTEOBLAST_DIFFERENTIATION | 109 | 0.316978 | 1.00337 | 0.475816 | 0.878069 |
| GOBP_BENZENE_CONTAINING_COMPOUND_METABOLIC_PROCESS | 16 | 0.426792 | 1.003262 | 0.461883 | 0.877848 |
| GOBP_POSITIVE_REGULATION_OF_GTPASE_ACTIVITY | 227 | 0.301701 | 1.003133 | 0.485232 | 0.877679 |
| GOBP_REGULATION_OF_MITOCHONDRIAL_MEMBRANE_POTENTIAL | 65 | 0.337473 | 1.002897 | 0.454436 | 0.877884 |
| GOBP_MITOCHONDRIAL_FISSION | 37 | 0.371397 | 1.002858 | 0.449409 | 0.87744 |
| GOBP_RNA_PHOSPHODIESTER_BOND_HYDROLYSIS_ENDONUCLEOLYTIC | 70 | 0.338246 | 1.002621 | 0.475212 | 0.877615 |
| GOBP_NEGATIVE_REGULATION_OF_CELL_PROJECTION_ORGANIZATION | 174 | 0.308173 | 1.00253 | 0.499465 | 0.877322 |
| GOBP_POSITIVE_REGULATION_OF_MITOTIC_NUCLEAR_DIVISION | 32 | 0.379075 | 1.002525 | 0.461538 | 0.876778 |
| GOBP_POSITIVE_REGULATION_OF_GLIOGENESIS | 52 | 0.351149 | 1.002155 | 0.494975 | 0.877375 |
| GOBP_POSITIVE_REGULATION_OF_ACTIN_FILAMENT_POLYMERIZATION | 45 | 0.358524 | 1.001815 | 0.471722 | 0.87788 |
| GOBP_AORTA_DEVELOPMENT | 54 | 0.350401 | 1.001699 | 0.478589 | 0.877669 |
| GOBP_XENOBIOTIC_TRANSPORT | 31 | 0.378613 | 1.001497 | 0.483871 | 0.87773 |

| | | | | | |
|---|---|---|---|---|---|
| GOBP_CELLULAR_RESPONSE_TO_NUTRIENT | 34 | 0.367173 | 1.001468 | 0.458498 | 0.877275 |
| GOBP_GASTRULATION_WITH_MOUTH_FORMING_SECOND | 25 | 0.39543 | 1.001269 | 0.482615 | 0.877319 |
| GOBP_REGULATION_OF_STRIATED_MUSCLE_CELL_DIFFERENTIATION | 68 | 0.336235 | 1.000961 | 0.461538 | 0.877732 |
| GOBP_POSITIVE_REGULATION_OF_PROTEIN_CONTAINING_COMPLEX_ASSEMBLY | 165 | 0.308516 | 1.000486 | 0.494635 | 0.878634 |
| GOBP_NEGATIVE_REGULATION_OF_NEURON_PROJECTION_DEVELOPMENT | 129 | 0.31134 | 1.000109 | 0.478842 | 0.879181 |
| GOBP_REGULATION_OF_DENDRITE_EXTENSION | 23 | 0.396348 | 0.999917 | 0.483146 | 0.879208 |
| GOBP_RESPONSE_TO_DSRNA | 38 | 0.367963 | 0.998716 | 0.470822 | 0.882378 |
| GOBP_VASCULAR_ENDOTHELIAL_CELL_PROLIFERATION | 18 | 0.419235 | 0.998489 | 0.469828 | 0.882473 |
| GOBP_NEUROEPITHELIAL_CELL_DIFFERENTIATION | 22 | 0.399583 | 0.998091 | 0.481375 | 0.883087 |
| GOBP_RENAL_SYSTEM_VASCULATURE_DEVELOPMENT | 23 | 0.396862 | 0.998082 | 0.462857 | 0.882554 |
| GOBP_THYMOCYTE_APOPTOTIC_PROCESS | 15 | 0.432616 | 0.997969 | 0.463679 | 0.882339 |
| GOBP_REGULATION_OF_SHORT_TERM_NEURONAL_SYNAPTIC_PLASTICITY | 15 | 0.443173 | 0.997468 | 0.491473 | 0.883269 |
| GOBP_EXTRINSIC_APOPTOTIC_SIGNALING_PATHWAY_VIA_DEATH_DOMAIN_RECEPTORS | 73 | 0.331929 | 0.99746 | 0.468523 | 0.882735 |
| GOBP_REGULATION_OF_NITRIC_OXIDE_SYNTHASE_ACTIVITY | 27 | 0.391599 | 0.99707 | 0.48711 | 0.883376 |
| GOBP_INTERLEUKIN_1_MEDIATED_SIGNALING_PATHWAY | 21 | 0.407178 | 0.996979 | 0.492898 | 0.883099 |
| GOBP_NEGATIVE_REGULATION_OF_MAP_KINASE_ACTIVITY | 49 | 0.346846 | 0.996671 | 0.492443 | 0.883431 |
| GOBP_REGULATION_OF_HETEROTYPIC_CELL_CELL_ADHESION | 16 | 0.434027 | 0.996594 | 0.483333 | 0.883117 |
| GOBP_REGULATION_OF_CALCINEURIN_MEDIATED_SIGNALING | 30 | 0.378997 | 0.996593 | 0.474114 | 0.882561 |
| GOBP_IMPORT_INTO_NUCLEUS | 145 | 0.309505 | 0.996547 | 0.495028 | 0.88215 |
| GOBP_RESPONSE_TO_RETINOIC_ACID | 86 | 0.327741 | 0.99562 | 0.478809 | 0.884447 |
| GOBP_ALCOHOL_CATABOLIC_PROCESS | 43 | 0.357282 | 0.995433 | 0.472329 | 0.884456 |
| GOBP_TRANSITION_METAL_ION_HOMEOSTASIS | 110 | 0.318931 | 0.995347 | 0.49093 | 0.884178 |
| GOBP_PEPTIDYL_TYROSINE_DEPHOSPHORYLATION | 92 | 0.318326 | 0.993686 | 0.487327 | 0.888697 |
| GOBP_GLYCEROPHOSPHOLIPID_METABOLIC_PROCESS | 273 | 0.294324 | 0.993527 | 0.516923 | 0.888616 |
| GOBP_HYALURONAN_METABOLIC_PROCESS | 21 | 0.401624 | 0.993465 | 0.482044 | 0.888245 |
| GOBP_AMMONIUM_ION_METABOLIC_PROCESS | 17 | 0.428202 | 0.993413 | 0.484627 | 0.887829 |
| GOBP_NEGATIVE_REGULATION_OF_CELL_SUBSTRATE_JUNCTION_ORGANIZATION | 16 | 0.437069 | 0.993046 | 0.467863 | 0.888375 |
| GOBP_POSITIVE_REGULATION_OF_RESPONSE_TO_CYTOKINE_STIMULUS | 51 | 0.344979 | 0.992955 | 0.47201 | 0.888101 |
| GOBP_VASCULAR_ASSOCIATED_SMOOTH_MUSCLE_CELL_DIFFERENTIATION | 23 | 0.394935 | 0.992801 | 0.477952 | 0.887999 |
| GOBP_TIGHT_JUNCTION_ORGANIZATION | 63 | 0.338999 | 0.992722 | 0.500617 | 0.887696 |
| GOBP_HYPOTHALAMUS_DEVELOPMENT | 19 | 0.418486 | 0.992621 | 0.485673 | 0.88742 |
| GOBP_MORPHOGENESIS_OF_AN_ENDOTHELIUM | 16 | 0.432413 | 0.992601 | 0.491803 | 0.886919 |
| GOBP_POSITIVE_REGULATION_OF_LIPID_KINASE_ACTIVITY | 31 | 0.380034 | 0.99255 | 0.483914 | 0.886507 |
| GOBP_ADRENAL_GLAND_DEVELOPMENT | 24 | 0.398547 | 0.991824 | 0.512409 | 0.888143 |
| GOBP_ACTIVATION_OF_INNATE_IMMUNE_RESPONSE | 42 | 0.363456 | 0.991739 | 0.5 | 0.887852 |
| GOBP_WOUND_HEALING_SPREADING_OF_EPIDERMAL_CELLS | 16 | 0.428356 | 0.991686 | 0.482456 | 0.88747 |
| GOBP_KERATINOCYTE_PROLIFERATION | 34 | 0.376002 | 0.99133 | 0.493188 | 0.887959 |
| GOBP_HEAD_MORPHOGENESIS | 32 | 0.372044 | 0.991216 | 0.485294 | 0.887744 |
| GOBP_PURINE_NUCLEOBASE_METABOLIC_PROCESS | 18 | 0.41938 | 0.991203 | 0.49705 | 0.887228 |
| GOBP_POST_ANAL_TAIL_MORPHOGENESIS | 16 | 0.422696 | 0.991202 | 0.5 | 0.886678 |
| GOBP_G_PROTEIN_COUPLED_RECEPTOR_SIGNALING_PATHWAY_COUPLED_TO_CYCLIC_ | 30 | 0.377446 | 0.990359 | 0.486933 | 0.888603 |
| GOBP_REGULATION_OF_PHOSPHATASE_ACTIVITY | 67 | 0.335019 | 0.990281 | 0.488429 | 0.888272 |
| GOBP_POSITIVE_REGULATION_OF_INSULIN_SECRETION_INVOLVED_IN_CELLULAR_RESPO | 23 | 0.390034 | 0.989932 | 0.472954 | 0.888756 |
| GOBP_NEPHRON_DEVELOPMENT | 120 | 0.310544 | 0.988038 | 0.502252 | 0.893914 |
| GOBP_REGULATION_OF_RECEPTOR_RECYCLING | 21 | 0.404737 | 0.987815 | 0.494444 | 0.894046 |
| GOBP_NEGATIVE_REGULATION_OF_DNA_BINDING_TRANSCRIPTION_FACTOR_ACTIVITY | 147 | 0.307207 | 0.987183 | 0.523344 | 0.895397 |
| GOBP_MICROTUBULE_BASED_MOVEMENT | 325 | 0.290164 | 0.987101 | 0.526749 | 0.895098 |
| GOBP_RESPONSE_TO_COPPER_ION | 31 | 0.378097 | 0.98708 | 0.475989 | 0.894595 |
| GOBP_ARGININE_METABOLIC_PROCESS | 16 | 0.419819 | 0.986584 | 0.47741 | 0.895511 |
| GOBP_REGULATION_OF_DIGESTIVE_SYSTEM_PROCESS | 23 | 0.393931 | 0.986425 | 0.493113 | 0.895417 |
| GOBP_REGULATION_OF_MYOBLAST_DIFFERENTIATION | 37 | 0.364431 | 0.986293 | 0.513298 | 0.895264 |
| GOBP_ORGANIC_HYDROXY_COMPOUND_TRANSPORT | 200 | 0.297509 | 0.986263 | 0.5228 | 0.894812 |
| GOBP_REGULATION_OF_ANATOMICAL_STRUCTURE_SIZE | 421 | 0.288479 | 0.985762 | 0.541248 | 0.895712 |
| GOBP_CONNECTIVE_TISSUE_DEVELOPMENT | 209 | 0.295216 | 0.985705 | 0.519669 | 0.895334 |
| GOBP_SIGNAL_TRANSDUCTION_BY_P53_CLASS_MEDIATOR | 158 | 0.304577 | 0.985195 | 0.499459 | 0.896262 |
| GOBP_LIPID_LOCALIZATION | 343 | 0.292419 | 0.985104 | 0.546476 | 0.895981 |
| GOBP_CYCLIC_NUCLEOTIDE_BIOSYNTHETIC_PROCESS | 18 | 0.417817 | 0.984991 | 0.487943 | 0.895767 |
| GOBP_REGULATION_OF_MEMBRANE_POTENTIAL | 350 | 0.287483 | 0.984667 | 0.5409 | 0.896167 |
| GOBP_REGULATION_OF_ENDOTHELIAL_CELL_DEVELOPMENT | 16 | 0.427927 | 0.984519 | 0.495652 | 0.896041 |
| GOBP_AUTONOMIC_NERVOUS_SYSTEM_DEVELOPMENT | 35 | 0.369267 | 0.984263 | 0.498708 | 0.896275 |
| GOBP_REGULATION_OF_NEUROTRANSMITTER_LEVELS | 188 | 0.300349 | 0.983288 | 0.534538 | 0.898625 |
| GOBP_CARDIAC_CONDUCTION | 79 | 0.321942 | 0.983137 | 0.514286 | 0.898498 |
| GOBP_NEURAL_CREST_CELL_MIGRATION | 54 | 0.337919 | 0.983064 | 0.513141 | 0.898172 |
| GOBP_PARASYMPATHETIC_NERVOUS_SYSTEM_DEVELOPMENT | 15 | 0.436753 | 0.981638 | 0.489583 | 0.901848 |
| GOBP_NEUROPEPTIDE_SIGNALING_PATHWAY | 68 | 0.33107 | 0.981363 | 0.513158 | 0.902096 |
| GOBP_ACTIVATION_OF_CYSTEINE_TYPE_ENDOPEPTIDASE_ACTIVITY | 18 | 0.406252 | 0.981097 | 0.49 | 0.902355 |
| GOBP_PLASMA_MEMBRANE_PHOSPHOLIPID_SCRAMBLING | 19 | 0.41487 | 0.98067 | 0.496434 | 0.902994 |
| GOBP_ORGANELLE_FISSION | 430 | 0.285875 | 0.979805 | 0.568687 | 0.90503 |
| GOBP_INTRINSIC_APOPTOTIC_SIGNALING_PATHWAY_BY_P53_CLASS_MEDIATOR | 74 | 0.323811 | 0.979772 | 0.504192 | 0.904567 |
| GOBP_POSITIVE_REGULATION_OF_DEPHOSPHORYLATION | 52 | 0.34036 | 0.979028 | 0.511222 | 0.906194 |
| GOBP_NEGATIVE_REGULATION_OF_CHEMOTAXIS | 50 | 0.340498 | 0.978872 | 0.498079 | 0.906143 |
| GOBP_PHASIC_SMOOTH_MUSCLE_CONTRACTION | 16 | 0.433983 | 0.978845 | 0.50511 | 0.905677 |
| GOBP_B_CELL_APOPTOTIC_PROCESS | 23 | 0.39118 | 0.978416 | 0.518414 | 0.90639 |
| GOBP_NEURON_MIGRATION | 144 | 0.302571 | 0.978302 | 0.512406 | 0.906181 |
| GOBP_CELL_CELL_JUNCTION_ORGANIZATION | 173 | 0.301107 | 0.978289 | 0.535181 | 0.905661 |
| GOBP_REGULATION_OF_SYNCYTIUM_FORMATION_BY_PLASMA_MEMBRANE_FUSION | 17 | 0.41771 | 0.978151 | 0.512445 | 0.905518 |
| GOBP_HIPPOCAMPUS_DEVELOPMENT | 66 | 0.326843 | 0.977868 | 0.498177 | 0.905801 |
| GOBP_FOCAL_ADHESION_ASSEMBLY | 80 | 0.319334 | 0.97776 | 0.515643 | 0.905572 |
| GOBP_GLIAL_CELL_MIGRATION | 41 | 0.352171 | 0.977438 | 0.517787 | 0.905993 |
| GOBP_REGULATION_OF_CHROMOSOME_SEGREGATION | 81 | 0.323546 | 0.977109 | 0.510059 | 0.906409 |
| GOBP_AMINOGLYCAN_METABOLIC_PROCESS | 109 | 0.310608 | 0.976918 | 0.516201 | 0.906413 |
| GOBP_AMYLOID_BETA_CLEARANCE | 30 | 0.36891 | 0.976918 | 0.513514 | 0.905863 |
| GOBP_INSULIN_SECRETION_INVOLVED_IN_CELLULAR_RESPONSE_TO_GLUCOSE_STIMUL | 52 | 0.344583 | 0.976527 | 0.509458 | 0.906442 |
| GOBP_PLASMA_MEMBRANE_ORGANIZATION | 122 | 0.310291 | 0.975941 | 0.518192 | 0.907575 |
| GOBP_ODONTOGENESIS | 96 | 0.315094 | 0.975682 | 0.528103 | 0.907801 |
| GOBP_DEFINITIVE_HEMOPOIESIS | 15 | 0.421661 | 0.975262 | 0.513393 | 0.908464 |
| GOBP_MODULATION_OF_EXCITATORY_POSTSYNAPTIC_POTENTIAL | 36 | 0.358352 | 0.975107 | 0.508152 | 0.908364 |
| GOBP_REGULATION_OF_EPIDERMIS_DEVELOPMENT | 52 | 0.339787 | 0.974623 | 0.495663 | 0.909228 |
| GOBP_POSITIVE_REGULATION_OF_CELL_CYCLE | 270 | 0.28703 | 0.974484 | 0.548354 | 0.909055 |
| GOBP_ACTOMYOSIN_STRUCTURE_ORGANIZATION | 176 | 0.297118 | 0.974461 | 0.541089 | 0.90856 |
| GOBP_NEGATIVE_REGULATION_OF_SMALL_GTPASE_MEDIATED_SIGNAL_TRANSDUCTION | 54 | 0.340043 | 0.974368 | 0.5025 | 0.908257 |
| GOBP_CELLULAR_IRON_ION_HOMEOSTASIS | 53 | 0.342648 | 0.974082 | 0.529915 | 0.908605 |
| GOBP_POSITIVE_REGULATION_OF_EPITHELIAL_CELL_PROLIFERATION | 150 | 0.300071 | 0.973865 | 0.557143 | 0.90872 |
| GOBP_DIGESTIVE_TRACT_MORPHOGENESIS | 34 | 0.367898 | 0.973763 | 0.515915 | 0.908474 |
| GOBP_PLACENTA_DEVELOPMENT | 130 | 0.300694 | 0.973425 | 0.538634 | 0.908916 |
| GOBP_RESPONSE_TO_ATP | 24 | 0.382808 | 0.973213 | 0.515775 | 0.908997 |
| GOBP_NEGATIVE_REGULATION_OF_PLATELET_ACTIVATION | 16 | 0.412798 | 0.972893 | 0.517037 | 0.90937 |
| GOBP_PROTEIN_TETRAMERIZATION | 75 | 0.322759 | 0.972527 | 0.519904 | 0.909904 |
| GOBP_EAR_DEVELOPMENT | 172 | 0.297512 | 0.972002 | 0.544102 | 0.910862 |
| GOBP_INTESTINAL_EPITHELIAL_CELL_DIFFERENTIATION | 18 | 0.413555 | 0.971836 | 0.49634 | 0.910804 |
| GOBP_HISTONE_H3_K14_ACETYLATION | 22 | 0.392783 | 0.971716 | 0.535354 | 0.910608 |
| GOBP_AMINE_CATABOLIC_PROCESS | 19 | 0.406643 | 0.970826 | 0.521866 | 0.912682 |
| GOBP_POSITIVE_REGULATION_OF_CELLULAR_RESPONSE_TO_INSULIN_STIMULUS | 19 | 0.411827 | 0.970503 | 0.525473 | 0.913056 |
| GOBP_REGULATION_OF_CELL_SUBSTRATE_JUNCTION_ORGANIZATION | 65 | 0.326747 | 0.970496 | 0.509047 | 0.912538 |
| GOBP_DIGESTIVE_SYSTEM_DEVELOPMENT | 103 | 0.313391 | 0.970326 | 0.545045 | 0.9125 |
| GOBP_POSITIVE_REGULATION_OF_SMOOTH_MUSCLE_CELL_MIGRATION | 25 | 0.389414 | 0.969993 | 0.535912 | 0.912941 |
| GOBP_RENAL_SYSTEM_PROCESS | 82 | 0.318877 | 0.969918 | 0.547847 | 0.91264 |
| GOBP_STEROID_METABOLIC_PROCESS | 232 | 0.290811 | 0.969294 | 0.560084 | 0.913968 |
| GOBP_IRON_ION_HOMEOSTASIS | 69 | 0.326871 | 0.968199 | 0.530393 | 0.916564 |
| GOBP_MESENCHYMAL_TO_EPITHELIAL_TRANSITION | 17 | 0.403532 | 0.968009 | 0.508321 | 0.916555 |
| GOBP_KIDNEY_EPITHELIUM_DEVELOPMENT | 118 | 0.306276 | 0.967658 | 0.534231 | 0.917029 |
| GOBP_PEPTIDYL_SERINE_MODIFICATION | 295 | 0.28625 | 0.967597 | 0.581706 | 0.916669 |
| GOBP_POSITIVE_REGULATION_OF_PROTEIN_KINASE_ACTIVITY | 336 | 0.285362 | 0.967535 | 0.592893 | 0.916284 |
| GOBP_REGULATION_OF_CELLULAR_RESPONSE_TO_TRANSFORMING_GROWTH_FACTOR_ | 112 | 0.308637 | 0.967491 | 0.561111 | 0.915849 |
| GOBP_POSITIVE_REGULATION_OF_TRANSLATION | 128 | 0.301268 | 0.967271 | 0.538117 | 0.915946 |
| GOBP_REGULATION_OF_ESTABLISHMENT_OF_PROTEIN_LOCALIZATION | 448 | 0.281418 | 0.967238 | 0.606244 | 0.915489 |
| GOBP_CALCIUM_INDEPENDENT_CELL_CELL_ADHESION_VIA_PLASMA_MEMBRANE_CELL_ | 16 | 0.416463 | 0.967013 | 0.517192 | 0.915594 |
| GOBP_PHOSPHOLIPID_METABOLIC_PROCESS | 337 | 0.28402 | 0.967 | 0.581276 | 0.915083 |
| GOBP_CYCLIC_NUCLEOTIDE_METABOLIC_PROCESS | 31 | 0.363589 | 0.966741 | 0.517711 | 0.915284 |
| GOBP_LIPID_TRANSLOCATION | 47 | 0.340176 | 0.966437 | 0.519901 | 0.915653 |
| GOBP_VASCULAR_ASSOCIATED_SMOOTH_MUSCLE_CONTRACTION | 20 | 0.400063 | 0.966248 | 0.524893 | 0.915652 |
| GOBP_EMBRYONIC_HEART_TUBE_MORPHOGENESIS | 59 | 0.329335 | 0.966137 | 0.510256 | 0.915428 |
| GOBP_MEMBRANE_REPOLARIZATION | 39 | 0.351793 | 0.9661 | 0.525132 | 0.914996 |
| GOBP_KERATINOCYTE_MIGRATION | 15 | 0.420657 | 0.965862 | 0.533137 | 0.915146 |
| GOBP_MORPHOGENESIS_OF_AN_EPITHELIAL_SHEET | 51 | 0.338957 | 0.965452 | 0.52381 | 0.915813 |
| GOBP_PROTEIN_MATURATION | 258 | 0.286006 | 0.965355 | 0.576883 | 0.91556 |
| GOBP_METANEPHRIC_NEPHRON_MORPHOGENESIS | 21 | 0.39397 | 0.965041 | 0.519553 | 0.915934 |
| GOBP_RESPONSE_TO_VITAMIN_D | 27 | 0.371908 | 0.964871 | 0.521858 | 0.915881 |
| GOBP_NUCLEAR_TRANSCRIBED_MRNA_POLY_A_TAIL_SHORTENING | 27 | 0.378041 | 0.964814 | 0.502116 | 0.915502 |
| GOBP_REGULATION_OF_GLIAL_CELL_DIFFERENTIATION | 59 | 0.330891 | 0.964517 | 0.542787 | 0.915843 |
| GOBP_POSITIVE_REGULATION_OF_PROTEIN_POLYUBIQUITINATION | 15 | 0.416213 | 0.963837 | 0.516871 | 0.91723 |
| GOBP_REGULATION_OF_ORGAN_GROWTH | 66 | 0.325453 | 0.963765 | 0.529483 | 0.916918 |
| GOBP_REGULATION_OF_MIRNA_TRANSCRIPTION | 54 | 0.330429 | 0.963527 | 0.522059 | 0.917096 |
| GOBP_ENDOCYTOSIS | 484 | 0.279084 | 0.963159 | 0.602616 | 0.917569 |
| GOBP_CELLULAR_RESPONSE_TO_ABIOTIC_STIMULUS | 272 | 0.285495 | 0.963158 | 0.586242 | 0.917036 |
| GOBP_REGULATION_OF_GLUTAMATE_SECRETION | 17 | 0.404431 | 0.963126 | 0.52381 | 0.916582 |
| GOBP_REGULATION_OF_PROTON_TRANSPORT | 22 | 0.392854 | 0.962887 | 0.525399 | 0.916718 |
| GOBP_CELL_DEATH_IN_RESPONSE_TO_HYDROGEN_PEROXIDE | 22 | 0.389867 | 0.962485 | 0.531034 | 0.917346 |
| GOBP_FC_RECEPTOR_SIGNALING_PATHWAY | 40 | 0.349367 | 0.962332 | 0.51933 | 0.91724 |
| GOBP_RESPONSE_TO_COLD | 46 | 0.346974 | 0.962282 | 0.51871 | 0.916826 |
| GOBP_POSITIVE_REGULATION_OF_STRESS_ACTIVATED_PROTEIN_KINASE_SIGNALING_CA | 111 | 0.30708 | 0.961449 | 0.535714 | 0.918669 |
| GOBP_ACTIVATION_OF_ADENYLATE_CYCLASE_ACTIVITY | 21 | 0.389982 | 0.961182 | 0.533333 | 0.918888 |
| GOBP_REGULATION_OF_PROTEIN_TARGETING_TO_MEMBRANE | 31 | 0.365079 | 0.960843 | 0.549525 | 0.919357 |
| GOBP_NUCLEIC_ACID_PHOSPHODIESTER_BOND_HYDROLYSIS | 233 | 0.286856 | 0.9608 | 0.584281 | 0.918934 |
| GOBP_CARDIAC_MYOFIBRIL_ASSEMBLY | 16 | 0.410868 | 0.960658 | 0.529582 | 0.918837 |
| GOBP_POSITIVE_REGULATION_OF_GROWTH | 206 | 0.289645 | 0.960572 | 0.580952 | 0.918534 |
| GOBP_INTERLEUKIN_4_PRODUCTION | 22 | 0.385715 | 0.959489 | 0.527007 | 0.92108 |
| GOBP_CELL_FATE_SPECIFICATION | 62 | 0.328262 | 0.959425 | 0.547708 | 0.920737 |
| GOBP_REGULATION_OF_VASCULAR_PERMEABILITY | 43 | 0.339875 | 0.959153 | 0.550065 | 0.920948 |
| GOBP_B_CELL_HOMEOSTASIS | 27 | 0.381499 | 0.959073 | 0.521505 | 0.920649 |
| GOBP_REGULATION_OF_DNA_METHYLATION | 21 | 0.395626 | 0.95875 | 0.527007 | 0.921001 |
| GOBP_VENTRICULAR_CARDIAC_MUSCLE_CELL_MEMBRANE_REPOLARIZATION | 21 | 0.399484 | 0.958556 | 0.538793 | 0.921003 |
| GOBP_SMALL_MOLECULE_CATABOLIC_PROCESS | 314 | 0.283708 | 0.958352 | 0.601842 | 0.921047 |
| GOBP_IN_UTERO_EMBRYONIC_DEVELOPMENT | 358 | 0.281982 | 0.957832 | 0.609756 | 0.921975 |
| GOBP_HISTONE_H3_K9_ACETYLATION | 16 | 0.416163 | 0.957579 | 0.515337 | 0.922127 |
| GOBP_JNK_CASCADE | 155 | 0.295218 | 0.957344 | 0.582876 | 0.922255 |
| GOBP_I_KAPPAB_KINASE_NF_KAPPAB_SIGNALING | 243 | 0.28214 | 0.95666 | 0.595016 | 0.923582 |

| | | | | | |
|---|---|---|---|---|---|
| GOBP_ANTIMICROBIAL_HUMORAL_IMMUNE_RESPONSE_MEDIATED_BY_ANTIMICROBIAL_P | 30 | 0.370243 | 0.956659 | 0.521678 | 0.923051 |
| GOBP_INTRINSIC_APOPTOTIC_SIGNALING_PATHWAY_IN_RESPONSE_TO_DNA_DAMAGE_B | 41 | 0.343584 | 0.954991 | 0.539153 | 0.927172 |
| GOBP_NEGATIVE_REGULATION_OF_RECEPTOR_SIGNALING_PATHWAY_VIA_STAT | 22 | 0.384987 | 0.954903 | 0.534188 | 0.926884 |
| GOBP_PHOSPHATIDYLINOSITOL_DEPHOSPHORYLATION | 29 | 0.368149 | 0.954665 | 0.539528 | 0.927 |
| GOBP_NEGATIVE_REGULATION_OF_KINASE_ACTIVITY | 206 | 0.288552 | 0.954378 | 0.603594 | 0.927251 |
| GOBP_PROTEIN_O_LINKED_GLYCOSYLATION | 81 | 0.314261 | 0.953846 | 0.560234 | 0.928216 |
| GOBP_SKIN_EPIDERMIS_DEVELOPMENT | 70 | 0.320215 | 0.953552 | 0.567308 | 0.928487 |
| GOBP_TRANSMEMBRANE_RECEPTOR_PROTEIN_SERINE_THREONINE_KINASE_SIGNALING | 309 | 0.283675 | 0.953279 | 0.613497 | 0.928721 |
| GOBP_NEGATIVE_REGULATION_OF_CELL_CYCLE_G1_S_PHASE_TRANSITION | 71 | 0.32092 | 0.953192 | 0.559855 | 0.928416 |
| GOBP_STEM_CELL_PROLIFERATION | 59 | 0.325465 | 0.953145 | 0.552096 | 0.928021 |
| GOBP_POSITIVE_REGULATION_OF_LIPID_CATABOLIC_PROCESS | 17 | 0.403332 | 0.952815 | 0.526316 | 0.928392 |
| GOBP_EMBRYONIC_MORPHOGENESIS | 489 | 0.277421 | 0.952766 | 0.641414 | 0.927994 |
| GOBP_NEGATIVE_REGULATION_OF_OSTEOCLAST_DIFFERENTIATION | 23 | 0.379944 | 0.952707 | 0.527007 | 0.927634 |
| GOBP_RESPONSE_TO_FLUID_SHEAR_STRESS | 31 | 0.361233 | 0.952395 | 0.550938 | 0.927993 |
| GOBP_VASOCONSTRICTION | 61 | 0.321285 | 0.952191 | 0.546126 | 0.928008 |
| GOBP_REGULATION_OF_LIPID_TRANSPORT | 101 | 0.305518 | 0.952076 | 0.55608 | 0.927801 |
| GOBP_RUFFLE_ORGANIZATION | 49 | 0.333899 | 0.951493 | 0.543883 | 0.928855 |
| GOBP_NEGATIVE_REGULATION_OF_CYCLIN_DEPENDENT_PROTEIN_SERINE_THREONINE_ | 27 | 0.373736 | 0.951466 | 0.529244 | 0.9284 |
| GOBP_COLUMNAR_CUBOIDAL_EPITHELIAL_CELL_DIFFERENTIATION | 67 | 0.321776 | 0.951029 | 0.568182 | 0.929029 |
| GOBP_BRANCHED_CHAIN_AMINO_ACID_METABOLIC_PROCESS | 23 | 0.378898 | 0.950995 | 0.529745 | 0.928596 |
| GOBP_REGULATION_OF_PEPTIDYL_LYSINE_ACETYLATION | 68 | 0.320503 | 0.950342 | 0.569544 | 0.929923 |
| GOBP_RNA_STABILIZATION | 56 | 0.328108 | 0.95017 | 0.566131 | 0.929846 |
| GOBP_NEGATIVE_REGULATION_OF_PROTEIN_SERINE_THREONINE_KINASE_ACTIVITY | 101 | 0.300912 | 0.949417 | 0.561582 | 0.931346 |
| GOBP_POSITIVE_REGULATION_OF_MUSCLE_CELL_APOPTOTIC_PROCESS | 19 | 0.395197 | 0.949303 | 0.573939 | 0.931126 |
| GOBP_NUCLEOSIDE_BISPHOSPHATE_METABOLIC_PROCESS | 101 | 0.306092 | 0.949219 | 0.581656 | 0.930842 |
| GOBP_POSITIVE_REGULATION_OF_MEIOTIC_CELL_CYCLE | 19 | 0.401624 | 0.94879 | 0.561047 | 0.931513 |
| GOBP_REGULATION_OF_MITOCHONDRIAL_MEMBRANE_PERMEABILITY_INVOLVED_IN_APO | 39 | 0.349957 | 0.948341 | 0.557545 | 0.932235 |
| GOBP_DSRNA_PROCESSING | 52 | 0.327762 | 0.947577 | 0.551425 | 0.933835 |
| GOBP_ORGAN_INDUCTION | 19 | 0.398823 | 0.947472 | 0.538813 | 0.93359 |
| GOBP_PROTEIN_LOCALIZATION_TO_PLASMA_MEMBRANE | 267 | 0.280294 | 0.945632 | 0.602669 | 0.938153 |
| GOBP_REGULATION_OF_SMAD_PROTEIN_SIGNAL_TRANSDUCTION | 28 | 0.361695 | 0.945197 | 0.5574 | 0.93878 |
| GOBP_POSITIVE_REGULATION_OF_CELL_DIVISION | 60 | 0.324027 | 0.944985 | 0.548662 | 0.938814 |
| GOBP_REGULATION_OF_MACROPHAGE_ACTIVATION | 35 | 0.354097 | 0.943348 | 0.550993 | 0.94265 |
| GOBP_VITAMIN_METABOLIC_PROCESS | 87 | 0.307447 | 0.943196 | 0.590012 | 0.942527 |
| GOBP_MECHANORECEPTOR_DIFFERENTIATION | 56 | 0.32417 | 0.943056 | 0.569136 | 0.942364 |
| GOBP_CELL_CELL_JUNCTION_MAINTENANCE | 17 | 0.404364 | 0.942892 | 0.555079 | 0.942284 |
| GOBP_CALCINEURIN_MEDIATED_SIGNALING | 40 | 0.339819 | 0.942384 | 0.534731 | 0.943144 |
| GOBP_REGULATION_OF_NEURAL_PRECURSOR_CELL_PROLIFERATION | 71 | 0.312383 | 0.941856 | 0.591335 | 0.944043 |
| GOBP_NEGATIVE_REGULATION_OF_MITOCHONDRION_ORGANIZATION | 45 | 0.337511 | 0.941847 | 0.562183 | 0.943532 |
| GOBP_REGULATION_OF_VASOCONSTRICTION | 47 | 0.335058 | 0.941648 | 0.563218 | 0.943579 |
| GOBP_REGULATION_OF_EPITHELIAL_TO_MESENCHYMAL_TRANSITION | 84 | 0.309185 | 0.941625 | 0.587059 | 0.943106 |
| GOBP_FC_EPSILON_RECEPTOR_SIGNALING_PATHWAY | 20 | 0.391849 | 0.941259 | 0.547826 | 0.943598 |
| GOBP_REGULATION_OF_TRANS_SYNAPTIC_SIGNALING | 371 | 0.27717 | 0.941015 | 0.64467 | 0.943717 |
| GOBP_RESPONSE_TO_ZINC_ION | 34 | 0.353048 | 0.940925 | 0.565274 | 0.943424 |
| GOBP_CAMP_METABOLIC_PROCESS | 18 | 0.390438 | 0.940923 | 0.539683 | 0.942896 |
| GOBP_POSITIVE_REGULATION_OF_STRIATED_MUSCLE_CELL_DIFFERENTIATION | 30 | 0.355832 | 0.940905 | 0.56016 | 0.942408 |
| GOBP_REGULATION_OF_ERYTHROCYTE_DIFFERENTIATION | 41 | 0.341723 | 0.940513 | 0.570871 | 0.942922 |
| GOBP_B_CELL_ACTIVATION_INVOLVED_IN_IMMUNE_RESPONSE | 65 | 0.321067 | 0.940385 | 0.551893 | 0.942738 |
| GOBP_NEGATIVE_REGULATION_OF_PHOSPHORYLATION | 332 | 0.275616 | 0.940211 | 0.656793 | 0.942686 |
| GOBP_APPENDAGE_DEVELOPMENT | 137 | 0.292722 | 0.939318 | 0.59736 | 0.944532 |
| GOBP_EMBRYONIC_PLACENTA_DEVELOPMENT | 78 | 0.312977 | 0.93904 | 0.589202 | 0.944743 |
| GOBP_POSITIVE_REGULATION_OF_SODIUM_ION_TRANSMEMBRANE_TRANSPORT | 20 | 0.390138 | 0.938501 | 0.568182 | 0.945628 |
| GOBP_POSITIVE_REGULATION_OF_TRIGLYCERIDE_METABOLIC_PROCESS | 18 | 0.385863 | 0.938023 | 0.584795 | 0.946394 |
| GOBP_APPENDAGE_MORPHOGENESIS | 107 | 0.299603 | 0.93795 | 0.582569 | 0.946067 |
| GOBP_BICARBONATE_TRANSPORT | 22 | 0.373187 | 0.937833 | 0.550992 | 0.945844 |
| GOBP_NEGATIVE_REGULATION_OF_AMINE_TRANSPORT | 23 | 0.37633 | 0.937497 | 0.568871 | 0.946235 |
| GOBP_NEGATIVE_REGULATION_OF_MRNA_METABOLIC_PROCESS | 83 | 0.307913 | 0.936967 | 0.59453 | 0.947126 |
| GOBP_POSITIVE_REGULATION_OF_DNA_RECOMBINATION | 62 | 0.31868 | 0.936876 | 0.57546 | 0.946844 |
| GOBP_NEGATIVE_REGULATION_OF_TYPE_I_INTERFERON_MEDIATED_SIGNALING_PATHWA | 16 | 0.404749 | 0.936237 | 0.540816 | 0.948015 |
| GOBP_REGULATION_OF_MUSCLE_ORGAN_DEVELOPMENT | 33 | 0.349933 | 0.935917 | 0.570248 | 0.948344 |
| GOBP_CELLULAR_RESPONSE_TO_VASCULAR_ENDOTHELIAL_GROWTH_FACTOR_STIMULU | 55 | 0.326665 | 0.935427 | 0.58486 | 0.949126 |
| GOBP_RESPONSE_TO_ACIDIC_PH | 20 | 0.390903 | 0.935301 | 0.567376 | 0.948931 |
| GOBP_RESPONSE_TO_UV | 133 | 0.287903 | 0.935258 | 0.603352 | 0.948507 |
| GOBP_STEROL_METABOLIC_PROCESS | 119 | 0.292641 | 0.934955 | 0.608743 | 0.948814 |
| GOBP_POSITIVE_REGULATION_OF_MYELINATION | 16 | 0.396628 | 0.934952 | 0.561086 | 0.948289 |
| GOBP_MITOCHONDRIAL_OUTER_MEMBRANE_PERMEABILIZATION_INVOLVED_IN_PROGRA | 36 | 0.34828 | 0.934853 | 0.550265 | 0.94804 |
| GOBP_SODIUM_ION_HOMEOSTASIS | 40 | 0.337605 | 0.934786 | 0.558704 | 0.947691 |
| GOBP_NEGATIVE_REGULATION_OF_HISTONE_MODIFICATION | 47 | 0.333563 | 0.934667 | 0.579214 | 0.9475 |
| GOBP_WATER_SOLUBLE_VITAMIN_METABOLIC_PROCESS | 56 | 0.323768 | 0.934469 | 0.583229 | 0.947505 |
| GOBP_PEPTIDYL_THREONINE_DEPHOSPHORYLATION | 16 | 0.411053 | 0.934404 | 0.56623 | 0.94715 |
| GOBP_REGULATION_OF_OXIDOREDUCTASE_ACTIVITY | 73 | 0.313667 | 0.934122 | 0.577515 | 0.947378 |
| GOBP_REGULATION_OF_ANOIKIS | 19 | 0.387122 | 0.933849 | 0.557522 | 0.947569 |
| GOBP_METANEPHRIC_NEPHRON_DEVELOPMENT | 30 | 0.355891 | 0.933771 | 0.544693 | 0.947236 |
| GOBP_VISUAL_BEHAVIOR | 42 | 0.334817 | 0.933665 | 0.567183 | 0.946991 |
| GOBP_CELLULAR_RESPONSE_TO_STEROID_HORMONE_STIMULUS | 170 | 0.28886 | 0.933629 | 0.636364 | 0.946574 |
| GOBP_POSITIVE_REGULATION_OF_NEURAL_PRECURSOR_CELL_PROLIFERATION | 42 | 0.338768 | 0.933586 | 0.584211 | 0.946155 |
| GOBP_POSITIVE_REGULATION_OF_REGULATED_SECRETORY_PATHWAY | 46 | 0.332793 | 0.933319 | 0.599744 | 0.94635 |
| GOBP_CELL_JUNCTION_ASSEMBLY | 365 | 0.275916 | 0.932218 | 0.691684 | 0.94875 |
| GOBP_CELL_PROLIFERATION_INVOLVED_IN_KIDNEY_DEVELOPMENT | 17 | 0.396908 | 0.932142 | 0.561482 | 0.948437 |
| GOBP_BONE_MINERALIZATION | 92 | 0.303389 | 0.93182 | 0.599068 | 0.94875 |
| GOBP_GLUTATHIONE_METABOLIC_PROCESS | 47 | 0.330242 | 0.931483 | 0.584693 | 0.949132 |
| GOBP_DOPAMINE_RECEPTOR_SIGNALING_PATHWAY | 36 | 0.346193 | 0.93037 | 0.572917 | 0.951513 |
| GOBP_SERTOLI_CELL_DIFFERENTIATION | 15 | 0.407565 | 0.930029 | 0.559347 | 0.951877 |
| GOBP_CATECHOLAMINE_SECRETION | 45 | 0.335688 | 0.930006 | 0.569427 | 0.951405 |
| GOBP_POSITIVE_REGULATION_OF_PROTEIN_LOCALIZATION_TO_MEMBRANE | 95 | 0.297638 | 0.929935 | 0.606236 | 0.951077 |
| GOBP_MORPHOGENESIS_OF_AN_EPITHELIUM | 432 | 0.27259 | 0.929573 | 0.714721 | 0.951508 |
| GOBP_POSITIVE_REGULATION_OF_MORPHOGENESIS_OF_AN_EPITHELIUM | 31 | 0.350996 | 0.929352 | 0.566443 | 0.951566 |
| GOBP_AMIDE_TRANSPORT | 228 | 0.281514 | 0.929256 | 0.654602 | 0.951296 |
| GOBP_NEGATIVE_REGULATION_OF_APOPTOTIC_SIGNALING_PATHWAY | 195 | 0.281749 | 0.928406 | 0.640474 | 0.953044 |
| GOBP_REGULATION_OF_CELL_DEVELOPMENT | 424 | 0.270532 | 0.927246 | 0.710977 | 0.955607 |
| GOBP_POSITIVE_REGULATION_OF_UBIQUITIN_DEPENDENT_PROTEIN_CATABOLIC_PROCE | 102 | 0.2943 | 0.926736 | 0.619048 | 0.956434 |
| GOBP_NEUROTRANSMITTER_RECEPTOR_INTERNALIZATION | 17 | 0.390115 | 0.926619 | 0.582744 | 0.95622 |
| GOBP_NEGATIVE_REGULATION_OF_JNK_CASCADE | 34 | 0.345763 | 0.926596 | 0.576503 | 0.955755 |
| GOBP_FIBROBLAST_GROWTH_FACTOR_RECEPTOR_SIGNALING_PATHWAY | 67 | 0.309895 | 0.926478 | 0.604141 | 0.955562 |
| GOBP_MULTICELLULAR_ORGANISMAL_SIGNALING | 127 | 0.288412 | 0.925814 | 0.635347 | 0.956731 |
| GOBP_ANTIGEN_PROCESSING_AND_PRESENTATION_OF_ENDOGENOUS_ANTIGEN | 21 | 0.380201 | 0.925763 | 0.566092 | 0.95634 |
| GOBP_SPERM_EGG_RECOGNITION | 24 | 0.364727 | 0.924871 | 0.575887 | 0.958179 |
| GOBP_ORGANELLE_MEMBRANE_FUSION | 99 | 0.293037 | 0.924633 | 0.616082 | 0.958269 |
| GOBP_CELLULAR_RESPONSE_TO_RADIATION | 156 | 0.288252 | 0.924572 | 0.644031 | 0.957904 |
| GOBP_ENDOCRINE_PROCESS | 62 | 0.313236 | 0.924396 | 0.607009 | 0.957861 |
| GOBP_CELLULAR_TRANSITION_METAL_ION_HOMEOSTASIS | 89 | 0.298423 | 0.924067 | 0.621622 | 0.958179 |
| GOBP_HEPATOCYTE_APOPTOTIC_PROCESS | 16 | 0.394673 | 0.923367 | 0.564029 | 0.959447 |
| GOBP_DORSAL_VENTRAL_PATTERN_FORMATION | 68 | 0.309693 | 0.921626 | 0.61639 | 0.963491 |
| GOBP_CELLULAR_RESPONSE_TO_PEPTIDE | 294 | 0.272437 | 0.920743 | 0.705102 | 0.965272 |
| GOBP_SYMPATHETIC_NERVOUS_SYSTEM_DEVELOPMENT | 17 | 0.398225 | 0.920661 | 0.581955 | 0.964962 |
| GOBP_INTRACELLULAR_RECEPTOR_SIGNALING_PATHWAY | 227 | 0.274249 | 0.920496 | 0.674004 | 0.964882 |
| GOBP_POLYOL_CATABOLIC_PROCESS | 26 | 0.354385 | 0.920391 | 0.579023 | 0.96463 |
| GOBP_REGULATION_OF_TRIGLYCERIDE_METABOLIC_PROCESS | 30 | 0.350199 | 0.92029 | 0.583445 | 0.964382 |
| GOBP_NEGATIVE_REGULATION_OF_PHOSPHORUS_METABOLIC_PROCESS | 381 | 0.270835 | 0.92028 | 0.716175 | 0.96388 |
| GOBP_NEGATIVE_REGULATION_OF_CELLULAR_CATABOLIC_PROCESS | 228 | 0.276172 | 0.920116 | 0.665971 | 0.963755 |
| GOBP_GLUCOSE_METABOLIC_PROCESS | 165 | 0.279954 | 0.920026 | 0.641328 | 0.96346 |
| GOBP_NEURAL_CREST_CELL_DIFFERENTIATION | 87 | 0.299958 | 0.919344 | 0.602579 | 0.964733 |
| GOBP_NEGATIVE_REGULATION_OF_PHOSPHOPROTEIN_PHOSPHATASE_ACTIVITY | 22 | 0.37102 | 0.917341 | 0.587912 | 0.969473 |
| GOBP_ANTIVIRAL_INNATE_IMMUNE_RESPONSE | 15 | 0.40495 | 0.917287 | 0.577485 | 0.969088 |
| GOBP_BRANCHED_CHAIN_AMINO_ACID_CATABOLIC_PROCESS | 19 | 0.3768 | 0.916301 | 0.599428 | 0.971119 |
| GOBP_POSITIVE_REGULATION_OF_KINASE_ACTIVITY | 405 | 0.268482 | 0.915936 | 0.734139 | 0.971544 |
| GOBP_THYMUS_DEVELOPMENT | 34 | 0.335148 | 0.915763 | 0.586071 | 0.971464 |
| GOBP_ORGANIC_CATION_TRANSPORT | 32 | 0.345622 | 0.915759 | 0.573856 | 0.970944 |
| GOBP_REGULATION_OF_CYCLASE_ACTIVITY | 44 | 0.326903 | 0.915662 | 0.597938 | 0.970689 |
| GOBP_REGULATION_OF_CELL_FATE_COMMITMENT | 17 | 0.390935 | 0.915213 | 0.584879 | 0.971293 |
| GOBP_CELLULAR_RESPONSE_TO_PEPTIDE_HORMONE_STIMULUS | 240 | 0.274147 | 0.91445 | 0.681913 | 0.972659 |
| GOBP_REGULATION_OF_CATECHOLAMINE_METABOLIC_PROCESS | 20 | 0.378173 | 0.914356 | 0.597475 | 0.972362 |
| GOBP_REGULATION_OF_RUFFLE_ASSEMBLY | 28 | 0.35275 | 0.914023 | 0.580381 | 0.972678 |
| GOBP_CELLULAR_RESPONSE_TO_CAMP | 41 | 0.330514 | 0.91378 | 0.6 | 0.972754 |
| GOBP_ENDOTHELIUM_DEVELOPMENT | 116 | 0.287719 | 0.913772 | 0.656983 | 0.972245 |
| GOBP_CARBOXYLIC_ACID_TRANSPORT | 204 | 0.272765 | 0.913719 | 0.680717 | 0.971855 |
| GOBP_TOLL_LIKE_RECEPTOR_SIGNALING_PATHWAY | 87 | 0.300523 | 0.913255 | 0.623705 | 0.972528 |
| GOBP_REGULATION_OF_MEMBRANE_DEPOLARIZATION | 38 | 0.339256 | 0.913164 | 0.604194 | 0.972239 |
| GOBP_NEGATIVE_REGULATION_OF_ENDOCYTOSIS | 47 | 0.324615 | 0.912915 | 0.593199 | 0.972351 |
| GOBP_REGULATION_OF_APPETITE | 15 | 0.393096 | 0.912537 | 0.569343 | 0.972801 |
| GOBP_EXCRETION | 44 | 0.323026 | 0.912438 | 0.594702 | 0.972528 |
| GOBP_RUFFLE_ASSEMBLY | 41 | 0.334258 | 0.912431 | 0.620735 | 0.97202 |
| GOBP_POSITIVE_REGULATION_OF_OSSIFICATION | 43 | 0.326918 | 0.912344 | 0.595579 | 0.971722 |
| GOBP_NEURON_APOPTOTIC_PROCESS | 210 | 0.272627 | 0.91197 | 0.676471 | 0.972178 |
| GOBP_REGULATION_OF_GLIOGENESIS | 78 | 0.302245 | 0.911944 | 0.627497 | 0.971724 |
| GOBP_ORGANIC_ANION_TRANSPORT | 277 | 0.270799 | 0.911185 | 0.703934 | 0.973065 |
| GOBP_OVULATION_CYCLE_PROCESS | 37 | 0.331641 | 0.911164 | 0.579576 | 0.972593 |
| GOBP_REGULATION_OF_MYELOID_CELL_DIFFERENTIATION | 163 | 0.279883 | 0.910943 | 0.668817 | 0.972616 |
| GOBP_NEGATIVE_REGULATION_OF_CATABOLIC_PROCESS | 278 | 0.270092 | 0.910745 | 0.712821 | 0.972596 |
| GOBP_POST_EMBRYONIC_DEVELOPMENT | 78 | 0.302056 | 0.910642 | 0.639766 | 0.972312 |
| GOBP_REGULATION_OF_PROTEIN_ACETYLATION | 76 | 0.301342 | 0.910448 | 0.618594 | 0.972247 |
| GOBP_CARTILAGE_DEVELOPMENT_INVOLVED_IN_ENDOCHONDRAL_BONE_MORPHOGENE | 25 | 0.354057 | 0.909397 | 0.59249 | 0.974369 |
| GOBP_EXTRACELLULAR_MATRIX_ASSEMBLY | 41 | 0.330365 | 0.909275 | 0.622865 | 0.974155 |
| GOBP_MONOAMINE_TRANSPORT | 65 | 0.30923 | 0.909265 | 0.631258 | 0.973658 |
| GOBP_ANION_TRANSPORT | 397 | 0.266235 | 0.90854 | 0.738648 | 0.975017 |
| GOBP_GMP_METABOLIC_PROCESS | 26 | 0.355706 | 0.908419 | 0.588721 | 0.974792 |
| GOBP_POSITIVE_REGULATION_OF_MICROTUBULE_POLYMERIZATION_OR_DEPOLYMERIZAT | 36 | 0.333479 | 0.908328 | 0.579156 | 0.974504 |
| GOBP_REGULATION_OF_CELL_SHAPE | 133 | 0.283663 | 0.908299 | 0.674082 | 0.974057 |
| GOBP_NEGATIVE_REGULATION_OF_DEPHOSPHORYLATION | 40 | 0.333137 | 0.908189 | 0.617876 | 0.973793 |
| GOBP_POSITIVE_REGULATION_OF_SMAD_PROTEIN_SIGNAL_TRANSDUCTION | 18 | 0.381989 | 0.907203 | 0.580552 | 0.975744 |
| GOBP_POSITIVE_REGULATION_OF_LIPASE_ACTIVITY | 44 | 0.318813 | 0.907159 | 0.594423 | 0.975355 |
| GOBP_LONG_TERM_SYNAPTIC_DEPRESSION | 29 | 0.35058 | 0.906936 | 0.604552 | 0.975391 |
| GOBP_REGULATION_OF_PHOSPHOLIPASE_C_ACTIVITY | 30 | 0.34552 | 0.906822 | 0.588315 | 0.975156 |
| GOBP_HORMONE_METABOLIC_PROCESS | 157 | 0.279311 | 0.906293 | 0.682119 | 0.976 |
| GOBP_REGULATION_OF_CATION_CHANNEL_ACTIVITY | 137 | 0.28317 | 0.905982 | 0.683333 | 0.976276 |
| GOBP_POSITIVE_REGULATION_OF_NERVOUS_SYSTEM_PROCESS | 29 | 0.35522 | 0.905981 | 0.617368 | 0.97576 |
| GOBP_CELLULAR_RESPONSE_TO_HORMONE_STIMULUS | 473 | 0.263859 | 0.904989 | 0.789579 | 0.977644 |

| | | | | | |
|---|---|---|---|---|---|
| GOBP_DNA_SYNTHESIS_INVOLVED_IN_DNA_REPAIR | 34 | 0.328425 | 0.904585 | 0.596838 | 0.978141 |
| GOBP_REGULATION_OF_MEMBRANE_LIPID_DISTRIBUTION | 53 | 0.314052 | 0.904101 | 0.636252 | 0.978847 |
| GOBP_REGULATION_OF_REACTIVE_OXYGEN_SPECIES_METABOLIC_PROCESS | 112 | 0.289465 | 0.904092 | 0.664773 | 0.978347 |
| GOBP_POSITIVE_REGULATION_OF_PROTEIN_MODIFICATION_BY_SMALL_PROTEIN_CONJU( | 128 | 0.282742 | 0.904013 | 0.655211 | 0.978019 |
| GOBP_ORGANIC_ACID_TRANSPORT | 240 | 0.269485 | 0.903973 | 0.697602 | 0.977594 |
| GOBP_NEGATIVE_REGULATION_OF_EPITHELIAL_CELL_APOPTOTIC_PROCESS | 39 | 0.330823 | 0.903847 | 0.607239 | 0.977389 |
| GOBP_AMYLOID_FIBRIL_FORMATION | 27 | 0.356251 | 0.903693 | 0.60411 | 0.977251 |
| GOBP_TERPENOID_METABOLIC_PROCESS | 64 | 0.306676 | 0.903365 | 0.640547 | 0.977556 |
| GOBP_MEGAKARYOCYTE_DIFFERENTIATION | 49 | 0.31593 | 0.903173 | 0.622525 | 0.977506 |
| GOBP_POSITIVE_REGULATION_OF_CYTOSKELETON_ORGANIZATION | 178 | 0.272989 | 0.902547 | 0.700107 | 0.978557 |
| GOBP_COENZYME_A_METABOLIC_PROCESS | 18 | 0.37856 | 0.902485 | 0.617816 | 0.978198 |
| GOBP_T_CELL_APOPTOTIC_PROCESS | 36 | 0.335739 | 0.90243 | 0.629482 | 0.977829 |
| GOBP_RELEASE_OF_SEQUESTERED_CALCIUM_ION_INTO_CYTOSOL_BY_ENDOPLASMIC_F | 26 | 0.350017 | 0.902412 | 0.595716 | 0.977363 |
| GOBP_ROOF_OF_MOUTH_DEVELOPMENT | 72 | 0.304146 | 0.902334 | 0.627359 | 0.977026 |
| GOBP_EMBRYONIC_APPENDAGE_MORPHOGENESIS | 88 | 0.294506 | 0.901819 | 0.658906 | 0.977774 |
| GOBP_POSITIVE_REGULATION_OF_NUCLEOCYTOPLASMIC_TRANSPORT | 55 | 0.312994 | 0.901723 | 0.651106 | 0.977479 |
| GOBP_NEGATIVE_CHEMOTAXIS | 45 | 0.317331 | 0.901559 | 0.611331 | 0.977341 |
| GOBP_NEGATIVE_REGULATION_OF_RECEPTOR_SIGNALING_PATHWAY_VIA_JAK_STAT | 20 | 0.368258 | 0.900972 | 0.608187 | 0.978238 |
| GOBP_SIGNAL_TRANSDUCTION_IN_RESPONSE_TO_DNA_DAMAGE | 165 | 0.273749 | 0.900495 | 0.685959 | 0.978889 |
| GOBP_POSITIVE_REGULATION_OF_SUPRAMOLECULAR_FIBER_ORGANIZATION | 161 | 0.27459 | 0.89933 | 0.69214 | 0.98121 |
| GOBP_MAMMARY_GLAND_EPITHELIAL_CELL_DIFFERENTIATION | 15 | 0.4005 | 0.899237 | 0.603631 | 0.980922 |
| GOBP_PROTEIN_TARGETING_TO_LYSOSOME | 23 | 0.362599 | 0.898957 | 0.61 | 0.981092 |
| GOBP_POSITIVE_REGULATION_OF_CELLULAR_AMIDE_METABOLIC_PROCESS | 151 | 0.274619 | 0.898097 | 0.67169 | 0.982649 |
| GOBP_REGULATION_OF_TOLL_LIKE_RECEPTOR_SIGNALING_PATHWAY | 52 | 0.318328 | 0.897785 | 0.645679 | 0.98286 |
| GOBP_REGULATION_OF_KERATINOCYTE_PROLIFERATION | 27 | 0.341755 | 0.897527 | 0.608635 | 0.982967 |
| GOBP_BRANCHING_MORPHOGENESIS_OF_AN_EPITHELIAL_TUBE | 121 | 0.284415 | 0.897039 | 0.667401 | 0.983593 |
| GOBP_NEGATIVE_REGULATION_OF_NUCLEOTIDE_METABOLIC_PROCESS | 21 | 0.363614 | 0.897008 | 0.608939 | 0.983154 |
| GOBP_ACTIN_FILAMENT_SEVERING | 16 | 0.382785 | 0.896872 | 0.575075 | 0.982966 |
| GOBP_ESTABLISHMENT_OF_ORGANELLE_LOCALIZATION | 367 | 0.266131 | 0.896836 | 0.76417 | 0.982528 |
| GOBP_HOMEOSTASIS_OF_NUMBER_OF_CELLS_WITHIN_A_TISSUE | 22 | 0.36684 | 0.896412 | 0.611111 | 0.983045 |
| GOBP_NEGATIVE_REGULATION_OF_NERVOUS_SYSTEM_DEVELOPMENT | 125 | 0.27838 | 0.896205 | 0.691128 | 0.983032 |
| GOBP_REGULATION_OF_INTRACELLULAR_PROTEIN_TRANSPORT | 206 | 0.268841 | 0.895938 | 0.712432 | 0.983182 |
| GOBP_PURINE_CONTAINING_COMPOUND_CATABOLIC_PROCESS | 46 | 0.315443 | 0.895052 | 0.632184 | 0.984814 |
| GOBP_RIBONUCLEOSIDE_MONOPHOSPHATE_METABOLIC_PROCESS | 55 | 0.30863 | 0.894645 | 0.631453 | 0.985308 |
| GOBP_ACTIN_FILAMENT_BUNDLE_ORGANIZATION | 152 | 0.277934 | 0.894351 | 0.712105 | 0.985532 |
| GOBP_REGULATION_OF_BLOOD_PRESSURE | 137 | 0.276554 | 0.894195 | 0.684795 | 0.985398 |
| GOBP_NUCLEOTIDE_SALVAGE | 16 | 0.382944 | 0.894172 | 0.611354 | 0.984943 |
| GOBP_REGULATION_OF_SYNAPTIC_PLASTICITY | 170 | 0.274526 | 0.89401 | 0.688507 | 0.984823 |
| GOBP_CARDIAC_MUSCLE_CELL_MEMBRANE_REPOLARIZATION | 30 | 0.34294 | 0.893701 | 0.637228 | 0.985048 |
| GOBP_NEGATIVE_REGULATION_OF_PROTEIN_POLYMERIZATION | 73 | 0.299659 | 0.893423 | 0.656398 | 0.985209 |
| GOBP_FACE_DEVELOPMENT | 46 | 0.320968 | 0.892526 | 0.650127 | 0.986859 |
| GOBP_REGULATION_OF_MONOOXYGENASE_ACTIVITY | 39 | 0.322769 | 0.892179 | 0.61823 | 0.987131 |
| GOBP_RECEPTOR_CATABOLIC_PROCESS | 27 | 0.346863 | 0.892128 | 0.64 | 0.986738 |
| GOBP_PLATELET_MORPHOGENESIS | 18 | 0.379149 | 0.892006 | 0.592431 | 0.986517 |
| GOBP_POSITIVE_REGULATION_OF_VIRAL_LIFE_CYCLE | 20 | 0.370046 | 0.891655 | 0.640114 | 0.986828 |
| GOBP_REGULATION_OF_EXTRINSIC_APOPTOTIC_SIGNALING_PATHWAY_VIA_DEATH_DOM/ | 41 | 0.319288 | 0.891443 | 0.650266 | 0.986821 |
| GOBP_NEGATIVE_REGULATION_OF_RESPONSE_TO_BIOTIC_STIMULUS | 86 | 0.291325 | 0.89105 | 0.665127 | 0.987262 |
| GOBP_REGULATION_OF_CELL_PROJECTION_ASSEMBLY | 200 | 0.267377 | 0.89104 | 0.710805 | 0.986776 |
| GOBP_REGULATION_OF_NUCLEOTIDE_METABOLIC_PROCESS | 71 | 0.295447 | 0.890835 | 0.644737 | 0.986758 |
| GOBP_POSITIVE_REGULATION_OF_CELL_DEVELOPMENT | 255 | 0.266356 | 0.89072 | 0.762055 | 0.986526 |
| GOBP_NEGATIVE_REGULATION_OF_GLYCOPROTEIN_METABOLIC_PROCESS | 15 | 0.386659 | 0.890674 | 0.614551 | 0.986134 |
| GOBP_NEGATIVE_REGULATION_OF_TUMOR_NECROSIS_FACTOR_SUPERFAMILY_CYTOKIN | 41 | 0.325139 | 0.890151 | 0.638853 | 0.986859 |
| GOBP_POSITIVE_REGULATION_OF_PROTEOLYSIS | 327 | 0.261581 | 0.889601 | 0.75567 | 0.987633 |
| GOBP_NUCLEAR_TRANSCRIBED_MRNA_CATABOLIC_PROCESS_DEADENYLATION_DEPENI | 60 | 0.30288 | 0.889039 | 0.638083 | 0.988454 |
| GOBP_SYNAPSE_MATURATION | 23 | 0.356078 | 0.88859 | 0.623776 | 0.988968 |
| GOBP_ACTION_POTENTIAL | 117 | 0.281325 | 0.88835 | 0.690157 | 0.989035 |
| GOBP_ORGANIC_ACID_CATABOLIC_PROCESS | 203 | 0.267916 | 0.888289 | 0.727368 | 0.988667 |
| GOBP_PROTEIN_HETEROOLIGOMERIZATION | 25 | 0.349571 | 0.888177 | 0.617978 | 0.988417 |
| GOBP_POSITIVE_REGULATION_OF_CELL_GROWTH | 144 | 0.273932 | 0.887213 | 0.703261 | 0.990175 |
| GOBP_FOREBRAIN_GENERATION_OF_NEURONS | 38 | 0.324061 | 0.88721 | 0.645806 | 0.989671 |
| GOBP_POSITIVE_REGULATION_OF_NEUROGENESIS | 196 | 0.268351 | 0.887168 | 0.728435 | 0.98926 |
| GOBP_REGULATION_OF_EPIDERMAL_GROWTH_FACTOR_ACTIVATED_RECEPTOR_ACTIVIT | 25 | 0.349516 | 0.887124 | 0.638268 | 0.988847 |
| GOBP_NITRIC_OXIDE_MEDIATED_SIGNAL_TRANSDUCTION | 18 | 0.365534 | 0.887016 | 0.638643 | 0.988585 |
| GOBP_AORTIC_VALVE_MORPHOGENESIS | 30 | 0.337995 | 0.88672 | 0.626359 | 0.988772 |
| GOBP_REGULATION_OF_APOPTOTIC_SIGNALING_PATHWAY | 315 | 0.260487 | 0.886347 | 0.772358 | 0.989166 |
| GOBP_RENAL_SODIUM_EXCRETION | 15 | 0.395431 | 0.886152 | 0.616208 | 0.989101 |
| GOBP_MAINTENANCE_OF_CELL_NUMBER | 127 | 0.275452 | 0.88547 | 0.691048 | 0.990186 |
| GOBP_MEMBRANE_LIPID_CATABOLIC_PROCESS | 32 | 0.331139 | 0.885365 | 0.635649 | 0.989919 |
| GOBP_REGULATION_OF_STEROID_METABOLIC_PROCESS | 75 | 0.292353 | 0.885031 | 0.656805 | 0.990212 |
| GOBP_NEPHRON_EPITHELIUM_DEVELOPMENT | 90 | 0.291059 | 0.88491 | 0.689376 | 0.990007 |
| GOBP_NOTOCHORD_DEVELOPMENT | 18 | 0.377063 | 0.884701 | 0.640244 | 0.990001 |
| GOBP_POST_TRANSLATIONAL_PROTEIN_MODIFICATION | 36 | 0.321622 | 0.884612 | 0.624 | 0.989708 |
| GOBP_FOREBRAIN_REGIONALIZATION | 16 | 0.382564 | 0.884334 | 0.636111 | 0.989877 |
| GOBP_CELLULAR_COMPONENT_DISASSEMBLY | 423 | 0.258902 | 0.883216 | 0.834511 | 0.992015 |
| GOBP_MITOTIC_SISTER_CHROMATID_COHESION | 27 | 0.344435 | 0.883215 | 0.636 | 0.991513 |
| GOBP_INORGANIC_ANION_TRANSPORT | 142 | 0.271857 | 0.882677 | 0.723077 | 0.992255 |
| GOBP_MITOCHONDRIAL_TRANSCRIPTION | 17 | 0.377194 | 0.882519 | 0.625899 | 0.992122 |
| GOBP_OSTEOBLAST_DIFFERENTIATION | 194 | 0.265469 | 0.881927 | 0.737844 | 0.992962 |
| GOBP_MYOBLAST_DIFFERENTIATION | 61 | 0.299319 | 0.881406 | 0.663772 | 0.993646 |
| GOBP_INTRINSIC_APOPTOTIC_SIGNALING_PATHWAY_IN_RESPONSE_TO_DNA_DAMAGE | 93 | 0.282513 | 0.881281 | 0.681818 | 0.993416 |
| GOBP_REGULATION_OF_PHOSPHATIDYLINOSITOL_3_KINASE_ACTIVITY | 47 | 0.312282 | 0.880378 | 0.669609 | 0.995047 |
| GOBP_INTERFERON_BETA_PRODUCTION | 50 | 0.312565 | 0.879687 | 0.668335 | 0.996121 |
| GOBP_REGULATION_OF_PROTEIN_LOCALIZATION_TO_MEMBRANE | 156 | 0.271177 | 0.879608 | 0.720395 | 0.995797 |
| GOBP_CELLULAR_MODIFIED_AMINO_ACID_METABOLIC_PROCESS | 147 | 0.270621 | 0.878712 | 0.720257 | 0.997405 |
| GOBP_REGULATION_OF_INTRINSIC_APOPTOTIC_SIGNALING_PATHWAY_BY_P53_CLASS_M | 29 | 0.336487 | 0.878687 | 0.640054 | 0.996954 |
| GOBP_CHONDROITIN_SULFATE_PROTEOGLYCAN_METABOLIC_PROCESS | 33 | 0.331039 | 0.878659 | 0.658999 | 0.996506 |
| GOBP_PROTEIN_LOCALIZATION_TO_CELL_PERIPHERY | 314 | 0.258392 | 0.878462 | 0.794688 | 0.996481 |
| GOBP_NON_MEMBRANE_BOUNDED_ORGANELLE_ASSEMBLY | 357 | 0.260576 | 0.878451 | 0.792857 | 0.996006 |
| GOBP_RESPIRATORY_SYSTEM_DEVELOPMENT | 164 | 0.26666 | 0.878402 | 0.739741 | 0.995606 |
| GOBP_PHAGOLYSOSOME_ASSEMBLY | 16 | 0.380359 | 0.87839 | 0.646809 | 0.995129 |
| GOBP_POSITIVE_REGULATION_OF_VIRAL_PROCESS | 57 | 0.303547 | 0.877804 | 0.681305 | 0.995925 |
| GOBP_GLYCOPROTEIN_CATABOLIC_PROCESS | 23 | 0.351362 | 0.877504 | 0.643388 | 0.996117 |
| GOBP_REGULATION_OF_PROTEIN_IMPORT | 55 | 0.301593 | 0.877493 | 0.682896 | 0.995639 |
| GOBP_REGULATION_OF_LIPID_CATABOLIC_PROCESS | 41 | 0.318504 | 0.877302 | 0.657588 | 0.995571 |
| GOBP_GLUCOCORTICOID_METABOLIC_PROCESS | 16 | 0.377686 | 0.877267 | 0.647149 | 0.995147 |
| GOBP_SIGNAL_TRANSDUCTION_IN_ABSENCE_OF_LIGAND | 53 | 0.307505 | 0.877181 | 0.660354 | 0.994834 |
| GOBP_POSITIVE_REGULATION_OF_PROTEOLYSIS_INVOLVED_IN_CELLULAR_PROTEIN_CA1 | 124 | 0.273597 | 0.877018 | 0.693584 | 0.994695 |
| GOBP_ODONTOGENESIS_OF_DENTIN_CONTAINING_TOOTH | 65 | 0.293925 | 0.875367 | 0.679222 | 0.997918 |
| GOBP_NEGATIVE_REGULATION_OF_DNA_BIOSYNTHETIC_PROCESS | 34 | 0.331559 | 0.874633 | 0.662585 | 0.999049 |
| GOBP_MITOCHONDRIAL_FUSION | 30 | 0.331438 | 0.874583 | 0.666216 | 0.998655 |
| GOBP_EPITHELIAL_CELL_MORPHOGENESIS | 30 | 0.332494 | 0.874475 | 0.661705 | 0.998407 |
| GOBP_CEREBRAL_CORTEX_DEVELOPMENT | 100 | 0.28007 | 0.873794 | 0.723898 | 0.99949 |
| GOBP_REGULATION_OF_T_CELL_APOPTOTIC_PROCESS | 22 | 0.354755 | 0.873363 | 0.646043 | 1 |
| GOBP_NEGATIVE_REGULATION_OF_PHAGOCYTOSIS | 18 | 0.362994 | 0.872885 | 0.650888 | 1 |
| GOBP_SYNAPTIC_TRANSMISSION_GABAERGIC | 46 | 0.307438 | 0.872141 | 0.65685 | 1 |
| GOBP_POSITIVE_REGULATION_OF_PROTEIN_LOCALIZATION | 398 | 0.255038 | 0.871931 | 0.827236 | 1 |
| GOBP_RESPONSE_TO_CAMP | 69 | 0.293498 | 0.871811 | 0.701583 | 1 |
| GOBP_REGULATION_OF_CELLULAR_RESPONSE_TO_INSULIN_STIMULUS | 52 | 0.305162 | 0.871739 | 0.690709 | 1 |
| GOBP_CELL_GROWTH | 434 | 0.253564 | 0.871632 | 0.845226 | 1 |
| GOBP_POSITIVE_REGULATION_OF_TRANSCRIPTION_FROM_RNA_POLYMERASE_II_PROMO | 22 | 0.351491 | 0.87143 | 0.660458 | 1 |
| GOBP_RNA_5_END_PROCESSING | 23 | 0.352041 | 0.871331 | 0.642959 | 1 |
| GOBP_NEGATIVE_REGULATION_OF_TRANSFERASE_ACTIVITY | 240 | 0.260468 | 0.871272 | 0.760331 | 1 |
| GOBP_EPITHELIAL_TUBE_MORPHOGENESIS | 283 | 0.257403 | 0.870287 | 0.801858 | 1 |
| GOBP_POSITIVE_REGULATION_OF_VASCULAR_PERMEABILITY | 17 | 0.371964 | 0.870279 | 0.627365 | 1 |
| GOBP_POSITIVE_REGULATION_OF_INTRACELLULAR_PROTEIN_TRANSPORT | 144 | 0.268184 | 0.869553 | 0.759912 | 1 |
| GOBP_VIRAL_PROCESS | 361 | 0.255181 | 0.869403 | 0.819096 | 1 |
| GOBP_POSITIVE_REGULATION_OF_GLUCOSE_METABOLIC_PROCESS | 33 | 0.326937 | 0.869289 | 0.6639 | 1 |
| GOBP_POSITIVE_REGULATION_OF_EXOCYTOSIS | 74 | 0.291738 | 0.869034 | 0.698113 | 1 |
| GOBP_POSITIVE_REGULATION_OF_VIRAL_GENOME_REPLICATION | 29 | 0.334638 | 0.868807 | 0.663989 | 1 |
| GOBP_AXONAL_TRANSPORT | 64 | 0.295786 | 0.868343 | 0.70529 | 1 |
| GOBP_SYNAPSE_ORGANIZATION | 374 | 0.255091 | 0.867944 | 0.832146 | 1 |
| GOBP_POSITIVE_REGULATION_OF_B_CELL_MEDIATED_IMMUNITY | 30 | 0.340517 | 0.867941 | 0.680822 | 1 |
| GOBP_REGULATION_OF_CELL_CYCLE_G1_S_PHASE_TRANSITION | 145 | 0.267083 | 0.867862 | 0.740332 | 1 |
| GOBP_REGULATION_OF_TUMOR_NECROSIS_FACTOR_MEDIATED_SIGNALING_PATHWAY | 46 | 0.310694 | 0.867816 | 0.682477 | 1 |
| GOBP_NEGATIVE_REGULATION_OF_I_KAPPAB_KINASE_NF_KAPPAB_SIGNALING | 40 | 0.314634 | 0.867667 | 0.661376 | 1 |
| GOBP_GLIAL_CELL_DEVELOPMENT | 92 | 0.283325 | 0.867489 | 0.707118 | 1 |
| GOBP_REGULATION_OF_GLUCOSE_TRANSMEMBRANE_TRANSPORT | 59 | 0.297284 | 0.867124 | 0.688889 | 1 |
| GOBP_PROTEIN_DESTABILIZATION | 43 | 0.311713 | 0.866749 | 0.648408 | 1 |
| GOBP_NEGATIVE_REGULATION_OF_MIRNA_TRANSCRIPTION | 17 | 0.373108 | 0.865918 | 0.66766 | 1 |
| GOBP_APOPTOTIC_MITOCHONDRIAL_CHANGES | 92 | 0.278794 | 0.865892 | 0.719037 | 1 |
| GOBP_PHOSPHATIDYLINOSITOL_3_KINASE_SIGNALING | 102 | 0.274492 | 0.865586 | 0.7319 | 1 |
| GOBP_STRESS_ACTIVATED_PROTEIN_KINASE_SIGNALING_CASCADE | 218 | 0.260731 | 0.865223 | 0.777083 | 1 |
| GOBP_NEPHRON_MORPHOGENESIS | 60 | 0.295386 | 0.864831 | 0.683761 | 1 |
| GOBP_ATRIAL_CARDIAC_MUSCLE_CELL_TO_AV_NODE_CELL_SIGNALING | 17 | 0.372783 | 0.864702 | 0.63901 | 1 |
| GOBP_ACTIN_FILAMENT_POLYMERIZATION | 149 | 0.267482 | 0.864161 | 0.738562 | 1 |
| GOBP_SPECIFICATION_OF_ANIMAL_ORGAN_IDENTITY | 28 | 0.334941 | 0.863568 | 0.68065 | 1 |
| GOBP_REGULATION_OF_SYNAPTIC_TRANSMISSION_GLUTAMATERGIC | 59 | 0.296872 | 0.863484 | 0.688199 | 1 |
| GOBP_PHOTORECEPTOR_CELL_MAINTENANCE | 35 | 0.317134 | 0.863459 | 0.669344 | 1 |
| GOBP_PURINE_NUCLEOSIDE_MONOPHOSPHATE_METABOLIC_PROCESS | 43 | 0.309649 | 0.861851 | 0.679739 | 1 |
| GOBP_REGULATION_OF_GLUCONEOGENESIS | 43 | 0.306828 | 0.861423 | 0.685972 | 1 |
| GOBP_MEMBRANE_LIPID_METABOLIC_PROCESS | 183 | 0.261969 | 0.861421 | 0.784103 | 1 |
| GOBP_REGULATION_OF_CELL_AGING | 42 | 0.313103 | 0.860662 | 0.692209 | 1 |
| GOBP_MITOCHONDRIAL_DEPOLARIZATION | 18 | 0.353477 | 0.860631 | 0.653147 | 1 |
| GOBP_REGULATION_OF_RESPONSE_TO_REACTIVE_OXYGEN_SPECIES | 28 | 0.324999 | 0.860048 | 0.66064 | 1 |
| GOBP_NEGATIVE_REGULATION_OF_PROTEIN_LOCALIZATION_TO_MEMBRANE | 28 | 0.334217 | 0.860026 | 0.658263 | 1 |
| GOBP_PEPTIDYL_PROLINE_MODIFICATION | 50 | 0.304299 | 0.858699 | 0.680958 | 1 |
| GOBP_GROWTH_PLATE_CARTILAGE_DEVELOPMENT | 16 | 0.366408 | 0.858332 | 0.675994 | 1 |
| GOBP_POSITIVE_REGULATION_OF_GLUCOSE_IMPORT | 30 | 0.327563 | 0.858127 | 0.680111 | 1 |
| GOBP_NEGATIVE_REGULATION_OF_CARBOHYDRATE_METABOLIC_PROCESS | 43 | 0.309569 | 0.858031 | 0.681579 | 1 |
| GOBP_PHARYNGEAL_SYSTEM_DEVELOPMENT | 25 | 0.338099 | 0.857948 | 0.674419 | 1 |
| GOBP_MITOCHONDRIAL_OUTER_MEMBRANE_PERMEABILIZATION | 31 | 0.327906 | 0.857895 | 0.673387 | 1 |
| GOBP_LACTATION | 39 | 0.314851 | 0.857792 | 0.68129 | 1 |
| GOBP_RESPONSE_TO_IMMOBILIZATION_STRESS | 16 | 0.373931 | 0.857698 | 0.680576 | 1 |
| GOBP_EMBRYONIC_SKELETAL_SYSTEM_DEVELOPMENT | 83 | 0.282538 | 0.857469 | 0.742243 | 1 |
| GOBP_CARDIAC_MUSCLE_CELL_ACTION_POTENTIAL_INVOLVED_IN_CONTRACTION | 45 | 0.302683 | 0.857362 | 0.699239 | 1 |
| GOBP_NEGATIVE_REGULATION_OF_FATTY_ACID_METABOLIC_PROCESS | 23 | 0.34706 | 0.857207 | 0.658263 | 1 |
| GOBP_RENAL_TUBULE_DEVELOPMENT | 76 | 0.284323 | 0.856826 | 0.692216 | 1 |

| | | | | | |
|---|---|---|---|---|---|
| GOBP_CELLULAR_RESPONSE_TO_STEROL_DEPLETION | 16 | 0.375304 | 0.856745 | 0.656012 | 1 |
| GOBP_LONG_CHAIN_FATTY_ACYL_COA_METABOLIC_PROCESS | 22 | 0.346582 | 0.856625 | 0.657593 | 1 |
| GOBP_FAT_SOLUBLE_VITAMIN_METABOLIC_PROCESS | 28 | 0.323979 | 0.856261 | 0.681511 | 1 |
| GOBP_ALPHA_AMINO_ACID_METABOLIC_PROCESS | 169 | 0.260882 | 0.856067 | 0.778256 | 1 |
| GOBP_REGULATION_OF_LYASE_ACTIVITY | 40 | 0.314519 | 0.855512 | 0.680958 | 1 |
| GOBP_CELL_RECOGNITION | 101 | 0.274939 | 0.853851 | 0.747486 | 1 |
| GOBP_GLYCEROPHOSPHOLIPID_BIOSYNTHETIC_PROCESS | 198 | 0.260286 | 0.853841 | 0.780149 | 1 |
| GOBP_PROTEIN_CONTAINING_COMPLEX_REMODELING | 20 | 0.356519 | 0.853421 | 0.663793 | 1 |
| GOBP_REGULATION_OF_GLUCAN_BIOSYNTHETIC_PROCESS | 23 | 0.346494 | 0.853377 | 0.671788 | 1 |
| GOBP_PIGMENTATION | 87 | 0.276728 | 0.85333 | 0.71967 | 1 |
| GOBP_STRIATED_MUSCLE_CELL_PROLIFERATION | 49 | 0.299129 | 0.853059 | 0.687254 | 1 |
| GOBP_REGULATION_OF_PROGRAMMED_NECROTIC_CELL_DEATH | 21 | 0.35479 | 0.852653 | 0.675824 | 1 |
| GOBP_REGULATION_OF_STRESS_ACTIVATED_PROTEIN_KINASE_SIGNALING_CASCADE | 172 | 0.260652 | 0.85253 | 0.778836 | 1 |
| GOBP_CELLULAR_RESPONSE_TO_INTERLEUKIN_1 | 66 | 0.290896 | 0.852512 | 0.723353 | 1 |
| GOBP_REGULATION_OF_DOUBLE_STRAND_BREAK_REPAIR_VIA_HOMOLOGOUS_RECOMB | 69 | 0.288392 | 0.852111 | 0.708738 | 1 |
| GOBP_POSITIVE_REGULATION_OF_BONE_MINERALIZATION | 35 | 0.317392 | 0.851976 | 0.666667 | 1 |
| GOBP_REGULATION_OF_CILIUM_MOVEMENT | 27 | 0.330193 | 0.850702 | 0.661705 | 1 |
| GOBP_TRANSEPITHELIAL_TRANSPORT | 28 | 0.331892 | 0.849948 | 0.694986 | 1 |
| GOBP_DENDRITE_EXTENSION | 33 | 0.323623 | 0.84985 | 0.69709 | 1 |
| GOBP_REGULATION_OF_INTRACELLULAR_TRANSPORT | 305 | 0.25143 | 0.849582 | 0.847959 | 1 |
| GOBP_POSITIVE_REGULATION_OF_SUBSTRATE_ADHESION_DEPENDENT_CELL_SPREADIN | 36 | 0.316352 | 0.847901 | 0.704486 | 1 |
| GOBP_REGULATION_OF_TOLL_LIKE_RECEPTOR_4_SIGNALING_PATHWAY | 17 | 0.366287 | 0.847489 | 0.677991 | 1 |
| GOBP_ORGANELLE_FUSION | 133 | 0.263513 | 0.847139 | 0.770563 | 1 |
| GOBP_RESPONSE_TO_GAMMA_RADIATION | 51 | 0.296949 | 0.847097 | 0.710127 | 1 |
| GOBP_RESPONSE_TO_HYPEROXIA | 17 | 0.365104 | 0.846993 | 0.637747 | 1 |
| GOBP_SPHINGOLIPID_METABOLIC_PROCESS | 139 | 0.264162 | 0.846639 | 0.772331 | 1 |
| GOBP_REGULATION_OF_ACTIN_FILAMENT_LENGTH | 141 | 0.262452 | 0.846339 | 0.775956 | 1 |
| GOBP_CELL_COMMUNICATION_INVOLVED_IN_CARDIAC_CONDUCTION | 52 | 0.296799 | 0.846081 | 0.720403 | 1 |
| GOBP_NEGATIVE_REGULATION_OF_PROTEIN_MODIFICATION_PROCESS | 458 | 0.24675 | 0.846027 | 0.884538 | 1 |
| GOBP_REGULATION_OF_SODIUM_ION_TRANSMEMBRANE_TRANSPORT | 57 | 0.28874 | 0.845651 | 0.720201 | 1 |
| GOBP_INTRINSIC_APOPTOTIC_SIGNALING_PATHWAY | 266 | 0.25003 | 0.845302 | 0.823347 | 1 |
| GOBP_REGULATION_OF_CENTRIOLE_REPLICATION | 21 | 0.343464 | 0.845035 | 0.698592 | 1 |
| GOBP_MEMBRANE_REPOLARIZATION_DURING_CARDIAC_MUSCLE_CELL_ACTION_POTENT | 17 | 0.355561 | 0.844408 | 0.678144 | 1 |
| GOBP_LONG_CHAIN_FATTY_ACID_BIOSYNTHETIC_PROCESS | 22 | 0.343113 | 0.844398 | 0.679487 | 1 |
| GOBP_POSITIVE_REGULATION_OF_ORGANELLE_ORGANIZATION | 436 | 0.246731 | 0.844169 | 0.889001 | 1 |
| GOBP_SYNAPTIC_VESICLE_PRIMING | 18 | 0.353499 | 0.843722 | 0.657933 | 1 |
| GOBP_REGULATION_OF_AUTOPHAGY_OF_MITOCHONDRION | 36 | 0.308222 | 0.842144 | 0.695883 | 1 |
| GOBP_TELENCEPHALON_DEVELOPMENT | 213 | 0.25352 | 0.841744 | 0.8 | 1 |
| GOBP_PROSTATE_GLAND_MORPHOGENESIS | 19 | 0.350224 | 0.841623 | 0.681948 | 1 |
| GOBP_NEGATIVE_REGULATION_OF_SMALL_MOLECULE_METABOLIC_PROCESS | 81 | 0.276324 | 0.841353 | 0.742206 | 1 |
| GOBP_REGULATION_OF_MEMBRANE_PERMEABILITY | 64 | 0.283909 | 0.841109 | 0.736397 | 1 |
| GOBP_NUCLEAR_CHROMOSOME_SEGREGATION | 261 | 0.251873 | 0.841102 | 0.836968 | 1 |
| GOBP_ACTIN_POLYMERIZATION_OR_DEPOLYMERIZATION | 177 | 0.253209 | 0.840845 | 0.796356 | 1 |
| GOBP_DEVELOPMENTAL_INDUCTION | 28 | 0.323839 | 0.840761 | 0.707349 | 1 |
| GOBP_POSITIVE_REGULATION_OF_NUCLEOTIDE_METABOLIC_PROCESS | 31 | 0.322249 | 0.840483 | 0.689891 | 1 |
| GOBP_PROTEIN_POLYUBIQUITINATION | 228 | 0.252182 | 0.839951 | 0.831579 | 1 |
| GOBP_NEGATIVE_REGULATION_OF_NEURON_DEATH | 182 | 0.255492 | 0.839619 | 0.812965 | 1 |
| GOBP_FOREBRAIN_DEVELOPMENT | 319 | 0.244536 | 0.839387 | 0.846389 | 1 |
| GOBP_REGULATION_OF_CARDIAC_MUSCLE_CELL_MEMBRANE_REPOLARIZATION | 21 | 0.347348 | 0.839004 | 0.694891 | 1 |
| GOBP_RESPONSE_TO_STEROL_DEPLETION | 18 | 0.353501 | 0.839002 | 0.676343 | 1 |
| GOBP_POSITIVE_REGULATION_OF_CHOLESTEROL_EFFLUX | 21 | 0.341158 | 0.838972 | 0.694329 | 1 |
| GOBP_SECONDARY_ALCOHOL_METABOLIC_PROCESS | 115 | 0.268703 | 0.838913 | 0.789001 | 1 |
| GOBP_PROTEOGLYCAN_METABOLIC_PROCESS | 74 | 0.275891 | 0.838896 | 0.734524 | 1 |
| GOBP_NEGATIVE_REGULATION_OF_AUTOPHAGY | 73 | 0.279691 | 0.838865 | 0.733491 | 1 |
| GOBP_RESPONSE_TO_FATTY_ACID | 49 | 0.295463 | 0.838624 | 0.70801 | 1 |
| GOBP_REGULATION_OF_ENDOTHELIAL_CELL_DIFFERENTIATION | 34 | 0.312413 | 0.838522 | 0.717362 | 1 |
| GOBP_KIDNEY_MORPHOGENESIS | 73 | 0.283497 | 0.838476 | 0.75179 | 1 |
| GOBP_POSITIVE_REGULATION_OF_NEURON_PROJECTION_DEVELOPMENT | 137 | 0.259505 | 0.838459 | 0.793066 | 1 |
| GOBP_CHROMOSOME_CONDENSATION | 40 | 0.305704 | 0.838025 | 0.708995 | 1 |
| GOBP_PHOTOTRANSDUCTION | 21 | 0.346414 | 0.837098 | 0.697872 | 1 |
| GOBP_INORGANIC_ANION_TRANSMEMBRANE_TRANSPORT | 72 | 0.279601 | 0.836237 | 0.728571 | 1 |
| GOBP_REGULATION_OF_PROTEIN_AUTOPHOSPHORYLATION | 39 | 0.305133 | 0.835534 | 0.701665 | 1 |
| GOBP_MICROVILLUS_ORGANIZATION | 28 | 0.326799 | 0.83519 | 0.707085 | 1 |
| GOBP_AMINO_ACID_IMPORT_ACROSS_PLASMA_MEMBRANE | 37 | 0.312191 | 0.834971 | 0.701299 | 1 |
| GOBP_PHAGOSOME_MATURATION | 24 | 0.329591 | 0.834883 | 0.710638 | 1 |
| GOBP_PROTEIN_AUTOUBIQUITINATION | 70 | 0.279393 | 0.834435 | 0.753641 | 1 |
| GOBP_REGULATION_OF_RETINOIC_ACID_RECEPTOR_SIGNALING_PATHWAY | 16 | 0.358301 | 0.833736 | 0.683212 | 1 |
| GOBP_POSITIVE_REGULATION_OF_MUSCLE_TISSUE_DEVELOPMENT | 16 | 0.36153 | 0.832939 | 0.676558 | 1 |
| GOBP_ORGANIC_CYCLIC_COMPOUND_CATABOLIC_PROCESS | 436 | 0.243395 | 0.832863 | 0.901408 | 1 |
| GOBP_AXO_DENDRITIC_TRANSPORT | 76 | 0.274937 | 0.832787 | 0.74821 | 1 |
| GOBP_REGULATION_OF_LONG_TERM_SYNAPTIC_POTENTIATION | 39 | 0.301451 | 0.832463 | 0.73038 | 1 |
| GOBP_SECONDARY_METABOLITE_BIOSYNTHETIC_PROCESS | 17 | 0.359997 | 0.832416 | 0.707951 | 1 |
| GOBP_LAMELLIPODIUM_ORGANIZATION | 81 | 0.275293 | 0.831897 | 0.763807 | 1 |
| GOBP_RESPONSE_TO_IONIZING_RADIATION | 131 | 0.257321 | 0.831477 | 0.801772 | 1 |
| GOBP_GLUCAN_BIOSYNTHETIC_PROCESS | 36 | 0.308327 | 0.831397 | 0.709986 | 1 |
| GOBP_NEGATIVE_REGULATION_OF_CELLULAR_SENESCENCE | 16 | 0.358451 | 0.830883 | 0.700292 | 1 |
| GOBP_POSITIVE_REGULATION_OF_P38MAPK_CASCADE | 25 | 0.325703 | 0.830778 | 0.70765 | 1 |
| GOBP_ENSHEATHMENT_OF_NEURONS | 126 | 0.26113 | 0.830497 | 0.786031 | 1 |
| GOBP_NEGATIVE_REGULATION_OF_PLASMA_MEMBRANE_BOUNDED_CELL_PROJECTION_ | 31 | 0.314061 | 0.830251 | 0.712329 | 1 |
| GOBP_REGULATION_OF_NEUROGENESIS | 315 | 0.245165 | 0.830154 | 0.877572 | 1 |
| GOBP_CELLULAR_RESPONSE_TO_GLUCOSE_STARVATION | 48 | 0.294898 | 0.830097 | 0.743261 | 1 |
| GOBP_REGULATION_OF_NEUROBLAST_PROLIFERATION | 24 | 0.330327 | 0.830018 | 0.697832 | 1 |
| GOBP_NEGATIVE_REGULATION_OF_AXONOGENESIS | 62 | 0.276961 | 0.829847 | 0.744214 | 1 |
| GOBP_GLIAL_CELL_DIFFERENTIATION | 178 | 0.254868 | 0.829388 | 0.833153 | 1 |
| GOBP_SARCOPLASMIC_RETICULUM_CALCIUM_ION_TRANSPORT | 29 | 0.315984 | 0.828896 | 0.707182 | 1 |
| GOBP_ANTEROGRADE_AXONAL_TRANSPORT | 48 | 0.295632 | 0.828609 | 0.741357 | 1 |
| GOBP_POSITIVE_REGULATION_OF_ANTIGEN_RECEPTOR_MEDIATED_SIGNALING_PATHWAY | 18 | 0.351514 | 0.828424 | 0.695522 | 1 |
| GOBP_MICROTUBULE_ORGANIZING_CENTER_LOCALIZATION | 32 | 0.30662 | 0.828336 | 0.717007 | 1 |
| GOBP_TELOMERE_MAINTENANCE_VIA_RECOMBINATION | 15 | 0.361029 | 0.828273 | 0.702381 | 1 |
| GOBP_REGULATION_OF_CALCIUM_ION_DEPENDENT_EXOCYTOSIS | 39 | 0.302439 | 0.828152 | 0.74173 | 1 |
| GOBP_HEMATOPOIETIC_STEM_CELL_PROLIFERATION | 23 | 0.336721 | 0.827457 | 0.704986 | 1 |
| GOBP_RRNA_METHYLATION | 25 | 0.325582 | 0.827394 | 0.719444 | 1 |
| GOBP_CGMP_MEDIATED_SIGNALING | 21 | 0.341517 | 0.827298 | 0.701977 | 1 |
| GOBP_VESICLE_CYTOSKELETAL_TRAFFICKING | 65 | 0.276123 | 0.827236 | 0.754902 | 1 |
| GOBP_NEGATIVE_REGULATION_OF_BIOMINERALIZATION | 22 | 0.331798 | 0.827226 | 0.694084 | 1 |
| GOBP_NEUROMUSCULAR_SYNAPTIC_TRANSMISSION | 20 | 0.342368 | 0.827172 | 0.705202 | 1 |
| GOBP_CELLULAR_AMINO_ACID_CATABOLIC_PROCESS | 90 | 0.270992 | 0.826397 | 0.774488 | 1 |
| GOBP_POSITIVE_REGULATION_OF_HEART_GROWTH | 21 | 0.339737 | 0.826344 | 0.738197 | 1 |
| GOBP_TORC1_SIGNALING | 50 | 0.289337 | 0.826031 | 0.75679 | 1 |
| GOBP_NEGATIVE_REGULATION_OF_CELL_CYCLE | 332 | 0.24411 | 0.825854 | 0.881994 | 1 |
| GOBP_REGULATION_OF_BIOMINERALIZATION | 74 | 0.277259 | 0.825416 | 0.761621 | 1 |
| GOBP_VASCULOGENESIS | 68 | 0.278964 | 0.825354 | 0.750302 | 1 |
| GOBP_REGULATION_OF_ENDOCYTOSIS | 181 | 0.251317 | 0.825222 | 0.832976 | 1 |
| GOBP_NEGATIVE_REGULATION_OF_PROTEIN_SECRETION | 52 | 0.291812 | 0.824379 | 0.765306 | 1 |
| GOBP_REGULATION_OF_CELLULAR_COMPONENT_SIZE | 322 | 0.242331 | 0.823677 | 0.87054 | 1 |
| GOBP_ZINC_ION_TRANSPORT | 23 | 0.332699 | 0.823215 | 0.726519 | 1 |
| GOBP_REGULATION_OF_DENDRITE_DEVELOPMENT | 92 | 0.265257 | 0.823179 | 0.781963 | 1 |
| GOBP_CARDIAC_ATRIUM_MORPHOGENESIS | 22 | 0.331439 | 0.823045 | 0.704708 | 1 |
| GOBP_DETECTION_OF_TEMPERATURE_STIMULUS | 17 | 0.353714 | 0.822171 | 0.734783 | 1 |
| GOBP_RESPONSE_TO_FIBROBLAST_GROWTH_FACTOR | 93 | 0.266405 | 0.821923 | 0.794755 | 1 |
| GOBP_NEURAL_PRECURSOR_CELL_PROLIFERATION | 123 | 0.256054 | 0.821337 | 0.809577 | 1 |
| GOBP_CELLULAR_RESPONSE_TO_EXTRACELLULAR_STIMULUS | 215 | 0.245566 | 0.821241 | 0.873431 | 1 |
| GOBP_REGULATION_OF_NEURON_PROJECTION_DEVELOPMENT | 400 | 0.241045 | 0.82083 | 0.913307 | 1 |
| GOBP_POSITIVE_REGULATION_OF_POTASSIUM_ION_TRANSMEMBRANE_TRANSPORTER_A | 21 | 0.33632 | 0.820804 | 0.721449 | 1 |
| GOBP_EMBRYONIC_DIGIT_MORPHOGENESIS | 46 | 0.285398 | 0.820625 | 0.751908 | 1 |
| GOBP_REGULATION_OF_NUCLEOCYTOPLASMIC_TRANSPORT | 97 | 0.265841 | 0.820521 | 0.794286 | 1 |
| GOBP_POSITIVE_REGULATION_OF_G1_S_TRANSITION_OF_MITOTIC_CELL_CYCLE | 40 | 0.299406 | 0.820448 | 0.732877 | 1 |
| GOBP_NEGATIVE_REGULATION_OF_CYTOKINE_PRODUCTION_INVOLVED_IN_INFLAMMATO | 15 | 0.35644 | 0.819555 | 0.70354 | 1 |
| GOBP_CELLULAR_GLUCAN_METABOLIC_PROCESS | 60 | 0.281621 | 0.819403 | 0.758874 | 1 |
| GOBP_POSITIVE_REGULATION_OF_OXIDOREDUCTASE_ACTIVITY | 45 | 0.294628 | 0.818902 | 0.719899 | 1 |
| GOBP_POSITIVE_REGULATION_OF_ACTIN_NUCLEATION | 16 | 0.356021 | 0.818405 | 0.709302 | 1 |
| GOBP_PURINE_NUCLEOTIDE_CATABOLIC_PROCESS | 41 | 0.296756 | 0.818232 | 0.732051 | 1 |
| GOBP_RESPONSE_TO_LEPTIN | 18 | 0.346471 | 0.81766 | 0.712271 | 1 |
| GOBP_REGULATION_OF_CELL_JUNCTION_ASSEMBLY | 176 | 0.251736 | 0.817634 | 0.821086 | 1 |
| GOBP_ORGANOPHOSPHATE_CATABOLIC_PROCESS | 138 | 0.256501 | 0.817522 | 0.821663 | 1 |
| GOBP_CHLORIDE_TRANSMEMBRANE_TRANSPORT | 50 | 0.292258 | 0.817099 | 0.761538 | 1 |
| GOBP_ASTROCYTE_DIFFERENTIATION | 59 | 0.279323 | 0.816961 | 0.778331 | 1 |
| GOBP_NEGATIVE_REGULATION_OF_CELL_DIVISION | 15 | 0.355364 | 0.816939 | 0.698813 | 1 |
| GOBP_REGULATION_OF_PROTEIN_SERINE_THREONINE_KINASE_ACTIVITY | 325 | 0.239939 | 0.816736 | 0.892638 | 1 |
| GOBP_PROTEIN_IMPORT | 191 | 0.245828 | 0.816181 | 0.862724 | 1 |
| GOBP_PURINE_CONTAINING_COMPOUND_METABOLIC_PROCESS | 350 | 0.23994 | 0.815957 | 0.903749 | 1 |
| GOBP_SYNAPTIC_TRANSMISSION_GLUTAMATERGIC | 80 | 0.267435 | 0.815698 | 0.779463 | 1 |
| GOBP_REGULATION_OF_VASCULOGENESIS | 15 | 0.348648 | 0.813923 | 0.720238 | 1 |
| GOBP_POSITIVE_REGULATION_OF_TRANSFERASE_ACTIVITY | 487 | 0.236557 | 0.812992 | 0.938816 | 1 |
| GOBP_CARTILAGE_DEVELOPMENT | 152 | 0.249603 | 0.812846 | 0.822703 | 1 |
| GOBP_NEGATIVE_REGULATION_OF_NF_KAPPAB_TRANSCRIPTION_FACTOR_ACTIVITY | 75 | 0.271887 | 0.812442 | 0.776596 | 1 |
| GOBP_ALPHA_AMINO_ACID_CATABOLIC_PROCESS | 74 | 0.269636 | 0.811031 | 0.797101 | 1 |
| GOBP_FEMALE_SEX_DIFFERENTIATION | 101 | 0.260062 | 0.81076 | 0.84 | 1 |
| GOBP_NEGATIVE_REGULATION_OF_HISTONE_METHYLATION | 23 | 0.325475 | 0.810701 | 0.721622 | 1 |
| GOBP_NEURAL_RETINA_DEVELOPMENT | 51 | 0.281714 | 0.809923 | 0.747837 | 1 |
| GOBP_REGULATION_OF_FATTY_ACID_BETA_OXIDATION | 16 | 0.349488 | 0.809332 | 0.732437 | 1 |
| GOBP_DETECTION_OF_STIMULUS_INVOLVED_IN_SENSORY_PERCEPTION_OF_PAIN | 22 | 0.330026 | 0.809096 | 0.736842 | 1 |
| GOBP_NEGATIVE_REGULATION_OF_EPITHELIAL_TO_MESENCHYMAL_TRANSITION | 31 | 0.305899 | 0.808926 | 0.752632 | 1 |
| GOBP_NEGATIVE_REGULATION_OF_MUSCLE_CONTRACTION | 21 | 0.328763 | 0.808393 | 0.732955 | 1 |
| GOBP_CELLULAR_AMINO_ACID_METABOLIC_PROCESS | 239 | 0.241803 | 0.807592 | 0.877466 | 1 |
| GOBP_CELLULAR_NITROGEN_COMPOUND_CATABOLIC_PROCESS | 407 | 0.236303 | 0.807183 | 0.93198 | 1 |
| GOBP_RETINA_DEVELOPMENT_IN_CAMERA_TYPE_EYE | 118 | 0.257593 | 0.806772 | 0.830168 | 1 |
| GOBP_POSITIVE_REGULATION_OF_BIOMINERALIZATION | 41 | 0.29095 | 0.806536 | 0.749344 | 1 |
| GOBP_NEUROTRANSMITTER_SECRETION | 130 | 0.253942 | 0.805551 | 0.833149 | 1 |
| GOBP_POSTSYNAPSE_ASSEMBLY | 27 | 0.310509 | 0.805482 | 0.734551 | 1 |
| GOBP_REGULATION_OF_HISTONE_MODIFICATION | 167 | 0.246451 | 0.805317 | 0.861141 | 1 |
| GOBP_NEGATIVE_REGULATION_OF_PROTEIN_CATABOLIC_PROCESS | 112 | 0.254977 | 0.805155 | 0.831461 | 1 |
| GOBP_NEURON_CELL_CELL_ADHESION | 16 | 0.344414 | 0.805035 | 0.711111 | 1 |
| GOBP_THYROID_GLAND_DEVELOPMENT | 15 | 0.351043 | 0.804489 | 0.735336 | 1 |
| GOBP_RNA_EXPORT_FROM_NUCLEUS | 75 | 0.27029 | 0.803759 | 0.78673 | 1 |
| GOBP_INTESTINAL_ABSORPTION | 22 | 0.322873 | 0.803425 | 0.729196 | 1 |
| GOBP_POSITIVE_REGULATION_OF_PHOSPHOLIPASE_ACTIVITY | 37 | 0.296912 | 0.8032 | 0.761097 | 1 |

| | | | | | |
|---|---|---|---|---|---|
| GOBP_RAC_PROTEIN_SIGNAL_TRANSDUCTION | 41 | 0.296937 | 0.802201 | 0.771392 | 1 |
| GOBP_SCHWANN_CELL_DEVELOPMENT | 29 | 0.313154 | 0.801804 | 0.742896 | 1 |
| GOBP_NEGATIVE_REGULATION_OF_BINDING | 137 | 0.249983 | 0.801608 | 0.848214 | 1 |
| GOBP_INOSITOL_LIPID_MEDIATED_SIGNALING | 132 | 0.249608 | 0.801451 | 0.841814 | 1 |
| GOBP_REGULATION_OF_GTP_BINDING | 15 | 0.351816 | 0.801361 | 0.725782 | 1 |
| GOBP_NEUROTRANSMITTER_TRANSPORT | 178 | 0.244933 | 0.801057 | 0.873932 | 1 |
| GOBP_PYRIDINE_NUCLEOTIDE_BIOSYNTHETIC_PROCESS | 21 | 0.325787 | 0.800868 | 0.719298 | 1 |
| GOBP_MITOTIC_DNA_INTEGRITY_CHECKPOINT_SIGNALING | 81 | 0.264102 | 0.800778 | 0.819477 | 1 |
| GOBP_RNA_DECAPPING | 18 | 0.341041 | 0.799958 | 0.758621 | 1 |
| GOBP_NEUROTRANSMITTER_REUPTAKE | 30 | 0.304563 | 0.799203 | 0.764626 | 1 |
| GOBP_P_BODY_ASSEMBLY | 21 | 0.329 | 0.798852 | 0.744681 | 1 |
| GOBP_INSULIN_LIKE_GROWTH_FACTOR_RECEPTOR_SIGNALING_PATHWAY | 27 | 0.307563 | 0.798309 | 0.752011 | 1 |
| GOBP_REGULATION_OF_NUCLEAR_TRANSCRIBED_MRNA_CATABOLIC_PROCESS_DEADEN | 24 | 0.311496 | 0.79766 | 0.755928 | 1 |
| GOBP_PATTERN_RECOGNITION_RECEPTOR_SIGNALING_PATHWAY | 132 | 0.249836 | 0.797555 | 0.843267 | 1 |
| GOBP_REGULATION_OF_EXTENT_OF_CELL_GROWTH | 104 | 0.254278 | 0.797536 | 0.828442 | 1 |
| GOBP_POSITIVE_REGULATION_OF_DNA_REPAIR | 91 | 0.258114 | 0.797426 | 0.82277 | 1 |
| GOBP_REACTIVE_OXYGEN_SPECIES_BIOSYNTHETIC_PROCESS | 43 | 0.285384 | 0.797388 | 0.788036 | 1 |
| GOBP_STEROID_HORMONE_SECRETION | 17 | 0.343786 | 0.79725 | 0.717037 | 1 |
| GOBP_ADAPTATION_OF_SIGNALING_PATHWAY | 19 | 0.330969 | 0.795955 | 0.726744 | 1 |
| GOBP_CELLULAR_RESPONSE_TO_ZINC_ION | 16 | 0.342348 | 0.795801 | 0.731003 | 1 |
| GOBP_REGULATION_OF_STEROL_TRANSPORT | 44 | 0.283776 | 0.795484 | 0.762626 | 1 |
| GOBP_CELL_CYCLE_CHECKPOINT_SIGNALING | 164 | 0.245827 | 0.794582 | 0.871767 | 1 |
| GOBP_ORGANOPHOSPHATE_BIOSYNTHETIC_PROCESS | 491 | 0.229654 | 0.794304 | 0.95674 | 1 |
| GOBP_RESPONSE_TO_AMINO_ACID_STARVATION | 48 | 0.280146 | 0.794163 | 0.775281 | 1 |
| GOBP_ESTABLISHMENT_OR_MAINTENANCE_OF_CELL_POLARITY | 198 | 0.239724 | 0.793581 | 0.872205 | 1 |
| GOBP_EXTRACELLULAR_TRANSPORT | 37 | 0.294471 | 0.793549 | 0.778772 | 1 |
| GOBP_REGULATION_OF_STEM_CELL_PROLIFERATION | 36 | 0.299148 | 0.793534 | 0.779507 | 1 |
| GOBP_RIBONUCLEOTIDE_CATABOLIC_PROCESS | 42 | 0.28562 | 0.793434 | 0.786822 | 1 |
| GOBP_VITAMIN_TRANSPORT | 35 | 0.29249 | 0.793424 | 0.758808 | 1 |
| GOBP_NEGATIVE_REGULATION_OF_NEURON_DIFFERENTIATION | 55 | 0.276315 | 0.793273 | 0.784339 | 1 |
| GOBP_SULFUR_COMPOUND_CATABOLIC_PROCESS | 26 | 0.313055 | 0.793043 | 0.747253 | 1 |
| GOBP_NEGATIVE_REGULATION_OF_EXTRINSIC_APOPTOTIC_SIGNALING_PATHWAY_VIA_DE | 24 | 0.314331 | 0.792926 | 0.753463 | 1 |
| GOBP_ANTIGEN_PROCESSING_AND_PRESENTATION_OF_ENDOGENOUS_PEPTIDE_ANTIGE | 18 | 0.336165 | 0.792835 | 0.724891 | 1 |
| GOBP_RECEPTOR_MEDIATED_ENDOCYTOSIS | 209 | 0.239355 | 0.791995 | 0.885472 | 1 |
| GOBP_PRIMARY_ALCOHOL_METABOLIC_PROCESS | 73 | 0.265323 | 0.791285 | 0.804734 | 1 |
| GOBP_REGULATION_OF_ENDOCRINE_PROCESS | 25 | 0.318495 | 0.790961 | 0.742446 | 1 |
| GOBP_ACTIVIN_RECEPTOR_SIGNALING_PATHWAY | 38 | 0.28792 | 0.790198 | 0.776062 | 1 |
| GOBP_POSITIVE_REGULATION_OF_NERVOUS_SYSTEM_DEVELOPMENT | 235 | 0.236794 | 0.789863 | 0.902618 | 1 |
| GOBP_RESPONSE_TO_AMYLOID_BETA | 39 | 0.289817 | 0.789532 | 0.774026 | 1 |
| GOBP_POSITIVE_REGULATION_OF_WNT_SIGNALING_PATHWAY | 126 | 0.249904 | 0.789421 | 0.859649 | 1 |
| GOBP_ENERGY_RESERVE_METABOLIC_PROCESS | 66 | 0.264071 | 0.789287 | 0.78773 | 1 |
| GOBP_LONG_TERM_MEMORY | 35 | 0.291878 | 0.789266 | 0.784676 | 1 |
| GOBP_REGULATION_OF_MEMBRANE_PROTEIN_ECTODOMAIN_PROTEOLYSIS | 20 | 0.323523 | 0.788624 | 0.738061 | 1 |
| GOBP_REGULATION_OF_STEM_CELL_POPULATION_MAINTENANCE | 29 | 0.297321 | 0.78858 | 0.772423 | 1 |
| GOBP_PHOSPHATIDYLINOSITOL_METABOLIC_PROCESS | 145 | 0.245158 | 0.788287 | 0.868048 | 1 |
| GOBP_REGULATION_OF_CELL_SIZE | 170 | 0.242538 | 0.787534 | 0.875796 | 1 |
| GOBP_INSULIN_RECEPTOR_SIGNALING_PATHWAY | 97 | 0.251703 | 0.787399 | 0.838934 | 1 |
| GOBP_NEGATIVE_REGULATION_OF_LYMPHOCYTE_APOPTOTIC_PROCESS | 26 | 0.306094 | 0.787246 | 0.770805 | 1 |
| GOBP_PHOSPHOLIPID_BIOSYNTHETIC_PROCESS | 234 | 0.233603 | 0.786581 | 0.91155 | 1 |
| GOBP_PROTEIN_POLYMERIZATION | 250 | 0.236076 | 0.78645 | 0.9207 | 1 |
| GOBP_REGULATION_OF_DNA_DAMAGE_RESPONSE_SIGNAL_TRANSDUCTION_BY_P53_CL | 34 | 0.289044 | 0.785875 | 0.784211 | 1 |
| GOBP_NEGATIVE_REGULATION_OF_FIBROBLAST_GROWTH_FACTOR_RECEPTOR_SIGNALIN | 19 | 0.322452 | 0.784287 | 0.75436 | 1 |
| GOBP_NOREPINEPHRINE_TRANSPORT | 16 | 0.332725 | 0.784079 | 0.743551 | 1 |
| GOBP_REGULATION_OF_BONE_MINERALIZATION | 60 | 0.270201 | 0.783514 | 0.811486 | 1 |
| GOBP_POSITIVE_REGULATION_OF_PROTEIN_LOCALIZATION_TO_NUCLEUS | 77 | 0.257324 | 0.783039 | 0.845499 | 1 |
| GOBP_CELLULAR_RESPONSE_TO_GAMMA_RADIATION | 28 | 0.302434 | 0.782554 | 0.772368 | 1 |
| GOBP_STEROID_CATABOLIC_PROCESS | 18 | 0.325706 | 0.782241 | 0.731118 | 1 |
| GOBP_REGULATION_OF_TRANSCRIPTION_REGULATORY_REGION_DNA_BINDING | 43 | 0.27979 | 0.782083 | 0.777349 | 1 |
| GOBP_POSITIVE_REGULATION_OF_CELL_CYCLE_PROCESS | 199 | 0.237551 | 0.781888 | 0.88959 | 1 |
| GOBP_DEVELOPMENTAL_CELL_GROWTH | 203 | 0.236747 | 0.781515 | 0.902413 | 1 |
| GOBP_CENTRAL_NERVOUS_SYSTEM_NEURON_DEVELOPMENT | 72 | 0.264871 | 0.781404 | 0.847418 | 1 |
| GOBP_POSITIVE_REGULATION_OF_CELL_PROJECTION_ORGANIZATION | 310 | 0.228875 | 0.781319 | 0.92718 | 1 |
| GOBP_NEGATIVE_REGULATION_OF_FIBROBLAST_PROLIFERATION | 27 | 0.307596 | 0.781127 | 0.798365 | 1 |
| GOBP_NEGATIVE_REGULATION_OF_PEPTIDYL_LYSINE_ACETYLATION | 18 | 0.326045 | 0.78108 | 0.736048 | 1 |
| GOBP_ESTABLISHMENT_OF_CELL_POLARITY | 130 | 0.244726 | 0.781004 | 0.861419 | 1 |
| GOBP_RESPONSE_TO_NICOTINE | 42 | 0.280783 | 0.780601 | 0.78329 | 1 |
| GOBP_REGULATION_OF_PROTEIN_LOCALIZATION_TO_CELL_PERIPHERY | 112 | 0.2478 | 0.780392 | 0.852478 | 1 |
| GOBP_POSITIVE_REGULATION_OF_AXON_EXTENSION | 38 | 0.288509 | 0.780203 | 0.778667 | 1 |
| GOBP_RIBONUCLEOSIDE_DIPHOSPHATE_METABOLIC_PROCESS | 100 | 0.248459 | 0.780071 | 0.836028 | 1 |
| GOBP_DNA_RECOMBINATION | 283 | 0.231787 | 0.779849 | 0.921509 | 1 |
| GOBP_REGULATION_OF_HISTONE_DEACETYLATION | 39 | 0.284254 | 0.779831 | 0.801825 | 1 |
| GOBP_REGULATION_OF_PROTEIN_POLYUBIQUITINATION | 25 | 0.310403 | 0.779312 | 0.7625 | 1 |
| GOBP_REGULATION_OF_PROTEIN_BINDING | 171 | 0.235448 | 0.779133 | 0.880645 | 1 |
| GOBP_DEOXYRIBONUCLEOSIDE_MONOPHOSPHATE_METABOLIC_PROCESS | 20 | 0.320686 | 0.779087 | 0.763121 | 1 |
| GOBP_REGULATION_OF_PROTEIN_LOCALIZATION_TO_PLASMA_MEMBRANE | 93 | 0.255615 | 0.778894 | 0.843607 | 1 |
| GOBP_HISTONE_MRNA_CATABOLIC_PROCESS | 15 | 0.334955 | 0.778427 | 0.72969 | 1 |
| GOBP_INTRACELLULAR_STEROL_TRANSPORT | 27 | 0.30281 | 0.778016 | 0.808451 | 1 |
| GOBP_REGULATION_OF_BICELLULAR_TIGHT_JUNCTION_ASSEMBLY | 17 | 0.335223 | 0.777892 | 0.7474 | 1 |
| GOBP_REGULATION_OF_CELL_DIVISION | 147 | 0.239126 | 0.777707 | 0.892857 | 1 |
| GOBP_RECEPTOR_CLUSTERING | 53 | 0.274053 | 0.777526 | 0.806818 | 1 |
| GOBP_NEGATIVE_REGULATION_OF_DNA_METABOLIC_PROCESS | 108 | 0.246276 | 0.777209 | 0.863892 | 1 |
| GOBP_AXONEMAL_DYNEIN_COMPLEX_ASSEMBLY | 29 | 0.298488 | 0.776914 | 0.802797 | 1 |
| GOBP_ADENYLATE_CYCLASE_INHIBITING_G_PROTEIN_COUPLED_RECEPTOR_SIGNALING_ | 57 | 0.268379 | 0.776282 | 0.807273 | 1 |
| GOBP_NUCLEOTIDE_SUGAR_BIOSYNTHETIC_PROCESS | 17 | 0.334198 | 0.776254 | 0.774524 | 1 |
| GOBP_NEGATIVE_REGULATION_OF_CELL_CYCLE_PROCESS | 261 | 0.229972 | 0.775805 | 0.93057 | 1 |
| GOBP_POSITIVE_REGULATION_OF_DNA_BINDING | 46 | 0.279003 | 0.775396 | 0.801567 | 1 |
| GOBP_NUCLEOBASE_CONTAINING_SMALL_MOLECULE_METABOLIC_PROCESS | 486 | 0.22387 | 0.774523 | 0.968844 | 1 |
| GOBP_REGULATION_OF_MORPHOGENESIS_OF_AN_EPITHELIUM | 48 | 0.274367 | 0.774382 | 0.79661 | 1 |
| GOBP_REGULATION_OF_PROTEIN_POLYMERIZATION | 181 | 0.23599 | 0.77426 | 0.898396 | 1 |
| GOBP_REGULATION_OF_SYNAPTIC_VESICLE_RECYCLING | 18 | 0.330173 | 0.773888 | 0.775 | 1 |
| GOBP_INOSITOL_PHOSPHATE_MEDIATED_SIGNALING | 47 | 0.276043 | 0.772897 | 0.799218 | 1 |
| GOBP_RESPONSE_TO_NERVE_GROWTH_FACTOR | 45 | 0.277802 | 0.772615 | 0.813452 | 1 |
| GOBP_POSITIVE_REGULATION_OF_TRANSCRIPTION_REGULATORY_REGION_DNA_BINDING | 18 | 0.327049 | 0.772586 | 0.759232 | 1 |
| GOBP_REGULATION_OF_HEART_RATE_BY_CARDIAC_CONDUCTION | 33 | 0.291662 | 0.772088 | 0.81586 | 1 |
| GOBP_CELLULAR_RESPONSE_TO_COPPER_ION | 18 | 0.324198 | 0.770281 | 0.792264 | 1 |
| GOBP_GLYCOPROTEIN_METABOLIC_PROCESS | 336 | 0.225775 | 0.770269 | 0.938588 | 1 |
| GOBP_LUNG_EPITHELIUM_DEVELOPMENT | 25 | 0.305465 | 0.769951 | 0.781719 | 1 |
| GOBP_VASCULAR_ENDOTHELIAL_GROWTH_FACTOR_SIGNALING_PATHWAY | 38 | 0.276935 | 0.76986 | 0.799228 | 1 |
| GOBP_RESPONSE_TO_X_RAY | 30 | 0.295682 | 0.769659 | 0.785034 | 1 |
| GOBP_REGULATION_OF_DNA_RECOMBINATION | 116 | 0.241941 | 0.76865 | 0.857948 | 1 |
| GOBP_NAD_METABOLIC_PROCESS | 25 | 0.302984 | 0.768626 | 0.797721 | 1 |
| GOBP_POSITIVE_REGULATION_OF_FOCAL_ADHESION_ASSEMBLY | 24 | 0.304744 | 0.768268 | 0.780387 | 1 |
| GOBP_PYRIDINE_CONTAINING_COMPOUND_BIOSYNTHETIC_PROCESS | 24 | 0.306548 | 0.767898 | 0.804408 | 1 |
| GOBP_NEURON_DEATH | 306 | 0.22653 | 0.767765 | 0.938081 | 1 |
| GOBP_POSITIVE_REGULATION_OF_CATABOLIC_PROCESS | 464 | 0.223773 | 0.767584 | 0.976884 | 1 |
| GOBP_STEROL_BIOSYNTHETIC_PROCESS | 55 | 0.26675 | 0.767508 | 0.821208 | 1 |
| GOBP_POSITIVE_REGULATION_OF_PROTEIN_SERINE_THREONINE_KINASE_ACTIVITY | 181 | 0.232868 | 0.767203 | 0.932828 | 1 |
| GOBP_INHIBITORY_SYNAPSE_ASSEMBLY | 15 | 0.338085 | 0.766262 | 0.752577 | 1 |
| GOBP_PHOSPHATIDYLINOSITOL_PHOSPHATE_BIOSYNTHETIC_PROCESS | 63 | 0.260453 | 0.766092 | 0.829028 | 1 |
| GOBP_ENERGY_HOMEOSTASIS | 35 | 0.283063 | 0.765946 | 0.806971 | 1 |
| GOBP_HISTONE_H3_K4_TRIMETHYLATION | 21 | 0.314642 | 0.76556 | 0.775281 | 1 |
| GOBP_REGULATION_OF_HEMATOPOIETIC_PROGENITOR_CELL_DIFFERENTIATION | 32 | 0.293427 | 0.765141 | 0.794667 | 1 |
| GOBP_KINETOCHORE_ORGANIZATION | 22 | 0.310597 | 0.764852 | 0.786957 | 1 |
| GOBP_CLATHRIN_COAT_ASSEMBLY | 18 | 0.318531 | 0.764103 | 0.785185 | 1 |
| GOBP_NEGATIVE_REGULATION_OF_ORGANIC_ACID_TRANSPORT | 19 | 0.319807 | 0.763446 | 0.788012 | 1 |
| GOBP_METAL_ION_EXPORT | 41 | 0.27542 | 0.763367 | 0.838422 | 1 |
| GOBP_RECEPTOR_RECYCLING | 31 | 0.287229 | 0.763243 | 0.815069 | 1 |
| GOBP_REGULATION_OF_MICROTUBULE_POLYMERIZATION | 53 | 0.268269 | 0.763094 | 0.821734 | 1 |
| GOBP_POSITIVE_REGULATION_OF_PROTEIN_TARGETING_TO_MEMBRANE | 27 | 0.299175 | 0.762998 | 0.790859 | 1 |
| GOBP_INTEGRATED_STRESS_RESPONSE_SIGNALING | 21 | 0.307555 | 0.762939 | 0.795745 | 1 |
| GOBP_POSITIVE_REGULATION_OF_ORGAN_GROWTH | 27 | 0.296852 | 0.762792 | 0.789826 | 1 |
| GOBP_POST_GOLGI_VESICLE_MEDIATED_TRANSPORT | 104 | 0.242496 | 0.762622 | 0.871707 | 1 |
| GOBP_MESONEPHRIC_TUBULE_MORPHOGENESIS | 49 | 0.266541 | 0.762534 | 0.826873 | 1 |
| GOBP_ORGANELLE_LOCALIZATION | 489 | 0.221438 | 0.761861 | 0.975782 | 1 |
| GOBP_CELLULAR_RESPONSE_TO_VITAMIN | 25 | 0.303588 | 0.761856 | 0.786816 | 1 |
| GOBP_REGULATION_OF_OSTEOCLAST_DIFFERENTIATION | 44 | 0.272765 | 0.761554 | 0.820546 | 1 |
| GOBP_REGULATION_OF_PROTEIN_CONTAINING_COMPLEX_ASSEMBLY | 348 | 0.224966 | 0.761323 | 0.964503 | 1 |
| GOBP_ACID_SECRETION | 34 | 0.286364 | 0.761155 | 0.810041 | 1 |
| GOBP_NEGATIVE_REGULATION_OF_AXON_EXTENSION | 42 | 0.275418 | 0.759912 | 0.816406 | 1 |
| GOBP_REGULATION_OF_SUBSTRATE_ADHESION_DEPENDENT_CELL_SPREADING | 51 | 0.265924 | 0.759827 | 0.843236 | 1 |
| GOBP_MEMBRANE_PROTEIN_ECTODOMAIN_PROTEOLYSIS | 41 | 0.278493 | 0.759769 | 0.820446 | 1 |
| GOBP_NEGATIVE_REGULATION_OF_TUMOR_NECROSIS_FACTOR_MEDIATED_SIGNALING_F | 20 | 0.314448 | 0.759136 | 0.759071 | 1 |
| GOBP_REGULATION_OF_LIPID_KINASE_ACTIVITY | 58 | 0.256739 | 0.759111 | 0.842041 | 1 |
| GOBP_HISTONE_METHYLATION | 148 | 0.235731 | 0.758729 | 0.910596 | 1 |
| GOBP_REGULATION_OF_INSULIN_RECEPTOR_SIGNALING_PATHWAY | 52 | 0.26453 | 0.758353 | 0.847799 | 1 |
| GOBP_MONOCARBOXYLIC_ACID_CATABOLIC_PROCESS | 105 | 0.242005 | 0.757792 | 0.887133 | 1 |
| GOBP_NUCLEOSIDE_MONOPHOSPHATE_METABOLIC_PROCESS | 72 | 0.25546 | 0.757234 | 0.877698 | 1 |
| GOBP_NEGATIVE_REGULATION_OF_STRESS_ACTIVATED_PROTEIN_KINASE_SIGNALING_C | 44 | 0.267429 | 0.757143 | 0.828066 | 1 |
| GOBP_POSITIVE_REGULATION_OF_HISTONE_MODIFICATION | 94 | 0.24552 | 0.756727 | 0.867797 | 1 |
| GOBP_POSITIVE_REGULATION_OF_CELLULAR_PROTEIN_CATABOLIC_PROCESS | 146 | 0.235091 | 0.756507 | 0.92053 | 1 |
| GOBP_SENSORY_PERCEPTION_OF_TEMPERATURE_STIMULUS | 20 | 0.308314 | 0.756487 | 0.791045 | 1 |
| GOBP_NEGATIVE_REGULATION_OF_TRANSLATIONAL_INITIATION | 19 | 0.314221 | 0.756466 | 0.79562 | 1 |
| GOBP_REGULATION_OF_CHOLESTEROL_METABOLIC_PROCESS | 28 | 0.288341 | 0.756068 | 0.80597 | 1 |
| GOBP_REGULATION_OF_AXONOGENESIS | 148 | 0.231934 | 0.754871 | 0.910675 | 1 |
| GOBP_CARDIAC_MUSCLE_CELL_ACTION_POTENTIAL | 65 | 0.252809 | 0.754595 | 0.864407 | 1 |
| GOBP_MUSCLE_CELL_APOPTOTIC_PROCESS | 57 | 0.260965 | 0.754289 | 0.850633 | 1 |
| GOBP_POSITIVE_REGULATION_OF_CANONICAL_WNT_SIGNALING_PATHWAY | 93 | 0.243418 | 0.754095 | 0.88734 | 1 |
| GOBP_POSITIVE_REGULATION_OF_AUTOPHAGY | 127 | 0.233908 | 0.753918 | 0.89769 | 1 |
| GOBP_POSITIVE_REGULATION_OF_PROTEIN_BINDING | 75 | 0.251975 | 0.753633 | 0.863799 | 1 |
| GOBP_REGULATION_OF_UBIQUITIN_DEPENDENT_PROTEIN_CATABOLIC_PROCESS | 154 | 0.23287 | 0.752602 | 0.915418 | 1 |
| GOBP_POSITIVE_REGULATION_OF_PROTEIN_LOCALIZATION_TO_CELL_PERIPHERY | 61 | 0.258475 | 0.752512 | 0.859085 | 1 |
| GOBP_REGULATION_OF_NEURON_MIGRATION | 42 | 0.271748 | 0.752498 | 0.846554 | 1 |
| GOBP_REGULATION_OF_HEART_GROWTH | 47 | 0.269489 | 0.75226 | 0.820802 | 1 |
| GOBP_CELLULAR_MONOVALENT_INORGANIC_CATION_HOMEOSTASIS | 92 | 0.244714 | 0.752016 | 0.863109 | 1 |
| GOBP_SPHINGOLIPID_BIOSYNTHETIC_PROCESS | 96 | 0.240407 | 0.751484 | 0.894495 | 1 |
| GOBP_SIGNALING_RECEPTOR_LIGAND_PRECURSOR_PROCESSING | 26 | 0.294499 | 0.75082 | 0.826516 | 1 |
| GOBP_EMBRYONIC_DIGESTIVE_TRACT_DEVELOPMENT | 21 | 0.30753 | 0.750587 | 0.789773 | 1 |

| | | | | | |
|---|---|---|---|---|---|
| GOBP_REGULATION_OF_PLASMA_LIPOPROTEIN_PARTICLE_LEVELS | 50 | 0.265451 | 0.750383 | 0.842893 | 1 |
| GOBP_PROTEIN_DEPHOSPHORYLATION | 259 | 0.22471 | 0.750053 | 0.948507 | 1 |
| GOBP_POSITIVE_REGULATION_OF_CYCLIN_DEPENDENT_PROTEIN_KINASE_ACTIVITY | 33 | 0.28145 | 0.749564 | 0.830873 | 1 |
| GOBP_PURINERGIC_NUCLEOTIDE_RECEPTOR_SIGNALING_PATHWAY | 20 | 0.316804 | 0.749277 | 0.806867 | 1 |
| GOBP_REGULATION_OF_RYANODINE_SENSITIVE_CALCIUM_RELEASE_CHANNEL_ACTIVITY | 24 | 0.295436 | 0.749269 | 0.810321 | 1 |
| GOBP_HYPEROSMOTIC_RESPONSE | 23 | 0.295928 | 0.747537 | 0.811698 | 1 |
| GOBP_MICROTUBULE_BASED_TRANSPORT | 188 | 0.226733 | 0.747515 | 0.941365 | 1 |
| GOBP_CALCIUM_ION_REGULATED_EXOCYTOSIS | 60 | 0.250625 | 0.747468 | 0.856622 | 1 |
| GOBP_GLYCOPROTEIN_BIOSYNTHETIC_PROCESS | 277 | 0.220508 | 0.746093 | 0.953464 | 1 |
| GOBP_POSITIVE_REGULATION_OF_STEROL_TRANSPORT | 28 | 0.289997 | 0.745631 | 0.843575 | 1 |
| GOBP_GAMMA_AMINOBUTYRIC_ACID_SIGNALING_PATHWAY | 22 | 0.30593 | 0.745228 | 0.815021 | 1 |
| GOBP_SYNAPSE_ASSEMBLY | 160 | 0.229475 | 0.744447 | 0.917559 | 1 |
| GOBP_REGULATION_OF_CYTOSKELETON_ORGANIZATION | 482 | 0.215652 | 0.743895 | 0.98783 | 1 |
| GOBP_POSITIVE_REGULATION_OF_TYPE_I_INTERFERON_PRODUCTION | 50 | 0.262648 | 0.743358 | 0.835459 | 1 |
| GOBP_NEGATIVE_REGULATION_OF_OXIDOREDUCTASE_ACTIVITY | 17 | 0.317541 | 0.743151 | 0.79795 | 1 |
| GOBP_NEGATIVE_REGULATION_OF_REACTIVE_OXYGEN_SPECIES_METABOLIC_PROCESS | 34 | 0.27986 | 0.742242 | 0.824219 | 1 |
| GOBP_REGULATION_OF_PATTERN_RECOGNITION_RECEPTOR_SIGNALING_PATHWAY | 79 | 0.245763 | 0.741605 | 0.880473 | 1 |
| GOBP_METANEPHRIC_EPITHELIUM_DEVELOPMENT | 16 | 0.315068 | 0.741431 | 0.792647 | 1 |
| GOBP_PEPTIDYL_GLUTAMIC_ACID_MODIFICATION | 24 | 0.291999 | 0.741279 | 0.833563 | 1 |
| GOBP_PEPTIDYL_LYSINE_METHYLATION | 133 | 0.231042 | 0.74082 | 0.9163 | 1 |
| GOBP_DEPHOSPHORYLATION | 371 | 0.21778 | 0.740646 | 0.972533 | 1 |
| GOBP_NEUROMUSCULAR_PROCESS_CONTROLLING_BALANCE | 39 | 0.26683 | 0.740447 | 0.841897 | 1 |
| GOBP_POSITIVE_REGULATION_OF_INTRACELLULAR_TRANSPORT | 181 | 0.226654 | 0.740411 | 0.942553 | 1 |
| GOBP_CERAMIDE_METABOLIC_PROCESS | 91 | 0.240296 | 0.739539 | 0.898225 | 1 |
| GOBP_THIOESTER_METABOLIC_PROCESS | 75 | 0.246342 | 0.739378 | 0.877672 | 1 |
| GOBP_REGULATION_OF_CHOLESTEROL_EFFLUX | 29 | 0.282938 | 0.739254 | 0.836927 | 1 |
| GOBP_LIPID_OXIDATION | 96 | 0.238867 | 0.738764 | 0.899207 | 1 |
| GOBP_CEREBRAL_CORTEX_RADIALLY_ORIENTED_CELL_MIGRATION | 30 | 0.283614 | 0.738755 | 0.833555 | 1 |
| GOBP_STEROID_BIOSYNTHETIC_PROCESS | 134 | 0.231459 | 0.73874 | 0.915385 | 1 |
| GOBP_DETERMINATION_OF_ADULT_LIFESPAN | 15 | 0.324378 | 0.738409 | 0.795827 | 1 |
| GOBP_NEGATIVE_REGULATION_OF_AXON_EXTENSION_INVOLVED_IN_AXON_GUIDANCE | 27 | 0.290489 | 0.737652 | 0.802503 | 1 |
| GOBP_PHOSPHATIDYLINOSITOL_BIOSYNTHETIC_PROCESS | 123 | 0.231062 | 0.737608 | 0.928251 | 1 |
| GOBP_CELLULAR_RESPONSE_TO_INSULIN_STIMULUS | 174 | 0.226651 | 0.737596 | 0.944625 | 1 |
| GOBP_DEVELOPMENT_OF_PRIMARY_FEMALE_SEXUAL_CHARACTERISTICS | 89 | 0.237799 | 0.737445 | 0.880233 | 1 |
| GOBP_REGULATION_OF_PROTEIN_DEACETYLATION | 54 | 0.255324 | 0.737299 | 0.85679 | 1 |
| GOBP_POSITIVE_REGULATION_OF_CELL_CYCLE_G2_M_PHASE_TRANSITION | 29 | 0.280919 | 0.737179 | 0.835635 | 1 |
| GOBP_CHROMOSOME_SEGREGATION | 319 | 0.218572 | 0.736734 | 0.968432 | 1 |
| GOBP_NEGATIVE_REGULATION_OF_LIPID_BIOSYNTHETIC_PROCESS | 47 | 0.263958 | 0.736518 | 0.859825 | 1 |
| GOBP_REGULATION_OF_SODIUM_ION_TRANSPORT | 80 | 0.241636 | 0.736402 | 0.887574 | 1 |
| GOBP_REGULATION_OF_RIG_I_SIGNALING_PATHWAY | 17 | 0.306293 | 0.735952 | 0.8125 | 1 |
| GOBP_MEMBRANE_PROTEIN_PROTEOLYSIS | 52 | 0.257063 | 0.735665 | 0.852273 | 1 |
| GOBP_NADP_METABOLIC_PROCESS | 35 | 0.274234 | 0.735548 | 0.849007 | 1 |
| GOBP_HEMATOPOIETIC_PROGENITOR_CELL_DIFFERENTIATION | 102 | 0.23686 | 0.735519 | 0.901136 | 1 |
| GOBP_CHONDROCYTE_DEVELOPMENT | 27 | 0.281792 | 0.735202 | 0.841313 | 1 |
| GOBP_REGULATION_OF_MAP_KINASE_ACTIVITY | 159 | 0.225578 | 0.735081 | 0.941998 | 1 |
| GOBP_TRANSLESION_SYNTHESIS | 24 | 0.293704 | 0.735048 | 0.820937 | 1 |
| GOBP_REGULATION_OF_DNA_METABOLIC_PROCESS | 368 | 0.215831 | 0.734869 | 0.964611 | 1 |
| GOBP_REGULATION_OF_BINDING | 321 | 0.216542 | 0.734656 | 0.973496 | 1 |
| GOBP_REGULATION_OF_CELL_MORPHOGENESIS | 276 | 0.219803 | 0.7345 | 0.972222 | 1 |
| GOBP_ADP_METABOLIC_PROCESS | 78 | 0.244117 | 0.733947 | 0.891355 | 1 |
| GOBP_SISTER_CHROMATID_SEGREGATION | 196 | 0.22088 | 0.733325 | 0.961945 | 1 |
| GOBP_CELLULAR_RESPONSE_TO_OSMOTIC_STRESS | 38 | 0.270523 | 0.732535 | 0.838542 | 1 |
| GOBP_POSITIVE_REGULATION_OF_CELL_SUBSTRATE_JUNCTION_ORGANIZATION | 29 | 0.280387 | 0.732516 | 0.846995 | 1 |
| GOBP_LAMELLIPODIUM_ASSEMBLY | 63 | 0.24955 | 0.732344 | 0.883495 | 1 |
| GOBP_NEGATIVE_REGULATION_OF_PROTEIN_DEPHOSPHORYLATION | 30 | 0.276791 | 0.731379 | 0.835408 | 1 |
| GOBP_FATTY_ACID_CATABOLIC_PROCESS | 92 | 0.234296 | 0.731024 | 0.917538 | 1 |
| GOBP_RESPONSE_TO_THYROID_HORMONE | 23 | 0.297907 | 0.730995 | 0.827035 | 1 |
| GOBP_REGULATION_OF_AEROBIC_RESPIRATION | 30 | 0.278694 | 0.730056 | 0.847594 | 1 |
| GOBP_NEGATIVE_REGULATION_OF_NEURAL_PRECURSOR_CELL_PROLIFERATION | 16 | 0.317276 | 0.72934 | 0.830384 | 1 |
| GOBP_ONE_CARBON_METABOLIC_PROCESS | 37 | 0.267938 | 0.728726 | 0.86193 | 1 |
| GOBP_REGULATION_OF_SUPRAMOLECULAR_FIBER_ORGANIZATION | 341 | 0.213717 | 0.728099 | 0.97665 | 1 |
| GOBP_POSITIVE_REGULATION_OF_PROTEIN_CATABOLIC_PROCESS | 212 | 0.218626 | 0.728045 | 0.963868 | 1 |
| GOBP_RECOMBINATIONAL_REPAIR | 158 | 0.222747 | 0.727716 | 0.93959 | 1 |
| GOBP_FIBROBLAST_MIGRATION | 46 | 0.260801 | 0.727689 | 0.875472 | 1 |
| GOBP_LUNG_MORPHOGENESIS | 39 | 0.267514 | 0.727507 | 0.850129 | 1 |
| GOBP_CELLULAR_RESPONSE_TO_FLUID_SHEAR_STRESS | 19 | 0.305606 | 0.727334 | 0.822511 | 1 |
| GOBP_DIOL_BIOSYNTHETIC_PROCESS | 22 | 0.292652 | 0.726854 | 0.835244 | 1 |
| GOBP_RESPONSE_TO_LEUKEMIA_INHIBITORY_FACTOR | 89 | 0.234014 | 0.726537 | 0.908881 | 1 |
| GOBP_CERAMIDE_BIOSYNTHETIC_PROCESS | 61 | 0.247486 | 0.726412 | 0.889697 | 1 |
| GOBP_MONOSACCHARIDE_METABOLIC_PROCESS | 222 | 0.21818 | 0.726388 | 0.959417 | 1 |
| GOBP_EXPORT_ACROSS_PLASMA_MEMBRANE | 55 | 0.247928 | 0.726316 | 0.897686 | 1 |
| GOBP_REGULATION_OF_PHOSPHOLIPID_METABOLIC_PROCESS | 29 | 0.27668 | 0.725812 | 0.859697 | 1 |
| GOBP_PROTEIN_TARGETING_TO_VACUOLE | 37 | 0.26654 | 0.724927 | 0.855782 | 1 |
| GOBP_NEGATIVE_REGULATION_OF_CELLULAR_RESPONSE_TO_INSULIN_STIMULUS | 32 | 0.27382 | 0.724646 | 0.851296 | 1 |
| GOBP_CELLULAR_RESPONSE_TO_VITAMIN_D | 18 | 0.307013 | 0.724489 | 0.829096 | 1 |
| GOBP_POSITIVE_REGULATION_OF_ERYTHROCYTE_DIFFERENTIATION | 28 | 0.278925 | 0.724263 | 0.838882 | 1 |
| GOBP_POSTREPLICATION_REPAIR | 34 | 0.266427 | 0.723389 | 0.856764 | 1 |
| GOBP_RIBOSE_PHOSPHATE_METABOLIC_PROCESS | 338 | 0.211229 | 0.723007 | 0.978395 | 1 |
| GOBP_RNA_MODIFICATION | 157 | 0.222377 | 0.722593 | 0.947484 | 1 |
| GOBP_NUCLEOBASE_METABOLIC_PROCESS | 37 | 0.263615 | 0.722217 | 0.845946 | 1 |
| GOBP_NEGATIVE_REGULATION_OF_PROTEIN_CONTAINING_COMPLEX_ASSEMBLY | 128 | 0.228302 | 0.721837 | 0.936455 | 1 |
| GOBP_POSITIVE_REGULATION_OF_LAMELLIPODIUM_ASSEMBLY | 24 | 0.287241 | 0.7216 | 0.847107 | 1 |
| GOBP_NEGATIVE_REGULATION_OF_PROTEIN_ACETYLATION | 20 | 0.298485 | 0.721459 | 0.835994 | 1 |
| GOBP_PROTEIN_METHYLATION | 187 | 0.218125 | 0.720933 | 0.964968 | 1 |
| GOBP_POLYAMINE_METABOLIC_PROCESS | 18 | 0.299208 | 0.720296 | 0.85044 | 1 |
| GOBP_REGULATION_OF_CELLULAR_AMINE_METABOLIC_PROCESS | 29 | 0.276104 | 0.720161 | 0.851296 | 1 |
| GOBP_REGULATION_OF_SYNAPTIC_TRANSMISSION_GABAERGIC | 30 | 0.281169 | 0.719746 | 0.867029 | 1 |
| GOBP_REGULATION_OF_OXIDATIVE_STRESS_INDUCED_CELL_DEATH | 56 | 0.247943 | 0.719661 | 0.891139 | 1 |
| GOBP_METANEPHRIC_TUBULE_DEVELOPMENT | 15 | 0.317776 | 0.719586 | 0.818874 | 1 |
| GOBP_NEURON_MATURATION | 40 | 0.258665 | 0.719161 | 0.882277 | 1 |
| GOBP_PERK_MEDIATED_UNFOLDED_PROTEIN_RESPONSE | 17 | 0.30185 | 0.717986 | 0.827011 | 1 |
| GOBP_POSITIVE_REGULATION_OF_STEROID_METABOLIC_PROCESS | 20 | 0.297552 | 0.717783 | 0.835949 | 1 |
| GOBP_PHOSPHOLIPID_DEPHOSPHORYLATION | 41 | 0.258459 | 0.717166 | 0.889594 | 1 |
| GOBP_MEMBRANE_LIPID_BIOSYNTHETIC_PROCESS | 132 | 0.225276 | 0.716672 | 0.946726 | 1 |
| GOBP_REGULATION_OF_CELL_CYCLE_PHASE_TRANSITION | 356 | 0.210472 | 0.715889 | 0.984741 | 1 |
| GOBP_NEGATIVE_REGULATION_OF_CELL_CYCLE_PHASE_TRANSITION | 221 | 0.216006 | 0.715779 | 0.954357 | 1 |
| GOBP_CELLULAR_HORMONE_METABOLIC_PROCESS | 89 | 0.233013 | 0.715757 | 0.922374 | 1 |
| GOBP_CELL_SURFACE_RECEPTOR_SIGNALING_PATHWAY_INVOLVED_IN_CELL_CELL_SIG | 479 | 0.206806 | 0.715449 | 0.993964 | 1 |
| GOBP_REGULATION_OF_SIGNALING_RECEPTOR_ACTIVITY | 127 | 0.226091 | 0.714742 | 0.939732 | 1 |
| GOBP_GLUTAMINE_FAMILY_AMINO_ACID_METABOLIC_PROCESS | 68 | 0.237215 | 0.713833 | 0.918093 | 1 |
| GOBP_REGULATION_OF_MITOTIC_NUCLEAR_DIVISION | 98 | 0.229248 | 0.713184 | 0.938286 | 1 |
| GOBP_POSITIVE_REGULATION_OF_INTERFERON_BETA_PRODUCTION | 34 | 0.261965 | 0.713138 | 0.870249 | 1 |
| GOBP_CELLULAR_METABOLIC_COMPOUND_SALVAGE | 26 | 0.283049 | 0.712111 | 0.858516 | 1 |
| GOBP_AXON_ENSHEATHMENT_IN_CENTRAL_NERVOUS_SYSTEM | 19 | 0.294039 | 0.712016 | 0.842795 | 1 |
| GOBP_NEGATIVE_REGULATION_OF_GTPASE_ACTIVITY | 28 | 0.273047 | 0.711622 | 0.857339 | 1 |
| GOBP_MRNA_CATABOLIC_PROCESS | 218 | 0.216249 | 0.71064 | 0.976017 | 1 |
| GOBP_REGULATION_OF_GLYCOPROTEIN_METABOLIC_PROCESS | 40 | 0.254235 | 0.710016 | 0.884817 | 1 |
| GOBP_MUCOPOLYSACCHARIDE_METABOLIC_PROCESS | 76 | 0.234738 | 0.709565 | 0.916468 | 1 |
| GOBP_REGULATION_OF_NEUROTRANSMITTER_RECEPTOR_ACTIVITY | 50 | 0.24884 | 0.709498 | 0.887484 | 1 |
| GOBP_POSITIVE_REGULATION_OF_EXTRINSIC_APOPTOTIC_SIGNALING_PATHWAY | 42 | 0.253126 | 0.709038 | 0.87808 | 1 |
| GOBP_CORTICAL_CYTOSKELETON_ORGANIZATION | 58 | 0.244066 | 0.708623 | 0.898137 | 1 |
| GOBP_DENDRITE_DEVELOPMENT | 217 | 0.210726 | 0.708412 | 0.966457 | 1 |
| GOBP_INTRACELLULAR_LIPID_TRANSPORT | 40 | 0.257741 | 0.707793 | 0.888214 | 1 |
| GOBP_POSITIVE_REGULATION_OF_AXONOGENESIS | 74 | 0.232455 | 0.707288 | 0.920428 | 1 |
| GOBP_REGULATION_OF_GENERATION_OF_PRECURSOR_METABOLITES_AND_ENERGY | 116 | 0.222717 | 0.706824 | 0.948603 | 1 |
| GOBP_GLYCOSYL_COMPOUND_BIOSYNTHETIC_PROCESS | 19 | 0.294848 | 0.706354 | 0.843705 | 1 |
| GOBP_CELLULAR_RESPONSE_TO_IONIZING_RADIATION | 69 | 0.237603 | 0.70617 | 0.906553 | 1 |
| GOBP_REGULATION_OF_PHOSPHOPROTEIN_PHOSPHATASE_ACTIVITY | 48 | 0.249641 | 0.705344 | 0.892308 | 1 |
| GOBP_POSITIVE_REGULATION_OF_STEROID_BIOSYNTHETIC_PROCESS | 16 | 0.299785 | 0.705172 | 0.847162 | 1 |
| GOBP_HISTONE_H3_K4_METHYLATION | 66 | 0.235428 | 0.704722 | 0.911007 | 1 |
| GOBP_CHOLESTEROL_STORAGE | 15 | 0.313818 | 0.70461 | 0.844118 | 1 |
| GOBP_NEURON_DEATH_IN_RESPONSE_TO_OXIDATIVE_STRESS | 27 | 0.276954 | 0.704302 | 0.872191 | 1 |
| GOBP_TYPE_I_INTERFERON_PRODUCTION | 81 | 0.230031 | 0.703371 | 0.926316 | 1 |
| GOBP_POSITIVE_REGULATION_OF_REACTIVE_OXYGEN_SPECIES_BIOSYNTHETIC_PROCES | 15 | 0.309311 | 0.703063 | 0.852725 | 1 |
| GOBP_MUSCLE_HYPERTROPHY_IN_RESPONSE_TO_STRESS | 20 | 0.29204 | 0.702983 | 0.85036 | 1 |
| GOBP_POSTSYNAPTIC_SPECIALIZATION_ASSEMBLY | 19 | 0.286838 | 0.702252 | 0.86963 | 1 |
| GOBP_REGULATION_OF_DEPHOSPHORYLATION | 110 | 0.223924 | 0.702212 | 0.946712 | 1 |
| GOBP_MEMBRANE_BIOGENESIS | 59 | 0.238283 | 0.70221 | 0.925428 | 1 |
| GOBP_NEGATIVE_REGULATION_OF_ERBB_SIGNALING_PATHWAY | 31 | 0.264555 | 0.70167 | 0.856746 | 1 |
| GOBP_POLYSACCHARIDE_METABOLIC_PROCESS | 89 | 0.225926 | 0.701557 | 0.946449 | 1 |
| GOBP_CELL_DIFFERENTIATION_IN_HINDBRAIN | 20 | 0.290597 | 0.701329 | 0.838983 | 1 |
| GOBP_POSITIVE_REGULATION_OF_CELL_CYCLE_G1_S_PHASE_TRANSITION | 51 | 0.244437 | 0.700776 | 0.892132 | 1 |
| GOBP_REGULATION_OF_DNA_TEMPLATED_TRANSCRIPTION_IN_RESPONSE_TO_STRESS | 49 | 0.247532 | 0.70075 | 0.902937 | 1 |
| GOBP_REGULATION_OF_TOR_SIGNALING | 98 | 0.225091 | 0.70054 | 0.936146 | 1 |
| GOBP_INCLUSION_BODY_ASSEMBLY | 22 | 0.280537 | 0.700359 | 0.864865 | 1 |
| GOBP_FATTY_ACYL_COA_METABOLIC_PROCESS | 33 | 0.264757 | 0.698888 | 0.896414 | 1 |
| GOBP_PLACENTA_BLOOD_VESSEL_DEVELOPMENT | 29 | 0.268108 | 0.698614 | 0.866758 | 1 |
| GOBP_POSITIVE_REGULATION_OF_MITOTIC_CELL_CYCLE_PHASE_TRANSITION | 83 | 0.228805 | 0.698602 | 0.934808 | 1 |
| GOBP_POSITIVE_REGULATION_OF_BINDING | 151 | 0.216722 | 0.696946 | 0.973713 | 1 |
| GOBP_KINETOCHORE_ASSEMBLY | 17 | 0.298677 | 0.696456 | 0.863568 | 1 |
| GOBP_POLYSACCHARIDE_BIOSYNTHETIC_PROCESS | 62 | 0.236951 | 0.695998 | 0.903423 | 1 |
| GOBP_NEGATIVE_REGULATION_OF_CELL_JUNCTION_ASSEMBLY | 27 | 0.270412 | 0.695287 | 0.875691 | 1 |
| GOBP_PURINE_CONTAINING_COMPOUND_TRANSMEMBRANE_TRANSPORT | 19 | 0.293229 | 0.69512 | 0.866097 | 1 |
| GOBP_RECEPTOR_INTERNALIZATION | 97 | 0.223643 | 0.695073 | 0.947674 | 1 |
| GOBP_HISTONE_MODIFICATION | 462 | 0.200403 | 0.694752 | 0.99496 | 1 |
| GOBP_REGULATION_OF_INCLUSION_BODY_ASSEMBLY | 16 | 0.295705 | 0.694343 | 0.87092 | 1 |
| GOBP_REGULATION_OF_HISTONE_H3_K9_METHYLATION | 23 | 0.280192 | 0.693386 | 0.877318 | 1 |
| GOBP_REGULATION_OF_POSTSYNAPTIC_NEUROTRANSMITTER_RECEPTOR_ACTIVITY | 17 | 0.296985 | 0.692839 | 0.874096 | 1 |
| GOBP_NON_CANONICAL_WNT_SIGNALING_PATHWAY | 68 | 0.232439 | 0.69214 | 0.93765 | 1 |
| GOBP_ESTABLISHMENT_OF_PIGMENT_GRANULE_LOCALIZATION | 22 | 0.278585 | 0.692026 | 0.858974 | 1 |
| GOBP_NEURON_RECOGNITION | 45 | 0.248182 | 0.691855 | 0.902089 | 1 |
| GOBP_ENTRAINMENT_OF_CIRCADIAN_CLOCK | 32 | 0.26358 | 0.691552 | 0.900277 | 1 |
| GOBP_CELLULAR_ALDEHYDE_METABOLIC_PROCESS | 54 | 0.239216 | 0.691398 | 0.898876 | 1 |
| GOBP_NUCLEOBASE_CONTAINING_SMALL_MOLECULE_BIOSYNTHETIC_PROCESS | 15 | 0.299418 | 0.690386 | 0.860429 | 1 |
| GOBP_REGULATION_OF_PROTEIN_LOCALIZATION_TO_CELL_SURFACE | 34 | 0.258561 | 0.690093 | 0.896325 | 1 |
| GOBP_LYTIC_VACUOLE_ORGANIZATION | 68 | 0.231279 | 0.689818 | 0.924849 | 1 |
| GOBP_NUCLEOTIDE_TRANSPORT | 33 | 0.252424 | 0.689736 | 0.899604 | 1 |

| | | | | | |
|---|---|---|---|---|---|
| GOBP_POSITIVE_REGULATION_OF_EXCITATORY_POSTSYNAPTIC_POTENTIAL | 24 | 0.276973 | 0.689586 | 0.863329 | 1 |
| GOBP_PURINE_RIBONUCLEOSIDE_METABOLIC_PROCESS | 19 | 0.287627 | 0.68957 | 0.875 | 1 |
| GOBP_CHROMOSOME_SEPARATION | 88 | 0.224853 | 0.689377 | 0.930994 | 1 |
| GOBP_NEGATIVE_REGULATION_OF_TRANSCRIPTION_REGULATORY_REGION_DNA_BINDIN | 17 | 0.293278 | 0.688584 | 0.865471 | 1 |
| GOBP_CARBOHYDRATE_TRANSMEMBRANE_TRANSPORT | 90 | 0.226638 | 0.688582 | 0.943794 | 1 |
| GOBP_NUCLEAR_TRANSPORT | 278 | 0.203444 | 0.688576 | 0.993834 | 1 |
| GOBP_MICROTUBULE_POLYMERIZATION | 81 | 0.223969 | 0.688163 | 0.930994 | 1 |
| GOBP_GOLGI_TO_PLASMA_MEMBRANE_TRANSPORT | 60 | 0.23619 | 0.687092 | 0.934227 | 1 |
| GOBP_NEGATIVE_REGULATION_OF_CELL_AGING | 21 | 0.280637 | 0.686601 | 0.855655 | 1 |
| GOBP_REGULATION_OF_CELLULAR_PROTEIN_CATABOLIC_PROCESS | 236 | 0.205615 | 0.685866 | 0.989594 | 1 |
| GOBP_CRISTAE_FORMATION | 16 | 0.29782 | 0.684861 | 0.878261 | 1 |
| GOBP_CYTOSKELETON_DEPENDENT_INTRACELLULAR_TRANSPORT | 188 | 0.207538 | 0.684713 | 0.985059 | 1 |
| GOBP_ER_NUCLEUS_SIGNALING_PATHWAY | 44 | 0.242401 | 0.684595 | 0.914174 | 1 |
| GOBP_REGULATION_OF_SYNAPTIC_VESICLE_EXOCYTOSIS | 49 | 0.239363 | 0.684579 | 0.922882 | 1 |
| GOBP_NEGATIVE_REGULATION_OF_MICROTUBULE_POLYMERIZATION_OR_DEPOLYMERIZA | 40 | 0.248329 | 0.684339 | 0.890339 | 1 |
| GOBP_GLUTAMINE_FAMILY_AMINO_ACID_CATABOLIC_PROCESS | 23 | 0.272281 | 0.683958 | 0.904432 | 1 |
| GOBP_REGULATION_OF_PROTEIN_CATABOLIC_PROCESS | 359 | 0.200838 | 0.683487 | 0.99495 | 1 |
| GOBP_POSTSYNAPTIC_MEMBRANE_ORGANIZATION | 34 | 0.256988 | 0.682921 | 0.896139 | 1 |
| GOBP_PROTEIN_LOCALIZATION_TO_LYSOSOME | 44 | 0.244838 | 0.681886 | 0.915723 | 1 |
| GOBP_PURINE_NUCLEOTIDE_TRANSPORT | 27 | 0.269493 | 0.681362 | 0.902703 | 1 |
| GOBP_CELLULAR_RESPONSE_TO_ALCOHOL | 73 | 0.224452 | 0.681092 | 0.940048 | 1 |
| GOBP_COPPER_ION_HOMEOSTASIS | 17 | 0.29101 | 0.680947 | 0.857357 | 1 |
| GOBP_MICROVILLUS_ASSEMBLY | 21 | 0.281177 | 0.680928 | 0.878999 | 1 |
| GOBP_CARBOHYDRATE_TRANSPORT | 117 | 0.214101 | 0.680249 | 0.968575 | 1 |
| GOBP_ASYMMETRIC_CELL_DIVISION | 17 | 0.287216 | 0.679683 | 0.868497 | 1 |
| GOBP_POSITIVE_REGULATION_OF_CARTILAGE_DEVELOPMENT | 27 | 0.264841 | 0.678973 | 0.893588 | 1 |
| GOBP_REGULATION_OF_UBIQUITIN_PROTEIN_LIGASE_ACTIVITY | 23 | 0.271352 | 0.678717 | 0.893352 | 1 |
| GOBP_INTERSTRAND_CROSS_LINK_REPAIR | 31 | 0.259147 | 0.67861 | 0.884932 | 1 |
| GOBP_MITOTIC_G2_DNA_DAMAGE_CHECKPOINT_SIGNALING | 35 | 0.254375 | 0.678319 | 0.907162 | 1 |
| GOBP_ENDOMEMBRANE_SYSTEM_ORGANIZATION | 475 | 0.196692 | 0.678223 | 0.998996 | 1 |
| GOBP_GLYCOSYLATION | 226 | 0.202867 | 0.678143 | 0.988566 | 1 |
| GOBP_NEGATIVE_REGULATION_OF_INTRINSIC_APOPTOTIC_SIGNALING_PATHWAY | 92 | 0.217912 | 0.677858 | 0.952711 | 1 |
| GOBP_INNER_EAR_RECEPTOR_CELL_STEREOCILIUM_ORGANIZATION | 27 | 0.266548 | 0.677803 | 0.891667 | 1 |
| GOBP_IMP_METABOLIC_PROCESS | 16 | 0.292298 | 0.677573 | 0.881737 | 1 |
| GOBP_REGULATION_OF_POLYSACCHARIDE_METABOLIC_PROCESS | 37 | 0.250072 | 0.676588 | 0.911227 | 1 |
| GOBP_MESENCHYMAL_CELL_PROLIFERATION | 30 | 0.260721 | 0.67518 | 0.904255 | 1 |
| GOBP_POSITIVE_REGULATION_OF_PROTEIN_POLYMERIZATION | 83 | 0.222084 | 0.674515 | 0.94921 | 1 |
| GOBP_RECEPTOR_METABOLIC_PROCESS | 147 | 0.20646 | 0.674259 | 0.971271 | 1 |
| GOBP_NEGATIVE_REGULATION_OF_MITOTIC_CELL_CYCLE | 201 | 0.204836 | 0.673627 | 0.978836 | 1 |
| GOBP_CRANIAL_NERVE_MORPHOGENESIS | 19 | 0.283613 | 0.673438 | 0.880171 | 1 |
| GOBP_PEPTIDYL_THREONINE_MODIFICATION | 114 | 0.211715 | 0.673362 | 0.972912 | 1 |
| GOBP_ENDOPLASMIC_RETICULUM_ORGANIZATION | 85 | 0.221261 | 0.672154 | 0.96256 | 1 |
| GOBP_PROTEIN_AUTOPHOSPHORYLATION | 206 | 0.205113 | 0.672031 | 0.978858 | 1 |
| GOBP_REGULATION_OF_SKELETAL_MUSCLE_TISSUE_DEVELOPMENT | 17 | 0.287142 | 0.671721 | 0.868538 | 1 |
| GOBP_RESPONSE_TO_STARVATION | 174 | 0.202814 | 0.671215 | 0.986111 | 1 |
| GOBP_RETINA_VASCULATURE_DEVELOPMENT_IN_CAMERA_TYPE_EYE | 17 | 0.284619 | 0.67099 | 0.865356 | 1 |
| GOBP_REGULATION_OF_MULTICELLULAR_ORGANISM_GROWTH | 55 | 0.230965 | 0.670568 | 0.945612 | 1 |
| GOBP_POSITIVE_REGULATION_OF_DNA_METABOLIC_PROCESS | 214 | 0.202943 | 0.670425 | 0.988397 | 1 |
| GOBP_REGULATION_OF_ACTIN_FILAMENT_ORGANIZATION | 243 | 0.20162 | 0.669089 | 0.991684 | 1 |
| GOBP_MONOSACCHARIDE_BIOSYNTHETIC_PROCESS | 72 | 0.222669 | 0.668268 | 0.94887 | 1 |
| GOBP_SCHWANN_CELL_DIFFERENTIATION | 35 | 0.250419 | 0.667167 | 0.920206 | 1 |
| GOBP_NEGATIVE_REGULATION_OF_PEPTIDYL_THREONINE_PHOSPHORYLATION | 15 | 0.299463 | 0.666373 | 0.880878 | 1 |
| GOBP_VOCALIZATION_BEHAVIOR | 15 | 0.294526 | 0.666342 | 0.875912 | 1 |
| GOBP_POSITIVE_REGULATION_OF_CALCIUM_MEDIATED_SIGNALING | 23 | 0.268209 | 0.666121 | 0.895184 | 1 |
| GOBP_POSITIVE_REGULATION_OF_RECEPTOR_MEDIATED_ENDOCYTOSIS | 44 | 0.241207 | 0.665695 | 0.936306 | 1 |
| GOBP_CELL_DEATH_IN_RESPONSE_TO_OXIDATIVE_STRESS | 78 | 0.222019 | 0.665604 | 0.9559 | 1 |
| GOBP_SNARE_COMPLEX_ASSEMBLY | 15 | 0.293218 | 0.665096 | 0.875562 | 1 |
| GOBP_REGULATION_OF_MITOTIC_CELL_CYCLE | 413 | 0.194264 | 0.664595 | 0.998992 | 1 |
| GOBP_ANDROGEN_RECEPTOR_SIGNALING_PATHWAY | 40 | 0.24273 | 0.664434 | 0.921569 | 1 |
| GOBP_POSITIVE_REGULATION_OF_ESTABLISHMENT_OF_PROTEIN_LOCALIZATION_TO_MIT | 34 | 0.251879 | 0.663705 | 0.949219 | 1 |
| GOBP_TRANSMISSION_OF_NERVE_IMPULSE | 53 | 0.230977 | 0.661808 | 0.938726 | 1 |
| GOBP_ENDOTHELIAL_CELL_DEVELOPMENT | 64 | 0.22682 | 0.661517 | 0.942683 | 1 |
| GOBP_TRANSPORT_ALONG_MICROTUBULE | 154 | 0.204407 | 0.661377 | 0.9814 | 1 |
| GOBP_REGULATION_OF_ENDOPLASMIC_RETICULUM_UNFOLDED_PROTEIN_RESPONSE | 28 | 0.25926 | 0.661061 | 0.922131 | 1 |
| GOBP_NEGATIVE_REGULATION_OF_ACTIN_FILAMENT_DEPOLYMERIZATION | 41 | 0.236803 | 0.659998 | 0.939394 | 1 |
| GOBP_POSITIVE_REGULATION_OF_CELLULAR_COMPONENT_BIOGENESIS | 435 | 0.192144 | 0.659209 | 0.996964 | 1 |
| GOBP_NEGATIVE_REGULATION_OF_SUPRAMOLECULAR_FIBER_ORGANIZATION | 149 | 0.205445 | 0.658939 | 0.984632 | 1 |
| GOBP_ACTIVATION_OF_PHOSPHOLIPASE_C_ACTIVITY | 18 | 0.278866 | 0.658647 | 0.88 | 1 |
| GOBP_RNA_PHOSPHODIESTER_BOND_HYDROLYSIS | 133 | 0.206736 | 0.658331 | 0.983498 | 1 |
| GOBP_BIOLOGICAL_PROCESS_INVOLVED_IN_INTERACTION_WITH_SYMBIONT | 68 | 0.220911 | 0.657736 | 0.947052 | 1 |
| GOBP_REGULATION_OF_CARDIAC_MUSCLE_CELL_PROLIFERATION | 29 | 0.247659 | 0.657618 | 0.912983 | 1 |
| GOBP_REGULATION_OF_WATER_LOSS_VIA_SKIN | 20 | 0.271835 | 0.656618 | 0.884298 | 1 |
| GOBP_ANTIGEN_PROCESSING_AND_PRESENTATION_OF_PEPTIDE_ANTIGEN_VIA_MHC_CL | 24 | 0.261745 | 0.656442 | 0.916103 | 1 |
| GOBP_MITOCHONDRIAL_RNA_METABOLIC_PROCESS | 49 | 0.227971 | 0.654949 | 0.939241 | 1 |
| GOBP_PEPTIDYL_LYSINE_MODIFICATION | 366 | 0.191172 | 0.654703 | 0.997972 | 1 |
| GOBP_REGULATION_OF_RESPONSE_TO_OXIDATIVE_STRESS | 78 | 0.216452 | 0.6545 | 0.968254 | 1 |
| GOBP_REGULATION_OF_PROTEIN_TYROSINE_KINASE_ACTIVITY | 76 | 0.213415 | 0.652836 | 0.957041 | 1 |
| GOBP_RNA_CATABOLIC_PROCESS | 261 | 0.193732 | 0.652675 | 0.992776 | 1 |
| GOBP_REGULATION_OF_OSSIFICATION | 94 | 0.212163 | 0.652526 | 0.964286 | 1 |
| GOBP_REGULATION_OF_CELLULAR_RESPONSE_TO_VASCULAR_ENDOTHELIAL_GROWTH_ | 20 | 0.266751 | 0.651962 | 0.897842 | 1 |
| GOBP_PYRIDINE_NUCLEOTIDE_METABOLIC_PROCESS | 25 | 0.258484 | 0.651799 | 0.898212 | 1 |
| GOBP_ESTABLISHMENT_OR_MAINTENANCE_OF_MONOPOLAR_CELL_POLARITY | 26 | 0.252479 | 0.651341 | 0.9125 | 1 |
| GOBP_PROTEIN_REFOLDING | 20 | 0.269259 | 0.649698 | 0.903361 | 1 |
| GOBP_STRESS_GRANULE_ASSEMBLY | 27 | 0.253458 | 0.6486 | 0.901685 | 1 |
| GOBP_SYNAPTIC_VESICLE_EXOCYTOSIS | 95 | 0.209226 | 0.648498 | 0.976959 | 1 |
| GOBP_REGULATION_OF_ERAD_PATHWAY | 20 | 0.268171 | 0.648296 | 0.919162 | 1 |
| GOBP_REGULATION_OF_DOUBLE_STRAND_BREAK_REPAIR | 108 | 0.208228 | 0.648008 | 0.976163 | 1 |
| GOBP_POSITIVE_REGULATION_OF_PROTEIN_DEPHOSPHORYLATION | 40 | 0.234216 | 0.647738 | 0.952866 | 1 |
| GOBP_PROTEIN_DEPOLYMERIZATION | 111 | 0.204681 | 0.647394 | 0.9819 | 1 |
| GOBP_REGULATION_OF_MICROTUBULE_BASED_PROCESS | 231 | 0.193259 | 0.645631 | 0.993776 | 1 |
| GOBP_VESICLE_ORGANIZATION | 278 | 0.190675 | 0.645371 | 0.996914 | 1 |
| GOBP_RNA_DESTABILIZATION | 88 | 0.211211 | 0.644887 | 0.971429 | 1 |
| GOBP_ACTIN_FILAMENT_DEPOLYMERIZATION | 54 | 0.22499 | 0.644824 | 0.94314 | 1 |
| GOBP_REGULATION_OF_CELLULAR_AMIDE_METABOLIC_PROCESS | 414 | 0.18827 | 0.644029 | 0.998991 | 1 |
| GOBP_TOR_SIGNALING | 120 | 0.202567 | 0.643912 | 0.988802 | 1 |
| GOBP_MAINTENANCE_OF_PROTEIN_LOCATION | 86 | 0.20753 | 0.643608 | 0.973472 | 1 |
| GOBP_GLUTAMATE_METABOLIC_PROCESS | 30 | 0.246367 | 0.641445 | 0.919463 | 1 |
| GOBP_TERTIARY_ALCOHOL_METABOLIC_PROCESS | 17 | 0.274101 | 0.640479 | 0.920705 | 1 |
| GOBP_CHAPERONE_MEDIATED_PROTEIN_COMPLEX_ASSEMBLY | 20 | 0.264089 | 0.63981 | 0.900277 | 1 |
| GOBP_REGULATION_OF_GLUCOSE_IMPORT | 46 | 0.226733 | 0.639542 | 0.947861 | 1 |
| GOBP_MODULATION_BY_HOST_OF_VIRAL_GENOME_REPLICATION | 17 | 0.277058 | 0.63852 | 0.894273 | 1 |
| GOBP_RESPONSE_TO_LECTIN | 15 | 0.275569 | 0.638195 | 0.910557 | 1 |
| GOBP_REGULATION_OF_PROTEIN_TARGETING | 73 | 0.213091 | 0.637453 | 0.958824 | 1 |
| GOBP_REGULATION_OF_PROTEIN_DEPOLYMERIZATION | 82 | 0.206902 | 0.636573 | 0.976608 | 1 |
| GOBP_MITOTIC_G2_M_TRANSITION_CHECKPOINT | 48 | 0.225512 | 0.636285 | 0.947631 | 1 |
| GOBP_LOW_DENSITY_LIPOPROTEIN_PARTICLE_CLEARANCE | 20 | 0.265376 | 0.635958 | 0.903319 | 1 |
| GOBP_SECRETORY_GRANULE_ORGANIZATION | 29 | 0.245637 | 0.634954 | 0.945205 | 1 |
| GOBP_EXOCYTIC_PROCESS | 68 | 0.215549 | 0.63486 | 0.968215 | 1 |
| GOBP_POSITIVE_REGULATION_OF_ANIMAL_ORGAN_MORPHOGENESIS | 29 | 0.247333 | 0.634615 | 0.929496 | 1 |
| GOBP_REGULATION_OF_INTRINSIC_APOPTOTIC_SIGNALING_PATHWAY | 151 | 0.193767 | 0.63414 | 0.990239 | 1 |
| GOBP_REGULATION_OF_WNT_SIGNALING_PATHWAY | 295 | 0.18705 | 0.634133 | 0.997953 | 1 |
| GOBP_POSITIVE_REGULATION_OF_INTERFERON_ALPHA_PRODUCTION | 17 | 0.269611 | 0.632357 | 0.906667 | 1 |
| GOBP_POSITIVE_REGULATION_OF_HISTONE_DEACETYLATION | 20 | 0.261391 | 0.632278 | 0.923513 | 1 |
| GOBP_RESPONSE_TO_INSULIN | 217 | 0.188477 | 0.630158 | 0.998941 | 1 |
| GOBP_REGULATION_OF_HISTONE_H3_K4_METHYLATION | 36 | 0.236079 | 0.629522 | 0.950327 | 1 |
| GOBP_ACTIVATION_OF_PROTEIN_KINASE_B_ACTIVITY | 25 | 0.248699 | 0.629316 | 0.928378 | 1 |
| GOBP_POSITIVE_REGULATION_OF_PLASMA_MEMBRANE_BOUNDED_CELL_PROJECTION_A | 93 | 0.20281 | 0.629093 | 0.97783 | 1 |
| GOBP_TOXIN_TRANSPORT | 34 | 0.232154 | 0.628775 | 0.948481 | 1 |
| GOBP_RESPONSE_TO_TESTOSTERONE | 32 | 0.237358 | 0.62801 | 0.948754 | 1 |
| GOBP_REGULATION_OF_AUTOPHAGY | 312 | 0.18507 | 0.627834 | 1 | 1 |
| GOBP_CELL_CYCLE_G1_S_PHASE_TRANSITION | 207 | 0.189298 | 0.627831 | 0.995812 | 1 |
| GOBP_MITOCHONDRIAL_GENOME_MAINTENANCE | 23 | 0.250981 | 0.627674 | 0.930791 | 1 |
| GOBP_MITOTIC_CELL_CYCLE_CHECKPOINT_SIGNALING | 125 | 0.195527 | 0.627514 | 0.993414 | 1 |
| GOBP_POSITIVE_REGULATION_OF_ACTIN_FILAMENT_BUNDLE_ASSEMBLY | 57 | 0.216422 | 0.627196 | 0.978022 | 1 |
| GOBP_INTERLEUKIN_13_PRODUCTION | 15 | 0.275844 | 0.626889 | 0.922039 | 1 |
| GOBP_LIPID_MODIFICATION | 178 | 0.191039 | 0.626475 | 0.996788 | 1 |
| GOBP_NEGATIVE_REGULATION_OF_CELLULAR_PROTEIN_CATABOLIC_PROCESS | 69 | 0.207902 | 0.626294 | 0.976526 | 1 |
| GOBP_RESPONSE_TO_PH | 33 | 0.233176 | 0.625583 | 0.947222 | 1 |
| GOBP_WATER_HOMEOSTASIS | 46 | 0.21996 | 0.625478 | 0.966794 | 1 |
| GOBP_EPITHELIAL_CELL_CELL_ADHESION | 16 | 0.268119 | 0.625289 | 0.91791 | 1 |
| GOBP_PEPTIDYL_LYSINE_DIMETHYLATION | 23 | 0.253981 | 0.62507 | 0.933518 | 1 |
| GOBP_POSITIVE_REGULATION_OF_LONG_TERM_SYNAPTIC_POTENTIATION | 21 | 0.256765 | 0.624956 | 0.92052 | 1 |
| GOBP_REGULATION_OF_CARDIAC_MUSCLE_CONTRACTION_BY_CALCIUM_ION_SIGNALING | 22 | 0.251828 | 0.624874 | 0.928673 | 1 |
| GOBP_METHIONINE_METABOLIC_PROCESS | 15 | 0.272945 | 0.62461 | 0.906627 | 1 |
| GOBP_CELLULAR_RESPONSE_TO_STARVATION | 145 | 0.191381 | 0.624147 | 0.993407 | 1 |
| GOBP_REGULATION_OF_PROTEIN_MODIFICATION_BY_SMALL_PROTEIN_CONJUGATION_OF | 228 | 0.187766 | 0.623883 | 1 | 1 |
| GOBP_POSITIVE_REGULATION_OF_MITOTIC_CELL_CYCLE | 109 | 0.196339 | 0.623309 | 0.985143 | 1 |
| GOBP_POSITIVE_REGULATION_OF_RESPONSE_TO_DNA_DAMAGE_STIMULUS | 124 | 0.194743 | 0.622356 | 0.992188 | 1 |
| GOBP_SULFUR_COMPOUND_METABOLIC_PROCESS | 283 | 0.183765 | 0.62124 | 1 | 1 |
| GOBP_TRICARBOXYLIC_ACID_CYCLE | 30 | 0.23889 | 0.621143 | 0.954787 | 1 |
| GOBP_NEGATIVE_REGULATION_OF_PROTEIN_LOCALIZATION | 169 | 0.188452 | 0.621089 | 0.996774 | 1 |
| GOBP_PROTEIN_PEPTIDYL_PROLYL_ISOMERIZATION | 35 | 0.231573 | 0.620643 | 0.941876 | 1 |
| GOBP_REGULATION_OF_ANDROGEN_RECEPTOR_SIGNALING_PATHWAY | 26 | 0.244171 | 0.620607 | 0.929059 | 1 |
| GOBP_STEROL_TRANSPORT | 89 | 0.198043 | 0.62002 | 0.986175 | 1 |
| GOBP_REGULATION_OF_ESTABLISHMENT_OR_MAINTENANCE_OF_CELL_POLARITY | 25 | 0.245006 | 0.619379 | 0.942513 | 1 |
| GOBP_REGULATION_OF_ESTABLISHMENT_OF_PLANAR_POLARITY | 55 | 0.212387 | 0.618827 | 0.96925 | 1 |
| GOBP_PIGMENT_GRANULE_LOCALIZATION | 24 | 0.248029 | 0.618791 | 0.951857 | 1 |
| GOBP_REGULATION_OF_T_CELL_DIFFERENTIATION_IN_THYMUS | 17 | 0.261816 | 0.618771 | 0.902857 | 1 |
| GOBP_PHOSPHATIDYLCHOLINE_BIOSYNTHETIC_PROCESS | 27 | 0.23962 | 0.618734 | 0.946629 | 1 |
| GOBP_PROTEIN_CATABOLIC_PROCESS_IN_THE_VACUOLE | 22 | 0.251934 | 0.618063 | 0.930935 | 1 |
| GOBP_GOLGI_TO_PLASMA_MEMBRANE_PROTEIN_TRANSPORT | 41 | 0.223549 | 0.617359 | 0.962963 | 1 |
| GOBP_SULFUR_AMINO_ACID_BIOSYNTHETIC_PROCESS | 16 | 0.269311 | 0.617331 | 0.921626 | 1 |
| GOBP_CELL_REDOX_HOMEOSTASIS | 34 | 0.226905 | 0.616427 | 0.955013 | 1 |
| GOBP_ESTABLISHMENT_OF_PROTEIN_LOCALIZATION_TO_VACUOLE | 48 | 0.21917 | 0.616338 | 0.964377 | 1 |
| GOBP_ESTABLISHMENT_OF_RNA_LOCALIZATION | 149 | 0.188346 | 0.61619 | 0.995647 | 1 |
| GOBP_REGULATION_OF_DNA_BINDING | 105 | 0.198486 | 0.61616 | 0.993159 | 1 |
| GOBP_CEREBRAL_CORTEX_CELL_MIGRATION | 41 | 0.219522 | 0.615655 | 0.966146 | 1 |

| | | | | | |
|---|---|---|---|---|---|
| GOBP_PROTEOGLYCAN_BIOSYNTHETIC_PROCESS | 53 | 0.21333 | 0.61369 | 0.969925 | 1 |
| GOBP_NEURONAL_ACTION_POTENTIAL | 27 | 0.238389 | 0.613426 | 0.950798 | 1 |
| GOBP_VESICLE_DOCKING | 53 | 0.214695 | 0.612804 | 0.976893 | 1 |
| GOBP_REGULATION_OF_ATP_BIOSYNTHETIC_PROCESS | 20 | 0.248784 | 0.612765 | 0.936288 | 1 |
| GOBP_LONG_CHAIN_FATTY_ACYL_COA_BIOSYNTHETIC_PROCESS | 16 | 0.26508 | 0.612453 | 0.929535 | 1 |
| GOBP_INNATE_IMMUNE_RESPONSE_ACTIVATING_SIGNAL_TRANSDUCTION | 15 | 0.266859 | 0.611708 | 0.924444 | 1 |
| GOBP_ORGANOPHOSPHATE_ESTER_TRANSPORT | 121 | 0.194953 | 0.610706 | 0.991091 | 1 |
| GOBP_NUCLEOTIDE_SUGAR_METABOLIC_PROCESS | 32 | 0.229021 | 0.61031 | 0.944444 | 1 |
| GOBP_NEGATIVE_REGULATION_OF_TYPE_I_INTERFERON_PRODUCTION | 24 | 0.241903 | 0.609731 | 0.936798 | 1 |
| GOBP_PTERIDINE_CONTAINING_COMPOUND_METABOLIC_PROCESS | 34 | 0.228935 | 0.609055 | 0.967302 | 1 |
| GOBP_CELL_JUNCTION_DISASSEMBLY | 16 | 0.264398 | 0.608723 | 0.931085 | 1 |
| GOBP_TETRAPYRROLE_METABOLIC_PROCESS | 52 | 0.211474 | 0.608621 | 0.972772 | 1 |
| GOBP_LUNG_CELL_DIFFERENTIATION | 15 | 0.264251 | 0.607616 | 0.923729 | 1 |
| GOBP_CELLULAR_RESPONSE_TO_ARSENIC_CONTAINING_SUBSTANCE | 21 | 0.253012 | 0.606778 | 0.93586 | 1 |
| GOBP_POSITIVE_REGULATION_OF_RESPONSE_TO_ENDOPLASMIC_RETICULUM_STRESS | 32 | 0.229867 | 0.606104 | 0.963585 | 1 |
| GOBP_POSITIVE_REGULATION_OF_MONOOXYGENASE_ACTIVITY | 24 | 0.241439 | 0.605452 | 0.943343 | 1 |
| GOBP_SNRNA_TRANSCRIPTION | 17 | 0.255817 | 0.604093 | 0.935438 | 1 |
| GOBP_NEGATIVE_REGULATION_OF_VASCULAR_PERMEABILITY | 17 | 0.255806 | 0.603425 | 0.93578 | 1 |
| GOBP_REGULATION_OF_MITOTIC_CELL_CYCLE_PHASE_TRANSITION | 274 | 0.178377 | 0.601141 | 0.998969 | 1 |
| GOBP_REGULATION_OF_STEROID_BIOSYNTHETIC_PROCESS | 56 | 0.204909 | 0.599727 | 0.979798 | 1 |
| GOBP_POSITIVE_REGULATION_OF_CELL_CYCLE_PHASE_TRANSITION | 103 | 0.190462 | 0.59946 | 0.996552 | 1 |
| GOBP_S_ADENOSYLMETHIONINE_METABOLIC_PROCESS | 15 | 0.256335 | 0.599334 | 0.922018 | 1 |
| GOBP_REGULATION_OF_MITOCHONDRIAL_MEMBRANE_PERMEABILITY | 54 | 0.207902 | 0.597796 | 0.979618 | 1 |
| GOBP_NEURONAL_STEM_CELL_POPULATION_MAINTENANCE | 21 | 0.244112 | 0.597751 | 0.958884 | 1 |
| GOBP_TRANSCRIPTION_BY_RNA_POLYMERASE_III | 53 | 0.208648 | 0.596106 | 0.964242 | 1 |
| GOBP_REGULATION_OF_P38MAPK_CASCADE | 36 | 0.220533 | 0.595674 | 0.963871 | 1 |
| GOBP_VESICLE_BUDDING_FROM_MEMBRANE | 61 | 0.203625 | 0.595662 | 0.978443 | 1 |
| GOBP_REGULATION_OF_MITOCHONDRIAL_OUTER_MEMBRANE_PERMEABILIZATION_INVOL | 20 | 0.247188 | 0.59477 | 0.945559 | 1 |
| GOBP_RESPONSE_TO_GLUCAGON | 15 | 0.265406 | 0.594147 | 0.928135 | 1 |
| GOBP_NEGATIVE_REGULATION_OF_MITOTIC_CELL_CYCLE_PHASE_TRANSITION | 156 | 0.182971 | 0.593889 | 1 | 1 |
| GOBP_REGULATION_OF_RAC_PROTEIN_SIGNAL_TRANSDUCTION | 19 | 0.243568 | 0.590977 | 0.940493 | 1 |
| GOBP_REGULATION_OF_RESPONSE_TO_DNA_DAMAGE_STIMULUS | 239 | 0.176312 | 0.590926 | 1 | 1 |
| GOBP_REGULATION_OF_RECEPTOR_MEDIATED_ENDOCYTOSIS | 93 | 0.19096 | 0.589047 | 0.990847 | 1 |
| GOBP_POSITIVE_REGULATION_OF_GLYCOPROTEIN_METABOLIC_PROCESS | 20 | 0.243295 | 0.588189 | 0.945245 | 1 |
| GOBP_AMP_METABOLIC_PROCESS | 23 | 0.234888 | 0.587861 | 0.966171 | 1 |
| GOBP_REGULATION_OF_PROTEIN_CONTAINING_COMPLEX_DISASSEMBLY | 117 | 0.181625 | 0.58635 | 0.995455 | 1 |
| GOBP_CYTOCHROME_COMPLEX_ASSEMBLY | 36 | 0.218961 | 0.586315 | 0.966844 | 1 |
| GOBP_POSITIVE_REGULATION_OF_INTRINSIC_APOPTOTIC_SIGNALING_PATHWAY | 52 | 0.206864 | 0.585378 | 0.988694 | 1 |
| GOBP_REGULATION_OF_CYCLIN_DEPENDENT_PROTEIN_KINASE_ACTIVITY | 91 | 0.188028 | 0.584309 | 0.992883 | 1 |
| GOBP_REGULATION_OF_CHROMATIN_ASSEMBLY_OR_DISASSEMBLY | 33 | 0.220108 | 0.583377 | 0.975477 | 1 |
| GOBP_AUTOPHAGY_OF_MITOCHONDRION | 86 | 0.189766 | 0.582914 | 0.996479 | 1 |
| GOBP_NEGATIVE_REGULATION_OF_ESTABLISHMENT_OF_PROTEIN_LOCALIZATION | 103 | 0.186545 | 0.582647 | 0.996536 | 1 |
| GOBP_RIBONUCLEOSIDE_MONOPHOSPHATE_BIOSYNTHETIC_PROCESS | 31 | 0.220353 | 0.582034 | 0.963315 | 1 |
| GOBP_REGULATION_OF_ACTIN_NUCLEATION | 31 | 0.21928 | 0.581282 | 0.95765 | 1 |
| GOBP_REGULATION_OF_PROTEIN_DEPHOSPHORYLATION | 79 | 0.192175 | 0.579461 | 0.988221 | 1 |
| GOBP_DEOXYRIBOSE_PHOSPHATE_CATABOLIC_PROCESS | 26 | 0.231939 | 0.579275 | 0.950495 | 1 |
| GOBP_CELLULAR_RESPONSE_TO_FATTY_ACID | 26 | 0.225993 | 0.578505 | 0.974428 | 1 |
| GOBP_REGULATION_OF_PROTEIN_EXPORT_FROM_NUCLEUS | 30 | 0.223673 | 0.578469 | 0.964674 | 1 |
| GOBP_NEGATIVE_REGULATION_OF_CELLULAR_MACROMOLECULE_BIOSYNTHETIC_PROCI | 239 | 0.172104 | 0.578317 | 1 | 1 |
| GOBP_VESICLE_LOCALIZATION | 177 | 0.175624 | 0.57796 | 1 | 1 |
| GOBP_RESPONSE_TO_MITOCHONDRIAL_DEPOLARISATION | 19 | 0.241345 | 0.577657 | 0.95791 | 1 |
| GOBP_GOLGI_ORGANIZATION | 141 | 0.177697 | 0.576876 | 0.998901 | 1 |
| GOBP_CELL_CYCLE_PHASE_TRANSITION | 472 | 0.168232 | 0.576216 | 1 | 1 |
| GOBP_CELLULAR_AMINO_ACID_BIOSYNTHETIC_PROCESS | 68 | 0.193004 | 0.575143 | 0.997579 | 1 |
| GOBP_REGULATION_OF_SIGNAL_TRANSDUCTION_BY_P53_CLASS_MEDIATOR | 97 | 0.185021 | 0.574004 | 0.997732 | 1 |
| GOBP_STEROID_HORMONE_MEDIATED_SIGNALING_PATHWAY | 113 | 0.180838 | 0.573967 | 1 | 1 |
| GOBP_PEPTIDYL_LYSINE_ACETYLATION | 176 | 0.175623 | 0.572946 | 0.997877 | 1 |
| GOBP_DNA_INTEGRITY_CHECKPOINT_SIGNALING | 118 | 0.179289 | 0.572623 | 0.998891 | 1 |
| GOBP_PLASMA_MEMBRANE_REPAIR | 23 | 0.230138 | 0.57106 | 0.954811 | 1 |
| GOBP_STRESS_FIBER_ASSEMBLY | 101 | 0.182272 | 0.57042 | 0.996591 | 1 |
| GOBP_REGULATION_OF_HISTONE_METHYLATION | 74 | 0.189791 | 0.569615 | 0.995221 | 1 |
| GOBP_NUCLEAR_EXPORT | 144 | 0.176031 | 0.56958 | 0.998925 | 1 |
| GOBP_RESPIRATORY_CHAIN_COMPLEX_IV_ASSEMBLY | 26 | 0.227193 | 0.568487 | 0.962861 | 1 |
| GOBP_RNA_MEDIATED_GENE_SILENCING_BY_INHIBITION_OF_TRANSLATION | 16 | 0.246784 | 0.567728 | 0.95421 | 1 |
| GOBP_HISTONE_H3_K9_TRIMETHYLATION | 16 | 0.245682 | 0.566039 | 0.958519 | 1 |
| GOBP_CELLULAR_RESPONSE_TO_UNFOLDED_PROTEIN | 92 | 0.183011 | 0.565256 | 0.995434 | 1 |
| GOBP_MEMBRANE_DOCKING | 79 | 0.186963 | 0.564233 | 0.99051 | 1 |
| GOBP_TELENCEPHALON_GLIAL_CELL_MIGRATION | 22 | 0.229082 | 0.56381 | 0.958101 | 1 |
| GOBP_MITOCHONDRIAL_DNA_METABOLIC_PROCESS | 15 | 0.248268 | 0.563551 | 0.94963 | 1 |
| GOBP_NEGATIVE_REGULATION_OF_PROTEIN_CONTAINING_COMPLEX_DISASSEMBLY | 76 | 0.18552 | 0.563418 | 0.995316 | 1 |
| GOBP_HORMONE_MEDIATED_SIGNALING_PATHWAY | 149 | 0.17543 | 0.563135 | 0.998921 | 1 |
| GOBP_PLASMA_LIPOPROTEIN_PARTICLE_CLEARANCE | 28 | 0.217454 | 0.563037 | 0.965007 | 1 |
| GOBP_CELLULAR_RESPONSE_TO_KETONE | 80 | 0.186886 | 0.562938 | 0.995311 | 1 |
| GOBP_PROTEIN_LOCALIZATION_TO_VACUOLE | 65 | 0.191158 | 0.562663 | 0.992754 | 1 |
| GOBP_MULTICELLULAR_ORGANISM_GROWTH | 124 | 0.176392 | 0.562283 | 0.99775 | 1 |
| GOBP_REGULATION_OF_NOTCH_SIGNALING_PATHWAY | 79 | 0.183494 | 0.558354 | 0.994005 | 1 |
| GOBP_PROTEIN_K48_LINKED_UBIQUITINATION | 66 | 0.187041 | 0.558142 | 0.986618 | 1 |
| GOBP_SYNAPTIC_VESICLE_LOCALIZATION | 51 | 0.198202 | 0.558075 | 0.99005 | 1 |
| GOBP_POSITIVE_REGULATION_OF_PROTEIN_DEACETYLATION | 24 | 0.221701 | 0.556953 | 0.969444 | 1 |
| GOBP_CHOLESTEROL_EFFLUX | 38 | 0.203533 | 0.55589 | 0.982574 | 1 |
| GOBP_ESTABLISHMENT_OF_ENDOTHELIAL_BARRIER | 49 | 0.196189 | 0.555466 | 0.993464 | 1 |
| GOBP_REGULATION_OF_NUCLEASE_ACTIVITY | 17 | 0.238215 | 0.555007 | 0.952035 | 1 |
| GOBP_PROTEIN_LOCALIZATION_TO_NUCLEUS | 269 | 0.164631 | 0.554646 | 1 | 1 |
| GOBP_CELLULAR_CARBOHYDRATE_CATABOLIC_PROCESS | 31 | 0.206833 | 0.554589 | 0.985392 | 1 |
| GOBP_ORGANISM_EMERGENCE_FROM_PROTECTIVE_STRUCTURE | 26 | 0.214932 | 0.553539 | 0.977685 | 1 |
| GOBP_REGULATION_OF_DNA_REPAIR | 148 | 0.170765 | 0.553314 | 0.998926 | 1 |
| GOBP_ACTIN_NUCLEATION | 50 | 0.192862 | 0.553236 | 0.991272 | 1 |
| GOBP_GENERATION_OF_PRECURSOR_METABOLITES_AND_ENERGY | 425 | 0.160232 | 0.550766 | 1 | 1 |
| GOBP_NUCLEOSIDE_PHOSPHATE_BIOSYNTHETIC_PROCESS | 224 | 0.165022 | 0.549268 | 1 | 1 |
| GOBP_PYRIDINE_CONTAINING_COMPOUND_METABOLIC_PROCESS | 32 | 0.208111 | 0.54861 | 0.979487 | 1 |
| GOBP_NEGATIVE_REGULATION_OF_PROTEIN_LOCALIZATION_TO_CELL_PERIPHERY | 24 | 0.223092 | 0.547573 | 0.969274 | 1 |
| GOBP_DNA_BIOSYNTHETIC_PROCESS | 182 | 0.165248 | 0.547519 | 1 | 1 |
| GOBP_TUBULIN_DEACETYLATION | 22 | 0.216008 | 0.546326 | 0.975183 | 1 |
| GOBP_MACROMOLECULE_DEACYLATION | 124 | 0.171798 | 0.545944 | 0.998895 | 1 |
| GOBP_RNA_METHYLATION | 81 | 0.17832 | 0.544639 | 0.9942 | 1 |
| GOBP_DEOXYRIBOSE_PHOSPHATE_METABOLIC_PROCESS | 41 | 0.199908 | 0.542469 | 0.996124 | 1 |
| GOBP_REGULATION_OF_CELL_CYCLE_G2_M_PHASE_TRANSITION | 101 | 0.173279 | 0.539822 | 0.996599 | 1 |
| GOBP_PURINE_CONTAINING_COMPOUND_BIOSYNTHETIC_PROCESS | 178 | 0.163883 | 0.5396 | 1 | 1 |
| GOBP_HISTONE_MRNA_METABOLIC_PROCESS | 24 | 0.209511 | 0.539083 | 0.980337 | 1 |
| GOBP_LYMPHOID_PROGENITOR_CELL_DIFFERENTIATION | 18 | 0.230433 | 0.537871 | 0.967262 | 1 |
| GOBP_ENDOSOME_TRANSPORT_VIA_MULTIVESICULAR_BODY_SORTING_PATHWAY | 31 | 0.202248 | 0.537832 | 0.989056 | 1 |
| GOBP_POSITIVE_REGULATION_OF_SODIUM_ION_TRANSPORT | 33 | 0.202658 | 0.537182 | 0.984993 | 1 |
| GOBP_ESTABLISHMENT_OF_PROTEIN_LOCALIZATION_TO_ORGANELLE | 400 | 0.158694 | 0.537103 | 1 | 1 |
| GOBP_ENERGY_DERIVATION_BY_OXIDATION_OF_ORGANIC_COMPOUNDS | 282 | 0.158331 | 0.537088 | 1 | 1 |
| GOBP_PROTEIN_ADP_RIBOSYLATION | 34 | 0.203649 | 0.536385 | 0.98679 | 1 |
| GOBP_MITOTIC_NUCLEAR_DIVISION | 276 | 0.158936 | 0.53503 | 1 | 1 |
| GOBP_PEPTIDYL_SERINE_DEPHOSPHORYLATION | 19 | 0.221657 | 0.53432 | 0.972543 | 1 |
| GOBP_NEGATIVE_REGULATION_OF_OXIDATIVE_STRESS_INDUCED_CELL_DEATH | 38 | 0.19617 | 0.532989 | 0.993515 | 1 |
| GOBP_ENDOPLASMIC_RETICULUM_UNFOLDED_PROTEIN_RESPONSE | 72 | 0.17698 | 0.532964 | 0.995249 | 1 |
| GOBP_CARBOHYDRATE_PHOSPHORYLATION | 24 | 0.210128 | 0.5322 | 0.972222 | 1 |
| GOBP_VESICLE_MEDIATED_TRANSPORT_TO_THE_PLASMA_MEMBRANE | 136 | 0.164313 | 0.530465 | 1 | 1 |
| GOBP_ORGANELLE_DISASSEMBLY | 119 | 0.16678 | 0.529513 | 1 | 1 |
| GOBP_MICROTUBULE_ORGANIZING_CENTER_ORGANIZATION | 140 | 0.163899 | 0.528297 | 1 | 1 |
| GOBP_PH_REDUCTION | 40 | 0.189954 | 0.525997 | 0.988417 | 1 |
| GOBP_REGULATION_OF_MRNA_METABOLIC_PROCESS | 272 | 0.155991 | 0.524415 | 1 | 1 |
| GOBP_MAINTENANCE_OF_PROTEIN_LOCATION_IN_NUCLEUS | 23 | 0.210893 | 0.524268 | 0.987143 | 1 |
| GOBP_ESTABLISHMENT_OF_PROTEIN_LOCALIZATION_TO_PLASMA_MEMBRANE | 61 | 0.177824 | 0.521404 | 0.996364 | 1 |
| GOBP_DOUBLE_STRAND_BREAK_REPAIR | 262 | 0.15606 | 0.520714 | 1 | 1 |
| GOBP_REGULATION_OF_RESPONSE_TO_ENDOPLASMIC_RETICULUM_STRESS | 75 | 0.172745 | 0.519816 | 0.997587 | 1 |
| GOBP_ACETYL_COA_METABOLIC_PROCESS | 28 | 0.202488 | 0.518367 | 0.98224 | 1 |
| GOBP_PIGMENT_METABOLIC_PROCESS | 68 | 0.176638 | 0.518302 | 0.997599 | 1 |
| GOBP_PROTEIN_ACETYLATION | 199 | 0.156152 | 0.517759 | 1 | 1 |
| GOBP_REGULATION_OF_DNA_REPLICATION | 122 | 0.163323 | 0.517484 | 1 | 1 |
| GOBP_NEGATIVE_REGULATION_OF_ORGANELLE_ORGANIZATION | 322 | 0.152632 | 0.517249 | 1 | 1 |
| GOBP_PHOTOPERIODISM | 29 | 0.198733 | 0.516179 | 0.984064 | 1 |
| GOBP_NUCLEOSIDE_MONOPHOSPHATE_BIOSYNTHETIC_PROCESS | 40 | 0.186682 | 0.516116 | 0.99466 | 1 |
| GOBP_NUCLEOBASE_CONTAINING_COMPOUND_TRANSPORT | 197 | 0.156828 | 0.514944 | 1 | 1 |
| GOBP_REGULATION_OF_INTRINSIC_APOPTOTIC_SIGNALING_PATHWAY_IN_RESPONSE_TO | 37 | 0.187423 | 0.514057 | 0.994716 | 1 |
| GOBP_NEGATIVE_REGULATION_OF_CELL_CYCLE_G2_M_PHASE_TRANSITION | 60 | 0.173105 | 0.512219 | 0.99757 | 1 |
| GOBP_PYRIMIDINE_NUCLEOSIDE_MONOPHOSPHATE_METABOLIC_PROCESS | 19 | 0.214746 | 0.511698 | 0.980226 | 1 |
| GOBP_NEGATIVE_REGULATION_OF_INTRINSIC_APOPTOTIC_SIGNALING_PATHWAY_IN_RES | 29 | 0.196242 | 0.511439 | 0.989404 | 1 |
| GOBP_REGULATION_OF_PROTEIN_LOCALIZATION_TO_NUCLEUS | 120 | 0.160162 | 0.509867 | 1 | 1 |
| GOBP_ESTABLISHMENT_OF_TISSUE_POLARITY | 67 | 0.170237 | 0.503322 | 0.998812 | 1 |
| GOBP_ENDOPLASMIC_RETICULUM_TUBULAR_NETWORK_ORGANIZATION | 18 | 0.20977 | 0.502527 | 0.977578 | 1 |
| GOBP_SPINDLE_ASSEMBLY | 126 | 0.156496 | 0.500329 | 0.998893 | 1 |
| GOBP_N_ACETYLGLUCOSAMINE_METABOLIC_PROCESS | 17 | 0.214371 | 0.500159 | 0.976642 | 1 |
| GOBP_ACTIVATION_OF_GTPASE_ACTIVITY | 96 | 0.16154 | 0.499996 | 0.998845 | 1 |
| GOBP_REGULATION_OF_CHROMATIN_ORGANIZATION | 42 | 0.18079 | 0.498652 | 0.988235 | 1 |
| GOBP_GLUCOSE_IMPORT | 60 | 0.170515 | 0.49819 | 0.998741 | 1 |
| GOBP_RRNA_CATABOLIC_PROCESS | 20 | 0.208617 | 0.496434 | 0.982249 | 1 |
| GOBP_CELLULAR_RESPONSE_TO_TOPOLOGICALLY_INCORRECT_PROTEIN | 111 | 0.157516 | 0.495764 | 1 | 1 |
| GOBP_REGULATION_OF_TRANSLATION_IN_RESPONSE_TO_STRESS | 20 | 0.205781 | 0.495364 | 0.99133 | 1 |
| GOBP_NUCLEOTIDE_TRANSMEMBRANE_TRANSPORT | 18 | 0.207662 | 0.493976 | 0.98557 | 1 |
| GOBP_PEPTIDYL_LYSINE_TRIMETHYLATION | 52 | 0.171933 | 0.492324 | 0.996341 | 1 |
| GOBP_RENAL_ABSORPTION | 17 | 0.2112 | 0.491782 | 0.975573 | 1 |
| GOBP_POSITIVE_REGULATION_OF_CELLULAR_RESPONSE_TO_TRANSFORMING_GROWTH_ | 29 | 0.186572 | 0.490303 | 0.995913 | 1 |
| GOBP_MUSCLE_CELL_CELLULAR_HOMEOSTASIS | 15 | 0.210508 | 0.487765 | 0.99689 | 1 |
| GOBP_REGULATION_OF_DNA_STRAND_ELONGATION | 16 | 0.210894 | 0.487265 | 0.979381 | 1 |
| GOBP_ACETYL_COA_BIOSYNTHETIC_PROCESS | 16 | 0.209559 | 0.486522 | 0.980597 | 1 |
| GOBP_N_TERMINAL_PROTEIN_AMINO_ACID_MODIFICATION | 27 | 0.188855 | 0.486072 | 0.994631 | 1 |
| GOBP_PURINE_NUCLEOSIDE_MONOPHOSPHATE_BIOSYNTHETIC_PROCESS | 21 | 0.197882 | 0.484184 | 0.989011 | 1 |
| GOBP_NEGATIVE_REGULATION_OF_CELLULAR_AMIDE_METABOLIC_PROCESS | 185 | 0.146424 | 0.483335 | 1 | 1 |
| GOBP_ATTACHMENT_OF_SPINDLE_MICROTUBULES_TO_KINETOCHORE | 36 | 0.179748 | 0.481151 | 0.997283 | 1 |
| GOBP_NUCLEOSIDE_BISPHOSPHATE_BIOSYNTHETIC_PROCESS | 54 | 0.166264 | 0.477334 | 1 | 1 |
| GOBP_REGULATION_OF_LIPOPOLYSACCHARIDE_MEDIATED_SIGNALING_PATHWAY | 15 | 0.207323 | 0.475213 | 0.989378 | 1 |
| GOBP_REGULATION_OF_EXOSOMAL_SECRETION | 18 | 0.199505 | 0.473932 | 0.991085 | 1 |
| GOBP_HETEROCHROMATIN_ORGANIZATION | 75 | 0.157258 | 0.472916 | 1 | 1 |

| | | | | | |
|---|---|---|---|---|---|
| GOBP_RESPONSE_TO_ARSENIC_CONTAINING_SUBSTANCE | 32 | 0.178831 | 0.470554 | 1 | 1 |
| GOBP_MITOCHONDRION_ORGANIZATION | 490 | 0.136405 | 0.470519 | 1 | 1 |
| GOBP_MAINTENANCE_OF_PROTEIN_LOCALIZATION_IN_ORGANELLE | 41 | 0.169548 | 0.467392 | 1 | 1 |
| GOBP_NEGATIVE_REGULATION_OF_RESPONSE_TO_ENDOPLASMIC_RETICULUM_STRESS | 40 | 0.17237 | 0.466409 | 0.998679 | 1 |
| GOBP_CELL_CYCLE_G2_M_PHASE_TRANSITION | 147 | 0.145568 | 0.465986 | 1 | 1 |
| GOBP_REGULATION_OF_INTRINSIC_APOPTOTIC_SIGNALING_PATHWAY_IN_RESPONSE_TO | 16 | 0.198507 | 0.46236 | 0.989811 | 1 |
| GOBP_NEGATIVE_REGULATION_OF_RELEASE_OF_CYTOCHROME_C_FROM_MITOCHONDR | 18 | 0.191586 | 0.461115 | 0.992775 | 1 |
| GOBP_REGULATION_OF_NEURON_PROJECTION_ARBORIZATION | 15 | 0.19844 | 0.459674 | 0.990881 | 1 |
| GOBP_NITRIC_OXIDE_SYNTHASE_BIOSYNTHETIC_PROCESS | 15 | 0.198305 | 0.457586 | 0.988423 | 1 |
| GOBP_MITOTIC_CELL_CYCLE_PHASE_TRANSITION | 387 | 0.133595 | 0.456131 | 1 | 1 |
| GOBP_MITOCHONDRIAL_CYTOCHROME_C_OXIDASE_ASSEMBLY | 22 | 0.184038 | 0.453264 | 0.988701 | 1 |
| GOBP_MORPHOGENESIS_OF_A_POLARIZED_EPITHELIUM | 90 | 0.147817 | 0.452913 | 1 | 1 |
| GOBP_ANOIKIS | 29 | 0.174446 | 0.452702 | 0.995913 | 1 |
| GOBP_ESTABLISHMENT_OF_MITOTIC_SPINDLE_LOCALIZATION | 32 | 0.173295 | 0.452231 | 0.998677 | 1 |
| GOBP_INTRINSIC_APOPTOTIC_SIGNALING_PATHWAY_IN_RESPONSE_TO_ENDOPLASMIC_F | 57 | 0.151819 | 0.44969 | 1 | 1 |
| GOBP_ESTABLISHMENT_OF_PROTEIN_LOCALIZATION_TO_MITOCHONDRIAL_MEMBRANE | 29 | 0.170248 | 0.448837 | 1 | 1 |
| GOBP_MITOTIC_INTRA_S_DNA_DAMAGE_CHECKPOINT_SIGNALING | 16 | 0.198796 | 0.446245 | 0.992593 | 1 |
| GOBP_PORPHYRIN_CONTAINING_COMPOUND_METABOLIC_PROCESS | 42 | 0.160541 | 0.445825 | 1 | 1 |
| GOBP_ATP_METABOLIC_PROCESS | 243 | 0.132662 | 0.445614 | 1 | 1 |
| GOBP_HISTONE_PHOSPHORYLATION | 43 | 0.15993 | 0.444706 | 0.997456 | 1 |
| GOBP_THIOESTER_BIOSYNTHETIC_PROCESS | 39 | 0.16219 | 0.443711 | 0.997446 | 1 |
| GOBP_MITOTIC_RECOMBINATION | 25 | 0.174627 | 0.4428 | 0.991655 | 1 |
| GOBP_HISTONE_DEACETYLATION | 92 | 0.142464 | 0.440941 | 0.997732 | 1 |
| GOBP_POSITIVE_REGULATION_OF_PROTEASOMAL_UBIQUITIN_DEPENDENT_PROTEIN_CAT | 88 | 0.142485 | 0.438014 | 1 | 1 |
| GOBP_ENDOSOMAL_TRANSPORT | 223 | 0.132791 | 0.43777 | 1 | 1 |
| GOBP_REGULATION_OF_VIRAL_INDUCED_CYTOPLASMIC_PATTERN_RECOGNITION_RECEF | 20 | 0.181655 | 0.436403 | 0.994398 | 1 |
| GOBP_PROTEIN_CONTAINING_COMPLEX_DISASSEMBLY | 232 | 0.130511 | 0.436218 | 1 | 1 |
| GOBP_REGULATION_OF_RRNA_PROCESSING | 16 | 0.187062 | 0.43576 | 0.995509 | 1 |
| GOBP_PEPTIDYL_ARGININE_METHYLATION | 15 | 0.192391 | 0.434638 | 0.992701 | 1 |
| GOBP_REGULATION_OF_CELL_CYCLE_CHECKPOINT | 32 | 0.163706 | 0.433399 | 0.997315 | 1 |
| GOBP_RNA_LOCALIZATION | 183 | 0.131546 | 0.429907 | 1 | 1 |
| GOBP_COPII_COATED_VESICLE_CARGO_LOADING | 15 | 0.183624 | 0.427076 | 0.995342 | 1 |
| GOBP_SNRNA_METABOLIC_PROCESS | 51 | 0.148259 | 0.425874 | 1 | 1 |
| GOBP_ESTROGEN_METABOLIC_PROCESS | 18 | 0.179353 | 0.425382 | 0.992978 | 1 |
| GOBP_PROTEIN_LOCALIZATION_TO_MICROTUBULE_ORGANIZING_CENTER | 35 | 0.159955 | 0.424966 | 0.997333 | 1 |
| GOBP_CYTOSOLIC_TRANSPORT | 168 | 0.128453 | 0.423806 | 1 | 1 |
| GOBP_CENTROSOME_DUPLICATION | 71 | 0.141707 | 0.423542 | 1 | 1 |
| GOBP_RRNA_MODIFICATION | 36 | 0.156753 | 0.421969 | 0.998718 | 1 |
| GOBP_REGULATION_OF_TRANSCRIPTION_BY_RNA_POLYMERASE_III | 29 | 0.160851 | 0.421139 | 0.998605 | 1 |
| GOBP_NEGATIVE_REGULATION_OF_PROTEASOMAL_PROTEIN_CATABOLIC_PROCESS | 43 | 0.15009 | 0.41799 | 1 | 1 |
| GOBP_RIG_I_SIGNALING_PATHWAY | 25 | 0.165178 | 0.417535 | 0.995902 | 1 |
| GOBP_REGULATION_OF_CELLULAR_RESPONSE_TO_HEAT | 17 | 0.176568 | 0.41263 | 0.997041 | 1 |
| GOBP_POSITIVE_REGULATION_OF_TRANSCRIPTION_ELONGATION_FROM_RNA_POLYMERA | 17 | 0.174658 | 0.412172 | 0.997089 | 1 |
| GOBP_LIPOPROTEIN_CATABOLIC_PROCESS | 16 | 0.178487 | 0.41056 | 0.995702 | 1 |
| GOBP_RESPONSE_TO_TOPOLOGICALLY_INCORRECT_PROTEIN | 149 | 0.124062 | 0.400339 | 1 | 1 |
| GOBP_ENDOSOME_ORGANIZATION | 81 | 0.131596 | 0.399876 | 1 | 1 |
| GOBP_NUCLEAR_TRANSCRIBED_MRNA_CATABOLIC_PROCESS | 118 | 0.123855 | 0.396457 | 1 | 1 |
| GOBP_POSITIVE_REGULATION_OF_TRANSCRIPTION_BY_RNA_POLYMERASE_III | 18 | 0.166476 | 0.396151 | 0.998521 | 1 |
| GOBP_PROTEIN_EXPORT_FROM_NUCLEUS | 56 | 0.136205 | 0.396077 | 1 | 1 |
| GOBP_ORGANELLE_INHERITANCE | 15 | 0.172746 | 0.394344 | 0.99696 | 1 |
| GOBP_MITOCHONDRIAL_MEMBRANE_ORGANIZATION | 109 | 0.123676 | 0.387297 | 1 | 1 |
| GOBP_CELLULAR_RESPIRATION | 212 | 0.114616 | 0.377977 | 1 | 1 |
| GOBP_NEGATIVE_REGULATION_OF_PROTEIN_MODIFICATION_BY_SMALL_PROTEIN_CONJU | 88 | 0.123027 | 0.374804 | 1 | 1 |
| GOBP_REGULATION_OF_CENTROSOME_CYCLE | 49 | 0.133003 | 0.374413 | 1 | 1 |
| GOBP_REGULATION_OF_PROTEASOMAL_UBIQUITIN_DEPENDENT_PROTEIN_CATABOLIC_P | 128 | 0.115105 | 0.368238 | 1 | 1 |
| GOBP_RIBOSOMAL_LARGE_SUBUNIT_ASSEMBLY | 26 | 0.143246 | 0.364114 | 1 | 1 |
| GOBP_FATTY_ACYL_COA_BIOSYNTHETIC_PROCESS | 21 | 0.150521 | 0.363453 | 1 | 1 |
| GOBP_NEGATIVE_REGULATION_OF_SMOOTHENED_SIGNALING_PATHWAY | 31 | 0.129338 | 0.341488 | 1 | 1 |
| GOBP_VESICLE_TARGETING_TO_FROM_OR_WITHIN_GOLGI | 21 | 0.139873 | 0.335893 | 1 | 1 |
| GOBP_AEROBIC_RESPIRATION | 175 | 0.100711 | 0.329915 | 1 | 1 |
| GOBP_MITOPHAGY | 34 | 0.117276 | 0.310774 | 1 | 1 |
| GOBP_PROTEIN_DEMANNOSYLATION | 17 | 0.129462 | 0.306665 | 1 | 1 |
| GOBP_COPII_COATED_VESICLE_BUDDING | 30 | 0.116346 | 0.304191 | 1 | 0.999989 |

**Table S5. GSEA of dysreguated genes in SF3B1$^{K700E}$ ES cells (K700E ES line 2).**

**GSEA Positive** Hematopoietic/Immune pathways are highlighted in red.

| Gene sets | SIZE | ES | NES | NOM p-val | FDR q-val |
|---|---|---|---|---|---|
| GOBP_HOMOLOGOUS_CHROMOSOME_PAIRING_AT_MEIOSIS | 38 | 0.719037 | 1.919143 | 0 | 0.110973 |
| GOBP_NEGATIVE_REGULATION_OF_REPRODUCTIVE_PROCESS | 45 | 0.685072 | 1.870529 | 0 | 0.186739 |
| GOBP_HOMOLOGOUS_CHROMOSOME_SEGREGATION | 47 | 0.680479 | 1.866505 | 0 | 0.137037 |
| GOBP_PIRNA_METABOLIC_PROCESS | 16 | 0.814969 | 1.864808 | 0 | 0.107483 |
| GOBP_ANTERIOR_POSTERIOR_AXIS_SPECIFICATION | 44 | 0.676067 | 1.835025 | 0 | 0.155346 |
| GOBP_SYNAPTONEMAL_COMPLEX_ORGANIZATION | 21 | 0.758895 | 1.826847 | 0.004292 | 0.147622 |
| GOBP_REGULATION_OF_HUMORAL_IMMUNE_RESPONSE | 23 | 0.746391 | 1.821481 | 0 | 0.140826 |
| GOBP_DEFENSE_RESPONSE_TO_FUNGUS | 17 | 0.792196 | 1.817904 | 0.001458 | 0.132137 |
| GOBP_AXIS_SPECIFICATION | 75 | 0.613292 | 1.817835 | 0 | 0.117675 |
| GOBP_GRANULOCYTE_CHEMOTAXIS | 73 | 0.624845 | 1.81625 | 0 | 0.109671 |
| GOBP_POSITIVE_REGULATION_OF_CD4_POSITIVE_ALPHA_BETA_T_CELL_ACTIVATION | 26 | 0.722998 | 1.812528 | 0.001376 | 0.106816 |
| GOBP_REGULATION_OF_TRANSPOSITION | 20 | 0.751154 | 1.807442 | 0 | 0.107248 |
| GOBP_GRANULOCYTE_ACTIVATION | 33 | 0.70331 | 1.805499 | 0.001304 | 0.103797 |
| GOBP_DNA_METHYLATION | 60 | 0.623766 | 1.801708 | 0.001256 | 0.103045 |
| GOBP_NEGATIVE_REGULATION_OF_IMMUNE_RESPONSE | 130 | 0.576921 | 1.798785 | 0 | 0.100927 |
| GOBP_NEGATIVE_REGULATION_OF_IMMUNE_EFFECTOR_PROCESS | 75 | 0.610512 | 1.793187 | 0 | 0.104582 |
| GOBP_SKELETAL_MUSCLE_ADAPTATION | 15 | 0.791539 | 1.786785 | 0.001536 | 0.111249 |
| GOBP_NEGATIVE_REGULATION_OF_CELL_KILLING | 16 | 0.768782 | 1.786252 | 0 | 0.106225 |
| GOBP_REGULATION_OF_T_CELL_MEDIATED_CYTOTOXICITY | 23 | 0.732109 | 1.786085 | 0.002793 | 0.101052 |
| GOBP_POSITIVE_REGULATION_OF_PROTEIN_DEACETYLATION | 26 | 0.714873 | 1.7777 | 0.001353 | 0.109511 |
| GOBP_GRANULOCYTE_MIGRATION | 91 | 0.589684 | 1.777065 | 0 | 0.105572 |
| GOBP_POSITIVE_REGULATION_OF_CD4_POSITIVE_ALPHA_BETA_T_CELL_DIFFERENTIA | 21 | 0.742379 | 1.770839 | 0 | 0.113827 |
| GOBP_NEGATIVE_REGULATION_OF_VIRAL_GENOME_REPLICATION | 45 | 0.64404 | 1.762063 | 0.001259 | 0.125499 |
| GOBP_DNA_METHYLATION_OR_DEMETHYLATION | 80 | 0.590682 | 1.7618 | 0 | 0.120642 |
| GOBP_RESPONSE_TO_GROWTH_HORMONE | 28 | 0.685077 | 1.755852 | 0.004127 | 0.127186 |
| GOBP_TISSUE_REGENERATION | 49 | 0.626691 | 1.754991 | 0.001247 | 0.123665 |
| GOBP_POSITIVE_REGULATION_OF_TOLL_LIKE_RECEPTOR_SIGNALING_PATHWAY | 21 | 0.72582 | 1.751884 | 0.004438 | 0.125506 |
| GOBP_NEGATIVE_REGULATION_OF_MYOTUBE_DIFFERENTIATION | 15 | 0.770273 | 1.74287 | 0.004491 | 0.139694 |
| GOBP_CHROMOSOME_ORGANIZATION_INVOLVED_IN_MEIOTIC_CELL_CYCLE | 55 | 0.629921 | 1.734791 | 0.001259 | 0.153015 |
| GOBP_INTERFERON_GAMMA_PRODUCTION | 66 | 0.603155 | 1.731949 | 0 | 0.15403 |
| GOBP_NEGATIVE_REGULATION_OF_LYMPHOCYTE_MEDIATED_IMMUNITY | 37 | 0.657432 | 1.730425 | 0 | 0.152986 |
| GOBP_POSITIVE_REGULATION_OF_INFLAMMATORY_RESPONSE | 92 | 0.574203 | 1.729003 | 0 | 0.151825 |
| GOBP_NEGATIVE_REGULATION_OF_INTERFERON_GAMMA_PRODUCTION | 24 | 0.702042 | 1.727621 | 0.001377 | 0.150409 |
| GOBP_REGULATION_OF_T_HELPER_1_TYPE_IMMUNE_RESPONSE | 16 | 0.761293 | 1.72737 | 0.001504 | 0.146539 |
| GOBP_PEPTIDE_CROSS_LINKING | 17 | 0.746036 | 1.72737 | 0.00428 | 0.142352 |
| GOBP_BONE_CELL_DEVELOPMENT | 30 | 0.672671 | 1.725656 | 0.002809 | 0.141889 |
| GOBP_MEIOSIS_I_CELL_CYCLE_PROCESS | 99 | 0.569954 | 1.722794 | 0 | 0.143884 |
| GOBP_LEUKOCYTE_CHEMOTAXIS | 148 | 0.545711 | 1.717031 | 0 | 0.152214 |
| GOBP_CELL_PROLIFERATION_INVOLVED_IN_HEART_MORPHOGENESIS | 16 | 0.741881 | 1.715751 | 0.004348 | 0.150951 |
| GOBP_OUTFLOW_TRACT_SEPTUM_MORPHOGENESIS | 22 | 0.71091 | 1.711338 | 0.004348 | 0.156119 |
| GOBP_HUMORAL_IMMUNE_RESPONSE | 102 | 0.559494 | 1.708737 | 0 | 0.158474 |
| GOBP_STRIATED_MUSCLE_CELL_DEVELOPMENT | 53 | 0.607476 | 1.706545 | 0.003713 | 0.159377 |
| GOBP_RESPONSE_TO_FUNGUS | 26 | 0.683977 | 1.706348 | 0.004231 | 0.156037 |
| GOBP_PEPTIDE_ANTIGEN_ASSEMBLY_WITH_MHC_PROTEIN_COMPLEX | 15 | 0.761594 | 1.704424 | 0.007519 | 0.156341 |
| GOBP_REGULATION_OF_IMMUNE_EFFECTOR_PROCESS | 236 | 0.522114 | 1.70184 | 0 | 0.158919 |
| GOBP_POSITIVE_REGULATION_OF_LEUKOCYTE_CELL_CELL_ADHESION | 167 | 0.535515 | 1.701642 | 0 | 0.155766 |
| GOBP_EMBRYONIC_AXIS_SPECIFICATION | 32 | 0.650711 | 1.701319 | 0.004144 | 0.153169 |
| GOBP_NUCLEUS_LOCALIZATION | 24 | 0.681741 | 1.700063 | 0.00142 | 0.152167 |
| GOBP_REGULATION_OF_T_CELL_MEDIATED_IMMUNITY | 56 | 0.598072 | 1.698203 | 0.002541 | 0.152943 |
| GOBP_T_HELPER_1_TYPE_IMMUNE_RESPONSE | 32 | 0.652368 | 1.696551 | 0.004093 | 0.153388 |
| GOBP_POSITIVE_REGULATION_OF_CHEMOKINE_PRODUCTION | 45 | 0.613665 | 1.690638 | 0.002541 | 0.162935 |
| GOBP_REGULATION_OF_LEUKOCYTE_DEGRANULATION | 38 | 0.646196 | 1.689559 | 0.00664 | 0.162355 |
| GOBP_ANTIGEN_PROCESSING_AND_PRESENTATION_OF_EXOGENOUS_PEPTIDE_ANTI | 20 | 0.708698 | 1.68907 | 0.009188 | 0.160095 |
| GOBP_POSITIVE_REGULATION_OF_DEFENSE_RESPONSE | 188 | 0.529379 | 1.689001 | 0 | 0.157204 |
| GOBP_NEGATIVE_REGULATION_OF_T_CELL_MEDIATED_IMMUNITY | 15 | 0.76292 | 1.688823 | 0.00744 | 0.154815 |
| GOBP_TRANSPOSITION | 24 | 0.691936 | 1.688769 | 0.005698 | 0.152104 |
| GOBP_NEGATIVE_REGULATION_OF_LEUKOCYTE_MEDIATED_IMMUNITY | 44 | 0.617938 | 1.687234 | 0.002532 | 0.15272 |
| GOBP_CARDIAC_CELL_DEVELOPMENT | 62 | 0.585411 | 1.685833 | 0.002503 | 0.15331 |
| GOBP_REGULATION_OF_MYOTUBE_DIFFERENTIATION | 34 | 0.651828 | 1.681448 | 0 | 0.159714 |
| GOBP_NEGATIVE_REGULATION_OF_IMMUNE_SYSTEM_PROCESS | 291 | 0.513844 | 1.680816 | 0 | 0.158307 |
| GOBP_RESPONSE_TO_ANGIOTENSIN | 23 | 0.696333 | 1.68 | 0.004317 | 0.157661 |
| GOBP_NUCLEOSIDE_CATABOLIC_PROCESS | 20 | 0.703438 | 1.677172 | 0.004342 | 0.161531 |
| GOBP_T_CELL_MEDIATED_IMMUNITY | 75 | 0.561515 | 1.675695 | 0.001189 | 0.162349 |
| GOBP_MYELOID_LEUKOCYTE_MIGRATION | 146 | 0.534755 | 1.673495 | 0 | 0.164444 |
| GOBP_MUSCLE_CELL_DEVELOPMENT | 142 | 0.530974 | 1.672753 | 0 | 0.163785 |
| GOBP_MYELOID_CELL_DEVELOPMENT | 58 | 0.583568 | 1.669748 | 0.002494 | 0.168375 |
| GOBP_POSITIVE_REGULATION_OF_EMBRYONIC_DEVELOPMENT | 20 | 0.696953 | 1.669306 | 0.005739 | 0.166674 |
| GOBP_ANTIGEN_PROCESSING_AND_PRESENTATION_OF_EXOGENOUS_ANTIGEN | 31 | 0.655321 | 1.669258 | 0.009576 | 0.16431 |
| GOBP_POSITIVE_REGULATION_OF_PATTERN_RECOGNITION_RECEPTOR_SIGNALING_F | 34 | 0.627743 | 1.668763 | 0.008075 | 0.163034 |
| GOBP_T_CELL_MEDIATED_CYTOTOXICITY | 31 | 0.65135 | 1.668494 | 0.005472 | 0.161242 |
| GOBP_REGULATION_OF_OSTEOBLAST_PROLIFERATION | 23 | 0.679077 | 1.666403 | 0.006868 | 0.163171 |
| GOBP_POSITIVE_REGULATION_OF_CELLULAR_CARBOHYDRATE_METABOLIC_PROCES | 45 | 0.613045 | 1.665273 | 0.006494 | 0.163286 |
| GOBP_POSITIVE_REGULATION_OF_IMMUNE_EFFECTOR_PROCESS | 163 | 0.523514 | 1.664849 | 0 | 0.161809 |
| GOBP_MALE_MEIOTIC_NUCLEAR_DIVISION | 36 | 0.628566 | 1.659748 | 0.00823 | 0.170923 |
| GOBP_NEUTROPHIL_ACTIVATION_INVOLVED_IN_IMMUNE_RESPONSE | 17 | 0.722419 | 1.658273 | 0.013453 | 0.172157 |
| GOBP_NEGATIVE_REGULATION_OF_VIRAL_PROCESS | 73 | 0.564621 | 1.656959 | 0.003667 | 0.172826 |
| GOBP_REGULATION_OF_MYELOID_LEUKOCYTE_MEDIATED_IMMUNITY | 46 | 0.599481 | 1.652658 | 0.007692 | 0.179986 |
| GOBP_POSITIVE_REGULATION_OF_CELL_KILLING | 33 | 0.636845 | 1.650552 | 0.006711 | 0.182761 |
| GOBP_OUTFLOW_TRACT_MORPHOGENESIS | 71 | 0.557961 | 1.649997 | 0.002398 | 0.181773 |
| GOBP_REGULATION_OF_ADAPTIVE_IMMUNE_RESPONSE | 132 | 0.530885 | 1.64764 | 0.001105 | 0.185022 |
| GOBP_POSITIVE_REGULATION_OF_CELL_CELL_ADHESION | 201 | 0.50808 | 1.647613 | 0 | 0.182799 |
| GOBP_NEGATIVE_REGULATION_OF_STRIATED_MUSCLE_CELL_DIFFERENTIATION | 32 | 0.637235 | 1.647271 | 0.009091 | 0.181138 |
| GOBP_SKELETAL_MUSCLE_CELL_DIFFERENTIATION | 53 | 0.585898 | 1.644937 | 0.003741 | 0.184195 |
| GOBP_REGULATION_OF_CD4_POSITIVE_ALPHA_BETA_T_CELL_ACTIVATION | 44 | 0.605829 | 1.642247 | 0.005038 | 0.188214 |
| GOBP_POSITIVE_REGULATION_OF_FATTY_ACID_METABOLIC_PROCESS | 27 | 0.654109 | 1.639312 | 0.005495 | 0.19271 |
| GOBP_NEGATIVE_REGULATION_OF_INNATE_IMMUNE_RESPONSE | 55 | 0.584869 | 1.639015 | 0.004963 | 0.191081 |
| GOBP_BONE_MORPHOGENESIS | 81 | 0.556962 | 1.635808 | 0 | 0.196111 |
| GOBP_STRIATED_MUSCLE_CELL_DIFFERENTIATION | 214 | 0.505891 | 1.635487 | 0 | 0.194547 |
| GOBP_POSITIVE_REGULATION_OF_HEMOPOIESIS | 117 | 0.524191 | 1.633231 | 0.00339 | 0.197801 |
| GOBP_NEUTROPHIL_MIGRATION | 73 | 0.557046 | 1.632769 | 0.003525 | 0.196606 |
| GOBP_REGULATION_OF_INFLAMMATORY_RESPONSE | 246 | 0.503556 | 1.63252 | 0 | 0.194935 |
| GOBP_POSITIVE_REGULATION_OF_IMMUNE_RESPONSE | 345 | 0.494765 | 1.631152 | 0 | 0.196325 |
| GOBP_DIGESTION | 73 | 0.559009 | 1.631025 | 0.002407 | 0.194523 |
| GOBP_NEUTROPHIL_CHEMOTAXIS | 57 | 0.583198 | 1.62855 | 0.004988 | 0.198146 |
| GOBP_MACROPHAGE_DIFFERENTIATION | 39 | 0.606309 | 1.625937 | 0.004 | 0.202221 |
| GOBP_ANTIGEN_PROCESSING_AND_PRESENTATION_OF_PEPTIDE_ANTIGEN | 46 | 0.592185 | 1.625216 | 0.006313 | 0.201848 |
| GOBP_ANTIGEN_PROCESSING_AND_PRESENTATION_OF_PEPTIDE_OR_POLYSACCHAR | 25 | 0.654632 | 1.624226 | 0.02069 | 0.202055 |
| GOBP_NEGATIVE_REGULATION_OF_ADAPTIVE_IMMUNE_RESPONSE | 40 | 0.609321 | 1.62381 | 0.007979 | 0.200854 |
| GOBP_ANTIGEN_PROCESSING_AND_PRESENTATION_OF_EXOGENOUS_PEPTIDE_ANTI | 27 | 0.650451 | 1.622672 | 0.009629 | 0.201387 |
| GOBP_MEIOTIC_CHROMOSOME_SEGREGATION | 73 | 0.56203 | 1.620699 | 0.00361 | 0.203999 |
| GOBP_SARCOMERE_ORGANIZATION | 33 | 0.628383 | 1.617808 | 0.011968 | 0.20862 |
| GOBP_POSITIVE_REGULATION_OF_T_CELL_MEDIATED_CYTOTOXICITY | 19 | 0.681473 | 1.612345 | 0.011976 | 0.220134 |
| GOBP_POSITIVE_REGULATION_OF_CELL_ACTIVATION | 247 | 0.490812 | 1.61205 | 0 | 0.218756 |
| GOBP_REGULATION_OF_FATTY_ACID_BIOSYNTHETIC_PROCESS | 33 | 0.611145 | 1.611728 | 0.01897 | 0.21765 |
| GOBP_IMMUNE_EFFECTOR_PROCESS | 393 | 0.481927 | 1.609305 | 0 | 0.222004 |
| GOBP_POSITIVE_REGULATION_OF_LYMPHOCYTE_DIFFERENTIATION | 77 | 0.546232 | 1.608911 | 0.00122 | 0.220936 |
| GOBP_MYOTUBE_DIFFERENTIATION | 94 | 0.537283 | 1.607965 | 0.001139 | 0.221278 |
| GOBP_POSITIVE_REGULATION_OF_CYTOKINE_PRODUCTION_INVOLVED_IN_IMMUNE_F | 49 | 0.569244 | 1.607314 | 0.015484 | 0.221017 |
| GOBP_NUCLEOBASE_CONTAINING_SMALL_MOLECULE_CATABOLIC_PROCESS | 25 | 0.64826 | 1.606059 | 0.016393 | 0.221943 |
| GOBP_POSITIVE_REGULATION_OF_LEUKOCYTE_APOPTOTIC_PROCESS | 18 | 0.687807 | 1.599064 | 0.02071 | 0.238319 |
| GOBP_NEGATIVE_REGULATION_OF_CARDIAC_MUSCLE_TISSUE_GROWTH | 21 | 0.65464 | 1.598479 | 0.018284 | 0.237813 |
| GOBP_GENE_SILENCING_BY_RNA | 118 | 0.517158 | 1.597823 | 0.002247 | 0.23744 |
| GOBP_NEUTROPHIL_MEDIATED_IMMUNITY | 29 | 0.624892 | 1.597225 | 0.017711 | 0.236776 |
| GOBP_ADAPTIVE_IMMUNE_RESPONSE | 274 | 0.487643 | 1.593701 | 0 | 0.24401 |
| GOBP_POSITIVE_REGULATION_OF_LEUKOCYTE_MIGRATION | 98 | 0.528368 | 1.593634 | 0.005708 | 0.242009 |
| GOBP_LEUKOCYTE_MIGRATION | 253 | 0.490326 | 1.593394 | 0 | 0.240623 |
| GOBP_KERATINIZATION | 16 | 0.699692 | 1.593303 | 0.01895 | 0.238854 |
| GOBP_BLASTODERM_SEGMENTATION | 20 | 0.6696 | 1.593031 | 0.022288 | 0.237527 |
| GOBP_ADAPTIVE_IMMUNE_RESPONSE_BASED_ON_SOMATIC_RECOMBINATION_OF_IM | 200 | 0.491563 | 1.592678 | 0 | 0.236313 |
| GOBP_RIBONUCLEOSIDE_CATABOLIC_PROCESS | 15 | 0.709259 | 1.591044 | 0.013235 | 0.238594 |
| GOBP_REGULATION_OF_CD4_POSITIVE_ALPHA_BETA_T_CELL_DIFFERENTIATION | 35 | 0.602495 | 1.590757 | 0.010512 | 0.237375 |
| GOBP_PLASMINOGEN_ACTIVATION | 19 | 0.681543 | 1.589975 | 0.021277 | 0.237589 |
| GOBP_REGULATION_OF_MYELOID_CELL_APOPTOTIC_PROCESS | 21 | 0.663069 | 1.586402 | 0.024251 | 0.244918 |
| GOBP_MYELOID_LEUKOCYTE_ACTIVATION | 155 | 0.503173 | 1.585653 | 0.001087 | 0.244756 |
| GOBP_EPITHELIAL_STRUCTURE_MAINTENANCE | 22 | 0.655013 | 1.584508 | 0.021583 | 0.245984 |
| GOBP_REGULATION_OF_HAIR_FOLLICLE_DEVELOPMENT | 15 | 0.69199 | 1.584305 | 0.017857 | 0.244456 |
| GOBP_TYPE_2_IMMUNE_RESPONSE | 24 | 0.643812 | 1.584177 | 0.019802 | 0.242859 |
| GOBP_ZYMOGEN_ACTIVATION | 48 | 0.57203 | 1.583664 | 0.003788 | 0.242299 |
| GOBP_NEGATIVE_REGULATION_OF_ALPHA_BETA_T_CELL_ACTIVATION | 31 | 0.615519 | 1.582713 | 0.014589 | 0.242648 |
| GOBP_AMELOGENESIS | 18 | 0.675634 | 1.580916 | 0.023669 | 0.245523 |
| GOBP_CELLULAR_RESPONSE_TO_EXOGENOUS_DSRNA | 16 | 0.691177 | 1.580754 | 0.023426 | 0.244111 |
| GOBP_TRIPARTITE_REGIONAL_SUBDIVISION | 16 | 0.697075 | 1.579741 | 0.023669 | 0.244723 |
| GOBP_T_CELL_ACTIVATION_INVOLVED_IN_IMMUNE_RESPONSE | 65 | 0.54371 | 1.579257 | 0.00738 | 0.244187 |
| GOBP_OSTEOBLAST_PROLIFERATION | 28 | 0.634467 | 1.578274 | 0.018056 | 0.245057 |
| GOBP_XENOBIOTIC_CATABOLIC_PROCESS | 18 | 0.702498 | 1.578264 | 0.008427 | 0.243249 |
| GOBP_DETECTION_OF_STIMULUS_INVOLVED_IN_SENSORY_PERCEPTION | 66 | 0.545725 | 1.577137 | 0.008696 | 0.244258 |
| GOBP_POSITIVE_REGULATION_OF_ERBB_SIGNALING_PATHWAY | 29 | 0.615015 | 1.577024 | 0.01752 | 0.242779 |
| GOBP_MONOCYTE_CHEMOTAXIS | 31 | 0.614555 | 1.575772 | 0.026316 | 0.244379 |
| GOBP_OSTEOCLAST_DEVELOPMENT | 15 | 0.698283 | 1.575592 | 0.041481 | 0.243034 |
| GOBP_INOSITOL_PHOSPHATE_BIOSYNTHETIC_PROCESS | 23 | 0.640649 | 1.575351 | 0.02381 | 0.241892 |
| GOBP_NEGATIVE_REGULATION_OF_HEMOPOIESIS | 78 | 0.53129 | 1.574147 | 0.008403 | 0.24342 |
| GOBP_RESPONSE_TO_VITAMIN | 62 | 0.543488 | 1.57363 | 0.008547 | 0.242884 |
| GOBP_TOOTH_MINERALIZATION | 17 | 0.6958 | 1.573564 | 0.022288 | 0.241276 |
| GOBP_REGULATION_OF_T_CELL_ACTIVATION | 233 | 0.48663 | 1.571375 | 0 | 0.245434 |
| GOBP_RESPIRATORY_BURST | 23 | 0.637775 | 1.571256 | 0.019608 | 0.244015 |
| GOBP_POSITIVE_REGULATION_OF_T_CELL_MEDIATED_IMMUNITY | 39 | 0.585147 | 1.569933 | 0.017219 | 0.245831 |
| GOBP_REGULATION_OF_LEUKOCYTE_DIFFERENTIATION | 207 | 0.486448 | 1.569682 | 0.001045 | 0.244596 |
| GOBP_PROTEIN_AUTOPROCESSING | 21 | 0.661207 | 1.569398 | 0.020173 | 0.243726 |

**GSEA Negative** Hematopoietic/Immune pathways are highlighted in green.

| Gene sets | SIZE | ES | NES | NOM p-val | FDR q-val |
|---|---|---|---|---|---|
| GOBP_DNA_REPLICATION_DEPENDENT_CHROMATIN_ORGANIZATION | 23 | -0.754451 | -2.18283 | 0 | 0.018784 |
| GOBP_DNA_REPLICATION_INDEPENDENT_CHROMATIN_ORGANIZATION | 28 | -0.642183 | -1.935541 | 0 | 0.42663 |
| GOBP_NUCLEOSOME_ASSEMBLY | 90 | -0.506826 | -1.91187 | 0 | 0.373467 |
| GOBP_N_GLYCAN_PROCESSING | 19 | -0.653017 | -1.863026 | 0 | 0.458716 |
| GOBP_NUCLEOSOME_ORGANIZATION | 135 | -0.446206 | -1.801062 | 0 | 0.673122 |
| GOBP_PROTEIN_DNA_COMPLEX_ASSEMBLY | 160 | -0.429746 | -1.796319 | 0 | 0.584526 |
| GOBP_TRACHEA_DEVELOPMENT | 15 | -0.655533 | -1.706257 | 0.021472 | 1 |
| GOBP_DNA_PACKAGING | 204 | -0.394261 | -1.696011 | 0 | 1 |
| GOBP_VIRAL_PROTEIN_PROCESSING | 30 | -0.556622 | -1.691215 | 0 | 0.954464 |
| GOBP_CHROMATIN_ASSEMBLY_OR_DISASSEMBLY | 190 | -0.402483 | -1.687932 | 0 | 0.878998 |
| GOBP_CELLULAR_RESPONSE_TO_COPPER_ION | 18 | -0.629611 | -1.659268 | 0.021672 | 0.992585 |
| GOBP_SIALYLATION | 17 | -0.600493 | -1.659158 | 0.012308 | 0.910472 |
| GOBP_L_GLUTAMATE_TRANSMEMBRANE_TRANSPORT | 24 | -0.55781 | -1.631334 | 0.020833 | 1 |
| GOBP_CHROMATIN_REMODELING | 219 | -0.38535 | -1.63086 | 0 | 0.954938 |
| GOBP_PROTEIN_DNA_COMPLEX_SUBUNIT_ORGANIZATION | 206 | -0.386009 | -1.624652 | 0 | 0.92977 |
| GOBP_DNA_STRAND_ELONGATION_INVOLVED_IN_DNA_REPLICATION | 15 | -0.639063 | -1.612846 | 0.020349 | 0.945015 |
| GOBP_POSITIVE_REGULATION_OF_RECEPTOR_INTERNALIZATION | 25 | -0.553887 | -1.601143 | 0.030717 | 0.964624 |
| GOBP_NEGATIVE_REGULATION_OF_CYTOKINE_PRODUCTION_INVOLVE | 15 | -0.615286 | -1.593007 | 0.037288 | 0.959304 |
| GOBP_DNA_CONFORMATION_CHANGE | 300 | -0.34307 | -1.587994 | 0 | 0.943298 |
| GOBP_RESPONSE_TO_SALT | 17 | -0.587927 | -1.573548 | 0.049669 | 0.986614 |
| GOBP_INTRACILIARY_TRANSPORT | 38 | -0.491845 | -1.566263 | 0.03252 | 0.98483 |
| GOBP_NEGATIVE_REGULATION_OF_INTERLEUKIN_1_BETA_PRODUCTI | 17 | -0.582668 | -1.562624 | 0.050296 | 0.96465 |
| GOBP_DNA_UNWINDING_INVOLVED_IN_DNA_REPLICATION | 22 | -0.544569 | -1.557988 | 0.025237 | 0.949889 |
| GOBP_NADPH_REGENERATION | 17 | -0.597508 | -1.552626 | 0.031153 | 0.942164 |
| GOBP_C21_STEROID_HORMONE_BIOSYNTHETIC_PROCESS | 16 | -0.573033 | -1.529511 | 0.057325 | 1 |
| GOBP_NEGATIVE_REGULATION_OF_PROTEIN_MATURATION | 20 | -0.54862 | -1.527885 | 0.052632 | 1 |
| GOBP_GLIAL_CELL_APOPTOTIC_PROCESS | 15 | -0.586001 | -1.523713 | 0.061584 | 1 |
| GOBP_REGULATION_OF_KETONE_BIOSYNTHETIC_PROCESS | 16 | -0.587877 | -1.522446 | 0.044068 | 0.978346 |
| GOBP_CELLULAR_RESPONSE_TO_ZINC_ION | 16 | -0.580002 | -1.519841 | 0.052786 | 0.959348 |
| GOBP_ADAPTATION_OF_SIGNALING_PATHWAY | 16 | -0.574731 | -1.5119 | 0.057878 | 0.97076 |
| GOBP_REGULATION_OF_SPINDLE_ORGANIZATION | 44 | -0.455644 | -1.504975 | 0.035088 | 0.979484 |
| GOBP_EMBRYONIC_PLACENTA_MORPHOGENESIS | 22 | -0.515717 | -1.496366 | 0.041379 | 1 |
| GOBP_MITOTIC_CHROMOSOME_CONDENSATION | 19 | -0.533658 | -1.484857 | 0.074324 | 1 |
| GOBP_MATING_BEHAVIOR | 16 | -0.542373 | -1.474092 | 0.052941 | 1 |
| GOBP_CALCIUM_DEPENDENT_CELL_CELL_ADHESION_VIA_PLASMA_M | 33 | -0.455568 | -1.459495 | 0.053498 | 1 |
| GOBP_RENAL_ABSORPTION | 17 | -0.539854 | -1.458766 | 0.052805 | 1 |
| GOBP_ZINC_ION_HOMEOSTASIS | 29 | -0.479322 | -1.450828 | 0.042857 | 1 |
| GOBP_PROTEIN_LOCALIZATION_TO_CILIUM | 69 | -0.404325 | -1.450541 | 0.02924 | 1 |
| GOBP_POSTTRANSLATIONAL_PROTEIN_TARGETING_TO_ENDOPLASMIC | 15 | -0.5665 | -1.444397 | 0.05949 | 1 |
| GOBP_CELL_CYCLE_DNA_REPLICATION | 41 | -0.428301 | -1.439217 | 0.022727 | 1 |
| GOBP_NEGATIVE_REGULATION_OF_NUCLEOCYTOPLASMIC_TRANSPOF | 20 | -0.498513 | -1.430883 | 0.064309 | 1 |
| GOBP_CYTOPLASMIC_TRANSLATIONAL_INITIATION | 27 | -0.459464 | -1.430432 | 0.053957 | 1 |
| GOBP_HISTONE_H2A_UBIQUITINATION | 26 | -0.476207 | -1.417661 | 0.082143 | 1 |
| GOBP_POSITIVE_REGULATION_OF_MRNA_SPLICING_VIA_SPLICEOSOMI | 20 | -0.504012 | -1.409788 | 0.099359 | 1 |
| GOBP_NEGATIVE_REGULATION_OF_RNA_SPLICING | 25 | -0.475175 | -1.407505 | 0.078689 | 1 |
| GOBP_REGULATION_OF_URINE_VOLUME | 15 | -0.558318 | -1.402545 | 0.083851 | 1 |
| GOBP_NEGATIVE_REGULATION_OF_INTERLEUKIN_1_PRODUCTION | 20 | -0.500453 | -1.399855 | 0.088889 | 1 |
| GOBP_POSITIVE_REGULATION_OF_HISTONE_H3_K4_METHYLATION | 25 | -0.465939 | -1.398513 | 0.088968 | 1 |
| GOBP_AXONEME_ASSEMBLY | 68 | -0.388206 | -1.390147 | 0.039548 | 1 |
| GOBP_REGULATION_OF_NON_CANONICAL_WNT_SIGNALING_PATHWAY | 22 | -0.479202 | -1.389443 | 0.105263 | 1 |
| GOBP_RESPONSE_TO_ZINC_ION | 32 | -0.452867 | -1.38849 | 0.084291 | 1 |
| GOBP_POSITIVE_REGULATION_OF_RNA_SPLICING | 34 | -0.43922 | -1.386715 | 0.070833 | 1 |
| GOBP_REGULATION_OF_MRNA_SPLICING_VIA_SPLICEOSOME | 97 | -0.360016 | -1.383596 | 0 | 1 |
| GOBP_OLEFINIC_COMPOUND_BIOSYNTHETIC_PROCESS | 18 | -0.508689 | -1.383329 | 0.080247 | 1 |
| GOBP_RESPONSE_TO_PLATELET_DERIVED_GROWTH_FACTOR | 23 | -0.470833 | -1.382857 | 0.07971 | 1 |
| GOBP_CYTOSKELETON_DEPENDENT_CYTOKINESIS | 93 | -0.354127 | -1.382427 | 0.044444 | 1 |
| GOBP_AUTOPHAGY_OF_NUCLEUS | 16 | -0.509185 | -1.379768 | 0.128931 | 1 |
| GOBP_DNA_TEMPLATED_TRANSCRIPTION_INITIATION | 117 | -0.343734 | -1.371389 | 0.018868 | 1 |
| GOBP_NUCLEOLUS_ORGANIZATION | 16 | -0.504757 | -1.365809 | 0.12963 | 1 |
| GOBP_SODIUM_ION_IMPORT_ACROSS_PLASMA_MEMBRANE | 18 | -0.492659 | -1.359926 | 0.127596 | 1 |
| GOBP_BASE_EXCISION_REPAIR | 43 | -0.410511 | -1.354772 | 0.084112 | 1 |
| GOBP_PROTEIN_INSERTION_INTO_ER_MEMBRANE | 22 | -0.470857 | -1.353611 | 0.102894 | 1 |
| GOBP_TAIL_ANCHORED_MEMBRANE_PROTEIN_INSERTION_INTO_ER_M | 17 | -0.515934 | -1.350776 | 0.133333 | 1 |
| GOBP_HISTONE_H4_K5_ACETYLATION | 16 | -0.509587 | -1.345997 | 0.11315 | 1 |
| GOBP_REGULATION_OF_ADENYLATE_CYCLASE_ACTIVITY | 27 | -0.447901 | -1.345293 | 0.129825 | 1 |
| GOBP_NEGATIVE_REGULATION_OF_MRNA_SPLICING_VIA_SPLICEOSOM | 20 | -0.47966 | -1.343486 | 0.142857 | 1 |
| GOBP_DNA_CATABOLIC_PROCESS_ENDONUCLEOLYTIC | 20 | -0.485645 | -1.341957 | 0.138037 | 1 |
| GOBP_CYTOPLASMIC_SEQUESTERING_OF_PROTEIN | 21 | -0.480314 | -1.341835 | 0.102474 | 1 |
| GOBP_NLS_BEARING_PROTEIN_IMPORT_INTO_NUCLEUS | 19 | -0.483033 | -1.340765 | 0.113372 | 1 |
| GOBP_POSITIVE_REGULATION_OF_DOUBLE_STRAND_BREAK_REPAIR_ | 16 | -0.506932 | -1.338791 | 0.138801 | 1 |
| GOBP_POSITIVE_REGULATION_OF_PROTEIN_AUTOPHOSPHORYLATION | 25 | -0.452455 | -1.338523 | 0.115672 | 1 |
| GOBP_INTERLEUKIN_1_PRODUCTION | 67 | -0.367742 | -1.338383 | 0.060109 | 1 |
| GOBP_GANGLIOSIDE_BIOSYNTHETIC_PROCESS | 15 | -0.521884 | -1.334461 | 0.135802 | 1 |
| GOBP_REGULATION_OF_DNA_DEPENDENT_DNA_REPLICATION | 49 | -0.395328 | -1.333819 | 0.076555 | 1 |
| GOBP_REGULATION_OF_MEGAKARYOCYTE_DIFFERENTIATION | 31 | -0.445498 | -1.33329 | 0.073593 | 1 |
| GOBP_REGULATION_OF_MRNA_PROCESSING | 133 | -0.329319 | -1.326872 | 0.018349 | 1 |
| GOBP_RESPONSE_TO_COPPER_ION | 31 | -0.432801 | -1.325762 | 0.120482 | 1 |
| GOBP_OUTER_DYNEIN_ARM_ASSEMBLY | 16 | -0.504462 | -1.322637 | 0.125806 | 1 |
| GOBP_REGULATION_OF_CYTOKINESIS | 78 | -0.362621 | -1.322141 | 0.056962 | 1 |
| GOBP_POSITIVE_REGULATION_OF_MRNA_PROCESSING | 29 | -0.432359 | -1.32106 | 0.130435 | 1 |
| GOBP_REGULATION_OF_EXIT_FROM_MITOSIS | 15 | -0.495804 | -1.32046 | 0.130303 | 1 |
| GOBP_REGULATION_OF_HORMONE_BIOSYNTHETIC_PROCESS | 20 | -0.457089 | -1.318906 | 0.148276 | 1 |
| GOBP_POSITIVE_REGULATION_OF_INTERLEUKIN_1_BETA_PRODUCTIO | 37 | -0.412811 | -1.318013 | 0.088983 | 1 |
| GOBP_POSITIVE_REGULATION_OF_AMYLOID_BETA_FORMATION | 16 | -0.492227 | -1.316682 | 0.136646 | 1 |
| GOBP_DNA_REPLICATION_INITIATION | 35 | -0.409763 | -1.316482 | 0.103004 | 1 |
| GOBP_MITOTIC_CYTOKINESIS | 69 | -0.355662 | -1.315328 | 0.069767 | 1 |
| GOBP_CYTOKINESIS | 155 | -0.318396 | -1.314542 | 0.044944 | 1 |
| GOBP_NEGATIVE_REGULATION_OF_UBIQUITIN_PROTEIN_TRANSFERAS | 22 | -0.458168 | -1.312007 | 0.149682 | 1 |
| GOBP_RNA_SPLICING_VIA_TRANSESTERIFICATION_REACTIONS | 281 | -0.306993 | -1.311709 | 0.047619 | 1 |
| GOBP_LABYRINTHINE_LAYER_MORPHOGENESIS | 18 | -0.477374 | -1.311569 | 0.146965 | 0.990061 |
| GOBP_FLUID_TRANSPORT | 19 | -0.467203 | -1.311392 | 0.120567 | 0.980069 |
| GOBP_HISTONE_UBIQUITINATION | 47 | -0.387426 | -1.307977 | 0.09589 | 0.986488 |
| GOBP_TRANSCRIPTION_PREINITIATION_COMPLEX_ASSEMBLY | 36 | -0.401999 | -1.307904 | 0.118367 | 0.976212 |
| GOBP_REGULATION_OF_EXECUTION_PHASE_OF_APOPTOSIS | 16 | -0.476082 | -1.301458 | 0.146341 | 0.998141 |
| GOBP_FACULTATIVE_HETEROCHROMATIN_ASSEMBLY | 30 | -0.424882 | -1.299858 | 0.137795 | 0.99577 |
| GOBP_REGULATION_OF_HORMONE_METABOLIC_PROCESS | 30 | -0.415919 | -1.292574 | 0.143939 | 1 |
| GOBP_ACIDIC_AMINO_ACID_TRANSPORT | 54 | -0.369798 | -1.283448 | 0.103825 | 1 |
| GOBP_GANGLIOSIDE_METABOLIC_PROCESS | 24 | -0.439065 | -1.278184 | 0.143333 | 1 |
| GOBP_GLUCOSE_6_PHOSPHATE_METABOLIC_PROCESS | 20 | -0.460205 | -1.273952 | 0.175497 | 1 |
| GOBP_ADENYLATE_CYCLASE_ACTIVATING_ADRENERGIC_RECEPTOR_ | 19 | -0.461345 | -1.272844 | 0.173633 | 1 |
| GOBP_OOCYTE_MATURATION | 24 | -0.433183 | -1.271599 | 0.142361 | 1 |
| GOBP_POSITIVE_REGULATION_OF_INTERLEUKIN_1_PRODUCTION | 43 | -0.370688 | -1.264553 | 0.117647 | 1 |
| GOBP_POSITIVE_REGULATION_OF_CYTOKINESIS | 30 | -0.413122 | -1.261805 | 0.164875 | 1 |
| GOBP_AXIS_ELONGATION | 22 | -0.437376 | -1.260384 | 0.161404 | 1 |
| GOBP_MITOTIC_METAPHASE_PLATE_CONGRESSION | 52 | -0.358358 | -1.259722 | 0.122549 | 1 |
| GOBP_CERAMIDE_TRANSPORT | 16 | -0.474494 | -1.259693 | 0.18018 | 1 |
| GOBP_CEREBRAL_CORTEX_NEURON_DIFFERENTIATION | 19 | -0.437774 | -1.258464 | 0.169231 | 1 |
| GOBP_HISTONE_DEUBIQUITINATION | 40 | -0.375892 | -1.256185 | 0.127119 | 1 |
| GOBP_C21_STEROID_HORMONE_METABOLIC_PROCESS | 27 | -0.405779 | -1.253131 | 0.165468 | 1 |
| GOBP_KETONE_BIOSYNTHETIC_PROCESS | 37 | -0.38281 | -1.250913 | 0.144444 | 1 |
| GOBP_MICROTUBULE_BUNDLE_FORMATION | 98 | -0.332359 | -1.246986 | 0.102041 | 1 |
| GOBP_MAINTENANCE_OF_SYNAPSE_STRUCTURE | 19 | -0.454554 | -1.245425 | 0.185668 | 1 |
| GOBP_NEGATIVE_REGULATION_OF_STEM_CELL_DIFFERENTIATION | 22 | -0.418659 | -1.244166 | 0.194346 | 1 |
| GOBP_RETINAL_GANGLION_CELL_AXON_GUIDANCE | 18 | -0.457362 | -1.242526 | 0.194888 | 1 |
| GOBP_SOMATIC_DIVERSIFICATION_OF_IMMUNE_RECEPTORS | 69 | -0.335409 | -1.24061 | 0.105882 | 1 |
| GOBP_EXIT_FROM_MITOSIS | 27 | -0.409362 | -1.240323 | 0.172932 | 1 |
| GOBP_SERINE_FAMILY_AMINO_ACID_METABOLIC_PROCESS | 33 | -0.390832 | -1.238226 | 0.159696 | 1 |
| GOBP_RESPONSE_TO_NICOTINE | 40 | -0.370743 | -1.234627 | 0.159292 | 1 |
| GOBP_MIDBODY_ABSCISSION | 17 | -0.457459 | -1.233062 | 0.210191 | 1 |
| GOBP_PROTEIN_LOCALIZATION_TO_ENDOSOME | 24 | -0.420635 | -1.231109 | 0.202614 | 1 |
| GOBP_TELOMERE_ORGANIZATION | 149 | -0.301781 | -1.225726 | 0.033708 | 1 |
| GOBP_POSITIVE_REGULATION_OF_DENDRITE_MORPHOGENESIS | 33 | -0.37316 | -1.225349 | 0.15444 | 1 |
| GOBP_PROTEIN_LOCALIZATION_TO_GOLGI_APPARATUS | 26 | -0.411251 | -1.224852 | 0.178832 | 1 |
| GOBP_NEGATIVE_REGULATION_OF_DNA_DEPENDENT_DNA_REPLICAT | 15 | -0.46707 | -1.222679 | 0.186275 | 1 |
| GOBP_NEGATIVE_REGULATION_OF_DNA_RECOMBINATION | 37 | -0.385363 | -1.219791 | 0.188462 | 1 |
| GOBP_PROTEIN_LOCALIZATION_TO_MICROTUBULE | 17 | -0.453029 | -1.218966 | 0.22327 | 1 |
| GOBP_REGULATION_OF_RECEPTOR_INTERNALIZATION | 52 | -0.345571 | -1.215418 | 0.126761 | 1 |
| GOBP_RNA_POLYMERASE_II_PREINITIATION_COMPLEX_ASSEMBLY | 24 | -0.425485 | -1.214416 | 0.193431 | 1 |
| GOBP_ENTEROENDOCRINE_CELL_DIFFERENTIATION | 20 | -0.429959 | -1.212021 | 0.205788 | 1 |
| GOBP_REGULATION_OF_SISTER_CHROMATID_COHESION | 21 | -0.426812 | -1.211001 | 0.200637 | 1 |
| GOBP_TRNA_WOBBLE_BASE_MODIFICATION | 17 | -0.463832 | -1.210288 | 0.222581 | 1 |
| GOBP_RENAL_SYSTEM_VASCULATURE_DEVELOPMENT | 22 | -0.414666 | -1.206146 | 0.206897 | 1 |
| GOBP_MOTILE_CILIUM_ASSEMBLY | 44 | -0.36036 | -1.201776 | 0.158654 | 1 |
| GOBP_C4_DICARBOXYLATE_TRANSPORT | 25 | -0.4068 | -1.19973 | 0.173469 | 1 |
| GOBP_POSITIVE_REGULATION_OF_DENDRITIC_SPINE_MORPHOGENES | 18 | -0.434684 | -1.194524 | 0.234177 | 1 |
| GOBP_PEPTIDYL_ASPARAGINE_MODIFICATION | 23 | -0.412392 | -1.193117 | 0.199313 | 1 |
| GOBP_REGULATION_OF_EARLY_ENDOSOME_TO_LATE_ENDOSOME_TF | 18 | -0.437625 | -1.191867 | 0.214953 | 1 |
| GOBP_DEOXYRIBONUCLEOSIDE_TRIPHOSPHATE_METABOLIC_PROCES | 15 | -0.457846 | -1.190844 | 0.227528 | 1 |
| GOBP_POSITIVE_REGULATION_OF_AMYLOID_PRECURSOR_PROTEIN_C | 21 | -0.42185 | -1.188692 | 0.255102 | 1 |
| GOBP_SOMITE_DEVELOPMENT | 66 | -0.331595 | -1.187707 | 0.167665 | 1 |
| GOBP_NUCLEAR_MEMBRANE_REASSEMBLY | 23 | -0.417067 | -1.186254 | 0.233202 | 1 |
| GOBP_SOMITOGENESIS | 50 | -0.345931 | -1.185894 | 0.175676 | 1 |
| GOBP_NEGATIVE_REGULATION_OF_GENE_EXPRESSION_EPIGENETIC | 75 | -0.315545 | -1.183369 | 0.158621 | 1 |
| GOBP_MATURATION_OF_SSU_RRNA_FROM_TRICISTRONIC_RRNA_TRA | 37 | -0.361503 | -1.182532 | 0.177273 | 1 |
| GOBP_RENAL_SODIUM_EXCRETION | 15 | -0.458845 | -1.178661 | 0.260736 | 1 |
| GOBP_NEGATIVE_REGULATION_OF_MRNA_PROCESSING | 29 | -0.398828 | -1.176861 | 0.213178 | 1 |
| GOBP_RNA_SPLICING | 388 | -0.263309 | -1.174695 | 0 | 1 |
| GOBP_MONOUBIQUITINATED_PROTEIN_DEUBIQUITINATION | 34 | -0.366315 | -1.173397 | 0.186992 | 1 |

| | | | | | |
|---|---|---|---|---|---|
| GOBP_SENSORY_PERCEPTION_OF_LIGHT_STIMULUS | 137 | 0.501738 | 1.569398 | 0.001098 | 0.24209 |
| GOBP_ANIMAL_ORGAN_REGENERATION | 49 | 0.571833 | 1.567289 | 0.009877 | 0.245825 |
| GOBP_CELL_CHEMOTAXIS | 198 | 0.49102 | 1.567279 | 0.001065 | 0.244224 |
| GOBP_REGULATION_OF_LYMPHOCYTE_DIFFERENTIATION | 130 | 0.499822 | 1.567277 | 0 | 0.242617 |
| GOBP_REGULATION_OF_VIRAL_GENOME_REPLICATION | 72 | 0.538402 | 1.567076 | 0.007067 | 0.241497 |
| GOBP_POSITIVE_REGULATION_OF_FATTY_ACID_BIOSYNTHETIC_PROCESS | 17 | 0.668962 | 1.566836 | 0.029326 | 0.240611 |
| GOBP_MYELOID_LEUKOCYTE_MEDIATED_IMMUNITY | 76 | 0.53656 | 1.566397 | 0.004711 | 0.240196 |
| GOBP_AMINE_BIOSYNTHETIC_PROCESS | 32 | 0.595985 | 1.566356 | 0.018568 | 0.238726 |
| GOBP_EMBRYONIC_PATTERN_SPECIFICATION | 55 | 0.559361 | 1.56331 | 0.007595 | 0.244336 |
| GOBP_ANTIGEN_PROCESSING_AND_PRESENTATION_OF_ENDOGENOUS_ANTIGEN | 21 | 0.647426 | 1.563111 | 0.018466 | 0.243221 |
| GOBP_REGULATION_OF_CELL_KILLING | 55 | 0.557286 | 1.562147 | 0.015385 | 0.244108 |
| GOBP_CALCIUM_ION_IMPORT_INTO_CYTOSOL | 20 | 0.661581 | 1.561678 | 0.026135 | 0.243877 |
| GOBP_REGULATION_OF_PLATELET_DERIVED_GROWTH_FACTOR_RECEPTOR_SIGNALI | 19 | 0.660648 | 1.561671 | 0.030043 | 0.242368 |
| GOBP_NEGATIVE_REGULATION_OF_CD4_POSITIVE_ALPHA_BETA_T_CELL_ACTIVATION | 23 | 0.638809 | 1.561251 | 0.01676 | 0.241959 |
| GOBP_REGULATION_OF_LEUKOCYTE_MIGRATION | 149 | 0.496119 | 1.561036 | 0 | 0.240998 |
| GOBP_TRIGLYCERIDE_BIOSYNTHETIC_PROCESS | 33 | 0.597806 | 1.560115 | 0.016043 | 0.241648 |
| GOBP_PROTEIN_HOMOTETRAMERIZATION | 48 | 0.560669 | 1.560106 | 0.015957 | 0.240214 |
| GOBP_T_CELL_DIFFERENTIATION | 184 | 0.483675 | 1.559356 | 0 | 0.240653 |
| GOBP_MYELOID_CELL_APOPTOTIC_PROCESS | 25 | 0.636077 | 1.558392 | 0.025105 | 0.241518 |
| GOBP_ENDOCHONDRAL_BONE_MORPHOGENESIS | 51 | 0.56213 | 1.556495 | 0.01938 | 0.244881 |
| GOBP_HETEROPHILIC_CELL_CELL_ADHESION_VIA_PLASMA_MEMBRANE_CELL_ADHE | 31 | 0.610252 | 1.555344 | 0.022606 | 0.246438 |
| GOBP_REGULATION_OF_T_HELPER_CELL_DIFFERENTIATION | 26 | 0.624135 | 1.555013 | 0.021246 | 0.245792 |
| GOBP_MUSCLE_ORGAN_MORPHOGENESIS | 63 | 0.535258 | 1.552517 | 0.006046 | 0.250976 |
| GOBP_INFLAMMATORY_CELL_APOPTOTIC_PROCESS | 17 | 0.669523 | 1.551895 | 0.027027 | 0.251112 |
| GOBP_CELL_MIGRATION_INVOLVED_IN_HEART_DEVELOPMENT | 16 | 0.674293 | 1.551719 | 0.032305 | 0.250186 |
| GOBP_REGULATION_OF_LYMPHOCYTE_ACTIVATION | 328 | 0.467757 | 1.551228 | 0 | 0.249971 |
| GOBP_REGULATION_OF_CELL_ACTIVATION | 422 | 0.466894 | 1.550954 | 0 | 0.249272 |
| GOBP_REGULATION_OF_MEIOTIC_NUCLEAR_DIVISION | 22 | 0.639885 | 1.54991 | 0.027548 | 0.250622 |
| GOBP_NEUTRAL_LIPID_BIOSYNTHETIC_PROCESS | 39 | 0.580506 | 1.547217 | 0.011842 | 0.255911 |
| GOBP_REGULATION_OF_NATURAL_KILLER_CELL_ACTIVATION | 27 | 0.615011 | 1.546909 | 0.025751 | 0.25523 |
| GOBP_INTERLEUKIN_12_PRODUCTION | 41 | 0.573862 | 1.546657 | 0.014304 | 0.254473 |
| GOBP_NEGATIVE_REGULATION_OF_LYMPHOCYTE_ACTIVATION | 108 | 0.504577 | 1.546509 | 0.009281 | 0.253456 |
| GOBP_POSITIVE_REGULATION_OF_REGULATORY_T_CELL_DIFFERENTIATION | 16 | 0.681609 | 1.546048 | 0.032624 | 0.253292 |
| GOBP_BONE_DEVELOPMENT | 180 | 0.484346 | 1.54529 | 0.00106 | 0.253707 |
| GOBP_REGULATION_OF_ERBB_SIGNALING_PATHWAY | 68 | 0.537055 | 1.545185 | 0.010949 | 0.252559 |
| GOBP_MYELOID_CELL_ACTIVATION_INVOLVED_IN_IMMUNE_RESPONSE | 67 | 0.536534 | 1.545158 | 0.008516 | 0.251251 |
| GOBP_CD8_POSITIVE_ALPHA_BETA_T_CELL_ACTIVATION | 16 | 0.690589 | 1.544102 | 0.031519 | 0.252688 |
| GOBP_DIGESTIVE_SYSTEM_PROCESS | 61 | 0.542783 | 1.543909 | 0.00978 | 0.251792 |
| GOBP_LEUKOCYTE_MEDIATED_CYTOTOXICITY | 75 | 0.526835 | 1.543823 | 0.006075 | 0.250647 |
| GOBP_PROTEIN_KINASE_C_SIGNALING | 27 | 0.605526 | 1.543187 | 0.025245 | 0.250978 |
| GOBP_HYDROGEN_PEROXIDE_CATABOLIC_PROCESS | 16 | 0.672677 | 1.542989 | 0.028316 | 0.250205 |
| GOBP_MAMMARY_GLAND_MORPHOGENESIS | 33 | 0.597166 | 1.54254 | 0.024032 | 0.249952 |
| GOBP_POSITIVE_REGULATION_OF_INTERFERON_GAMMA_PRODUCTION | 43 | 0.574107 | 1.542344 | 0.016497 | 0.249105 |
| GOBP_REGULATION_OF_BIOLOGICAL_PROCESS_INVOLVED_IN_SYMBIOTIC_INTERACTI | 46 | 0.563999 | 1.540617 | 0.016861 | 0.252135 |
| GOBP_REGULATION_OF_DEFENSE_RESPONSE | 436 | 0.461063 | 1.540345 | 0 | 0.251487 |
| GOBP_DETECTION_OF_STIMULUS | 145 | 0.491153 | 1.539787 | 0.00442 | 0.251631 |
| GOBP_POSITIVE_REGULATION_OF_RESPONSE_TO_EXTERNAL_STIMULUS | 296 | 0.470493 | 1.539584 | 0 | 0.250854 |
| GOBP_POSITIVE_REGULATION_OF_LEUKOCYTE_PROLIFERATION | 98 | 0.510945 | 1.538539 | 0.00346 | 0.252015 |
| GOBP_REGULATION_OF_T_CELL_DIFFERENTIATION | 105 | 0.501412 | 1.537832 | 0.00454 | 0.252532 |
| GOBP_CARDIOCYTE_DIFFERENTIATION | 112 | 0.495775 | 1.537141 | 0.005688 | 0.252919 |
| GOBP_TRANSFORMING_GROWTH_FACTOR_BETA_PRODUCTION | 34 | 0.579841 | 1.536675 | 0.027778 | 0.252902 |
| GOBP_MALE_MEIOSIS_I | 19 | 0.656676 | 1.535451 | 0.033141 | 0.254742 |
| GOBP_REGULATION_OF_FATTY_ACID_METABOLIC_PROCESS | 65 | 0.539943 | 1.53463 | 0.008495 | 0.255417 |
| GOBP_REGENERATION | 141 | 0.488707 | 1.534262 | 0.004444 | 0.255021 |
| GOBP_ACUTE_INFLAMMATORY_RESPONSE | 63 | 0.539078 | 1.533706 | 0.015951 | 0.255223 |
| GOBP_TISSUE_REMODELING | 122 | 0.493893 | 1.53367 | 0.003472 | 0.254064 |
| GOBP_DNA_MODIFICATION | 99 | 0.506671 | 1.533294 | 0.00348 | 0.25382 |
| GOBP_REGULATION_OF_LEUKOCYTE_MEDIATED_IMMUNITY | 162 | 0.48451 | 1.53305 | 0 | 0.253175 |
| GOBP_REGULATION_OF_LEUKOCYTE_APOPTOTIC_PROCESS | 62 | 0.535092 | 1.532113 | 0.008578 | 0.254267 |
| GOBP_CARDIAC_MUSCLE_TISSUE_DEVELOPMENT | 175 | 0.480897 | 1.531458 | 0.00106 | 0.254645 |
| GOBP_REGULATION_OF_REPRODUCTIVE_PROCESS | 124 | 0.493414 | 1.5303 | 0.0044 | 0.256256 |
| GOBP_LEUKOCYTE_DIFFERENTIATION | 403 | 0.460925 | 1.529319 | 0 | 0.257826 |
| GOBP_BINDING_OF_SPERM_TO_ZONA_PELLUCIDA | 22 | 0.629107 | 1.528976 | 0.03709 | 0.257421 |
| GOBP_SIGNAL_TRANSDUCTION_INVOLVED_IN_REGULATION_OF_GENE_EXPRESSION | 15 | 0.691899 | 1.528736 | 0.034024 | 0.256796 |
| GOBP_POSITIVE_REGULATION_OF_INTERLEUKIN_12_PRODUCTION | 28 | 0.598857 | 1.528154 | 0.028767 | 0.256942 |
| GOBP_REGULATION_OF_COAGULATION | 52 | 0.539027 | 1.527941 | 0.019455 | 0.256288 |
| GOBP_POSITIVE_REGULATION_OF_CELL_MIGRATION_INVOLVED_IN_SPROUTING_ANGI | 17 | 0.662114 | 1.527603 | 0.037202 | 0.255949 |
| GOBP_MEGAKARYOCYTE_DEVELOPMENT | 15 | 0.683991 | 1.527131 | 0.046579 | 0.25591 |
| GOBP_EXTRACELLULAR_MATRIX_DISASSEMBLY | 44 | 0.55732 | 1.526322 | 0.015326 | 0.256922 |
| GOBP_ENDOCRINE_SYSTEM_DEVELOPMENT | 93 | 0.511187 | 1.526283 | 0.010502 | 0.25583 |
| GOBP_TOLL_LIKE_RECEPTOR_4_SIGNALING_PATHWAY | 29 | 0.598705 | 1.52624 | 0.028494 | 0.254779 |
| GOBP_REGULATION_OF_LEUKOCYTE_PROLIFERATION | 162 | 0.478065 | 1.52576 | 0.003226 | 0.254765 |
| GOBP_NEUROINFLAMMATORY_RESPONSE | 21 | 0.640598 | 1.525255 | 0.035868 | 0.254706 |
| GOBP_POSITIVE_REGULATION_OF_ALPHA_BETA_T_CELL_DIFFERENTIATION | 35 | 0.57396 | 1.52415 | 0.029987 | 0.256213 |
| GOBP_REGULATION_OF_NITRIC_OXIDE_METABOLIC_PROCESS | 41 | 0.567381 | 1.523231 | 0.016949 | 0.257293 |
| GOBP_INOSITOL_PHOSPHATE_METABOLIC_PROCESS | 49 | 0.545684 | 1.52285 | 0.020592 | 0.25716 |
| GOBP_LEUKOCYTE_MEDIATED_IMMUNITY | 254 | 0.465796 | 1.522011 | 0 | 0.258082 |
| GOBP_INTERLEUKIN_8_PRODUCTION | 53 | 0.557445 | 1.52032 | 0.021223 | 0.260981 |
| GOBP_CYTOKINE_PRODUCTION_INVOLVED_IN_IMMUNE_RESPONSE | 70 | 0.521377 | 1.520075 | 0.012165 | 0.260433 |
| GOBP_ACTIVATION_OF_IMMUNE_RESPONSE | 213 | 0.469849 | 1.519893 | 0.001048 | 0.259677 |
| GOBP_B_CELL_PROLIFERATION | 52 | 0.53966 | 1.519557 | 0.02457 | 0.259343 |
| GOBP_NEGATIVE_REGULATION_OF_RESPONSE_TO_BIOTIC_STIMULUS | 89 | 0.512362 | 1.519426 | 0.013809 | 0.258547 |
| GOBP_POSITIVE_REGULATION_OF_RESPONSE_TO_BIOTIC_STIMULUS | 115 | 0.496762 | 1.519421 | 0.004525 | 0.257445 |
| GOBP_CELL_PROLIFERATION_IN_FOREBRAIN | 18 | 0.657775 | 1.518995 | 0.043668 | 0.257403 |
| GOBP_NEGATIVE_REGULATION_OF_ATP_DEPENDENT_ACTIVITY | 18 | 0.649758 | 1.51893 | 0.04271 | 0.25643 |
| GOBP_POSITIVE_REGULATION_OF_CELL_ADHESION | 334 | 0.463014 | 1.518661 | 0 | 0.255987 |
| GOBP_T_CELL_DIFFERENTIATION_INVOLVED_IN_IMMUNE_RESPONSE | 51 | 0.548662 | 1.517186 | 0.015019 | 0.258335 |
| GOBP_INTEGRIN_MEDIATED_SIGNALING_PATHWAY | 91 | 0.506968 | 1.51707 | 0.011614 | 0.25756 |
| GOBP_CELLULAR_RESPONSE_TO_DSRNA | 20 | 0.641178 | 1.515785 | 0.029369 | 0.259506 |
| GOBP_ACUTE_PHASE_RESPONSE | 26 | 0.607242 | 1.515351 | 0.0427 | 0.259378 |
| GOBP_LEUKOCYTE_PROLIFERATION | 211 | 0.470872 | 1.51483 | 0 | 0.259585 |
| GOBP_REGULATION_OF_T_CELL_PROLIFERATION | 112 | 0.496508 | 1.514677 | 0.004556 | 0.25887 |
| GOBP_FEMALE_GAMETE_GENERATION | 124 | 0.488824 | 1.514472 | 0.008879 | 0.25819 |
| GOBP_REGULATION_OF_LEUKOCYTE_MEDIATED_CYTOTOXICITY | 46 | 0.551447 | 1.514337 | 0.021384 | 0.25743 |
| GOBP_LEUKOCYTE_CELL_CELL_ADHESION | 254 | 0.464145 | 1.513198 | 0 | 0.259168 |
| GOBP_MAMMARY_GLAND_DUCT_MORPHOGENESIS | 23 | 0.624638 | 1.512869 | 0.041311 | 0.258861 |
| GOBP_CELL_COMMUNICATION_BY_ELECTRICAL_COUPLING | 30 | 0.603936 | 1.512598 | 0.03155 | 0.258434 |
| GOBP_NEGATIVE_REGULATION_OF_POTASSIUM_ION_TRANSPORT | 20 | 0.646565 | 1.512207 | 0.046647 | 0.258281 |
| GOBP_CELL_DIFFERENTIATION_INVOLVED_IN_EMBRYONIC_PLACENTA_DEVELOPMEN | 23 | 0.611821 | 1.512136 | 0.044034 | 0.257352 |
| GOBP_REGULATION_OF_VIRAL_PROCESS | 134 | 0.487616 | 1.511347 | 0.008919 | 0.258328 |
| GOBP_HEMOSTASIS | 171 | 0.473083 | 1.510236 | 0 | 0.259968 |
| GOBP_NEGATIVE_REGULATION_OF_PHOSPHATASE_ACTIVITY | 31 | 0.585708 | 1.510196 | 0.036536 | 0.259043 |
| GOBP_POSITIVE_REGULATION_OF_LEUKOCYTE_ADHESION_TO_VASCULAR_ENDOTHE | 15 | 0.668319 | 1.509661 | 0.050536 | 0.259222 |
| GOBP_NEGATIVE_REGULATION_OF_POTASSIUM_ION_TRANSMEMBRANE_TRANSPORT | 16 | 0.6556 | 1.508535 | 0.049563 | 0.260883 |
| GOBP_REGULATION_OF_HAIR_CYCLE | 23 | 0.61513 | 1.507241 | 0.040936 | 0.26298 |
| GOBP_REGULATION_OF_T_CELL_DIFFERENTIATION_IN_THYMUS | 18 | 0.656674 | 1.506849 | 0.051775 | 0.262846 |
| GOBP_CD4_POSITIVE_ALPHA_BETA_T_CELL_ACTIVATION | 69 | 0.518468 | 1.506389 | 0.011779 | 0.262922 |
| GOBP_REGULATION_OF_CARDIAC_CONDUCTION | 20 | 0.634496 | 1.505923 | 0.036657 | 0.26318 |
| GOBP_CALCIUM_MEDIATED_SIGNALING_USING_INTRACELLULAR_CALCIUM_SOURCE | 15 | 0.67124 | 1.504006 | 0.04048 | 0.266796 |
| GOBP_NEGATIVE_REGULATION_OF_ORGAN_GROWTH | 27 | 0.598415 | 1.503358 | 0.025503 | 0.267399 |
| GOBP_POSITIVE_REGULATION_OF_PROTEIN_DEPOLYMERIZATION | 17 | 0.653492 | 1.502438 | 0.046269 | 0.268601 |
| GOBP_MONONUCLEAR_CELL_DIFFERENTIATION | 308 | 0.455319 | 1.501077 | 0 | 0.270985 |
| GOBP_CELL_FATE_DETERMINATION | 25 | 0.608395 | 1.501014 | 0.038889 | 0.270076 |
| GOBP_PROGRAMMED_NECROTIC_CELL_DEATH | 37 | 0.570489 | 1.499337 | 0.031507 | 0.273305 |
| GOBP_REGULATION_OF_CELL_MIGRATION_INVOLVED_IN_SPROUTING_ANGIOGENESIS | 32 | 0.574871 | 1.499178 | 0.044655 | 0.272665 |
| GOBP_PYRIMIDINE_NUCLEOSIDE_METABOLIC_PROCESS | 18 | 0.648584 | 1.498565 | 0.046687 | 0.27325 |
| GOBP_NEGATIVE_REGULATION_OF_T_CELL_RECEPTOR_SIGNALING_PATHWAY | 17 | 0.658169 | 1.498191 | 0.048175 | 0.273141 |
| GOBP_CARDIAC_MUSCLE_CELL_DIFFERENTIATION | 86 | 0.503824 | 1.49811 | 0.009346 | 0.27233 |
| GOBP_CHEMOKINE_PRODUCTION | 61 | 0.524252 | 1.498067 | 0.016414 | 0.271399 |
| GOBP_INTERLEUKIN_2_PRODUCTION | 45 | 0.551236 | 1.4976 | 0.035851 | 0.27151 |
| GOBP_DIENCEPHALON_DEVELOPMENT | 55 | 0.527605 | 1.497536 | 0.024631 | 0.270644 |
| GOBP_DEFENSE_RESPONSE_TO_GRAM_NEGATIVE_BACTERIUM | 31 | 0.577022 | 1.49734 | 0.04336 | 0.27017 |
| GOBP_CELLULAR_RESPONSE_TO_ALKALOID | 29 | 0.590946 | 1.49629 | 0.034993 | 0.271568 |
| GOBP_SINGLE_STRANDED_VIRAL_RNA_REPLICATION_VIA_DOUBLE_STRANDED_DNA_ | 17 | 0.655517 | 1.496267 | 0.040698 | 0.270621 |
| GOBP_POSITIVE_REGULATION_OF_ION_TRANSMEMBRANE_TRANSPORT | 128 | 0.478779 | 1.49599 | 0.013544 | 0.270265 |
| GOBP_POSITIVE_REGULATION_OF_CALCIUM_ION_TRANSMEMBRANE_TRANSPORT | 56 | 0.524913 | 1.495656 | 0.024361 | 0.270052 |
| GOBP_SYNAPTIC_TRANSMISSION_CHOLINERGIC | 23 | 0.608019 | 1.495146 | 0.050754 | 0.270288 |
| GOBP_GLYCEROL_ETHER_METABOLIC_PROCESS | 21 | 0.61189 | 1.494861 | 0.051724 | 0.269947 |
| GOBP_MYELOID_LEUKOCYTE_DIFFERENTIATION | 164 | 0.474421 | 1.494762 | 0.003205 | 0.269166 |
| GOBP_SKELETAL_MUSCLE_TISSUE_REGENERATION | 27 | 0.594658 | 1.49467 | 0.048518 | 0.268422 |
| GOBP_PROTEIN_HOMOOLIGOMERIZATION | 158 | 0.46965 | 1.493142 | 0.012048 | 0.271178 |
| GOBP_DEFENSE_RESPONSE_TO_BACTERIUM | 126 | 0.486081 | 1.492815 | 0.007778 | 0.271023 |
| GOBP_REGULATION_OF_TOLL_LIKE_RECEPTOR_4_SIGNALING_PATHWAY | 18 | 0.656397 | 1.492718 | 0.030568 | 0.270256 |
| GOBP_LEUKOCYTE_DEGRANULATION | 57 | 0.53286 | 1.492336 | 0.023256 | 0.270221 |
| GOBP_COMPLEMENT_ACTIVATION | 28 | 0.597571 | 1.492125 | 0.038621 | 0.269784 |
| GOBP_T_CELL_ACTIVATION | 343 | 0.449918 | 1.490675 | 0 | 0.272357 |
| GOBP_ETHER_METABOLIC_PROCESS | 26 | 0.594944 | 1.489752 | 0.041609 | 0.273587 |
| GOBP_ACROSOME_REACTION | 26 | 0.6002 | 1.489477 | 0.041429 | 0.273389 |
| GOBP_POSITIVE_REGULATION_OF_DOUBLE_STRAND_BREAK_REPAIR_VIA_HOMOLOG | 36 | 0.562187 | 1.488908 | 0.052846 | 0.27373 |
| GOBP_CARDIAC_MUSCLE_TISSUE_MORPHOGENESIS | 48 | 0.540113 | 1.488036 | 0.029948 | 0.274881 |
| GOBP_REGULATION_OF_DEFENSE_RESPONSE_TO_BACTERIUM | 15 | 0.668014 | 1.487951 | 0.05303 | 0.274153 |
| GOBP_POSITIVE_REGULATION_OF_NEUTROPHIL_MIGRATION | 25 | 0.59933 | 1.487817 | 0.052999 | 0.273535 |
| GOBP_ARTERY_MORPHOGENESIS | 59 | 0.524007 | 1.487425 | 0.021684 | 0.273518 |
| GOBP_CELL_KILLING | 96 | 0.496202 | 1.486787 | 0.010345 | 0.274138 |
| GOBP_MUSCLE_TISSUE_DEVELOPMENT | 311 | 0.452382 | 1.485741 | 0 | 0.275645 |
| GOBP_CELL_AGGREGATION | 15 | 0.664874 | 1.483845 | 0.054381 | 0.279377 |
| GOBP_INTERLEUKIN_17_PRODUCTION | 24 | 0.618043 | 1.483462 | 0.024182 | 0.279384 |
| GOBP_POSITIVE_REGULATION_OF_T_CELL_PROLIFERATION | 61 | 0.516229 | 1.483159 | 0.026928 | 0.27911 |
| GOBP_MUSCLE_ORGAN_DEVELOPMENT | 268 | 0.453443 | 1.480678 | 0.004149 | 0.284278 |
| GOBP_REGULATION_OF_LEUKOCYTE_CHEMOTAXIS | 83 | 0.490652 | 1.48063 | 0.015606 | 0.283434 |
| GOBP_COLLAGEN_FIBRIL_ORGANIZATION | 56 | 0.524754 | 1.480219 | 0.024938 | 0.283473 |

| | | | | | |
|---|---|---|---|---|---|
| GOBP_L_GLUTAMATE_IMPORT_ACROSS_PLASMA_MEMBRANE | 15 | -0.448748 | -1.173368 | 0.243976 | 1 |
| GOBP_WALKING_BEHAVIOR | 26 | -0.394751 | -1.172936 | 0.233216 | 1 |
| GOBP_BENZENE_CONTAINING_COMPOUND_METABOLIC_PROCESS | 15 | -0.44944 | -1.171292 | 0.257862 | 1 |
| GOBP_HISTONE_H2A_MONOUBIQUITINATION | 17 | -0.436057 | -1.167409 | 0.275316 | 1 |
| GOBP_REGULATION_OF_TRANSCRIPTION_BY_RNA_POLYMERASE_I | 33 | -0.375098 | -1.166963 | 0.203704 | 1 |
| GOBP_TRANSCRIPTION_BY_RNA_POLYMERASE_I | 53 | -0.334837 | -1.165898 | 0.21 | 1 |
| GOBP_PROTEIN_REFOLDING | 20 | -0.413965 | -1.165066 | 0.231788 | 1 |
| GOBP_MEMBRANE_DEPOLARIZATION_DURING_ACTION_POTENTIAL | 33 | -0.37325 | -1.164692 | 0.239382 | 1 |
| GOBP_REGULATION_OF_INCLUSION_BODY_ASSEMBLY | 16 | -0.428737 | -1.164422 | 0.267913 | 1 |
| GOBP_CELLULAR_RESPONSE_TO_CADMIUM_ION | 29 | -0.379774 | -1.164184 | 0.238267 | 1 |
| GOBP_ATP_BIOSYNTHETIC_PROCESS | 52 | -0.334786 | -1.163906 | 0.180617 | 1 |
| GOBP_REGULATION_OF_REACTIVE_OXYGEN_SPECIES_BIOSYNTHETIC_ | 34 | -0.372302 | -1.161569 | 0.226496 | 1 |
| GOBP_V_D_J_RECOMBINATION | 18 | -0.433443 | -1.161461 | 0.26087 | 1 |
| GOBP_OLFACTORY_LOBE_DEVELOPMENT | 26 | -0.391061 | -1.158904 | 0.274131 | 1 |
| GOBP_NUCLEOLAR_LARGE_RRNA_TRANSCRIPTION_BY_RNA_POLYME | 18 | -0.419184 | -1.158157 | 0.263804 | 1 |
| GOBP_GLANDULAR_EPITHELIAL_CELL_DEVELOPMENT | 23 | -0.39907 | -1.152763 | 0.267974 | 1 |
| GOBP_RESPONSE_TO_COCAINE | 40 | -0.348975 | -1.152714 | 0.205479 | 1 |
| GOBP_DNA_TEMPLATED_TRANSCRIPTION_ELONGATION | 74 | -0.316909 | -1.15265 | 0.172662 | 1 |
| GOBP_CLEAVAGE_INVOLVED_IN_RRNA_PROCESSING | 27 | -0.380869 | -1.151176 | 0.256506 | 1 |
| GOBP_POSITIVE_REGULATION_OF_TRANSCRIPTION_BY_RNA_POLYME | 23 | -0.399034 | -1.150706 | 0.228956 | 1 |
| GOBP_STEROID_HORMONE_BIOSYNTHETIC_PROCESS | 28 | -0.374916 | -1.148244 | 0.247967 | 1 |
| GOBP_REGULATION_OF_RNA_SPLICING | 146 | -0.288101 | -1.146596 | 0.202247 | 1 |
| GOBP_TRANSLATIONAL_ELONGATION | 52 | -0.331145 | -1.146539 | 0.212963 | 1 |
| GOBP_DNA_STRAND_ELONGATION | 36 | -0.359671 | -1.145988 | 0.274194 | 1 |
| GOBP_RESPONSE_TO_CAMP | 68 | -0.313379 | -1.145557 | 0.233918 | 1 |
| GOBP_NEGATIVE_REGULATION_OF_CARTILAGE_DEVELOPMENT | 23 | -0.392169 | -1.144645 | 0.255591 | 1 |
| GOBP_RESPIRATORY_SYSTEM_PROCESS | 33 | -0.353485 | -1.142062 | 0.255814 | 1 |
| GOBP_NUCLEAR_TRANSCRIBED_MRNA_CATABOLIC_PROCESS_EXONU | 19 | -0.412178 | -1.140438 | 0.296296 | 1 |
| GOBP_HISTONE_H3_DEACETYLATION | 17 | -0.428246 | -1.140092 | 0.271293 | 1 |
| GOBP_TELOMERASE_RNA_LOCALIZATION | 19 | -0.414035 | -1.13975 | 0.277778 | 1 |
| GOBP_NUCLEOSIDE_TRIPHOSPHATE_METABOLIC_PROCESS | 101 | -0.292798 | -1.137431 | 0.178571 | 1 |
| GOBP_REGULATION_OF_SPINDLE_ASSEMBLY | 30 | -0.366557 | -1.137263 | 0.243728 | 1 |
| GOBP_MEGAKARYOCYTE_DIFFERENTIATION | 48 | -0.339749 | -1.134902 | 0.28 | 1 |
| GOBP_OLIGOSACCHARIDE_BIOSYNTHETIC_PROCESS | 18 | -0.405716 | -1.13414 | 0.257053 | 1 |
| GOBP_PARAXIAL_MESODERM_DEVELOPMENT | 16 | -0.426663 | -1.132945 | 0.286164 | 1 |
| GOBP_CILIUM_ORGANIZATION | 358 | -0.258547 | -1.13096 | 0.058824 | 1 |
| GOBP_POSITIVE_REGULATION_OF_TORC1_SIGNALING | 20 | -0.39479 | -1.129101 | 0.272109 | 1 |
| GOBP_SRP_DEPENDENT_COTRANSLATIONAL_PROTEIN_TARGETING_T | 18 | -0.412237 | -1.128191 | 0.28479 | 1 |
| GOBP_NEGATIVE_REGULATION_OF_ADENYLATE_CYCLASE_ACTIVITY | 16 | -0.420364 | -1.126221 | 0.283019 | 1 |
| GOBP_REGULATION_OF_TRANSLATIONAL_ELONGATION | 18 | -0.412588 | -1.12593 | 0.280936 | 1 |
| GOBP_NUCLEOTIDE_PHOSPHORYLATION | 84 | -0.297803 | -1.125802 | 0.20979 | 1 |
| GOBP_PROTEIN_FOLDING | 186 | -0.26566 | -1.124774 | 0.12 | 1 |
| GOBP_ADHERENS_JUNCTION_ORGANIZATION | 46 | -0.326985 | -1.1225 | 0.268908 | 1 |
| GOBP_REGULATION_OF_SYSTEMIC_ARTERIAL_BLOOD_PRESSURE | 69 | -0.310529 | -1.122457 | 0.216049 | 1 |
| GOBP_PYRUVATE_METABOLIC_PROCESS | 84 | -0.297641 | -1.121845 | 0.201299 | 1 |
| GOBP_POSITIVE_REGULATION_OF_CILIUM_ASSEMBLY | 24 | -0.374791 | -1.119577 | 0.307432 | 1 |
| GOBP_PROTEIN_TARGETING_TO_PEROXISOME | 18 | -0.415548 | -1.118427 | 0.308176 | 1 |
| GOBP_NEGATIVE_REGULATION_OF_OXIDATIVE_STRESS_INDUCED_NE | 17 | -0.410521 | -1.117611 | 0.311953 | 1 |
| GOBP_MATURATION_OF_5_8S_RRNA_FROM_TRICISTRONIC_RRNA_TR | 24 | -0.379052 | -1.117605 | 0.268707 | 1 |
| GOBP_ADRENERGIC_RECEPTOR_SIGNALING_PATHWAY | 24 | -0.37711 | -1.117413 | 0.296167 | 1 |
| GOBP_REGULATION_OF_SYSTEMIC_ARTERIAL_BLOOD_PRESSURE_BY | 25 | -0.382445 | -1.114879 | 0.308475 | 1 |
| GOBP_ASPARTATE_FAMILY_AMINO_ACID_BIOSYNTHETIC_PROCESS | 19 | -0.402235 | -1.112799 | 0.324841 | 1 |
| GOBP_ENDONUCLEOLYTIC_CLEAVAGE_INVOLVED_IN_RRNA_PROCES | 15 | -0.430176 | -1.112315 | 0.317221 | 1 |
| GOBP_VIRAL_GENE_EXPRESSION | 95 | -0.288661 | -1.112097 | 0.226563 | 1 |
| GOBP_GLOMERULAR_EPITHELIAL_CELL_DIFFERENTIATION | 19 | -0.394792 | -1.112004 | 0.307453 | 1 |
| GOBP_HISTONE_H4_K16_ACETYLATION | 27 | -0.367587 | -1.111919 | 0.292135 | 1 |
| GOBP_LUNG_ALVEOLUS_DEVELOPMENT | 32 | -0.356054 | -1.111531 | 0.301653 | 1 |
| GOBP_DNA_LIGATION | 15 | -0.427304 | -1.111132 | 0.316265 | 1 |
| GOBP_CELLULAR_CARBOHYDRATE_BIOSYNTHETIC_PROCESS | 63 | -0.309368 | -1.10816 | 0.246305 | 1 |
| GOBP_DOUBLE_STRAND_BREAK_REPAIR_VIA_NONHOMOLOGOUS_EN | 68 | -0.302648 | -1.105922 | 0.258427 | 1 |
| GOBP_CYTOKINE_PRODUCTION_INVOLVED_IN_INFLAMMATORY_RESP | 35 | -0.347411 | -1.105293 | 0.292308 | 1 |
| GOBP_PROTEIN_TRANSMEMBRANE_TRANSPORT | 63 | -0.308133 | -1.101576 | 0.247191 | 1 |
| GOBP_RNA_SURVEILLANCE | 15 | -0.412938 | -1.099191 | 0.333333 | 1 |
| GOBP_POSITIVE_REGULATION_OF_ISOTYPE_SWITCHING | 23 | -0.375787 | -1.098594 | 0.311258 | 1 |
| GOBP_GLOMERULUS_DEVELOPMENT | 54 | -0.31393 | -1.097286 | 0.277512 | 1 |
| GOBP_ENDOPLASMIC_RETICULUM_TO_GOLGI_VESICLE_MEDIATED_TR | 121 | -0.274484 | -1.09675 | 0.196721 | 1 |
| GOBP_REGULATION_OF_MITOTIC_SPINDLE_ASSEMBLY | 23 | -0.372115 | -1.096399 | 0.305136 | 1 |
| GOBP_COPULATION | 15 | -0.404885 | -1.096071 | 0.324759 | 1 |
| GOBP_NON_MOTILE_CILIUM_ASSEMBLY | 59 | -0.305165 | -1.095661 | 0.282353 | 1 |
| GOBP_CARBOHYDRATE_CATABOLIC_PROCESS | 123 | -0.272903 | -1.09444 | 0.252174 | 1 |
| GOBP_STEM_CELL_DIVISION | 24 | -0.382214 | -1.0944 | 0.300341 | 1 |
| GOBP_GLYCOSPHINGOLIPID_BIOSYNTHETIC_PROCESS | 29 | -0.36317 | -1.093852 | 0.31746 | 1 |
| GOBP_REGULATION_OF_EXCRETION | 17 | -0.403373 | -1.092831 | 0.333333 | 1 |
| GOBP_PROTEIN_IMPORT_INTO_MITOCHONDRIAL_MATRIX | 20 | -0.388431 | -1.092131 | 0.345912 | 1 |
| GOBP_NEGATIVE_REGULATION_OF_NF_KAPPAB_TRANSCRIPTION_FAC | 75 | -0.286369 | -1.092104 | 0.269006 | 1 |
| GOBP_ENDOCRINE_PROCESS | 61 | -0.303972 | -1.08982 | 0.287958 | 1 |
| GOBP_GLYCOSPHINGOLIPID_METABOLIC_PROCESS | 53 | -0.319177 | -1.089692 | 0.310345 | 1 |
| GOBP_REGULATION_OF_SODIUM_ION_TRANSPORT | 76 | -0.298822 | -1.089063 | 0.24026 | 1 |
| GOBP_RESPONSE_TO_AUDITORY_STIMULUS | 21 | -0.388566 | -1.088755 | 0.315457 | 1 |
| GOBP_RRNA_TRANSCRIPTION | 34 | -0.341701 | -1.088747 | 0.293907 | 1 |
| GOBP_BILE_ACID_METABOLIC_PROCESS | 36 | -0.330946 | -1.088033 | 0.278884 | 1 |
| GOBP_REGULATION_OF_SYSTEMIC_ARTERIAL_BLOOD_PRESSURE_ME | 32 | -0.361642 | -1.086849 | 0.334711 | 1 |
| GOBP_ADP_METABOLIC_PROCESS | 75 | -0.304711 | -1.083804 | 0.280702 | 1 |
| GOBP_INTRACELLULAR_PROTEIN_TRANSMEMBRANE_TRANSPORT | 52 | -0.317539 | -1.081937 | 0.285047 | 1 |
| GOBP_PROSTATE_GLAND_MORPHOGENESIS | 18 | -0.397392 | -1.080997 | 0.333333 | 1 |
| GOBP_PEROXISOMAL_TRANSPORT | 21 | -0.392833 | -1.080571 | 0.353571 | 1 |
| GOBP_MONOVALENT_INORGANIC_CATION_HOMEOSTASIS | 121 | -0.266597 | -1.080187 | 0.295082 | 1 |
| GOBP_PROTEIN_INSERTION_INTO_MEMBRANE | 53 | -0.309866 | -1.078725 | 0.31068 | 1 |
| GOBP_MICROTUBULE_NUCLEATION | 35 | -0.341676 | -1.07642 | 0.334783 | 1 |
| GOBP_INCLUSION_BODY_ASSEMBLY | 22 | -0.378097 | -1.072389 | 0.334405 | 1 |
| GOBP_GLYCOSIDE_METABOLIC_PROCESS | 17 | -0.400128 | -1.072232 | 0.385075 | 1 |
| GOBP_NEGATIVE_REGULATION_OF_CILIUM_ASSEMBLY | 17 | -0.394841 | -1.070919 | 0.346154 | 1 |
| GOBP_MATING | 30 | -0.348775 | -1.070664 | 0.345238 | 1 |
| GOBP_PROTEIN_NEDDYLATION | 17 | -0.394601 | -1.070143 | 0.347682 | 1 |
| GOBP_PROTEIN_ADP_RIBOSYLATION | 34 | -0.32896 | -1.069917 | 0.338235 | 1 |
| GOBP_REGULATION_OF_LYASE_ACTIVITY | 38 | -0.32681 | -1.069412 | 0.360731 | 1 |
| GOBP_NADP_METABOLIC_PROCESS | 34 | -0.337937 | -1.069319 | 0.315353 | 1 |
| GOBP_EPITHELIAL_TUBE_BRANCHING_INVOLVED_IN_LUNG_MORPHOG | 20 | -0.38854 | -1.067627 | 0.375427 | 1 |
| GOBP_PEPTIDYL_L_CYSTEINE_S_PALMITOYLATION | 23 | -0.375481 | -1.066081 | 0.380952 | 1 |
| GOBP_NEGATIVE_REGULATION_OF_PEPTIDE_SECRETION | 32 | -0.338836 | -1.064684 | 0.328302 | 1 |
| GOBP_MATURATION_OF_SSU_RRNA | 50 | -0.300245 | -1.063441 | 0.358491 | 1 |
| GOBP_POSITIVE_REGULATION_OF_SYNAPSE_ASSEMBLY | 51 | -0.311864 | -1.060651 | 0.357143 | 1 |
| GOBP_MEMBRANE_DEPOLARIZATION_DURING_CARDIAC_MUSCLE_CE | 20 | -0.370738 | -1.056947 | 0.359259 | 1 |
| GOBP_NEGATIVE_REGULATION_OF_DOUBLE_STRAND_BREAK_REPAIR | 19 | -0.377599 | -1.056326 | 0.321767 | 1 |
| GOBP_NUCLEOSIDE_DIPHOSPHATE_METABOLIC_PROCESS | 114 | -0.26225 | -1.055919 | 0.319149 | 1 |
| GOBP_REGULATION_OF_PH | 76 | -0.282517 | -1.055807 | 0.3125 | 1 |
| GOBP_NEGATIVE_REGULATION_OF_DNA_REPAIR | 34 | -0.334353 | -1.055426 | 0.374502 | 1 |
| GOBP_TYPE_B_PANCREATIC_CELL_DEVELOPMENT | 18 | -0.390895 | -1.055095 | 0.39322 | 1 |
| GOBP_REGULATION_OF_PRESYNAPSE_ORGANIZATION | 29 | -0.351299 | -1.052504 | 0.378906 | 1 |
| GOBP_PYRIMIDINE_NUCLEOSIDE_TRIPHOSPHATE_METABOLIC_PROCE | 24 | -0.348648 | -1.052301 | 0.372881 | 1 |
| GOBP_METAPHASE_PLATE_CONGRESSION | 65 | -0.296056 | -1.051695 | 0.341463 | 1 |
| GOBP_MATURATION_OF_LSU_RRNA_FROM_TRICISTRONIC_RRNA_TRA | 15 | -0.393069 | -1.051129 | 0.368902 | 1 |
| GOBP_MRNA_PROCESSING | 441 | -0.232121 | -1.05048 | 0.272727 | 1 |
| GOBP_NEGATIVE_REGULATION_OF_LYASE_ACTIVITY | 20 | -0.383249 | -1.047545 | 0.401254 | 1 |
| GOBP_SOMATIC_DIVERSIFICATION_OF_IMMUNOGLOBULINS | 57 | -0.306359 | -1.046683 | 0.347826 | 1 |
| GOBP_ARP2_3_COMPLEX_MEDIATED_ACTIN_NUCLEATION | 38 | -0.32204 | -1.046315 | 0.365546 | 1 |
| GOBP_CELLULAR_RESPONSE_TO_PROSTAGLANDIN_STIMULUS | 17 | -0.404159 | -1.045249 | 0.39604 | 1 |
| GOBP_ALTERNATIVE_MRNA_SPLICING_VIA_SPLICEOSOME | 72 | -0.283762 | -1.044577 | 0.309942 | 1 |
| GOBP_PROTEIN_TRANSMEMBRANE_IMPORT_INTO_INTRACELLULAR_O | 36 | -0.33575 | -1.043397 | 0.390438 | 1 |
| GOBP_PHOSPHATIDYLINOSITOL_3_PHOSPHATE_BIOSYNTHETIC_PROC | 19 | -0.376375 | -1.042195 | 0.385382 | 1 |
| GOBP_METAPHASE_ANAPHASE_TRANSITION_OF_CELL_CYCLE | 61 | -0.290548 | -1.041885 | 0.350877 | 1 |
| GOBP_OLIGOSACCHARIDE_METABOLIC_PROCESS | 46 | -0.317076 | -1.041591 | 0.39196 | 1 |
| GOBP_RESPONSE_TO_ATP | 23 | -0.348487 | -1.038897 | 0.367133 | 1 |
| GOBP_HINDBRAIN_DEVELOPMENT | 126 | -0.268534 | -1.038214 | 0.369369 | 1 |
| GOBP_REGULATION_OF_CLATHRIN_DEPENDENT_ENDOCYTOSIS | 18 | -0.379111 | -1.037619 | 0.403279 | 1 |
| GOBP_SOMATIC_DIVERSIFICATION_OF_IMMUNOGLOBULINS_INVOLVED | 42 | -0.313768 | -1.03746 | 0.365854 | 1 |
| GOBP_CHROMOSOME_LOCALIZATION | 78 | -0.276449 | -1.035939 | 0.322917 | 1 |
| GOBP_L_AMINO_ACID_TRANSPORT | 55 | -0.290219 | -1.03578 | 0.403226 | 1 |
| GOBP_MATURATION_OF_5_8S_RRNA | 35 | -0.320183 | -1.034165 | 0.395918 | 1 |
| GOBP_ANAPHASE_PROMOTING_COMPLEX_DEPENDENT_CATABOLIC_P | 16 | -0.391133 | -1.034023 | 0.408408 | 1 |
| GOBP_DICARBOXYLIC_ACID_TRANSPORT | 72 | -0.282431 | -1.033876 | 0.339744 | 1 |
| GOBP_COCHLEA_DEVELOPMENT | 40 | -0.321109 | -1.033745 | 0.44186 | 1 |
| GOBP_HISTONE_MONOUBIQUITINATION | 29 | -0.333715 | -1.032079 | 0.406015 | 1 |
| GOBP_NEGATIVE_REGULATION_OF_CALCIUM_ION_TRANSMEMBRANE_ | 29 | -0.336644 | -1.029217 | 0.420635 | 1 |
| GOBP_HORMONE_BIOSYNTHETIC_PROCESS | 47 | -0.297019 | -1.029204 | 0.384615 | 1 |
| GOBP_ASPARTATE_FAMILY_AMINO_ACID_CATABOLIC_PROCESS | 15 | -0.401644 | -1.028365 | 0.410405 | 1 |
| GOBP_PHOTOPERIODISM | 29 | -0.338907 | -1.0273 | 0.36965 | 1 |
| GOBP_RENAL_TUBULAR_SECRETION | 17 | -0.387524 | -1.023862 | 0.39939 | 1 |
| GOBP_REGULATION_OF_FIBROBLAST_MIGRATION | 33 | -0.325284 | -1.022989 | 0.390977 | 1 |
| GOBP_REGULATION_OF_DNA_TEMPLATED_TRANSCRIPTION_ELONGAT | 52 | -0.299657 | -1.019909 | 0.408867 | 1 |
| GOBP_NUCLEAR_TRANSCRIBED_MRNA_CATABOLIC_PROCESS_NONSE | 38 | -0.309901 | -1.019726 | 0.406504 | 1 |
| GOBP_DE_NOVO_PROTEIN_FOLDING | 41 | -0.314087 | -1.019251 | 0.410256 | 1 |
| GOBP_CELLULAR_RESPONSE_TO_INTERLEUKIN_1 | 65 | -0.283059 | -1.019248 | 0.365591 | 1 |
| GOBP_SODIUM_ION_TRANSPORT | 190 | -0.239331 | -1.018667 | 0.380282 | 1 |
| GOBP_HEMATOPOIETIC_STEM_CELL_DIFFERENTIATION | 24 | -0.347192 | -1.018174 | 0.40678 | 1 |
| GOBP_PORE_COMPLEX_ASSEMBLY | 20 | -0.383314 | -1.017631 | 0.444805 | 1 |
| GOBP_MRNA_CLEAVAGE | 18 | -0.378315 | -1.016928 | 0.40566 | 1 |
| GOBP_INNERVATION | 24 | -0.344644 | -1.01595 | 0.421053 | 1 |
| GOBP_GLOMERULAR_EPITHELIUM_DEVELOPMENT | 21 | -0.360606 | -1.015919 | 0.407407 | 1 |
| GOBP_PROTEIN_MODIFICATION_BY_SMALL_PROTEIN_REMOVAL | 146 | -0.242987 | -1.015353 | 0.428571 | 1 |
| GOBP_RESPONSE_TO_CADMIUM_ION | 47 | -0.300023 | -1.015348 | 0.383621 | 1 |

| | | | | | |
|---|---|---|---|---|---|
| GOBP_ENDODERM_DEVELOPMENT | 68 | 0.514142 | 1.479962 | 0.021557 | 0.283149 |
| GOBP_RIBONUCLEOSIDE_METABOLIC_PROCESS | 29 | 0.586487 | 1.478821 | 0.045822 | 0.285183 |
| GOBP_OVULATION | 19 | 0.631573 | 1.478003 | 0.040673 | 0.286467 |
| GOBP_ENDODERMAL_CELL_DIFFERENTIATION | 40 | 0.550398 | 1.477926 | 0.033378 | 0.285717 |
| GOBP_ICOSANOID_TRANSPORT | 34 | 0.571393 | 1.477807 | 0.034211 | 0.285057 |
| GOBP_REGULATION_OF_WOUND_HEALING | 98 | 0.493503 | 1.476951 | 0.017084 | 0.28626 |
| GOBP_REGULATION_OF_POTASSIUM_ION_TRANSPORT | 80 | 0.502866 | 1.474237 | 0.019231 | 0.292351 |
| GOBP_LIPOPOLYSACCHARIDE_MEDIATED_SIGNALING_PATHWAY | 44 | 0.544482 | 1.471695 | 0.035809 | 0.297632 |
| GOBP_DENDRITIC_CELL_DIFFERENTIATION | 34 | 0.569117 | 1.471259 | 0.038043 | 0.297764 |
| GOBP_HYPOTHALAMUS_DEVELOPMENT | 17 | 0.642591 | 1.470717 | 0.05772 | 0.298131 |
| GOBP_NEGATIVE_REGULATION_OF_DEVELOPMENTAL_GROWTH | 97 | 0.477534 | 1.469036 | 0.020737 | 0.301532 |
| GOBP_NEGATIVE_REGULATION_OF_MUSCLE_CELL_DIFFERENTIATION | 50 | 0.538094 | 1.468832 | 0.02372 | 0.301138 |
| GOBP_BLOOD_VESSEL_REMODELING | 32 | 0.55933 | 1.467735 | 0.05 | 0.30302 |
| GOBP_POSITIVE_REGULATION_OF_LEUKOCYTE_CHEMOTAXIS | 63 | 0.511578 | 1.467681 | 0.026994 | 0.302163 |
| GOBP_NEGATIVE_REGULATION_OF_CELL_ACTIVATION | 143 | 0.468502 | 1.467491 | 0.008696 | 0.301686 |
| GOBP_STRIATED_MUSCLE_ADAPTATION | 33 | 0.569899 | 1.467462 | 0.049159 | 0.300797 |
| GOBP_NEGATIVE_REGULATION_OF_MYELOID_LEUKOCYTE_DIFFERENTIATION | 37 | 0.553317 | 1.467384 | 0.038615 | 0.300046 |
| GOBP_REGULATION_OF_INNATE_IMMUNE_RESPONSE | 163 | 0.458491 | 1.466516 | 0.002139 | 0.301177 |
| GOBP_POSITIVE_REGULATION_OF_TRANSMEMBRANE_TRANSPORT | 168 | 0.461834 | 1.46623 | 0.005453 | 0.300984 |
| GOBP_POSITIVE_REGULATION_OF_INNATE_IMMUNE_RESPONSE | 93 | 0.485726 | 1.466152 | 0.017321 | 0.300242 |
| GOBP_MUSCLE_SYSTEM_PROCESS | 337 | 0.439567 | 1.464919 | 0 | 0.302486 |
| GOBP_REGULATION_OF_TISSUE_REMODELING | 52 | 0.529932 | 1.46459 | 0.030864 | 0.30233 |
| GOBP_POSITIVE_REGULATION_OF_ACUTE_INFLAMMATORY_RESPONSE | 16 | 0.640663 | 1.463989 | 0.060345 | 0.302939 |
| GOBP_CYTOKINE_MEDIATED_SIGNALING_PATHWAY | 309 | 0.442802 | 1.463748 | 0 | 0.302621 |
| GOBP_CELLULAR_RESPONSE_TO_INTERFERON_GAMMA | 76 | 0.495514 | 1.463445 | 0.018226 | 0.302485 |
| GOBP_POSITIVE_REGULATION_OF_SMALL_MOLECULE_METABOLIC_PROCESS | 107 | 0.479796 | 1.462776 | 0.019144 | 0.303259 |
| GOBP_ENDODERM_FORMATION | 48 | 0.530092 | 1.462299 | 0.031526 | 0.303659 |
| GOBP_POSITIVE_REGULATION_OF_PEPTIDYL_TYROSINE_PHOSPHORYLATION | 129 | 0.468517 | 1.461564 | 0.01559 | 0.304495 |
| GOBP_POSITIVE_REGULATION_OF_SMOOTH_MUSCLE_CELL_PROLIFERATION | 66 | 0.497664 | 1.461308 | 0.041045 | 0.30424 |
| GOBP_CELLULAR_RESPONSE_TO_GROWTH_HORMONE_STIMULUS | 20 | 0.621514 | 1.461045 | 0.053371 | 0.304025 |
| GOBP_GLYCOSYL_COMPOUND_CATABOLIC_PROCESS | 29 | 0.578765 | 1.460875 | 0.046089 | 0.303557 |
| GOBP_B_CELL_MEDIATED_IMMUNITY | 96 | 0.484522 | 1.46073 | 0.01837 | 0.303059 |
| GOBP_POSITIVE_REGULATION_OF_STEM_CELL_PROLIFERATION | 21 | 0.609164 | 1.460169 | 0.060393 | 0.303546 |
| GOBP_REGULATION_OF_RESPONSE_TO_BIOTIC_STIMULUS | 253 | 0.448088 | 1.45993 | 0.00309 | 0.303185 |
| GOBP_ANTIMICROBIAL_HUMORAL_RESPONSE | 45 | 0.537011 | 1.458895 | 0.036411 | 0.304941 |
| GOBP_REGULATION_OF_HISTONE_H3_K4_METHYLATION | 37 | 0.552968 | 1.458809 | 0.042691 | 0.304226 |
| GOBP_NEGATIVE_REGULATION_OF_CYSTEINE_TYPE_ENDOPEPTIDASE_ACTIVITY | 67 | 0.507332 | 1.456155 | 0.019536 | 0.310123 |
| GOBP_MUSCLE_ADAPTATION | 80 | 0.490666 | 1.455943 | 0.028369 | 0.309658 |
| GOBP_PITUITARY_GLAND_DEVELOPMENT | 30 | 0.577459 | 1.455604 | 0.055782 | 0.309606 |
| GOBP_RESPONSE_TO_DSRNA | 37 | 0.548085 | 1.455527 | 0.044156 | 0.308862 |
| GOBP_SMOOTH_MUSCLE_CELL_PROLIFERATION | 115 | 0.474474 | 1.455179 | 0.020619 | 0.308778 |
| GOBP_TOLL_LIKE_RECEPTOR_3_SIGNALING_PATHWAY | 15 | 0.657506 | 1.45443 | 0.067919 | 0.309828 |
| GOBP_ACTIVATION_OF_NF_KAPPAB_INDUCING_KINASE_ACTIVITY | 16 | 0.637729 | 1.452443 | 0.057949 | 0.314095 |
| GOBP_POSITIVE_REGULATION_OF_INTERLEUKIN_6_PRODUCTION | 63 | 0.50446 | 1.452151 | 0.0369 | 0.31399 |
| GOBP_NEGATIVE_REGULATION_OF_DEFENSE_RESPONSE | 176 | 0.453486 | 1.452122 | 0.007551 | 0.313164 |
| GOBP_POSITIVE_REGULATION_OF_LEUKOCYTE_MEDIATED_IMMUNITY | 90 | 0.486035 | 1.451602 | 0.011521 | 0.313488 |
| GOBP_REGULATION_OF_ALPHA_BETA_T_CELL_ACTIVATION | 71 | 0.492018 | 1.451211 | 0.029762 | 0.313624 |
| GOBP_ARACHIDONIC_ACID_METABOLIC_PROCESS | 39 | 0.537301 | 1.449936 | 0.043984 | 0.315916 |
| GOBP_BLOOD_VESSEL_ENDOTHELIAL_CELL_PROLIFERATION_INVOLVED_IN_SPROUTI | 21 | 0.606341 | 1.448885 | 0.055085 | 0.31771 |
| GOBP_PHOSPHATIDYLETHANOLAMINE_METABOLIC_PROCESS | 24 | 0.596068 | 1.448646 | 0.055944 | 0.317455 |
| GOBP_BIOMINERALIZATION | 124 | 0.466678 | 1.447791 | 0.016685 | 0.318766 |
| GOBP_ANTIGEN_PROCESSING_AND_PRESENTATION | 77 | 0.492938 | 1.446948 | 0.020359 | 0.320019 |
| GOBP_MUSCLE_CELL_DIFFERENTIATION | 294 | 0.444492 | 1.445997 | 0 | 0.321575 |
| GOBP_POSITIVE_REGULATION_OF_CARBOHYDRATE_METABOLIC_PROCESS | 62 | 0.511827 | 1.445684 | 0.027611 | 0.321525 |
| GOBP_NEGATIVE_REGULATION_OF_NECROTIC_CELL_DEATH | 15 | 0.65053 | 1.445455 | 0.060921 | 0.321221 |
| GOBP_MUSCLE_CONTRACTION | 270 | 0.440034 | 1.444838 | 0 | 0.321962 |
| GOBP_NEGATIVE_REGULATION_OF_TRANSMEMBRANE_RECEPTOR_PROTEIN_SERINE_ | 115 | 0.462892 | 1.444681 | 0.021041 | 0.321484 |
| GOBP_MEIOTIC_CELL_CYCLE | 207 | 0.44841 | 1.443119 | 0.003188 | 0.324756 |
| GOBP_HOMOLOGOUS_RECOMBINATION | 49 | 0.521923 | 1.442039 | 0.041667 | 0.326664 |
| GOBP_LYMPHOCYTE_MEDIATED_IMMUNITY | 184 | 0.451134 | 1.441779 | 0.008584 | 0.32649 |
| GOBP_EPIDERMIS_DEVELOPMENT | 222 | 0.443781 | 1.44139 | 0.005274 | 0.326657 |
| GOBP_POSITIVE_REGULATION_OF_TYROSINE_PHOSPHORYLATION_OF_STAT_PROTEIN | 36 | 0.55543 | 1.440788 | 0.048748 | 0.327286 |
| GOBP_SKELETAL_MUSCLE_ORGAN_DEVELOPMENT | 134 | 0.458878 | 1.440689 | 0.016593 | 0.326667 |
| GOBP_NEGATIVE_REGULATION_OF_OSTEOCLAST_DIFFERENTIATION | 23 | 0.585258 | 1.440453 | 0.067432 | 0.326411 |
| GOBP_CALCIUM_ION_TRANSPORT | 325 | 0.436307 | 1.439645 | 0.003046 | 0.327647 |
| GOBP_NEGATIVE_REGULATION_OF_INTERLEUKIN_2_PRODUCTION | 20 | 0.610933 | 1.439543 | 0.051799 | 0.326942 |
| GOBP_MEIOTIC_CELL_CYCLE_PROCESS | 158 | 0.461856 | 1.438292 | 0.009804 | 0.329304 |
| GOBP_AORTA_MORPHOGENESIS | 30 | 0.568041 | 1.437966 | 0.062241 | 0.329292 |
| GOBP_RESPONSE_TO_BACTERIUM | 374 | 0.43262 | 1.437578 | 0.002037 | 0.329386 |
| GOBP_LEUKOCYTE_APOPTOTIC_PROCESS | 83 | 0.479719 | 1.437013 | 0.022864 | 0.329962 |
| GOBP_INTERMEDIATE_FILAMENT_BASED_PROCESS | 38 | 0.541332 | 1.435222 | 0.047814 | 0.333584 |
| GOBP_POLYAMINE_METABOLIC_PROCESS | 19 | 0.60629 | 1.435011 | 0.066474 | 0.333293 |
| GOBP_NEGATIVE_REGULATION_OF_EXOCYTOSIS | 27 | 0.577951 | 1.434981 | 0.054775 | 0.332493 |
| GOBP_POSITIVE_REGULATION_OF_INTERLEUKIN_2_PRODUCTION | 24 | 0.589298 | 1.43477 | 0.076389 | 0.332157 |
| GOBP_G_PROTEIN_COUPLED_RECEPTOR_SIGNALING_PATHWAY_COUPLED_TO_CYCL | 31 | 0.556479 | 1.434704 | 0.0631 | 0.331438 |
| GOBP_OOGENESIS | 80 | 0.488078 | 1.434511 | 0.030697 | 0.331067 |
| GOBP_POSITIVE_REGULATION_OF_MYELOID_LEUKOCYTE_MEDIATED_IMMUNITY | 17 | 0.625511 | 1.434496 | 0.074074 | 0.330223 |
| GOBP_NEGATIVE_REGULATION_OF_MYOBLAST_DIFFERENTIATION | 19 | 0.616197 | 1.433805 | 0.085837 | 0.331258 |
| GOBP_SKELETAL_MUSCLE_CONTRACTION | 32 | 0.55576 | 1.433216 | 0.058108 | 0.331896 |
| GOBP_PARASYMPATHETIC_NERVOUS_SYSTEM_DEVELOPMENT | 15 | 0.631847 | 1.432771 | 0.065967 | 0.332118 |
| GOBP_NEGATIVE_REGULATION_OF_ENDOTHELIAL_CELL_PROLIFERATION | 34 | 0.545477 | 1.431905 | 0.060686 | 0.333584 |
| GOBP_POSITIVE_REGULATION_OF_ADAPTIVE_IMMUNE_RESPONSE | 80 | 0.482612 | 1.431567 | 0.027611 | 0.333598 |
| GOBP_NEGATIVE_REGULATION_OF_B_CELL_ACTIVATION | 27 | 0.575589 | 1.431566 | 0.059155 | 0.332724 |
| GOBP_BROWN_FAT_CELL_DIFFERENTIATION | 42 | 0.533355 | 1.430403 | 0.057441 | 0.33491 |
| GOBP_REGULATION_OF_FILOPODIUM_ASSEMBLY | 46 | 0.521429 | 1.430226 | 0.05168 | 0.334508 |
| GOBP_SPERM_EGG_RECOGNITION | 25 | 0.584615 | 1.429504 | 0.047076 | 0.335514 |
| GOBP_MACROPHAGE_ACTIVATION | 65 | 0.501526 | 1.42946 | 0.035496 | 0.334742 |
| GOBP_INFLAMMATORY_RESPONSE | 493 | 0.425741 | 1.429183 | 0.001009 | 0.334591 |
| GOBP_REGULATION_OF_ACTIVIN_RECEPTOR_SIGNALING_PATHWAY | 23 | 0.592872 | 1.429164 | 0.06347 | 0.333769 |
| GOBP_HUMORAL_IMMUNE_RESPONSE_MEDIATED_BY_CIRCULATING_IMMUNOGLOBUL | 26 | 0.565007 | 1.429037 | 0.078797 | 0.333247 |
| GOBP_NEGATIVE_REGULATION_OF_LEUKOCYTE_CELL_CELL_ADHESION | 88 | 0.479122 | 1.428831 | 0.023838 | 0.332939 |
| GOBP_CELLULAR_RESPONSE_TO_NUTRIENT | 35 | 0.546251 | 1.428764 | 0.074324 | 0.332262 |
| GOBP_FERTILIZATION | 118 | 0.469602 | 1.42843 | 0.022396 | 0.332216 |
| GOBP_SENSORY_PERCEPTION | 371 | 0.427853 | 1.428281 | 0.001009 | 0.331754 |
| GOBP_POLYAMINE_BIOSYNTHETIC_PROCESS | 15 | 0.639432 | 1.427733 | 0.083333 | 0.332303 |
| GOBP_CARDIAC_SEPTUM_MORPHOGENESIS | 62 | 0.498653 | 1.426991 | 0.045728 | 0.333465 |
| GOBP_REGULATION_OF_MACROPHAGE_DERIVED_FOAM_CELL_DIFFERENTIATION | 21 | 0.610485 | 1.426536 | 0.077372 | 0.333862 |
| GOBP_NECROTIC_CELL_DEATH | 49 | 0.512339 | 1.425327 | 0.040764 | 0.336232 |
| GOBP_POSITIVE_REGULATION_OF_ALCOHOL_BIOSYNTHETIC_PROCESS | 20 | 0.602062 | 1.425269 | 0.082489 | 0.335544 |
| GOBP_SPECIFICATION_OF_SYMMETRY | 113 | 0.45826 | 1.425138 | 0.014756 | 0.335033 |
| GOBP_EXTERNAL_ENCAPSULATING_STRUCTURE_ORGANIZATION | 249 | 0.439475 | 1.424893 | 0.008316 | 0.334899 |
| GOBP_CELLULAR_RESPONSE_TO_XENOBIOTIC_STIMULUS | 112 | 0.466736 | 1.424128 | 0.027964 | 0.33599 |
| GOBP_POSITIVE_REGULATION_OF_ALPHA_BETA_T_CELL_ACTIVATION | 47 | 0.516125 | 1.423924 | 0.050064 | 0.335693 |
| GOBP_ENDOTHELIAL_CELL_CHEMOTAXIS | 25 | 0.570037 | 1.42371 | 0.071331 | 0.335418 |
| GOBP_POSITIVE_REGULATION_OF_NATURAL_KILLER_CELL_ACTIVATION | 20 | 0.607209 | 1.423445 | 0.078762 | 0.335318 |
| GOBP_CELLULAR_RESPONSE_TO_TOXIC_SUBSTANCE | 93 | 0.47594 | 1.423121 | 0.025 | 0.335305 |
| GOBP_NUCLEOSIDE_METABOLIC_PROCESS | 54 | 0.505969 | 1.421671 | 0.032911 | 0.338606 |
| GOBP_T_CELL_CYTOKINE_PRODUCTION | 25 | 0.576552 | 1.421448 | 0.070621 | 0.338285 |
| GOBP_GLYCEROPHOSPHOLIPID_CATABOLIC_PROCESS | 28 | 0.566955 | 1.420743 | 0.069541 | 0.339331 |
| GOBP_ACTIN_CYTOSKELETON_REORGANIZATION | 97 | 0.463999 | 1.420612 | 0.037887 | 0.338855 |
| GOBP_NEGATIVE_REGULATION_OF_BMP_SIGNALING_PATHWAY | 50 | 0.511467 | 1.420308 | 0.050891 | 0.338786 |
| GOBP_RESPONSE_TO_STEROL | 23 | 0.578756 | 1.420073 | 0.066761 | 0.338539 |
| GOBP_ORGANIC_CATION_TRANSPORT | 33 | 0.545766 | 1.419763 | 0.062069 | 0.338594 |
| GOBP_DETECTION_OF_VISIBLE_LIGHT | 23 | 0.577831 | 1.419743 | 0.088023 | 0.337803 |
| GOBP_REGULATION_OF_MYELOID_LEUKOCYTE_DIFFERENTIATION | 86 | 0.482314 | 1.41898 | 0.033526 | 0.338975 |
| GOBP_NEGATIVE_REGULATION_OF_LYMPHOCYTE_DIFFERENTIATION | 40 | 0.5301 | 1.418695 | 0.061278 | 0.338984 |
| GOBP_CD4_POSITIVE_ALPHA_BETA_T_CELL_DIFFERENTIATION | 57 | 0.505491 | 1.41757 | 0.0475 | 0.341131 |
| GOBP_CHONDROCYTE_PROLIFERATION | 19 | 0.605312 | 1.417477 | 0.090511 | 0.340529 |
| GOBP_CORONARY_VASCULATURE_MORPHOGENESIS | 16 | 0.61776 | 1.415854 | 0.07511 | 0.344018 |
| GOBP_CELLULAR_PROCESS_INVOLVED_IN_REPRODUCTION_IN_MULTICELLULAR_OR | 284 | 0.428583 | 1.415773 | 0.006154 | 0.343377 |
| GOBP_INNATE_IMMUNE_RESPONSE | 498 | 0.422337 | 1.415388 | 0 | 0.343568 |
| GOBP_MAMMARY_GLAND_EPITHELIUM_DEVELOPMENT | 57 | 0.505395 | 1.415359 | 0.04914 | 0.342801 |
| GOBP_EMBRYONIC_FORELIMB_MORPHOGENESIS | 15 | 0.637566 | 1.415034 | 0.076462 | 0.342886 |
| GOBP_POLYOL_BIOSYNTHETIC_PROCESS | 54 | 0.499376 | 1.414747 | 0.055147 | 0.342846 |
| GOBP_MALE_SEX_DIFFERENTIATION | 127 | 0.455412 | 1.412396 | 0.018038 | 0.348339 |
| GOBP_CARDIAC_RIGHT_VENTRICLE_MORPHOGENESIS | 16 | 0.62714 | 1.412394 | 0.084465 | 0.347522 |
| GOBP_DOPAMINE_TRANSPORT | 43 | 0.521662 | 1.412223 | 0.062104 | 0.347192 |
| GOBP_REGULATION_OF_STRIATED_MUSCLE_CELL_DIFFERENTIATION | 67 | 0.491287 | 1.41133 | 0.03912 | 0.348769 |
| GOBP_NUCLEAR_MIGRATION | 21 | 0.592642 | 1.41124 | 0.088889 | 0.348173 |
| GOBP_POSITIVE_REGULATION_OF_TYPE_I_INTERFERON_MEDIATED_SIGNALING_PATH | 16 | 0.623178 | 1.410119 | 0.09538 | 0.350292 |
| GOBP_MICROGLIAL_CELL_ACTIVATION | 27 | 0.560965 | 1.409898 | 0.079561 | 0.3501 |
| GOBP_REGULATION_OF_LEUKOCYTE_ADHESION_TO_VASCULAR_ENDOTHELIAL_CELL | 20 | 0.593096 | 1.409891 | 0.081967 | 0.349306 |
| GOBP_RESPONSE_TO_CHEMOKINE | 45 | 0.519658 | 1.40968 | 0.04557 | 0.349066 |
| GOBP_REGULATION_OF_ALPHA_BETA_T_CELL_DIFFERENTIATION | 47 | 0.511106 | 1.409633 | 0.05243 | 0.348391 |
| GOBP_MUSCLE_CELL_PROLIFERATION | 154 | 0.447335 | 1.407307 | 0.015487 | 0.353632 |
| GOBP_NEGATIVE_REGULATION_OF_INFLAMMATORY_RESPONSE | 106 | 0.460501 | 1.40727 | 0.029246 | 0.352913 |
| GOBP_NEGATIVE_REGULATION_OF_BLOOD_VESSEL_ENDOTHELIAL_CELL_MIGRATION | 28 | 0.562892 | 1.406844 | 0.06469 | 0.353262 |
| GOBP_POSITIVE_REGULATION_OF_PROTEIN_CONTAINING_COMPLEX_DISASSEMBLY | 31 | 0.543782 | 1.406201 | 0.070573 | 0.35416 |
| GOBP_REGULATION_OF_LYMPHOCYTE_MEDIATED_IMMUNITY | 112 | 0.457541 | 1.406072 | 0.031008 | 0.353738 |
| GOBP_XENOBIOTIC_METABOLIC_PROCESS | 62 | 0.490091 | 1.405926 | 0.045067 | 0.353347 |
| GOBP_REGULATION_OF_MUSCLE_ADAPTATION | 64 | 0.490221 | 1.405858 | 0.045838 | 0.35272 |
| GOBP_SERTOLI_CELL_DIFFERENTIATION | 15 | 0.629357 | 1.405763 | 0.099561 | 0.352186 |
| GOBP_HEAD_MORPHOGENESIS | 30 | 0.553405 | 1.405488 | 0.066841 | 0.352106 |
| GOBP_B_CELL_ACTIVATION | 194 | 0.430924 | 1.405045 | 0.014815 | 0.352573 |
| GOBP_POSITIVE_REGULATION_OF_CHEMOTAXIS | 103 | 0.464604 | 1.404564 | 0.032672 | 0.353044 |
| GOBP_REGULATION_OF_ACUTE_INFLAMMATORY_RESPONSE | 28 | 0.558006 | 1.404529 | 0.080711 | 0.352335 |
| GOBP_TYROSINE_PHOSPHORYLATION_OF_STAT_PROTEIN | 49 | 0.51002 | 1.404284 | 0.061481 | 0.352245 |
| GOBP_PURINE_CONTAINING_COMPOUND_TRANSMEMBRANE_TRANSPORT | 20 | 0.580027 | 1.403733 | 0.086957 | 0.352939 |
| GOBP_POSITIVE_REGULATION_OF_RHO_PROTEIN_SIGNAL_TRANSDUCTION | 24 | 0.567613 | 1.403311 | 0.080622 | 0.353301 |
| GOBP_RESPONSE_TO_INTERFERON_ALPHA | 17 | 0.610087 | 1.403295 | 0.09299 | 0.352549 |
| GOBP_NEGATIVE_REGULATION_OF_TUMOR_NECROSIS_FACTOR_SUPERFAMILY_CYTO | 41 | 0.521334 | 1.403096 | 0.057428 | 0.352322 |

| | | | | | |
|---|---|---|---|---|---|
| GOBP_ESTABLISHMENT_OF_EPITHELIAL_CELL_POLARITY | 31 | -0.321291 | -1.013058 | 0.417323 | 1 |
| GOBP_RIBONUCLEOSIDE_DIPHOSPHATE_METABOLIC_PROCESS | 97 | -0.263202 | -1.0128 | 0.433566 | 1 |
| GOBP_DNA_METHYLATION_DEPENDENT_HETEROCHROMATIN_ASSEME | 22 | -0.349326 | -1.012657 | 0.410526 | 1 |
| GOBP_MICROTUBULE_CYTOSKELETON_ORGANIZATION_INVOLVED_IN_ | 149 | -0.248596 | -1.010977 | 0.4 | 1 |
| GOBP_ESTABLISHMENT_OF_PLANAR_POLARITY_OF_EMBRYONIC_EPIT | 15 | -0.378077 | -1.010904 | 0.392523 | 1 |
| GOBP_REGULATION_OF_NEURON_PROJECTION_REGENERATION | 26 | -0.329572 | -1.009822 | 0.421818 | 1 |
| GOBP_REGULATION_OF_ALTERNATIVE_MRNA_SPLICING_VIA_SPLICEO | 57 | -0.288624 | -1.008031 | 0.406091 | 1 |
| GOBP_REGULATION_OF_SODIUM_ION_TRANSMEMBRANE_TRANSPORT | 54 | -0.287328 | -1.007089 | 0.426606 | 1 |
| GOBP_REGULATION_OF_CHROMOSOME_SEPARATION | 66 | -0.273374 | -1.007003 | 0.446701 | 1 |
| GOBP_REGULATION_OF_CELLULAR_PH | 70 | -0.276996 | -1.006929 | 0.473373 | 1 |
| GOBP_LABYRINTHINE_LAYER_DEVELOPMENT | 40 | -0.305736 | -1.005524 | 0.413953 | 1 |
| GOBP_REGULATION_OF_BLOOD_PRESSURE | 136 | -0.243781 | -1.005072 | 0.425743 | 1 |
| GOBP_BILE_ACID_BIOSYNTHETIC_PROCESS | 27 | -0.331705 | -1.0037 | 0.416084 | 1 |
| GOBP_MITOTIC_SISTER_CHROMATID_SEGREGATION | 166 | -0.243845 | -1.003588 | 0.441176 | 1 |
| GOBP_POSITIVE_REGULATION_OF_CYSTEINE_TYPE_ENDOPEPTIDASE_ | 122 | -0.246518 | -0.998571 | 0.489362 | 1 |
| GOBP_DORSAL_VENTRAL_NEURAL_TUBE_PATTERNING | 19 | -0.366847 | -0.998488 | 0.439024 | 1 |
| GOBP_POSITIVE_REGULATION_OF_HEART_RATE | 23 | -0.342476 | -0.995495 | 0.472789 | 1 |
| GOBP_CHAPERONE_MEDIATED_PROTEIN_FOLDING | 67 | -0.278163 | -0.993797 | 0.450777 | 1 |
| GOBP_ASPARTATE_FAMILY_AMINO_ACID_METABOLIC_PROCESS | 44 | -0.295368 | -0.993009 | 0.453782 | 1 |
| GOBP_HISTONE_H4_ACETYLATION | 72 | -0.266153 | -0.992009 | 0.472727 | 1 |
| GOBP_NEGATIVE_REGULATION_OF_ESTABLISHMENT_OF_PROTEIN_LO | 101 | -0.253208 | -0.991717 | 0.47541 | 1 |
| GOBP_PYRIMIDINE_DEOXYRIBONUCLEOTIDE_METABOLIC_PROCESS | 22 | -0.340499 | -0.991549 | 0.437956 | 1 |
| GOBP_2_OXOGLUTARATE_METABOLIC_PROCESS | 16 | -0.37568 | -0.991328 | 0.468085 | 1 |
| GOBP_PEPTIDYL_GLUTAMIC_ACID_MODIFICATION | 24 | -0.342846 | -0.989453 | 0.465035 | 1 |
| GOBP_REGULATION_OF_ARP2_3_COMPLEX_MEDIATED_ACTIN_NUCLE | 21 | -0.346826 | -0.98889 | 0.484849 | 1 |
| GOBP_ESTABLISHMENT_OF_PROTEIN_LOCALIZATION_TO_CHROMOSON | 25 | -0.326816 | -0.987767 | 0.464789 | 1 |
| GOBP_NEGATIVE_REGULATION_OF_DNA_REPLICATION | 27 | -0.329113 | -0.987073 | 0.454545 | 1 |
| GOBP_REGULATION_OF_AMYLOID_PRECURSOR_PROTEIN_CATABOLIC | 37 | -0.312811 | -0.986594 | 0.502222 | 1 |
| GOBP_ORGANIC_ACID_TRANSMEMBRANE_TRANSPORT | 121 | -0.239088 | -0.985735 | 0.454545 | 1 |
| GOBP_SULFUR_COMPOUND_BIOSYNTHETIC_PROCESS | 130 | -0.247472 | -0.985378 | 0.509615 | 1 |
| GOBP_NEGATIVE_REGULATION_OF_DNA_TEMPLATED_TRANSCRIPTION | 16 | -0.372272 | -0.98428 | 0.425532 | 1 |
| GOBP_POSTSYNAPTIC_SPECIALIZATION_ORGANIZATION | 31 | -0.316555 | -0.981378 | 0.460967 | 1 |
| GOBP_SPERM_FLAGELLUM_ASSEMBLY | 24 | -0.33864 | -0.981119 | 0.477663 | 1 |
| GOBP_LIPOPROTEIN_BIOSYNTHETIC_PROCESS | 94 | -0.261533 | -0.979091 | 0.495935 | 1 |
| GOBP_NUCLEOSIDE_TRIPHOSPHATE_BIOSYNTHETIC_PROCESS | 79 | -0.261582 | -0.978775 | 0.496689 | 1 |
| GOBP_DICARBOXYLIC_ACID_CATABOLIC_PROCESS | 17 | -0.367991 | -0.976817 | 0.516418 | 1 |
| GOBP_COLUMNAR_CUBOIDAL_EPITHELIAL_CELL_DEVELOPMENT | 35 | -0.297301 | -0.976299 | 0.45815 | 1 |
| GOBP_REGULATION_OF_PROTEIN_MATURATION | 52 | -0.286119 | -0.975717 | 0.5 | 1 |
| GOBP_CHAPERONE_COFACTOR_DEPENDENT_PROTEIN_REFOLDING | 32 | -0.311396 | -0.975575 | 0.470339 | 1 |
| GOBP_REGULATION_OF_ISOTYPE_SWITCHING | 32 | -0.312353 | -0.970397 | 0.514184 | 1 |
| GOBP_REGULATION_OF_UBIQUITIN_PROTEIN_TRANSFERASE_ACTIVITY | 52 | -0.283398 | -0.970336 | 0.518519 | 1 |
| GOBP_REGULATION_OF_GLYCOLYTIC_PROCESS | 37 | -0.308863 | -0.969303 | 0.484127 | 1 |
| GOBP_REGULATION_OF_TRANSCRIPTION_OF_NUCLEOLAR_LARGE_RR | 15 | -0.363326 | -0.969214 | 0.501511 | 1 |
| GOBP_REGULATION_OF_ENDOCRINE_PROCESS | 23 | -0.342822 | -0.969102 | 0.504886 | 1 |
| GOBP_RIBONUCLEOSIDE_TRIPHOSPHATE_METABOLIC_PROCESS | 82 | -0.256641 | -0.966272 | 0.522293 | 1 |
| GOBP_AMINO_ACID_TRANSMEMBRANE_TRANSPORT | 77 | -0.261122 | -0.965376 | 0.551136 | 1 |
| GOBP_REGULATION_OF_INTRACELLULAR_ESTROGEN_RECEPTOR_SIG | 29 | -0.313578 | -0.964331 | 0.514286 | 1 |
| GOBP_PEROXISOME_ORGANIZATION | 33 | -0.305691 | -0.961208 | 0.512295 | 1 |
| GOBP_REGULATION_OF_PROTEIN_SUMOYLATION | 21 | -0.335728 | -0.958675 | 0.538732 | 1 |
| GOBP_CENTRIOLE_ASSEMBLY | 42 | -0.28492 | -0.957452 | 0.528455 | 1 |
| GOBP_NEGATIVE_REGULATION_OF_G_PROTEIN_COUPLED_RECEPTOR | 40 | -0.287376 | -0.954374 | 0.508696 | 1 |
| GOBP_DOPAMINE_RECEPTOR_SIGNALING_PATHWAY | 34 | -0.304244 | -0.95323 | 0.5 | 1 |
| GOBP_EXCITATORY_SYNAPSE_ASSEMBLY | 23 | -0.328424 | -0.951087 | 0.519856 | 1 |
| GOBP_REGULATION_OF_MYELINATION | 40 | -0.298321 | -0.950005 | 0.50885 | 1 |
| GOBP_PROTEIN_LOCALIZATION_TO_CYTOSKELETON | 56 | -0.276216 | -0.948631 | 0.581633 | 1 |
| GOBP_RENAL_SYSTEM_PROCESS | 80 | -0.255471 | -0.946198 | 0.582857 | 1 |
| GOBP_ATTACHMENT_OF_SPINDLE_MICROTUBULES_TO_KINETOCHORE | 36 | -0.302453 | -0.945902 | 0.552419 | 1 |
| GOBP_POLYSACCHARIDE_BIOSYNTHETIC_PROCESS | 62 | -0.268297 | -0.945881 | 0.52907 | 1 |
| GOBP_MITOTIC_CYTOKINETIC_PROCESS | 23 | -0.336599 | -0.945358 | 0.522807 | 1 |
| GOBP_ACID_SECRETION | 31 | -0.306577 | -0.944929 | 0.525692 | 1 |
| GOBP_TRANSCRIPTION_INITIATION_FROM_RNA_POLYMERASE_II_PROM | 72 | -0.257267 | -0.944314 | 0.583851 | 1 |
| GOBP_NEGATIVE_REGULATION_OF_CHROMOSOME_ORGANIZATION | 82 | -0.244797 | -0.943894 | 0.6 | 1 |
| GOBP_NEGATIVE_REGULATION_OF_INTRACELLULAR_PROTEIN_TRANS | 37 | -0.289355 | -0.943108 | 0.552 | 1 |
| GOBP_DNA_CATABOLIC_PROCESS | 25 | -0.319336 | -0.942753 | 0.507634 | 1 |
| GOBP_N_TERMINAL_PROTEIN_AMINO_ACID_ACETYLATION | 15 | -0.364828 | -0.942403 | 0.547231 | 1 |
| GOBP_POSITIVE_REGULATION_OF_ACTIN_NUCLEATION | 16 | -0.352951 | -0.942365 | 0.566879 | 1 |
| GOBP_REGULATION_OF_CHONDROCYTE_DIFFERENTIATION | 44 | -0.277901 | -0.942288 | 0.579208 | 1 |
| GOBP_MITOTIC_SPINDLE_ORGANIZATION | 125 | -0.234792 | -0.942025 | 0.666667 | 1 |
| GOBP_MORPHOGENESIS_OF_AN_EPITHELIAL_FOLD | 17 | -0.36103 | -0.94174 | 0.527607 | 1 |
| GOBP_LATE_ENDOSOME_TO_VACUOLE_TRANSPORT_VIA_MULTIVESIC | 17 | -0.344375 | -0.941716 | 0.508306 | 1 |
| GOBP_REGULATION_OF_CHROMOSOME_ORGANIZATION | 201 | -0.217894 | -0.941131 | 0.719298 | 1 |
| GOBP_PROTEIN_STABILIZATION | 165 | -0.236273 | -0.940099 | 0.642857 | 1 |
| GOBP_POSTSYNAPTIC_SIGNAL_TRANSDUCTION | 32 | -0.294091 | -0.939547 | 0.578125 | 1 |
| GOBP_POSITIVE_REGULATION_OF_SMOOTHENED_SIGNALING_PATHWA | 29 | -0.311052 | -0.938908 | 0.592593 | 1 |
| GOBP_REGULATION_OF_HEART_RATE | 81 | -0.250729 | -0.936743 | 0.596491 | 1 |
| GOBP_CELL_MIGRATION_IN_HINDBRAIN | 16 | -0.354523 | -0.936288 | 0.524096 | 1 |
| GOBP_PROTEIN_CONTAINING_COMPLEX_LOCALIZATION | 148 | -0.229441 | -0.936138 | 0.689655 | 1 |
| GOBP_FATTY_ACID_TRANSMEMBRANE_TRANSPORT | 15 | -0.361629 | -0.936002 | 0.552632 | 1 |
| GOBP_PROTEIN_PALMITOYLATION | 31 | -0.301048 | -0.933831 | 0.57037 | 1 |
| GOBP_RIBONUCLEOSIDE_TRIPHOSPHATE_BIOSYNTHETIC_PROCESS | 69 | -0.260581 | -0.933315 | 0.61017 | 1 |
| GOBP_NUCLEOSOME_MOBILIZATION | 20 | -0.33226 | -0.933233 | 0.553055 | 1 |
| GOBP_CELLULAR_RESPONSE_TO_KETONE | 80 | -0.251095 | -0.929921 | 0.651163 | 1 |
| GOBP_PRESYNAPSE_ORGANIZATION | 46 | -0.275122 | -0.929831 | 0.564593 | 1 |
| GOBP_POSITIVE_REGULATION_OF_LONG_TERM_SYNAPTIC_POTENTIAT | 21 | -0.332671 | -0.929427 | 0.562092 | 1 |
| GOBP_SERINE_FAMILY_AMINO_ACID_BIOSYNTHETIC_PROCESS | 18 | -0.345431 | -0.929106 | 0.545455 | 1 |
| GOBP_HISTONE_H3_K4_TRIMETHYLATION | 19 | -0.337465 | -0.927161 | 0.524096 | 1 |
| GOBP_REGULATION_OF_MORPHOGENESIS_OF_A_BRANCHING_STRUC | 35 | -0.29043 | -0.927121 | 0.583333 | 1 |
| GOBP_MULTIVESICULAR_BODY_ORGANIZATION | 31 | -0.297829 | -0.926892 | 0.584559 | 1 |
| GOBP_MUCOPOLYSACCHARIDE_METABOLIC_PROCESS | 76 | -0.246998 | -0.926685 | 0.645714 | 1 |
| GOBP_NEURON_PROJECTION_ORGANIZATION | 80 | -0.24702 | -0.926433 | 0.664474 | 1 |
| GOBP_G_PROTEIN_COUPLED_ACETYLCHOLINE_RECEPTOR_SIGNALIN | 15 | -0.355036 | -0.922071 | 0.536443 | 1 |
| GOBP_NONRIBOSOMAL_PEPTIDE_BIOSYNTHETIC_PROCESS | 15 | -0.351575 | -0.918139 | 0.563218 | 1 |
| GOBP_REGULATION_OF_RENAL_SYSTEM_PROCESS | 25 | -0.308222 | -0.917857 | 0.593985 | 1 |
| GOBP_LYMPHANGIOGENESIS | 16 | -0.34307 | -0.917513 | 0.552795 | 1 |
| GOBP_ENDOPLASMIC_RETICULUM_TO_CYTOSOL_TRANSPORT | 28 | -0.301742 | -0.917265 | 0.583658 | 1 |
| GOBP_POLYSACCHARIDE_METABOLIC_PROCESS | 89 | -0.24121 | -0.916734 | 0.681482 | 1 |
| GOBP_PRIMARY_ALCOHOL_METABOLIC_PROCESS | 72 | -0.246388 | -0.915889 | 0.635359 | 1 |
| GOBP_POSITIVE_REGULATION_OF_SMAD_PROTEIN_SIGNAL_TRANSDU | 18 | -0.326262 | -0.914997 | 0.535836 | 1 |
| GOBP_NEURAL_TUBE_FORMATION | 101 | -0.237215 | -0.913654 | 0.678571 | 1 |
| GOBP_RHYTHMIC_BEHAVIOR | 34 | -0.287544 | -0.913077 | 0.587302 | 1 |
| GOBP_TRANSLATIONAL_INITIATION | 111 | -0.229254 | -0.912593 | 0.644628 | 1 |
| GOBP_SPINDLE_ORGANIZATION | 190 | -0.221739 | -0.911653 | 0.779412 | 1 |
| GOBP_REGULATION_OF_GENE_EXPRESSION_EPIGENETIC | 126 | -0.230078 | -0.911238 | 0.651786 | 1 |
| GOBP_POSTSYNAPSE_ORGANIZATION | 145 | -0.221682 | -0.910369 | 0.731707 | 1 |
| GOBP_RETINA_VASCULATURE_DEVELOPMENT_IN_CAMERA_TYPE_EYE | 17 | -0.333568 | -0.908016 | 0.610592 | 1 |
| GOBP_PEPTIDYL_CYSTEINE_MODIFICATION | 44 | -0.266816 | -0.907919 | 0.682028 | 1 |
| GOBP_LIGAND_GATED_ION_CHANNEL_SIGNALING_PATHWAY | 18 | -0.340463 | -0.907764 | 0.586093 | 1 |
| GOBP_CLATHRIN_DEPENDENT_ENDOCYTOSIS | 45 | -0.272733 | -0.907748 | 0.624413 | 1 |
| GOBP_NEGATIVE_REGULATION_OF_INTRACELLULAR_TRANSPORT | 49 | -0.269733 | -0.905577 | 0.642241 | 1 |
| GOBP_AUDITORY_RECEPTOR_CELL_DEVELOPMENT | 18 | -0.33143 | -0.904828 | 0.567213 | 1 |
| GOBP_REGULATION_OF_VASCULAR_PERMEABILITY | 43 | -0.274552 | -0.904528 | 0.64 | 1 |
| GOBP_GAMMA_AMINOBUTYRIC_ACID_TRANSPORT | 15 | -0.340884 | -0.902866 | 0.633117 | 1 |
| GOBP_ASYMMETRIC_CELL_DIVISION | 17 | -0.333569 | -0.902443 | 0.573248 | 1 |
| GOBP_CHONDROCYTE_DEVELOPMENT | 28 | -0.298484 | -0.902279 | 0.599265 | 1 |
| GOBP_REGULATION_OF_VOLTAGE_GATED_CALCIUM_CHANNEL_ACTIV | 27 | -0.30016 | -0.899614 | 0.610236 | 1 |
| GOBP_DENDRITIC_SPINE_MORPHOGENESIS | 55 | -0.253601 | -0.896736 | 0.659898 | 1 |
| GOBP_MITOTIC_NUCLEAR_DIVISION | 274 | -0.202656 | -0.896306 | 0.888889 | 1 |
| GOBP_NEURON_PROJECTION_REGENERATION | 49 | -0.258994 | -0.895962 | 0.638191 | 1 |
| GOBP_REGULATION_OF_DNA_METHYLATION_DEPENDENT_HETEROCH | 16 | -0.334802 | -0.894758 | 0.594059 | 1 |
| GOBP_VESICLE_MEDIATED_TRANSPORT_BETWEEN_ENDOSOMAL_COM | 45 | -0.264281 | -0.893364 | 0.652542 | 1 |
| GOBP_NEURAL_TUBE_DEVELOPMENT | 141 | -0.216316 | -0.891264 | 0.755814 | 1 |
| GOBP_TRABECULA_FORMATION | 20 | -0.31837 | -0.890574 | 0.634304 | 1 |
| GOBP_REPRODUCTIVE_BEHAVIOR | 22 | -0.315904 | -0.890478 | 0.616129 | 1 |
| GOBP_QUINONE_METABOLIC_PROCESS | 31 | -0.287576 | -0.889647 | 0.664207 | 1 |
| GOBP_METENCEPHALON_DEVELOPMENT | 95 | -0.233814 | -0.887956 | 0.729167 | 1 |
| GOBP_VACUOLAR_ACIDIFICATION | 21 | -0.31791 | -0.887614 | 0.580328 | 1 |
| GOBP_VENTRICULAR_SYSTEM_DEVELOPMENT | 28 | -0.293421 | -0.887406 | 0.623077 | 1 |
| GOBP_REGULATION_OF_MRNA_POLYADENYLATION | 20 | -0.317332 | -0.886006 | 0.608392 | 1 |
| GOBP_SENSORY_PERCEPTION_OF_TASTE | 21 | -0.32067 | -0.885134 | 0.646209 | 1 |
| GOBP_IRON_SULFUR_CLUSTER_ASSEMBLY | 25 | -0.296882 | -0.88443 | 0.619231 | 1 |
| GOBP_CELLULAR_ALDEHYDE_METABOLIC_PROCESS | 50 | -0.257032 | -0.883511 | 0.699531 | 1 |
| GOBP_REGULATION_OF_STEM_CELL_POPULATION_MAINTENANCE | 30 | -0.289209 | -0.8835 | 0.643123 | 1 |
| GOBP_DIACYLGLYCEROL_METABOLIC_PROCESS | 23 | -0.298315 | -0.882154 | 0.607261 | 1 |
| GOBP_CELLULAR_CARBOHYDRATE_CATABOLIC_PROCESS | 32 | -0.284151 | -0.881687 | 0.601449 | 1 |
| GOBP_INTRACELLULAR_ESTROGEN_RECEPTOR_SIGNALING_PATHWAY | 43 | -0.268204 | -0.88039 | 0.638767 | 1 |
| GOBP_NEGATIVE_REGULATION_OF_TOLL_LIKE_RECEPTOR_SIGNALING | 27 | -0.296535 | -0.878744 | 0.626923 | 1 |
| GOBP_HYALURONAN_METABOLIC_PROCESS | 21 | -0.311956 | -0.878535 | 0.64 | 1 |
| GOBP_SODIUM_ION_TRANSMEMBRANE_TRANSPORT | 137 | -0.216849 | -0.878469 | 0.829787 | 1 |
| GOBP_SODIUM_ION_HOMEOSTASIS | 40 | -0.26712 | -0.877434 | 0.637097 | 1 |
| GOBP_POSITIVE_REGULATION_OF_CHROMATIN_ORGANIZATION | 15 | -0.33348 | -0.87729 | 0.6 | 1 |
| GOBP_CHROMOSOME_CONDENSATION | 37 | -0.267384 | -0.877098 | 0.706827 | 1 |
| GOBP_ACTIVATION_OF_ADENYLATE_CYCLASE_ACTIVITY | 20 | -0.316889 | -0.877004 | 0.61794 | 1 |
| GOBP_OLIGOPEPTIDE_TRANSPORT | 15 | -0.340927 | -0.875525 | 0.598837 | 1 |
| GOBP_CELL_JUNCTION_MAINTENANCE | 38 | -0.269415 | -0.87495 | 0.705394 | 1 |
| GOBP_REGULATION_OF_CARTILAGE_DEVELOPMENT | 59 | -0.250794 | -0.874651 | 0.723301 | 1 |
| GOBP_POSITIVE_REGULATION_OF_TELOMERASE_RNA_LOCALIZATION_ | 15 | -0.338145 | -0.872938 | 0.637363 | 1 |
| GOBP_PROTEIN_LOCALIZATION_TO_ENDOPLASMIC_RETICULUM | 66 | -0.238593 | -0.871372 | 0.768786 | 1 |
| GOBP_THIOESTER_BIOSYNTHETIC_PROCESS | 40 | -0.26439 | -0.869324 | 0.647321 | 1 |
| GOBP_REGULATION_OF_MITOTIC_SISTER_CHROMATID_SEGREGATION | 42 | -0.253775 | -0.86781 | 0.701195 | 1 |
| GOBP_POSITIVE_REGULATION_OF_VASCULAR_PERMEABILITY | 16 | -0.320236 | -0.86775 | 0.638095 | 1 |
| GOBP_POSTSYNAPTIC_MEMBRANE_ORGANIZATION | 33 | -0.276687 | -0.865815 | 0.670543 | 1 |

| | | | | | |
|---|---|---|---|---|---|
| GOBP_REGULATION_OF_NEUTROPHIL_MIGRATION | 35 | 0.537445 | 1.402055 | 0.064474 | 0.354313 |
| GOBP_REGULATION_OF_CELL_CELL_ADHESION | 320 | 0.425411 | 1.401994 | 0.002041 | 0.353687 |
| GOBP_PROXIMAL_DISTAL_PATTERN_FORMATION | 17 | 0.608532 | 1.401689 | 0.101796 | 0.353746 |
| GOBP_POSITIVE_REGULATION_OF_CALCIUM_ION_TRANSPORT | 90 | 0.464161 | 1.399811 | 0.029851 | 0.35813 |
| GOBP_REGULATION_OF_VESICLE_FUSION | 20 | 0.585078 | 1.399571 | 0.094286 | 0.357999 |
| GOBP_TOLERANCE_INDUCTION | 19 | 0.581295 | 1.399118 | 0.098431 | 0.358402 |
| GOBP_POSITIVE_REGULATION_OF_GLUCOSE_METABOLIC_PROCESS | 34 | 0.540294 | 1.39834 | 0.069241 | 0.359552 |
| GOBP_REGULATION_OF_LIPOPOLYSACCHARIDE_MEDIATED_SIGNALING_PATHWAY | 16 | 0.612608 | 1.398189 | 0.092754 | 0.359182 |
| GOBP_ANATOMICAL_STRUCTURE_HOMEOSTASIS | 238 | 0.42889 | 1.397503 | 0.006211 | 0.360258 |
| GOBP_REGULATION_OF_CHEMOTAXIS | 166 | 0.436995 | 1.397484 | 0.017335 | 0.359528 |
| GOBP_INFLAMMATORY_RESPONSE_TO_ANTIGENIC_STIMULUS | 41 | 0.513263 | 1.397375 | 0.059289 | 0.359039 |
| GOBP_XENOBIOTIC_TRANSPORT | 32 | 0.541218 | 1.396501 | 0.065104 | 0.360587 |
| GOBP_NEGATIVE_REGULATION_OF_MUSCLE_HYPERTROPHY | 24 | 0.57204 | 1.39583 | 0.079498 | 0.361667 |
| GOBP_RESPONSE_TO_NUTRIENT | 114 | 0.452941 | 1.395588 | 0.035714 | 0.361545 |
| GOBP_RESPONSE_TO_OXYGEN_RADICAL | 24 | 0.566877 | 1.39547 | 0.081514 | 0.361085 |
| GOBP_T_CELL_PROLIFERATION | 138 | 0.44879 | 1.395417 | 0.021882 | 0.360464 |
| GOBP_REGULATION_OF_CELLULAR_CARBOHYDRATE_METABOLIC_PROCESS | 126 | 0.451578 | 1.394798 | 0.020994 | 0.361353 |
| GOBP_PHOSPHATIDYLCHOLINE_METABOLIC_PROCESS | 62 | 0.492533 | 1.394452 | 0.065271 | 0.361546 |
| GOBP_THYROID_GLAND_DEVELOPMENT | 15 | 0.631485 | 1.394301 | 0.082822 | 0.361193 |
| GOBP_SKIN_DEVELOPMENT | 184 | 0.436274 | 1.394241 | 0.017076 | 0.360599 |
| GOBP_ACTIVATED_T_CELL_PROLIFERATION | 32 | 0.540833 | 1.393834 | 0.079088 | 0.360952 |
| GOBP_CALCIUM_ION_TRANSMEMBRANE_TRANSPORT | 244 | 0.429802 | 1.393596 | 0.003125 | 0.360829 |
| GOBP_REGULATION_OF_PEPTIDYL_TYROSINE_PHOSPHORYLATION | 189 | 0.431829 | 1.393428 | 0.013874 | 0.360529 |
| GOBP_EPIDERMAL_CELL_DIFFERENTIATION | 131 | 0.452786 | 1.393126 | 0.025814 | 0.360629 |
| GOBP_REGULATION_OF_EPIDERMAL_GROWTH_FACTOR_ACTIVATED_RECEPTOR_ACT | 26 | 0.561472 | 1.392791 | 0.077033 | 0.360789 |
| GOBP_REGULATION_OF_MACROPHAGE_DIFFERENTIATION | 20 | 0.599647 | 1.391898 | 0.067893 | 0.362397 |
| GOBP_POSITIVE_REGULATION_OF_G_PROTEIN_COUPLED_RECEPTOR_SIGNALING_PA | 21 | 0.584608 | 1.39151 | 0.082153 | 0.362674 |
| GOBP_LYMPHOCYTE_APOPTOTIC_PROCESS | 57 | 0.488151 | 1.390462 | 0.050369 | 0.364733 |
| GOBP_DENDRITIC_CELL_MIGRATION | 20 | 0.587607 | 1.39041 | 0.093838 | 0.364114 |
| GOBP_SKELETAL_SYSTEM_MORPHOGENESIS | 165 | 0.437806 | 1.390164 | 0.019737 | 0.364058 |
| GOBP_DETECTION_OF_ABIOTIC_STIMULUS | 90 | 0.464557 | 1.389124 | 0.048235 | 0.366003 |
| GOBP_T_CELL_DIFFERENTIATION_IN_THYMUS | 57 | 0.49246 | 1.388891 | 0.064039 | 0.365885 |
| GOBP_CELLULAR_COMPONENT_ASSEMBLY_INVOLVED_IN_MORPHOGENESIS | 90 | 0.467309 | 1.388608 | 0.037383 | 0.365845 |
| GOBP_SEROTONIN_TRANSPORT | 15 | 0.627882 | 1.388017 | 0.109034 | 0.366676 |
| GOBP_CELL_MATRIX_ADHESION | 196 | 0.430595 | 1.387208 | 0.012752 | 0.368095 |
| GOBP_WOUND_HEALING | 338 | 0.419511 | 1.386878 | 0.003067 | 0.368163 |
| GOBP_KERATINOCYTE_DIFFERENTIATION | 79 | 0.465945 | 1.38614 | 0.055229 | 0.3694 |
| GOBP_PEPTIDYL_ARGININE_MODIFICATION | 23 | 0.571458 | 1.385863 | 0.080851 | 0.369408 |
| GOBP_PRODUCTION_OF_MOLECULAR_MEDIATOR_OF_IMMUNE_RESPONSE | 154 | 0.438341 | 1.385845 | 0.028139 | 0.368718 |
| GOBP_NEGATIVE_REGULATION_OF_SMOOTH_MUSCLE_CELL_PROLIFERATION | 42 | 0.5088 | 1.385629 | 0.067183 | 0.368526 |
| GOBP_TISSUE_MIGRATION | 270 | 0.417961 | 1.385595 | 0.007322 | 0.367874 |
| GOBP_POSITIVE_REGULATION_OF_NIK_NF_KAPPAB_SIGNALING | 50 | 0.506719 | 1.384831 | 0.069498 | 0.369124 |
| GOBP_VENTRICULAR_CARDIAC_MUSCLE_TISSUE_DEVELOPMENT | 46 | 0.502817 | 1.384809 | 0.079487 | 0.368438 |
| GOBP_REGULATION_OF_LYMPHOCYTE_APOPTOTIC_PROCESS | 41 | 0.522213 | 1.383566 | 0.088507 | 0.371037 |
| GOBP_ANTIGEN_PROCESSING_AND_PRESENTATION_OF_ENDOGENOUS_PEPTIDE_AN | 18 | 0.607323 | 1.383326 | 0.104197 | 0.370936 |
| GOBP_PROTEIN_COMPLEX_OLIGOMERIZATION | 211 | 0.425043 | 1.383165 | 0.012685 | 0.370627 |
| GOBP_CELL_ACTIVATION_INVOLVED_IN_IMMUNE_RESPONSE | 194 | 0.430811 | 1.383158 | 0.011752 | 0.369901 |
| GOBP_CORONARY_VASCULATURE_DEVELOPMENT | 41 | 0.515264 | 1.382906 | 0.065891 | 0.369855 |
| GOBP_POSITIVE_REGULATION_OF_GLUCONEOGENESIS | 15 | 0.610084 | 1.382851 | 0.105263 | 0.36924 |
| GOBP_BRANCHING_INVOLVED_IN_MAMMARY_GLAND_DUCT_MORPHOGENESIS | 18 | 0.606074 | 1.382096 | 0.087537 | 0.370517 |
| GOBP_SMOOTH_MUSCLE_CONTRACTION | 91 | 0.461971 | 1.381091 | 0.034443 | 0.372552 |
| GOBP_ANTERIOR_POSTERIOR_PATTERN_SPECIFICATION | 145 | 0.438699 | 1.38078 | 0.023991 | 0.372604 |
| GOBP_CARDIAC_MUSCLE_CELL_PROLIFERATION | 37 | 0.517119 | 1.380746 | 0.079948 | 0.371927 |
| GOBP_CELL_ADHESION_MEDIATED_BY_INTEGRIN | 67 | 0.482067 | 1.380518 | 0.058458 | 0.371807 |
| GOBP_VENTRICULAR_SEPTUM_MORPHOGENESIS | 41 | 0.515763 | 1.380154 | 0.084635 | 0.37205 |
| GOBP_RESPONSE_TO_TUMOR_CELL | 28 | 0.5429 | 1.379458 | 0.097394 | 0.373187 |
| GOBP_EMBRYONIC_CAMERA_TYPE_EYE_MORPHOGENESIS | 17 | 0.598203 | 1.378663 | 0.096439 | 0.374606 |
| GOBP_VASCULAR_ASSOCIATED_SMOOTH_MUSCLE_CELL_PROLIFERATION | 46 | 0.50451 | 1.378556 | 0.079801 | 0.374167 |
| GOBP_LYMPHOCYTE_COSTIMULATION | 26 | 0.545019 | 1.378418 | 0.098291 | 0.373814 |
| GOBP_NEGATIVE_REGULATION_OF_CD4_POSITIVE_ALPHA_BETA_T_CELL_DIFFERENTI | 18 | 0.604824 | 1.378011 | 0.105263 | 0.374254 |
| GOBP_POSITIVE_REGULATION_OF_BONE_MINERALIZATION | 35 | 0.529402 | 1.377951 | 0.090909 | 0.373678 |
| GOBP_RECEPTOR_SIGNALING_PATHWAY_VIA_STAT | 105 | 0.449756 | 1.377888 | 0.04065 | 0.373134 |
| GOBP_MODULATION_BY_HOST_OF_VIRAL_PROCESS | 26 | 0.54853 | 1.377757 | 0.105624 | 0.372763 |
| GOBP_INTERFERON_GAMMA_MEDIATED_SIGNALING_PATHWAY | 20 | 0.590707 | 1.377632 | 0.090517 | 0.372376 |
| GOBP_BODY_MORPHOGENESIS | 40 | 0.518335 | 1.377384 | 0.086957 | 0.372326 |
| GOBP_NEGATIVE_REGULATION_OF_HYDROLASE_ACTIVITY | 247 | 0.419983 | 1.376726 | 0.010438 | 0.373405 |
| GOBP_TUMOR_NECROSIS_FACTOR_SUPERFAMILY_CYTOKINE_PRODUCTION | 112 | 0.449212 | 1.375822 | 0.03303 | 0.375137 |
| GOBP_POSITIVE_REGULATION_OF_RELEASE_OF_SEQUESTERED_CALCIUM_ION_INTO_ | 28 | 0.549457 | 1.374118 | 0.083916 | 0.379119 |
| GOBP_AORTA_DEVELOPMENT | 55 | 0.489589 | 1.373744 | 0.066919 | 0.379389 |
| GOBP_NEGATIVE_REGULATION_OF_DNA_BIOSYNTHETIC_PROCESS | 35 | 0.530383 | 1.37356 | 0.080491 | 0.37916 |
| GOBP_RESPONSE_TO_TYPE_I_INTERFERON | 53 | 0.489265 | 1.373133 | 0.073801 | 0.379648 |
| GOBP_VENTRICULAR_SEPTUM_DEVELOPMENT | 67 | 0.476543 | 1.373107 | 0.06357 | 0.378985 |
| GOBP_REGULATION_OF_B_CELL_ACTIVATION | 97 | 0.454315 | 1.373056 | 0.042286 | 0.378387 |
| GOBP_DNA_DOUBLE_STRAND_BREAK_PROCESSING | 16 | 0.603535 | 1.372855 | 0.100147 | 0.378192 |
| GOBP_IRON_ION_TRANSPORT | 46 | 0.505418 | 1.371696 | 0.08976 | 0.380664 |
| GOBP_POSITIVE_REGULATION_OF_LYMPHOCYTE_MEDIATED_IMMUNITY | 73 | 0.470283 | 1.370863 | 0.06401 | 0.382197 |
| GOBP_UNSATURATED_FATTY_ACID_METABOLIC_PROCESS | 83 | 0.460427 | 1.370448 | 0.042254 | 0.382585 |
| GOBP_INTERLEUKIN_10_PRODUCTION | 36 | 0.511557 | 1.370066 | 0.104712 | 0.382893 |
| GOBP_ACYLGLYCEROL_HOMEOSTASIS | 21 | 0.562053 | 1.369855 | 0.114243 | 0.382725 |
| GOBP_NEGATIVE_REGULATION_OF_T_CELL_DIFFERENTIATION | 32 | 0.522252 | 1.369618 | 0.100551 | 0.382667 |
| GOBP_SENSORY_PERCEPTION_OF_PAIN | 76 | 0.465453 | 1.366989 | 0.058264 | 0.389302 |
| GOBP_CATECHOL_CONTAINING_COMPOUND_BIOSYNTHETIC_PROCESS | 15 | 0.605388 | 1.366484 | 0.125382 | 0.389955 |
| GOBP_PHARYNGEAL_SYSTEM_DEVELOPMENT | 24 | 0.557606 | 1.365987 | 0.106354 | 0.390653 |
| GOBP_VENTRICULAR_CARDIAC_MUSCLE_TISSUE_MORPHOGENESIS | 39 | 0.500758 | 1.36587 | 0.081737 | 0.390272 |
| GOBP_IMMUNE_RESPONSE_REGULATING_SIGNALING_PATHWAY | 293 | 0.413431 | 1.365407 | 0.005165 | 0.390848 |
| GOBP_ENDOCARDIAL_CUSHION_FORMATION | 28 | 0.544939 | 1.364887 | 0.102817 | 0.391592 |
| GOBP_NEGATIVE_REGULATION_OF_PEPTIDASE_ACTIVITY | 158 | 0.435318 | 1.364145 | 0.02954 | 0.393117 |
| GOBP_PHAGOCYTOSIS | 175 | 0.428211 | 1.363835 | 0.021345 | 0.393314 |
| GOBP_ENDOTHELIAL_CELL_MIGRATION | 192 | 0.422805 | 1.363726 | 0.021186 | 0.392875 |
| GOBP_TAXIS | 449 | 0.409112 | 1.363688 | 0.003027 | 0.392237 |
| GOBP_TRANSFORMING_GROWTH_FACTOR_BETA_RECEPTOR_SIGNALING_PATHWAY | 175 | 0.428485 | 1.363537 | 0.026567 | 0.391939 |
| GOBP_POSITIVE_REGULATION_OF_ION_TRANSPORT | 214 | 0.423712 | 1.363121 | 0.017857 | 0.392401 |
| GOBP_MYOTUBE_CELL_DEVELOPMENT | 29 | 0.546813 | 1.36246 | 0.113976 | 0.393507 |
| GOBP_IMMUNE_RESPONSE_REGULATING_CELL_SURFACE_RECEPTOR_SIGNALING_P/ | 173 | 0.43105 | 1.36235 | 0.025532 | 0.393067 |
| GOBP_REGULATION_OF_GRANULOCYTE_CHEMOTAXIS | 36 | 0.522541 | 1.362006 | 0.081333 | 0.393348 |
| GOBP_PULMONARY_VALVE_MORPHOGENESIS | 16 | 0.598325 | 1.36183 | 0.123648 | 0.39315 |
| GOBP_REGIONALIZATION | 258 | 0.416562 | 1.360744 | 0.011294 | 0.395428 |
| GOBP_RESPONSE_TO_VITAMIN_D | 28 | 0.537324 | 1.360677 | 0.111565 | 0.394866 |
| GOBP_CELL_COMMUNICATION_BY_ELECTRICAL_COUPLING_INVOLVED_IN_CARDIAC_( | 26 | 0.540809 | 1.360145 | 0.105769 | 0.395535 |
| GOBP_REGULATION_OF_TYPE_I_INTERFERON_MEDIATED_SIGNALING_PATHWAY | 36 | 0.518966 | 1.359238 | 0.098143 | 0.397312 |
| GOBP_NEGATIVE_REGULATION_OF_RESPONSE_TO_EXTERNAL_STIMULUS | 295 | 0.414665 | 1.358789 | 0.009202 | 0.397907 |
| GOBP_OVULATION_CYCLE_PROCESS | 37 | 0.515913 | 1.358053 | 0.085676 | 0.399277 |
| GOBP_ARTERY_DEVELOPMENT | 84 | 0.45336 | 1.357913 | 0.05421 | 0.398956 |
| GOBP_MEIOTIC_CHROMOSOME_SEPARATION | 21 | 0.555697 | 1.357711 | 0.123167 | 0.398824 |
| GOBP_PODOSOME_ASSEMBLY | 16 | 0.608421 | 1.357277 | 0.127273 | 0.399287 |
| GOBP_POSITIVE_REGULATION_OF_MITOCHONDRION_ORGANIZATION | 66 | 0.470588 | 1.35701 | 0.081049 | 0.399304 |
| GOBP_VIRAL_GENOME_REPLICATION | 112 | 0.444587 | 1.356928 | 0.044674 | 0.39881 |
| GOBP_POSITIVE_REGULATION_OF_BIOMINERALIZATION | 41 | 0.503302 | 1.356871 | 0.077734 | 0.398251 |
| GOBP_NEGATIVE_REGULATION_OF_ALPHA_BETA_T_CELL_DIFFERENTIATION | 20 | 0.577133 | 1.35648 | 0.123386 | 0.398715 |
| GOBP_REGULATION_OF_EXOCYTOSIS | 176 | 0.424244 | 1.356293 | 0.02358 | 0.398465 |
| GOBP_POSITIVE_REGULATION_OF_SMALL_GTPASE_MEDIATED_SIGNAL_TRANSDUCTIC | 54 | 0.482579 | 1.355322 | 0.07875 | 0.400474 |
| GOBP_ICOSANOID_METABOLIC_PROCESS | 76 | 0.461544 | 1.355314 | 0.067776 | 0.399786 |
| GOBP_RESPONSE_TO_TRANSFORMING_GROWTH_FACTOR_BETA | 222 | 0.42007 | 1.355142 | 0.01797 | 0.399541 |
| GOBP_DETECTION_OF_LIGHT_STIMULUS | 34 | 0.518213 | 1.355013 | 0.106101 | 0.399201 |
| GOBP_REGULATION_OF_EPITHELIAL_CELL_DIFFERENTIATION | 114 | 0.447011 | 1.354776 | 0.04675 | 0.399216 |
| GOBP_MYELIN_MAINTENANCE | 16 | 0.597155 | 1.354589 | 0.122239 | 0.399059 |
| GOBP_MONONUCLEAR_CELL_MIGRATION | 120 | 0.436941 | 1.354239 | 0.042601 | 0.399384 |
| GOBP_MEMBRANE_REPOLARIZATION_DURING_ACTION_POTENTIAL | 22 | 0.565242 | 1.354048 | 0.102857 | 0.399238 |
| GOBP_REPLICATIVE_SENESCENCE | 15 | 0.609313 | 1.35351 | 0.123145 | 0.399955 |
| GOBP_COLLAGEN_METABOLIC_PROCESS | 69 | 0.469513 | 1.352691 | 0.077016 | 0.401574 |
| GOBP_CELLULAR_OXIDANT_DETOXIFICATION | 74 | 0.460352 | 1.351897 | 0.062127 | 0.403015 |
| GOBP_REGULATION_OF_PRODUCTION_OF_MOLECULAR_MEDIATOR_OF_IMMUNE_RES | 117 | 0.439359 | 1.351887 | 0.057955 | 0.402354 |
| GOBP_POSITIVE_REGULATION_OF_BLOOD_VESSEL_ENDOTHELIAL_CELL_MIGRATION | 49 | 0.487051 | 1.350066 | 0.085185 | 0.406689 |
| GOBP_POSITIVE_REGULATION_OF_COAGULATION | 18 | 0.591696 | 1.349681 | 0.115031 | 0.407046 |
| GOBP_ANTIGEN_RECEPTOR_MEDIATED_SIGNALING_PATHWAY | 128 | 0.435029 | 1.349615 | 0.037862 | 0.406496 |
| GOBP_REGULATION_OF_SKELETAL_MUSCLE_CELL_DIFFERENTIATION | 17 | 0.594184 | 1.349291 | 0.128012 | 0.406699 |
| GOBP_MALE_GAMETE_GENERATION | 396 | 0.404444 | 1.348534 | 0.00404 | 0.408101 |
| GOBP_POSITIVE_REGULATION_OF_SMOOTH_MUSCLE_CONTRACTION | 27 | 0.535049 | 1.348484 | 0.112033 | 0.407512 |
| GOBP_POSITIVE_REGULATION_OF_MUSCLE_CONTRACTION | 37 | 0.507464 | 1.348166 | 0.08399 | 0.407685 |
| GOBP_ICOSANOID_BIOSYNTHETIC_PROCESS | 33 | 0.5206 | 1.348077 | 0.108219 | 0.407214 |
| GOBP_PULMONARY_VALVE_DEVELOPMENT | 20 | 0.572078 | 1.347813 | 0.121481 | 0.407265 |
| GOBP_POSITIVE_REGULATION_OF_LIPID_BIOSYNTHETIC_PROCESS | 66 | 0.468337 | 1.347716 | 0.058752 | 0.406802 |
| GOBP_ENDOCRINE_PANCREAS_DEVELOPMENT | 31 | 0.520888 | 1.347156 | 0.101551 | 0.407565 |
| GOBP_FOAM_CELL_DIFFERENTIATION | 27 | 0.546001 | 1.347154 | 0.122392 | 0.406885 |
| GOBP_CELLULAR_SODIUM_ION_HOMEOSTASIS | 20 | 0.560996 | 1.346836 | 0.114493 | 0.407019 |
| GOBP_PROTEIN_TETRAMERIZATION | 75 | 0.46196 | 1.346698 | 0.075334 | 0.406744 |
| GOBP_CELL_MATURATION | 134 | 0.4317 | 1.346411 | 0.037445 | 0.406862 |
| GOBP_POSITIVE_REGULATION_OF_PHOSPHATIDYLINOSITOL_3_KINASE_SIGNALING | 54 | 0.473955 | 1.346402 | 0.084383 | 0.406202 |
| GOBP_CARDIAC_EPITHELIAL_TO_MESENCHYMAL_TRANSITION | 33 | 0.522559 | 1.346391 | 0.07947 | 0.405539 |
| GOBP_POSITIVE_REGULATION_OF_MACROPHAGE_DIFFERENTIATION | 15 | 0.592265 | 1.346329 | 0.159317 | 0.404998 |
| GOBP_RESPONSE_TO_INTERFERON_GAMMA | 92 | 0.448456 | 1.346191 | 0.0625 | 0.404732 |
| GOBP_REGULATION_OF_TOLL_LIKE_RECEPTOR_SIGNALING_PATHWAY | 53 | 0.483675 | 1.346121 | 0.095785 | 0.404251 |
| GOBP_FACE_DEVELOPMENT | 43 | 0.500063 | 1.346048 | 0.101023 | 0.403743 |
| GOBP_MONOAMINE_TRANSPORT | 63 | 0.471027 | 1.346018 | 0.065163 | 0.40314 |
| GOBP_POSITIVE_REGULATION_OF_RECEPTOR_SIGNALING_PATHWAY_VIA_STAT | 25 | 0.540518 | 1.345845 | 0.116312 | 0.402973 |
| GOBP_POSITIVE_REGULATION_OF_INTERLEUKIN_8_PRODUCTION | 40 | 0.504934 | 1.345755 | 0.099738 | 0.402567 |
| GOBP_RESPONSE_TO_MECHANICAL_STIMULUS | 164 | 0.421517 | 1.345637 | 0.033441 | 0.402255 |
| GOBP_POSITIVE_REGULATION_OF_I_KAPPAB_KINASE_NF_KAPPAB_SIGNALING | 163 | 0.421854 | 1.345434 | 0.031049 | 0.402137 |
| GOBP_OVULATION_CYCLE | 57 | 0.480011 | 1.344159 | 0.088378 | 0.405146 |
| GOBP_REGULATION_OF_RESPONSE_TO_WOUNDING | 126 | 0.433803 | 1.342555 | 0.050279 | 0.409023 |
| GOBP_LIMBIC_SYSTEM_DEVELOPMENT | 88 | 0.452732 | 1.342431 | 0.062061 | 0.408715 |

| | | | | | |
|---|---|---|---|---|---|
| GOBP_INNER_EAR_AUDITORY_RECEPTOR_CELL_DIFFERENTIATION | 31 | -0.281019 | -0.864529 | 0.686131 | 1 |
| GOBP_RESPONSE_TO_IRON_ION | 24 | -0.302013 | -0.864414 | 0.669014 | 1 |
| GOBP_PROTEIN_DNA_COMPLEX_DISASSEMBLY | 19 | -0.314401 | -0.861773 | 0.634551 | 1 |
| GOBP_OVARIAN_FOLLICLE_DEVELOPMENT | 46 | -0.247189 | -0.860828 | 0.717949 | 1 |
| GOBP_CELLULAR_MONOVALENT_INORGANIC_CATION_HOMEOSTASIS | 91 | -0.231636 | -0.860379 | 0.791667 | 1 |
| GOBP_LIPOSACCHARIDE_METABOLIC_PROCESS | 94 | -0.222777 | -0.859465 | 0.801418 | 1 |
| GOBP_OLIGODENDROCYTE_DEVELOPMENT | 33 | -0.272488 | -0.859394 | 0.69084 | 1 |
| GOBP_MITOTIC_SISTER_CHROMATID_COHESION | 27 | -0.286922 | -0.859307 | 0.733068 | 1 |
| GOBP_CELLULAR_RESPONSE_TO_ALCOHOL | 73 | -0.237934 | -0.859186 | 0.762712 | 1 |
| GOBP_RESPONSE_TO_INTERLEUKIN_4 | 29 | -0.267887 | -0.857834 | 0.669261 | 1 |
| GOBP_AMINOGLYCAN_BIOSYNTHETIC_PROCESS | 75 | -0.236453 | -0.857417 | 0.776471 | 1 |
| GOBP_ESTABLISHMENT_OF_MITOTIC_SPINDLE_LOCALIZATION | 32 | -0.2802 | -0.855805 | 0.687747 | 1 |
| GOBP_VIRAL_TRANSLATION | 18 | -0.311304 | -0.854515 | 0.663609 | 1 |
| GOBP_POSITIVE_REGULATION_OF_CHONDROCYTE_DIFFERENTIATION | 17 | -0.307941 | -0.853815 | 0.681388 | 1 |
| GOBP_DENDRITE_MORPHOGENESIS | 132 | -0.212108 | -0.852911 | 0.921348 | 1 |
| GOBP_REGULATION_OF_POSTSYNAPSE_ORGANIZATION | 72 | -0.233304 | -0.852771 | 0.837989 | 1 |
| GOBP_NEGATIVE_REGULATION_OF_CELL_CYCLE_G2_M_PHASE_TRAN | 60 | -0.232697 | -0.852405 | 0.754011 | 1 |
| GOBP_REGULATION_OF_CARBOHYDRATE_CATABOLIC_PROCESS | 46 | -0.253429 | -0.850564 | 0.740909 | 1 |
| GOBP_REGULATION_OF_TRANSCRIPTION_INVOLVED_IN_G1_S_TRANSI | 20 | -0.300193 | -0.849642 | 0.706625 | 1 |
| GOBP_REGULATION_OF_PROTEIN_LOCALIZATION_TO_SYNAPSE | 18 | -0.30281 | -0.849502 | 0.6875 | 1 |
| GOBP_POSITIVE_REGULATION_OF_CYTOSOLIC_CALCIUM_ION_CONCEI | 18 | -0.316913 | -0.849377 | 0.664653 | 1 |
| GOBP_NEGATIVE_REGULATION_OF_METAPHASE_ANAPHASE_TRANSITI | 39 | -0.265826 | -0.848696 | 0.723141 | 1 |
| GOBP_DENDRITIC_SPINE_MAINTENANCE | 15 | -0.32573 | -0.847778 | 0.652819 | 1 |
| GOBP_RECEPTOR_LOCALIZATION_TO_SYNAPSE | 51 | -0.240963 | -0.847498 | 0.733333 | 1 |
| GOBP_NEGATIVE_REGULATION_OF_PROTEIN_LOCALIZATION_TO_NUCL | 29 | -0.270398 | -0.84633 | 0.686792 | 1 |
| GOBP_NEGATIVE_REGULATION_OF_PROTEIN_LOCALIZATION | 166 | -0.202018 | -0.846248 | 0.936508 | 1 |
| GOBP_DEOXYRIBONUCLEOTIDE_BIOSYNTHETIC_PROCESS | 15 | -0.330203 | -0.845381 | 0.666667 | 1 |
| GOBP_RETINA_MORPHOGENESIS_IN_CAMERA_TYPE_EYE | 43 | -0.253662 | -0.844882 | 0.767635 | 1 |
| GOBP_PROTEIN_LOCALIZATION_TO_CELL_SURFACE | 58 | -0.24404 | -0.844708 | 0.771277 | 1 |
| GOBP_KERATAN_SULFATE_METABOLIC_PROCESS | 17 | -0.309489 | -0.844605 | 0.706949 | 1 |
| GOBP_ACETYL_COA_BIOSYNTHETIC_PROCESS | 16 | -0.317761 | -0.843461 | 0.693252 | 1 |
| GOBP_FATTY_ACYL_COA_BIOSYNTHETIC_PROCESS | 22 | -0.301903 | -0.842729 | 0.697819 | 1 |
| GOBP_POSITIVE_REGULATION_OF_PROTEIN_EXPORT_FROM_NUCLEUS | 16 | -0.317295 | -0.842126 | 0.667622 | 1 |
| GOBP_RNA_MEDIATED_GENE_SILENCING_BY_INHIBITION_OF_TRANSL/ | 16 | -0.31663 | -0.839879 | 0.678571 | 1 |
| GOBP_PROTEIN_MANNOSYLATION | 22 | -0.291294 | -0.839235 | 0.671378 | 1 |
| GOBP_SULFUR_AMINO_ACID_METABOLIC_PROCESS | 31 | -0.271911 | -0.837159 | 0.73606 | 1 |
| GOBP_MITOCHONDRIAL_TRANSMEMBRANE_TRANSPORT | 99 | -0.213274 | -0.836607 | 0.865672 | 1 |
| GOBP_CELLULAR_COMPONENT_DISASSEMBLY_INVOLVED_IN_EXECUT | 24 | -0.281694 | -0.834517 | 0.707143 | 1 |
| GOBP_NEGATIVE_REGULATION_OF_LIPID_CATABOLIC_PROCESS | 15 | -0.307641 | -0.834218 | 0.682779 | 1 |
| GOBP_PYRIMIDINE_NUCLEOTIDE_CATABOLIC_PROCESS | 18 | -0.304474 | -0.830858 | 0.692308 | 1 |
| GOBP_REGULATION_OF_CILIUM_ASSEMBLY | 82 | -0.224425 | -0.830605 | 0.837662 | 1 |
| GOBP_STRIATED_MUSCLE_CELL_APOPTOTIC_PROCESS | 32 | -0.263279 | -0.830362 | 0.723881 | 1 |
| GOBP_REGULATION_OF_TRANSLATIONAL_INITIATION | 73 | -0.223548 | -0.829643 | 0.865854 | 1 |
| GOBP_MODIFIED_AMINO_ACID_TRANSPORT | 25 | -0.285724 | -0.829408 | 0.716418 | 1 |
| GOBP_REGULATION_OF_PROTEIN_DEPHOSPHORYLATION | 79 | -0.219846 | -0.828816 | 0.887006 | 1 |
| GOBP_GLYCOLIPID_BIOSYNTHETIC_PROCESS | 65 | -0.230627 | -0.825969 | 0.810976 | 1 |
| GOBP_POSITIVE_REGULATION_OF_HISTONE_METHYLATION | 44 | -0.243575 | -0.825747 | 0.760684 | 1 |
| GOBP_RNA_INTERFERENCE | 17 | -0.306312 | -0.822024 | 0.72524 | 1 |
| GOBP_REGULATION_OF_ACTIN_NUCLEATION | 31 | -0.266275 | -0.821886 | 0.738197 | 1 |
| GOBP_MODULATION_OF_EXCITATORY_POSTSYNAPTIC_POTENTIAL | 36 | -0.257804 | -0.821513 | 0.785408 | 1 |
| GOBP_ESTROGEN_METABOLIC_PROCESS | 19 | -0.294274 | -0.820356 | 0.731928 | 1 |
| GOBP_NEGATIVE_REGULATION_OF_TRANSLATIONAL_INITIATION | 18 | -0.301697 | -0.818161 | 0.722561 | 1 |
| GOBP_POSITIVE_REGULATION_OF_CELL_JUNCTION_ASSEMBLY | 87 | -0.212006 | -0.817739 | 0.875 | 1 |
| GOBP_REGULATION_OF_CYTOPLASMIC_TRANSPORT | 27 | -0.266734 | -0.816028 | 0.740876 | 1 |
| GOBP_AMYLOID_BETA_FORMATION | 39 | -0.252538 | -0.815789 | 0.767241 | 1 |
| GOBP_STRIATUM_DEVELOPMENT | 16 | -0.31876 | -0.81494 | 0.716012 | 1 |
| GOBP_SCF_DEPENDENT_PROTEASOMAL_UBIQUITIN_DEPENDENT_PRO | 47 | -0.245101 | -0.814831 | 0.78341 | 1 |
| GOBP_POSITIVE_REGULATION_OF_REACTIVE_OXYGEN_SPECIES_BIOS | 15 | -0.315526 | -0.814739 | 0.717718 | 1 |
| GOBP_REGULATION_OF_DOUBLE_STRAND_BREAK_REPAIR_VIA_NONH | 31 | -0.266098 | -0.814235 | 0.766423 | 1 |
| GOBP_PROTEIN_K11_LINKED_UBIQUITINATION | 29 | -0.264456 | -0.812338 | 0.785441 | 1 |
| GOBP_BLASTOCYST_GROWTH | 19 | -0.294833 | -0.812304 | 0.723333 | 1 |
| GOBP_POLARIZED_EPITHELIAL_CELL_DIFFERENTIATION | 24 | -0.279863 | -0.811671 | 0.71987 | 1 |
| GOBP_POSITIVE_REGULATION_OF_MUSCLE_HYPERTROPHY | 20 | -0.28665 | -0.810975 | 0.70684 | 1 |
| GOBP_NEGATIVE_REGULATION_OF_CELL_JUNCTION_ASSEMBLY | 26 | -0.267613 | -0.810814 | 0.770833 | 1 |
| GOBP_POSITIVE_REGULATION_OF_ENDOCYTOSIS | 80 | -0.217333 | -0.810718 | 0.865772 | 1 |
| GOBP_DNA_GEOMETRIC_CHANGE | 94 | -0.212481 | -0.809585 | 0.876812 | 1 |
| GOBP_SOMATIC_STEM_CELL_POPULATION_MAINTENANCE | 35 | -0.261021 | -0.808963 | 0.794355 | 1 |
| GOBP_RIBOSOMAL_SMALL_SUBUNIT_BIOGENESIS | 73 | -0.220461 | -0.807026 | 0.901734 | 1 |
| GOBP_NEGATIVE_REGULATION_OF_PROTEIN_SECRETION | 50 | -0.235059 | -0.806876 | 0.835821 | 1 |
| GOBP_REGULATION_OF_PROTEIN_EXIT_FROM_ENDOPLASMIC_RETICU | 24 | -0.27182 | -0.805993 | 0.792593 | 1 |
| GOBP_ESTABLISHMENT_OF_ENDOTHELIAL_BARRIER | 49 | -0.23736 | -0.80565 | 0.827103 | 1 |
| GOBP_POSITIVE_REGULATION_OF_EXCITATORY_POSTSYNAPTIC_POTE | 24 | -0.277868 | -0.804838 | 0.726667 | 1 |
| GOBP_REGULATION_OF_TELOMERASE_ACTIVITY | 49 | -0.244666 | -0.803533 | 0.805263 | 1 |
| GOBP_ESTABLISHMENT_OF_PROTEIN_LOCALIZATION_TO_ENDOPLASM | 42 | -0.234045 | -0.802854 | 0.819005 | 1 |
| GOBP_REGULATION_OF_MICROTUBULE_CYTOSKELETON_ORGANIZATI( | 148 | -0.194417 | -0.801949 | 0.95122 | 1 |
| GOBP_POSITIVE_REGULATION_OF_LYASE_ACTIVITY | 16 | -0.298388 | -0.80188 | 0.762712 | 1 |
| GOBP_DNA_REPLICATION_CHECKPOINT_SIGNALING | 16 | -0.307249 | -0.801575 | 0.772871 | 1 |
| GOBP_REGULATION_OF_FIBROBLAST_GROWTH_FACTOR_RECEPTOR_$ | 30 | -0.254827 | -0.7995 | 0.76824 | 1 |
| GOBP_NEGATIVE_REGULATION_OF_DEFENSE_RESPONSE_TO_VIRUS | 25 | -0.271277 | -0.798969 | 0.783333 | 1 |
| GOBP_POSITIVE_REGULATION_OF_CYCLASE_ACTIVITY | 16 | -0.302738 | -0.798281 | 0.758621 | 1 |
| GOBP_PYRIMIDINE_NUCLEOSIDE_TRIPHOSPHATE_BIOSYNTHETIC_PRO | 19 | -0.284739 | -0.79794 | 0.777385 | 1 |
| GOBP_REGULATION_OF_FEEDING_BEHAVIOR | 16 | -0.307165 | -0.796954 | 0.733333 | 1 |
| GOBP_MITOTIC_SPINDLE_ASSEMBLY | 70 | -0.218936 | -0.795988 | 0.865922 | 1 |
| GOBP_PH_REDUCTION | 40 | -0.246832 | -0.795494 | 0.828571 | 1 |
| GOBP_NEGATIVE_REGULATION_OF_TELOMERE_MAINTENANCE_VIA_TE | 19 | -0.28569 | -0.79524 | 0.755853 | 1 |
| GOBP_PROTEIN_LOCALIZATION_TO_CHROMOSOME_TELOMERIC_REGI( | 30 | -0.254405 | -0.794731 | 0.816327 | 1 |
| GOBP_POSITIVE_REGULATION_OF_BLOOD_PRESSURE | 29 | -0.26002 | -0.79438 | 0.807547 | 1 |
| GOBP_PROTEIN_TARGETING_TO_MITOCHONDRION | 99 | -0.202709 | -0.794297 | 0.925676 | 1 |
| GOBP_NEGATIVE_REGULATION_OF_OXIDATIVE_STRESS_INDUCED_CEI | 37 | -0.24657 | -0.7921 | 0.769231 | 1 |
| GOBP_NUCLEUS_ORGANIZATION | 126 | -0.200613 | -0.792002 | 0.944954 | 1 |
| GOBP_RIBONUCLEOPROTEIN_COMPLEX_SUBUNIT_ORGANIZATION | 182 | -0.194948 | -0.790449 | 1 | 1 |
| GOBP_TETRAHYDROFOLATE_METABOLIC_PROCESS | 20 | -0.290206 | -0.789199 | 0.774295 | 1 |
| GOBP_REGULATION_OF_MORPHOGENESIS_OF_AN_EPITHELIUM | 46 | -0.234263 | -0.788789 | 0.831858 | 1 |
| GOBP_CYTOKINETIC_PROCESS | 39 | -0.239884 | -0.788721 | 0.834821 | 1 |
| GOBP_ALCOHOL_CATABOLIC_PROCESS | 42 | -0.235198 | -0.788082 | 0.879167 | 1 |
| GOBP_REGULATION_OF_ACTIN_FILAMENT_BUNDLE_ASSEMBLY | 95 | -0.204431 | -0.787529 | 0.914063 | 1 |
| GOBP_SPHINGOMYELIN_METABOLIC_PROCESS | 17 | -0.291943 | -0.786571 | 0.798246 | 1 |
| GOBP_CENTROSOME_DUPLICATION | 70 | -0.213834 | -0.785375 | 0.883333 | 1 |
| GOBP_REGULATION_OF_ORGANELLE_ASSEMBLY | 211 | -0.186223 | -0.785259 | 0.979167 | 1 |
| GOBP_POSITIVE_REGULATION_BY_HOST_OF_VIRAL_TRANSCRIPTION | 17 | -0.285487 | -0.78469 | 0.806557 | 1 |
| GOBP_LONG_TERM_SYNAPTIC_DEPRESSION | 27 | -0.261628 | -0.784558 | 0.818182 | 1 |
| GOBP_NUCLEAR_BODY_ORGANIZATION | 15 | -0.299184 | -0.783324 | 0.752322 | 1 |
| GOBP_MITOCHONDRIAL_DNA_REPLICATION | 15 | -0.299618 | -0.781795 | 0.791789 | 1 |
| GOBP_PROTEIN_MONOUBIQUITINATION | 78 | -0.209638 | -0.780673 | 0.908537 | 1 |
| GOBP_UBIQUITIN_DEPENDENT_ERAD_PATHWAY | 84 | -0.207788 | -0.780291 | 0.958621 | 1 |
| GOBP_CELLULAR_COMPONENT_MAINTENANCE | 57 | -0.225808 | -0.78022 | 0.866667 | 1 |
| GOBP_GOLGI_TO_ENDOSOME_TRANSPORT | 17 | -0.295821 | -0.77943 | 0.773639 | 1 |
| GOBP_DNA_DEPENDENT_DNA_REPLICATION | 152 | -0.191471 | -0.778668 | 0.988889 | 1 |
| GOBP_NEUROTROPHIN_SIGNALING_PATHWAY | 34 | -0.248495 | -0.778069 | 0.828358 | 1 |
| GOBP_SMOOTHENED_SIGNALING_PATHWAY | 120 | -0.193418 | -0.777546 | 0.961538 | 1 |
| GOBP_CELL_PROLIFERATION_INVOLVED_IN_KIDNEY_DEVELOPMENT | 17 | -0.288124 | -0.777424 | 0.772308 | 1 |
| GOBP_MODIFICATION_OF_SYNAPTIC_STRUCTURE | 19 | -0.282704 | -0.775589 | 0.833876 | 1 |
| GOBP_REGULATION_OF_DENDRITIC_SPINE_MORPHOGENESIS | 42 | -0.229696 | -0.775559 | 0.847291 | 1 |
| GOBP_PROTEIN_N_LINKED_GLYCOSYLATION | 65 | -0.217405 | -0.775087 | 0.93956 | 1 |
| GOBP_NAD_METABOLIC_PROCESS | 24 | -0.267046 | -0.775057 | 0.8 | 1 |
| GOBP_REGULATION_OF_MRNA_3_END_PROCESSING | 28 | -0.253085 | -0.774168 | 0.832776 | 1 |
| GOBP_EPHRIN_RECEPTOR_SIGNALING_PATHWAY | 51 | -0.227762 | -0.773634 | 0.851163 | 1 |
| GOBP_POSITIVE_REGULATION_OF_PROTEIN_DEPHOSPHORYLATION | 40 | -0.236161 | -0.772926 | 0.884774 | 1 |
| GOBP_GLANDULAR_EPITHELIAL_CELL_DIFFERENTIATION | 34 | -0.241693 | -0.772352 | 0.852713 | 1 |
| GOBP_REGULATION_OF_AMPA_RECEPTOR_ACTIVITY | 23 | -0.270747 | -0.772257 | 0.804124 | 1 |
| GOBP_SNARE_COMPLEX_ASSEMBLY | 15 | -0.300229 | -0.771628 | 0.803571 | 1 |
| GOBP_AMYLOID_PRECURSOR_PROTEIN_METABOLIC_PROCESS | 66 | -0.212598 | -0.770766 | 0.92 | 1 |
| GOBP_ACETYL_COA_METABOLIC_PROCESS | 28 | -0.246305 | -0.768824 | 0.840149 | 1 |
| GOBP_REGULATION_OF_STEM_CELL_DIFFERENTIATION | 54 | -0.213999 | -0.767997 | 0.899083 | 1 |
| GOBP_COTRANSLATIONAL_PROTEIN_TARGETING_TO_MEMBRANE | 24 | -0.258761 | -0.766896 | 0.834984 | 1 |
| GOBP_PROTEIN_LOCALIZATION_TO_CHROMOSOME | 90 | -0.199909 | -0.764648 | 0.946667 | 1 |
| GOBP_TRANSCRIPTION_ELONGATION_FROM_RNA_POLYMERASE_II_PF | 56 | -0.215247 | -0.764347 | 0.921951 | 1 |
| GOBP_SPINDLE_ASSEMBLY | 124 | -0.190955 | -0.764144 | 1 | 1 |
| GOBP_FOLIC_ACID_CONTAINING_COMPOUND_METABOLIC_PROCESS | 27 | -0.253241 | -0.763599 | 0.848709 | 1 |
| GOBP_LABYRINTHINE_LAYER_BLOOD_VESSEL_DEVELOPMENT | 18 | -0.282999 | -0.761608 | 0.825806 | 1 |
| GOBP_NADH_METABOLIC_PROCESS | 28 | -0.241552 | -0.761009 | 0.839695 | 1 |
| GOBP_PROTEIN_QUALITY_CONTROL_FOR_MISFOLDED_OR_INCOMPLE | 27 | -0.243444 | -0.75977 | 0.851724 | 1 |
| GOBP_G1_TO_G0_TRANSITION | 17 | -0.287078 | -0.759455 | 0.814103 | 1 |
| GOBP_NEGATIVE_REGULATION_OF_VASCULAR_PERMEABILITY | 17 | -0.286058 | -0.759347 | 0.791908 | 1 |
| GOBP_DNA_REPLICATION | 266 | -0.177993 | -0.759055 | 1 | 1 |
| GOBP_ENDOTHELIAL_CELL_DEVELOPMENT | 64 | -0.207573 | -0.757088 | 0.946309 | 1 |
| GOBP_REGULATION_OF_DNA_REPLICATION | 121 | -0.191488 | -0.753992 | 0.981651 | 1 |
| GOBP_RESPONSE_TO_INSULIN | 214 | -0.174202 | -0.753485 | 0.948718 | 1 |
| GOBP_NEURON_PROJECTION_ARBORIZATION | 29 | -0.245326 | -0.752983 | 0.854902 | 1 |
| GOBP_POSTSYNAPSE_ASSEMBLY | 26 | -0.250356 | -0.752674 | 0.830325 | 1 |
| GOBP_STEROID_CATABOLIC_PROCESS | 19 | -0.271196 | -0.75234 | 0.833866 | 1 |
| GOBP_NEGATIVE_REGULATION_OF_RESPONSE_TO_OXIDATIVE_STRES | 16 | -0.286307 | -0.751022 | 0.781759 | 1 |
| GOBP_NUCLEAR_ENVELOPE_ORGANIZATION | 55 | -0.216992 | -0.75088 | 0.918919 | 1 |
| GOBP_INTERMEMBRANE_LIPID_TRANSFER | 44 | -0.222122 | -0.750449 | 0.883621 | 1 |
| GOBP_REGULATION_OF_CHROMATIN_ASSEMBLY | 29 | -0.24514 | -0.75042 | 0.858238 | 1 |
| GOBP_SPLICEOSOMAL_COMPLEX_ASSEMBLY | 58 | -0.214088 | -0.749719 | 0.921053 | 1 |
| GOBP_3_UTR_MEDIATED_MRNA_STABILIZATION | 19 | -0.282929 | -0.749547 | 0.778125 | 1 |
| GOBP_INTERLEUKIN_1_MEDIATED_SIGNALING_PATHWAY | 20 | -0.265554 | -0.749416 | 0.813953 | 1 |
| GOBP_SNO_S_RNA_METABOLIC_PROCESS | 16 | -0.299513 | -0.749208 | 0.793594 | 1 |
| GOBP_ANDROGEN_METABOLIC_PROCESS | 17 | -0.272361 | -0.749173 | 0.789644 | 1 |
| GOBP_GPI_ANCHOR_METABOLIC_PROCESS | 30 | -0.240314 | -0.747554 | 0.854546 | 1 |

| | | | | | |
|---|---|---|---|---|---|
| GOBP_REGULATORY_T_CELL_DIFFERENTIATION | 25 | 0.551571 | 1.342357 | 0.122881 | 0.408258 |
| GOBP_ALPHA_BETA_T_CELL_ACTIVATION | 106 | 0.44309 | 1.342149 | 0.050459 | 0.408187 |
| GOBP_NEGATIVE_REGULATION_OF_EPITHELIAL_CELL_DIFFERENTIATION | 34 | 0.520052 | 1.342099 | 0.101583 | 0.407642 |
| GOBP_CATECHOLAMINE_UPTAKE | 16 | 0.595081 | 1.342081 | 0.144172 | 0.407019 |
| GOBP_REGULATION_OF_POSITIVE_CHEMOTAXIS | 21 | 0.561372 | 1.341948 | 0.130374 | 0.406706 |
| GOBP_REGULATION_OF_HEMOPOIESIS | 280 | 0.409405 | 1.341913 | 0.018424 | 0.406121 |
| GOBP_POSITIVE_CHEMOTAXIS | 42 | 0.498413 | 1.34189 | 0.10992 | 0.405512 |
| GOBP_CELL_SUBSTRATE_ADHESION | 312 | 0.407061 | 1.341784 | 0.006141 | 0.405088 |
| GOBP_POSITIVE_REGULATION_OF_POTASSIUM_ION_TRANSPORT | 41 | 0.501448 | 1.341625 | 0.098143 | 0.40483 |
| GOBP_POSITIVE_REGULATION_OF_MYELOID_CELL_DIFFERENTIATION | 77 | 0.452388 | 1.341066 | 0.060096 | 0.405858 |
| GOBP_REGULATION_OF_PROTEIN_DEACETYLATION | 55 | 0.48469 | 1.340859 | 0.093252 | 0.40575 |
| GOBP_INTERLEUKIN_6_PRODUCTION | 99 | 0.443709 | 1.340742 | 0.065728 | 0.405379 |
| GOBP_EPIDERMIS_MORPHOGENESIS | 22 | 0.548581 | 1.340688 | 0.123404 | 0.40488 |
| GOBP_POTASSIUM_ION_TRANSPORT | 181 | 0.418401 | 1.339474 | 0.039615 | 0.407515 |
| GOBP_REGULATION_OF_MUSCLE_CELL_DIFFERENTIATION | 106 | 0.43436 | 1.339222 | 0.055046 | 0.407624 |
| GOBP_NEGATIVE_REGULATION_OF_WOUND_HEALING | 52 | 0.478095 | 1.339136 | 0.099751 | 0.407183 |
| GOBP_SUPEROXIDE_METABOLIC_PROCESS | 51 | 0.484766 | 1.338764 | 0.09068 | 0.407476 |
| GOBP_AUTONOMIC_NERVOUS_SYSTEM_DEVELOPMENT | 35 | 0.503541 | 1.338652 | 0.102426 | 0.407156 |
| GOBP_REGULATION_OF_BIOMINERALIZATION | 74 | 0.460098 | 1.338501 | 0.071856 | 0.406909 |
| GOBP_RESPONSE_TO_CHOLESTEROL | 19 | 0.567755 | 1.33791 | 0.132668 | 0.407877 |
| GOBP_REGULATION_OF_MYOBLAST_DIFFERENTIATION | 37 | 0.505464 | 1.33687 | 0.109043 | 0.410034 |
| GOBP_REGULATION_OF_ENDOTHELIAL_CELL_CHEMOTAXIS | 19 | 0.570419 | 1.336549 | 0.121641 | 0.410281 |
| GOBP_CARDIAC_SEPTUM_DEVELOPMENT | 94 | 0.448985 | 1.335781 | 0.067352 | 0.41174 |
| GOBP_HEART_VALVE_DEVELOPMENT | 59 | 0.468382 | 1.335254 | 0.07375 | 0.412456 |
| GOBP_BLASTOCYST_DEVELOPMENT | 102 | 0.442716 | 1.33494 | 0.058957 | 0.412687 |
| GOBP_CALCIUM_ION_IMPORT | 71 | 0.458882 | 1.334883 | 0.08313 | 0.412206 |
| GOBP_REGULATION_OF_REGULATED_SECRETORY_PATHWAY | 118 | 0.42995 | 1.334817 | 0.054608 | 0.411734 |
| GOBP_ANATOMICAL_STRUCTURE_MATURATION | 186 | 0.419525 | 1.334758 | 0.030851 | 0.411238 |
| GOBP_UTERUS_DEVELOPMENT | 16 | 0.586216 | 1.334293 | 0.151976 | 0.411882 |
| GOBP_REGULATION_OF_MEIOTIC_CELL_CYCLE | 38 | 0.500257 | 1.334195 | 0.097911 | 0.411481 |
| GOBP_MITOCHONDRIAL_CALCIUM_ION_TRANSMEMBRANE_TRANSPORT | 18 | 0.569525 | 1.332863 | 0.132159 | 0.414682 |
| GOBP_PATTERN_SPECIFICATION_PROCESS | 343 | 0.403603 | 1.332257 | 0.007202 | 0.415736 |
| GOBP_POSITIVE_REGULATION_OF_LEUKOCYTE_DEGRANULATION | 21 | 0.555767 | 1.332126 | 0.117898 | 0.415436 |
| GOBP_REGULATION_OF_HEART_GROWTH | 47 | 0.483401 | 1.332007 | 0.082031 | 0.415116 |
| GOBP_PROSTANOID_BIOSYNTHETIC_PROCESS | 23 | 0.542749 | 1.331925 | 0.13205 | 0.414706 |
| GOBP_REGULATION_OF_NEUTROPHIL_CHEMOTAXIS | 25 | 0.529675 | 1.331706 | 0.12301 | 0.414594 |
| GOBP_REGULATION_OF_BONE_MINERALIZATION | 60 | 0.462409 | 1.331222 | 0.094059 | 0.415268 |
| GOBP_CARDIAC_VENTRICLE_DEVELOPMENT | 107 | 0.440513 | 1.330495 | 0.069635 | 0.416656 |
| GOBP_POSITIVE_REGULATION_OF_CHOLESTEROL_EFFLUX | 21 | 0.559024 | 1.330393 | 0.132312 | 0.41633 |
| GOBP_CELL_FATE_SPECIFICATION | 63 | 0.464203 | 1.329552 | 0.078358 | 0.418026 |
| GOBP_MITOTIC_G1_S_TRANSITION_CHECKPOINT_SIGNALING | 27 | 0.537659 | 1.328944 | 0.117241 | 0.41906 |
| GOBP_POSITIVE_REGULATION_OF_TYPE_I_INTERFERON_PRODUCTION | 50 | 0.477764 | 1.328164 | 0.096612 | 0.42063 |
| GOBP_DEFENSE_RESPONSE_TO_SYMBIONT | 212 | 0.413688 | 1.32816 | 0.028481 | 0.419986 |
| GOBP_MESENCHYME_MORPHOGENESIS | 48 | 0.480837 | 1.328154 | 0.099237 | 0.419348 |
| GOBP_POSITIVE_REGULATION_OF_COLD_INDUCED_THERMOGENESIS | 80 | 0.445681 | 1.328121 | 0.07377 | 0.418801 |
| GOBP_POSITIVE_REGULATION_OF_LIPID_LOCALIZATION | 81 | 0.446412 | 1.327178 | 0.080529 | 0.420815 |
| GOBP_GLYCEROLIPID_CATABOLIC_PROCESS | 52 | 0.482597 | 1.327136 | 0.089901 | 0.420287 |
| GOBP_CELLULAR_RESPONSE_TO_MOLECULE_OF_BACTERIAL_ORIGIN | 133 | 0.423479 | 1.326597 | 0.056167 | 0.420487 |
| GOBP_POSITIVE_REGULATION_OF_B_CELL_PROLIFERATION | 26 | 0.534281 | 1.326597 | 0.12763 | 0.421135 |
| GOBP_DNA_DEALKYLATION | 26 | 0.540715 | 1.326133 | 0.124118 | 0.421039 |
| GOBP_RESPONSE_TO_MUSCLE_STRETCH | 21 | 0.560312 | 1.325682 | 0.134181 | 0.42161 |
| GOBP_REGULATION_OF_CELL_ADHESION_MEDIATED_BY_INTEGRIN | 39 | 0.493504 | 1.325594 | 0.106579 | 0.421193 |
| GOBP_REGULATION_OF_FATTY_ACID_TRANSPORT | 25 | 0.537904 | 1.325581 | 0.138965 | 0.420583 |
| GOBP_POSITIVE_REGULATION_OF_PRODUCTION_OF_MOLECULAR_MEDIATOR_OF_IMM | 84 | 0.448002 | 1.325525 | 0.061758 | 0.420095 |
| GOBP_T_CELL_APOPTOTIC_PROCESS | 37 | 0.507779 | 1.325129 | 0.103947 | 0.420533 |
| GOBP_REGULATION_OF_ANTIGEN_RECEPTOR_MEDIATED_SIGNALING_PATHWAY | 45 | 0.477313 | 1.32509 | 0.122555 | 0.419997 |
| GOBP_REGULATION_OF_CARDIAC_MUSCLE_CELL_PROLIFERATION | 29 | 0.532045 | 1.324951 | 0.141873 | 0.419739 |
| GOBP_ALPHA_BETA_T_CELL_PROLIFERATION | 29 | 0.526875 | 1.324923 | 0.131171 | 0.419182 |
| GOBP_CELLULAR_RESPONSE_TO_MECHANICAL_STIMULUS | 60 | 0.467333 | 1.324803 | 0.083636 | 0.418854 |
| GOBP_REGULATION_OF_COLLAGEN_METABOLIC_PROCESS | 28 | 0.51468 | 1.324578 | 0.125529 | 0.418824 |
| GOBP_HEART_MORPHOGENESIS | 216 | 0.409741 | 1.323359 | 0.029289 | 0.42162 |
| GOBP_SINGLE_FERTILIZATION | 86 | 0.4462 | 1.322446 | 0.074799 | 0.423515 |
| GOBP_ODONTOGENESIS | 94 | 0.440365 | 1.321997 | 0.076655 | 0.424186 |
| GOBP_REGULATION_OF_CILIUM_DEPENDENT_CELL_MOTILITY | 18 | 0.579443 | 1.321991 | 0.147541 | 0.423576 |
| GOBP_SENSORY_PERCEPTION_OF_SMELL | 27 | 0.509713 | 1.321304 | 0.143654 | 0.424843 |
| GOBP_REGULATION_OF_VIRAL_ENTRY_INTO_HOST_CELL | 37 | 0.499823 | 1.319851 | 0.128342 | 0.428198 |
| GOBP_LIPID_DROPLET_ORGANIZATION | 22 | 0.548014 | 1.319587 | 0.129888 | 0.428348 |
| GOBP_CYTOSOLIC_CALCIUM_ION_TRANSPORT | 143 | 0.417435 | 1.319053 | 0.048405 | 0.429256 |
| GOBP_NEGATIVE_REGULATION_OF_RHO_PROTEIN_SIGNAL_TRANSDUCTION | 21 | 0.549675 | 1.318525 | 0.152327 | 0.430205 |
| GOBP_APOPTOTIC_PROCESS_INVOLVED_IN_MORPHOGENESIS | 23 | 0.5294 | 1.318397 | 0.120055 | 0.429943 |
| GOBP_POSITIVE_REGULATION_OF_PROTEIN_SECRETION | 103 | 0.432954 | 1.318233 | 0.057604 | 0.429766 |
| GOBP_RESPONSE_TO_MOLECULE_OF_BACTERIAL_ORIGIN | 217 | 0.406281 | 1.317876 | 0.030558 | 0.430073 |
| GOBP_RESPONSE_TO_REACTIVE_OXYGEN_SPECIES | 169 | 0.416817 | 1.317789 | 0.042105 | 0.429702 |
| GOBP_PHENOL_CONTAINING_COMPOUND_BIOSYNTHETIC_PROCESS | 31 | 0.506022 | 1.316977 | 0.129921 | 0.43124 |
| GOBP_REPLACEMENT_OSSIFICATION | 26 | 0.534391 | 1.316804 | 0.118812 | 0.431059 |
| GOBP_RESPONSE_TO_ESTROGEN | 53 | 0.476054 | 1.31648 | 0.105781 | 0.431338 |
| GOBP_POSITIVE_REGULATION_OF_DEFENSE_RESPONSE_TO_VIRUS_BY_HOST | 24 | 0.541211 | 1.315584 | 0.129445 | 0.433251 |
| GOBP_SEGMENTATION | 79 | 0.441601 | 1.314653 | 0.080702 | 0.43521 |
| GOBP_DNA_DAMAGE_RESPONSE_SIGNAL_TRANSDUCTION_BY_P53_CLASS_MEDIATO | 16 | 0.573025 | 1.314291 | 0.144838 | 0.435616 |
| GOBP_CELLULAR_RESPONSE_TO_OXYGEN_RADICAL | 20 | 0.563109 | 1.314088 | 0.131994 | 0.435546 |
| GOBP_REGULATION_OF_PROGRAMMED_NECROTIC_CELL_DEATH | 21 | 0.541541 | 1.313951 | 0.15156 | 0.435276 |
| GOBP_REGULATION_OF_ATP_DEPENDENT_ACTIVITY | 62 | 0.457136 | 1.313923 | 0.108344 | 0.434716 |
| GOBP_HEART_GROWTH | 62 | 0.457393 | 1.313666 | 0.094458 | 0.434807 |
| GOBP_CALCIUM_ION_TRANSPORT_INTO_CYTOSOL | 124 | 0.422013 | 1.313248 | 0.061867 | 0.435356 |
| GOBP_MULTICELLULAR_ORGANISMAL_MOVEMENT | 42 | 0.499761 | 1.312749 | 0.102902 | 0.436153 |
| GOBP_CELLULAR_RESPONSE_TO_VITAMIN_D | 19 | 0.557915 | 1.312611 | 0.147982 | 0.43586 |
| GOBP_POSITIVE_REGULATION_OF_LIPID_TRANSPORT | 64 | 0.449829 | 1.312358 | 0.092707 | 0.435991 |
| GOBP_REGULATION_OF_T_CELL_APOPTOTIC_PROCESS | 23 | 0.532803 | 1.312205 | 0.141304 | 0.435807 |
| GOBP_REACTIVE_NITROGEN_SPECIES_METABOLIC_PROCESS | 59 | 0.461045 | 1.312071 | 0.103194 | 0.435527 |
| GOBP_REGULATION_OF_BODY_FLUID_LEVELS | 293 | 0.400025 | 1.312012 | 0.029744 | 0.435075 |
| GOBP_HINDLIMB_MORPHOGENESIS | 21 | 0.554236 | 1.310581 | 0.156385 | 0.438505 |
| GOBP_POSITIVE_REGULATION_OF_GRANULOCYTE_CHEMOTAXIS | 19 | 0.562318 | 1.309216 | 0.160237 | 0.441671 |
| GOBP_GERM_CELL_DEVELOPMENT | 210 | 0.405876 | 1.30782 | 0.027197 | 0.445102 |
| GOBP_DEVELOPMENTAL_MATURATION | 234 | 0.403822 | 1.307691 | 0.033229 | 0.444818 |
| GOBP_HYDROGEN_PEROXIDE_METABOLIC_PROCESS | 34 | 0.505833 | 1.307624 | 0.124166 | 0.444384 |
| GOBP_REGULATION_OF_CALCIUM_MEDIATED_SIGNALING | 56 | 0.464782 | 1.307503 | 0.11344 | 0.4441 |
| GOBP_PHYSIOLOGICAL_CARDIAC_MUSCLE_HYPERTROPHY | 15 | 0.584014 | 1.307356 | 0.151563 | 0.443888 |
| GOBP_POSITIVE_REGULATION_OF_LOCOMOTION | 461 | 0.389119 | 1.306466 | 0.003003 | 0.445821 |
| GOBP_POSITIVE_REGULATION_OF_EPITHELIAL_CELL_MIGRATION | 136 | 0.41958 | 1.306377 | 0.057627 | 0.445473 |
| GOBP_TUMOR_NECROSIS_FACTOR_MEDIATED_SIGNALING_PATHWAY | 86 | 0.434684 | 1.304436 | 0.090376 | 0.450401 |
| GOBP_REGULATION_OF_EMBRYONIC_DEVELOPMENT | 57 | 0.464728 | 1.304403 | 0.093168 | 0.449841 |
| GOBP_NEGATIVE_REGULATION_OF_EPITHELIAL_CELL_PROLIFERATION | 106 | 0.428219 | 1.304333 | 0.075145 | 0.449393 |
| GOBP_NEGATIVE_REGULATION_OF_CELL_GROWTH | 165 | 0.411392 | 1.30409 | 0.052061 | 0.449481 |
| GOBP_RESPONSE_TO_WOUNDING | 443 | 0.390851 | 1.303215 | 0.006067 | 0.45133 |
| GOBP_ALPHA_BETA_T_CELL_DIFFERENTIATION | 77 | 0.44695 | 1.30299 | 0.100242 | 0.451375 |
| GOBP_SKELETAL_SYSTEM_DEVELOPMENT | 402 | 0.392681 | 1.302955 | 0.006067 | 0.45083 |
| GOBP_INFLAMMASOME_COMPLEX_ASSEMBLY | 16 | 0.571776 | 1.302305 | 0.172012 | 0.452112 |
| GOBP_MEMORY | 109 | 0.426245 | 1.302269 | 0.087401 | 0.451571 |
| GOBP_POSITIVE_REGULATION_OF_OXIDOREDUCTASE_ACTIVITY | 45 | 0.480598 | 1.301447 | 0.120155 | 0.453273 |
| GOBP_NOREPINEPHRINE_TRANSPORT | 17 | 0.573175 | 1.301325 | 0.171386 | 0.452936 |
| GOBP_REGULATION_OF_CARBOHYDRATE_BIOSYNTHETIC_PROCESS | 84 | 0.429266 | 1.300021 | 0.097122 | 0.456229 |
| GOBP_REGULATION_OF_DEFENSE_RESPONSE_TO_VIRUS_BY_HOST | 32 | 0.503637 | 1.299933 | 0.129959 | 0.455845 |
| GOBP_REGULATION_OF_MAST_CELL_ACTIVATION | 30 | 0.509286 | 1.299564 | 0.150815 | 0.456269 |
| GOBP_CELL_AGING | 104 | 0.429162 | 1.29929 | 0.077361 | 0.456369 |
| GOBP_RETINOIC_ACID_RECEPTOR_SIGNALING_PATHWAY | 27 | 0.52034 | 1.299201 | 0.130194 | 0.455958 |
| GOBP_REGULATION_OF_T_CELL_RECEPTOR_SIGNALING_PATHWAY | 32 | 0.504309 | 1.298243 | 0.123179 | 0.45803 |
| GOBP_CELLULAR_RESPONSE_TO_VITAMIN | 26 | 0.524086 | 1.298178 | 0.125 | 0.457604 |
| GOBP_POSITIVE_REGULATION_OF_WOUND_HEALING | 42 | 0.485891 | 1.29758 | 0.138562 | 0.458655 |
| GOBP_AMEBOIDAL_TYPE_CELL_MIGRATION | 372 | 0.393029 | 1.297309 | 0.015259 | 0.458786 |
| GOBP_NODAL_SIGNALING_PATHWAY | 16 | 0.564045 | 1.297113 | 0.163934 | 0.458718 |
| GOBP_MATERNAL_PLACENTA_DEVELOPMENT | 29 | 0.510633 | 1.296946 | 0.146309 | 0.458581 |
| GOBP_REGULATION_OF_ACTION_POTENTIAL | 45 | 0.476177 | 1.296671 | 0.119481 | 0.458773 |
| GOBP_INORGANIC_ION_IMPORT_ACROSS_PLASMA_MEMBRANE | 82 | 0.440274 | 1.295741 | 0.089848 | 0.460892 |
| GOBP_REGULATION_OF_TRANSMEMBRANE_RECEPTOR_PROTEIN_SERINE_THREONIN | 224 | 0.3961 | 1.294622 | 0.048117 | 0.463447 |
| GOBP_BRANCHING_INVOLVED_IN_SALIVARY_GLAND_MORPHOGENESIS | 17 | 0.556635 | 1.293491 | 0.161194 | 0.466091 |
| GOBP_REGULATION_OF_CELLULAR_RESPIRATION | 42 | 0.475139 | 1.292796 | 0.133333 | 0.467386 |
| GOBP_UNSATURATED_FATTY_ACID_BIOSYNTHETIC_PROCESS | 41 | 0.472474 | 1.292182 | 0.143596 | 0.468531 |
| GOBP_CELLULAR_SENESCENCE | 73 | 0.441141 | 1.291451 | 0.116364 | 0.470027 |
| GOBP_NEGATIVE_REGULATION_OF_CALCIUM_MEDIATED_SIGNALING | 17 | 0.556754 | 1.290801 | 0.143695 | 0.471279 |
| GOBP_REGULATION_OF_CALCIUM_ION_TRANSPORT | 186 | 0.402692 | 1.290713 | 0.048649 | 0.470904 |
| GOBP_ENDOCARDIAL_CUSHION_MORPHOGENESIS | 35 | 0.498288 | 1.290607 | 0.132353 | 0.470572 |
| GOBP_REGULATION_OF_LIPID_LOCALIZATION | 121 | 0.413911 | 1.290216 | 0.082207 | 0.471092 |
| GOBP_SPHINGOID_METABOLIC_PROCESS | 22 | 0.524336 | 1.290174 | 0.182203 | 0.470582 |
| GOBP_RESPONSE_TO_EXTRACELLULAR_STIMULUS | 389 | 0.3878 | 1.289915 | 0.017241 | 0.470692 |
| GOBP_POSITIVE_REGULATION_OF_NITRIC_OXIDE_METABOLIC_PROCESS | 26 | 0.513177 | 1.289865 | 0.165527 | 0.470217 |
| GOBP_POSITIVE_REGULATION_OF_INTERLEUKIN_10_PRODUCTION | 26 | 0.517936 | 1.28923 | 0.152144 | 0.471428 |
| GOBP_FATTY_ACID_BIOSYNTHETIC_PROCESS | 120 | 0.415893 | 1.287368 | 0.080682 | 0.476271 |
| GOBP_CELLULAR_RESPONSE_TO_HEAT | 61 | 0.447368 | 1.287283 | 0.106796 | 0.475893 |
| GOBP_PROSTANOID_METABOLIC_PROCESS | 37 | 0.48545 | 1.287243 | 0.138705 | 0.475361 |
| GOBP_REGULATION_OF_ACTIN_FILAMENT_BASED_MOVEMENT | 35 | 0.494254 | 1.286879 | 0.156785 | 0.475772 |
| GOBP_NUCLEOSIDE_MONOPHOSPHATE_CATABOLIC_PROCESS | 16 | 0.567663 | 1.286436 | 0.186186 | 0.476398 |
| GOBP_FATTY_ACID_METABOLIC_PROCESS | 302 | 0.391527 | 1.286258 | 0.026639 | 0.476259 |
| GOBP_APOPTOTIC_CELL_CLEARANCE | 40 | 0.476626 | 1.286128 | 0.140921 | 0.475995 |
| GOBP_3_PHOSPHOADENOSINE_5_PHOSPHOSULFATE_METABOLIC_PROCESS | 15 | 0.569989 | 1.285113 | 0.164653 | 0.478309 |
| GOBP_CARDIAC_VENTRICLE_MORPHOGENESIS | 59 | 0.449043 | 1.284669 | 0.106464 | 0.478965 |
| GOBP_PANCREAS_DEVELOPMENT | 54 | 0.460269 | 1.284368 | 0.12979 | 0.479222 |
| GOBP_OSTEOCLAST_DIFFERENTIATION | 71 | 0.449301 | 1.28408 | 0.106459 | 0.479411 |
| GOBP_INTERFERON_ALPHA_PRODUCTION | 20 | 0.54508 | 1.28402 | 0.177941 | 0.478929 |
| GOBP_INTERMEDIATE_FILAMENT_ORGANIZATION | 18 | 0.543556 | 1.282035 | 0.168588 | 0.484319 |
| GOBP_BONE_MINERALIZATION | 92 | 0.432377 | 1.282028 | 0.101499 | 0.483691 |
| GOBP_REGULATION_OF_CARDIOCYTE_DIFFERENTIATION | 22 | 0.537874 | 1.281891 | 0.158621 | 0.483468 |

| | | | | | |
|---|---|---|---|---|---|
| GOBP_ACTIN_NUCLEATION | 49 | -0.218468 | -0.746946 | 0.903382 | 1 |
| GOBP_INDOLE_CONTAINING_COMPOUND_METABOLIC_PROCESS | 15 | -0.290147 | -0.745633 | 0.829721 | 1 |
| GOBP_REGULATION_OF_DNA_DAMAGE_CHECKPOINT | 16 | -0.278571 | -0.745379 | 0.794798 | 1 |
| GOBP_NUCLEOTIDE_BINDING_DOMAIN_LEUCINE_RICH_REPEAT_CONT | 22 | -0.267499 | -0.745158 | 0.838951 | 1 |
| GOBP_NEGATIVE_REGULATION_OF_ORGANELLE_ASSEMBLY | 41 | -0.223956 | -0.74502 | 0.912134 | 1 |
| GOBP_ERAD_PATHWAY | 106 | -0.193846 | -0.744693 | 0.97037 | 1 |
| GOBP_MONOSACCHARIDE_CATABOLIC_PROCESS | 34 | -0.2349 | -0.744638 | 0.880165 | 1 |
| GOBP_AMYLOID_PRECURSOR_PROTEIN_CATABOLIC_PROCESS | 49 | -0.220184 | -0.744563 | 0.896226 | 1 |
| GOBP_CAMERA_TYPE_EYE_PHOTORECEPTOR_CELL_DIFFERENTIATION | 20 | -0.261339 | -0.744489 | 0.810496 | 1 |
| GOBP_RESPONSE_TO_INCREASED_OXYGEN_LEVELS | 24 | -0.257465 | -0.744301 | 0.837607 | 1 |
| GOBP_REGULATION_OF_DENDRITE_MORPHOGENESIS | 60 | -0.213616 | -0.743363 | 0.964103 | 1 |
| GOBP_HISTONE_MRNA_METABOLIC_PROCESS | 24 | -0.252176 | -0.743192 | 0.858974 | 1 |
| GOBP_IRE1_MEDIATED_UNFOLDED_PROTEIN_RESPONSE | 18 | -0.272426 | -0.741608 | 0.841122 | 1 |
| GOBP_NUCLEOSIDE_BISPHOSPHATE_BIOSYNTHETIC_PROCESS | 55 | -0.208748 | -0.738788 | 0.927184 | 1 |
| GOBP_ENTRAINMENT_OF_CIRCADIAN_CLOCK | 31 | -0.228792 | -0.738247 | 0.881679 | 1 |
| GOBP_SULFUR_AMINO_ACID_BIOSYNTHETIC_PROCESS | 16 | -0.275196 | -0.73802 | 0.800633 | 1 |
| GOBP_NEGATIVE_REGULATION_OF_NOTCH_SIGNALING_PATHWAY | 32 | -0.236241 | -0.737871 | 0.890977 | 1 |
| GOBP_POSITIVE_REGULATION_OF_OLIGODENDROCYTE_DIFFERENTIAT | 18 | -0.275369 | -0.737623 | 0.802899 | 1 |
| GOBP_TRNA_5_END_PROCESSING | 16 | -0.275751 | -0.737131 | 0.84058 | 1 |
| GOBP_TUBULIN_DEACETYLATION | 22 | -0.251227 | -0.736757 | 0.863014 | 1 |
| GOBP_GOLGI_VESICLE_TRANSPORT | 280 | -0.166335 | -0.736703 | 1 | 1 |
| GOBP_NEGATIVE_REGULATION_OF_LIPID_METABOLIC_PROCESS | 73 | -0.200582 | -0.735397 | 0.9375 | 1 |
| GOBP_GLUTAMINE_METABOLIC_PROCESS | 23 | -0.252541 | -0.734387 | 0.858108 | 1 |
| GOBP_POSITIVE_REGULATION_OF_PEPTIDYL_THREONINE_PHOSPHOR | 25 | -0.248736 | -0.733342 | 0.876405 | 1 |
| GOBP_MITOCHONDRIAL_DNA_METABOLIC_PROCESS | 15 | -0.280163 | -0.732427 | 0.820189 | 1 |
| GOBP_RIBONUCLEOPROTEIN_COMPLEX_BIOGENESIS | 418 | -0.163685 | -0.732057 | 1 | 1 |
| GOBP_POSITIVE_REGULATION_OF_IMMUNOGLOBULIN_PRODUCTION | 36 | -0.227358 | -0.732022 | 0.896396 | 1 |
| GOBP_REGULATION_OF_UBIQUITIN_PROTEIN_LIGASE_ACTIVITY | 23 | -0.254404 | -0.731829 | 0.850174 | 1 |
| GOBP_FAT_SOLUBLE_VITAMIN_METABOLIC_PROCESS | 26 | -0.247629 | -0.730414 | 0.889706 | 1 |
| GOBP_DIGESTIVE_TRACT_MORPHOGENESIS | 35 | -0.232299 | -0.72997 | 0.890977 | 1 |
| GOBP_HEART_TRABECULA_MORPHOGENESIS | 28 | -0.246525 | -0.728186 | 0.87108 | 1 |
| GOBP_NEGATIVE_REGULATION_OF_FIBROBLAST_GROWTH_FACTOR_R | 19 | -0.264138 | -0.727527 | 0.860465 | 1 |
| GOBP_POSITIVE_REGULATION_OF_EPITHELIAL_TO_MESENCHYMAL_TR | 42 | -0.214077 | -0.727404 | 0.918803 | 1 |
| GOBP_REGULATION_OF_DNA_TEMPLATED_TRANSCRIPTION_INITIATION | 51 | -0.21682 | -0.726993 | 0.928571 | 1 |
| GOBP_MRNA_TRANSCRIPTION | 39 | -0.222973 | -0.724219 | 0.892 | 1 |
| GOBP_PROTEIN_DEGLYCOSYLATION | 25 | -0.251901 | -0.724154 | 0.897887 | 1 |
| GOBP_POSITIVE_REGULATION_OF_STRESS_FIBER_ASSEMBLY | 48 | -0.213459 | -0.723751 | 0.910377 | 1 |
| GOBP_REGULATION_OF_CYCLIN_DEPENDENT_PROTEIN_KINASE_ACTI | 91 | -0.19414 | -0.723589 | 0.992593 | 1 |
| GOBP_VIRAL_BUDDING | 27 | -0.237506 | -0.723108 | 0.861487 | 1 |
| GOBP_ESTABLISHMENT_OF_PROTEIN_LOCALIZATION_TO_TELOMERE | 17 | -0.269343 | -0.722268 | 0.846154 | 1 |
| GOBP_SNRNA_PROCESSING | 25 | -0.240191 | -0.721782 | 0.884058 | 1 |
| GOBP_NEURAL_TUBE_PATTERNING | 29 | -0.237871 | -0.721507 | 0.880309 | 1 |
| GOBP_POSITIVE_REGULATION_OF_METAPHASE_ANAPHASE_TRANSITIO | 15 | -0.284463 | -0.721489 | 0.826087 | 1 |
| GOBP_NEUROTROPHIN_TRK_RECEPTOR_SIGNALING_PATHWAY | 22 | -0.257239 | -0.720492 | 0.834375 | 1 |
| GOBP_PROTON_TRANSMEMBRANE_TRANSPORT | 128 | -0.181122 | -0.720459 | 0.99115 | 1 |
| GOBP_HEART_FORMATION | 25 | -0.238786 | -0.719829 | 0.901754 | 1 |
| GOBP_FIBROBLAST_APOPTOTIC_PROCESS | 18 | -0.256259 | -0.716513 | 0.849231 | 1 |
| GOBP_NEGATIVE_REGULATION_OF_REACTIVE_OXYGEN_SPECIES_MET | 34 | -0.222849 | -0.715883 | 0.901288 | 1 |
| GOBP_REGULATION_OF_PROTEIN_AUTOPHOSPHORYLATION | 39 | -0.218197 | -0.715232 | 0.945098 | 1 |
| GOBP_MRNA_MODIFICATION | 25 | -0.23948 | -0.714877 | 0.869258 | 1 |
| GOBP_POSITIVE_REGULATION_OF_PROTEIN_POLYMERIZATION | 83 | -0.189296 | -0.714719 | 0.97973 | 1 |
| GOBP_PHOSPHOLIPID_TRANSPORT | 81 | -0.188149 | -0.714304 | 0.964789 | 1 |
| GOBP_CYTOPLASMIC_TRANSLATION | 146 | -0.171518 | -0.714107 | 0.970588 | 1 |
| GOBP_ESTABLISHMENT_OF_PROTEIN_LOCALIZATION_TO_ORGANELLE | 399 | -0.157623 | -0.712884 | 1 | 1 |
| GOBP_MICROTUBULE_ANCHORING | 24 | -0.244634 | -0.712544 | 0.873134 | 1 |
| GOBP_PROTEIN_O_LINKED_MANNOSYLATION | 17 | -0.267068 | -0.712521 | 0.888889 | 1 |
| GOBP_POSITIVE_REGULATION_OF_DENDRITIC_SPINE_DEVELOPMENT | 34 | -0.222046 | -0.712497 | 0.92 | 1 |
| GOBP_SPLICEOSOMAL_SNRNP_ASSEMBLY | 37 | -0.227119 | -0.711412 | 0.905512 | 1 |
| GOBP_MYELIN_ASSEMBLY | 22 | -0.250437 | -0.710622 | 0.894915 | 1 |
| GOBP_REGULATION_OF_CARDIAC_MUSCLE_CONTRACTION_BY_CALCI | 22 | -0.252242 | -0.708922 | 0.887097 | 1 |
| GOBP_PURINE_CONTAINING_COMPOUND_BIOSYNTHETIC_PROCESS | 177 | -0.16452 | -0.708736 | 0.987013 | 1 |
| GOBP_NEGATIVE_REGULATION_OF_UBIQUITIN_DEPENDENT_PROTEIN_ | 44 | -0.210477 | -0.707719 | 0.956332 | 1 |
| GOBP_MAINTENANCE_OF_CELL_POLARITY | 18 | -0.260233 | -0.707195 | 0.868512 | 1 |
| GOBP_PROTEIN_EXIT_FROM_ENDOPLASMIC_RETICULUM | 48 | -0.206529 | -0.706575 | 0.951456 | 1 |
| GOBP_BARBED_END_ACTIN_FILAMENT_CAPPING | 25 | -0.240132 | -0.706419 | 0.886207 | 1 |
| GOBP_RETROGRADE_VESICLE_MEDIATED_TRANSPORT_GOLGI_TO_EN | 49 | -0.206296 | -0.706138 | 0.930876 | 1 |
| GOBP_MIRNA_METABOLIC_PROCESS | 27 | -0.237442 | -0.705438 | 0.916981 | 1 |
| GOBP_B_CELL_ACTIVATION_INVOLVED_IN_IMMUNE_RESPONSE | 64 | -0.194563 | -0.704098 | 0.966102 | 1 |
| GOBP_REGULATION_OF_OXIDATIVE_STRESS_INDUCED_INTRINSIC_AP | 26 | -0.232802 | -0.704014 | 0.893617 | 1 |
| GOBP_PROTEIN_TARGETING | 295 | -0.163988 | -0.703271 | 1 | 1 |
| GOBP_ESTABLISHMENT_OF_EPITHELIAL_CELL_APICAL_BASAL_POLAR | 18 | -0.258104 | -0.702764 | 0.859813 | 1 |
| GOBP_PROTEIN_LOCALIZATION_TO_MITOCHONDRION | 121 | -0.178705 | -0.70276 | 1 | 1 |
| GOBP_REGULATION_OF_ESTABLISHMENT_OR_MAINTENANCE_OF_CEL | 25 | -0.235159 | -0.702403 | 0.908425 | 1 |
| GOBP_NUCLEOBASE_BIOSYNTHETIC_PROCESS | 19 | -0.255727 | -0.701796 | 0.866883 | 1 |
| GOBP_NEGATIVE_REGULATION_OF_TRANSCRIPTION_REGULATORY_R | 17 | -0.256752 | -0.70016 | 0.857616 | 1 |
| GOBP_POSITIVE_REGULATION_OF_MRNA_METABOLIC_PROCESS | 115 | -0.180563 | -0.69952 | 1 | 1 |
| GOBP_ESTABLISHMENT_OF_PIGMENT_GRANULE_LOCALIZATION | 22 | -0.24678 | -0.697745 | 0.869969 | 1 |
| GOBP_CELLULAR_RESPONSE_TO_INSULIN_STIMULUS | 172 | -0.162528 | -0.697202 | 1 | 1 |
| GOBP_PIGMENT_GRANULE_LOCALIZATION | 24 | -0.238939 | -0.697073 | 0.893471 | 1 |
| GOBP_CENTROMERE_COMPLEX_ASSEMBLY | 30 | -0.227109 | -0.696081 | 0.903915 | 1 |
| GOBP_HISTONE_H3_ACETYLATION | 74 | -0.189293 | -0.69559 | 0.977273 | 1 |
| GOBP_NEGATIVE_REGULATION_OF_RELEASE_OF_CYTOCHROME_C_FR | 18 | -0.266847 | -0.693486 | 0.856667 | 1 |
| GOBP_REGULATION_OF_RESPONSE_TO_OXIDATIVE_STRESS | 77 | -0.186546 | -0.691744 | 0.975758 | 1 |
| GOBP_REGULATION_OF_PEPTIDYL_THREONINE_PHOSPHORYLATION | 37 | -0.210909 | -0.689172 | 0.931624 | 1 |
| GOBP_MRNA_EXPORT_FROM_NUCLEUS | 54 | -0.199877 | -0.688042 | 0.977401 | 1 |
| GOBP_RNA_POLYADENYLATION | 47 | -0.208792 | -0.688003 | 0.942222 | 1 |
| GOBP_RIBOSE_PHOSPHATE_BIOSYNTHETIC_PROCESS | 173 | -0.162156 | -0.687913 | 1 | 1 |
| GOBP_VASCULAR_ASSOCIATED_SMOOTH_MUSCLE_CONTRACTION | 19 | -0.249089 | -0.68751 | 0.909091 | 1 |
| GOBP_BRAIN_MORPHOGENESIS | 32 | -0.224937 | -0.687345 | 0.933852 | 1 |
| GOBP_PEPTIDYL_LYSINE_TRIMETHYLATION | 50 | -0.202397 | -0.68705 | 0.927273 | 1 |
| GOBP_UBIQUINONE_METABOLIC_PROCESS | 18 | -0.25106 | -0.682754 | 0.866667 | 1 |
| GOBP_SUBSTANTIA_NIGRA_DEVELOPMENT | 40 | -0.212832 | -0.682355 | 0.952381 | 1 |
| GOBP_REGULATION_OF_CHROMATIN_ORGANIZATION | 42 | -0.209143 | -0.676894 | 0.961373 | 1 |
| GOBP_GTP_METABOLIC_PROCESS | 23 | -0.227519 | -0.676065 | 0.923913 | 1 |
| GOBP_REGULATION_OF_PROTEIN_STABILITY | 271 | -0.157144 | -0.672349 | 1 | 1 |
| GOBP_MRNA_METHYLATION | 16 | -0.258933 | -0.672327 | 0.904908 | 1 |
| GOBP_REGULATION_OF_CARDIAC_MUSCLE_CONTRACTION_BY_REGU | 18 | -0.241739 | -0.671627 | 0.915663 | 1 |
| GOBP_REGULATION_OF_ACTOMYOSIN_STRUCTURE_ORGANIZATION | 89 | -0.174457 | -0.671145 | 1 | 1 |
| GOBP_REGULATION_OF_TRANSLATION_IN_RESPONSE_TO_STRESS | 20 | -0.234919 | -0.668942 | 0.912621 | 1 |
| GOBP_REGULATION_OF_RRNA_PROCESSING | 16 | -0.254554 | -0.668753 | 0.904192 | 1 |
| GOBP_HISTONE_MRNA_CATABOLIC_PROCESS | 15 | -0.252045 | -0.664473 | 0.915254 | 1 |
| GOBP_MITOCHONDRIAL_TRANSLATION | 70 | -0.185351 | -0.664237 | 0.982857 | 1 |
| GOBP_AMYLOID_BETA_METABOLIC_PROCESS | 46 | -0.195303 | -0.661819 | 0.965812 | 1 |
| GOBP_INTRINSIC_APOPTOTIC_SIGNALING_PATHWAY_IN_RESPONSE_T | 42 | -0.199946 | -0.659852 | 0.968889 | 1 |
| GOBP_REGULATION_OF_POLYSACCHARIDE_METABOLIC_PROCESS | 37 | -0.200143 | -0.655791 | 0.958506 | 1 |
| GOBP_PROTEIN_LOCALIZATION_TO_MICROTUBULE_ORGANIZING_CEN | 35 | -0.201352 | -0.654938 | 0.977273 | 1 |
| GOBP_POSITIVE_REGULATION_OF_TELOMERASE_ACTIVITY | 34 | -0.203115 | -0.653583 | 0.983936 | 1 |
| GOBP_REGULATION_OF_RECEPTOR_LOCALIZATION_TO_SYNAPSE | 16 | -0.246371 | -0.653109 | 0.907692 | 1 |
| GOBP_LAMELLIPODIUM_ASSEMBLY | 63 | -0.180595 | -0.648102 | 0.989247 | 1 |
| GOBP_REGULATION_OF_INTRACELLULAR_STEROID_HORMONE_RECE | 64 | -0.18124 | -0.647754 | 0.988372 | 1 |
| GOBP_VIRAL_TRANSCRIPTION | 48 | -0.193807 | -0.647428 | 0.982143 | 1 |
| GOBP_PHOSPHATIDYLINOSITOL_PHOSPHATE_BIOSYNTHETIC_PROCES | 62 | -0.183298 | -0.646999 | 0.985646 | 1 |
| GOBP_NEGATIVE_REGULATION_OF_TELOMERE_MAINTENANCE_VIA_TE | 24 | -0.219155 | -0.646691 | 0.944828 | 1 |
| GOBP_CELLULAR_RESPONSE_TO_AMYLOID_BETA | 32 | -0.20558 | -0.64574 | 0.954064 | 1 |
| GOBP_REGULATION_OF_LONG_TERM_NEURONAL_SYNAPTIC_PLASTIC | 21 | -0.231611 | -0.645301 | 0.916944 | 1 |
| GOBP_MRNA_3_END_PROCESSING | 61 | -0.179957 | -0.64324 | 0.995349 | 1 |
| GOBP_MICROTUBULE_ORGANIZING_CENTER_ORGANIZATION | 139 | -0.158027 | -0.642438 | 1 | 1 |
| GOBP_VERY_LONG_CHAIN_FATTY_ACID_METABOLIC_PROCESS | 31 | -0.203487 | -0.637837 | 0.955466 | 1 |
| GOBP_ENDOCYTIC_RECYCLING | 74 | -0.170472 | -0.636215 | 0.988827 | 1 |
| GOBP_MITOCHONDRIAL_RNA_PROCESSING | 20 | -0.22827 | -0.634324 | 0.959707 | 1 |
| GOBP_REGULATION_OF_PHOSPHOLIPID_METABOLIC_PROCESS | 29 | -0.206367 | -0.632279 | 0.965116 | 1 |
| GOBP_PROTEIN_LOCALIZATION_TO_CHROMATIN | 30 | -0.204213 | -0.631042 | 0.953571 | 1 |
| GOBP_ESTABLISHMENT_OF_PROTEIN_LOCALIZATION_TO_MEMBRANE | 236 | -0.146518 | -0.629533 | 1 | 1 |
| GOBP_RRNA_METABOLIC_PROCESS | 258 | -0.141165 | -0.629419 | 1 | 1 |
| GOBP_PROTEIN_K63_LINKED_DEUBIQUITINATION | 33 | -0.201989 | -0.629217 | 0.968 | 1 |
| GOBP_REGULATION_OF_MITOCHONDRIAL_GENE_EXPRESSION | 28 | -0.203614 | -0.628443 | 0.974359 | 1 |
| GOBP_ESTABLISHMENT_OR_MAINTENANCE_OF_BIPOLAR_CELL_POLA | 48 | -0.184403 | -0.627402 | 0.990783 | 1 |
| GOBP_LATE_ENDOSOME_TO_VACUOLE_TRANSPORT | 24 | -0.209037 | -0.625365 | 0.952703 | 1 |
| GOBP_EXTRACELLULAR_VESICLE_BIOGENESIS | 24 | -0.219556 | -0.625227 | 0.97153 | 1 |
| GOBP_POSITIVE_REGULATION_OF_DNA_TEMPLATED_TRANSCRIPTION_ | 42 | -0.190456 | -0.624247 | 0.983539 | 1 |
| GOBP_RNA_DEPENDENT_DNA_BIOSYNTHETIC_PROCESS | 68 | -0.170317 | -0.623228 | 0.994286 | 1 |
| GOBP_DNA_REPAIR | 498 | -0.136134 | -0.623164 | 1 | 1 |
| GOBP_SULFUR_COMPOUND_TRANSPORT | 43 | -0.187593 | -0.620781 | 0.982609 | 1 |
| GOBP_REGULATION_OF_CELLULAR_RESPONSE_TO_HEAT | 17 | -0.229974 | -0.619881 | 0.913793 | 1 |
| GOBP_PROTEIN_LOCALIZATION_TO_CHROMOSOME_CENTROMERIC_R | 25 | -0.212754 | -0.619445 | 0.947368 | 1 |
| GOBP_N_TERMINAL_PROTEIN_AMINO_ACID_MODIFICATION | 27 | -0.210045 | -0.617712 | 0.944251 | 1 |
| GOBP_REGULATION_OF_CYTOPLASMIC_TRANSLATION | 27 | -0.200343 | -0.617076 | 0.972603 | 1 |
| GOBP_PROTEIN_LOCALIZATION_TO_POSTSYNAPSE | 31 | -0.191957 | -0.616412 | 0.984496 | 1 |
| GOBP_TERTIARY_ALCOHOL_METABOLIC_PROCESS | 17 | -0.216979 | -0.614475 | 0.951299 | 1 |
| GOBP_VIRION_ASSEMBLY | 36 | -0.186375 | -0.607599 | 0.961864 | 1 |
| GOBP_HISTONE_H2A_ACETYLATION | 23 | -0.207078 | -0.601691 | 0.967857 | 1 |
| GOBP_REGULATION_OF_ENDOCYTIC_RECYCLING | 17 | -0.223964 | -0.600804 | 0.948328 | 1 |
| GOBP_GOLGI_ORGANIZATION | 139 | -0.14744 | -0.600681 | 1 | 1 |
| GOBP_NEGATIVE_REGULATION_OF_PEPTIDYL_THREONINE_PHOSPHO | 15 | -0.228188 | -0.595959 | 0.955357 | 1 |
| GOBP_DOSAGE_COMPENSATION | 20 | -0.214528 | -0.595131 | 0.950495 | 1 |
| GOBP_TORC1_SIGNALING | 48 | -0.173435 | -0.593386 | 1 | 1 |
| GOBP_MISMATCH_REPAIR | 30 | -0.188479 | -0.592434 | 0.988189 | 1 |
| GOBP_NEGATIVE_REGULATION_OF_TORC1_SIGNALING | 17 | -0.22713 | -0.591391 | 0.960245 | 1 |
| GOBP_MICROVILLUS_ASSEMBLY | 21 | -0.205595 | -0.589324 | 0.960133 | 1 |
| GOBP_POSITIVE_REGULATION_OF_CHROMOSOME_ORGANIZATION | 85 | -0.152601 | -0.588227 | 1 | 1 |
| GOBP_MITOCHONDRIAL_TRANSPORT | 190 | -0.141734 | -0.587907 | 1 | 1 |

| | | | | | |
|---|---|---|---|---|---|
| GOBP_PYRIDINE_NUCLEOTIDE_METABOLIC_PROCESS | 26 | 0.501326 | 1.281882 | 0.142458 | 0.482849 |
| GOBP_FOREBRAIN_NEURON_DIFFERENTIATION | 32 | 0.488779 | 1.281795 | 0.155196 | 0.482488 |
| GOBP_MOLTING_CYCLE | 81 | 0.432921 | 1.281696 | 0.116686 | 0.482149 |
| GOBP_B_CELL_RECEPTOR_SIGNALING_PATHWAY | 39 | 0.475974 | 1.281067 | 0.147287 | 0.483398 |
| GOBP_REGULATION_OF_CALCINEURIN_MEDIATED_SIGNALING | 30 | 0.498146 | 1.280903 | 0.165113 | 0.483246 |
| GOBP_CELLULAR_RESPONSE_TO_BIOTIC_STIMULUS | 159 | 0.407898 | 1.28045 | 0.066955 | 0.483853 |
| GOBP_AMINOGLYCAN_CATABOLIC_PROCESS | 21 | 0.538339 | 1.279803 | 0.173427 | 0.485096 |
| GOBP_RESPONSE_TO_GONADOTROPIN | 21 | 0.534438 | 1.279054 | 0.187408 | 0.486628 |
| GOBP_REGULATION_OF_ORGAN_GROWTH | 65 | 0.450563 | 1.278631 | 0.10688 | 0.487324 |
| GOBP_REGULATION_OF_BMP_SIGNALING_PATHWAY | 87 | 0.432755 | 1.278377 | 0.110979 | 0.487418 |
| GOBP_POLYOL_METABOLIC_PROCESS | 101 | 0.418364 | 1.278086 | 0.095183 | 0.487637 |
| GOBP_POSITIVE_REGULATION_OF_MYELOID_LEUKOCYTE_DIFFERENTIATION | 40 | 0.482819 | 1.277912 | 0.147395 | 0.487454 |
| GOBP_GLAND_DEVELOPMENT | 338 | 0.386505 | 1.277614 | 0.021385 | 0.487707 |
| GOBP_HEART_VALVE_MORPHOGENESIS | 50 | 0.459276 | 1.277455 | 0.134443 | 0.487536 |
| GOBP_POSITIVE_REGULATION_OF_REPRODUCTIVE_PROCESS | 63 | 0.446501 | 1.277047 | 0.118582 | 0.488112 |
| GOBP_TRANSITION_METAL_ION_TRANSPORT | 82 | 0.431333 | 1.276791 | 0.110159 | 0.48828 |
| GOBP_REGULATION_OF_MEMBRANE_REPOLARIZATION | 27 | 0.514879 | 1.276714 | 0.162162 | 0.487867 |
| GOBP_POSITIVE_REGULATION_OF_CELL_SUBSTRATE_ADHESION | 104 | 0.424989 | 1.276017 | 0.09289 | 0.48932 |
| GOBP_SISTER_CHROMATID_COHESION | 58 | 0.451567 | 1.275794 | 0.122961 | 0.489308 |
| GOBP_REGULATION_OF_MUSCLE_SYSTEM_PROCESS | 183 | 0.4003 | 1.27462 | 0.076099 | 0.492204 |
| GOBP_ECTODERM_DEVELOPMENT | 18 | 0.536885 | 1.274111 | 0.172566 | 0.493153 |
| GOBP_MITOCHONDRION_LOCALIZATION | 48 | 0.459333 | 1.273776 | 0.137887 | 0.493516 |
| GOBP_LENS_FIBER_CELL_DIFFERENTIATION | 33 | 0.491798 | 1.273565 | 0.150611 | 0.493538 |
| GOBP_NEGATIVE_REGULATION_OF_TYPE_I_INTERFERON_MEDIATED_SIGNALING_PATH | 16 | 0.561467 | 1.272863 | 0.187959 | 0.495095 |
| GOBP_MAST_CELL_ACTIVATION | 46 | 0.461959 | 1.272023 | 0.147541 | 0.497051 |
| GOBP_MULTICELLULAR_ORGANISMAL_HOMEOSTASIS | 399 | 0.379042 | 1.272013 | 0.023185 | 0.49645 |
| GOBP_REGULATION_OF_PHOSPHATIDYLINOSITOL_3_KINASE_SIGNALING | 78 | 0.429737 | 1.271982 | 0.108619 | 0.495918 |
| GOBP_REGULATION_OF_MUSCLE_CONTRACTION | 132 | 0.406367 | 1.271596 | 0.09434 | 0.49647 |
| GOBP_REGULATION_OF_CARBOHYDRATE_METABOLIC_PROCESS | 154 | 0.400441 | 1.271593 | 0.093989 | 0.495837 |
| GOBP_POSITIVE_REGULATION_OF_FATTY_ACID_TRANSPORT | 17 | 0.551603 | 1.270016 | 0.192197 | 0.499895 |
| GOBP_COLLAGEN_CATABOLIC_PROCESS | 27 | 0.509015 | 1.269905 | 0.166209 | 0.499647 |
| GOBP_NEGATIVE_REGULATION_OF_COAGULATION | 35 | 0.489989 | 1.269597 | 0.143229 | 0.499921 |
| GOBP_AMINE_METABOLIC_PROCESS | 95 | 0.423078 | 1.269571 | 0.100793 | 0.499355 |
| GOBP_METANEPHROS_MORPHOGENESIS | 26 | 0.50891 | 1.269558 | 0.175 | 0.498746 |
| GOBP_NEGATIVE_REGULATION_OF_SIGNAL_TRANSDUCTION_IN_ABSENCE_OF_LIGANI | 25 | 0.502527 | 1.268948 | 0.193906 | 0.499938 |
| GOBP_SUBSTRATE_DEPENDENT_CELL_MIGRATION | 23 | 0.522361 | 1.268912 | 0.185083 | 0.499444 |
| GOBP_REGULATION_OF_CYTOSOLIC_CALCIUM_ION_CONCENTRATION | 240 | 0.394077 | 1.268286 | 0.05919 | 0.500753 |
| GOBP_REGULATION_OF_CELL_SUBSTRATE_ADHESION | 183 | 0.392582 | 1.268116 | 0.074231 | 0.500605 |
| GOBP_NEGATIVE_REGULATION_OF_VASCULAR_ASSOCIATED_SMOOTH_MUSCLE_CELI | 21 | 0.523328 | 1.268086 | 0.172012 | 0.500069 |
| GOBP_POSITIVE_REGULATION_OF_LIPID_METABOLIC_PROCESS | 116 | 0.417648 | 1.267927 | 0.096045 | 0.4999 |
| GOBP_CARDIAC_CHAMBER_MORPHOGENESIS | 105 | 0.415428 | 1.267879 | 0.101714 | 0.499398 |
| GOBP_VIRAL_LIFE_CYCLE | 270 | 0.387779 | 1.267466 | 0.039378 | 0.50003 |
| GOBP_VIRAL_RNA_GENOME_REPLICATION | 32 | 0.487277 | 1.26724 | 0.171582 | 0.500097 |
| GOBP_POSITIVE_REGULATION_OF_TRANSFORMING_GROWTH_FACTOR_BETA_PRODUC | 17 | 0.547985 | 1.26696 | 0.176558 | 0.500339 |
| GOBP_POSITIVE_REGULATION_OF_OSSIFICATION | 43 | 0.470318 | 1.266486 | 0.164508 | 0.501167 |
| GOBP_RESPONSE_TO_BMP | 143 | 0.402618 | 1.26639 | 0.084337 | 0.500809 |
| GOBP_POSITIVE_REGULATION_OF_ENDOTHELIAL_CELL_MIGRATION | 98 | 0.421087 | 1.266272 | 0.10842 | 0.50054 |
| GOBP_ANTIMICROBIAL_HUMORAL_IMMUNE_RESPONSE_MEDIATED_BY_ANTIMICROBIA | 32 | 0.491425 | 1.265894 | 0.173853 | 0.501018 |
| GOBP_SPROUTING_ANGIOGENESIS | 111 | 0.409812 | 1.265328 | 0.10333 | 0.502086 |
| GOBP_SEX_DIFFERENTIATION | 214 | 0.391548 | 1.265256 | 0.068565 | 0.50168 |
| GOBP_GLAND_MORPHOGENESIS | 95 | 0.419174 | 1.265235 | 0.101281 | 0.501128 |
| GOBP_MESENCHYME_DEVELOPMENT | 246 | 0.389379 | 1.265012 | 0.058273 | 0.501123 |
| GOBP_WATER_SOLUBLE_VITAMIN_METABOLIC_PROCESS | 57 | 0.447125 | 1.264923 | 0.140881 | 0.500733 |
| GOBP_RESPONSE_TO_VIRUS | 290 | 0.385776 | 1.264437 | 0.04918 | 0.50163 |
| GOBP_REGULATION_OF_RHO_PROTEIN_SIGNAL_TRANSDUCTION | 75 | 0.432283 | 1.264035 | 0.140995 | 0.502208 |
| GOBP_REGULATION_OF_PROTEIN_TYROSINE_KINASE_ACTIVITY | 76 | 0.430988 | 1.263553 | 0.12485 | 0.503011 |
| GOBP_METHYLATION | 339 | 0.379306 | 1.263099 | 0.033503 | 0.503793 |
| GOBP_RESPONSE_TO_XENOBIOTIC_STIMULUS | 304 | 0.382202 | 1.263063 | 0.039796 | 0.503292 |
| GOBP_NEGATIVE_REGULATION_OF_LEUKOCYTE_APOPTOTIC_PROCESS | 38 | 0.478906 | 1.262336 | 0.162667 | 0.504855 |
| GOBP_NEUROTRANSMITTER_METABOLIC_PROCESS | 22 | 0.522885 | 1.262321 | 0.182997 | 0.50427 |
| GOBP_HOMEOSTASIS_OF_NUMBER_OF_CELLS | 233 | 0.385247 | 1.2617 | 0.05625 | 0.505537 |
| GOBP_FORELIMB_MORPHOGENESIS | 19 | 0.531924 | 1.261606 | 0.206847 | 0.505208 |
| GOBP_REGULATION_OF_ANION_TRANSPORT | 69 | 0.432762 | 1.261511 | 0.124402 | 0.50487 |
| GOBP_NEGATIVE_REGULATION_OF_CELLULAR_RESPONSE_TO_GROWTH_FACTOR_ST | 89 | 0.4205 | 1.261191 | 0.112161 | 0.50518 |
| GOBP_SEMI_LUNAR_VALVE_DEVELOPMENT | 37 | 0.476239 | 1.259784 | 0.157182 | 0.508904 |
| GOBP_RENAL_FILTRATION | 22 | 0.52394 | 1.259472 | 0.187234 | 0.509229 |
| GOBP_REPLICATION_FORK_PROCESSING | 38 | 0.46835 | 1.259337 | 0.171613 | 0.509033 |
| GOBP_STRIATED_MUSCLE_CONTRACTION | 135 | 0.402633 | 1.259014 | 0.09476 | 0.509422 |
| GOBP_REGULATION_OF_MAST_CELL_ACTIVATION_INVOLVED_IN_IMMUNE_RESPONSE | 26 | 0.494691 | 1.258125 | 0.183311 | 0.511589 |
| GOBP_CELLULAR_RESPONSE_TO_REACTIVE_OXYGEN_SPECIES | 121 | 0.409305 | 1.258043 | 0.117845 | 0.511228 |
| GOBP_PEPTIDYL_TYROSINE_MODIFICATION | 294 | 0.380764 | 1.257875 | 0.045596 | 0.511129 |
| GOBP_POSITIVE_REGULATION_OF_VASCULAR_ASSOCIATED_SMOOTH_MUSCLE_CELL | 24 | 0.514508 | 1.257549 | 0.18637 | 0.511506 |
| GOBP_ORGAN_GROWTH | 123 | 0.406425 | 1.256737 | 0.103837 | 0.513369 |
| GOBP_T_CELL_HOMEOSTASIS | 29 | 0.4963 | 1.256726 | 0.179104 | 0.512792 |
| GOBP_T_CELL_RECEPTOR_SIGNALING_PATHWAY | 100 | 0.414033 | 1.25667 | 0.116763 | 0.512351 |
| GOBP_POSITIVE_REGULATION_OF_HISTONE_DEACETYLATION | 21 | 0.522583 | 1.256389 | 0.192416 | 0.512573 |
| GOBP_REGULATION_OF_ION_TRANSMEMBRANE_TRANSPORT | 372 | 0.379657 | 1.256219 | 0.030426 | 0.512456 |
| GOBP_KERATINOCYTE_PROLIFERATION | 35 | 0.47706 | 1.255998 | 0.160995 | 0.512502 |
| GOBP_PROTEIN_CONTAINING_COMPLEX_REMODELING | 21 | 0.521954 | 1.255952 | 0.200849 | 0.512017 |
| GOBP_MYOBLAST_DIFFERENTIATION | 61 | 0.43736 | 1.255756 | 0.14951 | 0.512005 |
| GOBP_POSITIVE_REGULATION_OF_TUMOR_NECROSIS_FACTOR_SUPERFAMILY_CYTOK | 70 | 0.429531 | 1.255705 | 0.160848 | 0.51153 |
| GOBP_MUSCLE_CELL_MIGRATION | 71 | 0.434987 | 1.255438 | 0.141646 | 0.511758 |
| GOBP_NEGATIVE_REGULATION_OF_CYCLIN_DEPENDENT_PROTEIN_SERINE_THREONI | 28 | 0.494661 | 1.25448 | 0.19209 | 0.514093 |
| GOBP_NEGATIVE_REGULATION_OF_DEPHOSPHORYLATION | 41 | 0.473972 | 1.254474 | 0.173228 | 0.513496 |
| GOBP_RELAXATION_OF_CARDIAC_MUSCLE | 16 | 0.556008 | 1.254315 | 0.211394 | 0.513404 |
| GOBP_REGULATION_OF_OSTEOCLAST_DIFFERENTIATION | 44 | 0.46374 | 1.254201 | 0.155583 | 0.513124 |
| GOBP_REGULATION_OF_CATION_TRANSMEMBRANE_TRANSPORT | 269 | 0.383288 | 1.254029 | 0.056936 | 0.513046 |
| GOBP_AXONAL_TRANSPORT_OF_MITOCHONDRION | 18 | 0.547943 | 1.253909 | 0.212214 | 0.512794 |
| GOBP_REGULATION_OF_EXTRINSIC_APOPTOTIC_SIGNALING_PATHWAY | 128 | 0.401495 | 1.253528 | 0.102679 | 0.513374 |
| GOBP_NEGATIVE_REGULATION_OF_ION_TRANSMEMBRANE_TRANSPORT | 66 | 0.433887 | 1.253356 | 0.151883 | 0.513284 |
| GOBP_REGULATION_OF_TRANSMEMBRANE_TRANSPORT | 442 | 0.376699 | 1.253294 | 0.024121 | 0.51285 |
| GOBP_GENITALIA_DEVELOPMENT | 29 | 0.501804 | 1.253287 | 0.185539 | 0.512261 |
| GOBP_RESPONSE_TO_TUMOR_NECROSIS_FACTOR | 177 | 0.392123 | 1.253188 | 0.089034 | 0.511966 |
| GOBP_DEVELOPMENT_OF_PRIMARY_SEXUAL_CHARACTERISTICS | 183 | 0.388587 | 1.252843 | 0.081741 | 0.512427 |
| GOBP_FILOPODIUM_ASSEMBLY | 62 | 0.439798 | 1.252807 | 0.143577 | 0.511912 |
| GOBP_POSITIVE_REGULATION_OF_COLLAGEN_METABOLIC_PROCESS | 19 | 0.545983 | 1.252081 | 0.208696 | 0.513488 |
| GOBP_T_HELPER_17_CELL_DIFFERENTIATION | 17 | 0.547101 | 1.251572 | 0.211852 | 0.514385 |
| GOBP_MEMBRANE_REPOLARIZATION | 39 | 0.464341 | 1.250958 | 0.180628 | 0.515654 |
| GOBP_NEGATIVE_REGULATION_OF_TRANSFORMING_GROWTH_FACTOR_BETA_RECEP | 70 | 0.427552 | 1.250953 | 0.14881 | 0.51506 |
| GOBP_POSITIVE_REGULATION_OF_CYTOKINE_PRODUCTION | 311 | 0.379878 | 1.250671 | 0.055328 | 0.515309 |
| GOBP_GLIOGENESIS | 234 | 0.385117 | 1.25063 | 0.070981 | 0.514824 |
| GOBP_EMBRYONIC_CAMERA_TYPE_EYE_DEVELOPMENT | 26 | 0.505054 | 1.250246 | 0.189116 | 0.515362 |
| GOBP_REGULATION_OF_MITOCHONDRIAL_FISSION | 28 | 0.503914 | 1.249985 | 0.170124 | 0.515515 |
| GOBP_PHOSPHATIDIC_ACID_METABOLIC_PROCESS | 33 | 0.473561 | 1.2494 | 0.176862 | 0.516661 |
| GOBP_GAMMA_AMINOBUTYRIC_ACID_SIGNALING_PATHWAY | 22 | 0.525767 | 1.24939 | 0.202837 | 0.516088 |
| GOBP_NEGATIVE_REGULATION_OF_T_CELL_PROLIFERATION | 43 | 0.453962 | 1.249262 | 0.166033 | 0.515895 |
| GOBP_PYRIDINE_NUCLEOTIDE_BIOSYNTHETIC_PROCESS | 22 | 0.508656 | 1.247979 | 0.175953 | 0.519226 |
| GOBP_REGULATION_OF_SMALL_MOLECULE_METABOLIC_PROCESS | 272 | 0.380569 | 1.247485 | 0.06095 | 0.520056 |
| GOBP_NEGATIVE_REGULATION_OF_AUTOPHAGY | 73 | 0.426821 | 1.247058 | 0.13866 | 0.520734 |
| GOBP_NEGATIVE_REGULATION_OF_GROWTH | 215 | 0.386921 | 1.246113 | 0.072862 | 0.522961 |
| GOBP_APOPTOTIC_PROCESS_INVOLVED_IN_DEVELOPMENT | 36 | 0.467545 | 1.245674 | 0.184032 | 0.523773 |
| GOBP_B_CELL_DIFFERENTIATION | 98 | 0.411625 | 1.245587 | 0.153488 | 0.523443 |
| GOBP_REGULATION_OF_LIPID_TRANSPORT | 101 | 0.409145 | 1.244675 | 0.121178 | 0.525725 |
| GOBP_REGULATED_EXOCYTOSIS | 190 | 0.387477 | 1.244657 | 0.070675 | 0.52518 |
| GOBP_POSITIVE_REGULATION_OF_SYNAPTIC_TRANSMISSION_GLUTAMATERGIC | 22 | 0.518357 | 1.244118 | 0.202279 | 0.526186 |
| GOBP_RETINA_HOMEOSTASIS | 50 | 0.448386 | 1.243466 | 0.161538 | 0.527599 |
| GOBP_CARDIAC_CHAMBER_DEVELOPMENT | 142 | 0.396409 | 1.243448 | 0.099671 | 0.527041 |
| GOBP_REGULATION_OF_NECROTIC_CELL_DEATH | 31 | 0.489906 | 1.243429 | 0.189873 | 0.526478 |
| GOBP_POSITIVE_REGULATION_OF_LAMELLIPODIUM_ORGANIZATION | 32 | 0.475232 | 1.242693 | 0.183267 | 0.52813 |
| GOBP_TEMPERATURE_HOMEOSTASIS | 146 | 0.396798 | 1.242619 | 0.097588 | 0.527756 |
| GOBP_PLATELET_DERIVED_GROWTH_FACTOR_RECEPTOR_SIGNALING_PATHWAY | 48 | 0.454729 | 1.242493 | 0.170732 | 0.527543 |
| GOBP_NEGATIVE_REGULATION_OF_PROTEOLYSIS | 228 | 0.383062 | 1.242446 | 0.084656 | 0.52709 |
| GOBP_NEUROTRANSMITTER_REUPTAKE | 29 | 0.491364 | 1.242163 | 0.213162 | 0.52731 |
| GOBP_REGULATION_OF_HISTONE_METHYLATION | 74 | 0.422937 | 1.242119 | 0.138922 | 0.526844 |
| GOBP_POSITIVE_REGULATION_OF_TRANSPORTER_ACTIVITY | 100 | 0.407844 | 1.241906 | 0.134189 | 0.526874 |
| GOBP_REGULATION_OF_SPROUTING_ANGIOGENESIS | 34 | 0.473282 | 1.241386 | 0.19085 | 0.527854 |
| GOBP_EPITHELIAL_CELL_DIFFERENTIATION | 484 | 0.369038 | 1.240612 | 0.023139 | 0.529651 |
| GOBP_REGULATION_OF_ANION_TRANSMEMBRANE_TRANSPORT | 19 | 0.532689 | 1.239963 | 0.197674 | 0.531038 |
| GOBP_SPERM_MOTILITY | 73 | 0.420552 | 1.239698 | 0.151775 | 0.531254 |
| GOBP_PROTEIN_KINASE_B_SIGNALING | 157 | 0.388958 | 1.239677 | 0.111597 | 0.530714 |
| GOBP_EXTRINSIC_APOPTOTIC_SIGNALING_PATHWAY | 188 | 0.386561 | 1.239654 | 0.084926 | 0.530167 |
| GOBP_REGULATION_OF_LIPASE_ACTIVITY | 62 | 0.426357 | 1.239598 | 0.136476 | 0.529727 |
| GOBP_ARACHIDONIC_ACID_SECRETION | 18 | 0.53862 | 1.239474 | 0.226879 | 0.529515 |
| GOBP_EMBRYONIC_HINDLIMB_MORPHOGENESIS | 18 | 0.527178 | 1.23936 | 0.205634 | 0.529268 |
| GOBP_REGULATION_OF_INFLAMMATORY_RESPONSE_TO_ANTIGENIC_STIMULUS | 31 | 0.482337 | 1.239021 | 0.179945 | 0.529684 |
| GOBP_DIOL_METABOLIC_PROCESS | 30 | 0.48558 | 1.238909 | 0.192256 | 0.529414 |
| GOBP_RESPONSE_TO_DIETARY_EXCESS | 24 | 0.497919 | 1.238432 | 0.192958 | 0.530194 |
| GOBP_EMBRYONIC_HEART_TUBE_DEVELOPMENT | 70 | 0.426473 | 1.238289 | 0.149466 | 0.530041 |
| GOBP_REGULATION_OF_STEROL_TRANSPORT | 45 | 0.457404 | 1.237819 | 0.171216 | 0.530884 |
| GOBP_EPITHELIAL_CELL_PROLIFERATION | 320 | 0.376072 | 1.237272 | 0.057201 | 0.53195 |
| GOBP_REGULATION_OF_ALCOHOL_BIOSYNTHETIC_PROCESS | 41 | 0.458476 | 1.237212 | 0.181462 | 0.531539 |
| GOBP_POSITIVE_REGULATION_OF_SYNAPTIC_TRANSMISSION | 126 | 0.399414 | 1.237131 | 0.113333 | 0.531196 |
| GOBP_RENAL_VESICLE_DEVELOPMENT | 16 | 0.551521 | 1.23705 | 0.229651 | 0.530823 |
| GOBP_NEGATIVE_REGULATION_OF_CYTOKINE_PRODUCTION | 189 | 0.386496 | 1.237032 | 0.095238 | 0.530272 |
| GOBP_TRIGLYCERIDE_METABOLIC_PROCESS | 73 | 0.421261 | 1.236916 | 0.160628 | 0.530032 |
| GOBP_REGULATION_OF_OSTEOBLAST_DIFFERENTIATION | 108 | 0.401283 | 1.236231 | 0.125287 | 0.531515 |
| GOBP_POSITIVE_REGULATION_OF_PATHWAY_RESTRICTED_SMAD_PROTEIN_PHOSPH | 41 | 0.463798 | 1.236168 | 0.189791 | 0.531115 |
| GOBP_LYMPH_VESSEL_DEVELOPMENT | 28 | 0.484435 | 1.23611 | 0.188679 | 0.530704 |
| GOBP_POSITIVE_REGULATION_OF_ALPHA_BETA_T_CELL_PROLIFERATION | 16 | 0.548403 | 1.235746 | 0.232061 | 0.531252 |
| GOBP_CELLULAR_RESPONSE_TO_VIRUS | 62 | 0.433074 | 1.235708 | 0.168317 | 0.530778 |
| GOBP_REGULATION_OF_STRIATED_MUSCLE_CONTRACTION | 71 | 0.42581 | 1.234878 | 0.161369 | 0.532727 |

| | | | | | |
|---|---|---|---|---|---|
| GOBP_NUCLEOTIDE_TRANSMEMBRANE_TRANSPORT | 18 | -0.213646 | -0.586071 | 0.955272 | 1 |
| GOBP_PROTEIN_LOCALIZATION_TO_PHAGOPHORE_ASSEMBLY_SITE | 17 | -0.212869 | -0.585453 | 0.951662 | 1 |
| GOBP_PLASMA_MEMBRANE_TUBULATION | 17 | -0.214887 | -0.584867 | 0.96 | 1 |
| GOBP_VESICLE_CARGO_LOADING | 21 | -0.20658 | -0.58079 | 0.961415 | 1 |
| GOBP_POSITIVE_REGULATION_OF_MITOCHONDRIAL_TRANSLATION | 15 | -0.230395 | -0.578285 | 0.954128 | 1 |
| GOBP_POSITIVE_REGULATION_OF_TRANSLATIONAL_INITIATION | 23 | -0.194306 | -0.576848 | 0.957055 | 1 |
| GOBP_HEPARAN_SULFATE_PROTEOGLYCAN_BIOSYNTHETIC_PROCESS | 24 | -0.198698 | -0.57644 | 0.971631 | 1 |
| GOBP_VESICLE_TARGETING | 40 | -0.176954 | -0.574882 | 0.996124 | 1 |
| GOBP_POSITIVE_REGULATION_OF_UBIQUITIN_PROTEIN_TRANSFERASE | 28 | -0.191106 | -0.573586 | 0.988764 | 1 |
| GOBP_RIBOSOME_ASSEMBLY | 60 | -0.161546 | -0.572988 | 1 | 1 |
| GOBP_NEGATIVE_REGULATION_OF_DNA_BINDING | 46 | -0.169749 | -0.572244 | 0.990338 | 1 |
| GOBP_RESPONSE_TO_TESTOSTERONE | 32 | -0.182364 | -0.571476 | 0.991936 | 1 |
| GOBP_HISTONE_DEACETYLATION | 92 | -0.148886 | -0.571406 | 1 | 1 |
| GOBP_RNA_3_END_PROCESSING | 104 | -0.145908 | -0.568051 | 1 | 1 |
| GOBP_P_BODY_ASSEMBLY | 20 | -0.205234 | -0.567876 | 0.974277 | 1 |
| GOBP_POSITIVE_REGULATION_OF_ORGANELLE_ASSEMBLY | 76 | -0.154068 | -0.567662 | 1 | 1 |
| GOBP_LIPID_IMPORT_INTO_CELL | 16 | -0.217748 | -0.567002 | 0.966197 | 1 |
| GOBP_REGULATION_OF_MACROAUTOPHAGY | 150 | -0.135907 | -0.566327 | 1 | 1 |
| GOBP_PROTEIN_LOCALIZATION_TO_NUCLEOLUS | 15 | -0.213374 | -0.559127 | 0.964169 | 1 |
| GOBP_AMINO_ACID_ACTIVATION | 45 | -0.16894 | -0.558188 | 1 | 1 |
| GOBP_NEGATIVE_REGULATION_OF_ACTIN_FILAMENT_BUNDLE_ASSEM | 29 | -0.183106 | -0.556967 | 0.988372 | 1 |
| GOBP_DNA_TEMPLATED_TRANSCRIPTION_TERMINATION | 21 | -0.197644 | -0.556644 | 0.987382 | 1 |
| GOBP_GLUCOSE_IMPORT | 59 | -0.159657 | -0.556383 | 0.995 | 1 |
| GOBP_PEPTIDYL_ARGININE_METHYLATION | 15 | -0.213501 | -0.555676 | 0.966667 | 1 |
| GOBP_NCRNA_3_END_PROCESSING | 39 | -0.171314 | -0.553731 | 0.995671 | 1 |
| GOBP_POSITIVE_REGULATION_OF_CHROMOSOME_SEPARATION | 17 | -0.208041 | -0.55302 | 0.965839 | 1 |
| GOBP_RIBOSOME_BIOGENESIS | 297 | -0.120837 | -0.550867 | 1 | 1 |
| GOBP_LONG_CHAIN_FATTY_ACYL_COA_BIOSYNTHETIC_PROCESS | 17 | -0.20322 | -0.548962 | 0.981595 | 1 |
| GOBP_ENDOSOME_ORGANIZATION | 81 | -0.143162 | -0.545916 | 1 | 1 |
| GOBP_REGULATION_OF_MITOTIC_NUCLEAR_DIVISION | 96 | -0.141777 | -0.540448 | 1 | 1 |
| GOBP_NUCLEAR_TRANSCRIBED_MRNA_CATABOLIC_PROCESS | 118 | -0.134333 | -0.535568 | 1 | 1 |
| GOBP_POSITIVE_REGULATION_OF_DNA_BIOSYNTHETIC_PROCESS | 70 | -0.146871 | -0.534606 | 1 | 1 |
| GOBP_RETROGRADE_TRANSPORT_ENDOSOME_TO_GOLGI | 92 | -0.138564 | -0.533864 | 1 | 1 |
| GOBP_COPII_COATED_VESICLE_CARGO_LOADING | 15 | -0.206509 | -0.533167 | 0.971098 | 1 |
| GOBP_PROTEIN_LOCALIZATION_TO_CILIARY_MEMBRANE | 17 | -0.198584 | -0.532994 | 0.978395 | 1 |
| GOBP_MITOCHONDRION_MORPHOGENESIS | 20 | -0.187536 | -0.528878 | 0.979381 | 1 |
| GOBP_REGULATION_OF_CENTROSOME_CYCLE | 49 | -0.157703 | -0.524865 | 1 | 1 |
| GOBP_POSITIVE_REGULATION_OF_TOR_SIGNALING | 43 | -0.157439 | -0.524209 | 1 | 1 |
| GOBP_MRNA_TRANSPORT | 116 | -0.135278 | -0.523752 | 1 | 1 |
| GOBP_PROTEIN_MATURATION_BY_IRON_SULFUR_CLUSTER_TRANSFE | 15 | -0.197055 | -0.522996 | 0.981982 | 1 |
| GOBP_LYSOSOMAL_TRANSPORT | 116 | -0.129673 | -0.521167 | 1 | 1 |
| GOBP_REGULATION_OF_DNA_STRAND_ELONGATION | 16 | -0.188827 | -0.514762 | 0.980263 | 1 |
| GOBP_PROTEIN_LOCALIZATION_TO_CONDENSED_CHROMOSOME | 19 | -0.18665 | -0.506894 | 0.994065 | 1 |
| GOBP_ENDOPLASMIC_RETICULUM_TUBULAR_NETWORK_ORGANIZATIO | 18 | -0.178812 | -0.497125 | 0.9967 | 1 |
| GOBP_POSITIVE_REGULATION_OF_TRANSCRIPTION_FROM_RNA_POLY | 22 | -0.178697 | -0.496658 | 1 | 1 |
| GOBP_PROTEIN_SUMOYLATION | 47 | -0.145281 | -0.496404 | 1 | 1 |
| GOBP_REGULATION_OF_MITOTIC_CELL_CYCLE_SPINDLE_ASSEMBLY_( | 15 | -0.190858 | -0.494662 | 0.984472 | 1 |
| GOBP_HEPARAN_SULFATE_PROTEOGLYCAN_METABOLIC_PROCESS | 30 | -0.1576 | -0.491196 | 0.996226 | 1 |
| GOBP_NEGATIVE_REGULATION_OF_ENDOPLASMIC_RETICULUM_STRES | 17 | -0.179234 | -0.487075 | 0.984568 | 1 |
| GOBP_COPII_COATED_VESICLE_BUDDING | 30 | -0.153357 | -0.485624 | 0.996403 | 1 |
| GOBP_REGULATION_OF_HISTONE_H3_K9_METHYLATION | 22 | -0.169987 | -0.48358 | 0.990291 | 1 |
| GOBP_TELOMERE_MAINTENANCE | 127 | -0.118022 | -0.480814 | 1 | 1 |
| GOBP_REGULATION_OF_CELL_CYCLE_CHECKPOINT | 32 | -0.156433 | -0.479564 | 1 | 1 |
| GOBP_NEGATIVE_REGULATION_OF_TELOMERE_MAINTENANCE | 32 | -0.152231 | -0.477945 | 1 | 1 |
| GOBP_RESPONSE_TO_ARSENIC_CONTAINING_SUBSTANCE | 32 | -0.149081 | -0.476009 | 1 | 1 |
| GOBP_TELOMERE_MAINTENANCE_VIA_TELOMERE_LENGTHENING | 76 | -0.126139 | -0.469148 | 1 | 1 |
| GOBP_UBIQUITIN_DEPENDENT_PROTEIN_CATABOLIC_PROCESS_VIA_T | 34 | -0.146283 | -0.464102 | 1 | 1 |
| GOBP_DSRNA_PROCESSING | 51 | -0.130041 | -0.449479 | 0.995556 | 1 |
| GOBP_POSITIVE_REGULATION_OF_NEURON_MIGRATION | 16 | -0.171076 | -0.448287 | 0.993333 | 1 |
| GOBP_KINETOCHORE_ASSEMBLY | 17 | -0.167779 | -0.444946 | 1 | 1 |
| GOBP_REGULATION_OF_NEURON_PROJECTION_ARBORIZATION | 15 | -0.170176 | -0.441815 | 0.997076 | 1 |
| GOBP_MUSCLE_CELL_CELLULAR_HOMEOSTASIS | 15 | -0.169093 | -0.436906 | 1 | 1 |
| GOBP_MITOCHONDRIAL_GENE_EXPRESSION | 102 | -0.111071 | -0.423526 | 1 | 1 |
| GOBP_NCRNA_PROCESSING | 373 | -0.096101 | -0.420171 | 1 | 1 |
| GOBP_REGULATION_OF_ENDOPLASMIC_RETICULUM_STRESS_INDUCE | 28 | -0.135203 | -0.409875 | 1 | 1 |
| GOBP_NADH_DEHYDROGENASE_COMPLEX_ASSEMBLY | 58 | -0.112349 | -0.394846 | 1 | 1 |
| GOBP_REGULATION_OF_TELOMERE_MAINTENANCE | 77 | -0.099983 | -0.380187 | 1 | 1 |
| GOBP_LYMPHOID_PROGENITOR_CELL_DIFFERENTIATION | 18 | -0.136459 | -0.373273 | 1 | 1 |
| GOBP_NCRNA_TRANSCRIPTION | 55 | -0.097738 | -0.336052 | 1 | 0.999981 |

| | | | | | |
|---|---|---|---|---|---|
| GOBP_MEMBRANE_REPOLARIZATION_DURING_CARDIAC_MUSCLE_CELL_ACTION_POT | 17 | 0.532165 | 1.234622 | 0.22061 | 0.532929 |
| GOBP_POSITIVE_REGULATION_OF_EPIDERMIS_DEVELOPMENT | 27 | 0.492295 | 1.234536 | 0.196133 | 0.532589 |
| GOBP_REGULATION_OF_CALCIUM_ION_TRANSMEMBRANE_TRANSPORT | 116 | 0.403683 | 1.234054 | 0.137853 | 0.533471 |
| GOBP_REPRODUCTIVE_SYSTEM_DEVELOPMENT | 354 | 0.373373 | 1.233342 | 0.043878 | 0.53502 |
| GOBP_INTERFERON_BETA_PRODUCTION | 50 | 0.447478 | 1.233102 | 0.174129 | 0.535151 |
| GOBP_MATERNAL_PROCESS_INVOLVED_IN_FEMALE_PREGNANCY | 45 | 0.450136 | 1.232316 | 0.184343 | 0.537 |
| GOBP_POSITIVE_REGULATION_OF_EPIDERMAL_CELL_DIFFERENTIATION | 23 | 0.509828 | 1.231234 | 0.199422 | 0.539747 |
| GOBP_POSITIVE_REGULATION_OF_MYELOID_LEUKOCYTE_CYTOKINE_PRODUCTION_IN | 20 | 0.513192 | 1.231133 | 0.222548 | 0.539474 |
| GOBP_POSITIVE_REGULATION_OF_CALCIUM_ION_TRANSPORT_INTO_CYTOSOL | 39 | 0.461827 | 1.230354 | 0.207301 | 0.541288 |
| GOBP_REGULATION_OF_METAL_ION_TRANSPORT | 303 | 0.376792 | 1.230285 | 0.056468 | 0.540934 |
| GOBP_INOSITOL_PHOSPHATE_CATABOLIC_PROCESS | 19 | 0.521821 | 1.230063 | 0.2066 | 0.540996 |
| GOBP_REGULATION_OF_SECRETION | 475 | 0.36822 | 1.229464 | 0.033133 | 0.542312 |
| GOBP_POSITIVE_REGULATION_OF_CALCIUM_ION_TRANSMEMBRANE_TRANSPORTER_ | 35 | 0.471604 | 1.229037 | 0.188 | 0.543018 |
| GOBP_NEGATIVE_REGULATION_OF_PROTEIN_KINASE_B_SIGNALING | 38 | 0.466156 | 1.22817 | 0.203187 | 0.545095 |
| GOBP_ERBB_SIGNALING_PATHWAY | 108 | 0.404224 | 1.228051 | 0.127252 | 0.544863 |
| GOBP_NEGATIVE_REGULATION_OF_TRANSMEMBRANE_TRANSPORT | 88 | 0.410661 | 1.227082 | 0.141204 | 0.547365 |
| GOBP_POSITIVE_REGULATION_OF_RESPONSE_TO_WOUNDING | 53 | 0.43722 | 1.226224 | 0.19401 | 0.549391 |
| GOBP_DNA_DAMAGE_RESPONSE_SIGNAL_TRANSDUCTION_BY_P53_CLASS_MEDIATO | 67 | 0.422509 | 1.226174 | 0.166268 | 0.548958 |
| GOBP_NEURON_MIGRATION | 140 | 0.394407 | 1.225214 | 0.127353 | 0.55136 |
| GOBP_HEART_DEVELOPMENT | 498 | 0.367576 | 1.224944 | 0.031219 | 0.551601 |
| GOBP_GRANULOCYTE_DIFFERENTIATION | 26 | 0.493833 | 1.224441 | 0.225313 | 0.55257 |
| GOBP_REGULATION_OF_MICROTUBULE_BASED_MOVEMENT | 40 | 0.452324 | 1.223727 | 0.203922 | 0.554113 |
| GOBP_RESPONSE_TO_PROSTAGLANDIN_E | 16 | 0.530342 | 1.223111 | 0.235988 | 0.555391 |
| GOBP_LYMPHOCYTE_ACTIVATION_INVOLVED_IN_IMMUNE_RESPONSE | 132 | 0.394708 | 1.222975 | 0.127374 | 0.555198 |
| GOBP_POSITIVE_REGULATION_OF_ENDOTHELIAL_CELL_CHEMOTAXIS | 15 | 0.549857 | 1.222542 | 0.228739 | 0.555884 |
| GOBP_POSITIVE_REGULATION_OF_VIRAL_LIFE_CYCLE | 20 | 0.513447 | 1.221831 | 0.211429 | 0.557506 |
| GOBP_REGULATION_OF_EPITHELIAL_CELL_MIGRATION | 202 | 0.381042 | 1.221181 | 0.112407 | 0.558953 |
| GOBP_MITOCHONDRIAL_CALCIUM_ION_HOMEOSTASIS | 24 | 0.497788 | 1.221122 | 0.212554 | 0.558533 |
| GOBP_TOXIN_TRANSPORT | 35 | 0.466627 | 1.220233 | 0.195538 | 0.560802 |
| GOBP_IMPORT_INTO_CELL | 183 | 0.381985 | 1.220197 | 0.109226 | 0.560302 |
| GOBP_POSITIVE_REGULATION_OF_BEHAVIOR | 15 | 0.540561 | 1.21986 | 0.224111 | 0.560723 |
| GOBP_DOPAMINE_SECRETION | 31 | 0.476995 | 1.219682 | 0.21067 | 0.560664 |
| GOBP_REGULATION_OF_DEVELOPMENTAL_GROWTH | 268 | 0.373404 | 1.21952 | 0.087269 | 0.560602 |
| GOBP_REGULATION_OF_RECEPTOR_BINDING | 18 | 0.528971 | 1.219335 | 0.218487 | 0.560534 |
| GOBP_REGULATION_OF_B_CELL_DIFFERENTIATION | 23 | 0.50813 | 1.218799 | 0.230137 | 0.561608 |
| GOBP_T_HELPER_17_TYPE_IMMUNE_RESPONSE | 27 | 0.487057 | 1.218474 | 0.222683 | 0.562088 |
| GOBP_POSITIVE_REGULATION_OF_PROTEIN_KINASE_B_SIGNALING | 89 | 0.405902 | 1.217833 | 0.137209 | 0.563463 |
| GOBP_NEGATIVE_REGULATION_OF_LYMPHOCYTE_APOPTOTIC_PROCESS | 27 | 0.478895 | 1.216726 | 0.20527 | 0.566317 |
| GOBP_REGULATION_OF_RECEPTOR_SIGNALING_PATHWAY_VIA_STAT | 59 | 0.423154 | 1.216433 | 0.180368 | 0.566586 |
| GOBP_SECRETION_BY_TISSUE | 31 | 0.46919 | 1.215835 | 0.236339 | 0.567822 |
| GOBP_LAMELLIPODIUM_MORPHOGENESIS | 17 | 0.530877 | 1.215388 | 0.225392 | 0.568729 |
| GOBP_SALIVARY_GLAND_DEVELOPMENT | 29 | 0.482751 | 1.215122 | 0.226902 | 0.568975 |
| GOBP_T_HELPER_1_CELL_DIFFERENTIATION | 17 | 0.524891 | 1.214729 | 0.237822 | 0.569622 |
| GOBP_BODY_FLUID_SECRETION | 74 | 0.420458 | 1.214706 | 0.148104 | 0.569104 |
| GOBP_REGULATION_OF_DIGESTIVE_SYSTEM_PROCESS | 23 | 0.493233 | 1.213602 | 0.230441 | 0.572061 |
| GOBP_NEGATIVE_REGULATION_OF_PEPTIDYL_SERINE_PHOSPHORYLATION | 25 | 0.490718 | 1.213364 | 0.239281 | 0.572196 |
| GOBP_INTRINSIC_APOPTOTIC_SIGNALING_PATHWAY_IN_RESPONSE_TO_DNA_DAMAG | 42 | 0.449332 | 1.213296 | 0.223087 | 0.571783 |
| GOBP_EXPLORATION_BEHAVIOR | 22 | 0.505917 | 1.212498 | 0.221289 | 0.573725 |
| GOBP_ADAPTIVE_THERMOGENESIS | 133 | 0.388993 | 1.212442 | 0.140505 | 0.573304 |
| GOBP_LEUKOCYTE_HOMEOSTASIS | 67 | 0.420923 | 1.212375 | 0.185319 | 0.572905 |
| GOBP_PURINE_NUCLEOSIDE_METABOLIC_PROCESS | 25 | 0.492385 | 1.211821 | 0.226075 | 0.574063 |
| GOBP_POSITIVE_REGULATION_OF_CARDIAC_MUSCLE_CELL_PROLIFERATION | 15 | 0.538233 | 1.211802 | 0.247706 | 0.573528 |
| GOBP_CARBOHYDRATE_DERIVATIVE_CATABOLIC_PROCESS | 142 | 0.388489 | 1.211452 | 0.13587 | 0.574025 |
| GOBP_CARDIAC_ATRIUM_DEVELOPMENT | 29 | 0.479193 | 1.211312 | 0.24343 | 0.573855 |
| GOBP_REGULATION_OF_SMOOTH_MUSCLE_CONTRACTION | 54 | 0.43418 | 1.210278 | 0.191411 | 0.576594 |
| GOBP_REGULATION_OF_ICOSANOID_SECRETION | 15 | 0.548909 | 1.209394 | 0.247006 | 0.578725 |
| GOBP_POSITIVE_REGULATION_OF_TRIGLYCERIDE_METABOLIC_PROCESS | 19 | 0.509163 | 1.20935 | 0.242424 | 0.57825 |
| GOBP_REGULATION_OF_PATTERN_RECOGNITION_RECEPTOR_SIGNALING_PATHWAY | 80 | 0.406934 | 1.209197 | 0.173145 | 0.578155 |
| GOBP_NEUROPEPTIDE_SIGNALING_PATHWAY | 67 | 0.419186 | 1.208215 | 0.183922 | 0.580613 |
| GOBP_MITOCHONDRIAL_TRANSCRIPTION | 17 | 0.523731 | 1.207978 | 0.250737 | 0.580787 |
| GOBP_POSITIVE_REGULATION_OF_JNK_CASCADE | 82 | 0.406027 | 1.207777 | 0.201935 | 0.580783 |
| GOBP_EMBRYONIC_EYE_MORPHOGENESIS | 25 | 0.480728 | 1.207736 | 0.217391 | 0.580314 |
| GOBP_SYNAPTIC_MEMBRANE_ADHESION | 25 | 0.493929 | 1.207366 | 0.234375 | 0.580847 |
| GOBP_SYNCYTIUM_FORMATION | 44 | 0.448958 | 1.207067 | 0.194771 | 0.581257 |
| GOBP_PHASIC_SMOOTH_MUSCLE_CONTRACTION | 15 | 0.540212 | 1.206679 | 0.248503 | 0.581956 |
| GOBP_REGULATION_OF_PHOSPHOLIPASE_ACTIVITY | 46 | 0.443403 | 1.206612 | 0.203125 | 0.58157 |
| GOBP_VASCULAR_WOUND_HEALING | 16 | 0.527278 | 1.206551 | 0.254873 | 0.58117 |
| GOBP_REGULATION_OF_G_PROTEIN_COUPLED_RECEPTOR_SIGNALING_PATHWAY | 110 | 0.395419 | 1.206524 | 0.150057 | 0.580647 |
| GOBP_METANEPHRIC_NEPHRON_MORPHOGENESIS | 21 | 0.505891 | 1.205133 | 0.257062 | 0.584523 |
| GOBP_RESPONSE_TO_CORTICOSTEROID | 118 | 0.394141 | 1.205037 | 0.162192 | 0.584204 |
| GOBP_POSITIVE_REGULATION_OF_EPITHELIAL_CELL_DIFFERENTIATION | 43 | 0.452047 | 1.204752 | 0.198454 | 0.584536 |
| GOBP_REGULATION_OF_ENDOTHELIAL_CELL_MIGRATION | 148 | 0.381499 | 1.204631 | 0.134929 | 0.584331 |
| GOBP_POSITIVE_REGULATION_OF_RELEASE_OF_CYTOCHROME_C_FROM_MITOCHONI | 24 | 0.490965 | 1.204075 | 0.235624 | 0.585438 |
| GOBP_NEGATIVE_REGULATION_OF_LIPID_STORAGE | 15 | 0.537461 | 1.203989 | 0.256598 | 0.585112 |
| GOBP_SEQUESTERING_OF_CALCIUM_ION | 94 | 0.402679 | 1.203304 | 0.176941 | 0.586625 |
| GOBP_POSITIVE_REGULATION_OF_PHOSPHATASE_ACTIVITY | 28 | 0.481445 | 1.202869 | 0.239554 | 0.587434 |
| GOBP_ENDOTHELIAL_CELL_PROLIFERATION | 129 | 0.389391 | 1.202828 | 0.158305 | 0.586944 |
| GOBP_NEGATIVE_REGULATION_OF_ENDOTHELIAL_CELL_APOPTOTIC_PROCESS | 25 | 0.483283 | 1.202342 | 0.242553 | 0.587883 |
| GOBP_BONE_GROWTH | 27 | 0.472897 | 1.202312 | 0.236413 | 0.587376 |
| GOBP_POSITIVE_REGULATION_OF_MAPK_CASCADE | 370 | 0.363534 | 1.201816 | 0.076375 | 0.588295 |
| GOBP_COPPER_ION_HOMEOSTASIS | 18 | 0.519457 | 1.201637 | 0.251479 | 0.588248 |
| GOBP_POSITIVE_REGULATION_OF_ERK1_AND_ERK2_CASCADE | 147 | 0.386281 | 1.201567 | 0.147346 | 0.58788 |
| GOBP_POSITIVE_REGULATION_OF_INTERFERON_BETA_PRODUCTION | 34 | 0.461961 | 1.200832 | 0.211286 | 0.589613 |
| GOBP_EXOCYTOSIS | 301 | 0.362917 | 1.200681 | 0.092593 | 0.589485 |
| GOBP_POSITIVE_REGULATION_OF_ORGAN_GROWTH | 27 | 0.470272 | 1.20033 | 0.220083 | 0.590012 |
| GOBP_IRON_ION_HOMEOSTASIS | 70 | 0.414226 | 1.200325 | 0.191566 | 0.589433 |
| GOBP_NEGATIVE_REGULATION_OF_VASCULATURE_DEVELOPMENT | 80 | 0.409313 | 1.200259 | 0.189635 | 0.58906 |
| GOBP_BIOLOGICAL_PROCESS_INVOLVED_IN_INTERACTION_WITH_SYMBIONT | 69 | 0.41534 | 1.200012 | 0.196078 | 0.589252 |
| GOBP_AMYLOID_BETA_CLEARANCE | 30 | 0.470371 | 1.199903 | 0.243775 | 0.589033 |
| GOBP_PATHWAY_RESTRICTED_SMAD_PROTEIN_PHOSPHORYLATION | 55 | 0.42411 | 1.199549 | 0.210859 | 0.589538 |
| GOBP_REGULATION_OF_TRIGLYCERIDE_BIOSYNTHETIC_PROCESS | 18 | 0.519869 | 1.199387 | 0.253256 | 0.589444 |
| GOBP_NIK_NF_KAPPAB_SIGNALING | 101 | 0.396918 | 1.198355 | 0.175377 | 0.592142 |
| GOBP_TROPHOBLAST_CELL_MIGRATION | 15 | 0.529365 | 1.198301 | 0.268562 | 0.591723 |
| GOBP_POSITIVE_REGULATION_OF_T_CELL_CYTOKINE_PRODUCTION | 18 | 0.524179 | 1.197544 | 0.258203 | 0.593503 |
| GOBP_RELAXATION_OF_MUSCLE | 27 | 0.473313 | 1.197449 | 0.256338 | 0.593194 |
| GOBP_REGULATION_OF_CHROMATIN_BINDING | 23 | 0.495418 | 1.197445 | 0.254261 | 0.592605 |
| GOBP_ADIPOSE_TISSUE_DEVELOPMENT | 41 | 0.446312 | 1.197311 | 0.218085 | 0.592445 |
| GOBP_RESPONSE_TO_LEAD_ION | 17 | 0.52201 | 1.196864 | 0.256677 | 0.593239 |
| GOBP_AMINE_TRANSPORT | 75 | 0.407773 | 1.1967 | 0.176471 | 0.593107 |
| GOBP_CELLULAR_COPPER_ION_HOMEOSTASIS | 15 | 0.534145 | 1.196496 | 0.269173 | 0.593197 |
| GOBP_MACROPHAGE_CYTOKINE_PRODUCTION | 15 | 0.5387 | 1.196453 | 0.251128 | 0.592733 |
| GOBP_REGULATION_OF_NEURON_MIGRATION | 41 | 0.43991 | 1.196227 | 0.226365 | 0.592831 |
| GOBP_NEGATIVE_REGULATION_OF_ANTIGEN_RECEPTOR_MEDIATED_SIGNALING_PATI | 22 | 0.495001 | 1.196162 | 0.267806 | 0.592459 |
| GOBP_CALCIUM_ION_REGULATED_EXOCYTOSIS_OF_NEUROTRANSMITTER | 15 | 0.534144 | 1.196077 | 0.260805 | 0.592132 |
| GOBP_CILIUM_OR_FLAGELLUM_DEPENDENT_CELL_MOTILITY | 92 | 0.403056 | 1.195738 | 0.185531 | 0.592649 |
| GOBP_ADENYLATE_CYCLASE_ACTIVATING_G_PROTEIN_COUPLED_RECEPTOR_SIGNA | 109 | 0.395106 | 1.195135 | 0.172336 | 0.593962 |
| GOBP_REGULATION_OF_POSTSYNAPTIC_MEMBRANE_POTENTIAL | 109 | 0.392412 | 1.194291 | 0.176267 | 0.596009 |
| GOBP_POSITIVE_REGULATION_OF_NEURAL_PRECURSOR_CELL_PROLIFERATION | 43 | 0.438337 | 1.193618 | 0.218421 | 0.597492 |
| GOBP_RESPONSE_TO_INTERFERON_BETA | 21 | 0.504598 | 1.193125 | 0.256522 | 0.598475 |
| GOBP_ARF_PROTEIN_SIGNAL_TRANSDUCTION | 19 | 0.511969 | 1.192963 | 0.24564 | 0.598437 |
| GOBP_LEUKOCYTE_ADHESION_TO_VASCULAR_ENDOTHELIAL_CELL | 31 | 0.458571 | 1.192957 | 0.237964 | 0.597866 |
| GOBP_INOSITOL_PHOSPHATE_MEDIATED_SIGNALING | 48 | 0.434055 | 1.192922 | 0.2225 | 0.597391 |
| GOBP_BLASTOCYST_FORMATION | 39 | 0.45058 | 1.192784 | 0.232 | 0.597229 |
| GOBP_EXOCRINE_SYSTEM_DEVELOPMENT | 37 | 0.449576 | 1.192177 | 0.234519 | 0.59862 |
| GOBP_POSITIVE_REGULATION_OF_MACROPHAGE_MIGRATION | 15 | 0.538285 | 1.192121 | 0.245455 | 0.598214 |
| GOBP_TRANSMEMBRANE_RECEPTOR_PROTEIN_SERINE_THREONINE_KINASE_SIGNAL | 311 | 0.362165 | 1.191998 | 0.103061 | 0.598058 |
| GOBP_FEMALE_SEX_DIFFERENTIATION | 101 | 0.39147 | 1.189652 | 0.156751 | 0.604895 |
| GOBP_PYRIDINE_CONTAINING_COMPOUND_BIOSYNTHETIC_PROCESS | 25 | 0.482552 | 1.189568 | 0.239281 | 0.604572 |
| GOBP_NEGATIVE_REGULATION_OF_EXTRINSIC_APOPTOTIC_SIGNALING_PATHWAY | 79 | 0.405172 | 1.189365 | 0.208766 | 0.604664 |
| GOBP_NEUROTRANSMITTER_RECEPTOR_INTERNALIZATION | 16 | 0.531987 | 1.189147 | 0.261398 | 0.604803 |
| GOBP_CELLULAR_RESPONSE_TO_RETINOIC_ACID | 52 | 0.425539 | 1.18872 | 0.210459 | 0.605587 |
| GOBP_SENSORY_ORGAN_MORPHOGENESIS | 212 | 0.368838 | 1.187671 | 0.134796 | 0.608343 |
| GOBP_AGING | 248 | 0.364811 | 1.187413 | 0.136601 | 0.608588 |
| GOBP_PLATELET_ACTIVATION | 100 | 0.393009 | 1.186828 | 0.174112 | 0.609881 |
| GOBP_NATURAL_KILLER_CELL_MEDIATED_IMMUNITY | 43 | 0.436621 | 1.186512 | 0.230065 | 0.610305 |
| GOBP_INHIBITORY_SYNAPSE_ASSEMBLY | 15 | 0.529731 | 1.185771 | 0.254815 | 0.612081 |
| GOBP_NEGATIVE_REGULATION_OF_PROTEIN_TYROSINE_KINASE_ACTIVITY | 29 | 0.460151 | 1.18547 | 0.251656 | 0.612437 |
| GOBP_PROTEIN_LOCALIZATION_TO_EXTRACELLULAR_REGION | 282 | 0.362444 | 1.185321 | 0.121399 | 0.612303 |
| GOBP_CARDIAC_MYOFIBRIL_ASSEMBLY | 16 | 0.524122 | 1.18528 | 0.275 | 0.61186 |
| GOBP_NEGATIVE_REGULATION_OF_REGULATED_SECRETORY_PATHWAY | 19 | 0.516527 | 1.18525 | 0.277299 | 0.611377 |
| GOBP_POSITIVE_REGULATION_OF_SECRETION | 241 | 0.366304 | 1.184269 | 0.129979 | 0.614014 |
| GOBP_POTASSIUM_ION_IMPORT_ACROSS_PLASMA_MEMBRANE | 38 | 0.441272 | 1.183897 | 0.231383 | 0.614649 |
| GOBP_REGULATION_OF_GLUCOSE_METABOLIC_PROCESS | 104 | 0.386843 | 1.183441 | 0.184807 | 0.615479 |
| GOBP_DECIDUALIZATION | 19 | 0.501244 | 1.18299 | 0.270195 | 0.616331 |
| GOBP_ENDOCARDIAL_CUSHION_DEVELOPMENT | 41 | 0.442996 | 1.182638 | 0.240885 | 0.616936 |
| GOBP_LIPID_EXPORT_FROM_CELL | 32 | 0.461159 | 1.182094 | 0.247283 | 0.618056 |
| GOBP_REGULATION_OF_PHOSPHATASE_ACTIVITY | 68 | 0.411782 | 1.182031 | 0.216867 | 0.617658 |
| GOBP_REGULATION_OF_GASTRULATION | 15 | 0.528534 | 1.181962 | 0.258493 | 0.617284 |
| GOBP_POSITIVE_REGULATION_OF_NEUROTRANSMITTER_SECRETION | 16 | 0.522176 | 1.181768 | 0.270423 | 0.61733 |
| GOBP_SKIN_EPIDERMIS_DEVELOPMENT | 69 | 0.406493 | 1.181608 | 0.216482 | 0.617245 |
| GOBP_REGULATION_OF_ORGANIC_ACID_TRANSPORT | 57 | 0.413757 | 1.181458 | 0.214377 | 0.61714 |
| GOBP_COLLATERAL_SPROUTING | 25 | 0.474054 | 1.181025 | 0.253219 | 0.617951 |
| GOBP_EXTRINSIC_APOPTOTIC_SIGNALING_PATHWAY_VIA_DEATH_DOMAIN_RECEPTOF | 73 | 0.403159 | 1.180805 | 0.22316 | 0.618067 |
| GOBP_NEGATIVE_REGULATION_OF_LEUKOCYTE_PROLIFERATION | 54 | 0.421587 | 1.180563 | 0.226533 | 0.6183 |
| GOBP_REGULATION_OF_PROTEIN_SECRETION | 195 | 0.368867 | 1.17997 | 0.158405 | 0.619528 |
| GOBP_BIOLOGICAL_PROCESS_INVOLVED_IN_INTRASPECIES_INTERACTION_BETWEEN | 42 | 0.432244 | 1.179913 | 0.234917 | 0.619127 |
| GOBP_CELL_FATE_COMMITMENT | 177 | 0.372338 | 1.179729 | 0.154749 | 0.619125 |
| GOBP_POSITIVE_REGULATION_OF_DNA_BINDING_TRANSCRIPTION_FACTOR_ACTIVITY | 216 | 0.365619 | 1.178879 | 0.160338 | 0.621253 |
| GOBP_NEUTRAL_LIPID_METABOLIC_PROCESS | 99 | 0.389891 | 1.178774 | 0.195205 | 0.621001 |
| GOBP_POSITIVE_REGULATION_OF_STRIATED_MUSCLE_CELL_DIFFERENTIATION | 30 | 0.459097 | 1.178652 | 0.230871 | 0.620801 |
| GOBP_ANTIVIRAL_INNATE_IMMUNE_RESPONSE | 15 | 0.521893 | 1.178017 | 0.280431 | 0.622275 |

| | | | | | |
|---|---|---|---|---|---|
| GOBP_POSITIVE_REGULATION_OF_FILOPODIUM_ASSEMBLY | 27 | 0.468299 | 1.177716 | 0.253425 | 0.622627 |
| GOBP_RESPONSE_TO_ESTRADIOL | 92 | 0.387809 | 1.177676 | 0.209602 | 0.622176 |
| GOBP_NEGATIVE_REGULATION_OF_RESPONSE_TO_WOUNDING | 67 | 0.408585 | 1.177558 | 0.224096 | 0.621961 |
| GOBP_METAL_ION_HOMEOSTASIS | 486 | 0.353648 | 1.177551 | 0.066869 | 0.621402 |
| GOBP_POSITIVE_REGULATION_OF_GLUCOSE_TRANSMEMBRANE_TRANSPORT | 35 | 0.457384 | 1.177389 | 0.253539 | 0.621339 |
| GOBP_DOPAMINERGIC_NEURON_DIFFERENTIATION | 27 | 0.478751 | 1.177038 | 0.276536 | 0.621929 |
| GOBP_MEMBRANE_FUSION | 133 | 0.380402 | 1.176799 | 0.189845 | 0.622121 |
| GOBP_REGULATION_OF_CALCIUM_ION_IMPORT | 30 | 0.466025 | 1.176393 | 0.257453 | 0.622835 |
| GOBP_HEMATOPOIETIC_STEM_CELL_HOMEOSTASIS | 18 | 0.509407 | 1.176275 | 0.28866 | 0.622606 |
| GOBP_REGULATION_OF_CELLULAR_KETONE_METABOLIC_PROCESS | 98 | 0.39105 | 1.17591 | 0.201595 | 0.623219 |
| GOBP_LONG_TERM_SYNAPTIC_POTENTIATION | 74 | 0.400815 | 1.175835 | 0.217497 | 0.622861 |
| GOBP_POSITIVE_REGULATION_OF_ATP_DEPENDENT_ACTIVITY | 38 | 0.445697 | 1.175703 | 0.248 | 0.622724 |
| GOBP_POSITIVE_REGULATION_OF_DENDRITE_DEVELOPMENT | 15 | 0.531634 | 1.175573 | 0.275362 | 0.622581 |
| GOBP_REGULATION_OF_OXIDOREDUCTASE_ACTIVITY | 72 | 0.400636 | 1.175377 | 0.213253 | 0.622612 |
| GOBP_GLIAL_CELL_PROLIFERATION | 32 | 0.455124 | 1.175345 | 0.249322 | 0.622139 |
| GOBP_REGULATION_OF_HORMONE_SECRETION | 180 | 0.366832 | 1.175096 | 0.159041 | 0.622348 |
| GOBP_EYE_MORPHOGENESIS | 127 | 0.380066 | 1.174856 | 0.189128 | 0.622572 |
| GOBP_BASEMENT_MEMBRANE_ORGANIZATION | 29 | 0.460161 | 1.174851 | 0.256342 | 0.622014 |
| GOBP_RESPONSE_TO_ISOQUINOLINE_ALKALOID | 26 | 0.469984 | 1.17482 | 0.264946 | 0.621537 |
| GOBP_REGULATION_OF_NEURAL_PRECURSOR_CELL_PROLIFERATION | 72 | 0.396519 | 1.174033 | 0.226619 | 0.623494 |
| GOBP_MYELOID_CELL_DIFFERENTIATION | 319 | 0.359095 | 1.173624 | 0.124873 | 0.624163 |
| GOBP_REGULATION_OF_NIK_NF_KAPPAB_SIGNALING | 79 | 0.399391 | 1.173279 | 0.220482 | 0.624739 |
| GOBP_IMMUNOGLOBULIN_PRODUCTION_INVOLVED_IN_IMMUNOGLOBULIN_MEDIATED | 56 | 0.412188 | 1.173262 | 0.236745 | 0.624203 |
| GOBP_REGULATION_OF_VASCULATURE_DEVELOPMENT | 227 | 0.360662 | 1.173162 | 0.162393 | 0.623935 |
| GOBP_SMOOTH_MUSCLE_TISSUE_DEVELOPMENT | 17 | 0.512905 | 1.172869 | 0.280753 | 0.624315 |
| GOBP_REGULATION_OF_NEUROTRANSMITTER_LEVELS | 183 | 0.361776 | 1.172666 | 0.181916 | 0.624371 |
| GOBP_ENTRY_INTO_HOST | 126 | 0.378124 | 1.172511 | 0.190581 | 0.624301 |
| GOBP_REGULATION_OF_LIPID_BIOSYNTHETIC_PROCESS | 136 | 0.37712 | 1.172416 | 0.193619 | 0.624058 |
| GOBP_REGULATION_OF_RELEASE_OF_SEQUESTERED_CALCIUM_ION_INTO_CYTOSOL | 61 | 0.406951 | 1.172042 | 0.233129 | 0.62462 |
| GOBP_MESENCHYMAL_TO_EPITHELIAL_TRANSITION | 17 | 0.510969 | 1.171347 | 0.268895 | 0.626301 |
| GOBP_CELLULAR_RESPONSE_TO_LIPID | 431 | 0.352013 | 1.171311 | 0.078471 | 0.625848 |
| GOBP_NEGATIVE_REGULATION_OF_CELL_CELL_ADHESION | 131 | 0.372772 | 1.171287 | 0.194164 | 0.625357 |
| GOBP_CELL_MIGRATION_INVOLVED_IN_SPROUTING_ANGIOGENESIS | 46 | 0.420002 | 1.171254 | 0.249017 | 0.624885 |
| GOBP_BEHAVIORAL_RESPONSE_TO_PAIN | 15 | 0.519364 | 1.170864 | 0.291233 | 0.625553 |
| GOBP_REGULATION_OF_NEUROTRANSMITTER_TRANSPORT | 85 | 0.39063 | 1.170723 | 0.230679 | 0.625411 |
| GOBP_SPLEEN_DEVELOPMENT | 25 | 0.472913 | 1.170596 | 0.264666 | 0.625223 |
| GOBP_NATURAL_KILLER_CELL_DIFFERENTIATION | 20 | 0.502963 | 1.170521 | 0.26383 | 0.624882 |
| GOBP_RESPONSE_TO_ACID_CHEMICAL | 107 | 0.380499 | 1.17008 | 0.212866 | 0.625711 |
| GOBP_VASCULAR_TRANSPORT | 78 | 0.395737 | 1.16928 | 0.234875 | 0.627645 |
| GOBP_PYRIDINE_CONTAINING_COMPOUND_METABOLIC_PROCESS | 33 | 0.449004 | 1.169115 | 0.265141 | 0.627608 |
| GOBP_EMBRYONIC_HEART_TUBE_MORPHOGENESIS | 58 | 0.415264 | 1.168966 | 0.23681 | 0.627483 |
| GOBP_CALCINEURIN_MEDIATED_SIGNALING | 40 | 0.438512 | 1.168864 | 0.2607 | 0.627231 |
| GOBP_RNA_PHOSPHODIESTER_BOND_HYDROLYSIS_ENDONUCLEOLYTIC | 70 | 0.401835 | 1.168845 | 0.23225 | 0.626725 |
| GOBP_NEURAL_PRECURSOR_CELL_PROLIFERATION | 123 | 0.377563 | 1.168225 | 0.185185 | 0.628077 |
| GOBP_NEGATIVE_REGULATION_OF_ION_TRANSPORT | 103 | 0.377575 | 1.168011 | 0.199313 | 0.628219 |
| GOBP_POSITIVE_REGULATION_OF_CELLULAR_EXTRAVASATION | 15 | 0.528398 | 1.167908 | 0.279817 | 0.627981 |
| GOBP_MORPHOGENESIS_OF_AN_ENDOTHELIUM | 15 | 0.510887 | 1.167807 | 0.293078 | 0.627738 |
| GOBP_CARDIAC_CONDUCTION | 79 | 0.393534 | 1.167707 | 0.222488 | 0.627494 |
| GOBP_CELLULAR_RESPONSE_TO_STEROL | 16 | 0.506323 | 1.167391 | 0.278184 | 0.627914 |
| GOBP_TRANSPORT_OF_VIRUS | 17 | 0.505144 | 1.16739 | 0.294205 | 0.627345 |
| GOBP_POSITIVE_REGULATION_OF_OSTEOBLAST_DIFFERENTIATION | 59 | 0.410722 | 1.167178 | 0.234858 | 0.627474 |
| GOBP_ERYTHROCYTE_DEVELOPMENT | 26 | 0.46989 | 1.166947 | 0.265647 | 0.627636 |
| GOBP_CAMERA_TYPE_EYE_MORPHOGENESIS | 99 | 0.386 | 1.166557 | 0.233065 | 0.628292 |
| GOBP_NEGATIVE_CHEMOTAXIS | 45 | 0.433885 | 1.166396 | 0.256881 | 0.628211 |
| GOBP_REGULATION_OF_MONONUCLEAR_CELL_MIGRATION | 77 | 0.400585 | 1.166097 | 0.207186 | 0.628599 |
| GOBP_CATECHOLAMINE_SECRETION | 43 | 0.43402 | 1.166048 | 0.25974 | 0.628183 |
| GOBP_CHOLESTEROL_EFFLUX | 39 | 0.433095 | 1.165779 | 0.26539 | 0.628501 |
| GOBP_PLATELET_AGGREGATION | 54 | 0.410963 | 1.165529 | 0.258065 | 0.628755 |
| GOBP_DETECTION_OF_BIOTIC_STIMULUS | 22 | 0.483736 | 1.165479 | 0.278689 | 0.628359 |
| GOBP_ORGAN_OR_TISSUE_SPECIFIC_IMMUNE_RESPONSE | 17 | 0.514188 | 1.165467 | 0.281818 | 0.627835 |
| GOBP_DIVALENT_INORGANIC_CATION_HOMEOSTASIS | 362 | 0.352939 | 1.165248 | 0.115736 | 0.627961 |
| GOBP_SPINDLE_LOCALIZATION | 48 | 0.428711 | 1.164562 | 0.242308 | 0.629548 |
| GOBP_MULTI_MULTICELLULAR_ORGANISM_PROCESS | 153 | 0.370806 | 1.163511 | 0.188184 | 0.632268 |
| GOBP_NEGATIVE_REGULATION_OF_CHEMOTAXIS | 51 | 0.4167 | 1.163482 | 0.24937 | 0.631793 |
| GOBP_REGULATION_OF_OSSIFICATION | 94 | 0.387269 | 1.163299 | 0.226852 | 0.631796 |
| GOBP_IMPORT_ACROSS_PLASMA_MEMBRANE | 141 | 0.374957 | 1.163253 | 0.190889 | 0.631375 |
| GOBP_TYPE_I_INTERFERON_PRODUCTION | 81 | 0.392921 | 1.162374 | 0.219429 | 0.63362 |
| GOBP_INTEGRIN_ACTIVATION | 20 | 0.495684 | 1.162286 | 0.284457 | 0.633373 |
| GOBP_REGULATION_OF_FAT_CELL_DIFFERENTIATION | 118 | 0.379237 | 1.161831 | 0.207719 | 0.634254 |
| GOBP_REGULATION_OF_NITRIC_OXIDE_SYNTHASE_ACTIVITY | 27 | 0.468414 | 1.161477 | 0.272853 | 0.634841 |
| GOBP_ACTIN_MEDIATED_CELL_CONTRACTION | 86 | 0.393225 | 1.161342 | 0.226779 | 0.634699 |
| GOBP_RETROGRADE_AXONAL_TRANSPORT | 21 | 0.483879 | 1.160005 | 0.289655 | 0.638426 |
| GOBP_SENSORY_PERCEPTION_OF_MECHANICAL_STIMULUS | 135 | 0.373197 | 1.159622 | 0.198676 | 0.639049 |
| GOBP_MESODERM_DEVELOPMENT | 107 | 0.378297 | 1.158954 | 0.229545 | 0.640638 |
| GOBP_METANEPHRIC_NEPHRON_DEVELOPMENT | 30 | 0.450718 | 1.15735 | 0.290634 | 0.645187 |
| GOBP_NEUTRAL_LIPID_CATABOLIC_PROCESS | 29 | 0.454564 | 1.156961 | 0.2875 | 0.645897 |
| GOBP_POSITIVE_REGULATION_OF_ACTIVATED_T_CELL_PROLIFERATION | 17 | 0.507309 | 1.155679 | 0.295559 | 0.649486 |
| GOBP_REGULATION_OF_COLLATERAL_SPROUTING | 19 | 0.497822 | 1.155619 | 0.314493 | 0.649131 |
| GOBP_PROTEIN_CATABOLIC_PROCESS_IN_THE_VACUOLE | 22 | 0.474218 | 1.155275 | 0.288703 | 0.649644 |
| GOBP_ORGANIC_HYDROXY_COMPOUND_TRANSPORT | 198 | 0.358996 | 1.154453 | 0.169831 | 0.651809 |
| GOBP_NEGATIVE_REGULATION_OF_CELL_SUBSTRATE_ADHESION | 51 | 0.414046 | 1.154371 | 0.249373 | 0.651471 |
| GOBP_INNER_EAR_MORPHOGENESIS | 79 | 0.390454 | 1.154368 | 0.224299 | 0.650907 |
| GOBP_BIOLOGICAL_PROCESS_INVOLVED_IN_SYMBIOTIC_INTERACTION | 227 | 0.355681 | 1.154029 | 0.171792 | 0.651376 |
| GOBP_EMBRYONIC_ORGAN_DEVELOPMENT | 351 | 0.351183 | 1.153867 | 0.14096 | 0.651328 |
| GOBP_REGULATION_OF_MITOCHONDRION_ORGANIZATION | 132 | 0.370965 | 1.153842 | 0.21 | 0.650843 |
| GOBP_NEPHRON_MORPHOGENESIS | 61 | 0.40216 | 1.153423 | 0.253416 | 0.651589 |
| GOBP_BLOOD_VESSEL_ENDOTHELIAL_CELL_MIGRATION | 101 | 0.377664 | 1.153357 | 0.212866 | 0.651254 |
| GOBP_APPENDAGE_MORPHOGENESIS | 105 | 0.375517 | 1.152793 | 0.213793 | 0.652417 |
| GOBP_POSITIVE_REGULATION_OF_SPROUTING_ANGIOGENESIS | 18 | 0.500182 | 1.152645 | 0.318182 | 0.652306 |
| GOBP_CARDIAC_MUSCLE_CELL_MEMBRANE_REPOLARIZATION | 30 | 0.447489 | 1.152281 | 0.270345 | 0.65297 |
| GOBP_HEART_PROCESS | 201 | 0.361734 | 1.151858 | 0.19641 | 0.653736 |
| GOBP_NEGATIVE_REGULATION_OF_CELL_ADHESION | 216 | 0.354079 | 1.151847 | 0.18896 | 0.653196 |
| GOBP_MAMMARY_GLAND_DEVELOPMENT | 114 | 0.378708 | 1.151302 | 0.225084 | 0.654457 |
| GOBP_NEGATIVE_REGULATION_OF_SMOOTH_MUSCLE_CELL_MIGRATION | 23 | 0.47366 | 1.151219 | 0.292717 | 0.654168 |
| GOBP_POSITIVE_REGULATION_OF_PROTEIN_TYROSINE_KINASE_ACTIVITY | 43 | 0.425154 | 1.15117 | 0.267442 | 0.653756 |
| GOBP_MAST_CELL_ACTIVATION_INVOLVED_IN_IMMUNE_RESPONSE | 40 | 0.42247 | 1.150764 | 0.282185 | 0.654443 |
| GOBP_REGULATION_OF_VASCULAR_ENDOTHELIAL_GROWTH_FACTOR_RECEPTOR_SI | 22 | 0.47393 | 1.150668 | 0.295198 | 0.654185 |
| GOBP_PHOTOTRANSDUCTION | 21 | 0.483998 | 1.150649 | 0.304167 | 0.653682 |
| GOBP_REGULATION_OF_BROWN_FAT_CELL_DIFFERENTIATION | 19 | 0.49441 | 1.150548 | 0.300709 | 0.653439 |
| GOBP_LENS_DEVELOPMENT_IN_CAMERA_TYPE_EYE | 71 | 0.390962 | 1.150297 | 0.240196 | 0.653689 |
| GOBP_PERIPHERAL_NERVOUS_SYSTEM_DEVELOPMENT | 69 | 0.389366 | 1.150271 | 0.248521 | 0.65321 |
| GOBP_SMOOTH_MUSCLE_CELL_DIFFERENTIATION | 48 | 0.41732 | 1.150037 | 0.275641 | 0.653392 |
| GOBP_NEUROMUSCULAR_PROCESS | 118 | 0.370639 | 1.149877 | 0.237136 | 0.653308 |
| GOBP_STEM_CELL_PROLIFERATION | 59 | 0.39918 | 1.149569 | 0.273632 | 0.653702 |
| GOBP_REGULATION_OF_PLASMA_LIPOPROTEIN_PARTICLE_LEVELS | 51 | 0.411622 | 1.149397 | 0.25827 | 0.653687 |
| GOBP_MYELOID_LEUKOCYTE_CYTOKINE_PRODUCTION | 27 | 0.456385 | 1.149073 | 0.295833 | 0.654199 |
| GOBP_REGULATION_OF_B_CELL_PROLIFERATION | 39 | 0.434957 | 1.148442 | 0.274026 | 0.65564 |
| GOBP_POSITIVE_REGULATION_OF_STRESS_ACTIVATED_PROTEIN_KINASE_SIGNALING | 112 | 0.373356 | 1.148158 | 0.238739 | 0.655988 |
| GOBP_AMINO_SUGAR_METABOLIC_PROCESS | 35 | 0.438366 | 1.147979 | 0.292 | 0.655986 |
| GOBP_SMOOTH_MUSCLE_CELL_MIGRATION | 59 | 0.402922 | 1.147879 | 0.273418 | 0.655743 |
| GOBP_REACTIVE_OXYGEN_SPECIES_METABOLIC_PROCESS | 170 | 0.361976 | 1.147874 | 0.213904 | 0.655197 |
| GOBP_PEPTIDYL_TYROSINE_AUTOPHOSPHORYLATION | 18 | 0.49261 | 1.147763 | 0.306122 | 0.654994 |
| GOBP_POSITIVE_REGULATION_OF_PROTEIN_MATURATION | 23 | 0.468698 | 1.147749 | 0.313031 | 0.654489 |
| GOBP_POSITIVE_REGULATION_OF_PEPTIDE_SECRETION | 69 | 0.396678 | 1.147487 | 0.260817 | 0.65479 |
| GOBP_REGULATION_OF_ERYTHROCYTE_DIFFERENTIATION | 41 | 0.422982 | 1.147221 | 0.27555 | 0.655085 |
| GOBP_REGULATION_OF_CILIUM_MOVEMENT | 26 | 0.455636 | 1.147211 | 0.291156 | 0.654557 |
| GOBP_POSITIVE_REGULATION_OF_INTERLEUKIN_4_PRODUCTION | 15 | 0.510855 | 1.147178 | 0.319826 | 0.654107 |
| GOBP_ATRIAL_SEPTUM_DEVELOPMENT | 19 | 0.483909 | 1.147053 | 0.304094 | 0.653953 |
| GOBP_POSITIVE_REGULATION_OF_LIPASE_ACTIVITY | 45 | 0.417638 | 1.146876 | 0.26087 | 0.653981 |
| GOBP_OOCYTE_DIFFERENTIATION | 48 | 0.416399 | 1.14687 | 0.276983 | 0.653435 |
| GOBP_CARTILAGE_DEVELOPMENT_INVOLVED_IN_ENDOCHONDRAL_BONE_MORPHOG | 26 | 0.468842 | 1.146592 | 0.293296 | 0.653747 |
| GOBP_CELLULAR_RESPONSE_TO_CORTICOSTEROID_STIMULUS | 48 | 0.419863 | 1.14621 | 0.264151 | 0.654447 |
| GOBP_SUBSTRATE_ADHESION_DEPENDENT_CELL_SPREADING | 99 | 0.381635 | 1.145994 | 0.240275 | 0.654585 |
| GOBP_NEUROTRANSMITTER_UPTAKE | 36 | 0.433828 | 1.145938 | 0.284367 | 0.654216 |
| GOBP_LYMPHOCYTE_HOMEOSTASIS | 48 | 0.420002 | 1.145923 | 0.273644 | 0.653706 |
| GOBP_POSITIVE_REGULATION_OF_SIGNAL_TRANSDUCTION_BY_P53_CLASS_MEDIATO | 26 | 0.47121 | 1.145387 | 0.307692 | 0.654815 |
| GOBP_SIGNAL_RELEASE | 364 | 0.345493 | 1.144218 | 0.157895 | 0.658081 |
| GOBP_LIPID_CATABOLIC_PROCESS | 244 | 0.353139 | 1.144209 | 0.191377 | 0.657552 |
| GOBP_CRANIAL_NERVE_MORPHOGENESIS | 19 | 0.479884 | 1.144138 | 0.305847 | 0.657227 |
| GOBP_NEGATIVE_REGULATION_OF_PRODUCTION_OF_MOLECULAR_MEDIATOR_OF_IM | 23 | 0.475062 | 1.144081 | 0.308451 | 0.656855 |
| GOBP_REGULATION_OF_MUSCLE_HYPERTROPHY | 45 | 0.416231 | 1.143931 | 0.265725 | 0.656829 |
| GOBP_HEMATOPOIETIC_STEM_CELL_PROLIFERATION | 23 | 0.47629 | 1.143551 | 0.297753 | 0.657468 |
| GOBP_ODONTOGENESIS_OF_DENTIN_CONTAINING_TOOTH | 65 | 0.39962 | 1.143532 | 0.262967 | 0.65696 |
| GOBP_VASCULAR_ENDOTHELIAL_GROWTH_FACTOR_RECEPTOR_SIGNALING_PATHWA | 47 | 0.418046 | 1.143407 | 0.259843 | 0.656786 |
| GOBP_NEURAL_CREST_CELL_MIGRATION | 53 | 0.413775 | 1.143076 | 0.299499 | 0.657235 |
| GOBP_APPENDAGE_DEVELOPMENT | 134 | 0.366584 | 1.142104 | 0.237458 | 0.659828 |
| GOBP_ORGANIC_HYDROXY_COMPOUND_BIOSYNTHETIC_PROCESS | 192 | 0.35474 | 1.142103 | 0.228145 | 0.659277 |
| GOBP_REGULATION_OF_B_CELL_MEDIATED_IMMUNITY | 47 | 0.412843 | 1.141574 | 0.276596 | 0.660453 |
| GOBP_ENDOTHELIAL_CELL_APOPTOTIC_PROCESS | 42 | 0.414122 | 1.141425 | 0.270833 | 0.660372 |
| GOBP_REGULATION_OF_CELLULAR_AMINE_METABOLIC_PROCESS | 28 | 0.452272 | 1.141161 | 0.294606 | 0.660622 |
| GOBP_BILE_ACID_AND_BILE_SALT_TRANSPORT | 16 | 0.493927 | 1.140939 | 0.304024 | 0.66078 |
| GOBP_REGULATION_OF_MONOCYTE_DIFFERENTIATION | 16 | 0.499803 | 1.14012 | 0.303354 | 0.662877 |
| GOBP_REGULATION_OF_CELL_MATRIX_ADHESION | 105 | 0.374012 | 1.13993 | 0.256587 | 0.662906 |
| GOBP_TRNA_METHYLATION | 41 | 0.42435 | 1.139815 | 0.283582 | 0.662702 |
| GOBP_POSITIVE_REGULATION_OF_VASOCONSTRICTION | 25 | 0.457819 | 1.139235 | 0.299861 | 0.663942 |
| GOBP_PURINE_DEOXYRIBONUCLEOTIDE_METABOLIC_PROCESS | 15 | 0.502457 | 1.139186 | 0.312977 | 0.663544 |
| GOBP_REGULATION_OF_MEMBRANE_POTENTIAL | 345 | 0.343479 | 1.138462 | 0.171429 | 0.665302 |
| GOBP_POSITIVE_REGULATION_OF_NF_KAPPAB_TRANSCRIPTION_FACTOR_ACTIVITY | 126 | 0.365086 | 1.137941 | 0.248053 | 0.666368 |
| GOBP_REGULATION_OF_HEART_CONTRACTION | 160 | 0.358537 | 1.137562 | 0.223185 | 0.667024 |
| GOBP_POSITIVE_REGULATION_OF_PROTEIN_IMPORT | 38 | 0.428651 | 1.137475 | 0.296248 | 0.666719 |
| GOBP_POSITIVE_REGULATION_OF_HISTONE_ACETYLATION | 29 | 0.450001 | 1.137433 | 0.305479 | 0.666303 |

| | | | | | |
|---|---|---|---|---|---|
| GOBP_NEUROMUSCULAR_SYNAPTIC_TRANSMISSION | 22 | 0.473527 | 1.137308 | 0.326118 | 0.66614 |
| GOBP_BONE_RESORPTION | 45 | 0.416503 | 1.137305 | 0.287565 | 0.665594 |
| GOBP_EPITHELIAL_CELL_APOPTOTIC_PROCESS | 85 | 0.37924 | 1.137038 | 0.262238 | 0.665899 |
| GOBP_NEGATIVE_REGULATION_OF_NUCLEAR_DIVISION | 51 | 0.413461 | 1.137021 | 0.277062 | 0.665397 |
| GOBP_NEGATIVE_REGULATION_OF_PROTEIN_BINDING | 79 | 0.38648 | 1.136982 | 0.249406 | 0.664995 |
| GOBP_NUCLEOTIDE_SALVAGE | 16 | 0.500113 | 1.13471 | 0.313411 | 0.671771 |
| GOBP_CELLULAR_RESPONSE_TO_ORGANIC_CYCLIC_COMPOUND | 457 | 0.34015 | 1.133699 | 0.157417 | 0.674527 |
| GOBP_REGULATION_OF_EXTRACELLULAR_MATRIX_ORGANIZATION | 41 | 0.419566 | 1.133673 | 0.28195 | 0.674064 |
| GOBP_CELLULAR_RESPONSE_TO_LIPOPROTEIN_PARTICLE_STIMULUS | 25 | 0.458857 | 1.133656 | 0.328338 | 0.673567 |
| GOBP_NEGATIVE_REGULATION_OF_CYTOKINE_PRODUCTION_INVOLVED_IN_IMMUNE_ | 16 | 0.5032 | 1.133645 | 0.319767 | 0.673044 |
| GOBP_REGULATION_OF_MULTICELLULAR_ORGANISM_GROWTH | 55 | 0.400571 | 1.133476 | 0.28875 | 0.673014 |
| GOBP_RESPONSE_TO_FOOD | 24 | 0.467899 | 1.132762 | 0.311634 | 0.67479 |
| GOBP_MUSCLE_HYPERTROPHY | 63 | 0.398983 | 1.13273 | 0.272616 | 0.674334 |
| GOBP_FOREBRAIN_REGIONALIZATION | 16 | 0.498338 | 1.132472 | 0.32641 | 0.674621 |
| GOBP_GLYCOSYL_COMPOUND_METABOLIC_PROCESS | 75 | 0.389606 | 1.132249 | 0.25956 | 0.674784 |
| GOBP_DEMETHYLATION | 59 | 0.400178 | 1.132059 | 0.276284 | 0.674828 |
| GOBP_CARDIAC_CONDUCTION_SYSTEM_DEVELOPMENT | 26 | 0.454217 | 1.131453 | 0.298898 | 0.676242 |
| GOBP_IRON_ION_TRANSMEMBRANE_TRANSPORT | 17 | 0.496533 | 1.130993 | 0.328035 | 0.677155 |
| GOBP_ESTABLISHMENT_OF_MITOCHONDRION_LOCALIZATION | 29 | 0.44539 | 1.130782 | 0.309524 | 0.677293 |
| GOBP_REGULATION_OF_CELLULAR_RESPONSE_TO_TRANSFORMING_GROWTH_FACT( | 115 | 0.37069 | 1.130374 | 0.264908 | 0.677988 |
| GOBP_ACTIVIN_RECEPTOR_SIGNALING_PATHWAY | 39 | 0.430308 | 1.1297 | 0.307285 | 0.679581 |
| GOBP_RESPONSE_TO_OXIDATIVE_STRESS | 370 | 0.34093 | 1.128844 | 0.163618 | 0.681857 |
| GOBP_ATP_TRANSPORT | 15 | 0.500175 | 1.128689 | 0.329193 | 0.681801 |
| GOBP_MONOCARBOXYLIC_ACID_TRANSPORT | 73 | 0.392674 | 1.128635 | 0.278302 | 0.681411 |
| GOBP_GLYCEROPHOSPHOLIPID_METABOLIC_PROCESS | 272 | 0.345862 | 1.12835 | 0.216272 | 0.68175 |
| GOBP_TRANSITION_METAL_ION_HOMEOSTASIS | 111 | 0.370701 | 1.128211 | 0.261117 | 0.681655 |
| GOBP_CELL_CELL_JUNCTION_ASSEMBLY | 117 | 0.36702 | 1.128031 | 0.255919 | 0.681691 |
| GOBP_POSITIVE_REGULATION_OF_DEVELOPMENTAL_GROWTH | 132 | 0.364208 | 1.127749 | 0.24605 | 0.682034 |
| GOBP_OSSIFICATION | 338 | 0.341036 | 1.127707 | 0.176892 | 0.681596 |
| GOBP_I_KAPPAB_KINASE_NF_KAPPAB_SIGNALING | 243 | 0.346745 | 1.127504 | 0.219792 | 0.681673 |
| GOBP_SENSORY_PERCEPTION_OF_CHEMICAL_STIMULUS | 53 | 0.39804 | 1.127268 | 0.292994 | 0.681871 |
| GOBP_COLLAGEN_BIOSYNTHETIC_PROCESS | 32 | 0.442781 | 1.126909 | 0.321038 | 0.682519 |
| GOBP_POSITIVE_REGULATION_OF_B_CELL_ACTIVATION | 58 | 0.392644 | 1.126733 | 0.297332 | 0.682564 |
| GOBP_POSITIVE_REGULATION_OF_ANION_TRANSPORT | 37 | 0.424406 | 1.126254 | 0.311497 | 0.683585 |
| GOBP_HAIR_CELL_DIFFERENTIATION | 39 | 0.420968 | 1.126078 | 0.296104 | 0.683625 |
| GOBP_REGULATION_OF_CHOLESTEROL_EFFLUX | 29 | 0.432821 | 1.125776 | 0.296 | 0.684045 |
| GOBP_SECOND_MESSENGER_MEDIATED_SIGNALING | 215 | 0.344956 | 1.125714 | 0.229885 | 0.683708 |
| GOBP_REGULATION_OF_VESICLE_MEDIATED_TRANSPORT | 447 | 0.33936 | 1.125518 | 0.159435 | 0.683778 |
| GOBP_PROSTATE_GLAND_DEVELOPMENT | 36 | 0.42853 | 1.125072 | 0.308497 | 0.684632 |
| GOBP_REGULATION_OF_ANOIKIS | 19 | 0.481376 | 1.124962 | 0.326211 | 0.684448 |
| GOBP_POSITIVE_REGULATION_OF_CELL_MATRIX_ADHESION | 47 | 0.414854 | 1.124672 | 0.299611 | 0.684805 |
| GOBP_ANTIGEN_PROCESSING_AND_PRESENTATION_OF_PEPTIDE_ANTIGEN_VIA_MHC | 24 | 0.454729 | 1.124405 | 0.303199 | 0.685106 |
| GOBP_CHLORIDE_TRANSMEMBRANE_TRANSPORT | 49 | 0.407062 | 1.123948 | 0.292776 | 0.686017 |
| GOBP_FC_GAMMA_RECEPTOR_SIGNALING_PATHWAY | 23 | 0.463131 | 1.123662 | 0.32656 | 0.686444 |
| GOBP_MEMBRANE_RAFT_ORGANIZATION | 19 | 0.481184 | 1.123549 | 0.310245 | 0.686243 |
| GOBP_EAR_DEVELOPMENT | 170 | 0.349999 | 1.123136 | 0.246502 | 0.686981 |
| GOBP_POSITIVE_REGULATION_OF_PHAGOCYTOSIS | 41 | 0.419562 | 1.123115 | 0.30916 | 0.686499 |
| GOBP_HISTONE_H3_K9_METHYLATION | 35 | 0.428894 | 1.122936 | 0.29585 | 0.686509 |
| GOBP_REGULATION_OF_CELLULAR_RESPONSE_TO_GROWTH_FACTOR_STIMULUS | 253 | 0.343754 | 1.122874 | 0.208551 | 0.686151 |
| GOBP_FATTY_ACID_TRANSPORT | 68 | 0.388995 | 1.122292 | 0.289157 | 0.687446 |
| GOBP_MYOBLAST_FUSION | 32 | 0.427414 | 1.122179 | 0.310811 | 0.687256 |
| GOBP_PROTEIN_KINASE_C_ACTIVATING_G_PROTEIN_COUPLED_RECEPTOR_SIGNALIN | 22 | 0.469353 | 1.122036 | 0.333809 | 0.687168 |
| GOBP_REGULATION_OF_LIPID_METABOLIC_PROCESS | 260 | 0.348161 | 1.121649 | 0.225309 | 0.687859 |
| GOBP_NEGATIVE_REGULATION_OF_NEURON_PROJECTION_DEVELOPMENT | 127 | 0.361853 | 1.121499 | 0.258489 | 0.687765 |
| GOBP_REGULATION_OF_CELLULAR_EXTRAVASATION | 27 | 0.449066 | 1.12146 | 0.315202 | 0.687344 |
| GOBP_NEGATIVE_REGULATION_OF_POSTTRANSCRIPTIONAL_GENE_SILENCING | 15 | 0.508305 | 1.120805 | 0.342688 | 0.688894 |
| GOBP_CELL_RECOGNITION | 100 | 0.374442 | 1.120769 | 0.285877 | 0.688457 |
| GOBP_POSITIVE_REGULATION_OF_MONONUCLEAR_CELL_MIGRATION | 42 | 0.414359 | 1.1206 | 0.320363 | 0.688455 |
| GOBP_MESODERM_MORPHOGENESIS | 60 | 0.398548 | 1.120397 | 0.301607 | 0.688519 |
| GOBP_RHO_PROTEIN_SIGNAL_TRANSDUCTION | 120 | 0.364757 | 1.120385 | 0.271091 | 0.688018 |
| GOBP_POSITIVE_REGULATION_OF_STEROL_TRANSPORT | 28 | 0.441641 | 1.120318 | 0.310014 | 0.687686 |
| GOBP_ALCOHOL_BIOSYNTHETIC_PROCESS | 119 | 0.365373 | 1.119891 | 0.274576 | 0.688471 |
| GOBP_NEGATIVE_REGULATION_OF_EPITHELIAL_CELL_APOPTOTIC_PROCESS | 39 | 0.422169 | 1.119817 | 0.322281 | 0.688194 |
| GOBP_TOLL_LIKE_RECEPTOR_SIGNALING_PATHWAY | 88 | 0.374522 | 1.119557 | 0.29186 | 0.688505 |
| GOBP_KIDNEY_MORPHOGENESIS | 74 | 0.388126 | 1.119398 | 0.272182 | 0.68845 |
| GOBP_MODULATION_BY_HOST_OF_SYMBIONT_PROCESS | 54 | 0.39433 | 1.119011 | 0.309024 | 0.689152 |
| GOBP_SMAD_PROTEIN_SIGNAL_TRANSDUCTION | 72 | 0.386273 | 1.118841 | 0.286578 | 0.689168 |
| GOBP_CELL_DIFFERENTIATION_IN_HINDBRAIN | 20 | 0.465729 | 1.118679 | 0.33477 | 0.689176 |
| GOBP_RESPONSE_TO_MURAMYL_DIPEPTIDE | 19 | 0.478568 | 1.118516 | 0.343478 | 0.689136 |
| GOBP_POSITIVE_REGULATION_OF_INSULIN_SECRETION | 55 | 0.397843 | 1.118444 | 0.308161 | 0.688816 |
| GOBP_NEGATIVE_REGULATION_OF_CELL_CYCLE_G1_S_PHASE_TRANSITION | 72 | 0.382089 | 1.11835 | 0.290516 | 0.688581 |
| GOBP_COENZYME_A_METABOLIC_PROCESS | 18 | 0.498723 | 1.118275 | 0.340974 | 0.688273 |
| GOBP_FORMATION_OF_PRIMARY_GERM_LAYER | 104 | 0.364057 | 1.117648 | 0.262683 | 0.689723 |
| GOBP_NEUROTRANSMITTER_TRANSPORT | 174 | 0.348986 | 1.117355 | 0.26455 | 0.690103 |
| GOBP_HISTONE_H3_K4_METHYLATION | 65 | 0.390591 | 1.11614 | 0.308354 | 0.693299 |
| GOBP_RESPONSE_TO_ANTIBIOTIC | 41 | 0.418171 | 1.116027 | 0.314322 | 0.693131 |
| GOBP_RESPONSE_TO_PEPTIDE_HORMONE | 322 | 0.338245 | 1.115881 | 0.219388 | 0.693046 |
| GOBP_REGULATION_OF_RESPONSE_TO_CYTOKINE_STIMULUS | 132 | 0.356445 | 1.115696 | 0.271224 | 0.693081 |
| GOBP_PROTEIN_LIPID_COMPLEX_SUBUNIT_ORGANIZATION | 34 | 0.425491 | 1.1153 | 0.318302 | 0.693793 |
| GOBP_CELLULAR_ION_HOMEOSTASIS | 498 | 0.332245 | 1.115266 | 0.181174 | 0.69337 |
| GOBP_CIRCULATORY_SYSTEM_PROCESS | 465 | 0.330903 | 1.115248 | 0.190524 | 0.692894 |
| GOBP_MAMMARY_GLAND_EPITHELIAL_CELL_DIFFERENTIATION | 15 | 0.48821 | 1.1145 | 0.352049 | 0.69472 |
| GOBP_INORGANIC_ANION_TRANSMEMBRANE_TRANSPORT | 70 | 0.387148 | 1.114456 | 0.303067 | 0.694336 |
| GOBP_INTRACELLULAR_RECEPTOR_SIGNALING_PATHWAY | 225 | 0.343145 | 1.114428 | 0.237552 | 0.693889 |
| GOBP_NEGATIVE_REGULATION_OF_PROTEIN_POLYMERIZATION | 73 | 0.381508 | 1.11433 | 0.29656 | 0.693654 |
| GOBP_CONNECTIVE_TISSUE_DEVELOPMENT | 209 | 0.342856 | 1.114176 | 0.248421 | 0.693609 |
| GOBP_REGULATION_OF_PEPTIDASE_ACTIVITY | 327 | 0.336851 | 1.113817 | 0.227366 | 0.694211 |
| GOBP_GLYCEROLIPID_METABOLIC_PROCESS | 334 | 0.337138 | 1.11359 | 0.210419 | 0.694415 |
| GOBP_PHOSPHOLIPID_CATABOLIC_PROCESS | 43 | 0.414309 | 1.113553 | 0.298969 | 0.694002 |
| GOBP_CELLULAR_EXTRAVASATION | 50 | 0.39267 | 1.113175 | 0.313548 | 0.694681 |
| GOBP_REGULATION_OF_PHAGOCYTOSIS | 67 | 0.385757 | 1.113003 | 0.302808 | 0.694666 |
| GOBP_ROOF_OF_MOUTH_DEVELOPMENT | 72 | 0.383879 | 1.112725 | 0.299394 | 0.695018 |
| GOBP_ATRIOVENTRICULAR_VALVE_DEVELOPMENT | 23 | 0.459325 | 1.112377 | 0.323691 | 0.695602 |
| GOBP_MONOCARBOXYLIC_ACID_BIOSYNTHETIC_PROCESS | 158 | 0.352933 | 1.112228 | 0.272336 | 0.695555 |
| GOBP_PROTEIN_PROCESSING | 188 | 0.347632 | 1.111767 | 0.264612 | 0.696511 |
| GOBP_CALCIUM_MEDIATED_SIGNALING | 141 | 0.355384 | 1.111702 | 0.269063 | 0.696201 |
| GOBP_REGULATION_OF_CELL_CYCLE_G1_S_PHASE_TRANSITION | 146 | 0.352589 | 1.111095 | 0.280435 | 0.697691 |
| GOBP_SYMPATHETIC_NERVOUS_SYSTEM_DEVELOPMENT | 17 | 0.484862 | 1.111051 | 0.348266 | 0.697289 |
| GOBP_ACTOMYOSIN_STRUCTURE_ORGANIZATION | 176 | 0.351959 | 1.111039 | 0.272436 | 0.696791 |
| GOBP_SENSORY_ORGAN_DEVELOPMENT | 440 | 0.333235 | 1.110768 | 0.185671 | 0.697104 |
| GOBP_EAR_MORPHOGENESIS | 93 | 0.371275 | 1.110608 | 0.291475 | 0.697111 |
| GOBP_POSITIVE_REGULATION_OF_MITOCHONDRIAL_FISSION | 20 | 0.470404 | 1.110225 | 0.349138 | 0.697801 |
| GOBP_PALLIUM_DEVELOPMENT | 144 | 0.352007 | 1.109283 | 0.267108 | 0.700301 |
| GOBP_CELLULAR_RESPONSE_TO_ACID_CHEMICAL | 71 | 0.379581 | 1.109044 | 0.309466 | 0.7005 |
| GOBP_LONG_CHAIN_FATTY_ACID_TRANSPORT | 47 | 0.399135 | 1.108947 | 0.314931 | 0.700269 |
| GOBP_NEGATIVE_REGULATION_OF_T_CELL_APOPTOTIC_PROCESS | 15 | 0.498896 | 1.108443 | 0.338171 | 0.701279 |
| GOBP_REGULATION_OF_EXTRINSIC_APOPTOTIC_SIGNALING_PATHWAY_IN_ABSENCE_ | 35 | 0.426738 | 1.108338 | 0.330779 | 0.701081 |
| GOBP_REGULATION_OF_MONOOXYGENASE_ACTIVITY | 39 | 0.409766 | 1.107472 | 0.326772 | 0.703293 |
| GOBP_POSITIVE_REGULATION_OF_CALCIUM_ION_IMPORT | 15 | 0.499332 | 1.107316 | 0.352308 | 0.703288 |
| GOBP_POSITIVE_REGULATION_OF_FAT_CELL_DIFFERENTIATION | 59 | 0.39269 | 1.107044 | 0.308824 | 0.703617 |
| GOBP_MESONEPHRIC_TUBULE_MORPHOGENESIS | 50 | 0.393467 | 1.106905 | 0.322663 | 0.703516 |
| GOBP_SYNAPTIC_TRANSMISSION_GABAERGIC | 45 | 0.400913 | 1.106364 | 0.3257 | 0.704666 |
| GOBP_MESENCHYMAL_CELL_DIFFERENTIATION | 205 | 0.340679 | 1.106099 | 0.268783 | 0.705007 |
| GOBP_REGULATION_OF_SMOOTH_MUSCLE_CELL_DIFFERENTIATION | 26 | 0.430399 | 1.105804 | 0.312325 | 0.705431 |
| GOBP_DETECTION_OF_TEMPERATURE_STIMULUS_INVOLVED_IN_SENSORY_PERCEPT | 15 | 0.489632 | 1.105576 | 0.358814 | 0.705606 |
| GOBP_REGULATION_OF_SENSORY_PERCEPTION | 24 | 0.449606 | 1.105509 | 0.332398 | 0.705281 |
| GOBP_PHOSPHOLIPASE_C_ACTIVATING_G_PROTEIN_COUPLED_RECEPTOR_SIGNALIN | 59 | 0.396661 | 1.105459 | 0.313896 | 0.704897 |
| GOBP_REGULATION_OF_TRIGLYCERIDE_METABOLIC_PROCESS | 31 | 0.430069 | 1.105183 | 0.324213 | 0.705254 |
| GOBP_NEGATIVE_REGULATION_OF_CELL_DEVELOPMENT | 148 | 0.35019 | 1.104948 | 0.298913 | 0.705453 |
| GOBP_FOREBRAIN_GENERATION_OF_NEURONS | 39 | 0.415872 | 1.104806 | 0.323569 | 0.705381 |
| GOBP_CHLORIDE_TRANSPORT | 81 | 0.373078 | 1.104745 | 0.306884 | 0.705047 |
| GOBP_REGULATION_OF_CALCIUM_ION_TRANSPORT_INTO_CYTOSOL | 78 | 0.377563 | 1.104698 | 0.320285 | 0.704655 |
| GOBP_POSITIVE_REGULATION_OF_P38MAPK_CASCADE | 26 | 0.443734 | 1.10431 | 0.345354 | 0.705295 |
| GOBP_POSITIVE_REGULATION_OF_MULTICELLULAR_ORGANISM_GROWTH | 25 | 0.445374 | 1.1042 | 0.32312 | 0.705096 |
| GOBP_VITAMIN_TRANSMEMBRANE_TRANSPORT | 15 | 0.498501 | 1.103521 | 0.356185 | 0.706732 |
| GOBP_ANGIOGENESIS_INVOLVED_IN_WOUND_HEALING | 25 | 0.443182 | 1.103433 | 0.342029 | 0.706462 |
| GOBP_NEGATIVE_REGULATION_OF_INFLAMMATORY_RESPONSE_TO_ANTIGENIC_STIM | 23 | 0.453799 | 1.10306 | 0.349515 | 0.707107 |
| GOBP_REGULATION_OF_JNK_CASCADE | 124 | 0.353638 | 1.102831 | 0.311732 | 0.707303 |
| GOBP_HISTONE_H3_K9_MODIFICATION | 49 | 0.400554 | 1.102735 | 0.322181 | 0.707081 |
| GOBP_CELLULAR_RESPONSE_TO_CHEMICAL_STRESS | 288 | 0.335495 | 1.102215 | 0.256198 | 0.708213 |
| GOBP_THYMOCYTE_APOPTOTIC_PROCESS | 15 | 0.490595 | 1.102141 | 0.364964 | 0.707927 |
| GOBP_ADENYLATE_CYCLASE_MODULATING_G_PROTEIN_COUPLED_RECEPTOR_SIGN | 174 | 0.345573 | 1.101108 | 0.305286 | 0.710673 |
| GOBP_NEGATIVE_REGULATION_OF_PEPTIDYL_TYROSINE_PHOSPHORYLATION | 51 | 0.39764 | 1.10072 | 0.318804 | 0.711311 |
| GOBP_ACTIN_FILAMENT_BASED_MOVEMENT | 113 | 0.358618 | 1.100488 | 0.322799 | 0.711484 |
| GOBP_BONE_MATURATION | 24 | 0.44916 | 1.1004 | 0.34903 | 0.711246 |
| GOBP_REGULATION_OF_DENDRITE_EXTENSION | 23 | 0.444653 | 1.099721 | 0.358042 | 0.712958 |
| GOBP_CELLULAR_CARBOHYDRATE_METABOLIC_PROCESS | 242 | 0.336462 | 1.09916 | 0.263267 | 0.714154 |
| GOBP_NEGATIVE_REGULATION_OF_INTRINSIC_APOPTOTIC_SIGNALING_PATHWAY_BY_ | 22 | 0.453787 | 1.098479 | 0.366959 | 0.715751 |
| GOBP_CARDIAC_MUSCLE_CONTRACTION | 108 | 0.358265 | 1.098434 | 0.331454 | 0.715367 |
| GOBP_DEFINITIVE_HEMOPOIESIS | 15 | 0.488926 | 1.098223 | 0.358118 | 0.715461 |
| GOBP_REGULATION_OF_PEPTIDE_TRANSPORT | 138 | 0.353367 | 1.098221 | 0.294972 | 0.714933 |
| GOBP_POSITIVE_REGULATION_OF_MUSCLE_CELL_DIFFERENTIATION | 45 | 0.395358 | 1.097345 | 0.34264 | 0.717172 |
| GOBP_MESENCHYMAL_CELL_DEVELOPMENT | 79 | 0.379284 | 1.097118 | 0.332151 | 0.717349 |
| GOBP_NEGATIVE_REGULATION_OF_CELL_PROJECTION_ORGANIZATION | 172 | 0.344981 | 1.097052 | 0.29118 | 0.717015 |
| GOBP_NEGATIVE_REGULATION_OF_CELL_MATRIX_ADHESION | 31 | 0.424794 | 1.097023 | 0.350068 | 0.716568 |
| GOBP_MULTICELLULAR_ORGANISMAL_SIGNALING | 127 | 0.352349 | 1.096067 | 0.306326 | 0.719097 |
| GOBP_SMOOTH_MUSCLE_CELL_APOPTOTIC_PROCESS | 18 | 0.470729 | 1.095746 | 0.354103 | 0.719599 |
| GOBP_DORSAL_VENTRAL_PATTERN_FORMATION | 69 | 0.382392 | 1.095329 | 0.353865 | 0.720416 |
| GOBP_REGULATION_OF_MACROPHAGE_MIGRATION | 24 | 0.442673 | 1.095172 | 0.359116 | 0.720333 |
| GOBP_CELLULAR_RESPONSE_TO_LOW_DENSITY_LIPOPROTEIN_PARTICLE_STIMULUS | 15 | 0.488242 | 1.095157 | 0.362298 | 0.719843 |
| GOBP_HORMONE_TRANSPORT | 228 | 0.342922 | 1.095035 | 0.277134 | 0.719695 |

| | | | | | |
|---|---|---|---|---|---|
| GOBP_PRODUCTION_OF_MOLECULAR_MEDIATOR_INVOLVED_IN_INFLAMMATORY_RES | 54 | 0.395635 | 1.094988 | 0.333333 | 0.719317 |
| GOBP_REGULATION_OF_SYNCYTIUM_FORMATION_BY_PLASMA_MEMBRANE_FUSION | 17 | 0.467947 | 1.094487 | 0.361319 | 0.720342 |
| GOBP_POSITIVE_REGULATION_OF_CELL_CYCLE | 268 | 0.330982 | 1.094243 | 0.266667 | 0.720552 |
| GOBP_MEMBRANE_PROTEIN_ECTODOMAIN_PROTEOLYSIS | 40 | 0.405049 | 1.09406 | 0.333766 | 0.72057 |
| GOBP_CELLULAR_RESPONSE_TO_EXTERNAL_STIMULUS | 269 | 0.335191 | 1.093654 | 0.269958 | 0.721308 |
| GOBP_NEGATIVE_REGULATION_OF_MONONUCLEAR_CELL_MIGRATION | 16 | 0.488107 | 1.093646 | 0.350548 | 0.720798 |
| GOBP_SUPEROXIDE_ANION_GENERATION | 29 | 0.426558 | 1.093467 | 0.344086 | 0.720835 |
| GOBP_RENAL_TUBULE_DEVELOPMENT | 77 | 0.375778 | 1.092572 | 0.348946 | 0.72314 |
| GOBP_STARTLE_RESPONSE | 19 | 0.463104 | 1.092335 | 0.379113 | 0.723349 |
| GOBP_INTERLEUKIN_13_PRODUCTION | 15 | 0.483847 | 1.092191 | 0.366032 | 0.723243 |
| GOBP_NEGATIVE_REGULATION_OF_AXON_EXTENSION | 42 | 0.409328 | 1.091635 | 0.34981 | 0.724503 |
| GOBP_POST_EMBRYONIC_ANIMAL_ORGAN_DEVELOPMENT | 16 | 0.477708 | 1.091321 | 0.370536 | 0.724969 |
| GOBP_PHENOL_CONTAINING_COMPOUND_METABOLIC_PROCESS | 79 | 0.368521 | 1.090655 | 0.328571 | 0.726515 |
| GOBP_NEGATIVE_REGULATION_OF_PROTEIN_SERINE_THREONINE_KINASE_ACTIVITY | 102 | 0.36136 | 1.090626 | 0.314451 | 0.726077 |
| GOBP_CELL_DIFFERENTIATION_INVOLVED_IN_METANEPHROS_DEVELOPMENT | 17 | 0.470512 | 1.090603 | 0.354412 | 0.725611 |
| GOBP_POSITIVE_REGULATION_OF_PROTEIN_KINASE_ACTIVITY | 329 | 0.328784 | 1.090576 | 0.268916 | 0.725174 |
| GOBP_FAT_CELL_DIFFERENTIATION | 203 | 0.338187 | 1.090469 | 0.294995 | 0.724974 |
| GOBP_ACTIVATION_OF_PROTEIN_KINASE_ACTIVITY | 119 | 0.355041 | 1.090332 | 0.314189 | 0.724871 |
| GOBP_POSITIVE_REGULATION_OF_VASCULATURE_DEVELOPMENT | 129 | 0.346772 | 1.090237 | 0.32167 | 0.724662 |
| GOBP_NEURAL_RETINA_DEVELOPMENT | 51 | 0.388358 | 1.090117 | 0.33715 | 0.724522 |
| GOBP_CELLULAR_RESPONSE_TO_ATP | 15 | 0.483851 | 1.088257 | 0.381306 | 0.729825 |
| GOBP_PATTERN_RECOGNITION_RECEPTOR_SIGNALING_PATHWAY | 133 | 0.350676 | 1.088092 | 0.335548 | 0.729847 |
| GOBP_SYNAPSE_MATURATION | 23 | 0.436843 | 1.088029 | 0.353846 | 0.729513 |
| GOBP_EPITHELIAL_TO_MESENCHYMAL_TRANSITION_INVOLVED_IN_ENDOCARDIAL_CU | 20 | 0.455219 | 1.088014 | 0.362702 | 0.729023 |
| GOBP_SENSORY_SYSTEM_DEVELOPMENT | 306 | 0.330383 | 1.087601 | 0.28777 | 0.729786 |
| GOBP_STRIATED_MUSCLE_CELL_PROLIFERATION | 48 | 0.389977 | 1.08636 | 0.336283 | 0.733125 |
| GOBP_NEGATIVE_REGULATION_OF_RESPONSE_TO_CYTOKINE_STIMULUS | 57 | 0.381943 | 1.085671 | 0.339695 | 0.734779 |
| GOBP_NEGATIVE_REGULATION_OF_GLYCOPROTEIN_METABOLIC_PROCESS | 15 | 0.48633 | 1.085442 | 0.380531 | 0.734928 |
| GOBP_REGULATION_OF_MYELOID_CELL_DIFFERENTIATION | 162 | 0.342195 | 1.085086 | 0.327234 | 0.735484 |
| GOBP_EPIBOLY | 33 | 0.422551 | 1.084253 | 0.359838 | 0.737639 |
| GOBP_NEURON_FATE_SPECIFICATION | 17 | 0.474463 | 1.084014 | 0.397929 | 0.737863 |
| GOBP_CELLULAR_RESPONSE_TO_INTERFERON_BETA | 15 | 0.480636 | 1.083991 | 0.370079 | 0.737423 |
| GOBP_ADULT_LOCOMOTORY_BEHAVIOR | 66 | 0.376536 | 1.083337 | 0.342432 | 0.738979 |
| GOBP_DEOXYRIBONUCLEOSIDE_MONOPHOSPHATE_METABOLIC_PROCESS | 20 | 0.463397 | 1.083091 | 0.390208 | 0.739222 |
| GOBP_NEGATIVE_REGULATION_OF_SIGNAL_TRANSDUCTION_BY_P53_CLASS_MEDIAT( | 32 | 0.423927 | 1.08239 | 0.364256 | 0.740895 |
| GOBP_RESPONSE_TO_TOXIC_SUBSTANCE | 184 | 0.338348 | 1.082204 | 0.312903 | 0.740953 |
| GOBP_CELL_CELL_RECOGNITION | 35 | 0.41834 | 1.081802 | 0.375505 | 0.741686 |
| GOBP_REGULATION_OF_LIPID_STORAGE | 39 | 0.405864 | 1.08156 | 0.382586 | 0.741911 |
| GOBP_RHYTHMIC_PROCESS | 251 | 0.328029 | 1.080976 | 0.311458 | 0.743224 |
| GOBP_DEVELOPMENT_OF_PRIMARY_FEMALE_SEXUAL_CHARACTERISTICS | 89 | 0.364497 | 1.080954 | 0.353216 | 0.742756 |
| GOBP_TRIGLYCERIDE_CATABOLIC_PROCESS | 21 | 0.456686 | 1.080739 | 0.366197 | 0.742909 |
| GOBP_POSITIVE_REGULATION_OF_PEPTIDYL_SERINE_PHOSPHORYLATION | 80 | 0.362669 | 1.080723 | 0.351572 | 0.742429 |
| GOBP_CARDIAC_ATRIUM_MORPHOGENESIS | 21 | 0.44167 | 1.080699 | 0.362845 | 0.741971 |
| GOBP_T_CELL_SELECTION | 30 | 0.420697 | 1.080135 | 0.364626 | 0.743228 |
| GOBP_BIOLOGICAL_PROCESS_INVOLVED_IN_INTERACTION_WITH_HOST | 163 | 0.33717 | 1.079835 | 0.334052 | 0.743669 |
| GOBP_INORGANIC_ANION_TRANSPORT | 138 | 0.346409 | 1.079813 | 0.329018 | 0.743216 |
| GOBP_REGULATION_OF_P38MAPK_CASCADE | 37 | 0.40029 | 1.078808 | 0.347601 | 0.745848 |
| GOBP_REGULATION_OF_BONE_RESORPTION | 27 | 0.435373 | 1.078667 | 0.378453 | 0.745794 |
| GOBP_DETECTION_OF_STIMULUS_INVOLVED_IN_SENSORY_PERCEPTION_OF_PAIN | 22 | 0.450337 | 1.076939 | 0.385257 | 0.750707 |
| GOBP_DETECTION_OF_MECHANICAL_STIMULUS | 40 | 0.40723 | 1.076829 | 0.364462 | 0.750521 |
| GOBP_NITRIC_OXIDE_MEDIATED_SIGNAL_TRANSDUCTION | 18 | 0.46159 | 1.076745 | 0.388102 | 0.750266 |
| GOBP_NEGATIVE_REGULATION_OF_AXON_EXTENSION_INVOLVED_IN_AXON_GUIDANC | 27 | 0.434189 | 1.076623 | 0.364005 | 0.750118 |
| GOBP_NEGATIVE_REGULATION_OF_TRANSPORT | 335 | 0.325623 | 1.07648 | 0.310204 | 0.750024 |
| GOBP_NEUROTRANSMITTER_SECRETION | 127 | 0.345664 | 1.075598 | 0.350112 | 0.752288 |
| GOBP_REGULATION_OF_POTASSIUM_ION_TRANSMEMBRANE_TRANSPORTER_ACTIVIT | 44 | 0.393804 | 1.075571 | 0.362338 | 0.751839 |
| GOBP_REGULATION_OF_CARDIAC_MUSCLE_CONTRACTION | 60 | 0.379268 | 1.075324 | 0.351154 | 0.752067 |
| GOBP_REGULATION_OF_MACROPHAGE_ACTIVATION | 34 | 0.414972 | 1.074242 | 0.384106 | 0.75494 |
| GOBP_POSITIVE_REGULATION_OF_HORMONE_SECRETION | 96 | 0.356214 | 1.073722 | 0.370455 | 0.756089 |
| GOBP_PROTEIN_KINASE_A_SIGNALING | 25 | 0.435696 | 1.07368 | 0.383754 | 0.755706 |
| GOBP_VASODILATION | 34 | 0.415755 | 1.073618 | 0.388736 | 0.75537 |
| GOBP_INSULIN_SECRETION | 145 | 0.335566 | 1.073158 | 0.347299 | 0.75627 |
| GOBP_DNA_DEPENDENT_DNA_REPLICATION_MAINTENANCE_OF_FIDELITY | 46 | 0.397014 | 1.072998 | 0.353786 | 0.756232 |
| GOBP_REGULATION_OF_RESPIRATORY_GASEOUS_EXCHANGE | 17 | 0.469412 | 1.072688 | 0.394387 | 0.756695 |
| GOBP_HISTONE_METHYLATION | 147 | 0.340993 | 1.07267 | 0.35691 | 0.756223 |
| GOBP_GLYCOSAMINOGLYCAN_CATABOLIC_PROCESS | 19 | 0.461639 | 1.072579 | 0.392496 | 0.755988 |
| GOBP_REGULATION_OF_SUPEROXIDE_ANION_GENERATION | 16 | 0.474834 | 1.072529 | 0.394118 | 0.75562 |
| GOBP_POSITIVE_REGULATION_OF_RESPONSE_TO_CYTOKINE_STIMULUS | 51 | 0.387841 | 1.071982 | 0.384049 | 0.756845 |
| GOBP_NEURON_CELLULAR_HOMEOSTASIS | 28 | 0.421208 | 1.071438 | 0.38 | 0.758036 |
| GOBP_LEUKOTRIENE_METABOLIC_PROCESS | 15 | 0.476855 | 1.070945 | 0.386297 | 0.759036 |
| GOBP_REGULATION_OF_LAMELLIPODIUM_ORGANIZATION | 46 | 0.388328 | 1.07071 | 0.378414 | 0.759217 |
| GOBP_REGULATION_OF_BLOOD_CIRCULATION | 197 | 0.331916 | 1.070647 | 0.350951 | 0.758886 |
| GOBP_GLIAL_CELL_MIGRATION | 41 | 0.396649 | 1.070349 | 0.382166 | 0.759301 |
| GOBP_CENTROSOME_SEPARATION | 15 | 0.476879 | 1.069553 | 0.397944 | 0.761292 |
| GOBP_COGNITION | 258 | 0.326814 | 1.069412 | 0.310559 | 0.76122 |
| GOBP_RESPONSE_TO_ETHANOL | 87 | 0.362298 | 1.069316 | 0.354386 | 0.760986 |
| GOBP_LENS_MORPHOGENESIS_IN_CAMERA_TYPE_EYE | 22 | 0.442406 | 1.069091 | 0.38081 | 0.761124 |
| GOBP_NEGATIVE_REGULATION_OF_LOCOMOTION | 277 | 0.326644 | 1.068795 | 0.323988 | 0.761542 |
| GOBP_TROPHECTODERMAL_CELL_DIFFERENTIATION | 16 | 0.480131 | 1.067626 | 0.39645 | 0.7647 |
| GOBP_ORGANELLE_MEMBRANE_FUSION | 98 | 0.35231 | 1.067382 | 0.348328 | 0.764966 |
| GOBP_CARTILAGE_DEVELOPMENT | 153 | 0.334777 | 1.067053 | 0.334778 | 0.765495 |
| GOBP_REGULATION_OF_RAS_PROTEIN_SIGNAL_TRANSDUCTION | 167 | 0.333278 | 1.067046 | 0.3452 | 0.764983 |
| GOBP_NUCLEOBASE_CONTAINING_SMALL_MOLECULE_BIOSYNTHETIC_PROCESS | 15 | 0.474507 | 1.067043 | 0.391768 | 0.764461 |
| GOBP_RESPONSE_TO_INORGANIC_SUBSTANCE | 429 | 0.320419 | 1.066921 | 0.314889 | 0.764326 |
| GOBP_POSITIVE_REGULATION_OF_KINASE_ACTIVITY | 398 | 0.321034 | 1.066723 | 0.322515 | 0.76443 |
| GOBP_POSITIVE_REGULATION_OF_DOUBLE_STRAND_BREAK_REPAIR | 63 | 0.375219 | 1.066061 | 0.379394 | 0.766015 |
| GOBP_TRANSCYTOSIS | 18 | 0.469968 | 1.065943 | 0.407725 | 0.76584 |
| GOBP_NEGATIVE_REGULATION_OF_APOPTOTIC_SIGNALING_PATHWAY | 195 | 0.331673 | 1.065932 | 0.353187 | 0.765345 |
| GOBP_REGULATION_OF_SIGNALING_RECEPTOR_ACTIVITY | 128 | 0.342937 | 1.065904 | 0.356347 | 0.764915 |
| GOBP_NEURON_PROJECTION_GUIDANCE | 206 | 0.33004 | 1.065758 | 0.338574 | 0.764862 |
| GOBP_REGULATION_OF_PHOSPHOLIPASE_C_ACTIVITY | 30 | 0.418788 | 1.065313 | 0.406593 | 0.765696 |
| GOBP_MORPHOGENESIS_OF_AN_EPITHELIAL_SHEET | 51 | 0.380846 | 1.064711 | 0.366584 | 0.767052 |
| GOBP_REGULATION_OF_CALCIUM_ION_DEPENDENT_EXOCYTOSIS | 38 | 0.400053 | 1.064642 | 0.397653 | 0.766741 |
| GOBP_REGULATION_OF_HORMONE_LEVELS | 365 | 0.31964 | 1.064556 | 0.325866 | 0.766482 |
| GOBP_VITAMIN_METABOLIC_PROCESS | 86 | 0.356288 | 1.064503 | 0.381287 | 0.766118 |
| GOBP_REGULATION_OF_CELL_AGING | 41 | 0.393178 | 1.064478 | 0.397933 | 0.765673 |
| GOBP_REGULATION_OF_CHOLESTEROL_METABOLIC_PROCESS | 28 | 0.415459 | 1.064199 | 0.390821 | 0.766031 |
| GOBP_FC_RECEPTOR_MEDIATED_STIMULATORY_SIGNALING_PATHWAY | 26 | 0.427105 | 1.063027 | 0.390604 | 0.769132 |
| GOBP_EMBRYONIC_ORGAN_MORPHOGENESIS | 214 | 0.327545 | 1.062364 | 0.349423 | 0.770609 |
| GOBP_NEGATIVE_REGULATION_OF_NIK_NF_KAPPAB_SIGNALING | 24 | 0.432215 | 1.062312 | 0.396384 | 0.770242 |
| GOBP_VITAMIN_BIOSYNTHETIC_PROCESS | 16 | 0.460454 | 1.062179 | 0.405732 | 0.770159 |
| GOBP_NEGATIVE_REGULATION_OF_LEUKOCYTE_MIGRATION | 33 | 0.416657 | 1.062016 | 0.404284 | 0.770129 |
| GOBP_REGULATION_OF_CHROMOSOME_SEGREGATION | 81 | 0.35934 | 1.061818 | 0.372781 | 0.770212 |
| GOBP_REGULATION_OF_BONE_REMODELING | 34 | 0.411592 | 1.061358 | 0.397864 | 0.771113 |
| GOBP_RELEASE_OF_CYTOCHROME_C_FROM_MITOCHONDRIA | 51 | 0.377443 | 1.061341 | 0.39 | 0.770642 |
| GOBP_NEGATIVE_REGULATION_OF_CATION_CHANNEL_ACTIVITY | 31 | 0.416828 | 1.061195 | 0.406803 | 0.770582 |
| GOBP_RESPONSE_TO_AXON_INJURY | 71 | 0.36556 | 1.060421 | 0.393902 | 0.77241 |
| GOBP_POSITIVE_REGULATION_OF_ESTABLISHMENT_OF_PROTEIN_LOCALIZATION | 271 | 0.323634 | 1.060295 | 0.335052 | 0.772279 |
| GOBP_ZINC_ION_TRANSPORT | 22 | 0.434727 | 1.060053 | 0.380682 | 0.772481 |
| GOBP_RESPONSE_TO_ACTIVITY | 43 | 0.391254 | 1.058456 | 0.388175 | 0.776843 |
| GOBP_ENDOCRINE_HORMONE_SECRETION | 35 | 0.397796 | 1.058305 | 0.395225 | 0.776798 |
| GOBP_POSITIVE_REGULATION_OF_TRANSMEMBRANE_RECEPTOR_PROTEIN_SERINE_T | 101 | 0.35183 | 1.058191 | 0.358277 | 0.776615 |
| GOBP_POSITIVE_REGULATION_OF_ERYTHROCYTE_DIFFERENTIATION | 28 | 0.417556 | 1.05818 | 0.38587 | 0.776125 |
| GOBP_MYELOID_CELL_HOMEOSTASIS | 140 | 0.337586 | 1.058128 | 0.360746 | 0.775768 |
| GOBP_INTERLEUKIN_4_PRODUCTION | 21 | 0.443505 | 1.057961 | 0.393713 | 0.775789 |
| GOBP_REGULATION_OF_CAMP_MEDIATED_SIGNALING | 24 | 0.432524 | 1.057858 | 0.41954 | 0.775586 |
| GOBP_NEGATIVE_REGULATION_OF_CALCIUM_ION_TRANSPORT_INTO_CYTOSOL | 16 | 0.467956 | 1.057718 | 0.422576 | 0.775497 |
| GOBP_FC_EPSILON_RECEPTOR_SIGNALING_PATHWAY | 20 | 0.449813 | 1.057511 | 0.412884 | 0.775615 |
| GOBP_POSITIVE_REGULATION_OF_POTASSIUM_ION_TRANSMEMBRANE_TRANSPORTE | 20 | 0.438079 | 1.056988 | 0.424198 | 0.776726 |
| GOBP_PROTEIN_DESTABILIZATION | 43 | 0.388191 | 1.0569 | 0.380155 | 0.776476 |
| GOBP_NEUROBLAST_PROLIFERATION | 49 | 0.383039 | 1.056432 | 0.380247 | 0.777444 |
| GOBP_GLUCOSAMINE_CONTAINING_COMPOUND_METABOLIC_PROCESS | 20 | 0.440942 | 1.056071 | 0.402032 | 0.777995 |
| GOBP_NEGATIVE_REGULATION_OF_CELLULAR_RESPONSE_TO_INSULIN_STIMULUS | 31 | 0.409989 | 1.055883 | 0.385559 | 0.778072 |
| GOBP_SPINAL_CORD_DEVELOPMENT | 76 | 0.364404 | 1.055131 | 0.373223 | 0.779868 |
| GOBP_VASCULAR_ENDOTHELIAL_GROWTH_FACTOR_PRODUCTION | 28 | 0.42058 | 1.055093 | 0.398417 | 0.77945 |
| GOBP_FOREBRAIN_DEVELOPMENT | 316 | 0.322174 | 1.054811 | 0.330964 | 0.779794 |
| GOBP_POSITIVE_REGULATION_OF_CELL_DEATH | 473 | 0.315338 | 1.054112 | 0.351406 | 0.781463 |
| GOBP_MAINTENANCE_OF_LOCATION_IN_CELL | 172 | 0.329754 | 1.053801 | 0.37897 | 0.781899 |
| GOBP_REGULATION_OF_EPIDERMIS_DEVELOPMENT | 52 | 0.374571 | 1.053742 | 0.398496 | 0.781553 |
| GOBP_FEAR_RESPONSE | 33 | 0.405603 | 1.053672 | 0.419619 | 0.781252 |
| GOBP_REGULATION_OF_ACTIN_CYTOSKELETON_REORGANIZATION | 34 | 0.411769 | 1.053542 | 0.410397 | 0.781104 |
| GOBP_ACTION_POTENTIAL | 116 | 0.342512 | 1.053362 | 0.391451 | 0.781137 |
| GOBP_MESONEPHROS_DEVELOPMENT | 80 | 0.353263 | 1.052901 | 0.381643 | 0.782072 |
| GOBP_FATTY_ACID_DERIVATIVE_METABOLIC_PROCESS | 52 | 0.377381 | 1.052436 | 0.397468 | 0.783019 |
| GOBP_FIBROBLAST_PROLIFERATION | 67 | 0.364319 | 1.052362 | 0.397576 | 0.782731 |
| GOBP_POSITIVE_REGULATION_OF_ENDOTHELIAL_CELL_PROLIFERATION | 79 | 0.354275 | 1.051215 | 0.395294 | 0.785783 |
| GOBP_REGULATION_OF_PEPTIDYL_SERINE_PHOSPHORYLATION | 110 | 0.342936 | 1.050275 | 0.390079 | 0.78819 |
| GOBP_SCHWANN_CELL_DEVELOPMENT | 29 | 0.414578 | 1.049712 | 0.396359 | 0.789458 |
| GOBP_REGULATION_OF_T_HELPER_17_TYPE_IMMUNE_RESPONSE | 17 | 0.462553 | 1.048053 | 0.42527 | 0.79409 |
| GOBP_PROTEIN_K63_LINKED_UBIQUITINATION | 54 | 0.373985 | 1.047833 | 0.389724 | 0.794235 |
| GOBP_POSITIVE_REGULATION_OF_MICROTUBULE_POLYMERIZATION_OR_DEPOLYMER | 36 | 0.387508 | 1.047641 | 0.410977 | 0.794315 |
| GOBP_REGULATION_OF_HIPPO_SIGNALING | 21 | 0.435978 | 1.047414 | 0.406026 | 0.794525 |
| GOBP_RESPONSE_TO_PURINE_CONTAINING_COMPOUND | 103 | 0.342786 | 1.047315 | 0.397059 | 0.794294 |
| GOBP_LIPID_STORAGE | 66 | 0.364989 | 1.047288 | 0.399757 | 0.793859 |
| GOBP_POSITIVE_REGULATION_OF_PROTEIN_MODIFICATION_BY_SMALL_PROTEIN_CON | 128 | 0.339995 | 1.04719 | 0.391892 | 0.793629 |
| GOBP_REGULATION_OF_DNA_DAMAGE_RESPONSE_SIGNAL_TRANSDUCTION_BY_P53 | 34 | 0.396122 | 1.047018 | 0.413747 | 0.793635 |
| GOBP_PYRIMIDINE_CONTAINING_COMPOUND_CATABOLIC_PROCESS | 35 | 0.40648 | 1.046997 | 0.416449 | 0.793175 |
| GOBP_PHOSPHATIDYLINOSITOL_3_KINASE_SIGNALING | 102 | 0.342871 | 1.046167 | 0.391405 | 0.795283 |
| GOBP_PHOSPHOLIPID_METABOLIC_PROCESS | 335 | 0.318188 | 1.045615 | 0.395112 | 0.79645 |
| GOBP_REGULATION_OF_THE_FORCE_OF_HEART_CONTRACTION | 19 | 0.441662 | 1.045545 | 0.411932 | 0.796142 |
| GOBP_SEROTONIN_RECEPTOR_SIGNALING_PATHWAY | 15 | 0.477303 | 1.045485 | 0.414878 | 0.795791 |
| GOBP_REGULATION_OF_HISTONE_MODIFICATION | 166 | 0.328096 | 1.045336 | 0.394822 | 0.795735 |

| | | | | | |
|---|---|---|---|---|---|
| GOBP_OSTEOBLAST_DIFFERENTIATION | 194 | 0.324081 | 1.045324 | 0.376858 | 0.795244 |
| GOBP_POSITIVE_REGULATION_OF_GROWTH | 204 | 0.324357 | 1.045094 | 0.379531 | 0.795388 |
| GOBP_REGULATION_OF_NEUROBLAST_PROLIFERATION | 24 | 0.431455 | 1.044977 | 0.419771 | 0.795236 |
| GOBP_RIBONUCLEOTIDE_CATABOLIC_PROCESS | 42 | 0.39188 | 1.044799 | 0.390836 | 0.795296 |
| GOBP_MONOCARBOXYLIC_ACID_METABOLIC_PROCESS | 448 | 0.312074 | 1.044755 | 0.364742 | 0.794911 |
| GOBP_NEGATIVE_REGULATION_OF_MUSCLE_CONTRACTION | 21 | 0.438175 | 1.044137 | 0.423898 | 0.796249 |
| GOBP_REGULATION_OF_NATURAL_KILLER_CELL_MEDIATED_IMMUNITY | 25 | 0.419751 | 1.043683 | 0.419805 | 0.797118 |
| GOBP_HISTONE_H3_K9_ACETYLATION | 16 | 0.466462 | 1.043259 | 0.421759 | 0.797881 |
| GOBP_REGULATION_OF_NUCLEOTIDE_BIOSYNTHETIC_PROCESS | 31 | 0.406123 | 1.043051 | 0.411279 | 0.798012 |
| GOBP_MORPHOGENESIS_OF_A_BRANCHING_STRUCTURE | 154 | 0.32532 | 1.042626 | 0.39404 | 0.798822 |
| GOBP_RESPONSE_TO_HEAT | 95 | 0.343788 | 1.042457 | 0.405814 | 0.798808 |
| GOBP_POSITIVE_REGULATION_OF_LIPID_CATABOLIC_PROCESS | 17 | 0.453564 | 1.042454 | 0.440678 | 0.79829 |
| GOBP_INTESTINAL_ABSORPTION | 22 | 0.42771 | 1.042256 | 0.417237 | 0.798352 |
| GOBP_CEREBELLAR_PURKINJE_CELL_LAYER_MORPHOGENESIS | 18 | 0.447423 | 1.042226 | 0.431686 | 0.797921 |
| GOBP_RESPONSE_TO_PEPTIDE | 390 | 0.313738 | 1.042144 | 0.384772 | 0.797656 |
| GOBP_SIGNAL_TRANSDUCTION_BY_P53_CLASS_MEDIATOR | 160 | 0.331234 | 1.041849 | 0.405983 | 0.798018 |
| GOBP_REGULATION_OF_SYSTEM_PROCESS | 431 | 0.313088 | 1.041848 | 0.388271 | 0.7975 |
| GOBP_REGULATION_OF_OXIDATIVE_PHOSPHORYLATION | 20 | 0.426437 | 1.041629 | 0.416422 | 0.797623 |
| GOBP_LEFT_RIGHT_PATTERN_FORMATION | 22 | 0.440871 | 1.041572 | 0.419399 | 0.797274 |
| GOBP_ANIMAL_ORGAN_MATURATION | 26 | 0.425289 | 1.040753 | 0.423803 | 0.799264 |
| GOBP_PLACENTA_DEVELOPMENT | 130 | 0.335537 | 1.040192 | 0.393805 | 0.800482 |
| GOBP_RESPONSE_TO_HYDROGEN_PEROXIDE | 103 | 0.34318 | 1.039542 | 0.408827 | 0.801945 |
| GOBP_RESPONSE_TO_CALCIUM_ION | 121 | 0.33792 | 1.039441 | 0.417808 | 0.801758 |
| GOBP_SULFATION | 16 | 0.458621 | 1.039269 | 0.435414 | 0.801768 |
| GOBP_GLYCOSYL_COMPOUND_BIOSYNTHETIC_PROCESS | 19 | 0.433832 | 1.038878 | 0.439266 | 0.802389 |
| GOBP_TELOMERE_MAINTENANCE_VIA_RECOMBINATION | 15 | 0.462953 | 1.038143 | 0.419355 | 0.804144 |
| GOBP_MITOCHONDRIAL_FISSION | 37 | 0.392139 | 1.038039 | 0.416449 | 0.803921 |
| GOBP_PEPTIDE_TRANSPORT | 195 | 0.327047 | 1.037974 | 0.417197 | 0.803597 |
| GOBP_ERK1_AND_ERK2_CASCADE | 246 | 0.319408 | 1.037905 | 0.404145 | 0.803276 |
| GOBP_VASCULAR_PROCESS_IN_CIRCULATORY_SYSTEM | 214 | 0.323026 | 1.037826 | 0.400831 | 0.803017 |
| GOBP_PURINERGIC_NUCLEOTIDE_RECEPTOR_SIGNALING_PATHWAY | 19 | 0.442805 | 1.03781 | 0.422741 | 0.802548 |
| GOBP_NEURAL_CREST_CELL_DIFFERENTIATION | 84 | 0.349407 | 1.037774 | 0.408115 | 0.802133 |
| GOBP_REGULATION_OF_STEM_CELL_PROLIFERATION | 36 | 0.391162 | 1.03758 | 0.407609 | 0.802201 |
| GOBP_RESPONSE_TO_PROSTAGLANDIN | 22 | 0.432887 | 1.037461 | 0.446176 | 0.802048 |
| GOBP_CATECHOL_CONTAINING_COMPOUND_METABOLIC_PROCESS | 46 | 0.377562 | 1.037351 | 0.423077 | 0.801861 |
| GOBP_RESPONSE_TO_TEMPERATURE_STIMULUS | 152 | 0.328065 | 1.037131 | 0.393514 | 0.802047 |
| GOBP_O_GLYCAN_PROCESSING | 36 | 0.393168 | 1.03618 | 0.41779 | 0.804412 |
| GOBP_MAINTENANCE_OF_LOCATION | 261 | 0.319425 | 1.036001 | 0.401235 | 0.804419 |
| GOBP_PHAGOCYTOSIS_RECOGNITION | 16 | 0.463204 | 1.035727 | 0.423581 | 0.804718 |
| GOBP_SPECIFICATION_OF_ANIMAL_ORGAN_IDENTITY | 27 | 0.413836 | 1.035119 | 0.430575 | 0.806081 |
| GOBP_MEMBRANE_DEPOLARIZATION | 71 | 0.356983 | 1.034895 | 0.441606 | 0.806255 |
| GOBP_RRNA_CATABOLIC_PROCESS | 20 | 0.442876 | 1.033733 | 0.422351 | 0.809236 |
| GOBP_POSITIVE_REGULATION_OF_G1_S_TRANSITION_OF_MITOTIC_CELL_CYCLE | 39 | 0.385019 | 1.033584 | 0.420699 | 0.809175 |
| GOBP_EMBRYO_IMPLANTATION | 48 | 0.381755 | 1.033429 | 0.445609 | 0.809107 |
| GOBP_MIDDLE_EAR_MORPHOGENESIS | 15 | 0.465538 | 1.033007 | 0.451807 | 0.809833 |
| GOBP_MACROPHAGE_CHEMOTAXIS | 24 | 0.41163 | 1.032679 | 0.424899 | 0.810301 |
| GOBP_REGULATION_OF_TRANSPORTER_ACTIVITY | 225 | 0.319126 | 1.032304 | 0.432914 | 0.810925 |
| GOBP_INOSITOL_LIPID_MEDIATED_SIGNALING | 132 | 0.3326 | 1.032157 | 0.413984 | 0.81083 |
| GOBP_UROGENITAL_SYSTEM_DEVELOPMENT | 292 | 0.314149 | 1.032153 | 0.410861 | 0.810322 |
| GOBP_GASTRULATION | 159 | 0.322707 | 1.032138 | 0.420022 | 0.809847 |
| GOBP_POSITIVE_REGULATION_OF_LIPID_KINASE_ACTIVITY | 31 | 0.399133 | 1.031829 | 0.440934 | 0.810233 |
| GOBP_DETOXIFICATION | 107 | 0.33869 | 1.031277 | 0.428733 | 0.811406 |
| GOBP_BONE_REMODELING | 66 | 0.360005 | 1.031257 | 0.43179 | 0.810949 |
| GOBP_GROWTH_PLATE_CARTILAGE_DEVELOPMENT | 17 | 0.449772 | 1.030935 | 0.433483 | 0.811459 |
| GOBP_MACROPHAGE_MIGRATION | 33 | 0.399453 | 1.030157 | 0.440318 | 0.813266 |
| GOBP_RESPONSE_TO_METAL_ION | 278 | 0.31517 | 1.028988 | 0.425 | 0.816232 |
| GOBP_MAMMARY_GLAND_EPITHELIAL_CELL_PROLIFERATION | 23 | 0.425119 | 1.02891 | 0.427577 | 0.815931 |
| GOBP_CARDIAC_MUSCLE_CELL_CONTRACTION | 63 | 0.359097 | 1.028596 | 0.428062 | 0.816377 |
| GOBP_POSITIVE_REGULATION_OF_ORGANIC_ACID_TRANSPORT | 34 | 0.391916 | 1.028152 | 0.420844 | 0.817186 |
| GOBP_EYE_PHOTORECEPTOR_CELL_DEVELOPMENT | 26 | 0.405469 | 1.02814 | 0.435211 | 0.816695 |
| GOBP_REGULATION_OF_CELL_DEVELOPMENT | 420 | 0.307026 | 1.028099 | 0.438066 | 0.816299 |
| GOBP_RESPONSE_TO_OXYGEN_LEVELS | 280 | 0.315353 | 1.028028 | 0.42872 | 0.815981 |
| GOBP_NEGATIVE_REGULATION_OF_KINASE_ACTIVITY | 207 | 0.318119 | 1.027806 | 0.413574 | 0.816108 |
| GOBP_REGULATION_OF_DNA_BINDING_TRANSCRIPTION_FACTOR_ACTIVITY | 363 | 0.309059 | 1.027747 | 0.43279 | 0.815772 |
| GOBP_REGULATION_OF_NUCLEAR_DIVISION | 115 | 0.334247 | 1.027693 | 0.429724 | 0.815424 |
| GOBP_HIPPO_SIGNALING | 40 | 0.386148 | 1.027001 | 0.431788 | 0.816966 |
| GOBP_REGULATION_OF_EXTRINSIC_APOPTOTIC_SIGNALING_PATHWAY_VIA_DEATH_D | 41 | 0.38129 | 1.025563 | 0.436997 | 0.820707 |
| GOBP_ENSHEATHMENT_OF_NEURONS | 125 | 0.331564 | 1.025094 | 0.429844 | 0.821626 |
| GOBP_ADULT_BEHAVIOR | 113 | 0.333778 | 1.025041 | 0.411561 | 0.821252 |
| GOBP_REGULATION_OF_ACTIN_FILAMENT_BASED_PROCESS | 350 | 0.306571 | 1.024446 | 0.43279 | 0.822555 |
| GOBP_SMALL_MOLECULE_BIOSYNTHETIC_PROCESS | 450 | 0.307564 | 1.024395 | 0.424823 | 0.822196 |
| GOBP_NEGATIVE_REGULATION_OF_MAPK_CASCADE | 146 | 0.327379 | 1.023606 | 0.428415 | 0.824072 |
| GOBP_REGULATION_OF_GLUCOSE_TRANSMEMBRANE_TRANSPORT | 59 | 0.358177 | 1.023227 | 0.431877 | 0.824738 |
| GOBP_ADRENAL_GLAND_DEVELOPMENT | 24 | 0.41321 | 1.022309 | 0.444444 | 0.826983 |
| GOBP_LYMPH_VESSEL_MORPHOGENESIS | 21 | 0.424758 | 1.022104 | 0.465418 | 0.827081 |
| GOBP_REGULATION_OF_BICELLULAR_TIGHT_JUNCTION_ASSEMBLY | 18 | 0.452523 | 1.021967 | 0.442105 | 0.826999 |
| GOBP_IMMUNOGLOBULIN_PRODUCTION | 91 | 0.339208 | 1.021923 | 0.445769 | 0.826603 |
| GOBP_NEGATIVE_REGULATION_OF_HISTONE_ACETYLATION | 15 | 0.454013 | 1.021545 | 0.457849 | 0.827216 |
| GOBP_CELL_CELL_ADHESION_VIA_PLASMA_MEMBRANE_ADHESION_MOLECULES | 222 | 0.314425 | 1.021199 | 0.461053 | 0.827724 |
| GOBP_PEPTIDYL_LYSINE_METHYLATION | 131 | 0.329135 | 1.020715 | 0.461366 | 0.828678 |
| GOBP_CELLULAR_RESPONSE_TO_ABIOTIC_STIMULUS | 273 | 0.309335 | 1.020157 | 0.433747 | 0.829847 |
| GOBP_EMBRYONIC_MORPHOGENESIS | 483 | 0.302852 | 1.019133 | 0.435176 | 0.832447 |
| GOBP_DIGESTIVE_SYSTEM_DEVELOPMENT | 102 | 0.333283 | 1.018534 | 0.437284 | 0.83371 |
| GOBP_POSITIVE_REGULATION_OF_EPITHELIAL_CELL_PROLIFERATION | 151 | 0.32293 | 1.018164 | 0.431718 | 0.834328 |
| GOBP_CELLULAR_IRON_ION_HOMEOSTASIS | 53 | 0.360446 | 1.018101 | 0.45614 | 0.833991 |
| GOBP_POSITIVE_REGULATION_OF_LIPID_STORAGE | 18 | 0.441238 | 1.017999 | 0.434191 | 0.833783 |
| GOBP_REGULATION_OF_SYNAPTIC_TRANSMISSION_GLUTAMATERGIC | 53 | 0.366664 | 1.01762 | 0.452088 | 0.834394 |
| GOBP_REGULATION_OF_TRANS_SYNAPTIC_SIGNALING | 363 | 0.305503 | 1.017398 | 0.452138 | 0.834537 |
| GOBP_VIRAL_PROCESS | 360 | 0.309164 | 1.017002 | 0.449746 | 0.835182 |
| GOBP_VENTRICULAR_CARDIAC_MUSCLE_CELL_MEMBRANE_REPOLARIZATION | 21 | 0.417401 | 1.016982 | 0.458453 | 0.834713 |
| GOBP_RESPONSE_TO_AMPHETAMINE | 26 | 0.402118 | 1.015863 | 0.443828 | 0.8376 |
| GOBP_DETECTION_OF_TEMPERATURE_STIMULUS | 17 | 0.443434 | 1.01562 | 0.449775 | 0.837742 |
| GOBP_REGULATION_OF_MACROPHAGE_CHEMOTAXIS | 17 | 0.446765 | 1.015154 | 0.45036 | 0.838649 |
| GOBP_CELLULAR_RESPONSE_TO_CAMP | 41 | 0.374091 | 1.01461 | 0.449198 | 0.83976 |
| GOBP_POSITIVE_REGULATION_OF_SUBSTRATE_ADHESION_DEPENDENT_CELL_SPREA | 36 | 0.382845 | 1.013937 | 0.438482 | 0.841273 |
| GOBP_REGULATION_OF_RELEASE_OF_CYTOCHROME_C_FROM_MITOCHONDRIA | 41 | 0.373778 | 1.013605 | 0.440945 | 0.841759 |
| GOBP_POSITIVE_REGULATION_OF_MIRNA_TRANSCRIPTION | 41 | 0.379709 | 1.013515 | 0.44591 | 0.841492 |
| GOBP_PYRIMIDINE_NUCLEOSIDE_MONOPHOSPHATE_METABOLIC_PROCESS | 19 | 0.428566 | 1.012962 | 0.469809 | 0.842604 |
| GOBP_THYMUS_DEVELOPMENT | 34 | 0.396269 | 1.012345 | 0.453425 | 0.843906 |
| GOBP_AORTIC_VALVE_MORPHOGENESIS | 28 | 0.395172 | 1.01229 | 0.452804 | 0.843542 |
| GOBP_ACTIVATION_OF_CYSTEINE_TYPE_ENDOPEPTIDASE_ACTIVITY_INVOLVED_IN_AF | 67 | 0.352019 | 1.012044 | 0.47136 | 0.843744 |
| GOBP_STEM_CELL_DIFFERENTIATION | 179 | 0.3185 | 1.011622 | 0.464324 | 0.844492 |
| GOBP_RAS_PROTEIN_SIGNAL_TRANSDUCTION | 313 | 0.305836 | 1.011606 | 0.444785 | 0.844011 |
| GOBP_NEGATIVE_REGULATION_OF_CELLULAR_CATABOLIC_PROCESS | 228 | 0.313213 | 1.011582 | 0.470772 | 0.843566 |
| GOBP_SYNAPTIC_VESICLE_PRIMING | 18 | 0.439037 | 1.011549 | 0.447059 | 0.843144 |
| GOBP_CELLULAR_KETONE_METABOLIC_PROCESS | 157 | 0.317765 | 1.011465 | 0.475 | 0.842864 |
| GOBP_RESPONSE_TO_ALCOHOL | 178 | 0.317821 | 1.011442 | 0.454252 | 0.842402 |
| GOBP_RESPONSE_TO_ELECTRICAL_STIMULUS | 31 | 0.396901 | 1.011236 | 0.461942 | 0.842487 |
| GOBP_POSITIVE_REGULATION_OF_INTERFERON_ALPHA_PRODUCTION | 17 | 0.43823 | 1.010945 | 0.455192 | 0.842843 |
| GOBP_NEGATIVE_REGULATION_OF_EPITHELIAL_CELL_MIGRATION | 56 | 0.354451 | 1.01062 | 0.449438 | 0.843269 |
| GOBP_REGULATION_OF_RETINOIC_ACID_RECEPTOR_SIGNALING_PATHWAY | 15 | 0.450581 | 1.010544 | 0.466267 | 0.842973 |
| GOBP_REGULATION_OF_TUBE_SIZE | 105 | 0.329365 | 1.010292 | 0.463218 | 0.843204 |
| GOBP_SENSORY_PERCEPTION_OF_TEMPERATURE_STIMULUS | 20 | 0.43397 | 1.010019 | 0.463989 | 0.843488 |
| GOBP_REGULATION_OF_STEROID_METABOLIC_PROCESS | 76 | 0.34039 | 1.00929 | 0.459394 | 0.845143 |
| GOBP_OLEFINIC_COMPOUND_METABOLIC_PROCESS | 108 | 0.328156 | 1.009086 | 0.450624 | 0.845227 |
| GOBP_ORGANELLE_FUSION | 132 | 0.324373 | 1.008818 | 0.467634 | 0.845486 |
| GOBP_JNK_CASCADE | 156 | 0.316992 | 1.00856 | 0.456568 | 0.845743 |
| GOBP_ERYTHROCYTE_HOMEOSTASIS | 117 | 0.324531 | 1.008317 | 0.473506 | 0.845921 |
| GOBP_ACTIN_FILAMENT_ORGANIZATION | 401 | 0.301704 | 1.008223 | 0.473313 | 0.845681 |
| GOBP_POSITIVE_REGULATION_OF_PEPTIDYL_LYSINE_ACETYLATION | 35 | 0.385459 | 1.007788 | 0.46462 | 0.846429 |
| GOBP_CHONDROCYTE_DIFFERENTIATION | 94 | 0.334264 | 1.007728 | 0.463387 | 0.846094 |
| GOBP_CRANIAL_SKELETAL_SYSTEM_DEVELOPMENT | 55 | 0.362339 | 1.00751 | 0.46088 | 0.846184 |
| GOBP_VASCULAR_ENDOTHELIAL_CELL_PROLIFERATION | 18 | 0.432116 | 1.007421 | 0.455752 | 0.845947 |
| GOBP_NEGATIVE_REGULATION_OF_AXONOGENESIS | 62 | 0.347054 | 1.007032 | 0.467153 | 0.846623 |
| GOBP_CELLULAR_TRANSITION_METAL_ION_HOMEOSTASIS | 89 | 0.334143 | 1.006937 | 0.45848 | 0.846372 |
| GOBP_REGULATION_OF_AMINO_ACID_TRANSPORT | 34 | 0.387168 | 1.006785 | 0.456671 | 0.846295 |
| GOBP_LIPID_LOCALIZATION | 334 | 0.305005 | 1.006708 | 0.467072 | 0.845995 |
| GOBP_POSITIVE_REGULATION_OF_VIRAL_PROCESS | 57 | 0.356075 | 1.006639 | 0.464596 | 0.845685 |
| GOBP_VASCULOGENESIS | 68 | 0.346391 | 1.006269 | 0.471306 | 0.846238 |
| GOBP_NEGATIVE_REGULATION_OF_MYELOID_CELL_DIFFERENTIATION | 70 | 0.34836 | 1.005713 | 0.491607 | 0.847394 |
| GOBP_REGULATION_OF_B_CELL_RECEPTOR_SIGNALING_PATHWAY | 15 | 0.450031 | 1.005574 | 0.476831 | 0.847278 |
| GOBP_ESTABLISHMENT_OF_SPINDLE_ORIENTATION | 32 | 0.385445 | 1.005263 | 0.450205 | 0.847713 |
| GOBP_VENTRICULAR_CARDIAC_MUSCLE_CELL_ACTION_POTENTIAL | 27 | 0.408237 | 1.00494 | 0.458108 | 0.848138 |
| GOBP_ANION_TRANSPORT | 389 | 0.302674 | 1.004231 | 0.492447 | 0.849721 |
| GOBP_NEGATIVE_REGULATION_OF_INTERLEUKIN_6_PRODUCTION | 30 | 0.387026 | 1.003999 | 0.469051 | 0.849933 |
| GOBP_NEGATIVE_REGULATION_OF_PROTEIN_CONTAINING_COMPLEX_ASSEMBLY | 128 | 0.323077 | 1.003069 | 0.484615 | 0.852179 |
| GOBP_POSITIVE_REGULATION_OF_PROTEIN_POLYUBIQUITINATION | 15 | 0.448954 | 1.002965 | 0.472264 | 0.851978 |
| GOBP_REGULATION_OF_RECEPTOR_RECYCLING | 21 | 0.422463 | 1.002947 | 0.457676 | 0.851512 |
| GOBP_GLYCEROLIPID_BIOSYNTHETIC_PROCESS | 225 | 0.305093 | 1.002925 | 0.488959 | 0.851067 |
| GOBP_HOMOTYPIC_CELL_CELL_ADHESION | 74 | 0.341498 | 1.001959 | 0.46247 | 0.853418 |
| GOBP_REGULATION_OF_RIG_I_SIGNALING_PATHWAY | 17 | 0.439289 | 1.001787 | 0.473054 | 0.853433 |
| GOBP_CALCIUM_INDEPENDENT_CELL_CELL_ADHESION_VIA_PLASMA_MEMBRANE_CE | 15 | 0.444071 | 1.001091 | 0.490196 | 0.854904 |
| GOBP_NEGATIVE_REGULATION_OF_MICROTUBULE_POLYMERIZATION_OR_DEPOLYME | 40 | 0.37259 | 1.001032 | 0.45756 | 0.854558 |
| GOBP_VASCULAR_ASSOCIATED_SMOOTH_MUSCLE_CELL_MIGRATION | 18 | 0.430402 | 1.000514 | 0.482558 | 0.855584 |
| GOBP_SPINAL_CORD_MOTOR_NEURON_DIFFERENTIATION | 20 | 0.42486 | 1.000217 | 0.47018 | 0.855979 |
| GOBP_METANEPHROS_DEVELOPMENT | 68 | 0.342434 | 0.999676 | 0.494611 | 0.857064 |
| GOBP_RESPONSE_TO_DOPAMINE | 61 | 0.347211 | 0.999208 | 0.462871 | 0.857881 |
| GOBP_INTRINSIC_APOPTOTIC_SIGNALING_PATHWAY_BY_P53_CLASS_MEDIATOR | 75 | 0.342478 | 0.998896 | 0.469965 | 0.858272 |
| GOBP_GLIAL_CELL_DIFFERENTIATION | 176 | 0.314854 | 0.998124 | 0.499465 | 0.860073 |
| GOBP_CILIUM_MOVEMENT | 123 | 0.32164 | 0.997756 | 0.472284 | 0.860616 |
| GOBP_PYRIMIDINE_RIBONUCLEOTIDE_METABOLIC_PROCESS | 28 | 0.401546 | 0.9977 | 0.504054 | 0.860273 |

| | | | | | |
|---|---|---|---|---|---|
| GOBP_CHOLESTEROL_STORAGE | 15 | 0.445414 | 0.997496 | 0.476744 | 0.860366 |
| GOBP_NEGATIVE_REGULATION_OF_MRNA_CATABOLIC_PROCESS | 55 | 0.352727 | 0.997363 | 0.476543 | 0.860239 |
| GOBP_RESPONSE_TO_STEROID_HORMONE | 261 | 0.307497 | 0.996979 | 0.508316 | 0.860879 |
| GOBP_CHEMICAL_SYNAPTIC_TRANSMISSION_POSTSYNAPTIC | 84 | 0.331962 | 0.996824 | 0.447779 | 0.860823 |
| GOBP_NEGATIVE_REGULATION_OF_DNA_METABOLIC_PROCESS | 104 | 0.325036 | 0.996694 | 0.480447 | 0.860699 |
| GOBP_REGULATION_OF_DOUBLE_STRAND_BREAK_REPAIR_VIA_HOMOLOGOUS_RECO | 69 | 0.344893 | 0.996609 | 0.476961 | 0.860442 |
| GOBP_ATRIAL_CARDIAC_MUSCLE_CELL_TO_AV_NODE_CELL_SIGNALING | 17 | 0.43733 | 0.996471 | 0.490358 | 0.86032 |
| GOBP_BEHAVIOR | 490 | 0.299144 | 0.994896 | 0.512538 | 0.864311 |
| GOBP_ORGANIC_ACID_TRANSPORT | 236 | 0.306319 | 0.994593 | 0.516913 | 0.864695 |
| GOBP_CELLULAR_RESPONSE_TO_CALCIUM_ION | 72 | 0.343237 | 0.994214 | 0.472859 | 0.865283 |
| GOBP_POSITIVE_REGULATION_OF_NUCLEAR_DIVISION | 40 | 0.371153 | 0.993822 | 0.490518 | 0.865909 |
| GOBP_POSITIVE_REGULATION_OF_PROTEOLYSIS_INVOLVED_IN_CELLULAR_PROTEIN_ | 124 | 0.319542 | 0.992134 | 0.500562 | 0.870258 |
| GOBP_PHOTORECEPTOR_CELL_DEVELOPMENT | 38 | 0.375849 | 0.992034 | 0.46789 | 0.87001 |
| GOBP_CARBOHYDRATE_BIOSYNTHETIC_PROCESS | 170 | 0.310443 | 0.991114 | 0.508091 | 0.872132 |
| GOBP_REGULATION_OF_INSULIN_SECRETION | 118 | 0.322606 | 0.991097 | 0.505543 | 0.871667 |
| GOBP_REGULATION_OF_EPITHELIAL_CELL_APOPTOTIC_PROCESS | 64 | 0.346768 | 0.990872 | 0.488971 | 0.871801 |
| GOBP_PRESYNAPTIC_ENDOCYTOSIS | 56 | 0.352781 | 0.990468 | 0.478315 | 0.872467 |
| GOBP_ORGANOPHOSPHATE_CATABOLIC_PROCESS | 139 | 0.31449 | 0.989673 | 0.513187 | 0.874263 |
| GOBP_REGULATION_OF_PROTEIN_IMPORT | 55 | 0.348916 | 0.988717 | 0.483313 | 0.87656 |
| GOBP_SYNAPTIC_VESICLE_EXOCYTOSIS | 92 | 0.327091 | 0.987825 | 0.486797 | 0.878633 |
| GOBP_NEGATIVE_REGULATION_OF_INTRACELLULAR_SIGNAL_TRANSDUCTION | 451 | 0.298684 | 0.987703 | 0.550959 | 0.878451 |
| GOBP_CHONDROITIN_SULFATE_PROTEOGLYCAN_METABOLIC_PROCESS | 33 | 0.384251 | 0.987525 | 0.490196 | 0.878438 |
| GOBP_NEGATIVE_REGULATION_OF_VIRAL_LIFE_CYCLE | 21 | 0.4157 | 0.987499 | 0.483405 | 0.877992 |
| GOBP_STEROL_TRANSPORT | 89 | 0.328362 | 0.986558 | 0.51373 | 0.88016 |
| GOBP_RESPONSE_TO_ACIDIC_PH | 20 | 0.409198 | 0.986328 | 0.476608 | 0.880292 |
| GOBP_SULFUR_COMPOUND_CATABOLIC_PROCESS | 26 | 0.400874 | 0.985796 | 0.502882 | 0.881366 |
| GOBP_REGULATION_OF_CELLULAR_RESPONSE_TO_INSULIN_STIMULUS | 51 | 0.354056 | 0.985702 | 0.493134 | 0.881093 |
| GOBP_NEGATIVE_REGULATION_OF_ENDOCYTOSIS | 47 | 0.357944 | 0.984637 | 0.513648 | 0.883672 |
| GOBP_REGULATION_OF_HEART_RATE_BY_CARDIAC_CONDUCTION | 33 | 0.377039 | 0.984614 | 0.502717 | 0.883206 |
| GOBP_REGULATION_OF_SMAD_PROTEIN_SIGNAL_TRANSDUCTION | 29 | 0.377288 | 0.984404 | 0.483916 | 0.883296 |
| GOBP_POSITIVE_REGULATION_OF_SIGNALING_RECEPTOR_ACTIVITY | 36 | 0.372852 | 0.984322 | 0.514801 | 0.882996 |
| GOBP_PURINE_CONTAINING_COMPOUND_CATABOLIC_PROCESS | 46 | 0.361348 | 0.984141 | 0.487959 | 0.88302 |
| GOBP_NEURON_RECOGNITION | 45 | 0.366133 | 0.98395 | 0.485825 | 0.883058 |
| GOBP_MULTICELLULAR_ORGANISM_GROWTH | 124 | 0.317195 | 0.983693 | 0.507743 | 0.883287 |
| GOBP_REGULATION_OF_STRESS_ACTIVATED_PROTEIN_KINASE_SIGNALING_CASCADE | 173 | 0.308697 | 0.983064 | 0.522581 | 0.884605 |
| GOBP_RIBONUCLEOSIDE_MONOPHOSPHATE_METABOLIC_PROCESS | 55 | 0.353015 | 0.982897 | 0.508366 | 0.884578 |
| GOBP_NEGATIVE_REGULATION_OF_CELL_CYCLE | 333 | 0.300439 | 0.982475 | 0.52863 | 0.885261 |
| GOBP_POSITIVE_REGULATION_OF_NUCLEOCYTOPLASMIC_TRANSPORT | 55 | 0.348871 | 0.98225 | 0.508537 | 0.885393 |
| GOBP_REGULATION_OF_MITOCHONDRIAL_MEMBRANE_PERMEABILITY_INVOLVED_IN_ | 39 | 0.367231 | 0.981304 | 0.502618 | 0.887619 |
| GOBP_REGULATION_OF_ESTABLISHMENT_OF_PROTEIN_LOCALIZATION | 444 | 0.294626 | 0.980699 | 0.566265 | 0.888868 |
| GOBP_EPITHELIAL_TO_MESENCHYMAL_TRANSITION | 137 | 0.312019 | 0.980281 | 0.518764 | 0.889541 |
| GOBP_REGULATION_OF_CARDIAC_MUSCLE_CELL_MEMBRANE_REPOLARIZATION | 21 | 0.410275 | 0.979978 | 0.502793 | 0.889906 |
| GOBP_SECONDARY_METABOLITE_BIOSYNTHETIC_PROCESS | 17 | 0.415269 | 0.979614 | 0.50295 | 0.890427 |
| GOBP_POSITIVE_REGULATION_OF_MONOOXYGENASE_ACTIVITY | 24 | 0.395422 | 0.97948 | 0.502801 | 0.890293 |
| GOBP_RESPONSE_TO_RADIATION | 365 | 0.295787 | 0.979339 | 0.56765 | 0.89016 |
| GOBP_PLASMA_MEMBRANE_ORGANIZATION | 118 | 0.321341 | 0.97929 | 0.507901 | 0.889798 |
| GOBP_RESPONSE_TO_KETONE | 151 | 0.310982 | 0.978628 | 0.536692 | 0.8912 |
| GOBP_AMMONIUM_ION_METABOLIC_PROCESS | 17 | 0.425995 | 0.97844 | 0.508824 | 0.891202 |
| GOBP_RESPONSE_TO_AMINE | 37 | 0.368586 | 0.977518 | 0.519634 | 0.893277 |
| GOBP_POSITIVE_REGULATION_OF_MEMBRANE_PERMEABILITY | 41 | 0.364865 | 0.977442 | 0.508431 | 0.892974 |
| GOBP_POSITIVE_REGULATION_OF_STEROID_METABOLIC_PROCESS | 21 | 0.405244 | 0.975972 | 0.500734 | 0.896622 |
| GOBP_PURINE_NUCLEOBASE_METABOLIC_PROCESS | 18 | 0.421955 | 0.975791 | 0.494286 | 0.896616 |
| GOBP_CELLULAR_RESPONSE_TO_VASCULAR_ENDOTHELIAL_GROWTH_FACTOR_STIM | 56 | 0.348631 | 0.975281 | 0.506158 | 0.897561 |
| GOBP_ORGANIC_HYDROXY_COMPOUND_METABOLIC_PROCESS | 424 | 0.293169 | 0.974776 | 0.570854 | 0.898391 |
| GOBP_PROTEIN_METHYLATION | 186 | 0.304007 | 0.974543 | 0.540741 | 0.89851 |
| GOBP_CELL_DIFFERENTIATION_IN_SPINAL_CORD | 35 | 0.369479 | 0.974497 | 0.507152 | 0.898112 |
| GOBP_CELL_COMMUNICATION_INVOLVED_IN_CARDIAC_CONDUCTION | 52 | 0.349619 | 0.974384 | 0.507692 | 0.897913 |
| GOBP_REGULATION_OF_AEROBIC_RESPIRATION | 29 | 0.37947 | 0.9743 | 0.501295 | 0.897636 |
| GOBP_CENTRAL_NERVOUS_SYSTEM_NEURON_DIFFERENTIATION | 134 | 0.311045 | 0.97401 | 0.551991 | 0.897958 |
| GOBP_RUFFLE_ORGANIZATION | 49 | 0.349646 | 0.973734 | 0.513241 | 0.898219 |
| GOBP_POSITIVE_REGULATION_OF_HEART_GROWTH | 21 | 0.411331 | 0.973624 | 0.516129 | 0.898004 |
| GOBP_POTASSIUM_ION_HOMEOSTASIS | 25 | 0.396167 | 0.973555 | 0.508966 | 0.897689 |
| GOBP_NEGATIVE_REGULATION_OF_ION_TRANSMEMBRANE_TRANSPORTER_ACTIVITY | 50 | 0.351311 | 0.973387 | 0.520779 | 0.897634 |
| GOBP_EXPORT_ACROSS_PLASMA_MEMBRANE | 53 | 0.345549 | 0.973178 | 0.518193 | 0.897728 |
| GOBP_INTRACELLULAR_STEROL_TRANSPORT | 27 | 0.391754 | 0.973109 | 0.52027 | 0.897404 |
| GOBP_CELLULAR_DEFENSE_RESPONSE | 22 | 0.409479 | 0.972935 | 0.516304 | 0.897389 |
| GOBP_CELLULAR_RESPONSE_TO_INORGANIC_SUBSTANCE | 177 | 0.30212 | 0.972895 | 0.522436 | 0.896986 |
| GOBP_NEGATIVE_REGULATION_OF_PHOSPHORYLATION | 330 | 0.29288 | 0.972376 | 0.574949 | 0.897965 |
| GOBP_LUNG_EPITHELIUM_DEVELOPMENT | 25 | 0.390485 | 0.972011 | 0.508403 | 0.898461 |
| GOBP_REGULATION_OF_CELL_SIZE | 167 | 0.309879 | 0.971105 | 0.537473 | 0.900466 |
| GOBP_REGULATION_OF_GLUCONEOGENESIS | 44 | 0.354583 | 0.970944 | 0.502532 | 0.900402 |
| GOBP_POSITIVE_REGULATION_OF_PHOSPHOLIPASE_ACTIVITY | 37 | 0.367359 | 0.970847 | 0.5 | 0.900149 |
| GOBP_HOMEOSTASIS_OF_NUMBER_OF_CELLS_WITHIN_A_TISSUE | 21 | 0.411226 | 0.969862 | 0.519006 | 0.902452 |
| GOBP_REGULATION_OF_LIPID_CATABOLIC_PROCESS | 40 | 0.370552 | 0.969852 | 0.502639 | 0.901961 |
| GOBP_REGULATION_OF_PLATELET_AGGREGATION | 20 | 0.399252 | 0.969784 | 0.508029 | 0.901631 |
| GOBP_POSITIVE_REGULATION_OF_EXTRINSIC_APOPTOTIC_SIGNALING_PATHWAY | 42 | 0.365832 | 0.969555 | 0.51871 | 0.901756 |
| GOBP_REGULATION_OF_DNA_RECOMBINATION | 114 | 0.314595 | 0.969448 | 0.536667 | 0.90152 |
| GOBP_HIPPOCAMPUS_DEVELOPMENT | 66 | 0.336403 | 0.969315 | 0.529769 | 0.901399 |
| GOBP_REGULATION_OF_APOPTOTIC_SIGNALING_PATHWAY | 315 | 0.295815 | 0.968809 | 0.567762 | 0.902328 |
| GOBP_GLUTAMINE_FAMILY_AMINO_ACID_BIOSYNTHETIC_PROCESS | 15 | 0.434567 | 0.968577 | 0.528529 | 0.902498 |
| GOBP_NEGATIVE_REGULATION_OF_SUPRAMOLECULAR_FIBER_ORGANIZATION | 149 | 0.303702 | 0.968318 | 0.544359 | 0.902673 |
| GOBP_POSITIVE_REGULATION_OF_T_CELL_MIGRATION | 21 | 0.400938 | 0.967533 | 0.505714 | 0.904381 |
| GOBP_REGULATION_OF_ANATOMICAL_STRUCTURE_SIZE | 417 | 0.291119 | 0.967466 | 0.593939 | 0.904043 |
| GOBP_ARGININE_METABOLIC_PROCESS | 15 | 0.42808 | 0.967428 | 0.520119 | 0.903629 |
| GOBP_ORGANIC_ANION_TRANSPORT | 272 | 0.295446 | 0.967321 | 0.565083 | 0.903416 |
| GOBP_ORGANIC_ACID_BIOSYNTHETIC_PROCESS | 237 | 0.29998 | 0.967297 | 0.572487 | 0.902948 |
| GOBP_POSITIVE_REGULATION_OF_NEUROTRANSMITTER_TRANSPORT | 21 | 0.398812 | 0.967145 | 0.516266 | 0.902869 |
| GOBP_IN_UTERO_EMBRYONIC_DEVELOPMENT | 357 | 0.291981 | 0.967082 | 0.594924 | 0.902536 |
| GOBP_STEROID_METABOLIC_PROCESS | 231 | 0.298437 | 0.966819 | 0.567368 | 0.902788 |
| GOBP_POSITIVE_REGULATION_OF_DNA_RECOMBINATION | 62 | 0.337576 | 0.966688 | 0.515403 | 0.90265 |
| GOBP_POSITIVE_REGULATION_OF_RESPONSE_TO_DNA_DAMAGE_STIMULUS | 125 | 0.314549 | 0.966635 | 0.56648 | 0.902288 |
| GOBP_NEGATIVE_REGULATION_OF_PHOSPHORUS_METABOLIC_PROCESS | 379 | 0.292618 | 0.966609 | 0.572581 | 0.901848 |
| GOBP_REGULATION_OF_SYNAPTIC_PLASTICITY | 167 | 0.305407 | 0.966564 | 0.56 | 0.90146 |
| GOBP_CAMERA_TYPE_EYE_DEVELOPMENT | 258 | 0.29836 | 0.966541 | 0.589506 | 0.901016 |
| GOBP_CELLULAR_COMPONENT_DISASSEMBLY | 418 | 0.294077 | 0.966173 | 0.583249 | 0.901531 |
| GOBP_PEROXISOME_PROLIFERATOR_ACTIVATED_RECEPTOR_SIGNALING_PATHWAY | 19 | 0.410945 | 0.965715 | 0.523546 | 0.902305 |
| GOBP_CARDIAC_MUSCLE_CELL_ACTION_POTENTIAL_INVOLVED_IN_CONTRACTION | 45 | 0.353846 | 0.965401 | 0.532383 | 0.902679 |
| GOBP_AXON_DEVELOPMENT | 431 | 0.291344 | 0.9653 | 0.589484 | 0.902469 |
| GOBP_RESPONSE_TO_ALKALOID | 84 | 0.322189 | 0.965275 | 0.544084 | 0.902034 |
| GOBP_AMINO_ACID_IMPORT_ACROSS_PLASMA_MEMBRANE | 37 | 0.367921 | 0.964593 | 0.523684 | 0.903404 |
| GOBP_CEREBELLAR_PURKINJE_CELL_LAYER_DEVELOPMENT | 26 | 0.392117 | 0.964466 | 0.539618 | 0.903236 |
| GOBP_DNA_DAMAGE_RESPONSE_SIGNAL_TRANSDUCTION_RESULTING_IN_TRANSCR | 17 | 0.422318 | 0.964168 | 0.527616 | 0.903547 |
| GOBP_LACTATION | 39 | 0.359803 | 0.963503 | 0.511749 | 0.904902 |
| GOBP_SMALL_MOLECULE_CATABOLIC_PROCESS | 307 | 0.294596 | 0.963435 | 0.592668 | 0.904579 |
| GOBP_REGULATION_OF_CYSTEINE_TYPE_ENDOPEPTIDASE_ACTIVITY | 190 | 0.301968 | 0.963397 | 0.567769 | 0.904174 |
| GOBP_VENTRAL_SPINAL_CORD_DEVELOPMENT | 33 | 0.374638 | 0.963176 | 0.534914 | 0.904278 |
| GOBP_GLUCOSE_METABOLIC_PROCESS | 164 | 0.302508 | 0.96276 | 0.559653 | 0.904895 |
| GOBP_NON_MEMBRANE_BOUNDED_ORGANELLE_ASSEMBLY | 354 | 0.291295 | 0.962013 | 0.598577 | 0.906484 |
| GOBP_REGULATION_OF_MIRNA_TRANSCRIPTION | 54 | 0.339466 | 0.960763 | 0.520807 | 0.909455 |
| GOBP_REGULATION_OF_INSULIN_SECRETION_INVOLVED_IN_CELLULAR_RESPONSE_ | 44 | 0.351659 | 0.960495 | 0.52459 | 0.909713 |
| GOBP_REACTIVE_OXYGEN_SPECIES_BIOSYNTHETIC_PROCESS | 44 | 0.354364 | 0.960341 | 0.543791 | 0.909662 |
| GOBP_AMIDE_TRANSPORT | 226 | 0.294544 | 0.960286 | 0.581081 | 0.909296 |
| GOBP_POSITIVE_REGULATION_OF_PROTEIN_LOCALIZATION_TO_MEMBRANE | 94 | 0.318094 | 0.959714 | 0.529895 | 0.910406 |
| GOBP_NEUROEPITHELIAL_CELL_DIFFERENTIATION | 22 | 0.393814 | 0.959697 | 0.537063 | 0.909938 |
| GOBP_CELLULAR_MODIFIED_AMINO_ACID_CATABOLIC_PROCESS | 22 | 0.403281 | 0.959403 | 0.519118 | 0.910208 |
| GOBP_REGULATION_OF_PROTEIN_BINDING | 171 | 0.303849 | 0.959226 | 0.56928 | 0.910186 |
| GOBP_NEGATIVE_REGULATION_OF_SECRETION | 122 | 0.310235 | 0.959133 | 0.556059 | 0.909932 |
| GOBP_TELENCEPHALON_DEVELOPMENT | 213 | 0.295958 | 0.959062 | 0.57492 | 0.909609 |
| GOBP_AMINE_CATABOLIC_PROCESS | 19 | 0.409813 | 0.958458 | 0.525148 | 0.910763 |
| GOBP_POSITIVE_REGULATION_OF_CALCIUM_ION_DEPENDENT_EXOCYTOSIS | 17 | 0.416965 | 0.958342 | 0.513006 | 0.910555 |
| GOBP_ORGANELLE_FISSION | 427 | 0.285665 | 0.958251 | 0.611895 | 0.910294 |
| GOBP_REGULATION_OF_MONOCYTE_CHEMOTAXIS | 16 | 0.423713 | 0.958103 | 0.558245 | 0.910186 |
| GOBP_NEGATIVE_REGULATION_OF_CATABOLIC_PROCESS | 276 | 0.294843 | 0.956787 | 0.59375 | 0.913229 |
| GOBP_REGULATION_OF_DEFENSE_RESPONSE_TO_VIRUS | 61 | 0.332885 | 0.956721 | 0.567073 | 0.912903 |
| GOBP_POSITIVE_REGULATION_OF_NEUROGENESIS | 193 | 0.298363 | 0.956496 | 0.585987 | 0.912991 |
| GOBP_CARBOXYLIC_ACID_TRANSPORT | 200 | 0.298393 | 0.956274 | 0.595388 | 0.913073 |
| GOBP_AMINOGLYCAN_METABOLIC_PROCESS | 109 | 0.312009 | 0.956196 | 0.531963 | 0.912763 |
| GOBP_REGULATION_OF_PEPTIDYL_LYSINE_ACETYLATION | 68 | 0.328253 | 0.955926 | 0.560588 | 0.913017 |
| GOBP_LEARNING | 127 | 0.30885 | 0.955895 | 0.564334 | 0.912595 |
| GOBP_NEGATIVE_REGULATION_OF_CALCIUM_ION_TRANSPORT | 44 | 0.349691 | 0.955814 | 0.571429 | 0.912292 |
| GOBP_NEGATIVE_REGULATION_OF_TRANSPORTER_ACTIVITY | 56 | 0.336275 | 0.955451 | 0.534418 | 0.912761 |
| GOBP_RESPONSE_TO_CATECHOLAMINE | 75 | 0.328065 | 0.955276 | 0.541417 | 0.91272 |
| GOBP_POSITIVE_REGULATION_OF_TRANSCRIPTION_FROM_RNA_POLYMERASE_II_PRO | 22 | 0.396101 | 0.954779 | 0.542466 | 0.913576 |
| GOBP_REGULATION_OF_EXTENT_OF_CELL_GROWTH | 103 | 0.313321 | 0.954514 | 0.558219 | 0.913807 |
| GOBP_POSITIVE_REGULATION_OF_NEURON_DEATH | 70 | 0.325186 | 0.954063 | 0.541667 | 0.914516 |
| GOBP_NEGATIVE_REGULATION_OF_TRANSFERASE_ACTIVITY | 241 | 0.292848 | 0.954001 | 0.592975 | 0.914191 |
| GOBP_METAL_ION_EXPORT | 39 | 0.360023 | 0.953704 | 0.55814 | 0.914473 |
| GOBP_BRANCHED_CHAIN_AMINO_ACID_CATABOLIC_PROCESS | 19 | 0.410267 | 0.953644 | 0.548991 | 0.914108 |
| GOBP_NEGATIVE_REGULATION_OF_BLOOD_PRESSURE | 34 | 0.367272 | 0.95364 | 0.559892 | 0.913611 |
| GOBP_POSITIVE_REGULATION_OF_REACTIVE_OXYGEN_SPECIES_METABOLIC_PROCES | 53 | 0.343602 | 0.953061 | 0.553059 | 0.914664 |
| GOBP_P38MAPK_CASCADE | 47 | 0.349237 | 0.95277 | 0.541087 | 0.914958 |
| GOBP_POSITIVE_REGULATION_OF_CELLULAR_RESPONSE_TO_TRANSFORMING_GROW | 29 | 0.369668 | 0.95248 | 0.535422 | 0.915245 |
| GOBP_PROTEIN_MATURATION | 255 | 0.290785 | 0.951951 | 0.593238 | 0.916218 |
| GOBP_CELL_GROWTH | 431 | 0.286498 | 0.951837 | 0.635081 | 0.916035 |
| GOBP_RESPONSE_TO_LEPTIN | 18 | 0.403529 | 0.951822 | 0.53343 | 0.915567 |
| GOBP_PURINE_NUCLEOTIDE_CATABOLIC_PROCESS | 41 | 0.357347 | 0.951794 | 0.53316 | 0.915137 |
| GOBP_ORGAN_INDUCTION | 18 | 0.4076 | 0.951522 | 0.543319 | 0.915356 |
| GOBP_POSITIVE_REGULATION_OF_NEURON_PROJECTION_DEVELOPMENT | 136 | 0.306175 | 0.95132 | 0.564215 | 0.915402 |
| GOBP_AMYLOID_FIBRIL_FORMATION | 27 | 0.380192 | 0.951197 | 0.521798 | 0.91523 |
| GOBP_EMBRYONIC_HEMOPOIESIS | 20 | 0.403097 | 0.951142 | 0.54623 | 0.914876 |
| GOBP_POSITIVE_REGULATION_OF_REGULATED_SECRETORY_PATHWAY | 46 | 0.347942 | 0.951041 | 0.552258 | 0.914627 |

| | | | | | |
|---|---|---|---|---|---|
| GOBP_RESPONSE_TO_MONOSACCHARIDE | 154 | 0.298922 | 0.950769 | 0.590217 | 0.914841 |
| GOBP_NUCLEIC_ACID_PHOSPHODIESTER_BOND_HYDROLYSIS | 234 | 0.292898 | 0.949887 | 0.616267 | 0.916698 |
| GOBP_SNRNA_TRANSCRIPTION | 18 | 0.40906 | 0.948433 | 0.549192 | 0.920148 |
| GOBP_EMBRYONIC_SKELETAL_SYSTEM_MORPHOGENESIS | 56 | 0.332877 | 0.94808 | 0.556401 | 0.920561 |
| GOBP_REGULATION_OF_CALCIUM_ION_TRANSMEMBRANE_TRANSPORTER_ACTIVITY | 71 | 0.325193 | 0.947669 | 0.57298 | 0.921159 |
| GOBP_REGULATION_OF_PROTEIN_SERINE_THREONINE_KINASE_ACTIVITY | 323 | 0.287047 | 0.947385 | 0.617587 | 0.921408 |
| GOBP_RELEASE_OF_SEQUESTERED_CALCIUM_ION_INTO_CYTOSOL_BY_ENDOPLASMI | 26 | 0.383897 | 0.946769 | 0.546196 | 0.922541 |
| GOBP_NEGATIVE_REGULATION_OF_PHOSPHOPROTEIN_PHOSPHATASE_ACTIVITY | 22 | 0.390258 | 0.946623 | 0.553009 | 0.922417 |
| GOBP_RESPONSE_TO_FATTY_ACID | 49 | 0.341264 | 0.945409 | 0.554293 | 0.925125 |
| GOBP_VITAMIN_TRANSPORT | 35 | 0.357878 | 0.945001 | 0.542092 | 0.925687 |
| GOBP_CELL_JUNCTION_ASSEMBLY | 362 | 0.28425 | 0.943343 | 0.636826 | 0.929583 |
| GOBP_EPITHELIAL_CELL_DEVELOPMENT | 174 | 0.29597 | 0.943254 | 0.613319 | 0.929316 |
| GOBP_POSITIVE_REGULATION_OF_FIBROBLAST_PROLIFERATION | 37 | 0.35868 | 0.942461 | 0.552318 | 0.930906 |
| GOBP_SEMAPHORIN_PLEXIN_SIGNALING_PATHWAY | 42 | 0.350291 | 0.942294 | 0.56383 | 0.930831 |
| GOBP_FEEDING_BEHAVIOR | 67 | 0.32709 | 0.941852 | 0.568127 | 0.931538 |
| GOBP_REGULATION_OF_NERVOUS_SYSTEM_PROCESS | 114 | 0.307439 | 0.941568 | 0.605145 | 0.931753 |
| GOBP_NEGATIVE_REGULATION_OF_HISTONE_MODIFICATION | 46 | 0.343179 | 0.941044 | 0.566624 | 0.932612 |
| GOBP_POSITIVE_REGULATION_OF_NUCLEOTIDE_BIOSYNTHETIC_PROCESS | 17 | 0.410578 | 0.940817 | 0.547789 | 0.932691 |
| GOBP_RESPONSE_TO_RETINOIC_ACID | 83 | 0.319987 | 0.940668 | 0.588517 | 0.932559 |
| GOBP_T_CELL_LINEAGE_COMMITMENT | 16 | 0.408376 | 0.940589 | 0.555716 | 0.932265 |
| GOBP_ENDOCYTOSIS | 477 | 0.281713 | 0.939692 | 0.671687 | 0.934081 |
| GOBP_POSITIVE_REGULATION_OF_PROTEIN_BINDING | 75 | 0.321259 | 0.93836 | 0.577778 | 0.937098 |
| GOBP_POSITIVE_REGULATION_OF_TRANSFERASE_ACTIVITY | 480 | 0.282165 | 0.938042 | 0.686808 | 0.937457 |
| GOBP_NEPHRON_EPITHELIUM_DEVELOPMENT | 90 | 0.317449 | 0.937896 | 0.585929 | 0.937333 |
| GOBP_REGULATION_OF_INTRINSIC_APOPTOTIC_SIGNALING_PATHWAY_BY_P53_CLASS | 30 | 0.369332 | 0.937807 | 0.570014 | 0.937061 |
| GOBP_REGULATION_OF_SMALL_GTPASE_MEDIATED_SIGNAL_TRANSDUCTION | 275 | 0.28589 | 0.93768 | 0.639835 | 0.936873 |
| GOBP_CIRCADIAN_RHYTHM | 173 | 0.290059 | 0.937597 | 0.604278 | 0.936585 |
| GOBP_ASTROCYTE_DEVELOPMENT | 29 | 0.365502 | 0.936926 | 0.55461 | 0.937896 |
| GOBP_NUCLEOTIDE_SUGAR_BIOSYNTHETIC_PROCESS | 17 | 0.405141 | 0.93598 | 0.556414 | 0.939872 |
| GOBP_AXON_ENSHEATHMENT_IN_CENTRAL_NERVOUS_SYSTEM | 19 | 0.393411 | 0.935766 | 0.56528 | 0.939931 |
| GOBP_BRANCHING_MORPHOGENESIS_OF_AN_EPITHELIAL_TUBE | 121 | 0.300729 | 0.934718 | 0.585118 | 0.942232 |
| GOBP_CRISTAE_FORMATION | 16 | 0.405469 | 0.934595 | 0.54964 | 0.942041 |
| GOBP_PURINE_NUCLEOTIDE_TRANSPORT | 27 | 0.376689 | 0.934461 | 0.57305 | 0.941896 |
| GOBP_REGULATION_OF_NEURON_PROJECTION_DEVELOPMENT | 397 | 0.283856 | 0.934419 | 0.679798 | 0.941506 |
| GOBP_EXTRACELLULAR_MATRIX_ASSEMBLY | 41 | 0.345639 | 0.934388 | 0.552835 | 0.941064 |
| GOBP_MECHANORECEPTOR_DIFFERENTIATION | 55 | 0.333463 | 0.934343 | 0.576485 | 0.940662 |
| GOBP_NEGATIVE_REGULATION_OF_SIGNALING_RECEPTOR_ACTIVITY | 39 | 0.352018 | 0.934029 | 0.577657 | 0.940952 |
| GOBP_REGULATION_OF_PROTEIN_TARGETING_TO_MEMBRANE | 31 | 0.364402 | 0.933495 | 0.581137 | 0.941836 |
| GOBP_POSITIVE_REGULATION_OF_CELL_DEVELOPMENT | 252 | 0.285262 | 0.933071 | 0.650155 | 0.942423 |
| GOBP_POSITIVE_REGULATION_OF_KERATINOCYTE_DIFFERENTIATION | 16 | 0.407895 | 0.932847 | 0.550933 | 0.942515 |
| GOBP_CENTRAL_NERVOUS_SYSTEM_NEURON_DEVELOPMENT | 70 | 0.32085 | 0.93269 | 0.578687 | 0.942449 |
| GOBP_REGULATION_OF_WATER_LOSS_VIA_SKIN | 20 | 0.397993 | 0.932527 | 0.555085 | 0.942363 |
| GOBP_RESPONSE_TO_PAIN | 26 | 0.374964 | 0.932448 | 0.579972 | 0.942056 |
| GOBP_POSITIVE_REGULATION_OF_VASCULAR_ENDOTHELIAL_GROWTH_FACTOR_PRO | 22 | 0.386682 | 0.931641 | 0.574344 | 0.943673 |
| GOBP_PROTEIN_AUTOUBIQUITINATION | 70 | 0.322774 | 0.931619 | 0.583738 | 0.943219 |
| GOBP_ACTIN_FILAMENT_DEPOLYMERIZATION | 54 | 0.334605 | 0.931156 | 0.597468 | 0.943902 |
| GOBP_MONOCYTE_DIFFERENTIATION | 29 | 0.36915 | 0.930892 | 0.575636 | 0.94412 |
| GOBP_CELLULAR_RESPONSE_TO_OXYGEN_LEVELS | 139 | 0.299461 | 0.93084 | 0.614618 | 0.943745 |
| GOBP_CELLULAR_RESPONSE_TO_EXTRACELLULAR_STIMULUS | 215 | 0.290013 | 0.930609 | 0.650743 | 0.943836 |
| GOBP_REGULATION_OF_DNA_METHYLATION | 20 | 0.389302 | 0.929918 | 0.564254 | 0.945071 |
| GOBP_RESPONSE_TO_MITOCHONDRIAL_DEPOLARISATION | 19 | 0.391589 | 0.92896 | 0.553672 | 0.947084 |
| GOBP_NEGATIVE_REGULATION_OF_PROTEIN_LOCALIZATION_TO_MEMBRANE | 28 | 0.366938 | 0.928938 | 0.566713 | 0.946633 |
| GOBP_CORTICAL_ACTIN_CYTOSKELETON_ORGANIZATION | 37 | 0.34983 | 0.928836 | 0.580603 | 0.94641 |
| GOBP_RIG_I_SIGNALING_PATHWAY | 25 | 0.376206 | 0.928673 | 0.580913 | 0.946338 |
| GOBP_NOTCH_SIGNALING_PATHWAY | 151 | 0.296604 | 0.928571 | 0.637761 | 0.946081 |
| GOBP_REGULATION_OF_CELL_JUNCTION_ASSEMBLY | 175 | 0.290571 | 0.927896 | 0.648649 | 0.947323 |
| GOBP_EPITHELIAL_CELL_DIFFERENTIATION_INVOLVED_IN_KIDNEY_DEVELOPMENT | 41 | 0.34239 | 0.927892 | 0.570513 | 0.946824 |
| GOBP_RNA_MODIFICATION | 157 | 0.290551 | 0.927416 | 0.633297 | 0.947593 |
| GOBP_MEMBRANE_PROTEIN_PROTEOLYSIS | 51 | 0.334519 | 0.92717 | 0.584833 | 0.947724 |
| GOBP_NEGATIVE_REGULATION_OF_NEURON_DIFFERENTIATION | 55 | 0.335554 | 0.926905 | 0.587039 | 0.947918 |
| GOBP_NEGATIVE_REGULATION_OF_NERVOUS_SYSTEM_DEVELOPMENT | 121 | 0.301147 | 0.926741 | 0.634119 | 0.947809 |
| GOBP_POSITIVE_REGULATION_OF_GTPASE_ACTIVITY | 227 | 0.283875 | 0.926295 | 0.649104 | 0.94846 |
| GOBP_NERVE_DEVELOPMENT | 64 | 0.323376 | 0.92614 | 0.604969 | 0.948342 |
| GOBP_SARCOPLASMIC_RETICULUM_CALCIUM_ION_TRANSPORT | 29 | 0.365418 | 0.926029 | 0.584958 | 0.948127 |
| GOBP_POSITIVE_REGULATION_OF_EPITHELIAL_CELL_APOPTOTIC_PROCESS | 22 | 0.378707 | 0.92554 | 0.550765 | 0.948896 |
| GOBP_POSITIVE_REGULATION_OF_PROTEIN_LOCALIZATION | 396 | 0.278692 | 0.925099 | 0.696538 | 0.949537 |
| GOBP_RETINA_DEVELOPMENT_IN_CAMERA_TYPE_EYE | 118 | 0.300985 | 0.924814 | 0.622517 | 0.949742 |
| GOBP_POSITIVE_REGULATION_OF_SMOOTH_MUSCLE_CELL_MIGRATION | 25 | 0.374589 | 0.924468 | 0.585573 | 0.95011 |
| GOBP_MORPHOGENESIS_OF_AN_EPITHELIUM | 425 | 0.277293 | 0.92422 | 0.721941 | 0.95024 |
| GOBP_REGULATION_OF_RUFFLE_ASSEMBLY | 28 | 0.359481 | 0.924013 | 0.568493 | 0.950276 |
| GOBP_NEGATIVE_REGULATION_OF_GTPASE_ACTIVITY | 29 | 0.36283 | 0.92374 | 0.590723 | 0.95048 |
| GOBP_REGULATION_OF_NEUROGENESIS | 311 | 0.280743 | 0.923578 | 0.672802 | 0.950395 |
| GOBP_PROTEIN_LOCALIZATION_TO_PLASMA_MEMBRANE | 264 | 0.283636 | 0.923533 | 0.680041 | 0.950012 |
| GOBP_REGULATION_OF_DENDRITE_DEVELOPMENT | 91 | 0.30839 | 0.923489 | 0.592506 | 0.949626 |
| GOBP_GLIAL_CELL_DEVELOPMENT | 90 | 0.30573 | 0.92344 | 0.607728 | 0.949258 |
| GOBP_REGULATION_OF_KERATINOCYTE_PROLIFERATION | 27 | 0.365138 | 0.923186 | 0.557103 | 0.949397 |
| GOBP_INSULIN_LIKE_GROWTH_FACTOR_RECEPTOR_SIGNALING_PATHWAY | 27 | 0.367261 | 0.922728 | 0.589333 | 0.950051 |
| GOBP_REGULATION_OF_CELL_CYCLE_PHASE_TRANSITION | 357 | 0.279389 | 0.922646 | 0.704197 | 0.949746 |
| GOBP_REGULATION_OF_ENDOCYTOSIS | 180 | 0.286601 | 0.922487 | 0.654467 | 0.949643 |
| GOBP_SYNAPTIC_TRANSMISSION_GLUTAMATERGIC | 74 | 0.31659 | 0.921976 | 0.614914 | 0.950477 |
| GOBP_DIOL_BIOSYNTHETIC_PROCESS | 22 | 0.378141 | 0.921911 | 0.587896 | 0.95014 |
| GOBP_KIDNEY_EPITHELIUM_DEVELOPMENT | 118 | 0.302454 | 0.921468 | 0.615215 | 0.950754 |
| GOBP_DNA_RECOMBINATION | 279 | 0.279514 | 0.920377 | 0.681115 | 0.952976 |
| GOBP_REGULATION_OF_BEHAVIOR | 47 | 0.332547 | 0.920353 | 0.600791 | 0.952525 |
| GOBP_RESPONSE_TO_INTERLEUKIN_6 | 29 | 0.357934 | 0.92027 | 0.554333 | 0.95221 |
| GOBP_NEGATIVE_REGULATION_OF_RECEPTOR_SIGNALING_PATHWAY_VIA_STAT | 22 | 0.376297 | 0.919916 | 0.571831 | 0.95263 |
| GOBP_PROTEIN_DEPOLYMERIZATION | 110 | 0.300107 | 0.919739 | 0.624714 | 0.952582 |
| GOBP_DEVELOPMENTAL_INDUCTION | 27 | 0.370925 | 0.919302 | 0.591892 | 0.953206 |
| GOBP_NEGATIVE_REGULATION_OF_NUCLEOTIDE_METABOLIC_PROCESS | 19 | 0.395636 | 0.918984 | 0.574048 | 0.953504 |
| GOBP_REGULATION_OF_GLIAL_CELL_DIFFERENTIATION | 57 | 0.324532 | 0.918766 | 0.601759 | 0.953549 |
| GOBP_ASTROCYTE_DIFFERENTIATION | 57 | 0.325732 | 0.918563 | 0.607721 | 0.953542 |
| GOBP_RESPONSE_TO_LITHIUM_ION | 17 | 0.400649 | 0.918281 | 0.564179 | 0.953774 |
| GOBP_NEGATIVE_REGULATION_OF_AMINE_TRANSPORT | 23 | 0.377775 | 0.918107 | 0.569863 | 0.953679 |
| GOBP_ALCOHOL_METABOLIC_PROCESS | 282 | 0.278924 | 0.918101 | 0.694444 | 0.953193 |
| GOBP_RESPONSE_TO_CARBOHYDRATE | 177 | 0.288324 | 0.917498 | 0.658432 | 0.954232 |
| GOBP_SCHWANN_CELL_DIFFERENTIATION | 35 | 0.349589 | 0.917419 | 0.592208 | 0.953947 |
| GOBP_POSITIVE_REGULATION_OF_PROTEIN_SERINE_THREONINE_KINASE_ACTIVITY | 178 | 0.289172 | 0.91732 | 0.661638 | 0.953692 |
| GOBP_REGULATION_OF_MITOCHONDRIAL_OUTER_MEMBRANE_PERMEABILIZATION_IN | 20 | 0.381046 | 0.91678 | 0.569715 | 0.954534 |
| GOBP_REGULATION_OF_B_CELL_APOPTOTIC_PROCESS | 18 | 0.391035 | 0.916467 | 0.578723 | 0.954796 |
| GOBP_POSITIVE_REGULATION_OF_B_CELL_MEDIATED_IMMUNITY | 30 | 0.360786 | 0.916186 | 0.591281 | 0.95503 |
| GOBP_POSITIVE_REGULATION_OF_PROTEIN_TARGETING_TO_MEMBRANE | 27 | 0.358656 | 0.916169 | 0.57438 | 0.954576 |
| GOBP_CELLULAR_LIPID_CATABOLIC_PROCESS | 180 | 0.288706 | 0.916088 | 0.673016 | 0.954283 |
| GOBP_NEGATIVE_REGULATION_OF_MAP_KINASE_ACTIVITY | 49 | 0.329453 | 0.915619 | 0.602302 | 0.954949 |
| GOBP_POSITIVE_REGULATION_OF_AMINE_TRANSPORT | 33 | 0.357827 | 0.915222 | 0.6 | 0.955507 |
| GOBP_NEGATIVE_REGULATION_OF_EXTRINSIC_APOPTOTIC_SIGNALING_PATHWAY_VIA | 24 | 0.371403 | 0.915006 | 0.599715 | 0.955549 |
| GOBP_REGULATION_OF_CELLULAR_COMPONENT_SIZE | 319 | 0.277979 | 0.914937 | 0.721193 | 0.955229 |
| GOBP_RUFFLE_ASSEMBLY | 41 | 0.34521 | 0.914113 | 0.589404 | 0.956763 |
| GOBP_EMBRYONIC_CRANIAL_SKELETON_MORPHOGENESIS | 37 | 0.342714 | 0.914068 | 0.580263 | 0.956383 |
| GOBP_STRESS_ACTIVATED_PROTEIN_KINASE_SIGNALING_CASCADE | 219 | 0.284516 | 0.914043 | 0.682489 | 0.95594 |
| GOBP_PEPTIDYL_TYROSINE_DEPHOSPHORYLATION | 92 | 0.303749 | 0.913788 | 0.631829 | 0.956101 |
| GOBP_VESICLE_MEDIATED_TRANSPORT_IN_SYNAPSE | 174 | 0.286917 | 0.913788 | 0.683422 | 0.955604 |
| GOBP_EMBRYONIC_APPENDAGE_MORPHOGENESIS | 85 | 0.306905 | 0.913491 | 0.607059 | 0.955892 |
| GOBP_CARBOHYDRATE_TRANSMEMBRANE_TRANSPORT | 89 | 0.308007 | 0.913465 | 0.642353 | 0.955459 |
| GOBP_RESPONSE_TO_INTERLEUKIN_1 | 87 | 0.304904 | 0.913158 | 0.623121 | 0.955705 |
| GOBP_SIGNAL_TRANSDUCTION_IN_ABSENCE_OF_LIGAND | 53 | 0.328572 | 0.912923 | 0.601504 | 0.955812 |
| GOBP_REGULATION_OF_NUCLEOTIDE_METABOLIC_PROCESS | 68 | 0.313304 | 0.912115 | 0.604567 | 0.957315 |
| GOBP_PIGMENT_METABOLIC_PROCESS | 68 | 0.314414 | 0.91131 | 0.635817 | 0.958855 |
| GOBP_MAINTENANCE_OF_BLOOD_BRAIN_BARRIER | 31 | 0.354911 | 0.910395 | 0.614173 | 0.960604 |
| GOBP_RESPIRATORY_CHAIN_COMPLEX_IV_ASSEMBLY | 26 | 0.363467 | 0.90986 | 0.598886 | 0.961403 |
| GOBP_VESICLE_DOCKING | 52 | 0.327856 | 0.909854 | 0.61519 | 0.960912 |
| GOBP_REGULATION_OF_LYMPHOCYTE_MIGRATION | 40 | 0.343943 | 0.909451 | 0.603627 | 0.961436 |
| GOBP_POSITIVE_REGULATION_OF_DNA_REPLICATION | 38 | 0.341788 | 0.909319 | 0.617268 | 0.961243 |
| GOBP_LIVER_REGENERATION | 23 | 0.375907 | 0.908764 | 0.6 | 0.962115 |
| GOBP_REGULATION_OF_PLATELET_ACTIVATION | 31 | 0.355909 | 0.908696 | 0.589333 | 0.961786 |
| GOBP_REGULATION_OF_ERAD_PATHWAY | 20 | 0.384662 | 0.908594 | 0.581633 | 0.961538 |
| GOBP_AMINO_ACID_IMPORT | 46 | 0.327549 | 0.908442 | 0.605634 | 0.961424 |
| GOBP_CRANIAL_NERVE_DEVELOPMENT | 38 | 0.338596 | 0.907996 | 0.603381 | 0.96205 |
| GOBP_RENAL_SYSTEM_PROCESS_INVOLVED_IN_REGULATION_OF_SYSTEMIC_ARTER | 15 | 0.401909 | 0.907589 | 0.571646 | 0.962546 |
| GOBP_POSITIVE_REGULATION_OF_EXTRACELLULAR_MATRIX_ORGANIZATION | 23 | 0.371805 | 0.907213 | 0.584507 | 0.962976 |
| GOBP_REGULATION_OF_PROTEIN_LOCALIZATION_TO_MEMBRANE | 155 | 0.286851 | 0.906648 | 0.650933 | 0.963913 |
| GOBP_PEPTIDYL_THREONINE_DEPHOSPHORYLATION | 16 | 0.391074 | 0.905983 | 0.570175 | 0.965083 |
| GOBP_NEPHRON_DEVELOPMENT | 120 | 0.291863 | 0.905887 | 0.644839 | 0.964828 |
| GOBP_MOTOR_NEURON_AXON_GUIDANCE | 28 | 0.356175 | 0.904737 | 0.588152 | 0.967106 |
| GOBP_REGULATION_OF_NUCLEAR_TRANSCRIBED_MRNA_POLY_A_TAIL_SHORTENING | 15 | 0.409871 | 0.904429 | 0.589231 | 0.967336 |
| GOBP_REGULATION_OF_RYANODINE_SENSITIVE_CALCIUM_RELEASE_CHANNEL_ACTI | 24 | 0.370774 | 0.904167 | 0.588889 | 0.967476 |
| GOBP_PURINE_RIBONUCLEOSIDE_METABOLIC_PROCESS | 20 | 0.376773 | 0.904144 | 0.625544 | 0.967039 |
| GOBP_NEURON_FATE_COMMITMENT | 37 | 0.345219 | 0.903396 | 0.622132 | 0.968395 |
| GOBP_POSITIVE_REGULATION_OF_UBIQUITIN_DEPENDENT_PROTEIN_CATABOLIC_PRO | 102 | 0.299721 | 0.902668 | 0.643023 | 0.9697 |
| GOBP_NUCLEOSIDE_BISPHOSPHATE_METABOLIC_PROCESS | 103 | 0.296461 | 0.902445 | 0.649366 | 0.969762 |
| GOBP_CELL_CELL_JUNCTION_MAINTENANCE | 17 | 0.38831 | 0.901972 | 0.587464 | 0.970398 |
| GOBP_EATING_BEHAVIOR | 25 | 0.359618 | 0.901645 | 0.591608 | 0.970705 |
| GOBP_LOCOMOTORY_BEHAVIOR | 164 | 0.284165 | 0.901435 | 0.673098 | 0.970732 |
| GOBP_NEGATIVE_REGULATION_OF_ERK1_AND_ERK2_CASCADE | 62 | 0.31298 | 0.901288 | 0.631579 | 0.970611 |
| GOBP_ANTIBACTERIAL_HUMORAL_RESPONSE | 22 | 0.378038 | 0.901237 | 0.596993 | 0.970233 |
| GOBP_PROTEIN_LOCALIZATION_TO_CELL_PERIPHERY | 311 | 0.27353 | 0.90076 | 0.734631 | 0.970898 |
| GOBP_RETINOIC_ACID_METABOLIC_PROCESS | 16 | 0.401862 | 0.900229 | 0.627193 | 0.971685 |
| GOBP_REGULATION_OF_SYNAPTIC_TRANSMISSION_GABAERGIC | 29 | 0.352969 | 0.900128 | 0.617978 | 0.971423 |
| GOBP_MONOSACCHARIDE_BIOSYNTHETIC_PROCESS | 71 | 0.306121 | 0.900103 | 0.643382 | 0.970982 |
| GOBP_REGULATION_OF_INTRINSIC_APOPTOTIC_SIGNALING_PATHWAY_IN_RESPONSE | 17 | 0.392587 | 0.899823 | 0.601744 | 0.971175 |
| GOBP_REGULATION_OF_CHOLESTEROL_BIOSYNTHETIC_PROCESS | 18 | 0.381778 | 0.899609 | 0.57855 | 0.971205 |
| GOBP_POSITIVE_REGULATION_OF_EXOCYTOSIS | 74 | 0.304038 | 0.899318 | 0.679087 | 0.971403 |

| | | | | | |
|---|---|---|---|---|---|
| GOBP_NEGATIVE_REGULATION_OF_INTRINSIC_APOPTOTIC_SIGNALING_PATHWAY_IN_ | 30 | 0.349383 | 0.898883 | 0.604749 | 0.971967 |
| GOBP_SUBPALLIUM_DEVELOPMENT | 18 | 0.380614 | 0.898849 | 0.61194 | 0.971549 |
| GOBP_REGULATION_OF_PROTEIN_DEPOLYMERIZATION | 82 | 0.301566 | 0.89853 | 0.65343 | 0.971827 |
| GOBP_REGULATION_OF_CATION_CHANNEL_ACTIVITY | 135 | 0.285419 | 0.898408 | 0.680435 | 0.971631 |
| GOBP_ORGANELLE_TRANSPORT_ALONG_MICROTUBULE | 83 | 0.301471 | 0.898111 | 0.646007 | 0.97185 |
| GOBP_CELLULAR_RESPONSE_TO_HYDROGEN_PEROXIDE | 70 | 0.306658 | 0.897765 | 0.660171 | 0.972209 |
| GOBP_INTRINSIC_APOPTOTIC_SIGNALING_PATHWAY_IN_RESPONSE_TO_DNA_DAMAG | 94 | 0.300469 | 0.897756 | 0.677201 | 0.971731 |
| GOBP_APICAL_JUNCTION_ASSEMBLY | 58 | 0.315054 | 0.897701 | 0.630086 | 0.971362 |
| GOBP_ACTIVATION_OF_INNATE_IMMUNE_RESPONSE | 40 | 0.336863 | 0.897649 | 0.601299 | 0.97099 |
| GOBP_RESPONSE_TO_HYPEROXIA | 17 | 0.38621 | 0.897471 | 0.59854 | 0.970928 |
| GOBP_GLUTAMATE_SECRETION | 22 | 0.377797 | 0.897349 | 0.587629 | 0.97073 |
| GOBP_SYNAPSE_ORGANIZATION | 369 | 0.270206 | 0.897311 | 0.748485 | 0.970323 |
| GOBP_PEPTIDE_CATABOLIC_PROCESS | 22 | 0.371807 | 0.897239 | 0.631215 | 0.970006 |
| GOBP_APOPTOTIC_MITOCHONDRIAL_CHANGES | 92 | 0.299753 | 0.897015 | 0.663934 | 0.970002 |
| GOBP_PROTEIN_HYDROXYLATION | 25 | 0.362599 | 0.896983 | 0.620112 | 0.969604 |
| GOBP_REGULATION_OF_PROTEIN_ACETYLATION | 76 | 0.30316 | 0.896753 | 0.647551 | 0.969673 |
| GOBP_NEURON_CELL_CELL_ADHESION | 15 | 0.391254 | 0.895903 | 0.609023 | 0.971213 |
| GOBP_MITOCHONDRIAL_DEPOLARIZATION | 17 | 0.391599 | 0.89557 | 0.624818 | 0.971548 |
| GOBP_HEPATICOBILIARY_SYSTEM_DEVELOPMENT | 111 | 0.289952 | 0.895522 | 0.65724 | 0.971177 |
| GOBP_RESPIRATORY_SYSTEM_DEVELOPMENT | 162 | 0.281886 | 0.895319 | 0.717063 | 0.971175 |
| GOBP_DEOXYRIBOSE_PHOSPHATE_CATABOLIC_PROCESS | 26 | 0.358299 | 0.895138 | 0.617403 | 0.971103 |
| GOBP_CALCIUM_ION_REGULATED_EXOCYTOSIS | 59 | 0.317433 | 0.894541 | 0.633971 | 0.972072 |
| GOBP_SYNAPSE_ASSEMBLY | 157 | 0.28111 | 0.894471 | 0.694624 | 0.971749 |
| GOBP_NEGATIVE_REGULATION_OF_ANION_TRANSPORT | 18 | 0.38868 | 0.893774 | 0.608947 | 0.972909 |
| GOBP_REGULATION_OF_SUPEROXIDE_METABOLIC_PROCESS | 23 | 0.365813 | 0.893413 | 0.623955 | 0.973273 |
| GOBP_HORMONE_METABOLIC_PROCESS | 153 | 0.281606 | 0.893396 | 0.695082 | 0.972828 |
| GOBP_NUCLEOTIDE_TRANSPORT | 32 | 0.344888 | 0.893215 | 0.603283 | 0.972758 |
| GOBP_RESPONSE_TO_PH | 33 | 0.342452 | 0.893149 | 0.629139 | 0.972431 |
| GOBP_POST_EMBRYONIC_DEVELOPMENT | 78 | 0.302608 | 0.893137 | 0.658879 | 0.97197 |
| GOBP_CELLULAR_MODIFIED_AMINO_ACID_METABOLIC_PROCESS | 144 | 0.286996 | 0.893079 | 0.691796 | 0.971616 |
| GOBP_MULTICELLULAR_ORGANISMAL_RESPONSE_TO_STRESS | 66 | 0.310655 | 0.892852 | 0.640491 | 0.971644 |
| GOBP_MITOCHONDRIAL_OUTER_MEMBRANE_PERMEABILIZATION_INVOLVED_IN_PROG | 36 | 0.334897 | 0.891113 | 0.629888 | 0.975282 |
| GOBP_RESPONSE_TO_IONIZING_RADIATION | 131 | 0.288144 | 0.89069 | 0.687704 | 0.975778 |
| GOBP_CELL_SUBSTRATE_JUNCTION_ORGANIZATION | 95 | 0.294255 | 0.890603 | 0.676368 | 0.975486 |
| GOBP_REGULATION_OF_ANIMAL_ORGAN_FORMATION | 21 | 0.371974 | 0.890283 | 0.622093 | 0.975778 |
| GOBP_GLUCAN_BIOSYNTHETIC_PROCESS | 36 | 0.337087 | 0.88962 | 0.628758 | 0.976862 |
| GOBP_CYTOCHROME_COMPLEX_ASSEMBLY | 36 | 0.332231 | 0.889492 | 0.622928 | 0.976679 |
| GOBP_REGULATION_OF_HISTONE_DEACETYLATION | 39 | 0.333111 | 0.88914 | 0.636483 | 0.977013 |
| GOBP_REGULATION_OF_GTPASE_ACTIVITY | 312 | 0.270253 | 0.888952 | 0.775385 | 0.97697 |
| GOBP_CELLULAR_METABOLIC_COMPOUND_SALVAGE | 26 | 0.354256 | 0.888418 | 0.622857 | 0.977739 |
| GOBP_B_CELL_HOMEOSTASIS | 27 | 0.358407 | 0.888153 | 0.622407 | 0.977891 |
| GOBP_SNRNA_TRANSCRIPTION_BY_RNA_POLYMERASE_II | 15 | 0.396522 | 0.887886 | 0.61062 | 0.978052 |
| GOBP_CORTICAL_CYTOSKELETON_ORGANIZATION | 58 | 0.316617 | 0.887473 | 0.643127 | 0.978534 |
| GOBP_PYRIMIDINE_CONTAINING_COMPOUND_METABOLIC_PROCESS | 78 | 0.298835 | 0.88745 | 0.677946 | 0.978106 |
| GOBP_MESODERMAL_CELL_DIFFERENTIATION | 22 | 0.368143 | 0.887401 | 0.610072 | 0.977734 |
| GOBP_NEGATIVE_REGULATION_OF_ERBB_SIGNALING_PATHWAY | 31 | 0.348786 | 0.887053 | 0.613889 | 0.978026 |
| GOBP_PROTEIN_HETEROOLIGOMERIZATION | 25 | 0.357907 | 0.886261 | 0.644258 | 0.979323 |
| GOBP_REGULATION_OF_T_CELL_MIGRATION | 28 | 0.351267 | 0.885965 | 0.637584 | 0.97952 |
| GOBP_NEGATIVE_REGULATION_OF_NEURON_APOPTOTIC_PROCESS | 125 | 0.285785 | 0.8857 | 0.716232 | 0.979659 |
| GOBP_SLEEP | 18 | 0.383235 | 0.885364 | 0.633188 | 0.979947 |
| GOBP_EPITHELIAL_TUBE_MORPHOGENESIS | 280 | 0.272494 | 0.885023 | 0.760083 | 0.980255 |
| GOBP_NEGATIVE_REGULATION_OF_OSTEOBLAST_DIFFERENTIATION | 39 | 0.331016 | 0.884025 | 0.657895 | 0.982092 |
| GOBP_POSITIVE_REGULATION_OF_ANTIGEN_RECEPTOR_MEDIATED_SIGNALING_PATH | 18 | 0.382179 | 0.883537 | 0.606695 | 0.982734 |
| GOBP_LONG_CHAIN_FATTY_ACID_METABOLIC_PROCESS | 85 | 0.299698 | 0.883457 | 0.670574 | 0.982434 |
| GOBP_SMALL_GTPASE_MEDIATED_SIGNAL_TRANSDUCTION | 460 | 0.261412 | 0.883122 | 0.813696 | 0.982738 |
| GOBP_NEGATIVE_REGULATION_OF_INTRINSIC_APOPTOTIC_SIGNALING_PATHWAY | 93 | 0.295148 | 0.882933 | 0.661955 | 0.982712 |
| GOBP_SPERMATID_DIFFERENTIATION | 119 | 0.282214 | 0.882803 | 0.704698 | 0.982507 |
| GOBP_DETECTION_OF_MECHANICAL_STIMULUS_INVOLVED_IN_SENSORY_PERCEPTIC | 28 | 0.349506 | 0.882469 | 0.648248 | 0.982784 |
| GOBP_PYRIMIDINE_RIBONUCLEOTIDE_BIOSYNTHETIC_PROCESS | 22 | 0.369013 | 0.882141 | 0.645658 | 0.983013 |
| GOBP_HISTONE_H3_K14_ACETYLATION | 22 | 0.3605 | 0.881725 | 0.600852 | 0.983497 |
| GOBP_POSITIVE_REGULATION_OF_HYDROLASE_ACTIVITY | 467 | 0.262788 | 0.881559 | 0.814329 | 0.983383 |
| GOBP_DNA_SYNTHESIS_INVOLVED_IN_DNA_REPAIR | 34 | 0.341778 | 0.881305 | 0.639148 | 0.983455 |
| GOBP_HISTONE_LYSINE_DEMETHYLATION | 24 | 0.358934 | 0.881196 | 0.620739 | 0.983218 |
| GOBP_NEGATIVE_REGULATION_OF_JNK_CASCADE | 34 | 0.338583 | 0.881149 | 0.637701 | 0.982844 |
| GOBP_NEUROMUSCULAR_JUNCTION_DEVELOPMENT | 38 | 0.332392 | 0.881142 | 0.686992 | 0.982375 |
| GOBP_POSITIVE_REGULATION_OF_PROTEIN_ACETYLATION | 44 | 0.326947 | 0.880802 | 0.664968 | 0.982654 |
| GOBP_RESPONSE_TO_FIBROBLAST_GROWTH_FACTOR | 93 | 0.29076 | 0.880617 | 0.69409 | 0.982553 |
| GOBP_NEGATIVE_REGULATION_OF_PHAGOCYTOSIS | 18 | 0.383601 | 0.880286 | 0.648094 | 0.982824 |
| GOBP_FOREBRAIN_NEURON_DEVELOPMENT | 22 | 0.364143 | 0.88005 | 0.641135 | 0.982845 |
| GOBP_REGULATION_OF_GTP_BINDING | 15 | 0.392256 | 0.879623 | 0.629742 | 0.983324 |
| GOBP_NEGATIVE_REGULATION_OF_HISTONE_METHYLATION | 23 | 0.355556 | 0.879535 | 0.640625 | 0.983041 |
| GOBP_FOCAL_ADHESION_ASSEMBLY | 80 | 0.295407 | 0.879497 | 0.718492 | 0.982652 |
| GOBP_REGULATION_OF_CARDIAC_MUSCLE_CELL_DIFFERENTIATION | 15 | 0.399767 | 0.879414 | 0.630564 | 0.982356 |
| GOBP_CAMP_MEDIATED_SIGNALING | 42 | 0.326977 | 0.878872 | 0.662371 | 0.983113 |
| GOBP_MICROTUBULE_BASED_MOVEMENT | 320 | 0.267046 | 0.878578 | 0.776758 | 0.983315 |
| GOBP_RAB_PROTEIN_SIGNAL_TRANSDUCTION | 21 | 0.367917 | 0.877568 | 0.631579 | 0.985127 |
| GOBP_GLYCOPROTEIN_CATABOLIC_PROCESS | 23 | 0.362575 | 0.877556 | 0.632514 | 0.984675 |
| GOBP_REGULATION_OF_MUSCLE_ORGAN_DEVELOPMENT | 32 | 0.337285 | 0.876783 | 0.647757 | 0.985886 |
| GOBP_RESPONSE_TO_DEXAMETHASONE | 33 | 0.343457 | 0.876725 | 0.652759 | 0.985533 |
| GOBP_NEGATIVE_REGULATION_OF_DNA_BINDING_TRANSCRIPTION_FACTOR_ACTIVITY | 146 | 0.277919 | 0.876656 | 0.733696 | 0.985204 |
| GOBP_NUCLEAR_CHROMOSOME_SEGREGATION | 259 | 0.269146 | 0.876205 | 0.765869 | 0.985757 |
| GOBP_MRNA_CIS_SPLICING_VIA_SPLICEOSOME | 25 | 0.353831 | 0.876167 | 0.629371 | 0.985362 |
| GOBP_CELL_CELL_JUNCTION_ORGANIZATION | 172 | 0.274566 | 0.875229 | 0.732899 | 0.98695 |
| GOBP_POSITIVE_REGULATION_OF_DNA_REPAIR | 91 | 0.292388 | 0.874815 | 0.687131 | 0.987396 |
| GOBP_CEREBRAL_CORTEX_DEVELOPMENT | 99 | 0.289531 | 0.874538 | 0.700113 | 0.987552 |
| GOBP_REGULATION_OF_AXONOGENESIS | 148 | 0.278395 | 0.874513 | 0.741512 | 0.987122 |
| GOBP_REGULATION_OF_PROTEIN_LOCALIZATION_TO_PLASMA_MEMBRANE | 92 | 0.29346 | 0.874479 | 0.707892 | 0.986722 |
| GOBP_PLASMA_LIPOPROTEIN_PARTICLE_CLEARANCE | 28 | 0.345536 | 0.87367 | 0.639161 | 0.988047 |
| GOBP_NEGATIVE_REGULATION_OF_RNA_CATABOLIC_PROCESS | 67 | 0.302452 | 0.873639 | 0.694444 | 0.987646 |
| GOBP_BRANCHED_CHAIN_AMINO_ACID_METABOLIC_PROCESS | 22 | 0.359405 | 0.872759 | 0.637394 | 0.989148 |
| GOBP_DOPAMINE_METABOLIC_PROCESS | 33 | 0.332903 | 0.871734 | 0.661705 | 0.990993 |
| GOBP_REGULATION_OF_INTRACELLULAR_PROTEIN_TRANSPORT | 207 | 0.271504 | 0.871321 | 0.754517 | 0.99146 |
| GOBP_POSITIVE_REGULATION_OF_CATION_CHANNEL_ACTIVITY | 57 | 0.304485 | 0.871095 | 0.679847 | 0.991488 |
| GOBP_REGULATION_OF_CELL_SHAPE | 133 | 0.279496 | 0.871073 | 0.745055 | 0.991051 |
| GOBP_DEFENSE_RESPONSE_TO_GRAM_POSITIVE_BACTERIUM | 45 | 0.316873 | 0.870947 | 0.670455 | 0.990838 |
| GOBP_RAC_PROTEIN_SIGNAL_TRANSDUCTION | 41 | 0.328345 | 0.870804 | 0.657895 | 0.990679 |
| GOBP_REGULATION_OF_CAMP_DEPENDENT_PROTEIN_KINASE_ACTIVITY | 18 | 0.372598 | 0.870711 | 0.617778 | 0.990391 |
| GOBP_EMBRYONIC_PLACENTA_DEVELOPMENT | 78 | 0.292635 | 0.870647 | 0.694146 | 0.990046 |
| GOBP_RESPONSE_TO_FLUID_SHEAR_STRESS | 29 | 0.34201 | 0.870147 | 0.666197 | 0.990659 |
| GOBP_CELLULAR_RESPONSE_TO_REACTIVE_NITROGEN_SPECIES | 15 | 0.381203 | 0.869858 | 0.637298 | 0.990809 |
| GOBP_REGULATION_OF_MRNA_CATABOLIC_PROCESS | 156 | 0.273407 | 0.869511 | 0.751357 | 0.991097 |
| GOBP_POSITIVE_REGULATION_OF_MITOTIC_NUCLEAR_DIVISION | 30 | 0.33874 | 0.86919 | 0.654939 | 0.991322 |
| GOBP_HISTONE_H3_K27_METHYLATION | 16 | 0.386645 | 0.868906 | 0.661218 | 0.99148 |
| GOBP_TRANSEPITHELIAL_TRANSPORT | 28 | 0.344249 | 0.868872 | 0.652482 | 0.991072 |
| GOBP_REGULATION_OF_CARDIAC_MUSCLE_CELL_ACTION_POTENTIAL | 25 | 0.346012 | 0.868281 | 0.649789 | 0.991967 |
| GOBP_REGULATION_OF_GENE_SILENCING_BY_RNA | 44 | 0.312387 | 0.866271 | 0.663212 | 0.995862 |
| GOBP_HOMOPHILIC_CELL_ADHESION_VIA_PLASMA_MEMBRANE_ADHESION_MOLECUL | 143 | 0.278038 | 0.865576 | 0.753019 | 0.996936 |
| GOBP_NEGATIVE_REGULATION_OF_CELL_CYCLE_PROCESS | 261 | 0.263274 | 0.864916 | 0.785417 | 0.997894 |
| GOBP_RESPONSE_TO_LIGHT_STIMULUS | 253 | 0.262993 | 0.864904 | 0.785047 | 0.997437 |
| GOBP_POSTTRANSCRIPTIONAL_REGULATION_OF_GENE_EXPRESSION | 491 | 0.258286 | 0.86469 | 0.86996 | 0.997413 |
| GOBP_POSITIVE_REGULATION_OF_CALCIUM_MEDIATED_SIGNALING | 22 | 0.358201 | 0.864549 | 0.64592 | 0.997226 |
| GOBP_PHOTORECEPTOR_CELL_DIFFERENTIATION | 48 | 0.310115 | 0.86361 | 0.673004 | 0.998803 |
| GOBP_NEGATIVE_REGULATION_OF_CELLULAR_SENESCENCE | 15 | 0.385788 | 0.863538 | 0.621329 | 0.998491 |
| GOBP_MITOCHONDRIAL_FUSION | 30 | 0.340866 | 0.862418 | 0.675134 | 1 |
| GOBP_GASTRULATION_WITH_MOUTH_FORMING_SECOND | 25 | 0.340541 | 0.862173 | 0.643454 | 1 |
| GOBP_RNA_METHYLATION | 82 | 0.292951 | 0.862065 | 0.709869 | 1 |
| GOBP_REGULATION_OF_VASOCONSTRICTION | 48 | 0.31364 | 0.862064 | 0.682603 | 0.999712 |
| GOBP_CELL_DIFFERENTIATION_INVOLVED_IN_KIDNEY_DEVELOPMENT | 48 | 0.310349 | 0.86189 | 0.679293 | 0.999616 |
| GOBP_RESPONSE_TO_COLD | 45 | 0.317199 | 0.861631 | 0.683354 | 0.999716 |
| GOBP_REGULATION_OF_NUCLEOCYTOPLASMIC_TRANSPORT | 97 | 0.283384 | 0.861367 | 0.716186 | 0.999808 |
| GOBP_DENDRITE_EXTENSION | 33 | 0.332966 | 0.861363 | 0.672775 | 0.999337 |
| GOBP_EMBRYONIC_SKELETAL_SYSTEM_DEVELOPMENT | 81 | 0.294068 | 0.860909 | 0.712264 | 0.999839 |
| GOBP_EMBRYONIC_DIGESTIVE_TRACT_DEVELOPMENT | 22 | 0.354643 | 0.860805 | 0.647143 | 0.999576 |
| GOBP_SIGNALING_RECEPTOR_LIGAND_PRECURSOR_PROCESSING | 26 | 0.345654 | 0.860748 | 0.657895 | 0.999232 |
| GOBP_POSITIVE_REGULATION_OF_BLOOD_CIRCULATION | 34 | 0.327929 | 0.860592 | 0.661438 | 0.999101 |
| GOBP_CELL_DEATH_IN_RESPONSE_TO_HYDROGEN_PEROXIDE | 22 | 0.357659 | 0.860321 | 0.677871 | 0.999223 |
| GOBP_REGULATION_OF_NEUROTRANSMITTER_RECEPTOR_ACTIVITY | 50 | 0.308975 | 0.86005 | 0.669261 | 0.999338 |
| GOBP_POSITIVE_REGULATION_OF_CELLULAR_PROTEIN_CATABOLIC_PROCESS | 146 | 0.274535 | 0.859702 | 0.752809 | 0.99957 |
| GOBP_SYNAPTIC_VESICLE_RECYCLING | 67 | 0.297701 | 0.859687 | 0.709001 | 0.999122 |
| GOBP_AXONAL_FASCICULATION | 21 | 0.364278 | 0.858462 | 0.667643 | 1 |
| GOBP_NUCLEOBASE_METABOLIC_PROCESS | 37 | 0.326125 | 0.858243 | 0.666218 | 1 |
| GOBP_POSITIVE_REGULATION_OF_CELL_CYCLE_G2_M_PHASE_TRANSITION | 29 | 0.336744 | 0.857185 | 0.654219 | 1 |
| GOBP_CELLULAR_AMINO_ACID_CATABOLIC_PROCESS | 87 | 0.287625 | 0.857085 | 0.717073 | 1 |
| GOBP_POSITIVE_REGULATION_OF_CELL_CYCLE_G1_S_PHASE_TRANSITION | 50 | 0.312711 | 0.856623 | 0.694696 | 1 |
| GOBP_NEGATIVE_REGULATION_OF_RECEPTOR_SIGNALING_PATHWAY_VIA_JAK_STAT | 20 | 0.365186 | 0.856555 | 0.653736 | 1 |
| GOBP_INSULIN_SECRETION_INVOLVED_IN_CELLULAR_RESPONSE_TO_GLUCOSE_STII | 53 | 0.310755 | 0.856462 | 0.70469 | 1 |
| GOBP_CELL_REDOX_HOMEOSTASIS | 34 | 0.327365 | 0.856408 | 0.689333 | 1 |
| GOBP_AMINO_ACID_TRANSPORT | 122 | 0.279815 | 0.856086 | 0.760135 | 1 |
| GOBP_AXO_DENDRITIC_TRANSPORT | 76 | 0.289422 | 0.855785 | 0.726415 | 1 |
| GOBP_POSITIVE_REGULATION_OF_CELL_CYCLE_PROCESS | 197 | 0.264162 | 0.85552 | 0.772967 | 1 |
| GOBP_REGULATION_OF_GLYCOPROTEIN_METABOLIC_PROCESS | 40 | 0.317139 | 0.855201 | 0.688172 | 1 |
| GOBP_MRNA_SPLICE_SITE_SELECTION | 32 | 0.332141 | 0.855096 | 0.662534 | 1 |
| GOBP_CELL_CYCLE_CHECKPOINT_SIGNALING | 164 | 0.268935 | 0.85473 | 0.776709 | 1 |
| GOBP_CELLULAR_RESPONSE_TO_DEXAMETHASONE_STIMULUS | 20 | 0.362217 | 0.853853 | 0.65896 | 1 |
| GOBP_REGULATION_OF_INTRACELLULAR_TRANSPORT | 305 | 0.26124 | 0.853621 | 0.835041 | 1 |
| GOBP_CERAMIDE_BIOSYNTHETIC_PROCESS | 60 | 0.298233 | 0.85338 | 0.702309 | 1 |
| GOBP_NEGATIVE_REGULATION_OF_CHONDROCYTE_DIFFERENTIATION | 18 | 0.365345 | 0.853344 | 0.641921 | 1 |
| GOBP_SECRETORY_GRANULE_ORGANIZATION | 29 | 0.335843 | 0.85313 | 0.6751 | 1 |
| GOBP_NEGATIVE_REGULATION_OF_WNT_SIGNALING_PATHWAY | 153 | 0.271284 | 0.852489 | 0.781182 | 1 |
| GOBP_NEGATIVE_REGULATION_OF_VIRAL_ENTRY_INTO_HOST_CELL | 18 | 0.372734 | 0.852333 | 0.679412 | 1 |
| GOBP_RESPONSE_TO_THYROID_HORMONE | 23 | 0.342359 | 0.851701 | 0.658708 | 1 |

| | | | | | |
|---|---|---|---|---|---|
| GOBP_POSITIVE_REGULATION_OF_NERVOUS_SYSTEM_DEVELOPMENT | 232 | 0.260212 | 0.850974 | 0.797706 | 1 |
| GOBP_NEGATIVE_REGULATION_OF_PROTEIN_CONTAINING_COMPLEX_DISASSEMBLY | 76 | 0.289395 | 0.850914 | 0.715152 | 1 |
| GOBP_NEGATIVE_REGULATION_OF_ACTIN_FILAMENT_POLYMERIZATION | 58 | 0.301393 | 0.850408 | 0.716584 | 1 |
| GOBP_IMPORT_INTO_NUCLEUS | 145 | 0.269322 | 0.849978 | 0.775751 | 1 |
| GOBP_CARBOHYDRATE_HOMEOSTASIS | 184 | 0.263816 | 0.849748 | 0.800215 | 1 |
| GOBP_CELLULAR_RESPONSE_TO_UV | 81 | 0.288951 | 0.849145 | 0.741218 | 1 |
| GOBP_POSITIVE_REGULATION_OF_RECEPTOR_MEDIATED_ENDOCYTOSIS | 45 | 0.317887 | 0.848999 | 0.707654 | 1 |
| GOBP_CELLULAR_RESPONSE_TO_RADIATION | 156 | 0.267022 | 0.848931 | 0.768489 | 1 |
| GOBP_LINOLEIC_ACID_METABOLIC_PROCESS | 17 | 0.370076 | 0.848855 | 0.669553 | 1 |
| GOBP_MAGNESIUM_ION_TRANSPORT | 16 | 0.37345 | 0.847926 | 0.674141 | 1 |
| GOBP_NEURON_DEATH_IN_RESPONSE_TO_OXIDATIVE_STRESS | 27 | 0.343224 | 0.847855 | 0.679831 | 1 |
| GOBP_NEGATIVE_REGULATION_OF_PLASMA_MEMBRANE_BOUNDED_CELL_PROJECTI | 31 | 0.32914 | 0.847733 | 0.695711 | 1 |
| GOBP_PORPHYRIN_CONTAINING_COMPOUND_METABOLIC_PROCESS | 42 | 0.309055 | 0.847507 | 0.719481 | 1 |
| GOBP_CELLULAR_RESPONSE_TO_IONIZING_RADIATION | 69 | 0.290408 | 0.846255 | 0.744604 | 1 |
| GOBP_MYELOID_DENDRITIC_CELL_ACTIVATION | 18 | 0.362458 | 0.845606 | 0.656433 | 1 |
| GOBP_POSITIVE_REGULATION_OF_PROTEIN_LOCALIZATION_TO_CELL_PERIPHERY | 60 | 0.296889 | 0.845483 | 0.724638 | 1 |
| GOBP_REGULATION_OF_LONG_TERM_SYNAPTIC_POTENTIATION | 39 | 0.319997 | 0.845072 | 0.706037 | 1 |
| GOBP_NEGATIVE_REGULATION_OF_CELL_CYCLE_PHASE_TRANSITION | 221 | 0.260311 | 0.845009 | 0.817419 | 1 |
| GOBP_NEGATIVE_REGULATION_OF_ACTIN_FILAMENT_DEPOLYMERIZATION | 41 | 0.314556 | 0.844675 | 0.70428 | 1 |
| GOBP_CELLULAR_GLUCOSE_HOMEOSTASIS | 122 | 0.272924 | 0.844383 | 0.766334 | 1 |
| GOBP_RESPONSE_TO_GAMMA_RADIATION | 51 | 0.305848 | 0.843314 | 0.721805 | 1 |
| GOBP_EYE_PHOTORECEPTOR_CELL_DIFFERENTIATION | 35 | 0.317707 | 0.842661 | 0.677333 | 1 |
| GOBP_NUCLEOSIDE_MONOPHOSPHATE_METABOLIC_PROCESS | 72 | 0.293793 | 0.842374 | 0.717445 | 1 |
| GOBP_GLUTATHIONE_METABOLIC_PROCESS | 46 | 0.309942 | 0.841967 | 0.720513 | 1 |
| GOBP_CIRCADIAN_REGULATION_OF_GENE_EXPRESSION | 63 | 0.293629 | 0.841398 | 0.738007 | 1 |
| GOBP_CELL_CELL_ADHESION_MEDIATED_BY_CADHERIN | 25 | 0.341533 | 0.841249 | 0.668532 | 1 |
| GOBP_REGULATION_OF_INTRINSIC_APOPTOTIC_SIGNALING_PATHWAY_IN_RESPONSE | 38 | 0.316778 | 0.841157 | 0.703851 | 1 |
| GOBP_ELECTRON_TRANSPORT_CHAIN | 150 | 0.270202 | 0.840703 | 0.785169 | 1 |
| GOBP_METANEPHRIC_EPITHELIUM_DEVELOPMENT | 16 | 0.371845 | 0.840677 | 0.692654 | 1 |
| GOBP_POSITIVE_REGULATION_OF_CELL_PROJECTION_ORGANIZATION | 309 | 0.255707 | 0.840645 | 0.840535 | 1 |
| GOBP_POSITIVE_REGULATION_OF_DEPHOSPHORYLATION | 52 | 0.302903 | 0.840275 | 0.716749 | 1 |
| GOBP_REGULATION_OF_GLIOGENESIS | 75 | 0.285473 | 0.840248 | 0.737226 | 1 |
| GOBP_REGULATION_OF_INTRINSIC_APOPTOTIC_SIGNALING_PATHWAY | 152 | 0.266393 | 0.839812 | 0.801724 | 1 |
| GOBP_NATURAL_KILLER_CELL_ACTIVATION | 54 | 0.29893 | 0.839436 | 0.728167 | 1 |
| GOBP_CELLULAR_RESPONSE_TO_LIGHT_STIMULUS | 95 | 0.276707 | 0.839278 | 0.75174 | 1 |
| GOBP_CYTOPLASMIC_PATTERN_RECOGNITION_RECEPTOR_SIGNALING_PATHWAY | 54 | 0.300842 | 0.838572 | 0.72705 | 1 |
| GOBP_POSITIVE_T_CELL_SELECTION | 20 | 0.356053 | 0.838517 | 0.688953 | 1 |
| GOBP_CELL_CYCLE_G1_S_PHASE_TRANSITION | 208 | 0.259328 | 0.838517 | 0.811975 | 1 |
| GOBP_REGULATION_OF_MEMBRANE_DEPOLARIZATION | 37 | 0.319748 | 0.838366 | 0.685452 | 1 |
| GOBP_NEGATIVE_REGULATION_OF_CELL_DIVISION | 15 | 0.376918 | 0.838058 | 0.665649 | 1 |
| GOBP_RESPONSE_TO_X_RAY | 30 | 0.325975 | 0.838011 | 0.692913 | 1 |
| GOBP_NEGATIVE_REGULATION_OF_BINDING | 137 | 0.267997 | 0.837788 | 0.798441 | 1 |
| GOBP_INTRINSIC_APOPTOTIC_SIGNALING_PATHWAY | 267 | 0.256293 | 0.837614 | 0.834532 | 1 |
| GOBP_REGULATION_OF_MAP_KINASE_ACTIVITY | 157 | 0.266556 | 0.837505 | 0.806067 | 1 |
| GOBP_POSITIVE_REGULATION_OF_STEM_CELL_DIFFERENTIATION | 15 | 0.375531 | 0.837408 | 0.69771 | 1 |
| GOBP_MITOCHONDRIAL_CYTOCHROME_C_OXIDASE_ASSEMBLY | 22 | 0.34785 | 0.837127 | 0.712644 | 1 |
| GOBP_REGULATION_OF_MITOCHONDRIAL_MEMBRANE_POTENTIAL | 62 | 0.293057 | 0.837003 | 0.733922 | 1 |
| GOBP_METANEPHRIC_TUBULE_DEVELOPMENT | 15 | 0.37686 | 0.836882 | 0.660661 | 1 |
| GOBP_HISTONE_MODIFICATION | 459 | 0.251651 | 0.836848 | 0.90091 | 1 |
| GOBP_LYMPHOCYTE_MIGRATION | 63 | 0.296587 | 0.836826 | 0.723247 | 1 |
| GOBP_CELLULAR_RESPONSE_TO_HORMONE_STIMULUS | 471 | 0.250109 | 0.835548 | 0.894472 | 1 |
| GOBP_POSITIVE_REGULATION_OF_OXIDATIVE_STRESS_INDUCED_CELL_DEATH | 15 | 0.374267 | 0.835236 | 0.699259 | 1 |
| GOBP_DEVELOPMENTAL_CELL_GROWTH | 201 | 0.257552 | 0.834992 | 0.822176 | 1 |
| GOBP_REGULATION_OF_OLIGODENDROCYTE_DIFFERENTIATION | 31 | 0.324953 | 0.834869 | 0.687254 | 1 |
| GOBP_ASSOCIATIVE_LEARNING | 69 | 0.284918 | 0.834337 | 0.751834 | 1 |
| GOBP_BICARBONATE_TRANSPORT | 21 | 0.348601 | 0.834235 | 0.682584 | 1 |
| GOBP_POSITIVE_REGULATION_OF_CREB_TRANSCRIPTION_FACTOR_ACTIVITY | 18 | 0.363803 | 0.8342 | 0.70015 | 1 |
| GOBP_TERPENOID_METABOLIC_PROCESS | 60 | 0.29032 | 0.834152 | 0.750925 | 1 |
| GOBP_NEGATIVE_REGULATION_OF_OSSIFICATION | 26 | 0.33627 | 0.834146 | 0.716956 | 1 |
| GOBP_EXCRETION | 44 | 0.307733 | 0.833746 | 0.72 | 1 |
| GOBP_POSITIVE_REGULATION_OF_NEURON_APOPTOTIC_PROCESS | 41 | 0.305262 | 0.833733 | 0.702128 | 1 |
| GOBP_REGULATION_OF_TUMOR_NECROSIS_FACTOR_MEDIATED_SIGNALING_PATHWA | 46 | 0.302607 | 0.833672 | 0.746497 | 1 |
| GOBP_TETRAPYRROLE_METABOLIC_PROCESS | 52 | 0.302565 | 0.833662 | 0.735404 | 1 |
| GOBP_GLYCEROPHOSPHOLIPID_BIOSYNTHETIC_PROCESS | 195 | 0.258959 | 0.832583 | 0.829787 | 1 |
| GOBP_RESPONSE_TO_MANGANESE_ION | 17 | 0.366554 | 0.832567 | 0.701807 | 1 |
| GOBP_REGULATION_OF_GLUCAN_BIOSYNTHETIC_PROCESS | 23 | 0.342185 | 0.832313 | 0.705964 | 1 |
| GOBP_REGULATION_OF_NERVOUS_SYSTEM_DEVELOPMENT | 375 | 0.249639 | 0.832311 | 0.883603 | 1 |
| GOBP_CARBOHYDRATE_TRANSPORT | 116 | 0.274544 | 0.832288 | 0.79865 | 1 |
| GOBP_POSITIVE_REGULATION_OF_JUN_KINASE_ACTIVITY | 37 | 0.321551 | 0.832171 | 0.717277 | 1 |
| GOBP_CELLULAR_RESPONSE_TO_GAMMA_RADIATION | 28 | 0.327126 | 0.832108 | 0.696133 | 1 |
| GOBP_POSITIVE_REGULATION_OF_MITOTIC_CELL_CYCLE_PHASE_TRANSITION | 82 | 0.280824 | 0.83209 | 0.78184 | 1 |
| GOBP_REGULATION_OF_PROTEIN_CONTAINING_COMPLEX_DISASSEMBLY | 116 | 0.271942 | 0.832082 | 0.793065 | 1 |
| GOBP_NEGATIVE_REGULATION_OF_PROTEIN_MODIFICATION_PROCESS | 456 | 0.248903 | 0.831964 | 0.900101 | 1 |
| GOBP_POSITIVE_REGULATION_OF_NERVOUS_SYSTEM_PROCESS | 28 | 0.329014 | 0.831851 | 0.712925 | 1 |
| GOBP_RESPONSE_TO_NERVE_GROWTH_FACTOR | 45 | 0.302786 | 0.83143 | 0.726463 | 1 |
| GOBP_NEURON_APOPTOTIC_PROCESS | 207 | 0.259751 | 0.830743 | 0.832258 | 1 |
| GOBP_REGULATION_OF_MITOTIC_CELL_CYCLE_PHASE_TRANSITION | 275 | 0.255414 | 0.829928 | 0.860996 | 1 |
| GOBP_MAINTENANCE_OF_PROTEIN_LOCATION_IN_NUCLEUS | 23 | 0.336869 | 0.829676 | 0.699029 | 1 |
| GOBP_REGULATION_OF_SUBSTRATE_ADHESION_DEPENDENT_CELL_SPREADING | 51 | 0.299173 | 0.829332 | 0.743462 | 1 |
| GOBP_ALDITOL_METABOLIC_PROCESS | 15 | 0.371075 | 0.828623 | 0.703297 | 1 |
| GOBP_PHOSPHATIDYLINOSITOL_DEPHOSPHORYLATION | 29 | 0.324827 | 0.82848 | 0.712162 | 1 |
| GOBP_MITOCHONDRIAL_GENOME_MAINTENANCE | 23 | 0.341492 | 0.827572 | 0.697222 | 1 |
| GOBP_TRANSMISSION_OF_NERVE_IMPULSE | 53 | 0.291691 | 0.827343 | 0.737172 | 1 |
| GOBP_STEROL_METABOLIC_PROCESS | 117 | 0.268875 | 0.82723 | 0.777156 | 1 |
| GOBP_EPITHELIAL_CELL_MORPHOGENESIS | 30 | 0.321527 | 0.827006 | 0.721245 | 1 |
| GOBP_NEGATIVE_REGULATION_OF_SMALL_MOLECULE_METABOLIC_PROCESS | 79 | 0.277727 | 0.826903 | 0.766628 | 1 |
| GOBP_ACTIVATION_OF_PHOSPHOLIPASE_C_ACTIVITY | 18 | 0.359822 | 0.826483 | 0.695588 | 1 |
| GOBP_REGULATION_OF_CATECHOLAMINE_METABOLIC_PROCESS | 19 | 0.342465 | 0.825551 | 0.693642 | 1 |
| GOBP_HYPEROSMOTIC_RESPONSE | 23 | 0.338589 | 0.825261 | 0.71049 | 1 |
| GOBP_RESPONSE_TO_ACETYLCHOLINE | 26 | 0.332436 | 0.825008 | 0.716788 | 1 |
| GOBP_VASCULAR_ASSOCIATED_SMOOTH_MUSCLE_CELL_DIFFERENTIATION | 23 | 0.342951 | 0.824866 | 0.722689 | 1 |
| GOBP_REGULATION_OF_GLUTAMATE_SECRETION | 17 | 0.363206 | 0.824626 | 0.682403 | 1 |
| GOBP_SECONDARY_PALATE_DEVELOPMENT | 22 | 0.339286 | 0.823931 | 0.707627 | 1 |
| GOBP_POSITIVE_REGULATION_OF_CELL_GROWTH | 142 | 0.260098 | 0.823653 | 0.817181 | 1 |
| GOBP_NUCLEOBASE_CONTAINING_SMALL_MOLECULE_METABOLIC_PROCESS | 484 | 0.247206 | 0.823379 | 0.918675 | 1 |
| GOBP_POSITIVE_REGULATION_OF_AUTOPHAGY | 127 | 0.267214 | 0.823103 | 0.799544 | 1 |
| GOBP_MATURE_B_CELL_DIFFERENTIATION | 22 | 0.343268 | 0.823026 | 0.721826 | 1 |
| GOBP_REGULATION_OF_CYCLASE_ACTIVITY | 42 | 0.302685 | 0.822237 | 0.727273 | 1 |
| GOBP_AROMATIC_AMINO_ACID_FAMILY_METABOLIC_PROCESS | 16 | 0.360951 | 0.822188 | 0.707646 | 1 |
| GOBP_MEMBRANE_INVAGINATION | 59 | 0.286622 | 0.822072 | 0.7597 | 1 |
| GOBP_NEGATIVE_REGULATION_OF_COLD_INDUCED_THERMOGENESIS | 40 | 0.309386 | 0.822028 | 0.732543 | 1 |
| GOBP_EXOCYTIC_PROCESS | 68 | 0.284321 | 0.821956 | 0.761194 | 1 |
| GOBP_LYMPHOCYTE_CHEMOTAXIS | 23 | 0.338236 | 0.821922 | 0.704225 | 1 |
| GOBP_NEGATIVE_REGULATION_OF_SYNAPTIC_TRANSMISSION | 59 | 0.292715 | 0.821453 | 0.772613 | 1 |
| GOBP_REGULATION_OF_MICROTUBULE_POLYMERIZATION | 53 | 0.293388 | 0.821349 | 0.743363 | 1 |
| GOBP_REGULATION_OF_SIGNAL_TRANSDUCTION_BY_P53_CLASS_MEDIATOR | 99 | 0.269569 | 0.820396 | 0.779838 | 1 |
| GOBP_REGULATION_OF_CELL_SUBSTRATE_JUNCTION_ORGANIZATION | 65 | 0.283651 | 0.820313 | 0.753659 | 1 |
| GOBP_PEPTIDYL_LYSINE_MODIFICATION | 364 | 0.249488 | 0.819993 | 0.911675 | 1 |
| GOBP_ORGANIC_ACID_CATABOLIC_PROCESS | 196 | 0.256908 | 0.819968 | 0.834728 | 1 |
| GOBP_CHROMOSOME_SEGREGATION | 317 | 0.246847 | 0.819749 | 0.880165 | 1 |
| GOBP_NEUTRAL_AMINO_ACID_TRANSPORT | 38 | 0.307041 | 0.819709 | 0.723958 | 1 |
| GOBP_REGULATION_OF_PROTEIN_LOCALIZATION_TO_CELL_PERIPHERY | 111 | 0.265985 | 0.819683 | 0.797282 | 1 |
| GOBP_DEPHOSPHORYLATION | 372 | 0.247107 | 0.819647 | 0.900305 | 1 |
| GOBP_REGULATION_OF_PROTEIN_POLYUBIQUITINATION | 25 | 0.327172 | 0.818463 | 0.712644 | 1 |
| GOBP_GLUCOCORTICOID_METABOLIC_PROCESS | 16 | 0.363073 | 0.818313 | 0.700447 | 1 |
| GOBP_ORGANOPHOSPHATE_BIOSYNTHETIC_PROCESS | 490 | 0.246981 | 0.81824 | 0.930653 | 1 |
| GOBP_REGULATION_OF_MICROTUBULE_DEPOLYMERIZATION | 31 | 0.318586 | 0.818218 | 0.716644 | 1 |
| GOBP_POSITIVE_REGULATION_OF_MUSCLE_CELL_APOPTOTIC_PROCESS | 19 | 0.342595 | 0.818205 | 0.717986 | 1 |
| GOBP_POSITIVE_REGULATION_OF_INTRACELLULAR_PROTEIN_TRANSPORT | 145 | 0.259849 | 0.818078 | 0.804324 | 1 |
| GOBP_REGULATION_OF_MICROTUBULE_BASED_PROCESS | 228 | 0.250504 | 0.817805 | 0.856243 | 1 |
| GOBP_PROTEIN_O_LINKED_GLYCOSYLATION | 81 | 0.27397 | 0.817608 | 0.775029 | 1 |
| GOBP_RECEPTOR_CATABOLIC_PROCESS | 27 | 0.330946 | 0.817149 | 0.735294 | 1 |
| GOBP_ISOPRENOID_METABOLIC_PROCESS | 77 | 0.27892 | 0.81711 | 0.780723 | 1 |
| GOBP_COCHLEA_MORPHOGENESIS | 17 | 0.358419 | 0.815047 | 0.71386 | 1 |
| GOBP_MACROMOLECULE_DEACYLATION | 125 | 0.260905 | 0.814532 | 0.812709 | 1 |
| GOBP_PURINE_NUCLEOSIDE_MONOPHOSPHATE_METABOLIC_PROCESS | 43 | 0.302282 | 0.814381 | 0.757813 | 1 |
| GOBP_REGULATION_OF_SYNAPTIC_VESICLE_EXOCYTOSIS | 47 | 0.294636 | 0.814274 | 0.757069 | 1 |
| GOBP_REGULATION_OF_BINDING | 320 | 0.248545 | 0.813862 | 0.89126 | 1 |
| GOBP_RESPONSE_TO_UV | 133 | 0.263629 | 0.81365 | 0.829508 | 1 |
| GOBP_CELLULAR_RESPIRATION | 210 | 0.252295 | 0.813512 | 0.845026 | 1 |
| GOBP_POSITIVE_REGULATION_OF_BINDING | 150 | 0.260797 | 0.813376 | 0.83446 | 1 |
| GOBP_CARDIAC_MUSCLE_CELL_ACTION_POTENTIAL | 65 | 0.279266 | 0.813141 | 0.780576 | 1 |
| GOBP_VESICLE_CYTOSKELETAL_TRAFFICKING | 64 | 0.284736 | 0.813094 | 0.779394 | 1 |
| GOBP_REGULATION_OF_INSULIN_RECEPTOR_SIGNALING_PATHWAY | 51 | 0.293271 | 0.812906 | 0.760248 | 1 |
| GOBP_GLUTAMINE_FAMILY_AMINO_ACID_METABOLIC_PROCESS | 67 | 0.27983 | 0.81285 | 0.773171 | 1 |
| GOBP_OLIGODENDROCYTE_DIFFERENTIATION | 77 | 0.278273 | 0.812833 | 0.762077 | 1 |
| GOBP_MULTICELLULAR_ORGANISM_AGING | 34 | 0.31241 | 0.812314 | 0.726908 | 1 |
| GOBP_NEGATIVE_REGULATION_OF_PROTEIN_CATABOLIC_PROCESS | 112 | 0.266674 | 0.812036 | 0.805936 | 1 |
| GOBP_RESPIRATORY_GASEOUS_EXCHANGE_BY_RESPIRATORY_SYSTEM | 56 | 0.289605 | 0.812002 | 0.7711 | 1 |
| GOBP_REGULATION_OF_GENERATION_OF_PRECURSOR_METABOLITES_AND_ENERGY | 113 | 0.264147 | 0.811216 | 0.827119 | 1 |
| GOBP_CERAMIDE_CATABOLIC_PROCESS | 18 | 0.351496 | 0.811169 | 0.733824 | 1 |
| GOBP_POSITIVE_REGULATION_OF_MITOTIC_CELL_CYCLE | 108 | 0.268831 | 0.811146 | 0.815789 | 1 |
| GOBP_ADENYLATE_CYCLASE_INHIBITING_G_PROTEIN_COUPLED_RECEPTOR_SIGNALI | 55 | 0.289392 | 0.811027 | 0.739777 | 1 |
| GOBP_EMBRYONIC_DIGIT_MORPHOGENESIS | 45 | 0.295811 | 0.810226 | 0.757106 | 1 |
| GOBP_REGULATION_OF_GLIAL_CELL_PROLIFERATION | 21 | 0.332173 | 0.809185 | 0.707736 | 1 |
| GOBP_POSITIVE_REGULATION_OF_INTRINSIC_APOPTOTIC_SIGNALING_PATHWAY | 52 | 0.288246 | 0.809064 | 0.764339 | 1 |
| GOBP_STEROL_HOMEOSTASIS | 62 | 0.279375 | 0.808475 | 0.754017 | 1 |
| GOBP_PROTEIN_DEMETHYLATION | 30 | 0.315729 | 0.808298 | 0.746649 | 1 |
| GOBP_REGULATION_OF_DEPHOSPHORYLATION | 111 | 0.268348 | 0.808286 | 0.815523 | 1 |
| GOBP_POSITIVE_REGULATION_OF_AMINO_ACID_TRANSPORT | 19 | 0.343587 | 0.808129 | 0.721739 | 1 |
| GOBP_PHOSPHOLIPID_BIOSYNTHETIC_PROCESS | 232 | 0.248865 | 0.807224 | 0.880802 | 1 |

| | | | | | |
|---|---|---|---|---|---|
| GOBP_ENGULFMENT_OF_APOPTOTIC_CELL | 17 | 0.353816 | 0.807074 | 0.716788 | 1 |
| GOBP_RRNA_METHYLATION | 25 | 0.330408 | 0.807027 | 0.731302 | 1 |
| GOBP_POSITIVE_REGULATION_OF_CELL_DIVISION | 60 | 0.285535 | 0.806892 | 0.782716 | 1 |
| GOBP_PHOSPHATIDYLINOSITOL_METABOLIC_PROCESS | 145 | 0.254005 | 0.806569 | 0.830601 | 1 |
| GOBP_LIPID_OXIDATION | 96 | 0.264056 | 0.806245 | 0.799771 | 1 |
| GOBP_NUCLEOSIDE_PHOSPHATE_CATABOLIC_PROCESS | 74 | 0.272924 | 0.805805 | 0.795372 | 1 |
| GOBP_ORGANELLE_LOCALIZATION | 488 | 0.241097 | 0.805804 | 0.930514 | 1 |
| GOBP_MICROVILLUS_ORGANIZATION | 28 | 0.31689 | 0.804814 | 0.747559 | 1 |
| GOBP_REGULATION_OF_DNA_BIOSYNTHETIC_PROCESS | 117 | 0.262764 | 0.803154 | 0.836712 | 1 |
| GOBP_CARBOHYDRATE_METABOLIC_PROCESS | 497 | 0.240849 | 0.802459 | 0.942829 | 1 |
| GOBP_ENERGY_DERIVATION_BY_OXIDATION_OF_ORGANIC_COMPOUNDS | 280 | 0.24541 | 0.801993 | 0.895769 | 1 |
| GOBP_I_KAPPAB_PHOSPHORYLATION | 16 | 0.352412 | 0.801668 | 0.717718 | 1 |
| GOBP_RECOMBINATIONAL_REPAIR | 157 | 0.255211 | 0.801003 | 0.855716 | 1 |
| GOBP_REGULATION_OF_SYNAPTIC_VESICLE_RECYCLING | 18 | 0.341149 | 0.800527 | 0.752174 | 1 |
| GOBP_GOLGI_TO_PLASMA_MEMBRANE_TRANSPORT | 59 | 0.281029 | 0.800294 | 0.800792 | 1 |
| GOBP_NEGATIVE_REGULATION_OF_FAT_CELL_DIFFERENTIATION | 46 | 0.293195 | 0.800252 | 0.76378 | 1 |
| GOBP_CELLULAR_RESPONSE_TO_OSMOTIC_STRESS | 37 | 0.304549 | 0.798954 | 0.765847 | 1 |
| GOBP_NEUROMUSCULAR_PROCESS_CONTROLLING_BALANCE | 38 | 0.300836 | 0.798641 | 0.792225 | 1 |
| GOBP_PROTEIN_CONTAINING_COMPLEX_DISASSEMBLY | 230 | 0.246066 | 0.797811 | 0.895178 | 1 |
| GOBP_HETEROTYPIC_CELL_CELL_ADHESION | 43 | 0.291869 | 0.797683 | 0.773946 | 1 |
| GOBP_NEGATIVE_REGULATION_OF_BIOMINERALIZATION | 21 | 0.33281 | 0.797037 | 0.746143 | 1 |
| GOBP_PROTEIN_LOCALIZATION_TO_CELL_CELL_JUNCTION | 20 | 0.335887 | 0.796994 | 0.742604 | 1 |
| GOBP_CEREBELLAR_CORTEX_FORMATION | 24 | 0.326944 | 0.796844 | 0.749648 | 1 |
| GOBP_OXIDATIVE_PHOSPHORYLATION | 128 | 0.256439 | 0.796265 | 0.84027 | 1 |
| GOBP_REGULATION_OF_ATP_BIOSYNTHETIC_PROCESS | 20 | 0.33527 | 0.796239 | 0.755928 | 1 |
| GOBP_RESPONSE_TO_STARVATION | 172 | 0.251058 | 0.79564 | 0.872727 | 1 |
| GOBP_POSITIVE_REGULATION_OF_PROTEIN_CATABOLIC_PROCESS | 212 | 0.246769 | 0.7956 | 0.884575 | 1 |
| GOBP_GOLGI_TO_VACUOLE_TRANSPORT | 19 | 0.335977 | 0.795445 | 0.734375 | 1 |
| GOBP_ESTABLISHMENT_OF_ORGANELLE_LOCALIZATION | 366 | 0.240632 | 0.794865 | 0.93667 | 1 |
| GOBP_NEGATIVE_REGULATION_OF_MITOCHONDRION_ORGANIZATION | 45 | 0.289887 | 0.794191 | 0.79079 | 1 |
| GOBP_MEMBRANE_LIPID_METABOLIC_PROCESS | 183 | 0.249518 | 0.793933 | 0.879144 | 1 |
| GOBP_POST_TRANSLATIONAL_PROTEIN_MODIFICATION | 36 | 0.303072 | 0.793795 | 0.770667 | 1 |
| GOBP_SEX_DETERMINATION | 16 | 0.350108 | 0.793728 | 0.732523 | 1 |
| GOBP_CEREBRAL_CORTEX_RADIALLY_ORIENTED_CELL_MIGRATION | 30 | 0.309176 | 0.793658 | 0.752703 | 1 |
| GOBP_NEGATIVE_REGULATION_OF_ATP_METABOLIC_PROCESS | 26 | 0.319946 | 0.79349 | 0.743412 | 1 |
| GOBP_NUCLEAR_TRANSCRIBED_MRNA_POLY_A_TAIL_SHORTENING | 28 | 0.315269 | 0.793473 | 0.762973 | 1 |
| GOBP_RETINA_LAYER_FORMATION | 19 | 0.335519 | 0.793459 | 0.74 | 1 |
| GOBP_POSITIVE_REGULATION_OF_INSULIN_SECRETION_INVOLVED_IN_CELLULAR_RE | 23 | 0.333575 | 0.793394 | 0.729844 | 1 |
| GOBP_CELLULAR_NITROGEN_COMPOUND_CATABOLIC_PROCESS | 405 | 0.238556 | 0.792926 | 0.940041 | 1 |
| GOBP_SIGNAL_TRANSDUCTION_IN_RESPONSE_TO_DNA_DAMAGE | 165 | 0.250682 | 0.792541 | 0.868817 | 1 |
| GOBP_POSITIVE_REGULATION_OF_LYMPHOCYTE_MIGRATION | 24 | 0.323625 | 0.792379 | 0.742978 | 1 |
| GOBP_EXTRACELLULAR_TRANSPORT | 37 | 0.301382 | 0.792272 | 0.769231 | 1 |
| GOBP_LONG_CHAIN_FATTY_ACID_BIOSYNTHETIC_PROCESS | 22 | 0.323648 | 0.79201 | 0.735616 | 1 |
| GOBP_POSITIVE_REGULATION_OF_MESENCHYMAL_CELL_PROLIFERATION | 16 | 0.353567 | 0.791293 | 0.761834 | 1 |
| GOBP_DENDRITE_DEVELOPMENT | 215 | 0.245343 | 0.791238 | 0.88125 | 1 |
| GOBP_REGULATION_OF_MEMBRANE_PROTEIN_ECTODOMAIN_PROTEOLYSIS | 20 | 0.336526 | 0.791021 | 0.734012 | 1 |
| GOBP_PLASMA_MEMBRANE_REPAIR | 23 | 0.32719 | 0.790861 | 0.756831 | 1 |
| GOBP_RNA_5_END_PROCESSING | 23 | 0.32538 | 0.790266 | 0.75 | 1 |
| GOBP_LONG_TERM_MEMORY | 35 | 0.302534 | 0.790205 | 0.765172 | 1 |
| GOBP_NEGATIVE_REGULATION_OF_LIPID_LOCALIZATION | 30 | 0.309075 | 0.790153 | 0.771768 | 1 |
| GOBP_WOUND_HEALING_SPREADING_OF_EPIDERMAL_CELLS | 16 | 0.344619 | 0.789973 | 0.73716 | 1 |
| GOBP_ANTEROGRADE_AXONAL_TRANSPORT | 48 | 0.290401 | 0.789836 | 0.789474 | 1 |
| GOBP_CELLULAR_RESPONSE_TO_PEPTIDE_HORMONE_STIMULUS | 237 | 0.242847 | 0.789794 | 0.909186 | 1 |
| GOBP_RNA_EXPORT_FROM_NUCLEUS | 75 | 0.268434 | 0.789704 | 0.811665 | 1 |
| GOBP_CYCLIC_NUCLEOTIDE_MEDIATED_SIGNALING | 59 | 0.27806 | 0.789263 | 0.80198 | 1 |
| GOBP_MUSCLE_CELL_APOPTOTIC_PROCESS | 56 | 0.280068 | 0.788727 | 0.811043 | 1 |
| GOBP_POSITIVE_REGULATION_OF_PROTEIN_LOCALIZATION_TO_NUCLEUS | 77 | 0.268819 | 0.788456 | 0.826969 | 1 |
| GOBP_ENDOPLASMIC_RETICULUM_ORGANIZATION | 85 | 0.263991 | 0.788027 | 0.811916 | 1 |
| GOBP_GLUCOSE_CATABOLIC_PROCESS | 17 | 0.343545 | 0.787867 | 0.734513 | 1 |
| GOBP_REGULATION_OF_DENDRITIC_SPINE_DEVELOPMENT | 52 | 0.279135 | 0.787831 | 0.774033 | 1 |
| GOBP_RESPONSE_TO_OSMOTIC_STRESS | 68 | 0.270595 | 0.787788 | 0.808894 | 1 |
| GOBP_NEURON_DEATH | 300 | 0.24032 | 0.787614 | 0.920782 | 1 |
| GOBP_NEGATIVE_REGULATION_OF_CELL_SUBSTRATE_JUNCTION_ORGANIZATION | 16 | 0.347734 | 0.787391 | 0.748181 | 1 |
| GOBP_FIBROBLAST_GROWTH_FACTOR_RECEPTOR_SIGNALING_PATHWAY | 67 | 0.273338 | 0.787326 | 0.815951 | 1 |
| GOBP_REGULATION_OF_AUTOPHAGY | 311 | 0.239788 | 0.787301 | 0.925281 | 1 |
| GOBP_MONOSACCHARIDE_METABOLIC_PROCESS | 221 | 0.242415 | 0.786841 | 0.907075 | 1 |
| GOBP_RETINOL_METABOLIC_PROCESS | 31 | 0.305676 | 0.786812 | 0.772358 | 1 |
| GOBP_REGULATION_OF_MEMBRANE_PERMEABILITY | 64 | 0.271724 | 0.786388 | 0.806412 | 1 |
| GOBP_CELLULAR_RESPONSE_TO_PEPTIDE | 289 | 0.238262 | 0.786278 | 0.925128 | 1 |
| GOBP_REGULATION_OF_DOUBLE_STRAND_BREAK_REPAIR | 107 | 0.259205 | 0.785762 | 0.851259 | 1 |
| GOBP_POSITIVE_REGULATION_OF_MUSCLE_TISSUE_DEVELOPMENT | 16 | 0.339579 | 0.785375 | 0.737609 | 1 |
| GOBP_NOTOCHORD_DEVELOPMENT | 18 | 0.347232 | 0.78535 | 0.734177 | 1 |
| GOBP_VASOCONSTRICTION | 62 | 0.273085 | 0.783692 | 0.810345 | 1 |
| GOBP_GLUTAMINE_FAMILY_AMINO_ACID_CATABOLIC_PROCESS | 23 | 0.318167 | 0.783664 | 0.741791 | 1 |
| GOBP_POSITIVE_REGULATION_OF_CELLULAR_RESPONSE_TO_INSULIN_STIMULUS | 19 | 0.339882 | 0.783253 | 0.7366 | 1 |
| GOBP_VISUAL_BEHAVIOR | 41 | 0.295373 | 0.782976 | 0.785135 | 1 |
| GOBP_CELL_JUNCTION_DISASSEMBLY | 15 | 0.342062 | 0.782331 | 0.757252 | 1 |
| GOBP_NEGATIVE_REGULATION_OF_PROTEIN_DEPHOSPHORYLATION | 30 | 0.304799 | 0.782164 | 0.784203 | 1 |
| GOBP_NEGATIVE_REGULATION_OF_TYPE_I_INTERFERON_PRODUCTION | 24 | 0.321326 | 0.782055 | 0.783784 | 1 |
| GOBP_ACTIN_POLYMERIZATION_OR_DEPOLYMERIZATION | 177 | 0.244506 | 0.781849 | 0.885562 | 1 |
| GOBP_POSITIVE_REGULATION_OF_ORGANELLE_ORGANIZATION | 434 | 0.234102 | 0.78174 | 0.957533 | 1 |
| GOBP_POSITIVE_REGULATION_OF_INTRACELLULAR_TRANSPORT | 182 | 0.247267 | 0.781477 | 0.893891 | 1 |
| GOBP_MEMBRANE_LIPID_BIOSYNTHETIC_PROCESS | 133 | 0.252194 | 0.781175 | 0.865639 | 1 |
| GOBP_POSITIVE_REGULATION_OF_CELL_CYCLE_PHASE_TRANSITION | 102 | 0.255402 | 0.780868 | 0.83802 | 1 |
| GOBP_GLUTAMATE_RECEPTOR_SIGNALING_PATHWAY | 45 | 0.282044 | 0.780663 | 0.796203 | 1 |
| GOBP_MAINTENANCE_OF_CELL_NUMBER | 127 | 0.251705 | 0.780511 | 0.86942 | 1 |
| GOBP_REGULATION_OF_POSTSYNAPTIC_NEUROTRANSMITTER_RECEPTOR_ACTIVITY | 17 | 0.339173 | 0.780231 | 0.749642 | 1 |
| GOBP_REGULATION_OF_CYSTEINE_TYPE_ENDOPEPTIDASE_ACTIVITY_INVOLVED_IN_A | 15 | 0.346128 | 0.779021 | 0.750725 | 1 |
| GOBP_REGULATION_OF_LAMELLIPODIUM_ASSEMBLY | 34 | 0.297158 | 0.778753 | 0.783959 | 1 |
| GOBP_RESPONSE_TO_IMMOBILIZATION_STRESS | 15 | 0.34753 | 0.778695 | 0.767267 | 1 |
| GOBP_NEGATIVE_REGULATION_OF_SMALL_GTPASE_MEDIATED_SIGNAL_TRANSDUCTI | 55 | 0.276928 | 0.778489 | 0.808271 | 1 |
| GOBP_ORGANIC_CYCLIC_COMPOUND_CATABOLIC_PROCESS | 434 | 0.235934 | 0.778453 | 0.953582 | 1 |
| GOBP_POSITIVE_REGULATION_OF_ANIMAL_ORGAN_MORPHOGENESIS | 28 | 0.307628 | 0.778417 | 0.773224 | 1 |
| GOBP_POSITIVE_REGULATION_OF_MEIOTIC_CELL_CYCLE | 19 | 0.332427 | 0.777778 | 0.75289 | 1 |
| GOBP_REGULATION_OF_NEURONAL_SYNAPTIC_PLASTICITY | 47 | 0.283736 | 0.777573 | 0.80916 | 1 |
| GOBP_REGULATION_OF_MITOTIC_CELL_CYCLE | 413 | 0.23424 | 0.776497 | 0.962245 | 1 |
| GOBP_ANOIKIS | 29 | 0.30336 | 0.775884 | 0.783265 | 1 |
| GOBP_HEMATOPOIETIC_PROGENITOR_CELL_DIFFERENTIATION | 102 | 0.256374 | 0.775467 | 0.858645 | 1 |
| GOBP_REGULATION_OF_MESENCHYMAL_CELL_PROLIFERATION | 22 | 0.319565 | 0.775351 | 0.784173 | 1 |
| GOBP_REGULATION_OF_CELL_FATE_COMMITMENT | 17 | 0.335017 | 0.775211 | 0.756168 | 1 |
| GOBP_ACTIN_FILAMENT_BUNDLE_ORGANIZATION | 152 | 0.242296 | 0.774645 | 0.886957 | 1 |
| GOBP_REGULATION_OF_RESPONSE_TO_ENDOPLASMIC_RETICULUM_STRESS | 75 | 0.262865 | 0.774379 | 0.837709 | 1 |
| GOBP_DEVELOPMENTAL_PIGMENTATION | 40 | 0.287843 | 0.774375 | 0.805951 | 1 |
| GOBP_REGULATION_OF_CELL_MORPHOGENESIS | 275 | 0.23655 | 0.774144 | 0.922919 | 1 |
| GOBP_GLUTAMATE_METABOLIC_PROCESS | 30 | 0.306144 | 0.773455 | 0.78841 | 1 |
| GOBP_TIGHT_JUNCTION_ORGANIZATION | 63 | 0.273067 | 0.773239 | 0.83891 | 1 |
| GOBP_DETERMINATION_OF_ADULT_LIFESPAN | 15 | 0.345528 | 0.772569 | 0.743202 | 1 |
| GOBP_CERAMIDE_METABOLIC_PROCESS | 90 | 0.25994 | 0.772539 | 0.843023 | 1 |
| GOBP_FATTY_ACID_DERIVATIVE_BIOSYNTHETIC_PROCESS | 35 | 0.292106 | 0.772437 | 0.811421 | 1 |
| GOBP_CHAPERONE_MEDIATED_AUTOPHAGY | 16 | 0.343886 | 0.771806 | 0.772464 | 1 |
| GOBP_SPHINGOLIPID_BIOSYNTHETIC_PROCESS | 97 | 0.255012 | 0.77175 | 0.85896 | 1 |
| GOBP_REGULATION_OF_VIRAL_INDUCED_CYTOPLASMIC_PATTERN_RECOGNITION_RE | 20 | 0.324243 | 0.771383 | 0.769671 | 1 |
| GOBP_POLYOL_CATABOLIC_PROCESS | 26 | 0.310809 | 0.7708 | 0.776389 | 1 |
| GOBP_REGULATION_OF_ASTROCYTE_DIFFERENTIATION | 23 | 0.310923 | 0.770459 | 0.778086 | 1 |
| GOBP_HETEROCHROMATIN_ORGANIZATION | 75 | 0.260162 | 0.770223 | 0.856639 | 1 |
| GOBP_TYPE_B_PANCREATIC_CELL_PROLIFERATION | 17 | 0.339368 | 0.770148 | 0.771852 | 1 |
| GOBP_REGULATION_OF_CELL_PROJECTION_ASSEMBLY | 198 | 0.240079 | 0.769658 | 0.91954 | 1 |
| GOBP_CELLULAR_RESPONSE_TO_FLUID_SHEAR_STRESS | 18 | 0.333285 | 0.769584 | 0.760402 | 1 |
| GOBP_REGULATION_OF_EXOSOMAL_SECRETION | 18 | 0.324851 | 0.769533 | 0.764886 | 1 |
| GOBP_REGULATION_OF_FATTY_ACID_OXIDATION | 24 | 0.310649 | 0.769406 | 0.777018 | 1 |
| GOBP_VASCULAR_ENDOTHELIAL_GROWTH_FACTOR_SIGNALING_PATHWAY | 39 | 0.289275 | 0.768832 | 0.802685 | 1 |
| GOBP_PEPTIDYL_SERINE_MODIFICATION | 291 | 0.232562 | 0.76863 | 0.949692 | 1 |
| GOBP_ACTIN_FILAMENT_BASED_TRANSPORT | 18 | 0.335127 | 0.768392 | 0.77266 | 1 |
| GOBP_REGULATION_OF_PROTEIN_CATABOLIC_PROCESS | 359 | 0.233384 | 0.767891 | 0.946356 | 1 |
| GOBP_NEGATIVE_REGULATION_OF_CANONICAL_WNT_SIGNALING_PATHWAY | 123 | 0.248696 | 0.766662 | 0.893617 | 1 |
| GOBP_REGULATION_OF_CELLULAR_PROTEIN_CATABOLIC_PROCESS | 236 | 0.234807 | 0.765705 | 0.92895 | 1 |
| GOBP_MAINTENANCE_OF_PROTEIN_LOCATION | 87 | 0.25678 | 0.765599 | 0.850897 | 1 |
| GOBP_REGULATION_OF_SYNAPSE_ASSEMBLY | 88 | 0.256208 | 0.765532 | 0.88242 | 1 |
| GOBP_RECEPTOR_MEDIATED_ENDOCYTOSIS | 207 | 0.238782 | 0.764622 | 0.928342 | 1 |
| GOBP_REGULATION_OF_DNA_METABOLIC_PROCESS | 363 | 0.230485 | 0.764216 | 0.957732 | 1 |
| GOBP_POSITIVE_REGULATION_OF_CANONICAL_WNT_SIGNALING_PATHWAY | 94 | 0.254259 | 0.764022 | 0.881257 | 1 |
| GOBP_ACTIN_FILAMENT_POLYMERIZATION | 149 | 0.242632 | 0.763942 | 0.873913 | 1 |
| GOBP_NEURAL_NUCLEUS_DEVELOPMENT | 53 | 0.273935 | 0.763935 | 0.842965 | 1 |
| GOBP_T_CELL_MIGRATION | 41 | 0.284408 | 0.763646 | 0.80914 | 1 |
| GOBP_TRNA_MODIFICATION | 86 | 0.256975 | 0.762641 | 0.869305 | 1 |
| GOBP_POSITIVE_REGULATION_OF_NUCLEOTIDE_METABOLIC_PROCESS | 30 | 0.302129 | 0.762601 | 0.805479 | 1 |
| GOBP_POSITIVE_REGULATION_OF_PROTEOLYSIS | 325 | 0.231266 | 0.762322 | 0.935517 | 1 |
| GOBP_L_ALPHA_AMINO_ACID_TRANSMEMBRANE_TRANSPORT | 54 | 0.271122 | 0.762256 | 0.847328 | 1 |
| GOBP_MELANOCYTE_DIFFERENTIATION | 23 | 0.311708 | 0.76186 | 0.788269 | 1 |
| GOBP_POSITIVE_REGULATION_OF_SUPRAMOLECULAR_FIBER_ORGANIZATION | 160 | 0.24146 | 0.760465 | 0.906696 | 1 |
| GOBP_REGULATION_OF_ANIMAL_ORGAN_MORPHOGENESIS | 107 | 0.249966 | 0.760315 | 0.872027 | 1 |
| GOBP_REGULATION_OF_PHOSPHATIDYLINOSITOL_3_KINASE_ACTIVITY | 46 | 0.279154 | 0.759891 | 0.811083 | 1 |
| GOBP_NITRIC_OXIDE_SYNTHASE_BIOSYNTHETIC_PROCESS | 15 | 0.335579 | 0.759556 | 0.780415 | 1 |
| GOBP_POSITIVE_REGULATION_OF_MYELINATION | 15 | 0.339511 | 0.759291 | 0.77266 | 1 |
| GOBP_REGULATION_OF_PROTEIN_POLYMERIZATION | 181 | 0.236251 | 0.759257 | 0.914104 | 1 |
| GOBP_REGULATION_OF_CIRCADIAN_RHYTHM | 102 | 0.251814 | 0.758945 | 0.870629 | 1 |
| GOBP_REGULATION_OF_RESPONSE_TO_DNA_DAMAGE_STIMULUS | 240 | 0.233729 | 0.758192 | 0.932008 | 1 |
| GOBP_LOW_DENSITY_LIPOPROTEIN_PARTICLE_CLEARANCE | 20 | 0.320899 | 0.75783 | 0.813711 | 1 |
| GOBP_REGULATION_OF_REACTIVE_OXYGEN_SPECIES_METABOLIC_PROCESS | 111 | 0.246173 | 0.757716 | 0.889273 | 1 |
| GOBP_ANION_TRANSMEMBRANE_TRANSPORT | 200 | 0.236359 | 0.75732 | 0.919492 | 1 |
| GOBP_CHROMOSOME_SEPARATION | 88 | 0.252413 | 0.756962 | 0.891892 | 1 |

| | | | | | |
|---|---|---|---|---|---|
| GOBP_MESENCHYMAL_CELL_PROLIFERATION | 30 | 0.297148 | 0.756524 | 0.821629 | 1 |
| GOBP_CELL_FATE_COMMITMENT_INVOLVED_IN_FORMATION_OF_PRIMARY_GERM_LAY | 25 | 0.302371 | 0.756348 | 0.779349 | 1 |
| GOBP_CELLULAR_RESPONSE_TO_CARBOHYDRATE_STIMULUS | 116 | 0.247885 | 0.756297 | 0.883295 | 1 |
| GOBP_SULFUR_COMPOUND_METABOLIC_PROCESS | 284 | 0.229045 | 0.755876 | 0.955852 | 1 |
| GOBP_CELL_SURFACE_RECEPTOR_SIGNALING_PATHWAY_INVOLVED_IN_HEART_DEV | 25 | 0.309469 | 0.755776 | 0.783032 | 1 |
| GOBP_REGULATION_OF_KERATINOCYTE_DIFFERENTIATION | 28 | 0.298082 | 0.755601 | 0.787755 | 1 |
| GOBP_CELLULAR_HORMONE_METABOLIC_PROCESS | 86 | 0.2535 | 0.755135 | 0.88019 | 1 |
| GOBP_VESICLE_DOCKING_INVOLVED_IN_EXOCYTOSIS | 34 | 0.293022 | 0.755093 | 0.832224 | 1 |
| GOBP_PHOTORECEPTOR_CELL_MAINTENANCE | 35 | 0.284358 | 0.754885 | 0.813299 | 1 |
| GOBP_LUNG_CELL_DIFFERENTIATION | 15 | 0.333637 | 0.754538 | 0.779087 | 1 |
| GOBP_STORE_OPERATED_CALCIUM_ENTRY | 18 | 0.330032 | 0.7543 | 0.786861 | 1 |
| GOBP_RIBONUCLEOSIDE_MONOPHOSPHATE_BIOSYNTHETIC_PROCESS | 31 | 0.29595 | 0.754057 | 0.816864 | 1 |
| GOBP_REGULATION_OF_HETEROTYPIC_CELL_CELL_ADHESION | 16 | 0.331878 | 0.753038 | 0.790977 | 1 |
| GOBP_REGULATION_OF_UBIQUITIN_DEPENDENT_PROTEIN_CATABOLIC_PROCESS | 154 | 0.239803 | 0.753033 | 0.913326 | 1 |
| GOBP_POSITIVE_REGULATION_OF_CATABOLIC_PROCESS | 462 | 0.224542 | 0.752615 | 0.981818 | 1 |
| GOBP_CELLULAR_RESPONSE_TO_STEROID_HORMONE_STIMULUS | 169 | 0.236894 | 0.752135 | 0.905681 | 1 |
| GOBP_MITOTIC_DNA_INTEGRITY_CHECKPOINT_SIGNALING | 81 | 0.25672 | 0.752024 | 0.865967 | 1 |
| GOBP_POSITIVE_REGULATION_OF_MORPHOGENESIS_OF_AN_EPITHELIUM | 29 | 0.292815 | 0.751583 | 0.813072 | 1 |
| GOBP_PROTEIN_POLYUBIQUITINATION | 228 | 0.233508 | 0.751437 | 0.93914 | 1 |
| GOBP_REGULATION_OF_BRANCHING_INVOLVED_IN_URETERIC_BUD_MORPHOGENESI | 16 | 0.331174 | 0.751384 | 0.775183 | 1 |
| GOBP_NEGATIVE_REGULATION_OF_CYTOSKELETON_ORGANIZATION | 149 | 0.238148 | 0.751132 | 0.918051 | 1 |
| GOBP_POSITIVE_REGULATION_OF_ATP_METABOLIC_PROCESS | 28 | 0.29263 | 0.750465 | 0.816576 | 1 |
| GOBP_THYROID_HORMONE_METABOLIC_PROCESS | 15 | 0.336758 | 0.750084 | 0.780715 | 1 |
| GOBP_CELLULAR_RESPONSE_TO_UNFOLDED_PROTEIN | 93 | 0.249079 | 0.750008 | 0.886705 | 1 |
| GOBP_REGULATION_OF_PROTEIN_MODIFICATION_BY_SMALL_PROTEIN_CONJUGATION | 228 | 0.232446 | 0.749879 | 0.946597 | 1 |
| GOBP_AXONAL_TRANSPORT | 64 | 0.262463 | 0.749566 | 0.865942 | 1 |
| GOBP_CEREBELLAR_CORTEX_MORPHOGENESIS | 33 | 0.29037 | 0.749468 | 0.832888 | 1 |
| GOBP_FEMALE_MEIOTIC_NUCLEAR_DIVISION | 29 | 0.290394 | 0.749204 | 0.805746 | 1 |
| GOBP_POSITIVE_REGULATION_OF_GLIOGENESIS | 50 | 0.270172 | 0.748929 | 0.86 | 1 |
| GOBP_CHAPERONE_MEDIATED_PROTEIN_COMPLEX_ASSEMBLY | 20 | 0.317776 | 0.74818 | 0.795714 | 1 |
| GOBP_REGULATION_OF_CYTOSKELETON_ORGANIZATION | 479 | 0.222965 | 0.748082 | 0.980808 | 1 |
| GOBP_PYRIMIDINE_RIBONUCLEOSIDE_TRIPHOSPHATE_BIOSYNTHETIC_PROCESS | 15 | 0.336643 | 0.747863 | 0.792285 | 1 |
| GOBP_POSITIVE_REGULATION_OF_NOTCH_SIGNALING_PATHWAY | 36 | 0.288835 | 0.747746 | 0.816733 | 1 |
| GOBP_SYNAPTIC_TRANSMISSION_DOPAMINERGIC | 20 | 0.314689 | 0.747598 | 0.790462 | 1 |
| GOBP_SUPPRESSION_OF_VIRAL_RELEASE_BY_HOST | 16 | 0.337042 | 0.74742 | 0.807175 | 1 |
| GOBP_PURINE_CONTAINING_COMPOUND_METABOLIC_PROCESS | 348 | 0.225256 | 0.747338 | 0.96945 | 1 |
| GOBP_CARBOHYDRATE_PHOSPHORYLATION | 23 | 0.307968 | 0.747094 | 0.803547 | 1 |
| GOBP_FC_RECEPTOR_SIGNALING_PATHWAY | 39 | 0.278883 | 0.74632 | 0.817577 | 1 |
| GOBP_VACUOLAR_LOCALIZATION | 68 | 0.256397 | 0.745818 | 0.862069 | 1 |
| GOBP_NEGATIVE_REGULATION_OF_ORGANIC_ACID_TRANSPORT | 19 | 0.321084 | 0.745519 | 0.817003 | 1 |
| GOBP_POSITIVE_REGULATION_OF_HISTONE_MODIFICATION | 94 | 0.24609 | 0.745241 | 0.871528 | 1 |
| GOBP_REGULATION_OF_KIDNEY_DEVELOPMENT | 24 | 0.297714 | 0.744753 | 0.808845 | 1 |
| GOBP_REGULATION_OF_PROTEIN_LOCALIZATION_TO_CELL_SURFACE | 34 | 0.285564 | 0.744574 | 0.844208 | 1 |
| GOBP_POSITIVE_REGULATION_OF_MAP_KINASE_ACTIVITY | 98 | 0.246381 | 0.743855 | 0.881473 | 1 |
| GOBP_CGMP_MEDIATED_SIGNALING | 19 | 0.316356 | 0.743742 | 0.793304 | 1 |
| GOBP_NEGATIVE_REGULATION_OF_HORMONE_SECRETION | 47 | 0.273846 | 0.743654 | 0.860969 | 1 |
| GOBP_PINOCYTOSIS | 21 | 0.315455 | 0.743067 | 0.796243 | 1 |
| GOBP_NUCLEOTIDE_SUGAR_METABOLIC_PROCESS | 32 | 0.286089 | 0.742803 | 0.833107 | 1 |
| GOBP_NEGATIVE_REGULATION_OF_STRIATED_MUSCLE_CELL_APOPTOTIC_PROCESS | 18 | 0.318025 | 0.742411 | 0.818444 | 1 |
| GOBP_NEGATIVE_REGULATION_OF_GLIAL_CELL_DIFFERENTIATION | 16 | 0.321923 | 0.742257 | 0.795888 | 1 |
| GOBP_MITOCHONDRIAL_ELECTRON_TRANSPORT_NADH_TO_UBIQUINONE | 48 | 0.268075 | 0.742164 | 0.859335 | 1 |
| GOBP_INTEGRATED_STRESS_RESPONSE_SIGNALING | 21 | 0.311826 | 0.742067 | 0.798592 | 1 |
| GOBP_POSITIVE_REGULATION_OF_TRANSLATION | 126 | 0.240607 | 0.741971 | 0.919214 | 1 |
| GOBP_NEGATIVE_REGULATION_OF_LIPID_BIOSYNTHETIC_PROCESS | 47 | 0.27029 | 0.741773 | 0.872587 | 1 |
| GOBP_PROTEIN_PEPTIDYL_PROLYL_ISOMERIZATION | 35 | 0.282525 | 0.741578 | 0.832902 | 1 |
| GOBP_VIRAL_RELEASE_FROM_HOST_CELL | 28 | 0.294689 | 0.741377 | 0.829016 | 1 |
| GOBP_RECEPTOR_RECYCLING | 31 | 0.2908 | 0.741331 | 0.816076 | 1 |
| GOBP_NEURON_MATURATION | 40 | 0.276737 | 0.74101 | 0.842313 | 1 |
| GOBP_LIPID_MODIFICATION | 180 | 0.232676 | 0.740776 | 0.933262 | 1 |
| GOBP_NEGATIVE_REGULATION_OF_FATTY_ACID_METABOLIC_PROCESS | 23 | 0.307038 | 0.739961 | 0.81882 | 1 |
| GOBP_NEGATIVE_REGULATION_OF_CELLULAR_PROTEIN_CATABOLIC_PROCESS | 69 | 0.253015 | 0.739765 | 0.860149 | 1 |
| GOBP_CYTOPLASMIC_PATTERN_RECOGNITION_RECEPTOR_SIGNALING_PATHWAY_IN_ | 31 | 0.28565 | 0.739651 | 0.814767 | 1 |
| GOBP_NEGATIVE_REGULATION_OF_GLIOGENESIS | 28 | 0.293406 | 0.739221 | 0.811798 | 1 |
| GOBP_STEROL_BIOSYNTHETIC_PROCESS | 54 | 0.262522 | 0.738692 | 0.867995 | 1 |
| GOBP_REGULATION_OF_RESPONSE_TO_EXTRACELLULAR_STIMULUS | 19 | 0.316485 | 0.738372 | 0.83915 | 1 |
| GOBP_NEGATIVE_REGULATION_OF_MUSCLE_CELL_APOPTOTIC_PROCESS | 30 | 0.288954 | 0.737928 | 0.81931 | 1 |
| GOBP_REGULATION_OF_JUN_KINASE_ACTIVITY | 50 | 0.267059 | 0.737879 | 0.855513 | 1 |
| GOBP_INTERLEUKIN_6_MEDIATED_SIGNALING_PATHWAY | 16 | 0.32192 | 0.73762 | 0.806867 | 1 |
| GOBP_LIPID_HOMEOSTASIS | 122 | 0.239322 | 0.736395 | 0.91535 | 1 |
| GOBP_REGULATION_OF_PROTEIN_CONTAINING_COMPLEX_ASSEMBLY | 347 | 0.222179 | 0.736199 | 0.974619 | 1 |
| GOBP_PROTEIN_POLYMERIZATION | 250 | 0.226451 | 0.736032 | 0.952381 | 1 |
| GOBP_CELL_SURFACE_RECEPTOR_SIGNALING_PATHWAY_INVOLVED_IN_CELL_CELL_ | 480 | 0.220445 | 0.736025 | 0.978852 | 1 |
| GOBP_CARBOHYDRATE_DERIVATIVE_TRANSPORT | 70 | 0.255911 | 0.735096 | 0.890663 | 1 |
| GOBP_ALPHA_AMINO_ACID_METABOLIC_PROCESS | 165 | 0.232802 | 0.734557 | 0.937902 | 1 |
| GOBP_RECEPTOR_CLUSTERING | 52 | 0.26498 | 0.7345 | 0.868726 | 1 |
| GOBP_STEROID_HORMONE_SECRETION | 16 | 0.327079 | 0.734396 | 0.802878 | 1 |
| GOBP_MITOCHONDRIAL_OUTER_MEMBRANE_PERMEABILIZATION | 31 | 0.287462 | 0.734122 | 0.856955 | 1 |
| GOBP_NEGATIVE_REGULATION_OF_STRESS_ACTIVATED_PROTEIN_KINASE_SIGNALING | 44 | 0.273652 | 0.732189 | 0.86606 | 1 |
| GOBP_POSITIVE_REGULATION_OF_GLUCOSE_IMPORT | 30 | 0.29112 | 0.731937 | 0.834455 | 1 |
| GOBP_MITOCHONDRIAL_RNA_METABOLIC_PROCESS | 49 | 0.261719 | 0.731863 | 0.864728 | 1 |
| GOBP_REGULATION_OF_RECEPTOR_MEDIATED_ENDOCYTOSIS | 93 | 0.242504 | 0.731666 | 0.900922 | 1 |
| GOBP_LYSOSOMAL_PROTEIN_CATABOLIC_PROCESS | 18 | 0.311802 | 0.731097 | 0.81994 | 1 |
| GOBP_COLUMNAR_CUBOIDAL_EPITHELIAL_CELL_DIFFERENTIATION | 66 | 0.253239 | 0.727922 | 0.885645 | 1 |
| GOBP_ORGANIC_HYDROXY_COMPOUND_CATABOLIC_PROCESS | 56 | 0.259467 | 0.727725 | 0.865747 | 1 |
| GOBP_TRANSCRIPTION_BY_RNA_POLYMERASE_III | 53 | 0.261764 | 0.727246 | 0.88404 | 1 |
| GOBP_NEGATIVE_REGULATION_OF_PROTEASOMAL_PROTEIN_CATABOLIC_PROCESS | 43 | 0.268627 | 0.726563 | 0.856397 | 1 |
| GOBP_POSITIVE_REGULATION_OF_CYTOSKELETON_ORGANIZATION | 176 | 0.230179 | 0.726389 | 0.939459 | 1 |
| GOBP_PEPTIDYL_LYSINE_ACETYLATION | 176 | 0.228638 | 0.726175 | 0.945279 | 1 |
| GOBP_NEGATIVE_REGULATION_OF_CELLULAR_MACROMOLECULE_BIOSYNTHETIC_PR | 239 | 0.22343 | 0.723901 | 0.973545 | 1 |
| GOBP_NEGATIVE_REGULATION_OF_NEURON_DEATH | 177 | 0.22776 | 0.723842 | 0.950802 | 1 |
| GOBP_PIGMENT_CELL_DIFFERENTIATION | 33 | 0.279903 | 0.723584 | 0.849734 | 1 |
| GOBP_REGULATION_OF_RESPONSE_TO_REACTIVE_OXYGEN_SPECIES | 28 | 0.283175 | 0.721864 | 0.862534 | 1 |
| GOBP_MEMBRANE_LIPID_CATABOLIC_PROCESS | 32 | 0.283882 | 0.721704 | 0.843137 | 1 |
| GOBP_GMP_METABOLIC_PROCESS | 26 | 0.292932 | 0.721576 | 0.840845 | 1 |
| GOBP_NEURON_PROJECTION_EXTENSION_INVOLVED_IN_NEURON_PROJECTION_GUI | 37 | 0.272369 | 0.720705 | 0.864829 | 1 |
| GOBP_CELL_CYCLE_PHASE_TRANSITION | 473 | 0.214783 | 0.719184 | 0.992915 | 1 |
| GOBP_REGULATION_OF_CANONICAL_WNT_SIGNALING_PATHWAY | 227 | 0.222078 | 0.719028 | 0.968421 | 1 |
| GOBP_RECEPTOR_INTERNALIZATION | 95 | 0.24221 | 0.718422 | 0.9383 | 1 |
| GOBP_RESPONSE_TO_PROGESTERONE | 31 | 0.277047 | 0.718345 | 0.846154 | 1 |
| GOBP_CELLULAR_AMINO_ACID_METABOLIC_PROCESS | 235 | 0.222616 | 0.718155 | 0.966667 | 1 |
| GOBP_RIBOSE_PHOSPHATE_METABOLIC_PROCESS | 336 | 0.217241 | 0.717717 | 0.97665 | 1 |
| GOBP_ALPHA_AMINO_ACID_CATABOLIC_PROCESS | 71 | 0.242306 | 0.716419 | 0.911271 | 1 |
| GOBP_NEGATIVE_REGULATION_OF_MACROAUTOPHAGY | 34 | 0.270991 | 0.716118 | 0.853437 | 1 |
| GOBP_MEMBRANE_DOCKING | 78 | 0.243595 | 0.715538 | 0.914561 | 1 |
| GOBP_PIGMENTATION | 87 | 0.239094 | 0.715469 | 0.91129 | 1 |
| GOBP_AXONEMAL_DYNEIN_COMPLEX_ASSEMBLY | 30 | 0.277801 | 0.71538 | 0.853964 | 1 |
| GOBP_INTESTINAL_EPITHELIAL_CELL_DIFFERENTIATION | 18 | 0.307982 | 0.715182 | 0.862267 | 1 |
| GOBP_REGULATION_OF_NEUROTRANSMITTER_UPTAKE | 15 | 0.321929 | 0.714413 | 0.802985 | 1 |
| GOBP_NEGATIVE_REGULATION_OF_CELL_AGING | 20 | 0.303569 | 0.713432 | 0.806078 | 1 |
| GOBP_PROTEOGLYCAN_METABOLIC_PROCESS | 73 | 0.244853 | 0.713353 | 0.914994 | 1 |
| GOBP_INTRACELLULAR_LIPID_TRANSPORT | 40 | 0.268101 | 0.713253 | 0.877124 | 1 |
| GOBP_REGULATION_OF_SUPRAMOLECULAR_FIBER_ORGANIZATION | 340 | 0.215916 | 0.713049 | 0.975535 | 1 |
| GOBP_PEPTIDYL_LYSINE_DIMETHYLATION | 23 | 0.294977 | 0.712969 | 0.84 | 1 |
| GOBP_REGULATION_OF_CELL_DIVISION | 147 | 0.22627 | 0.711859 | 0.954644 | 1 |
| GOBP_N_ACETYLGLUCOSAMINE_METABOLIC_PROCESS | 17 | 0.311501 | 0.710749 | 0.847162 | 1 |
| GOBP_SISTER_CHROMATID_SEGREGATION | 195 | 0.221161 | 0.710713 | 0.969506 | 1 |
| GOBP_B_CELL_APOPTOTIC_PROCESS | 23 | 0.292998 | 0.710539 | 0.862915 | 1 |
| GOBP_REGULATION_OF_NUCLEASE_ACTIVITY | 17 | 0.316635 | 0.710519 | 0.840299 | 1 |
| GOBP_NEGATIVE_REGULATION_OF_PEPTIDYL_LYSINE_ACETYLATION | 18 | 0.309525 | 0.709872 | 0.824638 | 1 |
| GOBP_REGULATION_OF_CENTRIOLE_REPLICATION | 21 | 0.294021 | 0.709856 | 0.84017 | 1 |
| GOBP_ENERGY_HOMEOSTASIS | 35 | 0.272025 | 0.709778 | 0.880658 | 1 |
| GOBP_REGULATION_OF_WNT_SIGNALING_PATHWAY | 296 | 0.214013 | 0.709661 | 0.976555 | 1 |
| GOBP_AEROBIC_RESPIRATION | 173 | 0.224036 | 0.709627 | 0.958155 | 1 |
| GOBP_PEPTIDYL_PROLINE_MODIFICATION | 50 | 0.259451 | 0.709588 | 0.883019 | 1 |
| GOBP_DEOXYRIBOSE_PHOSPHATE_METABOLIC_PROCESS | 41 | 0.261843 | 0.709522 | 0.889032 | 1 |
| GOBP_NUCLEOSIDE_MONOPHOSPHATE_BIOSYNTHETIC_PROCESS | 40 | 0.260417 | 0.709145 | 0.883721 | 1 |
| GOBP_ISOPRENOID_BIOSYNTHETIC_PROCESS | 22 | 0.296668 | 0.708695 | 0.856941 | 1 |
| GOBP_SPHINGOLIPID_METABOLIC_PROCESS | 139 | 0.226177 | 0.708459 | 0.951034 | 1 |
| GOBP_REGULATION_OF_NOTCH_SIGNALING_PATHWAY | 79 | 0.238127 | 0.707965 | 0.919048 | 1 |
| GOBP_POSITIVE_REGULATION_OF_AXONOGENESIS | 74 | 0.243808 | 0.706782 | 0.914286 | 1 |
| GOBP_NEGATIVE_REGULATION_OF_CARBOHYDRATE_METABOLIC_PROCESS | 42 | 0.258855 | 0.706501 | 0.885638 | 1 |
| GOBP_NEGATIVE_REGULATION_OF_PROTEIN_LOCALIZATION_TO_CELL_PERIPHERY | 24 | 0.285479 | 0.70625 | 0.837607 | 1 |
| GOBP_POSITIVE_REGULATION_OF_CELLULAR_AMIDE_METABOLIC_PROCESS | 149 | 0.222564 | 0.706114 | 0.957424 | 1 |
| GOBP_ORGANOPHOSPHATE_ESTER_TRANSPORT | 119 | 0.228532 | 0.705168 | 0.940909 | 1 |
| GOBP_CANONICAL_WNT_SIGNALING_PATHWAY | 271 | 0.214963 | 0.70496 | 0.983471 | 1 |
| GOBP_VESICLE_ORGANIZATION | 276 | 0.215907 | 0.704751 | 0.974174 | 1 |
| GOBP_REGULATION_OF_EPITHELIAL_TO_MESENCHYMAL_TRANSITION | 82 | 0.237398 | 0.704371 | 0.924617 | 1 |
| GOBP_HORMONE_MEDIATED_SIGNALING_PATHWAY | 149 | 0.224398 | 0.704352 | 0.951193 | 1 |
| GOBP_SECONDARY_ALCOHOL_METABOLIC_PROCESS | 112 | 0.229463 | 0.70409 | 0.948023 | 1 |
| GOBP_HINDBRAIN_MORPHOGENESIS | 40 | 0.266166 | 0.703608 | 0.889481 | 1 |
| GOBP_PIGMENT_BIOSYNTHETIC_PROCESS | 57 | 0.248631 | 0.703352 | 0.896679 | 1 |
| GOBP_ONE_CARBON_METABOLIC_PROCESS | 37 | 0.266831 | 0.702325 | 0.860743 | 1 |
| GOBP_ACTIVATION_OF_PROTEIN_KINASE_B_ACTIVITY | 25 | 0.284911 | 0.702233 | 0.86326 | 1 |
| GOBP_ANION_HOMEOSTASIS | 30 | 0.27311 | 0.702224 | 0.865465 | 1 |
| GOBP_ANDROGEN_RECEPTOR_SIGNALING_PATHWAY | 40 | 0.257796 | 0.701583 | 0.886544 | 1 |
| GOBP_MICROTUBULE_DEPOLYMERIZATION | 44 | 0.258024 | 0.701343 | 0.910156 | 1 |
| GOBP_MUSCLE_HYPERTROPHY_IN_RESPONSE_TO_STRESS | 19 | 0.297798 | 0.70134 | 0.863965 | 1 |
| GOBP_PYRIMIDINE_NUCLEOBASE_METABOLIC_PROCESS | 20 | 0.293048 | 0.701037 | 0.880882 | 1 |
| GOBP_NEGATIVE_REGULATION_OF_PROTEIN_ACETYLATION | 20 | 0.295892 | 0.70053 | 0.833333 | 1 |
| GOBP_REGULATION_OF_ACTIN_FILAMENT_LENGTH | 141 | 0.224891 | 0.700268 | 0.963934 | 1 |
| GOBP_NEGATIVE_REGULATION_OF_PLATELET_ACTIVATION | 16 | 0.308807 | 0.697971 | 0.833085 | 1 |

| | | | | | |
|---|---|---|---|---|---|
| GOBP_NEGATIVE_REGULATION_OF_RECEPTOR_INTERNALIZATION | 15 | 0.31379 | 0.697506 | 0.83179 | 1 |
| GOBP_PROTEIN_ACETYLATION | 199 | 0.219282 | 0.697485 | 0.968085 | 1 |
| GOBP_POSITIVE_REGULATION_OF_ACTIN_FILAMENT_BUNDLE_ASSEMBLY | 56 | 0.248422 | 0.696996 | 0.907268 | 1 |
| GOBP_POSITIVE_REGULATION_OF_LAMELLIPODIUM_ASSEMBLY | 24 | 0.279717 | 0.696969 | 0.847978 | 1 |
| GOBP_PYRIMIDINE_RIBONUCLEOSIDE_TRIPHOSPHATE_METABOLIC_PROCESS | 18 | 0.302499 | 0.696724 | 0.853067 | 1 |
| GOBP_ENDOMEMBRANE_SYSTEM_ORGANIZATION | 469 | 0.20847 | 0.695768 | 0.995996 | 1 |
| GOBP_POSITIVE_REGULATION_OF_PROTEIN_CONTAINING_COMPLEX_ASSEMBLY | 164 | 0.217267 | 0.69575 | 0.972013 | 1 |
| GOBP_CELLULAR_RESPONSE_TO_GLUCOSE_STARVATION | 48 | 0.253429 | 0.694969 | 0.910336 | 1 |
| GOBP_PROTEIN_DEPHOSPHORYLATION | 259 | 0.2123 | 0.693872 | 0.985386 | 1 |
| GOBP_POSITIVE_REGULATION_OF_CARTILAGE_DEVELOPMENT | 27 | 0.270865 | 0.693772 | 0.856738 | 1 |
| GOBP_POSITIVE_REGULATION_OF_TELOMERE_CAPPING | 17 | 0.302714 | 0.693574 | 0.854626 | 1 |
| GOBP_RESPONSE_TO_ISCHEMIA | 43 | 0.257406 | 0.693144 | 0.897172 | 1 |
| GOBP_ENDOTHELIUM_DEVELOPMENT | 114 | 0.225582 | 0.692605 | 0.936652 | 1 |
| GOBP_PTERIDINE_CONTAINING_COMPOUND_METABOLIC_PROCESS | 34 | 0.2632 | 0.691549 | 0.887582 | 1 |
| GOBP_ANIMAL_ORGAN_FORMATION | 46 | 0.249514 | 0.691458 | 0.894133 | 1 |
| GOBP_FOLIC_ACID_METABOLIC_PROCESS | 15 | 0.31318 | 0.691285 | 0.875 | 1 |
| GOBP_CYCLIC_NUCLEOTIDE_METABOLIC_PROCESS | 30 | 0.267174 | 0.691077 | 0.887363 | 1 |
| GOBP_NEGATIVE_REGULATION_OF_OXIDOREDUCTASE_ACTIVITY | 16 | 0.307459 | 0.690705 | 0.86963 | 1 |
| GOBP_MITOTIC_INTRA_S_DNA_DAMAGE_CHECKPOINT_SIGNALING | 16 | 0.304593 | 0.68827 | 0.847826 | 1 |
| GOBP_TUBE_FORMATION | 139 | 0.219414 | 0.68823 | 0.952222 | 1 |
| GOBP_NEGATIVE_REGULATION_OF_MRNA_METABOLIC_PROCESS | 83 | 0.230702 | 0.687911 | 0.94925 | 1 |
| GOBP_REGULATION_OF_NEURON_DIFFERENTIATION | 156 | 0.219018 | 0.687692 | 0.973422 | 1 |
| GOBP_REGULATION_OF_RAC_PROTEIN_SIGNAL_TRANSDUCTION | 19 | 0.292295 | 0.687373 | 0.848529 | 1 |
| GOBP_GENERATION_OF_PRECURSOR_METABOLITES_AND_ENERGY | 423 | 0.205379 | 0.686757 | 0.99498 | 1 |
| GOBP_ER_NUCLEUS_SIGNALING_PATHWAY | 44 | 0.250606 | 0.68673 | 0.89467 | 1 |
| GOBP_AMP_METABOLIC_PROCESS | 23 | 0.2833 | 0.686611 | 0.87482 | 1 |
| GOBP_NEURONAL_ACTION_POTENTIAL | 27 | 0.279767 | 0.686378 | 0.878702 | 1 |
| GOBP_EXECUTION_PHASE_OF_APOPTOSIS | 62 | 0.23929 | 0.685611 | 0.91769 | 1 |
| GOBP_PROTEIN_IMPORT | 191 | 0.213308 | 0.685349 | 0.980912 | 1 |
| GOBP_CELLULAR_GLUCAN_METABOLIC_PROCESS | 60 | 0.242802 | 0.684545 | 0.931457 | 1 |
| GOBP_DEVELOPMENTAL_GROWTH_INVOLVED_IN_MORPHOGENESIS | 213 | 0.212493 | 0.684421 | 0.977754 | 1 |
| GOBP_INSULIN_RECEPTOR_SIGNALING_PATHWAY | 95 | 0.2275 | 0.684299 | 0.950116 | 1 |
| GOBP_MEMBRANE_BIOGENESIS | 58 | 0.242954 | 0.683739 | 0.909434 | 1 |
| GOBP_LAMELLIPODIUM_ORGANIZATION | 81 | 0.231295 | 0.683446 | 0.948417 | 1 |
| GOBP_POSITIVE_REGULATION_OF_ACTIN_CYTOSKELETON_REORGANIZATION | 16 | 0.300291 | 0.683167 | 0.866959 | 1 |
| GOBP_REGULATION_OF_MICROTUBULE_POLYMERIZATION_OR_DEPOLYMERIZATION | 83 | 0.23092 | 0.68287 | 0.951364 | 1 |
| GOBP_FIBROBLAST_MIGRATION | 45 | 0.253658 | 0.682854 | 0.925974 | 1 |
| GOBP_NEGATIVE_REGULATION_OF_TUMOR_NECROSIS_FACTOR_MEDIATED_SIGNALIN | 20 | 0.284108 | 0.682394 | 0.877874 | 1 |
| GOBP_NUCLEOBASE_CONTAINING_COMPOUND_TRANSPORT | 196 | 0.212708 | 0.682058 | 0.981582 | 1 |
| GOBP_NEGATIVE_REGULATION_OF_NEURAL_PRECURSOR_CELL_PROLIFERATION | 16 | 0.298993 | 0.681628 | 0.851429 | 1 |
| GOBP_LEUKOCYTE_TETHERING_OR_ROLLING | 19 | 0.289284 | 0.681308 | 0.884451 | 1 |
| GOBP_NEGATIVE_REGULATION_OF_TOR_SIGNALING | 41 | 0.251993 | 0.680524 | 0.913613 | 1 |
| GOBP_MORPHOGENESIS_OF_EMBRYONIC_EPITHELIUM | 133 | 0.218552 | 0.680306 | 0.966231 | 1 |
| GOBP_PERK_MEDIATED_UNFOLDED_PROTEIN_RESPONSE | 17 | 0.297935 | 0.679724 | 0.873418 | 1 |
| GOBP_MITOTIC_G2_M_TRANSITION_CHECKPOINT | 48 | 0.244631 | 0.679199 | 0.902937 | 1 |
| GOBP_POSITIVE_REGULATION_OF_GLYCOPROTEIN_METABOLIC_PROCESS | 20 | 0.284702 | 0.678186 | 0.870229 | 1 |
| GOBP_NEGATIVE_REGULATION_OF_ORGANELLE_ORGANIZATION | 322 | 0.205488 | 0.678136 | 0.994903 | 1 |
| GOBP_NUCLEAR_TRANSCRIBED_MRNA_CATABOLIC_PROCESS_DEADENYLATION_DEF | 60 | 0.240449 | 0.677848 | 0.941992 | 1 |
| GOBP_NEGATIVE_REGULATION_OF_FIBROBLAST_PROLIFERATION | 27 | 0.27245 | 0.677539 | 0.90946 | 1 |
| GOBP_REGULATION_OF_PROTEIN_TARGETING | 73 | 0.23056 | 0.676476 | 0.955224 | 1 |
| GOBP_NEURON_PROJECTION_EXTENSION | 160 | 0.212156 | 0.675757 | 0.982514 | 1 |
| GOBP_RESPIRATORY_ELECTRON_TRANSPORT_CHAIN | 105 | 0.219924 | 0.675599 | 0.960182 | 1 |
| GOBP_REGULATION_OF_DNA_BINDING | 105 | 0.223237 | 0.675552 | 0.954861 | 1 |
| GOBP_CENTRAL_NERVOUS_SYSTEM_PROJECTION_NEURON_AXONOGENESIS | 25 | 0.276906 | 0.675525 | 0.89847 | 1 |
| GOBP_NUCLEOSIDE_PHOSPHATE_BIOSYNTHETIC_PROCESS | 224 | 0.208795 | 0.675243 | 0.989551 | 1 |
| GOBP_RNA_PHOSPHODIESTER_BOND_HYDROLYSIS | 134 | 0.216567 | 0.674061 | 0.967956 | 1 |
| GOBP_LIPOPROTEIN_CATABOLIC_PROCESS | 16 | 0.297786 | 0.673926 | 0.863976 | 1 |
| GOBP_METHIONINE_METABOLIC_PROCESS | 15 | 0.302733 | 0.672775 | 0.858462 | 1 |
| GOBP_MITOCHONDRION_ORGANIZATION | 487 | 0.202501 | 0.672072 | 0.998991 | 1 |
| GOBP_CEREBRAL_CORTEX_CELL_MIGRATION | 40 | 0.248679 | 0.671295 | 0.910156 | 1 |
| GOBP_SECONDARY_METABOLIC_PROCESS | 36 | 0.259687 | 0.671273 | 0.894244 | 1 |
| GOBP_REGULATION_OF_CELL_CYCLE_G2_M_PHASE_TRANSITION | 101 | 0.219634 | 0.671146 | 0.965599 | 1 |
| GOBP_REGULATION_OF_SYNAPSE_STRUCTURE_OR_ACTIVITY | 184 | 0.208985 | 0.670852 | 0.984076 | 1 |
| GOBP_TRABECULA_MORPHOGENESIS | 40 | 0.251862 | 0.670845 | 0.90276 | 1 |
| GOBP_POSITIVE_REGULATION_OF_APOPTOTIC_SIGNALING_PATHWAY | 112 | 0.216066 | 0.669232 | 0.969942 | 1 |
| GOBP_REGULATION_OF_VASCULOGENESIS | 15 | 0.296552 | 0.668988 | 0.890208 | 1 |
| GOBP_CHONDROITIN_SULFATE_PROTEOGLYCAN_BIOSYNTHETIC_PROCESS | 23 | 0.270168 | 0.668181 | 0.905109 | 1 |
| GOBP_CELLULAR_RESPONSE_TO_TOPOLOGICALLY_INCORRECT_PROTEIN | 112 | 0.218087 | 0.667279 | 0.975834 | 1 |
| GOBP_PLATELET_MORPHOGENESIS | 18 | 0.29119 | 0.667278 | 0.893895 | 1 |
| GOBP_SYNAPTIC_VESICLE_LOCALIZATION | 51 | 0.237511 | 0.665595 | 0.923674 | 1 |
| GOBP_REGULATION_OF_LIPID_KINASE_ACTIVITY | 57 | 0.235763 | 0.665196 | 0.933667 | 1 |
| GOBP_NEGATIVE_REGULATION_OF_MITOTIC_CELL_CYCLE | 201 | 0.205569 | 0.664784 | 0.990486 | 1 |
| GOBP_POSITIVE_REGULATION_OF_GLIAL_CELL_DIFFERENTIATION | 35 | 0.252 | 0.664761 | 0.921196 | 1 |
| GOBP_RESPONSE_TO_AMYLOID_BETA | 38 | 0.24731 | 0.662614 | 0.935083 | 1 |
| GOBP_POSITIVE_REGULATION_OF_PEPTIDASE_ACTIVITY | 164 | 0.210411 | 0.662248 | 0.980645 | 1 |
| GOBP_POSITIVE_REGULATION_OF_STEROID_BIOSYNTHETIC_PROCESS | 16 | 0.292847 | 0.661404 | 0.871907 | 1 |
| GOBP_RNA_STABILIZATION | 56 | 0.237593 | 0.66131 | 0.94875 | 1 |
| GOBP_NUCLEAR_TRANSPORT | 279 | 0.202231 | 0.660642 | 0.996914 | 1 |
| GOBP_CEREBELLAR_CORTEX_DEVELOPMENT | 47 | 0.242946 | 0.660617 | 0.929293 | 1 |
| GOBP_NEGATIVE_REGULATION_OF_RECEPTOR_MEDIATED_ENDOCYTOSIS | 28 | 0.264546 | 0.660157 | 0.916318 | 1 |
| GOBP_DNA_BIOSYNTHETIC_PROCESS | 182 | 0.207103 | 0.659722 | 0.983211 | 1 |
| GOBP_PROTEIN_LOCALIZATION_TO_LYSOSOME | 44 | 0.246978 | 0.659072 | 0.931035 | 1 |
| GOBP_ESTABLISHMENT_OF_CELL_POLARITY | 132 | 0.20994 | 0.658726 | 0.981278 | 1 |
| GOBP_PLACENTA_BLOOD_VESSEL_DEVELOPMENT | 29 | 0.261875 | 0.657525 | 0.907357 | 1 |
| GOBP_MYD88_DEPENDENT_TOLL_LIKE_RECEPTOR_SIGNALING_PATHWAY | 17 | 0.292669 | 0.657383 | 0.893275 | 1 |
| GOBP_RESPONSE_TO_SALT_STRESS | 20 | 0.279652 | 0.65648 | 0.889362 | 1 |
| GOBP_REGULATION_OF_MITOCHONDRIAL_MEMBRANE_PERMEABILITY | 54 | 0.236336 | 0.656173 | 0.949174 | 1 |
| GOBP_POLY_A_PLUS_MRNA_EXPORT_FROM_NUCLEUS | 15 | 0.294004 | 0.655356 | 0.86931 | 1 |
| GOBP_CELLULAR_RESPONSE_TO_FATTY_ACID | 26 | 0.265218 | 0.65383 | 0.915849 | 1 |
| GOBP_POSITIVE_REGULATION_OF_NEURON_DIFFERENTIATION | 68 | 0.225653 | 0.653808 | 0.959085 | 1 |
| GOBP_SEMAPHORIN_PLEXIN_SIGNALING_PATHWAY_INVOLVED_IN_NEURON_PROJEC | 16 | 0.294209 | 0.653681 | 0.880117 | 1 |
| GOBP_POSITIVE_REGULATION_OF_VIRAL_GENOME_REPLICATION | 29 | 0.259653 | 0.65303 | 0.924451 | 1 |
| GOBP_NEGATIVE_REGULATION_OF_MITOTIC_CELL_CYCLE_PHASE_TRANSITION | 156 | 0.208076 | 0.652734 | 0.992416 | 1 |
| GOBP_NCRNA_CATABOLIC_PROCESS | 43 | 0.244107 | 0.652477 | 0.947781 | 1 |
| GOBP_REGULATION_OF_ATP_METABOLIC_PROCESS | 74 | 0.222971 | 0.652468 | 0.964286 | 1 |
| GOBP_POSITIVE_REGULATION_OF_PROTEASOMAL_UBIQUITIN_DEPENDENT_PROTEIN_ | 88 | 0.217044 | 0.650291 | 0.976879 | 1 |
| GOBP_THIOESTER_METABOLIC_PROCESS | 77 | 0.223708 | 0.650244 | 0.969231 | 1 |
| GOBP_REGULATION_OF_ENDOTHELIAL_CELL_DEVELOPMENT | 16 | 0.287347 | 0.64971 | 0.890625 | 1 |
| GOBP_RESPONSE_TO_MINERALOCORTICOID | 25 | 0.259605 | 0.649022 | 0.911302 | 1 |
| GOBP_ACTIVATION_OF_CYSTEINE_TYPE_ENDOPEPTIDASE_ACTIVITY | 17 | 0.278956 | 0.648922 | 0.912072 | 1 |
| GOBP_NEGATIVE_REGULATION_OF_EPITHELIAL_TO_MESENCHYMAL_TRANSITION | 31 | 0.257473 | 0.648914 | 0.916553 | 1 |
| GOBP_POSITIVE_REGULATION_OF_MACROAUTOPHAGY | 69 | 0.222665 | 0.648713 | 0.951632 | 1 |
| GOBP_INNER_EAR_RECEPTOR_CELL_DEVELOPMENT | 37 | 0.242228 | 0.648068 | 0.944892 | 1 |
| GOBP_RECEPTOR_METABOLIC_PROCESS | 145 | 0.204946 | 0.647643 | 0.986726 | 1 |
| GOBP_VESICLE_FUSION_TO_PLASMA_MEMBRANE | 18 | 0.283499 | 0.645838 | 0.906158 | 1 |
| GOBP_REGULATION_OF_IMMUNOGLOBULIN_PRODUCTION | 52 | 0.231094 | 0.645393 | 0.953283 | 1 |
| GOBP_MIDBRAIN_DEVELOPMENT | 75 | 0.220943 | 0.644953 | 0.979369 | 1 |
| GOBP_CYCLIC_NUCLEOTIDE_BIOSYNTHETIC_PROCESS | 17 | 0.280093 | 0.64461 | 0.893275 | 1 |
| GOBP_ENERGY_RESERVE_METABOLIC_PROCESS | 66 | 0.223707 | 0.644359 | 0.963636 | 1 |
| GOBP_SYNAPTIC_VESICLE_MEMBRANE_ORGANIZATION | 23 | 0.262109 | 0.644306 | 0.917251 | 1 |
| GOBP_NEGATIVE_REGULATION_OF_MIRNA_TRANSCRIPTION | 17 | 0.281479 | 0.641926 | 0.912536 | 1 |
| GOBP_SYNAPTIC_VESICLE_CYTOSKELETAL_TRANSPORT | 19 | 0.273226 | 0.641922 | 0.930931 | 1 |
| GOBP_REGULATION_OF_HOMOTYPIC_CELL_CELL_ADHESION | 26 | 0.255474 | 0.641582 | 0.929478 | 1 |
| GOBP_REGULATION_OF_TRANSCRIPTION_REGULATORY_REGION_DNA_BINDING | 42 | 0.238486 | 0.641445 | 0.946753 | 1 |
| GOBP_POSITIVE_REGULATION_OF_WNT_SIGNALING_PATHWAY | 125 | 0.206672 | 0.641267 | 0.984073 | 1 |
| GOBP_RESPONSE_TO_AMINO_ACID_STARVATION | 48 | 0.235291 | 0.640762 | 0.960459 | 1 |
| GOBP_NEGATIVE_REGULATION_OF_BLOOD_CIRCULATION | 19 | 0.269577 | 0.640762 | 0.910791 | 1 |
| GOBP_PYRIMIDINE_CONTAINING_COMPOUND_BIOSYNTHETIC_PROCESS | 35 | 0.243715 | 0.640589 | 0.934297 | 1 |
| GOBP_PHOSPHATIDYLGLYCEROL_METABOLIC_PROCESS | 28 | 0.250068 | 0.640449 | 0.926554 | 1 |
| GOBP_POSITIVE_REGULATION_OF_ACTIN_FILAMENT_POLYMERIZATION | 45 | 0.233201 | 0.640428 | 0.946633 | 1 |
| GOBP_NEGATIVE_REGULATION_OF_SODIUM_ION_TRANSPORT | 16 | 0.280357 | 0.640111 | 0.917293 | 1 |
| GOBP_AXON_EXTENSION | 114 | 0.208391 | 0.638855 | 0.981714 | 1 |
| GOBP_NEGATIVE_REGULATION_OF_I_KAPPAB_KINASE_NF_KAPPAB_SIGNALING | 40 | 0.240122 | 0.638393 | 0.96066 | 1 |
| GOBP_PHOSPHATIDYLCHOLINE_BIOSYNTHETIC_PROCESS | 27 | 0.25744 | 0.637887 | 0.9362 | 1 |
| GOBP_CENTRAL_NERVOUS_SYSTEM_NEURON_AXONOGENESIS | 33 | 0.246202 | 0.637447 | 0.933159 | 1 |
| GOBP_MITOTIC_CELL_CYCLE_CHECKPOINT_SIGNALING | 125 | 0.205795 | 0.637257 | 0.990919 | 1 |
| GOBP_ATP_SYNTHESIS_COUPLED_ELECTRON_TRANSPORT | 84 | 0.21433 | 0.636102 | 0.97494 | 1 |
| GOBP_ENDOPLASMIC_RETICULUM_UNFOLDED_PROTEIN_RESPONSE | 73 | 0.215737 | 0.635933 | 0.973366 | 1 |
| GOBP_CYTOPLASMIC_MICROTUBULE_ORGANIZATION | 55 | 0.227637 | 0.635856 | 0.952678 | 1 |
| GOBP_CELLULAR_RESPONSE_TO_ESTRADIOL_STIMULUS | 32 | 0.245656 | 0.635738 | 0.924033 | 1 |
| GOBP_REGULATION_OF_PROTEIN_LOCALIZATION_TO_NUCLEUS | 119 | 0.20836 | 0.635487 | 0.988739 | 1 |
| GOBP_REGULATION_OF_CELL_CELL_ADHESION_MEDIATED_BY_CADHERIN | 18 | 0.27104 | 0.63523 | 0.892012 | 1 |
| GOBP_LIVER_MORPHOGENESIS | 19 | 0.270926 | 0.633153 | 0.911977 | 1 |
| GOBP_FATTY_ACYL_COA_METABOLIC_PROCESS | 34 | 0.24342 | 0.633137 | 0.92905 | 1 |
| GOBP_CELLULAR_MODIFIED_AMINO_ACID_BIOSYNTHETIC_PROCESS | 42 | 0.231105 | 0.63311 | 0.94852 | 1 |
| GOBP_MITOTIC_RECOMBINATION | 25 | 0.25355 | 0.632327 | 0.935754 | 1 |
| GOBP_REGULATION_OF_GLUCOSE_IMPORT | 46 | 0.232101 | 0.632038 | 0.962773 | 1 |
| GOBP_REGULATION_OF_PRODUCTION_OF_SMALL_RNA_INVOLVED_IN_GENE_SILENCI | 19 | 0.265156 | 0.631929 | 0.917417 | 1 |
| GOBP_NEGATIVE_REGULATION_OF_RESPONSE_TO_ENDOPLASMIC_RETICULUM_STRE | 40 | 0.23463 | 0.631924 | 0.950318 | 1 |
| GOBP_CELL_CELL_SIGNALING_INVOLVED_IN_CARDIAC_CONDUCTION | 27 | 0.252586 | 0.631136 | 0.941722 | 1 |
| GOBP_PHOSPHOLIPID_DEPHOSPHORYLATION | 42 | 0.234451 | 0.629797 | 0.952196 | 1 |
| GOBP_GLYCOPROTEIN_METABOLIC_PROCESS | 334 | 0.190505 | 0.62927 | 0.998982 | 1 |
| GOBP_ACTIN_FILAMENT_SEVERING | 16 | 0.274857 | 0.629181 | 0.913947 | 1 |
| GOBP_NON_CANONICAL_WNT_SIGNALING_PATHWAY | 66 | 0.21799 | 0.627001 | 0.974202 | 1 |
| GOBP_POSITIVE_REGULATION_OF_AXON_EXTENSION | 38 | 0.240113 | 0.626239 | 0.943079 | 1 |
| GOBP_RIBOSOMAL_LARGE_SUBUNIT_ASSEMBLY | 26 | 0.248398 | 0.625206 | 0.930041 | 1 |
| GOBP_MICROTUBULE_POLYMERIZATION | 81 | 0.20954 | 0.625151 | 0.986553 | 1 |
| GOBP_POSTSYNAPTIC_SPECIALIZATION_ASSEMBLY | 19 | 0.264214 | 0.624526 | 0.931133 | 1 |
| GOBP_REGULATION_OF_DNA_REPAIR | 147 | 0.198426 | 0.62441 | 0.99026 | 1 |
| GOBP_EPITHELIAL_TUBE_FORMATION | 127 | 0.202095 | 0.624286 | 0.991081 | 1 |
| GOBP_REGULATION_OF_HEMATOPOIETIC_PROGENITOR_CELL_DIFFERENTIATION | 32 | 0.240101 | 0.621409 | 0.943925 | 1 |

| | | | | | |
|---|---|---|---|---|---|
| GOBP_LYTIC_VACUOLE_ORGANIZATION | 66 | 0.214509 | 0.621322 | 0.976998 | 1 |
| GOBP_CELLULAR_RESPONSE_TO_STARVATION | 144 | 0.196841 | 0.621259 | 0.994595 | 1 |
| GOBP_POSITIVE_REGULATION_OF_FOCAL_ADHESION_ASSEMBLY | 24 | 0.252586 | 0.62092 | 0.928775 | 1 |
| GOBP_GLYCOSYLATION | 225 | 0.191228 | 0.620887 | 0.998945 | 1 |
| GOBP_NUCLEAR_MEMBRANE_ORGANIZATION | 42 | 0.228775 | 0.620362 | 0.961892 | 1 |
| GOBP_POSITIVE_REGULATION_OF_RESPONSE_TO_ENDOPLASMIC_RETICULUM_STRES | 32 | 0.240612 | 0.62023 | 0.962213 | 1 |
| GOBP_POSITIVE_REGULATION_OF_SODIUM_ION_TRANSMEMBRANE_TRANSPORTER_A | 15 | 0.280102 | 0.619963 | 0.93553 | 1 |
| GOBP_REGULATION_OF_ENDOTHELIAL_CELL_DIFFERENTIATION | 34 | 0.235785 | 0.619559 | 0.945726 | 1 |
| GOBP_RESPONSE_TO_TOPOLOGICALLY_INCORRECT_PROTEIN | 150 | 0.195573 | 0.617956 | 0.997807 | 1 |
| GOBP_MICROTUBULE_BASED_TRANSPORT | 186 | 0.193213 | 0.617838 | 0.998943 | 1 |
| GOBP_POSITIVE_REGULATION_OF_TRANSCRIPTION_ELONGATION_FROM_RNA_POLYM | 18 | 0.268376 | 0.617558 | 0.928675 | 1 |
| GOBP_POSITIVE_REGULATION_OF_DNA_METABOLIC_PROCESS | 212 | 0.190944 | 0.61617 | 0.998948 | 1 |
| GOBP_ORGANELLE_INHERITANCE | 15 | 0.281126 | 0.61612 | 0.922156 | 1 |
| GOBP_PROTEIN_LIPID_COMPLEX_ASSEMBLY | 23 | 0.251774 | 0.615259 | 0.922316 | 1 |
| GOBP_NEGATIVE_REGULATION_OF_VIRAL_TRANSCRIPTION | 15 | 0.27544 | 0.6152 | 0.919505 | 1 |
| GOBP_TELENCEPHALON_GLIAL_CELL_MIGRATION | 22 | 0.262134 | 0.614499 | 0.929191 | 1 |
| GOBP_TRNA_PROCESSING | 122 | 0.19671 | 0.613853 | 0.993392 | 1 |
| GOBP_PROTEIN_K48_LINKED_UBIQUITINATION | 66 | 0.213141 | 0.612639 | 0.968867 | 1 |
| GOBP_CELLULAR_RESPONSE_TO_PH | 21 | 0.254479 | 0.612548 | 0.921283 | 1 |
| GOBP_GOLGI_TO_PLASMA_MEMBRANE_PROTEIN_TRANSPORT | 40 | 0.231096 | 0.612159 | 0.969003 | 1 |
| GOBP_CAMP_METABOLIC_PROCESS | 18 | 0.265223 | 0.611644 | 0.913433 | 1 |
| GOBP_REGULATION_OF_MEMBRANE_LIPID_DISTRIBUTION | 50 | 0.217961 | 0.611641 | 0.969212 | 1 |
| GOBP_TETRAPYRROLE_BIOSYNTHETIC_PROCESS | 31 | 0.239988 | 0.610939 | 0.945551 | 1 |
| GOBP_POSITIVE_REGULATION_OF_ESTABLISHMENT_OF_PROTEIN_LOCALIZATION_TO_ | 34 | 0.232227 | 0.6102 | 0.938974 | 1 |
| GOBP_CELLULAR_AMINO_ACID_BIOSYNTHETIC_PROCESS | 66 | 0.206835 | 0.60827 | 0.988903 | 1 |
| GOBP_POSITIVE_REGULATION_OF_DNA_BINDING | 45 | 0.221286 | 0.607811 | 0.963303 | 1 |
| GOBP_REGULATION_OF_ENDOPLASMIC_RETICULUM_UNFOLDED_PROTEIN_RESPONS | 28 | 0.238654 | 0.607607 | 0.94504 | 1 |
| GOBP_DICARBOXYLIC_ACID_METABOLIC_PROCESS | 82 | 0.205131 | 0.607563 | 0.985899 | 1 |
| GOBP_MITOTIC_G2_DNA_DAMAGE_CHECKPOINT_SIGNALING | 35 | 0.233912 | 0.607297 | 0.952894 | 1 |
| GOBP_ESTABLISHMENT_OF_RNA_LOCALIZATION | 149 | 0.192332 | 0.60608 | 0.997807 | 1 |
| GOBP_NEGATIVE_REGULATION_OF_LIPID_TRANSPORT | 19 | 0.26148 | 0.605586 | 0.94152 | 1 |
| GOBP_VESICLE_MEDIATED_TRANSPORT_TO_THE_PLASMA_MEMBRANE | 135 | 0.192362 | 0.605474 | 0.993378 | 1 |
| GOBP_GLYCOPROTEIN_BIOSYNTHETIC_PROCESS | 275 | 0.18418 | 0.604351 | 1 | 1 |
| GOBP_ENDOPLASMIC_RETICULUM_CALCIUM_ION_HOMEOSTASIS | 25 | 0.241059 | 0.604307 | 0.953425 | 1 |
| GOBP_LIPID_TRANSLOCATION | 44 | 0.218348 | 0.603409 | 0.963109 | 1 |
| GOBP_ORGANELLE_DISASSEMBLY | 119 | 0.194792 | 0.602923 | 0.992152 | 1 |
| GOBP_POSITIVE_REGULATION_OF_CELLULAR_COMPONENT_BIOGENESIS | 432 | 0.181923 | 0.60267 | 1 | 1 |
| GOBP_PROTEIN_TARGETING_TO_LYSOSOME | 23 | 0.249763 | 0.602539 | 0.937073 | 1 |
| GOBP_PYRIMIDINE_NUCLEOTIDE_METABOLIC_PROCESS | 48 | 0.218746 | 0.602498 | 0.963338 | 1 |
| GOBP_MONOCARBOXYLIC_ACID_CATABOLIC_PROCESS | 100 | 0.195865 | 0.600994 | 0.993166 | 1 |
| GOBP_POST_GOLGI_VESICLE_MEDIATED_TRANSPORT | 103 | 0.19737 | 0.600841 | 0.990919 | 1 |
| GOBP_VESICLE_LOCALIZATION | 176 | 0.186992 | 0.600715 | 1 | 1 |
| GOBP_POSITIVE_REGULATION_OF_SODIUM_ION_TRANSPORT | 31 | 0.232088 | 0.600386 | 0.94751 | 1 |
| GOBP_POSITIVE_REGULATION_OF_NITRIC_OXIDE_SYNTHASE_ACTIVITY | 16 | 0.262531 | 0.599175 | 0.931686 | 1 |
| GOBP_FOREBRAIN_CELL_MIGRATION | 54 | 0.212292 | 0.599073 | 0.976687 | 1 |
| GOBP_REGULATION_OF_SKELETAL_MUSCLE_TISSUE_DEVELOPMENT | 16 | 0.267473 | 0.598681 | 0.930781 | 1 |
| GOBP_CELL_CELL_SIGNALING_BY_WNT | 399 | 0.179896 | 0.597997 | 1 | 1 |
| GOBP_REGULATION_OF_ACTIN_FILAMENT_ORGANIZATION | 242 | 0.183943 | 0.597882 | 1 | 1 |
| GOBP_RIBOSOMAL_SMALL_SUBUNIT_ASSEMBLY | 19 | 0.257754 | 0.597111 | 0.933333 | 1 |
| GOBP_BRANCHING_INVOLVED_IN_BLOOD_VESSEL_MORPHOGENESIS | 28 | 0.23427 | 0.595959 | 0.954967 | 1 |
| GOBP_MITOTIC_CELL_CYCLE_PHASE_TRANSITION | 388 | 0.178171 | 0.595048 | 1 | 1 |
| GOBP_PHOSPHATIDYLINOSITOL_BIOSYNTHETIC_PROCESS | 122 | 0.190992 | 0.594455 | 0.997725 | 1 |
| GOBP_LUNG_MORPHOGENESIS | 39 | 0.220488 | 0.593745 | 0.980315 | 1 |
| GOBP_NEURONAL_STEM_CELL_POPULATION_MAINTENANCE | 21 | 0.245389 | 0.593618 | 0.950355 | 1 |
| GOBP_REGULATION_OF_POSTSYNAPTIC_MEMBRANE_NEUROTRANSMITTER_RECEPTO | 51 | 0.214182 | 0.593337 | 0.979381 | 1 |
| GOBP_REGULATION_OF_CHROMATIN_ASSEMBLY_OR_DISASSEMBLY | 33 | 0.226755 | 0.5932 | 0.971925 | 1 |
| GOBP_REGULATION_OF_STEROID_BIOSYNTHETIC_PROCESS | 56 | 0.211029 | 0.593024 | 0.982968 | 1 |
| GOBP_WATER_HOMEOSTASIS | 45 | 0.215676 | 0.59211 | 0.978372 | 1 |
| GOBP_REGULATION_OF_PROTEIN_EXPORT_FROM_NUCLEUS | 30 | 0.232978 | 0.590424 | 0.975066 | 1 |
| GOBP_RNA_CATABOLIC_PROCESS | 261 | 0.182377 | 0.590331 | 1 | 1 |
| GOBP_HISTONE_PHOSPHORYLATION | 42 | 0.220076 | 0.589824 | 0.97047 | 1 |
| GOBP_INNER_MITOCHONDRIAL_MEMBRANE_ORGANIZATION | 40 | 0.214703 | 0.588152 | 0.983204 | 1 |
| GOBP_DOUBLE_STRAND_BREAK_REPAIR | 260 | 0.178536 | 0.587863 | 1 | 1 |
| GOBP_PROTEIN_LOCALIZATION_TO_NUCLEUS | 268 | 0.179807 | 0.587714 | 1 | 1 |
| GOBP_POSITIVE_REGULATION_OF_PLASMA_MEMBRANE_BOUNDED_CELL_PROJECTIO | 93 | 0.19566 | 0.587475 | 0.992933 | 1 |
| GOBP_PROTEIN_ACYLATION | 238 | 0.179515 | 0.584618 | 1 | 1 |
| GOBP_LIPOPROTEIN_METABOLIC_PROCESS | 121 | 0.189807 | 0.584086 | 1 | 1 |
| GOBP_PLASMA_MEMBRANE_PHOSPHOLIPID_SCRAMBLING | 16 | 0.256468 | 0.58339 | 0.951039 | 1 |
| GOBP_CELL_VOLUME_HOMEOSTASIS | 27 | 0.229979 | 0.583142 | 0.963064 | 1 |
| GOBP_ESTABLISHMENT_OR_MAINTENANCE_OF_MONOPOLAR_CELL_POLARITY | 26 | 0.232397 | 0.58215 | 0.965763 | 1 |
| GOBP_POSITIVE_REGULATION_OF_VACUOLE_ORGANIZATION | 21 | 0.246145 | 0.581773 | 0.960284 | 1 |
| GOBP_IMP_METABOLIC_PROCESS | 16 | 0.258122 | 0.581218 | 0.931232 | 1 |
| GOBP_MICROTUBULE_ORGANIZING_CENTER_LOCALIZATION | 32 | 0.226928 | 0.580842 | 0.965517 | 1 |
| GOBP_MITOCHONDRIAL_MEMBRANE_ORGANIZATION | 109 | 0.189465 | 0.58081 | 0.997748 | 1 |
| GOBP_REGULATION_OF_ANDROGEN_RECEPTOR_SIGNALING_PATHWAY | 26 | 0.233459 | 0.580503 | 0.965422 | 1 |
| GOBP_RESPONSE_TO_LEUKEMIA_INHIBITORY_FACTOR | 90 | 0.194191 | 0.578884 | 0.993111 | 1 |
| GOBP_MITOCHONDRIAL_ATP_SYNTHESIS_COUPLED_PROTON_TRANSPORT | 18 | 0.252353 | 0.576698 | 0.938067 | 1 |
| GOBP_ESTABLISHMENT_OR_MAINTENANCE_OF_CELL_POLARITY | 201 | 0.179928 | 0.573941 | 0.99894 | 1 |
| GOBP_STEROID_BIOSYNTHETIC_PROCESS | 133 | 0.184602 | 0.573763 | 0.998878 | 1 |
| GOBP_KERATINOCYTE_MIGRATION | 15 | 0.254279 | 0.573762 | 0.945865 | 1 |
| GOBP_REGULATION_OF_CELLULAR_RESPONSE_TO_VASCULAR_ENDOTHELIAL_GROW | 20 | 0.243004 | 0.573475 | 0.969208 | 1 |
| GOBP_RESPONSE_TO_HEPATOCYTE_GROWTH_FACTOR | 17 | 0.25348 | 0.573117 | 0.944024 | 1 |
| GOBP_NEGATIVE_REGULATION_OF_STEROID_METABOLIC_PROCESS | 22 | 0.237583 | 0.571323 | 0.963989 | 1 |
| GOBP_REGULATION_OF_GLYCOGEN_METABOLIC_PROCESS | 29 | 0.226346 | 0.571318 | 0.967877 | 1 |
| GOBP_RESPONSE_TO_MISFOLDED_PROTEIN | 24 | 0.231341 | 0.569198 | 0.962209 | 1 |
| GOBP_HEART_FIELD_SPECIFICATION | 15 | 0.252397 | 0.568865 | 0.944012 | 1 |
| GOBP_PROTEASOMAL_PROTEIN_CATABOLIC_PROCESS | 464 | 0.170991 | 0.568646 | 1 | 1 |
| GOBP_MRNA_CATABOLIC_PROCESS | 218 | 0.177864 | 0.568626 | 1 | 1 |
| GOBP_REGULATION_OF_TELOMERE_CAPPING | 26 | 0.232741 | 0.567979 | 0.97415 | 1 |
| GOBP_REGULATION_OF_CELLULAR_AMIDE_METABOLIC_PROCESS | 410 | 0.170765 | 0.567499 | 1 | 1 |
| GOBP_PROTEIN_LOCALIZATION_TO_VACUOLE | 65 | 0.197869 | 0.56749 | 0.991484 | 1 |
| GOBP_PROTEIN_LOCALIZATION_TO_SYNAPSE | 59 | 0.197302 | 0.565199 | 0.991369 | 1 |
| GOBP_PHAGOSOME_MATURATION | 22 | 0.237037 | 0.564749 | 0.958453 | 1 |
| GOBP_REGULATION_OF_TRANSLATIONAL_FIDELITY | 21 | 0.228199 | 0.562372 | 0.978022 | 1 |
| GOBP_POSITIVE_REGULATION_OF_NEUROBLAST_PROLIFERATION | 18 | 0.247054 | 0.561919 | 0.964392 | 1 |
| GOBP_POSITIVE_REGULATION_OF_BMP_SIGNALING_PATHWAY | 32 | 0.217908 | 0.560318 | 0.97139 | 1 |
| GOBP_TRANSLESION_SYNTHESIS | 24 | 0.225904 | 0.560055 | 0.964789 | 1 |
| GOBP_SELECTIVE_AUTOPHAGY | 77 | 0.190289 | 0.558667 | 0.993858 | 1 |
| GOBP_PROTEIN_LOCALIZATION_TO_CELL_JUNCTION | 87 | 0.186659 | 0.558255 | 0.996536 | 1 |
| GOBP_ORGANISM_EMERGENCE_FROM_PROTECTIVE_STRUCTURE | 25 | 0.222685 | 0.556224 | 0.968927 | 1 |
| GOBP_LONG_CHAIN_FATTY_ACYL_COA_METABOLIC_PROCESS | 23 | 0.228371 | 0.555929 | 0.973202 | 1 |
| GOBP_NUCLEAR_EXPORT | 144 | 0.177626 | 0.555583 | 0.997797 | 1 |
| GOBP_PROTEIN_AUTOPHOSPHORYLATION | 207 | 0.172601 | 0.555407 | 1 | 1 |
| GOBP_ACTIVATION_OF_GTPASE_ACTIVITY | 96 | 0.182994 | 0.555138 | 0.994138 | 1 |
| GOBP_CYTOSKELETON_DEPENDENT_INTRACELLULAR_TRANSPORT | 186 | 0.173278 | 0.554876 | 1 | 1 |
| GOBP_REGULATION_OF_PROTEASOMAL_UBIQUITIN_DEPENDENT_PROTEIN_CATABOLI | 128 | 0.178736 | 0.554382 | 1 | 1 |
| GOBP_REGULATION_OF_NMDA_RECEPTOR_ACTIVITY | 17 | 0.239538 | 0.553771 | 0.964497 | 1 |
| GOBP_EPITHELIAL_CELL_CELL_ADHESION | 16 | 0.243977 | 0.553353 | 0.966463 | 1 |
| GOBP_ATP_METABOLIC_PROCESS | 238 | 0.171508 | 0.55226 | 1 | 1 |
| GOBP_NEGATIVE_REGULATION_OF_RESPONSE_TO_DNA_DAMAGE_STIMULUS | 80 | 0.186186 | 0.552248 | 0.996471 | 1 |
| GOBP_MAINTENANCE_OF_PROTEIN_LOCATION_IN_CELL | 60 | 0.192737 | 0.552004 | 0.987745 | 1 |
| GOBP_MICROTUBULE_POLYMERIZATION_OR_DEPOLYMERIZATION | 117 | 0.180141 | 0.551673 | 0.997768 | 1 |
| GOBP_CELL_CYCLE_G2_M_PHASE_TRANSITION | 147 | 0.176067 | 0.551534 | 1 | 1 |
| GOBP_PROTEIN_TARGETING_TO_VACUOLE | 37 | 0.207247 | 0.551376 | 0.985372 | 1 |
| GOBP_CHROMATIN_DISASSEMBLY | 19 | 0.230037 | 0.549009 | 0.973538 | 1 |
| GOBP_KINETOCHORE_ORGANIZATION | 22 | 0.226256 | 0.547926 | 0.973865 | 1 |
| GOBP_RNA_DESTABILIZATION | 88 | 0.185906 | 0.547351 | 0.994279 | 1 |
| GOBP_REGULATION_OF_ESTABLISHMENT_OF_PLANAR_POLARITY | 54 | 0.194792 | 0.547289 | 0.995068 | 1 |
| GOBP_POSITIVE_REGULATION_OF_CYCLIN_DEPENDENT_PROTEIN_KINASE_ACTIVITY | 32 | 0.210208 | 0.546116 | 0.991892 | 1 |
| GOBP_FATTY_ACID_CATABOLIC_PROCESS | 90 | 0.179569 | 0.545602 | 0.998812 | 1 |
| GOBP_REGULATION_OF_PROTEIN_KINASE_C_SIGNALING | 17 | 0.232975 | 0.542948 | 0.958395 | 1 |
| GOBP_REGULATION_OF_PHOSPHOPROTEIN_PHOSPHATASE_ACTIVITY | 48 | 0.195732 | 0.54169 | 0.99125 | 1 |
| GOBP_INTRA_GOLGI_VESICLE_MEDIATED_TRANSPORT | 30 | 0.213845 | 0.541369 | 0.985115 | 1 |
| GOBP_REGULATION_OF_AUTOPHAGY_OF_MITOCHONDRION | 36 | 0.205313 | 0.540212 | 0.987903 | 1 |
| GOBP_POSITIVE_REGULATION_OF_CELL_SUBSTRATE_JUNCTION_ORGANIZATION | 29 | 0.209274 | 0.540185 | 0.983516 | 1 |
| GOBP_CELL_DEATH_IN_RESPONSE_TO_OXIDATIVE_STRESS | 77 | 0.18115 | 0.539051 | 0.996475 | 1 |
| GOBP_PROTEASOME_MEDIATED_UBIQUITIN_DEPENDENT_PROTEIN_CATABOLIC_PROC | 394 | 0.162102 | 0.538725 | 1 | 1 |
| GOBP_DNA_INTEGRITY_CHECKPOINT_SIGNALING | 118 | 0.17465 | 0.53789 | 0.997768 | 1 |
| GOBP_RNA_DECAPPING | 17 | 0.231569 | 0.536422 | 0.973875 | 1 |
| GOBP_REGULATION_OF_VIRAL_TRANSCRIPTION | 19 | 0.230546 | 0.534839 | 0.982609 | 1 |
| GOBP_REGULATION_OF_SMOOTHENED_SIGNALING_PATHWAY | 71 | 0.182412 | 0.533691 | 1 | 1 |
| GOBP_REGULATION_OF_PROTEASOMAL_PROTEIN_CATABOLIC_PROCESS | 173 | 0.16877 | 0.532508 | 1 | 1 |
| GOBP_MAINTENANCE_OF_PROTEIN_LOCALIZATION_IN_ORGANELLE | 41 | 0.198036 | 0.532347 | 0.990968 | 1 |
| GOBP_CELLULAR_RESPONSE_TO_STEROL_DEPLETION | 15 | 0.238724 | 0.531704 | 0.962264 | 1 |
| GOBP_DENDRITIC_SPINE_DEVELOPMENT | 86 | 0.179022 | 0.531654 | 0.997706 | 1 |
| GOBP_TRANSPORT_ALONG_MICROTUBULE | 152 | 0.168596 | 0.528978 | 1 | 1 |
| GOBP_STEROID_HORMONE_MEDIATED_SIGNALING_PATHWAY | 113 | 0.172118 | 0.528886 | 1 | 1 |
| GOBP_NEGATIVE_REGULATION_OF_PROTEOLYSIS_INVOLVED_IN_CELLULAR_PROTEIN | 57 | 0.188301 | 0.528058 | 0.996283 | 1 |
| GOBP_INNER_EAR_RECEPTOR_CELL_STEREOCILIUM_ORGANIZATION | 26 | 0.211375 | 0.527699 | 0.982167 | 1 |
| GOBP_MITOPHAGY | 34 | 0.203447 | 0.527067 | 0.989362 | 1 |
| GOBP_NEGATIVE_REGULATION_OF_INTRACELLULAR_STEROID_HORMONE_RECEPTOR | 33 | 0.203089 | 0.526683 | 0.982361 | 1 |
| GOBP_AUTOPHAGOSOME_MATURATION | 57 | 0.184005 | 0.524093 | 0.997456 | 1 |
| GOBP_PURINE_NUCLEOSIDE_MONOPHOSPHATE_BIOSYNTHETIC_PROCESS | 21 | 0.223261 | 0.523245 | 0.974286 | 1 |
| GOBP_INTRINSIC_APOPTOTIC_SIGNALING_PATHWAY_IN_RESPONSE_TO_ENDOPLASMI | 57 | 0.186202 | 0.522347 | 0.996231 | 1 |
| GOBP_REGULATION_OF_DNA_TEMPLATED_TRANSCRIPTION_IN_RESPONSE_TO_STRES | 49 | 0.188805 | 0.521318 | 0.994878 | 1 |
| GOBP_ESTABLISHMENT_OF_TISSUE_POLARITY | 66 | 0.179156 | 0.520268 | 0.997555 | 1 |
| GOBP_NEGATIVE_REGULATION_OF_PROTEASOMAL_UBIQUITIN_DEPENDENT_PROTEIN | 32 | 0.202754 | 0.51962 | 0.982713 | 1 |
| GOBP_TRNA_METABOLIC_PROCESS | 182 | 0.162694 | 0.518542 | 0.998925 | 1 |
| GOBP_CHONDROITIN_SULFATE_BIOSYNTHETIC_PROCESS | 18 | 0.222941 | 0.517664 | 0.976296 | 1 |
| GOBP_3_UTR_MEDIATED_MRNA_DESTABILIZATION | 15 | 0.231156 | 0.516796 | 0.974589 | 1 |
| GOBP_CARDIOLIPIN_METABOLIC_PROCESS | 15 | 0.229607 | 0.516638 | 0.965167 | 1 |

| | | | | | |
|---|---|---|---|---|---|
| GOBP_NEGATIVE_REGULATION_OF_CELLULAR_AMIDE_METABOLIC_PROCESS | 184 | 0.161204 | 0.516549 | 1 | 1 |
| GOBP_PIGMENT_GRANULE_ORGANIZATION | 27 | 0.207767 | 0.515778 | 0.980769 | 1 |
| GOBP_VESICLE_BUDDING_FROM_MEMBRANE | 60 | 0.177718 | 0.512985 | 0.998767 | 1 |
| GOBP_ESTABLISHMENT_OF_PROTEIN_LOCALIZATION_TO_PLASMA_MEMBRANE | 60 | 0.179888 | 0.511781 | 0.995012 | 1 |
| GOBP_RNA_LOCALIZATION | 183 | 0.157412 | 0.509997 | 1 | 1 |
| GOBP_S_ADENOSYLMETHIONINE_METABOLIC_PROCESS | 15 | 0.229757 | 0.509263 | 0.979259 | 1 |
| GOBP_NEGATIVE_REGULATION_OF_SMOOTHENED_SIGNALING_PATHWAY | 31 | 0.197598 | 0.508996 | 0.990057 | 1 |
| GOBP_PHOSPHATE_ION_TRANSPORT | 22 | 0.214994 | 0.507879 | 0.982118 | 1 |
| GOBP_REGULATION_OF_OXIDATIVE_STRESS_INDUCED_CELL_DEATH | 55 | 0.179201 | 0.507451 | 0.996207 | 1 |
| GOBP_PEPTIDYL_THREONINE_MODIFICATION | 114 | 0.162473 | 0.505784 | 0.998865 | 1 |
| GOBP_MORPHOGENESIS_OF_A_POLARIZED_EPITHELIUM | 89 | 0.1685 | 0.505093 | 0.997685 | 1 |
| GOBP_MULTIVESICULAR_BODY_SORTING_PATHWAY | 37 | 0.192092 | 0.50404 | 0.998679 | 1 |
| GOBP_TRANSLATIONAL_TERMINATION | 18 | 0.216698 | 0.504006 | 0.973837 | 1 |
| GOBP_RRNA_MODIFICATION | 36 | 0.190651 | 0.503772 | 0.992042 | 1 |
| GOBP_HISTONE_H3_K9_TRIMETHYLATION | 16 | 0.216469 | 0.501994 | 0.968703 | 1 |
| GOBP_PROTEIN_DEMANNOSYLATION | 17 | 0.219745 | 0.50145 | 0.980422 | 1 |
| GOBP_POSITIVE_REGULATION_OF_TRANSCRIPTION_REGULATORY_REGION_DNA_BIND | 17 | 0.217918 | 0.500463 | 0.986938 | 1 |
| GOBP_REGULATION_OF_TOR_SIGNALING | 96 | 0.165329 | 0.500173 | 0.998835 | 1 |
| GOBP_VESICLE_TARGETING_TO_FROM_OR_WITHIN_GOLGI | 21 | 0.208984 | 0.499918 | 0.987395 | 1 |
| GOBP_POSTREPLICATION_REPAIR | 34 | 0.190494 | 0.499828 | 0.985215 | 1 |
| GOBP_POSITIVE_REGULATION_OF_CELL_MORPHOGENESIS_INVOLVED_IN_DIFFERENT | 71 | 0.171393 | 0.49798 | 1 | 1 |
| GOBP_CELLULAR_RESPONSE_TO_ARSENIC_CONTAINING_SUBSTANCE | 21 | 0.214049 | 0.496509 | 0.98468 | 1 |
| GOBP_REGULATION_OF_CELL_MORPHOGENESIS_INVOLVED_IN_DIFFERENTIATION | 87 | 0.16637 | 0.496375 | 0.998835 | 1 |
| GOBP_RESPONSE_TO_EPIDERMAL_GROWTH_FACTOR | 44 | 0.180329 | 0.495758 | 0.996129 | 1 |
| GOBP_REGULATION_OF_PROTON_TRANSPORT | 20 | 0.208517 | 0.494745 | 0.976845 | 1 |
| GOBP_REGULATION_OF_FATTY_ACID_BETA_OXIDATION | 15 | 0.219866 | 0.493919 | 0.977444 | 1 |
| GOBP_RESPONSE_TO_STEROL_DEPLETION | 17 | 0.214297 | 0.491691 | 0.983264 | 1 |
| GOBP_RESPONSE_TO_ENDOPLASMIC_RETICULUM_STRESS | 241 | 0.15123 | 0.489622 | 1 | 1 |
| GOBP_NEGATIVE_REGULATION_OF_PROTEIN_MODIFICATION_BY_SMALL_PROTEIN_CO | 88 | 0.162856 | 0.488968 | 1 | 1 |
| GOBP_STRESS_GRANULE_ASSEMBLY | 27 | 0.193141 | 0.488607 | 0.99726 | 1 |
| GOBP_POSITIVE_REGULATION_OF_SODIUM_ION_TRANSMEMBRANE_TRANSPORT | 18 | 0.208282 | 0.488446 | 0.986726 | 1 |
| GOBP_AUTOPHAGY_OF_MITOCHONDRION | 86 | 0.163301 | 0.48842 | 1 | 1 |
| GOBP_MITOCHONDRIAL_RESPIRATORY_CHAIN_COMPLEX_ASSEMBLY | 94 | 0.162619 | 0.486671 | 1 | 1 |
| GOBP_ATP_SYNTHESIS_COUPLED_PROTON_TRANSPORT | 24 | 0.199037 | 0.486634 | 0.992878 | 1 |
| GOBP_REGULATION_OF_VACUOLE_ORGANIZATION | 49 | 0.177805 | 0.484501 | 0.99493 | 1 |
| GOBP_RESPONSE_TO_NITRIC_OXIDE | 15 | 0.215976 | 0.483091 | 0.989329 | 1 |
| GOBP_NEUROTRANSMITTER_RECEPTOR_TRANSPORT_TO_PLASMA_MEMBRANE | 16 | 0.214874 | 0.483074 | 0.984701 | 1 |
| GOBP_CYTOSOLIC_TRANSPORT | 168 | 0.151624 | 0.481699 | 1 | 1 |
| GOBP_PTERIDINE_CONTAINING_COMPOUND_BIOSYNTHETIC_PROCESS | 15 | 0.210874 | 0.480973 | 0.99393 | 1 |
| GOBP_REGULATION_OF_LIPOPROTEIN_PARTICLE_CLEARANCE | 15 | 0.218268 | 0.480855 | 0.989231 | 1 |
| GOBP_ENDOSOME_TRANSPORT_VIA_MULTIVESICULAR_BODY_SORTING_PATHWAY | 31 | 0.185455 | 0.475029 | 0.992074 | 1 |
| GOBP_PEPTIDYL_SERINE_DEPHOSPHORYLATION | 19 | 0.201593 | 0.473602 | 0.988588 | 1 |
| GOBP_PROTEIN_TARGETING_TO_MEMBRANE | 114 | 0.153066 | 0.471569 | 1 | 1 |
| GOBP_VACUOLE_ORGANIZATION | 174 | 0.147934 | 0.469999 | 1 | 1 |
| GOBP_MANNOSYLATION | 35 | 0.180253 | 0.469571 | 0.997382 | 1 |
| GOBP_INTERSTRAND_CROSS_LINK_REPAIR | 30 | 0.184644 | 0.467222 | 0.99187 | 1 |
| GOBP_VESICLE_TRANSPORT_ALONG_MICROTUBULE | 44 | 0.170548 | 0.461511 | 1 | 1 |
| GOBP_HISTONE_EXCHANGE | 17 | 0.200757 | 0.4613 | 0.989706 | 1 |
| GOBP_REGULATION_OF_MRNA_METABOLIC_PROCESS | 271 | 0.14124 | 0.460375 | 1 | 1 |
| GOBP_PROTEIN_K48_LINKED_DEUBIQUITINATION | 31 | 0.179887 | 0.459679 | 0.998665 | 1 |
| GOBP_REGULATION_OF_NUCLEAR_TRANSCRIBED_MRNA_CATABOLIC_PROCESS_DEA | 24 | 0.186139 | 0.458968 | 0.991632 | 1 |
| GOBP_ESTABLISHMENT_OF_PROTEIN_LOCALIZATION_TO_MITOCHONDRIAL_MEMBRAN | 29 | 0.179971 | 0.458302 | 0.997268 | 1 |
| GOBP_POSITIVE_REGULATION_OF_PHOSPHOPROTEIN_PHOSPHATASE_ACTIVITY | 17 | 0.199356 | 0.455248 | 0.990071 | 1 |
| GOBP_POSITIVE_REGULATION_OF_POSTTRANSCRIPTIONAL_GENE_SILENCING | 30 | 0.17847 | 0.453894 | 0.997245 | 1 |
| GOBP_LATE_ENDOSOME_TO_LYSOSOME_TRANSPORT | 18 | 0.198712 | 0.45309 | 0.992604 | 1 |
| GOBP_MATURATION_OF_LSU_RRNA | 28 | 0.180309 | 0.451675 | 0.99863 | 1 |
| GOBP_ENDOSOME_TO_LYSOSOME_TRANSPORT | 58 | 0.157435 | 0.450659 | 0.998755 | 1 |
| GOBP_EMBRYONIC_BRAIN_DEVELOPMENT | 15 | 0.199907 | 0.450176 | 0.990923 | 1 |
| GOBP_TRICARBOXYLIC_ACID_CYCLE | 30 | 0.174475 | 0.448716 | 0.997249 | 1 |
| GOBP_PHOSPHATIDYLSERINE_METABOLIC_PROCESS | 21 | 0.18439 | 0.444454 | 0.995833 | 1 |
| GOBP_STRESS_FIBER_ASSEMBLY | 100 | 0.143809 | 0.436824 | 1 | 1 |
| GOBP_TOR_SIGNALING | 118 | 0.140861 | 0.436397 | 1 | 1 |
| GOBP_NEGATIVE_REGULATION_OF_OXIDATIVE_STRESS_INDUCED_INTRINSIC_APOPTO | 18 | 0.188944 | 0.435012 | 0.994195 | 1 |
| GOBP_FATTY_ACID_BETA_OXIDATION | 67 | 0.151554 | 0.43306 | 1 | 1 |
| GOBP_CELLULAR_PIGMENTATION | 50 | 0.156405 | 0.432724 | 1 | 1 |
| GOBP_PROTEOGLYCAN_BIOSYNTHETIC_PROCESS | 52 | 0.15321 | 0.431385 | 1 | 1 |
| GOBP_AUTOPHAGOSOME_ORGANIZATION | 104 | 0.141261 | 0.430932 | 1 | 1 |
| GOBP_REGULATION_OF_ESTABLISHMENT_OF_PROTEIN_LOCALIZATION_TO_MITOCHON | 47 | 0.155715 | 0.427369 | 0.998744 | 1 |
| GOBP_RNA_PHOSPHODIESTER_BOND_HYDROLYSIS_EXONUCLEOLYTIC | 42 | 0.154924 | 0.418799 | 0.998713 | 1 |
| GOBP_NEGATIVE_REGULATION_OF_AMYLOID_PRECURSOR_PROTEIN_CATABOLIC_PR | 19 | 0.179072 | 0.418617 | 0.994236 | 1 |
| GOBP_MODULATION_BY_HOST_OF_VIRAL_GENOME_REPLICATION | 17 | 0.179913 | 0.416736 | 0.992571 | 1 |
| GOBP_POSITIVE_REGULATION_OF_DNA_TEMPLATED_TRANSCRIPTION_ELONGATION | 28 | 0.1652 | 0.412632 | 0.998641 | 1 |
| GOBP_HEPATOCYTE_APOPTOTIC_PROCESS | 16 | 0.17848 | 0.409906 | 0.997085 | 1 |
| GOBP_SYNAPTIC_VESICLE_TRANSPORT | 39 | 0.150348 | 0.404689 | 0.998724 | 1 |
| GOBP_OLIGOSACCHARIDE_LIPID_INTERMEDIATE_BIOSYNTHETIC_PROCESS | 19 | 0.169892 | 0.404413 | 0.998578 | 1 |
| GOBP_REGULATION_OF_TRANSCRIPTION_BY_RNA_POLYMERASE_III | 29 | 0.156528 | 0.400084 | 1 | 1 |
| GOBP_NEUROTRANSMITTER_RECEPTOR_TRANSPORT | 22 | 0.165506 | 0.398663 | 0.997046 | 1 |
| GOBP_POSITIVE_REGULATION_OF_TELOMERE_MAINTENANCE | 50 | 0.145667 | 0.397797 | 1 | 1 |
| GOBP_ESTABLISHMENT_OF_PROTEIN_LOCALIZATION_TO_VACUOLE | 48 | 0.143773 | 0.397137 | 1 | 1 |
| GOBP_SNRNA_METABOLIC_PROCESS | 52 | 0.140181 | 0.390657 | 1 | 1 |
| GOBP_NUCLEOTIDE_EXCISION_REPAIR | 56 | 0.137574 | 0.388174 | 1 | 1 |
| GOBP_CLATHRIN_COAT_ASSEMBLY | 18 | 0.165468 | 0.385742 | 1 | 1 |
| GOBP_ENDOSOMAL_TRANSPORT | 224 | 0.11897 | 0.383127 | 1 | 1 |
| GOBP_REGULATION_OF_TRANSCRIPTION_ELONGATION_FROM_RNA_POLYMERASE_II_ | 37 | 0.143635 | 0.380708 | 1 | 1 |
| GOBP_MACROAUTOPHAGY | 297 | 0.113891 | 0.375981 | 1 | 1 |
| GOBP_PROTEIN_EXPORT_FROM_NUCLEUS | 56 | 0.131465 | 0.375479 | 1 | 1 |
| GOBP_PSEUDOURIDINE_SYNTHESIS | 18 | 0.159427 | 0.374826 | 0.997089 | 1 |
| GOBP_POSITIVE_REGULATION_OF_TRANSCRIPTION_BY_RNA_POLYMERASE_III | 18 | 0.156069 | 0.356443 | 1 | 1 |
| GOBP_VACUOLAR_TRANSPORT | 148 | 0.113045 | 0.354036 | 1 | 1 |
| GOBP_POSITIVE_REGULATION_OF_TELOMERE_MAINTENANCE_VIA_TELOMERE_LENGT | 37 | 0.122394 | 0.327662 | 1 | 1 |
| GOBP_TELOMERE_CAPPING | 39 | 0.11845 | 0.318737 | 1 | 1 |
| GOBP_REGULATION_OF_GOLGI_ORGANIZATION | 18 | 0.134754 | 0.31693 | 1 | 1 |
| GOBP_RIBOSOMAL_LARGE_SUBUNIT_BIOGENESIS | 71 | 0.100913 | 0.297293 | 1 | 1 |
| GOBP_REGULATION_OF_TELOMERE_MAINTENANCE_VIA_TELOMERE_LENGTHENING | 58 | 0.091726 | 0.258485 | 1 | 1 |

**Table S5. GSEA of dysreguated genes in SF3B1$^{K700E}$ ES cells (K700E ES line 3).**

**GSEA Positive** Hematopoietic/Immune pathways are highlighted in red.

| Gene sets | SIZE | ES | NES | NOM p-val | FDR q-val |
|---|---|---|---|---|---|
| GOBP_OUTFLOW_TRACT_MORPHOGENESIS | 72 | 0.621518 | 1.984914 | 0 | 0.069237 |
| GOBP_APOPTOTIC_PROCESS_INVOLVED_IN_DEVELOPMENT | 38 | 0.691302 | 1.959832 | 0 | 0.066609 |
| GOBP_PIRNA_METABOLIC_PROCESS | 15 | 0.840825 | 1.958939 | 0 | 0.045428 |
| GOBP_POSITIVE_REGULATION_OF_FATTY_ACID_METABOLIC_PROCESS | 27 | 0.734415 | 1.95274 | 0 | 0.03845 |
| GOBP_POSITIVE_REGULATION_OF_FATTY_ACID_BIOSYNTHETIC_PROCESS | 16 | 0.796885 | 1.94607 | 0 | 0.037573 |
| GOBP_AXIS_SPECIFICATION | 74 | 0.610302 | 1.943905 | 0 | 0.032171 |
| GOBP_NEGATIVE_REGULATION_OF_INTERFERON_GAMMA_PRODUCTION | 23 | 0.746813 | 1.922396 | 0.001575 | 0.041247 |
| GOBP_ANTERIOR_POSTERIOR_AXIS_SPECIFICATION | 43 | 0.656234 | 1.912552 | 0 | 0.043284 |
| GOBP_GRANULOCYTE_ACTIVATION | 32 | 0.685221 | 1.911117 | 0 | 0.039396 |
| GOBP_CARDIOCYTE_DIFFERENTIATION | 114 | 0.56201 | 1.905318 | 0 | 0.040718 |
| GOBP_CARDIAC_SEPTUM_MORPHOGENESIS | 63 | 0.60398 | 1.88406 | 0 | 0.054735 |
| GOBP_POSITIVE_REGULATION_OF_BLOOD_VESSEL_ENDOTHELIAL_CELL_MIGRATION | 50 | 0.621425 | 1.866469 | 0 | 0.071817 |
| GOBP_POSITIVE_REGULATION_OF_CELL_MIGRATION_INVOLVED_IN_SPROUTING_ANGIOG | 18 | 0.745459 | 1.860647 | 0 | 0.074074 |
| GOBP_CARDIAC_MUSCLE_TISSUE_DEVELOPMENT | 178 | 0.525002 | 1.860224 | 0 | 0.069443 |
| GOBP_REGULATION_OF_CELL_MIGRATION_INVOLVED_IN_SPROUTING_ANGIOGENESIS | 33 | 0.675502 | 1.847959 | 0 | 0.079099 |
| GOBP_OUTFLOW_TRACT_SEPTUM_MORPHOGENESIS | 22 | 0.724834 | 1.847542 | 0 | 0.074609 |
| GOBP_CARDIAC_MUSCLE_CELL_DIFFERENTIATION | 87 | 0.562652 | 1.841755 | 0 | 0.076402 |
| GOBP_REGULATION_OF_MEIOTIC_NUCLEAR_DIVISION | 22 | 0.7121 | 1.838064 | 0.003241 | 0.076398 |
| GOBP_CARDIAC_SEPTUM_DEVELOPMENT | 95 | 0.5523 | 1.832799 | 0 | 0.078175 |
| GOBP_BLASTODERM_SEGMENTATION | 20 | 0.71857 | 1.82684 | 0.001553 | 0.081119 |
| GOBP_CARDIAC_VENTRICLE_DEVELOPMENT | 109 | 0.539461 | 1.818384 | 0 | 0.08885 |
| GOBP_CELL_ADHESION_MEDIATED_BY_INTEGRIN | 67 | 0.574201 | 1.815772 | 0 | 0.08837 |
| GOBP_PROXIMAL_DISTAL_PATTERN_FORMATION | 17 | 0.739214 | 1.812638 | 0.003096 | 0.088429 |
| GOBP_GRANULOCYTE_CHEMOTAXIS | 71 | 0.575782 | 1.810578 | 0 | 0.087409 |
| GOBP_STRIATED_MUSCLE_CELL_DEVELOPMENT | 52 | 0.598716 | 1.808945 | 0 | 0.086095 |
| GOBP_POSITIVE_REGULATION_OF_NEUTROPHIL_MIGRATION | 25 | 0.694831 | 1.806978 | 0.003175 | 0.085198 |
| GOBP_MESENCHYME_MORPHOGENESIS | 51 | 0.609495 | 1.80166 | 0 | 0.088797 |
| GOBP_CARDIAC_CHAMBER_MORPHOGENESIS | 107 | 0.538055 | 1.801525 | 0 | 0.085772 |
| GOBP_CARDIAC_CELL_DEVELOPMENT | 64 | 0.590328 | 1.797088 | 0 | 0.088619 |
| GOBP_REGULATION_OF_GRANULOCYTE_CHEMOTAXIS | 36 | 0.635216 | 1.794629 | 0.001499 | 0.08937 |
| GOBP_POSITIVE_REGULATION_OF_INFLAMMATORY_RESPONSE | 93 | 0.550313 | 1.792907 | 0 | 0.088118 |
| GOBP_BLOOD_VESSEL_REMODELING | 33 | 0.641557 | 1.792116 | 0.001425 | 0.086527 |
| GOBP_DNA_METHYLATION | 59 | 0.586763 | 1.791963 | 0 | 0.08406 |
| GOBP_CELL_MIGRATION_INVOLVED_IN_HEART_DEVELOPMENT | 17 | 0.73902 | 1.791514 | 0.001597 | 0.082042 |
| GOBP_HEART_VALVE_DEVELOPMENT | 61 | 0.575752 | 1.790691 | 0 | 0.080816 |
| GOBP_TRIPARTITE_REGIONAL_SUBDIVISION | 16 | 0.739234 | 1.790636 | 0.003205 | 0.078629 |
| GOBP_APOPTOTIC_PROCESS_INVOLVED_IN_MORPHOGENESIS | 24 | 0.684355 | 1.787702 | 0.001534 | 0.079873 |
| GOBP_STRIATED_MUSCLE_CELL_DIFFERENTIATION | 212 | 0.497976 | 1.783697 | 0 | 0.082409 |
| GOBP_REGULATION_OF_CARDIOCYTE_DIFFERENTIATION | 23 | 0.689083 | 1.782679 | 0.006163 | 0.081405 |
| GOBP_NUCLEOSIDE_CATABOLIC_PROCESS | 20 | 0.703445 | 1.779723 | 0.00321 | 0.083104 |
| GOBP_SKELETAL_MUSCLE_CELL_DIFFERENTIATION | 53 | 0.58981 | 1.775501 | 0 | 0.085655 |
| GOBP_HEART_VALVE_MORPHOGENESIS | 52 | 0.583962 | 1.774299 | 0 | 0.08506 |
| GOBP_PEPTIDE_CROSS_LINKING | 16 | 0.752916 | 1.769794 | 0.003407 | 0.088976 |
| GOBP_NEGATIVE_REGULATION_OF_REPRODUCTIVE_PROCESS | 44 | 0.602167 | 1.769496 | 0.001433 | 0.087328 |
| GOBP_CARDIAC_RIGHT_VENTRICLE_MORPHOGENESIS | 17 | 0.718955 | 1.76939 | 0.003205 | 0.085524 |
| GOBP_REGULATION_OF_NEUTROPHIL_MIGRATION | 35 | 0.633786 | 1.768791 | 0.004505 | 0.084313 |
| GOBP_EMBRYONIC_AXIS_SPECIFICATION | 32 | 0.64004 | 1.767629 | 0.004823 | 0.083897 |
| GOBP_PEPTIDYL_ARGININE_MODIFICATION | 23 | 0.688873 | 1.767093 | 0.003195 | 0.083093 |
| GOBP_REGULATION_OF_NEUTROPHIL_CHEMOTAXIS | 25 | 0.663479 | 1.765848 | 0.003101 | 0.082912 |
| GOBP_REGULATION_OF_CELL_SUBSTRATE_ADHESION | 182 | 0.501146 | 1.764558 | 0 | 0.082698 |
| GOBP_DNA_METHYLATION_OR_DEMETHYLATION | 79 | 0.546797 | 1.757662 | 0 | 0.088833 |
| GOBP_CELL_FATE_COMMITMENT | 179 | 0.501198 | 1.756938 | 0 | 0.087957 |
| GOBP_MAMMARY_GLAND_MORPHOGENESIS | 32 | 0.636129 | 1.75425 | 0.00149 | 0.089412 |
| GOBP_VENTRICULAR_CARDIAC_MUSCLE_TISSUE_DEVELOPMENT | 48 | 0.581366 | 1.752068 | 0 | 0.090485 |
| GOBP_ENDOCRINE_SYSTEM_DEVELOPMENT | 92 | 0.537137 | 1.749939 | 0.002611 | 0.091818 |
| GOBP_EMBRYONIC_HEART_TUBE_DEVELOPMENT | 72 | 0.549821 | 1.748112 | 0 | 0.092459 |
| GOBP_POSITIVE_REGULATION_OF_LEUKOCYTE_APOPTOTIC_PROCESS | 19 | 0.697362 | 1.746686 | 0.006279 | 0.092281 |
| GOBP_CARDIAC_ATRIUM_DEVELOPMENT | 30 | 0.639313 | 1.745769 | 0.005848 | 0.091847 |
| GOBP_ATRIOVENTRICULAR_VALVE_DEVELOPMENT | 25 | 0.657264 | 1.744339 | 0.00624 | 0.09193 |
| GOBP_MUSCLE_TISSUE_DEVELOPMENT | 313 | 0.469078 | 1.742075 | 0 | 0.093042 |
| GOBP_CARDIAC_CHAMBER_DEVELOPMENT | 144 | 0.503445 | 1.740774 | 0 | 0.092969 |
| GOBP_CELL_FATE_DETERMINATION | 26 | 0.654291 | 1.740464 | 0.001534 | 0.091834 |
| GOBP_CELL_MATRIX_ADHESION | 196 | 0.483296 | 1.737208 | 0 | 0.094338 |
| GOBP_REGULATION_OF_TRANSPOSITION | 20 | 0.705246 | 1.735956 | 0.011129 | 0.094571 |
| GOBP_VENTRICULAR_SEPTUM_DEVELOPMENT | 67 | 0.550547 | 1.73393 | 0 | 0.096149 |
| GOBP_ATRIAL_SEPTUM_DEVELOPMENT | 20 | 0.693027 | 1.73263 | 0.001524 | 0.096381 |
| GOBP_CARDIAC_VENTRICLE_MORPHOGENESIS | 61 | 0.569688 | 1.731055 | 0.001362 | 0.096652 |
| GOBP_ENDODERMAL_CELL_DIFFERENTIATION | 40 | 0.596671 | 1.729674 | 0.007163 | 0.097354 |
| GOBP_ARTERY_MORPHOGENESIS | 59 | 0.560768 | 1.724954 | 0.00137 | 0.101889 |
| GOBP_RESPONSE_TO_MUSCLE_STRETCH | 21 | 0.672064 | 1.72163 | 0.010903 | 0.104544 |
| GOBP_CARDIAC_MUSCLE_TISSUE_MORPHOGENESIS | 51 | 0.569795 | 1.716604 | 0.001374 | 0.109297 |
| GOBP_INTERFERON_GAMMA_PRODUCTION | 67 | 0.54754 | 1.715644 | 0.002663 | 0.109369 |
| GOBP_MUSCLE_ORGAN_DEVELOPMENT | 272 | 0.467104 | 1.714689 | 0 | 0.109169 |
| GOBP_AORTIC_VALVE_MORPHOGENESIS | 30 | 0.620905 | 1.710902 | 0.001497 | 0.112609 |
| GOBP_REGIONALIZATION | 257 | 0.472631 | 1.710257 | 0 | 0.112027 |
| GOBP_SYNAPTONEMAL_COMPLEX_ORGANIZATION | 20 | 0.683724 | 1.706908 | 0.004792 | 0.115202 |
| GOBP_NEGATIVE_REGULATION_OF_EXOCYTOSIS | 27 | 0.639114 | 1.706188 | 0.009231 | 0.114737 |
| GOBP_EMBRYONIC_CRANIAL_SKELETON_MORPHOGENESIS | 39 | 0.587281 | 1.704403 | 0.004458 | 0.115948 |
| GOBP_VENTRICULAR_CARDIAC_MUSCLE_TISSUE_MORPHOGENESIS | 41 | 0.582198 | 1.703257 | 0.00438 | 0.116242 |
| GOBP_SARCOMERE_ORGANIZATION | 32 | 0.627609 | 1.702668 | 0.00321 | 0.115663 |
| GOBP_NEGATIVE_REGULATION_OF_MYOBLAST_DIFFERENTIATION | 18 | 0.680256 | 1.701911 | 0.011551 | 0.115127 |
| GOBP_NUCLEOBASE_CONTAINING_SMALL_MOLECULE_CATABOLIC_PROCESS | 25 | 0.649903 | 1.69672 | 0.009479 | 0.121368 |
| GOBP_TISSUE_REGENERATION | 48 | 0.56802 | 1.696636 | 0.002911 | 0.119993 |
| GOBP_MUSCLE_CELL_DIFFERENTIATION | 291 | 0.461009 | 1.69625 | 0 | 0.119054 |
| GOBP_GRANULOCYTE_MIGRATION | 89 | 0.521486 | 1.696132 | 0.001252 | 0.117861 |
| GOBP_NEGATIVE_REGULATION_OF_STRIATED_MUSCLE_CELL_APOPTOTIC_PROCESS | 19 | 0.685563 | 1.694838 | 0.006369 | 0.118325 |
| GOBP_REGULATION_OF_CELL_MATRIX_ADHESION | 105 | 0.511525 | 1.693243 | 0 | 0.119323 |
| GOBP_VASCULAR_ASSOCIATED_SMOOTH_MUSCLE_CELL_PROLIFERATION | 46 | 0.577358 | 1.69254 | 0.004286 | 0.119043 |
| GOBP_SKELETAL_SYSTEM_MORPHOGENESIS | 165 | 0.477763 | 1.688878 | 0 | 0.123339 |
| GOBP_RIBONUCLEOSIDE_CATABOLIC_PROCESS | 15 | 0.728141 | 1.688177 | 0.008547 | 0.123092 |
| GOBP_POSITIVE_REGULATION_OF_FILOPODIUM_ASSEMBLY | 28 | 0.62478 | 1.687752 | 0.010622 | 0.122439 |
| GOBP_LEUKOCYTE_CHEMOTAXIS | 144 | 0.487061 | 1.686546 | 0 | 0.123029 |
| GOBP_MATERNAL_PROCESS_INVOLVED_IN_FEMALE_PREGNANCY | 45 | 0.57385 | 1.686041 | 0.004348 | 0.122483 |
| GOBP_TISSUE_MIGRATION | 268 | 0.458947 | 1.684814 | 0 | 0.122944 |
| GOBP_ENDOCARDIAL_CUSHION_DEVELOPMENT | 44 | 0.569159 | 1.684403 | 0 | 0.122387 |
| GOBP_MUSCLE_CELL_DEVELOPMENT | 142 | 0.491271 | 1.684123 | 0 | 0.121543 |
| GOBP_POSITIVE_REGULATION_OF_CD4_POSITIVE_ALPHA_BETA_T_CELL_ACTIVATION | 26 | 0.636776 | 1.683136 | 0.012857 | 0.121914 |
| GOBP_REGULATION_OF_INFLAMMATORY_RESPONSE_TO_ANTIGENIC_STIMULUS | 32 | 0.602153 | 1.682591 | 0.007508 | 0.121511 |
| GOBP_INFLAMMATORY_CELL_APOPTOTIC_PROCESS | 17 | 0.691486 | 1.681444 | 0.012442 | 0.121958 |
| GOBP_SEMI_LUNAR_VALVE_DEVELOPMENT | 39 | 0.588339 | 1.675389 | 0.001479 | 0.130049 |
| GOBP_POSITIVE_REGULATION_OF_GRANULOCYTE_CHEMOTAXIS | 19 | 0.671877 | 1.674052 | 0.009677 | 0.130331 |
| GOBP_REGULATION_OF_SKELETAL_MUSCLE_CELL_DIFFERENTIATION | 17 | 0.699589 | 1.672696 | 0.009569 | 0.130964 |
| GOBP_REGULATION_OF_REPRODUCTIVE_PROCESS | 117 | 0.487594 | 1.672631 | 0.002481 | 0.129813 |
| GOBP_SPECIFICATION_OF_SYMMETRY | 112 | 0.496552 | 1.669195 | 0.001267 | 0.133739 |
| GOBP_EMBRYONIC_FORELIMB_MORPHOGENESIS | 16 | 0.708595 | 1.665795 | 0.008143 | 0.138185 |
| GOBP_ENDOCARDIAL_CUSHION_MORPHOGENESIS | 37 | 0.59608 | 1.664299 | 0.007364 | 0.139156 |
| GOBP_CARDIAC_CONDUCTION_SYSTEM_DEVELOPMENT | 28 | 0.617015 | 1.664026 | 0.00744 | 0.138146 |
| GOBP_MYOTUBE_DIFFERENTIATION | 93 | 0.504898 | 1.662858 | 0.001326 | 0.138574 |
| GOBP_POSITIVE_REGULATION_OF_CELL_SUBSTRATE_ADHESION | 103 | 0.503076 | 1.662511 | 0.001241 | 0.137822 |
| GOBP_SPLEEN_DEVELOPMENT | 26 | 0.629137 | 1.661568 | 0.007849 | 0.138209 |
| GOBP_PHARYNGEAL_SYSTEM_DEVELOPMENT | 25 | 0.624218 | 1.660262 | 0.007463 | 0.138999 |
| GOBP_PROTEIN_KINASE_C_SIGNALING | 26 | 0.628416 | 1.658604 | 0.016591 | 0.140169 |
| GOBP_HEART_MORPHOGENESIS | 219 | 0.460856 | 1.657823 | 0 | 0.140078 |
| GOBP_PATTERN_SPECIFICATION_PROCESS | 341 | 0.44398 | 1.65694 | 0 | 0.140224 |
| GOBP_STRIATED_MUSCLE_CELL_APOPTOTIC_PROCESS | 32 | 0.599532 | 1.656814 | 0.00936 | 0.139193 |
| GOBP_REGULATION_OF_MYOTUBE_DIFFERENTIATION | 33 | 0.611167 | 1.655009 | 0.009105 | 0.140836 |
| GOBP_LEFT_RIGHT_PATTERN_FORMATION | 22 | 0.649224 | 1.653012 | 0.009804 | 0.142415 |
| GOBP_CELL_PROLIFERATION_INVOLVED_IN_HEART_MORPHOGENESIS | 17 | 0.684494 | 1.651907 | 0.011765 | 0.142892 |
| GOBP_ANTERIOR_POSTERIOR_PATTERN_SPECIFICATION | 145 | 0.477466 | 1.651857 | 0.001186 | 0.141761 |
| GOBP_POSITIVE_REGULATION_OF_ALPHA_BETA_T_CELL_PROLIFERATION | 16 | 0.697647 | 1.65182 | 0.007949 | 0.140648 |
| GOBP_CORONARY_VASCULATURE_DEVELOPMENT | 41 | 0.56222 | 1.651505 | 0.005789 | 0.140005 |
| GOBP_MAMMARY_GLAND_DUCT_MORPHOGENESIS | 22 | 0.660801 | 1.651244 | 0.006601 | 0.139247 |
| GOBP_VASCULAR_ENDOTHELIAL_GROWTH_FACTOR_PRODUCTION | 29 | 0.604838 | 1.650539 | 0.009494 | 0.139279 |
| GOBP_SMOOTH_MUSCLE_CELL_DIFFERENTIATION | 47 | 0.564393 | 1.647728 | 0.011189 | 0.14246 |
| GOBP_REGULATION_OF_EPITHELIAL_CELL_MIGRATION | 202 | 0.457765 | 1.645517 | 0.001157 | 0.144703 |
| GOBP_DENDRITIC_CELL_MIGRATION | 21 | 0.642985 | 1.644793 | 0.024834 | 0.144536 |
| GOBP_POSITIVE_REGULATION_OF_HEMOPOIESIS | 116 | 0.486496 | 1.64355 | 0.001259 | 0.145142 |
| GOBP_REGULATION_OF_FATTY_ACID_BIOSYNTHETIC_PROCESS | 31 | 0.601508 | 1.642746 | 0.010189 | 0.145193 |
| GOBP_POSITIVE_REGULATION_OF_COLD_INDUCED_THERMOGENESIS | 79 | 0.511306 | 1.642344 | 0.00672 | 0.144683 |
| GOBP_NEUTROPHIL_ACTIVATION_INVOLVED_IN_IMMUNE_RESPONSE | 16 | 0.688312 | 1.642102 | 0.004992 | 0.143912 |
| GOBP_REGULATION_OF_CELL_ADHESION_MEDIATED_BY_INTEGRIN | 39 | 0.574246 | 1.641195 | 0.014514 | 0.144251 |
| GOBP_DNA_DAMAGE_RESPONSE_SIGNAL_TRANSDUCTION_BY_P53_CLASS_MEDIATOR | 68 | 0.518092 | 1.640781 | 0.005413 | 0.143955 |
| GOBP_MALE_MEIOTIC_NUCLEAR_DIVISION | 36 | 0.592238 | 1.640733 | 0.01059 | 0.142934 |
| GOBP_POSITIVE_REGULATION_OF_VASCULATURE_DEVELOPMENT | 131 | 0.48242 | 1.63928 | 0 | 0.144098 |
| GOBP_REGULATION_OF_SPROUTING_ANGIOGENESIS | 33 | 0.585688 | 1.637866 | 0.008746 | 0.145128 |
| GOBP_T_HELPER_1_TYPE_IMMUNE_RESPONSE | 31 | 0.592744 | 1.636599 | 0.010511 | 0.145881 |
| GOBP_AMEBOIDAL_TYPE_CELL_MIGRATION | 372 | 0.436793 | 1.635844 | 0 | 0.14593 |
| GOBP_MONONUCLEAR_CELL_MIGRATION | 119 | 0.480875 | 1.63552 | 0 | 0.145388 |
| GOBP_HETEROPHILIC_CELL_CELL_ADHESION_VIA_PLASMA_MEMBRANE_CELL_ADHESIO | 31 | 0.603747 | 1.63536 | 0.006061 | 0.144564 |
| GOBP_VENTRICULAR_SEPTUM_MORPHOGENESIS | 40 | 0.572942 | 1.634471 | 0.010294 | 0.144931 |
| GOBP_CARDIAC_NEURAL_CREST_CELL_DIFFERENTIATION_INVOLVED_IN_HEART_DEVEL | 15 | 0.681873 | 1.634114 | 0.014493 | 0.144472 |
| GOBP_INTERLEUKIN_12_PRODUCTION | 42 | 0.564976 | 1.633588 | 0.008633 | 0.144333 |
| GOBP_MAMMARY_GLAND_EPITHELIUM_DEVELOPMENT | 56 | 0.531014 | 1.633078 | 0.005319 | 0.144173 |
| GOBP_ENDOTHELIAL_CELL_MIGRATION | 190 | 0.458451 | 1.633075 | 0 | 0.143172 |
| GOBP_BRANCHING_INVOLVED_IN_MAMMARY_GLAND_DUCT_MORPHOGENESIS | 18 | 0.665283 | 1.631435 | 0.009836 | 0.144735 |
| GOBP_ENDODERM_DEVELOPMENT | 70 | 0.516177 | 1.629875 | 0.002674 | 0.14611 |
| GOBP_SKELETAL_MUSCLE_TISSUE_REGENERATION | 28 | 0.603885 | 1.629142 | 0.01072 | 0.146288 |
| GOBP_REGULATION_OF_STRIATED_MUSCLE_CELL_DIFFERENTIATION | 66 | 0.521064 | 1.628813 | 0.00542 | 0.145822 |
| GOBP_POSITIVE_REGULATION_OF_CELL_ADHESION | 332 | 0.438853 | 1.627576 | 0 | 0.146733 |
| GOBP_POSITIVE_REGULATION_OF_INTERLEUKIN_4_PRODUCTION | 16 | 0.685845 | 1.626681 | 0.014041 | 0.147204 |
| GOBP_TRANSPOSITION | 24 | 0.627866 | 1.625817 | 0.021407 | 0.147547 |
| GOBP_POSITIVE_REGULATION_OF_CD4_POSITIVE_ALPHA_BETA_T_CELL_DIFFERENTIATIC | 21 | 0.638644 | 1.625019 | 0.016207 | 0.14781 |
| GOBP_EMBRYONIC_CAMERA_TYPE_EYE_MORPHOGENESIS | 18 | 0.67653 | 1.624345 | 0.004762 | 0.147909 |

**GSEA Negative** Hematopoietic/Immune pathways are highlighted in green.

| Gene sets | SIZE | ES | NES | NOM p-val | FDR q-val |
|---|---|---|---|---|---|
| GOBP_DNA_REPLICATION_DEPENDENT_CHROMATIN_ORGANIZATION | 21 | -0.807205 | -2.278853 | 0 | 9.23E-04 |
| GOBP_NUCLEOSOME_ASSEMBLY | 88 | -0.543113 | -2.108989 | 0 | 0.016182 |
| GOBP_NUCLEOSOME_ORGANIZATION | 133 | -0.497426 | -2.056483 | 0 | 0.029023 |
| GOBP_DNA_REPLICATION_INDEPENDENT_CHROMATIN_ORGANIZATION | 28 | -0.638257 | -1.956092 | 0 | 0.097741 |
| GOBP_PROTEIN_DNA_COMPLEX_ASSEMBLY | 158 | -0.463837 | -1.923578 | 0 | 0.122298 |
| GOBP_PROTEIN_DNA_COMPLEX_SUBUNIT_ORGANIZATION | 204 | -0.443144 | -1.920495 | 0 | 0.105467 |
| GOBP_L_GLUTAMATE_TRANSMEMBRANE_TRANSPORT | 24 | -0.647468 | -1.902091 | 0.003115 | 0.112665 |
| GOBP_CHROMATIN_ASSEMBLY_OR_DISASSEMBLY | 188 | -0.431188 | -1.81844 | 0 | 0.267777 |
| GOBP_REGULATION_OF_INTRINSIC_APOPTOTIC_SIGNALING_PATHWAY_IN_RESPONSE_T | 16 | -0.647037 | -1.759945 | 0.010363 | 0.42438 |
| GOBP_CHROMATIN_REMODELING | 217 | -0.403155 | -1.750548 | 0 | 0.416457 |
| GOBP_DNA_STRAND_ELONGATION_INVOLVED_IN_DNA_REPLICATION | 15 | -0.660472 | -1.737741 | 0.005128 | 0.429143 |
| GOBP_DNA_PACKAGING | 201 | -0.402442 | -1.736467 | 0 | 0.397547 |
| GOBP_ANTIBACTERIAL_HUMORAL_RESPONSE | 21 | -0.590945 | -1.703955 | 0.020408 | 0.493805 |
| GOBP_RNA_SPLICING_VIA_TRANSESTERIFICATION_REACTIONS | 280 | -0.374834 | -1.693858 | 0 | 0.505162 |
| GOBP_OLIGODENDROCYTE_DEVELOPMENT | 33 | -0.530794 | -1.67592 | 0.020057 | 0.551638 |
| GOBP_POSITIVE_REGULATION_OF_AMYLOID_BETA_FORMATION | 16 | -0.638395 | -1.667095 | 0.030471 | 0.555856 |
| GOBP_CHLORIDE_TRANSMEMBRANE_TRANSPORT | 48 | -0.483141 | -1.662747 | 0 | 0.543052 |
| GOBP_NEGATIVE_REGULATION_OF_INTRINSIC_APOPTOTIC_SIGNALING_PATHWAY_IN_RE | 29 | -0.536063 | -1.653206 | 0.016949 | 0.552731 |
| GOBP_CLEAVAGE_INVOLVED_IN_RRNA_PROCESSING | 27 | -0.548347 | -1.652604 | 0.008043 | 0.526375 |
| GOBP_ACID_SECRETION | 31 | -0.530691 | -1.651645 | 0.017192 | 0.504122 |
| GOBP_MATURATION_OF_5_8S_RRNA | 35 | -0.508048 | -1.649151 | 0.009259 | 0.48936 |
| GOBP_MATURATION_OF_SSU_RRNA_FROM_TRICISTRONIC_RRNA_TRANSCRIPT_SSU_RF | 37 | -0.503967 | -1.646281 | 0.009646 | 0.477963 |
| GOBP_DNA_UNWINDING_INVOLVED_IN_DNA_REPLICATION | 22 | -0.55945 | -1.642924 | 0.008086 | 0.470281 |
| GOBP_DNA_REPLICATION_INITIATION | 35 | -0.516867 | -1.640736 | 0.006431 | 0.459336 |
| GOBP_POSITIVE_REGULATION_OF_MRNA_SPLICING_VIA_SPLICEOSOME | 20 | -0.590668 | -1.640145 | 0.017493 | 0.443057 |
| GOBP_SYNAPTIC_VESICLE_MEMBRANE_ORGANIZATION | 23 | -0.555946 | -1.63973 | 0.014793 | 0.427851 |
| GOBP_GAMMA_AMINOBUTYRIC_ACID_SIGNALING_PATHWAY | 21 | -0.570317 | -1.635189 | 0.008174 | 0.427016 |
| GOBP_PYRIMIDINE_NUCLEOSIDE_TRIPHOSPHATE_METABOLIC_PROCESS | 24 | -0.542451 | -1.626558 | 0.011461 | 0.44231 |
| GOBP_THIOESTER_BIOSYNTHETIC_PROCESS | 40 | -0.491762 | -1.619547 | 0.0125 | 0.44937 |
| GOBP_C4_DICARBOXYLATE_TRANSPORT | 25 | -0.551191 | -1.614421 | 0.027027 | 0.452113 |
| GOBP_L_GLUTAMATE_IMPORT_ACROSS_PLASMA_MEMBRANE | 15 | -0.620499 | -1.602496 | 0.023316 | 0.481424 |
| GOBP_MATURATION_OF_SSU_RRNA | 50 | -0.453852 | -1.601183 | 0.013201 | 0.471572 |
| GOBP_CELL_CYCLE_DNA_REPLICATION | 41 | -0.474055 | -1.601043 | 0.016835 | 0.457812 |
| GOBP_POSITIVE_REGULATION_OF_METAPHASE_ANAPHASE_TRANSITION_OF_CELL_CYC | 15 | -0.617931 | -1.59927 | 0.023316 | 0.450658 |
| GOBP_OLEFINIC_COMPOUND_BIOSYNTHETIC_PROCESS | 17 | -0.583272 | -1.596823 | 0.023256 | 0.445425 |
| GOBP_ORGAN_OR_TISSUE_SPECIFIC_IMMUNE_RESPONSE | 16 | -0.597719 | -1.590627 | 0.038278 | 0.455126 |
| GOBP_INORGANIC_ANION_TRANSMEMBRANE_TRANSPORT | 69 | -0.429922 | -1.588439 | 0.003953 | 0.450233 |
| GOBP_NEUROTRANSMITTER_REUPTAKE | 26 | -0.533115 | -1.5868 | 0.019943 | 0.443876 |
| GOBP_POSITIVE_REGULATION_OF_AMYLOID_PRECURSOR_PROTEIN_CATABOLIC_PROCE | 21 | -0.568577 | -1.583555 | 0.020997 | 0.443389 |
| GOBP_REGULATION_OF_INTRINSIC_APOPTOTIC_SIGNALING_PATHWAY_IN_RESPONSE_T | 37 | -0.482241 | -1.582676 | 0.020057 | 0.435263 |
| GOBP_PROTEIN_LOCALIZATION_TO_MICROTUBULE | 17 | -0.584809 | -1.582085 | 0.036745 | 0.426876 |
| GOBP_ENDONUCLEOLYTIC_CLEAVAGE_INVOLVED_IN_RRNA_PROCESSING | 15 | -0.612414 | -1.570903 | 0.027027 | 0.455193 |
| GOBP_DE_NOVO_PROTEIN_FOLDING | 41 | -0.466223 | -1.570148 | 0.016181 | 0.44706 |
| GOBP_DNA_CONFORMATION_CHANGE | 297 | -0.348402 | -1.56631 | 0 | 0.449638 |
| GOBP_ANION_TRANSMEMBRANE_TRANSPORT | 196 | -0.358614 | -1.56445 | 0 | 0.445699 |
| GOBP_MODIFIED_AMINO_ACID_TRANSPORT | 25 | -0.537589 | -1.556059 | 0.043103 | 0.464271 |
| GOBP_CHAPERONE_COFACTOR_DEPENDENT_PROTEIN_REFOLDING | 32 | -0.497773 | -1.556046 | 0.026239 | 0.454413 |
| GOBP_SPLICEOSOMAL_SNRNP_ASSEMBLY | 37 | -0.470095 | -1.551197 | 0.021472 | 0.461301 |
| GOBP_CALCIUM_DEPENDENT_CELL_CELL_ADHESION_VIA_PLASMA_MEMBRANE_CELL_A | 33 | -0.48965 | -1.539113 | 0.025714 | 0.493702 |
| GOBP_POSITIVE_REGULATION_OF_CYTOKINESIS | 30 | -0.484079 | -1.538647 | 0.020896 | 0.485641 |
| GOBP_NEUROTRANSMITTER_UPTAKE | 33 | -0.493148 | -1.537168 | 0.029155 | 0.481077 |
| GOBP_G1_TO_G0_TRANSITION | 17 | -0.560707 | -1.52761 | 0.029333 | 0.504992 |
| GOBP_CHLORIDE_TRANSPORT | 81 | -0.404391 | -1.525511 | 0.012987 | 0.502979 |
| GOBP_REGULATION_OF_TRANSCRIPTION_BY_RNA_POLYMERASE_I | 33 | -0.483339 | -1.524195 | 0.02439 | 0.498284 |
| GOBP_WALKING_BEHAVIOR | 26 | -0.511876 | -1.523962 | 0.040936 | 0.489913 |
| GOBP_SRP_DEPENDENT_COTRANSLATIONAL_PROTEIN_TARGETING_TO_MEMBRANE | 18 | -0.554725 | -1.52247 | 0.042781 | 0.486532 |
| GOBP_RESPONSE_TO_IMMOBILIZATION_STRESS | 15 | -0.570572 | -1.522094 | 0.033419 | 0.479153 |
| GOBP_REGULATION_OF_LONG_TERM_NEURONAL_SYNAPTIC_PLASTICITY | 21 | -0.53603 | -1.522016 | 0.02957 | 0.471163 |
| GOBP_RESPONSE_TO_DOPAMINE | 59 | -0.427948 | -1.519985 | 0.015094 | 0.469826 |
| GOBP_POSITIVE_REGULATION_OF_DENDRITIC_SPINE_MORPHOGENESIS | 19 | -0.54121 | -1.518979 | 0.040404 | 0.465494 |
| GOBP_TELOMERE_ORGANIZATION | 146 | -0.362357 | -1.508404 | 0 | 0.492138 |
| GOBP_POSITIVE_REGULATION_OF_MRNA_PROCESSING | 29 | -0.49417 | -1.50598 | 0.021739 | 0.492733 |
| GOBP_DORSAL_VENTRAL_NEURAL_TUBE_PATTERNING | 19 | -0.520637 | -1.50594 | 0.036827 | 0.485087 |
| GOBP_REGULATION_OF_RECEPTOR_INTERNALIZATION | 51 | -0.435167 | -1.505379 | 0.017182 | 0.479327 |
| GOBP_VESICLE_FUSION_TO_PLASMA_MEMBRANE | 18 | -0.554999 | -1.505065 | 0.05042 | 0.473 |
| GOBP_VERY_LONG_CHAIN_FATTY_ACID_METABOLIC_PROCESS | 31 | -0.486617 | -1.502587 | 0.038889 | 0.474102 |
| GOBP_ACETYL_COA_BIOSYNTHETIC_PROCESS | 16 | -0.570103 | -1.501247 | 0.052356 | 0.471043 |
| GOBP_PROTEIN_LIPID_COMPLEX_ASSEMBLY | 22 | -0.530037 | -1.50024 | 0.029155 | 0.467284 |
| GOBP_INORGANIC_ANION_TRANSPORT | 137 | -0.368031 | -1.499541 | 0 | 0.462742 |
| GOBP_ANAPHASE_PROMOTING_COMPLEX_DEPENDENT_CATABOLIC_PROCESS | 16 | -0.570025 | -1.498248 | 0.052632 | 0.460196 |
| GOBP_MATURATION_OF_5_8S_RRNA_FROM_TRICISTRONIC_RRNA_TRANSCRIPT_SSU_R | 24 | -0.512722 | -1.483405 | 0.036827 | 0.501901 |
| GOBP_C21_STEROID_HORMONE_BIOSYNTHETIC_PROCESS | 16 | -0.550456 | -1.482408 | 0.047478 | 0.498155 |
| GOBP_L_AMINO_ACID_TRANSPORT | 54 | -0.416441 | -1.481719 | 0.026403 | 0.493566 |
| GOBP_BILE_ACID_BIOSYNTHETIC_PROCESS | 26 | -0.499971 | -1.474417 | 0.041209 | 0.511624 |
| GOBP_AXON_ENSHEATHMENT_IN_CENTRAL_NERVOUS_SYSTEM | 19 | -0.532053 | -1.473828 | 0.05168 | 0.506674 |
| GOBP_SODIUM_ION_HOMEOSTASIS | 36 | -0.449163 | -1.465382 | 0.042813 | 0.528173 |
| GOBP_PYRIMIDINE_NUCLEOSIDE_TRIPHOSPHATE_BIOSYNTHETIC_PROCESS | 19 | -0.538214 | -1.463399 | 0.068493 | 0.528316 |
| GOBP_SPHINGOMYELIN_METABOLIC_PROCESS | 16 | -0.540154 | -1.45534 | 0.082474 | 0.548499 |
| GOBP_POSITIVE_REGULATION_OF_TRANSCRIPTION_BY_RNA_POLYMERASE_I | 23 | -0.489705 | -1.449801 | 0.076923 | 0.561477 |
| GOBP_DNA_REPLICATION_CHECKPOINT_SIGNALING | 15 | -0.544378 | -1.437774 | 0.084577 | 0.598003 |
| GOBP_REGULATION_OF_NEURONAL_SYNAPTIC_PLASTICITY | 46 | -0.422774 | -1.436451 | 0.041096 | 0.595633 |
| GOBP_DOPAMINE_RECEPTOR_SIGNALING_PATHWAY | 34 | -0.445644 | -1.435475 | 0.066465 | 0.592344 |
| GOBP_REGULATION_OF_MEGAKARYOCYTE_DIFFERENTIATION | 31 | -0.456314 | -1.42001 | 0.070988 | 0.644647 |
| GOBP_INTERMEMBRANE_LIPID_TRANSFER | 43 | -0.424392 | -1.419657 | 0.052459 | 0.638351 |
| GOBP_RESPONSE_TO_CATECHOLAMINE | 72 | -0.381679 | -1.419389 | 0.02682 | 0.631843 |
| GOBP_FATTY_ACYL_COA_BIOSYNTHETIC_PROCESS | 22 | -0.493617 | -1.418717 | 0.072581 | 0.627065 |
| GOBP_TRACHEA_DEVELOPMENT | 15 | -0.533586 | -1.418083 | 0.079365 | 0.62213 |
| GOBP_INDOLE_CONTAINING_COMPOUND_METABOLIC_PROCESS | 15 | -0.560397 | -1.416951 | 0.09 | 0.619733 |
| GOBP_RESPONSE_TO_ACETYLCHOLINE | 25 | -0.474359 | -1.414916 | 0.088949 | 0.620591 |
| GOBP_OLIGOPEPTIDE_TRANSPORT | 15 | -0.544946 | -1.413444 | 0.094629 | 0.619381 |
| GOBP_REGULATION_OF_POSTSYNAPTIC_MEMBRANE_NEUROTRANSMITTER_RECEPTOR | 50 | -0.415554 | -1.411224 | 0.042105 | 0.620791 |
| GOBP_REGULATION_OF_SYSTEMIC_ARTERIAL_BLOOD_PRESSURE_BY_HORMONE | 23 | -0.485604 | -1.409835 | 0.065104 | 0.618921 |
| GOBP_POSITIVE_REGULATION_OF_DENDRITE_MORPHOGENESIS | 34 | -0.450262 | -1.407261 | 0.06192 | 0.621605 |
| GOBP_MONOVALENT_INORGANIC_CATION_HOMEOSTASIS | 115 | -0.353224 | -1.404286 | 0.015152 | 0.625722 |
| GOBP_POSTTRANSLATIONAL_PROTEIN_TARGETING_TO_ENDOPLASMIC_RETICULUM_ME | 15 | -0.541255 | -1.402051 | 0.115068 | 0.628032 |
| GOBP_POSITIVE_REGULATION_OF_DOUBLE_STRAND_BREAK_REPAIR_VIA_NONHOMOLO | 16 | -0.524383 | -1.396359 | 0.094629 | 0.642122 |
| GOBP_RESPONSE_TO_COCAINE | 40 | -0.418733 | -1.394523 | 0.068966 | 0.642415 |
| GOBP_RIBONUCLEOPROTEIN_COMPLEX_BIOGENESIS | 417 | -0.301166 | -1.393071 | 0 | 0.64139 |
| GOBP_DEFENSE_RESPONSE_TO_GRAM_POSITIVE_BACTERIUM | 44 | -0.411683 | -1.392685 | 0.042553 | 0.636401 |
| GOBP_CHAPERONE_MEDIATED_PROTEIN_FOLDING | 67 | -0.385382 | -1.389078 | 0.031496 | 0.643827 |
| GOBP_NEGATIVE_REGULATION_OF_DOUBLE_STRAND_BREAK_REPAIR_VIA_HOMOLOGOU | 19 | -0.489653 | -1.388296 | 0.107895 | 0.640538 |
| GOBP_PROTEIN_NEDDYLATION | 17 | -0.510393 | -1.387593 | 0.113065 | 0.636614 |
| GOBP_SPLICEOSOMAL_COMPLEX_ASSEMBLY | 57 | -0.397997 | -1.385992 | 0.036913 | 0.636478 |
| GOBP_NEGATIVE_REGULATION_OF_DNA_TEMPLATED_TRANSCRIPTION_ELONGATION | 16 | -0.521354 | -1.385555 | 0.087432 | 0.632118 |
| GOBP_TRANSCRIPTION_BY_RNA_POLYMERASE_I | 53 | -0.393523 | -1.384044 | 0.040816 | 0.631288 |
| GOBP_NUCLEOLAR_LARGE_RRNA_TRANSCRIPTION_BY_RNA_POLYMERASE_I | 18 | -0.508495 | -1.382703 | 0.07772 | 0.630332 |
| GOBP_POSTSYNAPTIC_SIGNAL_TRANSDUCTION | 31 | -0.44448 | -1.381789 | 0.067278 | 0.627867 |
| GOBP_ACIDIC_AMINO_ACID_TRANSPORT | 56 | -0.37919 | -1.377151 | 0.033557 | 0.6394 |
| GOBP_REGULATION_OF_MRNA_SPLICING_VIA_SPLICEOSOME | 97 | -0.348112 | -1.375522 | 0.019231 | 0.639571 |
| GOBP_POSITIVE_REGULATION_OF_RECEPTOR_INTERNALIZATION | 25 | -0.45436 | -1.369059 | 0.090141 | 0.657909 |
| GOBP_RIBOSOMAL_SMALL_SUBUNIT_BIOGENESIS | 73 | -0.366086 | -1.361906 | 0.054393 | 0.679399 |
| GOBP_RRNA_TRANSCRIPTION | 34 | -0.426546 | -1.361589 | 0.092199 | 0.674451 |
| GOBP_ACETYL_COA_METABOLIC_PROCESS | 28 | -0.44663 | -1.360862 | 0.081159 | 0.671421 |
| GOBP_MITOCHONDRIAL_RNA_PROCESSING | 20 | -0.477351 | -1.358677 | 0.105263 | 0.673291 |
| GOBP_RENAL_TUBULAR_SECRETION | 15 | -0.497898 | -1.356666 | 0.12 | 0.67495 |
| GOBP_NEGATIVE_REGULATION_OF_DNA_RECOMBINATION | 39 | -0.403275 | -1.35178 | 0.079511 | 0.68846 |
| GOBP_AMINO_ACID_TRANSMEMBRANE_TRANSPORT | 76 | -0.354986 | -1.351288 | 0.037594 | 0.684453 |
| GOBP_DNA_STRAND_ELONGATION | 36 | -0.423903 | -1.350462 | 0.075235 | 0.68195 |
| GOBP_SYNAPTIC_TRANSMISSION_GABAERGIC | 44 | -0.404242 | -1.349364 | 0.073955 | 0.680264 |
| GOBP_REGULATION_OF_EXCRETION | 15 | -0.515735 | -1.349033 | 0.11655 | 0.675859 |
| GOBP_PROTEIN_DNA_COMPLEX_DISASSEMBLY | 19 | -0.489277 | -1.348087 | 0.11054 | 0.674253 |
| GOBP_POSITIVE_REGULATION_OF_CHROMOSOME_SEPARATION | 17 | -0.499418 | -1.348039 | 0.125348 | 0.668877 |
| GOBP_ADENYLATE_CYCLASE_ACTIVATING_ADRENERGIC_RECEPTOR_SIGNALING_PATH\ | 19 | -0.483213 | -1.344738 | 0.110512 | 0.676572 |
| GOBP_NEGATIVE_REGULATION_OF_RNA_SPLICING | 25 | -0.449255 | -1.344622 | 0.107438 | 0.671554 |
| GOBP_ANION_HOMEOSTASIS | 30 | -0.439382 | -1.344515 | 0.103825 | 0.666762 |
| GOBP_REGULATION_OF_BEHAVIOR | 45 | -0.391608 | -1.344255 | 0.086262 | 0.662398 |
| GOBP_CELL_VOLUME_HOMEOSTASIS | 28 | -0.439414 | -1.343638 | 0.093939 | 0.659497 |
| GOBP_NEGATIVE_REGULATION_OF_MRNA_SPLICING_VIA_SPLICEOSOME | 20 | -0.474641 | -1.341043 | 0.129534 | 0.664058 |
| GOBP_TRNA_5_END_PROCESSING | 16 | -0.502142 | -1.338437 | 0.1175 | 0.668055 |
| GOBP_RHYTHMIC_BEHAVIOR | 33 | -0.420219 | -1.33678 | 0.069971 | 0.669224 |
| GOBP_PHOSPHATE_ION_TRANSPORT | 22 | -0.45767 | -1.335929 | 0.126316 | 0.667182 |
| GOBP_RRNA_METABOLIC_PROCESS | 258 | -0.301022 | -1.334967 | 0 | 0.665561 |
| GOBP_REGULATION_OF_DNA_DEPENDENT_DNA_REPLICATION | 49 | -0.384538 | -1.334647 | 0.075 | 0.661768 |
| GOBP_DICARBOXYLIC_ACID_TRANSPORT | 74 | -0.350684 | -1.334632 | 0.03937 | 0.656898 |
| GOBP_REGULATION_OF_SYSTEMIC_ARTERIAL_BLOOD_PRESSURE_MEDIATED_BY_A_CH | 30 | -0.424126 | -1.329788 | 0.105263 | 0.669588 |
| GOBP_KETONE_BIOSYNTHETIC_PROCESS | 36 | -0.416184 | -1.328787 | 0.075529 | 0.668099 |
| GOBP_POSITIVE_REGULATION_OF_DENDRITIC_SPINE_DEVELOPMENT | 35 | -0.414285 | -1.328452 | 0.108025 | 0.664525 |
| GOBP_RNA_SPLICING | 387 | -0.292148 | -1.3265 | 0.013699 | 0.667132 |
| GOBP_RIBOSOME_BIOGENESIS | 297 | -0.296972 | -1.326488 | 0.011765 | 0.662379 |
| GOBP_MONOAMINE_TRANSPORT | 61 | -0.367773 | -1.323399 | 0.052632 | 0.668737 |
| GOBP_NUCLEOLUS_ORGANIZATION | 16 | -0.482812 | -1.321543 | 0.106742 | 0.670899 |
| GOBP_MRNA_PROCESSING | 440 | -0.279851 | -1.320665 | 0 | 0.669449 |
| GOBP_NEGATIVE_REGULATION_OF_CALCIUM_ION_TRANSMEMBRANE_TRANSPORT | 30 | -0.427217 | -1.320339 | 0.126126 | 0.665892 |
| GOBP_SULFUR_COMPOUND_BIOSYNTHETIC_PROCESS | 130 | -0.326374 | -1.318251 | 0.02765 | 0.669163 |
| GOBP_POSITIVE_REGULATION_OF_TORC1_SIGNALING | 20 | -0.463353 | -1.315477 | 0.140921 | 0.674562 |
| GOBP_CYTOPLASMIC_SEQUESTERING_OF_PROTEIN | 21 | -0.467396 | -1.313489 | 0.147399 | 0.677518 |
| GOBP_SNO_S_RNA_METABOLIC_PROCESS | 16 | -0.491454 | -1.312555 | 0.15285 | 0.676236 |
| GOBP_REGULATION_OF_TRANSCRIPTION_OF_NUCLEOLAR_LARGE_RRNA_BY_RNA_POL | 15 | -0.489878 | -1.312439 | 0.131367 | 0.672073 |
| GOBP_ANION_TRANSPORT | 388 | -0.289727 | -1.311097 | 0 | 0.672395 |
| GOBP_NEGATIVE_REGULATION_OF_TOLL_LIKE_RECEPTOR_SIGNALING_PATHWAY | 27 | -0.443186 | -1.311035 | 0.13467 | 0.668128 |
| GOBP_MITOCHONDRION_MORPHOGENESIS | 20 | -0.461951 | -1.309621 | 0.12234 | 0.668728 |
| GOBP_STEROID_HORMONE_BIOSYNTHETIC_PROCESS | 27 | -0.425579 | -1.30853 | 0.127479 | 0.668349 |
| GOBP_REGULATION_OF_SYNAPSE_STRUCTURE_OR_ACTIVITY | 186 | -0.307303 | -1.307888 | 0.020833 | 0.666219 |

| | | | | | |
|---|---|---|---|---|---|
| GOBP_INTERLEUKIN_8_PRODUCTION | 52 | 0.535761 | 1.62272 | 0.013908 | 0.149587 |
| GOBP_TISSUE_REMODELING | 123 | 0.479002 | 1.622234 | 0 | 0.149295 |
| GOBP_CARDIAC_EPITHELIAL_TO_MESENCHYMAL_TRANSITION | 34 | 0.585231 | 1.620706 | 0.013554 | 0.150929 |
| GOBP_NEGATIVE_REGULATION_OF_MUSCLE_CELL_DIFFERENTIATION | 48 | 0.549001 | 1.620395 | 0.010101 | 0.150545 |
| GOBP_CELL_SUBSTRATE_ADHESION | 309 | 0.436044 | 1.61864 | 0 | 0.152404 |
| GOBP_REGULATION_OF_MUSCLE_CELL_DIFFERENTIATION | 104 | 0.491148 | 1.615612 | 0.002564 | 0.156486 |
| GOBP_REGULATION_OF_LEUKOCYTE_APOPTOTIC_PROCESS | 62 | 0.524769 | 1.615031 | 0.010695 | 0.156331 |
| GOBP_CARDIAC_ATRIUM_MORPHOGENESIS | 23 | 0.633958 | 1.614579 | 0.009419 | 0.155995 |
| GOBP_VASCULAR_WOUND_HEALING | 17 | 0.661075 | 1.614211 | 0.023372 | 0.15561 |
| GOBP_MUSCLE_CELL_PROLIFERATION | 153 | 0.462594 | 1.613959 | 0.001222 | 0.154958 |
| GOBP_SMOOTH_MUSCLE_CELL_PROLIFERATION | 114 | 0.476905 | 1.613623 | 0.008696 | 0.154617 |
| GOBP_BRANCHING_INVOLVED_IN_SALIVARY_GLAND_MORPHOGENESIS | 17 | 0.665175 | 1.613473 | 0.017828 | 0.153929 |
| GOBP_BONE_MORPHOGENESIS | 80 | 0.497727 | 1.612538 | 0.005376 | 0.154443 |
| GOBP_LYMPH_VESSEL_DEVELOPMENT | 29 | 0.601793 | 1.612369 | 0.016272 | 0.153796 |
| GOBP_INTERLEUKIN_4_PRODUCTION | 22 | 0.634818 | 1.612024 | 0.026563 | 0.153339 |
| GOBP_NEGATIVE_REGULATION_OF_STRIATED_MUSCLE_CELL_DIFFERENTIATION | 31 | 0.595626 | 1.611744 | 0.012195 | 0.152926 |
| GOBP_PHYSIOLOGICAL_CARDIAC_MUSCLE_HYPERTROPHY | 16 | 0.676399 | 1.611084 | 0.020124 | 0.152894 |
| GOBP_REGULATION_OF_INFLAMMATORY_RESPONSE | 243 | 0.444618 | 1.610617 | 0 | 0.152778 |
| GOBP_NODAL_SIGNALING_PATHWAY | 15 | 0.694465 | 1.609743 | 0.019512 | 0.153233 |
| GOBP_THYROID_HORMONE_METABOLIC_PROCESS | 15 | 0.689668 | 1.607887 | 0.031148 | 0.155301 |
| GOBP_ENDOCRINE_PANCREAS_DEVELOPMENT | 30 | 0.584846 | 1.607195 | 0.011182 | 0.155558 |
| GOBP_HEART_GROWTH | 63 | 0.517681 | 1.605872 | 0.004115 | 0.1567 |
| GOBP_CELL_MIGRATION_INVOLVED_IN_SPROUTING_ANGIOGENESIS | 47 | 0.539299 | 1.605429 | 0.009763 | 0.156547 |
| GOBP_POSITIVE_REGULATION_OF_EPITHELIAL_CELL_MIGRATION | 136 | 0.46899 | 1.605142 | 0 | 0.156087 |
| GOBP_MUSCLE_ORGAN_MORPHOGENESIS | 65 | 0.515962 | 1.605048 | 0.006614 | 0.155321 |
| GOBP_ALPHA_BETA_T_CELL_PROLIFERATION | 29 | 0.588816 | 1.604283 | 0.019939 | 0.155656 |
| GOBP_INFLAMMATORY_RESPONSE_TO_ANTIGENIC_STIMULUS | 41 | 0.561696 | 1.603929 | 0.009929 | 0.155467 |
| GOBP_POSITIVE_REGULATION_OF_INTERLEUKIN_12_PRODUCTION | 29 | 0.603247 | 1.603625 | 0.018576 | 0.155155 |
| GOBP_POSITIVE_REGULATION_OF_MACROPHAGE_DIFFERENTIATION | 15 | 0.67678 | 1.603378 | 0.019324 | 0.154607 |
| GOBP_POSITIVE_REGULATION_OF_RELEASE_OF_SEQUESTERED_CALCIUM_ION_INTO_CY | 28 | 0.600471 | 1.603009 | 0.023659 | 0.154405 |
| GOBP_LEUKOCYTE_APOPTOTIC_PROCESS | 83 | 0.495437 | 1.601915 | 0.004071 | 0.155179 |
| GOBP_POSITIVE_REGULATION_OF_CELL_CELL_ADHESION | 199 | 0.445739 | 1.60171 | 0 | 0.154585 |
| GOBP_POSITIVE_REGULATION_OF_CELLULAR_EXTRAVASATION | 15 | 0.688254 | 1.599787 | 0.018425 | 0.156702 |
| GOBP_REGULATION_OF_MYELOID_CELL_APOPTOTIC_PROCESS | 21 | 0.623708 | 1.599609 | 0.032159 | 0.156167 |
| GOBP_POSITIVE_REGULATION_OF_RHO_PROTEIN_SIGNAL_TRANSDUCTION | 23 | 0.626523 | 1.599232 | 0.026074 | 0.155983 |
| GOBP_MYELOID_LEUKOCYTE_MIGRATION | 141 | 0.466641 | 1.597948 | 0.00365 | 0.157014 |
| GOBP_POSITIVE_REGULATION_OF_LYMPHOCYTE_DIFFERENTIATION | 76 | 0.494494 | 1.594704 | 0.004121 | 0.161373 |
| GOBP_REGULATION_OF_RHO_PROTEIN_SIGNAL_TRANSDUCTION | 74 | 0.496606 | 1.593711 | 0.005457 | 0.161948 |
| GOBP_ENDOCARDIAL_CUSHION_FORMATION | 28 | 0.594426 | 1.593582 | 0.013534 | 0.161292 |
| GOBP_CRANIAL_SKELETAL_SYSTEM_DEVELOPMENT | 57 | 0.524321 | 1.593289 | 0.012712 | 0.160948 |
| GOBP_REGULATION_OF_PHOSPHATIDYLINOSITOL_3_KINASE_SIGNALING | 79 | 0.499483 | 1.592946 | 0.006545 | 0.160659 |
| GOBP_REGULATION_OF_CD4_POSITIVE_ALPHA_BETA_T_CELL_ACTIVATION | 44 | 0.548823 | 1.592664 | 0.019971 | 0.160305 |
| GOBP_POSITIVE_REGULATION_OF_VASCULAR_ENDOTHELIAL_GROWTH_FACTOR_PRODU | 23 | 0.631049 | 1.592443 | 0.025039 | 0.159698 |
| GOBP_POSITIVE_REGULATION_OF_LEUKOCYTE_CELL_CELL_ADHESION | 166 | 0.456511 | 1.591717 | 0.001203 | 0.159987 |
| GOBP_POSITIVE_REGULATION_OF_PATHWAY_RESTRICTED_SMAD_PROTEIN_PHOSPHORY | 39 | 0.558203 | 1.59129 | 0.008646 | 0.159887 |
| GOBP_VASODILATION | 34 | 0.567751 | 1.590058 | 0.009346 | 0.16099 |
| GOBP_REGULATION_OF_FILOPODIUM_ASSEMBLY | 47 | 0.539389 | 1.588859 | 0.014815 | 0.161958 |
| GOBP_REGULATION_OF_FATTY_ACID_METABOLIC_PROCESS | 64 | 0.5129 | 1.588689 | 0.005457 | 0.161367 |
| GOBP_BLOOD_VESSEL_ENDOTHELIAL_CELL_MIGRATION | 102 | 0.479678 | 1.587682 | 0.003856 | 0.162227 |
| GOBP_DNA_DAMAGE_RESPONSE_SIGNAL_TRANSDUCTION_BY_P53_CLASS_MEDIATOR_F | 16 | 0.659219 | 1.587608 | 0.022617 | 0.161555 |
| GOBP_REGULATION_OF_MUSCLE_CONTRACTION | 131 | 0.460835 | 1.587289 | 0.002454 | 0.1614 |
| GOBP_POSITIVE_REGULATION_OF_RESPONSE_TO_WOUNDING | 55 | 0.520056 | 1.586828 | 0.006944 | 0.161255 |
| GOBP_POSITIVE_REGULATION_OF_ALPHA_BETA_T_CELL_ACTIVATION | 47 | 0.538769 | 1.586298 | 0.014025 | 0.161339 |
| GOBP_TYPE_2_IMMUNE_RESPONSE | 24 | 0.598047 | 1.583548 | 0.028708 | 0.164906 |
| GOBP_MYELOID_CELL_APOPTOTIC_PROCESS | 25 | 0.605583 | 1.583039 | 0.027523 | 0.164866 |
| GOBP_NEGATIVE_REGULATION_OF_CYSTEINE_TYPE_ENDOPEPTIDASE_ACTIVITY | 65 | 0.509561 | 1.581708 | 0.005202 | 0.166079 |
| GOBP_MATERNAL_PLACENTA_DEVELOPMENT | 29 | 0.582869 | 1.580014 | 0.025373 | 0.16811 |
| GOBP_POSITIVE_REGULATION_OF_SMOOTH_MUSCLE_CELL_PROLIFERATION | 66 | 0.503074 | 1.57643 | 0.00955 | 0.173373 |
| GOBP_T_CELL_SELECTION | 31 | 0.575848 | 1.575851 | 0.026899 | 0.173464 |
| GOBP_RESPONSE_TO_GROWTH_HORMONE | 27 | 0.599251 | 1.57492 | 0.027735 | 0.174133 |
| GOBP_MESONEPHROS_DEVELOPMENT | 80 | 0.483422 | 1.573681 | 0.006596 | 0.175323 |
| GOBP_POSITIVE_REGULATION_OF_WOUND_HEALING | 44 | 0.542812 | 1.572937 | 0.015363 | 0.175635 |
| GOBP_MUSCLE_CONTRACTION | 269 | 0.428745 | 1.57179 | 0 | 0.176629 |
| GOBP_EPIDERMAL_CELL_DIFFERENTIATION | 131 | 0.455589 | 1.571243 | 0 | 0.176775 |
| GOBP_NEGATIVE_REGULATION_OF_MONONUCLEAR_CELL_MIGRATION | 15 | 0.659255 | 1.570863 | 0.027375 | 0.17655 |
| GOBP_ETHER_METABOLIC_PROCESS | 26 | 0.592279 | 1.570208 | 0.025037 | 0.176804 |
| GOBP_CELLULAR_RESPONSE_TO_MECHANICAL_STIMULUS | 60 | 0.510057 | 1.569155 | 0.009986 | 0.177771 |
| GOBP_NEGATIVE_REGULATION_OF_EPITHELIAL_CELL_MIGRATION | 56 | 0.512523 | 1.568601 | 0.00955 | 0.177941 |
| GOBP_ANGIOGENESIS_INVOLVED_IN_WOUND_HEALING | 25 | 0.595928 | 1.568223 | 0.02765 | 0.177706 |
| GOBP_HEART_DEVELOPMENT | 497 | 0.414244 | 1.567974 | 0 | 0.177358 |
| GOBP_POSITIVE_REGULATION_OF_LOCOMOTION | 460 | 0.414134 | 1.567701 | 0 | 0.177014 |
| GOBP_NEGATIVE_REGULATION_OF_MUSCLE_CELL_APOPTOTIC_PROCESS | 31 | 0.573465 | 1.56704 | 0.01372 | 0.177254 |
| GOBP_POSITIVE_REGULATION_OF_T_CELL_PROLIFERATION | 62 | 0.499004 | 1.564182 | 0.015559 | 0.181253 |
| GOBP_ADAPTIVE_THERMOGENESIS | 133 | 0.456725 | 1.563897 | 0.004914 | 0.180904 |
| GOBP_REGENERATION | 139 | 0.450351 | 1.563003 | 0 | 0.181478 |
| GOBP_REGULATION_OF_MYOBLAST_DIFFERENTIATION | 36 | 0.55369 | 1.562006 | 0.017021 | 0.182291 |
| GOBP_REGULATION_OF_T_HELPER_1_TYPE_IMMUNE_RESPONSE | 15 | 0.676502 | 1.561858 | 0.019802 | 0.181683 |
| GOBP_ARTERY_DEVELOPMENT | 84 | 0.480206 | 1.559538 | 0.007916 | 0.184866 |
| GOBP_NEUTROPHIL_CHEMOTAXIS | 56 | 0.50675 | 1.556377 | 0.012346 | 0.189445 |
| GOBP_REGULATION_OF_WOUND_HEALING | 99 | 0.468481 | 1.555739 | 0.011765 | 0.189747 |
| GOBP_POSITIVE_REGULATION_OF_ACUTE_INFLAMMATORY_RESPONSE | 17 | 0.643372 | 1.554042 | 0.029412 | 0.19201 |
| GOBP_MONOCYTE_CHEMOTAXIS | 30 | 0.565973 | 1.553385 | 0.027496 | 0.192313 |
| GOBP_BRANCHING_INVOLVED_IN_BLOOD_VESSEL_MORPHOGENESIS | 29 | 0.572236 | 1.553305 | 0.033333 | 0.191655 |
| GOBP_LEUKOCYTE_MIGRATION | 248 | 0.425069 | 1.552261 | 0.00112 | 0.192726 |
| GOBP_EMBRYONIC_PATTERN_SPECIFICATION | 54 | 0.515706 | 1.551511 | 0.022284 | 0.193176 |
| GOBP_REGULATION_OF_ENDOTHELIAL_CELL_MIGRATION | 148 | 0.445619 | 1.548131 | 0.00358 | 0.198289 |
| GOBP_REGULATION_OF_VASCULATURE_DEVELOPMENT | 229 | 0.427774 | 1.548029 | 0.001124 | 0.197648 |
| GOBP_REGULATION_OF_HAIR_FOLLICLE_DEVELOPMENT | 15 | 0.650496 | 1.54797 | 0.030151 | 0.196896 |
| GOBP_HINDLIMB_MORPHOGENESIS | 21 | 0.605222 | 1.547224 | 0.041475 | 0.19748 |
| GOBP_GLYCOSYL_COMPOUND_CATABOLIC_PROCESS | 29 | 0.585025 | 1.546346 | 0.019461 | 0.198229 |
| GOBP_NEGATIVE_REGULATION_OF_BLOOD_VESSEL_ENDOTHELIAL_CELL_MIGRATION | 28 | 0.585284 | 1.546237 | 0.022762 | 0.197632 |
| GOBP_CHONDROCYTE_DEVELOPMENT | 28 | 0.565591 | 1.543854 | 0.041159 | 0.200895 |
| GOBP_NEGATIVE_REGULATION_OF_EPITHELIAL_CELL_APOPTOTIC_PROCESS | 39 | 0.529591 | 1.542383 | 0.029197 | 0.20274 |
| GOBP_METANEPHROS_DEVELOPMENT | 69 | 0.488013 | 1.541114 | 0.010695 | 0.204234 |
| GOBP_EMBRYO_IMPLANTATION | 48 | 0.518109 | 1.540781 | 0.017417 | 0.204021 |
| GOBP_FORELIMB_MORPHOGENESIS | 20 | 0.612722 | 1.540295 | 0.026856 | 0.204071 |
| GOBP_EMBRYONIC_SKELETAL_SYSTEM_MORPHOGENESIS | 58 | 0.50324 | 1.53953 | 0.008208 | 0.204605 |
| GOBP_TOOTH_MINERALIZATION | 17 | 0.638614 | 1.539208 | 0.028391 | 0.204455 |
| GOBP_CELL_CHEMOTAXIS | 194 | 0.432665 | 1.538151 | 0.001183 | 0.205699 |
| GOBP_MYELOID_LEUKOCYTE_ACTIVATION | 151 | 0.442557 | 1.537492 | 0.002463 | 0.206064 |
| GOBP_LYMPHOCYTE_COSTIMULATION | 28 | 0.563545 | 1.535082 | 0.029688 | 0.209638 |
| GOBP_GLYCEROL_ETHER_METABOLIC_PROCESS | 21 | 0.609636 | 1.535054 | 0.037879 | 0.208872 |
| GOBP_MACROPHAGE_DIFFERENTIATION | 38 | 0.546485 | 1.534517 | 0.026034 | 0.209027 |
| GOBP_POSITIVE_REGULATION_OF_TRANSMEMBRANE_RECEPTOR_PROTEIN_SERINE_THI | 100 | 0.46288 | 1.53401 | 0.010336 | 0.209136 |
| GOBP_POSITIVE_REGULATION_OF_MUSCLE_CONTRACTION | 34 | 0.552079 | 1.533416 | 0.024169 | 0.209477 |
| GOBP_CARDIAC_MUSCLE_CELL_PROLIFERATION | 37 | 0.545177 | 1.533089 | 0.025797 | 0.209297 |
| GOBP_NEGATIVE_REGULATION_OF_CD4_POSITIVE_ALPHA_BETA_T_CELL_ACTIVATION | 22 | 0.598478 | 1.53258 | 0.03876 | 0.209444 |
| GOBP_CARDIOBLAST_DIFFERENTIATION | 16 | 0.632652 | 1.531562 | 0.034976 | 0.210497 |
| GOBP_NEGATIVE_REGULATION_OF_EPITHELIAL_CELL_DIFFERENTIATION | 34 | 0.535537 | 1.531147 | 0.023022 | 0.210492 |
| GOBP_POSITIVE_REGULATION_OF_ENDOTHELIAL_CELL_MIGRATION | 98 | 0.460261 | 1.530114 | 0.006477 | 0.211518 |
| GOBP_MUSCLE_CELL_APOPTOTIC_PROCESS | 56 | 0.499236 | 1.528607 | 0.020173 | 0.213372 |
| GOBP_LEUKOCYTE_MEDIATED_CYTOTOXICITY | 74 | 0.483654 | 1.527858 | 0.011811 | 0.214052 |
| GOBP_MYELOID_LEUKOCYTE_DIFFERENTIATION | 163 | 0.43498 | 1.527441 | 0.001195 | 0.213996 |
| GOBP_EPIBOLY | 33 | 0.556108 | 1.525791 | 0.033794 | 0.216085 |
| GOBP_KERATINOCYTE_DIFFERENTIATION | 79 | 0.475368 | 1.524407 | 0.01043 | 0.217763 |
| GOBP_PATHWAY_RESTRICTED_SMAD_PROTEIN_PHOSPHORYLATION | 53 | 0.511804 | 1.524306 | 0.017784 | 0.217215 |
| GOBP_NEUTROPHIL_MIGRATION | 72 | 0.476593 | 1.524272 | 0.01626 | 0.216471 |
| GOBP_BONE_DEVELOPMENT | 177 | 0.438172 | 1.524213 | 0.004684 | 0.215745 |
| GOBP_HOMOLOGOUS_CHROMOSOME_PAIRING_AT_MEIOSIS | 37 | 0.531345 | 1.523053 | 0.020115 | 0.217099 |
| GOBP_REGULATION_OF_STRIATED_MUSCLE_CONTRACTION | 73 | 0.477113 | 1.522909 | 0.014608 | 0.216576 |
| GOBP_INFLAMMASOME_COMPLEX_ASSEMBLY | 16 | 0.640253 | 1.521889 | 0.045959 | 0.217485 |
| GOBP_SEQUESTERING_OF_CALCIUM_ION | 96 | 0.461452 | 1.521822 | 0.011222 | 0.216825 |
| GOBP_MUSCLE_CELL_MIGRATION | 68 | 0.490767 | 1.521736 | 0.009736 | 0.216181 |
| GOBP_VASCULAR_ASSOCIATED_SMOOTH_MUSCLE_CELL_MIGRATION | 18 | 0.617224 | 1.521704 | 0.043902 | 0.215453 |
| GOBP_POSITIVE_REGULATION_OF_VASCULAR_ASSOCIATED_SMOOTH_MUSCLE_CELL_PI | 24 | 0.579635 | 1.520483 | 0.037398 | 0.217033 |
| GOBP_RESPONSE_TO_MECHANICAL_STIMULUS | 163 | 0.439479 | 1.520312 | 0.004745 | 0.216595 |
| GOBP_REPLICATIVE_SENESCENCE | 15 | 0.649158 | 1.519999 | 0.046281 | 0.216388 |
| GOBP_MESODERM_DEVELOPMENT | 104 | 0.454491 | 1.519236 | 0.010403 | 0.217114 |
| GOBP_CELL_PROLIFERATION_IN_FOREBRAIN | 18 | 0.61789 | 1.51807 | 0.042139 | 0.218603 |
| GOBP_PLASMINOGEN_ACTIVATION | 19 | 0.613714 | 1.517948 | 0.034596 | 0.218093 |
| GOBP_ENDOTHELIAL_CELL_APOPTOTIC_PROCESS | 42 | 0.535871 | 1.517564 | 0.017467 | 0.218025 |
| GOBP_NEGATIVE_REGULATION_OF_RHO_PROTEIN_SIGNAL_TRANSDUCTION | 21 | 0.60125 | 1.517529 | 0.040584 | 0.217336 |
| GOBP_REGULATION_OF_ALPHA_BETA_T_CELL_DIFFERENTIATION | 47 | 0.508827 | 1.517214 | 0.022099 | 0.217146 |
| GOBP_CELLULAR_COMPONENT_ASSEMBLY_INVOLVED_IN_MORPHOGENESIS | 89 | 0.464575 | 1.517022 | 0.011921 | 0.216766 |
| GOBP_NEGATIVE_REGULATION_OF_ALPHA_BETA_T_CELL_ACTIVATION | 30 | 0.556011 | 1.516414 | 0.047112 | 0.217098 |
| GOBP_CALCIUM_ION_TRANSPORT_INTO_CYTOSOL | 126 | 0.443582 | 1.515611 | 0.008861 | 0.217984 |
| GOBP_REGULATION_OF_CD4_POSITIVE_ALPHA_BETA_T_CELL_DIFFERENTIATION | 35 | 0.537083 | 1.515169 | 0.022796 | 0.218086 |
| GOBP_REGULATION_OF_PLATELET_DERIVED_GROWTH_FACTOR_RECEPTOR_SIGNALING | 20 | 0.598475 | 1.514728 | 0.034161 | 0.2181 |
| GOBP_NEUROEPITHELIAL_CELL_DIFFERENTIATION | 22 | 0.598289 | 1.513851 | 0.030488 | 0.218985 |
| GOBP_POSITIVE_REGULATION_OF_OXIDATIVE_STRESS_INDUCED_CELL_DEATH | 15 | 0.642761 | 1.51369 | 0.032573 | 0.218533 |
| GOBP_REGULATION_OF_MUSCLE_SYSTEM_PROCESS | 183 | 0.427397 | 1.513623 | 0.003509 | 0.217979 |
| GOBP_TROPHOBLAST_CELL_MIGRATION | 15 | 0.630372 | 1.512685 | 0.048465 | 0.21891 |
| GOBP_RESPONSE_TO_CHEMOKINE | 46 | 0.516413 | 1.512424 | 0.030568 | 0.218675 |
| GOBP_NUCLEOSIDE_MONOPHOSPHATE_CATABOLIC_PROCESS | 16 | 0.644335 | 1.512409 | 0.034036 | 0.217966 |
| GOBP_EMBRYONIC_CAMERA_TYPE_EYE_DEVELOPMENT | 27 | 0.572367 | 1.510825 | 0.042722 | 0.220229 |
| GOBP_TROPHECTODERMAL_CELL_DIFFERENTIATION | 16 | 0.618613 | 1.510782 | 0.065327 | 0.219576 |
| GOBP_POSITIVE_REGULATION_OF_ALPHA_BETA_T_CELL_DIFFERENTIATION | 35 | 0.537193 | 1.510527 | 0.035276 | 0.219345 |
| GOBP_REGULATION_OF_ORGAN_GROWTH | 64 | 0.47817 | 1.508745 | 0.015299 | 0.222103 |
| GOBP_REGULATION_OF_VIRAL_ENTRY_INTO_HOST_CELL | 37 | 0.536071 | 1.50874 | 0.029915 | 0.221371 |
| GOBP_NEGATIVE_REGULATION_OF_VASCULAR_ASSOCIATED_SMOOTH_MUSCLE_CELL_F | 21 | 0.602861 | 1.508225 | 0.032949 | 0.221511 |
| GOBP_EPITHELIAL_CELL_APOPTOTIC_PROCESS | 85 | 0.471136 | 1.507868 | 0.010884 | 0.221512 |
| GOBP_EPITHELIAL_CELL_DIFFERENTIATION | 483 | 0.396008 | 1.507523 | 0 | 0.221427 |
| GOBP_MUSCLE_SYSTEM_PROCESS | 336 | 0.40657 | 1.507238 | 0 | 0.221209 |
| GOBP_POSITIVE_REGULATION_OF_REGULATORY_T_CELL_DIFFERENTIATION | 16 | 0.631207 | 1.506267 | 0.042925 | 0.222284 |
| GOBP_NEUTROPHIL_MEDIATED_IMMUNITY | 28 | 0.567501 | 1.506261 | 0.038519 | 0.221566 |
| GOBP_REGULATION_OF_BIOLOGICAL_PROCESS_INVOLVED_IN_SYMBIOTIC_INTERACTION | 45 | 0.508517 | 1.505362 | 0.030792 | 0.222514 |
| GOBP_REGULATION_OF_ALPHA_BETA_T_CELL_ACTIVATION | 71 | 0.480853 | 1.50509 | 0.016438 | 0.222302 |

| | | | | | |
|---|---|---|---|---|---|
| GOBP_POSITIVE_REGULATION_OF_SYNAPSE_ASSEMBLY | 52 | -0.37233 | -1.304819 | 0.098039 | 0.672817 |
| GOBP_SODIUM_ION_TRANSMEMBRANE_TRANSPORT | 134 | -0.319001 | -1.303121 | 0.020408 | 0.675003 |
| GOBP_EXCITATORY_SYNAPSE_ASSEMBLY | 23 | -0.453268 | -1.302236 | 0.129199 | 0.67392 |
| GOBP_LONG_CHAIN_FATTY_ACYL_COA_BIOSYNTHETIC_PROCESS | 17 | -0.491907 | -1.300857 | 0.139651 | 0.674521 |
| GOBP_CELL_MIGRATION_IN_HINDBRAIN | 16 | -0.494362 | -1.298371 | 0.150418 | 0.679242 |
| GOBP_VASOCONSTRICTION | 59 | -0.364888 | -1.297234 | 0.078358 | 0.679053 |
| GOBP_MITOCHONDRIAL_TRANSMEMBRANE_TRANSPORT | 98 | -0.328305 | -1.296421 | 0.050459 | 0.678022 |
| GOBP_POSTSYNAPTIC_SPECIALIZATION_ORGANIZATION | 31 | -0.422691 | -1.294884 | 0.103125 | 0.679404 |
| GOBP_VISUAL_BEHAVIOR | 40 | -0.382859 | -1.291726 | 0.097484 | 0.686899 |
| GOBP_SODIUM_ION_TRANSPORT | 188 | -0.302651 | -1.290734 | 0.013699 | 0.686419 |
| GOBP_REGULATION_OF_MRNA_PROCESSING | 133 | -0.318316 | -1.28951 | 0.055556 | 0.686704 |
| GOBP_EPITHELIAL_TUBE_BRANCHING_INVOLVED_IN_LUNG_MORPHOGENESIS | 19 | -0.461823 | -1.289318 | 0.150273 | 0.683204 |
| GOBP_DNA_TEMPLATED_TRANSCRIPTION_ELONGATION | 73 | -0.3434 | -1.288693 | 0.078358 | 0.681431 |
| GOBP_N_GLYCAN_PROCESSING | 19 | -0.475178 | -1.286212 | 0.141791 | 0.686297 |
| GOBP_REGULATION_OF_ASTROCYTE_DIFFERENTIATION | 23 | -0.438265 | -1.284809 | 0.142458 | 0.686923 |
| GOBP_BILE_ACID_METABOLIC_PROCESS | 35 | -0.4051 | -1.282746 | 0.132075 | 0.690603 |
| GOBP_CERAMIDE_TRANSPORT | 16 | -0.489073 | -1.27913 | 0.169054 | 0.700452 |
| GOBP_COCHLEA_MORPHOGENESIS | 17 | -0.471421 | -1.277462 | 0.173469 | 0.702632 |
| GOBP_REGULATION_OF_AMPA_RECEPTOR_ACTIVITY | 23 | -0.432761 | -1.274561 | 0.158038 | 0.709296 |
| GOBP_ENDOPLASMIC_RETICULUM_TUBULAR_NETWORK_ORGANIZATION | 18 | -0.468481 | -1.272703 | 0.179028 | 0.711988 |
| GOBP_COCHLEA_DEVELOPMENT | 40 | -0.386811 | -1.269297 | 0.136364 | 0.720871 |
| GOBP_PYRIMIDINE_RIBONUCLEOSIDE_TRIPHOSPHATE_BIOSYNTHETIC_PROCESS | 15 | -0.481671 | -1.269092 | 0.176923 | 0.71753 |
| GOBP_DNA_TEMPLATED_TRANSCRIPTION_INITIATION | 118 | -0.311782 | -1.266209 | 0.058201 | 0.724326 |
| GOBP_NEGATIVE_REGULATION_OF_RESPONSE_TO_DNA_DAMAGE_STIMULUS | 80 | -0.335266 | -1.264429 | 0.064103 | 0.727317 |
| GOBP_SLEEP | 16 | -0.478976 | -1.259779 | 0.191257 | 0.741178 |
| GOBP_RECEPTOR_INTERNALIZATION | 95 | -0.319595 | -1.257262 | 0.065502 | 0.747218 |
| GOBP_MATING_BEHAVIOR | 16 | -0.475179 | -1.255831 | 0.196286 | 0.748663 |
| GOBP_CELLULAR_RESPONSE_TO_PROSTAGLANDIN_STIMULUS | 16 | -0.472217 | -1.254652 | 0.171149 | 0.74904 |
| GOBP_MRNA_MODIFICATION | 25 | -0.418923 | -1.254308 | 0.173333 | 0.746157 |
| GOBP_NUCLEOSIDE_BISPHOSPHATE_BIOSYNTHETIC_PROCESS | 55 | -0.342221 | -1.253575 | 0.10687 | 0.745043 |
| GOBP_REGULATION_OF_SYSTEMIC_ARTERIAL_BLOOD_PRESSURE | 66 | -0.335505 | -1.251405 | 0.074803 | 0.749331 |
| GOBP_ADENYLATE_CYCLASE_INHIBITING_G_PROTEIN_COUPLED_RECEPTOR_SIGNALIN( | 54 | -0.361657 | -1.25134 | 0.131313 | 0.745551 |
| GOBP_MRNA_CIS_SPLICING_VIA_SPLICEOSOME | 24 | -0.418636 | -1.250843 | 0.162162 | 0.743487 |
| GOBP_ASPARTATE_FAMILY_AMINO_ACID_METABOLIC_PROCESS | 44 | -0.370926 | -1.250629 | 0.130742 | 0.740385 |
| GOBP_REGULATION_OF_LONG_TERM_SYNAPTIC_POTENTIATION | 39 | -0.377527 | -1.249567 | 0.137705 | 0.740553 |
| GOBP_LIPID_HOMEOSTASIS | 120 | -0.310818 | -1.249419 | 0.044335 | 0.737178 |
| GOBP_EXPORT_ACROSS_PLASMA_MEMBRANE | 54 | -0.355098 | -1.249311 | 0.130742 | 0.733667 |
| GOBP_BASE_EXCISION_REPAIR | 43 | -0.385688 | -1.246609 | 0.139683 | 0.740127 |
| GOBP_REGULATION_OF_PROTEIN_SUMOYLATION | 21 | -0.431577 | -1.243155 | 0.179837 | 0.749613 |
| GOBP_POTASSIUM_ION_IMPORT_ACROSS_PLASMA_MEMBRANE | 38 | -0.377972 | -1.242772 | 0.15655 | 0.747211 |
| GOBP_MOTOR_NEURON_AXON_GUIDANCE | 28 | -0.402517 | -1.240006 | 0.152299 | 0.753825 |
| GOBP_L_ALPHA_AMINO_ACID_TRANSMEMBRANE_TRANSPORT | 53 | -0.355063 | -1.238152 | 0.161074 | 0.757204 |
| GOBP_REGULATION_OF_SYNAPTIC_PLASTICITY | 165 | -0.288754 | -1.238012 | 0.033333 | 0.753872 |
| GOBP_PROTEIN_LIPID_COMPLEX_SUBUNIT_ORGANIZATION | 32 | -0.396688 | -1.237112 | 0.16185 | 0.753561 |
| GOBP_REGULATION_OF_ADENYLATE_CYCLASE_ACTIVITY | 27 | -0.41056 | -1.235538 | 0.166667 | 0.755916 |
| GOBP_REGULATION_OF_VOLTAGE_GATED_CALCIUM_CHANNEL_ACTIVITY | 28 | -0.409465 | -1.233984 | 0.160112 | 0.758005 |
| GOBP_REGULATION_OF_POSTSYNAPSE_ORGANIZATION | 72 | -0.329482 | -1.233863 | 0.110236 | 0.754696 |
| GOBP_THIOESTER_METABOLIC_PROCESS | 76 | -0.333172 | -1.230579 | 0.087302 | 0.763325 |
| GOBP_REGULATION_OF_MACROPHAGE_ACTIVATION | 34 | -0.393566 | -1.230324 | 0.162577 | 0.760431 |
| GOBP_VENTRICULAR_SYSTEM_DEVELOPMENT | 28 | -0.394919 | -1.229963 | 0.193252 | 0.758106 |
| GOBP_PYRIMIDINE_NUCLEOTIDE_METABOLIC_PROCESS | 48 | -0.362195 | -1.226535 | 0.159322 | 0.767648 |
| GOBP_HISTONE_H2A_UBIQUITINATION | 26 | -0.405069 | -1.226454 | 0.174033 | 0.76421 |
| GOBP_NEGATIVE_REGULATION_OF_RECEPTOR_INTERNALIZATION | 15 | -0.462225 | -1.223548 | 0.198433 | 0.771848 |
| GOBP_MATURATION_OF_LSU_RRNA | 28 | -0.3955 | -1.223312 | 0.178886 | 0.76897 |
| GOBP_ADHERENS_JUNCTION_ORGANIZATION | 46 | -0.360408 | -1.222697 | 0.143836 | 0.767631 |
| GOBP_ADENYLATE_CYCLASE_MODULATING_G_PROTEIN_COUPLED_RECEPTOR_SIGNAL | 172 | -0.285608 | -1.221676 | 0.051282 | 0.767991 |
| GOBP_PYRIMIDINE_RIBONUCLEOSIDE_TRIPHOSPHATE_METABOLIC_PROCESS | 18 | -0.430833 | -1.22131 | 0.164384 | 0.765795 |
| GOBP_NEGATIVE_REGULATION_OF_INTRINSIC_APOPTOTIC_SIGNALING_PATHWAY_BY_P5 | 21 | -0.427842 | -1.220045 | 0.218659 | 0.766941 |
| GOBP_NEGATIVE_REGULATION_OF_VIRAL_TRANSCRIPTION | 15 | -0.467203 | -1.219738 | 0.227041 | 0.764461 |
| GOBP_POSITIVE_REGULATION_OF_FATTY_ACID_TRANSPORT | 18 | -0.454723 | -1.219342 | 0.2 | 0.762454 |
| GOBP_DENDRITIC_SPINE_MORPHOGENESIS | 56 | -0.343995 | -1.218592 | 0.144981 | 0.761694 |
| GOBP_ACTIVATION_OF_ADENYLATE_CYCLASE_ACTIVITY | 19 | -0.441168 | -1.218194 | 0.204244 | 0.759787 |
| GOBP_EXPLORATION_BEHAVIOR | 21 | -0.426343 | -1.217678 | 0.184211 | 0.758107 |
| GOBP_POTASSIUM_ION_HOMEOSTASIS | 25 | -0.422572 | -1.214235 | 0.203438 | 0.767681 |
| GOBP_NCRNA_PROCESSING | 372 | -0.263947 | -1.214216 | 0 | 0.764219 |
| GOBP_ORGANIC_ACID_TRANSMEMBRANE_TRANSPORT | 120 | -0.303366 | -1.213367 | 0.080214 | 0.76399 |
| GOBP_PROTEIN_TRANSMEMBRANE_TRANSPORT | 62 | -0.338446 | -1.213318 | 0.12 | 0.760651 |
| GOBP_KILLING_OF_CELLS_OF_ANOTHER_ORGANISM | 15 | -0.463548 | -1.212867 | 0.221411 | 0.758897 |
| GOBP_REGULATION_OF_DENDRITIC_SPINE_MORPHOGENESIS | 43 | -0.363165 | -1.212515 | 0.190625 | 0.756809 |
| GOBP_ASSOCIATIVE_LEARNING | 67 | -0.332424 | -1.211828 | 0.126437 | 0.756035 |
| GOBP_MUSCLE_HYPERTROPHY_IN_RESPONSE_TO_STRESS | 20 | -0.424182 | -1.208033 | 0.205333 | 0.76704 |
| GOBP_NEGATIVE_REGULATION_OF_ENDOCYTOSIS | 46 | -0.347315 | -1.207802 | 0.146789 | 0.764457 |
| GOBP_DEOXYRIBONUCLEOSIDE_TRIPHOSPHATE_METABOLIC_PROCESS | 15 | -0.477003 | -1.203978 | 0.211055 | 0.775845 |
| GOBP_RESPONSE_TO_SALT | 17 | -0.447114 | -1.20397 | 0.230769 | 0.772464 |
| GOBP_GPI_ANCHOR_METABOLIC_PROCESS | 30 | -0.391144 | -1.203732 | 0.193084 | 0.76993 |
| GOBP_REGULATION_OF_KETONE_BIOSYNTHETIC_PROCESS | 16 | -0.447238 | -1.203662 | 0.209974 | 0.766814 |
| GOBP_HISTONE_DEUBIQUITINATION | 40 | -0.369034 | -1.202763 | 0.175325 | 0.766951 |
| GOBP_REGULATION_OF_CELLULAR_AMINE_METABOLIC_PROCESS | 27 | -0.39448 | -1.202238 | 0.1875 | 0.765713 |
| GOBP_INTRACELLULAR_PROTEIN_TRANSMEMBRANE_TRANSPORT | 51 | -0.345871 | -1.200857 | 0.149153 | 0.767599 |
| GOBP_REGULATION_OF_PRESYNAPSE_ORGANIZATION | 29 | -0.392135 | -1.199997 | 0.195402 | 0.767475 |
| GOBP_NEGATIVE_REGULATION_OF_MRNA_PROCESSING | 29 | -0.384032 | -1.197739 | 0.21875 | 0.772811 |
| GOBP_TRNA_WOBBLE_BASE_MODIFICATION | 17 | -0.442335 | -1.195052 | 0.225888 | 0.780041 |
| GOBP_GAMMA_AMINOBUTYRIC_ACID_TRANSPORT | 16 | -0.457049 | -1.193041 | 0.233596 | 0.784693 |
| GOBP_RESPONSE_TO_AUDITORY_STIMULUS | 20 | -0.423171 | -1.192177 | 0.206304 | 0.784744 |
| GOBP_PYRIMIDINE_DEOXYRIBONUCLEOTIDE_METABOLIC_PROCESS | 22 | -0.41527 | -1.190158 | 0.215116 | 0.789253 |
| GOBP_MEMBRANE_DEPOLARIZATION_DURING_CARDIAC_MUSCLE_CELL_ACTION_POTEI | 20 | -0.428661 | -1.188737 | 0.246851 | 0.791493 |
| GOBP_STEROL_HOMEOSTASIS | 61 | -0.332034 | -1.187793 | 0.160279 | 0.791734 |
| GOBP_POSITIVE_REGULATION_OF_LONG_TERM_SYNAPTIC_POTENTIATION | 21 | -0.420433 | -1.186535 | 0.191136 | 0.793095 |
| GOBP_REGULATION_OF_SISTER_CHROMATID_COHESION | 21 | -0.421806 | -1.184648 | 0.230321 | 0.797146 |
| GOBP_VIRAL_PROTEIN_PROCESSING | 30 | -0.3795 | -1.182479 | 0.229651 | 0.802446 |
| GOBP_NEUROTRANSMITTER_TRANSPORT | 170 | -0.279655 | -1.182281 | 0.115152 | 0.799909 |
| GOBP_LIPOPROTEIN_BIOSYNTHETIC_PROCESS | 95 | -0.303267 | -1.180892 | 0.142857 | 0.802 |
| GOBP_DNA_LIGATION | 15 | -0.469201 | -1.179608 | 0.276316 | 0.803935 |
| GOBP_PROTEIN_INSERTION_INTO_ER_MEMBRANE | 22 | -0.413698 | -1.178392 | 0.241667 | 0.805165 |
| GOBP_REGULATION_OF_B_CELL_RECEPTOR_SIGNALING_PATHWAY | 15 | -0.452437 | -1.178321 | 0.235294 | 0.80222 |
| GOBP_NEGATIVE_REGULATION_OF_PHAGOCYTOSIS | 18 | -0.438501 | -1.176191 | 0.242105 | 0.807131 |
| GOBP_POSITIVE_REGULATION_OF_BLOOD_PRESSURE | 27 | -0.381917 | -1.175135 | 0.213068 | 0.808159 |
| GOBP_POSTSYNAPTIC_SPECIALIZATION_ASSEMBLY | 19 | -0.422052 | -1.174762 | 0.246684 | 0.80643 |
| GOBP_RECEPTOR_RECYCLING | 31 | -0.378754 | -1.172458 | 0.239521 | 0.812418 |
| GOBP_NEGATIVE_REGULATION_OF_CALCIUM_ION_TRANSPORT | 45 | -0.343622 | -1.171932 | 0.193443 | 0.811405 |
| GOBP_NEGATIVE_REGULATION_OF_BLOOD_CIRCULATION | 19 | -0.428671 | -1.171666 | 0.266839 | 0.809253 |
| GOBP_FACULTATIVE_HETEROCHROMATIN_ASSEMBLY | 30 | -0.378567 | -1.168448 | 0.237537 | 0.818886 |
| GOBP_HEPARAN_SULFATE_PROTEOGLYCAN_BIOSYNTHETIC_PROCESS | 24 | -0.397623 | -1.167824 | 0.26513 | 0.818183 |
| GOBP_LIPID_EXPORT_FROM_CELL | 31 | -0.371638 | -1.167401 | 0.255255 | 0.816721 |
| GOBP_REGULATION_OF_MEMBRANE_DEPOLARIZATION | 38 | -0.362327 | -1.166932 | 0.201166 | 0.815392 |
| GOBP_NEUROTRANSMITTER_RECEPTOR_INTERNALIZATION | 15 | -0.450805 | -1.166658 | 0.251309 | 0.813286 |
| GOBP_ALTERNATIVE_MRNA_SPLICING_VIA_SPLICEOSOME | 72 | -0.317307 | -1.166598 | 0.166008 | 0.810357 |
| GOBP_PROTEIN_IMPORT_INTO_MITOCHONDRIAL_MATRIX | 19 | -0.421757 | -1.165728 | 0.266491 | 0.810708 |
| GOBP_RIBONUCLEOPROTEIN_COMPLEX_SUBUNIT_ORGANIZATION | 181 | -0.267188 | -1.165125 | 0.107595 | 0.809979 |
| GOBP_HINDBRAIN_DEVELOPMENT | 126 | -0.281867 | -1.165122 | 0.12 | 0.806913 |
| GOBP_G_PROTEIN_COUPLED_RECEPTOR_SIGNALING_PATHWAY_COUPLED_TO_CYCLIC | 30 | -0.388234 | -1.163708 | 0.244318 | 0.809534 |
| GOBP_LEARNING | 126 | -0.289017 | -1.163661 | 0.135135 | 0.806643 |
| GOBP_CEREBRAL_CORTEX_NEURON_DIFFERENTIATION | 19 | -0.43139 | -1.163461 | 0.23743 | 0.804335 |
| GOBP_MEMBRANE_DEPOLARIZATION_DURING_ACTION_POTENTIAL | 31 | -0.379208 | -1.162807 | 0.237681 | 0.803922 |
| GOBP_NON_MOTILE_CILIUM_ASSEMBLY | 59 | -0.328877 | -1.162461 | 0.173285 | 0.802316 |
| GOBP_REGULATION_OF_ICOSANOID_SECRETION | 16 | -0.434993 | -1.161614 | 0.258794 | 0.802581 |
| GOBP_REGULATION_OF_RENAL_SYSTEM_PROCESS | 23 | -0.400732 | -1.161316 | 0.261838 | 0.800848 |
| GOBP_DOPAMINE_METABOLIC_PROCESS | 33 | -0.372566 | -1.160474 | 0.238095 | 0.801213 |
| GOBP_NEGATIVE_REGULATION_OF_DNA_REPLICATION | 27 | -0.382438 | -1.160116 | 0.244318 | 0.799583 |
| GOBP_RESPIRATORY_SYSTEM_PROCESS | 33 | -0.357262 | -1.159722 | 0.238636 | 0.798142 |
| GOBP_NEGATIVE_REGULATION_OF_MYELOID_CELL_DIFFERENTIATION | 67 | -0.320095 | -1.159597 | 0.190661 | 0.795742 |
| GOBP_REGULATION_OF_VASOCONSTRICTION | 45 | -0.339715 | -1.159218 | 0.23301 | 0.794342 |
| GOBP_ESTROGEN_METABOLIC_PROCESS | 18 | -0.424878 | -1.158532 | 0.257218 | 0.794033 |
| GOBP_DNA_REPLICATION | 265 | -0.259859 | -1.158403 | 0.072917 | 0.791607 |
| GOBP_POTASSIUM_ION_TRANSPORT | 184 | -0.273292 | -1.158036 | 0.140127 | 0.790144 |
| GOBP_METAPHASE_ANAPHASE_TRANSITION_OF_CELL_CYCLE | 61 | -0.327823 | -1.157854 | 0.168919 | 0.787959 |
| GOBP_ACUTE_PHASE_RESPONSE | 24 | -0.398624 | -1.156838 | 0.232493 | 0.788928 |
| GOBP_INTRACILIARY_TRANSPORT | 38 | -0.354216 | -1.15419 | 0.223022 | 0.796181 |
| GOBP_MITOCHONDRIAL_TRANSLATION | 70 | -0.312345 | -1.153386 | 0.199275 | 0.796486 |
| GOBP_DEOXYRIBONUCLEOTIDE_BIOSYNTHETIC_PROCESS | 15 | -0.429109 | -1.151753 | 0.267813 | 0.799663 |
| GOBP_FEAR_RESPONSE | 32 | -0.353426 | -1.151455 | 0.233333 | 0.798072 |
| GOBP_NONRIBOSOMAL_PEPTIDE_BIOSYNTHETIC_PROCESS | 15 | -0.433852 | -1.147195 | 0.285714 | 0.811475 |
| GOBP_REGULATION_OF_MYELINATION | 41 | -0.353351 | -1.146309 | 0.223602 | 0.812166 |
| GOBP_ORGANIC_HYDROXY_COMPOUND_TRANSPORT | 195 | -0.260694 | -1.145398 | 0.067227 | 0.812993 |
| GOBP_REGULATION_OF_DOUBLE_STRAND_BREAK_REPAIR_VIA_NONHOMOLOGOUS_ENI | 31 | -0.367535 | -1.145156 | 0.253165 | 0.81111 |
| GOBP_PROTEIN_LOCALIZATION_TO_SYNAPSE | 59 | -0.319928 | -1.14438 | 0.226721 | 0.81133 |
| GOBP_REGULATION_OF_SYNAPTIC_TRANSMISSION_GLUTAMATERGIC | 53 | -0.325255 | -1.143642 | 0.213483 | 0.811334 |
| GOBP_ESTABLISHMENT_OF_PLANAR_POLARITY_OF_EMBRYONIC_EPITHELIUM | 15 | -0.444991 | -1.142897 | 0.290244 | 0.811472 |
| GOBP_STEROID_BIOSYNTHETIC_PROCESS | 132 | -0.279532 | -1.142469 | 0.174359 | 0.810417 |
| GOBP_SULFUR_AMINO_ACID_METABOLIC_PROCESS | 31 | -0.362818 | -1.141254 | 0.263314 | 0.81229 |
| GOBP_PRESYNAPSE_ORGANIZATION | 46 | -0.326201 | -1.139096 | 0.248408 | 0.817916 |
| GOBP_STEROID_HORMONE_SECRETION | 15 | -0.434604 | -1.137815 | 0.275184 | 0.819961 |
| GOBP_MYELIN_ASSEMBLY | 22 | -0.393261 | -1.136949 | 0.269341 | 0.820742 |
| GOBP_PROTEIN_LOCALIZATION_TO_POSTSYNAPSE | 31 | -0.35796 | -1.136811 | 0.257962 | 0.818506 |
| GOBP_ORGANIC_ANION_TRANSPORT | 271 | -0.253103 | -1.136153 | 0.09009 | 0.818345 |
| GOBP_TAIL_ANCHORED_MEMBRANE_PROTEIN_INSERTION_INTO_ER_MEMBRANE | 17 | -0.424886 | -1.132958 | 0.274809 | 0.828312 |
| GOBP_AMMONIUM_ION_METABOLIC_PROCESS | 17 | -0.419914 | -1.131918 | 0.28125 | 0.829637 |
| GOBP_SYNAPTIC_VESICLE_CYTOSKELETAL_TRANSPORT | 19 | -0.409689 | -1.131172 | 0.307895 | 0.829792 |
| GOBP_AXONEME_ASSEMBLY | 66 | -0.30988 | -1.13097 | 0.223938 | 0.82776 |
| GOBP_DNA_DEPENDENT_DNA_REPLICATION | 152 | -0.270445 | -1.130759 | 0.174497 | 0.825814 |
| GOBP_NEGATIVE_REGULATION_OF_STEROID_METABOLIC_PROCESS | 22 | -0.392114 | -1.127316 | 0.297158 | 0.836486 |
| GOBP_RECEPTOR_METABOLIC_PROCESS | 145 | -0.26875 | -1.126101 | 0.159509 | 0.838604 |
| GOBP_TRANSCRIPTION_ELONGATION_FROM_RNA_POLYMERASE_II_PROMOTER | 55 | -0.316802 | -1.125169 | 0.227106 | 0.839514 |
| GOBP_REGULATION_OF_SYNAPSE_ASSEMBLY | 89 | -0.289992 | -1.124257 | 0.210084 | 0.840308 |
| GOBP_POSTSYNAPSE_ASSEMBLY | 27 | -0.377762 | -1.123673 | 0.284884 | 0.839826 |
| GOBP_REGULATION_OF_CHOLESTEROL_BIOSYNTHETIC_PROCESS | 18 | -0.412561 | -1.121993 | 0.305913 | 0.843925 |
| GOBP_POSITIVE_REGULATION_OF_SMOOTHENED_SIGNALING_PATHWAY | 30 | -0.35855 | -1.121209 | 0.295652 | 0.844168 |

| | | | | | |
|---|---|---|---|---|---|
| GOBP_EMBRYONIC_EYE_MORPHOGENESIS | 24 | 0.575967 | 1.50243 | 0.030303 | 0.226754 |
| GOBP_POSITIVE_REGULATION_OF_LEUKOCYTE_MIGRATION | 96 | 0.456941 | 1.500325 | 0.007864 | 0.229984 |
| GOBP_REGULATION_OF_EPITHELIAL_CELL_DIFFERENTIATION | 113 | 0.444757 | 1.500075 | 0.011321 | 0.229707 |
| GOBP_REGULATION_OF_ACTIVIN_RECEPTOR_SIGNALING_PATHWAY | 22 | 0.584864 | 1.498735 | 0.040359 | 0.231751 |
| GOBP_SKELETAL_MUSCLE_ORGAN_DEVELOPMENT | 134 | 0.436523 | 1.497747 | 0.006061 | 0.233015 |
| GOBP_CELLULAR_RESPONSE_TO_GROWTH_HORMONE_STIMULUS | 19 | 0.598805 | 1.497369 | 0.049834 | 0.232986 |
| GOBP_REGULATION_OF_RESPONSE_TO_WOUNDING | 127 | 0.436808 | 1.49723 | 0.010013 | 0.23255 |
| GOBP_REGULATION_OF_LEUKOCYTE_MIGRATION | 145 | 0.433793 | 1.49672 | 0.006024 | 0.232682 |
| GOBP_POSITIVE_REGULATION_OF_CALCIUM_ION_TRANSMEMBRANE_TRANSPORT | 56 | 0.492009 | 1.496212 | 0.025745 | 0.232896 |
| GOBP_BONE_GROWTH | 26 | 0.562658 | 1.495496 | 0.052308 | 0.233524 |
| GOBP_MORPHOGENESIS_OF_A_BRANCHING_STRUCTURE | 154 | 0.43307 | 1.494701 | 0.002404 | 0.234346 |
| GOBP_REGULATION_OF_CARDIAC_MUSCLE_CONTRACTION | 61 | 0.488727 | 1.494646 | 0.021186 | 0.233727 |
| GOBP_FEMALE_GAMETE_GENERATION | 126 | 0.438582 | 1.494208 | 0.006227 | 0.233836 |
| GOBP_NEGATIVE_REGULATION_OF_T_CELL_RECEPTOR_SIGNALING_PATHWAY | 17 | 0.609141 | 1.493317 | 0.0608 | 0.234866 |
| GOBP_NEGATIVE_REGULATION_OF_PROTEIN_MATURATION | 20 | 0.599078 | 1.492734 | 0.043411 | 0.235205 |
| GOBP_INTERLEUKIN_2_PRODUCTION | 46 | 0.504733 | 1.492532 | 0.030086 | 0.234911 |
| GOBP_MESENCHYME_DEVELOPMENT | 245 | 0.412872 | 1.491176 | 0.001111 | 0.236783 |
| GOBP_REGULATION_OF_NUCLEAR_TRANSCRIBED_MRNA_POLY_A_TAIL_SHORTENING | 15 | 0.629945 | 1.490037 | 0.061224 | 0.238163 |
| GOBP_PHOSPHATIDYLINOSITOL_3_KINASE_SIGNALING | 102 | 0.448136 | 1.488285 | 0.011378 | 0.240919 |
| GOBP_MESODERM_MORPHOGENESIS | 60 | 0.48693 | 1.487842 | 0.02849 | 0.241048 |
| GOBP_REGULATION_OF_MITOCHONDRIAL_FISSION | 28 | 0.545898 | 1.48761 | 0.062802 | 0.240845 |
| GOBP_EMBRYONIC_HINDLIMB_MORPHOGENESIS | 18 | 0.59435 | 1.486439 | 0.064873 | 0.242454 |
| GOBP_SPROUTING_ANGIOGENESIS | 110 | 0.439058 | 1.485877 | 0.007528 | 0.242835 |
| GOBP_REGULATION_OF_T_CELL_RECEPTOR_SIGNALING_PATHWAY | 33 | 0.534595 | 1.485661 | 0.044643 | 0.242559 |
| GOBP_LEUKOCYTE_DIFFERENTIATION | 401 | 0.394785 | 1.484246 | 0 | 0.244565 |
| GOBP_REGULATION_OF_LEUKOCYTE_CHEMOTAXIS | 80 | 0.458832 | 1.483454 | 0.01706 | 0.245346 |
| GOBP_REGULATION_OF_MONONUCLEAR_CELL_MIGRATION | 76 | 0.461392 | 1.482773 | 0.018135 | 0.245985 |
| GOBP_CARDIAC_MYOFIBRIL_ASSEMBLY | 16 | 0.619173 | 1.482485 | 0.064725 | 0.245865 |
| GOBP_ORGAN_GROWTH | 121 | 0.444453 | 1.482191 | 0.014851 | 0.245763 |
| GOBP_POSITIVE_REGULATION_OF_HISTONE_ACETYLATION | 30 | 0.550276 | 1.481648 | 0.03655 | 0.246104 |
| GOBP_EMBRYONIC_ORGAN_MORPHOGENESIS | 215 | 0.408763 | 1.481635 | 0.001186 | 0.245407 |
| GOBP_RESPONSE_TO_VITAMIN | 60 | 0.478459 | 1.480755 | 0.031856 | 0.246426 |
| GOBP_POSITIVE_REGULATION_OF_PHOSPHATIDYLINOSITOL_3_KINASE_SIGNALING | 54 | 0.491 | 1.479914 | 0.019746 | 0.247384 |
| GOBP_POSITIVE_REGULATION_OF_LEUKOCYTE_CHEMOTAXIS | 62 | 0.47579 | 1.479691 | 0.033784 | 0.247139 |
| GOBP_NEGATIVE_REGULATION_OF_ENDOTHELIAL_CELL_APOPTOTIC_PROCESS | 25 | 0.558933 | 1.479378 | 0.041538 | 0.247012 |
| GOBP_POSITIVE_REGULATION_OF_MUSCLE_CELL_DIFFERENTIATION | 45 | 0.505916 | 1.479316 | 0.033566 | 0.246399 |
| GOBP_TEMPERATURE_HOMEOSTASIS | 146 | 0.426387 | 1.478797 | 0.008485 | 0.246741 |
| GOBP_POSITIVE_REGULATION_OF_DEFENSE_RESPONSE | 188 | 0.415325 | 1.478247 | 0.001178 | 0.247108 |
| GOBP_DNA_MODIFICATION | 98 | 0.451064 | 1.477702 | 0.012804 | 0.247525 |
| GOBP_REGULATION_OF_SMOOTH_MUSCLE_CONTRACTION | 51 | 0.492821 | 1.476648 | 0.034819 | 0.248842 |
| GOBP_MESENCHYMAL_CELL_DEVELOPMENT | 81 | 0.451928 | 1.47623 | 0.012837 | 0.249008 |
| GOBP_SENSORY_ORGAN_MORPHOGENESIS | 211 | 0.410395 | 1.475901 | 0.004678 | 0.248944 |
| GOBP_CELL_FATE_SPECIFICATION | 64 | 0.477723 | 1.475603 | 0.020298 | 0.248777 |
| GOBP_SEGMENTATION | 80 | 0.460058 | 1.472972 | 0.01061 | 0.253563 |
| GOBP_RESPONSE_TO_BMP | 140 | 0.425974 | 1.472744 | 0.010843 | 0.253267 |
| GOBP_POSITIVE_REGULATION_OF_CHEMOTAXIS | 102 | 0.446762 | 1.472011 | 0.016839 | 0.254053 |
| GOBP_OVULATION | 18 | 0.604787 | 1.471225 | 0.049423 | 0.255007 |
| GOBP_HUMORAL_IMMUNE_RESPONSE | 100 | 0.437973 | 1.470829 | 0.023499 | 0.255087 |
| GOBP_REGULATION_OF_EPITHELIAL_CELL_APOPTOTIC_PROCESS | 64 | 0.47405 | 1.469988 | 0.029851 | 0.2561 |
| GOBP_NEGATIVE_REGULATION_OF_CELL_MATRIX_ADHESION | 31 | 0.552609 | 1.468842 | 0.047913 | 0.257761 |
| GOBP_POSITIVE_REGULATION_OF_LEUKOCYTE_PROLIFERATION | 98 | 0.446764 | 1.468441 | 0.014323 | 0.257921 |
| GOBP_NEGATIVE_REGULATION_OF_NECROTIC_CELL_DEATH | 15 | 0.61525 | 1.46762 | 0.076423 | 0.258946 |
| GOBP_POSITIVE_REGULATION_OF_MITOCHONDRION_ORGANIZATION | 66 | 0.46809 | 1.467297 | 0.027174 | 0.258846 |
| GOBP_EMBRYONIC_HEART_TUBE_MORPHOGENESIS | 59 | 0.47836 | 1.46714 | 0.040616 | 0.258461 |
| GOBP_RESPONSE_TO_DIETARY_EXCESS | 24 | 0.555558 | 1.467043 | 0.051829 | 0.257953 |
| GOBP_SKELETAL_MUSCLE_ADAPTATION | 16 | 0.61463 | 1.466958 | 0.060359 | 0.257477 |
| GOBP_SMOOTH_MUSCLE_CONTRACTION | 88 | 0.443005 | 1.465378 | 0.022368 | 0.260023 |
| GOBP_T_CELL_DIFFERENTIATION | 184 | 0.420691 | 1.465338 | 0.005834 | 0.259423 |
| GOBP_MITOTIC_G1_S_TRANSITION_CHECKPOINT_SIGNALING | 27 | 0.546834 | 1.464203 | 0.057632 | 0.260958 |
| GOBP_LEUKOCYTE_CELL_CELL_ADHESION | 254 | 0.395865 | 1.463589 | 0.003371 | 0.261532 |
| GOBP_NUCLEOTIDE_BINDING_OLIGOMERIZATION_DOMAIN_CONTAINING_2_SIGNALING_P | 15 | 0.623895 | 1.463571 | 0.055556 | 0.260849 |
| GOBP_POSITIVE_REGULATION_OF_STEM_CELL_PROLIFERATION | 21 | 0.575105 | 1.460383 | 0.0752 | 0.266941 |
| GOBP_REGULATION_OF_TRANSMEMBRANE_RECEPTOR_PROTEIN_SERINE_THREONINE_ | 216 | 0.404771 | 1.459776 | 0 | 0.267494 |
| GOBP_DENDRITIC_CELL_DIFFERENTIATION | 35 | 0.517915 | 1.45915 | 0.061562 | 0.268077 |
| GOBP_PHOSPHATIDYLETHANOLAMINE_METABOLIC_PROCESS | 24 | 0.566103 | 1.458998 | 0.057453 | 0.26775 |
| GOBP_ENDODERM_FORMATION | 48 | 0.49154 | 1.458819 | 0.038067 | 0.267375 |
| GOBP_BRANCHING_MORPHOGENESIS_OF_AN_EPITHELIAL_TUBE | 121 | 0.432651 | 1.458428 | 0.013819 | 0.267513 |
| GOBP_CELLULAR_DEFENSE_RESPONSE | 22 | 0.570452 | 1.458339 | 0.064915 | 0.266972 |
| GOBP_EPIDERMIS_DEVELOPMENT | 221 | 0.4072 | 1.458106 | 0.003497 | 0.266755 |
| GOBP_POSITIVE_REGULATION_OF_CELL_ACTIVATION | 246 | 0.40066 | 1.458085 | 0.003375 | 0.266107 |
| GOBP_PURINE_DEOXYRIBONUCLEOTIDE_METABOLIC_PROCESS | 15 | 0.622756 | 1.456986 | 0.076539 | 0.267778 |
| GOBP_MYOBLAST_DIFFERENTIATION | 61 | 0.477249 | 1.456411 | 0.032213 | 0.268316 |
| GOBP_REGULATION_OF_ACTION_POTENTIAL | 45 | 0.503629 | 1.455878 | 0.044248 | 0.268756 |
| GOBP_MALE_SEX_DIFFERENTIATION | 125 | 0.430436 | 1.454721 | 0.01118 | 0.270313 |
| GOBP_NATURAL_KILLER_CELL_MEDIATED_IMMUNITY | 42 | 0.499049 | 1.451545 | 0.045872 | 0.276522 |
| GOBP_POSITIVE_REGULATION_OF_SMALL_GTPASE_MEDIATED_SIGNAL_TRANSDUCTION | 53 | 0.480305 | 1.451079 | 0.043056 | 0.276826 |
| GOBP_SUBSTRATE_DEPENDENT_CELL_MIGRATION | 23 | 0.564648 | 1.451069 | 0.074016 | 0.276118 |
| GOBP_POSITIVE_REGULATION_OF_STRIATED_MUSCLE_CELL_DIFFERENTIATION | 30 | 0.542081 | 1.449903 | 0.04142 | 0.277765 |
| GOBP_ENTRY_INTO_HOST | 126 | 0.419574 | 1.449042 | 0.014981 | 0.278816 |
| GOBP_BROWN_FAT_CELL_DIFFERENTIATION | 42 | 0.495646 | 1.449015 | 0.036873 | 0.278153 |
| GOBP_REGULATION_OF_DEVELOPMENTAL_GROWTH | 267 | 0.395184 | 1.448742 | 0.005543 | 0.278003 |
| GOBP_POSITIVE_REGULATION_OF_LIPID_BIOSYNTHETIC_PROCESS | 65 | 0.468542 | 1.448694 | 0.034301 | 0.277394 |
| GOBP_CELL_CELL_RECOGNITION | 35 | 0.513277 | 1.447619 | 0.052632 | 0.278995 |
| GOBP_NEGATIVE_REGULATION_OF_SMOOTH_MUSCLE_CELL_PROLIFERATION | 43 | 0.491971 | 1.447445 | 0.05132 | 0.278621 |
| GOBP_INTERLEUKIN_1_PRODUCTION | 69 | 0.46344 | 1.447093 | 0.029536 | 0.278707 |
| GOBP_RESPONSE_TO_TYPE_I_INTERFERON | 54 | 0.477346 | 1.445061 | 0.041551 | 0.282228 |
| GOBP_PURINE_NUCLEOBASE_METABOLIC_PROCESS | 18 | 0.587352 | 1.444736 | 0.066775 | 0.282217 |
| GOBP_MONONUCLEAR_CELL_DIFFERENTIATION | 307 | 0.38946 | 1.444655 | 0.001098 | 0.281666 |
| GOBP_AUTONOMIC_NERVOUS_SYSTEM_DEVELOPMENT | 34 | 0.513866 | 1.4417 | 0.057269 | 0.287368 |
| GOBP_LEUKOCYTE_ADHESION_TO_VASCULAR_ENDOTHELIAL_CELL | 31 | 0.531056 | 1.441345 | 0.049254 | 0.287388 |
| GOBP_REGULATION_OF_CYTOSOLIC_CALCIUM_ION_CONCENTRATION | 238 | 0.39103 | 1.441344 | 0.006719 | 0.286671 |
| GOBP_NEGATIVE_REGULATION_OF_INTERLEUKIN_2_PRODUCTION | 20 | 0.57546 | 1.440141 | 0.0672 | 0.288519 |
| GOBP_POSITIVE_REGULATION_OF_NIK_NF_KAPPAB_SIGNALING | 51 | 0.485435 | 1.439394 | 0.02969 | 0.289259 |
| GOBP_NEGATIVE_REGULATION_OF_HISTONE_ACETYLATION | 15 | 0.603382 | 1.43708 | 0.08752 | 0.293706 |
| GOBP_REGULATION_OF_CELL_ACTIVATION | 418 | 0.378264 | 1.437054 | 0 | 0.29302 |
| GOBP_PULMONARY_VALVE_MORPHOGENESIS | 16 | 0.608444 | 1.436916 | 0.097479 | 0.292564 |
| GOBP_CELL_KILLING | 95 | 0.434927 | 1.435261 | 0.032425 | 0.29543 |
| GOBP_KERATINIZATION | 16 | 0.602995 | 1.434841 | 0.077419 | 0.295676 |
| GOBP_CALCIUM_MEDIATED_SIGNALING | 143 | 0.415097 | 1.434435 | 0.022975 | 0.295839 |
| GOBP_REGULATION_OF_MEIOTIC_CELL_CYCLE | 38 | 0.499307 | 1.434071 | 0.070073 | 0.29601 |
| GOBP_METANEPHRIC_NEPHRON_DEVELOPMENT | 30 | 0.529092 | 1.433786 | 0.069697 | 0.295917 |
| GOBP_CYTOSOLIC_CALCIUM_ION_TRANSPORT | 145 | 0.415555 | 1.433532 | 0.012019 | 0.295847 |
| GOBP_RESPONSE_TO_LEAD_ION | 17 | 0.606608 | 1.433311 | 0.068562 | 0.29561 |
| GOBP_CELL_FATE_COMMITMENT_INVOLVED_IN_FORMATION_OF_PRIMARY_GERM_LAYEI | 26 | 0.540562 | 1.433309 | 0.054348 | 0.294906 |
| GOBP_SKELETAL_SYSTEM_DEVELOPMENT | 397 | 0.382024 | 1.433115 | 0 | 0.29466 |
| GOBP_REGULATION_OF_LEUKOCYTE_PROLIFERATION | 161 | 0.414053 | 1.432587 | 0.013142 | 0.295042 |
| GOBP_SYMPATHETIC_NERVOUS_SYSTEM_DEVELOPMENT | 17 | 0.586129 | 1.432465 | 0.079077 | 0.294641 |
| GOBP_INTERLEUKIN_17_PRODUCTION | 24 | 0.548999 | 1.431218 | 0.074695 | 0.296592 |
| GOBP_NUCLEAR_TRANSCRIBED_MRNA_POLY_A_TAIL_SHORTENING | 28 | 0.543281 | 1.430325 | 0.045455 | 0.297732 |
| GOBP_REGULATION_OF_SMOOTH_MUSCLE_CELL_DIFFERENTIATION | 25 | 0.55084 | 1.42951 | 0.096423 | 0.298841 |
| GOBP_DNA_DEALKYLATION | 26 | 0.540757 | 1.428661 | 0.078947 | 0.299859 |
| GOBP_NEGATIVE_REGULATION_OF_LYMPHOCYTE_MEDIATED_IMMUNITY | 35 | 0.512512 | 1.428419 | 0.069424 | 0.299712 |
| GOBP_RESPONSE_TO_TUMOR_CELL | 28 | 0.531265 | 1.428354 | 0.077961 | 0.299164 |
| GOBP_PULMONARY_VALVE_DEVELOPMENT | 20 | 0.578934 | 1.427519 | 0.077778 | 0.300452 |
| GOBP_POSITIVE_REGULATION_OF_SMOOTH_MUSCLE_CONTRACTION | 24 | 0.559381 | 1.427058 | 0.05627 | 0.300796 |
| GOBP_POSITIVE_REGULATION_OF_RESPONSE_TO_EXTERNAL_STIMULUS | 294 | 0.383967 | 1.42594 | 0.003367 | 0.302618 |
| GOBP_RESPONSE_TO_MANGANESE_ION | 17 | 0.602548 | 1.425819 | 0.094703 | 0.30218 |
| GOBP_CORONARY_VASCULATURE_MORPHOGENESIS | 16 | 0.595943 | 1.425479 | 0.084261 | 0.302237 |
| GOBP_NEGATIVE_REGULATION_OF_LYMPHOCYTE_ACTIVATION | 107 | 0.425543 | 1.425074 | 0.023407 | 0.302419 |
| GOBP_ZYMOGEN_ACTIVATION | 47 | 0.483646 | 1.42393 | 0.043042 | 0.304176 |
| GOBP_T_CELL_ACTIVATION_INVOLVED_IN_IMMUNE_RESPONSE | 66 | 0.460298 | 1.423677 | 0.047753 | 0.304062 |
| GOBP_RESPONSE_TO_INTERFERON_ALPHA | 17 | 0.594125 | 1.423585 | 0.072785 | 0.303545 |
| GOBP_GLAND_MORPHOGENESIS | 94 | 0.437634 | 1.423292 | 0.031373 | 0.303542 |
| GOBP_REGULATION_OF_CELLULAR_EXTRAVASATION | 27 | 0.533301 | 1.422854 | 0.069337 | 0.303699 |
| GOBP_HEMATOPOIETIC_STEM_CELL_HOMEOSTASIS | 18 | 0.586721 | 1.422708 | 0.085484 | 0.303326 |
| GOBP_REGULATION_OF_ACTIN_FILAMENT_BASED_MOVEMENT | 36 | 0.51091 | 1.422587 | 0.06657 | 0.302906 |
| GOBP_RESPONSE_TO_DEXAMETHASONE | 34 | 0.512765 | 1.422565 | 0.063328 | 0.302258 |
| GOBP_EMBRYONIC_ORGAN_DEVELOPMENT | 349 | 0.382273 | 1.42245 | 0.00324 | 0.301829 |
| GOBP_REGULATION_OF_LEUKOCYTE_ADHESION_TO_VASCULAR_ENDOTHELIAL_CELL | 20 | 0.569375 | 1.421758 | 0.064567 | 0.30269 |
| GOBP_NEURON_MIGRATION | 144 | 0.409084 | 1.421264 | 0.017178 | 0.303166 |
| GOBP_EAR_MORPHOGENESIS | 95 | 0.431216 | 1.420674 | 0.022368 | 0.303801 |
| GOBP_REGULATION_OF_T_CELL_ACTIVATION | 232 | 0.392658 | 1.420394 | 0.011236 | 0.303719 |
| GOBP_REGULATION_OF_PROTEIN_AUTOPHOSPHORYLATION | 39 | 0.502205 | 1.419953 | 0.061047 | 0.303983 |
| GOBP_LYMPH_VESSEL_MORPHOGENESIS | 22 | 0.558952 | 1.419274 | 0.076324 | 0.304748 |
| GOBP_VASCULOGENESIS | 68 | 0.458204 | 1.418916 | 0.041841 | 0.304818 |
| GOBP_NEGATIVE_REGULATION_OF_VASCULATURE_DEVELOPMENT | 79 | 0.447379 | 1.418808 | 0.035912 | 0.304378 |
| GOBP_GRANULOCYTE_DIFFERENTIATION | 26 | 0.530989 | 1.418352 | 0.077273 | 0.304776 |
| GOBP_PANCREAS_DEVELOPMENT | 53 | 0.482048 | 1.418274 | 0.048815 | 0.304278 |
| GOBP_SYNAPTIC_TRANSMISSION_CHOLINERGIC | 20 | 0.5769 | 1.417991 | 0.074797 | 0.304183 |
| GOBP_WOUND_HEALING | 334 | 0.380796 | 1.417958 | 0.00547 | 0.303576 |
| GOBP_POSITIVE_REGULATION_OF_GLYCOPROTEIN_METABOLIC_PROCESS | 21 | 0.556853 | 1.417523 | 0.088468 | 0.303925 |
| GOBP_EPITHELIAL_CELL_DEVELOPMENT | 174 | 0.397717 | 1.416788 | 0.010539 | 0.304922 |
| GOBP_RESPONSE_TO_NITRIC_OXIDE | 16 | 0.598043 | 1.416014 | 0.089577 | 0.306055 |
| GOBP_NEGATIVE_REGULATION_OF_IMMUNE_SYSTEM_PROCESS | 283 | 0.385192 | 1.415585 | 0.008919 | 0.306369 |
| GOBP_RESPONSE_TO_TRANSFORMING_GROWTH_FACTOR_BETA | 216 | 0.390335 | 1.414658 | 0.013667 | 0.307842 |
| GOBP_INFLAMMATORY_RESPONSE | 486 | 0.372958 | 1.414449 | 0.001044 | 0.307617 |
| GOBP_MORPHOGENESIS_OF_AN_EPITHELIAL_SHEET | 51 | 0.470352 | 1.414269 | 0.052705 | 0.307361 |
| GOBP_PLATELET_DERIVED_GROWTH_FACTOR_RECEPTOR_SIGNALING_PATHWAY | 49 | 0.478812 | 1.413297 | 0.054393 | 0.308893 |
| GOBP_GLAND_DEVELOPMENT | 332 | 0.381924 | 1.412061 | 0.002225 | 0.310984 |
| GOBP_FOREBRAIN_NEURON_DEVELOPMENT | 22 | 0.548878 | 1.411615 | 0.073668 | 0.311352 |
| GOBP_REGULATION_OF_CARDIAC_CONDUCTION | 20 | 0.566297 | 1.411375 | 0.083072 | 0.311175 |
| GOBP_COLLATERAL_SPROUTING | 25 | 0.549609 | 1.41116 | 0.081761 | 0.311019 |
| GOBP_NEGATIVE_REGULATION_OF_EPITHELIAL_CELL_PROLIFERATION | 106 | 0.423237 | 1.410579 | 0.02125 | 0.311572 |
| GOBP_REGULATION_OF_OSTEOBLAST_PROLIFERATION | 22 | 0.538707 | 1.410363 | 0.102073 | 0.311455 |
| GOBP_SINGLE_STRANDED_VIRAL_RNA_REPLICATION_VIA_DOUBLE_STRANDED_DNA_INT | 17 | 0.578418 | 1.410138 | 0.081917 | 0.311329 |
| GOBP_ALPHA_BETA_T_CELL_ACTIVATION | 106 | 0.429456 | 1.409928 | 0.035847 | 0.311117 |
| GOBP_NEGATIVE_REGULATION_OF_CELL_ACTIVATION | 142 | 0.407267 | 1.409004 | 0.027027 | 0.312519 |

| | | | | | |
|---|---|---|---|---|---|
| GOBP_SOMITE_DEVELOPMENT | 67 | -0.308248 | -1.120955 | 0.209924 | 0.842556 |
| GOBP_ANTEROGRADE_AXONAL_TRANSPORT | 47 | -0.327189 | -1.120508 | 0.252459 | 0.841621 |
| GOBP_REGULATION_OF_GLIOGENESIS | 75 | -0.299808 | -1.119975 | 0.230216 | 0.841185 |
| GOBP_MEGAKARYOCYTE_DIFFERENTIATION | 47 | -0.325794 | -1.119814 | 0.242525 | 0.839093 |
| GOBP_RESPONSE_TO_AMPHETAMINE | 26 | -0.372494 | -1.119804 | 0.28453 | 0.836452 |
| GOBP_FATTY_ACYL_COA_METABOLIC_PROCESS | 34 | -0.345318 | -1.118838 | 0.283951 | 0.83753 |
| GOBP_ANTIMICROBIAL_HUMORAL_IMMUNE_RESPONSE_MEDIATED_BY_ANTIMICROBIAL_ | 32 | -0.361162 | -1.118083 | 0.283951 | 0.837955 |
| GOBP_PEPTIDYL_GLUTAMIC_ACID_MODIFICATION | 23 | -0.384564 | -1.117158 | 0.298667 | 0.838909 |
| GOBP_PEPTIDYL_CYSTEINE_MODIFICATION | 44 | -0.326346 | -1.115992 | 0.285266 | 0.840784 |
| GOBP_NEGATIVE_REGULATION_OF_INTERLEUKIN_6_PRODUCTION | 29 | -0.366119 | -1.113672 | 0.317221 | 0.846873 |
| GOBP_PHOSPHOLIPID_TRANSPORT | 82 | -0.290471 | -1.112863 | 0.22179 | 0.847352 |
| GOBP_TRNA_PROCESSING | 121 | -0.27453 | -1.112344 | 0.225275 | 0.846708 |
| GOBP_NCRNA_3_END_PROCESSING | 39 | -0.331858 | -1.11234 | 0.282799 | 0.844104 |
| GOBP_MATURATION_OF_LSU_RRNA_FROM_TRICISTRONIC_RRNA_TRANSCRIPT_SSU_RF | 15 | -0.422312 | -1.112131 | 0.335038 | 0.842291 |
| GOBP_CARBOXYLIC_ACID_TRANSPORT | 202 | -0.254945 | -1.110207 | 0.176871 | 0.847136 |
| GOBP_PYRIMIDINE_CONTAINING_COMPOUND_BIOSYNTHETIC_PROCESS | 35 | -0.343041 | -1.109619 | 0.277259 | 0.846715 |
| GOBP_REGULATION_OF_MICROTUBULE_BASED_MOVEMENT | 37 | -0.341289 | -1.10957 | 0.259887 | 0.844308 |
| GOBP_METENCEPHALON_DEVELOPMENT | 95 | -0.287777 | -1.108452 | 0.207965 | 0.846228 |
| GOBP_NEGATIVE_REGULATION_OF_NUCLEOCYTOPLASMIC_TRANSPORT | 20 | -0.397047 | -1.107975 | 0.300275 | 0.845559 |
| GOBP_MRNA_SPLICE_SITE_SELECTION | 31 | -0.35933 | -1.107755 | 0.296875 | 0.843803 |
| GOBP_MAINTENANCE_OF_CELL_POLARITY | 18 | -0.410595 | -1.106961 | 0.303258 | 0.844504 |
| GOBP_PROTEIN_LOCALIZATION_TO_CILIUM | 68 | -0.304837 | -1.106375 | 0.264706 | 0.844262 |
| GOBP_PROTEIN_REFOLDING | 20 | -0.388952 | -1.105938 | 0.302703 | 0.843432 |
| GOBP_REGULATION_OF_AMYLOID_PRECURSOR_PROTEIN_CATABOLIC_PROCESS | 38 | -0.336576 | -1.105808 | 0.253049 | 0.841378 |
| GOBP_MONOUBIQUITINATED_PROTEIN_DEUBIQUITINATION | 34 | -0.347935 | -1.105789 | 0.295265 | 0.83893 |
| GOBP_NUCLEOSOME_MOBILIZATION | 20 | -0.39106 | -1.103728 | 0.329923 | 0.844604 |
| GOBP_OOCYTE_MATURATION | 25 | -0.366121 | -1.103717 | 0.304843 | 0.842127 |
| GOBP_COTRANSLATIONAL_PROTEIN_TARGETING_TO_MEMBRANE | 24 | -0.368491 | -1.10361 | 0.311111 | 0.840094 |
| GOBP_REGULATION_OF_POSTSYNAPTIC_MEMBRANE_POTENTIAL | 106 | -0.282234 | -1.10352 | 0.208531 | 0.837963 |
| GOBP_REGULATION_OF_STEROID_BIOSYNTHETIC_PROCESS | 56 | -0.315419 | -1.102091 | 0.283154 | 0.841102 |
| GOBP_LOCOMOTORY_BEHAVIOR | 160 | -0.262095 | -1.10159 | 0.215909 | 0.840526 |
| GOBP_AMINO_ACID_TRANSPORT | 123 | -0.273425 | -1.101317 | 0.235849 | 0.839033 |
| GOBP_MOTILE_CILIUM_ASSEMBLY | 43 | -0.332366 | -1.100887 | 0.270968 | 0.838267 |
| GOBP_DENDRITIC_SPINE_DEVELOPMENT | 87 | -0.285713 | -1.100364 | 0.264591 | 0.837905 |
| GOBP_REGULATION_OF_DENDRITIC_SPINE_DEVELOPMENT | 53 | -0.307672 | -1.098349 | 0.257627 | 0.843266 |
| GOBP_NUCLEOSIDE_TRIPHOSPHATE_METABOLIC_PROCESS | 101 | -0.281768 | -1.098008 | 0.251337 | 0.842263 |
| GOBP_RESPONSE_TO_ISOQUINOLINE_ALKALOID | 24 | -0.374419 | -1.096128 | 0.336927 | 0.847204 |
| GOBP_NEURON_PROJECTION_ORGANIZATION | 81 | -0.287424 | -1.095272 | 0.260536 | 0.848101 |
| GOBP_DOPAMINE_TRANSPORT | 42 | -0.334959 | -1.094237 | 0.305031 | 0.849627 |
| GOBP_MICROTUBULE_BUNDLE_FORMATION | 96 | -0.277372 | -1.093752 | 0.27686 | 0.849215 |
| GOBP_PROTEIN_LOCALIZATION_TO_CELL_SURFACE | 58 | -0.306443 | -1.093629 | 0.294118 | 0.847303 |
| GOBP_HISTONE_UBIQUITINATION | 47 | -0.314675 | -1.092968 | 0.314685 | 0.847324 |
| GOBP_VASCULAR_TRANSPORT | 76 | -0.29322 | -1.092448 | 0.262712 | 0.846795 |
| GOBP_IMPORT_INTO_CELL | 181 | -0.257545 | -1.08975 | 0.2125 | 0.854848 |
| GOBP_REGULATION_OF_INTRINSIC_APOPTOTIC_SIGNALING_PATHWAY_BY_P53_CLASS_ | 29 | -0.357493 | -1.08806 | 0.327103 | 0.859058 |
| GOBP_NADPH_REGENERATION | 17 | -0.405084 | -1.086659 | 0.325301 | 0.861963 |
| GOBP_ADRENERGIC_RECEPTOR_SIGNALING_PATHWAY | 24 | -0.37506 | -1.085991 | 0.322581 | 0.862234 |
| GOBP_PORE_COMPLEX_ASSEMBLY | 21 | -0.384531 | -1.084734 | 0.324675 | 0.864803 |
| GOBP_NEGATIVE_REGULATION_OF_INTRACELLULAR_TRANSPORT | 49 | -0.313345 | -1.084182 | 0.295139 | 0.864688 |
| GOBP_POSITIVE_REGULATION_OF_MUSCLE_HYPERTROPHY | 20 | -0.395342 | -1.082237 | 0.355438 | 0.869892 |
| GOBP_REGULATION_OF_VIRAL_TRANSCRIPTION | 19 | -0.389999 | -1.081952 | 0.345109 | 0.868572 |
| GOBP_DOUBLE_STRAND_BREAK_REPAIR_VIA_NONHOMOLOGOUS_END_JOINING | 68 | -0.294722 | -1.081419 | 0.283088 | 0.868298 |
| GOBP_PROTEIN_LOCALIZATION_TO_NUCLEOLUS | 15 | -0.411316 | -1.077209 | 0.340483 | 0.883163 |
| GOBP_REGULATION_OF_DNA_REPLICATION | 120 | -0.267156 | -1.073813 | 0.310526 | 0.894211 |
| GOBP_REGULATION_OF_CATION_CHANNEL_ACTIVITY | 136 | -0.263518 | -1.07359 | 0.277487 | 0.892665 |
| GOBP_REGULATION_OF_HORMONE_BIOSYNTHETIC_PROCESS | 20 | -0.380771 | -1.073551 | 0.37464 | 0.890395 |
| GOBP_ZINC_ION_TRANSPORT | 22 | -0.376685 | -1.071682 | 0.348703 | 0.895447 |
| GOBP_ORGANIC_ACID_TRANSPORT | 238 | -0.239584 | -1.071597 | 0.241071 | 0.893368 |
| GOBP_PROTEIN_FOLDING | 184 | -0.250941 | -1.071351 | 0.272727 | 0.891863 |
| GOBP_REGULATION_OF_CATECHOLAMINE_METABOLIC_PROCESS | 19 | -0.390436 | -1.071272 | 0.327824 | 0.889756 |
| GOBP_NEGATIVE_REGULATION_OF_NERVOUS_SYSTEM_DEVELOPMENT | 121 | -0.262146 | -1.070467 | 0.267677 | 0.890666 |
| GOBP_RIBOSOMAL_LARGE_SUBUNIT_BIOGENESIS | 71 | -0.285255 | -1.068501 | 0.332 | 0.896106 |
| GOBP_NCRNA_TRANSCRIPTION | 54 | -0.303471 | -1.06778 | 0.338346 | 0.896634 |
| GOBP_GLUCOSE_6_PHOSPHATE_METABOLIC_PROCESS | 20 | -0.381939 | -1.067097 | 0.362857 | 0.897061 |
| GOBP_HISTONE_MONOUBIQUITINATION | 29 | -0.347459 | -1.064856 | 0.331361 | 0.903766 |
| GOBP_GLUTATHIONE_METABOLIC_PROCESS | 45 | -0.324092 | -1.062623 | 0.368771 | 0.910397 |
| GOBP_AMIDE_TRANSPORT | 219 | -0.241645 | -1.062573 | 0.259843 | 0.90816 |
| GOBP_REGULATION_OF_FATTY_ACID_TRANSPORT | 26 | -0.351972 | -1.06238 | 0.359043 | 0.906511 |
| GOBP_PHAGOSOME_MATURATION | 23 | -0.352894 | -1.062017 | 0.363897 | 0.905619 |
| GOBP_NEGATIVE_REGULATION_OF_LIPID_CATABOLIC_PROCESS | 15 | -0.410849 | -1.060627 | 0.386301 | 0.908895 |
| GOBP_SULFUR_AMINO_ACID_BIOSYNTHETIC_PROCESS | 16 | -0.389517 | -1.06044 | 0.409548 | 0.907233 |
| GOBP_POSITIVE_REGULATION_OF_CHROMATIN_ORGANIZATION | 15 | -0.403466 | -1.059941 | 0.359694 | 0.906932 |
| GOBP_V_D_J_RECOMBINATION | 18 | -0.388909 | -1.059268 | 0.342618 | 0.907375 |
| GOBP_ISOPRENOID_BIOSYNTHETIC_PROCESS | 22 | -0.368698 | -1.058834 | 0.365994 | 0.906649 |
| GOBP_REGULATION_OF_MICROTUBULE_CYTOSKELETON_ORGANIZATION | 149 | -0.253792 | -1.058512 | 0.314286 | 0.905573 |
| GOBP_SYNAPTIC_TRANSMISSION_GLUTAMATERGIC | 74 | -0.28767 | -1.05835 | 0.328125 | 0.90387 |
| GOBP_SECONDARY_PALATE_DEVELOPMENT | 22 | -0.367932 | -1.057012 | 0.367403 | 0.906861 |
| GOBP_POSITIVE_REGULATION_OF_HISTONE_H3_K4_METHYLATION | 25 | -0.360493 | -1.056102 | 0.372093 | 0.908085 |
| GOBP_REGULATION_OF_SODIUM_ION_TRANSMEMBRANE_TRANSPORT | 55 | -0.295601 | -1.054736 | 0.35743 | 0.911393 |
| GOBP_AMINO_ACID_IMPORT | 45 | -0.31038 | -1.054274 | 0.341936 | 0.910734 |
| GOBP_NEGATIVE_REGULATION_OF_GLIOGENESIS | 28 | -0.349763 | -1.051028 | 0.379747 | 0.9217 |
| GOBP_FEMALE_MEIOTIC_NUCLEAR_DIVISION | 29 | -0.34658 | -1.050396 | 0.354467 | 0.921895 |
| GOBP_POSITIVE_REGULATION_OF_CELL_JUNCTION_ASSEMBLY | 88 | -0.272808 | -1.049758 | 0.336364 | 0.922082 |
| GOBP_METAL_ION_EXPORT | 41 | -0.315941 | -1.049503 | 0.364486 | 0.920729 |
| GOBP_ESTABLISHMENT_OF_PROTEIN_LOCALIZATION_TO_CHROMOSOME | 26 | -0.345507 | -1.047144 | 0.385224 | 0.927836 |
| GOBP_POSITIVE_REGULATION_OF_CALCIUM_ION_DEPENDENT_EXOCYTOSIS | 17 | -0.389028 | -1.046324 | 0.400474 | 0.928796 |
| GOBP_NEGATIVE_REGULATION_OF_GENE_EXPRESSION_EPIGENETIC | 75 | -0.282722 | -1.045808 | 0.340081 | 0.928482 |
| GOBP_EMBRYONIC_PLACENTA_MORPHOGENESIS | 22 | -0.36542 | -1.04536 | 0.408602 | 0.928001 |
| GOBP_SIALYLATION | 17 | -0.379234 | -1.045254 | 0.388325 | 0.92617 |
| GOBP_DOPAMINE_SECRETION | 32 | -0.332601 | -1.043058 | 0.357349 | 0.932628 |
| GOBP_RESPONSE_TO_ALKALOID | 81 | -0.272048 | -1.042138 | 0.337255 | 0.934084 |
| GOBP_RESPONSE_TO_AMINE | 37 | -0.326925 | -1.04125 | 0.392638 | 0.935416 |
| GOBP_REGULATION_OF_NEUROTRANSMITTER_LEVELS | 180 | -0.249248 | -1.041181 | 0.384615 | 0.933336 |
| GOBP_REGULATION_OF_LIPID_KINASE_ACTIVITY | 58 | -0.290786 | -1.04042 | 0.377778 | 0.934174 |
| GOBP_ONE_CARBON_METABOLIC_PROCESS | 37 | -0.315902 | -1.039046 | 0.366762 | 0.93737 |
| GOBP_MICROTUBULE_DEPOLYMERIZATION | 45 | -0.304471 | -1.03878 | 0.394904 | 0.93613 |
| GOBP_CHONDROITIN_SULFATE_PROTEOGLYCAN_BIOSYNTHETIC_PROCESS | 23 | -0.35416 | -1.037297 | 0.407008 | 0.939771 |
| GOBP_PROSTAGLANDIN_TRANSPORT | 15 | -0.391714 | -1.036417 | 0.409574 | 0.941068 |
| GOBP_AMYLOID_BETA_FORMATION | 40 | -0.314231 | -1.035435 | 0.38255 | 0.942726 |
| GOBP_REGULATION_OF_ENDOCRINE_PROCESS | 19 | -0.366231 | -1.034619 | 0.393617 | 0.94371 |
| GOBP_REGULATION_OF_LYASE_ACTIVITY | 38 | -0.324318 | -1.034026 | 0.380814 | 0.943838 |
| GOBP_EMBRYONIC_DIGESTIVE_TRACT_DEVELOPMENT | 20 | -0.361427 | -1.031323 | 0.388021 | 0.952648 |
| GOBP_NEUROTRANSMITTER_RECEPTOR_TRANSPORT | 22 | -0.366902 | -1.030787 | 0.414493 | 0.952519 |
| GOBP_OLIGODENDROCYTE_DIFFERENTIATION | 77 | -0.272382 | -1.028828 | 0.401709 | 0.958228 |
| GOBP_RESPONSE_TO_ATP | 24 | -0.340244 | -1.028278 | 0.415978 | 0.958125 |
| GOBP_NEGATIVE_REGULATION_OF_RECEPTOR_MEDIATED_ENDOCYTOSIS | 27 | -0.339764 | -1.027597 | 0.404558 | 0.958629 |
| GOBP_AUTOPHAGY_OF_NUCLEUS | 16 | -0.377638 | -1.027582 | 0.409214 | 0.956419 |
| GOBP_SULFUR_COMPOUND_METABOLIC_PROCESS | 282 | -0.226212 | -1.027314 | 0.430108 | 0.955243 |
| GOBP_NEGATIVE_REGULATION_OF_AXON_EXTENSION | 42 | -0.301847 | -1.026972 | 0.408946 | 0.954311 |
| GOBP_VACUOLAR_ACIDIFICATION | 21 | -0.347695 | -1.025001 | 0.391429 | 0.960119 |
| GOBP_REGULATION_OF_DENDRITE_MORPHOGENESIS | 61 | -0.287237 | -1.023801 | 0.403571 | 0.962922 |
| GOBP_BICARBONATE_TRANSPORT | 19 | -0.373383 | -1.023776 | 0.446475 | 0.960732 |
| GOBP_AMINE_TRANSPORT | 75 | -0.273517 | -1.023704 | 0.37551 | 0.958722 |
| GOBP_AMYLOID_PRECURSOR_PROTEIN_CATABOLIC_PROCESS | 50 | -0.29145 | -1.023283 | 0.398649 | 0.958147 |
| GOBP_NOTOCHORD_DEVELOPMENT | 18 | -0.382125 | -1.023222 | 0.41129 | 0.956104 |
| GOBP_REGULATION_OF_CHROMOSOME_SEPARATION | 66 | -0.280652 | -1.022658 | 0.409894 | 0.956093 |
| GOBP_REGULATION_OF_NEUROTRANSMITTER_RECEPTOR_ACTIVITY | 50 | -0.297111 | -1.022096 | 0.403636 | 0.956122 |
| GOBP_REGULATION_OF_BLOOD_PRESSURE | 133 | -0.252513 | -1.020918 | 0.410256 | 0.958623 |
| GOBP_PROTEIN_CONTAINING_COMPLEX_REMODELING | 19 | -0.363995 | -1.018454 | 0.427822 | 0.96655 |
| GOBP_REGULATION_OF_TOLL_LIKE_RECEPTOR_SIGNALING_PATHWAY | 52 | -0.293626 | -1.018318 | 0.417219 | 0.964864 |
| GOBP_RECEPTOR_LOCALIZATION_TO_SYNAPSE | 51 | -0.293871 | -1.017797 | 0.426621 | 0.964758 |
| GOBP_SNRNA_PROCESSING | 25 | -0.35041 | -1.015989 | 0.414286 | 0.969873 |
| GOBP_REGULATION_OF_CYCLASE_ACTIVITY | 42 | -0.300392 | -1.014383 | 0.416357 | 0.974142 |
| GOBP_POSITIVE_REGULATION_OF_OLIGODENDROCYTE_DIFFERENTIATION | 18 | -0.368556 | -1.014062 | 0.409207 | 0.973177 |
| GOBP_MODIFICATION_OF_SYNAPTIC_STRUCTURE | 19 | -0.373541 | -1.014006 | 0.430168 | 0.971172 |
| GOBP_SULFUR_COMPOUND_TRANSPORT | 43 | -0.303465 | -1.01384 | 0.440895 | 0.96962 |
| GOBP_REGULATION_OF_POSTSYNAPTIC_NEUROTRANSMITTER_RECEPTOR_ACTIVITY | 17 | -0.382649 | -1.013053 | 0.438961 | 0.970622 |
| GOBP_REGULATION_OF_ALTERNATIVE_MRNA_SPLICING_VIA_SPLICEOSOME | 57 | -0.284398 | -1.012625 | 0.426282 | 0.970128 |
| GOBP_SENSORY_PERCEPTION_OF_SMELL | 26 | -0.340086 | -1.011246 | 0.423077 | 0.973421 |
| GOBP_LABYRINTHINE_LAYER_MORPHOGENESIS | 18 | -0.384182 | -1.011203 | 0.438692 | 0.971387 |
| GOBP_NON_CANONICAL_WNT_SIGNALING_PATHWAY | 67 | -0.273548 | -1.010992 | 0.415094 | 0.970024 |
| GOBP_POSITIVE_REGULATION_OF_CATION_CHANNEL_ACTIVITY | 57 | -0.281766 | -1.0097 | 0.423913 | 0.972999 |
| GOBP_RIBOSOME_ASSEMBLY | 60 | -0.278356 | -1.008871 | 0.390805 | 0.974193 |
| GOBP_SPERMATID_DIFFERENTIATION | 118 | -0.247669 | -1.008576 | 0.453202 | 0.973229 |
| GOBP_LIGAND_GATED_ION_CHANNEL_SIGNALING_PATHWAY | 18 | -0.36601 | -1.00851 | 0.454294 | 0.971308 |
| GOBP_SENSORY_PERCEPTION_OF_CHEMICAL_STIMULUS | 51 | -0.291365 | -1.007996 | 0.430868 | 0.971142 |
| GOBP_ANTIGEN_PROCESSING_AND_PRESENTATION_OF_PEPTIDE_ANTIGEN_VIA_MHC_C | 24 | -0.33831 | -1.007819 | 0.429363 | 0.969675 |
| GOBP_TRANSITION_METAL_ION_TRANSPORT | 81 | -0.267833 | -1.007809 | 0.401515 | 0.967552 |
| GOBP_METHIONINE_METABOLIC_PROCESS | 15 | -0.389359 | -1.007617 | 0.454094 | 0.966157 |
| GOBP_CYTOKINESIS | 155 | -0.243131 | -1.007357 | 0.43787 | 0.965028 |
| GOBP_VIRAL_GENE_EXPRESSION | 95 | -0.256806 | -1.006045 | 0.427313 | 0.968161 |
| GOBP_REGULATION_OF_TELOMERASE_ACTIVITY | 49 | -0.296572 | -1.004375 | 0.448161 | 0.972591 |
| GOBP_NUCLEAR_TRANSCRIBED_MRNA_CATABOLIC_PROCESS_EXONUCLEOLYTIC | 19 | -0.367864 | -1.004243 | 0.454545 | 0.970974 |
| GOBP_REGULATION_OF_CHROMOSOME_ORGANIZATION | 201 | -0.230297 | -1.003272 | 0.397059 | 0.97274 |
| GOBP_CELL_JUNCTION_MAINTENANCE | 38 | -0.303642 | -1.001338 | 0.434251 | 0.978538 |
| GOBP_REGULATION_OF_PH | 74 | -0.268839 | -1.000901 | 0.454545 | 0.97814 |
| GOBP_MITOCHONDRIAL_GENE_EXPRESSION | 102 | -0.253595 | -1.00075 | 0.455882 | 0.976566 |
| GOBP_STEM_CELL_DIVISION | 24 | -0.338181 | -0.999404 | 0.44507 | 0.979822 |
| GOBP_RENAL_SYSTEM_PROCESS | 77 | -0.269526 | -0.999221 | 0.463415 | 0.978395 |
| GOBP_CATECHOLAMINE_SECRETION | 44 | -0.302169 | -0.99873 | 0.442815 | 0.978303 |
| GOBP_REGULATION_OF_EARLY_ENDOSOME_TO_LATE_ENDOSOME_TRANSPORT | 18 | -0.366139 | -0.997934 | 0.469496 | 0.979404 |
| GOBP_TRNA_MODIFICATION | 85 | -0.263718 | -0.996804 | 0.454955 | 0.981728 |
| GOBP_NEGATIVE_REGULATION_OF_AMINE_TRANSPORT | 23 | -0.338356 | -0.996144 | 0.467236 | 0.98225 |
| GOBP_CHAPERONE_MEDIATED_PROTEIN_COMPLEX_ASSEMBLY | 20 | -0.347225 | -0.995963 | 0.442049 | 0.980877 |
| GOBP_POSTSYNAPSE_ORGANIZATION | 146 | -0.237894 | -0.995516 | 0.483696 | 0.980606 |
| GOBP_NADH_METABOLIC_PROCESS | 28 | -0.3185 | -0.995308 | 0.439227 | 0.979341 |
| GOBP_POSITIVE_REGULATION_OF_CHROMOSOME_ORGANIZATION | 85 | -0.261248 | -0.995035 | 0.442308 | 0.978321 |

| | | | | | |
|---|---|---|---|---|---|
| GOBP_DEFENSE_RESPONSE_TO_GRAM_NEGATIVE_BACTERIUM | 30 | 0.517393 | 1.408943 | 0.072086 | 0.311968 |
| GOBP_NEGATIVE_REGULATION_OF_IMMUNE_RESPONSE | 125 | 0.410568 | 1.408584 | 0.030979 | 0.312074 |
| GOBP_CELLULAR_PROCESS_INVOLVED_IN_REPRODUCTION_IN_MULTICELLULAR_ORGA | 282 | 0.385907 | 1.408171 | 0.004464 | 0.312304 |
| GOBP_REGULATION_OF_CELL_KILLING | 54 | 0.463983 | 1.407502 | 0.049645 | 0.313126 |
| GOBP_REGULATION_OF_HAIR_CYCLE | 23 | 0.541869 | 1.407329 | 0.084375 | 0.312853 |
| GOBP_REGULATION_OF_CELL_CELL_ADHESION | 318 | 0.380204 | 1.406895 | 0.0033 | 0.313099 |
| GOBP_OOGENESIS | 81 | 0.442148 | 1.406827 | 0.043825 | 0.312558 |
| GOBP_T_CELL_ACTIVATION | 343 | 0.38024 | 1.406212 | 0.004367 | 0.313266 |
| GOBP_ENDOCHONDRAL_BONE_MORPHOGENESIS | 49 | 0.47479 | 1.406114 | 0.064842 | 0.31283 |
| GOBP_MESODERMAL_CELL_DIFFERENTIATION | 22 | 0.551861 | 1.405602 | 0.088415 | 0.31332 |
| GOBP_KIDNEY_EPITHELIUM_DEVELOPMENT | 118 | 0.411882 | 1.405439 | 0.028061 | 0.313068 |
| GOBP_REGULATION_OF_T_CELL_PROLIFERATION | 112 | 0.419783 | 1.405422 | 0.029262 | 0.312447 |
| GOBP_BIOLOGICAL_PROCESS_INVOLVED_IN_SYMBIOTIC_INTERACTION | 225 | 0.38827 | 1.405154 | 0.006873 | 0.312431 |
| GOBP_NEGATIVE_REGULATION_OF_ALPHA_BETA_T_CELL_DIFFERENTIATION | 19 | 0.56827 | 1.404922 | 0.075829 | 0.312295 |
| GOBP_REGULATION_OF_ACUTE_INFLAMMATORY_RESPONSE | 29 | 0.521965 | 1.404188 | 0.069173 | 0.313225 |
| GOBP_DETECTION_OF_STIMULUS_INVOLVED_IN_SENSORY_PERCEPTION | 65 | 0.450656 | 1.403349 | 0.046479 | 0.314455 |
| GOBP_PLATELET_ACTIVATION | 98 | 0.42494 | 1.402469 | 0.037383 | 0.315767 |
| GOBP_REGULATION_OF_HEART_GROWTH | 46 | 0.479405 | 1.402061 | 0.060563 | 0.315961 |
| GOBP_POSITIVE_REGULATION_OF_CELL_DEATH | 471 | 0.368188 | 1.400879 | 0.001058 | 0.317935 |
| GOBP_REGULATION_OF_NATURAL_KILLER_CELL_ACTIVATION | 27 | 0.526943 | 1.399816 | 0.083082 | 0.319713 |
| GOBP_POSITIVE_REGULATION_OF_P38MAPK_CASCADE | 25 | 0.525784 | 1.399769 | 0.073132 | 0.319148 |
| GOBP_POSITIVE_REGULATION_OF_DNA_BINDING | 47 | 0.474138 | 1.399707 | 0.068017 | 0.318626 |
| GOBP_REGULATION_OF_DEFENSE_RESPONSE | 430 | 0.370117 | 1.399435 | 0.006424 | 0.318662 |
| GOBP_DEOXYRIBONUCLEOSIDE_MONOPHOSPHATE_METABOLIC_PROCESS | 20 | 0.566565 | 1.398777 | 0.085537 | 0.319483 |
| GOBP_SMOOTH_MUSCLE_CELL_MIGRATION | 56 | 0.465432 | 1.398661 | 0.050754 | 0.319095 |
| GOBP_CD8_POSITIVE_ALPHA_BETA_T_CELL_ACTIVATION | 16 | 0.570886 | 1.396489 | 0.10832 | 0.323295 |
| GOBP_MALE_MEIOSIS_I | 19 | 0.56236 | 1.395726 | 0.090202 | 0.32447 |
| GOBP_POSITIVE_REGULATION_OF_PEPTIDYL_LYSINE_ACETYLATION | 36 | 0.49596 | 1.394458 | 0.076812 | 0.326742 |
| GOBP_REPRODUCTIVE_SYSTEM_DEVELOPMENT | 349 | 0.373864 | 1.393332 | 0.005417 | 0.32874 |
| GOBP_NECROTIC_CELL_DEATH | 49 | 0.468503 | 1.39328 | 0.076056 | 0.328173 |
| GOBP_SCHWANN_CELL_DEVELOPMENT | 30 | 0.506789 | 1.392693 | 0.075734 | 0.32884 |
| GOBP_RETINOIC_ACID_RECEPTOR_SIGNALING_PATHWAY | 24 | 0.533939 | 1.392197 | 0.086567 | 0.329393 |
| GOBP_MEIOSIS_I_CELL_CYCLE_PROCESS | 98 | 0.426462 | 1.391166 | 0.033163 | 0.331049 |
| GOBP_HOMOLOGOUS_CHROMOSOME_SEGREGATION | 46 | 0.472181 | 1.390918 | 0.070201 | 0.33094 |
| GOBP_POSITIVE_REGULATION_OF_LEUKOCYTE_ADHESION_TO_VASCULAR_ENDOTHELIA | 15 | 0.59682 | 1.39063 | 0.102773 | 0.33092 |
| GOBP_REGULATION_OF_JNK_CASCADE | 123 | 0.407796 | 1.389831 | 0.037221 | 0.332149 |
| GOBP_REGULATION_OF_COLLATERAL_SPROUTING | 19 | 0.571109 | 1.389022 | 0.107843 | 0.333288 |
| GOBP_POSITIVE_REGULATION_OF_STEM_CELL_DIFFERENTIATION | 16 | 0.591699 | 1.38862 | 0.10859 | 0.333505 |
| GOBP_NEGATIVE_REGULATION_OF_INTERLEUKIN_1_PRODUCTION | 19 | 0.560119 | 1.388521 | 0.08769 | 0.333103 |
| GOBP_NEGATIVE_REGULATION_OF_CELLULAR_RESPONSE_TO_GROWTH_FACTOR_STIMU | 85 | 0.430298 | 1.388206 | 0.060052 | 0.333184 |
| GOBP_POSITIVE_REGULATION_OF_REACTIVE_OXYGEN_SPECIES_METABOLIC_PROCESS | 53 | 0.455131 | 1.387865 | 0.069832 | 0.333291 |
| GOBP_TRANSFORMING_GROWTH_FACTOR_BETA_RECEPTOR_SIGNALING_PATHWAY | 170 | 0.395187 | 1.387001 | 0.030374 | 0.334577 |
| GOBP_NEGATIVE_REGULATION_OF_IMMUNE_EFFECTOR_PROCESS | 71 | 0.442956 | 1.386674 | 0.04698 | 0.334742 |
| GOBP_AORTA_MORPHOGENESIS | 29 | 0.518831 | 1.386076 | 0.069219 | 0.335515 |
| GOBP_FOREBRAIN_NEURON_DIFFERENTIATION | 32 | 0.498354 | 1.385867 | 0.087021 | 0.335353 |
| GOBP_REGULATION_OF_LEUKOCYTE_DIFFERENTIATION | 204 | 0.385974 | 1.384668 | 0.007143 | 0.33764 |
| GOBP_POSITIVE_REGULATION_OF_SPROUTING_ANGIOGENESIS | 17 | 0.580605 | 1.38439 | 0.080592 | 0.337667 |
| GOBP_INNER_EAR_MORPHOGENESIS | 80 | 0.42901 | 1.384338 | 0.046392 | 0.337137 |
| GOBP_REGULATION_OF_CYSTEINE_TYPE_ENDOPEPTIDASE_ACTIVITY | 186 | 0.386776 | 1.382179 | 0.01993 | 0.341608 |
| GOBP_RESPONSE_TO_OXYGEN_RADICAL | 23 | 0.534433 | 1.381772 | 0.089623 | 0.341921 |
| GOBP_PURINE_NUCLEOSIDE_METABOLIC_PROCESS | 24 | 0.53159 | 1.381646 | 0.089419 | 0.34158 |
| GOBP_REGULATION_OF_BMP_SIGNALING_PATHWAY | 84 | 0.426437 | 1.381028 | 0.043767 | 0.342533 |
| GOBP_NEGATIVE_REGULATION_OF_CYTOKINE_PRODUCTION | 186 | 0.387101 | 1.380906 | 0.022962 | 0.342204 |
| GOBP_CELLULAR_RESPONSE_TO_INTERFERON_GAMMA | 76 | 0.436884 | 1.380882 | 0.069241 | 0.341602 |
| GOBP_GLYCEROLIPID_CATABOLIC_PROCESS | 49 | 0.471459 | 1.380679 | 0.070809 | 0.341434 |
| GOBP_MUSCLE_HYPERTROPHY | 65 | 0.440577 | 1.380327 | 0.0631 | 0.341649 |
| GOBP_VASCULAR_ASSOCIATED_SMOOTH_MUSCLE_CELL_DIFFERENTIATION | 23 | 0.534942 | 1.380287 | 0.111285 | 0.341086 |
| GOBP_STRIATED_MUSCLE_CELL_PROLIFERATION | 48 | 0.463331 | 1.379928 | 0.077465 | 0.341243 |
| GOBP_T_CELL_DIFFERENTIATION_IN_THYMUS | 57 | 0.445713 | 1.379599 | 0.071823 | 0.341413 |
| GOBP_CD4_POSITIVE_ALPHA_BETA_T_CELL_ACTIVATION | 69 | 0.438998 | 1.379372 | 0.047354 | 0.341235 |
| GOBP_NEURAL_CREST_CELL_DIFFERENTIATION | 86 | 0.423491 | 1.378415 | 0.056921 | 0.342716 |
| GOBP_RENAL_VESICLE_DEVELOPMENT | 16 | 0.590636 | 1.378334 | 0.096405 | 0.342236 |
| GOBP_NEURON_DEATH_IN_RESPONSE_TO_OXIDATIVE_STRESS | 27 | 0.529614 | 1.378085 | 0.095016 | 0.34212 |
| GOBP_REACTIVE_OXYGEN_SPECIES_METABOLIC_PROCESS | 170 | 0.396331 | 1.377981 | 0.023697 | 0.341743 |
| GOBP_HEART_PROCESS | 205 | 0.384507 | 1.377003 | 0.014994 | 0.343471 |
| GOBP_NEGATIVE_REGULATION_OF_REGULATED_SECRETORY_PATHWAY | 19 | 0.565062 | 1.376959 | 0.100629 | 0.34292 |
| GOBP_PROGRAMMED_NECROTIC_CELL_DEATH | 37 | 0.490642 | 1.376819 | 0.086331 | 0.342621 |
| GOBP_UROGENITAL_SYSTEM_DEVELOPMENT | 290 | 0.375114 | 1.376602 | 0.011148 | 0.342485 |
| GOBP_TRIGLYCERIDE_CATABOLIC_PROCESS | 20 | 0.5629 | 1.376552 | 0.10017 | 0.341954 |
| GOBP_ICOSANOID_METABOLIC_PROCESS | 73 | 0.433812 | 1.375545 | 0.055777 | 0.343744 |
| GOBP_REGULATION_OF_PEPTIDYL_THREONINE_PHOSPHORYLATION | 38 | 0.475019 | 1.375263 | 0.093093 | 0.343732 |
| GOBP_EPITHELIAL_TUBE_MORPHOGENESIS | 282 | 0.371121 | 1.374964 | 0.008999 | 0.343741 |
| GOBP_EXTRINSIC_APOPTOTIC_SIGNALING_PATHWAY | 189 | 0.384167 | 1.374244 | 0.017564 | 0.344839 |
| GOBP_REGULATION_OF_FATTY_ACID_BETA_OXIDATION | 16 | 0.571447 | 1.374072 | 0.112769 | 0.344539 |
| GOBP_NEGATIVE_REGULATION_OF_CYCLIN_DEPENDENT_PROTEIN_SERINE_THREONINE | 27 | 0.524873 | 1.37388 | 0.095665 | 0.344376 |
| GOBP_PYRIMIDINE_NUCLEOSIDE_METABOLIC_PROCESS | 18 | 0.563508 | 1.373555 | 0.11093 | 0.344526 |
| GOBP_REGULATION_OF_HETEROTYPIC_CELL_CELL_ADHESION | 15 | 0.586715 | 1.373085 | 0.108108 | 0.345051 |
| GOBP_POSITIVE_REGULATION_OF_MITOCHONDRIAL_FISSION | 20 | 0.543089 | 1.373023 | 0.086287 | 0.344577 |
| GOBP_POSITIVE_REGULATION_OF_NITRIC_OXIDE_METABOLIC_PROCESS | 26 | 0.508506 | 1.372816 | 0.094868 | 0.34443 |
| GOBP_MULTI_MULTICELLULAR_ORGANISM_PROCESS | 150 | 0.392122 | 1.371748 | 0.032622 | 0.346311 |
| GOBP_O_GLYCAN_PROCESSING | 35 | 0.496829 | 1.371687 | 0.090909 | 0.345794 |
| GOBP_POSITIVE_REGULATION_OF_INTERLEUKIN_6_PRODUCTION | 66 | 0.439964 | 1.371666 | 0.071023 | 0.345213 |
| GOBP_AMELOGENESIS | 18 | 0.564935 | 1.371564 | 0.092622 | 0.344842 |
| GOBP_STRIATED_MUSCLE_CONTRACTION | 138 | 0.40067 | 1.371523 | 0.03222 | 0.344302 |
| GOBP_NEPHRON_MORPHOGENESIS | 61 | 0.444614 | 1.371377 | 0.072193 | 0.344005 |
| GOBP_REGULATION_OF_RAS_PROTEIN_SIGNAL_TRANSDUCTION | 166 | 0.394552 | 1.370749 | 0.021951 | 0.344964 |
| GOBP_RESPONSE_TO_BACTERIUM | 373 | 0.364521 | 1.370637 | 0.005342 | 0.344596 |
| GOBP_RIBONUCLEOSIDE_METABOLIC_PROCESS | 28 | 0.511983 | 1.369982 | 0.092262 | 0.345585 |
| GOBP_PITUITARY_GLAND_DEVELOPMENT | 30 | 0.505553 | 1.369266 | 0.072508 | 0.346613 |
| GOBP_STEM_CELL_DIFFERENTIATION | 180 | 0.385468 | 1.369059 | 0.031395 | 0.346509 |
| GOBP_EMBRYONIC_HEMOPOIESIS | 20 | 0.549594 | 1.368838 | 0.114729 | 0.346383 |
| GOBP_NEUTRAL_LIPID_CATABOLIC_PROCESS | 28 | 0.514152 | 1.367754 | 0.109034 | 0.348234 |
| GOBP_GLYCEROPHOSPHOLIPID_CATABOLIC_PROCESS | 26 | 0.514683 | 1.367375 | 0.098131 | 0.348446 |
| GOBP_POSITIVE_REGULATION_OF_TRANSCRIPTION_REGULATORY_REGION_DNA_BINDIN | 18 | 0.554984 | 1.366642 | 0.1216 | 0.349537 |
| GOBP_POSITIVE_REGULATION_OF_PROTEIN_SECRETION | 103 | 0.412296 | 1.366165 | 0.038119 | 0.350022 |
| GOBP_REGULATION_OF_STRESS_ACTIVATED_PROTEIN_KINASE_SIGNALING_CASCADE | 173 | 0.387362 | 1.365768 | 0.01432 | 0.350303 |
| GOBP_CATECHOL_CONTAINING_COMPOUND_BIOSYNTHETIC_PROCESS | 15 | 0.592189 | 1.365163 | 0.125604 | 0.351071 |
| GOBP_RESPONSE_TO_FATTY_ACID | 49 | 0.45804 | 1.363015 | 0.088643 | 0.355507 |
| GOBP_PROTEIN_LOCALIZATION_TO_CELL_CELL_JUNCTION | 20 | 0.547062 | 1.362737 | 0.118936 | 0.355592 |
| GOBP_TOLERANCE_INDUCTION | 19 | 0.549127 | 1.362359 | 0.111455 | 0.355863 |
| GOBP_REGULATION_OF_MACROPHAGE_DIFFERENTIATION | 19 | 0.55642 | 1.361756 | 0.119163 | 0.356619 |
| GOBP_REGULATION_OF_LYMPHOCYTE_ACTIVATION | 325 | 0.367377 | 1.361508 | 0.009901 | 0.356595 |
| GOBP_VASCULAR_ENDOTHELIAL_GROWTH_FACTOR_RECEPTOR_SIGNALING_PATHWAY | 46 | 0.46321 | 1.361425 | 0.09621 | 0.356165 |
| GOBP_REGULATION_OF_T_CELL_DIFFERENTIATION | 104 | 0.40559 | 1.361031 | 0.055341 | 0.356525 |
| GOBP_UTERUS_DEVELOPMENT | 15 | 0.579081 | 1.360544 | 0.121212 | 0.357074 |
| GOBP_SECOND_MESSENGER_MEDIATED_SIGNALING | 217 | 0.374294 | 1.360494 | 0.021991 | 0.356564 |
| GOBP_POSITIVE_REGULATION_OF_PEPTIDYL_SERINE_PHOSPHORYLATION | 82 | 0.415439 | 1.360404 | 0.049745 | 0.35616 |
| GOBP_POSITIVE_REGULATION_OF_INTERLEUKIN_8_PRODUCTION | 40 | 0.47163 | 1.360237 | 0.090103 | 0.355903 |
| GOBP_RESPONSE_TO_FUNGUS | 25 | 0.518096 | 1.360009 | 0.11938 | 0.355838 |
| GOBP_BIOLOGICAL_PROCESS_INVOLVED_IN_INTERACTION_WITH_HOST | 163 | 0.382679 | 1.358832 | 0.037081 | 0.357928 |
| GOBP_CALCIUM_ION_TRANSMEMBRANE_TRANSPORT | 246 | 0.37524 | 1.358556 | 0.017897 | 0.357942 |
| GOBP_TRIGLYCERIDE_BIOSYNTHETIC_PROCESS | 32 | 0.494153 | 1.3564 | 0.093514 | 0.362431 |
| GOBP_PLATELET_AGGREGATION | 53 | 0.451049 | 1.356044 | 0.0699 | 0.3627 |
| GOBP_THYROID_GLAND_DEVELOPMENT | 15 | 0.582991 | 1.354921 | 0.117455 | 0.364948 |
| GOBP_REGULATION_OF_EXTRINSIC_APOPTOTIC_SIGNALING_PATHWAY | 128 | 0.395607 | 1.354655 | 0.053284 | 0.36504 |
| GOBP_REGULATION_OF_RELEASE_OF_SEQUESTERED_CALCIUM_ION_INTO_CYTOSOL | 61 | 0.437775 | 1.354526 | 0.071038 | 0.364743 |
| GOBP_POSITIVE_REGULATION_OF_ANTIGEN_RECEPTOR_MEDIATED_SIGNALING_PATHWA | 19 | 0.541979 | 1.354475 | 0.127419 | 0.364238 |
| GOBP_NEURAL_CREST_CELL_MIGRATION | 55 | 0.453147 | 1.354137 | 0.069638 | 0.364368 |
| GOBP_POSITIVE_REGULATION_OF_DEVELOPMENTAL_GROWTH | 129 | 0.398824 | 1.354108 | 0.051597 | 0.363806 |
| GOBP_CALCIUM_ION_TRANSPORT | 326 | 0.363812 | 1.354055 | 0.009804 | 0.363309 |
| GOBP_GENE_SILENCING_BY_RNA | 112 | 0.402462 | 1.353438 | 0.048995 | 0.364096 |
| GOBP_EPITHELIAL_CELL_PROLIFERATION | 312 | 0.364476 | 1.353317 | 0.014069 | 0.363754 |
| GOBP_REGULATION_OF_LYMPHOCYTE_DIFFERENTIATION | 128 | 0.395516 | 1.352933 | 0.025445 | 0.364015 |
| GOBP_REGULATION_OF_ANTIGEN_RECEPTOR_MEDIATED_SIGNALING_PATHWAY | 46 | 0.46473 | 1.352669 | 0.076056 | 0.364051 |
| GOBP_NEGATIVE_REGULATION_OF_DEVELOPMENTAL_GROWTH | 97 | 0.404537 | 1.351691 | 0.054638 | 0.365804 |
| GOBP_POSITIVE_REGULATION_OF_JNK_CASCADE | 81 | 0.422513 | 1.351394 | 0.060686 | 0.365863 |
| GOBP_POSITIVE_REGULATION_OF_TRANSCRIPTION_FROM_RNA_POLYMERASE_II_PROMO | 22 | 0.536554 | 1.351375 | 0.115016 | 0.36529 |
| GOBP_NEUROTRANSMITTER_METABOLIC_PROCESS | 22 | 0.532421 | 1.35136 | 0.104688 | 0.364704 |
| GOBP_BILE_ACID_AND_BILE_SALT_TRANSPORT | 17 | 0.549589 | 1.350736 | 0.118211 | 0.365509 |
| GOBP_MUSCLE_ADAPTATION | 82 | 0.410173 | 1.349403 | 0.065041 | 0.368142 |
| GOBP_POSITIVE_REGULATION_OF_PROTEIN_CONTAINING_COMPLEX_DISASSEMBLY | 32 | 0.490488 | 1.348517 | 0.100152 | 0.369683 |
| GOBP_REGULATION_OF_ESTABLISHMENT_OR_MAINTENANCE_OF_CELL_POLARITY | 25 | 0.516994 | 1.348273 | 0.121805 | 0.369697 |
| GOBP_HEMOSTASIS | 164 | 0.383827 | 1.348226 | 0.035838 | 0.369194 |
| GOBP_NEGATIVE_REGULATION_OF_ENDOTHELIAL_CELL_PROLIFERATION | 33 | 0.483484 | 1.348099 | 0.097826 | 0.368894 |
| GOBP_POSITIVE_REGULATION_OF_MIRNA_TRANSCRIPTION | 41 | 0.46832 | 1.34794 | 0.1 | 0.368709 |
| GOBP_RESPONSE_TO_REACTIVE_OXYGEN_SPECIES | 165 | 0.385738 | 1.347504 | 0.036557 | 0.369221 |
| GOBP_SMAD_PROTEIN_SIGNAL_TRANSDUCTION | 68 | 0.428841 | 1.347 | 0.071831 | 0.369803 |
| GOBP_POSITIVE_REGULATION_OF_RELEASE_OF_CYTOCHROME_C_FROM_MITOCHONDRI | 24 | 0.521552 | 1.34679 | 0.118729 | 0.369758 |
| GOBP_BLASTOCYST_FORMATION | 39 | 0.473101 | 1.345996 | 0.097953 | 0.371061 |
| GOBP_PHOSPHATIDIC_ACID_METABOLIC_PROCESS | 33 | 0.492887 | 1.345868 | 0.100295 | 0.370732 |
| GOBP_CELL_COMMUNICATION_BY_ELECTRICAL_COUPLING | 31 | 0.498872 | 1.345737 | 0.093502 | 0.37039 |
| GOBP_RESPONSE_TO_WOUNDING | 438 | 0.356017 | 1.345476 | 0.009464 | 0.370468 |
| GOBP_MORPHOGENESIS_OF_AN_EPITHELIUM | 428 | 0.355216 | 1.345442 | 0.007471 | 0.369955 |
| GOBP_RESPONSE_TO_HYPEROXIA | 17 | 0.567837 | 1.345138 | 0.107317 | 0.370085 |
| GOBP_ICOSANOID_TRANSPORT | 36 | 0.469538 | 1.344652 | 0.105581 | 0.370612 |
| GOBP_CAMERA_TYPE_EYE_MORPHOGENESIS | 100 | 0.405924 | 1.34465 | 0.053247 | 0.37001 |
| GOBP_KIDNEY_MORPHOGENESIS | 74 | 0.424677 | 1.343915 | 0.066396 | 0.371109 |
| GOBP_PROTEIN_KINASE_B_SIGNALING | 160 | 0.387026 | 1.343642 | 0.037736 | 0.371177 |
| GOBP_INTERFERON_GAMMA_MEDIATED_SIGNALING_PATHWAY | 20 | 0.536814 | 1.343169 | 0.111111 | 0.37174 |
| GOBP_ALPHA_BETA_T_CELL_DIFFERENTIATION | 77 | 0.419727 | 1.343106 | 0.074278 | 0.37132 |
| GOBP_ACTIVIN_RECEPTOR_SIGNALING_PATHWAY | 36 | 0.482267 | 1.342954 | 0.099554 | 0.371091 |
| GOBP_OOCYTE_DIFFERENTIATION | 49 | 0.452585 | 1.342647 | 0.086471 | 0.371186 |
| GOBP_REGULATION_OF_PEPTIDYL_LYSINE_ACETYLATION | 69 | 0.433272 | 1.342355 | 0.085482 | 0.371302 |
| GOBP_ACTOMYOSIN_STRUCTURE_ORGANIZATION | 174 | 0.386365 | 1.342336 | 0.038694 | 0.370747 |
| GOBP_REGULATION_OF_ANOIKIS | 20 | 0.531591 | 1.341595 | 0.129338 | 0.371894 |
| GOBP_POSITIVE_REGULATION_OF_NUCLEAR_DIVISION | 40 | 0.471886 | 1.341431 | 0.09887 | 0.371736 |
| GOBP_NEGATIVE_REGULATION_OF_CELL_SUBSTRATE_ADHESION | 51 | 0.450553 | 1.341131 | 0.084892 | 0.371811 |

| | | | | | |
|---|---|---|---|---|---|
| GOBP_POSITIVE_REGULATION_OF_NEUROBLAST_PROLIFERATION | 18 | -0.351822 | -0.993703 | 0.498615 | 0.981545 |
| GOBP_POSITIVE_REGULATION_OF_POTASSIUM_ION_TRANSMEMBRANE_TRANSPORTER_ | 20 | -0.355142 | -0.992351 | 0.456989 | 0.984765 |
| GOBP_ANTIGEN_PROCESSING_AND_PRESENTATION_OF_ENDOGENOUS_PEPTIDE_ANTIG | 18 | -0.355401 | -0.991304 | 0.479224 | 0.986752 |
| GOBP_LONG_CHAIN_FATTY_ACID_METABOLIC_PROCESS | 82 | -0.261633 | -0.990706 | 0.505338 | 0.987021 |
| GOBP_REGULATION_OF_DNA_TEMPLATED_TRANSCRIPTION_ELONGATION | 51 | -0.290291 | -0.990621 | 0.461538 | 0.985267 |
| GOBP_POSITIVE_REGULATION_OF_MITOCHONDRIAL_TRANSLATION | 15 | -0.380623 | -0.990385 | 0.455729 | 0.984107 |
| GOBP_CILIUM_ORGANIZATION | 357 | -0.219993 | -0.990136 | 0.47619 | 0.983058 |
| GOBP_DOPAMINERGIC_NEURON_DIFFERENTIATION | 25 | -0.334375 | -0.989815 | 0.448087 | 0.982236 |
| GOBP_CELLULAR_RESPONSE_TO_ESTRADIOL_STIMULUS | 32 | -0.318327 | -0.986964 | 0.43465 | 0.991388 |
| GOBP_POSITIVE_REGULATION_OF_CYCLIN_DEPENDENT_PROTEIN_KINASE_ACTIVITY | 32 | -0.30809 | -0.986272 | 0.493939 | 0.992061 |
| GOBP_POSITIVE_REGULATION_OF_NEURAL_PRECURSOR_CELL_PROLIFERATION | 42 | -0.297203 | -0.985623 | 0.478827 | 0.992502 |
| GOBP_AMYLOID_PRECURSOR_PROTEIN_METABOLIC_PROCESS | 67 | -0.267174 | -0.98526 | 0.501887 | 0.991933 |
| GOBP_REGULATION_OF_GLIAL_CELL_DIFFERENTIATION | 57 | -0.279972 | -0.985242 | 0.467857 | 0.989936 |
| GOBP_SMOOTH_MUSCLE_TISSUE_DEVELOPMENT | 16 | -0.368482 | -0.983941 | 0.459064 | 0.993032 |
| GOBP_SNARE_COMPLEX_ASSEMBLY | 15 | -0.379562 | -0.9836 | 0.498695 | 0.992244 |
| GOBP_NEGATIVE_REGULATION_OF_PROTEIN_LOCALIZATION_TO_NUCLEUS | 29 | -0.320309 | -0.982711 | 0.477478 | 0.993677 |
| GOBP_REGULATION_OF_MICROTUBULE_BASED_PROCESS | 226 | -0.224425 | -0.981583 | 0.483871 | 0.996087 |
| GOBP_ANTIGEN_PROCESSING_AND_PRESENTATION_OF_ENDOGENOUS_ANTIGEN | 21 | -0.344172 | -0.979825 | 0.472222 | 1 |
| GOBP_POSITIVE_REGULATION_OF_PROTEIN_EXPORT_FROM_NUCLEUS | 16 | -0.367576 | -0.979539 | 0.468421 | 1 |
| GOBP_GTP_METABOLIC_PROCESS | 23 | -0.337452 | -0.979315 | 0.465374 | 0.998939 |
| GOBP_MAINTENANCE_OF_SYNAPSE_STRUCTURE | 19 | -0.356881 | -0.97856 | 0.497268 | 0.999968 |
| GOBP_NUCLEOSIDE_TRIPHOSPHATE_BIOSYNTHETIC_PROCESS | 79 | -0.260155 | -0.978246 | 0.501873 | 0.99923 |
| GOBP_MICROTUBULE_POLYMERIZATION_OR_DEPOLYMERIZATION | 118 | -0.247832 | -0.977832 | 0.483871 | 0.998762 |
| GOBP_HORMONE_BIOSYNTHETIC_PROCESS | 46 | -0.282763 | -0.977823 | 0.488525 | 0.996773 |
| GOBP_POSITIVE_REGULATION_OF_LIPID_KINASE_ACTIVITY | 32 | -0.313003 | -0.977531 | 0.512821 | 0.995899 |
| GOBP_RESPONSE_TO_HEPATOCYTE_GROWTH_FACTOR | 17 | -0.362071 | -0.976989 | 0.463476 | 0.995999 |
| GOBP_NEGATIVE_REGULATION_OF_SIGNALING_RECEPTOR_ACTIVITY | 38 | -0.29426 | -0.975829 | 0.486405 | 0.998374 |
| GOBP_BARBED_END_ACTIN_FILAMENT_CAPPING | 25 | -0.329969 | -0.974895 | 0.501377 | 0.99988 |
| GOBP_NEGATIVE_REGULATION_OF_ION_TRANSMEMBRANE_TRANSPORTER_ACTIVITY | 51 | -0.277695 | -0.974739 | 0.480916 | 0.998499 |
| GOBP_NEGATIVE_REGULATION_OF_NEURAL_PRECURSOR_CELL_PROLIFERATION | 16 | -0.365 | -0.971981 | 0.496164 | 1 |
| GOBP_TRANSCRIPTION_PREINITIATION_COMPLEX_ASSEMBLY | 36 | -0.296468 | -0.971938 | 0.467532 | 1 |
| GOBP_DNA_TEMPLATED_TRANSCRIPTION_TERMINATION | 21 | -0.345871 | -0.971905 | 0.486559 | 1 |
| GOBP_PROTEIN_PALMITOYLATION | 31 | -0.30777 | -0.9703 | 0.469027 | 1 |
| GOBP_REGULATION_OF_CELLULAR_RESPONSE_TO_VASCULAR_ENDOTHELIAL_GROWTH | 20 | -0.350495 | -0.969976 | 0.487047 | 1 |
| GOBP_REGULATION_OF_MICROTUBULE_POLYMERIZATION_OR_DEPOLYMERIZATION | 84 | -0.254664 | -0.969857 | 0.493507 | 1 |
| GOBP_TRANSLATIONAL_TERMINATION | 18 | -0.34631 | -0.96947 | 0.479769 | 1 |
| GOBP_CHONDROITIN_SULFATE_BIOSYNTHETIC_PROCESS | 18 | -0.352985 | -0.968605 | 0.509859 | 1 |
| GOBP_RNA_POLYMERASE_II_PREINITIATION_COMPLEX_ASSEMBLY | 24 | -0.323407 | -0.9683 | 0.498471 | 1 |
| GOBP_REGULATION_OF_CALCIUM_ION_DEPENDENT_EXOCYTOSIS | 39 | -0.290935 | -0.968021 | 0.473988 | 1 |
| GOBP_REGULATION_OF_MICROTUBULE_DEPOLYMERIZATION | 31 | -0.305974 | -0.967837 | 0.519878 | 1 |
| GOBP_AMYLOID_BETA_METABOLIC_PROCESS | 47 | -0.28885 | -0.967552 | 0.508591 | 1 |
| GOBP_REPRODUCTIVE_BEHAVIOR | 22 | -0.33745 | -0.967202 | 0.513441 | 1 |
| GOBP_NEGATIVE_REGULATION_OF_DNA_REPAIR | 35 | -0.304922 | -0.96684 | 0.517799 | 1 |
| GOBP_LIPOPROTEIN_METABOLIC_PROCESS | 121 | -0.237258 | -0.966794 | 0.578378 | 0.998894 |
| GOBP_PROTEIN_CONTAINING_COMPLEX_LOCALIZATION | 146 | -0.235408 | -0.966546 | 0.560976 | 0.997867 |
| GOBP_REGULATION_OF_MITOTIC_CELL_CYCLE_SPINDLE_ASSEMBLY_CHECKPOINT | 15 | -0.378852 | -0.966282 | 0.505236 | 0.996926 |
| GOBP_REGULATION_OF_PROTEIN_LOCALIZATION_TO_CELL_SURFACE | 34 | -0.306978 | -0.965958 | 0.514881 | 0.996183 |
| GOBP_VITAMIN_TRANSMEMBRANE_TRANSPORT | 15 | -0.369616 | -0.964945 | 0.506562 | 0.998098 |
| GOBP_LONG_CHAIN_FATTY_ACYL_COA_METABOLIC_PROCESS | 23 | -0.338159 | -0.964464 | 0.497006 | 0.998068 |
| GOBP_ACTIVATION_OF_PHOSPHOLIPASE_C_ACTIVITY | 18 | -0.349208 | -0.963746 | 0.497355 | 0.99882 |
| GOBP_CELLULAR_IRON_ION_HOMEOSTASIS | 52 | -0.272619 | -0.963744 | 0.525362 | 0.996895 |
| GOBP_PYRIMIDINE_CONTAINING_COMPOUND_METABOLIC_PROCESS | 78 | -0.252915 | -0.962331 | 0.537594 | 1 |
| GOBP_MITOTIC_CHROMOSOME_CONDENSATION | 19 | -0.339549 | -0.961921 | 0.495957 | 1 |
| GOBP_POSITIVE_REGULATION_OF_GLIAL_CELL_DIFFERENTIATION | 35 | -0.29731 | -0.960151 | 0.504274 | 1 |
| GOBP_MRNA_METHYLATION | 16 | -0.371717 | -0.957668 | 0.509138 | 1 |
| GOBP_REGULATION_OF_CYTOKINESIS | 78 | -0.253909 | -0.955663 | 0.556522 | 1 |
| GOBP_NUCLEOTIDE_TRANSMEMBRANE_TRANSPORT | 18 | -0.348655 | -0.95532 | 0.520513 | 1 |
| GOBP_NEGATIVE_REGULATION_OF_CHROMOSOME_ORGANIZATION | 82 | -0.250849 | -0.955011 | 0.522088 | 1 |
| GOBP_NEGATIVE_REGULATION_OF_METAPHASE_ANAPHASE_TRANSITION_OF_CELL_CYC | 39 | -0.29037 | -0.954625 | 0.518293 | 1 |
| GOBP_AMINO_ACID_IMPORT_ACROSS_PLASMA_MEMBRANE | 36 | -0.299456 | -0.95414 | 0.521084 | 1 |
| GOBP_PTERIDINE_CONTAINING_COMPOUND_BIOSYNTHETIC_PROCESS | 15 | -0.362827 | -0.952632 | 0.524725 | 1 |
| GOBP_NEURON_PROJECTION_EXTENSION_INVOLVED_IN_NEURON_PROJECTION_GUIDA | 37 | -0.288669 | -0.952612 | 0.570552 | 1 |
| GOBP_REGULATION_OF_RECEPTOR_MEDIATED_ENDOCYTOSIS | 91 | -0.249432 | -0.952533 | 0.571429 | 1 |
| GOBP_NEGATIVE_REGULATION_OF_CARTILAGE_DEVELOPMENT | 22 | -0.332718 | -0.952145 | 0.527415 | 1 |
| GOBP_CHROMATIN_DISASSEMBLY | 19 | -0.337237 | -0.951412 | 0.48433 | 1 |
| GOBP_NEGATIVE_REGULATION_OF_AXON_EXTENSION_INVOLVED_IN_AXON_GUIDANCE | 27 | -0.316859 | -0.951394 | 0.540323 | 1 |
| GOBP_NEGATIVE_REGULATION_OF_DNA_DEPENDENT_DNA_REPLICATION | 15 | -0.361681 | -0.950697 | 0.542105 | 1 |
| GOBP_CIRCADIAN_RHYTHM | 174 | -0.223067 | -0.95039 | 0.623288 | 1 |
| GOBP_CELLULAR_RESPONSE_TO_PH | 21 | -0.336914 | -0.949719 | 0.528571 | 1 |
| GOBP_HISTONE_H2A_MONOUBIQUITINATION | 17 | -0.354692 | -0.949349 | 0.52815 | 1 |
| GOBP_REGULATION_OF_LIPOPROTEIN_PARTICLE_CLEARANCE | 15 | -0.363116 | -0.948874 | 0.506562 | 1 |
| GOBP_REGULATION_OF_PLASMA_LIPOPROTEIN_PARTICLE_LEVELS | 49 | -0.271429 | -0.948549 | 0.581882 | 1 |
| GOBP_NAD_METABOLIC_PROCESS | 24 | -0.32021 | -0.947361 | 0.542522 | 1 |
| GOBP_POSITIVE_REGULATION_OF_CYTOSOLIC_CALCIUM_ION_CONCENTRATION_INVOLV | 19 | -0.347768 | -0.946918 | 0.537275 | 1 |
| GOBP_C21_STEROID_HORMONE_METABOLIC_PROCESS | 27 | -0.314291 | -0.94685 | 0.487324 | 1 |
| GOBP_NEUROTRANSMITTER_SECRETION | 127 | -0.231826 | -0.946709 | 0.60396 | 1 |
| GOBP_REGULATION_OF_NON_CANONICAL_WNT_SIGNALING_PATHWAY | 23 | -0.335118 | -0.94557 | 0.508287 | 1 |
| GOBP_CYTOPLASMIC_PATTERN_RECOGNITION_RECEPTOR_SIGNALING_PATHWAY | 55 | -0.268924 | -0.94546 | 0.561837 | 1 |
| GOBP_TOXIN_TRANSPORT | 33 | -0.300015 | -0.945443 | 0.53869 | 1 |
| GOBP_ORGANOPHOSPHATE_ESTER_TRANSPORT | 121 | -0.236599 | -0.944813 | 0.585 | 1 |
| GOBP_PROTEIN_LOCALIZATION_TO_CHROMATIN | 31 | -0.305092 | -0.944806 | 0.535385 | 1 |
| GOBP_DNA_GEOMETRIC_CHANGE | 94 | -0.24001 | -0.944404 | 0.613208 | 1 |
| GOBP_PROTEIN_TRANSMEMBRANE_IMPORT_INTO_INTRACELLULAR_ORGANELLE | 35 | -0.286618 | -0.943955 | 0.555172 | 1 |
| GOBP_3_UTR_MEDIATED_MRNA_STABILIZATION | 19 | -0.337273 | -0.943903 | 0.528061 | 1 |
| GOBP_PHAGOCYTOSIS_RECOGNITION | 16 | -0.35612 | -0.943477 | 0.54026 | 1 |
| GOBP_NEGATIVE_REGULATION_OF_ESTABLISHMENT_OF_PROTEIN_LOCALIZATION | 98 | -0.240947 | -0.9427 | 0.606482 | 1 |
| GOBP_TRNA_METABOLIC_PROCESS | 181 | -0.221078 | -0.941726 | 0.616883 | 1 |
| GOBP_NUCLEAR_BODY_ORGANIZATION | 15 | -0.366874 | -0.94139 | 0.530769 | 1 |
| GOBP_REGULATION_OF_LIPID_TRANSPORT | 98 | -0.239509 | -0.941222 | 0.607143 | 1 |
| GOBP_ESTABLISHMENT_OF_PROTEIN_LOCALIZATION_TO_ENDOPLASMIC_RETICULUM | 42 | -0.284712 | -0.939577 | 0.554455 | 1 |
| GOBP_REGULATION_OF_RNA_SPLICING | 146 | -0.227701 | -0.93931 | 0.67284 | 1 |
| GOBP_PROTEIN_MATURATION_BY_IRON_SULFUR_CLUSTER_TRANSFER | 15 | -0.359893 | -0.938192 | 0.546135 | 1 |
| GOBP_SOMATIC_DIVERSIFICATION_OF_IMMUNE_RECEPTORS | 69 | -0.256131 | -0.937228 | 0.594595 | 1 |
| GOBP_POSITIVE_REGULATION_OF_LIPID_TRANSPORT | 63 | -0.25755 | -0.935458 | 0.599222 | 1 |
| GOBP_PATTERN_RECOGNITION_RECEPTOR_SIGNALING_PATHWAY | 133 | -0.228472 | -0.935294 | 0.645349 | 1 |
| GOBP_REGULATION_OF_DEFENSE_RESPONSE_TO_BACTERIUM | 15 | -0.357035 | -0.934811 | 0.532688 | 1 |
| GOBP_NEGATIVE_REGULATION_OF_INTRINSIC_APOPTOTIC_SIGNALING_PATHWAY | 92 | -0.243406 | -0.934788 | 0.618182 | 1 |
| GOBP_HORMONE_MEDIATED_SIGNALING_PATHWAY | 148 | -0.22049 | -0.934748 | 0.655556 | 1 |
| GOBP_NEGATIVE_REGULATION_OF_PROTEIN_CONTAINING_COMPLEX_DISASSEMBLY | 76 | -0.250063 | -0.93357 | 0.586207 | 1 |
| GOBP_POSITIVE_REGULATION_OF_TELOMERE_CAPPING | 17 | -0.347691 | -0.93338 | 0.560976 | 1 |
| GOBP_POSITIVE_REGULATION_OF_CILIUM_ASSEMBLY | 23 | -0.316961 | -0.932638 | 0.571802 | 1 |
| GOBP_REGULATION_OF_SPINDLE_ORGANIZATION | 44 | -0.27171 | -0.932536 | 0.559375 | 1 |
| GOBP_PLASMA_MEMBRANE_REPAIR | 23 | -0.314016 | -0.932052 | 0.559078 | 1 |
| GOBP_VOCALIZATION_BEHAVIOR | 15 | -0.353967 | -0.928267 | 0.517711 | 1 |
| GOBP_EPHRIN_RECEPTOR_SIGNALING_PATHWAY | 50 | -0.271745 | -0.927979 | 0.602606 | 1 |
| GOBP_RESPONSE_TO_NICOTINE | 38 | -0.285933 | -0.927255 | 0.565217 | 1 |
| GOBP_CELLULAR_TRANSITION_METAL_ION_HOMEOSTASIS | 87 | -0.241916 | -0.92592 | 0.644195 | 1 |
| GOBP_NEGATIVE_REGULATION_OF_MICROTUBULE_POLYMERIZATION_OR_DEPOLYMERIZ | 40 | -0.278243 | -0.925691 | 0.607143 | 1 |
| GOBP_POSITIVE_REGULATION_OF_T_CELL_MEDIATED_CYTOTOXICITY | 20 | -0.33316 | -0.925599 | 0.571023 | 1 |
| GOBP_ASYMMETRIC_CELL_DIVISION | 17 | -0.337403 | -0.92513 | 0.544555 | 1 |
| GOBP_ESTABLISHMENT_OR_MAINTENANCE_OF_BIPOLAR_CELL_POLARITY | 48 | -0.270702 | -0.925039 | 0.587189 | 1 |
| GOBP_REGULATION_OF_MITOCHONDRIAL_GENE_EXPRESSION | 28 | -0.308286 | -0.92335 | 0.560694 | 1 |
| GOBP_TELOMERASE_RNA_LOCALIZATION | 19 | -0.3372 | -0.921272 | 0.569482 | 1 |
| GOBP_REGULATION_OF_CELLULAR_PH | 68 | -0.247295 | -0.92034 | 0.645161 | 1 |
| GOBP_NEURON_PROJECTION_ARBORIZATION | 29 | -0.292221 | -0.919796 | 0.550432 | 1 |
| GOBP_ESTABLISHMENT_OF_PROTEIN_LOCALIZATION_TO_TELOMERE | 17 | -0.331058 | -0.919324 | 0.570732 | 1 |
| GOBP_REGULATION_OF_PROTEIN_DEPOLYMERIZATION | 82 | -0.245384 | -0.919151 | 0.618257 | 1 |
| GOBP_NEGATIVE_REGULATION_OF_INTRACELLULAR_PROTEIN_TRANSPORT | 37 | -0.279706 | -0.917107 | 0.591331 | 1 |
| GOBP_RIBONUCLEOSIDE_TRIPHOSPHATE_BIOSYNTHETIC_PROCESS | 69 | -0.249167 | -0.915769 | 0.642066 | 1 |
| GOBP_NEUROTRANSMITTER_RECEPTOR_TRANSPORT_TO_PLASMA_MEMBRANE | 16 | -0.340813 | -0.915475 | 0.570652 | 1 |
| GOBP_SERINE_FAMILY_AMINO_ACID_METABOLIC_PROCESS | 32 | -0.297787 | -0.915107 | 0.603659 | 1 |
| GOBP_REGULATION_OF_STEROID_METABOLIC_PROCESS | 75 | -0.24525 | -0.915043 | 0.645283 | 1 |
| GOBP_RESPONSE_TO_PH | 32 | -0.293588 | -0.913274 | 0.60061 | 1 |
| GOBP_NEGATIVE_REGULATION_OF_GLIAL_CELL_DIFFERENTIATION | 16 | -0.346547 | -0.913238 | 0.592896 | 1 |
| GOBP_PURINERGIC_NUCLEOTIDE_RECEPTOR_SIGNALING_PATHWAY | 20 | -0.327997 | -0.913216 | 0.549479 | 1 |
| GOBP_NUCLEOSIDE_PHOSPHATE_BIOSYNTHETIC_PROCESS | 224 | -0.208742 | -0.912231 | 0.758621 | 1 |
| GOBP_REGULATION_OF_PATTERN_RECOGNITION_RECEPTOR_SIGNALING_PATHWAY | 79 | -0.240223 | -0.910539 | 0.653226 | 1 |
| GOBP_REGULATION_OF_GENE_EXPRESSION_EPIGENETIC | 124 | -0.224065 | -0.910143 | 0.719807 | 1 |
| GOBP_SMOOTHENED_SIGNALING_PATHWAY | 120 | -0.228463 | -0.90991 | 0.712871 | 1 |
| GOBP_CYTOPLASMIC_PATTERN_RECOGNITION_RECEPTOR_SIGNALING_PATHWAY_IN_R | 31 | -0.288647 | -0.909494 | 0.628834 | 1 |
| GOBP_REGULATION_OF_ESTABLISHMENT_OF_PLANAR_POLARITY | 54 | -0.257288 | -0.90925 | 0.616197 | 1 |
| GOBP_MYD88_DEPENDENT_TOLL_LIKE_RECEPTOR_SIGNALING_PATHWAY | 17 | -0.33968 | -0.90917 | 0.572864 | 1 |
| GOBP_NEURAL_NUCLEUS_DEVELOPMENT | 52 | -0.26148 | -0.907917 | 0.627586 | 1 |
| GOBP_SECONDARY_ALCOHOL_METABOLIC_PROCESS | 113 | -0.226191 | -0.907804 | 0.742358 | 1 |
| GOBP_HUMORAL_IMMUNE_RESPONSE_MEDIATED_BY_CIRCULATING_IMMUNOGLOBULIN | 25 | -0.307243 | -0.905217 | 0.594241 | 1 |
| GOBP_VIRAL_TRANSCRIPTION | 48 | -0.259856 | -0.904736 | 0.651724 | 1 |
| GOBP_REGULATION_OF_INTRACELLULAR_STEROID_HORMONE_RECEPTOR_SIGNALING_ | 65 | -0.244964 | -0.904552 | 0.719368 | 1 |
| GOBP_TRNA_METHYLATION | 40 | -0.272334 | -0.902424 | 0.646497 | 1 |
| GOBP_CALCIUM_INDEPENDENT_CELL_CELL_ADHESION_VIA_PLASMA_MEMBRANE_CELL | 15 | -0.353831 | -0.90207 | 0.580153 | 1 |
| GOBP_AXON_EXTENSION | 114 | -0.225716 | -0.901404 | 0.73991 | 1 |
| GOBP_MIDBRAIN_DEVELOPMENT | 75 | -0.241857 | -0.899372 | 0.694215 | 1 |
| GOBP_PURINE_CONTAINING_COMPOUND_BIOSYNTHETIC_PROCESS | 178 | -0.211657 | -0.899204 | 0.741259 | 1 |
| GOBP_POSITIVE_REGULATION_OF_ISOTYPE_SWITCHING | 23 | -0.311703 | -0.89918 | 0.578212 | 1 |
| GOBP_PYRIMIDINE_RIBONUCLEOTIDE_BIOSYNTHETIC_PROCESS | 22 | -0.321303 | -0.898817 | 0.609065 | 1 |
| GOBP_LIPOSACCHARIDE_METABOLIC_PROCESS | 94 | -0.23144 | -0.898485 | 0.700855 | 1 |
| GOBP_REGULATION_OF_PROTEIN_DEPHOSPHORYLATION | 79 | -0.228865 | -0.898214 | 0.708 | 1 |
| GOBP_RESPONSE_TO_INTERLEUKIN_4 | 29 | -0.294844 | -0.896849 | 0.615385 | 1 |
| GOBP_POSITIVE_REGULATION_OF_GLYCOLYTIC_PROCESS | 15 | -0.343508 | -0.896481 | 0.586466 | 1 |
| GOBP_POSITIVE_REGULATION_OF_TOR_SIGNALING | 43 | -0.26868 | -0.896449 | 0.627063 | 1 |
| GOBP_DOUBLE_STRAND_BREAK_REPAIR | 260 | -0.200232 | -0.895058 | 0.902913 | 1 |
| GOBP_PROTEIN_DEPOLYMERIZATION | 111 | -0.224571 | -0.895018 | 0.753138 | 1 |
| GOBP_RIBONUCLEOSIDE_TRIPHOSPHATE_METABOLIC_PROCESS | 82 | -0.235107 | -0.894999 | 0.726496 | 1 |
| GOBP_CELLULAR_MODIFIED_AMINO_ACID_BIOSYNTHETIC_PROCESS | 42 | -0.262765 | -0.893006 | 0.625749 | 1 |
| GOBP_SEMAPHORIN_PLEXIN_SIGNALING_PATHWAY | 42 | -0.270959 | -0.890659 | 0.648562 | 1 |
| GOBP_REGULATION_OF_PHOSPHATIDYLINOSITOL_3_KINASE_ACTIVITY | 47 | -0.261527 | -0.890579 | 0.683168 | 1 |
| GOBP_NEGATIVE_REGULATION_OF_BLOOD_PRESSURE | 34 | -0.283022 | -0.889092 | 0.640449 | 1 |
| GOBP_REGULATION_OF_HORMONE_METABOLIC_PROCESS | 30 | -0.285078 | -0.888949 | 0.661721 | 1 |

| | | | | | |
|---|---|---|---|---|---|
| GOBP_BIOMINERALIZATION | 124 | 0.393909 | 1.340827 | 0.042945 | 0.371953 |
| GOBP_REGULATION_OF_ANION_TRANSMEMBRANE_TRANSPORT | 19 | 0.552274 | 1.340222 | 0.123577 | 0.372881 |
| GOBP_DIVALENT_INORGANIC_CATION_HOMEOSTASIS | 360 | 0.359451 | 1.340216 | 0.005382 | 0.372296 |
| GOBP_DORSAL_VENTRAL_PATTERN_FORMATION | 67 | 0.425454 | 1.339842 | 0.074526 | 0.37259 |
| GOBP_METANEPHRIC_NEPHRON_MORPHOGENESIS | 21 | 0.537683 | 1.339473 | 0.134647 | 0.372889 |
| GOBP_NEGATIVE_REGULATION_OF_PEPTIDASE_ACTIVITY | 150 | 0.382857 | 1.337345 | 0.050245 | 0.377661 |
| GOBP_ENDOTHELIAL_CELL_PROLIFERATION | 126 | 0.390472 | 1.336667 | 0.05335 | 0.378735 |
| GOBP_EYE_MORPHOGENESIS | 125 | 0.400071 | 1.335879 | 0.049383 | 0.380032 |
| GOBP_EPITHELIAL_CELL_DIFFERENTIATION_INVOLVED_IN_KIDNEY_DEVELOPMENT | 42 | 0.466326 | 1.335772 | 0.104634 | 0.37969 |
| GOBP_PROSTANOID_METABOLIC_PROCESS | 37 | 0.479864 | 1.335623 | 0.10917 | 0.379442 |
| GOBP_NEGATIVE_REGULATION_OF_LEUKOCYTE_MEDIATED_IMMUNITY | 42 | 0.46593 | 1.33464 | 0.09375 | 0.381328 |
| GOBP_PLACENTA_DEVELOPMENT | 129 | 0.393458 | 1.334404 | 0.049305 | 0.381271 |
| GOBP_POSITIVE_REGULATION_OF_I_KAPPAB_KINASE_NF_KAPPAB_SIGNALING | 161 | 0.37987 | 1.334234 | 0.043217 | 0.381121 |
| GOBP_NEGATIVE_REGULATION_OF_LEUKOCYTE_CELL_CELL_ADHESION | 88 | 0.410865 | 1.333943 | 0.074413 | 0.381277 |
| GOBP_3_PHOSPHOADENOSINE_5_PHOSPHOSULFATE_METABOLIC_PROCESS | 15 | 0.559496 | 1.333909 | 0.145867 | 0.380764 |
| GOBP_NEGATIVE_REGULATION_OF_TORC1_SIGNALING | 18 | 0.538208 | 1.33221 | 0.139571 | 0.384219 |
| GOBP_TAXIS | 445 | 0.353236 | 1.331733 | 0.006403 | 0.384801 |
| GOBP_PHOSPHATIDYLCHOLINE_METABOLIC_PROCESS | 61 | 0.425881 | 1.331588 | 0.077139 | 0.384593 |
| GOBP_REGULATION_OF_FAT_CELL_DIFFERENTIATION | 117 | 0.395151 | 1.331242 | 0.059406 | 0.384833 |
| GOBP_INTEGRIN_MEDIATED_SIGNALING_PATHWAY | 90 | 0.406621 | 1.331142 | 0.070968 | 0.384504 |
| GOBP_COLLAGEN_FIBRIL_ORGANIZATION | 55 | 0.437339 | 1.330923 | 0.09887 | 0.384472 |
| GOBP_REGULATION_OF_NATURAL_KILLER_CELL_MEDIATED_IMMUNITY | 24 | 0.519569 | 1.330653 | 0.137771 | 0.384539 |
| GOBP_FILOPODIUM_ASSEMBLY | 63 | 0.430349 | 1.330499 | 0.069209 | 0.384334 |
| GOBP_TRANSFORMING_GROWTH_FACTOR_BETA_PRODUCTION | 33 | 0.48453 | 1.330254 | 0.113846 | 0.384342 |
| GOBP_REGULATION_OF_FATTY_ACID_OXIDATION | 25 | 0.501091 | 1.329584 | 0.123457 | 0.385482 |
| GOBP_REGULATION_OF_DNA_BINDING_TRANSCRIPTION_FACTOR_ACTIVITY | 361 | 0.355098 | 1.329467 | 0.01087 | 0.385159 |
| GOBP_CYTOKINE_MEDIATED_SIGNALING_PATHWAY | 308 | 0.356978 | 1.328647 | 0.024472 | 0.386562 |
| GOBP_GASTRULATION | 159 | 0.380834 | 1.32802 | 0.046595 | 0.38751 |
| GOBP_FERTILIZATION | 113 | 0.395869 | 1.327936 | 0.050188 | 0.387097 |
| GOBP_ARACHIDONIC_ACID_METABOLIC_PROCESS | 36 | 0.466613 | 1.326903 | 0.10229 | 0.389015 |
| GOBP_REGULATION_OF_CARBOHYDRATE_BIOSYNTHETIC_PROCESS | 81 | 0.411639 | 1.326015 | 0.075916 | 0.390605 |
| GOBP_POSITIVE_REGULATION_OF_SMALL_MOLECULE_METABOLIC_PROCESS | 105 | 0.40059 | 1.325939 | 0.06727 | 0.390207 |
| GOBP_EMBRYONIC_SKELETAL_SYSTEM_DEVELOPMENT | 83 | 0.420555 | 1.325876 | 0.068182 | 0.389773 |
| GOBP_ESTABLISHMENT_OF_CELL_POLARITY | 132 | 0.389485 | 1.325568 | 0.05314 | 0.38991 |
| GOBP_POSITIVE_REGULATION_OF_HEART_GROWTH | 20 | 0.53371 | 1.325156 | 0.138756 | 0.390392 |
| GOBP_CARDIAC_MUSCLE_CONTRACTION | 111 | 0.400813 | 1.324789 | 0.05569 | 0.390642 |
| GOBP_POSITIVE_REGULATION_OF_MYELOID_LEUKOCYTE_DIFFERENTIATION | 40 | 0.459419 | 1.323776 | 0.10961 | 0.392638 |
| GOBP_T_CELL_CYTOKINE_PRODUCTION | 25 | 0.51037 | 1.323078 | 0.138346 | 0.393759 |
| GOBP_NEGATIVE_REGULATION_OF_POTASSIUM_ION_TRANSPORT | 20 | 0.531387 | 1.321747 | 0.153199 | 0.396343 |
| GOBP_EMBRYONIC_MORPHOGENESIS | 483 | 0.346743 | 1.321404 | 0.009524 | 0.39662 |
| GOBP_NEGATIVE_REGULATION_OF_CD4_POSITIVE_ALPHA_BETA_T_CELL_DIFFERENTIATI | 17 | 0.561327 | 1.320962 | 0.155698 | 0.397025 |
| GOBP_SENSORY_PERCEPTION | 367 | 0.353371 | 1.320784 | 0.017094 | 0.396889 |
| GOBP_POSITIVE_REGULATION_OF_DOUBLE_STRAND_BREAK_REPAIR_VIA_HOMOLOGOUS | 36 | 0.469969 | 1.320469 | 0.115265 | 0.397057 |
| GOBP_SALIVARY_GLAND_DEVELOPMENT | 29 | 0.488719 | 1.319544 | 0.135338 | 0.398786 |
| GOBP_REGULATION_OF_PROTEIN_MATURATION | 52 | 0.441927 | 1.3192 | 0.114731 | 0.399098 |
| GOBP_CALCIUM_ION_IMPORT_INTO_CYTOSOL | 20 | 0.535724 | 1.31831 | 0.135433 | 0.400697 |
| GOBP_REGULATION_OF_CELL_SUBSTRATE_JUNCTION_ORGANIZATION | 65 | 0.425566 | 1.318082 | 0.082321 | 0.400711 |
| GOBP_NEGATIVE_REGULATION_OF_COLD_INDUCED_THERMOGENESIS | 41 | 0.453851 | 1.317822 | 0.106881 | 0.400756 |
| GOBP_CHROMOSOME_ORGANIZATION_INVOLVED_IN_MEIOTIC_CELL_CYCLE | 54 | 0.434464 | 1.317441 | 0.097765 | 0.401135 |
| GOBP_REGULATION_OF_COAGULATION | 51 | 0.441181 | 1.317332 | 0.102899 | 0.400814 |
| GOBP_SYNCYTIUM_FORMATION | 43 | 0.453896 | 1.31677 | 0.098837 | 0.401674 |
| GOBP_COLUMNAR_CUBOIDAL_EPITHELIAL_CELL_DIFFERENTIATION | 66 | 0.417069 | 1.316697 | 0.091281 | 0.401267 |
| GOBP_REGULATION_OF_PEPTIDYL_SERINE_PHOSPHORYLATION | 112 | 0.389058 | 1.316638 | 0.071887 | 0.400811 |
| GOBP_ACTIVATION_OF_CYSTEINE_TYPE_ENDOPEPTIDASE_ACTIVITY_INVOLVED_IN_APOF | 65 | 0.421963 | 1.315975 | 0.090411 | 0.401912 |
| GOBP_POSITIVE_REGULATION_OF_CALCIUM_ION_TRANSMEMBRANE_TRANSPORTER_AC | 35 | 0.464102 | 1.315433 | 0.112426 | 0.402697 |
| GOBP_TRANSMEMBRANE_RECEPTOR_PROTEIN_SERINE_THREONINE_KINASE_SIGNALIN( | 301 | 0.356182 | 1.314846 | 0.018952 | 0.403616 |
| GOBP_REGULATION_OF_TISSUE_REMODELING | 52 | 0.436626 | 1.31465 | 0.097458 | 0.403526 |
| GOBP_POSITIVE_REGULATION_OF_PROTEIN_KINASE_B_SIGNALING | 90 | 0.402812 | 1.314557 | 0.077419 | 0.403203 |
| GOBP_OVULATION_CYCLE | 56 | 0.423779 | 1.314369 | 0.088276 | 0.403121 |
| GOBP_RUFFLE_ORGANIZATION | 50 | 0.437863 | 1.313848 | 0.115057 | 0.403882 |
| GOBP_REGULATION_OF_B_CELL_PROLIFERATION | 39 | 0.452477 | 1.311326 | 0.123881 | 0.409828 |
| GOBP_PROSTANOID_BIOSYNTHETIC_PROCESS | 23 | 0.499957 | 1.309639 | 0.153846 | 0.413619 |
| GOBP_REGULATION_OF_HUMORAL_IMMUNE_RESPONSE | 22 | 0.510989 | 1.30951 | 0.150794 | 0.413314 |
| GOBP_POSITIVE_REGULATION_OF_ION_TRANSMEMBRANE_TRANSPORT | 130 | 0.383808 | 1.309354 | 0.060226 | 0.413125 |
| GOBP_FEMALE_SEX_DIFFERENTIATION | 97 | 0.399289 | 1.308908 | 0.086229 | 0.413716 |
| GOBP_POSITIVE_REGULATION_OF_CALCIUM_ION_TRANSPORT | 91 | 0.39941 | 1.308516 | 0.088919 | 0.414154 |
| GOBP_PERIPHERAL_NERVOUS_SYSTEM_DEVELOPMENT | 71 | 0.410471 | 1.307533 | 0.090175 | 0.415984 |
| GOBP_REGULATION_OF_SENSORY_PERCEPTION | 23 | 0.50499 | 1.307352 | 0.131833 | 0.415869 |
| GOBP_REGULATION_OF_RELEASE_OF_CYTOCHROME_C_FROM_MITOCHONDRIA | 41 | 0.457044 | 1.306918 | 0.125 | 0.416417 |
| GOBP_NEGATIVE_REGULATION_OF_POTASSIUM_ION_TRANSMEMBRANE_TRANSPORT | 16 | 0.560029 | 1.306551 | 0.15873 | 0.416713 |
| GOBP_BIOLOGICAL_PROCESS_INVOLVED_IN_INTERACTION_WITH_SYMBIONT | 68 | 0.416313 | 1.306291 | 0.103448 | 0.41681 |
| GOBP_EXOCRINE_SYSTEM_DEVELOPMENT | 37 | 0.46337 | 1.305105 | 0.13037 | 0.419219 |
| GOBP_SKELETAL_MUSCLE_CONTRACTION | 33 | 0.474013 | 1.303519 | 0.12029 | 0.422647 |
| GOBP_EXTERNAL_ENCAPSULATING_STRUCTURE_ORGANIZATION | 246 | 0.355243 | 1.30309 | 0.048 | 0.423104 |
| GOBP_CELLULAR_EXTRAVASATION | 50 | 0.436994 | 1.302735 | 0.107093 | 0.423371 |
| GOBP_LEUKOCYTE_PROLIFERATION | 210 | 0.363844 | 1.302144 | 0.039863 | 0.424321 |
| GOBP_T_CELL_PROLIFERATION | 139 | 0.375833 | 1.300995 | 0.056763 | 0.426631 |
| GOBP_POSITIVE_REGULATION_OF_ORGAN_GROWTH | 25 | 0.505011 | 1.300613 | 0.153481 | 0.427014 |
| GOBP_REGULATION_OF_HEART_CONTRACTION | 164 | 0.372232 | 1.300284 | 0.054348 | 0.427323 |
| GOBP_PHENOL_CONTAINING_COMPOUND_BIOSYNTHETIC_PROCESS | 31 | 0.486635 | 1.299742 | 0.136434 | 0.428091 |
| GOBP_SEX_DIFFERENTIATION | 211 | 0.359297 | 1.299315 | 0.04084 | 0.428666 |
| GOBP_RESPONSE_TO_PAIN | 25 | 0.488662 | 1.299257 | 0.138187 | 0.428222 |
| GOBP_AORTA_DEVELOPMENT | 54 | 0.428331 | 1.299003 | 0.10101 | 0.428324 |
| GOBP_RELAXATION_OF_CARDIAC_MUSCLE | 16 | 0.548412 | 1.298756 | 0.158155 | 0.428371 |
| GOBP_NEGATIVE_REGULATION_OF_TRANSFORMING_GROWTH_FACTOR_BETA_RECEPTO | 67 | 0.415026 | 1.298541 | 0.093315 | 0.42835 |
| GOBP_T_CELL_DIFFERENTIATION_INVOLVED_IN_IMMUNE_RESPONSE | 51 | 0.424729 | 1.298295 | 0.120991 | 0.428339 |
| GOBP_CELL_DIFFERENTIATION_INVOLVED_IN_KIDNEY_DEVELOPMENT | 49 | 0.444337 | 1.298285 | 0.114613 | 0.427757 |
| GOBP_REGULATION_OF_PROTEIN_ACETYLATION | 77 | 0.40234 | 1.298078 | 0.110667 | 0.427686 |
| GOBP_REGULATION_OF_NIK_NF_KAPPAB_SIGNALING | 79 | 0.4059 | 1.298036 | 0.10582 | 0.427215 |
| GOBP_REGULATION_OF_CHEMOTAXIS | 163 | 0.366708 | 1.297124 | 0.059032 | 0.429036 |
| GOBP_HYALURONAN_METABOLIC_PROCESS | 21 | 0.51723 | 1.296333 | 0.153226 | 0.43057 |
| GOBP_REGULATION_OF_TRANSCRIPTION_REGULATORY_REGION_DNA_BINDING | 43 | 0.454953 | 1.295579 | 0.119597 | 0.431921 |
| GOBP_CELLULAR_RESPONSE_TO_CORTICOSTEROID_STIMULUS | 50 | 0.431405 | 1.295249 | 0.122781 | 0.432259 |
| GOBP_POSITIVE_REGULATION_OF_IMMUNE_RESPONSE | 342 | 0.346415 | 1.292669 | 0.019334 | 0.438412 |
| GOBP_REGULATION_OF_CELLULAR_RESPONSE_TO_GROWTH_FACTOR_STIMULUS | 246 | 0.349353 | 1.292658 | 0.041387 | 0.437834 |
| GOBP_POSITIVE_REGULATION_OF_CELL_MATRIX_ADHESION | 47 | 0.439768 | 1.291861 | 0.116379 | 0.439341 |
| GOBP_REGULATION_OF_NECROTIC_CELL_DEATH | 31 | 0.469459 | 1.291575 | 0.146667 | 0.439448 |
| GOBP_REGULATION_OF_LEUKOCYTE_MEDIATED_CYTOTOXICITY | 45 | 0.447053 | 1.291464 | 0.128242 | 0.439104 |
| GOBP_REGULATION_OF_REACTIVE_OXYGEN_SPECIES_METABOLIC_PROCESS | 113 | 0.382895 | 1.291397 | 0.080808 | 0.438682 |
| GOBP_CELL_DIFFERENTIATION_INVOLVED_IN_EMBRYONIC_PLACENTA_DEVELOPMENT | 23 | 0.499032 | 1.290724 | 0.151235 | 0.439788 |
| GOBP_RESPONSE_TO_CORTICOSTEROID | 121 | 0.381763 | 1.290483 | 0.078261 | 0.439788 |
| GOBP_SPERM_EGG_RECOGNITION | 24 | 0.494096 | 1.290332 | 0.162826 | 0.439556 |
| GOBP_NEURON_FATE_SPECIFICATION | 19 | 0.518464 | 1.290224 | 0.155763 | 0.439224 |
| GOBP_ACTIVATED_T_CELL_PROLIFERATION | 32 | 0.464285 | 1.289922 | 0.144131 | 0.4395 |
| GOBP_CYTOKINE_PRODUCTION_INVOLVED_IN_IMMUNE_RESPONSE | 70 | 0.412475 | 1.289469 | 0.119565 | 0.440075 |
| GOBP_SENSORY_ORGAN_DEVELOPMENT | 438 | 0.342477 | 1.288936 | 0.012474 | 0.440926 |
| GOBP_EAR_DEVELOPMENT | 172 | 0.357734 | 1.288369 | 0.058968 | 0.441861 |
| GOBP_EPIDERMIS_MORPHOGENESIS | 21 | 0.504107 | 1.288366 | 0.156958 | 0.441266 |
| GOBP_DETECTION_OF_STIMULUS | 142 | 0.37375 | 1.288078 | 0.068627 | 0.441425 |
| GOBP_MAINTENANCE_OF_LOCATION_IN_CELL | 174 | 0.366789 | 1.287544 | 0.063306 | 0.442246 |
| GOBP_POSITIVE_REGULATION_OF_EPIDERMAL_CELL_DIFFERENTIATION | 22 | 0.50508 | 1.286621 | 0.178797 | 0.444202 |
| GOBP_POSITIVE_REGULATION_OF_NEURON_APOPTOTIC_PROCESS | 41 | 0.442535 | 1.286323 | 0.130246 | 0.444362 |
| GOBP_FORMATION_OF_PRIMARY_GERM_LAYER | 104 | 0.387143 | 1.285804 | 0.096734 | 0.445165 |
| GOBP_CELLULAR_RESPONSE_TO_REACTIVE_OXYGEN_SPECIES | 118 | 0.381707 | 1.285464 | 0.095238 | 0.445465 |
| GOBP_AMYLOID_BETA_CLEARANCE | 28 | 0.471925 | 1.285432 | 0.141985 | 0.444964 |
| GOBP_NEGATIVE_REGULATION_OF_DNA_BINDING_TRANSCRIPTION_FACTOR_ACTIVITY | 146 | 0.370793 | 1.28543 | 0.068069 | 0.44437 |
| GOBP_MYELOID_CELL_ACTIVATION_INVOLVED_IN_IMMUNE_RESPONSE | 65 | 0.411353 | 1.28538 | 0.110067 | 0.4439 |
| GOBP_APOPTOTIC_CELL_CLEARANCE | 40 | 0.444626 | 1.283613 | 0.143288 | 0.448111 |
| GOBP_RESPONSE_TO_INTERFERON_GAMMA | 92 | 0.394848 | 1.283515 | 0.087179 | 0.447776 |
| GOBP_ARF_PROTEIN_SIGNAL_TRANSDUCTION | 19 | 0.520297 | 1.283503 | 0.163166 | 0.447217 |
| GOBP_NEPHRON_DEVELOPMENT | 121 | 0.38437 | 1.283217 | 0.083127 | 0.447387 |
| GOBP_APPENDAGE_MORPHOGENESIS | 104 | 0.38215 | 1.282669 | 0.100503 | 0.44827 |
| GOBP_POSITIVE_REGULATION_OF_MEIOTIC_CELL_CYCLE | 19 | 0.515543 | 1.281555 | 0.193443 | 0.45069 |
| GOBP_NUCLEOSIDE_METABOLIC_PROCESS | 53 | 0.418388 | 1.281251 | 0.127874 | 0.45092 |
| GOBP_INTERLEUKIN_10_PRODUCTION | 36 | 0.451705 | 1.280512 | 0.160294 | 0.45232 |
| GOBP_REGULATION_OF_MITOCHONDRION_ORGANIZATION | 132 | 0.373079 | 1.280497 | 0.075795 | 0.451766 |
| GOBP_REGULATION_OF_T_HELPER_CELL_DIFFERENTIATION | 26 | 0.49096 | 1.279575 | 0.148588 | 0.453631 |
| GOBP_FAT_CELL_DIFFERENTIATION | 201 | 0.354692 | 1.27928 | 0.054374 | 0.453737 |
| GOBP_SENSORY_PERCEPTION_OF_MECHANICAL_STIMULUS | 136 | 0.372971 | 1.278944 | 0.082512 | 0.454025 |
| GOBP_REGULATION_OF_CARDIAC_MUSCLE_CELL_DIFFERENTIATION | 15 | 0.54163 | 1.278524 | 0.165049 | 0.454513 |
| GOBP_EPITHELIAL_TO_MESENCHYMAL_TRANSITION_INVOLVED_IN_ENDOCARDIAL_CUSH | 20 | 0.509606 | 1.278186 | 0.140823 | 0.454808 |
| GOBP_CD4_POSITIVE_ALPHA_BETA_T_CELL_DIFFERENTIATION | 57 | 0.419567 | 1.278064 | 0.118852 | 0.454541 |
| GOBP_NEURON_APOPTOTIC_PROCESS | 211 | 0.356077 | 1.278061 | 0.06489 | 0.453954 |
| GOBP_POSITIVE_REGULATION_OF_STRESS_ACTIVATED_PROTEIN_KINASE_SIGNALING_C/ | 111 | 0.385669 | 1.27796 | 0.074906 | 0.453677 |
| GOBP_NEGATIVE_REGULATION_OF_INFLAMMATORY_RESPONSE | 104 | 0.386192 | 1.27743 | 0.094194 | 0.454522 |
| GOBP_REGULATION_OF_CALCIUM_ION_TRANSPORT | 188 | 0.35922 | 1.277075 | 0.068639 | 0.454888 |
| GOBP_SENSORY_SYSTEM_DEVELOPMENT | 304 | 0.346648 | 1.27706 | 0.042763 | 0.454333 |
| GOBP_NEGATIVE_REGULATION_OF_LEUKOCYTE_PROLIFERATION | 55 | 0.421932 | 1.275545 | 0.135802 | 0.457885 |
| GOBP_NEGATIVE_REGULATION_OF_LOCOMOTION | 272 | 0.347416 | 1.275438 | 0.044893 | 0.45757 |
| GOBP_MORPHOGENESIS_OF_AN_ENDOTHELIUM | 16 | 0.531574 | 1.275052 | 0.177236 | 0.458115 |
| GOBP_POSITIVE_REGULATION_OF_AXON_EXTENSION | 38 | 0.451326 | 1.274886 | 0.150568 | 0.457936 |
| GOBP_MULTICELLULAR_ORGANISMAL_HOMEOSTASIS | 397 | 0.338065 | 1.274846 | 0.025586 | 0.457454 |
| GOBP_POSITIVE_REGULATION_OF_PROTEIN_ACETYLATION | 45 | 0.431316 | 1.274808 | 0.138728 | 0.456964 |
| GOBP_REGULATION_OF_CYSTEINE_TYPE_ENDOPEPTIDASE_ACTIVITY_INVOLVED_IN_APC | 15 | 0.547394 | 1.274801 | 0.193333 | 0.45639 |
| GOBP_COPPER_ION_HOMEOSTASIS | 17 | 0.52534 | 1.274529 | 0.20098 | 0.45653 |
| GOBP_CONNECTIVE_TISSUE_DEVELOPMENT | 206 | 0.355214 | 1.274348 | 0.069767 | 0.456432 |
| GOBP_MYOBLAST_FUSION | 32 | 0.459959 | 1.274161 | 0.143072 | 0.456381 |
| GOBP_SKIN_DEVELOPMENT | 181 | 0.36002 | 1.274148 | 0.072551 | 0.455813 |
| GOBP_RETINAL_GANGLION_CELL_AXON_GUIDANCE | 19 | 0.513203 | 1.274083 | 0.192 | 0.455409 |
| GOBP_GERM_CELL_DEVELOPMENT | 210 | 0.354106 | 1.273556 | 0.048724 | 0.456188 |
| GOBP_AMINOGLYCAN_CATABOLIC_PROCESS | 21 | 0.502226 | 1.273194 | 0.161392 | 0.456554 |
| GOBP_DNA_DAMAGE_RESPONSE_SIGNAL_TRANSDUCTION_RESULTING_IN_TRANSCRIPT | 17 | 0.524528 | 1.272964 | 0.181115 | 0.45657 |
| GOBP_BINDING_OF_SPERM_TO_ZONA_PELLUCIDA | 21 | 0.496001 | 1.27228 | 0.166932 | 0.457766 |
| GOBP_PODOSOME_ASSEMBLY | 16 | 0.533377 | 1.272055 | 0.169518 | 0.457758 |
| GOBP_PHAGOCYTOSIS | 175 | 0.358675 | 1.271976 | 0.06974 | 0.457411 |
| GOBP_POSITIVE_REGULATION_OF_CYTOKINE_PRODUCTION | 314 | 0.342177 | 1.271004 | 0.043382 | 0.45948 |
| GOBP_MYELOID_LEUKOCYTE_MEDIATED_IMMUNITY | 74 | 0.399748 | 1.270805 | 0.099455 | 0.459466 |

| | | | | | |
|---|---|---|---|---|---|
| GOBP_NEGATIVE_REGULATION_OF_INTRACELLULAR_STEROID_HORMONE_RECEPTOR_S | 34 | -0.280257 | -0.885096 | 0.658754 | 1 |
| GOBP_CALCIUM_ION_REGULATED_EXOCYTOSIS | 60 | -0.250513 | -0.885057 | 0.702797 | 1 |
| GOBP_SCF_DEPENDENT_PROTEASOMAL_UBIQUITIN_DEPENDENT_PROTEIN_CATABOLIC | 47 | -0.254985 | -0.884965 | 0.642173 | 1 |
| GOBP_LIPID_MODIFICATION | 177 | -0.205362 | -0.882817 | 0.808383 | 1 |
| GOBP_RNA_SURVEILLANCE | 15 | -0.33908 | -0.880383 | 0.636592 | 1 |
| GOBP_POSITIVE_REGULATION_OF_TRANSLATIONAL_INITIATION | 23 | -0.297032 | -0.880165 | 0.609065 | 1 |
| GOBP_POSITIVE_REGULATION_OF_TELOMERASE_RNA_LOCALIZATION_TO_CAJAL_BODY | 15 | -0.333409 | -0.878999 | 0.645251 | 1 |
| GOBP_VESICLE_MEDIATED_TRANSPORT_IN_SYNAPSE | 175 | -0.208377 | -0.878597 | 0.848684 | 1 |
| GOBP_TETRAPYRROLE_BIOSYNTHETIC_PROCESS | 31 | -0.277142 | -0.877521 | 0.657224 | 1 |
| GOBP_SYNAPTIC_VESICLE_EXOCYTOSIS | 93 | -0.22662 | -0.877147 | 0.795455 | 1 |
| GOBP_ENSHEATHMENT_OF_NEURONS | 127 | -0.216066 | -0.876343 | 0.791667 | 1 |
| GOBP_DENDRITE_MORPHOGENESIS | 134 | -0.215761 | -0.875238 | 0.791045 | 1 |
| GOBP_REGULATION_OF_CELLULAR_RESPONSE_TO_HEAT | 17 | -0.319018 | -0.875026 | 0.63807 | 1 |
| GOBP_RESPONSE_TO_PROGESTERONE | 31 | -0.28053 | -0.874881 | 0.656442 | 1 |
| GOBP_REGULATION_OF_RECEPTOR_BINDING | 18 | -0.316602 | -0.873829 | 0.661333 | 1 |
| GOBP_PSEUDOURIDINE_SYNTHESIS | 18 | -0.321305 | -0.873538 | 0.639896 | 1 |
| GOBP_NEGATIVE_REGULATION_OF_STEM_CELL_DIFFERENTIATION | 22 | -0.307942 | -0.873344 | 0.619303 | 1 |
| GOBP_MISMATCH_REPAIR | 30 | -0.280415 | -0.872858 | 0.657895 | 1 |
| GOBP_LINOLEIC_ACID_METABOLIC_PROCESS | 16 | -0.328345 | -0.872223 | 0.643374 | 1 |
| GOBP_LONG_CHAIN_FATTY_ACID_BIOSYNTHETIC_PROCESS | 21 | -0.310577 | -0.871482 | 0.646341 | 1 |
| GOBP_PROTEIN_TARGETING_TO_PEROXISOME | 18 | -0.322703 | -0.870684 | 0.641873 | 1 |
| GOBP_PURINE_CONTAINING_COMPOUND_TRANSMEMBRANE_TRANSPORT | 18 | -0.32541 | -0.870285 | 0.620513 | 1 |
| GOBP_MRNA_EXPORT_FROM_NUCLEUS | 54 | -0.249069 | -0.869957 | 0.688172 | 1 |
| GOBP_CELL_CELL_JUNCTION_MAINTENANCE | 17 | -0.325178 | -0.869442 | 0.630319 | 1 |
| GOBP_VESICLE_DOCKING_INVOLVED_IN_EXOCYTOSIS | 34 | -0.275761 | -0.8691 | 0.69906 | 1 |
| GOBP_POSITIVE_REGULATION_OF_RECEPTOR_MEDIATED_ENDOCYTOSIS | 44 | -0.255194 | -0.868735 | 0.713826 | 1 |
| GOBP_REGULATION_OF_RECEPTOR_RECYCLING | 21 | -0.302447 | -0.868356 | 0.664103 | 1 |
| GOBP_AXIS_ELONGATION | 22 | -0.299651 | -0.86691 | 0.682997 | 1 |
| GOBP_PORPHYRIN_CONTAINING_COMPOUND_METABOLIC_PROCESS | 41 | -0.259306 | -0.866367 | 0.716867 | 1 |
| GOBP_ASPARTATE_FAMILY_AMINO_ACID_BIOSYNTHETIC_PROCESS | 19 | -0.305664 | -0.86463 | 0.657609 | 1 |
| GOBP_NUCLEOSIDE_BISPHOSPHATE_METABOLIC_PROCESS | 102 | -0.216239 | -0.862783 | 0.790393 | 1 |
| GOBP_POSITIVE_REGULATION_OF_INSULIN_SECRETION_INVOLVED_IN_CELLULAR_RESF | 22 | -0.298921 | -0.862353 | 0.655738 | 1 |
| GOBP_SOMATIC_DIVERSIFICATION_OF_IMMUNOGLOBULINS_INVOLVED_IN_IMMUNE_RES | 42 | -0.257719 | -0.859514 | 0.701299 | 1 |
| GOBP_PROTON_TRANSMEMBRANE_TRANSPORT | 129 | -0.212222 | -0.859215 | 0.857924 | 1 |
| GOBP_MITOTIC_SISTER_CHROMATID_SEGREGATION | 165 | -0.204791 | -0.858009 | 0.911765 | 1 |
| GOBP_NEGATIVE_REGULATION_OF_ADENYLATE_CYCLASE_ACTIVITY | 16 | -0.317532 | -0.857827 | 0.665012 | 1 |
| GOBP_REGULATION_OF_CIRCADIAN_RHYTHM | 102 | -0.217587 | -0.857161 | 0.843602 | 1 |
| GOBP_DIACYLGLYCEROL_METABOLIC_PROCESS | 23 | -0.295068 | -0.856946 | 0.633238 | 1 |
| GOBP_SUBSTANTIA_NIGRA_DEVELOPMENT | 40 | -0.262321 | -0.856178 | 0.699367 | 1 |
| GOBP_EXCRETION | 40 | -0.264611 | -0.855462 | 0.666667 | 1 |
| GOBP_DNA_REPAIR | 497 | -0.181431 | -0.855375 | 1 | 1 |
| GOBP_RENAL_SYSTEM_PROCESS_INVOLVED_IN_REGULATION_OF_SYSTEMIC_ARTERIAL | 15 | -0.332406 | -0.854606 | 0.682039 | 1 |
| GOBP_PEROXISOMAL_TRANSPORT | 21 | -0.302835 | -0.854311 | 0.665738 | 1 |
| GOBP_RIBOSE_PHOSPHATE_BIOSYNTHETIC_PROCESS | 174 | -0.201028 | -0.853082 | 0.909774 | 1 |
| GOBP_CYTOPLASMIC_MICROTUBULE_ORGANIZATION | 55 | -0.242932 | -0.850364 | 0.772727 | 1 |
| GOBP_UBIQUINONE_METABOLIC_PROCESS | 18 | -0.315315 | -0.84968 | 0.677686 | 1 |
| GOBP_REGULATION_OF_CHROMATIN_ASSEMBLY | 29 | -0.277819 | -0.847838 | 0.738028 | 1 |
| GOBP_HIPPOCAMPUS_DEVELOPMENT | 64 | -0.236849 | -0.846466 | 0.781132 | 1 |
| GOBP_STEROL_TRANSPORT | 89 | -0.218523 | -0.846307 | 0.832636 | 1 |
| GOBP_REGULATION_OF_ENDOCYTOSIS | 178 | -0.199982 | -0.845635 | 0.918129 | 1 |
| GOBP_POSITIVE_REGULATION_OF_GLIOGENESIS | 50 | -0.248079 | -0.845169 | 0.786713 | 1 |
| GOBP_REGULATION_OF_DNA_REPAIR | 148 | -0.20154 | -0.844522 | 0.887641 | 1 |
| GOBP_REGULATION_OF_NMDA_RECEPTOR_ACTIVITY | 17 | -0.319772 | -0.843942 | 0.681013 | 1 |
| GOBP_PHOTOPERIODISM | 29 | -0.277473 | -0.843079 | 0.720238 | 1 |
| GOBP_NEGATIVE_REGULATION_OF_PROTEIN_LOCALIZATION | 163 | -0.196948 | -0.841525 | 0.922078 | 1 |
| GOBP_REGULATION_OF_RESPONSE_TO_DNA_DAMAGE_STIMULUS | 240 | -0.193139 | -0.841025 | 0.936364 | 1 |
| GOBP_NEGATIVE_REGULATION_OF_PEPTIDE_SECRETION | 29 | -0.271137 | -0.83976 | 0.743516 | 1 |
| GOBP_GLYCOLIPID_BIOSYNTHETIC_PROCESS | 65 | -0.228391 | -0.838842 | 0.785978 | 1 |
| GOBP_PROTEIN_LOCALIZATION_TO_CHROMOSOME | 91 | -0.218041 | -0.83778 | 0.864865 | 1 |
| GOBP_PYRIMIDINE_RIBONUCLEOTIDE_METABOLIC_PROCESS | 28 | -0.276821 | -0.837225 | 0.71978 | 1 |
| GOBP_REGULATION_OF_CELL_CYCLE_CHECKPOINT | 32 | -0.264455 | -0.836972 | 0.728358 | 1 |
| GOBP_REGULATION_OF_DOUBLE_STRAND_BREAK_REPAIR | 108 | -0.211787 | -0.836572 | 0.843434 | 1 |
| GOBP_AXONAL_FASCICULATION | 21 | -0.302493 | -0.835125 | 0.723684 | 1 |
| GOBP_PLASMA_MEMBRANE_TUBULATION | 17 | -0.309552 | -0.835107 | 0.710256 | 1 |
| GOBP_CYTOSKELETON_DEPENDENT_CYTOKINESIS | 93 | -0.213034 | -0.834955 | 0.836538 | 1 |
| GOBP_LAMELLIPODIUM_ASSEMBLY | 63 | -0.234741 | -0.834629 | 0.813793 | 1 |
| GOBP_PROTEIN_MANNOSYLATION | 22 | -0.292248 | -0.833949 | 0.731507 | 1 |
| GOBP_IRON_SULFUR_CLUSTER_ASSEMBLY | 25 | -0.281211 | -0.833334 | 0.698508 | 1 |
| GOBP_HISTONE_H3_K27_METHYLATION | 16 | -0.309855 | -0.833305 | 0.665706 | 1 |
| GOBP_ATP_SYNTHESIS_COUPLED_PROTON_TRANSPORT | 24 | -0.288372 | -0.832413 | 0.724044 | 1 |
| GOBP_CELLULAR_COMPONENT_MAINTENANCE | 57 | -0.233919 | -0.832364 | 0.82971 | 1 |
| GOBP_HISTONE_H2A_ACETYLATION | 23 | -0.28918 | -0.83156 | 0.710306 | 1 |
| GOBP_SOMATIC_DIVERSIFICATION_OF_IMMUNOGLOBULINS | 57 | -0.240153 | -0.831501 | 0.767528 | 1 |
| GOBP_REGULATION_OF_DNA_DAMAGE_CHECKPOINT | 16 | -0.316051 | -0.830703 | 0.694087 | 1 |
| GOBP_REGULATION_OF_SMOOTHENED_SIGNALING_PATHWAY | 71 | -0.220435 | -0.830498 | 0.838951 | 1 |
| GOBP_PROTEIN_LOCALIZATION_TO_ENDOSOME | 24 | -0.281955 | -0.828678 | 0.720461 | 1 |
| GOBP_NEGATIVE_REGULATION_OF_CYTOSKELETON_ORGANIZATION | 149 | -0.199986 | -0.827757 | 0.94382 | 1 |
| GOBP_REGULATION_OF_MITOTIC_SISTER_CHROMATID_SEGREGATION | 42 | -0.251805 | -0.826732 | 0.786378 | 1 |
| GOBP_AXONAL_TRANSPORT | 63 | -0.230717 | -0.826651 | 0.825175 | 1 |
| GOBP_PROTEIN_SUMOYLATION | 47 | -0.243791 | -0.825955 | 0.78 | 1 |
| GOBP_ASPARTATE_FAMILY_AMINO_ACID_CATABOLIC_PROCESS | 15 | -0.317642 | -0.825845 | 0.700957 | 1 |
| GOBP_NEURAL_TUBE_DEVELOPMENT | 142 | -0.198089 | -0.825227 | 0.92823 | 1 |
| GOBP_REGULATION_OF_TRANSLATIONAL_FIDELITY | 21 | -0.289732 | -0.824811 | 0.715818 | 1 |
| GOBP_RESPONSE_TO_ISCHEMIA | 45 | -0.24021 | -0.824172 | 0.805825 | 1 |
| GOBP_PYRIMIDINE_NUCLEOTIDE_CATABOLIC_PROCESS | 18 | -0.301778 | -0.821524 | 0.727794 | 1 |
| GOBP_IRON_ION_HOMEOSTASIS | 68 | -0.220733 | -0.820522 | 0.828467 | 1 |
| GOBP_REGULATION_OF_ACTIN_FILAMENT_BUNDLE_ASSEMBLY | 93 | -0.212774 | -0.820316 | 0.87444 | 1 |
| GOBP_INTERLEUKIN_1_MEDIATED_SIGNALING_PATHWAY | 20 | -0.295122 | -0.816015 | 0.735955 | 1 |
| GOBP_REGULATION_OF_OXIDATIVE_STRESS_INDUCED_INTRINSIC_APOPTOTIC_SIGNALII | 26 | -0.272733 | -0.81595 | 0.762032 | 1 |
| GOBP_REGULATION_OF_RESPONSE_TO_EXTRACELLULAR_STIMULUS | 18 | -0.302445 | -0.815619 | 0.753846 | 1 |
| GOBP_POSITIVE_REGULATION_OF_RNA_SPLICING | 34 | -0.25915 | -0.815361 | 0.773585 | 1 |
| GOBP_REGULATION_OF_IMMUNOGLOBULIN_PRODUCTION | 52 | -0.229491 | -0.812492 | 0.86755 | 1 |
| GOBP_REGULATION_OF_T_CELL_DIFFERENTIATION_IN_THYMUS | 17 | -0.30538 | -0.812095 | 0.731183 | 1 |
| GOBP_POSITIVE_REGULATION_OF_MULTICELLULAR_ORGANISM_GROWTH | 24 | -0.277621 | -0.810924 | 0.758333 | 1 |
| GOBP_NEGATIVE_REGULATION_OF_SUPRAMOLECULAR_FIBER_ORGANIZATION | 149 | -0.196313 | -0.809978 | 0.912568 | 1 |
| GOBP_REGULATION_OF_MITOTIC_SPINDLE_ASSEMBLY | 23 | -0.276285 | -0.809198 | 0.785515 | 1 |
| GOBP_ESTABLISHMENT_OF_TISSUE_POLARITY | 66 | -0.225584 | -0.808697 | 0.847584 | 1 |
| GOBP_RESPONSE_TO_LIGHT_STIMULUS | 252 | -0.180829 | -0.808686 | 0.983607 | 1 |
| GOBP_SNRNA_METABOLIC_PROCESS | 51 | -0.23346 | -0.807391 | 0.810631 | 1 |
| GOBP_REGULATION_OF_SYNAPTIC_TRANSMISSION_GABAERGIC | 28 | -0.266815 | -0.806655 | 0.795107 | 1 |
| GOBP_POSITIVE_REGULATION_OF_CELL_CYCLE_PHASE_TRANSITION | 102 | -0.200604 | -0.806609 | 0.940594 | 1 |
| GOBP_AROMATIC_AMINO_ACID_FAMILY_METABOLIC_PROCESS | 16 | -0.302115 | -0.80623 | 0.753086 | 1 |
| GOBP_REGULATION_OF_CELL_FATE_COMMITMENT | 16 | -0.296708 | -0.805418 | 0.735941 | 1 |
| GOBP_ESTABLISHMENT_OF_PROTEIN_LOCALIZATION_TO_MEMBRANE | 235 | -0.181132 | -0.804968 | 0.983051 | 1 |
| GOBP_NEGATIVE_REGULATION_OF_OXIDATIVE_STRESS_INDUCED_INTRINSIC_APOPTOTI | 18 | -0.294124 | -0.803085 | 0.774011 | 1 |
| GOBP_REGULATION_OF_CYTOPLASMIC_TRANSPORT | 27 | -0.266318 | -0.802925 | 0.751497 | 1 |
| GOBP_HEMATOPOIETIC_STEM_CELL_DIFFERENTIATION | 24 | -0.2679 | -0.800891 | 0.766382 | 1 |
| GOBP_PROTEIN_TARGETING_TO_MEMBRANE | 113 | -0.200145 | -0.799158 | 0.915888 | 1 |
| GOBP_INHIBITORY_SYNAPSE_ASSEMBLY | 15 | -0.30525 | -0.798718 | 0.765432 | 1 |
| GOBP_NEGATIVE_REGULATION_OF_LYASE_ACTIVITY | 20 | -0.292425 | -0.798519 | 0.755435 | 1 |
| GOBP_FATTY_ACID_DERIVATIVE_BIOSYNTHETIC_PROCESS | 35 | -0.244009 | -0.797688 | 0.754491 | 1 |
| GOBP_RRNA_CATABOLIC_PROCESS | 20 | -0.285834 | -0.797667 | 0.777778 | 1 |
| GOBP_HINDBRAIN_MORPHOGENESIS | 40 | -0.236972 | -0.796113 | 0.833333 | 1 |
| GOBP_ESTABLISHMENT_OF_EPITHELIAL_CELL_POLARITY | 31 | -0.257941 | -0.795902 | 0.827273 | 1 |
| GOBP_DENDRITIC_SPINE_MAINTENANCE | 15 | -0.302209 | -0.795314 | 0.730198 | 1 |
| GOBP_VACUOLAR_LOCALIZATION | 67 | -0.217562 | -0.794897 | 0.866142 | 1 |
| GOBP_POSITIVE_REGULATION_OF_HISTONE_METHYLATION | 45 | -0.235396 | -0.794305 | 0.853741 | 1 |
| GOBP_VASCULAR_ENDOTHELIAL_GROWTH_FACTOR_SIGNALING_PATHWAY | 38 | -0.24022 | -0.789677 | 0.815789 | 1 |
| GOBP_OUTER_DYNEIN_ARM_ASSEMBLY | 15 | -0.302939 | -0.78921 | 0.727979 | 1 |
| GOBP_I_KAPPAB_PHOSPHORYLATION | 16 | -0.300851 | -0.788514 | 0.792929 | 1 |
| GOBP_RECOMBINATIONAL_REPAIR | 156 | -0.189482 | -0.788399 | 0.94186 | 1 |
| GOBP_DNA_RECOMBINATION | 278 | -0.177399 | -0.786158 | 1 | 1 |
| GOBP_DICARBOXYLIC_ACID_METABOLIC_PROCESS | 82 | -0.20866 | -0.785959 | 0.931452 | 1 |
| GOBP_RNA_3_END_PROCESSING | 104 | -0.201744 | -0.785891 | 0.952153 | 1 |
| GOBP_NUCLEUS_ORGANIZATION | 125 | -0.193698 | -0.785099 | 0.95098 | 1 |
| GOBP_PROTEIN_O_LINKED_MANNOSYLATION | 17 | -0.29057 | -0.784177 | 0.780051 | 1 |
| GOBP_LABYRINTHINE_LAYER_DEVELOPMENT | 40 | -0.242338 | -0.784044 | 0.798271 | 1 |
| GOBP_S_ADENOSYLMETHIONINE_METABOLIC_PROCESS | 15 | -0.302518 | -0.784018 | 0.743142 | 1 |
| GOBP_POSITIVE_REGULATION_OF_IMMUNOGLOBULIN_PRODUCTION | 36 | -0.240347 | -0.783399 | 0.830619 | 1 |
| GOBP_CEREBELLAR_CORTEX_DEVELOPMENT | 47 | -0.231689 | -0.783398 | 0.864516 | 1 |
| GOBP_PROTEIN_LOCALIZATION_TO_CELL_JUNCTION | 87 | -0.20757 | -0.782341 | 0.914634 | 1 |
| GOBP_NUCLEOSIDE_DIPHOSPHATE_METABOLIC_PROCESS | 116 | -0.196745 | -0.782008 | 0.943925 | 1 |
| GOBP_DETECTION_OF_TEMPERATURE_STIMULUS | 17 | -0.297577 | -0.77937 | 0.76 | 1 |
| GOBP_REGULATION_OF_CILIUM_MOVEMENT | 23 | -0.274153 | -0.777646 | 0.811989 | 1 |
| GOBP_NUCLEAR_TRANSCRIBED_MRNA_CATABOLIC_PROCESS_NONSENSE_MEDIATED_ | 38 | -0.233913 | -0.776552 | 0.829341 | 1 |
| GOBP_REGULATION_OF_ORGANELLE_ASSEMBLY | 211 | -0.177923 | -0.776204 | 0.986667 | 1 |
| GOBP_PEPTIDYL_L_CYSTEINE_S_PALMITOYLATION | 23 | -0.263513 | -0.775327 | 0.802597 | 1 |
| GOBP_REGULATION_OF_ISOTYPE_SWITCHING | 32 | -0.249381 | -0.77486 | 0.82243 | 1 |
| GOBP_CYTOPLASMIC_TRANSLATIONAL_INITIATION | 27 | -0.26078 | -0.774148 | 0.82235 | 1 |
| GOBP_PHOTOTRANSDUCTION | 22 | -0.271896 | -0.773095 | 0.808 | 1 |
| GOBP_NEGATIVE_REGULATION_OF_ACTIN_FILAMENT_DEPOLYMERIZATION | 41 | -0.231456 | -0.772422 | 0.854599 | 1 |
| GOBP_DEVELOPMENTAL_PIGMENTATION | 39 | -0.234128 | -0.770855 | 0.834891 | 1 |
| GOBP_CELLULAR_RESPONSE_TO_NUTRIENT | 32 | -0.245732 | -0.769077 | 0.84273 | 1 |
| GOBP_LAMELLIPODIUM_ORGANIZATION | 81 | -0.203156 | -0.768989 | 0.927419 | 1 |
| GOBP_MULTICELLULAR_ORGANISM_AGING | 33 | -0.23866 | -0.768719 | 0.847026 | 1 |
| GOBP_REGULATION_OF_CHOLESTEROL_METABOLIC_PROCESS | 28 | -0.252393 | -0.768016 | 0.844311 | 1 |
| GOBP_OLIGOSACCHARIDE_BIOSYNTHETIC_PROCESS | 18 | -0.284821 | -0.766705 | 0.794416 | 1 |
| GOBP_NEURAL_TUBE_FORMATION | 102 | -0.195564 | -0.765995 | 0.966667 | 1 |
| GOBP_PROTEIN_LOCALIZATION_TO_CHROMOSOME_TELOMERIC_REGION | 30 | -0.246875 | -0.765432 | 0.835329 | 1 |
| GOBP_DNA_METHYLATION_DEPENDENT_HETEROCHROMATIN_ASSEMBLY | 22 | -0.268676 | -0.765177 | 0.832861 | 1 |
| GOBP_MELANOCYTE_DIFFERENTIATION | 22 | -0.262487 | -0.764316 | 0.828255 | 1 |
| GOBP_PROTEIN_INSERTION_INTO_MEMBRANE | 53 | -0.216219 | -0.76062 | 0.910394 | 1 |
| GOBP_POSITIVE_REGULATION_OF_STEROL_TRANSPORT | 28 | -0.249462 | -0.75681 | 0.816384 | 1 |
| GOBP_NCRNA_CATABOLIC_PROCESS | 43 | -0.223499 | -0.755192 | 0.902913 | 1 |
| GOBP_REGULATION_OF_FEEDING_BEHAVIOR | 15 | -0.282131 | -0.754972 | 0.803192 | 1 |
| GOBP_ALCOHOL_BIOSYNTHETIC_PROCESS | 117 | -0.186758 | -0.754716 | 0.981132 | 1 |
| GOBP_PROSTATE_GLAND_MORPHOGENESIS | 18 | -0.2822 | -0.753203 | 0.826816 | 1 |

| | | | | | |
|---|---|---|---|---|---|
| GOBP_POSITIVE_REGULATION_OF_MYELOID_LEUKOCYTE_MEDIATED_IMMUNITY | 17 | 0.530115 | 1.27025 | 0.188312 | 0.46042 |
| GOBP_NEUROMUSCULAR_PROCESS | 118 | 0.374439 | 1.269834 | 0.105131 | 0.460969 |
| GOBP_INTESTINAL_ABSORPTION | 22 | 0.491179 | 1.269696 | 0.17101 | 0.460786 |
| GOBP_POSITIVE_REGULATION_OF_EMBRYONIC_DEVELOPMENT | 19 | 0.51066 | 1.268793 | 0.168053 | 0.462583 |
| GOBP_REGULATION_OF_MEMBRANE_REPOLARIZATION | 27 | 0.489081 | 1.268559 | 0.173979 | 0.462591 |
| GOBP_LYMPHOCYTE_MIGRATION | 65 | 0.412321 | 1.26805 | 0.12517 | 0.463351 |
| GOBP_REGULATION_OF_PEPTIDASE_ACTIVITY | 317 | 0.343191 | 1.267567 | 0.043094 | 0.464106 |
| GOBP_STARTLE_RESPONSE | 18 | 0.51544 | 1.267143 | 0.180685 | 0.464767 |
| GOBP_KERATINOCYTE_MIGRATION | 15 | 0.546456 | 1.267019 | 0.176752 | 0.464503 |
| GOBP_CALCIUM_MEDIATED_SIGNALING_USING_INTRACELLULAR_CALCIUM_SOURCE | 15 | 0.530746 | 1.265487 | 0.183007 | 0.467987 |
| GOBP_CELL_SUBSTRATE_JUNCTION_ORGANIZATION | 95 | 0.385695 | 1.265354 | 0.113466 | 0.467755 |
| GOBP_NEUTRAL_LIPID_BIOSYNTHETIC_PROCESS | 38 | 0.444163 | 1.264899 | 0.15303 | 0.468406 |
| GOBP_REGULATION_OF_OXIDATIVE_STRESS_INDUCED_CELL_DEATH | 57 | 0.412803 | 1.264745 | 0.130374 | 0.468216 |
| GOBP_HOMEOSTASIS_OF_NUMBER_OF_CELLS | 232 | 0.3515 | 1.264713 | 0.061364 | 0.467699 |
| GOBP_POSITIVE_REGULATION_OF_AXONOGENESIS | 75 | 0.398033 | 1.264573 | 0.119363 | 0.467534 |
| GOBP_NIK_NF_KAPPAB_SIGNALING | 99 | 0.383702 | 1.263547 | 0.105867 | 0.469698 |
| GOBP_POSITIVE_REGULATION_OF_EPIDERMIS_DEVELOPMENT | 26 | 0.47612 | 1.263416 | 0.18865 | 0.469489 |
| GOBP_POSITIVE_REGULATION_OF_MAPK_CASCADE | 368 | 0.340024 | 1.2631 | 0.039474 | 0.469783 |
| GOBP_NEGATIVE_REGULATION_OF_BMP_SIGNALING_PATHWAY | 46 | 0.428829 | 1.262889 | 0.160615 | 0.469735 |
| GOBP_REGULATION_OF_ENDOTHELIAL_CELL_CHEMOTAXIS | 19 | 0.51084 | 1.262417 | 0.173844 | 0.470451 |
| GOBP_NEGATIVE_REGULATION_OF_EXTRINSIC_APOPTOTIC_SIGNALING_PATHWAY | 79 | 0.397347 | 1.262041 | 0.107191 | 0.470901 |
| GOBP_PEPTIDYL_PROLINE_HYDROXYLATION | 15 | 0.5365 | 1.261575 | 0.214286 | 0.471606 |
| GOBP_NEGATIVE_REGULATION_OF_SIGNAL_TRANSDUCTION_IN_ABSENCE_OF_LIGAND | 25 | 0.482207 | 1.261371 | 0.190549 | 0.471521 |
| GOBP_REGULATION_OF_COLLAGEN_METABOLIC_PROCESS | 28 | 0.468074 | 1.261364 | 0.172524 | 0.470963 |
| GOBP_UNSATURATED_FATTY_ACID_METABOLIC_PROCESS | 80 | 0.394821 | 1.260146 | 0.129167 | 0.473641 |
| GOBP_POST_EMBRYONIC_ANIMAL_ORGAN_DEVELOPMENT | 16 | 0.533355 | 1.260095 | 0.196621 | 0.473193 |
| GOBP_VIRAL_LIFE_CYCLE | 268 | 0.34428 | 1.259802 | 0.055619 | 0.473351 |
| GOBP_APPENDAGE_DEVELOPMENT | 134 | 0.368697 | 1.259799 | 0.097356 | 0.472768 |
| GOBP_TRIGLYCERIDE_METABOLIC_PROCESS | 71 | 0.401528 | 1.259611 | 0.142282 | 0.472708 |
| GOBP_MYELIN_MAINTENANCE | 16 | 0.520032 | 1.259243 | 0.194929 | 0.47313 |
| GOBP_MODULATION_BY_HOST_OF_VIRAL_GENOME_REPLICATION | 17 | 0.526507 | 1.259009 | 0.191318 | 0.47322 |
| GOBP_DEVELOPMENT_OF_PRIMARY_FEMALE_SEXUAL_CHARACTERISTICS | 85 | 0.395338 | 1.258819 | 0.110193 | 0.47314 |
| GOBP_NEGATIVE_REGULATION_OF_SMALL_GTPASE_MEDIATED_SIGNAL_TRANSDUCTION | 55 | 0.417051 | 1.258814 | 0.147887 | 0.472581 |
| GOBP_ACUTE_INFLAMMATORY_RESPONSE | 61 | 0.407901 | 1.2587 | 0.128276 | 0.472307 |
| GOBP_MYELOID_CELL_HOMEOSTASIS | 138 | 0.361187 | 1.25867 | 0.085575 | 0.471815 |
| GOBP_DECIDUALIZATION | 19 | 0.510273 | 1.258603 | 0.192126 | 0.471422 |
| GOBP_INOSITOL_PHOSPHATE_BIOSYNTHETIC_PROCESS | 21 | 0.50453 | 1.258488 | 0.184375 | 0.471138 |
| GOBP_MESENCHYMAL_TO_EPITHELIAL_TRANSITION | 17 | 0.530299 | 1.258047 | 0.178752 | 0.471786 |
| GOBP_LIPID_CATABOLIC_PROCESS | 242 | 0.347019 | 1.255349 | 0.070707 | 0.478567 |
| GOBP_ERK1_AND_ERK2_CASCADE | 246 | 0.343638 | 1.255348 | 0.062937 | 0.477988 |
| GOBP_REGULATION_OF_CARDIAC_MUSCLE_CELL_ACTION_POTENTIAL | 25 | 0.481035 | 1.255315 | 0.193291 | 0.47749 |
| GOBP_MESENCHYMAL_CELL_DIFFERENTIATION | 205 | 0.353251 | 1.254611 | 0.069364 | 0.478897 |
| GOBP_RENAL_TUBULE_DEVELOPMENT | 77 | 0.390407 | 1.254517 | 0.129032 | 0.478546 |
| GOBP_POSITIVE_REGULATION_OF_INTERLEUKIN_2_PRODUCTION | 25 | 0.483384 | 1.254363 | 0.168 | 0.478433 |
| GOBP_HISTONE_LYSINE_DEMETHYLATION | 24 | 0.479755 | 1.254199 | 0.172303 | 0.478294 |
| GOBP_SIGNAL_TRANSDUCTION_BY_P53_CLASS_MEDIATOR | 159 | 0.361851 | 1.253901 | 0.082725 | 0.478522 |
| GOBP_DEVELOPMENT_OF_PRIMARY_SEXUAL_CHARACTERISTICS | 180 | 0.354916 | 1.253881 | 0.071179 | 0.477978 |
| GOBP_POSITIVE_REGULATION_OF_ERK1_AND_ERK2_CASCADE | 147 | 0.36459 | 1.253515 | 0.104738 | 0.478406 |
| GOBP_DETECTION_OF_MECHANICAL_STIMULUS | 40 | 0.440246 | 1.253163 | 0.17673 | 0.478833 |
| GOBP_REGULATION_OF_BIOMINERALIZATION | 74 | 0.395651 | 1.252692 | 0.119737 | 0.47962 |
| GOBP_POSITIVE_REGULATION_OF_RESPONSE_TO_ENDOPLASMIC_RETICULUM_STRESS | 32 | 0.459446 | 1.251743 | 0.182622 | 0.481608 |
| GOBP_EXTRINSIC_APOPTOTIC_SIGNALING_PATHWAY_VIA_DEATH_DOMAIN_RECEPTORS | 73 | 0.391097 | 1.251617 | 0.132199 | 0.481378 |
| GOBP_JNK_CASCADE | 155 | 0.358044 | 1.25137 | 0.096471 | 0.481529 |
| GOBP_CELLULAR_RESPONSE_TO_MOLECULE_OF_BACTERIAL_ORIGIN | 134 | 0.364236 | 1.251287 | 0.114532 | 0.481183 |
| GOBP_NEGATIVE_REGULATION_OF_SMOOTH_MUSCLE_CELL_MIGRATION | 23 | 0.492801 | 1.25086 | 0.182663 | 0.481814 |
| GOBP_SPHINGOID_METABOLIC_PROCESS | 21 | 0.496322 | 1.250624 | 0.186901 | 0.481895 |
| GOBP_MEIOTIC_CELL_CYCLE_PROCESS | 156 | 0.356818 | 1.250554 | 0.091995 | 0.481528 |
| GOBP_REGULATION_OF_SYNCYTIUM_FORMATION_BY_PLASMA_MEMBRANE_FUSION | 17 | 0.512174 | 1.249926 | 0.2032 | 0.482739 |
| GOBP_CELLULAR_RESPONSE_TO_CADMIUM_ION | 28 | 0.464525 | 1.249145 | 0.183056 | 0.484388 |
| GOBP_RESPONSE_TO_EXTRACELLULAR_STIMULUS | 384 | 0.328819 | 1.24867 | 0.039615 | 0.485179 |
| GOBP_IMMUNE_EFFECTOR_PROCESS | 386 | 0.336318 | 1.248101 | 0.046908 | 0.486231 |
| GOBP_OSTEOBLAST_PROLIFERATION | 27 | 0.472066 | 1.247955 | 0.183673 | 0.486054 |
| GOBP_FOREBRAIN_REGIONALIZATION | 16 | 0.516896 | 1.247604 | 0.214953 | 0.486466 |
| GOBP_GOLGI_TO_VACUOLE_TRANSPORT | 19 | 0.505233 | 1.247446 | 0.195122 | 0.486309 |
| GOBP_POSITIVE_REGULATION_OF_LIPID_METABOLIC_PROCESS | 117 | 0.371176 | 1.247152 | 0.107006 | 0.486518 |
| GOBP_DIENCEPHALON_DEVELOPMENT | 56 | 0.411027 | 1.24666 | 0.165981 | 0.487255 |
| GOBP_RESPONSE_TO_CHOLESTEROL | 18 | 0.511101 | 1.246655 | 0.210273 | 0.486701 |
| GOBP_REGULATION_OF_MIRNA_TRANSCRIPTION | 54 | 0.413475 | 1.245969 | 0.147564 | 0.488102 |
| GOBP_PROTEIN_HYDROXYLATION | 26 | 0.470248 | 1.245945 | 0.197411 | 0.487618 |
| GOBP_POSITIVE_REGULATION_OF_ALCOHOL_BIOSYNTHETIC_PROCESS | 18 | 0.505431 | 1.245895 | 0.192496 | 0.487191 |
| GOBP_POSITIVE_REGULATION_OF_TRANSMEMBRANE_TRANSPORT | 169 | 0.353783 | 1.24589 | 0.096154 | 0.486626 |
| GOBP_CRANIAL_NERVE_MORPHOGENESIS | 19 | 0.50834 | 1.245277 | 0.203175 | 0.487774 |
| GOBP_POSITIVE_REGULATION_OF_ESTABLISHMENT_OF_PROTEIN_LOCALIZATION | 270 | 0.338342 | 1.244764 | 0.069767 | 0.488581 |
| GOBP_CELL_MATURATION | 136 | 0.365361 | 1.244676 | 0.102117 | 0.488241 |
| GOBP_NEGATIVE_REGULATION_OF_HYDROLASE_ACTIVITY | 238 | 0.342678 | 1.244583 | 0.060364 | 0.487921 |
| GOBP_REGULATION_OF_NITRIC_OXIDE_METABOLIC_PROCESS | 41 | 0.426721 | 1.244169 | 0.182596 | 0.488505 |
| GOBP_POSITIVE_REGULATION_OF_DNA_BINDING_TRANSCRIPTION_FACTOR_ACTIVITY | 214 | 0.345031 | 1.243946 | 0.078795 | 0.488536 |
| GOBP_CELLULAR_RESPONSE_TO_BIOTIC_STIMULUS | 160 | 0.355355 | 1.243648 | 0.090244 | 0.488714 |
| GOBP_CIRCULATORY_SYSTEM_PROCESS | 461 | 0.327362 | 1.242446 | 0.03481 | 0.491554 |
| GOBP_POSITIVE_REGULATION_OF_CHEMOKINE_PRODUCTION | 45 | 0.42672 | 1.242268 | 0.168813 | 0.491482 |
| GOBP_POSITIVE_REGULATION_OF_TRANSFORMING_GROWTH_FACTOR_BETA_PRODUCTI | 17 | 0.511692 | 1.241964 | 0.202552 | 0.491676 |
| GOBP_NEGATIVE_REGULATION_OF_FAT_CELL_DIFFERENTIATION | 45 | 0.430446 | 1.241721 | 0.163447 | 0.491817 |
| GOBP_REGULATION_OF_CELLULAR_RESPONSE_TO_TRANSFORMING_GROWTH_FACTOR_ | 112 | 0.371237 | 1.241207 | 0.127341 | 0.492639 |
| GOBP_THYMUS_DEVELOPMENT | 34 | 0.443994 | 1.241037 | 0.196221 | 0.492499 |
| GOBP_INTESTINAL_EPITHELIAL_CELL_DIFFERENTIATION | 19 | 0.50473 | 1.23998 | 0.200647 | 0.494882 |
| GOBP_NEGATIVE_REGULATION_OF_CELL_ADHESION | 216 | 0.343414 | 1.239546 | 0.072748 | 0.495531 |
| GOBP_EXECUTION_PHASE_OF_APOPTOSIS | 63 | 0.397946 | 1.239207 | 0.168555 | 0.495869 |
| GOBP_REPLACEMENT_OSSIFICATION | 26 | 0.466757 | 1.239185 | 0.187595 | 0.495357 |
| GOBP_POSITIVE_REGULATION_OF_COLLAGEN_METABOLIC_PROCESS | 19 | 0.511335 | 1.239062 | 0.20625 | 0.49513 |
| GOBP_ENGULFMENT_OF_APOPTOTIC_CELL | 18 | 0.504553 | 1.23886 | 0.202899 | 0.495105 |
| GOBP_GLUTAMATE_SECRETION | 24 | 0.473535 | 1.237899 | 0.194234 | 0.497393 |
| GOBP_REGULATION_OF_MUSCLE_ADAPTATION | 66 | 0.404524 | 1.237629 | 0.150611 | 0.497568 |
| GOBP_VIRAL_RNA_GENOME_REPLICATION | 32 | 0.450385 | 1.237032 | 0.189313 | 0.498664 |
| GOBP_RELEASE_OF_CYTOCHROME_C_FROM_MITOCHONDRIA | 51 | 0.415054 | 1.236613 | 0.153409 | 0.499182 |
| GOBP_HISTONE_H3_K9_ACETYLATION | 16 | 0.522185 | 1.23661 | 0.230643 | 0.498622 |
| GOBP_TOLL_LIKE_RECEPTOR_4_SIGNALING_PATHWAY | 28 | 0.462382 | 1.236558 | 0.199081 | 0.498204 |
| GOBP_DEFENSE_RESPONSE_TO_FUNGUS | 16 | 0.529359 | 1.236183 | 0.224165 | 0.498724 |
| GOBP_REGULATION_OF_RESPONSE_TO_ENDOPLASMIC_RETICULUM_STRESS | 75 | 0.391775 | 1.235888 | 0.149333 | 0.498984 |
| GOBP_PEPTIDE_ANTIGEN_ASSEMBLY_WITH_MHC_PROTEIN_COMPLEX | 15 | 0.531435 | 1.235698 | 0.219156 | 0.498922 |
| GOBP_NEGATIVE_REGULATION_OF_JNK_CASCADE | 34 | 0.445733 | 1.23486 | 0.182109 | 0.500698 |
| GOBP_REGULATION_OF_CELLULAR_KETONE_METABOLIC_PROCESS | 97 | 0.37331 | 1.234837 | 0.153747 | 0.50019 |
| GOBP_REGULATION_OF_PLATELET_ACTIVATION | 31 | 0.458876 | 1.233755 | 0.178964 | 0.502655 |
| GOBP_GMP_METABOLIC_PROCESS | 26 | 0.465764 | 1.233616 | 0.215289 | 0.502478 |
| GOBP_RESPONSE_TO_LITHIUM_ION | 17 | 0.507921 | 1.233225 | 0.200315 | 0.503006 |
| GOBP_DEFENSE_RESPONSE_TO_BACTERIUM | 124 | 0.363258 | 1.232721 | 0.129397 | 0.503784 |
| GOBP_CGMP_MEDIATED_SIGNALING | 21 | 0.486299 | 1.231942 | 0.225859 | 0.5055 |
| GOBP_REGULATION_OF_DNA_DAMAGE_RESPONSE_SIGNAL_TRANSDUCTION_BY_P53_CL | 35 | 0.437113 | 1.231686 | 0.201481 | 0.505629 |
| GOBP_WATER_SOLUBLE_VITAMIN_METABOLIC_PROCESS | 56 | 0.402828 | 1.231579 | 0.158485 | 0.505341 |
| GOBP_PRODUCTION_OF_MOLECULAR_MEDIATOR_INVOLVED_IN_INFLAMMATORY_RESPO | 53 | 0.411476 | 1.231525 | 0.168033 | 0.504928 |
| GOBP_POSITIVE_REGULATION_OF_G_PROTEIN_COUPLED_RECEPTOR_SIGNALING_PATHW | 21 | 0.475704 | 1.231092 | 0.19906 | 0.50566 |
| GOBP_REGULATION_OF_P38MAPK_CASCADE | 36 | 0.44196 | 1.231065 | 0.211127 | 0.505157 |
| GOBP_RESPONSE_TO_LEPTIN | 17 | 0.512522 | 1.230436 | 0.216718 | 0.506322 |
| GOBP_REGULATION_OF_OSTEOBLAST_DIFFERENTIATION | 108 | 0.367795 | 1.229694 | 0.135377 | 0.507861 |
| GOBP_EMBRYONIC_APPENDAGE_MORPHOGENESIS | 85 | 0.376444 | 1.228944 | 0.14752 | 0.509456 |
| GOBP_RESPONSE_TO_MITOCHONDRIAL_DEPOLARISATION | 19 | 0.491871 | 1.22889 | 0.220986 | 0.509025 |
| GOBP_NEGATIVE_REGULATION_OF_INNATE_IMMUNE_RESPONSE | 53 | 0.414513 | 1.228756 | 0.172891 | 0.508827 |
| GOBP_ANTIGEN_PROCESSING_AND_PRESENTATION_OF_PEPTIDE_OR_POLYSACCHARIDI | 24 | 0.47482 | 1.228737 | 0.204871 | 0.508305 |
| GOBP_INTEGRATED_STRESS_RESPONSE_SIGNALING | 21 | 0.486755 | 1.228653 | 0.213636 | 0.507957 |
| GOBP_RESPONSE_TO_TUMOR_NECROSIS_FACTOR | 176 | 0.349002 | 1.228527 | 0.117085 | 0.507743 |
| GOBP_HOMOTYPIC_CELL_CELL_ADHESION | 73 | 0.386767 | 1.22836 | 0.144385 | 0.507682 |
| GOBP_POSITIVE_REGULATION_OF_REPRODUCTIVE_PROCESS | 58 | 0.401253 | 1.22836 | 0.16129 | 0.507119 |
| GOBP_MOLTING_CYCLE | 81 | 0.383089 | 1.227813 | 0.150066 | 0.508088 |
| GOBP_ACTIN_FILAMENT_BASED_MOVEMENT | 115 | 0.361552 | 1.227684 | 0.139303 | 0.507911 |
| GOBP_CELLULAR_RESPONSE_TO_GLUCOSE_STARVATION | 48 | 0.412839 | 1.22688 | 0.169631 | 0.509531 |
| GOBP_NEPHRON_EPITHELIUM_DEVELOPMENT | 91 | 0.374471 | 1.226756 | 0.146405 | 0.509328 |
| GOBP_POSITIVE_REGULATION_OF_MICROTUBULE_POLYMERIZATION_OR_DEPOLYMERIZA | 36 | 0.442883 | 1.226623 | 0.197368 | 0.509111 |
| GOBP_REGULATION_OF_TUBE_SIZE | 103 | 0.367115 | 1.226302 | 0.144501 | 0.509445 |
| GOBP_EMBRYONIC_DIGIT_MORPHOGENESIS | 46 | 0.414742 | 1.226212 | 0.185185 | 0.509171 |
| GOBP_POSITIVE_REGULATION_OF_CARBOHYDRATE_METABOLIC_PROCESS | 60 | 0.39996 | 1.225923 | 0.178763 | 0.509453 |
| GOBP_POSITIVE_REGULATION_OF_CELLULAR_RESPONSE_TO_TRANSFORMING_GROWTH | 29 | 0.451245 | 1.225576 | 0.205793 | 0.50984 |
| GOBP_ACTIVATION_OF_IMMUNE_RESPONSE | 209 | 0.341591 | 1.225458 | 0.086364 | 0.509638 |
| GOBP_FOCAL_ADHESION_ASSEMBLY | 80 | 0.381968 | 1.225051 | 0.14654 | 0.510231 |
| GOBP_NEGATIVE_REGULATION_OF_PEPTIDYL_LYSINE_ACETYLATION | 18 | 0.500449 | 1.224936 | 0.225266 | 0.509953 |
| GOBP_ACTIN_MEDIATED_CELL_CONTRACTION | 89 | 0.37605 | 1.224909 | 0.171315 | 0.509492 |
| GOBP_POSITIVE_REGULATION_OF_EPITHELIAL_CELL_DIFFERENTIATION | 42 | 0.425726 | 1.224742 | 0.197917 | 0.509397 |
| GOBP_NEGATIVE_REGULATION_OF_PEPTIDYL_SERINE_PHOSPHORYLATION | 25 | 0.474373 | 1.224611 | 0.210526 | 0.509182 |
| GOBP_POSITIVE_REGULATION_OF_MONONUCLEAR_CELL_MIGRATION | 43 | 0.429677 | 1.22416 | 0.177905 | 0.509938 |
| GOBP_REGULATION_OF_MYELOID_LEUKOCYTE_DIFFERENTIATION | 84 | 0.382556 | 1.224139 | 0.170051 | 0.50944 |
| GOBP_NEGATIVE_REGULATION_OF_FIBROBLAST_GROWTH_FACTOR_RECEPTOR_SIGNAL | 19 | 0.496735 | 1.223988 | 0.23871 | 0.509287 |
| GOBP_NEGATIVE_REGULATION_OF_INTERLEUKIN_1_BETA_PRODUCTION | 16 | 0.517111 | 1.223542 | 0.232172 | 0.509927 |
| GOBP_FOREBRAIN_GENERATION_OF_NEURONS | 39 | 0.425099 | 1.223476 | 0.175887 | 0.509526 |
| GOBP_NEGATIVE_REGULATION_OF_TOR_SIGNALING | 42 | 0.419518 | 1.222815 | 0.183309 | 0.510817 |
| GOBP_LYMPHANGIOGENESIS | 17 | 0.51301 | 1.222715 | 0.221138 | 0.510542 |
| GOBP_REGULATION_OF_T_CELL_MIGRATION | 28 | 0.452065 | 1.222503 | 0.202786 | 0.510565 |
| GOBP_REGULATION_OF_THE_FORCE_OF_HEART_CONTRACTION | 20 | 0.495052 | 1.222316 | 0.214063 | 0.510527 |
| GOBP_HEAD_MORPHOGENESIS | 31 | 0.448628 | 1.222105 | 0.202683 | 0.510574 |
| GOBP_REGULATORY_T_CELL_DIFFERENTIATION | 25 | 0.465499 | 1.221886 | 0.208333 | 0.510608 |
| GOBP_POSITIVE_REGULATION_OF_NEUROTRANSMITTER_SECRETION | 15 | 0.520563 | 1.221816 | 0.214984 | 0.51024 |
| GOBP_ENDOCRINE_HORMONE_SECRETION | 31 | 0.451423 | 1.220741 | 0.190096 | 0.512816 |
| GOBP_ACTIN_CYTOSKELETON_REORGANIZATION | 97 | 0.374625 | 1.220639 | 0.139386 | 0.51254 |
| GOBP_POSITIVE_REGULATION_OF_PATTERN_RECOGNITION_RECEPTOR_SIGNALING_PAT | 33 | 0.433161 | 1.219138 | 0.184132 | 0.516183 |
| GOBP_NEGATIVE_REGULATION_OF_TRANSMEMBRANE_RECEPTOR_PROTEIN_SERINE_TH | 108 | 0.365917 | 1.218434 | 0.140464 | 0.517628 |
| GOBP_ODONTOGENESIS_OF_DENTIN_CONTAINING_TOOTH | 64 | 0.390105 | 1.217263 | 0.172554 | 0.520279 |
| GOBP_METANEPHROS_MORPHOGENESIS | 26 | 0.45602 | 1.217078 | 0.216939 | 0.520262 |
| GOBP_REGULATION_OF_LYMPHOCYTE_MIGRATION | 40 | 0.422844 | 1.216892 | 0.204348 | 0.520231 |
| GOBP_I_KAPPAB_KINASE_NF_KAPPAB_SIGNALING | 240 | 0.331947 | 1.216727 | 0.086758 | 0.520135 |
| GOBP_MULTICELLULAR_ORGANISMAL_SIGNALING | 128 | 0.355565 | 1.216469 | 0.131936 | 0.52028 |

| | | | | | |
|---|---|---|---|---|---|
| GOBP_CARBOHYDRATE_DERIVATIVE_TRANSPORT | 70 | -0.202384 | -0.752654 | 0.936759 | 1 |
| GOBP_REGULATION_OF_TRANSLATIONAL_ELONGATION | 18 | -0.276942 | -0.75245 | 0.780899 | 1 |
| GOBP_CHOLESTEROL_EFFLUX | 38 | -0.230394 | -0.751839 | 0.888545 | 1 |
| GOBP_POSITIVE_REGULATION_OF_DNA_REPLICATION | 37 | -0.235675 | -0.749712 | 0.888554 | 1 |
| GOBP_RESPONSE_TO_EPIDERMAL_GROWTH_FACTOR | 44 | -0.219949 | -0.74437 | 0.907217 | 1 |
| GOBP_REGULATION_OF_PHOSPHOPROTEIN_PHOSPHATASE_ACTIVITY | 48 | -0.216456 | -0.743927 | 0.909385 | 1 |
| GOBP_GANGLIOSIDE_BIOSYNTHETIC_PROCESS | 15 | -0.283776 | -0.743272 | 0.784 | 1 |
| GOBP_REGULATION_OF_RECEPTOR_LOCALIZATION_TO_SYNAPSE | 16 | -0.285166 | -0.743006 | 0.798005 | 1 |
| GOBP_SISTER_CHROMATID_SEGREGATION | 194 | -0.174224 | -0.742691 | 0.992424 | 1 |
| GOBP_HEPARAN_SULFATE_PROTEOGLYCAN_METABOLIC_PROCESS | 30 | -0.23985 | -0.742507 | 0.872521 | 1 |
| GOBP_NUCLEOTIDE_PHOSPHORYLATION | 86 | -0.195475 | -0.739315 | 0.965368 | 1 |
| GOBP_SYNAPSE_MATURATION | 23 | -0.252141 | -0.737741 | 0.877384 | 1 |
| GOBP_RIBOSOMAL_SMALL_SUBUNIT_ASSEMBLY | 19 | -0.269889 | -0.737281 | 0.834734 | 1 |
| GOBP_LIPID_IMPORT_INTO_CELL | 16 | -0.27178 | -0.736337 | 0.818182 | 1 |
| GOBP_BENZENE_CONTAINING_COMPOUND_METABOLIC_PROCESS | 15 | -0.275838 | -0.735611 | 0.825243 | 1 |
| GOBP_NADP_METABOLIC_PROCESS | 34 | -0.230009 | -0.734728 | 0.886154 | 1 |
| GOBP_PIGMENT_METABOLIC_PROCESS | 67 | -0.197038 | -0.734564 | 0.929134 | 1 |
| GOBP_PROTEIN_ADP_RIBOSYLATION | 33 | -0.232068 | -0.733515 | 0.893372 | 1 |
| GOBP_REGULATION_OF_PROTEIN_CONTAINING_COMPLEX_DISASSEMBLY | 117 | -0.182028 | -0.733248 | 0.994737 | 1 |
| GOBP_FOREBRAIN_CELL_MIGRATION | 54 | -0.207608 | -0.733231 | 0.954545 | 1 |
| GOBP_DETECTION_OF_STIMULUS_INVOLVED_IN_SENSORY_PERCEPTION_OF_PAIN | 22 | -0.255292 | -0.73306 | 0.846797 | 1 |
| GOBP_REGULATION_OF_STEROL_TRANSPORT | 44 | -0.215009 | -0.732204 | 0.913333 | 1 |
| GOBP_MITOTIC_SISTER_CHROMATID_COHESION | 26 | -0.243851 | -0.731145 | 0.858859 | 1 |
| GOBP_TELOMERE_MAINTENANCE_VIA_TELOMERE_LENGTHENING | 76 | -0.193078 | -0.731058 | 0.946721 | 1 |
| GOBP_SOMATIC_STEM_CELL_POPULATION_MAINTENANCE | 35 | -0.226386 | -0.730201 | 0.908228 | 1 |
| GOBP_POSITIVE_REGULATION_OF_INTERFERON_BETA_PRODUCTION | 34 | -0.23093 | -0.730116 | 0.906149 | 1 |
| GOBP_REGULATION_OF_CELL_CELL_ADHESION_MEDIATED_BY_CADHERIN | 17 | -0.270734 | -0.729885 | 0.830601 | 1 |
| GOBP_TETRAHYDROFOLATE_METABOLIC_PROCESS | 20 | -0.252175 | -0.726679 | 0.861789 | 1 |
| GOBP_REGULATION_OF_DEPHOSPHORYLATION | 110 | -0.181007 | -0.726489 | 1 | 1 |
| GOBP_TETRAPYRROLE_METABOLIC_PROCESS | 51 | -0.20353 | -0.724636 | 0.925424 | 1 |
| GOBP_CYTOSKELETON_DEPENDENT_INTRACELLULAR_TRANSPORT | 184 | -0.171371 | -0.722872 | 0.993103 | 1 |
| GOBP_REGULATION_OF_VACUOLE_ORGANIZATION | 49 | -0.210264 | -0.719362 | 0.93662 | 1 |
| GOBP_REGULATION_OF_GLIAL_CELL_PROLIFERATION | 20 | -0.256114 | -0.718935 | 0.870466 | 1 |
| GOBP_POSITIVE_REGULATION_OF_CELL_CYCLE_G2_M_PHASE_TRANSITION | 29 | -0.228718 | -0.716384 | 0.882883 | 1 |
| GOBP_VESICLE_CARGO_LOADING | 21 | -0.250298 | -0.715745 | 0.860606 | 1 |
| GOBP_SYNAPTIC_VESICLE_TRANSPORT | 39 | -0.217134 | -0.715223 | 0.920578 | 1 |
| GOBP_B_CELL_ACTIVATION_INVOLVED_IN_IMMUNE_RESPONSE | 64 | -0.195455 | -0.709334 | 0.973282 | 1 |
| GOBP_TRANSLATIONAL_ELONGATION | 52 | -0.199876 | -0.708556 | 0.940141 | 1 |
| GOBP_TELOMERE_CAPPING | 39 | -0.210935 | -0.708042 | 0.933121 | 1 |
| GOBP_PROTEIN_CONTAINING_COMPLEX_DISASSEMBLY | 232 | -0.164216 | -0.707897 | 1 | 1 |
| GOBP_INTRA_GOLGI_VESICLE_MEDIATED_TRANSPORT | 29 | -0.225839 | -0.707782 | 0.91018 | 1 |
| GOBP_PEPTIDYL_ASPARAGINE_MODIFICATION | 23 | -0.244293 | -0.707534 | 0.888298 | 1 |
| GOBP_VACUOLE_ORGANIZATION | 176 | -0.165002 | -0.707479 | 0.994048 | 1 |
| GOBP_REGULATION_OF_SPINDLE_ASSEMBLY | 30 | -0.232151 | -0.706794 | 0.884726 | 1 |
| GOBP_NEGATIVE_REGULATION_OF_TELOMERE_MAINTENANCE_VIA_TELOMERASE | 19 | -0.254671 | -0.706609 | 0.894737 | 1 |
| GOBP_CELLULAR_METABOLIC_COMPOUND_SALVAGE | 26 | -0.236239 | -0.706439 | 0.889488 | 1 |
| GOBP_MITOTIC_INTRA_S_DNA_DAMAGE_CHECKPOINT_SIGNALING | 15 | -0.273513 | -0.704028 | 0.85641 | 1 |
| GOBP_NEGATIVE_REGULATION_OF_LYMPHOCYTE_APOPTOTIC_PROCESS | 26 | -0.235042 | -0.703439 | 0.915205 | 1 |
| GOBP_HISTONE_H3_DEACETYLATION | 17 | -0.256265 | -0.702117 | 0.886487 | 1 |
| GOBP_SERTOLI_CELL_DIFFERENTIATION | 15 | -0.269583 | -0.701949 | 0.865707 | 1 |
| GOBP_PROTEIN_LOCALIZATION_TO_CYTOSKELETON | 56 | -0.196731 | -0.699454 | 0.962712 | 1 |
| GOBP_MITOCHONDRIAL_TRANSPORT | 189 | -0.160815 | -0.698357 | 1 | 1 |
| GOBP_TELOMERE_MAINTENANCE | 126 | -0.176489 | -0.696737 | 1 | 1 |
| GOBP_ORGANISM_EMERGENCE_FROM_PROTECTIVE_STRUCTURE | 24 | -0.23339 | -0.693979 | 0.918605 | 1 |
| GOBP_REGULATION_OF_CHROMATIN_ORGANIZATION | 42 | -0.21024 | -0.69355 | 0.953416 | 1 |
| GOBP_NEGATIVE_REGULATION_OF_PROTEASOMAL_UBIQUITIN_DEPENDENT_PROTEIN_C | 32 | -0.22036 | -0.692705 | 0.927711 | 1 |
| GOBP_GOLGI_TO_PLASMA_MEMBRANE_PROTEIN_TRANSPORT | 40 | -0.203976 | -0.692611 | 0.938312 | 1 |
| GOBP_REGULATION_OF_DNA_METHYLATION_DEPENDENT_HETEROCHROMATIN_ASSEMI | 16 | -0.265324 | -0.692183 | 0.862797 | 1 |
| GOBP_PIGMENT_BIOSYNTHETIC_PROCESS | 57 | -0.195621 | -0.691615 | 0.954386 | 1 |
| GOBP_REGULATION_OF_CELL_MORPHOGENESIS_INVOLVED_IN_DIFFERENTIATION | 88 | -0.17977 | -0.688356 | 0.991597 | 1 |
| GOBP_CELLULAR_RESPONSE_TO_OSMOTIC_STRESS | 37 | -0.208593 | -0.686465 | 0.950617 | 1 |
| GOBP_NEGATIVE_REGULATION_OF_PROTEIN_LOCALIZATION_TO_CELL_PERIPHERY | 24 | -0.232002 | -0.684852 | 0.914729 | 1 |
| GOBP_NEGATIVE_REGULATION_OF_LIPID_TRANSPORT | 18 | -0.248651 | -0.683942 | 0.897019 | 1 |
| GOBP_POSITIVE_REGULATION_OF_CELL_MORPHOGENESIS_INVOLVED_IN_DIFFERENTIAT | 72 | -0.187633 | -0.682825 | 0.992509 | 1 |
| GOBP_PYRUVATE_METABOLIC_PROCESS | 87 | -0.176796 | -0.682194 | 0.987603 | 1 |
| GOBP_RESPONSE_TO_FOOD | 22 | -0.239734 | -0.681962 | 0.909621 | 1 |
| GOBP_PEROXISOME_ORGANIZATION | 33 | -0.215891 | -0.681062 | 0.943953 | 1 |
| GOBP_INNERVATION | 25 | -0.234404 | -0.678286 | 0.896359 | 1 |
| GOBP_REGULATION_OF_DNA_RECOMBINATION | 115 | -0.168806 | -0.676368 | 0.995516 | 1 |
| GOBP_REGULATION_OF_CILIUM_ASSEMBLY | 82 | -0.180198 | -0.674762 | 0.992537 | 1 |
| GOBP_NEGATIVE_REGULATION_OF_ORGANELLE_ASSEMBLY | 41 | -0.202526 | -0.672139 | 0.972028 | 1 |
| GOBP_MITOCHONDRIAL_DNA_REPLICATION | 15 | -0.25778 | -0.671801 | 0.894089 | 1 |
| GOBP_VESICLE_MEDIATED_TRANSPORT_BETWEEN_ENDOSOMAL_COMPARTMENTS | 45 | -0.192982 | -0.670978 | 0.969492 | 1 |
| GOBP_PIGMENT_GRANULE_LOCALIZATION | 24 | -0.226686 | -0.669787 | 0.93586 | 1 |
| GOBP_REGULATION_OF_NUCLEASE_ACTIVITY | 17 | -0.249258 | -0.66959 | 0.906566 | 1 |
| GOBP_ORGANIC_CATION_TRANSPORT | 30 | -0.212081 | -0.666583 | 0.944444 | 1 |
| GOBP_PROTEIN_MONOUBIQUITINATION | 78 | -0.18107 | -0.665538 | 0.982609 | 1 |
| GOBP_HISTONE_DEACETYLATION | 91 | -0.174029 | -0.665461 | 0.982979 | 1 |
| GOBP_PROTEIN_AUTOPROCESSING | 19 | -0.242108 | -0.664975 | 0.890585 | 1 |
| GOBP_NEGATIVE_REGULATION_OF_ACTIN_FILAMENT_BUNDLE_ASSEMBLY | 29 | -0.215326 | -0.664631 | 0.951515 | 1 |
| GOBP_REGULATION_OF_MEMBRANE_PROTEIN_ECTODOMAIN_PROTEOLYSIS | 20 | -0.238228 | -0.664615 | 0.910082 | 1 |
| GOBP_POSITIVE_REGULATION_OF_PROTEIN_DEPHOSPHORYLATION | 40 | -0.200521 | -0.664219 | 0.980392 | 1 |
| GOBP_RESPONSE_TO_DSRNA | 38 | -0.200227 | -0.663616 | 0.964052 | 1 |
| GOBP_POSITIVE_REGULATION_OF_TELOMERE_MAINTENANCE_VIA_TELOMERE_LENGTHE | 37 | -0.203127 | -0.662386 | 0.970968 | 1 |
| GOBP_EMBRYONIC_BRAIN_DEVELOPMENT | 15 | -0.255646 | -0.66166 | 0.895782 | 1 |
| GOBP_PROTEIN_LOCALIZATION_TO_ENDOPLASMIC_RETICULUM | 66 | -0.184451 | -0.660784 | 0.975 | 1 |
| GOBP_GOLGI_TO_PLASMA_MEMBRANE_TRANSPORT | 59 | -0.183184 | -0.660019 | 0.984733 | 1 |
| GOBP_PH_REDUCTION | 40 | -0.201226 | -0.658106 | 0.97205 | 1 |
| GOBP_GLUTAMINE_FAMILY_AMINO_ACID_BIOSYNTHETIC_PROCESS | 15 | -0.254595 | -0.657501 | 0.883544 | 1 |
| GOBP_ENDOPLASMIC_RETICULUM_TO_CYTOSOL_TRANSPORT | 28 | -0.215537 | -0.654903 | 0.953297 | 1 |
| GOBP_HISTONE_H4_ACETYLATION | 72 | -0.17719 | -0.653329 | 0.988764 | 1 |
| GOBP_NEGATIVE_REGULATION_OF_CILIUM_ASSEMBLY | 17 | -0.242857 | -0.652689 | 0.895442 | 1 |
| GOBP_MORPHOGENESIS_OF_A_POLARIZED_EPITHELIUM | 89 | -0.169436 | -0.652023 | 0.991936 | 1 |
| GOBP_PROTEIN_TARGETING_TO_MITOCHONDRION | 98 | -0.16729 | -0.650461 | 0.991968 | 1 |
| GOBP_POSITIVE_REGULATION_OF_SIGNALING_RECEPTOR_ACTIVITY | 35 | -0.20535 | -0.650245 | 0.960961 | 1 |
| GOBP_REGULATION_OF_PROTEIN_LOCALIZATION_TO_SYNAPSE | 18 | -0.237047 | -0.649987 | 0.914439 | 1 |
| GOBP_RESPONSE_TO_IRON_ION | 25 | -0.215812 | -0.649693 | 0.929155 | 1 |
| GOBP_GANGLIOSIDE_METABOLIC_PROCESS | 24 | -0.222425 | -0.649114 | 0.944134 | 1 |
| GOBP_REGULATION_OF_TRANSCRIPTION_ELONGATION_FROM_RNA_POLYMERASE_II_PF | 36 | -0.200621 | -0.64814 | 0.965409 | 1 |
| GOBP_REGULATION_OF_GOLGI_ORGANIZATION | 18 | -0.236484 | -0.647938 | 0.945545 | 1 |
| GOBP_ACTIN_NUCLEATION | 50 | -0.184668 | -0.647417 | 0.992647 | 1 |
| GOBP_CELLULAR_RESPONSE_TO_VIRUS | 62 | -0.178441 | -0.64704 | 1 | 1 |
| GOBP_MANNOSYLATION | 35 | -0.201821 | -0.646739 | 0.954268 | 1 |
| GOBP_MITOCHONDRIAL_CALCIUM_ION_TRANSMEMBRANE_TRANSPORT | 18 | -0.234692 | -0.645313 | 0.916667 | 1 |
| GOBP_MITOCHONDRIAL_ATP_SYNTHESIS_COUPLED_PROTON_TRANSPORT | 18 | -0.241339 | -0.64463 | 0.917706 | 1 |
| GOBP_RIG_I_SIGNALING_PATHWAY | 25 | -0.217336 | -0.64449 | 0.956897 | 1 |
| GOBP_UBIQUITIN_DEPENDENT_ERAD_PATHWAY | 84 | -0.167739 | -0.643445 | 0.995536 | 1 |
| GOBP_ENDOSOME_TO_LYSOSOME_TRANSPORT | 58 | -0.180982 | -0.643432 | 0.992806 | 1 |
| GOBP_VESICLE_TRANSPORT_ALONG_MICROTUBULE | 44 | -0.187922 | -0.640877 | 0.975779 | 1 |
| GOBP_NADH_DEHYDROGENASE_COMPLEX_ASSEMBLY | 58 | -0.178143 | -0.637146 | 0.996552 | 1 |
| GOBP_REGULATION_OF_LIPOPOLYSACCHARIDE_MEDIATED_SIGNALING_PATHWAY | 15 | -0.243084 | -0.636577 | 0.915167 | 1 |
| GOBP_TRANSPORT_ALONG_MICROTUBULE | 151 | -0.153237 | -0.635504 | 1 | 1 |
| GOBP_REGULATION_OF_EXIT_FROM_MITOSIS | 15 | -0.246572 | -0.634348 | 0.935135 | 1 |
| GOBP_ENDOPLASMIC_RETICULUM_TO_GOLGI_VESICLE_MEDIATED_TRANSPORT | 122 | -0.155886 | -0.633271 | 0.995074 | 1 |
| GOBP_REGULATION_OF_PROTEIN_KINASE_C_SIGNALING | 17 | -0.234615 | -0.629678 | 0.938719 | 1 |
| GOBP_RNA_PHOSPHODIESTER_BOND_HYDROLYSIS | 134 | -0.152901 | -0.62846 | 1 | 1 |
| GOBP_REGULATION_OF_INTRACELLULAR_ESTROGEN_RECEPTOR_SIGNALING_PATHWAY | 30 | -0.202806 | -0.627855 | 0.968571 | 1 |
| GOBP_MRNA_TRANSPORT | 115 | -0.155279 | -0.626253 | 1 | 1 |
| GOBP_ACTIN_FILAMENT_DEPOLYMERIZATION | 54 | -0.181883 | -0.624699 | 0.988327 | 1 |
| GOBP_VIRAL_TRANSLATION | 18 | -0.232575 | -0.622796 | 0.934211 | 1 |
| GOBP_MRNA_CLEAVAGE | 18 | -0.229031 | -0.622144 | 0.922886 | 1 |
| GOBP_REGULATION_OF_CLATHRIN_DEPENDENT_ENDOCYTOSIS | 18 | -0.226251 | -0.622034 | 0.945455 | 1 |
| GOBP_GLYCOSPHINGOLIPID_METABOLIC_PROCESS | 53 | -0.175631 | -0.620033 | 0.993443 | 1 |
| GOBP_RETROGRADE_VESICLE_MEDIATED_TRANSPORT_GOLGI_TO_ENDOPLASMIC_RETI | 48 | -0.178847 | -0.619767 | 0.986254 | 1 |
| GOBP_REGULATION_OF_MRNA_METABOLIC_PROCESS | 271 | -0.137859 | -0.61896 | 1 | 1 |
| GOBP_SULFUR_COMPOUND_CATABOLIC_PROCESS | 26 | -0.203405 | -0.618364 | 0.966759 | 1 |
| GOBP_NOREPINEPHRINE_TRANSPORT | 15 | -0.236547 | -0.61629 | 0.936288 | 1 |
| GOBP_NEGATIVE_REGULATION_OF_TELOMERE_MAINTENANCE | 32 | -0.191406 | -0.611719 | 0.981482 | 1 |
| GOBP_ESTABLISHMENT_OF_EPITHELIAL_CELL_APICAL_BASAL_POLARITY | 18 | -0.223279 | -0.610822 | 0.960733 | 1 |
| GOBP_MITOCHONDRIAL_DEPOLARIZATION | 18 | -0.223824 | -0.609466 | 0.937158 | 1 |
| GOBP_NEGATIVE_REGULATION_OF_LIPID_LOCALIZATION | 29 | -0.201843 | -0.607431 | 0.965333 | 1 |
| GOBP_HETEROCHROMATIN_ORGANIZATION | 75 | -0.165622 | -0.607187 | 0.995763 | 1 |
| GOBP_EXIT_FROM_MITOSIS | 27 | -0.201271 | -0.604283 | 0.968668 | 1 |
| GOBP_GLIAL_CELL_APOPTOTIC_PROCESS | 15 | -0.233685 | -0.604133 | 0.94686 | 1 |
| GOBP_TRANSLATIONAL_INITIATION | 111 | -0.153002 | -0.602619 | 1 | 1 |
| GOBP_REGULATION_OF_ENDOCYTIC_RECYCLING | 17 | -0.222304 | -0.602218 | 0.949333 | 1 |
| GOBP_ESTABLISHMENT_OF_PIGMENT_GRANULE_LOCALIZATION | 22 | -0.206742 | -0.597666 | 0.967391 | 1 |
| GOBP_GLUTAMATE_METABOLIC_PROCESS | 29 | -0.199448 | -0.596395 | 0.959375 | 1 |
| GOBP_PROTEIN_K63_LINKED_DEUBIQUITINATION | 33 | -0.185416 | -0.595145 | 0.982405 | 1 |
| GOBP_ATP_SYNTHESIS_COUPLED_ELECTRON_TRANSPORT | 84 | -0.152347 | -0.592752 | 1 | 1 |
| GOBP_INTRACELLULAR_ESTROGEN_RECEPTOR_SIGNALING_PATHWAY | 44 | -0.177074 | -0.59051 | 0.986928 | 1 |
| GOBP_REGULATION_OF_PROTEIN_STABILITY | 273 | -0.129804 | -0.582721 | 1 | 1 |
| GOBP_POSITIVE_REGULATION_OF_NITRIC_OXIDE_SYNTHASE_ACTIVITY | 16 | -0.223519 | -0.578313 | 0.949735 | 1 |
| GOBP_REGULATION_OF_TELOMERE_MAINTENANCE_VIA_TELOMERE_LENGTHENING | 58 | -0.160062 | -0.578304 | 0.993056 | 1 |
| GOBP_RRNA_MODIFICATION | 36 | -0.174038 | -0.576964 | 0.993691 | 1 |
| GOBP_VIRAL_RELEASE_FROM_HOST_CELL | 28 | -0.189021 | -0.576883 | 0.976261 | 1 |
| GOBP_HISTONE_H4_K16_ACETYLATION | 27 | -0.196146 | -0.576171 | 0.991304 | 1 |
| GOBP_CENTROMERE_COMPLEX_ASSEMBLY | 30 | -0.18806 | -0.575619 | 0.991525 | 1 |
| GOBP_PROTEIN_LOCALIZATION_TO_PHAGOPHORE_ASSEMBLY_SITE | 17 | -0.216187 | -0.574393 | 0.967005 | 1 |
| GOBP_ACTIN_FILAMENT_SEVERING | 16 | -0.217373 | -0.5728 | 0.94958 | 1 |
| GOBP_REGULATION_OF_UBIQUITIN_PROTEIN_LIGASE_ACTIVITY | 23 | -0.201224 | -0.571354 | 0.988304 | 1 |
| GOBP_POLARIZED_EPITHELIAL_CELL_DIFFERENTIATION | 24 | -0.191589 | -0.569892 | 0.97281 | 1 |
| GOBP_ELECTRON_TRANSPORT_CHAIN | 148 | -0.138664 | -0.569756 | 1 | 1 |
| GOBP_PROTEIN_EXIT_FROM_ENDOPLASMIC_RETICULUM | 48 | -0.163254 | -0.567723 | 0.996795 | 1 |
| GOBP_GLUCOSE_CATABOLIC_PROCESS | 17 | -0.213084 | -0.565208 | 0.974619 | 1 |
| GOBP_CYTOCHROME_COMPLEX_ASSEMBLY | 35 | -0.176696 | -0.56267 | 0.984663 | 1 |
| GOBP_AXONAL_TRANSPORT_OF_MITOCHONDRION | 17 | -0.209336 | -0.561062 | 0.96477 | 1 |

| | | | | | |
|---|---|---|---|---|---|
| GOBP_REGULATION_OF_LEUKOCYTE_DEGRANULATION | 37 | 0.433513 | 1.21643 | 0.205479 | 0.519843 |
| GOBP_CELL_DIFFERENTIATION_INVOLVED_IN_METANEPHROS_DEVELOPMENT | 17 | 0.508239 | 1.215945 | 0.244168 | 0.520685 |
| GOBP_INOSITOL_PHOSPHATE_METABOLIC_PROCESS | 47 | 0.409837 | 1.215832 | 0.184211 | 0.520476 |
| GOBP_CELLULAR_RESPONSE_TO_HYDROGEN_PEROXIDE | 70 | 0.385121 | 1.214822 | 0.173677 | 0.522841 |
| GOBP_REGULATION_OF_CARDIAC_MUSCLE_CELL_PROLIFERATION | 28 | 0.462602 | 1.214488 | 0.223101 | 0.523185 |
| GOBP_B_CELL_PROLIFERATION | 51 | 0.413231 | 1.214212 | 0.193966 | 0.523413 |
| GOBP_TOLL_LIKE_RECEPTOR_3_SIGNALING_PATHWAY | 15 | 0.523334 | 1.213659 | 0.239243 | 0.52443 |
| GOBP_CELL_ACTIVATION_INVOLVED_IN_IMMUNE_RESPONSE | 193 | 0.338039 | 1.213613 | 0.10984 | 0.524004 |
| GOBP_RHO_PROTEIN_SIGNAL_TRANSDUCTION | 119 | 0.359078 | 1.213215 | 0.140881 | 0.524541 |
| GOBP_T_CELL_MEDIATED_IMMUNITY | 74 | 0.38238 | 1.212968 | 0.175657 | 0.524762 |
| GOBP_RESPONSE_TO_PEPTIDE_HORMONE | 320 | 0.326834 | 1.212293 | 0.078947 | 0.526053 |
| GOBP_POSITIVE_REGULATION_OF_MORPHOGENESIS_OF_AN_EPITHELIUM | 31 | 0.437416 | 1.211948 | 0.211009 | 0.526488 |
| GOBP_INTERFERON_ALPHA_PRODUCTION | 20 | 0.493194 | 1.211759 | 0.244733 | 0.526482 |
| GOBP_ADAPTIVE_IMMUNE_RESPONSE | 271 | 0.332253 | 1.211497 | 0.091319 | 0.526618 |
| GOBP_ANTIGEN_RECEPTOR_MEDIATED_SIGNALING_PATHWAY | 127 | 0.353163 | 1.211246 | 0.145161 | 0.526766 |
| GOBP_REGULATION_OF_BONE_MINERALIZATION | 60 | 0.397857 | 1.211136 | 0.183424 | 0.526531 |
| GOBP_MACROPHAGE_CYTOKINE_PRODUCTION | 15 | 0.520399 | 1.210876 | 0.23126 | 0.526674 |
| GOBP_REGULATION_OF_BROWN_FAT_CELL_DIFFERENTIATION | 19 | 0.495894 | 1.210778 | 0.25 | 0.526389 |
| GOBP_POSITIVE_REGULATION_OF_CALCIUM_ION_TRANSPORT_INTO_CYTOSOL | 40 | 0.420144 | 1.210661 | 0.203812 | 0.526166 |
| GOBP_REGULATION_OF_ADAPTIVE_IMMUNE_RESPONSE | 129 | 0.350931 | 1.210641 | 0.128485 | 0.525669 |
| GOBP_EMBRYONIC_PLACENTA_DEVELOPMENT | 77 | 0.37832 | 1.210197 | 0.174667 | 0.526378 |
| GOBP_LEUKOCYTE_HOMEOSTASIS | 68 | 0.389047 | 1.210145 | 0.17374 | 0.525964 |
| GOBP_REGULATION_OF_CALCIUM_ION_TRANSMEMBRANE_TRANSPORT | 117 | 0.359515 | 1.210034 | 0.142683 | 0.525716 |
| GOBP_NEGATIVE_REGULATION_OF_DEFENSE_RESPONSE | 172 | 0.345031 | 1.20975 | 0.116307 | 0.525947 |
| GOBP_ADIPOSE_TISSUE_DEVELOPMENT | 40 | 0.410453 | 1.209698 | 0.207715 | 0.52555 |
| GOBP_RESPONSE_TO_NUTRIENT | 110 | 0.361452 | 1.209519 | 0.139842 | 0.525504 |
| GOBP_REGULATION_OF_VASCULAR_ENDOTHELIAL_GROWTH_FACTOR_RECEPTOR_SIGN | 21 | 0.478192 | 1.208889 | 0.248834 | 0.526714 |
| GOBP_REGULATION_OF_METAL_ION_TRANSPORT | 307 | 0.324139 | 1.208716 | 0.089617 | 0.526677 |
| GOBP_AMINE_BIOSYNTHETIC_PROCESS | 31 | 0.451211 | 1.20862 | 0.225291 | 0.526423 |
| GOBP_ANIMAL_ORGAN_REGENERATION | 47 | 0.409843 | 1.207594 | 0.197991 | 0.528802 |
| GOBP_POSITIVE_REGULATION_OF_FIBROBLAST_PROLIFERATION | 37 | 0.426697 | 1.206844 | 0.221068 | 0.530465 |
| GOBP_ICOSANOID_BIOSYNTHETIC_PROCESS | 33 | 0.434233 | 1.206626 | 0.227003 | 0.530552 |
| GOBP_CELL_COMMUNICATION_BY_ELECTRICAL_COUPLING_INVOLVED_IN_CARDIAC_CO | 26 | 0.469046 | 1.206016 | 0.214058 | 0.531684 |
| GOBP_MACROPHAGE_CHEMOTAXIS | 22 | 0.461857 | 1.205785 | 0.242857 | 0.53179 |
| GOBP_CHONDROCYTE_DIFFERENTIATION | 91 | 0.368805 | 1.205702 | 0.181818 | 0.531481 |
| GOBP_REGULATION_OF_IMMUNE_EFFECTOR_PROCESS | 231 | 0.331424 | 1.205599 | 0.112628 | 0.531204 |
| GOBP_NEGATIVE_REGULATION_OF_LEUKOCYTE_APOPTOTIC_PROCESS | 38 | 0.423347 | 1.20532 | 0.197674 | 0.53152 |
| GOBP_RESPONSE_TO_CADMIUM_ION | 46 | 0.415991 | 1.204639 | 0.204444 | 0.532932 |
| GOBP_NEGATIVE_REGULATION_OF_WOUND_HEALING | 52 | 0.400532 | 1.204337 | 0.191919 | 0.533265 |
| GOBP_CAMERA_TYPE_EYE_DEVELOPMENT | 258 | 0.328973 | 1.203844 | 0.093288 | 0.534081 |
| GOBP_RESPONSE_TO_PEPTIDE | 388 | 0.318404 | 1.203676 | 0.075372 | 0.533961 |
| GOBP_IMMUNE_RESPONSE_REGULATING_CELL_SURFACE_RECEPTOR_SIGNALING_PATH | 170 | 0.340331 | 1.202981 | 0.127078 | 0.535414 |
| GOBP_CELLULAR_RESPONSE_TO_OXYGEN_LEVELS | 140 | 0.347869 | 1.20298 | 0.151551 | 0.534876 |
| GOBP_DEFENSE_RESPONSE_TO_SYMBIONT | 212 | 0.335028 | 1.202637 | 0.122286 | 0.535286 |
| GOBP_REGULATION_OF_VIRAL_PROCESS | 132 | 0.351543 | 1.202409 | 0.140722 | 0.535359 |
| GOBP_HETEROTYPIC_CELL_CELL_ADHESION | 42 | 0.424587 | 1.202231 | 0.197368 | 0.535283 |
| GOBP_B_CELL_RECEPTOR_SIGNALING_PATHWAY | 39 | 0.418737 | 1.201935 | 0.224751 | 0.535544 |
| GOBP_REGULATION_OF_EXTRINSIC_APOPTOTIC_SIGNALING_PATHWAY_VIA_DEATH_DOM | 41 | 0.419427 | 1.201847 | 0.21944 | 0.535239 |
| GOBP_REGULATION_OF_VESICLE_FUSION | 20 | 0.492698 | 1.201803 | 0.24649 | 0.534814 |
| GOBP_MIDDLE_EAR_MORPHOGENESIS | 15 | 0.512027 | 1.201367 | 0.231156 | 0.535436 |
| GOBP_POSITIVE_REGULATION_OF_MYELOID_CELL_DIFFERENTIATION | 77 | 0.374352 | 1.201088 | 0.179866 | 0.535649 |
| GOBP_STRESS_ACTIVATED_PROTEIN_KINASE_SIGNALING_CASCADE | 219 | 0.329389 | 1.200676 | 0.1097 | 0.536326 |
| GOBP_EPITHELIAL_CELL_CELL_ADHESION | 15 | 0.507559 | 1.200504 | 0.256778 | 0.536268 |
| GOBP_METAL_ION_HOMEOSTASIS | 478 | 0.317687 | 1.200216 | 0.060988 | 0.53654 |
| GOBP_RESPONSE_TO_GONADOTROPIN | 20 | 0.474938 | 1.199885 | 0.248387 | 0.536916 |
| GOBP_RESPONSE_TO_MINERALOCORTICOID | 25 | 0.458228 | 1.199847 | 0.24424 | 0.536486 |
| GOBP_NEGATIVE_REGULATION_OF_MUSCLE_HYPERTROPHY | 24 | 0.462818 | 1.199746 | 0.241327 | 0.53621 |
| GOBP_POSITIVE_REGULATION_OF_CELLULAR_CARBOHYDRATE_METABOLIC_PROCESS | 42 | 0.408331 | 1.199741 | 0.208092 | 0.535682 |
| GOBP_REGULATION_OF_POSITIVE_CHEMOTAXIS | 20 | 0.472256 | 1.199421 | 0.237013 | 0.536047 |
| GOBP_REGULATION_OF_HEART_RATE_BY_CARDIAC_CONDUCTION | 35 | 0.427834 | 1.199168 | 0.216296 | 0.536186 |
| GOBP_NEGATIVE_REGULATION_OF_CHONDROCYTE_DIFFERENTIATION | 17 | 0.493399 | 1.198783 | 0.244592 | 0.536763 |
| GOBP_MALE_GAMETE_GENERATION | 393 | 0.318502 | 1.198379 | 0.091675 | 0.53736 |
| GOBP_FACE_DEVELOPMENT | 44 | 0.409262 | 1.197998 | 0.203366 | 0.537965 |
| GOBP_REGULATION_OF_STEM_CELL_PROLIFERATION | 36 | 0.4178 | 1.197761 | 0.218524 | 0.538062 |
| GOBP_LEUKOCYTE_DEGRANULATION | 55 | 0.391697 | 1.197291 | 0.19146 | 0.538816 |
| GOBP_REGULATION_OF_BLOOD_CIRCULATION | 197 | 0.333261 | 1.197101 | 0.123557 | 0.538801 |
| GOBP_CELLULAR_RESPONSE_TO_LIPOPROTEIN_PARTICLE_STIMULUS | 25 | 0.455439 | 1.197077 | 0.222059 | 0.53833 |
| GOBP_REGULATION_OF_LIPID_BIOSYNTHETIC_PROCESS | 134 | 0.346944 | 1.196377 | 0.154037 | 0.539757 |
| GOBP_LYMPHOCYTE_CHEMOTAXIS | 24 | 0.457447 | 1.196316 | 0.235562 | 0.539425 |
| GOBP_EXTRACELLULAR_MATRIX_DISASSEMBLY | 44 | 0.413708 | 1.196298 | 0.203008 | 0.538946 |
| GOBP_NEGATIVE_REGULATION_OF_T_CELL_PROLIFERATION | 44 | 0.409602 | 1.195538 | 0.203338 | 0.540454 |
| GOBP_NEGATIVE_REGULATION_OF_CELL_CELL_ADHESION | 131 | 0.351288 | 1.195329 | 0.169697 | 0.540544 |
| GOBP_NEGATIVE_REGULATION_OF_MITOCHONDRION_ORGANIZATION | 45 | 0.407803 | 1.194488 | 0.221226 | 0.542389 |
| GOBP_INTERMEDIATE_FILAMENT_BASED_PROCESS | 38 | 0.42493 | 1.193843 | 0.218524 | 0.543712 |
| GOBP_POSITIVE_REGULATION_OF_NEURON_DEATH | 70 | 0.375615 | 1.193715 | 0.171896 | 0.543543 |
| GOBP_REGULATION_OF_PROGRAMMED_NECROTIC_CELL_DEATH | 21 | 0.477672 | 1.19361 | 0.239683 | 0.543304 |
| GOBP_COLLAGEN_CATABOLIC_PROCESS | 27 | 0.451588 | 1.1929 | 0.24375 | 0.544761 |
| GOBP_ACTIN_FILAMENT_BASED_TRANSPORT | 17 | 0.49697 | 1.192852 | 0.263238 | 0.544369 |
| GOBP_POSITIVE_REGULATION_OF_PROTEIN_LOCALIZATION_TO_MEMBRANE | 94 | 0.367239 | 1.192839 | 0.174745 | 0.54387 |
| GOBP_REGULATION_OF_DNA_BINDING | 106 | 0.359162 | 1.192224 | 0.182171 | 0.545069 |
| GOBP_MULTICELLULAR_ORGANISMAL_MOVEMENT | 43 | 0.413068 | 1.192158 | 0.210219 | 0.544734 |
| GOBP_POSITIVE_REGULATION_OF_BIOMINERALIZATION | 41 | 0.408573 | 1.191617 | 0.203008 | 0.545773 |
| GOBP_REGULATION_OF_MONOCYTE_CHEMOTAXIS | 15 | 0.504937 | 1.191121 | 0.263682 | 0.546667 |
| GOBP_NUCLEAR_TRANSCRIBED_MRNA_CATABOLIC_PROCESS_DEADENYLATION_DEPEN | 60 | 0.392412 | 1.190393 | 0.206044 | 0.548142 |
| GOBP_ODONTOGENESIS | 95 | 0.365908 | 1.190329 | 0.179124 | 0.547788 |
| GOBP_NEURON_FATE_COMMITMENT | 39 | 0.417734 | 1.189533 | 0.224888 | 0.549591 |
| GOBP_POSITIVE_REGULATION_OF_B_CELL_PROLIFERATION | 26 | 0.456088 | 1.188964 | 0.244444 | 0.550701 |
| GOBP_NEGATIVE_REGULATION_OF_CELLULAR_SENESCENCE | 16 | 0.498529 | 1.188922 | 0.259319 | 0.55029 |
| GOBP_RESPONSE_TO_COPPER_ION | 31 | 0.435733 | 1.188838 | 0.240909 | 0.549995 |
| GOBP_RESPONSE_TO_OXYGEN_LEVELS | 279 | 0.323465 | 1.1888 | 0.110245 | 0.549555 |
| GOBP_NEGATIVE_REGULATION_OF_COAGULATION | 35 | 0.42636 | 1.188602 | 0.223583 | 0.549575 |
| GOBP_CARDIAC_CONDUCTION | 81 | 0.367832 | 1.188004 | 0.201571 | 0.550756 |
| GOBP_MYELOID_DENDRITIC_CELL_ACTIVATION | 18 | 0.491317 | 1.187969 | 0.250401 | 0.550334 |
| GOBP_ARACHIDONIC_ACID_SECRETION | 18 | 0.483456 | 1.187265 | 0.262658 | 0.551807 |
| GOBP_CELLULAR_RESPONSE_TO_FATTY_ACID | 26 | 0.451424 | 1.187172 | 0.239329 | 0.551569 |
| GOBP_DETECTION_OF_ABIOTIC_STIMULUS | 89 | 0.364648 | 1.187119 | 0.196053 | 0.551183 |
| GOBP_NEGATIVE_REGULATION_OF_NF_KAPPAB_TRANSCRIPTION_FACTOR_ACTIVITY | 75 | 0.374827 | 1.186749 | 0.188187 | 0.551708 |
| GOBP_POSITIVE_REGULATION_OF_IMMUNE_EFFECTOR_PROCESS | 163 | 0.338133 | 1.186526 | 0.16088 | 0.551804 |
| GOBP_MEIOTIC_CELL_CYCLE | 202 | 0.331733 | 1.186436 | 0.123843 | 0.551515 |
| GOBP_CELLULAR_RESPONSE_TO_DEXAMETHASONE_STIMULUS | 21 | 0.468419 | 1.185993 | 0.262739 | 0.552291 |
| GOBP_PROTEIN_K63_LINKED_UBIQUITINATION | 54 | 0.397623 | 1.18578 | 0.194051 | 0.552386 |
| GOBP_NEGATIVE_REGULATION_OF_B_CELL_ACTIVATION | 26 | 0.4504 | 1.185032 | 0.249608 | 0.554049 |
| GOBP_NEGATIVE_REGULATION_OF_NEURON_APOPTOTIC_PROCESS | 129 | 0.344662 | 1.184977 | 0.180488 | 0.553685 |
| GOBP_LIPID_DROPLET_ORGANIZATION | 22 | 0.470862 | 1.184911 | 0.2589 | 0.553324 |
| GOBP_REGULATION_OF_CHONDROCYTE_DIFFERENTIATION | 43 | 0.41127 | 1.184453 | 0.234722 | 0.554098 |
| GOBP_RELAXATION_OF_MUSCLE | 27 | 0.44945 | 1.18411 | 0.219243 | 0.554516 |
| GOBP_GROWTH_PLATE_CARTILAGE_DEVELOPMENT | 16 | 0.492533 | 1.183837 | 0.268997 | 0.554719 |
| GOBP_MACROPHAGE_ACTIVATION | 63 | 0.388635 | 1.183803 | 0.217391 | 0.554279 |
| GOBP_RENAL_SYSTEM_VASCULATURE_DEVELOPMENT | 23 | 0.459667 | 1.183602 | 0.264753 | 0.55433 |
| GOBP_DEMETHYLATION | 59 | 0.385799 | 1.183416 | 0.209877 | 0.554333 |
| GOBP_POSITIVE_REGULATION_OF_BMP_SIGNALING_PATHWAY | 33 | 0.436571 | 1.183377 | 0.226244 | 0.553917 |
| GOBP_POSITIVE_REGULATION_OF_PEPTIDYL_THREONINE_PHOSPHORYLATION | 26 | 0.455888 | 1.183088 | 0.25873 | 0.554197 |
| GOBP_REGULATION_OF_MACROPHAGE_DERIVED_FOAM_CELL_DIFFERENTIATION | 19 | 0.48326 | 1.182429 | 0.260163 | 0.555555 |
| GOBP_MODULATION_BY_HOST_OF_VIRAL_PROCESS | 25 | 0.450199 | 1.181926 | 0.259146 | 0.556451 |
| GOBP_POSITIVE_REGULATION_OF_COAGULATION | 18 | 0.485672 | 1.181008 | 0.244094 | 0.558547 |
| GOBP_REGULATION_OF_ION_TRANSMEMBRANE_TRANSPORT | 375 | 0.314647 | 1.180277 | 0.101293 | 0.560074 |
| GOBP_PHOTORECEPTOR_CELL_DIFFERENTIATION | 48 | 0.398367 | 1.180118 | 0.226496 | 0.559981 |
| GOBP_REGULATION_OF_PROTEIN_SECRETION | 191 | 0.330584 | 1.179639 | 0.149184 | 0.56074 |
| GOBP_RESPONSE_TO_OXIDATIVE_STRESS | 367 | 0.314206 | 1.179007 | 0.115756 | 0.562042 |
| GOBP_BONE_MINERALIZATION | 92 | 0.357219 | 1.178816 | 0.195059 | 0.562075 |
| GOBP_GLYCOSYL_COMPOUND_METABOLIC_PROCESS | 74 | 0.369441 | 1.178773 | 0.201351 | 0.56165 |
| GOBP_REGULATION_OF_CHROMATIN_BINDING | 23 | 0.446265 | 1.178447 | 0.246246 | 0.562021 |
| GOBP_INOSITOL_LIPID_MEDIATED_SIGNALING | 132 | 0.345499 | 1.178399 | 0.176179 | 0.56163 |
| GOBP_CELLULAR_RESPONSE_TO_LIPID | 428 | 0.311828 | 1.178021 | 0.090713 | 0.562142 |
| GOBP_NEGATIVE_REGULATION_OF_MUSCLE_CONTRACTION | 21 | 0.467346 | 1.177835 | 0.255054 | 0.562141 |
| GOBP_REGULATION_OF_CATION_TRANSMEMBRANE_TRANSPORT | 273 | 0.321414 | 1.177242 | 0.129032 | 0.563282 |
| GOBP_ENDOTHELIUM_DEVELOPMENT | 117 | 0.349493 | 1.177223 | 0.172121 | 0.562813 |
| GOBP_ESTABLISHMENT_OR_MAINTENANCE_OF_CELL_POLARITY | 200 | 0.329949 | 1.177043 | 0.162679 | 0.562831 |
| GOBP_POLYOL_METABOLIC_PROCESS | 97 | 0.355637 | 1.176903 | 0.17601 | 0.562715 |
| GOBP_ACTION_POTENTIAL | 116 | 0.347837 | 1.176632 | 0.189937 | 0.563002 |
| GOBP_MITOCHONDRIAL_FISSION | 37 | 0.415953 | 1.176493 | 0.219585 | 0.562843 |
| GOBP_ANTIMICROBIAL_HUMORAL_RESPONSE | 44 | 0.400929 | 1.176441 | 0.238849 | 0.562477 |
| GOBP_TYROSINE_PHOSPHORYLATION_OF_STAT_PROTEIN | 49 | 0.391957 | 1.175482 | 0.216292 | 0.564721 |
| GOBP_MAINTENANCE_OF_LOCATION | 261 | 0.322536 | 1.175109 | 0.131461 | 0.565286 |
| GOBP_CELLULAR_RESPONSE_TO_CHEMICAL_STRESS | 287 | 0.323215 | 1.174902 | 0.122222 | 0.565338 |
| GOBP_POSITIVE_REGULATION_OF_CYTOKINE_PRODUCTION_INVOLVED_IN_IMMUNE_RES | 49 | 0.408844 | 1.174011 | 0.224324 | 0.567402 |
| GOBP_REGULATION_OF_AEROBIC_RESPIRATION | 29 | 0.431639 | 1.173495 | 0.252396 | 0.568364 |
| GOBP_B_CELL_DIFFERENTIATION | 97 | 0.355965 | 1.173212 | 0.175416 | 0.568656 |
| GOBP_GENITALIA_DEVELOPMENT | 28 | 0.438754 | 1.173172 | 0.257919 | 0.568267 |
| GOBP_DIOL_METABOLIC_PROCESS | 29 | 0.432132 | 1.172514 | 0.262346 | 0.569594 |
| GOBP_CILIUM_OR_FLAGELLUM_DEPENDENT_CELL_MOTILITY | 91 | 0.356477 | 1.172095 | 0.207398 | 0.570253 |
| GOBP_PARAXIAL_MESODERM_DEVELOPMENT | 17 | 0.48322 | 1.171494 | 0.257319 | 0.571412 |
| GOBP_RESPONSE_TO_HYDROGEN_PEROXIDE | 103 | 0.353333 | 1.171364 | 0.200508 | 0.571265 |
| GOBP_POSITIVE_REGULATION_OF_BLOOD_CIRCULATION | 33 | 0.421229 | 1.170624 | 0.247093 | 0.572839 |
| GOBP_CELL_AGING | 105 | 0.349536 | 1.17023 | 0.191763 | 0.573448 |
| GOBP_COLLAGEN_METABOLIC_PROCESS | 70 | 0.368758 | 1.169903 | 0.228849 | 0.573853 |
| GOBP_REGULATION_OF_RESPONSE_TO_OXIDATIVE_STRESS | 79 | 0.361579 | 1.169392 | 0.21224 | 0.574794 |
| GOBP_NEURAL_RETINA_DEVELOPMENT | 52 | 0.386647 | 1.169314 | 0.225806 | 0.574481 |
| GOBP_REGULATION_OF_GLUCAN_BIOSYNTHETIC_PROCESS | 23 | 0.447224 | 1.169314 | 0.2672 | 0.573954 |
| GOBP_CELLULAR_RESPONSE_TO_INTERFERON_BETA | 16 | 0.490699 | 1.169204 | 0.271104 | 0.573723 |
| GOBP_RESPONSE_TO_INORGANIC_SUBSTANCE | 427 | 0.310668 | 1.16881 | 0.098501 | 0.574359 |
| GOBP_CELLULAR_SENESCENCE | 74 | 0.368759 | 1.168683 | 0.202997 | 0.574216 |
| GOBP_POSITIVE_REGULATION_OF_PHOSPHATASE_ACTIVITY | 28 | 0.436354 | 1.168344 | 0.24031 | 0.574632 |
| GOBP_AMINO_SUGAR_METABOLIC_PROCESS | 35 | 0.417235 | 1.167729 | 0.256223 | 0.575897 |
| GOBP_POSITIVE_REGULATION_OF_RESPONSE_TO_CYTOKINE_STIMULUS | 51 | 0.389028 | 1.16766 | 0.245364 | 0.575561 |
| GOBP_INTERLEUKIN_6_PRODUCTION | 101 | 0.352136 | 1.167042 | 0.214103 | 0.576833 |
| GOBP_NEGATIVE_REGULATION_OF_SYNAPTIC_TRANSMISSION | 59 | 0.375076 | 1.166989 | 0.23319 | 0.576479 |

| | | | | | |
|---|---|---|---|---|---|
| GOBP_MITOCHONDRIAL_RESPIRATORY_CHAIN_COMPLEX_ASSEMBLY | 93 | -0.147248 | -0.559714 | 1 | 1 |
| GOBP_VESICLE_MEDIATED_TRANSPORT_TO_THE_PLASMA_MEMBRANE | 135 | -0.13573 | -0.559208 | 1 | 1 |
| GOBP_RNA_DEPENDENT_DNA_BIOSYNTHETIC_PROCESS | 68 | -0.151035 | -0.556906 | 0.996198 | 1 |
| GOBP_NUCLEOBASE_CONTAINING_SMALL_MOLECULE_BIOSYNTHETIC_PROCESS | 15 | -0.215298 | -0.554423 | 0.969697 | 1 |
| GOBP_MITOTIC_NUCLEAR_DIVISION | 273 | -0.125135 | -0.553636 | 1 | 1 |
| GOBP_RNA_MODIFICATION | 156 | -0.13283 | -0.552711 | 1 | 1 |
| GOBP_REGULATION_OF_UBIQUITIN_PROTEIN_TRANSFERASE_ACTIVITY | 52 | -0.160375 | -0.552371 | 0.996575 | 1 |
| GOBP_ESTABLISHMENT_OF_PROTEIN_LOCALIZATION_TO_PLASMA_MEMBRANE | 60 | -0.152106 | -0.544434 | 0.996587 | 1 |
| GOBP_CELLULAR_RESPONSE_TO_STEROL_DEPLETION | 16 | -0.202323 | -0.541144 | 0.97449 | 1 |
| GOBP_FOLIC_ACID_CONTAINING_COMPOUND_METABOLIC_PROCESS | 27 | -0.179648 | -0.536709 | 0.988539 | 1 |
| GOBP_REGULATION_OF_TELOMERE_CAPPING | 26 | -0.176846 | -0.532111 | 0.989362 | 1 |
| GOBP_RNA_METHYLATION | 81 | -0.139434 | -0.530003 | 1 | 1 |
| GOBP_ATP_BIOSYNTHETIC_PROCESS | 52 | -0.152552 | -0.529975 | 1 | 1 |
| GOBP_MITOCHONDRIAL_CYTOCHROME_C_OXIDASE_ASSEMBLY | 21 | -0.182969 | -0.524487 | 0.988981 | 1 |
| GOBP_RESPIRATORY_CHAIN_COMPLEX_IV_ASSEMBLY | 25 | -0.176592 | -0.523977 | 0.985755 | 1 |
| GOBP_RNA_INTERFERENCE | 17 | -0.19186 | -0.519489 | 0.986945 | 1 |
| GOBP_RESPIRATORY_ELECTRON_TRANSPORT_CHAIN | 105 | -0.131594 | -0.517108 | 1 | 1 |
| GOBP_MITOCHONDRIAL_ELECTRON_TRANSPORT_NADH_TO_UBIQUINONE | 48 | -0.152024 | -0.515401 | 1 | 1 |
| GOBP_GLYCOSYL_COMPOUND_BIOSYNTHETIC_PROCESS | 19 | -0.186302 | -0.514137 | 0.99022 | 1 |
| GOBP_PROTEIN_LOCALIZATION_TO_CONDENSED_CHROMOSOME | 19 | -0.183962 | -0.513334 | 0.979592 | 1 |
| GOBP_PHAGOLYSOSOME_ASSEMBLY | 15 | -0.19968 | -0.513106 | 0.97549 | 1 |
| GOBP_NUCLEOBASE_BIOSYNTHETIC_PROCESS | 19 | -0.186694 | -0.512596 | 0.971649 | 1 |
| GOBP_REGULATION_OF_DNA_STRAND_ELONGATION | 16 | -0.189443 | -0.511821 | 0.980769 | 1 |
| GOBP_AUDITORY_RECEPTOR_CELL_MORPHOGENESIS | 15 | -0.191278 | -0.510604 | 0.973753 | 1 |
| GOBP_MITOTIC_SPINDLE_ASSEMBLY | 70 | -0.137116 | -0.509164 | 1 | 1 |
| GOBP_HISTONE_H3_K9_TRIMETHYLATION | 16 | -0.188908 | -0.501809 | 0.997362 | 1 |
| GOBP_MITOTIC_METAPHASE_PLATE_CONGRESSION | 52 | -0.143514 | -0.500737 | 1 | 1 |
| GOBP_POSITIVE_REGULATION_OF_TELOMERE_MAINTENANCE | 50 | -0.141521 | -0.497228 | 1 | 1 |
| GOBP_PROTEIN_QUALITY_CONTROL_FOR_MISFOLDED_OR_INCOMPLETELY_SYNTHESIZE | 27 | -0.164088 | -0.49591 | 1 | 1 |
| GOBP_HISTONE_EXCHANGE | 17 | -0.187914 | -0.495673 | 0.992105 | 1 |
| GOBP_ERAD_PATHWAY | 106 | -0.123429 | -0.489694 | 1 | 1 |
| GOBP_HISTONE_H4_K5_ACETYLATION | 16 | -0.182728 | -0.488355 | 0.994845 | 1 |
| GOBP_RESPONSE_TO_STEROL_DEPLETION | 18 | -0.181979 | -0.486297 | 1 | 1 |
| GOBP_AMINO_ACID_ACTIVATION | 45 | -0.143749 | -0.484208 | 1 | 1 |
| GOBP_NLS_BEARING_PROTEIN_IMPORT_INTO_NUCLEUS | 19 | -0.172325 | -0.482843 | 0.994832 | 1 |
| GOBP_MRNA_3_END_PROCESSING | 61 | -0.13215 | -0.47755 | 1 | 1 |
| GOBP_HISTONE_MRNA_METABOLIC_PROCESS | 24 | -0.161106 | -0.472474 | 0.997326 | 1 |
| GOBP_REGULATION_OF_MRNA_POLYADENYLATION | 20 | -0.168891 | -0.467686 | 0.985755 | 1 |
| GOBP_POSITIVE_REGULATION_OF_PROTEIN_DEACETYLATION | 24 | -0.158492 | -0.465111 | 1 | 1 |
| GOBP_DICARBOXYLIC_ACID_CATABOLIC_PROCESS | 17 | -0.167803 | -0.461856 | 0.997409 | 1 |
| GOBP_CHROMOSOME_LOCALIZATION | 78 | -0.122836 | -0.461229 | 1 | 1 |
| GOBP_CLATHRIN_COAT_ASSEMBLY | 18 | -0.168108 | -0.459764 | 0.992611 | 1 |
| GOBP_ARP2_3_COMPLEX_MEDIATED_ACTIN_NUCLEATION | 39 | -0.142049 | -0.458959 | 1 | 1 |
| GOBP_PEPTIDYL_ARGININE_METHYLATION | 15 | -0.173869 | -0.455144 | 0.989717 | 1 |
| GOBP_POSITIVE_REGULATION_OF_UBIQUITIN_PROTEIN_TRANSFERASE_ACTIVITY | 28 | -0.153364 | -0.455091 | 1 | 1 |
| GOBP_MITOCHONDRIAL_RNA_METABOLIC_PROCESS | 49 | -0.13176 | -0.454375 | 1 | 1 |
| GOBP_ENDOCYTIC_RECYCLING | 74 | -0.120957 | -0.45014 | 1 | 1 |
| GOBP_REGULATION_OF_MRNA_3_END_PROCESSING | 28 | -0.148837 | -0.449754 | 1 | 1 |
| GOBP_NEGATIVE_REGULATION_OF_UBIQUITIN_DEPENDENT_PROTEIN_CATABOLIC_PRO( | 44 | -0.131322 | -0.446438 | 1 | 1 |
| GOBP_POSITIVE_REGULATION_OF_TRANSCRIPTION_BY_RNA_POLYMERASE_III | 18 | -0.158757 | -0.438304 | 0.997403 | 1 |
| GOBP_REGULATION_OF_TELOMERE_MAINTENANCE | 77 | -0.113955 | -0.421627 | 1 | 1 |
| GOBP_REGULATION_OF_PROTEIN_EXIT_FROM_ENDOPLASMIC_RETICULUM | 24 | -0.145082 | -0.4216 | 1 | 1 |
| GOBP_POLY_A_PLUS_MRNA_EXPORT_FROM_NUCLEUS | 15 | -0.158733 | -0.408437 | 0.997333 | 1 |
| GOBP_MITOTIC_CYTOKINETIC_PROCESS | 23 | -0.142514 | -0.40356 | 1 | 1 |
| GOBP_NEGATIVE_REGULATION_OF_DEFENSE_RESPONSE_TO_VIRUS | 25 | -0.125746 | -0.370564 | 1 | 1 |
| GOBP_PEPTIDYL_LYSINE_TRIMETHYLATION | 51 | -0.103093 | -0.353759 | 1 | 1 |
| GOBP_NUCLEOSIDE_MONOPHOSPHATE_BIOSYNTHETIC_PROCESS | 40 | -0.098753 | -0.330998 | 1 | 0.999985 |

| | | | | | |
|---|---|---|---|---|---|
| GOBP_PHOSPHOLIPASE_C_ACTIVATING_G_PROTEIN_COUPLED_RECEPTOR_SIGNALING_ | 56 | 0.3803 | 1.166988 | 0.232194 | 0.575955 |
| GOBP_PROTEIN_LOCALIZATION_TO_EXTRACELLULAR_REGION | 276 | 0.318371 | 1.166821 | 0.146861 | 0.575958 |
| GOBP_RESPONSE_TO_FLUID_SHEAR_STRESS | 30 | 0.425272 | 1.166574 | 0.254815 | 0.576158 |
| GOBP_PHOTORECEPTOR_CELL_DEVELOPMENT | 38 | 0.414562 | 1.166068 | 0.268398 | 0.577113 |
| GOBP_ANATOMICAL_STRUCTURE_MATURATION | 187 | 0.331865 | 1.165478 | 0.159905 | 0.578256 |
| GOBP_POSITIVE_REGULATION_OF_AMINO_ACID_TRANSPORT | 20 | 0.460897 | 1.165362 | 0.25641 | 0.57805 |
| GOBP_POSITIVE_REGULATION_OF_ION_TRANSPORT | 214 | 0.322269 | 1.165263 | 0.159039 | 0.577821 |
| GOBP_ROOF_OF_MOUTH_DEVELOPMENT | 72 | 0.365583 | 1.16522 | 0.230769 | 0.577437 |
| GOBP_REGULATION_OF_TRANSMEMBRANE_TRANSPORT | 444 | 0.308024 | 1.165074 | 0.113924 | 0.577361 |
| GOBP_LEUKOCYTE_MEDIATED_IMMUNITY | 249 | 0.319183 | 1.164955 | 0.156841 | 0.577162 |
| GOBP_ENDOTHELIAL_CELL_CHEMOTAXIS | 24 | 0.443225 | 1.16324 | 0.275809 | 0.581612 |
| GOBP_NOTCH_SIGNALING_PATHWAY | 151 | 0.333469 | 1.162783 | 0.184783 | 0.582418 |
| GOBP_CELLULAR_RESPONSE_TO_ACID_CHEMICAL | 72 | 0.367578 | 1.162684 | 0.225806 | 0.582174 |
| GOBP_BLOOD_VESSEL_ENDOTHELIAL_CELL_PROLIFERATION_INVOLVED_IN_SPROUTING | 21 | 0.46117 | 1.162526 | 0.268174 | 0.582114 |
| GOBP_GLOMERULAR_EPITHELIAL_CELL_DIFFERENTIATION | 20 | 0.462056 | 1.162445 | 0.28614 | 0.581829 |
| GOBP_CARTILAGE_DEVELOPMENT_INVOLVED_IN_ENDOCHONDRAL_BONE_MORPHOGEN | 25 | 0.446887 | 1.162438 | 0.26489 | 0.581322 |
| GOBP_POSITIVE_REGULATION_OF_BONE_MINERALIZATION | 35 | 0.417796 | 1.16219 | 0.257732 | 0.581506 |
| GOBP_TELOMERE_MAINTENANCE_VIA_RECOMBINATION | 15 | 0.503702 | 1.16201 | 0.286424 | 0.581505 |
| GOBP_ACTIN_FILAMENT_ORGANIZATION | 399 | 0.309455 | 1.161774 | 0.134043 | 0.581662 |
| GOBP_REGULATION_OF_LEUKOCYTE_MEDIATED_IMMUNITY | 159 | 0.329717 | 1.160739 | 0.200713 | 0.584051 |
| GOBP_SUBSTRATE_ADHESION_DEPENDENT_CELL_SPREADING | 97 | 0.356157 | 1.160627 | 0.219512 | 0.583818 |
| GOBP_SCHWANN_CELL_DIFFERENTIATION | 36 | 0.41373 | 1.160425 | 0.257598 | 0.583898 |
| GOBP_RESPONSE_TO_INTERFERON_BETA | 22 | 0.452821 | 1.160354 | 0.258786 | 0.583579 |
| GOBP_REGULATION_OF_AUTOPHAGOSOME_MATURATION | 15 | 0.504679 | 1.160126 | 0.303777 | 0.583706 |
| GOBP_POSITIVE_REGULATION_OF_OSTEOBLAST_DIFFERENTIATION | 58 | 0.380969 | 1.159973 | 0.236915 | 0.583637 |
| GOBP_NEGATIVE_REGULATION_OF_PEPTIDYL_THREONINE_PHOSPHORYLATION | 15 | 0.499993 | 1.159952 | 0.292562 | 0.583166 |
| GOBP_REGULATION_OF_ERBB_SIGNALING_PATHWAY | 67 | 0.372676 | 1.159845 | 0.252022 | 0.58297 |
| GOBP_CELL_PROLIFERATION_INVOLVED_IN_KIDNEY_DEVELOPMENT | 17 | 0.483004 | 1.159025 | 0.276973 | 0.584782 |
| GOBP_DETECTION_OF_MECHANICAL_STIMULUS_INVOLVED_IN_SENSORY_PERCEPTION | 29 | 0.423436 | 1.158883 | 0.262918 | 0.584668 |
| GOBP_PHENOL_CONTAINING_COMPOUND_METABOLIC_PROCESS | 79 | 0.367154 | 1.158541 | 0.230971 | 0.585119 |
| GOBP_INNATE_IMMUNE_RESPONSE | 495 | 0.304108 | 1.15833 | 0.112853 | 0.585229 |
| GOBP_REGULATION_OF_SMALL_GTPASE_MEDIATED_SIGNAL_TRANSDUCTION | 274 | 0.314808 | 1.158119 | 0.171558 | 0.58535 |
| GOBP_REGULATION_OF_RUFFLE_ASSEMBLY | 28 | 0.426136 | 1.156788 | 0.256098 | 0.588589 |
| GOBP_RAS_PROTEIN_SIGNAL_TRANSDUCTION | 312 | 0.314517 | 1.156677 | 0.164835 | 0.5884 |
| GOBP_SPERM_MOTILITY | 71 | 0.36702 | 1.156148 | 0.252387 | 0.589417 |
| GOBP_TRABECULA_MORPHOGENESIS | 41 | 0.399902 | 1.155917 | 0.240059 | 0.589582 |
| GOBP_NEGATIVE_REGULATION_OF_INFLAMMATORY_RESPONSE_TO_ANTIGENIC_STIMUL | 23 | 0.447508 | 1.154226 | 0.27735 | 0.594042 |
| GOBP_CARDIAC_MUSCLE_CELL_CONTRACTION | 65 | 0.376247 | 1.154221 | 0.263158 | 0.593538 |
| GOBP_NEUTRAL_LIPID_METABOLIC_PROCESS | 97 | 0.347564 | 1.154049 | 0.232258 | 0.593525 |
| GOBP_POSITIVE_REGULATION_OF_AUTOPHAGY | 128 | 0.338696 | 1.153443 | 0.208763 | 0.594812 |
| GOBP_POSITIVE_REGULATION_OF_TUMOR_NECROSIS_FACTOR_SUPERFAMILY_CYTOKIN | 73 | 0.36058 | 1.153291 | 0.230769 | 0.594746 |
| GOBP_NEGATIVE_REGULATION_OF_PROTEIN_ACETYLATION | 20 | 0.460662 | 1.152487 | 0.266559 | 0.596597 |
| GOBP_CELL_CELL_ADHESION_VIA_PLASMA_MEMBRANE_ADHESION_MOLECULES | 222 | 0.319486 | 1.152369 | 0.170507 | 0.596416 |
| GOBP_ADAPTIVE_IMMUNE_RESPONSE_BASED_ON_SOMATIC_RECOMBINATION_OF_IMMU | 197 | 0.32515 | 1.151835 | 0.183234 | 0.597528 |
| GOBP_POLYOL_BIOSYNTHETIC_PROCESS | 52 | 0.379059 | 1.151174 | 0.266382 | 0.598882 |
| GOBP_REGULATION_OF_ACTIN_FILAMENT_BASED_PROCESS | 349 | 0.308512 | 1.150572 | 0.152104 | 0.600109 |
| GOBP_ANATOMICAL_STRUCTURE_HOMEOSTASIS | 235 | 0.316476 | 1.149926 | 0.181922 | 0.601422 |
| GOBP_RESPONSE_TO_ESTRADIOL | 90 | 0.353583 | 1.149712 | 0.232349 | 0.601527 |
| GOBP_REGULATION_OF_T_CELL_MEDIATED_IMMUNITY | 56 | 0.37865 | 1.149674 | 0.247911 | 0.601107 |
| GOBP_REGULATION_OF_NUCLEAR_DIVISION | 115 | 0.34135 | 1.149516 | 0.227848 | 0.60103 |
| GOBP_POSITIVE_REGULATION_OF_ENDOTHELIAL_CELL_PROLIFERATION | 76 | 0.360418 | 1.148877 | 0.242894 | 0.602374 |
| GOBP_HAIR_CELL_DIFFERENTIATION | 39 | 0.412572 | 1.148529 | 0.25289 | 0.602921 |
| GOBP_LENS_FIBER_CELL_DIFFERENTIATION | 32 | 0.421896 | 1.147958 | 0.267752 | 0.604049 |
| GOBP_REGULATION_OF_ENDOTHELIAL_CELL_DEVELOPMENT | 16 | 0.488347 | 1.147869 | 0.279743 | 0.603796 |
| GOBP_RESPONSE_TO_MOLECULE_OF_BACTERIAL_ORIGIN | 220 | 0.317812 | 1.147828 | 0.193699 | 0.603385 |
| GOBP_NATURAL_KILLER_CELL_DIFFERENTIATION | 20 | 0.464556 | 1.147542 | 0.298869 | 0.603689 |
| GOBP_RESPONSE_TO_ALCOHOL | 174 | 0.325642 | 1.147529 | 0.178241 | 0.603197 |
| GOBP_CELLULAR_RESPONSE_TO_INORGANIC_SUBSTANCE | 177 | 0.322916 | 1.14723 | 0.186019 | 0.603565 |
| GOBP_CELLULAR_RESPONSE_TO_UNFOLDED_PROTEIN | 93 | 0.350297 | 1.14692 | 0.247765 | 0.603924 |
| GOBP_REGULATION_OF_CALCIUM_ION_TRANSPORT_INTO_CYTOSOL | 79 | 0.360333 | 1.146719 | 0.228336 | 0.604002 |
| GOBP_NUCLEOBASE_METABOLIC_PROCESS | 37 | 0.405244 | 1.146622 | 0.279211 | 0.60378 |
| GOBP_REGULATION_OF_POTASSIUM_ION_TRANSPORT | 81 | 0.35048 | 1.146398 | 0.239305 | 0.603927 |
| GOBP_NEGATIVE_REGULATION_OF_PROTEOLYSIS | 220 | 0.314978 | 1.145936 | 0.188439 | 0.60476 |
| GOBP_ATRIAL_CARDIAC_MUSCLE_CELL_TO_AV_NODE_CELL_SIGNALING | 18 | 0.471124 | 1.145924 | 0.296 | 0.604279 |
| GOBP_REGULATION_OF_PROTEIN_TARGETING_TO_MEMBRANE | 31 | 0.415635 | 1.145723 | 0.271642 | 0.604355 |
| GOBP_NEUROPEPTIDE_SIGNALING_PATHWAY | 64 | 0.371081 | 1.145645 | 0.257618 | 0.604084 |
| GOBP_NEGATIVE_REGULATION_OF_MYELOID_LEUKOCYTE_DIFFERENTIATION | 35 | 0.407546 | 1.145585 | 0.294372 | 0.603716 |
| GOBP_SENSORY_PERCEPTION_OF_LIGHT_STIMULUS | 136 | 0.331482 | 1.145482 | 0.210657 | 0.603509 |
| GOBP_POSITIVE_REGULATION_OF_TYROSINE_PHOSPHORYLATION_OF_STAT_PROTEIN | 36 | 0.410719 | 1.14531 | 0.281481 | 0.603487 |
| GOBP_REGULATION_OF_HEMOPOIESIS | 275 | 0.311685 | 1.145219 | 0.173289 | 0.603211 |
| GOBP_CELLULAR_ION_HOMEOSTASIS | 491 | 0.301182 | 1.145212 | 0.131027 | 0.602711 |
| GOBP_POSITIVE_CHEMOTAXIS | 41 | 0.395747 | 1.144746 | 0.271955 | 0.60354 |
| GOBP_PARASYMPATHETIC_NERVOUS_SYSTEM_DEVELOPMENT | 15 | 0.490081 | 1.1443 | 0.295567 | 0.604345 |
| GOBP_AGING | 247 | 0.311964 | 1.144063 | 0.182127 | 0.604538 |
| GOBP_NEGATIVE_REGULATION_OF_SODIUM_ION_TRANSPORT | 16 | 0.482666 | 1.143993 | 0.308824 | 0.604218 |
| GOBP_NEGATIVE_REGULATION_OF_CELL_CYCLE_G1_S_PHASE_TRANSITION | 71 | 0.364138 | 1.143847 | 0.240216 | 0.60411 |
| GOBP_EPITHELIAL_TO_MESENCHYMAL_TRANSITION | 137 | 0.333575 | 1.143616 | 0.225885 | 0.604274 |
| GOBP_THYMOCYTE_APOPTOTIC_PROCESS | 15 | 0.479158 | 1.142493 | 0.3086 | 0.606973 |
| GOBP_PROTEIN_DESTABILIZATION | 44 | 0.386221 | 1.1423 | 0.263083 | 0.607051 |
| GOBP_NEGATIVE_REGULATION_OF_RESPONSE_TO_WOUNDING | 67 | 0.366351 | 1.142114 | 0.267038 | 0.607091 |
| GOBP_BONE_CELL_DEVELOPMENT | 28 | 0.430993 | 1.142018 | 0.279321 | 0.606827 |
| GOBP_PEPTIDYL_TYROSINE_DEPHOSPHORYLATION | 91 | 0.350121 | 1.141892 | 0.245478 | 0.606669 |
| GOBP_CHONDROCYTE_PROLIFERATION | 17 | 0.468474 | 1.141612 | 0.319328 | 0.60699 |
| GOBP_NEGATIVE_REGULATION_OF_MAPK_CASCADE | 147 | 0.32808 | 1.141556 | 0.224969 | 0.606629 |
| GOBP_NEGATIVE_REGULATION_OF_INTRACELLULAR_SIGNAL_TRANSDUCTION | 453 | 0.301044 | 1.141248 | 0.154886 | 0.60699 |
| GOBP_DIGESTION | 67 | 0.365156 | 1.141111 | 0.251374 | 0.606896 |
| GOBP_NEGATIVE_REGULATION_OF_FATTY_ACID_METABOLIC_PROCESS | 23 | 0.439794 | 1.140911 | 0.305085 | 0.606986 |
| GOBP_RESPONSE_TO_METAL_ION | 278 | 0.308879 | 1.140709 | 0.190949 | 0.60707 |
| GOBP_REGULATION_OF_INTRACELLULAR_PROTEIN_TRANSPORT | 206 | 0.318432 | 1.140291 | 0.218097 | 0.607789 |
| GOBP_SIGNAL_TRANSDUCTION_IN_RESPONSE_TO_DNA_DAMAGE | 165 | 0.324475 | 1.140146 | 0.214806 | 0.607683 |
| GOBP_INTRINSIC_APOPTOTIC_SIGNALING_PATHWAY_IN_RESPONSE_TO_ENDOPLASMIC_ | 57 | 0.372361 | 1.139769 | 0.273232 | 0.608243 |
| GOBP_NEGATIVE_REGULATION_OF_RESPONSE_TO_EXTERNAL_STIMULUS | 290 | 0.310582 | 1.139525 | 0.180401 | 0.608415 |
| GOBP_MAMMARY_GLAND_EPITHELIAL_CELL_PROLIFERATION | 23 | 0.441815 | 1.13943 | 0.311912 | 0.608173 |
| GOBP_NEGATIVE_REGULATION_OF_GROWTH | 211 | 0.316263 | 1.13881 | 0.186992 | 0.609487 |
| GOBP_CELLULAR_RESPONSE_TO_FLUID_SHEAR_STRESS | 18 | 0.462991 | 1.138666 | 0.287284 | 0.609396 |
| GOBP_POSITIVE_REGULATION_OF_PROTEIN_DEPOLYMERIZATION | 17 | 0.473343 | 1.138339 | 0.311667 | 0.60986 |
| GOBP_NEGATIVE_REGULATION_OF_ANTIGEN_RECEPTOR_MEDIATED_SIGNALING_PATHW | 22 | 0.445458 | 1.138249 | 0.299689 | 0.6096 |
| GOBP_REGULATION_OF_POLYSACCHARIDE_METABOLIC_PROCESS | 37 | 0.401575 | 1.13796 | 0.268895 | 0.609933 |
| GOBP_GLOMERULUS_DEVELOPMENT | 55 | 0.377975 | 1.137766 | 0.269553 | 0.609976 |
| GOBP_MAMMARY_GLAND_DEVELOPMENT | 112 | 0.33808 | 1.13753 | 0.248424 | 0.61017 |
| GOBP_NEGATIVE_REGULATION_OF_NIK_NF_KAPPAB_SIGNALING | 23 | 0.442388 | 1.137411 | 0.302432 | 0.610013 |
| GOBP_PHOSPHOLIPID_CATABOLIC_PROCESS | 41 | 0.395404 | 1.136699 | 0.285094 | 0.611522 |
| GOBP_LENS_DEVELOPMENT_IN_CAMERA_TYPE_EYE | 68 | 0.36636 | 1.136585 | 0.269601 | 0.61134 |
| GOBP_REGULATION_OF_CENTRIOLE_REPLICATION | 21 | 0.450397 | 1.13644 | 0.297872 | 0.611261 |
| GOBP_NUCLEUS_LOCALIZATION | 24 | 0.431255 | 1.136379 | 0.297214 | 0.610915 |
| GOBP_RELEASE_OF_SEQUESTERED_CALCIUM_ION_INTO_CYTOSOL_BY_ENDOPLASMIC_ | 26 | 0.424306 | 1.136367 | 0.3 | 0.610434 |
| GOBP_NEURONAL_ACTION_POTENTIAL | 26 | 0.434897 | 1.135891 | 0.2912 | 0.611291 |
| GOBP_CELLULAR_RESPONSE_TO_RETINOIC_ACID | 50 | 0.385951 | 1.135691 | 0.26257 | 0.611397 |
| GOBP_BLASTOCYST_DEVELOPMENT | 100 | 0.341251 | 1.134836 | 0.246415 | 0.613357 |
| GOBP_TRANSPORT_OF_VIRUS | 17 | 0.473844 | 1.13464 | 0.313115 | 0.613429 |
| GOBP_PEPTIDYL_THREONINE_DEPHOSPHORYLATION | 16 | 0.476913 | 1.134556 | 0.306397 | 0.613174 |
| GOBP_T_CELL_APOPTOTIC_PROCESS | 36 | 0.408697 | 1.134349 | 0.291982 | 0.6133 |
| GOBP_CELLULAR_RESPONSE_TO_LOW_DENSITY_LIPOPROTEIN_PARTICLE_STIMULUS | 15 | 0.478156 | 1.134343 | 0.319328 | 0.612808 |
| GOBP_NEGATIVE_REGULATION_OF_CYTOKINE_PRODUCTION_INVOLVED_IN_IMMUNE_RE | 16 | 0.47997 | 1.134142 | 0.3312 | 0.612859 |
| GOBP_PERK_MEDIATED_UNFOLDED_PROTEIN_RESPONSE | 17 | 0.470849 | 1.134097 | 0.321951 | 0.61249 |
| GOBP_ENDOPLASMIC_RETICULUM_UNFOLDED_PROTEIN_RESPONSE | 73 | 0.361413 | 1.133682 | 0.261518 | 0.613208 |
| GOBP_REGULATION_OF_EMBRYONIC_DEVELOPMENT | 56 | 0.372619 | 1.133297 | 0.267525 | 0.613794 |
| GOBP_LYMPHOCYTE_APOPTOTIC_PROCESS | 56 | 0.370049 | 1.133295 | 0.276006 | 0.613293 |
| GOBP_RESPONSE_TO_COLD | 46 | 0.381477 | 1.132842 | 0.298222 | 0.614073 |
| GOBP_SENSORY_PERCEPTION_OF_PAIN | 73 | 0.357214 | 1.131131 | 0.284553 | 0.618484 |
| GOBP_HISTONE_H3_K14_ACETYLATION | 22 | 0.448244 | 1.13106 | 0.30032 | 0.61816 |
| GOBP_REGULATION_OF_GLYCOPROTEIN_METABOLIC_PROCESS | 41 | 0.384034 | 1.130932 | 0.303468 | 0.618027 |
| GOBP_PLASMA_MEMBRANE_PHOSPHOLIPID_SCRAMBLING | 18 | 0.463609 | 1.130169 | 0.319287 | 0.619768 |
| GOBP_MITOCHONDRIAL_FUSION | 30 | 0.417667 | 1.130047 | 0.304878 | 0.619622 |
| GOBP_RESPONSE_TO_INTERLEUKIN_1 | 86 | 0.347527 | 1.130029 | 0.250326 | 0.619155 |
| GOBP_SKIN_EPIDERMIS_DEVELOPMENT | 69 | 0.363125 | 1.129762 | 0.268493 | 0.619483 |
| GOBP_RESPONSE_TO_ACID_CHEMICAL | 107 | 0.339151 | 1.128351 | 0.265122 | 0.623142 |
| GOBP_EATING_BEHAVIOR | 23 | 0.435304 | 1.12823 | 0.310289 | 0.622978 |
| GOBP_NEGATIVE_REGULATION_OF_PHOSPHORYLATION | 327 | 0.303651 | 1.127659 | 0.189749 | 0.624172 |
| GOBP_IN_UTERO_EMBRYONIC_DEVELOPMENT | 353 | 0.300477 | 1.127463 | 0.187432 | 0.624272 |
| GOBP_NEURON_DEATH | 304 | 0.306524 | 1.12723 | 0.20966 | 0.624436 |
| GOBP_POSITIVE_REGULATION_OF_INSULIN_SECRETION | 54 | 0.371782 | 1.126741 | 0.295058 | 0.625395 |
| GOBP_REGULATION_OF_KERATINOCYTE_PROLIFERATION | 27 | 0.422353 | 1.126325 | 0.302181 | 0.626128 |
| GOBP_NEGATIVE_REGULATION_OF_KINASE_ACTIVITY | 205 | 0.318826 | 1.125738 | 0.222477 | 0.627304 |
| GOBP_REGULATION_OF_HORMONE_LEVELS | 356 | 0.301903 | 1.125264 | 0.199571 | 0.628176 |
| GOBP_POST_EMBRYONIC_DEVELOPMENT | 78 | 0.347016 | 1.124869 | 0.240885 | 0.628826 |
| GOBP_CELLULAR_RESPONSE_TO_TOXIC_SUBSTANCE | 91 | 0.342289 | 1.124284 | 0.265013 | 0.630078 |
| GOBP_MEMBRANE_REPOLARIZATION_DURING_ACTION_POTENTIAL | 23 | 0.434254 | 1.124275 | 0.325688 | 0.629587 |
| GOBP_VENTRICULAR_CARDIAC_MUSCLE_CELL_ACTION_POTENTIAL | 28 | 0.418249 | 1.123968 | 0.31194 | 0.629996 |
| GOBP_DEVELOPMENTAL_CELL_GROWTH | 202 | 0.316 | 1.123725 | 0.232719 | 0.630161 |
| GOBP_SUPEROXIDE_METABOLIC_PROCESS | 49 | 0.377573 | 1.123538 | 0.285307 | 0.630196 |
| GOBP_OVULATION_CYCLE_PROCESS | 36 | 0.394547 | 1.123262 | 0.313343 | 0.630505 |
| GOBP_POSITIVE_REGULATION_OF_CYSTEINE_TYPE_ENDOPEPTIDASE_ACTIVITY | 120 | 0.329656 | 1.122659 | 0.244085 | 0.631792 |
| GOBP_RETINA_LAYER_FORMATION | 19 | 0.453863 | 1.12255 | 0.320786 | 0.631603 |
| GOBP_REGULATION_OF_RYANODINE_SENSITIVE_CALCIUM_RELEASE_CHANNEL_ACTIVIT | 24 | 0.434028 | 1.121758 | 0.301794 | 0.633384 |
| GOBP_POSITIVE_REGULATION_OF_ACTIN_FILAMENT_POLYMERIZATION | 46 | 0.376056 | 1.120621 | 0.301449 | 0.636302 |
| GOBP_NEGATIVE_REGULATION_OF_DNA_DAMAGE_RESPONSE_SIGNAL_TRANSDUCTION_ | 15 | 0.476777 | 1.119632 | 0.329618 | 0.638757 |
| GOBP_POSITIVE_REGULATION_OF_ORGANELLE_ORGANIZATION | 434 | 0.296604 | 1.119171 | 0.189968 | 0.639633 |
| GOBP_POSITIVE_REGULATION_OF_TOLL_LIKE_RECEPTOR_SIGNALING_PATHWAY | 20 | 0.446313 | 1.119089 | 0.338762 | 0.639361 |
| GOBP_NEGATIVE_REGULATION_OF_TRANSCRIPTION_REGULATORY_REGION_DNA_BINDI | 17 | 0.460962 | 1.119011 | 0.318699 | 0.639077 |
| GOBP_STEM_CELL_PROLIFERATION | 59 | 0.362675 | 1.118093 | 0.284741 | 0.64132 |
| GOBP_NEGATIVE_REGULATION_OF_OXIDATIVE_STRESS_INDUCED_CELL_DEATH | 39 | 0.389712 | 1.117983 | 0.302326 | 0.641127 |
| GOBP_CELLULAR_RESPONSE_TO_OXYGEN_RADICAL | 19 | 0.447759 | 1.117385 | 0.316535 | 0.642411 |
| GOBP_EYE_PHOTORECEPTOR_CELL_DEVELOPMENT | 26 | 0.431712 | 1.116832 | 0.316358 | 0.643588 |
| GOBP_REGULATION_OF_MUSCLE_ORGAN_DEVELOPMENT | 33 | 0.409671 | 1.116681 | 0.329446 | 0.643548 |
| GOBP_NEGATIVE_REGULATION_OF_OSTEOCLAST_DIFFERENTIATION | 22 | 0.442383 | 1.115963 | 0.333333 | 0.645188 |
| GOBP_BODY_MORPHOGENESIS | 41 | 0.385493 | 1.115679 | 0.323486 | 0.645525 |

| | | | | | |
|---|---|---|---|---|---|
| GOBP_GLYCEROLIPID_METABOLIC_PROCESS | 330 | 0.302749 | 1.115662 | 0.202586 | 0.645054 |
| GOBP_DIGESTIVE_SYSTEM_DEVELOPMENT | 100 | 0.339179 | 1.11559 | 0.280566 | 0.644729 |
| GOBP_POSITIVE_REGULATION_OF_INTERFERON_GAMMA_PRODUCTION | 44 | 0.381502 | 1.114658 | 0.288732 | 0.646887 |
| GOBP_REGULATION_OF_NERVOUS_SYSTEM_PROCESS | 114 | 0.332563 | 1.1141 | 0.283767 | 0.647931 |
| GOBP_REGULATION_OF_LYMPHOCYTE_MEDIATED_IMMUNITY | 110 | 0.32801 | 1.113616 | 0.258512 | 0.648847 |
| GOBP_PROTEIN_TARGETING_TO_VACUOLE | 37 | 0.391907 | 1.113128 | 0.286957 | 0.649707 |
| GOBP_REGULATION_OF_SYSTEM_PROCESS | 429 | 0.295843 | 1.112855 | 0.2 | 0.649977 |
| GOBP_ACTIVATION_OF_PROTEIN_KINASE_ACTIVITY | 116 | 0.332989 | 1.112788 | 0.27182 | 0.649645 |
| GOBP_RESPONSE_TO_KETONE | 149 | 0.32409 | 1.112548 | 0.270205 | 0.649831 |
| GOBP_REGULATION_OF_BODY_FLUID_LEVELS | 282 | 0.304479 | 1.112416 | 0.232923 | 0.649738 |
| GOBP_RESPONSE_TO_TOXIC_SUBSTANCE | 179 | 0.314416 | 1.112181 | 0.245327 | 0.6499 |
| GOBP_CELLULAR_LIPID_CATABOLIC_PROCESS | 179 | 0.311299 | 1.111213 | 0.261484 | 0.652232 |
| GOBP_CELLULAR_RESPONSE_TO_EXTERNAL_STIMULUS | 266 | 0.30571 | 1.110859 | 0.241031 | 0.652724 |
| GOBP_T_CELL_RECEPTOR_SIGNALING_PATHWAY | 99 | 0.334072 | 1.110503 | 0.281046 | 0.653257 |
| GOBP_MESONEPHRIC_TUBULE_MORPHOGENESIS | 50 | 0.379192 | 1.110214 | 0.299857 | 0.653598 |
| GOBP_POSITIVE_REGULATION_OF_NATURAL_KILLER_CELL_ACTIVATION | 20 | 0.449474 | 1.110027 | 0.330203 | 0.653603 |
| GOBP_OVARIAN_FOLLICLE_DEVELOPMENT | 45 | 0.379658 | 1.109688 | 0.31686 | 0.654067 |
| GOBP_FOAM_CELL_DIFFERENTIATION | 25 | 0.422258 | 1.10898 | 0.323124 | 0.655657 |
| GOBP_MICROTUBULE_ORGANIZING_CENTER_LOCALIZATION | 32 | 0.401082 | 1.107767 | 0.319572 | 0.658749 |
| GOBP_RESPONSE_TO_ANTIBIOTIC | 42 | 0.381232 | 1.107624 | 0.321429 | 0.658631 |
| GOBP_REGULATION_OF_ACTIN_CYTOSKELETON_REORGANIZATION | 34 | 0.401896 | 1.107477 | 0.3125 | 0.658537 |
| GOBP_CENTRAL_NERVOUS_SYSTEM_NEURON_DEVELOPMENT | 70 | 0.350537 | 1.107307 | 0.30163 | 0.658536 |
| GOBP_REGULATION_OF_G_PROTEIN_COUPLED_RECEPTOR_SIGNALING_PATHWAY | 112 | 0.330384 | 1.10708 | 0.302149 | 0.658699 |
| GOBP_BRANCHED_CHAIN_AMINO_ACID_CATABOLIC_PROCESS | 20 | 0.452249 | 1.106807 | 0.339434 | 0.659008 |
| GOBP_RESPONSE_TO_ETHANOL | 85 | 0.339795 | 1.10661 | 0.295276 | 0.659063 |
| GOBP_T_HELPER_1_CELL_DIFFERENTIATION | 16 | 0.457979 | 1.104702 | 0.344209 | 0.66413 |
| GOBP_CELLULAR_RESPONSE_TO_ATP | 16 | 0.466197 | 1.104666 | 0.343089 | 0.663739 |
| GOBP_MAGNESIUM_ION_TRANSPORT | 17 | 0.46604 | 1.10427 | 0.349259 | 0.664392 |
| GOBP_POSITIVE_REGULATION_OF_PROTEIN_IMPORT | 38 | 0.393111 | 1.104096 | 0.329463 | 0.664387 |
| GOBP_PURINE_NUCLEOSIDE_MONOPHOSPHATE_METABOLIC_PROCESS | 43 | 0.381735 | 1.103485 | 0.326501 | 0.665689 |
| GOBP_SMALL_MOLECULE_CATABOLIC_PROCESS | 308 | 0.297884 | 1.102725 | 0.25247 | 0.667442 |
| GOBP_POSITIVE_REGULATION_OF_CELL_SUBSTRATE_JUNCTION_ORGANIZATION | 29 | 0.407196 | 1.10217 | 0.329134 | 0.668602 |
| GOBP_MEMBRANE_REPOLARIZATION | 40 | 0.372784 | 1.102091 | 0.321075 | 0.668307 |
| GOBP_ANTIGEN_PROCESSING_AND_PRESENTATION | 77 | 0.344482 | 1.101198 | 0.316438 | 0.67038 |
| GOBP_NEGATIVE_REGULATION_OF_PHOSPHORUS_METABOLIC_PROCESS | 375 | 0.292722 | 1.101044 | 0.229614 | 0.670302 |
| GOBP_POSITIVE_REGULATION_OF_PROTEIN_POLYUBIQUITINATION | 15 | 0.470099 | 1.100896 | 0.333898 | 0.670234 |
| GOBP_NEGATIVE_REGULATION_OF_ADAPTIVE_IMMUNE_RESPONSE | 39 | 0.386307 | 1.100675 | 0.335277 | 0.670351 |
| GOBP_FIBROBLAST_GROWTH_FACTOR_RECEPTOR_SIGNALING_PATHWAY | 66 | 0.358785 | 1.100649 | 0.317373 | 0.669905 |
| GOBP_PALLIUM_DEVELOPMENT | 141 | 0.319494 | 1.100357 | 0.283111 | 0.670246 |
| GOBP_WOUND_HEALING_SPREADING_OF_EPIDERMAL_CELLS | 16 | 0.460117 | 1.100059 | 0.361809 | 0.670647 |
| GOBP_REGULATION_OF_CELL_JUNCTION_ASSEMBLY | 175 | 0.310343 | 1.099913 | 0.295537 | 0.670542 |
| GOBP_NEGATIVE_CHEMOTAXIS | 44 | 0.374606 | 1.099881 | 0.307365 | 0.670124 |
| GOBP_REGULATION_OF_CELLULAR_CARBOHYDRATE_METABOLIC_PROCESS | 123 | 0.324922 | 1.099011 | 0.280788 | 0.672154 |
| GOBP_MACROPHAGE_MIGRATION | 31 | 0.409147 | 1.098821 | 0.355556 | 0.672169 |
| GOBP_REGULATION_OF_B_CELL_ACTIVATION | 96 | 0.332183 | 1.09847 | 0.297845 | 0.672667 |
| GOBP_RESPONSE_TO_NERVE_GROWTH_FACTOR | 46 | 0.377717 | 1.098462 | 0.315714 | 0.67217 |
| GOBP_INTRINSIC_APOPTOTIC_SIGNALING_PATHWAY_BY_P53_CLASS_MEDIATOR | 74 | 0.345472 | 1.098401 | 0.296875 | 0.671856 |
| GOBP_NEGATIVE_REGULATION_OF_PROTEIN_SERINE_THREONINE_KINASE_ACTIVITY | 101 | 0.332197 | 1.098221 | 0.300512 | 0.671864 |
| GOBP_REGULATION_OF_TRANSPORTER_ACTIVITY | 228 | 0.30096 | 1.097053 | 0.261067 | 0.6749 |
| GOBP_NEGATIVE_REGULATION_OF_DNA_BIOSYNTHETIC_PROCESS | 35 | 0.387225 | 1.096692 | 0.340491 | 0.675452 |
| GOBP_FATTY_ACID_METABOLIC_PROCESS | 295 | 0.298057 | 1.096637 | 0.260382 | 0.675113 |
| GOBP_PEPTIDYL_TYROSINE_AUTOPHOSPHORYLATION | 18 | 0.44971 | 1.096023 | 0.341463 | 0.676426 |
| GOBP_POSITIVE_REGULATION_OF_MEMBRANE_PERMEABILITY | 41 | 0.380197 | 1.096004 | 0.314244 | 0.675963 |
| GOBP_TRABECULA_FORMATION | 21 | 0.435944 | 1.095552 | 0.342146 | 0.676843 |
| GOBP_RESPONSE_TO_VIRUS | 290 | 0.297655 | 1.095346 | 0.257709 | 0.67693 |
| GOBP_CELL_DEATH_IN_RESPONSE_TO_OXIDATIVE_STRESS | 79 | 0.336819 | 1.095224 | 0.31444 | 0.676801 |
| GOBP_RESPONSE_TO_LEUKEMIA_INHIBITORY_FACTOR | 90 | 0.333194 | 1.094864 | 0.299223 | 0.677383 |
| GOBP_AMYLOID_FIBRIL_FORMATION | 27 | 0.413426 | 1.09468 | 0.339422 | 0.677414 |
| GOBP_HORMONE_TRANSPORT | 219 | 0.304417 | 1.093449 | 0.261124 | 0.680613 |
| GOBP_CARDIAC_MUSCLE_CELL_MEMBRANE_REPOLARIZATION | 31 | 0.400777 | 1.0934 | 0.352679 | 0.680243 |
| GOBP_LYMPHOCYTE_ACTIVATION_INVOLVED_IN_IMMUNE_RESPONSE | 133 | 0.320522 | 1.093388 | 0.285894 | 0.679764 |
| GOBP_REGULATION_OF_LAMELLIPODIUM_ASSEMBLY | 34 | 0.396196 | 1.093386 | 0.338438 | 0.679252 |
| GOBP_REGULATION_OF_AUTOPHAGY | 313 | 0.296194 | 1.093057 | 0.269945 | 0.679765 |
| GOBP_REGULATION_OF_ESTABLISHMENT_OF_PROTEIN_LOCALIZATION | 439 | 0.287876 | 1.093001 | 0.234177 | 0.679437 |
| GOBP_REGULATION_OF_EPIDERMIS_DEVELOPMENT | 51 | 0.360748 | 1.092743 | 0.335294 | 0.679685 |
| GOBP_REGULATION_OF_AUTOPHAGY_OF_MITOCHONDRION | 36 | 0.38646 | 1.092329 | 0.345534 | 0.68045 |
| GOBP_REGULATION_OF_CELLULAR_RESPIRATION | 42 | 0.382077 | 1.092214 | 0.34963 | 0.680272 |
| GOBP_VESICLE_TARGETING_TO_FROM_OR_WITHIN_GOLGI | 21 | 0.430264 | 1.092206 | 0.363495 | 0.679783 |
| GOBP_ERBB_SIGNALING_PATHWAY | 107 | 0.32629 | 1.092125 | 0.300885 | 0.679515 |
| GOBP_NEGATIVE_REGULATION_OF_VIRAL_ENTRY_INTO_HOST_CELL | 18 | 0.450296 | 1.09143 | 0.360374 | 0.681025 |
| GOBP_VASCULAR_ENDOTHELIAL_CELL_PROLIFERATION | 17 | 0.457106 | 1.091347 | 0.372852 | 0.680744 |
| GOBP_POSITIVE_REGULATION_OF_INTRACELLULAR_PROTEIN_TRANSPORT | 144 | 0.319207 | 1.091142 | 0.318681 | 0.680804 |
| GOBP_GLYCEROPHOSPHOLIPID_METABOLIC_PROCESS | 269 | 0.299304 | 1.090694 | 0.272216 | 0.681642 |
| GOBP_PROTEIN_TARGETING_TO_LYSOSOME | 23 | 0.419766 | 1.090178 | 0.350467 | 0.682715 |
| GOBP_POSITIVE_REGULATION_OF_T_CELL_MIGRATION | 21 | 0.421676 | 1.090131 | 0.34551 | 0.682327 |
| GOBP_RESPONSE_TO_ESTROGEN | 52 | 0.362573 | 1.089543 | 0.334262 | 0.683573 |
| GOBP_REGULATION_OF_RESPONSE_TO_REACTIVE_OXYGEN_SPECIES | 28 | 0.402023 | 1.089324 | 0.343548 | 0.683732 |
| GOBP_POSITIVE_REGULATION_OF_PEPTIDASE_ACTIVITY | 162 | 0.308785 | 1.088816 | 0.28487 | 0.684723 |
| GOBP_NEGATIVE_REGULATION_OF_STRESS_ACTIVATED_PROTEIN_KINASE_SIGNALING_C | 44 | 0.377475 | 1.087658 | 0.35044 | 0.687621 |
| GOBP_NATURAL_KILLER_CELL_ACTIVATION | 54 | 0.360838 | 1.087406 | 0.328488 | 0.687861 |
| GOBP_MEIOTIC_CHROMOSOME_SEGREGATION | 72 | 0.344456 | 1.087263 | 0.320548 | 0.687793 |
| GOBP_B_CELL_ACTIVATION | 193 | 0.302862 | 1.086941 | 0.28284 | 0.688189 |
| GOBP_REGULATION_OF_SMALL_MOLECULE_METABOLIC_PROCESS | 269 | 0.297551 | 1.086912 | 0.282998 | 0.687733 |
| GOBP_AMINE_METABOLIC_PROCESS | 94 | 0.330518 | 1.086886 | 0.319574 | 0.687292 |
| GOBP_REGULATION_OF_PEPTIDYL_TYROSINE_PHOSPHORYLATION | 186 | 0.306667 | 1.086711 | 0.298472 | 0.687298 |
| GOBP_POSITIVE_REGULATION_OF_CARTILAGE_DEVELOPMENT | 27 | 0.402219 | 1.086232 | 0.360429 | 0.688214 |
| GOBP_NEGATIVE_REGULATION_OF_LEUKOCYTE_MIGRATION | 31 | 0.401292 | 1.086157 | 0.358824 | 0.68792 |
| GOBP_HEPATICOBILIARY_SYSTEM_DEVELOPMENT | 108 | 0.32297 | 1.08586 | 0.323155 | 0.688289 |
| GOBP_PROTEIN_AUTOPHOSPHORYLATION | 207 | 0.299987 | 1.085831 | 0.282201 | 0.687863 |
| GOBP_MYELOID_CELL_DIFFERENTIATION | 316 | 0.292124 | 1.085497 | 0.283649 | 0.688353 |
| GOBP_REGULATION_OF_ANION_TRANSPORT | 70 | 0.339451 | 1.084985 | 0.315291 | 0.689322 |
| GOBP_PEPTIDYL_TYROSINE_MODIFICATION | 289 | 0.29458 | 1.084716 | 0.275901 | 0.689658 |
| GOBP_BIOLOGICAL_PROCESS_INVOLVED_IN_INTRASPECIES_INTERACTION_BETWEEN_OI | 42 | 0.374841 | 1.084295 | 0.351003 | 0.690329 |
| GOBP_MEMBRANE_INVAGINATION | 60 | 0.354192 | 1.083587 | 0.329184 | 0.691905 |
| GOBP_POSITIVE_REGULATION_OF_ENDOTHELIAL_CELL_CHEMOTAXIS | 15 | 0.457599 | 1.08349 | 0.368601 | 0.691662 |
| GOBP_FC_RECEPTOR_MEDIATED_STIMULATORY_SIGNALING_PATHWAY | 25 | 0.41104 | 1.08213 | 0.363636 | 0.695146 |
| GOBP_SARCOPLASMIC_RETICULUM_CALCIUM_ION_TRANSPORT | 29 | 0.393385 | 1.081897 | 0.352665 | 0.695316 |
| GOBP_POSITIVE_REGULATION_OF_INTERLEUKIN_1_PRODUCTION | 45 | 0.365179 | 1.081622 | 0.360176 | 0.695642 |
| GOBP_RIBONUCLEOSIDE_MONOPHOSPHATE_METABOLIC_PROCESS | 55 | 0.356899 | 1.08156 | 0.361227 | 0.695298 |
| GOBP_REGULATION_OF_ENDOPLASMIC_RETICULUM_STRESS_INDUCED_INTRINSIC_APO | 28 | 0.397634 | 1.081065 | 0.348653 | 0.696293 |
| GOBP_RESPONSE_TO_STEROID_HORMONE | 263 | 0.296056 | 1.081003 | 0.310155 | 0.695982 |
| GOBP_DEVELOPMENTAL_MATURATION | 235 | 0.299685 | 1.080922 | 0.3 | 0.695712 |
| GOBP_POSITIVE_REGULATION_OF_ANION_TRANSPORT | 38 | 0.380555 | 1.080919 | 0.362687 | 0.695205 |
| GOBP_REGULATION_OF_OSSIFICATION | 92 | 0.333401 | 1.08047 | 0.333763 | 0.695992 |
| GOBP_CHEMOKINE_PRODUCTION | 60 | 0.3519 | 1.080403 | 0.351678 | 0.695704 |
| GOBP_NEGATIVE_REGULATION_OF_VIRAL_LIFE_CYCLE | 21 | 0.429562 | 1.080253 | 0.381459 | 0.695651 |
| GOBP_ADRENAL_GLAND_DEVELOPMENT | 24 | 0.414617 | 1.079381 | 0.353752 | 0.697766 |
| GOBP_REGULATION_OF_PROTEIN_LOCALIZATION_TO_MEMBRANE | 155 | 0.311467 | 1.079366 | 0.320293 | 0.697293 |
| GOBP_MUCOPOLYSACCHARIDE_METABOLIC_PROCESS | 76 | 0.336364 | 1.079245 | 0.344 | 0.697118 |
| GOBP_HOMOPHILIC_CELL_ADHESION_VIA_PLASMA_MEMBRANE_ADHESION_MOLECULES | 143 | 0.310635 | 1.078934 | 0.313559 | 0.697552 |
| GOBP_BONE_RESORPTION | 45 | 0.368136 | 1.078874 | 0.344477 | 0.69721 |
| GOBP_POSITIVE_REGULATION_OF_DNA_TEMPLATED_TRANSCRIPTION_INITIATION | 43 | 0.373986 | 1.078786 | 0.346211 | 0.696967 |
| GOBP_NEUROINFLAMMATORY_RESPONSE | 21 | 0.423612 | 1.078395 | 0.356589 | 0.69756 |
| GOBP_CYCLIC_NUCLEOTIDE_MEDIATED_SIGNALING | 60 | 0.351664 | 1.078124 | 0.338192 | 0.697834 |
| GOBP_REGULATION_OF_HEMATOPOIETIC_PROGENITOR_CELL_DIFFERENTIATION | 32 | 0.390292 | 1.078098 | 0.342647 | 0.697388 |
| GOBP_ZINC_ION_HOMEOSTASIS | 29 | 0.396237 | 1.077898 | 0.362406 | 0.697452 |
| GOBP_REGULATION_OF_ERYTHROCYTE_DIFFERENTIATION | 41 | 0.373414 | 1.077709 | 0.350224 | 0.697514 |
| GOBP_KERATINOCYTE_PROLIFERATION | 34 | 0.385815 | 1.077219 | 0.365892 | 0.698426 |
| GOBP_REGULATION_OF_CILIUM_DEPENDENT_CELL_MOTILITY | 15 | 0.465358 | 1.076499 | 0.401284 | 0.700031 |
| GOBP_REGULATION_OF_MACROPHAGE_CHEMOTAXIS | 15 | 0.461245 | 1.076472 | 0.392799 | 0.699588 |
| GOBP_CELL_DIFFERENTIATION_IN_SPINAL_CORD | 37 | 0.386263 | 1.076354 | 0.347181 | 0.699428 |
| GOBP_VITAMIN_METABOLIC_PROCESS | 83 | 0.335986 | 1.075964 | 0.336461 | 0.700018 |
| GOBP_VENTRAL_SPINAL_CORD_DEVELOPMENT | 35 | 0.383255 | 1.075919 | 0.379161 | 0.699643 |
| GOBP_REACTIVE_NITROGEN_SPECIES_METABOLIC_PROCESS | 59 | 0.355545 | 1.075209 | 0.369081 | 0.701274 |
| GOBP_LYMPHOCYTE_MEDIATED_IMMUNITY | 181 | 0.30355 | 1.075132 | 0.322543 | 0.700981 |
| GOBP_MEMBRANE_REPOLARIZATION_DURING_CARDIAC_MUSCLE_CELL_ACTION_POTEN | 18 | 0.45495 | 1.075081 | 0.388084 | 0.700617 |
| GOBP_NEGATIVE_REGULATION_OF_TISSUE_REMODELING | 15 | 0.463962 | 1.074517 | 0.375817 | 0.701712 |
| GOBP_REGULATION_OF_PHOSPHOLIPASE_ACTIVITY | 46 | 0.365122 | 1.07398 | 0.377168 | 0.702795 |
| GOBP_INTERMEDIATE_FILAMENT_ORGANIZATION | 18 | 0.437198 | 1.073605 | 0.363339 | 0.703391 |
| GOBP_RESPONSE_TO_STEROL | 22 | 0.427886 | 1.073497 | 0.409583 | 0.703215 |
| GOBP_CENTRAL_NERVOUS_SYSTEM_NEURON_DIFFERENTIATION | 136 | 0.308959 | 1.072594 | 0.328767 | 0.705377 |
| GOBP_NEGATIVE_REGULATION_OF_HEMOPOIESIS | 75 | 0.33703 | 1.072364 | 0.345382 | 0.705518 |
| GOBP_REGULATION_OF_LIPID_METABOLIC_PROCESS | 260 | 0.293389 | 1.071654 | 0.317621 | 0.707156 |
| GOBP_REGULATION_OF_AMINO_ACID_TRANSPORT | 35 | 0.385879 | 1.071647 | 0.38253 | 0.706671 |
| GOBP_PROTEIN_DEMETHYLATION | 30 | 0.396081 | 1.071217 | 0.390318 | 0.707452 |
| GOBP_INORGANIC_ION_IMPORT_ACROSS_PLASMA_MEMBRANE | 82 | 0.333302 | 1.070965 | 0.348285 | 0.707719 |
| GOBP_POSITIVE_REGULATION_OF_MYELOID_LEUKOCYTE_CYTOKINE_PRODUCTION_INVO | 20 | 0.424247 | 1.069942 | 0.368341 | 0.710279 |
| GOBP_RETINA_DEVELOPMENT_IN_CAMERA_TYPE_EYE | 118 | 0.315851 | 1.069825 | 0.340102 | 0.710147 |
| GOBP_POSITIVE_REGULATION_OF_EXTRINSIC_APOPTOTIC_SIGNALING_PATHWAY | 42 | 0.370146 | 1.069095 | 0.369723 | 0.711868 |
| GOBP_CARBOHYDRATE_HOMEOSTASIS | 182 | 0.302903 | 1.068968 | 0.350236 | 0.71171 |
| GOBP_REGULATION_OF_CARBOHYDRATE_METABOLIC_PROCESS | 152 | 0.307957 | 1.068862 | 0.324029 | 0.711512 |
| GOBP_PROTEIN_KINASE_A_SIGNALING | 25 | 0.403958 | 1.067323 | 0.372855 | 0.715553 |
| GOBP_POSITIVE_REGULATION_OF_GROWTH | 202 | 0.297333 | 1.067179 | 0.317857 | 0.715492 |
| GOBP_POSITIVE_REGULATION_OF_PROTEIN_TARGETING_TO_MEMBRANE | 27 | 0.404672 | 1.067151 | 0.370717 | 0.715064 |
| GOBP_POSITIVE_REGULATION_OF_VIRAL_GENOME_REPLICATION | 28 | 0.400596 | 1.067002 | 0.358712 | 0.714985 |
| GOBP_NEGATIVE_REGULATION_OF_LIPID_BIOSYNTHETIC_PROCESS | 47 | 0.36365 | 1.066833 | 0.376281 | 0.714968 |
| GOBP_POSITIVE_REGULATION_OF_EPITHELIAL_CELL_PROLIFERATION | 145 | 0.30992 | 1.065887 | 0.329723 | 0.717336 |
| GOBP_REGULATION_OF_NITRIC_OXIDE_SYNTHASE_ACTIVITY | 27 | 0.396216 | 1.065858 | 0.390625 | 0.716907 |
| GOBP_CEREBRAL_CORTEX_DEVELOPMENT | 98 | 0.322976 | 1.065425 | 0.334184 | 0.717658 |
| GOBP_REGULATION_OF_TRIGLYCERIDE_BIOSYNTHETIC_PROCESS | 18 | 0.431128 | 1.065244 | 0.381538 | 0.717644 |
| GOBP_LONG_CHAIN_FATTY_ACID_TRANSPORT | 47 | 0.358935 | 1.065046 | 0.359238 | 0.717729 |
| GOBP_ALPHA_AMINO_ACID_CATABOLIC_PROCESS | 73 | 0.341186 | 1.064875 | 0.357527 | 0.717701 |
| GOBP_REGULATION_OF_RESPONSE_TO_CYTOKINE_STIMULUS | 131 | 0.309532 | 1.064849 | 0.324937 | 0.717254 |
| GOBP_CELLULAR_RESPONSE_TO_ZINC_ION | 16 | 0.449298 | 1.06458 | 0.408547 | 0.717566 |
| GOBP_REGULATION_OF_DENDRITE_EXTENSION | 23 | 0.410873 | 1.063994 | 0.364366 | 0.718848 |
| GOBP_CARTILAGE_DEVELOPMENT | 149 | 0.308684 | 1.063845 | 0.346348 | 0.718764 |
| GOBP_REGULATION_OF_EXTRACELLULAR_MATRIX_ORGANIZATION | 41 | 0.372405 | 1.063452 | 0.352855 | 0.719451 |

| | | | | | |
|---|---|---|---|---|---|
| GOBP_MEMORY | 105 | 0.318813 | 1.063257 | 0.358876 | 0.719528 |
| GOBP_REGULATION_OF_APOPTOTIC_SIGNALING_PATHWAY | 314 | 0.289138 | 1.062627 | 0.324973 | 0.720845 |
| GOBP_REGULATION_OF_LAMELLIPODIUM_ORGANIZATION | 46 | 0.363725 | 1.062445 | 0.373027 | 0.720896 |
| GOBP_CELLULAR_RESPONSE_TO_IONIZING_RADIATION | 69 | 0.337291 | 1.062241 | 0.369919 | 0.721025 |
| GOBP_POSITIVE_REGULATION_OF_INTRINSIC_APOPTOTIC_SIGNALING_PATHWAY | 52 | 0.356016 | 1.062126 | 0.349794 | 0.720848 |
| GOBP_CELLULAR_AMINO_ACID_CATABOLIC_PROCESS | 88 | 0.326319 | 1.061786 | 0.37751 | 0.721335 |
| GOBP_PHOSPHATIDYLSERINE_METABOLIC_PROCESS | 21 | 0.423165 | 1.061445 | 0.401305 | 0.721874 |
| GOBP_CELL_SURFACE_RECEPTOR_SIGNALING_PATHWAY_INVOLVED_IN_HEART_DEVEL( | 25 | 0.406206 | 1.06126 | 0.398074 | 0.72191 |
| GOBP_PROSTATE_GLAND_DEVELOPMENT | 34 | 0.378239 | 1.061037 | 0.391104 | 0.722055 |
| GOBP_RESPONSE_TO_FIBROBLAST_GROWTH_FACTOR | 92 | 0.325074 | 1.061003 | 0.349677 | 0.721633 |
| GOBP_PLACENTA_BLOOD_VESSEL_DEVELOPMENT | 29 | 0.401806 | 1.060938 | 0.382883 | 0.721334 |
| GOBP_POSITIVE_REGULATION_OF_NEUROTRANSMITTER_TRANSPORT | 20 | 0.431203 | 1.060916 | 0.392744 | 0.720877 |
| GOBP_SECONDARY_METABOLIC_PROCESS | 36 | 0.371619 | 1.060644 | 0.377219 | 0.721212 |
| GOBP_INSULIN_LIKE_GROWTH_FACTOR_RECEPTOR_SIGNALING_PATHWAY | 27 | 0.402927 | 1.060245 | 0.390769 | 0.721879 |
| GOBP_CELLULAR_MODIFIED_AMINO_ACID_CATABOLIC_PROCESS | 23 | 0.404455 | 1.06006 | 0.388037 | 0.721954 |
| GOBP_POSITIVE_REGULATION_OF_FOCAL_ADHESION_ASSEMBLY | 24 | 0.404363 | 1.060054 | 0.403756 | 0.721464 |
| GOBP_IMMUNE_RESPONSE_REGULATING_SIGNALING_PATHWAY | 290 | 0.285705 | 1.05992 | 0.328918 | 0.721355 |
| GOBP_NEGATIVE_REGULATION_OF_ION_TRANSMEMBRANE_TRANSPORT | 68 | 0.339867 | 1.059905 | 0.371812 | 0.720891 |
| GOBP_TRANSMISSION_OF_NERVE_IMPULSE | 52 | 0.352461 | 1.059592 | 0.377168 | 0.721342 |
| GOBP_DEOXYRIBOSE_PHOSPHATE_CATABOLIC_PROCESS | 26 | 0.39565 | 1.058645 | 0.394984 | 0.723639 |
| GOBP_SPINAL_CORD_MOTOR_NEURON_DIFFERENTIATION | 22 | 0.413913 | 1.058275 | 0.377236 | 0.724252 |
| GOBP_NEGATIVE_REGULATION_OF_TRANSMEMBRANE_TRANSPORT | 90 | 0.327075 | 1.058047 | 0.368153 | 0.72444 |
| GOBP_REGULATION_OF_EXTRINSIC_APOPTOTIC_SIGNALING_PATHWAY_IN_ABSENCE_OF | 35 | 0.386119 | 1.058047 | 0.369208 | 0.723934 |
| GOBP_T_CELL_LINEAGE_COMMITMENT | 16 | 0.442352 | 1.057151 | 0.40536 | 0.726003 |
| GOBP_POSITIVE_REGULATION_OF_PROTEIN_KINASE_ACTIVITY | 326 | 0.279926 | 1.056759 | 0.339499 | 0.726638 |
| GOBP_OSTEOBLAST_DIFFERENTIATION | 192 | 0.296826 | 1.056698 | 0.362664 | 0.726298 |
| GOBP_DIGESTIVE_SYSTEM_PROCESS | 58 | 0.343421 | 1.05614 | 0.362984 | 0.727474 |
| GOBP_REGULATION_OF_MORPHOGENESIS_OF_AN_EPITHELIUM | 48 | 0.357538 | 1.056069 | 0.393419 | 0.727201 |
| GOBP_GLUCAN_BIOSYNTHETIC_PROCESS | 36 | 0.370829 | 1.056043 | 0.395044 | 0.726773 |
| GOBP_POSITIVE_REGULATION_OF_ERBB_SIGNALING_PATHWAY | 28 | 0.390933 | 1.055298 | 0.394495 | 0.72842 |
| GOBP_REGULATION_OF_CENTROSOME_CYCLE | 49 | 0.348664 | 1.055128 | 0.365782 | 0.728431 |
| GOBP_NEGATIVE_REGULATION_OF_REACTIVE_OXYGEN_SPECIES_METABOLIC_PROCESS | 35 | 0.369169 | 1.054318 | 0.388724 | 0.730312 |
| GOBP_FIBROBLAST_PROLIFERATION | 66 | 0.335941 | 1.053903 | 0.366667 | 0.730998 |
| GOBP_RESPONSE_TO_MONOSACCHARIDE | 151 | 0.30393 | 1.053572 | 0.374241 | 0.731446 |
| GOBP_REGULATION_OF_EXOCYTOSIS | 176 | 0.295137 | 1.052899 | 0.348193 | 0.732953 |
| GOBP_REGULATION_OF_MACROPHAGE_MIGRATION | 22 | 0.415999 | 1.052828 | 0.398773 | 0.732637 |
| GOBP_POSITIVE_REGULATION_OF_FAT_CELL_DIFFERENTIATION | 59 | 0.338498 | 1.052219 | 0.371669 | 0.733959 |
| GOBP_LIMBIC_SYSTEM_DEVELOPMENT | 87 | 0.323464 | 1.052082 | 0.359008 | 0.733842 |
| GOBP_NUCLEOTIDE_SUGAR_BIOSYNTHETIC_PROCESS | 17 | 0.431234 | 1.051932 | 0.397833 | 0.733755 |
| GOBP_NEGATIVE_REGULATION_OF_AMYLOID_PRECURSOR_PROTEIN_CATABOLIC_PROC | 20 | 0.420256 | 1.051827 | 0.392465 | 0.733545 |
| GOBP_RESPONSE_TO_XENOBIOTIC_STIMULUS | 303 | 0.286547 | 1.051497 | 0.355221 | 0.733982 |
| GOBP_PROTEIN_O_LINKED_GLYCOSYLATION | 80 | 0.325721 | 1.050666 | 0.382393 | 0.735956 |
| GOBP_POSITIVE_REGULATION_OF_SECRETION | 236 | 0.289373 | 1.050665 | 0.362471 | 0.73545 |
| GOBP_LYMPHOCYTE_HOMEOSTASIS | 49 | 0.352266 | 1.050527 | 0.411189 | 0.735374 |
| GOBP_NEGATIVE_REGULATION_OF_PROTEIN_MODIFICATION_PROCESS | 454 | 0.277378 | 1.049996 | 0.341799 | 0.736455 |
| GOBP_RESPONSE_TO_TEMPERATURE_STIMULUS | 152 | 0.300728 | 1.049969 | 0.37006 | 0.736029 |
| GOBP_GLYCEROLIPID_BIOSYNTHETIC_PROCESS | 224 | 0.28837 | 1.049822 | 0.365604 | 0.735973 |
| GOBP_BRANCHED_CHAIN_AMINO_ACID_METABOLIC_PROCESS | 23 | 0.402038 | 1.049798 | 0.416933 | 0.735544 |
| GOBP_POSITIVE_REGULATION_OF_DENDRITE_DEVELOPMENT | 15 | 0.450493 | 1.049752 | 0.410802 | 0.73518 |
| GOBP_NEGATIVE_REGULATION_OF_ERK1_AND_ERK2_CASCADE | 62 | 0.341838 | 1.049699 | 0.380753 | 0.734842 |
| GOBP_MEIOTIC_CHROMOSOME_SEPARATION | 21 | 0.415851 | 1.048112 | 0.419811 | 0.73904 |
| GOBP_REGULATION_OF_VIRAL_GENOME_REPLICATION | 70 | 0.33092 | 1.04781 | 0.373315 | 0.739446 |
| GOBP_MESENCHYMAL_CELL_PROLIFERATION | 30 | 0.390143 | 1.047775 | 0.416794 | 0.739041 |
| GOBP_NEGATIVE_REGULATION_OF_TRANSFERASE_ACTIVITY | 239 | 0.288309 | 1.047738 | 0.352413 | 0.738662 |
| GOBP_POSITIVE_REGULATION_OF_SUPRAMOLECULAR_FIBER_ORGANIZATION | 159 | 0.298429 | 1.04687 | 0.357227 | 0.740761 |
| GOBP_REGULATION_OF_MONOOXYGENASE_ACTIVITY | 39 | 0.364747 | 1.04609 | 0.392749 | 0.742541 |
| GOBP_ANTIGEN_PROCESSING_AND_PRESENTATION_OF_EXOGENOUS_PEPTIDE_ANTIGEI | 19 | 0.426304 | 1.045773 | 0.424765 | 0.742957 |
| GOBP_NEGATIVE_REGULATION_OF_VIRAL_PROCESS | 72 | 0.328199 | 1.04535 | 0.381018 | 0.743711 |
| GOBP_ANTIGEN_PROCESSING_AND_PRESENTATION_OF_EXOGENOUS_ANTIGEN | 30 | 0.380011 | 1.045214 | 0.413686 | 0.743615 |
| GOBP_POSITIVE_REGULATION_OF_CYTOSKELETON_ORGANIZATION | 176 | 0.296587 | 1.044525 | 0.383626 | 0.745124 |
| GOBP_CARDIAC_MUSCLE_CELL_ACTION_POTENTIAL_INVOLVED_IN_CONTRACTION | 46 | 0.352575 | 1.044521 | 0.412463 | 0.744635 |
| GOBP_EPITHELIAL_STRUCTURE_MAINTENANCE | 22 | 0.410109 | 1.044465 | 0.407524 | 0.744286 |
| GOBP_REGULATION_OF_MYELOID_LEUKOCYTE_MEDIATED_IMMUNITY | 45 | 0.356826 | 1.044386 | 0.380117 | 0.744022 |
| GOBP_BASEMENT_MEMBRANE_ORGANIZATION | 29 | 0.38705 | 1.043948 | 0.405819 | 0.744844 |
| GOBP_TUMOR_NECROSIS_FACTOR_SUPERFAMILY_CYTOKINE_PRODUCTION | 113 | 0.308074 | 1.043447 | 0.377113 | 0.745742 |
| GOBP_POSITIVE_REGULATION_OF_SIGNAL_TRANSDUCTION_BY_P53_CLASS_MEDIATOR | 26 | 0.396819 | 1.043243 | 0.413433 | 0.74585 |
| GOBP_DEVELOPMENTAL_INDUCTION | 27 | 0.38838 | 1.043002 | 0.402124 | 0.74602 |
| GOBP_POSITIVE_REGULATION_OF_NUCLEOTIDE_BIOSYNTHETIC_PROCESS | 17 | 0.43886 | 1.042533 | 0.409023 | 0.746893 |
| GOBP_REGULATION_OF_POTASSIUM_ION_TRANSMEMBRANE_TRANSPORTER_ACTIVITY | 44 | 0.355214 | 1.042523 | 0.392655 | 0.746414 |
| GOBP_REGULATION_OF_GTPASE_ACTIVITY | 313 | 0.277242 | 1.042179 | 0.380435 | 0.746886 |
| GOBP_POSITIVE_REGULATION_OF_PEPTIDYL_TYROSINE_PHOSPHORYLATION | 126 | 0.303845 | 1.042157 | 0.37218 | 0.746447 |
| GOBP_INSULIN_SECRETION | 140 | 0.303634 | 1.041397 | 0.384328 | 0.74818 |
| GOBP_RESPONSE_TO_STARVATION | 172 | 0.292991 | 1.041256 | 0.383255 | 0.748053 |
| GOBP_CELLULAR_RESPONSE_TO_XENOBIOTIC_STIMULUS | 112 | 0.310949 | 1.040947 | 0.375949 | 0.748491 |
| GOBP_TERTIARY_ALCOHOL_METABOLIC_PROCESS | 17 | 0.430221 | 1.04092 | 0.397611 | 0.748059 |
| GOBP_NEUROMUSCULAR_PROCESS_CONTROLLING_BALANCE | 38 | 0.364655 | 1.040426 | 0.398507 | 0.749036 |
| GOBP_INTERLEUKIN_13_PRODUCTION | 15 | 0.440396 | 1.039956 | 0.422371 | 0.749853 |
| GOBP_NON_MEMBRANE_BOUNDED_ORGANELLE_ASSEMBLY | 356 | 0.278821 | 1.039805 | 0.372737 | 0.749809 |
| GOBP_ANTIGEN_PROCESSING_AND_PRESENTATION_OF_EXOGENOUS_PEPTIDE_ANTIGEI | 26 | 0.389377 | 1.039782 | 0.401216 | 0.749372 |
| GOBP_CELL_JUNCTION_ASSEMBLY | 364 | 0.27812 | 1.038828 | 0.37897 | 0.751722 |
| GOBP_MAINTENANCE_OF_BLOOD_BRAIN_BARRIER | 31 | 0.377014 | 1.038657 | 0.418182 | 0.75173 |
| GOBP_REGULATION_OF_ORGANIC_ACID_TRANSPORT | 58 | 0.341227 | 1.038455 | 0.429986 | 0.751803 |
| GOBP_REGULATION_OF_CELL_CYCLE_G1_S_PHASE_TRANSITION | 144 | 0.300629 | 1.037914 | 0.389769 | 0.752882 |
| GOBP_HYDROGEN_PEROXIDE_METABOLIC_PROCESS | 32 | 0.375136 | 1.037702 | 0.409861 | 0.753003 |
| GOBP_TUMOR_NECROSIS_FACTOR_MEDIATED_SIGNALING_PATHWAY | 85 | 0.319501 | 1.037669 | 0.387728 | 0.752589 |
| GOBP_REGULATION_OF_ENDOPLASMIC_RETICULUM_UNFOLDED_PROTEIN_RESPONSE | 28 | 0.388662 | 1.037026 | 0.42812 | 0.753925 |
| GOBP_REGULATION_OF_TUMOR_NECROSIS_FACTOR_MEDIATED_SIGNALING_PATHWAY | 45 | 0.351892 | 1.036509 | 0.416667 | 0.754912 |
| GOBP_NEGATIVE_REGULATION_OF_NEURON_DIFFERENTIATION | 54 | 0.341348 | 1.036177 | 0.407353 | 0.755373 |
| GOBP_CATECHOL_CONTAINING_COMPOUND_METABOLIC_PROCESS | 46 | 0.353605 | 1.0357 | 0.406907 | 0.756281 |
| GOBP_NEGATIVE_REGULATION_OF_OXIDATIVE_STRESS_INDUCED_NEURON_DEATH | 17 | 0.436036 | 1.035615 | 0.427621 | 0.756022 |
| GOBP_POSITIVE_REGULATION_OF_SUBSTRATE_ADHESION_DEPENDENT_CELL_SPREADI | 36 | 0.369318 | 1.035484 | 0.400307 | 0.755926 |
| GOBP_REGULATION_OF_TRIGLYCERIDE_METABOLIC_PROCESS | 30 | 0.38207 | 1.034892 | 0.410769 | 0.757163 |
| GOBP_CELLULAR_KETONE_METABOLIC_PROCESS | 155 | 0.296732 | 1.034687 | 0.405172 | 0.757247 |
| GOBP_POSITIVE_REGULATION_OF_LEUKOCYTE_MEDIATED_IMMUNITY | 90 | 0.316374 | 1.034454 | 0.385027 | 0.757452 |
| GOBP_VIRAL_GENOME_REPLICATION | 110 | 0.308306 | 1.034343 | 0.407022 | 0.757309 |
| GOBP_MICROGLIAL_CELL_ACTIVATION | 26 | 0.386791 | 1.034197 | 0.410016 | 0.757211 |
| GOBP_HIPPO_SIGNALING | 39 | 0.367297 | 1.034014 | 0.428358 | 0.757247 |
| GOBP_CARBOHYDRATE_DERIVATIVE_CATABOLIC_PROCESS | 142 | 0.297421 | 1.033899 | 0.404177 | 0.757063 |
| GOBP_TRANSCYTOSIS | 18 | 0.421407 | 1.03386 | 0.418985 | 0.756669 |
| GOBP_FC_EPSILON_RECEPTOR_SIGNALING_PATHWAY | 20 | 0.416676 | 1.032919 | 0.422297 | 0.758934 |
| GOBP_POSITIVE_REGULATION_OF_CHONDROCYTE_DIFFERENTIATION | 17 | 0.422463 | 1.032842 | 0.414286 | 0.758643 |
| GOBP_NEGATIVE_REGULATION_OF_PROTEIN_KINASE_B_SIGNALING | 38 | 0.362679 | 1.032618 | 0.399711 | 0.758777 |
| GOBP_POSITIVE_REGULATION_OF_CELL_KILLING | 34 | 0.37663 | 1.032012 | 0.442922 | 0.759992 |
| GOBP_REGULATION_OF_MUSCLE_HYPERTROPHY | 45 | 0.349436 | 1.031676 | 0.425591 | 0.760481 |
| GOBP_VASCULAR_ASSOCIATED_SMOOTH_MUSCLE_CONTRACTION | 19 | 0.422416 | 1.031296 | 0.439542 | 0.761101 |
| GOBP_EYE_PHOTORECEPTOR_CELL_DIFFERENTIATION | 35 | 0.377274 | 1.030853 | 0.419847 | 0.761934 |
| GOBP_HORMONE_METABOLIC_PROCESS | 152 | 0.2948 | 1.030836 | 0.396806 | 0.761497 |
| GOBP_CRANIOFACIAL_SUTURE_MORPHOGENESIS | 15 | 0.438587 | 1.030483 | 0.443182 | 0.76203 |
| GOBP_NEGATIVE_REGULATION_OF_OSTEOBLAST_DIFFERENTIATION | 40 | 0.357471 | 1.030321 | 0.423358 | 0.761994 |
| GOBP_REGULATION_OF_CELL_PROJECTION_ASSEMBLY | 200 | 0.288432 | 1.030175 | 0.416374 | 0.761936 |
| GOBP_REGULATION_OF_KIDNEY_DEVELOPMENT | 24 | 0.394225 | 1.02981 | 0.43622 | 0.762495 |
| GOBP_REGULATION_OF_SKELETAL_MUSCLE_TISSUE_DEVELOPMENT | 17 | 0.426511 | 1.029732 | 0.409619 | 0.762231 |
| GOBP_REGULATION_OF_CARDIAC_MUSCLE_CONTRACTION_BY_CALCIUM_ION_SIGNALIN | 22 | 0.408137 | 1.029603 | 0.447619 | 0.762118 |
| GOBP_POSITIVE_REGULATION_OF_RESPONSE_TO_BIOTIC_STIMULUS | 115 | 0.306069 | 1.029457 | 0.40708 | 0.762019 |
| GOBP_MONOCYTE_DIFFERENTIATION | 29 | 0.373611 | 1.029267 | 0.430124 | 0.762105 |
| GOBP_POSITIVE_REGULATION_OF_PROTEOLYSIS_INVOLVED_IN_CELLULAR_PROTEIN_CA | 123 | 0.304657 | 1.029256 | 0.4 | 0.761633 |
| GOBP_REGULATION_OF_B_CELL_DIFFERENTIATION | 22 | 0.395001 | 1.028722 | 0.440061 | 0.762775 |
| GOBP_SULFATION | 16 | 0.436107 | 1.02779 | 0.443548 | 0.764975 |
| GOBP_POSITIVE_REGULATION_OF_BINDING | 153 | 0.29628 | 1.027739 | 0.397163 | 0.764627 |
| GOBP_CELLULAR_RESPONSE_TO_ABIOTIC_STIMULUS | 274 | 0.27917 | 1.027548 | 0.424036 | 0.764693 |
| GOBP_CELL_COMMUNICATION_INVOLVED_IN_CARDIAC_CONDUCTION | 53 | 0.337558 | 1.027364 | 0.408901 | 0.764753 |
| GOBP_SENSORY_PERCEPTION_OF_TASTE | 21 | 0.411271 | 1.027307 | 0.437294 | 0.764434 |
| GOBP_CARDIAC_MUSCLE_CELL_ACTION_POTENTIAL | 66 | 0.328782 | 1.026849 | 0.448895 | 0.765234 |
| GOBP_REGULATION_OF_LIPID_CATABOLIC_PROCESS | 40 | 0.355979 | 1.026142 | 0.415698 | 0.766771 |
| GOBP_RESPONSE_TO_AMYLOID_BETA | 38 | 0.360892 | 1.025842 | 0.434844 | 0.76717 |
| GOBP_POSITIVE_REGULATION_OF_VIRAL_LIFE_CYCLE | 20 | 0.403706 | 1.025783 | 0.450156 | 0.766845 |
| GOBP_POSITIVE_REGULATION_OF_CELL_CYCLE | 267 | 0.283457 | 1.02563 | 0.416667 | 0.766802 |
| GOBP_NEGATIVE_REGULATION_OF_PHOSPHATASE_ACTIVITY | 30 | 0.368169 | 1.025291 | 0.440242 | 0.767311 |
| GOBP_ENDOCRINE_PROCESS | 55 | 0.337727 | 1.025234 | 0.428966 | 0.766968 |
| GOBP_NEGATIVE_REGULATION_OF_VIRAL_GENOME_REPLICATION | 44 | 0.357035 | 1.025204 | 0.427338 | 0.76654 |
| GOBP_REGULATION_OF_HEART_RATE | 83 | 0.31484 | 1.025017 | 0.42027 | 0.7666 |
| GOBP_CELLULAR_GLUCOSE_HOMEOSTASIS | 120 | 0.303769 | 1.024399 | 0.415 | 0.767873 |
| GOBP_POSITIVE_REGULATION_OF_TRANSCRIPTION_FROM_RNA_POLYMERASE_II_PROMO | 22 | 0.401888 | 1.023965 | 0.468153 | 0.76861 |
| GOBP_MONOCARBOXYLIC_ACID_TRANSPORT | 75 | 0.321849 | 1.023662 | 0.423435 | 0.768964 |
| GOBP_POSITIVE_REGULATION_OF_SMOOTH_MUSCLE_CELL_MIGRATION | 25 | 0.395134 | 1.02355 | 0.431221 | 0.768797 |
| GOBP_MEMBRANE_FUSION | 133 | 0.299879 | 1.023457 | 0.425031 | 0.768564 |
| GOBP_POSITIVE_REGULATION_OF_T_CELL_MEDIATED_IMMUNITY | 40 | 0.367367 | 1.023426 | 0.437143 | 0.768161 |
| GOBP_ASTROCYTE_DEVELOPMENT | 28 | 0.38536 | 1.023007 | 0.444954 | 0.768863 |
| GOBP_CELL_GROWTH | 430 | 0.272617 | 1.022805 | 0.424211 | 0.768938 |
| GOBP_CELL_RECOGNITION | 100 | 0.306843 | 1.022474 | 0.412516 | 0.769422 |
| GOBP_POSITIVE_REGULATION_OF_SYNAPTIC_TRANSMISSION | 124 | 0.298297 | 1.022436 | 0.410564 | 0.769022 |
| GOBP_COLUMNAR_CUBOIDAL_EPITHELIAL_CELL_DEVELOPMENT | 34 | 0.364284 | 1.022383 | 0.416296 | 0.76868 |
| GOBP_METHYLATION | 336 | 0.275663 | 1.02211 | 0.434311 | 0.768941 |
| GOBP_RESPONSE_TO_RETINOIC_ACID | 82 | 0.317862 | 1.021823 | 0.431635 | 0.769286 |
| GOBP_NEGATIVE_REGULATION_OF_AUTOPHAGY | 74 | 0.322611 | 1.021603 | 0.402667 | 0.769437 |
| GOBP_POSITIVE_REGULATION_OF_PROTEIN_LOCALIZATION | 395 | 0.271789 | 1.021328 | 0.437768 | 0.769715 |
| GOBP_REGULATION_OF_SUBSTRATE_ADHESION_DEPENDENT_CELL_SPREADING | 51 | 0.341839 | 1.02119 | 0.440341 | 0.769611 |
| GOBP_MITOPHAGY | 34 | 0.365766 | 1.021123 | 0.440299 | 0.769311 |
| GOBP_RESPONSE_TO_GAMMA_RADIATION | 51 | 0.337665 | 1.020954 | 0.408696 | 0.769304 |
| GOBP_CHEMICAL_SYNAPTIC_TRANSMISSION_POSTSYNAPTIC | 83 | 0.314827 | 1.020602 | 0.424516 | 0.76985 |
| GOBP_POSITIVE_REGULATION_OF_CATABOLIC_PROCESS | 462 | 0.269698 | 1.020403 | 0.419422 | 0.769916 |
| GOBP_FOREBRAIN_DEVELOPMENT | 312 | 0.276206 | 1.020233 | 0.43798 | 0.769903 |
| GOBP_POSITIVE_REGULATION_OF_T_CELL_CYTOKINE_PRODUCTION | 18 | 0.417098 | 1.020179 | 0.465625 | 0.769559 |
| GOBP_RESPONSE_TO_TOPOLOGICALLY_INCORRECT_PROTEIN | 150 | 0.2967 | 1.020164 | 0.437055 | 0.769111 |
| GOBP_REGULATION_OF_PLATELET_AGGREGATION | 20 | 0.404555 | 1.020158 | 0.433962 | 0.768632 |
| GOBP_POSITIVE_REGULATION_OF_EXTRACELLULAR_MATRIX_ORGANIZATION | 23 | 0.395447 | 1.020118 | 0.44288 | 0.768248 |
| GOBP_RESPONSE_TO_INCREASED_OXYGEN_LEVELS | 24 | 0.390918 | 1.019759 | 0.433283 | 0.768801 |

| | | | | | |
|---|---|---|---|---|---|
| GOBP_OSSIFICATION | 335 | 0.276558 | 1.019634 | 0.42246 | 0.768668 |
| GOBP_NEGATIVE_REGULATION_OF_DNA_METABOLIC_PROCESS | 107 | 0.303416 | 1.019464 | 0.431447 | 0.768678 |
| GOBP_DIGESTIVE_TRACT_MORPHOGENESIS | 35 | 0.363968 | 1.019326 | 0.430303 | 0.768576 |
| GOBP_FATTY_ACID_BIOSYNTHETIC_PROCESS | 117 | 0.30076 | 1.018542 | 0.450439 | 0.770278 |
| GOBP_REGULATION_OF_MESENCHYMAL_CELL_PROLIFERATION | 22 | 0.399829 | 1.018509 | 0.445483 | 0.769883 |
| GOBP_PRODUCTION_OF_MOLECULAR_MEDIATOR_OF_IMMUNE_RESPONSE | 154 | 0.292114 | 1.017612 | 0.441916 | 0.771985 |
| GOBP_SMALL_GTPASE_MEDIATED_SIGNAL_TRANSDUCTION | 459 | 0.268054 | 1.017159 | 0.435058 | 0.7728 |
| GOBP_BEHAVIOR | 483 | 0.266608 | 1.016697 | 0.432203 | 0.773624 |
| GOBP_REGULATION_OF_PRODUCTION_OF_MOLECULAR_MEDIATOR_OF_IMMUNE_RESPO | 117 | 0.301301 | 1.016159 | 0.45512 | 0.774701 |
| GOBP_CELLULAR_RESPONSE_TO_CALCIUM_ION | 72 | 0.316118 | 1.016145 | 0.427408 | 0.774254 |
| GOBP_COMPLEMENT_ACTIVATION | 27 | 0.392403 | 1.015537 | 0.449231 | 0.775486 |
| GOBP_GLOMERULAR_EPITHELIUM_DEVELOPMENT | 22 | 0.391618 | 1.015172 | 0.437991 | 0.776036 |
| GOBP_CELLULAR_RESPONSE_TO_INTERLEUKIN_1 | 64 | 0.33035 | 1.014943 | 0.458839 | 0.776172 |
| GOBP_NEGATIVE_REGULATION_OF_CELLULAR_AMIDE_METABOLIC_PROCESS | 185 | 0.285031 | 1.014332 | 0.441142 | 0.777503 |
| GOBP_REGULATION_OF_ANIMAL_ORGAN_FORMATION | 21 | 0.405583 | 1.013669 | 0.437002 | 0.778977 |
| GOBP_POSITIVE_REGULATION_OF_PROTEIN_CONTAINING_COMPLEX_ASSEMBLY | 164 | 0.289841 | 1.013597 | 0.44335 | 0.778686 |
| GOBP_POSITIVE_REGULATION_OF_TRANSPORTER_ACTIVITY | 100 | 0.30768 | 1.013464 | 0.458498 | 0.778567 |
| GOBP_POSITIVE_REGULATION_OF_TRIGLYCERIDE_METABOLIC_PROCESS | 18 | 0.410751 | 1.01307 | 0.44795 | 0.779166 |
| GOBP_POSITIVE_REGULATION_OF_OSSIFICATION | 42 | 0.353619 | 1.012394 | 0.456738 | 0.780596 |
| GOBP_REGULATION_OF_PROTEIN_SERINE_THREONINE_KINASE_ACTIVITY | 323 | 0.274364 | 1.011924 | 0.444812 | 0.781472 |
| GOBP_NEGATIVE_REGULATION_OF_RESPONSE_TO_BIOTIC_STIMULUS | 86 | 0.321646 | 1.011658 | 0.441799 | 0.781738 |
| GOBP_RESPONSE_TO_IONIZING_RADIATION | 131 | 0.297179 | 1.010535 | 0.448101 | 0.784406 |
| GOBP_MEMBRANE_RAFT_ORGANIZATION | 19 | 0.393573 | 1.009621 | 0.459069 | 0.786547 |
| GOBP_NEGATIVE_REGULATION_OF_G_PROTEIN_COUPLED_RECEPTOR_SIGNALING_PATH | 41 | 0.352482 | 1.009243 | 0.46583 | 0.787168 |
| GOBP_REGULATION_OF_STEM_CELL_DIFFERENTIATION | 55 | 0.34084 | 1.00908 | 0.460114 | 0.787124 |
| GOBP_LEUKOCYTE_TETHERING_OR_ROLLING | 19 | 0.406097 | 1.008756 | 0.476115 | 0.787543 |
| GOBP_CELLULAR_RESPONSE_TO_TOPOLOGICALLY_INCORRECT_PROTEIN | 112 | 0.301755 | 1.00865 | 0.466584 | 0.787347 |
| GOBP_POLYSACCHARIDE_BIOSYNTHETIC_PROCESS | 62 | 0.32829 | 1.008535 | 0.456241 | 0.787174 |
| GOBP_HYPOTHALAMUS_DEVELOPMENT | 18 | 0.418813 | 1.007862 | 0.446483 | 0.788604 |
| GOBP_REGULATION_OF_LIPASE_ACTIVITY | 61 | 0.329716 | 1.007789 | 0.457182 | 0.788326 |
| GOBP_CELLULAR_COMPONENT_DISASSEMBLY_INVOLVED_IN_EXECUTION_PHASE_OF_A | 24 | 0.389555 | 1.007644 | 0.468118 | 0.788269 |
| GOBP_RESPONSE_TO_MURAMYL_DIPEPTIDE | 19 | 0.395353 | 1.007268 | 0.463054 | 0.788873 |
| GOBP_CELLULAR_RESPONSE_TO_LIGHT_STIMULUS | 96 | 0.307024 | 1.006889 | 0.441558 | 0.789437 |
| GOBP_VIRAL_PROCESS | 358 | 0.272136 | 1.006574 | 0.464635 | 0.789858 |
| GOBP_RESPONSE_TO_HEAT | 94 | 0.308028 | 1.006528 | 0.441678 | 0.789494 |
| GOBP_FC_GAMMA_RECEPTOR_SIGNALING_PATHWAY | 22 | 0.398739 | 1.006429 | 0.469291 | 0.789299 |
| GOBP_CELLULAR_RESPONSE_TO_ORGANIC_CYCLIC_COMPOUND | 454 | 0.267333 | 1.005852 | 0.480588 | 0.790434 |
| GOBP_NUCLEAR_MIGRATION | 21 | 0.393649 | 1.005667 | 0.443439 | 0.790454 |
| GOBP_HISTONE_H3_K9_MODIFICATION | 50 | 0.33936 | 1.005554 | 0.452693 | 0.790291 |
| GOBP_FEEDING_BEHAVIOR | 65 | 0.32542 | 1.00494 | 0.477454 | 0.791525 |
| GOBP_RESPONSE_TO_VITAMIN_D | 26 | 0.379846 | 1.004198 | 0.458589 | 0.793143 |
| GOBP_POSITIVE_REGULATION_OF_APOPTOTIC_SIGNALING_PATHWAY | 111 | 0.299297 | 1.003961 | 0.454662 | 0.793302 |
| GOBP_SYNAPSE_ASSEMBLY | 159 | 0.287441 | 1.003071 | 0.45 | 0.795256 |
| GOBP_RESPONSE_TO_CARBOHYDRATE | 174 | 0.284639 | 1.002542 | 0.460796 | 0.796264 |
| GOBP_POSITIVE_REGULATION_OF_INTRACELLULAR_TRANSPORT | 181 | 0.281258 | 1.002397 | 0.471412 | 0.796191 |
| GOBP_REGULATION_OF_HISTONE_MODIFICATION | 166 | 0.282005 | 1.00234 | 0.466744 | 0.795859 |
| GOBP_CELLULAR_RESPONSE_TO_UV | 81 | 0.307387 | 1.002195 | 0.448871 | 0.795788 |
| GOBP_POSITIVE_REGULATION_OF_HYDROLASE_ACTIVITY | 465 | 0.265207 | 1.001886 | 0.477152 | 0.796188 |
| GOBP_CERAMIDE_CATABOLIC_PROCESS | 18 | 0.402972 | 1.00133 | 0.433931 | 0.797275 |
| GOBP_POSITIVE_REGULATION_OF_SMAD_PROTEIN_SIGNAL_TRANSDUCTION | 18 | 0.412065 | 1.001084 | 0.463884 | 0.797434 |
| GOBP_POSITIVE_REGULATION_OF_LIPID_CATABOLIC_PROCESS | 17 | 0.408088 | 1.000919 | 0.490066 | 0.797423 |
| GOBP_POSITIVE_REGULATION_OF_MESENCHYMAL_CELL_PROLIFERATION | 16 | 0.424796 | 1.000781 | 0.473344 | 0.797328 |
| GOBP_HOMOLOGOUS_RECOMBINATION | 48 | 0.340491 | 1.000508 | 0.486607 | 0.797618 |
| GOBP_NEURON_RECOGNITION | 45 | 0.338426 | 1.00041 | 0.47654 | 0.79742 |
| GOBP_ORGANIC_ACID_CATABOLIC_PROCESS | 198 | 0.279294 | 1.000396 | 0.442375 | 0.796963 |
| GOBP_RUFFLE_ASSEMBLY | 41 | 0.342371 | 0.999492 | 0.45911 | 0.799041 |
| GOBP_REGULATION_OF_BONE_REMODELING | 34 | 0.360499 | 0.998745 | 0.461774 | 0.800642 |
| GOBP_REGULATION_OF_CALCIUM_ION_TRANSMEMBRANE_TRANSPORTER_ACTIVITY | 72 | 0.311731 | 0.998671 | 0.471218 | 0.800376 |
| GOBP_T_CELL_MIGRATION | 41 | 0.348741 | 0.998423 | 0.482861 | 0.80055 |
| GOBP_PROTEIN_HOMOOLIGOMERIZATION | 156 | 0.284698 | 0.997997 | 0.45728 | 0.801271 |
| GOBP_POSITIVE_REGULATION_OF_SODIUM_ION_TRANSPORT | 32 | 0.361871 | 0.997788 | 0.482456 | 0.801354 |
| GOBP_ORGANIC_HYDROXY_COMPOUND_METABOLIC_PROCESS | 414 | 0.264164 | 0.99756 | 0.512766 | 0.801501 |
| GOBP_NEGATIVE_REGULATION_OF_POSTTRANSCRIPTIONAL_GENE_SILENCING | 15 | 0.423081 | 0.997159 | 0.46549 | 0.802122 |
| GOBP_IMPORT_INTO_NUCLEUS | 144 | 0.28794 | 0.996536 | 0.467073 | 0.803435 |
| GOBP_NEGATIVE_REGULATION_OF_HISTONE_MODIFICATION | 45 | 0.337446 | 0.996503 | 0.474359 | 0.80304 |
| GOBP_SPINAL_CORD_DEVELOPMENT | 78 | 0.310511 | 0.996272 | 0.471448 | 0.803216 |
| GOBP_POSITIVE_REGULATION_OF_KINASE_ACTIVITY | 395 | 0.263792 | 0.996256 | 0.488347 | 0.80277 |
| GOBP_NEGATIVE_REGULATION_OF_WNT_SIGNALING_PATHWAY | 152 | 0.287077 | 0.996014 | 0.477356 | 0.802969 |
| GOBP_NEGATIVE_REGULATION_OF_CARDIAC_MUSCLE_TISSUE_GROWTH | 21 | 0.384468 | 0.995501 | 0.470209 | 0.803916 |
| GOBP_VESICLE_TARGETING | 40 | 0.349139 | 0.994645 | 0.486676 | 0.805827 |
| GOBP_NEGATIVE_REGULATION_OF_EXTRINSIC_APOPTOTIC_SIGNALING_PATHWAY_VIA_D | 24 | 0.380825 | 0.994173 | 0.484227 | 0.8066 |
| GOBP_ACYLGLYCEROL_HOMEOSTASIS | 19 | 0.401236 | 0.993896 | 0.487725 | 0.806834 |
| GOBP_PYRIDINE_CONTAINING_COMPOUND_BIOSYNTHETIC_PROCESS | 24 | 0.380338 | 0.99334 | 0.481947 | 0.807887 |
| GOBP_ANIMAL_ORGAN_FORMATION | 47 | 0.334084 | 0.992471 | 0.475317 | 0.809865 |
| GOBP_REGULATION_OF_DEFENSE_RESPONSE_TO_VIRUS_BY_HOST | 32 | 0.359555 | 0.992184 | 0.47619 | 0.81018 |
| GOBP_PROTEIN_PROCESSING | 187 | 0.278246 | 0.992052 | 0.5 | 0.810039 |
| GOBP_RHYTHMIC_PROCESS | 252 | 0.269659 | 0.99175 | 0.488215 | 0.810436 |
| GOBP_NEGATIVE_REGULATION_OF_CATION_CHANNEL_ACTIVITY | 32 | 0.355989 | 0.991426 | 0.468606 | 0.810888 |
| GOBP_RESPONSE_TO_PLATELET_DERIVED_GROWTH_FACTOR | 23 | 0.391336 | 0.99072 | 0.490415 | 0.812361 |
| GOBP_HEART_TRABECULA_MORPHOGENESIS | 28 | 0.37301 | 0.990395 | 0.496894 | 0.812767 |
| GOBP_AMINOGLYCAN_METABOLIC_PROCESS | 107 | 0.299044 | 0.990309 | 0.463878 | 0.812514 |
| GOBP_REGULATION_OF_MULTICELLULAR_ORGANISM_GROWTH | 54 | 0.33168 | 0.989444 | 0.477961 | 0.814441 |
| GOBP_SIGNAL_TRANSDUCTION_IN_ABSENCE_OF_LIGAND | 53 | 0.331054 | 0.989315 | 0.463768 | 0.814298 |
| GOBP_POSITIVE_REGULATION_OF_LEUKOCYTE_DEGRANULATION | 21 | 0.390583 | 0.988986 | 0.50077 | 0.814724 |
| GOBP_P38MAPK_CASCADE | 46 | 0.340279 | 0.988954 | 0.455752 | 0.814324 |
| GOBP_POSITIVE_REGULATION_OF_NF_KAPPAB_TRANSCRIPTION_FACTOR_ACTIVITY | 125 | 0.291724 | 0.988777 | 0.487867 | 0.814287 |
| GOBP_REGULATION_OF_SODIUM_ION_TRANSPORT | 78 | 0.307444 | 0.988578 | 0.468338 | 0.814388 |
| GOBP_CELLULAR_MODIFIED_AMINO_ACID_METABOLIC_PROCESS | 144 | 0.28417 | 0.988466 | 0.488169 | 0.814216 |
| GOBP_INTRACELLULAR_RECEPTOR_SIGNALING_PATHWAY | 223 | 0.275157 | 0.988164 | 0.491972 | 0.814548 |
| GOBP_CELLULAR_RESPONSE_TO_CARBOHYDRATE_STIMULUS | 114 | 0.292187 | 0.986921 | 0.484549 | 0.817474 |
| GOBP_RETINOIC_ACID_METABOLIC_PROCESS | 17 | 0.412032 | 0.986833 | 0.508972 | 0.81724 |
| GOBP_PEPTIDYL_SERINE_MODIFICATION | 292 | 0.267078 | 0.986779 | 0.498904 | 0.816903 |
| GOBP_REGULATION_OF_VASCULAR_PERMEABILITY | 40 | 0.347741 | 0.986629 | 0.515015 | 0.816873 |
| GOBP_REGULATION_OF_DNA_TEMPLATED_TRANSCRIPTION_INITIATION | 52 | 0.328932 | 0.985366 | 0.495833 | 0.819826 |
| GOBP_VASCULAR_PROCESS_IN_CIRCULATORY_SYSTEM | 209 | 0.27671 | 0.985194 | 0.50464 | 0.819821 |
| GOBP_FATTY_ACID_TRANSPORT | 69 | 0.313522 | 0.985036 | 0.499299 | 0.819759 |
| GOBP_REGULATION_OF_CARTILAGE_DEVELOPMENT | 58 | 0.32129 | 0.984908 | 0.469444 | 0.819632 |
| GOBP_RECEPTOR_SIGNALING_PATHWAY_VIA_STAT | 105 | 0.295528 | 0.984817 | 0.489744 | 0.819395 |
| GOBP_REGULATION_OF_GENE_SILENCING_BY_RNA | 44 | 0.339021 | 0.984768 | 0.500734 | 0.819028 |
| GOBP_POSITIVE_REGULATION_OF_INTERLEUKIN_1_BETA_PRODUCTION | 39 | 0.348287 | 0.983454 | 0.47181 | 0.822229 |
| GOBP_POSITIVE_REGULATION_OF_TYPE_I_INTERFERON_MEDIATED_SIGNALING_PATHWA | 16 | 0.420107 | 0.982593 | 0.470588 | 0.824104 |
| GOBP_CELL_CELL_JUNCTION_ASSEMBLY | 117 | 0.293511 | 0.982529 | 0.49375 | 0.823796 |
| GOBP_XENOBIOTIC_METABOLIC_PROCESS | 62 | 0.315117 | 0.982303 | 0.503477 | 0.823944 |
| GOBP_RESPONSE_TO_AMINO_ACID_STARVATION | 48 | 0.329312 | 0.982124 | 0.508646 | 0.823964 |
| GOBP_POSITIVE_REGULATION_OF_DEFENSE_RESPONSE_TO_VIRUS_BY_HOST | 24 | 0.385475 | 0.98162 | 0.476408 | 0.82488 |
| GOBP_REGULATION_OF_INSULIN_SECRETION | 113 | 0.293282 | 0.981602 | 0.505013 | 0.824436 |
| GOBP_EXOCYTOSIS | 299 | 0.26715 | 0.981359 | 0.537694 | 0.82461 |
| GOBP_NEGATIVE_REGULATION_OF_CARBOHYDRATE_METABOLIC_PROCESS | 41 | 0.334727 | 0.981246 | 0.463557 | 0.824431 |
| GOBP_INCLUSION_BODY_ASSEMBLY | 22 | 0.384223 | 0.981149 | 0.486614 | 0.824207 |
| GOBP_NEGATIVE_REGULATION_OF_PLASMA_MEMBRANE_BOUNDED_CELL_PROJECTION | 31 | 0.361336 | 0.98095 | 0.48605 | 0.824265 |
| GOBP_REGULATION_OF_EXTENT_OF_CELL_GROWTH | 103 | 0.294698 | 0.980564 | 0.489873 | 0.824875 |
| GOBP_POSITIVE_REGULATION_OF_CELL_GROWTH | 143 | 0.284489 | 0.980312 | 0.520988 | 0.825149 |
| GOBP_IRON_ION_TRANSMEMBRANE_TRANSPORT | 18 | 0.401312 | 0.980127 | 0.503195 | 0.825183 |
| GOBP_NEGATIVE_REGULATION_OF_ION_TRANSPORT | 105 | 0.295716 | 0.980005 | 0.494764 | 0.825027 |
| GOBP_ENERGY_RESERVE_METABOLIC_PROCESS | 66 | 0.313892 | 0.979755 | 0.504965 | 0.825254 |
| GOBP_RNA_DESTABILIZATION | 88 | 0.303123 | 0.979655 | 0.524766 | 0.825047 |
| GOBP_ANOIKIS | 30 | 0.35997 | 0.979303 | 0.488131 | 0.825548 |
| GOBP_NEUROTROPHIN_SIGNALING_PATHWAY | 35 | 0.353411 | 0.979239 | 0.502276 | 0.825247 |
| GOBP_GLYCOPROTEIN_BIOSYNTHETIC_PROCESS | 276 | 0.263321 | 0.978618 | 0.526024 | 0.826436 |
| GOBP_RESPIRATORY_GASEOUS_EXCHANGE_BY_RESPIRATORY_SYSTEM | 56 | 0.32154 | 0.978476 | 0.496444 | 0.826358 |
| GOBP_POSITIVE_REGULATION_OF_PROTEOLYSIS | 322 | 0.264352 | 0.978442 | 0.527808 | 0.825983 |
| GOBP_POSITIVE_REGULATION_OF_CANONICAL_WNT_SIGNALING_PATHWAY | 94 | 0.297347 | 0.978316 | 0.503251 | 0.825827 |
| GOBP_CELLULAR_RESPONSE_TO_CAMP | 39 | 0.337892 | 0.977873 | 0.498551 | 0.826547 |
| GOBP_REGULATION_OF_INNATE_IMMUNE_RESPONSE | 160 | 0.280267 | 0.977685 | 0.516207 | 0.826544 |
| GOBP_REGULATION_OF_CALCIUM_MEDIATED_SIGNALING | 56 | 0.320521 | 0.977601 | 0.501393 | 0.826283 |
| GOBP_POSITIVE_REGULATION_OF_CELL_PROJECTION_ORGANIZATION | 311 | 0.265435 | 0.977493 | 0.537528 | 0.826111 |
| GOBP_CELLULAR_RESPONSE_TO_PEPTIDE | 288 | 0.264537 | 0.977239 | 0.524917 | 0.826341 |
| GOBP_CELLULAR_RESPONSE_TO_VASCULAR_ENDOTHELIAL_GROWTH_FACTOR_STIMUL | 55 | 0.324688 | 0.976843 | 0.478992 | 0.826984 |
| GOBP_MYELOID_CELL_DEVELOPMENT | 56 | 0.321996 | 0.976608 | 0.51314 | 0.827139 |
| GOBP_REGULATION_OF_BONE_RESORPTION | 27 | 0.366564 | 0.976505 | 0.508423 | 0.826926 |
| GOBP_CELLULAR_RESPONSE_TO_PEPTIDE_HORMONE_STIMULUS | 236 | 0.268107 | 0.976447 | 0.524831 | 0.826609 |
| GOBP_MULTICELLULAR_ORGANISMAL_RESPONSE_TO_STRESS | 64 | 0.311561 | 0.976276 | 0.515193 | 0.826605 |
| GOBP_P_BODY_ASSEMBLY | 20 | 0.381958 | 0.976066 | 0.519048 | 0.826731 |
| GOBP_POSITIVE_REGULATION_OF_CELL_DEVELOPMENT | 253 | 0.266158 | 0.975381 | 0.516892 | 0.828205 |
| GOBP_OLEFINIC_COMPOUND_METABOLIC_PROCESS | 104 | 0.293445 | 0.974879 | 0.501971 | 0.829088 |
| GOBP_CELLULAR_RESPONSE_TO_COPPER_ION | 18 | 0.398406 | 0.974578 | 0.513029 | 0.829417 |
| GOBP_POSITIVE_REGULATION_OF_EPITHELIAL_CELL_APOPTOTIC_PROCESS | 22 | 0.388096 | 0.9743 | 0.502385 | 0.829712 |
| GOBP_DETOXIFICATION | 105 | 0.289693 | 0.974047 | 0.51178 | 0.829908 |
| GOBP_NEGATIVE_REGULATION_OF_CELL_CYCLE | 327 | 0.260493 | 0.973856 | 0.550599 | 0.829953 |
| GOBP_POSITIVE_REGULATION_OF_UBIQUITIN_DEPENDENT_PROTEIN_CATABOLIC_PROCE | 101 | 0.29492 | 0.972859 | 0.516005 | 0.832246 |
| GOBP_DETECTION_OF_VISIBLE_LIGHT | 23 | 0.375004 | 0.972638 | 0.496835 | 0.832373 |
| GOBP_CELLULAR_RESPONSE_TO_RADIATION | 157 | 0.277635 | 0.971921 | 0.534442 | 0.833893 |
| GOBP_POSITIVE_REGULATION_OF_ORGANIC_ACID_TRANSPORT | 35 | 0.343371 | 0.971908 | 0.496263 | 0.833441 |
| GOBP_NEURAL_PRECURSOR_CELL_PROLIFERATION | 121 | 0.2852 | 0.9717 | 0.525935 | 0.833514 |
| GOBP_REGULATION_OF_DNA_METHYLATION | 20 | 0.389677 | 0.971249 | 0.4928 | 0.834299 |
| GOBP_NEGATIVE_REGULATION_OF_APOPTOTIC_SIGNALING_PATHWAY | 195 | 0.273155 | 0.971236 | 0.540216 | 0.833848 |
| GOBP_NEGATIVE_REGULATION_OF_CELL_AGING | 21 | 0.375355 | 0.971047 | 0.506977 | 0.833907 |
| GOBP_CELLULAR_RESPONSE_TO_HORMONE_STIMULUS | 468 | 0.255733 | 0.970691 | 0.564756 | 0.834402 |
| GOBP_NEGATIVE_REGULATION_OF_RESPONSE_TO_CYTOKINE_STIMULUS | 56 | 0.316589 | 0.970519 | 0.527778 | 0.834393 |
| GOBP_ACROSOME_REACTION | 23 | 0.379713 | 0.969907 | 0.517611 | 0.835571 |
| GOBP_REGULATION_OF_NUCLEAR_TRANSCRIBED_MRNA_CATABOLIC_PROCESS_DEADE | 24 | 0.373303 | 0.969754 | 0.524738 | 0.835514 |
| GOBP_SIGNAL_RELEASE | 357 | 0.260121 | 0.969581 | 0.557112 | 0.835516 |
| GOBP_PRESYNAPTIC_ENDOCYTOSIS | 57 | 0.318697 | 0.969278 | 0.514685 | 0.835903 |
| GOBP_ESTABLISHMENT_OF_SPINDLE_ORIENTATION | 32 | 0.354775 | 0.968621 | 0.496274 | 0.837172 |
| GOBP_PROTEIN_HOMOTETRAMERIZATION | 46 | 0.32957 | 0.968202 | 0.508571 | 0.837833 |
| GOBP_IMPORT_ACROSS_PLASMA_MEMBRANE | 140 | 0.2816 | 0.967714 | 0.519901 | 0.838649 |
| GOBP_LIPOPOLYSACCHARIDE_MEDIATED_SIGNALING_PATHWAY | 43 | 0.338981 | 0.967701 | 0.533333 | 0.838197 |

| | | | | | |
|---|---|---|---|---|---|
| GOBP_REGULATION_OF_MEMBRANE_POTENTIAL | 343 | 0.260932 | 0.967245 | 0.554054 | 0.838889 |
| GOBP_REGULATION_OF_OSTEOCLAST_DIFFERENTIATION | 43 | 0.332968 | 0.967028 | 0.5218 | 0.838993 |
| GOBP_B_CELL_APOPTOTIC_PROCESS | 23 | 0.380126 | 0.966576 | 0.548438 | 0.839741 |
| GOBP_UNSATURATED_FATTY_ACID_BIOSYNTHETIC_PROCESS | 40 | 0.337721 | 0.966532 | 0.529674 | 0.839368 |
| GOBP_REGULATION_OF_NEUROBLAST_PROLIFERATION | 24 | 0.359254 | 0.965373 | 0.522013 | 0.842043 |
| GOBP_GLUCOSAMINE_CONTAINING_COMPOUND_METABOLIC_PROCESS | 20 | 0.383587 | 0.965072 | 0.493691 | 0.842431 |
| GOBP_POSITIVE_REGULATION_OF_WNT_SIGNALING_PATHWAY | 126 | 0.283592 | 0.965041 | 0.551171 | 0.842017 |
| GOBP_CELLULAR_OXIDANT_DETOXIFICATION | 72 | 0.309187 | 0.964969 | 0.52611 | 0.841723 |
| GOBP_ORGANELLE_FISSION | 425 | 0.256634 | 0.964695 | 0.572785 | 0.841975 |
| GOBP_POSITIVE_REGULATION_OF_INTERLEUKIN_10_PRODUCTION | 26 | 0.369833 | 0.964396 | 0.523365 | 0.842265 |
| GOBP_NEGATIVE_REGULATION_OF_PRODUCTION_OF_MOLECULAR_MEDIATOR_OF_IMMU | 23 | 0.376001 | 0.964282 | 0.501618 | 0.84208 |
| GOBP_NEGATIVE_REGULATION_OF_I_KAPPAB_KINASE_NF_KAPPAB_SIGNALING | 40 | 0.342121 | 0.963853 | 0.525074 | 0.842822 |
| GOBP_INTRINSIC_APOPTOTIC_SIGNALING_PATHWAY_IN_RESPONSE_TO_DNA_DAMAGE_E | 41 | 0.334474 | 0.96385 | 0.520059 | 0.842347 |
| GOBP_REGULATION_OF_PROTEIN_TARGETING | 73 | 0.306892 | 0.963423 | 0.539267 | 0.843011 |
| GOBP_T_CELL_MEDIATED_CYTOTOXICITY | 31 | 0.353749 | 0.962896 | 0.512668 | 0.843964 |
| GOBP_FATTY_ACID_TRANSMEMBRANE_TRANSPORT | 15 | 0.416603 | 0.962839 | 0.520129 | 0.843641 |
| GOBP_CELLULAR_RESPONSE_TO_ALCOHOL | 71 | 0.303095 | 0.962772 | 0.535475 | 0.843331 |
| GOBP_REGULATION_OF_ANATOMICAL_STRUCTURE_SIZE | 416 | 0.25524 | 0.962597 | 0.582279 | 0.843313 |
| GOBP_RNA_MEDIATED_GENE_SILENCING_BY_INHIBITION_OF_TRANSLATION | 16 | 0.402983 | 0.962136 | 0.513957 | 0.84411 |
| GOBP_B_CELL_HOMEOSTASIS | 27 | 0.363335 | 0.961907 | 0.509203 | 0.844253 |
| GOBP_REGULATION_OF_HORMONE_SECRETION | 171 | 0.272025 | 0.961521 | 0.56401 | 0.844785 |
| GOBP_REGULATION_OF_AXONOGENESIS | 149 | 0.275546 | 0.961247 | 0.546442 | 0.845059 |
| GOBP_REGULATION_OF_CAMP_MEDIATED_SIGNALING | 25 | 0.364497 | 0.960998 | 0.525424 | 0.845241 |
| GOBP_REGULATION_OF_ANIMAL_ORGAN_MORPHOGENESIS | 108 | 0.288887 | 0.960591 | 0.50706 | 0.845831 |
| GOBP_NEGATIVE_REGULATION_OF_NEURON_DEATH | 181 | 0.271965 | 0.960146 | 0.56615 | 0.846547 |
| GOBP_POSITIVE_REGULATION_OF_AMINE_TRANSPORT | 33 | 0.344863 | 0.959886 | 0.520424 | 0.846758 |
| GOBP_POSITIVE_REGULATION_OF_PROTEIN_CATABOLIC_PROCESS | 210 | 0.266212 | 0.95971 | 0.561707 | 0.846773 |
| GOBP_REGULATION_OF_TRANSLATION_IN_RESPONSE_TO_STRESS | 20 | 0.389388 | 0.959072 | 0.507813 | 0.847995 |
| GOBP_POSITIVE_REGULATION_OF_ACTIVATED_T_CELL_PROLIFERATION | 17 | 0.396626 | 0.95901 | 0.509375 | 0.847683 |
| GOBP_CILIUM_MOVEMENT | 120 | 0.282078 | 0.958444 | 0.553325 | 0.848716 |
| GOBP_SINGLE_FERTILIZATION | 82 | 0.292318 | 0.957521 | 0.509934 | 0.850663 |
| GOBP_POSITIVE_REGULATION_OF_G1_S_TRANSITION_OF_MITOTIC_CELL_CYCLE | 39 | 0.33354 | 0.957226 | 0.530192 | 0.85099 |
| GOBP_ORGANELLE_INHERITANCE | 15 | 0.406654 | 0.956683 | 0.504132 | 0.851963 |
| GOBP_GLYCOSAMINOGLYCAN_CATABOLIC_PROCESS | 19 | 0.388583 | 0.955816 | 0.534733 | 0.853883 |
| GOBP_REGULATION_OF_SUPEROXIDE_ANION_GENERATION | 16 | 0.393196 | 0.955766 | 0.515625 | 0.853545 |
| GOBP_POSITIVE_REGULATION_OF_MUSCLE_TISSUE_DEVELOPMENT | 16 | 0.396403 | 0.955744 | 0.518283 | 0.853121 |
| GOBP_GLIOGENESIS | 231 | 0.259876 | 0.955696 | 0.565119 | 0.852785 |
| GOBP_PROTEIN_HETEROOLIGOMERIZATION | 25 | 0.358633 | 0.955448 | 0.515447 | 0.852985 |
| GOBP_CYTOKINE_PRODUCTION_INVOLVED_IN_INFLAMMATORY_RESPONSE | 35 | 0.342041 | 0.954333 | 0.525714 | 0.855548 |
| GOBP_REGULATION_OF_OXIDOREDUCTASE_ACTIVITY | 72 | 0.300618 | 0.954322 | 0.530184 | 0.855091 |
| GOBP_NEGATIVE_REGULATION_OF_PEPTIDYL_TYROSINE_PHOSPHORYLATION | 51 | 0.319253 | 0.954104 | 0.54089 | 0.855189 |
| GOBP_POSITIVE_REGULATION_OF_POTASSIUM_ION_TRANSPORT | 41 | 0.326781 | 0.95352 | 0.555556 | 0.856284 |
| GOBP_RIBONUCLEOTIDE_CATABOLIC_PROCESS | 42 | 0.327666 | 0.953362 | 0.522828 | 0.856206 |
| GOBP_DNA_CATABOLIC_PROCESS | 26 | 0.36164 | 0.953309 | 0.5 | 0.855871 |
| GOBP_POSITIVE_REGULATION_OF_EPITHELIAL_TO_MESENCHYMAL_TRANSITION | 44 | 0.328943 | 0.952684 | 0.543964 | 0.857106 |
| GOBP_PROTEIN_DEMANNOSYLATION | 17 | 0.391023 | 0.951917 | 0.519071 | 0.858706 |
| GOBP_SMOOTH_MUSCLE_CELL_APOPTOTIC_PROCESS | 18 | 0.38959 | 0.951456 | 0.533442 | 0.859471 |
| GOBP_POSITIVE_REGULATION_OF_ADAPTIVE_IMMUNE_RESPONSE | 79 | 0.294775 | 0.951436 | 0.555708 | 0.859042 |
| GOBP_NEURON_PROJECTION_GUIDANCE | 207 | 0.263938 | 0.951298 | 0.57346 | 0.858937 |
| GOBP_REGULATION_OF_OXIDATIVE_PHOSPHORYLATION | 20 | 0.38822 | 0.95106 | 0.528939 | 0.859068 |
| GOBP_ADENYLATE_CYCLASE_ACTIVATING_G_PROTEIN_COUPLED_RECEPTOR_SIGNALIN | 109 | 0.285451 | 0.950834 | 0.557867 | 0.859167 |
| GOBP_CRISTAE_FORMATION | 16 | 0.407553 | 0.950696 | 0.537438 | 0.859057 |
| GOBP_NEGATIVE_REGULATION_OF_PLATELET_ACTIVATION | 16 | 0.400398 | 0.950516 | 0.522109 | 0.859047 |
| GOBP_REGULATION_OF_SECRETION | 466 | 0.251099 | 0.950196 | 0.639535 | 0.85944 |
| GOBP_NEGATIVE_REGULATION_OF_ENDOPLASMIC_RETICULUM_STRESS_INDUCED_INTR | 17 | 0.401548 | 0.950012 | 0.540902 | 0.859462 |
| GOBP_RESPONSE_TO_CALCIUM_ION | 121 | 0.280705 | 0.94995 | 0.587654 | 0.859156 |
| GOBP_PEPTIDYL_THREONINE_MODIFICATION | 114 | 0.282581 | 0.949735 | 0.569288 | 0.85924 |
| GOBP_OSTEOCLAST_DIFFERENTIATION | 70 | 0.304553 | 0.949386 | 0.549862 | 0.859651 |
| GOBP_INOSITOL_PHOSPHATE_CATABOLIC_PROCESS | 19 | 0.386849 | 0.949116 | 0.528391 | 0.859879 |
| GOBP_CEREBELLAR_PURKINJE_CELL_LAYER_MORPHOGENESIS | 18 | 0.388016 | 0.948788 | 0.529118 | 0.860264 |
| GOBP_REGULATION_OF_RESPONSE_TO_BIOTIC_STIMULUS | 249 | 0.260115 | 0.948286 | 0.578231 | 0.861105 |
| GOBP_ECTODERM_DEVELOPMENT | 18 | 0.394898 | 0.947953 | 0.545315 | 0.861478 |
| GOBP_NEGATIVE_REGULATION_OF_RESPONSE_TO_ENDOPLASMIC_RETICULUM_STRESS | 40 | 0.328965 | 0.947587 | 0.538787 | 0.861942 |
| GOBP_NEGATIVE_REGULATION_OF_PROTEIN_TYROSINE_KINASE_ACTIVITY | 29 | 0.354214 | 0.94739 | 0.544901 | 0.86197 |
| GOBP_POSITIVE_REGULATION_OF_TRANSFERASE_ACTIVITY | 477 | 0.249762 | 0.94688 | 0.633612 | 0.862833 |
| GOBP_NEGATIVE_REGULATION_OF_CELL_GROWTH | 162 | 0.267861 | 0.94646 | 0.57783 | 0.863447 |
| GOBP_SECRETORY_GRANULE_ORGANIZATION | 29 | 0.341284 | 0.946258 | 0.558912 | 0.863478 |
| GOBP_RESPONSE_TO_ZINC_ION | 33 | 0.344475 | 0.946215 | 0.562404 | 0.863117 |
| GOBP_ACTIVATION_OF_GTPASE_ACTIVITY | 97 | 0.286144 | 0.946192 | 0.561224 | 0.862694 |
| GOBP_CIRCADIAN_REGULATION_OF_GENE_EXPRESSION | 63 | 0.308339 | 0.946079 | 0.536755 | 0.862505 |
| GOBP_ENERGY_HOMEOSTASIS | 35 | 0.342582 | 0.945851 | 0.562408 | 0.862613 |
| GOBP_REPLICATION_FORK_PROCESSING | 38 | 0.332165 | 0.94577 | 0.548435 | 0.862336 |
| GOBP_NEGATIVE_REGULATION_OF_T_CELL_DIFFERENTIATION | 31 | 0.344368 | 0.945527 | 0.538231 | 0.862492 |
| GOBP_RESPONSE_TO_PURINE_CONTAINING_COMPOUND | 102 | 0.285108 | 0.94531 | 0.578145 | 0.862597 |
| GOBP_REGULATION_OF_STEM_CELL_POPULATION_MAINTENANCE | 30 | 0.349596 | 0.944562 | 0.548387 | 0.864088 |
| GOBP_REGULATION_OF_CELL_DEVELOPMENT | 419 | 0.249596 | 0.94442 | 0.647184 | 0.863981 |
| GOBP_REGULATION_OF_DNA_TEMPLATED_TRANSCRIPTION_IN_RESPONSE_TO_STRESS | 49 | 0.31762 | 0.944277 | 0.574468 | 0.863864 |
| GOBP_MITOTIC_DNA_INTEGRITY_CHECKPOINT_SIGNALING | 80 | 0.293415 | 0.944169 | 0.559572 | 0.863664 |
| GOBP_RECEPTOR_CATABOLIC_PROCESS | 27 | 0.348438 | 0.943787 | 0.531546 | 0.864197 |
| GOBP_POSITIVE_REGULATION_OF_MITOTIC_NUCLEAR_DIVISION | 30 | 0.344774 | 0.943288 | 0.56 | 0.864964 |
| GOBP_NEGATIVE_REGULATION_OF_BINDING | 136 | 0.276992 | 0.94191 | 0.603194 | 0.86815 |
| GOBP_NEGATIVE_REGULATION_OF_MIRNA_TRANSCRIPTION | 17 | 0.389456 | 0.941725 | 0.543444 | 0.868151 |
| GOBP_NEGATIVE_REGULATION_OF_NUCLEOTIDE_METABOLIC_PROCESS | 19 | 0.376384 | 0.941216 | 0.530303 | 0.869075 |
| GOBP_PLATELET_MORPHOGENESIS | 18 | 0.385126 | 0.940092 | 0.552381 | 0.871586 |
| GOBP_REGULATION_OF_T_CELL_APOPTOTIC_PROCESS | 22 | 0.373605 | 0.939729 | 0.548092 | 0.872033 |
| GOBP_CELLULAR_GLUCAN_METABOLIC_PROCESS | 60 | 0.306323 | 0.938307 | 0.578363 | 0.875291 |
| GOBP_VENTRICULAR_CARDIAC_MUSCLE_CELL_MEMBRANE_REPOLARIZATION | 22 | 0.379185 | 0.937461 | 0.581731 | 0.877034 |
| GOBP_REGULATION_OF_PEPTIDE_TRANSPORT | 133 | 0.272827 | 0.93703 | 0.574879 | 0.877676 |
| GOBP_POSITIVE_REGULATION_OF_SODIUM_ION_TRANSMEMBRANE_TRANSPORTER_ACT | 15 | 0.398223 | 0.936876 | 0.531546 | 0.877623 |
| GOBP_COLLAGEN_BIOSYNTHETIC_PROCESS | 32 | 0.339266 | 0.936801 | 0.552239 | 0.877325 |
| GOBP_MONOCARBOXYLIC_ACID_METABOLIC_PROCESS | 444 | 0.246358 | 0.936574 | 0.667728 | 0.877406 |
| GOBP_POSITIVE_REGULATION_OF_NUCLEOCYTOPLASMIC_TRANSPORT | 55 | 0.312271 | 0.936335 | 0.556474 | 0.877556 |
| GOBP_REGULATION_OF_LYMPHOCYTE_APOPTOTIC_PROCESS | 40 | 0.326945 | 0.93631 | 0.558824 | 0.877132 |
| GOBP_POSITIVE_REGULATION_OF_NEUROGENESIS | 194 | 0.261755 | 0.936282 | 0.612903 | 0.87672 |
| GOBP_BONE_REMODELING | 66 | 0.299286 | 0.936114 | 0.57754 | 0.876694 |
| GOBP_POSITIVE_REGULATION_OF_GTPASE_ACTIVITY | 228 | 0.256338 | 0.93576 | 0.620968 | 0.877123 |
| GOBP_NITRIC_OXIDE_MEDIATED_SIGNAL_TRANSDUCTION | 18 | 0.380439 | 0.93553 | 0.531915 | 0.877233 |
| GOBP_INTRINSIC_APOPTOTIC_SIGNALING_PATHWAY | 266 | 0.254336 | 0.933726 | 0.639509 | 0.881498 |
| GOBP_MAINTENANCE_OF_PROTEIN_LOCATION_IN_NUCLEUS | 23 | 0.360778 | 0.933709 | 0.55521 | 0.881055 |
| GOBP_ENDOPLASMIC_RETICULUM_CALCIUM_ION_HOMEOSTASIS | 25 | 0.357926 | 0.93327 | 0.566564 | 0.8817 |
| GOBP_LIPID_STORAGE | 65 | 0.297977 | 0.932661 | 0.573677 | 0.882806 |
| GOBP_T_HELPER_17_TYPE_IMMUNE_RESPONSE | 27 | 0.353899 | 0.932503 | 0.578538 | 0.882738 |
| GOBP_NEGATIVE_REGULATION_OF_ANION_TRANSPORT | 18 | 0.376661 | 0.931896 | 0.545603 | 0.883849 |
| GOBP_REGULATION_OF_REGULATED_SECRETORY_PATHWAY | 118 | 0.273084 | 0.931842 | 0.598991 | 0.883503 |
| GOBP_REGULATION_OF_T_CELL_MEDIATED_CYTOTOXICITY | 23 | 0.362526 | 0.931483 | 0.558735 | 0.883926 |
| GOBP_SPECIFICATION_OF_ANIMAL_ORGAN_IDENTITY | 28 | 0.349626 | 0.931432 | 0.573939 | 0.883583 |
| GOBP_POSITIVE_REGULATION_OF_PRODUCTION_OF_MOLECULAR_MEDIATOR_OF_IMMUN | 84 | 0.289566 | 0.930937 | 0.587106 | 0.884367 |
| GOBP_REGULATION_OF_MONOCYTE_DIFFERENTIATION | 16 | 0.39394 | 0.930909 | 0.5469 | 0.883963 |
| GOBP_REGULATION_OF_BINDING | 322 | 0.251116 | 0.930457 | 0.640437 | 0.884633 |
| GOBP_HISTONE_MRNA_CATABOLIC_PROCESS | 15 | 0.396933 | 0.930374 | 0.561688 | 0.884338 |
| GOBP_POSITIVE_REGULATION_OF_PROTEIN_LOCALIZATION_TO_NUCLEUS | 77 | 0.293972 | 0.929931 | 0.588158 | 0.885027 |
| GOBP_PROTEIN_AUTOUBIQUITINATION | 70 | 0.295792 | 0.929144 | 0.592298 | 0.886567 |
| GOBP_COGNITION | 256 | 0.254869 | 0.929126 | 0.662934 | 0.886129 |
| GOBP_CELLULAR_RESPONSE_TO_EXOGENOUS_DSRNA | 16 | 0.385 | 0.928242 | 0.569106 | 0.887898 |
| GOBP_POSITIVE_REGULATION_OF_PROTEIN_SERINE_THREONINE_KINASE_ACTIVITY | 179 | 0.262797 | 0.928219 | 0.625296 | 0.887471 |
| GOBP_MATING | 29 | 0.346399 | 0.928055 | 0.563077 | 0.887402 |
| GOBP_PYRIDINE_NUCLEOTIDE_BIOSYNTHETIC_PROCESS | 21 | 0.370492 | 0.927723 | 0.553991 | 0.887723 |
| GOBP_ERYTHROCYTE_HOMEOSTASIS | 115 | 0.277943 | 0.927403 | 0.614124 | 0.888076 |
| GOBP_VESICLE_BUDDING_FROM_MEMBRANE | 61 | 0.295041 | 0.927352 | 0.590971 | 0.88774 |
| GOBP_CERAMIDE_METABOLIC_PROCESS | 90 | 0.283905 | 0.927326 | 0.616402 | 0.88732 |
| GOBP_ALCOHOL_METABOLIC_PROCESS | 278 | 0.250484 | 0.927325 | 0.643333 | 0.886841 |
| GOBP_POSITIVE_T_CELL_SELECTION | 20 | 0.36734 | 0.927168 | 0.550725 | 0.88677 |
| GOBP_MECHANORECEPTOR_DIFFERENTIATION | 56 | 0.306986 | 0.926863 | 0.568214 | 0.887067 |
| GOBP_NEGATIVE_REGULATION_OF_TRANSPORT | 330 | 0.247922 | 0.926831 | 0.663821 | 0.886666 |
| GOBP_MYELOID_LEUKOCYTE_CYTOKINE_PRODUCTION | 27 | 0.346468 | 0.926326 | 0.550225 | 0.887481 |
| GOBP_TRANSITION_METAL_ION_HOMEOSTASIS | 108 | 0.276208 | 0.925793 | 0.618205 | 0.8884 |
| GOBP_DENDRITE_EXTENSION | 33 | 0.335525 | 0.924445 | 0.574815 | 0.8914 |
| GOBP_MICROVILLUS_ORGANIZATION | 28 | 0.344523 | 0.92335 | 0.567073 | 0.893747 |
| GOBP_ACTIN_FILAMENT_BUNDLE_ORGANIZATION | 149 | 0.263671 | 0.922188 | 0.633854 | 0.896278 |
| GOBP_POST_ANAL_TAIL_MORPHOGENESIS | 15 | 0.395324 | 0.922136 | 0.576737 | 0.895918 |
| GOBP_CALCINEURIN_MEDIATED_SIGNALING | 40 | 0.322588 | 0.921678 | 0.592988 | 0.896599 |
| GOBP_PROTEIN_COMPLEX_OLIGOMERIZATION | 209 | 0.257951 | 0.921522 | 0.661972 | 0.896518 |
| GOBP_CELL_SURFACE_RECEPTOR_SIGNALING_PATHWAY_INVOLVED_IN_CELL_CELL_SIG | 481 | 0.243826 | 0.921406 | 0.715645 | 0.896321 |
| GOBP_REGULATION_OF_SMAD_PROTEIN_SIGNAL_TRANSDUCTION | 27 | 0.342851 | 0.920988 | 0.580745 | 0.896913 |
| GOBP_REGULATION_OF_CARDIAC_MUSCLE_CONTRACTION_BY_REGULATION_OF_THE_R | 18 | 0.379918 | 0.920793 | 0.583748 | 0.896936 |
| GOBP_RECEPTOR_CLUSTERING | 53 | 0.303523 | 0.920493 | 0.603129 | 0.897236 |
| GOBP_CELL_REDOX_HOMEOSTASIS | 34 | 0.330097 | 0.920202 | 0.59375 | 0.897522 |
| GOBP_REGULATION_OF_MICROTUBULE_POLYMERIZATION | 53 | 0.307977 | 0.920172 | 0.602322 | 0.897119 |
| GOBP_FLUID_TRANSPORT | 19 | 0.372905 | 0.920076 | 0.592477 | 0.896882 |
| GOBP_CELL_JUNCTION_DISASSEMBLY | 15 | 0.398803 | 0.919314 | 0.565978 | 0.898305 |
| GOBP_CELLULAR_CARBOHYDRATE_METABOLIC_PROCESS | 237 | 0.252552 | 0.918923 | 0.648492 | 0.898819 |
| GOBP_REGULATION_OF_EXOSOMAL_SECRETION | 18 | 0.377921 | 0.918901 | 0.583199 | 0.898393 |
| GOBP_CORTICAL_CYTOSKELETON_ORGANIZATION | 58 | 0.29564 | 0.918009 | 0.596465 | 0.900203 |
| GOBP_CARBOHYDRATE_BIOSYNTHETIC_PROCESS | 168 | 0.257939 | 0.91792 | 0.643623 | 0.899957 |
| GOBP_NEURONAL_STEM_CELL_POPULATION_MAINTENANCE | 21 | 0.358887 | 0.9179 | 0.576737 | 0.899524 |
| GOBP_POSITIVE_REGULATION_OF_INNATE_IMMUNE_RESPONSE | 93 | 0.277038 | 0.917531 | 0.629032 | 0.899986 |
| GOBP_CAMP_MEDIATED_SIGNALING | 42 | 0.316115 | 0.917244 | 0.622585 | 0.900213 |
| GOBP_VESICLE_CYTOSKELETAL_TRAFFICKING | 63 | 0.30011 | 0.916456 | 0.598938 | 0.901764 |
| GOBP_RESPONSE_TO_ELECTRICAL_STIMULUS | 29 | 0.345481 | 0.916172 | 0.592126 | 0.902015 |
| GOBP_HEMATOPOIETIC_STEM_CELL_PROLIFERATION | 23 | 0.354701 | 0.916145 | 0.567188 | 0.90159 |
| GOBP_DNA_CATABOLIC_PROCESS_ENDONUCLEOLYTIC | 21 | 0.363844 | 0.915952 | 0.571429 | 0.90159 |
| GOBP_POSITIVE_REGULATION_OF_CREB_TRANSCRIPTION_FACTOR_ACTIVITY | 17 | 0.382062 | 0.915699 | 0.57561 | 0.901746 |
| GOBP_POSITIVE_REGULATION_OF_CELLULAR_PROTEIN_CATABOLIC_PROCESS | 145 | 0.262666 | 0.915549 | 0.631391 | 0.901666 |
| GOBP_REGULATION_OF_PROTEIN_CONTAINING_COMPLEX_ASSEMBLY | 348 | 0.243569 | 0.91511 | 0.706392 | 0.90232 |
| GOBP_DIOL_BIOSYNTHETIC_PROCESS | 22 | 0.360637 | 0.914837 | 0.577778 | 0.902553 |
| GOBP_REGULATION_OF_SUPEROXIDE_METABOLIC_PROCESS | 23 | 0.351599 | 0.914685 | 0.583832 | 0.902454 |
| GOBP_POSITIVE_REGULATION_OF_EXOCYTOSIS | 74 | 0.292076 | 0.914638 | 0.61831 | 0.902089 |

| | | | | | |
|---|---|---|---|---|---|
| GOBP_REGULATION_OF_NEURON_DIFFERENTIATION | 153 | 0.262905 | 0.914534 | 0.65659 | 0.901876 |
| GOBP_LONG_TERM_SYNAPTIC_POTENTIATION | 74 | 0.288675 | 0.914069 | 0.617219 | 0.902558 |
| GOBP_NEGATIVE_REGULATION_OF_MRNA_CATABOLIC_PROCESS | 55 | 0.30372 | 0.913904 | 0.591978 | 0.902508 |
| GOBP_NEGATIVE_REGULATION_OF_CELL_SUBSTRATE_JUNCTION_ORGANIZATION | 16 | 0.376098 | 0.913747 | 0.565495 | 0.902441 |
| GOBP_REACTIVE_OXYGEN_SPECIES_BIOSYNTHETIC_PROCESS | 43 | 0.313527 | 0.913665 | 0.588064 | 0.902185 |
| GOBP_MAINTENANCE_OF_CELL_NUMBER | 128 | 0.268294 | 0.912949 | 0.651422 | 0.903507 |
| GOBP_CELLULAR_RESPONSE_TO_AMYLOID_BETA | 32 | 0.33835 | 0.912755 | 0.581845 | 0.903509 |
| GOBP_NUCLEOSIDE_MONOPHOSPHATE_METABOLIC_PROCESS | 72 | 0.288071 | 0.912694 | 0.619624 | 0.903184 |
| GOBP_NUCLEOTIDE_BINDING_DOMAIN_LEUCINE_RICH_REPEAT_CONTAINING_RECEPTOR | 23 | 0.349958 | 0.912656 | 0.57504 | 0.902801 |
| GOBP_CELLULAR_RESPONSE_TO_STARVATION | 143 | 0.267581 | 0.912525 | 0.654888 | 0.902651 |
| GOBP_POSITIVE_REGULATION_OF_LYMPHOCYTE_MEDIATED_IMMUNITY | 73 | 0.289019 | 0.911085 | 0.630229 | 0.90582 |
| GOBP_GLANDULAR_EPITHELIAL_CELL_DIFFERENTIATION | 33 | 0.328697 | 0.910783 | 0.591118 | 0.906104 |
| GOBP_SECRETION_BY_TISSUE | 28 | 0.339242 | 0.91072 | 0.597037 | 0.90578 |
| GOBP_NEGATIVE_REGULATION_OF_GTPASE_ACTIVITY | 29 | 0.335333 | 0.91057 | 0.598507 | 0.905667 |
| GOBP_REGULATION_OF_WNT_SIGNALING_PATHWAY | 298 | 0.247645 | 0.910373 | 0.705171 | 0.905709 |
| GOBP_REGULATION_OF_DENDRITE_DEVELOPMENT | 92 | 0.2755 | 0.910296 | 0.634241 | 0.905414 |
| GOBP_RETINA_HOMEOSTASIS | 48 | 0.31373 | 0.909587 | 0.59322 | 0.906661 |
| GOBP_CRANIAL_NERVE_DEVELOPMENT | 39 | 0.31697 | 0.909269 | 0.605531 | 0.906964 |
| GOBP_SYNAPTIC_MEMBRANE_ADHESION | 25 | 0.35156 | 0.908688 | 0.598746 | 0.907889 |
| GOBP_MORPHOGENESIS_OF_EMBRYONIC_EPITHELIUM | 134 | 0.267046 | 0.908608 | 0.660025 | 0.907634 |
| GOBP_TELENCEPHALON_DEVELOPMENT | 208 | 0.251627 | 0.908462 | 0.679954 | 0.907517 |
| GOBP_FIBROBLAST_MIGRATION | 44 | 0.309155 | 0.90831 | 0.608631 | 0.907435 |
| GOBP_REGULATION_OF_NEURAL_PRECURSOR_CELL_PROLIFERATION | 71 | 0.290225 | 0.908206 | 0.622691 | 0.907199 |
| GOBP_NEGATIVE_REGULATION_OF_CATABOLIC_PROCESS | 277 | 0.247855 | 0.90772 | 0.714124 | 0.907957 |
| GOBP_REGULATION_OF_EPIDERMAL_GROWTH_FACTOR_ACTIVATED_RECEPTOR_ACTIVIT | 25 | 0.349408 | 0.90717 | 0.597826 | 0.908837 |
| GOBP_LONG_TERM_SYNAPTIC_DEPRESSION | 27 | 0.344977 | 0.90713 | 0.589896 | 0.908456 |
| GOBP_REGULATION_OF_INTRACELLULAR_TRANSPORT | 304 | 0.246713 | 0.907054 | 0.711432 | 0.908169 |
| GOBP_PEPTIDE_CATABOLIC_PROCESS | 23 | 0.349394 | 0.907038 | 0.596434 | 0.907733 |
| GOBP_POSITIVE_REGULATION_OF_HORMONE_SECRETION | 91 | 0.283291 | 0.906814 | 0.614891 | 0.907822 |
| GOBP_REGULATION_OF_BICELLULAR_TIGHT_JUNCTION_ASSEMBLY | 17 | 0.380749 | 0.906518 | 0.610853 | 0.908071 |
| GOBP_NEGATIVE_REGULATION_OF_SECRETION | 116 | 0.267508 | 0.906427 | 0.641154 | 0.907826 |
| GOBP_REGULATION_OF_NEURON_PROJECTION_DEVELOPMENT | 398 | 0.242557 | 0.906191 | 0.727949 | 0.907952 |
| GOBP_REGULATION_OF_FIBROBLAST_MIGRATION | 33 | 0.329181 | 0.905628 | 0.582222 | 0.90887 |
| GOBP_REGULATION_OF_PROTEIN_IMPORT | 55 | 0.298968 | 0.905427 | 0.607578 | 0.908879 |
| GOBP_POSITIVE_REGULATION_OF_PROTEIN_BINDING | 76 | 0.284598 | 0.905427 | 0.626234 | 0.908405 |
| GOBP_NEGATIVE_REGULATION_OF_CELLULAR_CATABOLIC_PROCESS | 229 | 0.252347 | 0.905255 | 0.664027 | 0.908327 |
| GOBP_POSITIVE_REGULATION_OF_PROTEASOMAL_UBIQUITIN_DEPENDENT_PROTEIN_CA | 88 | 0.275022 | 0.904901 | 0.657289 | 0.90872 |
| GOBP_NEGATIVE_REGULATION_OF_OSSIFICATION | 25 | 0.350407 | 0.903842 | 0.600299 | 0.910872 |
| GOBP_POSITIVE_REGULATION_OF_VIRAL_PROCESS | 56 | 0.295894 | 0.903774 | 0.621697 | 0.910577 |
| GOBP_RESPONSE_TO_GLUCAGON | 16 | 0.377276 | 0.903466 | 0.603015 | 0.91086 |
| GOBP_STEROID_METABOLIC_PROCESS | 227 | 0.248467 | 0.903394 | 0.6916 | 0.910561 |
| GOBP_NEGATIVE_REGULATION_OF_NUCLEAR_DIVISION | 51 | 0.299832 | 0.90331 | 0.627778 | 0.910293 |
| GOBP_REGULATION_OF_MAP_KINASE_ACTIVITY | 158 | 0.259581 | 0.902662 | 0.669065 | 0.911414 |
| GOBP_GLUTAMATE_RECEPTOR_SIGNALING_PATHWAY | 45 | 0.311549 | 0.902005 | 0.615836 | 0.912541 |
| GOBP_EXTRACELLULAR_TRANSPORT | 38 | 0.3196 | 0.901967 | 0.611833 | 0.912156 |
| GOBP_MYOTUBE_CELL_DEVELOPMENT | 29 | 0.335378 | 0.901787 | 0.609984 | 0.912106 |
| GOBP_RESPONSE_TO_PROSTAGLANDIN_E | 16 | 0.378731 | 0.901279 | 0.606109 | 0.912828 |
| GOBP_POSITIVE_REGULATION_OF_PEPTIDE_SECRETION | 68 | 0.290178 | 0.90114 | 0.630854 | 0.91268 |
| GOBP_REGULATION_OF_CALCINEURIN_MEDIATED_SIGNALING | 30 | 0.331635 | 0.900766 | 0.60597 | 0.913169 |
| GOBP_NEGATIVE_REGULATION_OF_TRANSPORTER_ACTIVITY | 58 | 0.294112 | 0.899485 | 0.639257 | 0.915891 |
| GOBP_CENTRAL_NERVOUS_SYSTEM_PROJECTION_NEURON_AXONOGENESIS | 25 | 0.340366 | 0.89885 | 0.597222 | 0.916996 |
| GOBP_POSITIVE_REGULATION_OF_DEPHOSPHORYLATION | 52 | 0.301378 | 0.898523 | 0.643678 | 0.917332 |
| GOBP_NEGATIVE_REGULATION_OF_CHEMOTAXIS | 49 | 0.30111 | 0.898407 | 0.646199 | 0.917145 |
| GOBP_MODULATION_BY_HOST_OF_SYMBIONT_PROCESS | 53 | 0.295631 | 0.898333 | 0.633286 | 0.916849 |
| GOBP_PEPTIDE_TRANSPORT | 189 | 0.254106 | 0.898322 | 0.685714 | 0.916405 |
| GOBP_FC_RECEPTOR_SIGNALING_PATHWAY | 38 | 0.316267 | 0.897915 | 0.616809 | 0.916908 |
| GOBP_REGULATION_OF_TRANS_SYNAPTIC_SIGNALING | 359 | 0.240368 | 0.897855 | 0.751921 | 0.916573 |
| GOBP_NEGATIVE_REGULATION_OF_DNA_BINDING | 45 | 0.308926 | 0.897784 | 0.615272 | 0.916274 |
| GOBP_ATP_TRANSPORT | 15 | 0.378818 | 0.896915 | 0.589905 | 0.917915 |
| GOBP_CENTROSOME_DUPLICATION | 71 | 0.281876 | 0.896269 | 0.63807 | 0.919014 |
| GOBP_REGULATION_OF_NUCLEOCYTOPLASMIC_TRANSPORT | 97 | 0.275183 | 0.896244 | 0.655128 | 0.918592 |
| GOBP_CEREBRAL_CORTEX_CELL_MIGRATION | 40 | 0.312112 | 0.896157 | 0.626292 | 0.918347 |
| GOBP_STRIATED_MUSCLE_ADAPTATION | 34 | 0.320703 | 0.895997 | 0.636914 | 0.918244 |
| GOBP_INTERLEUKIN_6_MEDIATED_SIGNALING_PATHWAY | 16 | 0.376553 | 0.895569 | 0.588133 | 0.918825 |
| GOBP_SUPEROXIDE_ANION_GENERATION | 28 | 0.33479 | 0.895525 | 0.606013 | 0.918463 |
| GOBP_PEPTIDYL_PROLINE_MODIFICATION | 51 | 0.303907 | 0.895278 | 0.654015 | 0.918585 |
| GOBP_CELLULAR_RESPONSE_TO_STEROID_HORMONE_STIMULUS | 171 | 0.254174 | 0.894293 | 0.699284 | 0.920498 |
| GOBP_REGULATION_OF_EPITHELIAL_TO_MESENCHYMAL_TRANSITION | 82 | 0.27726 | 0.893352 | 0.661784 | 0.922284 |
| GOBP_ORGANIC_HYDROXY_COMPOUND_CATABOLIC_PROCESS | 55 | 0.297518 | 0.893314 | 0.636608 | 0.921889 |
| GOBP_PURINE_CONTAINING_COMPOUND_CATABOLIC_PROCESS | 46 | 0.306917 | 0.892659 | 0.643875 | 0.922977 |
| GOBP_NEGATIVE_REGULATION_OF_OXIDOREDUCTASE_ACTIVITY | 16 | 0.376927 | 0.892297 | 0.589018 | 0.923375 |
| GOBP_GLYCOPROTEIN_METABOLIC_PROCESS | 335 | 0.238883 | 0.891902 | 0.752759 | 0.923883 |
| GOBP_CELLULAR_ALDEHYDE_METABOLIC_PROCESS | 51 | 0.297735 | 0.891824 | 0.626016 | 0.923594 |
| GOBP_REGULATION_OF_DEFENSE_RESPONSE_TO_VIRUS | 61 | 0.289206 | 0.891598 | 0.644793 | 0.923654 |
| GOBP_NEGATIVE_REGULATION_OF_SIGNAL_TRANSDUCTION_BY_P53_CLASS_MEDIATOR | 32 | 0.323917 | 0.891528 | 0.635793 | 0.923348 |
| GOBP_GLYCEROPHOSPHOLIPID_BIOSYNTHETIC_PROCESS | 195 | 0.251549 | 0.891411 | 0.711449 | 0.923172 |
| GOBP_CEREBELLAR_PURKINJE_CELL_LAYER_DEVELOPMENT | 26 | 0.335045 | 0.891073 | 0.608964 | 0.923541 |
| GOBP_RESPONSE_TO_ENDOPLASMIC_RETICULUM_STRESS | 241 | 0.243839 | 0.890662 | 0.723906 | 0.924042 |
| GOBP_NEGATIVE_REGULATION_OF_CELL_PROJECTION_ORGANIZATION | 172 | 0.252181 | 0.889755 | 0.705113 | 0.925713 |
| GOBP_POSITIVE_REGULATION_OF_PROTEIN_AUTOPHOSPHORYLATION | 25 | 0.335494 | 0.889591 | 0.59403 | 0.92564 |
| GOBP_SPHINGOLIPID_BIOSYNTHETIC_PROCESS | 96 | 0.268611 | 0.887827 | 0.670984 | 0.929432 |
| GOBP_CERAMIDE_BIOSYNTHETIC_PROCESS | 60 | 0.291236 | 0.887591 | 0.648383 | 0.929511 |
| GOBP_NEGATIVE_REGULATION_OF_CELLULAR_MACROMOLECULE_BIOSYNTHETIC_PROC | 239 | 0.244643 | 0.887465 | 0.724524 | 0.929348 |
| GOBP_POLYSACCHARIDE_METABOLIC_PROCESS | 89 | 0.274113 | 0.887465 | 0.679688 | 0.928876 |
| GOBP_POSITIVE_REGULATION_OF_LIPASE_ACTIVITY | 44 | 0.30266 | 0.887457 | 0.626609 | 0.928424 |
| GOBP_SPINDLE_LOCALIZATION | 48 | 0.297315 | 0.887138 | 0.628895 | 0.928705 |
| GOBP_POSITIVE_REGULATION_OF_PROTEIN_MATURATION | 23 | 0.342495 | 0.886963 | 0.617694 | 0.928658 |
| GOBP_POST_TRANSLATIONAL_PROTEIN_MODIFICATION | 36 | 0.312679 | 0.886896 | 0.627068 | 0.928347 |
| GOBP_SPERM_FLAGELLUM_ASSEMBLY | 24 | 0.345901 | 0.886802 | 0.640127 | 0.928111 |
| GOBP_NEGATIVE_REGULATION_OF_MAP_KINASE_ACTIVITY | 49 | 0.296767 | 0.886641 | 0.645954 | 0.928024 |
| GOBP_POSITIVE_REGULATION_OF_TRANSLATION | 126 | 0.263904 | 0.886475 | 0.686104 | 0.927921 |
| GOBP_NEGATIVE_REGULATION_OF_PROTEIN_BINDING | 78 | 0.276804 | 0.886455 | 0.651132 | 0.927503 |
| GOBP_REGULATION_OF_PHOSPHOLIPID_METABOLIC_PROCESS | 29 | 0.334617 | 0.886115 | 0.647239 | 0.927827 |
| GOBP_REGULATION_OF_RAC_PROTEIN_SIGNAL_TRANSDUCTION | 19 | 0.356311 | 0.885976 | 0.603804 | 0.927684 |
| GOBP_NEGATIVE_REGULATION_OF_ORGANIC_ACID_TRANSPORT | 19 | 0.3567 | 0.885269 | 0.61646 | 0.928862 |
| GOBP_POSITIVE_REGULATION_OF_LYMPHOCYTE_MIGRATION | 24 | 0.337104 | 0.884639 | 0.616541 | 0.929876 |
| GOBP_CENTRAL_NERVOUS_SYSTEM_NEURON_AXONOGENESIS | 33 | 0.320839 | 0.884291 | 0.634361 | 0.930229 |
| GOBP_POSITIVE_REGULATION_OF_NUCLEOTIDE_METABOLIC_PROCESS | 31 | 0.323283 | 0.88343 | 0.630178 | 0.931808 |
| GOBP_ADAPTATION_OF_SIGNALING_PATHWAY | 17 | 0.371695 | 0.882988 | 0.622517 | 0.932353 |
| GOBP_POSITIVE_REGULATION_OF_PROTEIN_POLYMERIZATION | 84 | 0.273495 | 0.882824 | 0.686828 | 0.932271 |
| GOBP_REGULATION_OF_FIBROBLAST_GROWTH_FACTOR_RECEPTOR_SIGNALING_PATHW | 29 | 0.334082 | 0.882675 | 0.656347 | 0.932167 |
| GOBP_PHOSPHATIDYLINOSITOL_DEPHOSPHORYLATION | 29 | 0.324941 | 0.882348 | 0.642415 | 0.932488 |
| GOBP_ENTRAINMENT_OF_CIRCADIAN_CLOCK | 32 | 0.32167 | 0.882345 | 0.629464 | 0.932023 |
| GOBP_POSITIVE_REGULATION_OF_SODIUM_ION_TRANSMEMBRANE_TRANSPORT | 19 | 0.351857 | 0.881742 | 0.64658 | 0.933 |
| GOBP_ENDOCYTOSIS | 477 | 0.231382 | 0.881575 | 0.793249 | 0.932921 |
| GOBP_POSITIVE_REGULATION_OF_REGULATED_SECRETORY_PATHWAY | 46 | 0.306022 | 0.880883 | 0.669054 | 0.934097 |
| GOBP_NEGATIVE_REGULATION_OF_NOTCH_SIGNALING_PATHWAY | 32 | 0.317468 | 0.880371 | 0.628912 | 0.934827 |
| GOBP_REGULATION_OF_DNA_BIOSYNTHETIC_PROCESS | 117 | 0.260843 | 0.880162 | 0.718078 | 0.934844 |
| GOBP_LIPID_LOCALIZATION | 333 | 0.235389 | 0.880049 | 0.779493 | 0.934667 |
| GOBP_COENZYME_A_METABOLIC_PROCESS | 18 | 0.36109 | 0.879349 | 0.633174 | 0.93582 |
| GOBP_POSITIVE_REGULATION_OF_B_CELL_ACTIVATION | 58 | 0.285715 | 0.878911 | 0.668022 | 0.936392 |
| GOBP_RESPONSE_TO_THYROID_HORMONE | 22 | 0.341838 | 0.878858 | 0.628225 | 0.93605 |
| GOBP_DEOXYRIBOSE_PHOSPHATE_METABOLIC_PROCESS | 41 | 0.30812 | 0.878716 | 0.644068 | 0.935928 |
| GOBP_HEMATOPOIETIC_PROGENITOR_CELL_DIFFERENTIATION | 102 | 0.26225 | 0.878707 | 0.694058 | 0.935478 |
| GOBP_BODY_FLUID_SECRETION | 70 | 0.277881 | 0.878545 | 0.664374 | 0.935395 |
| GOBP_NEURON_PROJECTION_REGENERATION | 50 | 0.292477 | 0.878197 | 0.658537 | 0.935723 |
| GOBP_VITAMIN_TRANSPORT | 34 | 0.31003 | 0.877469 | 0.640483 | 0.936968 |
| GOBP_ESTABLISHMENT_OF_ENDOTHELIAL_BARRIER | 49 | 0.295862 | 0.877132 | 0.65368 | 0.937247 |
| GOBP_SYNAPSE_ORGANIZATION | 371 | 0.234238 | 0.876953 | 0.814973 | 0.937225 |
| GOBP_POSITIVE_REGULATION_OF_KERATINOCYTE_DIFFERENTIATION | 15 | 0.37714 | 0.876694 | 0.648829 | 0.937382 |
| GOBP_RNA_DECAPPING | 17 | 0.362287 | 0.875539 | 0.629325 | 0.939615 |
| GOBP_INNER_EAR_AUDITORY_RECEPTOR_CELL_DIFFERENTIATION | 32 | 0.317746 | 0.875155 | 0.645833 | 0.940079 |
| GOBP_POSITIVE_REGULATION_OF_ANIMAL_ORGAN_MORPHOGENESIS | 28 | 0.324472 | 0.874024 | 0.665102 | 0.942225 |
| GOBP_REGULATION_OF_ENDOTHELIAL_CELL_DIFFERENTIATION | 34 | 0.313729 | 0.873264 | 0.650655 | 0.943461 |
| GOBP_POSITIVE_REGULATION_OF_EXCITATORY_POSTSYNAPTIC_POTENTIAL | 24 | 0.333845 | 0.873067 | 0.636785 | 0.94344 |
| GOBP_DNA_DEPENDENT_DNA_REPLICATION_MAINTENANCE_OF_FIDELITY | 46 | 0.298135 | 0.873066 | 0.686012 | 0.942971 |
| GOBP_CELL_CELL_JUNCTION_ORGANIZATION | 173 | 0.248349 | 0.872982 | 0.724138 | 0.942694 |
| GOBP_REGULATION_OF_PROTEIN_POLYUBIQUITINATION | 25 | 0.33278 | 0.872937 | 0.643879 | 0.942338 |
| GOBP_PLASMA_MEMBRANE_ORGANIZATION | 119 | 0.256824 | 0.872899 | 0.725641 | 0.941957 |
| GOBP_RESPONSE_TO_X_RAY | 30 | 0.31999 | 0.872484 | 0.632813 | 0.942448 |
| GOBP_REGULATION_OF_SIGNALING_RECEPTOR_ACTIVITY | 126 | 0.256588 | 0.872407 | 0.737745 | 0.942151 |
| GOBP_GOLGI_ORGANIZATION | 141 | 0.253771 | 0.872341 | 0.7164 | 0.941819 |
| GOBP_DNA_SYNTHESIS_INVOLVED_IN_DNA_REPAIR | 33 | 0.313249 | 0.871956 | 0.647761 | 0.942248 |
| GOBP_T_CELL_HOMEOSTASIS | 29 | 0.321828 | 0.871404 | 0.627943 | 0.943116 |
| GOBP_IRON_ION_TRANSPORT | 45 | 0.295354 | 0.870894 | 0.656077 | 0.943812 |
| GOBP_POSITIVE_REGULATION_OF_MONOOXYGENASE_ACTIVITY | 24 | 0.331569 | 0.870736 | 0.639319 | 0.943714 |
| GOBP_CELLULAR_RESPONSE_TO_ARSENIC_CONTAINING_SUBSTANCE | 21 | 0.349258 | 0.870725 | 0.642857 | 0.943273 |
| GOBP_AUDITORY_RECEPTOR_CELL_DEVELOPMENT | 19 | 0.349499 | 0.870478 | 0.628059 | 0.943382 |
| GOBP_AXON_DEVELOPMENT | 432 | 0.230478 | 0.870002 | 0.832101 | 0.943984 |
| GOBP_MORPHOGENESIS_OF_AN_EPITHELIAL_FOLD | 16 | 0.361336 | 0.869866 | 0.634993 | 0.943822 |
| GOBP_RESPONSE_TO_AXON_INJURY | 71 | 0.275489 | 0.869405 | 0.690638 | 0.944407 |
| GOBP_B_CELL_MEDIATED_IMMUNITY | 95 | 0.264193 | 0.869338 | 0.712951 | 0.944085 |
| GOBP_NEGATIVE_REGULATION_OF_TRANSLATIONAL_INITIATION | 18 | 0.358653 | 0.869235 | 0.634494 | 0.943865 |
| GOBP_SUBPALLIUM_DEVELOPMENT | 16 | 0.368906 | 0.868731 | 0.646362 | 0.944548 |
| GOBP_STEROL_METABOLIC_PROCESS | 116 | 0.258623 | 0.86811 | 0.717029 | 0.945486 |
| GOBP_NEGATIVE_REGULATION_OF_PROTEIN_CONTAINING_COMPLEX_ASSEMBLY | 128 | 0.251522 | 0.867009 | 0.718205 | 0.947599 |
| GOBP_PHOSPHATIDYLCHOLINE_BIOSYNTHETIC_PROCESS | 27 | 0.320287 | 0.865974 | 0.67134 | 0.949487 |
| GOBP_PROTEIN_MATURATION | 254 | 0.236519 | 0.865451 | 0.781963 | 0.950243 |
| GOBP_CELLULAR_COMPONENT_DISASSEMBLY | 420 | 0.230404 | 0.865366 | 0.832808 | 0.949965 |
| GOBP_NUCLEIC_ACID_PHOSPHODIESTER_BOND_HYDROLYSIS | 233 | 0.240511 | 0.865086 | 0.784027 | 0.950151 |
| GOBP_REGULATION_OF_CELL_SHAPE | 133 | 0.252151 | 0.864907 | 0.719902 | 0.950088 |
| GOBP_POSITIVE_REGULATION_OF_PHAGOCYTOSIS | 41 | 0.298233 | 0.864665 | 0.654357 | 0.950178 |
| GOBP_PHOSPHATIDYLINOSITOL_METABOLIC_PROCESS | 144 | 0.251148 | 0.86453 | 0.750605 | 0.950013 |
| GOBP_REGULATION_OF_KERATINOCYTE_DIFFERENTIATION | 27 | 0.325546 | 0.864209 | 0.646967 | 0.950266 |
| GOBP_CELL_DEATH_IN_RESPONSE_TO_HYDROGEN_PEROXIDE | 22 | 0.339452 | 0.863677 | 0.645802 | 0.950986 |
| GOBP_POSITIVE_REGULATION_OF_LAMELLIPODIUM_ORGANIZATION | 32 | 0.315618 | 0.863641 | 0.682635 | 0.950598 |
| GOBP_REGULATION_OF_NEUROGENESIS | 312 | 0.233772 | 0.863625 | 0.819475 | 0.950167 |

| | | | | | |
|---|---|---|---|---|---|
| GOBP_SODIUM_ION_IMPORT_ACROSS_PLASMA_MEMBRANE | 18 | 0.355438 | 0.863547 | 0.647727 | 0.949885 |
| GOBP_RESPIRATORY_SYSTEM_DEVELOPMENT | 159 | 0.246106 | 0.863424 | 0.753864 | 0.949694 |
| GOBP_NEGATIVE_REGULATION_OF_TUMOR_NECROSIS_FACTOR_SUPERFAMILY_CYTOKI | 40 | 0.29633 | 0.863058 | 0.676991 | 0.950052 |
| GOBP_REGULATION_OF_B_CELL_APOPTOTIC_PROCESS | 18 | 0.350628 | 0.862445 | 0.63711 | 0.950959 |
| GOBP_ADULT_LOCOMOTORY_BEHAVIOR | 65 | 0.27991 | 0.861589 | 0.689655 | 0.952407 |
| GOBP_CELLULAR_RESPONSE_TO_EXTRACELLULAR_STIMULUS | 212 | 0.237331 | 0.861165 | 0.771991 | 0.952904 |
| GOBP_POSITIVE_REGULATION_OF_STEROID_METABOLIC_PROCESS | 20 | 0.340822 | 0.860886 | 0.651125 | 0.953035 |
| GOBP_APICAL_JUNCTION_ASSEMBLY | 57 | 0.276388 | 0.860816 | 0.68306 | 0.952735 |
| GOBP_REGULATION_OF_GLUCONEOGENESIS | 43 | 0.294831 | 0.860792 | 0.696148 | 0.952321 |
| GOBP_NEGATIVE_REGULATION_OF_CANONICAL_WNT_SIGNALING_PATHWAY | 122 | 0.254065 | 0.860576 | 0.7325 | 0.952332 |
| GOBP_POSITIVE_REGULATION_OF_PROTEIN_LOCALIZATION_TO_CELL_PERIPHERY | 60 | 0.281585 | 0.860554 | 0.707317 | 0.951921 |
| GOBP_POSITIVE_REGULATION_OF_DOUBLE_STRAND_BREAK_REPAIR | 63 | 0.275511 | 0.859564 | 0.687237 | 0.953611 |
| GOBP_MICROTUBULE_BASED_MOVEMENT | 317 | 0.230221 | 0.859553 | 0.82988 | 0.953169 |
| GOBP_PURINE_NUCLEOTIDE_CATABOLIC_PROCESS | 41 | 0.29701 | 0.859192 | 0.671449 | 0.953538 |
| GOBP_CELLULAR_RESPONSE_TO_DSRNA | 21 | 0.340163 | 0.858977 | 0.636081 | 0.95357 |
| GOBP_CALCIUM_ION_REGULATED_EXOCYTOSIS_OF_NEUROTRANSMITTER | 15 | 0.372001 | 0.858914 | 0.656786 | 0.953253 |
| GOBP_REGULATION_OF_DNA_METABOLIC_PROCESS | 365 | 0.229214 | 0.858469 | 0.820043 | 0.953755 |
| GOBP_RESPONSE_TO_ACTIVITY | 42 | 0.29705 | 0.857988 | 0.690377 | 0.954374 |
| GOBP_REGULATION_OF_PROTEIN_CATABOLIC_PROCESS | 357 | 0.228745 | 0.857896 | 0.838608 | 0.954108 |
| GOBP_GLIAL_CELL_MIGRATION | 41 | 0.298466 | 0.857699 | 0.679274 | 0.954079 |
| GOBP_REGULATION_OF_T_HELPER_17_TYPE_IMMUNE_RESPONSE | 17 | 0.360123 | 0.857594 | 0.64377 | 0.95385 |
| GOBP_POSITIVE_REGULATION_OF_ACTIN_NUCLEATION | 16 | 0.368192 | 0.856673 | 0.661765 | 0.955458 |
| GOBP_HEPATOCYTE_APOPTOTIC_PROCESS | 16 | 0.362812 | 0.856345 | 0.649266 | 0.955737 |
| GOBP_TRANSEPITHELIAL_TRANSPORT | 28 | 0.324328 | 0.856187 | 0.672986 | 0.955632 |
| GOBP_POSITIVE_REGULATION_OF_PLASMA_MEMBRANE_BOUNDED_CELL_PROJECTION_ | 94 | 0.258231 | 0.855522 | 0.728682 | 0.956672 |
| GOBP_REGULATION_OF_CYTOSKELETON_ORGANIZATION | 479 | 0.225837 | 0.855496 | 0.864638 | 0.956265 |
| GOBP_REGULATED_EXOCYTOSIS | 190 | 0.239196 | 0.854863 | 0.794725 | 0.957207 |
| GOBP_HEART_FIELD_SPECIFICATION | 16 | 0.352461 | 0.854857 | 0.634551 | 0.956752 |
| GOBP_NEGATIVE_REGULATION_OF_RECEPTOR_SIGNALING_PATHWAY_VIA_STAT | 22 | 0.335673 | 0.854633 | 0.661585 | 0.956818 |
| GOBP_DEPHOSPHORYLATION | 370 | 0.226485 | 0.854278 | 0.852052 | 0.957143 |
| GOBP_POSITIVE_REGULATION_OF_INTERFERON_ALPHA_PRODUCTION | 17 | 0.360415 | 0.853717 | 0.645 | 0.957919 |
| GOBP_DNA_DOUBLE_STRAND_BREAK_PROCESSING | 16 | 0.353529 | 0.852899 | 0.655116 | 0.959247 |
| GOBP_POSITIVE_REGULATION_OF_REACTIVE_OXYGEN_SPECIES_BIOSYNTHETIC_PROCE | 15 | 0.369921 | 0.852825 | 0.668277 | 0.958947 |
| GOBP_MITOCHONDRIAL_CALCIUM_ION_HOMEOSTASIS | 24 | 0.329974 | 0.852471 | 0.678288 | 0.959259 |
| GOBP_T_HELPER_17_CELL_DIFFERENTIATION | 17 | 0.360736 | 0.852214 | 0.684729 | 0.959346 |
| GOBP_REGULATION_OF_NEURON_MIGRATION | 41 | 0.297667 | 0.851138 | 0.669004 | 0.9612 |
| GOBP_APOPTOTIC_MITOCHONDRIAL_CHANGES | 92 | 0.259593 | 0.850338 | 0.746154 | 0.962512 |
| GOBP_NEGATIVE_REGULATION_OF_ORGAN_GROWTH | 27 | 0.322924 | 0.84958 | 0.677177 | 0.963669 |
| GOBP_REGULATION_OF_JUN_KINASE_ACTIVITY | 50 | 0.286266 | 0.849363 | 0.70963 | 0.963673 |
| GOBP_PROTEIN_IMPORT | 189 | 0.236615 | 0.849263 | 0.796231 | 0.963419 |
| GOBP_ALPHA_AMINO_ACID_METABOLIC_PROCESS | 165 | 0.242149 | 0.849028 | 0.79078 | 0.963461 |
| GOBP_POSITIVE_REGULATION_OF_HISTONE_MODIFICATION | 95 | 0.25957 | 0.848691 | 0.744681 | 0.963734 |
| GOBP_ORGANOPHOSPHATE_CATABOLIC_PROCESS | 137 | 0.246058 | 0.848567 | 0.755528 | 0.963546 |
| GOBP_REGULATION_OF_REACTIVE_OXYGEN_SPECIES_BIOSYNTHETIC_PROCESS | 35 | 0.297094 | 0.846571 | 0.694529 | 0.967415 |
| GOBP_PHOSPHOLIPID_METABOLIC_PROCESS | 331 | 0.228472 | 0.846528 | 0.837134 | 0.967041 |
| GOBP_POSITIVE_REGULATION_OF_CALCIUM_ION_IMPORT | 15 | 0.358141 | 0.846464 | 0.665079 | 0.966714 |
| GOBP_NEGATIVE_REGULATION_OF_CALCIUM_ION_TRANSPORT_INTO_CYTOSOL | 16 | 0.354837 | 0.846006 | 0.65534 | 0.967209 |
| GOBP_NERVE_DEVELOPMENT | 65 | 0.267714 | 0.845286 | 0.746055 | 0.968305 |
| GOBP_PHOSPHATIDYLINOSITOL_PHOSPHATE_BIOSYNTHETIC_PROCESS | 62 | 0.277195 | 0.844967 | 0.732782 | 0.968531 |
| GOBP_IMMUNOGLOBULIN_PRODUCTION_INVOLVED_IN_IMMUNOGLOBULIN_MEDIATED_IM | 56 | 0.277151 | 0.844962 | 0.705219 | 0.968076 |
| GOBP_REGULATION_OF_SIGNAL_TRANSDUCTION_BY_P53_CLASS_MEDIATOR | 98 | 0.255662 | 0.844738 | 0.764783 | 0.968065 |
| GOBP_RNA_EXPORT_FROM_NUCLEUS | 74 | 0.264403 | 0.844533 | 0.735571 | 0.968053 |
| GOBP_NEGATIVE_REGULATION_OF_CYTOKINE_PRODUCTION_INVOLVED_IN_INFLAMMATO | 15 | 0.353191 | 0.844405 | 0.674497 | 0.967865 |
| GOBP_GLIAL_CELL_DEVELOPMENT | 90 | 0.259958 | 0.844352 | 0.742782 | 0.967525 |
| GOBP_INTERFERON_BETA_PRODUCTION | 50 | 0.284597 | 0.844181 | 0.728571 | 0.967426 |
| GOBP_MODULATION_OF_EXCITATORY_POSTSYNAPTIC_POTENTIAL | 36 | 0.297598 | 0.843257 | 0.689708 | 0.96898 |
| GOBP_ALCOHOL_CATABOLIC_PROCESS | 42 | 0.286974 | 0.842863 | 0.712209 | 0.969377 |
| GOBP_SOMITOGENESIS | 51 | 0.281213 | 0.842327 | 0.707071 | 0.970058 |
| GOBP_REGULATION_OF_NEUROTRANSMITTER_TRANSPORT | 83 | 0.261478 | 0.84103 | 0.762943 | 0.972385 |
| GOBP_NEGATIVE_REGULATION_OF_PHOSPHOPROTEIN_PHOSPHATASE_ACTIVITY | 22 | 0.32391 | 0.840092 | 0.678196 | 0.973906 |
| GOBP_NEGATIVE_REGULATION_OF_SMALL_MOLECULE_METABOLIC_PROCESS | 79 | 0.261558 | 0.840031 | 0.74829 | 0.973569 |
| GOBP_RESPONSE_TO_SALT_STRESS | 20 | 0.337252 | 0.839664 | 0.683465 | 0.97388 |
| GOBP_PROTEIN_DEPHOSPHORYLATION | 258 | 0.230672 | 0.839309 | 0.854586 | 0.974179 |
| GOBP_POSITIVE_REGULATION_OF_ATP_METABOLIC_PROCESS | 29 | 0.305181 | 0.838683 | 0.676471 | 0.975002 |
| GOBP_NEURON_CELLULAR_HOMEOSTASIS | 28 | 0.304486 | 0.838423 | 0.700475 | 0.975105 |
| GOBP_CELLULAR_RESPONSE_TO_HEAT | 61 | 0.272964 | 0.838025 | 0.719274 | 0.97549 |
| GOBP_REGULATION_OF_MYELOID_CELL_DIFFERENTIATION | 159 | 0.238854 | 0.83753 | 0.792176 | 0.976056 |
| GOBP_PHOSPHOLIPID_BIOSYNTHETIC_PROCESS | 231 | 0.231604 | 0.837503 | 0.837868 | 0.97565 |
| GOBP_CEREBELLAR_CORTEX_FORMATION | 24 | 0.324071 | 0.836844 | 0.660465 | 0.976577 |
| GOBP_REGULATION_OF_CARDIAC_MUSCLE_CELL_MEMBRANE_REPOLARIZATION | 21 | 0.331442 | 0.836734 | 0.706169 | 0.97634 |
| GOBP_NEGATIVE_REGULATION_OF_RELEASE_OF_CYTOCHROME_C_FROM_MITOCHONDR | 18 | 0.339182 | 0.836633 | 0.694772 | 0.976097 |
| GOBP_CELLULAR_RESPONSE_TO_GAMMA_RADIATION | 28 | 0.312469 | 0.835001 | 0.71518 | 0.979037 |
| GOBP_NEGATIVE_REGULATION_OF_RNA_CATABOLIC_PROCESS | 67 | 0.265444 | 0.834938 | 0.744741 | 0.978686 |
| GOBP_MICROTUBULE_POLYMERIZATION | 81 | 0.261497 | 0.834828 | 0.77047 | 0.978444 |
| GOBP_AUTOPHAGY_OF_MITOCHONDRION | 86 | 0.253239 | 0.834796 | 0.765873 | 0.978052 |
| GOBP_VITAMIN_BIOSYNTHETIC_PROCESS | 16 | 0.348787 | 0.834764 | 0.686084 | 0.977654 |
| GOBP_CELLULAR_RESPONSE_TO_REACTIVE_NITROGEN_SPECIES | 15 | 0.363496 | 0.834625 | 0.708126 | 0.977471 |
| GOBP_MITOCHONDRIAL_OUTER_MEMBRANE_PERMEABILIZATION_INVOLVED_IN_PROGRA | 36 | 0.297299 | 0.83462 | 0.69037 | 0.977017 |
| GOBP_REGULATION_OF_PHOSPHATASE_ACTIVITY | 67 | 0.267324 | 0.834506 | 0.735772 | 0.976792 |
| GOBP_DEVELOPMENTAL_GROWTH_INVOLVED_IN_MORPHOGENESIS | 213 | 0.23152 | 0.834368 | 0.83882 | 0.976625 |
| GOBP_PROTEIN_POLYUBIQUITINATION | 228 | 0.228957 | 0.833681 | 0.847802 | 0.977597 |
| GOBP_REGULATION_OF_NUCLEOTIDE_METABOLIC_PROCESS | 69 | 0.264289 | 0.83368 | 0.748299 | 0.977137 |
| GOBP_POSITIVE_REGULATION_OF_MUSCLE_CELL_APOPTOTIC_PROCESS | 18 | 0.344374 | 0.833389 | 0.682566 | 0.977248 |
| GOBP_ANIMAL_ORGAN_MATURATION | 25 | 0.321308 | 0.833117 | 0.722675 | 0.977383 |
| GOBP_CELLULAR_SODIUM_ION_HOMEOSTASIS | 19 | 0.33896 | 0.832875 | 0.689764 | 0.977407 |
| GOBP_SERINE_FAMILY_AMINO_ACID_BIOSYNTHETIC_PROCESS | 18 | 0.335412 | 0.832532 | 0.679487 | 0.977678 |
| GOBP_ANTIVIRAL_INNATE_IMMUNE_RESPONSE | 15 | 0.350191 | 0.83126 | 0.679117 | 0.9798 |
| GOBP_EXTRACELLULAR_MATRIX_ASSEMBLY | 40 | 0.288488 | 0.831005 | 0.697194 | 0.979855 |
| GOBP_SYNAPTIC_VESICLE_RECYCLING | 68 | 0.262551 | 0.830731 | 0.779706 | 0.979959 |
| GOBP_POSITIVE_REGULATION_OF_ERYTHROCYTE_DIFFERENTIATION | 28 | 0.306308 | 0.830692 | 0.721078 | 0.979574 |
| GOBP_REGULATION_OF_SUPRAMOLECULAR_FIBER_ORGANIZATION | 339 | 0.222935 | 0.830214 | 0.863044 | 0.980082 |
| GOBP_CELL_CYCLE_G1_S_PHASE_TRANSITION | 206 | 0.230718 | 0.829951 | 0.849078 | 0.980155 |
| GOBP_SISTER_CHROMATID_COHESION | 57 | 0.277015 | 0.829434 | 0.745902 | 0.980758 |
| GOBP_REGULATION_OF_GLYCOGEN_METABOLIC_PROCESS | 29 | 0.30359 | 0.829367 | 0.70679 | 0.980437 |
| GOBP_POSTTRANSCRIPTIONAL_REGULATION_OF_GENE_EXPRESSION | 485 | 0.218004 | 0.829003 | 0.923475 | 0.980729 |
| GOBP_REGULATION_OF_MITOTIC_CELL_CYCLE | 409 | 0.221596 | 0.828626 | 0.893089 | 0.981062 |
| GOBP_INOSITOL_PHOSPHATE_MEDIATED_SIGNALING | 47 | 0.283102 | 0.828145 | 0.741667 | 0.981582 |
| GOBP_POSITIVE_REGULATION_OF_MYELINATION | 16 | 0.344276 | 0.828047 | 0.696486 | 0.981325 |
| GOBP_ER_NUCLEUS_SIGNALING_PATHWAY | 44 | 0.283718 | 0.82802 | 0.765557 | 0.980919 |
| GOBP_RESPONSE_TO_PROSTAGLANDIN | 21 | 0.331403 | 0.826923 | 0.701967 | 0.982686 |
| GOBP_REGULATION_OF_MEMBRANE_PERMEABILITY | 64 | 0.264009 | 0.826726 | 0.760753 | 0.982597 |
| GOBP_REGULATION_OF_GLUCOSE_METABOLIC_PROCESS | 103 | 0.247599 | 0.826078 | 0.813385 | 0.983443 |
| GOBP_ORGAN_INDUCTION | 18 | 0.336699 | 0.825962 | 0.710611 | 0.983209 |
| GOBP_REGULATION_OF_MAST_CELL_ACTIVATION | 30 | 0.304458 | 0.825753 | 0.732733 | 0.983163 |
| GOBP_MONOCARBOXYLIC_ACID_BIOSYNTHETIC_PROCESS | 155 | 0.237752 | 0.825588 | 0.831541 | 0.983039 |
| GOBP_NITRIC_OXIDE_SYNTHASE_BIOSYNTHETIC_PROCESS | 15 | 0.349943 | 0.825393 | 0.68438 | 0.982994 |
| GOBP_NEGATIVE_REGULATION_OF_VASCULAR_PERMEABILITY | 17 | 0.343008 | 0.825052 | 0.698854 | 0.983241 |
| GOBP_REGULATION_OF_CELLULAR_PROTEIN_CATABOLIC_PROCESS | 235 | 0.226848 | 0.824988 | 0.862344 | 0.982911 |
| GOBP_NEGATIVE_REGULATION_OF_CELL_JUNCTION_ASSEMBLY | 26 | 0.309018 | 0.824789 | 0.728814 | 0.982835 |
| GOBP_ACTIVATION_OF_PROTEIN_KINASE_B_ACTIVITY | 25 | 0.310391 | 0.824711 | 0.733124 | 0.982522 |
| GOBP_POSITIVE_REGULATION_OF_POSTTRANSCRIPTIONAL_GENE_SILENCING | 30 | 0.304938 | 0.824393 | 0.714286 | 0.982712 |
| GOBP_CALCIUM_ION_IMPORT | 70 | 0.260501 | 0.824297 | 0.756831 | 0.982453 |
| GOBP_RIBONUCLEOSIDE_DIPHOSPHATE_METABOLIC_PROCESS | 99 | 0.247878 | 0.823829 | 0.793333 | 0.982938 |
| GOBP_PROTEIN_KINASE_C_ACTIVATING_G_PROTEIN_COUPLED_RECEPTOR_SIGNALING_ | 21 | 0.328767 | 0.823331 | 0.726283 | 0.983483 |
| GOBP_ORGANIC_HYDROXY_COMPOUND_BIOSYNTHETIC_PROCESS | 188 | 0.230921 | 0.823226 | 0.847302 | 0.983235 |
| GOBP_RESPONSE_TO_TESTOSTERONE | 32 | 0.300511 | 0.822743 | 0.726048 | 0.983809 |
| GOBP_MEMBRANE_DEPOLARIZATION | 70 | 0.257937 | 0.82251 | 0.771784 | 0.983832 |
| GOBP_POSITIVE_REGULATION_OF_PROTEIN_MODIFICATION_BY_SMALL_PROTEIN_CONJU | 127 | 0.241594 | 0.822489 | 0.81273 | 0.983418 |
| GOBP_PHOSPHATIDYLINOSITOL_BIOSYNTHETIC_PROCESS | 122 | 0.241374 | 0.822408 | 0.822816 | 0.983126 |
| GOBP_REGULATION_OF_PROTEIN_LOCALIZATION_TO_CELL_PERIPHERY | 111 | 0.246717 | 0.821931 | 0.817136 | 0.983651 |
| GOBP_OLFACTORY_LOBE_DEVELOPMENT | 26 | 0.312218 | 0.821583 | 0.703416 | 0.983866 |
| GOBP_EPITHELIAL_CELL_MORPHOGENESIS | 30 | 0.304158 | 0.821567 | 0.70219 | 0.983446 |
| GOBP_ORGANIC_ACID_BIOSYNTHETIC_PROCESS | 234 | 0.226324 | 0.821566 | 0.870307 | 0.982993 |
| GOBP_ORGANELLE_FUSION | 131 | 0.238903 | 0.821385 | 0.802454 | 0.98288 |
| GOBP_SUPPRESSION_OF_VIRAL_RELEASE_BY_HOST | 16 | 0.347 | 0.821192 | 0.685938 | 0.982833 |
| GOBP_REGULATION_OF_NERVOUS_SYSTEM_DEVELOPMENT | 376 | 0.219052 | 0.82041 | 0.889366 | 0.983915 |
| GOBP_CAMERA_TYPE_EYE_PHOTORECEPTOR_CELL_DIFFERENTIATION | 21 | 0.327894 | 0.820259 | 0.712264 | 0.983753 |
| GOBP_REGULATION_OF_MRNA_CATABOLIC_PROCESS | 156 | 0.234478 | 0.819569 | 0.840476 | 0.984658 |
| GOBP_CARBOHYDRATE_METABOLIC_PROCESS | 492 | 0.216165 | 0.819307 | 0.920585 | 0.984718 |
| GOBP_NEGATIVE_REGULATION_OF_HORMONE_SECRETION | 42 | 0.284988 | 0.818795 | 0.714286 | 0.985303 |
| GOBP_CELL_CELL_SIGNALING_BY_WNT | 401 | 0.216244 | 0.818756 | 0.918259 | 0.984929 |
| GOBP_REGULATION_OF_CAMP_DEPENDENT_PROTEIN_KINASE_ACTIVITY | 18 | 0.33088 | 0.818641 | 0.703377 | 0.984702 |
| GOBP_PEROXISOME_PROLIFERATOR_ACTIVATED_RECEPTOR_SIGNALING_PATHWAY | 19 | 0.327147 | 0.818201 | 0.696401 | 0.985107 |
| GOBP_REGULATION_OF_B_CELL_MEDIATED_IMMUNITY | 46 | 0.279067 | 0.818134 | 0.75469 | 0.984772 |
| GOBP_POSITIVE_REGULATION_OF_DNA_RECOMBINATION | 62 | 0.269567 | 0.818006 | 0.775862 | 0.984569 |
| GOBP_PRIMARY_ALCOHOL_METABOLIC_PROCESS | 71 | 0.257715 | 0.817652 | 0.775266 | 0.984802 |
| GOBP_MITOTIC_G2_DNA_DAMAGE_CHECKPOINT_SIGNALING | 35 | 0.289148 | 0.817629 | 0.712782 | 0.984402 |
| GOBP_TYPE_B_PANCREATIC_CELL_DEVELOPMENT | 18 | 0.337774 | 0.817469 | 0.707463 | 0.984246 |
| GOBP_ESTABLISHMENT_OF_PROTEIN_LOCALIZATION_TO_VACUOLE | 48 | 0.275498 | 0.817208 | 0.762108 | 0.984297 |
| GOBP_AMINE_CATABOLIC_PROCESS | 19 | 0.331285 | 0.817097 | 0.722311 | 0.984062 |
| GOBP_REGULATION_OF_PROTEIN_LOCALIZATION_TO_PLASMA_MEMBRANE | 92 | 0.24848 | 0.816906 | 0.811989 | 0.983986 |
| GOBP_TORC1_SIGNALING | 49 | 0.273919 | 0.816851 | 0.753644 | 0.983642 |
| GOBP_SYNAPTIC_TRANSMISSION_DOPAMINERGIC | 20 | 0.321715 | 0.816181 | 0.706731 | 0.984522 |
| GOBP_NEGATIVE_REGULATION_OF_LYMPHOCYTE_DIFFERENTIATION | 38 | 0.288363 | 0.816062 | 0.758621 | 0.984292 |
| GOBP_LYMPHOID_PROGENITOR_CELL_DIFFERENTIATION | 18 | 0.338854 | 0.815794 | 0.713592 | 0.984344 |
| GOBP_GLYCOSYLATION | 224 | 0.228069 | 0.815711 | 0.857981 | 0.984069 |
| GOBP_POSITIVE_REGULATION_OF_NEURON_DIFFERENTIATION | 67 | 0.257692 | 0.815373 | 0.802168 | 0.984278 |
| GOBP_ESTABLISHMENT_OF_ORGANELLE_LOCALIZATION | 363 | 0.218736 | 0.814959 | 0.906623 | 0.984588 |
| GOBP_MONOCARBOXYLIC_ACID_CATABOLIC_PROCESS | 102 | 0.243506 | 0.814845 | 0.829237 | 0.984363 |
| GOBP_RECEPTOR_MEDIATED_ENDOCYTOSIS | 207 | 0.229048 | 0.814755 | 0.860165 | 0.984082 |
| GOBP_REGULATION_OF_PROTEIN_TYROSINE_KINASE_ACTIVITY | 74 | 0.260816 | 0.81465 | 0.798141 | 0.983836 |
| GOBP_INTEGRIN_ACTIVATION | 19 | 0.330977 | 0.814272 | 0.732484 | 0.984128 |
| GOBP_MICROVILLUS_ASSEMBLY | 21 | 0.32344 | 0.814022 | 0.719685 | 0.984142 |
| GOBP_GASTRULATION_WITH_MOUTH_FORMING_SECOND | 24 | 0.314288 | 0.81308 | 0.728571 | 0.985463 |
| GOBP_INSULIN_SECRETION_INVOLVED_IN_CELLULAR_RESPONSE_TO_GLUCOSE_STIMUI | 50 | 0.269157 | 0.812782 | 0.788099 | 0.985554 |
| GOBP_NEGATIVE_REGULATION_OF_RECEPTOR_SIGNALING_PATHWAY_VIA_JAK_STAT | 20 | 0.323727 | 0.812415 | 0.704969 | 0.985786 |
| GOBP_REGULATION_OF_RESPIRATORY_GASEOUS_EXCHANGE | 17 | 0.338943 | 0.812052 | 0.739269 | 0.986033 |

| | | | | | |
|---|---|---|---|---|---|
| GOBP_RETINOL_METABOLIC_PROCESS | 30 | 0.302707 | 0.81171 | 0.721649 | 0.986245 |
| GOBP_CELL_DIFFERENTIATION_IN_HINDBRAIN | 20 | 0.314387 | 0.811347 | 0.740219 | 0.986484 |
| GOBP_STEROID_CATABOLIC_PROCESS | 17 | 0.335117 | 0.81132 | 0.724409 | 0.986082 |
| GOBP_REGULATION_OF_VASCULOGENESIS | 15 | 0.345191 | 0.810303 | 0.708333 | 0.987518 |
| GOBP_POSITIVE_REGULATION_OF_MAP_KINASE_ACTIVITY | 99 | 0.244916 | 0.809795 | 0.809886 | 0.988011 |
| GOBP_REGULATION_OF_ALCOHOL_BIOSYNTHETIC_PROCESS | 39 | 0.278903 | 0.809597 | 0.753228 | 0.987924 |
| GOBP_REGULATION_OF_PROTEIN_BINDING | 171 | 0.229371 | 0.80959 | 0.866272 | 0.987488 |
| GOBP_NEGATIVE_REGULATION_OF_CELL_CYCLE_PROCESS | 259 | 0.220258 | 0.809569 | 0.894972 | 0.987082 |
| GOBP_POSITIVE_REGULATION_OF_NERVOUS_SYSTEM_DEVELOPMENT | 232 | 0.223989 | 0.809055 | 0.866817 | 0.987602 |
| GOBP_CELLULAR_HORMONE_METABOLIC_PROCESS | 85 | 0.249175 | 0.808212 | 0.817945 | 0.98871 |
| GOBP_TIGHT_JUNCTION_ORGANIZATION | 62 | 0.261926 | 0.808164 | 0.777778 | 0.988352 |
| GOBP_REGULATION_OF_TOLL_LIKE_RECEPTOR_4_SIGNALING_PATHWAY | 17 | 0.340192 | 0.806731 | 0.73132 | 0.990591 |
| GOBP_RESPONSE_TO_INSULIN | 215 | 0.223815 | 0.806149 | 0.90281 | 0.991241 |
| GOBP_REGULATION_OF_CELL_MORPHOGENESIS | 276 | 0.221503 | 0.806031 | 0.899669 | 0.991021 |
| GOBP_POSITIVE_REGULATION_OF_CELL_CYCLE_PROCESS | 196 | 0.224657 | 0.80491 | 0.87949 | 0.992634 |
| GOBP_NEGATIVE_REGULATION_OF_LIPID_METABOLIC_PROCESS | 73 | 0.252694 | 0.804522 | 0.808684 | 0.992883 |
| GOBP_ESTABLISHMENT_OF_PROTEIN_LOCALIZATION_TO_MITOCHONDRIAL_MEMBRANE | 29 | 0.299344 | 0.804283 | 0.734139 | 0.992869 |
| GOBP_POSITIVE_REGULATION_OF_ACTIN_CYTOSKELETON_REORGANIZATION | 16 | 0.334409 | 0.803941 | 0.715928 | 0.993039 |
| GOBP_EPITHELIAL_TUBE_FORMATION | 128 | 0.233607 | 0.80331 | 0.849624 | 0.993735 |
| GOBP_POSITIVE_REGULATION_OF_RESPONSE_TO_DNA_DAMAGE_STIMULUS | 125 | 0.236596 | 0.803266 | 0.848293 | 0.993362 |
| GOBP_DNA_BIOSYNTHETIC_PROCESS | 181 | 0.227278 | 0.803073 | 0.86215 | 0.993259 |
| GOBP_ADP_METABOLIC_PROCESS | 77 | 0.250901 | 0.800257 | 0.818425 | 0.997983 |
| GOBP_SPHINGOLIPID_METABOLIC_PROCESS | 138 | 0.231389 | 0.799659 | 0.86618 | 0.998653 |
| GOBP_CELLULAR_AMINO_ACID_BIOSYNTHETIC_PROCESS | 66 | 0.257435 | 0.79927 | 0.800554 | 0.998917 |
| GOBP_PEPTIDYL_LYSINE_ACETYLATION | 176 | 0.227153 | 0.798612 | 0.897129 | 0.999629 |
| GOBP_POSTSYNAPTIC_MEMBRANE_ORGANIZATION | 34 | 0.289703 | 0.798308 | 0.77266 | 0.999737 |
| GOBP_CENTRIOLE_ASSEMBLY | 43 | 0.273095 | 0.798232 | 0.783668 | 0.999435 |
| GOBP_DENDRITE_DEVELOPMENT | 216 | 0.221102 | 0.797689 | 0.901136 | 0.999971 |
| GOBP_POSITIVE_REGULATION_OF_NERVOUS_SYSTEM_PROCESS | 29 | 0.295332 | 0.79766 | 0.751479 | 0.999572 |
| GOBP_REGULATION_OF_NUCLEOTIDE_BIOSYNTHETIC_PROCESS | 31 | 0.287011 | 0.796911 | 0.76006 | 1 |
| GOBP_NUCLEOTIDE_SALVAGE | 16 | 0.32732 | 0.796727 | 0.755556 | 1 |
| GOBP_PROTEIN_TETRAMERIZATION | 73 | 0.250387 | 0.796258 | 0.80764 | 1 |
| GOBP_REGULATION_OF_CELLULAR_COMPONENT_SIZE | 320 | 0.216214 | 0.795966 | 0.934211 | 1 |
| GOBP_POSITIVE_REGULATION_OF_JUN_KINASE_ACTIVITY | 37 | 0.278874 | 0.795508 | 0.761976 | 1 |
| GOBP_CELLULAR_RESPONSE_TO_KETONE | 80 | 0.246642 | 0.795325 | 0.821622 | 1 |
| GOBP_SELECTIVE_AUTOPHAGY | 77 | 0.246383 | 0.794594 | 0.840054 | 1 |
| GOBP_NEGATIVE_REGULATION_OF_MITOTIC_CELL_CYCLE | 198 | 0.222611 | 0.794132 | 0.896119 | 1 |
| GOBP_RESPONSE_TO_OSMOTIC_STRESS | 68 | 0.254648 | 0.793715 | 0.809052 | 1 |
| GOBP_MAST_CELL_ACTIVATION | 45 | 0.269936 | 0.793393 | 0.786744 | 1 |
| GOBP_PROTEIN_K48_LINKED_UBIQUITINATION | 66 | 0.255042 | 0.793266 | 0.809257 | 1 |
| GOBP_WATER_HOMEOSTASIS | 45 | 0.269991 | 0.792797 | 0.777612 | 1 |
| GOBP_RESPONSE_TO_CAMP | 67 | 0.249429 | 0.7926 | 0.800536 | 1 |
| GOBP_MULTICELLULAR_ORGANISM_GROWTH | 121 | 0.235604 | 0.792008 | 0.874378 | 1 |
| GOBP_POSITIVE_REGULATION_OF_CELL_CYCLE_G1_S_PHASE_TRANSITION | 50 | 0.26479 | 0.791913 | 0.808067 | 1 |
| GOBP_RETROGRADE_AXONAL_TRANSPORT | 20 | 0.318811 | 0.791695 | 0.737261 | 1 |
| GOBP_ANTIGEN_PROCESSING_AND_PRESENTATION_OF_PEPTIDE_ANTIGEN | 45 | 0.270456 | 0.791247 | 0.778255 | 1 |
| GOBP_REGULATION_OF_CALCIUM_ION_IMPORT | 30 | 0.29161 | 0.791224 | 0.753012 | 1 |
| GOBP_XENOBIOTIC_TRANSPORT | 31 | 0.289565 | 0.790309 | 0.786585 | 1 |
| GOBP_NEUROTROPHIN_TRK_RECEPTOR_SIGNALING_PATHWAY | 22 | 0.312007 | 0.789895 | 0.759554 | 1 |
| GOBP_REGULATION_OF_LIPID_LOCALIZATION | 118 | 0.230568 | 0.789723 | 0.877193 | 1 |
| GOBP_CANONICAL_WNT_SIGNALING_PATHWAY | 270 | 0.21405 | 0.78956 | 0.931461 | 1 |
| GOBP_REGULATION_OF_PHAGOCYTOSIS | 67 | 0.252261 | 0.789257 | 0.830111 | 1 |
| GOBP_REGULATION_OF_ERAD_PATHWAY | 20 | 0.320653 | 0.78851 | 0.756211 | 1 |
| GOBP_MEMBRANE_BIOGENESIS | 59 | 0.260375 | 0.788218 | 0.811453 | 1 |
| GOBP_REGULATION_OF_ACTIN_FILAMENT_LENGTH | 142 | 0.229558 | 0.787811 | 0.871481 | 1 |
| GOBP_REGULATION_OF_CELL_CYCLE_PHASE_TRANSITION | 354 | 0.212339 | 0.787761 | 0.946623 | 1 |
| GOBP_XENOBIOTIC_CATABOLIC_PROCESS | 16 | 0.333146 | 0.787443 | 0.738989 | 1 |
| GOBP_POSITIVE_REGULATION_OF_LIPID_LOCALIZATION | 80 | 0.247522 | 0.786839 | 0.839189 | 1 |
| GOBP_REGULATION_OF_PROTEIN_POLYMERIZATION | 182 | 0.221947 | 0.786106 | 0.912515 | 1 |
| GOBP_PROTEIN_LOCALIZATION_TO_LYSOSOME | 44 | 0.269225 | 0.785162 | 0.801136 | 1 |
| GOBP_LIPID_OXIDATION | 95 | 0.236052 | 0.78415 | 0.878628 | 1 |
| GOBP_ADULT_BEHAVIOR | 111 | 0.234006 | 0.784038 | 0.861147 | 1 |
| GOBP_REGULATION_OF_ACTIN_FILAMENT_ORGANIZATION | 241 | 0.216539 | 0.783312 | 0.917885 | 1 |
| GOBP_PROTEIN_LOCALIZATION_TO_MICROTUBULE_ORGANIZING_CENTER | 35 | 0.283428 | 0.782307 | 0.758671 | 1 |
| GOBP_PROTEIN_ACETYLATION | 199 | 0.217897 | 0.782105 | 0.909091 | 1 |
| GOBP_GLUTAMINE_FAMILY_AMINO_ACID_CATABOLIC_PROCESS | 23 | 0.303686 | 0.781811 | 0.785714 | 1 |
| GOBP_HISTONE_H3_K9_METHYLATION | 36 | 0.278847 | 0.781674 | 0.797015 | 1 |
| GOBP_CELL_CELL_ADHESION_MEDIATED_BY_CADHERIN | 24 | 0.299753 | 0.781197 | 0.785605 | 1 |
| GOBP_REGULATION_OF_HOMOTYPIC_CELL_CELL_ADHESION | 26 | 0.295944 | 0.781145 | 0.776744 | 1 |
| GOBP_NUCLEAR_EXPORT | 143 | 0.227455 | 0.780857 | 0.902985 | 1 |
| GOBP_REGULATION_OF_VESICLE_MEDIATED_TRANSPORT | 445 | 0.207371 | 0.780308 | 0.958019 | 1 |
| GOBP_NEGATIVE_REGULATION_OF_PROTEIN_CATABOLIC_PROCESS | 112 | 0.232573 | 0.780213 | 0.873434 | 1 |
| GOBP_MAINTENANCE_OF_PROTEIN_LOCATION | 86 | 0.239331 | 0.779775 | 0.852564 | 1 |
| GOBP_MULTIVESICULAR_BODY_SORTING_PATHWAY | 37 | 0.277272 | 0.779526 | 0.796584 | 1 |
| GOBP_REGULATION_OF_PROTON_TRANSPORT | 21 | 0.305298 | 0.778737 | 0.769475 | 1 |
| GOBP_LYSOSOMAL_TRANSPORT | 116 | 0.234445 | 0.778393 | 0.876106 | 1 |
| GOBP_GLUCOSE_METABOLIC_PROCESS | 162 | 0.22167 | 0.778337 | 0.910736 | 1 |
| GOBP_RENAL_FILTRATION | 22 | 0.306748 | 0.778167 | 0.767145 | 1 |
| GOBP_NEGATIVE_REGULATION_OF_FIBROBLAST_PROLIFERATION | 27 | 0.29488 | 0.777785 | 0.785395 | 1 |
| GOBP_NUCLEAR_TRANSPORT | 277 | 0.211749 | 0.77749 | 0.948889 | 1 |
| GOBP_STEROL_BIOSYNTHETIC_PROCESS | 55 | 0.258112 | 0.77743 | 0.826503 | 1 |
| GOBP_POSITIVE_REGULATION_OF_NEURON_PROJECTION_DEVELOPMENT | 137 | 0.227051 | 0.777251 | 0.899003 | 1 |
| GOBP_POSITIVE_REGULATION_OF_OXIDOREDUCTASE_ACTIVITY | 45 | 0.267072 | 0.777076 | 0.811239 | 1 |
| GOBP_REGULATION_OF_PRODUCTION_OF_SMALL_RNA_INVOLVED_IN_GENE_SILENCING | 19 | 0.315406 | 0.776682 | 0.770313 | 1 |
| GOBP_POSITIVE_REGULATION_OF_VASOCONSTRICTION | 22 | 0.309029 | 0.776593 | 0.766458 | 1 |
| GOBP_REGULATION_OF_DIGESTIVE_SYSTEM_PROCESS | 20 | 0.315122 | 0.776224 | 0.776509 | 1 |
| GOBP_RNA_PHOSPHODIESTER_BOND_HYDROLYSIS_ENDONUCLEOLYTIC | 70 | 0.249969 | 0.776202 | 0.848285 | 1 |
| GOBP_3_UTR_MEDIATED_MRNA_DESTABILIZATION | 15 | 0.322697 | 0.776115 | 0.749186 | 1 |
| GOBP_REGULATION_OF_INSULIN_SECRETION_INVOLVED_IN_CELLULAR_RESPONSE_TO_ | 41 | 0.272027 | 0.775866 | 0.820059 | 1 |
| GOBP_MEMBRANE_LIPID_METABOLIC_PROCESS | 182 | 0.220388 | 0.775103 | 0.909518 | 1 |
| GOBP_PROTEOGLYCAN_METABOLIC_PROCESS | 74 | 0.246374 | 0.774947 | 0.834912 | 1 |
| GOBP_HISTONE_H3_K4_METHYLATION | 65 | 0.249074 | 0.774755 | 0.843191 | 1 |
| GOBP_POST_GOLGI_VESICLE_MEDIATED_TRANSPORT | 103 | 0.234199 | 0.774737 | 0.859849 | 1 |
| GOBP_CELLULAR_CARBOHYDRATE_BIOSYNTHETIC_PROCESS | 63 | 0.251735 | 0.774157 | 0.832394 | 1 |
| GOBP_NEGATIVE_REGULATION_OF_CELL_DEVELOPMENT | 146 | 0.222511 | 0.77401 | 0.902644 | 1 |
| GOBP_REGULATION_OF_PROTEASOMAL_PROTEIN_CATABOLIC_PROCESS | 173 | 0.219351 | 0.774 | 0.917857 | 1 |
| GOBP_NUCLEOTIDE_EXCISION_REPAIR | 55 | 0.261504 | 0.773299 | 0.808081 | 1 |
| GOBP_POSITIVE_REGULATION_OF_MACROAUTOPHAGY | 70 | 0.24689 | 0.772949 | 0.858667 | 1 |
| GOBP_MICROTUBULE_ORGANIZING_CENTER_ORGANIZATION | 140 | 0.226339 | 0.772801 | 0.894932 | 1 |
| GOBP_RETINA_MORPHOGENESIS_IN_CAMERA_TYPE_EYE | 44 | 0.269531 | 0.772621 | 0.807069 | 1 |
| GOBP_NEGATIVE_REGULATION_OF_BIOMINERALIZATION | 21 | 0.308873 | 0.772453 | 0.780063 | 1 |
| GOBP_NEURON_PROJECTION_EXTENSION | 160 | 0.220246 | 0.772238 | 0.920482 | 1 |
| GOBP_VACUOLAR_TRANSPORT | 148 | 0.223826 | 0.772046 | 0.903941 | 1 |
| GOBP_RESPONSE_TO_UV | 133 | 0.224737 | 0.771835 | 0.898263 | 1 |
| GOBP_AXONEMAL_DYNEIN_COMPLEX_ASSEMBLY | 29 | 0.28948 | 0.771643 | 0.798206 | 1 |
| GOBP_CELLULAR_RESPONSE_TO_ALKALOID | 28 | 0.282433 | 0.771422 | 0.797256 | 1 |
| GOBP_RAC_PROTEIN_SIGNAL_TRANSDUCTION | 41 | 0.26818 | 0.770798 | 0.818978 | 1 |
| GOBP_RESPONSE_TO_ACIDIC_PH | 19 | 0.309109 | 0.769994 | 0.7792 | 1 |
| GOBP_PURINE_CONTAINING_COMPOUND_METABOLIC_PROCESS | 349 | 0.206254 | 0.769807 | 0.959651 | 1 |
| GOBP_RESPONSE_TO_RADIATION | 362 | 0.207398 | 0.76957 | 0.963402 | 1 |
| GOBP_RESPONSE_TO_ANGIOTENSIN | 21 | 0.308948 | 0.769446 | 0.794953 | 1 |
| GOBP_PROTEIN_LOCALIZATION_TO_VACUOLE | 65 | 0.247623 | 0.769362 | 0.842818 | 1 |
| GOBP_NEGATIVE_REGULATION_OF_NEURON_PROJECTION_DEVELOPMENT | 127 | 0.223728 | 0.769229 | 0.907035 | 1 |
| GOBP_POSITIVE_REGULATION_OF_GLUCOSE_METABOLIC_PROCESS | 33 | 0.278364 | 0.769025 | 0.796407 | 1 |
| GOBP_INSULIN_RECEPTOR_SIGNALING_PATHWAY | 96 | 0.234357 | 0.768302 | 0.888305 | 1 |
| GOBP_REGULATION_OF_INTRINSIC_APOPTOTIC_SIGNALING_PATHWAY | 151 | 0.221022 | 0.768289 | 0.908434 | 1 |
| GOBP_CELLULAR_RESPONSE_TO_VITAMIN_D | 17 | 0.31282 | 0.768156 | 0.768025 | 1 |
| GOBP_NEGATIVE_REGULATION_OF_PROTEIN_POLYMERIZATION | 73 | 0.236995 | 0.76802 | 0.846154 | 1 |
| GOBP_REGULATION_OF_ATP_DEPENDENT_ACTIVITY | 61 | 0.249936 | 0.767835 | 0.840483 | 1 |
| GOBP_TUBE_FORMATION | 140 | 0.224079 | 0.767627 | 0.903776 | 1 |
| GOBP_CEREBRAL_CORTEX_RADIALLY_ORIENTED_CELL_MIGRATION | 30 | 0.28326 | 0.767235 | 0.810893 | 1 |
| GOBP_REGULATION_OF_CELL_SIZE | 167 | 0.219502 | 0.766823 | 0.920768 | 1 |
| GOBP_VESICLE_LOCALIZATION | 175 | 0.216027 | 0.766477 | 0.920804 | 1 |
| GOBP_POSITIVE_REGULATION_OF_PHOSPHOLIPASE_ACTIVITY | 37 | 0.266285 | 0.766393 | 0.803681 | 1 |
| GOBP_ACTIVATION_OF_CYSTEINE_TYPE_ENDOPEPTIDASE_ACTIVITY | 17 | 0.315078 | 0.76615 | 0.76183 | 1 |
| GOBP_LIPID_TRANSLOCATION | 46 | 0.258865 | 0.765491 | 0.82948 | 1 |
| GOBP_REGULATION_OF_MITOTIC_CELL_CYCLE_PHASE_TRANSITION | 272 | 0.209701 | 0.765101 | 0.953125 | 1 |
| GOBP_DETECTION_OF_TEMPERATURE_STIMULUS_INVOLVED_IN_SENSORY_PERCEPTION | 15 | 0.329801 | 0.764884 | 0.781906 | 1 |
| GOBP_SENSORY_PERCEPTION_OF_TEMPERATURE_STIMULUS | 20 | 0.304183 | 0.764656 | 0.788217 | 1 |
| GOBP_POSITIVE_REGULATION_OF_SYNAPTIC_TRANSMISSION_GLUTAMATERGIC | 22 | 0.299215 | 0.764312 | 0.775835 | 1 |
| GOBP_MEMBRANE_LIPID_CATABOLIC_PROCESS | 32 | 0.278892 | 0.763521 | 0.797704 | 1 |
| GOBP_ENDOTHELIAL_CELL_DEVELOPMENT | 64 | 0.24476 | 0.76313 | 0.846365 | 1 |
| GOBP_CYCLIC_NUCLEOTIDE_BIOSYNTHETIC_PROCESS | 18 | 0.307978 | 0.762016 | 0.773246 | 1 |
| GOBP_NEGATIVE_REGULATION_OF_ATP_DEPENDENT_ACTIVITY | 17 | 0.314084 | 0.761882 | 0.783522 | 1 |
| GOBP_NEGATIVE_REGULATION_OF_ATP_METABOLIC_PROCESS | 26 | 0.290388 | 0.761841 | 0.82641 | 1 |
| GOBP_REGULATION_OF_LIPID_STORAGE | 38 | 0.269317 | 0.761312 | 0.830104 | 1 |
| GOBP_ORGANIC_CYCLIC_COMPOUND_CATABOLIC_PROCESS | 432 | 0.201706 | 0.760295 | 0.976621 | 1 |
| GOBP_ASTROCYTE_DIFFERENTIATION | 56 | 0.246406 | 0.760179 | 0.852368 | 1 |
| GOBP_MITOCHONDRIAL_TRANSCRIPTION | 17 | 0.316833 | 0.760071 | 0.783094 | 1 |
| GOBP_SMALL_MOLECULE_BIOSYNTHETIC_PROCESS | 445 | 0.199727 | 0.758522 | 0.981152 | 1 |
| GOBP_POSITIVE_REGULATION_OF_CELLULAR_COMPONENT_BIOGENESIS | 431 | 0.201963 | 0.758465 | 0.976645 | 1 |
| GOBP_POSITIVE_REGULATION_OF_LIPID_STORAGE | 18 | 0.31599 | 0.758311 | 0.788732 | 1 |
| GOBP_REGULATION_OF_ESTABLISHMENT_OF_PROTEIN_LOCALIZATION_TO_MITOCHONDR | 47 | 0.256072 | 0.757493 | 0.823789 | 1 |
| GOBP_REGULATION_OF_CANONICAL_WNT_SIGNALING_PATHWAY | 226 | 0.210206 | 0.757092 | 0.961712 | 1 |
| GOBP_CELL_CELL_SIGNALING_INVOLVED_IN_CARDIAC_CONDUCTION | 28 | 0.277032 | 0.75681 | 0.797297 | 1 |
| GOBP_PROTEIN_POLYMERIZATION | 251 | 0.209204 | 0.756384 | 0.947846 | 1 |
| GOBP_ORGANELLE_LOCALIZATION | 484 | 0.199923 | 0.75637 | 0.98431 | 1 |
| GOBP_POSITIVE_REGULATION_OF_PHOSPHOPROTEIN_PHOSPHATASE_ACTIVITY | 17 | 0.311194 | 0.756101 | 0.786408 | 1 |
| GOBP_MONOSACCHARIDE_METABOLIC_PROCESS | 219 | 0.210122 | 0.754109 | 0.950461 | 1 |
| GOBP_NEGATIVE_REGULATION_OF_MITOTIC_CELL_CYCLE_PHASE_TRANSITION | 154 | 0.215774 | 0.754071 | 0.924298 | 1 |
| GOBP_BRAIN_MORPHOGENESIS | 32 | 0.271602 | 0.753785 | 0.82194 | 1 |
| GOBP_REGULATION_OF_MITOCHONDRIAL_MEMBRANE_POTENTIAL | 64 | 0.245314 | 0.753616 | 0.85346 | 1 |
| GOBP_POSITIVE_REGULATION_OF_MITOTIC_CELL_CYCLE_PHASE_TRANSITION | 82 | 0.236599 | 0.753565 | 0.886212 | 1 |
| GOBP_CENTROSOME_SEPARATION | 15 | 0.320076 | 0.753041 | 0.789389 | 1 |
| GOBP_PURINE_NUCLEOTIDE_TRANSPORT | 28 | 0.285569 | 0.752469 | 0.786248 | 1 |
| GOBP_MITOCHONDRIAL_GENOME_MAINTENANCE | 23 | 0.294981 | 0.752432 | 0.803487 | 1 |
| GOBP_REGULATION_OF_UBIQUITIN_DEPENDENT_PROTEIN_CATABOLIC_PROCESS | 153 | 0.216969 | 0.751942 | 0.933171 | 1 |

| | | | | | |
|---|---|---|---|---|---|
| GOBP_TERPENOID_METABOLIC_PROCESS | 60 | 0.244052 | 0.75174 | 0.881543 | 1 |
| GOBP_ORGANELLE_DISASSEMBLY | 119 | 0.222323 | 0.751454 | 0.921519 | 1 |
| GOBP_NUCLEOTIDE_TRANSPORT | 34 | 0.265636 | 0.750515 | 0.815634 | 1 |
| GOBP_GLIAL_CELL_DIFFERENTIATION | 175 | 0.212981 | 0.74997 | 0.929245 | 1 |
| GOBP_STRESS_FIBER_ASSEMBLY | 98 | 0.226539 | 0.749426 | 0.898936 | 1 |
| GOBP_NUCLEOTIDE_SUGAR_METABOLIC_PROCESS | 32 | 0.274971 | 0.749231 | 0.833583 | 1 |
| GOBP_MITOCHONDRIAL_OUTER_MEMBRANE_PERMEABILIZATION | 31 | 0.271474 | 0.749064 | 0.824666 | 1 |
| GOBP_NEGATIVE_REGULATION_OF_AXONOGENESIS | 62 | 0.243904 | 0.748273 | 0.866391 | 1 |
| GOBP_DETECTION_OF_LIGHT_STIMULUS | 34 | 0.268807 | 0.748153 | 0.845468 | 1 |
| GOBP_RNA_5_END_PROCESSING | 23 | 0.294182 | 0.747772 | 0.80721 | 1 |
| GOBP_GLUCOSE_IMPORT | 59 | 0.243885 | 0.74754 | 0.895461 | 1 |
| GOBP_POSITIVE_REGULATION_OF_CALCIUM_MEDIATED_SIGNALING | 22 | 0.291611 | 0.747439 | 0.827103 | 1 |
| GOBP_POSITIVE_REGULATION_OF_CELLULAR_AMIDE_METABOLIC_PROCESS | 149 | 0.215408 | 0.747417 | 0.925532 | 1 |
| GOBP_PROTEIN_LOCALIZATION_TO_PLASMA_MEMBRANE | 263 | 0.203603 | 0.747161 | 0.973003 | 1 |
| GOBP_NEURON_CELL_CELL_ADHESION | 15 | 0.310349 | 0.746753 | 0.802698 | 1 |
| GOBP_CELLULAR_RESPONSE_TO_INSULIN_STIMULUS | 172 | 0.209245 | 0.746744 | 0.944771 | 1 |
| GOBP_REGULATION_OF_DOUBLE_STRAND_BREAK_REPAIR_VIA_HOMOLOGOUS_RECOME | 69 | 0.239644 | 0.746634 | 0.878621 | 1 |
| GOBP_REGULATION_OF_PROTEIN_EXPORT_FROM_NUCLEUS | 30 | 0.273327 | 0.745408 | 0.827744 | 1 |
| GOBP_LONG_TERM_MEMORY | 34 | 0.272158 | 0.74493 | 0.838806 | 1 |
| GOBP_POSITIVE_REGULATION_BY_HOST_OF_VIRAL_TRANSCRIPTION | 17 | 0.30796 | 0.744596 | 0.798669 | 1 |
| GOBP_POSITIVE_REGULATION_OF_ENDOCYTOSIS | 79 | 0.234172 | 0.744239 | 0.887715 | 1 |
| GOBP_REGULATION_OF_HISTONE_H3_K4_METHYLATION | 36 | 0.263677 | 0.743126 | 0.83105 | 1 |
| GOBP_MAMMARY_GLAND_EPITHELIAL_CELL_DIFFERENTIATION | 15 | 0.320801 | 0.743047 | 0.802326 | 1 |
| GOBP_REGULATION_OF_CELL_DIVISION | 146 | 0.213721 | 0.742809 | 0.944645 | 1 |
| GOBP_POSITIVE_REGULATION_OF_RECEPTOR_SIGNALING_PATHWAY_VIA_STAT | 25 | 0.282251 | 0.742786 | 0.813846 | 1 |
| GOBP_POSITIVE_REGULATION_OF_CHOLESTEROL_EFFLUX | 21 | 0.294184 | 0.742567 | 0.779528 | 1 |
| GOBP_POSITIVE_REGULATION_OF_HEART_RATE | 23 | 0.288457 | 0.741824 | 0.815047 | 1 |
| GOBP_CARBOHYDRATE_CATABOLIC_PROCESS | 124 | 0.21603 | 0.741557 | 0.935443 | 1 |
| GOBP_PROTEIN_LOCALIZATION_TO_CELL_PERIPHERY | 310 | 0.200931 | 0.741274 | 0.981319 | 1 |
| GOBP_TRANSCRIPTION_BY_RNA_POLYMERASE_III | 53 | 0.242659 | 0.741054 | 0.857546 | 1 |
| GOBP_SECONDARY_METABOLITE_BIOSYNTHETIC_PROCESS | 17 | 0.306644 | 0.740968 | 0.822153 | 1 |
| GOBP_INTRINSIC_APOPTOTIC_SIGNALING_PATHWAY_IN_RESPONSE_TO_DNA_DAMAGE | 93 | 0.226954 | 0.740965 | 0.908189 | 1 |
| GOBP_RENAL_ABSORPTION | 17 | 0.301726 | 0.740178 | 0.797997 | 1 |
| GOBP_REGULATION_OF_CYTOPLASMIC_TRANSLATION | 27 | 0.274049 | 0.739871 | 0.849921 | 1 |
| GOBP_SEMAPHORIN_PLEXIN_SIGNALING_PATHWAY_INVOLVED_IN_NEURON_PROJECTIO | 16 | 0.308931 | 0.739057 | 0.838915 | 1 |
| GOBP_NUCLEAR_TRANSCRIBED_MRNA_CATABOLIC_PROCESS | 118 | 0.218452 | 0.7379 | 0.93617 | 1 |
| GOBP_REGULATION_OF_GENERATION_OF_PRECURSOR_METABOLITES_AND_ENERGY | 114 | 0.216022 | 0.737466 | 0.94802 | 1 |
| GOBP_VESICLE_DOCKING | 52 | 0.246306 | 0.737276 | 0.888889 | 1 |
| GOBP_MITOTIC_CELL_CYCLE_CHECKPOINT_SIGNALING | 124 | 0.213923 | 0.736561 | 0.936275 | 1 |
| GOBP_ACTIN_POLYMERIZATION_OR_DEPOLYMERIZATION | 177 | 0.205698 | 0.735721 | 0.96483 | 1 |
| GOBP_HYPEROSMOTIC_RESPONSE | 23 | 0.286907 | 0.735619 | 0.827586 | 1 |
| GOBP_RAB_PROTEIN_SIGNAL_TRANSDUCTION | 21 | 0.291872 | 0.735536 | 0.853211 | 1 |
| GOBP_TRANSLESION_SYNTHESIS | 24 | 0.280301 | 0.735479 | 0.81129 | 1 |
| GOBP_FATTY_ACID_DERIVATIVE_METABOLIC_PROCESS | 52 | 0.24705 | 0.734645 | 0.860882 | 1 |
| GOBP_N_TERMINAL_PROTEIN_AMINO_ACID_MODIFICATION | 27 | 0.28081 | 0.733397 | 0.843567 | 1 |
| GOBP_LATE_ENDOSOME_TO_LYSOSOME_TRANSPORT | 18 | 0.290288 | 0.731829 | 0.808581 | 1 |
| GOBP_POSITIVE_REGULATION_OF_B_CELL_MEDIATED_IMMUNITY | 29 | 0.272161 | 0.731121 | 0.84816 | 1 |
| GOBP_PIGMENTATION | 86 | 0.226616 | 0.730477 | 0.925099 | 1 |
| GOBP_PYRIMIDINE_CONTAINING_COMPOUND_CATABOLIC_PROCESS | 35 | 0.262531 | 0.73032 | 0.840804 | 1 |
| GOBP_POSITIVE_REGULATION_OF_STRESS_FIBER_ASSEMBLY | 46 | 0.2501 | 0.729323 | 0.869873 | 1 |
| GOBP_LEUKOTRIENE_METABOLIC_PROCESS | 15 | 0.311038 | 0.729251 | 0.820097 | 1 |
| GOBP_REGULATION_OF_CELL_AGING | 42 | 0.250028 | 0.728945 | 0.866485 | 1 |
| GOBP_POSITIVE_REGULATION_OF_ESTABLISHMENT_OF_PROTEIN_LOCALIZATION_TO_MIT | 34 | 0.263455 | 0.728731 | 0.845468 | 1 |
| GOBP_LENS_MORPHOGENESIS_IN_CAMERA_TYPE_EYE | 21 | 0.28663 | 0.728126 | 0.817784 | 1 |
| GOBP_POSITIVE_REGULATION_OF_ATP_DEPENDENT_ACTIVITY | 38 | 0.25502 | 0.727808 | 0.871201 | 1 |
| GOBP_REGULATION_OF_MORPHOGENESIS_OF_A_BRANCHING_STRUCTURE | 35 | 0.25597 | 0.727619 | 0.844347 | 1 |
| GOBP_REGULATION_OF_GLUTAMATE_SECRETION | 18 | 0.297115 | 0.727234 | 0.813312 | 1 |
| GOBP_NEGATIVE_REGULATION_OF_TYPE_I_INTERFERON_PRODUCTION | 24 | 0.27939 | 0.72715 | 0.85842 | 1 |
| GOBP_LABYRINTHINE_LAYER_BLOOD_VESSEL_DEVELOPMENT | 18 | 0.295479 | 0.726822 | 0.83689 | 1 |
| GOBP_REGULATION_OF_ANDROGEN_RECEPTOR_SIGNALING_PATHWAY | 26 | 0.277455 | 0.726009 | 0.847662 | 1 |
| GOBP_REGULATION_OF_SYNAPTIC_VESICLE_RECYCLING | 18 | 0.293136 | 0.724846 | 0.815534 | 1 |
| GOBP_CHOLESTEROL_STORAGE | 15 | 0.30421 | 0.724276 | 0.843072 | 1 |
| GOBP_GOLGI_TO_ENDOSOME_TRANSPORT | 17 | 0.298809 | 0.723672 | 0.812401 | 1 |
| GOBP_CHONDROITIN_SULFATE_PROTEOGLYCAN_METABOLIC_PROCESS | 33 | 0.261857 | 0.723387 | 0.860465 | 1 |
| GOBP_PLASMA_LIPOPROTEIN_PARTICLE_CLEARANCE | 28 | 0.269645 | 0.722665 | 0.857798 | 1 |
| GOBP_NEGATIVE_REGULATION_OF_ERBB_SIGNALING_PATHWAY | 31 | 0.266164 | 0.72242 | 0.860465 | 1 |
| GOBP_POSITIVE_REGULATION_OF_MITOTIC_CELL_CYCLE | 108 | 0.216323 | 0.722281 | 0.954944 | 1 |
| GOBP_NEGATIVE_REGULATION_OF_CELL_DIVISION | 15 | 0.305222 | 0.722265 | 0.810767 | 1 |
| GOBP_REGULATION_OF_CHROMOSOME_SEGREGATION | 81 | 0.22533 | 0.721913 | 0.923377 | 1 |
| GOBP_SYNAPTIC_VESICLE_PRIMING | 18 | 0.297389 | 0.721835 | 0.824074 | 1 |
| GOBP_CELL_CYCLE_CHECKPOINT_SIGNALING | 163 | 0.206617 | 0.7217 | 0.963139 | 1 |
| GOBP_TYPE_B_PANCREATIC_CELL_PROLIFERATION | 15 | 0.305848 | 0.721584 | 0.812298 | 1 |
| GOBP_LIVER_REGENERATION | 22 | 0.28536 | 0.721219 | 0.828201 | 1 |
| GOBP_EXOCYTIC_PROCESS | 68 | 0.228247 | 0.721054 | 0.912517 | 1 |
| GOBP_MEMBRANE_LIPID_BIOSYNTHETIC_PROCESS | 132 | 0.208555 | 0.720603 | 0.941992 | 1 |
| GOBP_CELLULAR_RESPONSE_TO_STEROL | 15 | 0.313636 | 0.720212 | 0.833871 | 1 |
| GOBP_GLYCOSIDE_METABOLIC_PROCESS | 17 | 0.305842 | 0.719716 | 0.825817 | 1 |
| GOBP_PURINE_RIBONUCLEOSIDE_METABOLIC_PROCESS | 19 | 0.291864 | 0.719483 | 0.816425 | 1 |
| GOBP_PROTEIN_LOCALIZATION_TO_NUCLEUS | 267 | 0.195243 | 0.718358 | 0.986425 | 1 |
| GOBP_POSITIVE_REGULATION_OF_CELL_DIVISION | 59 | 0.233325 | 0.718198 | 0.910112 | 1 |
| GOBP_ACTIN_FILAMENT_POLYMERIZATION | 150 | 0.208117 | 0.717897 | 0.968862 | 1 |
| GOBP_REGULATION_OF_CYCLIN_DEPENDENT_PROTEIN_KINASE_ACTIVITY | 90 | 0.219233 | 0.717573 | 0.944444 | 1 |
| GOBP_REGULATION_OF_SYNAPTIC_VESICLE_EXOCYTOSIS | 47 | 0.240545 | 0.717419 | 0.908062 | 1 |
| GOBP_AMINOGLYCAN_BIOSYNTHETIC_PROCESS | 73 | 0.227419 | 0.717263 | 0.930171 | 1 |
| GOBP_LUNG_MORPHOGENESIS | 38 | 0.250961 | 0.717096 | 0.878832 | 1 |
| GOBP_ORGANELLE_MEMBRANE_FUSION | 98 | 0.216571 | 0.716318 | 0.934447 | 1 |
| GOBP_NEGATIVE_REGULATION_OF_CELL_CYCLE_PHASE_TRANSITION | 219 | 0.19789 | 0.715535 | 0.981777 | 1 |
| GOBP_CARBOHYDRATE_TRANSMEMBRANE_TRANSPORT | 88 | 0.220435 | 0.715304 | 0.937662 | 1 |
| GOBP_MITOCHONDRION_LOCALIZATION | 46 | 0.242316 | 0.715268 | 0.909613 | 1 |
| GOBP_PROTEIN_K11_LINKED_UBIQUITINATION | 29 | 0.269912 | 0.715027 | 0.868343 | 1 |
| GOBP_HEART_FORMATION | 26 | 0.272812 | 0.714757 | 0.852535 | 1 |
| GOBP_CYCLIC_NUCLEOTIDE_METABOLIC_PROCESS | 31 | 0.264006 | 0.714643 | 0.864583 | 1 |
| GOBP_HISTONE_MODIFICATION | 458 | 0.188942 | 0.71453 | 0.99787 | 1 |
| GOBP_NEGATIVE_REGULATION_OF_CELLULAR_RESPONSE_TO_INSULIN_STIMULUS | 31 | 0.2577 | 0.714297 | 0.871065 | 1 |
| GOBP_GLUTAMINE_FAMILY_AMINO_ACID_METABOLIC_PROCESS | 66 | 0.230093 | 0.713994 | 0.90068 | 1 |
| GOBP_REGULATION_OF_MITOCHONDRIAL_MEMBRANE_PERMEABILITY_INVOLVED_IN_AP( | 39 | 0.253201 | 0.713878 | 0.87426 | 1 |
| GOBP_ESTABLISHMENT_OF_PROTEIN_LOCALIZATION_TO_ORGANELLE | 399 | 0.189402 | 0.713218 | 0.99467 | 1 |
| GOBP_SEX_DETERMINATION | 15 | 0.306104 | 0.712662 | 0.82504 | 1 |
| GOBP_CELLULAR_AMINO_ACID_METABOLIC_PROCESS | 235 | 0.196564 | 0.711945 | 0.986207 | 1 |
| GOBP_TOLL_LIKE_RECEPTOR_SIGNALING_PATHWAY | 87 | 0.217098 | 0.711736 | 0.953519 | 1 |
| GOBP_RETINA_VASCULATURE_DEVELOPMENT_IN_CAMERA_TYPE_EYE | 17 | 0.292093 | 0.711239 | 0.855118 | 1 |
| GOBP_PYRIDINE_NUCLEOTIDE_METABOLIC_PROCESS | 25 | 0.27464 | 0.710849 | 0.845917 | 1 |
| GOBP_POLYOL_CATABOLIC_PROCESS | 26 | 0.266517 | 0.709677 | 0.850863 | 1 |
| GOBP_N_TERMINAL_PROTEIN_AMINO_ACID_ACETYLATION | 15 | 0.301621 | 0.708047 | 0.842697 | 1 |
| GOBP_NUCLEOBASE_CONTAINING_SMALL_MOLECULE_METABOLIC_PROCESS | 484 | 0.185805 | 0.707974 | 0.995833 | 1 |
| GOBP_NEURAL_TUBE_PATTERNING | 29 | 0.262505 | 0.705399 | 0.869628 | 1 |
| GOBP_MEMBRANE_DOCKING | 78 | 0.219109 | 0.704497 | 0.954856 | 1 |
| GOBP_SYNAPTIC_VESICLE_LOCALIZATION | 51 | 0.23543 | 0.704362 | 0.904432 | 1 |
| GOBP_ENDOMEMBRANE_SYSTEM_ORGANIZATION | 472 | 0.184811 | 0.704226 | 1 | 1 |
| GOBP_RESPONSE_TO_ARSENIC_CONTAINING_SUBSTANCE | 32 | 0.252933 | 0.702949 | 0.872671 | 1 |
| GOBP_REGULATION_OF_CELLULAR_AMIDE_METABOLIC_PROCESS | 410 | 0.187357 | 0.701175 | 0.998953 | 1 |
| GOBP_NEGATIVE_REGULATION_OF_PROTEIN_SECRETION | 47 | 0.239038 | 0.700919 | 0.9125 | 1 |
| GOBP_MAINTENANCE_OF_PROTEIN_LOCATION_IN_CELL | 60 | 0.228233 | 0.700871 | 0.920765 | 1 |
| GOBP_PROTEIN_LOCALIZATION_TO_MITOCHONDRION | 120 | 0.205847 | 0.7004 | 0.965693 | 1 |
| GOBP_CAMP_METABOLIC_PROCESS | 18 | 0.28675 | 0.700355 | 0.843296 | 1 |
| GOBP_TOR_SIGNALING | 119 | 0.205661 | 0.698855 | 0.974874 | 1 |
| GOBP_POSITIVE_REGULATION_OF_STEROID_BIOSYNTHETIC_PROCESS | 16 | 0.2956 | 0.698401 | 0.867395 | 1 |
| GOBP_ORGANOPHOSPHATE_BIOSYNTHETIC_PROCESS | 487 | 0.183089 | 0.698074 | 0.998958 | 1 |
| GOBP_POSITIVE_REGULATION_OF_LAMELLIPODIUM_ASSEMBLY | 24 | 0.271836 | 0.69804 | 0.879518 | 1 |
| GOBP_POSITIVE_REGULATION_OF_DNA_BIOSYNTHETIC_PROCESS | 70 | 0.224644 | 0.697825 | 0.93842 | 1 |
| GOBP_HOMEOSTASIS_OF_NUMBER_OF_CELLS_WITHIN_A_TISSUE | 22 | 0.272663 | 0.697717 | 0.869852 | 1 |
| GOBP_QUINONE_METABOLIC_PROCESS | 31 | 0.251522 | 0.697323 | 0.884731 | 1 |
| GOBP_REGULATION_OF_BRANCHING_INVOLVED_IN_URETERIC_BUD_MORPHOGENESIS | 16 | 0.290412 | 0.696996 | 0.855241 | 1 |
| GOBP_REGULATION_OF_TOR_SIGNALING | 97 | 0.211268 | 0.696413 | 0.951031 | 1 |
| GOBP_MAINTENANCE_OF_PROTEIN_LOCALIZATION_IN_ORGANELLE | 41 | 0.245406 | 0.696328 | 0.897472 | 1 |
| GOBP_REGULATION_OF_OLIGODENDROCYTE_DIFFERENTIATION | 31 | 0.252171 | 0.695902 | 0.869242 | 1 |
| GOBP_REGULATION_OF_MAST_CELL_ACTIVATION_INVOLVED_IN_IMMUNE_RESPONSE | 26 | 0.261661 | 0.695646 | 0.892308 | 1 |
| GOBP_LUNG_EPITHELIUM_DEVELOPMENT | 25 | 0.26732 | 0.695176 | 0.872894 | 1 |
| GOBP_NEUTRAL_AMINO_ACID_TRANSPORT | 38 | 0.242739 | 0.693813 | 0.907353 | 1 |
| GOBP_HISTONE_METHYLATION | 146 | 0.198643 | 0.69361 | 0.977833 | 1 |
| GOBP_AUTOPHAGOSOME_MATURATION | 58 | 0.22258 | 0.693501 | 0.929293 | 1 |
| GOBP_FATTY_ACID_BETA_OXIDATION | 68 | 0.217479 | 0.692521 | 0.94702 | 1 |
| GOBP_REGULATION_OF_HISTONE_H3_K9_METHYLATION | 23 | 0.266861 | 0.692486 | 0.899514 | 1 |
| GOBP_BLASTOCYST_GROWTH | 19 | 0.27801 | 0.692027 | 0.864434 | 1 |
| GOBP_NEUROMUSCULAR_JUNCTION_DEVELOPMENT | 38 | 0.243653 | 0.69192 | 0.896403 | 1 |
| GOBP_CEREBELLAR_CORTEX_MORPHOGENESIS | 33 | 0.248888 | 0.690613 | 0.901471 | 1 |
| GOBP_POSITIVE_REGULATION_OF_CYCLASE_ACTIVITY | 16 | 0.288618 | 0.689594 | 0.857143 | 1 |
| GOBP_REGULATION_OF_ACTOMYOSIN_STRUCTURE_ORGANIZATION | 87 | 0.212078 | 0.689121 | 0.963612 | 1 |
| GOBP_TRICARBOXYLIC_ACID_CYCLE | 30 | 0.256239 | 0.688886 | 0.888071 | 1 |
| GOBP_NEGATIVE_REGULATION_OF_HISTONE_METHYLATION | 22 | 0.274908 | 0.688825 | 0.874618 | 1 |
| GOBP_REGULATION_OF_MEMBRANE_LIPID_DISTRIBUTION | 52 | 0.229098 | 0.688568 | 0.933518 | 1 |
| GOBP_HISTONE_H3_ACETYLATION | 74 | 0.217766 | 0.687602 | 0.946565 | 1 |
| GOBP_MONOSACCHARIDE_BIOSYNTHETIC_PROCESS | 71 | 0.217867 | 0.687317 | 0.957333 | 1 |
| GOBP_DSRNA_PROCESSING | 51 | 0.228009 | 0.686108 | 0.921196 | 1 |
| GOBP_IMMUNOGLOBULIN_PRODUCTION | 91 | 0.208816 | 0.685978 | 0.967532 | 1 |
| GOBP_LUNG_ALVEOLUS_DEVELOPMENT | 32 | 0.250008 | 0.685943 | 0.901813 | 1 |
| GOBP_NUCLEAR_MEMBRANE_ORGANIZATION | 42 | 0.236487 | 0.683912 | 0.924107 | 1 |
| GOBP_REGULATION_OF_ATP_METABOLIC_PROCESS | 75 | 0.214816 | 0.683578 | 0.953519 | 1 |
| GOBP_ESTABLISHMENT_OF_MITOTIC_SPINDLE_LOCALIZATION | 32 | 0.24844 | 0.68354 | 0.893455 | 1 |
| GOBP_ATTACHMENT_OF_SPINDLE_MICROTUBULES_TO_KINETOCHORE | 36 | 0.243165 | 0.682838 | 0.897929 | 1 |
| GOBP_RNA_STABILIZATION | 56 | 0.225979 | 0.682455 | 0.923944 | 1 |
| GOBP_LIVER_MORPHOGENESIS | 19 | 0.27068 | 0.681807 | 0.875 | 1 |
| GOBP_REGULATION_OF_CARBOHYDRATE_CATABOLIC_PROCESS | 47 | 0.233755 | 0.679637 | 0.942775 | 1 |
| GOBP_REGULATION_OF_EXECUTION_PHASE_OF_APOPTOSIS | 16 | 0.282305 | 0.678494 | 0.884127 | 1 |
| GOBP_INTRACELLULAR_STEROL_TRANSPORT | 27 | 0.258736 | 0.678339 | 0.904468 | 1 |
| GOBP_REGULATION_OF_RECEPTOR_SIGNALING_PATHWAY_VIA_STAT | 59 | 0.219981 | 0.678319 | 0.950617 | 1 |
| GOBP_NEUROMUSCULAR_SYNAPTIC_TRANSMISSION | 20 | 0.268697 | 0.67821 | 0.877934 | 1 |

| | | | | | |
|---|---|---|---|---|---|
| GOBP_RESPONSE_TO_INTERLEUKIN_6 | 28 | 0.256054 | 0.67734 | 0.921296 | 1 |
| GOBP_NEUROBLAST_PROLIFERATION | 48 | 0.229083 | 0.677148 | 0.932011 | 1 |
| GOBP_CELLULAR_MONOVALENT_INORGANIC_CATION_HOMEOSTASIS | 88 | 0.209044 | 0.676604 | 0.971622 | 1 |
| GOBP_CELLULAR_NITROGEN_COMPOUND_CATABOLIC_PROCESS | 406 | 0.178689 | 0.676591 | 1 | 1 |
| GOBP_NEGATIVE_REGULATION_OF_PROTEASOMAL_PROTEIN_CATABOLIC_PROCESS | 43 | 0.230939 | 0.675748 | 0.939794 | 1 |
| GOBP_MITOTIC_G2_M_TRANSITION_CHECKPOINT | 47 | 0.227935 | 0.675143 | 0.940751 | 1 |
| GOBP_RESPIRATORY_BURST | 21 | 0.271428 | 0.674427 | 0.894819 | 1 |
| GOBP_RIBOSE_PHOSPHATE_METABOLIC_PROCESS | 338 | 0.181636 | 0.67413 | 0.998902 | 1 |
| GOBP_METANEPHRIC_EPITHELIUM_DEVELOPMENT | 16 | 0.28256 | 0.674065 | 0.872517 | 1 |
| GOBP_PHOSPHATIDYLINOSITOL_3_PHOSPHATE_BIOSYNTHETIC_PROCESS | 19 | 0.273812 | 0.673743 | 0.899514 | 1 |
| GOBP_ANDROGEN_RECEPTOR_SIGNALING_PATHWAY | 40 | 0.231461 | 0.67309 | 0.899851 | 1 |
| GOBP_NEURON_MATURATION | 40 | 0.228411 | 0.671891 | 0.920792 | 1 |
| GOBP_REGULATION_OF_NOTCH_SIGNALING_PATHWAY | 78 | 0.210961 | 0.671804 | 0.969292 | 1 |
| GOBP_POSITIVE_REGULATION_OF_VACUOLE_ORGANIZATION | 21 | 0.265592 | 0.671473 | 0.886943 | 1 |
| GOBP_POSITIVE_REGULATION_OF_DNA_METABOLIC_PROCESS | 212 | 0.18629 | 0.671195 | 0.993119 | 1 |
| GOBP_ENERGY_DERIVATION_BY_OXIDATION_OF_ORGANIC_COMPOUNDS | 280 | 0.1819 | 0.669966 | 0.992126 | 1 |
| GOBP_REGULATION_OF_PHOSPHOLIPASE_C_ACTIVITY | 30 | 0.245221 | 0.669724 | 0.917683 | 1 |
| GOBP_CORTICAL_ACTIN_CYTOSKELETON_ORGANIZATION | 37 | 0.238463 | 0.669713 | 0.925 | 1 |
| GOBP_PROTEIN_ACYLATION | 238 | 0.184578 | 0.669539 | 0.995402 | 1 |
| GOBP_PROTEOGLYCAN_BIOSYNTHETIC_PROCESS | 53 | 0.220839 | 0.669498 | 0.947149 | 1 |
| GOBP_NEGATIVE_REGULATION_OF_TUMOR_NECROSIS_FACTOR_MEDIATED_SIGNALING_ | 19 | 0.270808 | 0.669383 | 0.903879 | 1 |
| GOBP_AXO_DENDRITIC_TRANSPORT | 75 | 0.208464 | 0.668975 | 0.973404 | 1 |
| GOBP_NEGATIVE_REGULATION_OF_DEPHOSPHORYLATION | 40 | 0.231026 | 0.668939 | 0.93037 | 1 |
| GOBP_REGULATION_OF_GTP_BINDING | 15 | 0.2883 | 0.668838 | 0.881579 | 1 |
| GOBP_GLANDULAR_EPITHELIAL_CELL_DEVELOPMENT | 22 | 0.263089 | 0.668373 | 0.910798 | 1 |
| GOBP_PHOSPHOLIPID_DEPHOSPHORYLATION | 41 | 0.232328 | 0.667978 | 0.919453 | 1 |
| GOBP_CELLULAR_RESPONSE_TO_VITAMIN | 24 | 0.261652 | 0.667547 | 0.911859 | 1 |
| GOBP_METANEPHRIC_TUBULE_DEVELOPMENT | 15 | 0.28939 | 0.666927 | 0.862969 | 1 |
| GOBP_GLYCOSPHINGOLIPID_BIOSYNTHETIC_PROCESS | 29 | 0.246638 | 0.666514 | 0.906344 | 1 |
| GOBP_PYRIMIDINE_NUCLEOSIDE_MONOPHOSPHATE_METABOLIC_PROCESS | 19 | 0.271888 | 0.666 | 0.871336 | 1 |
| GOBP_MITOTIC_RECOMBINATION | 25 | 0.255348 | 0.6658 | 0.881356 | 1 |
| GOBP_POSITIVE_REGULATION_OF_DNA_REPAIR | 91 | 0.202468 | 0.665263 | 0.976285 | 1 |
| GOBP_ENDOSOMAL_TRANSPORT | 223 | 0.18385 | 0.663466 | 0.995429 | 1 |
| GOBP_PHOTORECEPTOR_CELL_MAINTENANCE | 34 | 0.239531 | 0.66313 | 0.926094 | 1 |
| GOBP_PEPTIDYL_LYSINE_DIMETHYLATION | 23 | 0.257787 | 0.662878 | 0.899371 | 1 |
| GOBP_MICROTUBULE_BASED_TRANSPORT | 186 | 0.184845 | 0.661646 | 0.991803 | 1 |
| GOBP_CELL_CYCLE_G2_M_PHASE_TRANSITION | 146 | 0.19082 | 0.661636 | 0.991453 | 1 |
| GOBP_STRESS_GRANULE_ASSEMBLY | 27 | 0.251086 | 0.660732 | 0.909798 | 1 |
| GOBP_GENERATION_OF_PRECURSOR_METABOLITES_AND_ENERGY | 424 | 0.174582 | 0.660037 | 1 | 1 |
| GOBP_2_OXOGLUTARATE_METABOLIC_PROCESS | 16 | 0.2784 | 0.659985 | 0.888889 | 1 |
| GOBP_POLYAMINE_METABOLIC_PROCESS | 18 | 0.268997 | 0.659476 | 0.881579 | 1 |
| GOBP_POSITIVE_REGULATION_OF_PROTEIN_TYROSINE_KINASE_ACTIVITY | 41 | 0.228702 | 0.659328 | 0.935532 | 1 |
| GOBP_STEROID_HORMONE_MEDIATED_SIGNALING_PATHWAY | 114 | 0.196112 | 0.657732 | 0.982323 | 1 |
| GOBP_POSITIVE_REGULATION_OF_LYASE_ACTIVITY | 16 | 0.272955 | 0.656932 | 0.882645 | 1 |
| GOBP_MICROTUBULE_ANCHORING | 24 | 0.254476 | 0.656724 | 0.908669 | 1 |
| GOBP_BONE_MATURATION | 23 | 0.252845 | 0.655929 | 0.904531 | 1 |
| GOBP_EXTRACELLULAR_VESICLE_BIOGENESIS | 24 | 0.259195 | 0.654676 | 0.916667 | 1 |
| GOBP_REGULATION_OF_PROTEASOMAL_UBIQUITIN_DEPENDENT_PROTEIN_CATABOLIC_ | 128 | 0.193039 | 0.652977 | 0.992443 | 1 |
| GOBP_ENTEROENDOCRINE_CELL_DIFFERENTIATION | 20 | 0.261669 | 0.65194 | 0.917722 | 1 |
| GOBP_ENDOSOME_TRANSPORT_VIA_MULTIVESICULAR_BODY_SORTING_PATHWAY | 31 | 0.243105 | 0.651589 | 0.920489 | 1 |
| GOBP_NEGATIVE_REGULATION_OF_PROTEOLYSIS_INVOLVED_IN_CELLULAR_PROTEIN_C | 57 | 0.210032 | 0.65102 | 0.951977 | 1 |
| GOBP_REGULATION_OF_MITOTIC_NUCLEAR_DIVISION | 96 | 0.196423 | 0.65067 | 0.982984 | 1 |
| GOBP_TYPE_I_INTERFERON_PRODUCTION | 81 | 0.20281 | 0.649422 | 0.974934 | 1 |
| GOBP_CLATHRIN_DEPENDENT_ENDOCYTOSIS | 45 | 0.219781 | 0.649401 | 0.955903 | 1 |
| GOBP_REGULATION_OF_GLYCOLYTIC_PROCESS | 38 | 0.227075 | 0.649099 | 0.948758 | 1 |
| GOBP_REGULATION_OF_NEURON_PROJECTION_REGENERATION | 26 | 0.241785 | 0.64834 | 0.910078 | 1 |
| GOBP_MRNA_CATABOLIC_PROCESS | 218 | 0.179744 | 0.648201 | 0.998837 | 1 |
| GOBP_FATTY_ACID_CATABOLIC_PROCESS | 91 | 0.198229 | 0.643791 | 0.988064 | 1 |
| GOBP_TELENCEPHALON_GLIAL_CELL_MIGRATION | 22 | 0.252481 | 0.64306 | 0.915309 | 1 |
| GOBP_GLYCOPROTEIN_CATABOLIC_PROCESS | 23 | 0.252519 | 0.643026 | 0.930159 | 1 |
| GOBP_SIGNALING_RECEPTOR_LIGAND_PRECURSOR_PROCESSING | 26 | 0.245239 | 0.642084 | 0.937405 | 1 |
| GOBP_PTERIDINE_CONTAINING_COMPOUND_METABOLIC_PROCESS | 34 | 0.233532 | 0.640669 | 0.938144 | 1 |
| GOBP_LUNG_CELL_DIFFERENTIATION | 15 | 0.269373 | 0.639986 | 0.9 | 1 |
| GOBP_DETERMINATION_OF_ADULT_LIFESPAN | 15 | 0.274185 | 0.639148 | 0.917355 | 1 |
| GOBP_REGULATION_OF_CHROMATIN_ASSEMBLY_OR_DISASSEMBLY | 33 | 0.232836 | 0.639044 | 0.95122 | 1 |
| GOBP_MITOCHONDRIAL_DNA_METABOLIC_PROCESS | 15 | 0.266527 | 0.637692 | 0.896147 | 1 |
| GOBP_NEGATIVE_REGULATION_OF_MRNA_METABOLIC_PROCESS | 83 | 0.197475 | 0.637252 | 0.988048 | 1 |
| GOBP_PYRIDINE_CONTAINING_COMPOUND_METABOLIC_PROCESS | 32 | 0.230952 | 0.636692 | 0.933131 | 1 |
| GOBP_POSITIVE_REGULATION_OF_GLUCOSE_IMPORT | 30 | 0.230278 | 0.635284 | 0.942857 | 1 |
| GOBP_PROTEIN_METHYLATION | 185 | 0.178769 | 0.635022 | 0.998804 | 1 |
| GOBP_NEGATIVE_REGULATION_OF_CELL_CYCLE_G2_M_PHASE_TRANSITION | 59 | 0.209593 | 0.634307 | 0.970549 | 1 |
| GOBP_ISOPRENOID_METABOLIC_PROCESS | 77 | 0.198861 | 0.633839 | 0.985564 | 1 |
| GOBP_REGULATION_OF_RRNA_PROCESSING | 16 | 0.267065 | 0.63267 | 0.905473 | 1 |
| GOBP_GLUTAMINE_METABOLIC_PROCESS | 23 | 0.245079 | 0.632378 | 0.929713 | 1 |
| GOBP_REGULATION_OF_CHOLESTEROL_EFFLUX | 29 | 0.233846 | 0.632072 | 0.944611 | 1 |
| GOBP_CELL_CYCLE_PHASE_TRANSITION | 470 | 0.166646 | 0.631902 | 1 | 1 |
| GOBP_CHROMOSOME_CONDENSATION | 39 | 0.223595 | 0.631868 | 0.942393 | 1 |
| GOBP_REGULATION_OF_ATP_BIOSYNTHETIC_PROCESS | 20 | 0.25585 | 0.631189 | 0.920584 | 1 |
| GOBP_PROTEIN_LOCALIZATION_TO_GOLGI_APPARATUS | 26 | 0.241432 | 0.630335 | 0.923077 | 1 |
| GOBP_REGULATION_OF_HISTONE_METHYLATION | 73 | 0.199112 | 0.628666 | 0.971774 | 1 |
| GOBP_OLIGOSACCHARIDE_LIPID_INTERMEDIATE_BIOSYNTHETIC_PROCESS | 19 | 0.256811 | 0.628129 | 0.92283 | 1 |
| GOBP_MIRNA_METABOLIC_PROCESS | 27 | 0.233989 | 0.62485 | 0.940639 | 1 |
| GOBP_REGULATION_OF_GLUCOSE_TRANSMEMBRANE_TRANSPORT | 58 | 0.206247 | 0.62469 | 0.979224 | 1 |
| GOBP_ESTABLISHMENT_OF_RNA_LOCALIZATION | 147 | 0.180377 | 0.624613 | 0.996416 | 1 |
| GOBP_POSITIVE_REGULATION_OF_ACTIN_FILAMENT_BUNDLE_ASSEMBLY | 54 | 0.205679 | 0.62357 | 0.966527 | 1 |
| GOBP_REGULATION_OF_TYPE_I_INTERFERON_MEDIATED_SIGNALING_PATHWAY | 36 | 0.219368 | 0.623535 | 0.959276 | 1 |
| GOBP_ESTABLISHMENT_OF_MITOCHONDRION_LOCALIZATION | 28 | 0.228497 | 0.623293 | 0.953148 | 1 |
| GOBP_PIGMENT_GRANULE_ORGANIZATION | 27 | 0.238883 | 0.622702 | 0.941177 | 1 |
| GOBP_POSTREPLICATION_REPAIR | 34 | 0.224252 | 0.621985 | 0.95 | 1 |
| GOBP_PROTEIN_CATABOLIC_PROCESS_IN_THE_VACUOLE | 21 | 0.250083 | 0.621341 | 0.935681 | 1 |
| GOBP_POSITIVE_REGULATION_OF_GLUCOSE_TRANSMEMBRANE_TRANSPORT | 34 | 0.224687 | 0.620912 | 0.963415 | 1 |
| GOBP_POSITIVE_REGULATION_OF_MRNA_METABOLIC_PROCESS | 115 | 0.183171 | 0.620763 | 0.997497 | 1 |
| GOBP_NEGATIVE_REGULATION_OF_CELLULAR_PROTEIN_CATABOLIC_PROCESS | 69 | 0.199538 | 0.619724 | 0.984911 | 1 |
| GOBP_SNRNA_TRANSCRIPTION | 17 | 0.252523 | 0.619496 | 0.933225 | 1 |
| GOBP_CELLULAR_CARBOHYDRATE_CATABOLIC_PROCESS | 31 | 0.223583 | 0.618683 | 0.939759 | 1 |
| GOBP_FAT_SOLUBLE_VITAMIN_METABOLIC_PROCESS | 24 | 0.239867 | 0.618486 | 0.931142 | 1 |
| GOBP_NEGATIVE_REGULATION_OF_ORGANELLE_ORGANIZATION | 322 | 0.1668 | 0.617945 | 0.998891 | 1 |
| GOBP_PEPTIDYL_LYSINE_METHYLATION | 131 | 0.179908 | 0.617588 | 0.995122 | 1 |
| GOBP_POSITIVE_REGULATION_OF_ORGANELLE_ASSEMBLY | 75 | 0.193815 | 0.61707 | 0.978581 | 1 |
| GOBP_REGULATION_OF_CELL_CYCLE_G2_M_PHASE_TRANSITION | 100 | 0.186535 | 0.617001 | 0.99493 | 1 |
| GOBP_DNA_INTEGRITY_CHECKPOINT_SIGNALING | 117 | 0.18262 | 0.616269 | 0.992366 | 1 |
| GOBP_CHROMOSOME_SEPARATION | 88 | 0.188876 | 0.615329 | 0.996063 | 1 |
| GOBP_PIGMENT_CELL_DIFFERENTIATION | 32 | 0.226589 | 0.614874 | 0.953632 | 1 |
| GOBP_VESICLE_ORGANIZATION | 277 | 0.166316 | 0.614582 | 1 | 1 |
| GOBP_ANDROGEN_METABOLIC_PROCESS | 16 | 0.256469 | 0.614352 | 0.916667 | 1 |
| GOBP_CHAPERONE_MEDIATED_AUTOPHAGY | 16 | 0.255075 | 0.614289 | 0.922581 | 1 |
| GOBP_NUCLEAR_ENVELOPE_ORGANIZATION | 55 | 0.203895 | 0.61425 | 0.968363 | 1 |
| GOBP_NEGATIVE_REGULATION_OF_EPITHELIAL_TO_MESENCHYMAL_TRANSITION | 30 | 0.22518 | 0.613972 | 0.961136 | 1 |
| GOBP_LYTIC_VACUOLE_ORGANIZATION | 67 | 0.193698 | 0.612636 | 0.988966 | 1 |
| GOBP_POSITIVE_REGULATION_OF_TELOMERASE_ACTIVITY | 34 | 0.221634 | 0.612613 | 0.964286 | 1 |
| GOBP_DETECTION_OF_BIOTIC_STIMULUS | 22 | 0.236626 | 0.610472 | 0.93049 | 1 |
| GOBP_NEGATIVE_REGULATION_OF_CALCIUM_MEDIATED_SIGNALING | 17 | 0.250964 | 0.608782 | 0.930448 | 1 |
| GOBP_REGULATION_OF_TRANSLATIONAL_INITIATION | 73 | 0.195207 | 0.608653 | 0.982644 | 1 |
| GOBP_FOLIC_ACID_METABOLIC_PROCESS | 15 | 0.260814 | 0.608119 | 0.930464 | 1 |
| GOBP_GLIAL_CELL_PROLIFERATION | 30 | 0.225822 | 0.607917 | 0.943452 | 1 |
| GOBP_REGULATION_OF_TRANSCRIPTION_INVOLVED_IN_G1_S_TRANSITION_OF_MITOTIC_ | 20 | 0.238637 | 0.606165 | 0.945425 | 1 |
| GOBP_REGULATION_OF_PROTEIN_LOCALIZATION_TO_NUCLEUS | 119 | 0.179158 | 0.605732 | 0.997512 | 1 |
| GOBP_NEGATIVE_REGULATION_OF_PROTEIN_LOCALIZATION_TO_MEMBRANE | 28 | 0.228202 | 0.605676 | 0.969697 | 1 |
| GOBP_MACROAUTOPHAGY | 298 | 0.163588 | 0.60563 | 1 | 1 |
| GOBP_NUCLEAR_CHROMOSOME_SEGREGATION | 257 | 0.16427 | 0.605356 | 1 | 1 |
| GOBP_IMP_METABOLIC_PROCESS | 16 | 0.255249 | 0.604645 | 0.941569 | 1 |
| GOBP_PROTEIN_TARGETING | 293 | 0.164015 | 0.604179 | 1 | 1 |
| GOBP_CELLULAR_PIGMENTATION | 50 | 0.199397 | 0.603438 | 0.98209 | 1 |
| GOBP_OLIGOSACCHARIDE_METABOLIC_PROCESS | 45 | 0.202609 | 0.602586 | 0.971469 | 1 |
| GOBP_CARBOHYDRATE_TRANSPORT | 115 | 0.177928 | 0.602537 | 0.996236 | 1 |
| GOBP_POSITIVE_REGULATION_OF_NOTCH_SIGNALING_PATHWAY | 35 | 0.211909 | 0.601601 | 0.971514 | 1 |
| GOBP_TRANSCRIPTION_INITIATION_FROM_RNA_POLYMERASE_II_PROMOTER | 72 | 0.1893 | 0.599549 | 0.992948 | 1 |
| GOBP_PEPTIDYL_SERINE_DEPHOSPHORYLATION | 19 | 0.245231 | 0.599145 | 0.941807 | 1 |
| GOBP_NUCLEOBASE_CONTAINING_COMPOUND_TRANSPORT | 196 | 0.166687 | 0.596655 | 1 | 1 |
| GOBP_REGULATION_OF_MITOCHONDRIAL_OUTER_MEMBRANE_PERMEABILIZATION_INVO | 20 | 0.238303 | 0.595468 | 0.951027 | 1 |
| GOBP_REGULATION_OF_TRANSCRIPTION_BY_RNA_POLYMERASE_III | 29 | 0.221449 | 0.595418 | 0.953383 | 1 |
| GOBP_VIRAL_BUDDING | 27 | 0.223901 | 0.594829 | 0.969925 | 1 |
| GOBP_PROTEIN_MODIFICATION_BY_SMALL_PROTEIN_REMOVAL | 147 | 0.173447 | 0.59475 | 0.998782 | 1 |
| GOBP_AMP_METABOLIC_PROCESS | 23 | 0.227112 | 0.593394 | 0.955906 | 1 |
| GOBP_POSITIVE_REGULATION_OF_TYPE_I_INTERFERON_PRODUCTION | 50 | 0.202186 | 0.592748 | 0.973901 | 1 |
| GOBP_PEPTIDYL_LYSINE_MODIFICATION | 364 | 0.159287 | 0.592166 | 1 | 1 |
| GOBP_LOW_DENSITY_LIPOPROTEIN_PARTICLE_CLEARANCE | 20 | 0.235852 | 0.591477 | 0.943038 | 1 |
| GOBP_SPINDLE_ASSEMBLY | 126 | 0.174859 | 0.590502 | 1 | 1 |
| GOBP_REGULATION_OF_PROTEIN_DEACETYLATION | 53 | 0.195674 | 0.588845 | 0.98042 | 1 |
| GOBP_NEGATIVE_REGULATION_OF_PROTEIN_DEPHOSPHORYLATION | 30 | 0.21617 | 0.588089 | 0.967017 | 1 |
| GOBP_NEGATIVE_REGULATION_OF_ACTIN_FILAMENT_POLYMERIZATION | 58 | 0.192242 | 0.587812 | 0.983264 | 1 |
| GOBP_MITOCHONDRION_ORGANIZATION | 485 | 0.15445 | 0.587742 | 1 | 1 |
| GOBP_STORE_OPERATED_CALCIUM_ENTRY | 18 | 0.239533 | 0.586374 | 0.952846 | 1 |
| GOBP_REGULATION_OF_INCLUSION_BODY_ASSEMBLY | 16 | 0.244857 | 0.586127 | 0.941463 | 1 |
| GOBP_LACTATION | 39 | 0.201258 | 0.581838 | 0.980854 | 1 |
| GOBP_MITOTIC_CELL_CYCLE_PHASE_TRANSITION | 385 | 0.15494 | 0.580939 | 1 | 1 |
| GOBP_REGULATION_OF_MACROAUTOPHAGY | 151 | 0.166883 | 0.58047 | 0.997567 | 1 |
| GOBP_MATURE_B_CELL_DIFFERENTIATION | 22 | 0.224245 | 0.577264 | 0.981191 | 1 |
| GOBP_ATP_METABOLIC_PROCESS | 241 | 0.159718 | 0.574901 | 1 | 1 |
| GOBP_AEROBIC_RESPIRATION | 173 | 0.162001 | 0.573776 | 1 | 1 |
| GOBP_REGULATION_OF_MITOCHONDRIAL_MEMBRANE_PERMEABILITY | 54 | 0.190377 | 0.573245 | 0.993036 | 1 |
| GOBP_MONOSACCHARIDE_CATABOLIC_PROCESS | 34 | 0.209918 | 0.573069 | 0.975073 | 1 |
| GOBP_MICROTUBULE_CYTOSKELETON_ORGANIZATION_INVOLVED_IN_MITOSIS | 149 | 0.165764 | 0.571882 | 1 | 1 |
| GOBP_PINOCYTOSIS | 21 | 0.227447 | 0.571181 | 0.959029 | 1 |
| GOBP_REGULATION_OF_PROTEIN_MODIFICATION_BY_SMALL_PROTEIN_CONJUGATION_C | 227 | 0.157843 | 0.569999 | 1 | 1 |
| GOBP_PROTEIN_EXPORT_FROM_NUCLEUS | 56 | 0.185105 | 0.569543 | 0.987569 | 1 |
| GOBP_LIPOPROTEIN_CATABOLIC_PROCESS | 16 | 0.23899 | 0.569473 | 0.954839 | 1 |

| | | | | | |
|---|---|---|---|---|---|
| GOBP_MITOCHONDRIAL_MEMBRANE_ORGANIZATION | 109 | 0.169257 | 0.569111 | 0.998703 | 1 |
| GOBP_INTRACELLULAR_LIPID_TRANSPORT | 40 | 0.199917 | 0.567824 | 0.981791 | 1 |
| GOBP_ORGANELLE_TRANSPORT_ALONG_MICROTUBULE | 82 | 0.17504 | 0.567301 | 1 | 1 |
| GOBP_NUCLEOSIDE_PHOSPHATE_CATABOLIC_PROCESS | 74 | 0.178204 | 0.565634 | 0.995995 | 1 |
| GOBP_MITOTIC_SPINDLE_ORGANIZATION | 125 | 0.167379 | 0.565539 | 0.998744 | 1 |
| GOBP_PROTEIN_K48_LINKED_DEUBIQUITINATION | 31 | 0.205857 | 0.565357 | 0.975073 | 1 |
| GOBP_MAST_CELL_ACTIVATION_INVOLVED_IN_IMMUNE_RESPONSE | 39 | 0.198296 | 0.561724 | 0.979798 | 1 |
| GOBP_REGULATION_OF_ARP2_3_COMPLEX_MEDIATED_ACTIN_NUCLEATION | 21 | 0.226068 | 0.560826 | 0.972222 | 1 |
| GOBP_POSITIVE_REGULATION_OF_CELLULAR_RESPONSE_TO_INSULIN_STIMULUS | 19 | 0.228941 | 0.560435 | 0.957211 | 1 |
| GOBP_PYRIMIDINE_NUCLEOBASE_METABOLIC_PROCESS | 20 | 0.224345 | 0.559847 | 0.968454 | 1 |
| GOBP_INTRINSIC_APOPTOTIC_SIGNALING_PATHWAY_IN_RESPONSE_TO_OXIDATIVE_STR | 42 | 0.194132 | 0.558989 | 0.991354 | 1 |
| GOBP_REGULATION_OF_RIG_I_SIGNALING_PATHWAY | 17 | 0.227164 | 0.556756 | 0.964467 | 1 |
| GOBP_AUTOPHAGOSOME_ORGANIZATION | 105 | 0.168528 | 0.553218 | 0.997413 | 1 |
| GOBP_ENDOSOME_ORGANIZATION | 81 | 0.172679 | 0.552295 | 0.998626 | 1 |
| GOBP_UBIQUITIN_DEPENDENT_PROTEIN_CATABOLIC_PROCESS_VIA_THE_MULTIVESICUL | 34 | 0.19885 | 0.551532 | 0.988113 | 1 |
| GOBP_REGULATION_OF_WATER_LOSS_VIA_SKIN | 20 | 0.22205 | 0.551305 | 0.973603 | 1 |
| GOBP_DOSAGE_COMPENSATION | 20 | 0.224586 | 0.550759 | 0.960876 | 1 |
| GOBP_ENDOPLASMIC_RETICULUM_ORGANIZATION | 85 | 0.170544 | 0.548953 | 1 | 1 |
| GOBP_REGULATION_OF_CELLULAR_RESPONSE_TO_INSULIN_STIMULUS | 51 | 0.183104 | 0.548739 | 0.989914 | 1 |
| GOBP_NEGATIVE_REGULATION_OF_PROTEIN_MODIFICATION_BY_SMALL_PROTEIN_CONJ | 88 | 0.168608 | 0.546137 | 0.998703 | 1 |
| GOBP_SPINDLE_ORGANIZATION | 192 | 0.15294 | 0.546004 | 1 | 1 |
| GOBP_PROTEASOMAL_PROTEIN_CATABOLIC_PROCESS | 464 | 0.142844 | 0.545132 | 1 | 1 |
| GOBP_LYSOSOMAL_PROTEIN_CATABOLIC_PROCESS | 18 | 0.220858 | 0.545104 | 0.977883 | 1 |
| GOBP_HISTONE_PHOSPHORYLATION | 42 | 0.186058 | 0.543353 | 0.989736 | 1 |
| GOBP_ESTABLISHMENT_OR_MAINTENANCE_OF_MONOPOLAR_CELL_POLARITY | 26 | 0.202063 | 0.539902 | 0.982866 | 1 |
| GOBP_REGULATION_OF_ACTIN_NUCLEATION | 31 | 0.192734 | 0.536025 | 0.982998 | 1 |
| GOBP_N_ACETYLGLUCOSAMINE_METABOLIC_PROCESS | 17 | 0.222938 | 0.534839 | 0.974643 | 1 |
| GOBP_TUBULIN_DEACETYLATION | 22 | 0.212143 | 0.534435 | 0.982622 | 1 |
| GOBP_CHROMOSOME_SEGREGATION | 315 | 0.143483 | 0.533689 | 1 | 1 |
| GOBP_NEGATIVE_REGULATION_OF_SMOOTHENED_SIGNALING_PATHWAY | 30 | 0.191816 | 0.53054 | 0.991111 | 1 |
| GOBP_LATE_ENDOSOME_TO_VACUOLE_TRANSPORT_VIA_MULTIVESICULAR_BODY_SORT | 17 | 0.218331 | 0.530493 | 0.967051 | 1 |
| GOBP_CARDIOLIPIN_METABOLIC_PROCESS | 15 | 0.226324 | 0.530133 | 0.974603 | 1 |
| GOBP_INNER_EAR_RECEPTOR_CELL_DEVELOPMENT | 38 | 0.18399 | 0.527316 | 0.994083 | 1 |
| GOBP_ARGININE_METABOLIC_PROCESS | 15 | 0.221656 | 0.527288 | 0.971714 | 1 |
| GOBP_CARBOHYDRATE_PHOSPHORYLATION | 24 | 0.202453 | 0.526776 | 0.987578 | 1 |
| GOBP_PROTEASOME_MEDIATED_UBIQUITIN_DEPENDENT_PROTEIN_CATABOLIC_PROCES | 394 | 0.139699 | 0.525686 | 1 | 1 |
| GOBP_INNER_EAR_RECEPTOR_CELL_STEREOCILIUM_ORGANIZATION | 27 | 0.197355 | 0.524542 | 0.981453 | 1 |
| GOBP_KERATAN_SULFATE_METABOLIC_PROCESS | 17 | 0.215809 | 0.524199 | 0.967791 | 1 |
| GOBP_MACROMOLECULE_DEACYLATION | 123 | 0.153464 | 0.523394 | 1 | 1 |
| GOBP_LAMELLIPODIUM_MORPHOGENESIS | 17 | 0.216327 | 0.522747 | 0.976351 | 1 |
| GOBP_NEGATIVE_REGULATION_OF_GLYCOPROTEIN_METABOLIC_PROCESS | 15 | 0.220138 | 0.522239 | 0.97054 | 1 |
| GOBP_HISTONE_H3_K4_TRIMETHYLATION | 20 | 0.206464 | 0.519633 | 0.97678 | 1 |
| GOBP_CYTOSOLIC_TRANSPORT | 167 | 0.14888 | 0.517763 | 1 | 1 |
| GOBP_REGULATION_OF_INSULIN_RECEPTOR_SIGNALING_PATHWAY | 51 | 0.17079 | 0.514447 | 0.995665 | 1 |
| GOBP_RNA_CATABOLIC_PROCESS | 261 | 0.141748 | 0.514194 | 1 | 1 |
| GOBP_RRNA_METHYLATION | 25 | 0.195261 | 0.513044 | 0.986154 | 1 |
| GOBP_CELLULAR_RESPIRATION | 210 | 0.14227 | 0.511615 | 1 | 1 |
| GOBP_MIDBODY_ABSCISSION | 17 | 0.215326 | 0.510731 | 0.976153 | 1 |
| GOBP_REGULATION_OF_GLUCOSE_IMPORT | 46 | 0.172978 | 0.510137 | 0.997155 | 1 |
| GOBP_PROTEIN_LOCALIZATION_TO_CILIARY_MEMBRANE | 17 | 0.213975 | 0.509641 | 0.982839 | 1 |
| GOBP_NEGATIVE_REGULATION_OF_MACROAUTOPHAGY | 34 | 0.182287 | 0.509554 | 0.990937 | 1 |
| GOBP_GLUCOCORTICOID_METABOLIC_PROCESS | 16 | 0.21336 | 0.508193 | 0.977383 | 1 |
| GOBP_INNER_MITOCHONDRIAL_MEMBRANE_ORGANIZATION | 40 | 0.177234 | 0.506082 | 0.998555 | 1 |
| GOBP_NEGATIVE_REGULATION_OF_TELOMERE_MAINTENANCE_VIA_TELOMERE_LENGTH | 24 | 0.193712 | 0.501516 | 0.995169 | 1 |
| GOBP_MICROTUBULE_NUCLEATION | 35 | 0.177583 | 0.49805 | 0.994228 | 1 |
| GOBP_RNA_LOCALIZATION | 181 | 0.140749 | 0.496875 | 1 | 1 |
| GOBP_PROTEIN_PEPTIDYL_PROLYL_ISOMERIZATION | 35 | 0.177167 | 0.496592 | 0.996928 | 1 |
| GOBP_COPII_COATED_VESICLE_CARGO_LOADING | 15 | 0.205422 | 0.493572 | 0.982877 | 1 |
| GOBP_MRNA_TRANSCRIPTION | 38 | 0.174029 | 0.493426 | 0.99393 | 1 |
| GOBP_PROTEIN_STABILIZATION | 167 | 0.139312 | 0.491951 | 1 | 1 |
| GOBP_REGULATION_OF_HISTONE_DEACETYLATION | 38 | 0.17361 | 0.489101 | 0.998529 | 1 |
| GOBP_MUSCLE_CELL_CELLULAR_HOMEOSTASIS | 15 | 0.205332 | 0.487903 | 0.981132 | 1 |
| GOBP_REGULATION_OF_NEURON_PROJECTION_ARBORIZATION | 15 | 0.210359 | 0.487152 | 0.987138 | 1 |
| GOBP_MEMBRANE_PROTEIN_PROTEOLYSIS | 52 | 0.160263 | 0.481473 | 0.997179 | 1 |
| GOBP_KINETOCHORE_ASSEMBLY | 17 | 0.196811 | 0.477464 | 0.995185 | 1 |
| GOBP_FIBROBLAST_APOPTOTIC_PROCESS | 18 | 0.195554 | 0.476705 | 0.993538 | 1 |
| GOBP_PROTEIN_DEGLYCOSYLATION | 25 | 0.179443 | 0.469839 | 0.998457 | 1 |
| GOBP_INTERSTRAND_CROSS_LINK_REPAIR | 30 | 0.171771 | 0.468709 | 0.993949 | 1 |
| GOBP_VIRION_ASSEMBLY | 36 | 0.166225 | 0.467758 | 1 | 1 |
| GOBP_COPII_COATED_VESICLE_BUDDING | 30 | 0.171233 | 0.461469 | 0.996974 | 1 |
| GOBP_POSITIVE_REGULATION_OF_NEURON_MIGRATION | 16 | 0.197346 | 0.45957 | 0.991817 | 1 |
| GOBP_PHOSPHATIDYLGLYCEROL_METABOLIC_PROCESS | 26 | 0.17282 | 0.453043 | 0.995461 | 1 |
| GOBP_POSITIVE_REGULATION_OF_TRANSCRIPTION_ELONGATION_FROM_RNA_POLYMER | 17 | 0.1885 | 0.452874 | 0.987768 | 1 |
| GOBP_RETROGRADE_TRANSPORT_ENDOSOME_TO_GOLGI | 91 | 0.136516 | 0.452653 | 1 | 1 |
| GOBP_MULTIVESICULAR_BODY_ORGANIZATION | 31 | 0.162448 | 0.450643 | 1 | 1 |
| GOBP_RESPONSE_TO_MISFOLDED_PROTEIN | 24 | 0.176738 | 0.4471 | 0.993507 | 1 |
| GOBP_MEMBRANE_PROTEIN_ECTODOMAIN_PROTEOLYSIS | 41 | 0.156534 | 0.44555 | 1 | 1 |
| GOBP_POSITIVE_REGULATION_OF_HISTONE_DEACETYLATION | 20 | 0.173167 | 0.442897 | 0.995305 | 1 |
| GOBP_RNA_PHOSPHODIESTER_BOND_HYDROLYSIS_EXONUCLEOLYTIC | 42 | 0.152442 | 0.439444 | 1 | 1 |
| GOBP_REGULATION_OF_HIPPO_SIGNALING | 20 | 0.172187 | 0.431465 | 0.998405 | 1 |
| GOBP_MITOTIC_CYTOKINESIS | 69 | 0.136454 | 0.430899 | 1 | 1 |
| GOBP_KINETOCHORE_ORGANIZATION | 22 | 0.168734 | 0.428406 | 0.996914 | 1 |
| GOBP_OXIDATIVE_PHOSPHORYLATION | 128 | 0.124235 | 0.42344 | 1 | 1 |
| GOBP_NEGATIVE_REGULATION_OF_TYPE_I_INTERFERON_MEDIATED_SIGNALING_PATHW | 16 | 0.175527 | 0.422952 | 0.995253 | 1 |
| GOBP_ERYTHROCYTE_DEVELOPMENT | 26 | 0.160674 | 0.422133 | 1 | 1 |
| GOBP_NEGATIVE_REGULATION_OF_UBIQUITIN_PROTEIN_TRANSFERASE_ACTIVITY | 22 | 0.165676 | 0.420104 | 0.995246 | 1 |
| GOBP_METAPHASE_PLATE_CONGRESSION | 65 | 0.134428 | 0.417181 | 1 | 1 |
| GOBP_CYTOKINETIC_PROCESS | 39 | 0.144048 | 0.416241 | 1 | 1 |
| GOBP_NEGATIVE_REGULATION_OF_RESPONSE_TO_OXIDATIVE_STRESS | 16 | 0.175674 | 0.41476 | 0.995246 | 1 |
| GOBP_NUCLEAR_MEMBRANE_REASSEMBLY | 23 | 0.1599 | 0.412014 | 1 | 1 |
| GOBP_POSITIVE_REGULATION_OF_DNA_TEMPLATED_TRANSCRIPTION_ELONGATION | 27 | 0.158669 | 0.41095 | 0.998512 | 1 |
| GOBP_PURINE_NUCLEOSIDE_MONOPHOSPHATE_BIOSYNTHETIC_PROCESS | 21 | 0.162137 | 0.409124 | 0.996909 | 1 |
| GOBP_RNA_POLYADENYLATION | 47 | 0.137108 | 0.402814 | 1 | 1 |
| GOBP_PROTEIN_N_LINKED_GLYCOSYLATION | 65 | 0.12827 | 0.3993 | 1 | 1 |
| GOBP_ACTIVATION_OF_INNATE_IMMUNE_RESPONSE | 40 | 0.1368 | 0.394462 | 1 | 1 |
| GOBP_GOLGI_VESICLE_TRANSPORT | 280 | 0.106565 | 0.389965 | 1 | 1 |
| GOBP_LATE_ENDOSOME_TO_VACUOLE_TRANSPORT | 24 | 0.146507 | 0.384697 | 0.998353 | 1 |
| GOBP_IRE1_MEDIATED_UNFOLDED_PROTEIN_RESPONSE | 18 | 0.155628 | 0.383834 | 0.99841 | 1 |
| GOBP_RIBONUCLEOSIDE_MONOPHOSPHATE_BIOSYNTHETIC_PROCESS | 31 | 0.139487 | 0.37763 | 1 | 1 |
| GOBP_PROTEIN_LOCALIZATION_TO_CHROMOSOME_CENTROMERIC_REGION | 25 | 0.143307 | 0.377422 | 1 | 1 |
| GOBP_RIBOSOMAL_LARGE_SUBUNIT_ASSEMBLY | 26 | 0.143127 | 0.373033 | 1 | 1 |
| GOBP_REGULATION_OF_VIRAL_INDUCED_CYTOPLASMIC_PATTERN_RECOGNITION_RECE | 20 | 0.141099 | 0.357806 | 0.99681 | 1 |
| GOBP_CYTOPLASMIC_TRANSLATION | 146 | 0.066497 | 0.231369 | 1 | 1 |
