## Supplementary material for "Chemokine Receptor 1 and its associated immune pathway are downregulated in SF3B1^MT^ blood and non-blood cancers": Table_S6

**Table S6. Shared immune gene sets in GSEA positive of SF3B1$^{K700E}$ ES lines.** NES (normalized enrichment score) are shown for three lines. Gene sets with p-values < 0.05 are listed and chemokine-related pathways are highlighted in red.

| Gene sets | NES | | |
|---|---|---|---|
| | SF3B1$^{K700E}$ ES line 1 | SF3B1$^{K700E}$ ES line 2 | SF3B1$^{K700E}$ ES line 3 |
| GOBP_GRANULOCYTE_ACTIVATION | 1.9353536 | 1.8054994 | 1.9111171 |
| GOBP_NEUTROPHIL_MEDIATED_IMMUNITY | 1.8026063 | 1.5972247 | 1.5062608 |
| GOBP_GRANULOCYTE_MIGRATION | 1.8010272 | 1.777065 | 1.6961323 |
| GOBP_GRANULOCYTE_CHEMOTAXIS | 1.800437 | 1.8162504 | 1.8105782 |
| GOBP_MONOCYTE_CHEMOTAXIS | 1.7978711 | 1.5757718 | 1.5533849 |
| GOBP_POSITIVE_REGULATION_OF_T_CELL_PROLIFERATION | 1.7638497 | 1.4831594 | 1.5641818 |
| GOBP_MYELOID_LEUKOCYTE_MIGRATION | 1.7554848 | 1.6734947 | 1.5979475 |
| GOBP_LEUKOCYTE_CHEMOTAXIS | 1.7545179 | 1.7170309 | 1.6865455 |
| GOBP_LEUKOCYTE_MIGRATION | 1.7444276 | 1.5933937 | 1.5522609 |
| GOBP_NEUTROPHIL_ACTIVATION_INVOLVED_IN_IMMUNE_RESPONSE | 1.7069777 | 1.6582732 | 1.6421021 |
| GOBP_POSITIVE_REGULATION_OF_LEUKOCYTE_PROLIFERATION | 1.7033479 | 1.5385394 | 1.4684414 |
| GOBP_INTERLEUKIN_8_PRODUCTION | 1.6962907 | 1.5203203 | 1.6227201 |
| GOBP_NEGATIVE_REGULATION_OF_INTERFERON_GAMMA_PRODUCTION | 1.6759768 | 1.7276211 | 1.9223958 |
| GOBP_POSITIVE_REGULATION_OF_LEUKOCYTE_CELL_CELL_ADHESION | 1.6699414 | 1.7016416 | 1.5917172 |
| GOBP_NEUTROPHIL_MIGRATION | 1.6615901 | 1.6327692 | 1.5242716 |
| GOBP_REGULATION_OF_LEUKOCYTE_MIGRATION | 1.6580168 | 1.5610356 | 1.4967196 |
| GOBP_MYELOID_LEUKOCYTE_ACTIVATION | 1.6536512 | 1.5856533 | 1.537492 |
| GOBP_REGULATION_OF_T_CELL_PROLIFERATION | 1.6490484 | 1.5146772 | 1.4054219 |
| GOBP_POSITIVE_REGULATION_OF_IMMUNE_RESPONSE | 1.6434894 | 1.6311518 | 1.2926693 |
| GOBP_REGULATION_OF_LEUKOCYTE_PROLIFERATION | 1.6272827 | 1.5257598 | 1.4325868 |
| GOBP_POSITIVE_REGULATION_OF_LEUKOCYTE_MIGRATION | 1.6258199 | 1.5936339 | 1.5003251 |
| GOBP_MACROPHAGE_DIFFERENTIATION | 1.619454 | 1.6259371 | 1.5345172 |
| GOBP_HUMORAL_IMMUNE_RESPONSE | 1.6063786 | 1.7087374 | 1.4708287 |
| GOBP_IMMUNE_EFFECTOR_PROCESS | 1.605233 | 1.6093051 | 1.2481014 |
| GOBP_POSITIVE_REGULATION_OF_INFLAMMATORY_RESPONSE | 1.6026944 | 1.7290034 | 1.7929065 |
| GOBP_LEUKOCYTE_CELL_CELL_ADHESION | 1.6001824 | 1.5131979 | 1.4635885 |
| GOBP_NEUTROPHIL_CHEMOTAXIS | 1.5993718 | 1.6285495 | 1.5563774 |
| GOBP_POSITIVE_REGULATION_OF_CD4_POSITIVE_ALPHA_BETA_T_CELL_ACTIVATIO | 1.5854201 | 1.812528 | 1.6831356 |
| GOBP_LEUKOCYTE_MEDIATED_CYTOTOXICITY | 1.5825696 | 1.5438228 | 1.5278581 |
| GOBP_REGULATION_OF_T_HELPER_1_TYPE_IMMUNE_RESPONSE | 1.580166 | 1.7273704 | 1.5618584 |
| GOBP_POSITIVE_REGULATION_OF_LEUKOCYTE_CHEMOTAXIS | 1.578578 | 1.467681 | 1.4796906 |
| GOBP_LEUKOCYTE_PROLIFERATION | 1.5760497 | 1.5148302 | 1.3021438 |
| GOBP_NEGATIVE_REGULATION_OF_ALPHA_BETA_T_CELL_ACTIVATION | 1.5612907 | 1.5827127 | 1.5164137 |
| GOBP_REGULATION_OF_INFLAMMATORY_RESPONSE | 1.5598786 | 1.6325203 | 1.6106169 |
| GOBP_NEGATIVE_REGULATION_OF_LYMPHOCYTE_ACTIVATION | 1.5592586 | 1.5465086 | 1.4250743 |
| GOBP_MYELOID_CELL_APOPTOTIC_PROCESS | 1.5584497 | 1.5583924 | 1.5830393 |
| GOBP_NEGATIVE_REGULATION_OF_IMMUNE_EFFECTOR_PROCESS | 1.5557295 | 1.7931865 | 1.3866743 |
| GOBP_T_CELL_ACTIVATION_INVOLVED_IN_IMMUNE_RESPONSE | 1.5518135 | 1.5792574 | 1.4236768 |
| GOBP_INFLAMMATORY_CELL_APOPTOTIC_PROCESS | 1.5509335 | 1.5518951 | 1.6814438 |
| GOBP_REGULATION_OF_ALPHA_BETA_T_CELL_ACTIVATION | 1.5508096 | 1.451211 | 1.50509 |
| GOBP_REGULATION_OF_T_CELL_ACTIVATION | 1.5506935 | 1.571375 | 1.4203935 |
| GOBP_TYPE_2_IMMUNE_RESPONSE | 1.5459174 | 1.5841765 | 1.5835477 |
| GOBP_RESPONSE_TO_CHEMOKINE | 1.5445112 | 1.4096801 | 1.5124242 |
| GOBP_POSITIVE_REGULATION_OF_CD4_POSITIVE_ALPHA_BETA_T_CELL_DIFFERENT | 1.5434407 | 1.7708387 | 1.6250192 |
| GOBP_INTERFERON_GAMMA_PRODUCTION | 1.534856 | 1.7319489 | 1.7156436 |
| GOBP_REGULATION_OF_LYMPHOCYTE_ACTIVATION | 1.5341733 | 1.5512276 | 1.361508 |
| GOBP_REGULATION_OF_LEUKOCYTE_CHEMOTAXIS | 1.5319239 | 1.4806296 | 1.4834539 |
| GOBP_T_CELL_ACTIVATION | 1.530798 | 1.4906747 | 1.4062117 |
| GOBP_NEGATIVE_REGULATION_OF_IMMUNE_RESPONSE | 1.5148497 | 1.798785 | 1.4085842 |
| GOBP_POSITIVE_REGULATION_OF_REGULATORY_T_CELL_DIFFERENTIATION | 1.5085077 | 1.5460477 | 1.5062668 |
| GOBP_LEUKOCYTE_APOPTOTIC_PROCESS | 1.480612 | 1.4370134 | 1.6019149 |
| GOBP_NEGATIVE_REGULATION_OF_IMMUNE_SYSTEM_PROCESS | 1.4786646 | 1.6808155 | 1.415585 |
| GOBP_T_CELL_DIFFERENTIATION | 1.4755586 | 1.5593555 | 1.4653376 |
| GOBP_CD4_POSITIVE_ALPHA_BETA_T_CELL_ACTIVATION | 1.4715511 | 1.5063893 | 1.3793724 |
| GOBP_LEUKOCYTE_DIFFERENTIATION | 1.4686122 | 1.5293185 | 1.4842457 |
| GOBP_T_HELPER_1_TYPE_IMMUNE_RESPONSE | 1.4551858 | 1.6965508 | 1.6365988 |
| GOBP_INFLAMMATORY_RESPONSE | 1.4495611 | 1.4291826 | 1.4144489 |
| GOBP_REGULATION_OF_LEUKOCYTE_APOPTOTIC_PROCESS | 1.4153107 | 1.5321132 | 1.6150314 |
| GOBP_POSITIVE_REGULATION_OF_LYMPHOCYTE_DIFFERENTIATION | 1.37709 | 1.608911 | 1.5947037 |
| GOBP_REGULATION_OF_LEUKOCYTE_DIFFERENTIATION | 1.3723793 | 1.5696824 | 1.384668 |
| GOBP_REGULATION_OF_LYMPHOCYTE_DIFFERENTIATION | 1.3638217 | 1.5672766 | 1.3529334 |
| GOBP_MYELOID_LEUKOCYTE_DIFFERENTIATION | 1.3580112 | 1.494762 | 1.5274414 |
| GOBP_CYTOKINE_MEDIATED_SIGNALING_PATHWAY | 1.3560992 | 1.4637483 | 1.3286468 |
