## Supplementary material for "Chemokine Receptor 1 and its associated immune pathway are downregulated in SF3B1^MT^ blood and non-blood cancers": Table_S7

| **Table S7. Shared upregulated immune genes in SF3B1$^{K700E}$ ES lines.** Genes with a fold change (FC) ≥1.5 and p-value < 0.05 are shown. In SF3B1$^{K700E}$ ES lines 1, 2 and 3, 23%, 22%, and 22% of significantly upregulated genes have immune functions, respectively. * indicates p-value ≥ 0.05 | | | |
|---|---|---|---|
| **Gene** | **FC** | | |
| | **SF3B1$^{K700E}$ ES line 1** | **SF3B1$^{K700E}$ ES line 2** | **SF3B1$^{K700E}$ ES line 3** |
| ABCC3 | 4.144606416 | 4.035394512 | 3.320833385 |
| ABI3 | 2.976852897 | 2.851838404 | 1.900236559 |
| ACP5 | 2.229572147 | 1.527463955 | 1.750826855 |
| ADCY7 | 2.030453203 | 1.833404984 | 1.582276474 |
| ADGRE5 | 3.876355128 | 4.27198281 | 2.238854186 |
| ADGRG3 | 4.148619677 | 4.027694198 | 5.00491191 |
| ALDH1A1 | 2.639997278 | 2.049827927 | 3.99221704 |
| ALOX5 | 2.916530289 | 2.962416002 | 3.251566832 |
| AMIGO2 | 2.007629233 | 2.3012872 | 2.277930964 |
| ANXA1 | 15.52221943 | 17.15139687 | 37.22306998 |
| ANXA2 | 2.26753128 | 2.419550358 | 1.508590584 |
| ANXA3 | 2.784334773 | 2.202469528 | 5.157401378 |
| APOBEC3F | 1.892340439 | 1.578068219 | 1.770477941 |
| ARID5B | 4.980696422 | 4.702435185 | 3.141418833 |
| ASB2 | 6.059505926 | 14.07358777 | 3.469715109 |
| ASH1L | 2.212456957 | 1.95431661 | 1.645519073 |
| ATXN1 | 3.143302734 | 2.706710608 | 2.854812412 |
| BANK1 | 3.751701136 | 3.366617334 | 4.121086201 |
| BTG2 | 4.940458901 | 3.892818028 | 4.73068249 |
| CAV1 | 3.793948001 | 4.424790399 | 2.69473194 |
| CAV2 | 2.696729285 | 2.264700178 | 1.994343368 |
| CDKN1A | 7.36834405 | 4.253749207 | 8.255602741 |
| CEACAM1 | 3.928235587 | 2.607874256 | 4.291379184 |
| CFD | 3.804318405 | 2.87710021 | 2.865612152 |
| CMKLR1 | 5.737160482 | 2.89925283 | 5.502954072 |
| COL8A1 | 12.51701916 | 9.766148723 | 14.27817944 |
| COLEC11 | 3.454717726 | 5.134002353 | 4.423670449 |
| CPEB4 | 2.314223529 | 2.12332013 | 2.085621592 |
| CR2 | 2.760676229 | 2.463679409 | 1.660960907 |
| CREB5 | 4.394442886 | 2.536332809 | 6.956928498 |
| CSF2RA | 1.861123563 | 1.857566787 | 1.900674477 |
| DGKH | 1.928757049 | 1.789493842 | 2.780974026 |
| DMTN | 3.400366661 | 1.760638061 | 1.869930238 |
| DUSP10 | 2.973044633 | 2.523125903 | 2.710154459 |
| EBF2 | 18.7156129 | 12.12813959 | 10.64063638 |
| EBI3 | 4.110176461 | 3.026516261 | 3.975962251 |
| EDA2R | 2.313416751 | 1.530202989 | 2.025414396 |
| EMILIN1 | 2.113749696 | 1.575314465 | 1.508307607 |
| ENPP2 | 2.898395386 | 2.52323278 | 1.838385606 |
| F10 | 2.687151049 | 2.878183112 | 2.229941041 |
| F7 | 16.99220691 | 34.13900622 | 7.17725697 |
| FAXDC2 | 2.437041629 | 2.207814088 | 2.285633665 |

| | | | |
|---|---|---|---|
| FES | 1.711034545 | 1.806995308 | 1.595366487 |
| FLI1 | 4.80924461 | 3.956869655 | 7.892197198 |
| FN1 | 1.693919987 | 1.787864371 | 2.448457426 |
| FZD5 | 2.256173634 | 2.077737333 | 2.042114503 |
| FZD6 | 1.556071236 | 1.476579512 | 1.606151841 |
| GADD45B | 1.674547855 | 1.507018647 | 3.86040917 |
| GADD45G | 2.007878284 | 1.973841435 | 2.386873338 |
| GALR1 | 7.289369911 | 3.977084962 | 12.652298 |
| GDF15 | 16.72575521 | 11.73576917 | 46.5822295 |
| GDF3 | 3.888660435 | 2.399842166 | 3.095712351 |
| HAPLN1 | 4.405716359 | 3.433943298 | 3.953228333 |
| HCLS1 | 6.679359978 | 4.028098566 | 10.4344593 |
| HLA-DQA1 | 5.467587357 | 6.713187656 | 3.021078318 |
| HLA-DQB1 | 3.208188508 | 3.526051527 | 2.037845087 |
| HLA-DRB1 | 1.761923878 | 1.965629837 | 1.845009673 |
| HLA-F | 1.791741424 | 1.946057278 | 1.810536607 |
| HPGD | 2.721096542 | 2.875902216 | 2.467608144 |
| HSPA2 | 3.895639502 | 3.923003606 | 2.900897389 |
| ICAM3 | 2.43571121 | 1.649758375 | 1.571247357 |
| IFI35 | 2.777020229 | 3.259364379 | 2.226631565 |
| IFIT1 | 2.38514683 | 1.962095087 | 1.739629641 |
| IFIT3 | 5.131966277 | 5.215861946 | 3.670916288 |
| IFITM1 | 1.910384208 | 1.710691366 | 2.039119346 |
| IL13RA1 | 1.805655484 | 1.417724955 | 1.635353128 |
| IL15 | 3.14386866 | 2.640061838 | 2.548214763 |
| IL6R | 3.205377795 | 3.008698914 | 2.256279116 |
| INPP5D | 2.565559831 | 1.795894823 | 3.334300753 |
| IRAK1 | 2.062292636 | 2.321456075 | 1.521462687 |
| ITGB2 | 2.865193556 | 2.413237378 | 2.404310072 |
| ITPR2 | 2.39775887 | 1.875717731 | 3.020497024 |
| JAK1 | 1.852372896 | 1.40192363 | 1.925231484 |
| JUN | 1.673785824 | 1.506294864 | 2.597773974 |
| LAMA2 | 4.285919703 | 3.098477609 | 4.57067965 |
| LAMP3 | 3.443346529 | 3.168990508 | 7.862689348 |
| LCP1 | 4.140863753 | 2.991608727 | 8.822476391 |
| LEFTY1 | 8.172798945 | 5.564626602 | 5.276539308 |
| LEFTY2 | 10.89780586 | 8.511686838 | 4.099297248 |
| LEPR | 2.358007779 | 2.55623047 | 2.805040687 |
| LGALS1 | 3.184079192 | 3.7082736 | 2.574285728 |
| LGALS12 | 38.31034971 | 17.74870692 | 29.81427232 |
| LIF | 2.50399159 | 2.024469145 | 1.931158823 |
| LIPA | 3.399564089 | 2.617484293 | 1.822981125 |
| LPL | 2.373983261 | 2.968679031 | 2.000059528 |
| LRFN5 | 2.769593897 | 2.825888494 | 7.109211554 |
| LTBR | 2.047766672 | 1.803114765 | 2.316601584 |
| MAFB | 2.648917038 | 2.126490588 | 5.027208596 |
| MAML2 | 2.160311284 | 1.59682286 | 2.601423502 |
| MASP1 | 1.531247237 | 2.181769632 | 2.662868731 |

| | | | |
|---|---|---|---|
| MECOM | 3.223347806 | 4.309059982 | 5.598486844 |
| MIR17HG | 2.109994373 | 1.7733603 | 1.530668161 |
| MMP9 | 5.345828833 | 2.749674997 | 4.427345084 |
| MUC12 | 1.937957762 | 2.058544477 | 2.404551344 |
| MX2 | 3.138015987 | 3.390441281 | 1.785248672 |
| MYL9 | 1.554484251 | 1.700763511 | 2.135720877 |
| MYO1F | 3.372393439 | 2.351622647 | 2.503781388 |
| NAV3 | 1.985763078 | 2.322540494 | 2.566907627 |
| NDRG1 | 2.63763366 | 2.130438901 | 2.718527173 |
| NEFH | 2.292918177 | 1.656581299 | 2.068909978 |
| NEK7 | 2.779120869 | 2.330200841 | 1.689185231 |
| NFAM1 | 1.981836497 | 1.942624661 | 1.804596708 |
| NLRP1 | 5.619491828 | 3.924604914 | 3.20458846 |
| NMI | 2.929651237 | 3.499881459 | 2.077354042 |
| NODAL | 5.644989154 | 2.860032625 | 6.78739855 |
| NOS3 | 3.445250267 | 2.650597689 | 2.488600045 |
| NRROS | 2.712366182 | 2.80571261 | 1.946962027 |
| NTN1 | 3.571246357 | 3.531758398 | 1.938687909 |
| P2RX4 | 1.725873541 | 1.630606163 | 1.841391555 |
| PADI2 | 2.382267578 | 1.978632758 | 3.071462072 |
| PDCD1 | 3.984231362 | 3.789353042 | 2.116362679 |
| PDGFRB | 1.771282869 | 1.668903712 | 1.954583406 |
| PF4 | 18.66081438 | 4.519650589 | 4.46741479 |
| PITX2 | 11.1279483 | 16.84283224 | 27.98354158 |
| PLA2G2A | 3.869527413 | 3.223351821 | 9.437446775 |
| PLA2G4C | 7.595558128 | 7.055302479 | 3.41310139 |
| PLA2G6 | 1.670577957 | 1.406703991 | 2.043470528 |
| PLAT | 3.37054795 | 3.418386601 | 1.850703347 |
| PLD1 | 2.308722011 | 1.798093442 | 1.983748686 |
| PLK2 | 3.756319121 | 3.231693218 | 5.441535367 |
| PRDM1 | 7.609229233 | 4.390984683 | 8.853153945 |
| PRNP | 1.578379902 | 1.651279849 | 1.93720661 |
| PRTN3 | 3.231981817 | 3.005210266 | 3.392001122 |
| PTAFR | 3.499578805 | 2.796512719 | 2.507012444 |
| PTPN6 | 2.119861275 | 1.850916658 | 1.672107259 |
| PTPRB | 9.875513855 | 3.415453018 | 12.298837 |
| PURA | 1.882408283 | 1.890407376 | 2.121994748 |
| QPCT | 2.613083421 | 1.772857464 | 2.067200159 |
| RAC2 | 3.370469421 | 2.947536515 | 1.912103466 |
| RBM15 | 2.150515566 | 1.967924718 | 2.321566405 |
| RGS2 | 2.14826144 | 2.445916307 | 1.899837455 |
| RGS4 | 9.715691949 | 6.341098356 | 10.77575921 |
| RORA | 1.983656712 | 1.584171653 | 1.641102921 |
| SCARF1 | 4.096482856 | 2.488011553 | 2.713982255 |
| SCG2 | 12.2012711 | 18.61382377 | 18.06781751 |
| SERPINB1 | 2.54178507 | 2.144110045 | 2.138126341 |
| SERPINB9 | 4.712902314 | 5.667520796 | 6.665060635 |
| SERPINE1 | 3.790088101 | 3.294852553 | 10.3141017 |

| | | | |
|---|---|---|---|
| SLFN5 | 1.929847271 | 1.731175294 | 1.659382659 |
| SOCS1 | 1.595793938 | 1.459483114 | 1.673668895 |
| SOX15 | 5.477805808 | 5.912687812 | 2.083639121 |
| SPINK5 | 2.007151322 | 1.579015161 | 1.5307248 |
| SPSB2 | 2.29663001 | 3.090865455 | 2.814745021 |
| STARD13 | 2.387729746 | 2.259621664 | 4.168678539 |
| STAT6 | 2.482605444 | 2.213881813 | 1.816150183 |
| TAPBPL | 2.442774349 | 2.373652847 | 2.260921448 |
| TCIRG1 | 1.919346216 | 1.853590174 | 1.486582172 |
| TGM2 | 2.93362188 | 2.969327324 | 2.408673661 |
| TNFAIP6 | 2.667305252 | 3.76773737 | 2.768292142 |
| TNFRSF10C | 5.84868336 | 3.667021151 | 4.938647089 |
| TNFSF9 | 3.424395446 | 3.501564705 | 4.144549016 |
| TRIM5 | 1.69704355 | 10.153516 | 1.772656119 |
| TRIM56 | 1.8048212 | 1.626626688 | 1.738671131 |
| TRPM2 | 2.033464757 | 3.934203114 | 3.578534487 |
| TRPV1 | 2.033464757 | 1.972460446 | 1.67173351 |
| TRPV4 | 2.07834965 | 1.875076569 | 2.359828407 |
| VAV1 | 11.70977174 | 9.116505065 | 5.307338113 |
| VIM | 2.4937278 | 2.870704662 | 2.743616913 |
| VSIR | 5.236173862 | 5.307391148 | 4.613562841 |
| WNT3 | 1.593494475 | 2.279380022 | 3.498929494 |
| ZDHHC1 | 1.844953152 | 1.584209345 | 5.276539308 |
| ZDHHC21 | 2.007187687 | 1.908886225 | 2.010827548 |
| ZNF91 | 1.976240903 | 1.602365576 | 2.36998201 |
| MEF2C * | 1.88288986 | 2.192980087 | 2.36286007 |

**Table S7. Upregulated immune genes in** SF3B1$^{K700E}$ ES line 1. Genes with a fold change (FC) ≥1.5 and p-value < 0.05 are shown.

| Gene | FC | p-value |
|---|---|---|
| PRAM1 | 90.95669163 | 1.44E-06 |
| CD48 | 52.63805037 | 2.07E-04 |
| IL6 | 49.85350505 | 1.38E-04 |
| MAS1 | 39.17631199 | 3.63E-06 |
| LGALS12 | 38.31034971 | 1.88E-08 |
| PLA2G2C | 24.30679503 | 2.79E-03 |
| ICAM2 | 20.10840914 | 4.51E-03 |
| EBF2 | 18.7156129 | 9.30E-04 |
| PF4 | 18.66081438 | 3.19E-06 |
| CH25H | 17.52212601 | 8.70E-03 |
| F7 | 16.99220691 | 8.92E-05 |
| GDF15 | 16.72575521 | 4.51E-133 |
| CHI3L1 | 16.3940263 | 1.16E-02 |
| AGBL4 | 16.18547136 | 1.76E-04 |
| ANXA1 | 15.52221943 | 4.33E-09 |
| TRIM58 | 14.7397322 | 9.51E-37 |
| COL8A1 | 12.51701916 | 1.55E-08 |
| ADTRP | 12.50984772 | 8.25E-03 |
| SCG2 | 12.2012711 | 1.35E-10 |
| VAV1 | 11.70977174 | 1.06E-57 |
| MPO | 11.50659437 | 1.21E-02 |
| PITX2 | 11.1279483 | 6.00E-10 |
| LCP2 | 10.99761829 | 1.53E-02 |
| LEFTY2 | 10.89780586 | 2.79E-63 |
| PTPRB | 9.875513855 | 6.16E-15 |
| RGS4 | 9.715691949 | 6.32E-05 |
| LAPTM5 | 8.569216181 | 7.97E-05 |
| LEFTY1 | 8.172798945 | 1.55E-154 |
| GZMM | 8.018260332 | 5.68E-04 |
| ALOX15B | 7.943671763 | 2.09E-04 |
| SLC16A3 | 7.911018463 | 1.27E-100 |
| PRDM1 | 7.609229233 | 6.36E-05 |
| PLA2G4C | 7.595558128 | 1.00E-24 |
| CDKN1A | 7.36834405 | 7.52E-180 |
| GALR1 | 7.289369911 | 2.65E-04 |
| HCLS1 | 6.679359978 | 1.57E-16 |
| CD163 | 6.59523614 | 4.52E-02 |
| IL1A | 6.116711302 | 3.99E-04 |
| ASB2 | 6.059505926 | 5.48E-04 |
| PTPRH | 5.981296225 | 2.51E-03 |

| | | |
|---|---|---|
| TNFRSF10C | 5.84868336 | 4.04E-17 |
| CMKLR1 | 5.737160482 | 1.36E-15 |
| NODAL | 5.644989154 | 1.78E-36 |
| NLRP1 | 5.619491828 | 1.79E-07 |
| FOXA2 | 5.480384292 | 1.60E-05 |
| SOX15 | 5.477805808 | 3.46E-24 |
| HLA-DQA1 | 5.467587357 | 2.74E-07 |
| SIGLEC15 | 5.427124562 | 3.32E-04 |
| MMP9 | 5.345828833 | 9.70E-44 |
| VSIR | 5.236173862 | 8.91E-04 |
| TRPM2 | 5.158912312 | 1.40E-04 |
| IFIT3 | 5.131966277 | 1.01E-03 |
| BIRC7 | 5.003362349 | 3.57E-02 |
| ARID5B | 4.980696422 | 6.68E-06 |
| BTG2 | 4.940458901 | 1.16E-34 |
| FLI1 | 4.80924461 | 8.53E-06 |
| TFF3 | 4.78469048 | 2.81E-08 |
| GATA4 | 4.780316859 | 4.80E-02 |
| CFHR3 | 4.750986168 | 1.09E-02 |
| TRIM4 | 4.735942469 | 1.21E-23 |
| SERPINB9 | 4.712902314 | 3.70E-104 |
| CD93 | 4.695247978 | 2.63E-02 |
| TGFB2 | 4.485100936 | 3.51E-04 |
| PTGS2 | 4.456995012 | 2.96E-02 |
| HAPLN1 | 4.405716359 | 1.05E-14 |
| CREB5 | 4.394442886 | 1.68E-06 |
| CD8A | 4.355055726 | 5.95E-03 |
| CSMD1 | 4.286994495 | 4.54E-04 |
| LAMA2 | 4.285919703 | 9.78E-26 |
| CR1L | 4.220654916 | 3.98E-07 |
| CFP | 4.205028632 | 8.42E-03 |
| RASAL3 | 4.201855145 | 1.30E-05 |
| ADGRG3 | 4.148619677 | 2.06E-02 |
| ABCC3 | 4.144606416 | 6.58E-03 |
| LCP1 | 4.140863753 | 1.15E-14 |
| EBI3 | 4.110176461 | 1.67E-04 |
| SCARF1 | 4.096482856 | 1.44E-36 |
| APOBEC3H | 4.039753058 | 2.55E-02 |
| PDE4B | 3.992518607 | 9.42E-59 |
| PDCD1 | 3.984231362 | 7.15E-10 |
| CXCL1 | 3.965159788 | 8.07E-03 |
| CEACAM1 | 3.928235587 | 2.79E-06 |
| WNT3 | 3.902490273 | 3.43E-35 |
| HSPA2 | 3.895639502 | 1.56E-140 |
| GDF3 | 3.888660435 | 3.46E-39 |

| | | |
|---|---|---|
| ADGRE5 | 3.876355128 | 1.42E-14 |
| PLA2G2A | 3.869527413 | 1.38E-13 |
| CFD | 3.804318405 | 4.87E-11 |
| CAV1 | 3.793948001 | 1.55E-30 |
| SERPINE1 | 3.790088101 | 4.62E-18 |
| PLK2 | 3.756319121 | 9.22E-66 |
| BANK1 | 3.751701136 | 8.37E-05 |
| GP1BA | 3.715607404 | 3.24E-03 |
| CD79B | 3.711552816 | 1.04E-04 |
| NTN1 | 3.571246357 | 1.51E-27 |
| RORC | 3.540871196 | 4.93E-04 |
| FOXJ1 | 3.533329874 | 5.92E-36 |
| PTAFR | 3.499578805 | 2.30E-12 |
| COLEC11 | 3.454717726 | 2.58E-03 |
| NOS3 | 3.445250267 | 3.93E-16 |
| LAMP3 | 3.443346529 | 1.96E-02 |
| TNFSF9 | 3.424395446 | 9.93E-14 |
| DMTN | 3.400366661 | 7.47E-08 |
| LIPA | 3.399564089 | 8.05E-90 |
| ADCY8 | 3.381410376 | 2.14E-11 |
| GREM2 | 3.372544365 | 2.74E-02 |
| MYO1F | 3.372393439 | 7.81E-22 |
| PLAT | 3.37054795 | 1.96E-34 |
| RAC2 | 3.370469421 | 9.29E-11 |
| CD19 | 3.334533266 | 4.62E-03 |
| MT1G | 3.324500393 | 5.24E-90 |
| DBH | 3.319059847 | 3.20E-04 |
| APOBEC3D | 3.303680602 | 7.99E-03 |
| CLC | 3.290254314 | 9.33E-04 |
| S100A4 | 3.282315451 | 6.26E-26 |
| LYST | 3.23921498 | 3.93E-02 |
| PRTN3 | 3.231981817 | 3.30E-02 |
| MECOM | 3.223347806 | 3.76E-03 |
| HLA-DQB1 | 3.208188508 | 1.28E-50 |
| IL6R | 3.205377795 | 1.37E-26 |
| LGALS1 | 3.184079192 | 1.01E-33 |
| TBC1D10C | 3.165141773 | 1.03E-03 |
| IL15 | 3.14386866 | 1.20E-03 |
| ATXN1 | 3.143302734 | 3.41E-07 |
| MX2 | 3.138015987 | 1.29E-13 |
| CD7 | 3.118157843 | 2.87E-04 |
| PSTPIP1 | 3.099168605 | 5.09E-03 |
| PTK6 | 3.022061441 | 1.75E-02 |
| ABI3 | 2.976852897 | 1.16E-08 |
| DUSP10 | 2.973044633 | 2.49E-04 |

| | | |
|---|---|---|
| GGT1 | 2.95819329 | 1.08E-05 |
| TGM2 | 2.93362188 | 4.34E-12 |
| NMI | 2.929651237 | 3.72E-08 |
| ALOX5 | 2.916530289 | 1.28E-07 |
| TSPAN32 | 2.899462824 | 3.31E-02 |
| ARRB1 | 2.898854853 | 3.57E-58 |
| ENPP2 | 2.898395386 | 1.00E-06 |
| C5AR2 | 2.897355573 | 2.54E-02 |
| ITGB2 | 2.865193556 | 4.70E-03 |
| MOG | 2.860686699 | 1.71E-02 |
| LCN2 | 2.836229489 | 7.32E-03 |
| ANXA3 | 2.784334773 | 5.94E-37 |
| NEK7 | 2.779120869 | 1.31E-23 |
| IFI35 | 2.777020229 | 3.61E-05 |
| LRFN5 | 2.769593897 | 7.38E-03 |
| CR2 | 2.760676229 | 5.47E-26 |
| APOBEC3G | 2.754275761 | 5.95E-05 |
| AIF1 | 2.752858317 | 2.51E-07 |
| HPGD | 2.721096542 | 1.11E-04 |
| AZU1 | 2.717348544 | 1.33E-02 |
| NRROS | 2.712366182 | 1.01E-14 |
| CAV2 | 2.696729285 | 1.95E-05 |
| F10 | 2.687151049 | 5.29E-08 |
| MASP1 | 2.667610333 | 2.79E-07 |
| TNFAIP6 | 2.667305252 | 4.40E-02 |
| FGF4 | 2.665712872 | 2.30E-05 |
| MAFB | 2.648917038 | 6.49E-07 |
| WFDC1 | 2.641527477 | 6.24E-03 |
| ALDH1A1 | 2.639997278 | 1.55E-04 |
| NDRG1 | 2.63763366 | 1.31E-08 |
| NTSR1 | 2.615797196 | 3.81E-04 |
| QPCT | 2.613083421 | 5.06E-12 |
| PROX1 | 2.58186139 | 4.08E-03 |
| INPP5D | 2.565559831 | 2.97E-35 |
| IRF2BP2 | 2.560970412 | 1.54E-51 |
| SERPINB1 | 2.54178507 | 1.85E-24 |
| TRAF1 | 2.53575668 | 1.45E-05 |
| S100A14 | 2.509708636 | 2.58E-04 |
| LIF | 2.50399159 | 1.66E-13 |
| WNT9B | 2.497735473 | 2.81E-02 |
| PDGFB | 2.49396885 | 1.69E-18 |
| VIM | 2.4937278 | 1.47E-23 |
| SAMSN1 | 2.483880647 | 3.94E-02 |
| STAT6 | 2.482605444 | 1.70E-88 |
| KCNN4 | 2.481548689 | 1.31E-02 |

| | | |
|---|---|---|
| KLK7 | 2.459449543 | 2.50E-03 |
| CSRP1 | 2.458512179 | 3.31E-02 |
| TAPBPL | 2.442774349 | 2.63E-02 |
| TLR3 | 2.437091898 | 6.29E-03 |
| FAXDC2 | 2.437041629 | 4.01E-06 |
| ICAM3 | 2.43571121 | 6.42E-10 |
| CD300A | 2.412200876 | 1.16E-02 |
| ITPR2 | 2.39775887 | 8.89E-54 |
| MXRA8 | 2.390043575 | 1.17E-21 |
| STARD13 | 2.387729746 | 2.32E-08 |
| IFIT1 | 2.38514683 | 1.40E-04 |
| PADI2 | 2.382267578 | 1.44E-15 |
| SULF1 | 2.377033292 | 4.70E-19 |
| LPL | 2.373983261 | 1.52E-14 |
| SRMS | 2.368876347 | 4.23E-02 |
| LEPR | 2.358007779 | 4.04E-05 |
| PAX5 | 2.354866873 | 8.86E-03 |
| TLR5 | 2.352306244 | 2.32E-07 |
| CRLF1 | 2.335834399 | 4.96E-15 |
| CPEB4 | 2.314223529 | 1.08E-17 |
| EDA2R | 2.313416751 | 1.32E-14 |
| TRIM22 | 2.311211436 | 3.09E-32 |
| PLD1 | 2.308722011 | 8.24E-04 |
| SPSB2 | 2.29663001 | 2.63E-12 |
| NEFH | 2.292918177 | 3.80E-13 |
| C1QTNF12 | 2.29096283 | 1.28E-03 |
| ANXA2 | 2.26753128 | 3.81E-72 |
| FZD5 | 2.256173634 | 2.20E-112 |
| ACP5 | 2.229572147 | 1.65E-09 |
| IL2RB | 2.216447112 | 7.93E-05 |
| ASH1L | 2.212456957 | 1.74E-30 |
| TRPV2 | 2.211950877 | 4.93E-04 |
| PLA2G4A | 2.191295584 | 4.77E-04 |
| RAG1 | 2.174148651 | 1.27E-02 |
| TIRAP | 2.165933644 | 7.26E-03 |
| MAML2 | 2.160311284 | 9.24E-10 |
| HSH2D | 2.15169061 | 4.95E-03 |
| RBM15 | 2.150515566 | 2.13E-28 |
| RGS2 | 2.14826144 | 1.59E-15 |
| PTPN6 | 2.119861275 | 3.95E-28 |
| GCSAM | 2.119162404 | 1.34E-02 |
| TRIML2 | 2.116717601 | 3.75E-32 |
| EMILIN1 | 2.113749696 | 1.92E-13 |
| APOBEC3B | 2.100902681 | 3.17E-08 |
| MC1R | 2.093006413 | 3.99E-02 |

| | | |
|---|---|---|
| DUOX1 | 2.080256485 | 5.97E-03 |
| TRPV4 | 2.07834965 | 1.20E-04 |
| IRAK1 | 2.062292636 | 1.00E-54 |
| LTBR | 2.047766672 | 2.08E-04 |
| LY75 | 2.046257909 | 2.49E-03 |
| SPIB | 2.043491492 | 1.80E-08 |
| TRPV1 | 2.033464757 | 1.99E-04 |
| SIT1 | 2.031493111 | 6.72E-06 |
| ADCY7 | 2.030453203 | 5.61E-08 |
| APOBEC3C | 2.027150545 | 1.62E-25 |
| SPN | 2.02630356 | 6.62E-04 |
| BATF2 | 2.016622249 | 3.26E-03 |
| CDH11 | 2.015793024 | 2.85E-15 |
| STAB1 | 2.01465548 | 1.53E-02 |
| GADD45G | 2.007878284 | 6.88E-13 |
| AMIGO2 | 2.007629233 | 2.71E-10 |
| ZDHHC21 | 2.007187687 | 2.86E-06 |
| SPINK5 | 2.007151322 | 9.05E-06 |
| PLD5 | 1.999929507 | 2.19E-05 |
| RAB20 | 1.997203873 | 1.39E-22 |
| NAV3 | 1.985763078 | 2.32E-03 |
| RORA | 1.983656712 | 2.89E-04 |
| NFAM1 | 1.981836497 | 1.03E-02 |
| ZNF91 | 1.976240903 | 1.95E-18 |
| C3 | 1.971439373 | 9.14E-17 |
| HLA-DPA1 | 1.958445145 | 1.58E-03 |
| HLA-DRA | 1.9478542 | 5.24E-04 |
| LACC1 | 1.943420562 | 2.58E-04 |
| MUC12 | 1.937957762 | 3.22E-02 |
| IL23A | 1.932415311 | 2.29E-02 |
| ZBTB7B | 1.932399195 | 5.96E-41 |
| SLFN5 | 1.929847271 | 4.10E-07 |
| DGKH | 1.928757049 | 7.66E-07 |
| TCIRG1 | 1.919346216 | 2.19E-17 |
| SCYL3 | 1.91372929 | 5.89E-08 |
| CCL26 | 1.911615293 | 1.04E-04 |
| IFITM1 | 1.910384208 | 2.87E-08 |
| CD46 | 1.908751974 | 5.90E-38 |
| OSCAR | 1.899633555 | 4.37E-04 |
| TRIM6 | 1.896614605 | 1.22E-13 |
| APOBEC3F | 1.892340439 | 1.72E-05 |
| HMGB3 | 1.887469418 | 6.43E-124 |
| PURA | 1.882408283 | 4.95E-06 |
| LRRK1 | 1.875983318 | 1.98E-05 |
| KITLG | 1.872427251 | 5.83E-06 |

| | | |
|---|---|---|
| GPSM3 | 1.871073727 | 3.92E-05 |
| ADGRE2 | 1.870535886 | 9.08E-03 |
| CSF2RA | 1.861123563 | 1.49E-02 |
| PDCD4 | 1.860104151 | 4.94E-14 |
| JAK1 | 1.852372896 | 1.15E-28 |
| MGST2 | 1.850938491 | 6.57E-05 |
| HLA-G | 1.850298616 | 7.20E-04 |
| ZDHHC1 | 1.844953152 | 4.85E-04 |
| CD74 | 1.84291999 | 1.29E-29 |
| BTN3A2 | 1.8348439 | 1.53E-04 |
| DOCK2 | 1.833640529 | 2.59E-07 |
| EFNA1 | 1.811983442 | 4.29E-04 |
| ARNT | 1.807028156 | 2.63E-32 |
| CD70 | 1.806692945 | 2.03E-02 |
| IL13RA1 | 1.805655484 | 1.56E-24 |
| TRIM5 | 1.8048212 | 1.14E-30 |
| TRIM56 | 1.8048212 | 1.14E-30 |
| DAGLA | 1.798009111 | 3.30E-11 |
| NLRP7 | 1.797941553 | 2.48E-02 |
| HLA-F | 1.791741424 | 2.35E-03 |
| SEMG1 | 1.791360106 | 1.98E-02 |
| IL17C | 1.79102467 | 3.04E-02 |
| VENTX | 1.788080886 | 1.73E-09 |
| CASP8 | 1.788028269 | 1.30E-03 |
| PDGFRB | 1.771282869 | 4.62E-13 |
| FZD9 | 1.770761143 | 1.07E-06 |
| HLA-DRB1 | 1.761923878 | 3.08E-11 |
| NOS2 | 1.758318887 | 2.71E-04 |
| TNFRSF1B | 1.747158697 | 1.37E-05 |
| IL1RAP | 1.743426383 | 1.77E-05 |
| SLC39A8 | 1.733387533 | 7.10E-20 |
| P2RY1 | 1.731617131 | 7.35E-18 |
| EPRS1 | 1.726723302 | 2.29E-78 |
| P2RX4 | 1.725873541 | 1.72E-05 |
| JUNB | 1.72102595 | 4.35E-05 |
| CGAS | 1.715843905 | 3.86E-03 |
| GAL | 1.714896777 | 7.21E-20 |
| FES | 1.711034545 | 2.35E-05 |
| POU2F2 | 1.709750534 | 4.40E-05 |
| BATF3 | 1.70745899 | 3.37E-03 |
| DYRK3 | 1.707000442 | 3.82E-09 |
| MAPK13 | 1.706814191 | 2.49E-11 |
| TIMP4 | 1.7060446 | 6.91E-04 |
| DDX60 | 1.699585236 | 5.90E-03 |
| REL | 1.698012677 | 7.06E-03 |

| | | |
|---|---|---|
| FN1 | 1.693919987 | 1.97E-26 |
| ELF3 | 1.689278802 | 2.14E-03 |
| GADD45B | 1.674547855 | 4.55E-05 |
| JUN | 1.673785824 | 9.26E-10 |
| HSPG2 | 1.672813229 | 3.18E-39 |
| PLA2G6 | 1.670577957 | 8.42E-06 |
| ITGB1 | 1.651690846 | 4.44E-55 |
| ZFP36 | 1.642307438 | 4.41E-04 |
| ZFP36 | 1.642307438 | 4.41E-04 |
| TFPI | 1.63922489 | 7.54E-04 |
| TFR2 | 1.635770071 | 4.09E-05 |
| IRAK4 | 1.629200964 | 2.88E-02 |
| PBXIP1 | 1.628887704 | 1.28E-18 |
| ATP7A | 1.62675548 | 9.01E-03 |
| CXCL5 | 1.625737016 | 4.05E-03 |
| RICTOR | 1.620831651 | 1.26E-06 |
| TRIM32 | 1.620454456 | 1.72E-11 |
| PTP4A3 | 1.606951129 | 1.77E-16 |
| TRAF3IP2 | 1.601275312 | 6.67E-14 |
| COLEC12 | 1.598000933 | 1.62E-02 |
| GAP43 | 1.597596935 | 3.32E-11 |
| ITPKB | 1.597223457 | 8.62E-09 |
| IGFBP2 | 1.596006571 | 1.81E-08 |
| SOCS1 | 1.595793938 | 3.91E-07 |
| PGF | 1.59312393 | 2.95E-04 |
| MCAM | 1.591994594 | 1.39E-14 |
| CD1D | 1.586857109 | 3.88E-02 |
| SH3PXD2A | 1.586266105 | 2.00E-15 |
| PRNP | 1.578379902 | 7.61E-12 |
| HLA-DRB5 | 1.574640128 | 1.60E-02 |
| TNFRSF10B | 1.573817451 | 4.46E-22 |
| FGF19 | 1.569281264 | 5.11E-10 |
| JAG2 | 1.565287962 | 1.12E-11 |
| MFNG | 1.564465981 | 4.33E-03 |
| RIPOR2 | 1.562656405 | 1.25E-14 |
| FZD6 | 1.556071236 | 1.53E-04 |
| MYL9 | 1.554484251 | 2.58E-27 |
| NCSTN | 1.548138219 | 8.64E-24 |
| INAVA | 1.534778331 | 1.12E-15 |
| SERPINF2 | 1.53199088 | 2.88E-02 |
| ICAM1 | 1.528420601 | 9.04E-03 |
| TRIM68 | 1.524344509 | 2.50E-04 |
| BCL11B | 1.52419812 | 1.02E-02 |
| PRKG1 | 1.518904372 | 1.16E-04 |
| PARP1 | 1.517101953 | 6.38E-62 |

| | | |
|---|---|---|
| IMPDH2 | 1.516637166 | 2.44E-39 |
| KDM6B | 1.513657182 | 2.77E-05 |
| CYBA | 1.510928061 | 1.79E-27 |
| IKBKE | 1.510810672 | 3.67E-06 |
| MYSM1 | 1.510132758 | 3.21E-14 |
| PYCARD | 1.508642888 | 2.38E-03 |
| SMAD3 | 1.507901682 | 2.52E-09 |
| FLNB | 1.507890754 | 3.86E-36 |
| XIAP | 1.507844309 | 1.15E-11 |
| COCH | 1.506122199 | 4.99E-05 |
| FNIP1 | 1.503469005 | 6.46E-04 |
| ABCB10 | 1.503287427 | 4.76E-06 |
| CX3CL1 | 1.500644354 | 3.23E-02 |

| **Table S7. Upregulated immune genes in SF3B1$^{K700E}$ ES line 2. Genes with a fold change (FC) ≥1.5 and p-value < 0.05 are shown.** | | |
|---|---|---|
| **Gene** | **FC** | **p-value** |
| LTF | 61.48610311 | 3.92E-05 |
| REG1A | 50.86404385 | 1.63E-04 |
| PRAM1 | 50.70598138 | 1.44E-04 |
| COL3A1 | 36.827441 | 3.81E-02 |
| F7 | 34.13900622 | 9.48E-08 |
| ICAM2 | 21.41808485 | 4.32E-03 |
| SCG2 | 18.61382377 | 2.27E-15 |
| LGALS12 | 17.74870692 | 7.99E-05 |
| ANXA1 | 17.15139687 | 2.80E-10 |
| MPO | 14.80759925 | 3.52E-03 |
| ASB2 | 14.07358777 | 1.11E-09 |
| EBF2 | 12.12813959 | 9.73E-03 |
| GDF15 | 11.73576917 | 6.94E-85 |
| PRSS3 | 10.95780214 | 1.45E-02 |
| MAS1 | 10.53496474 | 1.72E-02 |
| AGBL4 | 10.32909765 | 3.79E-03 |
| TRIM58 | 10.153516 | 1.35E-23 |
| F5 | 10.10937974 | 2.32E-02 |
| COL8A1 | 9.766148723 | 1.46E-06 |
| VAV1 | 9.116505065 | 7.47E-39 |
| NCF1 | 8.993524381 | 2.83E-02 |
| LEFTY2 | 8.511686838 | 1.91E-120 |
| PLA2G4C | 7.055302479 | 1.47E-21 |
| HLA-DQA1 | 6.713187656 | 2.41E-09 |
| LAIR1 | 6.449128245 | 1.79E-02 |
| RGS4 | 6.341098356 | 5.03E-03 |
| SOX15 | 5.912687812 | 2.77E-27 |
| SERPINB9 | 5.667520796 | 2.47E-126 |
| LEFTY1 | 5.564626602 | 4.82E-89 |
| VSIR | 5.307391148 | 8.16E-04 |
| IFIT3 | 5.215861946 | 1.05E-03 |
| COLEC11 | 5.134002353 | 1.69E-05 |
| NLRP10 | 4.909609772 | 2.73E-02 |
| CSRP1 | 4.888322825 | 2.38E-05 |
| GPER1 | 4.850280423 | 4.80E-02 |
| CFHR3 | 4.84665037 | 1.39E-02 |
| ARID5B | 4.702435185 | 2.35E-05 |
| PF4 | 4.519650589 | 5.17E-03 |
| CAV1 | 4.424790399 | 1.39E-39 |
| TGFB2 | 4.398145677 | 9.75E-04 |

| | | |
|---|---|---|
| PRDM1 | 4.390984683 | 1.18E-02 |
| RORC | 4.371676095 | 2.03E-05 |
| IL18 | 4.35806654 | 9.10E-03 |
| IL1A | 4.329064037 | 1.13E-02 |
| MECOM | 4.309059982 | 2.71E-04 |
| ADGRE5 | 4.27198281 | 8.90E-16 |
| CDKN1A | 4.253749207 | 1.28E-81 |
| TRIM4 | 4.191211743 | 1.62E-18 |
| ABCC3 | 4.035394512 | 8.58E-03 |
| HCLS1 | 4.028098566 | 1.91E-07 |
| ADGRG3 | 4.027694198 | 3.09E-02 |
| GALR1 | 3.977084962 | 2.46E-02 |
| FLI1 | 3.956869655 | 2.43E-04 |
| TRPM2 | 3.934203114 | 5.22E-03 |
| NLRP1 | 3.924604914 | 2.74E-04 |
| HSPA2 | 3.923003606 | 3.04E-129 |
| BTG2 | 3.892818028 | 1.45E-23 |
| TRADD | 3.824079864 | 9.53E-03 |
| BMPER | 3.806201658 | 3.27E-02 |
| PSTPIP1 | 3.800923683 | 4.33E-04 |
| PDCD1 | 3.789353042 | 2.57E-09 |
| SIGLEC15 | 3.769116153 | 9.74E-03 |
| TNFAIP6 | 3.76773737 | 1.89E-03 |
| LGALS1 | 3.7082736 | 2.14E-41 |
| TNFRSF10C | 3.667021151 | 1.23E-07 |
| ZP3 | 3.623490765 | 3.63E-03 |
| ADGRE2 | 3.600967034 | 2.84E-10 |
| NTN1 | 3.531758398 | 2.46E-26 |
| HLA-DQB1 | 3.526051527 | 2.54E-49 |
| PDE4B | 3.522776816 | 9.44E-42 |
| CD79B | 3.50400378 | 1.75E-04 |
| TNFSF9 | 3.501564705 | 1.62E-13 |
| NMI | 3.499881459 | 5.83E-12 |
| ABCC2 | 3.456728569 | 1.77E-03 |
| HAPLN1 | 3.433943298 | 3.86E-09 |
| PLAT | 3.418386601 | 5.24E-34 |
| TFF3 | 3.415520508 | 2.07E-04 |
| PTPRB | 3.415453018 | 1.64E-03 |
| MX2 | 3.390441281 | 6.55E-15 |
| BANK1 | 3.366617334 | 2.51E-03 |
| HLA-DMB | 3.33127114 | 3.11E-03 |
| ALPK1 | 3.330220601 | 2.11E-02 |
| THPO | 3.308838065 | 9.39E-03 |
| SERPINE1 | 3.294852553 | 7.01E-12 |
| IFI35 | 3.259364379 | 6.07E-07 |

| | | |
|---|---|---|
| MXRA8 | 3.233924664 | 3.59E-42 |
| PLK2 | 3.231693218 | 1.57E-47 |
| PLA2G2A | 3.223351821 | 3.84E-10 |
| LAMP3 | 3.168990508 | 3.88E-02 |
| LAMA2 | 3.098477609 | 3.35E-14 |
| SPSB2 | 3.090865455 | 6.95E-23 |
| EBI3 | 3.026516261 | 5.08E-03 |
| IL6R | 3.008698914 | 2.90E-22 |
| PRTN3 | 3.005210266 | 5.00E-02 |
| HLA-DRA | 2.994862572 | 3.56E-10 |
| LCP1 | 2.991608727 | 5.11E-08 |
| TGM2 | 2.969327324 | 6.71E-12 |
| LPL | 2.968679031 | 7.68E-24 |
| ALOX5 | 2.962416002 | 1.38E-07 |
| RAC2 | 2.947536515 | 7.46E-08 |
| CMKLR1 | 2.89925283 | 1.13E-04 |
| FOXA2 | 2.896064354 | 2.23E-02 |
| PLD5 | 2.880146954 | 1.45E-10 |
| F10 | 2.878183112 | 7.50E-09 |
| CFD | 2.87710021 | 1.42E-06 |
| HPGD | 2.875902216 | 3.38E-05 |
| VIM | 2.870704662 | 1.08E-32 |
| NODAL | 2.860032625 | 1.73E-10 |
| ABI3 | 2.851838404 | 2.20E-08 |
| CD19 | 2.843959763 | 1.99E-02 |
| LRFN5 | 2.825888494 | 7.79E-03 |
| NRROS | 2.80571261 | 6.02E-15 |
| PTAFR | 2.796512719 | 2.06E-07 |
| TBC1D10C | 2.765818555 | 1.10E-02 |
| MMP9 | 2.749674997 | 1.69E-11 |
| ATXN1 | 2.706710608 | 3.35E-05 |
| S100A4 | 2.667697395 | 1.29E-15 |
| PTGER3 | 2.660023126 | 7.60E-03 |
| NOS3 | 2.650597689 | 3.57E-09 |
| TLR3 | 2.645053614 | 3.22E-03 |
| IL15 | 2.640061838 | 1.34E-02 |
| FOXJ1 | 2.620461543 | 8.65E-18 |
| LIPA | 2.617484293 | 9.51E-49 |
| CEACAM1 | 2.607874256 | 3.78E-03 |
| KCNN4 | 2.605843176 | 1.46E-02 |
| GGT1 | 2.597360647 | 3.23E-04 |
| LEPR | 2.55623047 | 7.24E-06 |
| KLK7 | 2.551601706 | 2.31E-03 |
| CREB5 | 2.536332809 | 8.11E-03 |
| ENPP2 | 2.52323278 | 9.66E-05 |

| | | |
|---|---|---|
| DUSP10 | 2.523125903 | 4.24E-03 |
| NOD2 | 2.521735116 | 7.97E-03 |
| APOBEC3B | 2.518342981 | 6.28E-12 |
| ELF3 | 2.491543483 | 2.58E-09 |
| SCARF1 | 2.488011553 | 1.59E-10 |
| WFDC1 | 2.469617101 | 9.63E-03 |
| CR2 | 2.463679409 | 2.30E-18 |
| RGS2 | 2.445916307 | 2.68E-21 |
| CD7 | 2.433462051 | 7.08E-03 |
| ANXA2 | 2.419550358 | 1.45E-69 |
| ITGB2 | 2.413237378 | 2.51E-02 |
| SPIB | 2.401522482 | 6.71E-11 |
| GDF3 | 2.399842166 | 1.08E-13 |
| IL2RB | 2.380502523 | 7.81E-06 |
| TAPBPL | 2.373652847 | 2.18E-02 |
| OAS3 | 2.370426417 | 1.79E-03 |
| CD1D | 2.369572067 | 1.40E-05 |
| F8 | 2.359086453 | 1.14E-04 |
| NOS2 | 2.358173061 | 4.86E-09 |
| MYO1F | 2.351622647 | 1.06E-09 |
| SIT1 | 2.332649457 | 1.85E-10 |
| NEK7 | 2.330200841 | 1.74E-14 |
| FGF4 | 2.327239855 | 4.74E-04 |
| NAV3 | 2.322540494 | 1.26E-04 |
| IRAK1 | 2.321456075 | 6.83E-73 |
| AMIGO2 | 2.3012872 | 1.94E-12 |
| WNT3 | 2.279380022 | 4.47E-10 |
| CAV2 | 2.264700178 | 9.94E-04 |
| APOBEC3G | 2.262406758 | 2.22E-03 |
| STARD13 | 2.259621664 | 3.19E-07 |
| STAT6 | 2.213881813 | 1.25E-52 |
| FAXDC2 | 2.207814088 | 4.19E-05 |
| DAGLA | 2.206568959 | 3.35E-22 |
| ANXA3 | 2.202469528 | 2.22E-19 |
| MEF2C | 2.192980087 | 1.94E-02 |
| MASP1 | 2.181769632 | 5.78E-05 |
| INHBA | 2.181179107 | 4.22E-02 |
| TIRAP | 2.180661438 | 8.41E-03 |
| NTSR1 | 2.180207741 | 4.46E-03 |
| SERPINB1 | 2.144110045 | 1.10E-16 |
| NDRG1 | 2.130438901 | 4.61E-05 |
| ZBTB7B | 2.129013316 | 7.35E-49 |
| MAFB | 2.126490588 | 5.84E-04 |
| IRF2BP2 | 2.125078116 | 9.98E-29 |
| CPEB4 | 2.12332013 | 3.47E-13 |

| | | |
|---|---|---|
| DYRK3 | 2.121280405 | 2.22E-17 |
| CD247 | 2.111290357 | 5.29E-04 |
| GCSAM | 2.085055152 | 1.85E-02 |
| FZD5 | 2.077737333 | 3.24E-64 |
| MUC12 | 2.058544477 | 2.07E-02 |
| ALDH1A1 | 2.049827927 | 7.04E-03 |
| CDH11 | 2.039438436 | 1.44E-14 |
| LIF | 2.024469145 | 9.89E-07 |
| CGAS | 2.010015241 | 2.33E-04 |
| HLA-G | 2.006273787 | 1.27E-04 |
| AIF1 | 2.006099514 | 9.50E-04 |
| PADI2 | 1.978632758 | 6.97E-09 |
| GADD45G | 1.973841435 | 1.06E-11 |
| TRPV1 | 1.972460446 | 8.40E-04 |
| HFE | 1.968124571 | 2.29E-02 |
| RBM15 | 1.967924718 | 4.87E-20 |
| HLA-DRB1 | 1.965629837 | 3.75E-15 |
| IFIT1 | 1.962095087 | 6.05E-03 |
| ASH1L | 1.95431661 | 2.79E-21 |
| RNF128 | 1.946265239 | 3.08E-03 |
| HLA-F | 1.946057278 | 7.74E-04 |
| NFAM1 | 1.942624661 | 1.33E-02 |
| PROS1 | 1.932169853 | 2.73E-02 |
| CD70 | 1.930270264 | 7.97E-03 |
| PLA2G4A | 1.915024167 | 5.89E-03 |
| ZDHHC21 | 1.908886225 | 5.94E-05 |
| PURA | 1.890407376 | 1.46E-05 |
| ITPR2 | 1.875717731 | 8.70E-21 |
| TRPV4 | 1.875076569 | 2.17E-03 |
| HMGB3 | 1.867323998 | 5.60E-90 |
| NLRC5 | 1.866983519 | 1.36E-02 |
| CSF2RA | 1.857566787 | 1.61E-02 |
| TCIRG1 | 1.853590174 | 1.97E-12 |
| GAP43 | 1.852385784 | 1.06E-17 |
| PTPN6 | 1.850916658 | 9.35E-17 |
| ADCY7 | 1.833404984 | 2.67E-05 |
| FES | 1.806995308 | 1.06E-05 |
| LTBR | 1.803114765 | 4.27E-03 |
| CASP8 | 1.79831321 | 1.57E-03 |
| PLD1 | 1.798093442 | 2.56E-02 |
| KLKB1 | 1.797644909 | 5.12E-07 |
| MST1R | 1.796458814 | 1.53E-03 |
| INPP5D | 1.795894823 | 8.87E-11 |
| DGKH | 1.789493842 | 3.71E-05 |
| FN1 | 1.787864371 | 2.59E-25 |

| | | |
|---|---|---|
| QPCT | 1.772857464 | 2.01E-04 |
| TNFRSF18 | 1.771738059 | 3.66E-02 |
| ACKR3 | 1.771061757 | 3.86E-02 |
| FZD9 | 1.770772722 | 2.06E-06 |
| DMTN | 1.760638061 | 3.92E-02 |
| HLA-DPA1 | 1.747988991 | 1.31E-02 |
| BTN3A2 | 1.744733498 | 2.30E-04 |
| SLFN5 | 1.731175294 | 7.08E-05 |
| ADAMTS1 | 1.720602866 | 1.02E-05 |
| IFITM1 | 1.710691366 | 6.71E-05 |
| CD74 | 1.707853483 | 7.02E-18 |
| DHX58 | 1.703413367 | 1.22E-03 |
| MYL9 | 1.700763511 | 5.49E-32 |
| ILDR2 | 1.700721468 | 4.99E-02 |
| CD55 | 1.681399363 | 8.55E-11 |
| CHRNB2 | 1.66926946 | 1.32E-05 |
| PDGFRB | 1.668903712 | 5.17E-10 |
| NEFH | 1.656581299 | 4.58E-05 |
| HYAL1 | 1.654781914 | 9.63E-04 |
| PRNP | 1.651279849 | 1.17E-12 |
| ICAM3 | 1.649758375 | 2.49E-03 |
| SH3PXD2A | 1.648759454 | 6.64E-16 |
| BTN3A1 | 1.648584811 | 2.70E-03 |
| SCYL3 | 1.640765053 | 9.82E-05 |
| TNFRSF1B | 1.639537633 | 2.95E-04 |
| TICAM1 | 1.636493372 | 4.69E-02 |
| COLEC12 | 1.633481424 | 1.28E-02 |
| P2RX4 | 1.630606163 | 2.50E-04 |
| SLC9B2 | 1.62740906 | 4.61E-03 |
| SYT11 | 1.626785958 | 7.41E-08 |
| TRIM56 | 1.626626688 | 6.37E-18 |
| TFR2 | 1.62208869 | 6.23E-05 |
| HLA-DRB5 | 1.61041045 | 9.32E-03 |
| CHRNA4 | 1.606722825 | 1.77E-03 |
| ZNF91 | 1.602365576 | 9.89E-09 |
| PTGES | 1.59716547 | 3.88E-03 |
| MAML2 | 1.59682286 | 6.27E-04 |
| ISG15 | 1.59627014 | 1.85E-09 |
| ZFP36 | 1.595198103 | 1.48E-03 |
| CLU | 1.594500574 | 1.39E-40 |
| CD47 | 1.594005262 | 8.73E-08 |
| GPR89A | 1.593410921 | 3.38E-04 |
| MAPK11 | 1.590470745 | 1.49E-02 |
| CX3CL1 | 1.587885207 | 1.75E-02 |
| NMB | 1.587553764 | 8.91E-06 |

| | | |
|---|---|---|
| ZDHHC1 | 1.584209345 | 1.01E-02 |
| RORA | 1.584171653 | 3.30E-02 |
| INAVA | 1.582440399 | 7.33E-17 |
| SPINK5 | 1.579015161 | 1.26E-02 |
| APOBEC3F | 1.578068219 | 4.60E-03 |
| ZNFX1 | 1.575397364 | 8.34E-09 |
| EMILIN1 | 1.575314465 | 7.94E-05 |
| C3 | 1.571468736 | 2.77E-07 |
| GAL | 1.569662942 | 2.97E-12 |
| CXCL5 | 1.569394002 | 1.60E-02 |
| PIAS3 | 1.566574476 | 6.40E-15 |
| PBXIP1 | 1.564738628 | 3.90E-15 |
| PSMB4 | 1.562539933 | 8.56E-39 |
| TRIM11 | 1.54925013 | 1.39E-11 |
| H19 | 1.549220019 | 2.92E-02 |
| LIFR | 1.546967845 | 1.95E-05 |
| SERPINF2 | 1.546180492 | 2.80E-02 |
| NUMBL | 1.544649135 | 1.98E-11 |
| RNF187 | 1.538172272 | 2.30E-26 |
| EPRS1 | 1.537964086 | 1.89E-33 |
| COCH | 1.536711722 | 1.26E-05 |
| LGALS3 | 1.532700078 | 3.07E-02 |
| APOBEC3C | 1.532215433 | 2.14E-08 |
| ADORA1 | 1.53167053 | 9.60E-04 |
| PROCR | 1.530622204 | 8.55E-04 |
| EDA2R | 1.530202989 | 1.80E-03 |
| ACP5 | 1.527463955 | 3.30E-03 |
| JAG2 | 1.527125181 | 5.20E-10 |
| ABCB10 | 1.526886146 | 2.26E-06 |
| ETS1 | 1.522972396 | 1.70E-18 |
| FZD7 | 1.520786809 | 1.58E-24 |
| MX1 | 1.519192341 | 1.33E-02 |
| ARNT | 1.518830956 | 6.59E-14 |
| TRIM6 | 1.517854931 | 1.20E-05 |
| CXCL3 | 1.508751794 | 3.81E-02 |
| GADD45B | 1.507018647 | 1.90E-03 |
| JUN | 1.506294864 | 5.91E-06 |
| PDE12 | 1.503244516 | 3.75E-07 |
| NFKBIZ | 1.502623317 | 1.10E-03 |
| IMPDH2 | 1.500766963 | 2.53E-27 |
| F12 | 1.500127116 | 4.07E-02 |

**Table S7. Upregulated immune genes in $SF3B1^{K700E}$ ES line 3. Genes with a fold change (FC) ≥1.5 and p-value < 0.05 are shown.**

| Gene | FC | p-value |
|---|---|---|
| GDF15 | 46.5822295 | 2.69E-279 |
| REG1A | 45.35711588 | 5.84E-04 |
| ANXA1 | 37.22306998 | 1.43E-18 |
| CH25H | 36.51399237 | 2.46E-04 |
| LGALS12 | 29.81427232 | 1.23E-06 |
| PITX2 | 27.98354158 | 2.97E-22 |
| CCR7 | 23.47792286 | 2.94E-03 |
| SCG2 | 18.06781751 | 1.56E-13 |
| COL8A1 | 14.27817944 | 3.05E-09 |
| GALR1 | 12.652298 | 1.19E-07 |
| PTPRB | 12.298837 | 3.50E-17 |
| RGS4 | 10.77575921 | 4.05E-05 |
| EBF2 | 10.64063638 | 2.37E-02 |
| HCLS1 | 10.4344593 | 2.41E-27 |
| SERPINE1 | 10.3141017 | 3.98E-69 |
| TRPC6 | 10.23641459 | 3.01E-09 |
| PLA2G2A | 9.437446775 | 7.84E-47 |
| PRDM1 | 8.853153945 | 1.46E-05 |
| LCP1 | 8.822476391 | 2.18E-39 |
| LAPTM5 | 8.641145268 | 1.28E-04 |
| NLRP10 | 8.308556076 | 5.74E-04 |
| CDKN1A | 8.255602741 | 1.12E-201 |
| FLI1 | 7.892197198 | 2.67E-10 |
| LAMP3 | 7.862689348 | 2.70E-06 |
| PTPRH | 7.811389132 | 6.72E-04 |
| F7 | 7.17725697 | 2.77E-02 |
| LRFN5 | 7.109211554 | 2.33E-10 |
| CREB5 | 6.956928498 | 6.56E-12 |
| NODAL | 6.78739855 | 1.28E-40 |
| SERPINB9 | 6.665060635 | 4.40E-134 |
| ARHGAP24 | 6.38624796 | 7.16E-06 |
| GPER1 | 5.955300098 | 1.95E-02 |
| MECOM | 5.598486844 | 2.03E-06 |
| CMKLR1 | 5.502954072 | 3.18E-13 |
| PLK2 | 5.441535367 | 3.91E-105 |
| VAV1 | 5.307338113 | 3.13E-16 |
| LEFTY1 | 5.276539308 | 1.17E-73 |
| ANXA3 | 5.157401378 | 8.79E-91 |
| MAFB | 5.027208596 | 8.22E-19 |
| GATA4 | 5.022426573 | 4.39E-02 |

| | | |
|---|---|---|
| ADGRG3 | 5.00491191 | 8.45E-03 |
| TNFRSF10C | 4.938647089 | 5.06E-12 |
| BTG2 | 4.73068249 | 1.44E-31 |
| PRKN | 4.697698791 | 1.34E-04 |
| CDKN2A | 4.672239307 | 1.92E-02 |
| SCNN1B | 4.634787999 | 8.17E-04 |
| VSIR | 4.613562841 | 4.50E-03 |
| LAMA2 | 4.57067965 | 1.88E-25 |
| TWIST1 | 4.567698968 | 9.73E-03 |
| PF4 | 4.46741479 | 8.98E-03 |
| MMP9 | 4.427345084 | 1.03E-25 |
| COLEC11 | 4.423670449 | 3.15E-04 |
| CSMD1 | 4.411954372 | 5.90E-04 |
| CEACAM1 | 4.291379184 | 9.21E-07 |
| STARD13 | 4.168678539 | 7.05E-25 |
| DDIT3 | 4.154392978 | 2.05E-11 |
| MT1G | 4.151744113 | 1.61E-95 |
| TNFSF9 | 4.144549016 | 7.01E-17 |
| PADI4 | 4.134246324 | 3.05E-02 |
| BANK1 | 4.121086201 | 3.66E-05 |
| LEFTY2 | 4.099297248 | 6.00E-34 |
| TAC1 | 4.088589001 | 2.55E-16 |
| PDGFB | 4.084082625 | 1.38E-40 |
| ALDH1A1 | 3.99221704 | 2.71E-09 |
| EBI3 | 3.975962251 | 2.66E-04 |
| HAPLN1 | 3.953228333 | 4.68E-11 |
| BMPER | 3.92137288 | 4.09E-02 |
| GADD45B | 3.86040917 | 6.76E-35 |
| NLRC3 | 3.685301657 | 1.94E-02 |
| IFIT3 | 3.670916288 | 2.03E-02 |
| PAX5 | 3.611728661 | 8.04E-05 |
| GP1BA | 3.589349989 | 7.47E-03 |
| NUPR1 | 3.588642707 | 9.10E-03 |
| TRPM2 | 3.578534487 | 9.22E-03 |
| WNT3 | 3.498929494 | 5.67E-25 |
| ASB2 | 3.469715109 | 4.53E-02 |
| ULBP1 | 3.457196265 | 4.12E-04 |
| ITGAM | 3.450490626 | 8.12E-03 |
| PLA2G4C | 3.41310139 | 2.24E-06 |
| CYP26B1 | 3.403895574 | 3.98E-03 |
| PRTN3 | 3.392001122 | 3.79E-02 |
| INPP5D | 3.334300753 | 4.86E-51 |
| ABCC3 | 3.320833385 | 3.64E-02 |
| CD8A | 3.27224726 | 4.95E-02 |
| ALOX5 | 3.251566832 | 1.24E-08 |

| | | |
|---|---|---|
| SLC16A3 | 3.2374421 | 6.85E-19 |
| TRADD | 3.209192241 | 3.78E-02 |
| NLRP1 | 3.20458846 | 4.04E-03 |
| AZU1 | 3.15372573 | 5.66E-03 |
| ARID5B | 3.141418833 | 7.08E-03 |
| ARHGAP25 | 3.134882086 | 3.93E-05 |
| GDF3 | 3.095712351 | 8.05E-20 |
| PADI2 | 3.071462072 | 1.96E-25 |
| HLA-DQA1 | 3.021078318 | 7.01E-03 |
| ITPR2 | 3.020497024 | 1.12E-54 |
| HSPA2 | 2.900897389 | 6.40E-57 |
| CFD | 2.865612152 | 7.81E-06 |
| ATXN1 | 2.854812412 | 1.19E-05 |
| CRLF1 | 2.847719896 | 6.63E-17 |
| SPSB2 | 2.814745021 | 3.32E-16 |
| LEPR | 2.805040687 | 1.19E-06 |
| ONECUT1 | 2.796870213 | 4.51E-05 |
| DGKH | 2.780974026 | 8.36E-13 |
| TNFAIP6 | 2.768292142 | 3.51E-02 |
| HAS2 | 2.757662631 | 1.14E-15 |
| ZP3 | 2.744026775 | 4.12E-02 |
| VIM | 2.743616913 | 1.35E-27 |
| NDRG1 | 2.718527173 | 6.49E-08 |
| SCARF1 | 2.713982255 | 1.08E-13 |
| DUSP10 | 2.710154459 | 2.27E-03 |
| CAV1 | 2.69473194 | 4.58E-13 |
| CDH5 | 2.689635732 | 4.44E-02 |
| KITLG | 2.665311592 | 1.18E-12 |
| MASP1 | 2.662868731 | 1.03E-06 |
| THBS1 | 2.616554474 | 4.60E-33 |
| MAML2 | 2.601423502 | 7.28E-15 |
| JUN | 2.597773974 | 8.33E-29 |
| LACC1 | 2.575136203 | 1.22E-07 |
| DHRS2 | 2.574917064 | 2.69E-05 |
| LGALS1 | 2.574285728 | 6.20E-18 |
| TRIM22 | 2.570139027 | 2.35E-25 |
| NAV3 | 2.566907627 | 9.61E-05 |
| IL15 | 2.548214763 | 1.71E-02 |
| PTAFR | 2.507012444 | 8.00E-06 |
| MYO1F | 2.503781388 | 1.58E-10 |
| NOS3 | 2.488600045 | 1.30E-07 |
| HPGD | 2.467608144 | 1.31E-03 |
| FN1 | 2.448457426 | 5.72E-37 |
| CR1L | 2.430966241 | 1.24E-02 |
| ARRB1 | 2.428077702 | 7.68E-23 |

| | | |
|---|---|---|
| SULF1 | 2.42698032 | 1.41E-16 |
| TGM2 | 2.408673661 | 3.01E-07 |
| MUC12 | 2.404551344 | 8.67E-03 |
| ITGB2 | 2.404310072 | 3.02E-02 |
| GADD45G | 2.386873338 | 1.83E-16 |
| ZNF91 | 2.36998201 | 3.88E-27 |
| MEF2C | 2.36286007 | 6.09E-03 |
| TRPV4 | 2.359828407 | 1.44E-05 |
| TRPV2 | 2.356371959 | 2.58E-04 |
| CXCR2 | 2.329934861 | 1.44E-02 |
| CLC | 2.327892183 | 3.95E-02 |
| RBM15 | 2.321566405 | 2.45E-20 |
| LTBR | 2.316601584 | 2.27E-05 |
| VENTX | 2.313552809 | 8.40E-18 |
| FAXDC2 | 2.285633665 | 7.25E-05 |
| AMIGO2 | 2.277930964 | 9.17E-13 |
| RIPOR2 | 2.269953008 | 1.45E-44 |
| TAPBPL | 2.260921448 | 3.57E-02 |
| SLIT2 | 2.26007117 | 9.12E-17 |
| IL6R | 2.256279116 | 1.89E-10 |
| IL20RB | 2.254092154 | 9.71E-04 |
| NLRP7 | 2.240466407 | 2.49E-03 |
| ADGRE5 | 2.238854186 | 3.22E-04 |
| KDM6B | 2.236191526 | 2.11E-16 |
| F10 | 2.229941041 | 6.48E-05 |
| IFI35 | 2.226631565 | 3.81E-03 |
| CCN2 | 2.218162808 | 1.53E-37 |
| SERPINB1 | 2.138126341 | 1.21E-14 |
| MYL9 | 2.135720877 | 6.26E-50 |
| TFPI2 | 2.124009275 | 1.25E-08 |
| PURA | 2.121994748 | 3.71E-07 |
| PDCD1 | 2.116362679 | 5.12E-03 |
| CPEB4 | 2.085621592 | 6.06E-12 |
| SOX15 | 2.083639121 | 1.31E-03 |
| NMI | 2.077354042 | 8.00E-04 |
| RUNX2 | 2.071275436 | 3.43E-02 |
| HSPG2 | 2.069302848 | 3.63E-24 |
| NEFH | 2.068909978 | 8.75E-09 |
| RAG1 | 2.068605445 | 2.94E-02 |
| QPCT | 2.067200159 | 2.71E-05 |
| AOC3 | 2.060061444 | 2.71E-03 |
| RND3 | 2.053359739 | 4.46E-31 |
| DUOX1 | 2.04688915 | 1.30E-02 |
| PLA2G6 | 2.043470528 | 6.61E-09 |
| FZD5 | 2.042114503 | 6.64E-47 |

| | | |
|---|---|---|
| IFITM1 | 2.039119346 | 2.03E-18 |
| HLA-DQB1 | 2.037845087 | 7.94E-14 |
| TMEM64 | 2.034230835 | 3.84E-21 |
| EDA2R | 2.025414396 | 1.58E-07 |
| ZDHHC21 | 2.010827548 | 6.07E-06 |
| LPL | 2.000059528 | 2.08E-08 |
| CAV2 | 1.994343368 | 9.24E-03 |
| PLD1 | 1.983748686 | 1.00E-02 |
| PRKG1 | 1.962884544 | 2.21E-09 |
| PLAU | 1.961477026 | 9.69E-12 |
| PDGFRB | 1.954583406 | 5.63E-15 |
| NRROS | 1.946962027 | 6.30E-06 |
| NTN1 | 1.938687909 | 7.61E-06 |
| PRNP | 1.93720661 | 2.64E-19 |
| DDX60 | 1.934046299 | 3.52E-04 |
| LIF | 1.931158823 | 1.34E-05 |
| JAK1 | 1.925231484 | 1.31E-22 |
| HLA-E | 1.918563062 | 2.04E-32 |
| RAC2 | 1.912103466 | 7.00E-03 |
| CSF2RA | 1.900674477 | 1.79E-02 |
| ABI3 | 1.900236559 | 3.33E-03 |
| RGS2 | 1.899837455 | 1.22E-09 |
| DMTN | 1.869930238 | 3.22E-02 |
| SPN | 1.863878613 | 6.39E-03 |
| ADCY8 | 1.858927435 | 7.05E-03 |
| PLAT | 1.850703347 | 5.72E-07 |
| HLA-DRB1 | 1.845009673 | 2.99E-11 |
| NR5A2 | 1.843325653 | 2.93E-07 |
| P2RX4 | 1.841391555 | 7.54E-06 |
| ENPP2 | 1.838385606 | 2.53E-02 |
| TRAF1 | 1.830024252 | 3.93E-02 |
| RAB20 | 1.82850154 | 5.34E-14 |
| LIPA | 1.822981125 | 3.71E-14 |
| SPP1 | 1.820546184 | 9.87E-09 |
| STAT6 | 1.816150183 | 2.58E-19 |
| HLA-F | 1.810536607 | 3.64E-03 |
| ATP7A | 1.805814812 | 2.49E-03 |
| NFAM1 | 1.804596708 | 3.80E-02 |
| NLRC5 | 1.801460596 | 2.67E-02 |
| FOXO3 | 1.801142408 | 2.33E-14 |
| MX2 | 1.785248672 | 2.62E-03 |
| TRIM5 | 1.772656119 | 7.42E-10 |
| TRIML2 | 1.771514659 | 4.17E-12 |
| FNIP1 | 1.770812114 | 3.94E-06 |
| APOBEC3F | 1.770477941 | 3.93E-04 |

| | | |
|---|---|---|
| PLCB1 | 1.761536229 | 1.23E-05 |
| FDXR | 1.757572591 | 9.22E-23 |
| ACP5 | 1.750826855 | 1.22E-04 |
| VCL | 1.744327426 | 9.96E-40 |
| IFIT1 | 1.739629641 | 4.08E-02 |
| TRIM56 | 1.738671131 | 1.15E-18 |
| BATF2 | 1.737673421 | 3.16E-02 |
| AKAP13 | 1.727707853 | 1.41E-10 |
| SERINC5 | 1.727092396 | 5.80E-20 |
| MPP1 | 1.721095503 | 6.79E-09 |
| TRIM68 | 1.704195466 | 1.52E-05 |
| XIAP | 1.69368021 | 6.83E-12 |
| WLS | 1.693631715 | 1.32E-02 |
| NFATC1 | 1.68979877 | 3.17E-05 |
| NEK7 | 1.689185231 | 1.12E-03 |
| PIK3R1 | 1.688878806 | 2.68E-07 |
| FLNB | 1.687533962 | 1.38E-21 |
| SLC9B2 | 1.68055045 | 3.48E-03 |
| SOCS1 | 1.673668895 | 2.18E-06 |
| PTPN6 | 1.672107259 | 1.61E-10 |
| TRPV1 | 1.67173351 | 2.88E-02 |
| TNFRSF10B | 1.663504147 | 1.50E-17 |
| CR2 | 1.660960907 | 2.12E-05 |
| SLFN5 | 1.659382659 | 1.03E-03 |
| SOX4 | 1.659375498 | 2.73E-34 |
| CASP10 | 1.657111931 | 1.55E-02 |
| ASH1L | 1.645519073 | 5.70E-09 |
| RORA | 1.641102921 | 1.61E-02 |
| IL13RA1 | 1.635353128 | 2.50E-12 |
| RRAS | 1.62600847 | 1.10E-07 |
| NFAT5 | 1.625800934 | 5.41E-07 |
| IL18BP | 1.621025285 | 4.69E-03 |
| ITGB1 | 1.618477081 | 7.89E-24 |
| P2RY1 | 1.617778115 | 5.59E-09 |
| SMAD7 | 1.614549374 | 5.99E-17 |
| TGFBR2 | 1.611948606 | 1.25E-06 |
| SERINC3 | 1.609184183 | 5.73E-14 |
| CTH | 1.607053334 | 4.74E-08 |
| FZD6 | 1.606151841 | 3.92E-05 |
| ID2 | 1.602242829 | 2.74E-07 |
| FES | 1.595366487 | 5.27E-04 |
| ICAM1 | 1.587745657 | 7.08E-03 |
| ADCY7 | 1.582276474 | 2.26E-03 |
| ITGA3 | 1.574585135 | 2.74E-15 |
| CDK6 | 1.573670106 | 3.38E-06 |

| | | |
|---|---|---|
| MAVS | 1.573606411 | 1.76E-17 |
| GCNT1 | 1.572549363 | 1.71E-05 |
| ICAM3 | 1.571247357 | 1.07E-02 |
| SHC3 | 1.56903187 | 1.55E-06 |
| IL1RAP | 1.564223409 | 8.05E-04 |
| DFFA | 1.561185838 | 4.79E-16 |
| ZDHHC1 | 1.56067581 | 1.89E-02 |
| RICTOR | 1.559743173 | 6.73E-05 |
| KIF13B | 1.558027206 | 8.00E-06 |
| APP | 1.548259912 | 7.72E-29 |
| EPHA2 | 1.548010502 | 6.77E-16 |
| TRAF3IP2 | 1.547260848 | 1.11E-09 |
| SMAD3 | 1.546766505 | 6.81E-08 |
| TFRC | 1.545703932 | 5.86E-17 |
| JUND | 1.540622971 | 2.02E-11 |
| LRRK1 | 1.535352937 | 8.24E-03 |
| DHRS7B | 1.532575355 | 9.87E-06 |
| SPINK5 | 1.5307248 | 2.65E-02 |
| LIFR | 1.530016607 | 4.21E-05 |
| BCL6 | 1.527371189 | 4.28E-02 |
| ACTG1 | 1.525758953 | 8.15E-30 |
| PLGRKT | 1.525729343 | 2.56E-04 |
| IRAK1 | 1.521462687 | 6.98E-15 |
| BLOC1S3 | 1.520882591 | 1.07E-03 |
| ANXA2 | 1.508590584 | 1.18E-11 |
| EMILIN1 | 1.508307607 | 6.20E-04 |
| CD99L2 | 1.507040101 | 2.12E-07 |
| HECTD1 | 1.506560269 | 3.61E-11 |
| TSC22D3 | 1.506096268 | 1.78E-09 |
| TFPI | 1.503055175 | 1.28E-02 |
| PJA2 | 1.502327658 | 1.04E-12 |
