## Supplementary material for "Chemokine Receptor 1 and its associated immune pathway are downregulated in SF3B1^MT^ blood and non-blood cancers": Table_S8

**Table S8. GSEA analysis in SF3B1$^{MT}$ and SF3B1$^{WT}$ MDS patients.** See Table S16 for patient information.

**GSEA Positive** Hematopoietic/Immune pathways are highlighted in red.

| Gene sets | SIZE | ES | NES | NOM p-val | FDR q-val |
|---|---|---|---|---|---|
| GOBP_RDNA_HETEROCHROMATIN_ASSEMBLY | 35 | 0.769441 | 2.677348 | 0 | 0 |
| GOBP_TETRAPYRROLE_BIOSYNTHETIC_PROCESS | 28 | 0.777877 | 2.567594 | 0 | 0 |
| GOBP_CENTROMERE_COMPLEX_ASSEMBLY | 55 | 0.68912 | 2.560799 | 0 | 0 |
| GOBP_CHROMATIN_REMODELING_AT_CENTROMERE | 46 | 0.701922 | 2.55141 | 0 | 0 |
| GOBP_DNA_REPLICATION_DEPENDENT_NUCLEOSOME_ORGANIZATION | 30 | 0.745594 | 2.543164 | 0 | 0 |
| GOBP_DNA_REPLICATION_INDEPENDENT_NUCLEOSOME_ORGANIZATION | 53 | 0.67298 | 2.526515 | 0 | 0 |
| GOBP_NUCLEOSOME_ASSEMBLY | 123 | 0.560226 | 2.476471 | 0 | 0 |
| GOBP_AMINO_ACID_ACTIVATION | 49 | 0.639171 | 2.399184 | 0 | 1.37E-04 |
| GOBP_HISTONE_EXCHANGE | 53 | 0.652692 | 2.398513 | 0 | 1.22E-04 |
| GOBP_TELOMERE_ORGANIZATION | 168 | 0.52534 | 2.376912 | 0 | 1.10E-04 |
| GOBP_MITOTIC_SISTER_CHROMATID_SEGREGATION | 161 | 0.523285 | 2.363227 | 0 | 9.97E-05 |
| GOBP_HEME_METABOLIC_PROCESS | 30 | 0.706762 | 2.342814 | 0 | 9.14E-05 |
| GOBP_NUCLEOSOME_ORGANIZATION | 156 | 0.520277 | 2.342786 | 0 | 8.44E-05 |
| GOBP_DNA_PACKAGING | 209 | 0.503348 | 2.342384 | 0 | 7.83E-05 |
| GOBP_MITOCHONDRIAL_TRANSLATION | 133 | 0.525694 | 2.328263 | 0 | 7.31E-05 |
| GOBP_CELL_CYCLE_DNA_REPLICATION | 64 | 0.592092 | 2.318444 | 0 | 6.85E-05 |
| GOBP_DNA_REPLICATION_INITIATION | 39 | 0.648511 | 2.311417 | 0 | 6.45E-05 |
| GOBP_PROTEIN_DNA_COMPLEX_SUBUNIT_ORGANIZATION | 252 | 0.496068 | 2.285123 | 0 | 1.84E-04 |
| GOBP_TELOMERE_MAINTENANCE_VIA_SEMI_CONSERVATIVE_REPLICATION | 27 | 0.710417 | 2.264334 | 0 | 2.35E-04 |
| GOBP_ATTACHMENT_OF_SPINDLE_MICROTUBULES_TO_KINETOCHORE | 35 | 0.635606 | 2.243511 | 0 | 4.49E-04 |
| GOBP_PROTEIN_LOCALIZATION_TO_CHROMOSOME | 90 | 0.537693 | 2.242175 | 0 | 4.27E-04 |
| GOBP_DNA_CONFORMATION_CHANGE | 320 | 0.45966 | 2.239972 | 0 | 4.08E-04 |
| GOBP_NEGATIVE_REGULATION_OF_MEGAKARYOCYTE_DIFFERENTIATION | 17 | 0.78212 | 2.237235 | 0 | 4.39E-04 |
| GOBP_DNA_DEPENDENT_DNA_REPLICATION | 155 | 0.500475 | 2.236061 | 0 | 4.21E-04 |
| GOBP_MITOCHONDRIAL_GENE_EXPRESSION | 164 | 0.493945 | 2.231504 | 0 | 4.04E-04 |
| GOBP_CHROMATIN_ASSEMBLY_OR_DISASSEMBLY | 196 | 0.482756 | 2.222691 | 0 | 4.31E-04 |
| GOBP_TRANSLATIONAL_TERMINATION | 104 | 0.512127 | 2.196239 | 0 | 7.06E-04 |
| GOBP_PROTEIN_LOCALIZATION_TO_CHROMOSOME_CENTROMERIC_REGION | 25 | 0.675585 | 2.177286 | 0 | 9.62E-04 |
| GOBP_PORPHYRIN_CONTAINING_COMPOUND_METABOLIC_PROCESS | 36 | 0.632193 | 2.166142 | 0 | 0.001162 |
| GOBP_CHROMOSOME_SEGREGATION | 322 | 0.447781 | 2.16513 | 0 | 0.001161 |
| GOBP_MITOCHONDRIAL_TRANSLATIONAL_TERMINATION | 88 | 0.531434 | 2.154679 | 0 | 0.001408 |
| GOBP_SISTER_CHROMATID_SEGREGATION | 193 | 0.468837 | 2.154424 | 0 | 0.001432 |
| GOBP_NUCLEAR_CHROMOSOME_SEGREGATION | 261 | 0.450356 | 2.152903 | 0 | 0.001389 |
| GOBP_KINETOCHORE_ORGANIZATION | 22 | 0.700809 | 2.150295 | 0 | 0.001381 |
| GOBP_DNA_STRAND_ELONGATION | 26 | 0.667097 | 2.143579 | 0 | 0.001501 |
| GOBP_NEGATIVE_REGULATION_OF_GENE_EXPRESSION_EPIGENETIC | 109 | 0.506051 | 2.140521 | 0 | 0.00149 |
| GOBP_CHROMOSOME_ORGANIZATION_INVOLVED_IN_MEIOTIC_CELL_CYCLE | 59 | 0.552697 | 2.136556 | 0 | 0.001541 |
| GOBP_ATP_DEPENDENT_CHROMATIN_REMODELING | 84 | 0.522997 | 2.126117 | 0 | 0.001882 |
| GOBP_PROTEIN_LOCALIZATION_TO_KINETOCHORE | 19 | 0.72204 | 2.12386 | 0 | 0.00189 |
| GOBP_CHROMOSOME_SEPARATION | 93 | 0.5007 | 2.115411 | 0 | 0.002092 |
| GOBP_HETEROCHROMATIN_ORGANIZATION | 75 | 0.531371 | 2.102608 | 0 | 0.002426 |
| GOBP_GAS_TRANSPORT | 16 | 0.735366 | 2.102399 | 0 | 0.002368 |
| GOBP_CRISTAE_FORMATION | 35 | 0.619017 | 2.101125 | 0 | 0.002365 |
| GOBP_CHROMATIN_ORGANIZATION_INVOLVED_IN_REGULATION_OF_TRANSCRIPTION | 137 | 0.477856 | 2.100607 | 0 | 0.002361 |
| GOBP_REGULATION_OF_MEGAKARYOCYTE_DIFFERENTIATION | 79 | 0.516109 | 2.098207 | 0 | 0.002357 |
| GOBP_REGULATION_OF_CHROMOSOME_SEGREGATION | 84 | 0.513957 | 2.08688 | 0 | 0.002724 |
| GOBP_REGULATION_OF_CHROMOSOME_SEPARATION | 69 | 0.521795 | 2.067488 | 0 | 0.003375 |
| GOBP_METAPHASE_ANAPHASE_TRANSITION_OF_CELL_CYCLE | 62 | 0.538543 | 2.062528 | 0 | 0.003491 |
| GOBP_CHROMOSOME_CONDENSATION | 39 | 0.591321 | 2.052171 | 0 | 0.003964 |
| GOBP_DNA_REPLICATION | 275 | 0.428679 | 2.031856 | 0 | 0.005118 |
| GOBP_DNA_STRAND_ELONGATION_INVOLVED_IN_DNA_REPLICATION | 19 | 0.682584 | 2.016594 | 0 | 0.006192 |
| GOBP_REGULATION_OF_CHROMATIN_SILENCING | 15 | 0.71461 | 2.014292 | 0 | 0.006138 |
| GOBP_TRNA_METABOLIC_PROCESS | 178 | 0.440645 | 2.011932 | 0 | 0.006212 |
| GOBP_PIGMENT_BIOSYNTHETIC_PROCESS | 51 | 0.541742 | 1.993676 | 0 | 0.007765 |
| GOBP_INNER_MITOCHONDRIAL_MEMBRANE_ORGANIZATION | 56 | 0.524268 | 1.986791 | 0 | 0.00815 |
| GOBP_INTERSTRAND_CROSS_LINK_REPAIR | 55 | 0.526378 | 1.978654 | 0 | 0.008722 |
| GOBP_IRON_ION_HOMEOSTASIS | 82 | 0.488962 | 1.969573 | 0 | 0.009815 |
| GOBP_REGULATION_OF_GENE_SILENCING | 133 | 0.450074 | 1.958077 | 0 | 0.011318 |
| GOBP_REGULATION_OF_DNA_DEPENDENT_DNA_REPLICATION | 47 | 0.52483 | 1.939666 | 0 | 0.013716 |
| GOBP_DOUBLE_STRAND_BREAK_REPAIR | 259 | 0.399544 | 1.934296 | 0 | 0.014063 |
| GOBP_POSITIVE_REGULATION_OF_TELOMERASE_RNA_LOCALIZATION_TO_CAJAL_BODY | 15 | 0.693157 | 1.934288 | 0 | 0.013832 |
| GOBP_NON_RECOMBINATIONAL_REPAIR | 103 | 0.462386 | 1.923823 | 0 | 0.015384 |
| GOBP_ANAPHASE_PROMOTING_COMPLEX_DEPENDENT_CATABOLIC_PROCESS | 80 | 0.479825 | 1.919522 | 0 | 0.015902 |
| GOBP_DNA_UNWINDING_INVOLVED_IN_DNA_REPLICATION | 16 | 0.697029 | 1.917168 | 0 | 0.016075 |
| GOBP_SERINE_FAMILY_AMINO_ACID_METABOLIC_PROCESS | 41 | 0.527757 | 1.899295 | 0 | 0.019209 |
| GOBP_PIGMENT_METABOLIC_PROCESS | 59 | 0.497482 | 1.897483 | 0 | 0.019376 |
| GOBP_MITOTIC_NUCLEAR_DIVISION | 286 | 0.394162 | 1.88763 | 0 | 0.020936 |
| GOBP_COPPER_ION_TRANSPORT | 16 | 0.649943 | 1.883895 | 0.008499 | 0.021533 |
| GOBP_NEGATIVE_REGULATION_OF_METAPHASE_ANAPHASE_TRANSITION_OF_CELL_CYCLE | 40 | 0.535067 | 1.876546 | 0 | 0.022745 |
| GOBP_KINETOCHORE_ASSEMBLY | 17 | 0.663378 | 1.868575 | 0.002632 | 0.024474 |
| GOBP_ATP_SYNTHESIS_COUPLED_ELECTRON_TRANSPORT | 97 | 0.444608 | 1.86808 | 0 | 0.024286 |
| GOBP_MITOTIC_RECOMBINATION | 24 | 0.589568 | 1.867463 | 0.002907 | 0.024106 |
| GOBP_MEIOTIC_CHROMOSOME_SEGREGATION | 80 | 0.460463 | 1.866347 | 0.003937 | 0.02405 |
| GOBP_MITOCHONDRIAL_ATP_SYNTHESIS_COUPLED_PROTON_TRANSPORT | 22 | 0.604677 | 1.861969 | 0 | 0.024677 |
| GOBP_CELLULAR_IRON_ION_HOMEOSTASIS | 66 | 0.479185 | 1.85987 | 0 | 0.024767 |
| GOBP_TELOMERASE_RNA_LOCALIZATION | 19 | 0.630548 | 1.855611 | 0.002725 | 0.025556 |
| GOBP_MEGAKARYOCYTE_DIFFERENTIATION | 96 | 0.442865 | 1.853283 | 0 | 0.025778 |
| GOBP_REGULATION_OF_MITOTIC_SISTER_CHROMATID_SEGREGATION | 45 | 0.517424 | 1.849413 | 0 | 0.026437 |
| GOBP_METAPHASE_PLATE_CONGRESSION | 63 | 0.474926 | 1.83648 | 0 | 0.030156 |
| GOBP_NONRIBOSOMAL_PEPTIDE_BIOSYNTHETIC_PROCESS | 17 | 0.631683 | 1.835467 | 0.002762 | 0.030016 |
| GOBP_REGULATION_OF_MITOCHONDRIAL_GENE_EXPRESSION | 29 | 0.55717 | 1.835238 | 0.003175 | 0.029742 |
| GOBP_HOMOLOGOUS_CHROMOSOME_SEGREGATION | 53 | 0.489281 | 1.829668 | 0 | 0.030895 |
| GOBP_TELOMERE_CAPPING | 52 | 0.490703 | 1.819281 | 0 | 0.034143 |
| GOBP_REGULATION_OF_GENE_EXPRESSION_EPIGENETIC | 188 | 0.3942 | 1.814559 | 0 | 0.035519 |
| GOBP_GLYCOSYLCERAMIDE_METABOLIC_PROCESS | 17 | 0.631218 | 1.812393 | 0.00578 | 0.035743 |
| GOBP_NEGATIVE_REGULATION_OF_CHROMOSOME_ORGANIZATION | 88 | 0.436546 | 1.809334 | 0 | 0.036341 |
| GOBP_NEGATIVE_REGULATION_OF_NUCLEAR_DIVISION | 51 | 0.485776 | 1.806598 | 0 | 0.036758 |
| GOBP_MITOCHONDRIAL_ELECTRON_TRANSPORT_NADH_TO_UBIQUINONE | 54 | 0.476777 | 1.80657 | 0 | 0.036352 |
| GOBP_VASCULAR_TRANSPORT | 83 | 0.436358 | 1.801007 | 0 | 0.038125 |
| GOBP_CHROMOSOME_LOCALIZATION | 76 | 0.451448 | 1.795266 | 0 | 0.040007 |
| GOBP_REGULATION_OF_GENE_SILENCING_BY_RNA | 117 | 0.416873 | 1.780149 | 0 | 0.045323 |
| GOBP_TETRAHYDROFOLATE_METABOLIC_PROCESS | 19 | 0.608076 | 1.777495 | 0.008746 | 0.045998 |
| GOBP_REGULATION_OF_CELL_CYCLE_G2_M_PHASE_TRANSITION | 213 | 0.381545 | 1.777362 | 0 | 0.045565 |
| GOBP_CHROMATIN_REMODELING | 197 | 0.386479 | 1.773029 | 0 | 0.046647 |
| GOBP_MICROTUBULE_CYTOSKELETON_ORGANIZATION_INVOLVED_IN_MITOSIS | 141 | 0.394052 | 1.766664 | 0 | 0.048982 |
| GOBP_DNA_REPLICATION_CHECKPOINT | 17 | 0.622775 | 1.765811 | 0.00266 | 0.048869 |
| GOBP_SYNAPTONEMAL_COMPLEX_ORGANIZATION | 22 | 0.577097 | 1.765626 | 0.013298 | 0.048482 |
| GOBP_ORGANELLE_FISSION | 459 | 0.347715 | 1.76499 | 0 | 0.04825 |
| GOBP_MICROTUBULE_ORGANIZING_CENTER_ORGANIZATION | 135 | 0.405213 | 1.759837 | 0 | 0.050514 |
| GOBP_CENTROSOME_DUPLICATION | 69 | 0.437877 | 1.751189 | 0 | 0.053832 |
| GOBP_PROTEIN_LOCALIZATION_TO_CHROMATIN | 29 | 0.539886 | 1.748103 | 0.005952 | 0.054782 |
| GOBP_DNA_GEOMETRIC_CHANGE | 113 | 0.404184 | 1.742166 | 0 | 0.057549 |
| GOBP_HYDROGEN_PEROXIDE_CATABOLIC_PROCESS | 29 | 0.534857 | 1.741675 | 0.002967 | 0.057186 |
| GOBP_MATURATION_OF_5_8S_RRNA_FROM_TRICISTRONIC_RRNA_TRANSCRIPT_SSU_RRNA_5_ | 24 | 0.548222 | 1.737354 | 0.005831 | 0.05889 |
| GOBP_MEIOSIS_I_CELL_CYCLE_PROCESS | 115 | 0.403624 | 1.733965 | 0 | 0.060244 |
| GOBP_PTERIDINE_CONTAINING_COMPOUND_METABOLIC_PROCESS | 33 | 0.503629 | 1.733889 | 0.002841 | 0.059738 |
| GOBP_MITOTIC_SISTER_CHROMATID_COHESION | 26 | 0.532425 | 1.730641 | 0.005571 | 0.060495 |
| GOBP_SHORT_CHAIN_FATTY_ACID_METABOLIC_PROCESS | 16 | 0.614156 | 1.729582 | 0.01039 | 0.060441 |
| GOBP_RECOMBINATIONAL_REPAIR | 137 | 0.396946 | 1.728112 | 0 | 0.060683 |
| GOBP_MITOCHONDRIAL_RNA_METABOLIC_PROCESS | 47 | 0.480263 | 1.726323 | 0 | 0.060856 |
| GOBP_ESTABLISHMENT_OF_PROTEIN_LOCALIZATION_TO_CHROMOSOME | 27 | 0.550445 | 1.724332 | 0.014493 | 0.061415 |
| GOBP_RESPIRATORY_ELECTRON_TRANSPORT_CHAIN | 114 | 0.400539 | 1.723869 | 0 | 0.061097 |
| GOBP_MITOTIC_DNA_REPLICATION | 16 | 0.617783 | 1.716387 | 0.010554 | 0.064516 |
| GOBP_MATURATION_OF_5_8S_RRNA | 35 | 0.497635 | 1.713179 | 0.009063 | 0.065577 |
| GOBP_MITOTIC_SPINDLE_ORGANIZATION | 117 | 0.391918 | 1.707757 | 0 | 0.067888 |
| GOBP_NUCLEOTIDE_EXCISION_REPAIR_DNA_GAP_FILLING | 23 | 0.539368 | 1.704208 | 0.005698 | 0.06926 |
| GOBP_VENTRICULAR_SYSTEM_DEVELOPMENT | 25 | 0.533056 | 1.702346 | 0.008671 | 0.06976 |
| GOBP_PSEUDOURIDINE_SYNTHESIS | 18 | 0.581903 | 1.700781 | 0.014164 | 0.070076 |
| GOBP_HOMOLOGOUS_RECOMBINATION | 57 | 0.453737 | 1.700108 | 0.003448 | 0.06991 |
| GOBP_NCRNA_3_END_PROCESSING | 40 | 0.485414 | 1.695921 | 0.002976 | 0.071504 |
| GOBP_BASE_EXCISION_REPAIR | 39 | 0.480592 | 1.683579 | 0.003125 | 0.078323 |
| GOBP_REGULATION_OF_GENE_EXPRESSION_BY_GENETIC_IMPRINTING | 16 | 0.589904 | 1.682667 | 0.024523 | 0.078359 |
| GOBP_REGULATION_OF_MITOTIC_NUCLEAR_DIVISION | 105 | 0.387017 | 1.682537 | 0 | 0.077784 |
| GOBP_CLEAVAGE_INVOLVED_IN_RRNA_PROCESSING | 27 | 0.517948 | 1.67988 | 0.005731 | 0.078616 |
| GOBP_ATTACHMENT_OF_MITOTIC_SPINDLE_MICROTUBULES_TO_KINETOCHORE | 15 | 0.596564 | 1.673922 | 0.011494 | 0.081626 |
| GOBP_TRANSITION_METAL_ION_HOMEOSTASIS | 126 | 0.389409 | 1.669886 | 0 | 0.083763 |
| GOBP_HOMOLOGOUS_CHROMOSOME_PAIRING_AT_MEIOSIS | 44 | 0.469038 | 1.669472 | 0.006993 | 0.083386 |
| GOBP_FOLIC_ACID_CONTAINING_COMPOUND_METABOLIC_PROCESS | 27 | 0.506421 | 1.666629 | 0.003086 | 0.08472 |
| GOBP_AMINO_ACID_IMPORT | 44 | 0.459188 | 1.65913 | 0.003311 | 0.089369 |
| GOBP_PYRIMIDINE_NUCLEOSIDE_TRIPHOSPHATE_BIOSYNTHETIC_PROCESS | 19 | 0.557679 | 1.658865 | 0.021798 | 0.088871 |
| GOBP_ONE_CARBON_METABOLIC_PROCESS | 38 | 0.469094 | 1.657805 | 0.002959 | 0.08886 |
| GOBP_NEGATIVE_REGULATION_OF_DNA_RECOMBINATION | 41 | 0.46052 | 1.655277 | 0.00627 | 0.090144 |
| GOBP_SULFUR_AMINO_ACID_BIOSYNTHETIC_PROCESS | 17 | 0.572369 | 1.653422 | 0.025568 | 0.090878 |
| GOBP_NADH_DEHYDROGENASE_COMPLEX_ASSEMBLY | 65 | 0.416791 | 1.650949 | 0.003521 | 0.091852 |
| GOBP_MITOTIC_CELL_CYCLE_CHECKPOINT | 156 | 0.361327 | 1.649402 | 0 | 0.092332 |
| GOBP_TELOMERE_MAINTENANCE_VIA_TELOMERE_LENGTHENING | 77 | 0.405689 | 1.649181 | 0 | 0.091791 |
| GOBP_TETRAPYRROLE_METABOLIC_PROCESS | 56 | 0.434755 | 1.648408 | 0 | 0.091703 |
| GOBP_NEGATIVE_REGULATION_OF_DNA_DEPENDENT_DNA_REPLICATION | 17 | 0.580848 | 1.641723 | 0.028646 | 0.096014 |
| GOBP_NCRNA_EXPORT_FROM_NUCLEUS | 38 | 0.480919 | 1.640163 | 0.006154 | 0.096505 |
| GOBP_RESOLUTION_OF_MEIOTIC_RECOMBINATION_INTERMEDIATES | 18 | 0.552657 | 1.638405 | 0.010753 | 0.097253 |
| GOBP_PYRIMIDINE_NUCLEOSIDE_TRIPHOSPHATE_METABOLIC_PROCESS | 24 | 0.518933 | 1.635036 | 0.015337 | 0.099034 |
| GOBP_TRANSLATIONAL_ELONGATION | 131 | 0.373298 | 1.630507 | 0 | 0.101392 |
| GOBP_POSITIVE_REGULATION_OF_MITOCHONDRIAL_TRANSLATION | 16 | 0.575473 | 1.624248 | 0.029101 | 0.105533 |
| GOBP_MITOTIC_METAPHASE_PLATE_CONGRESSION | 50 | 0.438029 | 1.622447 | 0.012158 | 0.106345 |
| GOBP_NUCLEOBASE_CONTAINING_SMALL_MOLECULE_INTERCONVERSION | 27 | 0.506672 | 1.617694 | 0.013263 | 0.10955 |
| GOBP_TRANSCRIPTION_COUPLED_NUCLEOTIDE_EXCISION_REPAIR | 73 | 0.403468 | 1.616948 | 0 | 0.10937 |
| GOBP_SISTER_CHROMATID_COHESION | 59 | 0.425794 | 1.613062 | 0.00365 | 0.111884 |
| GOBP_MITOTIC_CHROMOSOME_CONDENSATION | 15 | 0.595199 | 1.611564 | 0.045455 | 0.112265 |
| GOBP_MEIOTIC_CELL_CYCLE | 224 | 0.342346 | 1.607847 | 0 | 0.114548 |
| GOBP_TRNA_PROCESSING | 129 | 0.372671 | 1.603321 | 0 | 0.117319 |
| GOBP_GLYCOSPHINGOLIPID_BIOSYNTHETIC_PROCESS | 25 | 0.511198 | 1.598614 | 0.008523 | 0.120764 |
| GOBP_NUCLEOTIDE_EXCISION_REPAIR | 106 | 0.377819 | 1.595519 | 0.004329 | 0.122502 |
| GOBP_OXIDATIVE_PHOSPHORYLATION | 141 | 0.356161 | 1.59324 | 0 | 0.123649 |
| GOBP_MEIOTIC_CELL_CYCLE_PROCESS | 170 | 0.351921 | 1.59161 | 0 | 0.124347 |
| GOBP_ATP_SYNTHESIS_COUPLED_PROTON_TRANSPORT | 27 | 0.502943 | 1.587863 | 0.017595 | 0.126998 |
| GOBP_ESTABLISHMENT_OF_MITOTIC_SPINDLE_LOCALIZATION | 31 | 0.472103 | 1.585752 | 0.024691 | 0.128241 |
| GOBP_MITOCHONDRIAL_RESPIRATORY_CHAIN_COMPLEX_ASSEMBLY | 101 | 0.381361 | 1.585051 | 0.004525 | 0.128171 |
| GOBP_ERROR_PRONE_TRANSLESION_SYNTHESIS | 21 | 0.52606 | 1.58452 | 0.02681 | 0.127848 |
| GOBP_DNA_RECOMBINATION | 283 | 0.32744 | 1.58159 | 0 | 0.12932 |
| GOBP_CELLULAR_TRANSITION_METAL_ION_HOMEOSTASIS | 105 | 0.376683 | 1.581197 | 0.004184 | 0.12896 |
| GOBP_ERYTHROCYTE_DEVELOPMENT | 32 | 0.474592 | 1.580083 | 0.003236 | 0.129406 |
| GOBP_INTRA_S_DNA_DAMAGE_CHECKPOINT | 17 | 0.543303 | 1.579894 | 0.031915 | 0.128772 |
| GOBP_SPINDLE_ORGANIZATION | 180 | 0.34522 | 1.578885 | 0 | 0.128923 |
| GOBP_CELLULAR_MODIFIED_AMINO_ACID_BIOSYNTHETIC_PROCESS | 43 | 0.432614 | 1.574042 | 0.013378 | 0.13242 |
| GOBP_NUCLEOBASE_BIOSYNTHETIC_PROCESS | 18 | 0.53683 | 1.572637 | 0.032258 | 0.132883 |

**GSEA Negative** Hematopoietic/Immune pathways are highlighted in green.

| Gene sets | SIZE | ES | NES | NOM p-val | FDR q-val |
|---|---|---|---|---|---|
| GOBP_ADENYLATE_CYCLASE_ACTIVATING_G_PROTEIN_COUPLED_RECEPTOR_SIGNALING_PATI | 109 | -0.557681 | -2.051381 | 0 | 0.022884 |
| GOBP_INTERFERON_GAMMA_MEDIATED_SIGNALING_PATHWAY | 88 | -0.56373 | -2.048416 | 0 | 0.012392 |
| GOBP_REGULATION_OF_ALPHA_BETA_T_CELL_DIFFERENTIATION | 66 | -0.595866 | -2.040144 | 0 | 0.010174 |
| GOBP_REGULATION_OF_ALPHA_BETA_T_CELL_ACTIVATION | 98 | -0.553647 | -2.020073 | 0 | 0.011675 |
| GOBP_POSITIVE_REGULATION_OF_CELL_KILLING | 62 | -0.60411 | -2.017937 | 0 | 0.009531 |
| GOBP_CELLULAR_RESPONSE_TO_MOLECULE_OF_BACTERIAL_ORIGIN | 190 | -0.509247 | -2.017034 | 0 | 0.008263 |
| GOBP_ADENYLATE_CYCLASE_MODULATING_G_PROTEIN_COUPLED_RECEPTOR_SIGNALING_PA | 175 | -0.51256 | -2.003464 | 0 | 0.009395 |
| GOBP_LEUKOCYTE_MEDIATED_CYTOTOXICITY | 107 | -0.539795 | -1.987522 | 0 | 0.010967 |
| GOBP_REGULATION_OF_LEUKOCYTE_MEDIATED_CYTOTOXICITY | 77 | -0.568563 | -1.983032 | 0 | 0.010278 |
| GOBP_INTERLEUKIN_6_PRODUCTION | 131 | -0.514518 | -1.973178 | 0 | 0.011732 |
| GOBP_CELLULAR_RESPONSE_TO_BIOTIC_STIMULUS | 214 | -0.491206 | -1.970822 | 0 | 0.011013 |
| GOBP_NEGATIVE_REGULATION_OF_CYTOKINE_PRODUCTION_INVOLVED_IN_IMMUNE_RESPONS | 25 | -0.694405 | -1.968472 | 0 | 0.010732 |
| GOBP_INTERLEUKIN_2_PRODUCTION | 58 | -0.592419 | -1.968458 | 0 | 0.009907 |
| GOBP_RESPONSE_TO_INTERFERON_GAMMA | 181 | -0.500664 | -1.965221 | 0 | 0.009677 |
| GOBP_INTERLEUKIN_12_PRODUCTION | 57 | -0.579609 | -1.958482 | 0 | 0.010618 |
| GOBP_ALPHA_BETA_T_CELL_ACTIVATION | 143 | -0.514034 | -1.95721 | 0 | 0.010252 |
| GOBP_ALPHA_BETA_T_CELL_DIFFERENTIATION | 104 | -0.530083 | -1.935752 | 0 | 0.014366 |
| GOBP_GLOMERULAR_EPITHELIUM_DEVELOPMENT | 23 | -0.685579 | -1.930031 | 0 | 0.014363 |
| GOBP_REGULATION_OF_CELL_KILLING | 95 | -0.526027 | -1.926126 | 0 | 0.014665 |
| GOBP_REGULATION_OF_CD4_POSITIVE_ALPHA_BETA_T_CELL_DIFFERENTIATION | 50 | -0.601203 | -1.925822 | 0 | 0.01398 |
| GOBP_POSITIVE_REGULATION_OF_PRI_MIRNA_TRANSCRIPTION_BY_RNA_POLYMERASE_II | 40 | -0.604802 | -1.903034 | 0 | 0.01963 |
| GOBP_REGULATION_OF_NATURAL_KILLER_CELL_MEDIATED_IMMUNITY | 45 | -0.59425 | -1.897089 | 0 | 0.020822 |
| GOBP_REGULATION_OF_CD4_POSITIVE_ALPHA_BETA_T_CELL_ACTIVATION | 64 | -0.558527 | -1.892928 | 0 | 0.021243 |
| GOBP_CYTOKINE_PRODUCTION_INVOLVED_IN_IMMUNE_RESPONSE | 91 | -0.523938 | -1.891391 | 0 | 0.020597 |
| GOBP_POSITIVE_REGULATION_OF_ALPHA_BETA_T_CELL_ACTIVATION | 64 | -0.550041 | -1.878402 | 0 | 0.023785 |
| GOBP_NEGATIVE_REGULATION_OF_INTERLEUKIN_2_PRODUCTION | 26 | -0.64148 | -1.874093 | 0 | 0.024562 |
| GOBP_REGULATION_OF_T_CELL_DIFFERENTIATION | 141 | -0.483874 | -1.872658 | 0 | 0.024218 |
| GOBP_POSITIVE_REGULATION_OF_LEUKOCYTE_MEDIATED_IMMUNITY | 122 | -0.495607 | -1.872278 | 0 | 0.023558 |
| GOBP_POSITIVE_REGULATION_OF_T_CELL_MEDIATED_CYTOTOXICITY | 29 | -0.628501 | -1.871236 | 0 | 0.023105 |
| GOBP_POSITIVE_REGULATION_OF_LEUKOCYTE_CELL_CELL_ADHESION | 227 | -0.466179 | -1.869985 | 0 | 0.022812 |
| GOBP_POSITIVE_REGULATION_OF_ALPHA_BETA_T_CELL_DIFFERENTIATION | 48 | -0.577982 | -1.867543 | 0 | 0.023 |
| GOBP_ANIMAL_ORGAN_FORMATION | 47 | -0.567291 | -1.862794 | 0 | 0.023982 |
| GOBP_RECEPTOR_SIGNALING_PATHWAY_VIA_STAT | 138 | -0.486559 | -1.862168 | 0 | 0.023429 |
| GOBP_REGULATION_OF_KILLING_OF_CELLS_OF_OTHER_ORGANISM | 15 | -0.746271 | -1.861321 | 0.001597 | 0.022993 |
| GOBP_RESPONSE_TO_MOLECULE_OF_BACTERIAL_ORIGIN | 310 | -0.449749 | -1.860288 | 0 | 0.022608 |
| GOBP_NATURAL_KILLER_CELL_MEDIATED_IMMUNITY | 63 | -0.548079 | -1.859358 | 0 | 0.022351 |
| GOBP_REGULATION_OF_T_HELPER_CELL_DIFFERENTIATION | 38 | -0.594761 | -1.856567 | 0.001431 | 0.022857 |
| GOBP_POSITIVE_REGULATION_OF_INTERFERON_GAMMA_PRODUCTION | 64 | -0.545903 | -1.855723 | 0 | 0.022584 |
| GOBP_T_CELL_MEDIATED_IMMUNITY | 99 | -0.510352 | -1.853911 | 0 | 0.022568 |
| GOBP_POSITIVE_REGULATION_OF_CELL_CELL_ADHESION | 261 | -0.455205 | -1.843184 | 0 | 0.026578 |
| GOBP_REGULATION_OF_T_CELL_ACTIVATION | 313 | -0.446519 | -1.834335 | 0 | 0.029794 |
| GOBP_POSITIVE_REGULATION_OF_RHO_PROTEIN_SIGNAL_TRANSDUCTION | 33 | -0.604872 | -1.827409 | 0 | 0.032742 |
| GOBP_GLOMERULAR_EPITHELIAL_CELL_DIFFERENTIATION | 20 | -0.688959 | -1.827345 | 0 | 0.032025 |
| GOBP_T_CELL_DIFFERENTIATION | 239 | -0.450272 | -1.821876 | 0 | 0.033618 |
| GOBP_POSITIVE_REGULATION_OF_NATURAL_KILLER_CELL_MEDIATED_IMMUNITY | 29 | -0.62533 | -1.818666 | 0 | 0.034441 |
| GOBP_REGULATION_OF_ANIMAL_ORGAN_FORMATION | 19 | -0.668262 | -1.81383 | 0.001531 | 0.036325 |
| GOBP_T_HELPER_17_TYPE_IMMUNE_RESPONSE | 31 | -0.609205 | -1.81164 | 0 | 0.036669 |
| GOBP_POSITIVE_REGULATION_OF_CD4_POSITIVE_ALPHA_BETA_T_CELL_ACTIVATION | 37 | -0.594054 | -1.810621 | 0 | 0.036481 |
| GOBP_REGULATION_OF_LYMPHOCYTE_DIFFERENTIATION | 169 | -0.461396 | -1.80948 | 0 | 0.03642 |
| GOBP_POSITIVE_REGULATION_OF_T_CELL_MEDIATED_IMMUNITY | 51 | -0.555624 | -1.808366 | 0.001418 | 0.036206 |
| GOBP_T_CELL_ACTIVATION | 443 | -0.426947 | -1.808132 | 0 | 0.035663 |
| GOBP_REGULATION_OF_LYMPHOCYTE_MEDIATED_IMMUNITY | 152 | -0.464489 | -1.806575 | 0 | 0.035691 |
| GOBP_CD4_POSITIVE_ALPHA_BETA_T_CELL_DIFFERENTIATION | 79 | -0.508191 | -1.805988 | 0 | 0.035486 |
| GOBP_RESPONSE_TO_MURAMYL_DIPEPTIDE | 20 | -0.655965 | -1.803732 | 0.001479 | 0.035855 |
| GOBP_NEGATIVE_REGULATION_OF_TYPE_I_INTERFERON_PRODUCTION | 46 | -0.556187 | -1.802786 | 0 | 0.035775 |
| GOBP_INTERLEUKIN_10_PRODUCTION | 57 | -0.535885 | -1.799668 | 0.001429 | 0.036502 |
| GOBP_PRI_MIRNA_TRANSCRIPTION_BY_RNA_POLYMERASE_II | 52 | -0.54516 | -1.794696 | 0 | 0.038255 |
| GOBP_POSITIVE_REGULATION_OF_INTERLEUKIN_6_PRODUCTION | 80 | -0.507293 | -1.794092 | 0 | 0.038006 |
| GOBP_REGULATION_OF_LEUKOCYTE_MEDIATED_IMMUNITY | 202 | -0.447264 | -1.793173 | 0 | 0.037848 |
| GOBP_POSITIVE_REGULATION_OF_INTERLEUKIN_12_PRODUCTION | 38 | -0.574642 | -1.792671 | 0.001427 | 0.037537 |
| GOBP_AMINE_TRANSPORT | 76 | -0.502656 | -1.792296 | 0 | 0.037093 |
| GOBP_REGULATION_OF_T_HELPER_17_TYPE_IMMUNE_RESPONSE | 20 | -0.660379 | -1.788288 | 0 | 0.038418 |
| GOBP_SPECIFICATION_OF_ANIMAL_ORGAN_IDENTITY | 26 | -0.617253 | -1.78812 | 0.001553 | 0.037899 |
| GOBP_POSITIVE_REGULATION_OF_ADAPTIVE_IMMUNE_RESPONSE | 101 | -0.484138 | -1.787943 | 0 | 0.037396 |
| GOBP_CD4_POSITIVE_ALPHA_BETA_T_CELL_ACTIVATION | 96 | -0.489276 | -1.787359 | 0 | 0.037232 |
| GOBP_ANTIGEN_PROCESSING_AND_PRESENTATION_OF_ENDOGENOUS_ANTIGEN | 26 | -0.622157 | -1.78692 | 0.001479 | 0.036827 |
| GOBP_REGULATION_OF_T_CELL_MEDIATED_CYTOTOXICITY | 38 | -0.575309 | -1.782328 | 0.001479 | 0.038716 |
| GOBP_POSITIVE_REGULATION_OF_IMMUNE_EFFECTOR_PROCESS | 204 | -0.445943 | -1.781686 | 0 | 0.038468 |
| GOBP_NEGATIVE_REGULATION_OF_LYMPHOCYTE_DIFFERENTIATION | 51 | -0.539656 | -1.781109 | 0.001391 | 0.038173 |
| GOBP_INTERFERON_GAMMA_PRODUCTION | 105 | -0.484392 | -1.777082 | 0.001302 | 0.039807 |
| GOBP_CELL_KILLING | 151 | -0.460442 | -1.776092 | 0 | 0.039878 |
| GOBP_MATURE_B_CELL_DIFFERENTIATION | 30 | -0.589658 | -1.774325 | 0.003035 | 0.040357 |
| GOBP_REGULATION_OF_T_CELL_MEDIATED_IMMUNITY | 76 | -0.51298 | -1.771074 | 0 | 0.041568 |
| GOBP_POSITIVE_REGULATION_OF_LEUKOCYTE_ADHESION_TO_VASCULAR_ENDOTHELIAL_CELI | 20 | -0.651057 | -1.769572 | 0.003067 | 0.041832 |
| GOBP_LEUKOCYTE_CELL_CELL_ADHESION | 344 | -0.425712 | -1.768753 | 0 | 0.041822 |
| GOBP_RESPONSE_TO_CORTICOSTERONE | 15 | -0.696102 | -1.767527 | 0.001527 | 0.041987 |
| GOBP_PROSTATE_GLAND_DEVELOPMENT | 34 | -0.587408 | -1.767136 | 0 | 0.041664 |
| GOBP_MONONUCLEAR_CELL_DIFFERENTIATION | 381 | -0.423356 | -1.764942 | 0 | 0.042429 |
| GOBP_NEGATIVE_REGULATION_OF_ALPHA_BETA_T_CELL_DIFFERENTIATION | 26 | -0.608212 | -1.763607 | 0.007485 | 0.042678 |
| GOBP_POSITIVE_REGULATION_OF_CD4_POSITIVE_ALPHA_BETA_T_CELL_DIFFERENTIATION | 31 | -0.60342 | -1.759598 | 0.005997 | 0.044588 |
| GOBP_DORSAL_VENTRAL_PATTERN_FORMATION | 55 | -0.532458 | -1.758935 | 0.001406 | 0.044508 |
| GOBP_REGULATORY_T_CELL_DIFFERENTIATION | 34 | -0.579859 | -1.758576 | 0.002886 | 0.044141 |
| GOBP_CRANIAL_NERVE_DEVELOPMENT | 36 | -0.570927 | -1.758407 | 0.002911 | 0.043735 |
| GOBP_LIPOPOLYSACCHARIDE_MEDIATED_SIGNALING_PATHWAY | 56 | -0.524486 | -1.757876 | 0.001427 | 0.04351 |
| GOBP_T_CELL_ACTIVATION_INVOLVED_IN_IMMUNE_RESPONSE | 95 | -0.482134 | -1.754862 | 0 | 0.044995 |
| GOBP_POSITIVE_REGULATION_OF_CELL_ACTIVATION | 384 | -0.418907 | -1.754692 | 0 | 0.04456 |
| GOBP_HEART_FIELD_SPECIFICATION | 15 | -0.690133 | -1.75401 | 0.003205 | 0.044454 |
| GOBP_REGULATION_OF_CELL_CELL_ADHESION | 405 | -0.416318 | -1.751066 | 0 | 0.045499 |
| GOBP_POSITIVE_REGULATION_OF_INTERLEUKIN_10_PRODUCTION | 37 | -0.572267 | -1.75086 | 0 | 0.045106 |
| GOBP_T_CELL_MEDIATED_CYTOTOXICITY | 46 | -0.540984 | -1.750668 | 0 | 0.0447 |
| GOBP_REGULATION_OF_LYMPHOCYTE_ACTIVATION | 477 | -0.411896 | -1.748427 | 0 | 0.04535 |
| GOBP_POSITIVE_REGULATION_OF_LYMPHOCYTE_DIFFERENTIATION | 99 | -0.478994 | -1.747062 | 0 | 0.045792 |
| GOBP_PROSTATE_GLAND_MORPHOGENESIS | 21 | -0.648309 | -1.746698 | 0.00624 | 0.045628 |
| GOBP_MATURE_B_CELL_DIFFERENTIATION_INVOLVED_IN_IMMUNE_RESPONSE | 25 | -0.618138 | -1.745326 | 0 | 0.045945 |
| GOBP_INTERLEUKIN_1_PRODUCTION | 100 | -0.48161 | -1.741247 | 0 | 0.047641 |
| GOBP_POSITIVE_REGULATION_OF_CYTOKINE_PRODUCTION_INVOLVED_IN_INFLAMMATORY_RES | 18 | -0.656175 | -1.740486 | 0.001605 | 0.047731 |
| GOBP_CD8_POSITIVE_ALPHA_BETA_T_CELL_ACTIVATION | 25 | -0.603347 | -1.740252 | 0.009302 | 0.047337 |
| GOBP_POSITIVE_REGULATION_OF_NATURAL_KILLER_CELL_MEDIATED_CYTOTOXICITY | 25 | -0.6184 | -1.738133 | 0 | 0.048177 |
| GOBP_REGULATION_OF_CELL_FATE_COMMITMENT | 26 | -0.604937 | -1.735407 | 0.006126 | 0.049618 |
| GOBP_LEUKOCYTE_PROLIFERATION | 282 | -0.422095 | -1.732874 | 0 | 0.050898 |
| GOBP_POSITIVE_REGULATION_OF_AMYLOID_PRECURSOR_PROTEIN_CATABOLIC_PROCESS | 22 | -0.622267 | -1.732776 | 0 | 0.050441 |
| GOBP_POSITIVE_REGULATION_OF_CYTOKINE_PRODUCTION | 408 | -0.4113 | -1.730509 | 0 | 0.051695 |
| GOBP_NEGATIVE_REGULATION_OF_CD4_POSITIVE_ALPHA_BETA_T_CELL_DIFFERENTIATION | 22 | -0.630514 | -1.72454 | 0.00152 | 0.054969 |
| GOBP_INTERLEUKIN_1_BETA_PRODUCTION | 88 | -0.477675 | -1.724465 | 0.001319 | 0.054486 |
| GOBP_MESONEPHROS_DEVELOPMENT | 76 | -0.485266 | -1.722577 | 0 | 0.055149 |
| GOBP_POSITIVE_REGULATION_OF_INTERLEUKIN_2_PRODUCTION | 30 | -0.568859 | -1.721909 | 0.002967 | 0.05498 |
| GOBP_REGULATION_OF_ICOSANOID_SECRETION | 15 | -0.677561 | -1.721349 | 0.007825 | 0.054804 |
| GOBP_KIDNEY_EPITHELIUM_DEVELOPMENT | 110 | -0.459605 | -1.718315 | 0 | 0.056242 |
| GOBP_TYPE_2_IMMUNE_RESPONSE | 35 | -0.573951 | -1.717747 | 0.00149 | 0.05604 |
| GOBP_T_HELPER_2_CELL_DIFFERENTIATION | 15 | -0.678443 | -1.717319 | 0.004762 | 0.0558 |
| GOBP_REGULATION_OF_LEUKOCYTE_PROLIFERATION | 226 | -0.426112 | -1.716484 | 0 | 0.056071 |
| GOBP_REGULATION_OF_PROTEIN_LOCALIZATION_TO_SYNAPSE | 27 | -0.603077 | -1.716456 | 0.007645 | 0.055579 |
| GOBP_CELL_SURFACE_RECEPTOR_SIGNALING_PATHWAY_INVOLVED_IN_HEART_DEVELOPMEN | 25 | -0.594714 | -1.713619 | 0.00312 | 0.056928 |
| GOBP_RESPONSE_TO_INTERFERON_BETA | 29 | -0.581494 | -1.709636 | 0.003012 | 0.059361 |
| GOBP_REGULATION_OF_CD8_POSITIVE_ALPHA_BETA_T_CELL_ACTIVATION | 19 | -0.640716 | -1.701936 | 0.006431 | 0.064896 |
| GOBP_REGULATION_OF_NITRIC_OXIDE_METABOLIC_PROCESS | 54 | -0.507697 | -1.701795 | 0.002789 | 0.064403 |
| GOBP_OUTFLOW_TRACT_MORPHOGENESIS | 62 | -0.503324 | -1.700939 | 0.002766 | 0.064488 |
| GOBP_PROTEIN_KINASE_C_SIGNALING | 28 | -0.581124 | -1.700285 | 0.007474 | 0.064411 |
| GOBP_NEGATIVE_REGULATION_OF_INTERLEUKIN_12_PRODUCTION | 17 | -0.637992 | -1.697924 | 0.007752 | 0.065923 |
| GOBP_REGULATION_OF_B_CELL_PROLIFERATION | 60 | -0.506336 | -1.69728 | 0 | 0.065995 |
| GOBP_RESPONSE_TO_AUDITORY_STIMULUS | 21 | -0.623217 | -1.693535 | 0.006182 | 0.068196 |
| GOBP_POSITIVE_REGULATION_OF_ACUTE_INFLAMMATORY_RESPONSE | 24 | -0.599207 | -1.692085 | 0.004673 | 0.068607 |
| GOBP_B_CELL_PROLIFERATION | 80 | -0.475848 | -1.691725 | 0 | 0.068414 |
| GOBP_RESPONSE_TO_PROGESTERONE | 42 | -0.528221 | -1.690511 | 0.002789 | 0.068702 |
| GOBP_PROTEIN_LOCALIZATION_TO_SYNAPSE | 73 | -0.480582 | -1.68842 | 0.004087 | 0.069862 |
| GOBP_CELL_DIFFERENTIATION_INVOLVED_IN_KIDNEY_DEVELOPMENT | 49 | -0.515167 | -1.685404 | 0.007153 | 0.071864 |
| GOBP_MYELOID_DENDRITIC_CELL_ACTIVATION | 27 | -0.588312 | -1.684179 | 0.003017 | 0.072407 |
| GOBP_RESPONSE_TO_ISCHEMIA | 48 | -0.519818 | -1.682549 | 0.001412 | 0.073242 |
| GOBP_RESPONSE_TO_TYPE_I_INTERFERON | 86 | -0.468293 | -1.680743 | 0.002577 | 0.074071 |
| GOBP_NOREPINEPHRINE_SECRETION | 15 | -0.675535 | -1.679991 | 0.008039 | 0.074222 |
| GOBP_ORGAN_INDUCTION | 15 | -0.650932 | -1.679907 | 0.011094 | 0.07375 |
| GOBP_REACTIVE_NITROGEN_SPECIES_METABOLIC_PROCESS | 70 | -0.48063 | -1.679232 | 0.001353 | 0.073777 |
| GOBP_GLOMERULUS_DEVELOPMENT | 55 | -0.502495 | -1.677802 | 0.001368 | 0.074478 |
| GOBP_REGULATION_OF_ADAPTIVE_IMMUNE_RESPONSE | 166 | -0.433092 | -1.677622 | 0 | 0.074036 |
| GOBP_POSITIVE_REGULATION_OF_REGULATORY_T_CELL_DIFFERENTIATION | 16 | -0.647942 | -1.675704 | 0.004673 | 0.075106 |
| GOBP_RESPONSE_TO_KETONE | 175 | -0.418618 | -1.675006 | 0.001222 | 0.075242 |
| GOBP_MESONEPHRIC_TUBULE_MORPHOGENESIS | 49 | -0.514538 | -1.674812 | 0.007022 | 0.074874 |
| GOBP_PEPTIDYL_THREONINE_DEPHOSPHORYLATION | 21 | -0.62242 | -1.673459 | 0.010989 | 0.075458 |
| GOBP_NEGATIVE_REGULATION_OF_CYTOKINE_PRODUCTION | 279 | -0.407269 | -1.672381 | 0 | 0.075782 |
| GOBP_DOPAMINERGIC_NEURON_DIFFERENTIATION | 25 | -0.59395 | -1.672307 | 0.005891 | 0.075289 |
| GOBP_NEGATIVE_REGULATION_OF_T_HELPER_CELL_DIFFERENTIATION | 18 | -0.628752 | -1.668714 | 0.008996 | 0.078029 |
| GOBP_REGULATION_OF_EPITHELIAL_TO_MESENCHYMAL_TRANSITION | 87 | -0.460989 | -1.667738 | 0.00266 | 0.078441 |
| GOBP_CYTOKINE_PRODUCTION_INVOLVED_IN_INFLAMMATORY_RESPONSE | 50 | -0.508615 | -1.666029 | 0.005517 | 0.07934 |
| GOBP_NEGATIVE_REGULATION_OF_LEUKOCYTE_CELL_CELL_ADHESION | 124 | -0.441143 | -1.663865 | 0 | 0.080804 |
| GOBP_HEART_FORMATION | 23 | -0.597358 | -1.663511 | 0.00624 | 0.080503 |
| GOBP_POSITIVE_REGULATION_OF_PHAGOCYTOSIS | 61 | -0.495247 | -1.662233 | 0.002813 | 0.081266 |
| GOBP_POSITIVE_REGULATION_OF_LEUKOCYTE_PROLIFERATION | 139 | -0.440927 | -1.661815 | 0.001221 | 0.081109 |
| GOBP_SECONDARY_PALATE_DEVELOPMENT | 20 | -0.616167 | -1.661429 | 0.011006 | 0.080941 |
| GOBP_B_CELL_HOMEOSTASIS | 28 | -0.570805 | -1.660743 | 0.007764 | 0.080917 |
| GOBP_NEGATIVE_REGULATION_OF_IMMUNE_EFFECTOR_PROCESS | 119 | -0.441744 | -1.660173 | 0.001282 | 0.080873 |
| GOBP_NEGATIVE_REGULATION_OF_ALPHA_BETA_T_CELL_ACTIVATION | 40 | -0.527839 | -1.657822 | 0.005952 | 0.082303 |
| GOBP_GLIAL_CELL_MIGRATION | 48 | -0.507407 | -1.656594 | 0.005814 | 0.082916 |
| GOBP_PRODUCTION_OF_MOLECULAR_MEDIATOR_OF_IMMUNE_RESPONSE | 266 | -0.407357 | -1.65372 | 0 | 0.085125 |
| GOBP_NEPHRON_EPITHELIUM_DEVELOPMENT | 89 | -0.459291 | -1.647626 | 0.001321 | 0.090626 |
| GOBP_REGULATION_OF_ANTIGEN_RECEPTOR_MEDIATED_SIGNALING_PATHWAY | 63 | -0.483408 | -1.647371 | 0 | 0.090214 |
| GOBP_POSITIVE_REGULATION_OF_MUSCLE_CELL_APOPTOTIC_PROCESS | 23 | -0.591462 | -1.647138 | 0.01037 | 0.089869 |
| GOBP_PROSTAGLANDIN_TRANSPORT | 16 | -0.639021 | -1.646029 | 0.008224 | 0.090324 |
| GOBP_REGULATION_OF_LIPID_CATABOLIC_PROCESS | 51 | -0.504837 | -1.642475 | 0.00289 | 0.093231 |
| GOBP_NEGATIVE_REGULATION_OF_LIPID_CATABOLIC_PROCESS | 22 | -0.597391 | -1.64216 | 0.005935 | 0.092909 |
| GOBP_MORPHOGENESIS_OF_AN_EPITHELIAL_FOLD | 18 | -0.635714 | -1.642061 | 0.011725 | 0.092484 |
| GOBP_NEGATIVE_REGULATION_OF_LYMPHOCYTE_ACTIVATION | 144 | -0.432207 | -1.641542 | 0 | 0.092437 |
| GOBP_NEGATIVE_REGULATION_OF_LYMPHOCYTE_MEDIATED_IMMUNITY | 43 | -0.509916 | -1.640717 | 0.005722 | 0.092821 |
| GOBP_NEGATIVE_REGULATION_OF_PRODUCTION_OF_MOLECULAR_MEDIATOR_OF_IMMUNE_RE | 37 | -0.538139 | -1.63964 | 0.008415 | 0.093274 |
| GOBP_REGULATION_OF_IMMUNE_EFFECTOR_PROCESS | 433 | -0.387583 | -1.63899 | 0 | 0.093479 |
| GOBP_EPITHELIAL_TUBE_BRANCHING_INVOLVED_IN_LUNG_MORPHOGENESIS | 22 | -0.588577 | -1.638068 | 0.018349 | 0.093688 |

| | | | | | |
|---|---|---|---|---|---|
| GOBP_NEGATIVE_REGULATION_OF_CELL_CYCLE_G2_M_PHASE_TRANSITION | 107 | 0.3694 | 1.572192 | 0 | 0.132486 |
| GOBP_RESPONSE_TO_NERVE_GROWTH_FACTOR | 44 | 0.430409 | 1.571436 | 0.019868 | 0.132415 |
| GOBP_REGULATION_OF_TRANSCRIPTION_INVOLVED_IN_G1_S_TRANSITION_OF_MITOTIC_CELL_ | 35 | 0.452298 | 1.568602 | 0.02349 | 0.134185 |
| GOBP_REGULATION_OF_UBIQUITIN_PROTEIN_LIGASE_ACTIVITY | 22 | 0.527688 | 1.562026 | 0.031161 | 0.1399 |
| GOBP_DNA_DAMAGE_RESPONSE_DETECTION_OF_DNA_DAMAGE | 38 | 0.458852 | 1.561223 | 0.030303 | 0.139841 |
| GOBP_CHROMATIN_SILENCING | 65 | 0.407687 | 1.559609 | 0.011029 | 0.140846 |
| GOBP_MICROTUBULE_ORGANIZING_CENTER_LOCALIZATION | 29 | 0.477825 | 1.558525 | 0.022346 | 0.141123 |
| GOBP_CILIARY_BASAL_BODY_PLASMA_MEMBRANE_DOCKING | 95 | 0.371162 | 1.558489 | 0.012397 | 0.140327 |
| GOBP_IRON_ION_TRANSPORT | 72 | 0.384405 | 1.555084 | 0.007194 | 0.143105 |
| GOBP_POSITIVE_REGULATION_OF_BMP_SIGNALING_PATHWAY | 30 | 0.472818 | 1.554715 | 0.027322 | 0.142645 |
| GOBP_ESTABLISHMENT_OF_PROTEIN_LOCALIZATION_TO_TELOMERE | 18 | 0.543967 | 1.553024 | 0.043597 | 0.143483 |
| GOBP_RESPONSE_TO_MITOCHONDRIAL_DEPOLARISATION | 20 | 0.52344 | 1.55111 | 0.048 | 0.144589 |
| GOBP_CELL_CYCLE_G2_M_PHASE_TRANSITION | 268 | 0.32123 | 1.547643 | 0 | 0.147348 |
| GOBP_ERROR_FREE_TRANSLESION_SYNTHESIS | 22 | 0.50717 | 1.547076 | 0.043353 | 0.147031 |
| GOBP_REGULATION_OF_CENTROSOME_CYCLE | 47 | 0.421524 | 1.542359 | 0.010309 | 0.151363 |
| GOBP_PTERIDINE_CONTAINING_COMPOUND_BIOSYNTHETIC_PROCESS | 15 | 0.554951 | 1.540536 | 0.042895 | 0.152411 |
| GOBP_CELL_CYCLE_CHECKPOINT | 202 | 0.335118 | 1.538787 | 0 | 0.153299 |
| GOBP_NEGATIVE_REGULATION_OF_DNA_REPLICATION | 32 | 0.464345 | 1.536458 | 0.017291 | 0.154735 |
| GOBP_TRNA_TRANSPORT | 37 | 0.441966 | 1.532652 | 0.027682 | 0.158091 |
| GOBP_GLYCOLIPID_BIOSYNTHETIC_PROCESS | 61 | 0.402068 | 1.532178 | 0.010949 | 0.157773 |
| GOBP_FOLIC_ACID_METABOLIC_PROCESS | 20 | 0.517625 | 1.528522 | 0.043836 | 0.160755 |
| GOBP_REGULATION_OF_CELLULAR_AMINO_ACID_METABOLIC_PROCESS | 57 | 0.391887 | 1.527239 | 0.007782 | 0.161236 |
| GOBP_REGULATION_OF_HEMATOPOIETIC_STEM_CELL_DIFFERENTIATION | 72 | 0.382497 | 1.52492 | 0.004016 | 0.163162 |
| GOBP_NUCLEOTIDE_EXCISION_REPAIR_DNA_INCISION | 39 | 0.438481 | 1.524242 | 0.040248 | 0.163117 |
| GOBP_CELLULAR_AMINO_ACID_METABOLIC_PROCESS | 309 | 0.313006 | 1.522644 | 0 | 0.164009 |
| GOBP_POSITIVE_REGULATION_OF_CELL_CYCLE_G2_M_PHASE_TRANSITION | 29 | 0.465015 | 1.522577 | 0.039773 | 0.163208 |
| GOBP_POSITIVE_REGULATION_OF_CHROMOSOME_SEPARATION | 17 | 0.533096 | 1.519004 | 0.050398 | 0.166227 |
| GOBP_NEGATIVE_REGULATION_OF_SINGLE_STRANDED_VIRAL_RNA_REPLICATION_VIA_DOUBL | 16 | 0.540617 | 1.517262 | 0.05291 | 0.167302 |
| GOBP_IRON_ION_TRANSMEMBRANE_TRANSPORT | 19 | 0.513457 | 1.516725 | 0.048387 | 0.167094 |
| GOBP_CELL_AGGREGATION | 16 | 0.530287 | 1.511293 | 0.050746 | 0.172415 |
| GOBP_GENETIC_IMPRINTING | 24 | 0.48246 | 1.508897 | 0.058824 | 0.174125 |
| GOBP_REGULATION_OF_CHOLESTEROL_BIOSYNTHETIC_PROCESS | 43 | 0.421504 | 1.506253 | 0.018116 | 0.176189 |
| GOBP_EYE_PHOTORECEPTOR_CELL_DEVELOPMENT | 23 | 0.495843 | 1.505622 | 0.042945 | 0.176023 |
| GOBP_COENZYME_A_METABOLIC_PROCESS | 15 | 0.551734 | 1.504045 | 0.063776 | 0.177054 |
| GOBP_TRANSFERRIN_TRANSPORT | 33 | 0.445797 | 1.501739 | 0.038835 | 0.178826 |
| GOBP_PROTEIN_LOCALIZATION_TO_CHROMOSOME_TELOMERIC_REGION | 30 | 0.45248 | 1.498125 | 0.036364 | 0.182064 |
| GOBP_MITOTIC_SPINDLE_ASSEMBLY | 64 | 0.389229 | 1.49663 | 0.023529 | 0.18283 |
| GOBP_MICROTUBULE_NUCLEATION | 30 | 0.457048 | 1.494377 | 0.046729 | 0.184795 |
| GOBP_DICARBOXYLIC_ACID_METABOLIC_PROCESS | 88 | 0.362659 | 1.49391 | 0.003937 | 0.184518 |
| GOBP_RNA_CAPPING | 34 | 0.432936 | 1.493164 | 0.030387 | 0.184544 |
| GOBP_RETINA_VASCULATURE_DEVELOPMENT_IN_CAMERA_TYPE_EYE | 16 | 0.529522 | 1.486898 | 0.07027 | 0.191808 |
| GOBP_COLLAGEN_FIBRIL_ORGANIZATION | 50 | 0.395644 | 1.481695 | 0.006757 | 0.197481 |
| GOBP_TRNA_5_END_PROCESSING | 16 | 0.524907 | 1.47091 | 0.055263 | 0.21038 |
| GOBP_NEGATIVE_REGULATION_OF_GENE_SILENCING | 31 | 0.440001 | 1.46754 | 0.040115 | 0.21401 |
| GOBP_GENE_SILENCING | 325 | 0.297799 | 1.46749 | 0 | 0.213092 |
| GOBP_MEMBRANE_DISASSEMBLY | 15 | 0.541428 | 1.464501 | 0.084507 | 0.216398 |
| GOBP_PROTEIN_KINASE_A_SIGNALING | 31 | 0.440248 | 1.464052 | 0.056548 | 0.215969 |
| GOBP_NEGATIVE_REGULATION_OF_VASCULAR_PERMEABILITY | 16 | 0.511625 | 1.463855 | 0.059659 | 0.215207 |
| GOBP_NEURON_PROJECTION_EXTENSION_INVOLVED_IN_NEURON_PROJECTION_GUIDANCE | 35 | 0.427097 | 1.45963 | 0.051724 | 0.220209 |
| GOBP_POSTREPLICATION_REPAIR | 52 | 0.386459 | 1.458642 | 0.026144 | 0.220499 |
| GOBP_CELLULAR_OXIDANT_DETOXIFICATION | 90 | 0.357214 | 1.458147 | 0.007968 | 0.220204 |
| GOBP_NEGATIVE_REGULATION_OF_MUSCLE_CELL_APOPTOTIC_PROCESS | 32 | 0.437477 | 1.45747 | 0.043228 | 0.220142 |
| GOBP_PROTEIN_REFOLDING | 23 | 0.475319 | 1.45486 | 0.088889 | 0.222818 |
| GOBP_EMBRYONIC_HEMOPOIESIS | 22 | 0.481702 | 1.454582 | 0.058309 | 0.222194 |
| GOBP_SCHWANN_CELL_DEVELOPMENT | 28 | 0.431813 | 1.453827 | 0.058824 | 0.222185 |
| GOBP_NEGATIVE_REGULATION_OF_GLUCONEOGENESIS | 16 | 0.50697 | 1.450043 | 0.075949 | 0.226703 |
| GOBP_CAMERA_TYPE_EYE_PHOTORECEPTOR_CELL_DIFFERENTIATION | 21 | 0.481664 | 1.4489 | 0.059946 | 0.227388 |
| GOBP_CELLULAR_RESPONSE_TO_TOXIC_SUBSTANCE | 107 | 0.339891 | 1.447531 | 0.021008 | 0.228353 |
| GOBP_CELLULAR_RESPONSE_TO_DEXAMETHASONE_STIMULUS | 28 | 0.457693 | 1.445846 | 0.042683 | 0.229813 |
| GOBP_STEROID_HORMONE_SECRETION | 16 | 0.496684 | 1.441441 | 0.069364 | 0.235554 |
| GOBP_HEPATOCYTE_GROWTH_FACTOR_RECEPTOR_SIGNALING_PATHWAY | 15 | 0.529784 | 1.440441 | 0.057637 | 0.235916 |
| GOBP_SPINDLE_ASSEMBLY | 113 | 0.336998 | 1.436783 | 0.0131 | 0.24042 |
| GOBP_DNA_DEPENDENT_DNA_REPLICATION_MAINTENANCE_OF_FIDELITY | 48 | 0.387239 | 1.435904 | 0.030508 | 0.240555 |
| GOBP_ALPHA_AMINO_ACID_METABOLIC_PROCESS | 174 | 0.317419 | 1.435887 | 0 | 0.239528 |
| GOBP_POSITIVE_REGULATION_OF_MITOCHONDRIAL_FISSION | 17 | 0.496301 | 1.432689 | 0.07438 | 0.243371 |
| GOBP_IRON_SULFUR_CLUSTER_ASSEMBLY | 24 | 0.464715 | 1.431728 | 0.050595 | 0.24394 |
| GOBP_RESPONSE_TO_ACID_CHEMICAL | 107 | 0.327696 | 1.431195 | 0.008511 | 0.243635 |
| GOBP_TELOMERE_MAINTENANCE_VIA_RECOMBINATION | 15 | 0.524757 | 1.430319 | 0.093995 | 0.243945 |
| GOBP_RNA_MODIFICATION | 166 | 0.313594 | 1.427986 | 0 | 0.246608 |
| GOBP_ESTABLISHMENT_OF_MITOTIC_SPINDLE_ORIENTATION | 26 | 0.45009 | 1.426807 | 0.052198 | 0.247166 |
| GOBP_NEGATIVE_REGULATION_OF_TORC1_SIGNALING | 17 | 0.485356 | 1.423889 | 0.05698 | 0.250423 |
| GOBP_G0_TO_G1_TRANSITION | 44 | 0.396137 | 1.422678 | 0.041801 | 0.251418 |
| GOBP_INTEGRIN_ACTIVATION | 24 | 0.454078 | 1.419403 | 0.063536 | 0.255693 |
| GOBP_POSITIVE_REGULATION_OF_GENE_EXPRESSION_EPIGENETIC | 55 | 0.367183 | 1.416634 | 0.026756 | 0.258979 |
| GOBP_ERYTHROCYTE_HOMEOSTASIS | 121 | 0.326552 | 1.412567 | 0.009804 | 0.264281 |
| GOBP_NEGATIVE_REGULATION_OF_PROTEIN_LOCALIZATION_TO_NUCLEUS | 30 | 0.422768 | 1.411204 | 0.061584 | 0.265465 |
| GOBP_POSITIVE_REGULATION_OF_METAPHASE_ANAPHASE_TRANSITION_OF_CELL_CYCLE | 15 | 0.51149 | 1.410831 | 0.060847 | 0.265041 |
| GOBP_POSITIVE_REGULATION_OF_CELL_CYCLE_PHASE_TRANSITION | 93 | 0.343075 | 1.410012 | 0.020747 | 0.26534 |
| GOBP_NEGATIVE_REGULATION_OF_CELL_CYCLE_PHASE_TRANSITION | 240 | 0.29682 | 1.405707 | 0.007937 | 0.271283 |
| GOBP_REGULATION_OF_SPINDLE_CHECKPOINT | 15 | 0.506698 | 1.405294 | 0.073569 | 0.270857 |
| GOBP_REGULATION_OF_CELL_CYCLE_PHASE_TRANSITION | 431 | 0.280687 | 1.402896 | 0 | 0.273736 |
| GOBP_NUCLEOTIDE_EXCISION_REPAIR_PREINCISION_COMPLEX_STABILIZATION | 21 | 0.464782 | 1.401804 | 0.067692 | 0.274595 |
| GOBP_MEIOTIC_CHROMOSOME_SEPARATION | 24 | 0.451016 | 1.400085 | 0.057637 | 0.276442 |
| GOBP_REGULATION_OF_CELL_CYCLE_CHECKPOINT | 30 | 0.424063 | 1.398718 | 0.059701 | 0.277469 |
| GOBP_REGULATION_OF_PROTEIN_KINASE_A_SIGNALING | 20 | 0.454731 | 1.397375 | 0.103245 | 0.278647 |
| GOBP_L_ALPHA_AMINO_ACID_TRANSMEMBRANE_TRANSPORT | 56 | 0.372431 | 1.394467 | 0.036496 | 0.282249 |
| GOBP_REGULATION_OF_NUCLEAR_DIVISION | 126 | 0.32116 | 1.391703 | 0.004975 | 0.285851 |
| GOBP_TRICARBOXYLIC_ACID_CYCLE | 33 | 0.413075 | 1.390879 | 0.056291 | 0.286279 |
| GOBP_HYDROGEN_PEROXIDE_METABOLIC_PROCESS | 51 | 0.36582 | 1.389707 | 0.041379 | 0.287173 |
| GOBP_OLIGOSACCHARIDE_LIPID_INTERMEDIATE_BIOSYNTHETIC_PROCESS | 21 | 0.456848 | 1.387494 | 0.091429 | 0.289965 |
| GOBP_ASPARTATE_FAMILY_AMINO_ACID_CATABOLIC_PROCESS | 22 | 0.454068 | 1.384787 | 0.110778 | 0.293667 |
| GOBP_CENTRIOLE_ASSEMBLY | 41 | 0.389448 | 1.382807 | 0.045603 | 0.296106 |
| GOBP_ASPARTATE_FAMILY_AMINO_ACID_BIOSYNTHETIC_PROCESS | 20 | 0.472113 | 1.382106 | 0.085227 | 0.296239 |
| GOBP_FEMALE_MEIOTIC_NUCLEAR_DIVISION | 25 | 0.441711 | 1.379693 | 0.081461 | 0.299465 |
| GOBP_FATTY_ACYL_COA_METABOLIC_PROCESS | 41 | 0.39823 | 1.37719 | 0.039344 | 0.302865 |
| GOBP_TRANSLESION_SYNTHESIS | 42 | 0.379772 | 1.372213 | 0.047619 | 0.310869 |
| GOBP_PHYSIOLOGICAL_CARDIAC_MUSCLE_HYPERTROPHY | 22 | 0.448496 | 1.370934 | 0.082153 | 0.311998 |
| GOBP_MALE_MEIOTIC_NUCLEAR_DIVISION | 38 | 0.396028 | 1.367568 | 0.070175 | 0.317292 |
| GOBP_NEURAL_RETINA_DEVELOPMENT | 53 | 0.369738 | 1.367196 | 0.05802 | 0.316835 |
| GOBP_NEGATIVE_REGULATION_OF_RHO_PROTEIN_SIGNAL_TRANSDUCTION | 20 | 0.456958 | 1.366813 | 0.106145 | 0.316328 |
| GOBP_CLATHRIN_DEPENDENT_ENDOCYTOSIS | 45 | 0.3808 | 1.357928 | 0.055556 | 0.331832 |
| GOBP_PROTEIN_LOCALIZATION_TO_MICROTUBULE_ORGANIZING_CENTER | 35 | 0.387494 | 1.357556 | 0.064103 | 0.331292 |
| GOBP_SNRNA_PROCESSING | 26 | 0.418568 | 1.35715 | 0.080495 | 0.330803 |
| GOBP_CELLULAR_AMINO_ACID_BIOSYNTHETIC_PROCESS | 71 | 0.334448 | 1.356916 | 0.028 | 0.330098 |
| GOBP_CELLULAR_PROTEIN_COMPLEX_DISASSEMBLY | 216 | 0.296847 | 1.354816 | 0 | 0.332921 |
| GOBP_DNA_INTEGRITY_CHECKPOINT | 152 | 0.306971 | 1.354652 | 0.011111 | 0.331977 |
| GOBP_REGULATION_OF_CHOLESTEROL_METABOLIC_PROCESS | 53 | 0.365294 | 1.354475 | 0.068404 | 0.331066 |
| GOBP_DNA_METHYLATION_INVOLVED_IN_GAMETE_GENERATION | 17 | 0.486794 | 1.353877 | 0.128947 | 0.330971 |
| GOBP_ENDONUCLEOLYTIC_CLEAVAGE_INVOLVED_IN_RRNA_PROCESSING | 15 | 0.48419 | 1.351942 | 0.127479 | 0.333661 |
| GOBP_NEUTRAL_AMINO_ACID_TRANSPORT | 37 | 0.384304 | 1.351705 | 0.070707 | 0.332842 |
| GOBP_TRANSITION_METAL_ION_TRANSPORT | 115 | 0.30883 | 1.351696 | 0.037657 | 0.331652 |
| GOBP_DNA_SYNTHESIS_INVOLVED_IN_DNA_REPAIR | 51 | 0.359226 | 1.349661 | 0.059233 | 0.334458 |
| GOBP_EYE_PHOTORECEPTOR_CELL_DIFFERENTIATION | 34 | 0.398797 | 1.347249 | 0.086053 | 0.337825 |
| GOBP_ASPARTATE_FAMILY_AMINO_ACID_METABOLIC_PROCESS | 48 | 0.37039 | 1.346789 | 0.067742 | 0.337557 |
| GOBP_REGULATION_OF_CHROMOSOME_ORGANIZATION | 262 | 0.286787 | 1.346137 | 0.014925 | 0.33764 |
| GOBP_MITOCHONDRIAL_DEPOLARIZATION | 21 | 0.451871 | 1.343455 | 0.093664 | 0.341678 |
| GOBP_CELLULAR_RESPIRATION | 181 | 0.294701 | 1.342367 | 0 | 0.342551 |
| GOBP_SPINDLE_LOCALIZATION | 47 | 0.369508 | 1.341247 | 0.073333 | 0.343484 |
| GOBP_DIOL_BIOSYNTHETIC_PROCESS | 22 | 0.424922 | 1.340587 | 0.100295 | 0.343686 |
| GOBP_RESPONSE_TO_PH | 34 | 0.392109 | 1.338759 | 0.072508 | 0.346268 |
| GOBP_NEGATIVE_REGULATION_OF_PEPTIDYL_LYSINE_ACETYLATION | 18 | 0.462154 | 1.337263 | 0.114058 | 0.347983 |
| GOBP_BASE_CONVERSION_OR_SUBSTITUTION_EDITING | 17 | 0.475824 | 1.336938 | 0.116667 | 0.347477 |
| GOBP_NEGATIVE_REGULATION_OF_CELL_CYCLE_PROCESS | 321 | 0.275419 | 1.334604 | 0.010101 | 0.351044 |
| GOBP_NEGATIVE_REGULATION_OF_CELL_DIVISION | 15 | 0.47784 | 1.331324 | 0.144385 | 0.356512 |
| GOBP_POSITIVE_REGULATION_OF_FOCAL_ADHESION_ASSEMBLY | 24 | 0.411963 | 1.330086 | 0.102564 | 0.357815 |
| GOBP_NEUROTRANSMITTER_METABOLIC_PROCESS | 20 | 0.446272 | 1.327699 | 0.117486 | 0.361474 |
| GOBP_NEGATIVE_REGULATION_OF_PROTEIN_ACETYLATION | 20 | 0.448355 | 1.327341 | 0.143713 | 0.361042 |
| GOBP_GLUTAMINE_FAMILY_AMINO_ACID_METABOLIC_PROCESS | 67 | 0.331986 | 1.327161 | 0.035587 | 0.360192 |
| GOBP_REGULATION_OF_CENTRIOLE_REPLICATION | 20 | 0.439198 | 1.323632 | 0.117166 | 0.366299 |
| GOBP_TRNA_MODIFICATION | 89 | 0.327738 | 1.32358 | 0.070833 | 0.365171 |
| GOBP_DNA_DOUBLE_STRAND_BREAK_PROCESSING | 23 | 0.423787 | 1.323421 | 0.115044 | 0.364205 |
| GOBP_REGULATION_OF_DNA_REPLICATION | 105 | 0.310439 | 1.322845 | 0.031746 | 0.36424 |
| GOBP_ALPHA_AMINO_ACID_BIOSYNTHETIC_PROCESS | 63 | 0.341744 | 1.320414 | 0.059701 | 0.367813 |
| GOBP_SULFUR_COMPOUND_TRANSPORT | 46 | 0.364869 | 1.317696 | 0.068729 | 0.372288 |
| GOBP_PREASSEMBLY_OF_GPI_ANCHOR_IN_ER_MEMBRANE | 16 | 0.474956 | 1.316585 | 0.140401 | 0.373402 |
| GOBP_ESTABLISHMENT_OF_SPINDLE_ORIENTATION | 30 | 0.406911 | 1.314851 | 0.097179 | 0.375786 |
| GOBP_AMINO_ACID_TRANSMEMBRANE_TRANSPORT | 81 | 0.318312 | 1.31409 | 0.035433 | 0.376165 |
| GOBP_NUCLEOTIDE_EXCISION_REPAIR_PREINCISION_COMPLEX_ASSEMBLY | 29 | 0.400906 | 1.31243 | 0.076023 | 0.37863 |
| GOBP_POSITIVE_REGULATION_OF_CELL_CYCLE_PROCESS | 269 | 0.270455 | 1.311829 | 0 | 0.37871 |
| GOBP_NCRNA_METABOLIC_PROCESS | 474 | 0.261993 | 1.310041 | 0 | 0.381296 |
| GOBP_RESPONSE_TO_GLUCAGON | 34 | 0.386909 | 1.307384 | 0.102804 | 0.38579 |
| GOBP_MITOCHONDRIAL_TRANSMEMBRANE_TRANSPORT | 105 | 0.309504 | 1.306549 | 0.049793 | 0.386461 |
| GOBP_SOMATIC_DIVERSIFICATION_OF_IMMUNE_RECEPTORS_VIA_SOMATIC_MUTATION | 16 | 0.466345 | 1.304999 | 0.156566 | 0.388643 |
| GOBP_VASCULAR_ENDOTHELIAL_GROWTH_FACTOR_SIGNALING_PATHWAY | 42 | 0.368457 | 1.302712 | 0.068627 | 0.392557 |
| GOBP_PLASMA_MEMBRANE_FUSION | 17 | 0.457205 | 1.302137 | 0.146132 | 0.392448 |
| GOBP_CELLULAR_RESPONSE_TO_ACID_CHEMICAL | 65 | 0.332445 | 1.30173 | 0.071161 | 0.392019 |
| GOBP_MATERNAL_PLACENTA_DEVELOPMENT | 27 | 0.410115 | 1.301649 | 0.152439 | 0.390964 |
| GOBP_PROTEIN_AUTOPROCESSING | 21 | 0.433201 | 1.299787 | 0.125348 | 0.393861 |
| GOBP_MISMATCH_REPAIR | 36 | 0.385838 | 1.29667 | 0.116352 | 0.399552 |
| GOBP_CELLULAR_RESPONSE_TO_PH | 20 | 0.444821 | 1.295543 | 0.128655 | 0.400922 |
| GOBP_NEGATIVE_REGULATION_OF_DNA_METABOLIC_PROCESS | 115 | 0.304844 | 1.293442 | 0.0553 | 0.404274 |
| GOBP_RNA_5_END_PROCESSING | 23 | 0.419177 | 1.293346 | 0.137536 | 0.403218 |
| GOBP_REGULATION_OF_CYTOKINESIS | 81 | 0.316388 | 1.291902 | 0.052209 | 0.405307 |
| GOBP_INTERLEUKIN_7_MEDIATED_SIGNALING_PATHWAY | 27 | 0.408718 | 1.290932 | 0.127329 | 0.406026 |
| GOBP_TRANSCRIPTION_ELONGATION_FROM_RNA_POLYMERASE_I_PROMOTER | 30 | 0.388609 | 1.290578 | 0.135542 | 0.405587 |
| GOBP_NUCLEOLAR_LARGE_RRNA_TRANSCRIPTION_BY_RNA_POLYMERASE_I | 17 | 0.442675 | 1.289329 | 0.141304 | 0.407111 |
| GOBP_REGULATION_OF_HEMATOPOIETIC_PROGENITOR_CELL_DIFFERENTIATION | 87 | 0.320308 | 1.2884 | 0.070175 | 0.407896 |
| GOBP_SNRNA_METABOLIC_PROCESS | 37 | 0.370108 | 1.2857 | 0.117825 | 0.412982 |
| GOBP_CHONDROCYTE_PROLIFERATION | 18 | 0.449 | 1.285246 | 0.156156 | 0.412776 |
| GOBP_ATRIAL_SEPTUM_DEVELOPMENT | 18 | 0.432292 | 1.283317 | 0.164265 | 0.415719 |
| GOBP_PROTEIN_DEPOLYMERIZATION | 108 | 0.299388 | 1.281409 | 0.070175 | 0.418707 |
| GOBP_NEGATIVE_REGULATION_OF_CHROMATIN_ORGANIZATION | 57 | 0.337435 | 1.277403 | 0.10989 | 0.426888 |
| GOBP_CELLULAR_CARBOHYDRATE_BIOSYNTHETIC_PROCESS | 77 | 0.313119 | 1.276891 | 0.068592 | 0.426777 |
| GOBP_ESTABLISHMENT_OF_PIGMENT_GRANULE_LOCALIZATION | 24 | 0.401569 | 1.275631 | 0.14377 | 0.428453 |
| GOBP_URETER_DEVELOPMENT | 15 | 0.459827 | 1.275588 | 0.176 | 0.427243 |
| GOBP_WALKING_BEHAVIOR | 20 | 0.422811 | 1.275479 | 0.169355 | 0.426194 |
| GOBP_MYELOID_CELL_HOMEOSTASIS | 145 | 0.284539 | 1.2729 | 0.034483 | 0.431138 |
| GOBP_INTERMEDIATE_FILAMENT_BASED_PROCESS | 44 | 0.355549 | 1.270888 | 0.096463 | 0.434411 |

| | | | | | |
|---|---|---|---|---|---|
| GOBP_NEGATIVE_REGULATION_OF_INTERLEUKIN_6_PRODUCTION | 42 | -0.518551 | -1.637902 | 0.002829 | 0.093285 |
| GOBP_P38MAPK_CASCADE | 41 | -0.519764 | -1.636399 | 0.007496 | 0.094326 |
| GOBP_POSITIVE_REGULATION_OF_LEUKOCYTE_APOPTOTIC_PROCESS | 23 | -0.588203 | -1.635709 | 0.010753 | 0.094394 |
| GOBP_ICOSANOID_TRANSPORT | 40 | -0.523315 | -1.635302 | 0.008547 | 0.094265 |
| GOBP_POSITIVE_REGULATION_OF_CHEMOKINE_PRODUCTION | 56 | -0.488122 | -1.635063 | 0.00569 | 0.09398 |
| GOBP_RESPONSE_TO_DSRNA | 39 | -0.517666 | -1.633906 | 0.012894 | 0.094714 |
| GOBP_REGULATION_OF_SIGNALING_RECEPTOR_ACTIVITY | 155 | -0.421239 | -1.633801 | 0 | 0.094263 |
| GOBP_RUFFLE_ORGANIZATION | 52 | -0.499585 | -1.633705 | 0.008451 | 0.093801 |
| GOBP_POSITIVE_REGULATION_OF_FATTY_ACID_BIOSYNTHETIC_PROCESS | 17 | -0.631308 | -1.633354 | 0.013534 | 0.093591 |
| GOBP_POSITIVE_REGULATION_OF_SIGNALING_RECEPTOR_ACTIVITY | 41 | -0.513001 | -1.632721 | 0.005908 | 0.093706 |
| GOBP_POSITIVE_REGULATION_OF_EPITHELIAL_TO_MESENCHYMAL_TRANSITION | 45 | -0.506844 | -1.632261 | 0.002894 | 0.093694 |
| GOBP_NEGATIVE_REGULATION_OF_B_CELL_ACTIVATION | 33 | -0.53408 | -1.632177 | 0.007353 | 0.093246 |
| GOBP_POSITIVE_REGULATION_OF_CYTOKINE_PRODUCTION_INVOLVED_IN_IMMUNE_RESPONSE | 52 | -0.503274 | -1.632103 | 0.011429 | 0.09277 |
| GOBP_KILLING_OF_CELLS_OF_OTHER_ORGANISM | 45 | -0.502847 | -1.631574 | 0.008811 | 0.092742 |
| GOBP_VITAMIN_D_METABOLIC_PROCESS | 18 | -0.631365 | -1.629922 | 0.01037 | 0.093855 |
| GOBP_POSITIVE_REGULATION_OF_EPIDERMAL_CELL_DIFFERENTIATION | 23 | -0.577054 | -1.628958 | 0.007429 | 0.094369 |
| GOBP_RESPONSE_TO_MINERALOCORTICOID | 26 | -0.557977 | -1.626865 | 0.009346 | 0.096074 |
| GOBP_T_CELL_PROLIFERATION | 188 | -0.411379 | -1.626798 | 0 | 0.095643 |
| GOBP_T_CELL_DIFFERENTIATION_INVOLVED_IN_IMMUNE_RESPONSE | 72 | -0.467773 | -1.62572 | 0.006623 | 0.09618 |
| GOBP_POSITIVE_REGULATION_OF_REACTIVE_OXYGEN_SPECIES_METABOLIC_PROCESS | 92 | -0.448504 | -1.62537 | 0.002639 | 0.096037 |
| GOBP_RESPONSE_TO_PROSTAGLANDIN_E | 22 | -0.583363 | -1.624654 | 0.006116 | 0.09624 |
| GOBP_RESPONSE_TO_CHEMOKINE | 88 | -0.449825 | -1.624318 | 0.004 | 0.096022 |
| GOBP_CELLULAR_RESPONSE_TO_LIPID | 496 | -0.382048 | -1.623406 | 0 | 0.09648 |
| GOBP_REGULATION_OF_THE_FORCE_OF_HEART_CONTRACTION | 23 | -0.581291 | -1.621924 | 0.007463 | 0.097452 |
| GOBP_EPITHELIAL_CELL_DIFFERENTIATION_INVOLVED_IN_KIDNEY_DEVELOPMENT | 41 | -0.519205 | -1.621175 | 0.005848 | 0.097768 |
| GOBP_ICOSANOID_SECRETION | 32 | -0.545217 | -1.620729 | 0.005988 | 0.097696 |
| GOBP_NEGATIVE_REGULATION_OF_INFLAMMATORY_RESPONSE | 118 | -0.433336 | -1.62018 | 0.001292 | 0.097758 |
| GOBP_MACROPHAGE_ACTIVATION | 92 | -0.450994 | -1.61964 | 0 | 0.097741 |
| GOBP_POSITIVE_REGULATION_OF_T_HELPER_CELL_DIFFERENTIATION | 21 | -0.591699 | -1.618909 | 0.009464 | 0.098069 |
| GOBP_NEGATIVE_REGULATION_OF_LIPID_LOCALIZATION | 45 | -0.5097 | -1.618871 | 0.008596 | 0.097586 |
| GOBP_POSITIVE_REGULATION_OF_EXTRINSIC_APOPTOTIC_SIGNALING_PATHWAY | 46 | -0.499665 | -1.61552 | 0.011611 | 0.100632 |
| GOBP_NEGATIVE_REGULATION_OF_CELL_ACTIVATION | 184 | -0.407216 | -1.614858 | 0 | 0.100852 |
| GOBP_INTERLEUKIN_6_MEDIATED_SIGNALING_PATHWAY | 22 | -0.585395 | -1.614323 | 0.009524 | 0.100878 |
| GOBP_LEUKOCYTE_DIFFERENTIATION | 486 | -0.379465 | -1.614053 | 0 | 0.100669 |
| GOBP_POSITIVE_REGULATION_OF_NITRIC_OXIDE_METABOLIC_PROCESS | 35 | -0.528983 | -1.610155 | 0.012784 | 0.104308 |
| GOBP_NEGATIVE_REGULATION_OF_INTERLEUKIN_1_PRODUCTION | 31 | -0.542491 | -1.609818 | 0.014556 | 0.104226 |
| GOBP_NEGATIVE_REGULATION_OF_IMMUNE_SYSTEM_PROCESS | 367 | -0.38235 | -1.609262 | 0 | 0.104348 |
| GOBP_CHEMOKINE_PRODUCTION | 88 | -0.449987 | -1.608338 | 0.005208 | 0.104797 |
| GOBP_MACROPHAGE_MIGRATION | 48 | -0.502593 | -1.607577 | 0.00561 | 0.105111 |
| GOBP_CATECHOLAMINE_SECRETION | 43 | -0.509334 | -1.606598 | 0.005874 | 0.105539 |
| GOBP_NEGATIVE_REGULATION_OF_AMINE_TRANSPORT | 21 | -0.593037 | -1.605473 | 0.013889 | 0.106169 |
| GOBP_NEGATIVE_REGULATION_OF_INTERLEUKIN_1_BETA_PRODUCTION | 26 | -0.546631 | -1.604514 | 0.023006 | 0.106634 |
| GOBP_REGULATION_OF_LEUKOCYTE_DIFFERENTIATION | 262 | -0.395074 | -1.602563 | 0 | 0.108185 |
| GOBP_LYMPHOCYTE_MEDIATED_IMMUNITY | 334 | -0.385737 | -1.600455 | 0 | 0.109964 |
| GOBP_NEUROINFLAMMATORY_RESPONSE | 33 | -0.527658 | -1.600107 | 0.008708 | 0.10979 |
| GOBP_FAT_SOLUBLE_VITAMIN_METABOLIC_PROCESS | 37 | -0.515772 | -1.598032 | 0.008392 | 0.111566 |
| GOBP_NEGATIVE_REGULATION_OF_HEMOPOIESIS | 98 | -0.440259 | -1.597237 | 0.002577 | 0.111841 |
| GOBP_POSITIVE_REGULATION_OF_AMYLOID_BETA_FORMATION | 17 | -0.613818 | -1.595964 | 0.029595 | 0.112705 |
| GOBP_REGULATION_OF_RECEPTOR_SIGNALING_PATHWAY_VIA_STAT | 80 | -0.453531 | -1.595318 | 0.007003 | 0.112878 |
| GOBP_POSITIVE_REGULATION_OF_ANIMAL_ORGAN_MORPHOGENESIS | 24 | -0.559396 | -1.594507 | 0.015408 | 0.113306 |
| GOBP_RESPONSE_TO_COLD | 44 | -0.499434 | -1.593555 | 0.002853 | 0.113864 |
| GOBP_POSITIVE_REGULATION_OF_EPIDERMIS_DEVELOPMENT | 28 | -0.550892 | -1.593509 | 0.011958 | 0.113414 |
| GOBP_POSITIVE_REGULATION_OF_HEMOPOIESIS | 146 | -0.412461 | -1.592533 | 0.001258 | 0.114011 |
| GOBP_LYMPHOCYTE_ACTIVATION_INVOLVED_IN_IMMUNE_RESPONSE | 171 | -0.409475 | -1.591855 | 0 | 0.114184 |
| GOBP_POSITIVE_REGULATION_OF_INTERLEUKIN_1_PRODUCTION | 57 | -0.474946 | -1.589086 | 0.006821 | 0.116953 |
| GOBP_NEGATIVE_REGULATION_OF_LEUKOCYTE_MEDIATED_IMMUNITY | 50 | -0.479735 | -1.588537 | 0.011852 | 0.116998 |
| GOBP_AORTA_DEVELOPMENT | 48 | -0.49586 | -1.587827 | 0.012894 | 0.117293 |
| GOBP_POSITIVE_REGULATION_OF_ERBB_SIGNALING_PATHWAY | 40 | -0.505785 | -1.585442 | 0.01122 | 0.11959 |
| GOBP_NEGATIVE_REGULATION_OF_CD4_POSITIVE_ALPHA_BETA_T_CELL_ACTIVATION | 31 | -0.522658 | -1.584264 | 0.01497 | 0.120492 |
| GOBP_PARAXIAL_MESODERM_DEVELOPMENT | 16 | -0.61834 | -1.582204 | 0.023148 | 0.12257 |
| GOBP_WNT_SIGNALING_PATHWAY_CALCIUM_MODULATING_PATHWAY | 38 | -0.505383 | -1.580274 | 0.016492 | 0.124485 |
| GOBP_RUFFLE_ASSEMBLY | 41 | -0.505239 | -1.579687 | 0.012857 | 0.124663 |
| GOBP_TUMOR_NECROSIS_FACTOR_SUPERFAMILY_CYTOKINE_PRODUCTION | 140 | -0.412802 | -1.577305 | 0 | 0.126891 |
| GOBP_REGULATION_OF_LEUKOCYTE_ADHESION_TO_VASCULAR_ENDOTHELIAL_CELL | 27 | -0.546321 | -1.577135 | 0.025148 | 0.126553 |
| GOBP_POSITIVE_REGULATION_OF_FATTY_ACID_METABOLIC_PROCESS | 31 | -0.529107 | -1.575571 | 0.013333 | 0.127828 |
| GOBP_POSITIVE_REGULATION_OF_MAST_CELL_ACTIVATION | 18 | -0.604338 | -1.575149 | 0.026357 | 0.127774 |
| GOBP_TYROSINE_PHOSPHORYLATION_OF_STAT_PROTEIN | 73 | -0.447941 | -1.574286 | 0.006702 | 0.128285 |
| GOBP_REGULATION_OF_ANTIGEN_PROCESSING_AND_PRESENTATION | 19 | -0.594611 | -1.573067 | 0.021311 | 0.129183 |
| GOBP_REGULATION_OF_B_CELL_ACTIVATION | 180 | -0.402394 | -1.571715 | 0 | 0.130328 |
| GOBP_RENAL_SYSTEM_VASCULATURE_DEVELOPMENT | 22 | -0.571035 | -1.570489 | 0.011527 | 0.131361 |
| GOBP_NEPHRON_DEVELOPMENT | 118 | -0.419625 | -1.569421 | 0.002571 | 0.132087 |
| GOBP_REGULATION_OF_VASOCONSTRICTION | 52 | -0.484157 | -1.568128 | 0.010264 | 0.133041 |
| GOBP_REGULATION_OF_MACROPHAGE_ACTIVATION | 52 | -0.475037 | -1.567555 | 0.009859 | 0.133227 |
| GOBP_REGULATION_OF_TYPE_2_IMMUNE_RESPONSE | 29 | -0.538114 | -1.567298 | 0.015152 | 0.133009 |
| GOBP_POSITIVE_REGULATION_OF_REACTIVE_OXYGEN_SPECIES_BIOSYNTHETIC_PROCESS | 48 | -0.480085 | -1.567145 | 0.010401 | 0.132618 |
| GOBP_REGULATION_OF_RECEPTOR_LOCALIZATION_TO_SYNAPSE | 18 | -0.603425 | -1.566787 | 0.027735 | 0.13257 |
| GOBP_POSITIVE_REGULATION_OF_TUMOR_NECROSIS_FACTOR_SUPERFAMILY_CYTOKINE_PRO | 84 | -0.445772 | -1.565884 | 0.004011 | 0.133256 |
| GOBP_NEGATIVE_REGULATION_OF_LEUKOCYTE_CHEMOTAXIS | 17 | -0.604385 | -1.564736 | 0.028614 | 0.134121 |
| GOBP_TOLERANCE_INDUCTION | 28 | -0.536909 | -1.5645 | 0.021841 | 0.13379 |
| GOBP_RESPONSE_TO_STIMULUS_INVOLVED_IN_REGULATION_OF_MUSCLE_ADAPTATION | 15 | -0.630296 | -1.563172 | 0.017742 | 0.134957 |
| GOBP_RESPONSE_TO_VIRUS | 322 | -0.375886 | -1.563135 | 0 | 0.134436 |
| GOBP_EXCITATORY_SYNAPSE_ASSEMBLY | 23 | -0.566373 | -1.562985 | 0.028571 | 0.134081 |
| GOBP_REGULATION_OF_INFLAMMATORY_RESPONSE | 303 | -0.380057 | -1.562533 | 0 | 0.134079 |
| GOBP_ALPHA_BETA_T_CELL_PROLIFERATION | 36 | -0.510739 | -1.561184 | 0.024355 | 0.135276 |
| GOBP_REGULATION_OF_B_CELL_DIFFERENTIATION | 30 | -0.530464 | -1.560531 | 0.017621 | 0.135513 |
| GOBP_NERVE_DEVELOPMENT | 55 | -0.465706 | -1.559509 | 0.012228 | 0.136182 |
| GOBP_DEFENSE_RESPONSE_TO_VIRUS | 234 | -0.386289 | -1.558647 | 0 | 0.136729 |
| GOBP_BILE_ACID_METABOLIC_PROCESS | 42 | -0.488222 | -1.558491 | 0.015714 | 0.136389 |
| GOBP_POSITIVE_REGULATION_OF_RESPONSE_TO_ENDOPLASMIC_RETICULUM_STRESS | 35 | -0.504599 | -1.55837 | 0.014347 | 0.135971 |
| GOBP_MEMBRANE_DEPOLARIZATION_DURING_CARDIAC_MUSCLE_CELL_ACTION_POTENTIAL | 19 | -0.586235 | -1.557445 | 0.027108 | 0.136646 |
| GOBP_POSITIVE_REGULATION_OF_T_CELL_PROLIFERATION | 96 | -0.428352 | -1.555971 | 0.003891 | 0.137954 |
| GOBP_REGULATION_OF_CYTOSOLIC_CALCIUM_ION_CONCENTRATION | 300 | -0.377595 | -1.554041 | 0 | 0.139987 |
| GOBP_I_KAPPAB_KINASE_NF_KAPPAB_SIGNALING | 271 | -0.378953 | -1.553418 | 0 | 0.140183 |
| GOBP_SPINAL_CORD_MOTOR_NEURON_DIFFERENTIATION | 22 | -0.563319 | -1.553067 | 0.028401 | 0.140036 |
| GOBP_REGULATION_OF_MONONUCLEAR_CELL_MIGRATION | 106 | -0.42648 | -1.552657 | 0.006614 | 0.140044 |
| GOBP_NEGATIVE_REGULATION_OF_ANION_TRANSMEMBRANE_TRANSPORT | 29 | -0.524025 | -1.551088 | 0.026786 | 0.141497 |
| GOBP_ARACHIDONIC_ACID_METABOLIC_PROCESS | 49 | -0.478232 | -1.549576 | 0.015449 | 0.142927 |
| GOBP_PRODUCTION_OF_MOLECULAR_MEDIATOR_INVOLVED_IN_INFLAMMATORY_RESPONSE | 67 | -0.449001 | -1.54947 | 0.005464 | 0.142547 |
| GOBP_POSITIVE_REGULATION_OF_DNA_BINDING_TRANSCRIPTION_FACTOR_ACTIVITY | 249 | -0.382214 | -1.548044 | 0 | 0.143858 |
| GOBP_POSITIVE_REGULATION_OF_DEFENSE_RESPONSE | 322 | -0.371772 | -1.546523 | 0 | 0.145286 |
| GOBP_CONVERGENT_EXTENSION | 16 | -0.597281 | -1.546415 | 0.03271 | 0.144876 |
| GOBP_CELLULAR_RESPONSE_TO_PROSTAGLANDIN_E_STIMULUS | 15 | -0.618824 | -1.544178 | 0.025682 | 0.147334 |
| GOBP_NEGATIVE_REGULATION_OF_BMP_SIGNALING_PATHWAY | 43 | -0.491866 | -1.542825 | 0.012587 | 0.148636 |
| GOBP_ADRENAL_GLAND_DEVELOPMENT | 18 | -0.580041 | -1.54076 | 0.035115 | 0.151001 |
| GOBP_POSITIVE_REGULATION_OF_PEPTIDYL_TYROSINE_PHOSPHORYLATION | 172 | -0.394947 | -1.539894 | 0.003695 | 0.151579 |
| GOBP_PRIMARY_NEURAL_TUBE_FORMATION | 84 | -0.427237 | -1.538069 | 0.005326 | 0.153746 |
| GOBP_OSTEOCLAST_DIFFERENTIATION | 86 | -0.433806 | -1.537277 | 0.004138 | 0.15437 |
| GOBP_NATURAL_KILLER_CELL_ACTIVATION | 67 | -0.446189 | -1.537142 | 0.005398 | 0.154018 |
| GOBP_POSITIVE_REGULATION_OF_TYROSINE_PHOSPHORYLATION_OF_STAT_PROTEIN | 58 | -0.459978 | -1.536272 | 0.012605 | 0.154585 |
| GOBP_NEGATIVE_REGULATION_OF_INTRINSIC_APOPTOTIC_SIGNALING_PATHWAY_BY_P53_CLAS | 20 | -0.562571 | -1.53563 | 0.025559 | 0.154988 |
| GOBP_POSITIVE_REGULATION_OF_CELL_ADHESION | 403 | -0.364753 | -1.534978 | 0 | 0.155425 |
| GOBP_POSITIVE_REGULATION_OF_LIPID_CATABOLIC_PROCESS | 21 | -0.551309 | -1.534815 | 0.029052 | 0.155074 |
| GOBP_RELAXATION_OF_CARDIAC_MUSCLE | 16 | -0.600592 | -1.534529 | 0.028754 | 0.154886 |
| GOBP_CELLULAR_RESPONSE_TO_INTERFERON_BETA | 20 | -0.569181 | -1.534112 | 0.031532 | 0.154967 |
| GOBP_POSITIVE_REGULATION_OF_NF_KAPPAB_TRANSCRIPTION_FACTOR_ACTIVITY | 155 | -0.397174 | -1.533981 | 0.002389 | 0.154612 |
| GOBP_RENAL_TUBULE_DEVELOPMENT | 74 | -0.444077 | -1.533453 | 0.009321 | 0.154791 |
| GOBP_LYMPHOID_PROGENITOR_CELL_DIFFERENTIATION | 20 | -0.57177 | -1.533211 | 0.023585 | 0.154577 |
| GOBP_REGULATION_OF_RESPONSE_TO_CYTOKINE_STIMULUS | 165 | -0.392587 | -1.532685 | 0.002454 | 0.154738 |
| GOBP_REGULATION_OF_PRODUCTION_OF_MOLECULAR_MEDIATOR_OF_IMMUNE_RESPONSE | 137 | -0.400588 | -1.532199 | 0.001238 | 0.154854 |
| GOBP_LIVER_MORPHOGENESIS | 19 | -0.566056 | -1.531733 | 0.023112 | 0.155008 |
| GOBP_REGULATION_OF_EXTRINSIC_APOPTOTIC_SIGNALING_PATHWAY | 145 | -0.399965 | -1.530962 | 0.001222 | 0.155496 |
| GOBP_NEURON_PROJECTION_ARBORIZATION | 28 | -0.517704 | -1.530687 | 0.022489 | 0.155324 |
| GOBP_MACROPHAGE_CHEMOTAXIS | 34 | -0.502138 | -1.529813 | 0.022489 | 0.156021 |
| GOBP_REGULATION_OF_GLIAL_CELL_MIGRATION | 19 | -0.574958 | -1.529795 | 0.029827 | 0.155514 |
| GOBP_HINDLIMB_MORPHOGENESIS | 28 | -0.524009 | -1.528751 | 0.024922 | 0.15647 |
| GOBP_REGULATION_OF_CALCIUM_ION_DEPENDENT_EXOCYTOSIS | 36 | -0.5002 | -1.528533 | 0.020408 | 0.156264 |
| GOBP_FIBRINOLYSIS | 19 | -0.565868 | -1.527949 | 0.044978 | 0.156523 |
| GOBP_POSITIVE_REGULATION_OF_B_CELL_PROLIFERATION | 39 | -0.4933 | -1.527814 | 0.019774 | 0.156171 |
| GOBP_REGULATION_OF_DEFENSE_RESPONSE_TO_VIRUS_BY_HOST | 39 | -0.491323 | -1.527594 | 0.020319 | 0.155952 |
| GOBP_T_CELL_CYTOKINE_PRODUCTION | 35 | -0.499686 | -1.527295 | 0.023324 | 0.155817 |
| GOBP_HOMOPHILIC_CELL_ADHESION_VIA_PLASMA_MEMBRANE_ADHESION_MOLECULES | 129 | -0.406054 | -1.526578 | 0.003672 | 0.156369 |
| GOBP_NEGATIVE_REGULATION_OF_CELL_CELL_ADHESION | 171 | -0.389903 | -1.526395 | 0.00235 | 0.156081 |
| GOBP_INTERLEUKIN_17_PRODUCTION | 33 | -0.502285 | -1.525731 | 0.022489 | 0.156464 |
| GOBP_T_CELL_SELECTION | 48 | -0.462874 | -1.525515 | 0.017518 | 0.156247 |
| GOBP_POSITIVE_REGULATION_OF_MACROPHAGE_MIGRATION | 23 | -0.556982 | -1.525211 | 0.029806 | 0.156146 |
| GOBP_POSITIVE_REGULATION_OF_ANTIGEN_RECEPTOR_MEDIATED_SIGNALING_PATHWAY | 21 | -0.552321 | -1.523259 | 0.026316 | 0.15836 |
| GOBP_SENSORY_PERCEPTION_OF_TASTE | 45 | -0.470661 | -1.523038 | 0.021398 | 0.158105 |
| GOBP_CELLULAR_RESPONSE_TO_ALCOHOL | 79 | -0.43665 | -1.522148 | 0.014805 | 0.158796 |
| GOBP_ARTERY_DEVELOPMENT | 78 | -0.430055 | -1.52155 | 0.012195 | 0.159109 |
| GOBP_LUNG_MORPHOGENESIS | 39 | -0.483299 | -1.520842 | 0.02819 | 0.159604 |
| GOBP_RESPONSE_TO_INTERLEUKIN_6 | 37 | -0.500738 | -1.519968 | 0.028902 | 0.16037 |
| GOBP_DIGESTION | 95 | -0.422628 | -1.519112 | 0.005215 | 0.161144 |
| GOBP_DENDRITIC_CELL_DIFFERENTIATION | 44 | -0.47541 | -1.518784 | 0.02 | 0.161073 |
| GOBP_REGULATION_OF_MONOCYTE_DIFFERENTIATION | 19 | -0.564368 | -1.518635 | 0.036566 | 0.160694 |
| GOBP_IMMUNE_RESPONSE_REGULATING_SIGNALING_PATHWAY | 484 | -0.35787 | -1.517907 | 0 | 0.161232 |
| GOBP_LYMPHOCYTE_COSTIMULATION | 58 | -0.453867 | -1.516952 | 0.015625 | 0.162116 |
| GOBP_CELLULAR_DEFENSE_RESPONSE | 50 | -0.459802 | -1.516918 | 0.025352 | 0.161639 |
| GOBP_EMBRYONIC_PATTERN_SPECIFICATION | 51 | -0.462166 | -1.516754 | 0.014493 | 0.16134 |
| GOBP_ANTIGEN_RECEPTOR_MEDIATED_SIGNALING_PATHWAY | 302 | -0.367541 | -1.516746 | 0 | 0.160835 |
| GOBP_UNSATURATED_FATTY_ACID_METABOLIC_PROCESS | 96 | -0.417179 | -1.516014 | 0.015789 | 0.161357 |
| GOBP_REGULATION_OF_PRESYNAPSE_ORGANIZATION | 22 | -0.547824 | -1.514804 | 0.02381 | 0.16258 |
| GOBP_REGULATION_OF_CHEMOTAXIS | 202 | -0.381159 | -1.51473 | 0 | 0.162131 |
| GOBP_POSITIVE_REGULATION_OF_KERATINOCYTE_DIFFERENTIATION | 16 | -0.585975 | -1.514093 | 0.043546 | 0.162535 |
| GOBP_DEVELOPMENTAL_INDUCTION | 24 | -0.531942 | -1.512085 | 0.032659 | 0.164891 |
| GOBP_SOMATIC_STEM_CELL_POPULATION_MAINTENANCE | 60 | -0.44678 | -1.511234 | 0.016438 | 0.165574 |
| GOBP_NEGATIVE_REGULATION_OF_SMOOTH_MUSCLE_CELL_PROLIFERATION | 43 | -0.475157 | -1.510629 | 0.02391 | 0.165989 |
| GOBP_REGULATION_OF_OSTEOCLAST_DIFFERENTIATION | 57 | -0.442662 | -1.510159 | 0.015515 | 0.166182 |
| GOBP_AXIS_ELONGATION | 23 | -0.54559 | -1.509512 | 0.033588 | 0.166654 |
| GOBP_ADAPTIVE_IMMUNE_RESPONSE_BASED_ON_SOMATIC_RECOMBINATION_OF_IMMUNE_RE | 343 | -0.363992 | -1.509481 | 0 | 0.166199 |
| GOBP_NEGATIVE_REGULATION_OF_DEFENSE_RESPONSE | 187 | -0.384716 | -1.50905 | 0.001185 | 0.166287 |
| GOBP_POSITIVE_REGULATION_OF_RECEPTOR_SIGNALING_PATHWAY_VIA_STAT | 43 | -0.470414 | -1.508936 | 0.029283 | 0.165935 |
| GOBP_POSITIVE_REGULATION_OF_RESPONSE_TO_CYTOKINE_STIMULUS | 55 | -0.448868 | -1.508504 | 0.012465 | 0.166025 |
| GOBP_CELLULAR_RESPONSE_TO_ZINC_ION | 15 | -0.608344 | -1.50842 | 0.043831 | 0.165627 |
| GOBP_PULMONARY_VALVE_DEVELOPMENT | 19 | -0.554576 | -1.508325 | 0.046296 | 0.165216 |
| GOBP_VENTRICULAR_SEPTUM_MORPHOGENESIS | 32 | -0.5097 | -1.508208 | 0.025678 | 0.16488 |
| GOBP_REGULATION_OF_MACROPHAGE_MIGRATION | 36 | -0.499418 | -1.508166 | 0.028107 | 0.164442 |
| GOBP_REGULATION_OF_CAMP_MEDIATED_SIGNALING | 32 | -0.504484 | -1.507374 | 0.017831 | 0.16512 |
| GOBP_INTERLEUKIN_8_PRODUCTION | 82 | -0.423204 | -1.506621 | 0.015027 | 0.165755 |

| | | | | | |
|---|---|---|---|---|---|
| GOBP_CELLULAR_ALDEHYDE_METABOLIC_PROCESS | 54 | 0.336846 | 1.269111 | 0.085911 | 0.437229 |
| GOBP_POSITIVE_REGULATION_OF_UBIQUITIN_PROTEIN_TRANSFERASE_ACTIVITY | 30 | 0.375006 | 1.268116 | 0.138365 | 0.438297 |
| GOBP_CLATHRIN_COAT_ASSEMBLY | 17 | 0.444528 | 1.267452 | 0.163934 | 0.43866 |
| GOBP_G2_DNA_DAMAGE_CHECKPOINT | 34 | 0.367834 | 1.266192 | 0.173633 | 0.440463 |
| GOBP_NADH_METABOLIC_PROCESS | 43 | 0.35417 | 1.265479 | 0.115756 | 0.440848 |
| GOBP_PIGMENT_GRANULE_LOCALIZATION | 26 | 0.395573 | 1.264802 | 0.110749 | 0.441206 |
| GOBP_MEMBRANE_FISSION | 15 | 0.458364 | 1.263949 | 0.180822 | 0.44201 |
| GOBP_FOREBRAIN_CELL_MIGRATION | 47 | 0.341001 | 1.263776 | 0.115756 | 0.441101 |
| GOBP_NCRNA_PROCESSING | 385 | 0.257611 | 1.263551 | 0.009434 | 0.440427 |
| GOBP_REGULATION_OF_KETONE_BIOSYNTHETIC_PROCESS | 16 | 0.45124 | 1.26339 | 0.155172 | 0.439486 |
| GOBP_RETINA_MORPHOGENESIS_IN_CAMERA_TYPE_EYE | 46 | 0.344687 | 1.262849 | 0.094406 | 0.439493 |
| GOBP_DNA_LIGATION | 15 | 0.462984 | 1.26245 | 0.188144 | 0.439292 |
| GOBP_SCF_DEPENDENT_PROTEASOMAL_UBIQUITIN_DEPENDENT_PROTEIN_CATABOLIC_PROC | 91 | 0.30592 | 1.261988 | 0.084 | 0.439103 |
| GOBP_GPI_ANCHOR_METABOLIC_PROCESS | 31 | 0.382868 | 1.261905 | 0.168067 | 0.438027 |
| GOBP_FATTY_ACYL_COA_BIOSYNTHETIC_PROCESS | 31 | 0.381736 | 1.259087 | 0.133127 | 0.44328 |
| GOBP_MITOCHONDRIAL_TRANSCRIPTION | 17 | 0.437606 | 1.258032 | 0.172414 | 0.444529 |
| GOBP_MITOTIC_DNA_INTEGRITY_CHECKPOINT | 104 | 0.298835 | 1.257922 | 0.08 | 0.443547 |
| GOBP_CELLULAR_MODIFIED_AMINO_ACID_CATABOLIC_PROCESS | 24 | 0.399127 | 1.254539 | 0.161677 | 0.450272 |
| GOBP_REGULATION_OF_GLUCAN_BIOSYNTHETIC_PROCESS | 27 | 0.387967 | 1.254097 | 0.14121 | 0.449985 |
| GOBP_CELLULAR_RESPONSE_TO_GROWTH_HORMONE_STIMULUS | 21 | 0.413953 | 1.25229 | 0.169139 | 0.45303 |
| GOBP_POSITIVE_REGULATION_OF_TRANSMEMBRANE_RECEPTOR_PROTEIN_SERINE_THREONIN | 94 | 0.305787 | 1.250466 | 0.088496 | 0.456453 |
| GOBP_METANEPHRIC_TUBULE_DEVELOPMENT | 16 | 0.448471 | 1.25039 | 0.191667 | 0.455353 |
| GOBP_DETOXIFICATION | 119 | 0.292166 | 1.249613 | 0.075758 | 0.45596 |
| GOBP_CELLULAR_MODIFIED_AMINO_ACID_METABOLIC_PROCESS | 166 | 0.276828 | 1.249502 | 0.03352 | 0.454946 |
| GOBP_RESPONSE_TO_OXYGEN_RADICAL | 24 | 0.408897 | 1.247117 | 0.175793 | 0.459412 |
| GOBP_MALE_GAMETE_GENERATION | 445 | 0.244111 | 1.246722 | 0.012346 | 0.459093 |
| GOBP_ANATOMICAL_STRUCTURE_HOMEOSTASIS | 423 | 0.250953 | 1.246489 | 0 | 0.458447 |
| GOBP_SPROUTING_ANGIOGENESIS | 121 | 0.286143 | 1.242734 | 0.059091 | 0.46633 |
| GOBP_DNA_BIOSYNTHETIC_PROCESS | 183 | 0.272642 | 1.241921 | 0.053892 | 0.467071 |
| GOBP_MRNA_MODIFICATION | 23 | 0.410023 | 1.241065 | 0.18984 | 0.468011 |
| GOBP_BASEMENT_MEMBRANE_ORGANIZATION | 29 | 0.382719 | 1.240953 | 0.178363 | 0.467054 |
| GOBP_NEGATIVE_REGULATION_OF_MITOTIC_CELL_CYCLE | 290 | 0.257945 | 1.23882 | 0.02381 | 0.471157 |
| GOBP_POSITIVE_REGULATION_OF_LONG_TERM_SYNAPTIC_POTENTIATION | 21 | 0.409272 | 1.238052 | 0.18 | 0.471741 |
| GOBP_RESPONSE_TO_INTERLEUKIN_7 | 37 | 0.355954 | 1.237304 | 0.14557 | 0.472317 |
| GOBP_RNA_DEPENDENT_DNA_BIOSYNTHETIC_PROCESS | 68 | 0.316695 | 1.237227 | 0.099602 | 0.471181 |
| GOBP_SIGNAL_TRANSDUCTION_INVOLVED_IN_CELL_CYCLE_CHECKPOINT | 72 | 0.313053 | 1.235914 | 0.118705 | 0.473198 |
| GOBP_RETROGRADE_VESICLE_MEDIATED_TRANSPORT_GOLGI_TO_ENDOPLASMIC_RETICULUM | 86 | 0.300726 | 1.235566 | 0.089069 | 0.472825 |
| GOBP_SMOOTH_MUSCLE_CELL_MIGRATION | 65 | 0.309746 | 1.233642 | 0.10989 | 0.476126 |
| GOBP_REPLICATION_FORK_PROCESSING | 39 | 0.350599 | 1.232721 | 0.149837 | 0.477213 |
| GOBP_WATER_TRANSPORT | 17 | 0.429335 | 1.230122 | 0.199475 | 0.482521 |
| GOBP_NUCLEIC_ACID_PHOSPHODIESTER_BOND_HYDROLYSIS | 282 | 0.257826 | 1.229407 | 0.054054 | 0.483037 |
| GOBP_SULFUR_COMPOUND_CATABOLIC_PROCESS | 47 | 0.340588 | 1.228229 | 0.136213 | 0.484771 |
| GOBP_RRNA_CATABOLIC_PROCESS | 20 | 0.411442 | 1.226324 | 0.186301 | 0.48822 |
| GOBP_SULFUR_AMINO_ACID_METABOLIC_PROCESS | 34 | 0.359311 | 1.223555 | 0.165714 | 0.493937 |
| GOBP_PURINE_CONTAINING_COMPOUND_TRANSMEMBRANE_TRANSPORT | 15 | 0.444013 | 1.223384 | 0.185185 | 0.493135 |
| GOBP_GLUTAMINE_FAMILY_AMINO_ACID_BIOSYNTHETIC_PROCESS | 16 | 0.439283 | 1.22325 | 0.203804 | 0.492176 |
| GOBP_ALDITOL_METABOLIC_PROCESS | 20 | 0.404985 | 1.220368 | 0.196676 | 0.49849 |
| GOBP_RNA_METHYLATION | 84 | 0.300094 | 1.219962 | 0.097458 | 0.498214 |
| GOBP_NEGATIVE_REGULATION_OF_SMOOTH_MUSCLE_CELL_MIGRATION | 20 | 0.404908 | 1.218631 | 0.205128 | 0.500354 |
| GOBP_POTASSIUM_ION_IMPORT_ACROSS_PLASMA_MEMBRANE | 41 | 0.345658 | 1.216574 | 0.170492 | 0.504488 |
| GOBP_POSITIVE_REGULATION_OF_MITOTIC_CELL_CYCLE | 107 | 0.288434 | 1.214283 | 0.092105 | 0.50924 |
| GOBP_BLOOD_VESSEL_REMODELING | 37 | 0.346037 | 1.211354 | 0.16358 | 0.515781 |
| GOBP_FATTY_ACID_DERIVATIVE_BIOSYNTHETIC_PROCESS | 53 | 0.320774 | 1.210992 | 0.158491 | 0.515413 |
| GOBP_POSITIVE_REGULATION_OF_STEM_CELL_DIFFERENTIATION | 16 | 0.433561 | 1.207399 | 0.238636 | 0.523738 |
| GOBP_REGULATION_OF_POLYSACCHARIDE_BIOSYNTHETIC_PROCESS | 35 | 0.350962 | 1.206596 | 0.193878 | 0.524522 |
| GOBP_PROTEIN_CONTAINING_COMPLEX_REMODELING | 25 | 0.381858 | 1.206425 | 0.202857 | 0.523599 |
| GOBP_GLYCINE_METABOLIC_PROCESS | 18 | 0.420671 | 1.206305 | 0.207865 | 0.5226 |
| GOBP_VASCULAR_ENDOTHELIAL_GROWTH_FACTOR_RECEPTOR_SIGNALING_PATHWAY | 88 | 0.294134 | 1.20397 | 0.102459 | 0.527586 |
| GOBP_DECIDUALIZATION | 19 | 0.405988 | 1.20367 | 0.202817 | 0.527059 |
| GOBP_REGULATION_OF_CLATHRIN_DEPENDENT_ENDOCYTOSIS | 18 | 0.413502 | 1.202198 | 0.207101 | 0.529756 |
| GOBP_TERTIARY_ALCOHOL_METABOLIC_PROCESS | 18 | 0.412864 | 1.200962 | 0.227273 | 0.531845 |
| GOBP_RESPONSE_TO_IRON_ION | 26 | 0.37834 | 1.200811 | 0.204334 | 0.53094 |
| GOBP_POSITIVE_REGULATION_OF_SMOOTH_MUSCLE_CELL_MIGRATION | 37 | 0.33492 | 1.200042 | 0.178218 | 0.531727 |
| GOBP_RESPONSE_TO_DEXAMETHASONE | 35 | 0.345576 | 1.199436 | 0.180124 | 0.532153 |
| GOBP_CELLULAR_RESPONSE_TO_VASCULAR_ENDOTHELIAL_GROWTH_FACTOR_STIMULUS | 60 | 0.317532 | 1.199201 | 0.164875 | 0.531538 |
| GOBP_POSITIVE_REGULATION_OF_CELL_CYCLE_ARREST | 77 | 0.300277 | 1.199139 | 0.152893 | 0.530399 |
| GOBP_REGULATION_OF_WATER_LOSS_VIA_SKIN | 23 | 0.391201 | 1.1991 | 0.219444 | 0.529193 |
| GOBP_REGULATION_OF_CELL_DIVISION | 151 | 0.261194 | 1.197731 | 0.089385 | 0.531541 |
| GOBP_POSITIVE_REGULATION_OF_GLYCOGEN_METABOLIC_PROCESS | 16 | 0.42295 | 1.196322 | 0.251366 | 0.534144 |
| GOBP_TRIGLYCERIDE_BIOSYNTHETIC_PROCESS | 37 | 0.343557 | 1.19502 | 0.160991 | 0.536275 |
| GOBP_V_D_J_RECOMBINATION | 16 | 0.416463 | 1.190909 | 0.229462 | 0.546588 |
| GOBP_AUTOPHAGY_OF_MITOCHONDRION | 74 | 0.295492 | 1.187824 | 0.15748 | 0.553606 |
| GOBP_PYRIMIDINE_NUCLEOSIDE_METABOLIC_PROCESS | 34 | 0.351228 | 1.187733 | 0.200637 | 0.552465 |
| GOBP_ORGANIC_CATION_TRANSPORT | 30 | 0.357825 | 1.184586 | 0.219136 | 0.559908 |
| GOBP_REGULATION_OF_MITOCHONDRIAL_MEMBRANE_POTENTIAL | 69 | 0.296567 | 1.184105 | 0.163823 | 0.56001 |
| GOBP_MUSCLE_HYPERTROPHY | 78 | 0.290909 | 1.183953 | 0.1417 | 0.559048 |
| GOBP_NUCLEOSIDE_BISPHOSPHATE_BIOSYNTHETIC_PROCESS | 66 | 0.304885 | 1.179233 | 0.180392 | 0.571242 |
| GOBP_POSITIVE_REGULATION_OF_GLUCOSE_METABOLIC_PROCESS | 36 | 0.340816 | 1.178132 | 0.194969 | 0.573335 |
| GOBP_NEGATIVE_REGULATION_OF_ACTIN_FILAMENT_DEPOLYMERIZATION | 44 | 0.327713 | 1.17696 | 0.203279 | 0.575383 |
| GOBP_LONG_CHAIN_FATTY_ACYL_COA_METABOLIC_PROCESS | 25 | 0.375762 | 1.175022 | 0.227811 | 0.579924 |
| GOBP_L_AMINO_ACID_TRANSPORT | 60 | 0.309616 | 1.173744 | 0.173432 | 0.582255 |
| GOBP_CARTILAGE_DEVELOPMENT | 162 | 0.258386 | 1.172039 | 0.084337 | 0.585911 |
| GOBP_ACTIN_FILAMENT_DEPOLYMERIZATION | 58 | 0.31131 | 1.171842 | 0.205298 | 0.585088 |
| GOBP_THIOESTER_BIOSYNTHETIC_PROCESS | 52 | 0.313467 | 1.171483 | 0.195035 | 0.584774 |
| GOBP_BRANCHING_INVOLVED_IN_BLOOD_VESSEL_MORPHOGENESIS | 29 | 0.35615 | 1.170862 | 0.214706 | 0.585305 |
| GOBP_MITOTIC_G2_M_TRANSITION_CHECKPOINT | 36 | 0.336186 | 1.170386 | 0.198083 | 0.585321 |
| GOBP_AEROBIC_RESPIRATION | 83 | 0.285467 | 1.166835 | 0.154812 | 0.594565 |
| GOBP_POSITIVE_REGULATION_OF_AXON_EXTENSION | 37 | 0.33681 | 1.164362 | 0.203593 | 0.600466 |
| GOBP_REGULATION_OF_TRANSLATIONAL_FIDELITY | 20 | 0.40112 | 1.162443 | 0.264151 | 0.604972 |
| GOBP_VASCULAR_PROCESS_IN_CIRCULATORY_SYSTEM | 219 | 0.249927 | 1.161443 | 0.096552 | 0.606611 |
| GOBP_IMPORT_ACROSS_PLASMA_MEMBRANE | 144 | 0.26516 | 1.161151 | 0.135417 | 0.606113 |
| GOBP_TRIGLYCERIDE_CATABOLIC_PROCESS | 24 | 0.374728 | 1.160194 | 0.25 | 0.607629 |
| GOBP_PROTEIN_LIPID_COMPLEX_SUBUNIT_ORGANIZATION | 44 | 0.317085 | 1.159574 | 0.20743 | 0.607974 |
| GOBP_VASCULAR_ASSOCIATED_SMOOTH_MUSCLE_CELL_MIGRATION | 21 | 0.373639 | 1.159378 | 0.25 | 0.607192 |
| GOBP_DNA_METHYLATION_DEPENDENT_HETEROCHROMATIN_ASSEMBLY | 19 | 0.393736 | 1.159148 | 0.236686 | 0.606494 |
| GOBP_PYRIMIDINE_RIBONUCLEOSIDE_TRIPHOSPHATE_BIOSYNTHETIC_PROCESS | 15 | 0.426953 | 1.156817 | 0.271739 | 0.612201 |
| GOBP_REGULATION_OF_LONG_TERM_SYNAPTIC_POTENTIATION | 41 | 0.330911 | 1.156418 | 0.202572 | 0.61208 |
| GOBP_NUCLEOSIDE_PHOSPHATE_BIOSYNTHETIC_PROCESS | 252 | 0.24473 | 1.156135 | 0.073171 | 0.611523 |
| GOBP_CELL_CYCLE_G1_S_PHASE_TRANSITION | 261 | 0.24264 | 1.154114 | 0.101563 | 0.616461 |
| GOBP_NUCLEOBASE_METABOLIC_PROCESS | 33 | 0.347366 | 1.152056 | 0.241935 | 0.62134 |
| GOBP_BRANCHED_CHAIN_AMINO_ACID_METABOLIC_PROCESS | 23 | 0.367679 | 1.15188 | 0.241279 | 0.620452 |
| GOBP_SINGLE_STRANDED_VIRAL_RNA_REPLICATION_VIA_DOUBLE_STRANDED_DNA_INTERME | 20 | 0.377597 | 1.151545 | 0.243323 | 0.620078 |
| GOBP_GLUTATHIONE_METABOLIC_PROCESS | 50 | 0.306966 | 1.150253 | 0.192547 | 0.622531 |
| GOBP_REGULATION_OF_INSULIN_LIKE_GROWTH_FACTOR_RECEPTOR_SIGNALING_PATHWAY | 18 | 0.401475 | 1.149203 | 0.258312 | 0.624359 |
| GOBP_RNA_SPLICING_VIA_ENDONUCLEOLYTIC_CLEAVAGE_AND_LIGATION | 16 | 0.395104 | 1.148451 | 0.295652 | 0.625224 |
| GOBP_HISTONE_PHOSPHORYLATION | 39 | 0.334023 | 1.148049 | 0.219672 | 0.625014 |
| GOBP_RETINA_DEVELOPMENT_IN_CAMERA_TYPE_EYE | 116 | 0.27023 | 1.147942 | 0.136986 | 0.623892 |
| GOBP_BICARBONATE_TRANSPORT | 40 | 0.331111 | 1.147827 | 0.220183 | 0.622879 |
| GOBP_RESPONSE_TO_TOXIC_SUBSTANCE | 214 | 0.244144 | 1.1477 | 0.10274 | 0.621855 |
| GOBP_POSITIVE_REGULATION_OF_CYTOKINESIS | 31 | 0.350341 | 1.146717 | 0.240854 | 0.623505 |
| GOBP_AXON_ENSHEATHMENT_IN_CENTRAL_NERVOUS_SYSTEM | 20 | 0.391387 | 1.146667 | 0.271955 | 0.622217 |
| GOBP_PYRIMIDINE_RIBONUCLEOSIDE_TRIPHOSPHATE_METABOLIC_PROCESS | 18 | 0.394931 | 1.146066 | 0.259669 | 0.622696 |
| GOBP_CELLULAR_RESPONSE_TO_CORTICOSTEROID_STIMULUS | 52 | 0.311285 | 1.145867 | 0.211221 | 0.621916 |
| GOBP_NEGATIVE_REGULATION_OF_PLATELET_ACTIVATION | 16 | 0.399367 | 1.14579 | 0.277457 | 0.620775 |
| GOBP_AMINO_ACID_TRANSPORT | 131 | 0.261848 | 1.145557 | 0.129353 | 0.620139 |
| GOBP_PYRIMIDINE_NUCLEOTIDE_BIOSYNTHETIC_PROCESS | 29 | 0.350749 | 1.145233 | 0.258258 | 0.619756 |
| GOBP_2_OXOGLUTARATE_METABOLIC_PROCESS | 16 | 0.408669 | 1.14509 | 0.276536 | 0.618817 |
| GOBP_MICROTUBULE_POLYMERIZATION_OR_DEPOLYMERIZATION | 109 | 0.268469 | 1.144882 | 0.170306 | 0.618068 |
| GOBP_MUSCLE_CELL_MIGRATION | 78 | 0.280146 | 1.144103 | 0.205776 | 0.618969 |
| GOBP_POSITIVE_REGULATION_OF_GLUCOSE_IMPORT | 32 | 0.338075 | 1.14402 | 0.257764 | 0.617812 |
| GOBP_NEGATIVE_REGULATION_OF_MYELOID_CELL_DIFFERENTIATION | 88 | 0.27558 | 1.142176 | 0.199115 | 0.621992 |
| GOBP_THIOESTER_METABOLIC_PROCESS | 100 | 0.272547 | 1.141962 | 0.16 | 0.621324 |
| GOBP_RETINA_LAYER_FORMATION | 20 | 0.377621 | 1.140354 | 0.255556 | 0.625109 |
| GOBP_TORC1_SIGNALING | 47 | 0.313132 | 1.139274 | 0.241935 | 0.627094 |
| GOBP_ALCOHOL_BIOSYNTHETIC_PROCESS | 143 | 0.259469 | 1.138948 | 0.132353 | 0.626731 |
| GOBP_RIBOSOME_BIOGENESIS | 303 | 0.235915 | 1.137289 | 0.125 | 0.630516 |
| GOBP_NEGATIVE_REGULATION_OF_STRIATED_MUSCLE_CELL_APOPTOTIC_PROCESS | 20 | 0.377153 | 1.136707 | 0.275072 | 0.630894 |
| GOBP_RESPONSE_TO_AXON_INJURY | 70 | 0.286078 | 1.13618 | 0.195572 | 0.631123 |
| GOBP_REGULATION_OF_INTEGRIN_ACTIVATION | 16 | 0.408549 | 1.135642 | 0.27809 | 0.631486 |
| GOBP_POSITIVE_REGULATION_OF_CELL_SUBSTRATE_JUNCTION_ORGANIZATION | 29 | 0.35154 | 1.135514 | 0.270517 | 0.630527 |
| GOBP_INSULIN_LIKE_GROWTH_FACTOR_RECEPTOR_SIGNALING_PATHWAY | 29 | 0.346269 | 1.133467 | 0.245098 | 0.635726 |
| GOBP_MITOTIC_CYTOKINESIS | 70 | 0.289862 | 1.132359 | 0.232 | 0.637856 |
| GOBP_PROXIMAL_DISTAL_PATTERN_FORMATION | 24 | 0.361577 | 1.131538 | 0.290323 | 0.639122 |
| GOBP_STEROL_BIOSYNTHETIC_PROCESS | 70 | 0.288142 | 1.131288 | 0.219231 | 0.638495 |
| GOBP_PURINE_CONTAINING_COMPOUND_BIOSYNTHETIC_PROCESS | 196 | 0.239977 | 1.130687 | 0.125 | 0.639152 |
| GOBP_REGULATION_OF_TORC1_SIGNALING | 39 | 0.315295 | 1.130452 | 0.271875 | 0.638576 |
| GOBP_PROTEIN_TARGETING_TO_MITOCHONDRION | 99 | 0.271288 | 1.129704 | 0.211009 | 0.639632 |
| GOBP_REGULATION_OF_CHROMATIN_ASSEMBLY_OR_DISASSEMBLY | 24 | 0.369548 | 1.128102 | 0.295322 | 0.643234 |
| GOBP_CORNIFICATION | 57 | 0.298289 | 1.128042 | 0.2397 | 0.642044 |
| GOBP_GLUTAMATE_METABOLIC_PROCESS | 29 | 0.342939 | 1.12774 | 0.28655 | 0.641631 |
| GOBP_IMPORT_INTO_CELL | 205 | 0.239917 | 1.124847 | 0.125786 | 0.649328 |
| GOBP_REGULATION_OF_CELLULAR_AMINE_METABOLIC_PROCESS | 69 | 0.283616 | 1.123756 | 0.244526 | 0.651526 |
| GOBP_VESICLE_MEDIATED_TRANSPORT_IN_SYNAPSE | 180 | 0.244494 | 1.122746 | 0.155556 | 0.653318 |
| GOBP_ISOPRENOID_BIOSYNTHETIC_PROCESS | 25 | 0.357095 | 1.122716 | 0.311111 | 0.652028 |
| GOBP_BONE_MORPHOGENESIS | 82 | 0.28002 | 1.122399 | 0.211155 | 0.651681 |
| GOBP_RRNA_MODIFICATION | 40 | 0.320496 | 1.121952 | 0.250814 | 0.651676 |
| GOBP_NEGATIVE_REGULATION_OF_SMALL_MOLECULE_METABOLIC_PROCESS | 82 | 0.274087 | 1.121772 | 0.240458 | 0.650879 |
| GOBP_BONE_MINERALIZATION | 103 | 0.267815 | 1.121623 | 0.198276 | 0.650018 |
| GOBP_RESPONSE_TO_HEPATOCYTE_GROWTH_FACTOR | 16 | 0.395839 | 1.120887 | 0.321244 | 0.650987 |
| GOBP_NEGATIVE_REGULATION_OF_OXIDATIVE_STRESS_INDUCED_CELL_DEATH | 39 | 0.313219 | 1.120397 | 0.272152 | 0.651233 |
| GOBP_HEMATOPOIETIC_STEM_CELL_DIFFERENTIATION | 86 | 0.268202 | 1.119973 | 0.207407 | 0.651227 |
| GOBP_CELLULAR_AMINO_ACID_CATABOLIC_PROCESS | 96 | 0.269442 | 1.118718 | 0.262009 | 0.653926 |
| GOBP_GLIAL_CELL_DIFFERENTIATION | 190 | 0.242575 | 1.118578 | 0.152047 | 0.653045 |
| GOBP_POSITIVE_REGULATION_OF_GLUCOSE_TRANSMEMBRANE_TRANSPORT | 37 | 0.322998 | 1.118207 | 0.27 | 0.652949 |
| GOBP_NAD_METABOLIC_PROCESS | 50 | 0.305553 | 1.117936 | 0.256757 | 0.652438 |
| GOBP_NEGATIVE_REGULATION_OF_CELL_DEVELOPMENT | 152 | 0.246198 | 1.115161 | 0.167539 | 0.65993 |
| GOBP_RIBOSE_PHOSPHATE_BIOSYNTHETIC_PROCESS | 190 | 0.243372 | 1.115047 | 0.159236 | 0.658944 |
| GOBP_ADAPTATION_OF_SIGNALING_PATHWAY | 20 | 0.369142 | 1.114223 | 0.292754 | 0.660305 |
| GOBP_SCHWANN_CELL_DIFFERENTIATION | 36 | 0.327027 | 1.113629 | 0.29653 | 0.660871 |
| GOBP_TRIGLYCERIDE_METABOLIC_PROCESS | 83 | 0.271923 | 1.113322 | 0.235521 | 0.66051 |
| GOBP_RESPONSE_TO_ACIDIC_PH | 21 | 0.362427 | 1.111958 | 0.30117 | 0.663673 |
| GOBP_GLOBAL_GENOME_NUCLEOTIDE_EXCISION_REPAIR | 26 | 0.347856 | 1.110706 | 0.30226 | 0.666305 |
| GOBP_RRNA_METABOLIC_PROCESS | 236 | 0.230877 | 1.110029 | 0.107143 | 0.667064 |
| GOBP_RESPONSE_TO_ATP | 31 | 0.328371 | 1.109905 | 0.295775 | 0.66616 |
| GOBP_PYRIMIDINE_CONTAINING_COMPOUND_METABOLIC_PROCESS | 82 | 0.268122 | 1.109067 | 0.234615 | 0.667654 |
| GOBP_LIPID_IMPORT_INTO_CELL | 34 | 0.321971 | 1.108762 | 0.257426 | 0.667348 |
| GOBP_ESTABLISHMENT_OR_MAINTENANCE_OF_CELL_POLARITY | 204 | 0.241017 | 1.108092 | 0.186667 | 0.668318 |
| GOBP_POSITIVE_REGULATION_OF_TELOMERE_MAINTENANCE_VIA_TELOMERE_LENGTHENING | 36 | 0.322158 | 1.107464 | 0.291429 | 0.669142 |

| | | | | | |
|---|---|---|---|---|---|
| GOBP_POSITIVE_REGULATION_OF_NEUROBLAST_PROLIFERATION | 18 | -0.565042 | -1.506595 | 0.048154 | 0.165288 |
| GOBP_POSITIVE_REGULATION_OF_RESPONSE_TO_EXTERNAL_STIMULUS | 458 | -0.353614 | -1.505914 | 0 | 0.165783 |
| GOBP_B_CELL_RECEPTOR_SIGNALING_PATHWAY | 119 | -0.401467 | -1.505664 | 0.008872 | 0.165676 |
| GOBP_GAMMA_DELTA_T_CELL_ACTIVATION | 19 | -0.566155 | -1.504006 | 0.028125 | 0.167537 |
| GOBP_BIOLOGICAL_PROCESS_INVOLVED_IN_INTRASPECIES_INTERACTION_BETWEEN_ORGANIS | 45 | -0.470465 | -1.503718 | 0.014265 | 0.167504 |
| GOBP_TOLL_LIKE_RECEPTOR_2_SIGNALING_PATHWAY | 17 | -0.566999 | -1.503548 | 0.027464 | 0.167232 |
| GOBP_CD4_POSITIVE_ALPHA_BETA_T_CELL_CYTOKINE_PRODUCTION | 17 | -0.570874 | -1.503357 | 0.043614 | 0.167012 |
| GOBP_REGULATION_OF_INNATE_IMMUNE_RESPONSE | 291 | -0.369588 | -1.502884 | 0.003468 | 0.167175 |
| GOBP_POSITIVE_REGULATION_OF_ENDOTHELIAL_CELL_APOPTOTIC_PROCESS | 17 | -0.573848 | -1.501438 | 0.038278 | 0.168678 |
| GOBP_POSITIVE_REGULATION_OF_SMALL_GTPASE_MEDIATED_SIGNAL_TRANSDUCTION | 74 | -0.428085 | -1.501164 | 0.007762 | 0.168603 |
| GOBP_BILE_ACID_BIOSYNTHETIC_PROCESS | 33 | -0.492969 | -1.500465 | 0.03003 | 0.169091 |
| GOBP_POSITIVE_REGULATION_OF_MACROPHAGE_ACTIVATION | 24 | -0.529196 | -1.499802 | 0.048921 | 0.169597 |
| GOBP_PROTEIN_LOCALIZATION_TO_CELL_JUNCTION | 102 | -0.4096 | -1.499617 | 0.011658 | 0.169388 |
| GOBP_REGULATION_OF_RESPONSE_TO_BIOTIC_STIMULUS | 383 | -0.356883 | -1.498987 | 0 | 0.169909 |
| GOBP_CELL_DIFFERENTIATION_IN_SPINAL_CORD | 32 | -0.50996 | -1.498604 | 0.026393 | 0.170041 |
| GOBP_REGULATION_OF_RESPONSE_TO_INTERFERON_GAMMA | 24 | -0.536551 | -1.496422 | 0.042618 | 0.172744 |
| GOBP_REGULATION_OF_RECEPTOR_INTERNALIZATION | 50 | -0.453675 | -1.496409 | 0.020862 | 0.172261 |
| GOBP_HORMONE_MEDIATED_SIGNALING_PATHWAY | 163 | -0.385483 | -1.49636 | 0.002454 | 0.171844 |
| GOBP_NEGATIVE_REGULATION_OF_EPITHELIAL_CELL_PROLIFERATION | 118 | -0.40038 | -1.495382 | 0.007585 | 0.172694 |
| GOBP_NEGATIVE_REGULATION_OF_RESPONSE_TO_EXTERNAL_STIMULUS | 320 | -0.360035 | -1.495293 | 0.001129 | 0.172337 |
| GOBP_LYMPH_NODE_DEVELOPMENT | 16 | -0.57591 | -1.495261 | 0.048077 | 0.171907 |
| GOBP_ICOSANOID_METABOLIC_PROCESS | 97 | -0.41057 | -1.495218 | 0.007958 | 0.171485 |
| GOBP_MONOCYTE_DIFFERENTIATION | 34 | -0.492674 | -1.493908 | 0.027859 | 0.172826 |
| GOBP_REGULATION_OF_PEPTIDYL_TYROSINE_PHOSPHORYLATION | 234 | -0.371534 | -1.49377 | 0 | 0.17255 |
| GOBP_EPITHELIAL_TO_MESENCHYMAL_TRANSITION | 136 | -0.394003 | -1.493675 | 0.007509 | 0.172183 |
| GOBP_GLAND_MORPHOGENESIS | 98 | -0.407504 | -1.492559 | 0.011858 | 0.173367 |
| GOBP_POSITIVE_REGULATION_OF_DEFENSE_RESPONSE_TO_VIRUS_BY_HOST | 31 | -0.49618 | -1.492448 | 0.042424 | 0.173029 |
| GOBP_POSITIVE_REGULATION_OF_WOUND_HEALING | 51 | -0.453198 | -1.491676 | 0.026798 | 0.173662 |
| GOBP_NEURAL_TUBE_FORMATION | 91 | -0.415428 | -1.490606 | 0.002729 | 0.174743 |
| GOBP_REGULATION_OF_PHAGOCYTOSIS | 87 | -0.414158 | -1.489737 | 0.006649 | 0.175682 |
| GOBP_NEGATIVE_REGULATION_OF_TRANSMEMBRANE_RECEPTOR_PROTEIN_SERINE_THREONI | 115 | -0.400217 | -1.488425 | 0.003812 | 0.177199 |
| GOBP_REGULATION_OF_LYMPHOCYTE_MIGRATION | 60 | -0.441422 | -1.488211 | 0.017833 | 0.177052 |
| GOBP_DORSAL_VENTRAL_NEURAL_TUBE_PATTERNING | 19 | -0.552577 | -1.485778 | 0.047544 | 0.180236 |
| GOBP_POSITIVE_REGULATION_OF_GLIOGENESIS | 57 | -0.435143 | -1.485622 | 0.015278 | 0.179997 |
| GOBP_COCHLEA_DEVELOPMENT | 36 | -0.48462 | -1.485521 | 0.031884 | 0.179664 |
| GOBP_TOLL_LIKE_RECEPTOR_4_SIGNALING_PATHWAY | 38 | -0.471165 | -1.485015 | 0.041958 | 0.179958 |
| GOBP_VASCULAR_ENDOTHELIAL_CELL_PROLIFERATION | 22 | -0.530475 | -1.484601 | 0.045732 | 0.180118 |
| GOBP_REGULATION_OF_B_CELL_RECEPTOR_SIGNALING_PATHWAY | 25 | -0.527204 | -1.484573 | 0.035821 | 0.179669 |
| GOBP_REGULATION_OF_MAST_CELL_ACTIVATION | 39 | -0.471409 | -1.482218 | 0.041899 | 0.182756 |
| GOBP_NEGATIVE_REGULATION_OF_NATURAL_KILLER_CELL_MEDIATED_IMMUNITY | 18 | -0.571722 | -1.480251 | 0.044554 | 0.185299 |
| GOBP_B_CELL_ACTIVATION | 294 | -0.3602 | -1.480218 | 0.001134 | 0.18485 |
| GOBP_NEGATIVE_REGULATION_OF_PEPTIDYL_SERINE_PHOSPHORYLATION | 27 | -0.510773 | -1.479983 | 0.035139 | 0.184738 |
| GOBP_POSITIVE_REGULATION_OF_INFLAMMATORY_RESPONSE | 116 | -0.398663 | -1.47994 | 0.011335 | 0.184321 |
| GOBP_NEPHRON_MORPHOGENESIS | 59 | -0.437142 | -1.478359 | 0.026316 | 0.186236 |
| GOBP_REGULATION_OF_SMOOTH_MUSCLE_CELL_DIFFERENTIATION | 30 | -0.500804 | -1.477957 | 0.027356 | 0.186324 |
| GOBP_PERICARDIUM_DEVELOPMENT | 15 | -0.583208 | -1.477831 | 0.054602 | 0.186021 |
| GOBP_POSITIVE_REGULATION_OF_ALPHA_BETA_T_CELL_PROLIFERATION | 21 | -0.538843 | -1.477639 | 0.049521 | 0.185853 |
| GOBP_DETECTION_OF_CHEMICAL_STIMULUS | 118 | -0.395152 | -1.477196 | 0.017566 | 0.186002 |
| GOBP_EXTRINSIC_APOPTOTIC_SIGNALING_PATHWAY | 206 | -0.368927 | -1.477088 | 0.002398 | 0.185697 |
| GOBP_NEGATIVE_REGULATION_OF_CARTILAGE_DEVELOPMENT | 21 | -0.537924 | -1.477038 | 0.050536 | 0.185277 |
| GOBP_VENTRAL_SPINAL_CORD_DEVELOPMENT | 30 | -0.503007 | -1.475962 | 0.032211 | 0.186432 |
| GOBP_PYROPTOSIS | 18 | -0.567898 | -1.475753 | 0.049005 | 0.186265 |
| GOBP_PROSTANOID_BIOSYNTHETIC_PROCESS | 26 | -0.513246 | -1.47536 | 0.03937 | 0.18639 |
| GOBP_REGULATION_OF_MICROGLIAL_CELL_ACTIVATION | 17 | -0.551634 | -1.47444 | 0.051724 | 0.187255 |
| GOBP_ANTIGEN_PROCESSING_AND_PRESENTATION_OF_ENDOGENOUS_PEPTIDE_ANTIGEN | 19 | -0.556249 | -1.473791 | 0.041854 | 0.187886 |
| GOBP_TRABECULA_FORMATION | 20 | -0.539468 | -1.471926 | 0.052067 | 0.19043 |
| GOBP_LEUKOCYTE_CHEMOTAXIS | 208 | -0.369032 | -1.471672 | 0.002389 | 0.190377 |
| GOBP_ESTABLISHMENT_OF_PROTEIN_LOCALIZATION_TO_ENDOPLASMIC_RETICULUM | 119 | -0.392728 | -1.470399 | 0.00885 | 0.191766 |
| GOBP_NEGATIVE_REGULATION_OF_T_CELL_PROLIFERATION | 63 | -0.433701 | -1.469022 | 0.024931 | 0.19351 |
| GOBP_REGULATION_OF_T_CELL_RECEPTOR_SIGNALING_PATHWAY | 40 | -0.471296 | -1.468665 | 0.035613 | 0.193615 |
| GOBP_REGULATION_OF_DNA_BINDING_TRANSCRIPTION_FACTOR_ACTIVITY | 396 | -0.348686 | -1.468533 | 0.002232 | 0.193332 |
| GOBP_REGULATION_OF_SKELETAL_MUSCLE_TISSUE_DEVELOPMENT | 21 | -0.529209 | -1.466548 | 0.052632 | 0.195966 |
| GOBP_CELL_CHEMOTAXIS | 274 | -0.358669 | -1.466299 | 0.001126 | 0.195952 |
| GOBP_NEGATIVE_REGULATION_OF_LIPASE_ACTIVITY | 16 | -0.565015 | -1.465928 | 0.0512 | 0.196069 |
| GOBP_ESTROGEN_METABOLIC_PROCESS | 26 | -0.507903 | -1.465838 | 0.045249 | 0.195696 |
| GOBP_NEURAL_TUBE_DEVELOPMENT | 135 | -0.384722 | -1.465806 | 0.008986 | 0.195263 |
| GOBP_PULMONARY_VALVE_MORPHOGENESIS | 15 | -0.583844 | -1.464991 | 0.059006 | 0.196145 |
| GOBP_NEGATIVE_REGULATION_OF_T_CELL_MEDIATED_IMMUNITY | 21 | -0.534429 | -1.464942 | 0.037538 | 0.195761 |
| GOBP_ADENYLATE_CYCLASE_ACTIVATING_ADRENERGIC_RECEPTOR_SIGNALING_PATHWAY | 18 | -0.557211 | -1.464633 | 0.047468 | 0.195765 |
| GOBP_REACTIVE_OXYGEN_SPECIES_BIOSYNTHETIC_PROCESS | 106 | -0.397758 | -1.464261 | 0.007624 | 0.195831 |
| GOBP_REGULATION_OF_MONOCYTE_CHEMOTAXIS | 24 | -0.514359 | -1.464172 | 0.043939 | 0.195459 |
| GOBP_POSTSYNAPTIC_SPECIALIZATION_ASSEMBLY | 17 | -0.573867 | -1.464152 | 0.058372 | 0.195026 |
| GOBP_UROGENITAL_SYSTEM_DEVELOPMENT | 281 | -0.355971 | -1.463292 | 0 | 0.195933 |
| GOBP_MYELOID_DENDRITIC_CELL_DIFFERENTIATION | 18 | -0.552702 | -1.463213 | 0.061611 | 0.195576 |
| GOBP_STORE_OPERATED_CALCIUM_ENTRY | 18 | -0.563904 | -1.462857 | 0.037707 | 0.195657 |
| GOBP_REGULATION_OF_LYMPHOCYTE_CHEMOTAXIS | 24 | -0.516568 | -1.460924 | 0.051622 | 0.198257 |
| GOBP_DEFENSE_RESPONSE_TO_BACTERIUM | 245 | -0.358466 | -1.459896 | 0.002281 | 0.199554 |
| GOBP_NEGATIVE_REGULATION_OF_CELLULAR_RESPONSE_TO_GROWTH_FACTOR_STIMULUS | 88 | -0.409276 | -1.45895 | 0.023873 | 0.200501 |
| GOBP_MYELOID_LEUKOCYTE_MIGRATION | 198 | -0.368938 | -1.457351 | 0.001183 | 0.202659 |
| GOBP_POSITIVE_REGULATION_OF_INTERLEUKIN_17_PRODUCTION | 17 | -0.560179 | -1.45692 | 0.068006 | 0.202873 |
| GOBP_NEGATIVE_REGULATION_OF_IMMUNE_RESPONSE | 133 | -0.385209 | -1.456699 | 0.005128 | 0.202795 |
| GOBP_RESPONSE_TO_PROTOZOAN | 25 | -0.515224 | -1.456445 | 0.036419 | 0.202788 |
| GOBP_REGULATION_OF_POSTSYNAPTIC_MEMBRANE_NEUROTRANSMITTER_RECEPTOR_LEVELS | 54 | -0.444401 | -1.455801 | 0.035568 | 0.203343 |
| GOBP_RECEPTOR_LOCALIZATION_TO_SYNAPSE | 47 | -0.452988 | -1.455578 | 0.040057 | 0.20327 |
| GOBP_B_CELL_DIFFERENTIATION | 120 | -0.389863 | -1.455463 | 0.009126 | 0.20295 |
| GOBP_MICROGLIAL_CELL_ACTIVATION | 43 | -0.457828 | -1.454883 | 0.034884 | 0.203352 |
| GOBP_CORONARY_VASCULATURE_DEVELOPMENT | 38 | -0.469963 | -1.453819 | 0.032164 | 0.204586 |
| GOBP_FEMALE_SEX_DIFFERENTIATION | 96 | -0.400358 | -1.453333 | 0.016021 | 0.204918 |
| GOBP_INSULIN_SECRETION | 183 | -0.364746 | -1.453266 | 0.006075 | 0.204518 |
| GOBP_RESPONSE_TO_TESTOSTERONE | 39 | -0.462698 | -1.453013 | 0.04271 | 0.204482 |
| GOBP_KIDNEY_MESENCHYME_DEVELOPMENT | 16 | -0.559563 | -1.452627 | 0.051037 | 0.204643 |
| GOBP_POSITIVE_REGULATION_OF_P38MAPK_CASCADE | 24 | -0.523812 | -1.452135 | 0.05296 | 0.20499 |
| GOBP_GENITALIA_DEVELOPMENT | 31 | -0.493263 | -1.44939 | 0.04893 | 0.209069 |
| GOBP_NEGATIVE_REGULATION_OF_CELL_JUNCTION_ASSEMBLY | 27 | -0.495784 | -1.448575 | 0.057143 | 0.209975 |
| GOBP_NEGATIVE_REGULATION_OF_PEPTIDASE_ACTIVITY | 186 | -0.368536 | -1.447753 | 0.0012 | 0.210834 |
| GOBP_NEUROTRANSMITTER_RECEPTOR_INTERNALIZATION | 21 | -0.522617 | -1.447605 | 0.06298 | 0.210598 |
| GOBP_NEGATIVE_REGULATION_OF_OSSIFICATION | 30 | -0.494683 | -1.446986 | 0.060241 | 0.211028 |
| GOBP_NEURAL_TUBE_PATTERNING | 31 | -0.486663 | -1.446312 | 0.059406 | 0.211637 |
| GOBP_POSITIVE_REGULATION_OF_SKELETAL_MUSCLE_TISSUE_DEVELOPMENT | 15 | -0.575998 | -1.446279 | 0.049689 | 0.211211 |
| GOBP_REGULATION_OF_WOUND_HEALING | 111 | -0.386538 | -1.445834 | 0.015345 | 0.211479 |
| GOBP_POSITIVE_REGULATION_OF_CHEMOTAXIS | 132 | -0.383061 | -1.445705 | 0.019704 | 0.211167 |
| GOBP_SMOOTH_MUSCLE_CELL_PROLIFERATION | 130 | -0.377893 | -1.445471 | 0.010013 | 0.211072 |
| GOBP_RESPONSE_TO_FLUID_SHEAR_STRESS | 30 | -0.47744 | -1.445096 | 0.055882 | 0.211193 |
| GOBP_EPITHELIAL_TUBE_FORMATION | 114 | -0.389916 | -1.444942 | 0.015267 | 0.210974 |
| GOBP_T_CELL_CHEMOTAXIS | 25 | -0.503049 | -1.444765 | 0.053571 | 0.210839 |
| GOBP_NEGATIVE_REGULATION_OF_EPITHELIAL_TO_MESENCHYMAL_TRANSITION | 33 | -0.476501 | -1.444734 | 0.041605 | 0.210398 |
| GOBP_NOREPINEPHRINE_TRANSPORT | 21 | -0.525908 | -1.444291 | 0.057437 | 0.210711 |
| GOBP_NEGATIVE_REGULATION_OF_VASCULATURE_DEVELOPMENT | 96 | -0.39266 | -1.443997 | 0.020633 | 0.210729 |
| GOBP_CALCIUM_ION_IMPORT_INTO_CYTOSOL | 18 | -0.545825 | -1.442921 | 0.056489 | 0.211972 |
| GOBP_SPLICEOSOMAL_TRI_SNRNP_COMPLEX_ASSEMBLY | 23 | -0.508581 | -1.442678 | 0.057453 | 0.211876 |
| GOBP_REGIONALIZATION | 259 | -0.352754 | -1.442496 | 0.001172 | 0.211688 |
| GOBP_SEX_DETERMINATION | 15 | -0.569015 | -1.442466 | 0.076312 | 0.211264 |
| GOBP_REGULATION_OF_NEUROTRANSMITTER_RECEPTOR_ACTIVITY | 69 | -0.416935 | -1.441316 | 0.041494 | 0.212695 |
| GOBP_NEGATIVE_REGULATION_OF_CHEMOKINE_PRODUCTION | 22 | -0.514641 | -1.440828 | 0.068249 | 0.212968 |
| GOBP_RESPONSE_TO_FUNGUS | 36 | -0.47028 | -1.440762 | 0.061765 | 0.212591 |
| GOBP_REGULATION_OF_EPIDERMAL_GROWTH_FACTOR_ACTIVATED_RECEPTOR_ACTIVITY | 29 | -0.497929 | -1.439017 | 0.04414 | 0.214967 |
| GOBP_NUCLEOTIDE_BINDING_DOMAIN_LEUCINE_RICH_REPEAT_CONTAINING_RECEPTOR_SIGN | 41 | -0.453439 | -1.438449 | 0.052933 | 0.215539 |
| GOBP_POSITIVE_CHEMOTAXIS | 54 | -0.436887 | -1.438411 | 0.030899 | 0.215118 |
| GOBP_REGULATION_OF_LEUKOCYTE_CHEMOTAXIS | 112 | -0.385703 | -1.437786 | 0.012887 | 0.215665 |
| GOBP_NEGATIVE_REGULATION_OF_LIPID_TRANSPORT | 29 | -0.499293 | -1.437738 | 0.067365 | 0.215284 |
| GOBP_PEPTIDYL_GLUTAMIC_ACID_MODIFICATION | 27 | -0.504136 | -1.436917 | 0.055031 | 0.216182 |
| GOBP_DETECTION_OF_BIOTIC_STIMULUS | 33 | -0.47342 | -1.436893 | 0.055046 | 0.215742 |
| GOBP_DIGESTIVE_TRACT_MORPHOGENESIS | 38 | -0.467881 | -1.436463 | 0.039695 | 0.216019 |
| GOBP_POSITIVE_REGULATION_OF_T_CELL_CYTOKINE_PRODUCTION | 20 | -0.523389 | -1.434781 | 0.076923 | 0.218342 |
| GOBP_ACTIVATION_OF_NF_KAPPAB_INDUCING_KINASE_ACTIVITY | 18 | -0.538114 | -1.434767 | 0.07109 | 0.217888 |
| GOBP_POSITIVE_REGULATION_OF_MONONUCLEAR_CELL_MIGRATION | 59 | -0.422795 | -1.434405 | 0.026316 | 0.218016 |
| GOBP_POSITIVE_REGULATION_OF_RESPONSE_TO_BIOTIC_STIMULUS | 231 | -0.353886 | -1.432399 | 0.004592 | 0.220914 |
| GOBP_UTERUS_DEVELOPMENT | 15 | -0.564627 | -1.432169 | 0.065916 | 0.220869 |
| GOBP_ADENYLATE_CYCLASE_INHIBITING_G_PROTEIN_COUPLED_RECEPTOR_SIGNALING_PATH | 64 | -0.430522 | -1.431477 | 0.040334 | 0.221636 |
| GOBP_TRABECULA_MORPHOGENESIS | 40 | -0.446354 | -1.431434 | 0.040404 | 0.221244 |
| GOBP_MAMMARY_GLAND_ALVEOLUS_DEVELOPMENT | 15 | -0.569325 | -1.430852 | 0.067103 | 0.221753 |
| GOBP_CALCIUM_ION_IMPORT | 72 | -0.408145 | -1.430277 | 0.028378 | 0.222273 |
| GOBP_MONOAMINE_TRANSPORT | 64 | -0.420234 | -1.430229 | 0.032653 | 0.22188 |
| GOBP_COTRANSLATIONAL_PROTEIN_TARGETING_TO_MEMBRANE | 105 | -0.387671 | -1.430149 | 0.023286 | 0.221544 |
| GOBP_SECRETORY_GRANULE_ORGANIZATION | 29 | -0.48533 | -1.429897 | 0.061765 | 0.221493 |
| GOBP_NUCLEOTIDE_BINDING_OLIGOMERIZATION_DOMAIN_CONTAINING_2_SIGNALING_PATHWA | 16 | -0.56346 | -1.429871 | 0.076433 | 0.221069 |
| GOBP_REGULATION_OF_ENDOTHELIAL_CELL_DIFFERENTIATION | 34 | -0.464524 | -1.429085 | 0.042489 | 0.221881 |
| GOBP_MACROPHAGE_DIFFERENTIATION | 45 | -0.441722 | -1.427561 | 0.03125 | 0.224162 |
| GOBP_REGULATION_OF_DEFENSE_RESPONSE_TO_VIRUS | 67 | -0.41311 | -1.427046 | 0.037552 | 0.224574 |
| GOBP_CELL_CELL_ADHESION_MEDIATED_BY_INTEGRIN | 17 | -0.54738 | -1.426911 | 0.06869 | 0.22437 |
| GOBP_NEGATIVE_REGULATION_OF_CALCIUM_MEDIATED_SIGNALING | 17 | -0.550199 | -1.426185 | 0.082942 | 0.225239 |
| GOBP_REGULATION_OF_LEUKOCYTE_MIGRATION | 193 | -0.359672 | -1.425542 | 0.008226 | 0.225837 |
| GOBP_INTERFERON_BETA_PRODUCTION | 52 | -0.437891 | -1.424958 | 0.035261 | 0.226398 |
| GOBP_MAST_CELL_ACTIVATION | 58 | -0.421916 | -1.423947 | 0.035714 | 0.227747 |
| GOBP_REGULATION_OF_CALCIUM_ION_IMPORT | 33 | -0.479028 | -1.423836 | 0.054993 | 0.22743 |
| GOBP_HEAT_GENERATION | 16 | -0.558224 | -1.423291 | 0.072327 | 0.227845 |
| GOBP_REGULATION_OF_TUBE_SIZE | 107 | -0.384203 | -1.423236 | 0.020305 | 0.22746 |
| GOBP_NEGATIVE_REGULATION_OF_LEUKOCYTE_MEDIATED_CYTOTOXICITY | 21 | -0.525625 | -1.423213 | 0.069021 | 0.227031 |
| GOBP_T_CELL_RECEPTOR_SIGNALING_PATHWAY | 198 | -0.357777 | -1.422696 | 0.005917 | 0.227488 |
| GOBP_SPINAL_CORD_DEVELOPMENT | 75 | -0.401778 | -1.421894 | 0.024096 | 0.228366 |
| GOBP_REGULATION_OF_NERVOUS_SYSTEM_PROCESS | 119 | -0.373131 | -1.420517 | 0.014121 | 0.230235 |
| GOBP_REGULATION_OF_EPITHELIAL_CELL_DIFFERENTIATION | 122 | -0.378469 | -1.419737 | 0.017544 | 0.231213 |
| GOBP_RETINAL_GANGLION_CELL_AXON_GUIDANCE | 15 | -0.555167 | -1.419455 | 0.091493 | 0.231214 |
| GOBP_TRANSFORMING_GROWTH_FACTOR_BETA_PRODUCTION | 39 | -0.452529 | -1.418775 | 0.049536 | 0.231927 |
| GOBP_AXIS_SPECIFICATION | 64 | -0.41283 | -1.418726 | 0.040595 | 0.231528 |
| GOBP_KIDNEY_MORPHOGENESIS | 73 | -0.403898 | -1.418 | 0.039326 | 0.232371 |
| GOBP_CALCIUM_MEDIATED_SIGNALING | 184 | -0.358225 | -1.417874 | 0.009581 | 0.232146 |
| GOBP_NEGATIVE_REGULATION_OF_CHEMOTAXIS | 53 | -0.42964 | -1.417092 | 0.036711 | 0.233016 |
| GOBP_POSITIVE_REGULATION_OF_COAGULATION | 21 | -0.514095 | -1.417041 | 0.072214 | 0.232634 |
| GOBP_POSITIVE_REGULATION_OF_TRANSFORMING_GROWTH_FACTOR_BETA_PRODUCTION | 19 | -0.526428 | -1.415228 | 0.070015 | 0.235288 |
| GOBP_POSITIVE_REGULATION_OF_LIPID_BIOSYNTHETIC_PROCESS | 71 | -0.412443 | -1.414277 | 0.033195 | 0.236501 |
| GOBP_POSITIVE_REGULATION_OF_EPITHELIAL_CELL_APOPTOTIC_PROCESS | 32 | -0.470521 | -1.413261 | 0.075301 | 0.237876 |
| GOBP_POSITIVE_REGULATION_OF_B_CELL_ACTIVATION | 137 | -0.372834 | -1.412211 | 0.017348 | 0.239333 |
| GOBP_REGULATION_OF_EPIDERMIS_DEVELOPMENT | 58 | -0.421118 | -1.411665 | 0.041609 | 0.239849 |
| GOBP_REGULATION_OF_NEURON_PROJECTION_ARBORIZATION | 15 | -0.563008 | -1.410815 | 0.090214 | 0.240938 |
| GOBP_MULTICELLULAR_ORGANISMAL_RESPONSE_TO_STRESS | 54 | -0.422912 | -1.4079 | 0.037139 | 0.245876 |
| GOBP_RESPONSE_TO_ALCOHOL | 210 | -0.352407 | -1.407254 | 0.007084 | 0.246559 |
| GOBP_MONONUCLEAR_CELL_MIGRATION | 175 | -0.359502 | -1.406794 | 0.008526 | 0.246922 |

| | | | | | |
|---|---|---|---|---|---|
| GOBP_DNA_TEMPLATED_TRANSCRIPTION_TERMINATION | 75 | 0.27972 | 1.106359 | 0.251969 | 0.671396 |
| GOBP_ANION_TRANSMEMBRANE_TRANSPORT | 476 | 0.218615 | 1.106315 | 0.042857 | 0.670198 |
| GOBP_NEGATIVE_REGULATION_OF_CELLULAR_AMIDE_METABOLIC_PROCESS | 166 | 0.245731 | 1.105839 | 0.205263 | 0.670481 |
| GOBP_PURINE_NUCLEOSIDE_MONOPHOSPHATE_BIOSYNTHETIC_PROCESS | 23 | 0.357502 | 1.105782 | 0.289086 | 0.669319 |
| GOBP_AMINO_ACID_BETAINE_METABOLIC_PROCESS | 15 | 0.401494 | 1.105109 | 0.31016 | 0.670179 |
| GOBP_MEMBRANE_REPOLARIZATION_DURING_ACTION_POTENTIAL | 25 | 0.34658 | 1.104918 | 0.291291 | 0.669432 |
| GOBP_SULFUR_COMPOUND_BIOSYNTHETIC_PROCESS | 176 | 0.243365 | 1.104886 | 0.225275 | 0.668211 |
| GOBP_POSITIVE_REGULATION_OF_CELL_CYCLE | 353 | 0.223451 | 1.104651 | 0.156627 | 0.667622 |
| GOBP_MYELIN_MAINTENANCE | 16 | 0.392215 | 1.104478 | 0.318302 | 0.666899 |
| GOBP_CELLULAR_RESPONSE_TO_OXYGEN_RADICAL | 20 | 0.374327 | 1.103581 | 0.320359 | 0.66853 |
| GOBP_REGULATION_OF_CHROMATIN_BINDING | 21 | 0.365376 | 1.102478 | 0.315493 | 0.670645 |
| GOBP_LACTATE_METABOLIC_PROCESS | 15 | 0.395511 | 1.101931 | 0.29397 | 0.671129 |
| GOBP_BILE_ACID_AND_BILE_SALT_TRANSPORT | 25 | 0.351437 | 1.101879 | 0.290909 | 0.669971 |
| GOBP_LIPOPROTEIN_CATABOLIC_PROCESS | 15 | 0.408275 | 1.101123 | 0.316527 | 0.671086 |
| GOBP_REGULATION_OF_AUTOPHAGY_OF_MITOCHONDRION | 31 | 0.330128 | 1.099203 | 0.294461 | 0.675981 |
| GOBP_ORGANIC_ACID_CATABOLIC_PROCESS | 241 | 0.234196 | 1.098685 | 0.159722 | 0.67634 |
| GOBP_REGULATION_OF_DNA_RECOMBINATION | 100 | 0.267255 | 1.098356 | 0.246575 | 0.67606 |
| GOBP_PHOTORECEPTOR_CELL_MAINTENANCE | 36 | 0.320903 | 1.097558 | 0.322785 | 0.677443 |
| GOBP_NEURON_PROJECTION_REGENERATION | 54 | 0.289608 | 1.097299 | 0.247423 | 0.676993 |
| GOBP_SERINE_FAMILY_AMINO_ACID_BIOSYNTHETIC_PROCESS | 20 | 0.370172 | 1.096806 | 0.32687 | 0.677211 |
| GOBP_INTERMEDIATE_FILAMENT_ORGANIZATION | 19 | 0.369762 | 1.095825 | 0.305638 | 0.679119 |
| GOBP_REGULATION_OF_MITOCHONDRIAL_FISSION | 27 | 0.35049 | 1.095618 | 0.310559 | 0.678503 |
| GOBP_POSITIVE_REGULATION_OF_CELL_DIVISION | 70 | 0.276342 | 1.095611 | 0.276515 | 0.677234 |
| GOBP_REGULATION_OF_MULTICELLULAR_ORGANISM_GROWTH | 52 | 0.296751 | 1.095453 | 0.292517 | 0.676451 |
| GOBP_WATER_SOLUBLE_VITAMIN_METABOLIC_PROCESS | 82 | 0.268518 | 1.09491 | 0.259259 | 0.676974 |
| GOBP_POSITIVE_REGULATION_OF_CELL_MATRIX_ADHESION | 52 | 0.284749 | 1.093829 | 0.313869 | 0.679134 |
| GOBP_CGMP_MEDIATED_SIGNALING | 22 | 0.351749 | 1.091383 | 0.339031 | 0.686069 |
| GOBP_PROTEIN_INSERTION_INTO_ER_MEMBRANE | 19 | 0.369648 | 1.090007 | 0.334262 | 0.689305 |
| GOBP_NUCLEOTIDE_EXCISION_REPAIR_DNA_DAMAGE_RECOGNITION | 23 | 0.348608 | 1.089437 | 0.303951 | 0.689984 |
| GOBP_PROTEIN_TETRAMERIZATION | 76 | 0.27214 | 1.088561 | 0.291188 | 0.691545 |
| GOBP_PURINE_NUCLEOBASE_METABOLIC_PROCESS | 18 | 0.37938 | 1.087295 | 0.321229 | 0.694403 |
| GOBP_FEEDING_BEHAVIOR | 67 | 0.274598 | 1.086956 | 0.283688 | 0.694265 |
| GOBP_PROTEIN_LOCALIZATION_TO_CYTOSKELETON | 58 | 0.283855 | 1.085437 | 0.262136 | 0.697942 |
| GOBP_POLYAMINE_METABOLIC_PROCESS | 17 | 0.365674 | 1.083724 | 0.353261 | 0.702429 |
| GOBP_BLASTOCYST_GROWTH | 19 | 0.365104 | 1.083218 | 0.34294 | 0.70279 |
| GOBP_APOPTOTIC_PROCESS_INVOLVED_IN_DEVELOPMENT | 28 | 0.33328 | 1.083112 | 0.33642 | 0.701815 |
| GOBP_GLYCOLYTIC_PROCESS_THROUGH_FRUCTOSE_6_PHOSPHATE | 29 | 0.326983 | 1.082843 | 0.32948 | 0.701368 |
| GOBP_REGULATION_OF_PROTEIN_DEPOLYMERIZATION | 83 | 0.267872 | 1.082652 | 0.268775 | 0.700662 |
| GOBP_REGULATION_OF_ALCOHOL_BIOSYNTHETIC_PROCESS | 68 | 0.275966 | 1.081884 | 0.31 | 0.701806 |
| GOBP_NEUTRAL_LIPID_BIOSYNTHETIC_PROCESS | 43 | 0.299081 | 1.081468 | 0.280936 | 0.701844 |
| GOBP_PHOTORECEPTOR_CELL_DEVELOPMENT | 35 | 0.317402 | 1.081349 | 0.302326 | 0.700908 |
| GOBP_CARDIAC_ATRIUM_DEVELOPMENT | 29 | 0.324661 | 1.079855 | 0.342857 | 0.704588 |
| GOBP_REGULATION_OF_TELOMERE_MAINTENANCE_VIA_TELOMERE_LENGTHENING | 58 | 0.28412 | 1.079583 | 0.329825 | 0.704208 |
| GOBP_CELLULAR_PROCESS_INVOLVED_IN_REPRODUCTION_IN_MULTICELLULAR_ORGANISM | 289 | 0.223476 | 1.079233 | 0.214953 | 0.70408 |
| GOBP_MITOTIC_G1_S_TRANSITION_CHECKPOINT | 62 | 0.281825 | 1.078893 | 0.295533 | 0.703994 |
| GOBP_EPITHELIAL_CELL_MORPHOGENESIS | 28 | 0.330525 | 1.078841 | 0.327434 | 0.702921 |
| GOBP_LONG_CHAIN_FATTY_ACYL_COA_BIOSYNTHETIC_PROCESS | 19 | 0.358518 | 1.078405 | 0.349296 | 0.703033 |
| GOBP_CELLULAR_GLUCOSE_HOMEOSTASIS | 127 | 0.250834 | 1.077525 | 0.276498 | 0.704727 |
| GOBP_BONE_GROWTH | 23 | 0.34856 | 1.077006 | 0.357349 | 0.705244 |
| GOBP_TERMINATION_OF_RNA_POLYMERASE_II_TRANSCRIPTION | 35 | 0.318971 | 1.076643 | 0.305389 | 0.705228 |
| GOBP_INORGANIC_ION_IMPORT_ACROSS_PLASMA_MEMBRANE | 85 | 0.263357 | 1.07574 | 0.275862 | 0.707039 |
| GOBP_NEGATIVE_REGULATION_OF_PROTEIN_CONTAINING_COMPLEX_DISASSEMBLY | 79 | 0.266515 | 1.075559 | 0.328413 | 0.70632 |
| GOBP_VIRAL_TRANSLATION | 16 | 0.377879 | 1.071697 | 0.347826 | 0.717847 |
| GOBP_AUTONOMIC_NERVOUS_SYSTEM_DEVELOPMENT | 36 | 0.314448 | 1.07085 | 0.344411 | 0.719359 |
| GOBP_DNA_CATABOLIC_PROCESS | 35 | 0.306769 | 1.070181 | 0.31761 | 0.720302 |
| GOBP_ENDOCARDIAL_CUSHION_DEVELOPMENT | 36 | 0.307144 | 1.070115 | 0.316923 | 0.71921 |
| GOBP_VENTRICULAR_CARDIAC_MUSCLE_CELL_MEMBRANE_REPOLARIZATION | 23 | 0.346042 | 1.069149 | 0.340782 | 0.721278 |
| GOBP_RENAL_ABSORPTION | 17 | 0.380866 | 1.068502 | 0.360215 | 0.722094 |
| GOBP_DNA_MODIFICATION | 109 | 0.254076 | 1.068239 | 0.3125 | 0.721722 |
| GOBP_REGULATION_OF_HISTONE_H3_K9_METHYLATION | 20 | 0.362529 | 1.067909 | 0.34072 | 0.721568 |
| GOBP_ATP_METABOLIC_PROCESS | 299 | 0.223356 | 1.067873 | 0.214953 | 0.720405 |
| GOBP_ATRIOVENTRICULAR_VALVE_DEVELOPMENT | 23 | 0.347847 | 1.067856 | 0.345133 | 0.71921 |
| GOBP_INNATE_IMMUNE_RESPONSE_IN_MUCOSA | 24 | 0.338078 | 1.067771 | 0.341246 | 0.718229 |
| GOBP_MITOCHONDRIAL_RNA_PROCESSING | 17 | 0.377436 | 1.067747 | 0.358974 | 0.717049 |
| GOBP_HISTONE_MRNA_METABOLIC_PROCESS | 24 | 0.342536 | 1.067517 | 0.341737 | 0.71651 |
| GOBP_RESPONSE_TO_MANGANESE_ION | 18 | 0.373807 | 1.066618 | 0.365169 | 0.718235 |
| GOBP_GLYCEROL_ETHER_METABOLIC_PROCESS | 21 | 0.350849 | 1.066132 | 0.350829 | 0.718608 |
| GOBP_POSITIVE_REGULATION_OF_PROTEASOMAL_UBIQUITIN_DEPENDENT_PROTEIN_CATABOL | 86 | 0.256727 | 1.065132 | 0.311024 | 0.720645 |
| GOBP_CALCIUM_INDEPENDENT_CELL_CELL_ADHESION_VIA_PLASMA_MEMBRANE_CELL_ADHE | 18 | 0.369996 | 1.06321 | 0.388571 | 0.725831 |
| GOBP_FEMALE_GAMETE_GENERATION | 112 | 0.249462 | 1.063209 | 0.333333 | 0.724566 |
| GOBP_REGULATION_OF_TELOMERE_MAINTENANCE | 77 | 0.261929 | 1.063115 | 0.281955 | 0.723595 |
| GOBP_OLIGODENDROCYTE_DIFFERENTIATION | 83 | 0.268321 | 1.06299 | 0.329502 | 0.722776 |
| GOBP_POSITIVE_REGULATION_OF_LIPASE_ACTIVITY | 61 | 0.275874 | 1.061269 | 0.342373 | 0.727306 |
| GOBP_TRNA_METHYLATION | 40 | 0.308108 | 1.06112 | 0.352542 | 0.726484 |
| GOBP_ORGANIC_ACID_TRANSMEMBRANE_TRANSPORT | 136 | 0.240953 | 1.061027 | 0.309524 | 0.72551 |
| GOBP_WATER_HOMEOSTASIS | 64 | 0.273222 | 1.06018 | 0.338182 | 0.72709 |
| GOBP_REGULATION_OF_GRANULOCYTE_DIFFERENTIATION | 15 | 0.37912 | 1.058107 | 0.361538 | 0.73289 |
| GOBP_MOLTING_CYCLE | 84 | 0.25271 | 1.057872 | 0.320988 | 0.73245 |
| GOBP_REGULATION_OF_SISTER_CHROMATID_COHESION | 22 | 0.34727 | 1.056367 | 0.351744 | 0.736098 |
| GOBP_DNA_METHYLATION_OR_DEMETHYLATION | 88 | 0.255444 | 1.055046 | 0.338645 | 0.739169 |
| GOBP_NEGATIVE_REGULATION_OF_LIPID_BIOSYNTHETIC_PROCESS | 45 | 0.290851 | 1.053976 | 0.327759 | 0.741601 |
| GOBP_DNA_DEMETHYLATION | 26 | 0.332726 | 1.053437 | 0.353293 | 0.742173 |
| GOBP_CARBOHYDRATE_BIOSYNTHETIC_PROCESS | 190 | 0.231658 | 1.052182 | 0.3 | 0.745171 |
| GOBP_DEVELOPMENTAL_CELL_GROWTH | 195 | 0.231793 | 1.051991 | 0.281879 | 0.744499 |
| GOBP_DNA_DEALKYLATION | 32 | 0.311658 | 1.050903 | 0.376855 | 0.746959 |
| GOBP_NUCLEOTIDE_EXCISION_REPAIR_DNA_DUPLEX_UNWINDING | 22 | 0.346695 | 1.050837 | 0.361878 | 0.745909 |
| GOBP_NEGATIVE_REGULATION_OF_DEVELOPMENTAL_GROWTH | 92 | 0.253471 | 1.049595 | 0.347458 | 0.749032 |
| GOBP_PROTEIN_CONTAINING_COMPLEX_DISASSEMBLY | 319 | 0.211166 | 1.048316 | 0.256881 | 0.752227 |
| GOBP_CARDIAC_CELL_DEVELOPMENT | 63 | 0.271249 | 1.04592 | 0.356115 | 0.759259 |
| GOBP_LIPID_DROPLET_ORGANIZATION | 26 | 0.328847 | 1.045288 | 0.387283 | 0.760256 |
| GOBP_TERMINATION_OF_RNA_POLYMERASE_I_TRANSCRIPTION | 31 | 0.31277 | 1.045199 | 0.38206 | 0.759259 |
| GOBP_RNA_3_END_PROCESSING | 141 | 0.232679 | 1.044545 | 0.359788 | 0.760348 |
| GOBP_REGULATION_OF_CELL_SUBSTRATE_JUNCTION_ORGANIZATION | 68 | 0.262318 | 1.043893 | 0.353383 | 0.761276 |
| GOBP_REGULATION_OF_CELL_CYCLE_ARREST | 98 | 0.249631 | 1.04358 | 0.325893 | 0.761091 |
| GOBP_HOMEOSTASIS_OF_NUMBER_OF_CELLS | 245 | 0.217768 | 1.042971 | 0.328244 | 0.761918 |
| GOBP_ALPHA_AMINO_ACID_CATABOLIC_PROCESS | 78 | 0.260049 | 1.042764 | 0.372 | 0.761379 |
| GOBP_SUBSTRATE_DEPENDENT_CELL_MIGRATION | 23 | 0.340248 | 1.04206 | 0.402266 | 0.762588 |
| GOBP_REGULATION_OF_STEM_CELL_DIFFERENTIATION | 104 | 0.24228 | 1.041886 | 0.323944 | 0.761942 |
| GOBP_POSITIVE_REGULATION_OF_MITOTIC_NUCLEAR_DIVISION | 40 | 0.298237 | 1.041515 | 0.385382 | 0.761905 |
| GOBP_REGULATION_OF_CAMP_DEPENDENT_PROTEIN_KINASE_ACTIVITY | 19 | 0.358623 | 1.040959 | 0.38253 | 0.76257 |
| GOBP_POSITIVE_REGULATION_OF_TELOMERE_MAINTENANCE | 50 | 0.282024 | 1.040053 | 0.351792 | 0.764323 |
| GOBP_OOCYTE_DIFFERENTIATION | 43 | 0.290738 | 1.038766 | 0.369775 | 0.767487 |
| GOBP_NUCLEOSIDE_TRIPHOSPHATE_BIOSYNTHETIC_PROCESS | 83 | 0.252883 | 1.03849 | 0.393574 | 0.767195 |
| GOBP_MICROTUBULE_DEPOLYMERIZATION | 42 | 0.292369 | 1.038021 | 0.357576 | 0.767419 |
| GOBP_SIGNALING_RECEPTOR_LIGAND_PRECURSOR_PROCESSING | 27 | 0.322177 | 1.037001 | 0.399351 | 0.76988 |
| GOBP_NEGATIVE_REGULATION_OF_ORGANELLE_ORGANIZATION | 329 | 0.213592 | 1.036786 | 0.299065 | 0.769301 |
| GOBP_NEUROTROPHIN_TRK_RECEPTOR_SIGNALING_PATHWAY | 27 | 0.323149 | 1.03613 | 0.379747 | 0.770329 |
| GOBP_POSITIVE_REGULATION_OF_ENDOTHELIAL_CELL_PROLIFERATION | 87 | 0.24922 | 1.035628 | 0.406504 | 0.770825 |
| GOBP_MEMBRANE_REPOLARIZATION | 40 | 0.289735 | 1.034858 | 0.398693 | 0.772369 |
| GOBP_MYELOID_CELL_DEVELOPMENT | 66 | 0.265811 | 1.034141 | 0.377289 | 0.773622 |
| GOBP_POSITIVE_REGULATION_OF_BONE_RESORPTION | 18 | 0.363221 | 1.033106 | 0.401099 | 0.776105 |
| GOBP_REGULATION_OF_VASCULAR_PERMEABILITY | 41 | 0.288813 | 1.033021 | 0.407767 | 0.775147 |
| GOBP_POSITIVE_REGULATION_OF_PATHWAY_RESTRICTED_SMAD_PROTEIN_PHOSPHORYLATIO | 42 | 0.292064 | 1.032893 | 0.40264 | 0.774339 |
| GOBP_NEGATIVE_REGULATION_OF_CELLULAR_SENESCENCE | 19 | 0.355199 | 1.032796 | 0.423497 | 0.773405 |
| GOBP_NEGATIVE_REGULATION_OF_COLD_INDUCED_THERMOGENESIS | 46 | 0.282778 | 1.03195 | 0.396226 | 0.775197 |
| GOBP_CELLULAR_RESPONSE_TO_GLUCAGON_STIMULUS | 23 | 0.32746 | 1.030706 | 0.396067 | 0.778161 |
| GOBP_MICROTUBULE_POLYMERIZATION | 74 | 0.257537 | 1.030539 | 0.384058 | 0.777521 |
| GOBP_RESPONSE_TO_AMYLOID_BETA | 41 | 0.295094 | 1.029681 | 0.412371 | 0.77922 |
| GOBP_MONOSACCHARIDE_BIOSYNTHETIC_PROCESS | 88 | 0.25494 | 1.028505 | 0.349442 | 0.782212 |
| GOBP_ORGAN_OR_TISSUE_SPECIFIC_IMMUNE_RESPONSE | 35 | 0.303829 | 1.02697 | 0.391437 | 0.786262 |
| GOBP_PHOTORECEPTOR_CELL_DIFFERENTIATION | 49 | 0.27922 | 1.025149 | 0.396166 | 0.791457 |
| GOBP_CELL_MATRIX_ADHESION | 214 | 0.222865 | 1.025106 | 0.356643 | 0.79034 |
| GOBP_REGENERATION | 173 | 0.224861 | 1.02477 | 0.374359 | 0.790225 |
| GOBP_GLUCOSE_CATABOLIC_PROCESS | 36 | 0.294097 | 1.022353 | 0.436893 | 0.797559 |
| GOBP_ADULT_LOCOMOTORY_BEHAVIOR | 49 | 0.279452 | 1.022153 | 0.412969 | 0.796985 |
| GOBP_CHONDROITIN_SULFATE_BIOSYNTHETIC_PROCESS | 24 | 0.328072 | 1.022128 | 0.376855 | 0.795803 |
| GOBP_ENSHEATHMENT_OF_NEURONS | 121 | 0.23485 | 1.020781 | 0.405941 | 0.799289 |
| GOBP_REGULATION_OF_DNA_REPAIR | 127 | 0.232691 | 1.02055 | 0.418605 | 0.798824 |
| GOBP_PLATELET_MORPHOGENESIS | 21 | 0.343765 | 1.019858 | 0.432277 | 0.799978 |
| GOBP_TOXIN_TRANSPORT | 38 | 0.285953 | 1.019733 | 0.393939 | 0.799142 |
| GOBP_CELL_DEATH_IN_RESPONSE_TO_HYDROGEN_PEROXIDE | 23 | 0.327294 | 1.019533 | 0.409471 | 0.798531 |
| GOBP_PROTEIN_K11_LINKED_UBIQUITINATION | 29 | 0.319445 | 1.018675 | 0.424856 | 0.800321 |
| GOBP_MODULATION_BY_HOST_OF_VIRAL_PROCESS | 29 | 0.306561 | 1.016084 | 0.403614 | 0.808642 |
| GOBP_REGULATION_OF_CARBOHYDRATE_BIOSYNTHETIC_PROCESS | 88 | 0.24445 | 1.015695 | 0.396624 | 0.808764 |
| GOBP_SKELETAL_SYSTEM_MORPHOGENESIS | 186 | 0.222895 | 1.015642 | 0.393064 | 0.807706 |
| GOBP_APOPTOTIC_PROCESS_INVOLVED_IN_MORPHOGENESIS | 19 | 0.351023 | 1.015497 | 0.436782 | 0.806961 |
| GOBP_SPERM_MOTILITY | 76 | 0.253929 | 1.014244 | 0.433198 | 0.810063 |
| GOBP_REGULATION_OF_CELL_MATRIX_ADHESION | 115 | 0.232213 | 1.013233 | 0.434783 | 0.812361 |
| GOBP_REGULATION_OF_MYELOID_CELL_DIFFERENTIATION | 243 | 0.213481 | 1.012411 | 0.463576 | 0.814004 |
| GOBP_REGULATION_OF_CHROMATIN_ORGANIZATION | 173 | 0.220573 | 1.011974 | 0.430233 | 0.814269 |
| GOBP_CELLULAR_RESPONSE_TO_PEPTIDE_HORMONE_STIMULUS | 293 | 0.211878 | 1.011702 | 0.377049 | 0.814057 |
| GOBP_HEMOGLOBIN_METABOLIC_PROCESS | 16 | 0.358718 | 1.010745 | 0.421717 | 0.816318 |
| GOBP_FERTILIZATION | 142 | 0.225721 | 1.010655 | 0.370558 | 0.815324 |
| GOBP_CELL_PROLIFERATION_IN_FOREBRAIN | 21 | 0.334546 | 1.010614 | 0.436747 | 0.814237 |
| GOBP_SYNAPSE_MATURATION | 24 | 0.319013 | 1.010511 | 0.416901 | 0.813366 |
| GOBP_CARDIAC_ATRIUM_MORPHOGENESIS | 23 | 0.328708 | 1.010043 | 0.431169 | 0.813739 |
| GOBP_AMMONIUM_ION_METABOLIC_PROCESS | 22 | 0.337711 | 1.009556 | 0.451613 | 0.814171 |
| GOBP_PROTEIN_LIPID_COMPLEX_ASSEMBLY | 29 | 0.302845 | 1.009177 | 0.39759 | 0.814312 |
| GOBP_MALE_MEIOSIS_I | 20 | 0.344286 | 1.008912 | 0.428994 | 0.814045 |
| GOBP_NEGATIVE_REGULATION_OF_SMOOTHENED_SIGNALING_PATHWAY | 30 | 0.307152 | 1.008774 | 0.448795 | 0.813322 |
| GOBP_RESPONSE_TO_EXTRACELLULAR_STIMULUS | 423 | 0.203126 | 1.008228 | 0.435294 | 0.813972 |
| GOBP_NEUTRAL_LIPID_CATABOLIC_PROCESS | 34 | 0.295828 | 1.007546 | 0.428571 | 0.815181 |
| GOBP_SLEEP | 27 | 0.306139 | 1.00754 | 0.457576 | 0.813955 |
| GOBP_BASIC_AMINO_ACID_TRANSPORT | 17 | 0.349484 | 1.007309 | 0.418919 | 0.813539 |
| GOBP_POLYSACCHARIDE_BIOSYNTHETIC_PROCESS | 71 | 0.253052 | 1.005945 | 0.447581 | 0.817095 |
| GOBP_MICROTUBULE_BASED_MOVEMENT | 332 | 0.204999 | 1.005426 | 0.398058 | 0.81764 |
| GOBP_NEGATIVE_REGULATION_OF_AXONOGENESIS | 61 | 0.259503 | 1.004573 | 0.458333 | 0.819354 |
| GOBP_NEURON_PROJECTION_EXTENSION | 151 | 0.227821 | 1.004404 | 0.444444 | 0.818704 |
| GOBP_MATURATION_OF_LSU_RRNA_FROM_TRICISTRONIC_RRNA_TRANSCRIPT_SSU_RRNA_5_ | 15 | 0.357427 | 1.004058 | 0.426273 | 0.818724 |
| GOBP_CYTOCHROME_COMPLEX_ASSEMBLY | 36 | 0.290161 | 1.003905 | 0.444444 | 0.818024 |
| GOBP_GRANULOCYTE_DIFFERENTIATION | 32 | 0.301469 | 1.002658 | 0.464832 | 0.82127 |
| GOBP_RESPONSE_TO_FATTY_ACID | 54 | 0.262523 | 1.002241 | 0.425373 | 0.821513 |
| GOBP_NEGATIVE_REGULATION_OF_CELL_CYCLE_G1_S_PHASE_TRANSITION | 100 | 0.23487 | 1.001938 | 0.461538 | 0.821315 |
| GOBP_REGULATION_OF_GLUCONEOGENESIS | 42 | 0.280596 | 1.00164 | 0.434084 | 0.821094 |
| GOBP_POSITIVE_REGULATION_OF_ANION_TRANSMEMBRANE_TRANSPORT | 53 | 0.266198 | 1.000875 | 0.430508 | 0.822592 |
| GOBP_NEGATIVE_REGULATION_OF_AXON_EXTENSION_INVOLVED_IN_AXON_GUIDANCE | 24 | 0.316459 | 1.000656 | 0.45858 | 0.822093 |
| GOBP_REGULATION_OF_HAIR_CYCLE | 25 | 0.315149 | 1.000585 | 0.440828 | 0.821095 |
| GOBP_LONG_TERM_SYNAPTIC_POTENTIATION | 75 | 0.248921 | 0.999958 | 0.452282 | 0.822046 |

| | | | | | |
|---|---|---|---|---|---|
| GOBP_SKELETAL_MUSCLE_ORGAN_DEVELOPMENT | 135 | -0.373013 | -1.406791 | 0.02104 | 0.24644 |
| GOBP_T_CELL_MIGRATION | 63 | -0.416524 | -1.406624 | 0.036415 | 0.246281 |
| GOBP_DIVALENT_INORGANIC_CATION_HOMEOSTASIS | 433 | -0.332803 | -1.406595 | 0.004334 | 0.245859 |
| GOBP_RAB_PROTEIN_SIGNAL_TRANSDUCTION | 21 | -0.519359 | -1.406479 | 0.076563 | 0.245554 |
| GOBP_DIGESTIVE_SYSTEM_DEVELOPMENT | 112 | -0.377171 | -1.406418 | 0.023747 | 0.245199 |
| GOBP_PEPTIDYL_SERINE_DEPHOSPHORYLATION | 19 | -0.530103 | -1.406372 | 0.083205 | 0.244813 |
| GOBP_REGULATION_OF_PEPTIDYL_THREONINE_PHOSPHORYLATION | 44 | -0.444432 | -1.406248 | 0.049157 | 0.244526 |
| GOBP_POSITIVE_REGULATION_OF_NUCLEOCYTOPLASMIC_TRANSPORT | 60 | -0.414686 | -1.405663 | 0.039548 | 0.245023 |
| GOBP_ACID_SECRETION | 34 | -0.46011 | -1.40559 | 0.059701 | 0.244694 |
| GOBP_STEROID_CATABOLIC_PROCESS | 22 | -0.512397 | -1.405051 | 0.071763 | 0.245264 |
| GOBP_REGULATION_OF_T_HELPER_1_TYPE_IMMUNE_RESPONSE | 26 | -0.490593 | -1.404336 | 0.078195 | 0.246081 |
| GOBP_REGULATION_OF_DENDRITE_EXTENSION | 22 | -0.502676 | -1.404054 | 0.071203 | 0.24609 |
| GOBP_NEGATIVE_REGULATION_OF_BLOOD_PRESSURE | 33 | -0.465162 | -1.403232 | 0.051128 | 0.247136 |
| GOBP_INFLAMMATORY_RESPONSE_TO_ANTIGENIC_STIMULUS | 50 | -0.428387 | -1.400533 | 0.066572 | 0.251682 |
| GOBP_EXTRACELLULAR_VESICLE_BIOGENESIS | 22 | -0.504072 | -1.400464 | 0.077778 | 0.251318 |
| GOBP_ENDOTHELIAL_CELL_APOPTOTIC_PROCESS | 52 | -0.423724 | -1.400194 | 0.052482 | 0.251319 |
| GOBP_SPHINGOLIPID_MEDIATED_SIGNALING_PATHWAY | 15 | -0.560531 | -1.399274 | 0.082935 | 0.252587 |
| GOBP_REGULATION_OF_PROTEIN_KINASE_C_SIGNALING | 15 | -0.552975 | -1.398766 | 0.091216 | 0.253072 |
| GOBP_METANEPHRIC_NEPHRON_DEVELOPMENT | 29 | -0.471776 | -1.398227 | 0.062682 | 0.253571 |
| GOBP_MYOBLAST_DIFFERENTIATION | 70 | -0.407609 | -1.396867 | 0.031788 | 0.255728 |
| GOBP_NEGATIVE_REGULATION_OF_LEUKOCYTE_PROLIFERATION | 83 | -0.386231 | -1.396748 | 0.036 | 0.255483 |
| GOBP_POSITIVE_REGULATION_OF_MEMBRANE_PROTEIN_ECTODOMAIN_PROTEOLYSIS | 15 | -0.556473 | -1.395501 | 0.087149 | 0.257259 |
| GOBP_PROSTANOID_METABOLIC_PROCESS | 40 | -0.446958 | -1.395315 | 0.043353 | 0.257097 |
| GOBP_G_PROTEIN_COUPLED_RECEPTOR_SIGNALING_PATHWAY_COUPLED_TO_CYCLIC_NUCLE | 35 | -0.458282 | -1.395189 | 0.080058 | 0.256862 |
| GOBP_SOMITE_DEVELOPMENT | 72 | -0.406053 | -1.394213 | 0.03877 | 0.258202 |
| GOBP_SECOND_MESSENGER_MEDIATED_SIGNALING | 273 | -0.339198 | -1.394164 | 0.004651 | 0.257801 |
| GOBP_MUSCLE_ORGAN_DEVELOPMENT | 275 | -0.340383 | -1.394141 | 0.00904 | 0.257352 |
| GOBP_HUMORAL_IMMUNE_RESPONSE | 290 | -0.342154 | -1.393905 | 0.004635 | 0.257287 |
| GOBP_POSITIVE_REGULATION_OF_SMALL_MOLECULE_METABOLIC_PROCESS | 120 | -0.375877 | -1.393402 | 0.020356 | 0.257806 |
| GOBP_NEGATIVE_REGULATION_OF_EPITHELIAL_CELL_DIFFERENTIATION | 35 | -0.454446 | -1.393288 | 0.064468 | 0.257568 |
| GOBP_VASOCONSTRICTION | 66 | -0.406253 | -1.392704 | 0.053597 | 0.258188 |
| GOBP_NEGATIVE_REGULATION_OF_TISSUE_REMODELING | 17 | -0.539743 | -1.392269 | 0.100156 | 0.258439 |
| GOBP_REGULATION_OF_RUFFLE_ASSEMBLY | 27 | -0.485024 | -1.392167 | 0.070073 | 0.25815 |
| GOBP_SENSORY_PERCEPTION_OF_CHEMICAL_STIMULUS | 129 | -0.369791 | -1.391672 | 0.02104 | 0.258711 |
| GOBP_MODULATION_BY_SYMBIONT_OF_ENTRY_INTO_HOST | 42 | -0.434545 | -1.391364 | 0.052709 | 0.258853 |
| GOBP_NEGATIVE_REGULATION_OF_CELL_KILLING | 24 | -0.491864 | -1.391133 | 0.076577 | 0.258799 |
| GOBP_REGULATION_OF_NMDA_RECEPTOR_ACTIVITY | 33 | -0.456538 | -1.389721 | 0.057101 | 0.261196 |
| GOBP_MUSCLE_CELL_PROLIFERATION | 176 | -0.354341 | -1.389583 | 0.013597 | 0.260993 |
| GOBP_IMMUNOGLOBULIN_PRODUCTION | 187 | -0.352417 | -1.388673 | 0.011806 | 0.262247 |
| GOBP_NEGATIVE_REGULATION_OF_INNATE_IMMUNE_RESPONSE | 58 | -0.416713 | -1.38814 | 0.055633 | 0.262812 |
| GOBP_TOOTH_MINERALIZATION | 18 | -0.534866 | -1.387981 | 0.08871 | 0.262667 |
| GOBP_REGULATION_OF_REACTIVE_OXYGEN_SPECIES_BIOSYNTHETIC_PROCESS | 85 | -0.389581 | -1.387304 | 0.038095 | 0.263538 |
| GOBP_DRUG_METABOLIC_PROCESS | 34 | -0.457479 | -1.38721 | 0.071327 | 0.263247 |
| GOBP_POSITIVE_REGULATION_OF_G_PROTEIN_COUPLED_RECEPTOR_SIGNALING_PATHWAY | 22 | -0.497261 | -1.386806 | 0.080916 | 0.263566 |
| GOBP_INNER_EAR_AUDITORY_RECEPTOR_CELL_DIFFERENTIATION | 31 | -0.460285 | -1.386573 | 0.089552 | 0.263562 |
| GOBP_PEPTIDYL_TYROSINE_MODIFICATION | 339 | -0.334167 | -1.386206 | 0.001112 | 0.263846 |
| GOBP_ESTABLISHMENT_OF_ENDOTHELIAL_BARRIER | 46 | -0.424009 | -1.386181 | 0.061885 | 0.263416 |
| GOBP_CELL_CELL_ADHESION_VIA_PLASMA_MEMBRANE_ADHESION_MOLECULES | 219 | -0.346996 | -1.38601 | 0.005952 | 0.263293 |
| GOBP_CHAPERONE_MEDIATED_AUTOPHAGY | 16 | -0.537653 | -1.384987 | 0.085308 | 0.26485 |
| GOBP_DENDRITIC_SPINE_MORPHOGENESIS | 60 | -0.415544 | -1.384984 | 0.033803 | 0.264377 |
| GOBP_NEGATIVE_REGULATION_OF_ORGANIC_ACID_TRANSPORT | 16 | -0.54 | -1.384937 | 0.105678 | 0.263977 |
| GOBP_NEGATIVE_REGULATION_OF_RECEPTOR_SIGNALING_PATHWAY_VIA_STAT | 23 | -0.492603 | -1.384742 | 0.08147 | 0.263897 |
| GOBP_POSITIVE_REGULATION_OF_PRODUCTION_OF_MOLECULAR_MEDIATOR_OF_IMMUNE_RES | 93 | -0.376858 | -1.383085 | 0.026008 | 0.266521 |
| GOBP_ADRENERGIC_RECEPTOR_SIGNALING_PATHWAY | 22 | -0.507459 | -1.382195 | 0.107544 | 0.267709 |
| GOBP_COLUMNAR_CUBOIDAL_EPITHELIAL_CELL_DIFFERENTIATION | 68 | -0.403163 | -1.381883 | 0.057024 | 0.267833 |
| GOBP_PHOSPHOLIPASE_C_ACTIVATING_G_PROTEIN_COUPLED_RECEPTOR_SIGNALING_PATHW | 80 | -0.393311 | -1.381605 | 0.042466 | 0.267923 |
| GOBP_REGULATION_OF_SYSTEM_PROCESS | 490 | -0.324373 | -1.381399 | 0.002134 | 0.267867 |
| GOBP_REGULATION_OF_BLOOD_CIRCULATION | 240 | -0.340617 | -1.380387 | 0.007109 | 0.269337 |
| GOBP_PEPTIDE_HORMONE_SECRETION | 215 | -0.345661 | -1.380189 | 0.016189 | 0.269239 |
| GOBP_REGULATION_OF_PEPTIDE_HORMONE_SECRETION | 181 | -0.350803 | -1.379867 | 0.009709 | 0.269434 |
| GOBP_POLARIZED_EPITHELIAL_CELL_DIFFERENTIATION | 20 | -0.515148 | -1.37967 | 0.085758 | 0.269349 |
| GOBP_LIPID_TRANSLOCATION | 45 | -0.42003 | -1.378555 | 0.060781 | 0.271122 |
| GOBP_POSITIVE_REGULATION_OF_I_KAPPAB_KINASE_NF_KAPPAB_SIGNALING | 178 | -0.352417 | -1.378402 | 0.015644 | 0.270943 |
| GOBP_EPITHELIAL_CELL_DEVELOPMENT | 187 | -0.350765 | -1.377976 | 0.012063 | 0.271298 |
| GOBP_NEGATIVE_REGULATION_OF_INTERLEUKIN_8_PRODUCTION | 20 | -0.515633 | -1.377884 | 0.080696 | 0.271022 |
| GOBP_SYNCYTIUM_FORMATION | 51 | -0.418271 | -1.375042 | 0.069241 | 0.276236 |
| GOBP_POSTSYNAPTIC_NEUROTRANSMITTER_RECEPTOR_INTERNALIZATION | 18 | -0.518096 | -1.374505 | 0.097403 | 0.276845 |
| GOBP_3_UTR_MEDIATED_MRNA_DESTABILIZATION | 17 | -0.518826 | -1.374314 | 0.090772 | 0.276787 |
| GOBP_CELLULAR_RESPONSE_TO_CALCIUM_ION | 73 | -0.393294 | -1.374198 | 0.048177 | 0.276521 |
| GOBP_UNSATURATED_FATTY_ACID_BIOSYNTHETIC_PROCESS | 45 | -0.426724 | -1.37393 | 0.069602 | 0.276563 |
| GOBP_IMMUNE_RESPONSE_TO_TUMOR_CELL | 20 | -0.510566 | -1.37388 | 0.084848 | 0.276164 |
| GOBP_NEGATIVE_REGULATION_OF_ADAPTIVE_IMMUNE_RESPONSE | 51 | -0.419009 | -1.372552 | 0.056604 | 0.278327 |
| GOBP_SOMITOGENESIS | 54 | -0.418407 | -1.371628 | 0.063739 | 0.279712 |
| GOBP_LEUKOCYTE_MIGRATION | 478 | -0.323348 | -1.371388 | 0.00107 | 0.279715 |
| GOBP_POSITIVE_REGULATION_OF_MYELOID_LEUKOCYTE_MEDIATED_IMMUNITY | 17 | -0.532019 | -1.370685 | 0.097072 | 0.28067 |
| GOBP_MYELOID_LEUKOCYTE_DIFFERENTIATION | 190 | -0.349987 | -1.370349 | 0.016726 | 0.280866 |
| GOBP_POSITIVE_REGULATION_OF_VASOCONSTRICTION | 25 | -0.475246 | -1.36995 | 0.085329 | 0.281134 |
| GOBP_NEGATIVE_REGULATION_OF_MACROPHAGE_ACTIVATION | 16 | -0.53768 | -1.369444 | 0.092767 | 0.281655 |
| GOBP_CYTOLYSIS | 21 | -0.501469 | -1.369229 | 0.091047 | 0.281624 |
| GOBP_POSITIVE_REGULATION_OF_INTERLEUKIN_8_PRODUCTION | 56 | -0.409871 | -1.368613 | 0.0625 | 0.282369 |
| GOBP_MYOTUBE_DIFFERENTIATION | 96 | -0.374499 | -1.368042 | 0.037284 | 0.283033 |
| GOBP_POSITIVE_REGULATION_OF_REGULATED_SECRETORY_PATHWAY | 47 | -0.420914 | -1.368018 | 0.073759 | 0.282599 |
| GOBP_REGULATION_OF_TRANSPORTER_ACTIVITY | 245 | -0.334171 | -1.367665 | 0.005794 | 0.282757 |
| GOBP_REGULATION_OF_EXTRINSIC_APOPTOTIC_SIGNALING_PATHWAY_VIA_DEATH_DOMAIN_RE | 56 | -0.404184 | -1.367241 | 0.056101 | 0.283067 |
| GOBP_PROTEIN_LOCALIZATION_TO_POSTSYNAPSE | 38 | -0.436504 | -1.366921 | 0.07215 | 0.283214 |
| GOBP_NEGATIVE_REGULATION_OF_ERK1_AND_ERK2_CASCADE | 68 | -0.397635 | -1.365076 | 0.053793 | 0.286426 |
| GOBP_EMBRYONIC_AXIS_SPECIFICATION | 28 | -0.474498 | -1.365038 | 0.09687 | 0.286031 |
| GOBP_REGULATION_OF_FATTY_ACID_BETA_OXIDATION | 19 | -0.508295 | -1.364917 | 0.093949 | 0.285809 |
| GOBP_REGULATION_OF_PROTEIN_LOCALIZATION_TO_CELL_PERIPHERY | 113 | -0.366155 | -1.364652 | 0.027273 | 0.285894 |
| GOBP_NUCLEAR_TRANSCRIBED_MRNA_CATABOLIC_PROCESS_NONSENSE_MEDIATED_DECAY | 120 | -0.362356 | -1.364223 | 0.033987 | 0.286305 |
| GOBP_NEGATIVE_REGULATION_OF_SECRETION | 137 | -0.360723 | -1.363802 | 0.034611 | 0.286721 |
| GOBP_NEGATIVE_REGULATION_OF_HORMONE_SECRETION | 55 | -0.408622 | -1.363718 | 0.067024 | 0.286418 |
| GOBP_REGULATION_OF_MEMBRANE_PROTEIN_ECTODOMAIN_PROTEOLYSIS | 23 | -0.489127 | -1.363264 | 0.08855 | 0.28685 |
| GOBP_REGULATION_OF_ENDOTHELIAL_CELL_DEVELOPMENT | 17 | -0.538629 | -1.363137 | 0.108597 | 0.286631 |
| GOBP_POSITIVE_REGULATION_OF_NON_CANONICAL_WNT_SIGNALING_PATHWAY | 15 | -0.540538 | -1.36256 | 0.093313 | 0.287363 |
| GOBP_CELLULAR_RESPONSE_TO_LOW_DENSITY_LIPOPROTEIN_PARTICLE_STIMULUS | 18 | -0.508089 | -1.362252 | 0.10443 | 0.287487 |
| GOBP_NEGATIVE_REGULATION_OF_LOCOMOTION | 293 | -0.330253 | -1.361965 | 0.008979 | 0.287637 |
| GOBP_MULTI_MULTICELLULAR_ORGANISM_PROCESS | 173 | -0.348082 | -1.360772 | 0.018634 | 0.289553 |
| GOBP_TUBE_FORMATION | 129 | -0.362907 | -1.360596 | 0.028607 | 0.28942 |
| GOBP_NEGATIVE_REGULATION_OF_CYCLIN_DEPENDENT_PROTEIN_KINASE_ACTIVITY | 29 | -0.464553 | -1.360202 | 0.102339 | 0.289734 |
| GOBP_POSITIVE_REGULATION_OF_CYTOSOLIC_CALCIUM_ION_CONCENTRATION_INVOLVED_IN_ | 30 | -0.464267 | -1.358961 | 0.095808 | 0.291836 |
| GOBP_MEMBRANE_DEPOLARIZATION_DURING_ACTION_POTENTIAL | 31 | -0.454648 | -1.358872 | 0.089872 | 0.291537 |
| GOBP_MEMBRANE_INVAGINATION | 130 | -0.358718 | -1.357765 | 0.03949 | 0.293345 |
| GOBP_CD40_SIGNALING_PATHWAY | 15 | -0.541821 | -1.356995 | 0.102439 | 0.294501 |
| GOBP_EMBRYONIC_HINDLIMB_MORPHOGENESIS | 25 | -0.476733 | -1.356859 | 0.095164 | 0.294286 |
| GOBP_SIGNAL_RELEASE | 414 | -0.324426 | -1.356553 | 0.005359 | 0.294448 |
| GOBP_NEGATIVE_REGULATION_OF_REPRODUCTIVE_PROCESS | 44 | -0.425356 | -1.355941 | 0.069869 | 0.295267 |
| GOBP_DEFENSE_RESPONSE_TO_FUNGUS | 23 | -0.489545 | -1.35552 | 0.112179 | 0.295698 |
| GOBP_LENS_FIBER_CELL_DIFFERENTIATION | 25 | -0.485128 | -1.355065 | 0.094595 | 0.296151 |
| GOBP_REGULATION_OF_MYOBLAST_DIFFERENTIATION | 42 | -0.428744 | -1.354954 | 0.064234 | 0.295888 |
| GOBP_POSITIVE_REGULATION_OF_RECEPTOR_INTERNALIZATION | 22 | -0.48736 | -1.354898 | 0.113069 | 0.295513 |
| GOBP_REGULATION_OF_ACUTE_INFLAMMATORY_RESPONSE | 43 | -0.421692 | -1.354328 | 0.069209 | 0.296146 |
| GOBP_ROOF_OF_MOUTH_DEVELOPMENT | 74 | -0.387642 | -1.353243 | 0.055777 | 0.297907 |
| GOBP_ANGIOGENESIS_INVOLVED_IN_WOUND_HEALING | 25 | -0.479636 | -1.352793 | 0.084877 | 0.298363 |
| GOBP_POSITIVE_REGULATION_OF_NEUROTRANSMITTER_SECRETION | 17 | -0.518525 | -1.350833 | 0.100775 | 0.302086 |
| GOBP_REGULATION_OF_NIK_NF_KAPPAB_SIGNALING | 92 | -0.371975 | -1.35077 | 0.041499 | 0.301721 |
| GOBP_SENSORY_PERCEPTION_OF_BITTER_TASTE | 25 | -0.475528 | -1.350735 | 0.107413 | 0.301292 |
| GOBP_NEUROTRANSMITTER_RECEPTOR_LOCALIZATION_TO_POSTSYNAPTIC_SPECIALIZATION_M | 15 | -0.535486 | -1.350611 | 0.11875 | 0.301102 |
| GOBP_NEGATIVE_REGULATION_OF_LEUKOCYTE_MIGRATION | 41 | -0.433377 | -1.349986 | 0.085457 | 0.301977 |
| GOBP_TYPE_I_INTERFERON_PRODUCTION | 126 | -0.356388 | -1.349612 | 0.051443 | 0.302228 |
| GOBP_B_CELL_ACTIVATION_INVOLVED_IN_IMMUNE_RESPONSE | 78 | -0.380294 | -1.349374 | 0.051007 | 0.302247 |
| GOBP_NEGATIVE_REGULATION_OF_LIPID_STORAGE | 20 | -0.501398 | -1.348625 | 0.119195 | 0.303315 |
| GOBP_B_CELL_APOPTOTIC_PROCESS | 25 | -0.474981 | -1.346108 | 0.093272 | 0.30816 |
| GOBP_REGULATION_OF_PROTEIN_LOCALIZATION_TO_PLASMA_MEMBRANE | 94 | -0.371425 | -1.346098 | 0.055772 | 0.307693 |
| GOBP_DEFENSE_RESPONSE_TO_GRAM_POSITIVE_BACTERIUM | 70 | -0.38326 | -1.34591 | 0.059392 | 0.307561 |
| GOBP_DETECTION_OF_OTHER_ORGANISM | 17 | -0.51195 | -1.345484 | 0.137318 | 0.307927 |
| GOBP_OUTFLOW_TRACT_SEPTUM_MORPHOGENESIS | 21 | -0.490093 | -1.345458 | 0.115385 | 0.307486 |
| GOBP_POSITIVE_REGULATION_OF_ORGANIC_ACID_TRANSPORT | 32 | -0.451892 | -1.345386 | 0.11226 | 0.307145 |
| GOBP_RESPONSE_TO_CALCIUM_ION | 128 | -0.354482 | -1.344813 | 0.042875 | 0.307883 |
| GOBP_REGULATION_OF_NUCLEOTIDE_BIOSYNTHETIC_PROCESS | 35 | -0.436436 | -1.34325 | 0.102489 | 0.31067 |
| GOBP_ENDOCRINE_PROCESS | 69 | -0.389621 | -1.341794 | 0.066291 | 0.313277 |
| GOBP_POSITIVE_REGULATION_OF_TYPE_2_IMMUNE_RESPONSE | 16 | -0.527086 | -1.341318 | 0.110938 | 0.31381 |
| GOBP_NEGATIVE_REGULATION_OF_FIBROBLAST_GROWTH_FACTOR_RECEPTOR_SIGNALING_PA | 18 | -0.50628 | -1.341069 | 0.117829 | 0.313851 |
| GOBP_REGULATION_OF_ORGANIC_ACID_TRANSPORT | 52 | -0.410171 | -1.341009 | 0.068279 | 0.313502 |
| GOBP_T_HELPER_1_TYPE_IMMUNE_RESPONSE | 40 | -0.434213 | -1.340937 | 0.086083 | 0.313192 |
| GOBP_REGULATION_OF_METAL_ION_TRANSPORT | 220 | -0.334619 | -1.340857 | 0.022145 | 0.312864 |
| GOBP_REGULATION_OF_MORPHOGENESIS_OF_AN_EPITHELIUM | 49 | -0.414693 | -1.340251 | 0.077247 | 0.313724 |
| GOBP_NEGATIVE_REGULATION_OF_MONONUCLEAR_CELL_MIGRATION | 23 | -0.481471 | -1.339162 | 0.104412 | 0.315584 |
| GOBP_FC_RECEPTOR_MEDIATED_STIMULATORY_SIGNALING_PATHWAY | 141 | -0.353476 | -1.339142 | 0.045283 | 0.315131 |
| GOBP_FIBROBLAST_MIGRATION | 43 | -0.421149 | -1.337601 | 0.084406 | 0.317947 |
| GOBP_NEGATIVE_REGULATION_OF_PHOSPHORYLATION | 373 | -0.31997 | -1.336102 | 0.008677 | 0.320786 |
| GOBP_CAMP_MEDIATED_SIGNALING | 54 | -0.408513 | -1.336003 | 0.082489 | 0.320512 |
| GOBP_RESPONSE_TO_INTERLEUKIN_1 | 187 | -0.3392 | -1.335993 | 0.031991 | 0.320035 |
| GOBP_POSTSYNAPSE_ASSEMBLY | 26 | -0.470302 | -1.335863 | 0.102446 | 0.319842 |
| GOBP_REGULATION_OF_MUSCLE_ORGAN_DEVELOPMENT | 35 | -0.438664 | -1.335495 | 0.091172 | 0.320161 |
| GOBP_ACTIVATION_OF_MAPKK_ACTIVITY | 48 | -0.411791 | -1.334745 | 0.091164 | 0.3213 |
| GOBP_REGULATION_OF_PEPTIDASE_ACTIVITY | 369 | -0.321274 | -1.334726 | 0.010056 | 0.320832 |
| GOBP_MONOCYTE_CHEMOTAXIS | 56 | -0.399199 | -1.333879 | 0.078621 | 0.322171 |
| GOBP_REGULATION_OF_LYMPHOCYTE_APOPTOTIC_PROCESS | 50 | -0.410249 | -1.333568 | 0.084821 | 0.322342 |
| GOBP_REGULATION_OF_CELLULAR_RESPONSE_TO_VASCULAR_ENDOTHELIAL_GROWTH_FACT | 22 | -0.481286 | -1.332974 | 0.120944 | 0.323234 |
| GOBP_NEGATIVE_REGULATION_OF_ADENYLATE_CYCLASE_ACTIVITY | 17 | -0.508457 | -1.332929 | 0.137008 | 0.322838 |
| GOBP_POST_EMBRYONIC_DEVELOPMENT | 73 | -0.381305 | -1.332902 | 0.071329 | 0.322395 |
| GOBP_POSITIVE_REGULATION_OF_OSTEOCLAST_DIFFERENTIATION | 23 | -0.472428 | -1.332288 | 0.118087 | 0.323227 |
| GOBP_MUSCLE_ORGAN_MORPHOGENESIS | 60 | -0.394085 | -1.33224 | 0.081492 | 0.322826 |
| GOBP_REGULATION_OF_B_CELL_APOPTOTIC_PROCESS | 19 | -0.496805 | -1.330897 | 0.115031 | 0.325319 |
| GOBP_FORELIMB_MORPHOGENESIS | 29 | -0.448171 | -1.330713 | 0.119938 | 0.325235 |
| GOBP_REGULATION_OF_G_PROTEIN_COUPLED_RECEPTOR_SIGNALING_PATHWAY | 125 | -0.355177 | -1.330556 | 0.047014 | 0.325033 |
| GOBP_REGULATION_OF_MORPHOGENESIS_OF_A_BRANCHING_STRUCTURE | 39 | -0.425501 | -1.329277 | 0.091298 | 0.32737 |
| GOBP_MYELOID_LEUKOCYTE_CYTOKINE_PRODUCTION | 26 | -0.474993 | -1.328994 | 0.109256 | 0.327485 |
| GOBP_NATURAL_KILLER_CELL_ACTIVATION_INVOLVED_IN_IMMUNE_RESPONSE | 15 | -0.523112 | -1.328183 | 0.116057 | 0.328713 |
| GOBP_RESPONSE_TO_SALT_STRESS | 23 | -0.467943 | -1.327882 | 0.12069 | 0.32884 |
| GOBP_ACUTE_INFLAMMATORY_RESPONSE | 91 | -0.366485 | -1.326887 | 0.065596 | 0.330538 |
| GOBP_DETECTION_OF_CHEMICAL_STIMULUS_INVOLVED_IN_SENSORY_PERCEPTION_OF_TASTE | 26 | -0.458619 | -1.324836 | 0.119822 | 0.334662 |
| GOBP_NEGATIVE_REGULATION_OF_G_PROTEIN_COUPLED_RECEPTOR_SIGNALING_PATHWAY | 45 | -0.41598 | -1.323131 | 0.090647 | 0.337972 |
| GOBP_GRANULOCYTE_MIGRATION | 134 | -0.342697 | -1.323128 | 0.048507 | 0.337466 |
| GOBP_POSITIVE_REGULATION_OF_MYOBLAST_DIFFERENTIATION | 15 | -0.527377 | -1.322743 | 0.128713 | 0.337777 |
| GOBP_POSITIVE_REGULATION_OF_LIPID_METABOLIC_PROCESS | 131 | -0.350662 | -1.32159 | 0.054264 | 0.339956 |
| GOBP_CELL_PROLIFERATION_INVOLVED_IN_KIDNEY_DEVELOPMENT | 18 | -0.503131 | -1.32145 | 0.132921 | 0.339743 |

| | | | | | |
|---|---|---|---|---|---|
| GOBP_SULFUR_COMPOUND_METABOLIC_PROCESS | 345 | 0.203503 | 0.99954 | 0.418367 | 0.822187 |
| GOBP_RESPONSE_TO_NITRIC_OXIDE | 16 | 0.345463 | 0.999532 | 0.470914 | 0.820985 |
| GOBP_SPERMATID_DIFFERENTIATION | 126 | 0.232466 | 0.999006 | 0.445415 | 0.821545 |
| GOBP_REGULATION_OF_MYOBLAST_FUSION | 16 | 0.35859 | 0.997964 | 0.474227 | 0.823987 |
| GOBP_POSITIVE_REGULATION_OF_AXONOGENESIS | 72 | 0.25375 | 0.997715 | 0.45098 | 0.823629 |
| GOBP_GTP_METABOLIC_PROCESS | 24 | 0.324079 | 0.997534 | 0.464497 | 0.823067 |
| GOBP_REGULATION_OF_CARDIOCYTE_DIFFERENTIATION | 26 | 0.308085 | 0.997318 | 0.434132 | 0.822596 |
| GOBP_POSITIVE_REGULATION_OF_DNA_REPAIR | 71 | 0.248199 | 0.997283 | 0.433962 | 0.821487 |
| GOBP_CGMP_METABOLIC_PROCESS | 16 | 0.355615 | 0.997046 | 0.4375 | 0.821135 |
| GOBP_DNA_CATABOLIC_PROCESS_ENDONUCLEOLYTIC | 27 | 0.307313 | 0.996856 | 0.420613 | 0.820603 |
| GOBP_PYRUVATE_METABOLIC_PROCESS | 142 | 0.223154 | 0.995943 | 0.456989 | 0.82244 |
| GOBP_REGULATION_OF_INCLUSION_BODY_ASSEMBLY | 15 | 0.35504 | 0.995462 | 0.443526 | 0.822915 |
| GOBP_BONE_DEVELOPMENT | 180 | 0.217933 | 0.995008 | 0.462428 | 0.823278 |
| GOBP_RETINA_HOMEOSTASIS | 64 | 0.260253 | 0.993867 | 0.435374 | 0.825963 |
| GOBP_NEGATIVE_REGULATION_OF_AMYLOID_PRECURSOR_PROTEIN_CATABOLIC_PROCESS | 23 | 0.327936 | 0.993444 | 0.464912 | 0.826279 |
| GOBP_REGULATION_OF_GLUCOSE_METABOLIC_PROCESS | 110 | 0.235822 | 0.992969 | 0.5 | 0.826777 |
| GOBP_REGULATION_OF_MACROPHAGE_DIFFERENTIATION | 20 | 0.333613 | 0.992433 | 0.45584 | 0.827401 |
| GOBP_CYTOKINESIS | 159 | 0.219405 | 0.991941 | 0.489796 | 0.827877 |
| GOBP_NEGATIVE_REGULATION_OF_CARBOHYDRATE_METABOLIC_PROCESS | 46 | 0.270863 | 0.991726 | 0.471761 | 0.827395 |
| GOBP_NUCLEOSIDE_METABOLIC_PROCESS | 101 | 0.238082 | 0.991101 | 0.446429 | 0.82831 |
| GOBP_GLUCOSE_METABOLIC_PROCESS | 192 | 0.219179 | 0.990966 | 0.462857 | 0.8276 |
| GOBP_MITOCHONDRIAL_FUSION | 28 | 0.309784 | 0.989427 | 0.478964 | 0.831827 |
| GOBP_FATTY_ACID_DERIVATIVE_METABOLIC_PROCESS | 70 | 0.247342 | 0.989394 | 0.493377 | 0.830726 |
| GOBP_NEGATIVE_REGULATION_OF_PROTEIN_POLYMERIZATION | 76 | 0.248004 | 0.989024 | 0.459144 | 0.830779 |
| GOBP_NEGATIVE_REGULATION_OF_TRANSLATIONAL_INITIATION | 19 | 0.342666 | 0.988735 | 0.446483 | 0.830526 |
| GOBP_CATECHOL_CONTAINING_COMPOUND_METABOLIC_PROCESS | 38 | 0.285105 | 0.988705 | 0.49495 | 0.829439 |
| GOBP_NUCLEOSIDE_MONOPHOSPHATE_METABOLIC_PROCESS | 72 | 0.249776 | 0.988574 | 0.470817 | 0.828665 |
| GOBP_RESPONSE_TO_HEAT | 155 | 0.217914 | 0.988066 | 0.487047 | 0.829258 |
| GOBP_NEGATIVE_REGULATION_OF_NOTCH_SIGNALING_PATHWAY | 36 | 0.288925 | 0.98801 | 0.455657 | 0.828257 |
| GOBP_DETECTION_OF_LIGHT_STIMULUS | 56 | 0.259554 | 0.98708 | 0.495082 | 0.830215 |
| GOBP_RESPONSE_TO_OXIDATIVE_STRESS | 411 | 0.199176 | 0.985795 | 0.505618 | 0.833443 |
| GOBP_CELLULAR_RESPONSE_TO_RADIATION | 171 | 0.215334 | 0.985513 | 0.473054 | 0.833234 |
| GOBP_DEFINITIVE_HEMOPOIESIS | 19 | 0.33242 | 0.985394 | 0.480556 | 0.832431 |
| GOBP_PYRIDINE_CONTAINING_COMPOUND_BIOSYNTHETIC_PROCESS | 33 | 0.288249 | 0.984246 | 0.444109 | 0.835303 |
| GOBP_NEURON_CELLULAR_HOMEOSTASIS | 31 | 0.294603 | 0.984037 | 0.46875 | 0.834813 |
| GOBP_DICARBOXYLIC_ACID_CATABOLIC_PROCESS | 16 | 0.341629 | 0.983679 | 0.470588 | 0.834871 |
| GOBP_PYRIMIDINE_NUCLEOSIDE_CATABOLIC_PROCESS | 21 | 0.330859 | 0.982777 | 0.482353 | 0.836698 |
| GOBP_NEGATIVE_REGULATION_OF_ATPASE_ACTIVITY | 17 | 0.347952 | 0.98267 | 0.498592 | 0.835887 |
| GOBP_GLYCOSYL_COMPOUND_METABOLIC_PROCESS | 124 | 0.225799 | 0.982459 | 0.491453 | 0.83545 |
| GOBP_STEM_CELL_DIFFERENTIATION | 227 | 0.208358 | 0.982287 | 0.507692 | 0.834867 |
| GOBP_CHAPERONE_MEDIATED_PROTEIN_COMPLEX_ASSEMBLY | 20 | 0.323749 | 0.982255 | 0.448925 | 0.833804 |
| GOBP_AMINE_METABOLIC_PROCESS | 144 | 0.22129 | 0.981902 | 0.482558 | 0.833908 |
| GOBP_FATTY_ACID_CATABOLIC_PROCESS | 102 | 0.234943 | 0.981554 | 0.484305 | 0.833895 |
| GOBP_REGULATION_OF_OXIDATIVE_STRESS_INDUCED_CELL_DEATH | 55 | 0.265483 | 0.981078 | 0.501695 | 0.834321 |
| GOBP_REGULATION_OF_DNA_METABOLIC_PROCESS | 334 | 0.200295 | 0.981078 | 0.533981 | 0.833154 |
| GOBP_ACTIN_CYTOSKELETON_REORGANIZATION | 98 | 0.231469 | 0.97933 | 0.485106 | 0.838119 |
| GOBP_LUNG_ALVEOLUS_DEVELOPMENT | 31 | 0.30161 | 0.978782 | 0.49375 | 0.838772 |
| GOBP_NUCLEOSIDE_TRIPHOSPHATE_METABOLIC_PROCESS | 107 | 0.230099 | 0.978187 | 0.479452 | 0.839683 |
| GOBP_REGULATION_OF_HORMONE_METABOLIC_PROCESS | 32 | 0.294987 | 0.977412 | 0.489297 | 0.841187 |
| GOBP_REGULATION_OF_PRESYNAPTIC_CYTOSOLIC_CALCIUM_ION_CONCENTRATION | 15 | 0.355394 | 0.976819 | 0.484536 | 0.842163 |
| GOBP_EMBRYONIC_PLACENTA_MORPHOGENESIS | 21 | 0.331632 | 0.975951 | 0.473837 | 0.843933 |
| GOBP_LIPOSACCHARIDE_METABOLIC_PROCESS | 104 | 0.232507 | 0.97559 | 0.488987 | 0.843997 |
| GOBP_POSITIVE_REGULATION_OF_STEM_CELL_PROLIFERATION | 20 | 0.333444 | 0.974068 | 0.449721 | 0.848194 |
| GOBP_PYRIDINE_CONTAINING_COMPOUND_METABOLIC_PROCESS | 39 | 0.27824 | 0.972874 | 0.49848 | 0.851179 |
| GOBP_CELLULAR_RESPONSE_TO_LIGHT_STIMULUS | 116 | 0.229305 | 0.972814 | 0.504587 | 0.85022 |
| GOBP_NEURON_MATURATION | 41 | 0.275411 | 0.970998 | 0.489097 | 0.855354 |
| GOBP_RIBOSOMAL_LARGE_SUBUNIT_BIOGENESIS | 71 | 0.247163 | 0.969141 | 0.512097 | 0.860634 |
| GOBP_INTESTINAL_EPITHELIAL_CELL_DIFFERENTIATION | 19 | 0.334222 | 0.968965 | 0.497093 | 0.86004 |
| GOBP_POTASSIUM_ION_HOMEOSTASIS | 24 | 0.313914 | 0.968753 | 0.48433 | 0.859591 |
| GOBP_REGULATION_OF_POLYSACCHARIDE_METABOLIC_PROCESS | 42 | 0.271914 | 0.968382 | 0.526846 | 0.859812 |
| GOBP_TAIL_ANCHORED_MEMBRANE_PROTEIN_INSERTION_INTO_ER_MEMBRANE | 16 | 0.336804 | 0.968323 | 0.513812 | 0.858842 |
| GOBP_REGULATION_OF_UBIQUITIN_PROTEIN_TRANSFERASE_ACTIVITY | 49 | 0.266806 | 0.967961 | 0.537415 | 0.858965 |
| GOBP_CERAMIDE_BIOSYNTHETIC_PROCESS | 65 | 0.245851 | 0.967928 | 0.520446 | 0.857913 |
| GOBP_ELECTRON_TRANSPORT_CHAIN | 169 | 0.212096 | 0.967301 | 0.541667 | 0.858866 |
| GOBP_CELLULAR_RESPONSE_TO_PEPTIDE | 353 | 0.198739 | 0.966732 | 0.537634 | 0.859624 |
| GOBP_CARDIAC_MUSCLE_CELL_MEMBRANE_REPOLARIZATION | 33 | 0.28511 | 0.96625 | 0.516923 | 0.860102 |
| GOBP_POSITIVE_REGULATION_OF_MULTICELLULAR_ORGANISM_GROWTH | 23 | 0.313455 | 0.966164 | 0.510638 | 0.859261 |
| GOBP_REGULATION_OF_PLASMA_LIPOPROTEIN_PARTICLE_LEVELS | 78 | 0.23973 | 0.965941 | 0.510121 | 0.858921 |
| GOBP_VIRAL_RNA_GENOME_REPLICATION | 34 | 0.283468 | 0.965891 | 0.488024 | 0.857951 |
| GOBP_PROTEIN_K48_LINKED_UBIQUITINATION | 61 | 0.254001 | 0.965773 | 0.509363 | 0.857219 |
| GOBP_REGULATION_OF_STEM_CELL_PROLIFERATION | 36 | 0.279821 | 0.965492 | 0.520767 | 0.856929 |
| GOBP_ATRIAL_CARDIAC_MUSCLE_CELL_TO_AV_NODE_CELL_SIGNALING | 18 | 0.327105 | 0.965261 | 0.496 | 0.856534 |
| GOBP_CELL_DEATH_IN_RESPONSE_TO_OXIDATIVE_STRESS | 76 | 0.241537 | 0.965027 | 0.539326 | 0.856175 |
| GOBP_CARBOHYDRATE_TRANSPORT | 133 | 0.219456 | 0.965002 | 0.538462 | 0.85511 |
| GOBP_REGULATION_OF_GLYCOGEN_METABOLIC_PROCESS | 34 | 0.284123 | 0.96493 | 0.509091 | 0.85421 |
| GOBP_PRIMARY_ALCOHOL_METABOLIC_PROCESS | 78 | 0.244851 | 0.964578 | 0.51046 | 0.854295 |
| GOBP_REGULATION_OF_CHROMATIN_ASSEMBLY | 19 | 0.327178 | 0.964239 | 0.494536 | 0.854232 |
| GOBP_NUCLEOSIDE_MONOPHOSPHATE_BIOSYNTHETIC_PROCESS | 41 | 0.272831 | 0.963754 | 0.533784 | 0.854757 |
| GOBP_NEGATIVE_REGULATION_OF_POTASSIUM_ION_TRANSPORT | 26 | 0.29321 | 0.963259 | 0.507418 | 0.855317 |
| GOBP_POSTSYNAPTIC_MEMBRANE_ORGANIZATION | 34 | 0.291902 | 0.96279 | 0.522152 | 0.855756 |
| GOBP_POSITIVE_REGULATION_OF_MRNA_PROCESSING | 30 | 0.293449 | 0.962427 | 0.45858 | 0.855849 |
| GOBP_STEROID_HORMONE_BIOSYNTHETIC_PROCESS | 39 | 0.273012 | 0.962323 | 0.505119 | 0.855096 |
| GOBP_CELLULAR_RESPONSE_TO_EXTRACELLULAR_STIMULUS | 220 | 0.206805 | 0.961569 | 0.583333 | 0.856527 |
| GOBP_MATERNAL_PROCESS_INVOLVED_IN_FEMALE_PREGNANCY | 52 | 0.258024 | 0.960421 | 0.518644 | 0.85922 |
| GOBP_CELLULAR_BIOGENIC_AMINE_METABOLIC_PROCESS | 89 | 0.234648 | 0.959588 | 0.542751 | 0.860931 |
| GOBP_DNA_METHYLATION | 65 | 0.246347 | 0.959404 | 0.526502 | 0.860396 |
| GOBP_G_PROTEIN_COUPLED_RECEPTOR_INTERNALIZATION | 15 | 0.353366 | 0.959393 | 0.502793 | 0.859284 |
| GOBP_POSITIVE_REGULATION_OF_CELLULAR_EXTRAVASATION | 16 | 0.343853 | 0.959275 | 0.508671 | 0.858531 |
| GOBP_GLYCOSPHINGOLIPID_METABOLIC_PROCESS | 63 | 0.246228 | 0.958712 | 0.542254 | 0.85928 |
| GOBP_PYRIMIDINE_CONTAINING_COMPOUND_BIOSYNTHETIC_PROCESS | 39 | 0.270666 | 0.958235 | 0.527508 | 0.859744 |
| GOBP_NUCLEOSIDE_BISPHOSPHATE_METABOLIC_PROCESS | 132 | 0.219528 | 0.95814 | 0.594059 | 0.85887 |
| GOBP_MITOCHONDRIAL_TRANSPORT | 264 | 0.198822 | 0.958104 | 0.627586 | 0.857849 |
| GOBP_PRESYNAPTIC_ENDOCYTOSIS | 57 | 0.251021 | 0.957483 | 0.527211 | 0.858731 |
| GOBP_POSITIVE_REGULATION_OF_HISTONE_DEACETYLATION | 16 | 0.33633 | 0.957479 | 0.508571 | 0.857637 |
| GOBP_ESTABLISHMENT_OF_CELL_POLARITY | 130 | 0.22078 | 0.956938 | 0.567164 | 0.858317 |
| GOBP_INTRA_GOLGI_VESICLE_MEDIATED_TRANSPORT | 34 | 0.278643 | 0.956817 | 0.501587 | 0.857589 |
| GOBP_POTASSIUM_ION_TRANSPORT | 180 | 0.212625 | 0.956446 | 0.598837 | 0.857642 |
| GOBP_DEOXYRIBONUCLEOSIDE_TRIPHOSPHATE_METABOLIC_PROCESS | 15 | 0.34231 | 0.954763 | 0.486559 | 0.861859 |
| GOBP_NUCLEAR_TRANSCRIBED_MRNA_CATABOLIC_PROCESS_EXONUCLEOLYTIC | 35 | 0.276387 | 0.954348 | 0.501558 | 0.862121 |
| GOBP_RIBOSOME_ASSEMBLY | 62 | 0.247067 | 0.954076 | 0.575758 | 0.861915 |
| GOBP_GLUCAN_BIOSYNTHETIC_PROCESS | 42 | 0.274057 | 0.95397 | 0.516892 | 0.861148 |
| GOBP_PROTEIN_INSERTION_INTO_MEMBRANE | 75 | 0.242023 | 0.953741 | 0.571429 | 0.860783 |
| GOBP_POSITIVE_REGULATION_OF_VASCULAR_ENDOTHELIAL_GROWTH_FACTOR_PRODUCTION | 25 | 0.296102 | 0.953108 | 0.485333 | 0.861796 |
| GOBP_PROTON_TRANSMEMBRANE_TRANSPORT | 141 | 0.215645 | 0.9529 | 0.549505 | 0.861386 |
| GOBP_VESICLE_MEDIATED_TRANSPORT_BETWEEN_ENDOSOMAL_COMPARTMENTS | 43 | 0.268444 | 0.952834 | 0.528701 | 0.860483 |
| GOBP_NEGATIVE_CHEMOTAXIS | 42 | 0.268132 | 0.951547 | 0.51875 | 0.863619 |
| GOBP_PYRIMIDINE_NUCLEOBASE_METABOLIC_PROCESS | 16 | 0.340121 | 0.951388 | 0.536023 | 0.863026 |
| GOBP_GLUCOCORTICOID_BIOSYNTHETIC_PROCESS | 15 | 0.344665 | 0.950512 | 0.534091 | 0.864798 |
| GOBP_SYNAPTIC_VESICLE_RECYCLING | 69 | 0.248358 | 0.949937 | 0.567669 | 0.865462 |
| GOBP_CAMERA_TYPE_EYE_MORPHOGENESIS | 91 | 0.229526 | 0.949713 | 0.602459 | 0.865069 |
| GOBP_POSITIVE_REGULATION_OF_CELL_SUBSTRATE_ADHESION | 113 | 0.220872 | 0.949669 | 0.565217 | 0.864135 |
| GOBP_PROTEIN_HOMOTETRAMERIZATION | 50 | 0.255107 | 0.949572 | 0.55137 | 0.863401 |
| GOBP_POSITIVE_REGULATION_OF_TORC1_SIGNALING | 17 | 0.325401 | 0.949242 | 0.513889 | 0.863379 |
| GOBP_N_ACETYLGLUCOSAMINE_METABOLIC_PROCESS | 17 | 0.327746 | 0.948622 | 0.507937 | 0.864335 |
| GOBP_REGULATION_OF_CELL_CYCLE_G1_S_PHASE_TRANSITION | 160 | 0.20473 | 0.946761 | 0.655172 | 0.869117 |
| GOBP_REGULATION_OF_MICROTUBULE_BASED_PROCESS | 214 | 0.200935 | 0.945972 | 0.647482 | 0.8706 |
| GOBP_MITOCHONDRION_LOCALIZATION | 45 | 0.266242 | 0.945764 | 0.558394 | 0.870161 |
| GOBP_CELLULAR_RESPONSE_TO_INSULIN_STIMULUS | 209 | 0.204261 | 0.9454 | 0.641892 | 0.870223 |
| GOBP_CELLULAR_RESPONSE_TO_THYROID_HORMONE_STIMULUS | 16 | 0.328148 | 0.945127 | 0.532967 | 0.870012 |
| GOBP_REGULATION_OF_DNA_DAMAGE_CHECKPOINT | 15 | 0.334384 | 0.944709 | 0.498534 | 0.87024 |
| GOBP_POSITIVE_REGULATION_OF_VASCULAR_ASSOCIATED_SMOOTH_MUSCLE_CELL_MIGRATI | 16 | 0.340166 | 0.943868 | 0.554622 | 0.871873 |
| GOBP_OLEFINIC_COMPOUND_BIOSYNTHETIC_PROCESS | 18 | 0.320006 | 0.943671 | 0.536932 | 0.871368 |
| GOBP_REGULATION_OF_MRNA_3_END_PROCESSING | 28 | 0.289322 | 0.943625 | 0.547468 | 0.870408 |
| GOBP_NEGATIVE_REGULATION_OF_ACTIN_FILAMENT_POLYMERIZATION | 62 | 0.247431 | 0.941916 | 0.541045 | 0.874718 |
| GOBP_REGULATION_OF_RESPONSE_TO_OXIDATIVE_STRESS | 77 | 0.235279 | 0.941649 | 0.576471 | 0.874478 |
| GOBP_RESPONSE_TO_IONIZING_RADIATION | 138 | 0.215495 | 0.941287 | 0.637931 | 0.874504 |
| GOBP_CELL_VOLUME_HOMEOSTASIS | 27 | 0.294706 | 0.940793 | 0.548961 | 0.874988 |
| GOBP_VENTRICULAR_CARDIAC_MUSCLE_CELL_ACTION_POTENTIAL | 32 | 0.27961 | 0.940512 | 0.553398 | 0.874759 |
| GOBP_PYRIMIDINE_NUCLEOTIDE_METABOLIC_PROCESS | 49 | 0.258341 | 0.940398 | 0.546667 | 0.874024 |
| GOBP_NEUROTROPHIN_SIGNALING_PATHWAY | 33 | 0.283141 | 0.939479 | 0.551515 | 0.875916 |
| GOBP_REGULATION_OF_DOUBLE_STRAND_BREAK_REPAIR_VIA_NONHOMOLOGOUS_END_JOIN | 28 | 0.292947 | 0.93834 | 0.536585 | 0.878426 |
| GOBP_ETHER_METABOLIC_PROCESS | 26 | 0.303258 | 0.937679 | 0.534954 | 0.87943 |
| GOBP_FOCAL_ADHESION_ASSEMBLY | 84 | 0.225242 | 0.937568 | 0.600775 | 0.878664 |
| GOBP_REGULATION_OF_CYCLIN_DEPENDENT_PROTEIN_KINASE_ACTIVITY | 96 | 0.227068 | 0.936381 | 0.598394 | 0.881332 |
| GOBP_SOMATIC_DIVERSIFICATION_OF_IMMUNE_RECEPTORS | 75 | 0.232967 | 0.935594 | 0.590038 | 0.882695 |
| GOBP_INNER_EAR_RECEPTOR_CELL_STEREOCILIUM_ORGANIZATION | 27 | 0.285917 | 0.933736 | 0.549254 | 0.887401 |
| GOBP_POSITIVE_REGULATION_OF_PROTEIN_MATURATION | 24 | 0.298071 | 0.933137 | 0.583333 | 0.888221 |
| GOBP_VACUOLAR_ACIDIFICATION | 22 | 0.305305 | 0.932956 | 0.565934 | 0.887694 |
| GOBP_POLYOL_BIOSYNTHETIC_PROCESS | 60 | 0.240044 | 0.932646 | 0.592466 | 0.887568 |
| GOBP_NEUROMUSCULAR_JUNCTION_DEVELOPMENT | 44 | 0.254391 | 0.930668 | 0.579114 | 0.892532 |
| GOBP_REGULATION_OF_SYNAPTIC_VESICLE_EXOCYTOSIS | 53 | 0.242735 | 0.929709 | 0.575862 | 0.894512 |
| GOBP_RENAL_WATER_HOMEOSTASIS | 30 | 0.274604 | 0.927907 | 0.591837 | 0.898879 |
| GOBP_TISSUE_REGENERATION | 61 | 0.23985 | 0.927491 | 0.619772 | 0.899057 |
| GOBP_SEROTONIN_TRANSPORT | 17 | 0.317606 | 0.926892 | 0.531328 | 0.899783 |
| GOBP_HISTONE_H2A_ACETYLATION | 17 | 0.328421 | 0.925245 | 0.579897 | 0.90377 |
| GOBP_VERY_LONG_CHAIN_FATTY_ACID_METABOLIC_PROCESS | 33 | 0.267639 | 0.924002 | 0.602484 | 0.906584 |
| GOBP_RESPIRATORY_SYSTEM_PROCESS | 29 | 0.280803 | 0.923976 | 0.596273 | 0.905543 |
| GOBP_NEGATIVE_REGULATION_OF_FATTY_ACID_METABOLIC_PROCESS | 24 | 0.29921 | 0.922569 | 0.579853 | 0.908793 |
| GOBP_CELLULAR_CARBOHYDRATE_METABOLIC_PROCESS | 264 | 0.191507 | 0.922166 | 0.784 | 0.908909 |
| GOBP_SIGNAL_TRANSDUCTION_BY_P53_CLASS_MEDIATOR | 258 | 0.191535 | 0.922075 | 0.77193 | 0.908087 |
| GOBP_NEGATIVE_REGULATION_OF_CELL_AGING | 25 | 0.291997 | 0.921605 | 0.570248 | 0.908541 |
| GOBP_PHOTOTRANSDUCTION | 42 | 0.263265 | 0.921189 | 0.57764 | 0.908747 |
| GOBP_PROTEIN_MATURATION | 260 | 0.191505 | 0.920555 | 0.840708 | 0.909586 |
| GOBP_NEGATIVE_REGULATION_OF_OXIDATIVE_STRESS_INDUCED_INTRINSIC_APOPTOTIC_SIGN | 19 | 0.316626 | 0.919912 | 0.560694 | 0.910484 |
| GOBP_RESPIRATORY_CHAIN_COMPLEX_IV_ASSEMBLY | 26 | 0.280394 | 0.918706 | 0.569733 | 0.912988 |
| GOBP_REGULATION_OF_NEURON_PROJECTION_REGENERATION | 30 | 0.282133 | 0.918628 | 0.583333 | 0.912147 |
| GOBP_SMOOTH_MUSCLE_CELL_APOPTOTIC_PROCESS | 23 | 0.29764 | 0.917516 | 0.588406 | 0.914378 |
| GOBP_NCRNA_CATABOLIC_PROCESS | 36 | 0.268187 | 0.917391 | 0.598706 | 0.913707 |
| GOBP_PEPTIDYL_TYROSINE_DEPHOSPHORYLATION | 96 | 0.218213 | 0.916937 | 0.680672 | 0.913972 |
| GOBP_ENERGY_DERIVATION_BY_OXIDATION_OF_ORGANIC_COMPOUNDS | 264 | 0.191778 | 0.916774 | 0.813008 | 0.913395 |
| GOBP_PURINE_CONTAINING_COMPOUND_METABOLIC_PROCESS | 431 | 0.183502 | 0.915971 | 0.864865 | 0.914638 |
| GOBP_REGULATION_OF_AXONOGENESIS | 141 | 0.209621 | 0.915726 | 0.686275 | 0.914296 |
| GOBP_C21_STEROID_HORMONE_METABOLIC_PROCESS | 34 | 0.267028 | 0.914278 | 0.60061 | 0.917673 |
| GOBP_GENERATION_OF_PRECURSOR_METABOLITES_AND_ENERGY | 496 | 0.180363 | 0.912258 | 0.901408 | 0.922639 |
| GOBP_NEGATIVE_REGULATION_OF_CYTOSKELETON_ORGANIZATION | 150 | 0.207603 | 0.910433 | 0.739583 | 0.926944 |
| GOBP_DE_NOVO_PROTEIN_FOLDING | 43 | 0.255439 | 0.910163 | 0.647975 | 0.926617 |
| GOBP_NEGATIVE_REGULATION_OF_STEROID_METABOLIC_PROCESS | 20 | 0.309414 | 0.909606 | 0.586103 | 0.927088 |
| GOBP_MITOCHONDRION_DISTRIBUTION | 15 | 0.324577 | 0.90838 | 0.571429 | 0.929638 |

| | | | | | |
|---|---|---|---|---|---|
| GOBP_MONOCYTE_CHEMOTACTIC_PROTEIN_1_PRODUCTION | 16 | -0.520517 | -1.321316 | 0.1168 | 0.339535 |
| GOBP_LYMPHOCYTE_MIGRATION | 108 | -0.355953 | -1.320257 | 0.059645 | 0.341547 |
| GOBP_MEMBRANE_DEPOLARIZATION | 76 | -0.375861 | -1.320156 | 0.077236 | 0.341275 |
| GOBP_DIGESTIVE_SYSTEM_PROCESS | 76 | -0.374576 | -1.320051 | 0.074425 | 0.341006 |
| GOBP_EXTRINSIC_APOPTOTIC_SIGNALING_PATHWAY_VIA_DEATH_DOMAIN_RECEPTORS | 80 | -0.38134 | -1.319818 | 0.074224 | 0.341055 |
| GOBP_MEMBRANE_PROTEIN_PROTEOLYSIS | 60 | -0.391631 | -1.319035 | 0.080221 | 0.34235 |
| GOBP_POSITIVE_REGULATION_OF_PEPTIDYL_THREONINE_PHOSPHORYLATION | 28 | -0.461642 | -1.318236 | 0.114114 | 0.343689 |
| GOBP_NEGATIVE_REGULATION_OF_SIGNAL_TRANSDUCTION_BY_P53_CLASS_MEDIATOR | 30 | -0.452776 | -1.318103 | 0.10767 | 0.34345 |
| GOBP_POSITIVE_REGULATION_OF_RESPONSE_TO_WOUNDING | 62 | -0.392554 | -1.31802 | 0.087379 | 0.34314 |
| GOBP_RESPONSE_TO_GONADOTROPIN | 21 | -0.478435 | -1.317909 | 0.118902 | 0.342891 |
| GOBP_NEGATIVE_REGULATION_OF_PEPTIDE_SECRETION | 66 | -0.383848 | -1.317529 | 0.09498 | 0.34323 |
| GOBP_CENTRAL_NERVOUS_SYSTEM_NEURON_DIFFERENTIATION | 140 | -0.344518 | -1.317412 | 0.039801 | 0.343012 |
| GOBP_VENTRICULAR_SEPTUM_DEVELOPMENT | 56 | -0.394526 | -1.317381 | 0.083676 | 0.342568 |
| GOBP_POSITIVE_REGULATION_OF_STRIATED_MUSCLE_CELL_DIFFERENTIATION | 36 | -0.42894 | -1.316021 | 0.113798 | 0.345177 |
| GOBP_NEURON_FATE_COMMITMENT | 38 | -0.432136 | -1.315744 | 0.108504 | 0.345286 |
| GOBP_ETHANOL_METABOLIC_PROCESS | 18 | -0.498247 | -1.315662 | 0.136923 | 0.344978 |
| GOBP_MECHANORECEPTOR_DIFFERENTIATION | 53 | -0.398128 | -1.315186 | 0.083099 | 0.345584 |
| GOBP_REGULATION_OF_T_CELL_APOPTOTIC_PROCESS | 30 | -0.445168 | -1.31501 | 0.121212 | 0.345478 |
| GOBP_ANTIMICROBIAL_HUMORAL_IMMUNE_RESPONSE_MEDIATED_BY_ANTIMICROBIAL_PEPTID | 51 | -0.401403 | -1.313846 | 0.081346 | 0.347615 |
| GOBP_FOREBRAIN_GENERATION_OF_NEURONS | 40 | -0.424463 | -1.3136 | 0.10981 | 0.347701 |
| GOBP_ANTIMICROBIAL_HUMORAL_RESPONSE | 94 | -0.3659 | -1.311878 | 0.061842 | 0.351286 |
| GOBP_LIPID_EXPORT_FROM_CELL | 72 | -0.374309 | -1.311817 | 0.072802 | 0.350939 |
| GOBP_SEGMENTATION | 80 | -0.369828 | -1.311291 | 0.082794 | 0.351648 |
| GOBP_PRESYNAPSE_ORGANIZATION | 39 | -0.419782 | -1.310934 | 0.107914 | 0.351954 |
| GOBP_INTERFERON_ALPHA_PRODUCTION | 27 | -0.4556 | -1.309912 | 0.121396 | 0.353885 |
| GOBP_GLANDULAR_EPITHELIAL_CELL_DIFFERENTIATION | 41 | -0.412455 | -1.308825 | 0.1 | 0.355868 |
| GOBP_LIPID_STORAGE | 77 | -0.371357 | -1.308204 | 0.082988 | 0.356842 |
| GOBP_EOSINOPHIL_CHEMOTAXIS | 15 | -0.524578 | -1.307887 | 0.164782 | 0.357023 |
| GOBP_GRANULOCYTE_CHEMOTAXIS | 113 | -0.354626 | -1.306929 | 0.056051 | 0.358755 |
| GOBP_NEGATIVE_REGULATION_OF_SIGNALING_RECEPTOR_ACTIVITY | 48 | -0.397272 | -1.306677 | 0.093931 | 0.358812 |
| GOBP_REGULATION_OF_HETEROTYPIC_CELL_CELL_ADHESION | 23 | -0.466397 | -1.306524 | 0.146789 | 0.35864 |
| GOBP_REGULATION_OF_INTRINSIC_APOPTOTIC_SIGNALING_PATHWAY_BY_P53_CLASS_MEDIAT | 27 | -0.450884 | -1.306519 | 0.117199 | 0.358145 |
| GOBP_CHEMOKINE_C_X_C_MOTIF_LIGAND_2_PRODUCTION | 15 | -0.519788 | -1.306482 | 0.15655 | 0.357724 |
| GOBP_DENDRITIC_SPINE_DEVELOPMENT | 98 | -0.358572 | -1.306476 | 0.058902 | 0.357227 |
| GOBP_REGULATION_OF_KIDNEY_DEVELOPMENT | 26 | -0.457454 | -1.306283 | 0.126718 | 0.357148 |
| GOBP_POSITIVE_REGULATION_OF_PROTEIN_TYROSINE_KINASE_ACTIVITY | 55 | -0.390868 | -1.305973 | 0.096419 | 0.357361 |
| GOBP_REGULATION_OF_CATION_TRANSMEMBRANE_TRANSPORT | 293 | -0.31648 | -1.305733 | 0.02381 | 0.357396 |
| GOBP_NOTOCHORD_DEVELOPMENT | 16 | -0.509805 | -1.305554 | 0.155449 | 0.357287 |
| GOBP_REGULATION_OF_CALCIUM_MEDIATED_SIGNALING | 60 | -0.388465 | -1.305397 | 0.103164 | 0.357168 |
| GOBP_REGULATION_OF_HORMONE_SECRETION | 221 | -0.325011 | -1.305347 | 0.029002 | 0.356804 |
| GOBP_RELEASE_OF_CYTOCHROME_C_FROM_MITOCHONDRIA | 54 | -0.397899 | -1.304391 | 0.088534 | 0.358451 |
| GOBP_CELL_FATE_SPECIFICATION | 64 | -0.379919 | -1.304237 | 0.091691 | 0.358298 |
| GOBP_CYTOPLASMIC_SEQUESTERING_OF_TRANSCRIPTION_FACTOR | 15 | -0.510875 | -1.303401 | 0.167712 | 0.359784 |
| GOBP_REGULATION_OF_FATTY_ACID_TRANSPORT | 41 | -0.410592 | -1.302571 | 0.106195 | 0.361295 |
| GOBP_DENDRITIC_CELL_MIGRATION | 28 | -0.440135 | -1.302224 | 0.128983 | 0.361611 |
| GOBP_EOSINOPHIL_MIGRATION | 19 | -0.495684 | -1.301961 | 0.139969 | 0.361748 |
| GOBP_POSITIVE_REGULATION_OF_INSULIN_SECRETION | 61 | -0.383184 | -1.301563 | 0.09517 | 0.362238 |
| GOBP_POSITIVE_REGULATION_OF_NUCLEOTIDE_BIOSYNTHETIC_PROCESS | 16 | -0.503114 | -1.30052 | 0.151274 | 0.364182 |
| GOBP_NEGATIVE_REGULATION_OF_PROTEOLYSIS | 270 | -0.318529 | -1.30046 | 0.02662 | 0.36382 |
| GOBP_DEVELOPMENT_OF_PRIMARY_FEMALE_SEXUAL_CHARACTERISTICS | 84 | -0.365889 | -1.299983 | 0.086551 | 0.364498 |
| GOBP_REGULATION_OF_VASCULATURE_DEVELOPMENT | 266 | -0.320159 | -1.299969 | 0.030752 | 0.36402 |
| GOBP_NEGATIVE_REGULATION_OF_ANION_TRANSPORT | 180 | -0.33293 | -1.299585 | 0.047158 | 0.364411 |
| GOBP_REGULATION_OF_LIPOPOLYSACCHARIDE_MEDIATED_SIGNALING_PATHWAY | 23 | -0.464893 | -1.298592 | 0.142636 | 0.36631 |
| GOBP_NEGATIVE_REGULATION_OF_DNA_BINDING_TRANSCRIPTION_FACTOR_ACTIVITY | 147 | -0.336753 | -1.298348 | 0.053549 | 0.366373 |
| GOBP_CALCIUM_ION_TRANSPORT | 361 | -0.310952 | -1.29828 | 0.018868 | 0.366023 |
| GOBP_SKELETAL_MUSCLE_CELL_DIFFERENTIATION | 51 | -0.401147 | -1.298261 | 0.099037 | 0.365566 |
| GOBP_REGULATION_OF_VOLTAGE_GATED_CALCIUM_CHANNEL_ACTIVITY | 31 | -0.434012 | -1.297922 | 0.141631 | 0.365864 |
| GOBP_RESPONSE_TO_HYPEROXIA | 21 | -0.480879 | -1.296931 | 0.144796 | 0.36762 |
| GOBP_RETINOIC_ACID_RECEPTOR_SIGNALING_PATHWAY | 29 | -0.445115 | -1.296751 | 0.126075 | 0.367535 |
| GOBP_NEGATIVE_REGULATION_OF_OSTEOCLAST_DIFFERENTIATION | 26 | -0.453384 | -1.295882 | 0.141745 | 0.3691 |
| GOBP_NEGATIVE_REGULATION_OF_HYDROLASE_ACTIVITY | 367 | -0.308239 | -1.295064 | 0.022002 | 0.37057 |
| GOBP_HORMONE_TRANSPORT | 267 | -0.318083 | -1.294987 | 0.02746 | 0.370238 |
| GOBP_ANTERIOR_POSTERIOR_AXIS_SPECIFICATION | 36 | -0.416203 | -1.294613 | 0.11791 | 0.370659 |
| GOBP_CELL_FATE_COMMITMENT | 195 | -0.326131 | -1.294593 | 0.032569 | 0.370205 |
| GOBP_VASODILATION | 30 | -0.438848 | -1.294516 | 0.118619 | 0.369866 |
| GOBP_NEGATIVE_REGULATION_OF_RECEPTOR_SIGNALING_PATHWAY_VIA_JAK_STAT | 22 | -0.468299 | -1.294329 | 0.156442 | 0.369796 |
| GOBP_NEGATIVE_REGULATION_OF_CELL_ADHESION | 265 | -0.319162 | -1.293849 | 0.036405 | 0.370416 |
| GOBP_POSITIVE_REGULATION_OF_NEURON_DEATH | 83 | -0.361926 | -1.293792 | 0.07651 | 0.370035 |
| GOBP_REGULATION_OF_PROTEIN_IMPORT | 57 | -0.385706 | -1.293637 | 0.091034 | 0.369883 |
| GOBP_AMELOGENESIS | 16 | -0.509578 | -1.292933 | 0.146646 | 0.371046 |
| GOBP_CD4_POSITIVE_OR_CD8_POSITIVE_ALPHA_BETA_T_CELL_LINEAGE_COMMITMENT | 22 | -0.471236 | -1.29277 | 0.139535 | 0.370929 |
| GOBP_DIENCEPHALON_DEVELOPMENT | 53 | -0.394811 | -1.292457 | 0.109065 | 0.371175 |
| GOBP_RELAXATION_OF_MUSCLE | 26 | -0.457233 | -1.292057 | 0.136637 | 0.371644 |
| GOBP_NEGATIVE_REGULATION_OF_RESPONSE_TO_BIOTIC_STIMULUS | 90 | -0.359478 | -1.291797 | 0.083784 | 0.371728 |
| GOBP_NEGATIVE_REGULATION_OF_VIRAL_TRANSCRIPTION | 15 | -0.513916 | -1.291758 | 0.168675 | 0.37135 |
| GOBP_POSITIVE_T_CELL_SELECTION | 36 | -0.419279 | -1.291744 | 0.098361 | 0.370891 |
| GOBP_POSITIVE_REGULATION_OF_PROTEIN_IMPORT | 39 | -0.417352 | -1.291455 | 0.121701 | 0.371086 |
| GOBP_MESENCHYMAL_CELL_DIFFERENTIATION | 204 | -0.326274 | -1.291383 | 0.046893 | 0.370778 |
| GOBP_OVARIAN_FOLLICLE_DEVELOPMENT | 46 | -0.401164 | -1.290891 | 0.11831 | 0.37145 |
| GOBP_TOLL_LIKE_RECEPTOR_3_SIGNALING_PATHWAY | 19 | -0.480905 | -1.290802 | 0.145161 | 0.37114 |
| GOBP_POSITIVE_REGULATION_OF_NEURAL_PRECURSOR_CELL_PROLIFERATION | 40 | -0.408246 | -1.288875 | 0.121849 | 0.375189 |
| GOBP_REGULATION_OF_AMYLOID_BETA_CLEARANCE | 15 | -0.509139 | -1.287134 | 0.155146 | 0.378881 |
| GOBP_MEMBRANE_PROTEIN_ECTODOMAIN_PROTEOLYSIS | 42 | -0.409864 | -1.287095 | 0.118143 | 0.378492 |
| GOBP_CARDIAC_VENTRICLE_DEVELOPMENT | 94 | -0.355311 | -1.286792 | 0.071244 | 0.378717 |
| GOBP_ACTIVATION_OF_ADENYLATE_CYCLASE_ACTIVITY | 30 | -0.436153 | -1.286537 | 0.122939 | 0.378818 |
| GOBP_MORPHOGENESIS_OF_AN_ENDOTHELIUM | 16 | -0.500164 | -1.286534 | 0.163492 | 0.378328 |
| GOBP_PATTERN_SPECIFICATION_PROCESS | 343 | -0.309601 | -1.286457 | 0.026461 | 0.378015 |
| GOBP_REGULATION_OF_TOLERANCE_INDUCTION | 18 | -0.482059 | -1.286419 | 0.15969 | 0.377598 |
| GOBP_DETECTION_OF_STIMULUS_INVOLVED_IN_SENSORY_PERCEPTION | 141 | -0.338646 | -1.286218 | 0.063354 | 0.377588 |
| GOBP_PHAGOCYTOSIS | 355 | -0.307479 | -1.286092 | 0.021064 | 0.377383 |
| GOBP_MYOTUBE_CELL_DEVELOPMENT | 30 | -0.43783 | -1.285748 | 0.129412 | 0.377738 |
| GOBP_EPITHELIAL_TUBE_MORPHOGENESIS | 269 | -0.314466 | -1.28554 | 0.032445 | 0.377737 |
| GOBP_REGULATION_OF_RESPONSE_TO_WOUNDING | 141 | -0.336173 | -1.285293 | 0.071611 | 0.377851 |
| GOBP_REGULATION_OF_ASTROCYTE_DIFFERENTIATION | 21 | -0.470199 | -1.285146 | 0.141337 | 0.377707 |
| GOBP_GLUTAMATE_SECRETION | 37 | -0.417789 | -1.284895 | 0.127721 | 0.377812 |
| GOBP_CELLULAR_RESPONSE_TO_KETONE | 86 | -0.362771 | -1.283544 | 0.095745 | 0.380522 |
| GOBP_REGULATION_OF_OSSIFICATION | 99 | -0.349472 | -1.283353 | 0.082368 | 0.380471 |
| GOBP_CRANIAL_NERVE_MORPHOGENESIS | 22 | -0.457489 | -1.282571 | 0.160363 | 0.381953 |
| GOBP_NEGATIVE_REGULATION_OF_ERBB_SIGNALING_PATHWAY | 51 | -0.388909 | -1.282545 | 0.128655 | 0.381509 |
| GOBP_SECRETION_BY_TISSUE | 24 | -0.457622 | -1.282093 | 0.150077 | 0.382136 |
| GOBP_NEGATIVE_REGULATION_OF_LYMPHOCYTE_MIGRATION | 15 | -0.516193 | -1.281942 | 0.164251 | 0.382004 |
| GOBP_REGULATION_OF_KERATINOCYTE_DIFFERENTIATION | 33 | -0.42298 | -1.281586 | 0.131241 | 0.382362 |
| GOBP_REGULATION_OF_ADENYLATE_CYCLASE_ACTIVITY | 26 | -0.442217 | -1.280986 | 0.152516 | 0.383324 |
| GOBP_MORPHOGENESIS_OF_EMBRYONIC_EPITHELIUM | 127 | -0.340065 | -1.280767 | 0.079427 | 0.383372 |
| GOBP_CALCIUM_ION_TRANSPORT_INTO_CYTOSOL | 134 | -0.33609 | -1.280692 | 0.068493 | 0.383082 |
| GOBP_DETECTION_OF_MECHANICAL_STIMULUS_INVOLVED_IN_SENSORY_PERCEPTION | 28 | -0.444462 | -1.2806 | 0.154688 | 0.382819 |
| GOBP_PHOSPHOLIPID_TRANSPORT | 84 | -0.362625 | -1.280348 | 0.096561 | 0.382928 |
| GOBP_NEGATIVE_REGULATION_OF_NF_KAPPAB_TRANSCRIPTION_FACTOR_ACTIVITY | 80 | -0.3622 | -1.280037 | 0.099723 | 0.383204 |
| GOBP_SMOOTH_MUSCLE_CELL_DIFFERENTIATION | 53 | -0.391176 | -1.279564 | 0.11544 | 0.383819 |
| GOBP_REGULATION_OF_POSITIVE_CHEMOTAXIS | 24 | -0.450045 | -1.278965 | 0.131661 | 0.384844 |
| GOBP_POSITIVE_REGULATION_OF_VASCULAR_ENDOTHELIAL_CELL_PROLIFERATION | 17 | -0.495371 | -1.278883 | 0.177953 | 0.384528 |
| GOBP_POSITIVE_REGULATION_OF_STRESS_ACTIVATED_PROTEIN_KINASE_SIGNALING_CASCADI | 110 | -0.34729 | -1.278798 | 0.069281 | 0.384234 |
| GOBP_MUSCLE_FILAMENT_SLIDING | 32 | -0.427106 | -1.278644 | 0.14095 | 0.384131 |
| GOBP_REGULATION_OF_NATURAL_KILLER_CELL_ACTIVATION | 32 | -0.424317 | -1.278514 | 0.144756 | 0.383951 |
| GOBP_DENDRITIC_CELL_CHEMOTAXIS | 23 | -0.464578 | -1.27767 | 0.162328 | 0.385499 |
| GOBP_POSITIVE_REGULATION_OF_INTERFERON_BETA_PRODUCTION | 33 | -0.429878 | -1.277285 | 0.145646 | 0.385972 |
| GOBP_GLUTATHIONE_DERIVATIVE_METABOLIC_PROCESS | 18 | -0.48485 | -1.276861 | 0.173502 | 0.386462 |
| GOBP_REGULATION_OF_REACTIVE_OXYGEN_SPECIES_METABOLIC_PROCESS | 171 | -0.327873 | -1.275764 | 0.05068 | 0.388623 |
| GOBP_RESPONSE_TO_ESTRADIOL | 111 | -0.338963 | -1.275572 | 0.06743 | 0.388589 |
| GOBP_NEGATIVE_REGULATION_OF_STEM_CELL_DIFFERENTIATION | 19 | -0.475376 | -1.275518 | 0.148773 | 0.388226 |
| GOBP_RESPONSE_TO_STEROL | 32 | -0.419684 | -1.274444 | 0.138763 | 0.390395 |
| GOBP_NEGATIVE_REGULATION_OF_ACTIN_FILAMENT_BUNDLE_ASSEMBLY | 30 | -0.428235 | -1.273553 | 0.160606 | 0.392057 |
| GOBP_NEGATIVE_REGULATION_OF_PHOSPHORUS_METABOLIC_PROCESS | 481 | -0.299661 | -1.273137 | 0.009544 | 0.392558 |
| GOBP_HUMORAL_IMMUNE_RESPONSE_MEDIATED_BY_CIRCULATING_IMMUNOGLOBULIN | 139 | -0.333197 | -1.272584 | 0.075094 | 0.393425 |
| GOBP_MULTICELLULAR_ORGANISMAL_SIGNALING | 170 | -0.326874 | -1.272519 | 0.067797 | 0.393075 |
| GOBP_NEGATIVE_REGULATION_OF_KINASE_ACTIVITY | 246 | -0.314777 | -1.272443 | 0.04388 | 0.392785 |
| GOBP_REGULATION_OF_OSTEOBLAST_PROLIFERATION | 23 | -0.459403 | -1.272408 | 0.149031 | 0.392389 |
| GOBP_REGULATION_OF_BLOOD_PRESSURE | 154 | -0.334118 | -1.272083 | 0.084975 | 0.39272 |
| GOBP_NEURON_RECOGNITION | 39 | -0.40189 | -1.272073 | 0.141593 | 0.392251 |
| GOBP_REGULATION_OF_CATION_CHANNEL_ACTIVITY | 149 | -0.332627 | -1.271362 | 0.073048 | 0.393538 |
| GOBP_REGULATION_OF_MEMBRANE_INVAGINATION | 16 | -0.505511 | -1.271035 | 0.164852 | 0.393881 |
| GOBP_POSITIVE_REGULATION_OF_MUSCLE_TISSUE_DEVELOPMENT | 19 | -0.472585 | -1.270732 | 0.190252 | 0.394137 |
| GOBP_REGULATION_OF_CYSTEINE_TYPE_ENDOPEPTIDASE_ACTIVITY_INVOLVED_IN_APOPTOTIC | 17 | -0.493488 | -1.270596 | 0.166403 | 0.393998 |
| GOBP_AROMATIC_AMINO_ACID_FAMILY_CATABOLIC_PROCESS | 19 | -0.475633 | -1.270544 | 0.166667 | 0.393626 |
| GOBP_RESPONSE_TO_FOOD | 29 | -0.433191 | -1.26927 | 0.153285 | 0.396257 |
| GOBP_SENSORY_PERCEPTION | 481 | -0.299239 | -1.268629 | 0.018028 | 0.3974 |
| GOBP_HAIR_CELL_DIFFERENTIATION | 38 | -0.412569 | -1.268316 | 0.132374 | 0.397696 |
| GOBP_NEGATIVE_REGULATION_OF_NITRIC_OXIDE_METABOLIC_PROCESS | 15 | -0.510856 | -1.267692 | 0.157978 | 0.398811 |
| GOBP_MHC_CLASS_II_BIOSYNTHETIC_PROCESS | 17 | -0.483481 | -1.266614 | 0.174383 | 0.401081 |
| GOBP_NEGATIVE_REGULATION_OF_OSTEOBLAST_DIFFERENTIATION | 40 | -0.405899 | -1.266253 | 0.150492 | 0.401509 |
| GOBP_EMBRYO_IMPLANTATION | 43 | -0.392433 | -1.26617 | 0.134393 | 0.401204 |
| GOBP_EPITHELIAL_CELL_CELL_ADHESION | 17 | -0.486 | -1.265328 | 0.18507 | 0.402886 |
| GOBP_OSTEOCLAST_DEVELOPMENT | 16 | -0.492323 | -1.264307 | 0.196594 | 0.405016 |
| GOBP_NEGATIVE_REGULATION_OF_NEURON_DIFFERENTIATION | 47 | -0.389862 | -1.26279 | 0.132231 | 0.408372 |
| GOBP_POSITIVE_REGULATION_OF_CYCLASE_ACTIVITY | 15 | -0.50319 | -1.262245 | 0.192192 | 0.40923 |
| GOBP_INFLAMMATORY_CELL_APOPTOTIC_PROCESS | 20 | -0.463125 | -1.262172 | 0.18003 | 0.408888 |
| GOBP_REGULATION_OF_PEPTIDE_SECRETION | 269 | -0.309213 | -1.261699 | 0.046857 | 0.4096 |
| GOBP_CELLULAR_RESPONSE_TO_ORGANIC_CYCLIC_COMPOUND | 494 | -0.296439 | -1.26145 | 0.02234 | 0.4097 |
| GOBP_SYNAPTIC_VESICLE_PRIMING | 17 | -0.487914 | -1.261435 | 0.199695 | 0.40923 |
| GOBP_MAINTENANCE_OF_PROTEIN_LOCATION_IN_NUCLEUS | 23 | -0.461325 | -1.261395 | 0.168421 | 0.408834 |
| GOBP_POSITIVE_REGULATION_OF_AMINE_TRANSPORT | 29 | -0.424586 | -1.261324 | 0.159399 | 0.408518 |
| GOBP_METANEPHRIC_NEPHRON_MORPHOGENESIS | 19 | -0.466301 | -1.260837 | 0.167421 | 0.409256 |
| GOBP_RESPONSE_TO_PAIN | 22 | -0.453913 | -1.260657 | 0.156204 | 0.409195 |
| GOBP_ADIPOSE_TISSUE_DEVELOPMENT | 35 | -0.414319 | -1.260213 | 0.151424 | 0.409829 |
| GOBP_DETECTION_OF_EXTERNAL_BIOTIC_STIMULUS | 22 | -0.4618 | -1.259681 | 0.187311 | 0.410716 |
| GOBP_CALCIUM_DEPENDENT_CELL_CELL_ADHESION_VIA_PLASMA_MEMBRANE_CELL_ADHESI | 34 | -0.410853 | -1.259599 | 0.153731 | 0.410427 |
| GOBP_REGULATION_OF_AMPA_RECEPTOR_ACTIVITY | 21 | -0.46474 | -1.25959 | 0.16208 | 0.409959 |
| GOBP_ATP_TRANSPORT | 16 | -0.492178 | -1.259409 | 0.171384 | 0.409911 |
| GOBP_ODONTOGENESIS_OF_DENTIN_CONTAINING_TOOTH | 66 | -0.367449 | -1.259342 | 0.11967 | 0.409604 |
| GOBP_CARDIAC_EPITHELIAL_TO_MESENCHYMAL_TRANSITION | 29 | -0.435507 | -1.259266 | 0.149927 | 0.409282 |
| GOBP_NEUTROPHIL_CHEMOTAXIS | 94 | -0.347243 | -1.259194 | 0.103038 | 0.408969 |
| GOBP_DETECTION_OF_STIMULUS | 287 | -0.305835 | -1.259171 | 0.042093 | 0.408542 |
| GOBP_PSEUDOPODIUM_ORGANIZATION | 16 | -0.496355 | -1.259068 | 0.174881 | 0.408338 |
| GOBP_REGULATION_OF_GLIOGENESIS | 82 | -0.35623 | -1.258842 | 0.086137 | 0.408416 |
| GOBP_CARDIAC_SEPTUM_DEVELOPMENT | 84 | -0.347163 | -1.258663 | 0.109948 | 0.408374 |
| GOBP_REGULATION_OF_SYNCYTIUM_FORMATION_BY_PLASMA_MEMBRANE_FUSION | 23 | -0.45062 | -1.258437 | 0.149691 | 0.408445 |
| GOBP_POSITIVE_REGULATION_OF_LYMPHOCYTE_CHEMOTAXIS | 18 | -0.488732 | -1.258307 | 0.171381 | 0.408266 |
| GOBP_RESPONSE_TO_MECHANICAL_STIMULUS | 189 | -0.321094 | -1.258191 | 0.077473 | 0.408095 |

| | | | | | |
|---|---|---|---|---|---|
| GOBP_CYTOSKELETON_DEPENDENT_CYTOKINESIS | 95 | 0.21631 | 0.907702 | 0.715517 | 0.930507 |
| GOBP_RIBOSOMAL_SMALL_SUBUNIT_BIOGENESIS | 75 | 0.226571 | 0.90747 | 0.636364 | 0.930071 |
| GOBP_MITOPHAGY | 24 | 0.287975 | 0.906843 | 0.608955 | 0.93076 |
| GOBP_MEMBRANE_LIPID_BIOSYNTHETIC_PROCESS | 142 | 0.205266 | 0.90666 | 0.791667 | 0.930217 |
| GOBP_LABYRINTHINE_LAYER_MORPHOGENESIS | 17 | 0.323259 | 0.906438 | 0.590529 | 0.929757 |
| GOBP_CELL_DIFFERENTIATION_INVOLVED_IN_EMBRYONIC_PLACENTA_DEVELOPMENT | 21 | 0.298079 | 0.905923 | 0.62029 | 0.93015 |
| GOBP_POSITIVE_REGULATION_OF_CYCLIN_DEPENDENT_PROTEIN_KINASE_ACTIVITY | 30 | 0.278142 | 0.905738 | 0.623794 | 0.929582 |
| GOBP_GLYCEROLIPID_BIOSYNTHETIC_PROCESS | 245 | 0.192312 | 0.905553 | 0.777778 | 0.929006 |
| GOBP_RESPONSE_TO_MUSCLE_ACTIVITY | 23 | 0.296684 | 0.905058 | 0.60221 | 0.929304 |
| GOBP_PROTEIN_LOCALIZATION_TO_MITOCHONDRION | 144 | 0.199204 | 0.904579 | 0.735135 | 0.92964 |
| GOBP_MITOCHONDRIAL_MEMBRANE_ORGANIZATION | 139 | 0.208439 | 0.903181 | 0.716895 | 0.93282 |
| GOBP_FLUID_TRANSPORT | 27 | 0.282084 | 0.902731 | 0.613208 | 0.933048 |
| GOBP_RNA_POLYMERASE_II_PREINITIATION_COMPLEX_ASSEMBLY | 24 | 0.290311 | 0.902243 | 0.60989 | 0.933417 |
| GOBP_RESPONSE_TO_RADIATION | 410 | 0.183451 | 0.901741 | 0.901235 | 0.933734 |
| GOBP_REGULATION_OF_PROTEASOMAL_PROTEIN_CATABOLIC_PROCESS | 177 | 0.199181 | 0.899922 | 0.770186 | 0.937858 |
| GOBP_ESTABLISHMENT_OF_MITOCHONDRION_LOCALIZATION | 24 | 0.291292 | 0.899661 | 0.612426 | 0.937546 |
| GOBP_CELLULAR_RESPONSE_TO_CHEMICAL_STRESS | 319 | 0.185545 | 0.899554 | 0.854167 | 0.936727 |
| GOBP_VISUAL_BEHAVIOR | 43 | 0.249568 | 0.897767 | 0.657321 | 0.940736 |
| GOBP_POSITIVE_REGULATION_OF_SODIUM_ION_TRANSMEMBRANE_TRANSPORT | 20 | 0.298246 | 0.895902 | 0.60452 | 0.944947 |
| GOBP_RESPONSE_TO_LITHIUM_ION | 18 | 0.306216 | 0.895681 | 0.616188 | 0.944454 |
| GOBP_POSITIVE_REGULATION_OF_STEROL_TRANSPORT | 30 | 0.271593 | 0.895372 | 0.598784 | 0.944187 |
| GOBP_NCRNA_TRANSCRIPTION | 107 | 0.207845 | 0.895231 | 0.729258 | 0.943493 |
| GOBP_CARTILAGE_DEVELOPMENT_INVOLVED_IN_ENDOCHONDRAL_BONE_MORPHOGENESIS | 24 | 0.281861 | 0.895225 | 0.613636 | 0.94241 |
| GOBP_DOPAMINE_METABOLIC_PROCESS | 27 | 0.27667 | 0.894652 | 0.619632 | 0.942972 |
| GOBP_RRNA_METHYLATION | 28 | 0.269238 | 0.894425 | 0.616766 | 0.942513 |
| GOBP_EXTRACELLULAR_TRANSPORT | 33 | 0.263366 | 0.894415 | 0.62069 | 0.94145 |
| GOBP_PURINE_NUCLEOSIDE_BIOSYNTHETIC_PROCESS | 22 | 0.291875 | 0.89408 | 0.633929 | 0.941273 |
| GOBP_RNA_INTERFERENCE | 16 | 0.312029 | 0.894015 | 0.5961 | 0.94035 |
| GOBP_CEREBRAL_CORTEX_DEVELOPMENT | 98 | 0.213964 | 0.893822 | 0.758755 | 0.939796 |
| GOBP_REGULATION_OF_APPETITE | 16 | 0.323211 | 0.893376 | 0.61017 | 0.939877 |
| GOBP_PIGMENTATION | 85 | 0.216906 | 0.893031 | 0.712551 | 0.939752 |
| GOBP_HISTONE_H3_K9_METHYLATION | 33 | 0.264475 | 0.890495 | 0.655882 | 0.94562 |
| GOBP_NEGATIVE_REGULATION_OF_CELL_MATRIX_ADHESION | 35 | 0.256966 | 0.890339 | 0.652997 | 0.944967 |
| GOBP_DNA_TEMPLATED_TRANSCRIPTION_INITIATION | 238 | 0.189086 | 0.889898 | 0.849673 | 0.945115 |
| GOBP_POSITIVE_REGULATION_OF_TRANSCRIPTION_BY_RNA_POLYMERASE_I | 22 | 0.289106 | 0.889638 | 0.609195 | 0.944749 |
| GOBP_MONOSACCHARIDE_METABOLIC_PROCESS | 250 | 0.186174 | 0.889583 | 0.903704 | 0.943825 |
| GOBP_REGULATION_OF_CELLULAR_RESPONSE_TO_HEAT | 79 | 0.216365 | 0.889384 | 0.715953 | 0.943296 |
| GOBP_REGULATION_OF_TRANSCRIPTION_FROM_RNA_POLYMERASE_II_PROMOTER_IN_RESPO | 76 | 0.220363 | 0.888856 | 0.715302 | 0.943622 |
| GOBP_AMINOGLYCAN_BIOSYNTHETIC_PROCESS | 108 | 0.209502 | 0.888828 | 0.777778 | 0.942625 |
| GOBP_RIBOSOMAL_SMALL_SUBUNIT_ASSEMBLY | 21 | 0.293777 | 0.88811 | 0.633766 | 0.943514 |
| GOBP_PEPTIDYL_PROLINE_HYDROXYLATION | 15 | 0.318153 | 0.888056 | 0.586667 | 0.942579 |
| GOBP_MATURATION_OF_LSU_RRNA | 28 | 0.269615 | 0.88654 | 0.6375 | 0.945606 |
| GOBP_ENDODERMAL_CELL_DIFFERENTIATION | 43 | 0.243193 | 0.886413 | 0.670034 | 0.944865 |
| GOBP_L_GLUTAMATE_IMPORT_ACROSS_PLASMA_MEMBRANE | 15 | 0.319003 | 0.885819 | 0.602564 | 0.945373 |
| GOBP_ORGANELLE_DISASSEMBLY | 106 | 0.210819 | 0.885485 | 0.75576 | 0.945193 |
| GOBP_GLUCOCORTICOID_METABOLIC_PROCESS | 21 | 0.297321 | 0.885026 | 0.627566 | 0.945385 |
| GOBP_REGULATION_OF_PROTEIN_CONTAINING_COMPLEX_DISASSEMBLY | 118 | 0.203098 | 0.884952 | 0.788288 | 0.944523 |
| GOBP_HYALURONAN_CATABOLIC_PROCESS | 16 | 0.311635 | 0.883898 | 0.624294 | 0.94629 |
| GOBP_PLASMA_LIPOPROTEIN_PARTICLE_CLEARANCE | 50 | 0.23472 | 0.882922 | 0.738019 | 0.947867 |
| GOBP_CELLULAR_RESPONSE_TO_REACTIVE_OXYGEN_SPECIES | 140 | 0.200596 | 0.882452 | 0.811881 | 0.94806 |
| GOBP_NEGATIVE_REGULATION_OF_NEURON_PROJECTION_DEVELOPMENT | 125 | 0.204607 | 0.879915 | 0.837104 | 0.953879 |
| GOBP_REGULATION_OF_OXIDATIVE_STRESS_INDUCED_INTRINSIC_APOPTOTIC_SIGNALING_PAT | 27 | 0.273283 | 0.878965 | 0.674157 | 0.955328 |
| GOBP_REGULATION_OF_PROTEIN_LOCALIZATION_TO_NUCLEUS | 127 | 0.203963 | 0.878026 | 0.777778 | 0.956729 |
| GOBP_ESTABLISHMENT_OF_ORGANELLE_LOCALIZATION | 413 | 0.176417 | 0.877842 | 0.934066 | 0.956113 |
| GOBP_NUCLEOBASE_CONTAINING_SMALL_MOLECULE_BIOSYNTHETIC_PROCESS | 111 | 0.20748 | 0.877371 | 0.773148 | 0.956311 |
| GOBP_POSITIVE_REGULATION_OF_PROTEIN_DEPOLYMERIZATION | 18 | 0.308204 | 0.876911 | 0.625749 | 0.956447 |
| GOBP_PROTEIN_MATURATION_BY_IRON_SULFUR_CLUSTER_TRANSFER | 16 | 0.309394 | 0.874995 | 0.637795 | 0.960366 |
| GOBP_NEGATIVE_REGULATION_OF_TOR_SIGNALING | 43 | 0.250751 | 0.874003 | 0.670808 | 0.961878 |
| GOBP_REGULATION_OF_SKELETAL_MUSCLE_CELL_DIFFERENTIATION | 17 | 0.305448 | 0.872136 | 0.664706 | 0.965624 |
| GOBP_MODULATION_BY_HOST_OF_VIRAL_GENOME_REPLICATION | 19 | 0.306402 | 0.872029 | 0.672365 | 0.964811 |
| GOBP_GLYCEROLIPID_METABOLIC_PROCESS | 387 | 0.178043 | 0.871215 | 0.928571 | 0.965798 |
| GOBP_REGULATION_OF_TOR_SIGNALING | 97 | 0.209567 | 0.870506 | 0.77381 | 0.96653 |
| GOBP_OOGENESIS | 73 | 0.216393 | 0.87033 | 0.721992 | 0.965925 |
| GOBP_SEMAPHORIN_PLEXIN_SIGNALING_PATHWAY | 37 | 0.242497 | 0.870264 | 0.660131 | 0.965013 |
| GOBP_NEGATIVE_REGULATION_OF_SUPRAMOLECULAR_FIBER_ORGANIZATION | 150 | 0.19877 | 0.870221 | 0.846939 | 0.964019 |
| GOBP_RRNA_CONTAINING_RIBONUCLEOPROTEIN_COMPLEX_EXPORT_FROM_NUCLEUS | 15 | 0.314309 | 0.870104 | 0.652291 | 0.963244 |
| GOBP_EMBRYONIC_DIGESTIVE_TRACT_DEVELOPMENT | 24 | 0.275168 | 0.866172 | 0.655556 | 0.97198 |
| GOBP_ACTIVATION_OF_PROTEIN_KINASE_A_ACTIVITY | 17 | 0.309966 | 0.865889 | 0.622535 | 0.971638 |
| GOBP_SUPEROXIDE_METABOLIC_PROCESS | 60 | 0.223183 | 0.865574 | 0.742647 | 0.97133 |
| GOBP_CELLULAR_RESPONSE_TO_STARVATION | 149 | 0.194285 | 0.863699 | 0.856436 | 0.974987 |
| GOBP_REGULATION_OF_GENERATION_OF_PRECURSOR_METABOLITES_AND_ENERGY | 153 | 0.192507 | 0.863364 | 0.83 | 0.974699 |
| GOBP_MUSCLE_CELL_CELLULAR_HOMEOSTASIS | 19 | 0.294582 | 0.862961 | 0.65404 | 0.974633 |
| GOBP_POSITIVE_REGULATION_OF_UBIQUITIN_DEPENDENT_PROTEIN_CATABOLIC_PROCESS | 100 | 0.204621 | 0.862553 | 0.809524 | 0.974614 |
| GOBP_CELLULAR_COMPONENT_DISASSEMBLY_INVOLVED_IN_EXECUTION_PHASE_OF_APOPTO | 31 | 0.258832 | 0.862369 | 0.70625 | 0.97401 |
| GOBP_LONG_CHAIN_FATTY_ACID_IMPORT_INTO_CELL | 17 | 0.299098 | 0.861446 | 0.667553 | 0.975243 |
| GOBP_CELLULAR_RESPONSE_TO_UV | 85 | 0.20794 | 0.860597 | 0.801587 | 0.97624 |
| GOBP_REGULATION_OF_TRANSCRIPTION_BY_RNA_POLYMERASE_I | 32 | 0.253957 | 0.859959 | 0.702614 | 0.976781 |
| GOBP_REGULATION_OF_EXTENT_OF_CELL_GROWTH | 94 | 0.206364 | 0.859871 | 0.783186 | 0.97592 |
| GOBP_REGULATION_OF_PHOSPHOLIPASE_C_ACTIVITY | 42 | 0.244046 | 0.859707 | 0.721894 | 0.975235 |
| GOBP_PHOTOTRANSDUCTION_VISIBLE_LIGHT | 26 | 0.267624 | 0.858205 | 0.677419 | 0.977773 |
| GOBP_NEGATIVE_REGULATION_OF_POTASSIUM_ION_TRANSMEMBRANE_TRANSPORT | 20 | 0.283084 | 0.857174 | 0.668622 | 0.979212 |
| GOBP_REGULATION_OF_HISTONE_H3_K4_METHYLATION | 26 | 0.271363 | 0.856005 | 0.682243 | 0.980952 |
| GOBP_KERATINOCYTE_MIGRATION | 16 | 0.304976 | 0.853709 | 0.662198 | 0.985212 |
| GOBP_XENOBIOTIC_TRANSPORT | 38 | 0.245993 | 0.853484 | 0.725166 | 0.984671 |
| GOBP_PURINE_NUCLEOSIDE_MONOPHOSPHATE_METABOLIC_PROCESS | 42 | 0.242045 | 0.852906 | 0.728435 | 0.984973 |
| GOBP_RIBONUCLEOSIDE_MONOPHOSPHATE_BIOSYNTHETIC_PROCESS | 33 | 0.249596 | 0.852732 | 0.698171 | 0.984308 |
| GOBP_REGULATION_OF_RIG_I_SIGNALING_PATHWAY | 18 | 0.284599 | 0.851954 | 0.665714 | 0.985026 |
| GOBP_AMYLOID_PRECURSOR_PROTEIN_BIOSYNTHETIC_PROCESS | 15 | 0.311432 | 0.850757 | 0.684066 | 0.986719 |
| GOBP_POSITIVE_REGULATION_OF_DENDRITIC_SPINE_MORPHOGENESIS | 19 | 0.28859 | 0.850747 | 0.678977 | 0.98568 |
| GOBP_SYNAPTIC_MEMBRANE_ADHESION | 21 | 0.281919 | 0.850193 | 0.685237 | 0.98581 |
| GOBP_REGULATION_OF_SUBSTRATE_ADHESION_DEPENDENT_CELL_SPREADING | 54 | 0.225644 | 0.84838 | 0.76324 | 0.988889 |
| GOBP_PROTEIN_K63_LINKED_UBIQUITINATION | 54 | 0.222363 | 0.845423 | 0.76 | 0.994636 |
| GOBP_REGULATION_OF_AUTOPHAGOSOME_ASSEMBLY | 39 | 0.239145 | 0.844673 | 0.71028 | 0.995254 |
| GOBP_MITOCHONDRIAL_CYTOCHROME_C_OXIDASE_ASSEMBLY | 22 | 0.278837 | 0.844586 | 0.685879 | 0.994372 |
| GOBP_NEGATIVE_REGULATION_OF_CATION_CHANNEL_ACTIVITY | 37 | 0.240092 | 0.843671 | 0.721212 | 0.995385 |
| GOBP_KETONE_BIOSYNTHETIC_PROCESS | 37 | 0.245408 | 0.843617 | 0.736487 | 0.994432 |
| GOBP_RESPONSE_TO_INSULIN | 256 | 0.177796 | 0.843405 | 0.944 | 0.993864 |
| GOBP_NEGATIVE_REGULATION_OF_ANOIKIS | 17 | 0.295568 | 0.843304 | 0.69382 | 0.993023 |
| GOBP_THYROID_HORMONE_GENERATION | 16 | 0.28856 | 0.843084 | 0.70195 | 0.992467 |
| GOBP_CALCIUM_MEDIATED_SIGNALING_USING_INTRACELLULAR_CALCIUM_SOURCE | 21 | 0.282349 | 0.842377 | 0.74359 | 0.992939 |
| GOBP_NEGATIVE_REGULATION_OF_AXON_EXTENSION | 40 | 0.239125 | 0.841676 | 0.753333 | 0.993458 |
| GOBP_RNA_SURVEILLANCE | 15 | 0.30099 | 0.840771 | 0.67688 | 0.994441 |
| GOBP_CILIUM_MOVEMENT | 131 | 0.192782 | 0.840371 | 0.855204 | 0.994236 |
| GOBP_ORGANELLE_INHERITANCE | 15 | 0.305053 | 0.84009 | 0.67624 | 0.993838 |
| GOBP_PHOSPHOLIPID_BIOSYNTHETIC_PROCESS | 253 | 0.175968 | 0.839751 | 0.942149 | 0.993549 |
| GOBP_NUCLEOSIDE_DIPHOSPHATE_METABOLIC_PROCESS | 152 | 0.187536 | 0.83911 | 0.912281 | 0.993889 |
| GOBP_POSITIVE_REGULATION_OF_CHROMOSOME_ORGANIZATION | 157 | 0.189185 | 0.839044 | 0.905473 | 0.992987 |
| GOBP_HISTONE_H3_K9_MODIFICATION | 44 | 0.232278 | 0.837056 | 0.785276 | 0.996332 |
| GOBP_NEGATIVE_REGULATION_OF_EPITHELIAL_CELL_APOPTOTIC_PROCESS | 40 | 0.236426 | 0.835842 | 0.772586 | 0.997929 |
| GOBP_PROTEASOMAL_PROTEIN_CATABOLIC_PROCESS | 469 | 0.164434 | 0.834231 | 0.968254 | 1 |
| GOBP_RNA_LOCALIZATION | 226 | 0.177358 | 0.834023 | 0.95946 | 0.999767 |
| GOBP_PROTEIN_FOLDING | 218 | 0.17928 | 0.833715 | 0.960526 | 0.999393 |
| GOBP_RIBONUCLEOSIDE_METABOLIC_PROCESS | 70 | 0.2133 | 0.833588 | 0.823045 | 0.998619 |
| GOBP_REGULATION_OF_SIGNAL_TRANSDUCTION_BY_P53_CLASS_MEDIATOR | 175 | 0.185871 | 0.833336 | 0.934641 | 0.998103 |
| GOBP_GLYCOSYL_COMPOUND_BIOSYNTHETIC_PROCESS | 42 | 0.239184 | 0.832898 | 0.728223 | 0.997997 |
| GOBP_POSITIVE_REGULATION_OF_DENDRITE_MORPHOGENESIS | 35 | 0.249043 | 0.832346 | 0.755776 | 0.998101 |
| GOBP_POSITIVE_REGULATION_OF_CANONICAL_WNT_SIGNALING_PATHWAY | 140 | 0.186205 | 0.832005 | 0.896175 | 0.997719 |
| GOBP_MRNA_3_END_PROCESSING | 98 | 0.198598 | 0.831731 | 0.857143 | 0.997239 |
| GOBP_MEMBRANE_DOCKING | 180 | 0.183934 | 0.830522 | 0.91358 | 0.998713 |
| GOBP_POSITIVE_REGULATION_OF_VIRAL_GENOME_REPLICATION | 31 | 0.24463 | 0.830113 | 0.789809 | 0.998535 |
| GOBP_LATE_ENDOSOME_TO_VACUOLE_TRANSPORT | 24 | 0.263949 | 0.829398 | 0.734756 | 0.999007 |
| GOBP_AXON_EXTENSION | 107 | 0.194397 | 0.82936 | 0.919598 | 0.998038 |
| GOBP_DIOL_METABOLIC_PROCESS | 29 | 0.252324 | 0.82921 | 0.730887 | 0.997332 |
| GOBP_CILIUM_ORGANIZATION | 385 | 0.163307 | 0.82906 | 1 | 0.996607 |
| GOBP_REGULATION_OF_CELL_MORPHOGENESIS_INVOLVED_IN_DIFFERENTIATION | 91 | 0.200966 | 0.828822 | 0.84375 | 0.99604 |
| GOBP_CYTOSOLIC_TRANSPORT | 164 | 0.183002 | 0.828626 | 0.933673 | 0.995414 |
| GOBP_ASSOCIATIVE_LEARNING | 62 | 0.21304 | 0.828371 | 0.825175 | 0.994882 |
| GOBP_VIRAL_RELEASE_FROM_HOST_CELL | 33 | 0.2472 | 0.827288 | 0.758018 | 0.996076 |
| GOBP_REGULATION_OF_DNA_BIOSYNTHETIC_PROCESS | 104 | 0.194693 | 0.826944 | 0.907563 | 0.995778 |
| GOBP_MITOTIC_G2_DNA_DAMAGE_CHECKPOINT | 23 | 0.264079 | 0.825694 | 0.731707 | 0.997187 |
| GOBP_SNRNA_TRANSCRIPTION | 74 | 0.202018 | 0.825461 | 0.854406 | 0.996659 |
| GOBP_RNA_PHOSPHODIESTER_BOND_HYDROLYSIS_ENDONUCLEOLYTIC | 75 | 0.207941 | 0.82522 | 0.816794 | 0.996152 |
| GOBP_RESPONSE_TO_LIGHT_STIMULUS | 286 | 0.169876 | 0.820589 | 0.991667 | 1 |
| GOBP_MODIFIED_AMINO_ACID_TRANSPORT | 28 | 0.25435 | 0.820177 | 0.754777 | 1 |
| GOBP_NADP_METABOLIC_PROCESS | 34 | 0.237093 | 0.820174 | 0.779553 | 1 |
| GOBP_ARGININE_METABOLIC_PROCESS | 17 | 0.288044 | 0.819943 | 0.736364 | 1 |
| GOBP_NEGATIVE_REGULATION_OF_ORGANELLE_ASSEMBLY | 40 | 0.232434 | 0.819149 | 0.78882 | 1 |
| GOBP_POSITIVE_REGULATION_OF_PROTEOLYSIS_INVOLVED_IN_CELLULAR_PROTEIN_CATABOL | 123 | 0.188355 | 0.817367 | 0.925581 | 1 |
| GOBP_PURINE_CONTAINING_COMPOUND_SALVAGE | 15 | 0.294786 | 0.816163 | 0.722372 | 1 |
| GOBP_AMINO_SUGAR_METABOLIC_PROCESS | 36 | 0.242038 | 0.815833 | 0.782875 | 1 |
| GOBP_FORMATION_OF_CYTOPLASMIC_TRANSLATION_INITIATION_COMPLEX | 16 | 0.289563 | 0.815311 | 0.691843 | 1 |
| GOBP_REGULATION_OF_ERAD_PATHWAY | 19 | 0.272276 | 0.814898 | 0.708215 | 1 |
| GOBP_NEGATIVE_REGULATION_OF_PROTEIN_CONTAINING_COMPLEX_ASSEMBLY | 130 | 0.186635 | 0.81298 | 0.926316 | 1 |
| GOBP_SOMATIC_DIVERSIFICATION_OF_IMMUNOGLOBULINS | 64 | 0.205557 | 0.812438 | 0.856061 | 1 |
| GOBP_POSITIVE_REGULATION_OF_CELL_CYCLE_G1_S_PHASE_TRANSITION | 45 | 0.226982 | 0.812336 | 0.798742 | 1 |
| GOBP_SPHINGOLIPID_BIOSYNTHETIC_PROCESS | 106 | 0.192148 | 0.811439 | 0.894737 | 1 |
| GOBP_FATTY_ACID_BETA_OXIDATION_USING_ACYL_COA_OXIDASE | 15 | 0.287615 | 0.810788 | 0.731183 | 1 |
| GOBP_NEGATIVE_REGULATION_OF_CELL_SUBSTRATE_JUNCTION_ORGANIZATION | 18 | 0.278324 | 0.810031 | 0.760989 | 1 |
| GOBP_REGULATION_OF_MICROTUBULE_POLYMERIZATION | 48 | 0.218544 | 0.809621 | 0.82866 | 1 |
| GOBP_CELLULAR_PIGMENTATION | 51 | 0.220283 | 0.809195 | 0.826625 | 1 |
| GOBP_RNA_EXPORT_FROM_NUCLEUS | 136 | 0.184306 | 0.809053 | 0.95 | 1 |
| GOBP_ACETYL_COA_METABOLIC_PROCESS | 35 | 0.241766 | 0.808147 | 0.766667 | 1 |
| GOBP_TYPE_B_PANCREATIC_CELL_PROLIFERATION | 17 | 0.278967 | 0.806166 | 0.740458 | 1 |
| GOBP_ACTIVATION_OF_CYSTEINE_TYPE_ENDOPEPTIDASE_ACTIVITY | 15 | 0.2983 | 0.805925 | 0.742931 | 1 |
| GOBP_ESTABLISHMENT_OF_TISSUE_POLARITY | 120 | 0.183665 | 0.805742 | 0.910569 | 1 |
| GOBP_REPLICATIVE_SENESCENCE | 16 | 0.287545 | 0.80543 | 0.738903 | 1 |
| GOBP_CELLULAR_LIPID_CATABOLIC_PROCESS | 205 | 0.171298 | 0.805342 | 0.975758 | 1 |
| GOBP_NUCLEAR_ENVELOPE_REASSEMBLY | 18 | 0.279221 | 0.805127 | 0.748649 | 1 |
| GOBP_LOW_DENSITY_LIPOPROTEIN_PARTICLE_CLEARANCE | 33 | 0.246278 | 0.80511 | 0.790274 | 1 |
| GOBP_MITOTIC_CYTOKINETIC_PROCESS | 23 | 0.263938 | 0.80356 | 0.772334 | 1 |
| GOBP_NEGATIVE_REGULATION_OF_SODIUM_ION_TRANSPORT | 18 | 0.273315 | 0.80332 | 0.74359 | 1 |
| GOBP_REGULATION_OF_PROTEASOMAL_UBIQUITIN_DEPENDENT_PROTEIN_CATABOLIC_PROCE | 128 | 0.183429 | 0.803234 | 0.922374 | 1 |
| GOBP_NEGATIVE_REGULATION_OF_TELOMERE_MAINTENANCE | 33 | 0.235788 | 0.803205 | 0.78209 | 1 |
| GOBP_NEGATIVE_REGULATION_OF_HISTONE_ACETYLATION | 15 | 0.287296 | 0.802406 | 0.742382 | 1 |
| GOBP_TOR_SIGNALING | 119 | 0.186582 | 0.800634 | 0.921296 | 1 |
| GOBP_RESPONSE_TO_UV | 140 | 0.182702 | 0.799578 | 0.973404 | 1 |
| GOBP_BARBED_END_ACTIN_FILAMENT_CAPPING | 25 | 0.253418 | 0.799423 | 0.78 | 1 |
| GOBP_REGULATION_OF_MICROTUBULE_POLYMERIZATION_OR_DEPOLYMERIZATION | 78 | 0.20401 | 0.799412 | 0.905882 | 1 |

| | | | | | |
|---|---|---|---|---|---|
| GOBP_RESPONSE_TO_FIBROBLAST_GROWTH_FACTOR | 125 | -0.333217 | -1.258099 | 0.099754 | 0.407867 |
| GOBP_REGULATION_OF_TOLL_LIKE_RECEPTOR_SIGNALING_PATHWAY | 70 | -0.361159 | -1.258012 | 0.111409 | 0.407583 |
| GOBP_NEGATIVE_REGULATION_OF_RESPONSE_TO_CYTOKINE_STIMULUS | 58 | -0.373908 | -1.257838 | 0.134286 | 0.40755 |
| GOBP_POSITIVE_REGULATION_OF_LYASE_ACTIVITY | 15 | -0.48642 | -1.257767 | 0.191759 | 0.407258 |
| GOBP_NEGATIVE_REGULATION_OF_VIRAL_GENOME_REPLICATION | 51 | -0.376631 | -1.257612 | 0.131542 | 0.407205 |
| GOBP_CELLULAR_RESPONSE_TO_VIRUS | 61 | -0.37037 | -1.257524 | 0.124498 | 0.406954 |
| GOBP_REGULATION_OF_AMINO_ACID_TRANSPORT | 27 | -0.4357 | -1.257331 | 0.176119 | 0.406989 |
| GOBP_LYMPHOCYTE_CHEMOTAXIS | 54 | -0.377688 | -1.256997 | 0.132333 | 0.407375 |
| GOBP_RESPONSE_TO_ELECTRICAL_STIMULUS | 37 | -0.407349 | -1.25673 | 0.146487 | 0.407594 |
| GOBP_NEGATIVE_REGULATION_OF_TRANSFERASE_ACTIVITY | 274 | -0.305867 | -1.255822 | 0.050171 | 0.40928 |
| GOBP_TRANSMISSION_OF_NERVE_IMPULSE | 51 | -0.382411 | -1.255468 | 0.115489 | 0.409679 |
| GOBP_REGULATION_OF_DIGESTIVE_SYSTEM_PROCESS | 30 | -0.427104 | -1.255276 | 0.164706 | 0.409679 |
| GOBP_REGULATION_OF_FIBROBLAST_GROWTH_FACTOR_RECEPTOR_SIGNALING_PATHWAY | 26 | -0.440257 | -1.255175 | 0.172876 | 0.409431 |
| GOBP_NEGATIVE_REGULATION_OF_B_CELL_PROLIFERATION | 17 | -0.487269 | -1.254753 | 0.187097 | 0.410018 |
| GOBP_REGULATION_OF_EXECUTION_PHASE_OF_APOPTOSIS | 20 | -0.462209 | -1.253809 | 0.167178 | 0.41182 |
| GOBP_VASCULAR_WOUND_HEALING | 17 | -0.485625 | -1.253439 | 0.176378 | 0.412269 |
| GOBP_POSITIVE_REGULATION_OF_EXCITATORY_POSTSYNAPTIC_POTENTIAL | 26 | -0.435235 | -1.253277 | 0.181679 | 0.412214 |
| GOBP_NEUROEPITHELIAL_CELL_DIFFERENTIATION | 16 | -0.484378 | -1.252291 | 0.171073 | 0.414128 |
| GOBP_SKELETAL_MUSCLE_CONTRACTION | 38 | -0.405396 | -1.252013 | 0.176387 | 0.41433 |
| GOBP_REGULATION_OF_NEURON_DIFFERENTIATION | 150 | -0.32471 | -1.251751 | 0.092138 | 0.414506 |
| GOBP_NEGATIVE_REGULATION_OF_ENDOTHELIAL_CELL_PROLIFERATION | 42 | -0.394595 | -1.251268 | 0.15495 | 0.415202 |
| GOBP_REGULATION_OF_CELL_MIGRATION_INVOLVED_IN_SPROUTING_ANGIOGENESIS | 39 | -0.403241 | -1.250365 | 0.1402 | 0.416916 |
| GOBP_RESPONSE_TO_RETINOIC_ACID | 84 | -0.34586 | -1.250348 | 0.099196 | 0.416471 |
| GOBP_REGULATION_OF_RAS_PROTEIN_SIGNAL_TRANSDUCTION | 183 | -0.315979 | -1.250346 | 0.0761 | 0.415992 |
| GOBP_REGULATION_OF_CELL_CELL_ADHESION_MEDIATED_BY_CADHERIN | 18 | -0.472561 | -1.250158 | 0.177524 | 0.415976 |
| GOBP_INNERVATION | 19 | -0.468972 | -1.249505 | 0.182663 | 0.417177 |
| GOBP_KERATINOCYTE_PROLIFERATION | 37 | -0.40677 | -1.249452 | 0.162996 | 0.416822 |
| GOBP_POSITIVE_REGULATION_OF_STEROID_METABOLIC_PROCESS | 22 | -0.44687 | -1.249359 | 0.151563 | 0.416578 |
| GOBP_NEGATIVE_REGULATION_OF_MAP_KINASE_ACTIVITY | 67 | -0.360736 | -1.249337 | 0.12763 | 0.416161 |
| GOBP_COGNITION | 236 | -0.310489 | -1.249333 | 0.068427 | 0.415694 |
| GOBP_CELLULAR_RESPONSE_TO_LIPOPROTEIN_PARTICLE_STIMULUS | 29 | -0.42772 | -1.248723 | 0.172107 | 0.416741 |
| GOBP_POSITIVE_REGULATION_OF_LEUKOCYTE_CHEMOTAXIS | 86 | -0.348189 | -1.248185 | 0.118825 | 0.417659 |
| GOBP_POSITIVE_REGULATION_OF_MUSCLE_CELL_DIFFERENTIATION | 64 | -0.369608 | -1.247701 | 0.13278 | 0.418406 |
| GOBP_I_KAPPAB_PHOSPHORYLATION | 19 | -0.467368 | -1.247692 | 0.195122 | 0.41795 |
| GOBP_REGULATION_OF_PEPTIDYL_SERINE_PHOSPHORYLATION | 116 | -0.334758 | -1.247267 | 0.101759 | 0.418577 |
| GOBP_MYD88_INDEPENDENT_TOLL_LIKE_RECEPTOR_SIGNALING_PATHWAY | 34 | -0.417389 | -1.247039 | 0.159292 | 0.418673 |
| GOBP_POSITIVE_REGULATION_OF_NOTCH_SIGNALING_PATHWAY | 50 | -0.384019 | -1.246984 | 0.145275 | 0.41834 |
| GOBP_INTERLEUKIN_4_PRODUCTION | 32 | -0.409491 | -1.246722 | 0.169839 | 0.418513 |
| GOBP_LEUKOCYTE_HOMEOSTASIS | 78 | -0.356532 | -1.246459 | 0.134511 | 0.418736 |
| GOBP_PHARYNGEAL_SYSTEM_DEVELOPMENT | 19 | -0.469572 | -1.246403 | 0.171875 | 0.418388 |
| GOBP_PEROXISOME_PROLIFERATOR_ACTIVATED_RECEPTOR_SIGNALING_PATHWAY | 20 | -0.456768 | -1.246105 | 0.16411 | 0.418665 |
| GOBP_PEPTIDYL_L_CYSTEINE_S_PALMITOYLATION | 23 | -0.438925 | -1.244754 | 0.164818 | 0.421588 |
| GOBP_EPIDERMAL_CELL_DIFFERENTIATION | 194 | -0.317039 | -1.244173 | 0.076651 | 0.422591 |
| GOBP_MESENCHYME_DEVELOPMENT | 245 | -0.307392 | -1.2434 | 0.068605 | 0.424099 |
| GOBP_HYDROGEN_PEROXIDE_BIOSYNTHETIC_PROCESS | 15 | -0.485679 | -1.243112 | 0.17817 | 0.424409 |
| GOBP_CARDIAC_SEPTUM_MORPHOGENESIS | 54 | -0.376218 | -1.243106 | 0.168345 | 0.42394 |
| GOBP_NEUROTRANSMITTER_RECEPTOR_TRANSPORT | 42 | -0.393903 | -1.242876 | 0.162554 | 0.424072 |
| GOBP_POSITIVE_REGULATION_OF_OSSIFICATION | 43 | -0.393609 | -1.242618 | 0.14936 | 0.424195 |
| GOBP_POSITIVE_REGULATION_OF_MONOCYTE_CHEMOTAXIS | 17 | -0.477871 | -1.242576 | 0.208775 | 0.423799 |
| GOBP_REGULATION_OF_LIPID_KINASE_ACTIVITY | 65 | -0.362421 | -1.242198 | 0.131469 | 0.424358 |
| GOBP_BODY_FLUID_SECRETION | 72 | -0.357047 | -1.24168 | 0.130674 | 0.425221 |
| GOBP_LYMPHOCYTE_APOPTOTIC_PROCESS | 67 | -0.363232 | -1.241291 | 0.144928 | 0.425721 |
| GOBP_DETECTION_OF_MECHANICAL_STIMULUS | 43 | -0.392187 | -1.241201 | 0.163889 | 0.425483 |
| GOBP_3_PHOSPHOADENOSINE_5_PHOSPHOSULFATE_METABOLIC_PROCESS | 22 | -0.439987 | -1.240561 | 0.17716 | 0.426682 |
| GOBP_REGULATION_OF_HEART_CONTRACTION | 202 | -0.3123 | -1.239309 | 0.081049 | 0.42942 |
| GOBP_MORPHOGENESIS_OF_AN_EPITHELIAL_SHEET | 52 | -0.374823 | -1.239259 | 0.13986 | 0.429058 |
| GOBP_ACTIVATION_OF_PROTEIN_KINASE_ACTIVITY | 306 | -0.300048 | -1.238515 | 0.051665 | 0.430484 |
| GOBP_INNER_EAR_RECEPTOR_CELL_DEVELOPMENT | 35 | -0.40807 | -1.238513 | 0.171171 | 0.43001 |
| GOBP_REGULATION_OF_SYSTEMIC_ARTERIAL_BLOOD_PRESSURE | 76 | -0.349372 | -1.238456 | 0.13459 | 0.429665 |
| GOBP_REGULATION_OF_ENDOPLASMIC_RETICULUM_STRESS_INDUCED_INTRINSIC_APOPTOTIC | 28 | -0.421675 | -1.238448 | 0.176737 | 0.429206 |
| GOBP_HETEROTYPIC_CELL_CELL_ADHESION | 56 | -0.372069 | -1.23826 | 0.149927 | 0.429177 |
| GOBP_REGULATION_OF_MESENCHYMAL_CELL_PROLIFERATION | 24 | -0.438109 | -1.238181 | 0.196262 | 0.428894 |
| GOBP_POSITIVE_REGULATION_OF_AMINO_ACID_TRANSPORT | 15 | -0.498228 | -1.238028 | 0.191626 | 0.428781 |
| GOBP_COBALAMIN_METABOLIC_PROCESS | 20 | -0.458187 | -1.237212 | 0.195686 | 0.430392 |
| GOBP_REGULATION_OF_ACTIN_FILAMENT_BUNDLE_ASSEMBLY | 95 | -0.337869 | -1.237152 | 0.124147 | 0.430078 |
| GOBP_LEUKOCYTE_ADHESION_TO_VASCULAR_ENDOTHELIAL_CELL | 44 | -0.38012 | -1.237049 | 0.150355 | 0.429851 |
| GOBP_HEART_MORPHOGENESIS | 205 | -0.307774 | -1.236944 | 0.072029 | 0.429628 |
| GOBP_ACUTE_INFLAMMATORY_RESPONSE_TO_ANTIGENIC_STIMULUS | 21 | -0.450254 | -1.236733 | 0.194745 | 0.429684 |
| GOBP_POSITIVE_REGULATION_OF_LYMPHOCYTE_MIGRATION | 35 | -0.399122 | -1.236261 | 0.168909 | 0.430452 |
| GOBP_PROTEIN_PALMITOYLATION | 31 | -0.420658 | -1.235573 | 0.179083 | 0.431863 |
| GOBP_METANEPHROS_DEVELOPMENT | 65 | -0.365695 | -1.235444 | 0.128 | 0.431721 |
| GOBP_REGULATION_OF_GLYCOPROTEIN_METABOLIC_PROCESS | 44 | -0.386362 | -1.23498 | 0.14556 | 0.432418 |
| GOBP_SENSORY_PERCEPTION_OF_MECHANICAL_STIMULUS | 139 | -0.324933 | -1.23445 | 0.110974 | 0.433341 |
| GOBP_INNER_EAR_MORPHOGENESIS | 73 | -0.352282 | -1.234029 | 0.132128 | 0.433957 |
| GOBP_REGULATION_OF_HORMONE_LEVELS | 423 | -0.2927 | -1.233815 | 0.024758 | 0.434061 |
| GOBP_RESPONSE_TO_PROSTAGLANDIN | 29 | -0.415786 | -1.233722 | 0.181684 | 0.433829 |
| GOBP_REGULATION_OF_NON_CANONICAL_WNT_SIGNALING_PATHWAY | 25 | -0.434175 | -1.233462 | 0.202808 | 0.433987 |
| GOBP_RESPONSE_TO_CORTICOSTEROID | 131 | -0.325809 | -1.232657 | 0.104193 | 0.43559 |
| GOBP_NIK_NF_KAPPAB_SIGNALING | 166 | -0.315338 | -1.232623 | 0.110169 | 0.435184 |
| GOBP_NEGATIVE_REGULATION_OF_CYSTEINE_TYPE_ENDOPEPTIDASE_ACTIVITY | 71 | -0.357722 | -1.231515 | 0.137466 | 0.437587 |
| GOBP_NEGATIVE_REGULATION_OF_ANTIGEN_RECEPTOR_MEDIATED_SIGNALING_PATHWAY | 29 | -0.418012 | -1.231065 | 0.181269 | 0.43829 |
| GOBP_REGULATION_OF_SYNAPSE_STRUCTURE_OR_ACTIVITY | 188 | -0.311727 | -1.230944 | 0.091017 | 0.438133 |
| GOBP_NEURON_PROJECTION_GUIDANCE | 242 | -0.303329 | -1.230806 | 0.069204 | 0.438045 |
| GOBP_PIGMENT_CELL_DIFFERENTIATION | 30 | -0.417416 | -1.23007 | 0.18006 | 0.439487 |
| GOBP_LENS_DEVELOPMENT_IN_CAMERA_TYPE_EYE | 61 | -0.364779 | -1.229886 | 0.134163 | 0.439508 |
| GOBP_RECEPTOR_INTERNALIZATION | 102 | -0.332246 | -1.229855 | 0.123016 | 0.439094 |
| GOBP_CYTOPLASMIC_PATTERN_RECOGNITION_RECEPTOR_SIGNALING_PATHWAY | 70 | -0.356654 | -1.229454 | 0.133425 | 0.439659 |
| GOBP_REGULATION_OF_NUCLEASE_ACTIVITY | 22 | -0.448159 | -1.228833 | 0.197269 | 0.440814 |
| GOBP_REGULATION_OF_INTRINSIC_APOPTOTIC_SIGNALING_PATHWAY_IN_RESPONSE_TO_DNA_ | 17 | -0.467095 | -1.228747 | 0.220472 | 0.440576 |
| GOBP_NEGATIVE_REGULATION_OF_MYELOID_LEUKOCYTE_DIFFERENTIATION | 45 | -0.382702 | -1.228687 | 0.167421 | 0.440267 |
| GOBP_REGULATION_OF_SYSTEMIC_ARTERIAL_BLOOD_PRESSURE_MEDIATED_BY_A_CHEMICAL | 38 | -0.395286 | -1.228149 | 0.168889 | 0.441197 |
| GOBP_REGULATION_OF_TISSUE_REMODELING | 67 | -0.356287 | -1.226844 | 0.144847 | 0.444047 |
| GOBP_REGULATION_OF_FAT_CELL_DIFFERENTIATION | 118 | -0.327109 | -1.226773 | 0.11457 | 0.443753 |
| GOBP_POSITIVE_REGULATION_OF_MACROPHAGE_CHEMOTAXIS | 16 | -0.477735 | -1.226507 | 0.222222 | 0.443967 |
| GOBP_REGULATION_OF_NUCLEOCYTOPLASMIC_TRANSPORT | 101 | -0.333228 | -1.226232 | 0.129458 | 0.444237 |
| GOBP_NEGATIVE_REGULATION_OF_EXTRINSIC_APOPTOTIC_SIGNALING_PATHWAY | 96 | -0.34105 | -1.226149 | 0.135208 | 0.443978 |
| GOBP_RESPONSE_TO_INCREASED_OXYGEN_LEVELS | 28 | -0.415971 | -1.225737 | 0.172876 | 0.444592 |
| GOBP_COPII_COATED_VESICLE_CARGO_LOADING | 15 | -0.489584 | -1.225663 | 0.22427 | 0.444311 |
| GOBP_RESPONSE_TO_TUMOR_NECROSIS_FACTOR | 287 | -0.295643 | -1.225454 | 0.066897 | 0.444393 |
| GOBP_MEMBRANE_PROTEIN_INTRACELLULAR_DOMAIN_PROTEOLYSIS | 20 | -0.447215 | -1.225332 | 0.198438 | 0.444252 |
| GOBP_CHONDROITIN_SULFATE_PROTEOGLYCAN_METABOLIC_PROCESS | 41 | -0.393922 | -1.22498 | 0.158501 | 0.444684 |
| GOBP_POSITIVE_REGULATION_OF_REPRODUCTIVE_PROCESS | 65 | -0.356121 | -1.224194 | 0.161379 | 0.446342 |
| GOBP_REGULATION_OF_CELLULAR_RESPIRATION | 21 | -0.443369 | -1.224151 | 0.193893 | 0.445975 |
| GOBP_DEVELOPMENTAL_PIGMENTATION | 39 | -0.391143 | -1.224055 | 0.19209 | 0.445748 |
| GOBP_REGULATION_OF_HEMOPOIESIS | 394 | -0.292173 | -1.223423 | 0.046002 | 0.446898 |
| GOBP_INTRACELLULAR_RECEPTOR_SIGNALING_PATHWAY | 255 | -0.301681 | -1.223382 | 0.08168 | 0.446523 |
| GOBP_POSITIVE_REGULATION_OF_GLYCOPROTEIN_METABOLIC_PROCESS | 23 | -0.439558 | -1.223373 | 0.193846 | 0.446075 |
| GOBP_PANCREAS_DEVELOPMENT | 56 | -0.366591 | -1.223329 | 0.156863 | 0.445716 |
| GOBP_ENDOTHELIUM_DEVELOPMENT | 118 | -0.32874 | -1.223134 | 0.120406 | 0.445755 |
| GOBP_REGULATION_OF_MACROPHAGE_CHEMOTAXIS | 25 | -0.431489 | -1.223132 | 0.193646 | 0.445293 |
| GOBP_POSITIVE_REGULATION_OF_BIOMINERALIZATION | 43 | -0.384488 | -1.22264 | 0.167862 | 0.446049 |
| GOBP_TOLL_LIKE_RECEPTOR_9_SIGNALING_PATHWAY | 25 | -0.423432 | -1.222591 | 0.199074 | 0.445695 |
| GOBP_NEURON_PROJECTION_ORGANIZATION | 88 | -0.341379 | -1.222442 | 0.132992 | 0.445635 |
| GOBP_VOCALIZATION_BEHAVIOR | 17 | -0.468783 | -1.22214 | 0.202853 | 0.445999 |
| GOBP_EPIDERMIS_DEVELOPMENT | 284 | -0.2982 | -1.222136 | 0.066589 | 0.445534 |
| GOBP_EPITHELIAL_CELL_PROLIFERATION | 347 | -0.295155 | -1.222043 | 0.059351 | 0.445342 |
| GOBP_REGULATION_OF_INTRACELLULAR_ESTROGEN_RECEPTOR_SIGNALING_PATHWAY | 37 | -0.39601 | -1.222022 | 0.188321 | 0.444932 |
| GOBP_CALCIUM_ION_TRANSMEMBRANE_TRANSPORT | 269 | -0.299401 | -1.221623 | 0.070358 | 0.445536 |
| GOBP_DIACYLGLYCEROL_METABOLIC_PROCESS | 23 | -0.438662 | -1.221462 | 0.212403 | 0.445482 |
| GOBP_BLASTODERM_SEGMENTATION | 15 | -0.486028 | -1.221089 | 0.198003 | 0.445997 |
| GOBP_REGULATION_OF_RELEASE_OF_CYTOCHROME_C_FROM_MITOCHONDRIA | 44 | -0.382651 | -1.220682 | 0.168639 | 0.446619 |
| GOBP_REGULATION_OF_APOPTOTIC_SIGNALING_PATHWAY | 328 | -0.29413 | -1.220566 | 0.056117 | 0.446442 |
| GOBP_PROTEIN_ACTIVATION_CASCADE | 21 | -0.447518 | -1.220392 | 0.206587 | 0.446445 |
| GOBP_POSITIVE_REGULATION_OF_NERVOUS_SYSTEM_DEVELOPMENT | 237 | -0.299799 | -1.220373 | 0.076477 | 0.446045 |
| GOBP_POSITIVE_REGULATION_OF_PROTEIN_KINASE_B_SIGNALING | 153 | -0.317503 | -1.219807 | 0.096894 | 0.447035 |
| GOBP_LAMELLIPODIUM_MORPHOGENESIS | 15 | -0.486866 | -1.219655 | 0.21732 | 0.446928 |
| GOBP_NEURON_CELL_CELL_ADHESION | 16 | -0.478264 | -1.218791 | 0.221705 | 0.448754 |
| GOBP_NEUTROPHIL_MIGRATION | 112 | -0.329412 | -1.218481 | 0.130208 | 0.449068 |
| GOBP_ACUTE_PHASE_RESPONSE | 34 | -0.401102 | -1.218336 | 0.192825 | 0.44896 |
| GOBP_REGULATION_OF_HOMOTYPIC_CELL_CELL_ADHESION | 25 | -0.429024 | -1.2181 | 0.207154 | 0.44907 |
| GOBP_RESPONSE_TO_STEROID_HORMONE | 295 | -0.295799 | -1.218059 | 0.070234 | 0.448715 |
| GOBP_CELL_FATE_DETERMINATION | 28 | -0.411798 | -1.217985 | 0.209231 | 0.448459 |
| GOBP_STRESS_FIBER_ASSEMBLY | 98 | -0.333561 | -1.217822 | 0.13272 | 0.448376 |
| GOBP_POSITIVE_REGULATION_OF_HEART_GROWTH | 33 | -0.395197 | -1.217693 | 0.169231 | 0.448223 |
| GOBP_REGULATION_OF_CYSTEINE_TYPE_ENDOPEPTIDASE_ACTIVITY | 210 | -0.306921 | -1.217642 | 0.083333 | 0.447887 |
| GOBP_EMBRYONIC_MORPHOGENESIS | 489 | -0.286028 | -1.21737 | 0.037274 | 0.448186 |
| GOBP_B_CELL_MEDIATED_IMMUNITY | 208 | -0.307973 | -1.217361 | 0.091014 | 0.447741 |
| GOBP_NEGATIVE_REGULATION_OF_INTRINSIC_APOPTOTIC_SIGNALING_PATHWAY_IN_RESPONS | 30 | -0.412058 | -1.216257 | 0.202399 | 0.450168 |
| GOBP_POSITIVE_REGULATION_OF_SMOOTH_MUSCLE_CONTRACTION | 22 | -0.448102 | -1.216228 | 0.229985 | 0.44978 |
| GOBP_ENERGY_HOMEOSTASIS | 35 | -0.405107 | -1.215975 | 0.20324 | 0.449973 |
| GOBP_REGULATION_OF_MEMBRANE_LIPID_DISTRIBUTION | 51 | -0.367465 | -1.215551 | 0.184211 | 0.450624 |
| GOBP_POSITIVE_REGULATION_OF_NITRIC_OXIDE_SYNTHASE_BIOSYNTHETIC_PROCESS | 15 | -0.479891 | -1.215432 | 0.221698 | 0.450459 |
| GOBP_POSITIVE_REGULATION_OF_MYELOID_LEUKOCYTE_CYTOKINE_PRODUCTION_INVOLVED_ | 16 | -0.472074 | -1.215308 | 0.21865 | 0.450315 |
| GOBP_MULTICELLULAR_ORGANISMAL_MOVEMENT | 47 | -0.377045 | -1.21506 | 0.182328 | 0.450503 |
| GOBP_REGULATION_OF_HEART_GROWTH | 57 | -0.362148 | -1.214957 | 0.174825 | 0.450315 |
| GOBP_POSITIVE_REGULATION_OF_PROTEIN_EXPORT_FROM_NUCLEUS | 20 | -0.447343 | -1.214542 | 0.231746 | 0.450977 |
| GOBP_NEGATIVE_REGULATION_OF_TUMOR_NECROSIS_FACTOR_SUPERFAMILY_CYTOKINE_PRO | 55 | -0.363549 | -1.214276 | 0.170596 | 0.451247 |
| GOBP_REGULATION_OF_HUMORAL_IMMUNE_RESPONSE | 119 | -0.322894 | -1.214171 | 0.14361 | 0.451085 |
| GOBP_CELLULAR_RESPONSE_TO_VITAMIN_D | 20 | -0.455989 | -1.21409 | 0.224888 | 0.450812 |
| GOBP_REGULATION_OF_LIPOPROTEIN_LIPASE_ACTIVITY | 18 | -0.461611 | -1.214039 | 0.223089 | 0.450481 |
| GOBP_REGULATION_OF_PROTEIN_LOCALIZATION_TO_CELL_SURFACE | 36 | -0.394883 | -1.214037 | 0.185507 | 0.450034 |
| GOBP_PEPTIDE_SECRETION | 352 | -0.290549 | -1.213934 | 0.059867 | 0.449852 |
| GOBP_PROTEIN_KINASE_B_SIGNALING | 233 | -0.298378 | -1.213753 | 0.100928 | 0.449883 |
| GOBP_NEGATIVE_REGULATION_OF_LYMPHOCYTE_APOPTOTIC_PROCESS | 30 | -0.408856 | -1.213507 | 0.204204 | 0.450112 |
| GOBP_NEGATIVE_REGULATION_OF_INTERFERON_GAMMA_PRODUCTION | 35 | -0.398497 | -1.213431 | 0.19 | 0.44984 |
| GOBP_SENSORY_ORGAN_DEVELOPMENT | 434 | -0.287186 | -1.213159 | 0.04476 | 0.450139 |
| GOBP_CAMERA_TYPE_EYE_DEVELOPMENT | 249 | -0.297621 | -1.212489 | 0.091124 | 0.45139 |
| GOBP_RENAL_SYSTEM_PROCESS_INVOLVED_IN_REGULATION_OF_SYSTEMIC_ARTERIAL_BLOO | 18 | -0.462156 | -1.212468 | 0.229955 | 0.450989 |
| GOBP_NEGATIVE_REGULATION_OF_PEPTIDE_HORMONE_SECRETION | 38 | -0.398185 | -1.211916 | 0.198006 | 0.451952 |
| GOBP_REGULATION_OF_JUN_KINASE_ACTIVITY | 80 | -0.338106 | -1.211645 | 0.158184 | 0.452157 |
| GOBP_BEHAVIOR | 449 | -0.286681 | -1.211275 | 0.054348 | 0.452627 |
| GOBP_REGULATION_OF_TRANSMEMBRANE_TRANSPORT | 478 | -0.286146 | -1.210269 | 0.046759 | 0.454758 |
| GOBP_POSITIVE_REGULATION_OF_GLIAL_CELL_DIFFERENTIATION | 37 | -0.394996 | -1.210003 | 0.199414 | 0.454975 |
| GOBP_INTERLEUKIN_5_PRODUCTION | 17 | -0.465118 | -1.209639 | 0.220635 | 0.455493 |
| GOBP_REGULATION_OF_MYOTUBE_DIFFERENTIATION | 31 | -0.409901 | -1.2094 | 0.204052 | 0.455602 |
| GOBP_POSITIVE_REGULATION_OF_GLIAL_CELL_PROLIFERATION | 17 | -0.468491 | -1.209231 | 0.237082 | 0.455592 |
| GOBP_GASTRULATION | 159 | -0.31254 | -1.208671 | 0.116995 | 0.456601 |
| GOBP_REGULATION_OF_FATTY_ACID_BIOSYNTHETIC_PROCESS | 39 | -0.38763 | -1.20864 | 0.216095 | 0.456227 |

| | | | | | |
|---|---|---|---|---|---|
| GOBP_REGULATION_OF_PROTEIN_AUTOPHOSPHORYLATION | 45 | 0.22105 | 0.798383 | 0.864238 | 1 |
| GOBP_SNO_S_RNA_METABOLIC_PROCESS | 15 | 0.284153 | 0.797908 | 0.761364 | 1 |
| GOBP_NUCLEOTIDE_PHOSPHORYLATION | 129 | 0.181579 | 0.797457 | 0.950739 | 1 |
| GOBP_CARDIAC_RIGHT_VENTRICLE_MORPHOGENESIS | 15 | 0.294106 | 0.797191 | 0.763587 | 1 |
| GOBP_RRNA_TRANSCRIPTION | 33 | 0.235186 | 0.796593 | 0.82866 | 1 |
| GOBP_PHOSPHATIDYLINOSITOL_BIOSYNTHETIC_PROCESS | 112 | 0.188156 | 0.795964 | 0.938272 | 1 |
| GOBP_PROTEIN_TRANSMEMBRANE_IMPORT_INTO_INTRACELLULAR_ORGANELLE | 36 | 0.231217 | 0.795578 | 0.796721 | 1 |
| GOBP_PROTEIN_QUALITY_CONTROL_FOR_MISFOLDED_OR_INCOMPLETELY_SYNTHESIZED_PRO | 28 | 0.242997 | 0.79419 | 0.807692 | 1 |
| GOBP_GLUTAMINE_FAMILY_AMINO_ACID_CATABOLIC_PROCESS | 22 | 0.250329 | 0.794083 | 0.790368 | 1 |
| GOBP_RIBONUCLEOSIDE_TRIPHOSPHATE_METABOLIC_PROCESS | 86 | 0.197161 | 0.793722 | 0.908696 | 1 |
| GOBP_RIBONUCLEOPROTEIN_COMPLEX_BIOGENESIS | 452 | 0.159483 | 0.791701 | 1 | 1 |
| GOBP_POSITIVE_REGULATION_OF_VASCULAR_ASSOCIATED_SMOOTH_MUSCLE_CELL_PROLIFE | 35 | 0.232029 | 0.790394 | 0.814925 | 1 |
| GOBP_MRNA_METHYLATION | 16 | 0.287665 | 0.790342 | 0.76781 | 1 |
| GOBP_POSITIVE_REGULATION_OF_DNA_METABOLIC_PROCESS | 192 | 0.171402 | 0.790009 | 0.980263 | 1 |
| GOBP_MULTI_ORGANISM_LOCALIZATION | 66 | 0.20023 | 0.788976 | 0.884892 | 1 |
| GOBP_REGULATION_OF_ATPASE_ACTIVITY | 78 | 0.193614 | 0.788135 | 0.941177 | 1 |
| GOBP_REGULATION_OF_HISTONE_METHYLATION | 59 | 0.202726 | 0.78628 | 0.900383 | 1 |
| GOBP_CELL_COMMUNICATION_BY_ELECTRICAL_COUPLING | 28 | 0.238436 | 0.786274 | 0.798701 | 1 |
| GOBP_MICROTUBULE_BASED_TRANSPORT | 185 | 0.170688 | 0.785416 | 1 | 1 |
| GOBP_NEGATIVE_REGULATION_OF_DNA_BIOSYNTHETIC_PROCESS | 37 | 0.231206 | 0.784894 | 0.816199 | 1 |
| GOBP_MITOCHONDRIAL_CALCIUM_ION_TRANSMEMBRANE_TRANSPORT | 25 | 0.251125 | 0.784405 | 0.798799 | 1 |
| GOBP_GOLGI_TO_VACUOLE_TRANSPORT | 19 | 0.259738 | 0.784325 | 0.764179 | 1 |
| GOBP_MITOCHONDRIAL_FISSION | 42 | 0.217757 | 0.783603 | 0.820896 | 1 |
| GOBP_POSITIVE_REGULATION_OF_CELLULAR_RESPONSE_TO_INSULIN_STIMULUS | 22 | 0.257122 | 0.782919 | 0.785276 | 1 |
| GOBP_RIBOSOMAL_LARGE_SUBUNIT_ASSEMBLY | 27 | 0.246693 | 0.782628 | 0.809384 | 1 |
| GOBP_DETECTION_OF_LIGHT_STIMULUS_INVOLVED_IN_SENSORY_PERCEPTION | 15 | 0.278611 | 0.782401 | 0.795918 | 1 |
| GOBP_PROTEIN_DESTABILIZATION | 46 | 0.216271 | 0.781942 | 0.860317 | 1 |
| GOBP_SUPEROXIDE_ANION_GENERATION | 31 | 0.235498 | 0.78182 | 0.823188 | 1 |
| GOBP_COPPER_ION_HOMEOSTASIS | 16 | 0.287521 | 0.7808 | 0.804094 | 1 |
| GOBP_REGULATION_OF_ACTIN_CYTOSKELETON_REORGANIZATION | 36 | 0.232115 | 0.780577 | 0.825688 | 1 |
| GOBP_MACROMOLECULE_METHYLATION | 287 | 0.161573 | 0.778748 | 0.991379 | 1 |
| GOBP_BLOOD_VESSEL_ENDOTHELIAL_CELL_PROLIFERATION_INVOLVED_IN_SPROUTING_ANGI | 25 | 0.249281 | 0.777294 | 0.821317 | 1 |
| GOBP_PYRIMIDINE_CONTAINING_COMPOUND_CATABOLIC_PROCESS | 37 | 0.229793 | 0.775502 | 0.859873 | 1 |
| GOBP_REGULATION_OF_MICROTUBULE_CYTOSKELETON_ORGANIZATION | 133 | 0.176196 | 0.774994 | 0.967593 | 1 |
| GOBP_REGULATION_OF_TRANSLATIONAL_ELONGATION | 19 | 0.264787 | 0.7748 | 0.80057 | 1 |
| GOBP_PROTEIN_MONOUBIQUITINATION | 66 | 0.196395 | 0.774673 | 0.923913 | 1 |
| GOBP_PROTEIN_IMPORT_INTO_MITOCHONDRIAL_MATRIX | 20 | 0.258688 | 0.772646 | 0.810127 | 1 |
| GOBP_MEGAKARYOCYTE_DEVELOPMENT | 16 | 0.274019 | 0.772338 | 0.776882 | 1 |
| GOBP_SPERM_EGG_RECOGNITION | 43 | 0.214744 | 0.771457 | 0.86121 | 1 |
| GOBP_REGULATION_OF_PROTEIN_TARGETING_TO_MITOCHONDRION | 43 | 0.216149 | 0.770563 | 0.85489 | 1 |
| GOBP_POSITIVE_REGULATION_OF_PROTEIN_TARGETING_TO_MITOCHONDRION | 31 | 0.232875 | 0.770209 | 0.845118 | 1 |
| GOBP_MITOCHONDRIAL_ELECTRON_TRANSPORT_CYTOCHROME_C_TO_OXYGEN | 19 | 0.261948 | 0.769566 | 0.790561 | 1 |
| GOBP_METHYLATION | 335 | 0.156994 | 0.768706 | 1 | 1 |
| GOBP_POSITIVE_REGULATION_OF_ATPASE_ACTIVITY | 55 | 0.198235 | 0.767831 | 0.874126 | 1 |
| GOBP_REGULATION_OF_GLUCOSE_IMPORT | 49 | 0.208537 | 0.767266 | 0.911972 | 1 |
| GOBP_PROTEIN_METHYLATION | 161 | 0.17166 | 0.767155 | 0.989796 | 1 |
| GOBP_POSITIVE_REGULATION_OF_TRANSCRIPTION_INITIATION_FROM_RNA_POLYMERASE_II_PI | 27 | 0.238169 | 0.766902 | 0.834783 | 1 |
| GOBP_REGULATION_OF_DOUBLE_STRAND_BREAK_REPAIR_VIA_HOMOLOGOUS_RECOMBINATIC | 47 | 0.20916 | 0.766535 | 0.896552 | 1 |
| GOBP_REGULATION_OF_TRANSCRIPTION_INITIATION_FROM_RNA_POLYMERASE_II_PROMOTER | 33 | 0.226189 | 0.766151 | 0.841463 | 1 |
| GOBP_GUANOSINE_CONTAINING_COMPOUND_METABOLIC_PROCESS | 40 | 0.221712 | 0.765877 | 0.893417 | 1 |
| GOBP_REGULATION_OF_PROTEIN_TARGETING | 78 | 0.190056 | 0.765248 | 0.939759 | 1 |
| GOBP_REGULATION_OF_MUSCLE_HYPERTROPHY | 53 | 0.201163 | 0.764721 | 0.896057 | 1 |
| GOBP_GLYCEROPHOSPHOLIPID_METABOLIC_PROCESS | 323 | 0.156017 | 0.763665 | 1 | 1 |
| GOBP_POSITIVE_REGULATION_OF_PROTEIN_MODIFICATION_BY_SMALL_PROTEIN_CONJUGATIO | 130 | 0.176496 | 0.762699 | 0.974249 | 1 |
| GOBP_REGULATION_OF_CELL_SIZE | 160 | 0.16931 | 0.760832 | 0.986425 | 1 |
| GOBP_PHOSPHOLIPID_METABOLIC_PROCESS | 387 | 0.155908 | 0.760677 | 1 | 1 |
| GOBP_REGULATION_OF_VIRAL_INDUCED_CYTOPLASMIC_PATTERN_RECOGNITION_RECEPTOR_ | 21 | 0.24995 | 0.760502 | 0.76161 | 1 |
| GOBP_NEGATIVE_REGULATION_OF_MICROTUBULE_POLYMERIZATION_OR_DEPOLYMERIZATION | 37 | 0.223739 | 0.759943 | 0.846395 | 1 |
| GOBP_RNA_POLYADENYLATION | 45 | 0.215223 | 0.759589 | 0.877483 | 1 |
| GOBP_RIBONUCLEOSIDE_TRIPHOSPHATE_BIOSYNTHETIC_PROCESS | 72 | 0.190886 | 0.759279 | 0.932836 | 1 |
| GOBP_EMBRYONIC_CAMERA_TYPE_EYE_DEVELOPMENT | 25 | 0.246381 | 0.758654 | 0.824451 | 1 |
| GOBP_CYTOPLASMIC_TRANSLATIONAL_INITIATION | 34 | 0.222731 | 0.758649 | 0.833333 | 1 |
| GOBP_MATURATION_OF_SSU_RRNA_FROM_TRICISTRONIC_RRNA_TRANSCRIPT_SSU_RRNA_5_ | 37 | 0.221457 | 0.758117 | 0.852941 | 1 |
| GOBP_AMYLOID_FIBRIL_FORMATION | 23 | 0.245214 | 0.75617 | 0.822157 | 1 |
| GOBP_REGULATION_OF_CYTOPLASMIC_TRANSLATION | 20 | 0.246716 | 0.753573 | 0.809942 | 1 |
| GOBP_EXPORT_ACROSS_PLASMA_MEMBRANE | 54 | 0.197617 | 0.751569 | 0.942761 | 1 |
| GOBP_OUTER_DYNEIN_ARM_ASSEMBLY | 18 | 0.264278 | 0.748849 | 0.827684 | 1 |
| GOBP_CHAPERONE_MEDIATED_PROTEIN_FOLDING | 61 | 0.19204 | 0.747321 | 0.940741 | 1 |
| GOBP_MIDBODY_ABSCISSION | 17 | 0.26387 | 0.747245 | 0.833333 | 1 |
| GOBP_MITOCHONDRIAL_CALCIUM_ION_HOMEOSTASIS | 25 | 0.245981 | 0.745153 | 0.863777 | 1 |
| GOBP_NEGATIVE_REGULATION_OF_DNA_REPAIR | 36 | 0.209539 | 0.744258 | 0.90303 | 1 |
| GOBP_LATE_ENDOSOME_TO_VACUOLE_TRANSPORT_VIA_MULTIVESICULAR_BODY_SORTING_P | 17 | 0.26387 | 0.743432 | 0.841808 | 1 |
| GOBP_RECEPTOR_CATABOLIC_PROCESS | 35 | 0.215574 | 0.743257 | 0.880342 | 1 |
| GOBP_CELLULAR_METABOLIC_COMPOUND_SALVAGE | 34 | 0.223002 | 0.742134 | 0.874636 | 1 |
| GOBP_TRANSCRIPTION_PREINITIATION_COMPLEX_ASSEMBLY | 36 | 0.216562 | 0.740767 | 0.917492 | 1 |
| GOBP_LIPID_MODIFICATION | 250 | 0.153209 | 0.740745 | 1 | 1 |
| GOBP_POSITIVE_REGULATION_OF_MICROTUBULE_POLYMERIZATION_OR_DEPOLYMERIZATION | 32 | 0.22503 | 0.739222 | 0.873156 | 1 |
| GOBP_ACETYL_COA_BIOSYNTHETIC_PROCESS | 21 | 0.246332 | 0.738647 | 0.849231 | 1 |
| GOBP_GLUCOSE_IMPORT | 63 | 0.194736 | 0.737055 | 0.926056 | 1 |
| GOBP_PYRIMIDINE_RIBONUCLEOTIDE_BIOSYNTHETIC_PROCESS | 22 | 0.239789 | 0.736024 | 0.845029 | 1 |
| GOBP_PHOSPHATIDIC_ACID_METABOLIC_PROCESS | 42 | 0.206992 | 0.73507 | 0.88961 | 1 |
| GOBP_PHOSPHOLIPID_DEPHOSPHORYLATION | 42 | 0.203371 | 0.735012 | 0.891525 | 1 |
| GOBP_POSITIVE_REGULATION_OF_ISOTYPE_SWITCHING | 26 | 0.228843 | 0.734326 | 0.864198 | 1 |
| GOBP_REGULATION_OF_DOUBLE_STRAND_BREAK_REPAIR | 84 | 0.182465 | 0.733832 | 0.95339 | 1 |
| GOBP_NUCLEAR_ENVELOPE_ORGANIZATION | 51 | 0.197142 | 0.729516 | 0.912281 | 1 |
| GOBP_REGULATION_OF_PROTEIN_TARGETING_TO_MEMBRANE | 34 | 0.213278 | 0.726736 | 0.908497 | 1 |
| GOBP_ORGANELLE_TRANSPORT_ALONG_MICROTUBULE | 80 | 0.177855 | 0.726529 | 0.978648 | 1 |
| GOBP_REGULATION_OF_TRANSLATION_NCRNA_MEDIATED | 25 | 0.229834 | 0.726513 | 0.872093 | 1 |
| GOBP_METAL_ION_EXPORT | 43 | 0.202139 | 0.72629 | 0.91815 | 1 |
| GOBP_NEUROMUSCULAR_SYNAPTIC_TRANSMISSION | 20 | 0.24415 | 0.726254 | 0.840708 | 1 |
| GOBP_PYRIMIDINE_DEOXYRIBONUCLEOTIDE_METABOLIC_PROCESS | 22 | 0.234251 | 0.725734 | 0.871345 | 1 |
| GOBP_REGULATION_OF_OXIDATIVE_PHOSPHORYLATION | 23 | 0.235379 | 0.723323 | 0.871951 | 1 |
| GOBP_RNA_PHOSPHODIESTER_BOND_HYDROLYSIS | 145 | 0.1641 | 0.722224 | 1 | 1 |
| GOBP_MICROTUBULE_ANCHORING | 27 | 0.222787 | 0.720501 | 0.883582 | 1 |
| GOBP_MITOCHONDRION_MORPHOGENESIS | 19 | 0.240789 | 0.719704 | 0.87069 | 1 |
| GOBP_POSITIVE_REGULATION_OF_RESPONSE_TO_DNA_DAMAGE_STIMULUS | 104 | 0.170527 | 0.717187 | 0.977578 | 1 |
| GOBP_REGULATION_OF_PROTEIN_STABILITY | 282 | 0.14933 | 0.715391 | 1 | 1 |
| GOBP_PIRNA_METABOLIC_PROCESS | 15 | 0.265256 | 0.715368 | 0.833795 | 1 |
| GOBP_SULFATE_TRANSPORT | 15 | 0.259627 | 0.714392 | 0.836461 | 1 |
| GOBP_REGULATION_OF_TRANSCRIPTION_REGULATORY_REGION_DNA_BINDING | 44 | 0.200081 | 0.714317 | 0.938983 | 1 |
| GOBP_TRANSPORT_ALONG_MICROTUBULE | 155 | 0.158639 | 0.714225 | 0.994819 | 1 |
| GOBP_HIPPO_SIGNALING | 44 | 0.199302 | 0.710941 | 0.938312 | 1 |
| GOBP_REGULATION_OF_SPINDLE_ORGANIZATION | 39 | 0.199242 | 0.709574 | 0.942771 | 1 |
| GOBP_REGULATION_OF_CELLULAR_AMIDE_METABOLIC_PROCESS | 413 | 0.142161 | 0.706334 | 1 | 1 |
| GOBP_POSITIVE_REGULATION_OF_TRANSCRIPTION_OF_NOTCH_RECEPTOR_TARGET | 18 | 0.244249 | 0.705866 | 0.848649 | 1 |
| GOBP_REGULATION_OF_RESPONSE_TO_DNA_DAMAGE_STIMULUS | 217 | 0.151269 | 0.703842 | 1 | 1 |
| GOBP_SYNAPTIC_TRANSMISSION_DOPAMINERGIC | 18 | 0.244446 | 0.702877 | 0.900293 | 1 |
| GOBP_REGULATION_OF_NUCLEOBASE_CONTAINING_COMPOUND_TRANSPORT | 15 | 0.256761 | 0.701904 | 0.86351 | 1 |
| GOBP_PROTEASOMAL_UBIQUITIN_INDEPENDENT_PROTEIN_CATABOLIC_PROCESS | 21 | 0.231149 | 0.70104 | 0.894879 | 1 |
| GOBP_NEGATIVE_REGULATION_OF_AUTOPHAGOSOME_ASSEMBLY | 15 | 0.249967 | 0.700035 | 0.859649 | 1 |
| GOBP_PROTEIN_NEDDYLATION | 18 | 0.243582 | 0.699954 | 0.861972 | 1 |
| GOBP_APOPTOTIC_DNA_FRAGMENTATION | 20 | 0.235384 | 0.699563 | 0.892442 | 1 |
| GOBP_CELLULAR_SODIUM_ION_HOMEOSTASIS | 15 | 0.261735 | 0.699059 | 0.846154 | 1 |
| GOBP_PEPTIDYL_LYSINE_METHYLATION | 118 | 0.162324 | 0.698734 | 0.995575 | 1 |
| GOBP_POSITIVE_REGULATION_OF_G1_S_TRANSITION_OF_MITOTIC_CELL_CYCLE | 36 | 0.203748 | 0.696395 | 0.936709 | 1 |
| GOBP_NUCLEUS_ORGANIZATION | 118 | 0.158499 | 0.696126 | 0.985714 | 1 |
| GOBP_MAGNESIUM_ION_TRANSPORT | 16 | 0.244279 | 0.694187 | 0.904899 | 1 |
| GOBP_POSITIVE_REGULATION_OF_MUSCLE_HYPERTROPHY | 27 | 0.214015 | 0.69363 | 0.918478 | 1 |
| GOBP_MATURATION_OF_SSU_RRNA | 50 | 0.188949 | 0.690042 | 0.950877 | 1 |
| GOBP_UBIQUINONE_METABOLIC_PROCESS | 18 | 0.236505 | 0.688718 | 0.886957 | 1 |
| GOBP_PEPTIDYL_PROLINE_MODIFICATION | 52 | 0.182871 | 0.687598 | 0.953901 | 1 |
| GOBP_POSITIVE_REGULATION_OF_ERYTHROCYTE_DIFFERENTIATION | 32 | 0.211555 | 0.687408 | 0.936364 | 1 |
| GOBP_SMOOTHENED_SIGNALING_PATHWAY | 122 | 0.156789 | 0.686043 | 1 | 1 |
| GOBP_POSITIVE_REGULATION_OF_DOUBLE_STRAND_BREAK_REPAIR_VIA_NONHOMOLOGOUS_ | 16 | 0.237892 | 0.685707 | 0.894879 | 1 |
| GOBP_REGULATION_OF_RECEPTOR_RECYCLING | 23 | 0.224654 | 0.682159 | 0.911175 | 1 |
| GOBP_PROTEIN_STABILIZATION | 182 | 0.148662 | 0.681635 | 1 | 1 |
| GOBP_REGULATION_OF_PROTON_TRANSPORT | 23 | 0.218934 | 0.681603 | 0.931694 | 1 |
| GOBP_PEPTIDE_CATABOLIC_PROCESS | 25 | 0.217507 | 0.681165 | 0.918919 | 1 |
| GOBP_SIALYLATION | 17 | 0.234863 | 0.678332 | 0.920118 | 1 |
| GOBP_REGULATION_OF_INTEGRIN_MEDIATED_SIGNALING_PATHWAY | 15 | 0.243153 | 0.676012 | 0.901042 | 1 |
| GOBP_CELL_REDOX_HOMEOSTASIS | 42 | 0.186948 | 0.67373 | 0.960854 | 1 |
| GOBP_POSITIVE_REGULATION_OF_HISTONE_METHYLATION | 34 | 0.201341 | 0.6728 | 0.929577 | 1 |
| GOBP_MITOCHONDRIAL_GENOME_MAINTENANCE | 21 | 0.22087 | 0.672646 | 0.94302 | 1 |
| GOBP_ESTABLISHMENT_OF_RNA_LOCALIZATION | 193 | 0.143268 | 0.67131 | 1 | 1 |
| GOBP_POSITIVE_REGULATION_OF_DNA_RECOMBINATION | 44 | 0.188093 | 0.668788 | 0.971246 | 1 |
| GOBP_LYMPHANGIOGENESIS | 15 | 0.242655 | 0.668422 | 0.903485 | 1 |
| GOBP_CELLULAR_RESPONSE_TO_GAMMA_RADIATION | 26 | 0.20988 | 0.667955 | 0.930514 | 1 |
| GOBP_CELL_COMMUNICATION_BY_ELECTRICAL_COUPLING_INVOLVED_IN_CARDIAC_CONDUCT | 23 | 0.212603 | 0.666862 | 0.929972 | 1 |
| GOBP_MICROVILLUS_ORGANIZATION | 23 | 0.216753 | 0.665895 | 0.926254 | 1 |
| GOBP_GOLGI_TO_ENDOSOME_TRANSPORT | 17 | 0.232921 | 0.663784 | 0.928962 | 1 |
| GOBP_REGULATION_OF_SPINDLE_ASSEMBLY | 25 | 0.213756 | 0.663351 | 0.945619 | 1 |
| GOBP_PLATELET_AGGREGATION | 56 | 0.176199 | 0.663061 | 0.972881 | 1 |
| GOBP_POSITIVE_REGULATION_OF_RELEASE_OF_SEQUESTERED_CALCIUM_ION_INTO_CYTOSOL | 35 | 0.195139 | 0.661506 | 0.973856 | 1 |
| GOBP_INTRACILIARY_TRANSPORT_INVOLVED_IN_CILIUM_ASSEMBLY | 40 | 0.187521 | 0.660905 | 0.973684 | 1 |
| GOBP_LIPOPROTEIN_METABOLIC_PROCESS | 125 | 0.150626 | 0.660423 | 1 | 1 |
| GOBP_POSITIVE_REGULATION_OF_MACROAUTOPHAGY | 59 | 0.175225 | 0.657178 | 0.989474 | 1 |
| GOBP_POSITIVE_REGULATION_OF_CALCIUM_ION_TRANSMEMBRANE_TRANSPORT | 62 | 0.169731 | 0.657039 | 0.992565 | 1 |
| GOBP_INCLUSION_BODY_ASSEMBLY | 21 | 0.21939 | 0.656232 | 0.92515 | 1 |
| GOBP_NEGATIVE_REGULATION_OF_RNA_CATABOLIC_PROCESS | 62 | 0.165233 | 0.653732 | 0.996032 | 1 |
| GOBP_CARBOHYDRATE_PHOSPHORYLATION | 24 | 0.212859 | 0.653163 | 0.922652 | 1 |
| GOBP_REGULATION_OF_MICROTUBULE_DEPOLYMERIZATION | 29 | 0.197183 | 0.652753 | 0.952681 | 1 |
| GOBP_REGULATION_OF_VACUOLE_ORGANIZATION | 47 | 0.176351 | 0.652495 | 0.976431 | 1 |
| GOBP_POSITIVE_REGULATION_OF_FILOPODIUM_ASSEMBLY | 28 | 0.197914 | 0.651711 | 0.934579 | 1 |
| GOBP_RIG_I_SIGNALING_PATHWAY | 24 | 0.212396 | 0.651467 | 0.946746 | 1 |
| GOBP_SPERM_AXONEME_ASSEMBLY | 20 | 0.218315 | 0.646265 | 0.921127 | 1 |
| GOBP_REGULATION_OF_TELOMERE_CAPPING | 26 | 0.202993 | 0.644878 | 0.95614 | 1 |
| GOBP_CYTOKINETIC_PROCESS | 38 | 0.180687 | 0.640725 | 0.986577 | 1 |
| GOBP_COVALENT_CHROMATIN_MODIFICATION | 437 | 0.127904 | 0.640622 | 1 | 1 |
| GOBP_CHROMATIN_DISASSEMBLY | 18 | 0.214942 | 0.638911 | 0.939891 | 1 |
| GOBP_HISTONE_METHYLATION | 123 | 0.148259 | 0.63759 | 1 | 1 |
| GOBP_POSITIVE_REGULATION_OF_CHROMATIN_ORGANIZATION | 95 | 0.153241 | 0.63746 | 0.995652 | 1 |
| GOBP_N_TERMINAL_PROTEIN_AMINO_ACID_MODIFICATION | 26 | 0.200131 | 0.637297 | 0.962963 | 1 |
| GOBP_PROTEIN_DNA_COMPLEX_DISASSEMBLY | 18 | 0.21553 | 0.636875 | 0.951483 | 1 |
| GOBP_REGULATION_OF_UBIQUITIN_DEPENDENT_PROTEIN_CATABOLIC_PROCESS | 155 | 0.142396 | 0.636634 | 1 | 1 |
| GOBP_NEGATIVE_REGULATION_OF_STRIATED_MUSCLE_CELL_DIFFERENTIATION | 32 | 0.191022 | 0.635995 | 0.963855 | 1 |
| GOBP_GOLGI_VESICLE_TRANSPORT | 361 | 0.128014 | 0.63314 | 1 | 1 |
| GOBP_POSITIVE_REGULATION_OF_RNA_SPLICING | 34 | 0.185195 | 0.630105 | 0.980456 | 1 |
| GOBP_MRNA_EXPORT_FROM_NUCLEUS | 110 | 0.14749 | 0.62914 | 1 | 1 |

| | | | | | |
|---|---|---|---|---|---|
| GOBP_REGULATION_OF_ERBB_SIGNALING_PATHWAY | 92 | -0.338305 | -1.208624 | 0.14433 | 0.455814 |
| GOBP_METAL_ION_TRANSPORT | 431 | -0.288269 | -1.207998 | 0.060086 | 0.456994 |
| GOBP_CALCIUM_ION_REGULATED_EXOCYTOSIS_OF_NEUROTRANSMITTER | 16 | -0.468763 | -1.207928 | 0.215655 | 0.456734 |
| GOBP_REGULATION_OF_MAP_KINASE_ACTIVITY | 285 | -0.293676 | -1.207672 | 0.074492 | 0.456969 |
| GOBP_FOREBRAIN_NEURON_DEVELOPMENT | 20 | -0.440113 | -1.207564 | 0.22292 | 0.456799 |
| GOBP_REGULATION_OF_GLIAL_CELL_PROLIFERATION | 29 | -0.404476 | -1.207141 | 0.209064 | 0.457416 |
| GOBP_REGULATION_OF_TRANSMEMBRANE_RECEPTOR_PROTEIN_SERINE_THREONINE_KINASE | 215 | -0.299039 | -1.20704 | 0.09319 | 0.457226 |
| GOBP_LYTIC_VACUOLE_ORGANIZATION | 67 | -0.347397 | -1.206874 | 0.180217 | 0.457214 |
| GOBP_REGULATION_OF_NEURAL_PRECURSOR_CELL_PROLIFERATION | 67 | -0.347498 | -1.206841 | 0.160221 | 0.456842 |
| GOBP_NEGATIVE_REGULATION_OF_T_CELL_APOPTOTIC_PROCESS | 19 | -0.453153 | -1.206185 | 0.241325 | 0.458132 |
| GOBP_POSITIVE_REGULATION_OF_CREB_TRANSCRIPTION_FACTOR_ACTIVITY | 17 | -0.464042 | -1.206078 | 0.227273 | 0.457966 |
| GOBP_CELLULAR_RESPONSE_TO_NUTRIENT | 38 | -0.38914 | -1.206075 | 0.206795 | 0.457526 |
| GOBP_REGULATION_OF_INFLAMMATORY_RESPONSE_TO_ANTIGENIC_STIMULUS | 27 | -0.415092 | -1.205835 | 0.204268 | 0.457686 |
| GOBP_POSITIVE_REGULATION_OF_NUCLEOTIDE_METABOLIC_PROCESS | 39 | -0.388575 | -1.205715 | 0.20292 | 0.457585 |
| GOBP_REGULATION_OF_CYCLASE_ACTIVITY | 34 | -0.400865 | -1.20546 | 0.201794 | 0.457778 |
| GOBP_GLIAL_CELL_PROLIFERATION | 44 | -0.376852 | -1.205315 | 0.207407 | 0.457699 |
| GOBP_INACTIVATION_OF_MAPK_ACTIVITY | 23 | -0.434757 | -1.205299 | 0.212766 | 0.457304 |
| GOBP_COCHLEA_MORPHOGENESIS | 18 | -0.458645 | -1.204958 | 0.216216 | 0.45779 |
| GOBP_REGULATION_OF_LEUKOCYTE_APOPTOTIC_PROCESS | 75 | -0.345402 | -1.204572 | 0.158184 | 0.458361 |
| GOBP_FUCOSYLATION | 16 | -0.4753 | -1.203596 | 0.214724 | 0.460506 |
| GOBP_REGULATION_OF_REGULATED_SECRETORY_PATHWAY | 127 | -0.321604 | -1.203312 | 0.131378 | 0.460772 |
| GOBP_REGULATION_OF_OSTEOBLAST_DIFFERENTIATION | 108 | -0.32646 | -1.203256 | 0.137662 | 0.460463 |
| GOBP_NATURAL_KILLER_CELL_DIFFERENTIATION | 17 | -0.463314 | -1.203107 | 0.221184 | 0.460389 |
| GOBP_POSITIVE_REGULATION_OF_OLIGODENDROCYTE_DIFFERENTIATION | 19 | -0.452214 | -1.202802 | 0.230415 | 0.460731 |
| GOBP_CIRCADIAN_SLEEP_WAKE_CYCLE | 19 | -0.449812 | -1.202561 | 0.231861 | 0.460893 |
| GOBP_NEGATIVE_REGULATION_OF_TRANSPORT | 379 | -0.28529 | -1.202232 | 0.077778 | 0.46133 |
| GOBP_RESPONSE_TO_ZINC_ION | 43 | -0.373179 | -1.20212 | 0.191679 | 0.461142 |
| GOBP_PROTEIN_CATABOLIC_PROCESS_IN_THE_VACUOLE | 16 | -0.471093 | -1.201968 | 0.229068 | 0.461097 |
| GOBP_REGULATION_OF_PHOSPHATIDYLINOSITOL_3_KINASE_ACTIVITY | 53 | -0.373284 | -1.201652 | 0.172714 | 0.461478 |
| GOBP_NEGATIVE_REGULATION_OF_MAPK_CASCADE | 158 | -0.307135 | -1.201598 | 0.148962 | 0.461157 |
| GOBP_CELLULAR_RESPONSE_TO_VITAMIN | 28 | -0.407247 | -1.20121 | 0.233687 | 0.461729 |
| GOBP_REGULATION_OF_MUSCLE_CELL_DIFFERENTIATION | 126 | -0.317237 | -1.200769 | 0.146944 | 0.462442 |
| GOBP_CARDIAC_CHAMBER_DEVELOPMENT | 127 | -0.317817 | -1.200358 | 0.151629 | 0.463081 |
| GOBP_CORPUS_CALLOSUM_DEVELOPMENT | 15 | -0.475517 | -1.200291 | 0.248387 | 0.462839 |
| GOBP_CELLULAR_RESPONSE_TO_MECHANICAL_STIMULUS | 73 | -0.343318 | -1.200231 | 0.166895 | 0.462535 |
| GOBP_SUBPALLIUM_DEVELOPMENT | 21 | -0.431354 | -1.199567 | 0.223473 | 0.46385 |
| GOBP_ENTEROENDOCRINE_CELL_DIFFERENTIATION | 23 | -0.438163 | -1.199375 | 0.201299 | 0.463926 |
| GOBP_REGULATION_OF_EXTRACELLULAR_MATRIX_DISASSEMBLY | 15 | -0.475478 | -1.199329 | 0.24037 | 0.463609 |
| GOBP_RESPIRATORY_SYSTEM_DEVELOPMENT | 165 | -0.309672 | -1.199301 | 0.119244 | 0.463257 |
| GOBP_FORMATION_OF_PRIMARY_GERM_LAYER | 100 | -0.326821 | -1.19918 | 0.154345 | 0.463117 |
| GOBP_POSITIVE_REGULATION_OF_SIGNAL_TRANSDUCTION_BY_P53_CLASS_MEDIATOR | 22 | -0.431997 | -1.199134 | 0.223765 | 0.462805 |
| GOBP_HEART_PROCESS | 241 | -0.298735 | -1.199079 | 0.089655 | 0.462509 |
| GOBP_REGULATION_OF_STRIATED_MUSCLE_CELL_DIFFERENTIATION | 73 | -0.343819 | -1.198923 | 0.184828 | 0.462514 |
| GOBP_REGULATION_OF_VESICLE_MEDIATED_TRANSPORT | 477 | -0.283086 | -1.198492 | 0.056204 | 0.463155 |
| GOBP_REGULATION_OF_COAGULATION | 62 | -0.351015 | -1.198467 | 0.178378 | 0.462774 |
| GOBP_NEGATIVE_REGULATION_OF_NIK_NF_KAPPAB_SIGNALING | 24 | -0.424917 | -1.197061 | 0.217778 | 0.466025 |
| GOBP_POSITIVE_REGULATION_OF_LEUKOCYTE_MIGRATION | 127 | -0.316852 | -1.196554 | 0.150617 | 0.466903 |
| GOBP_RAS_PROTEIN_SIGNAL_TRANSDUCTION | 328 | -0.288707 | -1.19567 | 0.092945 | 0.468839 |
| GOBP_ORGANOPHOSPHATE_ESTER_TRANSPORT | 120 | -0.317136 | -1.195633 | 0.15679 | 0.468488 |
| GOBP_REGULATION_OF_MYELOID_LEUKOCYTE_DIFFERENTIATION | 107 | -0.327089 | -1.195383 | 0.147906 | 0.468684 |
| GOBP_ENDOCRINE_HORMONE_SECRETION | 39 | -0.381041 | -1.195195 | 0.201646 | 0.468747 |
| GOBP_POSITIVE_REGULATION_OF_DNA_DAMAGE_RESPONSE_SIGNAL_TRANSDUCTION_BY_P53 | 15 | -0.470327 | -1.195121 | 0.235835 | 0.468493 |
| GOBP_POSITIVE_REGULATION_OF_CALCIUM_MEDIATED_SIGNALING | 27 | -0.415782 | -1.194402 | 0.25 | 0.469956 |
| GOBP_REGULATION_OF_MULTI_ORGANISM_PROCESS | 41 | -0.383261 | -1.194367 | 0.204142 | 0.469603 |
| GOBP_ARACHIDONIC_ACID_SECRETION | 20 | -0.438048 | -1.192767 | 0.238547 | 0.473441 |
| GOBP_STRIATUM_DEVELOPMENT | 17 | -0.468536 | -1.192162 | 0.234206 | 0.474642 |
| GOBP_SENSORY_PERCEPTION_OF_SMELL | 80 | -0.338231 | -1.19213 | 0.199187 | 0.474284 |
| GOBP_POSITIVE_REGULATION_OF_TRIGLYCERIDE_METABOLIC_PROCESS | 17 | -0.451258 | -1.191679 | 0.229483 | 0.475073 |
| GOBP_EAR_DEVELOPMENT | 168 | -0.304312 | -1.191654 | 0.13848 | 0.474697 |
| GOBP_LYMPHOCYTE_HOMEOSTASIS | 56 | -0.358839 | -1.191377 | 0.198582 | 0.474994 |
| GOBP_REGULATION_OF_TOLL_LIKE_RECEPTOR_4_SIGNALING_PATHWAY | 22 | -0.430985 | -1.191129 | 0.229135 | 0.475228 |
| GOBP_NEGATIVE_REGULATION_OF_GLYCOPROTEIN_METABOLIC_PROCESS | 16 | -0.465299 | -1.191071 | 0.222405 | 0.474931 |
| GOBP_POSITIVE_REGULATION_OF_NIK_NF_KAPPAB_SIGNALING | 66 | -0.34669 | -1.191006 | 0.17688 | 0.474678 |
| GOBP_NEGATIVE_REGULATION_OF_SMOOTH_MUSCLE_CELL_DIFFERENTIATION | 17 | -0.460613 | -1.19091 | 0.241706 | 0.474502 |
| GOBP_POSTSYNAPTIC_SPECIALIZATION_ORGANIZATION | 27 | -0.41152 | -1.190855 | 0.237609 | 0.474222 |
| GOBP_MULTICELLULAR_ORGANISM_AGING | 26 | -0.412012 | -1.19058 | 0.226225 | 0.474466 |
| GOBP_STARTLE_RESPONSE | 18 | -0.463453 | -1.190406 | 0.24961 | 0.474486 |
| GOBP_CARDIAC_CONDUCTION | 126 | -0.319059 | -1.189996 | 0.160804 | 0.475156 |
| GOBP_AMYLOID_BETA_CLEARANCE | 34 | -0.393363 | -1.189707 | 0.22576 | 0.475482 |
| GOBP_REGULATION_OF_NEURONAL_SYNAPTIC_PLASTICITY | 41 | -0.378626 | -1.189365 | 0.213043 | 0.4759 |
| GOBP_HEART_TRABECULA_MORPHOGENESIS | 27 | -0.414177 | -1.188689 | 0.237189 | 0.477314 |
| GOBP_T_CELL_DIFFERENTIATION_IN_THYMUS | 72 | -0.339691 | -1.18803 | 0.173442 | 0.478682 |
| GOBP_REGULATION_OF_PROTEIN_SUMOYLATION | 22 | -0.434664 | -1.187095 | 0.257669 | 0.480655 |
| GOBP_CARDIAC_CHAMBER_MORPHOGENESIS | 95 | -0.329592 | -1.186617 | 0.183862 | 0.481554 |
| GOBP_SYNAPSE_ASSEMBLY | 145 | -0.309038 | -1.186323 | 0.154862 | 0.481876 |
| GOBP_CALCIUM_ION_REGULATED_EXOCYTOSIS | 55 | -0.360425 | -1.186128 | 0.198006 | 0.48197 |
| GOBP_STEROL_HOMEOSTASIS | 76 | -0.332896 | -1.185875 | 0.180519 | 0.482218 |
| GOBP_POSITIVE_REGULATION_OF_BEHAVIOR | 19 | -0.436774 | -1.18363 | 0.254658 | 0.487734 |
| GOBP_POSITIVE_REGULATION_OF_DNA_REPLICATION | 37 | -0.380986 | -1.183341 | 0.218391 | 0.48805 |
| GOBP_HINDBRAIN_MORPHOGENESIS | 36 | -0.387858 | -1.18261 | 0.221429 | 0.489553 |
| GOBP_MONOCARBOXYLIC_ACID_TRANSPORT | 149 | -0.306236 | -1.1824 | 0.14557 | 0.489653 |
| GOBP_ACTIN_FILAMENT_BASED_TRANSPORT | 21 | -0.428167 | -1.182253 | 0.242199 | 0.489615 |
| GOBP_REGULATION_OF_CELLULAR_RESPONSE_TO_TRANSFORMING_GROWTH_FACTOR_BETA_ | 113 | -0.321941 | -1.182125 | 0.179819 | 0.489516 |
| GOBP_INOSITOL_PHOSPHATE_BIOSYNTHETIC_PROCESS | 24 | -0.417913 | -1.182005 | 0.241486 | 0.489415 |
| GOBP_RIBONUCLEOTIDE_CATABOLIC_PROCESS | 38 | -0.380833 | -1.181967 | 0.21398 | 0.489083 |
| GOBP_REGULATION_OF_POSTSYNAPTIC_MEMBRANE_POTENTIAL | 108 | -0.320753 | -1.181789 | 0.178244 | 0.48912 |
| GOBP_BRANCHING_MORPHOGENESIS_OF_AN_EPITHELIAL_TUBE | 120 | -0.315968 | -1.181662 | 0.160714 | 0.489032 |
| GOBP_REGULATION_OF_RHO_PROTEIN_SIGNAL_TRANSDUCTION | 80 | -0.335751 | -1.181543 | 0.176144 | 0.488912 |
| GOBP_POSITIVE_REGULATION_OF_SECRETION | 262 | -0.290957 | -1.181476 | 0.121387 | 0.488619 |
| GOBP_LEFT_RIGHT_PATTERN_FORMATION | 19 | -0.453938 | -1.18085 | 0.250755 | 0.489905 |
| GOBP_POSITIVE_REGULATION_OF_MAPK_CASCADE | 459 | -0.278411 | -1.180756 | 0.095289 | 0.489723 |
| GOBP_POSITIVE_REGULATION_OF_TRANSCRIPTION_FROM_RNA_POLYMERASE_II_PROMOTER_II | 23 | -0.426371 | -1.180543 | 0.227414 | 0.489831 |
| GOBP_TUMOR_NECROSIS_FACTOR_MEDIATED_SIGNALING_PATHWAY | 164 | -0.303018 | -1.180046 | 0.153563 | 0.490663 |
| GOBP_POSITIVE_REGULATION_OF_CELL_DEVELOPMENT | 265 | -0.290019 | -1.179319 | 0.137656 | 0.492163 |
| GOBP_REGULATION_OF_PROTEIN_TYROSINE_KINASE_ACTIVITY | 88 | -0.327084 | -1.17914 | 0.191744 | 0.492159 |
| GOBP_METANEPHROS_MORPHOGENESIS | 23 | -0.419445 | -1.179037 | 0.246835 | 0.491995 |
| GOBP_POSITIVE_REGULATION_OF_INTERFERON_ALPHA_PRODUCTION | 21 | -0.421684 | -1.178901 | 0.253543 | 0.491915 |
| GOBP_ENDOCRINE_SYSTEM_DEVELOPMENT | 95 | -0.325108 | -1.17824 | 0.186755 | 0.493282 |
| GOBP_REGULATION_OF_SMOOTH_MUSCLE_CONTRACTION | 47 | -0.364914 | -1.178174 | 0.208333 | 0.49301 |
| GOBP_REGULATION_OF_NEUROBLAST_PROLIFERATION | 24 | -0.423082 | -1.177999 | 0.233494 | 0.493043 |
| GOBP_REGULATION_OF_HEART_RATE_BY_CARDIAC_CONDUCTION | 37 | -0.38116 | -1.177717 | 0.242009 | 0.493382 |
| GOBP_SPLEEN_DEVELOPMENT | 27 | -0.403135 | -1.177168 | 0.234421 | 0.494379 |
| GOBP_SENSORY_SYSTEM_DEVELOPMENT | 299 | -0.287716 | -1.177131 | 0.105562 | 0.494036 |
| GOBP_RESPONSE_TO_LEPTIN | 22 | -0.430856 | -1.17677 | 0.252412 | 0.494511 |
| GOBP_REGULATION_OF_FIBROBLAST_MIGRATION | 34 | -0.389491 | -1.176455 | 0.220211 | 0.494898 |
| GOBP_REGULATION_OF_RESPONSE_TO_ENDOPLASMIC_RETICULUM_STRESS | 79 | -0.331754 | -1.176323 | 0.188859 | 0.494793 |
| GOBP_ANIMAL_ORGAN_REGENERATION | 69 | -0.342171 | -1.176026 | 0.192819 | 0.495178 |
| GOBP_MONOCARBOXYLIC_ACID_BIOSYNTHETIC_PROCESS | 194 | -0.295029 | -1.175992 | 0.135514 | 0.494802 |
| GOBP_REGULATION_OF_T_CELL_MIGRATION | 40 | -0.372041 | -1.175948 | 0.22734 | 0.494467 |
| GOBP_REGULATION_OF_VASCULOGENESIS | 15 | -0.469331 | -1.175669 | 0.277778 | 0.49477 |
| GOBP_REGULATION_OF_REPRODUCTIVE_PROCESS | 128 | -0.308887 | -1.17558 | 0.180812 | 0.494568 |
| GOBP_FATTY_ACID_TRANSPORT | 130 | -0.312426 | -1.175214 | 0.175505 | 0.495075 |
| GOBP_PROTEIN_TARGETING_TO_MEMBRANE | 200 | -0.293539 | -1.174731 | 0.132629 | 0.495943 |
| GOBP_REGULATION_OF_CELLULAR_RESPONSE_TO_GROWTH_FACTOR_STIMULUS | 242 | -0.291889 | -1.174692 | 0.122066 | 0.495611 |
| GOBP_CELLULAR_COMPONENT_ASSEMBLY_INVOLVED_IN_MORPHOGENESIS | 85 | -0.329152 | -1.174654 | 0.178244 | 0.495289 |
| GOBP_AUDITORY_RECEPTOR_CELL_DEVELOPMENT | 18 | -0.448757 | -1.174467 | 0.27673 | 0.495368 |
| GOBP_MESODERM_DEVELOPMENT | 102 | -0.325378 | -1.174378 | 0.192802 | 0.495188 |
| GOBP_RESPONSE_TO_CAMP | 85 | -0.332108 | -1.174297 | 0.185864 | 0.494973 |
| GOBP_RESPONSE_TO_INTERLEUKIN_4 | 33 | -0.388287 | -1.17382 | 0.244648 | 0.495797 |
| GOBP_T_HELPER_CELL_LINEAGE_COMMITMENT | 17 | -0.457524 | -1.173237 | 0.283699 | 0.496912 |
| GOBP_FAT_CELL_DIFFERENTIATION | 197 | -0.296083 | -1.172245 | 0.149644 | 0.499145 |
| GOBP_NEGATIVE_REGULATION_OF_ION_TRANSPORT | 270 | -0.289046 | -1.172165 | 0.12704 | 0.498917 |
| GOBP_REGULATION_OF_MUSCLE_CONTRACTION | 131 | -0.311002 | -1.171924 | 0.186567 | 0.499147 |
| GOBP_REGULATION_OF_BONE_REMODELING | 46 | -0.36308 | -1.171884 | 0.231198 | 0.498818 |
| GOBP_ENDOTHELIAL_CELL_PROLIFERATION | 143 | -0.303139 | -1.17182 | 0.151961 | 0.498539 |
| GOBP_CYTOSOLIC_CALCIUM_ION_TRANSPORT | 155 | -0.302734 | -1.171718 | 0.153382 | 0.498386 |
| GOBP_CELL_JUNCTION_ASSEMBLY | 373 | -0.279978 | -1.171671 | 0.123922 | 0.498068 |
| GOBP_POSITIVE_REGULATION_OF_PROTEIN_LOCALIZATION_TO_CELL_SURFACE | 15 | -0.463244 | -1.170778 | 0.268608 | 0.500079 |
| GOBP_POSITIVE_REGULATION_OF_BLOOD_VESSEL_ENDOTHELIAL_CELL_MIGRATION | 56 | -0.348077 | -1.17033 | 0.229282 | 0.500841 |
| GOBP_NEGATIVE_REGULATION_OF_VIRAL_PROCESS | 82 | -0.332611 | -1.169953 | 0.196477 | 0.501414 |
| GOBP_SYNAPSE_ORGANIZATION | 365 | -0.279881 | -1.169514 | 0.096452 | 0.502216 |
| GOBP_REGULATION_OF_NEURON_MIGRATION | 34 | -0.379646 | -1.168435 | 0.221745 | 0.504684 |
| GOBP_CYTOPLASMIC_SEQUESTERING_OF_PROTEIN | 22 | -0.417317 | -1.168177 | 0.261062 | 0.504943 |
| GOBP_EMBRYONIC_ORGAN_MORPHOGENESIS | 229 | -0.289588 | -1.167851 | 0.153488 | 0.505393 |
| GOBP_NEGATIVE_REGULATION_OF_NUCLEOTIDE_METABOLIC_PROCESS | 20 | -0.427451 | -1.16781 | 0.263804 | 0.505067 |
| GOBP_POSITIVE_REGULATION_OF_DNA_BINDING | 48 | -0.363683 | -1.167706 | 0.24507 | 0.504899 |
| GOBP_REGULATION_OF_CALCIUM_ION_TRANSMEMBRANE_TRANSPORTER_ACTIVITY | 72 | -0.335942 | -1.1677 | 0.208614 | 0.504471 |
| GOBP_EPIBOLY | 33 | -0.383432 | -1.166878 | 0.226879 | 0.506309 |
| GOBP_PURINERGIC_NUCLEOTIDE_RECEPTOR_SIGNALING_PATHWAY | 31 | -0.390846 | -1.166011 | 0.238026 | 0.508225 |
| GOBP_HORMONE_METABOLIC_PROCESS | 173 | -0.298097 | -1.165921 | 0.166262 | 0.508032 |
| GOBP_DENDRITE_SELF_AVOIDANCE | 15 | -0.474867 | -1.165715 | 0.272025 | 0.508145 |
| GOBP_POSITIVE_REGULATION_OF_PROTEIN_KINASE_ACTIVITY | 490 | -0.273521 | -1.164967 | 0.099476 | 0.509809 |
| GOBP_ERK1_AND_ERK2_CASCADE | 268 | -0.287767 | -1.164852 | 0.147936 | 0.509689 |
| GOBP_AXONAL_FASCICULATION | 18 | -0.448817 | -1.164735 | 0.264659 | 0.509542 |
| GOBP_NEGATIVE_REGULATION_OF_MYOBLAST_DIFFERENTIATION | 22 | -0.420486 | -1.164628 | 0.273994 | 0.509387 |
| GOBP_HEPARAN_SULFATE_PROTEOGLYCAN_BIOSYNTHETIC_PROCESS | 23 | -0.414094 | -1.164257 | 0.26087 | 0.509941 |
| GOBP_POSITIVE_REGULATION_OF_CALCIUM_ION_TRANSPORT | 107 | -0.315565 | -1.164089 | 0.194236 | 0.509975 |
| GOBP_POSITIVE_REGULATION_OF_ACTIN_FILAMENT_POLYMERIZATION | 93 | -0.320786 | -1.16383 | 0.19709 | 0.510256 |
| GOBP_NEGATIVE_REGULATION_OF_CELL_MIGRATION_INVOLVED_IN_SPROUTING_ANGIOGENESI | 17 | -0.446695 | -1.162832 | 0.266876 | 0.51256 |
| GOBP_RECEPTOR_METABOLIC_PROCESS | 158 | -0.298216 | -1.162754 | 0.169365 | 0.512346 |
| GOBP_REGULATION_OF_CALCIUM_ION_TRANSMEMBRANE_TRANSPORT | 125 | -0.309567 | -1.161783 | 0.19546 | 0.514518 |
| GOBP_G_PROTEIN_COUPLED_ACETYLCHOLINE_RECEPTOR_SIGNALING_PATHWAY | 16 | -0.453549 | -1.161432 | 0.283912 | 0.515005 |
| GOBP_REGULATION_OF_CELL_JUNCTION_ASSEMBLY | 174 | -0.293843 | -1.161347 | 0.18399 | 0.514781 |
| GOBP_NEGATIVE_REGULATION_OF_APOPTOTIC_SIGNALING_PATHWAY | 211 | -0.291825 | -1.160987 | 0.167268 | 0.515363 |
| GOBP_CELLULAR_RESPONSE_TO_DSRNA | 22 | -0.419463 | -1.160787 | 0.266768 | 0.515505 |
| GOBP_NEGATIVE_REGULATION_OF_PROTEIN_SERINE_THREONINE_KINASE_ACTIVITY | 127 | -0.307561 | -1.160786 | 0.190955 | 0.515066 |
| GOBP_POSITIVE_REGULATION_OF_BINDING | 158 | -0.299537 | -1.160678 | 0.201923 | 0.514917 |
| GOBP_NEGATIVE_REGULATION_OF_TOLL_LIKE_RECEPTOR_SIGNALING_PATHWAY | 38 | -0.376805 | -1.160089 | 0.243478 | 0.51615 |
| GOBP_MACROPHAGE_ACTIVATION_INVOLVED_IN_IMMUNE_RESPONSE | 17 | -0.444273 | -1.160029 | 0.276827 | 0.515855 |
| GOBP_INTERLEUKIN_1_MEDIATED_SIGNALING_PATHWAY | 99 | -0.31586 | -1.15996 | 0.180978 | 0.515587 |
| GOBP_DEOXYRIBOSE_PHOSPHATE_CATABOLIC_PROCESS | 26 | -0.402145 | -1.159809 | 0.252941 | 0.515628 |
| GOBP_PLATELET_DERIVED_GROWTH_FACTOR_RECEPTOR_SIGNALING_PATHWAY | 54 | -0.34802 | -1.159703 | 0.242507 | 0.515486 |
| GOBP_RESPONSE_TO_XENOBIOTIC_STIMULUS | 103 | -0.315335 | -1.159688 | 0.214192 | 0.515089 |
| GOBP_POSITIVE_REGULATION_OF_PROTEIN_LOCALIZATION_TO_CELL_PERIPHERY | 63 | -0.344284 | -1.159495 | 0.223448 | 0.515179 |
| GOBP_AUDITORY_RECEPTOR_CELL_MORPHOGENESIS | 15 | -0.459466 | -1.158752 | 0.273885 | 0.516798 |
| GOBP_NEGATIVE_REGULATION_OF_TRANSFORMING_GROWTH_FACTOR_BETA_RECEPTOR_SIGN | 76 | -0.333008 | -1.158534 | 0.215506 | 0.516947 |
| GOBP_RNA_DESTABILIZATION | 35 | -0.381595 | -1.1584 | 0.244807 | 0.516867 |
| GOBP_REPRODUCTIVE_SYSTEM_DEVELOPMENT | 348 | -0.280642 | -1.158261 | 0.114537 | 0.516833 |

| | | | | | |
|---|---|---|---|---|---|
| GOBP_POLYOL_CATABOLIC_PROCESS | 25 | 0.203147 | 0.62912 | 0.952077 | 1 |
| GOBP_PROTEIN_PEPTIDYL_PROLYL_ISOMERIZATION | 36 | 0.183038 | 0.627681 | 0.987461 | 1 |
| GOBP_PYRIMIDINE_RIBONUCLEOTIDE_METABOLIC_PROCESS | 29 | 0.192697 | 0.626355 | 0.956772 | 1 |
| GOBP_PROTEIN_AUTOUBIQUITINATION | 69 | 0.159827 | 0.622454 | 0.992933 | 1 |
| GOBP_HISTONE_H3_K4_METHYLATION | 50 | 0.165693 | 0.61956 | 0.986755 | 1 |
| GOBP_RESPONSE_TO_EPINEPHRINE | 15 | 0.230068 | 0.617106 | 0.916442 | 1 |
| GOBP_RETROGRADE_TRANSPORT_ENDOSOME_TO_GOLGI | 90 | 0.152813 | 0.617001 | 1 | 1 |
| GOBP_POSITIVE_REGULATION_OF_CALCIUM_ION_TRANSMEMBRANE_TRANSPORTER_ACTIVITY | 35 | 0.175438 | 0.614289 | 0.963087 | 1 |
| GOBP_PROTEIN_HYDROXYLATION | 27 | 0.190084 | 0.605353 | 0.962145 | 1 |
| GOBP_LIPOPROTEIN_BIOSYNTHETIC_PROCESS | 90 | 0.145747 | 0.604582 | 0.995984 | 1 |
| GOBP_POSITIVE_REGULATION_OF_TELOMERE_CAPPING | 17 | 0.207867 | 0.599059 | 0.965517 | 1 |
| GOBP_PEPTIDYL_LYSINE_MODIFICATION | 372 | 0.118479 | 0.596778 | 1 | 1 |
| GOBP_REGULATION_OF_MRNA_POLYADENYLATION | 17 | 0.209259 | 0.596229 | 0.957507 | 1 |
| GOBP_SELECTIVE_AUTOPHAGY | 62 | 0.154855 | 0.592824 | 1 | 1 |
| GOBP_POSITIVE_REGULATION_OF_ESTABLISHMENT_OF_PROTEIN_LOCALIZATION_TO_MITOCHO | 57 | 0.155045 | 0.591093 | 0.996337 | 1 |
| GOBP_POSITIVE_REGULATION_OF_PROTEIN_DEACETYLATION | 21 | 0.192622 | 0.5904 | 0.982659 | 1 |
| GOBP_REGULATION_OF_HYDROGEN_PEROXIDE_METABOLIC_PROCESS | 18 | 0.201463 | 0.581653 | 0.958333 | 1 |
| GOBP_TRANSCRIPTION_BY_RNA_POLYMERASE_I | 67 | 0.15023 | 0.580012 | 1 | 1 |
| GOBP_REGULATION_OF_CARDIAC_MUSCLE_CONTRACTION_BY_REGULATION_OF_THE_RELEAS | 18 | 0.203221 | 0.576751 | 0.96875 | 1 |
| GOBP_REGULATION_OF_HISTONE_DEACETYLATION | 25 | 0.185239 | 0.576326 | 0.97486 | 1 |
| GOBP_POSITIVE_REGULATION_OF_MRNA_SPLICING_VIA_SPLICEOSOME | 20 | 0.19737 | 0.573702 | 0.968116 | 1 |
| GOBP_NUCLEOBASE_CONTAINING_COMPOUND_TRANSPORT | 241 | 0.119569 | 0.571144 | 1 | 1 |
| GOBP_REGULATION_OF_FILOPODIUM_ASSEMBLY | 43 | 0.158524 | 0.570133 | 0.993378 | 1 |
| GOBP_INTRACILIARY_TRANSPORT | 53 | 0.151782 | 0.567054 | 0.989011 | 1 |
| GOBP_LYMPH_VESSEL_MORPHOGENESIS | 19 | 0.191974 | 0.565167 | 0.965944 | 1 |
| GOBP_METHIONINE_METABOLIC_PROCESS | 15 | 0.212818 | 0.563871 | 0.976945 | 1 |
| GOBP_LONG_TERM_SYNAPTIC_DEPRESSION | 24 | 0.181871 | 0.561489 | 0.997333 | 1 |
| GOBP_POSITIVE_REGULATION_OF_DOUBLE_STRAND_BREAK_REPAIR | 39 | 0.15881 | 0.560484 | 1 | 1 |
| GOBP_ENDOPLASMIC_RETICULUM_TUBULAR_NETWORK_ORGANIZATION | 18 | 0.194832 | 0.558498 | 0.966154 | 1 |
| GOBP_PROTEIN_N_LINKED_GLYCOSYLATION | 71 | 0.137856 | 0.544232 | 1 | 1 |
| GOBP_REGULATION_OF_MRNA_CATABOLIC_PROCESS | 199 | 0.118865 | 0.543096 | 1 | 1 |
| GOBP_HISTONE_H4_K16_ACETYLATION | 21 | 0.175049 | 0.540704 | 0.97654 | 1 |
| GOBP_RESPONSE_TO_EPIDERMAL_GROWTH_FACTOR | 42 | 0.150373 | 0.535508 | 0.996855 | 1 |
| GOBP_MRNA_TRANSPORT | 147 | 0.120744 | 0.533124 | 1 | 1 |
| GOBP_NEGATIVE_REGULATION_OF_MACROAUTOPHAGY | 35 | 0.154078 | 0.529403 | 0.997015 | 1 |
| GOBP_REGULATION_OF_STEM_CELL_POPULATION_MAINTENANCE | 26 | 0.158402 | 0.522004 | 0.993846 | 1 |
| GOBP_NUCLEAR_EXPORT | 195 | 0.112426 | 0.520771 | 1 | 1 |
| GOBP_HISTONE_MONOUBIQUITINATION | 29 | 0.157618 | 0.520591 | 0.994152 | 1 |
| GOBP_REGULATION_OF_DNA_TEMPLATED_TRANSCRIPTION_IN_RESPONSE_TO_STRESS | 113 | 0.120213 | 0.51634 | 1 | 1 |
| GOBP_ENDOPLASMIC_RETICULUM_TO_CYTOSOL_TRANSPORT | 28 | 0.156118 | 0.507237 | 0.994135 | 1 |
| GOBP_GLUCOSE_6_PHOSPHATE_METABOLIC_PROCESS | 22 | 0.162231 | 0.505092 | 0.994253 | 1 |
| GOBP_RNA_PHOSPHODIESTER_BOND_HYDROLYSIS_EXONUCLEOLYTIC | 43 | 0.141023 | 0.505031 | 1 | 1 |
| GOBP_REGULATION_OF_MITOTIC_SPINDLE_ASSEMBLY | 18 | 0.167111 | 0.491078 | 0.994286 | 1 |
| GOBP_NEGATIVE_REGULATION_OF_TELOMERE_MAINTENANCE_VIA_TELOMERE_LENGTHENING | 25 | 0.158115 | 0.490025 | 1 | 1 |
| GOBP_PROTEIN_TRANSMEMBRANE_TRANSPORT | 60 | 0.128119 | 0.487403 | 1 | 1 |
| GOBP_ENDOSOME_TRANSPORT_VIA_MULTIVESICULAR_BODY_SORTING_PATHWAY | 31 | 0.145482 | 0.484831 | 1 | 1 |
| GOBP_OXIDATIVE_DEMETHYLATION | 17 | 0.168208 | 0.482985 | 0.99726 | 1 |
| GOBP_EXIT_FROM_MITOSIS | 26 | 0.146227 | 0.467295 | 0.993976 | 1 |
| GOBP_RIBONUCLEOPROTEIN_COMPLEX_SUBUNIT_ORGANIZATION | 213 | 0.100104 | 0.46622 | 1 | 1 |
| GOBP_MULTIVESICULAR_BODY_SORTING_PATHWAY | 37 | 0.13073 | 0.453243 | 1 | 1 |
| GOBP_MODULATION_BY_VIRUS_OF_HOST_PROCESS | 28 | 0.14171 | 0.44652 | 0.997015 | 1 |
| GOBP_TRNA_WOBBLE_BASE_MODIFICATION | 21 | 0.129089 | 0.38709 | 1 | 1 |
| GOBP_RNA_SPLICING_VIA_TRANSESTERIFICATION_REACTIONS | 357 | 0.062834 | 0.313801 | 1 | 0.999997 |

| | | | | | |
|---|---|---|---|---|---|
| GOBP_NEGATIVE_REGULATION_OF_NEURAL_PRECURSOR_CELL_PROLIFERATION | 18 | -0.436411 | -1.157788 | 0.295707 | 0.517726 |
| GOBP_MACROPHAGE_CYTOKINE_PRODUCTION | 15 | -0.456615 | -1.156327 | 0.28481 | 0.521395 |
| GOBP_OVULATION_CYCLE | 55 | -0.342967 | -1.156225 | 0.220506 | 0.521231 |
| GOBP_BETA_CATENIN_DESTRUCTION_COMPLEX_DISASSEMBLY | 19 | -0.43161 | -1.155767 | 0.256329 | 0.522091 |
| GOBP_CELLULAR_RESPONSE_TO_COPPER_ION | 20 | -0.4296 | -1.155652 | 0.281669 | 0.521972 |
| GOBP_APPENDAGE_MORPHOGENESIS | 112 | -0.312316 | -1.155123 | 0.217005 | 0.522936 |
| GOBP_PATTERN_RECOGNITION_RECEPTOR_SIGNALING_PATHWAY | 194 | -0.287818 | -1.155109 | 0.163484 | 0.522533 |
| GOBP_ENDOTHELIAL_CELL_DEVELOPMENT | 62 | -0.340334 | -1.15486 | 0.242091 | 0.522785 |
| GOBP_STRIATED_MUSCLE_CELL_DEVELOPMENT | 88 | -0.322396 | -1.154149 | 0.212766 | 0.524221 |
| GOBP_ANTIGEN_PROCESSING_AND_PRESENTATION | 227 | -0.284899 | -1.154046 | 0.165476 | 0.524067 |
| GOBP_DEVELOPMENT_OF_PRIMARY_SEXUAL_CHARACTERISTICS | 185 | -0.290783 | -1.154003 | 0.178998 | 0.523747 |
| GOBP_DOPAMINE_SECRETION | 25 | -0.408063 | -1.153977 | 0.258258 | 0.523379 |
| GOBP_REGULATION_OF_ENDOPLASMIC_RETICULUM_UNFOLDED_PROTEIN_RESPONSE | 29 | -0.388075 | -1.153322 | 0.246512 | 0.524739 |
| GOBP_T_CELL_APOPTOTIC_PROCESS | 45 | -0.360527 | -1.153115 | 0.244666 | 0.524872 |
| GOBP_RETINOIC_ACID_METABOLIC_PROCESS | 25 | -0.403041 | -1.152963 | 0.273684 | 0.52487 |
| GOBP_COLLAGEN_CATABOLIC_PROCESS | 41 | -0.364188 | -1.152615 | 0.258493 | 0.525335 |
| GOBP_ENDOCRINE_PANCREAS_DEVELOPMENT | 36 | -0.375877 | -1.152598 | 0.248097 | 0.524933 |
| GOBP_POSITIVE_REGULATION_OF_ERK1_AND_ERK2_CASCADE | 171 | -0.293797 | -1.152278 | 0.183374 | 0.525388 |
| GOBP_FATTY_ACID_BIOSYNTHETIC_PROCESS | 144 | -0.298801 | -1.152275 | 0.203727 | 0.524951 |
| GOBP_POSITIVE_REGULATION_OF_MAP_KINASE_ACTIVITY | 216 | -0.28769 | -1.152186 | 0.177647 | 0.52475 |
| GOBP_RESPONSE_TO_VITAMIN_D | 29 | -0.388971 | -1.152186 | 0.266272 | 0.524311 |
| GOBP_IMMUNOLOGICAL_MEMORY_PROCESS | 17 | -0.443443 | -1.152172 | 0.284566 | 0.523908 |
| GOBP_INNER_DYNEIN_ARM_ASSEMBLY | 15 | -0.453287 | -1.151652 | 0.292079 | 0.524958 |
| GOBP_NEGATIVE_REGULATION_OF_WOUND_HEALING | 60 | -0.340494 | -1.151301 | 0.237964 | 0.525425 |
| GOBP_RESPONSE_TO_ETHANOL | 114 | -0.306817 | -1.151142 | 0.204342 | 0.525432 |
| GOBP_REGULATION_OF_PROTEIN_BINDING | 187 | -0.294411 | -1.150932 | 0.189061 | 0.52558 |
| GOBP_NEGATIVE_REGULATION_OF_MULTI_ORGANISM_PROCESS | 15 | -0.458782 | -1.150782 | 0.31068 | 0.525554 |
| GOBP_POSITIVE_REGULATION_OF_TYPE_I_INTERFERON_PRODUCTION | 76 | -0.326318 | -1.150567 | 0.240766 | 0.525746 |
| GOBP_REGULATION_OF_RAC_PROTEIN_SIGNAL_TRANSDUCTION | 18 | -0.430398 | -1.150525 | 0.261756 | 0.525431 |
| GOBP_INORGANIC_ANION_TRANSPORT | 138 | -0.303819 | -1.150263 | 0.189422 | 0.525724 |
| GOBP_POSITIVE_REGULATION_OF_HORMONE_SECRETION | 104 | -0.313443 | -1.150103 | 0.204993 | 0.525739 |
| GOBP_FEAR_RESPONSE | 30 | -0.389238 | -1.150093 | 0.276853 | 0.525333 |
| GOBP_ACTIVATED_T_CELL_PROLIFERATION | 43 | -0.35735 | -1.149904 | 0.245014 | 0.525444 |
| GOBP_MODULATION_OF_PROCESS_OF_OTHER_ORGANISM | 112 | -0.310345 | -1.14981 | 0.207332 | 0.525268 |
| GOBP_PLACENTA_BLOOD_VESSEL_DEVELOPMENT | 26 | -0.396563 | -1.149674 | 0.252266 | 0.525229 |
| GOBP_REGULATION_OF_ANIMAL_ORGAN_MORPHOGENESIS | 162 | -0.297121 | -1.14957 | 0.184242 | 0.525091 |
| GOBP_ARF_PROTEIN_SIGNAL_TRANSDUCTION | 19 | -0.429333 | -1.148971 | 0.296998 | 0.52634 |
| GOBP_CELL_CELL_SIGNALING_INVOLVED_IN_CARDIAC_CONDUCTION | 28 | -0.392035 | -1.148755 | 0.266471 | 0.526478 |
| GOBP_SYNAPTIC_TRANSMISSION_CHOLINERGIC | 24 | -0.412502 | -1.148729 | 0.26161 | 0.526111 |
| GOBP_REGULATION_OF_SYNAPSE_ASSEMBLY | 79 | -0.32472 | -1.148691 | 0.216802 | 0.525781 |
| GOBP_LEUKOCYTE_APOPTOTIC_PROCESS | 97 | -0.315346 | -1.148451 | 0.224227 | 0.5261 |
| GOBP_EAR_MORPHOGENESIS | 92 | -0.319606 | -1.148097 | 0.224516 | 0.526611 |
| GOBP_SENSORY_ORGAN_MORPHOGENESIS | 199 | -0.286923 | -1.148037 | 0.182783 | 0.52634 |
| GOBP_POSITIVE_REGULATION_OF_INTERLEUKIN_4_PRODUCTION | 24 | -0.407373 | -1.147976 | 0.282675 | 0.526078 |
| GOBP_REGULATION_OF_STRESS_ACTIVATED_PROTEIN_KINASE_SIGNALING_CASCADE | 168 | -0.297501 | -1.147557 | 0.2 | 0.526823 |
| GOBP_CHLORIDE_TRANSPORT | 84 | -0.323077 | -1.147253 | 0.210383 | 0.52722 |
| GOBP_GASTRIC_ACID_SECRETION | 16 | -0.44951 | -1.147131 | 0.288644 | 0.527133 |
| GOBP_HOMEOSTASIS_OF_NUMBER_OF_CELLS_WITHIN_A_TISSUE | 24 | -0.402534 | -1.14683 | 0.27454 | 0.527565 |
| GOBP_COPULATION | 17 | -0.445548 | -1.146361 | 0.297508 | 0.528408 |
| GOBP_REGULATION_OF_BODY_FLUID_LEVELS | 440 | -0.27051 | -1.14506 | 0.13113 | 0.531658 |
| GOBP_REGULATION_OF_CIRCADIAN_SLEEP_WAKE_CYCLE | 15 | -0.456364 | -1.145038 | 0.298246 | 0.53129 |
| GOBP_EPIDERMIS_MORPHOGENESIS | 24 | -0.409606 | -1.144581 | 0.300151 | 0.532113 |
| GOBP_POSITIVE_REGULATION_OF_B_CELL_DIFFERENTIATION | 15 | -0.460732 | -1.144566 | 0.302251 | 0.531718 |
| GOBP_REGULATION_OF_ACTOMYOSIN_STRUCTURE_ORGANIZATION | 89 | -0.318177 | -1.14431 | 0.247354 | 0.532009 |
| GOBP_INTRINSIC_APOPTOTIC_SIGNALING_PATHWAY_IN_RESPONSE_TO_DNA_DAMAGE_BY_P53 | 44 | -0.359613 | -1.143954 | 0.248596 | 0.532529 |
| GOBP_CHEMICAL_SYNAPTIC_TRANSMISSION_POSTSYNAPTIC | 91 | -0.314981 | -1.143943 | 0.218509 | 0.532125 |
| GOBP_CONNECTIVE_TISSUE_DEVELOPMENT | 214 | -0.285209 | -1.143691 | 0.178404 | 0.532379 |
| GOBP_REGULATION_OF_KERATINOCYTE_PROLIFERATION | 28 | -0.389388 | -1.143512 | 0.274336 | 0.532417 |
| GOBP_POSITIVE_REGULATION_OF_RECEPTOR_MEDIATED_ENDOCYTOSIS | 45 | -0.358599 | -1.143504 | 0.245588 | 0.532008 |
| GOBP_ZINC_ION_HOMEOSTASIS | 32 | -0.380005 | -1.142627 | 0.259542 | 0.533997 |
| GOBP_NEGATIVE_REGULATION_OF_BLOOD_VESSEL_ENDOTHELIAL_CELL_MIGRATION | 35 | -0.374317 | -1.142531 | 0.266667 | 0.533836 |
| GOBP_NEGATIVE_REGULATION_OF_MUSCLE_CONTRACTION | 19 | -0.423213 | -1.142526 | 0.277244 | 0.533421 |
| GOBP_REGULATION_OF_PROTEIN_EXIT_FROM_ENDOPLASMIC_RETICULUM | 26 | -0.401143 | -1.142069 | 0.272588 | 0.534251 |
| GOBP_KERATINOCYTE_DIFFERENTIATION | 141 | -0.295019 | -1.141836 | 0.210526 | 0.534448 |
| GOBP_CEREBRAL_CORTEX_RADIALLY_ORIENTED_CELL_MIGRATION | 26 | -0.40577 | -1.141809 | 0.276973 | 0.534087 |
| GOBP_POSITIVE_REGULATION_OF_VASCULATURE_DEVELOPMENT | 148 | -0.295923 | -1.140827 | 0.202424 | 0.53642 |
| GOBP_RESPONSE_TO_TUMOR_CELL | 27 | -0.390605 | -1.140478 | 0.301685 | 0.536941 |
| GOBP_BLOOD_VESSEL_ENDOTHELIAL_CELL_MIGRATION | 116 | -0.306534 | -1.140254 | 0.209738 | 0.537124 |
| GOBP_CELL_RECOGNITION | 188 | -0.290689 | -1.139812 | 0.20936 | 0.5379 |
| GOBP_SMOOTH_MUSCLE_CONTRACTION | 83 | -0.321172 | -1.139557 | 0.253012 | 0.538194 |
| GOBP_REGULATION_OF_MEMBRANE_POTENTIAL | 365 | -0.273879 | -1.139294 | 0.146421 | 0.53849 |
| GOBP_RNA_DECAPPING | 16 | -0.436745 | -1.139167 | 0.292049 | 0.538422 |
| GOBP_NEGATIVE_REGULATION_OF_BONE_MINERALIZATION | 16 | -0.435992 | -1.139115 | 0.303079 | 0.538133 |
| GOBP_O_GLYCAN_PROCESSING | 51 | -0.344752 | -1.137681 | 0.258741 | 0.541638 |
| GOBP_POSITIVE_REGULATION_OF_MESENCHYMAL_CELL_PROLIFERATION | 19 | -0.417482 | -1.137252 | 0.274478 | 0.542363 |
| GOBP_ENDOTHELIAL_CELL_MIGRATION | 211 | -0.283476 | -1.136968 | 0.200469 | 0.542773 |
| GOBP_REGULATION_OF_EPITHELIAL_CELL_APOPTOTIC_PROCESS | 76 | -0.326261 | -1.136705 | 0.23427 | 0.543017 |
| GOBP_NITRIC_OXIDE_SYNTHASE_BIOSYNTHETIC_PROCESS | 19 | -0.419733 | -1.136544 | 0.283721 | 0.543021 |
| GOBP_CELLULAR_RESPONSE_TO_CADMIUM_ION | 31 | -0.382308 | -1.136326 | 0.28529 | 0.543202 |
| GOBP_PROTEOGLYCAN_METABOLIC_PROCESS | 81 | -0.318368 | -1.135596 | 0.235135 | 0.544778 |
| GOBP_POSITIVE_REGULATION_OF_CALCIUM_ION_DEPENDENT_EXOCYTOSIS | 16 | -0.439044 | -1.135504 | 0.299517 | 0.544609 |
| GOBP_TISSUE_MIGRATION | 289 | -0.277498 | -1.135011 | 0.173469 | 0.545502 |
| GOBP_COPII_COATED_VESICLE_BUDDING | 72 | -0.329112 | -1.134409 | 0.248634 | 0.546701 |
| GOBP_POSITIVE_REGULATION_OF_NEUROGENESIS | 198 | -0.28478 | -1.134302 | 0.200234 | 0.546575 |
| GOBP_REGULATION_OF_FATTY_ACID_METABOLIC_PROCESS | 79 | -0.318666 | -1.134234 | 0.248996 | 0.546368 |
| GOBP_MYD88_DEPENDENT_TOLL_LIKE_RECEPTOR_SIGNALING_PATHWAY | 34 | -0.374142 | -1.134212 | 0.279762 | 0.546001 |
| GOBP_SODIUM_ION_TRANSMEMBRANE_TRANSPORT | 144 | -0.293947 | -1.13416 | 0.228571 | 0.545716 |
| GOBP_ENGULFMENT_OF_APOPTOTIC_CELL | 18 | -0.428426 | -1.13414 | 0.295752 | 0.54535 |
| GOBP_MORPHOGENESIS_OF_A_BRANCHING_STRUCTURE | 156 | -0.292759 | -1.133871 | 0.211098 | 0.545605 |
| GOBP_POSITIVE_REGULATION_OF_MONOOXYGENASE_ACTIVITY | 29 | -0.3853 | -1.133783 | 0.289318 | 0.545401 |
| GOBP_REGULATION_OF_ENDOTHELIAL_CELL_CHEMOTAXIS | 18 | -0.436491 | -1.13371 | 0.301092 | 0.545161 |
| GOBP_OVULATION_CYCLE_PROCESS | 37 | -0.364886 | -1.133574 | 0.279539 | 0.545126 |
| GOBP_RESPONSE_TO_METAL_ION | 307 | -0.275791 | -1.133376 | 0.167598 | 0.545227 |
| GOBP_GLIOGENESIS | 256 | -0.278971 | -1.133319 | 0.199768 | 0.544959 |
| GOBP_PROTEIN_LOCALIZATION_TO_ENDOPLASMIC_RETICULUM | 147 | -0.295974 | -1.13328 | 0.223313 | 0.544643 |
| GOBP_PEPTIDYL_THREONINE_MODIFICATION | 120 | -0.299706 | -1.132891 | 0.23192 | 0.545294 |
| GOBP_GLAND_DEVELOPMENT | 360 | -0.272766 | -1.132393 | 0.175615 | 0.546242 |
| GOBP_MUSCLE_CONTRACTION | 300 | -0.273815 | -1.132377 | 0.177928 | 0.545858 |
| GOBP_MAINTENANCE_OF_GASTROINTESTINAL_EPITHELIUM | 15 | -0.451901 | -1.132371 | 0.317263 | 0.545455 |
| GOBP_POSITIVE_REGULATION_OF_PEPTIDE_HORMONE_SECRETION | 79 | -0.317601 | -1.132226 | 0.265013 | 0.545464 |
| GOBP_RENAL_FILTRATION | 24 | -0.393728 | -1.132075 | 0.284375 | 0.545445 |
| GOBP_INORGANIC_ANION_TRANSMEMBRANE_TRANSPORT | 93 | -0.314991 | -1.130455 | 0.233023 | 0.549498 |
| GOBP_REGULATION_OF_MUSCLE_SYSTEM_PROCESS | 196 | -0.283028 | -1.130177 | 0.209964 | 0.549845 |
| GOBP_CELLULAR_RESPONSE_TO_INORGANIC_SUBSTANCE | 187 | -0.286502 | -1.129774 | 0.217654 | 0.550533 |
| GOBP_POSITIVE_REGULATION_OF_SMOOTHENED_SIGNALING_PATHWAY | 34 | -0.374304 | -1.129741 | 0.292683 | 0.550195 |
| GOBP_REGULATION_OF_CARTILAGE_DEVELOPMENT | 58 | -0.334494 | -1.129697 | 0.271468 | 0.549885 |
| GOBP_RESPONSE_TO_ESTROGEN | 68 | -0.327804 | -1.129624 | 0.255659 | 0.549663 |
| GOBP_CELLULAR_EXTRAVASATION | 67 | -0.328313 | -1.129208 | 0.253388 | 0.55041 |
| GOBP_FUCOSE_METABOLIC_PROCESS | 15 | -0.452368 | -1.128828 | 0.319115 | 0.551018 |
| GOBP_REGULATION_OF_RENAL_SYSTEM_PROCESS | 29 | -0.380062 | -1.12881 | 0.300305 | 0.55064 |
| GOBP_MUSCLE_FIBER_DEVELOPMENT | 47 | -0.339606 | -1.128196 | 0.259259 | 0.551975 |
| GOBP_POSITIVE_REGULATION_OF_JNK_CASCADE | 81 | -0.316607 | -1.12816 | 0.252312 | 0.551646 |
| GOBP_MAMMARY_GLAND_DEVELOPMENT | 118 | -0.303645 | -1.127885 | 0.240949 | 0.551962 |
| GOBP_ODONTOGENESIS | 104 | -0.308296 | -1.127776 | 0.234839 | 0.551848 |
| GOBP_ESTABLISHMENT_OR_MAINTENANCE_OF_MONOPOLAR_CELL_POLARITY | 21 | -0.417169 | -1.12726 | 0.298762 | 0.552865 |
| GOBP_HINDBRAIN_DEVELOPMENT | 117 | -0.301385 | -1.127121 | 0.239637 | 0.552808 |
| GOBP_POSITIVE_REGULATION_OF_NEUROTRANSMITTER_TRANSPORT | 22 | -0.410469 | -1.126886 | 0.306502 | 0.553003 |
| GOBP_POSITIVE_REGULATION_OF_MEIOTIC_CELL_CYCLE | 18 | -0.422719 | -1.126863 | 0.326563 | 0.552636 |
| GOBP_POSITIVE_REGULATION_OF_MUSCLE_CONTRACTION | 33 | -0.380764 | -1.126834 | 0.302395 | 0.552298 |
| GOBP_POSITIVE_REGULATION_OF_MYELOID_LEUKOCYTE_DIFFERENTIATION | 52 | -0.341688 | -1.126602 | 0.277778 | 0.55247 |
| GOBP_DEFENSE_RESPONSE_TO_GRAM_NEGATIVE_BACTERIUM | 57 | -0.335321 | -1.126569 | 0.283668 | 0.552135 |
| GOBP_SEX_DIFFERENTIATION | 218 | -0.280598 | -1.126391 | 0.203529 | 0.55222 |
| GOBP_STRESS_ACTIVATED_PROTEIN_KINASE_SIGNALING_CASCADE | 256 | -0.277066 | -1.126203 | 0.18358 | 0.552358 |
| GOBP_MESODERM_MORPHOGENESIS | 57 | -0.338621 | -1.126119 | 0.279494 | 0.552168 |
| GOBP_REGULATION_OF_SYSTEMIC_ARTERIAL_BLOOD_PRESSURE_BY_HORMONE | 31 | -0.385612 | -1.126022 | 0.310185 | 0.552017 |
| GOBP_TRANSMEMBRANE_RECEPTOR_PROTEIN_SERINE_THREONINE_KINASE_SIGNALING_PATH | 308 | -0.272581 | -1.12507 | 0.19909 | 0.554276 |
| GOBP_POSITIVE_REGULATION_OF_SYNAPSE_ASSEMBLY | 50 | -0.346648 | -1.124959 | 0.264624 | 0.554153 |
| GOBP_SULFATION | 19 | -0.428061 | -1.124915 | 0.31135 | 0.553847 |
| GOBP_SMALL_GTPASE_MEDIATED_SIGNAL_TRANSDUCTION | 488 | -0.263924 | -1.124692 | 0.1606 | 0.554037 |
| GOBP_REGULATION_OF_BINDING | 333 | -0.272358 | -1.124499 | 0.194413 | 0.554145 |
| GOBP_AMEBOIDAL_TYPE_CELL_MIGRATION | 389 | -0.26789 | -1.124269 | 0.17121 | 0.554373 |
| GOBP_ESTABLISHMENT_OF_PROTEIN_LOCALIZATION_TO_POSTSYNAPTIC_MEMBRANE | 15 | -0.436147 | -1.12403 | 0.3056 | 0.554587 |
| GOBP_POSITIVE_REGULATION_OF_CYSTEINE_TYPE_ENDOPEPTIDASE_ACTIVITY | 141 | -0.292782 | -1.123949 | 0.250927 | 0.554387 |
| GOBP_DICARBOXYLIC_ACID_TRANSPORT | 84 | -0.318087 | -1.123664 | 0.282967 | 0.554754 |
| GOBP_DEOXYRIBONUCLEOSIDE_MONOPHOSPHATE_METABOLIC_PROCESS | 16 | -0.4294 | -1.12364 | 0.343354 | 0.554389 |
| GOBP_NEGATIVE_REGULATION_OF_RESPONSE_TO_REACTIVE_OXYGEN_SPECIES | 15 | -0.438284 | -1.123295 | 0.325843 | 0.554927 |
| GOBP_EPHRIN_RECEPTOR_SIGNALING_PATHWAY | 81 | -0.319849 | -1.123258 | 0.258065 | 0.554594 |
| GOBP_RECEPTOR_MEDIATED_ENDOCYTOSIS | 304 | -0.270941 | -1.123209 | 0.192865 | 0.554302 |
| GOBP_REGULATION_OF_AMYLOID_PRECURSOR_PROTEIN_CATABOLIC_PROCESS | 42 | -0.35473 | -1.123178 | 0.288406 | 0.553961 |
| GOBP_REGULATION_OF_PROTEIN_LOCALIZATION_TO_MEMBRANE | 184 | -0.283511 | -1.123087 | 0.223971 | 0.553788 |
| GOBP_WOUND_HEALING | 473 | -0.265505 | -1.122169 | 0.171067 | 0.55595 |
| GOBP_STRIATED_MUSCLE_CELL_DIFFERENTIATION | 227 | -0.278969 | -1.121725 | 0.21028 | 0.556728 |
| GOBP_CENTRAL_NERVOUS_SYSTEM_PROJECTION_NEURON_AXONOGENESIS | 25 | -0.393413 | -1.120866 | 0.313253 | 0.558668 |
| GOBP_VITAMIN_BIOSYNTHETIC_PROCESS | 18 | -0.430384 | -1.12054 | 0.326498 | 0.559151 |
| GOBP_REGULATION_OF_EXOCYTOSIS | 184 | -0.282563 | -1.120361 | 0.235995 | 0.559203 |
| GOBP_SMOOTH_MUSCLE_TISSUE_DEVELOPMENT | 18 | -0.418255 | -1.120329 | 0.312789 | 0.558877 |
| GOBP_ARTERY_MORPHOGENESIS | 54 | -0.338996 | -1.118646 | 0.277383 | 0.563155 |
| GOBP_MAMMARY_GLAND_DUCT_MORPHOGENESIS | 25 | -0.390039 | -1.117067 | 0.310448 | 0.567178 |
| GOBP_POSITIVE_REGULATION_OF_PROTEIN_BINDING | 81 | -0.320149 | -1.117041 | 0.246338 | 0.566815 |
| GOBP_ASTROCYTE_ACTIVATION | 22 | -0.400569 | -1.116702 | 0.321767 | 0.567348 |
| GOBP_POSITIVE_REGULATION_OF_PHOSPHATIDYLINOSITOL_3_KINASE_SIGNALING | 83 | -0.317414 | -1.116368 | 0.280362 | 0.567878 |
| GOBP_POSITIVE_REGULATION_OF_EPITHELIAL_CELL_MIGRATION | 148 | -0.288517 | -1.116334 | 0.245509 | 0.567548 |
| GOBP_TOLL_LIKE_RECEPTOR_SIGNALING_PATHWAY | 144 | -0.291399 | -1.11596 | 0.250923 | 0.568112 |
| GOBP_ALCOHOL_CATABOLIC_PROCESS | 46 | -0.344884 | -1.115656 | 0.28734 | 0.56851 |
| GOBP_POSITIVE_REGULATION_OF_EPITHELIAL_CELL_DIFFERENTIATION | 45 | -0.349519 | -1.115271 | 0.319061 | 0.569163 |
| GOBP_POSITIVE_REGULATION_OF_APOPTOTIC_SIGNALING_PATHWAY | 119 | -0.299873 | -1.115074 | 0.270166 | 0.569297 |
| GOBP_PATHWAY_RESTRICTED_SMAD_PROTEIN_PHOSPHORYLATION | 58 | -0.338334 | -1.114877 | 0.283476 | 0.569396 |
| GOBP_PROTEIN_LOCALIZATION_TO_CELL_SURFACE | 62 | -0.325485 | -1.114793 | 0.272984 | 0.569204 |
| GOBP_NEGATIVE_REGULATION_OF_BIOMINERALIZATION | 25 | -0.390888 | -1.114234 | 0.321656 | 0.57036 |
| GOBP_RESPONSE_TO_TRANSFORMING_GROWTH_FACTOR_BETA | 228 | -0.276654 | -1.11418 | 0.220183 | 0.570067 |
| GOBP_REGULATION_OF_EXCRETION | 20 | -0.409184 | -1.113378 | 0.321317 | 0.571896 |
| GOBP_REGULATION_OF_NITRIC_OXIDE_SYNTHASE_ACTIVITY | 44 | -0.347855 | -1.11259 | 0.300423 | 0.57371 |
| GOBP_EMBRYONIC_SKELETAL_SYSTEM_DEVELOPMENT | 102 | -0.303031 | -1.112294 | 0.258732 | 0.574154 |
| GOBP_REGULATION_OF_PLATELET_DERIVED_GROWTH_FACTOR_RECEPTOR_SIGNALING_PATHW | 21 | -0.406014 | -1.111394 | 0.334906 | 0.576241 |
| GOBP_MUSCLE_CELL_DIFFERENTIATION | 303 | -0.269763 | -1.111316 | 0.21768 | 0.576029 |
| GOBP_REGULATION_OF_BIOMINERALIZATION | 83 | -0.309015 | -1.111044 | 0.278884 | 0.576338 |
| GOBP_P_BODY_ASSEMBLY | 21 | -0.412879 | -1.111012 | 0.31129 | 0.575999 |
| GOBP_RESPONSE_TO_CADMIUM_ION | 53 | -0.334466 | -1.111006 | 0.299584 | 0.575583 |

| | | | | | |
|---|---|---|---|---|---|
| GOBP_CELLULAR_RESPONSE_TO_FLUID_SHEAR_STRESS | 17 | -0.424237 | -1.110864 | 0.308917 | 0.575575 |
| GOBP_SODIUM_ION_TRANSPORT | 195 | -0.28126 | -1.11056 | 0.243081 | 0.575976 |
| GOBP_REGULATION_OF_ACTIN_FILAMENT_ORGANIZATION | 258 | -0.272592 | -1.110387 | 0.229432 | 0.576018 |
| GOBP_MAMMARY_GLAND_EPITHELIUM_DEVELOPMENT | 56 | -0.332063 | -1.110365 | 0.308451 | 0.575657 |
| GOBP_REGULATION_OF_GLIAL_CELL_DIFFERENTIATION | 60 | -0.329434 | -1.109722 | 0.297297 | 0.577062 |
| GOBP_CIRCADIAN_SLEEP_WAKE_CYCLE_SLEEP | 16 | -0.42709 | -1.109349 | 0.330261 | 0.577688 |
| GOBP_EMBRYONIC_EYE_MORPHOGENESIS | 24 | -0.392007 | -1.108453 | 0.323657 | 0.57982 |
| GOBP_PHAGOSOME_MATURATION | 42 | -0.350178 | -1.108429 | 0.302521 | 0.579466 |
| GOBP_OLEFINIC_COMPOUND_METABOLIC_PROCESS | 99 | -0.305646 | -1.108083 | 0.279528 | 0.580014 |
| GOBP_C4_DICARBOXYLATE_TRANSPORT | 23 | -0.390187 | -1.108078 | 0.334361 | 0.579596 |
| GOBP_EXTRACELLULAR_MATRIX_DISASSEMBLY | 68 | -0.320058 | -1.10781 | 0.292683 | 0.579945 |
| GOBP_ENTRAINMENT_OF_CIRCADIAN_CLOCK | 30 | -0.371079 | -1.107624 | 0.325826 | 0.58002 |
| GOBP_NEGATIVE_REGULATION_OF_INTRACELLULAR_SIGNAL_TRANSDUCTION | 464 | -0.261937 | -1.10751 | 0.192182 | 0.579924 |
| GOBP_MYOBLAST_FUSION | 37 | -0.358682 | -1.107449 | 0.289318 | 0.579663 |
| GOBP_STEROID_METABOLIC_PROCESS | 276 | -0.270427 | -1.10682 | 0.224352 | 0.580979 |
| GOBP_MAINTENANCE_OF_LOCATION | 291 | -0.271541 | -1.106614 | 0.221854 | 0.581113 |
| GOBP_CEREBELLAR_CORTEX_MORPHOGENESIS | 27 | -0.383814 | -1.106199 | 0.334849 | 0.581835 |
| GOBP_REGULATION_OF_PATTERN_RECOGNITION_RECEPTOR_SIGNALING_PATHWAY | 97 | -0.302884 | -1.105796 | 0.281818 | 0.582525 |
| GOBP_MUSCLE_SYSTEM_PROCESS | 378 | -0.264629 | -1.105546 | 0.193443 | 0.58278 |
| GOBP_POSITIVE_REGULATION_OF_PHOSPHOPROTEIN_PHOSPHATASE_ACTIVITY | 20 | -0.410046 | -1.10523 | 0.308917 | 0.583262 |
| GOBP_REGULATION_OF_SODIUM_ION_TRANSMEMBRANE_TRANSPORTER_ACTIVITY | 50 | -0.339946 | -1.105029 | 0.298077 | 0.583411 |
| GOBP_ANTERIOR_POSTERIOR_PATTERN_SPECIFICATION | 164 | -0.286151 | -1.104607 | 0.259615 | 0.584179 |
| GOBP_REGULATION_OF_ATP_BIOSYNTHETIC_PROCESS | 20 | -0.407072 | -1.104481 | 0.346645 | 0.584114 |
| GOBP_MUSCLE_TISSUE_DEVELOPMENT | 311 | -0.267415 | -1.10409 | 0.224581 | 0.584803 |
| GOBP_MUCOPOLYSACCHARIDE_METABOLIC_PROCESS | 108 | -0.300124 | -1.103853 | 0.26242 | 0.585082 |
| GOBP_MAMMARY_GLAND_MORPHOGENESIS | 36 | -0.364896 | -1.103625 | 0.296852 | 0.585306 |
| GOBP_POSITIVE_REGULATION_OF_DEVELOPMENTAL_GROWTH | 144 | -0.284506 | -1.103505 | 0.269521 | 0.58518 |
| GOBP_CENTRAL_NERVOUS_SYSTEM_NEURON_DEVELOPMENT | 70 | -0.322604 | -1.102717 | 0.293056 | 0.586958 |
| GOBP_CELL_ADHESION_MEDIATED_BY_INTEGRIN | 67 | -0.323707 | -1.102474 | 0.283401 | 0.587229 |
| GOBP_REGULATION_OF_PROTEIN_EXPORT_FROM_NUCLEUS | 32 | -0.368663 | -1.102361 | 0.313546 | 0.587097 |
| GOBP_ACROSOME_REACTION | 29 | -0.370641 | -1.102348 | 0.327189 | 0.586701 |
| GOBP_POSITIVE_REGULATION_OF_SYNAPTIC_TRANSMISSION_GLUTAMATERGIC | 24 | -0.389264 | -1.101656 | 0.33627 | 0.588235 |
| GOBP_NUCLEAR_TRANSCRIBED_MRNA_CATABOLIC_PROCESS | 206 | -0.275249 | -1.101628 | 0.23804 | 0.587883 |
| GOBP_INTEGRATED_STRESS_RESPONSE_SIGNALING | 26 | -0.376599 | -1.101612 | 0.336991 | 0.587505 |
| GOBP_REGULATION_OF_B_CELL_MEDIATED_IMMUNITY | 54 | -0.336921 | -1.101319 | 0.323611 | 0.587899 |
| GOBP_NEURONAL_ACTION_POTENTIAL | 26 | -0.383119 | -1.101291 | 0.310496 | 0.587543 |
| GOBP_BIOMINERALIZATION | 134 | -0.292384 | -1.101137 | 0.272258 | 0.587577 |
| GOBP_MESENCHYME_MORPHOGENESIS | 41 | -0.346493 | -1.100763 | 0.317251 | 0.588201 |
| GOBP_ACTIN_FILAMENT_BUNDLE_ORGANIZATION | 148 | -0.289021 | -1.100624 | 0.279245 | 0.58817 |
| GOBP_REGULATION_OF_MONOOXYGENASE_ACTIVITY | 53 | -0.334584 | -1.100384 | 0.30109 | 0.588431 |
| GOBP_PROTEIN_LOCALIZATION_TO_PLASMA_MEMBRANE | 263 | -0.268039 | -1.100161 | 0.257471 | 0.588616 |
| GOBP_TRANSFORMING_GROWTH_FACTOR_BETA_RECEPTOR_SIGNALING_PATHWAY | 178 | -0.277995 | -1.099046 | 0.262195 | 0.59131 |
| GOBP_OSTEOBLAST_PROLIFERATION | 28 | -0.379828 | -1.098811 | 0.33985 | 0.591517 |
| GOBP_NEGATIVE_REGULATION_OF_TRANSMEMBRANE_TRANSPORT | 103 | -0.299663 | -1.098505 | 0.30125 | 0.591953 |
| GOBP_ICOSANOID_BIOSYNTHETIC_PROCESS | 41 | -0.353583 | -1.098364 | 0.33474 | 0.591934 |
| GOBP_CELLULAR_HORMONE_METABOLIC_PROCESS | 105 | -0.300448 | -1.098149 | 0.275381 | 0.592116 |
| GOBP_VESICLE_ORGANIZATION | 326 | -0.266842 | -1.097475 | 0.24345 | 0.593605 |
| GOBP_SENSORY_PERCEPTION_OF_PAIN | 78 | -0.310846 | -1.097181 | 0.297222 | 0.594017 |
| GOBP_HEART_DEVELOPMENT | 476 | -0.256502 | -1.096853 | 0.217063 | 0.594509 |
| GOBP_REGULATION_OF_SODIUM_ION_TRANSPORT | 78 | -0.312631 | -1.096825 | 0.289093 | 0.594154 |
| GOBP_ACTIVATION_OF_CYSTEINE_TYPE_ENDOPEPTIDASE_ACTIVITY_INVOLVED_IN_APOPTOTIC_ | 82 | -0.306511 | -1.096796 | 0.295455 | 0.59381 |
| GOBP_POSITIVE_REGULATION_OF_CALCIUM_ION_IMPORT | 16 | -0.430025 | -1.096482 | 0.340062 | 0.594301 |
| GOBP_ACTIVATION_OF_MAPK_ACTIVITY | 143 | -0.287894 | -1.09566 | 0.294494 | 0.596185 |
| GOBP_HYALURONAN_METABOLIC_PROCESS | 36 | -0.355565 | -1.095644 | 0.329562 | 0.595813 |
| GOBP_REACTIVE_OXYGEN_SPECIES_METABOLIC_PROCESS | 253 | -0.27191 | -1.095374 | 0.260023 | 0.596157 |
| GOBP_ENDODERM_FORMATION | 50 | -0.339878 | -1.094647 | 0.341259 | 0.597782 |
| GOBP_POSITIVE_REGULATION_OF_MORPHOGENESIS_OF_AN_EPITHELIUM | 29 | -0.375524 | -1.094462 | 0.360119 | 0.597867 |
| GOBP_NEGATIVE_REGULATION_OF_DNA_DAMAGE_RESPONSE_SIGNAL_TRANSDUCTION_BY_P5 | 15 | -0.438149 | -1.094282 | 0.355556 | 0.597884 |
| GOBP_ORGANIC_HYDROXY_COMPOUND_CATABOLIC_PROCESS | 64 | -0.322243 | -1.094244 | 0.304288 | 0.597577 |
| GOBP_DENDRITE_MORPHOGENESIS | 138 | -0.285207 | -1.093845 | 0.278562 | 0.598272 |
| GOBP_TELENCEPHALON_GLIAL_CELL_MIGRATION | 20 | -0.407283 | -1.093818 | 0.369159 | 0.59793 |
| GOBP_ORGANIC_ANION_TRANSPORT | 318 | -0.266456 | -1.093798 | 0.243902 | 0.597562 |
| GOBP_RHYTHMIC_BEHAVIOR | 36 | -0.355949 | -1.093151 | 0.324286 | 0.598951 |
| GOBP_NEGATIVE_REGULATION_OF_INTERLEUKIN_10_PRODUCTION | 19 | -0.409563 | -1.092158 | 0.333333 | 0.601343 |
| GOBP_REGULATION_OF_TYPE_I_INTERFERON_MEDIATED_SIGNALING_PATHWAY | 32 | -0.359993 | -1.092043 | 0.33888 | 0.601266 |
| GOBP_CANONICAL_WNT_SIGNALING_PATHWAY | 308 | -0.262435 | -1.091736 | 0.256236 | 0.601703 |
| GOBP_THYMIC_T_CELL_SELECTION | 22 | -0.400143 | -1.091501 | 0.347826 | 0.601957 |
| GOBP_MORPHOGENESIS_OF_AN_EPITHELIUM | 472 | -0.255465 | -1.091408 | 0.234914 | 0.601815 |
| GOBP_POSITIVE_REGULATION_OF_PHOSPHATASE_ACTIVITY | 31 | -0.377041 | -1.090945 | 0.346264 | 0.602678 |
| GOBP_SEQUESTERING_OF_CALCIUM_ION | 106 | -0.294754 | -1.090242 | 0.299606 | 0.604257 |
| GOBP_FOREBRAIN_DEVELOPMENT | 302 | -0.263092 | -1.089806 | 0.259259 | 0.605069 |
| GOBP_VASCULAR_ASSOCIATED_SMOOTH_MUSCLE_CELL_DIFFERENTIATION | 25 | -0.383446 | -1.089231 | 0.352496 | 0.606252 |
| GOBP_REGULATION_OF_VIRAL_GENOME_REPLICATION | 80 | -0.309747 | -1.088916 | 0.298222 | 0.606701 |
| GOBP_INTRACELLULAR_ESTROGEN_RECEPTOR_SIGNALING_PATHWAY | 54 | -0.328389 | -1.08882 | 0.319776 | 0.606557 |
| GOBP_CELLULAR_RESPONSE_TO_RETINOIC_ACID | 51 | -0.331412 | -1.088609 | 0.322314 | 0.606724 |
| GOBP_RESPONSE_TO_MUSCLE_STRETCH | 21 | -0.392813 | -1.088085 | 0.362121 | 0.607805 |
| GOBP_HEART_VALVE_DEVELOPMENT | 57 | -0.326391 | -1.087985 | 0.339056 | 0.607675 |
| GOBP_REGULATION_OF_NERVOUS_SYSTEM_DEVELOPMENT | 375 | -0.260666 | -1.087816 | 0.237785 | 0.607745 |
| GOBP_REGULATION_OF_RESPONSE_TO_TUMOR_CELL | 16 | -0.425925 | -1.087464 | 0.384735 | 0.60832 |
| GOBP_SENSORY_PERCEPTION_OF_LIGHT_STIMULUS | 160 | -0.280361 | -1.08713 | 0.304562 | 0.608833 |
| GOBP_AMINOGLYCAN_METABOLIC_PROCESS | 160 | -0.278383 | -1.087032 | 0.306846 | 0.608674 |
| GOBP_POSITIVE_REGULATION_OF_MYELOID_CELL_DIFFERENTIATION | 96 | -0.296944 | -1.086917 | 0.278205 | 0.608578 |
| GOBP_POSITIVE_REGULATION_OF_SMOOTH_MUSCLE_CELL_PROLIFERATION | 83 | -0.303821 | -1.08678 | 0.325397 | 0.60854 |
| GOBP_POSITIVE_REGULATION_OF_EXOCYTOSIS | 79 | -0.307199 | -1.086774 | 0.302989 | 0.608131 |
| GOBP_THYMOCYTE_APOPTOTIC_PROCESS | 16 | -0.42781 | -1.086761 | 0.347334 | 0.607736 |
| GOBP_REGULATION_OF_BMP_SIGNALING_PATHWAY | 78 | -0.310635 | -1.086391 | 0.31516 | 0.608332 |
| GOBP_RESPONSE_TO_VITAMIN | 80 | -0.307215 | -1.085486 | 0.329624 | 0.610473 |
| GOBP_REGULATION_OF_LIPID_LOCALIZATION | 152 | -0.280468 | -1.085304 | 0.291463 | 0.610559 |
| GOBP_REGULATION_OF_POSTSYNAPSE_ORGANIZATION | 79 | -0.309303 | -1.085161 | 0.328767 | 0.610555 |
| GOBP_HEMATOPOIETIC_STEM_CELL_PROLIFERATION | 25 | -0.379645 | -1.084642 | 0.369661 | 0.611577 |
| GOBP_RESPONSE_TO_INTERLEUKIN_12 | 49 | -0.332426 | -1.084494 | 0.337535 | 0.611558 |
| GOBP_ANDROGEN_METABOLIC_PROCESS | 24 | -0.385767 | -1.083797 | 0.344985 | 0.613147 |
| GOBP_CARDIAC_MUSCLE_CELL_ACTION_POTENTIAL_INVOLVED_IN_CONTRACTION | 49 | -0.332256 | -1.083041 | 0.323864 | 0.614922 |
| GOBP_REGULATION_OF_SYSTEMIC_ARTERIAL_BLOOD_PRESSURE_BY_CIRCULATORY_RENIN_A | 15 | -0.427659 | -1.083029 | 0.367284 | 0.614524 |
| GOBP_CYCLIC_NUCLEOTIDE_MEDIATED_SIGNALING | 74 | -0.307378 | -1.082829 | 0.300401 | 0.614652 |
| GOBP_REGULATION_OF_PHOSPHATIDYLINOSITOL_3_KINASE_SIGNALING | 116 | -0.291719 | -1.082754 | 0.317647 | 0.614448 |
| GOBP_NEGATIVE_REGULATION_OF_GROWTH | 223 | -0.269331 | -1.082097 | 0.298349 | 0.615869 |
| GOBP_REPLACEMENT_OSSIFICATION | 28 | -0.376432 | -1.081864 | 0.361991 | 0.616099 |
| GOBP_REGULATION_OF_EPITHELIAL_CELL_MIGRATION | 224 | -0.268922 | -1.081694 | 0.293567 | 0.616149 |
| GOBP_REGULATION_OF_DNA_DAMAGE_RESPONSE_SIGNAL_TRANSDUCTION_BY_P53_CLASS_M | 34 | -0.359748 | -1.081613 | 0.32106 | 0.615935 |
| GOBP_POSITIVE_REGULATION_OF_GLYCOLYTIC_PROCESS | 23 | -0.393796 | -1.081417 | 0.362013 | 0.61602 |
| GOBP_CARBOXYLIC_ACID_TRANSPORT | 234 | -0.268559 | -1.081381 | 0.276256 | 0.615709 |
| GOBP_REGULATION_OF_ANATOMICAL_STRUCTURE_SIZE | 448 | -0.255605 | -1.081337 | 0.247588 | 0.615418 |
| GOBP_REGULATION_OF_BEHAVIOR | 51 | -0.335416 | -1.080977 | 0.331909 | 0.616026 |
| GOBP_MATING | 31 | -0.359982 | -1.079227 | 0.351515 | 0.620478 |
| GOBP_POSITIVE_REGULATION_OF_CELL_JUNCTION_ASSEMBLY | 85 | -0.300435 | -1.078746 | 0.318653 | 0.621408 |
| GOBP_EMBRYONIC_PLACENTA_DEVELOPMENT | 71 | -0.30713 | -1.078605 | 0.334711 | 0.621387 |
| GOBP_LONG_CHAIN_FATTY_ACID_TRANSPORT | 61 | -0.317326 | -1.078406 | 0.33662 | 0.621531 |
| GOBP_REGULATION_OF_INTRINSIC_APOPTOTIC_SIGNALING_PATHWAY_IN_RESPONSE_TO_DNA_ | 37 | -0.349725 | -1.078267 | 0.348228 | 0.621485 |
| GOBP_NEGATIVE_REGULATION_OF_VIRAL_ENTRY_INTO_HOST_CELL | 18 | -0.406615 | -1.078038 | 0.371473 | 0.621732 |
| GOBP_REGULATION_OF_VIRAL_LIFE_CYCLE | 138 | -0.282479 | -1.076943 | 0.321519 | 0.624493 |
| GOBP_STEM_CELL_PROLIFERATION | 58 | -0.315611 | -1.076941 | 0.34874 | 0.624081 |
| GOBP_REGULATION_OF_LIPID_TRANSPORT | 125 | -0.286527 | -1.076313 | 0.334171 | 0.625442 |
| GOBP_NEGATIVE_REGULATION_OF_CALCIUM_ION_TRANSPORT | 48 | -0.331838 | -1.075905 | 0.337111 | 0.62614 |
| GOBP_CELLULAR_RESPONSE_TO_EXOGENOUS_DSRNA | 18 | -0.412042 | -1.075777 | 0.3712 | 0.626089 |
| GOBP_PROTEIN_LOCALIZATION_TO_CELL_PERIPHERY | 310 | -0.258794 | -1.075149 | 0.284414 | 0.627506 |
| GOBP_CELLULAR_RESPONSE_TO_GLUCOSE_STARVATION | 48 | -0.329094 | -1.074777 | 0.35014 | 0.628166 |
| GOBP_REGULATION_OF_COLLAGEN_METABOLIC_PROCESS | 39 | -0.346569 | -1.074668 | 0.354074 | 0.62804 |
| GOBP_REGULATION_OF_EXTRACELLULAR_MATRIX_ORGANIZATION | 44 | -0.336106 | -1.074462 | 0.332386 | 0.628182 |
| GOBP_BONE_REMODELING | 81 | -0.303027 | -1.073812 | 0.32636 | 0.629606 |
| GOBP_MAINTENANCE_OF_CELL_NUMBER | 134 | -0.281495 | -1.073151 | 0.306122 | 0.631094 |
| GOBP_CELL_FATE_COMMITMENT_INVOLVED_IN_FORMATION_OF_PRIMARY_GERM_LAYER | 25 | -0.372418 | -1.072931 | 0.358407 | 0.631329 |
| GOBP_VESICLE_FUSION_TO_PLASMA_MEMBRANE | 20 | -0.404122 | -1.072717 | 0.363499 | 0.631491 |
| GOBP_POSITIVE_REGULATION_OF_NEURON_APOPTOTIC_PROCESS | 46 | -0.327274 | -1.072708 | 0.346705 | 0.631088 |
| GOBP_NEUROTRANSMITTER_SECRETION | 142 | -0.280308 | -1.07212 | 0.306072 | 0.632359 |
| GOBP_REGULATION_OF_CELLULAR_EXTRAVASATION | 31 | -0.356623 | -1.071953 | 0.369048 | 0.632387 |
| GOBP_REGULATION_OF_OXIDOREDUCTASE_ACTIVITY | 92 | -0.296422 | -1.071671 | 0.337255 | 0.632726 |
| GOBP_POSITIVE_REGULATION_OF_OSTEOBLAST_DIFFERENTIATION | 57 | -0.317554 | -1.071631 | 0.349794 | 0.632401 |
| GOBP_FC_RECEPTOR_SIGNALING_PATHWAY | 234 | -0.267032 | -1.070895 | 0.308943 | 0.634112 |
| GOBP_POSITIVE_REGULATION_OF_TRANSLATIONAL_INITIATION | 25 | -0.372997 | -1.070702 | 0.367868 | 0.634232 |
| GOBP_REGULATION_OF_STRIATED_MUSCLE_CONTRACTION | 75 | -0.303488 | -1.069875 | 0.336054 | 0.636093 |
| GOBP_NEURON_MIGRATION | 118 | -0.285859 | -1.069833 | 0.324967 | 0.635775 |
| GOBP_INSULIN_SECRETION_INVOLVED_IN_CELLULAR_RESPONSE_TO_GLUCOSE_STIMULUS | 58 | -0.318762 | -1.06945 | 0.357735 | 0.636431 |
| GOBP_POSITIVE_REGULATION_OF_CATION_CHANNEL_ACTIVITY | 60 | -0.313275 | -1.068813 | 0.358904 | 0.637881 |
| GOBP_REGULATION_OF_EXTRINSIC_APOPTOTIC_SIGNALING_PATHWAY_IN_ABSENCE_OF_LIGAN | 38 | -0.343163 | -1.068709 | 0.367128 | 0.637741 |
| GOBP_POSITIVE_REGULATION_OF_ACTIN_FILAMENT_BUNDLE_ASSEMBLY | 59 | -0.318267 | -1.06867 | 0.361644 | 0.637421 |
| GOBP_COAGULATION | 309 | -0.257516 | -1.068454 | 0.304972 | 0.6376 |
| GOBP_NEGATIVE_REGULATION_OF_I_KAPPAB_KINASE_NF_KAPPAB_SIGNALING | 43 | -0.33626 | -1.068122 | 0.353474 | 0.638094 |
| GOBP_POSITIVE_REGULATION_OF_FAT_CELL_DIFFERENTIATION | 57 | -0.321786 | -1.068091 | 0.356241 | 0.637752 |
| GOBP_AXON_DEVELOPMENT | 453 | -0.251728 | -1.067997 | 0.28169 | 0.637593 |
| GOBP_PRODUCTION_OF_SMALL_RNA_INVOLVED_IN_GENE_SILENCING_BY_RNA | 53 | -0.328705 | -1.067743 | 0.367832 | 0.637879 |
| GOBP_CARDIAC_MUSCLE_TISSUE_MORPHOGENESIS | 47 | -0.327934 | -1.067668 | 0.365357 | 0.63766 |
| GOBP_SEQUESTERING_OF_TRIGLYCERIDE | 15 | -0.422194 | -1.067566 | 0.382445 | 0.637536 |
| GOBP_REGULATION_OF_TUMOR_NECROSIS_FACTOR_MEDIATED_SIGNALING_PATHWAY | 57 | -0.319389 | -1.067501 | 0.337931 | 0.637283 |
| GOBP_NECROTIC_CELL_DEATH | 56 | -0.321168 | -1.067494 | 0.362379 | 0.636882 |
| GOBP_NEGATIVE_REGULATION_OF_PEPTIDYL_THREONINE_PHOSPHORYLATION | 19 | -0.397106 | -1.066721 | 0.403481 | 0.638691 |
| GOBP_MITOTIC_CELL_CYCLE_ARREST | 16 | -0.412501 | -1.065816 | 0.374224 | 0.640827 |
| GOBP_NEGATIVE_REGULATION_OF_NEURON_APOPTOTIC_PROCESS | 110 | -0.288154 | -1.065807 | 0.340996 | 0.640423 |
| GOBP_POSITIVE_REGULATION_OF_CARTILAGE_DEVELOPMENT | 27 | -0.366138 | -1.065115 | 0.37386 | 0.641981 |
| GOBP_POSITIVE_REGULATION_OF_RELEASE_OF_CYTOCHROME_C_FROM_MITOCHONDRIA | 27 | -0.365529 | -1.064284 | 0.35905 | 0.643903 |
| GOBP_LONG_CHAIN_FATTY_ACID_BIOSYNTHETIC_PROCESS | 31 | -0.35569 | -1.063664 | 0.373887 | 0.645258 |
| GOBP_REGULATION_OF_RESPIRATORY_GASEOUS_EXCHANGE | 17 | -0.402809 | -1.063587 | 0.378086 | 0.645048 |
| GOBP_POSITIVE_REGULATION_OF_CARBOHYDRATE_METABOLIC_PROCESS | 72 | -0.313347 | -1.06333 | 0.356643 | 0.645341 |
| GOBP_PLACENTA_DEVELOPMENT | 118 | -0.28341 | -1.063224 | 0.340426 | 0.645211 |
| GOBP_HEART_GROWTH | 75 | -0.303338 | -1.06241 | 0.342355 | 0.647114 |
| GOBP_REGULATION_OF_MUSCLE_ADAPTATION | 77 | -0.305345 | -1.061589 | 0.367429 | 0.648993 |
| GOBP_POLYSACCHARIDE_CATABOLIC_PROCESS | 24 | -0.379892 | -1.061127 | 0.392424 | 0.649929 |
| GOBP_UDP_N_ACETYLGLUCOSAMINE_METABOLIC_PROCESS | 15 | -0.414255 | -1.061045 | 0.387521 | 0.649753 |
| GOBP_POSITIVE_REGULATION_OF_CELL_ADHESION_MEDIATED_BY_INTEGRIN | 20 | -0.392612 | -1.06099 | 0.388626 | 0.649474 |
| GOBP_INTRINSIC_APOPTOTIC_SIGNALING_PATHWAY_IN_RESPONSE_TO_DNA_DAMAGE | 100 | -0.290826 | -1.060863 | 0.347313 | 0.649378 |
| GOBP_NEGATIVE_REGULATION_OF_INTRINSIC_APOPTOTIC_SIGNALING_PATHWAY | 89 | -0.302181 | -1.060682 | 0.360324 | 0.649434 |
| GOBP_APOPTOTIC_CELL_CLEARANCE | 49 | -0.320086 | -1.060566 | 0.373156 | 0.649331 |
| GOBP_NEGATIVE_REGULATION_OF_JNK_CASCADE | 34 | -0.353845 | -1.059548 | 0.360632 | 0.65183 |
| GOBP_REPRODUCTIVE_BEHAVIOR | 22 | -0.383713 | -1.059463 | 0.406107 | 0.651643 |
| GOBP_BONE_RESORPTION | 56 | -0.31581 | -1.058795 | 0.384397 | 0.653111 |
| GOBP_REGULATION_OF_TRANS_SYNAPTIC_SIGNALING | 358 | -0.254691 | -1.058663 | 0.316757 | 0.653062 |
| GOBP_REGULATION_OF_RESPIRATORY_BURST | 15 | -0.425462 | -1.058227 | 0.385671 | 0.653865 |
| GOBP_EPIDERMAL_GROWTH_FACTOR_RECEPTOR_SIGNALING_PATHWAY | 118 | -0.282519 | -1.058022 | 0.362681 | 0.654022 |

| | | | | | |
|---|---|---|---|---|---|
| GOBP_RESPONSE_TO_UV_B | 15 | -0.419282 | -1.057639 | 0.413953 | 0.654644 |
| GOBP_CARDIAC_VENTRICLE_MORPHOGENESIS | 55 | -0.317988 | -1.056965 | 0.375867 | 0.656109 |
| GOBP_VITAMIN_METABOLIC_PROCESS | 120 | -0.283784 | -1.056797 | 0.356436 | 0.656156 |
| GOBP_DENDRITE_DEVELOPMENT | 226 | -0.263443 | -1.056592 | 0.355713 | 0.656317 |
| GOBP_PROTEIN_COMPLEX_OLIGOMERIZATION | 201 | -0.267572 | -1.056518 | 0.321719 | 0.656089 |
| GOBP_CELL_CELL_ADHESION_MEDIATED_BY_CADHERIN | 25 | -0.363466 | -1.056412 | 0.370044 | 0.655939 |
| GOBP_PROTEIN_LOCALIZATION_TO_LYSOSOME | 41 | -0.333852 | -1.0564 | 0.376093 | 0.655535 |
| GOBP_REGULATION_OF_TRIGLYCERIDE_BIOSYNTHETIC_PROCESS | 18 | -0.405088 | -1.056383 | 0.398792 | 0.655149 |
| GOBP_LONG_CHAIN_FATTY_ACID_METABOLIC_PROCESS | 106 | -0.287248 | -1.056275 | 0.363522 | 0.655029 |
| GOBP_NEURAL_CREST_CELL_DIFFERENTIATION | 78 | -0.300313 | -1.056143 | 0.357333 | 0.654973 |
| GOBP_REGULATION_OF_SENSORY_PERCEPTION | 29 | -0.363954 | -1.056002 | 0.404624 | 0.654975 |
| GOBP_POSITIVE_REGULATION_OF_LAMELLIPODIUM_ORGANIZATION | 36 | -0.343404 | -1.055865 | 0.362963 | 0.654951 |
| GOBP_AMYLOID_PRECURSOR_PROTEIN_CATABOLIC_PROCESS | 54 | -0.31745 | -1.055698 | 0.382734 | 0.655008 |
| GOBP_COLUMNAR_CUBOIDAL_EPITHELIAL_CELL_DEVELOPMENT | 34 | -0.347429 | -1.055657 | 0.393539 | 0.654698 |
| GOBP_PROTEOGLYCAN_BIOSYNTHETIC_PROCESS | 58 | -0.312316 | -1.05543 | 0.368644 | 0.654899 |
| GOBP_REGULATION_OF_LIPID_STORAGE | 48 | -0.328068 | -1.054784 | 0.374819 | 0.656255 |
| GOBP_EXCRETION | 48 | -0.326191 | -1.054707 | 0.373427 | 0.656057 |
| GOBP_WOUND_HEALING_SPREADING_OF_EPIDERMAL_CELLS | 18 | -0.402791 | -1.054581 | 0.374618 | 0.655972 |
| GOBP_REGULATION_OF_MEMBRANE_DEPOLARIZATION | 36 | -0.340893 | -1.054468 | 0.374269 | 0.655881 |
| GOBP_RESPONSE_TO_VITAMIN_A | 18 | -0.397869 | -1.054453 | 0.410769 | 0.6555 |
| GOBP_NEGATIVE_REGULATION_OF_NEURON_DEATH | 166 | -0.271591 | -1.054283 | 0.355721 | 0.655546 |
| GOBP_NECROPTOTIC_PROCESS | 37 | -0.344916 | -1.053533 | 0.362857 | 0.657254 |
| GOBP_METANEPHRIC_EPITHELIUM_DEVELOPMENT | 19 | -0.392537 | -1.053508 | 0.397554 | 0.656901 |
| GOBP_RELEASE_OF_SEQUESTERED_CALCIUM_ION_INTO_CYTOSOL_BY_ENDOPLASMIC_RETICU | 29 | -0.353564 | -1.053279 | 0.380734 | 0.657131 |
| GOBP_OVULATION | 15 | -0.424404 | -1.053196 | 0.411765 | 0.65694 |
| GOBP_PROTEIN_O_LINKED_GLYCOSYLATION | 95 | -0.291806 | -1.053152 | 0.357326 | 0.656631 |
| GOBP_NEGATIVE_REGULATION_OF_VASCULAR_ASSOCIATED_SMOOTH_MUSCLE_CELL_PROLIFE | 20 | -0.39108 | -1.052925 | 0.38653 | 0.656849 |
| GOBP_GLUTAMATE_RECEPTOR_SIGNALING_PATHWAY | 41 | -0.335192 | -1.052883 | 0.384173 | 0.656537 |
| GOBP_DETECTION_OF_STIMULUS_INVOLVED_IN_SENSORY_PERCEPTION_OF_PAIN | 21 | -0.384507 | -1.052675 | 0.40412 | 0.656688 |
| GOBP_MIDBRAIN_DEVELOPMENT | 74 | -0.299753 | -1.052426 | 0.37535 | 0.65696 |
| GOBP_EPITHELIAL_CELL_APOPTOTIC_PROCESS | 97 | -0.291886 | -1.052332 | 0.370656 | 0.656795 |
| GOBP_EMBRYONIC_ORGAN_DEVELOPMENT | 352 | -0.253241 | -1.052017 | 0.346239 | 0.657274 |
| GOBP_NEGATIVE_REGULATION_OF_WNT_SIGNALING_PATHWAY | 198 | -0.266565 | -1.051889 | 0.352872 | 0.657201 |
| GOBP_REGULATION_OF_LYASE_ACTIVITY | 37 | -0.335732 | -1.051817 | 0.382646 | 0.65699 |
| GOBP_CELL_MIGRATION_INVOLVED_IN_SPROUTING_ANGIOGENESIS | 54 | -0.314684 | -1.051749 | 0.376405 | 0.656762 |
| GOBP_POSITIVE_REGULATION_OF_ANION_TRANSPORT | 439 | -0.248684 | -1.051691 | 0.329694 | 0.656504 |
| GOBP_APOPTOTIC_MITOCHONDRIAL_CHANGES | 111 | -0.28139 | -1.051225 | 0.379224 | 0.657394 |
| GOBP_REGULATION_OF_RECEPTOR_BINDING | 23 | -0.378117 | -1.050955 | 0.397882 | 0.657726 |
| GOBP_POSITIVE_REGULATION_OF_SYNAPTIC_TRANSMISSION | 126 | -0.277198 | -1.050492 | 0.34596 | 0.658621 |
| GOBP_MELANOCYTE_DIFFERENTIATION | 20 | -0.385881 | -1.050409 | 0.391026 | 0.658412 |
| GOBP_ENDOCYTIC_RECYCLING | 48 | -0.320431 | -1.04956 | 0.388966 | 0.660336 |
| GOBP_RAC_PROTEIN_SIGNAL_TRANSDUCTION | 39 | -0.335213 | -1.049295 | 0.385174 | 0.660645 |
| GOBP_RHO_PROTEIN_SIGNAL_TRANSDUCTION | 128 | -0.278447 | -1.049098 | 0.360303 | 0.660787 |
| GOBP_ACIDIC_AMINO_ACID_TRANSPORT | 62 | -0.309942 | -1.048764 | 0.374663 | 0.661372 |
| GOBP_NITRIC_OXIDE_MEDIATED_SIGNAL_TRANSDUCTION | 20 | -0.387722 | -1.048691 | 0.416159 | 0.661154 |
| GOBP_RESPONSE_TO_GROWTH_HORMONE | 35 | -0.340926 | -1.048401 | 0.377551 | 0.66153 |
| GOBP_INTRINSIC_APOPTOTIC_SIGNALING_PATHWAY_BY_P53_CLASS_MEDIATOR | 73 | -0.29821 | -1.048245 | 0.369128 | 0.661546 |
| GOBP_THYROID_GLAND_DEVELOPMENT | 18 | -0.405982 | -1.047902 | 0.399689 | 0.662092 |
| GOBP_SEMI_LUNAR_VALVE_DEVELOPMENT | 38 | -0.337555 | -1.047725 | 0.389127 | 0.662174 |
| GOBP_REGULATION_OF_JNK_CASCADE | 122 | -0.280604 | -1.047206 | 0.372215 | 0.663195 |
| GOBP_REGULATION_OF_VASCULAR_ENDOTHELIAL_GROWTH_FACTOR_RECEPTOR_SIGNALING_ | 25 | -0.368908 | -1.047114 | 0.396605 | 0.663053 |
| GOBP_POSITIVE_REGULATION_OF_B_CELL_MEDIATED_IMMUNITY | 38 | -0.343059 | -1.046992 | 0.394161 | 0.662955 |
| GOBP_MEMBRANE_FUSION | 159 | -0.270107 | -1.046843 | 0.358025 | 0.662944 |
| GOBP_NEGATIVE_REGULATION_OF_T_CELL_RECEPTOR_SIGNALING_PATHWAY | 22 | -0.380416 | -1.04681 | 0.380665 | 0.662593 |
| GOBP_NEGATIVE_REGULATION_OF_PROTEIN_MODIFICATION_BY_SMALL_PROTEIN_CONJUGATIO | 85 | -0.29352 | -1.046221 | 0.378162 | 0.663847 |
| GOBP_SPERM_CAPACITATION | 19 | -0.391942 | -1.046056 | 0.396226 | 0.663871 |
| GOBP_ANTIGEN_PROCESSING_AND_PRESENTATION_OF_PEPTIDE_OR_POLYSACCHARIDE_ANTI | 100 | -0.283613 | -1.04585 | 0.371681 | 0.664014 |
| GOBP_POSITIVE_REGULATION_OF_SPROUTING_ANGIOGENESIS | 20 | -0.384268 | -1.045701 | 0.390551 | 0.663991 |
| GOBP_EXTERNAL_ENCAPSULATING_STRUCTURE_ORGANIZATION | 354 | -0.252795 | -1.045548 | 0.365449 | 0.663986 |
| GOBP_REGULATION_OF_NEUROTRANSMITTER_TRANSPORT | 90 | -0.291848 | -1.045035 | 0.378129 | 0.665014 |
| GOBP_POSITIVE_REGULATION_OF_PEPTIDYL_SERINE_PHOSPHORYLATION | 84 | -0.29365 | -1.045012 | 0.383202 | 0.664652 |
| GOBP_POSITIVE_REGULATION_OF_CHONDROCYTE_DIFFERENTIATION | 15 | -0.41398 | -1.044891 | 0.418462 | 0.664568 |
| GOBP_SARCOPLASMIC_RETICULUM_CALCIUM_ION_TRANSPORT | 32 | -0.345951 | -1.044708 | 0.394203 | 0.664661 |
| GOBP_SARCOMERE_ORGANIZATION | 31 | -0.350979 | -1.044366 | 0.395349 | 0.665206 |
| GOBP_LIPID_CATABOLIC_PROCESS | 294 | -0.253323 | -1.044357 | 0.38339 | 0.664806 |
| GOBP_POSITIVE_REGULATION_OF_GROWTH | 220 | -0.261109 | -1.043966 | 0.36197 | 0.665499 |
| GOBP_RESPONSE_TO_COPPER_ION | 34 | -0.349167 | -1.043634 | 0.399417 | 0.665993 |
| GOBP_ENTRY_INTO_HOST | 143 | -0.270269 | -1.043562 | 0.383648 | 0.665758 |
| GOBP_REGULATION_OF_OLIGODENDROCYTE_DIFFERENTIATION | 33 | -0.347973 | -1.042861 | 0.36903 | 0.667322 |
| GOBP_ACTOMYOSIN_STRUCTURE_ORGANIZATION | 174 | -0.266019 | -1.042156 | 0.380838 | 0.668917 |
| GOBP_OSSIFICATION | 358 | -0.248778 | -1.041234 | 0.363234 | 0.671083 |
| GOBP_REGULATION_OF_CILIUM_MOVEMENT | 27 | -0.364842 | -1.041217 | 0.423781 | 0.670703 |
| GOBP_LIPID_LOCALIZATION | 438 | -0.24474 | -1.041207 | 0.347267 | 0.670304 |
| GOBP_RESPONSE_TO_ORGANOPHOSPHORUS | 122 | -0.278944 | -1.040839 | 0.395109 | 0.670925 |
| GOBP_MEMBRANE_BIOGENESIS | 39 | -0.330805 | -1.040727 | 0.387931 | 0.670823 |
| GOBP_EXOCYTIC_PROCESS | 74 | -0.299181 | -1.040641 | 0.395473 | 0.670634 |
| GOBP_NEGATIVE_REGULATION_OF_ATP_METABOLIC_PROCESS | 27 | -0.355676 | -1.040451 | 0.409985 | 0.670739 |
| GOBP_REGULATION_OF_BIOLOGICAL_PROCESS_INVOLVED_IN_SYMBIOTIC_INTERACTION | 184 | -0.262158 | -1.040408 | 0.380895 | 0.670441 |
| GOBP_POSITIVE_REGULATION_OF_ALCOHOL_BIOSYNTHETIC_PROCESS | 18 | -0.397687 | -1.04035 | 0.403427 | 0.670179 |
| GOBP_POSITIVE_REGULATION_OF_POTASSIUM_ION_TRANSPORT | 38 | -0.336325 | -1.040281 | 0.403509 | 0.66995 |
| GOBP_REGULATION_OF_LIPID_METABOLIC_PROCESS | 353 | -0.250508 | -1.040172 | 0.378349 | 0.669836 |
| GOBP_PHOSPHATIDYLGLYCEROL_METABOLIC_PROCESS | 31 | -0.348947 | -1.039997 | 0.400881 | 0.669888 |
| GOBP_VESICLE_CARGO_LOADING | 25 | -0.362903 | -1.039949 | 0.395245 | 0.669602 |
| GOBP_MAST_CELL_ACTIVATION_INVOLVED_IN_IMMUNE_RESPONSE | 45 | -0.321933 | -1.039834 | 0.416176 | 0.669508 |
| GOBP_RESPONSE_TO_INORGANIC_SUBSTANCE | 474 | -0.243842 | -1.039634 | 0.356838 | 0.669661 |
| GOBP_REGULATION_OF_ACTION_POTENTIAL | 46 | -0.326367 | -1.039262 | 0.419591 | 0.670319 |
| GOBP_POSITIVE_REGULATION_OF_MYELINATION | 17 | -0.398036 | -1.038969 | 0.406492 | 0.670677 |
| GOBP_PROTEIN_O_LINKED_MANNOSYLATION | 18 | -0.391597 | -1.038251 | 0.444262 | 0.672229 |
| GOBP_REGULATION_OF_CELL_MORPHOGENESIS | 294 | -0.25297 | -1.038205 | 0.377551 | 0.671941 |
| GOBP_MULTICELLULAR_ORGANISMAL_HOMEOSTASIS | 459 | -0.245375 | -1.03817 | 0.367679 | 0.671622 |
| GOBP_REGULATION_OF_FATTY_ACID_OXIDATION | 31 | -0.348421 | -1.03813 | 0.401515 | 0.671315 |
| GOBP_INOSITOL_PHOSPHATE_METABOLIC_PROCESS | 66 | -0.303544 | -1.037924 | 0.369771 | 0.671465 |
| GOBP_EPITHELIAL_TO_MESENCHYMAL_TRANSITION_INVOLVED_IN_ENDOCARDIAL_CUSHION_FO | 16 | -0.393936 | -1.037814 | 0.419512 | 0.671369 |
| GOBP_REGULATION_OF_CARDIAC_MUSCLE_CELL_PROLIFERATION | 35 | -0.336896 | -1.037802 | 0.401471 | 0.670978 |
| GOBP_ACTIN_FILAMENT_ORGANIZATION | 409 | -0.248699 | -1.037268 | 0.359002 | 0.67206 |
| GOBP_NEGATIVE_REGULATION_OF_GLIAL_CELL_DIFFERENTIATION | 19 | -0.385459 | -1.037221 | 0.426357 | 0.671775 |
| GOBP_POSITIVE_REGULATION_OF_ATP_METABOLIC_PROCESS | 39 | -0.327794 | -1.037208 | 0.409289 | 0.671393 |
| GOBP_RESPONSE_TO_ALKALOID | 85 | -0.29153 | -1.037202 | 0.40239 | 0.670995 |
| GOBP_STEROID_BIOSYNTHETIC_PROCESS | 163 | -0.266105 | -1.036704 | 0.388134 | 0.671933 |
| GOBP_FOAM_CELL_DIFFERENTIATION | 34 | -0.337215 | -1.036459 | 0.404011 | 0.672262 |
| GOBP_REGULATION_OF_SYNAPTIC_VESICLE_ENDOCYTOSIS | 15 | -0.415223 | -1.036311 | 0.422619 | 0.672254 |
| GOBP_NEURAL_PRECURSOR_CELL_PROLIFERATION | 119 | -0.27462 | -1.035521 | 0.383929 | 0.674082 |
| GOBP_NEGATIVE_REGULATION_OF_PEPTIDYL_TYROSINE_PHOSPHORYLATION | 52 | -0.316021 | -1.035177 | 0.41523 | 0.674657 |
| GOBP_NEGATIVE_REGULATION_OF_PHAGOCYTOSIS | 21 | -0.37818 | -1.035078 | 0.431988 | 0.674496 |
| GOBP_POSITIVE_REGULATION_OF_ACTIVATED_T_CELL_PROLIFERATION | 24 | -0.363325 | -1.034934 | 0.413686 | 0.674468 |
| GOBP_ANION_HOMEOSTASIS | 27 | -0.35724 | -1.03441 | 0.425339 | 0.675531 |
| GOBP_POSITIVE_REGULATION_OF_BONE_MINERALIZATION | 36 | -0.340243 | -1.034405 | 0.415584 | 0.675126 |
| GOBP_POSITIVE_REGULATION_OF_JUN_KINASE_ACTIVITY | 66 | -0.302445 | -1.033878 | 0.393056 | 0.676173 |
| GOBP_CELL_CELL_JUNCTION_ASSEMBLY | 125 | -0.274627 | -1.03382 | 0.412776 | 0.675917 |
| GOBP_ORGANIC_HYDROXY_COMPOUND_TRANSPORT | 220 | -0.255792 | -1.033747 | 0.390995 | 0.675713 |
| GOBP_ORGAN_GROWTH | 135 | -0.273383 | -1.033604 | 0.402967 | 0.675715 |
| GOBP_LACTATION | 42 | -0.326129 | -1.032964 | 0.412371 | 0.677142 |
| GOBP_POSITIVE_REGULATION_OF_PEPTIDE_SECRETION | 132 | -0.271175 | -1.032802 | 0.409439 | 0.677153 |
| GOBP_REGULATION_OF_CELL_DEVELOPMENT | 434 | -0.243469 | -1.032617 | 0.390086 | 0.67726 |
| GOBP_POSITIVE_REGULATION_OF_CYTOSKELETON_ORGANIZATION | 209 | -0.256877 | -1.032533 | 0.389732 | 0.677089 |
| GOBP_REGULATION_OF_MICROTUBULE_BASED_MOVEMENT | 42 | -0.322565 | -1.032202 | 0.408955 | 0.677593 |
| GOBP_CEREBRAL_CORTEX_NEURON_DIFFERENTIATION | 16 | -0.403966 | -1.03212 | 0.425926 | 0.677399 |
| GOBP_INNATE_IMMUNE_RESPONSE_ACTIVATING_SIGNAL_TRANSDUCTION | 109 | -0.279983 | -1.031506 | 0.399471 | 0.678704 |
| GOBP_NEGATIVE_REGULATION_OF_RELEASE_OF_CYTOCHROME_C_FROM_MITOCHONDRIA | 17 | -0.396432 | -1.031469 | 0.420402 | 0.678382 |
| GOBP_PLATELET_ACTIVATION | 146 | -0.270034 | -1.03146 | 0.395782 | 0.67799 |
| GOBP_ACTION_POTENTIAL | 116 | -0.277642 | -1.031366 | 0.417323 | 0.677828 |
| GOBP_VESICLE_BUDDING_FROM_MEMBRANE | 107 | -0.280676 | -1.031229 | 0.417596 | 0.677777 |
| GOBP_DETECTION_OF_TEMPERATURE_STIMULUS | 18 | -0.38496 | -1.030699 | 0.424242 | 0.678768 |
| GOBP_STRIATED_MUSCLE_CONTRACTION | 151 | -0.268402 | -1.029932 | 0.399504 | 0.680457 |
| GOBP_CEREBELLAR_CORTEX_DEVELOPMENT | 37 | -0.335448 | -1.029841 | 0.404444 | 0.680275 |
| GOBP_SIGNAL_TRANSDUCTION_IN_ABSENCE_OF_LIGAND | 62 | -0.298794 | -1.029691 | 0.422192 | 0.68027 |
| GOBP_CELL_SUBSTRATE_ADHESION | 334 | -0.24695 | -1.029536 | 0.399333 | 0.680292 |
| GOBP_ENDOTHELIAL_CELL_CHEMOTAXIS | 26 | -0.357641 | -1.029484 | 0.414861 | 0.680019 |
| GOBP_REGULATION_OF_LIPASE_ACTIVITY | 84 | -0.289955 | -1.029331 | 0.411126 | 0.680037 |
| GOBP_RESPONSE_TO_ARSENIC_CONTAINING_SUBSTANCE | 30 | -0.344462 | -1.029199 | 0.426647 | 0.680003 |
| GOBP_POSITIVE_REGULATION_OF_PEPTIDASE_ACTIVITY | 183 | -0.263729 | -1.0289 | 0.42629 | 0.680412 |
| GOBP_MUSCLE_ADAPTATION | 95 | -0.283785 | -1.028776 | 0.397098 | 0.680337 |
| GOBP_REGULATION_OF_INTRINSIC_APOPTOTIC_SIGNALING_PATHWAY | 150 | -0.266229 | -1.027571 | 0.407767 | 0.683267 |
| GOBP_REGULATION_OF_SODIUM_ION_TRANSMEMBRANE_TRANSPORT | 58 | -0.305005 | -1.027246 | 0.414634 | 0.683769 |
| GOBP_POSITIVE_REGULATION_OF_POTASSIUM_ION_TRANSMEMBRANE_TRANSPORT | 32 | -0.343991 | -1.027047 | 0.447717 | 0.683907 |
| GOBP_RENAL_SYSTEM_PROCESS | 104 | -0.278236 | -1.027025 | 0.386928 | 0.683554 |
| GOBP_RESPONSE_TO_DOPAMINE | 63 | -0.305681 | -1.026987 | 0.430575 | 0.683239 |
| GOBP_REGULATION_OF_CARDIAC_MUSCLE_CELL_DIFFERENTIATION | 17 | -0.400574 | -1.026754 | 0.408321 | 0.683495 |
| GOBP_MEMBRANE_LIPID_CATABOLIC_PROCESS | 36 | -0.333043 | -1.025098 | 0.41411 | 0.687737 |
| GOBP_CENTRAL_NERVOUS_SYSTEM_NEURON_AXONOGENESIS | 33 | -0.342159 | -1.02505 | 0.436519 | 0.687454 |
| GOBP_CELLULAR_RESPONSE_TO_PROSTAGLANDIN_STIMULUS | 20 | -0.384919 | -1.024759 | 0.431988 | 0.687849 |
| GOBP_REGULATION_OF_RECEPTOR_MEDIATED_ENDOCYTOSIS | 94 | -0.282114 | -1.024618 | 0.404639 | 0.687832 |
| GOBP_NEGATIVE_REGULATION_OF_EPITHELIAL_CELL_MIGRATION | 68 | -0.294411 | -1.024479 | 0.41555 | 0.687782 |
| GOBP_POSITIVE_REGULATION_OF_STRESS_FIBER_ASSEMBLY | 50 | -0.316166 | -1.023924 | 0.43553 | 0.688936 |
| GOBP_ACTIN_MEDIATED_CELL_CONTRACTION | 106 | -0.276891 | -1.023709 | 0.418182 | 0.689143 |
| GOBP_GLYCOPROTEIN_METABOLIC_PROCESS | 375 | -0.243341 | -1.023557 | 0.408791 | 0.689138 |
| GOBP_CEREBELLAR_CORTEX_FORMATION | 18 | -0.379655 | -1.023402 | 0.433588 | 0.689165 |
| GOBP_NEUROTRANSMITTER_TRANSPORT | 182 | -0.259064 | -1.02275 | 0.4 | 0.690551 |
| GOBP_DRUG_CATABOLIC_PROCESS | 17 | -0.390877 | -1.022647 | 0.424806 | 0.69042 |
| GOBP_LOCALIZATION_WITHIN_MEMBRANE | 68 | -0.295571 | -1.022152 | 0.415869 | 0.691392 |
| GOBP_AORTA_MORPHOGENESIS | 27 | -0.354524 | -1.022138 | 0.413146 | 0.691017 |
| GOBP_NEGATIVE_REGULATION_OF_PROTEIN_LOCALIZATION_TO_CELL_PERIPHERY | 22 | -0.373954 | -1.02203 | 0.448889 | 0.690884 |
| GOBP_POSITIVE_REGULATION_OF_MRNA_CATABOLIC_PROCESS | 51 | -0.316674 | -1.022 | 0.41791 | 0.690553 |
| GOBP_NEGATIVE_REGULATION_OF_CYTOKINE_PRODUCTION_INVOLVED_IN_INFLAMMATORY_RE | 23 | -0.366255 | -1.021288 | 0.44288 | 0.692127 |
| GOBP_REGULATION_OF_GRANULOCYTE_CHEMOTAXIS | 45 | -0.319166 | -1.021195 | 0.449721 | 0.691956 |
| GOBP_NEGATIVE_REGULATION_OF_CELLULAR_RESPONSE_TO_INSULIN_STIMULUS | 36 | -0.334251 | -1.02072 | 0.428571 | 0.69284 |
| GOBP_PEPTIDYL_SERINE_MODIFICATION | 302 | -0.246795 | -1.020505 | 0.425989 | 0.693028 |
| GOBP_REGULATION_OF_MYELOID_LEUKOCYTE_MEDIATED_IMMUNITY | 50 | -0.311201 | -1.019908 | 0.433718 | 0.694321 |
| GOBP_REGULATION_OF_INTRACELLULAR_STEROID_HORMONE_RECEPTOR_SIGNALING_PATHW | 68 | -0.292208 | -1.019534 | 0.417687 | 0.694976 |
| GOBP_PIGMENT_GRANULE_ORGANIZATION | 27 | -0.353203 | -1.019491 | 0.430454 | 0.694673 |
| GOBP_REGULATION_OF_CARDIAC_CONDUCTION | 50 | -0.312845 | -1.019319 | 0.426778 | 0.694708 |
| GOBP_ASTROCYTE_DIFFERENTIATION | 67 | -0.29799 | -1.019137 | 0.43364 | 0.694785 |
| GOBP_MUSCLE_CELL_DEVELOPMENT | 138 | -0.268293 | -1.018851 | 0.4325 | 0.69517 |
| GOBP_DOPAMINE_TRANSPORT | 36 | -0.327475 | -1.018109 | 0.418021 | 0.696822 |
| GOBP_SPECIFICATION_OF_SYMMETRY | 106 | -0.274511 | -1.018094 | 0.449219 | 0.696444 |
| GOBP_POSITIVE_REGULATION_OF_PROTEIN_POLYMERIZATION | 126 | -0.273675 | -1.018071 | 0.407862 | 0.696087 |
| GOBP_REGULATION_OF_NUCLEOTIDE_METABOLIC_PROCESS | 111 | -0.272012 | -1.017899 | 0.435733 | 0.69614 |
| GOBP_CELLULAR_RESPONSE_TO_CAMP | 50 | -0.313408 | -1.016818 | 0.441958 | 0.698708 |

| | | | | | |
|---|---|---|---|---|---|
| GOBP_REGULATION_OF_SMALL_GTPASE_MEDIATED_SIGNAL_TRANSDUCTION | 310 | -0.246915 | -1.016419 | 0.430958 | 0.699396 |
| GOBP_POSITIVE_REGULATION_OF_OXIDOREDUCTASE_ACTIVITY | 51 | -0.311509 | -1.016253 | 0.431259 | 0.699455 |
| GOBP_TISSUE_REMODELING | 148 | -0.263282 | -1.016134 | 0.43494 | 0.699371 |
| GOBP_NLS_BEARING_PROTEIN_IMPORT_INTO_NUCLEUS | 18 | -0.385212 | -1.016125 | 0.433437 | 0.698983 |
| GOBP_NEGATIVE_REGULATION_OF_RESPONSE_TO_OXIDATIVE_STRESS | 19 | -0.383735 | -1.015845 | 0.442748 | 0.699348 |
| GOBP_RESPONSE_TO_THYROID_HORMONE | 23 | -0.363018 | -1.015656 | 0.439324 | 0.699467 |
| GOBP_CELL_JUNCTION_DISASSEMBLY | 20 | -0.379223 | -1.015631 | 0.461774 | 0.699121 |
| GOBP_MYOFIBRIL_ASSEMBLY | 49 | -0.311832 | -1.015573 | 0.451117 | 0.698877 |
| GOBP_ANATOMICAL_STRUCTURE_MATURATION | 189 | -0.257162 | -1.015445 | 0.426859 | 0.698809 |
| GOBP_REGULATION_OF_STEROL_TRANSPORT | 52 | -0.306588 | -1.01538 | 0.415746 | 0.698559 |
| GOBP_REGULATION_OF_DEVELOPMENTAL_GROWTH | 272 | -0.248074 | -1.015322 | 0.439498 | 0.698291 |
| GOBP_VASCULOGENESIS | 67 | -0.296122 | -1.014986 | 0.445215 | 0.698814 |
| GOBP_POSITIVE_REGULATION_OF_ENDOCYTOSIS | 91 | -0.278458 | -1.014772 | 0.445019 | 0.698999 |
| GOBP_FIBROBLAST_GROWTH_FACTOR_RECEPTOR_SIGNALING_PATHWAY | 94 | -0.28145 | -1.0143 | 0.438551 | 0.699943 |
| GOBP_ER_NUCLEUS_SIGNALING_PATHWAY | 52 | -0.306983 | -1.014277 | 0.434462 | 0.699596 |
| GOBP_OOCYTE_MATURATION | 23 | -0.365316 | -1.013595 | 0.454121 | 0.701095 |
| GOBP_SPLICEOSOMAL_SNRNP_ASSEMBLY | 50 | -0.310512 | -1.013204 | 0.432277 | 0.701757 |
| GOBP_RESPONSE_TO_ISOQUINOLINE_ALKALOID | 23 | -0.35727 | -1.012941 | 0.449102 | 0.702062 |
| GOBP_PHASIC_SMOOTH_MUSCLE_CONTRACTION | 18 | -0.381905 | -1.012428 | 0.446677 | 0.70309 |
| GOBP_RESPONSE_TO_PURINE_CONTAINING_COMPOUND | 135 | -0.265176 | -1.010633 | 0.435897 | 0.707675 |
| GOBP_RESPONSE_TO_OSMOTIC_STRESS | 76 | -0.288326 | -1.010326 | 0.456908 | 0.708122 |
| GOBP_POSITIVE_REGULATION_OF_PEPTIDYL_LYSINE_ACETYLATION | 32 | -0.339761 | -1.010266 | 0.451709 | 0.707865 |
| GOBP_NEGATIVE_REGULATION_OF_LIPID_METABOLIC_PROCESS | 80 | -0.285301 | -1.010003 | 0.434663 | 0.708188 |
| GOBP_POSITIVE_REGULATION_OF_DEPHOSPHORYLATION | 56 | -0.301751 | -1.009544 | 0.440225 | 0.709037 |
| GOBP_REGULATION_OF_FEEDING_BEHAVIOR | 19 | -0.384663 | -1.009345 | 0.452229 | 0.709156 |
| GOBP_MOTOR_NEURON_AXON_GUIDANCE | 28 | -0.34367 | -1.009153 | 0.432927 | 0.709243 |
| GOBP_REGULATION_OF_DNA_METHYLATION | 20 | -0.374807 | -1.009141 | 0.440495 | 0.708869 |
| GOBP_NEGATIVE_REGULATION_OF_SIGNAL_TRANSDUCTION_IN_ABSENCE_OF_LIGAND | 29 | -0.344616 | -1.0091 | 0.451613 | 0.708564 |
| GOBP_REGULATION_OF_ENDOTHELIAL_CELL_MIGRATION | 165 | -0.259958 | -1.008928 | 0.449398 | 0.708642 |
| GOBP_CELLULAR_RESPONSE_TO_ESTRADIOL_STIMULUS | 31 | -0.340302 | -1.008635 | 0.461427 | 0.709013 |
| GOBP_RESPONSE_TO_INTERFERON_ALPHA | 21 | -0.374678 | -1.00858 | 0.448171 | 0.708746 |
| GOBP_GAMMA_AMINOBUTYRIC_ACID_SIGNALING_PATHWAY | 17 | -0.384856 | -1.008382 | 0.436066 | 0.708894 |
| GOBP_POSITIVE_REGULATION_OF_MACROPHAGE_DERIVED_FOAM_CELL_DIFFERENTIATION | 16 | -0.403442 | -1.008308 | 0.470588 | 0.708689 |
| GOBP_REGULATION_OF_INSULIN_RECEPTOR_SIGNALING_PATHWAY | 63 | -0.293509 | -1.008277 | 0.464828 | 0.708378 |
| GOBP_BODY_MORPHOGENESIS | 45 | -0.317019 | -1.008229 | 0.439655 | 0.708093 |
| GOBP_NEGATIVE_REGULATION_OF_LYASE_ACTIVITY | 22 | -0.369836 | -1.008054 | 0.436137 | 0.70815 |
| GOBP_PHOSPHATIDYLCHOLINE_METABOLIC_PROCESS | 79 | -0.285893 | -1.007994 | 0.437677 | 0.707902 |
| GOBP_POSITIVE_REGULATION_OF_BLOOD_CIRCULATION | 32 | -0.335978 | -1.007993 | 0.459168 | 0.707496 |
| GOBP_NEGATIVE_REGULATION_OF_COAGULATION | 44 | -0.314497 | -1.007867 | 0.452586 | 0.70744 |
| GOBP_ORGANISM_EMERGENCE_FROM_PROTECTIVE_STRUCTURE | 21 | -0.371735 | -1.007824 | 0.452915 | 0.707149 |
| GOBP_NEGATIVE_REGULATION_OF_LEUKOCYTE_APOPTOTIC_PROCESS | 44 | -0.315919 | -1.007604 | 0.446198 | 0.707367 |
| GOBP_CARBOHYDRATE_DERIVATIVE_TRANSPORT | 77 | -0.287016 | -1.007591 | 0.44474 | 0.706986 |
| GOBP_MRNA_SPLICE_SITE_SELECTION | 39 | -0.322615 | -1.007365 | 0.460641 | 0.707179 |
| GOBP_KILLING_OF_CELLS_IN_OTHER_ORGANISM_INVOLVED_IN_SYMBIOTIC_INTERACTION | 17 | -0.396722 | -1.006991 | 0.462145 | 0.707777 |
| GOBP_RESPONSE_TO_PEPTIDE | 467 | -0.236162 | -1.006868 | 0.43871 | 0.707716 |
| GOBP_POSITIVE_REGULATION_OF_LIPID_TRANSPORT | 78 | -0.284066 | -1.005818 | 0.444444 | 0.710216 |
| GOBP_TRANSCYTOSIS | 19 | -0.37551 | -1.004817 | 0.448855 | 0.712616 |
| GOBP_ORGANIC_ACID_TRANSPORT | 273 | -0.245478 | -1.004742 | 0.479452 | 0.712411 |
| GOBP_MATING_BEHAVIOR | 16 | -0.390695 | -1.004666 | 0.453271 | 0.712199 |
| GOBP_CELLULAR_RESPONSE_TO_STEROL | 22 | -0.362656 | -1.004571 | 0.452239 | 0.712042 |
| GOBP_ACTIVIN_RECEPTOR_SIGNALING_PATHWAY | 36 | -0.322831 | -1.004563 | 0.46176 | 0.711651 |
| GOBP_SKIN_DEVELOPMENT | 244 | -0.249276 | -1.004543 | 0.443801 | 0.711297 |
| GOBP_FATTY_ACID_DERIVATIVE_CATABOLIC_PROCESS | 15 | -0.40094 | -1.004537 | 0.465927 | 0.710897 |
| GOBP_CELL_COMMUNICATION_INVOLVED_IN_CARDIAC_CONDUCTION | 50 | -0.30765 | -1.00434 | 0.450139 | 0.711011 |
| GOBP_CAMP_METABOLIC_PROCESS | 18 | -0.375856 | -1.004008 | 0.452381 | 0.711502 |
| GOBP_JNK_CASCADE | 181 | -0.255836 | -1.003051 | 0.463768 | 0.713685 |
| GOBP_DOPAMINE_RECEPTOR_SIGNALING_PATHWAY | 35 | -0.325523 | -1.002777 | 0.456456 | 0.714052 |
| GOBP_MEMORY | 88 | -0.280822 | -1.00227 | 0.465426 | 0.715048 |
| GOBP_MAMMARY_GLAND_EPITHELIAL_CELL_PROLIFERATION | 26 | -0.346745 | -1.002098 | 0.466767 | 0.7151 |
| GOBP_NEGATIVE_REGULATION_OF_CHONDROCYTE_DIFFERENTIATION | 17 | -0.380535 | -1.002092 | 0.465753 | 0.7147 |
| GOBP_VITAMIN_TRANSPORT | 36 | -0.331385 | -1.001539 | 0.480463 | 0.715824 |
| GOBP_REGULATION_OF_ACTIN_FILAMENT_BASED_PROCESS | 362 | -0.241315 | -1.001512 | 0.467706 | 0.715481 |
| GOBP_EMBRYONIC_FORELIMB_MORPHOGENESIS | 25 | -0.353089 | -1.001258 | 0.464768 | 0.715788 |
| GOBP_VESICLE_TARGETING | 94 | -0.275947 | -1.001171 | 0.464052 | 0.715622 |
| GOBP_PODOSOME_ASSEMBLY | 16 | -0.385062 | -1.001144 | 0.464789 | 0.715284 |
| GOBP_POSITIVE_REGULATION_OF_ESTABLISHMENT_OF_PROTEIN_LOCALIZATION | 309 | -0.243768 | -1.001011 | 0.476244 | 0.715212 |
| GOBP_LAMELLIPODIUM_ORGANIZATION | 84 | -0.280509 | -1.000841 | 0.466408 | 0.715251 |
| GOBP_DETECTION_OF_ABIOTIC_STIMULUS | 114 | -0.26798 | -1.000707 | 0.447402 | 0.715187 |
| GOBP_SEROTONIN_RECEPTOR_SIGNALING_PATHWAY | 17 | -0.389741 | -0.99995 | 0.459752 | 0.716894 |
| GOBP_RHYTHMIC_PROCESS | 262 | -0.248201 | -0.999772 | 0.478507 | 0.71692 |
| GOBP_CARDIOCYTE_DIFFERENTIATION | 111 | -0.270037 | -0.99969 | 0.459948 | 0.716729 |
| GOBP_NEGATIVE_REGULATION_OF_CELL_SUBSTRATE_ADHESION | 58 | -0.297048 | -0.999263 | 0.463415 | 0.717482 |
| GOBP_MODULATION_OF_EXCITATORY_POSTSYNAPTIC_POTENTIAL | 40 | -0.314537 | -0.998919 | 0.478382 | 0.717965 |
| GOBP_RESPONSE_TO_HYDROGEN_PEROXIDE | 123 | -0.266867 | -0.998745 | 0.466667 | 0.71804 |
| GOBP_RESPONSE_TO_ANGIOTENSIN | 22 | -0.361314 | -0.998012 | 0.459337 | 0.719597 |
| GOBP_NEUROBLAST_PROLIFERATION | 45 | -0.310295 | -0.997847 | 0.468443 | 0.71964 |
| GOBP_TEMPERATURE_HOMEOSTASIS | 161 | -0.260396 | -0.99764 | 0.473236 | 0.719798 |
| GOBP_THYMUS_DEVELOPMENT | 40 | -0.318904 | -0.997579 | 0.458993 | 0.719569 |
| GOBP_STRIATED_MUSCLE_CELL_APOPTOTIC_PROCESS | 31 | -0.335678 | -0.997496 | 0.459736 | 0.719378 |
| GOBP_REGULATION_OF_WNT_SIGNALING_PATHWAY | 343 | -0.241147 | -0.996984 | 0.489083 | 0.72038 |
| GOBP_HEPARAN_SULFATE_PROTEOGLYCAN_METABOLIC_PROCESS | 29 | -0.339667 | -0.996764 | 0.464392 | 0.72058 |
| GOBP_ACTIN_FILAMENT_BASED_MOVEMENT | 136 | -0.263034 | -0.996029 | 0.48289 | 0.722143 |
| GOBP_REGULATION_OF_LONG_TERM_NEURONAL_SYNAPTIC_PLASTICITY | 19 | -0.366874 | -0.996004 | 0.467615 | 0.721799 |
| GOBP_VIRAL_GENOME_REPLICATION | 125 | -0.263425 | -0.995143 | 0.472329 | 0.723723 |
| GOBP_REGULATION_OF_STEROID_METABOLIC_PROCESS | 101 | -0.26724 | -0.994901 | 0.470052 | 0.723991 |
| GOBP_3_UTR_MEDIATED_MRNA_STABILIZATION | 19 | -0.366874 | -0.994884 | 0.47438 | 0.723627 |
| GOBP_POSTSYNAPSE_ORGANIZATION | 148 | -0.260167 | -0.994733 | 0.471769 | 0.723616 |
| GOBP_POSITIVE_REGULATION_OF_EPITHELIAL_CELL_PROLIFERATION | 167 | -0.256257 | -0.994533 | 0.493268 | 0.723734 |
| GOBP_FACE_DEVELOPMENT | 42 | -0.312855 | -0.994508 | 0.460432 | 0.723389 |
| GOBP_REGULATION_OF_LIPID_BIOSYNTHETIC_PROCESS | 173 | -0.255113 | -0.993896 | 0.4914 | 0.72461 |
| GOBP_EYE_MORPHOGENESIS | 114 | -0.268253 | -0.993339 | 0.486284 | 0.725736 |
| GOBP_RESPONSE_TO_CATECHOLAMINE | 81 | -0.278963 | -0.99314 | 0.478969 | 0.725885 |
| GOBP_ORGANIC_ACID_BIOSYNTHETIC_PROCESS | 291 | -0.24118 | -0.992815 | 0.482143 | 0.726369 |
| GOBP_RESPONSE_TO_LEUKEMIA_INHIBITORY_FACTOR | 77 | -0.282081 | -0.991966 | 0.464286 | 0.72831 |
| GOBP_ANTIBACTERIAL_HUMORAL_RESPONSE | 44 | -0.314667 | -0.991882 | 0.453297 | 0.72812 |
| GOBP_REGULATION_OF_PHOSPHOLIPID_METABOLIC_PROCESS | 32 | -0.327376 | -0.991405 | 0.474702 | 0.729025 |
| GOBP_TERPENOID_METABOLIC_PROCESS | 92 | -0.269494 | -0.991378 | 0.470669 | 0.728695 |
| GOBP_CELLULAR_RESPONSE_TO_ARSENIC_CONTAINING_SUBSTANCE | 18 | -0.381086 | -0.991162 | 0.462848 | 0.72884 |
| GOBP_POSITIVE_REGULATION_OF_GRANULOCYTE_CHEMOTAXIS | 25 | -0.347104 | -0.990946 | 0.474398 | 0.729007 |
| GOBP_CELL_JUNCTION_MAINTENANCE | 29 | -0.337953 | -0.990649 | 0.469512 | 0.729388 |
| GOBP_BRANCHING_INVOLVED_IN_SALIVARY_GLAND_MORPHOGENESIS | 17 | -0.374651 | -0.990348 | 0.474016 | 0.729776 |
| GOBP_NEGATIVE_REGULATION_OF_INTRACELLULAR_STEROID_HORMONE_RECEPTOR_SIGNALI | 32 | -0.324691 | -0.990114 | 0.468795 | 0.730008 |
| GOBP_NEURON_DEATH | 302 | -0.239964 | -0.989991 | 0.493348 | 0.729933 |
| GOBP_BONE_CELL_DEVELOPMENT | 32 | -0.327482 | -0.989931 | 0.485496 | 0.729688 |
| GOBP_REGULATION_OF_LEUKOCYTE_DEGRANULATION | 41 | -0.316497 | -0.989737 | 0.478832 | 0.729814 |
| GOBP_REGULATION_OF_CALCIUM_ION_TRANSPORT_INTO_CYTOSOL | 83 | -0.274346 | -0.989682 | 0.484646 | 0.72954 |
| GOBP_CARDIAC_MUSCLE_CELL_ACTION_POTENTIAL | 68 | -0.28375 | -0.989298 | 0.486072 | 0.730177 |
| GOBP_LABYRINTHINE_LAYER_DEVELOPMENT | 37 | -0.327801 | -0.989236 | 0.483357 | 0.729946 |
| GOBP_POSITIVE_REGULATION_OF_ENDOTHELIAL_CELL_MIGRATION | 107 | -0.266277 | -0.98871 | 0.479221 | 0.730982 |
| GOBP_CELL_CELL_SIGNALING_BY_WNT | 484 | -0.231499 | -0.988253 | 0.52099 | 0.731794 |
| GOBP_POSITIVE_REGULATION_OF_INSULIN_SECRETION_INVOLVED_IN_CELLULAR_RESPONSE_ | 30 | -0.338013 | -0.987834 | 0.479876 | 0.732524 |
| GOBP_MULTI_ORGANISM_CELLULAR_PROCESS | 16 | -0.384371 | -0.987286 | 0.490385 | 0.73364 |
| GOBP_REGULATION_OF_ENDOCYTOSIS | 191 | -0.251117 | -0.987119 | 0.483986 | 0.733664 |
| GOBP_ENDODERM_DEVELOPMENT | 69 | -0.290478 | -0.9871 | 0.47019 | 0.733314 |
| GOBP_ORGANELLE_MEMBRANE_FUSION | 106 | -0.266467 | -0.987091 | 0.469755 | 0.732931 |
| GOBP_GLYCOPROTEIN_CATABOLIC_PROCESS | 23 | -0.357175 | -0.9869 | 0.482143 | 0.733028 |
| GOBP_INTERLEUKIN_13_PRODUCTION | 19 | -0.368642 | -0.986828 | 0.487961 | 0.732816 |
| GOBP_REGULATION_OF_EMBRYONIC_DEVELOPMENT | 54 | -0.295387 | -0.986487 | 0.503526 | 0.733309 |
| GOBP_IMPORT_INTO_NUCLEUS | 155 | -0.254461 | -0.986368 | 0.483051 | 0.733222 |
| GOBP_REGULATION_OF_MAST_CELL_ACTIVATION_INVOLVED_IN_IMMUNE_RESPONSE | 28 | -0.343078 | -0.986249 | 0.464122 | 0.733116 |
| GOBP_RESPONSE_TO_MONOSACCHARIDE | 173 | -0.252403 | -0.986044 | 0.500602 | 0.733251 |
| GOBP_CELLULAR_RESPONSE_TO_CHOLESTEROL | 17 | -0.377166 | -0.986035 | 0.493691 | 0.732872 |
| GOBP_PHOSPHATIDYLETHANOLAMINE_ACYL_CHAIN_REMODELING | 19 | -0.370975 | -0.986019 | 0.49403 | 0.73251 |
| GOBP_RESPONSE_TO_DRUG | 320 | -0.240112 | -0.985892 | 0.510734 | 0.732439 |
| GOBP_REGULATION_OF_SYNAPTIC_PLASTICITY | 157 | -0.260688 | -0.985778 | 0.496124 | 0.732321 |
| GOBP_CARDIAC_MUSCLE_CELL_PROLIFERATION | 44 | -0.303051 | -0.985575 | 0.483595 | 0.732479 |
| GOBP_REGULATION_OF_BONE_MINERALIZATION | 68 | -0.285119 | -0.984834 | 0.49858 | 0.734048 |
| GOBP_RESPONSE_TO_BMP | 142 | -0.257824 | -0.98481 | 0.503106 | 0.733708 |
| GOBP_TRANSEPITHELIAL_TRANSPORT | 28 | -0.339489 | -0.984663 | 0.487062 | 0.733706 |
| GOBP_REGULATION_OF_HEART_RATE | 80 | -0.281134 | -0.984572 | 0.488801 | 0.73355 |
| GOBP_PLASMINOGEN_ACTIVATION | 18 | -0.376791 | -0.984083 | 0.485531 | 0.734461 |
| GOBP_REGULATION_OF_TRANSCRIPTION_ELONGATION_FROM_RNA_POLYMERASE_II_PROMOTE | 30 | -0.336679 | -0.9835 | 0.472347 | 0.735598 |
| GOBP_NEGATIVE_REGULATION_OF_ESTABLISHMENT_OF_PROTEIN_LOCALIZATION | 114 | -0.26451 | -0.983133 | 0.491294 | 0.736154 |
| GOBP_NEGATIVE_REGULATION_OF_GTPASE_ACTIVITY | 43 | -0.308554 | -0.982816 | 0.48927 | 0.736583 |
| GOBP_POSITIVE_REGULATION_OF_LIPID_KINASE_ACTIVITY | 35 | -0.324455 | -0.982725 | 0.482249 | 0.736421 |
| GOBP_PYRIMIDINE_NUCLEOSIDE_MONOPHOSPHATE_METABOLIC_PROCESS | 18 | -0.369045 | -0.982626 | 0.486312 | 0.736284 |
| GOBP_POSTSYNAPTIC_SIGNAL_TRANSDUCTION | 28 | -0.34198 | -0.982488 | 0.494799 | 0.736254 |
| GOBP_POSITIVE_REGULATION_OF_INTRACELLULAR_PROTEIN_TRANSPORT | 174 | -0.249595 | -0.982465 | 0.526253 | 0.735911 |
| GOBP_BENZENE_CONTAINING_COMPOUND_METABOLIC_PROCESS | 22 | -0.355106 | -0.982151 | 0.495549 | 0.736363 |
| GOBP_POSITIVE_REGULATION_OF_LIPID_LOCALIZATION | 98 | -0.271342 | -0.981846 | 0.497436 | 0.736791 |
| GOBP_CERAMIDE_CATABOLIC_PROCESS | 20 | -0.365014 | -0.981252 | 0.485075 | 0.738005 |
| GOBP_REGULATION_OF_NEUROTRANSMITTER_LEVELS | 189 | -0.248103 | -0.981207 | 0.513889 | 0.737726 |
| GOBP_VESICLE_TARGETING_TO_FROM_OR_WITHIN_GOLGI | 73 | -0.277741 | -0.981196 | 0.480641 | 0.737353 |
| GOBP_INOSITOL_PHOSPHATE_MEDIATED_SIGNALING | 47 | -0.298209 | -0.980882 | 0.487069 | 0.737759 |
| GOBP_PROGRAMMED_NECROTIC_CELL_DEATH | 45 | -0.305037 | -0.980764 | 0.516987 | 0.737656 |
| GOBP_CELL_CELL_JUNCTION_ORGANIZATION | 186 | -0.250859 | -0.980694 | 0.509047 | 0.737444 |
| GOBP_NEGATIVE_REGULATION_OF_RESPONSE_TO_WOUNDING | 75 | -0.28008 | -0.980542 | 0.478854 | 0.73742 |
| GOBP_REGULATION_OF_PROTEIN_SERINE_THREONINE_KINASE_ACTIVITY | 455 | -0.229932 | -0.980236 | 0.550586 | 0.737809 |
| GOBP_ASTROCYTE_DEVELOPMENT | 38 | -0.310188 | -0.979986 | 0.491971 | 0.738094 |
| GOBP_OSTEOBLAST_DIFFERENTIATION | 198 | -0.245309 | -0.97973 | 0.5 | 0.738411 |
| GOBP_RESPONSE_TO_LEAD_ION | 20 | -0.357483 | -0.979571 | 0.49923 | 0.738419 |
| GOBP_NEGATIVE_REGULATION_OF_GLIOGENESIS | 30 | -0.336382 | -0.979098 | 0.475038 | 0.739275 |
| GOBP_DEVELOPMENTAL_MATURATION | 237 | -0.243805 | -0.978982 | 0.54748 | 0.739168 |
| GOBP_LEUKOTRIENE_METABOLIC_PROCESS | 27 | -0.333806 | -0.978942 | 0.507396 | 0.738863 |
| GOBP_PROTEIN_HOMOOLIGOMERIZATION | 151 | -0.252032 | -0.978644 | 0.507519 | 0.739225 |
| GOBP_POSITIVE_REGULATION_OF_T_CELL_MIGRATION | 28 | -0.339465 | -0.977512 | 0.500734 | 0.741823 |
| GOBP_NEGATIVE_REGULATION_OF_CARDIAC_MUSCLE_TISSUE_GROWTH | 22 | -0.358244 | -0.977295 | 0.494118 | 0.742021 |
| GOBP_RESPIRATORY_GASEOUS_EXCHANGE_BY_RESPIRATORY_SYSTEM | 58 | -0.288119 | -0.977233 | 0.502052 | 0.741774 |
| GOBP_TISSUE_HOMEOSTASIS | 225 | -0.241525 | -0.976922 | 0.518343 | 0.742184 |
| GOBP_TELENCEPHALON_DEVELOPMENT | 199 | -0.24576 | -0.976121 | 0.519048 | 0.743914 |
| GOBP_ACTIVATION_OF_INNATE_IMMUNE_RESPONSE | 137 | -0.256118 | -0.975996 | 0.511628 | 0.743835 |
| GOBP_VACUOLE_ORGANIZATION | 169 | -0.247965 | -0.975223 | 0.537241 | 0.745562 |
| GOBP_REGULATION_OF_SUPRAMOLECULAR_FIBER_ORGANIZATION | 348 | -0.232757 | -0.974822 | 0.552141 | 0.746237 |
| GOBP_CELLULAR_RESPONSE_TO_STEROID_HORMONE_STIMULUS | 182 | -0.245559 | -0.974211 | 0.510638 | 0.74744 |
| GOBP_NEGATIVE_REGULATION_OF_FIBROBLAST_PROLIFERATION | 28 | -0.337172 | -0.974038 | 0.514593 | 0.747504 |
| GOBP_PEPTIDE_CROSS_LINKING | 27 | -0.33705 | -0.973119 | 0.516535 | 0.749527 |
| GOBP_STRIATED_MUSCLE_CELL_PROLIFERATION | 57 | -0.289484 | -0.973037 | 0.495225 | 0.749338 |
| GOBP_POSITIVE_REGULATION_OF_NATURAL_KILLER_CELL_ACTIVATION | 20 | -0.362582 | -0.972539 | 0.505935 | 0.750229 |

| | | | | | |
|---|---|---|---|---|---|
| GOBP_AROMATIC_AMINO_ACID_FAMILY_METABOLIC_PROCESS | 28 | -0.336481 | -0.971754 | 0.486446 | 0.751852 |
| GOBP_METENCEPHALON_DEVELOPMENT | 85 | -0.273082 | -0.970706 | 0.523179 | 0.754196 |
| GOBP_POSITIVE_REGULATION_OF_INTRACELLULAR_TRANSPORT | 211 | -0.245693 | -0.970558 | 0.532622 | 0.754194 |
| GOBP_SYNAPTIC_VESICLE_CYTOSKELETAL_TRANSPORT | 19 | -0.360792 | -0.970169 | 0.506231 | 0.754831 |
| GOBP_MALE_SEX_DIFFERENTIATION | 130 | -0.255592 | -0.969399 | 0.53 | 0.756445 |
| GOBP_HEAD_MORPHOGENESIS | 33 | -0.316884 | -0.969154 | 0.524664 | 0.756682 |
| GOBP_EMBRYONIC_HEART_TUBE_DEVELOPMENT | 65 | -0.28053 | -0.967419 | 0.521622 | 0.760851 |
| GOBP_REGULATION_OF_DENDRITIC_SPINE_MORPHOGENESIS | 45 | -0.301364 | -0.967209 | 0.495714 | 0.760994 |
| GOBP_ACTIN_POLYMERIZATION_OR_DEPOLYMERIZATION | 207 | -0.241378 | -0.966753 | 0.5549 | 0.761829 |
| GOBP_ENDOCARDIAL_CUSHION_FORMATION | 21 | -0.353113 | -0.966634 | 0.496124 | 0.761715 |
| GOBP_REGULATION_OF_ACTIN_FILAMENT_LENGTH | 178 | -0.246794 | -0.966163 | 0.552381 | 0.762554 |
| GOBP_SKELETAL_SYSTEM_DEVELOPMENT | 429 | -0.229075 | -0.966024 | 0.594391 | 0.762512 |
| GOBP_DOSAGE_COMPENSATION | 17 | -0.373363 | -0.965771 | 0.515106 | 0.762764 |
| GOBP_GLIAL_CELL_DEVELOPMENT | 104 | -0.259728 | -0.964797 | 0.544061 | 0.764917 |
| GOBP_ACYLGLYCEROL_HOMEOSTASIS | 27 | -0.329529 | -0.964193 | 0.498507 | 0.76612 |
| GOBP_REGULATION_OF_PH | 98 | -0.265372 | -0.963571 | 0.51774 | 0.767346 |
| GOBP_EATING_BEHAVIOR | 22 | -0.346747 | -0.963563 | 0.514925 | 0.766953 |
| GOBP_MODIFICATION_OF_SYNAPTIC_STRUCTURE | 19 | -0.363311 | -0.963419 | 0.509091 | 0.766908 |
| GOBP_CELLULAR_RESPONSE_TO_CARBOHYDRATE_STIMULUS | 122 | -0.254283 | -0.962763 | 0.541087 | 0.768231 |
| GOBP_POSITIVE_REGULATION_OF_ORGAN_GROWTH | 43 | -0.301564 | -0.96192 | 0.530172 | 0.770089 |
| GOBP_POSITIVE_REGULATION_OF_CELL_GROWTH | 146 | -0.249816 | -0.961766 | 0.556242 | 0.770075 |
| GOBP_REGULATION_OF_ACTIVIN_RECEPTOR_SIGNALING_PATHWAY | 21 | -0.349596 | -0.9617 | 0.527439 | 0.769825 |
| GOBP_ESTABLISHMENT_OR_MAINTENANCE_OF_BIPOLAR_CELL_POLARITY | 43 | -0.301514 | -0.961141 | 0.518678 | 0.770851 |
| GOBP_NEGATIVE_REGULATION_OF_ENDOTHELIAL_CELL_APOPTOTIC_PROCESS | 28 | -0.332042 | -0.960759 | 0.51194 | 0.771423 |
| GOBP_NEGATIVE_REGULATION_OF_CANONICAL_WNT_SIGNALING_PATHWAY | 166 | -0.245541 | -0.959792 | 0.563791 | 0.773533 |
| GOBP_FIBROBLAST_PROLIFERATION | 73 | -0.276261 | -0.959306 | 0.537063 | 0.77439 |
| GOBP_VIRAL_GENE_EXPRESSION | 198 | -0.240369 | -0.958237 | 0.586826 | 0.776754 |
| GOBP_RESPONSE_TO_AMPHETAMINE | 27 | -0.330296 | -0.95651 | 0.531479 | 0.780983 |
| GOBP_ENDOPLASMIC_RETICULUM_UNFOLDED_PROTEIN_RESPONSE | 123 | -0.255314 | -0.95615 | 0.557908 | 0.781507 |
| GOBP_INOSITOL_LIPID_MEDIATED_SIGNALING | 179 | -0.242673 | -0.955883 | 0.557298 | 0.781776 |
| GOBP_CELLULAR_COMPONENT_MAINTENANCE | 55 | -0.2855 | -0.955423 | 0.539919 | 0.782567 |
| GOBP_POSITIVE_REGULATION_OF_LEUKOCYTE_DEGRANULATION | 22 | -0.347254 | -0.95536 | 0.5125 | 0.782313 |
| GOBP_REGULATION_OF_GTPASE_ACTIVITY | 456 | -0.225758 | -0.954794 | 0.61828 | 0.783371 |
| GOBP_PROTEIN_HETEROOLIGOMERIZATION | 22 | -0.341937 | -0.954284 | 0.52795 | 0.78434 |
| GOBP_MICROTUBULE_BUNDLE_FORMATION | 97 | -0.258755 | -0.954283 | 0.55291 | 0.783926 |
| GOBP_EMBRYONIC_APPENDAGE_MORPHOGENESIS | 94 | -0.267136 | -0.953631 | 0.559413 | 0.785218 |
| GOBP_CELL_CELL_RECOGNITION | 60 | -0.285523 | -0.953302 | 0.536111 | 0.785619 |
| GOBP_LIPID_HOMEOSTASIS | 134 | -0.251894 | -0.953254 | 0.575377 | 0.785333 |
| GOBP_PHOSPHATIDYLSERINE_ACYL_CHAIN_REMODELING | 16 | -0.36764 | -0.953137 | 0.530488 | 0.785248 |
| GOBP_REGULATION_OF_COMPLEMENT_ACTIVATION | 103 | -0.258615 | -0.952647 | 0.560265 | 0.7861 |
| GOBP_NEGATIVE_REGULATION_OF_CELL_GROWTH | 171 | -0.243056 | -0.952531 | 0.56401 | 0.785967 |
| GOBP_PHAGOCYTOSIS_RECOGNITION | 81 | -0.270706 | -0.952397 | 0.55496 | 0.785903 |
| GOBP_REGULATION_OF_CELL_SUBSTRATE_ADHESION | 198 | -0.239126 | -0.952365 | 0.592899 | 0.785563 |
| GOBP_POLYOL_METABOLIC_PROCESS | 122 | -0.25244 | -0.952201 | 0.557478 | 0.785584 |
| GOBP_ENERGY_RESERVE_METABOLIC_PROCESS | 79 | -0.267553 | -0.952055 | 0.534301 | 0.785567 |
| GOBP_PINOCYTOSIS | 22 | -0.346918 | -0.95194 | 0.5117 | 0.785463 |
| GOBP_POSITIVE_REGULATION_OF_PROTEIN_LOCALIZATION_TO_MEMBRANE | 122 | -0.253756 | -0.950931 | 0.554278 | 0.787679 |
| GOBP_ACTIN_NUCLEATION | 50 | -0.294348 | -0.950574 | 0.554131 | 0.788212 |
| GOBP_IONOTROPIC_GLUTAMATE_RECEPTOR_SIGNALING_PATHWAY | 21 | -0.345934 | -0.950304 | 0.533011 | 0.788489 |
| GOBP_DNA_DAMAGE_RESPONSE_SIGNAL_TRANSDUCTION_RESULTING_IN_TRANSCRIPTION | 18 | -0.35437 | -0.950169 | 0.553159 | 0.788406 |
| GOBP_REGULATION_OF_IMMUNOGLOBULIN_PRODUCTION | 66 | -0.279505 | -0.950132 | 0.544966 | 0.788091 |
| GOBP_POSITIVE_REGULATION_OF_DNA_TEMPLATED_TRANSCRIPTION_ELONGATION | 26 | -0.333964 | -0.949534 | 0.547511 | 0.789252 |
| GOBP_REGULATION_OF_NEUROGENESIS | 311 | -0.230084 | -0.949242 | 0.615642 | 0.789591 |
| GOBP_PHOSPHATIDYLETHANOLAMINE_METABOLIC_PROCESS | 28 | -0.325466 | -0.94922 | 0.533752 | 0.789241 |
| GOBP_COLLAGEN_METABOLIC_PROCESS | 94 | -0.25997 | -0.949136 | 0.55102 | 0.789032 |
| GOBP_REGULATION_OF_CARDIAC_MUSCLE_CELL_ACTION_POTENTIAL | 24 | -0.335995 | -0.949111 | 0.523881 | 0.788689 |
| GOBP_REGULATION_OF_BROWN_FAT_CELL_DIFFERENTIATION | 19 | -0.358686 | -0.948356 | 0.548686 | 0.790286 |
| GOBP_AORTIC_VALVE_MORPHOGENESIS | 28 | -0.327673 | -0.947613 | 0.528975 | 0.791793 |
| GOBP_MESENCHYMAL_CELL_PROLIFERATION | 33 | -0.31225 | -0.94711 | 0.550445 | 0.792622 |
| GOBP_DERMATAN_SULFATE_PROTEOGLYCAN_METABOLIC_PROCESS | 16 | -0.36921 | -0.946749 | 0.543779 | 0.793129 |
| GOBP_CARDIAC_MUSCLE_CONTRACTION | 117 | -0.254581 | -0.946702 | 0.579767 | 0.792826 |
| GOBP_INTRINSIC_APOPTOTIC_SIGNALING_PATHWAY | 271 | -0.234166 | -0.946624 | 0.628604 | 0.79262 |
| GOBP_REGULATION_OF_ORGAN_GROWTH | 82 | -0.266545 | -0.945614 | 0.569191 | 0.794841 |
| GOBP_POSITIVE_REGULATION_OF_TRANSPORTER_ACTIVITY | 101 | -0.261669 | -0.94553 | 0.585752 | 0.794643 |
| GOBP_POSITIVE_REGULATION_OF_GTPASE_ACTIVITY | 383 | -0.22456 | -0.945447 | 0.634973 | 0.794428 |
| GOBP_CELL_DIFFERENTIATION_INVOLVED_IN_METANEPHROS_DEVELOPMENT | 17 | -0.365926 | -0.945256 | 0.549317 | 0.794489 |
| GOBP_NEGATIVE_REGULATION_OF_PROTEIN_DEPHOSPHORYLATION | 61 | -0.280322 | -0.945149 | 0.568871 | 0.79435 |
| GOBP_REGULATION_OF_CHOLESTEROL_EFFLUX | 32 | -0.313101 | -0.945083 | 0.545588 | 0.794108 |
| GOBP_NOTCH_SIGNALING_PATHWAY | 171 | -0.243585 | -0.944875 | 0.622892 | 0.794237 |
| GOBP_NEGATIVE_REGULATION_OF_NERVOUS_SYSTEM_DEVELOPMENT | 120 | -0.252445 | -0.944747 | 0.580124 | 0.794158 |
| GOBP_FATTY_ACID_METABOLIC_PROCESS | 344 | -0.226492 | -0.943827 | 0.660419 | 0.796098 |
| GOBP_REGULATION_OF_SYNAPTIC_TRANSMISSION_GLUTAMATERGIC | 55 | -0.278599 | -0.943595 | 0.551375 | 0.796269 |
| GOBP_FC_EPSILON_RECEPTOR_SIGNALING_PATHWAY | 165 | -0.243064 | -0.942918 | 0.602941 | 0.797588 |
| GOBP_NEGATIVE_REGULATION_OF_SMALL_GTPASE_MEDIATED_SIGNAL_TRANSDUCTION | 55 | -0.280979 | -0.942466 | 0.558172 | 0.798366 |
| GOBP_BETA_CATENIN_TCF_COMPLEX_ASSEMBLY | 31 | -0.316908 | -0.941623 | 0.562054 | 0.80016 |
| GOBP_ESTABLISHMENT_OF_PROTEIN_LOCALIZATION_TO_MEMBRANE | 342 | -0.226492 | -0.940609 | 0.637569 | 0.802327 |
| GOBP_PROTEIN_LOCALIZATION_TO_VACUOLE | 65 | -0.277876 | -0.940244 | 0.567204 | 0.802861 |
| GOBP_FILOPODIUM_ASSEMBLY | 60 | -0.279158 | -0.939983 | 0.588811 | 0.803103 |
| GOBP_POSITIVE_REGULATION_OF_NUCLEAR_DIVISION | 51 | -0.28749 | -0.939692 | 0.54963 | 0.803431 |
| GOBP_EMBRYONIC_HEART_TUBE_MORPHOGENESIS | 58 | -0.282622 | -0.939612 | 0.574101 | 0.803219 |
| GOBP_CHONDROCYTE_DEVELOPMENT | 26 | -0.329961 | -0.939536 | 0.564886 | 0.80299 |
| GOBP_PROTEIN_IMPORT | 186 | -0.236799 | -0.939081 | 0.622412 | 0.803726 |
| GOBP_LONG_TERM_MEMORY | 26 | -0.324406 | -0.938608 | 0.558245 | 0.804514 |
| GOBP_CHRONIC_INFLAMMATORY_RESPONSE | 18 | -0.353208 | -0.938274 | 0.55521 | 0.804947 |
| GOBP_DNA_DAMAGE_RESPONSE_SIGNAL_TRANSDUCTION_BY_P53_CLASS_MEDIATOR | 105 | -0.254018 | -0.93787 | 0.586873 | 0.805568 |
| GOBP_COMPLEMENT_ACTIVATION | 152 | -0.246509 | -0.937855 | 0.605867 | 0.805183 |
| GOBP_EMBRYONIC_SKELETAL_SYSTEM_MORPHOGENESIS | 78 | -0.267424 | -0.937851 | 0.582432 | 0.804774 |
| GOBP_CELLULAR_RESPONSE_TO_LEPTIN_STIMULUS | 17 | -0.355813 | -0.936814 | 0.550077 | 0.807004 |
| GOBP_REGULATION_OF_CHONDROCYTE_DIFFERENTIATION | 42 | -0.296039 | -0.93656 | 0.570815 | 0.807244 |
| GOBP_NEGATIVE_REGULATION_OF_VIRAL_LIFE_CYCLE | 21 | -0.340037 | -0.935357 | 0.577035 | 0.809935 |
| GOBP_PHOSPHATIDYLINOSITOL_3_KINASE_SIGNALING | 141 | -0.244766 | -0.934566 | 0.616625 | 0.811563 |
| GOBP_POSITIVE_REGULATION_OF_MEMBRANE_POTENTIAL | 15 | -0.372149 | -0.934205 | 0.539658 | 0.81209 |
| GOBP_RAP_PROTEIN_SIGNAL_TRANSDUCTION | 15 | -0.37096 | -0.934093 | 0.5488 | 0.811963 |
| GOBP_APPENDAGE_DEVELOPMENT | 140 | -0.244178 | -0.933655 | 0.609125 | 0.81267 |
| GOBP_RESPONSE_TO_DIETARY_EXCESS | 18 | -0.352894 | -0.933573 | 0.550388 | 0.812477 |
| GOBP_DEVELOPMENTAL_GROWTH_INVOLVED_IN_MORPHOGENESIS | 201 | -0.236288 | -0.933503 | 0.632775 | 0.81223 |
| GOBP_SINGLE_FERTILIZATION | 112 | -0.251531 | -0.933273 | 0.605459 | 0.812377 |
| GOBP_HORMONE_BIOSYNTHETIC_PROCESS | 57 | -0.28039 | -0.93301 | 0.587017 | 0.812595 |
| GOBP_POSITIVE_REGULATION_OF_PROTEIN_SERINE_THREONINE_KINASE_ACTIVITY | 292 | -0.225377 | -0.932672 | 0.655133 | 0.813052 |
| GOBP_REGULATION_OF_PHOSPHOPROTEIN_PHOSPHATASE_ACTIVITY | 114 | -0.252252 | -0.932513 | 0.592122 | 0.813062 |
| GOBP_POSITIVE_REGULATION_OF_TRANSCRIPTION_REGULATORY_REGION_DNA_BINDING | 21 | -0.34362 | -0.931316 | 0.586207 | 0.81573 |
| GOBP_EXECUTION_PHASE_OF_APOPTOSIS | 80 | -0.262054 | -0.9312 | 0.592391 | 0.815596 |
| GOBP_ISOPRENOID_METABOLIC_PROCESS | 110 | -0.25105 | -0.930873 | 0.627907 | 0.816014 |
| GOBP_ENDOCHONDRAL_BONE_MORPHOGENESIS | 50 | -0.280963 | -0.93042 | 0.585091 | 0.816735 |
| GOBP_MIDDLE_EAR_MORPHOGENESIS | 16 | -0.360356 | -0.930304 | 0.563718 | 0.816607 |
| GOBP_VENTRICULAR_CARDIAC_MUSCLE_TISSUE_DEVELOPMENT | 43 | -0.287991 | -0.929725 | 0.564655 | 0.817617 |
| GOBP_NEGATIVE_REGULATION_OF_BLOOD_CIRCULATION | 18 | -0.344501 | -0.928841 | 0.571429 | 0.819435 |
| GOBP_REGULATION_OF_RESPONSE_TO_EXTRACELLULAR_STIMULUS | 22 | -0.33732 | -0.928665 | 0.553707 | 0.819469 |
| GOBP_RESPONSE_TO_TEMPERATURE_STIMULUS | 216 | -0.231845 | -0.928572 | 0.644311 | 0.819285 |
| GOBP_REGULATION_OF_PRODUCTION_OF_SMALL_RNA_INVOLVED_IN_GENE_SILENCING_BY_RI | 23 | -0.335067 | -0.927967 | 0.569069 | 0.820435 |
| GOBP_PROTEIN_MANNOSYLATION | 23 | -0.335615 | -0.92781 | 0.573416 | 0.820403 |
| GOBP_STEROID_HORMONE_MEDIATED_SIGNALING_PATHWAY | 122 | -0.24877 | -0.927436 | 0.620988 | 0.820956 |
| GOBP_CARDIAC_MUSCLE_CELL_CONTRACTION | 66 | -0.270568 | -0.927385 | 0.61178 | 0.820662 |
| GOBP_CIRCADIAN_RHYTHM | 187 | -0.23616 | -0.92688 | 0.644769 | 0.821525 |
| GOBP_SODIUM_ION_IMPORT_ACROSS_PLASMA_MEMBRANE | 17 | -0.35547 | -0.926595 | 0.587253 | 0.821813 |
| GOBP_REGULATION_OF_SPROUTING_ANGIOGENESIS | 37 | -0.297709 | -0.925649 | 0.599138 | 0.823796 |
| GOBP_OLIGODENDROCYTE_DEVELOPMENT | 39 | -0.298366 | -0.92562 | 0.568768 | 0.823456 |
| GOBP_OLFACTORY_LOBE_DEVELOPMENT | 22 | -0.343452 | -0.925567 | 0.576923 | 0.823166 |
| GOBP_CHONDROITIN_SULFATE_PROTEOGLYCAN_BIOSYNTHETIC_PROCESS | 29 | -0.315321 | -0.925463 | 0.567143 | 0.822998 |
| GOBP_POSITIVE_REGULATION_OF_CILIUM_ASSEMBLY | 23 | -0.332363 | -0.925177 | 0.575261 | 0.823292 |
| GOBP_NEGATIVE_REGULATION_OF_EXTRINSIC_APOPTOTIC_SIGNALING_PATHWAY_VIA_DEATH_ | 38 | -0.301363 | -0.925125 | 0.586402 | 0.823007 |
| GOBP_POSITIVE_REGULATION_BY_HOST_OF_VIRAL_TRANSCRIPTION | 16 | -0.35524 | -0.925007 | 0.560372 | 0.822885 |
| GOBP_PHENOL_CONTAINING_COMPOUND_BIOSYNTHETIC_PROCESS | 30 | -0.314476 | -0.92464 | 0.591185 | 0.823397 |
| GOBP_MEMBRANE_RAFT_ORGANIZATION | 23 | -0.329214 | -0.924589 | 0.570325 | 0.823112 |
| GOBP_PLATELET_DEGRANULATION | 119 | -0.24709 | -0.924509 | 0.627426 | 0.822879 |
| GOBP_REGULATION_OF_NOTCH_SIGNALING_PATHWAY | 98 | -0.25543 | -0.924389 | 0.63871 | 0.822779 |
| GOBP_POSITIVE_REGULATION_OF_AUTOPHAGY | 117 | -0.248955 | -0.924277 | 0.619171 | 0.822667 |
| GOBP_PERIPHERAL_NERVOUS_SYSTEM_DEVELOPMENT | 64 | -0.271894 | -0.924228 | 0.604167 | 0.822383 |
| GOBP_REGULATION_OF_LAMELLIPODIUM_ORGANIZATION | 48 | -0.283517 | -0.923728 | 0.587464 | 0.823233 |
| GOBP_POSITIVE_REGULATION_OF_POSTTRANSCRIPTIONAL_GENE_SILENCING | 29 | -0.311051 | -0.92366 | 0.590015 | 0.82299 |
| GOBP_SYNAPTIC_TRANSMISSION_GABAERGIC | 33 | -0.312334 | -0.923287 | 0.591615 | 0.823516 |
| GOBP_RESPONSE_TO_ACETYLCHOLINE | 24 | -0.326327 | -0.922872 | 0.585438 | 0.824161 |
| GOBP_INTERMEMBRANE_LIPID_TRANSFER | 44 | -0.291327 | -0.922846 | 0.570783 | 0.823807 |
| GOBP_STEROL_TRANSPORT | 102 | -0.249633 | -0.922407 | 0.61811 | 0.8245 |
| GOBP_SODIUM_ION_HOMEOSTASIS | 43 | -0.293258 | -0.922386 | 0.583573 | 0.824133 |
| GOBP_ACTIVATION_OF_JUN_KINASE_ACTIVITY | 39 | -0.296144 | -0.921961 | 0.601744 | 0.824793 |
| GOBP_PROTEIN_INSERTION_INTO_MITOCHONDRIAL_MEMBRANE_INVOLVED_IN_APOPTOTIC_SIG | 30 | -0.314649 | -0.921114 | 0.598551 | 0.826504 |
| GOBP_NEGATIVE_REGULATION_OF_RECEPTOR_MEDIATED_ENDOCYTOSIS | 28 | -0.311974 | -0.920953 | 0.57855 | 0.826477 |
| GOBP_NEGATIVE_REGULATION_OF_INTRACELLULAR_TRANSPORT | 53 | -0.278757 | -0.920357 | 0.581062 | 0.82754 |
| GOBP_POSITIVE_REGULATION_OF_ENDOTHELIAL_CELL_CHEMOTAXIS | 15 | -0.364543 | -0.919567 | 0.571429 | 0.829078 |
| GOBP_ADULT_BEHAVIOR | 98 | -0.249727 | -0.919168 | 0.632199 | 0.829678 |
| GOBP_FATTY_ACID_TRANSMEMBRANE_TRANSPORT | 48 | -0.286011 | -0.918785 | 0.622126 | 0.830216 |
| GOBP_MESODERMAL_CELL_DIFFERENTIATION | 26 | -0.31807 | -0.918719 | 0.566964 | 0.829948 |
| GOBP_RESPONSE_TO_CARBOHYDRATE | 196 | -0.231979 | -0.918125 | 0.674941 | 0.831003 |
| GOBP_SYNAPTIC_VESICLE_EXOCYTOSIS | 97 | -0.254238 | -0.917968 | 0.638889 | 0.830987 |
| GOBP_MYELIN_ASSEMBLY | 19 | -0.343404 | -0.917846 | 0.579845 | 0.830887 |
| GOBP_REGULATION_OF_ANOIKIS | 24 | -0.323479 | -0.917537 | 0.606725 | 0.831258 |
| GOBP_NEURONAL_STEM_CELL_POPULATION_MAINTENANCE | 17 | -0.348155 | -0.917517 | 0.586524 | 0.830894 |
| GOBP_RENAL_SODIUM_EXCRETION | 18 | -0.347997 | -0.91733 | 0.605634 | 0.830958 |
| GOBP_INTESTINAL_LIPID_ABSORPTION | 15 | -0.36887 | -0.917326 | 0.560387 | 0.830553 |
| GOBP_REGULATION_OF_DNA_TEMPLATED_TRANSCRIPTION_ELONGATION | 50 | -0.28041 | -0.917181 | 0.611111 | 0.830511 |
| GOBP_REGULATION_OF_ENDOCRINE_PROCESS | 27 | -0.321038 | -0.917093 | 0.577982 | 0.83032 |
| GOBP_BINDING_OF_SPERM_TO_ZONA_PELLUCIDA | 30 | -0.304405 | -0.916184 | 0.584337 | 0.832122 |
| GOBP_RESPONSE_TO_IMMOBILIZATION_STRESS | 17 | -0.347975 | -0.915694 | 0.54893 | 0.832886 |
| GOBP_LEUKOCYTE_MIGRATION_INVOLVED_IN_INFLAMMATORY_RESPONSE | 18 | -0.345683 | -0.915684 | 0.573209 | 0.8325 |
| GOBP_INDOLALKYLAMINE_METABOLIC_PROCESS | 16 | -0.353414 | -0.915348 | 0.601942 | 0.832931 |
| GOBP_SENSORY_PERCEPTION_OF_TEMPERATURE_STIMULUS | 19 | -0.343868 | -0.915057 | 0.563863 | 0.833256 |
| GOBP_AMINOGLYCAN_CATABOLIC_PROCESS | 63 | -0.268129 | -0.914644 | 0.610193 | 0.833861 |
| GOBP_POSITIVE_REGULATION_OF_NEURON_DIFFERENTIATION | 67 | -0.260532 | -0.914624 | 0.631436 | 0.833493 |
| GOBP_REGULATION_OF_VIRAL_TRANSCRIPTION | 42 | -0.290096 | -0.914288 | 0.574815 | 0.833901 |
| GOBP_CALCINEURIN_MEDIATED_SIGNALING | 40 | -0.294282 | -0.913907 | 0.612335 | 0.834412 |
| GOBP_SYNAPTIC_TRANSMISSION_GLUTAMATERGIC | 70 | -0.265031 | -0.91383 | 0.622132 | 0.834197 |
| GOBP_POSITIVE_REGULATION_OF_BLOOD_PRESSURE | 25 | -0.323075 | -0.913334 | 0.576453 | 0.835005 |
| GOBP_RESPONSE_TO_GAMMA_RADIATION | 51 | -0.277929 | -0.913127 | 0.616901 | 0.835115 |
| GOBP_HOMOTYPIC_CELL_CELL_ADHESION | 79 | -0.255703 | -0.913122 | 0.615283 | 0.834712 |
| GOBP_ESTABLISHMENT_OF_EPITHELIAL_CELL_POLARITY | 27 | -0.31392 | -0.912796 | 0.596626 | 0.835096 |
| GOBP_POSITIVE_REGULATION_OF_LAMELLIPODIUM_ASSEMBLY | 28 | -0.319965 | -0.912524 | 0.592424 | 0.835295 |
| GOBP_REGULATION_OF_ANION_TRANSMEMBRANE_TRANSPORT | 96 | -0.253226 | -0.912084 | 0.65088 | 0.835902 |

| | | | | | |
|---|---|---|---|---|---|
| GOBP_PHOSPHATIDYLCHOLINE_BIOSYNTHETIC_PROCESS | 38 | -0.296166 | -0.911124 | 0.600571 | 0.837772 |
| GOBP_LEUKOCYTE_TETHERING_OR_ROLLING | 28 | -0.313124 | -0.910654 | 0.605463 | 0.838497 |
| GOBP_GLUCOSAMINE_CONTAINING_COMPOUND_METABOLIC_PROCESS | 23 | -0.32771 | -0.910549 | 0.582712 | 0.838335 |
| GOBP_REGULATION_OF_MACROPHAGE_DERIVED_FOAM_CELL_DIFFERENTIATION | 28 | -0.308273 | -0.910543 | 0.593509 | 0.837938 |
| GOBP_T_CELL_LINEAGE_COMMITMENT | 27 | -0.313665 | -0.910236 | 0.608896 | 0.838302 |
| GOBP_CELLULAR_RESPONSE_TO_ALKALOID | 27 | -0.315138 | -0.909944 | 0.602699 | 0.838595 |
| GOBP_RESPONSE_TO_NUTRIENT | 150 | -0.2372 | -0.909668 | 0.653113 | 0.838838 |
| GOBP_VESICLE_CYTOSKELETAL_TRAFFICKING | 70 | -0.26306 | -0.9093 | 0.606061 | 0.839351 |
| GOBP_NEUROMUSCULAR_PROCESS | 71 | -0.262565 | -0.909264 | 0.655944 | 0.839036 |
| GOBP_CELL_MATURATION | 143 | -0.238506 | -0.909172 | 0.667892 | 0.838827 |
| GOBP_NEUROPEPTIDE_SIGNALING_PATHWAY | 64 | -0.268875 | -0.908829 | 0.635117 | 0.839255 |
| GOBP_EPITHELIAL_STRUCTURE_MAINTENANCE | 21 | -0.333498 | -0.908492 | 0.594595 | 0.83968 |
| GOBP_NEGATIVE_REGULATION_OF_HISTONE_METHYLATION | 21 | -0.333751 | -0.908064 | 0.591049 | 0.840316 |
| GOBP_LIPID_DIGESTION | 16 | -0.355468 | -0.907845 | 0.604938 | 0.840419 |
| GOBP_CILIUM_OR_FLAGELLUM_DEPENDENT_CELL_MOTILITY | 101 | -0.250296 | -0.90774 | 0.663185 | 0.840258 |
| GOBP_NEGATIVE_REGULATION_OF_BINDING | 146 | -0.237949 | -0.907727 | 0.662129 | 0.839874 |
| GOBP_REGULATION_OF_CELL_ADHESION_MEDIATED_BY_INTEGRIN | 45 | -0.283689 | -0.907454 | 0.608511 | 0.840111 |
| GOBP_GMP_METABOLIC_PROCESS | 15 | -0.350347 | -0.907436 | 0.588618 | 0.839748 |
| GOBP_LAMELLIPODIUM_ASSEMBLY | 68 | -0.262513 | -0.907354 | 0.649425 | 0.839523 |
| GOBP_REGULATION_OF_SYNAPTIC_TRANSMISSION_GABAERGIC | 25 | -0.322141 | -0.9073 | 0.600619 | 0.839241 |
| GOBP_POLY_A_PLUS_MRNA_EXPORT_FROM_NUCLEUS | 20 | -0.333398 | -0.906992 | 0.581184 | 0.839582 |
| GOBP_NEUROTRANSMITTER_UPTAKE | 32 | -0.307696 | -0.906489 | 0.62029 | 0.840387 |
| GOBP_PLASMA_MEMBRANE_ORGANIZATION | 107 | -0.245023 | -0.90628 | 0.653165 | 0.840466 |
| GOBP_INTRACELLULAR_STEROL_TRANSPORT | 24 | -0.31976 | -0.90574 | 0.611367 | 0.841366 |
| GOBP_REGULATION_OF_MEIOTIC_CELL_CYCLE | 39 | -0.288865 | -0.905318 | 0.600865 | 0.841959 |
| GOBP_NEGATIVE_REGULATION_OF_CATABOLIC_PROCESS | 287 | -0.21939 | -0.904681 | 0.704598 | 0.843093 |
| GOBP_REGULATION_OF_TRANSPOSITION | 23 | -0.319279 | -0.90411 | 0.592537 | 0.844052 |
| GOBP_NEGATIVE_REGULATION_OF_HISTONE_MODIFICATION | 42 | -0.28949 | -0.903938 | 0.627874 | 0.844056 |
| GOBP_CHONDROCYTE_DIFFERENTIATION | 94 | -0.249809 | -0.903715 | 0.665803 | 0.844176 |
| GOBP_ENDOSOME_ORGANIZATION | 80 | -0.252721 | -0.902068 | 0.642091 | 0.847726 |
| GOBP_REGULATION_OF_HORMONE_BIOSYNTHETIC_PROCESS | 20 | -0.330936 | -0.901197 | 0.626959 | 0.849417 |
| GOBP_INDOLE_CONTAINING_COMPOUND_METABOLIC_PROCESS | 22 | -0.326409 | -0.900401 | 0.636364 | 0.850925 |
| GOBP_REGULATION_OF_CARDIAC_MUSCLE_CONTRACTION | 62 | -0.261354 | -0.900192 | 0.627424 | 0.851028 |
| GOBP_POSITIVE_REGULATION_OF_NITRIC_OXIDE_SYNTHASE_ACTIVITY | 22 | -0.326166 | -0.899975 | 0.613333 | 0.851112 |
| GOBP_REGULATION_OF_BRANCHING_INVOLVED_IN_URETERIC_BUD_MORPHOGENESIS | 16 | -0.350443 | -0.899757 | 0.611285 | 0.851211 |
| GOBP_POSITIVE_REGULATION_OF_PROTEIN_ACETYLATION | 41 | -0.285518 | -0.899642 | 0.640173 | 0.851076 |
| GOBP_CELLULAR_RESPONSE_TO_EXTERNAL_STIMULUS | 287 | -0.21965 | -0.8991 | 0.735955 | 0.851978 |
| GOBP_CHOLESTEROL_EFFLUX | 48 | -0.274831 | -0.898719 | 0.65123 | 0.852487 |
| GOBP_REGULATION_OF_TRANSCRIPTION_BY_RNA_POLYMERASE_III | 23 | -0.321044 | -0.898493 | 0.638889 | 0.852608 |
| GOBP_GASTRULATION_WITH_MOUTH_FORMING_SECOND | 19 | -0.333734 | -0.897614 | 0.630468 | 0.854288 |
| GOBP_CRANIAL_SKELETAL_SYSTEM_DEVELOPMENT | 57 | -0.268494 | -0.89742 | 0.637241 | 0.85433 |
| GOBP_GANGLIOSIDE_METABOLIC_PROCESS | 21 | -0.333604 | -0.897417 | 0.609907 | 0.853921 |
| GOBP_LIPID_OXIDATION | 104 | -0.24476 | -0.897319 | 0.682243 | 0.85375 |
| GOBP_ANDROGEN_RECEPTOR_SIGNALING_PATHWAY | 36 | -0.288638 | -0.897063 | 0.620839 | 0.85395 |
| GOBP_CARDIAC_MUSCLE_TISSUE_DEVELOPMENT | 169 | -0.231607 | -0.896454 | 0.70632 | 0.854947 |
| GOBP_CARBOHYDRATE_TRANSMEMBRANE_TRANSPORT | 101 | -0.242785 | -0.896363 | 0.681638 | 0.854738 |
| GOBP_NEURON_APOPTOTIC_PROCESS | 196 | -0.227727 | -0.896093 | 0.729117 | 0.854955 |
| GOBP_GLYCOPROTEIN_BIOSYNTHETIC_PROCESS | 308 | -0.219001 | -0.896046 | 0.748879 | 0.854649 |
| GOBP_REGULATION_OF_MITOCHONDRIAL_MEMBRANE_PERMEABILITY | 70 | -0.260798 | -0.895962 | 0.651882 | 0.854436 |
| GOBP_SNARE_COMPLEX_ASSEMBLY | 18 | -0.342725 | -0.89573 | 0.599384 | 0.854549 |
| GOBP_POSITIVE_REGULATION_OF_CELLULAR_CARBOHYDRATE_METABOLIC_PROCESS | 48 | -0.274103 | -0.895511 | 0.661041 | 0.854646 |
| GOBP_REGULATION_OF_RESPONSE_TO_FOOD | 15 | -0.353876 | -0.895337 | 0.59845 | 0.854638 |
| GOBP_RESPONSE_TO_X_RAY | 32 | -0.300469 | -0.894824 | 0.632504 | 0.855405 |
| GOBP_BLASTOCYST_FORMATION | 32 | -0.29462 | -0.894353 | 0.63893 | 0.856102 |
| GOBP_RESPONSE_TO_NICOTINE | 37 | -0.287737 | -0.894053 | 0.637594 | 0.856408 |
| GOBP_REGULATION_OF_WNT_SIGNALING_PATHWAY_PLANAR_CELL_POLARITY_PATHWAY | 15 | -0.345433 | -0.893541 | 0.613893 | 0.857195 |
| GOBP_REGULATION_OF_ER_TO_GOLGI_VESICLE_MEDIATED_TRANSPORT | 15 | -0.350229 | -0.893041 | 0.605723 | 0.857945 |
| GOBP_POSITIVE_REGULATION_OF_COLLAGEN_METABOLIC_PROCESS | 25 | -0.315072 | -0.892877 | 0.618012 | 0.857919 |
| GOBP_SYNAPTIC_VESICLE_TRANSPORT | 42 | -0.283609 | -0.892686 | 0.637313 | 0.857951 |
| GOBP_MAINTENANCE_OF_LOCATION_IN_CELL | 191 | -0.224922 | -0.891826 | 0.720958 | 0.859492 |
| GOBP_CELLULAR_RESPONSE_TO_ATP | 17 | -0.338496 | -0.89156 | 0.613235 | 0.859695 |
| GOBP_NEGATIVE_REGULATION_OF_STRESS_ACTIVATED_PROTEIN_KINASE_SIGNALING_CASCAD | 44 | -0.27889 | -0.890805 | 0.6411 | 0.861109 |
| GOBP_ENDOSOME_TO_LYSOSOME_TRANSPORT | 52 | -0.2699 | -0.890208 | 0.65113 | 0.86207 |
| GOBP_REGULATION_OF_OXIDATIVE_STRESS_INDUCED_NEURON_DEATH | 22 | -0.322413 | -0.889935 | 0.626524 | 0.862258 |
| GOBP_REGULATION_OF_CELL_PROJECTION_ASSEMBLY | 175 | -0.226331 | -0.889288 | 0.71705 | 0.863341 |
| GOBP_POSITIVE_REGULATION_OF_PROTEOLYSIS | 344 | -0.214663 | -0.889083 | 0.775419 | 0.863366 |
| GOBP_REGULATION_OF_ACTIN_NUCLEATION | 28 | -0.300555 | -0.888249 | 0.630137 | 0.864893 |
| GOBP_ESTABLISHMENT_OF_PROTEIN_LOCALIZATION_TO_VACUOLE | 48 | -0.2687 | -0.887733 | 0.646648 | 0.865634 |
| GOBP_MODULATION_BY_SYMBIONT_OF_HOST_CELLULAR_PROCESS | 23 | -0.318712 | -0.887641 | 0.626292 | 0.865436 |
| GOBP_MOVEMENT_IN_HOST_ENVIRONMENT | 165 | -0.228303 | -0.886837 | 0.72705 | 0.866865 |
| GOBP_POSITIVE_REGULATION_OF_CATION_TRANSMEMBRANE_TRANSPORT | 123 | -0.235896 | -0.88648 | 0.698842 | 0.867272 |
| GOBP_REGULATION_OF_NEUTROPHIL_MIGRATION | 40 | -0.279404 | -0.885086 | 0.635432 | 0.870098 |
| GOBP_SMAD_PROTEIN_SIGNAL_TRANSDUCTION | 66 | -0.258546 | -0.885033 | 0.671979 | 0.869803 |
| GOBP_NEGATIVE_REGULATION_OF_PROTEIN_LOCALIZATION_TO_MEMBRANE | 27 | -0.308589 | -0.884936 | 0.655539 | 0.869601 |
| GOBP_REGULATION_OF_SUPEROXIDE_METABOLIC_PROCESS | 27 | -0.30894 | -0.883971 | 0.633634 | 0.871371 |
| GOBP_RESPONSE_TO_AMINE | 38 | -0.283849 | -0.883964 | 0.634006 | 0.87097 |
| GOBP_NEGATIVE_REGULATION_OF_FAT_CELL_DIFFERENTIATION | 46 | -0.274896 | -0.883806 | 0.660538 | 0.870921 |
| GOBP_REGULATION_OF_POTASSIUM_ION_TRANSPORT | 79 | -0.249345 | -0.883607 | 0.692308 | 0.870936 |
| GOBP_REGULATION_OF_NECROTIC_CELL_DEATH | 32 | -0.293161 | -0.883395 | 0.645833 | 0.870999 |
| GOBP_KERATINIZATION | 73 | -0.252024 | -0.883212 | 0.669388 | 0.871023 |
| GOBP_HEPATICOBILIARY_SYSTEM_DEVELOPMENT | 128 | -0.236122 | -0.882317 | 0.706983 | 0.872685 |
| GOBP_POSITIVE_REGULATION_OF_PLASMA_MEMBRANE_BOUNDED_CELL_PROJECTION_ASSEM | 101 | -0.241438 | -0.882229 | 0.699219 | 0.872475 |
| GOBP_NEGATIVE_REGULATION_OF_TRANSPORTER_ACTIVITY | 69 | -0.255821 | -0.881968 | 0.693243 | 0.872658 |
| GOBP_REGULATION_OF_T_CELL_DIFFERENTIATION_IN_THYMUS | 22 | -0.315157 | -0.881461 | 0.653673 | 0.873444 |
| GOBP_POSITIVE_REGULATION_OF_CATABOLIC_PROCESS | 431 | -0.208522 | -0.881272 | 0.820847 | 0.873459 |
| GOBP_NEGATIVE_REGULATION_OF_PROTEIN_BINDING | 90 | -0.245075 | -0.881185 | 0.696335 | 0.87324 |
| GOBP_RESPONSE_TO_PEPTIDE_HORMONE | 391 | -0.208978 | -0.881126 | 0.810364 | 0.872965 |
| GOBP_CIRCADIAN_REGULATION_OF_GENE_EXPRESSION | 63 | -0.26033 | -0.880917 | 0.663073 | 0.873023 |
| GOBP_REGULATION_OF_DELAYED_RECTIFIER_POTASSIUM_CHANNEL_ACTIVITY | 17 | -0.343097 | -0.879667 | 0.645768 | 0.875401 |
| GOBP_PITUITARY_GLAND_DEVELOPMENT | 28 | -0.307759 | -0.879659 | 0.650298 | 0.875006 |
| GOBP_RHODOPSIN_MEDIATED_SIGNALING_PATHWAY | 22 | -0.318497 | -0.878082 | 0.633234 | 0.878289 |
| GOBP_REGULATION_OF_POTASSIUM_ION_TRANSMEMBRANE_TRANSPORTER_ACTIVITY | 48 | -0.27091 | -0.878052 | 0.676136 | 0.87794 |
| GOBP_CARDIAC_MUSCLE_CELL_DIFFERENTIATION | 88 | -0.245746 | -0.877807 | 0.694848 | 0.878061 |
| GOBP_ANTIGEN_PROCESSING_AND_PRESENTATION_OF_PEPTIDE_ANTIGEN | 189 | -0.22322 | -0.876189 | 0.754177 | 0.88127 |
| GOBP_CELLULAR_KETONE_METABOLIC_PROCESS | 221 | -0.217994 | -0.876028 | 0.764706 | 0.881219 |
| GOBP_CYCLIC_NUCLEOTIDE_METABOLIC_PROCESS | 35 | -0.290468 | -0.875958 | 0.659056 | 0.880964 |
| GOBP_REGULATION_OF_CELLULAR_CARBOHYDRATE_METABOLIC_PROCESS | 133 | -0.229277 | -0.875633 | 0.749054 | 0.881294 |
| GOBP_POSITIVE_REGULATION_OF_CARDIAC_MUSCLE_CELL_PROLIFERATION | 22 | -0.318646 | -0.875456 | 0.657012 | 0.881279 |
| GOBP_CARBOHYDRATE_CATABOLIC_PROCESS | 186 | -0.223569 | -0.875267 | 0.756333 | 0.881276 |
| GOBP_NEGATIVE_REGULATION_OF_PROTEIN_TYROSINE_KINASE_ACTIVITY | 29 | -0.304932 | -0.875253 | 0.652504 | 0.880885 |
| GOBP_REGULATION_OF_PLATELET_ACTIVATION | 31 | -0.293929 | -0.875055 | 0.665663 | 0.880943 |
| GOBP_ESTABLISHMENT_OF_PROTEIN_LOCALIZATION_TO_PLASMA_MEMBRANE | 58 | -0.260321 | -0.875012 | 0.694294 | 0.880621 |
| GOBP_T_HELPER_1_CELL_DIFFERENTIATION | 18 | -0.334372 | -0.874837 | 0.649351 | 0.880618 |
| GOBP_STEM_CELL_DIVISION | 23 | -0.311945 | -0.874106 | 0.635528 | 0.881856 |
| GOBP_C21_STEROID_HORMONE_BIOSYNTHETIC_PROCESS | 23 | -0.316198 | -0.87391 | 0.641369 | 0.881888 |
| GOBP_CELLULAR_RESPONSE_TO_AMYLOID_BETA | 35 | -0.286425 | -0.873056 | 0.655639 | 0.883376 |
| GOBP_HISTONE_H3_K4_TRIMETHYLATION | 16 | -0.338823 | -0.872688 | 0.624204 | 0.883782 |
| GOBP_VASCULAR_ASSOCIATED_SMOOTH_MUSCLE_CONTRACTION | 22 | -0.319897 | -0.872652 | 0.636364 | 0.883441 |
| GOBP_REGULATION_OF_EXTRACELLULAR_MATRIX_ASSEMBLY | 15 | -0.350648 | -0.872614 | 0.677165 | 0.883103 |
| GOBP_SUBSTRATE_ADHESION_DEPENDENT_CELL_SPREADING | 104 | -0.237115 | -0.872032 | 0.703226 | 0.883962 |
| GOBP_NEUROTRANSMITTER_REUPTAKE | 26 | -0.302651 | -0.871819 | 0.659226 | 0.88402 |
| GOBP_POSITIVE_REGULATION_OF_SMAD_PROTEIN_SIGNAL_TRANSDUCTION | 15 | -0.345807 | -0.870843 | 0.635671 | 0.885786 |
| GOBP_NEGATIVE_REGULATION_OF_DEFENSE_RESPONSE_TO_VIRUS | 24 | -0.311406 | -0.870754 | 0.660551 | 0.885574 |
| GOBP_REGULATION_OF_CYTOSKELETON_ORGANIZATION | 481 | -0.204797 | -0.870435 | 0.84952 | 0.885885 |
| GOBP_ACTIVATION_OF_PROTEIN_KINASE_B_ACTIVITY | 29 | -0.300228 | -0.870306 | 0.642314 | 0.885759 |
| GOBP_POSITIVE_REGULATION_OF_FIBROBLAST_PROLIFERATION | 43 | -0.271795 | -0.870299 | 0.662824 | 0.885359 |
| GOBP_REGULATION_OF_CARBOHYDRATE_METABOLIC_PROCESS | 194 | -0.219065 | -0.870267 | 0.772462 | 0.885017 |
| GOBP_PEROXISOMAL_MEMBRANE_TRANSPORT | 20 | -0.329018 | -0.870152 | 0.661994 | 0.884854 |
| GOBP_POSITIVE_REGULATION_OF_INTRINSIC_APOPTOTIC_SIGNALING_PATHWAY | 56 | -0.258429 | -0.869756 | 0.722067 | 0.88532 |
| GOBP_POSITIVE_REGULATION_OF_CELL_MIGRATION_INVOLVED_IN_SPROUTING_ANGIOGENESIS | 20 | -0.318053 | -0.868973 | 0.657277 | 0.886672 |
| GOBP_ANTEROGRADE_AXONAL_TRANSPORT | 46 | -0.266051 | -0.86896 | 0.689415 | 0.886283 |
| GOBP_PROTEIN_TRIMERIZATION | 17 | -0.336036 | -0.868736 | 0.659476 | 0.886373 |
| GOBP_REGULATION_OF_STEROID_BIOSYNTHETIC_PROCESS | 78 | -0.243403 | -0.868054 | 0.729655 | 0.887452 |
| GOBP_REGULATION_OF_DENDRITE_DEVELOPMENT | 94 | -0.240334 | -0.867068 | 0.746719 | 0.889174 |
| GOBP_ORGANIC_HYDROXY_COMPOUND_METABOLIC_PROCESS | 479 | -0.204109 | -0.866863 | 0.854856 | 0.88922 |
| GOBP_POSITIVE_REGULATION_OF_MULTI_ORGANISM_PROCESS | 17 | -0.334243 | -0.865927 | 0.630724 | 0.890846 |
| GOBP_REGULATION_OF_PROTEIN_DEACETYLATION | 40 | -0.274404 | -0.865709 | 0.680516 | 0.890901 |
| GOBP_MONOCARBOXYLIC_ACID_CATABOLIC_PROCESS | 122 | -0.228664 | -0.86552 | 0.745247 | 0.89087 |
| GOBP_ERBB_SIGNALING_PATHWAY | 138 | -0.229254 | -0.865454 | 0.746173 | 0.890608 |
| GOBP_REGULATION_OF_MYELINATION | 40 | -0.275767 | -0.865151 | 0.682927 | 0.890864 |
| GOBP_REGULATION_OF_CELL_SHAPE | 147 | -0.225508 | -0.864965 | 0.776529 | 0.890873 |
| GOBP_REGULATION_OF_NEURON_PROJECTION_DEVELOPMENT | 403 | -0.205355 | -0.864777 | 0.84222 | 0.890883 |
| GOBP_REGULATION_OF_URINE_VOLUME | 19 | -0.324648 | -0.864739 | 0.648062 | 0.890547 |
| GOBP_GLANDULAR_EPITHELIAL_CELL_DEVELOPMENT | 23 | -0.307694 | -0.864082 | 0.668189 | 0.89157 |
| GOBP_POSITIVE_REGULATION_OF_COLD_INDUCED_THERMOGENESIS | 90 | -0.236764 | -0.863491 | 0.739187 | 0.892451 |
| GOBP_CORTICAL_CYTOSKELETON_ORGANIZATION | 58 | -0.25853 | -0.863265 | 0.710564 | 0.892525 |
| GOBP_REGULATION_OF_MYELOID_CELL_APOPTOTIC_PROCESS | 22 | -0.317015 | -0.863054 | 0.663623 | 0.892568 |
| GOBP_MYELOID_CELL_DIFFERENTIATION | 401 | -0.207496 | -0.863053 | 0.861322 | 0.892155 |
| GOBP_REGULATION_OF_RELEASE_OF_SEQUESTERED_CALCIUM_ION_INTO_CYTOSOL | 65 | -0.251978 | -0.862996 | 0.715076 | 0.891876 |
| GOBP_REGULATION_OF_CELLULAR_RESPONSE_TO_INSULIN_STIMULUS | 60 | -0.25245 | -0.862917 | 0.71108 | 0.891638 |
| GOBP_LUNG_EPITHELIUM_DEVELOPMENT | 28 | -0.293523 | -0.862653 | 0.675676 | 0.891799 |
| GOBP_T_CELL_HOMEOSTASIS | 35 | -0.28204 | -0.862393 | 0.684583 | 0.891951 |
| GOBP_HISTONE_H3_DEACETYLATION | 19 | -0.321156 | -0.86223 | 0.657771 | 0.891913 |
| GOBP_REGULATION_OF_RESPONSE_TO_REACTIVE_OXYGEN_SPECIES | 28 | -0.294796 | -0.861816 | 0.679245 | 0.892374 |
| GOBP_CARBOHYDRATE_HOMEOSTASIS | 200 | -0.21721 | -0.86072 | 0.788716 | 0.894283 |
| GOBP_POSITIVE_REGULATION_OF_LIPID_STORAGE | 22 | -0.312416 | -0.860709 | 0.654867 | 0.893892 |
| GOBP_PEPTIDYL_CYSTEINE_MODIFICATION | 44 | -0.26924 | -0.860392 | 0.706314 | 0.894156 |
| GOBP_REGULATION_OF_PROTEIN_POLYMERIZATION | 214 | -0.216076 | -0.859855 | 0.807916 | 0.894887 |
| GOBP_REGULATION_OF_PROTEIN_DEPHOSPHORYLATION | 137 | -0.224217 | -0.859726 | 0.771536 | 0.894744 |
| GOBP_RETROGRADE_TRANSPORT_ENDOSOME_TO_PLASMA_MEMBRANE | 30 | -0.289661 | -0.859537 | 0.688172 | 0.894738 |
| GOBP_CHOLESTEROL_STORAGE | 18 | -0.325197 | -0.85907 | 0.664063 | 0.895341 |
| GOBP_POSITIVE_REGULATION_OF_CELLULAR_COMPONENT_BIOGENESIS | 499 | -0.201434 | -0.858712 | 0.883103 | 0.895687 |
| GOBP_MAINTENANCE_OF_PROTEIN_LOCATION | 90 | -0.238207 | -0.858362 | 0.748656 | 0.896076 |
| GOBP_IMMUNOGLOBULIN_PRODUCTION_INVOLVED_IN_IMMUNOGLOBULIN_MEDIATED_IMMUNE_ | 54 | -0.260516 | -0.858208 | 0.713287 | 0.895996 |
| GOBP_POSITIVE_REGULATION_OF_POTASSIUM_ION_TRANSMEMBRANE_TRANSPORTER_ACTIVIT | 19 | -0.328653 | -0.857927 | 0.655864 | 0.896182 |
| GOBP_POSITIVE_REGULATION_OF_CHOLESTEROL_EFFLUX | 21 | -0.31511 | -0.857617 | 0.670245 | 0.896429 |
| GOBP_PHOSPHATIDYLSERINE_METABOLIC_PROCESS | 30 | -0.287799 | -0.857555 | 0.665179 | 0.896143 |
| GOBP_SMALL_MOLECULE_CATABOLIC_PROCESS | 402 | -0.20425 | -0.857335 | 0.847661 | 0.896217 |
| GOBP_CARBOHYDRATE_DERIVATIVE_CATABOLIC_PROCESS | 182 | -0.218948 | -0.85665 | 0.778182 | 0.897238 |
| GOBP_NAD_BIOSYNTHETIC_PROCESS | 27 | -0.296349 | -0.855613 | 0.709531 | 0.899051 |
| GOBP_REGULATION_OF_VESICLE_FUSION | 22 | -0.308425 | -0.854999 | 0.673716 | 0.899925 |
| GOBP_REGULATION_OF_TRIGLYCERIDE_METABOLIC_PROCESS | 29 | -0.288513 | -0.854435 | 0.67971 | 0.9007 |
| GOBP_CYTOPLASMIC_MICROTUBULE_ORGANIZATION | 61 | -0.253649 | -0.853848 | 0.734143 | 0.901502 |
| GOBP_NEURAL_CREST_CELL_MIGRATION | 45 | -0.265706 | -0.853015 | 0.705799 | 0.902823 |
| GOBP_PLASMA_MEMBRANE_PHOSPHOLIPID_SCRAMBLING | 16 | -0.331222 | -0.852998 | 0.658647 | 0.902445 |
| GOBP_LIMBIC_SYSTEM_DEVELOPMENT | 78 | -0.241857 | -0.852783 | 0.737063 | 0.90249 |
| GOBP_AMYLOID_PRECURSOR_PROTEIN_METABOLIC_PROCESS | 71 | -0.241949 | -0.852479 | 0.713528 | 0.902743 |
| GOBP_REGULATION_OF_RYANODINE_SENSITIVE_CALCIUM_RELEASE_CHANNEL_ACTIVITY | 22 | -0.31036 | -0.85241 | 0.673653 | 0.90247 |

| | | | | | |
|---|---|---|---|---|---|
| GOBP_ADAPTIVE_THERMOGENESIS | 144 | -0.222602 | -0.852212 | 0.78882 | 0.902475 |
| GOBP_CELL_CYCLE_ARREST | 216 | -0.212825 | -0.851717 | 0.817422 | 0.903116 |
| GOBP_NUCLEOSIDE_SALVAGE | 16 | -0.333072 | -0.850424 | 0.664099 | 0.905372 |
| GOBP_PROTEIN_KINASE_C_ACTIVATING_G_PROTEIN_COUPLED_RECEPTOR_SIGNALING_PATHW | 23 | -0.303302 | -0.849869 | 0.677372 | 0.906124 |
| GOBP_PURINE_NUCLEOTIDE_TRANSPORT | 24 | -0.306548 | -0.849753 | 0.678179 | 0.905952 |
| GOBP_REGULATION_OF_INTRACELLULAR_TRANSPORT | 336 | -0.204808 | -0.849503 | 0.86674 | 0.906074 |
| GOBP_MONOVALENT_INORGANIC_CATION_HOMEOSTASIS | 146 | -0.223256 | -0.849467 | 0.797739 | 0.905732 |
| GOBP_REGULATION_OF_VASCULAR_ASSOCIATED_SMOOTH_MUSCLE_CELL_PROLIFERATION | 54 | -0.257977 | -0.849166 | 0.750712 | 0.905943 |
| GOBP_PHOTOPERIODISM | 30 | -0.285905 | -0.84837 | 0.69697 | 0.907174 |
| GOBP_REGULATION_OF_SMALL_MOLECULE_METABOLIC_PROCESS | 392 | -0.201823 | -0.847697 | 0.859155 | 0.908141 |
| GOBP_SYMPATHETIC_NERVOUS_SYSTEM_DEVELOPMENT | 15 | -0.338533 | -0.847423 | 0.671667 | 0.908313 |
| GOBP_POSITIVE_REGULATION_OF_CELLULAR_PROTEIN_LOCALIZATION | 284 | -0.205411 | -0.846614 | 0.857143 | 0.909606 |
| GOBP_PEPTIDYL_LYSINE_DIMETHYLATION | 21 | -0.313382 | -0.846557 | 0.681607 | 0.909318 |
| GOBP_PROTEIN_AUTOPHOSPHORYLATION | 219 | -0.211691 | -0.846032 | 0.825983 | 0.909984 |
| GOBP_REGULATION_OF_DNA_BINDING | 107 | -0.229427 | -0.845305 | 0.777195 | 0.911093 |
| GOBP_HIPPOCAMPUS_DEVELOPMENT | 59 | -0.246634 | -0.84367 | 0.735795 | 0.913975 |
| GOBP_HEMATOPOIETIC_STEM_CELL_HOMEOSTASIS | 15 | -0.335855 | -0.843635 | 0.671362 | 0.913638 |
| GOBP_VITAMIN_TRANSMEMBRANE_TRANSPORT | 15 | -0.334449 | -0.843601 | 0.703526 | 0.913301 |
| GOBP_LYSOSOMAL_TRANSPORT | 106 | -0.228749 | -0.843166 | 0.795165 | 0.913758 |
| GOBP_CELLULAR_RESPONSE_TO_OSMOTIC_STRESS | 38 | -0.268826 | -0.842976 | 0.723837 | 0.913736 |
| GOBP_PH_REDUCTION | 53 | -0.254483 | -0.842721 | 0.75321 | 0.913844 |
| GOBP_GERM_CELL_DEVELOPMENT | 211 | -0.212851 | -0.842465 | 0.842657 | 0.913936 |
| GOBP_REGULATION_OF_BICELLULAR_TIGHT_JUNCTION_ASSEMBLY | 22 | -0.306509 | -0.841554 | 0.701997 | 0.915327 |
| GOBP_REGULATION_OF_CELLULAR_COMPONENT_SIZE | 348 | -0.201398 | -0.841509 | 0.884828 | 0.915 |
| GOBP_POSITIVE_REGULATION_OF_SODIUM_ION_TRANSPORT | 33 | -0.275616 | -0.841202 | 0.71 | 0.915198 |
| GOBP_INTESTINAL_ABSORPTION | 31 | -0.282851 | -0.841193 | 0.721574 | 0.914801 |
| GOBP_MYELOID_CELL_APOPTOTIC_PROCESS | 27 | -0.289645 | -0.841186 | 0.707281 | 0.9144 |
| GOBP_REGULATION_OF_SYSTEMIC_ARTERIAL_BLOOD_PRESSURE_BY_RENIN_ANGIOTENSIN | 19 | -0.315449 | -0.840604 | 0.670927 | 0.915205 |
| GOBP_EMBRYONIC_DIGIT_MORPHOGENESIS | 45 | -0.263502 | -0.83964 | 0.706569 | 0.916699 |
| GOBP_ADHERENS_JUNCTION_ORGANIZATION | 60 | -0.248861 | -0.838966 | 0.75657 | 0.917697 |
| GOBP_POSITIVE_REGULATION_OF_TOR_SIGNALING | 42 | -0.264568 | -0.838865 | 0.730827 | 0.917482 |
| GOBP_RESPONSE_TO_REACTIVE_OXYGEN_SPECIES | 203 | -0.210253 | -0.838807 | 0.841795 | 0.917182 |
| GOBP_POLYSACCHARIDE_METABOLIC_PROCESS | 103 | -0.227247 | -0.837764 | 0.775726 | 0.91882 |
| GOBP_SPHINGOID_METABOLIC_PROCESS | 22 | -0.305481 | -0.837585 | 0.701378 | 0.918759 |
| GOBP_LOCOMOTORY_BEHAVIOR | 129 | -0.223932 | -0.836385 | 0.80129 | 0.920767 |
| GOBP_GLYCEROLIPID_CATABOLIC_PROCESS | 56 | -0.252127 | -0.836216 | 0.767218 | 0.920689 |
| GOBP_ANIMAL_ORGAN_MATURATION | 23 | -0.29755 | -0.83517 | 0.718845 | 0.922384 |
| GOBP_NEGATIVE_REGULATION_OF_REGULATED_SECRETORY_PATHWAY | 20 | -0.30714 | -0.835031 | 0.704334 | 0.922255 |
| GOBP_DENDRITE_EXTENSION | 31 | -0.279607 | -0.834624 | 0.734838 | 0.922647 |
| GOBP_POSITIVE_REGULATION_OF_CELLULAR_AMIDE_METABOLIC_PROCESS | 151 | -0.216989 | -0.834594 | 0.815094 | 0.922287 |
| GOBP_NEURAL_NUCLEUS_DEVELOPMENT | 52 | -0.254328 | -0.834472 | 0.725968 | 0.922112 |
| GOBP_NEGATIVE_REGULATION_OF_CALCIUM_ION_TRANSMEMBRANE_TRANSPORT | 33 | -0.279487 | -0.834423 | 0.707463 | 0.921791 |
| GOBP_REGULATION_OF_PHOSPHATASE_ACTIVITY | 165 | -0.211631 | -0.834045 | 0.843862 | 0.922146 |
| GOBP_SIGNAL_TRANSDUCTION_IN_RESPONSE_TO_DNA_DAMAGE | 128 | -0.219495 | -0.833196 | 0.851385 | 0.923406 |
| GOBP_POSITIVE_REGULATION_OF_EXTRACELLULAR_MATRIX_ORGANIZATION | 22 | -0.302978 | -0.833196 | 0.69906 | 0.922991 |
| GOBP_CELLULAR_RESPONSE_TO_HEAT | 119 | -0.22224 | -0.832943 | 0.787995 | 0.923072 |
| GOBP_PLASMA_MEMBRANE_TUBULATION | 20 | -0.302196 | -0.831807 | 0.694888 | 0.924894 |
| GOBP_NEGATIVE_REGULATION_OF_SYNAPTIC_TRANSMISSION | 54 | -0.251652 | -0.831438 | 0.76681 | 0.925232 |
| GOBP_REGULATION_OF_DEFENSE_RESPONSE_TO_VIRUS_BY_VIRUS | 29 | -0.283191 | -0.831104 | 0.739583 | 0.925499 |
| GOBP_NEGATIVE_REGULATION_OF_ENDOPLASMIC_RETICULUM_STRESS_INDUCED_INTRINSIC_ | 16 | -0.330685 | -0.829053 | 0.708333 | 0.929082 |
| GOBP_BRAIN_MORPHOGENESIS | 32 | -0.277401 | -0.828995 | 0.737089 | 0.928779 |
| GOBP_HETEROPHILIC_CELL_CELL_ADHESION_VIA_PLASMA_MEMBRANE_CELL_ADHESION_MOL | 41 | -0.262896 | -0.82861 | 0.737374 | 0.929127 |
| GOBP_CORTICAL_ACTIN_CYTOSKELETON_ORGANIZATION | 38 | -0.268562 | -0.828562 | 0.75 | 0.928796 |
| GOBP_REGULATION_OF_CELLULAR_KETONE_METABOLIC_PROCESS | 159 | -0.211627 | -0.828216 | 0.836759 | 0.929052 |
| GOBP_SALIVARY_GLAND_DEVELOPMENT | 28 | -0.283557 | -0.827594 | 0.722059 | 0.929826 |
| GOBP_REGULATION_OF_EXOSOMAL_SECRETION | 16 | -0.322791 | -0.826988 | 0.718995 | 0.930612 |
| GOBP_CELLULAR_RESPONSE_TO_FATTY_ACID | 32 | -0.279864 | -0.826447 | 0.733906 | 0.931244 |
| GOBP_POSITIVE_REGULATION_OF_NEUTROPHIL_MIGRATION | 28 | -0.28379 | -0.826439 | 0.730253 | 0.930841 |
| GOBP_CELL_GROWTH | 430 | -0.196057 | -0.826197 | 0.918015 | 0.930882 |
| GOBP_NON_CANONICAL_WNT_SIGNALING_PATHWAY | 150 | -0.213645 | -0.826167 | 0.821159 | 0.930525 |
| GOBP_REGULATION_OF_INTRACELLULAR_PROTEIN_TRANSPORT | 241 | -0.203565 | -0.826065 | 0.872538 | 0.9303 |
| GOBP_AMYLOID_BETA_METABOLIC_PROCESS | 54 | -0.247511 | -0.825568 | 0.760773 | 0.93089 |
| GOBP_POSITIVE_REGULATION_OF_TRANSMEMBRANE_TRANSPORT | 181 | -0.209205 | -0.825305 | 0.841727 | 0.930972 |
| GOBP_REGULATION_OF_SMAD_PROTEIN_SIGNAL_TRANSDUCTION | 23 | -0.29048 | -0.824282 | 0.733835 | 0.932527 |
| GOBP_APICAL_JUNCTION_ASSEMBLY | 73 | -0.233669 | -0.824215 | 0.763052 | 0.932243 |
| GOBP_PURINE_CONTAINING_COMPOUND_CATABOLIC_PROCESS | 48 | -0.250964 | -0.823938 | 0.769014 | 0.932367 |
| GOBP_N_TERMINAL_PROTEIN_AMINO_ACID_ACETYLATION | 16 | -0.319704 | -0.823846 | 0.705882 | 0.932131 |
| GOBP_ENDOMEMBRANE_SYSTEM_ORGANIZATION | 458 | -0.194997 | -0.823336 | 0.938232 | 0.932676 |
| GOBP_FIBROBLAST_APOPTOTIC_PROCESS | 20 | -0.304885 | -0.821185 | 0.742726 | 0.936292 |
| GOBP_REGULATION_OF_MAMMARY_GLAND_EPITHELIAL_CELL_PROLIFERATION | 17 | -0.315931 | -0.821169 | 0.716952 | 0.935903 |
| GOBP_CELLULAR_RESPONSE_TO_DRUG | 59 | -0.242714 | -0.820539 | 0.764543 | 0.936655 |
| GOBP_REGULATION_OF_CYTOPLASMIC_TRANSPORT | 27 | -0.287402 | -0.820528 | 0.724551 | 0.936259 |
| GOBP_MUSCLE_CELL_APOPTOTIC_PROCESS | 60 | -0.242711 | -0.820514 | 0.780172 | 0.935869 |
| GOBP_NUCLEUS_LOCALIZATION | 22 | -0.299573 | -0.819859 | 0.72943 | 0.936708 |
| GOBP_SPHINGOLIPID_METABOLIC_PROCESS | 161 | -0.208076 | -0.819857 | 0.868358 | 0.936297 |
| GOBP_REGULATION_OF_DENDRITIC_SPINE_DEVELOPMENT | 60 | -0.243438 | -0.819252 | 0.801145 | 0.93703 |
| GOBP_PHOSPHATIDYLINOSITOL_DEPHOSPHORYLATION | 30 | -0.27956 | -0.818831 | 0.741456 | 0.937381 |
| GOBP_NUCLEOTIDE_TRANSPORT | 30 | -0.274508 | -0.818809 | 0.757991 | 0.937011 |
| GOBP_ENDOCARDIAL_CUSHION_MORPHOGENESIS | 29 | -0.276365 | -0.817678 | 0.744557 | 0.938668 |
| GOBP_HISTONE_H2A_UBIQUITINATION | 26 | -0.286429 | -0.817235 | 0.75188 | 0.939069 |
| GOBP_REGULATION_OF_VENTRICULAR_CARDIAC_MUSCLE_CELL_MEMBRANE_REPOLARIZATIO | 19 | -0.307814 | -0.816834 | 0.731438 | 0.939376 |
| GOBP_RESPONSE_TO_ANTIBIOTIC | 42 | -0.264189 | -0.815923 | 0.774815 | 0.940638 |
| GOBP_REGULATION_OF_SPHINGOLIPID_BIOSYNTHETIC_PROCESS | 18 | -0.313572 | -0.81562 | 0.723781 | 0.940767 |
| GOBP_OLIGOSACCHARIDE_METABOLIC_PROCESS | 46 | -0.257063 | -0.815556 | 0.79561 | 0.940474 |
| GOBP_NEGATIVE_REGULATION_OF_RESPONSE_TO_DNA_DAMAGE_STIMULUS | 82 | -0.231831 | -0.815136 | 0.793478 | 0.940833 |
| GOBP_RESPONSE_TO_PLATELET_DERIVED_GROWTH_FACTOR | 19 | -0.301059 | -0.814695 | 0.71248 | 0.941266 |
| GOBP_MULTICELLULAR_ORGANISM_GROWTH | 117 | -0.218354 | -0.814577 | 0.840796 | 0.941073 |
| GOBP_CERAMIDE_METABOLIC_PROCESS | 98 | -0.222781 | -0.814196 | 0.805951 | 0.941323 |
| GOBP_POSITIVE_REGULATION_OF_VIRAL_LIFE_CYCLE | 31 | -0.270344 | -0.813444 | 0.736607 | 0.942287 |
| GOBP_NEGATIVE_REGULATION_OF_OXIDATIVE_STRESS_INDUCED_NEURON_DEATH | 16 | -0.323567 | -0.813067 | 0.737705 | 0.942551 |
| GOBP_RECEPTOR_CLUSTERING | 49 | -0.249967 | -0.812886 | 0.780112 | 0.942476 |
| GOBP_REGULATION_OF_ATP_METABOLIC_PROCESS | 114 | -0.219156 | -0.812126 | 0.83268 | 0.943435 |
| GOBP_POSITIVE_REGULATION_OF_TOLL_LIKE_RECEPTOR_SIGNALING_PATHWAY | 27 | -0.278481 | -0.811943 | 0.747006 | 0.943356 |
| GOBP_NEGATIVE_REGULATION_OF_OXIDOREDUCTASE_ACTIVITY | 21 | -0.297329 | -0.811763 | 0.743119 | 0.94326 |
| GOBP_PROTEIN_DEMETHYLATION | 33 | -0.268301 | -0.811268 | 0.752907 | 0.943747 |
| GOBP_NEGATIVE_REGULATION_OF_MUSCLE_CELL_DIFFERENTIATION | 53 | -0.244879 | -0.810939 | 0.780347 | 0.94391 |
| GOBP_PROTEIN_LOCALIZATION_TO_GOLGI_APPARATUS | 28 | -0.278848 | -0.810421 | 0.746224 | 0.94443 |
| GOBP_NEGATIVE_REGULATION_OF_EXOCYTOSIS | 28 | -0.279377 | -0.810375 | 0.741353 | 0.944107 |
| GOBP_RESPIRATORY_BURST | 35 | -0.265887 | -0.810326 | 0.775988 | 0.943773 |
| GOBP_VASCULAR_ENDOTHELIAL_GROWTH_FACTOR_PRODUCTION | 37 | -0.26203 | -0.810289 | 0.756598 | 0.943424 |
| GOBP_CELL_SUBSTRATE_JUNCTION_ORGANIZATION | 104 | -0.220371 | -0.809813 | 0.861075 | 0.943832 |
| GOBP_MAINTENANCE_OF_BLOOD_BRAIN_BARRIER | 32 | -0.269783 | -0.80979 | 0.751914 | 0.943458 |
| GOBP_POSITIVE_REGULATION_OF_WNT_SIGNALING_PATHWAY | 171 | -0.208665 | -0.808786 | 0.893333 | 0.944842 |
| GOBP_REGULATION_OF_PHOSPHOLIPASE_ACTIVITY | 63 | -0.236411 | -0.808737 | 0.801939 | 0.94452 |
| GOBP_NEGATIVE_REGULATION_OF_TUMOR_NECROSIS_FACTOR_MEDIATED_SIGNALING_PATHW | 18 | -0.31301 | -0.808229 | 0.734884 | 0.945023 |
| GOBP_SKIN_EPIDERMIS_DEVELOPMENT | 73 | -0.230931 | -0.808119 | 0.796053 | 0.944789 |
| GOBP_RESPONSE_TO_MISFOLDED_PROTEIN | 24 | -0.286088 | -0.807117 | 0.759434 | 0.946127 |
| GOBP_EXPLORATION_BEHAVIOR | 16 | -0.314204 | -0.806662 | 0.734824 | 0.946543 |
| GOBP_ANTIGEN_PROCESSING_AND_PRESENTATION_OF_EXOGENOUS_PEPTIDE_ANTIGEN_VIA_ | 79 | -0.225331 | -0.806378 | 0.809722 | 0.946633 |
| GOBP_LEARNING | 117 | -0.217424 | -0.80626 | 0.860902 | 0.946441 |
| GOBP_REGULATION_OF_CILIUM_ASSEMBLY | 55 | -0.241795 | -0.806197 | 0.797241 | 0.946128 |
| GOBP_NEGATIVE_REGULATION_OF_REACTIVE_OXYGEN_SPECIES_BIOSYNTHETIC_PROCESS | 28 | -0.275944 | -0.805939 | 0.788732 | 0.946182 |
| GOBP_PYRIMIDINE_NUCLEOTIDE_CATABOLIC_PROCESS | 19 | -0.301231 | -0.805528 | 0.751515 | 0.946475 |
| GOBP_POSITIVE_REGULATION_OF_PROTEIN_CATABOLIC_PROCESS | 218 | -0.200462 | -0.805519 | 0.898734 | 0.946077 |
| GOBP_BLASTOCYST_DEVELOPMENT | 87 | -0.223965 | -0.80531 | 0.839065 | 0.946051 |
| GOBP_BRANCHING_INVOLVED_IN_MAMMARY_GLAND_DUCT_MORPHOGENESIS | 18 | -0.306852 | -0.805104 | 0.741486 | 0.945997 |
| GOBP_NEGATIVE_REGULATION_OF_CELLULAR_PROTEIN_LOCALIZATION | 108 | -0.218779 | -0.804975 | 0.847545 | 0.945814 |
| GOBP_NUCLEOBASE_CONTAINING_SMALL_MOLECULE_CATABOLIC_PROCESS | 48 | -0.248468 | -0.804668 | 0.78863 | 0.945923 |
| GOBP_LYSOSOME_LOCALIZATION | 72 | -0.228225 | -0.803661 | 0.827913 | 0.94727 |
| GOBP_RIBONUCLEOSIDE_CATABOLIC_PROCESS | 23 | -0.286614 | -0.803638 | 0.753823 | 0.946897 |
| GOBP_PROTEIN_LOCALIZATION_TO_NUCLEUS | 265 | -0.195357 | -0.803005 | 0.916856 | 0.94762 |
| GOBP_POSITIVE_REGULATION_OF_NEURON_MIGRATION | 15 | -0.317006 | -0.802946 | 0.764706 | 0.947309 |
| GOBP_BROWN_FAT_CELL_DIFFERENTIATION | 40 | -0.256008 | -0.802588 | 0.762115 | 0.947509 |
| GOBP_CARDIOLIPIN_METABOLIC_PROCESS | 17 | -0.305451 | -0.801077 | 0.741325 | 0.949716 |
| GOBP_CEREBELLAR_PURKINJE_CELL_LAYER_DEVELOPMENT | 20 | -0.29563 | -0.801062 | 0.748092 | 0.949327 |
| GOBP_NEGATIVE_REGULATION_OF_CALCIUM_ION_TRANSMEMBRANE_TRANSPORTER_ACTIVITY | 28 | -0.27597 | -0.800597 | 0.740795 | 0.949688 |
| GOBP_BONE_MATURATION | 20 | -0.293255 | -0.799027 | 0.737557 | 0.951949 |
| GOBP_REGULATION_OF_PROTEIN_MATURATION | 56 | -0.237929 | -0.79899 | 0.823529 | 0.951603 |
| GOBP_MOTILE_CILIUM_ASSEMBLY | 18 | -0.302363 | -0.79836 | 0.739198 | 0.952243 |
| GOBP_REGULATION_OF_MEMBRANE_PERMEABILITY | 80 | -0.226924 | -0.798106 | 0.834225 | 0.952256 |
| GOBP_REGULATION_OF_RETINOIC_ACID_RECEPTOR_SIGNALING_PATHWAY | 15 | -0.315218 | -0.797983 | 0.743961 | 0.952039 |
| GOBP_NUCLEOSIDE_PHOSPHATE_CATABOLIC_PROCESS | 79 | -0.225705 | -0.797422 | 0.837057 | 0.952554 |
| GOBP_NEGATIVE_REGULATION_OF_REACTIVE_OXYGEN_SPECIES_METABOLIC_PROCESS | 52 | -0.242703 | -0.796959 | 0.817558 | 0.95291 |
| GOBP_ANOIKIS | 34 | -0.262633 | -0.796493 | 0.78125 | 0.953249 |
| GOBP_ORGANIC_HYDROXY_COMPOUND_BIOSYNTHETIC_PROCESS | 218 | -0.198721 | -0.796135 | 0.925837 | 0.953422 |
| GOBP_STRIATED_MUSCLE_ADAPTATION | 40 | -0.251321 | -0.795784 | 0.788462 | 0.95359 |
| GOBP_ARP2_3_COMPLEX_MEDIATED_ACTIN_NUCLEATION | 37 | -0.260416 | -0.795312 | 0.783582 | 0.953985 |
| GOBP_REGULATION_OF_ISOTYPE_SWITCHING | 35 | -0.259423 | -0.795249 | 0.775988 | 0.953691 |
| GOBP_POSITIVE_REGULATION_OF_PROTEIN_CONTAINING_COMPLEX_ASSEMBLY | 245 | -0.196381 | -0.795173 | 0.947066 | 0.953394 |
| GOBP_L_GLUTAMATE_TRANSMEMBRANE_TRANSPORT | 23 | -0.284606 | -0.794793 | 0.754545 | 0.95361 |
| GOBP_NEGATIVE_REGULATION_OF_RESPONSE_TO_ENDOPLASMIC_RETICULUM_STRESS | 42 | -0.251816 | -0.79393 | 0.808542 | 0.954632 |
| GOBP_POSITIVE_REGULATION_OF_PROTEIN_AUTOPHOSPHORYLATION | 26 | -0.277794 | -0.792577 | 0.78626 | 0.956503 |
| GOBP_HISTONE_H3_K27_METHYLATION | 17 | -0.307878 | -0.792467 | 0.758842 | 0.956269 |
| GOBP_INTRINSIC_APOPTOTIC_SIGNALING_PATHWAY_IN_RESPONSE_TO_ENDOPLASMIC_RETICU | 58 | -0.235865 | -0.791903 | 0.829609 | 0.956809 |
| GOBP_PHOSPHATE_ION_TRANSPORT | 23 | -0.281915 | -0.791838 | 0.773292 | 0.956502 |
| GOBP_PROTEIN_POLYMERIZATION | 280 | -0.193249 | -0.791289 | 0.942463 | 0.957012 |
| GOBP_ORGANELLE_FUSION | 141 | -0.205634 | -0.791167 | 0.897883 | 0.956782 |
| GOBP_POSITIVE_REGULATION_OF_HEART_RATE | 20 | -0.294792 | -0.790913 | 0.758779 | 0.956786 |
| GOBP_CHAPERONE_COFACTOR_DEPENDENT_PROTEIN_REFOLDING | 34 | -0.259869 | -0.790274 | 0.794948 | 0.957449 |
| GOBP_POSITIVE_REGULATION_OF_MITOCHONDRION_ORGANIZATION | 83 | -0.220742 | -0.789028 | 0.86326 | 0.959033 |
| GOBP_LIVER_REGENERATION | 29 | -0.271843 | -0.7888 | 0.778281 | 0.958984 |
| GOBP_REGULATION_OF_METALLOPEPTIDASE_ACTIVITY | 15 | -0.309841 | -0.786439 | 0.770079 | 0.962248 |
| GOBP_POSITIVE_REGULATION_OF_NERVOUS_SYSTEM_PROCESS | 31 | -0.267881 | -0.786327 | 0.789877 | 0.962021 |
| GOBP_PROTEIN_LOCALIZATION_TO_ENDOSOME | 25 | -0.273887 | -0.78596 | 0.777266 | 0.962172 |
| GOBP_POSITIVE_REGULATION_OF_PROTEIN_DEPHOSPHORYLATION | 44 | -0.246863 | -0.785412 | 0.834294 | 0.962626 |
| GOBP_NEGATIVE_REGULATION_OF_ORGAN_GROWTH | 29 | -0.26641 | -0.784705 | 0.791667 | 0.963306 |
| GOBP_VIRAL_BUDDING_VIA_HOST_ESCRT_COMPLEX | 21 | -0.287456 | -0.783909 | 0.752294 | 0.964141 |
| GOBP_MRNA_CLEAVAGE | 22 | -0.282202 | -0.78288 | 0.783866 | 0.965323 |
| GOBP_REGULATION_OF_ACTIN_FILAMENT_BASED_MOVEMENT | 34 | -0.261108 | -0.782682 | 0.809816 | 0.965218 |
| GOBP_REGULATION_OF_NEUTROPHIL_CHEMOTAXIS | 30 | -0.264344 | -0.782443 | 0.791429 | 0.965177 |
| GOBP_AGING | 262 | -0.191592 | -0.78223 | 0.951835 | 0.965075 |
| GOBP_REGULATION_OF_VASCULAR_ASSOCIATED_SMOOTH_MUSCLE_CELL_DIFFERENTIATION | 15 | -0.314856 | -0.782103 | 0.775701 | 0.964868 |
| GOBP_NEGATIVE_REGULATION_OF_PROTEIN_KINASE_B_SIGNALING | 47 | -0.240188 | -0.781814 | 0.846821 | 0.964886 |
| GOBP_NUCLEAR_MIGRATION | 19 | -0.290685 | -0.781759 | 0.77812 | 0.964566 |
| GOBP_NEUROMUSCULAR_PROCESS_CONTROLLING_BALANCE | 32 | -0.261851 | -0.781691 | 0.792109 | 0.964244 |
| GOBP_TRANSPOSITION | 26 | -0.275273 | -0.78131 | 0.778802 | 0.964427 |
| GOBP_POSITIVE_REGULATION_OF_ACTIN_CYTOSKELETON_REORGANIZATION | 18 | -0.289472 | -0.780582 | 0.760736 | 0.965131 |
| GOBP_RESPONSE_TO_HYDROPEROXIDE | 16 | -0.307619 | -0.780571 | 0.774092 | 0.964736 |

| | | | | | |
|---|---|---|---|---|---|
| GOBP_NEGATIVE_REGULATION_OF_MUSCLE_HYPERTROPHY | 27 | -0.270296 | -0.779713 | 0.802691 | 0.965625 |
| GOBP_POSITIVE_REGULATION_OF_NEURON_PROJECTION_DEVELOPMENT | 144 | -0.20352 | -0.779616 | 0.900645 | 0.965371 |
| GOBP_BIOLOGICAL_PROCESS_INVOLVED_IN_INTERACTION_WITH_SYMBIONT | 79 | -0.219726 | -0.778834 | 0.854993 | 0.966094 |
| GOBP_NEGATIVE_REGULATION_OF_MITOCHONDRION_ORGANIZATION | 45 | -0.241013 | -0.778382 | 0.826516 | 0.966357 |
| GOBP_REGULATION_OF_DENDRITE_MORPHOGENESIS | 65 | -0.227365 | -0.778128 | 0.884507 | 0.966333 |
| GOBP_REGULATION_OF_DEPHOSPHORYLATION | 199 | -0.195131 | -0.777908 | 0.927907 | 0.96627 |
| GOBP_ANTIGEN_PROCESSING_AND_PRESENTATION_OF_PEPTIDE_ANTIGEN_VIA_MHC_CLASS_I | 98 | -0.215225 | -0.777899 | 0.884467 | 0.965873 |
| GOBP_POSITIVE_REGULATION_OF_CELL_PROJECTION_ORGANIZATION | 323 | -0.187353 | -0.777588 | 0.958011 | 0.965913 |
| GOBP_ERBB2_SIGNALING_PATHWAY | 30 | -0.263013 | -0.77646 | 0.802651 | 0.967232 |
| GOBP_NEGATIVE_REGULATION_OF_AUTOPHAGY | 79 | -0.220062 | -0.776377 | 0.856574 | 0.966944 |
| GOBP_REGULATION_OF_MITOCHONDRION_ORGANIZATION | 149 | -0.204097 | -0.776119 | 0.919699 | 0.966916 |
| GOBP_RETINOL_METABOLIC_PROCESS | 35 | -0.253358 | -0.774036 | 0.821483 | 0.969613 |
| GOBP_INTEGRIN_MEDIATED_SIGNALING_PATHWAY | 103 | -0.208103 | -0.773351 | 0.888149 | 0.970185 |
| GOBP_NUCLEOSIDE_CATABOLIC_PROCESS | 32 | -0.261413 | -0.773259 | 0.796784 | 0.969907 |
| GOBP_PROTEIN_PROCESSING | 193 | -0.195017 | -0.772952 | 0.935407 | 0.969949 |
| GOBP_VIRAL_LIFE_CYCLE | 332 | -0.186368 | -0.770948 | 0.964286 | 0.972371 |
| GOBP_PROTEIN_PHOSPHOPANTETHEINYLATION | 296 | -0.186462 | -0.769813 | 0.968218 | 0.973608 |
| GOBP_PEPTIDYL_ASPARAGINE_MODIFICATION | 30 | -0.260465 | -0.769539 | 0.821752 | 0.973588 |
| GOBP_CELLULAR_PROTEIN_CONTAINING_COMPLEX_LOCALIZATION | 21 | -0.281299 | -0.76939 | 0.797806 | 0.97339 |
| GOBP_REGULATION_OF_CIRCADIAN_RHYTHM | 103 | -0.209583 | -0.76927 | 0.917969 | 0.973146 |
| GOBP_VIRAL_BUDDING | 26 | -0.265318 | -0.768887 | 0.829085 | 0.973266 |
| GOBP_REGULATION_OF_PROTEIN_CONTAINING_COMPLEX_ASSEMBLY | 423 | -0.182562 | -0.768847 | 0.982869 | 0.972908 |
| GOBP_DETERMINATION_OF_ADULT_LIFESPAN | 16 | -0.304705 | -0.768771 | 0.771791 | 0.972611 |
| GOBP_ZYMOGEN_ACTIVATION | 50 | -0.231411 | -0.768332 | 0.834522 | 0.972848 |
| GOBP_REGULATION_OF_GLUCOSE_TRANSMEMBRANE_TRANSPORT | 64 | -0.227519 | -0.76831 | 0.834925 | 0.972467 |
| GOBP_REGULATION_OF_MEMBRANE_REPOLARIZATION | 30 | -0.260725 | -0.767827 | 0.800298 | 0.972745 |
| GOBP_NEUTRAL_LIPID_METABOLIC_PROCESS | 108 | -0.209749 | -0.767252 | 0.912698 | 0.973155 |
| GOBP_HISTONE_H4_K5_ACETYLATION | 16 | -0.303565 | -0.76676 | 0.785047 | 0.973441 |
| GOBP_REGULATION_OF_CARDIAC_MUSCLE_CELL_MEMBRANE_REPOLARIZATION | 24 | -0.273847 | -0.766504 | 0.794207 | 0.973395 |
| GOBP_POSITIVE_REGULATION_OF_VASCULAR_PERMEABILITY | 15 | -0.302549 | -0.765844 | 0.768212 | 0.973918 |
| GOBP_UBIQUITIN_DEPENDENT_PROTEIN_CATABOLIC_PROCESS_VIA_THE_MULTIVESICULAR_BO | 23 | -0.275341 | -0.764971 | 0.808576 | 0.974675 |
| GOBP_POSITIVE_REGULATION_OF_TELOMERASE_ACTIVITY | 32 | -0.255629 | -0.764019 | 0.827834 | 0.975561 |
| GOBP_AXONEME_ASSEMBLY | 66 | -0.222745 | -0.763721 | 0.866131 | 0.975551 |
| GOBP_CEREBRAL_CORTEX_CELL_MIGRATION | 35 | -0.249214 | -0.762657 | 0.848752 | 0.976591 |
| GOBP_REGULATION_OF_PLATELET_AGGREGATION | 17 | -0.295145 | -0.76168 | 0.791332 | 0.977528 |
| GOBP_MIRNA_METABOLIC_PROCESS | 28 | -0.261814 | -0.761599 | 0.844136 | 0.977218 |
| GOBP_EXOCRINE_SYSTEM_DEVELOPMENT | 36 | -0.247107 | -0.761403 | 0.810931 | 0.977083 |
| GOBP_MACROMOLECULE_DEACYLATION | 105 | -0.20697 | -0.761369 | 0.925099 | 0.976725 |
| GOBP_TYPE_B_PANCREATIC_CELL_DEVELOPMENT | 18 | -0.289697 | -0.760884 | 0.806154 | 0.976968 |
| GOBP_POSITIVE_REGULATION_OF_IMMUNOGLOBULIN_PRODUCTION | 44 | -0.236485 | -0.760621 | 0.859155 | 0.976906 |
| GOBP_SECONDARY_METABOLITE_BIOSYNTHETIC_PROCESS | 17 | -0.293682 | -0.76044 | 0.814815 | 0.97675 |
| GOBP_MEMBRANE_LIPID_METABOLIC_PROCESS | 205 | -0.190304 | -0.75921 | 0.961538 | 0.977948 |
| GOBP_IN_UTERO_EMBRYONIC_DEVELOPMENT | 311 | -0.185114 | -0.758572 | 0.974501 | 0.978376 |
| GOBP_N_GLYCAN_PROCESSING | 17 | -0.288459 | -0.758437 | 0.791461 | 0.978123 |
| GOBP_LUNG_CELL_DIFFERENTIATION | 17 | -0.29001 | -0.757681 | 0.805281 | 0.978687 |
| GOBP_REGULATION_OF_TELOMERASE_ACTIVITY | 45 | -0.234824 | -0.757417 | 0.868876 | 0.978624 |
| GOBP_BIOLOGICAL_PROCESS_INVOLVED_IN_INTERACTION_WITH_HOST | 207 | -0.190556 | -0.756285 | 0.957678 | 0.979729 |
| GOBP_HEMATOPOIETIC_PROGENITOR_CELL_DIFFERENTIATION | 156 | -0.196642 | -0.754557 | 0.941392 | 0.981531 |
| GOBP_NEGATIVE_REGULATION_OF_PROTEIN_MATURATION | 24 | -0.26889 | -0.754413 | 0.813534 | 0.981303 |
| GOBP_REGULATION_OF_SMOOTHENED_SIGNALING_PATHWAY | 73 | -0.217722 | -0.754044 | 0.895095 | 0.981367 |
| GOBP_CELLULAR_RESPONSE_TO_ABIOTIC_STIMULUS | 303 | -0.182782 | -0.754033 | 0.978555 | 0.980972 |
| GOBP_SECONDARY_METABOLIC_PROCESS | 42 | -0.238673 | -0.752371 | 0.859216 | 0.98276 |
| GOBP_MAINTENANCE_OF_PROTEIN_LOCALIZATION_IN_ORGANELLE | 42 | -0.235905 | -0.752148 | 0.861671 | 0.98263 |
| GOBP_NEGATIVE_REGULATION_OF_NECROTIC_CELL_DEATH | 17 | -0.285527 | -0.751112 | 0.802198 | 0.983561 |
| GOBP_NEGATIVE_REGULATION_OF_CATION_TRANSMEMBRANE_TRANSPORT | 75 | -0.214957 | -0.750334 | 0.896505 | 0.984122 |
| GOBP_GLYCEROPHOSPHOLIPID_CATABOLIC_PROCESS | 27 | -0.259867 | -0.749955 | 0.830104 | 0.984162 |
| GOBP_OSTEOBLAST_DEVELOPMENT | 16 | -0.298406 | -0.749733 | 0.820827 | 0.984027 |
| GOBP_SPERMATID_NUCLEUS_DIFFERENTIATION | 15 | -0.298464 | -0.749695 | 0.808065 | 0.983662 |
| GOBP_PHOSPHATIDYLINOSITOL_METABOLIC_PROCESS | 173 | -0.191534 | -0.74962 | 0.961353 | 0.983351 |
| GOBP_AUTOPHAGOSOME_MATURATION | 41 | -0.236606 | -0.748795 | 0.865672 | 0.983949 |
| GOBP_ALCOHOL_METABOLIC_PROCESS | 332 | -0.18078 | -0.748045 | 0.983642 | 0.984493 |
| GOBP_LIPOXYGENASE_PATHWAY | 15 | -0.296374 | -0.747571 | 0.80303 | 0.984643 |
| GOBP_NEGATIVE_REGULATION_OF_CELL_PROJECTION_ORGANIZATION | 166 | -0.190471 | -0.747228 | 0.964029 | 0.984642 |
| GOBP_NEGATIVE_REGULATION_OF_PROTEIN_CATABOLIC_PROCESS | 113 | -0.199766 | -0.747003 | 0.939744 | 0.984505 |
| GOBP_NEGATIVE_REGULATION_OF_INTRACELLULAR_PROTEIN_TRANSPORT | 40 | -0.235073 | -0.746505 | 0.859575 | 0.984666 |
| GOBP_CYCLIC_NUCLEOTIDE_BIOSYNTHETIC_PROCESS | 20 | -0.271607 | -0.746414 | 0.797771 | 0.984372 |
| GOBP_LYMPH_VESSEL_DEVELOPMENT | 24 | -0.267235 | -0.746026 | 0.814307 | 0.984455 |
| GOBP_POSITIVE_REGULATION_OF_ORGANELLE_ASSEMBLY | 65 | -0.217467 | -0.745351 | 0.883008 | 0.98488 |
| GOBP_EMBRYONIC_CRANIAL_SKELETON_MORPHOGENESIS | 42 | -0.232743 | -0.745259 | 0.870588 | 0.984585 |
| GOBP_INTRACELLULAR_STEROID_HORMONE_RECEPTOR_SIGNALING_PATHWAY | 104 | -0.202068 | -0.74521 | 0.915617 | 0.984238 |
| GOBP_FATTY_ACID_BETA_OXIDATION | 73 | -0.214683 | -0.745023 | 0.915323 | 0.98405 |
| GOBP_REGULATION_OF_ARP2_3_COMPLEX_MEDIATED_ACTIN_NUCLEATION | 20 | -0.272452 | -0.744884 | 0.811178 | 0.983813 |
| GOBP_NEGATIVE_REGULATION_OF_PLASMA_MEMBRANE_BOUNDED_CELL_PROJECTION_ASSEM | 27 | -0.259204 | -0.743556 | 0.831845 | 0.984986 |
| GOBP_PROTEIN_TARGETING_TO_LYSOSOME | 23 | -0.26373 | -0.742784 | 0.826667 | 0.985477 |
| GOBP_QUINONE_METABOLIC_PROCESS | 32 | -0.249343 | -0.742514 | 0.848787 | 0.985376 |
| GOBP_REGULATION_OF_CELL_MATURATION | 18 | -0.284222 | -0.74244 | 0.829653 | 0.985056 |
| GOBP_ZINC_ION_TRANSPORT | 24 | -0.263268 | -0.741736 | 0.851796 | 0.985497 |
| GOBP_NEUROTRANSMITTER_RECEPTOR_TRANSPORT_TO_PLASMA_MEMBRANE | 16 | -0.285223 | -0.74105 | 0.806748 | 0.985901 |
| GOBP_REGULATION_OF_PROTEIN_CATABOLIC_PROCESS | 368 | -0.178971 | -0.740399 | 0.991021 | 0.986247 |
| GOBP_LIPID_PHOSPHORYLATION | 68 | -0.213552 | -0.739196 | 0.892807 | 0.987236 |
| GOBP_SYNAPTIC_VESICLE_LOCALIZATION | 53 | -0.225937 | -0.738059 | 0.885269 | 0.988146 |
| GOBP_CELLULAR_GLUCAN_METABOLIC_PROCESS | 70 | -0.212102 | -0.737461 | 0.901639 | 0.988443 |
| GOBP_POSITIVE_REGULATION_OF_PROTEIN_LOCALIZATION_TO_NUCLEUS | 83 | -0.207658 | -0.737058 | 0.91363 | 0.988509 |
| GOBP_NEGATIVE_REGULATION_OF_MRNA_SPLICING_VIA_SPLICEOSOME | 20 | -0.275075 | -0.736924 | 0.845577 | 0.988242 |
| GOBP_SKELETAL_MUSCLE_ADAPTATION | 21 | -0.27133 | -0.735467 | 0.841555 | 0.989486 |
| GOBP_NEGATIVE_REGULATION_OF_POSTTRANSCRIPTIONAL_GENE_SILENCING | 20 | -0.269802 | -0.734901 | 0.830218 | 0.98972 |
| GOBP_PHAGOSOME_ACIDIFICATION | 25 | -0.255722 | -0.734894 | 0.853169 | 0.989319 |
| GOBP_GOLGI_ORGANIZATION | 142 | -0.192479 | -0.734444 | 0.951128 | 0.989436 |
| GOBP_GLYCOSYLATION | 242 | -0.181746 | -0.732829 | 0.984866 | 0.990824 |
| GOBP_REGULATION_OF_CILIUM_DEPENDENT_CELL_MOTILITY | 17 | -0.282972 | -0.732468 | 0.841615 | 0.990787 |
| GOBP_NEGATIVE_REGULATION_OF_RNA_SPLICING | 25 | -0.256781 | -0.731045 | 0.863924 | 0.991933 |
| GOBP_NUCLEAR_TRANSPORT | 330 | -0.175154 | -0.729517 | 0.995628 | 0.993147 |
| GOBP_IRE1_MEDIATED_UNFOLDED_PROTEIN_RESPONSE | 66 | -0.212829 | -0.728613 | 0.903401 | 0.993694 |
| GOBP_GLUTAMINE_METABOLIC_PROCESS | 22 | -0.264826 | -0.728378 | 0.862654 | 0.993542 |
| GOBP_CELLULAR_RESPONSE_TO_HYDROGEN_PEROXIDE | 79 | -0.206383 | -0.726957 | 0.926997 | 0.994588 |
| GOBP_RESPONSE_TO_STARVATION | 182 | -0.185958 | -0.726523 | 0.963743 | 0.994642 |
| GOBP_REGULATION_OF_SUPEROXIDE_ANION_GENERATION | 19 | -0.273896 | -0.726404 | 0.825959 | 0.994357 |
| GOBP_SPLICEOSOMAL_COMPLEX_ASSEMBLY | 65 | -0.214829 | -0.726113 | 0.905687 | 0.994256 |
| GOBP_POSITIVE_REGULATION_OF_CALCIUM_ION_TRANSPORT_INTO_CYTOSOL | 47 | -0.223837 | -0.72607 | 0.897959 | 0.993888 |
| GOBP_REGULATION_OF_MEIOTIC_NUCLEAR_DIVISION | 25 | -0.253102 | -0.725765 | 0.856269 | 0.99381 |
| GOBP_AMINE_CATABOLIC_PROCESS | 29 | -0.242176 | -0.725634 | 0.862745 | 0.993538 |
| GOBP_SUBSTANTIA_NIGRA_DEVELOPMENT | 41 | -0.230653 | -0.72552 | 0.880419 | 0.99325 |
| GOBP_LABYRINTHINE_LAYER_BLOOD_VESSEL_DEVELOPMENT | 16 | -0.282459 | -0.724621 | 0.821372 | 0.993771 |
| GOBP_PROTEIN_LOCALIZATION_TO_MICROTUBULE | 18 | -0.272969 | -0.722454 | 0.84019 | 0.99558 |
| GOBP_HISTONE_H2A_MONOUBIQUITINATION | 17 | -0.274892 | -0.722382 | 0.840376 | 0.99524 |
| GOBP_POSITIVE_REGULATION_OF_PROTEIN_CONTAINING_COMPLEX_DISASSEMBLY | 34 | -0.237905 | -0.721843 | 0.884558 | 0.995367 |
| GOBP_INTRACELLULAR_LIPID_TRANSPORT | 44 | -0.228122 | -0.72178 | 0.896011 | 0.995023 |
| GOBP_REGULATION_OF_NUCLEAR_TRANSCRIBED_MRNA_CATABOLIC_PROCESS_DEADENYLAT | 16 | -0.281252 | -0.721598 | 0.845295 | 0.994804 |
| GOBP_PHENOL_CONTAINING_COMPOUND_METABOLIC_PROCESS | 81 | -0.200558 | -0.72148 | 0.937008 | 0.994511 |
| GOBP_CELLULAR_MONOVALENT_INORGANIC_CATION_HOMEOSTASIS | 106 | -0.195756 | -0.721297 | 0.944966 | 0.994288 |
| GOBP_COLLAGEN_BIOSYNTHETIC_PROCESS | 43 | -0.223412 | -0.72105 | 0.892962 | 0.994127 |
| GOBP_NLRP3_INFLAMMASOME_COMPLEX_ASSEMBLY | 15 | -0.291268 | -0.720846 | 0.807154 | 0.993937 |
| GOBP_REGULATION_OF_AUTOPHAGY | 319 | -0.173638 | -0.719122 | 0.992135 | 0.995217 |
| GOBP_RESPONSE_TO_OXYGEN_LEVELS | 355 | -0.173098 | -0.718563 | 0.995556 | 0.995358 |
| GOBP_DENDRITIC_SPINE_MAINTENANCE | 17 | -0.273163 | -0.718417 | 0.856467 | 0.995088 |
| GOBP_DEOXYRIBONUCLEOTIDE_METABOLIC_PROCESS | 39 | -0.231684 | -0.717722 | 0.894203 | 0.995357 |
| GOBP_NEGATIVE_REGULATION_OF_ENDOCYTOSIS | 47 | -0.222932 | -0.717383 | 0.907381 | 0.995279 |
| GOBP_NEGATIVE_REGULATION_OF_CELLULAR_CATABOLIC_PROCESS | 234 | -0.176865 | -0.717118 | 0.989631 | 0.995137 |
| GOBP_NEGATIVE_REGULATION_OF_DNA_BINDING | 48 | -0.220525 | -0.717088 | 0.909884 | 0.994765 |
| GOBP_STEROL_METABOLIC_PROCESS | 147 | -0.185097 | -0.716273 | 0.971189 | 0.995154 |
| GOBP_ORGANOPHOSPHATE_CATABOLIC_PROCESS | 144 | -0.188882 | -0.71599 | 0.969287 | 0.995033 |
| GOBP_PEPTIDYL_ARGININE_MODIFICATION | 22 | -0.253439 | -0.714244 | 0.869301 | 0.996279 |
| GOBP_NEGATIVE_REGULATION_OF_DEPHOSPHORYLATION | 102 | -0.192447 | -0.713206 | 0.965649 | 0.996822 |
| GOBP_LINOLEIC_ACID_METABOLIC_PROCESS | 18 | -0.271138 | -0.710439 | 0.867089 | 0.999068 |
| GOBP_PROTEIN_LOCALIZATION_TO_CELL_CELL_JUNCTION | 21 | -0.259736 | -0.710414 | 0.86015 | 0.998692 |
| GOBP_REGULATION_OF_TRANSLATIONAL_INITIATION_IN_RESPONSE_TO_STRESS | 15 | -0.286184 | -0.710385 | 0.855306 | 0.99831 |
| GOBP_POSITIVE_REGULATION_OF_CELLULAR_RESPONSE_TO_TRANSFORMING_GROWTH_FACT | 25 | -0.253548 | -0.707682 | 0.888227 | 1 |
| GOBP_PROTEIN_DEPHOSPHORYLATION | 308 | -0.17135 | -0.707572 | 0.9967 | 1 |
| GOBP_PROTEIN_TARGETING_TO_VACUOLE | 40 | -0.225597 | -0.707483 | 0.90113 | 0.999708 |
| GOBP_PHOSPHORYLATED_CARBOHYDRATE_DEPHOSPHORYLATION | 16 | -0.276938 | -0.707398 | 0.855814 | 0.999375 |
| GOBP_POSITIVE_REGULATION_OF_SODIUM_ION_TRANSMEMBRANE_TRANSPORTER_ACTIVITY | 16 | -0.27045 | -0.705898 | 0.837359 | 1 |
| GOBP_PALLIUM_DEVELOPMENT | 135 | -0.185501 | -0.705518 | 0.97449 | 1 |
| GOBP_REGULATION_OF_PROGRAMMED_NECROTIC_CELL_DEATH | 22 | -0.25657 | -0.705057 | 0.863976 | 1 |
| GOBP_POSITIVE_REGULATION_OF_DENDRITIC_SPINE_DEVELOPMENT | 42 | -0.22695 | -0.703954 | 0.915835 | 1 |
| GOBP_PEPTIDYL_TYROSINE_AUTOPHOSPHORYLATION | 20 | -0.262883 | -0.703238 | 0.866987 | 1 |
| GOBP_NEGATIVE_REGULATION_OF_NUCLEOCYTOPLASMIC_TRANSPORT | 20 | -0.262288 | -0.702986 | 0.874807 | 1 |
| GOBP_POSITIVE_REGULATION_OF_SUBSTRATE_ADHESION_DEPENDENT_CELL_SPREADING | 39 | -0.225365 | -0.702725 | 0.916074 | 1 |
| GOBP_MONOSACCHARIDE_CATABOLIC_PROCESS | 60 | -0.205482 | -0.701368 | 0.941512 | 1 |
| GOBP_PROTEIN_TRANSPORT_ALONG_MICROTUBULE | 68 | -0.202503 | -0.701353 | 0.936022 | 1 |
| GOBP_EXTRACELLULAR_MATRIX_ASSEMBLY | 43 | -0.223059 | -0.698519 | 0.936391 | 1 |
| GOBP_POSITIVE_REGULATION_OF_PROTEIN_TARGETING_TO_MEMBRANE | 29 | -0.237075 | -0.698189 | 0.902798 | 1 |
| GOBP_REGULATION_OF_CARBOHYDRATE_CATABOLIC_PROCESS | 87 | -0.196208 | -0.69733 | 0.967655 | 1 |
| GOBP_REGULATION_OF_ERYTHROCYTE_DIFFERENTIATION | 47 | -0.217106 | -0.697157 | 0.921238 | 1 |
| GOBP_AXONEMAL_DYNEIN_COMPLEX_ASSEMBLY | 32 | -0.232217 | -0.696674 | 0.885542 | 1 |
| GOBP_ENDOPLASMIC_RETICULUM_CALCIUM_ION_HOMEOSTASIS | 25 | -0.239793 | -0.692862 | 0.909091 | 1 |
| GOBP_VESICLE_MEDIATED_TRANSPORT_TO_THE_PLASMA_MEMBRANE | 90 | -0.192122 | -0.689999 | 0.962766 | 1 |
| GOBP_PROTEIN_K63_LINKED_DEUBIQUITINATION | 34 | -0.232286 | -0.68905 | 0.914119 | 1 |
| GOBP_REGULATION_OF_NUCLEAR_TRANSCRIBED_MRNA_POLY_A_TAIL_SHORTENING | 15 | -0.273201 | -0.688288 | 0.891547 | 1 |
| GOBP_PHOSPHATIDYLCHOLINE_ACYL_CHAIN_REMODELING | 23 | -0.250424 | -0.687187 | 0.880419 | 1 |
| GOBP_PROTEIN_CONTAINING_COMPLEX_LOCALIZATION | 274 | -0.168106 | -0.685313 | 0.996599 | 1 |
| GOBP_GLYCOSYL_COMPOUND_CATABOLIC_PROCESS | 42 | -0.216066 | -0.68246 | 0.929799 | 1 |
| GOBP_DEPHOSPHORYLATION | 459 | -0.161717 | -0.68225 | 1 | 1 |
| GOBP_MANNOSYLATION | 36 | -0.219854 | -0.681853 | 0.941423 | 1 |
| GOBP_POSITIVE_REGULATION_OF_MITOCHONDRIAL_OUTER_MEMBRANE_PERMEABILIZATION_IN | 35 | -0.224147 | -0.681527 | 0.931133 | 1 |
| GOBP_POSITIVE_REGULATION_OF_MRNA_METABOLIC_PROCESS | 79 | -0.190945 | -0.681016 | 0.969274 | 1 |
| GOBP_TONGUE_DEVELOPMENT | 16 | -0.259929 | -0.680876 | 0.88254 | 1 |
| GOBP_REGULATION_OF_DNA_TEMPLATED_TRANSCRIPTION_INITIATION | 41 | -0.218632 | -0.680774 | 0.929379 | 1 |
| GOBP_RESPONSE_TO_ACTIVITY | 65 | -0.201629 | -0.68043 | 0.9573 | 1 |
| GOBP_NEGATIVE_REGULATION_OF_CELLULAR_PROTEIN_CATABOLIC_PROCESS | 72 | -0.198285 | -0.680148 | 0.956464 | 1 |
| GOBP_RESPONSE_TO_ENDOPLASMIC_RETICULUM_STRESS | 284 | -0.16467 | -0.679852 | 0.998828 | 1 |
| GOBP_SKELETAL_MUSCLE_TISSUE_REGENERATION | 34 | -0.224576 | -0.678504 | 0.918155 | 1 |
| GOBP_POSITIVE_REGULATION_OF_MEMBRANE_PERMEABILITY | 63 | -0.196727 | -0.678234 | 0.95423 | 1 |
| GOBP_NUCLEAR_TRANSCRIBED_MRNA_POLY_A_TAIL_SHORTENING | 34 | -0.221799 | -0.676273 | 0.953556 | 1 |
| GOBP_ACTIN_FILAMENT_SEVERING | 16 | -0.266165 | -0.675451 | 0.909836 | 1 |
| GOBP_HISTONE_DEUBIQUITINATION | 23 | -0.24194 | -0.6752 | 0.920852 | 1 |
| GOBP_REGULATION_OF_HAIR_FOLLICLE_DEVELOPMENT | 17 | -0.254721 | -0.67476 | 0.920382 | 1 |
| GOBP_AXO_DENDRITIC_TRANSPORT | 68 | -0.194129 | -0.674359 | 0.961003 | 1 |
| GOBP_KERATAN_SULFATE_METABOLIC_PROCESS | 31 | -0.229246 | -0.674224 | 0.924088 | 1 |

| | | | | | |
|---|---|---|---|---|---|
| GOBP_CELLULAR_RESPONSE_TO_OXYGEN_LEVELS | 209 | -0.169606 | -0.673317 | 0.997683 | 1 |
| GOBP_KERATAN_SULFATE_BIOSYNTHETIC_PROCESS | 26 | -0.232853 | -0.672801 | 0.911894 | 1 |
| GOBP_REGULATION_OF_LAMELLIPODIUM_ASSEMBLY | 37 | -0.217324 | -0.672711 | 0.935484 | 1 |
| GOBP_CELLULAR_RESPONSE_TO_TOPOLOGICALLY_INCORRECT_PROTEIN | 163 | -0.172222 | -0.672331 | 0.993873 | 1 |
| GOBP_INOSITOL_PHOSPHATE_CATABOLIC_PROCESS | 18 | -0.254831 | -0.671585 | 0.887279 | 1 |
| GOBP_NADPH_REGENERATION | 17 | -0.255378 | -0.670005 | 0.903614 | 1 |
| GOBP_VESICLE_DOCKING | 64 | -0.196158 | -0.669422 | 0.956044 | 1 |
| GOBP_ENDOPLASMIC_RETICULUM_ORGANIZATION | 85 | -0.188572 | -0.668355 | 0.975275 | 1 |
| GOBP_MAINTENANCE_OF_CELL_POLARITY | 18 | -0.255013 | -0.667551 | 0.905426 | 1 |
| GOBP_POSITIVE_REGULATION_OF_DNA_TEMPLATED_TRANSCRIPTION_INITIATION | 34 | -0.219915 | -0.666978 | 0.936872 | 1 |
| GOBP_PROTEIN_TARGETING | 423 | -0.158782 | -0.666712 | 0.998911 | 1 |
| GOBP_REGULATION_OF_RNA_SPLICING | 136 | -0.174346 | -0.66667 | 0.994845 | 1 |
| GOBP_REGULATION_OF_CELL_GROWTH_INVOLVED_IN_CARDIAC_MUSCLE_CELL_DEVELOPMEN | 17 | -0.256917 | -0.665031 | 0.90755 | 1 |
| GOBP_RESPONSE_TO_TOPOLOGICALLY_INCORRECT_PROTEIN | 197 | -0.167978 | -0.664906 | 0.998824 | 1 |
| GOBP_PROTEIN_ACYLATION | 224 | -0.166193 | -0.663518 | 0.998833 | 1 |
| GOBP_CELLULAR_RESPONSE_TO_IONIZING_RADIATION | 62 | -0.194285 | -0.66278 | 0.970954 | 1 |
| GOBP_MICROVILLUS_ASSEMBLY | 16 | -0.254332 | -0.66248 | 0.911179 | 1 |
| GOBP_REGULATION_OF_SYNAPTIC_VESICLE_RECYCLING | 23 | -0.237463 | -0.661615 | 0.927711 | 1 |
| GOBP_GLYCOSIDE_METABOLIC_PROCESS | 19 | -0.24954 | -0.66139 | 0.898714 | 1 |
| GOBP_MITOCHONDRIAL_OUTER_MEMBRANE_PERMEABILIZATION | 55 | -0.196634 | -0.661292 | 0.95904 | 1 |
| GOBP_CYTOPLASMIC_PATTERN_RECOGNITION_RECEPTOR_SIGNALING_PATHWAY_IN_RESPONS | 30 | -0.227142 | -0.660697 | 0.93617 | 1 |
| GOBP_TRANSLATIONAL_INITIATION | 188 | -0.166246 | -0.658619 | 0.996399 | 1 |
| GOBP_POST_TRANSLATIONAL_PROTEIN_MODIFICATION | 332 | -0.159456 | -0.656967 | 0.998893 | 1 |
| GOBP_INSULIN_RECEPTOR_SIGNALING_PATHWAY | 131 | -0.173043 | -0.655014 | 0.992376 | 1 |
| GOBP_POTASSIUM_ION_EXPORT_ACROSS_PLASMA_MEMBRANE | 16 | -0.256255 | -0.655005 | 0.895082 | 1 |
| GOBP_POSITIVE_REGULATION_OF_CELLULAR_PROTEIN_CATABOLIC_PROCESS | 145 | -0.16821 | -0.652634 | 0.995037 | 1 |
| GOBP_MORPHOGENESIS_OF_A_POLARIZED_EPITHELIUM | 139 | -0.172195 | -0.652619 | 0.994943 | 1 |
| GOBP_MODULATION_BY_HOST_OF_SYMBIONT_PROCESS | 60 | -0.196372 | -0.652557 | 0.964937 | 1 |
| GOBP_HYPEROSMOTIC_RESPONSE | 23 | -0.231158 | -0.651721 | 0.934251 | 1 |
| GOBP_VESICLE_TRANSPORT_ALONG_MICROTUBULE | 46 | -0.203531 | -0.651505 | 0.958884 | 1 |
| GOBP_CELLULAR_SENESCENCE | 67 | -0.191018 | -0.651233 | 0.973721 | 1 |
| GOBP_ACTIVATION_OF_PHOSPHOLIPASE_C_ACTIVITY | 29 | -0.222245 | -0.650707 | 0.935143 | 1 |
| GOBP_REGULATION_OF_HIPPO_SIGNALING | 18 | -0.242942 | -0.650651 | 0.915994 | 1 |
| GOBP_POSITIVE_REGULATION_OF_CELL_MORPHOGENESIS_INVOLVED_IN_DIFFERENTIATION | 76 | -0.182777 | -0.648996 | 0.982827 | 1 |
| GOBP_REGULATION_OF_ESTABLISHMENT_OR_MAINTENANCE_OF_CELL_POLARITY | 25 | -0.229188 | -0.647974 | 0.926145 | 1 |
| GOBP_POSITIVE_REGULATION_OF_PATTERN_RECOGNITION_RECEPTOR_SIGNALING_PATHWAY | 41 | -0.204067 | -0.645322 | 0.960756 | 1 |
| GOBP_REGULATION_OF_EARLY_ENDOSOME_TO_LATE_ENDOSOME_TRANSPORT | 18 | -0.245128 | -0.643992 | 0.929448 | 1 |
| GOBP_NEGATIVE_REGULATION_OF_PROTEASOMAL_UBIQUITIN_DEPENDENT_PROTEIN_CATABOL | 34 | -0.211645 | -0.643926 | 0.966818 | 1 |
| GOBP_POSITIVE_REGULATION_OF_DNA_BIOSYNTHETIC_PROCESS | 64 | -0.185816 | -0.642992 | 0.97486 | 1 |
| GOBP_REGULATION_OF_ALTERNATIVE_MRNA_SPLICING_VIA_SPLICEOSOME | 53 | -0.19466 | -0.642609 | 0.962861 | 1 |
| GOBP_PHOSPHOLIPID_CATABOLIC_PROCESS | 43 | -0.202751 | -0.642064 | 0.965418 | 1 |
| GOBP_MAINTENANCE_OF_PROTEIN_LOCATION_IN_CELL | 63 | -0.188788 | -0.640521 | 0.977901 | 1 |
| GOBP_REGULATION_OF_PEPTIDYL_LYSINE_ACETYLATION | 58 | -0.191439 | -0.639864 | 0.973721 | 1 |
| GOBP_THYROID_HORMONE_METABOLIC_PROCESS | 20 | -0.234955 | -0.639181 | 0.926563 | 1 |
| GOBP_SOMATIC_RECOMBINATION_OF_IMMUNOGLOBULIN_GENE_SEGMENTS | 54 | -0.192854 | -0.638399 | 0.975291 | 1 |
| GOBP_EMBRYONIC_CAMERA_TYPE_EYE_MORPHOGENESIS | 17 | -0.244046 | -0.638024 | 0.91875 | 1 |
| GOBP_GOLGI_TO_PLASMA_MEMBRANE_TRANSPORT | 57 | -0.188283 | -0.635608 | 0.977992 | 1 |
| GOBP_POSITIVE_REGULATION_OF_CELL_AGING | 15 | -0.250967 | -0.635281 | 0.925566 | 1 |
| GOBP_NEURON_DEATH_IN_RESPONSE_TO_OXIDATIVE_STRESS | 26 | -0.223729 | -0.634038 | 0.955023 | 1 |
| GOBP_REGULATION_OF_LIPOPROTEIN_PARTICLE_CLEARANCE | 17 | -0.242532 | -0.633768 | 0.931035 | 1 |
| GOBP_POSITIVE_REGULATION_OF_TRANSCRIPTION_FROM_RNA_POLYMERASE_II_PROMOTER_II | 21 | -0.228272 | -0.63307 | 0.926791 | 1 |
| GOBP_RESPONSE_TO_STEROL_DEPLETION | 18 | -0.238588 | -0.630619 | 0.934351 | 1 |
| GOBP_MRNA_CIS_SPLICING_VIA_SPLICEOSOME | 29 | -0.215637 | -0.629471 | 0.953593 | 1 |
| GOBP_POSITIVE_REGULATION_OF_TRANSLATION | 123 | -0.168436 | -0.628205 | 0.996119 | 1 |
| GOBP_TRANSCRIPTION_INITIATION_FROM_RNA_POLYMERASE_II_PROMOTER | 178 | -0.161155 | -0.626728 | 1 | 1 |
| GOBP_NEGATIVE_REGULATION_OF_PROGRAMMED_NECROTIC_CELL_DEATH | 15 | -0.24689 | -0.62634 | 0.92 | 1 |
| GOBP_CYTOSKELETON_DEPENDENT_INTRACELLULAR_TRANSPORT | 192 | -0.158678 | -0.6257 | 1 | 1 |
| GOBP_PROTEIN_SUMOYLATION | 78 | -0.179245 | -0.625626 | 0.98545 | 1 |
| GOBP_REGULATION_OF_PROTEIN_MODIFICATION_BY_SMALL_PROTEIN_CONJUGATION_OR_REM | 226 | -0.155973 | -0.62456 | 1 | 1 |
| GOBP_AUTOPHAGOSOME_ORGANIZATION | 97 | -0.170446 | -0.624063 | 0.990716 | 1 |
| GOBP_NEGATIVE_REGULATION_OF_DOUBLE_STRAND_BREAK_REPAIR_VIA_HOMOLOGOUS_REC | 19 | -0.236776 | -0.623101 | 0.936728 | 1 |
| GOBP_PURINE_NUCLEOSIDE_METABOLIC_PROCESS | 60 | -0.183397 | -0.623063 | 0.977273 | 1 |
| GOBP_RIBONUCLEOSIDE_MONOPHOSPHATE_METABOLIC_PROCESS | 55 | -0.185642 | -0.619624 | 0.973126 | 1 |
| GOBP_POST_GOLGI_VESICLE_MEDIATED_TRANSPORT | 102 | -0.169646 | -0.619413 | 0.993548 | 1 |
| GOBP_INTRINSIC_APOPTOTIC_SIGNALING_PATHWAY_IN_RESPONSE_TO_OXIDATIVE_STRESS | 43 | -0.194367 | -0.618596 | 0.974063 | 1 |
| GOBP_VESICLE_LOCALIZATION | 220 | -0.153828 | -0.61791 | 0.998812 | 1 |
| GOBP_NEGATIVE_REGULATION_OF_CELL_CYCLE_ARREST | 19 | -0.229469 | -0.616539 | 0.925466 | 1 |
| GOBP_POSITIVE_REGULATION_OF_EMBRYONIC_DEVELOPMENT | 16 | -0.238067 | -0.616012 | 0.936508 | 1 |
| GOBP_CELL_AGING | 101 | -0.16726 | -0.615525 | 0.990979 | 1 |
| GOBP_PHOSPHATIDYLINOSITOL_PHOSPHORYLATION | 54 | -0.185015 | -0.614043 | 0.990057 | 1 |
| GOBP_MODULATION_OF_PROCESS_OF_OTHER_ORGANISM_INVOLVED_IN_SYMBIOTIC_INTERACT | 95 | -0.168781 | -0.614021 | 1 | 1 |
| GOBP_ATP_BIOSYNTHETIC_PROCESS | 55 | -0.182762 | -0.613053 | 0.984529 | 1 |
| GOBP_REGULATION_OF_MRNA_SPLICING_VIA_SPLICEOSOME | 93 | -0.166958 | -0.610429 | 0.992136 | 1 |
| GOBP_MYOBLAST_PROLIFERATION | 17 | -0.235081 | -0.606517 | 0.928918 | 1 |
| GOBP_REGULATION_OF_CELL_AGING | 49 | -0.185176 | -0.60609 | 0.977044 | 1 |
| GOBP_GOLGI_TO_PLASMA_MEMBRANE_PROTEIN_TRANSPORT | 38 | -0.196146 | -0.605879 | 0.971429 | 1 |
| GOBP_PROTEIN_LOCALIZATION_TO_CILIUM | 58 | -0.17739 | -0.602917 | 0.983146 | 1 |
| GOBP_MULTIVESICULAR_BODY_ORGANIZATION | 31 | -0.201649 | -0.602782 | 0.960294 | 1 |
| GOBP_NEGATIVE_REGULATION_OF_UBIQUITIN_DEPENDENT_PROTEIN_CATABOLIC_PROCESS | 47 | -0.184509 | -0.597805 | 0.990113 | 1 |
| GOBP_REGULATION_OF_TRANSLATION_IN_RESPONSE_TO_STRESS | 21 | -0.219673 | -0.596554 | 0.955658 | 1 |
| GOBP_REGULATION_OF_CALCINEURIN_MEDIATED_SIGNALING | 28 | -0.200816 | -0.592192 | 0.970724 | 1 |
| GOBP_REGULATION_OF_HISTONE_MODIFICATION | 133 | -0.15525 | -0.590156 | 0.998761 | 1 |
| GOBP_REGULATION_OF_CARDIAC_MUSCLE_CONTRACTION_BY_CALCIUM_ION_SIGNALING | 23 | -0.213141 | -0.586695 | 0.970543 | 1 |
| GOBP_PROTEIN_DEMANNOSYLATION | 20 | -0.215866 | -0.586396 | 0.94582 | 1 |
| GOBP_MUSCLE_HYPERTROPHY_IN_RESPONSE_TO_STRESS | 21 | -0.214961 | -0.586248 | 0.956587 | 1 |
| GOBP_RECEPTOR_RECYCLING | 32 | -0.194404 | -0.585459 | 0.969559 | 1 |
| GOBP_PROTEIN_EXIT_FROM_ENDOPLASMIC_RETICULUM | 46 | -0.1794 | -0.585385 | 0.989736 | 1 |
| GOBP_REGULATION_OF_ESTABLISHMENT_OF_PROTEIN_LOCALIZATION_TO_MITOCHONDRION | 71 | -0.166778 | -0.583053 | 0.992997 | 1 |
| GOBP_CELLULAR_RESPONSE_TO_STEROL_DEPLETION | 16 | -0.227736 | -0.583002 | 0.94795 | 1 |
| GOBP_AMINE_BIOSYNTHETIC_PROCESS | 31 | -0.196911 | -0.580891 | 0.980854 | 1 |
| GOBP_TRANSCRIPTION_BY_RNA_POLYMERASE_III | 47 | -0.180777 | -0.579461 | 0.99169 | 1 |
| GOBP_REGULATION_OF_MITOCHONDRIAL_OUTER_MEMBRANE_PERMEABILIZATION_INVOLVED_I | 44 | -0.182713 | -0.579187 | 0.986706 | 1 |
| GOBP_REGULATION_OF_PROTEIN_ACETYLATION | 68 | -0.166335 | -0.576965 | 0.997234 | 1 |
| GOBP_VIRION_ASSEMBLY | 40 | -0.181712 | -0.57689 | 0.991202 | 1 |
| GOBP_HISTONE_DEACETYLATION | 73 | -0.164705 | -0.573803 | 0.992042 | 1 |
| GOBP_NEGATIVE_REGULATION_OF_PROTEASOMAL_PROTEIN_CATABOLIC_PROCESS | 47 | -0.178707 | -0.572812 | 0.983051 | 1 |
| GOBP_CYTOPLASMIC_TRANSLATION | 101 | -0.155065 | -0.571543 | 0.998728 | 1 |
| GOBP_RESPONSE_TO_COCAINE | 38 | -0.18451 | -0.57151 | 0.982456 | 1 |
| GOBP_RETROGRADE_AXONAL_TRANSPORT | 20 | -0.211328 | -0.570149 | 0.964286 | 1 |
| GOBP_NEGATIVE_REGULATION_OF_PROTEOLYSIS_INVOLVED_IN_CELLULAR_PROTEIN_CATABO | 62 | -0.169595 | -0.570146 | 0.99863 | 1 |
| GOBP_NUCLEOTIDE_SUGAR_BIOSYNTHETIC_PROCESS | 23 | -0.201177 | -0.569199 | 0.96423 | 1 |
| GOBP_NEGATIVE_REGULATION_OF_MRNA_METABOLIC_PROCESS | 80 | -0.161243 | -0.568285 | 0.998626 | 1 |
| GOBP_REGULATION_OF_CELLULAR_PROTEIN_CATABOLIC_PROCESS | 241 | -0.140163 | -0.566946 | 1 | 1 |
| GOBP_NUCLEAR_TRANSCRIBED_MRNA_CATABOLIC_PROCESS_DEADENYLATION_DEPENDENT_ | 75 | -0.163938 | -0.565806 | 0.997355 | 1 |
| GOBP_NEGATIVE_REGULATION_OF_DNA_TEMPLATED_TRANSCRIPTION_ELONGATION | 17 | -0.217222 | -0.565069 | 0.966074 | 1 |
| GOBP_ESTABLISHMENT_OF_PROTEIN_LOCALIZATION_TO_MITOCHONDRIAL_MEMBRANE | 53 | -0.171529 | -0.564806 | 0.995798 | 1 |
| GOBP_ALTERNATIVE_MRNA_SPLICING_VIA_SPLICEOSOME | 67 | -0.164903 | -0.563938 | 0.997279 | 1 |
| GOBP_STRESS_GRANULE_ASSEMBLY | 23 | -0.203107 | -0.563892 | 0.976708 | 1 |
| GOBP_RESPONSE_TO_CAFFEINE | 16 | -0.219182 | -0.563542 | 0.970405 | 1 |
| GOBP_NEGATIVE_REGULATION_OF_MRNA_PROCESSING | 29 | -0.192981 | -0.562112 | 0.978134 | 1 |
| GOBP_NEGATIVE_REGULATION_OF_MRNA_CATABOLIC_PROCESS | 51 | -0.17294 | -0.561414 | 0.994195 | 1 |
| GOBP_PYRIMIDINE_NUCLEOSIDE_BIOSYNTHETIC_PROCESS | 16 | -0.218525 | -0.560064 | 0.964401 | 1 |
| GOBP_PORE_COMPLEX_ASSEMBLY | 20 | -0.207732 | -0.558776 | 0.974961 | 1 |
| GOBP_NON_MOTILE_CILIUM_ASSEMBLY | 53 | -0.167707 | -0.553165 | 0.993084 | 1 |
| GOBP_ENDOSOMAL_TRANSPORT | 237 | -0.138013 | -0.552797 | 1 | 1 |
| GOBP_VESICLE_DOCKING_INVOLVED_IN_EXOCYTOSIS | 43 | -0.174152 | -0.551301 | 0.994334 | 1 |
| GOBP_PROTEIN_ACETYLATION | 184 | -0.13987 | -0.551223 | 1 | 1 |
| GOBP_PEPTIDYL_LYSINE_TRIMETHYLATION | 43 | -0.17402 | -0.55122 | 0.989928 | 1 |
| GOBP_DEMETHYLATION | 68 | -0.160599 | -0.547228 | 1 | 1 |
| GOBP_ACTIVATION_OF_GTPASE_ACTIVITY | 104 | -0.147855 | -0.546281 | 0.99871 | 1 |
| GOBP_PEPTIDYL_LYSINE_ACETYLATION | 158 | -0.14002 | -0.543587 | 1 | 1 |
| GOBP_REGULATION_OF_MACROAUTOPHAGY | 158 | -0.137542 | -0.539434 | 1 | 1 |
| GOBP_POSITIVE_REGULATION_OF_HISTONE_H3_K4_METHYLATION | 16 | -0.209122 | -0.537777 | 0.973186 | 1 |
| GOBP_CELLULAR_CARBOHYDRATE_CATABOLIC_PROCESS | 43 | -0.165123 | -0.532707 | 0.995601 | 1 |
| GOBP_PROTEIN_DEGLYCOSYLATION | 28 | -0.183128 | -0.532379 | 0.995569 | 1 |
| GOBP_RNA_CATABOLIC_PROCESS | 401 | -0.124234 | -0.525922 | 1 | 1 |
| GOBP_MODULATION_BY_VIRUS_OF_HOST_CELLULAR_PROCESS | 17 | -0.198669 | -0.524451 | 0.987539 | 1 |
| GOBP_PEROXISOME_ORGANIZATION | 80 | -0.145851 | -0.522064 | 1 | 1 |
| GOBP_VACUOLAR_TRANSPORT | 148 | -0.13586 | -0.521974 | 1 | 1 |
| GOBP_NEGATIVE_REGULATION_OF_TRANSCRIPTION_REGULATORY_REGION_DNA_BINDING | 17 | -0.20139 | -0.518702 | 0.976084 | 1 |
| GOBP_RESPONSE_TO_AMINO_ACID_STARVATION | 49 | -0.160812 | -0.517945 | 0.998607 | 1 |
| GOBP_REGULATION_OF_ORGANELLE_ASSEMBLY | 177 | -0.131639 | -0.517715 | 1 | 1 |
| GOBP_HISTONE_H3_ACETYLATION | 59 | -0.153326 | -0.517157 | 0.995856 | 1 |
| GOBP_REGULATION_OF_TRANSLATIONAL_INITIATION | 75 | -0.147627 | -0.516214 | 0.998661 | 1 |
| GOBP_PROTEIN_POLYUBIQUITINATION | 324 | -0.123459 | -0.509516 | 1 | 1 |
| GOBP_ENDOPLASMIC_RETICULUM_MANNOSE_TRIMMING | 16 | -0.198302 | -0.50766 | 0.987539 | 1 |
| GOBP_UBIQUITIN_DEPENDENT_ERAD_PATHWAY | 77 | -0.144032 | -0.506554 | 1 | 1 |
| GOBP_SPHINGOMYELIN_METABOLIC_PROCESS | 15 | -0.199956 | -0.506207 | 0.986025 | 1 |
| GOBP_POSITIVE_REGULATION_OF_VIRAL_TRANSCRIPTION | 26 | -0.177307 | -0.505279 | 0.992582 | 1 |
| GOBP_TRANSCRIPTION_INITIATION_FROM_RNA_POLYMERASE_I_PROMOTER | 38 | -0.162219 | -0.502389 | 0.995745 | 1 |
| GOBP_NEGATIVE_REGULATION_OF_TELOMERE_MAINTENANCE_VIA_TELOMERASE | 19 | -0.189128 | -0.500049 | 0.990066 | 1 |
| GOBP_AMP_METABOLIC_PROCESS | 16 | -0.193329 | -0.494704 | 0.993779 | 1 |
| GOBP_PROTEIN_ADP_RIBOSYLATION | 33 | -0.160897 | -0.487404 | 0.988287 | 1 |
| GOBP_POSITIVE_REGULATION_OF_VIRAL_PROCESS | 90 | -0.134007 | -0.484453 | 1 | 1 |
| GOBP_NUCLEAR_MEMBRANE_ORGANIZATION | 17 | -0.180973 | -0.474531 | 0.995169 | 1 |
| GOBP_NEGATIVE_REGULATION_OF_UBIQUITIN_PROTEIN_TRANSFERASE_ACTIVITY | 16 | -0.188426 | -0.471662 | 0.99682 | 1 |
| GOBP_MODULATION_BY_SYMBIONT_OF_HOST_PROCESS | 36 | -0.155584 | -0.470202 | 0.994253 | 1 |
| GOBP_REGULATION_OF_GOLGI_ORGANIZATION | 17 | -0.180761 | -0.468091 | 0.992175 | 1 |
| GOBP_NUCLEOTIDE_SUGAR_METABOLIC_PROCESS | 36 | -0.154397 | -0.467025 | 1 | 1 |
| GOBP_HISTONE_UBIQUITINATION | 46 | -0.140787 | -0.453816 | 1 | 1 |
| GOBP_PROTEIN_MODIFICATION_BY_SMALL_PROTEIN_REMOVAL | 272 | -0.109653 | -0.446378 | 1 | 1 |
| GOBP_REGULATION_OF_ANDROGEN_RECEPTOR_SIGNALING_PATHWAY | 24 | -0.157243 | -0.444415 | 0.997063 | 1 |
| GOBP_ERAD_PATHWAY | 99 | -0.118954 | -0.437033 | 1 | 1 |
| GOBP_MACROAUTOPHAGY | 306 | -0.104483 | -0.43618 | 1 | 1 |
| GOBP_DNA_TEMPLATED_TRANSCRIPTION_ELONGATION | 113 | -0.116387 | -0.433171 | 1 | 1 |
| GOBP_ENDOPLASMIC_RETICULUM_TO_GOLGI_VESICLE_MEDIATED_TRANSPORT | 199 | -0.109184 | -0.43305 | 1 | 1 |
| GOBP_TRANSCRIPTION_ELONGATION_FROM_RNA_POLYMERASE_II_PROMOTER | 84 | -0.121017 | -0.432382 | 1 | 1 |
| GOBP_COLLATERAL_SPROUTING | 19 | -0.160598 | -0.431778 | 0.996942 | 1 |
| GOBP_REGULATION_OF_LYSOSOMAL_LUMEN_PH | 15 | -0.171404 | -0.429386 | 0.9968 | 1 |
| GOBP_REGULATION_OF_MRNA_PROCESSING | 132 | -0.11239 | -0.424236 | 1 | 1 |
| GOBP_HISTONE_H4_ACETYLATION | 67 | -0.122256 | -0.419443 | 1 | 1 |
| GOBP_ASYMMETRIC_CELL_DIVISION | 16 | -0.163721 | -0.414065 | 0.99688 | 1 |
| GOBP_PEROXISOMAL_TRANSPORT | 68 | -0.116789 | -0.403098 | 1 | 1 |
| GOBP_PROTEIN_K48_LINKED_DEUBIQUITINATION | 33 | -0.130658 | -0.392896 | 1 | 1 |
| GOBP_REGULATION_OF_MRNA_METABOLIC_PROCESS | 315 | -0.09166 | -0.379027 | 1 | 1 |
| GOBP_REGULATION_OF_PROTEIN_POLYUBIQUITINATION | 23 | -0.135281 | -0.372382 | 1 | 1 |
| GOBP_RNA_SPLICING | 446 | -0.069524 | -0.292349 | 1 | 0.999998 |
