## Supplementary material for "Chemokine Receptor 1 and its associated immune pathway are downregulated in SF3B1^MT^ blood and non-blood cancers": Table_S9

| Table S9. Enriched pathways in module eigengenes, related to Figure 4. |
|---|
| **M1_green** |
| green "SRP-dependent cotranslational protein targeting to membrane" |
| green "cotranslational protein targeting to membrane" |
| green "establishment of protein localization to endoplasmic reticulum" |
| green "protein targeting to ER" |
| green "cytosolic ribosome" |
| green "protein localization to endoplasmic reticulum" |
| green "nuclear-transcribed mRNA catabolic process |
| green "structural constituent of ribosome" |
| green "translational initiation" |
| green "protein targeting to membrane" |
| **M2_red** |
| red "mitochondrial inner membrane" |
| red "organelle inner membrane" |
| red "mitochondrial envelope" |
| red "mitochondrial ribosome" |
| red "mitochondrial translational elongation" |
| red "mitochondrial translational termination" |
| red "mitochondrial membrane" |
| red "translational termination" |
| red "mitochondrion" |
| red "translational elongation" |
| **M3_lightcyan** |
| lightcyan "positive regulation of immunoglobulin production" |
| lightcyan "apical plasma membrane" |
| lightcyan "regulation of B cell proliferation" |
| lightcyan "positive regulation of immunoglobulin biosynthetic process" |
| lightcyan "cellular lactam catabolic process" |
| lightcyan "regulation of immunoglobulin production" |
| lightcyan "apical part of cell" |
| lightcyan "B cell proliferation" |
| lightcyan "positive regulation of T cell mediated immune response to tumor cell" |
| lightcyan "cellular amide catabolic process" |
| **M4_black** |
| black "extracellular region" |
| black "extracellular matrix" |
| black "extracellular space" |
| black "collagen-containing extracellular matrix" |
| black "extracellular matrix structural constituent" |
| black "anatomical structure morphogenesis" |
| black "extracellular matrix organization" |
| black "extracellular structure organization" |
| black "multicellular organism development" |
| black "cell migration" |

| M5_blue |
| --- |
| blue "mitotic cell cycle" |
| blue "cell cycle" |
| blue "DNA replication" |
| blue "condensed chromosome" |
| blue "cell cycle phase transition" |
| blue "mitotic cell cycle phase transition" |
| blue "chromosome organization" |
| blue "regulation of cell cycle process" |
| blue "regulation of mitotic cell cycle phase transition" |
| blue "chromosome" |
| **M6_purple** |
| purple "MHC class II protein complex" |
| purple "cell periphery" |
| purple "plasma membrane" |
| purple "MHC class II protein complex binding" |
| purple "response to stimulus" |
| purple "positive regulation of T cell proliferation" |
| purple "positive regulation of lymphocyte proliferation" |
| purple "positive regulation of mononuclear cell proliferation" |
| purple "adaptive immune response" |
| purple "positive regulation of leukocyte proliferation" |
| **M7_salmon** |
| salmon "B cell proliferation" |
| salmon "lymphocyte activation" |
| salmon "regulation of lymphocyte activation" |
| salmon "regulation of immunoglobulin secretion" |
| salmon "B cell receptor signaling pathway" |
| salmon "immunoglobulin secretion" |
| salmon "T cell activation" |
| salmon "regulation of cell activation" |
| salmon "regulation of B cell receptor signaling pathway" |
| salmon "lymphocyte differentiation" |
| **M8_brown** |
| brown "external side of plasma membrane" |
| brown "plasma membrane" |
| brown "T cell receptor complex" |
| brown "antigen receptor-mediated signaling pathway" |
| brown "cell periphery" |
| brown "homophilic cell adhesion via plasma membrane adhesion molecules" |
| brown "integral component of plasma membrane" |
| brown "intrinsic component of plasma membrane" |
| brown "signaling receptor activity" |
| brown "cell killing" |
| **M9_turquoise** |

| |
|---|
| turquoise "regulation of RNA metabolic process" |
| turquoise "DNA-binding transcription factor activity" |
| turquoise "RNA biosynthetic process" |
| turquoise "DNA-binding transcription factor activity |
| turquoise "transcription |
| turquoise "nucleic acid binding" |
| turquoise "RNA metabolic process" |
| turquoise "DNA binding" |
| turquoise "regulation of nucleic acid-templated transcription" |
| turquoise "regulation of RNA biosynthetic process" |
| **M10_midnightblue** |
| midnightblue "cell activation" |
| midnightblue "leukocyte activation" |
| midnightblue "leukocyte activation involved in immune response" |
| midnightblue "cell activation involved in immune response" |
| midnightblue "neutrophil degranulation" |
| midnightblue "neutrophil activation involved in immune response" |
| midnightblue "neutrophil mediated immunity" |
| midnightblue "neutrophil activation" |
| midnightblue "leukocyte degranulation" |
| midnightblue "myeloid leukocyte activation" |
| **M11_tan** |
| tan "neutrophil degranulation" |
| tan "neutrophil activation involved in immune response" |
| tan "secretory granule membrane" |
| tan "neutrophil mediated immunity" |
| tan "neutrophil activation" |
| tan "specific granule membrane" |
| tan "vesicle-mediated transport" |
| tan "cellular lipid metabolic process" |
| tan "leukocyte degranulation" |
| tan "exocytosis" |
| **M12_greenyellow** |
| greenyellow "leukocyte activation involved in immune response" |
| greenyellow "cell activation involved in immune response" |
| greenyellow "leukocyte mediated immunity" |
| greenyellow "leukocyte degranulation" |
| greenyellow "neutrophil activation" |
| greenyellow "myeloid leukocyte activation" |
| greenyellow "cell activation" |
| greenyellow "regulated exocytosis" |
| greenyellow "neutrophil mediated immunity" |
| greenyellow "neutrophil activation involved in immune response" |
| **M13_magenta** |
| magenta "neutrophil degranulation" |

| |
|---|
| magenta "neutrophil activation involved in immune response" |
| magenta "neutrophil mediated immunity" |
| magenta "neutrophil activation" |
| magenta "leukocyte degranulation" |
| magenta "specific granule" |
| magenta "secretory vesicle" |
| magenta "secretory granule" |
| magenta "regulated exocytosis" |
| magenta "myeloid leukocyte activation" |
| **M14_cyan** |
| cyan "glutamate dehydrogenase (NAD+) activity" |
| cyan "glutamate dehydrogenase [NAD(P)+] activity" |
| cyan "actin cortical patch assembly" |
| cyan "glutamate biosynthetic process" |
| cyan "ribonuclease MRP complex" |
| cyan "glutamate catabolic process" |
| cyan "actin cortical patch localization" |
| cyan "leucine binding" |
| cyan "negative regulation of lipopolysaccharide-mediated signaling pathway" |
| cyan "multimeric ribonuclease P complex" |
| **M15_pink** |
| pink "nucleoplasm" |
| pink "nuclear lumen" |
| pink "nucleus" |
| pink "transcription coactivator activity" |
| pink "intracellular membrane-bounded organelle" |
| pink "intracellular organelle" |
| pink "intracellular" |
| pink "nitrogen compound metabolic process" |
| pink "nucleocytoplasmic transport" |
| pink "positive regulation of nitrogen compound metabolic process" |
| **M16_yellow** |
| yellow "intracellular organelle" |
| yellow "intracellular membrane-bounded organelle" |
| yellow "membrane-bounded organelle" |
| yellow "vesicle organization" |
| yellow "intracellular" |
| yellow "Golgi membrane" |
| yellow "Golgi vesicle transport" |
| yellow "GTPase activity" |
| yellow "endoplasmic reticulum to Golgi vesicle-mediated transport" |
| yellow "endomembrane system organization" |
| **Gray module contains unrelated members** |
| grey "receptor antagonist activity" |
| grey "startle response" |

| |
|---|
| grey "motile cilium" |
| grey "drinking behavior" |
| grey "regulation of polarized epithelial cell differentiation" |
| grey "receptor inhibitor activity" |
| grey "L-aspartate transmembrane transport" |
| grey "thyroid hormone metabolic process" |
| grey "homophilic cell adhesion via plasma membrane adhesion molecules" |
| grey "non-motile cilium" |
