## Supplementary material for "Chemokine Receptor 1 and its associated immune pathway are downregulated in SF3B1^MT^ blood and non-blood cancers": Table_S10

| **Table S10. Gene interaction in M6_purple module**. Weight columns refers to the connection strength between two nodes (genes). 100 interactions with the highest weights are shown. This package of WGCNA is limited to undirected networks. | | | | | |
|---|---|---|---|---|---|
| **From gene name** | **To gene name** | **Weight** | **Direction** | **From node** | **To node** |
| AMIGO2 | DYRK2 | 0.217490795 | undirected | ENSG00000139211 | ENSG00000127334 |
| AKNA | NLRP1 | 0.215596426 | undirected | ENSG00000106948 | ENSG00000091592 |
| SERPINB9 | DYRK2 | 0.210338906 | undirected | ENSG00000170542 | ENSG00000127334 |
| FUT11 | BRIP1 | 0.208507453 | undirected | ENSG00000196968 | ENSG00000136492 |
| DYRK2 | MORN3 | 0.207973203 | undirected | ENSG00000127334 | ENSG00000139714 |
| STAB1 | TRIO | 0.206879284 | undirected | ENSG00000010327 | ENSG00000038382 |
| SPTBN5 | NLRP1 | 0.206166303 | undirected | ENSG00000137877 | ENSG00000091592 |
| HIVEP3 | HAVCR2 | 0.206131615 | undirected | ENSG00000127124 | ENSG00000135077 |
| SCML2 | NLRP1 | 0.206093669 | undirected | ENSG00000102098 | ENSG00000091592 |
| TRIO | SECTM1 | 0.205845628 | undirected | ENSG00000038382 | ENSG00000141574 |
| PLD2 | NLRP1 | 0.205648954 | undirected | ENSG00000129219 | ENSG00000091592 |
| HIVEP3 | DOK2 | 0.205387444 | undirected | ENSG00000127124 | ENSG00000147443 |
| NLRP1 | CYTH4 | 0.204687726 | undirected | ENSG00000091592 | ENSG00000100055 |
| DYRK2 | RALGAPA1 | 0.204658346 | undirected | ENSG00000127334 | ENSG00000174373 |
| GSTCD | NLRP1 | 0.204312962 | undirected | ENSG00000138780 | ENSG00000091592 |
| HIVEP3 | MAP3K14 | 0.204244324 | undirected | ENSG00000127124 | ENSG00000006062 |
| HIVEP3 | CD300E | 0.204223331 | undirected | ENSG00000127124 | ENSG00000186407 |
| NBPF9 | TRIO | 0.203931787 | undirected | ENSG00000269713 | ENSG00000038382 |
| TRIO | ANO8 | 0.203847484 | undirected | ENSG00000038382 | ENSG00000074855 |
| DOK2 | NLRP1 | 0.203818676 | undirected | ENSG00000147443 | ENSG00000091592 |
| TRIO | PCSK5 | 0.20354535 | undirected | ENSG00000038382 | ENSG00000099139 |
| TRIO | SLC2A6 | 0.202991768 | undirected | ENSG00000038382 | ENSG00000160326 |
| HIVEP3 | TRIO | 0.202939852 | undirected | ENSG00000127124 | ENSG00000038382 |
| HIVEP3 | NBPF9 | 0.202830081 | undirected | ENSG00000127124 | ENSG00000269713 |
| HIVEP3 | SERPINB9 | 0.202367948 | undirected | ENSG00000127124 | ENSG00000170542 |
| DOK2 | DYRK2 | 0.202011702 | undirected | ENSG00000147443 | ENSG00000127334 |
| HIVEP3 | DYRK2 | 0.201922822 | undirected | ENSG00000127124 | ENSG00000127334 |
| DYRK2 | APOL6 | 0.201909879 | undirected | ENSG00000127334 | ENSG00000221963 |
| TRIO | ADGRB1 | 0.201870232 | undirected | ENSG00000038382 | ENSG00000181790 |
| AAK1 | NLRP1 | 0.201693693 | undirected | ENSG00000115977 | ENSG00000091592 |
| HIVEP3 | AMIGO2 | 0.20162174 | undirected | ENSG00000127124 | ENSG00000139211 |
| ADGRB1 | NLRP1 | 0.201475933 | undirected | ENSG00000181790 | ENSG00000091592 |
| APBB1 | DYRK2 | 0.200818749 | undirected | ENSG00000166313 | ENSG00000127334 |
| FUT11 | TIPIN | 0.200770984 | undirected | ENSG00000196968 | ENSG00000075131 |
| DYRK2 | BRIP1 | 0.200595645 | undirected | ENSG00000127334 | ENSG00000136492 |
| RASA3 | NLRP1 | 0.20043176 | undirected | ENSG00000185989 | ENSG00000091592 |
| TNFSF8 | NLRP1 | 0.200201974 | undirected | ENSG00000106952 | ENSG00000091592 |
| MED8 | NLRP1 | 0.200194066 | undirected | ENSG00000159479 | ENSG00000091592 |
| TRIO | IQSEC2 | 0.200002234 | undirected | ENSG00000038382 | ENSG00000124313 |
| HIVEP3 | MED8 | 0.199831935 | undirected | ENSG00000127124 | ENSG00000159479 |
| HIVEP3 | NLRP1 | 0.199228561 | undirected | ENSG00000127124 | ENSG00000091592 |
| NLRP1 | JAK3 | 0.199122795 | undirected | ENSG00000091592 | ENSG00000105639 |
| TRIO | GIT1 | 0.199055678 | undirected | ENSG00000038382 | ENSG00000108262 |
| TRIO | RAB39A | 0.19896966 | undirected | ENSG00000038382 | ENSG00000179331 |
| HIVEP3 | PLD2 | 0.19877232 | undirected | ENSG00000127124 | ENSG00000129219 |
| FUT11 | JAK3 | 0.198732974 | undirected | ENSG00000196968 | ENSG00000105639 |
| FAM131B | NLRP1 | 0.198657264 | undirected | ENSG00000159784 | ENSG00000091592 |
| SERPINB9 | FUT11 | 0.198638268 | undirected | ENSG00000170542 | ENSG00000196968 |

| | | | | | |
|---|---|---|---|---|---|
| TRIO | PANX1 | 0.198428721 | undirected | ENSG00000038382 | ENSG00000110218 |
| NLRP1 | SECTM1 | 0.198423382 | undirected | ENSG00000091592 | ENSG00000141574 |
| SCG5 | NLRP1 | 0.198407058 | undirected | ENSG00000166922 | ENSG00000091592 |
| ARHGEF10L | TRIO | 0.198307238 | undirected | ENSG00000074964 | ENSG00000038382 |
| ADGRB1 | FUT11 | 0.198094721 | undirected | ENSG00000181790 | ENSG00000196968 |
| TRIO | TMEM268 | 0.198001307 | undirected | ENSG00000038382 | ENSG00000157693 |
| FUT11 | DYRK2 | 0.197646727 | undirected | ENSG00000196968 | ENSG00000127334 |
| HIVEP3 | TMEM268 | 0.197475978 | undirected | ENSG00000127124 | ENSG00000157693 |
| HIVEP3 | TIPIN | 0.197447582 | undirected | ENSG00000127124 | ENSG00000075131 |
| DYRK2 | TIPIN | 0.197335134 | undirected | ENSG00000127334 | ENSG00000075131 |
| SETD7 | TRIO | 0.197045773 | undirected | ENSG00000145391 | ENSG00000038382 |
| FIGNL1 | NLRP1 | 0.197023097 | undirected | ENSG00000132436 | ENSG00000091592 |
| TOGARAM2 | NLRP1 | 0.196949531 | undirected | ENSG00000189350 | ENSG00000091592 |
| TRIO | DOK2 | 0.196907196 | undirected | ENSG00000038382 | ENSG00000147443 |
| TIPIN | NLRP1 | 0.196838454 | undirected | ENSG00000075131 | ENSG00000091592 |
| HIVEP3 | SETD7 | 0.196797612 | undirected | ENSG00000127124 | ENSG00000145391 |
| HIVEP3 | PLXNB2 | 0.196557 | undirected | ENSG00000127124 | ENSG00000196576 |
| DYRK2 | RASA3 | 0.196499362 | undirected | ENSG00000127334 | ENSG00000185989 |
| PLS1 | FUT11 | 0.196370375 | undirected | ENSG00000120756 | ENSG00000196968 |
| USP48 | DYRK2 | 0.196335666 | undirected | ENSG00000090686 | ENSG00000127334 |
| CCDC88C | NLRP1 | 0.195988595 | undirected | ENSG00000015133 | ENSG00000091592 |
| TRIO | ADAP2 | 0.195825122 | undirected | ENSG00000038382 | ENSG00000184060 |
| FUT11 | AMIGO2 | 0.195753408 | undirected | ENSG00000196968 | ENSG00000139211 |
| TRIO | C20orf194 | 0.195741898 | undirected | ENSG00000038382 | ENSG00000088854 |
| HIVEP3 | SECTM1 | 0.195624025 | undirected | ENSG00000127124 | ENSG00000141574 |
| SERPINB9 | NLRP1 | 0.195615796 | undirected | ENSG00000170542 | ENSG00000091592 |
| DYRK2 | RNF125 | 0.195559836 | undirected | ENSG00000127334 | ENSG00000101695 |
| SLC2A6 | NLRP1 | 0.195541531 | undirected | ENSG00000160326 | ENSG00000091592 |
| HIVEP3 | APOL6 | 0.195518047 | undirected | ENSG00000127124 | ENSG00000221963 |
| DNER | TRIO | 0.195512887 | undirected | ENSG00000187957 | ENSG00000038382 |
| TRIO | PLXNB2 | 0.195438296 | undirected | ENSG00000038382 | ENSG00000196576 |
| TRIO | LINC01503 | 0.195212135 | undirected | ENSG00000038382 | ENSG00000233901 |
| TRIO | PYGB | 0.195202104 | undirected | ENSG00000038382 | ENSG00000100994 |
| SCML2 | FUT11 | 0.195134062 | undirected | ENSG00000102098 | ENSG00000196968 |
| TRIO | NLRP1 | 0.194977576 | undirected | ENSG00000038382 | ENSG00000091592 |
| TRIO | CD4 | 0.194929133 | undirected | ENSG00000038382 | ENSG00000010610 |
| HIVEP3 | RTL5 | 0.194521333 | undirected | ENSG00000127124 | ENSG00000242732 |
| TRIO | TIPIN | 0.194435311 | undirected | ENSG00000038382 | ENSG00000075131 |
| HIVEP3 | FUT11 | 0.194275696 | undirected | ENSG00000127124 | ENSG00000196968 |
| TRIO | BRIP1 | 0.194184044 | undirected | ENSG00000038382 | ENSG00000136492 |
| HAVCR2 | DYRK2 | 0.194123954 | undirected | ENSG00000135077 | ENSG00000127334 |
| HIVEP3 | CCR1 | 0.193919873 | undirected | ENSG00000127124 | ENSG00000163823 |
| CCDC150 | DYRK2 | 0.19378695 | undirected | ENSG00000144395 | ENSG00000127334 |
| XPNPEP2 | DYRK2 | 0.193773499 | undirected | ENSG00000122121 | ENSG00000127334 |
| TRIO | PLD2 | 0.193730278 | undirected | ENSG00000038382 | ENSG00000129219 |
| FUT11 | APOL6 | 0.193718448 | undirected | ENSG00000196968 | ENSG00000221963 |
| HIVEP3 | RASA3 | 0.19365397 | undirected | ENSG00000127124 | ENSG00000185989 |
| TRIO | FAM131B | 0.193645536 | undirected | ENSG00000038382 | ENSG00000159784 |
| FUT11 | MORN3 | 0.193613674 | undirected | ENSG00000196968 | ENSG00000139714 |
| TRIO | LGALS2 | 0.193570143 | undirected | ENSG00000038382 | ENSG00000100079 |
| HIVEP3 | APBB1 | 0.193569165 | undirected | ENSG00000127124 | ENSG00000166313 |
| HIVEP3 | PLEKHO1 | 0.193543908 | undirected | ENSG00000127124 | ENSG00000023902 |

| **Table S10. Gene interaction in M7_salmon module.** Weight columns refer to the connection strength between two nodes (genes). 100 interactions with the highest weights are shown. This package of WGCNA is limited to undirected networks. | | | | | |
|---|---|---|---|---|---|
| **From gene name** | **To gene name** | **Weight** | **Direction** | **From node** | **To node** |
| CYSTM1 | MCU | 0.19893189 | undirected | ENSG00000120306 | ENSG00000156026 |
| CYSTM1 | MTSS1 | 0.193627274 | undirected | ENSG00000120306 | ENSG00000170873 |
| CYSTM1 | CD72 | 0.193054385 | undirected | ENSG00000120306 | ENSG00000137101 |
| LNPK | CYSTM1 | 0.187213885 | undirected | ENSG00000144320 | ENSG00000120306 |
| CYSTM1 | SUN1 | 0.186817012 | undirected | ENSG00000120306 | ENSG00000164828 |
| CYSTM1 | BACH2 | 0.186019573 | undirected | ENSG00000120306 | ENSG00000112182 |
| CYSTM1 | CD40LG | 0.184911797 | undirected | ENSG00000120306 | ENSG00000102245 |
| ABLIM2 | CYSTM1 | 0.183837736 | undirected | ENSG00000163995 | ENSG00000120306 |
| MTSS1 | MCU | 0.182374554 | undirected | ENSG00000170873 | ENSG00000156026 |
| CYSTM1 | LAG3 | 0.18228815 | undirected | ENSG00000120306 | ENSG00000089692 |
| SUN1 | MCU | 0.182087154 | undirected | ENSG00000164828 | ENSG00000156026 |
| CELSR2 | CYSTM1 | 0.181776949 | undirected | ENSG00000143126 | ENSG00000120306 |
| MCU | CRTC3 | 0.180643569 | undirected | ENSG00000156026 | ENSG00000140577 |
| CYSTM1 | KIR2DL1 | 0.179599648 | undirected | ENSG00000120306 | ENSG00000125498 |
| CYSTM1 | LINC00278 | 0.179250341 | undirected | ENSG00000120306 | ENSG00000231535 |
| CYSTM1 | CRTC3 | 0.178504418 | undirected | ENSG00000120306 | ENSG00000140577 |
| XRCC5 | CYSTM1 | 0.178062689 | undirected | ENSG00000079246 | ENSG00000120306 |
| CYSTM1 | KLC2 | 0.177423331 | undirected | ENSG00000120306 | ENSG00000174996 |
| CYSTM1 | PLXNA3 | 0.177367843 | undirected | ENSG00000120306 | ENSG00000130827 |
| CD72 | MCU | 0.177058282 | undirected | ENSG00000137101 | ENSG00000156026 |
| CYSTM1 | ZNF503-AS1 | 0.176323832 | undirected | ENSG00000120306 | ENSG00000226051 |
| CDCA7L | MCU | 0.175890213 | undirected | ENSG00000164649 | ENSG00000156026 |
| CYSTM1 | DDX3Y | 0.175834704 | undirected | ENSG00000120306 | ENSG00000067048 |
| CD40LG | LPAR5 | 0.175069665 | undirected | ENSG00000102245 | ENSG00000184574 |
| CYSTM1 | RNF217 | 0.175062326 | undirected | ENSG00000120306 | ENSG00000146373 |
| MCU | CD40 | 0.174393246 | undirected | ENSG00000156026 | ENSG00000101017 |
| NFU1 | CYSTM1 | 0.174311296 | undirected | ENSG00000169599 | ENSG00000120306 |
| BACH2 | MCU | 0.174194285 | undirected | ENSG00000112182 | ENSG00000156026 |
| HSPA4L | MCU | 0.173703161 | undirected | ENSG00000164070 | ENSG00000156026 |
| CYSTM1 | PIMREG | 0.173452829 | undirected | ENSG00000120306 | ENSG00000129195 |
| CYSTM1 | ELFN2 | 0.173211784 | undirected | ENSG00000120306 | ENSG00000166897 |
| ABLIM2 | MCU | 0.172658321 | undirected | ENSG00000163995 | ENSG00000156026 |
| MCU | ATP23 | 0.172027834 | undirected | ENSG00000156026 | ENSG00000166896 |
| CYSTM1 | ATP23 | 0.171348503 | undirected | ENSG00000120306 | ENSG00000166896 |
| MCU | SUOX | 0.171070179 | undirected | ENSG00000156026 | ENSG00000139531 |
| LY9 | MCU | 0.170869167 | undirected | ENSG00000122224 | ENSG00000156026 |
| CELSR2 | MCU | 0.170606239 | undirected | ENSG00000143126 | ENSG00000156026 |
| CYSTM1 | KIR3DL2 | 0.170569819 | undirected | ENSG00000120306 | ENSG00000240403 |
| LY9 | CYSTM1 | 0.170542492 | undirected | ENSG00000122224 | ENSG00000120306 |
| CYSTM1 | CD40 | 0.170258585 | undirected | ENSG00000120306 | ENSG00000101017 |
| CYSTM1 | RGS9 | 0.17011638 | undirected | ENSG00000120306 | ENSG00000108370 |
| CYSTM1 | ZBTB4 | 0.170103817 | undirected | ENSG00000120306 | ENSG00000174282 |
| NFU1 | LAG3 | 0.169865844 | undirected | ENSG00000169599 | ENSG00000089692 |
| PIK3C2B | MCU | 0.16968119 | undirected | ENSG00000133056 | ENSG00000156026 |
| PPP3CC | MCU | 0.169263431 | undirected | ENSG00000120910 | ENSG00000156026 |
| CYSTM1 | UTY | 0.169070711 | undirected | ENSG00000120306 | ENSG00000183878 |
| CYSTM1 | ZFY | 0.168920307 | undirected | ENSG00000120306 | ENSG00000067646 |
| CYSTM1 | LPAR5 | 0.168639687 | undirected | ENSG00000120306 | ENSG00000184574 |

| | | | | | |
|---|---|---|---|---|---|
| CYSTM1 | EBF4 | 0.168401513 | undirected | ENSG00000120306 | ENSG00000088881 |
| MCU | RGS9 | 0.168155878 | undirected | ENSG00000156026 | ENSG00000108370 |
| MCU | LAG3 | 0.167786372 | undirected | ENSG00000156026 | ENSG00000089692 |
| SUCLG1 | CYSTM1 | 0.167750288 | undirected | ENSG00000163541 | ENSG00000120306 |
| CYSTM1 | KIFC3 | 0.167475937 | undirected | ENSG00000120306 | ENSG00000140859 |
| PIK3C2B | CYSTM1 | 0.167236485 | undirected | ENSG00000133056 | ENSG00000120306 |
| LNPK | MCU | 0.167199642 | undirected | ENSG00000144320 | ENSG00000156026 |
| CYSTM1 | SLC29A2 | 0.167125597 | undirected | ENSG00000120306 | ENSG00000174669 |
| CYSTM1 | XIST | 0.167071797 | undirected | ENSG00000120306 | ENSG00000229807 |
| HSPA4L | CYSTM1 | 0.166916537 | undirected | ENSG00000164070 | ENSG00000120306 |
| TRAF1 | GATA3 | 0.166738553 | undirected | ENSG00000056558 | ENSG00000107485 |
| MCU | KIR2DL1 | 0.16627588 | undirected | ENSG00000156026 | ENSG00000125498 |
| MCU | CD22 | 0.166150319 | undirected | ENSG00000156026 | ENSG00000012124 |
| PLEKHA6 | CYSTM1 | 0.166051085 | undirected | ENSG00000143850 | ENSG00000120306 |
| CD40LG | DIP2C | 0.165743786 | undirected | ENSG00000102245 | ENSG00000151240 |
| MCU | KIR3DL2 | 0.165720214 | undirected | ENSG00000156026 | ENSG00000240403 |
| DIP2C | LPAR5 | 0.165716767 | undirected | ENSG00000151240 | ENSG00000184574 |
| CYSTM1 | SUOX | 0.165147703 | undirected | ENSG00000120306 | ENSG00000139531 |
| PLXNA3 | MCU | 0.164913317 | undirected | ENSG00000130827 | ENSG00000156026 |
| CYSTM1 | CDCA7L | 0.164587249 | undirected | ENSG00000120306 | ENSG00000164649 |
| CYSTM1 | RPS4Y1 | 0.164577802 | undirected | ENSG00000120306 | ENSG00000129824 |
| BACH2 | LPAR5 | 0.164257627 | undirected | ENSG00000112182 | ENSG00000184574 |
| NFU1 | XRCC5 | 0.163977449 | undirected | ENSG00000169599 | ENSG00000079246 |
| CYSTM1 | KDM5D | 0.163673659 | undirected | ENSG00000120306 | ENSG00000012817 |
| MCU | ELFN2 | 0.163613237 | undirected | ENSG00000156026 | ENSG00000166897 |
| MCU | PIMREG | 0.163320118 | undirected | ENSG00000156026 | ENSG00000129195 |
| CYSTM1 | MAPK11 | 0.163180853 | undirected | ENSG00000120306 | ENSG00000185386 |
| KLC2 | MCU | 0.163148678 | undirected | ENSG00000174996 | ENSG00000156026 |
| KLC2 | LPAR5 | 0.162866309 | undirected | ENSG00000174996 | ENSG00000184574 |
| CYSTM1 | RORA | 0.162763581 | undirected | ENSG00000120306 | ENSG00000069667 |
| CYSTM1 | SFMBT2 | 0.162674963 | undirected | ENSG00000120306 | ENSG00000198879 |
| PLXNA3 | LARGE2 | 0.16249801 | undirected | ENSG00000130827 | ENSG00000165905 |
| TRAF1 | PGGHG | 0.162463413 | undirected | ENSG00000056558 | ENSG00000142102 |
| TRAF1 | SPOCK2 | 0.162081835 | undirected | ENSG00000056558 | ENSG00000107742 |
| NFU1 | PLXNA3 | 0.162050119 | undirected | ENSG00000169599 | ENSG00000130827 |
| MCU | RORA | 0.161923221 | undirected | ENSG00000156026 | ENSG00000069667 |
| SLC29A2 | DIP2C | 0.161830302 | undirected | ENSG00000174669 | ENSG00000151240 |
| CYSTM1 | CD22 | 0.161806439 | undirected | ENSG00000120306 | ENSG00000012124 |
| TRAF1 | CD7 | 0.161615868 | undirected | ENSG00000056558 | ENSG00000173762 |
| CYSTM1 | TTTY14 | 0.16126084 | undirected | ENSG00000120306 | ENSG00000176728 |
| MCU | EBF4 | 0.161100503 | undirected | ENSG00000156026 | ENSG00000088881 |
| RNF217 | MCU | 0.160925297 | undirected | ENSG00000146373 | ENSG00000156026 |
| LPAR5 | KIR2DL1 | 0.160684162 | undirected | ENSG00000184574 | ENSG00000125498 |
| DUSP7 | CYSTM1 | 0.160375661 | undirected | ENSG00000164086 | ENSG00000120306 |
| MCU | KIFC3 | 0.160221528 | undirected | ENSG00000156026 | ENSG00000140859 |
| ARID5B | MCU | 0.160164886 | undirected | ENSG00000150347 | ENSG00000156026 |
| CD72 | LPAR5 | 0.159835309 | undirected | ENSG00000137101 | ENSG00000184574 |
| CYSTM1 | TMSB4Y | 0.159464756 | undirected | ENSG00000120306 | ENSG00000154620 |
| LY9 | LPAR5 | 0.15922032 | undirected | ENSG00000122224 | ENSG00000184574 |
| CD40LG | SLC29A2 | 0.159151191 | undirected | ENSG00000102245 | ENSG00000174669 |
| MCU | TMEM241 | 0.158900262 | undirected | ENSG00000156026 | ENSG00000134490 |
| CYSTM1 | PPP3CC | 0.158847132 | undirected | ENSG00000120306 | ENSG00000120910 |
